# MAFB drives differentiation by permitting WT1 binding to podocyte specific promoters

**DOI:** 10.1101/2023.09.06.555670

**Authors:** Filippo M. Massa, Fariba Jian-Motamedi, Marijus Šerys, Amelie Tison, Agnès Loubat, Sandra Lacas-Gervais, Luc Martin, Hassiba Belahbib, Sandrine Sarrazin, Michael H. Sieweke, Andreas Schedl

**Affiliations:** Université Côte d’Azur, Inserm, CNRS, iBV, Nice, 06108, France; Université Côte d’Azur, CCMA, Nice, F-06108, France; Center for Regenerative Therapies Dresden (CRTD), Technische Universität Dresden, Germany; Aix Marseille University, CNRS, INSERM, CIML, Marseille, France

## Abstract

Podocytes are highly specialized cells, but their chromatin status and the precise molecular events leading to their differentiation remain poorly defined. Here we used ChIP-Seq analysis for H3K4me3, H3K4me1 and H3K27me3 to establish the histone methylation map in adult mouse podocytes. Our data demonstrate open chromatin across podocyte specific genes and reveals that genes expressed in the mesoderm lineage become actively repressed upon podocyte differentiation. To better understand the transcriptional control of podocyte differentiation, we studied the role of transcription factor MAFB. ChIP-Seq experiments and functional analysis in conditional knockout mice identified a set of direct MAFB targets including *Nphs1*, *Nphs2, Vegfa* and *Tcf21*. Loss of *MafB* led to the deposition of extracellular matrix, progressive foot process effacement, and kidney disease. ChIP experiments in knockout animals revealed that during development MAFB is essential for H3K4me3 methylation and the recruitment of WT1 to the promoters of the podocyte specific genes *Nphs1* and *Nphs2*. Taken together our data reveal the crucial function of MAFB by permitting chromatin accessibility at podocyte-specific genes during development and maintaining terminal differentiation in adults.

## Introduction

Cellular differentiation relies on the combined action of transcriptional regulators that activate specific genes to give a cell type its characteristic traits. Epigenetic modifications, such as histone methylations modulate chromatin structure and thus strongly impact the ability of transcription factors to bind to DNA and activate downstream target genes. Depending on the type of protein modifications, histone marks can be activating (H3K4me3), enhancing (H3K4me1, H3K27ac) or suppressing (H3K27me3) ^1^. Epigenetic marks are not static, but are shaped by developmental processes and cellular differentiation. Deposition of the repressive mark H3K27me3 involves the polycomb repressive complex 2 (PRC2) that has been shown to be essential for early embryonic development^2^. Removal of suppressive histone marks is achieved through specific enzymes that work together with so called pioneering factors that are able to access closed chromatin.

Podocytes are highly specialised cells that are crucial components of the glomerular filter. Differentiation of progenitors involves a number of transcription factors including members of the FOXC gene family ^3–7^, LMX1B ^4,8–10^, TCF21 ^11^, the Wilms’ tumor homologue WT1 ^12–18^ and MAFB ^19,20^. WT1 in particular has been shown to be essential for podocyte differentiation and as many as 50% of podocyte specific genes appear to be direct WT1 targets. Indeed, deletion of WT1 in adult mice leads to a rapid loss of podocyte specific gene expression and, as a consequence renal failure ^14,22^. While WT1 is essential for the expression of podocyte specific genes such as nephrin (*Nphs1*) ^23^, it is not sufficient for their activation. How WT1 can access enhancers and promoters and activate podocyte specific genes is presently poorly understood.

MAFB belongs to the family of large MAF proteins that are characterised by a basic domain involved in DNA binding, a leucine zipper structure that permits hetero and homo-dimerization and an acidic domain involved in transactivation of gene expression ^24,25^. MAFB has originally been identified as the gene underlying the Kreisler mutation in mice ^26^, a spontaneous mutant that results in defects of segmentation, brain patterning, pancreas development and macrophage differentiation (for review see ^27^). Further analyses have revealed an important function for podocyte development and function ^20,28,19^, and heterozygous point mutations in *MAFB* have been identified in patients suffering from FSGS and Duane retraction syndrome ^21^, as well as in cases of multicentric carpotarsal osteolysis^29^, a syndrome that is associated with a high risk of developing chronic kidney disease. How MAFB contributes to podocyte differentiation and maintenance, as well as its direct targets in this cell type remain poorly defined.

Here we set out to perform an unbiased study of the epigenetic state of the fully mature podocytes and identify the genome wide targets of MAFB. By employing constitutive and inducible podocyte specific knockout animals, we show that MAFB is essential for the activation of the podocyte specific gene program, by opening chromatin and permitting WT1 to bind to promoters of podocyte specific genes such as *Nphs1* and *Nphs2*.

## Materials and Methods

### Mice

Mice were housed in a conventional animal facility with free access to food and water. Deletion of *MafB* in fully differentiated podocytes was performed using the inducible *Wt1:Cre^ERT^*^2^ line ^30^ in combination with the conditional *Mafb^LoxP/LoxP^* ^31^ allele and the GFP reporter *Rosa26R^mTmG^* (Ref. ^30^). *Mafb* deletion was induced in adult male mice (8-10 weeks) with two consecutive days intraperitoneal injection of Tamoxifen (SIGMA) at 40mg/kg dissolved in corn oil. The developmental analysis was performed on E18.5 embryonic kidneys obtained by mating animals carrying the *Mafb*^GFP/+^ constitutive knockin allele ^32^. All experiments were performed according to national guidelines and approved by the French Minister of Higher Education (APAFIS#22471-2019011513265550v8).

### Chromatin Immune Precipitation

Adult podocytes were isolated as previously described^33^ with minor modifications. Briefly, glomeruli were isolated by paramagnetic beads perfusion and podocytes released by further digestion. Single cell suspension was then crosslinked with methanol-free Formaldehyde 1% (Thermofisher) for 10 minutes at RT before quenching, permeabilization and staining with anti-Podocin (SIGMA). Isolated podocytes were aliquoted in sonication buffer (50 mM HEPES pH 7.4, 1 mM EDTA, 150 mM NaCl, 0.4 % SDS, 1% Triton ×100, 1x protease inhibitor) and immediately sonicated using the sonicator Covaris S220 (peak incidence 150W, duty factor 10%, duration 20 minutes). For the epigenetic analysis during kidney development, whole kidneys were isolated from E18.5 embryos, crosslinked, mechanically dissociated in sonication buffer and sonicated with the previous parameters. Chromatin immuno-precipitation was performed using 200000 cells per condition (adult podocytes) or 3µg of chromatin per condition (immature podocytes) in combination with the following kit, reactives and antibodies from Diagenode, if not otherwise specified: True MicroChip-seq kit (ref. C01010132), anti-H3K4me3 (ref. C15410030), anti-H3K4me1 (ref. C15410037), anti-H3K27me3 (ref. C15410195), rabbit IgG (C15410206), anti WT1 (SantaCruz, ref. sc192), anti MAFB, (SIGMA, ref. HPA005653). Libraries were synthetized from the obtained chromatin using the MicroPlex Library Preparation Kit v2 (ref. C05010013) and sequenced.

Bionformatic analysis was performed on two independent biological replicates for each epigenetic mark or transcription factor using Galaxy based approaches (http://usegalaxy.org). Trimmed, groomed reads were aligned to the reference genome mm10 and the peaks were called using the MACS2 algorithm (FDR<0.05). WT1 and MAFB ChIP peaks were further refined using IDR<0.01 and IDR<0.05, respectively, as a threshold. The datasets obtained were further polished with the exclusion of blacklisted loci recently published. For histone marks no IDR analysis was performed but enriched regions consistent between both datasets were considered as peaks. MEME analysis ^34^ (http://meme-suite.org) was carried out to define *de novo* consensus binding regions and search for other TF binding sites using *in silico* databases. GREAT algorithm ^35^ (http://bejerano.stanford.edu/great/public/html/) was used to analyze the gene ontology terms correlated with the regions bound by the transcription factors or the modified histones. Heat maps and genomic tracks were generated using the Easeq software ^36^.

H3K4me3 enrichment in embryonic kidneys was assessed by RT-qPCR. Immunoprecipitated chromatin was purified and the enrichment for *Nphs1*, *Nphs2* and *Synpo* promoters was compared against IgG ChIP experiments and regions distant from the discovered peaks. Primers sequences used for the RT-qPCR are listed in Suppl. Table 5.

### Gene expression analysis

RNA-seq analysis was performed on FACS-sorted isolated podocytes. Kidneys were harvested from 8 weeks old mice, 2 days after tamoxifen induction. Podocytes were isolated using previously described protocols^33^ and taking advantage of the *Rosa26R* mTmG reporter. In brief, GFP+ (recombined) podocytes were sorted by FACS from dissociated kidneys and isolated cells immediately processed with RNeasy micro kit (Qiagen) to obtain purified total RNA. A total of 5 samples for control (*Mafb^LoxP/+^; WT1 Cre ^ERT^*^2^*; Rosa26^mTmG^*) and KO (*Mafb^LoxP/LoxP^; WT1 Cre ^ERT^*^2^*; Rosa26^mTmG^*) were selected. RNAs were quality assessed (Agilent 2100 Bioanalyzer, RIN>9) and RNA libraries were prepared using NEBNext® Ultra TM RNA Library Prep Kit for Illumina® (NEB, USA) following manufacturer’s protocols. The clustering of the index-coded samples was performed by Novogene on a cBot Cluster Generation System using PE Cluster Kit cBot-HS (Illumina). Paired-end clean reads were mapped to the reference genome using HISAT2 software and for samples with biological replicates, differential expression analysis of two conditions/groups was performed using the DESeq2 R package.

Single cell analysis in Fig. 4B was performed using the fetal nephron dataset from ^37^ in the publicly available database (https://cellxgene.cziscience.com/e/08073b32-d389-41f4-a4fd-616de76915ab.cxg/).

### Tissue analysis, histology, EM analysis

Immune-fluorescence and histological analyses were performed on renal tissue, embedded in paraffine after fixation ON at 4°C with PFA 4%, and sectioned at 5um thickness. Primary antibodies used in the study included: anti-WT1 (Dako, ref M3561, clone 6F-H2), anti-MAFB (BETHYL Laboratories, ref IHC-00351), anti-NPHS1 (R&D Systems, ref AF3159), anti-NPHS2 (Sigma, ref P0372), anti-SYNPO (PROGEN Biotechnik, ref 65194).

Transmission electron microscopy (TEM) and scanning electron microscopy (SEM) analyses were performed essentially as described ^12,38^. For the quantification of foot process effacement and glomerular basal membrane expansion, 3 animals per genotype were analyzed, using 20 images per animal containing an average of 1.33 basal membrane transversal sections. The basal membrane thickness was measured every two micrometers along each section, whereas the diameters of the foot processes was assessed for each event present along each segment.

### Urine analysis

Spot urines were collected at the same hour of the day (10 am) and centrifuged to eliminate aggregates and impurity. 5ul of the urines for each sample and time point was loaded in denaturing/reducing buffer (1x Laemmli) in 10% acrylamide/bis acrylamide gel and resolved by electric field. The gel was then stained by Blue Coomassie solution for 3 hours and destained until proper contrast was achieved.

## Results

### Determining the histone profile in adult mouse podocytes

Epigenetic marks are highly cell type specific and it is therefore essential to start with pure cell populations to map histone modifications. To achieve this goal we isolated glomeruli from perfused adult mice, dissociated them into single cells and FACS sorted them based on the expression of podocin ^33^ (Fig. 1A). ChIP-Seq experiments (GEO GSE233911) for H3K4me3 (active), H3K4me1 (primed) and H3K27me3 (repressed) were carried out to determine the genome wide chromatin status in podocytes under homeostatic conditions. Biological duplicates showed highly reproducible peaks (Suppl. Figure 1A&B) and all further analyses were therefore carried out with a merged data set. Heat maps across a set of podocyte-specific genes (Suppl. Table 1: GSE119531) ^39^ showed the expected histone methylation patterns with strong H3K4me3 at the transcription start site (TSS), enrichment of H3K4me1 in the gene body and depletion of H3K27me3 throughout expressed loci (Fig. 1B-C). H3K4me3 enriched regions mostly cluster around the TSS and relate to GO biological processes involved in ‘*nucleosome assembly’* and ‘*protein folding*’ (Fig. 1D). Consistent with the podocyte specific signature, the GO-term ‘Absent podocyte slit diaphragm’ featured in the *Mouse Phenotype* category (Suppl. Table 1 worksheet 2; FDR q-value= 0.000119119.) Indeed, close inspection of the methylation pattern around podocyte-specific genes confirmed strong H3K4me3 peaks at TSS, enrichment of H3K4me1 throughout out the gene body and depletion of the H3K27me3 repressive mark as exemplified by *Wt1, Nphs1, Nphs2* and *Tdrd5* (Fig. 1F-G; Suppl. Figure 1A).

**Figure 1:**
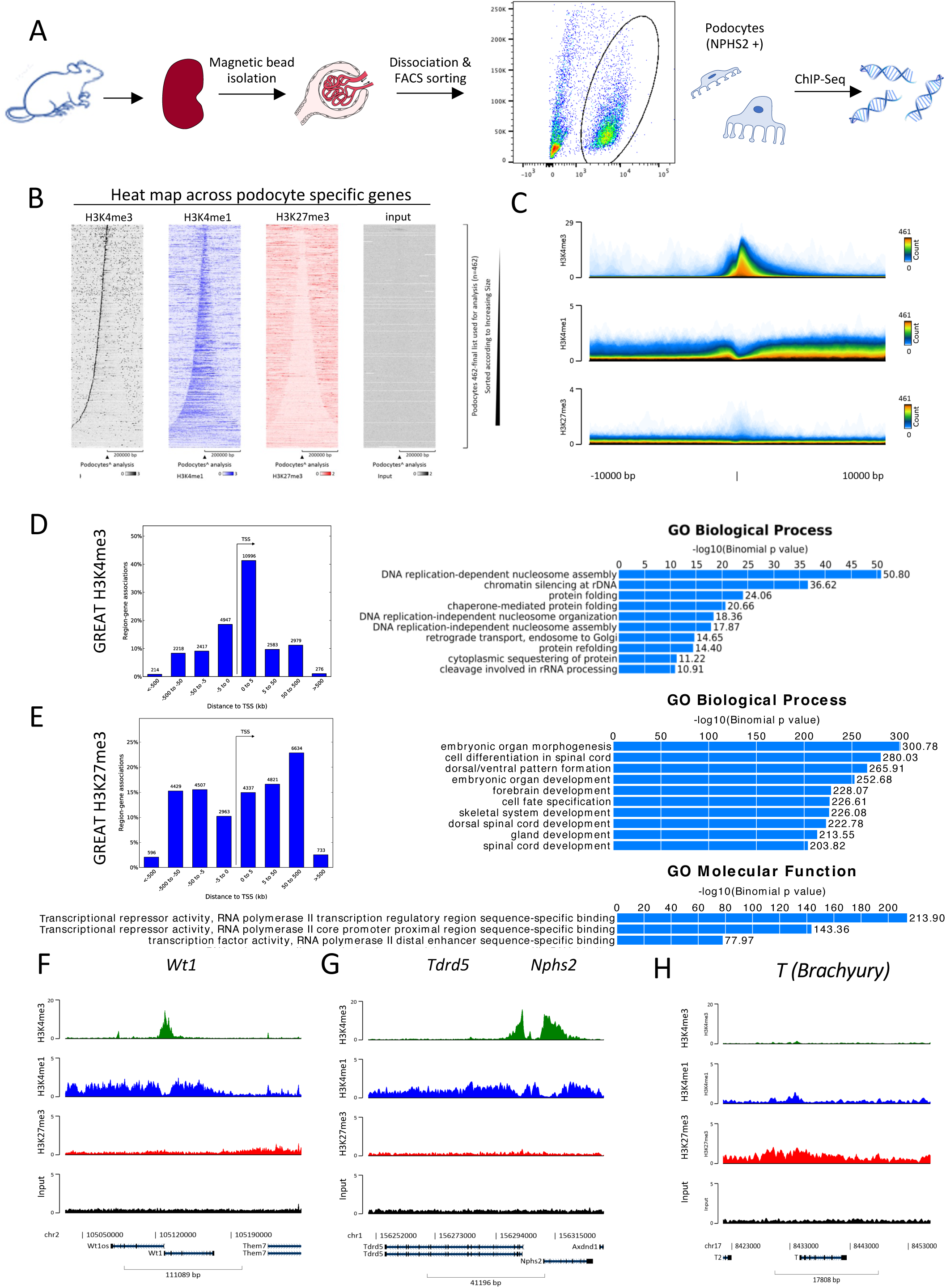
The genome wide histone methylation patterns in adult mouse podocytes. A) Schematic illustration of the podocyte isolation procedure. B) Heatmap across a set of 462 podocyte specific genes (Suppl. Table 4) ^39^ sorted according to gene size. H3K4me3 enrichment was found at the TSS, gene bodies were enriched for H3K4me1 and depleted for H3K27me3. C) Overlay of peaks at the TSS of podocyte specific genes confirmed a strong enrichment of H3K4me3 at the TSS. D) GREAT analysis for H3K4me3 peaks identified peak enrichment at TSS and enriched GO biological processes related to ‘*nucleosome assembly’* and ‘*protein folding*’. E) H3K27me3 peaks are spread further upstream and downstream of the TSS and showed a strong GO term enrichment for ‘*transcription factors’* and ‘d*evelopmental abnormalities*’. F-H) Examples of methylation tracks across the podocyte specific genes *Wt1* (F), *Nphs2* (G) and the mesoderm regulator *Brachyury* (H).

By contrast H3K27me3 peaks were broader, and GREAT analysis identified GO biological terms that relate to embryonic development and differentiation processes (Fig. 1E). Examples included the early mesoderm marker T (Brachyury) (Fig. 1H), the paraxial mesoderm markers *paraxis* (Suppl. Figure 1C), the lateral plate markers *Gata4* (Suppl. Figure 1D), the gonadal marker *Dmrt1* (Suppl. Figure 1E), and the kidney progenitor marker *Six2* (Suppl. Figure 1F). This H3K27me3 methylation pattern is consistent with an active suppression of genes that orchestrate development into alternative developmental linages or genes that are expressed in early stages of kidney progenitors (such as *Six2*), but that are switched off during terminal differentiation. Interestingly, *Hox* gene clusters were split with the *Hoxc10/11/12* paralogues being actively repressed, whereas downstream *Hox* genes (*Hoxd9*-*Hoxd1*) were devoid of H3K27me3 marks (Suppl. Figure 1G), which is in agreement with their expression in adult podocytes ^40^. Taken together this data set provides a blueprint of the methylation status of podocytes under homeostatic conditions.

### Identification of direct MAFB targets in adult podocytes

To gain further insights into podocyte-specific gene regulation, we performed ChIP-Seq analysis for the transcription factors WT1 and MAFB on isolated glomeruli of adult mice (GEO GSE233910). Consistent with published data ^13,41^, WT1 bound a large number of genomic regions (17097 peaks; IDR <0.01) mapping not only close to genes, but also to intergenic regions, which is in agreement with the somewhat non-classical action of WT1 (Suppl. Figure 2B). Comparison of our data with previously published findings ^13^ confirmed the WT1-binding motif (Suppl. Figure 2C) and an overlap of 87% of previously identified peaks (14946 peaks out of 17098 total peaks, data summarized in Suppl. Fig. 2D). As expected, GREAT analysis of the 17097 peaks showed an enrichment of podocyte-related GO terms (Suppl. Figure 2E-F).

Bioinformatic analysis of MAFB ChIP-Seq data from adult glomeruli identified 347 consistent peaks (IDR<0.05). MEME analysis (http://meme-suite.org) confirmed the presence of the MAFB core binding sequence that has been previously identified *in silico* (jaspar.genereg.net) (Fig. 2A). The analysis revealed a high recurrence of palindromic MAFB sites, a finding that is consistent with binding of this transcriptional regulator as a homodimer ^42^. 57% of MAFB binding sites were found within 3kb of the transcriptions start sites (Fig. 2B). GREAT analysis of peak-associated genes showed GO term enrichment for cell matrix adhesion (q-value=0.004) and the cellular components of the podosome (q-value=0.01). In the mouse phenotype category, the most enriched GO terms related to abnormal podocyte physiology, including abnormal/expanded mesangial matrix morphology (q-value=0.0125 and q-value=0.0107), podocyte foot process effacement (q-value= 0.0161) and abnormal slit junction morphology (q-value=0.0164) (Fig. 2C and Suppl. Table 2, Worksheet2). Comparison of the list of genes associated with MAFB binding peaks with a list of 462 podocyte specific genes^39^ showed that MAFB binds to 42 genes (Suppl. Table 2; worksheet2).

**Figure 2:**
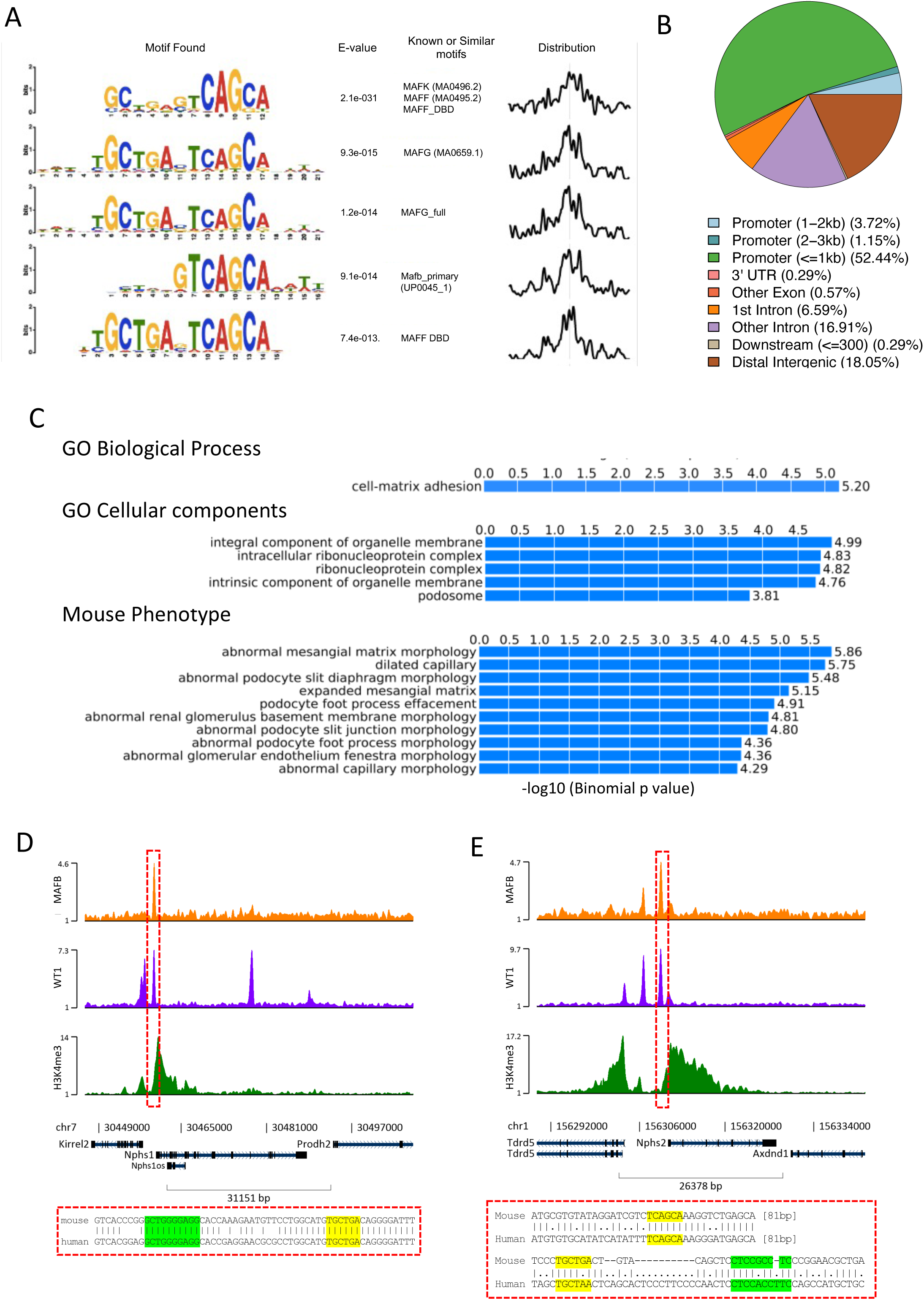
MAFB ChIP-Seq analysis. A) MEME analysis of the peak centred enriched regions reveals MAFB monomeric and dimeric binding sequences. B) MAFB binding site distribution shows strong enrichment at promoter regions. C) GREAT analysis reveals binding at genes involved in cell-matrix adhesion and podocytes pathological processes. D-E) MAFB (orange), WT1 (violet) and H3K4me3 (green) tracks across the podocyte specific genes *Nphs1* and *Nphs2* and the conservation of binding sites in the mouse and human genome.

Genome wide comparison between WT1 and MAFB IDR peaks revealed 256 chromosomal regions that were bound by both transcription factors (Fig. 2D-E and Suppl. Table 2 worksheet 5). Closer inspection of these showed no precise spacing between the TF binding sites, which may suggest that the two TFs bind to podocyte specific genes in an independent manner. Consistent with this hypothesis, pull-down experiment in co-transfection experiments (HEK293 cells) did not show direct interaction between WT1 and MAFB (data not shown).

### MAFB is required to maintain adult podocytes fully differentiated

A recent study has shown that MAFB is not only important during development, but also for podocyte maintenance in adult animals^19^. To obtain further insights into the molecular mechanisms driven by MAFB, we employed a tamoxifen inducible *Wt1-CreERT2* driver^30^ (*Mafb^flox/flox^; Wt1-CreERT* from now on called *Mafb^cko^*). Despite very efficient deletion of *Mafb* as early as 2 days after tamoxifen injection (Fig. 3A&B), loss of kidney function was slow, developing into overt proteinuria only 8 weeks after induction (Fig. 3C). Scanning and transmission electron microscopy showed a diffuse fusion of podocyte foot processes and thickening of the basement membrane (Fig. 3D I to IV). Histological analysis confirmed focal segmental glomerulosclerosis with excessive deposition of collagen matrix (Fig. 3D V and VI). Interestingly, qPCR analysis revealed only a mild reduction of podocyte specific genes even 30 days after deletion (e.g. *THSD7a*, *Synpo, Podxl, Nphs2;* Fig. 3F).

**Figure 3:**
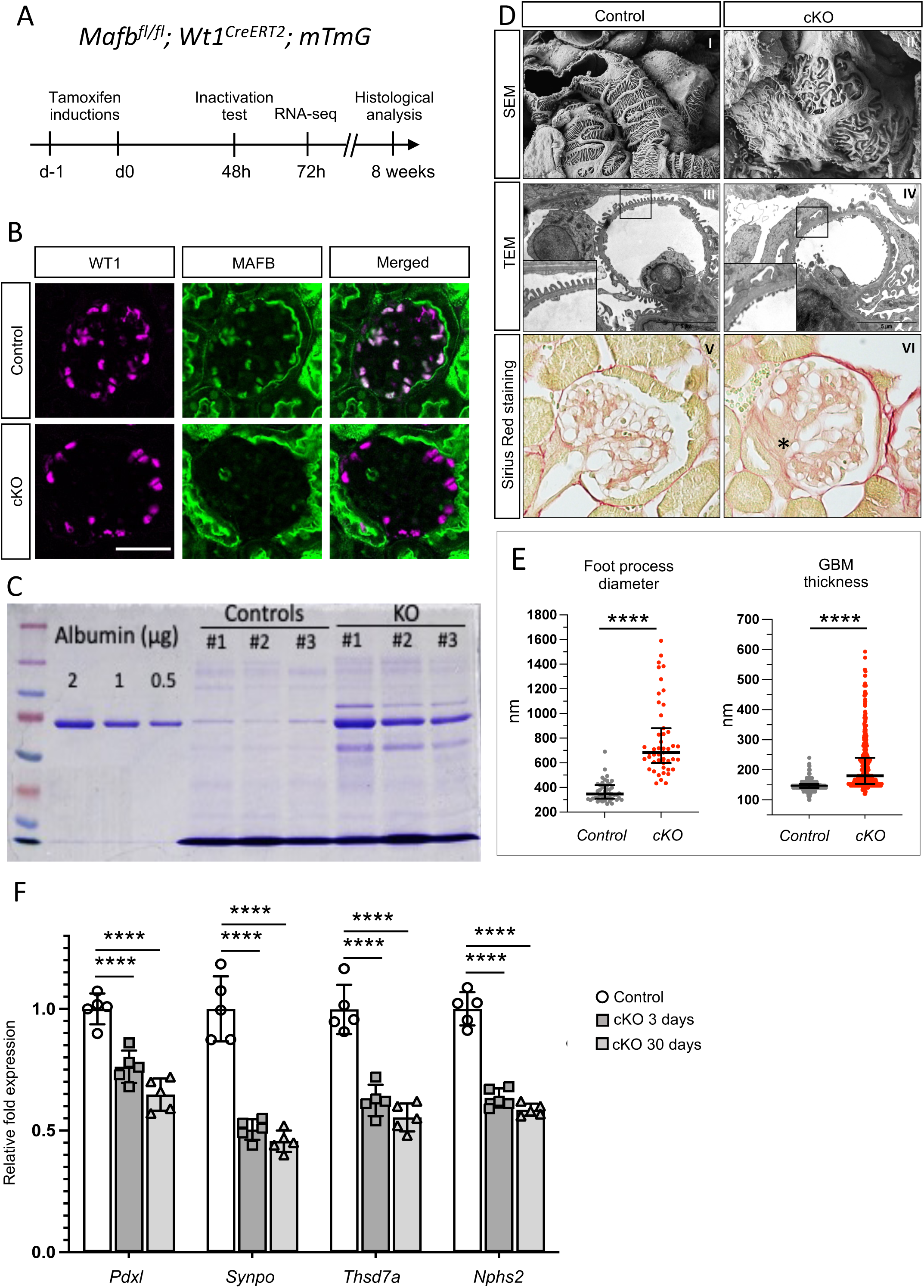
Phenotypic analysis in conditional *Mafb* knockout mice reveals progressive kidney disease with podocyte effacement. A) Strategy for the conditional knockout in *Mafb^cko^ (Wt1^CreERT^*; *Mafb^flox/flox^; mTmG*) mice. B) Confirmation of efficient loss of MAFB protein at 72h after induction (note that non-nuclear staining with the MAFB antibody is background) (scale bar: 50µm). C) Electrophoresis of urine samples showed proteinuria at 8 weeks after induction, accompanied by (D) abnormal footprocesses (SEM, I-II), thickening of the basement membrane (TEM, III-IV) and segmental sclerosis (Sirius RED, V-VI). E) Quantification of foot processes and basement membrane thickness. (n= 3 animals per condition, 20 images per animal, **** p<0.001). F) Expression levels of different podocytes specific genes 3 and 30 days after TAM induction. (n=5, **** p<0.001). Abbreviations: SEM=scanning electron microscope; TEM=Transmission electron microscope. GBM=glomerular basement membrane.

To obtain an unbiased view of genes that directly depend on MAFB in adult podocytes, we next performed RNA-Seq analysis (GEO GSE233912) on glomeruli isolated two days after tamoxifen induction, a time point when the expression of *MafB* was lost (Fig. 3B), *Nphs2* (a known target of MAFB) was reduced (Suppl. Figure 3A), but no overt phenotypic abnormalities were present. Pearson and PCA analysis confirmed clustering of wildtype and knockout samples into two different groups (Suppl. Figure 3B&C). Differential expression analysis identified 1109 (693) and 1178 (724) genes to be decreased and increased in *Mafb^cko^* animals at an adjusted p-Value of p_adj_ <0.05 (p_adj_ <0.01), respectively (Fig. 4A and Suppl. Table 3). *Enrichr* analysis of downregulated genes identified GO-terms including ‘*Genes controlling nephrogenesis’* (p=1.6×10^-6^), nephrotic syndrome (p=2.0×10^-5^) and ‘*VEGFA-VEGFR2 signaling’* (p=2.0×10^-5^) (Fig. 4B). GO-terms for upregulated genes included ‘*Tgf-β regulation of extracellular matrix’* (p=9×10^-6^), ‘*nephrin interactions’* (p=1×10^-4^) and ‘*Rac1/Rho1 motility signaling pathways’* (p=1×10^-3^) (Fig. 4C), which probably reflects the cellular response of the podocyte to the reduction of key components of the slit diaphragm such as *Nphs1* and *Nphs2*. GSEA analysis with a list of podocyte-specific differentiation markers ^40^ showed a significant loss of podocyte specific genes (Fig. 4D; enrichment score = 0.65) and a return to a precursor phenotype (Fig. 4E; enrichment score=-0.4).

**Figure 4:**
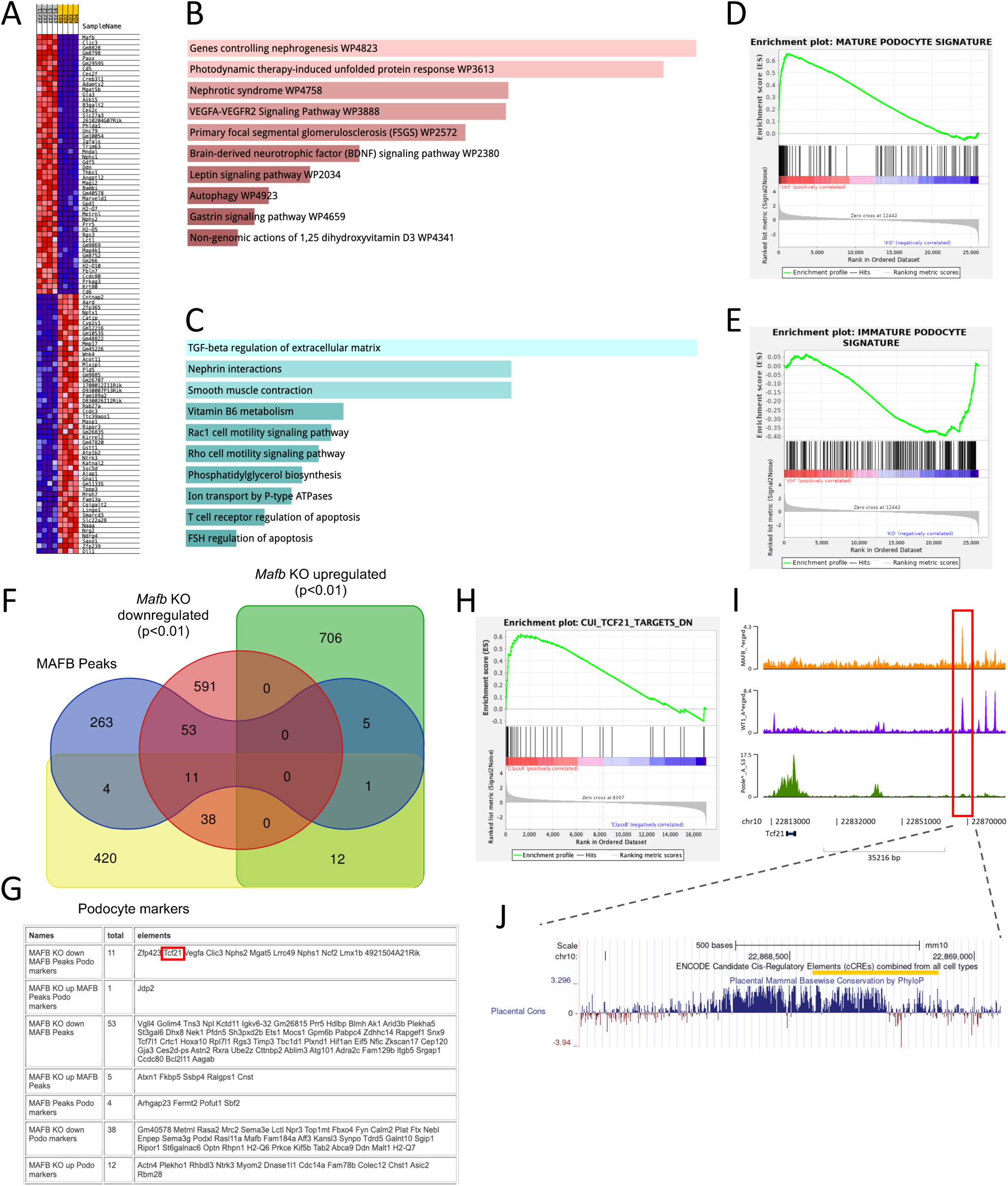
RNAseq analysis in *Mafb^cko^* podocytes. A) Heat map of deregulated genes three days after tamoxifen induction. B) *Enrichr* analysis of 693 downregulated genes (p_adj_<0.01) reveals GO-term enrichment for ‘*podocyte dysfunction*’ and ‘*VEGFA/VEGFR2 signalling’.* C) Go Term enrichment analysis for upregulated genes highlighted ‘*Tgfb regulation of extracellular matrix’* and ‘*Nephrin interactions’*. D-E) GSEA analysis of the deregulated genes indicate a rapid reversion of podocytes to a more undifferentiated state after *Mafb* inactivation. F) Venn diagram and G) gene list of the intersections between the different datasets (*Mafb^cko^* upregulated genes, *Mafb^cko^* downregulated genes, MAFB ChIP peaks, Podocyte specific genes) identified direct and indirect MAFB targets. H) GSEA analysis of the RNA-seq data highlighted ‘*TCF21/POD1 targets’* to be affected in *Mafb^cko^* animals. I) ChIP-Seq tracks reveal peaks for MAFB and WT1 approximately 49.5kb upstream of the *Tcf21* promoter. J) Inspection of the corresponding chromosomal region reveals a high degree of evolutionary conservation.

Comparison of significantly deregulated transcripts with genes that map close to MAFB ChIP-Seq peaks indicates 64 genes to be directly regulated by this transcription factor, 11 of which appeared to be podocyte specific (Fig. 4 F&G). The much higher number of significantly downregulated genes in absence of MAFB (p<0.01, n=693) compared to the relatively low number of direct targets detected by ChIP-Seq may suggest secondary effects already at this early stage after deletion. Interestingly, we found *Pod1/Tcf21*, a gene with important functions in glomerular development ^8–11^, to be reduced upon *MafB* deletion (p_adj_=2.0×10^-7^; 1.8fold; Suppl. Table 3). Furthermore, GSEA analysis revealed ‘*Tcf21 targets’* as a key GO-term enriched in control over knockout samples (enrichment score 0.6; Fig. 4H). Examination of the genomic locus revealed peaks for MAFB and WT1 48.5kb upstream of the *Tcf21* transcription start site (Fig. 4I&J), a region that is evolutionary conserved in placental animals and has been identified as a candidate cis-regulatory region (Encode project). We conclude that MAFB achieves its podocyte-specific function by directly regulating a subset of podocyte specific genes including other transcription factors required for terminal differentiation, such as *Tcf21*.

### MAFB is required for chromatin accessibility of podocyte specific promoters

To gain further insights into the molecular mechanisms underlying podocyte differentiation we turned our attention to development. Comparison of the expression pattern on mouse tissues revealed that MAFB expression occurs after WT1, but slightly before NPHS1 and NPHS2 (Suppl. Figure 4A). Analysis of a publicly available single cell database (https://cellxgene.cziscience.com/e/08073b32-d389-41f4-a4fd-616de76915ab.cxg/) ^37^ confirmed a similar expression profile in human nephron development (Suppl. Figure 4B). Mice homozygous for the *Mafb^GFP^* knock-in that represents a loss-of-function allele completely lacked NPHS1 and NPHS2 despite the persistence of WT1 expression (Fig. 5A). We therefore hypothesized that MAFB may be involved in chromatin opening thus allowing other transcription factors such as WT1 to access promoter regions of podocyte specific genes. To test this model, we carried out ChIP experiments with antibodies against the active chromatin mark H3K4me3 on E17.5 wildtype and *Mafb^Gfp/Gfp^* kidneys. As expected, *Nphs1*, *Nphs2*, *Synpo* promoter regions were enriched for H3K4me3 modification implying an open chromatin structure in wildtype samples (Fig. 5B). Regions downstream of these genes were not enriched and served as negative controls. By contrast, in *Mafb^Gfp/Gfp^* samples chromatin at the *Nphs1* and *Nphs2* promoter failed to be precipitated with H3K4me3 antibodies. Of note, the *Synpo* promoter remained open as demonstrated by maintenance of the enrichment and the residual expression level observed in the IF. To test if changes in the methylation pattern would directly impact binding of other podocyte specific transcription factors, we evaluated the binding of WT1 to these regions. Strikingly, WT1 enrichment was completely lost at *Nphs1/Nphs2* promoters, but only slightly reduced at the regulatory region of *Synpo* (Fig. 5B), a gene that is not bound by MAFB (Suppl. Table 2, Worksheet 1). We conclude that MAFB is crucial to permit chromatin accessibility at its targets, thus allowing transcription factors, such as WT1, to bind and activate key genes of the podocyte.

**Figure 5:**
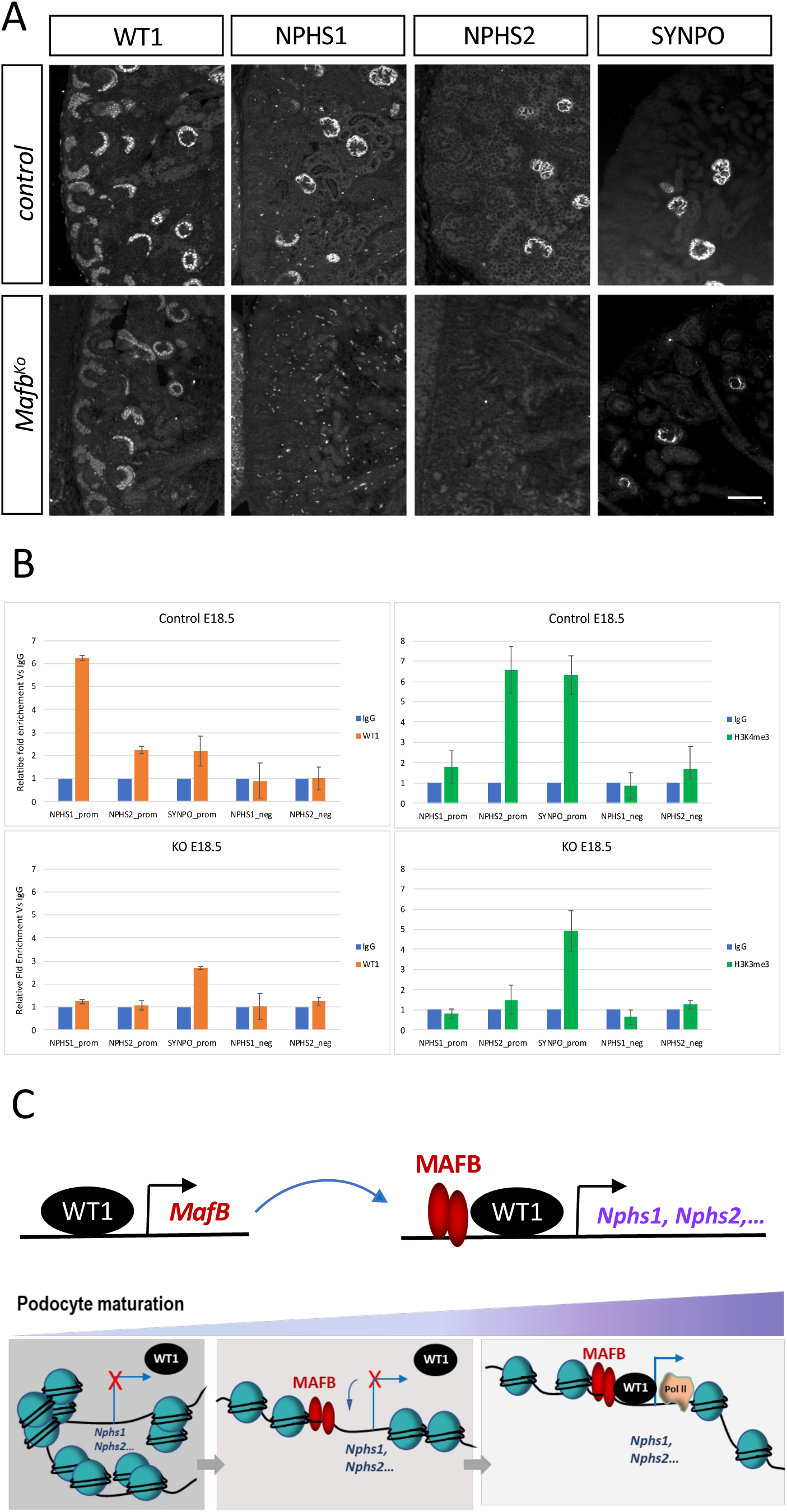
ChIP-qPCR analysis of chromatin conformation in *Mafb* KO embryonic kidney. **A)** Immuno-florescence analysis in E18.5 homozygous *Mafb* knockout animals (*Mafb^GFP/GFP^*) shows complete loss of the direct targets NPHS1 and NPHS2, whereas WT1 and synaptopodin (SYNPO) remain expressed (scale bar: 100µm). B) qPCR analysis of replicated immunoprecipitated chromatin with WT1 (yellow bars) and H3K4me3 (green bars) antibodies in E18.5 embryonic kidneys reveals loss of histone methylation and WT1 binding at *Nphs1* and *Nphs2* promoters. C) Schematic illustration of the role of MAFB in podocyte development. *Mafb* is a target of WT1, but is also required to permit access of WT1 to other podocyte-specific genes.

## Discussion

The transcriptional profile of a cell is shaped by the epigenetic landscape that permits or interferes with binding of TFs and thus the activation of cell type specific genes. In this study we determined the map of histone marks in the adult podocyte, which provides important insights into specific gene regulation and will serve as a resource for future analysis. Not surprisingly, strong H3K4me3 peaks were found at transcription start sites of a large number of genes, with podocyte specific genes often showing very broad peaks. H3K4me3 peak breadth has been shown previously to be highly dynamic and directly associated with transcriptional activity ^43^. In contrast to H3K4me3 modifications, the repressive H3K27me3 mark was predominantly found covering developmental genes. In particular, genes encoding transcription factors that are active at an earlier time point (e.gi. *T, Gata4, Six2*) were covered with repressive marks. This is consistent with the known role of the PRC2 complex that actively represses genes to lock in developmental decisions ^44^. Hox genes are peculiar as they show collinearity of gene expression during early stages of development and progressive silencing of members within a cluster is accompanied by histone methylation via the PRC complex ^45^. In podocytes, H3K27me3 methylation followed this pattern with only the silent portion of each cluster being CpG-marked.

Active suppression by H3K27me3 is important as podocyte specific deletion of *Ezh2*, a crucial member of the PRC2 complex, sensitizes mice to glomerular disease^46^. On the molecular level this has been suggested to be due to an increase of Notch signalling and in particular a loss of *Jag1* H3K27me3 methylation. In our data we did not find strong H3K27me3 methylation at the *Jag1* locus, which may suggest that demethylation at this locus is not a primary cause of dysfunction in *Ezh2* mutant podocytes. In the future it will be interesting to investigate in detail epigenetic changes on a genome wide scale in glomerular diseases and link them with the observed changes in gene expression.

We also describe the genome wide binding sites of MAFB in podocytes *in vivo*. Compared to WT1, which shows very promiscuous binding (17098 peaks), the number of MAFB peaks with an IDR <0.05 was relatively low (349 peaks), suggesting a more selective target site selection. This is consistent with a recent study that showed enrichment for WT1 and FOXC2, but not MAFB in glomeruli specific super-enhancers^3^. Previous studies in other organ systems have identified a larger number of MAFB targets, which may be due to a less stringent IDR analysis in their studies ^47,48^. The binding motif detected precisely fits the expected MAFB recognition sequence, with a large proportion of sites showing palindromic sites indicating binding as a homodimer. 256 of the identified MAFB regions (including *Nphs1* and *Nphs2* promoters) appear to be also bound by WT1, which may suggest that these two transcription factors cooperate in the activation of downstream targets. The absence of a precise spacing between the binding sites and the lack of co-precipitation after co-transfection into HEK293 cells suggests this cooperation not to involve physical interaction. Instead we propose that MAFB is required to allow WT1 factors to access chromatin and thus permit activation of a subset of podocyte specific genes (Fig. 5C).

While this study was in progress, a manuscript reported that deletion of MAFB with the *Nphs2-CreERT2* driver in adult podocytes leads to glomerular defects 8 weeks after induction^19^. Our own data using a different Cre driver (*Wt1-CreERT2*) that induces >95% deletion of *Mafb* at protein level three days after induction confirms this relatively slow, but progressive kidney disease. In contrast to the previously published study that reported transcriptomic analysis at a rather late time point (8 weeks after induction), our RNA-Seq analysis was performed only two days after tamoxifen induction, which allowed us to look at primary events. Despite the rapid loss of MAFB protein after inactivation, we observed only a relatively mild reduction of a broad spectrum of genes, including podocyte marker proteins. The intersection of RNA-seq and ChIPseq data allowed us to elaborate a transcriptional cascade. Indeed, *Lmx1b* and *TCF21/Pod1*, are directly regulated by MAFB, whereas the key regulator *WT1* remains normally expressed in absence of MAFB. MAFB can therefore be considered in an intermediate hierarchic position, downstream of WT1 (by which it is activated), but responsible for high *TCF21/Pod1* and *Lmx1b* expression levels and, as a consequence, their downstream effector genes. Contrary to published *in vitro* experiments^19^, we did not find significant enrichment at the *Tcf21* start site in *our in vivo* ChIP-Seq experiments. Instead we identified an evolutionary conserved element 49.5kb upstream of *Tcf21* that is bound by both WT1 and MAFB and has been identified as a potential cis-regulatory element by the ENCODE project. Given the important function of TCF21 in podocyte differentiation, it will be interesting to test this region for mutations in patients with unexplained cases of glomerulopathies.

In contrast to the relatively mild kidney phenotype after tamoxifen induced deletion of *Mafb* in adults, constitutive deletion abolishes terminal podocyte differentiation. Our data point to a crucial role for MAFB as a pioneer transcription factor that allows chromatin to open and other factors to bind ^49^. Expression of MAFB has been shown in the epidermis to be both necessary and sufficient to exit cell cycle and promote progenitor differentiation ^50^. Once chromatin has been opened, MAFB may no longer be essential to permit continuous transcription of podocyte specific genes. This model is supported by our ChIP studies in developmental kidneys which demonstrate that loss of MAFB leads to a lack of open chromatin and strongly reduced binding of WT1 at the *Nphs1* and *Nphs2* loci.

Taken together our study describes the epigenetic landscape of mouse podocytes, identified genome wide binding sites for MAFB and discovered an important role of MAFB during differentiation in opening chromatin and other TFs expression and binding to promoters of podocyte specific genes.

## Acknowledgements

We would like to thank the entire staff of the iBV animal facility for their dedication. We are grateful to William Pu for providing us with the *Wt1^CreERT^*^2^ strain. This study was supported by institutional grants from TU Dresden, the Institut National de la Santé et de la Recherche Médicale, Centre National de la Recherche Scientifique, University Cote d’Azur and Aix-Marseille University and grants to A.S. from the Fondation pour la Recherche Médicale (ING20160435020), ANR (ANR-12-BSV 1-0033-02; ANR-11-LABX-0028-01), the European Commission (EURenOmics Grant agreement 305608), Conseil Régional Sud Est PACA”, the “Conseil Départemental 06”, to M.H. Sieweke from Fondation pour la Recherche Médicale (DEQ. 20110421320), the ‘Agence Nationale de la Recherche’ (ANR-11-BSV3-026-01, ANR-17-CE15-0007-01 and ANR-18-CE12-0019-03), INCA (13-10/405/AB-LC-HS), Fondation ARC pour la Recherche sur le Cancer (PGA1 RF20170205515), an INSERM-Helmholtz cooperation and the European Research Council (ERC) under the European Union’s Horizon 2020 research and innovation program (grant agreement number 695093 MacAge). M.H. Sieweke is an Alexander von Humboldt Professor at TU Dresden. The authors acknowledge the University’s CCMA, Electron Microscopy facility (Centre Commun de Microscopie Appliquée, Université Côte d’Azur) and MICA Imaging platform Côte d’Azur supported by Université Côte d’Azur, the “Conseil Régional Sud Est PACA”, the “Conseil Départemental 06 “ and Gis Ibisa, and Christelle Boscagli for her technical help.

---

**Suppl. Figure 1:**
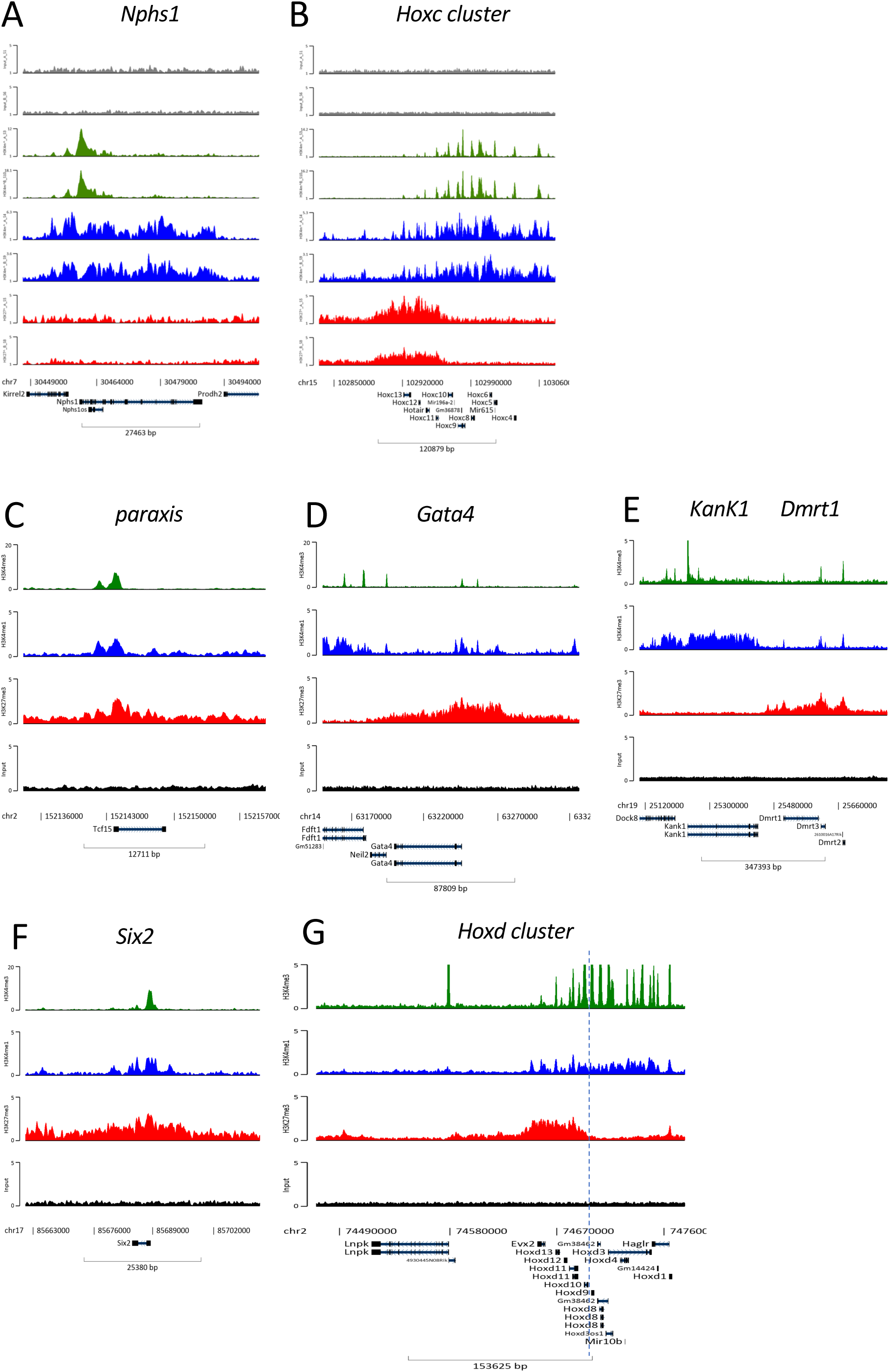
Representation of ChIPseq tracks. **A-B)** Side-by side comparison of two independent ChIP-seq experiments at the *Nphs1* (podocytes specific) and *Hoxc* cluster showed highly reproducible results. All further analyses were therefore performed with merged datasets. C-G) Examples of actively repressed developmental genes: *paraxis*, *Gata4*, *Dmrt1*, *Six2* and *Hoxd* genes cluster, dotted line in G indicate the separation of active and inactive genes of the cluster. Colour scheme: H3K4m3 (green), H3K4me1 (blue,) H3K27me3 (red) and the Input (raw sonicated chromatin, grey).

**Suppl. Figure 2:**
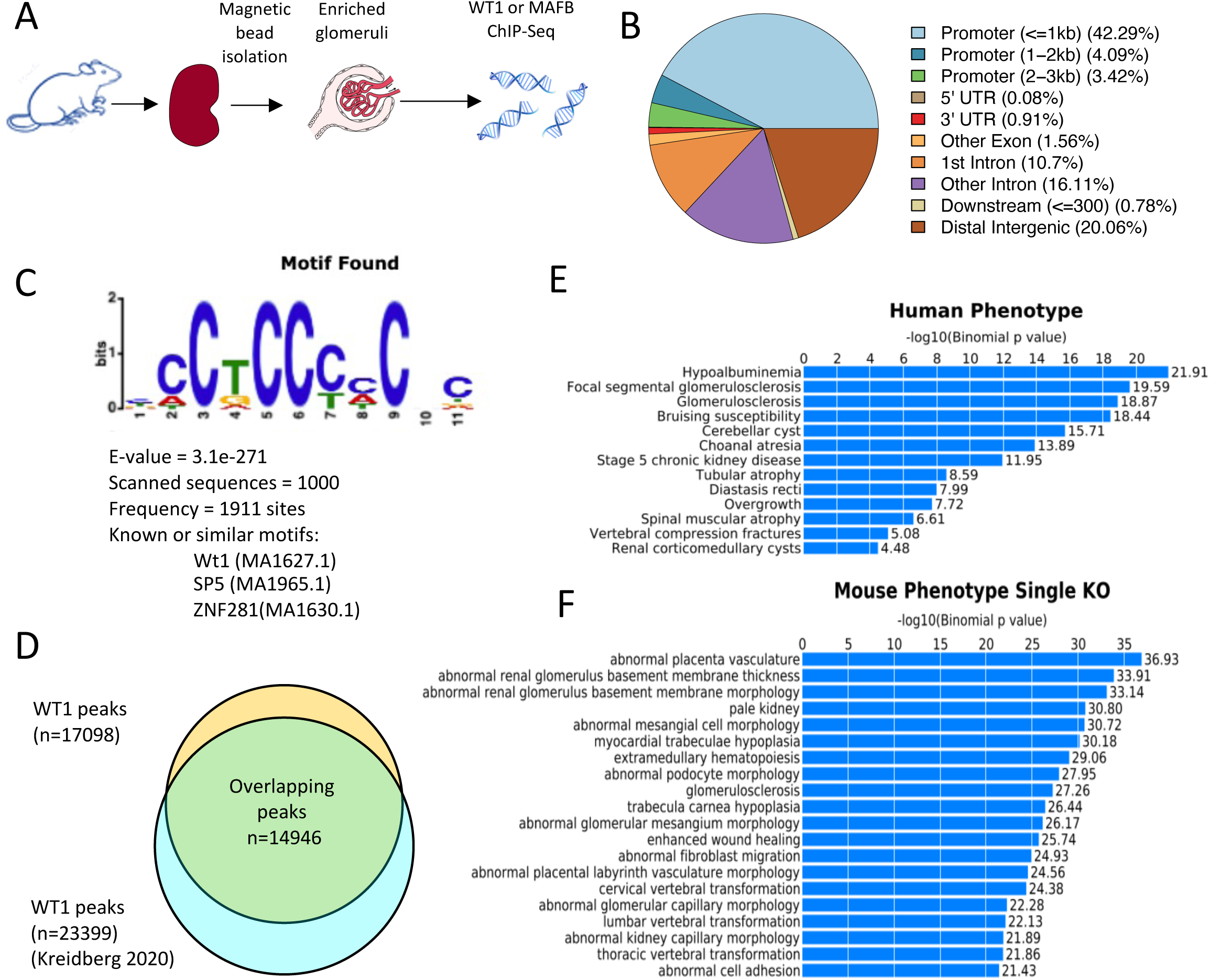
WT1 ChIP-seq analysis. **A**) Schematic illustration of the glomerular isolation procedure used for ChIP experiments. B) Distribution of WT1 peaks shows a large proportion (42.29%) of peaks to map to promoter regions (<1kb). C) *De novo* MEME analysis of the top 1000 enriched regions identified a core sequence highly similar to the previously identified consensus sequence for WT1 binding. D) Overlap between newly identified and previously published WT1 binding regions. E-F) GREAT analysis identifies ‘*focal segmental glomerulosclerosis*’ and ‘*abnormal renal glomerulus basement membrane thickness’* as highly enriched GO terms.

**Suppl. Figure 3:**
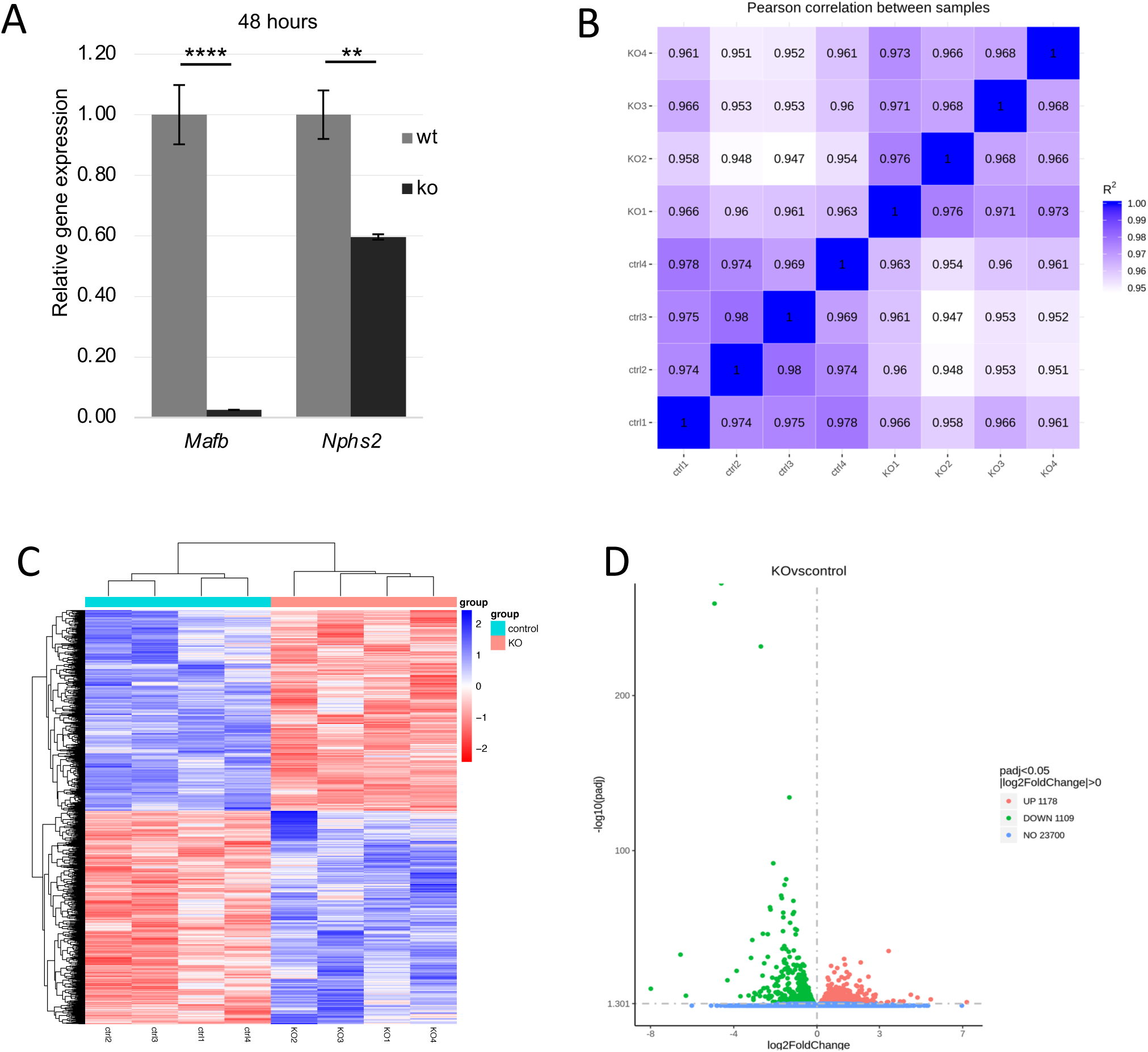
Validation of the RNA-seq analysis. A) qPCR analysis showed absence of *Mafb* and significant reduction of *Nphs2* expression 48hours after tamoxifen induction (n=5, ****p<0.001, **p<0.01). B) Pearson correlation and C) hierarchical clustering of samples. D) Volcano plot revealing the spread of down-and up-regulated genes in *Mafb^cko^* glomeruli.

**Suppl. Figure 4:**
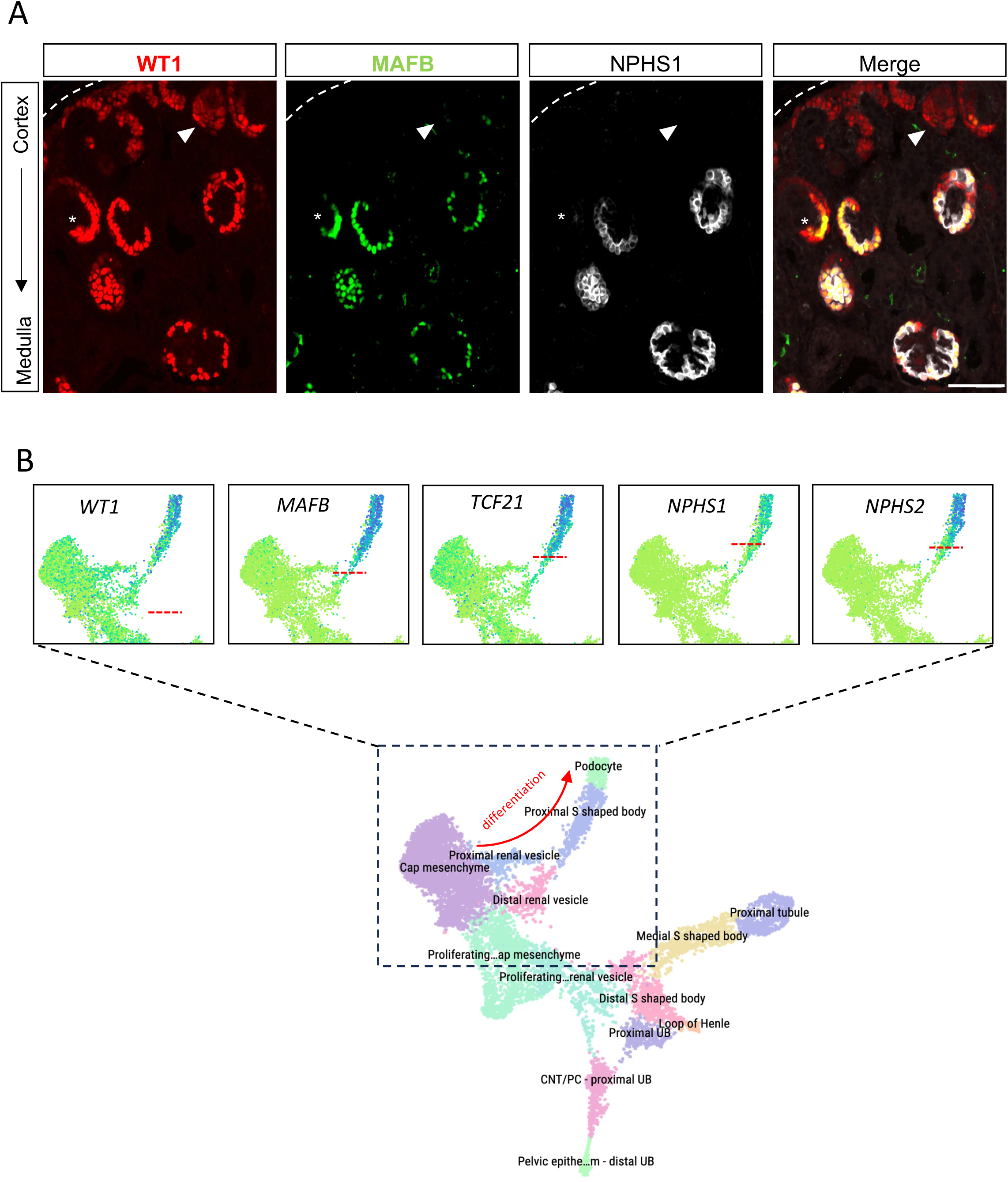
MAFB expression pattern during podocyte maturation. A) Immunofluorescence staining shows that MAFB is not detectable at the early stage of nephrogenesis (vesicle stage, arrowhead), but it appears at the very first steps of podocyte differentiation (comma shaped bodies/precapillary loop stage, asterisk) and remains expressed in adult podocytes. Scale bar: 100µm. B) Single cell analysis in developing human kidneys. Dotted lines identify the early expression of *MAFB* in proximal S shaped bodies, comparable with the one of *WT1* and slightly earlier than *TCF21* compared with the appearance of late podocytes markers *NPHS1* and *NPHS1*

**Suppl. Table 1:** Histone methylation peaks detected by ChIP-Seq analysis

**Suppl. Table 2:** ChIP-Seq peaks for MAFB and WT1

**Suppl. Table 3:** RNA-Seq analysis

**Suppl. Table 4:** List of Primers

## References

1. Jambhekar A, Dhall A, Shi Y: Roles and regulation of histone methylation in animal development [Internet]. Nat. Rev. Mol. Cell Biol. 20: 625–641, 2019 Available from: https://pubmed-ncbi-nlm-nih-gov.proxy.insermbiblio.inist.fr/31267065/ [cited 2021 May 11]

2. O’Carroll D, Erhardt S, Pagani M, Barton SC, Surani MA, Jenuwein T: The Polycomb -Group Gene Ezh2 Is Required for Early Mouse Development. Mol. Cell. Biol. [Internet] 21: 4330–4336, 2001 Available from: https://journals.asm.org/doi/10.1128/MCB.21.13.4330-4336.2001 [cited 2022 Dec 2]

3. Yang J, Zhang D, Motojima M, Kume T, Hou Q, Pan Y, Duan A, Zhang M, Jiang S, Hou J, Shi J, Qin Z, Liu Z: Super-Enhancer-Associated Transcription Factors Maintain Transcriptional Regulation in Mature Podocytes. J. Am. Soc. Nephrol. [Internet] ASN.2020081177, 2021 Available from: https://pubmed.ncbi.nlm.nih.gov/33771836/ [cited 2021 May 11]

4. He B, Ebarasi L, Zhao Z, Guo J, Ojala JR, Hultenby K, De Val S, Betsholtz C, Tryggvason K: Lmx1b and FoxC combinatorially regulate podocin expression in podocytes. J Am Soc Nephrol [Internet] 25: 2764– 2777, 2014 Available from: https://www.ncbi.nlm.nih.gov/pubmed/24854274

5. O’Brien LL, Grimaldi M, Kostun Z, Wingert RA, Selleck R, Davidson AJ: Wt1a, Foxc1a, and the Notch mediator Rbpj physically interact and regulate the formation of podocytes in zebrafish. Dev Biol [Internet] 358: 318–330, 2011 Available from: http://www.ncbi.nlm.nih.gov/pubmed/21871448

6. White JT, Zhang B, Cerqueira DM, Tran U, Wessely O: Notch signaling, wt1 and foxc2 are key regulators of the podocyte gene regulatory network in Xenopus. Development [Internet] 137: 1863–1873, 2010 Available from: https://www.ncbi.nlm.nih.gov/pubmed/20431116

7. Motojima M, Kume T, Matsusaka T: Foxc1 and Foxc2 are necessary to maintain glomerular podocytes. Exp. Cell Res. 352: 265–272, 2017

8. Morello R, Zhou G, Dreyer SD, Harvey SJ, Ninomiya Y, Thorner PS, Miner JH, Cole W, Winterpacht A, Zabel B, Oberg KC, Lee B: Regulation of glomerular basement membrane collagen expression by LMX1B contributes to renal disease in nail patella syndrome. Nat Genet [Internet] 27: 205–208, 2001 Available from: https://www.ncbi.nlm.nih.gov/pubmed/11175791

9. Rohr C, Prestel J, Heidet L, Hosser H, Kriz W, Johnson RL, Antignac C, Witzgall R: The LIM-homeodomain transcription factor Lmx1b plays a crucial role in podocytes. J. Clin. Invest. 109: 1073–1082, 2002

10. Miner JH, Morello R, Andrews KL, Li C, Antignac C, Shaw AS, Lee B: Transcriptional induction of slit diaphragm genes by Lmx1b is required in podocyte differentiation. J. Clin. Invest. [Internet] 109: 1065– 1072, 2002 Available from: https://pubmed.ncbi.nlm.nih.gov/11956244/ [cited 2021 May 11]

11. Maezawa Y, Onay T, Scott RP, Keir LS, Dimke H, Li C, Eremina V, Maezawa Y, Jeansson M, Shan J, Binnie M, Lewin M, Ghosh A, Miner JH, Vainio SJ, Quaggin SE: Loss of the podocyte-expressed transcription factor Tcf21/Pod1 Results in podocyte differentiation defects and FSGS. J. Am. Soc. Nephrol. 25: 2459– 2470, 2014

12. Lefebvre J, Clarkson M, Massa F, Bradford ST, Charlet A, Buske F, Lacas-Gervais S, Schulz H, Gimpel C, Hata Y, Schaefer F, Schedl A: Alternatively spliced isoforms of WT1 control podocyte-specific gene expression. Kidney Int. 2015

13. Kann M, Ettou S, Jung YL, Lenz MO, Taglienti ME, Park PJ, Schermer B, Benzing T, Kreidberg JA: Genome-Wide Analysis of Wilms’ Tumor 1-Controlled Gene Expression in Podocytes Reveals Key Regulatory Mechanisms. J Am Soc Nephrol [Internet] 2015 Available from: http://www.ncbi.nlm.nih.gov/pubmed/25636411

14. Dong L, Pietsch S, Tan Z, Perner B, Sierig R, Kruspe D, Groth M, Witzgall R, Grone HJ, Platzer M, Englert C: Integration of Cistromic and Transcriptomic Analyses Identifies Nphs2, Mafb, and Magi2 as Wilms’ Tumor 1 Target Genes in Podocyte Differentiation and Maintenance. J Am Soc Nephrol [Internet] 2015 Available from: http://www.ncbi.nlm.nih.gov/pubmed/25556170

15. Barbaux S, Niaudet P, Gubler MC, Grunfeld JP, Jaubert F, Kuttenn F, Fekete CN, Souleyreau Therville N, Thibaud E, Fellous M, McElreavey K: Donor splice-site mutations in WT1 are responsible for Frasier syndrome. Nat Genet 17: 467–470, 1997

16. Klamt B, Koziell A, Poulat F, Wieacker P, Scambler P, Berta P, Gessler M: Frasier syndrome is caused by defective alternative splicing of WT1 leading to an altered ratio of WT1 +/-KTS splice isoforms. Hum Mol Genet. 7: 709–714, 1998

17. Li FP, Breslow NE, Morgan JM, Ghahremani M, Miller GA, Grundy PE, Green DM, Diller LR, Pelletier J: Germline WT1 mutations in Wilms’ tumor patients: preliminary results. Med.Pediatr.Oncol. 27: 404– 407, 1996

18. Pelletier J, Bruening W, Kashtan CE, Mauer SM, Manivel JC, Striegel JE, Houghton DC, Junien C, Habib R, Fouser L, et al: Germline mutations in the Wilms’ tumor suppressor gene are associated with abnormal urogenital development in Denys-Drash syndrome. Cell. 67: 437–447, 1991

19. Usui T, Morito N, Shawki HH, Sato Y, Tsukaguchi H, Hamada M, Jeon H, Yadav MK, Kuno A, Tsunakawa Y, Okada R, Ojima T, Kanai M, Asano K, Imamura Y, Koshida R, Yoh K, Usui J, Yokoi H, Kasahara M, Yoshimura A, Muratani M, Kudo T, Oishi H, Yamagata K, Takahashi S: Transcription factor MafB in podocytes protects against the development of focal segmental glomerulosclerosis. Kidney Int. [Internet] 98: 391–403, 2020 Available from: https://pubmed.ncbi.nlm.nih.gov/32622525/ [cited 2021 May 11]

20. Moriguchi T, Hamada M, Morito N, Terunuma T, Hasegawa K, Zhang C, Yokomizo T, Esaki R, Kuroda E, Yoh K, Kudo T, Nagata M, Greaves DR, Engel JD, Yamamoto M, Takahashi S: MafB is essential for renal development and F4/80 expression in macrophages. Mol Cell Biol [Internet] 26: 5715–5727, 2006 Available from: http://www.ncbi.nlm.nih.gov/pubmed/16847325

21. Sato Y, Tsukaguchi H, Morita H, Higasa K, Tran MTN, Hamada M, Usui T, Morito N, Horita S, Hayashi T, Takagi J, Yamaguchi I, Nguyen HT, Harada M, Inui K, Maruta Y, Inoue Y, Koiwa F, Sato H, Matsuda F, Ayabe S, Mizuno S, Sugiyama F, Takahashi S, Yoshimura A: A mutation in transcription factor MAFB causes Focal Segmental Glomerulosclerosis with Duane Retraction Syndrome. Kidney Int. [Internet] 94: 396–407, 2018 Available from: https://pubmed.ncbi.nlm.nih.gov/29779709/ [cited 2021 May 11]

22. Chau YY, Brownstein D, Mjoseng H, Lee WC, Buza-Vidas N, Nerlov C, Jacobsen SE, Perry P, Berry R, Thornburn A, Sexton D, Morton N, Hohenstein P, Freyer E, Samuel K, Van’t Hof R, Hastie N: Acute multiple organ failure in adult mice deleted for the developmental regulator wt1. PLoS Genet [Internet] 7: e1002404, 2011 Available from: http://www.ncbi.nlm.nih.gov/entrez/query.fcgi?cmd=Retrieve&db=PubMed&dopt=Citation&list_uids=22216009

23. Wagner N, Wagner KD, Xing Y, Scholz H, Schedl A: The major podocyte protein nephrin is transcriptionally activated by the Wilms’ tumor suppressor WT1. J. Am. Soc. Nephrol. [Internet] 15: 3044–3051, 2004 Available from: https://pubmed.ncbi.nlm.nih.gov/15579507/ [cited 2021 May 11]

24. Blank V, Andrews NC: The Maf transcription factors: Regulators of differentiation [Internet]. Trends Biochem. Sci. 22: 437–441, 1997 Available from: https://pubmed.ncbi.nlm.nih.gov/9397686/ [cited 2021 May 11]

25. Motohashi H, Shavit JA, Igarashi K, Yamamoto M, Engel JD: The world according to Maf. Nucleic Acids Res. [Internet] 25: 2953–2959, 1997 Available from: https://pubmed.ncbi.nlm.nih.gov/9224592/ [cited 2021 May 11]

26. Cordes SP, Barsh GS: The mouse segmentation gene kr encodes a novel basic domain-leucine zipper transcription factor. Cell [Internet] 79: 1025–1034, 1994 Available from: https://pubmed.ncbi.nlm.nih.gov/8001130/ [cited 2021 May 11]

27. Takahashi S: Functional analysis of large MAF transcription factors and elucidation of their relationships with human diseases. Exp. Anim. 70: 264–271, 2021

28. Sadl VS, Jin F, Yu J, Cui S, Holmyard D, Quaggin SE, Barsh GS, Cordes SP: The mouse Kreisler (Krml1/MafB) segmentation gene is required for differentiation of glomerular visceral epithelial cells. Dev. Biol. [Internet] 249: 16–29, 2002 Available from: https://pubmed.ncbi.nlm.nih.gov/12217315/[cited 2021 May 11]

29. Zankl A, Duncan EL, Leo PJ, Clark GR, Glazov EA, Addor MC, Herlin T, Kim CA, Leheup BP, McGill J, McTaggart S, Mittas S, Mitchell AL, Mortier GR, Robertson SP, Schroeder M, Terhal P, Brown MA: Multicentric carpotarsal osteolysis is caused by mutations clustering in the amino-terminal transcriptional activation domain of MAFB. Am. J. Hum. Genet. [Internet] 90: 494–501, 2012 Available from: https://pubmed.ncbi.nlm.nih.gov/22387013/ [cited 2023 Jan 16]

30. Zhou B, Ma Q, Rajagopal S, Wu SM, Domian I, Rivera-Feliciano J, Jiang D, von Gise A, Ikeda S, Chien KR, Pu WT: Epicardial progenitors contribute to the cardiomyocyte lineage in the developing heart. Nature [Internet] 454: 109–113, 2008 Available from: http://www.ncbi.nlm.nih.gov/entrez/query.fcgi?cmd=Retrieve&db=PubMed&dopt=Citation&list_uids=18568026

31. Matcovitch-Natan O, Winter DR, Giladi A, Aguilar SV, Spinrad A, Sarrazin S, Ben-Yehuda H, David E, González FZ, Perrin P, Keren-Shaul H, Gury M, Lara-Astaiso D, Thaiss CA, Cohen M, Halpern KB, Baruch K, Deczkowska A, Lorenzo-Vivas E, Itzkovitz S, Elinav E, Sieweke MH, Schwartz M, Amit I: Microglia development follows a stepwise program to regulate brain homeostasis. Science [Internet] 353: 2016 Available from: https://pubmed-ncbi-nlm-nih-gov.proxy.insermbiblio.inist.fr/27338705/ [cited 2023 Jun 8]

32. Blanchi B, Kelly LM, Viemari JC, Lafon I, Burnet H, Bevengut M, Tillmanns S, Daniel L, Graf T, Hilaire G, Sieweke MH: MafB deficiency causes defective respiratory rhythmogenesis and fatal central apnea at birth. Nat Neurosci [Internet] 6: 1091–1100, 2003 Available from: https://www.ncbi.nlm.nih.gov/pubmed/14513037

33. Boerries M, Grahammer F, Eiselein S, Buck M, Meyer C, Goedel M, Bechtel W, Zschiedrich S, Pfeifer D, Laloë D, Arrondel C, Gonçalves S, Krüger M, Harvey SJ, Busch H, Dengjel J, Huber TB: Molecular fingerprinting of the podocyte reveals novel gene and protein regulatory networks. Kidney Int. [Internet] 83: 1052–1064, 2013 Available from: https://pubmed.ncbi.nlm.nih.gov/23364521/ [cited 2023 Jun 9]

34. Bailey TL, Johnson J, Grant CE, Noble WS: The MEME Suite. Nucleic Acids Res. [Internet] 43: W39–W49, 2015 Available from: https://pubmed.ncbi.nlm.nih.gov/25953851/ [cited 2023 Jun 9]

35. McLean CY, Bristor D, Hiller M, Clarke SL, Schaar BT, Lowe CB, Wenger AM, Bejerano G: GREAT improves functional interpretation of cis-regulatory regions. Nat Biotechnol [Internet] 28: 495–501, 2010 Available from: http://www.ncbi.nlm.nih.gov/pubmed/20436461

36. Lerdrup M, Johansen JV, Agrawal-Singh S, Hansen K: An interactive environment for agile analysis and visualization of ChIP-sequencing data. Nat. Struct. Mol. Biol. [Internet] 23: 349–357, 2016 Available from: https://pubmed.ncbi.nlm.nih.gov/26926434/ [cited 2023 Jun 9]

37. Stewart BJ, Ferdinand JR, Young MD, Mitchell TJ, Loudon KW, Riding AM, Richoz N, Frazer GL, Staniforth JUL, Braga FAV, Botting RA, Popescu DM, Vento-Tormo R, Stephenson E, Cagan A, Farndon SJ, Polanski K, Efremova M, Green K, Velasco-Herrera MDC, Guzzo C, Collord G, Mamanova L, Aho T, Armitage JN, Riddick ACP, Mushtaq I, Farrell S, Rampling D, Nicholson J, Filby A, Burge J, Lisgo S, Lindsay S, Bajenoff M, Warren AY, Stewart GD, Sebire N, Coleman N, Haniffa M, Teichmann SA, Behjati S, Clatworthy MR: Spatiotemporal immune zonation of the human kidney. Science [Internet] 365: 1461–1466, 2019 Available from: https://pubmed.ncbi.nlm.nih.gov/31604275/ [cited 2023 Jun 7]

38. Courtney M, Gjernes E, Druelle N, Ravaud C, Vieira A, Ben-Othman N, Pfeifer A, Avolio F, Leuckx G, Lacas-Gervais S, Burel-Vandenbos F, Ambrosetti D, Hecksher-Sorensen J, Ravassard P, Heimberg H, Mansouri A, Collombat P: The inactivation of Arx in pancreatic alpha-cells triggers their neogenesis and conversion into functional beta-like cells. PLoS Genet [Internet] 9: e1003934, 2013 Available from: http://www.ncbi.nlm.nih.gov/pubmed/24204325

39. Kirita Y, Wu H, Uchimura K, Wilson PC, Humphreys BD: Cell profiling of mouse acute kidney injury reveals conserved cellular responses to injury. Proc. Natl. Acad. Sci. U. S. A. [Internet] 117: 15874– 15883, 2020 Available from: https://pubmed.ncbi.nlm.nih.gov/32571916/ [cited 2023 May 1]

40. Brunskill EW, Georgas K, Rumballe B, Little MH, Potter SS: Defining the molecular character of the developing and adult kidney podocyte. PLoS One [Internet] 6: e24640, 2011 Available from: http://www.ncbi.nlm.nih.gov/pubmed/21931791

41. Motamedi FJ, Badro DA, Clarkson M, Lecca MR, Bradford ST, Buske FA, Saar K, Hübner N, Brändli AW, Schedl A: WT1 controls antagonistic FGF and BMP-pSMAD pathways in early renal progenitors. Nat. Commun. 5: 2014

42. Pogenberg V, Consani Textor L, Vanhille L, Holton SJ, Sieweke MH, Wilmanns M: Design of a bZip transcription factor with homo/heterodimer-induced DNA-binding preference. Structure [Internet] 22: 466–477, 2014 Available from: https://pubmed.ncbi.nlm.nih.gov/24530283/ [cited 2023 Jun 9]

43. Liu X, Wang C, Liu W, Li J, Li C, Kou X, Chen J, Zhao Y, Gao H, Wang H, Zhang Y, Gao Y, Gao S: Distinct features of H3K4me3 and H3K27me3 chromatin domains in pre-implantation embryos. Nat. 2016 5377621 [Internet] 537: 558–562, 2016 Available from: https://www.nature.com/articles/nature19362 [cited 2021 Aug 20]

44. Margueron R, Reinberg D: The Polycomb complex PRC2 and its mark in life. Nature [Internet] 469: 343–349, 2011 Available from: https://pubmed.ncbi.nlm.nih.gov/21248841/ [cited 2023 Jun 9]

45. Soshnikova N, Duboule D: Epigenetic Temporal Control of Mouse Hox Genes in Vivo. Science (80-.). [Internet] 324: 1320–1323, 2009 Available from: https://science.sciencemag.org/content/324/5932/1320 [cited 2021 Aug 20]

46. Majumder S, Thieme K, Batchu SN, Alghamdi TA, Bowskill BB, Golam Kabir M, Liu Y, Advani SL, White KE, Geldenhuys L, Tennankore KK, Poyah P, Siddiqi FS, Advani A: Shifts in podocyte histone H3K27me3 regulate mouse and human glomerular disease. J. Clin. Invest. [Internet] 128: 483–499, 2018 Available from: https://pubmed.ncbi.nlm.nih.gov/29227285/ [cited 2023 Jun 9]

47. Jeong H-W, Hernández-Rodríguez B, Kim J, Kim K-P, Enriquez-Gasca R, Yoon J, Adams S, Schöler HR, Vaquerizas JM, Adams RH: Transcriptional regulation of endothelial cell behavior during sprouting angiogenesis. Nat. Commun. 2017 81 [Internet] 8: 1–14, 2017 Available from: https://www.nature.com/articles/s41467-017-00738-7 [cited 2021 Aug 23]

48. Pasquali L, Gaulton KJ, Rodríguez-Seguí SA, Mularoni L, Miguel-Escalada I, Akerman İ, Tena JJ, Morán I, Gómez-Marín C, Bunt M van de Ponsa-Cobas J, Castro N, Nammo T, Cebola I, García-Hurtado J, Maestro MA, Pattou F, Piemonti L, Berney T, Gloyn AL, Ravassard P, Skarmeta JLG, Müller F, McCarthy MI, Ferrer J: Pancreatic islet enhancer clusters enriched in type 2 diabetes risk–associated variants. Nat. Genet. [Internet] 46: 136, 2014 Available from: /pmc/articles/PMC3935450/ [cited 2021 Aug 23]

49. Zaret KS: Pioneer Transcription Factors Initiating Gene Network Changes. Annu. Rev. Genet. [Internet] 54: 367, 2020 Available from: /pmc/articles/PMC7900943/ [cited 2023 Jan 19]

50. Lopez-Pajares V, Qu K, Zhang J, Webster DE, Barajas BC, Siprashvili Z, Zarnegar BJ, Boxer LD, Rios EJ, Tao S, Kretz M, Khavari PA: A LncRNA-MAF:MAFB Transcription Factor Network Regulates Epidermal Differentiation. Dev. Cell 32: 693–706, 2015

