## Supplementary Table 1 for "MAFB drives differentiation by permitting WT1 binding to podocyte specific promoters"

**Analysis Pipeline**

1. Fastq alignment with mm10 -> BAM files
2. MACS2 peakcalling with Input as background (broad option for H3K27me3)
3. Intersect peaks from replicate A and B
4. Intersect common peaks with mm10 blacklist (-v option)
5. Final list for GREAT analysis

| Chrom | Start | End | -LOG10(pvalue) | fold | -LOG10(q-value) | annotation | geneChr | geneStart | geneEnd | geneLength | geneStrand | transcriptId | distanceToTS | geneName | geneId |
|---|---|---|---|---|---|---|---|---|---|---|---|---|---|---|---|
| chr17 | 4626080 | 4626682 | 79.22768 | 354.44785 | 346.18814 | Distal Interge | 17 | 4678020 | 4678126 | 107 | 1 | ENSMUST00 | -51338 | Gm23186 | ENSMUSG00000096200.1 |
| chr8 | 20296814 | 20298046 | 42.50412 | 269.88489 | 262.64221 | Promoter | 8 | 20268286 | 20297432 | 29147 | 2 | ENSMUST00 | 0 | 6820431F20 | ENSMUSG00000071796.8 |
| chr8 | 19784244 | 19785701 | 32.82438 | 269.4866 | 262.2467 | Distal Interge | 8 | 19789798 | 19798548 | 8751 | 1 | ENSMUST00 | -4097 | Gm26853 | ENSMUSG00000097230.1 |
| chr1 | 88276574 | 88278131 | 25.44072 | 266.78333 | 259.59549 | Promoter | 1 | 88277661 | 88308073 | 30413 | 1 | ENSMUST00 | 0 | Trpm8 | ENSMUSG00000036251.11 |
| chr17 | 35393440 | 35395937 | 55.63129 | 266.09174 | 258.92075 | Promoter | 17 | 35394126 | 35395632 | 1507 | 1 | ENSMUST00 | 0 | H2-Q5 | ENSMUSG00000055413.12 |
| chr1 | 178334667 | 178338600 | 62.49637 | 223.48865 | 216.8802 | Promoter | 1 | 178324343 | 178337797 | 13455 | 2 | ENSMUST00 | 0 | Hnrnpu | ENSMUSG00000039630.9 |
| chr17 | 15040898 | 15041535 | 58.36058 | 205.03804 | 198.55309 | Promoter | 17 | 14995787 | 15041537 | 45751 | 2 | ENSMUST00 | 2 | Gm3448 | ENSMUSG00000079710.4 |
| chr17 | 6475994 | 6477493 | 45.40116 | 200.98293 | 194.51932 | Intron (ENSM | 17 | 6441402 | 6467022 | 25621 | 2 | ENSMUST00 | -8973 | Tmem181b- | ENSMUSG00000096780.2 |
| chr8 | 20549763 | 20550962 | 32.37673 | 198.61781 | 192.18805 | Promoter | 8 | 20550331 | 20581999 | 31669 | 1 | ENSMUST00 | 0 | Gm21092 | ENSMUSG00000095403.2 |
| chr17 | 15009455 | 15011012 | 43.8213 | 195.4646 | 189.06622 | Promoter | 17 | 15010266 | 15022448 | 12183 | 1 | ENSMUST00 | 0 | Ermard | ENSMUSG00000036552.11 |
| chr5 | 4191986 | 4193509 | 43.81247 | 194.26276 | 187.87305 | Promoter | 5 | 4192367 | 4197651 | 5285 | 1 | ENSMUST00 | 0 | Mterf1b | ENSMUSG00000053178.5 |
| chr17 | 14960475 | 14962497 | 54.36761 | 193.18201 | 186.79927 | Promoter | 17 | 14945009 | 14961273 | 16265 | 2 | ENSMUST00 | 0 | Phf10 | ENSMUSG00000023883.8 |
| chr14 | 27038953 | 27040019 | 54.02456 | 188.93643 | 182.61591 | Promoter | 14 | 27039001 | 27107286 | 68286 | 1 | ENSMUST00 | 0 | Il17rd | ENSMUSG00000040717.5 |
| chr4 | 126254944 | 126257391 | 48.06229 | 188.02159 | 181.71353 | Promoter | 4 | 126232167 | 126256277 | 24111 | 2 | ENSMUST00 | 0 | Map7d1 | ENSMUSG00000028849.12 |
| chr16 | 96081146 | 96083699 | 52.79799 | 184.38852 | 178.14023 | Promoter | 16 | 95992092 | 96082428 | 90337 | 2 | ENSMUST00 | 0 | Brwd1 | ENSMUSG00000022914.10 |
| chr13 | 98924893 | 98927111 | 50.0437 | 183.17024 | 176.94609 | Promoter | 13 | 98839019 | 98926384 | 87366 | 2 | ENSMUST00 | 0 | Tnpo1 | ENSMUSG00000009470.11 |
| chr3 | 58524418 | 58526697 | 43.8714 | 181.39459 | 175.20454 | Promoter | 3 | 58521969 | 58525873 | 3905 | 2 | ENSMUST00 | 0 | Serp1 | ENSMUSG00000027808.7 |
| chr18 | 31910082 | 31911842 | 50.16549 | 180.98639 | 174.80148 | Promoter | 18 | 31909094 | 31911903 | 2810 | 2 | ENSMUST00 | 61 | Sft2d3 | ENSMUSG00000044982.6 |
| chr16 | 62845942 | 62847074 | 52.11642 | 180.80226 | 174.62196 | Promoter | 16 | 62793685 | 62847040 | 53356 | 2 | ENSMUST00 | 0 | Arl13b | ENSMUSG00000022911.10 |
| chr9 | 104262073 | 104263571 | 50.59046 | 179.33908 | 173.18294 | Promoter | 9 | 104151282 | 104262930 | 111649 | 2 | ENSMUST00 | 0 | Dnajc13 | ENSMUSG00000032560.8 |
| chr9 | 92541328 | 92546074 | 51.17007 | 179.15596 | 173.00638 | Promoter | 9 | 92542223 | 92608428 | 66206 | 1 | ENSMUST00 | 0 | Plod2 | ENSMUSG00000032374.9 |
| chr1 | 178187528 | 178188717 | 46.18486 | 178.62439 | 172.4859 | Promoter | 1 | 178187532 | 178252598 | 65067 | 1 | ENSMUST00 | 0 | Desi2 | ENSMUSG00000026502.8 |
| chr12 | 118300522 | 118302892 | 46.18486 | 178.62439 | 172.4859 | Promoter | 12 | 118234933 | 118301440 | 66508 | 2 | ENSMUST00 | 0 | Sp4 | ENSMUSG00000025323.9 |
| chr16 | 84834034 | 84836397 | 49.45498 | 178.53914 | 172.40457 | Promoter | 16 | 84834925 | 84861315 | 26391 | 1 | ENSMUST00 | 0 | Gabpa | ENSMUSG00000008976.11 |
| chr15 | 94588792 | 94589913 | 51.45567 | 178.45879 | 172.32768 | Promoter | 15 | 94577951 | 94589879 | 11929 | 2 | ENSMUST00 | 0 | Twf1 | ENSMUSG00000022451.7 |
| chr7 | 19663940 | 19666077 | 50.52507 | 176.83885 | 170.7393 | Promoter | 7 | 19631581 | 19665030 | 33450 | 2 | ENSMUST00 | 0 | Clptm1 | ENSMUSG00000002981.9 |
| chr10 | 7779970 | 7781381 | 51.61714 | 176.35683 | 170.26894 | Promoter | 10 | 7767947 | 7780917 | 12971 | 2 | ENSMUST00 | 0 | Ginm1 | ENSMUSG00000040006.8 |
| chr11 | 94211421 | 94215068 | 50.8003 | 176.12823 | 170.0448 | Promoter | 11 | 94211454 | 94215495 | 4042 | 1 | ENSMUST00 | 0 | Tob1 | ENSMUSG00000037573.5 |
| chr18 | 6134705 | 6136106 | 49.92314 | 175.78799 | 169.70897 | Promoter | 18 | 6024785 | 6135907 | 111123 | 2 | ENSMUST00 | 0 | Arhgap12 | ENSMUSG00000041225.11 |
| chr15 | 11994777 | 11996183 | 50.38949 | 175.41335 | 169.3387 | Promoter | 15 | 11981339 | 11996058 | 14720 | 2 | ENSMUST00 | 0 | Sub1 | ENSMUSG00000022205.10 |
| chr13 | 37992943 | 37994742 | 50.25226 | 173.98802 | 167.94014 | Promoter | 13 | 37971401 | 37994190 | 22790 | 2 | ENSMUST00 | 0 | Ssr1 | ENSMUSG00000021427.9 |
| chr2 | 148874090 | 148875697 | 49.19598 | 173.32986 | 167.2899 | Promoter | 2 | 148871722 | 148875576 | 3855 | 2 | ENSMUST00 | 0 | Cst3 | ENSMUSG00000027447.6 |
| chr17 | 14992064 | 14992914 | 51.00806 | 172.93202 | 166.90143 | Downstream | 17 | 14978864 | 14991577 | 12714 | 1 | ENSMUST00 | 13201 | Ermard | ENSMUSG00000036552.11 |
| chr4 | 149424565 | 149427122 | 48.19765 | 172.75977 | 166.73389 | Promoter | 4 | 149328416 | 149426749 | 98334 | 2 | ENSMUST00 | 0 | Ube4b | ENSMUSG00000028960.13 |
| chr18 | 34330397 | 34332264 | 50.11335 | 172.5629 | 166.54207 | Promoter | 18 | 34330847 | 34336345 | 5499 | 1 | ENSMUST00 | 0 | Srp19 | ENSMUSG00000014504.11 |
| chr2 | 164070106 | 164071514 | 50.11335 | 172.5629 | 166.54207 | Promoter | 2 | 164070322 | 164086591 | 16270 | 1 | ENSMUST00 | 0 | Stk4 | ENSMUSG00000018209.10 |
| chr7 | 113207453 | 113208961 | 48.0577 | 171.33241 | 165.33356 | Promoter | 7 | 113207465 | 113314122 | 106658 | 1 | ENSMUST00 | 0 | Arntl | ENSMUSG00000055116.7 |
| chr13 | 59822193 | 59823730 | 50.54853 | 170.95712 | 164.9677 | Promoter | 13 | 59771879 | 59823147 | 51269 | 2 | ENSMUST00 | 0 | Zcchc6 | ENSMUSG00000035248.8 |
| chr17 | 80289467 | 80290712 | 50.54853 | 170.95712 | 164.9677 | Promoter | 17 | 80238304 | 80290476 | 52173 | 2 | ENSMUST00 | 0 | Dhx57 | ENSMUSG00000035051.9 |
| chr4 | 152325753 | 152326885 | 50.54853 | 170.95712 | 164.9677 | Promoter | 4 | 152325853 | 152332480 | 6628 | 1 | ENSMUST00 | 0 | Rpl22 | ENSMUSG00000028936.10 |
| chr8 | 111094098 | 111095587 | 50.54853 | 170.95712 | 164.9677 | Promoter | 8 | 111094630 | 111112579 | 17950 | 1 | ENSMUST00 | 0 | Pdpr | ENSMUSG00000033624.5 |
| chr1 | 93753617 | 93755649 | 47.77854 | 170.84077 | 164.85675 | Promoter | 1 | 93701935 | 93754780 | 52846 | 2 | ENSMUST00 | 0 | Thap4 | ENSMUSG00000026279.9 |
| chr2 | 156990769 | 156992419 | 47.77854 | 170.84077 | 164.85675 | Promoter | 2 | 156927345 | 156992043 | 64699 | 2 | ENSMUST00 | 0 | Ndrg3 | ENSMUSG00000027634.9 |
| chr15 | 98868485 | 98872894 | 47.14861 | 169.72867 | 163.76503 | Promoter | 15 | 98831669 | 98871183 | 39515 | 2 | ENSMUST00 | 0 | Kmt2d | ENSMUSG00000048154.11 |
| chr8 | 41239069 | 41241158 | 48.76767 | 169.04231 | 163.0952 | Promoter | 8 | 41239759 | 41334086 | 94328 | 1 | ENSMUST00 | 0 | Pcm1 | ENSMUSG00000031592.9 |
| chr11 | 121353162 | 121355254 | 47.35943 | 168.9256 | 162.98769 | Promoter | 11 | 121327224 | 121354445 | 27222 | 2 | ENSMUST00 | 0 | Wdr45b | ENSMUSG00000025173.8 |
| chr12 | 73907239 | 73909233 | 47.35943 | 168.9256 | 162.98769 | Promoter | 12 | 73907904 | 73933076 | 25173 | 1 | ENSMUST00 | 0 | Hif1a | ENSMUSG00000021109.8 |
| chr6 | 128760132 | 128761444 | 47.35943 | 168.9256 | 162.98769 | Distal Interge | 6 | 128780043 | 128788486 | 8444 | 2 | ENSMUST00 | 27042 | Klrb1c | ENSMUSG00000030325.11 |
| chr8 | 85071472 | 85072711 | 44.60098 | 167.51549 | 161.60988 | Promoter | 8 | 85071757 | 85075162 | 3406 | 1 | ENSMUST00 | 0 | Dhps | ENSMUSG00000060038.9 |
| chr9 | 37488179 | 37490348 | 44.60098 | 167.51549 | 161.60988 | Promoter | 9 | 37489321 | 37524148 | 34828 | 1 | ENSMUST00 | 0 | Msantd2 | ENSMUSG00000042138.6 |
| chr10 | 79668844 | 79670401 | 49.17356 | 167.40045 | 161.50195 | Promoter | 10 | 79669410 | 79676126 | 6717 | 1 | ENSMUST00 | 0 | Tpgs1 | ENSMUSG00000020308.5 |
| chr11 | 5706711 | 5707980 | 49.62947 | 167.0191 | 161.12611 | Promoter | 11 | 5703983 | 5707687 | 3705 | 2 | ENSMUST00 | 0 | Mrps24 | ENSMUSG00000020477.5 |
| chr13 | 24761042 | 24762251 | 49.62947 | 167.0191 | 161.12611 | Promoter | 13 | 24751845 | 24761861 | 10017 | 2 | ENSMUST00 | 0 | Gmnn | ENSMUSG00000006715.6 |
| chr6 | 122308371 | 122309857 | 46.94032 | 167.01425 | 161.12611 | Promoter | 6 | 122308720 | 122317680 | 8961 | 1 | ENSMUST00 | 0 | M6pr | ENSMUSG00000007458.9 |
| chr17 | 6269849 | 6271411 | 38.84867 | 166.99442 | 161.11731 | Promoter | 17 | 6270475 | 6305783 | 35309 | 1 | ENSMUST00 | 0 | Tmem181a | ENSMUSG00000038141.8 |
| chr6 | 108783011 | 108784750 | 48.77102 | 166.70177 | 160.83063 | Promoter | 6 | 108783059 | 108823723 | 40665 | 1 | ENSMUST00 | 0 | Arl8b | ENSMUSG00000030105.6 |
| chr7 | 143548070 | 143549784 | 45.42985 | 166.67271 | 160.80592 | Promoter | 7 | 143513579 | 143549090 | 35512 | 2 | ENSMUST00 | 0 | Nap1l4 | ENSMUSG00000059119.8 |
| chr4 | 141444174 | 141445618 | 48.03872 | 166.59781 | 160.73184 | Promoter | 4 | 141444654 | 141467225 | 22572 | 1 | ENSMUST00 | 0 | Zbtb17 | ENSMUSG00000006215.7 |
| chr11 | 75461573 | 75463115 | 48.67142 | 166.52875 | 160.66557 | Promoter | 11 | 75461715 | 75469663 | 7949 | 1 | ENSMUST00 | 0 | Tlcd2 | ENSMUSG00000038217.8 |
| chr8 | 34964185 | 34966282 | 48.57222 | 166.35635 | 160.49539 | Promoter | 8 | 34829179 | 34965690 | 136512 | 2 | ENSMUST00 | 0 | Tnks | ENSMUSG00000031529.4 |
| chr17 | 73107245 | 73108937 | 43.83018 | 166.11598 | 160.25778 | Promoter | 17 | 73107985 | 73161991 | 54007 | 1 | ENSMUST00 | 0 | Lclat1 | ENSMUSG00000054469.8 |
| chr16 | 38521905 | 38523495 | 47.75257 | 166.09818 | 160.24055 | Promoter | 16 | 38498347 | 38522663 | 24317 | 2 | ENSMUST00 | 0 | Timmdc1 | ENSMUSG00000002846.9 |
| chr16 | 36875051 | 36876088 | 48.37502 | 166.01349 | 160.15747 | Promoter | 16 | 36875140 | 36887598 | 12459 | 1 | ENSMUST00 | 0 | Golgb1 | ENSMUSG00000034243.11 |
| chr7 | 100120657 | 100122085 | 46.23398 | 165.7672 | 159.91937 | Promoter | 7 | 100082111 | 100121565 | 39455 | 2 | ENSMUST00 | 0 | Pold3 | ENSMUSG00000030726.10 |
| chr19 | 44930334 | 44932028 | 46.79302 | 165.58527 | 159.74297 | Promoter | 19 | 44931119 | 44983787 | 52669 | 1 | ENSMUST00 | 0 | Fam178a | ENSMUSG00000036097.7 |
| chr10 | 78412065 | 78413345 | 47.28315 | 165.27718 | 159.4395 | Promoter | 10 | 78400362 | 78413043 | 12682 | 2 | ENSMUST00 | 0 | Rrp1 | ENSMUSG00000061032.8 |
| chr19 | 56821099 | 56822749 | 44.61662 | 165.21439 | 159.37868 | Promoter | 19 | 56787481 | 56822190 | 34710 | 2 | ENSMUST00 | 0 | A630007B06 | ENSMUSG00000035173.9 |
| chr16 | 10784188 | 10786523 | 46.52121 | 165.10674 | 159.2771 | Promoter | 16 | 10783808 | 10785536 | 1729 | 2 | ENSMUST00 | 0 | Socs1 | ENSMUSG00000038037.4 |
| chr17 | 45571510 | 45574755 | 48.52219 | 165.10075 | 159.2771 | Promoter | 17 | 45567775 | 45573271 | 5497 | 2 | ENSMUST00 | 0 | Hsp90ab1 | ENSMUSG00000023944.9 |
| chr18 | 34758740 | 34760539 | 46.52121 | 165.10674 | 159.2771 | Promoter | 18 | 34758906 | 34773760 | 14855 | 1 | ENSMUST00 | 0 | Fam53c | ENSMUSG00000034300.11 |
| chr8 | 66859836 | 66861417 | 46.52121 | 165.10674 | 159.2771 | Promoter | 8 | 66860217 | 66890564 | 30348 | 1 | ENSMUST00 | 0 | Naf1 | ENSMUSG00000014907.9 |
| chr2 | 33467578 | 33468998 | 49.16994 | 165.05611 | 159.23541 | Promoter | 2 | 33450287 | 33468532 | 18246 | 2 | ENSMUST00 | 0 | Zbtb43 | ENSMUSG00000026788.8 |
| chr4 | 33246794 | 33249051 | 49.16994 | 165.05611 | 159.23541 | Promoter | 4 | 33245423 | 33248510 | 3088 | 2 | ENSMUST00 | 0 | Pnrc1 | ENSMUSG00000040128.9 |
| chr6 | 39376973 | 39378071 | 48.32358 | 164.75595 | 158.94608 | Promoter | 6 | 39334770 | 39377707 | 42938 | 2 | ENSMUST00 | 0 | Slc37a3 | ENSMUSG00000029924.9 |
| chr13 | 23738302 | 23740845 | 31.6659 | 164.62325 | 158.82181 | Promoter | 13 | 23738808 | 23740367 | 1560 | 1 | ENSMUST00 | 0 | Hist1h1c | ENSMUSG00000036181.2 |
| chr11 | 58167749 | 58168721 | 48.1266 | 164.41373 | 158.61591 | Promoter | 11 | 58120002 | 58168539 | 48538 | 2 | ENSMUST00 | 0 | Gemin5 | ENSMUSG00000037275.9 |
| chr16 | 91009983 | 91011976 | 43.18097 | 163.77325 | 157.99008 | Promoter | 16 | 90936092 | 91011308 | 75217 | 2 | ENSMUST00 | 0 | Synj1 | ENSMUSG00000022973.12 |
| chr9 | 62979447 | 62981327 | 43.18097 | 163.77325 | 157.99008 | Promoter | 9 | 62880077 | 62980879 | 100803 | 2 | ENSMUST00 | 0 | Pias1 | ENSMUSG00000032405.9 |
| chr12 | 71015870 | 71017300 | 46.37146 | 163.67729 | 157.89816 | Promoter | 12 | 71015990 | 71098592 | 82603 | 1 | ENSMUST00 | 0 | Arid4a | ENSMUSG00000048118.12 |
| chr2 | 59611415 | 59614138 | 45.5631 | 163.41409 | 157.64189 | Promoter | 2 | 59612042 | 59845717 | 233676 | 1 | ENSMUST00 | 0 | Tanc1 | ENSMUSG00000035168.11 |
| chr11 | 61173977 | 61175124 | 40.47052 | 163.32245 | 157.55278 | Promoter | 11 | 61151785 | 61175055 | 23271 | 2 | ENSMUST00 | 0 | Usp22 | ENSMUSG00000042506.10 |
| chr15 | 77837742 | 77842188 | 46.1021 | 163.20312 | 157.4384 | Promoter | 15 | 77760587 | 77842175 | 81589 | 2 | ENSMUST00 | 0 | Myh9 | ENSMUSG00000022443.11 |
| chr15 | 97246325 | 97248785 | 46.1021 | 163.20312 | 157.4384 | Promoter | 15 | 97247107 | 97385689 | 138583 | 1 | ENSMUST00 | 0 | Pced1b | ENSMUSG00000044250.7 |
| chr9 | 46012067 | 46014680 | 46.1021 | 163.20312 | 157.4384 | Promoter | 9 | 46012820 | 46224194 | 211375 | 1 | ENSMUST00 | 0 | Sik3 | ENSMUSG00000034135.9 |
| chr3 | 129830284 | 129831977 | 48.7104 | 163.09714 | 157.34039 | Promoter | 3 | 129824912 | 129831396 | 6485 | 2 | ENSMUST00 | 0 | Gar1 | ENSMUSG00000028010.11 |
| chr6 | 120835572 | 120837055 | 48.7104 | 163.09714 | 157.34039 | Promoter | 6 | 120836230 | 120892842 | 56613 | 1 | ENSMUST00 | 0 | Bcl2l13 | ENSMUSG00000009112.2 |
| chr5 | 25497212 | 25498824 | 47.35447 | 163.06985 | 157.31989 | Promoter | 5 | 25273609 | 25498745 | 225137 | 2 | ENSMUST00 | 0 | Kmt2c | ENSMUSG00000038056.10 |
| chr8 | 20122172 | 20123103 | 16.79035 | 163.0587 | 157.31015 | Promoter | 8 | 20122365 | 20126624 | 4260 | 1 | ENSMUST00 | 0 | Gm21769 | ENSMUSG00000094614.1 |
| chr14 | 47297691 | 47299602 | 42.71415 | 162.91933 | 157.17317 | Promoter | 14 | 47298314 | 47323091 | 24778 | 1 | ENSMUST00 | 0 | Mapk1ip1l | ENSMUSG00000021840.11 |
| chr7 | 141901297 | 141902486 | 47.16529 | 162.73997 | 156.99878 | Promoter | 7 | 141874813 | 141902415 | 27603 | 2 | ENSMUST00 | 0 | Tollip | ENSMUSG00000025139.8 |
| chr4 | 11485225 | 11487018 | 47.07127 | 162.57588 | 156.84026 | Promoter | 4 | 11485958 | 11527621 | 41664 | 1 | ENSMUST00 | 0 | 1110037F02 | ENSMUSG00000040720.10 |
| chr5 | 145280391 | 145281483 | 47.584 | 162.30655 | 156.57771 | Distal Interge | 5 | 145283343 | 145291469 | 8127 | 1 | ENSMUST00 | -1860 | Zscan25 | ENSMUSG00000070420.3 |
| chr2 | 166905495 | 166907259 | 47.48741 | 162.13861 | 156.41418 | Promoter | 2 | 166906040 | 166944550 | 38511 | 1 | ENSMUST00 | 0 | Cse1l | ENSMUSG00000002718.9 |
| chr4 | 134930934 | 134931820 | 45.49427 | 162.13052 | 156.40686 | Promoter | 4 | 134931010 | 134936078 | 5069 | 1 | ENSMUST00 | 0 | Syf2 | ENSMUSG00000028821.3 |
| chr7 | 47050135 | 47051334 | 45.49427 | 162.13052 | 156.40686 | Promoter | 7 | 47050640 | 47054776 | 4137 | 1 | ENSMUST00 | 0 | Tmem86a | ENSMUSG00000010307.5 |
| chr2 | 39225225 | 39226736 | 28.88857 | 161.91336 | 156.20052 | Promoter | 2 | 39194354 | 39226451 | 32098 | 2 | ENSMUST00 | 0 | Ppp6c | ENSMUSG00000026753.3 |
| chr17 | 35438644 | 35441380 | 29.97639 | 161.56351 | 155.85881 | Promoter | 17 | 35439155 | 35443643 | 4489 | 1 | ENSMUST00 | 0 | H2-Q7 | ENSMUSG00000060550.10 |
| chr10 | 42501601 | 42503518 | 47.82016 | 161.55559 | 155.85097 | Promoter | 10 | 42502030 | 42534787 | 32758 | 1 | ENSMUST00 | 0 | Snx3 | ENSMUSG00000019804.7 |
| chr10 | 93160225 | 93162546 | 40.13043 | 161.53474 | 155.83125 | Promoter | 10 | 93160876 | 93241342 | 80467 | 1 | ENSMUST00 | 0 | Cdk17 | ENSMUSG00000020015.9 |
| chr15 | 80255173 | 80257302 | 45.68299 | 161.30347 | 155.60802 | Promoter | 15 | 80255184 | 80257540 | 2357 | 1 | ENSMUST00 | 0 | Atf4 | ENSMUSG00000042406.7 |
| chr17 | 12317546 | 12319360 | 45.68299 | 161.30347 | 155.60802 | Promoter | 17 | 12227621 | 12318660 | 91040 | 2 | ENSMUST00 | 0 | Map3k4 | ENSMUSG00000014426.8 |
| chr5 | 23431644 | 23437822 | 48.25087 | 161.14223 | 155.45619 | Promoter | 5 | 23434441 | 23504235 | 69795 | 1 | ENSMUST00 | 0 | Kmt2e | ENSMUSG00000029004.10 |
| chr9 | 48984533 | 48986205 | 48.25087 | 161.14223 | 155.45619 | Promoter | 9 | 48985385 | 49042517 | 57133 | 1 | ENSMUST00 | 0 | Usp28 | ENSMUSG00000032267.7 |
| chr6 | 35538743 | 35539878 | 44.23433 | 161.04973 | 155.37192 | Promoter | 6 | 35508906 | 35539888 | 30983 | 2 | ENSMUST00 | 10 | Mtpn | ENSMUSG00000029840.5 |
| chr8 | 20424027 | 20425282 | 27.72379 | 160.87808 | 155.20201 | Promoter | 8 | 20385782 | 20424814 | 39033 | 2 | ENSMUST00 | 0 | 2610005L07 | ENSMUSG00000071793.7 |
| chr7 | 141130877 | 141132105 | 47.4287 | 160.87634 | 155.20065 | Promoter | 7 | 141131286 | 141157021 | 25736 | 1 | ENSMUST00 | 0 | Ptdss2 | ENSMUSG00000025495.9 |
| chr4 | 59625627 | 59627522 | 46.54264 | 160.49254 | 154.82993 | Promoter | 4 | 59626211 | 59761439 | 135229 | 1 | ENSMUST00 | 0 | E130308A19 | ENSMUSG00000045071.8 |
| chr2 | 131351759 | 131353433 | 45.08069 | 160.24072 | 154.5827 | Promoter | 2 | 131298064 | 131352881 | 54818 | 2 | ENSMUST00 | 0 | Rnf24 | ENSMUSG00000048911.10 |
| chr12 | 110600951 | 110602880 | 42.42999 | 160.09743 | 154.44568 | Promoter | 12 | 110601452 | 110666945 | 65494 | 1 | ENSMUST00 | 0 | Dync1h1 | ENSMUSG00000018707.8 |
| chr16 | 57120751 | 57122572 | 42.42999 | 160.09743 | 154.44568 | Promoter | 16 | 57121714 | 57154530 | 32817 | 1 | ENSMUST00 | 0 | Tomm70a | ENSMUSG00000022752.8 |
| chr19 | 29100369 | 29102323 | 42.42999 | 160.09743 | 154.44568 | Promoter | 19 | 29101375 | 29143843 | 42469 | 1 | ENSMUST00 | 0 | Rcl1 | ENSMUSG00000024785.6 |
| chr7 | 28179405 | 28180796 | 42.42999 | 160.09743 | 154.44568 | Promoter | 7 | 28179469 | 28187294 | 7826 | 1 | ENSMUST00 | 0 | Dyrk1b | ENSMUSG00000002409.12 |
| chr7 | 99979459 | 99980922 | 42.42999 | 160.09743 | 154.44568 | Promoter | 7 | 99920257 | 99980458 | 60202 | 2 | ENSMUST00 | 0 | Rnf169 | ENSMUSG00000058761.4 |
| chr18 | 52465053 | 52466791 | 46.9483 | 160.04147 | 154.39299 | Promoter | 18 | 52465693 | 52490738 | 25046 | 1 | ENSMUST00 | 0 | Srfbp1 | ENSMUSG00000024528.7 |
| chr17 | 33289126 | 33290195 | 37.60551 | 160.01111 | 154.36346 | Promoter | 17 | 33289649 | 33303196 | 13548 | 1 | ENSMUST00 | 0 | Zfp955b | ENSMUSG00000096910.1 |
| chr7 | 126976071 | 126977438 | 37.60551 | 160.01111 | 154.36346 | Promoter | 7 | 126976349 | 126980499 | 4151 | 1 | ENSMUST00 | 0 | Cdipt | ENSMUSG00000030682.3 |
| chr11 | 113649012 | 113650540 | 45.52835 | 159.87318 | 154.22893 | Promoter | 11 | 113649169 | 113662947 | 13779 | 1 | ENSMUST00 | 0 | Cog1 | ENSMUSG00000018661.12 |
| chr18 | 77937879 | 77939097 | 46.75885 | 159.71181 | 154.07388 | Promoter | 18 | 77938467 | 78035027 | 96561 | 1 | ENSMUST00 | 0 | Epg5 | ENSMUSG00000039840.7 |
| chr2 | 155276013 | 155277317 | 47.36903 | 159.61537 | 153.98001 | Promoter | 2 | 155276297 | 155278073 | 1777 | 1 | ENSMUST00 | 0 | Map1lc3a | ENSMUSG00000027602.4 |
| chr9 | 35420777 | 35422586 | 47.36903 | 159.61537 | 153.98001 | Promoter | 9 | 35421128 | 35455365 | 34238 | 1 | ENSMUST00 | 0 | Cdon | ENSMUSG00000038119.9 |
| chr14 | 121878075 | 121879743 | 45.99361 | 159.53282 | 153.89999 | Promoter | 14 | 121878606 | 122021034 | 142429 | 1 | ENSMUST00 | 0 | Ubac2 | ENSMUSG00000041765.5 |
| chr4 | 149098516 | 149100205 | 45.26388 | 159.40776 | 153.7765 | Promoter | 4 | 148960535 | 149099876 | 139342 | 2 | ENSMUST00 | 0 | Pex14 | ENSMUSG00000028975.11 |
| chr9 | 63602084 | 63603798 | 45.26388 | 159.40776 | 153.7765 | Promoter | 9 | 63602655 | 63641889 | 39235 | 1 | ENSMUST00 | 0 | Aagab | ENSMUSG00000037257.7 |
| chr5 | 117356598 | 117359461 | 46.57093 | 159.38457 | 153.76053 | Promoter | 5 | 117357304 | 117378601 | 21298 | 1 | ENSMUST00 | 0 | Wsb2 | ENSMUSG00000029364.10 |
| chr2 | 72977775 | 72981250 | 47.79134 | 159.19145 | 153.57301 | Promoter | 2 | 72936427 | 72980446 | 44020 | 2 | ENSMUST00 | 0 | Sp3 | ENSMUSG00000027109.11 |
| chr19 | 46761481 | 46763325 | 44.24375 | 158.75748 | 153.15265 | Promoter | 19 | 46761596 | 46878795 | 117200 | 1 | ENSMUST00 | 0 | Cnnm2 | ENSMUSG00000064105.7 |
| chr10 | 62448773 | 62450405 | 46.01611 | 158.41685 | 152.81914 | Promoter | 10 | 62429209 | 62449738 | 20530 | 2 | ENSMUST00 | 0 | Supv3l1 | ENSMUSG00000020079.10 |

| | | | | | | | | | | | | | | | |
|---|---|---|---|---|---|---|---|---|---|---|---|---|---|---|---|
| chr11 | 50376642 | 50379439 | 45.10679 | 157.97711 | 152.39525 | Promoter | 11 | 50376990 | 50386528 | 9539 | 1 | ENSMUST00 | 0 | Hnrnph1 | ENSMUSG00000007850.11 |
| chr11 | 61929327 | 61931046 | 45.10679 | 157.97711 | 152.39525 | Promoter | 11 | 61871307 | 61930252 | 58946 | 2 | ENSMUST00 | 0 | Akap10 | ENSMUSG00000047804.10 |
| chr9 | 96118703 | 96120086 | 45.10679 | 157.97711 | 152.39525 | Promoter | 9 | 96119362 | 96180190 | 60829 | 1 | ENSMUST00 | 0 | Gk5 | ENSMUSG00000041440.13 |
| chr7 | 55961612 | 55963013 | 46.41137 | 157.95216 | 152.37489 | Promoter | 7 | 55931287 | 55962245 | 30959 | 2 | ENSMUST00 | 0 | Nipa2 | ENSMUSG00000030452.11 |
| chr12 | 3425990 | 3428222 | 44.33166 | 157.76141 | 152.18738 | Promoter | 12 | 3426857 | 3506847 | 79991 | 1 | ENSMUST00 | 0 | Asxl2 | ENSMUSG00000037486.12 |
| chr10 | 81135087 | 81137270 | 45.56376 | 157.62607 | 152.05525 | Promoter | 10 | 81135220 | 81144496 | 9277 | 1 | ENSMUST00 | 0 | Zbtb7a | ENSMUSG00000035011.10 |
| chr9 | 13748500 | 13750046 | 44.84737 | 157.52063 | 151.9514 | Promoter | 9 | 13749181 | 13806481 | 57301 | 1 | ENSMUST00 | 0 | Mtmr2 | ENSMUSG00000031918.11 |
| chr7 | 131409343 | 131411661 | 44.84477 | 157.51604 | 151.94687 | Promoter | 7 | 131410601 | 131446211 | 35611 | 1 | ENSMUST00 | 0 | Acadsb | ENSMUSG00000030861.9 |
| chr11 | 54785734 | 54788212 | 43.43732 | 157.3205 | 151.76292 | Promoter | 11 | 54717456 | 54787675 | 70220 | 2 | ENSMUST00 | 0 | Cdc42se2 | ENSMUSG00000052298.7 |
| chr18 | 34953189 | 34955577 | 43.43732 | 157.3205 | 151.76292 | Promoter | 18 | 34937414 | 34954357 | 16944 | 2 | ENSMUST00 | 0 | Hspa9 | ENSMUSG00000024359.8 |
| chr1 | 86064354 | 86065837 | 46.03907 | 157.30389 | 151.74776 | Promoter | 1 | 86064387 | 86070776 | 6390 | 1 | ENSMUST00 | 0 | Psmd1 | ENSMUSG00000026229.12 |
| chr11 | 32454239 | 32456649 | 46.03907 | 157.30389 | 151.74776 | Promoter | 11 | 32455370 | 32516687 | 61318 | 1 | ENSMUST00 | 0 | Ubtd2 | ENSMUSG00000044949.4 |
| chr2 | 73910284 | 73911845 | 46.03907 | 157.30389 | 151.74776 | Promoter | 2 | 73908447 | 73911325 | 2879 | 2 | ENSMUST00 | 0 | Atp5g3 | ENSMUSG00000018770.4 |
| chr12 | 29937557 | 29939537 | 47.33181 | 157.24483 | 151.69102 | Promoter | 12 | 29938036 | 30017658 | 79623 | 1 | ENSMUST00 | 0 | Pxdn | ENSMUSG00000020674.11 |
| chr8 | 109736843 | 109738166 | 47.33181 | 157.24483 | 151.69102 | Promoter | 8 | 109726451 | 109737739 | 11289 | 2 | ENSMUST00 | 0 | Atxn1l | ENSMUSG00000069895.3 |
| chr3 | 90462429 | 90466691 | 46.53382 | 157.01303 | 151.46753 | Promoter | 3 | 90450591 | 90465866 | 15276 | 2 | ENSMUST00 | 0 | Npr1 | ENSMUSG00000027931.7 |
| chr7 | 108934283 | 108935797 | 46.53382 | 157.01303 | 151.46753 | Promoter | 7 | 108934417 | 108942951 | 8535 | 1 | ENSMUST00 | 0 | Eif3f | ENSMUSG00000031029.6 |
| chr17 | 63498560 | 63500998 | 43.83784 | 156.88544 | 151.34334 | Promoter | 17 | 63057452 | 63500017 | 442566 | 2 | ENSMUST00 | 0 | Fbxl17 | ENSMUSG00000023965.8 |
| chr4 | 129740674 | 129743306 | 43.83784 | 156.88544 | 151.34334 | Promoter | 4 | 129703164 | 129742297 | 39134 | 2 | ENSMUST00 | 0 | Khdrbs1 | ENSMUSG00000028790.8 |
| chr14 | 31336155 | 31337880 | 45.12022 | 156.84895 | 151.30879 | Promoter | 14 | 31336638 | 31371984 | 35347 | 1 | ENSMUST00 | 0 | Capn7 | ENSMUSG00000021893.8 |
| chr11 | 117114277 | 117117694 | 46.43878 | 156.84802 | 151.30803 | Promoter | 11 | 117115168 | 117152185 | 37018 | 1 | ENSMUST00 | 0 | Sec14l1 | ENSMUSG00000020823.11 |
| chr2 | 69722273 | 69724225 | 44.33739 | 156.62123 | 151.08617 | Promoter | 2 | 69722545 | 69723789 | 1245 | 1 | ENSMUST00 | 0 | Ppig | ENSMUSG00000042133.11 |
| chr8 | 26118269 | 26120274 | 44.25352 | 156.47305 | 150.94255 | Promoter | 8 | 26119368 | 26143871 | 24504 | 1 | ENSMUST00 | 0 | Rnf170 | ENSMUSG00000013878.12 |
| chr1 | 87755464 | 87757081 | 41.67902 | 156.4369 | 150.90947 | Promoter | 1 | 87755870 | 87771398 | 15529 | 1 | ENSMUST00 | 0 | Atg16l1 | ENSMUSG00000026289.10 |
| chr11 | 94320406 | 94322646 | 41.67902 | 156.4369 | 150.90947 | Promoter | 11 | 94287890 | 94321958 | 34069 | 2 | ENSMUST00 | 0 | Luc7l3 | ENSMUSG00000020863.10 |
| chr7 | 63935484 | 63939001 | 41.67902 | 156.4369 | 150.90947 | Promoter | 7 | 63886347 | 63938915 | 52569 | 2 | ENSMUST00 | 0 | Klf13 | ENSMUSG00000052040.9 |
| chr14 | 64652037 | 64653671 | 42.29403 | 156.41156 | 150.88818 | Promoter | 14 | 64652531 | 64806296 | 153766 | 1 | ENSMUST00 | 0 | Kif13b | ENSMUSG00000060012.7 |
| chr16 | 33250592 | 33252470 | 45.49167 | 156.34886 | 150.8266 | Promoter | 16 | 33251456 | 33299562 | 48107 | 1 | ENSMUST00 | 0 | Snx4 | ENSMUSG00000022808.4 |
| chr10 | 81320912 | 81322806 | 46.06248 | 156.19411 | 150.67332 | Promoter | 10 | 81321103 | 81326251 | 5149 | 1 | ENSMUST00 | 0 | Cactin | ENSMUSG00000034889.7 |
| chr19 | 45781513 | 45783689 | 39.11016 | 156.19371 | 150.67332 | Promoter | 19 | 45750261 | 45783520 | 33260 | 2 | ENSMUST00 | 0 | Mgea5 | ENSMUSG00000025220.7 |
| chr4 | 121016006 | 121017644 | 46.06248 | 156.19411 | 150.67332 | Promoter | 4 | 120968317 | 121017247 | 48931 | 2 | ENSMUST00 | 0 | Smap2 | ENSMUSG00000032870.8 |
| chr6 | 88083933 | 88085458 | 39.11016 | 156.19371 | 150.67332 | Promoter | 6 | 88084473 | 88105304 | 20832 | 1 | ENSMUST00 | 0 | Rpn1 | ENSMUSG00000030062.6 |
| chr13 | 69611916 | 69612758 | 44.68523 | 156.08508 | 150.56929 | Promoter | 13 | 69612016 | 69634334 | 22319 | 1 | ENSMUST00 | 0 | Nsun2 | ENSMUSG00000021595.13 |
| chr7 | 128744460 | 128746030 | 44.68523 | 156.08508 | 150.56929 | Promoter | 7 | 128744870 | 128784836 | 39967 | 1 | ENSMUST00 | 0 | Sec23ip | ENSMUSG00000055319.7 |
| chr17 | 35163257 | 35164931 | 45.22282 | 155.87894 | 150.36807 | Promoter | 17 | 35149076 | 35164897 | 15822 | 2 | ENSMUST00 | 0 | Prrc2a | ENSMUSG00000024393.9 |
| chr2 | 91729790 | 91731263 | 45.22282 | 155.87894 | 150.36807 | Promoter | 2 | 91730134 | 91918849 | 188716 | 1 | ENSMUST00 | 0 | Ambra1 | ENSMUSG00000040506.11 |
| chr8 | 40510828 | 40512559 | 43.03881 | 155.46181 | 149.96747 | Promoter | 8 | 40511783 | 40551134 | 39352 | 1 | ENSMUST00 | 0 | Vps37a | ENSMUSG00000031600.10 |
| chr12 | 85110097 | 85111989 | 45.50933 | 155.23103 | 149.75105 | Promoter | 12 | 85110833 | 85134091 | 23259 | 1 | ENSMUST00 | 0 | Dlst | ENSMUSG00000004789.9 |
| chr19 | 53142680 | 53143596 | 41.5631 | 155.08545 | 149.60895 | Promoter | 19 | 53142756 | 53247399 | 104644 | 1 | ENSMUST00 | 0 | Add3 | ENSMUSG00000025026.9 |
| chr8 | 83697311 | 83699766 | 46.69001 | 154.98724 | 149.51399 | Promoter | 8 | 83669762 | 83699179 | 29418 | 2 | ENSMUST00 | 0 | Pkn1 | ENSMUSG00000057672.10 |
| chr1 | 131910423 | 131911726 | 46.59246 | 154.81853 | 149.34816 | Promoter | 1 | 131910534 | 131932679 | 22146 | 1 | ENSMUST00 | 0 | Nucks1 | ENSMUSG00000026434.7 |
| chr18 | 44659797 | 44662967 | 44.60771 | 154.80135 | 149.33388 | Promoter | 18 | 44425061 | 44662266 | 237206 | 2 | ENSMUST00 | 0 | Mcc | ENSMUSG00000071856.5 |
| chr11 | 105126417 | 105127420 | 46.49532 | 154.65041 | 149.18355 | Promoter | 11 | 105126425 | 105140394 | 13970 | 1 | ENSMUST00 | 0 | Mettl2 | ENSMUSG00000020691.8 |
| chr8 | 111939834 | 111941588 | 37.1464 | 154.63618 | 149.16937 | Promoter | 8 | 111940703 | 111953612 | 12910 | 1 | ENSMUST00 | 0 | Gabarapl2 | ENSMUSG00000031950.7 |
| chr1 | 189006835 | 189008219 | 41.30354 | 154.61249 | 149.14572 | Promoter | 1 | 188971098 | 189007840 | 36743 | 2 | ENSMUST00 | 0 | Kctd3 | ENSMUSG00000026608.7 |
| chr15 | 80710752 | 80712434 | 41.30354 | 154.61249 | 149.14572 | Promoter | 15 | 80711319 | 80941086 | 229768 | 1 | ENSMUST00 | 0 | Tnrc6b | ENSMUSG00000047888.9 |
| chr19 | 5490396 | 5492187 | 41.30354 | 154.61249 | 149.14572 | Promoter | 19 | 5490485 | 5494029 | 3545 | 1 | ENSMUST00 | 0 | Cfl1 | ENSMUSG00000056201.6 |
| chr4 | 3938776 | 3939740 | 41.30354 | 154.61249 | 149.14572 | Promoter | 4 | 3938888 | 3943523 | 4636 | 1 | ENSMUST00 | 0 | Chchd7 | ENSMUSG00000042198.7 |
| chr4 | 132535479 | 132536616 | 41.30354 | 154.61249 | 149.14572 | Promoter | 4 | 132535550 | 132546646 | 11097 | 1 | ENSMUST00 | 0 | Dnajc8 | ENSMUSG00000054405.9 |
| chr1 | 182408150 | 182410996 | 46.39859 | 154.48299 | 149.02768 | Promoter | 1 | 182409172 | 182462432 | 53261 | 1 | ENSMUST00 | 0 | Trp53bp2 | ENSMUSG00000026510.9 |
| chr17 | 80726713 | 80728878 | 38.77008 | 154.42084 | 148.9682 | Promoter | 17 | 80580512 | 80728093 | 147582 | 2 | ENSMUST00 | 0 | Map4k3 | ENSMUSG00000024242.9 |
| chr4 | 127243709 | 127244760 | 38.77008 | 154.42084 | 148.9682 | Promoter | 4 | 127243784 | 127247809 | 4026 | 1 | ENSMUST00 | 0 | Smim12 | ENSMUSG00000042380.8 |
| chr8 | 94837445 | 94839091 | 38.77008 | 154.42084 | 148.9682 | Promoter | 8 | 94819818 | 94838358 | 18541 | 2 | ENSMUST00 | 0 | Ciapin1 | ENSMUSG00000031781.8 |
| chr18 | 76059035 | 76063179 | 44.34918 | 154.34741 | 148.89743 | Promoter | 18 | 76059458 | 76146443 | 86986 | 1 | ENSMUST00 | 0 | Zbtb7c | ENSMUSG00000044646.9 |
| chr2 | 125858259 | 125859691 | 44.9693 | 154.28859 | 148.83946 | Promoter | 2 | 125830304 | 125859139 | 28836 | 2 | ENSMUST00 | 0 | Cops2 | ENSMUSG00000027206.8 |
| chr16 | 55973101 | 55975647 | 44.26367 | 154.19713 | 148.75002 | Promoter | 16 | 55973883 | 56008913 | 35031 | 1 | ENSMUST00 | 0 | Zbtb11 | ENSMUSG00000022601.11 |
| chr16 | 38088967 | 38093101 | 44.26367 | 154.19713 | 148.75002 | Promoter | 16 | 38089001 | 38246084 | 157084 | 1 | ENSMUST00 | 0 | Gsk3b | ENSMUSG00000022812.8 |
| chr4 | 94555818 | 94557256 | 44.26367 | 154.19713 | 148.75002 | Promoter | 4 | 94500081 | 94556796 | 56716 | 2 | ENSMUST00 | 0 | Caap1 | ENSMUSG00000028578.8 |
| chr2 | 75702478 | 75705502 | 45.52735 | 154.11624 | 148.67334 | Promoter | 2 | 75675513 | 75704641 | 29129 | 2 | ENSMUST00 | 0 | Nfe2l2 | ENSMUSG00000015839.6 |
| chr7 | 16815903 | 16817616 | 45.52735 | 154.11624 | 148.67334 | Promoter | 7 | 16816317 | 16840927 | 24611 | 1 | ENSMUST00 | 0 | Strn4 | ENSMUSG00000030374.10 |
| chr6 | 113739771 | 113741011 | 45.43549 | 153.95641 | 148.5168 | Promoter | 6 | 113728067 | 113740743 | 12677 | 2 | ENSMUST00 | 0 | Sec13 | ENSMUSG00000030298.5 |
| chr14 | 57998533 | 57999620 | 44.70407 | 153.82483 | 148.38618 | Promoter | 14 | 57916280 | 57999262 | 82983 | 2 | ENSMUST00 | 0 | Micu2 | ENSMUSG00000021973.8 |
| chr10 | 128034885 | 128037186 | 46.01562 | 153.81966 | 148.38319 | Promoter | 10 | 128035346 | 128048637 | 13292 | 1 | ENSMUST00 | 0 | Naca | ENSMUSG00000061315.8 |
| chr5 | 134175699 | 134176820 | 46.01562 | 153.81966 | 148.38319 | Promoter | 5 | 134148054 | 134176774 | 28721 | 2 | ENSMUST00 | 0 | Wbscr16 | ENSMUSG00000061979.3 |
| chr14 | 26579063 | 26580603 | 44.00655 | 153.74478 | 148.311 | Promoter | 14 | 26579578 | 26633394 | 53817 | 1 | ENSMUST00 | 0 | Dennd6a | ENSMUSG00000040818.7 |
| chr3 | 137917963 | 137919376 | 44.00655 | 153.74478 | 148.311 | Promoter | 3 | 137918526 | 137928765 | 10240 | 1 | ENSMUST00 | 0 | Lamtor3 | ENSMUSG00000091512.1 |
| chr12 | 87146697 | 87148637 | 42.6403 | 153.60713 | 148.18196 | Promoter | 12 | 87147707 | 87164721 | 17015 | 1 | ENSMUST00 | 0 | Gstz1 | ENSMUSG00000021033.10 |
| chr4 | 12088959 | 12090339 | 32.87366 | 153.46529 | 148.04802 | Promoter | 4 | 12039645 | 12090020 | 50376 | 2 | ENSMUST00 | 0 | Tmem67 | ENSMUSG00000049488.9 |
| chr16 | 44138136 | 44141263 | 46.41274 | 153.3642 | 147.94775 | Promoter | 16 | 44139830 | 44162474 | 22645 | 1 | ENSMUST00 | 0 | Naa50 | ENSMUSG00000022698.12 |
| chr12 | 108178081 | 108179348 | 43.6742 | 153.15909 | 147.75397 | Promoter | 12 | 108106431 | 108179314 | 72884 | 2 | ENSMUST00 | 0 | Setd3 | ENSMUSG00000056770.10 |
| chr2 | 30966825 | 30968309 | 43.02603 | 153.1534 | 147.74835 | Promoter | 2 | 30960627 | 30967933 | 7307 | 2 | ENSMUST00 | 0 | Tor1a | ENSMUSG00000026849.13 |
| chr7 | 105735728 | 105738289 | 44.26889 | 153.06256 | 147.66058 | Promoter | 7 | 105732207 | 105736781 | 4575 | 2 | ENSMUST00 | 0 | Rrp8 | ENSMUSG00000030888.8 |
| chr10 | 95514318 | 95516303 | 45.54573 | 153.00453 | 147.60376 | Promoter | 10 | 95515162 | 95545658 | 30497 | 1 | ENSMUST00 | 0 | Ube2n | ENSMUSG00000074781.4 |
| chr14 | 54360471 | 54361828 | 45.54573 | 153.00453 | 147.60376 | Promoter | 14 | 54360841 | 54369669 | 8829 | 1 | ENSMUST00 | 0 | Oxa1l | ENSMUSG00000000959.5 |
| chr13 | 55104952 | 55106061 | 40.92805 | 152.79198 | 147.3963 | Promoter | 13 | 55105311 | 55122478 | 17168 | 1 | ENSMUST00 | 0 | Zfp346 | ENSMUSG00000021481.6 |
| chr3 | 27708580 | 27711670 | 41.52505 | 152.7413 | 147.35226 | Promoter | 3 | 27416162 | 27710439 | 294278 | 2 | ENSMUST00 | 0 | Fndc3b | ENSMUSG00000039286.7 |
| chr11 | 64978947 | 64980027 | 43.42635 | 152.72153 | 147.3335 | Promoter | 11 | 64979038 | 65002069 | 23032 | 1 | ENSMUST00 | 0 | Elac2 | ENSMUSG00000020549.9 |
| chr4 | 148590542 | 148592352 | 43.42635 | 152.72153 | 147.3335 | Promoter | 4 | 148591503 | 148594619 | 3117 | 1 | ENSMUST00 | 0 | Srm | ENSMUSG00000006442.10 |
| chr3 | 19162104 | 19163829 | 44.71377 | 152.69855 | 147.31396 | Promoter | 3 | 19132146 | 19163065 | 30920 | 2 | ENSMUST00 | 0 | Armc1 | ENSMUSG00000027599.8 |
| chr9 | 72984801 | 72986380 | 44.71377 | 152.69855 | 147.31396 | Promoter | 9 | 72985429 | 73016340 | 30912 | 1 | ENSMUST00 | 0 | Ccpg1 | ENSMUSG00000034563.11 |
| chr6 | 38533951 | 38535632 | 38.42999 | 152.65175 | 147.26898 | Promoter | 6 | 38534823 | 38539449 | 4627 | 1 | ENSMUST00 | 0 | 1110001J03R | ENSMUSG00000019689.4 |
| chr7 | 100607615 | 100609354 | 38.42999 | 152.65175 | 147.26898 | Promoter | 7 | 100607690 | 100641268 | 33579 | 1 | ENSMUST00 | 0 | Rab6a | ENSMUSG00000030704.9 |
| chr7 | 97578774 | 97581491 | 38.42999 | 152.65175 | 147.26898 | Promoter | 7 | 97579896 | 97692782 | 112887 | 1 | ENSMUST00 | 0 | Rsf1 | ENSMUSG00000035623.9 |
| chr8 | 111144711 | 111146229 | 38.42999 | 152.65175 | 147.26898 | Promoter | 8 | 111128886 | 111145459 | 16574 | 2 | ENSMUST00 | 0 | 9430091E24I | ENSMUSG00000084808.2 |
| chr11 | 5520607 | 5522588 | 43.84211 | 152.31328 | 146.93933 | Promoter | 11 | 5520659 | 5525893 | 5235 | 1 | ENSMUST00 | 0 | Xbp1 | ENSMUSG00000020484.13 |
| chr10 | 80560439 | 80562186 | 45.75331 | 152.2233 | 146.85388 | Promoter | 10 | 80540925 | 80561560 | 20636 | 2 | ENSMUST00 | 0 | Rexo1 | ENSMUSG00000047417.12 |
| chr14 | 101653775 | 101654926 | 45.08533 | 152.20454 | 146.8354 | Promoter | 14 | 101653967 | 101696124 | 42158 | 1 | ENSMUST00 | 0 | Uchl3 | ENSMUSG00000022111.8 |
| chr17 | 15499114 | 15500699 | 38.08887 | 152.00154 | 146.63557 | Promoter | 17 | 15499888 | 15504605 | 4718 | 1 | ENSMUST00 | 0 | Tbp | ENSMUSG00000014767.11 |
| chr8 | 111299240 | 111300356 | 40.48558 | 151.98326 | 146.61765 | Promoter | 8 | 111270944 | 111300222 | 29279 | 2 | ENSMUST00 | 0 | Rfwd3 | ENSMUSG00000033596.4 |
| chr10 | 71237106 | 71238605 | 44.27422 | 151.93042 | 146.56525 | Promoter | 10 | 71225464 | 71238280 | 12817 | 2 | ENSMUST00 | 0 | Tfam | ENSMUSG00000003923.9 |
| chr2 | 28640968 | 28642143 | 44.90377 | 151.88861 | 146.52586 | Promoter | 2 | 28641228 | 28672507 | 31280 | 1 | ENSMUST00 | 0 | Tsc1 | ENSMUSG00000026812.11 |
| chr1 | 24677760 | 24679595 | 43.58744 | 151.86528 | 146.50356 | Promoter | 1 | 24678630 | 24765448 | 86819 | 1 | ENSMUST00 | 0 | Lmbrd1 | ENSMUSG00000073725.3 |
| chr11 | 69088066 | 69090646 | 43.58744 | 151.86528 | 146.50356 | Promoter | 11 | 69088490 | 69092384 | 3895 | 1 | ENSMUST00 | 0 | Vamp2 | ENSMUSG00000020894.11 |
| chr12 | 51690328 | 51693137 | 43.58744 | 151.86528 | 146.50356 | Promoter | 12 | 51608541 | 51691914 | 83374 | 2 | ENSMUST00 | 0 | Strn3 | ENSMUSG00000020954.10 |
| chr13 | 54686089 | 54688786 | 43.58744 | 151.86528 | 146.50356 | Promoter | 13 | 54679399 | 54687808 | 8410 | 2 | ENSMUST00 | 0 | Rnf44 | ENSMUSG00000034928.11 |
| chr17 | 28690674 | 28692802 | 43.58744 | 151.86528 | 146.50356 | Promoter | 17 | 28691342 | 28741164 | 49823 | 1 | ENSMUST00 | 0 | Mapk14 | ENSMUSG00000053436.9 |
| chr2 | 164878547 | 164879851 | 43.58744 | 151.86528 | 146.50356 | Promoter | 2 | 164879304 | 164883199 | 3896 | 1 | ENSMUST00 | 0 | Pcif1 | ENSMUSG00000039849.5 |
| chr3 | 152209361 | 152212419 | 43.58744 | 151.86528 | 146.50356 | Promoter | 3 | 152210458 | 152236830 | 26373 | 1 | ENSMUST00 | 0 | Fubp1 | ENSMUSG00000028034.10 |
| chr4 | 132074095 | 132075741 | 43.58744 | 151.86528 | 146.50356 | Promoter | 4 | 131923413 | 132075292 | 151880 | 2 | ENSMUST00 | 0 | Epb4.1 | ENSMUSG00000028906.10 |
| chr5 | 125388036 | 125390303 | 44.18734 | 151.7784 | 146.42473 | Promoter | 5 | 125385965 | 125390014 | 4050 | 2 | ENSMUST00 | 0 | Ubc | ENSMUSG00000008348.9 |
| chr2 | 174326582 | 174332166 | 45.47067 | 151.73354 | 146.38181 | Promoter | 2 | 174327570 | 174345241 | 17672 | 1 | ENSMUST00 | 0 | Gnas | ENSMUSG00000027523.14 |
| chr4 | 114904677 | 114911326 | 45.47067 | 151.73354 | 146.38181 | Promoter | 4 | 114906282 | 114908873 | 2592 | 2 | ENSMUST00 | 0 | Foxd2 | ENSMUSG00000055210.4 |
| chr9 | 66349776 | 66351501 | 45.47067 | 151.73354 | 146.38181 | Promoter | 9 | 66350450 | 66374070 | 23621 | 1 | ENSMUST00 | 0 | Herc1 | ENSMUSG00000038664.11 |
| chr16 | 13671401 | 13672923 | 43.50657 | 151.72289 | 146.37334 | Promoter | 16 | 13671858 | 13703612 | 31755 | 1 | ENSMUST00 | 0 | Bfar | ENSMUSG00000022684.9 |
| chr2 | 37451563 | 37453152 | 43.50657 | 151.72289 | 146.37334 | Promoter | 2 | 37452245 | 37475363 | 23119 | 1 | ENSMUST00 | 0 | Rabgap1 | ENSMUSG00000035437.10 |
| chr14 | 26532901 | 26535352 | 44.72366 | 151.575 | 146.22792 | Promoter | 14 | 26413168 | 26534931 | 121764 | 2 | ENSMUST00 | 0 | Slmap | ENSMUSG00000021870.12 |
| chr16 | 17927209 | 17928907 | 44.72366 | 151.575 | 146.22792 | Promoter | 16 | 17925211 | 17928219 | 3009 | 2 | ENSMUST00 | 0 | Slc25a1 | ENSMUSG00000003528.9 |
| chr6 | 56714320 | 56716114 | 44.72366 | 151.575 | 146.22792 | Promoter | 6 | 56714927 | 56761911 | 46985 | 1 | ENSMUST00 | 0 | Avl9 | ENSMUSG00000029787.8 |
| chr6 | 117906713 | 117908804 | 44.72366 | 151.575 | 146.22792 | Promoter | 6 | 117906790 | 117923798 | 17009 | 1 | ENSMUST00 | 0 | Hnrnpf | ENSMUSG00000042079.8 |
| chr12 | 55229467 | 55231445 | 45.95321 | 151.4303 | 146.08812 | Distal Interge | 12 | 55239326 | 55263020 | 23695 | 1 | ENSMUST00 | -7881 | Srp54b | ENSMUSG00000079108.5 |
| chr17 | 47592999 | 47595153 | 45.95321 | 151.4303 | 146.08812 | Promoter | 17 | 47593436 | 47598174 | 4739 | 1 | ENSMUST00 | 0 | Ccnd3 | ENSMUSG00000034165.11 |
| chr17 | 15023482 | 15025305 | 45.95321 | 151.4303 | 146.08812 | Intron (ENSM | 17 | 15010266 | 15022448 | 12183 | 1 | ENSMUST00 | 13217 | Ermard | ENSMUSG00000036552.11 |
| chr8 | 81013302 | 81014990 | 45.95321 | 151.4303 | 146.08812 | Promoter | 8 | 81013576 | 81038584 | 25009 | 1 | ENSMUST00 | 0 | Gm9725 | ENSMUSG00000037982.10 |
| chr11 | 118247199 | 118248609 | 42.62012 | 151.29346 | 145.9586 | Promoter | 11 | 118132019 | 118248502 | 116484 | 2 | ENSMUST00 | 0 | Cyth1 | ENSMUSG00000017132.12 |
| chr3 | 139073799 | 139075230 | 42.62012 | 151.29346 | 145.9586 | Promoter | 3 | 138925906 | 139075199 | 149294 | 2 | ENSMUST00 | 0 | Rap1gds1 | ENSMUSG00000028149.7 |
| chr4 | 98382241 | 98383502 | 45.87056 | 151.28743 | 145.95461 | Promoter | 4 | 98355370 | 98383060 | 27691 | 2 | ENSMUST00 | 0 | Tm2d1 | ENSMUSG00000028563.11 |
| chr17 | 25013319 | 25015876 | 45.1915 | 151.24933 | 145.91733 | Promoter | 17 | 24961228 | 25015230 | 54003 | 2 | ENSMUST00 | 0 | Cramp1l | ENSMUSG00000038002.8 |
| chr13 | 95374547 | 95375851 | 43.84323 | 151.17558 | 145.84569 | Promoter | 13 | 95350683 | 95375352 | 24670 | 2 | ENSMUST00 | 0 | Aggf1 | ENSMUSG00000021681.8 |
| chr8 | 25784255 | 25786297 | 44.4562 | 151.10883 | 145.78059 | Promoter | 8 | 25785591 | 25803975 | 18385 | 1 | ENSMUST00 | 0 | Lsm1 | ENSMUSG00000037296.6 |
| chr1 | 185362319 | 185364185 | 40.55256 | 150.97545 | 145.64915 | Promoter | 1 | 185363095 | 185428355 | 65261 | 1 | ENSMUST00 | 0 | Eprs | ENSMUSG00000026615.9 |
| chr11 | 97439863 | 97443213 | 40.55256 | 150.97545 | 145.64915 | Intron (ENSM | 11 | 97436285 | 97444228 | 7944 | 1 | ENSMUST00 | 3579 | Arhgap23 | ENSMUSG00000049807.11 |
| chr17 | 55944460 | 55945971 | 40.55256 | 150.97545 | 145.64915 | Promoter | 17 | 55938381 | 55945259 | 6879 | 2 | ENSMUST00 | 0 | Zfp119b | ENSMUSG00000062101.7 |
| chr9 | 108516286 | 108518500 | 45.67889 | 150.95599 | 145.63583 | Promoter | 9 | 108517087 | 108560167 | 43081 | 1 | ENSMUST00 | 0 | Qrich1 | ENSMUSG00000006673.9 |
| chr5 | 115237429 | 115238970 | 44.36775 | 150.95453 | 145.63559 | Promoter | 5 | 115237979 | 115240242 | 2264 | 1 | ENSMUST00 | 0 | Pop5 | ENSMUSG00000060152.9 |
| chr15 | 102278807 | 102280361 | 41.14056 | 150.91219 | 145.59421 | Promoter | 15 | 102279456 | 102281744 | 2289 | 1 | ENSMUST00 | 0 | Mfsd5 | ENSMUSG00000045665.4 |
| chr8 | 69715689 | 69716993 | 38.0899 | 150.88651 | 145.56995 | Distal Interge | 8 | 69706332 | 69710724 | 4393 | 2 | ENSMUST00 | -4966 | Zfp869 | ENSMUSG00000054648.7 |
| chr1 | 170884514 | 170885772 | 43.01277 | 150.85193 | 145.53938 | Promoter | 1 | 170873498 | 170885457 | 11960 | 2 | ENSMUST00 | 0 | Dusp12 | ENSMUSG00000026659.8 |
| chr10 | 81349311 | 81350520 | 43.01277 | 150.85193 | 145.53938 | Promoter | 10 | 81346048 | 81350480 | 4433 | 2 | ENSMUST00 | 0 | Hmg20b | ENSMUSG00000020232.12 |

| | | | | | | | | | | | | | | | |
|---|---|---|---|---|---|---|---|---|---|---|---|---|---|---|---|
| chr18 | 39488384 | 39492046 | 43.01277 | 150.85193 | 145.53938 | Promoter | 18 | 39410545 | 39490674 | 80130 | 2 | ENSMUST00 | 0 | Nr3c1 | ENSMUSG00000024431.9 |
| chr10 | 128083259 | 128084528 | 44.27966 | 150.80078 | 145.49101 | Promoter | 10 | 128083273 | 128090391 | 7119 | 1 | ENSMUST00 | 0 | Atp5b | ENSMUSG00000025393.7 |
| chr5 | 41706910 | 41708875 | 45.58365 | 150.79124 | 145.48296 | Promoter | 5 | 41624976 | 41708155 | 83180 | 2 | ENSMUST00 | 0 | Rab28 | ENSMUSG00000029128.9 |
| chr1 | 171037112 | 171038718 | 44.91574 | 150.77055 | 145.46295 | Distal Interge | 1 | 171027965 | 171028129 | 165 | 1 | ENSMUST00 | 9148 | Gm26110 | ENSMUSG00000065905.1 |
| chr2 | 122117578 | 122118852 | 39.80928 | 150.74051 | 145.43391 | Promoter | 2 | 122053520 | 122118386 | 64867 | 2 | ENSMUST00 | 0 | Spg11 | ENSMUSG00000033396.8 |
| chr9 | 108001749 | 108004053 | 42.93156 | 150.70844 | 145.40213 | Promoter | 9 | 108002501 | 108048780 | 46280 | 1 | ENSMUST00 | 0 | Ip6k1 | ENSMUSG00000032594.6 |
| chr2 | 34869918 | 34871751 | 44.10451 | 150.49493 | 145.19151 | Promoter | 2 | 34852088 | 34870939 | 18852 | 2 | ENSMUST00 | 0 | Psmd5 | ENSMUSG00000026869.7 |
| chr1 | 43444845 | 43448373 | 43.42055 | 150.43358 | 145.13264 | Promoter | 1 | 43445751 | 43462194 | 16444 | 1 | ENSMUST00 | 0 | Nck2 | ENSMUSG00000066877.6 |
| chr11 | 95309751 | 95310300 | 43.42055 | 150.43358 | 145.13264 | Promoter | 11 | 95274259 | 95310201 | 35943 | 2 | ENSMUST00 | 0 | Kat7 | ENSMUSG00000038909.11 |
| chr17 | 42874504 | 42877282 | 43.42055 | 150.43358 | 145.13264 | Promoter | 17 | 42792951 | 42876424 | 83474 | 2 | ENSMUST00 | 0 | Cd2ap | ENSMUSG00000061665.6 |
| chr2 | 170511215 | 170512373 | 43.42055 | 150.43358 | 145.13264 | Promoter | 2 | 170511235 | 170519123 | 7889 | 1 | ENSMUST00 | 0 | Pfdn4 | ENSMUSG00000052033.8 |
| chr2 | 94405707 | 94407604 | 44.01745 | 150.34283 | 145.04555 | Promoter | 2 | 94300767 | 94406679 | 105913 | 2 | ENSMUST00 | 0 | Ttc17 | ENSMUSG00000027194.11 |
| chr15 | 34237454 | 34239100 | 45.30031 | 150.3008 | 145.00395 | Promoter | 15 | 34238026 | 34284295 | 46270 | 1 | ENSMUST00 | 0 | Laptm4b | ENSMUSG00000022257.3 |
| chr17 | 6429668 | 6430969 | 29.58566 | 150.20636 | 144.911 | Promoter | 17 | 6430112 | 6436293 | 6182 | 1 | ENSMUST00 | 0 | Dynlt1b | ENSMUSG00000096255.1 |
| chr16 | 20097199 | 20098659 | 32.34344 | 150.1674 | 144.87335 | Promoter | 16 | 20097554 | 20127744 | 30191 | 1 | ENSMUST00 | 0 | Klhl24 | ENSMUSG00000062901.2 |
| chr19 | 12793084 | 12795521 | 45.20664 | 150.13857 | 144.8454 | Promoter | 19 | 12763660 | 12796126 | 32467 | 2 | ENSMUST00 | 605 | Zfp91 | ENSMUSG00000024695.10 |
| chr1 | 10231713 | 10233939 | 43.16833 | 149.98996 | 144.69951 | Promoter | 1 | 10137571 | 10232670 | 95100 | 2 | ENSMUST00 | 0 | Arfgef1 | ENSMUSG00000067851.6 |
| chr17 | 6777892 | 6783808 | 43.16833 | 149.98996 | 144.69951 | Promoter | 17 | 6738041 | 6782784 | 44744 | 2 | ENSMUST00 | 0 | Ezr | ENSMUSG00000052397.8 |
| chr6 | 88517663 | 88519629 | 44.46354 | 149.9843 | 144.69951 | Promoter | 6 | 88503601 | 88518905 | 15305 | 2 | ENSMUST00 | 0 | Sec61a1 | ENSMUSG00000030082.9 |
| chr7 | 81762203 | 81763860 | 43.16833 | 149.98996 | 144.69951 | Promoter | 7 | 81762925 | 81769491 | 6567 | 1 | ENSMUST00 | 0 | Fam103a1 | ENSMUSG00000038646.8 |
| chr15 | 79670476 | 79671513 | 41.84328 | 149.90991 | 144.6302 | Promoter | 15 | 79670861 | 79672862 | 2002 | 1 | ENSMUST00 | 0 | Tomm22 | ENSMUSG00000022427.3 |
| chr3 | 66985288 | 66986374 | 45.49368 | 149.5007 | 144.23251 | Promoter | 3 | 66985670 | 66996473 | 10804 | 1 | ENSMUST00 | 0 | Rsrc1 | ENSMUSG00000034544.12 |
| chr11 | 87126333 | 87127988 | 43.50226 | 149.44124 | 144.17897 | Promoter | 11 | 87127077 | 87220683 | 93607 | 1 | ENSMUST00 | 0 | Trim37 | ENSMUSG00000018548.10 |
| chr11 | 90637418 | 90639067 | 42.21422 | 149.43762 | 144.17581 | Promoter | 11 | 90638173 | 90644931 | 6759 | 1 | ENSMUST00 | 0 | Cox11 | ENSMUSG00000020544.9 |
| chr7 | 75455049 | 75456716 | 42.21422 | 149.43762 | 144.17581 | Promoter | 7 | 75455534 | 75754609 | 299076 | 1 | ENSMUST00 | 0 | Akap13 | ENSMUSG00000066406.9 |
| chr4 | 6190582 | 6191771 | 44.10828 | 149.36552 | 144.10564 | Promoter | 4 | 6191098 | 6221688 | 30591 | 1 | ENSMUST00 | 0 | Ubxn2b | ENSMUSG00000028243.5 |
| chr3 | 35753595 | 35755263 | 44.74406 | 149.33661 | 144.07773 | Promoter | 3 | 35754134 | 35829688 | 75555 | 1 | ENSMUST00 | 0 | Atp11b | ENSMUSG00000037400.11 |
| chr12 | 85373644 | 85375468 | 40.17707 | 149.16289 | 143.90889 | Promoter | 12 | 85340614 | 85374717 | 34104 | 2 | ENSMUST00 | 0 | Tmed10 | ENSMUSG00000021248.8 |
| chr16 | 21422763 | 21423824 | 40.17707 | 149.16289 | 143.90889 | Promoter | 16 | 21423118 | 21579247 | 156130 | 1 | ENSMUST00 | 0 | Vps8 | ENSMUSG00000033653.12 |
| chr19 | 8819501 | 8822631 | 40.17707 | 149.16289 | 143.90889 | Promoter | 19 | 8819401 | 8834142 | 14742 | 1 | ENSMUST00 | 101 | Hnrnpul2 | ENSMUSG00000071659.3 |
| chr11 | 53298936 | 53301316 | 37.74981 | 149.12509 | 143.87827 | Promoter | 11 | 53259814 | 53300457 | 40644 | 2 | ENSMUST00 | 0 | Hspa4 | ENSMUSG00000020361.8 |
| chr15 | 36790381 | 36794895 | 37.74981 | 149.12509 | 143.87827 | Promoter | 15 | 36770770 | 36794538 | 23769 | 2 | ENSMUST00 | 0 | Ywhaz | ENSMUSG00000022285.11 |
| chr18 | 38992277 | 38994637 | 37.74981 | 149.12509 | 143.87827 | Promoter | 18 | 38993145 | 39100149 | 107005 | 1 | ENSMUST00 | 0 | Arhgap26 | ENSMUSG00000036452.12 |
| chr18 | 62547837 | 62549716 | 37.74981 | 149.12509 | 143.87827 | Promoter | 18 | 62504069 | 62548743 | 44675 | 2 | ENSMUST00 | 0 | Fbxo38 | ENSMUSG00000042211.6 |
| chr3 | 122274316 | 122275023 | 37.74981 | 149.12509 | 143.87827 | Promoter | 3 | 122274414 | 122285269 | 10856 | 1 | ENSMUST00 | 0 | Dnttip2 | ENSMUSG00000039756.8 |
| chr11 | 116851455 | 116854638 | 40.75607 | 149.08708 | 143.84373 | Promoter | 11 | 116852440 | 116874712 | 22273 | 1 | ENSMUST00 | 0 | Mfsd11 | ENSMUSG00000020818.11 |
| chr9 | 36796370 | 36797563 | 43.93277 | 149.05946 | 143.81693 | Promoter | 9 | 36779159 | 36797273 | 18115 | 2 | ENSMUST00 | 0 | Ei24 | ENSMUSG00000062762.10 |
| chr12 | 69183312 | 69185321 | 42.59918 | 148.98648 | 143.74643 | Promoter | 12 | 69184157 | 69186770 | 2614 | 1 | ENSMUST00 | 0 | Mgat2 | ENSMUSG00000043998.10 |
| chr5 | 73254020 | 73256650 | 42.59918 | 148.98648 | 143.74643 | Promoter | 5 | 73020322 | 73256618 | 236297 | 2 | ENSMUST00 | 0 | Fryl | ENSMUSG00000070733.8 |
| chr8 | 91133202 | 91134090 | 42.59918 | 148.98648 | 143.74643 | Promoter | 8 | 91123499 | 91133942 | 10444 | 2 | ENSMUST00 | 0 | Aktip | ENSMUSG00000031667.10 |
| chr9 | 120491999 | 120493744 | 43.84554 | 148.90724 | 143.66989 | Promoter | 9 | 120492606 | 120495327 | 2722 | 1 | ENSMUST00 | 0 | Eif1b | ENSMUSG00000006941.4 |
| chr9 | 27028962 | 27031181 | 42.51875 | 148.84439 | 143.61023 | Promoter | 9 | 27004503 | 27030094 | 25592 | 2 | ENSMUST00 | 0 | Vps26b | ENSMUSG00000031988.9 |
| chr4 | 129599730 | 129601428 | 43.75865 | 148.75557 | 143.52182 | Promoter | 4 | 129600668 | 129607558 | 6891 | 1 | ENSMUST00 | 0 | Tmem234 | ENSMUSG00000028797.15 |
| chr10 | 81863536 | 81864659 | 34.16928 | 148.64955 | 143.4183 | Distal Interge | 10 | 81821127 | 81823360 | 2234 | 1 | ENSMUST00 | 42410 | Gm4767 | ENSMUSG00000094076.2 |
| chr3 | 40799381 | 40800896 | 44.94042 | 148.54384 | 143.31813 | Promoter | 3 | 40800019 | 40816883 | 16865 | 1 | ENSMUST00 | 0 | Plk4 | ENSMUSG00000025758.6 |
| chr4 | 155991804 | 155994301 | 44.94042 | 148.54384 | 143.31813 | Promoter | 4 | 155992872 | 156010210 | 17339 | 1 | ENSMUST00 | 0 | Sdf4 | ENSMUSG00000029076.9 |
| chr9 | 108460183 | 108462029 | 43.58591 | 148.45386 | 143.22935 | Promoter | 9 | 108460518 | 108465945 | 5428 | 1 | ENSMUST00 | 0 | Ccdc71 | ENSMUSG00000049305.5 |
| chr8 | 94601927 | 94603405 | 42.91624 | 148.41264 | 143.18958 | Promoter | 8 | 94601955 | 94660275 | 58321 | 1 | ENSMUST00 | 0 | Rspry1 | ENSMUSG00000050079.9 |
| chr11 | 52360043 | 52362840 | 43.41453 | 148.15424 | 142.93486 | Promoter | 11 | 52360860 | 52389397 | 28538 | 1 | ENSMUST00 | 0 | Vdac1 | ENSMUSG00000020402.6 |
| chr2 | 180589025 | 180590552 | 43.41452 | 148.15424 | 142.93486 | Promoter | 2 | 180589245 | 180595836 | 6592 | 1 | ENSMUST00 | 0 | Ogfr | ENSMUSG00000049401.10 |
| chr4 | 129188679 | 129191045 | 43.41452 | 148.15424 | 142.93486 | Promoter | 4 | 129189760 | 129219607 | 29848 | 1 | ENSMUST00 | 0 | Yars | ENSMUSG00000028811.6 |
| chr5 | 114567128 | 114570164 | 43.41453 | 148.15424 | 142.93486 | Promoter | 5 | 114568016 | 114613220 | 45205 | 1 | ENSMUST00 | 0 | Fam222a | ENSMUSG00000041930.7 |
| chr8 | 11634291 | 11636731 | 43.41453 | 148.15424 | 142.93486 | Promoter | 8 | 11611583 | 11635754 | 24172 | 2 | ENSMUST00 | 0 | Ankrd10 | ENSMUSG00000031508.8 |
| chr9 | 85748545 | 85750182 | 43.41453 | 148.15424 | 142.93486 | Promoter | 9 | 85687360 | 85749334 | 61975 | 2 | ENSMUST00 | 0 | Ibtk | ENSMUSG00000035941.10 |
| chr1 | 152765791 | 152768258 | 42.74922 | 148.11885 | 142.90198 | Promoter | 1 | 152766542 | 152775597 | 9056 | 1 | ENSMUST00 | 0 | Arpc5 | ENSMUSG00000008475.8 |
| chr1 | 93342383 | 93344516 | 42.74922 | 148.11885 | 142.90198 | Promoter | 1 | 93308778 | 93342789 | 34012 | 2 | ENSMUST00 | 0 | Pask | ENSMUSG00000026274.6 |
| chr10 | 63022070 | 63025156 | 42.74922 | 148.11885 | 142.90198 | Promoter | 10 | 63014664 | 63023894 | 9231 | 2 | ENSMUST00 | 0 | Hnrnph3 | ENSMUSG00000020069.11 |
| chr11 | 108920064 | 108922921 | 42.74922 | 148.11885 | 142.90198 | Promoter | 11 | 108920349 | 108950783 | 30435 | 1 | ENSMUST00 | 0 | Axin2 | ENSMUSG00000000142.10 |
| chr11 | 115602764 | 115604989 | 42.74922 | 148.11885 | 142.90198 | Promoter | 11 | 115584255 | 115604051 | 19797 | 2 | ENSMUST00 | 0 | Gga3 | ENSMUSG00000020740.8 |
| chr14 | 20929354 | 20930865 | 42.74922 | 148.11885 | 142.90198 | Promoter | 14 | 20929433 | 21033673 | 104241 | 1 | ENSMUST00 | 0 | Vcl | ENSMUSG00000021823.9 |
| chr15 | 35937043 | 35938280 | 42.74922 | 148.11885 | 142.90198 | Promoter | 15 | 35931976 | 35938246 | 6271 | 2 | ENSMUST00 | 0 | Cox6c | ENSMUSG00000014313.9 |
| chr15 | 38661292 | 38662773 | 42.74922 | 148.11885 | 142.90198 | Promoter | 15 | 38661904 | 38692443 | 30540 | 1 | ENSMUST00 | 0 | Atp6v1c1 | ENSMUSG00000022295.7 |
| chr15 | 97784332 | 97785489 | 42.74922 | 148.11885 | 142.90198 | Promoter | 15 | 97784355 | 97792692 | 8338 | 1 | ENSMUST00 | 0 | Slc48a1 | ENSMUSG00000081534.2 |
| chr17 | 12991036 | 12993295 | 42.74922 | 148.11885 | 142.90198 | Promoter | 17 | 12966796 | 12992497 | 25702 | 2 | ENSMUST00 | 0 | Wtap | ENSMUSG00000060475.7 |
| chr2 | 121473103 | 121474739 | 42.74922 | 148.11885 | 142.90198 | Promoter | 2 | 121460235 | 121474067 | 13833 | 2 | ENSMUST00 | 0 | Mfap1b | ENSMUSG00000048222.3 |
| chr4 | 139309988 | 139311917 | 42.74922 | 148.11885 | 142.90198 | Promoter | 4 | 139310744 | 139318426 | 7683 | 1 | ENSMUST00 | 0 | Akr7a5 | ENSMUSG00000028743.7 |
| chr4 | 151057007 | 151058085 | 42.74922 | 148.11885 | 142.90198 | Promoter | 4 | 151047300 | 151057953 | 10654 | 2 | ENSMUST00 | 0 | Vamp3 | ENSMUSG00000028955.3 |
| chr6 | 83914249 | 83915341 | 42.74922 | 148.11885 | 142.90198 | Promoter | 6 | 83914353 | 83986869 | 72517 | 1 | ENSMUST00 | 0 | Zfml | ENSMUSG00000030016.9 |
| chr7 | 28311136 | 28312482 | 42.74922 | 148.11885 | 142.90198 | Promoter | 7 | 28305516 | 28312072 | 6557 | 2 | ENSMUST00 | 0 | Timm50 | ENSMUSG00000003438.12 |
| chr13 | 21530204 | 21531455 | 41.44478 | 148.06744 | 142.8609 | Promoter | 13 | 21513222 | 21531120 | 17899 | 2 | ENSMUST00 | 0 | Zkscan8 | ENSMUSG00000063894.9 |
| chr14 | 46787422 | 46788855 | 41.44478 | 148.06744 | 142.8609 | Promoter | 14 | 46775582 | 46788411 | 12830 | 2 | ENSMUST00 | 0 | Cnih1 | ENSMUSG00000015759.5 |
| chr11 | 102554733 | 102556749 | 43.93484 | 147.93027 | 142.7272 | Promoter | 11 | 102475915 | 102556199 | 80285 | 2 | ENSMUST00 | 0 | Gpatch8 | ENSMUSG00000034621.9 |
| chr8 | 117157074 | 117158922 | 44.57026 | 147.90271 | 142.70154 | Promoter | 8 | 117158135 | 117205186 | 47052 | 1 | ENSMUST00 | 0 | Gan | ENSMUSG00000052557.6 |
| chr15 | 83171457 | 83172969 | 43.84673 | 147.77686 | 142.57693 | Promoter | 15 | 83153494 | 83172592 | 19099 | 2 | ENSMUST00 | 0 | Cyb5r3 | ENSMUSG00000018042.12 |
| chr1 | 55130651 | 55132400 | 44.47867 | 147.74396 | 142.5466 | Promoter | 1 | 55131231 | 55153938 | 22708 | 1 | ENSMUST00 | 0 | Mob4 | ENSMUSG00000025979.8 |
| chr15 | 5184679 | 5186578 | 41.88607 | 147.724 | 142.52785 | Promoter | 15 | 5185525 | 5218336 | 32812 | 1 | ENSMUST00 | 0 | Ttc33 | ENSMUSG00000022151.11 |
| chr1 | 170174111 | 170175592 | 43.75897 | 147.62399 | 142.42812 | Promoter | 1 | 170141938 | 170174957 | 33020 | 2 | ENSMUST00 | 0 | Uap1 | ENSMUSG00000026670.10 |
| chr4 | 122884994 | 122885972 | 43.75897 | 147.62399 | 142.42812 | Promoter | 4 | 122859047 | 122885911 | 26865 | 2 | ENSMUST00 | 0 | Cap1 | ENSMUSG00000028656.9 |
| chr6 | 38432966 | 38435255 | 41.80831 | 147.58597 | 142.39091 | Promoter | 6 | 38433950 | 38512763 | 78814 | 1 | ENSMUST00 | 0 | Ubn2 | ENSMUSG00000038538.12 |
| chr15 | 99971934 | 99974376 | 45.03415 | 147.57545 | 142.38377 | Promoter | 15 | 99972780 | 100016351 | 43572 | 1 | ENSMUST00 | 0 | Larp4 | ENSMUSG00000023025.10 |
| chr2 | 119476247 | 119478890 | 45.03415 | 147.57545 | 142.38377 | Promoter | 2 | 119373042 | 119477687 | 104646 | 2 | ENSMUST00 | 0 | Ino80 | ENSMUSG00000034154.10 |
| chr3 | 38483727 | 38484902 | 45.03415 | 147.57545 | 142.38377 | Promoter | 3 | 38449261 | 38484844 | 35584 | 2 | ENSMUST00 | 0 | Ankrd50 | ENSMUSG00000044864.11 |
| chr6 | 47672087 | 47673247 | 11.30501 | 147.47621 | 142.28978 | Distal Interge | 6 | 47637143 | 47637219 | 77 | 1 | ENSMUST00 | 34945 | Gm25283 | ENSMUSG00000087897.1 |
| chr4 | 136020557 | 136021737 | 44.95315 | 147.43549 | 142.24948 | Promoter | 4 | 136003368 | 136021763 | 18396 | 2 | ENSMUST00 | 26 | Tceb3 | ENSMUSG00000028668.5 |
| chr10 | 77515129 | 77516528 | 37.40972 | 147.3676 | 142.18408 | Promoter | 10 | 77486656 | 77515813 | 29158 | 2 | ENSMUST00 | 0 | Fam207a | ENSMUSG00000032977.8 |
| chr11 | 93994683 | 93997734 | 37.40972 | 147.3676 | 142.18408 | Promoter | 11 | 93996091 | 94043628 | 47538 | 1 | ENSMUST00 | 0 | Spag9 | ENSMUSG00000020859.11 |
| chr12 | 110695420 | 110696379 | 37.40972 | 147.3676 | 142.18408 | Promoter | 12 | 110691028 | 110696289 | 5262 | 2 | ENSMUST00 | 0 | Hsp90aa1 | ENSMUSG00000021270.8 |
| chr17 | 72836567 | 72838139 | 37.40972 | 147.3676 | 142.18408 | Promoter | 17 | 72836704 | 72851195 | 14492 | 1 | ENSMUST00 | 0 | Ypel5 | ENSMUSG00000039770.5 |
| chr6 | 34316487 | 34318211 | 37.40972 | 147.3676 | 142.18408 | Promoter | 6 | 34303934 | 34317497 | 13564 | 2 | ENSMUST00 | 0 | Akr1b3 | ENSMUSG00000001642.13 |
| chr13 | 37825344 | 37828365 | 39.80159 | 147.3544 | 142.17488 | Promoter | 13 | 37825975 | 37893826 | 67852 | 1 | ENSMUST00 | 0 | Rreb1 | ENSMUSG00000039087.11 |
| chr17 | 71615666 | 71617723 | 39.80159 | 147.3544 | 142.17488 | Promoter | 17 | 71616215 | 71659031 | 42817 | 1 | ENSMUST00 | 0 | Wdr43 | ENSMUSG00000041057.9 |
| chr18 | 10180415 | 10182911 | 39.80159 | 147.3544 | 142.17488 | Promoter | 18 | 10064401 | 10181792 | 117392 | 2 | ENSMUST00 | 0 | Rock1 | ENSMUSG00000024290.8 |
| chr19 | 6363840 | 6364939 | 39.80159 | 147.3544 | 142.17488 | Promoter | 19 | 6363922 | 6365955 | 2034 | 1 | ENSMUST00 | 0 | Sf1 | ENSMUSG00000024949.12 |
| chr2 | 31688201 | 31689640 | 39.80159 | 147.3544 | 142.17488 | Promoter | 2 | 31688376 | 31698336 | 9961 | 1 | ENSMUST00 | 0 | Abl1 | ENSMUSG00000026842.11 |
| chr4 | 119195282 | 119196013 | 39.80159 | 147.3544 | 142.17488 | Promoter | 4 | 119195306 | 119201298 | 5993 | 1 | ENSMUST00 | 0 | Ccdc23 | ENSMUSG00000028643.5 |
| chr5 | 35574339 | 35575582 | 39.80159 | 147.3544 | 142.17488 | Promoter | 5 | 35557206 | 35575070 | 17865 | 2 | ENSMUST00 | 0 | Trmt44 | ENSMUSG00000029097.7 |
| chr9 | 108306306 | 108308038 | 39.80159 | 147.3544 | 142.17488 | Promoter | 9 | 108306205 | 108337939 | 31735 | 1 | ENSMUST00 | 102 | Rhoa | ENSMUSG00000007815.8 |
| chr9 | 113740556 | 113742818 | 39.80159 | 147.3544 | 142.17488 | Distal Interge | 9 | 113651744 | 113708259 | 56516 | 2 | ENSMUST00 | -32298 | Pdcd6ip | ENSMUSG00000032504.10 |
| chr11 | 82930300 | 82933972 | 44.20615 | 147.27122 | 142.09979 | Promoter | 11 | 82933602 | 82937757 | 4156 | 1 | ENSMUST00 | 0 | Unc45b | ENSMUSG00000018845.9 |
| chr1 | 161249724 | 161251227 | 43.4978 | 147.16869 | 141.99896 | Promoter | 1 | 161240112 | 161251219 | 11108 | 2 | ENSMUST00 | 0 | Prdx6 | ENSMUSG00000026701.10 |
| chr13 | 30544221 | 30545909 | 42.1856 | 147.12523 | 141.95732 | Promoter | 13 | 30540308 | 30545362 | 5055 | 2 | ENSMUST00 | 0 | Uqcrfs1 | ENSMUSG00000038462.3 |
| chr17 | 24669204 | 24670761 | 42.1856 | 147.12523 | 141.95732 | Promoter | 17 | 24669752 | 24682015 | 12264 | 1 | ENSMUST00 | 0 | Zfp598 | ENSMUSG00000041130.9 |
| chr7 | 126501677 | 126504111 | 42.82515 | 147.12178 | 141.95673 | Promoter | 7 | 126491708 | 126503437 | 11730 | 2 | ENSMUST00 | 0 | Atxn2l | ENSMUSG00000032637.10 |
| chr10 | 117062841 | 117063868 | 42.74226 | 146.9763 | 141.81439 | Promoter | 10 | 117050998 | 117063814 | 12817 | 2 | ENSMUST00 | 0 | Cct2 | ENSMUSG00000034024.7 |
| chr8 | 25596721 | 25597950 | 42.0266 | 146.8443 | 141.68535 | Promoter | 8 | 25578492 | 25597487 | 18996 | 2 | ENSMUST00 | 0 | Letm2 | ENSMUSG00000037363.8 |
| chr19 | 7216506 | 7217703 | 43.93695 | 146.80379 | 141.64571 | Promoter | 19 | 7215158 | 7217616 | 2459 | 2 | ENSMUST00 | 0 | Cox8a | ENSMUSG00000035885.4 |
| chr17 | 33999048 | 34000816 | 41.34773 | 146.7668 | 141.60938 | Promoter | 17 | 33996017 | 34000289 | 4273 | 2 | ENSMUST00 | 0 | H2-K1 | ENSMUSG00000061232.10 |
| chr3 | 148986735 | 148991143 | 40.70288 | 146.73875 | 141.58145 | Promoter | 3 | 148888941 | 148989316 | 100376 | 2 | ENSMUST00 | 0 | Lphn2 | ENSMUSG00000028184.8 |
| chr11 | 78164224 | 78166188 | 42.57743 | 146.68684 | 141.53157 | Promoter | 11 | 78158499 | 78165589 | 7091 | 2 | ENSMUST00 | 0 | Traf4 | ENSMUSG00000017386.5 |
| chr3 | 122983401 | 122985518 | 42.57743 | 146.68684 | 141.53157 | Promoter | 3 | 122933601 | 122984447 | 50847 | 2 | ENSMUST00 | 0 | Usp53 | ENSMUSG00000039701.7 |
| chr19 | 32711296 | 32713053 | 44.48648 | 146.62837 | 141.47598 | Promoter | 19 | 32672563 | 32712298 | 39736 | 2 | ENSMUST00 | 0 | Atad1 | ENSMUSG00000013662.5 |
| chr2 | 173658896 | 173660911 | 43.15437 | 146.56914 | 141.41742 | Promoter | 2 | 173659760 | 173707315 | 47556 | 1 | ENSMUST00 | 0 | Rab22a | ENSMUSG00000027519.5 |
| chr15 | 55089912 | 55091131 | 39.3713 | 146.56828 | 141.41737 | Promoter | 15 | 55076099 | 55090491 | 14393 | 2 | ENSMUST00 | 0 | Dscc1 | ENSMUSG00000022422.8 |
| chr6 | 71830473 | 71832268 | 39.3713 | 146.56828 | 141.41737 | Promoter | 6 | 71831331 | 71875244 | 43914 | 1 | ENSMUST00 | 0 | Immt | ENSMUSG00000052337.10 |
| chr18 | 65430914 | 65432485 | 41.86879 | 146.56517 | 141.41492 | Promoter | 18 | 65430963 | 65478823 | 47861 | 1 | ENSMUST00 | 0 | Malt1 | ENSMUSG00000032688.7 |
| chr13 | 100743765 | 100745190 | 44.3943 | 146.46883 | 141.31972 | Promoter | 13 | 100735955 | 100744659 | 8705 | 2 | ENSMUST00 | 0 | Mrps36 | ENSMUSG00000061474.6 |
| chr14 | 30653838 | 30655023 | 44.3943 | 146.46883 | 141.31972 | Promoter | 14 | 30654360 | 30691317 | 36958 | 1 | ENSMUST00 | 0 | Rft1 | ENSMUSG00000052395.10 |
| chr4 | 141872917 | 141875442 | 44.3943 | 146.46883 | 141.31972 | Promoter | 4 | 141858142 | 141874920 | 16779 | 2 | ENSMUST00 | 0 | Efhd2 | ENSMUSG00000040659.3 |
| chr8 | 69972761 | 69974894 | 43.06936 | 146.42058 | 141.27246 | Promoter | 8 | 69907069 | 69974383 | 67315 | 2 | ENSMUST00 | 0 | Gatad2a | ENSMUSG00000036180.9 |
| chr13 | 100124304 | 100125912 | 42.98468 | 146.27251 | 141.12608 | Promoter | 13 | 100124852 | 100137690 | 12839 | 1 | ENSMUST00 | 0 | Smn1 | ENSMUSG00000021645.11 |
| chr7 | 130865745 | 130867110 | 42.98468 | 146.27251 | 141.12608 | Promoter | 7 | 130865756 | 130913312 | 47557 | 1 | ENSMUST00 | 0 | Plekha1 | ENSMUSG00000040268.11 |
| chr11 | 120484051 | 120485543 | 42.33011 | 146.25195 | 141.10883 | Promoter | 11 | 120484613 | 120489065 | 4453 | 1 | ENSMUST00 | 0 | Mrpl12 | ENSMUSG00000039640.7 |
| chr13 | 58214535 | 58216489 | 42.33011 | 146.25195 | 141.10883 | Promoter | 13 | 58176156 | 58215653 | 39498 | 2 | ENSMUST00 | 0 | Ubqln1 | ENSMUSG00000005312.9 |
| chr13 | 12394669 | 12396294 | 42.33011 | 146.25195 | 141.10883 | Promoter | 13 | 12395398 | 12438677 | 43280 | 1 | ENSMUST00 | 0 | Heatr1 | ENSMUSG00000050244.8 |
| chr16 | 11174891 | 11177068 | 42.33011 | 146.25195 | 141.10883 | Promoter | 16 | 11136592 | 11176393 | 39802 | 2 | ENSMUST00 | 0 | Zc3h7a | ENSMUSG00000037965.9 |
| chr2 | 26932319 | 26934235 | 42.33011 | 146.25195 | 141.10883 | Promoter | 2 | 26920040 | 26933928 | 13889 | 2 | ENSMUST00 | 0 | Surf4 | ENSMUSG00000014867.4 |
| chr2 | 162930504 | 162932824 | 42.33011 | 146.25195 | 141.10883 | Promoter | 2 | 162931528 | 162937121 | 5594 | 1 | ENSMUST00 | 0 | Srsf6 | ENSMUSG00000016921.9 |
| chr11 | 59838681 | 59840380 | 41.04627 | 146.22916 | 141.09622 | Promoter | 11 | 59817795 | 59839838 | 22044 | 2 | ENSMUST00 | 0 | Cops3 | ENSMUSG00000019373.6 |

| | | | | | | | | | | | | | | | |
|---|---|---|---|---|---|---|---|---|---|---|---|---|---|---|---|
| chr15 | 8168292 | 8169857 | 41.04627 | 146.22916 | 141.09622 | Promoter | 15 | 8169106 | 8271158 | 102053 | 1 | ENSMUST00 | 0 | 2410089E03I | ENSMUSG00000039801.6 |
| chr6 | 31557739 | 31563996 | 41.04627 | 146.22916 | 141.09622 | Promoter | 6 | 31519488 | 31563981 | 44494 | 2 | ENSMUST00 | 0 | Podxl | ENSMUSG00000025608.9 |
| chr8 | 64946606 | 64948077 | 41.04627 | 146.22916 | 141.09622 | Promoter | 8 | 64947185 | 64969034 | 21850 | 1 | ENSMUST00 | 0 | Tmem192 | ENSMUSG00000025521.9 |
| chr1 | 180640589 | 180642144 | 35.11919 | 146.21799 | 141.08704 | Promoter | 1 | 180641150 | 180690518 | 49369 | 1 | ENSMUST00 | 0 | Lin9 | ENSMUSG00000058729.8 |
| chr17 | 29031776 | 29035290 | 39.67387 | 146.00111 | 140.87399 | Promoter | 17 | 29032673 | 29042243 | 9571 | 1 | ENSMUST00 | 0 | Srsf3 | ENSMUSG00000071172.7 |
| chr19 | 5271956 | 5273622 | 41.4024 | 145.73851 | 140.61644 | Promoter | 19 | 5133688 | 5273119 | 139432 | 2 | ENSMUST00 | 0 | Pacs1 | ENSMUSG00000024855.9 |
| chr2 | 157565403 | 157567312 | 41.4024 | 145.73851 | 140.61644 | Promoter | 2 | 157556362 | 157566337 | 9976 | 2 | ENSMUST00 | 0 | Blcap | ENSMUSG00000067787.5 |
| chr4 | 129671587 | 129673549 | 41.4024 | 145.73851 | 140.61644 | Promoter | 4 | 129643980 | 129672767 | 28788 | 2 | ENSMUST00 | 0 | Kpna6 | ENSMUSG00000003731.8 |
| chr6 | 146576879 | 146579175 | 41.4024 | 145.73851 | 140.61644 | Promoter | 6 | 146577203 | 146592865 | 15663 | 1 | ENSMUST00 | 0 | Fgfr1op2 | ENSMUSG00000040242.9 |
| chr8 | 85026831 | 85028363 | 41.4024 | 145.73851 | 140.61644 | Promoter | 8 | 85026833 | 85030285 | 3453 | 1 | ENSMUST00 | 0 | 2310036O22 | ENSMUSG00000041203.8 |
| chr1 | 65185759 | 65187604 | 42.64928 | 145.68546 | 140.56778 | Promoter | 1 | 65186683 | 65205706 | 19024 | 1 | ENSMUST00 | 0 | Pikfyve | ENSMUSG00000025949.10 |
| chr11 | 46436468 | 46438045 | 42.64928 | 145.68546 | 140.56778 | Promoter | 11 | 46436925 | 46440858 | 3934 | 1 | ENSMUST00 | 0 | Med7 | ENSMUSG00000020397.10 |
| chr1 | 193129321 | 193130888 | 44.57462 | 145.65463 | 140.53799 | Promoter | 1 | 193104403 | 193130251 | 25849 | 2 | ENSMUST00 | 0 | Diexf | ENSMUSG00000016181.4 |
| chr10 | 75763631 | 75765317 | 44.57462 | 145.65463 | 140.53799 | Promoter | 10 | 75646110 | 75764357 | 118248 | 2 | ENSMUST00 | 0 | Cabin1 | ENSMUSG00000020196.9 |
| chr10 | 120897818 | 120899446 | 44.57462 | 145.65463 | 140.53799 | Promoter | 10 | 120781096 | 120899101 | 118006 | 2 | ENSMUST00 | 0 | Msrb3 | ENSMUSG00000051236.8 |
| chr13 | 91806936 | 91808633 | 44.57462 | 145.65463 | 140.53799 | Promoter | 13 | 91796533 | 91807705 | 11173 | 2 | ENSMUST00 | 0 | Zcchc9 | ENSMUSG00000021621.10 |
| chr13 | 9093707 | 9095228 | 44.57462 | 145.65463 | 140.53799 | Promoter | 13 | 9093881 | 9171104 | 77224 | 1 | ENSMUST00 | 0 | Larp4b | ENSMUSG00000033499.8 |
| chr17 | 84789802 | 84791615 | 44.57462 | 145.65463 | 140.53799 | Promoter | 17 | 84705247 | 84790558 | 85312 | 2 | ENSMUST00 | 0 | Lrpprc | ENSMUSG00000024120.7 |
| chr18 | 12941134 | 12942141 | 44.57462 | 145.65463 | 140.53799 | Promoter | 18 | 12755314 | 12941841 | 186528 | 2 | ENSMUST00 | 0 | Osbpl1a | ENSMUSG00000044252.12 |
| chr19 | 28009765 | 28012498 | 44.57462 | 145.65463 | 140.53799 | Promoter | 19 | 27766441 | 28011048 | 244608 | 2 | ENSMUST00 | 0 | Rfx3 | ENSMUSG00000040929.11 |
| chr18 | 34505681 | 34507604 | 43.23485 | 145.58217 | 140.47385 | Promoter | 18 | 34442352 | 34506823 | 64472 | 2 | ENSMUST00 | 0 | Fam13b | ENSMUSG00000036501.6 |
| chr5 | 93205257 | 93207249 | 43.23485 | 145.58217 | 140.47385 | Promoter | 5 | 93206518 | 93213475 | 6958 | 1 | ENSMUST00 | 0 | 2010109A12I | ENSMUSG00000029384.4 |
| chr17 | 80060963 | 80063060 | 39.4261 | 145.54997 | 140.44281 | Promoter | 17 | 80029487 | 80062227 | 32741 | 2 | ENSMUST00 | 0 | Hnrnpll | ENSMUSG00000024095.10 |
| chr17 | 56583240 | 56584610 | 39.4261 | 145.54997 | 140.44281 | Promoter | 17 | 56562939 | 56584512 | 21574 | 2 | ENSMUST00 | 0 | Safb2 | ENSMUSG00000042625.11 |
| chr18 | 64886807 | 64889297 | 39.4261 | 145.54997 | 140.44281 | Promoter | 18 | 64887756 | 65217826 | 330071 | 1 | ENSMUST00 | 0 | Nedd4l | ENSMUSG00000024589.10 |
| chr4 | 111719377 | 111720955 | 39.4261 | 145.54997 | 140.44281 | Promoter | 4 | 111719984 | 111829142 | 109159 | 1 | ENSMUST00 | 0 | Spata6 | ENSMUSG00000034401.11 |
| chr5 | 31526093 | 31527959 | 39.4261 | 145.54997 | 140.44281 | Promoter | 5 | 31526995 | 31529405 | 2411 | 1 | ENSMUST00 | 0 | Slc4a1ap | ENSMUSG00000029141.11 |
| chr9 | 65293303 | 65295315 | 39.4261 | 145.54997 | 140.44281 | Promoter | 9 | 65294260 | 65330657 | 36398 | 1 | ENSMUST00 | 0 | Clpx | ENSMUSG00000015357.5 |
| chr9 | 118039935 | 118041951 | 39.4261 | 145.54997 | 140.44281 | Promoter | 9 | 118012821 | 118040259 | 27439 | 2 | ENSMUST00 | 0 | Zcwpw2 | ENSMUSG00000032443.10 |
| chr11 | 86755991 | 86758872 | 42.56624 | 145.53995 | 140.43991 | Promoter | 11 | 86694351 | 86757565 | 63215 | 2 | ENSMUST00 | 0 | Cltc | ENSMUSG00000047126.12 |
| chr15 | 8442340 | 8446554 | 43.84918 | 145.52415 | 140.42462 | Promoter | 15 | 8291225 | 8444463 | 153239 | 2 | ENSMUST00 | 0 | Nipbl | ENSMUSG00000022141.6 |
| chr3 | 97900742 | 97902162 | 43.84918 | 145.52415 | 140.42462 | Promoter | 3 | 97901190 | 97923276 | 22087 | 1 | ENSMUST00 | 0 | Sec22b | ENSMUSG00000027879.4 |
| chr3 | 121425851 | 121428534 | 43.84918 | 145.52415 | 140.42462 | Promoter | 3 | 121426541 | 121458205 | 31665 | 1 | ENSMUST00 | 0 | Cnn3 | ENSMUSG00000053931.7 |
| chr9 | 74952562 | 74953836 | 37.5617 | 145.4381 | 140.34033 | Promoter | 9 | 74953053 | 75032468 | 79416 | 1 | ENSMUST00 | 0 | Fam214a | ENSMUSG00000034858.10 |
| chr5 | 32853439 | 32854321 | 43.14866 | 145.43195 | 140.33446 | Promoter | 5 | 32789218 | 32854256 | 65039 | 2 | ENSMUST00 | 0 | Prr14l | ENSMUSG00000054280.9 |
| chr7 | 127478104 | 127480347 | 38.71362 | 145.36029 | 140.26395 | Distal Interge | 7 | 127475501 | 127476757 | 1257 | 1 | ENSMUST00 | 2604 | Prr14 | ENSMUSG00000030822.10 |
| chr1 | 160905985 | 160907996 | 41.77201 | 145.26822 | 140.17451 | Promoter | 1 | 160906418 | 160974978 | 68561 | 1 | ENSMUST00 | 0 | Rc3h1 | ENSMUSG00000040423.5 |
| chr11 | 101464825 | 101466732 | 41.77201 | 145.26822 | 140.17451 | Promoter | 11 | 101464999 | 101471853 | 6855 | 1 | ENSMUST00 | 0 | Rnd2 | ENSMUSG00000001313.7 |
| chr14 | 20694576 | 20696294 | 41.77201 | 145.26822 | 140.17451 | Promoter | 14 | 20694968 | 20700197 | 5230 | 1 | ENSMUST00 | 0 | Fut11 | ENSMUSG00000039357.1 |
| chr4 | 108878856 | 108880715 | 41.77201 | 145.26822 | 140.17451 | Promoter | 4 | 108879063 | 108943324 | 64262 | 1 | ENSMUST00 | 0 | Rab3b | ENSMUSG00000003411.5 |
| chr8 | 40307383 | 40308874 | 41.77201 | 145.26822 | 140.17451 | Promoter | 8 | 40279725 | 40308331 | 28607 | 2 | ENSMUST00 | 0 | Fgf20 | ENSMUSG00000031603.9 |
| chr19 | 10041037 | 10042840 | 43.67043 | 145.21399 | 140.12416 | Promoter | 19 | 10041548 | 10059671 | 18124 | 1 | ENSMUST00 | 0 | Fads3 | ENSMUSG00000024664.4 |
| chr11 | 59201710 | 59203125 | 42.97731 | 145.13309 | 140.04448 | Promoter | 11 | 59202486 | 59205927 | 3442 | 1 | ENSMUST00 | 0 | Mrpl55 | ENSMUSG00000036860.9 |
| chr2 | 59881834 | 59882942 | 43.58161 | 145.05977 | 139.97232 | Promoter | 2 | 59852364 | 59882584 | 30221 | 2 | ENSMUST00 | 0 | Wdsub1 | ENSMUSG00000026988.10 |
| chr9 | 79792262 | 79793432 | 40.31151 | 144.91344 | 139.82919 | Promoter | 9 | 79768943 | 79793430 | 24488 | 2 | ENSMUST00 | 0 | Tmem30a | ENSMUSG00000032328.7 |
| chr8 | 94857164 | 94858232 | 43.49314 | 144.90613 | 139.82271 | Promoter | 8 | 94857450 | 94864242 | 6793 | 1 | ENSMUST00 | 0 | Polr2c | ENSMUSG00000031783.9 |
| chr12 | 85338342 | 85339915 | 42.15588 | 144.81987 | 139.73779 | Promoter | 12 | 85299514 | 85339362 | 39849 | 2 | ENSMUST00 | 0 | Nek9 | ENSMUSG00000034290.9 |
| chr17 | 21961677 | 21963474 | 42.15588 | 144.81987 | 139.73779 | Promoter | 17 | 21962552 | 21994366 | 31815 | 1 | ENSMUST00 | 0 | Zfp943 | ENSMUSG00000053347.9 |
| chr18 | 61555228 | 61560694 | 42.15588 | 144.81987 | 139.73779 | Promoter | 18 | 61555274 | 61590061 | 34788 | 1 | ENSMUST00 | 0 | Csnk1a1 | ENSMUSG00000024576.9 |
| chr19 | 47089681 | 47091451 | 42.15588 | 144.81987 | 139.73779 | Promoter | 19 | 47090766 | 47131143 | 40378 | 1 | ENSMUST00 | 0 | Pdcd11 | ENSMUSG00000025047.8 |
| chr2 | 14054882 | 14056514 | 42.15588 | 144.81987 | 139.73779 | Promoter | 2 | 13975563 | 14056020 | 80458 | 2 | ENSMUST00 | 0 | Ptpla | ENSMUSG00000063275.10 |
| chr3 | 88501420 | 88503310 | 37.81545 | 144.80544 | 139.72641 | Promoter | 3 | 88483205 | 88503307 | 20103 | 2 | ENSMUST00 | 0 | Lmna | ENSMUSG00000028063.10 |
| chr12 | 59066132 | 59067965 | 38.99987 | 144.77138 | 139.69241 | Promoter | 12 | 59066919 | 59074017 | 7099 | 1 | ENSMUST00 | 0 | Pnn | ENSMUSG00000020994.4 |
| chr2 | 181519666 | 181521722 | 38.99987 | 144.77138 | 139.69241 | Promoter | 2 | 181520485 | 181555133 | 34649 | 1 | ENSMUST00 | 0 | Dnajc5 | ENSMUSG00000000826.11 |
| chr3 | 107331120 | 107333142 | 38.99987 | 144.77138 | 139.69241 | Promoter | 3 | 107326110 | 107333289 | 7180 | 2 | ENSMUST00 | 147 | Rbm15 | ENSMUSG00000048109.9 |
| chr2 | 3284114 | 3285676 | 43.40503 | 144.75304 | 139.67491 | Promoter | 2 | 3284212 | 3328877 | 44666 | 1 | ENSMUST00 | 0 | Nmt2 | ENSMUSG00000026643.11 |
| chr7 | 113368620 | 113370289 | 43.40503 | 144.75304 | 139.67491 | Promoter | 7 | 113315626 | 113369339 | 53714 | 2 | ENSMUST00 | 0 | Btbd10 | ENSMUSG00000038187.9 |
| chr8 | 109778088 | 109780170 | 43.40503 | 144.75304 | 139.67491 | Promoter | 8 | 109778554 | 109864204 | 85651 | 1 | ENSMUST00 | 0 | Ap1g1 | ENSMUSG00000031731.11 |
| chr7 | 17026986 | 17028377 | 42.72282 | 144.68874 | 139.61232 | Promoter | 7 | 17004640 | 17027916 | 23277 | 2 | ENSMUST00 | 0 | Ppp5c | ENSMUSG00000003099.6 |
| chr11 | 105180525 | 105182894 | 40.64776 | 144.39505 | 139.32506 | Promoter | 11 | 105180746 | 105209855 | 29110 | 1 | ENSMUST00 | 0 | Tlk2 | ENSMUSG00000020694.11 |
| chr15 | 59075869 | 59083046 | 40.64776 | 144.39505 | 139.32506 | Promoter | 15 | 58941244 | 59082026 | 140783 | 2 | ENSMUST00 | 0 | Mtss1 | ENSMUSG00000022353.9 |
| chr15 | 82046399 | 82048156 | 40.64776 | 144.39505 | 139.32506 | Promoter | 15 | 82041319 | 82047598 | 6280 | 2 | ENSMUST00 | 0 | Nhp2l1 | ENSMUSG00000063480.7 |
| chr2 | 5951360 | 5952315 | 42.55484 | 144.39514 | 139.32506 | Promoter | 2 | 5951469 | 6056703 | 105235 | 1 | ENSMUST00 | 0 | Upf2 | ENSMUSG00000043241.9 |
| chr3 | 90079128 | 90080759 | 41.91343 | 144.39362 | 139.32506 | Promoter | 3 | 90079540 | 90100901 | 21362 | 1 | ENSMUST00 | 0 | Tpm3 | ENSMUSG00000027940.13 |
| chr1 | 57377189 | 57378473 | 41.911 | 144.38934 | 139.32462 | Promoter | 1 | 57377620 | 57385423 | 7804 | 1 | ENSMUST00 | 0 | 1700066M21 | ENSMUSG00000038323.3 |
| chr18 | 80200011 | 80201112 | 41.911 | 144.38934 | 139.32462 | Promoter | 18 | 80192265 | 80200658 | 8394 | 2 | ENSMUST00 | 0 | Rbfa | ENSMUSG00000024570.6 |
| chr19 | 53389502 | 53391452 | 41.911 | 144.38934 | 139.32462 | Promoter | 19 | 53379214 | 53390573 | 11360 | 2 | ENSMUST00 | 0 | Smndc1 | ENSMUSG00000025024.6 |
| chr2 | 52072429 | 52073853 | 41.911 | 144.38934 | 139.32462 | Promoter | 2 | 52072837 | 52122381 | 49545 | 1 | ENSMUST00 | 0 | Rif1 | ENSMUSG00000036202.10 |
| chr2 | 128966468 | 128968509 | 41.911 | 144.38934 | 139.32462 | Promoter | 2 | 128967402 | 129018563 | 51162 | 1 | ENSMUST00 | 0 | Zc3h6 | ENSMUSG00000042851.12 |
| chr4 | 155704220 | 155706443 | 41.911 | 144.38934 | 139.32462 | Promoter | 4 | 155704800 | 155733879 | 29080 | 1 | ENSMUST00 | 0 | Ssu72 | ENSMUSG00000029038.4 |
| chr4 | 11321290 | 11322745 | 41.911 | 144.38934 | 139.32462 | Promoter | 4 | 11261315 | 11322137 | 60823 | 2 | ENSMUST00 | 0 | Dpy19l4 | ENSMUSG00000045205.11 |
| chr7 | 44815266 | 44817335 | 41.911 | 144.38934 | 139.32462 | Promoter | 7 | 44812257 | 44815658 | 3402 | 2 | ENSMUST00 | 0 | Atf5 | ENSMUSG00000038539.9 |
| chr9 | 120127671 | 120128492 | 41.911 | 144.38934 | 139.32462 | Promoter | 9 | 120127766 | 120132369 | 4604 | 1 | ENSMUST00 | 0 | Rpsa | ENSMUSG00000032518.5 |
| chr14 | 14119872 | 14121052 | 41.83324 | 144.2525 | 139.19977 | Promoter | 14 | 14112184 | 14120984 | 8801 | 2 | ENSMUST00 | 0 | Psmd6 | ENSMUSG00000021737.3 |
| chr5 | 21784084 | 21785577 | 42.47133 | 144.2491 | 139.19682 | Promoter | 5 | 21785283 | 21803787 | 18505 | 1 | ENSMUST00 | 0 | Psmc2 | ENSMUSG00000028932.7 |
| chr1 | 92472809 | 92474597 | 39.29602 | 144.19646 | 139.14511 | Promoter | 1 | 92439010 | 92473860 | 34851 | 2 | ENSMUST00 | 0 | Ndufa10 | ENSMUSG00000026260.7 |
| chr10 | 127210787 | 127212119 | 39.29602 | 144.19646 | 139.14511 | Promoter | 10 | 127197067 | 127211622 | 14556 | 2 | ENSMUST00 | 0 | Pip4k2c | ENSMUSG00000025417.7 |
| chr7 | 89403160 | 89405751 | 38.07755 | 144.18416 | 139.13362 | Promoter | 7 | 89156448 | 89404222 | 247775 | 2 | ENSMUST00 | 0 | Tmem135 | ENSMUSG00000039428.5 |
| chr4 | 140961260 | 140962243 | 41.75335 | 144.11185 | 139.06203 | Promoter | 4 | 140961401 | 140976796 | 15396 | 1 | ENSMUST00 | 0 | Sdhb | ENSMUSG00000009863.9 |
| chr12 | 80642771 | 80645369 | 42.38816 | 144.10358 | 139.05386 | Promoter | 12 | 80634022 | 80644341 | 10320 | 2 | ENSMUST00 | 0 | Erh | ENSMUSG00000021131.8 |
| chr8 | 95433852 | 95434912 | 42.38816 | 144.10358 | 139.05386 | Promoter | 8 | 95420252 | 95434869 | 14618 | 2 | ENSMUST00 | 0 | Gtl3 | ENSMUSG00000031796.6 |
| chr1 | 106795662 | 106797288 | 42.9698 | 143.99599 | 138.94749 | Promoter | 1 | 106764071 | 106796667 | 32597 | 2 | ENSMUST00 | 0 | Vps4b | ENSMUSG00000009907.12 |
| chr10 | 45577641 | 45578826 | 42.9698 | 143.99599 | 138.94749 | Promoter | 10 | 45577829 | 45712345 | 134517 | 1 | ENSMUST00 | 0 | Hace1 | ENSMUSG00000038822.10 |
| chr2 | 179975983 | 179977691 | 42.9698 | 143.99599 | 138.94749 | Promoter | 2 | 179912146 | 179976646 | 64501 | 2 | ENSMUST00 | 0 | Taf4a | ENSMUSG00000039117.11 |
| chr10 | 115250494 | 115252670 | 40.9965 | 143.89526 | 138.84959 | Promoter | 10 | 115197877 | 115251493 | 53617 | 2 | ENSMUST00 | 0 | Tbc1d15 | ENSMUSG00000020130.8 |
| chr10 | 128458402 | 128460755 | 40.9965 | 143.89526 | 138.84959 | Promoter | 10 | 128459236 | 128490173 | 30938 | 1 | ENSMUST00 | 0 | Smarcc2 | ENSMUSG00000025369.9 |
| chr13 | 55634627 | 55635987 | 40.9965 | 143.89526 | 138.84959 | Promoter | 13 | 55635027 | 55681256 | 46230 | 1 | ENSMUST00 | 0 | Ddx46 | ENSMUSG00000021500.10 |
| chr14 | 46820959 | 46822650 | 40.9965 | 143.89526 | 138.84959 | Promoter | 14 | 46808149 | 46822242 | 14094 | 2 | ENSMUST00 | 0 | Gmfb | ENSMUSG00000062014.7 |
| chr4 | 40721673 | 40724055 | 32.90589 | 143.88075 | 138.83896 | Promoter | 4 | 40722150 | 40723795 | 1646 | 1 | ENSMUST00 | 0 | Dnaja1 | ENSMUSG00000028410.8 |
| chr1 | 89070287 | 89072598 | 36.72955 | 143.86444 | 138.82292 | Promoter | 1 | 89070415 | 89155068 | 84654 | 1 | ENSMUST00 | 0 | Sh3bp4 | ENSMUSG00000036206.7 |
| chr10 | 63243852 | 63245055 | 36.72955 | 143.86444 | 138.82292 | Promoter | 10 | 63243068 | 63244153 | 1086 | 2 | ENSMUST00 | 0 | 1700120B22I | ENSMUSG00000045391.4 |
| chr10 | 42273937 | 42278830 | 36.72955 | 143.86444 | 138.82292 | Promoter | 10 | 42181841 | 42276752 | 94912 | 2 | ENSMUST00 | 0 | Foxo3 | ENSMUSG00000048756.6 |
| chr16 | 31273924 | 31275287 | 36.72955 | 143.86444 | 138.82292 | Promoter | 16 | 31251537 | 31275277 | 23741 | 2 | ENSMUST00 | 0 | Ppp1r2 | ENSMUSG00000047714.9 |
| chr9 | 78107503 | 78109725 | 36.72955 | 143.86444 | 138.82292 | Promoter | 9 | 78081499 | 78108656 | 27158 | 2 | ENSMUST00 | 0 | Fbxo9 | ENSMUSG00000001366.9 |
| chr8 | 111536021 | 111538448 | 42.22279 | 143.81401 | 138.77701 | Promoter | 8 | 111536097 | 111625720 | 89624 | 1 | ENSMUST00 | 0 | Znrf1 | ENSMUSG00000033545.9 |
| chr9 | 58253191 | 58253958 | 42.22279 | 143.81401 | 138.77701 | Promoter | 9 | 58253164 | 58262523 | 9360 | 1 | ENSMUST00 | 28 | Stoml1 | ENSMUSG00000032333.5 |
| chr10 | 84575899 | 84577895 | 39.05061 | 143.74966 | 138.71384 | Promoter | 10 | 84576626 | 84614359 | 37734 | 1 | ENSMUST00 | 0 | Tcp11l2 | ENSMUSG00000020034.7 |
| chr10 | 62577526 | 62578495 | 39.05061 | 143.74966 | 138.71384 | Promoter | 10 | 62538626 | 62578440 | 39815 | 2 | ENSMUST00 | 0 | 2510003E04I | ENSMUSG00000036955.10 |
| chr13 | 54468153 | 54469196 | 39.05061 | 143.74966 | 138.71384 | Promoter | 13 | 54458837 | 54468849 | 10013 | 2 | ENSMUST00 | 0 | Thoc3 | ENSMUSG00000025872.5 |
| chr13 | 106935938 | 106937353 | 39.05061 | 143.74966 | 138.71384 | Promoter | 13 | 106794439 | 106936958 | 142520 | 2 | ENSMUST00 | 0 | Ipo11 | ENSMUSG00000042590.11 |
| chr14 | 40965844 | 40966967 | 39.05061 | 143.74966 | 138.71384 | Promoter | 14 | 40906445 | 40966807 | 60363 | 2 | ENSMUST00 | 0 | Tspan14 | ENSMUSG00000037824.5 |
| chr6 | 13069130 | 13070822 | 39.05061 | 143.74966 | 138.71384 | Promoter | 6 | 13069759 | 13075094 | 5336 | 1 | ENSMUST00 | 0 | Tmem106b | ENSMUSG00000029571.9 |
| chr6 | 39204806 | 39207527 | 39.05061 | 143.74966 | 138.71384 | Promoter | 6 | 39136623 | 39206789 | 70167 | 2 | ENSMUST00 | 0 | Kdm7a | ENSMUSG00000042599.8 |
| chr7 | 45813398 | 45814380 | 39.05061 | 143.74966 | 138.71384 | Promoter | 7 | 45806637 | 45814316 | 7680 | 2 | ENSMUST00 | 0 | Cyth2 | ENSMUSG00000003269.9 |
| chr9 | 25251253 | 25253817 | 39.05061 | 143.74966 | 138.71384 | Promoter | 9 | 25252439 | 25308571 | 56133 | 1 | ENSMUST00 | 0 | Sep-07 | ENSMUSG00000001833.11 |
| chr11 | 117986409 | 117987584 | 44.11508 | 143.73827 | 138.70853 | Promoter | 11 | 117986885 | 118024011 | 37127 | 1 | ENSMUST00 | 0 | Pgs1 | ENSMUSG00000017715.6 |
| chr13 | 34873812 | 34876956 | 44.11508 | 143.73827 | 138.70853 | Promoter | 13 | 34875494 | 34902877 | 27384 | 1 | ENSMUST00 | 0 | Prpf4b | ENSMUSG00000021413.8 |
| chr19 | 46396798 | 46397572 | 44.11508 | 143.73827 | 138.70853 | Promoter | 19 | 46396896 | 46488804 | 91909 | 1 | ENSMUST00 | 0 | Sufu | ENSMUSG00000025231.11 |
| chr4 | 47352092 | 47354375 | 44.11508 | 143.73827 | 138.70853 | Promoter | 4 | 47353222 | 47414931 | 61710 | 1 | ENSMUST00 | 0 | Tgfbr1 | ENSMUSG00000007613.10 |
| chr6 | 119847023 | 119849337 | 44.11508 | 143.73827 | 138.70853 | Promoter | 6 | 119570796 | 119848150 | 277355 | 2 | ENSMUST00 | 0 | Erc1 | ENSMUSG00000030172.10 |
| chr7 | 19457183 | 19459389 | 44.11508 | 143.73827 | 138.70853 | Promoter | 7 | 19426075 | 19458494 | 32420 | 2 | ENSMUST00 | 0 | Mark4 | ENSMUSG00000030397.9 |
| chr19 | 36553796 | 36555516 | 42.71282 | 143.54831 | 138.52826 | Promoter | 19 | 36554639 | 36621132 | 66494 | 1 | ENSMUST00 | 0 | Hectd2 | ENSMUSG00000041180.8 |
| chr7 | 101107784 | 101110527 | 43.94354 | 143.44185 | 138.423 | Promoter | 7 | 101108775 | 101284382 | 175608 | 1 | ENSMUST00 | 0 | Fchsd2 | ENSMUSG00000030691.9 |
| chr1 | 86525464 | 86529269 | 41.35843 | 143.41548 | 138.39708 | Promoter | 1 | 86526726 | 86530712 | 3987 | 1 | ENSMUST00 | 0 | Ptma | ENSMUSG00000026238.9 |
| chr17 | 75005050 | 75006665 | 41.35843 | 143.41548 | 138.39708 | Promoter | 17 | 75005568 | 75391767 | 386200 | 1 | ENSMUST00 | 0 | Ltbp1 | ENSMUSG00000001870.10 |
| chr2 | 152829469 | 152831763 | 41.35843 | 143.41548 | 138.39708 | Promoter | 2 | 152780668 | 152831728 | 51061 | 2 | ENSMUST00 | 0 | Bcl2l1 | ENSMUSG00000007659.13 |
| chr3 | 54734296 | 54736530 | 41.35843 | 143.41548 | 138.39708 | Promoter | 3 | 54728678 | 54735393 | 6716 | 2 | ENSMUST00 | 0 | Exosc8 | ENSMUSG00000027752.10 |
| chr6 | 113237289 | 113238881 | 41.35843 | 143.41548 | 138.39708 | Promoter | 6 | 113237843 | 113281202 | 43360 | 1 | ENSMUST00 | 0 | Mtmr14 | ENSMUSG00000030269.9 |
| chr6 | 83053281 | 83056016 | 41.35843 | 143.41548 | 138.39708 | Promoter | 6 | 83054653 | 83057680 | 3028 | 1 | ENSMUST00 | 0 | Aup1 | ENSMUSG00000068328.7 |
| chr8 | 119344417 | 119345409 | 41.35843 | 143.41548 | 138.39708 | Promoter | 8 | 119344538 | 119348927 | 4390 | 1 | ENSMUST00 | 0 | Hsbp1 | ENSMUSG00000031839.6 |
| chr19 | 4395327 | 4397114 | 41.97713 | 143.38342 | 138.36984 | Promoter | 19 | 4316170 | 4397077 | 80908 | 2 | ENSMUST00 | 0 | Kdm2a | ENSMUSG00000054611.11 |
| chr15 | 100227015 | 100229191 | 43.22482 | 143.31757 | 138.30418 | Promoter | 15 | 100227819 | 100261244 | 33426 | 1 | ENSMUST00 | 0 | Atf1 | ENSMUSG00000023027.7 |
| chr2 | 74711584 | 74713681 | 43.85173 | 143.28317 | 138.27043 | Promoter | 2 | 74703248 | 74712054 | 8807 | 2 | ENSMUST00 | 0 | Hoxd3os1 | ENSMUSG00000052371.5 |
| chr17 | 29614281 | 29615934 | 43.13689 | 143.16493 | 138.15439 | Promoter | 17 | 29614790 | 29641664 | 26875 | 1 | ENSMUST00 | 0 | Rnf8 | ENSMUSG00000090083.5 |
| chr16 | 58669867 | 58671062 | 42.45889 | 143.10539 | 138.09682 | Promoter | 16 | 58670208 | 58680391 | 10184 | 1 | ENSMUST00 | 0 | Cpox | ENSMUSG00000022742.5 |
| chr12 | 33429503 | 33430420 | 39.92014 | 143.09229 | 138.08496 | Promoter | 12 | 33429624 | 33439380 | 9757 | 1 | ENSMUST00 | 0 | Twistnb | ENSMUSG00000020561.7 |
| chr5 | 90223166 | 90224759 | 43.04933 | 143.01288 | 138.00682 | Promoter | 5 | 90214724 | 90223972 | 9249 | 2 | ENSMUST00 | 0 | Cox18 | ENSMUSG00000035505.9 |
| chr10 | 34206373 | 34207924 | 41.12505 | 143.00313 | 137.99791 | Promoter | 10 | 34151394 | 34207551 | 56158 | 2 | ENSMUST00 | 0 | Dse | ENSMUSG00000039497.7 |

| | | | | | | | | | | | | | | | |
|---|---|---|---|---|---|---|---|---|---|---|---|---|---|---|---|
| chr7 | 127587645 | 127589549 | 43.66926 | 142.96762 | 137.96356 | Promoter | 7 | 127588698 | 127603605 | 14908 | 1 | ENSMUST00 | 0 | Rnf40 | ENSMUSG00000030816.7 |
| chr13 | 47104757 | 47106907 | 41.73432 | 142.95721 | 137.9541 | Promoter | 13 | 47084782 | 47106190 | 21409 | 2 | ENSMUST00 | 0 | Dek | ENSMUSG00000021377.9 |
| chr6 | 95117804 | 95118814 | 41.73432 | 142.95721 | 137.9541 | Promoter | 6 | 95117880 | 95129718 | 11839 | 1 | ENSMUST00 | 0 | Kbtbd8 | ENSMUSG00000030031.9 |
| chr6 | 66895983 | 66897619 | 41.73432 | 142.95721 | 137.9541 | Promoter | 6 | 66896397 | 67021350 | 124954 | 1 | ENSMUST00 | 0 | Gng12 | ENSMUSG00000036402.8 |
| chr12 | 31948692 | 31950317 | 42.96212 | 142.86137 | 137.8616 | Promoter | 12 | 31929383 | 31950223 | 20841 | 2 | ENSMUST00 | 0 | Hbp1 | ENSMUSG00000002996.12 |
| chr10 | 80100966 | 80102919 | 34.49762 | 142.80731 | 137.81027 | Promoter | 10 | 80057416 | 80102698 | 45283 | 2 | ENSMUST00 | 0 | Sbno2 | ENSMUSG00000035673.9 |
| chr5 | 117976342 | 117977742 | 34.49762 | 142.80731 | 137.81027 | Promoter | 5 | 117976770 | 118010191 | 33422 | 1 | ENSMUST00 | 0 | Fbxo21 | ENSMUSG00000032898.6 |
| chr8 | 69790270 | 69792190 | 34.49762 | 142.80731 | 137.81027 | Promoter | 8 | 69791163 | 69807749 | 16587 | 1 | ENSMUST00 | 0 | Atp13a1 | ENSMUSG00000031862.10 |
| chr16 | 17619013 | 17620255 | 42.87527 | 142.71045 | 137.71519 | Promoter | 16 | 17619354 | 17644828 | 25475 | 1 | ENSMUST00 | 0 | Smpd4 | ENSMUSG00000005899.7 |
| chr3 | 58414128 | 58418695 | 40.32275 | 142.69632 | 137.70172 | Promoter | 3 | 58415689 | 58466787 | 51099 | 1 | ENSMUST00 | 0 | Tsc22d2 | ENSMUSG00000027806.5 |
| chr13 | 13589771 | 13591179 | 43.4883 | 142.6545 | 137.66066 | Promoter | 13 | 13590409 | 13777440 | 187032 | 1 | ENSMUST00 | 0 | Lyst | ENSMUSG00000019726.10 |
| chr12 | 69296264 | 69297928 | 40.24926 | 142.56519 | 137.57257 | Promoter | 12 | 69296681 | 69301117 | 4437 | 1 | ENSMUST00 | 0 | Klhdc2 | ENSMUSG00000020978.5 |
| chr7 | 109751569 | 109753100 | 40.24926 | 142.56519 | 137.57257 | Promoter | 7 | 109735834 | 109752279 | 16446 | 2 | ENSMUST00 | 0 | Tmem9b | ENSMUSG00000031021.8 |
| chr3 | 87905898 | 87907578 | 42.78876 | 142.56003 | 137.56947 | Promoter | 3 | 87906321 | 87916132 | 9812 | 1 | ENSMUST00 | 0 | Hdgf | ENSMUSG00000004897.11 |
| chr9 | 60793797 | 60796563 | 42.78876 | 142.56003 | 137.56947 | Promoter | 9 | 60794548 | 60880370 | 85823 | 1 | ENSMUST00 | 0 | Uaca | ENSMUSG00000034485.9 |
| chr12 | 82168802 | 82171414 | 42.12499 | 142.52214 | 137.54099 | Promoter | 12 | 82170016 | 82451782 | 281767 | 1 | ENSMUST00 | 0 | Sipa1l1 | ENSMUSG00000042700.10 |
| chr15 | 98762094 | 98763689 | 41.49189 | 142.53105 | 137.54099 | Promoter | 15 | 98737621 | 98763211 | 25591 | 2 | ENSMUST00 | 0 | Arf3 | ENSMUSG00000051853.8 |
| chr17 | 48408332 | 48409762 | 41.49189 | 142.53105 | 137.54099 | Promoter | 17 | 48389081 | 48409730 | 20650 | 2 | ENSMUST00 | 0 | Nfya | ENSMUSG00000023994.8 |
| chr18 | 63691141 | 63692990 | 42.12499 | 142.52214 | 137.54099 | Promoter | 18 | 63662801 | 63692359 | 29559 | 2 | ENSMUST00 | 0 | Txnl1 | ENSMUSG00000024583.2 |
| chr5 | 32610381 | 32612364 | 41.49189 | 142.53105 | 137.54099 | Promoter | 5 | 32611171 | 32687066 | 75896 | 1 | ENSMUST00 | 0 | Yes1 | ENSMUSG00000014932.10 |
| chr5 | 86171817 | 86173491 | 41.49189 | 142.53105 | 137.54099 | Promoter | 5 | 86110720 | 86172803 | 62084 | 2 | ENSMUST00 | 0 | Uba6 | ENSMUSG00000035898.8 |
| chr7 | 16921011 | 16924086 | 41.49189 | 142.53105 | 137.54099 | Promoter | 7 | 16917144 | 16923829 | 6686 | 2 | ENSMUST00 | 0 | Calm3 | ENSMUSG00000019370.10 |
| chr7 | 127903936 | 127904880 | 41.49189 | 142.53105 | 137.54099 | Promoter | 7 | 127904082 | 127909664 | 5583 | 1 | ENSMUST00 | 0 | Bckdk | ENSMUSG00000030802.9 |
| chr8 | 13757380 | 13758515 | 42.12499 | 142.52214 | 137.54099 | Promoter | 8 | 13757676 | 13781938 | 24263 | 1 | ENSMUST00 | 0 | Cdc16 | ENSMUSG00000038416.9 |
| chr3 | 116421128 | 116425134 | 42.70261 | 142.41019 | 137.43451 | Promoter | 3 | 116272553 | 116423986 | 151434 | 2 | ENSMUST00 | 0 | Cdc14a | ENSMUSG00000033502.9 |
| chr8 | 80877775 | 80881139 | 42.61679 | 142.26089 | 137.289 | Promoter | 8 | 80764434 | 80880479 | 116046 | 2 | ENSMUST00 | 0 | Gab1 | ENSMUSG00000031714.8 |
| chr1 | 162569974 | 162571928 | 32.61975 | 142.21701 | 137.24615 | Promoter | 1 | 162570515 | 162599084 | 28570 | 1 | ENSMUST00 | 0 | Vamp4 | ENSMUSG00000026696.10 |
| chr16 | 8830049 | 8831883 | 32.61975 | 142.21701 | 137.24615 | Promoter | 16 | 8830100 | 8858916 | 28817 | 1 | ENSMUST00 | 0 | 1810013L24F | ENSMUSG00000022507.5 |
| chr4 | 134533990 | 134535992 | 32.61975 | 142.21701 | 137.24615 | Promoter | 4 | 134525550 | 134535268 | 9719 | 2 | ENSMUST00 | 0 | Mtfr1l | ENSMUSG00000046671.13 |
| chr5 | 143622170 | 143624398 | 40.66609 | 142.19028 | 137.22005 | Promoter | 5 | 143622447 | 143710250 | 87804 | 1 | ENSMUST00 | 0 | Cyth3 | ENSMUSG00000018001.13 |
| chr11 | 72606702 | 72608877 | 36.38946 | 142.11884 | 137.14919 | Promoter | 11 | 72607283 | 72686481 | 79199 | 1 | ENSMUST00 | 0 | Ube2g1 | ENSMUSG00000020794.8 |
| chr13 | 48545980 | 48546574 | 36.38946 | 142.11884 | 137.14919 | Distal Interge | 13 | 48538171 | 48538270 | 100 | 2 | ENSMUST00 | -7711 | Mirlet7a-1 | ENSMUSG00000065421.2 |
| chr14 | 122106300 | 122108494 | 36.38946 | 142.11884 | 137.14919 | Promoter | 14 | 122107038 | 122159604 | 52567 | 1 | ENSMUST00 | 0 | Tm9sf2 | ENSMUSG00000025544.8 |
| chr15 | 78796787 | 78803936 | 36.38946 | 142.11884 | 137.14919 | Promoter | 15 | 78775138 | 78803042 | 27905 | 2 | ENSMUST00 | 0 | Card10 | ENSMUSG00000033170.9 |
| chr2 | 10079446 | 10081412 | 36.38946 | 142.11884 | 137.14919 | Promoter | 2 | 10056016 | 10080427 | 24412 | 2 | ENSMUST00 | 0 | Atp5c1 | ENSMUSG00000025781.9 |
| chr7 | 29155320 | 29156385 | 36.38946 | 142.11884 | 137.14919 | Promoter | 7 | 29152511 | 29156210 | 3700 | 2 | ENSMUST00 | 0 | Fam98c | ENSMUSG00000030590.9 |
| chr12 | 111375721 | 111378772 | 42.53133 | 142.11215 | 137.14622 | Promoter | 12 | 111292972 | 111377718 | 84747 | 2 | ENSMUST00 | 0 | Cdc42bpb | ENSMUSG00000021279.4 |
| chr4 | 135946033 | 135947539 | 41.87798 | 142.08998 | 137.12619 | Promoter | 4 | 135946448 | 135962617 | 16170 | 1 | ENSMUST00 | 0 | Hmgcl | ENSMUSG00000028672.8 |
| chr14 | 50930207 | 50930957 | 40.5906 | 142.05632 | 137.09286 | Promoter | 14 | 50926070 | 50930849 | 4780 | 2 | ENSMUST00 | 0 | Tmem55b | ENSMUSG00000035953.8 |
| chr15 | 58075166 | 58077523 | 40.5906 | 142.05632 | 137.09286 | Promoter | 15 | 58047003 | 58076245 | 29243 | 2 | ENSMUST00 | 0 | Zhx1 | ENSMUSG00000022361.9 |
| chr4 | 148625679 | 148627458 | 40.5906 | 142.05632 | 137.09286 | Promoter | 4 | 148612382 | 148626996 | 14615 | 2 | ENSMUST00 | 0 | Tardbp | ENSMUSG00000041459.10 |
| chr1 | 160211644 | 160212976 | 38.67513 | 141.95352 | 136.99454 | Promoter | 1 | 160202367 | 160212892 | 10526 | 2 | ENSMUST00 | 0 | Cacybp | ENSMUSG00000014226.5 |
| chr1 | 191905836 | 191908156 | 38.67513 | 141.95352 | 136.99454 | Promoter | 1 | 191906767 | 191913247 | 6481 | 1 | ENSMUST00 | 0 | Slc30a1 | ENSMUSG00000037434.7 |
| chr12 | 59217895 | 59220396 | 38.67513 | 141.95352 | 136.99454 | Promoter | 12 | 59200655 | 59219725 | 19071 | 2 | ENSMUST00 | 0 | Fbxo33 | ENSMUSG00000035329.7 |
| chr14 | 63543062 | 63544610 | 38.67513 | 141.95352 | 136.99454 | Promoter | 14 | 63523616 | 63543953 | 20338 | 2 | ENSMUST00 | 0 | Mtmr9 | ENSMUSG00000035078.5 |
| chr5 | 147859955 | 147861658 | 38.67513 | 141.95352 | 136.99454 | Promoter | 5 | 147860628 | 147875778 | 15151 | 1 | ENSMUST00 | 0 | Pomp | ENSMUSG00000029649.7 |
| chr8 | 39005471 | 39006885 | 38.67513 | 141.95352 | 136.99454 | Promoter | 8 | 39005867 | 39165112 | 159246 | 1 | ENSMUST00 | 0 | Tusc3 | ENSMUSG00000039530.11 |
| chr9 | 69989429 | 69990676 | 39.2181 | 141.82785 | 136.87613 | Promoter | 9 | 69989466 | 70008317 | 18852 | 1 | ENSMUST00 | 0 | Bnip2 | ENSMUSG00000011958.12 |
| chr9 | 106890890 | 106892979 | 39.2181 | 141.82785 | 136.87613 | Promoter | 9 | 106838312 | 106891872 | 53561 | 2 | ENSMUST00 | 0 | Manf | ENSMUSG00000032575.11 |
| chr11 | 21369063 | 21371817 | 43.65555 | 141.82639 | 136.87535 | Promoter | 11 | 21321138 | 21370477 | 49340 | 2 | ENSMUST00 | 0 | Ugp2 | ENSMUSG00000001891.11 |
| chr11 | 70562192 | 70564153 | 43.65555 | 141.82639 | 136.87535 | Promoter | 11 | 70562881 | 70614482 | 51602 | 1 | ENSMUST00 | 0 | Mink1 | ENSMUSG00000020827.13 |
| chr4 | 133010598 | 133013973 | 43.65555 | 141.82639 | 136.87535 | Promoter | 4 | 133011260 | 133060820 | 49561 | 1 | ENSMUST00 | 0 | Ahdc1 | ENSMUSG00000037692.9 |
| chr8 | 71542828 | 71543684 | 43.65555 | 141.82639 | 136.87535 | Promoter | 8 | 71542930 | 71548024 | 5095 | 1 | ENSMUST00 | 0 | Mvb12a | ENSMUSG00000031813.7 |
| chr1 | 191395099 | 191398253 | 42.9543 | 141.72932 | 136.78545 | Promoter | 1 | 191351981 | 191397041 | 45061 | 2 | ENSMUST00 | 0 | Ppp2r5a | ENSMUSG00000026626.6 |
| chr4 | 3874596 | 3875649 | 42.9543 | 141.72932 | 136.78545 | Distal Interge | 4 | 3870657 | 3872105 | 1449 | 2 | ENSMUST00 | -2492 | Mos | ENSMUSG00000078365.5 |
| chr10 | 89685331 | 89686796 | 42.27697 | 141.66908 | 136.72713 | Promoter | 10 | 89638721 | 89686285 | 47565 | 2 | ENSMUST00 | 0 | Scyl2 | ENSMUSG00000069539.6 |
| chr17 | 71551587 | 71552821 | 40.94485 | 141.56706 | 136.62593 | Promoter | 17 | 71552061 | 71578700 | 26640 | 1 | ENSMUST00 | 0 | Spdya | ENSMUSG00000052525.9 |
| chr18 | 12127847 | 12129840 | 40.94485 | 141.56706 | 136.62593 | Promoter | 18 | 12128850 | 12157367 | 28518 | 1 | ENSMUST00 | 0 | Riok3 | ENSMUSG00000024404.6 |
| chr2 | 25607393 | 25608770 | 40.94485 | 141.56706 | 136.62593 | Promoter | 2 | 25583018 | 25608521 | 25504 | 2 | ENSMUST00 | 0 | Rabl6 | ENSMUSG00000015087.9 |
| chr5 | 48371961 | 48373134 | 40.94485 | 141.56706 | 136.62593 | Promoter | 5 | 48372392 | 48388756 | 16365 | 1 | ENSMUST00 | 0 | Pacrgl | ENSMUSG00000029089.4 |
| chr17 | 28232691 | 28233753 | 43.4858 | 141.53311 | 136.59532 | Promoter | 17 | 28232754 | 28301469 | 68716 | 1 | ENSMUST00 | 0 | Ppard | ENSMUSG00000002250.10 |
| chr8 | 88360901 | 88363027 | 43.4858 | 141.53311 | 136.59532 | Promoter | 8 | 88331059 | 88362194 | 31136 | 2 | ENSMUST00 | 0 | Brd7 | ENSMUSG00000031660.8 |
| chr17 | 23770889 | 23771715 | 42.19286 | 141.52246 | 136.58501 | Promoter | 17 | 23755423 | 23771591 | 16169 | 2 | ENSMUST00 | 0 | Flywch1 | ENSMUSG00000040097.10 |
| chr5 | 143403208 | 143404946 | 40.86754 | 141.43057 | 136.49426 | Promoter | 5 | 143403820 | 143421900 | 18081 | 1 | ENSMUST00 | 0 | Kdelr2 | ENSMUSG00000079111.2 |
| chr1 | 53296545 | 53297894 | 43.39494 | 141.3761 | 136.44113 | Promoter | 1 | 53297095 | 53301953 | 4859 | 1 | ENSMUST00 | 0 | Ormdl1 | ENSMUSG00000026097.4 |
| chr16 | 17200175 | 17201755 | 43.39494 | 141.3761 | 136.44113 | Promoter | 16 | 17152015 | 17201492 | 49478 | 2 | ENSMUST00 | 0 | Ube2l3 | ENSMUSG00000038965.10 |
| chr2 | 152331574 | 152333091 | 43.39494 | 141.3761 | 136.44113 | Promoter | 2 | 152316334 | 152332412 | 16079 | 2 | ENSMUST00 | 0 | Rbck1 | ENSMUSG00000027466.10 |
| chr11 | 59947795 | 59948892 | 39.52876 | 141.27539 | 136.34271 | Promoter | 11 | 59948206 | 59962205 | 14000 | 1 | ENSMUST00 | 0 | Med9 | ENSMUSG00000061650.6 |
| chr17 | 25271619 | 25274673 | 39.52876 | 141.27539 | 136.34271 | Promoter | 17 | 25261916 | 25274310 | 12395 | 2 | ENSMUST00 | 0 | Ube2i | ENSMUSG00000015120.10 |
| chr5 | 23787616 | 23788615 | 39.52876 | 141.27539 | 136.34271 | Promoter | 5 | 23787711 | 23790003 | 2293 | 1 | ENSMUST00 | 0 | Rint1 | ENSMUSG00000028999.10 |
| chr14 | 49065401 | 49067299 | 40.14342 | 141.26135 | 136.33099 | Promoter | 14 | 49066495 | 49087723 | 21229 | 1 | ENSMUST00 | 0 | Ap5m1 | ENSMUSG00000036291.5 |
| chr13 | 34001889 | 34003343 | 38.25702 | 141.19 | 136.26103 | Promoter | 13 | 34002363 | 34033677 | 31315 | 1 | ENSMUST00 | 0 | Ripk1 | ENSMUSG00000021408.9 |
| chr15 | 79108405 | 79110109 | 42.60553 | 141.12407 | 136.19626 | Promoter | 15 | 79108972 | 79136897 | 27926 | 1 | ENSMUST00 | 0 | Micall1 | ENSMUSG00000033039.9 |
| chr10 | 99106403 | 99109000 | 34.18682 | 141.10779 | 136.18079 | Promoter | 10 | 99107036 | 99197988 | 90953 | 1 | ENSMUST00 | 0 | Poc1b | ENSMUSG00000019952.9 |
| chr5 | 129019359 | 129021396 | 34.18682 | 141.10779 | 136.18079 | Promoter | 5 | 129020069 | 129024323 | 4255 | 1 | ENSMUST00 | 0 | Ran | ENSMUSG00000029430.8 |
| chr12 | 116280328 | 116282437 | 41.31276 | 141.09892 | 136.17311 | Promoter | 12 | 116281222 | 116373096 | 91875 | 1 | ENSMUST00 | 0 | Esyt2 | ENSMUSG00000021171.7 |
| chr15 | 68360668 | 68363243 | 41.31276 | 141.09892 | 136.17311 | Promoter | 15 | 68340983 | 68363176 | 22194 | 2 | ENSMUST00 | 0 | Gm20732 | ENSMUSG00000093387.1 |
| chr16 | 5048737 | 5051282 | 41.31276 | 141.09892 | 136.17311 | Promoter | 16 | 5013906 | 5049910 | 36005 | 2 | ENSMUST00 | 0 | Glyr1 | ENSMUSG00000022536.8 |
| chr18 | 52767566 | 52770026 | 41.31276 | 141.09892 | 136.17311 | Promoter | 18 | 52767709 | 52915864 | 148156 | 1 | ENSMUST00 | 0 | Sncaip | ENSMUSG00000024534.10 |
| chr5 | 21737047 | 21737944 | 41.31276 | 141.09892 | 136.17311 | Promoter | 5 | 21737141 | 21757152 | 20012 | 1 | ENSMUST00 | 0 | Pmpcb | ENSMUSG00000029017.8 |
| chr7 | 97695965 | 97697507 | 41.31276 | 141.09892 | 136.17311 | Promoter | 7 | 97696634 | 97718879 | 22246 | 1 | ENSMUST00 | 0 | Clns1a | ENSMUSG00000025439.4 |
| chr8 | 119861809 | 119862974 | 41.31276 | 141.09892 | 136.17311 | Promoter | 8 | 119862305 | 119876988 | 14684 | 1 | ENSMUST00 | 0 | Klhl36 | ENSMUSG00000031828.8 |
| chr18 | 76929562 | 76931051 | 41.94252 | 141.08568 | 136.16306 | Promoter | 18 | 76930126 | 76971449 | 41324 | 1 | ENSMUST00 | 0 | Hdhd2 | ENSMUSG00000025421.9 |
| chr12 | 108422545 | 108423952 | 42.51921 | 140.97417 | 136.05247 | Promoter | 12 | 108422816 | 108539564 | 116749 | 1 | ENSMUST00 | 0 | Eml1 | ENSMUSG00000058070.9 |
| chr16 | 4878216 | 4880538 | 42.51921 | 140.97417 | 136.05247 | Promoter | 16 | 4874778 | 4880315 | 5538 | 2 | ENSMUST00 | 0 | Ubald1 | ENSMUSG00000039568.6 |
| chr4 | 114405755 | 114407430 | 42.51921 | 140.97417 | 136.05247 | Promoter | 4 | 114406724 | 114615098 | 208375 | 1 | ENSMUST00 | 0 | Trabd2b | ENSMUSG00000070867.4 |
| chr17 | 65883962 | 65886522 | 41.23326 | 140.95923 | 136.03836 | Promoter | 17 | 65848433 | 65884923 | 36491 | 2 | ENSMUST00 | 0 | Ralbp1 | ENSMUSG00000024096.11 |
| chr19 | 6234264 | 6236344 | 43.03529 | 140.75403 | 135.83646 | Promoter | 19 | 6227765 | 6235872 | 8108 | 2 | ENSMUST00 | 0 | Ppp2r5b | ENSMUSG00000024777.9 |
| chr11 | 70652869 | 70656609 | 41.07278 | 140.67711 | 135.76277 | Promoter | 11 | 70651850 | 70654644 | 2795 | 2 | ENSMUST00 | 0 | Pfn1 | ENSMUSG00000018293.4 |
| chr11 | 97361882 | 97364182 | 41.07278 | 140.67711 | 135.76277 | Promoter | 11 | 97362551 | 97398542 | 35992 | 1 | ENSMUST00 | 0 | Socs7 | ENSMUSG00000038485.5 |
| chr11 | 98794305 | 98797777 | 41.07278 | 140.67711 | 135.76277 | Promoter | 11 | 98795516 | 98807859 | 12344 | 1 | ENSMUST00 | 0 | Msl1 | ENSMUSG00000052915.9 |
| chr12 | 75733833 | 75736200 | 41.07278 | 140.67711 | 135.76277 | Promoter | 12 | 75714248 | 75735729 | 21482 | 2 | ENSMUST00 | 0 | Sgpp1 | ENSMUSG00000021054.5 |
| chr13 | 74639074 | 74640621 | 41.07278 | 140.67711 | 135.76277 | Promoter | 13 | 74639872 | 74691875 | 52004 | 1 | ENSMUST00 | 0 | Erap1 | ENSMUSG00000021583.6 |
| chr14 | 21051549 | 21053380 | 41.07278 | 140.67711 | 135.76277 | Promoter | 14 | 21031442 | 21052450 | 21009 | 2 | ENSMUST00 | 0 | Ap3m1 | ENSMUSG00000021824.7 |
| chr14 | 118937104 | 118938956 | 41.07278 | 140.67711 | 135.76277 | Promoter | 14 | 118937932 | 118981702 | 43771 | 1 | ENSMUST00 | 0 | Dnajc3 | ENSMUSG00000022136.7 |
| chr15 | 25842720 | 25844970 | 41.07278 | 140.67711 | 135.76277 | Promoter | 15 | 25843264 | 25973687 | 130424 | 1 | ENSMUST00 | 0 | Fam134b | ENSMUSG00000022270.10 |
| chr16 | 64850498 | 64851808 | 42.34764 | 140.67596 | 135.76277 | Promoter | 16 | 64851996 | 64859507 | 7512 | 1 | ENSMUST00 | -188 | Cggbp1 | ENSMUSG00000054604.7 |
| chr18 | 61398143 | 61401028 | 41.07278 | 140.67711 | 135.76277 | Promoter | 18 | 61298136 | 61400431 | 102296 | 2 | ENSMUST00 | 0 | Ppargc1b | ENSMUSG00000033871.9 |
| chr19 | 53902548 | 53904667 | 41.07278 | 140.67711 | 135.76277 | Promoter | 19 | 53903351 | 53929861 | 26511 | 1 | ENSMUST00 | 0 | Pdcd4 | ENSMUSG00000024975.7 |
| chr19 | 60225670 | 60226851 | 41.07278 | 140.67711 | 135.76277 | Promoter | 19 | 60226334 | 60227678 | 1345 | 1 | ENSMUST00 | 0 | 4933412A08I | ENSMUSG00000043623.7 |
| chr2 | 166995030 | 166996893 | 41.07278 | 140.67711 | 135.76277 | Promoter | 2 | 166947549 | 166996278 | 48730 | 2 | ENSMUST00 | 0 | Stau1 | ENSMUSG00000039536.11 |
| chr4 | 24897370 | 24898737 | 41.07278 | 140.67711 | 135.76277 | Promoter | 4 | 24898083 | 24905001 | 6919 | 1 | ENSMUST00 | 0 | Ndufaf4 | ENSMUSG00000028261.8 |
| chr4 | 117155140 | 117156629 | 41.07278 | 140.67711 | 135.76277 | Promoter | 4 | 117153827 | 117156243 | 2417 | 2 | ENSMUST00 | 0 | Rps8 | ENSMUSG00000047675.10 |
| chr6 | 7844040 | 7846005 | 41.07278 | 140.67711 | 135.76277 | Promoter | 6 | 7844842 | 7872047 | 27206 | 1 | ENSMUST00 | 0 | C1galt1 | ENSMUSG00000042460.4 |
| chr7 | 75847679 | 75849583 | 41.07278 | 140.67711 | 135.76277 | Promoter | 7 | 75848338 | 75874130 | 25793 | 1 | ENSMUST00 | 0 | Klhl25 | ENSMUSG00000055652.9 |
| chr8 | 84831548 | 84833112 | 41.07278 | 140.67711 | 135.76277 | Promoter | 8 | 84831522 | 84835482 | 3961 | 1 | ENSMUST00 | 27 | Gadd45gip1 | ENSMUSG00000033751.5 |
| chr8 | 108703474 | 108705192 | 41.07278 | 140.67711 | 135.76277 | Promoter | 8 | 108703100 | 108800270 | 97171 | 1 | ENSMUST00 | 375 | Zfhx3 | ENSMUSG00000038872.9 |
| chr9 | 54863122 | 54864695 | 41.07278 | 140.67711 | 135.76277 | Promoter | 9 | 54863755 | 54912534 | 48780 | 1 | ENSMUST00 | 0 | Ireb2 | ENSMUSG00000032293.7 |
| chr6 | 52227601 | 52235632 | 41.69514 | 140.65352 | 135.74945 | Promoter | 6 | 52231197 | 52234939 | 3743 | 2 | ENSMUST00 | 0 | Hoxa10 | ENSMUSG00000000938.11 |
| chr12 | 70110700 | 70112394 | 38.54033 | 140.59962 | 135.69896 | Promoter | 12 | 70018740 | 70111925 | 93186 | 2 | ENSMUST00 | 0 | Nin | ENSMUSG00000021068.10 |
| chr10 | 108161189 | 108163933 | 32.33361 | 140.55707 | 135.65718 | Promoter | 10 | 108162400 | 108277575 | 115176 | 1 | ENSMUST00 | 0 | Ppp1r12a | ENSMUSG00000019907.9 |
| chr2 | 39189954 | 39190808 | 32.33361 | 140.55707 | 135.65718 | Promoter | 2 | 39066214 | 39190730 | 124517 | 2 | ENSMUST00 | 0 | Scai | ENSMUSG00000035236.12 |
| chr1 | 135146032 | 135147917 | 42.26237 | 140.52768 | 135.6286 | Promoter | 1 | 135146824 | 135156269 | 9446 | 1 | ENSMUST00 | 0 | Arl8a | ENSMUSG00000026426.5 |
| chr6 | 115037211 | 115038032 | 42.26237 | 140.52768 | 135.6286 | Promoter | 6 | 115004381 | 115037876 | 33496 | 2 | ENSMUST00 | 0 | Tamm41 | ENSMUSG00000030316.8 |
| chr4 | 119538635 | 119541529 | 41.61333 | 140.51047 | 135.61182 | Promoter | 4 | 119539661 | 119629119 | 89459 | 1 | ENSMUST00 | 0 | Foxj3 | ENSMUSG00000032998.11 |
| chr12 | 79171474 | 79173188 | 35.51886 | 140.45512 | 135.55692 | Promoter | 12 | 79156017 | 79172667 | 16651 | 2 | ENSMUST00 | 0 | Vti1b | ENSMUSG00000021124.8 |
| chr2 | 120849487 | 120850506 | 42.17744 | 140.37996 | 135.48283 | Promoter | 2 | 120718186 | 120850128 | 131943 | 2 | ENSMUST00 | 0 | Cdan1 | ENSMUSG00000027284.11 |
| chr1 | 182340569 | 182342123 | 36.04937 | 140.37729 | 135.48044 | Promoter | 1 | 182313102 | 182341606 | 28505 | 2 | ENSMUST00 | 0 | Fbxo28 | ENSMUSG00000047539.4 |
| chr13 | 49187428 | 49188528 | 36.04937 | 140.37729 | 135.48044 | Promoter | 13 | 49187485 | 49196244 | 8760 | 1 | ENSMUST00 | 0 | Ninj1 | ENSMUSG00000037966.10 |
| chr16 | 22263924 | 22266550 | 36.04937 | 140.37729 | 135.48044 | Promoter | 16 | 22244549 | 22266005 | 21457 | 2 | ENSMUST00 | 0 | Tra2b | ENSMUSG00000022858.10 |
| chr7 | 92560717 | 92562437 | 36.04937 | 140.37729 | 135.48044 | Promoter | 7 | 92561149 | 92564720 | 3572 | 1 | ENSMUST00 | 0 | Ccdc90b | ENSMUSG00000030613.8 |
| chr10 | 81105337 | 81106833 | 41.53183 | 140.36792 | 135.47412 | Promoter | 10 | 81105915 | 81124697 | 18783 | 1 | ENSMUST00 | 0 | Map2k2 | ENSMUSG00000035027.12 |
| chr19 | 60889254 | 60892116 | 41.53183 | 140.36792 | 135.47412 | Promoter | 19 | 60889749 | 61092553 | 202805 | 1 | ENSMUST00 | 0 | Grk5 | ENSMUSG00000003228.8 |
| chr11 | 29171793 | 29174277 | 42.09286 | 140.23274 | 135.34074 | Promoter | 11 | 29172890 | 29209641 | 36752 | 1 | ENSMUST00 | 0 | Smek2 | ENSMUSG00000020463.10 |
| chr4 | 59805261 | 59807196 | 42.09286 | 140.23274 | 135.34074 | Promoter | 4 | 59805840 | 59904737 | 98898 | 1 | ENSMUST00 | 0 | Snx30 | ENSMUSG00000028385.9 |
| chr9 | 59616681 | 59618061 | 42.09286 | 140.23274 | 135.34074 | Promoter | 9 | 59617284 | 59650290 | 33007 | 1 | ENSMUST00 | 0 | Parp6 | ENSMUSG00000025237.10 |
| chr7 | 110060246 | 110061386 | 39.56231 | 140.2242 | 135.33536 | Promoter | 7 | 110057987 | 110061325 | 3339 | 2 | ENSMUST00 | 0 | AA474408 | ENSMUSG00000073867.2 |
| chr1 | 153486180 | 153488090 | 40.18469 | 140.22169 | 135.33289 | Promoter | 1 | 153455758 | 153487608 | 31851 | 2 | ENSMUST00 | 0 | Dhx9 | ENSMUSG00000042699.6 |

| | | | | | | | | | | | | | | | |
|---|---|---|---|---|---|---|---|---|---|---|---|---|---|---|---|
| chr4 | 44755579 | 44757654 | 40.18469 | 140.22169 | 135.33289 | Promoter | 4 | 44755877 | 44761971 | 6095 | 1 | ENSMUST00 | 0 | Zcchc7 | ENSMUSG00000035649.12 |
| chr11 | 86682913 | 86684911 | 38.29964 | 140.16156 | 135.27679 | Promoter | 11 | 86583865 | 86683836 | 99972 | 2 | ENSMUST00 | 0 | Vmp1 | ENSMUSG00000018171.4 |
| chr14 | 73379381 | 73385386 | 38.29964 | 140.16156 | 135.27679 | Promoter | 14 | 73362231 | 73385271 | 23041 | 2 | ENSMUST00 | 0 | Itm2b | ENSMUSG00000022108.7 |
| chr17 | 20945375 | 20947082 | 38.29964 | 140.16156 | 135.27679 | Promoter | 17 | 20945487 | 20958784 | 13298 | 1 | ENSMUST00 | 0 | Ppp2r1a | ENSMUSG00000007564.9 |
| chr2 | 118925995 | 118927451 | 38.29964 | 140.16156 | 135.27679 | Promoter | 2 | 118926496 | 118928585 | 2090 | 1 | ENSMUST00 | 0 | Chst14 | ENSMUSG00000074916.5 |
| chr3 | 146811927 | 146813727 | 38.29964 | 140.16156 | 135.27679 | Promoter | 3 | 146729579 | 146812960 | 83382 | 2 | ENSMUST00 | 0 | Prkacb | ENSMUSG00000005034.10 |
| chr4 | 139192872 | 139194254 | 38.29964 | 140.16156 | 135.27679 | Promoter | 4 | 139192899 | 139291818 | 98920 | 1 | ENSMUST00 | 0 | Capzb | ENSMUSG00000028745.13 |
| chr4 | 148803997 | 148806784 | 38.29964 | 140.16156 | 135.27679 | Promoter | 4 | 148804429 | 148945129 | 140701 | 1 | ENSMUST00 | 0 | Casz1 | ENSMUSG00000028977.11 |
| chr7 | 44553090 | 44554471 | 38.29964 | 140.16156 | 135.27679 | Promoter | 7 | 44549616 | 44553686 | 4071 | 2 | ENSMUST00 | 0 | Nr1h2 | ENSMUSG00000060601.7 |
| chr9 | 53666273 | 53667755 | 38.29964 | 140.16156 | 135.27679 | Promoter | 9 | 53614582 | 53667507 | 52926 | 2 | ENSMUST00 | 0 | Cul5 | ENSMUSG00000032030.11 |
| chr18 | 7867839 | 7871023 | 42.68155 | 140.14148 | 135.26276 | Promoter | 18 | 7868832 | 7926505 | 57674 | 1 | ENSMUST00 | 0 | Wac | ENSMUSG00000024283.9 |
| chr6 | 49262330 | 49264348 | 38.83361 | 140.02353 | 135.14601 | Promoter | 6 | 49243921 | 49264052 | 20132 | 2 | ENSMUST00 | 0 | Tra2a | ENSMUSG00000029817.9 |
| chr9 | 50856535 | 50857714 | 38.83361 | 140.02353 | 135.14601 | Promoter | 9 | 50856924 | 50892797 | 35874 | 1 | ENSMUST00 | 0 | Ppp2r1b | ENSMUSG00000032058.9 |
| chr9 | 36766509 | 36767731 | 38.83361 | 140.02353 | 135.14601 | Promoter | 9 | 36729344 | 36767679 | 38336 | 2 | ENSMUST00 | 0 | Stt3a | ENSMUSG00000032116.12 |
| chr5 | 129500513 | 129502418 | 37.61795 | 140.01674 | 135.14008 | Promoter | 5 | 129501231 | 129571383 | 70153 | 1 | ENSMUST00 | 0 | Sfswap | ENSMUSG00000029439.10 |
| chr12 | 98900319 | 98901839 | 41.28926 | 139.94322 | 135.06778 | Promoter | 12 | 98786604 | 98901484 | 114881 | 2 | ENSMUST00 | 0 | Eml5 | ENSMUSG00000051166.9 |
| chr10 | 79704012 | 79705228 | 43.19602 | 139.91907 | 135.04468 | Promoter | 10 | 79704491 | 79711969 | 7479 | 1 | ENSMUST00 | 0 | Bsg | ENSMUSG00000023175.10 |
| chr14 | 57889078 | 57890767 | 43.19602 | 139.91907 | 135.04468 | Promoter | 14 | 57832702 | 57890262 | 57561 | 2 | ENSMUST00 | 0 | Zdhhc20 | ENSMUSG00000021969.7 |
| chr16 | 20535449 | 20536335 | 43.19602 | 139.91907 | 135.04468 | Promoter | 16 | 20535478 | 20544056 | 8579 | 1 | ENSMUST00 | 0 | Ap2m1 | ENSMUSG00000022841.9 |
| chr8 | 121590065 | 121591628 | 43.19602 | 139.91907 | 135.04468 | Promoter | 8 | 121590361 | 121598760 | 8400 | 1 | ENSMUST00 | 0 | Map1lc3b | ENSMUSG00000031812.11 |
| chr1 | 155557631 | 155559183 | 42.50686 | 139.83865 | 134.96973 | Promoter | 1 | 155558120 | 155572288 | 14169 | 1 | ENSMUST00 | 0 | Acbd6 | ENSMUSG00000033701.8 |
| chr10 | 81543738 | 81545170 | 42.50686 | 139.83865 | 134.96973 | Promoter | 10 | 81528724 | 81545190 | 16467 | 2 | ENSMUST00 | 20 | Gna11 | ENSMUSG00000034781.5 |
| chr12 | 70974273 | 70975788 | 42.50686 | 139.83865 | 134.96973 | Promoter | 12 | 70974621 | 70996347 | 21727 | 1 | ENSMUST00 | 0 | Psma3 | ENSMUSG00000060073.4 |
| chr6 | 47835563 | 47836836 | 41.84113 | 139.79433 | 134.92708 | Promoter | 6 | 47835661 | 47843647 | 7987 | 1 | ENSMUST00 | 0 | Zfp398 | ENSMUSG00000062519.8 |
| chr11 | 52098650 | 52100712 | 40.53126 | 139.72299 | 134.85652 | Promoter | 11 | 52098681 | 52127778 | 29098 | 1 | ENSMUST00 | 0 | Ppp2ca | ENSMUSG00000020349.4 |
| chr12 | 84316044 | 84317963 | 40.53126 | 139.72299 | 134.85652 | Promoter | 12 | 84316859 | 84343830 | 26972 | 1 | ENSMUST00 | 0 | Zfp410 | ENSMUSG00000042472.10 |
| chr13 | 76097951 | 76099536 | 40.53126 | 139.72299 | 134.85652 | Promoter | 13 | 76060422 | 76098648 | 38227 | 2 | ENSMUST00 | 0 | Arsk | ENSMUSG00000021592.10 |
| chr4 | 155890767 | 155893165 | 40.53126 | 139.72299 | 134.85652 | Promoter | 4 | 155891822 | 155907250 | 15429 | 1 | ENSMUST00 | 0 | Acap3 | ENSMUSG00000029033.11 |
| chr9 | 118149159 | 118150425 | 40.53126 | 139.72299 | 134.85652 | Promoter | 9 | 118064529 | 118150196 | 85668 | 2 | ENSMUST00 | 0 | Cmc1 | ENSMUSG00000039163.9 |
| chr15 | 93517856 | 93520341 | 41.67497 | 139.50464 | 134.64323 | Promoter | 15 | 93499114 | 93519523 | 20410 | 2 | ENSMUST00 | 0 | Prickle1 | ENSMUSG00000036158.7 |
| chr11 | 50130903 | 50132247 | 42.93816 | 139.47357 | 134.61256 | Promoter | 11 | 50131396 | 50172030 | 40635 | 1 | ENSMUST00 | 0 | Tbc1d9b | ENSMUSG00000036644.10 |
| chr3 | 51408086 | 51409036 | 42.93816 | 139.47357 | 134.61256 | Promoter | 3 | 51404677 | 51408988 | 4312 | 2 | ENSMUST00 | 0 | Ndufc1 | ENSMUSG00000037152.6 |
| chr10 | 121625236 | 121627023 | 39.13739 | 139.46275 | 134.60301 | Promoter | 10 | 121587380 | 121626316 | 38937 | 2 | ENSMUST00 | 0 | Xpot | ENSMUSG00000034667.7 |
| chr10 | 85101964 | 85103897 | 40.3785 | 139.45323 | 134.59462 | Promoter | 10 | 85097019 | 85102495 | 5477 | 2 | ENSMUST00 | 0 | Fhl4 | ENSMUSG00000050035.6 |
| chr4 | 63544737 | 63545610 | 33.87604 | 139.4122 | 134.55379 | Promoter | 4 | 63544772 | 63550750 | 5979 | 1 | ENSMUST00 | 0 | Atp6v1g1 | ENSMUSG00000039105.6 |
| chr12 | 81859409 | 81861509 | 37.88559 | 139.40556 | 134.54842 | Promoter | 12 | 81860030 | 82000924 | 140895 | 1 | ENSMUST00 | 0 | Pcnx | ENSMUSG00000021140.8 |
| chr11 | 77982697 | 77985582 | 42.24747 | 139.38866 | 134.53271 | Promoter | 11 | 77982754 | 78026214 | 43461 | 1 | ENSMUST00 | 0 | Phf12 | ENSMUSG00000037791.10 |
| chr12 | 3571809 | 3573315 | 41.59238 | 139.36057 | 134.50562 | Promoter | 12 | 3572381 | 3781796 | 209416 | 1 | ENSMUST00 | 0 | Dtnb | ENSMUSG00000071454.8 |
| chr10 | 91122573 | 91124044 | 40.8912 | 139.24503 | 134.39108 | Promoter | 10 | 91116791 | 91123957 | 7167 | 2 | ENSMUST00 | 0 | Slc25a3 | ENSMUSG00000061904.7 |
| chr12 | 102469150 | 102470031 | 40.8912 | 139.24503 | 134.39108 | Promoter | 12 | 102469910 | 102497902 | 27993 | 1 | ENSMUST00 | 0 | Golga5 | ENSMUSG00000021192.10 |
| chr6 | 90369051 | 90371189 | 40.8912 | 139.24503 | 134.39108 | Promoter | 6 | 90369492 | 90385084 | 15593 | 1 | ENSMUST00 | 0 | Zxdc | ENSMUSG00000034430.11 |
| chr9 | 104002614 | 104003433 | 40.8912 | 139.24503 | 134.39108 | Promoter | 9 | 104002654 | 104153483 | 150830 | 1 | ENSMUST00 | 0 | Nphp3 | ENSMUSG00000032558.13 |
| chr11 | 115419229 | 115421314 | 42.16172 | 139.23978 | 134.38916 | Promoter | 11 | 115420128 | 115431274 | 11147 | 1 | ENSMUST00 | 0 | Kctd2 | ENSMUSG00000016940.13 |
| chr9 | 99242651 | 99244141 | 42.16172 | 139.23978 | 134.38916 | Promoter | 9 | 99243367 | 99263762 | 20396 | 1 | ENSMUST00 | 0 | Cep70 | ENSMUSG00000056267.9 |
| chr11 | 63918689 | 63922332 | 41.51012 | 139.21701 | 134.36754 | Promoter | 11 | 63885792 | 63922290 | 36499 | 2 | ENSMUST00 | 0 | Hs3st3b1 | ENSMUSG00000070407.5 |
| chr5 | 72914347 | 72917697 | 41.51012 | 139.21701 | 134.36754 | Promoter | 5 | 72914418 | 72918485 | 4068 | 1 | ENSMUST00 | 0 | Slain2 | ENSMUSG00000036087.13 |
| chr2 | 122368187 | 122369883 | 42.75948 | 139.16466 | 134.31596 | Promoter | 2 | 122348893 | 122368918 | 20026 | 2 | ENSMUST00 | 0 | Shf | ENSMUSG00000033256.9 |
| chr7 | 44868285 | 44870536 | 42.75948 | 139.16466 | 134.31596 | Promoter | 7 | 44863068 | 44869788 | 6721 | 2 | ENSMUST00 | 0 | Ptov1 | ENSMUSG00000038502.10 |
| chr1 | 43933120 | 43935537 | 42.07631 | 139.09143 | 134.24335 | Promoter | 1 | 43933647 | 43949405 | 15759 | 1 | ENSMUST00 | 0 | Tpp2 | ENSMUSG00000041763.9 |
| chr16 | 3883780 | 3885605 | 42.07631 | 139.09143 | 134.24335 | Promoter | 16 | 3884619 | 3904780 | 20162 | 1 | ENSMUST00 | 0 | Naa60 | ENSMUSG00000005982.8 |
| chr18 | 36560556 | 36561910 | 42.07631 | 139.09143 | 134.24335 | Promoter | 18 | 36560611 | 36658908 | 98298 | 1 | ENSMUST00 | 0 | Ankhd1 | ENSMUSG00000024483.13 |
| chr1 | 90603053 | 90604104 | 41.42819 | 139.07397 | 134.22679 | Promoter | 1 | 90603476 | 90613124 | 9649 | 1 | ENSMUST00 | 0 | Cops8 | ENSMUSG00000034432.6 |
| chr1 | 75191232 | 75192989 | 39.45224 | 138.91832 | 134.07426 | Promoter | 1 | 75192151 | 75195084 | 2934 | 1 | ENSMUST00 | 0 | Ankzf1 | ENSMUSG00000026199.12 |
| chr1 | 170866958 | 170869093 | 39.45224 | 138.91832 | 134.07426 | Promoter | 1 | 170704674 | 170867771 | 163098 | 2 | ENSMUST00 | 0 | Atf6 | ENSMUSG00000026663.6 |
| chr12 | 16810615 | 16811935 | 39.45224 | 138.91832 | 134.07426 | Promoter | 12 | 16810965 | 16826743 | 15779 | 1 | ENSMUST00 | 0 | E2f6 | ENSMUSG00000057469.7 |
| chr9 | 15279108 | 15280707 | 39.45224 | 138.91832 | 134.07426 | Promoter | 9 | 15260352 | 15279867 | 19516 | 2 | ENSMUST00 | 0 | Med17 | ENSMUSG00000031935.8 |
| chr1 | 185328457 | 185329651 | 32.04747 | 138.90092 | 134.05882 | Promoter | 1 | 185284726 | 185329396 | 44671 | 2 | ENSMUST00 | 0 | Iars2 | ENSMUSG00000026618.6 |
| chr13 | 54692600 | 54693974 | 32.04747 | 138.90092 | 134.05882 | Promoter | 13 | 54679410 | 54693847 | 14438 | 2 | ENSMUST00 | 0 | Rnf44 | ENSMUSG00000034928.11 |
| chr17 | 63862964 | 63864912 | 32.04747 | 138.90092 | 134.05882 | Promoter | 17 | 63863300 | 63863791 | 492 | 2 | ENSMUST00 | 0 | A930002H24 | ENSMUSG00000045506.3 |
| chr13 | 21749989 | 21750947 | 25.81531 | 138.84267 | 134.0024 | Promoter | 13 | 21750194 | 21750505 | 312 | 2 | ENSMUST00 | 0 | Hist1h4k | ENSMUSG00000064288.4 |
| chr8 | 107055713 | 107057716 | 40.65603 | 138.83176 | 133.99156 | Promoter | 8 | 107046289 | 107056689 | 10401 | 2 | ENSMUST00 | 0 | Cog8 | ENSMUSG00000031916.12 |
| chr11 | 88046872 | 88048999 | 40.65367 | 138.82761 | 133.98773 | Promoter | 11 | 88047373 | 88053755 | 6383 | 1 | ENSMUST00 | 0 | Srsf1 | ENSMUSG00000018379.12 |
| chr11 | 106215465 | 106217940 | 40.65367 | 138.82761 | 133.98773 | Promoter | 11 | 106216926 | 106249139 | 32214 | 1 | ENSMUST00 | 0 | Ddx42 | ENSMUSG00000020705.6 |
| chr12 | 54794178 | 54796105 | 40.65367 | 138.82761 | 133.98773 | Promoter | 12 | 54746349 | 54795703 | 49355 | 2 | ENSMUST00 | 0 | Snx6 | ENSMUSG00000005656.8 |
| chr12 | 40444767 | 40448342 | 40.65367 | 138.82761 | 133.98773 | Promoter | 12 | 40446053 | 40846486 | 400434 | 1 | ENSMUST00 | 0 | Dock4 | ENSMUSG00000035954.9 |
| chr2 | 130628866 | 130630406 | 40.65367 | 138.82761 | 133.98773 | Promoter | 2 | 130590002 | 130630038 | 40037 | 2 | ENSMUST00 | 0 | Ubox5 | ENSMUSG00000027300.9 |
| chr3 | 34019437 | 34021363 | 40.65367 | 138.82761 | 133.98773 | Promoter | 3 | 34020079 | 34069342 | 49264 | 1 | ENSMUST00 | 0 | Fxr1 | ENSMUSG00000027680.10 |
| chr3 | 67429814 | 67430976 | 40.65367 | 138.82761 | 133.98773 | Promoter | 3 | 67430096 | 67476529 | 46434 | 1 | ENSMUST00 | 0 | Gfm1 | ENSMUSG00000027774.11 |
| chr4 | 132883493 | 132885139 | 40.65367 | 138.82761 | 133.98773 | Promoter | 4 | 132853501 | 132884509 | 31009 | 2 | ENSMUST00 | 0 | Stx12 | ENSMUSG00000028879.4 |
| chr5 | 142902951 | 142907768 | 40.65367 | 138.82761 | 133.98773 | Promoter | 5 | 142903115 | 142906754 | 3640 | 2 | ENSMUST00 | 0 | Actb | ENSMUSG00000029580.9 |
| chr8 | 47673181 | 47676469 | 40.65367 | 138.82761 | 133.98773 | Promoter | 8 | 47667178 | 47673689 | 6512 | 2 | ENSMUST00 | 0 | Ing2 | ENSMUSG00000063049.4 |
| chr15 | 84191749 | 84192877 | 38.16249 | 138.80753 | 133.97699 | Promoter | 15 | 84192233 | 84214303 | 22071 | 1 | ENSMUST00 | 0 | Samm50 | ENSMUSG00000022437.6 |
| chr11 | 75678797 | 75680664 | 41.2653 | 138.78937 | 133.96024 | Promoter | 11 | 75679259 | 75706065 | 26807 | 1 | ENSMUST00 | 0 | Crk | ENSMUSG00000017776.10 |
| chr15 | 55071338 | 55072780 | 41.2653 | 138.78937 | 133.96024 | Promoter | 15 | 55015129 | 55072152 | 57024 | 2 | ENSMUST00 | 0 | Taf2 | ENSMUSG00000037343.7 |
| chr16 | 16302532 | 16303955 | 41.2653 | 138.78937 | 133.96024 | Promoter | 16 | 16302965 | 16308942 | 5978 | 1 | ENSMUST00 | 0 | Yars2 | ENSMUSG00000022792.11 |
| chr4 | 141301143 | 141303243 | 41.2653 | 138.78937 | 133.96024 | Promoter | 4 | 141301240 | 141329384 | 28145 | 1 | ENSMUST00 | 0 | Epha2 | ENSMUSG00000006445.3 |
| chr9 | 20951839 | 20953287 | 41.26529 | 138.78937 | 133.96024 | Promoter | 9 | 20907206 | 20952623 | 45418 | 2 | ENSMUST00 | 0 | Dnmt1 | ENSMUSG00000004099.10 |
| chr9 | 80164358 | 80166802 | 41.26529 | 138.78937 | 133.96024 | Promoter | 9 | 80165031 | 80311729 | 146699 | 1 | ENSMUST00 | 0 | Myo6 | ENSMUSG00000033577.13 |
| chr2 | 165992265 | 165994229 | 41.82214 | 138.64967 | 133.82387 | Promoter | 2 | 165992636 | 166071470 | 78835 | 1 | ENSMUST00 | 0 | Ncoa3 | ENSMUSG00000027678.12 |
| chr14 | 69731620 | 69733869 | 41.18432 | 138.64781 | 133.82259 | Promoter | 14 | 69716979 | 69732570 | 15592 | 2 | ENSMUST00 | 0 | Chmp7 | ENSMUSG00000034190.8 |
| chr1 | 166002909 | 166003759 | 35.70928 | 138.63979 | 133.81488 | Promoter | 1 | 166002960 | 166038256 | 35297 | 1 | ENSMUST00 | 0 | Gm26665 | ENSMUSG00000097196.2 |
| chr2 | 152413063 | 152415644 | 35.70928 | 138.63979 | 133.81488 | Promoter | 2 | 152415587 | 152444330 | 28744 | 1 | ENSMUST00 | 0 | 6820408C15I | ENSMUSG00000032680.6 |
| chr6 | 56703816 | 56704805 | 35.70928 | 138.63979 | 133.81488 | Promoter | 6 | 56701063 | 56704692 | 3630 | 2 | ENSMUST00 | 0 | Lsm5 | ENSMUSG00000091625.3 |
| chr14 | 25693771 | 25695225 | 41.10367 | 138.50676 | 133.686 | Promoter | 14 | 25694154 | 25700464 | 6311 | 1 | ENSMUST00 | 0 | Ppif | ENSMUSG00000021868.6 |
| chr2 | 32606837 | 32611447 | 41.10367 | 138.50676 | 133.686 | Promoter | 2 | 32606961 | 32620804 | 13844 | 1 | ENSMUST00 | 0 | St6galnac6 | ENSMUSG00000026811.13 |
| chr4 | 136309751 | 136312569 | 36.17052 | 138.42075 | 133.60089 | Promoter | 4 | 136310942 | 136340681 | 29740 | 1 | ENSMUST00 | 0 | Hnrnpr | ENSMUSG00000066037.9 |
| chr9 | 107562702 | 107564101 | 42.31924 | 138.40276 | 133.58318 | Promoter | 9 | 107563255 | 107566108 | 2854 | 1 | ENSMUST00 | 0 | Tusc2 | ENSMUSG00000010054.3 |
| chr1 | 97768601 | 97771049 | 39.77878 | 138.39142 | 133.5723 | Promoter | 1 | 97770172 | 97792969 | 22798 | 1 | ENSMUST00 | 0 | Gin1 | ENSMUSG00000026333.9 |
| chr16 | 48992858 | 48994613 | 39.77878 | 138.39142 | 133.5723 | Promoter | 16 | 48994185 | 49019705 | 25521 | 1 | ENSMUST00 | 0 | C330027C09I | ENSMUSG00000033031.8 |
| chr1 | 91412402 | 91414604 | 37.92415 | 138.37384 | 133.55817 | Promoter | 1 | 91411483 | 91413208 | 1726 | 2 | ENSMUST00 | 0 | Hes6 | ENSMUSG00000067071.3 |
| chr11 | 17158587 | 17160727 | 37.92415 | 138.37384 | 133.55817 | Promoter | 11 | 17159263 | 17200375 | 41113 | 1 | ENSMUST00 | 0 | Ppp3r1 | ENSMUSG00000033953.10 |
| chr12 | 87386808 | 87388351 | 37.92415 | 138.37384 | 133.55817 | Promoter | 12 | 87346734 | 87388241 | 41508 | 2 | ENSMUST00 | 0 | Sptlc2 | ENSMUSG00000021036.5 |
| chr14 | 26667928 | 26670575 | 37.92415 | 138.37384 | 133.55817 | Promoter | 14 | 26659958 | 26669883 | 9926 | 2 | ENSMUST00 | 0 | Pde12 | ENSMUSG00000043702.9 |
| chr4 | 116555293 | 116559172 | 37.92415 | 138.37384 | 133.55817 | Promoter | 4 | 116557658 | 116593882 | 36225 | 1 | ENSMUST00 | 0 | Gpbp1l1 | ENSMUSG00000034042.11 |
| chr5 | 21053536 | 21057043 | 37.92415 | 138.37384 | 133.55817 | Promoter | 5 | 20986645 | 21055911 | 69267 | 2 | ENSMUST00 | 0 | Ptpn12 | ENSMUSG00000028771.8 |
| chr8 | 71380023 | 71382831 | 37.92415 | 138.37384 | 133.55817 | Promoter | 8 | 71374123 | 71381906 | 7784 | 2 | ENSMUST00 | 0 | Nr2f6 | ENSMUSG00000002393.9 |
| chr8 | 107030699 | 107032235 | 37.92415 | 138.37384 | 133.55817 | Promoter | 8 | 107031222 | 107045755 | 14534 | 1 | ENSMUST00 | 0 | Vps4a | ENSMUSG00000031913.9 |
| chr9 | 113706732 | 113708611 | 37.92415 | 138.37384 | 133.55817 | Promoter | 9 | 113651744 | 113708259 | 56516 | 2 | ENSMUST00 | 0 | Pdcd6ip | ENSMUSG00000032504.10 |
| chr9 | 65435657 | 65436522 | 37.92415 | 138.37384 | 133.55817 | Promoter | 9 | 65435782 | 65453054 | 17273 | 1 | ENSMUST00 | 0 | Mtfmt | ENSMUSG00000059183.7 |
| chr3 | 19310411 | 19312172 | 41.65439 | 138.35785 | 133.54871 | Promoter | 3 | 19223108 | 19311322 | 88215 | 2 | ENSMUST00 | 0 | Pde7a | ENSMUSG00000069094.7 |
| chr3 | 53862964 | 53863991 | 42.23227 | 138.25212 | 133.44542 | Promoter | 3 | 53855212 | 53863830 | 8619 | 2 | ENSMUST00 | 0 | Ufm1 | ENSMUSG00000027746.8 |
| chr8 | 79638419 | 79640775 | 37.25272 | 138.2439 | 133.43781 | Promoter | 8 | 79639618 | 79677724 | 38107 | 1 | ENSMUST00 | 0 | Otud4 | ENSMUSG00000036990.8 |
| chr1 | 59236516 | 59238149 | 38.44912 | 138.22351 | 133.41835 | Promoter | 1 | 59162926 | 59237221 | 74306 | 2 | ENSMUST00 | 0 | Als2 | ENSMUSG00000026024.9 |
| chr11 | 98026005 | 98027843 | 38.44912 | 138.22351 | 133.41835 | Promoter | 11 | 98026710 | 98030492 | 3783 | 1 | ENSMUST00 | 0 | Rpl19 | ENSMUSG00000017404.7 |
| chr16 | 10544783 | 10546131 | 38.44912 | 138.22351 | 133.41835 | Promoter | 16 | 10545339 | 10744878 | 199540 | 1 | ENSMUST00 | 0 | Clec16a | ENSMUSG00000068663.8 |
| chr2 | 38286358 | 38287858 | 41.48799 | 138.06813 | 133.26578 | Promoter | 2 | 37798991 | 38287375 | 488385 | 2 | ENSMUST00 | 0 | Dennd1a | ENSMUSG00000035392.12 |
| chr4 | 116626405 | 116628009 | 41.48799 | 138.06813 | 133.26578 | Promoter | 4 | 116601052 | 116627497 | 26446 | 2 | ENSMUST00 | 0 | Nasp | ENSMUSG00000028693.10 |
| chr1 | 128243734 | 128245254 | 42.73649 | 138.01634 | 133.21471 | Promoter | 1 | 128243964 | 128259535 | 15572 | 1 | ENSMUST00 | 0 | Ubxn4 | ENSMUSG00000026353.4 |
| chr14 | 54463669 | 54465005 | 42.73649 | 138.01634 | 133.21471 | Promoter | 14 | 54464164 | 54471497 | 7334 | 1 | ENSMUST00 | 0 | Lrp10 | ENSMUSG00000022175.8 |
| chr17 | 47923048 | 47925384 | 42.73649 | 138.01634 | 133.21471 | Promoter | 17 | 47867133 | 47924645 | 57513 | 2 | ENSMUST00 | 0 | Foxp4 | ENSMUSG00000023991.11 |
| chr5 | 108628398 | 108630468 | 42.73649 | 138.01634 | 133.21471 | Promoter | 5 | 108629770 | 108644030 | 14261 | 1 | ENSMUST00 | 0 | Tmem175 | ENSMUSG00000013495.10 |
| chr6 | 47920226 | 47921724 | 42.73649 | 138.01634 | 133.21471 | Promoter | 6 | 47920476 | 47932639 | 12164 | 1 | ENSMUST00 | 0 | Zfp212 | ENSMUSG00000052763.7 |
| chr11 | 76845296 | 76847512 | 42.05941 | 137.95251 | 133.15681 | Promoter | 11 | 76778424 | 76847018 | 68595 | 2 | ENSMUST00 | 0 | Cpd | ENSMUSG00000020841.5 |
| chr11 | 102316341 | 102321029 | 40.78419 | 137.94737 | 133.15321 | Promoter | 11 | 102304560 | 102319096 | 14537 | 2 | ENSMUST00 | 0 | Ubtf | ENSMUSG00000020923.12 |
| chr17 | 79353864 | 79356423 | 40.11768 | 137.88335 | 133.09009 | Promoter | 17 | 79334025 | 79355091 | 21067 | 2 | ENSMUST00 | 0 | Cdc42ep3 | ENSMUSG00000036533.8 |
| chr19 | 29804622 | 29807012 | 40.11768 | 137.88335 | 133.09009 | Promoter | 19 | 29714402 | 29805507 | 91106 | 2 | ENSMUST00 | 0 | 9930021J03R | ENSMUSG00000046138.9 |
| chr9 | 64736169 | 64738075 | 40.11768 | 137.88335 | 133.09009 | Promoter | 9 | 64715299 | 64737758 | 22460 | 2 | ENSMUST00 | 0 | Rab11a | ENSMUSG00000004771.7 |
| chr16 | 31663033 | 31664931 | 42.5703 | 137.72931 | 132.9411 | Promoter | 16 | 31663443 | 31873350 | 209908 | 1 | ENSMUST00 | 0 | Dlg1 | ENSMUSG00000022770.11 |
| chr11 | 86992502 | 86994164 | 33.56525 | 137.72058 | 132.93262 | Promoter | 11 | 86936425 | 86993707 | 57283 | 2 | ENSMUST00 | 0 | Ypel2 | ENSMUSG00000018427.7 |
| chr3 | 108590867 | 108592394 | 33.56525 | 137.72058 | 132.93262 | Promoter | 3 | 108591279 | 108645719 | 54441 | 1 | ENSMUST00 | 0 | Wdr47 | ENSMUSG00000040389.10 |
| chr10 | 94687776 | 94689607 | 40.62631 | 137.67056 | 132.88402 | Promoter | 10 | 94688790 | 94790336 | 101547 | 1 | ENSMUST00 | 0 | Cep83 | ENSMUSG00000020024.6 |
| chr1 | 165459002 | 165461051 | 38.74602 | 137.65443 | 132.86864 | Promoter | 1 | 165461037 | 165481214 | 20178 | 1 | ENSMUST00 | 0 | Mpc2 | ENSMUSG00000026568.5 |
| chr6 | 148211039 | 148212835 | 38.74602 | 137.65443 | 132.86864 | Promoter | 6 | 148179079 | 148212374 | 33296 | 2 | ENSMUST00 | 0 | Ergic2 | ENSMUSG00000030304.6 |
| chr3 | 102734254 | 102736336 | 41.24086 | 137.63742 | 132.85336 | Promoter | 3 | 102734529 | 102772292 | 37764 | 1 | ENSMUST00 | 0 | Tspan2 | ENSMUSG00000027858.8 |
| chr5 | 108460692 | 108462315 | 41.24086 | 137.63742 | 132.85336 | Promoter | 5 | 108461232 | 108489224 | 27993 | 1 | ENSMUST00 | 0 | Pcgf3 | ENSMUSG00000033623.8 |
| chr16 | 33185017 | 33186455 | 37.51416 | 137.62538 | 132.8419 | Promoter | 16 | 33185071 | 33243311 | 58241 | 1 | ENSMUST00 | 0 | Osbpl11 | ENSMUSG00000022807.8 |
| chr18 | 75018570 | 75019764 | 37.51416 | 137.62538 | 132.8419 | Promoter | 18 | 75018772 | 75286966 | 268195 | 1 | ENSMUST00 | 0 | Dym | ENSMUSG00000035765.9 |
| chr18 | 31759113 | 31761179 | 42.48137 | 137.57564 | 132.7934 | Promoter | 18 | 31759824 | 31784071 | 24248 | 1 | ENSMUST00 | 0 | Ammecr1l | ENSMUSG00000041915.8 |

| | | | | | | | | | | | | | | | |
|---|---|---|---|---|---|---|---|---|---|---|---|---|---|---|---|
| chr10 | 128096543 | 128098298 | 40.54782 | 137.53284 | 132.75229 | Intron (ENSM | 10 | 128092783 | 128129303 | 36521 | 1 | ENSMUST00 | 3761 | Baz2a | ENSMUSG00000040054.10 |
| chr10 | 80754650 | 80756360 | 41.80276 | 137.50731 | 132.72684 | Promoter | 10 | 80755206 | 80795461 | 40256 | 1 | ENSMUST00 | 0 | Dot1l | ENSMUSG00000061589.9 |
| chr11 | 103411928 | 103413358 | 41.80276 | 137.50731 | 132.72684 | Promoter | 11 | 103364275 | 103412687 | 48413 | 2 | ENSMUST00 | 0 | Plekhm1 | ENSMUSG00000034247.9 |
| chr14 | 31435079 | 31436535 | 41.80276 | 137.50731 | 132.72684 | Promoter | 14 | 31359880 | 31436078 | 76199 | 2 | ENSMUST00 | 0 | Sh3bp5 | ENSMUSG00000021892.9 |
| chr16 | 96191509 | 96192644 | 40.46964 | 137.39561 | 132.61725 | Promoter | 16 | 96158407 | 96192250 | 33844 | 2 | ENSMUST00 | 0 | Lca5l | ENSMUSG00000045275.10 |
| chr2 | 70660882 | 70662957 | 40.46964 | 137.39561 | 132.61725 | Promoter | 2 | 70661576 | 70691725 | 30150 | 1 | ENSMUST00 | 0 | Gorasp2 | ENSMUSG00000014959.7 |
| chr7 | 12921403 | 12923067 | 40.46964 | 137.39561 | 132.61725 | Promoter | 7 | 12922290 | 12926685 | 4396 | 1 | ENSMUST00 | 0 | Rps5 | ENSMUSG00000012848.9 |
| chr4 | 45010792 | 45012460 | 41.07773 | 137.35284 | 132.57864 | Promoter | 4 | 44991242 | 45012394 | 21153 | 2 | ENSMUST00 | 0 | Zbtb5 | ENSMUSG00000049657.9 |
| chr6 | 94499787 | 94501059 | 41.07773 | 137.35284 | 132.57864 | Promoter | 6 | 94500314 | 94604648 | 104335 | 1 | ENSMUST00 | 0 | Slc25a26 | ENSMUSG00000045100.9 |
| chr8 | 47712794 | 47714015 | 31.76134 | 137.24866 | 132.47624 | Promoter | 8 | 47709344 | 47713931 | 4588 | 2 | ENSMUST00 | 0 | Cdkn2aip | ENSMUSG00000038069.5 |
| chr10 | 61694995 | 61697126 | 39.05373 | 137.10143 | 132.33189 | Promoter | 10 | 61695514 | 61702773 | 7260 | 1 | ENSMUST00 | 0 | Tysnd1 | ENSMUSG00000020087.4 |
| chr11 | 98681882 | 98683642 | 39.05373 | 137.10143 | 132.33189 | Promoter | 11 | 98682554 | 98695979 | 13426 | 1 | ENSMUST00 | 0 | Psmd3 | ENSMUSG00000017221.9 |
| chr15 | 58888090 | 58890604 | 39.05373 | 137.10143 | 132.33189 | Promoter | 15 | 58889229 | 58902390 | 13162 | 1 | ENSMUST00 | 0 | Rnf139 | ENSMUSG00000037075.6 |
| chr16 | 91930944 | 91932195 | 39.05373 | 137.10143 | 132.33189 | Promoter | 16 | 91925214 | 91931687 | 6474 | 2 | ENSMUST00 | 0 | Atp5o | ENSMUSG00000022956.5 |
| chr2 | 23038901 | 23040223 | 39.05373 | 137.10143 | 132.33189 | Promoter | 2 | 22940073 | 23040195 | 100123 | 2 | ENSMUST00 | 0 | Abi1 | ENSMUSG00000058835.9 |
| chr4 | 141147189 | 141149189 | 39.05373 | 137.10143 | 132.33189 | Promoter | 4 | 141147913 | 141204062 | 56150 | 1 | ENSMUST00 | 0 | Fbxo42 | ENSMUSG00000028920.9 |
| chr4 | 106909808 | 106912767 | 41.54923 | 137.06703 | 132.30061 | Promoter | 4 | 106910701 | 107031069 | 120369 | 1 | ENSMUST00 | 0 | Ssbp3 | ENSMUSG00000061887.9 |
| chr1 | 57995547 | 57996485 | 37.78464 | 137.01973 | 132.25398 | Promoter | 1 | 57995971 | 58025899 | 29929 | 1 | ENSMUST00 | 0 | Sgol2 | ENSMUSG00000026039.9 |
| chr14 | 46831260 | 46833192 | 37.78464 | 137.01973 | 132.25398 | Promoter | 14 | 46832134 | 46854193 | 22060 | 1 | ENSMUST00 | 0 | Cgrrf1 | ENSMUSG00000055128.9 |
| chr8 | 72161256 | 72162149 | 37.78464 | 137.01973 | 132.25398 | Promoter | 8 | 72161313 | 72175863 | 14551 | 1 | ENSMUST00 | 0 | Rab8a | ENSMUSG00000003037.11 |
| chr9 | 88438145 | 88439793 | 37.78464 | 137.01973 | 132.25398 | Promoter | 9 | 88376750 | 88438951 | 62202 | 2 | ENSMUST00 | 0 | Snx14 | ENSMUSG00000032422.12 |
| chr10 | 7210033 | 7213358 | 40.23456 | 136.98251 | 132.21786 | Promoter | 10 | 7119063 | 7212237 | 93175 | 2 | ENSMUST00 | 0 | Cnksr3 | ENSMUSG00000015202.8 |
| chr11 | 95073478 | 95078046 | 40.23456 | 136.98251 | 132.21786 | Promoter | 11 | 95044474 | 95076714 | 32241 | 2 | ENSMUST00 | 0 | Itga3 | ENSMUSG00000001507.11 |
| chr11 | 44518262 | 44520961 | 40.23456 | 136.98251 | 132.21786 | Promoter | 11 | 44518964 | 44565520 | 46557 | 1 | ENSMUST00 | 0 | Rnf145 | ENSMUSG00000019189.8 |
| chr13 | 14612429 | 14614120 | 40.23456 | 136.98251 | 132.21786 | Promoter | 13 | 14613242 | 14625671 | 12430 | 1 | ENSMUST00 | 0 | Psma2 | ENSMUSG00000015671.10 |
| chr19 | 4877826 | 4879348 | 40.23456 | 136.98251 | 132.21786 | Promoter | 19 | 4861223 | 4877909 | 16687 | 2 | ENSMUST00 | 0 | Actn3 | ENSMUSG00000006457.3 |
| chr2 | 71054773 | 71056569 | 40.23456 | 136.98251 | 132.21786 | Promoter | 2 | 71055328 | 71086618 | 31291 | 1 | ENSMUST00 | 0 | Dcaf17 | ENSMUSG00000041966.13 |
| chr5 | 74194700 | 74196223 | 40.23456 | 136.98251 | 132.21786 | Promoter | 5 | 74195288 | 74199481 | 4194 | 1 | ENSMUST00 | 0 | Rasl11b | ENSMUSG00000049907.7 |
| chr6 | 88445685 | 88446905 | 40.23456 | 136.98251 | 132.21786 | Promoter | 6 | 88257334 | 88446539 | 189206 | 2 | ENSMUST00 | 0 | Eefsec | ENSMUSG00000033216.7 |
| chr6 | 85332329 | 85333980 | 40.23456 | 136.98251 | 132.21786 | Promoter | 6 | 85213051 | 85333422 | 120372 | 2 | ENSMUST00 | 0 | Sfxn5 | ENSMUSG00000033720.7 |
| chr2 | 164074051 | 164075217 | 42.12929 | 136.96686 | 132.21103 | Promoter | 2 | 164074138 | 164155524 | 81387 | 1 | ENSMUST00 | 0 | Stk4 | ENSMUSG00000018209.10 |
| chr8 | 95805980 | 95808104 | 42.12929 | 136.96686 | 132.21103 | Promoter | 8 | 95719451 | 95807462 | 88012 | 2 | ENSMUST00 | 0 | Cnot1 | ENSMUSG00000036550.10 |
| chr1 | 36709036 | 36710077 | 40.83545 | 136.9297 | 132.17467 | Promoter | 1 | 36698116 | 36710008 | 11893 | 2 | ENSMUST00 | 0 | Actr1b | ENSMUSG00000037351.4 |
| chr1 | 64956081 | 64957136 | 40.83545 | 136.9297 | 132.17467 | Promoter | 1 | 64785983 | 64956824 | 170842 | 2 | ENSMUST00 | 0 | Plekhm3 | ENSMUSG00000051344.8 |
| chr18 | 10608994 | 10610706 | 40.83545 | 136.9297 | 132.17467 | Promoter | 18 | 10566611 | 10610129 | 43519 | 2 | ENSMUST00 | 0 | Esco1 | ENSMUSG00000024293.10 |
| chr5 | 103910526 | 103911538 | 40.83545 | 136.9297 | 132.17467 | Promoter | 5 | 103862050 | 103911259 | 49210 | 2 | ENSMUST00 | 0 | Klhl8 | ENSMUSG00000029312.7 |
| chr8 | 57510779 | 57511644 | 40.83545 | 136.9297 | 132.17467 | Promoter | 8 | 57511843 | 57515999 | 4157 | 1 | ENSMUST00 | -199 | Hmgb2 | ENSMUSG00000054717.6 |
| chr1 | 105662921 | 105664954 | 35.36919 | 136.9064 | 132.15405 | Promoter | 1 | 105663861 | 105687141 | 23281 | 1 | ENSMUST00 | 0 | 2310035C23I | ENSMUSG00000026319.8 |
| chr19 | 7493707 | 7495706 | 35.36919 | 136.9064 | 132.15405 | Promoter | 19 | 7494040 | 7538608 | 44569 | 1 | ENSMUST00 | 0 | Atl3 | ENSMUSG00000024759.8 |
| chr3 | 116594187 | 116595616 | 35.36919 | 136.9064 | 132.15405 | Promoter | 3 | 116595008 | 116630984 | 35977 | 1 | ENSMUST00 | 0 | Sass6 | ENSMUSG00000027959.9 |
| chr5 | 103424449 | 103426532 | 35.36919 | 136.9064 | 132.15405 | Promoter | 5 | 103425192 | 103598359 | 173168 | 1 | ENSMUST00 | 0 | Ptpn13 | ENSMUSG00000034573.10 |
| chr7 | 96951008 | 96954739 | 35.36919 | 136.9064 | 132.15405 | Promoter | 7 | 96951505 | 97064761 | 113257 | 1 | ENSMUST00 | 0 | Nars2 | ENSMUSG00000018995.7 |
| chr7 | 78895371 | 78896901 | 35.36919 | 136.9064 | 132.15405 | Promoter | 7 | 78895854 | 78911209 | 15356 | 1 | ENSMUST00 | 0 | Aen | ENSMUSG00000030609.13 |
| chr7 | 25753297 | 25756211 | 35.36919 | 136.9064 | 132.15405 | Promoter | 7 | 25721498 | 25754730 | 33233 | 2 | ENSMUST00 | 0 | Hnrnpul1 | ENSMUSG00000040725.8 |
| chr8 | 11497061 | 11498571 | 35.36919 | 136.9064 | 132.15405 | Promoter | 8 | 11497506 | 11513287 | 15782 | 1 | ENSMUST00 | 0 | Carkd | ENSMUSG00000031505.5 |
| chr13 | 86045854 | 86046886 | 40.15991 | 136.85123 | 132.10168 | Promoter | 13 | 86044816 | 86046460 | 1645 | 2 | ENSMUST00 | 0 | Cox7c | ENSMUSG00000017778.9 |
| chr6 | 119478571 | 119481610 | 42.04218 | 136.8161 | 132.06688 | Promoter | 6 | 119479668 | 119482630 | 2963 | 1 | ENSMUST00 | 0 | Fbxl14 | ENSMUSG00000030019.6 |
| chr10 | 85956815 | 85958197 | 40.75532 | 136.78966 | 132.04059 | Promoter | 10 | 85938637 | 85957823 | 19187 | 2 | ENSMUST00 | 0 | Rtcb | ENSMUSG00000001783.3 |
| chr7 | 102249024 | 102251102 | 40.75532 | 136.78966 | 132.04059 | Promoter | 7 | 102239123 | 102250088 | 10966 | 2 | ENSMUST00 | 0 | Rhog | ENSMUSG00000073982.5 |
| chr10 | 77621807 | 77623491 | 41.3819 | 136.77621 | 132.02756 | Promoter | 10 | 77622275 | 77645993 | 23719 | 1 | ENSMUST00 | 0 | Ube2g2 | ENSMUSG00000009293.12 |
| chr8 | 72722964 | 72724301 | 35.82273 | 136.67671 | 131.92941 | Promoter | 8 | 72723288 | 72739910 | 16623 | 1 | ENSMUST00 | 0 | Sin3b | ENSMUSG00000031622.10 |
| chr4 | 83524743 | 83526364 | 41.95543 | 136.66595 | 131.91884 | Promoter | 4 | 83525545 | 83864670 | 339126 | 1 | ENSMUST00 | 0 | Ccdc171 | ENSMUSG00000052407.11 |
| chr4 | 114986035 | 114987681 | 41.29875 | 136.63161 | 131.88535 | Promoter | 4 | 114959336 | 114987241 | 27906 | 2 | ENSMUST00 | 0 | Cmpk1 | ENSMUSG00000028719.8 |
| chr11 | 96976345 | 96977744 | 37.54867 | 136.59039 | 131.84451 | Promoter | 11 | 96953340 | 96977688 | 24349 | 2 | ENSMUST00 | 0 | Sp2 | ENSMUSG00000018678.7 |
| chr14 | 54863467 | 54864678 | 37.54867 | 136.59039 | 131.84451 | Promoter | 14 | 54853787 | 54864158 | 10372 | 2 | ENSMUST00 | 0 | Homez | ENSMUSG00000057156.10 |
| chr19 | 60789055 | 60791515 | 37.54867 | 136.59039 | 131.84451 | Promoter | 19 | 60761116 | 60790693 | 29578 | 2 | ENSMUST00 | 0 | Eif3a | ENSMUSG00000024991.7 |
| chr2 | 163642816 | 163645495 | 37.54867 | 136.59039 | 131.84451 | Promoter | 2 | 163623272 | 163645131 | 21860 | 2 | ENSMUST00 | 0 | Serinc3 | ENSMUSG00000017707.9 |
| chr3 | 40744587 | 40746633 | 37.54867 | 136.59039 | 131.84451 | Promoter | 3 | 40745382 | 40781598 | 36217 | 1 | ENSMUST00 | 0 | Hspa4l | ENSMUSG00000025757.8 |
| chr4 | 126320878 | 126322123 | 37.54867 | 136.59039 | 131.84451 | Promoter | 4 | 126316047 | 126321369 | 5323 | 2 | ENSMUST00 | 0 | Adprhl2 | ENSMUSG00000042558.11 |
| chr4 | 116994964 | 116996323 | 37.54867 | 136.59039 | 131.84451 | Promoter | 4 | 116995317 | 117005277 | 9961 | 1 | ENSMUST00 | 0 | Hectd3 | ENSMUSG00000046861.9 |
| chr5 | 74530768 | 74532217 | 37.54867 | 136.59039 | 131.84451 | Promoter | 5 | 74204816 | 74531742 | 326927 | 2 | ENSMUST00 | 0 | Scfd2 | ENSMUSG00000062110.9 |
| chr6 | 143099152 | 143100613 | 37.54867 | 136.59039 | 131.84451 | Promoter | 6 | 143010920 | 143100107 | 89188 | 2 | ENSMUST00 | 0 | C2cd5 | ENSMUSG00000030279.10 |
| chr7 | 97081213 | 97083356 | 37.54867 | 136.59039 | 131.84451 | Promoter | 7 | 97081751 | 97308940 | 227190 | 1 | ENSMUST00 | 0 | Gab2 | ENSMUSG00000004508.5 |
| chr14 | 70577252 | 70578497 | 39.37288 | 136.56555 | 131.82602 | Promoter | 14 | 70577847 | 70582571 | 4725 | 1 | ENSMUST00 | 0 | Nudt18 | ENSMUSG00000045211.4 |
| chr7 | 120634318 | 120636330 | 39.37288 | 136.56555 | 131.82602 | Promoter | 7 | 120635176 | 120659478 | 24303 | 1 | ENSMUST00 | 0 | Uqcrc2 | ENSMUSG00000030884.6 |
| chr17 | 32388391 | 32389436 | 41.21593 | 136.4875 | 131.75212 | Promoter | 17 | 32354055 | 32388644 | 34590 | 2 | ENSMUST00 | 0 | Wiz | ENSMUSG00000024050.10 |
| chr18 | 80206157 | 80207976 | 38.06463 | 136.4278 | 131.69504 | Promoter | 18 | 80206765 | 80211493 | 4729 | 1 | ENSMUST00 | 0 | Gm16286 | ENSMUSG00000024571.9 |
| chr2 | 104027441 | 104029300 | 40.51682 | 136.37238 | 131.6409 | Promoter | 2 | 104027721 | 104063240 | 35520 | 1 | ENSMUST00 | 0 | Fbxo3 | ENSMUSG00000027180.7 |
| chr17 | 28484845 | 28486181 | 41.78299 | 136.36734 | 131.63638 | Promoter | 17 | 28399095 | 28486149 | 87055 | 2 | ENSMUST00 | 0 | Fkbp5 | ENSMUSG00000024222.11 |
| chr8 | 109705066 | 109706517 | 40.43793 | 136.23427 | 131.50453 | Promoter | 8 | 109705549 | 109724932 | 19384 | 1 | ENSMUST00 | 0 | Zfp821 | ENSMUSG00000031728.9 |
| chr10 | 3739900 | 3742294 | 41.69731 | 136.21887 | 131.48979 | Promoter | 10 | 3740364 | 3967303 | 226940 | 1 | ENSMUST00 | 0 | Plekhg1 | ENSMUSG00000040624.12 |
| chr5 | 113907486 | 113909403 | 41.69731 | 136.21887 | 131.48979 | Promoter | 5 | 113842436 | 113908705 | 66270 | 2 | ENSMUST00 | 0 | Coro1c | ENSMUSG00000004530.7 |
| chr1 | 66815744 | 66818353 | 39.77934 | 136.18091 | 131.45303 | Promoter | 1 | 66719363 | 66817546 | 98184 | 2 | ENSMUST00 | 0 | Kansl1l | ENSMUSG00000026004.10 |
| chr13 | 34162481 | 34163721 | 42.27695 | 136.11826 | 131.39047 | Promoter | 13 | 34162964 | 34178168 | 15205 | 1 | ENSMUST00 | 0 | Psmg4 | ENSMUSG00000071451.4 |
| chr4 | 125029594 | 125030624 | 42.27695 | 136.11826 | 131.39047 | Promoter | 4 | 125029992 | 125055376 | 25385 | 1 | ENSMUST00 | 0 | Gnl2 | ENSMUSG00000028869.8 |
| chr4 | 124849422 | 124852147 | 42.27695 | 136.11826 | 131.39047 | Promoter | 4 | 124850679 | 124855242 | 4564 | 1 | ENSMUST00 | 0 | Yrdc | ENSMUSG00000028889.12 |
| chr4 | 140986613 | 140987704 | 42.27695 | 136.11826 | 131.39047 | Promoter | 4 | 140986873 | 141005087 | 18215 | 1 | ENSMUST00 | 0 | Atp13a2 | ENSMUSG00000036622.10 |
| chr4 | 147940819 | 147942100 | 42.27695 | 136.11826 | 131.39047 | Promoter | 4 | 147940895 | 147947314 | 6420 | 1 | ENSMUST00 | 0 | 2510039O18 | ENSMUSG00000044496.6 |
| chr6 | 147042302 | 147043788 | 42.27695 | 136.11826 | 131.39047 | Promoter | 6 | 147042764 | 147050580 | 7817 | 1 | ENSMUST00 | 0 | Mrps35 | ENSMUSG00000040112.7 |
| chr9 | 108346953 | 108348997 | 42.27695 | 136.11826 | 131.39047 | Promoter | 9 | 108347831 | 108392529 | 44699 | 1 | ENSMUST00 | 0 | Usp4 | ENSMUSG00000032612.9 |
| chr10 | 80432209 | 80434035 | 41.61197 | 136.07095 | 131.34828 | Promoter | 10 | 80409514 | 80433637 | 24124 | 2 | ENSMUST00 | 0 | Tcf3 | ENSMUSG00000020167.9 |
| chr9 | 108648312 | 108650207 | 41.61197 | 136.07095 | 131.34828 | Promoter | 9 | 108602942 | 108649380 | 46439 | 2 | ENSMUST00 | 0 | Arih2 | ENSMUSG00000064145.7 |
| chr4 | 133129620 | 133132418 | 40.96944 | 136.05835 | 131.33708 | Promoter | 4 | 133130505 | 133199756 | 69252 | 1 | ENSMUST00 | 0 | Wasf2 | ENSMUSG00000028868.8 |
| chr17 | 87671934 | 87673378 | 39.70409 | 136.04816 | 131.32747 | Promoter | 17 | 87672330 | 87723713 | 51384 | 1 | ENSMUST00 | 0 | Msh2 | ENSMUSG00000024151.8 |
| chr4 | 21847275 | 21849107 | 39.70409 | 136.04816 | 131.32747 | Promoter | 4 | 21847583 | 21876475 | 28893 | 1 | ENSMUST00 | 0 | Pnisr | ENSMUSG00000028248.10 |
| chr4 | 48278566 | 48280414 | 39.70409 | 136.04816 | 131.32747 | Promoter | 4 | 48193323 | 48279558 | 86236 | 2 | ENSMUST00 | 0 | Erp44 | ENSMUSG00000028343.5 |
| chr1 | 56971937 | 56973997 | 33.25446 | 136.03296 | 131.3158 | Promoter | 1 | 56793981 | 56972502 | 178522 | 2 | ENSMUST00 | 0 | Satb2 | ENSMUSG00000038331.10 |
| chr2 | 25254741 | 25256479 | 33.25446 | 136.03296 | 131.3158 | Promoter | 2 | 25244821 | 25255383 | 10563 | 2 | ENSMUST00 | 0 | Ndor1 | ENSMUSG00000006471.12 |
| chr1 | 43097729 | 43099549 | 40.28109 | 135.95943 | 131.24442 | Promoter | 1 | 43098710 | 43115947 | 17238 | 1 | ENSMUST00 | 0 | AI597479 | ENSMUSG00000010290.7 |
| chr11 | 120783522 | 120785265 | 37.14274 | 135.84946 | 131.13702 | Promoter | 11 | 120784272 | 120786333 | 2062 | 1 | ENSMUST00 | 0 | Gps1 | ENSMUSG00000025156.11 |
| chr3 | 30599675 | 30603392 | 38.35464 | 135.85051 | 131.13702 | Promoter | 3 | 30602087 | 30618492 | 16406 | 1 | ENSMUST00 | 0 | Mynn | ENSMUSG00000037730.8 |
| chr5 | 112325411 | 112327353 | 38.35464 | 135.85051 | 131.13702 | Promoter | 5 | 112326358 | 112338073 | 11716 | 1 | ENSMUST00 | 0 | Tfip11 | ENSMUSG00000029345.8 |
| chr7 | 35584486 | 35587154 | 38.35464 | 135.85051 | 131.13702 | Promoter | 7 | 35586244 | 35639226 | 52983 | 1 | ENSMUST00 | 0 | Ankrd27 | ENSMUSG00000034867.11 |
| chr5 | 43232302 | 43236237 | 40.20312 | 135.82269 | 131.11238 | Promoter | 5 | 43233463 | 43289724 | 56262 | 1 | ENSMUST00 | 0 | Cpeb2 | ENSMUSG00000039782.9 |
| chr8 | 72134816 | 72136882 | 41.44235 | 135.77681 | 131.06697 | Promoter | 8 | 72134975 | 72144709 | 9735 | 1 | ENSMUST00 | 0 | Tpm4 | ENSMUSG00000031799.9 |
| chr13 | 51202561 | 51203102 | 31.4752 | 135.60028 | 130.8949 | Promoter | 13 | 51202688 | 51203065 | 378 | 2 | ENSMUST00 | 0 | Hist1h2al | ENSMUSG00000091383.1 |
| chr14 | 79480316 | 79483475 | 31.4752 | 135.60028 | 130.8949 | Promoter | 14 | 79459937 | 79481268 | 21332 | 2 | ENSMUST00 | 0 | Wbp4 | ENSMUSG00000022023.5 |
| chr7 | 110018185 | 110019429 | 31.4752 | 135.60028 | 130.8949 | Promoter | 7 | 110018425 | 110055114 | 36690 | 1 | ENSMUST00 | 0 | Ipo7 | ENSMUSG00000066232.3 |
| chr14 | 30624866 | 30626772 | 40.04808 | 135.55066 | 130.84587 | Promoter | 14 | 30595356 | 30626208 | 30853 | 2 | ENSMUST00 | 0 | Prkcd | ENSMUSG00000021948.10 |
| chr14 | 33446132 | 33447281 | 40.64533 | 135.49321 | 130.79207 | Promoter | 14 | 33377898 | 33447158 | 69261 | 2 | ENSMUST00 | 0 | Mapk8 | ENSMUSG00000021936.8 |
| chr11 | 69058617 | 69061280 | 41.8497 | 135.3801 | 130.68082 | Promoter | 11 | 69059717 | 69061578 | 1862 | 1 | ENSMUST00 | 0 | 2310047M10 | ENSMUSG00000045176.3 |
| chr9 | 119339312 | 119342200 | 36.28143 | 135.35555 | 130.65697 | Promoter | 9 | 119335934 | 119340041 | 4108 | 2 | ENSMUST00 | 0 | Myd88 | ENSMUSG00000032508.8 |
| chr5 | 67306320 | 67308135 | 40.56511 | 135.35318 | 130.65489 | Promoter | 5 | 67306955 | 67355336 | 48382 | 1 | ENSMUST00 | 0 | Slc30a9 | ENSMUSG00000029221.10 |
| chr11 | 93884540 | 93887391 | 41.19049 | 135.33968 | 130.64169 | Promoter | 11 | 93859243 | 93885766 | 26524 | 2 | ENSMUST00 | 0 | Utp18 | ENSMUSG00000054079.7 |
| chr16 | 87454436 | 87456031 | 38.65522 | 135.28891 | 130.59163 | Promoter | 16 | 87454703 | 87483517 | 28815 | 1 | ENSMUST00 | 0 | Usp16 | ENSMUSG00000025616.9 |
| chr9 | 110131232 | 110132839 | 38.65522 | 135.28891 | 130.59163 | Promoter | 9 | 110132037 | 110184303 | 52267 | 1 | ENSMUST00 | 0 | Smarcc1 | ENSMUSG00000032481.11 |
| chr5 | 107900520 | 107901721 | 37.4068 | 135.23624 | 130.54147 | Promoter | 5 | 107900572 | 107904123 | 3552 | 1 | ENSMUST00 | 0 | Rpl5 | ENSMUSG00000058558.7 |
| chr3 | 102203071 | 102204781 | 41.76281 | 135.22986 | 130.53618 | Promoter | 3 | 102156701 | 102204693 | 47993 | 2 | ENSMUST00 | 0 | Vangl1 | ENSMUSG00000027860.10 |
| chr4 | 131872840 | 131874986 | 41.76281 | 135.22986 | 130.53618 | Promoter | 4 | 131873617 | 131901706 | 28090 | 1 | ENSMUST00 | 0 | Srsf4 | ENSMUSG00000028911.11 |
| chr8 | 47989082 | 47991874 | 41.76281 | 135.22986 | 130.53618 | Promoter | 8 | 47826083 | 47990543 | 164461 | 2 | ENSMUST00 | 0 | Wwc2 | ENSMUSG00000031563.7 |
| chr11 | 23496813 | 23498021 | 41.10722 | 135.19504 | 130.50221 | Promoter | 11 | 23487882 | 23497930 | 10049 | 2 | ENSMUST00 | 0 | Ahsa2 | ENSMUSG00000020288.8 |
| chr11 | 51605527 | 51607486 | 35.02911 | 135.1772 | 130.48503 | Promoter | 11 | 51600100 | 51606847 | 6748 | 2 | ENSMUST00 | 0 | Hnrnpab | ENSMUSG00000020358.12 |
| chr11 | 74722916 | 74725014 | 35.02911 | 135.1772 | 130.48503 | Promoter | 11 | 74673949 | 74724335 | 50387 | 2 | ENSMUST00 | 0 | Pafah1b1 | ENSMUSG00000020745.10 |
| chr12 | 24650599 | 24652468 | 35.02911 | 135.1772 | 130.48503 | Promoter | 12 | 24651371 | 24662774 | 11404 | 1 | ENSMUST00 | 0 | Klf11 | ENSMUSG00000020653.6 |
| chr12 | 76368766 | 76371973 | 35.02911 | 135.1772 | 130.48503 | Promoter | 12 | 76370266 | 76396950 | 26685 | 1 | ENSMUST00 | 0 | Zbtb1 | ENSMUSG00000033454.6 |
| chr13 | 17802516 | 17806006 | 35.02911 | 135.1772 | 130.48503 | Promoter | 13 | 17715962 | 17805097 | 89136 | 2 | ENSMUST00 | 0 | Cdk13 | ENSMUSG00000041297.7 |
| chr15 | 100494362 | 100495672 | 35.02911 | 135.1772 | 130.48503 | Promoter | 15 | 100479570 | 100495239 | 15670 | 2 | ENSMUST00 | 0 | Csrnp2 | ENSMUSG00000044636.5 |
| chr4 | 57955210 | 57956381 | 35.02911 | 135.1772 | 130.48503 | Promoter | 4 | 57943373 | 57956411 | 13039 | 2 | ENSMUST00 | 30 | Txn1 | ENSMUSG00000028367.5 |
| chr6 | 108827844 | 108830493 | 35.02911 | 135.1772 | 130.48503 | Promoter | 6 | 108828641 | 108859356 | 30716 | 1 | ENSMUST00 | 0 | Edem1 | ENSMUSG00000030104.7 |
| chr9 | 78480141 | 78481717 | 35.02911 | 135.1772 | 130.48503 | Promoter | 9 | 78479278 | 78480688 | 1411 | 2 | ENSMUST00 | 0 | Eef1a1 | ENSMUSG00000037742.9 |
| chr1 | 155416315 | 155417702 | 39.81545 | 135.14195 | 130.45284 | Promoter | 1 | 155275701 | 155417309 | 141609 | 2 | ENSMUST00 | 0 | Xpr1 | ENSMUSG00000026469.9 |
| chr11 | 17952987 | 17954262 | 39.81545 | 135.14195 | 130.45284 | Promoter | 11 | 17938756 | 17953875 | 15120 | 2 | ENSMUST00 | 0 | Etaa1 | ENSMUSG00000016984.7 |
| chr11 | 117076313 | 117077570 | 39.81545 | 135.14195 | 130.45284 | Promoter | 11 | 117076790 | 117077457 | 668 | 1 | ENSMUST00 | 0 | 2810008D09I | ENSMUSG00000086859.3 |
| chr18 | 23954036 | 23955938 | 39.81545 | 135.14195 | 130.45284 | Promoter | 18 | 23954688 | 23964671 | 9984 | 1 | ENSMUST00 | 0 | Zfp397 | ENSMUSG00000024276.6 |
| chr19 | 55314944 | 55317042 | 39.81545 | 135.14195 | 130.45284 | Promoter | 19 | 55298301 | 55316032 | 17732 | 2 | ENSMUST00 | 0 | Zdhhc6 | ENSMUSG00000024982.9 |
| chr2 | 163471692 | 163472926 | 39.81545 | 135.14195 | 130.45284 | Promoter | 2 | 163466379 | 163472629 | 6251 | 2 | ENSMUST00 | 0 | Fitm2 | ENSMUSG00000048486.6 |
| chr4 | 8689673 | 8692342 | 39.81545 | 135.14195 | 130.45284 | Promoter | 4 | 8690406 | 8755906 | 65501 | 1 | ENSMUST00 | 0 | Chd7 | ENSMUSG00000041235.6 |
| chr7 | 127448218 | 127449677 | 39.81545 | 135.14195 | 130.45284 | Promoter | 7 | 127443752 | 127449158 | 5407 | 2 | ENSMUST00 | 0 | Zfp689 | ENSMUSG00000048921.6 |

| | | | | | | | | | | | | | | | |
|---|---|---|---|---|---|---|---|---|---|---|---|---|---|---|---|
| chr9 | 123851001 | 123851932 | 39.81545 | 135.14195 | 130.45284 | Promoter | 9 | 123789510 | 123851899 | 62390 | 2 | ENSMUST00 | 0 | Fyco1 | ENSMUSG00000025241.10 |
| chr13 | 85189018 | 85190300 | 41.67628 | 135.08018 | 130.40039 | Promoter | 13 | 85189408 | 85213723 | 24316 | 1 | ENSMUST00 | 0 | Ccnh | ENSMUSG00000021548.5 |
| chr11 | 106271426 | 106273693 | 40.4056 | 135.07462 | 130.39536 | Promoter | 11 | 106263179 | 106272972 | 9794 | 2 | ENSMUST00 | 0 | Smarcd2 | ENSMUSG00000078619.5 |
| chr18 | 53175802 | 53177208 | 35.47494 | 134.93686 | 130.2605 | Promoter | 18 | 53176365 | 53220860 | 44496 | 1 | ENSMUST00 | 0 | Snx2 | ENSMUSG00000034484.7 |
| chr8 | 13158659 | 13160275 | 41.59011 | 134.93111 | 130.25542 | Promoter | 8 | 13159135 | 13175338 | 16204 | 1 | ENSMUST00 | 0 | Lamp1 | ENSMUSG00000031447.6 |
| chr7 | 80992503 | 80994889 | 39.66568 | 134.87851 | 130.20418 | Promoter | 7 | 80993681 | 81045164 | 51484 | 1 | ENSMUST00 | 0 | Zfp592 | ENSMUSG00000005621.11 |
| chr13 | 54564424 | 54565700 | 39.03945 | 134.87268 | 130.19839 | Promoter | 13 | 54551218 | 54565435 | 14218 | 2 | ENSMUST00 | 0 | 4833439L19R | ENSMUSG00000025871.13 |
| chr1 | 16664920 | 16666201 | 37.17318 | 134.81126 | 130.13722 | Promoter | 1 | 16665207 | 16678195 | 12989 | 1 | ENSMUST00 | 0 | Tmem70 | ENSMUSG00000025940.6 |
| chr1 | 36938405 | 36940540 | 37.17318 | 134.81126 | 130.13722 | Promoter | 1 | 36792194 | 36939527 | 147334 | 2 | ENSMUST00 | 0 | Tmem131 | ENSMUSG00000026116.6 |
| chr10 | 98913573 | 98916645 | 37.17318 | 134.81126 | 130.13722 | Promoter | 10 | 98915152 | 99026143 | 110992 | 1 | ENSMUST00 | 0 | Atp2b1 | ENSMUSG00000019943.9 |
| chr10 | 62650076 | 62651299 | 37.17318 | 134.81126 | 130.13722 | Promoter | 10 | 62616023 | 62651218 | 35196 | 2 | ENSMUST00 | 0 | Ddx50 | ENSMUSG00000020076.6 |
| chr11 | 96826971 | 96829996 | 37.17318 | 134.81126 | 130.13722 | Promoter | 11 | 96817414 | 96829502 | 12089 | 2 | ENSMUST00 | 0 | Nfe2l1 | ENSMUSG00000038615.12 |
| chr13 | 55361595 | 55363242 | 37.17318 | 134.81126 | 130.13722 | Promoter | 13 | 55343833 | 55362783 | 18951 | 2 | ENSMUST00 | 0 | Lman2 | ENSMUSG00000021484.7 |
| chr14 | 31250459 | 31252425 | 37.17318 | 134.81126 | 130.13722 | Promoter | 14 | 31251450 | 31259944 | 8495 | 1 | ENSMUST00 | 0 | Bap1 | ENSMUSG00000021901.5 |
| chr15 | 7128546 | 7131106 | 37.17318 | 134.81126 | 130.13722 | Promoter | 15 | 7129559 | 7191743 | 62185 | 1 | ENSMUST00 | 0 | Lifr | ENSMUSG00000054263.6 |
| chr16 | 17890469 | 17892118 | 37.17318 | 134.81126 | 130.13722 | Promoter | 16 | 17839482 | 17891654 | 52173 | 2 | ENSMUST00 | 0 | Dgcr2 | ENSMUSG00000003166.14 |
| chr16 | 13902166 | 13904013 | 37.17318 | 134.81126 | 130.13722 | Promoter | 16 | 13903161 | 13949619 | 46459 | 1 | ENSMUST00 | 0 | Mpv17l | ENSMUSG00000022679.8 |
| chr17 | 34605337 | 34606830 | 37.17318 | 134.81126 | 130.13722 | Promoter | 17 | 34605861 | 34612443 | 6583 | 1 | ENSMUST00 | 0 | Agpat1 | ENSMUSG00000034254.12 |
| chr2 | 121413138 | 121415144 | 37.17318 | 134.81126 | 130.13722 | Promoter | 2 | 121413775 | 121438687 | 24913 | 1 | ENSMUST00 | 0 | Pdia3 | ENSMUSG00000027248.8 |
| chr4 | 149484616 | 149486348 | 37.17318 | 134.81126 | 130.13722 | Promoter | 4 | 149485229 | 149488122 | 2894 | 1 | ENSMUST00 | 0 | Lzic | ENSMUSG00000028990.8 |
| chr4 | 16162201 | 16163614 | 37.17318 | 134.81126 | 130.13722 | Promoter | 4 | 16122741 | 16163643 | 40903 | 2 | ENSMUST00 | 29 | Ripk2 | ENSMUSG00000041135.10 |
| chr7 | 28958851 | 28962304 | 37.17318 | 134.81126 | 130.13722 | Promoter | 7 | 28898620 | 28962223 | 63604 | 2 | ENSMUST00 | 0 | Actn4 | ENSMUSG00000054808.8 |
| chr9 | 57763942 | 57766341 | 37.17318 | 134.81126 | 130.13722 | Promoter | 9 | 57750711 | 57765860 | 15150 | 2 | ENSMUST00 | 0 | Clk3 | ENSMUSG00000032316.7 |
| chr11 | 50290597 | 50293020 | 40.24734 | 134.79797 | 130.13031 | Promoter | 11 | 50255634 | 50291804 | 36171 | 2 | ENSMUST00 | 0 | Maml1 | ENSMUSG00000050567.11 |
| chr15 | 76008984 | 76010319 | 41.50429 | 134.78256 | 130.11549 | Promoter | 15 | 76001093 | 76009498 | 8406 | 2 | ENSMUST00 | 0 | Fam83h | ENSMUSG00000046761.7 |
| chr6 | 142386389 | 142388158 | 41.50429 | 134.78256 | 130.11549 | Promoter | 6 | 142350342 | 142387036 | 36695 | 2 | ENSMUST00 | 0 | Recql | ENSMUSG00000030243.11 |
| chr2 | 91709479 | 91711625 | 38.96697 | 134.74414 | 130.07794 | Promoter | 2 | 91710852 | 91712327 | 1476 | 1 | ENSMUST00 | 0 | Harbi1 | ENSMUSG00000027243.8 |
| chr4 | 148448508 | 148449485 | 38.96697 | 134.74414 | 130.07794 | Promoter | 4 | 148448611 | 148454969 | 6359 | 1 | ENSMUST00 | 0 | Mtor | ENSMUSG00000028991.10 |
| chr8 | 11006639 | 11010129 | 40.16868 | 134.66035 | 129.99733 | Promoter | 8 | 10986980 | 11008458 | 21479 | 2 | ENSMUST00 | 0 | Irs2 | ENSMUSG00000038894.6 |
| chr1 | 55405130 | 55407992 | 37.68014 | 134.63641 | 129.97372 | Promoter | 1 | 55405921 | 55754285 | 348365 | 1 | ENSMUST00 | 0 | Plcl1 | ENSMUSG00000038349.10 |
| chr14 | 54685456 | 54686709 | 37.68014 | 134.63641 | 129.97372 | Promoter | 14 | 54642162 | 54686616 | 44455 | 2 | ENSMUST00 | 0 | Acin1 | ENSMUSG00000022185.13 |
| chr7 | 55841802 | 55843400 | 37.68014 | 134.63641 | 129.97372 | Promoter | 7 | 55842071 | 55930700 | 88630 | 1 | ENSMUST00 | 0 | Cyfip1 | ENSMUSG00000030447.10 |
| chr6 | 120821692 | 120822848 | 41.41883 | 134.6346 | 129.97316 | Promoter | 6 | 120795245 | 120822685 | 27441 | 2 | ENSMUST00 | 0 | Atp6v1e1 | ENSMUSG00000019210.8 |
| chr11 | 70844185 | 70845713 | 40.77746 | 134.62177 | 129.96082 | Promoter | 11 | 70844778 | 70943105 | 98328 | 1 | ENSMUST00 | 0 | Rabep1 | ENSMUSG00000020817.11 |
| chr12 | 87470984 | 87472395 | 40.77746 | 134.62177 | 129.96082 | Promoter | 12 | 87449936 | 87472299 | 22364 | 2 | ENSMUST00 | 0 | Snw1 | ENSMUSG00000021039.8 |
| chr19 | 6980136 | 6981173 | 40.77746 | 134.62177 | 129.96082 | Promoter | 19 | 6977741 | 6980440 | 2700 | 2 | ENSMUST00 | 0 | Fkbp2 | ENSMUSG00000056629.10 |
| chr4 | 46137489 | 46139715 | 40.77746 | 134.62177 | 129.96082 | Promoter | 4 | 46138613 | 46172403 | 33791 | 1 | ENSMUST00 | 0 | Ncbp1 | ENSMUSG00000028330.8 |
| chr9 | 41156374 | 41158065 | 40.77746 | 134.62177 | 129.96082 | Promoter | 9 | 41011098 | 41158062 | 146965 | 2 | ENSMUST00 | 0 | Ubash3b | ENSMUSG00000032020.10 |
| chr18 | 38418542 | 38420768 | 41.33371 | 134.48721 | 129.82906 | Promoter | 18 | 38418975 | 38464399 | 45425 | 1 | ENSMUST00 | 0 | Ndfip1 | ENSMUSG00000024425.3 |
| chr8 | 120100411 | 120102247 | 40.69584 | 134.47972 | 129.82225 | Promoter | 8 | 120081095 | 120101482 | 20388 | 2 | ENSMUST00 | 0 | Zdhhc7 | ENSMUSG00000031823.8 |
| chr8 | 13104180 | 13106753 | 40.01227 | 134.38658 | 129.72945 | Promoter | 8 | 13105621 | 13147940 | 42320 | 1 | ENSMUST00 | 0 | Cul4a | ENSMUSG00000031446.9 |
| chr1 | 58421831 | 58424072 | 32.94367 | 134.34938 | 129.69266 | Promoter | 1 | 58410189 | 58424066 | 13878 | 2 | ENSMUST00 | 0 | Clk1 | ENSMUSG00000026034.12 |
| chr12 | 70937304 | 70938812 | 32.94367 | 134.34938 | 129.69266 | Promoter | 12 | 70937857 | 70964718 | 26862 | 1 | ENSMUST00 | 0 | Actr10 | ENSMUSG00000021076.5 |
| chr6 | 83774759 | 83776337 | 32.94367 | 134.34938 | 129.69266 | Promoter | 6 | 83768885 | 83775813 | 6929 | 2 | ENSMUST00 | 0 | Tex261 | ENSMUSG00000014748.8 |
| chr3 | 10439271 | 10440425 | 41.24895 | 134.34035 | 129.68512 | Promoter | 3 | 10417819 | 10440124 | 22306 | 2 | ENSMUST00 | 0 | Snx16 | ENSMUSG00000027534.11 |
| chr10 | 13008406 | 13009372 | 40.61456 | 134.33821 | 129.68364 | 5' UTR | 10 | 13005343 | 13009183 | 3841 | 1 | ENSMUST00 | 3064 | Sf3b5 | ENSMUSG00000078348.4 |
| chr15 | 102073793 | 102075076 | 40.61456 | 134.33821 | 129.68364 | Promoter | 15 | 102073773 | 102097172 | 23400 | 1 | ENSMUST00 | 21 | Eif4b | ENSMUSG00000058655.8 |
| chr17 | 46855436 | 46858855 | 40.61456 | 134.33821 | 129.68364 | Distal Interge | 17 | 46798116 | 46831413 | 33298 | 2 | ENSMUST00 | -24024 | Gltscr1l | ENSMUSG00000036568.6 |
| chr6 | 86732345 | 86734030 | 35.69496 | 134.26468 | 129.61096 | Promoter | 6 | 86691768 | 86733378 | 41611 | 2 | ENSMUST00 | 0 | Gmcl1 | ENSMUSG00000001157.8 |
| chr10 | 58446089 | 58447838 | 41.81742 | 134.22485 | 129.57179 | Promoter | 10 | 58446852 | 58494155 | 47304 | 1 | ENSMUST00 | 0 | Ranbp2 | ENSMUSG00000003226.6 |
| chr13 | 111806971 | 111810084 | 41.81742 | 134.22485 | 129.57179 | Promoter | 13 | 111746428 | 111808993 | 62566 | 2 | ENSMUST00 | 0 | Map3k1 | ENSMUSG00000021754.12 |
| chr14 | 98164311 | 98170276 | 41.81742 | 134.22485 | 129.57179 | Promoter | 14 | 97786853 | 98169543 | 382691 | 2 | ENSMUST00 | 0 | Dach1 | ENSMUSG00000055639.11 |
| chr17 | 8282625 | 8284867 | 41.81742 | 134.22485 | 129.57179 | Promoter | 17 | 8282904 | 8296885 | 13982 | 1 | ENSMUST00 | 0 | Mpc1 | ENSMUSG00000023861.12 |
| chr9 | 72531033 | 72533138 | 41.81742 | 134.22485 | 129.57179 | Promoter | 9 | 72532240 | 72622829 | 90590 | 1 | ENSMUST00 | 0 | Rfx7 | ENSMUSG00000037674.10 |
| chr10 | 22730228 | 22732192 | 39.29051 | 134.21745 | 129.56854 | Promoter | 10 | 22703879 | 22731447 | 27569 | 2 | ENSMUST00 | 0 | Tbpl1 | ENSMUSG00000071359.8 |
| chr19 | 57361054 | 57363157 | 39.29051 | 134.21745 | 129.56854 | Promoter | 19 | 57361009 | 57389594 | 28586 | 1 | ENSMUST00 | 46 | Fam160b1 | ENSMUSG00000033478.4 |
| chr19 | 34877357 | 34879837 | 39.29051 | 134.21745 | 129.56854 | Promoter | 19 | 34810894 | 34879455 | 68562 | 2 | ENSMUST00 | 0 | Pank1 | ENSMUSG00000033610.10 |
| chr5 | 122421531 | 122423402 | 39.29051 | 134.21745 | 129.56854 | Promoter | 5 | 122421693 | 122444912 | 23220 | 1 | ENSMUST00 | 0 | Anapc7 | ENSMUSG00000029466.6 |
| chr6 | 72344520 | 72345612 | 39.29051 | 134.21745 | 129.56854 | Promoter | 6 | 72318678 | 72345184 | 26507 | 2 | ENSMUST00 | 0 | Usp39 | ENSMUSG00000056305.3 |
| chr7 | 122066447 | 122068300 | 39.29051 | 134.21745 | 129.56854 | Promoter | 7 | 122067175 | 122082194 | 15020 | 1 | ENSMUST00 | 0 | Ubfd1 | ENSMUSG00000030870.7 |
| chr17 | 78735419 | 78737563 | 40.53359 | 134.1972 | 129.55179 | Promoter | 17 | 78649913 | 78737196 | 87284 | 2 | ENSMUST00 | 0 | Strn | ENSMUSG00000024077.10 |
| chr6 | 18847967 | 18849308 | 41.16454 | 134.19406 | 129.54916 | Promoter | 6 | 18848579 | 18854053 | 5475 | 1 | ENSMUST00 | 0 | Lsm8 | ENSMUSG00000044155.8 |
| chr6 | 72437281 | 72440865 | 28.40054 | 134.09712 | 129.45372 | Promoter | 6 | 72432799 | 72439558 | 6760 | 2 | ENSMUST00 | 0 | Mat2a | ENSMUSG00000053907.9 |
| chr2 | 10047371 | 10049480 | 36.77131 | 134.07788 | 129.4352 | Promoter | 2 | 9914552 | 10048596 | 134045 | 2 | ENSMUST00 | 0 | Taf3 | ENSMUSG00000025782.7 |
| chr3 | 106546888 | 106548859 | 36.77131 | 134.07788 | 129.4352 | Promoter | 3 | 106547693 | 106573930 | 26238 | 1 | ENSMUST00 | 0 | Dram2 | ENSMUSG00000027900.10 |
| chr5 | 33781894 | 33783416 | 36.77131 | 134.07788 | 129.4352 | Promoter | 5 | 33739673 | 33782817 | 43145 | 2 | ENSMUST00 | 0 | Letm1 | ENSMUSG00000005299.5 |
| chr11 | 95039880 | 95041440 | 40.45295 | 134.0567 | 129.41484 | Promoter | 11 | 95026258 | 95041354 | 15097 | 2 | ENSMUST00 | 0 | Pdk2 | ENSMUSG00000038967.8 |
| chr16 | 31199908 | 31201268 | 37.96327 | 134.05096 | 129.40988 | Promoter | 16 | 31092413 | 31201238 | 108826 | 2 | ENSMUST00 | 0 | Acap2 | ENSMUSG00000049076.9 |
| chr4 | 130360126 | 130360954 | 41.08046 | 134.04831 | 129.40907 | Promoter | 4 | 130360132 | 130390026 | 29895 | 1 | ENSMUST00 | 0 | Snrnp40 | ENSMUSG00000074088.6 |
| chr12 | 21111284 | 21113355 | 41.65482 | 133.94409 | 129.3063 | Promoter | 12 | 21111748 | 21270171 | 158424 | 1 | ENSMUST00 | 0 | Asap2 | ENSMUSG00000052632.10 |
| chr17 | 34980890 | 34982714 | 40.37263 | 133.9167 | 129.27921 | Promoter | 17 | 34981862 | 34983981 | 2120 | 1 | ENSMUST00 | 0 | Lsm2 | ENSMUSG00000007050.12 |
| chr8 | 23237067 | 23237778 | 40.37263 | 133.9167 | 129.27921 | Promoter | 8 | 23226610 | 23237668 | 11059 | 2 | ENSMUST00 | 0 | Gins4 | ENSMUSG00000031546.5 |
| chr13 | 53376577 | 53377871 | 40.99673 | 133.90312 | 129.26613 | Promoter | 13 | 53332716 | 53377365 | 44650 | 2 | ENSMUST00 | 0 | Sptlc1 | ENSMUSG00000021468.7 |
| chr5 | 113770528 | 113773818 | 34.91946 | 133.88905 | 129.25287 | Promoter | 5 | 113742446 | 113771649 | 29204 | 2 | ENSMUST00 | 0 | Sart3 | ENSMUSG00000018974.3 |
| chr10 | 117147475 | 117149206 | 41.56779 | 133.79375 | 129.15814 | Promoter | 10 | 117070129 | 117148474 | 78346 | 2 | ENSMUST00 | 0 | Frs2 | ENSMUSG00000020170.3 |
| chr17 | 35262477 | 35264430 | 41.56779 | 133.79375 | 129.15814 | Promoter | 17 | 35262730 | 35267499 | 4770 | 1 | ENSMUST00 | 0 | H2-D1 | ENSMUSG00000073411.6 |
| chr6 | 58831048 | 58832945 | 41.56779 | 133.79375 | 129.15814 | Promoter | 6 | 58831465 | 58843728 | 12264 | 1 | ENSMUST00 | 0 | Herc3 | ENSMUSG00000029804.11 |
| chr7 | 83882272 | 83884889 | 41.56779 | 133.79375 | 129.15814 | Promoter | 7 | 83884466 | 83897940 | 13475 | 1 | ENSMUST00 | 0 | Mesdc2 | ENSMUSG00000038503.9 |
| chr17 | 23563495 | 23564961 | 40.29262 | 133.77719 | 129.14264 | Promoter | 17 | 23556769 | 23564226 | 7458 | 2 | ENSMUST00 | 0 | Zfp213 | ENSMUSG00000071256.3 |
| chr4 | 134766368 | 134768617 | 40.29262 | 133.77719 | 129.14264 | Promoter | 4 | 134744735 | 134768024 | 23290 | 2 | ENSMUST00 | 0 | Ldlrap1 | ENSMUSG00000037295.6 |
| chr4 | 59782788 | 59783948 | 40.29262 | 133.77719 | 129.14264 | Promoter | 4 | 59769637 | 59783866 | 14230 | 2 | ENSMUST00 | 0 | Inip | ENSMUSG00000038544.9 |
| chr17 | 64330658 | 64331974 | 40.91335 | 133.75845 | 129.12437 | Promoter | 17 | 64281005 | 64331916 | 50912 | 2 | ENSMUST00 | 0 | Pja2 | ENSMUSG00000024083.10 |
| chr19 | 56547547 | 56549649 | 39.62653 | 133.71027 | 129.07751 | Promoter | 19 | 56529167 | 56548147 | 18981 | 2 | ENSMUST00 | 0 | Dclre1a | ENSMUSG00000025077.8 |
| chr3 | 33799366 | 33800882 | 39.62653 | 133.71027 | 129.07751 | Promoter | 3 | 33799955 | 33814860 | 14906 | 1 | ENSMUST00 | 0 | Ttc14 | ENSMUSG00000027677.12 |
| chr8 | 104247668 | 104248919 | 39.62653 | 133.71027 | 129.07751 | Promoter | 8 | 104226691 | 104248558 | 21868 | 2 | ENSMUST00 | 0 | Tk2 | ENSMUSG00000035824.6 |
| chr9 | 78447873 | 78449175 | 39.62653 | 133.71027 | 129.07751 | Promoter | 9 | 78448208 | 78474152 | 25945 | 1 | ENSMUST00 | 0 | Mto1 | ENSMUSG00000032342.8 |
| chr4 | 56862824 | 56865430 | 41.48112 | 133.644 | 129.01402 | Promoter | 4 | 56810935 | 56865188 | 54254 | 2 | ENSMUST00 | 0 | Ctnnal1 | ENSMUSG00000038816.9 |
| chr6 | 38551430 | 38552912 | 41.48112 | 133.644 | 129.01402 | Promoter | 6 | 38551465 | 38606367 | 54903 | 1 | ENSMUST00 | 0 | Luc7l2 | ENSMUSG00000029823.11 |
| chr15 | 38517684 | 38520107 | 40.21293 | 133.6382 | 129.0087 | Promoter | 15 | 38487430 | 38519266 | 31837 | 2 | ENSMUST00 | 0 | Azin1 | ENSMUSG00000037458.9 |
| chr7 | 99139179 | 99141230 | 40.8303 | 133.61433 | 128.98549 | Promoter | 7 | 98886743 | 99141144 | 254402 | 2 | ENSMUST00 | 0 | Uvrag | ENSMUSG00000035354.8 |
| chr10 | 119238719 | 119241052 | 38.25672 | 133.48087 | 128.85442 | Promoter | 10 | 119199255 | 119240055 | 40801 | 2 | ENSMUST00 | 0 | Cand1 | ENSMUSG00000020114.7 |
| chr5 | 115118586 | 115119761 | 38.25672 | 133.48087 | 128.85442 | Promoter | 5 | 115110299 | 115119287 | 8989 | 2 | ENSMUST00 | 0 | Acads | ENSMUSG00000029545.8 |
| chr6 | 145745878 | 145749136 | 38.25672 | 133.48087 | 128.85442 | Promoter | 6 | 145746748 | 145821079 | 74332 | 1 | ENSMUST00 | 0 | Rassf8 | ENSMUSG00000030259.7 |
| chr9 | 22467532 | 22468784 | 38.25672 | 133.48087 | 128.85442 | Promoter | 9 | 22448311 | 22468356 | 20046 | 2 | ENSMUST00 | 0 | Rp9 | ENSMUSG00000032239.9 |
| chr2 | 174415534 | 174416555 | 37.02895 | 133.45709 | 128.83354 | Promoter | 2 | 174415804 | 174427502 | 11699 | 1 | ENSMUST00 | 0 | Nelfcd | ENSMUSG00000016253.7 |
| chr11 | 30266216 | 30269045 | 34.68901 | 133.45216 | 128.82967 | Promoter | 11 | 30099395 | 30268169 | 168775 | 2 | ENSMUST00 | 0 | Sptbn1 | ENSMUSG00000020315.13 |
| chr11 | 115162609 | 115165876 | 34.68901 | 133.45216 | 128.82967 | Promoter | 11 | 115163341 | 115181181 | 17841 | 1 | ENSMUST00 | 0 | Slc9a3r1 | ENSMUSG00000020733.3 |
| chr12 | 77237711 | 77240432 | 34.68901 | 133.45216 | 128.82967 | Promoter | 12 | 77238104 | 77475996 | 237893 | 1 | ENSMUST00 | 0 | Fut8 | ENSMUSG00000021065.10 |
| chr16 | 91546579 | 91548069 | 34.68901 | 133.45216 | 128.82967 | Promoter | 16 | 91547072 | 91565169 | 18098 | 1 | ENSMUST00 | 0 | Ifngr2 | ENSMUSG00000022965.7 |
| chr18 | 77712518 | 77714741 | 34.68901 | 133.45216 | 128.82967 | Promoter | 18 | 77633281 | 77714010 | 80730 | 2 | ENSMUST00 | 0 | 8030462N17 | ENSMUSG00000047466.9 |
| chr3 | 75555872 | 75556922 | 34.68901 | 133.45216 | 128.82967 | Promoter | 3 | 75516490 | 75556856 | 40367 | 2 | ENSMUST00 | 0 | Pdcd10 | ENSMUSG00000027835.6 |
| chr4 | 117550274 | 117551211 | 34.68901 | 133.45216 | 128.82967 | Promoter | 4 | 117550365 | 117674297 | 123933 | 1 | ENSMUST00 | 0 | Eri3 | ENSMUSG00000033423.11 |
| chr4 | 141546127 | 141547460 | 34.68901 | 133.45216 | 128.82967 | Promoter | 4 | 141546189 | 141547892 | 1704 | 1 | ENSMUST00 | 0 | B330016D10 | ENSMUSG00000048406.5 |
| chr5 | 93267238 | 93268862 | 34.68901 | 133.45216 | 128.82967 | Promoter | 5 | 93267257 | 93276231 | 8975 | 1 | ENSMUST00 | 0 | Ccng2 | ENSMUSG00000029385.9 |
| chr5 | 67260051 | 67261484 | 34.68901 | 133.45216 | 128.82967 | Promoter | 5 | 67260565 | 67291461 | 30897 | 1 | ENSMUST00 | 0 | Tmem33 | ENSMUSG00000037720.11 |
| chr9 | 44603277 | 44606301 | 40.0545 | 133.36163 | 128.74257 | Promoter | 9 | 44604892 | 44640731 | 35840 | 1 | ENSMUST00 | 0 | Ddx6 | ENSMUSG00000032097.9 |
| chr18 | 25167893 | 25169670 | 40.6652 | 133.3277 | 128.70952 | Promoter | 18 | 25127223 | 25168694 | 41472 | 2 | ENSMUST00 | 0 | Tpgs2 | ENSMUSG00000024269.6 |
| chr10 | 75920566 | 75922292 | 39.39634 | 133.30592 | 128.68834 | Promoter | 10 | 75896769 | 75921617 | 24849 | 2 | ENSMUST00 | 0 | Smarcb1 | ENSMUSG00000000902.8 |
| chr10 | 80740869 | 80742306 | 39.39634 | 133.30592 | 128.68834 | Promoter | 10 | 80706978 | 80742211 | 35234 | 2 | ENSMUST00 | 0 | Ap3d1 | ENSMUSG00000020198.7 |
| chr12 | 17690029 | 17692705 | 39.39634 | 133.30592 | 128.68834 | Promoter | 12 | 17690814 | 17791926 | 101113 | 1 | ENSMUST00 | 0 | Hpcal1 | ENSMUSG00000071379.1 |
| chr13 | 45962952 | 45966515 | 39.39634 | 133.30592 | 128.68834 | Promoter | 13 | 45965253 | 45966508 | 1256 | 1 | ENSMUST00 | 0 | 5033430I15R | ENSMUSG00000038152.4 |
| chr16 | 44723737 | 44725506 | 39.39634 | 133.30592 | 128.68834 | Promoter | 16 | 44724301 | 44737284 | 12984 | 1 | ENSMUST00 | 0 | BC027231 | ENSMUSG00000036208.8 |
| chr17 | 45563840 | 45564964 | 39.39634 | 133.30592 | 128.68834 | Promoter | 17 | 45563964 | 45567669 | 3706 | 1 | ENSMUST00 | 0 | Slc35b2 | ENSMUSG00000037089.9 |
| chr4 | 94978064 | 94979668 | 39.39634 | 133.30592 | 128.68834 | Promoter | 4 | 94942040 | 94979100 | 37061 | 2 | ENSMUST00 | 0 | Mysm1 | ENSMUSG00000062627.4 |
| chr5 | 107437510 | 107438662 | 39.39634 | 133.30592 | 128.68834 | Promoter | 5 | 107437997 | 107491628 | 53632 | 1 | ENSMUST00 | 0 | Btbd8 | ENSMUSG00000070632.3 |
| chr8 | 11554880 | 11558049 | 39.39634 | 133.30592 | 128.68834 | Promoter | 8 | 11556066 | 11563250 | 7185 | 1 | ENSMUST00 | 0 | Ing1 | ENSMUSG00000045969.7 |
| chr9 | 70541924 | 70543925 | 39.39634 | 133.30592 | 128.68834 | Promoter | 9 | 70542920 | 70592237 | 49318 | 1 | ENSMUST00 | 0 | Sltm | ENSMUSG00000032212.9 |
| chr1 | 91397475 | 91399330 | 39.97575 | 133.22409 | 128.61581 | Promoter | 1 | 91373861 | 91398700 | 24840 | 2 | ENSMUST00 | 0 | Ilkap | ENSMUSG00000026309.9 |
| chr10 | 126978573 | 126984050 | 39.97575 | 133.22411 | 128.61581 | Promoter | 10 | 126978717 | 126999975 | 21259 | 1 | ENSMUST00 | 0 | Ctdsp2 | ENSMUSG00000078429.3 |
| chr18 | 90509491 | 90510945 | 39.97575 | 133.22411 | 128.61581 | Promoter | 18 | 90510154 | 90543267 | 33114 | 1 | ENSMUST00 | 0 | Tmx3 | ENSMUSG00000024614.6 |
| chr4 | 4792040 | 4793755 | 39.97575 | 133.22411 | 128.61581 | Promoter | 4 | 4762484 | 4793355 | 30872 | 2 | ENSMUST00 | 0 | Impad1 | ENSMUSG00000066324.2 |
| chr5 | 130143888 | 130145604 | 39.97575 | 133.22411 | 128.61581 | Promoter | 5 | 130144861 | 130155806 | 10946 | 1 | ENSMUST00 | 0 | Kctd7 | ENSMUSG00000034110.3 |
| chr5 | 92434120 | 92435750 | 39.97575 | 133.22411 | 128.61581 | Promoter | 5 | 92415540 | 92435219 | 19680 | 2 | ENSMUST00 | 0 | Nup54 | ENSMUSG00000034826.9 |
| chr5 | 140648001 | 140649046 | 39.97575 | 133.22409 | 128.61581 | Promoter | 5 | 140620586 | 140649031 | 28446 | 2 | ENSMUST00 | 0 | Ttyh3 | ENSMUSG00000036565.3 |
| chr6 | 85450706 | 85452985 | 39.97575 | 133.22411 | 128.61581 | Promoter | 6 | 85451505 | 85468477 | 16973 | 1 | ENSMUST00 | 0 | Cct7 | ENSMUSG00000030007.6 |
| chr1 | 10037131 | 10040542 | 35.12714 | 133.20123 | 128.59534 | Promoter | 1 | 10024601 | 10038127 | 13527 | 2 | ENSMUST00 | 0 | Cops5 | ENSMUSG00000025917.4 |
| chr19 | 10603045 | 10606602 | 41.22328 | 133.19821 | 128.59248 | Promoter | 19 | 10605625 | 10629813 | 24189 | 1 | ENSMUST00 | 0 | Ddb1 | ENSMUSG00000024740.9 |
| chr11 | 74896187 | 74898875 | 40.58316 | 133.18515 | 128.57994 | Promoter | 11 | 74898071 | 74904940 | 6870 | 1 | ENSMUST00 | 0 | Tsr1 | ENSMUSG00000038335.8 |

| | | | | | | | | | | | | | | | |
|---|---|---|---|---|---|---|---|---|---|---|---|---|---|---|---|
| chr17 | 45685260 | 45687208 | 40.58316 | 133.18515 | 128.57994 | Promoter | 17 | 45686322 | 45698495 | 12174 | 1 | ENSMUST00 | 0 | Mrpl14 | ENSMUSG00000023939.7 |
| chr16 | 18836265 | 18837323 | 41.13805 | 133.05075 | 128.44682 | Promoter | 16 | 18836578 | 18840160 | 3583 | 1 | ENSMUST00 | 0 | 2510002D24I | ENSMUSG00000071632.5 |
| chr7 | 98655540 | 98657644 | 39.24815 | 133.0453 | 128.44171 | Promoter | 7 | 98590615 | 98656569 | 65955 | 2 | ENSMUST00 | 0 | 2210018M11 | ENSMUSG00000035401.8 |
| chr1 | 93444164 | 93446315 | 36.7977 | 133.03651 | 128.43381 | Promoter | 1 | 93407762 | 93445686 | 37925 | 2 | ENSMUST00 | 0 | Hdlbp | ENSMUSG00000034088.10 |
| chr11 | 120823409 | 120825174 | 36.7977 | 133.03651 | 128.43381 | Promoter | 11 | 120805940 | 120824178 | 18239 | 2 | ENSMUST00 | 0 | Fasn | ENSMUSG00000025153.8 |
| chr11 | 70687182 | 70688680 | 36.7977 | 133.03651 | 128.43381 | Promoter | 11 | 70669463 | 70687943 | 18481 | 2 | ENSMUST00 | 0 | Camta2 | ENSMUSG00000040712.11 |
| chr12 | 55835937 | 55837231 | 36.7977 | 133.03651 | 128.43381 | Promoter | 12 | 55836366 | 55869735 | 33370 | 1 | ENSMUST00 | 0 | Brms1l | ENSMUSG00000012076.7 |
| chr13 | 38658335 | 38659414 | 36.7977 | 133.03651 | 128.43381 | Promoter | 13 | 38645698 | 38659028 | 13331 | 2 | ENSMUST00 | 0 | Eef1e1 | ENSMUSG00000001707.7 |
| chr15 | 99473692 | 99475027 | 36.7977 | 133.03651 | 128.43381 | Promoter | 15 | 99470084 | 99474730 | 4647 | 2 | ENSMUST00 | 0 | Bcdin3d | ENSMUSG00000037525.4 |
| chr16 | 4726000 | 4727245 | 36.7977 | 133.03651 | 128.43381 | Promoter | 16 | 4726361 | 4764848 | 38488 | 1 | ENSMUST00 | 0 | Hmox2 | ENSMUSG00000004070.10 |
| chr16 | 52030893 | 52033595 | 36.7977 | 133.03651 | 128.43381 | Promoter | 16 | 52031549 | 52208047 | 176499 | 1 | ENSMUST00 | 0 | Cblb | ENSMUSG00000022637.10 |
| chr18 | 10725216 | 10727065 | 36.7977 | 133.03651 | 128.43381 | Promoter | 18 | 10725548 | 10818578 | 93031 | 1 | ENSMUST00 | 0 | Mib1 | ENSMUSG00000024294.8 |
| chr2 | 173021539 | 173022981 | 36.7977 | 133.03651 | 128.43381 | Promoter | 2 | 173021899 | 173033913 | 12015 | 1 | ENSMUST00 | 0 | Rbm38 | ENSMUSG00000027510.12 |
| chr2 | 34801899 | 34803367 | 36.7977 | 133.03651 | 128.43381 | Downstream | 2 | 34777556 | 34799912 | 22357 | 2 | ENSMUST00 | -1988 | Rabepk | ENSMUSG00000070953.8 |
| chr4 | 138215850 | 138218184 | 36.7977 | 133.03651 | 128.43381 | Promoter | 4 | 138216296 | 138244683 | 28388 | 1 | ENSMUST00 | 0 | Hp1bp3 | ENSMUSG00000028759.8 |
| chr4 | 132921541 | 132922574 | 36.7977 | 133.03651 | 128.43381 | Promoter | 4 | 132899213 | 132922558 | 23346 | 2 | ENSMUST00 | 0 | Fam76a | ENSMUSG00000028878.6 |
| chr4 | 120824626 | 120826112 | 36.7977 | 133.03651 | 128.43381 | Promoter | 4 | 120757438 | 120825706 | 68269 | 2 | ENSMUST00 | 0 | Nfyc | ENSMUSG00000032897.12 |
| chr6 | 13607283 | 13608435 | 36.7977 | 133.03651 | 128.43381 | Promoter | 6 | 13580687 | 13608100 | 27414 | 2 | ENSMUST00 | 0 | Tmem168 | ENSMUSG00000029569.4 |
| chr6 | 125096062 | 125099104 | 36.7977 | 133.03651 | 128.43381 | Promoter | 6 | 125096162 | 125130591 | 34430 | 1 | ENSMUST00 | 0 | Chd4 | ENSMUSG00000063870.7 |
| chr8 | 123948488 | 123950292 | 36.7977 | 133.03651 | 128.43381 | Promoter | 8 | 123897123 | 123949265 | 52143 | 2 | ENSMUST00 | 0 | Nup133 | ENSMUSG00000039509.7 |
| chr1 | 134559887 | 134561761 | 32.2491 | 132.9626 | 128.36652 | Promoter | 1 | 134560171 | 134635285 | 75115 | 1 | ENSMUST00 | 0 | Kdm5b | ENSMUSG00000042207.12 |
| chr17 | 46201994 | 46203778 | 36.15705 | 132.95093 | 128.35489 | Promoter | 17 | 46172004 | 46202625 | 30622 | 2 | ENSMUST00 | 0 | Polh | ENSMUSG00000023953.8 |
| chr10 | 61451227 | 61452888 | 38.56107 | 132.92723 | 128.33237 | Promoter | 10 | 61432497 | 61452669 | 20173 | 2 | ENSMUST00 | 0 | Eif4ebp2 | ENSMUSG00000020091.9 |
| chr18 | 61176314 | 61177815 | 34.3811 | 132.86635 | 128.27481 | Promoter | 18 | 61131279 | 61177050 | 45772 | 2 | ENSMUST00 | 0 | Hmgxb3 | ENSMUSG00000024622.7 |
| chr5 | 65390336 | 65392150 | 37.29565 | 132.84946 | 128.25806 | Promoter | 5 | 65391497 | 65410693 | 19197 | 1 | ENSMUST00 | 0 | Lias | ENSMUSG00000029199.6 |
| chr11 | 88098491 | 88100196 | 39.74135 | 132.8143 | 128.22414 | Promoter | 11 | 88099146 | 88194140 | 94995 | 1 | ENSMUST00 | 0 | Cuedc1 | ENSMUSG00000018378.8 |
| chr16 | 32608478 | 32609750 | 39.74135 | 132.8143 | 128.22414 | Promoter | 16 | 32608920 | 32632794 | 23875 | 1 | ENSMUST00 | 0 | Tfrc | ENSMUSG00000022797.10 |
| chr11 | 79143897 | 79146881 | 38.4896 | 132.80048 | 128.21092 | Promoter | 11 | 79013440 | 79146407 | 132968 | 2 | ENSMUST00 | 0 | Ksr1 | ENSMUSG00000018334.11 |
| chr11 | 51755431 | 51757268 | 40.33899 | 132.76065 | 128.17128 | Promoter | 11 | 51692264 | 51756618 | 64355 | 2 | ENSMUST00 | 0 | Sec24a | ENSMUSG00000036391.11 |
| chr12 | 64964474 | 64968476 | 40.33899 | 132.76065 | 128.17128 | Promoter | 12 | 64942440 | 64965536 | 23097 | 2 | ENSMUST00 | 0 | Klhl28 | ENSMUSG00000020948.8 |
| chr2 | 92023121 | 92024855 | 40.33899 | 132.76065 | 128.17128 | Promoter | 2 | 91982328 | 92024502 | 42175 | 2 | ENSMUST00 | 0 | Creb3l1 | ENSMUSG00000027230.9 |
| chr5 | 92137086 | 92138804 | 40.33899 | 132.76065 | 128.17128 | Promoter | 5 | 92137938 | 92202794 | 64857 | 1 | ENSMUST00 | 0 | Uso1 | ENSMUSG00000029407.7 |
| chr11 | 79992377 | 79994546 | 39.66383 | 132.67865 | 128.09186 | Promoter | 11 | 79993106 | 80034122 | 41017 | 1 | ENSMUST00 | 0 | Suz12 | ENSMUSG00000017548.10 |
| chr2 | 122737814 | 122739246 | 39.66383 | 132.67865 | 128.09186 | Promoter | 2 | 122738503 | 122749475 | 10973 | 1 | ENSMUST00 | 0 | Bloc1s6 | ENSMUSG00000005804.9 |
| chr8 | 104441803 | 104443201 | 39.66383 | 132.67865 | 128.09186 | Promoter | 8 | 104417674 | 104443047 | 25374 | 2 | ENSMUST00 | 0 | Dync1li2 | ENSMUSG00000035770.7 |
| chr10 | 128524149 | 128526021 | 32.63288 | 132.66988 | 128.08362 | Promoter | 10 | 128510248 | 128525859 | 15612 | 2 | ENSMUST00 | 0 | Esyt1 | ENSMUSG00000025366.7 |
| chr11 | 4593858 | 4594834 | 32.63288 | 132.66988 | 128.08362 | Promoter | 11 | 4480868 | 4594815 | 113948 | 2 | ENSMUST00 | 0 | Mtmr3 | ENSMUSG00000034354.12 |
| chr14 | 56885267 | 56886353 | 32.63288 | 132.66988 | 128.08362 | Distal Interge | 14 | 56887795 | 56962579 | 74785 | 1 | ENSMUST00 | -1442 | Zmym2 | ENSMUSG00000021945.8 |
| chr2 | 180041455 | 180043558 | 32.63288 | 132.66988 | 128.08362 | Promoter | 2 | 180036374 | 180042395 | 6022 | 2 | ENSMUST00 | 0 | Psma7 | ENSMUSG00000027566.9 |
| chr9 | 95954057 | 95955654 | 32.63288 | 132.66988 | 128.08362 | Promoter | 9 | 95954760 | 96053027 | 98268 | 1 | ENSMUST00 | 0 | Xrn1 | ENSMUSG00000032410.8 |
| chr17 | 8310558 | 8311862 | 39.02457 | 132.65163 | 128.06667 | Promoter | 17 | 8311102 | 8327442 | 16341 | 1 | ENSMUST00 | 0 | Sft2d1 | ENSMUSG00000073468.6 |
| chr17 | 34836196 | 34838013 | 40.25826 | 132.62018 | 128.03537 | Promoter | 17 | 34837019 | 34838648 | 1630 | 1 | ENSMUST00 | 0 | Dxo | ENSMUSG00000040482.10 |
| chr2 | 52857275 | 52859971 | 40.88443 | 132.61177 | 128.02734 | Promoter | 2 | 52857860 | 53133804 | 275945 | 1 | ENSMUST00 | 0 | Fmnl2 | ENSMUSG00000036053.11 |
| chr7 | 45394890 | 45395811 | 40.88443 | 132.61177 | 128.02734 | Promoter | 7 | 45376462 | 45395692 | 19231 | 2 | ENSMUST00 | 0 | Snrnp70 | ENSMUSG00000063511.8 |
| chr17 | 71474211 | 71476282 | 40.80059 | 132.46654 | 127.88422 | Promoter | 17 | 71344489 | 71475343 | 130855 | 2 | ENSMUST00 | 0 | Smchd1 | ENSMUSG00000024054.8 |
| chr4 | 120950859 | 120952375 | 40.80059 | 132.46654 | 127.88422 | Promoter | 4 | 120930137 | 120951699 | 21563 | 2 | ENSMUST00 | 0 | Zfp69 | ENSMUSG00000064141.9 |
| chr4 | 35225022 | 35226833 | 39.50969 | 132.40687 | 127.82707 | Promoter | 4 | 35191285 | 35226123 | 34839 | 2 | ENSMUST00 | 0 | 3110043O21 | ENSMUSG00000028300.9 |
| chr1 | 177441041 | 177442001 | 38.87692 | 132.39133 | 127.80991 | Intron (ENSM | 1 | 177429047 | 177443270 | 14224 | 2 | ENSMUST00 | 1269 | Gm26801 | ENSMUSG00000097892.1 |
| chr10 | 91081827 | 91083688 | 38.87692 | 132.39133 | 127.80991 | Promoter | 10 | 91082940 | 91102607 | 19668 | 1 | ENSMUST00 | 0 | Ikbip | ENSMUSG00000019975.6 |
| chr11 | 50915587 | 50916437 | 38.87692 | 132.39133 | 127.80991 | Promoter | 11 | 50898712 | 50916142 | 17431 | 2 | ENSMUST00 | 0 | Zfp2 | ENSMUSG00000049321.12 |
| chr12 | 26468972 | 26470438 | 38.87692 | 132.39133 | 127.80991 | Promoter | 12 | 26469204 | 26479835 | 10632 | 1 | ENSMUST00 | 0 | Cmpk2 | ENSMUSG00000020638.7 |
| chr19 | 10947643 | 10949342 | 38.87692 | 132.39133 | 127.80991 | Promoter | 19 | 10941481 | 10949266 | 7786 | 2 | ENSMUST00 | 0 | Ccdc86 | ENSMUSG00000024732.7 |
| chr4 | 25280968 | 25282172 | 38.87692 | 132.39133 | 127.80991 | Promoter | 4 | 25248600 | 25281760 | 33161 | 2 | ENSMUST00 | 0 | Ufl1 | ENSMUSG00000040359.9 |
| chr5 | 45855601 | 45858324 | 38.87692 | 132.39133 | 127.80991 | Promoter | 5 | 45697181 | 45857615 | 160435 | 2 | ENSMUST00 | 0 | Lcorl | ENSMUSG00000015882.12 |
| chr6 | 120665084 | 120668136 | 38.87692 | 132.39133 | 127.80991 | Promoter | 6 | 120666369 | 120724920 | 58552 | 1 | ENSMUST00 | 0 | Cecr2 | ENSMUSG00000071226.6 |
| chr8 | 91069467 | 91071360 | 38.87692 | 132.39133 | 127.80991 | Promoter | 8 | 91070057 | 91123844 | 53788 | 1 | ENSMUST00 | 0 | Rbl2 | ENSMUSG00000031666.9 |
| chr1 | 7397106 | 7398006 | 41.35789 | 132.3362 | 127.75856 | Promoter | 1 | 7349406 | 7397801 | 48396 | 2 | ENSMUST00 | 0 | Gm26901 | ENSMUSG00000097797.1 |
| chr11 | 98936314 | 98938742 | 41.35789 | 132.3362 | 127.75856 | Promoter | 11 | 98937698 | 98974162 | 36465 | 1 | ENSMUST00 | 0 | Rara | ENSMUSG00000037992.11 |
| chr11 | 20740526 | 20742745 | 41.35789 | 132.3362 | 127.75856 | Promoter | 11 | 20685084 | 20741589 | 56506 | 2 | ENSMUST00 | 0 | Aftph | ENSMUSG00000049659.10 |
| chr12 | 79006402 | 79007629 | 41.35789 | 132.3362 | 127.75856 | Promoter | 12 | 78961795 | 79007627 | 45833 | 2 | ENSMUST00 | 0 | Tmem229b | ENSMUSG00000046157.8 |
| chr2 | 112239017 | 112240610 | 41.35789 | 132.3362 | 127.75856 | Promoter | 2 | 112239468 | 112247111 | 7644 | 1 | ENSMUST00 | 0 | Lpcat4 | ENSMUSG00000027134.4 |
| chr7 | 140137230 | 140138492 | 41.35789 | 132.3362 | 127.75856 | Promoter | 7 | 140137564 | 140150786 | 13223 | 1 | ENSMUST00 | 0 | Mtg1 | ENSMUSG00000039018.9 |
| chr2 | 73891600 | 73892667 | 40.71709 | 132.32185 | 127.74851 | Promoter | 2 | 73816509 | 73892536 | 76028 | 2 | ENSMUST00 | 0 | Atf2 | ENSMUSG00000027104.13 |
| chr9 | 51007800 | 51009147 | 40.71709 | 132.32185 | 127.74851 | Promoter | 9 | 50892801 | 51008936 | 116136 | 2 | ENSMUST00 | 0 | Sik2 | ENSMUSG00000037112.11 |
| chr3 | 142880321 | 142883208 | 30.90292 | 132.31535 | 127.74332 | Promoter | 3 | 142790902 | 142882004 | 91103 | 2 | ENSMUST00 | 0 | Pkn2 | ENSMUSG00000004591.11 |
| chr2 | 154791040 | 154792992 | 39.43306 | 132.27451 | 127.70399 | Promoter | 2 | 154791096 | 154867245 | 76150 | 1 | ENSMUST00 | 0 | Raly | ENSMUSG00000027593.10 |
| chr9 | 57130158 | 57131721 | 37.5719 | 132.25589 | 127.68574 | Promoter | 9 | 57130690 | 57134155 | 3466 | 1 | ENSMUST00 | 0 | Man2c1 | ENSMUSG00000032295.10 |
| chr7 | 120917145 | 120918777 | 41.2835 | 132.20779 | 127.63918 | Promoter | 7 | 120917744 | 120947432 | 29689 | 1 | ENSMUST00 | 0 | Polr3e | ENSMUSG00000030880.8 |
| chr4 | 109279957 | 109281494 | 39.93851 | 132.06328 | 127.49693 | Promoter | 4 | 109280268 | 109387817 | 107550 | 1 | ENSMUST00 | 0 | Eps15 | ENSMUSG00000028552.8 |
| chr15 | 89314018 | 89315977 | 41.19707 | 132.05855 | 127.49266 | Promoter | 15 | 89288236 | 89315311 | 27076 | 2 | ENSMUST00 | 0 | Sbf1 | ENSMUSG00000036529.12 |
| chr5 | 120502797 | 120503656 | 40.55112 | 132.03412 | 127.46851 | Promoter | 5 | 120483282 | 120503625 | 20344 | 2 | ENSMUST00 | 0 | Plbd2 | ENSMUSG00000029598.7 |
| chr6 | 38874171 | 38877070 | 40.55112 | 132.03412 | 127.46851 | Promoter | 6 | 38875404 | 38952973 | 77570 | 1 | ENSMUST00 | 0 | Tbxas1 | ENSMUSG00000029925.8 |
| chr7 | 92667825 | 92669960 | 39.85937 | 131.92529 | 127.36071 | Promoter | 7 | 92643712 | 92669933 | 26222 | 2 | ENSMUST00 | 0 | Pcf11 | ENSMUSG00000041328.10 |
| chr17 | 45506779 | 45507331 | 41.111 | 131.90991 | 127.34579 | Promoter | 17 | 45506841 | 45520842 | 14002 | 1 | ENSMUST00 | 0 | Aars2 | ENSMUSG00000023938.7 |
| chr17 | 26932571 | 26934332 | 41.111 | 131.90991 | 127.34579 | Promoter | 17 | 26933127 | 26937890 | 4764 | 1 | ENSMUST00 | 0 | Phf1 | ENSMUSG00000024193.7 |
| chr1 | 125559921 | 125561892 | 40.46864 | 131.89105 | 127.3278 | Promoter | 1 | 125560595 | 125568573 | 7979 | 1 | ENSMUST00 | 0 | Slc35f5 | ENSMUSG00000026342.5 |
| chr15 | 57692968 | 57696625 | 40.46864 | 131.89105 | 127.3278 | Promoter | 15 | 57694665 | 57839832 | 145168 | 1 | ENSMUST00 | 0 | Zhx2 | ENSMUSG00000071757.5 |
| chr1 | 181210182 | 181212891 | 39.20496 | 131.87447 | 127.31225 | Promoter | 1 | 181173228 | 181211552 | 38325 | 2 | ENSMUST00 | 0 | Wdr26 | ENSMUSG00000038733.8 |
| chr3 | 141464561 | 141466743 | 39.20496 | 131.87447 | 127.31225 | Promoter | 3 | 141465564 | 141834924 | 369361 | 1 | ENSMUST00 | 0 | Unc5c | ENSMUSG00000059921.10 |
| chr4 | 45319754 | 45321065 | 39.20496 | 131.87447 | 127.31225 | Promoter | 4 | 45316613 | 45320629 | 4017 | 2 | ENSMUST00 | 0 | Exosc3 | ENSMUSG00000028322.6 |
| chr5 | 25759438 | 25760604 | 39.20496 | 131.87447 | 127.31225 | Promoter | 5 | 25759997 | 25850342 | 90346 | 1 | ENSMUST00 | 0 | Actr3b | ENSMUSG00000056367.9 |
| chr8 | 121828734 | 121830237 | 39.20496 | 131.87447 | 127.31225 | Promoter | 8 | 121796313 | 121829569 | 33257 | 2 | ENSMUST00 | 0 | Klhdc4 | ENSMUSG00000040263.11 |
| chr9 | 46282506 | 46283758 | 39.20496 | 131.87447 | 127.31225 | Promoter | 9 | 46283008 | 46294099 | 11092 | 1 | ENSMUST00 | 0 | Bud13 | ENSMUSG00000032077.4 |
| chr9 | 58582230 | 58583686 | 39.20496 | 131.87447 | 127.31225 | Promoter | 9 | 58582240 | 58651318 | 69079 | 1 | ENSMUST00 | 0 | Nptn | ENSMUSG00000032336.12 |
| chr2 | 60879546 | 60881897 | 39.78054 | 131.78781 | 127.22884 | Promoter | 2 | 60751953 | 60881438 | 129486 | 2 | ENSMUST00 | 0 | Rbms1 | ENSMUSG00000026970.11 |
| chr9 | 51962656 | 51963991 | 40.38649 | 131.74852 | 127.19042 | Promoter | 9 | 51943307 | 51963533 | 20227 | 2 | ENSMUST00 | 0 | Fdx1 | ENSMUSG00000032051.8 |
| chr1 | 166379287 | 166380486 | 34.34893 | 131.73137 | 127.17394 | Promoter | 1 | 166379317 | 166388928 | 9612 | 1 | ENSMUST00 | 0 | Tada1 | ENSMUSG00000026563.8 |
| chr10 | 54074002 | 54076827 | 34.34893 | 131.73137 | 127.17394 | Promoter | 10 | 53904788 | 54075796 | 171009 | 2 | ENSMUST00 | 0 | Man1a | ENSMUSG00000003746.10 |
| chr12 | 101957223 | 101958769 | 34.34893 | 131.73137 | 127.17394 | Promoter | 12 | 101918901 | 101958246 | 39346 | 2 | ENSMUST00 | 0 | Atxn3 | ENSMUSG00000021189.6 |
| chr15 | 76326793 | 76328235 | 34.34893 | 131.73137 | 127.17394 | Promoter | 15 | 76306487 | 76328015 | 21529 | 2 | ENSMUST00 | 0 | Oplah | ENSMUSG00000022562.8 |
| chr15 | 84680404 | 84684554 | 34.34893 | 131.73137 | 127.17394 | Promoter | 15 | 84681002 | 84703672 | 22671 | 1 | ENSMUST00 | 0 | Prr5 | ENSMUSG00000036106.10 |
| chr18 | 30266866 | 30268583 | 34.34893 | 131.73137 | 127.17394 | Distal Interge | 18 | 30272747 | 30348126 | 75380 | 1 | ENSMUST00 | -4164 | Pik3c3 | ENSMUSG00000033628.10 |
| chr3 | 133090816 | 133092747 | 34.34893 | 131.73137 | 127.17394 | Promoter | 3 | 133091840 | 133110988 | 19149 | 1 | ENSMUST00 | 0 | Ints12 | ENSMUSG00000028016.8 |
| chr5 | 136986283 | 136987996 | 34.34893 | 131.73137 | 127.17394 | Promoter | 5 | 136987019 | 136996648 | 9630 | 1 | ENSMUST00 | 0 | Plod3 | ENSMUSG00000004846.5 |
| chr6 | 47593688 | 47596080 | 34.34893 | 131.73137 | 127.17394 | Promoter | 6 | 47530274 | 47595030 | 64757 | 2 | ENSMUST00 | 0 | Ezh2 | ENSMUSG00000029687.11 |
| chr6 | 124711365 | 124713309 | 36.6511 | 131.68239 | 127.12846 | Promoter | 6 | 124712336 | 124713068 | 733 | 1 | ENSMUST00 | 0 | Phb2 | ENSMUSG00000004264.12 |
| chr9 | 65020724 | 65022246 | 36.6511 | 131.68239 | 127.12846 | Promoter | 9 | 64986985 | 65021717 | 34733 | 2 | ENSMUST00 | 0 | Ptplad1 | ENSMUSG00000033629.5 |
| chr1 | 134954938 | 134956311 | 37.85821 | 131.67731 | 127.12434 | Promoter | 1 | 134754658 | 134955940 | 201283 | 2 | ENSMUST00 | 0 | Ppp1r12b | ENSMUSG00000073557.6 |
| chr3 | 89772697 | 89775534 | 37.85821 | 131.67731 | 127.12434 | Promoter | 3 | 89773609 | 89783997 | 10389 | 1 | ENSMUST00 | 0 | Ube2q1 | ENSMUSG00000042572.8 |
| chr5 | 123523316 | 123524882 | 37.85821 | 131.67731 | 127.12434 | Promoter | 5 | 123509765 | 123524143 | 14379 | 2 | ENSMUST00 | 0 | Diablo | ENSMUSG00000029433.9 |
| chr9 | 104061947 | 104064222 | 37.85821 | 131.67731 | 127.12434 | Promoter | 9 | 104046599 | 104063134 | 16536 | 2 | ENSMUST00 | 0 | Uba5 | ENSMUSG00000032557.6 |
| chr15 | 100422204 | 100424245 | 39.70201 | 131.65079 | 127.10004 | Promoter | 15 | 100387898 | 100423055 | 35158 | 2 | ENSMUST00 | 0 | Slc11a2 | ENSMUSG00000023030.11 |
| chr14 | 121034923 | 121037682 | 40.93993 | 131.61429 | 127.06391 | Promoter | 14 | 121035200 | 121235089 | 199890 | 1 | ENSMUST00 | 0 | Farp1 | ENSMUSG00000025555.9 |
| chr16 | 96126477 | 96128557 | 40.93993 | 131.61429 | 127.06391 | Promoter | 16 | 96120618 | 96127729 | 7112 | 2 | ENSMUST00 | 0 | Hmgn1 | ENSMUSG00000040681.9 |
| chr5 | 121660016 | 121661566 | 39.62381 | 131.51428 | 126.96502 | Promoter | 5 | 121621026 | 121660479 | 39454 | 2 | ENSMUST00 | 0 | Acad10 | ENSMUSG00000029456.7 |
| chr10 | 41809166 | 41812188 | 38.97949 | 131.47847 | 126.9297 | Promoter | 10 | 41718841 | 41809613 | 90773 | 2 | ENSMUST00 | 0 | Cep57l1 | ENSMUSG00000019813.10 |
| chr11 | 6199324 | 6200851 | 38.97949 | 131.47847 | 126.9297 | Promoter | 11 | 6105691 | 6200415 | 94725 | 2 | ENSMUST00 | 0 | Nudcd3 | ENSMUSG00000053838.9 |
| chr11 | 61493320 | 61495096 | 38.97723 | 131.47449 | 126.92584 | Promoter | 11 | 61488812 | 61494266 | 5455 | 2 | ENSMUST00 | 0 | Mapk7 | ENSMUSG00000001034.12 |
| chr11 | 50208222 | 50211414 | 38.97723 | 131.47449 | 126.92584 | Promoter | 11 | 50199366 | 50210791 | 11426 | 2 | ENSMUST00 | 0 | Sqstm1 | ENSMUSG00000015837.10 |
| chr11 | 115611872 | 115613594 | 40.85493 | 131.46736 | 126.92584 | Promoter | 11 | 115607918 | 115612503 | 4586 | 2 | ENSMUST00 | 0 | Mif4gd | ENSMUSG00000020743.10 |
| chr12 | 31654187 | 31655804 | 38.97723 | 131.47449 | 126.92584 | Promoter | 12 | 31640055 | 31654826 | 14772 | 2 | ENSMUST00 | 0 | Dus4l | ENSMUSG00000020648.2 |
| chr14 | 34673302 | 34675036 | 38.97723 | 131.47449 | 126.92584 | Promoter | 14 | 34673928 | 34747983 | 74056 | 1 | ENSMUST00 | 0 | Wapal | ENSMUSG00000041408.11 |
| chr16 | 17721985 | 17723021 | 38.97723 | 131.47449 | 126.92584 | Promoter | 16 | 17651221 | 17722930 | 71710 | 2 | ENSMUST00 | 0 | Med15 | ENSMUSG00000012114.11 |
| chr16 | 5233175 | 5234300 | 38.97723 | 131.47449 | 126.92584 | Promoter | 16 | 5233621 | 5244903 | 11283 | 1 | ENSMUST00 | 0 | Alg1 | ENSMUSG00000039427.9 |
| chr18 | 3507865 | 3510534 | 38.97723 | 131.47449 | 126.92584 | Promoter | 18 | 3507957 | 3516404 | 8448 | 1 | ENSMUST00 | 0 | Bambi | ENSMUSG00000024232.1 |
| chr2 | 22894736 | 22896484 | 38.97723 | 131.47449 | 126.92584 | Promoter | 2 | 22895522 | 22912699 | 17178 | 1 | ENSMUST00 | 0 | Pdss1 | ENSMUSG00000026784.9 |
| chr3 | 19894342 | 19895800 | 38.97723 | 131.47449 | 126.92584 | Promoter | 3 | 19894886 | 19895962 | 1077 | 1 | ENSMUST00 | 0 | 4632415L05R | ENSMUSG00000048106.4 |
| chr4 | 126103908 | 126104979 | 38.97723 | 131.47449 | 126.92584 | Promoter | 4 | 126103957 | 126141028 | 37072 | 1 | ENSMUST00 | 0 | Stk40 | ENSMUSG00000042608.10 |
| chr5 | 21424867 | 21425843 | 38.97723 | 131.47449 | 126.92584 | Promoter | 5 | 21424958 | 21482124 | 57167 | 1 | ENSMUST00 | 0 | Fam185a | ENSMUSG00000047221.5 |
| chr6 | 83316883 | 83317776 | 38.97723 | 131.47449 | 126.92584 | Promoter | 6 | 83305691 | 83317606 | 11916 | 2 | ENSMUST00 | 0 | Mthfd2 | ENSMUSG00000005667.6 |
| chr6 | 143166144 | 143168273 | 38.97723 | 131.47449 | 126.92584 | Promoter | 6 | 143167230 | 143208542 | 41313 | 1 | ENSMUST00 | 0 | Etnk1 | ENSMUSG00000030275.4 |
| chr7 | 39449007 | 39450319 | 38.97723 | 131.47449 | 126.92584 | Promoter | 7 | 39449518 | 39477413 | 27896 | 1 | ENSMUST00 | 0 | Zfp939 | ENSMUSG00000030424.10 |
| chr9 | 50774680 | 50776188 | 38.97723 | 131.47449 | 126.92584 | Promoter | 9 | 50775019 | 50843528 | 68510 | 1 | ENSMUST00 | 0 | Alg9 | ENSMUSG00000032059.8 |
| chr9 | 83547863 | 83550185 | 38.97723 | 131.47449 | 126.92584 | Promoter | 9 | 83548327 | 83638801 | 90475 | 1 | ENSMUST00 | 0 | Sh3bgrl2 | ENSMUSG00000032261.11 |
| chr9 | 57644457 | 57645948 | 38.97723 | 131.47449 | 126.92584 | Promoter | 9 | 57626647 | 57645653 | 19007 | 2 | ENSMUST00 | 0 | Csk | ENSMUSG00000032312.6 |
| chr11 | 120989882 | 120991796 | 26.812 | 131.41335 | 126.87386 | Promoter | 11 | 120961749 | 120991092 | 29344 | 2 | ENSMUST00 | 0 | Csnk1d | ENSMUSG00000025162.12 |
| chr11 | 115764905 | 115766491 | 39.54591 | 131.37822 | 126.83877 | Promoter | 11 | 115765433 | 115799366 | 33934 | 1 | ENSMUST00 | 0 | 2310067B10 | ENSMUSG00000020747.12 |
| chr15 | 58133824 | 58135588 | 39.54591 | 131.3782 | 126.83877 | Promoter | 15 | 58094047 | 58135082 | 41036 | 2 | ENSMUST00 | 0 | Atad2 | ENSMUSG00000022360.7 |

| | | | | | | | | | | | | | | | |
|---|---|---|---|---|---|---|---|---|---|---|---|---|---|---|---|
| chr2 | 26388432 | 26390237 | 39.54591 | 131.37822 | 126.83877 | Promoter | 2 | 26389339 | 26397122 | 7784 | 1 | ENSMUST00 | 0 | Pmpca | ENSMUSG00000026926.9 |
| chr3 | 90508236 | 90509564 | 39.54591 | 131.3782 | 126.83877 | Promoter | 3 | 90498956 | 90509450 | 10495 | 2 | ENSMUST00 | 0 | Chtop | ENSMUSG00000001017.10 |
| chr4 | 46649573 | 46650981 | 39.54591 | 131.37822 | 126.83877 | Promoter | 4 | 46604390 | 46650209 | 45820 | 2 | ENSMUST00 | 0 | Tbc1d2 | ENSMUSG00000039813.9 |
| chr7 | 4657404 | 4659752 | 39.54591 | 131.3782 | 126.83877 | Promoter | 7 | 4631501 | 4658950 | 27450 | 2 | ENSMUST00 | 0 | Ppp6r1 | ENSMUSG00000052296.6 |
| chr1 | 130715318 | 130718138 | 36.42221 | 131.26616 | 126.73016 | Promoter | 1 | 130689183 | 130715734 | 26552 | 2 | ENSMUST00 | 0 | Pfkfb2 | ENSMUSG00000026409.9 |
| chr11 | 35833557 | 35834665 | 36.42221 | 131.26616 | 126.73016 | Promoter | 11 | 35808381 | 35834506 | 26126 | 2 | ENSMUST00 | 0 | Rars | ENSMUSG00000018848.4 |
| chr11 | 98148294 | 98150721 | 36.42221 | 131.26616 | 126.73016 | Promoter | 11 | 98082556 | 98149612 | 67057 | 2 | ENSMUST00 | 0 | Fbxl20 | ENSMUSG00000020883.13 |
| chr11 | 120624082 | 120625528 | 36.42221 | 131.26616 | 126.73016 | Promoter | 11 | 120618372 | 120624969 | 6598 | 2 | ENSMUST00 | 0 | Sirt7 | ENSMUSG00000025138.9 |
| chr11 | 52231184 | 52233477 | 36.42221 | 131.26616 | 126.73016 | Promoter | 11 | 52231995 | 52245094 | 13100 | 1 | ENSMUST00 | 0 | Skp1a | ENSMUSG00000036309.9 |
| chr12 | 52502712 | 52505830 | 36.42221 | 131.26616 | 126.73016 | Distal Interge | 12 | 52516077 | 52567852 | 51776 | 1 | ENSMUST00 | -10247 | Arhgap5 | ENSMUSG00000035133.8 |
| chr13 | 58400187 | 58403692 | 36.42221 | 131.26616 | 126.73016 | Promoter | 13 | 58391142 | 58403343 | 12202 | 2 | ENSMUST00 | 0 | Hnrnpk | ENSMUSG00000021546.12 |
| chr19 | 38480630 | 38483282 | 36.42221 | 131.26616 | 126.73016 | Promoter | 19 | 38481109 | 38524066 | 42958 | 1 | ENSMUST00 | 0 | Plce1 | ENSMUSG00000024998.12 |
| chr19 | 47067164 | 47068977 | 36.42221 | 131.26616 | 126.73016 | Promoter | 19 | 47067748 | 47083479 | 15732 | 1 | ENSMUST00 | 0 | Taf5 | ENSMUSG00000025049.5 |
| chr3 | 51275540 | 51278105 | 36.42221 | 131.26616 | 126.73016 | Promoter | 3 | 51255665 | 51277514 | 21850 | 2 | ENSMUST00 | 0 | Elf2 | ENSMUSG00000037174.13 |
| chr4 | 33309868 | 33311407 | 36.42221 | 131.26616 | 126.73016 | Promoter | 4 | 33310311 | 33502614 | 192304 | 1 | ENSMUST00 | 0 | Rngtt | ENSMUSG00000028274.12 |
| chr5 | 110769828 | 110770769 | 36.42221 | 131.26616 | 126.73016 | Promoter | 5 | 110664373 | 110770527 | 106155 | 2 | ENSMUST00 | 0 | Ep400 | ENSMUSG00000029505.12 |
| chr5 | 65491649 | 65493977 | 36.42221 | 131.26616 | 126.73016 | Promoter | 5 | 65446844 | 65493051 | 46208 | 2 | ENSMUST00 | 0 | Smim14 | ENSMUSG00000037822.7 |
| chr5 | 115156953 | 115159326 | 36.42221 | 131.26616 | 126.73016 | Promoter | 5 | 115142981 | 115158179 | 15199 | 2 | ENSMUST00 | 0 | Mlec | ENSMUSG00000048578.8 |
| chr6 | 39573285 | 39574769 | 36.42221 | 131.26616 | 126.73016 | Promoter | 6 | 39573873 | 39588769 | 14897 | 1 | ENSMUST00 | 0 | Adck2 | ENSMUSG00000046947.7 |
| chr9 | 57157469 | 57158754 | 36.42221 | 131.26616 | 126.73016 | Promoter | 9 | 57155041 | 57158299 | 3259 | 2 | ENSMUST00 | 0 | Commd4 | ENSMUSG00000032299.9 |
| chr9 | 67839796 | 67841312 | 36.42221 | 131.26616 | 126.73016 | Promoter | 9 | 67840396 | 67995634 | 155239 | 1 | ENSMUST00 | 0 | Vps13c | ENSMUSG00000035284.9 |
| chr2 | 30827493 | 30828386 | 35.79183 | 131.19528 | 126.66647 | Promoter | 2 | 30824470 | 30828265 | 3796 | 2 | ENSMUST00 | 0 | Asb6 | ENSMUSG00000039483.5 |
| chr6 | 8208860 | 8210167 | 35.79183 | 131.19528 | 126.66647 | Promoter | 6 | 8209222 | 8236274 | 27053 | 1 | ENSMUST00 | 0 | Mios | ENSMUSG00000042447.8 |
| chr7 | 34167607 | 34168979 | 35.79183 | 131.19528 | 126.66647 | Promoter | 7 | 34140689 | 34168575 | 27887 | 2 | ENSMUST00 | 0 | Uba2 | ENSMUSG00000052997.10 |
| chr12 | 21285266 | 21287173 | 38.15516 | 131.11487 | 126.58785 | Promoter | 12 | 21286297 | 21315056 | 28760 | 1 | ENSMUST00 | 0 | Cpsf3 | ENSMUSG00000054309.8 |
| chr13 | 17695034 | 17696297 | 38.15516 | 131.11487 | 126.58785 | Promoter | 13 | 17695413 | 17699105 | 3693 | 1 | ENSMUST00 | 0 | Mplkip | ENSMUSG00000012429.8 |
| chr14 | 61556004 | 61557669 | 38.15516 | 131.11487 | 126.58785 | Promoter | 14 | 61531993 | 61556886 | 24894 | 2 | ENSMUST00 | 0 | Spryd7 | ENSMUSG00000021930.9 |
| chr18 | 61725550 | 61726987 | 38.15516 | 131.11487 | 126.58785 | Promoter | 18 | 61712440 | 61726331 | 13892 | 2 | ENSMUST00 | 0 | Grpel2 | ENSMUSG00000024580.7 |
| chr9 | 70678949 | 70680437 | 38.15516 | 131.11487 | 126.58785 | Promoter | 9 | 70678997 | 70778382 | 99386 | 1 | ENSMUST00 | 0 | Adam10 | ENSMUSG00000054693.9 |
| chr14 | 73324628 | 73326699 | 39.39101 | 131.10754 | 126.58253 | Promoter | 14 | 73195502 | 73325822 | 130321 | 2 | ENSMUST00 | 0 | Rb1 | ENSMUSG00000022105.6 |
| chr17 | 46710999 | 46712571 | 39.39101 | 131.10754 | 126.58253 | Promoter | 17 | 46711463 | 46725541 | 14079 | 1 | ENSMUST00 | 0 | Pex6 | ENSMUSG00000002763.11 |
| chr2 | 127269433 | 127271238 | 36.91116 | 131.06699 | 126.54268 | Promoter | 2 | 127270218 | 127284584 | 14367 | 1 | ENSMUST00 | 0 | Stard7 | ENSMUSG00000027367.11 |
| chr2 | 122630073 | 122631776 | 36.91116 | 131.06699 | 126.54268 | Promoter | 2 | 122630625 | 122632704 | 2080 | 1 | ENSMUST00 | 0 | Spata5l1 | ENSMUSG00000074876.5 |
| chr7 | 109518375 | 109520221 | 36.91116 | 131.06699 | 126.54268 | Promoter | 7 | 109519147 | 109522367 | 3221 | 1 | ENSMUST00 | 0 | Rpl27a | ENSMUSG00000046364.9 |
| chr9 | 44919668 | 44921597 | 36.91116 | 131.06699 | 126.54268 | Promoter | 9 | 44913250 | 44920742 | 7493 | 2 | ENSMUST00 | 0 | Atp5l | ENSMUSG00000038717.7 |
| chr6 | 147475681 | 147477054 | 40.60202 | 131.02992 | 126.5071 | Promoter | 6 | 147475871 | 147632612 | 156742 | 1 | ENSMUST00 | 0 | Ccdc91 | ENSMUSG00000030301.12 |
| chr2 | 132263073 | 132264530 | 32.32209 | 130.99449 | 126.47215 | Promoter | 2 | 132263148 | 132296193 | 33046 | 1 | ENSMUST00 | 0 | Cds2 | ENSMUSG00000058793.9 |
| chr2 | 29964945 | 29967047 | 38.08445 | 130.98944 | 126.4684 | Promoter | 2 | 29965560 | 30031448 | 65889 | 1 | ENSMUST00 | 0 | Sptan1 | ENSMUSG00000057738.8 |
| chr9 | 88718912 | 88720593 | 27.90661 | 130.94769 | 126.42739 | Promoter | 9 | 88719629 | 88806264 | 86636 | 1 | ENSMUST00 | 0 | Gm10634 | ENSMUSG00000074149.4 |
| chr10 | 80167723 | 80169106 | 39.90052 | 130.90422 | 126.38404 | Promoter | 10 | 80167778 | 80171655 | 3878 | 1 | ENSMUST00 | 0 | Cirbp | ENSMUSG00000045193.8 |
| chr11 | 103131357 | 103132961 | 39.90052 | 130.90422 | 126.38404 | Promoter | 11 | 103132429 | 103138498 | 6070 | 1 | ENSMUST00 | 0 | Hexim2 | ENSMUSG00000043372.7 |
| chr13 | 95890627 | 95892671 | 39.90052 | 130.90422 | 126.38404 | Promoter | 13 | 95627177 | 95891922 | 264746 | 2 | ENSMUST00 | 0 | Iqgap2 | ENSMUSG00000021676.9 |
| chr8 | 83715135 | 83716233 | 39.90052 | 130.90422 | 126.38404 | Promoter | 8 | 83715177 | 83723348 | 8172 | 1 | ENSMUST00 | 0 | Ddx39 | ENSMUSG00000005481.12 |
| chr3 | 53462704 | 53465185 | 39.23734 | 130.83873 | 126.32064 | Promoter | 3 | 53448588 | 53463291 | 14704 | 2 | ENSMUST00 | 0 | Nhlrc3 | ENSMUSG00000042997.8 |
| chr18 | 32163027 | 32164275 | 38.60941 | 130.82727 | 126.30969 | Promoter | 18 | 32163089 | 32236751 | 73663 | 1 | ENSMUST00 | 0 | Map3k2 | ENSMUSG00000024383.8 |
| chr5 | 33935251 | 33936336 | 34.98813 | 130.7849 | 126.26756 | Promoter | 5 | 33919125 | 33936250 | 17126 | 2 | ENSMUST00 | 0 | Nelfa | ENSMUSG00000029111.6 |
| chr5 | 114813303 | 114814331 | 34.98813 | 130.7849 | 126.26756 | Promoter | 5 | 114808196 | 114813976 | 5781 | 2 | ENSMUST00 | 0 | 1500011B03I | ENSMUSG00000072694.7 |
| chr1 | 180725507 | 180727314 | 39.82066 | 130.7653 | 126.24818 | Promoter | 1 | 180726043 | 180754203 | 28161 | 1 | ENSMUST00 | 0 | Acbd3 | ENSMUSG00000026499.4 |
| chr10 | 95562613 | 95564657 | 39.82066 | 130.7653 | 126.24818 | Promoter | 10 | 95547007 | 95564167 | 17161 | 2 | ENSMUST00 | 0 | Nudt4 | ENSMUSG00000020029.5 |
| chr6 | 12745333 | 12750108 | 39.82066 | 130.7653 | 126.24818 | Promoter | 6 | 12311610 | 12749253 | 437644 | 2 | ENSMUST00 | 0 | Thsd7a | ENSMUSG00000032625.9 |
| chr9 | 111271774 | 111273023 | 39.74112 | 130.62688 | 126.11213 | Promoter | 9 | 111271929 | 111279089 | 7161 | 1 | ENSMUST00 | 0 | Epm2aip1 | ENSMUSG00000046785.7 |
| chr8 | 111742412 | 111744224 | 40.35223 | 130.59746 | 126.08324 | Promoter | 8 | 111710485 | 111743849 | 33365 | 2 | ENSMUST00 | 0 | Bcar1 | ENSMUSG00000031955.9 |
| chr10 | 80576172 | 80578703 | 38.46334 | 130.56975 | 126.05653 | Promoter | 10 | 80567124 | 80577321 | 10198 | 2 | ENSMUST00 | 0 | Klf16 | ENSMUSG00000035397.8 |
| chr11 | 104231437 | 104234756 | 38.46334 | 130.56975 | 126.05653 | Promoter | 11 | 104231479 | 104295317 | 63839 | 1 | ENSMUST00 | 0 | Mapt | ENSMUSG00000018411.12 |
| chr11 | 17210870 | 17212623 | 38.46334 | 130.56975 | 126.05653 | Promoter | 11 | 17203198 | 17211568 | 8371 | 2 | ENSMUST00 | 0 | Pno1 | ENSMUSG00000020116.7 |
| chr11 | 20248487 | 20250058 | 38.46334 | 130.56975 | 126.05653 | Promoter | 11 | 20227037 | 20249418 | 22382 | 2 | ENSMUST00 | 0 | Cep68 | ENSMUSG00000044066.9 |
| chr2 | 18996496 | 18998878 | 38.46334 | 130.56975 | 126.05653 | Promoter | 2 | 18998343 | 18999801 | 1459 | 1 | ENSMUST00 | 0 | 4930426L09I | ENSMUSG00000026736.1 |
| chr2 | 163418188 | 163419609 | 38.46334 | 130.56975 | 126.05653 | Promoter | 2 | 163405822 | 163419528 | 13707 | 2 | ENSMUST00 | 0 | Oser1 | ENSMUSG00000035399.7 |
| chr2 | 29249202 | 29253779 | 38.46334 | 130.56975 | 126.05653 | Promoter | 2 | 29203822 | 29253005 | 49184 | 2 | ENSMUST00 | 0 | Ntng2 | ENSMUSG00000035513.14 |
| chr4 | 41722397 | 41723573 | 38.46334 | 130.56975 | 126.05653 | Promoter | 4 | 41714798 | 41723163 | 8366 | 2 | ENSMUST00 | 0 | Dctn3 | ENSMUSG00000028447.6 |
| chr5 | 115271649 | 115273337 | 38.46334 | 130.56975 | 126.05653 | Promoter | 5 | 115241412 | 115272898 | 31487 | 2 | ENSMUST00 | 0 | Rnf10 | ENSMUSG00000041740.11 |
| chr7 | 130935422 | 130939180 | 38.46334 | 130.56975 | 126.05653 | Promoter | 7 | 130936111 | 130979934 | 43824 | 1 | ENSMUST00 | 0 | Htra1 | ENSMUSG00000006205.8 |
| chr10 | 115383165 | 115387078 | 36.02846 | 130.54782 | 126.03754 | Promoter | 10 | 115369966 | 115384435 | 14470 | 2 | ENSMUST00 | 0 | Thap2 | ENSMUSG00000020137.4 |
| chr13 | 23530050 | 23532457 | 25.63199 | 130.5271 | 126.01785 | Promoter | 13 | 23531050 | 23531522 | 473 | 1 | ENSMUST00 | 0 | Hist1h4h | ENSMUSG00000060981.6 |
| chr2 | 130177981 | 130180043 | 37.18052 | 130.46529 | 125.95659 | Promoter | 2 | 130171414 | 130179303 | 7890 | 2 | ENSMUST00 | 0 | Snrpb | ENSMUSG00000027404.10 |
| chr9 | 121710227 | 121711199 | 37.18052 | 130.46529 | 125.95659 | Promoter | 9 | 121710389 | 121712921 | 2533 | 1 | ENSMUST00 | 0 | Deb1 | ENSMUSG00000032526.10 |
| chr4 | 115737296 | 115738872 | 40.26965 | 130.45439 | 125.94711 | Promoter | 4 | 115737744 | 115774435 | 36692 | 1 | ENSMUST00 | 0 | Efcab14 | ENSMUSG00000034210.8 |
| chr5 | 148928039 | 148929495 | 40.26965 | 130.45439 | 125.94711 | Promoter | 5 | 148874332 | 148928647 | 54316 | 2 | ENSMUST00 | 0 | Katnal1 | ENSMUSG00000041298.10 |
| chr1 | 153899280 | 153901286 | 40.89836 | 130.45236 | 125.94646 | Promoter | 1 | 153899944 | 153909723 | 9780 | 1 | ENSMUST00 | 0 | Glul | ENSMUSG00000026473.10 |
| chr10 | 79926277 | 79928478 | 40.89836 | 130.45236 | 125.94646 | Promoter | 10 | 79927043 | 79955018 | 27976 | 1 | ENSMUST00 | 0 | Arid3a | ENSMUSG00000019564.7 |
| chr12 | 72760286 | 72762586 | 40.89836 | 130.45236 | 125.94646 | Promoter | 12 | 72761211 | 72794940 | 33730 | 1 | ENSMUST00 | 0 | Ppm1a | ENSMUSG00000021096.10 |
| chr15 | 82146565 | 82148559 | 40.89836 | 130.45236 | 125.94646 | Promoter | 15 | 82147266 | 82205376 | 58111 | 1 | ENSMUST00 | 0 | Srebf2 | ENSMUSG00000022463.7 |
| chr17 | 87106372 | 87108483 | 40.89836 | 130.45236 | 125.94646 | Promoter | 17 | 87107679 | 87137583 | 29905 | 1 | ENSMUST00 | 0 | Socs5 | ENSMUSG00000037104.8 |
| chr3 | 108415035 | 108416402 | 40.89836 | 130.45236 | 125.94646 | Promoter | 3 | 108390851 | 108415552 | 24702 | 2 | ENSMUST00 | 0 | Celsr2 | ENSMUSG00000068740.8 |
| chr4 | 128993195 | 128994389 | 40.89836 | 130.45236 | 125.94646 | Promoter | 4 | 128993224 | 129008401 | 15178 | 1 | ENSMUST00 | 0 | Ak2 | ENSMUSG00000028792.9 |
| chr4 | 139091610 | 139093306 | 40.89836 | 130.45236 | 125.94646 | Promoter | 4 | 139082284 | 139092993 | 10710 | 2 | ENSMUST00 | 0 | Nbl1 | ENSMUSG00000041120.6 |
| chr9 | 120110272 | 120111312 | 40.89836 | 130.45236 | 125.94646 | Promoter | 9 | 120110374 | 120124504 | 14131 | 1 | ENSMUST00 | 0 | Slc25a38 | ENSMUSG00000032519.6 |
| chr5 | 110176136 | 110178000 | 38.39072 | 130.44165 | 125.93981 | Promoter | 5 | 110176701 | 110226470 | 49770 | 1 | ENSMUST00 | 0 | Golga3 | ENSMUSG00000029502.9 |
| chr13 | 14063753 | 14065567 | 39.00905 | 130.43898 | 125.93717 | Promoter | 13 | 14063784 | 14169960 | 106177 | 1 | ENSMUST00 | 0 | Arid4b | ENSMUSG00000039219.12 |
| chr11 | 43433369 | 43434819 | 39.58299 | 130.35152 | 125.85003 | Promoter | 11 | 43433744 | 43438276 | 4533 | 1 | ENSMUST00 | 0 | Slu7 | ENSMUSG00000020409.5 |
| chr13 | 59585249 | 59586287 | 40.18741 | 130.31186 | 125.8112 | Promoter | 13 | 59585259 | 59599790 | 14532 | 1 | ENSMUST00 | 0 | Naa35 | ENSMUSG00000021555.9 |
| chr15 | 39005741 | 39007041 | 40.18741 | 130.31186 | 125.8112 | Promoter | 15 | 39006280 | 39038186 | 31907 | 1 | ENSMUST00 | 0 | Fzd6 | ENSMUSG00000022297.9 |
| chr16 | 44745467 | 44746436 | 39.5044 | 130.21455 | 125.71494 | Promoter | 16 | 44736768 | 44746363 | 9596 | 2 | ENSMUST00 | 0 | Gtpbp8 | ENSMUSG00000022668.5 |
| chr5 | 76183382 | 76185050 | 40.73933 | 130.17783 | 125.67887 | Promoter | 5 | 76183880 | 76209245 | 25366 | 1 | ENSMUST00 | 0 | Tmem165 | ENSMUSG00000029234.8 |
| chr13 | 3609837 | 3611659 | 38.85833 | 130.17477 | 125.676 | Promoter | 13 | 3566036 | 3611108 | 45073 | 2 | ENSMUST00 | 0 | Fam208b | ENSMUSG00000033799.9 |
| chr10 | 77258691 | 77260213 | 40.1055 | 130.16986 | 125.67132 | Promoter | 10 | 77259300 | 77269586 | 10287 | 1 | ENSMUST00 | 0 | Pofut2 | ENSMUSG00000020260.8 |
| chr7 | 118704843 | 118706115 | 38.78341 | 130.04332 | 125.54643 | Promoter | 7 | 118688545 | 118705778 | 17234 | 2 | ENSMUST00 | 0 | Gde1 | ENSMUSG00000033917.9 |
| chr14 | 30548537 | 30550051 | 40.65421 | 130.03084 | 125.53634 | Promoter | 14 | 30549131 | 30574722 | 25592 | 1 | ENSMUST00 | 0 | Tkt | ENSMUSG00000021957.6 |
| chr11 | 98201924 | 98205269 | 34.00884 | 130.01486 | 125.52227 | Promoter | 11 | 98203059 | 98251564 | 48506 | 1 | ENSMUST00 | 0 | Cdk12 | ENSMUSG00000003119.9 |
| chr12 | 24830727 | 24832387 | 34.00884 | 130.01486 | 125.52227 | Promoter | 12 | 24831599 | 24960301 | 128703 | 1 | ENSMUST00 | 0 | Mboat2 | ENSMUSG00000020646.11 |
| chr13 | 56134430 | 56136012 | 34.00884 | 130.01486 | 125.52227 | Promoter | 13 | 56073619 | 56135631 | 62013 | 2 | ENSMUST00 | 0 | H2afy | ENSMUSG00000015937.9 |
| chr16 | 17451315 | 17453886 | 34.00884 | 130.01486 | 125.52227 | Promoter | 16 | 17451987 | 17486255 | 34269 | 1 | ENSMUST00 | 0 | Crkl | ENSMUSG00000006134.3 |
| chr16 | 37867771 | 37870669 | 34.00884 | 130.01486 | 125.52227 | Promoter | 16 | 37868389 | 37888858 | 20470 | 1 | ENSMUST00 | 0 | Lrrc58 | ENSMUSG00000034158.9 |
| chr17 | 31511650 | 31512835 | 34.00884 | 130.01486 | 125.52227 | Promoter | 17 | 31494323 | 31512268 | 17946 | 2 | ENSMUST00 | 0 | Wdr4 | ENSMUSG00000024037.7 |
| chr17 | 81064431 | 81065310 | 34.00884 | 130.01486 | 125.52227 | Promoter | 17 | 81026327 | 81065085 | 38759 | 2 | ENSMUST00 | 0 | Thumpd2 | ENSMUSG00000024246.8 |
| chr18 | 84588660 | 84590582 | 34.00884 | 130.01486 | 125.52227 | Promoter | 18 | 84207704 | 84589504 | 381801 | 2 | ENSMUST00 | 0 | Zfp407 | ENSMUSG00000048410.11 |
| chr19 | 10688136 | 10689836 | 34.00884 | 130.01486 | 125.52227 | Promoter | 19 | 10688815 | 10714419 | 25605 | 1 | ENSMUST00 | 0 | Vps37c | ENSMUSG00000048832.8 |
| chr2 | 90745429 | 90746834 | 34.00884 | 130.01486 | 125.52227 | Promoter | 2 | 90745604 | 90776475 | 30872 | 1 | ENSMUST00 | 0 | Fnbp4 | ENSMUSG00000008200.9 |
| chr3 | 103913684 | 103916036 | 34.00884 | 130.01486 | 125.52227 | Promoter | 3 | 103914120 | 103966613 | 52494 | 1 | ENSMUST00 | 0 | Rsbn1 | ENSMUSG00000044098.9 |
| chr5 | 31219428 | 31221550 | 34.00884 | 130.01486 | 125.52227 | Promoter | 5 | 31202668 | 31220545 | 17878 | 2 | ENSMUST00 | 0 | Ppm1g | ENSMUSG00000029147.7 |
| chr5 | 123075871 | 123077894 | 34.00884 | 130.01486 | 125.52227 | Promoter | 5 | 123076275 | 123117749 | 41475 | 1 | ENSMUST00 | 0 | Tmem120b | ENSMUSG00000054434.9 |
| chr6 | 85068481 | 85069756 | 34.00884 | 130.01486 | 125.52227 | Promoter | 6 | 84618487 | 85069513 | 451027 | 2 | ENSMUST00 | 0 | Exoc6b | ENSMUSG00000033769.11 |
| chr7 | 19603486 | 19604804 | 34.00884 | 130.01486 | 125.52227 | Promoter | 7 | 19581044 | 19604468 | 23425 | 2 | ENSMUST00 | 0 | Clasrp | ENSMUSG00000061028.6 |
| chr5 | 30869281 | 30870597 | 39.34814 | 129.94209 | 125.45322 | Promoter | 5 | 30869579 | 30879191 | 9613 | 1 | ENSMUST00 | 0 | Tmem214 | ENSMUSG00000038828.8 |
| chr5 | 134550699 | 134552810 | 36.27325 | 129.91212 | 125.42401 | Promoter | 5 | 134489386 | 134552434 | 63049 | 2 | ENSMUST00 | 0 | Clip2 | ENSMUSG00000063146.6 |
| chr6 | 128437487 | 128439261 | 36.27325 | 129.91212 | 125.42401 | Promoter | 6 | 128429735 | 128438677 | 8943 | 2 | ENSMUST00 | 0 | Fkbp4 | ENSMUSG00000030357.5 |
| chr9 | 57520719 | 57522380 | 36.27325 | 129.91212 | 125.42401 | Promoter | 9 | 57521232 | 57532424 | 11193 | 1 | ENSMUST00 | 0 | Cox5a | ENSMUSG00000000088.6 |
| chr10 | 14704375 | 14705862 | 37.4597 | 129.87831 | 125.39164 | Promoter | 10 | 14705591 | 14766771 | 61181 | 1 | ENSMUST00 | 0 | Nmbr | ENSMUSG00000019865.4 |
| chr11 | 87085908 | 87087315 | 37.4597 | 129.87831 | 125.39164 | Promoter | 11 | 87077732 | 87086774 | 9043 | 2 | ENSMUST00 | 0 | Smg8 | ENSMUSG00000020495.8 |
| chr11 | 52396013 | 52397309 | 37.4597 | 129.87831 | 125.39164 | Promoter | 11 | 52396428 | 52404242 | 7815 | 1 | ENSMUST00 | 0 | 9530068E07I | ENSMUSG00000036275.8 |
| chr12 | 8497962 | 8501028 | 37.4597 | 129.87831 | 125.39164 | Promoter | 12 | 8497763 | 8499985 | 2223 | 2 | ENSMUST00 | 0 | Rhob | ENSMUSG00000054364.4 |
| chr3 | 104637906 | 104639646 | 37.4597 | 129.87831 | 125.39164 | Promoter | 3 | 104638668 | 104658462 | 19795 | 1 | ENSMUST00 | 0 | Slc16a1 | ENSMUSG00000032902.1 |
| chr5 | 150522183 | 150523236 | 37.4597 | 129.87831 | 125.39164 | Promoter | 5 | 150522630 | 150569746 | 47117 | 1 | ENSMUST00 | 0 | Brca2 | ENSMUSG00000041147.7 |
| chr7 | 126199284 | 126201146 | 37.4597 | 129.87831 | 125.39164 | Promoter | 7 | 126101715 | 126200501 | 98787 | 2 | ENSMUST00 | 0 | Xpo6 | ENSMUSG00000000131.10 |
| chr18 | 34930321 | 34933202 | 39.27047 | 129.80659 | 125.32217 | Promoter | 18 | 34902785 | 34932007 | 29223 | 2 | ENSMUST00 | 0 | Etf1 | ENSMUSG00000024360.7 |
| chr9 | 102716871 | 102721558 | 32.23346 | 129.76328 | 125.27943 | Promoter | 9 | 102717289 | 102720733 | 3445 | 1 | ENSMUST00 | 0 | Amotl2 | ENSMUSG00000032531.10 |
| chr17 | 35860920 | 35861831 | 39.86176 | 129.74701 | 125.26318 | Promoter | 17 | 35861318 | 35865402 | 4085 | 1 | ENSMUST00 | 0 | Nrm | ENSMUSG00000059791.9 |
| chr19 | 41980347 | 41982830 | 34.43156 | 129.74289 | 125.25954 | Promoter | 19 | 41941086 | 41981114 | 40029 | 2 | ENSMUST00 | 0 | Mms19 | ENSMUSG00000025159.3 |
| chr5 | 76529226 | 76530180 | 39.19312 | 129.67159 | 125.18892 | Promoter | 5 | 76529311 | 76570294 | 40984 | 1 | ENSMUST00 | 0 | Exoc1 | ENSMUSG00000036435.8 |
| chr1 | 106171618 | 106174296 | 38.55812 | 129.64772 | 125.16568 | Promoter | 1 | 106171752 | 106394250 | 222499 | 1 | ENSMUST00 | 0 | Phlpp1 | ENSMUSG00000044340.7 |
| chr12 | 104750854 | 104752922 | 38.55812 | 129.64772 | 125.16568 | Promoter | 12 | 104687742 | 104751952 | 64211 | 2 | ENSMUST00 | 0 | Dicer1 | ENSMUSG00000041415.9 |
| chr14 | 48119853 | 48122326 | 38.55812 | 129.64772 | 125.16568 | Promoter | 14 | 48120869 | 48260883 | 140015 | 1 | ENSMUST00 | 0 | Peli2 | ENSMUSG00000021846.8 |
| chr18 | 67448219 | 67449289 | 38.55812 | 129.64772 | 125.16568 | Promoter | 18 | 67404767 | 67449166 | 44400 | 2 | ENSMUST00 | 0 | Afg3l2 | ENSMUSG00000024527.9 |
| chr2 | 48948386 | 48950830 | 38.55812 | 129.64772 | 125.16568 | Promoter | 2 | 48902824 | 48949276 | 46453 | 2 | ENSMUST00 | 0 | Orc4 | ENSMUSG00000026761.7 |
| chr2 | 29059962 | 29061201 | 38.55812 | 129.64772 | 125.16568 | Promoter | 2 | 29060262 | 29087656 | 27395 | 1 | ENSMUST00 | 0 | Ttf1 | ENSMUSG00000026803.12 |
| chr3 | 110249623 | 110251413 | 38.55812 | 129.64772 | 125.16568 | Promoter | 3 | 110246109 | 110250998 | 4890 | 2 | ENSMUST00 | 0 | Prmt6 | ENSMUSG00000049300.11 |
| chr6 | 23838567 | 23840477 | 38.55812 | 129.64772 | 125.16568 | Promoter | 6 | 23262773 | 23839137 | 576365 | 2 | ENSMUST00 | 0 | Cadps2 | ENSMUSG00000017978.13 |
| chr6 | 119902226 | 119903655 | 38.55812 | 129.64772 | 125.16568 | Promoter | 6 | 119902698 | 119915335 | 12638 | 1 | ENSMUST00 | 0 | Rad52 | ENSMUSG00000030166.9 |

| | | | | | | | | | | | | | | | |
|---|---|---|---|---|---|---|---|---|---|---|---|---|---|---|---|
| chr7 | 67950399 | 67954090 | 38.55812 | 129.64772 | 125.16568 | Promoter | 7 | 67952859 | 68226780 | 273922 | 1 | ENSMUST00 | 0 | Igf1r | ENSMUSG00000005533.9 |
| chr7 | 28810797 | 28812960 | 38.55812 | 129.64772 | 125.16568 | Promoter | 7 | 28810890 | 28822266 | 11377 | 1 | ENSMUST00 | 0 | Hnrnpl | ENSMUSG00000015165.11 |
| chr7 | 128297368 | 128298548 | 38.55812 | 129.64772 | 125.16568 | Promoter | 7 | 128271379 | 128298170 | 26792 | 2 | ENSMUST00 | 0 | BC017158 | ENSMUSG00000030780.10 |
| chr8 | 94385758 | 94388107 | 38.55812 | 129.64772 | 125.16568 | Promoter | 8 | 94386438 | 94395377 | 8940 | 1 | ENSMUST00 | 0 | Herpud1 | ENSMUSG00000031770.10 |
| chr9 | 62340409 | 62345269 | 38.55812 | 129.64772 | 125.16568 | Promoter | 9 | 62341293 | 62377701 | 36409 | 1 | ENSMUST00 | 0 | Anp32a | ENSMUSG00000032249.9 |
| chr9 | 53247237 | 53248682 | 38.55812 | 129.64772 | 125.16568 | Promoter | 9 | 53098454 | 53248112 | 149659 | 2 | ENSMUST00 | 0 | Ddx10 | ENSMUSG00000053289.7 |
| chr10 | 111972697 | 111973969 | 39.78118 | 129.60712 | 125.13316 | Promoter | 10 | 111972757 | 111976515 | 3759 | 1 | ENSMUST00 | 0 | Krr1 | ENSMUSG00000063334.11 |
| chr11 | 82780387 | 82782393 | 39.11606 | 129.53703 | 125.06399 | Promoter | 11 | 82781108 | 82802862 | 21755 | 1 | ENSMUST00 | 0 | Lig3 | ENSMUSG00000020697.11 |
| chr12 | 106009676 | 106011099 | 39.11606 | 129.53703 | 125.06399 | Promoter | 12 | 106010263 | 106077410 | 67148 | 1 | ENSMUST00 | 0 | Vrk1 | ENSMUSG00000021115.9 |
| chr14 | 12283936 | 12285129 | 39.11606 | 129.53703 | 125.06399 | Promoter | 14 | 12284209 | 12303231 | 19023 | 1 | ENSMUST00 | 0 | 3830406C13I | ENSMUSG00000033111.10 |
| chr2 | 156007603 | 156008754 | 39.11606 | 129.53703 | 125.06399 | Promoter | 2 | 156008045 | 156018279 | 10235 | 1 | ENSMUST00 | 0 | Ergic3 | ENSMUSG00000005881.8 |
| chr8 | 71591371 | 71593108 | 39.11606 | 129.53703 | 125.06399 | Promoter | 8 | 71592176 | 71596269 | 4094 | 1 | ENSMUST00 | 0 | Pgls | ENSMUSG00000031807.5 |
| chr8 | 105170151 | 105172219 | 39.11606 | 129.53703 | 125.06399 | Promoter | 8 | 105170674 | 105217989 | 47316 | 1 | ENSMUST00 | 0 | Cbfb | ENSMUSG00000031885.8 |
| chr1 | 121326874 | 121328605 | 36.04672 | 129.50026 | 125.02953 | Promoter | 1 | 121304353 | 121328024 | 23672 | 2 | ENSMUST00 | 0 | Insig2 | ENSMUSG00000003721.9 |
| chr1 | 138617343 | 138620436 | 36.04672 | 129.50026 | 125.02953 | Promoter | 1 | 138483837 | 138619749 | 135913 | 2 | ENSMUST00 | 0 | Nek7 | ENSMUSG00000026393.5 |
| chr10 | 62979812 | 62981157 | 36.04672 | 129.50026 | 125.02953 | Promoter | 10 | 62980223 | 62990293 | 10071 | 1 | ENSMUST00 | 0 | Rufy2 | ENSMUSG00000020070.13 |
| chr10 | 53376609 | 53380307 | 36.04672 | 129.50026 | 125.02953 | Promoter | 10 | 53278081 | 53379851 | 101771 | 2 | ENSMUST00 | 0 | Cep85l | ENSMUSG00000038594.8 |
| chr11 | 70409194 | 70410542 | 36.04672 | 129.50026 | 125.02953 | Promoter | 11 | 70392883 | 70410028 | 17146 | 2 | ENSMUST00 | 0 | Pelp1 | ENSMUSG00000018921.6 |
| chr12 | 83920493 | 83921919 | 36.04672 | 129.50026 | 125.02953 | Promoter | 12 | 83794071 | 83921844 | 127774 | 2 | ENSMUST00 | 0 | Numb | ENSMUSG00000021224.9 |
| chr14 | 62759434 | 62761182 | 36.04672 | 129.50026 | 125.02953 | Promoter | 14 | 62676330 | 62761112 | 84783 | 2 | ENSMUST00 | 0 | Ints6 | ENSMUSG00000035161.6 |
| chr16 | 91043343 | 91045245 | 36.04672 | 129.50026 | 125.02953 | Promoter | 16 | 91014037 | 91044543 | 30507 | 2 | ENSMUST00 | 0 | Paxbp1 | ENSMUSG00000022974.7 |
| chr18 | 14681165 | 14683465 | 36.04672 | 129.50026 | 125.02953 | Promoter | 18 | 14624198 | 14682766 | 58569 | 2 | ENSMUST00 | 0 | Ss18 | ENSMUSG00000037013.10 |
| chr2 | 79635281 | 79639748 | 36.04672 | 129.50026 | 125.02953 | Promoter | 2 | 79635352 | 79672966 | 37615 | 1 | ENSMUST00 | 0 | Ssfa2 | ENSMUSG00000027007.11 |
| chr2 | 177956754 | 177957411 | 36.04672 | 129.50026 | 125.02953 | Promoter | 2 | 177935993 | 177957288 | 21296 | 2 | ENSMUST00 | 0 | Gm14326 | ENSMUSG00000078862.5 |
| chr4 | 127021239 | 127023612 | 36.04672 | 129.50026 | 125.02953 | Promoter | 4 | 127021324 | 127031236 | 9913 | 1 | ENSMUST00 | 0 | Sfpq | ENSMUSG00000028820.8 |
| chr4 | 93334516 | 93335970 | 36.04672 | 129.50026 | 125.02953 | Promoter | 4 | 93334138 | 93335511 | 1374 | 2 | ENSMUST00 | 0 | Tusc1 | ENSMUSG00000054000.5 |
| chr5 | 100820155 | 100821412 | 36.04672 | 129.50026 | 125.02953 | Promoter | 5 | 100805192 | 100820940 | 15749 | 2 | ENSMUST00 | 0 | Fam175a | ENSMUSG00000035234.13 |
| chr5 | 124860882 | 124863843 | 36.04672 | 129.50026 | 125.02953 | Promoter | 5 | 124834418 | 124861885 | 27468 | 2 | ENSMUST00 | 0 | Ccdc92 | ENSMUSG00000037979.8 |
| chr7 | 118490421 | 118492334 | 36.04672 | 129.50026 | 125.02953 | Promoter | 7 | 118485114 | 118491975 | 6862 | 2 | ENSMUST00 | 0 | Itpripl2 | ENSMUSG00000095115.1 |
| chr9 | 44482529 | 44484376 | 36.04672 | 129.50026 | 125.02953 | Distal Interg | 9 | 44478778 | 44482180 | 3403 | 1 | ENSMUST00 | 3752 | C030014I23R | ENSMUSG00000060380.2 |
| chr11 | 60931193 | 60933311 | 39.70091 | 129.46771 | 125.00398 | Promoter | 11 | 60932033 | 60952811 | 20779 | 1 | ENSMUST00 | 0 | Map2k3 | ENSMUSG00000018932.4 |
| chr14 | 8172240 | 8173816 | 39.70091 | 129.46771 | 125.00398 | Promoter | 14 | 8165991 | 8172992 | 7002 | 2 | ENSMUST00 | 0 | Pdhb | ENSMUSG00000021748.8 |
| chr5 | 107868027 | 107869684 | 40.31728 | 129.44856 | 124.98527 | Promoter | 5 | 107788287 | 107869168 | 80882 | 2 | ENSMUST00 | 0 | Evi5 | ENSMUSG00000011831.11 |
| chr7 | 29517714 | 29519538 | 35.42661 | 129.44405 | 124.98113 | Promoter | 7 | 29400762 | 29518612 | 117851 | 2 | ENSMUST00 | 0 | Sipa1l3 | ENSMUSG00000030583.11 |
| chr7 | 123123418 | 123125182 | 35.42661 | 129.44405 | 124.98113 | Promoter | 7 | 123123885 | 123195288 | 71404 | 1 | ENSMUST00 | 0 | Tnrc6a | ENSMUSG00000052707.7 |
| chr7 | 116333174 | 116334905 | 39.03931 | 129.40295 | 124.94073 | Promoter | 7 | 116333171 | 116333257 | 87 | 2 | ENSMUST00 | 0 | Snord14a | ENSMUSG00000064741.1 |
| chr13 | 55692609 | 55694450 | 39.62098 | 129.32881 | 124.86707 | Promoter | 13 | 55693124 | 55703499 | 10376 | 1 | ENSMUST00 | 0 | B230219D22 | ENSMUSG00000045767.8 |
| chr2 | 156861938 | 156864135 | 39.62098 | 129.32881 | 124.86707 | Promoter | 2 | 156863128 | 156873563 | 10436 | 1 | ENSMUST00 | 0 | 1110008F13I | ENSMUSG00000027637.3 |
| chr11 | 100937693 | 100939519 | 32.0113 | 129.32326 | 124.86235 | Promoter | 11 | 100885098 | 100939404 | 54307 | 2 | ENSMUST00 | 0 | Stat3 | ENSMUSG00000004040.11 |
| chr11 | 80428159 | 80429904 | 32.0113 | 129.32326 | 124.86235 | Promoter | 11 | 80428615 | 80434538 | 5924 | 1 | ENSMUST00 | 0 | Psmd11 | ENSMUSG00000017428.11 |
| chr11 | 97798972 | 97802073 | 32.0113 | 129.32326 | 124.86235 | Promoter | 11 | 97799000 | 97806839 | 7840 | 1 | ENSMUST00 | 0 | Lasp1 | ENSMUSG00000038366.10 |
| chr14 | 75753380 | 75754982 | 32.0113 | 129.32326 | 124.86235 | Promoter | 14 | 75702351 | 75754493 | 52143 | 2 | ENSMUST00 | 0 | Cog3 | ENSMUSG00000034893.7 |
| chr18 | 35770990 | 35772691 | 32.0113 | 129.32326 | 124.86235 | Promoter | 18 | 35771559 | 35807172 | 35614 | 1 | ENSMUST00 | 0 | Ube2d2a | ENSMUSG00000091896.2 |
| chr2 | 121449316 | 121450775 | 32.0113 | 129.32326 | 124.86235 | Promoter | 2 | 121449368 | 121453005 | 3638 | 1 | ENSMUST00 | 0 | Serf2 | ENSMUSG00000074884.7 |
| chr4 | 107177443 | 107180020 | 32.0113 | 129.32326 | 124.86235 | Promoter | 4 | 107178399 | 107200996 | 22598 | 1 | ENSMUST00 | 0 | Tmem59 | ENSMUSG00000028618.6 |
| chr5 | 124424344 | 124426339 | 32.0113 | 129.32326 | 124.86235 | Promoter | 5 | 124368702 | 124425914 | 57213 | 2 | ENSMUST00 | 0 | Sbno1 | ENSMUSG00000038095.10 |
| chr5 | 114772533 | 114775921 | 32.0113 | 129.32326 | 124.86235 | Promoter | 5 | 114774677 | 114805822 | 31146 | 1 | ENSMUST00 | 0 | Ankrd13a | ENSMUSG00000041870.11 |
| chr6 | 117716372 | 117717125 | 32.0113 | 129.32326 | 124.86235 | Promoter | 6 | 117646694 | 117717263 | 70570 | 2 | ENSMUST00 | 138 | RP24-479J14 | ENSMUSG00000099869.1 |
| chr1 | 170588872 | 170590414 | 37.74925 | 129.3071 | 124.84747 | Promoter | 1 | 170317459 | 170589861 | 272403 | 2 | ENSMUST00 | 0 | Nos1ap | ENSMUSG00000038473.9 |
| chr10 | 41518482 | 41521441 | 37.74925 | 129.3071 | 124.84747 | Promoter | 10 | 41519414 | 41531048 | 11635 | 1 | ENSMUST00 | 0 | Cd164 | ENSMUSG00000019818.10 |
| chr10 | 111008979 | 111011604 | 37.74925 | 129.3071 | 124.84747 | Promoter | 10 | 110941782 | 111010066 | 68285 | 2 | ENSMUST00 | 0 | Zdhhc17 | ENSMUSG00000035798.8 |
| chr3 | 108256390 | 108258058 | 37.74925 | 129.3071 | 124.84747 | Promoter | 3 | 108256926 | 108279974 | 23049 | 1 | ENSMUST00 | 0 | Psma5 | ENSMUSG00000068749.5 |
| chr8 | 123373094 | 123374909 | 40.23391 | 129.30441 | 124.84723 | Promoter | 8 | 123373824 | 123401892 | 28069 | 1 | ENSMUST00 | 0 | Tcf25 | ENSMUSG00000001472.11 |
| chr8 | 121082669 | 121085168 | 40.23391 | 129.30441 | 124.84723 | Promoter | 8 | 121084386 | 121088144 | 3759 | 1 | ENSMUST00 | 0 | Foxf1 | ENSMUSG00000042812.5 |
| chr1 | 87326615 | 87328066 | 36.52666 | 129.28897 | 124.83229 | Promoter | 1 | 87326998 | 87355722 | 28725 | 1 | ENSMUST00 | 0 | Gigyf2 | ENSMUSG00000048000.10 |
| chr15 | 76476684 | 76478589 | 39.54136 | 129.19044 | 124.73553 | Promoter | 15 | 76477445 | 76500972 | 23528 | 1 | ENSMUST00 | 0 | Hsf1 | ENSMUSG00000022556.9 |
| chr8 | 83566640 | 83567815 | 39.54136 | 129.19044 | 124.73553 | Promoter | 8 | 83566671 | 83571626 | 4956 | 1 | ENSMUST00 | 0 | Ndufb7 | ENSMUSG00000033938.5 |
| chr15 | 100615584 | 100616898 | 37.6793 | 129.18304 | 124.72858 | Promoter | 15 | 100615628 | 100620731 | 5104 | 1 | ENSMUST00 | 0 | Dazap2 | ENSMUSG00000000346.8 |
| chr8 | 88137475 | 88138849 | 37.6793 | 129.18304 | 124.72858 | Promoter | 8 | 88137855 | 88172027 | 34173 | 1 | ENSMUST00 | 0 | Heatr3 | ENSMUSG00000031657.11 |
| chr2 | 30171290 | 30172407 | 40.15089 | 129.16078 | 124.70641 | Promoter | 2 | 30171493 | 30174069 | 2577 | 1 | ENSMUST00 | 0 | Endog | ENSMUSG00000015337.5 |
| chr4 | 124692638 | 124694006 | 40.15089 | 129.16078 | 124.70641 | Promoter | 4 | 124678160 | 124693600 | 15441 | 2 | ENSMUST00 | 0 | Utp11l | ENSMUSG00000028907.7 |
| chr12 | 105683809 | 105686329 | 38.8867 | 129.13618 | 124.68224 | Promoter | 12 | 105613539 | 105685241 | 71703 | 2 | ENSMUST00 | 0 | Atg2b | ENSMUSG00000041341.9 |
| chr15 | 78982979 | 78984171 | 28.89701 | 129.08754 | 124.63428 | Promoter | 15 | 78983041 | 79005864 | 22824 | 1 | ENSMUST00 | 0 | Triobp | ENSMUSG00000033088.13 |
| chr6 | 149308882 | 149310424 | 28.89701 | 129.08754 | 124.63428 | Promoter | 6 | 149309414 | 149326671 | 17258 | 1 | ENSMUST00 | 0 | 2810474O19 | ENSMUSG00000032712.11 |
| chr17 | 25918691 | 25919790 | 39.46206 | 129.05257 | 124.59962 | Promoter | 17 | 25882114 | 25919556 | 37443 | 2 | ENSMUST00 | 0 | Rab40c | ENSMUSG00000025730.9 |
| chr2 | 90940019 | 90941495 | 34.63471 | 129.05151 | 124.59962 | Promoter | 2 | 90940382 | 90987754 | 47373 | 1 | ENSMUST00 | 0 | Celf1 | ENSMUSG00000005506.11 |
| chr11 | 78511525 | 78512886 | 30.33064 | 129.04645 | 124.59567 | Promoter | 11 | 78512193 | 78522736 | 10544 | 1 | ENSMUST00 | 0 | Poldip2 | ENSMUSG00000001100.5 |
| chr19 | 55098118 | 55100331 | 30.33064 | 129.04645 | 124.59567 | Promoter | 19 | 55069734 | 55099451 | 29718 | 2 | ENSMUST00 | 0 | Gpam | ENSMUSG00000024978.10 |
| chr2 | 52422513 | 52425752 | 30.33064 | 129.04645 | 124.59567 | Promoter | 2 | 52397951 | 52424901 | 26951 | 2 | ENSMUST00 | 0 | Arl5a | ENSMUSG00000036093.7 |
| chr2 | 18693722 | 18694863 | 39.38308 | 128.91518 | 124.46609 | Promoter | 2 | 18694032 | 18698884 | 4853 | 1 | ENSMUST00 | 0 | Gm13334 | ENSMUSG00000087182.2 |
| chr6 | 33059175 | 33060356 | 39.38308 | 128.91518 | 124.46609 | Promoter | 6 | 32792217 | 33060260 | 268044 | 2 | ENSMUST00 | 0 | Chchd3 | ENSMUSG00000053768.8 |
| chr11 | 23632854 | 23634167 | 39.98587 | 128.8752 | 124.42683 | Promoter | 11 | 23564961 | 23633549 | 68589 | 2 | ENSMUST00 | 0 | 0610010F05I | ENSMUSG00000042208.10 |
| chr12 | 86678281 | 86680383 | 39.98587 | 128.8752 | 124.42683 | Promoter | 12 | 86678700 | 86692091 | 13392 | 1 | ENSMUST00 | 0 | Vash1 | ENSMUSG00000021256.5 |
| chr15 | 100038564 | 100039854 | 39.98587 | 128.8752 | 124.42683 | Promoter | 15 | 100038664 | 100219473 | 180810 | 1 | ENSMUST00 | 0 | Dip2b | ENSMUSG00000023026.10 |
| chr17 | 27622163 | 27624145 | 39.98587 | 128.8752 | 124.42683 | Promoter | 17 | 27579382 | 27623465 | 44084 | 2 | ENSMUST00 | 0 | Nudt3 | ENSMUSG00000024213.8 |
| chr16 | 32098925 | 32100392 | 38.73528 | 128.87126 | 124.42339 | Promoter | 16 | 32084418 | 32099727 | 15310 | 2 | ENSMUST00 | 0 | Pigx | ENSMUSG00000023791.9 |
| chr4 | 134923576 | 134925935 | 35.65703 | 128.78946 | 124.34206 | Promoter | 4 | 134923592 | 134927671 | 4080 | 1 | ENSMUST00 | 0 | Rsrp1 | ENSMUSG00000037266.13 |
| chr1 | 134454842 | 134456279 | 39.30441 | 128.77829 | 124.33173 | Promoter | 1 | 134455531 | 134466570 | 11040 | 1 | ENSMUST00 | 0 | Klhl12 | ENSMUSG00000026455.9 |
| chr1 | 183368584 | 183370028 | 38.04976 | 128.75285 | 124.30675 | Promoter | 1 | 183326725 | 183369529 | 42805 | 2 | ENSMUST00 | 0 | Mia3 | ENSMUSG00000056050.7 |
| chr15 | 86058221 | 86059708 | 38.04976 | 128.75285 | 124.30675 | Promoter | 15 | 86058727 | 86137618 | 78892 | 1 | ENSMUST00 | 0 | Gramd4 | ENSMUSG00000035900.13 |
| chr17 | 15703965 | 15706505 | 38.04976 | 128.75285 | 124.30675 | Promoter | 17 | 15704967 | 15772610 | 67644 | 1 | ENSMUST00 | 0 | Chd1 | ENSMUSG00000023852.8 |
| chr17 | 34849538 | 34851078 | 38.04976 | 128.75285 | 124.30675 | Promoter | 17 | 34850391 | 34856369 | 5979 | 1 | ENSMUST00 | 0 | Nelfe | ENSMUSG00000024369.11 |
| chr2 | 147012182 | 147014400 | 38.04976 | 128.75285 | 124.30675 | Promoter | 2 | 147012996 | 147078000 | 65005 | 1 | ENSMUST00 | 0 | Xrn2 | ENSMUSG00000027433.5 |
| chr5 | 33334893 | 33336925 | 38.04976 | 128.75285 | 124.30675 | Promoter | 5 | 33335572 | 33373294 | 37723 | 1 | ENSMUST00 | 0 | Maea | ENSMUSG00000079562.3 |
| chr7 | 126791558 | 126792941 | 38.04976 | 128.75285 | 124.30675 | Promoter | 7 | 126785907 | 126792471 | 6565 | 2 | ENSMUST00 | 0 | Ppp4c | ENSMUSG00000030697.6 |
| chr7 | 125706384 | 125708627 | 38.04976 | 128.75285 | 124.30675 | Promoter | 7 | 125707915 | 125761832 | 53918 | 1 | ENSMUST00 | 0 | D430042O09 | ENSMUSG00000032743.10 |
| chr2 | 31140653 | 31142585 | 38.66002 | 128.73949 | 124.2961 | Promoter | 2 | 31026206 | 31141802 | 115597 | 2 | ENSMUST00 | 0 | Fnbp1 | ENSMUSG00000075415.8 |
| chr12 | 55199105 | 55200451 | 36.20828 | 128.71358 | 124.27102 | Promoter | 12 | 55199533 | 55214076 | 14544 | 1 | ENSMUST00 | 0 | 1700047I17R | ENSMUSG00000094103.1 |
| chr11 | 82870035 | 82871276 | 36.78915 | 128.67926 | 124.23671 | Promoter | 11 | 82803822 | 82870744 | 66923 | 2 | ENSMUST00 | 0 | Rffl | ENSMUSG00000020696.13 |
| chr12 | 40221511 | 40223282 | 36.78915 | 128.67926 | 124.23671 | Promoter | 12 | 40201567 | 40223189 | 21623 | 2 | ENSMUST00 | 0 | Ifrd1 | ENSMUSG00000001627.6 |
| chr6 | 29735189 | 29737881 | 36.78915 | 128.67926 | 124.23671 | Promoter | 6 | 29735694 | 29761365 | 25672 | 1 | ENSMUST00 | 0 | Smo | ENSMUSG00000001761.6 |
| chr4 | 43058535 | 43059823 | 39.22606 | 128.64191 | 124.20071 | Promoter | 4 | 43058953 | 43166122 | 107170 | 1 | ENSMUST00 | 0 | Unc13b | ENSMUSG00000028456.12 |
| chr2 | 28467819 | 28468909 | 37.97791 | 128.62614 | 124.18539 | Promoter | 2 | 28468066 | 28471178 | 3113 | 1 | ENSMUST00 | 0 | Mrps2 | ENSMUSG00000035772.6 |
| chr1 | 53351756 | 53352941 | 39.82221 | 128.59171 | 124.15125 | Promoter | 1 | 53344617 | 53352752 | 8136 | 2 | ENSMUST00 | 0 | Asnsd1 | ENSMUSG00000026095.10 |
| chr12 | 57230001 | 57231414 | 39.82221 | 128.59171 | 124.15125 | Promoter | 12 | 57230427 | 57308695 | 78269 | 1 | ENSMUST00 | 0 | Mipol1 | ENSMUSG00000047022.13 |
| chr15 | 100598613 | 100600666 | 39.82221 | 128.59171 | 124.15125 | Promoter | 15 | 100575318 | 100599984 | 24667 | 2 | ENSMUST00 | 0 | Pou6f1 | ENSMUSG00000009739.11 |
| chr17 | 34119478 | 34123860 | 39.82221 | 128.59171 | 124.15125 | Promoter | 17 | 34112023 | 34121747 | 9725 | 2 | ENSMUST00 | 0 | Brd2 | ENSMUSG00000024335.14 |
| chr6 | 88842418 | 88843127 | 39.82221 | 128.59171 | 124.15125 | Promoter | 6 | 88842854 | 88847274 | 4421 | 1 | ENSMUST00 | 0 | Gm15612 | ENSMUSG00000085828.1 |
| chr12 | 54694977 | 54696798 | 40.43883 | 128.57333 | 124.13428 | Promoter | 12 | 54670340 | 54695881 | 25542 | 2 | ENSMUST00 | 0 | Eapp | ENSMUSG00000054302.10 |
| chr15 | 89428885 | 89430631 | 40.43883 | 128.57333 | 124.13428 | Promoter | 15 | 89416405 | 89429696 | 13292 | 2 | ENSMUST00 | 0 | Chkb | ENSMUSG00000022617.9 |
| chr17 | 29490723 | 29492947 | 40.43883 | 128.57333 | 124.13428 | Promoter | 17 | 29490812 | 29496111 | 5300 | 1 | ENSMUST00 | 0 | Pim1 | ENSMUSG00000024014.7 |
| chr17 | 26252486 | 26254060 | 40.43883 | 128.57333 | 124.13428 | Promoter | 17 | 26252910 | 26280719 | 27810 | 1 | ENSMUST00 | 0 | Luc7l | ENSMUSG00000024188.10 |
| chr17 | 33928712 | 33931164 | 40.43883 | 128.57333 | 124.13428 | Promoter | 17 | 33929543 | 33930495 | 953 | 1 | ENSMUST00 | 0 | Rgl2 | ENSMUSG00000041354.10 |
| chr2 | 3474048 | 3475781 | 40.43883 | 128.57333 | 124.13428 | Promoter | 2 | 3455815 | 3475031 | 19217 | 2 | ENSMUST00 | 0 | Suv39h2 | ENSMUSG00000026646.11 |
| chr17 | 35379510 | 35380703 | 40.36609 | 128.44781 | 124.01388 | Promoter | 17 | 35379617 | 35384674 | 5058 | 1 | ENSMUST00 | 0 | H2-Q4 | ENSMUSG00000035929.11 |
| chr5 | 52741243 | 52742932 | 39.07029 | 128.37054 | 123.93713 | Promoter | 5 | 52741587 | 52769344 | 27758 | 1 | ENSMUST00 | 0 | Pi4k2b | ENSMUSG00000029186.7 |
| chr4 | 94602271 | 94604038 | 34.84454 | 128.36943 | 123.93644 | Promoter | 4 | 94569167 | 94603244 | 34078 | 2 | ENSMUST00 | 0 | Plaa | ENSMUSG00000028577.8 |
| chr6 | 115601705 | 115602741 | 34.84454 | 128.36943 | 123.93644 | Promoter | 6 | 115601938 | 115618670 | 16733 | 1 | ENSMUST00 | 0 | Mkrn2 | ENSMUSG00000000439.8 |
| chr19 | 5456222 | 5458183 | 38.43596 | 128.34685 | 123.91427 | Promoter | 19 | 5456789 | 5457397 | 609 | 2 | ENSMUST00 | 0 | Ccdc85b | ENSMUSG00000095098.1 |
| chr11 | 73175882 | 73178809 | 33.66875 | 128.30269 | 123.87101 | Promoter | 11 | 73177083 | 73182152 | 5070 | 1 | ENSMUST00 | 0 | Tax1bp3 | ENSMUSG00000040158.7 |
| chr13 | 69809455 | 69810921 | 33.66875 | 128.30269 | 123.87101 | Promoter | 13 | 69809882 | 69816094 | 6213 | 1 | ENSMUST00 | 0 | Med10 | ENSMUSG00000021598.8 |
| chr2 | 57123281 | 57124500 | 33.66875 | 128.30269 | 123.87101 | Promoter | 2 | 57106830 | 57124003 | 17174 | 2 | ENSMUST00 | 0 | Nr4a2 | ENSMUSG00000026826.8 |
| chr2 | 24934249 | 24935881 | 33.66875 | 128.30269 | 123.87101 | Promoter | 2 | 24925352 | 24935163 | 9812 | 2 | ENSMUST00 | 0 | Arrdc1 | ENSMUSG00000026972.10 |
| chr3 | 69003957 | 69006802 | 33.66875 | 128.30269 | 123.87101 | Promoter | 3 | 68892499 | 69004543 | 112045 | 2 | ENSMUST00 | 0 | Ift80 | ENSMUSG00000027778.10 |
| chr4 | 133338191 | 133339666 | 33.66875 | 128.30269 | 123.87101 | Promoter | 4 | 133292459 | 133339324 | 46866 | 2 | ENSMUST00 | 0 | Wdtc1 | ENSMUSG00000037622.9 |
| chr5 | 124528429 | 124531965 | 33.66875 | 128.30269 | 123.87101 | Promoter | 5 | 124493082 | 124531391 | 38310 | 2 | ENSMUST00 | 0 | Rilpl1 | ENSMUSG00000029392.8 |
| chr9 | 55937295 | 55938834 | 33.66875 | 128.30269 | 123.87101 | Distal Interg | 9 | 55549883 | 55919605 | 369723 | 2 | ENSMUST00 | -17691 | Scaper | ENSMUSG00000034007.9 |
| chr9 | 103759904 | 103762409 | 33.66875 | 128.30269 | 123.87101 | Promoter | 9 | 103483778 | 103761837 | 278060 | 2 | ENSMUST00 | 0 | Tmem108 | ENSMUSG00000042757.10 |
| chr14 | 36967727 | 36969299 | 38.99286 | 128.2356 | 123.80784 | Promoter | 14 | 36874936 | 36968687 | 93752 | 2 | ENSMUST00 | 0 | Ccser2 | ENSMUSG00000058690.9 |
| chr2 | 167931436 | 167933867 | 38.36185 | 128.21687 | 123.78947 | Promoter | 2 | 167932057 | 167974737 | 42681 | 1 | ENSMUST00 | 0 | Ptpn1 | ENSMUSG00000027540.8 |
| chr4 | 116074263 | 116077477 | 38.36185 | 128.21687 | 123.78947 | Promoter | 4 | 116075269 | 116097043 | 21775 | 1 | ENSMUST00 | 0 | Lrrc41 | ENSMUSG00000028703.9 |
| chr6 | 88901581 | 88903846 | 38.36185 | 128.21687 | 123.78947 | Promoter | 6 | 88902251 | 88912238 | 9988 | 1 | ENSMUST00 | 0 | Tpra1 | ENSMUSG00000002871.9 |
| chr2 | 4880757 | 4883313 | 40.19742 | 128.15663 | 123.73233 | Promoter | 2 | 4881564 | 4910557 | 28994 | 1 | ENSMUST00 | 0 | Sephs1 | ENSMUSG00000026662.8 |
| chr9 | 88480664 | 88483536 | 40.19742 | 128.15663 | 123.73233 | Promoter | 9 | 88449731 | 88482248 | 32518 | 2 | ENSMUST00 | 0 | Syncrip | ENSMUSG00000032423.7 |
| chr1 | 93634478 | 93635780 | 35.89541 | 128.14636 | 123.72295 | Promoter | 1 | 93619751 | 93635722 | 15972 | 2 | ENSMUST00 | 0 | Stk25 | ENSMUSG00000026277.8 |
| chr5 | 110807512 | 110810111 | 35.89541 | 128.14636 | 123.72295 | Promoter | 5 | 110784488 | 110810097 | 25610 | 2 | ENSMUST00 | 0 | Ulk1 | ENSMUSG00000029512.7 |
| chr11 | 69995731 | 69997499 | 38.91574 | 128.10115 | 123.6785 | Promoter | 11 | 69995777 | 70000009 | 4233 | 1 | ENSMUST00 | 0 | Phf23 | ENSMUSG00000018572.5 |
| chr2 | 44927335 | 44928141 | 38.28802 | 128.08731 | 123.66512 | Promoter | 2 | 44635264 | 44927657 | 292394 | 2 | ENSMUST00 | 0 | Gtdc1 | ENSMUSG00000036890.8 |

| | | | | | | | | | | | | | | | |
|---|---|---|---|---|---|---|---|---|---|---|---|---|---|---|---|
| chr10 | 92721740 | 92722810 | 37.0612 | 128.08391 | 123.66181 | Promoter | 10 | 92684746 | 92722420 | 37675 | 2 | ENSMUST00 | 0 | Nedd1 | ENSMUSG00000019988.6 |
| chr14 | 34548457 | 34549653 | 37.0612 | 128.08391 | 123.66181 | Intron (ENSM | 14 | 34511619 | 34522801 | 11183 | 2 | ENSMUST00 | -25657 | 9230112D13 | ENSMUSG00000021797.1 |
| chr18 | 21001889 | 21003402 | 37.0612 | 128.08391 | 123.66181 | Promoter | 18 | 21001341 | 21028223 | 26883 | 1 | ENSMUST00 | 549 | Rnf138 | ENSMUSG00000024317.9 |
| chr19 | 5023330 | 5024767 | 37.0612 | 128.08391 | 123.66181 | Promoter | 19 | 5024006 | 5031971 | 7966 | 1 | ENSMUST00 | 0 | Slc29a2 | ENSMUSG00000024891.5 |
| chr4 | 41313584 | 41315385 | 37.0612 | 128.08391 | 123.66181 | Promoter | 4 | 41291300 | 41314889 | 23590 | 2 | ENSMUST00 | 0 | Dcaf12 | ENSMUSG00000028436.8 |
| chr5 | 45519885 | 45521123 | 37.0612 | 128.08391 | 123.66181 | Promoter | 5 | 45520221 | 45529276 | 9056 | 1 | ENSMUST00 | 0 | Med28 | ENSMUSG00000015804.9 |
| chr9 | 70206932 | 70210342 | 37.0612 | 128.08391 | 123.66181 | Promoter | 9 | 70207350 | 70400067 | 192718 | 1 | ENSMUST00 | 0 | Myo1e | ENSMUSG00000032220.9 |
| chr14 | 76109940 | 76111691 | 34.08376 | 128.02023 | 123.60056 | Promoter | 14 | 76086232 | 76110815 | 24584 | 2 | ENSMUST00 | 0 | Gpalpp1 | ENSMUSG00000022008.3 |
| chr2 | 153291021 | 153292117 | 34.08376 | 128.02023 | 123.60056 | Promoter | 2 | 153291413 | 153333390 | 41978 | 1 | ENSMUST00 | 0 | Kif3b | ENSMUSG00000027475.9 |
| chr11 | 53519402 | 53521185 | 40.11361 | 128.01187 | 123.59241 | Promoter | 11 | 53519746 | 53544095 | 24350 | 1 | ENSMUST00 | 0 | Sep-08 | ENSMUSG00000018398.13 |
| chr4 | 136601489 | 136602829 | 37.62274 | 127.9987 | 123.57949 | Promoter | 4 | 136550540 | 136602723 | 52184 | 2 | ENSMUST00 | 0 | Kdm1a | ENSMUSG00000036940.10 |
| chr2 | 109890222 | 109892197 | 38.83893 | 127.96716 | 123.54797 | Promoter | 2 | 109890853 | 109901003 | 10151 | 1 | ENSMUST00 | 0 | Lin7c | ENSMUSG00000027162.7 |
| chr8 | 82863357 | 82865470 | 38.83893 | 127.96716 | 123.54797 | Promoter | 8 | 82863356 | 83091271 | 227916 | 1 | ENSMUST00 | 2 | Rnf150 | ENSMUSG00000047747.9 |
| chr10 | 76468166 | 76469686 | 39.41886 | 127.89232 | 123.47375 | Promoter | 10 | 76468971 | 76515856 | 46886 | 1 | ENSMUST00 | 0 | Mcm3ap | ENSMUSG00000001150.6 |
| chr15 | 94543018 | 94544299 | 39.41886 | 127.89232 | 123.47375 | Promoter | 15 | 94522688 | 94543547 | 20860 | 2 | ENSMUST00 | 0 | Pus7l | ENSMUSG00000033356.3 |
| chr15 | 81585366 | 81588048 | 40.03016 | 127.86769 | 123.44967 | Promoter | 15 | 81586220 | 81624218 | 37999 | 1 | ENSMUST00 | 0 | Ep300 | ENSMUSG00000055024.7 |
| chr8 | 46470894 | 46472109 | 40.03016 | 127.86769 | 123.44967 | Promoter | 8 | 46471041 | 46536051 | 65011 | 1 | ENSMUST00 | 0 | Acsl1 | ENSMUSG00000018796.7 |
| chr19 | 30029826 | 30031769 | 38.76242 | 127.83364 | 123.41602 | Promoter | 19 | 30030513 | 30058812 | 28300 | 1 | ENSMUST00 | 0 | Uhrf2 | ENSMUSG00000024817.8 |
| chr1 | 75140886 | 75143697 | 38.13901 | 127.82562 | 123.40861 | Promoter | 1 | 75134554 | 75142638 | 8085 | 2 | ENSMUST00 | 0 | Cnppd1 | ENSMUSG00000033159.9 |
| chr10 | 80115705 | 80117886 | 38.13901 | 127.82562 | 123.40861 | Promoter | 10 | 80115803 | 80130469 | 14667 | 1 | ENSMUST00 | 0 | Stk11 | ENSMUSG00000003068.10 |
| chr10 | 120977509 | 120980049 | 38.13901 | 127.82562 | 123.40861 | Promoter | 10 | 120923413 | 120979332 | 55920 | 2 | ENSMUST00 | 0 | Lemd3 | ENSMUSG00000048661.9 |
| chr11 | 97743491 | 97745609 | 38.13901 | 127.82562 | 123.40861 | Promoter | 11 | 97715157 | 97744704 | 29548 | 2 | ENSMUST00 | 0 | Pip4k2b | ENSMUSG00000018547.8 |
| chr11 | 58104039 | 58105114 | 38.13901 | 127.82562 | 123.40861 | Promoter | 11 | 58104153 | 58118594 | 14442 | 1 | ENSMUST00 | 0 | Cnot8 | ENSMUSG00000020515.8 |
| chr13 | 92354035 | 92355747 | 38.13901 | 127.82562 | 123.40861 | Promoter | 13 | 92211872 | 92354995 | 143124 | 2 | ENSMUST00 | 0 | Msh3 | ENSMUSG00000014850.10 |
| chr14 | 54426315 | 54427561 | 38.13901 | 127.82562 | 123.40861 | Promoter | 14 | 54426909 | 54429750 | 2842 | 1 | ENSMUST00 | 0 | Mrpl52 | ENSMUSG00000010406.7 |
| chr16 | 22009422 | 22010175 | 38.13901 | 127.82562 | 123.40861 | Promoter | 16 | 22009484 | 22049269 | 39786 | 1 | ENSMUST00 | 0 | Senp2 | ENSMUSG00000022855.8 |
| chr2 | 73774661 | 73776103 | 38.13901 | 127.82562 | 123.40861 | Promoter | 2 | 73610660 | 73775346 | 164687 | 2 | ENSMUST00 | 0 | Chn1 | ENSMUSG00000056486.12 |
| chr3 | 104510945 | 104512658 | 38.13901 | 127.82562 | 123.40861 | Promoter | 3 | 104453984 | 104511856 | 57873 | 2 | ENSMUST00 | 0 | Lrig2 | ENSMUSG00000032913.9 |
| chr4 | 123410776 | 123412702 | 38.13901 | 127.82562 | 123.40861 | Intron (ENSM | 4 | 123397275 | 123407515 | 10241 | 2 | ENSMUST00 | -3262 | Macf1 | ENSMUSG00000028649.12 |
| chr4 | 141535704 | 141538627 | 38.13901 | 127.82562 | 123.40861 | Promoter | 4 | 141467890 | 141538449 | 70560 | 2 | ENSMUST00 | 0 | Spen | ENSMUSG00000040761.11 |
| chr4 | 139574658 | 139577815 | 38.13901 | 127.82562 | 123.40861 | Promoter | 4 | 139574720 | 139620382 | 45663 | 1 | ENSMUST00 | 0 | Iffo2 | ENSMUSG00000041025.10 |
| chr5 | 122613796 | 122614751 | 38.13901 | 127.82562 | 123.40861 | Promoter | 5 | 122550204 | 122614518 | 64315 | 2 | ENSMUST00 | 0 | Ift81 | ENSMUSG00000029469.9 |
| chr5 | 136116279 | 136118675 | 38.13901 | 127.82562 | 123.40861 | Promoter | 5 | 136116631 | 136122723 | 6093 | 1 | ENSMUST00 | 0 | Polr2j | ENSMUSG00000039771.8 |
| chr9 | 78112889 | 78114174 | 38.13901 | 127.82562 | 123.40861 | Promoter | 9 | 78113241 | 78134119 | 20879 | 1 | ENSMUST00 | 0 | Ick | ENSMUSG00000009828.10 |
| chr10 | 52416804 | 52418699 | 39.33916 | 127.75399 | 123.34521 | Promoter | 10 | 52417547 | 52440192 | 22646 | 1 | ENSMUST00 | 0 | Nus1 | ENSMUSG00000023068.9 |
| chr9 | 92249860 | 92252220 | 39.33916 | 127.75399 | 123.34521 | Promoter | 9 | 92250342 | 92271975 | 21634 | 1 | ENSMUST00 | 0 | Plscr1 | ENSMUSG00000032369.8 |
| chr19 | 17393476 | 17395235 | 37.48252 | 127.75053 | 123.3422 | Promoter | 19 | 17394043 | 17401349 | 7307 | 1 | ENSMUST00 | 0 | Rfk | ENSMUSG00000024712.4 |
| chr7 | 112022814 | 112025018 | 37.48252 | 127.75053 | 123.3422 | Promoter | 7 | 112023506 | 112111386 | 87881 | 1 | ENSMUST00 | 0 | Usp47 | ENSMUSG00000059263.7 |
| chr10 | 78008931 | 78010334 | 35.67123 | 127.73884 | 123.33056 | Promoter | 10 | 77986947 | 78009807 | 22861 | 2 | ENSMUST00 | 0 | Pfkl | ENSMUSG00000020277.9 |
| chr11 | 74648841 | 74650947 | 35.67123 | 127.73884 | 123.33056 | Promoter | 11 | 74649495 | 74656551 | 7057 | 1 | ENSMUST00 | 0 | Cluh | ENSMUSG00000020741.12 |
| chr11 | 45943874 | 45945010 | 35.67123 | 127.73884 | 123.33056 | Promoter | 11 | 45928269 | 45944935 | 16667 | 2 | ENSMUST00 | 0 | Lsm11 | ENSMUSG00000044847.8 |
| chr15 | 93274138 | 93275225 | 35.67123 | 127.73884 | 123.33056 | Promoter | 15 | 93239742 | 93275161 | 35420 | 2 | ENSMUST00 | 0 | Gxylt1 | ENSMUSG00000036197.10 |
| chr16 | 11252796 | 11255094 | 35.67123 | 127.73884 | 123.33056 | Promoter | 16 | 11219292 | 11254270 | 34979 | 2 | ENSMUST00 | 0 | Gspt1 | ENSMUSG00000062203.9 |
| chr18 | 21299173 | 21300235 | 35.67123 | 127.73884 | 123.33056 | Promoter | 18 | 21127201 | 21300138 | 172938 | 2 | ENSMUST00 | 0 | Garem | ENSMUSG00000042680.7 |
| chr2 | 180708615 | 180711658 | 35.67123 | 127.73884 | 123.33056 | Promoter | 2 | 180710117 | 180718733 | 8617 | 1 | ENSMUST00 | 0 | Gid8 | ENSMUSG00000027573.6 |
| chr4 | 11075579 | 11076683 | 35.67123 | 127.73884 | 123.33056 | Promoter | 4 | 11051045 | 11076204 | 25160 | 2 | ENSMUST00 | 0 | Ndufaf6 | ENSMUSG00000050323.9 |
| chr5 | 33819761 | 33821890 | 35.67123 | 127.73884 | 123.33056 | Promoter | 5 | 33820725 | 33897966 | 77242 | 1 | ENSMUST00 | 0 | Whsc1 | ENSMUSG00000057406.11 |
| chr7 | 30649422 | 30650247 | 35.67123 | 127.73884 | 123.33056 | Promoter | 7 | 30640999 | 30650144 | 9146 | 2 | ENSMUST00 | 0 | Rbm42 | ENSMUSG00000036733.10 |
| chr7 | 143052294 | 143056592 | 35.67123 | 127.73884 | 123.33056 | Promoter | 7 | 143052739 | 143067934 | 15196 | 1 | ENSMUST00 | 0 | Cd81 | ENSMUSG00000037706.11 |
| chr7 | 24547077 | 24547597 | 39.94704 | 127.72406 | 123.32267 | Promoter | 7 | 24547150 | 24573438 | 26289 | 1 | ENSMUST00 | 0 | Xrcc1 | ENSMUSG00000051768.8 |
| chr1 | 184844456 | 184846734 | 38.68621 | 127.7006 | 123.29971 | Promoter | 1 | 184813068 | 184845881 | 32814 | 2 | ENSMUST00 | 0 | Mar-02 | ENSMUSG00000073481.4 |
| chr11 | 120795477 | 120796856 | 38.68621 | 127.7006 | 123.29971 | Promoter | 11 | 120789201 | 120796403 | 7203 | 2 | ENSMUST00 | 0 | Dus1l | ENSMUSG00000025155.10 |
| chr15 | 5239457 | 5245131 | 38.68621 | 127.7006 | 123.29971 | Promoter | 15 | 5206661 | 5242710 | 36050 | 2 | ENSMUST00 | 0 | Ptger4 | ENSMUSG00000039942.10 |
| chr13 | 85287580 | 85289440 | 35.06139 | 127.69727 | 123.29834 | Promoter | 13 | 85214780 | 85289029 | 74250 | 2 | ENSMUST00 | 0 | Rasa1 | ENSMUSG00000021549.8 |
| chr1 | 37299838 | 37301548 | 31.70051 | 127.6562 | 123.25787 | Promoter | 1 | 37299865 | 37348512 | 48648 | 1 | ENSMUST00 | 0 | Inpp4a | ENSMUSG00000026113.12 |
| chr1 | 57774051 | 57776163 | 31.70051 | 127.6562 | 123.25787 | Promoter | 1 | 57774162 | 57885768 | 111607 | 1 | ENSMUST00 | 0 | Spats2l | ENSMUSG00000038305.9 |
| chr10 | 116580190 | 116583053 | 31.70051 | 127.6562 | 123.25787 | Promoter | 10 | 116485161 | 116581511 | 96351 | 2 | ENSMUST00 | 0 | Cnot2 | ENSMUSG00000020166.9 |
| chr14 | 72709140 | 72711049 | 31.70051 | 127.6562 | 123.25787 | Promoter | 14 | 72537946 | 72710003 | 172058 | 2 | ENSMUST00 | 0 | Fndc3a | ENSMUSG00000033487.9 |
| chr17 | 34647120 | 34647999 | 31.70051 | 127.6562 | 123.25787 | Promoter | 17 | 34647146 | 34655074 | 7929 | 1 | ENSMUST00 | 0 | Atf6b | ENSMUSG00000015461.10 |
| chr19 | 16779373 | 16781820 | 31.70051 | 127.6562 | 123.25787 | Promoter | 19 | 16616763 | 16780822 | 164060 | 2 | ENSMUST00 | 0 | Vps13a | ENSMUSG00000046230.9 |
| chr3 | 54806800 | 54808512 | 31.70051 | 127.6562 | 123.25787 | Promoter | 3 | 54803115 | 54807791 | 4677 | 2 | ENSMUST00 | 0 | Rfxap | ENSMUSG00000036615.5 |
| chr4 | 141789947 | 141791329 | 31.70051 | 127.6562 | 123.25787 | Promoter | 4 | 141760189 | 141790891 | 30703 | 2 | ENSMUST00 | 0 | Dnajc16 | ENSMUSG00000040697.5 |
| chr7 | 30587480 | 30589984 | 31.70051 | 127.6562 | 123.25787 | Promoter | 7 | 30568858 | 30588717 | 19860 | 2 | ENSMUST00 | 0 | Kmt2b | ENSMUSG00000006307.11 |
| chr9 | 54586037 | 54587439 | 31.70051 | 127.6562 | 123.25787 | Promoter | 9 | 54586511 | 54604661 | 18151 | 1 | ENSMUST00 | 0 | Idh3a | ENSMUSG00000032279.10 |
| chr9 | 56159942 | 56161222 | 31.70051 | 127.6562 | 123.25787 | Promoter | 9 | 56135885 | 56161070 | 25186 | 2 | ENSMUST00 | 0 | Tspan3 | ENSMUSG00000032324.10 |
| chr14 | 45317289 | 45319128 | 39.25979 | 127.61615 | 123.21935 | Promoter | 14 | 45283092 | 45318572 | 35481 | 2 | ENSMUST00 | 0 | Ero1l | ENSMUSG00000021831.8 |
| chr17 | 14942533 | 14944424 | 37.99554 | 127.57345 | 123.17763 | Promoter | 17 | 14943184 | 14945936 | 2753 | 1 | ENSMUST00 | 0 | 1600012H06 | ENSMUSG00000050088.9 |
| chr4 | 126676980 | 126678614 | 38.6103 | 127.56802 | 123.17235 | Promoter | 4 | 126677630 | 126709714 | 32085 | 1 | ENSMUST00 | 0 | Psmb2 | ENSMUSG00000028837.3 |
| chr10 | 112927882 | 112929555 | 36.14217 | 127.51552 | 123.12013 | Promoter | 10 | 112925430 | 112928995 | 3566 | 2 | ENSMUST00 | 0 | Atxn7l3b | ENSMUSG00000074748.3 |
| chr13 | 30973358 | 30974593 | 36.14217 | 127.51552 | 123.12013 | Promoter | 13 | 30813919 | 30974047 | 160129 | 2 | ENSMUST00 | 0 | Exoc2 | ENSMUSG00000021357.8 |
| chr4 | 135352017 | 135353251 | 36.14217 | 127.51552 | 123.12013 | Promoter | 4 | 135320491 | 135353244 | 32754 | 2 | ENSMUST00 | 0 | Srrm1 | ENSMUSG00000028809.11 |
| chr7 | 101301299 | 101302819 | 36.14217 | 127.51552 | 123.12013 | Promoter | 7 | 101289392 | 101302027 | 12636 | 2 | ENSMUST00 | 0 | Atg16l2 | ENSMUSG00000047767.11 |
| chr15 | 101173412 | 101176072 | 37.34335 | 127.50397 | 123.10989 | Promoter | 15 | 101174067 | 101213679 | 39613 | 1 | ENSMUST00 | 0 | Acvr1b | ENSMUSG00000000532.10 |
| chr16 | 87495038 | 87496706 | 37.34335 | 127.50397 | 123.10989 | Promoter | 16 | 87483326 | 87495873 | 12548 | 2 | ENSMUST00 | 0 | Cct8 | ENSMUSG00000025613.8 |
| chr3 | 116661343 | 116663117 | 37.34335 | 127.50397 | 123.10989 | Promoter | 3 | 116631164 | 116662677 | 31514 | 2 | ENSMUST00 | 0 | Hiat1 | ENSMUSG00000089911.2 |
| chr6 | 51467769 | 51469887 | 37.34335 | 127.50397 | 123.10989 | Promoter | 6 | 51461124 | 51469845 | 8722 | 2 | ENSMUST00 | 0 | Hnrnpa2b1 | ENSMUSG00000004980.11 |
| chr7 | 90442171 | 90445003 | 37.34335 | 127.50397 | 123.10989 | Promoter | 7 | 90442729 | 90446104 | 3376 | 1 | ENSMUST00 | 0 | Crebzf | ENSMUSG00000051451.5 |
| chr8 | 95715278 | 95716679 | 37.34335 | 127.50397 | 123.10989 | Promoter | 8 | 95715915 | 95716781 | 867 | 1 | ENSMUST00 | 0 | Setd6 | ENSMUSG00000031671.5 |
| chr17 | 66100653 | 66102325 | 39.18074 | 127.47883 | 123.08725 | Promoter | 17 | 66078795 | 66101559 | 22765 | 2 | ENSMUST00 | 0 | Ndufv2 | ENSMUSG00000024099.9 |
| chr12 | 84361285 | 84363000 | 38.53469 | 127.4359 | 123.04581 | Promoter | 12 | 84345309 | 84361833 | 16525 | 2 | ENSMUST00 | 0 | Fam161b | ENSMUSG00000021234.4 |
| chr2 | 150903888 | 150905232 | 38.53469 | 127.4359 | 123.04581 | Promoter | 2 | 150832493 | 150904741 | 72249 | 2 | ENSMUST00 | 0 | Abhd12 | ENSMUSG00000032046.10 |
| chr3 | 87970233 | 87975133 | 30.04451 | 127.41808 | 123.02824 | Promoter | 3 | 87971078 | 87973391 | 2314 | 1 | ENSMUST00 | 0 | Nes | ENSMUSG00000004891.11 |
| chr8 | 123253289 | 123255503 | 39.10201 | 127.34201 | 122.95288 | Promoter | 8 | 123242356 | 123254176 | 11821 | 2 | ENSMUST00 | 0 | Vps9d1 | ENSMUSG00000001062.12 |
| chr11 | 121259204 | 121261148 | 37.85098 | 127.31907 | 122.93079 | Promoter | 11 | 121259990 | 121309896 | 49907 | 1 | ENSMUST00 | 0 | Foxk2 | ENSMUSG00000039275.9 |
| chr1 | 190911038 | 190912446 | 39.02359 | 127.20569 | 122.81842 | Promoter | 1 | 190700202 | 190911747 | 211546 | 2 | ENSMUST00 | 0 | Rps6kc1 | ENSMUSG00000089872.5 |
| chr11 | 21238334 | 21240491 | 39.02359 | 127.20569 | 122.81842 | Promoter | 11 | 21239281 | 21321122 | 81842 | 1 | ENSMUST00 | 0 | Vps54 | ENSMUSG00000020128.10 |
| chr17 | 78834676 | 78836245 | 39.02359 | 127.20569 | 122.81842 | Promoter | 17 | 78752906 | 78835381 | 82476 | 2 | ENSMUST00 | 0 | Heatr5b | ENSMUSG00000039414.9 |
| chr4 | 101263825 | 101265786 | 39.02359 | 127.20569 | 122.81842 | Promoter | 4 | 101152367 | 101265282 | 112916 | 2 | ENSMUST00 | 0 | Jak1 | ENSMUSG00000028530.9 |
| chr7 | 19628045 | 19629963 | 39.02359 | 127.20569 | 122.81842 | Promoter | 7 | 19606217 | 19629438 | 23222 | 2 | ENSMUST00 | 0 | Relb | ENSMUSG00000002983.11 |
| chr8 | 104640850 | 104642429 | 39.02359 | 127.20569 | 122.81842 | Promoter | 8 | 104641807 | 104658587 | 16781 | 1 | ENSMUST00 | 0 | 1700082M22 | ENSMUSG00000023737.8 |
| chr3 | 95929228 | 95930742 | 39.61801 | 127.15503 | 122.77008 | Promoter | 3 | 95929246 | 95947390 | 18145 | 1 | ENSMUST00 | 0 | Anp32e | ENSMUSG00000015749.7 |
| chr3 | 69315733 | 69318906 | 39.61801 | 127.15503 | 122.77008 | Promoter | 3 | 69316861 | 69560802 | 243942 | 1 | ENSMUST00 | 0 | Ppm1l | ENSMUSG00000027784.9 |
| chr5 | 140419171 | 140420345 | 38.94548 | 127.06984 | 122.68537 | Promoter | 5 | 140419305 | 140443358 | 24054 | 1 | ENSMUST00 | 0 | Eif3b | ENSMUSG00000056076.9 |
| chr11 | 5260004 | 5262317 | 38.30962 | 127.04226 | 122.65842 | Promoter | 11 | 5191552 | 5261558 | 70007 | 2 | ENSMUST00 | 0 | Kremen1 | ENSMUSG00000020393.11 |
| chr18 | 59174453 | 59176473 | 38.30962 | 127.04226 | 122.65842 | Promoter | 18 | 59175401 | 59410446 | 235046 | 1 | ENSMUST00 | 0 | Chsy3 | ENSMUSG00000058152.7 |
| chr1 | 121430723 | 121431968 | 35.2856 | 127.03558 | 122.65217 | Promoter | 1 | 121431049 | 121506460 | 75412 | 1 | ENSMUST00 | 0 | Ccdc93 | ENSMUSG00000026339.13 |
| chr10 | 117709420 | 117711415 | 35.2856 | 127.03558 | 122.65217 | Promoter | 10 | 117688888 | 117710758 | 21871 | 2 | ENSMUST00 | 0 | Mdm2 | ENSMUSG00000020184.9 |
| chr10 | 20311327 | 20313751 | 35.2856 | 127.03558 | 122.65217 | Promoter | 10 | 20312469 | 20342501 | 30033 | 1 | ENSMUST00 | 0 | Bclaf1 | ENSMUSG00000037608.10 |
| chr10 | 63428590 | 63430584 | 35.2856 | 127.03558 | 122.65217 | Promoter | 10 | 63430098 | 65002965 | 1572868 | 1 | ENSMUST00 | 0 | Ctnna3 | ENSMUSG00000060843.6 |
| chr2 | 129800009 | 129802353 | 35.2856 | 127.03558 | 122.65217 | Promoter | 2 | 129800517 | 129828974 | 28458 | 1 | ENSMUST00 | 0 | Stk35 | ENSMUSG00000037885.13 |
| chr2 | 152226184 | 152227818 | 35.2856 | 127.03558 | 122.65217 | Promoter | 2 | 152226839 | 152281852 | 55014 | 1 | ENSMUST00 | 0 | Csnk2a1 | ENSMUSG00000074698.5 |
| chr1 | 58504109 | 58505616 | 37.63617 | 126.94061 | 122.55849 | Promoter | 1 | 58462771 | 58505109 | 42339 | 2 | ENSMUST00 | 0 | Orc2 | ENSMUSG00000026037.9 |
| chr1 | 9746906 | 9748499 | 37.63617 | 126.94061 | 122.55849 | Promoter | 1 | 9718622 | 9748382 | 29761 | 2 | ENSMUST00 | 0 | Vcpip1 | ENSMUSG00000045210.8 |
| chr10 | 117223735 | 117225006 | 37.63617 | 126.94061 | 122.55849 | Promoter | 10 | 117215142 | 117224507 | 9366 | 2 | ENSMUST00 | 0 | Yeats4 | ENSMUSG00000020171.7 |
| chr2 | 32450723 | 32451991 | 37.63617 | 126.94061 | 122.55849 | Promoter | 2 | 32414487 | 32451422 | 36936 | 2 | ENSMUST00 | 0 | Slc25a25 | ENSMUSG00000026819.10 |
| chr2 | 155728761 | 155729774 | 37.63617 | 126.94061 | 122.55849 | Promoter | 2 | 155701677 | 155729475 | 27799 | 2 | ENSMUST00 | 0 | Edem2 | ENSMUSG00000038312.5 |
| chr2 | 29619788 | 29621117 | 37.63617 | 126.94061 | 122.55849 | Promoter | 2 | 29619910 | 29738327 | 118418 | 1 | ENSMUST00 | 0 | Rapgef1 | ENSMUSG00000039844.14 |
| chr4 | 98922058 | 98924971 | 37.63617 | 126.94061 | 122.55849 | Promoter | 4 | 98923810 | 98935543 | 11734 | 1 | ENSMUST00 | 0 | Usp1 | ENSMUSG00000028560.6 |
| chr6 | 8508635 | 8513604 | 37.63617 | 126.94061 | 122.55849 | Promoter | 6 | 8509600 | 8597548 | 87949 | 1 | ENSMUST00 | 0 | Glcci1 | ENSMUSG00000029638.12 |
| chr8 | 83331567 | 83333039 | 37.63617 | 126.94061 | 122.55849 | Promoter | 8 | 83312632 | 83332486 | 19855 | 2 | ENSMUST00 | 0 | Elmod2 | ENSMUSG00000035151.7 |
| chr18 | 31633629 | 31635801 | 38.86769 | 126.9345 | 122.55549 | Promoter | 18 | 31634371 | 31723061 | 88691 | 1 | ENSMUST00 | 0 | Sap130 | ENSMUSG00000024260.8 |
| chr4 | 139379645 | 139381792 | 38.86769 | 126.9345 | 122.55549 | Promoter | 4 | 139380659 | 139489530 | 108872 | 1 | ENSMUST00 | 0 | Ubr4 | ENSMUSG00000066036.9 |
| chr9 | 20652043 | 20652744 | 38.86769 | 126.9345 | 122.55549 | Promoter | 9 | 20652130 | 20666584 | 14455 | 1 | ENSMUST00 | 0 | Pin1 | ENSMUSG00000032171.6 |
| chr10 | 80264951 | 80266229 | 38.23518 | 126.91195 | 122.53338 | Promoter | 10 | 80264984 | 80288408 | 23425 | 1 | ENSMUST00 | 0 | Dazap1 | ENSMUSG00000069565.7 |
| chr19 | 46152470 | 46153499 | 38.23518 | 126.91195 | 122.53338 | Promoter | 19 | 46152509 | 46286510 | 134002 | 1 | ENSMUST00 | 0 | Gbf1 | ENSMUSG00000025224.9 |
| chr2 | 140066120 | 140067385 | 36.39777 | 126.8978 | 122.51947 | Promoter | 2 | 139833480 | 140066760 | 233281 | 2 | ENSMUST00 | 0 | Tasp1 | ENSMUSG00000039033.6 |
| chr3 | 88685742 | 88687601 | 36.39777 | 126.8978 | 122.51947 | Promoter | 3 | 88685794 | 88712933 | 27140 | 1 | ENSMUST00 | 0 | 2810403A07 | ENSMUSG00000028060.10 |
| chr1 | 131527827 | 131528977 | 37.56511 | 126.8153 | 122.43879 | Promoter | 1 | 131527903 | 131539872 | 11970 | 1 | ENSMUST00 | 0 | Fam72a | ENSMUSG00000055184.4 |
| chr5 | 3801956 | 3804579 | 38.79021 | 126.79964 | 122.42325 | Promoter | 5 | 3803165 | 3844515 | 41351 | 1 | ENSMUST00 | 0 | Krit1 | ENSMUSG00000000600.10 |
| chr4 | 115784604 | 115785862 | 38.16103 | 126.78209 | 122.40619 | Promoter | 4 | 115784812 | 115798006 | 13195 | 1 | ENSMUST00 | 0 | Atpaf1 | ENSMUSG00000028710.11 |
| chr2 | 94009838 | 94011480 | 39.37477 | 126.73389 | 122.35824 | Promoter | 2 | 93980550 | 94010807 | 30258 | 2 | ENSMUST00 | 0 | Alkbh3 | ENSMUSG00000040174.9 |
| chr3 | 41081402 | 41083110 | 39.37477 | 126.73389 | 122.35824 | Promoter | 3 | 41066326 | 41083046 | 16721 | 2 | ENSMUST00 | 0 | Pgrmc2 | ENSMUSG00000049940.7 |
| chr4 | 149166032 | 149167740 | 39.37477 | 126.73389 | 122.35824 | Promoter | 4 | 149149991 | 149166707 | 16717 | 2 | ENSMUST00 | 0 | Pgd | ENSMUSG00000028961.10 |
| chr5 | 28071277 | 28072315 | 39.37477 | 126.73389 | 122.35824 | Promoter | 5 | 28071363 | 28078662 | 7300 | 1 | ENSMUST00 | 0 | Insig1 | ENSMUSG00000045294.10 |
| chr12 | 65073017 | 65076367 | 39.97929 | 126.69925 | 122.32521 | Promoter | 12 | 65062436 | 65073944 | 11509 | 2 | ENSMUST00 | 0 | Fkbp3 | ENSMUSG00000020949.8 |
| chr18 | 42511414 | 42512593 | 39.97929 | 126.69925 | 122.32521 | Promoter | 18 | 42511510 | 42575551 | 64042 | 1 | ENSMUST00 | 0 | Tcerg1 | ENSMUSG00000024498.10 |
| chr5 | 124551849 | 124553504 | 39.97929 | 126.69925 | 122.32521 | Promoter | 5 | 124552864 | 124569660 | 16797 | 1 | ENSMUST00 | 0 | Ddx55 | ENSMUSG00000029389.12 |
| chr6 | 124915360 | 124918752 | 39.97929 | 126.69925 | 122.32521 | Promoter | 6 | 124913681 | 124917946 | 4266 | 2 | ENSMUST00 | 0 | Ptms | ENSMUSG00000030122.10 |

| | | | | | | | | | | | | | | | |
|---|---|---|---|---|---|---|---|---|---|---|---|---|---|---|---|
| chr2 | 138256276 | 138258341 | 37.49432 | 126.6904 | 122.3202 | Promoter | 2 | 138256565 | 138287422 | 30858 | 1 | ENSMUST00 | 0 | Btbd3 | ENSMUSG00000062098.6 |
| chr4 | 155942945 | 155944933 | 38.71304 | 126.66526 | 122.2951 | Promoter | 4 | 155943831 | 155957711 | 13881 | 1 | ENSMUST00 | 0 | Ube2j2 | ENSMUSG00000023286.11 |
| chr8 | 106892093 | 106894289 | 38.08716 | 126.65268 | 122.28292 | Promoter | 8 | 106893640 | 106923093 | 29454 | 1 | ENSMUST00 | 0 | Cirh1a | ENSMUSG00000041438.7 |
| chr1 | 176813017 | 176815457 | 33.32866 | 126.59486 | 122.2257 | Promoter | 1 | 176814660 | 176845497 | 30838 | 1 | ENSMUST00 | 0 | Sdccag8 | ENSMUSG00000026504.12 |
| chr1 | 160791788 | 160793644 | 39.29435 | 126.59459 | 122.2257 | Promoter | 1 | 160256572 | 160792938 | 536367 | 2 | ENSMUST00 | 0 | Rabgap1l | ENSMUSG00000026721.10 |
| chr13 | 23620451 | 23623409 | 33.32866 | 126.59486 | 122.2257 | Promoter | 13 | 23583742 | 23621094 | 37353 | 2 | ENSMUST00 | 0 | Hist1h2be | ENSMUSG00000047246.6 |
| chr13 | 108043523 | 108045394 | 33.32866 | 126.59486 | 122.2257 | Distal Interge | 13 | 108046424 | 108049146 | 2723 | 1 | ENSMUST00 | -1030 | Smim15 | ENSMUSG00000071180.4 |
| chr19 | 55741009 | 55745168 | 33.32866 | 126.59486 | 122.2257 | Promoter | 19 | 55741810 | 55933654 | 191845 | 1 | ENSMUST00 | 0 | Tcf7l2 | ENSMUSG00000024985.13 |
| chr4 | 150236391 | 150238373 | 33.32866 | 126.59486 | 122.2257 | Promoter | 4 | 150236721 | 150244289 | 7569 | 1 | ENSMUST00 | 0 | Eno1 | ENSMUSG00000063524.8 |
| chr5 | 122391528 | 122392929 | 33.32866 | 126.59486 | 122.2257 | Promoter | 5 | 122391878 | 122406179 | 14302 | 1 | ENSMUST00 | 0 | Arpc3 | ENSMUSG00000029465.9 |
| chr6 | 85373844 | 85375332 | 33.32866 | 126.59486 | 122.2257 | Promoter | 6 | 85334963 | 85374634 | 39672 | 2 | ENSMUST00 | 0 | Rab11fip5 | ENSMUSG00000051343.9 |
| chr7 | 3645204 | 3646176 | 33.32866 | 126.59486 | 122.2257 | Promoter | 7 | 3645290 | 3661108 | 15819 | 1 | ENSMUST00 | 0 | Cnot3 | ENSMUSG00000035632.11 |
| chr12 | 79089061 | 79090014 | 39.90739 | 126.57519 | 122.21001 | Promoter | 12 | 79080673 | 79089670 | 8998 | 2 | ENSMUST00 | 0 | Pigh | ENSMUSG00000021120.10 |
| chr14 | 18237831 | 18239850 | 37.42379 | 126.5659 | 122.20106 | Promoter | 14 | 18204054 | 18239127 | 35074 | 2 | ENSMUST00 | 0 | Nr1d2 | ENSMUSG00000021775.10 |
| chr1 | 37996353 | 37999132 | 38.63617 | 126.53137 | 122.16663 | Promoter | 1 | 37998010 | 38055579 | 57570 | 1 | ENSMUST00 | 0 | Eif5b | ENSMUSG00000026083.7 |
| chr11 | 23255903 | 23258180 | 38.01358 | 126.5237 | 122.15934 | Promoter | 11 | 23256041 | 23297597 | 41557 | 1 | ENSMUST00 | 0 | Xpo1 | ENSMUSG00000020290.9 |
| chr11 | 72689595 | 72691265 | 37.94029 | 126.39516 | 122.03232 | Promoter | 11 | 72690006 | 72725180 | 35175 | 1 | ENSMUST00 | 0 | Ankfy1 | ENSMUSG00000020790.9 |
| chr13 | 55599565 | 55600944 | 37.94029 | 126.39516 | 122.03232 | Promoter | 13 | 55599896 | 55610443 | 10548 | 1 | ENSMUST00 | 0 | B4galt7 | ENSMUSG00000021504.9 |
| chr4 | 43380841 | 43383763 | 37.94029 | 126.39516 | 122.03232 | Promoter | 4 | 43381979 | 43427088 | 45110 | 1 | ENSMUST00 | 0 | Rusc2 | ENSMUSG00000035969.10 |
| chr3 | 130729348 | 130730993 | 35.51756 | 126.38514 | 122.02445 | Promoter | 3 | 130726831 | 130730337 | 3507 | 2 | ENSMUST00 | 0 | Rpl34 | ENSMUSG00000062006.7 |
| chr7 | 119719932 | 119721266 | 35.51756 | 126.38514 | 122.02445 | Promoter | 7 | 119715096 | 119720765 | 5670 | 2 | ENSMUST00 | 0 | Thumpd1 | ENSMUSG00000030942.7 |
| chr10 | 85916118 | 85917629 | 39.1345 | 126.3175 | 121.95772 | Promoter | 10 | 85891970 | 85916945 | 24976 | 2 | ENSMUST00 | 0 | Prdm4 | ENSMUSG00000035529.9 |
| chr5 | 3646616 | 3648093 | 39.1345 | 126.3175 | 121.95772 | Promoter | 5 | 3639961 | 3647934 | 7974 | 2 | ENSMUST00 | 0 | Gatad1 | ENSMUSG00000007415.10 |
| chr15 | 43281778 | 43283136 | 33.73597 | 126.30199 | 121.9427 | Promoter | 15 | 43250040 | 43282736 | 32697 | 2 | ENSMUST00 | 0 | Eif3e | ENSMUSG00000022336.2 |
| chr9 | 66945369 | 66946939 | 33.73597 | 126.30199 | 121.9427 | Promoter | 9 | 66946086 | 66949512 | 3427 | 1 | ENSMUST00 | 0 | Rps27l | ENSMUSG00000036781.8 |
| chr10 | 40348278 | 40350333 | 36.66269 | 126.29414 | 121.93508 | Promoter | 10 | 40349308 | 40483818 | 134511 | 1 | ENSMUST00 | 0 | Cdk19 | ENSMUSG00000038481.7 |
| chr11 | 72795110 | 72797378 | 36.66269 | 126.29414 | 121.93508 | Promoter | 11 | 72796226 | 72851380 | 55155 | 1 | ENSMUST00 | 0 | Zzef1 | ENSMUSG00000055670.8 |
| chr13 | 55444309 | 55446120 | 36.66269 | 126.29414 | 121.93508 | Promoter | 13 | 55445334 | 55459577 | 14244 | 1 | ENSMUST00 | 0 | Grk6 | ENSMUSG00000074886.5 |
| chr4 | 141077406 | 141078403 | 36.66269 | 126.29414 | 121.93508 | Promoter | 4 | 141066512 | 141078342 | 11831 | 2 | ENSMUST00 | 0 | Necap2 | ENSMUSG00000028923.9 |
| chr5 | 104046126 | 104048099 | 36.66269 | 126.29414 | 121.93508 | Promoter | 5 | 104046306 | 104058202 | 11897 | 1 | ENSMUST00 | 0 | Nudt9 | ENSMUSG00000029310.8 |
| chr6 | 115807458 | 115808995 | 36.66269 | 126.29414 | 121.93508 | Promoter | 6 | 115805505 | 115808747 | 3243 | 2 | ENSMUST00 | 0 | Rpl32 | ENSMUSG00000057841.3 |
| chr8 | 35494364 | 35496178 | 36.66269 | 126.29414 | 121.93508 | Promoter | 8 | 35465265 | 35495533 | 30269 | 2 | ENSMUST00 | 0 | Eri1 | ENSMUSG00000031527.6 |
| chr14 | 59597024 | 59598554 | 39.74063 | 126.28735 | 121.93008 | Promoter | 14 | 59560896 | 59597836 | 36941 | 2 | ENSMUST00 | 0 | Cdadc1 | ENSMUSG00000021982.8 |
| chr19 | 6940978 | 6943193 | 39.74063 | 126.28735 | 121.93008 | Promoter | 19 | 6938069 | 6941507 | 3439 | 2 | ENSMUST00 | 0 | Gpr137 | ENSMUSG00000024958.7 |
| chr9 | 35199293 | 35200483 | 39.74063 | 126.28735 | 121.93008 | Promoter | 9 | 35184551 | 35200291 | 15741 | 2 | ENSMUST00 | 0 | Tirap | ENSMUSG00000032041.10 |
| chr2 | 137114213 | 137117889 | 37.86728 | 126.26704 | 121.91066 | Promoter | 2 | 137081456 | 137116644 | 35189 | 2 | ENSMUST00 | 0 | Jag1 | ENSMUSG00000027276.7 |
| chr4 | 124802003 | 124803469 | 38.48335 | 126.265 | 121.90878 | Promoter | 4 | 124802104 | 124849379 | 47276 | 1 | ENSMUST00 | 0 | Mtf1 | ENSMUSG00000028890.8 |
| chr9 | 59035272 | 59037558 | 38.48335 | 126.265 | 121.90878 | Promoter | 9 | 58874679 | 59036441 | 161763 | 2 | ENSMUST00 | 0 | Neo1 | ENSMUSG00000032340.7 |
| chr15 | 82353227 | 82354298 | 27.16573 | 126.25231 | 121.89658 | Promoter | 15 | 82350139 | 82354291 | 4153 | 2 | ENSMUST00 | 0 | Ndufa6 | ENSMUSG00000022450.5 |
| chr1 | 156299255 | 156304171 | 39.05507 | 126.17973 | 121.82412 | Promoter | 1 | 156255296 | 156303664 | 48369 | 2 | ENSMUST00 | 0 | Tdrd5 | ENSMUSG00000060985.10 |
| chr11 | 120548775 | 120550285 | 39.05507 | 126.17973 | 121.82412 | Promoter | 11 | 120542888 | 120549701 | 6814 | 2 | ENSMUST00 | 0 | Fam195b | ENSMUSG00000061111.8 |
| chr11 | 45851500 | 45852960 | 39.65778 | 126.14424 | 121.78909 | Promoter | 11 | 45852051 | 45910625 | 58575 | 1 | ENSMUST00 | 0 | Clint1 | ENSMUSG00000006169.14 |
| chr6 | 89361182 | 89362641 | 38.97595 | 126.04247 | 121.68831 | Promoter | 6 | 89316316 | 89362613 | 46298 | 2 | ENSMUST00 | 0 | Plxna1 | ENSMUSG00000030084.6 |
| chr2 | 27425224 | 27427512 | 33.03282 | 126.03245 | 121.67878 | Promoter | 2 | 27262104 | 27426912 | 164809 | 2 | ENSMUST00 | 0 | Vav2 | ENSMUSG00000009621.13 |
| chr4 | 34549956 | 34552534 | 33.03282 | 126.03245 | 121.67878 | Promoter | 4 | 34550937 | 34566908 | 15972 | 1 | ENSMUST00 | 0 | Akirin2 | ENSMUSG00000028291.7 |
| chr5 | 105698985 | 105701609 | 37.72209 | 126.01213 | 121.65854 | Promoter | 5 | 105699969 | 105815215 | 115247 | 1 | ENSMUST00 | 0 | Lrrc8d | ENSMUSG00000046079.11 |
| chr1 | 183344773 | 183346260 | 37.7199 | 126.00829 | 121.65485 | Promoter | 1 | 183326725 | 183345296 | 18572 | 2 | ENSMUST00 | 0 | Mia3 | ENSMUSG00000056050.7 |
| chr10 | 31608164 | 31610207 | 37.7199 | 126.00829 | 121.65485 | Promoter | 10 | 31493193 | 31609184 | 115992 | 2 | ENSMUST00 | 0 | Rnf217 | ENSMUSG00000063760.9 |
| chr11 | 87986201 | 87988020 | 37.7199 | 126.00829 | 121.65485 | Promoter | 11 | 87979525 | 87986585 | 7061 | 2 | ENSMUST00 | 0 | Dynll2 | ENSMUSG00000020483.9 |
| chr12 | 73964146 | 73965410 | 37.7199 | 126.00829 | 121.65485 | Promoter | 12 | 73964530 | 73984820 | 20291 | 1 | ENSMUST00 | 0 | Snapc1 | ENSMUSG00000021113.4 |
| chr12 | 69680563 | 69681199 | 37.7199 | 126.00829 | 121.65485 | Promoter | 12 | 69583766 | 69681852 | 98087 | 2 | ENSMUST00 | 0 | Sos2 | ENSMUSG00000034801.8 |
| chr12 | 54202081 | 54203966 | 37.7199 | 126.00829 | 121.65485 | Promoter | 12 | 54178981 | 54203860 | 24880 | 2 | ENSMUST00 | 0 | Egln3 | ENSMUSG00000035105.5 |
| chr13 | 74316462 | 74317945 | 37.7199 | 126.00829 | 121.65485 | Promoter | 13 | 74303121 | 74317326 | 14206 | 2 | ENSMUST00 | 0 | Pdcd6 | ENSMUSG00000021576.5 |
| chr13 | 100650333 | 100652423 | 37.7199 | 126.00829 | 121.65485 | Promoter | 13 | 100617165 | 100651059 | 33895 | 2 | ENSMUST00 | 0 | Rad17 | ENSMUSG00000021635.8 |
| chr13 | 95324447 | 95327308 | 37.7199 | 126.00829 | 121.65485 | Promoter | 13 | 95325237 | 95337841 | 12605 | 1 | ENSMUST00 | 0 | Zbed3 | ENSMUSG00000041995.7 |
| chr14 | 70442608 | 70444060 | 37.7199 | 126.00829 | 121.65485 | Promoter | 14 | 70438748 | 70443227 | 4480 | 2 | ENSMUST00 | 0 | Polr3d | ENSMUSG00000000776.7 |
| chr14 | 60265179 | 60266233 | 37.7199 | 126.00829 | 121.65485 | Promoter | 14 | 60265205 | 60302370 | 37166 | 1 | ENSMUST00 | 0 | Mtmr6 | ENSMUSG00000021987.8 |
| chr14 | 49171516 | 49174414 | 37.7199 | 126.00829 | 121.65485 | Promoter | 14 | 49172226 | 49190866 | 18641 | 1 | ENSMUST00 | 0 | Naa30 | ENSMUSG00000036282.9 |
| chr16 | 10991534 | 10993175 | 37.7199 | 126.00829 | 121.65485 | Promoter | 16 | 10959275 | 10993121 | 33847 | 2 | ENSMUST00 | 0 | Litaf | ENSMUSG00000022500.9 |
| chr2 | 37442474 | 37444150 | 37.7199 | 126.00829 | 121.65485 | Promoter | 2 | 37443279 | 37495589 | 52311 | 1 | ENSMUST00 | 0 | Rabgap1 | ENSMUSG00000035437.10 |
| chr2 | 32150787 | 32152576 | 37.7199 | 126.00829 | 121.65485 | Promoter | 2 | 32151082 | 32234537 | 83456 | 1 | ENSMUST00 | 0 | Prrc2b | ENSMUSG00000039262.11 |
| chr2 | 167630879 | 167632622 | 37.7199 | 126.00829 | 121.65485 | Promoter | 2 | 167607638 | 167632095 | 24458 | 2 | ENSMUST00 | 0 | Ube2v1 | ENSMUSG00000078923.5 |
| chr4 | 154159759 | 154161545 | 37.7199 | 126.00829 | 121.65485 | Promoter | 4 | 154157485 | 154160666 | 3182 | 2 | ENSMUST00 | 0 | Tprgl | ENSMUSG00000029030.9 |
| chr5 | 108268150 | 108270837 | 37.7199 | 126.00829 | 121.65485 | Promoter | 5 | 108268897 | 108280526 | 11630 | 1 | ENSMUST00 | 0 | Dr1 | ENSMUSG00000029265.4 |
| chr5 | 100036831 | 100040800 | 37.7199 | 126.00829 | 121.65485 | Promoter | 5 | 100039994 | 100068762 | 28769 | 1 | ENSMUST00 | 0 | Enoph1 | ENSMUSG00000029326.8 |
| chr5 | 134637596 | 134639924 | 37.7199 | 126.00829 | 121.65485 | Promoter | 5 | 134619880 | 134639348 | 19469 | 2 | ENSMUST00 | 0 | Eif4h | ENSMUSG00000040731.10 |
| chr6 | 91683631 | 91686902 | 37.7199 | 126.00829 | 121.65485 | Promoter | 6 | 91684067 | 91759063 | 74997 | 1 | ENSMUST00 | 0 | Slc6a6 | ENSMUSG00000030096.7 |
| chr8 | 41416158 | 41417536 | 37.7199 | 126.00829 | 121.65485 | Promoter | 8 | 41397457 | 41417118 | 19662 | 2 | ENSMUST00 | 0 | Frg1 | ENSMUSG00000031590.7 |
| chr9 | 44766600 | 44768050 | 37.7199 | 126.00829 | 121.65485 | Promoter | 9 | 44767908 | 44782233 | 14326 | 1 | ENSMUST00 | 0 | Ift46 | ENSMUSG00000002031.6 |
| chr9 | 44318922 | 44320981 | 39.57526 | 126.00172 | 121.65485 | Promoter | 9 | 44309237 | 44320282 | 11046 | 2 | ENSMUST00 | 0 | C2cd2l | ENSMUSG00000032120.10 |
| chr11 | 80079550 | 80081430 | 31.38972 | 125.99339 | 121.64841 | Promoter | 11 | 80046493 | 80080938 | 34446 | 2 | ENSMUST00 | 0 | Crlf3 | ENSMUSG00000017561.11 |
| chr12 | 86241512 | 86242916 | 31.38972 | 125.99339 | 121.64841 | Promoter | 12 | 86241890 | 86291414 | 49525 | 1 | ENSMUST00 | 0 | Gpatch2l | ENSMUSG00000021254.8 |
| chr14 | 65836958 | 65838434 | 31.38972 | 125.99339 | 121.64841 | Promoter | 14 | 65837302 | 65866604 | 29303 | 1 | ENSMUST00 | 0 | Ccdc25 | ENSMUSG00000022035.5 |
| chr19 | 18630087 | 18632373 | 31.38972 | 125.99339 | 121.64841 | Promoter | 19 | 18579328 | 18631789 | 52462 | 2 | ENSMUST00 | 0 | Ostf1 | ENSMUSG00000024725.7 |
| chr19 | 3767123 | 3769600 | 24.98033 | 125.99245 | 121.64841 | Promoter | 19 | 3767421 | 3808829 | 41409 | 1 | ENSMUST00 | 0 | Suv420h1 | ENSMUSG00000045098.13 |
| chr2 | 83643993 | 83645302 | 31.38972 | 125.99339 | 121.64841 | Promoter | 2 | 83644435 | 83664622 | 20188 | 1 | ENSMUST00 | 0 | Zc3h15 | ENSMUSG00000027091.9 |
| chr5 | 88720019 | 88722250 | 31.38972 | 125.99339 | 121.64841 | Promoter | 5 | 88720855 | 88747350 | 26496 | 1 | ENSMUST00 | 0 | Mob1b | ENSMUSG00000006262.10 |
| chr9 | 75556512 | 75559968 | 31.38972 | 125.99339 | 121.64841 | Promoter | 9 | 75497796 | 75559657 | 61862 | 2 | ENSMUST00 | 0 | Tmod3 | ENSMUSG00000058587.7 |
| chr1 | 179802281 | 179804474 | 35.29575 | 125.98199 | 121.63858 | Promoter | 1 | 179744894 | 179803680 | 58787 | 2 | ENSMUST00 | 0 | Ahctf1 | ENSMUSG00000026491.8 |
| chr1 | 78657393 | 78659137 | 35.29575 | 125.98199 | 121.63858 | Promoter | 1 | 78657825 | 78707743 | 49919 | 1 | ENSMUST00 | 0 | Acsl3 | ENSMUSG00000032883.10 |
| chr1 | 151755243 | 151756505 | 35.29575 | 125.98199 | 121.63858 | Promoter | 1 | 151755371 | 151818935 | 63565 | 1 | ENSMUST00 | 0 | Edem3 | ENSMUSG00000043019.7 |
| chr10 | 45066499 | 45067924 | 35.29575 | 125.98199 | 121.63858 | Promoter | 10 | 45067206 | 45158997 | 91792 | 1 | ENSMUST00 | 0 | Prep | ENSMUSG00000019849.10 |
| chr10 | 11148950 | 11150431 | 35.29575 | 125.98199 | 121.63858 | Promoter | 10 | 11149427 | 11188150 | 38724 | 1 | ENSMUST00 | 0 | Shprh | ENSMUSG00000090112.3 |
| chr11 | 88067386 | 88069722 | 35.29575 | 125.98199 | 121.63858 | Promoter | 11 | 88068279 | 88084729 | 16451 | 1 | ENSMUST00 | 0 | Vezf1 | ENSMUSG00000018377.5 |
| chr11 | 5787587 | 5789480 | 35.29575 | 125.98199 | 121.63858 | Promoter | 11 | 5788488 | 5800629 | 12142 | 1 | ENSMUST00 | 0 | Dbnl | ENSMUSG00000020476.9 |
| chr12 | 4475646 | 4477790 | 35.29575 | 125.98199 | 121.63858 | Promoter | 12 | 4247363 | 4477182 | 229820 | 2 | ENSMUST00 | 0 | Ncoa1 | ENSMUSG00000020647.9 |
| chr14 | 70765491 | 70767927 | 35.29575 | 125.98199 | 121.63858 | Promoter | 14 | 70654246 | 70766628 | 112383 | 2 | ENSMUST00 | 0 | Xpo7 | ENSMUSG00000022100.8 |
| chr14 | 52102822 | 52104066 | 35.29575 | 125.98199 | 121.63858 | Promoter | 14 | 52073380 | 52104028 | 30649 | 2 | ENSMUST00 | 0 | Hnrnpc | ENSMUSG00000060373.9 |
| chr15 | 25622489 | 25624944 | 35.29575 | 125.98199 | 121.63858 | Promoter | 15 | 25622525 | 25813673 | 191149 | 1 | ENSMUST00 | 0 | Myo10 | ENSMUSG00000022272.12 |
| chr17 | 85089519 | 85091531 | 35.29575 | 125.98199 | 121.63858 | Promoter | 17 | 85063477 | 85090267 | 26791 | 2 | ENSMUST00 | 0 | Prepl | ENSMUSG00000024127.9 |
| chr17 | 46772099 | 46773542 | 35.29575 | 125.98199 | 121.63858 | Promoter | 17 | 46772635 | 46773407 | 773 | 1 | ENSMUST00 | 0 | 2310039H08 | ENSMUSG00000062619.5 |
| chr18 | 77773316 | 77775134 | 35.29575 | 125.98199 | 121.63858 | Promoter | 18 | 77773729 | 77782869 | 9141 | 1 | ENSMUST00 | 0 | Atp5a1 | ENSMUSG00000025428.10 |
| chr18 | 84084710 | 84090178 | 35.29575 | 125.98199 | 121.63858 | Promoter | 18 | 84011627 | 84086404 | 74778 | 2 | ENSMUST00 | 0 | Tshz1 | ENSMUSG00000046982.10 |
| chr2 | 160361390 | 160367044 | 35.29575 | 125.98199 | 121.63858 | Promoter | 2 | 160363703 | 160367065 | 3363 | 2 | ENSMUST00 | 21 | Mafb | ENSMUSG00000074622.4 |
| chr3 | 9002549 | 9004984 | 35.29575 | 125.98199 | 121.63858 | Promoter | 3 | 8928626 | 9004478 | 75853 | 2 | ENSMUST00 | 0 | Tpd52 | ENSMUSG00000027506.10 |
| chr4 | 103213425 | 103215747 | 35.29575 | 125.98199 | 121.63858 | Promoter | 4 | 103170649 | 103215164 | 44516 | 2 | ENSMUST00 | 0 | Slc35d1 | ENSMUSG00000028521.12 |
| chr4 | 32656653 | 32658078 | 35.29575 | 125.98199 | 121.63858 | Promoter | 4 | 32657119 | 32775207 | 118089 | 1 | ENSMUST00 | 0 | Mdn1 | ENSMUSG00000058006.7 |
| chr9 | 109931060 | 109933251 | 35.29575 | 125.98199 | 121.63858 | Promoter | 9 | 109931460 | 110083947 | 152488 | 1 | ENSMUST00 | 0 | Map4 | ENSMUSG00000032479.10 |
| chr9 | 108560313 | 108561124 | 38.89715 | 125.90571 | 121.56978 | Promoter | 9 | 108560534 | 108565519 | 4986 | 1 | ENSMUST00 | 0 | Impdh2 | ENSMUSG00000062867.8 |
| chr1 | 172147589 | 172149840 | 28.36679 | 125.90208 | 121.5666 | Promoter | 1 | 172148015 | 172196393 | 48379 | 1 | ENSMUST00 | 0 | Dcaf8 | ENSMUSG00000026554.10 |
| chr11 | 70982268 | 70983069 | 37.64991 | 125.88531 | 121.55012 | Promoter | 11 | 70977836 | 70983026 | 5191 | 2 | ENSMUST00 | 0 | C1qbp | ENSMUSG00000018446.8 |
| chr13 | 110394346 | 110396681 | 37.64991 | 125.88531 | 121.55012 | Promoter | 13 | 110395044 | 110400844 | 5801 | 1 | ENSMUST00 | 0 | Plk2 | ENSMUSG00000021701.7 |
| chr17 | 28313508 | 28314368 | 38.25637 | 125.86897 | 121.53396 | Promoter | 17 | 28313530 | 28326567 | 13038 | 1 | ENSMUST00 | 0 | Fance | ENSMUSG00000007570.10 |
| chr17 | 24886605 | 24887644 | 38.25637 | 125.86897 | 121.53396 | Promoter | 17 | 24886643 | 24892149 | 5507 | 1 | ENSMUST00 | 0 | Spsb3 | ENSMUSG00000024160.11 |
| chr5 | 135149595 | 135150436 | 38.25636 | 125.86897 | 121.53396 | Promoter | 5 | 135149657 | 135163873 | 14217 | 1 | ENSMUST00 | 0 | Tbl2 | ENSMUSG00000005374.8 |
| chr3 | 89213228 | 89214738 | 39.4931 | 125.85976 | 121.52685 | Promoter | 3 | 89209081 | 89214430 | 5350 | 2 | ENSMUST00 | 0 | Mtx1 | ENSMUSG00000064068.8 |
| chr12 | 69371247 | 69375626 | 29.75837 | 125.79383 | 121.46131 | Promoter | 12 | 69372150 | 69375979 | 3830 | 1 | ENSMUST00 | 0 | Arf6 | ENSMUSG00000044147.7 |
| chr16 | 90282811 | 90285323 | 29.75837 | 125.79383 | 121.46131 | Promoter | 16 | 90228957 | 90284308 | 55352 | 2 | ENSMUST00 | 0 | Scaf4 | ENSMUSG00000022983.10 |
| chr19 | 42518305 | 42519945 | 29.75837 | 125.79383 | 121.46131 | Promoter | 19 | 42518759 | 42592343 | 73585 | 1 | ENSMUST00 | 0 | R3hcc1l | ENSMUSG00000025184.5 |
| chr3 | 152713711 | 152714806 | 29.75837 | 125.79383 | 121.46131 | Promoter | 3 | 152714100 | 152788898 | 74799 | 1 | ENSMUST00 | 0 | Pigk | ENSMUSG00000039047.12 |
| chr16 | 5202695 | 5204408 | 35.75768 | 125.74667 | 121.41521 | Promoter | 16 | 5195289 | 5204002 | 8714 | 2 | ENSMUST00 | 0 | Nagpa | ENSMUSG00000023143.5 |
| chr4 | 135971920 | 135973034 | 35.75768 | 125.74667 | 121.41521 | Promoter | 4 | 135968224 | 135971929 | 3706 | 2 | ENSMUST00 | 0 | Lypla2 | ENSMUSG00000028670.9 |
| chr9 | 31210441 | 31212694 | 35.75768 | 125.74667 | 121.41521 | Promoter | 9 | 31149557 | 31211815 | 62259 | 2 | ENSMUST00 | 0 | Aplp2 | ENSMUSG00000031996.11 |
| chr16 | 22856742 | 22858785 | 38.1813 | 125.73788 | 121.40766 | Promoter | 16 | 22857845 | 22872388 | 14544 | 1 | ENSMUST00 | 0 | Dnajb11 | ENSMUSG00000004460.11 |
| chr5 | 130221509 | 130223056 | 38.1813 | 125.73788 | 121.40766 | Promoter | 5 | 130222335 | 130243765 | 21431 | 1 | ENSMUST00 | 0 | Tmem248 | ENSMUSG00000053094.10 |
| chr13 | 38036426 | 38038278 | 39.41127 | 125.71832 | 121.38846 | Promoter | 13 | 38036989 | 38061433 | 24445 | 1 | ENSMUST00 | 0 | Riok1 | ENSMUSG00000021428.8 |
| chr7 | 127911928 | 127913051 | 39.41127 | 125.71832 | 121.38846 | Promoter | 7 | 127912517 | 127925832 | 13316 | 1 | ENSMUST00 | 0 | Kat8 | ENSMUSG00000030801.9 |
| chr11 | 62602203 | 62603985 | 36.93744 | 125.70555 | 121.37619 | Promoter | 11 | 62603079 | 62603150 | 72 | 1 | ENSMUST00 | 0 | Snord49b | ENSMUSG00000064437.1 |
| chr12 | 87199298 | 87200635 | 36.93744 | 125.70555 | 121.37619 | Promoter | 12 | 87166242 | 87200229 | 33988 | 2 | ENSMUST00 | 0 | Tmed8 | ENSMUSG00000034111.5 |
| chr14 | 74944710 | 74948380 | 36.93744 | 125.70555 | 121.37619 | Promoter | 14 | 74754674 | 74947877 | 193204 | 2 | ENSMUST00 | 0 | Lrch1 | ENSMUSG00000068015.6 |
| chr15 | 83594176 | 83595449 | 36.93744 | 125.70555 | 121.37619 | Promoter | 15 | 83575119 | 83595157 | 20039 | 2 | ENSMUST00 | 0 | Ttll12 | ENSMUSG00000016757.5 |
| chr4 | 24495942 | 24497269 | 36.93744 | 125.70555 | 121.37619 | Promoter | 4 | 24496451 | 24581338 | 84888 | 1 | ENSMUST00 | 0 | Mms22l | ENSMUSG00000045751.11 |
| chr4 | 45107082 | 45108494 | 36.93744 | 125.70555 | 121.37619 | Promoter | 4 | 45105208 | 45108114 | 2907 | 2 | ENSMUST00 | 0 | Tomm5 | ENSMUSG00000078713.3 |
| chr9 | 122922617 | 122923865 | 36.93744 | 125.70555 | 121.37619 | Promoter | 9 | 122923072 | 122931028 | 7957 | 1 | ENSMUST00 | 0 | Zfp105 | ENSMUSG00000057895.6 |
| chr11 | 70000485 | 70001538 | 38.74051 | 125.63367 | 121.30666 | Promoter | 11 | 70000595 | 70010111 | 9517 | 1 | ENSMUST00 | 0 | Dvl2 | ENSMUSG00000020888.11 |
| chr8 | 120666950 | 120669119 | 38.74051 | 125.63367 | 121.30666 | Promoter | 8 | 120668222 | 120674207 | 5986 | 1 | ENSMUST00 | 0 | Cox4i1 | ENSMUSG00000031818.7 |
| chr6 | 92090752 | 92093151 | 37.50639 | 125.63292 | 121.30651 | Promoter | 6 | 92091390 | 92173057 | 81668 | 1 | ENSMUST00 | 0 | Nr2c2 | ENSMUSG00000005893.9 |
| chr19 | 57609779 | 57612310 | 33.92788 | 125.59804 | 121.27206 | Promoter | 19 | 57611034 | 58133338 | 522305 | 1 | ENSMUST00 | 0 | Atrnl1 | ENSMUSG00000054843.8 |

| | | | | | | | | | | | | | | | |
|---|---|---|---|---|---|---|---|---|---|---|---|---|---|---|---|
| chr11 | 101225447 | 101226781 | 39.32978 | 125.57745 | 121.25191 | Promoter | 11 | 101191115 | 101226463 | 35349 | 2 | ENSMUST00 | 0 | Ezh1 | ENSMUSG00000006920.9 |
| chr10 | 81704406 | 81705593 | 31.16153 | 125.53995 | 121.21483 | Distal Interge | 10 | 81693718 | 81694185 | 468 | 2 | ENSMUST00 | -10222 | Gm8112 | ENSMUSG00000096623.1 |
| chr10 | 19014041 | 19016117 | 37.43503 | 125.50735 | 121.18225 | Promoter | 10 | 19000910 | 19015410 | 14501 | 2 | ENSMUST00 | 0 | Tnfaip3 | ENSMUSG00000019850.6 |
| chr14 | 77873889 | 77875294 | 38.66266 | 125.49842 | 121.17354 | Promoter | 14 | 77826217 | 77874917 | 48701 | 2 | ENSMUST00 | 0 | Dnajc15 | ENSMUSG00000022013.3 |
| chr9 | 64172434 | 64174358 | 36.8008 | 125.46325 | 121.13905 | Promoter | 9 | 64173387 | 64178562 | 5176 | 1 | ENSMUST00 | 0 | Rpl4 | ENSMUSG00000032399.7 |
| chr14 | 31640285 | 31641710 | 39.24862 | 125.4371 | 121.11293 | Promoter | 14 | 31552905 | 31640878 | 87974 | 2 | ENSMUST00 | 0 | Hacl1 | ENSMUSG00000021884.12 |
| chr2 | 164910073 | 164912468 | 39.24862 | 125.4371 | 121.11293 | Promoter | 2 | 164891882 | 164911757 | 19876 | 2 | ENSMUST00 | 0 | Zfp335 | ENSMUSG00000039834.12 |
| chr3 | 9249532 | 9253181 | 39.24862 | 125.4371 | 121.11293 | Promoter | 3 | 9250602 | 9285333 | 34732 | 1 | ENSMUST00 | 0 | Zbtb10 | ENSMUSG00000069114.8 |
| chr1 | 36444527 | 36445300 | 38.58512 | 125.36365 | 121.04008 | Promoter | 1 | 36422065 | 36445271 | 23207 | 2 | ENSMUST00 | 0 | Lman2l | ENSMUSG00000001143.8 |
| chr1 | 66700375 | 66701781 | 38.58512 | 125.36365 | 121.04008 | Promoter | 1 | 66700831 | 66715194 | 14364 | 1 | ENSMUST00 | 0 | Rpe | ENSMUSG00000026005.10 |
| chr13 | 67813698 | 67814919 | 38.58512 | 125.36365 | 121.04008 | Promoter | 13 | 67813740 | 67827002 | 13263 | 1 | ENSMUST00 | 0 | Zfp273 | ENSMUSG00000030446.12 |
| chr14 | 61438917 | 61440050 | 38.58512 | 125.36365 | 121.04008 | Promoter | 14 | 61365186 | 61439947 | 74762 | 2 | ENSMUST00 | 0 | Kpna3 | ENSMUSG00000021929.8 |
| chr17 | 78417090 | 78418392 | 38.58512 | 125.36365 | 121.04008 | Promoter | 17 | 78377885 | 78418131 | 40247 | 2 | ENSMUST00 | 0 | Fez2 | ENSMUSG00000056121.10 |
| chr6 | 113342483 | 113344245 | 38.58512 | 125.36365 | 121.04008 | Promoter | 6 | 113334124 | 113343984 | 9861 | 2 | ENSMUST00 | 0 | Camk1 | ENSMUSG00000030272.10 |
| chr13 | 93497758 | 93500424 | 37.95786 | 125.34732 | 121.02549 | Promoter | 13 | 93430097 | 93499808 | 69712 | 2 | ENSMUST00 | 0 | Jmy | ENSMUSG00000021690.9 |
| chr2 | 28839265 | 28841606 | 36.73285 | 125.34267 | 121.02122 | Promoter | 2 | 28840406 | 28847511 | 7106 | 1 | ENSMUST00 | 0 | Ddx31 | ENSMUSG00000026806.10 |
| chr10 | 117792041 | 117792773 | 33.20772 | 125.30651 | 120.98507 | Promoter | 10 | 117750624 | 117792698 | 42075 | 2 | ENSMUST00 | 0 | Nup107 | ENSMUSG00000052798.7 |
| chr10 | 127266048 | 127267362 | 34.91417 | 125.28625 | 120.96531 | Promoter | 10 | 127266262 | 127281959 | 15698 | 1 | ENSMUST00 | 0 | Dctn2 | ENSMUSG00000025410.9 |
| chr14 | 16363963 | 16366999 | 34.91417 | 125.28625 | 120.96531 | Promoter | 14 | 16365179 | 16430787 | 65609 | 1 | ENSMUST00 | 0 | Top2b | ENSMUSG00000017485.5 |
| chr3 | 51415428 | 51417724 | 34.91417 | 125.28625 | 120.96531 | Promoter | 3 | 51416016 | 51475985 | 59970 | 1 | ENSMUST00 | 0 | Naa15 | ENSMUSG00000063273.6 |
| chr1 | 128416488 | 128417732 | 25.88745 | 125.26566 | 120.94565 | Promoter | 1 | 128363707 | 128417359 | 53653 | 2 | ENSMUST00 | 0 | Dars | ENSMUSG00000026356.10 |
| chr17 | 26505964 | 26508522 | 39.08731 | 125.158 | 120.83892 | Promoter | 17 | 26505590 | 26508519 | 2930 | 2 | ENSMUST00 | 0 | Dusp1 | ENSMUSG00000024190.6 |
| chr2 | 167659265 | 167661579 | 39.08731 | 125.158 | 120.83892 | Promoter | 2 | 167607850 | 167661496 | 53647 | 2 | ENSMUST00 | 0 | Gm20431 | ENSMUSG00000089739.2 |
| chr5 | 29377672 | 29379069 | 39.08731 | 125.158 | 120.83892 | Promoter | 5 | 29229814 | 29378390 | 148577 | 2 | ENSMUST00 | 0 | Lmbr1 | ENSMUSG00000010721.10 |
| chr14 | 99045187 | 99047254 | 37.22259 | 125.13314 | 120.81451 | Promoter | 14 | 99046427 | 99074537 | 28111 | 1 | ENSMUST00 | 0 | Bora | ENSMUSG00000022070.6 |
| chr16 | 59600114 | 59601432 | 37.22259 | 125.13314 | 120.81451 | Promoter | 16 | 59558013 | 59600886 | 42874 | 2 | ENSMUST00 | 0 | Gm9581 | ENSMUSG00000091562.3 |
| chr17 | 80478875 | 80481064 | 37.22259 | 125.13314 | 120.81451 | Promoter | 17 | 80393752 | 80480453 | 86702 | 2 | ENSMUST00 | 0 | Sos1 | ENSMUSG00000024241.6 |
| chr18 | 36280325 | 36282308 | 37.22259 | 125.13314 | 120.81451 | Promoter | 18 | 36281097 | 36301521 | 20425 | 1 | ENSMUST00 | 0 | Pura | ENSMUSG00000043991.3 |
| chr3 | 90253884 | 90255447 | 37.22259 | 125.13314 | 120.81451 | Promoter | 3 | 90254163 | 90264125 | 9963 | 1 | ENSMUST00 | 0 | Crtc2 | ENSMUSG00000027936.13 |
| chr7 | 45015107 | 45016269 | 37.22259 | 125.13314 | 120.81451 | Promoter | 7 | 45010260 | 45016131 | 5872 | 2 | ENSMUST00 | 0 | Scaf1 | ENSMUSG00000038406.10 |
| chr8 | 18594764 | 18596012 | 37.22259 | 125.13314 | 120.81451 | Promoter | 8 | 18595131 | 18803189 | 208059 | 1 | ENSMUST00 | 0 | Mcph1 | ENSMUSG00000039842.10 |
| chr2 | 120729861 | 120732120 | 36.0064 | 125.12101 | 120.80552 | Promoter | 2 | 120716154 | 120731517 | 15364 | 2 | ENSMUST00 | 0 | Cdan1 | ENSMUSG00000027284.11 |
| chr7 | 105810243 | 105811081 | 36.0064 | 125.12101 | 120.80552 | Promoter | 7 | 105809657 | 105810958 | 1302 | 2 | ENSMUST00 | 0 | Mrpl17 | ENSMUSG00000030879.9 |
| chr4 | 115838693 | 115839945 | 38.43098 | 125.09557 | 120.78131 | Promoter | 4 | 115839198 | 115879250 | 40053 | 1 | ENSMUST00 | 0 | Mknk1 | ENSMUSG00000028708.11 |
| chr2 | 160645955 | 160647510 | 32.53129 | 125.07367 | 120.7602 | Promoter | 2 | 160645980 | 160650107 | 4128 | 1 | ENSMUST00 | 0 | Top1 | ENSMUSG00000070544.6 |
| chr12 | 70825459 | 70828098 | 39.00716 | 125.01923 | 120.70579 | Promoter | 12 | 70825514 | 70902232 | 76719 | 1 | ENSMUST00 | 0 | Frmd6 | ENSMUSG00000048285.8 |
| chr1 | 155097721 | 155099716 | 34.1261 | 124.90291 | 120.59063 | Promoter | 1 | 155096361 | 155099636 | 3276 | 2 | ENSMUST00 | 0 | Ier5 | ENSMUSG00000056708.5 |
| chr2 | 29934966 | 29936160 | 34.1261 | 124.90291 | 120.59063 | Promoter | 2 | 29935426 | 29960371 | 24946 | 1 | ENSMUST00 | 0 | Gle1 | ENSMUSG00000019715.9 |
| chr1 | 39535231 | 39536703 | 32.98857 | 124.89146 | 120.57959 | Promoter | 1 | 39535802 | 39546873 | 11072 | 1 | ENSMUST00 | 0 | Cnot11 | ENSMUSG00000003135.10 |
| chr1 | 177794901 | 177796574 | 32.98857 | 124.89146 | 120.57959 | Promoter | 1 | 177762962 | 177796511 | 33550 | 2 | ENSMUST00 | 0 | Adss | ENSMUSG00000015961.8 |
| chr10 | 88730430 | 88732564 | 32.98857 | 124.89146 | 120.57959 | Promoter | 10 | 88730858 | 88735911 | 5054 | 1 | ENSMUST00 | 0 | Arl1 | ENSMUSG00000060904.9 |
| chr11 | 40754366 | 40755681 | 32.98857 | 124.89146 | 120.57959 | Promoter | 11 | 40748552 | 40755311 | 6760 | 2 | ENSMUST00 | 0 | Ccng1 | ENSMUSG00000020326.7 |
| chr13 | 42051845 | 42053677 | 32.98857 | 124.89146 | 120.57959 | Promoter | 13 | 42052021 | 42185026 | 133006 | 1 | ENSMUST00 | 0 | Hivep1 | ENSMUSG00000021366.7 |
| chr14 | 21750151 | 21752138 | 32.98857 | 124.89146 | 120.57959 | Promoter | 14 | 21744874 | 21751181 | 6308 | 2 | ENSMUST00 | 0 | Dusp13 | ENSMUSG00000021768.10 |
| chr15 | 51989574 | 51991790 | 32.98857 | 124.89146 | 120.57959 | Promoter | 15 | 51962605 | 51991760 | 29156 | 2 | ENSMUST00 | 0 | Rad21 | ENSMUSG00000022314.9 |
| chr15 | 82015070 | 82016938 | 32.98857 | 124.89146 | 120.57959 | Promoter | 15 | 82015519 | 82040085 | 24567 | 1 | ENSMUST00 | 0 | Xrcc6 | ENSMUSG00000022471.8 |
| chr16 | 32430226 | 32431794 | 32.98857 | 124.89146 | 120.57959 | Promoter | 16 | 32430921 | 32474704 | 43784 | 1 | ENSMUST00 | 0 | Pcyt1a | ENSMUSG00000005615.9 |
| chr2 | 167502904 | 167504857 | 32.98857 | 124.89146 | 120.57959 | Promoter | 2 | 167503089 | 167516173 | 13085 | 1 | ENSMUST00 | 0 | Rnf114 | ENSMUSG00000006418.12 |
| chr2 | 37702046 | 37704173 | 32.98857 | 124.89146 | 120.57959 | Promoter | 2 | 37583156 | 37703275 | 120120 | 2 | ENSMUST00 | 0 | Strbp | ENSMUSG00000026915.11 |
| chr2 | 121505826 | 121507746 | 32.98857 | 124.89146 | 120.57959 | Promoter | 2 | 121506723 | 121543404 | 36682 | 1 | ENSMUST00 | 0 | Wdr76 | ENSMUSG00000027242.9 |
| chr4 | 74251001 | 74253232 | 32.98857 | 124.89146 | 120.57959 | Promoter | 4 | 74251936 | 74405860 | 153925 | 1 | ENSMUST00 | 0 | Kdm4c | ENSMUSG00000028397.8 |
| chr4 | 116720168 | 116722603 | 32.98857 | 124.89146 | 120.57959 | Promoter | 4 | 116720948 | 116804242 | 83295 | 1 | ENSMUST00 | 0 | Tesk2 | ENSMUSG00000033985.12 |
| chr4 | 151994455 | 151997533 | 32.98857 | 124.89146 | 120.57959 | Promoter | 4 | 151989633 | 151996258 | 6626 | 2 | ENSMUST00 | 0 | Phf13 | ENSMUSG00000047777.9 |
| chr5 | 30154081 | 30156417 | 32.98857 | 124.89146 | 120.57959 | Promoter | 5 | 30118301 | 30155162 | 36862 | 2 | ENSMUST00 | 0 | Hadha | ENSMUSG00000025745.6 |
| chr6 | 39419347 | 39420676 | 32.98857 | 124.89146 | 120.57959 | Promoter | 6 | 39397804 | 39419776 | 21973 | 2 | ENSMUST00 | 0 | Mkrn1 | ENSMUSG00000029922.10 |
| chr8 | 13972915 | 13975381 | 32.98857 | 124.89146 | 120.57959 | Promoter | 8 | 13952009 | 13974777 | 22769 | 2 | ENSMUST00 | 0 | Tdrp | ENSMUSG00000050052.6 |
| chr10 | 8517801 | 8519623 | 39.51976 | 124.83012 | 120.52397 | Promoter | 10 | 8204753 | 8518825 | 314073 | 2 | ENSMUST00 | 0 | Ust | ENSMUSG00000047712.7 |
| chr11 | 108342513 | 108344600 | 38.27806 | 124.82941 | 120.52397 | Promoter | 11 | 108343354 | 108395905 | 52552 | 1 | ENSMUST00 | 0 | Apoh | ENSMUSG00000000049.6 |
| chr11 | 61853941 | 61855671 | 39.51976 | 124.83012 | 120.52397 | Promoter | 11 | 61775649 | 61855073 | 79425 | 2 | ENSMUST00 | 0 | Ulk2 | ENSMUSG00000004798.9 |
| chr16 | 20672338 | 20674211 | 39.51976 | 124.83012 | 120.52397 | Promoter | 16 | 20672723 | 20692884 | 20162 | 1 | ENSMUST00 | 0 | Eif4g1 | ENSMUSG00000045983.10 |
| chr16 | 50528359 | 50530576 | 39.51976 | 124.83012 | 120.52397 | Promoter | 16 | 50528502 | 50558681 | 30180 | 1 | ENSMUST00 | 0 | G730013B05 | ENSMUSG00000097694.2 |
| chr17 | 43016512 | 43018370 | 39.51976 | 124.83012 | 120.52397 | Promoter | 17 | 43016555 | 43089188 | 72634 | 1 | ENSMUST00 | 0 | Tnfrsf21 | ENSMUSG00000023915.4 |
| chr17 | 46159755 | 46161963 | 39.51976 | 124.83012 | 120.52397 | Promoter | 17 | 46161032 | 46169368 | 8337 | 1 | ENSMUST00 | 0 | Gtpbp2 | ENSMUSG00000023952.8 |
| chr19 | 14595042 | 14598151 | 39.51976 | 124.83012 | 120.52397 | Promoter | 19 | 14448072 | 14597983 | 149912 | 2 | ENSMUST00 | 0 | Tle4 | ENSMUSG00000024642.11 |
| chr5 | 115340384 | 115342249 | 38.27806 | 124.82941 | 120.52397 | Promoter | 5 | 115341225 | 115343569 | 2345 | 1 | ENSMUST00 | 0 | Triap1 | ENSMUSG00000029535.4 |
| chr5 | 115348143 | 115349660 | 39.51976 | 124.83012 | 120.52397 | Promoter | 5 | 115345642 | 115348981 | 3340 | 2 | ENSMUST00 | 0 | Cox6a1 | ENSMUSG00000041697.8 |
| chr6 | 99665745 | 99667267 | 38.27806 | 124.82941 | 120.52397 | Promoter | 6 | 99257484 | 99666797 | 409314 | 2 | ENSMUST00 | 0 | Gm20696 | ENSMUSG00000030068.3 |
| chr7 | 24506592 | 24508247 | 39.51976 | 124.83012 | 120.52397 | Promoter | 7 | 24507006 | 24515419 | 8414 | 1 | ENSMUST00 | 0 | Zfp428 | ENSMUSG00000064264.9 |
| chr8 | 124520671 | 124521927 | 39.51976 | 124.83012 | 120.52397 | Promoter | 8 | 124520767 | 124552008 | 31242 | 1 | ENSMUST00 | 0 | Cog2 | ENSMUSG00000031979.12 |
| chr13 | 21179205 | 21181506 | 37.01254 | 124.7626 | 120.46037 | Promoter | 13 | 21179445 | 21180416 | 972 | 1 | ENSMUST00 | 0 | Trim27 | ENSMUSG00000021326.6 |
| chr17 | 5491522 | 5494381 | 31.29462 | 124.76181 | 120.45966 | Promoter | 17 | 5492600 | 5753891 | 261292 | 1 | ENSMUST00 | 0 | Zdhhc14 | ENSMUSG00000034265.7 |
| chr15 | 98144731 | 98146076 | 38.84783 | 124.74329 | 120.44118 | Promoter | 15 | 98135714 | 98145659 | 9946 | 2 | ENSMUST00 | 0 | Asb8 | ENSMUSG00000048175.8 |
| chr5 | 33273539 | 33275535 | 38.84783 | 124.74329 | 120.44118 | Promoter | 5 | 33247724 | 33275004 | 27281 | 2 | ENSMUST00 | 0 | Ctbp1 | ENSMUSG00000037373.9 |
| chr6 | 47654721 | 47655533 | 15.44916 | 124.7347 | 120.43309 | Distal Interge | 6 | 47637143 | 47637219 | 77 | 1 | ENSMUST00 | 17579 | Gm25283 | ENSMUSG00000087897.1 |
| chr15 | 89210792 | 89212473 | 39.44868 | 124.7075 | 120.4059 | Exon (ENSMU | 15 | 89211559 | 89215012 | 3454 | 2 | ENSMUST00 | 2539 | Gm26798 | ENSMUSG00000097578.1 |
| chr17 | 71182601 | 71187348 | 37.59121 | 124.70534 | 120.4041 | Promoter | 17 | 71183556 | 71232306 | 48751 | 1 | ENSMUST00 | 0 | Lpin2 | ENSMUSG00000024052.12 |
| chr11 | 70237200 | 70240661 | 38.20206 | 124.69703 | 120.39601 | Promoter | 11 | 70235206 | 70237853 | 2648 | 2 | ENSMUST00 | 0 | 0610010K14 | ENSMUSG00000020831.13 |
| chr15 | 81398585 | 81400981 | 38.20206 | 124.69703 | 120.39601 | Promoter | 15 | 81400138 | 81456807 | 56670 | 1 | ENSMUST00 | 0 | Xpnpep3 | ENSMUSG00000022401.6 |
| chr10 | 121739395 | 121740639 | 35.13972 | 124.62851 | 120.32814 | Promoter | 10 | 121739937 | 121752865 | 12929 | 1 | ENSMUST00 | 0 | BC048403 | ENSMUSG00000053684.8 |
| chr12 | 83687692 | 83689481 | 35.13972 | 124.62851 | 120.32814 | Promoter | 12 | 83688203 | 83734113 | 45911 | 1 | ENSMUST00 | 0 | Psen1 | ENSMUSG00000019969.8 |
| chr10 | 41449557 | 41451045 | 38.76865 | 124.60608 | 120.30663 | Promoter | 10 | 41450396 | 41465571 | 15176 | 1 | ENSMUST00 | 0 | Zbtb24 | ENSMUSG00000019826.10 |
| chr1 | 171287191 | 171288223 | 33.38818 | 124.5882 | 120.28912 | Promoter | 1 | 171281945 | 171287991 | 6047 | 2 | ENSMUST00 | 0 | Usp21 | ENSMUSG00000053483.8 |
| chr12 | 110737110 | 110739434 | 33.38818 | 124.5882 | 120.28912 | Promoter | 12 | 110737949 | 110795028 | 57080 | 1 | ENSMUST00 | 0 | Wdr20 | ENSMUSG00000037957.9 |
| chr4 | 3834635 | 3836092 | 33.38818 | 124.5882 | 120.28912 | Promoter | 4 | 3831334 | 3835665 | 4332 | 2 | ENSMUST00 | 0 | Rps20 | ENSMUSG00000028234.6 |
| chr11 | 80300381 | 80302828 | 37.51873 | 124.57824 | 120.27934 | Promoter | 11 | 80300912 | 80355955 | 55044 | 1 | ENSMUST00 | 0 | Rhbdl3 | ENSMUSG00000017692.8 |
| chr11 | 84879481 | 84881173 | 37.51873 | 124.57824 | 120.27934 | Promoter | 11 | 84880148 | 84911226 | 31079 | 1 | ENSMUST00 | 0 | Myo19 | ENSMUSG00000020527.9 |
| chr14 | 20706964 | 20709035 | 37.51873 | 124.57824 | 120.27934 | Promoter | 14 | 20707552 | 20723618 | 16067 | 1 | ENSMUST00 | 0 | Zswim8 | ENSMUSG00000021819.10 |
| chr2 | 157006297 | 157007564 | 37.51873 | 124.57824 | 120.27934 | Promoter | 2 | 156995265 | 157007075 | 11811 | 2 | ENSMUST00 | 0 | Dsn1 | ENSMUSG00000027635.10 |
| chr4 | 126428459 | 126430348 | 37.51873 | 124.57824 | 120.27934 | Promoter | 4 | 126332732 | 126429556 | 96825 | 2 | ENSMUST00 | 0 | Ago3 | ENSMUSG00000028842.10 |
| chr5 | 34335513 | 34337600 | 37.51873 | 124.57824 | 120.27934 | Promoter | 5 | 34336289 | 34353436 | 17148 | 1 | ENSMUST00 | 0 | Rnf4 | ENSMUSG00000029110.8 |
| chr7 | 79464733 | 79467421 | 37.51873 | 124.57824 | 120.27934 | Promoter | 7 | 79448791 | 79466362 | 17572 | 2 | ENSMUST00 | 0 | Polg | ENSMUSG00000039176.12 |
| chr17 | 24894213 | 24897261 | 38.12635 | 124.56513 | 120.26845 | Promoter | 17 | 24895116 | 24896289 | 1174 | 1 | ENSMUST00 | 0 | Mrps34 | ENSMUSG00000038880.8 |
| chr7 | 44996606 | 44999011 | 39.36609 | 124.56497 | 120.26845 | Promoter | 7 | 44997648 | 45002848 | 5201 | 1 | ENSMUST00 | 0 | Irf3 | ENSMUSG00000003184.9 |
| chr11 | 107479008 | 107480788 | 36.26418 | 124.50906 | 120.21295 | Promoter | 11 | 107479484 | 107498173 | 18690 | 1 | ENSMUST00 | 0 | Psmd12 | ENSMUSG00000020720.8 |
| chr11 | 121701795 | 121706690 | 36.26418 | 124.50906 | 120.21295 | Promoter | 11 | 121702411 | 121716306 | 13896 | 1 | ENSMUST00 | 0 | Metrnl | ENSMUSG00000039208.10 |
| chr11 | 98412075 | 98414297 | 36.26418 | 124.50906 | 120.21295 | Promoter | 11 | 98412470 | 98437716 | 25247 | 1 | ENSMUST00 | 0 | Erbb2 | ENSMUSG00000062312.5 |
| chr12 | 80944805 | 80947053 | 36.26418 | 124.50906 | 120.21295 | Promoter | 12 | 80945504 | 80950507 | 5004 | 1 | ENSMUST00 | 0 | Srsf5 | ENSMUSG00000021134.12 |
| chr3 | 108910150 | 108912387 | 36.26418 | 124.50906 | 120.21295 | Promoter | 3 | 108902667 | 108911696 | 9030 | 2 | ENSMUST00 | 0 | Prpf38b | ENSMUSG00000027881.9 |
| chr16 | 35154215 | 35157004 | 38.68979 | 124.4694 | 120.17514 | Promoter | 16 | 35154877 | 35305737 | 150861 | 1 | ENSMUST00 | 0 | Adcy5 | ENSMUSG00000022840.8 |
| chr10 | 59322839 | 59324728 | 37.44653 | 124.45157 | 120.15771 | Promoter | 10 | 59323296 | 59373304 | 50009 | 1 | ENSMUST00 | 0 | P4ha1 | ENSMUSG00000019916.9 |
| chr13 | 12460437 | 12462111 | 39.28384 | 124.42301 | 120.12974 | Promoter | 13 | 12439417 | 12461738 | 22322 | 2 | ENSMUST00 | 0 | Lgals8 | ENSMUSG00000057554.8 |
| chr18 | 80255462 | 80257280 | 39.28384 | 124.42301 | 120.12974 | Promoter | 18 | 80255836 | 80284071 | 28236 | 1 | ENSMUST00 | 0 | Pqlc1 | ENSMUSG00000034006.11 |
| chr3 | 5575250 | 5576976 | 39.28384 | 124.42301 | 120.12974 | Promoter | 3 | 5560188 | 5576150 | 15963 | 2 | ENSMUST00 | 0 | Pex2 | ENSMUSG00000040374.8 |
| chr4 | 32982548 | 32984479 | 39.28384 | 124.42301 | 120.12974 | Promoter | 4 | 32983037 | 33022180 | 39144 | 1 | ENSMUST00 | 0 | Rragd | ENSMUSG00000028278.9 |
| chr11 | 71018459 | 71019368 | 31.07893 | 124.33488 | 120.0426 | Promoter | 11 | 71007164 | 71019242 | 12079 | 2 | ENSMUST00 | 0 | Derl2 | ENSMUSG00000018442.8 |
| chr12 | 101081809 | 101084598 | 31.07893 | 124.33488 | 120.0426 | Promoter | 12 | 101039409 | 101083702 | 44294 | 2 | ENSMUST00 | 0 | Smek1 | ENSMUSG00000041846.9 |
| chr13 | 93771425 | 93772881 | 31.07893 | 124.33488 | 120.0426 | Promoter | 13 | 93771679 | 93943016 | 171338 | 1 | ENSMUST00 | 0 | Arsb | ENSMUSG00000042082.6 |
| chr14 | 20733546 | 20735008 | 31.07893 | 124.33488 | 120.0426 | 3' UTR | 14 | 20723739 | 20730327 | 6589 | 2 | ENSMUST00 | -3220 | Ndst2 | ENSMUSG00000039308.5 |
| chr2 | 34772049 | 34773448 | 31.07893 | 124.33488 | 120.0426 | Promoter | 2 | 34772092 | 34776531 | 4440 | 1 | ENSMUST00 | 0 | Hspa5 | ENSMUSG00000026864.8 |
| chr3 | 96696049 | 96697777 | 31.07893 | 124.33488 | 120.0426 | Promoter | 3 | 96696384 | 96705962 | 9579 | 1 | ENSMUST00 | 0 | Pias3 | ENSMUSG00000028101.13 |
| chr3 | 130708277 | 130709489 | 31.07893 | 124.33488 | 120.0426 | Promoter | 3 | 130695921 | 130709444 | 13524 | 2 | ENSMUST00 | 0 | Ostc | ENSMUSG00000041084.7 |
| chr3 | 90264901 | 90266211 | 38.61126 | 124.33321 | 120.0426 | Promoter | 3 | 90265185 | 90280669 | 15485 | 1 | ENSMUST00 | 0 | Dennd4b | ENSMUSG00000042404.11 |
| chr5 | 76974036 | 76975655 | 31.07893 | 124.33488 | 120.0426 | Promoter | 5 | 76974683 | 76999937 | 25255 | 1 | ENSMUST00 | 0 | Srp72 | ENSMUSG00000036323.9 |
| chr2 | 144009741 | 144011296 | 37.3746 | 124.32532 | 120.03539 | Promoter | 2 | 143947395 | 144011263 | 63869 | 2 | ENSMUST00 | 0 | Rrbp1 | ENSMUSG00000027422.10 |
| chr16 | 23106844 | 23109861 | 28.10168 | 124.31538 | 120.02569 | Promoter | 16 | 23107444 | 23113947 | 6504 | 1 | ENSMUST00 | 0 | Eif4a2 | ENSMUSG00000022884.9 |
| chr5 | 92504372 | 92506287 | 28.10168 | 124.31538 | 120.02569 | Promoter | 5 | 92443873 | 92505608 | 61736 | 2 | ENSMUST00 | 0 | Scarb2 | ENSMUSG00000029426.7 |
| chr5 | 23850105 | 23851328 | 28.10168 | 124.31538 | 120.02569 | Promoter | 5 | 23850597 | 23854924 | 4328 | 1 | ENSMUST00 | 0 | 2700038G22 | ENSMUSG00000097180.1 |
| chr2 | 71211651 | 71212884 | 39.20194 | 124.28158 | 119.99248 | Promoter | 2 | 71211706 | 71263298 | 51593 | 1 | ENSMUST00 | 0 | Dync1i2 | ENSMUSG00000027012.10 |
| chr1 | 59912260 | 59915684 | 34.92026 | 124.22972 | 119.94103 | Promoter | 1 | 59913006 | 59985346 | 72341 | 1 | ENSMUST00 | 0 | Fam117b | ENSMUSG00000041040.7 |
| chr10 | 53629220 | 53630057 | 34.92026 | 124.22972 | 119.94103 | Promoter | 10 | 53536329 | 53630439 | 94111 | 2 | ENSMUST00 | 382 | Mcm9 | ENSMUSG00000058298.7 |
| chr11 | 100968093 | 100971128 | 34.92026 | 124.22972 | 119.94103 | Promoter | 11 | 100956733 | 100970887 | 14155 | 2 | ENSMUST00 | 0 | Ptrf | ENSMUSG00000004044.9 |
| chr11 | 20631080 | 20633425 | 34.92026 | 124.22972 | 119.94103 | Promoter | 11 | 20631979 | 20653021 | 21043 | 1 | ENSMUST00 | 0 | Sertad2 | ENSMUSG00000049800.8 |
| chr11 | 101315586 | 101317002 | 34.92026 | 124.22972 | 119.94103 | Promoter | 11 | 101316213 | 101320008 | 3796 | 1 | ENSMUST00 | 0 | Psme3 | ENSMUSG00000078652.4 |
| chr12 | 28634869 | 28636511 | 34.92026 | 124.22972 | 119.94103 | Promoter | 12 | 28630847 | 28635953 | 5107 | 2 | ENSMUST00 | 0 | Rps7 | ENSMUSG00000061477.3 |
| chr13 | 94284370 | 94286213 | 34.92026 | 124.22972 | 119.94103 | Promoter | 13 | 94201310 | 94285829 | 84520 | 2 | ENSMUST00 | 0 | Scamp1 | ENSMUSG00000021687.9 |
| chr18 | 33794397 | 33795757 | 34.92026 | 124.22972 | 119.94103 | Promoter | 18 | 33794892 | 33795986 | 1095 | 1 | ENSMUST00 | 0 | 2410004N09 | ENSMUSG00000087590.2 |
| chr2 | 157278838 | 157280074 | 34.92026 | 124.22972 | 119.94103 | Promoter | 2 | 157279017 | 157289561 | 10545 | 1 | ENSMUST00 | 0 | Rpn2 | ENSMUSG00000027642.10 |
| chr3 | 94886069 | 94887318 | 34.92026 | 124.22972 | 119.94103 | Promoter | 3 | 94884093 | 94886961 | 2869 | 2 | ENSMUST00 | 0 | Psmb4 | ENSMUSG00000005779.7 |

| | | | | | | | | | | | | | | | |
|---|---|---|---|---|---|---|---|---|---|---|---|---|---|---|---|
| chr4 | 135920683 | 135922132 | 34.92026 | 124.22972 | 119.94103 | Promoter | 4 | 135920735 | 135940311 | 19577 | 1 | ENSMUST00 | 0 | Fuca1 | ENSMUSG00000028673.5 |
| chr4 | 151933681 | 151934863 | 34.92026 | 124.22972 | 119.94103 | Promoter | 4 | 151933691 | 151982137 | 48447 | 1 | ENSMUST00 | 0 | Dnajc11 | ENSMUSG00000039768.11 |
| chr5 | 110230299 | 110231981 | 34.92026 | 124.22972 | 119.94103 | Promoter | 5 | 110231004 | 110256651 | 25648 | 1 | ENSMUST00 | 0 | Ankle2 | ENSMUSG00000029501.9 |
| chr5 | 115506647 | 115510205 | 34.92026 | 124.22972 | 119.94103 | Promoter | 5 | 115506676 | 115555987 | 49312 | 1 | ENSMUST00 | 0 | Pxn | ENSMUSG00000029528.12 |
| chr7 | 80231843 | 80233760 | 34.92026 | 124.22972 | 119.94103 | Promoter | 7 | 80227156 | 80232813 | 5658 | 2 | ENSMUST00 | 0 | Cib1 | ENSMUSG00000030538.10 |
| chr1 | 172375815 | 172377599 | 34.33094 | 124.21722 | 119.93477 | Promoter | 1 | 172376531 | 172384099 | 7569 | 1 | ENSMUST00 | 0 | Pigm | ENSMUSG00000050229.2 |
| chr2 | 167491749 | 167493590 | 34.33094 | 124.21722 | 119.93477 | Promoter | 2 | 167492645 | 167516166 | 23522 | 1 | ENSMUST00 | 0 | Rnf114 | ENSMUSG00000006418.12 |
| chr6 | 136827277 | 136828231 | 34.33094 | 124.21722 | 119.93477 | Promoter | 6 | 136819728 | 136828143 | 8416 | 2 | ENSMUST00 | 0 | Wbp11 | ENSMUSG00000030216.10 |
| chr7 | 133636998 | 133638678 | 34.33094 | 124.21722 | 119.93477 | Promoter | 7 | 133637543 | 133672971 | 35429 | 1 | ENSMUST00 | 0 | Edrf1 | ENSMUSG00000039990.10 |
| chr11 | 69670578 | 69672426 | 37.30296 | 124.19952 | 119.9176 | Promoter | 11 | 69666936 | 69672317 | 5382 | 2 | ENSMUST00 | 0 | Eif4a1 | ENSMUSG00000059796.11 |
| chr9 | 52167458 | 52169267 | 37.30296 | 124.19952 | 119.9176 | Promoter | 9 | 52111985 | 52168111 | 56127 | 2 | ENSMUST00 | 0 | Zc3h12c | ENSMUSG00000035164.7 |
| chr10 | 121474826 | 121476326 | 37.30079 | 124.19571 | 119.91465 | Promoter | 10 | 121410350 | 121476250 | 65901 | 2 | ENSMUST00 | 0 | Rassf3 | ENSMUSG00000025795.7 |
| chr10 | 22814781 | 22820826 | 37.30079 | 124.19571 | 119.91465 | Promoter | 10 | 22817279 | 22820128 | 2850 | 2 | ENSMUST00 | 0 | Tcf21 | ENSMUSG00000045680.7 |
| chr11 | 29692865 | 29694637 | 37.30079 | 124.19571 | 119.91465 | Promoter | 11 | 29692947 | 29742916 | 49970 | 1 | ENSMUST00 | 0 | Rtn4 | ENSMUSG00000020458.11 |
| chr11 | 84828291 | 84829675 | 37.30079 | 124.19571 | 119.91465 | Promoter | 11 | 84820856 | 84828964 | 8109 | 2 | ENSMUST00 | 0 | Dhrs11 | ENSMUSG00000034449.7 |
| chr12 | 91848313 | 91849226 | 37.30079 | 124.19571 | 119.91465 | Promoter | 12 | 91806043 | 91849157 | 43115 | 2 | ENSMUST00 | 0 | Sel1l | ENSMUSG00000020964.9 |
| chr12 | 101717845 | 101719039 | 37.30079 | 124.19571 | 119.91465 | Promoter | 12 | 101645443 | 101718523 | 73081 | 2 | ENSMUST00 | 0 | Tc2n | ENSMUSG00000021187.9 |
| chr13 | 48966128 | 48969320 | 37.30079 | 124.19571 | 119.91465 | Promoter | 13 | 48879219 | 48968017 | 88799 | 2 | ENSMUST00 | 0 | Fam120a | ENSMUSG00000038014.7 |
| chr13 | 20089916 | 20091932 | 37.30079 | 124.19571 | 119.91465 | Promoter | 13 | 20090596 | 20264007 | 173412 | 1 | ENSMUST00 | 0 | Elmo1 | ENSMUSG00000041112.10 |
| chr14 | 27428244 | 27430149 | 37.30079 | 124.19571 | 119.91465 | Promoter | 14 | 27428847 | 27483552 | 54706 | 1 | ENSMUST00 | 0 | Fam208a | ENSMUSG00000040651.8 |
| chr15 | 97764504 | 97768671 | 37.30079 | 124.19571 | 119.91465 | Promoter | 15 | 97744770 | 97767846 | 23077 | 2 | ENSMUST00 | 0 | Rapgef3 | ENSMUSG00000022469.11 |
| chr15 | 99701792 | 99702839 | 37.30079 | 124.19571 | 119.91465 | Promoter | 15 | 99702287 | 99713995 | 11709 | 1 | ENSMUST00 | 0 | Smarcd1 | ENSMUSG00000023018.4 |
| chr17 | 28621359 | 28622908 | 37.30079 | 124.19571 | 119.91465 | Promoter | 17 | 28587648 | 28622521 | 34874 | 2 | ENSMUST00 | 0 | Srpk1 | ENSMUSG00000004865.10 |
| chr17 | 32829461 | 32830595 | 37.30079 | 124.19571 | 119.91465 | Distal Interge | 17 | 32815455 | 32822200 | 6746 | 2 | ENSMUST00 | -7262 | Zfp799 | ENSMUSG00000095253.1 |
| chr18 | 22342934 | 22345013 | 37.30079 | 124.19571 | 119.91465 | Promoter | 18 | 22344883 | 22378950 | 34068 | 1 | ENSMUST00 | 0 | Asxl3 | ENSMUSG00000045215.11 |
| chr3 | 51339237 | 51342406 | 37.30079 | 124.19571 | 119.91465 | Promoter | 3 | 51255665 | 51340566 | 84902 | 2 | ENSMUST00 | 0 | Elf2 | ENSMUSG00000037174.13 |
| chr4 | 138304724 | 138305652 | 37.30079 | 124.19571 | 119.91465 | Promoter | 4 | 138304730 | 138312628 | 7899 | 1 | ENSMUST00 | 0 | Ddost | ENSMUSG00000028757.4 |
| chr5 | 127616053 | 127617842 | 37.30079 | 124.19571 | 119.91465 | Promoter | 5 | 127595664 | 127617416 | 21753 | 2 | ENSMUST00 | 0 | Slc15a4 | ENSMUSG00000029416.12 |
| chr5 | 143234870 | 143236127 | 37.30079 | 124.19571 | 119.91465 | Promoter | 5 | 143235163 | 143247948 | 12786 | 1 | ENSMUST00 | 0 | Zfp12 | ENSMUSG00000029587.11 |
| chr5 | 118154837 | 118155756 | 37.30079 | 124.19571 | 119.91465 | Promoter | 5 | 118064981 | 118155458 | 90478 | 2 | ENSMUST00 | 0 | Fbxw8 | ENSMUSG00000032867.7 |
| chr5 | 110447113 | 110449296 | 37.30079 | 124.19571 | 119.91465 | Promoter | 5 | 110361754 | 110448503 | 86750 | 2 | ENSMUST00 | 0 | Fbrsl1 | ENSMUSG00000043323.11 |
| chr7 | 25686448 | 25689783 | 37.30079 | 124.19571 | 119.91465 | Promoter | 7 | 25687002 | 25705077 | 18076 | 1 | ENSMUST00 | 0 | Tgfb1 | ENSMUSG00000002603.10 |
| chr7 | 43312052 | 43313787 | 37.30079 | 124.19571 | 119.91465 | Promoter | 7 | 43296197 | 43313247 | 17051 | 2 | ENSMUST00 | 0 | Zfp715 | ENSMUSG00000012640.11 |
| chr7 | 121980991 | 121982510 | 37.30079 | 124.19571 | 119.91465 | Promoter | 7 | 121922839 | 121981711 | 58873 | 2 | ENSMUST00 | 0 | Cog7 | ENSMUSG00000034951.9 |
| chr7 | 25077166 | 25078317 | 37.30079 | 124.19571 | 119.91465 | Promoter | 7 | 25077205 | 25083492 | 6288 | 1 | ENSMUST00 | 0 | Zfp574 | ENSMUSG00000045252.10 |
| chr9 | 57835728 | 57836991 | 37.30079 | 124.19571 | 119.91465 | Promoter | 9 | 57790353 | 57836706 | 46354 | 2 | ENSMUST00 | 0 | Arid3b | ENSMUSG00000004661.10 |
| chr9 | 45905634 | 45907785 | 37.30079 | 124.19571 | 119.91465 | Promoter | 9 | 45906569 | 45929692 | 23124 | 1 | ENSMUST00 | 0 | Pcsk7 | ENSMUSG00000035382.8 |
| chr2 | 29888930 | 29890840 | 29.47223 | 124.17374 | 119.90009 | Promoter | 2 | 29889221 | 29901107 | 11887 | 1 | ENSMUST00 | 0 | Odf2 | ENSMUSG00000026790.14 |
| chr3 | 105051756 | 105053687 | 29.47223 | 124.17374 | 119.90009 | Promoter | 3 | 105001915 | 105052953 | 51039 | 2 | ENSMUST00 | 0 | Cttnbp2nl | ENSMUSG00000062127.6 |
| chr12 | 55398093 | 55400049 | 37.90103 | 124.17223 | 119.89915 | Promoter | 12 | 55398775 | 55418454 | 19680 | 1 | ENSMUST00 | 0 | Psma6 | ENSMUSG00000021024.9 |
| chr14 | 67716051 | 67717209 | 37.90103 | 124.17223 | 119.89915 | Promoter | 14 | 67716098 | 67742310 | 26213 | 1 | ENSMUST00 | 0 | Kctd9 | ENSMUSG00000034327.12 |
| chr12 | 84996250 | 84997380 | 39.12038 | 124.14072 | 119.86815 | Promoter | 12 | 84996321 | 85070492 | 74172 | 1 | ENSMUST00 | 0 | Ylpm1 | ENSMUSG00000021244.10 |
| chr19 | 7381970 | 7383712 | 39.12038 | 124.14072 | 119.86815 | Promoter | 19 | 7356464 | 7383026 | 26563 | 2 | ENSMUST00 | 0 | AI846148 | ENSMUSG00000024970.4 |
| chr5 | 138618861 | 138619827 | 38.45514 | 124.06234 | 119.79023 | Promoter | 5 | 138604616 | 138619761 | 15146 | 2 | ENSMUST00 | 0 | Zfp68 | ENSMUSG00000058291.9 |
| chr1 | 136017641 | 136019273 | 37.82652 | 124.04217 | 119.77048 | Promoter | 1 | 136017967 | 136035024 | 17058 | 1 | ENSMUST00 | 0 | Tmem9 | ENSMUSG00000026411.12 |
| chr11 | 106786098 | 106790164 | 37.82652 | 124.04217 | 119.77048 | Promoter | 11 | 106789252 | 106818860 | 29609 | 1 | ENSMUST00 | 0 | Cep95 | ENSMUSG00000018372.8 |
| chr13 | 106946275 | 106947899 | 37.82652 | 124.04217 | 119.77048 | Promoter | 13 | 106947129 | 106960224 | 13096 | 1 | ENSMUST00 | 0 | Dimt1 | ENSMUSG00000021692.8 |
| chr17 | 87281464 | 87284423 | 37.82652 | 124.04217 | 119.77048 | Promoter | 17 | 87282886 | 87381769 | 98884 | 1 | ENSMUST00 | 0 | Ttc7 | ENSMUSG00000036918.10 |
| chr7 | 127272769 | 127274480 | 37.82652 | 124.04217 | 119.77048 | Promoter | 7 | 127272561 | 127273919 | 1359 | 2 | ENSMUST00 | 0 | Sephs2 | ENSMUSG00000049091.5 |
| chr2 | 39007436 | 39009098 | 36.59948 | 124.03223 | 119.76238 | Promoter | 2 | 39008076 | 39015877 | 7802 | 1 | ENSMUST00 | 0 | Arpc5l | ENSMUSG00000026755.9 |
| chr1 | 156717889 | 156719597 | 39.03915 | 124.0004 | 119.73064 | Promoter | 1 | 156678532 | 156719072 | 40541 | 2 | ENSMUST00 | 0 | Fam20b | ENSMUSG00000033557.12 |
| chr17 | 56716333 | 56718467 | 39.03915 | 124.0004 | 119.73064 | Promoter | 17 | 56717762 | 56724248 | 6487 | 1 | ENSMUST00 | 0 | Ndufa11 | ENSMUSG00000002379.6 |
| chr1 | 55236585 | 55238734 | 35.37319 | 123.98246 | 119.71299 | Promoter | 1 | 55237177 | 55248470 | 11294 | 1 | ENSMUST00 | 0 | mars-02 | ENSMUSG00000046994.9 |
| chr19 | 36347832 | 36349860 | 35.37319 | 123.98246 | 119.71299 | Distal Interge | 19 | 36379067 | 36456192 | 77126 | 1 | ENSMUST00 | -29207 | Pcgf5 | ENSMUSG00000024805.10 |
| chr7 | 128611271 | 128612373 | 35.37319 | 123.98246 | 119.71299 | Promoter | 7 | 128611328 | 128671622 | 60295 | 1 | ENSMUST00 | 0 | Inpp5f | ENSMUSG00000042105.13 |
| chr19 | 6060656 | 6062281 | 37.16048 | 123.94915 | 119.68073 | Promoter | 19 | 6061192 | 6062472 | 1281 | 1 | ENSMUST00 | 0 | Znhit2 | ENSMUSG00000075227.6 |
| chr11 | 22286225 | 22287137 | 38.37755 | 123.92766 | 119.65938 | Promoter | 11 | 22005828 | 22286795 | 280968 | 2 | ENSMUST00 | 0 | Ehbp1 | ENSMUSG00000042302.9 |
| chr3 | 30995072 | 30997340 | 37.7523 | 123.91259 | 119.6447 | Promoter | 3 | 30995747 | 31052741 | 56995 | 1 | ENSMUST00 | 0 | Prkci | ENSMUSG00000037643.9 |
| chr1 | 167283561 | 167285860 | 36.53154 | 123.91184 | 119.64436 | Promoter | 1 | 167226090 | 167285127 | 59038 | 2 | ENSMUST00 | 0 | Uck2 | ENSMUSG00000026558.8 |
| chr1 | 157411007 | 157413312 | 36.53154 | 123.91184 | 119.64436 | Promoter | 1 | 157412352 | 157420236 | 7885 | 1 | ENSMUST00 | 0 | 2810025M15 | ENSMUSG00000049881.8 |
| chr14 | 79389734 | 79391248 | 36.53154 | 123.91184 | 119.64436 | Promoter | 14 | 79334507 | 79390778 | 56272 | 2 | ENSMUST00 | 0 | Naa16 | ENSMUSG00000022020.9 |
| chr14 | 54875917 | 54877905 | 36.53154 | 123.91184 | 119.64436 | Promoter | 14 | 54872681 | 54877550 | 4870 | 2 | ENSMUST00 | 0 | Ppp1r3e | ENSMUSG00000072494.6 |
| chr15 | 99369582 | 99370569 | 36.53154 | 123.91184 | 119.64436 | Promoter | 15 | 99317225 | 99370449 | 53225 | 2 | ENSMUST00 | 0 | Fmnl3 | ENSMUSG00000023008.13 |
| chr17 | 35978627 | 35981131 | 36.53154 | 123.91184 | 119.64436 | Promoter | 17 | 35979851 | 35989462 | 9612 | 1 | ENSMUST00 | 0 | Gnl1 | ENSMUSG00000024429.9 |
| chr19 | 21777956 | 21779995 | 36.53154 | 123.91184 | 119.64436 | Promoter | 19 | 21778342 | 21858327 | 79986 | 1 | ENSMUST00 | 0 | Tmem2 | ENSMUSG00000024754.7 |
| chr6 | 120035822 | 120038688 | 36.53154 | 123.91184 | 119.64436 | Promoter | 6 | 119923969 | 120038672 | 114704 | 2 | ENSMUST00 | 0 | Wnk1 | ENSMUSG00000045962.11 |
| chr8 | 12671452 | 12672797 | 36.53154 | 123.91184 | 119.64436 | Promoter | 8 | 12614277 | 12672099 | 57823 | 2 | ENSMUST00 | 0 | Tubgcp3 | ENSMUSG00000000759.9 |
| chr9 | 111270886 | 111271633 | 36.53154 | 123.91184 | 119.64436 | Promoter | 9 | 111228228 | 111271546 | 43319 | 2 | ENSMUST00 | 0 | Mlh1 | ENSMUSG00000032498.4 |
| chr4 | 88720960 | 88722819 | 33.57447 | 123.878 | 119.61288 | Promoter | 4 | 88718292 | 88722465 | 4174 | 2 | ENSMUST00 | 0 | Klhl9 | ENSMUSG00000070923.4 |
| chr9 | 105053051 | 105054049 | 33.57447 | 123.878 | 119.61288 | Promoter | 9 | 105053239 | 105076472 | 23234 | 1 | ENSMUST00 | 0 | Mrpl3 | ENSMUSG00000032563.10 |
| chr2 | 154656983 | 154658473 | 38.95827 | 123.86062 | 119.59576 | Promoter | 2 | 154657018 | 154694785 | 37768 | 1 | ENSMUST00 | 0 | Chmp4b | ENSMUSG00000038467.10 |
| chr16 | 91688128 | 91689446 | 38.30027 | 123.79345 | 119.52906 | Promoter | 16 | 91679265 | 91688765 | 9501 | 2 | ENSMUST00 | 0 | Donson | ENSMUSG00000022960.7 |
| chr15 | 31223848 | 31225303 | 37.67836 | 123.78344 | 119.51971 | Promoter | 15 | 31224314 | 31274341 | 50028 | 1 | ENSMUST00 | 0 | Dap | ENSMUSG00000039168.10 |
| chr13 | 118386114 | 118387875 | 25.65631 | 123.7383 | 119.47491 | Promoter | 13 | 118380110 | 118387252 | 7143 | 2 | ENSMUST00 | 0 | Mrps30 | ENSMUSG00000021731.9 |
| chr11 | 6415138 | 6417437 | 36.39639 | 123.67226 | 119.40948 | Promoter | 11 | 6415443 | 6419817 | 4375 | 1 | ENSMUST00 | 0 | Ppia | ENSMUSG00000071866.6 |
| chr2 | 4717210 | 4718922 | 37.60472 | 123.65472 | 119.39239 | Promoter | 2 | 4717831 | 4800563 | 82733 | 1 | ENSMUST00 | 0 | Bend7 | ENSMUSG00000048186.9 |
| chr7 | 80801785 | 80803839 | 36.94879 | 123.57673 | 119.31536 | Promoter | 7 | 80711583 | 80803331 | 91749 | 2 | ENSMUST00 | 0 | Iqgap1 | ENSMUSG00000030536.9 |
| chr1 | 133130586 | 133135428 | 34.54275 | 123.54154 | 119.28033 | Promoter | 1 | 133131143 | 133139783 | 8641 | 1 | ENSMUST00 | 0 | Ppp1r15b | ENSMUSG00000046062.5 |
| chr5 | 115631117 | 115632856 | 34.54275 | 123.54154 | 119.28033 | Promoter | 5 | 115631929 | 115647736 | 15808 | 1 | ENSMUST00 | 0 | Rab35 | ENSMUSG00000029518.10 |
| chr14 | 59200711 | 59202259 | 38.14666 | 123.52654 | 119.2661 | Promoter | 14 | 59201209 | 59223214 | 22006 | 1 | ENSMUST00 | 0 | Rcbtb1 | ENSMUSG00000035469.12 |
| chr17 | 32283246 | 32285341 | 38.14666 | 123.52654 | 119.2661 | Promoter | 17 | 32196274 | 32284097 | 87824 | 2 | ENSMUST00 | 0 | Brd4 | ENSMUSG00000024002.11 |
| chr8 | 105496272 | 105497231 | 38.14666 | 123.52654 | 119.2661 | Distal Interge | 8 | 105472481 | 105484350 | 11870 | 2 | ENSMUST00 | -11923 | Zdhhc1 | ENSMUSG00000039199.5 |
| chr9 | 98562946 | 98564432 | 37.53136 | 123.5265 | 119.2661 | Promoter | 9 | 98563721 | 98567316 | 3596 | 1 | ENSMUST00 | 0 | Copb2 | ENSMUSG00000032458.6 |
| chr6 | 146501332 | 146502915 | 36.87877 | 123.45338 | 119.19501 | Promoter | 6 | 146108299 | 146502099 | 393801 | 2 | ENSMUST00 | 0 | Itpr2 | ENSMUSG00000030287.10 |
| chr1 | 119836203 | 119837955 | 38.7176 | 123.44448 | 119.18625 | Promoter | 1 | 119652467 | 119837613 | 185147 | 2 | ENSMUST00 | 0 | Ptpn4 | ENSMUSG00000026384.8 |
| chr13 | 9276148 | 9278135 | 38.7176 | 123.44448 | 119.18625 | Promoter | 13 | 9276528 | 9668928 | 392401 | 1 | ENSMUST00 | 0 | Dip2c | ENSMUSG00000048264.10 |
| chr15 | 11902214 | 11907460 | 38.7176 | 123.44448 | 119.18625 | Promoter | 15 | 11839896 | 11905674 | 65779 | 2 | ENSMUST00 | 0 | Npr3 | ENSMUSG00000022206.6 |
| chr16 | 4963351 | 4965084 | 37.45829 | 123.39867 | 119.14091 | Promoter | 16 | 4941415 | 4964237 | 22823 | 2 | ENSMUST00 | 0 | Anks3 | ENSMUSG00000022515.5 |
| chr1 | 52630408 | 52631322 | 38.0703 | 123.3938 | 119.13641 | Promoter | 1 | 52630703 | 52651919 | 21217 | 1 | ENSMUST00 | 0 | Tmem194b | ENSMUSG00000043015.10 |
| chr10 | 19591951 | 19594959 | 38.0703 | 123.3938 | 119.13641 | Promoter | 10 | 19591958 | 19597851 | 5894 | 1 | ENSMUST00 | 0 | Ifngr1 | ENSMUSG00000020009.7 |
| chr15 | 10712738 | 10715620 | 38.0703 | 123.3938 | 119.13641 | Promoter | 15 | 10568979 | 10713540 | 144562 | 2 | ENSMUST00 | 0 | Rai14 | ENSMUSG00000022246.8 |
| chr5 | 74535063 | 74536477 | 38.0703 | 123.3938 | 119.13641 | Promoter | 5 | 74535449 | 74598790 | 63342 | 1 | ENSMUST00 | 0 | Fip1l1 | ENSMUSG00000029227.10 |
| chr11 | 61577769 | 61579775 | 35.61502 | 123.34891 | 119.09213 | Promoter | 11 | 61517249 | 61579669 | 62421 | 2 | ENSMUST00 | 0 | Epn2 | ENSMUSG00000001036.12 |
| chr16 | 31947500 | 31949536 | 35.61502 | 123.34891 | 119.09213 | Promoter | 16 | 31948513 | 31958472 | 9960 | 1 | ENSMUST00 | 0 | Ncbp2 | ENSMUSG00000022774.7 |
| chr16 | 55820908 | 55822916 | 35.61502 | 123.34891 | 119.09213 | Promoter | 16 | 55811375 | 55822138 | 10764 | 2 | ENSMUST00 | 0 | Nfkbiz | ENSMUSG00000035356.11 |
| chr18 | 3382779 | 3384086 | 35.61502 | 123.34891 | 119.09213 | Promoter | 18 | 3382988 | 3436374 | 53387 | 1 | ENSMUST00 | 0 | Cul2 | ENSMUSG00000024231.9 |
| chr7 | 64871935 | 64873471 | 35.61502 | 123.34891 | 119.09213 | Promoter | 7 | 64867052 | 64872997 | 5946 | 2 | ENSMUST00 | 0 | Ndnl2 | ENSMUSG00000070520.2 |
| chr1 | 58972637 | 58974577 | 36.809 | 123.33047 | 119.07505 | Promoter | 1 | 58973522 | 58981475 | 7954 | 1 | ENSMUST00 | 0 | Stradb | ENSMUSG00000026027.8 |
| chr13 | 24830659 | 24832491 | 36.809 | 123.33047 | 119.07505 | Promoter | 13 | 24817948 | 24831517 | 13570 | 2 | ENSMUST00 | 0 | Acot13 | ENSMUSG00000006717.6 |
| chr17 | 24072662 | 24073555 | 36.809 | 123.33047 | 119.07505 | Promoter | 17 | 24047734 | 24073485 | 25752 | 2 | ENSMUST00 | 0 | Kctd5 | ENSMUSG00000016946.5 |
| chr18 | 6514144 | 6516887 | 36.809 | 123.33047 | 119.07505 | Promoter | 18 | 6435951 | 6516108 | 80158 | 2 | ENSMUST00 | 0 | Epc1 | ENSMUSG00000024240.8 |
| chr19 | 8756088 | 8757919 | 36.809 | 123.33047 | 119.07505 | Promoter | 19 | 8757073 | 8770922 | 13850 | 1 | ENSMUST00 | 0 | Nxf1 | ENSMUSG00000010097.8 |
| chr3 | 103057815 | 103059145 | 36.809 | 123.33047 | 119.07505 | Promoter | 3 | 103058339 | 103067914 | 9576 | 1 | ENSMUST00 | 0 | Nras | ENSMUSG00000027852.9 |
| chr5 | 124111237 | 124113178 | 36.809 | 123.33047 | 119.07505 | Promoter | 5 | 124112297 | 124115483 | 3187 | 1 | ENSMUST00 | 0 | Ogfod2 | ENSMUSG00000023707.8 |
| chr6 | 28423162 | 28424631 | 36.809 | 123.33047 | 119.07505 | Promoter | 6 | 28423560 | 28426602 | 3043 | 1 | ENSMUST00 | 0 | Arf5 | ENSMUSG00000020440.8 |
| chr6 | 136808205 | 136809562 | 36.809 | 123.33047 | 119.07505 | Promoter | 6 | 136808244 | 136810070 | 1827 | 1 | ENSMUST00 | 0 | H2afj | ENSMUSG00000060032.5 |
| chr9 | 105494013 | 105496076 | 36.809 | 123.33047 | 119.07505 | Promoter | 9 | 105403546 | 105494420 | 90875 | 2 | ENSMUST00 | 0 | Atp2c1 | ENSMUSG00000032570.11 |
| chr4 | 155760283 | 155761996 | 38.63803 | 123.30683 | 119.05459 | Promoter | 4 | 155740641 | 155761093 | 20453 | 2 | ENSMUST00 | 0 | Atad3a | ENSMUSG00000029036.13 |
| chr4 | 101068633 | 101070848 | 38.63803 | 123.30683 | 119.05459 | Promoter | 4 | 101068983 | 101135356 | 66374 | 1 | ENSMUST00 | 0 | Raver2 | ENSMUSG00000035275.9 |
| chr1 | 132416868 | 132418544 | 37.38551 | 123.27131 | 119.01954 | Promoter | 1 | 132417555 | 132466958 | 49404 | 1 | ENSMUST00 | 0 | Dstyk | ENSMUSG00000042046.10 |
| chr12 | 40036276 | 40038537 | 36.73951 | 123.20795 | 118.95689 | Promoter | 12 | 40005447 | 40037367 | 31921 | 2 | ENSMUST00 | 0 | Arl4a | ENSMUSG00000047446.13 |
| chr15 | 9139812 | 9141459 | 36.73951 | 123.20795 | 118.95689 | Promoter | 15 | 9140621 | 9198318 | 57698 | 1 | ENSMUST00 | 0 | Lmbrd2 | ENSMUSG00000039704.6 |
| chr1 | 151427619 | 151429673 | 32.64849 | 123.19247 | 118.94151 | Promoter | 1 | 151367699 | 151428455 | 60757 | 2 | ENSMUST00 | 0 | Swt1 | ENSMUSG00000052748.9 |
| chr10 | 91170021 | 91172319 | 32.64849 | 123.19247 | 118.94151 | Promoter | 10 | 91147571 | 91171582 | 24012 | 2 | ENSMUST00 | 0 | Tmpo | ENSMUSG00000019961.11 |
| chr11 | 102406444 | 102408601 | 32.64849 | 123.19247 | 118.94151 | Promoter | 11 | 102402985 | 102407489 | 4505 | 2 | ENSMUST00 | 0 | Slc25a39 | ENSMUSG00000018677.4 |
| chr17 | 66111173 | 66112438 | 32.64849 | 123.19247 | 118.94151 | Promoter | 17 | 66111546 | 66120503 | 8958 | 1 | ENSMUST00 | 0 | Wash | ENSMUSG00000024101.9 |
| chr2 | 74825369 | 74826551 | 32.64849 | 123.19247 | 118.94151 | Promoter | 2 | 74825803 | 74878431 | 52629 | 1 | ENSMUST00 | 0 | Mtx2 | ENSMUSG00000027099.9 |
| chr2 | 128125321 | 128128418 | 32.64849 | 123.19247 | 118.94151 | Promoter | 2 | 128126038 | 128130015 | 3978 | 1 | ENSMUST00 | 0 | Bcl2l11 | ENSMUSG00000027381.11 |
| chr3 | 105686766 | 105688240 | 32.64849 | 123.19247 | 118.94151 | Promoter | 3 | 105678270 | 105687574 | 9305 | 2 | ENSMUST00 | 0 | Ddx20 | ENSMUSG00000027905.11 |
| chr3 | 84813640 | 84816188 | 32.64849 | 123.19247 | 118.94151 | Promoter | 3 | 84815268 | 84979198 | 163931 | 1 | ENSMUST00 | 0 | Fbxw7 | ENSMUSG00000028086.9 |
| chr4 | 120404779 | 120406505 | 32.64849 | 123.19247 | 118.94151 | Promoter | 4 | 120405281 | 120454515 | 49235 | 1 | ENSMUST00 | 0 | Scmh1 | ENSMUSG00000000085.11 |
| chr4 | 19569105 | 19570425 | 32.64849 | 123.19247 | 118.94151 | Promoter | 4 | 19519254 | 19570108 | 50855 | 2 | ENSMUST00 | 0 | Cpne3 | ENSMUSG00000028228.5 |
| chr4 | 108779751 | 108781324 | 32.64849 | 123.19247 | 118.94151 | Promoter | 4 | 108637466 | 108780503 | 143038 | 2 | ENSMUST00 | 0 | Zfyve9 | ENSMUSG00000034557.9 |
| chr6 | 113377011 | 113379381 | 32.64849 | 123.19247 | 118.94151 | Promoter | 6 | 113366025 | 113377883 | 11859 | 2 | ENSMUST00 | 0 | Tada3 | ENSMUSG00000048930.7 |
| chr7 | 78846364 | 78847461 | 32.64849 | 123.19247 | 118.94151 | Promoter | 7 | 78827474 | 78847236 | 19763 | 2 | ENSMUST00 | 0 | Det1 | ENSMUSG00000030610.8 |
| chr8 | 70476054 | 70477789 | 32.64849 | 123.19247 | 118.94151 | Promoter | 8 | 70450228 | 70476943 | 26716 | 2 | ENSMUST00 | 0 | Klhl26 | ENSMUSG00000055707.7 |
| chr11 | 5955184 | 5956586 | 33.76688 | 123.17645 | 118.92934 | Promoter | 11 | 5955693 | 5967780 | 12088 | 1 | ENSMUST00 | 0 | Ykt6 | ENSMUSG00000002741.9 |
| chr3 | 137863702 | 137865762 | 33.76688 | 123.17645 | 118.92934 | Promoter | 3 | 137864487 | 137866922 | 2436 | 1 | ENSMUST00 | 0 | H2afz | ENSMUSG00000037894.8 |

| | | | | | | | | | | | | | | | |
|---|---|---|---|---|---|---|---|---|---|---|---|---|---|---|---|
| chr15 | 96459601 | 96460930 | 38.5588 | 123.16969 | 118.923 | Promoter | 15 | 96411698 | 96460843 | 49146 | 2 | ENSMUST00 | 0 | Scaf11 | ENSMUSG00000033228.7 |
| chr12 | 111712442 | 111714088 | 37.91852 | 123.12976 | 118.88376 | Promoter | 12 | 111713261 | 111734220 | 20960 | 1 | ENSMUST00 | 0 | Apopt1 | ENSMUSG00000037787.8 |
| chr16 | 97850536 | 97852480 | 37.91852 | 123.12976 | 118.88376 | Promoter | 16 | 97791467 | 97851227 | 59761 | 2 | ENSMUST00 | 0 | Prdm15 | ENSMUSG00000014039.12 |
| chr1 | 191325662 | 191326896 | 38.47989 | 123.03305 | 118.78763 | Promoter | 1 | 191325912 | 191350914 | 25003 | 1 | ENSMUST00 | 0 | Tmem206 | ENSMUSG00000026627.7 |
| chr15 | 44619120 | 44620360 | 38.47989 | 123.03305 | 118.78763 | Promoter | 15 | 44619641 | 44641024 | 21384 | 1 | ENSMUST00 | 0 | Ebag9 | ENSMUSG00000022339.8 |
| chr2 | 181457277 | 181460692 | 38.47989 | 123.03305 | 118.78763 | Promoter | 2 | 181387762 | 181459426 | 71665 | 2 | ENSMUST00 | 0 | Zbtb46 | ENSMUSG00000027583.8 |
| chr8 | 78820304 | 78821690 | 38.47989 | 123.03305 | 118.78763 | Promoter | 8 | 78804865 | 78821140 | 16276 | 2 | ENSMUST00 | 0 | Lsm6 | ENSMUSG00000031683.11 |
| chr5 | 44225630 | 44227675 | 37.24078 | 123.01789 | 118.77389 | Promoter | 5 | 44175164 | 44226606 | 51443 | 2 | ENSMUST00 | 0 | Tapt1 | ENSMUSG00000046985.7 |
| chr1 | 151499350 | 151501598 | 39.06023 | 122.966 | 118.72274 | Promoter | 1 | 151469405 | 151500811 | 31407 | 2 | ENSMUST00 | 0 | Rnf2 | ENSMUSG00000026484.8 |
| chr10 | 94034845 | 94036859 | 39.06023 | 122.966 | 118.72274 | Promoter | 10 | 94036001 | 94145339 | 109339 | 1 | ENSMUST00 | 0 | Fgd6 | ENSMUSG00000020021.4 |
| chr16 | 78575560 | 78577511 | 39.06023 | 122.966 | 118.72274 | Promoter | 16 | 78544012 | 78576657 | 32646 | 2 | ENSMUST00 | 0 | D16Ertd472e | ENSMUSG00000022864.8 |
| chr17 | 27856364 | 27857489 | 39.06023 | 122.966 | 118.72274 | Promoter | 17 | 27856490 | 27900040 | 43551 | 1 | ENSMUST00 | 0 | Uhrf1bp1 | ENSMUSG00000039512.11 |
| chr18 | 46523340 | 46525949 | 39.06023 | 122.966 | 118.72274 | Promoter | 18 | 46501746 | 46525971 | 24226 | 2 | ENSMUST00 | 22 | Fem1c | ENSMUSG00000033319.5 |
| chr3 | 89089368 | 89091624 | 39.06023 | 122.966 | 118.72274 | Promoter | 3 | 89083981 | 89090031 | 6051 | 2 | ENSMUST00 | 0 | Rusc1 | ENSMUSG00000041263.9 |
| chr5 | 123682686 | 123685179 | 39.06023 | 122.966 | 118.72274 | Promoter | 5 | 123577795 | 123684322 | 106528 | 2 | ENSMUST00 | 0 | Clip1 | ENSMUSG00000049550.12 |
| chr6 | 48593384 | 48594992 | 39.06023 | 122.966 | 118.72274 | Promoter | 6 | 48593883 | 48597387 | 3505 | 1 | ENSMUST00 | 0 | Repin1 | ENSMUSG00000052751.11 |
| chr8 | 4166452 | 4167108 | 39.06023 | 122.966 | 118.72274 | Promoter | 8 | 4166567 | 4186880 | 20314 | 1 | ENSMUST00 | 0 | Evi5l | ENSMUSG00000011832.10 |
| chr8 | 94214485 | 94215240 | 39.06023 | 122.966 | 118.72274 | Promoter | 8 | 94214597 | 94314857 | 100261 | 1 | ENSMUST00 | 0 | Nup93 | ENSMUSG00000032939.8 |
| chr9 | 98600851 | 98602075 | 39.06023 | 122.966 | 118.72274 | Promoter | 9 | 98588730 | 98601660 | 12931 | 2 | ENSMUST00 | 0 | Mrps22 | ENSMUSG00000032459.9 |
| chr8 | 33384518 | 33386849 | 38.4013 | 122.89693 | 118.65718 | Promoter | 8 | 33234384 | 33385527 | 151144 | 2 | ENSMUST00 | 0 | Wrn | ENSMUSG00000031583.7 |
| chr11 | 79710752 | 79712376 | 34.76187 | 122.87653 | 118.63768 | Promoter | 11 | 79711969 | 79712034 | 66 | 1 | ENSMUST00 | 0 | Mir193a | ENSMUSG00000065395.1 |
| chr14 | 57743310 | 57746899 | 34.76187 | 122.87653 | 118.63768 | Promoter | 14 | 57642077 | 57746103 | 104027 | 2 | ENSMUST00 | 0 | Lats2 | ENSMUSG00000021959.10 |
| chr13 | 51099595 | 51102016 | 37.76794 | 122.86759 | 118.62959 | Promoter | 13 | 51100880 | 51152546 | 51667 | 1 | ENSMUST00 | 0 | Spin1 | ENSMUSG00000021395.10 |
| chr7 | 128739046 | 128740883 | 37.76794 | 122.86759 | 118.62959 | Promoter | 7 | 128696441 | 128740495 | 44055 | 2 | ENSMUST00 | 0 | Mcmbp | ENSMUSG00000048170.9 |
| chr9 | 48494451 | 48496186 | 36.53257 | 122.84278 | 118.60545 | Promoter | 9 | 48495345 | 48495964 | 620 | 1 | ENSMUST00 | 0 | Gm5617 | ENSMUSG00000042293.7 |
| chr1 | 97660633 | 97662645 | 37.09717 | 122.76616 | 118.52905 | Promoter | 1 | 97606318 | 97662074 | 55757 | 2 | ENSMUST00 | 0 | D1Ertd622e | ENSMUSG00000044768.11 |
| chr10 | 43540096 | 43541068 | 37.09717 | 122.76616 | 118.52905 | Promoter | 10 | 43525121 | 43540994 | 15874 | 2 | ENSMUST00 | 0 | 1700021F05I | ENSMUSG00000019797.5 |
| chr13 | 49420540 | 49422122 | 37.09717 | 122.76616 | 118.52905 | Promoter | 13 | 49421311 | 49462990 | 41680 | 1 | ENSMUST00 | 0 | Ippk | ENSMUSG00000021385.9 |
| chr16 | 4683437 | 4684929 | 37.09717 | 122.76616 | 118.52905 | Promoter | 16 | 4684070 | 4707695 | 23626 | 1 | ENSMUST00 | 0 | Dnaja3 | ENSMUSG00000004069.11 |
| chr16 | 87353948 | 87354949 | 37.09717 | 122.76616 | 118.52905 | Promoter | 16 | 87354185 | 87368742 | 14558 | 1 | ENSMUST00 | 0 | N6amt1 | ENSMUSG00000044442.6 |
| chr18 | 4634673 | 4636017 | 37.09717 | 122.76616 | 118.52905 | Promoter | 18 | 4634929 | 4682868 | 47940 | 1 | ENSMUST00 | 0 | 9430020K01I | ENSMUSG00000033960.5 |
| chr6 | 125312582 | 125314359 | 37.09717 | 122.76616 | 118.52905 | Promoter | 6 | 125306571 | 125313885 | 7315 | 2 | ENSMUST00 | 0 | Ltbr | ENSMUSG00000030339.7 |
| chr8 | 25975808 | 25977366 | 37.09717 | 122.76616 | 118.52905 | Promoter | 8 | 25944459 | 25976744 | 32286 | 2 | ENSMUST00 | 0 | Hgsnat | ENSMUSG00000037260.7 |
| chr11 | 5541261 | 5542456 | 38.32303 | 122.76134 | 118.52611 | Promoter | 11 | 5528887 | 5542187 | 13301 | 2 | ENSMUST00 | 0 | Ccdc117 | ENSMUSG00000020482.4 |
| chr2 | 94156989 | 94157992 | 38.32303 | 122.76134 | 118.52611 | Promoter | 2 | 94032689 | 94157964 | 125276 | 2 | ENSMUST00 | 0 | Hsd17b12 | ENSMUSG00000027195.5 |
| chr14 | 105176084 | 105177955 | 37.6931 | 122.73721 | 118.50226 | Promoter | 14 | 105114965 | 105176860 | 61896 | 2 | ENSMUST00 | 0 | Rbm26 | ENSMUSG00000022119.10 |
| chr15 | 51864544 | 51865978 | 37.6931 | 122.73721 | 118.50226 | Promoter | 15 | 51786558 | 51865481 | 78924 | 2 | ENSMUST00 | 0 | Eif3h | ENSMUSG00000022312.10 |
| chr4 | 118427573 | 118429131 | 37.6931 | 122.73721 | 118.50226 | Promoter | 4 | 118428093 | 118432953 | 4861 | 1 | ENSMUST00 | 0 | Elovl1 | ENSMUSG00000006390.10 |
| chr5 | 72558763 | 72559731 | 37.6931 | 122.73721 | 118.50226 | Promoter | 5 | 72513301 | 72559684 | 46384 | 2 | ENSMUST00 | 0 | Nfxl1 | ENSMUSG00000072889.4 |
| chr14 | 21499033 | 21500514 | 27.83657 | 122.73275 | 118.49821 | Promoter | 14 | 21499770 | 21672071 | 172302 | 1 | ENSMUST00 | 0 | Kat6b | ENSMUSG00000021767.11 |
| chr15 | 52712029 | 52713383 | 27.83657 | 122.73275 | 118.49821 | Promoter | 15 | 52712445 | 52730431 | 17987 | 1 | ENSMUST00 | 0 | Med30 | ENSMUSG00000038622.7 |
| chr2 | 27474588 | 27476587 | 35.86567 | 122.72874 | 118.49439 | Promoter | 2 | 27445579 | 27475398 | 29820 | 2 | ENSMUST00 | 0 | Brd3 | ENSMUSG00000026918.11 |
| chr4 | 49631400 | 49632901 | 35.86567 | 122.72874 | 118.49439 | Promoter | 4 | 49632006 | 49638702 | 6697 | 1 | ENSMUST00 | 0 | Rnf20 | ENSMUSG00000028309.9 |
| chr15 | 51877054 | 51878261 | 30.76814 | 122.68065 | 118.44827 | Promoter | 15 | 51877429 | 51884614 | 7186 | 1 | ENSMUST00 | 0 | Utp23 | ENSMUSG00000022313.10 |
| chr18 | 35561702 | 35564147 | 30.76814 | 122.68065 | 118.44827 | Promoter | 18 | 35562146 | 35572490 | 10345 | 1 | ENSMUST00 | 0 | Matr3 | ENSMUSG00000037236.9 |
| chr6 | 115674958 | 115677277 | 30.76814 | 122.68065 | 118.44827 | Promoter | 6 | 115618569 | 115675964 | 57396 | 2 | ENSMUST00 | 0 | Raf1 | ENSMUSG00000000441.12 |
| chr7 | 90456547 | 90458058 | 30.76814 | 122.68065 | 118.44827 | Promoter | 7 | 90450700 | 90457208 | 6509 | 2 | ENSMUST00 | 0 | Tmem126a | ENSMUSG00000030615.6 |
| chr9 | 99567865 | 99569427 | 30.76814 | 122.68065 | 118.44827 | Promoter | 9 | 99478372 | 99568899 | 90528 | 2 | ENSMUST00 | 0 | Armc8 | ENSMUSG00000032468.6 |
| chr9 | 55511381 | 55512691 | 32.35868 | 122.64178 | 118.41108 | Promoter | 9 | 55454508 | 55512243 | 57736 | 2 | ENSMUST00 | 0 | Etfa | ENSMUSG00000032314.9 |
| chr17 | 6079019 | 6080497 | 38.82705 | 122.56369 | 118.33408 | Promoter | 17 | 6042196 | 6079741 | 37546 | 2 | ENSMUST00 | 0 | Serac1 | ENSMUSG00000015659.7 |
| chr18 | 84950665 | 84952385 | 38.82705 | 122.56369 | 118.33408 | Promoter | 18 | 84947294 | 84951524 | 4231 | 2 | ENSMUST00 | 0 | Timm21 | ENSMUSG00000024645.4 |
| chr5 | 115279443 | 115280437 | 38.82705 | 122.56369 | 118.33408 | Promoter | 5 | 115279666 | 115296972 | 17307 | 1 | ENSMUST00 | 0 | Coq5 | ENSMUSG00000041733.5 |
| chr19 | 37206366 | 37209314 | 29.18609 | 122.55783 | 118.32927 | Promoter | 19 | 37207543 | 37222139 | 14597 | 1 | ENSMUST00 | 0 | Mar-05 | ENSMUSG00000023307.8 |
| chr5 | 66617073 | 66618806 | 29.18609 | 122.55783 | 118.32927 | Promoter | 5 | 66301872 | 66618663 | 316792 | 2 | ENSMUST00 | 0 | Apbb2 | ENSMUSG00000029207.11 |
| chr9 | 54500833 | 54502265 | 29.18609 | 122.55783 | 118.32927 | Promoter | 9 | 54365158 | 54501626 | 136469 | 2 | ENSMUST00 | 0 | Dmxl2 | ENSMUSG00000041268.11 |
| chr17 | 46247345 | 46249077 | 38.16745 | 122.49164 | 118.26377 | Promoter | 17 | 46243920 | 46248054 | 4135 | 2 | ENSMUST00 | 0 | Polr1c | ENSMUSG00000067148.10 |
| chr11 | 106486553 | 106488525 | 33.96572 | 122.48407 | 118.2566 | Promoter | 11 | 106394650 | 106487796 | 93147 | 2 | ENSMUST00 | 0 | Ern1 | ENSMUSG00000020715.4 |
| chr3 | 63962339 | 63965122 | 33.96572 | 122.48407 | 118.2566 | Promoter | 3 | 63933507 | 63964666 | 31160 | 2 | ENSMUST00 | 0 | Slc33a1 | ENSMUSG00000027822.11 |
| chr1 | 74235907 | 74238302 | 34.54477 | 122.48211 | 118.25522 | Promoter | 1 | 74236084 | 74236888 | 805 | 1 | ENSMUST00 | 0 | Arpc2 | ENSMUSG00000006304.9 |
| chr10 | 57531168 | 57532829 | 34.54477 | 122.48211 | 118.25522 | Promoter | 10 | 57515774 | 57532530 | 16757 | 2 | ENSMUST00 | 0 | Serinc1 | ENSMUSG00000019877.5 |
| chr10 | 42678492 | 42680079 | 34.54477 | 122.48211 | 118.25522 | Promoter | 10 | 42678916 | 42702459 | 23544 | 1 | ENSMUST00 | 0 | Ostm1 | ENSMUSG00000038280.7 |
| chr11 | 100394893 | 100397863 | 34.54477 | 122.48211 | 118.25522 | Promoter | 11 | 100370619 | 100397749 | 27131 | 2 | ENSMUST00 | 0 | Jup | ENSMUSG00000001552.9 |
| chr11 | 57644082 | 57647951 | 34.54477 | 122.48211 | 118.25522 | Promoter | 11 | 57645442 | 57787514 | 142073 | 1 | ENSMUST00 | 0 | Galnt10 | ENSMUSG00000020520.9 |
| chr11 | 106084543 | 106085881 | 34.54477 | 122.48211 | 118.25522 | Promoter | 11 | 106084613 | 106155446 | 70834 | 1 | ENSMUST00 | 0 | Map3k3 | ENSMUSG00000020700.11 |
| chr11 | 77685741 | 77688196 | 34.54477 | 122.48211 | 118.25522 | Promoter | 11 | 77686155 | 77712741 | 26587 | 1 | ENSMUST00 | 0 | Nufip2 | ENSMUSG00000037857.11 |
| chr12 | 32820328 | 32822070 | 34.54477 | 122.48211 | 118.25522 | Promoter | 12 | 32820335 | 32853369 | 33035 | 1 | ENSMUST00 | 0 | Nampt | ENSMUSG00000020572.7 |
| chr12 | 102438988 | 102439747 | 34.54477 | 122.48211 | 118.25522 | Promoter | 12 | 102394098 | 102439813 | 45716 | 2 | ENSMUST00 | 66 | Lgmn | ENSMUSG00000021190.9 |
| chr13 | 32801439 | 32803311 | 34.54477 | 122.48211 | 118.25522 | Promoter | 13 | 32802038 | 32822609 | 20572 | 1 | ENSMUST00 | 0 | Wrnip1 | ENSMUSG00000021400.6 |
| chr13 | 94357953 | 94359860 | 37.5443 | 122.47784 | 118.25522 | Promoter | 13 | 94358960 | 94566316 | 207357 | 1 | ENSMUST00 | 0 | Ap3b1 | ENSMUSG00000021686.4 |
| chr15 | 82898547 | 82900341 | 34.54477 | 122.48211 | 118.25522 | Promoter | 15 | 82899080 | 82905818 | 6739 | 1 | ENSMUST00 | 0 | RP23-223M2 | ENSMUSG00000100199.1 |
| chr16 | 65561439 | 65563288 | 34.54477 | 122.48211 | 118.25522 | Promoter | 16 | 65539133 | 65562697 | 23565 | 2 | ENSMUST00 | 0 | Chmp2b | ENSMUSG00000004843.6 |
| chr16 | 93711890 | 93712743 | 34.54477 | 122.48211 | 118.25522 | Promoter | 16 | 93711907 | 93810585 | 98679 | 1 | ENSMUST00 | 0 | Dopey2 | ENSMUSG00000022946.8 |
| chr17 | 24648245 | 24650753 | 34.54477 | 122.48211 | 118.25522 | Promoter | 17 | 24639282 | 24650305 | 11024 | 2 | ENSMUST00 | 0 | Slc9a3r2 | ENSMUSG00000002504.9 |
| chr2 | 25223627 | 25224712 | 34.54477 | 122.48211 | 118.25522 | Promoter | 2 | 25222160 | 25224702 | 2543 | 2 | ENSMUST00 | 0 | Tubb4b | ENSMUSG00000036752.4 |
| chr2 | 142900680 | 142902004 | 37.5443 | 122.47784 | 118.25522 | Promoter | 2 | 142617474 | 142901531 | 284058 | 2 | ENSMUST00 | 0 | Kif16b | ENSMUSG00000038844.8 |
| chr2 | 69896922 | 69899033 | 34.54477 | 122.48211 | 118.25522 | Promoter | 2 | 69897246 | 70024013 | 126768 | 1 | ENSMUST00 | 0 | Ubr3 | ENSMUSG00000044308.12 |
| chr3 | 96904606 | 96905822 | 34.54477 | 122.48211 | 118.25522 | Promoter | 3 | 96868281 | 96905346 | 37066 | 2 | ENSMUST00 | 0 | Gpr89 | ENSMUSG00000028096.8 |
| chr3 | 138741744 | 138743902 | 34.54477 | 122.48211 | 118.25522 | Promoter | 3 | 138742195 | 138904433 | 162239 | 1 | ENSMUST00 | 0 | Tspan5 | ENSMUSG00000028152.5 |
| chr4 | 116463048 | 116464210 | 34.54477 | 122.48211 | 118.25522 | Promoter | 4 | 116306762 | 116463751 | 156990 | 2 | ENSMUST00 | 0 | Mast2 | ENSMUSG00000003810.7 |
| chr4 | 133583753 | 133585184 | 34.54477 | 122.48211 | 118.25522 | Promoter | 4 | 133584362 | 133591735 | 7374 | 1 | ENSMUST00 | 0 | Gpn2 | ENSMUSG00000028848.8 |
| chr4 | 155668118 | 155669937 | 34.54477 | 122.48211 | 118.25522 | Promoter | 4 | 155654682 | 155669198 | 14517 | 2 | ENSMUST00 | 0 | Mib2 | ENSMUSG00000029060.12 |
| chr4 | 108300058 | 108301693 | 34.54477 | 122.48211 | 118.25522 | Promoter | 4 | 108229724 | 108301096 | 71373 | 2 | ENSMUST00 | 0 | Zyg11b | ENSMUSG00000034636.9 |
| chr4 | 58910802 | 58913238 | 34.54477 | 122.48211 | 118.25522 | Promoter | 4 | 58798911 | 58912749 | 113839 | 2 | ENSMUST00 | 0 | AI314180 | ENSMUSG00000050812.13 |
| chr5 | 105414998 | 105416445 | 34.54477 | 122.48211 | 118.25522 | Promoter | 5 | 105415775 | 105486189 | 70415 | 1 | ENSMUST00 | 0 | Lrrc8b | ENSMUSG00000070639.4 |
| chr6 | 39724148 | 39725926 | 34.54477 | 122.48211 | 118.25522 | Promoter | 6 | 39603237 | 39725463 | 122227 | 2 | ENSMUST00 | 0 | Braf | ENSMUSG00000002413.10 |
| chr7 | 13022963 | 13025610 | 34.54477 | 122.48211 | 118.25522 | Promoter | 7 | 13024152 | 13031035 | 6884 | 1 | ENSMUST00 | 0 | Trim28 | ENSMUSG00000005566.8 |
| chr7 | 121033856 | 121035911 | 34.54477 | 122.48211 | 118.25522 | Promoter | 7 | 121034445 | 121076834 | 42390 | 1 | ENSMUST00 | 0 | Mettl9 | ENSMUSG00000030876.6 |
| chr7 | 120101899 | 120103846 | 34.54477 | 122.48211 | 118.25522 | Promoter | 7 | 120102376 | 120120986 | 18611 | 1 | ENSMUST00 | 0 | Tmem159 | ENSMUSG00000030917.7 |
| chr7 | 16613597 | 16615423 | 36.32796 | 122.48113 | 118.25522 | Promoter | 7 | 16493719 | 16614993 | 121275 | 2 | ENSMUST00 | 0 | Arhgap35 | ENSMUSG00000058230.7 |
| chr8 | 60982289 | 60984001 | 34.54477 | 122.48211 | 118.25522 | Promoter | 8 | 60910389 | 60983300 | 72912 | 2 | ENSMUST00 | 0 | Clcn3 | ENSMUSG00000004319.10 |
| chr8 | 88118680 | 88119684 | 34.54477 | 122.48211 | 118.25522 | Promoter | 8 | 88118769 | 88135106 | 16338 | 1 | ENSMUST00 | 0 | Cnep1r1 | ENSMUSG00000036810.11 |
| chr9 | 118505823 | 118507445 | 34.54477 | 122.48211 | 118.25522 | Promoter | 9 | 118506318 | 118582522 | 76205 | 1 | ENSMUST00 | 0 | Golga4 | ENSMUSG00000038708.9 |
| chr1 | 16103566 | 16106735 | 36.88168 | 122.38801 | 118.16834 | Promoter | 1 | 16105774 | 16133734 | 27961 | 1 | ENSMUST00 | 0 | Rdh10 | ENSMUSG00000025921.7 |
| chr1 | 30872553 | 30874500 | 36.88168 | 122.38801 | 118.16834 | Promoter | 1 | 30802339 | 30873581 | 71243 | 2 | ENSMUST00 | 0 | Phf3 | ENSMUSG00000048874.10 |
| chr10 | 34018380 | 34019734 | 36.88168 | 122.38801 | 118.16834 | Promoter | 10 | 33996555 | 34019624 | 23070 | 2 | ENSMUST00 | 0 | Rwdd1 | ENSMUSG00000019782.9 |
| chr12 | 75594986 | 75597036 | 36.88168 | 122.38801 | 118.16834 | Promoter | 12 | 75450883 | 75596200 | 145318 | 2 | ENSMUST00 | 0 | Ppp2r5e | ENSMUSG00000021051.9 |
| chr12 | 76961239 | 76962304 | 36.88168 | 122.38801 | 118.16834 | Promoter | 12 | 76937269 | 76962248 | 24980 | 2 | ENSMUST00 | 0 | Max | ENSMUSG00000059436.7 |
| chr15 | 95790185 | 95792142 | 36.88168 | 122.38801 | 118.16834 | Promoter | 15 | 95790843 | 95975464 | 184622 | 1 | ENSMUST00 | 0 | Ano6 | ENSMUSG00000064210.6 |
| chr17 | 7170029 | 7171203 | 36.88168 | 122.38801 | 118.16834 | Promoter | 17 | 7170115 | 7303313 | 133199 | 1 | ENSMUST00 | 0 | Rps6ka2 | ENSMUSG00000023809.9 |
| chr18 | 65800046 | 65801424 | 36.88168 | 122.38801 | 118.16834 | Promoter | 18 | 65800549 | 65817665 | 17117 | 1 | ENSMUST00 | 0 | Sec11c | ENSMUSG00000024516.7 |
| chr3 | 154596276 | 154597735 | 36.88168 | 122.38801 | 118.16834 | Promoter | 3 | 154596711 | 154623182 | 26472 | 1 | ENSMUST00 | 0 | Cryz | ENSMUSG00000028199.13 |
| chr3 | 101923665 | 101924883 | 36.88168 | 122.38801 | 118.16834 | Promoter | 3 | 101858930 | 101924449 | 65520 | 2 | ENSMUST00 | 0 | Slc22a15 | ENSMUSG00000033147.11 |
| chr4 | 126046577 | 126047763 | 36.88168 | 122.38801 | 118.16834 | Promoter | 4 | 126046925 | 126055532 | 8608 | 1 | ENSMUST00 | 0 | Mrps15 | ENSMUSG00000028861.8 |
| chr4 | 152039099 | 152040303 | 36.88168 | 122.38801 | 118.16834 | Promoter | 4 | 152039321 | 152052229 | 12909 | 1 | ENSMUST00 | 0 | Nol9 | ENSMUSG00000028948.11 |
| chr4 | 155811762 | 155813813 | 36.88168 | 122.38801 | 118.16834 | Promoter | 4 | 155812489 | 155824543 | 12055 | 1 | ENSMUST00 | 0 | Ccnl2 | ENSMUSG00000029068.11 |
| chr6 | 4600584 | 4601859 | 36.88168 | 122.38801 | 118.16834 | Promoter | 6 | 4600840 | 4643381 | 42542 | 1 | ENSMUST00 | 0 | Casd1 | ENSMUSG00000015189.7 |
| chr6 | 72787705 | 72789802 | 36.88168 | 122.38801 | 118.16834 | Promoter | 6 | 72626378 | 72789254 | 162877 | 2 | ENSMUST00 | 0 | Tcf7l1 | ENSMUSG00000055799.8 |
| chr7 | 29983214 | 29984733 | 36.88168 | 122.38801 | 118.16834 | Promoter | 7 | 29983955 | 30025111 | 41157 | 1 | ENSMUST00 | 0 | Zfp568 | ENSMUSG00000074221.6 |
| chr8 | 57486536 | 57488606 | 36.88168 | 122.38801 | 118.16834 | Promoter | 8 | 57482707 | 57487860 | 5154 | 2 | ENSMUST00 | 0 | Sap30 | ENSMUSG00000031609.3 |
| chr2 | 51146410 | 51149200 | 38.09013 | 122.35755 | 118.14518 | Promoter | 2 | 51130438 | 51149111 | 18674 | 2 | ENSMUST00 | 0 | Rnd3 | ENSMUSG00000017144.3 |
| chr12 | 35046630 | 35048578 | 37.47034 | 122.34885 | 118.13707 | Promoter | 12 | 35047189 | 35147469 | 100281 | 1 | ENSMUST00 | 0 | Snx13 | ENSMUSG00000020590.10 |
| chr8 | 34154121 | 34155652 | 38.66549 | 122.28474 | 118.07326 | Promoter | 8 | 34154563 | 34170845 | 16283 | 1 | ENSMUST00 | 0 | Tmem66 | ENSMUSG00000031532.5 |
| chr13 | 45389204 | 45392361 | 36.81325 | 122.26781 | 118.05663 | Promoter | 13 | 45389742 | 45412019 | 22278 | 1 | ENSMUST00 | 0 | Mylip | ENSMUSG00000038175.9 |
| chr8 | 22592554 | 22594510 | 36.81325 | 122.26781 | 118.05663 | Promoter | 8 | 22577075 | 22593813 | 16739 | 2 | ENSMUST00 | 0 | Vdac3 | ENSMUSG00000008892.6 |
| chr3 | 90388356 | 90390337 | 38.01312 | 122.22395 | 118.01301 | Promoter | 3 | 90385239 | 90389927 | 4689 | 2 | ENSMUST00 | 0 | Slc27a3 | ENSMUSG00000027932.9 |
| chr6 | 72956611 | 72959041 | 38.01312 | 122.22395 | 118.01301 | Promoter | 6 | 72957347 | 72958488 | 1142 | 2 | ENSMUST00 | 0 | Tmsb10 | ENSMUSG00000079523.3 |
| chr12 | 98257958 | 98259946 | 34.9887 | 122.22292 | 118.01233 | Promoter | 12 | 98202304 | 98259459 | 57156 | 2 | ENSMUST00 | 0 | Galc | ENSMUSG00000021003.9 |
| chr19 | 10524504 | 10526311 | 34.9887 | 122.22292 | 118.01233 | Promoter | 19 | 10500513 | 10525209 | 24697 | 2 | ENSMUST00 | 0 | Sdhaf2 | ENSMUSG00000024668.7 |
| chr5 | 38219326 | 38221199 | 34.9887 | 122.22292 | 118.01233 | Promoter | 5 | 38200079 | 38220457 | 20379 | 2 | ENSMUST00 | 0 | Zbtb49 | ENSMUSG00000029127.10 |
| chr2 | 6322201 | 6324135 | 37.39668 | 122.22031 | 118.01075 | Promoter | 2 | 6322667 | 6446390 | 123724 | 1 | ENSMUST00 | 0 | Usp6nl | ENSMUSG00000039046.10 |
| chr4 | 47473550 | 47475815 | 37.39667 | 122.2203 | 118.01075 | Promoter | 4 | 47469833 | 47474333 | 4501 | 2 | ENSMUST00 | 0 | Alg2 | ENSMUSG00000039740.6 |
| chr7 | 65366546 | 65371948 | 37.39668 | 122.22031 | 118.01075 | Promoter | 7 | 65296165 | 65371239 | 75075 | 2 | ENSMUST00 | 0 | Tjp1 | ENSMUSG00000030516.9 |
| chr1 | 152385821 | 152386981 | 33.22105 | 122.16254 | 117.95486 | Promoter | 1 | 152370735 | 152386682 | 15948 | 2 | ENSMUST00 | 0 | Tsen15 | ENSMUSG00000014980.9 |
| chr4 | 86747876 | 86749925 | 33.22105 | 122.16254 | 117.95486 | Promoter | 4 | 86748555 | 86830167 | 81613 | 1 | ENSMUST00 | 0 | Dennd4c | ENSMUSG00000038024.12 |
| chr11 | 4186221 | 4188124 | 38.58521 | 122.14608 | 117.93867 | Promoter | 11 | 4186789 | 4214885 | 28097 | 1 | ENSMUST00 | 0 | Tbc1d10a | ENSMUSG00000034412.8 |
| chr9 | 122886523 | 122887634 | 36.74294 | 122.14426 | 117.93718 | Promoter | 9 | 122888471 | 122896124 | 7654 | 1 | ENSMUST00 | -837 | Zkscan7 | ENSMUSG00000063488.5 |
| chr17 | 35896499 | 35897924 | 36.12563 | 122.12295 | 117.91594 | Promoter | 17 | 35892677 | 35897120 | 4444 | 2 | ENSMUST00 | 0 | 2310061I04R | ENSMUSG00000050705.10 |
| chr3 | 129878062 | 129879464 | 36.12563 | 122.12295 | 117.91594 | Promoter | 3 | 129878622 | 129895821 | 17200 | 1 | ENSMUST00 | 0 | Pla2g12a | ENSMUSG00000027999.10 |

| | | | | | | | | | | | | | | | |
|---|---|---|---|---|---|---|---|---|---|---|---|---|---|---|---|
| chr4 | 133966850 | 133968680 | 36.12563 | 122.12295 | 117.91594 | Promoter | 4 | 133964738 | 133968001 | 3264 | 2 | ENSMUST00 | 0 | Hmgn2 | ENSMUSG00000003038.10 |
| chr4 | 43655687 | 43657198 | 36.12563 | 122.12295 | 117.91594 | Promoter | 4 | 43654227 | 43656443 | 2217 | 2 | ENSMUST00 | 0 | Hint2 | ENSMUSG00000028470.5 |
| chr4 | 44071698 | 44073354 | 36.12563 | 122.12295 | 117.91594 | Promoter | 4 | 44036826 | 44072778 | 35953 | 2 | ENSMUST00 | 0 | Gne | ENSMUSG00000028479.13 |
| chr5 | 135543812 | 135545408 | 36.12563 | 122.12295 | 117.91594 | Promoter | 5 | 135406523 | 135545122 | 138600 | 2 | ENSMUST00 | 0 | Hip1 | ENSMUSG00000039959.9 |
| chr5 | 23674847 | 23676543 | 36.12563 | 122.12295 | 117.91594 | Promoter | 5 | 23674289 | 23674381 | 93 | 2 | ENSMUST00 | -467 | Gm24009 | ENSMUSG00000088359.1 |
| chr6 | 140423533 | 140426226 | 36.12563 | 122.12295 | 117.91594 | Promoter | 6 | 140424099 | 140594906 | 170808 | 1 | ENSMUST00 | 0 | Plekha5 | ENSMUSG00000030231.8 |
| chr6 | 57824019 | 57825531 | 36.12563 | 122.12295 | 117.91594 | Promoter | 6 | 57727807 | 57825136 | 97330 | 2 | ENSMUST00 | 0 | Vopp1 | ENSMUSG00000037788.9 |
| chr7 | 34312738 | 34314229 | 36.12563 | 122.12295 | 117.91594 | Promoter | 7 | 34236707 | 34313551 | 76845 | 2 | ENSMUST00 | 0 | 4931406P16 | ENSMUSG00000066571.8 |
| chr11 | 60911793 | 60914427 | 37.32329 | 122.0922 | 117.88763 | Promoter | 11 | 60902246 | 60913812 | 11567 | 2 | ENSMUST00 | 0 | Gm16515 | ENSMUSG00000018931.3 |
| chr3 | 107695484 | 107696643 | 36.67291 | 122.02113 | 117.81725 | Promoter | 3 | 107663120 | 107696560 | 33441 | 2 | ENSMUST00 | 0 | Ahcyl1 | ENSMUSG00000027893.9 |
| chr4 | 11005934 | 11008487 | 38.50526 | 122.00793 | 117.8042 | Promoter | 4 | 10988662 | 11007676 | 19015 | 2 | ENSMUST00 | 0 | Plekhf2 | ENSMUSG00000049969.3 |
| chr15 | 9070331 | 9072400 | 37.86004 | 121.9582 | 117.75514 | Promoter | 15 | 9071260 | 9110487 | 39228 | 1 | ENSMUST00 | 0 | Nadk2 | ENSMUSG00000022253.9 |
| chr7 | 140035620 | 140037192 | 37.86004 | 121.9582 | 117.75514 | Promoter | 7 | 140036391 | 140040605 | 4215 | 1 | ENSMUST00 | 0 | Zfp511 | ENSMUSG00000025470.10 |
| chr11 | 68851734 | 68853148 | 35.99199 | 121.88605 | 117.68387 | Promoter | 11 | 68821434 | 68853135 | 31702 | 2 | ENSMUST00 | 0 | Ndel1 | ENSMUSG00000018736.9 |
| chr2 | 155691354 | 155692393 | 37.17739 | 121.83733 | 117.63558 | Promoter | 2 | 155634271 | 155692384 | 58114 | 2 | ENSMUST00 | 0 | Trpc4ap | ENSMUSG00000038324.8 |
| chr6 | 128299157 | 128301648 | 37.17739 | 121.83733 | 117.63558 | Promoter | 6 | 128227143 | 128300813 | 73671 | 2 | ENSMUST00 | 0 | Tead4 | ENSMUSG00000030353.10 |
| chr1 | 139454422 | 139455665 | 37.78396 | 121.82604 | 117.62474 | Promoter | 1 | 139454773 | 139474334 | 19562 | 1 | ENSMUST00 | 0 | Aspm | ENSMUSG00000033952.9 |
| chr5 | 121710758 | 121713663 | 37.78396 | 121.82604 | 117.62474 | Promoter | 5 | 121711337 | 121749183 | 37847 | 1 | ENSMUST00 | 0 | Atxn2 | ENSMUSG00000042605.10 |
| chr16 | 57230448 | 57231979 | 34.17132 | 121.80145 | 117.60049 | Promoter | 16 | 57168866 | 57231466 | 62601 | 2 | ENSMUST00 | 0 | Tbc1d23 | ENSMUSG00000022749.6 |
| chr18 | 56707031 | 56709440 | 34.17132 | 121.80145 | 117.60049 | Promoter | 18 | 56707813 | 56753424 | 45612 | 1 | ENSMUST00 | 0 | Lmnb1 | ENSMUSG00000024590.8 |
| chr2 | 168080973 | 168086113 | 34.17132 | 121.80145 | 117.60049 | Promoter | 2 | 168081004 | 168101203 | 20200 | 1 | ENSMUST00 | 0 | Pard6b | ENSMUSG00000044641.7 |
| chr17 | 34024085 | 34025483 | 38.34636 | 121.73327 | 117.53308 | Promoter | 17 | 34020792 | 34024680 | 3889 | 2 | ENSMUST00 | 0 | Ring1 | ENSMUSG00000024325.8 |
| chr19 | 32755517 | 32759967 | 38.34636 | 121.73327 | 117.53308 | Promoter | 19 | 32757497 | 32826160 | 68664 | 1 | ENSMUST00 | 0 | Pten | ENSMUSG00000013663.7 |
| chr18 | 5591062 | 5593921 | 37.10487 | 121.71056 | 117.5108 | Promoter | 18 | 5591860 | 5775467 | 183608 | 1 | ENSMUST00 | 0 | Zeb1 | ENSMUSG00000024238.8 |
| chr1 | 91539734 | 91541534 | 37.70819 | 121.69438 | 117.49508 | Promoter | 1 | 91540544 | 91556088 | 15545 | 1 | ENSMUST00 | 0 | Asb1 | ENSMUSG00000026311.11 |
| chr15 | 96697901 | 96700652 | 37.70819 | 121.69438 | 117.49508 | Promoter | 15 | 96687394 | 96699698 | 12305 | 2 | ENSMUST00 | 0 | Slc38a2 | ENSMUSG00000022462.6 |
| chr16 | 32331774 | 32333666 | 37.70819 | 121.69438 | 117.49508 | Promoter | 16 | 32332252 | 32393747 | 61496 | 1 | ENSMUST00 | 0 | Ubxn7 | ENSMUSG00000053774.8 |
| chr3 | 108653419 | 108654919 | 37.70819 | 121.69438 | 117.49508 | Promoter | 3 | 108653913 | 108678840 | 24928 | 1 | ENSMUST00 | 0 | Clcc1 | ENSMUSG00000027884.11 |
| chr6 | 117840572 | 117842150 | 37.70819 | 121.69438 | 117.49508 | Promoter | 6 | 117841180 | 117844518 | 3339 | 1 | ENSMUST00 | 0 | Zfp637 | ENSMUSG00000059689.9 |
| chr11 | 120571973 | 120572955 | 36.4644 | 121.65416 | 117.45652 | Promoter | 11 | 120560945 | 120572841 | 11897 | 2 | ENSMUST00 | 0 | P4hb | ENSMUSG00000025130.6 |
| chr5 | 8422049 | 8423680 | 36.4644 | 121.65416 | 117.45652 | Promoter | 5 | 8396973 | 8422680 | 25708 | 2 | ENSMUST00 | 0 | Dbf4 | ENSMUSG00000002297.10 |
| chr10 | 88501897 | 88504448 | 38.2674 | 121.59672 | 117.39927 | Promoter | 10 | 88469514 | 88504073 | 34560 | 2 | ENSMUST00 | 0 | Chpt1 | ENSMUSG00000060002.9 |
| chr18 | 60560230 | 60561617 | 38.2674 | 121.59672 | 117.39927 | Promoter | 18 | 60560736 | 60572810 | 12075 | 1 | ENSMUST00 | 0 | Rbm22 | ENSMUSG00000024604.7 |
| chr4 | 41134685 | 41137061 | 38.2674 | 121.59672 | 117.39927 | Promoter | 4 | 41135743 | 41193380 | 57638 | 1 | ENSMUST00 | 0 | Ube2r2 | ENSMUSG00000036241.3 |
| chr19 | 5387442 | 5390053 | 26.42485 | 121.59232 | 117.39527 | Promoter | 19 | 5377523 | 5388703 | 11181 | 2 | ENSMUST00 | 0 | Sart1 | ENSMUSG00000039148.4 |
| chr2 | 32395195 | 32397243 | 37.03263 | 121.58421 | 117.38725 | Promoter | 2 | 32395896 | 32402739 | 6844 | 1 | ENSMUST00 | 0 | Ptges2 | ENSMUSG00000026820.4 |
| chr16 | 32913705 | 32915472 | 35.22365 | 121.58154 | 117.38489 | Promoter | 16 | 32914100 | 33015647 | 101548 | 1 | ENSMUST00 | 0 | Lrch3 | ENSMUSG00000022801.8 |
| chr3 | 98012900 | 98014942 | 35.22365 | 121.58154 | 117.38489 | Promoter | 3 | 98013538 | 98150367 | 136830 | 1 | ENSMUST00 | 0 | Notch2 | ENSMUSG00000027878.10 |
| chr3 | 145937966 | 145938964 | 35.22365 | 121.58154 | 117.38489 | Promoter | 3 | 145938014 | 145944829 | 6816 | 1 | ENSMUST00 | 0 | 2410004B18 | ENSMUSG00000036873.8 |
| chr10 | 67126347 | 67127904 | 37.63272 | 121.56319 | 117.36774 | Promoter | 10 | 67127258 | 67256318 | 129061 | 1 | ENSMUST00 | 0 | Jmjd1c | ENSMUSG00000037876.11 |
| chr4 | 121214004 | 121215785 | 37.63272 | 121.56319 | 117.36774 | Promoter | 4 | 121145885 | 121215084 | 69200 | 2 | ENSMUST00 | 0 | Rlf | ENSMUSG00000049878.7 |
| chr5 | 123227842 | 123228997 | 37.63272 | 121.56319 | 117.36774 | Promoter | 5 | 123228190 | 123245268 | 17079 | 1 | ENSMUST00 | 0 | Psmd9 | ENSMUSG00000029440.11 |
| chr11 | 44469697 | 44471607 | 35.79337 | 121.53349 | 117.33853 | Promoter | 11 | 44454571 | 44470498 | 15928 | 2 | ENSMUST00 | 0 | Ublcp1 | ENSMUSG00000041231.10 |
| chr10 | 21992730 | 21996734 | 36.39542 | 121.53264 | 117.33772 | Promoter | 10 | 21993890 | 21999891 | 6002 | 1 | ENSMUST00 | 0 | Sgk1 | ENSMUSG00000019970.10 |
| chr11 | 115512932 | 115514445 | 36.39542 | 121.53264 | 117.33772 | Promoter | 11 | 115497353 | 115514387 | 17035 | 2 | ENSMUST00 | 0 | Hn1 | ENSMUSG00000020737.7 |
| chr11 | 50826888 | 50828109 | 36.39542 | 121.53264 | 117.33772 | Promoter | 11 | 50811086 | 50827690 | 16605 | 2 | ENSMUST00 | 0 | Zfp354c | ENSMUSG00000044807.8 |
| chr11 | 19923664 | 19926262 | 36.39542 | 121.53264 | 117.33772 | Promoter | 11 | 19924375 | 20024026 | 99652 | 1 | ENSMUST00 | 0 | Spred2 | ENSMUSG00000045671.12 |
| chr12 | 4906536 | 4908244 | 36.39542 | 121.53264 | 117.33772 | Promoter | 12 | 4879032 | 4907705 | 28674 | 2 | ENSMUST00 | 0 | Ubxn2a | ENSMUSG00000020634.7 |
| chr14 | 60250596 | 60252614 | 36.39542 | 121.53264 | 117.33772 | Promoter | 14 | 60219204 | 60251507 | 32304 | 2 | ENSMUST00 | 0 | Nupl1 | ENSMUSG00000063895.6 |
| chr15 | 99392917 | 99394158 | 36.39542 | 121.53264 | 117.33772 | Promoter | 15 | 99392986 | 99408536 | 15551 | 1 | ENSMUST00 | 0 | Tmbim6 | ENSMUSG00000023010.9 |
| chr17 | 49428325 | 49430731 | 36.39542 | 121.53264 | 117.33772 | Promoter | 17 | 49428362 | 49437941 | 9580 | 1 | ENSMUST00 | 0 | Mocs1 | ENSMUSG00000064120.8 |
| chr18 | 24602821 | 24604600 | 36.39542 | 121.53264 | 117.33772 | Promoter | 18 | 24579881 | 24603817 | 23937 | 2 | ENSMUST00 | 0 | Slc39a6 | ENSMUSG00000024270.6 |
| chr1 | 180255386 | 180256935 | 32.3084 | 121.498 | 117.30614 | Promoter | 1 | 180227004 | 180256300 | 29297 | 2 | ENSMUST00 | 0 | Psen2 | ENSMUSG00000010609.10 |
| chr10 | 89744559 | 89746078 | 32.3084 | 121.498 | 117.30614 | Promoter | 10 | 89744991 | 89819869 | 74879 | 1 | ENSMUST00 | 0 | Uhrf1bp1l | ENSMUSG00000019951.9 |
| chr11 | 107546712 | 107548785 | 32.3084 | 121.498 | 117.30614 | Promoter | 11 | 107547930 | 107578156 | 30227 | 1 | ENSMUST00 | 0 | Helz | ENSMUSG00000020721.11 |
| chr12 | 111538117 | 111539637 | 32.3084 | 121.498 | 117.30614 | Promoter | 12 | 111538853 | 111544923 | 6071 | 1 | ENSMUST00 | 0 | Eif5 | ENSMUSG00000021282.11 |
| chr12 | 71831028 | 71832533 | 32.3084 | 121.498 | 117.30614 | Promoter | 12 | 71831078 | 71992367 | 161290 | 1 | ENSMUST00 | 0 | Daam1 | ENSMUSG00000034574.9 |
| chr12 | 103424964 | 103425929 | 32.3084 | 121.498 | 117.30614 | Promoter | 12 | 103407976 | 103425780 | 17805 | 2 | ENSMUST00 | 0 | Ddx24 | ENSMUSG00000041645.8 |
| chr13 | 96541915 | 96544264 | 32.3084 | 121.498 | 117.30614 | Promoter | 13 | 96480689 | 96542485 | 61797 | 2 | ENSMUST00 | 0 | Polk | ENSMUSG00000021668.9 |
| chr13 | 104108731 | 104110687 | 32.3084 | 121.498 | 117.30614 | Promoter | 13 | 104023439 | 104109614 | 86176 | 2 | ENSMUST00 | 0 | Nln | ENSMUSG00000021710.10 |
| chr16 | 11133497 | 11135404 | 32.3084 | 121.498 | 117.30614 | Promoter | 16 | 11074911 | 11134650 | 59740 | 2 | ENSMUST00 | 0 | Txndc11 | ENSMUSG00000022498.11 |
| chr17 | 17622986 | 17626193 | 32.3084 | 121.498 | 117.30614 | Promoter | 17 | 17527723 | 17624489 | 96767 | 2 | ENSMUST00 | 0 | Lnpep | ENSMUSG00000023845.6 |
| chr17 | 24959729 | 24961229 | 32.3084 | 121.498 | 117.30614 | Promoter | 17 | 24942156 | 24960636 | 18481 | 2 | ENSMUST00 | 0 | Hn1l | ENSMUSG00000024165.8 |
| chr17 | 35969007 | 35970467 | 32.3084 | 121.498 | 117.30614 | Promoter | 17 | 35956819 | 35969761 | 12943 | 2 | ENSMUST00 | 0 | Abcf1 | ENSMUSG00000038762.10 |
| chr19 | 5770509 | 5771473 | 32.3084 | 121.498 | 117.30614 | Promoter | 19 | 5758427 | 5771401 | 12975 | 2 | ENSMUST00 | 0 | Scyl1 | ENSMUSG00000024941.8 |
| chr2 | 163994559 | 163996605 | 32.3084 | 121.498 | 117.30614 | Promoter | 2 | 163994960 | 164018588 | 23629 | 1 | ENSMUST00 | 0 | Ywhab | ENSMUSG00000018326.4 |
| chr2 | 32875288 | 32879031 | 32.3084 | 121.498 | 117.30614 | Promoter | 2 | 32876114 | 32925254 | 49141 | 1 | ENSMUST00 | 0 | Fam129b | ENSMUSG00000026796.11 |
| chr2 | 151493919 | 151495065 | 32.3084 | 121.498 | 117.30614 | Promoter | 2 | 151494182 | 151511310 | 17129 | 1 | ENSMUST00 | 0 | Nsfl1c | ENSMUSG00000027455.11 |
| chr4 | 123916660 | 123919069 | 32.3084 | 121.498 | 117.30614 | Promoter | 4 | 123917446 | 123936997 | 19552 | 1 | ENSMUST00 | 0 | Rragc | ENSMUSG00000028646.11 |
| chr4 | 11965470 | 11967183 | 32.3084 | 121.498 | 117.30614 | Promoter | 4 | 11958184 | 11966452 | 8269 | 2 | ENSMUST00 | 0 | Pdp1 | ENSMUSG00000049225.9 |
| chr5 | 139791432 | 139793535 | 32.3084 | 121.498 | 117.30614 | Promoter | 5 | 139791513 | 139802653 | 11141 | 1 | ENSMUST00 | 0 | Mafk | ENSMUSG00000018143.5 |
| chr5 | 63967664 | 63969562 | 32.3084 | 121.498 | 117.30614 | Promoter | 5 | 63908896 | 63968905 | 60010 | 2 | ENSMUST00 | 0 | Rell1 | ENSMUSG00000047881.9 |
| chr7 | 12478018 | 12479279 | 32.3084 | 121.498 | 117.30614 | Promoter | 7 | 12478293 | 12496235 | 17943 | 1 | ENSMUST00 | 0 | Zfp606 | ENSMUSG00000030386.12 |
| chr9 | 66974580 | 66975855 | 32.3084 | 121.498 | 117.30614 | Promoter | 9 | 66955393 | 66975484 | 20092 | 2 | ENSMUST00 | 0 | Lactb | ENSMUSG00000032370.7 |
| chr9 | 21424870 | 21426244 | 32.3084 | 121.498 | 117.30614 | Promoter | 9 | 21424908 | 21507754 | 82847 | 1 | ENSMUST00 | 0 | Dnm2 | ENSMUSG00000033335.12 |
| chr9 | 70011985 | 70013521 | 32.3084 | 121.498 | 117.30614 | Promoter | 9 | 70012550 | 70022866 | 10317 | 1 | ENSMUST00 | 0 | Gtf2a2 | ENSMUSG00000033543.12 |
| chr9 | 114400674 | 114402241 | 32.3084 | 121.498 | 117.30614 | Promoter | 9 | 114401076 | 114474376 | 73301 | 1 | ENSMUST00 | 0 | Glb1 | ENSMUSG00000045594.9 |
| chr10 | 80053524 | 80055167 | 38.18876 | 121.46067 | 117.27263 | Promoter | 10 | 80054035 | 80055107 | 1073 | 1 | ENSMUST00 | 0 | Gpx4 | ENSMUSG00000075706.5 |
| chr8 | 71611041 | 71612143 | 38.18876 | 121.46067 | 117.27263 | Promoter | 8 | 71611024 | 71624909 | 13886 | 1 | ENSMUST00 | 18 | Colgalt1 | ENSMUSG00000034807.8 |
| chr8 | 124230883 | 124232575 | 38.18876 | 121.46067 | 117.27263 | Promoter | 8 | 124231391 | 124345722 | 114332 | 1 | ENSMUST00 | 0 | Galnt2 | ENSMUSG00000089704.4 |
| chr1 | 86581764 | 86583368 | 36.96067 | 121.45832 | 117.27072 | Promoter | 1 | 86542994 | 86582547 | 39554 | 2 | ENSMUST00 | 0 | Pde6d | ENSMUSG00000026239.9 |
| chr11 | 60776300 | 60778387 | 36.96067 | 121.45832 | 117.27072 | Promoter | 11 | 60740058 | 60777365 | 37308 | 2 | ENSMUST00 | 0 | Top3a | ENSMUSG00000002814.10 |
| chr12 | 51970357 | 51971857 | 33.40766 | 121.45458 | 117.2672 | Promoter | 12 | 51875873 | 51971321 | 95449 | 2 | ENSMUST00 | 0 | Heatr5a | ENSMUSG00000035181.5 |
| chr7 | 80113906 | 80115785 | 33.40766 | 121.45458 | 117.2672 | Promoter | 7 | 80094846 | 80115392 | 20547 | 2 | ENSMUST00 | 0 | Idh2 | ENSMUSG00000030541.11 |
| chr9 | 97109890 | 97111659 | 30.11944 | 121.38741 | 117.20103 | Promoter | 9 | 97074961 | 97111157 | 36197 | 2 | ENSMUST00 | 0 | Slc25a36 | ENSMUSG00000032449.8 |
| chr11 | 80377345 | 80378452 | 36.88899 | 121.33285 | 117.14651 | Promoter | 11 | 80360492 | 80378015 | 17524 | 2 | ENSMUST00 | 0 | 5730455P16 | ENSMUSG00000057181.10 |
| chr13 | 58273089 | 58275023 | 36.88899 | 121.33285 | 117.14651 | Promoter | 13 | 58233351 | 58274188 | 40838 | 2 | ENSMUST00 | 0 | Gkap1 | ENSMUSG00000021552.6 |
| chr3 | 16182684 | 16184590 | 38.11044 | 121.32513 | 117.13898 | Promoter | 3 | 16183183 | 16217035 | 33853 | 1 | ENSMUST00 | 0 | Ythdf3 | ENSMUSG00000047213.9 |
| chr14 | 55721408 | 55722416 | 37.48268 | 121.30223 | 117.11643 | Promoter | 14 | 55715873 | 55722198 | 6326 | 2 | ENSMUST00 | 0 | Rabggta | ENSMUSG00000040472.9 |
| chr8 | 85554247 | 85556254 | 36.81759 | 121.2078 | 117.02259 | Promoter | 8 | 85537640 | 85555271 | 17632 | 2 | ENSMUST00 | 0 | Dnaja2 | ENSMUSG00000031701.5 |
| chr19 | 3707741 | 3709052 | 38.03245 | 121.19012 | 117.00504 | Promoter | 19 | 3708333 | 3717881 | 9549 | 1 | ENSMUST00 | 0 | 1810055G02 | ENSMUSG00000035372.1 |
| chr5 | 65334890 | 65336115 | 38.03245 | 121.19012 | 117.00504 | Promoter | 5 | 65261852 | 65335670 | 73819 | 2 | ENSMUST00 | 0 | Rfc1 | ENSMUSG00000029191.11 |
| chr7 | 30444157 | 30445874 | 38.03245 | 121.19012 | 117.00504 | Promoter | 7 | 30434981 | 30445582 | 10602 | 2 | ENSMUST00 | 0 | Aplp1 | ENSMUSG00000006651.7 |
| chr8 | 27173868 | 27175786 | 38.03245 | 121.19012 | 117.00504 | Promoter | 8 | 27138773 | 27174646 | 35874 | 2 | ENSMUST00 | 0 | Rab11fip1 | ENSMUSG00000031488.8 |
| chr8 | 77517035 | 77518647 | 38.03245 | 121.19012 | 117.00504 | Promoter | 8 | 77250366 | 77517907 | 267542 | 2 | ENSMUST00 | 0 | Arhgap10 | ENSMUSG00000037148.7 |
| chr9 | 109094798 | 109099441 | 38.03245 | 121.19012 | 117.00504 | Promoter | 9 | 109093844 | 109097796 | 3953 | 2 | ENSMUST00 | 0 | Gm7628 | ENSMUSG00000025644.10 |
| chr11 | 69913419 | 69914903 | 32.69259 | 121.17415 | 116.99028 | Promoter | 11 | 69913888 | 69916576 | 2689 | 1 | ENSMUST00 | 0 | Gps2 | ENSMUSG00000023170.9 |
| chr3 | 93554542 | 93557324 | 37.4081 | 121.17245 | 116.98877 | Promoter | 3 | 93555080 | 93564643 | 9564 | 1 | ENSMUST00 | 0 | S100a10 | ENSMUSG00000041959.9 |
| chr8 | 72491599 | 72492739 | 36.19004 | 121.17046 | 116.98717 | Promoter | 8 | 72480641 | 72492614 | 11974 | 2 | ENSMUST00 | 0 | Slc35e1 | ENSMUSG00000019731.8 |
| chr17 | 87974423 | 87976176 | 27.57146 | 121.15424 | 116.97103 | Promoter | 17 | 87975050 | 87990883 | 15834 | 1 | ENSMUST00 | 0 | Msh6 | ENSMUSG00000005370.3 |
| chr10 | 24869515 | 24870614 | 34.38402 | 121.12925 | 116.94628 | Promoter | 10 | 24869986 | 24913681 | 43696 | 1 | ENSMUST00 | 0 | Med23 | ENSMUSG00000019984.11 |
| chr14 | 31829329 | 31831818 | 34.38402 | 121.12925 | 116.94628 | Promoter | 14 | 31700015 | 31830415 | 130401 | 2 | ENSMUST00 | 0 | Ankrd28 | ENSMUSG00000014496.7 |
| chr6 | 124414215 | 124416051 | 34.38402 | 121.12925 | 116.94628 | Promoter | 6 | 124396816 | 124415067 | 18252 | 2 | ENSMUST00 | 0 | Pex5 | ENSMUSG00000005069.7 |
| chr7 | 144581347 | 144582972 | 34.38402 | 121.12925 | 116.94628 | Promoter | 7 | 144577318 | 144582463 | 5146 | 2 | ENSMUST00 | 0 | Fadd | ENSMUSG00000031077.7 |
| chr1 | 36307691 | 36309096 | 38.6007 | 121.10697 | 116.92481 | Promoter | 1 | 36307733 | 36319203 | 11471 | 1 | ENSMUST00 | 0 | Arid5a | ENSMUSG00000037447.11 |
| chr10 | 77581692 | 77584244 | 38.6007 | 121.10697 | 116.92481 | Promoter | 10 | 77581720 | 77598732 | 17013 | 1 | ENSMUST00 | 0 | Pttg1ip | ENSMUSG00000009291.7 |
| chr12 | 65035953 | 65037435 | 38.6007 | 121.10697 | 116.92481 | Promoter | 12 | 65036333 | 65063386 | 27054 | 1 | ENSMUST00 | 0 | Prpf39 | ENSMUSG00000035597.13 |
| chr13 | 54610388 | 54611912 | 38.6007 | 121.10697 | 116.92481 | Promoter | 13 | 54592401 | 54611344 | 18944 | 2 | ENSMUST00 | 0 | Cltb | ENSMUSG00000047547.9 |
| chr15 | 93594446 | 93596861 | 38.6007 | 121.10697 | 116.92481 | Promoter | 15 | 93499114 | 93595891 | 96778 | 2 | ENSMUST00 | 0 | Prickle1 | ENSMUSG00000036158.7 |
| chr4 | 117133149 | 117134433 | 38.6007 | 121.10697 | 116.92481 | Promoter | 4 | 117128655 | 117133944 | 5290 | 2 | ENSMUST00 | 0 | Plk3 | ENSMUSG00000028680.9 |
| chr4 | 59034915 | 59035880 | 38.6007 | 121.10697 | 116.92481 | Promoter | 4 | 59035088 | 59041903 | 6816 | 1 | ENSMUST00 | 0 | Gng10 | ENSMUSG00000038607.7 |
| chr4 | 32950135 | 32951389 | 38.6007 | 121.10697 | 116.92481 | Promoter | 4 | 32804035 | 32950826 | 146792 | 2 | ENSMUST00 | 0 | Ankrd6 | ENSMUSG00000040183.8 |
| chr7 | 80860265 | 80861546 | 38.6007 | 121.10697 | 116.92481 | Promoter | 7 | 80860920 | 80874969 | 14050 | 1 | ENSMUST00 | 0 | Zscan2 | ENSMUSG00000038797.8 |
| chr8 | 117256201 | 117258587 | 38.6007 | 121.10697 | 116.92481 | Promoter | 8 | 117257019 | 117461503 | 204485 | 1 | ENSMUST00 | 0 | Cmip | ENSMUSG00000034390.10 |
| chr8 | 36247653 | 36249429 | 38.6007 | 121.10697 | 116.92481 | Promoter | 8 | 36216064 | 36249516 | 33453 | 2 | ENSMUST00 | 87 | Lonrf1 | ENSMUSG00000039633.6 |
| chr16 | 13779904 | 13781578 | 37.95477 | 121.05561 | 116.87667 | Promoter | 16 | 13780699 | 13814840 | 34142 | 1 | ENSMUST00 | 0 | Rrn3 | ENSMUSG00000022682.7 |
| chr9 | 87021424 | 87022785 | 37.33383 | 121.04314 | 116.86485 | Promoter | 9 | 87022014 | 87077774 | 55761 | 1 | ENSMUST00 | 0 | Cyb5r4 | ENSMUSG00000032872.11 |
| chr1 | 13659369 | 13661089 | 30.45735 | 121.03082 | 116.85291 | Promoter | 1 | 13623330 | 13660546 | 37217 | 2 | ENSMUST00 | 0 | Lactb2 | ENSMUSG00000025937.6 |
| chr12 | 101974869 | 101976971 | 30.45735 | 121.03082 | 116.85291 | Promoter | 12 | 101975974 | 102005993 | 30020 | 1 | ENSMUST00 | 0 | Cpsf2 | ENSMUSG00000041781.8 |
| chr15 | 11398921 | 11400293 | 30.45735 | 121.03082 | 116.85291 | Promoter | 15 | 11383663 | 11399658 | 15996 | 2 | ENSMUST00 | 0 | Tars | ENSMUSG00000022241.5 |
| chr17 | 36270565 | 36272542 | 30.45735 | 121.03082 | 116.85291 | Promoter | 17 | 36258873 | 36271567 | 12695 | 2 | ENSMUST00 | 0 | Trim39 | ENSMUSG00000045409.11 |
| chr2 | 130838998 | 130840815 | 30.45735 | 121.03082 | 116.85291 | Promoter | 2 | 130706200 | 130840134 | 133935 | 2 | ENSMUST00 | 0 | 4930402H24 | ENSMUSG00000027309.13 |
| chr2 | 153442889 | 153444629 | 30.45735 | 121.03082 | 116.85291 | Promoter | 2 | 153407466 | 153444479 | 37014 | 2 | ENSMUST00 | 0 | Nol4l | ENSMUSG00000061411.7 |
| chr3 | 9830234 | 9834375 | 30.45735 | 121.03082 | 116.85291 | Promoter | 3 | 9687479 | 9833679 | 146201 | 2 | ENSMUST00 | 0 | Pag1 | ENSMUSG00000027508.10 |
| chr6 | 52157936 | 52159956 | 30.45735 | 121.03082 | 116.85291 | Promoter | 6 | 52155590 | 52158317 | 2728 | 2 | ENSMUST00 | 0 | Hoxa1 | ENSMUSG00000029844.9 |
| chr6 | 82652157 | 82653322 | 30.45735 | 121.03082 | 116.85291 | Promoter | 6 | 82621852 | 82652870 | 31019 | 2 | ENSMUST00 | 0 | Pole4 | ENSMUSG00000030042.10 |
| chr13 | 46726280 | 46728669 | 36.67561 | 120.95901 | 116.78313 | Promoter | 13 | 46679905 | 46727940 | 48036 | 2 | ENSMUST00 | 0 | Nup153 | ENSMUSG00000021374.9 |
| chr2 | 144599540 | 144600703 | 36.67561 | 120.95901 | 116.78313 | Promoter | 2 | 144599897 | 144768758 | 168862 | 1 | ENSMUST00 | 0 | Dtd1 | ENSMUSG00000027430.7 |

| | | | | | | | | | | | | | | | |
|---|---|---|---|---|---|---|---|---|---|---|---|---|---|---|---|
| chr4 | 62359649 | 62361552 | 36.67561 | 120.95901 | 116.78313 | Promoter | 4 | 62360727 | 62391769 | 31043 | 1 | ENSMUST00 | 0 | Slc31a1 | ENSMUSG00000066150.7 |
| chr9 | 44232995 | 44234966 | 36.67561 | 120.95901 | 116.78313 | Promoter | 9 | 44149262 | 44234046 | 84785 | 2 | ENSMUST00 | 0 | Cbl | ENSMUSG00000034342.8 |
| chr13 | 31805773 | 31812441 | 35.46717 | 120.9532 | 116.77927 | Promoter | 13 | 31806691 | 31810643 | 3953 | 1 | ENSMUST00 | 0 | Foxc1 | ENSMUSG00000050295.3 |
| chr3 | 95314707 | 95316369 | 35.46717 | 120.9532 | 116.77927 | Promoter | 3 | 95314791 | 95321880 | 7090 | 1 | ENSMUST00 | 0 | Cers2 | ENSMUSG00000015714.6 |
| chr4 | 49596702 | 49598204 | 35.46717 | 120.9532 | 116.77927 | Promoter | 4 | 49584506 | 49597876 | 13371 | 2 | ENSMUST00 | 0 | Tmem246 | ENSMUSG00000039611.2 |
| chr1 | 43826816 | 43828457 | 28.89995 | 120.94616 | 116.77392 | Promoter | 1 | 43746966 | 43827800 | 80835 | 2 | ENSMUST00 | 0 | Uxs1 | ENSMUSG00000057363.7 |
| chr12 | 87442576 | 87444803 | 28.89995 | 120.94616 | 116.77392 | Promoter | 12 | 87443896 | 87452206 | 8311 | 1 | ENSMUST00 | 0 | Slirp | ENSMUSG00000021040.10 |
| chr13 | 111488368 | 111489932 | 28.89995 | 120.94616 | 116.77392 | Promoter | 13 | 111425932 | 111490028 | 64097 | 2 | ENSMUST00 | 96 | Gpbp1 | ENSMUSG00000032745.12 |
| chr17 | 35424438 | 35426991 | 28.89995 | 120.94616 | 116.77392 | Promoter | 17 | 35424850 | 35430055 | 5206 | 1 | ENSMUST00 | 0 | H2-Q6 | ENSMUSG00000073409.7 |
| chr17 | 83213509 | 83220055 | 37.87741 | 120.9216 | 116.74998 | Promoter | 17 | 83215292 | 83225070 | 9779 | 1 | ENSMUST00 | 0 | Pkdcc | ENSMUSG00000024247.9 |
| chr11 | 75486703 | 75488165 | 38.4506 | 120.84806 | 116.67712 | Promoter | 11 | 75486816 | 75509447 | 22632 | 1 | ENSMUST00 | 0 | Prpf8 | ENSMUSG00000020850.9 |
| chr1 | 9943023 | 9944627 | 37.18615 | 120.78587 | 116.61589 | Promoter | 1 | 9941959 | 9944118 | 2160 | 2 | ENSMUST00 | 0 | Snhg6 | ENSMUSG00000098234.2 |
| chr19 | 29812174 | 29813688 | 37.18615 | 120.78587 | 116.61589 | Promoter | 19 | 29808108 | 29812974 | 4867 | 2 | ENSMUST00 | 0 | Ranbp6 | ENSMUSG00000074909.3 |
| chr4 | 32238283 | 32240386 | 37.18615 | 120.78587 | 116.61589 | Promoter | 4 | 32238804 | 32586104 | 347301 | 1 | ENSMUST00 | 0 | Bach2 | ENSMUSG00000040270.11 |
| chr10 | 71284153 | 71285709 | 34.16929 | 120.7392 | 116.5701 | Promoter | 10 | 71254980 | 71285262 | 30283 | 2 | ENSMUST00 | 0 | Ube2d1 | ENSMUSG00000019927.6 |
| chr11 | 83753232 | 83754816 | 34.16929 | 120.7392 | 116.5701 | Promoter | 11 | 83753696 | 83783754 | 30059 | 1 | ENSMUST00 | 0 | Heatr6 | ENSMUSG00000000976.8 |
| chr12 | 54861687 | 54863386 | 34.16929 | 120.7392 | 116.5701 | Promoter | 12 | 54858817 | 54862877 | 4061 | 2 | ENSMUST00 | 0 | Cfl2 | ENSMUSG00000062929.7 |
| chr16 | 10411593 | 10412611 | 34.16929 | 120.7392 | 116.5701 | Promoter | 16 | 10411948 | 10424428 | 12481 | 1 | ENSMUST00 | 0 | Nubp1 | ENSMUSG00000022503.5 |
| chr16 | 35363020 | 35364493 | 34.16929 | 120.7392 | 116.5701 | Promoter | 16 | 35311131 | 35363918 | 52788 | 2 | ENSMUST00 | 0 | Sec22a | ENSMUSG00000034473.8 |
| chr17 | 24454317 | 24455384 | 34.16929 | 120.7392 | 116.5701 | Promoter | 17 | 24443792 | 24455311 | 11520 | 2 | ENSMUST00 | 0 | E4f1 | ENSMUSG00000024137.8 |
| chr18 | 42274785 | 42276250 | 34.16929 | 120.7392 | 116.5701 | Promoter | 18 | 42275353 | 42341542 | 66190 | 1 | ENSMUST00 | 0 | Rbm27 | ENSMUSG00000024491.10 |
| chr18 | 66001779 | 66003151 | 34.16929 | 120.7392 | 116.5701 | Promoter | 18 | 65980754 | 66002637 | 21884 | 2 | ENSMUST00 | 0 | Lman1 | ENSMUSG00000041891.10 |
| chr18 | 67774259 | 67776254 | 34.16929 | 120.7392 | 116.5701 | Promoter | 18 | 67774876 | 67795461 | 20586 | 1 | ENSMUST00 | 0 | Seh1l | ENSMUSG00000079614.3 |
| chr19 | 6839670 | 6841212 | 34.16929 | 120.7392 | 116.5701 | Promoter | 19 | 6829085 | 6840601 | 11517 | 2 | ENSMUST00 | 0 | Rps6ka4 | ENSMUSG00000024952.7 |
| chr19 | 11911493 | 11913356 | 34.16929 | 120.7392 | 116.5701 | Promoter | 19 | 11912399 | 11945096 | 32698 | 1 | ENSMUST00 | 0 | Patl1 | ENSMUSG00000046139.7 |
| chr2 | 39064853 | 39066259 | 34.16929 | 120.7392 | 116.5701 | Promoter | 2 | 39016261 | 39065488 | 49228 | 2 | ENSMUST00 | 0 | Golga1 | ENSMUSG00000026754.11 |
| chr2 | 156178007 | 156180253 | 34.16929 | 120.7392 | 116.5701 | Promoter | 2 | 156147239 | 156180238 | 33000 | 2 | ENSMUST00 | 0 | Rbm39 | ENSMUSG00000027620.11 |
| chr3 | 79590859 | 79592433 | 34.16929 | 120.7392 | 116.5701 | Promoter | 3 | 79591342 | 79603276 | 11935 | 1 | ENSMUST00 | 0 | Ppid | ENSMUSG00000027804.8 |
| chr3 | 104863463 | 104866019 | 34.16929 | 120.7392 | 116.5701 | Promoter | 3 | 104864503 | 104926860 | 62358 | 1 | ENSMUST00 | 0 | St7l | ENSMUSG00000045576.11 |
| chr3 | 135438324 | 135440830 | 34.16929 | 120.7392 | 116.5701 | Promoter | 3 | 135438759 | 135467178 | 28420 | 1 | ENSMUST00 | 0 | Ube2d3 | ENSMUSG00000078578.4 |
| chr4 | 119733173 | 119734497 | 34.16929 | 120.7392 | 116.5701 | Promoter | 4 | 119733784 | 119814748 | 80965 | 1 | ENSMUST00 | 0 | Hivep3 | ENSMUSG00000028634.11 |
| chr4 | 119232006 | 119234210 | 34.16929 | 120.7392 | 116.5701 | Promoter | 4 | 119232915 | 119248972 | 16058 | 1 | ENSMUST00 | 0 | Lepre1 | ENSMUSG00000028641.11 |
| chr4 | 150280873 | 150282960 | 34.16929 | 120.7392 | 116.5701 | Promoter | 4 | 150281646 | 150621966 | 340321 | 1 | ENSMUST00 | 0 | Rere | ENSMUSG00000039852.11 |
| chr5 | 9100104 | 9101841 | 34.16929 | 120.7392 | 116.5701 | Promoter | 5 | 9100737 | 9118983 | 18247 | 1 | ENSMUST00 | 0 | Tmem243 | ENSMUSG00000079659.1 |
| chr6 | 47453518 | 47455734 | 34.16929 | 120.7392 | 116.5701 | Promoter | 6 | 47453576 | 47485193 | 31618 | 1 | ENSMUST00 | 0 | Cul1 | ENSMUSG00000029686.10 |
| chr6 | 52221266 | 52227348 | 34.16929 | 120.7392 | 116.5701 | Promoter | 6 | 52224121 | 52226189 | 2069 | 2 | ENSMUST00 | 0 | Hoxa9 | ENSMUSG00000038227.10 |
| chr7 | 142532699 | 142533937 | 34.16929 | 120.7392 | 116.5701 | Promoter | 7 | 142532799 | 142540737 | 7939 | 1 | ENSMUST00 | 0 | Mrpl23 | ENSMUSG00000037772.7 |
| chr8 | 25015978 | 25017012 | 34.16929 | 120.7392 | 116.5701 | Promoter | 8 | 24949611 | 25016922 | 67312 | 2 | ENSMUST00 | 0 | Adam9 | ENSMUSG00000031555.8 |
| chr8 | 57511815 | 57513277 | 34.16929 | 120.7392 | 116.5701 | Promoter | 8 | 57511843 | 57515999 | 4157 | 1 | ENSMUST00 | 0 | Hmgb2 | ENSMUSG00000054717.6 |
| chr6 | 86523216 | 86527818 | 38.37027 | 120.70945 | 116.54665 | Promoter | 6 | 86524492 | 86526321 | 1830 | 2 | ENSMUST00 | 0 | Pcbp1 | ENSMUSG00000051695.6 |
| chr19 | 59343941 | 59346591 | 37.72364 | 120.6551 | 116.49378 | Promoter | 19 | 59296084 | 59345780 | 49697 | 2 | ENSMUST00 | 0 | Pdzd8 | ENSMUSG00000074746.2 |
| chr1 | 36243287 | 36244763 | 36.46257 | 120.58523 | 116.42456 | Promoter | 1 | 36141530 | 36244305 | 102776 | 2 | ENSMUST00 | 0 | Uggt1 | ENSMUSG00000037470.9 |
| chr11 | 85310340 | 85313068 | 36.46257 | 120.58523 | 116.42456 | Promoter | 11 | 85311244 | 85347066 | 35823 | 1 | ENSMUST00 | 0 | Ppm1d | ENSMUSG00000020525.12 |
| chr12 | 111573966 | 111576269 | 36.46257 | 120.58523 | 116.42456 | Promoter | 12 | 111574510 | 111656227 | 81718 | 1 | ENSMUST00 | 0 | Mark3 | ENSMUSG00000007411.10 |
| chr13 | 59556269 | 59557388 | 36.46257 | 120.58523 | 116.42456 | Promoter | 13 | 59449538 | 59557289 | 107752 | 2 | ENSMUST00 | 0 | Agtpbp1 | ENSMUSG00000021557.9 |
| chr13 | 65240352 | 65242591 | 36.46257 | 120.58523 | 116.42456 | Distal Interge | 13 | 65259539 | 65260826 | 1288 | 1 | ENSMUST00 | -16948 | Gm10775 | ENSMUSG00000074847.2 |
| chr14 | 48446036 | 48447259 | 36.46257 | 120.58523 | 116.42456 | Promoter | 14 | 48446128 | 48512790 | 66663 | 1 | ENSMUST00 | 0 | Tmem260 | ENSMUSG00000036339.12 |
| chr15 | 79544943 | 79546662 | 36.46257 | 120.58523 | 116.42456 | Promoter | 15 | 79527707 | 79546741 | 19035 | 2 | ENSMUST00 | 79 | Ddx17 | ENSMUSG00000055065.6 |
| chr17 | 26113050 | 26114059 | 36.46257 | 120.58523 | 116.42456 | Promoter | 17 | 26113316 | 26123254 | 9939 | 1 | ENSMUST00 | 0 | Tmem8 | ENSMUSG00000024180.8 |
| chr18 | 35498166 | 35499383 | 36.46257 | 120.58523 | 116.42456 | Promoter | 18 | 35266400 | 35498925 | 232526 | 2 | ENSMUST00 | 0 | Sil1 | ENSMUSG00000024357.9 |
| chr18 | 60774063 | 60775805 | 36.46257 | 120.58523 | 116.42456 | Promoter | 18 | 60774510 | 60778546 | 4037 | 1 | ENSMUST00 | 0 | Rps14 | ENSMUSG00000024608.5 |
| chr19 | 46327814 | 46329602 | 36.46257 | 120.58523 | 116.42456 | Promoter | 19 | 46328184 | 46330446 | 2263 | 1 | ENSMUST00 | 0 | Fbxl15 | ENSMUSG00000025226.6 |
| chr2 | 128697969 | 128703570 | 36.46257 | 120.58523 | 116.42456 | Promoter | 2 | 128698956 | 128802894 | 103939 | 1 | ENSMUST00 | 0 | Mertk | ENSMUSG00000014361.5 |
| chr2 | 112367215 | 112368751 | 36.46257 | 120.58523 | 116.42456 | Promoter | 2 | 112363011 | 112368027 | 5017 | 2 | ENSMUST00 | 0 | Emc4 | ENSMUSG00000027131.3 |
| chr2 | 172345482 | 172346421 | 36.46257 | 120.58523 | 116.42456 | Promoter | 2 | 172345565 | 172355749 | 10185 | 1 | ENSMUST00 | 0 | Fam210b | ENSMUSG00000027495.4 |
| chr2 | 3331893 | 3333526 | 36.46257 | 120.58523 | 116.42456 | Promoter | 2 | 3328949 | 3332643 | 3695 | 2 | ENSMUST00 | 0 | Rpp38 | ENSMUSG00000049950.6 |
| chr4 | 70533328 | 70535844 | 36.46257 | 120.58523 | 116.42456 | Promoter | 4 | 70427065 | 70534928 | 107864 | 2 | ENSMUST00 | 0 | Megf9 | ENSMUSG00000039270.4 |
| chr5 | 76303304 | 76305577 | 36.46257 | 120.58523 | 116.42456 | Promoter | 5 | 76212177 | 76304548 | 92372 | 2 | ENSMUST00 | 0 | Clock | ENSMUSG00000029238.8 |
| chr5 | 136565019 | 136569262 | 36.46257 | 120.58523 | 116.42456 | Promoter | 5 | 136248135 | 136567248 | 319114 | 2 | ENSMUST00 | 0 | Cux1 | ENSMUSG00000029705.12 |
| chr6 | 120363210 | 120365573 | 36.46257 | 120.58523 | 116.42456 | Promoter | 6 | 120324323 | 120364374 | 40052 | 2 | ENSMUST00 | 0 | Ccdc77 | ENSMUSG00000030177.9 |
| chr7 | 126395961 | 126396874 | 36.46257 | 120.58523 | 116.42456 | Promoter | 7 | 126382854 | 126396737 | 13884 | 2 | ENSMUST00 | 0 | Nfatc2ip | ENSMUSG00000030722.7 |
| chr7 | 99381440 | 99384522 | 36.46257 | 120.58523 | 116.42456 | Promoter | 7 | 99381549 | 99460983 | 79435 | 1 | ENSMUST00 | 0 | Gdpd5 | ENSMUSG00000035314.8 |
| chr8 | 124661452 | 124663780 | 36.46257 | 120.58523 | 116.42456 | Promoter | 8 | 124635766 | 124663369 | 27604 | 2 | ENSMUST00 | 0 | 2310022B05 | ENSMUSG00000031983.6 |
| chr9 | 46272634 | 46274141 | 36.46257 | 120.58523 | 116.42456 | Promoter | 9 | 46273064 | 46282643 | 9580 | 1 | ENSMUST00 | 0 | Zpr1 | ENSMUSG00000032078.11 |
| chr9 | 72110445 | 72112518 | 36.46257 | 120.58523 | 116.42456 | Promoter | 9 | 71842688 | 72111847 | 269160 | 2 | ENSMUST00 | 0 | Tcf12 | ENSMUSG00000032228.11 |
| chr7 | 97788076 | 97789484 | 38.29027 | 120.57137 | 116.41801 | Distal Interge | 7 | 97726379 | 97738247 | 11869 | 2 | ENSMUST00 | -49830 | Aqp11 | ENSMUSG00000042797.8 |
| chr3 | 94658090 | 94659611 | 37.03965 | 120.53043 | 116.37734 | Promoter | 3 | 94612757 | 94658871 | 46115 | 2 | ENSMUST00 | 0 | Tuft1 | ENSMUSG00000005968.9 |
| chr15 | 77928467 | 77929042 | 37.64722 | 120.52259 | 116.3698 | Promoter | 15 | 77915047 | 77929006 | 13960 | 2 | ENSMUST00 | 0 | Txn2 | ENSMUSG00000005354.10 |
| chr17 | 33890057 | 33890723 | 37.64722 | 120.52259 | 116.3698 | Promoter | 17 | 33875666 | 33890661 | 14996 | 2 | ENSMUST00 | 0 | Kifc1 | ENSMUSG00000079553.5 |
| chr14 | 99098902 | 99100541 | 34.60421 | 120.46815 | 116.3158 | Promoter | 14 | 99099433 | 99254493 | 155061 | 1 | ENSMUST00 | 0 | Pibf1 | ENSMUSG00000022064.9 |
| chr14 | 45388277 | 45389470 | 34.60421 | 120.46815 | 116.3158 | Promoter | 14 | 45376334 | 45388802 | 12469 | 2 | ENSMUST00 | 0 | Gnpnat1 | ENSMUSG00000037722.7 |
| chr5 | 104459201 | 104460760 | 34.60421 | 120.46815 | 116.3158 | Promoter | 5 | 104459450 | 104505819 | 46370 | 1 | ENSMUST00 | 0 | Pkd2 | ENSMUSG00000034462.9 |
| chr5 | 100415482 | 100416869 | 34.60421 | 120.46815 | 116.3158 | Promoter | 5 | 100361649 | 100416234 | 54586 | 2 | ENSMUST00 | 0 | Sec31a | ENSMUSG00000035325.11 |
| chr8 | 119574208 | 119575594 | 34.60421 | 120.46815 | 116.3158 | Promoter | 8 | 119575235 | 119598454 | 23220 | 1 | ENSMUST00 | 0 | Dnaaf1 | ENSMUSG00000031831.6 |
| chr18 | 20895173 | 20897177 | 32.86763 | 120.45166 | 116.30058 | Promoter | 18 | 20817224 | 20896078 | 78855 | 2 | ENSMUST00 | 0 | Trappc8 | ENSMUSG00000033382.8 |
| chr8 | 114439605 | 114440402 | 38.2106 | 120.43382 | 116.28301 | Promoter | 8 | 114439655 | 115352708 | 913054 | 1 | ENSMUST00 | 0 | Wwox | ENSMUSG00000004637.10 |
| chr8 | 120113193 | 120115008 | 38.2106 | 120.43382 | 116.28301 | Promoter | 8 | 120114152 | 120154574 | 40423 | 1 | ENSMUST00 | 0 | 6430548M08 | ENSMUSG00000031824.9 |
| chr11 | 58329516 | 58331743 | 36.96683 | 120.40339 | 116.25284 | Promoter | 11 | 58330724 | 58347728 | 17005 | 1 | ENSMUST00 | 0 | Sh3bp5l | ENSMUSG00000013646.12 |
| chr7 | 120980868 | 120983246 | 36.96683 | 120.40339 | 116.25284 | Promoter | 7 | 120982509 | 121014787 | 32279 | 1 | ENSMUST00 | 0 | 4933427G17 | ENSMUSG00000030877.5 |
| chr8 | 95330917 | 95332916 | 36.96683 | 120.40339 | 116.25284 | Promoter | 8 | 95332292 | 95347507 | 15216 | 1 | ENSMUST00 | 0 | Usb1 | ENSMUSG00000031792.9 |
| chr7 | 45127341 | 45128827 | 37.57111 | 120.39056 | 116.24171 | Promoter | 7 | 45127382 | 45127464 | 83 | 2 | ENSMUST00 | 0 | Snord32a | ENSMUSG00000065219.1 |
| chr8 | 26014771 | 26015921 | 37.57111 | 120.39056 | 116.24171 | Promoter | 8 | 25998729 | 26015601 | 16873 | 2 | ENSMUST00 | 0 | Fnta | ENSMUSG00000015994.9 |
| chr9 | 77542586 | 77546045 | 36.32541 | 120.3443 | 116.1959 | Promoter | 9 | 77430823 | 77544870 | 114048 | 2 | ENSMUST00 | 0 | Lrrc1 | ENSMUSG00000032352.10 |
| chr14 | 55884665 | 55886221 | 35.71972 | 120.33891 | 116.19067 | Promoter | 14 | 55884969 | 55896781 | 11813 | 1 | ENSMUST00 | 0 | Khnyn | ENSMUSG00000047153.3 |
| chr15 | 103365373 | 103367168 | 35.71972 | 120.33891 | 116.19067 | Promoter | 15 | 103344289 | 103366763 | 22475 | 2 | ENSMUST00 | 0 | Itga5 | ENSMUSG00000000555.6 |
| chr2 | 151631861 | 151632838 | 35.71972 | 120.33891 | 116.19067 | Promoter | 2 | 151590549 | 151632549 | 42001 | 2 | ENSMUST00 | 0 | Snph | ENSMUSG00000027457.10 |
| chr5 | 33018827 | 33020735 | 35.71972 | 120.33891 | 116.19067 | Promoter | 5 | 33019761 | 33026976 | 7216 | 1 | ENSMUST00 | 0 | Ywhah | ENSMUSG00000018965.10 |
| chr5 | 30814249 | 30815503 | 35.71972 | 120.33891 | 116.19067 | Promoter | 5 | 30814641 | 30866106 | 51466 | 1 | ENSMUST00 | 0 | Mapre3 | ENSMUSG00000029166.9 |
| chr12 | 110445884 | 110447835 | 36.89429 | 120.27677 | 116.13108 | Distal Interge | 12 | 110485739 | 110583061 | 97323 | 1 | ENSMUST00 | -37904 | Ppp2r5c | ENSMUSG00000017843.8 |
| chr16 | 28928458 | 28930569 | 36.89429 | 120.27677 | 116.13108 | Promoter | 16 | 28826176 | 28929698 | 103523 | 2 | ENSMUST00 | 0 | Mb21d2 | ENSMUSG00000051065.8 |
| chr2 | 11776733 | 11779311 | 36.89429 | 120.27677 | 116.13108 | Promoter | 2 | 11777876 | 11778859 | 984 | 1 | ENSMUST00 | 0 | Ankrd16 | ENSMUSG00000047909.6 |
| chr6 | 113045546 | 113047223 | 37.49531 | 120.259 | 116.11364 | Promoter | 6 | 113046225 | 113068273 | 22049 | 1 | ENSMUST00 | 0 | Thumpd3 | ENSMUSG00000030264.9 |
| chr3 | 67582348 | 67583528 | 32.18068 | 120.21006 | 116.06535 | Promoter | 3 | 67582768 | 67604231 | 21464 | 1 | ENSMUST00 | 0 | Mfsd1 | ENSMUSG00000027775.8 |
| chr7 | 16098562 | 16099601 | 32.18068 | 120.21006 | 116.06535 | Promoter | 7 | 16098608 | 16113535 | 14928 | 1 | ENSMUST00 | 0 | Napa | ENSMUSG00000006024.7 |
| chr14 | 60784529 | 60785401 | 38.05226 | 120.16034 | 116.01576 | Promoter | 14 | 60784566 | 60903610 | 119045 | 1 | ENSMUST00 | 0 | Mipep | ENSMUSG00000021993.9 |
| chr18 | 62977412 | 62979177 | 38.05226 | 120.16034 | 116.01576 | Promoter | 18 | 62977836 | 62999450 | 21615 | 1 | ENSMUST00 | 0 | Napg | ENSMUSG00000024581.7 |
| chr4 | 149305850 | 149308579 | 36.82204 | 120.15061 | 116.00636 | Promoter | 4 | 149176319 | 149307692 | 131374 | 2 | ENSMUST00 | 0 | Kif1b | ENSMUSG00000063077.9 |
| chr4 | 132421420 | 132422946 | 33.79989 | 120.06606 | 115.92274 | Promoter | 4 | 132355923 | 132422489 | 66567 | 2 | ENSMUST00 | 0 | Phactr4 | ENSMUSG00000066043.8 |
| chr10 | 39369298 | 39371780 | 26.17789 | 120.04698 | 115.90433 | Promoter | 10 | 39369764 | 39565381 | 195618 | 1 | ENSMUST00 | 0 | Fyn | ENSMUSG00000019843.9 |
| chr7 | 19561027 | 19563379 | 26.17789 | 120.04698 | 115.90433 | Promoter | 7 | 19530967 | 19562398 | 31432 | 2 | ENSMUST00 | 0 | Ppp1r37 | ENSMUSG00000051403.8 |
| chr19 | 3282042 | 3283633 | 37.97358 | 120.02438 | 115.8822 | Promoter | 19 | 3282901 | 3292837 | 9937 | 1 | ENSMUST00 | 0 | Mrpl21 | ENSMUSG00000024829.7 |
| chr2 | 156311610 | 156313091 | 37.34461 | 119.99735 | 115.85562 | Promoter | 2 | 156312299 | 156375635 | 63337 | 1 | ENSMUST00 | 0 | Cnbd2 | ENSMUSG00000038085.8 |
| chr9 | 114842973 | 114844728 | 37.34461 | 119.99735 | 115.85562 | Promoter | 9 | 114789345 | 114844152 | 54808 | 2 | ENSMUST00 | 0 | Cmtm8 | ENSMUSG00000041012.3 |
| chr8 | 72546955 | 72549355 | 36.11848 | 119.98036 | 115.83894 | Promoter | 8 | 72494558 | 72548310 | 53753 | 2 | ENSMUST00 | 0 | Med26 | ENSMUSG00000045248.5 |
| chr1 | 133659960 | 133661826 | 36.67838 | 119.89957 | 115.75835 | Promoter | 1 | 133619871 | 133661380 | 41510 | 2 | ENSMUST00 | 0 | Zbed6 | ENSMUSG00000094410.2 |
| chr1 | 127773212 | 127775687 | 37.89522 | 119.88892 | 115.74806 | Promoter | 1 | 127774164 | 127793589 | 19426 | 1 | ENSMUST00 | 0 | Ccnt2 | ENSMUSG00000026349.9 |
| chr2 | 30236663 | 30238701 | 37.89522 | 119.88892 | 115.74806 | Promoter | 2 | 30237715 | 30263785 | 26071 | 1 | ENSMUST00 | 0 | Lrrc8a | ENSMUSG00000007476.13 |
| chr2 | 152395767 | 152399538 | 37.89522 | 119.88892 | 115.74806 | Promoter | 2 | 152393611 | 152398063 | 4453 | 2 | ENSMUST00 | 0 | Sox12 | ENSMUSG00000051817.8 |
| chr16 | 10834773 | 10835534 | 37.26972 | 119.86725 | 115.72688 | Promoter | 16 | 10835059 | 10843238 | 8180 | 1 | ENSMUST00 | 0 | Rmi2 | ENSMUSG00000037991.9 |
| chr16 | 20140561 | 20141914 | 37.26972 | 119.86725 | 115.72688 | Promoter | 16 | 20141063 | 20232573 | 91511 | 1 | ENSMUST00 | 0 | Yeats2 | ENSMUSG00000041215.10 |
| chr2 | 5061506 | 5064203 | 37.26972 | 119.86725 | 115.72688 | Promoter | 2 | 5020642 | 5064045 | 43404 | 2 | ENSMUST00 | 0 | Optn | ENSMUSG00000026672.6 |
| chr3 | 138442368 | 138443783 | 37.26972 | 119.86725 | 115.72688 | Promoter | 3 | 138443093 | 138455499 | 12407 | 1 | ENSMUST00 | 0 | Adh5 | ENSMUSG00000028138.3 |
| chr5 | 145083439 | 145084861 | 37.26972 | 119.86725 | 115.72688 | Promoter | 5 | 145083830 | 145108761 | 24932 | 1 | ENSMUST00 | 0 | Arpc1a | ENSMUSG00000029621.9 |
| chr5 | 20950039 | 20952699 | 37.26972 | 119.86725 | 115.72688 | Promoter | 5 | 20893028 | 20951822 | 58795 | 2 | ENSMUST00 | 0 | Rsbn1l | ENSMUSG00000039968.5 |
| chr9 | 21287215 | 21288457 | 37.26972 | 119.86725 | 115.72688 | Promoter | 9 | 21273457 | 21287969 | 14513 | 2 | ENSMUST00 | 0 | Kri1 | ENSMUSG00000035047.4 |
| chr5 | 110106963 | 110108282 | 34.83228 | 119.81895 | 115.68042 | Promoter | 5 | 110103975 | 110108197 | 4223 | 2 | ENSMUST00 | 0 | Gtpbp6 | ENSMUSG00000033434.10 |
| chr6 | 86403561 | 86405415 | 34.83228 | 119.81895 | 115.68042 | Promoter | 6 | 86404219 | 86431201 | 26983 | 1 | ENSMUST00 | 0 | Tia1 | ENSMUSG00000071337.6 |
| chr7 | 140125241 | 140126186 | 34.83228 | 119.81895 | 115.68042 | Promoter | 7 | 140125657 | 140134334 | 8678 | 1 | ENSMUST00 | 0 | Paox | ENSMUSG00000025464.9 |
| chr10 | 128922352 | 128924170 | 31.96831 | 119.80809 | 115.67081 | Promoter | 10 | 128918420 | 128922888 | 4469 | 2 | ENSMUST00 | 0 | Rdh5 | ENSMUSG00000025350.10 |
| chr10 | 18234526 | 18235785 | 31.96831 | 119.80809 | 115.67081 | Promoter | 10 | 18213676 | 18234996 | 21321 | 2 | ENSMUST00 | 0 | Ccdc28a | ENSMUSG00000059554.7 |
| chr11 | 84868751 | 84871497 | 31.96831 | 119.80809 | 115.67081 | Promoter | 11 | 84832361 | 84870738 | 38378 | 2 | ENSMUST00 | 0 | Ggnbp2 | ENSMUSG00000020530.9 |
| chr11 | 70699759 | 70701628 | 31.96831 | 119.80809 | 115.67081 | Promoter | 11 | 70700548 | 70707568 | 7021 | 1 | ENSMUST00 | 0 | Kif1c | ENSMUSG00000020821.12 |
| chr13 | 100832248 | 100833839 | 31.96831 | 119.80809 | 115.67081 | Promoter | 13 | 100802649 | 100833427 | 30779 | 2 | ENSMUST00 | 0 | Slc30a5 | ENSMUSG00000021629.9 |
| chr15 | 89195732 | 89197334 | 31.96831 | 119.80809 | 115.67081 | Promoter | 15 | 89182217 | 89196474 | 14258 | 2 | ENSMUST00 | 0 | Dennd6b | ENSMUSG00000015377.9 |
| chr18 | 49755061 | 49756385 | 31.96831 | 119.80809 | 115.67081 | Promoter | 18 | 49696145 | 49755601 | 59457 | 2 | ENSMUST00 | 0 | Dtwd2 | ENSMUSG00000024505.10 |
| chr18 | 32836880 | 32838394 | 31.96831 | 119.80809 | 115.67081 | Promoter | 18 | 32837225 | 32866350 | 29126 | 1 | ENSMUST00 | 0 | Wdr36 | ENSMUSG00000038299.10 |
| chr2 | 160872139 | 160873587 | 31.96831 | 119.80809 | 115.67081 | Promoter | 2 | 160748708 | 160872998 | 124291 | 2 | ENSMUST00 | 0 | Zhx3 | ENSMUSG00000035877.12 |
| chr4 | 59002771 | 59003983 | 31.96831 | 119.80809 | 115.67081 | Promoter | 4 | 59003172 | 59025573 | 22402 | 1 | ENSMUST00 | 0 | Dnajc25 | ENSMUSG00000070972.8 |

| | | | | | | | | | | | | | | | |
|---|---|---|---|---|---|---|---|---|---|---|---|---|---|---|---|
| chr5 | 38560225 | 38562421 | 31.96831 | 119.80809 | 115.67081 | Promoter | 5 | 38526814 | 38561666 | 34853 | 2 | ENSMUST00 | 0 | Wdr1 | ENSMUSG00000005103.9 |
| chr5 | 34186146 | 34187794 | 31.96831 | 119.80809 | 115.67081 | Promoter | 5 | 34173887 | 34187710 | 13824 | 2 | ENSMUST00 | 0 | Mxd4 | ENSMUSG00000037235.8 |
| chr5 | 33628562 | 33630518 | 31.96831 | 119.80809 | 115.67081 | Promoter | 5 | 33600351 | 33629635 | 29285 | 2 | ENSMUST00 | 0 | Fam53a | ENSMUSG00000037339.12 |
| chr7 | 5079795 | 5081124 | 31.96831 | 119.80809 | 115.67081 | Promoter | 7 | 5080235 | 5098178 | 17944 | 1 | ENSMUST00 | 0 | Epn1 | ENSMUSG00000035203.10 |
| chr8 | 84845023 | 84846923 | 31.96831 | 119.80809 | 115.67081 | Promoter | 8 | 84845437 | 84846830 | 1394 | 2 | ENSMUST00 | 0 | Calr | ENSMUSG00000003814.8 |
| chr8 | 105859948 | 105861575 | 31.96831 | 119.80809 | 115.67081 | Promoter | 8 | 105860634 | 105880402 | 19769 | 1 | ENSMUST00 | 0 | Nutf2 | ENSMUSG00000008450.7 |
| chr8 | 25601119 | 25603913 | 31.96831 | 119.80809 | 115.67081 | Promoter | 8 | 25601601 | 25677214 | 75614 | 1 | ENSMUST00 | 0 | Whsc1l1 | ENSMUSG00000054823.10 |
| chr9 | 66123546 | 66125254 | 31.96831 | 119.80809 | 115.67081 | Promoter | 9 | 66088133 | 66124886 | 36754 | 2 | ENSMUST00 | 0 | Snx1 | ENSMUSG00000032382.9 |
| chr9 | 98985853 | 98987475 | 31.96831 | 119.80809 | 115.67081 | Promoter | 9 | 98986373 | 99002021 | 15649 | 1 | ENSMUST00 | 0 | Faim | ENSMUSG00000032463.5 |
| chr5 | 52782729 | 52783710 | 36.60697 | 119.77473 | 115.64114 | Promoter | 5 | 52783055 | 52823571 | 40517 | 1 | ENSMUST00 | 0 | Zcchc4 | ENSMUSG00000029179.9 |
| chr10 | 53596886 | 53598164 | 37.19513 | 119.7376 | 115.60696 | Promoter | 10 | 53596961 | 53609213 | 12253 | 1 | ENSMUST00 | 0 | Asf1a | ENSMUSG00000019857.6 |
| chr10 | 75516392 | 75518670 | 35.98183 | 119.73972 | 115.60696 | Promoter | 10 | 75517551 | 75537373 | 19823 | 1 | ENSMUST00 | 0 | Snrpd3 | ENSMUSG00000020180.9 |
| chr11 | 116489203 | 116491230 | 35.98183 | 119.73972 | 115.60696 | Promoter | 11 | 116470845 | 116490314 | 19470 | 2 | ENSMUST00 | 0 | Prpsap1 | ENSMUSG00000015869.11 |
| chr11 | 120229779 | 120231698 | 35.98183 | 119.73972 | 115.60696 | Promoter | 11 | 120229802 | 120231585 | 1784 | 2 | ENSMUST00 | 0 | 2900052L18F | ENSMUSG00000043993.6 |
| chr12 | 86725747 | 86726552 | 33.04844 | 119.73734 | 115.60696 | Promoter | 12 | 86700502 | 86726460 | 25959 | 2 | ENSMUST00 | 0 | Angel1 | ENSMUSG00000021257.9 |
| chr13 | 94788578 | 94789846 | 37.19513 | 119.7376 | 115.60696 | Promoter | 13 | 94788943 | 94842899 | 53957 | 1 | ENSMUST00 | 0 | Tbca | ENSMUSG00000042043.5 |
| chr14 | 34310717 | 34312662 | 35.98183 | 119.73972 | 115.60696 | Promoter | 14 | 34310727 | 34345144 | 34418 | 1 | ENSMUST00 | 0 | Glud1 | ENSMUSG00000021794.10 |
| chr14 | 73552312 | 73553765 | 35.98183 | 119.73972 | 115.60696 | Promoter | 14 | 73552632 | 73556827 | 4196 | 1 | ENSMUST00 | 0 | Sucla2 | ENSMUSG00000022110.8 |
| chr15 | 89075128 | 89077118 | 35.98183 | 119.73972 | 115.60696 | Promoter | 15 | 89076013 | 89087077 | 11065 | 1 | ENSMUST00 | 0 | Trabd | ENSMUSG00000015363.8 |
| chr18 | 64658246 | 64661826 | 35.98183 | 119.73972 | 115.60696 | Promoter | 18 | 64528979 | 64661000 | 132022 | 2 | ENSMUST00 | 0 | Atp8b1 | ENSMUSG00000039529.8 |
| chr2 | 69646793 | 69647959 | 35.98183 | 119.73972 | 115.60696 | Promoter | 2 | 69647171 | 69667318 | 20148 | 1 | ENSMUST00 | 0 | Bbs5 | ENSMUSG00000063145.5 |
| chr4 | 9668334 | 9669818 | 35.98183 | 119.73972 | 115.60696 | Promoter | 4 | 9449079 | 9669162 | 220084 | 2 | ENSMUST00 | 0 | Asph | ENSMUSG00000028207.13 |
| chr4 | 155563720 | 155564892 | 35.98183 | 119.73972 | 115.60696 | Promoter | 4 | 155563826 | 155577096 | 13271 | 1 | ENSMUST00 | 0 | Nadk | ENSMUSG00000029063.11 |
| chr9 | 65032063 | 65033438 | 35.98183 | 119.73972 | 115.60696 | Promoter | 9 | 65032458 | 65082650 | 50193 | 1 | ENSMUST00 | 0 | Dpp8 | ENSMUSG00000032393.9 |
| chr9 | 108261324 | 108263997 | 35.98183 | 119.73972 | 115.60696 | Promoter | 9 | 108205955 | 108263918 | 57964 | 2 | ENSMUST00 | 0 | Dag1 | ENSMUSG00000039952.10 |
| chr9 | 123529641 | 123531067 | 36.53584 | 119.65029 | 115.52113 | Promoter | 9 | 123529882 | 123592598 | 62717 | 1 | ENSMUST00 | 0 | Sacm1l | ENSMUSG00000025240.8 |
| chr11 | 105291956 | 105293037 | 37.73948 | 119.61958 | 115.49072 | Promoter | 11 | 105292643 | 105326891 | 34249 | 1 | ENSMUST00 | 0 | Mrc2 | ENSMUSG00000020695.9 |
| chr13 | 96924242 | 96925782 | 37.73948 | 119.61958 | 115.49072 | Promoter | 13 | 96924689 | 96950912 | 26224 | 1 | ENSMUST00 | 0 | Gcnt4 | ENSMUSG00000091387.1 |
| chr17 | 32363372 | 32365041 | 37.73948 | 119.61958 | 115.49072 | Promoter | 17 | 32361675 | 32364463 | 2789 | 2 | ENSMUST00 | 0 | Wiz | ENSMUSG00000024050.10 |
| chr13 | 101766190 | 101769111 | 27.30634 | 119.5799 | 115.45193 | Promoter | 13 | 101680563 | 101768217 | 87655 | 2 | ENSMUST00 | 0 | Pik3r1 | ENSMUSG00000041417.10 |
| chr3 | 37419826 | 37421635 | 27.30634 | 119.5799 | 115.45193 | Promoter | 3 | 37419896 | 37579096 | 159201 | 1 | ENSMUST00 | 0 | Spata5 | ENSMUSG00000027722.9 |
| chr3 | 126595914 | 126598682 | 36.46498 | 119.52629 | 115.39857 | Promoter | 3 | 126596302 | 126684688 | 88387 | 1 | ENSMUST00 | 0 | Camk2d | ENSMUSG00000053819.11 |
| chr19 | 42170194 | 42171469 | 35.84622 | 119.50067 | 115.37318 | Promoter | 19 | 42170567 | 42194590 | 24024 | 1 | ENSMUST00 | 0 | Zfyve27 | ENSMUSG00000018820.8 |
| chr6 | 81965172 | 81966163 | 35.84622 | 119.50067 | 115.37318 | Promoter | 6 | 81957851 | 81965958 | 8108 | 2 | ENSMUST00 | 0 | Mrpl19 | ENSMUSG00000030045.10 |
| chr2 | 33129759 | 33132001 | 37.04683 | 119.47973 | 115.35273 | Promoter | 2 | 33054158 | 33131436 | 77279 | 2 | ENSMUST00 | 0 | Garnl3 | ENSMUSG00000038860.10 |
| chr7 | 138845531 | 138847675 | 37.04683 | 119.47973 | 115.35273 | Promoter | 7 | 138846079 | 138871702 | 25624 | 1 | ENSMUST00 | 0 | Ppp2r2d | ENSMUSG00000041769.8 |
| chr11 | 120347215 | 120350938 | 34.00618 | 119.38669 | 115.26049 | Promoter | 11 | 120348678 | 120351190 | 2513 | 1 | ENSMUST00 | 0 | 0610009L18F | ENSMUSG00000043644.4 |
| chr12 | 44208849 | 44210745 | 34.00618 | 119.38669 | 115.26049 | Promoter | 12 | 44205897 | 44210068 | 4172 | 2 | ENSMUST00 | 0 | Dnajb9 | ENSMUSG00000014905.3 |
| chr9 | 90112697 | 90115394 | 34.00618 | 119.38669 | 115.26049 | Promoter | 9 | 90091665 | 90114715 | 23051 | 2 | ENSMUST00 | 0 | Morf4l1 | ENSMUSG00000062270.8 |
| chr1 | 55025654 | 55027961 | 30.14656 | 119.38538 | 115.2599 | Promoter | 1 | 54985169 | 55027481 | 42313 | 2 | ENSMUST00 | 0 | Sf3b1 | ENSMUSG00000025982.8 |
| chr11 | 5740331 | 5741629 | 30.14656 | 119.38538 | 115.2599 | Promoter | 11 | 5713417 | 5741150 | 27734 | 2 | ENSMUST00 | 0 | Urgcp | ENSMUSG00000049680.12 |
| chr12 | 11265266 | 11267287 | 30.14656 | 119.38538 | 115.2599 | Promoter | 12 | 11265886 | 11319785 | 53900 | 1 | ENSMUST00 | 0 | Smc6 | ENSMUSG00000020608.6 |
| chr13 | 69532246 | 69535657 | 30.14656 | 119.38538 | 115.2599 | Promoter | 13 | 69533746 | 69615468 | 81723 | 1 | ENSMUST00 | 0 | Nsun2 | ENSMUSG00000021595.13 |
| chr17 | 74528308 | 74529934 | 30.14656 | 119.38538 | 115.2599 | Promoter | 17 | 74528468 | 74702383 | 173916 | 1 | ENSMUST00 | 0 | Birc6 | ENSMUSG00000024073.9 |
| chr18 | 56924506 | 56926376 | 30.14656 | 119.38538 | 115.2599 | Promoter | 18 | 56761716 | 56925548 | 163833 | 2 | ENSMUST00 | 0 | Mar-03 | ENSMUSG00000032656.9 |
| chr2 | 151038378 | 151040035 | 30.14656 | 119.38538 | 115.2599 | Promoter | 2 | 151029685 | 151039398 | 9714 | 2 | ENSMUST00 | 0 | Nanp | ENSMUSG00000053916.4 |
| chr5 | 100797920 | 100799510 | 30.14656 | 119.38538 | 115.2599 | Promoter | 5 | 100798627 | 100804471 | 5845 | 1 | ENSMUST00 | 0 | Mrps18c | ENSMUSG00000016833.9 |
| chr5 | 31240267 | 31242202 | 30.14656 | 119.38538 | 115.2599 | Promoter | 5 | 31240864 | 31251560 | 10697 | 1 | ENSMUST00 | 0 | Nrbp1 | ENSMUSG00000029148.10 |
| chr6 | 120530741 | 120532312 | 30.14656 | 119.38538 | 115.2599 | Promoter | 6 | 120509494 | 120531319 | 21826 | 2 | ENSMUST00 | 0 | Cecr5 | ENSMUSG00000058979.6 |
| chr8 | 84686292 | 84688224 | 30.14656 | 119.38538 | 115.2599 | Promoter | 8 | 84686307 | 84690936 | 4630 | 1 | ENSMUST00 | 0 | Trmt1 | ENSMUSG00000001909.10 |
| chr8 | 125012518 | 125013834 | 30.14656 | 119.38538 | 115.2599 | Promoter | 8 | 125012997 | 125034192 | 21196 | 1 | ENSMUST00 | 0 | Tsnax | ENSMUSG00000056820.6 |
| chr9 | 115307644 | 115311336 | 30.14656 | 119.38538 | 115.2599 | Promoter | 9 | 115242607 | 115310421 | 67815 | 2 | ENSMUST00 | 0 | Stt3b | ENSMUSG00000032437.9 |
| chr1 | 45794798 | 45796261 | 37.58501 | 119.35227 | 115.22867 | Promoter | 1 | 45795166 | 45823619 | 28454 | 1 | ENSMUST00 | 0 | Wdr75 | ENSMUSG00000025995.11 |
| chr16 | 20548593 | 20549523 | 37.58501 | 119.35227 | 115.22867 | Promoter | 16 | 20548577 | 20561379 | 12803 | 1 | ENSMUST00 | 17 | Abcf3 | ENSMUSG00000003234.10 |
| chr7 | 80197944 | 80199464 | 37.58501 | 119.35227 | 115.22867 | Promoter | 7 | 80198538 | 80214065 | 15528 | 1 | ENSMUST00 | 0 | Sema4b | ENSMUSG00000030539.8 |
| chr1 | 155972112 | 155973264 | 28.61382 | 119.33878 | 115.2169 | Promoter | 1 | 155844966 | 155972887 | 127922 | 2 | ENSMUST00 | 0 | Cep350 | ENSMUSG00000033671.12 |
| chr15 | 79402108 | 79404130 | 28.61382 | 119.33878 | 115.2169 | Promoter | 15 | 79360684 | 79402919 | 42236 | 2 | ENSMUST00 | 0 | Tmem184b | ENSMUSG00000009035.8 |
| chr18 | 80468317 | 80470727 | 28.61382 | 119.33878 | 115.2169 | Promoter | 18 | 80407959 | 80469695 | 61737 | 2 | ENSMUST00 | 0 | Ctdp1 | ENSMUSG00000033323.8 |
| chr2 | 117249432 | 117250793 | 28.61382 | 119.33878 | 115.2169 | Promoter | 2 | 117249739 | 117260243 | 10505 | 1 | ENSMUST00 | 0 | Fam98b | ENSMUSG00000027349.5 |
| chr3 | 96246039 | 96246579 | 28.61382 | 119.33878 | 115.2169 | Promoter | 3 | 96243550 | 96246380 | 2831 | 2 | ENSMUST00 | 0 | Gm20633 | ENSMUSG00000093553.1 |
| chr4 | 11155635 | 11158102 | 28.61382 | 119.33878 | 115.2169 | Promoter | 4 | 11156431 | 11174379 | 17949 | 1 | ENSMUST00 | 0 | Trp53inp1 | ENSMUSG00000028211.6 |
| chr1 | 74390907 | 74393162 | 38.14117 | 119.25307 | 115.13219 | Promoter | 1 | 74391509 | 74397285 | 5777 | 1 | ENSMUST00 | 0 | Ctdsp1 | ENSMUSG00000026176.8 |
| chr1 | 191317309 | 191318274 | 38.14117 | 119.25307 | 115.13219 | Promoter | 1 | 191306789 | 191318194 | 11406 | 2 | ENSMUST00 | 0 | Nenf | ENSMUSG00000037499.9 |
| chr11 | 114668534 | 114670048 | 38.14117 | 119.25307 | 115.13219 | Promoter | 11 | 114668548 | 114672331 | 3784 | 1 | ENSMUST00 | 0 | Rpl38 | ENSMUSG00000057322.7 |
| chr11 | 106500875 | 106501735 | 38.14117 | 119.25307 | 115.13219 | Promoter | 11 | 106500993 | 106501062 | 70 | 1 | ENSMUST00 | 0 | Snord104 | ENSMUSG00000065126.1 |
| chr14 | 24244111 | 24246301 | 38.14117 | 119.25307 | 115.13219 | Promoter | 14 | 24133953 | 24245920 | 111968 | 2 | ENSMUST00 | 0 | Dlg5 | ENSMUSG00000021782.9 |
| chr16 | 33828900 | 33834465 | 38.14117 | 119.25307 | 115.13219 | Promoter | 16 | 33829665 | 33949146 | 119482 | 1 | ENSMUST00 | 0 | Itgb5 | ENSMUSG00000022817.9 |
| chr3 | 96487928 | 96489306 | 38.14117 | 119.25307 | 115.13219 | Promoter | 3 | 96488729 | 96488893 | 165 | 1 | ENSMUST00 | 0 | Gm22614 | ENSMUSG00000094812.1 |
| chr4 | 147936057 | 147937085 | 38.14117 | 119.25307 | 115.13219 | Promoter | 4 | 147909753 | 147936767 | 27015 | 2 | ENSMUST00 | 0 | Plod1 | ENSMUSG00000019055.10 |
| chr4 | 153956707 | 153958174 | 38.14117 | 119.25307 | 115.13219 | Promoter | 4 | 153957237 | 153961918 | 4682 | 1 | ENSMUST00 | 0 | A430005L14F | ENSMUSG00000047613.5 |
| chr5 | 125441176 | 125442448 | 38.14117 | 119.25307 | 115.13219 | Promoter | 5 | 125441568 | 125460872 | 19305 | 1 | ENSMUST00 | 0 | Bri3bp | ENSMUSG00000037905.9 |
| chr5 | 9160571 | 9162609 | 38.14117 | 119.25307 | 115.13219 | Promoter | 5 | 9118801 | 9161776 | 42976 | 2 | ENSMUST00 | 0 | Dmtf1 | ENSMUSG00000042508.11 |
| chr7 | 44928662 | 44929560 | 38.14117 | 119.25307 | 115.13219 | Promoter | 7 | 44900373 | 44929496 | 29124 | 2 | ENSMUST00 | 0 | Ap2a1 | ENSMUSG00000060279.8 |
| chr8 | 69808372 | 69809481 | 38.14117 | 119.25307 | 115.13219 | Promoter | 8 | 69808679 | 69821870 | 13192 | 1 | ENSMUST00 | 0 | Gmip | ENSMUSG00000036246.9 |
| chr9 | 62838121 | 62839384 | 38.14117 | 119.25307 | 115.13219 | Promoter | 9 | 62838785 | 62852006 | 13222 | 1 | ENSMUST00 | 0 | Cln6 | ENSMUSG00000032245.7 |
| chr2 | 26236459 | 26237867 | 36.89971 | 119.22369 | 115.10591 | Promoter | 2 | 26209125 | 26237525 | 28401 | 2 | ENSMUST00 | 0 | Qsox2 | ENSMUSG00000036327.13 |
| chr1 | 36511402 | 36513251 | 37.50825 | 119.21938 | 115.10198 | Promoter | 1 | 36511867 | 36528237 | 16371 | 1 | ENSMUST00 | 0 | Cnnm3 | ENSMUSG00000001138.8 |
| chr11 | 119602379 | 119604058 | 35.06866 | 119.18253 | 115.06551 | Promoter | 11 | 119602905 | 119823176 | 220272 | 1 | ENSMUST00 | 0 | Rptor | ENSMUSG00000025583.10 |
| chr16 | 8671377 | 8672582 | 35.06866 | 119.18253 | 115.06551 | Promoter | 16 | 8658587 | 8672153 | 13567 | 2 | ENSMUST00 | 0 | Carhsp1 | ENSMUSG00000008393.8 |
| chr19 | 7205120 | 7206836 | 35.06866 | 119.18253 | 115.06551 | Promoter | 19 | 7198202 | 7206316 | 8115 | 2 | ENSMUST00 | 0 | Otub1 | ENSMUSG00000024767.6 |
| chr3 | 146520338 | 146522001 | 35.06866 | 119.18253 | 115.06551 | Promoter | 3 | 146506348 | 146521423 | 15076 | 2 | ENSMUST00 | 0 | Rpf1 | ENSMUSG00000028187.6 |
| chr3 | 122292329 | 122294466 | 35.06866 | 119.18253 | 115.06551 | Promoter | 3 | 122255958 | 122293547 | 37590 | 2 | ENSMUST00 | 0 | Gm17494 | ENSMUSG00000057359.6 |
| chr4 | 11190318 | 11192533 | 35.06866 | 119.18253 | 115.06551 | Promoter | 4 | 11191351 | 11193961 | 2611 | 1 | ENSMUST00 | 0 | Ccne2 | ENSMUSG00000028212.11 |
| chr4 | 15957819 | 15958996 | 35.06866 | 119.18253 | 115.06551 | Promoter | 4 | 15957925 | 15992589 | 34665 | 1 | ENSMUST00 | 0 | Nbn | ENSMUSG00000028224.9 |
| chr10 | 7680225 | 7682548 | 36.25406 | 119.15681 | 115.04134 | Promoter | 10 | 7644332 | 7681136 | 36805 | 2 | ENSMUST00 | 0 | Pcmt1 | ENSMUSG00000019795.12 |
| chr11 | 100849708 | 100851276 | 36.25406 | 119.15681 | 115.04134 | Promoter | 11 | 100780731 | 100850724 | 69994 | 2 | ENSMUST00 | 0 | Stat5b | ENSMUSG00000020919.6 |
| chr12 | 100109768 | 100111167 | 36.25406 | 119.15681 | 115.04134 | Promoter | 12 | 100110186 | 100123364 | 13179 | 1 | ENSMUST00 | 0 | Psmc1 | ENSMUSG00000021178.8 |
| chr14 | 56575064 | 56576957 | 36.25406 | 119.15681 | 115.04134 | Promoter | 14 | 56575619 | 56659794 | 84176 | 1 | ENSMUST00 | 0 | Parp4 | ENSMUSG00000054509.7 |
| chr17 | 34027371 | 34028281 | 36.25406 | 119.15681 | 115.04134 | Promoter | 17 | 34026033 | 34028056 | 2024 | 2 | ENSMUST00 | 0 | H2-Ke6 | ENSMUSG00000073422.5 |
| chr7 | 100159002 | 100160115 | 36.25406 | 119.15681 | 115.04134 | Promoter | 7 | 100159277 | 100161365 | 2089 | 1 | ENSMUST00 | 0 | Lipt2 | ENSMUSG00000030725.3 |
| chr8 | 25518487 | 25520589 | 36.25406 | 119.15681 | 115.04134 | Promoter | 8 | 25518772 | 25575714 | 56943 | 1 | ENSMUST00 | 0 | Fgfr1 | ENSMUSG00000031565.12 |
| chr13 | 35740437 | 35743017 | 37.4318 | 119.08698 | 114.97395 | Promoter | 13 | 35741313 | 35874058 | 132746 | 1 | ENSMUST00 | 0 | Cdyl | ENSMUSG00000059288.8 |
| chr17 | 25874471 | 25876274 | 33.23528 | 119.03175 | 114.9192 | Promoter | 17 | 25875464 | 25877160 | 1697 | 1 | ENSMUST00 | 0 | 0610011F06F | ENSMUSG00000025731.10 |
| chr10 | 80322860 | 80323983 | 37.99286 | 118.9973 | 114.88904 | Promoter | 10 | 80321287 | 80323126 | 1840 | 2 | ENSMUST00 | 0 | Pcsk4 | ENSMUSG00000020131.9 |
| chr10 | 80398121 | 80399546 | 33.7938 | 119.00099 | 114.88904 | Promoter | 10 | 80392839 | 80399422 | 6584 | 2 | ENSMUST00 | 0 | Mbd3 | ENSMUSG00000035478.9 |
| chr11 | 97279259 | 97281276 | 33.7938 | 119.00099 | 114.88904 | Promoter | 11 | 97205842 | 97280486 | 74645 | 2 | ENSMUST00 | 0 | Npepps | ENSMUSG00000001441.8 |
| chr11 | 97698478 | 97701285 | 33.7938 | 119.00099 | 114.88904 | Promoter | 11 | 97688823 | 97699384 | 10562 | 2 | ENSMUST00 | 0 | Pcgf2 | ENSMUSG00000018537.11 |
| chr12 | 80789969 | 80791597 | 33.7938 | 119.00099 | 114.88904 | Promoter | 12 | 80790532 | 80880832 | 90301 | 1 | ENSMUST00 | 0 | 4933426M11 | ENSMUSG00000021133.8 |
| chr12 | 108792910 | 108795425 | 33.7938 | 119.00099 | 114.88904 | Promoter | 12 | 108792973 | 108816632 | 23660 | 1 | ENSMUST00 | 0 | Yy1 | ENSMUSG00000021264.11 |
| chr14 | 76151945 | 76153435 | 33.7938 | 119.00099 | 114.88904 | Distal Interge | 14 | 76110891 | 76137379 | 26489 | 1 | ENSMUST00 | 41055 | Nufip1 | ENSMUSG00000022009.1 |
| chr14 | 29967742 | 29969586 | 33.7938 | 119.00099 | 114.88904 | Promoter | 14 | 29968308 | 29975074 | 6767 | 1 | ENSMUST00 | 0 | Selk | ENSMUSG00000042682.8 |
| chr16 | 4076892 | 4078331 | 33.7938 | 119.00099 | 114.88904 | Promoter | 16 | 4039971 | 4077827 | 37857 | 2 | ENSMUST00 | 0 | Trap1 | ENSMUSG00000005981.11 |
| chr17 | 24350879 | 24352896 | 33.7938 | 119.00099 | 114.88904 | Promoter | 17 | 24351950 | 24397783 | 45834 | 1 | ENSMUST00 | 0 | Abca3 | ENSMUSG00000024130.10 |
| chr17 | 27202960 | 27204531 | 33.7938 | 119.00099 | 114.88904 | Promoter | 17 | 27189601 | 27204438 | 14838 | 2 | ENSMUST00 | 0 | Lemd2 | ENSMUSG00000044857.7 |
| chr18 | 44379865 | 44381364 | 33.7938 | 119.00099 | 114.88904 | Promoter | 18 | 44380500 | 44424969 | 44470 | 1 | ENSMUST00 | 0 | Dcp2 | ENSMUSG00000024472.8 |
| chr19 | 36833456 | 36836298 | 33.7938 | 119.00099 | 114.88904 | Promoter | 19 | 36834232 | 36893477 | 59246 | 1 | ENSMUST00 | 0 | Tnks2 | ENSMUSG00000024811.6 |
| chr19 | 8941466 | 8943320 | 33.7938 | 119.00099 | 114.88904 | Promoter | 19 | 8941875 | 8952303 | 10429 | 1 | ENSMUST00 | 0 | Mta2 | ENSMUSG00000071646.4 |
| chr2 | 35199491 | 35201101 | 33.7938 | 119.00099 | 114.88904 | Promoter | 2 | 35189900 | 35200789 | 10890 | 2 | ENSMUST00 | 0 | Rab14 | ENSMUSG00000026878.11 |
| chr2 | 80579951 | 80581955 | 33.7938 | 119.00099 | 114.88904 | Promoter | 2 | 80500512 | 80581264 | 80753 | 2 | ENSMUST00 | 0 | Nckap1 | ENSMUSG00000027002.8 |
| chr2 | 84678347 | 84679231 | 33.7938 | 119.00099 | 114.88904 | Promoter | 2 | 84678402 | 84688215 | 9814 | 1 | ENSMUST00 | 0 | Med19 | ENSMUSG00000027080.6 |
| chr3 | 52267261 | 52270933 | 33.7938 | 119.00099 | 114.88904 | Promoter | 3 | 52268337 | 52350109 | 81773 | 1 | ENSMUST00 | 0 | Foxo1 | ENSMUSG00000044167.5 |
| chr4 | 123904277 | 123905645 | 33.7938 | 119.00099 | 114.88904 | Promoter | 4 | 123904832 | 123912269 | 7438 | 1 | ENSMUST00 | 0 | Mycbp | ENSMUSG00000028647.8 |
| chr4 | 108458983 | 108461770 | 33.7938 | 119.00099 | 114.88904 | Promoter | 4 | 108459426 | 108479153 | 19728 | 1 | ENSMUST00 | 0 | Zcchc11 | ENSMUSG00000034610.9 |
| chr5 | 138441396 | 138442611 | 33.7938 | 119.00099 | 114.88904 | Promoter | 5 | 138441468 | 138460694 | 19227 | 1 | ENSMUST00 | 0 | Zfp157 | ENSMUSG00000036898.11 |
| chr5 | 88554397 | 88555369 | 33.7938 | 119.00099 | 114.88904 | Promoter | 5 | 88554483 | 88556083 | 1601 | 1 | ENSMUST00 | 0 | Utp3 | ENSMUSG00000070697.3 |
| chr6 | 55037488 | 55039171 | 33.7938 | 119.00099 | 114.88904 | Promoter | 6 | 55038001 | 55079504 | 41504 | 1 | ENSMUST00 | 0 | Gars | ENSMUSG00000029777.9 |
| chr7 | 46918979 | 46920516 | 33.7938 | 119.00099 | 114.88904 | Promoter | 7 | 46888949 | 46919969 | 31021 | 2 | ENSMUST00 | 0 | Tsg101 | ENSMUSG00000014402.9 |
| chr7 | 28350054 | 28351837 | 33.7938 | 119.00099 | 114.88904 | Promoter | 7 | 28350652 | 28352696 | 2045 | 1 | ENSMUST00 | 0 | Rps16 | ENSMUSG00000037563.9 |
| chr7 | 89337799 | 89339259 | 33.7938 | 119.00099 | 114.88904 | Promoter | 7 | 89139723 | 89338787 | 199065 | 2 | ENSMUST00 | 0 | Tmem135 | ENSMUSG00000039428.5 |
| chr8 | 61223517 | 61225207 | 33.7938 | 119.00099 | 114.88904 | Promoter | 8 | 61224171 | 61396072 | 171902 | 1 | ENSMUST00 | 0 | Sh3rf1 | ENSMUSG00000031642.8 |
| chr8 | 123982163 | 123983624 | 33.7938 | 119.00099 | 114.88904 | Promoter | 8 | 123952459 | 123983122 | 30664 | 2 | ENSMUST00 | 0 | Abcb10 | ENSMUSG00000031974.8 |
| chr9 | 64252324 | 64253690 | 33.7938 | 119.00099 | 114.88904 | Promoter | 9 | 64185770 | 64253631 | 67862 | 2 | ENSMUST00 | 0 | Map2k1 | ENSMUSG00000004936.8 |
| chr9 | 21616122 | 21617444 | 33.7938 | 119.00099 | 114.88904 | Promoter | 9 | 21616169 | 21704217 | 88049 | 1 | ENSMUST00 | 0 | Smarca4 | ENSMUSG00000032187.11 |
| chr9 | 70502669 | 70504229 | 33.7938 | 119.00099 | 114.88904 | Promoter | 9 | 70425429 | 70503725 | 78297 | 2 | ENSMUST00 | 0 | Rnf111 | ENSMUSG00000032217.7 |
| chr6 | 30693171 | 30694240 | 36.75376 | 118.96947 | 114.86417 | Promoter | 6 | 30653457 | 30693749 | 40293 | 2 | ENSMUST00 | 0 | Cep41 | ENSMUSG00000029790.11 |
| chr3 | 51223991 | 51225879 | 37.35566 | 118.95506 | 114.85009 | Promoter | 3 | 51224447 | 51251644 | 27198 | 1 | ENSMUST00 | 0 | Ccrn4l | ENSMUSG00000023087.10 |
| chr6 | 86371039 | 86372324 | 37.35566 | 118.95506 | 114.85009 | Promoter | 6 | 86371583 | 86378902 | 7320 | 1 | ENSMUST00 | 0 | Snrpg | ENSMUSG00000057278.6 |
| chr15 | 28023999 | 28025987 | 36.11478 | 118.91257 | 114.80801 | Promoter | 15 | 27730652 | 28025848 | 295197 | 2 | ENSMUST00 | 0 | Trio | ENSMUSG00000022263.9 |

| | | | | | | | | | | | | | | | |
|---|---|---|---|---|---|---|---|---|---|---|---|---|---|---|---|
| chr1 | 52231711 | 52233894 | 37.91348 | 118.86035 | 114.75604 | Promoter | 1 | 52163448 | 52233223 | 69776 | 2 | ENSMUST00 | 0 | Gls | ENSMUSG00000026103.9 |
| chr15 | 66577170 | 66578557 | 37.91348 | 118.86035 | 114.75604 | Promoter | 15 | 66577572 | 66645255 | 67684 | 1 | ENSMUST00 | 0 | Phf20l1 | ENSMUSG00000072501.4 |
| chr2 | 104492244 | 104495149 | 37.91348 | 118.86035 | 114.75604 | Promoter | 2 | 104426481 | 104494446 | 67966 | 2 | ENSMUST00 | 0 | Hipk3 | ENSMUSG00000027177.9 |
| chr4 | 40851570 | 40854097 | 37.91348 | 118.86035 | 114.75604 | Promoter | 4 | 40804602 | 40854005 | 49404 | 2 | ENSMUST00 | 0 | B4galt1 | ENSMUSG00000028413.8 |
| chr10 | 123195574 | 123197730 | 37.27983 | 118.82364 | 114.72062 | Promoter | 10 | 123113244 | 123196935 | 83692 | 2 | ENSMUST00 | 0 | Usp15 | ENSMUSG00000020124.9 |
| chr2 | 35255869 | 35258912 | 37.27983 | 118.82364 | 114.72062 | Distal Interge | 2 | 35279663 | 35293567 | 13905 | 1 | ENSMUST00 | -20751 | Gsn | ENSMUSG00000026879.9 |
| chr2 | 84826391 | 84827862 | 37.27983 | 118.82364 | 114.72062 | Promoter | 2 | 84826997 | 84830213 | 3217 | 1 | ENSMUST00 | 0 | Timm10 | ENSMUSG00000027076.5 |
| chr10 | 79908302 | 79909152 | 36.04346 | 118.78738 | 114.68494 | Promoter | 10 | 79894708 | 79908921 | 14214 | 2 | ENSMUST00 | 0 | Med16 | ENSMUSG00000013833.10 |
| chr10 | 19907213 | 19907743 | 36.04346 | 118.78738 | 114.68494 | Promoter | 10 | 19859929 | 19907689 | 47761 | 2 | ENSMUST00 | 0 | Pex7 | ENSMUSG00000020003.10 |
| chr11 | 21091018 | 21093123 | 36.04346 | 118.78738 | 114.68494 | Promoter | 11 | 21091291 | 21143005 | 51715 | 1 | ENSMUST00 | 0 | Peli1 | ENSMUSG00000020134.8 |
| chr11 | 105589174 | 105590737 | 36.04346 | 118.78738 | 114.68494 | Promoter | 11 | 105589986 | 105776834 | 186849 | 1 | ENSMUST00 | 0 | Tanc2 | ENSMUSG00000053580.10 |
| chr12 | 84407946 | 84409513 | 36.04346 | 118.78738 | 114.68494 | Promoter | 12 | 84373875 | 84409029 | 35155 | 2 | ENSMUST00 | 0 | Entpd5 | ENSMUSG00000021236.11 |
| chr13 | 19394935 | 19396398 | 36.04346 | 118.78738 | 114.68494 | Promoter | 13 | 19357677 | 19395752 | 38076 | 2 | ENSMUST00 | 0 | Stard3nl | ENSMUSG00000003062.8 |
| chr14 | 70519471 | 70520854 | 36.04346 | 118.78738 | 114.68494 | Promoter | 14 | 70474557 | 70520260 | 45704 | 2 | ENSMUST00 | 0 | Bmp1 | ENSMUSG00000022098.9 |
| chr15 | 12117175 | 12119324 | 36.04346 | 118.78738 | 114.68494 | Promoter | 15 | 12117831 | 12185683 | 67853 | 1 | ENSMUST00 | 0 | Zfr | ENSMUSG00000022201.10 |
| chr15 | 64059457 | 64060826 | 36.04346 | 118.78738 | 114.68494 | Promoter | 15 | 63929097 | 64060448 | 131352 | 2 | ENSMUST00 | 0 | Fam49b | ENSMUSG00000022378.8 |
| chr17 | 88063293 | 88066770 | 36.04346 | 118.78738 | 114.68494 | Promoter | 17 | 87990859 | 88065028 | 74170 | 2 | ENSMUST00 | 0 | Fbxo11 | ENSMUSG00000005371.9 |
| chr17 | 80205681 | 80207946 | 36.04346 | 118.78738 | 114.68494 | Promoter | 17 | 80200089 | 80207305 | 7217 | 2 | ENSMUST00 | 0 | Srsf7 | ENSMUSG00000024097.10 |
| chr18 | 67932777 | 67934639 | 36.04346 | 118.78738 | 114.68494 | Promoter | 18 | 67933257 | 68255549 | 322293 | 1 | ENSMUST00 | 0 | Ldlrad4 | ENSMUSG00000024544.8 |
| chr2 | 29346771 | 29347866 | 36.04346 | 118.78738 | 114.68494 | Promoter | 2 | 29346819 | 29524793 | 177975 | 1 | ENSMUST00 | 0 | Med27 | ENSMUSG00000026799.10 |
| chr2 | 180024135 | 180026031 | 36.04346 | 118.78738 | 114.68494 | Promoter | 2 | 180024987 | 180035461 | 10475 | 1 | ENSMUST00 | 0 | Lsm14b | ENSMUSG00000039108.9 |
| chr4 | 40757058 | 40758347 | 36.04346 | 118.78738 | 114.68494 | Promoter | 4 | 40736542 | 40757923 | 21382 | 2 | ENSMUST00 | 0 | Smu1 | ENSMUSG00000028409.6 |
| chr4 | 134852051 | 134854505 | 36.04346 | 118.78738 | 114.68494 | Promoter | 4 | 134802759 | 134853345 | 50587 | 2 | ENSMUST00 | 0 | Tmem57 | ENSMUSG00000028826.9 |
| chr4 | 132210818 | 132212608 | 36.04346 | 118.78738 | 114.68494 | Promoter | 4 | 132184912 | 132212303 | 27392 | 2 | ENSMUST00 | 0 | Ythdf2 | ENSMUSG00000040025.12 |
| chr5 | 143314251 | 143315661 | 36.04346 | 118.78738 | 114.68494 | Promoter | 5 | 143301195 | 143315064 | 13870 | 2 | ENSMUST00 | 0 | E130309D02 | ENSMUSG00000039244.5 |
| chr6 | 148944401 | 148947082 | 36.04346 | 118.78738 | 114.68494 | Promoter | 6 | 148921035 | 148946467 | 25433 | 2 | ENSMUST00 | 0 | Fam60a | ENSMUSG00000039985.9 |
| chr6 | 119174737 | 119176198 | 36.04346 | 118.78738 | 114.68494 | Promoter | 6 | 119175253 | 119221616 | 46364 | 1 | ENSMUST00 | 0 | Dcp1b | ENSMUSG00000041477.9 |
| chr6 | 71631634 | 71633659 | 36.04346 | 118.78738 | 114.68494 | Promoter | 6 | 71588974 | 71632686 | 43713 | 2 | ENSMUST00 | 0 | Kdm3a | ENSMUSG00000053470.8 |
| chr7 | 12965377 | 12966527 | 36.04346 | 118.78738 | 114.68494 | Promoter | 7 | 12965838 | 12973822 | 7985 | 1 | ENSMUST00 | 0 | Zfp324 | ENSMUSG00000004500.8 |
| chr7 | 48880592 | 48882262 | 36.04346 | 118.78738 | 114.68494 | Promoter | 7 | 48866429 | 48881041 | 14613 | 2 | ENSMUST00 | 0 | E2f8 | ENSMUSG00000046179.12 |
| chr8 | 94666142 | 94667466 | 36.04346 | 118.78738 | 114.68494 | Promoter | 8 | 94666755 | 94674417 | 7663 | 1 | ENSMUST00 | 0 | Arl2bp | ENSMUSG00000031776.11 |
| chr9 | 44343310 | 44344479 | 36.04346 | 118.78738 | 114.68494 | Promoter | 9 | 44336350 | 44344228 | 7879 | 2 | ENSMUST00 | 0 | Hmbs | ENSMUSG00000032126.10 |
| chr9 | 96195685 | 96197584 | 36.04346 | 118.78738 | 114.68494 | Promoter | 9 | 96196275 | 96318878 | 122604 | 1 | ENSMUST00 | 0 | Tfdp2 | ENSMUSG00000032411.10 |
| chr6 | 146642099 | 146643520 | 32.51422 | 118.74544 | 114.65017 | Promoter | 6 | 146642547 | 146650602 | 8056 | 1 | ENSMUST00 | 0 | Med21 | ENSMUSG00000030291.7 |
| chr8 | 95633371 | 95634393 | 32.51422 | 118.74544 | 114.65017 | Promoter | 8 | 95633559 | 95645059 | 11501 | 1 | ENSMUST00 | 0 | Gins3 | ENSMUSG00000031669.9 |
| chr17 | 30003745 | 30005950 | 37.83443 | 118.72395 | 114.62898 | Promoter | 17 | 30005087 | 30210019 | 204933 | 1 | ENSMUST00 | 0 | Zfand3 | ENSMUSG00000044477.10 |
| chr1 | 44147255 | 44148910 | 34.21972 | 118.71816 | 114.6235 | Promoter | 1 | 44147744 | 44149522 | 1779 | 1 | ENSMUST00 | 0 | Ercc5 | ENSMUSG00000026048.11 |
| chr16 | 26102904 | 26106511 | 34.21972 | 118.71816 | 114.6235 | Promoter | 16 | 25959288 | 26105784 | 146497 | 2 | ENSMUST00 | 0 | Leprel1 | ENSMUSG00000038168.5 |
| chr19 | 46500135 | 46506359 | 34.21972 | 118.71816 | 114.6235 | Promoter | 19 | 46501648 | 46516444 | 14797 | 1 | ENSMUST00 | 0 | Trim8 | ENSMUSG00000025034.8 |
| chr7 | 46794979 | 46796929 | 34.21972 | 118.71816 | 114.6235 | Promoter | 7 | 46796103 | 46823800 | 27698 | 1 | ENSMUST00 | 0 | Gtf2h1 | ENSMUSG00000006599.8 |
| chr4 | 103118846 | 103120943 | 35.3795 | 118.67607 | 114.58318 | Promoter | 4 | 103119286 | 103165752 | 46467 | 1 | ENSMUST00 | 0 | Mier1 | ENSMUSG00000028522.11 |
| chr3 | 30792690 | 30794526 | 35.97658 | 118.66996 | 114.57709 | Promoter | 3 | 30792876 | 30821263 | 28388 | 1 | ENSMUST00 | 0 | Sec62 | ENSMUSG00000027706.7 |
| chr5 | 150664539 | 150665702 | 35.97658 | 118.66996 | 114.57709 | Promoter | 5 | 150635972 | 150665612 | 29641 | 2 | ENSMUST00 | 0 | N4bp2l2 | ENSMUSG00000029655.12 |
| chr10 | 78168944 | 78170313 | 36.53698 | 118.59147 | 114.49885 | Promoter | 10 | 78162067 | 78169768 | 7702 | 2 | ENSMUST00 | 0 | D10Jhu81e | ENSMUSG00000053329.6 |
| chr11 | 23305515 | 23307804 | 36.53698 | 118.59147 | 114.49885 | Promoter | 11 | 23306895 | 23490560 | 183666 | 1 | ENSMUST00 | 0 | Usp34 | ENSMUSG00000056342.11 |
| chr17 | 57246907 | 57247770 | 36.53698 | 118.59147 | 114.49885 | Promoter | 17 | 57234914 | 57247641 | 12728 | 2 | ENSMUST00 | 0 | Gpr108 | ENSMUSG00000005823.8 |
| chr14 | 21984527 | 21987918 | 37.75571 | 118.58805 | 114.49699 | 5' UTR | 14 | 21983962 | 21989601 | 5640 | 2 | ENSMUST00 | 1683 | Zfp503 | ENSMUSG00000039081.9 |
| chr17 | 32035826 | 32037064 | 37.1291 | 118.56225 | 114.47147 | Promoter | 17 | 32036100 | 32062865 | 26766 | 1 | ENSMUST00 | 0 | Rrp1b | ENSMUSG00000058392.7 |
| chr13 | 64151825 | 64153338 | 35.31382 | 118.55977 | 114.46947 | Promoter | 13 | 64133022 | 64153202 | 20181 | 2 | ENSMUST00 | 0 | Zfp367 | ENSMUSG00000044934.7 |
| chr16 | 78301158 | 78303001 | 35.31382 | 118.55977 | 114.46947 | Promoter | 16 | 78301496 | 78340752 | 39257 | 1 | ENSMUST00 | 0 | Cxadr | ENSMUSG00000022865.8 |
| chr5 | 52833457 | 52835017 | 35.31382 | 118.55977 | 114.46947 | Promoter | 5 | 52834015 | 52867797 | 33783 | 1 | ENSMUST00 | 0 | Anapc4 | ENSMUSG00000029176.8 |
| chr5 | 116023224 | 116024959 | 35.31382 | 118.55977 | 114.46947 | Promoter | 5 | 116013586 | 116024462 | 10877 | 2 | ENSMUST00 | 0 | Prkab1 | ENSMUSG00000029513.9 |
| chr5 | 86803694 | 86805523 | 35.31382 | 118.55977 | 114.46947 | Promoter | 5 | 86804221 | 86836654 | 32434 | 1 | ENSMUST00 | 0 | Ythdc1 | ENSMUSG00000035851.9 |
| chr1 | 64615346 | 64618004 | 35.90788 | 118.54927 | 114.46114 | Promoter | 1 | 64606473 | 64617242 | 10770 | 2 | ENSMUST00 | 0 | Mettl21a | ENSMUSG00000025956.5 |
| chr9 | 77916929 | 77918711 | 35.90788 | 118.54927 | 114.46114 | Promoter | 9 | 77917364 | 77984519 | 67156 | 1 | ENSMUST00 | 0 | Elovl5 | ENSMUSG00000032349.8 |
| chr3 | 38884547 | 38888528 | 37.05418 | 118.43227 | 114.34519 | Promoter | 3 | 38886940 | 39011985 | 125046 | 1 | ENSMUST00 | 0 | Fat4 | ENSMUSG00000046743.6 |
| chr6 | 133104654 | 133106469 | 37.05418 | 118.43227 | 114.34519 | Promoter | 6 | 133105239 | 133107750 | 2512 | 1 | ENSMUST00 | 0 | 2700089E24 | ENSMUSG00000072704.5 |
| chr15 | 59373437 | 59375064 | 35.83944 | 118.42897 | 114.34225 | Promoter | 15 | 59331998 | 59374167 | 42170 | 2 | ENSMUST00 | 0 | E430025E21 | ENSMUSG00000022350.6 |
| chr15 | 80233528 | 80235026 | 31.20348 | 118.35023 | 114.26361 | Promoter | 15 | 80234080 | 80253366 | 19287 | 1 | ENSMUST00 | 0 | Mief1 | ENSMUSG00000022412.7 |
| chr4 | 149737028 | 149738543 | 36.39388 | 118.3417 | 114.25511 | Promoter | 4 | 149715375 | 149738044 | 22670 | 2 | ENSMUST00 | 0 | Tmem201 | ENSMUSG00000044700.10 |
| chr15 | 31364914 | 31369386 | 33.42846 | 118.3354 | 114.24911 | Promoter | 15 | 31291479 | 31367759 | 76281 | 2 | ENSMUST00 | 0 | Ankrd33b | ENSMUSG00000022237.11 |
| chr13 | 114387066 | 114388440 | 37.59926 | 118.31785 | 114.23224 | Promoter | 13 | 114287795 | 114388094 | 100300 | 2 | ENSMUST00 | 0 | Ndufs4 | ENSMUSG00000021764.6 |
| chr13 | 43479386 | 43482222 | 37.59926 | 118.31785 | 114.23224 | Promoter | 13 | 43402673 | 43480973 | 78301 | 2 | ENSMUST00 | 0 | Ranbp9 | ENSMUSG00000038546.8 |
| chr12 | 24707869 | 24708986 | 36.97958 | 118.30276 | 114.21766 | Promoter | 12 | 24708241 | 24714146 | 5906 | 1 | ENSMUST00 | 0 | Rrm2 | ENSMUSG00000020649.6 |
| chr16 | 32245965 | 32247998 | 36.97958 | 118.30276 | 114.21766 | Promoter | 16 | 32247227 | 32256753 | 9527 | 1 | ENSMUST00 | 0 | Wdr53 | ENSMUSG00000022787.5 |
| chr3 | 127836661 | 127838171 | 36.97958 | 118.30276 | 114.21766 | Promoter | 3 | 127807264 | 127837492 | 30229 | 2 | ENSMUST00 | 0 | Ap1ar | ENSMUSG00000074238.2 |
| chr19 | 53600047 | 53601711 | 37.52151 | 118.18354 | 114.0993 | Promoter | 19 | 53600398 | 53645833 | 45436 | 1 | ENSMUST00 | 0 | Smc3 | ENSMUSG00000024974.10 |
| chr10 | 43477903 | 43480632 | 36.90527 | 118.17374 | 114.08985 | Promoter | 10 | 43479140 | 43515396 | 36257 | 1 | ENSMUST00 | 0 | Bend3 | ENSMUSG00000038214.9 |
| chr14 | 52256664 | 52258350 | 36.90527 | 118.17374 | 114.08985 | Distal Interge | 14 | 52261769 | 52269106 | 7338 | 2 | ENSMUST00 | 10756 | Rab2b | ENSMUSG00000022159.11 |
| chr7 | 102209316 | 102210236 | 36.90527 | 118.17374 | 114.08985 | Promoter | 7 | 102210208 | 102223483 | 13276 | 1 | ENSMUST00 | 0 | Pgap2 | ENSMUSG00000030990.12 |
| chr1 | 55052067 | 55054172 | 31.62822 | 118.12274 | 114.03909 | Promoter | 1 | 55052770 | 55071953 | 19184 | 1 | ENSMUST00 | 0 | Coq10b | ENSMUSG00000025981.8 |
| chr1 | 53784111 | 53785238 | 31.62822 | 118.12274 | 114.03909 | Promoter | 1 | 53755506 | 53785224 | 29719 | 2 | ENSMUST00 | 0 | Stk17b | ENSMUSG00000026094.9 |
| chr10 | 76344402 | 76346573 | 31.62822 | 118.12274 | 114.03909 | Promoter | 10 | 76262736 | 76345291 | 82556 | 2 | ENSMUST00 | 0 | Dip2a | ENSMUSG00000020231.10 |
| chr11 | 99229408 | 99231335 | 31.62822 | 118.12274 | 114.03909 | Promoter | 11 | 99209047 | 99231017 | 21971 | 2 | ENSMUST00 | 0 | Smarce1 | ENSMUSG00000037935.11 |
| chr13 | 112463544 | 112465442 | 31.62822 | 118.12274 | 114.03909 | Promoter | 13 | 112464070 | 112510086 | 46017 | 1 | ENSMUST00 | 0 | Il6st | ENSMUSG00000021756.7 |
| chr14 | 50784823 | 50785681 | 31.62822 | 118.12274 | 114.03909 | Promoter | 14 | 50765415 | 50785518 | 20104 | 2 | ENSMUST00 | 0 | Ttc5 | ENSMUSG00000006288.7 |
| chr15 | 73511913 | 73514307 | 31.62822 | 118.12274 | 114.03909 | Promoter | 15 | 73512560 | 73572242 | 59683 | 1 | ENSMUST00 | 0 | Dennd3 | ENSMUSG00000036661.8 |
| chr15 | 81190019 | 81190846 | 31.62822 | 118.12274 | 114.03909 | Promoter | 15 | 81012281 | 81190733 | 178453 | 2 | ENSMUST00 | 0 | Mkl1 | ENSMUSG00000042292.11 |
| chr16 | 45157695 | 45160217 | 31.62822 | 118.12274 | 114.03909 | Promoter | 16 | 45158785 | 45188538 | 29754 | 1 | ENSMUST00 | 0 | Atg3 | ENSMUSG00000022663.3 |
| chr17 | 56035704 | 56036685 | 31.62822 | 118.12274 | 114.03909 | Promoter | 17 | 56016753 | 56036637 | 19885 | 2 | ENSMUST00 | 0 | Sh3gl1 | ENSMUSG00000003200.9 |
| chr17 | 27839197 | 27840871 | 31.62822 | 118.12274 | 114.03909 | Promoter | 17 | 27839974 | 27851964 | 11991 | 1 | ENSMUST00 | 0 | Snrpc | ENSMUSG00000024217.9 |
| chr18 | 58658862 | 58660585 | 31.62822 | 118.12274 | 114.03909 | Promoter | 18 | 58659482 | 58679570 | 20089 | 1 | ENSMUST00 | 0 | Isoc1 | ENSMUSG00000024601.8 |
| chr19 | 8870791 | 8872513 | 31.62822 | 118.12274 | 114.03909 | Promoter | 19 | 8871559 | 8875655 | 4097 | 1 | ENSMUST00 | 0 | Ubxn1 | ENSMUSG00000071655.6 |
| chr2 | 18391673 | 18393132 | 31.62822 | 118.12274 | 114.03909 | Promoter | 2 | 18206333 | 18392811 | 186479 | 2 | ENSMUST00 | 0 | Dnajc1 | ENSMUSG00000026740.7 |
| chr5 | 110652774 | 110654437 | 31.62822 | 118.12274 | 114.03909 | Promoter | 5 | 110653451 | 110660496 | 7046 | 1 | ENSMUST00 | 0 | Ddx51 | ENSMUSG00000029504.5 |
| chr5 | 142628759 | 142630558 | 31.62822 | 118.12274 | 114.03909 | Promoter | 5 | 142629542 | 142670588 | 41047 | 1 | ENSMUST00 | 0 | Wipi2 | ENSMUSG00000029578.10 |
| chr5 | 137778297 | 137779860 | 31.62822 | 118.12274 | 114.03909 | Promoter | 5 | 137778849 | 137779762 | 914 | 1 | ENSMUST00 | 0 | Ppp1r35 | ENSMUSG00000029725.5 |
| chr6 | 97148008 | 97149516 | 31.62822 | 118.12274 | 114.03909 | Promoter | 6 | 97110024 | 97149182 | 39159 | 2 | ENSMUST00 | 0 | Eogt | ENSMUSG00000035245.7 |
| chr8 | 106058803 | 106061197 | 31.62822 | 118.12274 | 114.03909 | Promoter | 8 | 106059603 | 106130537 | 70935 | 1 | ENSMUST00 | 0 | Nfatc3 | ENSMUSG00000031902.9 |
| chr8 | 124896500 | 124898851 | 31.62822 | 118.12274 | 114.03909 | Promoter | 8 | 124897886 | 124903813 | 5928 | 1 | ENSMUST00 | 0 | Sprtn | ENSMUSG00000031986.5 |
| chr8 | 123042552 | 123043579 | 31.62822 | 118.12274 | 114.03909 | Promoter | 8 | 123042466 | 123044602 | 2137 | 1 | ENSMUST00 | 87 | RP23-460F2 | ENSMUSG00000099881.1 |
| chr5 | 121190294 | 121191978 | 36.25188 | 118.09364 | 114.014 | Promoter | 5 | 121130533 | 121191397 | 60865 | 2 | ENSMUST00 | 0 | Ptpn11 | ENSMUSG00000043733.9 |
| chr1 | 134414933 | 134416522 | 34.4409 | 118.06123 | 113.98219 | Promoter | 1 | 134415378 | 134433351 | 17974 | 1 | ENSMUST00 | 0 | Adipor1 | ENSMUSG00000026457.9 |
| chr17 | 35821005 | 35823894 | 34.4409 | 118.06123 | 113.98219 | Promoter | 17 | 35821684 | 35822923 | 1240 | 1 | ENSMUST00 | 0 | Ier3 | ENSMUSG00000003541.6 |
| chr6 | 140622736 | 140625497 | 34.4409 | 118.06123 | 113.98219 | Promoter | 6 | 140623499 | 140652826 | 29328 | 1 | ENSMUST00 | 0 | Aebp2 | ENSMUSG00000030232.12 |
| chr6 | 149187700 | 149189217 | 34.4409 | 118.06123 | 113.98219 | Promoter | 6 | 149157147 | 149188712 | 31566 | 2 | ENSMUST00 | 0 | Amn1 | ENSMUSG00000068250.8 |
| chr7 | 28808053 | 28810551 | 34.4409 | 118.06123 | 113.98219 | Promoter | 7 | 28808541 | 28822266 | 13726 | 1 | ENSMUST00 | 0 | Hnrnpl | ENSMUSG00000015165.11 |
| chr12 | 8673672 | 8675883 | 37.44409 | 118.04973 | 113.97176 | Promoter | 12 | 8674134 | 8751454 | 77321 | 1 | ENSMUST00 | 0 | Pum2 | ENSMUSG00000020594.9 |
| chr12 | 112588574 | 112590424 | 36.83125 | 118.04518 | 113.96772 | Promoter | 12 | 112588784 | 112615556 | 26773 | 1 | ENSMUST00 | 0 | Inf2 | ENSMUSG00000037679.8 |
| chr13 | 21467932 | 21469104 | 36.83125 | 118.04518 | 113.96772 | Promoter | 13 | 21467047 | 21468509 | 1463 | 2 | ENSMUST00 | 0 | Nkapl | ENSMUSG00000059395.4 |
| chr2 | 83723513 | 83726191 | 36.83125 | 118.04518 | 113.96772 | Promoter | 2 | 83724397 | 83756797 | 32401 | 1 | ENSMUST00 | 0 | Itgav | ENSMUSG00000027087.6 |
| chr6 | 125131400 | 125132564 | 36.83125 | 118.04518 | 113.96772 | Promoter | 6 | 125131883 | 125144753 | 12871 | 1 | ENSMUST00 | 0 | Nop2 | ENSMUSG00000038279.8 |
| chr7 | 19269289 | 19272839 | 36.83125 | 118.04518 | 113.96772 | Promoter | 7 | 19256929 | 19271817 | 14889 | 2 | ENSMUST00 | 0 | Vasp | ENSMUSG00000030403.8 |
| chr2 | 30414758 | 30416971 | 32.68921 | 118.02481 | 113.94904 | Promoter | 2 | 30400471 | 30415771 | 15301 | 2 | ENSMUST00 | 0 | Crat | ENSMUSG00000026853.10 |
| chr10 | 128695458 | 128696358 | 27.04123 | 118.00974 | 113.93439 | Promoter | 10 | 128677193 | 128696268 | 19076 | 2 | ENSMUST00 | 0 | Rab5b | ENSMUSG00000000711.2 |
| chr12 | 110696524 | 110697214 | 36.18131 | 117.97025 | 113.89517 | Promoter | 12 | 110690605 | 110696395 | 5791 | 2 | ENSMUST00 | -130 | Hsp90aa1 | ENSMUSG00000021270.8 |
| chr11 | 58977051 | 58978959 | 35.56825 | 117.95176 | 113.87694 | Promoter | 11 | 58978093 | 58991455 | 13363 | 1 | ENSMUST00 | 0 | Trim11 | ENSMUSG00000020455.11 |
| chr11 | 3647807 | 3650043 | 35.56825 | 117.95176 | 113.87694 | Promoter | 11 | 3649494 | 3690370 | 40877 | 1 | ENSMUST00 | 0 | Morc2a | ENSMUSG00000034543.9 |
| chr11 | 40691371 | 40693824 | 35.56825 | 117.95176 | 113.87694 | Promoter | 11 | 40679314 | 40692774 | 13461 | 2 | ENSMUST00 | 0 | Mat2b | ENSMUSG00000042032.8 |
| chr12 | 8207616 | 8208862 | 35.56825 | 117.95176 | 113.87694 | Promoter | 12 | 8208107 | 8285758 | 77652 | 1 | ENSMUST00 | 0 | 1110057K04 | ENSMUSG00000037669.9 |
| chr15 | 75841123 | 75842571 | 35.56825 | 117.95176 | 113.87694 | Promoter | 15 | 75754432 | 75841915 | 87484 | 2 | ENSMUST00 | 0 | Zc3h3 | ENSMUSG00000075600.2 |
| chr18 | 74441849 | 74443301 | 35.56825 | 117.95176 | 113.87694 | Promoter | 18 | 74442537 | 74771493 | 328957 | 1 | ENSMUST00 | 0 | Myo5b | ENSMUSG00000025885.13 |
| chr19 | 8773568 | 8775578 | 35.56825 | 117.95176 | 113.87694 | Promoter | 19 | 8772522 | 8774467 | 1946 | 2 | ENSMUST00 | 0 | Taf6l | ENSMUSG00000003680.10 |
| chr2 | 132846572 | 132848126 | 35.56825 | 117.95176 | 113.87694 | Promoter | 2 | 132846666 | 132866785 | 20120 | 1 | ENSMUST00 | 0 | Crls1 | ENSMUSG00000027357.11 |
| chr2 | 38925035 | 38926515 | 35.56825 | 117.95176 | 113.87694 | Promoter | 2 | 38723374 | 38926217 | 202844 | 2 | ENSMUST00 | 0 | Nr6a1 | ENSMUSG00000063972.8 |
| chr3 | 142393844 | 142396226 | 35.56825 | 117.95176 | 113.87694 | Promoter | 3 | 142239586 | 142395696 | 156111 | 2 | ENSMUST00 | 0 | Pdlim5 | ENSMUSG00000028273.10 |
| chr6 | 67034307 | 67037493 | 35.56825 | 117.95176 | 113.87694 | Promoter | 6 | 67035096 | 67037406 | 2311 | 2 | ENSMUST00 | 0 | Gadd45a | ENSMUSG00000036390.6 |
| chr7 | 5019648 | 5021260 | 35.56825 | 117.95176 | 113.87694 | Promoter | 7 | 5020376 | 5033138 | 12763 | 1 | ENSMUST00 | 0 | Zfp865 | ENSMUSG00000074405.5 |
| chr8 | 25099212 | 25103419 | 35.56825 | 117.95176 | 113.87694 | Promoter | 8 | 25039144 | 25102194 | 63051 | 2 | ENSMUST00 | 0 | Plekha2 | ENSMUSG00000031557.10 |
| chr8 | 111056253 | 111057313 | 35.56825 | 117.95176 | 113.87694 | 3' UTR | 8 | 111047912 | 111050010 | 2099 | 1 | ENSMUST00 | 8342 | Aars | ENSMUSG00000031960.9 |
| chr17 | 74716981 | 74718307 | 36.75754 | 117.9171 | 113.84505 | Promoter | 17 | 74717750 | 74863570 | 145821 | 1 | ENSMUST00 | 0 | Ttc27 | ENSMUSG00000024078.6 |
| chr2 | 18063641 | 18065839 | 37.36699 | 117.91644 | 113.84495 | Promoter | 2 | 18064394 | 18122348 | 57955 | 1 | ENSMUST00 | 0 | Mllt10 | ENSMUSG00000026743.11 |
| chr17 | 46649997 | 46651455 | 36.111 | 117.84728 | 113.77644 | Promoter | 17 | 46650337 | 46664364 | 14028 | 1 | ENSMUST00 | 0 | Cul7 | ENSMUSG00000038545.8 |
| chr2 | 172999755 | 173001100 | 31.997 | 117.77848 | 113.70881 | Promoter | 2 | 173000117 | 173015739 | 15623 | 1 | ENSMUST00 | 0 | Rae1 | ENSMUSG00000027509.6 |
| chr2 | 158794749 | 158795388 | 31.997 | 117.77848 | 113.70881 | Promoter | 2 | 158794807 | 158858214 | 63408 | 1 | ENSMUST00 | 0 | Dhx35 | ENSMUSG00000027655.9 |
| chr4 | 10873211 | 10875421 | 31.997 | 117.77848 | 113.70881 | Promoter | 4 | 10874498 | 10899421 | 24924 | 1 | ENSMUST00 | 0 | 2610301B20 | ENSMUSG00000059482.10 |
| chr1 | 16617928 | 16619784 | 29.83578 | 117.74437 | 113.67492 | Promoter | 1 | 16540790 | 16619312 | 78523 | 2 | ENSMUST00 | 0 | Ube2w | ENSMUSG00000025939.12 |
| chr11 | 117967602 | 117970051 | 29.83578 | 117.74437 | 113.67492 | Promoter | 11 | 117966079 | 117969186 | 3108 | 2 | ENSMUST00 | 0 | Socs3 | ENSMUSG00000053113.3 |

| | | | | | | | | | | | | | | | |
|---|---|---|---|---|---|---|---|---|---|---|---|---|---|---|---|
| chr12 | 69158171 | 69159793 | 29.83578 | 117.74437 | 113.67492 | Promoter | 12 | 69157722 | 69159186 | 1465 | 2 | ENSMUST00 | 0 | Rps29 | ENSMUSG00000034892.8 |
| chr13 | 98315862 | 98317835 | 29.83578 | 117.74437 | 113.67492 | Promoter | 13 | 98309896 | 98316967 | 7072 | 2 | ENSMUST00 | 0 | Btf3 | ENSMUSG00000021660.9 |
| chr3 | 82357582 | 82358686 | 29.83578 | 117.74437 | 113.67492 | Promoter | 3 | 82358072 | 82395268 | 37197 | 1 | ENSMUST00 | 0 | Map9 | ENSMUSG00000033900.8 |
| chr13 | 41000191 | 41002089 | 28.32768 | 117.73571 | 113.66808 | Promoter | 13 | 41001023 | 41013015 | 11993 | 1 | ENSMUST00 | 0 | Pak1ip1 | ENSMUSG00000038683.10 |
| chr17 | 87024579 | 87026295 | 28.32768 | 117.73571 | 113.66808 | Promoter | 17 | 86997256 | 87025406 | 28151 | 2 | ENSMUST00 | 0 | Pigf | ENSMUSG00000024145.6 |
| chr19 | 60579332 | 60581756 | 28.32768 | 117.73571 | 113.66808 | Promoter | 19 | 60524696 | 60581023 | 56328 | 2 | ENSMUST00 | 0 | Cacul1 | ENSMUSG00000033417.10 |
| chr3 | 86223104 | 86225747 | 28.32768 | 117.73571 | 113.66808 | Promoter | 3 | 86224690 | 86782693 | 558004 | 1 | ENSMUST00 | 0 | Lrba | ENSMUSG00000028080.10 |
| chr1 | 177256972 | 177259264 | 37.21373 | 117.65135 | 113.58522 | Promoter | 1 | 177022116 | 177258203 | 236088 | 2 | ENSMUST00 | 0 | Akt3 | ENSMUSG00000019699.10 |
| chr1 | 87213120 | 87215128 | 37.21373 | 117.65135 | 113.58522 | Promoter | 1 | 87213914 | 87226607 | 12694 | 1 | ENSMUST00 | 0 | Eif4e2 | ENSMUSG00000026254.12 |
| chr3 | 139075336 | 139076229 | 33.62833 | 117.64893 | 113.58318 | Promoter | 3 | 138925906 | 139075199 | 149294 | 2 | ENSMUST00 | -138 | Rap1gds1 | ENSMUSG00000028149.7 |
| chr4 | 43492637 | 43494474 | 33.62833 | 117.64893 | 113.58318 | Promoter | 4 | 43492900 | 43494205 | 1306 | 1 | ENSMUST00 | 0 | Ccdc107 | ENSMUSG00000028461.7 |
| chr10 | 111164281 | 111166238 | 36.53815 | 117.53558 | 113.47088 | Promoter | 10 | 111164802 | 111297247 | 132446 | 1 | ENSMUST00 | 0 | Osbpl8 | ENSMUSG00000020189.8 |
| chr11 | 103966649 | 103967830 | 37.13757 | 117.51956 | 113.45509 | Promoter | 11 | 103966739 | 103985337 | 18599 | 1 | ENSMUST00 | 0 | Arf2 | ENSMUSG00000062421.8 |
| chr19 | 46055978 | 46057815 | 37.13757 | 117.51956 | 113.45509 | Promoter | 19 | 46056539 | 46072911 | 16373 | 1 | ENSMUST00 | 0 | Pprc1 | ENSMUSG00000055491.8 |
| chr2 | 84714015 | 84716660 | 37.13757 | 117.51956 | 113.45509 | Promoter | 2 | 84687970 | 84715180 | 27211 | 2 | ENSMUST00 | 0 | Zdhhc5 | ENSMUSG00000034075.10 |
| chr6 | 72096911 | 72098643 | 37.13757 | 117.51956 | 113.45509 | Promoter | 6 | 72097592 | 72154571 | 56980 | 1 | ENSMUST00 | 0 | St3gal5 | ENSMUSG00000056091.7 |
| chr14 | 47567643 | 47568983 | 34.67015 | 117.41676 | 113.35385 | Promoter | 14 | 47540894 | 47568434 | 27541 | 2 | ENSMUST00 | 0 | Atg14 | ENSMUSG00000037526.6 |
| chr19 | 3321491 | 3326282 | 34.67015 | 117.41676 | 113.35385 | Promoter | 19 | 3323301 | 3385732 | 62432 | 1 | ENSMUST00 | 0 | Cpt1a | ENSMUSG00000024900.4 |
| chr4 | 150568860 | 150571152 | 34.67015 | 117.41676 | 113.35385 | Promoter | 4 | 150569588 | 150570495 | 908 | 1 | ENSMUST00 | 0 | Rere | ENSMUSG00000039852.11 |
| chr9 | 88547470 | 88549208 | 34.67015 | 117.41676 | 113.35385 | Promoter | 9 | 88548020 | 88571061 | 23042 | 1 | ENSMUST00 | 0 | Zfp949 | ENSMUSG00000032425.10 |
| chr13 | 43398310 | 43399139 | 37.68163 | 117.40437 | 113.3432 | Promoter | 13 | 43398376 | 43402858 | 4483 | 1 | ENSMUST00 | 0 | Nol7 | ENSMUSG00000063200.3 |
| chr14 | 105258237 | 105259618 | 37.68163 | 117.40437 | 113.3432 | Promoter | 14 | 105258573 | 105309298 | 50726 | 1 | ENSMUST00 | 0 | Ndfip2 | ENSMUSG00000053253.11 |
| chr16 | 38549386 | 38550580 | 37.68163 | 117.40437 | 113.3432 | Promoter | 16 | 38525137 | 38550258 | 25122 | 2 | ENSMUST00 | 0 | Poglut1 | ENSMUSG00000034064.9 |
| chr16 | 97961578 | 97963139 | 37.68163 | 117.40437 | 113.3432 | Promoter | 16 | 97947435 | 97962621 | 15187 | 2 | ENSMUST00 | 0 | Zbtb21 | ENSMUSG00000046962.10 |
| chr19 | 28963503 | 28964799 | 37.68163 | 117.40437 | 113.3432 | Promoter | 19 | 28963953 | 28966811 | 2859 | 1 | ENSMUST00 | 0 | Ppapdc2 | ENSMUSG00000040105.3 |
| chr19 | 46003066 | 46004412 | 37.68163 | 117.40437 | 113.3432 | Promoter | 19 | 46003478 | 46006173 | 2696 | 1 | ENSMUST00 | 0 | Hps6 | ENSMUSG00000074811.2 |
| chr5 | 100497647 | 100499112 | 37.68163 | 117.40437 | 113.3432 | Promoter | 5 | 100441928 | 100498591 | 56664 | 2 | ENSMUST00 | 0 | Lin54 | ENSMUSG00000035310.10 |
| chr6 | 89643634 | 89645154 | 37.68163 | 117.40437 | 113.3432 | Promoter | 6 | 89643988 | 89675529 | 31542 | 1 | ENSMUST00 | 0 | Txnrd3 | ENSMUSG00000000811.8 |
| chr7 | 100705978 | 100707534 | 37.68163 | 117.40437 | 113.3432 | Promoter | 7 | 100706651 | 100841313 | 134663 | 1 | ENSMUST00 | 0 | Fam168a | ENSMUSG00000029461.10 |
| chr7 | 45052080 | 45054223 | 37.68163 | 117.40437 | 113.3432 | Promoter | 7 | 45027707 | 45052881 | 25175 | 2 | ENSMUST00 | 0 | Prr12 | ENSMUSG00000046574.7 |
| chr9 | 43742978 | 43745680 | 37.68163 | 117.40437 | 113.3432 | Promoter | 9 | 43744576 | 43807454 | 62879 | 1 | ENSMUST00 | 0 | Pvrl1 | ENSMUSG00000032012.7 |
| chrX | 94796903 | 94798710 | 37.68163 | 117.40437 | 113.3432 | Promoter | 20 | 94795563 | 94798183 | 2621 | 2 | ENSMUST00 | 0 | Zxda | ENSMUSG00000073060.4 |
| chr5 | 117120034 | 117121290 | 37.06172 | 117.38827 | 113.32996 | Promoter | 5 | 117120129 | 117275217 | 155089 | 1 | ENSMUST00 | 0 | Taok3 | ENSMUSG00000061288.8 |
| chr10 | 122678407 | 122680961 | 35.8325 | 117.35963 | 113.30182 | Promoter | 10 | 122678762 | 122880333 | 201572 | 1 | ENSMUST00 | 0 | Ppm1h | ENSMUSG00000034613.7 |
| chr15 | 12204978 | 12206098 | 35.8325 | 117.35963 | 113.30182 | Promoter | 15 | 12205028 | 12272240 | 67213 | 1 | ENSMUST00 | 0 | Mtmr12 | ENSMUSG00000039458.9 |
| chr2 | 60208946 | 60211053 | 35.8325 | 117.35963 | 113.30182 | Promoter | 2 | 60209887 | 60249211 | 39325 | 1 | ENSMUST00 | 0 | Mar-07 | ENSMUSG00000026977.12 |
| chr2 | 80314850 | 80316280 | 35.8325 | 117.35963 | 113.30182 | Promoter | 2 | 80315466 | 80324788 | 9323 | 1 | ENSMUST00 | 0 | Dnajc10 | ENSMUSG00000027006.8 |
| chr7 | 65692880 | 65694150 | 35.8325 | 117.35963 | 113.30182 | Promoter | 7 | 65693417 | 65701902 | 8486 | 1 | ENSMUST00 | 0 | Tm2d3 | ENSMUSG00000078681.5 |
| chr3 | 108145284 | 108146677 | 29.61888 | 117.31387 | 113.25776 | Promoter | 3 | 108107280 | 108146146 | 38867 | 2 | ENSMUST00 | 0 | Gnai3 | ENSMUSG00000000001.4 |
| chr2 | 112378870 | 112380430 | 32.87005 | 117.31267 | 113.25658 | Promoter | 2 | 112379211 | 112414240 | 35030 | 1 | ENSMUST00 | 0 | Katnbl1 | ENSMUSG00000027132.3 |
| chr9 | 121856654 | 121858574 | 32.87005 | 117.31267 | 113.25658 | Promoter | 9 | 121848560 | 121858000 | 9441 | 2 | ENSMUST00 | 0 | Higd1a | ENSMUSG00000038412.7 |
| chr8 | 84249054 | 84250048 | 37.61386 | 117.28751 | 113.23196 | Promoter | 8 | 84246237 | 84249761 | 3525 | 2 | ENSMUST00 | 0 | D8Ertd738e | ENSMUSG00000019362.7 |
| chr1 | 119646090 | 119649158 | 33.41831 | 117.26761 | 113.21259 | Promoter | 1 | 119545037 | 119648936 | 103900 | 2 | ENSMUST00 | 0 | Epb4.1l5 | ENSMUSG00000026383.9 |
| chr1 | 170214386 | 170215718 | 33.41831 | 117.26761 | 113.21259 | Promoter | 1 | 170193420 | 170214734 | 21315 | 2 | ENSMUST00 | 0 | Uhmk1 | ENSMUSG00000026667.9 |
| chr1 | 165763380 | 165764596 | 33.41831 | 117.26761 | 113.21259 | Promoter | 1 | 165763746 | 165775308 | 11563 | 1 | ENSMUST00 | 0 | Creg1 | ENSMUSG00000040713.7 |
| chr1 | 171149870 | 171151259 | 33.41831 | 117.26761 | 113.21259 | Promoter | 1 | 171150711 | 171161130 | 10420 | 1 | ENSMUST00 | 0 | Mpz | ENSMUSG00000056569.5 |
| chr10 | 17721687 | 17725607 | 33.41831 | 117.26761 | 113.21259 | Promoter | 10 | 17723228 | 17725673 | 2446 | 1 | ENSMUST00 | 0 | Cited2 | ENSMUSG00000039910.9 |
| chr10 | 128490711 | 128495254 | 33.41831 | 117.26761 | 113.21259 | Promoter | 10 | 128490860 | 128493875 | 3016 | 2 | ENSMUST00 | 0 | Myl6 | ENSMUSG00000090841.1 |
| chr11 | 74924900 | 74926913 | 33.41831 | 117.26761 | 113.21259 | Promoter | 11 | 74906359 | 74925798 | 19440 | 2 | ENSMUST00 | 0 | Srr | ENSMUSG00000001323.12 |
| chr11 | 32641800 | 32643992 | 33.41831 | 117.26761 | 113.21259 | Promoter | 11 | 32642724 | 32746569 | 103846 | 1 | ENSMUST00 | 0 | Fbxw11 | ENSMUSG00000020271.12 |
| chr11 | 50324465 | 50326594 | 33.41831 | 117.26761 | 113.21259 | Promoter | 11 | 50293961 | 50325673 | 31713 | 2 | ENSMUST00 | 0 | Canx | ENSMUSG00000020368.10 |
| chr11 | 54955487 | 54956498 | 33.41831 | 117.26761 | 113.21259 | Promoter | 11 | 54910785 | 54956122 | 45338 | 2 | ENSMUST00 | 0 | Tnip1 | ENSMUSG00000020400.12 |
| chr13 | 63814553 | 63816042 | 33.41831 | 117.26761 | 113.21259 | Promoter | 13 | 63815240 | 63853810 | 38571 | 1 | ENSMUST00 | 0 | Ercc6l2 | ENSMUSG00000021470.12 |
| chr13 | 38634179 | 38635474 | 33.41831 | 117.26761 | 113.21259 | Promoter | 13 | 38602709 | 38635109 | 32401 | 2 | ENSMUST00 | 0 | Bloc1s5 | ENSMUSG00000038982.9 |
| chr13 | 13953612 | 13955626 | 33.41831 | 117.26761 | 113.21259 | Promoter | 13 | 13954674 | 13999068 | 44395 | 1 | ENSMUST00 | 0 | B3galnt2 | ENSMUSG00000039242.9 |
| chr14 | 55642637 | 55644415 | 33.41831 | 117.26761 | 113.21259 | Promoter | 14 | 55635965 | 55643251 | 7287 | 2 | ENSMUST00 | 0 | Tm9sf1 | ENSMUSG00000002320.10 |
| chr14 | 19822763 | 19824633 | 33.41831 | 117.26761 | 113.21259 | Promoter | 14 | 19811402 | 19823823 | 12422 | 2 | ENSMUST00 | 0 | 2700060E02I | ENSMUSG00000021807.5 |
| chr14 | 20347149 | 20349125 | 33.41831 | 117.26761 | 113.21259 | Promoter | 14 | 20319859 | 20348121 | 28263 | 2 | ENSMUST00 | 0 | Ecd | ENSMUSG00000021810.2 |
| chr14 | 78535987 | 78537459 | 33.41831 | 117.26761 | 113.21259 | Promoter | 14 | 78492246 | 78536808 | 44563 | 2 | ENSMUST00 | 0 | Akap11 | ENSMUSG00000022016.10 |
| chr14 | 65357764 | 65359579 | 33.41831 | 117.26761 | 113.21259 | Promoter | 14 | 65358676 | 65398930 | 40255 | 1 | ENSMUST00 | 0 | Zfp395 | ENSMUSG00000034522.9 |
| chr16 | 13255775 | 13257573 | 33.41831 | 117.26761 | 113.21259 | Promoter | 16 | 13256481 | 13415063 | 158583 | 1 | ENSMUST00 | 0 | Mkl2 | ENSMUSG00000009569.9 |
| chr16 | 4938754 | 4940192 | 33.41831 | 117.26761 | 113.21259 | Promoter | 16 | 4939111 | 4941020 | 1910 | 1 | ENSMUST00 | 0 | Nudt16l1 | ENSMUSG00000022516.9 |
| chr16 | 87431822 | 87432918 | 33.41831 | 117.26761 | 113.21259 | Promoter | 16 | 87376651 | 87432606 | 55956 | 2 | ENSMUST00 | 0 | Ltn1 | ENSMUSG00000052299.8 |
| chr17 | 31519590 | 31521063 | 33.41831 | 117.26761 | 113.21259 | Promoter | 17 | 31503526 | 31519965 | 16440 | 2 | ENSMUST00 | 0 | Wdr4 | ENSMUSG00000024037.7 |
| chr17 | 83513267 | 83515007 | 33.41831 | 117.26761 | 113.21259 | Promoter | 17 | 83501919 | 83514333 | 12415 | 2 | ENSMUST00 | 0 | Cox7a2l | ENSMUSG00000024248.8 |
| chr17 | 10319435 | 10321332 | 33.41831 | 117.26761 | 113.21259 | Promoter | 17 | 10319945 | 10320229 | 285 | 2 | ENSMUST00 | 0 | Qk | ENSMUSG00000044407.6 |
| chr18 | 70567903 | 70569818 | 33.41831 | 117.26761 | 113.21259 | Promoter | 18 | 70568189 | 70626131 | 57943 | 1 | ENSMUST00 | 0 | Mbd2 | ENSMUSG00000024513.11 |
| chr18 | 74216143 | 74217076 | 33.41831 | 117.26761 | 113.21259 | Promoter | 18 | 74216131 | 74221491 | 5361 | 1 | ENSMUST00 | 13 | Cxxc1 | ENSMUSG00000024560.6 |
| chr18 | 34859236 | 34862643 | 33.41831 | 117.26761 | 113.21259 | Promoter | 18 | 34859823 | 34864984 | 5162 | 1 | ENSMUST00 | 0 | Egr1 | ENSMUSG00000038418.7 |
| chr18 | 36285948 | 36288201 | 33.41831 | 117.26761 | 113.21259 | 5' UTR | 18 | 36281097 | 36301521 | 20425 | 1 | ENSMUST00 | 4852 | Pura | ENSMUSG00000043991.3 |
| chr19 | 21651908 | 21654841 | 33.41831 | 117.26761 | 113.21259 | Promoter | 19 | 21581202 | 21652728 | 71527 | 2 | ENSMUST00 | 0 | 1110059E24I | ENSMUSG00000035171.4 |
| chr19 | 8988723 | 8990788 | 33.41831 | 117.26761 | 113.21259 | Promoter | 19 | 8989284 | 9019195 | 29912 | 1 | ENSMUST00 | 0 | Ahnak | ENSMUSG00000069833.7 |
| chr2 | 128686446 | 128687786 | 33.41831 | 117.26761 | 113.21259 | Promoter | 2 | 128610104 | 128687391 | 77288 | 2 | ENSMUST00 | 0 | Anapc1 | ENSMUSG00000014355.5 |
| chr2 | 112265293 | 112267278 | 33.41831 | 117.26761 | 113.21259 | Promoter | 2 | 112265825 | 112359127 | 93303 | 1 | ENSMUST00 | 0 | Slc12a6 | ENSMUSG00000027130.10 |
| chr2 | 172369956 | 172371698 | 33.41831 | 117.26761 | 113.21259 | Promoter | 2 | 172356190 | 172370535 | 14346 | 2 | ENSMUST00 | 0 | Aurka | ENSMUSG00000027496.10 |
| chr3 | 119782091 | 119783812 | 33.41831 | 117.26761 | 113.21259 | Promoter | 3 | 119718742 | 119783388 | 64647 | 2 | ENSMUST00 | 0 | Ptbp2 | ENSMUSG00000028134.7 |
| chr3 | 87617020 | 87619659 | 33.41831 | 117.26761 | 113.21259 | Promoter | 3 | 87617559 | 87699145 | 81587 | 1 | ENSMUST00 | 0 | Arhgef11 | ENSMUSG00000041977.12 |
| chr3 | 96171376 | 96173402 | 33.41831 | 117.26761 | 113.21259 | Promoter | 3 | 96172332 | 96177564 | 5233 | 1 | ENSMUST00 | 0 | Sf3b4 | ENSMUSG00000068856.3 |
| chr5 | 99251538 | 99253454 | 33.41831 | 117.26761 | 113.21259 | Promoter | 5 | 99217426 | 99252927 | 35502 | 2 | ENSMUST00 | 0 | Rasgef1b | ENSMUSG00000029333.9 |
| chr5 | 137629072 | 137631603 | 33.41831 | 117.26761 | 113.21259 | Promoter | 5 | 137629175 | 137642899 | 13725 | 1 | ENSMUST00 | 0 | Gm20605 | ENSMUSG00000029720.9 |
| chr5 | 144357066 | 144358946 | 33.41831 | 117.26761 | 113.21259 | Promoter | 5 | 144264526 | 144358112 | 93587 | 2 | ENSMUST00 | 0 | Baiap2l1 | ENSMUSG00000038859.7 |
| chr6 | 116337597 | 116338886 | 33.41831 | 117.26761 | 113.21259 | Promoter | 6 | 116338024 | 116409540 | 71517 | 1 | ENSMUST00 | 0 | Mar-08 | ENSMUSG00000025702.10 |
| chr7 | 110121226 | 110123539 | 33.41831 | 117.26761 | 113.21259 | Promoter | 7 | 110122350 | 110142112 | 19763 | 1 | ENSMUST00 | 0 | Wee1 | ENSMUSG00000031016.8 |
| chr7 | 35802007 | 35803236 | 33.41831 | 117.26761 | 113.21259 | Promoter | 7 | 35772343 | 35802989 | 30647 | 2 | ENSMUST00 | 0 | Zfp507 | ENSMUSG00000044452.5 |
| chr8 | 83608165 | 83609506 | 33.41831 | 117.26761 | 113.21259 | Promoter | 8 | 83608175 | 83611902 | 3728 | 1 | ENSMUST00 | 0 | Dnajb1 | ENSMUSG00000005483.9 |
| chr8 | 124721276 | 124722857 | 33.41831 | 117.26761 | 113.21259 | Promoter | 8 | 124722139 | 124734123 | 11985 | 1 | ENSMUST00 | 0 | Arv1 | ENSMUSG00000031982.4 |
| chr9 | 62810356 | 62812424 | 33.41831 | 117.26761 | 113.21259 | Promoter | 9 | 62791832 | 62811648 | 19817 | 2 | ENSMUST00 | 0 | Fem1b | ENSMUSG00000032244.8 |
| chr9 | 92457272 | 92458605 | 33.41831 | 117.26761 | 113.21259 | Promoter | 9 | 92457373 | 92492456 | 35084 | 1 | ENSMUST00 | 0 | Plscr4 | ENSMUSG00000032377.8 |
| chr9 | 99138922 | 99141004 | 33.41831 | 117.26761 | 113.21259 | Promoter | 9 | 99036654 | 99140087 | 103434 | 2 | ENSMUST00 | 0 | Pik3cb | ENSMUSG00000032462.9 |
| chr9 | 86467107 | 86468011 | 33.41831 | 117.26761 | 113.21259 | Promoter | 9 | 86467154 | 86555923 | 88770 | 1 | ENSMUST00 | 0 | Dopey1 | ENSMUSG00000034973.11 |
| chr9 | 110531641 | 110533828 | 33.41831 | 117.26761 | 113.21259 | Promoter | 9 | 110532597 | 110618633 | 86037 | 1 | ENSMUST00 | 0 | Setd2 | ENSMUSG00000044791.11 |
| chr9 | 101073983 | 101079109 | 33.41831 | 117.26761 | 113.21259 | Promoter | 9 | 101074101 | 101101785 | 27685 | 1 | ENSMUST00 | 0 | Msl2 | ENSMUSG00000066415.4 |
| chr17 | 3113719 | 3116459 | 36.98618 | 117.25745 | 113.20857 | Promoter | 17 | 3114972 | 3198855 | 83884 | 1 | ENSMUST00 | 0 | Scaf8 | ENSMUSG00000046201.8 |
| chr3 | 103171443 | 103172541 | 35.16908 | 117.24756 | 113.199 | Promoter | 3 | 103171655 | 103179152 | 7498 | 1 | ENSMUST00 | 0 | Bcas2 | ENSMUSG00000005687.9 |
| chr17 | 65612263 | 65614741 | 35.76354 | 117.23875 | 113.19022 | Promoter | 17 | 65580056 | 65613555 | 33500 | 2 | ENSMUST00 | 0 | Vapa | ENSMUSG00000024091.8 |
| chr11 | 95712399 | 95714519 | 37.53511 | 117.15172 | 113.10373 | Promoter | 11 | 95712673 | 95772353 | 59681 | 1 | ENSMUST00 | 0 | Zfp652 | ENSMUSG00000075595.4 |
| chr10 | 7954510 | 7957213 | 36.91095 | 117.12714 | 113.07945 | Promoter | 10 | 7905653 | 7956230 | 50578 | 2 | ENSMUST00 | 0 | Tab2 | ENSMUSG00000015755.11 |
| chr8 | 25017207 | 25018218 | 36.91095 | 117.12714 | 113.07945 | Promoter | 8 | 25017211 | 25023259 | 6049 | 1 | ENSMUST00 | 0 | Tm2d2 | ENSMUSG00000031556.5 |
| chr3 | 88335235 | 88337205 | 37.45669 | 117.01644 | 112.96991 | Promoter | 3 | 88336260 | 88362337 | 26078 | 1 | ENSMUST00 | 0 | Smg5 | ENSMUSG00000001415.5 |
| chr6 | 71271644 | 71273367 | 37.45669 | 117.01644 | 112.96991 | Promoter | 6 | 71272019 | 71285318 | 13300 | 1 | ENSMUST00 | 0 | Krcc1 | ENSMUSG00000053012.12 |
| chr1 | 152901741 | 152904742 | 36.83603 | 116.99731 | 112.9521 | Promoter | 1 | 152836995 | 152902622 | 65628 | 2 | ENSMUST00 | 0 | Smg7 | ENSMUSG00000042772.10 |
| chr15 | 36607571 | 36609023 | 36.83603 | 116.99731 | 112.9521 | Promoter | 15 | 36595661 | 36608973 | 13313 | 2 | ENSMUST00 | 0 | Pabpc1 | ENSMUSG00000022283.9 |
| chr10 | 81357080 | 81358122 | 35.62435 | 116.99458 | 112.94973 | Promoter | 10 | 81357491 | 81366225 | 8735 | 1 | ENSMUST00 | 0 | Mfsd12 | ENSMUSG00000034854.8 |
| chr11 | 98809154 | 98810753 | 35.62435 | 116.99458 | 112.94973 | Promoter | 11 | 98797707 | 98810087 | 12381 | 2 | ENSMUST00 | 0 | Gm12359 | ENSMUSG00000075576.7 |
| chr12 | 36155299 | 36158688 | 35.62435 | 116.99458 | 112.94973 | Promoter | 12 | 36091846 | 36156825 | 64980 | 2 | ENSMUST00 | 0 | Bzw2 | ENSMUSG00000020547.4 |
| chr14 | 66296421 | 66298807 | 35.62435 | 116.99458 | 112.94973 | Promoter | 14 | 66297031 | 66311424 | 14394 | 1 | ENSMUST00 | 0 | Trim35 | ENSMUSG00000022043.13 |
| chr18 | 34776612 | 34778537 | 35.62435 | 116.99458 | 112.94973 | Promoter | 18 | 34777008 | 34839370 | 62363 | 1 | ENSMUST00 | 0 | Kdm3b | ENSMUSG00000038773.10 |
| chr2 | 181600688 | 181602190 | 35.62435 | 116.99458 | 112.94973 | Promoter | 2 | 181601316 | 181655651 | 54336 | 1 | ENSMUST00 | 0 | Prpf6 | ENSMUSG00000002455.9 |
| chr2 | 173274005 | 173277330 | 35.62435 | 116.99458 | 112.94973 | Promoter | 2 | 173224458 | 173276533 | 52076 | 2 | ENSMUST00 | 0 | Pmepa1 | ENSMUSG00000038400.10 |
| chr4 | 119292585 | 119295348 | 35.62435 | 116.99458 | 112.94973 | Promoter | 4 | 119277981 | 119294604 | 16624 | 2 | ENSMUST00 | 0 | Ybx1 | ENSMUSG00000028639.9 |
| chr5 | 101764380 | 101766811 | 35.62435 | 116.99458 | 112.94973 | Promoter | 5 | 101765130 | 101823858 | 58729 | 1 | ENSMUST00 | 0 | Cds1 | ENSMUSG00000029330.8 |
| chr5 | 117114778 | 117116319 | 35.62435 | 116.99458 | 112.94973 | Promoter | 5 | 117091682 | 117115993 | 24312 | 2 | ENSMUST00 | 0 | Suds3 | ENSMUSG00000066900.6 |
| chr6 | 28260448 | 28262923 | 35.62435 | 116.99458 | 112.94973 | Promoter | 6 | 28239931 | 28261911 | 21981 | 2 | ENSMUST00 | 0 | Zfp800 | ENSMUSG00000039841.9 |
| chr8 | 122676506 | 122679041 | 35.62435 | 116.99458 | 112.94973 | Promoter | 8 | 122629463 | 122678072 | 48610 | 2 | ENSMUST00 | 0 | Cbfa2t3 | ENSMUSG00000006362.10 |
| chr8 | 70041755 | 70043748 | 35.62435 | 116.99458 | 112.94973 | Promoter | 8 | 70042813 | 70071953 | 29141 | 1 | ENSMUST00 | 0 | Sugp1 | ENSMUSG00000011306.6 |
| chr9 | 110116637 | 110118293 | 35.62435 | 116.99458 | 112.94973 | Promoter | 9 | 110084320 | 110117422 | 33103 | 2 | ENSMUST00 | 0 | Dhx30 | ENSMUSG00000032480.12 |
| chr9 | 80066534 | 80068862 | 35.62435 | 116.99458 | 112.94973 | Promoter | 9 | 80066903 | 80094484 | 27582 | 1 | ENSMUST00 | 0 | Senp6 | ENSMUSG00000034252.9 |
| chr9 | 121296988 | 121298937 | 35.62435 | 116.99458 | 112.94973 | Distal Interge | 9 | 121102544 | 121277197 | 174654 | 2 | ENSMUST00 | -19792 | Ulk4 | ENSMUSG00000040936.9 |
| chr11 | 5097235 | 5100631 | 33.83522 | 116.97303 | 112.93475 | Promoter | 11 | 5069689 | 5099266 | 29578 | 2 | ENSMUST00 | 0 | Ewsr1 | ENSMUSG00000009079.11 |
| chr3 | 116711370 | 116713116 | 33.83522 | 116.97303 | 112.93475 | Promoter | 3 | 116669469 | 116712650 | 43182 | 2 | ENSMUST00 | 0 | Slc35a3 | ENSMUSG00000027957.8 |
| chr3 | 62338524 | 62341788 | 33.83522 | 116.97303 | 112.93475 | Promoter | 3 | 62338999 | 62379719 | 40721 | 1 | ENSMUST00 | 0 | Arhgef26 | ENSMUSG00000036885.9 |
| chr4 | 95048614 | 95052213 | 33.83522 | 116.97303 | 112.93475 | Promoter | 4 | 95051534 | 95067422 | 15889 | 1 | ENSMUST00 | 0 | Junos | ENSMUSG00000087366.2 |
| chr9 | 119894501 | 119895905 | 33.83522 | 116.97303 | 112.93475 | Promoter | 9 | 119894895 | 119926579 | 31685 | 1 | ENSMUST00 | 0 | Wdr48 | ENSMUSG00000032512.6 |
| chr17 | 32349208 | 32350688 | 36.17826 | 116.9087 | 112.87152 | Promoter | 17 | 32321425 | 32350581 | 29157 | 2 | ENSMUST00 | 0 | Akap8l | ENSMUSG00000002625.9 |
| chr10 | 81429715 | 81431421 | 37.37859 | 116.88169 | 112.84478 | Promoter | 10 | 81400953 | 81431005 | 30053 | 2 | ENSMUST00 | 0 | Nfic | ENSMUSG00000055053.12 |
| chr15 | 79074546 | 79075935 | 37.37859 | 116.88169 | 112.84478 | Promoter | 15 | 79075223 | 79094400 | 19178 | 1 | ENSMUST00 | 0 | Eif3l | ENSMUSG00000033047.4 |
| chr17 | 15824815 | 15827519 | 37.37859 | 116.88169 | 112.84478 | Promoter | 17 | 15806253 | 15826586 | 20334 | 2 | ENSMUST00 | 0 | Rgmb | ENSMUSG00000048027.7 |
| chr15 | 80977670 | 80978584 | 36.76141 | 116.86796 | 112.83147 | Promoter | 15 | 80977765 | 81012289 | 34525 | 1 | ENSMUST00 | 0 | Sgsm3 | ENSMUSG00000042303.11 |

| | | | | | | | | | | | | | | | |
|---|---|---|---|---|---|---|---|---|---|---|---|---|---|---|---|
| chr18 | 10617431 | 10618746 | 36.76141 | 116.86796 | 112.83147 | Promoter | 18 | 10617796 | 10628230 | 10435 | 1 | ENSMUST00 | 0 | Snrpd1 | ENSMUSG00000002477.4 |
| chr4 | 148900254 | 148902977 | 36.76141 | 116.86796 | 112.83147 | Promoter | 4 | 148901149 | 148932896 | 31748 | 1 | ENSMUST00 | 0 | Casz1 | ENSMUSG00000028977.11 |
| chr8 | 90907503 | 90909657 | 36.76141 | 116.86796 | 112.83147 | Promoter | 8 | 90907939 | 90908415 | 477 | 2 | ENSMUST00 | 0 | Gm6658 | ENSMUSG00000074171.1 |
| chr15 | 39942756 | 39944821 | 31.49598 | 116.8353 | 112.79929 | Promoter | 15 | 39870603 | 39943788 | 73186 | 2 | ENSMUST00 | 0 | Lrp12 | ENSMUSG00000022305.7 |
| chr1 | 134361026 | 134362594 | 36.10714 | 116.78466 | 112.7497 | Promoter | 1 | 134346097 | 134361979 | 15883 | 2 | ENSMUST00 | 0 | Tmem183a | ENSMUSG00000042305.7 |
| chr10 | 4386149 | 4389234 | 34.90791 | 116.7856 | 112.7497 | Promoter | 10 | 4367074 | 4388108 | 21035 | 2 | ENSMUST00 | 0 | Zbtb2 | ENSMUSG00000075327.4 |
| chr12 | 86881379 | 86885421 | 34.90791 | 116.7856 | 112.7497 | Promoter | 12 | 86880703 | 86884814 | 4112 | 2 | ENSMUST00 | 0 | Irf2bpl | ENSMUSG00000034168.6 |
| chr17 | 74315512 | 74317085 | 36.10714 | 116.78466 | 112.7497 | Promoter | 17 | 74299474 | 74316394 | 16921 | 2 | ENSMUST00 | 0 | Dpy30 | ENSMUSG00000024067.8 |
| chr2 | 163016704 | 163018201 | 34.90791 | 116.7856 | 112.7497 | Promoter | 2 | 163017354 | 163046141 | 28788 | 1 | ENSMUST00 | 0 | Ift52 | ENSMUSG00000017858.7 |
| chr2 | 178022973 | 178024115 | 36.10714 | 116.78466 | 112.7497 | Promoter | 2 | 178023284 | 178032287 | 9004 | 1 | ENSMUST00 | 0 | Etohi1 | ENSMUSG00000074519.6 |
| chr3 | 57846610 | 57848374 | 36.10713 | 116.78466 | 112.7497 | Promoter | 3 | 57841895 | 57847564 | 5670 | 2 | ENSMUST00 | 0 | Pfn2 | ENSMUSG00000027805.11 |
| chr4 | 8535506 | 8537034 | 36.10713 | 116.78466 | 112.7497 | Promoter | 4 | 8535644 | 8607778 | 72135 | 1 | ENSMUST00 | 0 | Rab2a | ENSMUSG00000047187.9 |
| chr7 | 38018689 | 38019816 | 34.90791 | 116.7856 | 112.7497 | Promoter | 7 | 37960002 | 38019542 | 59541 | 2 | ENSMUST00 | 0 | Uri1 | ENSMUSG00000030421.8 |
| chr9 | 69453593 | 69455744 | 36.10714 | 116.78466 | 112.7497 | Promoter | 9 | 69453620 | 69491795 | 38176 | 1 | ENSMUST00 | 0 | Anxa2 | ENSMUSG00000032231.9 |
| chr5 | 111330131 | 111331640 | 35.49035 | 116.75928 | 112.72716 | Promoter | 5 | 111330763 | 111388359 | 57597 | 1 | ENSMUST00 | 0 | Pitpnb | ENSMUSG00000050017.7 |
| chr3 | 138488534 | 138490299 | 37.30082 | 116.74746 | 112.71554 | Promoter | 3 | 138458960 | 138489382 | 30423 | 2 | ENSMUST00 | 0 | Metap1 | ENSMUSG00000005813.8 |
| chr11 | 106998544 | 106999580 | 36.03628 | 116.66104 | 112.62998 | Promoter | 11 | 106988629 | 106999440 | 10812 | 2 | ENSMUST00 | 0 | Kpna2 | ENSMUSG00000018362.9 |
| chr6 | 108184884 | 108185916 | 36.61306 | 116.61066 | 112.58018 | Promoter | 6 | 108107028 | 108185582 | 78555 | 2 | ENSMUST00 | 0 | Sumf1 | ENSMUSG00000030101.10 |
| chr12 | 52602865 | 52605494 | 33.05704 | 116.60957 | 112.57934 | Promoter | 12 | 52604485 | 52604591 | 107 | 2 | ENSMUST00 | 0 | Gm23296 | ENSMUSG00000064489.1 |
| chr2 | 180919906 | 180921040 | 33.05704 | 116.60957 | 112.57934 | Promoter | 2 | 180904377 | 180920949 | 16573 | 2 | ENSMUST00 | 0 | Ythdf1 | ENSMUSG00000038848.9 |
| chr2 | 26444037 | 26445254 | 34.77877 | 116.55683 | 112.52724 | Promoter | 2 | 26409431 | 26445214 | 35784 | 2 | ENSMUST00 | 0 | Sec16a | ENSMUSG00000026924.12 |
| chr17 | 78936228 | 78937796 | 35.3553 | 116.52192 | 112.49273 | Promoter | 17 | 78919006 | 78937070 | 18065 | 2 | ENSMUST00 | 0 | Cebpz | ENSMUSG00000024081.8 |
| chr2 | 73311841 | 73313736 | 29.72989 | 116.51302 | 112.48396 | Promoter | 2 | 73312601 | 73337818 | 25218 | 1 | ENSMUST00 | 0 | Scrn3 | ENSMUSG00000008226.9 |
| chr4 | 53439450 | 53441541 | 36.53934 | 116.48274 | 112.45372 | Promoter | 4 | 53440413 | 53550200 | 109788 | 1 | ENSMUST00 | 0 | Slc44a1 | ENSMUSG00000028412.12 |
| chr7 | 122132388 | 122133812 | 36.53934 | 116.48274 | 112.45372 | Promoter | 7 | 122133041 | 122149044 | 16004 | 1 | ENSMUST00 | 0 | Dctn5 | ENSMUSG00000030868.6 |
| chr9 | 50602958 | 50604667 | 36.53934 | 116.48274 | 112.45372 | Promoter | 9 | 50596338 | 50603849 | 7512 | 2 | ENSMUST00 | 0 | Sdhd | ENSMUSG00000000171.4 |
| chr16 | 55933701 | 55935186 | 37.14626 | 116.48058 | 112.45192 | Promoter | 16 | 55899888 | 55934855 | 34968 | 2 | ENSMUST00 | 0 | Cep97 | ENSMUSG00000022604.13 |
| chr3 | 52102288 | 52105222 | 37.14626 | 116.48058 | 112.45192 | Promoter | 3 | 51687611 | 52104891 | 417281 | 2 | ENSMUST00 | 0 | Maml3 | ENSMUSG00000061143.10 |
| chr4 | 134286315 | 134288340 | 26.77612 | 116.44383 | 112.41551 | Promoter | 4 | 134275002 | 134287239 | 12238 | 2 | ENSMUST00 | 0 | Pdik1l | ENSMUSG00000050890.10 |
| chr7 | 141228433 | 141229762 | 26.77612 | 116.44383 | 112.41551 | Promoter | 7 | 141228784 | 141262742 | 33959 | 1 | ENSMUST00 | 0 | Phrf1 | ENSMUSG00000038611.11 |
| chr11 | 109649925 | 109652561 | 31.28813 | 116.44205 | 112.41405 | Promoter | 11 | 109650575 | 109661753 | 11179 | 1 | ENSMUST00 | 0 | Prkar1a | ENSMUSG00000020612.11 |
| chr12 | 69356386 | 69357823 | 31.28813 | 116.44205 | 112.41405 | Promoter | 12 | 69311544 | 69357176 | 45633 | 2 | ENSMUST00 | 0 | Nemf | ENSMUSG00000020982.8 |
| chr13 | 93303134 | 93306100 | 31.28813 | 116.44205 | 112.41405 | Promoter | 13 | 93303757 | 93386744 | 82988 | 1 | ENSMUST00 | 0 | Homer1 | ENSMUSG00000007617.12 |
| chr15 | 65786665 | 65788013 | 31.28813 | 116.44205 | 112.41405 | Promoter | 15 | 65787034 | 65791351 | 4318 | 1 | ENSMUST00 | 0 | Efr3a | ENSMUSG00000015002.11 |
| chr16 | 20301284 | 20302649 | 31.28813 | 116.44205 | 112.41405 | Promoter | 16 | 20279820 | 20302345 | 22526 | 2 | ENSMUST00 | 0 | Parl | ENSMUSG00000033918.10 |
| chr16 | 21693648 | 21695637 | 31.28813 | 116.44205 | 112.41405 | Promoter | 16 | 21649045 | 21694665 | 45621 | 2 | ENSMUST00 | 0 | 2510009E07I | ENSMUSG00000043391.9 |
| chr17 | 74489143 | 74490468 | 31.28813 | 116.44205 | 112.41405 | Promoter | 17 | 74489493 | 74500277 | 10785 | 1 | ENSMUST00 | 0 | Yipf4 | ENSMUSG00000024072.8 |
| chr18 | 36348253 | 36349847 | 31.28813 | 116.44205 | 112.41405 | Promoter | 18 | 36348697 | 36402980 | 54284 | 1 | ENSMUST00 | 0 | Cystm1 | ENSMUSG00000046727.7 |
| chr19 | 7481900 | 7483910 | 31.28813 | 116.44205 | 112.41405 | Promoter | 19 | 7425901 | 7483281 | 57381 | 2 | ENSMUST00 | 0 | Rtn3 | ENSMUSG00000024758.9 |
| chr2 | 3511999 | 3513696 | 31.28813 | 116.44205 | 112.41405 | Promoter | 2 | 3513030 | 3523822 | 10793 | 1 | ENSMUST00 | 0 | Cdnf | ENSMUSG00000039496.3 |
| chr2 | 161107862 | 161109129 | 31.28813 | 116.44205 | 112.41405 | Promoter | 2 | 160946978 | 161109022 | 162045 | 2 | ENSMUST00 | 0 | Chd6 | ENSMUSG00000057133.9 |
| chr4 | 137356219 | 137358176 | 31.28813 | 116.44205 | 112.41405 | Promoter | 4 | 137319696 | 137357720 | 38025 | 2 | ENSMUST00 | 0 | Cdc42 | ENSMUSG00000006699.12 |
| chr4 | 108832647 | 108835582 | 31.28813 | 116.44205 | 112.41405 | Promoter | 4 | 108834601 | 108862127 | 27527 | 1 | ENSMUST00 | 0 | Txndc12 | ENSMUSG00000028567.8 |
| chr4 | 156196002 | 156198318 | 31.28813 | 116.44205 | 112.41405 | Promoter | 4 | 156165778 | 156197488 | 31711 | 2 | ENSMUST00 | 0 | Agrn | ENSMUSG00000041936.13 |
| chr4 | 15265209 | 15266911 | 31.28813 | 116.44205 | 112.41405 | Promoter | 4 | 15265831 | 15286753 | 20923 | 1 | ENSMUST00 | 0 | Tmem64 | ENSMUSG00000043252.8 |
| chr9 | 22306872 | 22308190 | 31.28813 | 116.44205 | 112.41405 | Promoter | 9 | 22276748 | 22307638 | 30891 | 2 | ENSMUST00 | 0 | Zfp810 | ENSMUSG00000066829.6 |
| chr11 | 116023135 | 116024573 | 35.28817 | 116.40383 | 112.37991 | Promoter | 11 | 116021961 | 116024504 | 2544 | 2 | ENSMUST00 | 0 | H3f3b | ENSMUSG00000016559.9 |
| chr11 | 77215383 | 77217053 | 35.28817 | 116.40383 | 112.37991 | Promoter | 11 | 77216287 | 77393153 | 176867 | 1 | ENSMUST00 | 0 | Ssh2 | ENSMUSG00000037926.10 |
| chr12 | 108834590 | 108837062 | 32.32999 | 116.31703 | 112.294 | Promoter | 12 | 108825873 | 108835883 | 10011 | 2 | ENSMUST00 | 0 | Slc25a29 | ENSMUSG00000021265.3 |
| chr18 | 34005244 | 34007563 | 32.32999 | 116.31703 | 112.294 | Promoter | 18 | 33796327 | 34007206 | 210880 | 2 | ENSMUST00 | 0 | Epb4.1l4a | ENSMUSG00000024376.6 |
| chr2 | 180272453 | 180273457 | 34.04953 | 116.30843 | 112.28568 | Promoter | 2 | 180262855 | 180273446 | 10592 | 2 | ENSMUST00 | 0 | Cables2 | ENSMUSG00000038990.9 |
| chr3 | 101602862 | 101605252 | 34.04953 | 116.30843 | 112.28568 | Promoter | 3 | 101576219 | 101604603 | 28385 | 2 | ENSMUST00 | 0 | Atp1a1 | ENSMUSG00000033161.9 |
| chr6 | 30400851 | 30402755 | 34.04953 | 116.30843 | 112.28568 | Promoter | 6 | 30401868 | 30453288 | 51421 | 1 | ENSMUST00 | 0 | Klhdc10 | ENSMUSG00000029775.9 |
| chr6 | 137734431 | 137736103 | 34.04953 | 116.30843 | 112.28568 | Promoter | 6 | 137735078 | 137751932 | 16855 | 1 | ENSMUST00 | 0 | Strap | ENSMUSG00000030224.10 |
| chr8 | 85839917 | 85841768 | 34.04953 | 116.30843 | 112.28568 | Promoter | 8 | 85717557 | 85840949 | 123393 | 2 | ENSMUST00 | 0 | Itfg1 | ENSMUSG00000031703.7 |
| chr9 | 70655957 | 70657918 | 34.04953 | 116.30843 | 112.28568 | Promoter | 9 | 70599014 | 70657174 | 58161 | 2 | ENSMUST00 | 0 | Fam63b | ENSMUSG00000042444.9 |
| chr7 | 131361788 | 131363493 | 35.82539 | 116.29276 | 112.27101 | Promoter | 7 | 131342718 | 131362698 | 19981 | 2 | ENSMUST00 | 0 | 2310057M21 | ENSMUSG00000040177.9 |
| chr8 | 4323747 | 4325926 | 35.82539 | 116.29276 | 112.27101 | Promoter | 8 | 4325210 | 4359998 | 34789 | 1 | ENSMUST00 | 0 | Ccl25 | ENSMUSG00000023235.8 |
| chr3 | 32817158 | 32819036 | 36.39278 | 116.22827 | 112.20707 | Promoter | 3 | 32817546 | 32938071 | 120526 | 1 | ENSMUST00 | 0 | Usp13 | ENSMUSG00000056900.8 |
| chr4 | 133545129 | 133546433 | 35.75565 | 116.17085 | 112.15157 | Promoter | 4 | 133532542 | 133545996 | 13455 | 2 | ENSMUST00 | 0 | Nudc | ENSMUSG00000028851.6 |
| chr8 | 68793896 | 68794703 | 35.75565 | 116.17085 | 112.15157 | Promoter | 8 | 68793929 | 68829410 | 35482 | 1 | ENSMUST00 | 0 | Ints10 | ENSMUSG00000031864.10 |
| chr1 | 44551388 | 44552745 | 35.15467 | 116.16882 | 112.14974 | Promoter | 1 | 44551511 | 44796838 | 245328 | 1 | ENSMUST00 | 0 | Gulp1 | ENSMUSG00000056870.4 |
| chr12 | 21316058 | 21317124 | 35.15467 | 116.16882 | 112.14974 | Promoter | 12 | 21316392 | 21323605 | 7214 | 1 | ENSMUST00 | 0 | Iah1 | ENSMUSG00000062054.6 |
| chr14 | 61646907 | 61648830 | 35.15467 | 116.16882 | 112.14974 | Promoter | 14 | 61603589 | 61648822 | 45234 | 2 | ENSMUST00 | 0 | Dleu2 | ENSMUSG00000097589.4 |
| chr18 | 46596626 | 46598838 | 35.15467 | 116.16882 | 112.14974 | Promoter | 18 | 46560235 | 46597314 | 37080 | 2 | ENSMUST00 | 0 | Tmed7 | ENSMUSG00000033184.9 |
| chr19 | 4624885 | 4626451 | 35.15467 | 116.16882 | 112.14974 | Promoter | 19 | 4622551 | 4625617 | 3067 | 2 | ENSMUST00 | 0 | Rce1 | ENSMUSG00000024889.3 |
| chr19 | 8953000 | 8954531 | 35.15467 | 116.16882 | 112.14974 | Promoter | 19 | 8953850 | 8966207 | 12358 | 1 | ENSMUST00 | 0 | Tut1 | ENSMUSG00000071645.5 |
| chr5 | 29434237 | 29435857 | 35.15467 | 116.16882 | 112.14974 | Promoter | 5 | 29434664 | 29453506 | 18843 | 1 | ENSMUST00 | 0 | Nom1 | ENSMUSG00000001569.9 |
| chr5 | 29735258 | 29737284 | 35.15467 | 116.16882 | 112.14974 | Promoter | 5 | 29735688 | 29786478 | 50791 | 1 | ENSMUST00 | 0 | Dnajb6 | ENSMUSG00000029131.9 |
| chr5 | 113735182 | 113736906 | 35.15467 | 116.16882 | 112.14974 | Promoter | 5 | 113735782 | 113740600 | 4819 | 1 | ENSMUST00 | 0 | Ficd | ENSMUSG00000053334.5 |
| chr6 | 81923433 | 81924504 | 35.15467 | 116.16882 | 112.14974 | Promoter | 6 | 81923669 | 81959915 | 36247 | 1 | ENSMUST00 | 0 | Gcfc2 | ENSMUSG00000035125.5 |
| chr8 | 69088367 | 69089628 | 35.15467 | 116.16882 | 112.14974 | Promoter | 8 | 69088646 | 69113711 | 25066 | 1 | ENSMUST00 | 0 | Atp6v1b2 | ENSMUSG00000006273.9 |
| chr8 | 22652683 | 22653856 | 35.15467 | 116.16882 | 112.14974 | Promoter | 8 | 22628119 | 22653437 | 25319 | 2 | ENSMUST00 | 0 | Polb | ENSMUSG00000031536.5 |
| chr13 | 119690292 | 119691178 | 28.04154 | 116.13699 | 112.1206 | Promoter | 13 | 119690462 | 119708086 | 17625 | 1 | ENSMUST00 | 0 | Hmgcs1 | ENSMUSG00000093930.1 |
| chr16 | 64769876 | 64771589 | 28.04154 | 116.13699 | 112.1206 | Promoter | 16 | 64762136 | 64771161 | 9026 | 2 | ENSMUST00 | 0 | 4930453N24 | ENSMUSG00000059920.9 |
| chr4 | 56801586 | 56803102 | 28.04154 | 116.13699 | 112.1206 | Promoter | 4 | 56749680 | 56802331 | 52652 | 2 | ENSMUST00 | 0 | Ikbkap | ENSMUSG00000028431.6 |
| chr7 | 140881281 | 140883193 | 28.04154 | 116.13699 | 112.1206 | Promoter | 7 | 140863670 | 140881869 | 18200 | 2 | ENSMUST00 | 0 | Sirt3 | ENSMUSG00000025486.11 |
| chr9 | 21797469 | 21799074 | 28.04154 | 116.13699 | 112.1206 | Promoter | 9 | 21766773 | 21798546 | 31774 | 2 | ENSMUST00 | 0 | Kank2 | ENSMUSG00000032194.8 |
| chr9 | 31029400 | 31031742 | 28.04154 | 116.13699 | 112.1206 | Promoter | 9 | 31030644 | 31075879 | 45236 | 1 | ENSMUST00 | 0 | Zbtb44 | ENSMUSG00000047412.10 |
| chr1 | 75316459 | 75317532 | 29.52499 | 116.10789 | 112.09218 | Promoter | 1 | 75307940 | 75316993 | 9054 | 2 | ENSMUST00 | 0 | Dnpep | ENSMUSG00000026209.10 |
| chr1 | 74661680 | 74662863 | 29.52499 | 116.10789 | 112.09218 | Promoter | 1 | 74661745 | 74703730 | 41986 | 1 | ENSMUST00 | 0 | Ttll4 | ENSMUSG00000033257.9 |
| chr11 | 69765911 | 69767941 | 29.52499 | 116.10789 | 112.09218 | Promoter | 11 | 69765912 | 69780209 | 14298 | 1 | ENSMUST00 | 0 | Zbtb4 | ENSMUSG00000018750.9 |
| chr14 | 34500567 | 34503606 | 29.52499 | 116.10789 | 112.09218 | Promoter | 14 | 34410734 | 34502687 | 91954 | 2 | ENSMUST00 | 0 | Bmpr1a | ENSMUSG00000021796.9 |
| chr15 | 34081757 | 34084754 | 29.52499 | 116.10789 | 112.09218 | Promoter | 15 | 34082694 | 34103345 | 20652 | 1 | ENSMUST00 | 0 | Mtdh | ENSMUSG00000022255.10 |
| chr15 | 39111953 | 39113834 | 29.52499 | 116.10789 | 112.09218 | Promoter | 15 | 39094191 | 39112716 | 18526 | 2 | ENSMUST00 | 0 | Slc25a32 | ENSMUSG00000022299.8 |
| chr2 | 53190411 | 53192920 | 29.52499 | 116.10789 | 112.09218 | Promoter | 2 | 53191726 | 53217534 | 25809 | 1 | ENSMUST00 | 0 | Arl6ip6 | ENSMUSG00000026960.6 |
| chr3 | 105958303 | 105960525 | 29.52499 | 116.10789 | 112.09218 | Promoter | 3 | 105959369 | 105969760 | 10392 | 1 | ENSMUST00 | 0 | Wdr77 | ENSMUSG00000000561.9 |
| chr4 | 76447761 | 76450636 | 29.52499 | 116.10789 | 112.09218 | Promoter | 4 | 75941238 | 76449980 | 508743 | 2 | ENSMUST00 | 0 | Ptprd | ENSMUSG00000028399.12 |
| chr5 | 145166174 | 145168160 | 29.52499 | 116.10789 | 112.09218 | Promoter | 5 | 145147514 | 145167108 | 19595 | 2 | ENSMUST00 | 0 | Ptcd1 | ENSMUSG00000029624.10 |
| chr8 | 105964976 | 105966946 | 29.52499 | 116.10789 | 112.09218 | Promoter | 8 | 105943590 | 105966044 | 22455 | 2 | ENSMUST00 | 0 | Slc12a4 | ENSMUSG00000017765.11 |
| chr8 | 86883811 | 86885338 | 29.52499 | 116.10789 | 112.09218 | Promoter | 8 | 86841139 | 86885258 | 44120 | 2 | ENSMUST00 | 0 | N4bp1 | ENSMUSG00000031652.10 |
| chr9 | 116174150 | 116176045 | 29.52499 | 116.10789 | 112.09218 | Promoter | 9 | 116087698 | 116175363 | 87666 | 2 | ENSMUST00 | 0 | Tgfbr2 | ENSMUSG00000032440.7 |
| chr12 | 80258838 | 80261410 | 36.31995 | 116.10173 | 112.08785 | Promoter | 12 | 80167542 | 80260371 | 92830 | 2 | ENSMUST00 | 0 | Actn1 | ENSMUSG00000015143.9 |
| chr13 | 64273883 | 64275903 | 36.31995 | 116.10173 | 112.08785 | Promoter | 13 | 64192545 | 64274973 | 82429 | 2 | ENSMUST00 | 0 | Cdc14b | ENSMUSG00000033102.9 |
| chr14 | 45528002 | 45530952 | 36.31995 | 116.10173 | 112.08785 | Promoter | 14 | 45458792 | 45530118 | 71327 | 2 | ENSMUST00 | 0 | Fermt2 | ENSMUSG00000037712.10 |
| chr19 | 4192114 | 4193553 | 36.31995 | 116.10173 | 112.08785 | Promoter | 19 | 4192158 | 4195419 | 3262 | 1 | ENSMUST00 | 0 | Ppp1ca | ENSMUSG00000040385.6 |
| chr7 | 133121202 | 133124729 | 36.31995 | 116.10173 | 112.08785 | Promoter | 7 | 130162451 | 133123350 | 2960900 | 2 | ENSMUST00 | 0 | Fgfr2 | ENSMUSG00000030849.12 |
| chr9 | 21955316 | 21956539 | 36.2474 | 115.97563 | 111.96324 | Promoter | 9 | 21955753 | 21958270 | 2518 | 1 | ENSMUST00 | 0 | Swsap1 | ENSMUSG00000051238.5 |
| chr6 | 17636985 | 17638327 | 36.84092 | 115.9529 | 111.94093 | Promoter | 6 | 17637035 | 17659609 | 22575 | 1 | ENSMUST00 | 0 | Capza2 | ENSMUSG00000015733.8 |
| chr17 | 44776597 | 44778038 | 33.25048 | 115.91603 | 111.90475 | Promoter | 17 | 44777152 | 45119290 | 342139 | 1 | ENSMUST00 | 0 | Supt3 | ENSMUSG00000038954.9 |
| chr4 | 108619225 | 108620804 | 33.25048 | 115.91603 | 111.90475 | Promoter | 4 | 108619940 | 108634123 | 14184 | 1 | ENSMUST00 | 0 | Cc2d1b | ENSMUSG00000028582.9 |
| chr19 | 29046371 | 29048801 | 36.17514 | 115.85 | 111.8395 | Promoter | 19 | 29020833 | 29047961 | 27129 | 2 | ENSMUST00 | 0 | Ak3 | ENSMUSG00000024782.7 |
| chr2 | 60961189 | 60963742 | 36.17514 | 115.85 | 111.8395 | Promoter | 2 | 60750193 | 60963192 | 213000 | 2 | ENSMUST00 | 0 | Rbms1 | ENSMUSG00000026970.11 |
| chr8 | 70538131 | 70541044 | 36.17514 | 115.85 | 111.8395 | Promoter | 8 | 70539675 | 70592858 | 53184 | 1 | ENSMUST00 | 0 | Ell | ENSMUSG00000070002.6 |
| chr14 | 55745042 | 55746841 | 36.76537 | 115.82222 | 111.81207 | Promoter | 14 | 55739020 | 55745599 | 6580 | 2 | ENSMUST00 | 0 | Dhrs1 | ENSMUSG00000002332.10 |
| chr2 | 119207787 | 119209563 | 36.76537 | 115.82222 | 111.81207 | Promoter | 2 | 119172500 | 119208795 | 36296 | 2 | ENSMUST00 | 0 | Dnajc17 | ENSMUSG00000034278.3 |
| chr9 | 108093689 | 108095691 | 36.76537 | 115.82222 | 111.81207 | Promoter | 9 | 108085417 | 108094480 | 9064 | 2 | ENSMUST00 | 0 | Apeh | ENSMUSG00000032590.9 |
| chr2 | 85036527 | 85038036 | 34.95629 | 115.81918 | 111.80935 | Promoter | 2 | 85037234 | 85047109 | 9876 | 1 | ENSMUST00 | 0 | Ssrp1 | ENSMUSG00000027067.11 |
| chr1 | 180811048 | 180813568 | 35.54802 | 115.80763 | 111.79785 | Promoter | 1 | 180802697 | 180813547 | 10851 | 2 | ENSMUST00 | 0 | H3f3a | ENSMUSG00000060743.7 |
| chr2 | 32534502 | 32536789 | 36.10317 | 115.72481 | 111.71513 | Promoter | 2 | 32535332 | 32569750 | 34419 | 1 | ENSMUST00 | 0 | Fam102a | ENSMUSG00000039157.7 |
| chr6 | 8258438 | 8260172 | 36.10317 | 115.72481 | 111.71513 | Promoter | 6 | 8255936 | 8259173 | 3238 | 2 | ENSMUST00 | 0 | Rpa3 | ENSMUSG00000012483.4 |
| chr12 | 112678302 | 112679730 | 36.69013 | 115.69205 | 111.68276 | Promoter | 12 | 112678828 | 112682747 | 3920 | 1 | ENSMUST00 | 0 | Zbtb42 | ENSMUSG00000037638.5 |
| chr17 | 4993664 | 4997887 | 36.69013 | 115.69205 | 111.68276 | Promoter | 17 | 4994332 | 5097733 | 103402 | 1 | ENSMUST00 | 0 | Arid1b | ENSMUSG00000069729.8 |
| chr6 | 134565101 | 134567582 | 36.69013 | 115.69205 | 111.68276 | Promoter | 6 | 134446476 | 134566913 | 120438 | 2 | ENSMUST00 | 0 | Lrp6 | ENSMUSG00000030201.10 |
| chr8 | 105224498 | 105225991 | 36.69013 | 115.69205 | 111.68276 | Promoter | 8 | 105225145 | 105253053 | 27909 | 1 | ENSMUST00 | 0 | D230025D16 | ENSMUSG00000031889.4 |
| chr8 | 70351451 | 70353355 | 35.47934 | 115.68738 | 111.67922 | Promoter | 8 | 70331522 | 70353273 | 21752 | 2 | ENSMUST00 | 0 | Upf1 | ENSMUSG00000058301.6 |
| chr1 | 39192818 | 39194810 | 34.27164 | 115.65593 | 111.648 | Promoter | 1 | 39193731 | 39287615 | 93885 | 1 | ENSMUST00 | 0 | Npas2 | ENSMUSG00000026077.10 |
| chr1 | 71102039 | 71103659 | 34.27164 | 115.65593 | 111.648 | Promoter | 1 | 71027498 | 71103146 | 75649 | 2 | ENSMUST00 | 0 | Bard1 | ENSMUSG00000026196.7 |
| chr10 | 100486525 | 100487414 | 34.27164 | 115.65593 | 111.648 | Promoter | 10 | 100443902 | 100487350 | 43449 | 2 | ENSMUST00 | 0 | Tmtc3 | ENSMUSG00000036676.9 |
| chr11 | 53890629 | 53891792 | 34.27164 | 115.65593 | 111.648 | Promoter | 11 | 53864542 | 53891660 | 27119 | 2 | ENSMUST00 | 0 | Slc22a5 | ENSMUSG00000018900.7 |
| chr11 | 3265618 | 3267287 | 34.27164 | 115.65593 | 111.648 | Promoter | 11 | 3249907 | 3266393 | 16487 | 2 | ENSMUST00 | 0 | Drg1 | ENSMUSG00000020457.8 |
| chr13 | 24801064 | 24803492 | 34.27164 | 115.65593 | 111.648 | Promoter | 13 | 24801657 | 24814013 | 12357 | 1 | ENSMUST00 | 0 | BC005537 | ENSMUSG00000019132.5 |
| chr14 | 45329739 | 45330690 | 34.27164 | 115.65593 | 111.648 | Promoter | 14 | 45329824 | 45349071 | 19248 | 1 | ENSMUST00 | 0 | Psmc6 | ENSMUSG00000021832.7 |
| chr19 | 43673418 | 43674960 | 34.27164 | 115.65593 | 111.648 | Promoter | 19 | 43663822 | 43674881 | 11060 | 2 | ENSMUST00 | 0 | Slc25a28 | ENSMUSG00000040414.7 |
| chr3 | 28804965 | 28806327 | 34.27164 | 115.65593 | 111.648 | Promoter | 3 | 28805436 | 28807352 | 1917 | 1 | ENSMUST00 | 0 | Rpl22l1 | ENSMUSG00000039221.5 |
| chr4 | 88094020 | 88095311 | 34.27164 | 115.65593 | 111.648 | Promoter | 4 | 88094629 | 88121812 | 27184 | 1 | ENSMUST00 | 0 | Focad | ENSMUSG00000038368.11 |
| chr8 | 126421748 | 126423577 | 34.27164 | 115.65593 | 111.648 | Promoter | 8 | 126422501 | 126425434 | 2934 | 1 | ENSMUST00 | 0 | Coa6 | ENSMUSG00000051671.6 |
| chr15 | 100550821 | 100552541 | 36.03148 | 115.60005 | 111.59352 | Promoter | 15 | 100502799 | 100552008 | 49210 | 2 | ENSMUST00 | 0 | Tfcp2 | ENSMUSG00000009733.8 |

| | | | | | | | | | | | | | | | |
|---|---|---|---|---|---|---|---|---|---|---|---|---|---|---|---|
| chr13 | 96388227 | 96389149 | 32.50483 | 115.59841 | 111.59217 | Promoter | 13 | 96388294 | 96415587 | 27294 | 1 | ENSMUST00 | 0 | Poc5 | ENSMUSG00000021671.8 |
| chr17 | 69382406 | 69384678 | 32.50483 | 115.59841 | 111.59217 | Promoter | 17 | 69383050 | 69390750 | 7701 | 1 | ENSMUST00 | 0 | Zbtb14 | ENSMUSG00000049672.9 |
| chr9 | 65460642 | 65461561 | 32.50483 | 115.59841 | 111.59217 | Promoter | 9 | 65460937 | 65488470 | 27534 | 1 | ENSMUST00 | 0 | Spg21 | ENSMUSG00000032388.6 |
| chr15 | 31529514 | 31531708 | 35.41093 | 115.56754 | 111.56183 | Promoter | 15 | 31455899 | 31531037 | 75139 | 2 | ENSMUST00 | 0 | Mar-06 | ENSMUSG00000039100.9 |
| chr17 | 56008496 | 56010059 | 35.41093 | 115.56754 | 111.56183 | Promoter | 17 | 56009201 | 56016790 | 7590 | 1 | ENSMUST00 | 0 | Mpnd | ENSMUSG00000003199.11 |
| chr19 | 32107593 | 32108438 | 35.41093 | 115.56754 | 111.56183 | Distal Interge | 19 | 31984654 | 32061469 | 76816 | 2 | ENSMUST00 | -46125 | Asah2 | ENSMUSG00000024887.9 |
| chr2 | 144269317 | 144271392 | 35.41093 | 115.56754 | 111.56183 | Promoter | 2 | 144250123 | 144270572 | 20450 | 2 | ENSMUST00 | 0 | Snx5 | ENSMUSG00000027423.10 |
| chr12 | 85219440 | 85220409 | 36.6152 | 115.56235 | 111.55841 | Promoter | 12 | 85219481 | 85226628 | 7148 | 1 | ENSMUST00 | 0 | Eif2b2 | ENSMUSG00000004788.6 |
| chr7 | 97331532 | 97333065 | 36.6152 | 115.56235 | 111.55841 | Promoter | 7 | 97309380 | 97332020 | 22641 | 2 | ENSMUST00 | 0 | Usp35 | ENSMUSG00000035713.10 |
| chr9 | 49055099 | 49056378 | 36.6152 | 115.56235 | 111.55841 | Promoter | 9 | 49055581 | 49078772 | 23192 | 1 | ENSMUST00 | 0 | Zw10 | ENSMUSG00000032264.8 |
| chr12 | 55490207 | 55493180 | 37.2221 | 115.56093 | 111.55734 | Promoter | 12 | 55489411 | 55492647 | 3237 | 2 | ENSMUST00 | 0 | Nfkbia | ENSMUSG00000021025.7 |
| chr12 | 80462742 | 80464239 | 37.2221 | 115.56093 | 111.55734 | Promoter | 12 | 80463095 | 80498135 | 35041 | 1 | ENSMUST00 | 0 | Exd2 | ENSMUSG00000032705.8 |
| chr12 | 20990013 | 20991467 | 37.2221 | 115.56093 | 111.55734 | Promoter | 12 | 20990459 | 21200919 | 210461 | 1 | ENSMUST00 | 0 | Gm21989 | ENSMUSG00000090370.3 |
| chr14 | 119098724 | 119100034 | 37.2221 | 115.56093 | 111.55734 | Promoter | 14 | 118985039 | 119099430 | 114392 | 2 | ENSMUST00 | 0 | Uggt2 | ENSMUSG00000042104.13 |
| chr17 | 88440015 | 88442495 | 37.2221 | 115.56093 | 111.55734 | Promoter | 17 | 88440711 | 88490533 | 49823 | 1 | ENSMUST00 | 0 | Foxn2 | ENSMUSG00000034998.13 |
| chr2 | 45108545 | 45111853 | 37.2221 | 115.56093 | 111.55734 | Promoter | 2 | 44987125 | 45110464 | 123340 | 2 | ENSMUST00 | 0 | Zeb2 | ENSMUSG00000026872.11 |
| chr6 | 72898418 | 72900908 | 37.2221 | 115.56093 | 111.55734 | Promoter | 6 | 72841114 | 72899979 | 58866 | 2 | ENSMUST00 | 0 | Kcmf1 | ENSMUSG00000055239.8 |
| chr9 | 68653168 | 68655850 | 37.2221 | 115.56093 | 111.55734 | Promoter | 9 | 68653786 | 68664967 | 11182 | 1 | ENSMUST00 | 0 | Rora | ENSMUSG00000032238.12 |
| chr1 | 67038121 | 67039355 | 33.04283 | 115.53907 | 111.53828 | Promoter | 1 | 67000517 | 67038865 | 38349 | 2 | ENSMUST00 | 0 | Lancl1 | ENSMUSG00000026000.11 |
| chr10 | 78243998 | 78245435 | 33.04283 | 115.53907 | 111.53828 | Promoter | 10 | 78186725 | 78244642 | 57918 | 2 | ENSMUST00 | 0 | Trappc10 | ENSMUSG00000000374.7 |
| chr10 | 79960056 | 79960967 | 33.04283 | 115.53907 | 111.53828 | Promoter | 10 | 79960152 | 79969246 | 9095 | 1 | ENSMUST00 | 0 | Wdr18 | ENSMUSG00000035754.7 |
| chr10 | 96616981 | 96620333 | 33.04283 | 115.53907 | 111.53828 | Promoter | 10 | 96617001 | 96622813 | 5813 | 1 | ENSMUST00 | 0 | Btg1 | ENSMUSG00000036478.7 |
| chr10 | 58254917 | 58256362 | 33.04283 | 115.53907 | 111.53828 | Promoter | 10 | 58255497 | 58305599 | 50103 | 1 | ENSMUST00 | 0 | Gcc2 | ENSMUSG00000038039.8 |
| chr11 | 49846235 | 49847738 | 33.04283 | 115.53907 | 111.53828 | Promoter | 11 | 49846751 | 49886421 | 39671 | 1 | ENSMUST00 | 0 | Mapk9 | ENSMUSG00000020366.13 |
| chr11 | 59505222 | 59507156 | 33.04283 | 115.53907 | 111.53828 | Promoter | 11 | 59485520 | 59506644 | 21125 | 2 | ENSMUST00 | 0 | Zkscan17 | ENSMUSG00000020472.9 |
| chr11 | 60877694 | 60879849 | 33.04283 | 115.53907 | 111.53828 | Promoter | 11 | 60864452 | 60879038 | 14587 | 2 | ENSMUST00 | 0 | Tmem11 | ENSMUSG00000043284.7 |
| chr12 | 17266519 | 17267741 | 33.04283 | 115.53907 | 111.53828 | Promoter | 12 | 17266545 | 17281810 | 15266 | 1 | ENSMUST00 | 0 | Pdia6 | ENSMUSG00000020571.7 |
| chr12 | 69196366 | 69198595 | 33.04283 | 115.53907 | 111.53828 | Promoter | 12 | 69189087 | 69198429 | 9343 | 2 | ENSMUST00 | 0 | Dnaaf2 | ENSMUSG00000020973.7 |
| chr12 | 59128963 | 59132130 | 33.04283 | 115.53907 | 111.53828 | Promoter | 12 | 59129720 | 59190176 | 60457 | 1 | ENSMUST00 | 0 | Ctage5 | ENSMUSG00000021000.10 |
| chr12 | 99561991 | 99564031 | 33.04283 | 115.53907 | 111.53828 | Promoter | 12 | 99563841 | 99620038 | 56198 | 1 | ENSMUST00 | 0 | Gm26839 | ENSMUSG00000097917.1 |
| chr13 | 21158768 | 21161864 | 33.04283 | 115.53907 | 111.53828 | Distal Interge | 13 | 21142026 | 21143081 | 1056 | 2 | ENSMUST00 | -15688 | Olfr1368 | ENSMUSG00000045474.4 |
| chr14 | 31000426 | 31002352 | 33.04283 | 115.53907 | 111.53828 | Promoter | 14 | 31001393 | 31012441 | 11049 | 1 | ENSMUST00 | 0 | Glt8d1 | ENSMUSG00000021916.10 |
| chr15 | 12320620 | 12323549 | 33.04283 | 115.53907 | 111.53828 | Promoter | 15 | 12321496 | 12351265 | 29770 | 1 | ENSMUST00 | 0 | Golph3 | ENSMUSG00000022200.7 |
| chr15 | 102135500 | 102136850 | 33.04283 | 115.53907 | 111.53828 | Promoter | 15 | 102116528 | 102136192 | 19665 | 2 | ENSMUST00 | 0 | Spryd3 | ENSMUSG00000036966.10 |
| chr16 | 18776124 | 18780444 | 33.04283 | 115.53907 | 111.53828 | Promoter | 16 | 18776847 | 18778260 | 1414 | 1 | ENSMUST00 | 0 | Cldn5 | ENSMUSG00000041378.1 |
| chr17 | 15496997 | 15499015 | 33.04283 | 115.53907 | 111.53828 | Promoter | 17 | 15475721 | 15498276 | 22556 | 2 | ENSMUST00 | 0 | Psmb1 | ENSMUSG00000014769.9 |
| chr17 | 24550016 | 24551137 | 33.04283 | 115.53907 | 111.53828 | Promoter | 17 | 24549950 | 24596514 | 46565 | 1 | ENSMUST00 | 67 | Pkd1 | ENSMUSG00000032855.5 |
| chr18 | 80707078 | 80708822 | 33.04283 | 115.53907 | 111.53828 | Promoter | 18 | 80606205 | 80708173 | 101969 | 2 | ENSMUST00 | 0 | Nfatc1 | ENSMUSG00000033016.10 |
| chr19 | 31082319 | 31083953 | 33.04283 | 115.53907 | 111.53828 | Promoter | 19 | 31082841 | 31086590 | 3750 | 1 | ENSMUST00 | 0 | Cstf2t | ENSMUSG00000053536.6 |
| chr3 | 104817345 | 104818906 | 33.04283 | 115.53907 | 111.53828 | Promoter | 3 | 104794836 | 104818242 | 23407 | 2 | ENSMUST00 | 0 | Mov10 | ENSMUSG00000002227.10 |
| chr3 | 62506047 | 62507456 | 33.04283 | 115.53907 | 111.53828 | Promoter | 3 | 62470039 | 62507004 | 36966 | 2 | ENSMUST00 | 0 | Dhx36 | ENSMUSG00000027770.4 |
| chr5 | 90364700 | 90367410 | 33.04283 | 115.53907 | 111.53828 | Promoter | 5 | 90367204 | 90368487 | 1284 | 1 | ENSMUST00 | 0 | Gm9958 | ENSMUSG00000054945.3 |
| chr6 | 30047101 | 30049195 | 33.04283 | 115.53907 | 111.53828 | Promoter | 6 | 30047988 | 30153458 | 105471 | 1 | ENSMUST00 | 0 | Nrf1 | ENSMUSG00000058440.9 |
| chr7 | 28765706 | 28767655 | 33.04283 | 115.53907 | 111.53828 | Promoter | 7 | 28766735 | 28788660 | 21926 | 1 | ENSMUST00 | 0 | Sirt2 | ENSMUSG00000015149.8 |
| chr7 | 113512797 | 113514889 | 33.04283 | 115.53907 | 111.53828 | Promoter | 7 | 113513834 | 113571511 | 57678 | 1 | ENSMUST00 | 0 | Far1 | ENSMUSG00000030759.11 |
| chr8 | 94011226 | 94013098 | 33.04283 | 115.53907 | 111.53828 | Promoter | 8 | 93971588 | 94012663 | 41076 | 2 | ENSMUST00 | 0 | Amfr | ENSMUSG00000031751.8 |
| chr8 | 25240215 | 25241431 | 33.04283 | 115.53907 | 111.53828 | Distal Interge | 8 | 25154552 | 25201449 | 46898 | 2 | ENSMUST00 | -38767 | Tacc1 | ENSMUSG00000065954.5 |
| chr9 | 14783999 | 14785418 | 33.04283 | 115.53907 | 111.53828 | Promoter | 9 | 14784654 | 14834646 | 49993 | 1 | ENSMUST00 | 0 | Mre11a | ENSMUSG00000031928.9 |
| chr9 | 56417092 | 56419722 | 33.04283 | 115.53907 | 111.53828 | Promoter | 9 | 56418644 | 56496884 | 78241 | 1 | ENSMUST00 | 0 | Hmg20a | ENSMUSG00000032329.3 |
| chr9 | 111117612 | 111119605 | 33.04283 | 115.53907 | 111.53828 | Promoter | 9 | 111118111 | 111225668 | 107558 | 1 | ENSMUST00 | 0 | Lrrfip2 | ENSMUSG00000032497.10 |
| chr11 | 6388725 | 6390322 | 35.96008 | 115.47577 | 111.48098 | Promoter | 11 | 6389074 | 6406158 | 17085 | 1 | ENSMUST00 | 0 | Zmiz2 | ENSMUSG00000041164.10 |
| chr2 | 119617910 | 119619334 | 35.34279 | 115.4481 | 111.45375 | Promoter | 2 | 119618298 | 119650160 | 31863 | 1 | ENSMUST00 | 0 | Nusap1 | ENSMUSG00000027306.10 |
| chr10 | 62791319 | 62793219 | 36.54057 | 115.43316 | 111.4393 | Promoter | 10 | 62743928 | 62792368 | 48441 | 2 | ENSMUST00 | 0 | Ccar1 | ENSMUSG00000020074.6 |
| chr11 | 118476842 | 118477691 | 36.54057 | 115.43316 | 111.4393 | Promoter | 11 | 118476941 | 118479176 | 2236 | 1 | ENSMUST00 | 0 | Engase | ENSMUSG00000033857.7 |
| chr9 | 20897701 | 20898916 | 36.54057 | 115.43316 | 111.4393 | Promoter | 9 | 20894352 | 20898600 | 4249 | 2 | ENSMUST00 | 0 | Eif3g | ENSMUSG00000070319.5 |
| chr12 | 112999391 | 113000799 | 35.88896 | 115.35191 | 111.35841 | Promoter | 12 | 112959862 | 113000621 | 40760 | 2 | ENSMUST00 | 0 | Brf1 | ENSMUSG00000011158.7 |
| chr13 | 17943115 | 17944986 | 35.88896 | 115.35191 | 111.35841 | Promoter | 13 | 17880575 | 17944239 | 63665 | 2 | ENSMUST00 | 0 | Rala | ENSMUSG00000008859.7 |
| chr16 | 97921637 | 97923023 | 31.80739 | 115.34708 | 111.35392 | Promoter | 16 | 97855210 | 97922633 | 67424 | 2 | ENSMUST00 | 0 | C2cd2 | ENSMUSG00000045975.7 |
| chr5 | 74093016 | 74093839 | 34.08526 | 115.32381 | 111.33112 | Promoter | 5 | 74093060 | 74094336 | 1277 | 1 | ENSMUST00 | 0 | Dancr | ENSMUSG00000053656.5 |
| chr2 | 26915357 | 26917174 | 32.90924 | 115.29609 | 111.30405 | Promoter | 2 | 26916367 | 26920183 | 3817 | 1 | ENSMUST00 | 0 | Surf2 | ENSMUSG00000014873.10 |
| chr13 | 49215165 | 49216478 | 33.45073 | 115.2328 | 111.2408 | Promoter | 13 | 49202951 | 49216026 | 13076 | 2 | ENSMUST00 | 0 | 1110007C09I | ENSMUSG00000037960.10 |
| chr14 | 54894103 | 54895521 | 33.45073 | 115.2328 | 111.2408 | Promoter | 14 | 54894148 | 54898156 | 4009 | 1 | ENSMUST00 | 0 | Pabpn1 | ENSMUSG00000022194.10 |
| chr5 | 108659935 | 108661207 | 33.45073 | 115.2328 | 111.2408 | Promoter | 5 | 108646693 | 108660811 | 14119 | 2 | ENSMUST00 | 0 | Dgkq | ENSMUSG00000004815.7 |
| chr5 | 31192419 | 31194275 | 33.45073 | 115.2328 | 111.2408 | Promoter | 5 | 31187558 | 31193430 | 5873 | 2 | ENSMUST00 | 0 | Eif2b4 | ENSMUSG00000029145.7 |
| chr6 | 121108833 | 121110689 | 33.45073 | 115.2328 | 111.2408 | Distal Interge | 6 | 121007241 | 121081609 | 74369 | 2 | ENSMUST00 | -27225 | Mical3 | ENSMUSG00000003178.7 |
| chr8 | 3392521 | 3397432 | 33.45073 | 115.2328 | 111.2408 | Promoter | 8 | 3393006 | 3450640 | 57635 | 1 | ENSMUST00 | 0 | Arhgef18 | ENSMUSG00000004568.7 |
| chr10 | 62485689 | 62487643 | 35.20524 | 115.20684 | 111.21615 | Promoter | 10 | 62454843 | 62486598 | 31756 | 2 | ENSMUST00 | 0 | Vps26a | ENSMUSG00000020078.10 |
| chr10 | 80826565 | 80828230 | 35.20524 | 115.20684 | 111.21615 | Promoter | 10 | 80826656 | 80829290 | 2635 | 1 | ENSMUST00 | 0 | Oaz1 | ENSMUSG00000035242.10 |
| chr10 | 40300536 | 40303093 | 35.20524 | 115.20684 | 111.21615 | Promoter | 10 | 40287458 | 40302188 | 14731 | 2 | ENSMUST00 | 0 | Amd1 | ENSMUSG00000075232.4 |
| chr11 | 101069577 | 101070879 | 35.20524 | 115.20684 | 111.21615 | Promoter | 11 | 101070012 | 101077672 | 7661 | 1 | ENSMUST00 | 0 | Naglu | ENSMUSG00000001751.9 |
| chr11 | 51762984 | 51764445 | 35.20524 | 115.20684 | 111.21615 | Promoter | 11 | 51763687 | 51791925 | 28239 | 1 | ENSMUST00 | 0 | Sar1b | ENSMUSG00000020386.5 |
| chr11 | 102184448 | 102185831 | 35.20524 | 115.20684 | 111.21615 | Promoter | 11 | 102162497 | 102185229 | 22733 | 2 | ENSMUST00 | 0 | Lsm12 | ENSMUSG00000020922.6 |
| chr11 | 69919331 | 69921345 | 35.20524 | 115.20684 | 111.21615 | Promoter | 11 | 69916714 | 69921085 | 4372 | 2 | ENSMUST00 | 0 | Eif5a | ENSMUSG00000078812.5 |
| chr13 | 58126701 | 58129768 | 35.20524 | 115.20684 | 111.21615 | Promoter | 13 | 58125879 | 58128556 | 2678 | 2 | ENSMUST00 | 0 | Hnrnpa0 | ENSMUSG00000007836.6 |
| chr14 | 17659199 | 17661639 | 35.20524 | 115.20684 | 111.21615 | Promoter | 14 | 17660960 | 18036559 | 375600 | 1 | ENSMUST00 | 0 | Thrb | ENSMUSG00000021779.11 |
| chr15 | 76816817 | 76818846 | 35.20524 | 115.20684 | 111.21615 | Promoter | 15 | 76723985 | 76818083 | 94099 | 2 | ENSMUST00 | 0 | Arhgap39 | ENSMUSG00000033697.10 |
| chr16 | 50429049 | 50432934 | 35.20524 | 115.20684 | 111.21615 | Promoter | 16 | 50191844 | 50432390 | 240547 | 2 | ENSMUST00 | 0 | Bbx | ENSMUSG00000022641.10 |
| chr17 | 56183230 | 56184385 | 35.20524 | 115.20684 | 111.21615 | Promoter | 17 | 56175744 | 56183920 | 8177 | 2 | ENSMUST00 | 0 | D17Wsu104e | ENSMUSG00000019579.8 |
| chr18 | 31803509 | 31805708 | 35.20524 | 115.20684 | 111.21615 | Promoter | 18 | 31804057 | 31841642 | 37586 | 1 | ENSMUST00 | 0 | Wdr33 | ENSMUSG00000024400.10 |
| chr2 | 58566062 | 58567838 | 35.20524 | 115.20684 | 111.21615 | Promoter | 2 | 58446441 | 58566826 | 120386 | 2 | ENSMUST00 | 0 | Acvr1 | ENSMUSG00000026836.10 |
| chr3 | 108561517 | 108563451 | 35.20524 | 115.20684 | 111.21615 | Promoter | 3 | 108556425 | 108562170 | 5746 | 2 | ENSMUST00 | 0 | Tmem167b | ENSMUSG00000068732.5 |
| chr4 | 6365096 | 6367078 | 35.20524 | 115.20684 | 111.21615 | Promoter | 4 | 6365650 | 6396284 | 30635 | 1 | ENSMUST00 | 0 | Sdcbp | ENSMUSG00000028249.10 |
| chr4 | 111414260 | 111415782 | 35.20524 | 115.20684 | 111.21615 | Promoter | 4 | 111415006 | 111460298 | 45293 | 1 | ENSMUST00 | 0 | Bend5 | ENSMUSG00000028545.8 |
| chr4 | 142237879 | 142239546 | 35.20524 | 115.20684 | 111.21615 | Promoter | 4 | 142139240 | 142239401 | 100162 | 2 | ENSMUST00 | 0 | Kazn | ENSMUSG00000040606.8 |
| chr5 | 135186308 | 135188508 | 35.20524 | 115.20684 | 111.21615 | Promoter | 5 | 135187264 | 135246129 | 58866 | 1 | ENSMUST00 | 0 | Baz1b | ENSMUSG00000002748.6 |
| chr6 | 57702226 | 57703887 | 35.20524 | 115.20684 | 111.21615 | Promoter | 6 | 57702601 | 57739438 | 36838 | 1 | ENSMUST00 | 0 | Lancl2 | ENSMUSG00000062190.7 |
| chr7 | 4445041 | 4445810 | 35.20524 | 115.20684 | 111.21615 | Promoter | 7 | 4424770 | 4445601 | 20832 | 2 | ENSMUST00 | 0 | Rdh13 | ENSMUSG00000008435.10 |
| chr7 | 116307045 | 116309157 | 35.20524 | 115.20684 | 111.21615 | Promoter | 7 | 116123493 | 116308376 | 184884 | 2 | ENSMUST00 | 0 | Plekha7 | ENSMUSG00000045659.12 |
| chr7 | 19747811 | 19749602 | 35.20524 | 115.20684 | 111.21615 | Promoter | 7 | 19716644 | 19749573 | 32930 | 2 | ENSMUST00 | 0 | Pvrl2 | ENSMUSG00000062300.8 |
| chr8 | 112010549 | 112012599 | 35.20524 | 115.20684 | 111.21615 | Promoter | 8 | 111993440 | 112011288 | 17849 | 2 | ENSMUST00 | 0 | Kars | ENSMUSG00000031948.8 |
| chr9 | 50658630 | 50660762 | 35.20524 | 115.20684 | 111.21615 | Promoter | 9 | 50634633 | 50659780 | 25148 | 2 | ENSMUST00 | 0 | Dlat | ENSMUSG00000000168.4 |
| chr1 | 42850828 | 42852273 | 36.9999 | 115.1778 | 111.19392 | Promoter | 1 | 42851233 | 42905682 | 54450 | 1 | ENSMUST00 | 0 | Mrps9 | ENSMUSG00000060679.9 |
| chr11 | 89059680 | 89061154 | 36.9999 | 115.1778 | 111.19392 | Promoter | 11 | 89037582 | 89060748 | 23167 | 2 | ENSMUST00 | 0 | Dgke | ENSMUSG00000000276.6 |
| chr12 | 111038168 | 111040621 | 36.9999 | 115.1778 | 111.19392 | Promoter | 12 | 111039351 | 111115901 | 76551 | 1 | ENSMUST00 | 0 | Rcor1 | ENSMUSG00000037896.12 |
| chr12 | 69241385 | 69242592 | 36.9999 | 115.1778 | 111.19392 | Promoter | 12 | 69241832 | 69284632 | 42801 | 1 | ENSMUST00 | 0 | Klhdc1 | ENSMUSG00000051890.8 |
| chr4 | 40970221 | 40972014 | 36.9999 | 115.1778 | 111.19392 | Promoter | 4 | 40970906 | 41012608 | 41703 | 1 | ENSMUST00 | 0 | Nfx1 | ENSMUSG00000028423.10 |
| chr11 | 95336380 | 95337867 | 36.39222 | 115.17619 | 111.19315 | Promoter | 11 | 95337018 | 95381872 | 44855 | 1 | ENSMUST00 | 0 | Fam117a | ENSMUSG00000038893.7 |
| chr7 | 110843466 | 110844572 | 36.39222 | 115.17619 | 111.19315 | Promoter | 7 | 110813531 | 110844457 | 30927 | 2 | ENSMUST00 | 0 | Rnf141 | ENSMUSG00000030788.10 |
| chr11 | 61407443 | 61409214 | 31.15363 | 115.15495 | 111.17228 | Distal Interge | 11 | 61416402 | 61416815 | 414 | 2 | ENSMUST00 | 7601 | Gm12271 | ENSMUSG00000082032.1 |
| chr9 | 14499824 | 14501389 | 34.56618 | 115.13003 | 111.14746 | Promoter | 9 | 14500617 | 14510577 | 9961 | 1 | ENSMUST00 | 0 | Cwc15 | ENSMUSG00000004096.8 |
| chr16 | 95701830 | 95704226 | 35.74757 | 115.10552 | 111.12301 | Promoter | 16 | 95702075 | 95721051 | 18977 | 1 | ENSMUST00 | 0 | Ets2 | ENSMUSG00000022895.9 |
| chr13 | 65111921 | 65113528 | 35.13992 | 115.09217 | 111.10991 | Promoter | 13 | 65064663 | 65112975 | 48313 | 2 | ENSMUST00 | 0 | Hiatl1 | ENSMUSG00000038212.10 |
| chr15 | 81729016 | 81729988 | 36.3185 | 115.04841 | 111.06631 | Promoter | 15 | 81704248 | 81729919 | 25672 | 2 | ENSMUST00 | 0 | Rangap1 | ENSMUSG00000022391.10 |
| chr3 | 121814226 | 121815863 | 36.3185 | 115.04841 | 111.06631 | Promoter | 3 | 121758909 | 121815302 | 56394 | 2 | ENSMUST00 | 0 | Abcd3 | ENSMUSG00000028127.5 |
| chr2 | 31151729 | 31153157 | 36.92276 | 115.04471 | 111.06307 | Promoter | 2 | 31152316 | 31218775 | 66460 | 1 | ENSMUST00 | 0 | Gpr107 | ENSMUSG00000000194.8 |
| chr14 | 20544831 | 20546595 | 34.50201 | 115.01646 | 111.03505 | Promoter | 14 | 20499364 | 20546573 | 47210 | 2 | ENSMUST00 | 0 | Ppp3cb | ENSMUSG00000021816.5 |
| chr15 | 76343160 | 76344618 | 34.50201 | 115.01646 | 111.03505 | Promoter | 15 | 76343523 | 76345934 | 2412 | 1 | ENSMUST00 | 0 | Cyc1 | ENSMUSG00000022551.7 |
| chr15 | 75704224 | 75711444 | 34.50201 | 115.01646 | 111.03505 | Promoter | 15 | 75704280 | 75714408 | 10129 | 1 | ENSMUST00 | 0 | Rhpn1 | ENSMUSG00000022580.7 |
| chr15 | 75909652 | 75911214 | 34.50201 | 115.01646 | 111.03505 | Promoter | 15 | 75909760 | 75912148 | 2389 | 1 | ENSMUST00 | 0 | Tigd5 | ENSMUSG00000047166.9 |
| chr1 | 79775655 | 79776899 | 35.67729 | 114.98298 | 111.00362 | Promoter | 1 | 79776018 | 79781445 | 5428 | 1 | ENSMUST00 | 0 | Mrpl44 | ENSMUSG00000026248.9 |
| chr11 | 59208056 | 59209347 | 35.67729 | 114.98298 | 111.00362 | Promoter | 11 | 59208321 | 59210738 | 2418 | 1 | ENSMUST00 | 0 | 2310033P09I | ENSMUSG00000020441.6 |
| chr8 | 34145755 | 34147509 | 35.67729 | 114.98298 | 111.00362 | Promoter | 8 | 34135572 | 34147033 | 11462 | 2 | ENSMUST00 | 0 | Leprotl1 | ENSMUSG00000031513.8 |
| chr7 | 122969333 | 122971749 | 35.07281 | 114.97435 | 110.99648 | Promoter | 7 | 122970564 | 123002557 | 31994 | 1 | ENSMUST00 | 0 | Rbbp6 | ENSMUSG00000030779.10 |
| chr19 | 42147079 | 42151271 | 36.24507 | 114.92111 | 110.94343 | Promoter | 19 | 42147400 | 42151703 | 4304 | 1 | ENSMUST00 | 0 | Marveld1 | ENSMUSG00000044345.9 |
| chr1 | 4857039 | 4858858 | 36.84594 | 114.91216 | 110.93493 | Promoter | 1 | 4857814 | 4897905 | 40092 | 1 | ENSMUST00 | 0 | Tcea1 | ENSMUSG00000033813.10 |
| chr2 | 65237620 | 65239624 | 36.84594 | 114.91216 | 110.93493 | Promoter | 2 | 65088339 | 65238626 | 150288 | 2 | ENSMUST00 | 0 | Cobll1 | ENSMUSG00000034903.13 |
| chr2 | 134643280 | 134644493 | 32.68561 | 114.88856 | 110.91166 | Promoter | 2 | 134594185 | 134644145 | 49961 | 2 | ENSMUST00 | 0 | Tmx4 | ENSMUSG00000034723.6 |
| chr7 | 127966431 | 127969481 | 32.68561 | 114.88856 | 110.91166 | Promoter | 7 | 127967457 | 127982368 | 14912 | 1 | ENSMUST00 | 0 | Fus | ENSMUSG00000030795.13 |
| chr17 | 17293826 | 17295206 | 26.51101 | 114.88218 | 110.90597 | Promoter | 17 | 17294443 | 17318104 | 23662 | 1 | ENSMUST00 | 0 | Gm6712 | ENSMUSG00000072761.5 |
| chr2 | 118900502 | 118903046 | 26.51101 | 114.88218 | 110.90597 | Promoter | 2 | 118901453 | 118924528 | 23076 | 1 | ENSMUST00 | 0 | Bahd1 | ENSMUSG00000040007.8 |
| chr11 | 78261654 | 78262508 | 35.60728 | 114.86086 | 110.88493 | Promoter | 11 | 78261752 | 78264032 | 2281 | 1 | ENSMUST00 | 0 | 2610507B11I | ENSMUSG00000010277.7 |
| chr3 | 63897829 | 63900101 | 35.60728 | 114.86086 | 110.88493 | Promoter | 3 | 63696731 | 63899472 | 202742 | 2 | ENSMUST00 | 0 | Plch1 | ENSMUSG00000036834.11 |
| chr10 | 44267494 | 44269302 | 35.00596 | 114.85688 | 110.88117 | Promoter | 10 | 44268358 | 44364291 | 95934 | 1 | ENSMUST00 | 0 | Atg5 | ENSMUSG00000038160.6 |
| chr17 | 87265120 | 87266517 | 35.00596 | 114.85688 | 110.88117 | Promoter | 17 | 87254658 | 87265935 | 11278 | 2 | ENSMUST00 | 0 | Mcfd2 | ENSMUSG00000024150.6 |
| chr3 | 153943430 | 153945325 | 35.00596 | 114.85688 | 110.88117 | Promoter | 3 | 153922357 | 153944632 | 22276 | 2 | ENSMUST00 | 0 | Acadm | ENSMUSG00000062908.7 |
| chr11 | 78696349 | 78697538 | 36.17194 | 114.79427 | 110.81874 | Promoter | 11 | 78567168 | 78697373 | 130206 | 2 | ENSMUST00 | 0 | Nlk | ENSMUSG00000017376.10 |
| chr17 | 86144536 | 86145235 | 36.17194 | 114.79427 | 110.81874 | Promoter | 17 | 85984665 | 86145175 | 160511 | 2 | ENSMUST00 | 0 | Srbd1 | ENSMUSG00000024135.9 |
| chr2 | 164485834 | 164487505 | 36.17194 | 114.79427 | 110.81874 | Promoter | 2 | 164486106 | 164490767 | 4662 | 1 | ENSMUST00 | 0 | Dbndd2 | ENSMUSG00000017734.10 |

| | | | | | | | | | | | | | | | |
|---|---|---|---|---|---|---|---|---|---|---|---|---|---|---|---|
| chr2 | 24949244 | 24950807 | 36.17194 | 114.79427 | 110.81874 | Promoter | 2 | 24949792 | 24962075 | 12284 | 1 | ENSMUST00 | 0 | Zmynd19 | ENSMUSG00000026974.6 |
| chr11 | 60104773 | 60106757 | 36.76944 | 114.78011 | 110.80492 | Promoter | 11 | 60105013 | 60199195 | 94183 | 1 | ENSMUST00 | 0 | Rai1 | ENSMUSG00000062115.10 |
| chr1 | 64735947 | 64737853 | 30.94804 | 114.76607 | 110.79114 | Promoter | 1 | 64730558 | 64737751 | 7194 | 2 | ENSMUST00 | 0 | Fzd5 | ENSMUSG00000045005.9 |
| chr10 | 42477676 | 42478837 | 30.94804 | 114.76607 | 110.79114 | Promoter | 10 | 42312585 | 42478280 | 165696 | 2 | ENSMUST00 | 0 | Lace1 | ENSMUSG00000038302.9 |
| chr10 | 34280801 | 34283989 | 30.94804 | 114.76607 | 110.79114 | Promoter | 10 | 34282190 | 34285275 | 3086 | 1 | ENSMUST00 | 0 | Tspyl1 | ENSMUSG00000047514.6 |
| chr12 | 15810380 | 15817601 | 30.94804 | 114.76607 | 110.79114 | Promoter | 12 | 15791727 | 15816785 | 25059 | 2 | ENSMUST00 | 0 | Trib2 | ENSMUSG00000020601.7 |
| chr13 | 34650232 | 34653634 | 30.94804 | 114.76607 | 110.79114 | Promoter | 13 | 34627840 | 34652671 | 24832 | 2 | ENSMUST00 | 0 | Pxdc1 | ENSMUSG00000021411.10 |
| chr14 | 21830875 | 21833127 | 30.94804 | 114.76607 | 110.79114 | Promoter | 14 | 21831269 | 21845879 | 14611 | 1 | ENSMUST00 | 0 | Vdac2 | ENSMUSG00000021771.8 |
| chr15 | 81871296 | 81873351 | 30.94804 | 114.76607 | 110.79114 | Promoter | 15 | 81872309 | 81915133 | 42825 | 1 | ENSMUST00 | 0 | Aco2 | ENSMUSG00000022477.7 |
| chr16 | 18392488 | 18395490 | 30.94804 | 114.76607 | 110.79114 | Promoter | 16 | 18392588 | 18407076 | 14489 | 1 | ENSMUST00 | 0 | Arvcf | ENSMUSG00000000325.10 |
| chr16 | 85171787 | 85174244 | 30.94804 | 114.76607 | 110.79114 | Promoter | 16 | 84954440 | 85173707 | 219268 | 2 | ENSMUST00 | 0 | App | ENSMUSG00000022892.10 |
| chr17 | 31854734 | 31855826 | 30.94804 | 114.76607 | 110.79114 | Promoter | 17 | 31844248 | 31855792 | 11545 | 2 | ENSMUST00 | 0 | Sik1 | ENSMUSG00000024042.6 |
| chr17 | 8164716 | 8166297 | 30.94804 | 114.76607 | 110.79114 | Promoter | 17 | 8165501 | 8196804 | 31304 | 1 | ENSMUST00 | 0 | Fgfr1op | ENSMUSG00000069135.6 |
| chr19 | 29250740 | 29253200 | 30.94804 | 114.76607 | 110.79114 | Promoter | 19 | 29251828 | 29313080 | 61253 | 1 | ENSMUST00 | 0 | Jak2 | ENSMUSG00000024789.7 |
| chr19 | 53037782 | 53039160 | 30.94804 | 114.76607 | 110.79114 | Promoter | 19 | 52943417 | 53038553 | 95137 | 2 | ENSMUST00 | 0 | Xpnpep1 | ENSMUSG00000025027.12 |
| chr2 | 32711568 | 32713700 | 30.94804 | 114.76607 | 110.79114 | Promoter | 2 | 32707438 | 32712784 | 5347 | 2 | ENSMUST00 | 0 | Cdk9 | ENSMUSG00000009555.11 |
| chr2 | 144526905 | 144528448 | 30.94804 | 114.76607 | 110.79114 | Promoter | 2 | 144527718 | 144541995 | 14278 | 1 | ENSMUST00 | 0 | Polr3f | ENSMUSG00000027427.8 |
| chr2 | 153014535 | 153015443 | 30.94804 | 114.76607 | 110.79114 | Promoter | 2 | 153008890 | 153015427 | 6538 | 2 | ENSMUST00 | 0 | Pdrg1 | ENSMUSG00000027472.9 |
| chr3 | 7503032 | 7504295 | 30.94804 | 114.76607 | 110.79114 | Promoter | 3 | 7503426 | 7553848 | 50423 | 1 | ENSMUST00 | 0 | Zc2hc1a | ENSMUSG00000043542.7 |
| chr4 | 53713868 | 53714963 | 30.94804 | 114.76607 | 110.79114 | Promoter | 4 | 53713998 | 53765785 | 51788 | 1 | ENSMUST00 | 0 | Fktn | ENSMUSG00000028414.11 |
| chr5 | 35892978 | 35895135 | 30.94804 | 114.76607 | 110.79114 | Promoter | 5 | 35893319 | 36003923 | 110605 | 1 | ENSMUST00 | 0 | Afap1 | ENSMUSG00000029094.6 |
| chr6 | 87849237 | 87851675 | 30.94804 | 114.76607 | 110.79114 | Promoter | 6 | 87842615 | 87851106 | 8492 | 2 | ENSMUST00 | 0 | Cnbp | ENSMUSG00000030057.10 |
| chr7 | 141561658 | 141563249 | 30.94804 | 114.76607 | 110.79114 | Promoter | 7 | 141562180 | 141633011 | 70832 | 1 | ENSMUST00 | 0 | Ap2a2 | ENSMUSG00000002957.8 |
| chr7 | 27227052 | 27229240 | 30.94804 | 114.76607 | 110.79114 | Promoter | 7 | 27207172 | 27228628 | 21457 | 2 | ENSMUST00 | 0 | Itpkc | ENSMUSG00000003752.8 |
| chr7 | 101835409 | 101838921 | 30.94804 | 114.76607 | 110.79114 | Promoter | 7 | 101822632 | 101838076 | 15445 | 2 | ENSMUST00 | 0 | Inppl1 | ENSMUSG00000032737.7 |
| chr7 | 92636301 | 92637898 | 30.94804 | 114.76607 | 110.79114 | Promoter | 7 | 92581723 | 92637102 | 55380 | 2 | ENSMUST00 | 0 | Ankrd42 | ENSMUSG00000041343.13 |
| chr7 | 44245631 | 44247708 | 30.94804 | 114.76607 | 110.79114 | Promoter | 7 | 44246722 | 44252319 | 5598 | 1 | ENSMUST00 | 0 | 2410002F23I | ENSMUSG00000045411.11 |
| chr8 | 83972900 | 83974112 | 30.94804 | 114.76607 | 110.79114 | Promoter | 8 | 83972978 | 83996445 | 23468 | 1 | ENSMUST00 | 0 | Prkaca | ENSMUSG00000005469.9 |
| chr9 | 44304800 | 44305745 | 30.94804 | 114.76607 | 110.79114 | Promoter | 9 | 44295673 | 44305671 | 9999 | 2 | ENSMUST00 | 0 | Hinfp | ENSMUSG00000032119.4 |
| chr9 | 31279611 | 31282029 | 30.94804 | 114.76607 | 110.79114 | Promoter | 9 | 31280538 | 31330618 | 50081 | 1 | ENSMUST00 | 0 | Prdm10 | ENSMUSG00000042496.12 |
| chr16 | 16212978 | 16214696 | 35.53755 | 114.73917 | 110.76805 | Promoter | 16 | 16213318 | 16227287 | 13970 | 1 | ENSMUST00 | 0 | Pkp2 | ENSMUSG00000041957.10 |
| chr10 | 13964189 | 13968566 | 36.09911 | 114.6679 | 110.69713 | Promoter | 10 | 13966075 | 14150589 | 184515 | 1 | ENSMUST00 | 0 | Hivep2 | ENSMUSG00000015501.5 |
| chr10 | 80147630 | 80154081 | 36.09911 | 114.6679 | 110.69713 | Promoter | 10 | 80148272 | 80158367 | 10096 | 1 | ENSMUST00 | 0 | Midn | ENSMUSG00000035621.8 |
| chr8 | 61589311 | 61591302 | 36.69325 | 114.64857 | 110.67819 | Promoter | 8 | 61513248 | 61591142 | 77895 | 2 | ENSMUST00 | 0 | Palld | ENSMUSG00000058056.10 |
| chr2 | 103795571 | 103798285 | 34.87302 | 114.62315 | 110.65308 | Promoter | 2 | 103762941 | 103797649 | 34709 | 2 | ENSMUST00 | 0 | Caprin1 | ENSMUSG00000027184.9 |
| chr13 | 77708286 | 77709324 | 35.46809 | 114.61787 | 110.64792 | Promoter | 13 | 77708690 | 78166238 | 457549 | 1 | ENSMUST00 | 0 | Fam172a | ENSMUSG00000064138.8 |
| chr13 | 99343881 | 99345450 | 31.97077 | 114.61404 | 110.64433 | Promoter | 13 | 99319649 | 99344678 | 25030 | 2 | ENSMUST00 | 0 | Ptcd2 | ENSMUSG00000021650.6 |
| chr1 | 191717578 | 191719447 | 33.65816 | 114.56059 | 110.59126 | Promoter | 1 | 191717834 | 191719398 | 1565 | 1 | ENSMUST00 | 0 | Lpgat1 | ENSMUSG00000026623.11 |
| chr15 | 4025971 | 4028201 | 33.65816 | 114.56059 | 110.59126 | Promoter | 15 | 4026383 | 4155344 | 128962 | 1 | ENSMUST00 | 0 | Oxct1 | ENSMUSG00000022186.9 |
| chr2 | 76369918 | 76370856 | 33.65816 | 114.56059 | 110.59126 | Promoter | 2 | 76369984 | 76383768 | 13785 | 1 | ENSMUST00 | 0 | Rbm45 | ENSMUSG00000042369.8 |
| chr3 | 96103895 | 96107098 | 33.65816 | 114.56059 | 110.59126 | Promoter | 3 | 96104527 | 96141518 | 36992 | 1 | ENSMUST00 | 0 | Otud7b | ENSMUSG00000038495.9 |
| chr5 | 143526334 | 143528725 | 33.65816 | 114.56059 | 110.59126 | Promoter | 5 | 143505478 | 143528036 | 22559 | 2 | ENSMUST00 | 0 | Rac1 | ENSMUSG00000001847.9 |
| chr5 | 147429490 | 147431889 | 33.65816 | 114.56059 | 110.59126 | Promoter | 5 | 147430161 | 147548502 | 118342 | 1 | ENSMUST00 | 0 | Pan3 | ENSMUSG00000029647.10 |
| chr8 | 109564705 | 109566641 | 33.65816 | 114.56059 | 110.59126 | Promoter | 8 | 109565892 | 109574051 | 8160 | 1 | ENSMUST00 | 0 | Txnl4b | ENSMUSG00000031723.3 |
| chr12 | 36041013 | 36042881 | 27.7554 | 114.54269 | 110.5744 | Promoter | 12 | 36014557 | 36042489 | 27933 | 2 | ENSMUST00 | 0 | Tspan13 | ENSMUSG00000020577.10 |
| chr16 | 35022017 | 35023359 | 27.7554 | 114.54269 | 110.5744 | Promoter | 16 | 35022421 | 35109175 | 86755 | 1 | ENSMUST00 | 0 | Ptplb | ENSMUSG00000035376.9 |
| chr2 | 155381057 | 155382543 | 27.7554 | 114.54269 | 110.5744 | Promoter | 2 | 155381059 | 155386057 | 4999 | 1 | ENSMUST00 | 0 | Trp53inp2 | ENSMUSG00000038375.10 |
| chr2 | 168229172 | 168231717 | 27.7554 | 114.54269 | 110.5744 | Promoter | 2 | 168230622 | 168232594 | 1973 | 1 | ENSMUST00 | 0 | Mocs3 | ENSMUSG00000074576.4 |
| chr6 | 52713330 | 52714872 | 27.7554 | 114.54269 | 110.5744 | Promoter | 6 | 52713729 | 52766780 | 53052 | 1 | ENSMUST00 | 0 | Tax1bp1 | ENSMUSG00000004535.7 |
| chr6 | 29347377 | 29349332 | 27.7554 | 114.54269 | 110.5744 | Promoter | 6 | 29348069 | 29374503 | 26435 | 1 | ENSMUST00 | 0 | Calu | ENSMUSG00000029767.11 |
| chr6 | 51470042 | 51471568 | 27.7554 | 114.54269 | 110.5744 | Promoter | 6 | 51470360 | 51483033 | 12674 | 1 | ENSMUST00 | 0 | Cbx3 | ENSMUSG00000029836.10 |
| chr19 | 11818037 | 11820164 | 36.61738 | 114.51753 | 110.55032 | Promoter | 19 | 11775119 | 11818893 | 43775 | 2 | ENSMUST00 | 0 | Stx3 | ENSMUSG00000041488.10 |
| chr2 | 181496321 | 181497904 | 35.3989 | 114.49703 | 110.53023 | Promoter | 2 | 181497142 | 181515115 | 17974 | 1 | ENSMUST00 | 0 | Tpd52l2 | ENSMUSG00000000827.13 |
| chr13 | 91223518 | 91224518 | 29.2142 | 114.47596 | 110.50954 | Promoter | 13 | 90935349 | 91223987 | 288639 | 2 | ENSMUST00 | 0 | Atg10 | ENSMUSG00000021619.5 |
| chr2 | 132252245 | 132253796 | 29.2142 | 114.47596 | 110.50954 | Promoter | 2 | 132249162 | 132253314 | 4153 | 2 | ENSMUST00 | 0 | Pcna | ENSMUSG00000027342.9 |
| chr4 | 44166984 | 44168121 | 29.2142 | 114.47596 | 110.50954 | Promoter | 4 | 44126210 | 44167608 | 41399 | 2 | ENSMUST00 | 0 | Rnf38 | ENSMUSG00000035696.10 |
| chr6 | 31215613 | 31219210 | 29.2142 | 114.47596 | 110.50954 | Promoter | 6 | 31093559 | 31218474 | 124916 | 2 | ENSMUST00 | 0 | Lincpint | ENSMUSG00000044471.6 |
| chr13 | 38204537 | 38205923 | 35.95432 | 114.41653 | 110.45183 | Promoter | 13 | 38204939 | 38227663 | 22725 | 1 | ENSMUST00 | 0 | Snrnp48 | ENSMUSG00000021431.9 |
| chr13 | 92609843 | 92612075 | 35.95432 | 114.41653 | 110.45183 | Promoter | 13 | 92611138 | 92711944 | 100807 | 1 | ENSMUST00 | 0 | Serinc5 | ENSMUSG00000021703.8 |
| chr15 | 98917164 | 98918235 | 35.95432 | 114.41653 | 110.45183 | Promoter | 15 | 98903921 | 98918231 | 14311 | 2 | ENSMUST00 | 0 | Lmbr1l | ENSMUSG00000022999.9 |
| chr5 | 115134185 | 115135180 | 35.95432 | 114.41653 | 110.45183 | Promoter | 5 | 115122550 | 115134975 | 12426 | 2 | ENSMUST00 | 0 | Unc119b | ENSMUSG00000046562.5 |
| chr6 | 24662156 | 24665827 | 35.95432 | 114.41653 | 110.45183 | Promoter | 6 | 24613805 | 24665009 | 51205 | 2 | ENSMUST00 | 0 | Wasl | ENSMUSG00000029684.9 |
| chr7 | 29767817 | 29769439 | 35.95432 | 114.41653 | 110.45183 | Promoter | 7 | 29768552 | 29779821 | 11270 | 1 | ENSMUST00 | 0 | Zfp84 | ENSMUSG00000046185.10 |
| chr8 | 126589312 | 126594549 | 35.95432 | 114.41653 | 110.45183 | Promoter | 8 | 126588296 | 126593986 | 5691 | 2 | ENSMUST00 | 0 | Irf2bp2 | ENSMUSG00000051495.7 |
| chr16 | 16870250 | 16871600 | 34.74108 | 114.39095 | 110.42815 | Promoter | 16 | 16870736 | 16885849 | 15114 | 1 | ENSMUST00 | 0 | Top3b | ENSMUSG00000022779.9 |
| chr16 | 15886653 | 15888845 | 34.74108 | 114.39095 | 110.42815 | Promoter | 16 | 15887286 | 15888466 | 1181 | 1 | ENSMUST00 | 0 | Cebpd | ENSMUSG00000071637.4 |
| chr17 | 26972386 | 26974099 | 34.74108 | 114.39095 | 110.42815 | Promoter | 17 | 26973217 | 27036378 | 63162 | 1 | ENSMUST00 | 0 | Ggnbp1 | ENSMUSG00000048731.9 |
| chr2 | 130283581 | 130285094 | 34.74108 | 114.39095 | 110.42815 | Promoter | 2 | 130279309 | 130284547 | 5239 | 2 | ENSMUST00 | 0 | Idh3b | ENSMUSG00000027406.7 |
| chr3 | 96726884 | 96728769 | 34.74108 | 114.39095 | 110.42815 | Promoter | 3 | 96727611 | 96791155 | 63545 | 1 | ENSMUST00 | 0 | Rnf115 | ENSMUSG00000028098.9 |
| chr5 | 24840760 | 24842707 | 34.74108 | 114.39095 | 110.42815 | Promoter | 5 | 24802823 | 24842624 | 39802 | 2 | ENSMUST00 | 0 | Rheb | ENSMUSG00000028945.8 |
| chr5 | 137306616 | 137308486 | 34.74108 | 114.39095 | 110.42815 | Promoter | 5 | 137295704 | 137307674 | 11971 | 2 | ENSMUST00 | 0 | Srrt | ENSMUSG00000037364.7 |
| chr6 | 47830002 | 47830883 | 34.74108 | 114.39095 | 110.42815 | Promoter | 6 | 47819266 | 47830867 | 11602 | 2 | ENSMUST00 | 0 | Zfp786 | ENSMUSG00000051499.6 |
| chr7 | 137437318 | 137438870 | 34.74108 | 114.39095 | 110.42815 | Promoter | 7 | 137437648 | 137468594 | 30947 | 1 | ENSMUST00 | 0 | Glrx3 | ENSMUSG00000031068.7 |
| chr8 | 126970328 | 126971599 | 34.74108 | 114.39095 | 110.42815 | Promoter | 8 | 126947175 | 126971079 | 23905 | 2 | ENSMUST00 | 0 | Rbm34 | ENSMUSG00000033931.8 |
| chr11 | 53567057 | 53568144 | 35.32998 | 114.37659 | 110.41663 | Promoter | 11 | 53567379 | 53601105 | 33727 | 1 | ENSMUST00 | 0 | Kif3a | ENSMUSG00000018395.14 |
| chr19 | 5460296 | 5461341 | 35.32998 | 114.37659 | 110.41663 | Promoter | 19 | 5460607 | 5465052 | 4446 | 1 | ENSMUST00 | 0 | Fibp | ENSMUSG00000024911.5 |
| chr8 | 125568205 | 125569883 | 35.88236 | 114.29153 | 110.33186 | Distal Interge | 8 | 125418063 | 125492710 | 74648 | 2 | ENSMUST00 | -75496 | Sipa1l2 | ENSMUSG00000001995.8 |
| chr4 | 133670509 | 133673626 | 34.67549 | 114.27542 | 110.31616 | Promoter | 4 | 133660387 | 133672647 | 12261 | 2 | ENSMUST00 | 0 | Pigv | ENSMUSG00000043257.10 |
| chr2 | 158027591 | 158029892 | 36.46658 | 114.25698 | 110.29781 | Promoter | 2 | 157981803 | 158028433 | 46631 | 2 | ENSMUST00 | 0 | Tti1 | ENSMUSG00000027650.7 |
| chr8 | 11311497 | 11314028 | 36.46658 | 114.25698 | 110.29781 | Promoter | 8 | 11198423 | 11312826 | 114404 | 2 | ENSMUST00 | 0 | Col4a1 | ENSMUSG00000031502.9 |
| chr16 | 18288146 | 18289955 | 30.67334 | 114.24463 | 110.28603 | Promoter | 16 | 18253948 | 18289246 | 35299 | 2 | ENSMUST00 | 0 | Dgcr8 | ENSMUSG00000022718.6 |
| chr17 | 26715594 | 26717210 | 32.87264 | 114.18805 | 110.22955 | Promoter | 17 | 26715650 | 26776635 | 60986 | 1 | ENSMUST00 | 0 | Crebrf | ENSMUSG00000048249.9 |
| chr3 | 90490305 | 90491371 | 32.87264 | 114.18805 | 110.22955 | Promoter | 3 | 90488026 | 90490428 | 2403 | 2 | ENSMUST00 | 0 | Snapin | ENSMUSG00000001018.10 |
| chr14 | 52279596 | 52280414 | 34.61014 | 114.16027 | 110.20278 | Promoter | 14 | 52279617 | 52280124 | 508 | 1 | ENSMUST00 | 0 | Tox4 | ENSMUSG00000016831.11 |
| chr4 | 44012051 | 44013901 | 30.08283 | 114.13763 | 110.18025 | Promoter | 4 | 44012638 | 44032846 | 20209 | 1 | ENSMUST00 | 0 | Clta | ENSMUSG00000028478.13 |
| chr8 | 22756921 | 22764179 | 35.19295 | 114.13696 | 110.1796 | Promoter | 8 | 22757744 | 22782847 | 25104 | 1 | ENSMUST00 | 0 | Plat | ENSMUSG00000031538.5 |
| chr10 | 79854526 | 79858632 | 33.99707 | 114.12061 | 110.16386 | Promoter | 10 | 79854605 | 79859712 | 5108 | 1 | ENSMUST00 | 0 | Ptbp1 | ENSMUSG00000006498.11 |
| chr3 | 85970752 | 85972621 | 36.31701 | 113.99837 | 110.04225 | Promoter | 3 | 85971109 | 86083538 | 112430 | 1 | ENSMUST00 | 0 | Sh3d19 | ENSMUSG00000028082.8 |
| chr4 | 143211838 | 143213341 | 36.31701 | 113.99837 | 110.04225 | Promoter | 4 | 143107391 | 143212705 | 105315 | 2 | ENSMUST00 | 0 | Prdm2 | ENSMUSG00000057637.8 |
| chr9 | 59750841 | 59752440 | 36.31701 | 113.99837 | 110.04225 | Promoter | 9 | 59750896 | 59928866 | 177971 | 1 | ENSMUST00 | 0 | Myo9a | ENSMUSG00000039585.10 |
| chr7 | 15945139 | 15946579 | 34.48018 | 113.93111 | 109.97533 | Promoter | 7 | 15937838 | 15946108 | 8271 | 2 | ENSMUST00 | 0 | Gltscr2 | ENSMUSG00000041560.8 |
| chr19 | 16131695 | 16134440 | 35.66819 | 113.91924 | 109.96352 | Promoter | 19 | 16132831 | 16387463 | 254633 | 1 | ENSMUST00 | 0 | Gnaq | ENSMUSG00000024639.5 |
| chr6 | 113076843 | 113079361 | 35.66819 | 113.91924 | 109.96352 | Promoter | 6 | 113077365 | 113107561 | 30197 | 1 | ENSMUST00 | 0 | Setd5 | ENSMUSG00000034269.7 |
| chr8 | 105854750 | 105856486 | 35.66819 | 113.91924 | 109.96352 | Promoter | 8 | 105855103 | 105856950 | 1848 | 1 | ENSMUST00 | 0 | Thap11 | ENSMUSG00000036442.7 |
| chr1 | 105989454 | 105992404 | 33.87313 | 113.90014 | 109.94483 | Promoter | 1 | 105990406 | 106034074 | 43669 | 1 | ENSMUST00 | 0 | Zcchc2 | ENSMUSG00000038866.10 |
| chr11 | 58322321 | 58324170 | 33.87313 | 113.90014 | 109.94483 | Promoter | 11 | 58315114 | 58323346 | 8233 | 2 | ENSMUST00 | 0 | Zfp672 | ENSMUSG00000049755.9 |
| chr13 | 8869857 | 8872185 | 33.87313 | 113.90014 | 109.94483 | Promoter | 13 | 8802968 | 8871730 | 68763 | 2 | ENSMUST00 | 0 | Wdr37 | ENSMUSG00000021147.11 |
| chr13 | 77134514 | 77136098 | 33.87313 | 113.90014 | 109.94483 | Promoter | 13 | 77043088 | 77135473 | 92386 | 2 | ENSMUST00 | 0 | Ankrd32 | ENSMUSG00000021597.11 |
| chr16 | 46495554 | 46498310 | 33.87313 | 113.90014 | 109.94483 | Promoter | 16 | 46387706 | 46496967 | 109262 | 2 | ENSMUST00 | 0 | Pvrl3 | ENSMUSG00000022656.10 |
| chr16 | 92465196 | 92466250 | 33.87313 | 113.90014 | 109.94483 | Promoter | 16 | 92391953 | 92466146 | 74194 | 2 | ENSMUST00 | 0 | Rcan1 | ENSMUSG00000022951.10 |
| chr16 | 29578898 | 29580266 | 35.05697 | 113.89898 | 109.94483 | Promoter | 16 | 29579334 | 29654884 | 75551 | 1 | ENSMUST00 | 0 | Opa1 | ENSMUSG00000038084.11 |
| chr4 | 62518215 | 62520467 | 33.87313 | 113.90014 | 109.94483 | Promoter | 4 | 62509169 | 62519881 | 10713 | 2 | ENSMUST00 | 0 | Alad | ENSMUSG00000028393.5 |
| chr6 | 17064773 | 17066140 | 33.87313 | 113.90014 | 109.94483 | Promoter | 6 | 17065149 | 17105823 | 40675 | 1 | ENSMUST00 | 0 | Tes | ENSMUSG00000029552.14 |
| chr6 | 72346936 | 72348379 | 33.87313 | 113.90014 | 109.94483 | Promoter | 6 | 72347317 | 72353158 | 5842 | 1 | ENSMUST00 | 0 | 0610030E20I | ENSMUSG00000058706.3 |
| chr13 | 64368915 | 64371185 | 32.13961 | 113.88897 | 109.9352 | Promoter | 13 | 64363214 | 64370306 | 7093 | 2 | ENSMUST00 | 0 | Ctsl | ENSMUSG00000021477.8 |
| chr13 | 49340856 | 49342745 | 32.13961 | 113.88897 | 109.9352 | Promoter | 13 | 49341549 | 49387025 | 45477 | 1 | ENSMUST00 | 0 | Bicd2 | ENSMUSG00000037933.10 |
| chr16 | 32867519 | 32868790 | 32.13961 | 113.88897 | 109.9352 | Promoter | 16 | 32821703 | 32868366 | 46664 | 2 | ENSMUST00 | 0 | 1700021K19I | ENSMUSG00000035629.12 |
| chr12 | 41023693 | 41024806 | 36.24269 | 113.86979 | 109.91653 | Promoter | 12 | 41024090 | 41955588 | 931499 | 1 | ENSMUST00 | 0 | Immp2l | ENSMUSG00000056899.5 |
| chr15 | 79140754 | 79141965 | 36.24269 | 113.86979 | 109.91653 | Promoter | 15 | 79141351 | 79151774 | 10424 | 1 | ENSMUST00 | 0 | Polr2f | ENSMUSG00000033020.6 |
| chr5 | 129583568 | 129586214 | 36.24269 | 113.86979 | 109.91653 | Promoter | 5 | 129584169 | 129608875 | 24707 | 1 | ENSMUST00 | 0 | Mmp17 | ENSMUSG00000029436.8 |
| chr8 | 107150149 | 107151371 | 36.24269 | 113.86979 | 109.91653 | Promoter | 8 | 107150661 | 107187470 | 36810 | 1 | ENSMUST00 | 0 | Cyb5b | ENSMUSG00000031924.3 |
| chr1 | 178405272 | 178406881 | 32.66734 | 113.81544 | 109.86345 | Promoter | 1 | 178405881 | 178483247 | 77367 | 1 | ENSMUST00 | 0 | Efcab2 | ENSMUSG00000026495.7 |
| chr10 | 41489428 | 41491002 | 32.66734 | 113.81544 | 109.86345 | Promoter | 10 | 41485642 | 41490330 | 4689 | 2 | ENSMUST00 | 0 | Smpd2 | ENSMUSG00000019822.7 |
| chr11 | 49711694 | 49713635 | 32.66734 | 113.81544 | 109.86345 | Promoter | 11 | 49671503 | 49712723 | 41221 | 2 | ENSMUST00 | 0 | Cnot6 | ENSMUSG00000020362.8 |
| chr11 | 50024682 | 50026322 | 32.66734 | 113.81544 | 109.86345 | Promoter | 11 | 50025346 | 50104758 | 79413 | 1 | ENSMUST00 | 0 | Rnf130 | ENSMUSG00000020376.11 |
| chr11 | 107468617 | 107471214 | 32.66734 | 113.81544 | 109.86345 | Promoter | 11 | 107207892 | 107470699 | 262808 | 2 | ENSMUST00 | 0 | Pitpnc1 | ENSMUSG00000040430.13 |
| chr11 | 98740244 | 98743094 | 32.66734 | 113.81544 | 109.86345 | Promoter | 11 | 98740638 | 98753379 | 12742 | 1 | ENSMUST00 | 0 | Thra | ENSMUSG00000058756.8 |
| chr12 | 110888374 | 110890178 | 32.66734 | 113.81544 | 109.86345 | Promoter | 12 | 110889264 | 110972392 | 83129 | 1 | ENSMUST00 | 0 | Tecpr2 | ENSMUSG00000021275.10 |
| chr12 | 28751304 | 28752397 | 32.66734 | 113.81544 | 109.86345 | Promoter | 12 | 28751828 | 28867491 | 115664 | 1 | ENSMUST00 | 0 | Tssc1 | ENSMUSG00000036613.7 |
| chr12 | 110840131 | 110841366 | 32.66734 | 113.81544 | 109.86345 | Promoter | 12 | 110807798 | 110840930 | 33133 | 2 | ENSMUST00 | 0 | Mok | ENSMUSG00000056458.11 |
| chr12 | 108272978 | 108276795 | 32.66734 | 113.81544 | 109.86345 | Promoter | 12 | 108206345 | 108275417 | 69073 | 2 | ENSMUST00 | 0 | Ccdc85c | ENSMUSG00000084883.1 |
| chr13 | 47043467 | 47044803 | 32.66734 | 113.81544 | 109.86345 | Promoter | 13 | 47025368 | 47044664 | 19297 | 2 | ENSMUST00 | 0 | Tpmt | ENSMUSG00000021376.5 |
| chr14 | 56811934 | 56813530 | 32.66734 | 113.81544 | 109.86345 | Promoter | 14 | 56811953 | 56815597 | 3645 | 1 | ENSMUST00 | 0 | RP23-170I24 | ENSMUSG00000100017.1 |
| chr15 | 76379922 | 76381414 | 32.66734 | 113.81544 | 109.86345 | Promoter | 15 | 76380261 | 76439744 | 59484 | 1 | ENSMUST00 | 0 | Mroh1 | ENSMUSG00000022558.10 |
| chr16 | 58520006 | 58524671 | 32.66734 | 113.81544 | 109.86345 | Promoter | 16 | 58468125 | 58523426 | 55302 | 2 | ENSMUST00 | 0 | St3gal6 | ENSMUSG00000022747.11 |
| chr16 | 57166539 | 57167829 | 32.66734 | 113.81544 | 109.86345 | Promoter | 16 | 57156671 | 57167332 | 10662 | 2 | ENSMUST00 | 0 | Nit2 | ENSMUSG00000022751.8 |
| chr17 | 46023564 | 46033809 | 32.66734 | 113.81544 | 109.86345 | Promoter | 17 | 46016993 | 46031815 | 14823 | 2 | ENSMUST00 | 0 | Vegfa | ENSMUSG00000023951.11 |
| chr17 | 25867687 | 25868978 | 32.66734 | 113.81544 | 109.86345 | Promoter | 17 | 25863698 | 25868738 | 5041 | 2 | ENSMUST00 | 0 | Fam195a | ENSMUSG00000025732.5 |

| | | | | | | | | | | | | | | | |
|---|---|---|---|---|---|---|---|---|---|---|---|---|---|---|---|
| chr17 | 17345857 | 17346432 | 32.66734 | 113.81544 | 109.86345 | Promoter | 17 | 17345456 | 17346140 | 685 | 2 | ENSMUST00 | 0 | Oaz1-ps | ENSMUSG00000043801.6 |
| chr19 | 27428956 | 27431040 | 32.66734 | 113.81544 | 109.86345 | Promoter | 19 | 27388702 | 27429820 | 41119 | 2 | ENSMUST00 | 0 | D19Bwg1357 | ENSMUSG00000041360.7 |
| chr2 | 27009437 | 27010847 | 32.66734 | 113.81544 | 109.86345 | Promoter | 2 | 27009926 | 27016849 | 6924 | 1 | ENSMUST00 | 0 | Cacfd1 | ENSMUSG00000015488.9 |
| chr2 | 165324516 | 165326561 | 32.66734 | 113.81544 | 109.86345 | Promoter | 2 | 165289155 | 165326479 | 37325 | 2 | ENSMUST00 | 0 | Elmo2 | ENSMUSG00000017670.11 |
| chr2 | 168205498 | 168207679 | 32.66734 | 113.81544 | 109.86345 | Promoter | 2 | 168180986 | 168206922 | 25937 | 2 | ENSMUST00 | 0 | Adnp | ENSMUSG00000051149.10 |
| chr4 | 86575183 | 86576814 | 32.66734 | 113.81544 | 109.86345 | Promoter | 4 | 86575668 | 86577285 | 1618 | 1 | ENSMUST00 | 0 | Rraga | ENSMUSG00000070934.6 |
| chr4 | 156235465 | 156237064 | 32.66734 | 113.81544 | 109.86345 | Promoter | 4 | 156236010 | 156247616 | 11607 | 1 | ENSMUST00 | 0 | Noc2l | ENSMUSG00000095567.2 |
| chr5 | 92082682 | 92084325 | 32.66734 | 113.81544 | 109.86345 | Promoter | 5 | 92052146 | 92082917 | 30772 | 2 | ENSMUST00 | 0 | G3bp2 | ENSMUSG00000029405.11 |
| chr5 | 125433367 | 125434198 | 32.66734 | 113.81544 | 109.86345 | Promoter | 5 | 125414406 | 125434048 | 19643 | 2 | ENSMUST00 | 0 | Dhx37 | ENSMUSG00000029480.9 |
| chr5 | 34660365 | 34661530 | 32.66734 | 113.81544 | 109.86345 | Promoter | 5 | 34660379 | 34701083 | 40705 | 1 | ENSMUST00 | 0 | Grk4 | ENSMUSG00000052783.11 |
| chr6 | 86396283 | 86397795 | 32.66734 | 113.81544 | 109.86345 | Promoter | 6 | 86386006 | 86397150 | 11145 | 2 | ENSMUST00 | 0 | Pcyox1 | ENSMUSG00000029998.9 |
| chr7 | 78782266 | 78783924 | 32.66734 | 113.81544 | 109.86345 | Promoter | 7 | 78783119 | 78792989 | 9871 | 1 | ENSMUST00 | 0 | Mrps11 | ENSMUSG00000030611.5 |
| chr7 | 6154864 | 6156928 | 32.66734 | 113.81544 | 109.86345 | Promoter | 7 | 6131489 | 6155971 | 24483 | 2 | ENSMUST00 | 0 | Zfp787 | ENSMUSG00000046792.7 |
| chr8 | 60631383 | 60633721 | 32.66734 | 113.81544 | 109.86345 | Promoter | 8 | 60632827 | 60674933 | 42107 | 1 | ENSMUST00 | 0 | Mfap3l | ENSMUSG00000031647.5 |
| chr8 | 110845795 | 110847888 | 32.66734 | 113.81544 | 109.86345 | Promoter | 8 | 110846600 | 110881895 | 35296 | 1 | ENSMUST00 | 0 | Cog4 | ENSMUSG00000031753.11 |
| chr8 | 87957648 | 87961172 | 32.66734 | 113.81544 | 109.86345 | Promoter | 8 | 87661810 | 87959595 | 297786 | 2 | ENSMUST00 | 0 | Zfp423 | ENSMUSG00000045333.10 |
| chr9 | 102625338 | 102627283 | 32.66734 | 113.81544 | 109.86345 | Promoter | 9 | 102586578 | 102626124 | 39547 | 2 | ENSMUST00 | 0 | Cep63 | ENSMUSG00000032534.11 |
| chr1 | 91494397 | 91495532 | 35.59737 | 113.79602 | 109.85027 | Promoter | 1 | 91494647 | 91527956 | 33310 | 1 | ENSMUST00 | 0 | Traf3ip1 | ENSMUSG00000034292.8 |
| chr14 | 20388139 | 20389535 | 34.98938 | 113.78057 | 109.83514 | Promoter | 14 | 20384638 | 20388910 | 4273 | 2 | ENSMUST00 | 0 | Dnajc9 | ENSMUSG00000021811.6 |
| chr17 | 56673291 | 56674310 | 34.98938 | 113.78057 | 109.83514 | Promoter | 17 | 56673225 | 56711769 | 38545 | 1 | ENSMUST00 | 67 | Ranbp3 | ENSMUSG00000002372.8 |
| chr17 | 29549466 | 29550457 | 34.98938 | 113.78057 | 109.83514 | Promoter | 17 | 29549788 | 29606895 | 57108 | 1 | ENSMUST00 | 0 | Tbc1d22b | ENSMUSG00000042203.7 |
| chr2 | 119757645 | 119759259 | 34.98938 | 113.78057 | 109.83514 | Promoter | 2 | 119751320 | 119758525 | 7206 | 2 | ENSMUST00 | 0 | Ltk | ENSMUSG00000027297.9 |
| chr5 | 123572247 | 123573243 | 34.98938 | 113.78057 | 109.83514 | Promoter | 5 | 123528764 | 123573015 | 44252 | 2 | ENSMUST00 | 0 | Vps33a | ENSMUSG00000029434.8 |
| chr15 | 35154948 | 35155944 | 36.16867 | 113.74171 | 109.7979 | Promoter | 15 | 34875496 | 35155806 | 280311 | 2 | ENSMUST00 | 0 | Stk3 | ENSMUSG00000022329.9 |
| chr9 | 15044502 | 15045972 | 36.16867 | 113.74171 | 109.7979 | Promoter | 9 | 15005161 | 15045478 | 40318 | 2 | ENSMUST00 | 0 | Panx1 | ENSMUSG00000031934.8 |
| chr1 | 39900215 | 39903023 | 36.76257 | 113.72284 | 109.77938 | Promoter | 1 | 39900913 | 40026310 | 125398 | 1 | ENSMUST00 | 0 | Map4k4 | ENSMUSG00000026074.9 |
| chr15 | 100468181 | 100469981 | 36.76257 | 113.72284 | 109.77938 | Promoter | 15 | 100469034 | 100479252 | 10219 | 1 | ENSMUST00 | 0 | Letmd1 | ENSMUSG00000037353.8 |
| chr15 | 100636116 | 100637647 | 36.76257 | 113.72284 | 109.77938 | Promoter | 15 | 100621342 | 100636855 | 15514 | 2 | ENSMUST00 | 0 | Smagp | ENSMUSG00000053559.7 |
| chr15 | 73089864 | 73091428 | 36.76257 | 113.72284 | 109.77938 | Promoter | 15 | 73090392 | 73094075 | 3684 | 1 | ENSMUST00 | 0 | Chrac1 | ENSMUSG00000068391.3 |
| chr2 | 167688507 | 167691530 | 36.76257 | 113.72284 | 109.77938 | Promoter | 2 | 167691177 | 167729172 | 37996 | 1 | ENSMUST00 | 0 | A530013C23I | ENSMUSG00000006462.6 |
| chr2 | 93044878 | 93046685 | 36.76257 | 113.72284 | 109.77938 | Promoter | 2 | 92965151 | 93045571 | 80421 | 2 | ENSMUST00 | 0 | Prdm11 | ENSMUSG00000075028.6 |
| chr3 | 32616014 | 32617147 | 36.76257 | 113.72284 | 109.77938 | Promoter | 3 | 32580332 | 32616585 | 36254 | 2 | ENSMUST00 | 0 | Gnb4 | ENSMUSG00000027669.9 |
| chr3 | 115887293 | 115888898 | 36.76257 | 113.72284 | 109.77938 | Promoter | 3 | 115887837 | 115929086 | 41250 | 1 | ENSMUST00 | 0 | Dph5 | ENSMUSG00000033554.12 |
| chr3 | 36612848 | 36613878 | 36.76257 | 113.72284 | 109.77938 | Promoter | 3 | 36573142 | 36613389 | 40248 | 2 | ENSMUST00 | 0 | Bbs7 | ENSMUSG00000037325.5 |
| chr4 | 46450413 | 46452821 | 36.76257 | 113.72284 | 109.77938 | Promoter | 4 | 46450902 | 46472657 | 21756 | 1 | ENSMUST00 | 0 | Anp32b | ENSMUSG00000028333.10 |
| chr5 | 122049044 | 122050724 | 36.76257 | 113.72284 | 109.77938 | Promoter | 5 | 121858387 | 122049851 | 191465 | 2 | ENSMUST00 | 0 | Cux2 | ENSMUSG00000042589.13 |
| chr5 | 149677956 | 149679210 | 36.76257 | 113.72284 | 109.77938 | Promoter | 5 | 149678230 | 149762599 | 84370 | 1 | ENSMUST00 | 0 | B3glct | ENSMUSG00000051950.8 |
| chr6 | 113696360 | 113698200 | 36.76257 | 113.72284 | 109.77938 | Promoter | 6 | 113697050 | 113711069 | 14020 | 1 | ENSMUST00 | 0 | Tatdn2 | ENSMUSG00000056952.8 |
| chr7 | 28391883 | 28393806 | 36.76257 | 113.72284 | 109.77938 | Promoter | 7 | 28392951 | 28399388 | 6438 | 1 | ENSMUST00 | 0 | Paf1 | ENSMUSG00000003437.9 |
| chr7 | 29783817 | 29784795 | 36.76257 | 113.72284 | 109.77938 | Promoter | 7 | 29783998 | 29794702 | 10705 | 1 | ENSMUST00 | 0 | Zfp30 | ENSMUSG00000047473.9 |
| chr7 | 112678205 | 112681471 | 36.76257 | 113.72284 | 109.77938 | Promoter | 7 | 112679318 | 112899459 | 220142 | 1 | ENSMUST00 | 0 | Tead1 | ENSMUSG00000055320.11 |
| chr4 | 105108502 | 105110562 | 33.18295 | 113.70766 | 109.76701 | Promoter | 4 | 105029874 | 105109890 | 80017 | 2 | ENSMUST00 | 0 | Prkaa2 | ENSMUSG00000028518.8 |
| chr1 | 33813492 | 33814814 | 34.3512 | 113.70341 | 109.76276 | Promoter | 1 | 33761544 | 33814595 | 53052 | 2 | ENSMUST00 | 0 | Zfp451 | ENSMUSG00000042197.8 |
| chr2 | 119156068 | 119157543 | 34.3512 | 113.70341 | 109.76276 | Promoter | 2 | 119137001 | 119157034 | 20034 | 2 | ENSMUST00 | 0 | Rmdn3 | ENSMUSG00000070730.6 |
| chr7 | 126776355 | 126777937 | 31.45397 | 113.65508 | 109.71495 | Promoter | 7 | 126776955 | 126780514 | 3560 | 1 | ENSMUST00 | 0 | Ypel3 | ENSMUSG00000042675.10 |
| chr10 | 122984594 | 122987014 | 36.09495 | 113.61411 | 109.67428 | Promoter | 10 | 122985634 | 122985723 | 90 | 2 | ENSMUST00 | 0 | Mirlet7i | ENSMUSG00000065406.2 |
| chr4 | 97776660 | 97779285 | 35.45657 | 113.55091 | 109.61168 | Promoter | 4 | 97777608 | 98118871 | 341264 | 1 | ENSMUST00 | 0 | Nfia | ENSMUSG00000028565.13 |
| chr2 | 152293204 | 152294766 | 34.85497 | 113.54498 | 109.60606 | Promoter | 2 | 152293828 | 152313996 | 20169 | 1 | ENSMUST00 | 0 | Tbc1d20 | ENSMUSG00000027465.8 |
| chr1 | 105993233 | 105994297 | 33.06624 | 113.49754 | 109.55873 | Intron (ENSM | 1 | 105990689 | 106034067 | 43379 | 1 | ENSMUST00 | 2545 | Zcchc2 | ENSMUSG00000038866.10 |
| chr12 | 31498404 | 31500658 | 33.06624 | 113.49754 | 109.55873 | Promoter | 12 | 31484829 | 31499616 | 14788 | 2 | ENSMUST00 | 0 | Cbll1 | ENSMUSG00000020659.10 |
| chr14 | 101607475 | 101609516 | 33.06624 | 113.49754 | 109.55873 | Promoter | 14 | 101442360 | 101609033 | 166674 | 2 | ENSMUST00 | 0 | Tbc1d4 | ENSMUSG00000033083.11 |
| chr16 | 32077855 | 32079646 | 33.06624 | 113.49754 | 109.55873 | Promoter | 16 | 32016290 | 32079342 | 63053 | 2 | ENSMUST00 | 0 | Pak2 | ENSMUSG00000022781.8 |
| chr17 | 25183813 | 25185433 | 33.06624 | 113.49754 | 109.55873 | Promoter | 17 | 25184561 | 25187661 | 3101 | 1 | ENSMUST00 | 0 | BC003965 | ENSMUSG00000067722.3 |
| chr4 | 132047947 | 132049501 | 33.06624 | 113.49754 | 109.55873 | Promoter | 4 | 131923429 | 132049078 | 125650 | 2 | ENSMUST00 | 0 | Epb4.1 | ENSMUSG00000028906.10 |
| chr7 | 79919930 | 79921230 | 33.06624 | 113.49754 | 109.55873 | Promoter | 7 | 79875332 | 79920640 | 45309 | 2 | ENSMUST00 | 0 | Ap3s2 | ENSMUSG00000063801.6 |
| chr8 | 94600788 | 94601772 | 33.06624 | 113.49754 | 109.55873 | Promoter | 8 | 94574943 | 94601726 | 26784 | 2 | ENSMUST00 | 0 | Fam192a | ENSMUSG00000031774.7 |
| chr5 | 53588566 | 53591937 | 30.81129 | 113.47932 | 109.54176 | Promoter | 5 | 53590215 | 53657445 | 67231 | 1 | ENSMUST00 | 0 | Rbpj | ENSMUSG00000039191.7 |
| chr1 | 39477826 | 39479423 | 34.78613 | 113.42423 | 109.48744 | Promoter | 1 | 39371499 | 39478747 | 107249 | 2 | ENSMUST00 | 0 | Tbc1d8 | ENSMUSG00000003134.5 |
| chr1 | 4807230 | 4808912 | 34.78613 | 113.42423 | 109.48744 | Promoter | 1 | 4807788 | 4848410 | 40623 | 1 | ENSMUST00 | 0 | Lypla1 | ENSMUSG00000025903.9 |
| chr1 | 60097058 | 60099158 | 34.78613 | 113.42423 | 109.48744 | Promoter | 1 | 60098247 | 60151032 | 52786 | 1 | ENSMUST00 | 0 | Carf | ENSMUSG00000026017.8 |
| chr1 | 53313115 | 53314391 | 34.78613 | 113.42423 | 109.48744 | Promoter | 1 | 53313624 | 53326343 | 12720 | 1 | ENSMUST00 | 0 | Osgepl1 | ENSMUSG00000026096.9 |
| chr10 | 26372988 | 26376464 | 34.78613 | 113.42423 | 109.48744 | Promoter | 10 | 26274468 | 26373956 | 99489 | 2 | ENSMUST00 | 0 | L3mbtl3 | ENSMUSG00000039089.9 |
| chr11 | 110336758 | 110338226 | 34.78613 | 113.42423 | 109.48744 | Promoter | 11 | 110269369 | 110337678 | 68310 | 2 | ENSMUST00 | 0 | Abca5 | ENSMUSG00000018800.9 |
| chr11 | 98021697 | 98023289 | 34.78613 | 113.42423 | 109.48744 | Promoter | 11 | 98001508 | 98022634 | 21127 | 2 | ENSMUST00 | 0 | Cacnb1 | ENSMUSG00000020882.12 |
| chr12 | 31350526 | 31351929 | 34.78613 | 113.42423 | 109.48744 | Promoter | 12 | 31331564 | 31351437 | 19874 | 2 | ENSMUST00 | 0 | Dld | ENSMUSG00000020664.9 |
| chr13 | 46928802 | 46931023 | 34.78613 | 113.42423 | 109.48744 | Promoter | 13 | 46749087 | 46929718 | 180632 | 2 | ENSMUST00 | 0 | Kif13a | ENSMUSG00000021375.9 |
| chr14 | 46882071 | 46885731 | 34.78613 | 113.42423 | 109.48744 | Promoter | 14 | 46882854 | 47103150 | 220297 | 1 | ENSMUST00 | 0 | Samd4 | ENSMUSG00000021838.11 |
| chr16 | 15593731 | 15594927 | 34.78613 | 113.42423 | 109.48744 | Promoter | 16 | 15552677 | 15594524 | 41848 | 2 | ENSMUST00 | 0 | Ube2v2 | ENSMUSG00000022674.9 |
| chr16 | 16357981 | 16359179 | 34.78613 | 113.42423 | 109.48744 | Promoter | 16 | 16312235 | 16359030 | 46796 | 2 | ENSMUST00 | 0 | Dnm1l | ENSMUSG00000022789.9 |
| chr17 | 47687260 | 47689501 | 34.78613 | 113.42423 | 109.48744 | Promoter | 17 | 47689030 | 47699551 | 10522 | 1 | ENSMUST00 | 0 | Frs3 | ENSMUSG00000023266.6 |
| chr17 | 78200179 | 78202236 | 34.78613 | 113.42423 | 109.48744 | Promoter | 17 | 78200248 | 78376592 | 176345 | 1 | ENSMUST00 | 0 | Crim1 | ENSMUSG00000024074.7 |
| chr18 | 60621048 | 60625338 | 34.78613 | 113.42423 | 109.48744 | Promoter | 18 | 60593973 | 60624336 | 30364 | 2 | ENSMUST00 | 0 | Synpo | ENSMUSG00000043079.12 |
| chr19 | 4305223 | 4306424 | 34.78613 | 113.42423 | 109.48744 | Promoter | 19 | 4286001 | 4305955 | 19955 | 2 | ENSMUST00 | 0 | Adrbk1 | ENSMUSG00000024858.8 |
| chr2 | 76405887 | 76407684 | 34.78613 | 113.42423 | 109.48744 | Promoter | 2 | 76406508 | 76407931 | 1424 | 1 | ENSMUST00 | 0 | Osbpl6 | ENSMUSG00000042359.13 |
| chr2 | 30285584 | 30287487 | 34.78613 | 113.42423 | 109.48744 | Promoter | 2 | 30286397 | 30307732 | 21336 | 1 | ENSMUST00 | 0 | Nup188 | ENSMUSG00000052533.9 |
| chr2 | 30065820 | 30067950 | 34.78613 | 113.42423 | 109.48744 | Promoter | 2 | 30066456 | 30072577 | 6122 | 1 | ENSMUST00 | 0 | Set | ENSMUSG00000054766.8 |
| chr2 | 156110584 | 156112392 | 34.78613 | 113.42423 | 109.48744 | Promoter | 2 | 156071842 | 156111935 | 40094 | 2 | ENSMUST00 | 0 | Cpne1 | ENSMUSG00000074643.7 |
| chr3 | 95281057 | 95284014 | 34.78613 | 113.42423 | 109.48744 | Promoter | 3 | 95253674 | 95282076 | 28403 | 2 | ENSMUST00 | 0 | Prune | ENSMUSG00000015711.8 |
| chr3 | 85886565 | 85888451 | 34.78613 | 113.42423 | 109.48744 | Promoter | 3 | 85801047 | 85887518 | 86472 | 2 | ENSMUST00 | 0 | Glt28d2 | ENSMUSG00000031286.5 |
| chr3 | 79566758 | 79568522 | 34.78613 | 113.42423 | 109.48744 | Promoter | 3 | 79455971 | 79567679 | 111709 | 2 | ENSMUST00 | 0 | Fnip2 | ENSMUSG00000061175.6 |
| chr4 | 123749157 | 123750996 | 34.78613 | 113.42423 | 109.48744 | Promoter | 4 | 123734559 | 123750345 | 15787 | 2 | ENSMUST00 | 0 | Akirin1 | ENSMUSG00000023075.9 |
| chr4 | 129461576 | 129462666 | 34.78613 | 113.42423 | 109.48744 | Promoter | 4 | 129461581 | 129488498 | 26918 | 1 | ENSMUST00 | 0 | Bsdc1 | ENSMUSG00000040859.9 |
| chr5 | 123720026 | 123721639 | 34.78613 | 113.42423 | 109.48744 | Promoter | 5 | 123698302 | 123721044 | 22743 | 2 | ENSMUST00 | 0 | Zcchc8 | ENSMUSG00000029427.7 |
| chr5 | 65107033 | 65108648 | 34.78613 | 113.42423 | 109.48744 | Promoter | 5 | 65107568 | 65168188 | 60621 | 1 | ENSMUST00 | 0 | Klhl5 | ENSMUSG00000054920.7 |
| chr5 | 142400816 | 142402701 | 34.78613 | 113.42423 | 109.48744 | Promoter | 5 | 142401497 | 142462014 | 60518 | 1 | ENSMUST00 | 0 | Foxk1 | ENSMUSG00000056493.8 |
| chr5 | 25498998 | 25500639 | 34.78613 | 113.42423 | 109.48744 | Promoter | 5 | 25499067 | 25501541 | 2475 | 1 | ENSMUST00 | 0 | 4831440E17I | ENSMUSG00000097236.1 |
| chr6 | 85960339 | 85962539 | 34.78613 | 113.42423 | 109.48744 | Promoter | 6 | 85942268 | 85961667 | 19400 | 2 | ENSMUST00 | 0 | Dusp11 | ENSMUSG00000030002.10 |
| chr7 | 66387468 | 66389353 | 34.78613 | 113.42423 | 109.48744 | Promoter | 7 | 66258745 | 66388350 | 129606 | 2 | ENSMUST00 | 0 | Lrrk1 | ENSMUSG00000015133.11 |
| chr7 | 120830240 | 120831642 | 34.78613 | 113.42423 | 109.48744 | Distal Interge | 7 | 120842831 | 120907219 | 64389 | 1 | ENSMUST00 | -11189 | Eef2k | ENSMUSG00000035064.11 |
| chr8 | 84856782 | 84857858 | 34.78613 | 113.42423 | 109.48744 | Promoter | 8 | 84856989 | 84869257 | 12269 | 1 | ENSMUST00 | 0 | Farsa | ENSMUSG00000003808.13 |
| chr8 | 88198612 | 88200888 | 34.78613 | 113.42423 | 109.48744 | Promoter | 8 | 88199213 | 88259716 | 60504 | 1 | ENSMUST00 | 0 | Papd5 | ENSMUSG00000036779.7 |
| chr8 | 70896375 | 70898212 | 34.78613 | 113.42423 | 109.48744 | Promoter | 8 | 70894722 | 70897443 | 2722 | 2 | ENSMUST00 | 0 | Rpl18a | ENSMUSG00000045128.8 |
| chr9 | 21237217 | 21239638 | 34.78613 | 113.42423 | 109.48744 | Promoter | 9 | 21229730 | 21239332 | 9603 | 2 | ENSMUST00 | 0 | Keap1 | ENSMUSG00000003308.9 |
| chr1 | 171345063 | 171346482 | 34.15952 | 113.36459 | 109.4342 | Promoter | 1 | 171345670 | 171359254 | 13585 | 1 | ENSMUST00 | 0 | Pfdn2 | ENSMUSG00000006412.5 |
| chr15 | 5143151 | 5144825 | 35.94841 | 113.36032 | 109.43 | Promoter | 15 | 5143861 | 5181899 | 38039 | 1 | ENSMUST00 | 0 | Prkaa1 | ENSMUSG00000050697.8 |
| chr10 | 128704299 | 128705622 | 36.54311 | 113.3445 | 109.41462 | Promoter | 10 | 128697939 | 128705051 | 7113 | 2 | ENSMUST00 | 0 | Cdk2 | ENSMUSG00000025358.10 |
| chr19 | 45997563 | 45998973 | 36.54311 | 113.3445 | 109.41462 | Promoter | 19 | 45818144 | 45998488 | 180345 | 2 | ENSMUST00 | 0 | 9130011E15I | ENSMUSG00000039901.9 |
| chr7 | 127392833 | 127394475 | 36.54311 | 113.3445 | 109.41462 | Promoter | 7 | 127389673 | 127393619 | 3947 | 2 | ENSMUST00 | 0 | E430018J23I | ENSMUSG00000078580.3 |
| chr12 | 71135727 | 71138170 | 26.2459 | 113.32486 | 109.39578 | Promoter | 12 | 71123173 | 71136675 | 13503 | 2 | ENSMUST00 | 0 | Timm9 | ENSMUSG00000021079.10 |
| chr15 | 8098513 | 8099303 | 26.2459 | 113.32486 | 109.39578 | Promoter | 15 | 7873055 | 8099209 | 226155 | 2 | ENSMUST00 | 0 | Wdr70 | ENSMUSG00000039828.6 |
| chr4 | 48124476 | 48125633 | 26.2459 | 113.32486 | 109.39578 | Promoter | 4 | 48124915 | 48185911 | 60997 | 1 | ENSMUST00 | 0 | Stx17 | ENSMUSG00000061455.8 |
| chr9 | 106788240 | 106789925 | 26.2459 | 113.32486 | 109.39578 | Promoter | 9 | 106688082 | 106789194 | 101113 | 2 | ENSMUST00 | 0 | Rad54l2 | ENSMUSG00000040661.6 |
| chr15 | 44426984 | 44429083 | 34.72158 | 113.31097 | 109.38217 | Promoter | 15 | 44428111 | 44437685 | 9575 | 1 | ENSMUST00 | 0 | Eny2 | ENSMUSG00000022338.6 |
| chr5 | 129786551 | 129788333 | 35.31687 | 113.30753 | 109.37892 | Promoter | 5 | 129765558 | 129787204 | 21647 | 2 | ENSMUST00 | 0 | Psph | ENSMUSG00000029446.9 |
| chr10 | 67284045 | 67285362 | 34.0961 | 113.25239 | 109.32404 | Promoter | 10 | 67266689 | 67285305 | 18617 | 2 | ENSMUST00 | 0 | Nrbf2 | ENSMUSG00000075000.6 |
| chr13 | 34037326 | 34038351 | 34.0961 | 113.25239 | 109.32404 | Promoter | 13 | 34037641 | 34074074 | 36434 | 1 | ENSMUST00 | 0 | Bphl | ENSMUSG00000038286.10 |
| chr17 | 32786896 | 32788757 | 34.0961 | 113.25239 | 109.32404 | Promoter | 17 | 32771236 | 32788287 | 17052 | 2 | ENSMUST00 | 0 | Zfp871 | ENSMUSG00000024298.9 |
| chr2 | 77945290 | 77946596 | 34.0961 | 113.25239 | 109.32404 | Promoter | 2 | 77881159 | 77946180 | 65022 | 2 | ENSMUST00 | 0 | Cwc22 | ENSMUSG00000027014.9 |
| chr2 | 130905723 | 130907477 | 34.0961 | 113.25239 | 109.32404 | Promoter | 2 | 130812866 | 130906406 | 93541 | 2 | ENSMUST00 | 0 | 4930402H24I | ENSMUSG00000027309.13 |
| chr7 | 126694880 | 126696774 | 34.0961 | 113.25239 | 109.32404 | Promoter | 7 | 126695401 | 126696687 | 1287 | 1 | ENSMUST00 | 0 | Bola2 | ENSMUSG00000047721.7 |
| chr9 | 77754467 | 77755628 | 35.87559 | 113.23412 | 109.30762 | Promoter | 9 | 77754535 | 77794485 | 39951 | 1 | ENSMUST00 | 0 | Gclc | ENSMUSG00000032350.8 |
| chr1 | 55086793 | 55088783 | 35.24744 | 113.18649 | 109.26083 | Promoter | 1 | 55077835 | 55088024 | 10190 | 2 | ENSMUST00 | 0 | Hspd1 | ENSMUSG00000025980.9 |
| chr19 | 32386186 | 32390069 | 35.24744 | 113.18649 | 109.26083 | Promoter | 19 | 32122727 | 32388512 | 265786 | 2 | ENSMUST00 | 0 | Sgms1 | ENSMUSG00000040451.12 |
| chr9 | 86571006 | 86572641 | 35.24744 | 113.18649 | 109.26083 | Promoter | 9 | 86571991 | 86574733 | 2743 | 1 | ENSMUST00 | 0 | Rwdd2a | ENSMUSG00000032417.5 |
| chr2 | 155133033 | 155134461 | 32.31418 | 113.17245 | 109.24831 | Promoter | 2 | 155133509 | 155139289 | 5781 | 1 | ENSMUST00 | 0 | Itch | ENSMUSG00000027598.11 |
| chr5 | 135743398 | 135744879 | 32.31418 | 113.17245 | 109.24831 | Promoter | 5 | 135735485 | 135744271 | 8787 | 2 | ENSMUST00 | 0 | Tmem120a | ENSMUSG00000039886.3 |
| chr5 | 144767881 | 144769477 | 32.31418 | 113.17245 | 109.24831 | Promoter | 5 | 144768092 | 144787907 | 19816 | 1 | ENSMUST00 | 0 | Trrap | ENSMUSG00000045482.10 |
| chr1 | 60565241 | 60567491 | 30.60796 | 113.09483 | 109.17188 | Promoter | 1 | 60482292 | 60566779 | 84488 | 2 | ENSMUST00 | 0 | Raph1 | ENSMUSG00000026014.10 |
| chr1 | 156034962 | 156037742 | 30.60796 | 113.09483 | 109.17188 | Promoter | 1 | 156004599 | 156036480 | 31882 | 2 | ENSMUST00 | 0 | Tor1aip1 | ENSMUSG00000026466.11 |
| chr10 | 30617599 | 30619020 | 30.60796 | 113.09483 | 109.17188 | Promoter | 10 | 30608141 | 30618479 | 10339 | 2 | ENSMUST00 | 0 | Hint3 | ENSMUSG00000019791.6 |
| chr11 | 51262442 | 51264681 | 30.60796 | 113.09483 | 109.17188 | Promoter | 11 | 51263114 | 51281764 | 18651 | 1 | ENSMUST00 | 0 | Clk4 | ENSMUSG00000020385.11 |
| chr11 | 101424134 | 101425924 | 30.60796 | 113.09483 | 109.17188 | Promoter | 11 | 101425085 | 101435673 | 10589 | 1 | ENSMUST00 | 0 | Rundc1 | ENSMUSG00000035007.5 |
| chr12 | 17544833 | 17546364 | 30.60796 | 113.09483 | 109.17188 | Promoter | 12 | 17544873 | 17551502 | 6630 | 1 | ENSMUST00 | 0 | Odc1 | ENSMUSG00000011179.7 |
| chr13 | 8995116 | 8996764 | 30.60796 | 113.09483 | 109.17188 | Promoter | 13 | 8972071 | 8996017 | 23947 | 2 | ENSMUST00 | 0 | Gtpbp4 | ENSMUSG00000021149.7 |
| chr13 | 48624908 | 48626279 | 30.60796 | 113.09483 | 109.17188 | Promoter | 13 | 48577873 | 48625573 | 47701 | 2 | ENSMUST00 | 0 | Ptpdc1 | ENSMUSG00000038042.9 |
| chr15 | 12823969 | 12825597 | 30.60796 | 113.09483 | 109.17188 | Promoter | 15 | 12824815 | 12935290 | 110476 | 1 | ENSMUST00 | 0 | Drosha | ENSMUSG00000022191.10 |
| chr15 | 58140916 | 58142176 | 30.60796 | 113.09483 | 109.17188 | Promoter | 15 | 58141436 | 58158652 | 17217 | 1 | ENSMUST00 | 0 | Wdyhv1 | ENSMUSG00000022359.9 |
| chr15 | 86184618 | 86186620 | 30.60796 | 113.09483 | 109.17188 | Promoter | 15 | 86139129 | 86186141 | 47013 | 2 | ENSMUST00 | 0 | Cerk | ENSMUSG00000035891.11 |

| | | | | | | | | | | | | | | | |
|---|---|---|---|---|---|---|---|---|---|---|---|---|---|---|---|
| chr16 | 18247796 | 18249956 | 30.60796 | 113.09483 | 109.17188 | Promoter | 16 | 18239784 | 18248732 | 8949 | 2 | ENSMUST00 | 0 | Ranbp1 | ENSMUSG00000005732.9 |
| chr16 | 64851908 | 64853636 | 30.60796 | 113.09483 | 109.17188 | Promoter | 16 | 64851996 | 64859507 | 7512 | 1 | ENSMUST00 | 0 | Cggbp1 | ENSMUSG00000054604.7 |
| chr18 | 61910130 | 61911876 | 30.60796 | 113.09483 | 109.17188 | Promoter | 18 | 61799395 | 61911824 | 112430 | 2 | ENSMUST00 | 0 | Ablim3 | ENSMUSG00000032735.9 |
| chr18 | 7625447 | 7627366 | 30.60796 | 113.09483 | 109.17188 | Promoter | 18 | 7347962 | 7626863 | 278902 | 2 | ENSMUST00 | 0 | Mpp7 | ENSMUSG00000057440.7 |
| chr5 | 130171056 | 130173066 | 30.60796 | 113.09483 | 109.17188 | Promoter | 5 | 130171798 | 130214342 | 42545 | 1 | ENSMUST00 | 0 | Rabgef1 | ENSMUSG00000025340.9 |
| chr5 | 121397352 | 121398901 | 30.60796 | 113.09483 | 109.17188 | Promoter | 5 | 121397936 | 121442409 | 44474 | 1 | ENSMUST00 | 0 | Naa25 | ENSMUSG00000042719.12 |
| chr6 | 97178114 | 97179837 | 30.60796 | 113.09483 | 109.17188 | Promoter | 6 | 97155266 | 97179037 | 23772 | 2 | ENSMUST00 | 0 | Tmf1 | ENSMUSG00000030059.10 |
| chr8 | 73351993 | 73354929 | 30.60796 | 113.09483 | 109.17188 | Promoter | 8 | 72814599 | 73352556 | 537958 | 2 | ENSMUST00 | 0 | Large | ENSMUSG00000004383.12 |
| chr8 | 22705691 | 22707007 | 30.60796 | 113.09483 | 109.17188 | Promoter | 8 | 22659212 | 22706578 | 47367 | 2 | ENSMUST00 | 0 | Ikbkb | ENSMUSG00000031537.10 |
| chr9 | 40191649 | 40193103 | 30.60796 | 113.09483 | 109.17188 | Promoter | 9 | 40192316 | 40213604 | 21289 | 1 | ENSMUST00 | 0 | Zfp202 | ENSMUSG00000025602.8 |
| chr8 | 13676275 | 13678189 | 36.39105 | 113.08218 | 109.16298 | Promoter | 8 | 13566950 | 13677603 | 110654 | 2 | ENSMUST00 | 0 | Rasa3 | ENSMUSG00000031453.10 |
| chr11 | 11808460 | 11809562 | 34.58922 | 113.07852 | 109.15961 | Promoter | 11 | 11787431 | 11808925 | 21495 | 2 | ENSMUST00 | 0 | Fignl1 | ENSMUSG00000035455.7 |
| chr19 | 44293182 | 44296580 | 35.73082 | 112.98314 | 109.06473 | Promoter | 19 | 44293676 | 44306864 | 13189 | 1 | ENSMUST00 | 0 | Scd2 | ENSMUSG00000025203.5 |
| chr4 | 46195548 | 46196735 | 34.52342 | 112.96288 | 109.04475 | Promoter | 4 | 46155347 | 46196311 | 40965 | 2 | ENSMUST00 | 0 | Xpa | ENSMUSG00000028329.8 |
| chr1 | 180941860 | 180943495 | 27.46926 | 112.95284 | 109.03484 | Promoter | 1 | 180942344 | 180952172 | 9829 | 1 | ENSMUST00 | 0 | Tmem63a | ENSMUSG00000026519.11 |
| chr13 | 93191112 | 93192929 | 27.46926 | 112.95284 | 109.03484 | Promoter | 13 | 93147400 | 93192283 | 44884 | 2 | ENSMUST00 | 0 | Papd4 | ENSMUSG00000042167.5 |
| chr14 | 70099533 | 70101030 | 27.46926 | 112.95284 | 109.03484 | Promoter | 14 | 70100145 | 70138191 | 38047 | 1 | ENSMUST00 | 0 | Bin3 | ENSMUSG00000022089.8 |
| chr15 | 64311537 | 64313724 | 27.46926 | 112.95284 | 109.03484 | Promoter | 15 | 64086857 | 64312673 | 225817 | 2 | ENSMUST00 | 0 | Asap1 | ENSMUSG00000022377.11 |
| chr17 | 56256022 | 56257981 | 27.46926 | 112.95284 | 109.03484 | Promoter | 17 | 56256793 | 56263608 | 6816 | 1 | ENSMUST00 | 0 | Fem1a | ENSMUSG00000043683.3 |
| chr2 | 119674410 | 119675809 | 27.46926 | 112.95284 | 109.03484 | Promoter | 2 | 119675068 | 119735407 | 60340 | 1 | ENSMUST00 | 0 | Rtf1 | ENSMUSG00000027304.9 |
| chr2 | 70823887 | 70826522 | 27.46926 | 112.95284 | 109.03484 | Promoter | 2 | 70712407 | 70825397 | 112991 | 2 | ENSMUST00 | 0 | Tlk1 | ENSMUSG00000041997.11 |
| chr1 | 69685182 | 69688560 | 36.31549 | 112.95179 | 109.03448 | Promoter | 1 | 69531214 | 69685821 | 154608 | 2 | ENSMUST00 | 0 | Ikzf2 | ENSMUSG00000025997.8 |
| chr1 | 179959117 | 179962379 | 36.31549 | 112.95179 | 109.03448 | Promoter | 1 | 179960472 | 180163156 | 202685 | 1 | ENSMUST00 | 0 | Cdc42bpa | ENSMUSG00000026490.12 |
| chr1 | 177983280 | 177983856 | 36.31549 | 112.95179 | 109.03448 | Distal Interge | 1 | 177991435 | 178048291 | 56857 | 1 | ENSMUST00 | -7579 | Gm16432 | ENSMUSG00000091476.1 |
| chr4 | 156109916 | 156110573 | 36.31549 | 112.95179 | 109.03448 | Promoter | 4 | 156109982 | 156127265 | 17284 | 1 | ENSMUST00 | 0 | 9430015G10 | ENSMUSG00000059939.8 |
| chr7 | 30955688 | 30958528 | 36.31549 | 112.95179 | 109.03448 | Promoter | 7 | 30945252 | 30956176 | 10925 | 2 | ENSMUST00 | 0 | Usf2 | ENSMUSG00000058239.8 |
| chr4 | 154011313 | 154012981 | 35.10938 | 112.94566 | 109.02863 | Promoter | 4 | 154011731 | 154021513 | 9783 | 1 | ENSMUST00 | 0 | Lrrc47 | ENSMUSG00000029028.9 |
| chr3 | 96635047 | 96636574 | 31.61155 | 112.91588 | 108.99915 | Promoter | 3 | 96635376 | 96645366 | 9991 | 1 | ENSMUST00 | 0 | Pex11b | ENSMUSG00000028102.10 |
| chr6 | 146633664 | 146634785 | 31.61155 | 112.91588 | 108.99915 | Promoter | 6 | 146602352 | 146634597 | 32246 | 2 | ENSMUST00 | 0 | Tm7sf3 | ENSMUSG00000040234.11 |
| chr7 | 64287101 | 64288643 | 31.61155 | 112.91588 | 108.99915 | Promoter | 7 | 64287653 | 64303172 | 15520 | 1 | ENSMUST00 | 0 | Mtmr10 | ENSMUSG00000030522.9 |
| chr7 | 127025218 | 127028304 | 35.65887 | 112.85833 | 108.94193 | Promoter | 7 | 127022143 | 127026479 | 4337 | 2 | ENSMUST00 | 0 | Maz | ENSMUSG00000030678.6 |
| chr1 | 131095659 | 131098154 | 28.90341 | 112.84859 | 108.93253 | Promoter | 1 | 131053700 | 131097826 | 44127 | 2 | ENSMUST00 | 0 | Mapkapk2 | ENSMUSG00000016528.5 |
| chr10 | 82082990 | 82083936 | 28.90341 | 112.84859 | 108.93253 | Distal Interge | 10 | 82058417 | 82061329 | 2913 | 1 | ENSMUST00 | 24574 | Zfp873 | ENSMUSG00000061371.6 |
| chr14 | 54574109 | 54578573 | 28.90341 | 112.84859 | 108.93253 | Promoter | 14 | 54567469 | 54577661 | 10193 | 2 | ENSMUST00 | 0 | Ajuba | ENSMUSG00000022178.10 |
| chr15 | 93335351 | 93337833 | 28.90341 | 112.84859 | 108.93253 | Promoter | 15 | 93284278 | 93336808 | 52531 | 2 | ENSMUST00 | 0 | Yaf2 | ENSMUSG00000022634.9 |
| chr2 | 31973976 | 31975444 | 28.90341 | 112.84859 | 108.93253 | Promoter | 2 | 31974436 | 32052371 | 77936 | 1 | ENSMUST00 | 0 | Nup214 | ENSMUSG00000001855.10 |
| chr5 | 139774931 | 139775984 | 28.90341 | 112.84859 | 108.93253 | Promoter | 5 | 139751282 | 139775678 | 24397 | 2 | ENSMUST00 | 0 | Ints1 | ENSMUSG00000029547.6 |
| chr9 | 107769446 | 107771471 | 28.90341 | 112.84859 | 108.93253 | Promoter | 9 | 107740371 | 107770996 | 30626 | 2 | ENSMUST00 | 0 | Rbm5 | ENSMUSG00000032580.7 |
| chr5 | 108065126 | 108066927 | 35.04076 | 112.82586 | 108.91161 | Promoter | 5 | 108065674 | 108107163 | 41490 | 1 | ENSMUST00 | 0 | Mtf2 | ENSMUSG00000029267.12 |
| chr6 | 145211095 | 145211950 | 35.04076 | 112.82586 | 108.91161 | Promoter | 6 | 145211134 | 145215791 | 4658 | 1 | ENSMUST00 | 0 | Lyrm5 | ENSMUSG00000040370.8 |
| chr11 | 20111757 | 20113819 | 36.24025 | 112.82189 | 108.90788 | Promoter | 11 | 20062304 | 20112913 | 50610 | 2 | ENSMUST00 | 0 | Actr2 | ENSMUSG00000020152.7 |
| chr19 | 38836126 | 38837723 | 33.26678 | 112.81776 | 108.90401 | Promoter | 19 | 38836579 | 38919923 | 83345 | 1 | ENSMUST00 | 0 | Tbc1d12 | ENSMUSG00000048720.7 |
| chr8 | 34107409 | 34109112 | 33.26678 | 112.81776 | 108.90401 | Promoter | 8 | 34090423 | 34108796 | 18374 | 2 | ENSMUST00 | 0 | Dctn6 | ENSMUSG00000031516.6 |
| chr11 | 78536348 | 78537492 | 34.39256 | 112.73273 | 108.82028 | Promoter | 11 | 78536361 | 78541737 | 5377 | 1 | ENSMUST00 | 0 | Ift20 | ENSMUSG00000001105.10 |
| chr12 | 72085706 | 72087055 | 34.3275 | 112.61823 | 108.70669 | Promoter | 12 | 72085789 | 72094015 | 8227 | 1 | ENSMUST00 | 0 | Jkamp | ENSMUSG00000005078.11 |
| chr13 | 43302084 | 43304284 | 34.3275 | 112.61823 | 108.70669 | Promoter | 13 | 43195519 | 43304172 | 108654 | 2 | ENSMUST00 | 0 | Gfod1 | ENSMUSG00000051335.5 |
| chr15 | 81855855 | 81859985 | 34.3275 | 112.61823 | 108.70669 | Promoter | 15 | 81848270 | 81858326 | 10057 | 2 | ENSMUST00 | 0 | Tob2 | ENSMUSG00000048546.7 |
| chr4 | 19574627 | 19575981 | 34.3275 | 112.61823 | 108.70669 | Promoter | 4 | 19575162 | 19598654 | 23493 | 1 | ENSMUST00 | 0 | Rmdn1 | ENSMUSG00000028229.6 |
| chr4 | 40142530 | 40144718 | 34.3275 | 112.61823 | 108.70669 | Promoter | 4 | 40143081 | 40198338 | 55258 | 1 | ENSMUST00 | 0 | Aco1 | ENSMUSG00000028405.9 |
| chr5 | 67606389 | 67611595 | 34.3275 | 112.61823 | 108.70669 | Promoter | 5 | 67607883 | 67613992 | 6110 | 1 | ENSMUST00 | 0 | Shisa3 | ENSMUSG00000050010.7 |
| chr6 | 106799187 | 106800432 | 34.3275 | 112.61823 | 108.70669 | Promoter | 6 | 106778249 | 106800069 | 21821 | 2 | ENSMUST00 | 0 | Crbn | ENSMUSG00000005362.9 |
| chr11 | 95384608 | 95385767 | 35.51585 | 112.61008 | 108.7009 | Promoter | 11 | 95384692 | 95391776 | 7085 | 1 | ENSMUST00 | 0 | Slc35b1 | ENSMUSG00000020873.8 |
| chr16 | 18875990 | 18878206 | 35.51585 | 112.61008 | 108.7009 | Promoter | 16 | 18877037 | 18928030 | 50994 | 1 | ENSMUST00 | 0 | Hira | ENSMUSG00000022702.10 |
| chr3 | 32708548 | 32709897 | 35.51585 | 112.61008 | 108.7009 | Promoter | 3 | 32709164 | 32718656 | 9493 | 1 | ENSMUST00 | 0 | Actl6a | ENSMUSG00000027671.9 |
| chr4 | 131843093 | 131844092 | 35.51585 | 112.61008 | 108.7009 | Promoter | 4 | 131843470 | 131867786 | 24317 | 1 | ENSMUST00 | 0 | Mecr | ENSMUSG00000028910.7 |
| chr6 | 90735442 | 90737552 | 35.51585 | 112.61008 | 108.7009 | Promoter | 6 | 90690662 | 90736687 | 46026 | 2 | ENSMUST00 | 0 | Iqsec1 | ENSMUSG00000034312.8 |
| chr8 | 128684852 | 128687886 | 35.51585 | 112.61008 | 108.7009 | Promoter | 8 | 128685654 | 128733200 | 47547 | 1 | ENSMUST00 | 0 | Itgb1 | ENSMUSG00000025809.10 |
| chr2 | 124609423 | 124611639 | 34.90432 | 112.58753 | 108.68004 | Promoter | 2 | 124610106 | 124662885 | 52780 | 1 | ENSMUST00 | 0 | Sema6d | ENSMUSG00000027200.12 |
| chr15 | 73060364 | 73061115 | 30.33627 | 112.57923 | 108.67202 | Promoter | 15 | 72589621 | 73061204 | 471584 | 2 | ENSMUST00 | 89 | Trappc9 | ENSMUSG00000047921.10 |
| chr4 | 138325647 | 138326702 | 30.33627 | 112.57923 | 108.67202 | Promoter | 4 | 138313412 | 138326307 | 12896 | 2 | ENSMUST00 | 0 | Pink1 | ENSMUSG00000028756.7 |
| chr2 | 51934510 | 51935549 | 36.09069 | 112.56361 | 108.65645 | Promoter | 2 | 51924448 | 51935007 | 10560 | 2 | ENSMUST00 | 0 | Rbm43 | ENSMUSG00000036249.11 |
| chr3 | 108283322 | 108284697 | 35.44477 | 112.48663 | 108.5799 | Promoter | 3 | 108284091 | 108361511 | 77421 | 1 | ENSMUST00 | 0 | Sort1 | ENSMUSG00000068747.9 |
| chr10 | 11280393 | 11282529 | 34.8365 | 112.46898 | 108.56265 | Promoter | 10 | 11281330 | 11298052 | 16723 | 1 | ENSMUST00 | 0 | Fbxo30 | ENSMUSG00000047648.8 |
| chr18 | 49831667 | 49833802 | 34.8365 | 112.46898 | 108.56265 | Promoter | 18 | 49832670 | 49964254 | 131585 | 1 | ENSMUST00 | 0 | Dmxl1 | ENSMUSG00000037416.7 |
| chr4 | 48472261 | 48473732 | 32.49479 | 112.46503 | 108.55889 | Promoter | 4 | 48430858 | 48473459 | 42602 | 2 | ENSMUST00 | 0 | Tex10 | ENSMUSG00000028345.10 |
| chr5 | 65763081 | 65765578 | 32.49479 | 112.46503 | 108.55889 | Promoter | 5 | 65763521 | 65830108 | 66588 | 1 | ENSMUST00 | 0 | N4bp2 | ENSMUSG00000037795.9 |
| chr10 | 75059656 | 75062040 | 36.01637 | 112.4352 | 108.52973 | Promoter | 10 | 75060896 | 75184923 | 124028 | 1 | ENSMUST00 | 0 | Bcr | ENSMUSG00000009681.9 |
| chr12 | 69580317 | 69583372 | 28.69329 | 112.43184 | 108.52676 | Promoter | 12 | 69577628 | 69583028 | 5401 | 2 | ENSMUST00 | 0 | Vcpkmt | ENSMUSG00000049882.9 |
| chr6 | 21949556 | 21950660 | 34.76894 | 112.35082 | 108.44631 | Promoter | 6 | 21949571 | 21976038 | 26468 | 1 | ENSMUST00 | 0 | Ing3 | ENSMUSG00000029670.7 |
| chr10 | 18055122 | 18057311 | 35.94236 | 112.30728 | 108.40293 | Promoter | 10 | 18055861 | 18125130 | 69270 | 1 | ENSMUST00 | 0 | Reps1 | ENSMUSG00000019854.11 |
| chr5 | 106963862 | 106965219 | 35.94236 | 112.30728 | 108.40293 | Promoter | 5 | 106964322 | 106984431 | 20110 | 1 | ENSMUST00 | 0 | Cdc7 | ENSMUSG00000029283.12 |
| chr17 | 23803067 | 23805197 | 35.30345 | 112.24108 | 108.33721 | Promoter | 17 | 23803187 | 23824736 | 21550 | 1 | ENSMUST00 | 0 | Srrm2 | ENSMUSG00000039218.11 |
| chr10 | 7662375 | 7663966 | 31.77438 | 112.18448 | 108.28101 | Promoter | 10 | 7629373 | 7663264 | 33892 | 2 | ENSMUST00 | 0 | Pcmt1 | ENSMUSG00000019795.12 |
| chr9 | 65345462 | 65347318 | 31.77438 | 112.18448 | 108.28101 | Promoter | 9 | 65346068 | 65359642 | 13575 | 1 | ENSMUST00 | 0 | Pdcd7 | ENSMUSG00000041837.3 |
| chr1 | 118310257 | 118311782 | 33.47463 | 112.14942 | 108.24687 | Promoter | 1 | 118294778 | 118311416 | 16639 | 2 | ENSMUST00 | 0 | Tsn | ENSMUSG00000026374.9 |
| chr13 | 80882761 | 80887175 | 33.47463 | 112.14942 | 108.24687 | Promoter | 13 | 80883384 | 80889643 | 6260 | 1 | ENSMUST00 | 0 | Arrdc3 | ENSMUSG00000074794.4 |
| chr14 | 37134341 | 37136021 | 33.47463 | 112.14942 | 108.24687 | Promoter | 14 | 37120444 | 37135139 | 14696 | 2 | ENSMUST00 | 0 | Ghitm | ENSMUSG00000041028.10 |
| chr17 | 24168379 | 24170164 | 33.47463 | 112.14942 | 108.24687 | Promoter | 17 | 24163866 | 24169461 | 5596 | 2 | ENSMUST00 | 0 | Atp6v0c | ENSMUSG00000024121.8 |
| chr19 | 5037979 | 5040018 | 33.47463 | 112.14942 | 108.24687 | Promoter | 19 | 5038826 | 5041134 | 2309 | 1 | ENSMUST00 | 0 | B3gnt1 | ENSMUSG00000047379.6 |
| chr2 | 181580738 | 181582236 | 33.47463 | 112.14942 | 108.24687 | Promoter | 2 | 181569149 | 181582008 | 12860 | 2 | ENSMUST00 | 0 | Uckl1 | ENSMUSG00000089917.2 |
| chr4 | 137593688 | 137595481 | 33.47463 | 112.14942 | 108.24687 | Promoter | 4 | 137593755 | 137605908 | 12154 | 1 | ENSMUST00 | 0 | Usp48 | ENSMUSG00000043411.10 |
| chr5 | 115299554 | 115300971 | 33.47463 | 112.14942 | 108.24687 | Promoter | 5 | 115298505 | 115300912 | 2408 | 2 | ENSMUST00 | 0 | Dynll1 | ENSMUSG00000009013.4 |
| chr9 | 66918618 | 66920405 | 33.47463 | 112.14942 | 108.24687 | Promoter | 9 | 66843672 | 66919705 | 76034 | 2 | ENSMUST00 | 0 | Rab8b | ENSMUSG00000036943.8 |
| chr10 | 108331582 | 108333603 | 35.23322 | 112.11895 | 108.21771 | Promoter | 10 | 108332189 | 108414391 | 82203 | 1 | ENSMUST00 | 0 | Pawr | ENSMUSG00000035873.7 |
| chr7 | 127243936 | 127245261 | 34.6346 | 112.11573 | 108.21484 | Promoter | 7 | 127244526 | 127254803 | 10278 | 1 | ENSMUST00 | 0 | Zfp771 | ENSMUSG00000054716.3 |
| chr1 | 59669650 | 59671580 | 32.29185 | 112.09676 | 108.19604 | Promoter | 1 | 59586558 | 59670792 | 84235 | 2 | ENSMUST00 | 0 | Sumo1 | ENSMUSG00000026021.10 |
| chr1 | 23921556 | 23922782 | 32.29185 | 112.09676 | 108.19604 | Promoter | 1 | 23845625 | 23922317 | 76693 | 2 | ENSMUST00 | 0 | Smap1 | ENSMUSG00000026155.8 |
| chr1 | 118321395 | 118322864 | 32.29185 | 112.09676 | 108.19604 | Promoter | 1 | 118321839 | 118333822 | 11984 | 1 | ENSMUST00 | 0 | Nifk | ENSMUSG00000026377.7 |
| chr1 | 167308081 | 167309546 | 32.29185 | 112.09676 | 108.19604 | Promoter | 1 | 167308670 | 167333978 | 25309 | 1 | ENSMUST00 | 0 | Tmco1 | ENSMUSG00000052428.5 |
| chr11 | 60416404 | 60418696 | 32.29185 | 112.09676 | 108.19604 | Promoter | 11 | 60417145 | 60450927 | 33783 | 1 | ENSMUST00 | 0 | Gid4 | ENSMUSG00000018415.9 |
| chr11 | 67051730 | 67053147 | 32.29185 | 112.09676 | 108.19604 | Promoter | 11 | 67037905 | 67052594 | 14690 | 2 | ENSMUST00 | 0 | Adprm | ENSMUSG00000020910.11 |
| chr12 | 16588648 | 16589858 | 32.29185 | 112.09676 | 108.19604 | Promoter | 12 | 16535669 | 16589770 | 54102 | 2 | ENSMUST00 | 0 | Lpin1 | ENSMUSG00000020593.9 |
| chr12 | 69986329 | 69987632 | 32.29185 | 112.09676 | 108.19604 | Promoter | 12 | 69965012 | 69987002 | 21991 | 2 | ENSMUST00 | 0 | Sav1 | ENSMUSG00000021067.7 |
| chr12 | 102704128 | 102705752 | 32.29185 | 112.09676 | 108.19604 | Promoter | 12 | 102568582 | 102704930 | 136349 | 2 | ENSMUST00 | 0 | Itpk1 | ENSMUSG00000057963.4 |
| chr16 | 17131439 | 17132792 | 32.29185 | 112.09676 | 108.19604 | Promoter | 16 | 17130138 | 17132383 | 2246 | 2 | ENSMUST00 | 0 | Sdf2l1 | ENSMUSG00000022769.7 |
| chr16 | 62786007 | 62787329 | 32.29185 | 112.09676 | 108.19604 | Promoter | 16 | 62732444 | 62786777 | 54334 | 2 | ENSMUST00 | 0 | Nsun3 | ENSMUSG00000050312.6 |
| chr17 | 22224926 | 22226140 | 32.29185 | 112.09676 | 108.19604 | Promoter | 17 | 22197265 | 22225614 | 28350 | 2 | ENSMUST00 | 0 | Gm4944 | ENSMUSG00000096433.1 |
| chr19 | 8897342 | 8899234 | 32.29185 | 112.09676 | 108.19604 | Promoter | 19 | 8898090 | 8916663 | 18574 | 1 | ENSMUST00 | 0 | Ganab | ENSMUSG00000071650.5 |
| chr2 | 180223494 | 180226303 | 32.29185 | 112.09676 | 108.19604 | Promoter | 2 | 180176373 | 180225859 | 49487 | 2 | ENSMUST00 | 0 | Lama5 | ENSMUSG00000015647.9 |
| chr2 | 130563252 | 130564759 | 32.29185 | 112.09676 | 108.19604 | Promoter | 2 | 130563742 | 130565394 | 1653 | 1 | ENSMUST00 | 0 | Mrps26 | ENSMUSG00000037740.8 |
| chr2 | 61592715 | 61594305 | 32.29185 | 112.09676 | 108.19604 | Promoter | 2 | 61593077 | 61654171 | 61095 | 1 | ENSMUST00 | 0 | Tank | ENSMUSG00000064289.10 |
| chr3 | 97609302 | 97610592 | 32.29185 | 112.09676 | 108.19604 | Promoter | 3 | 97560742 | 97610203 | 49462 | 2 | ENSMUST00 | 0 | Chd1l | ENSMUSG00000028089.4 |
| chr5 | 121544415 | 121546654 | 32.29185 | 112.09676 | 108.19604 | Promoter | 5 | 121518613 | 121545482 | 26870 | 2 | ENSMUST00 | 0 | Mapkapk5 | ENSMUSG00000029454.10 |
| chr5 | 137785173 | 137786774 | 32.29185 | 112.09676 | 108.19604 | Promoter | 5 | 137781906 | 137786663 | 4758 | 2 | ENSMUST00 | 0 | Mepce | ENSMUSG00000029726.11 |
| chr5 | 67845830 | 67848432 | 32.29185 | 112.09676 | 108.19604 | Promoter | 5 | 67618140 | 67847432 | 229293 | 2 | ENSMUST00 | 0 | Atp8a1 | ENSMUSG00000037685.10 |
| chr7 | 118854934 | 118856478 | 32.29185 | 112.09676 | 108.19604 | Promoter | 7 | 118842222 | 118855998 | 13777 | 2 | ENSMUST00 | 0 | Knop1 | ENSMUSG00000030980.11 |
| chr7 | 81570859 | 81572804 | 32.29185 | 112.09676 | 108.19604 | Promoter | 7 | 81571292 | 81596836 | 25545 | 1 | ENSMUST00 | 0 | Whamm | ENSMUSG00000045795.9 |
| chr8 | 84269409 | 84270481 | 32.29185 | 112.09676 | 108.19604 | Promoter | 8 | 84257795 | 84270380 | 12586 | 2 | ENSMUST00 | 0 | Ccdc130 | ENSMUSG00000004994.7 |
| chr11 | 98585990 | 98587917 | 35.79525 | 112.05288 | 108.1582 | Promoter | 11 | 98581256 | 98587368 | 6113 | 2 | ENSMUST00 | 0 | Ormdl3 | ENSMUSG00000038150.7 |
| chr6 | 11925352 | 11926813 | 35.79525 | 112.05288 | 108.1582 | Promoter | 6 | 11925881 | 12008013 | 82133 | 1 | ENSMUST00 | 0 | Phf14 | ENSMUSG00000029629.11 |
| chr7 | 44985691 | 44986485 | 35.79525 | 112.05288 | 108.1582 | Promoter | 7 | 44976727 | 44986356 | 9630 | 2 | ENSMUST00 | 0 | Prmt1 | ENSMUSG00000052429.12 |
| chr16 | 17110329 | 17111347 | 34.56782 | 111.9988 | 108.10535 | Promoter | 16 | 17086555 | 17111252 | 24698 | 2 | ENSMUST00 | 0 | Ppil2 | ENSMUSG00000022771.10 |
| chr3 | 87929380 | 87931180 | 34.56782 | 111.9988 | 108.10535 | Promoter | 3 | 87922601 | 87930127 | 7527 | 2 | ENSMUST00 | 0 | Rrnad1 | ENSMUSG00000004896.11 |
| chr3 | 51483418 | 51485063 | 34.56782 | 111.9988 | 108.10535 | Promoter | 3 | 51483966 | 51495905 | 11940 | 1 | ENSMUST00 | 0 | Rab33b | ENSMUSG00000027739.8 |
| chr6 | 83794348 | 83795957 | 35.16326 | 111.99725 | 108.10535 | Promoter | 6 | 83794982 | 83802556 | 7575 | 1 | ENSMUST00 | 0 | Nagk | ENSMUSG00000034744.8 |
| chr8 | 77609924 | 77611314 | 34.56782 | 111.9988 | 108.10535 | Promoter | 8 | 77595982 | 77610653 | 14672 | 2 | ENSMUST00 | 0 | Tmem184c | ENSMUSG00000031617.11 |
| chr8 | 4612522 | 4614038 | 34.56782 | 111.9988 | 108.10535 | Distal Interge | 8 | 4625840 | 4629380 | 3541 | 1 | ENSMUST00 | -11802 | Zfp958 | ENSMUSG00000058748.5 |
| chr1 | 155158626 | 155159615 | 31.10056 | 111.96791 | 108.07632 | Promoter | 1 | 155158703 | 155203517 | 44815 | 1 | ENSMUST00 | 0 | Stx6 | ENSMUSG00000026470.9 |
| chr1 | 98094581 | 98096244 | 36.30304 | 111.89011 | 107.99921 | Promoter | 1 | 97821094 | 98095621 | 274528 | 2 | ENSMUST00 | 0 | Pam | ENSMUSG00000026335.11 |
| chr11 | 76406837 | 76407845 | 36.30304 | 111.89011 | 107.99921 | Promoter | 11 | 76406952 | 76414329 | 7378 | 1 | ENSMUST00 | 0 | Timm22 | ENSMUSG00000020843.10 |
| chr12 | 3364808 | 3366248 | 36.30304 | 111.89011 | 107.99921 | Promoter | 12 | 3365132 | 3406494 | 41363 | 1 | ENSMUST00 | 0 | Kif3c | ENSMUSG00000020668.9 |
| chr2 | 173737113 | 173738489 | 36.30304 | 111.89011 | 107.99921 | Promoter | 2 | 173737511 | 173784336 | 46826 | 1 | ENSMUST00 | 0 | Vapb | ENSMUSG00000054455.4 |
| chr6 | 134926609 | 134929329 | 36.30304 | 111.89011 | 107.99921 | Promoter | 6 | 134929092 | 134951577 | 22486 | 1 | ENSMUST00 | 0 | 1190002F15I | ENSMUSG00000098318.2 |
| chr15 | 77305678 | 77307532 | 34.5013 | 111.88224 | 107.9941 | Promoter | 15 | 77078990 | 77307053 | 228064 | 2 | ENSMUST00 | 0 | Rbfox2 | ENSMUSG00000033565.11 |
| chr8 | 27259721 | 27261776 | 30.46894 | 111.80851 | 107.92088 | Promoter | 8 | 27260327 | 27275656 | 15330 | 1 | ENSMUST00 | 0 | Eif4ebp1 | ENSMUSG00000031490.5 |
| chr5 | 142894253 | 142895809 | 35.64933 | 111.80038 | 107.9129 | Promoter | 5 | 142866946 | 142895421 | 28476 | 2 | ENSMUST00 | 0 | Fbxl18 | ENSMUSG00000066640.6 |
| chr1 | 6213191 | 6216341 | 25.98079 | 111.7719 | 107.88497 | Promoter | 1 | 6214645 | 6276648 | 62004 | 1 | ENSMUST00 | 0 | Rb1cc1 | ENSMUSG00000025907.9 |
| chr10 | 121364588 | 121366171 | 25.98079 | 111.7719 | 107.88497 | Promoter | 10 | 121365090 | 121397245 | 32156 | 1 | ENSMUST00 | 0 | Gns | ENSMUSG00000034707.6 |

| | | | | | | | | | | | | | | | |
|---|---|---|---|---|---|---|---|---|---|---|---|---|---|---|---|
| chr2 | 177839449 | 177840645 | 25.98079 | 111.7719 | 107.88497 | Promoter | 2 | 177831791 | 177840318 | 8528 | 2 | ENSMUST00 | 0 | Gm14325 | ENSMUSG00000095362.2 |
| chr15 | 79347265 | 79348573 | 32.68175 | 111.76729 | 107.88066 | Promoter | 15 | 79347541 | 79359072 | 11532 | 1 | ENSMUST00 | 0 | Maff | ENSMUSG00000042622.8 |
| chr3 | 136669097 | 136672052 | 32.68175 | 111.76729 | 107.88066 | Promoter | 3 | 136670124 | 136797838 | 127715 | 1 | ENSMUST00 | 0 | Ppp3ca | ENSMUSG00000028161.12 |
| chr4 | 108118102 | 108118887 | 32.68175 | 111.76729 | 107.88066 | Promoter | 4 | 108043839 | 108118532 | 74694 | 2 | ENSMUST00 | 0 | Scp2 | ENSMUSG00000028603.10 |
| chr5 | 139826060 | 139827232 | 32.68175 | 111.76729 | 107.88066 | Promoter | 5 | 139823592 | 139826885 | 3294 | 2 | ENSMUST00 | 0 | Psmg3 | ENSMUSG00000029551.8 |
| chr16 | 58407933 | 58410191 | 35.02417 | 111.75519 | 107.8695 | Promoter | 16 | 58408443 | 58469727 | 61285 | 1 | ENSMUST00 | 0 | Dcbld2 | ENSMUSG00000035107.8 |
| chr2 | 90846506 | 90848212 | 35.02417 | 111.75519 | 107.8695 | Promoter | 2 | 90847155 | 90866810 | 19656 | 1 | ENSMUST00 | 0 | Mtch2 | ENSMUSG00000027282.12 |
| chr3 | 144568873 | 144570248 | 35.02417 | 111.75519 | 107.8695 | Promoter | 3 | 144431103 | 144570181 | 139079 | 2 | ENSMUST00 | 0 | Hs2st1 | ENSMUSG00000040151.8 |
| chr5 | 64969293 | 64970870 | 35.57682 | 111.67485 | 107.78959 | Promoter | 5 | 64970071 | 65041874 | 71804 | 1 | ENSMUST00 | 0 | Fam114a1 | ENSMUSG00000029185.9 |
| chr10 | 105840876 | 105842352 | 34.36702 | 111.64683 | 107.76208 | Promoter | 10 | 105841067 | 105847833 | 6767 | 1 | ENSMUST00 | 0 | Ccdc59 | ENSMUSG00000019897.8 |
| chr10 | 61145939 | 61148106 | 34.36702 | 111.64683 | 107.76208 | Promoter | 10 | 61098642 | 61147644 | 49003 | 2 | ENSMUST00 | 0 | Sgpl1 | ENSMUSG00000020097.9 |
| chr10 | 62228042 | 62231601 | 34.36702 | 111.64683 | 107.76208 | Promoter | 10 | 62185396 | 62231218 | 45823 | 2 | ENSMUST00 | 0 | Tspan15 | ENSMUSG00000037031.9 |
| chr11 | 69979551 | 69981936 | 34.36702 | 111.64683 | 107.76208 | Promoter | 11 | 69981156 | 69990601 | 9446 | 1 | ENSMUST00 | 0 | Ctdnep1 | ENSMUSG00000018559.11 |
| chr11 | 121387488 | 121388799 | 34.36702 | 111.64683 | 107.76208 | Promoter | 11 | 121356125 | 121388251 | 32127 | 2 | ENSMUST00 | 0 | Rab40b | ENSMUSG00000025170.8 |
| chr11 | 29373114 | 29374752 | 34.36702 | 111.64683 | 107.76208 | Promoter | 11 | 29373658 | 29510808 | 137151 | 1 | ENSMUST00 | 0 | Ccdc88a | ENSMUSG00000032740.11 |
| chr12 | 85279704 | 85280762 | 34.36702 | 111.64683 | 107.76208 | Promoter | 12 | 85272398 | 85280435 | 8038 | 2 | ENSMUST00 | 0 | Acyp1 | ENSMUSG00000008822.10 |
| chr12 | 32953340 | 32955472 | 34.36702 | 111.64683 | 107.76208 | Promoter | 12 | 32953891 | 32979050 | 25160 | 1 | ENSMUST00 | 0 | Sypl | ENSMUSG00000020570.9 |
| chr13 | 44729311 | 44730489 | 36.16187 | 111.64684 | 107.76208 | Promoter | 13 | 44729474 | 44731574 | 2101 | 1 | ENSMUST00 | 0 | Jarid2 | ENSMUSG00000038518.10 |
| chr14 | 57423453 | 57424648 | 34.36702 | 111.64683 | 107.76208 | Promoter | 14 | 57424062 | 57517936 | 93875 | 1 | ENSMUST00 | 0 | Ift88 | ENSMUSG00000040040.11 |
| chr15 | 79061997 | 79063003 | 34.36702 | 111.64683 | 107.76208 | Promoter | 15 | 79053094 | 79062885 | 9792 | 2 | ENSMUST00 | 0 | Ankrd54 | ENSMUSG00000033055.5 |
| chr16 | 77013187 | 77016626 | 34.36702 | 111.64683 | 107.76208 | Promoter | 16 | 77014069 | 77116779 | 102711 | 1 | ENSMUST00 | 0 | Usp25 | ENSMUSG00000022867.9 |
| chr17 | 15395635 | 15396981 | 34.36702 | 111.64683 | 107.76208 | Promoter | 17 | 15396246 | 15433580 | 37335 | 1 | ENSMUST00 | 0 | Fam120b | ENSMUSG00000014763.7 |
| chr17 | 44105080 | 44106469 | 34.36702 | 111.64683 | 107.76208 | Promoter | 17 | 44096308 | 44105809 | 9502 | 2 | ENSMUST00 | 0 | Enpp4 | ENSMUSG00000023961.11 |
| chr2 | 117119728 | 117123489 | 34.36702 | 111.64683 | 107.76208 | Promoter | 2 | 117121374 | 117182279 | 60906 | 1 | ENSMUST00 | 0 | Spred1 | ENSMUSG00000027351.9 |
| chr2 | 72475659 | 72477136 | 34.36702 | 111.64683 | 107.76208 | Promoter | 2 | 72476159 | 72486893 | 10735 | 1 | ENSMUST00 | 0 | Cdca7 | ENSMUSG00000055612.10 |
| chr4 | 3936695 | 3938445 | 34.36702 | 111.64683 | 107.76208 | Promoter | 4 | 3900996 | 3938382 | 37387 | 2 | ENSMUST00 | 0 | Plag1 | ENSMUSG00000003282.4 |
| chr4 | 152086343 | 152087761 | 34.36702 | 111.64683 | 107.76208 | Promoter | 4 | 152086858 | 152115390 | 28533 | 1 | ENSMUST00 | 0 | Plekhg5 | ENSMUSG00000039713.11 |
| chr4 | 26345607 | 26347297 | 34.36702 | 111.64683 | 107.76208 | Promoter | 4 | 26324506 | 26346891 | 22386 | 2 | ENSMUST00 | 0 | Manea | ENSMUSG00000040520.7 |
| chr5 | 65536471 | 65538620 | 34.36702 | 111.64683 | 107.76208 | Promoter | 5 | 65537233 | 65598988 | 61756 | 1 | ENSMUST00 | 0 | Ube2k | ENSMUSG00000029203.11 |
| chr5 | 77265048 | 77267299 | 34.36702 | 111.64683 | 107.76208 | Promoter | 5 | 77265491 | 77286432 | 20942 | 1 | ENSMUST00 | 0 | Rest | ENSMUSG00000029249.10 |
| chr5 | 140321083 | 140322497 | 34.36702 | 111.64683 | 107.76208 | Promoter | 5 | 140008689 | 140321550 | 312862 | 2 | ENSMUST00 | 0 | Mad1l1 | ENSMUSG00000029554.10 |
| chr5 | 24164403 | 24165642 | 34.36702 | 111.64683 | 107.76208 | Promoter | 5 | 24164963 | 24184013 | 19051 | 1 | ENSMUST00 | 0 | Nupl2 | ENSMUSG00000048439.10 |
| chr6 | 125191150 | 125193125 | 34.36702 | 111.64683 | 107.76208 | Promoter | 6 | 125191801 | 125193558 | 1758 | 1 | ENSMUST00 | 0 | Mrpl51 | ENSMUSG00000030335.5 |
| chr6 | 88044096 | 88045316 | 34.36702 | 111.64683 | 107.76208 | Promoter | 6 | 87999106 | 88045270 | 46165 | 2 | ENSMUST00 | 0 | Rab7 | ENSMUSG00000079477.4 |
| chr7 | 126376523 | 126378139 | 34.36702 | 111.64683 | 107.76208 | Promoter | 7 | 126370060 | 126377450 | 7391 | 2 | ENSMUST00 | 0 | Spns1 | ENSMUSG00000030741.10 |
| chr7 | 84150810 | 84152691 | 34.36702 | 111.64683 | 107.76208 | Promoter | 7 | 84109356 | 84151893 | 42538 | 2 | ENSMUST00 | 0 | Abhd17c | ENSMUSG00000038459.9 |
| chr7 | 90129152 | 90132776 | 34.36702 | 111.64683 | 107.76208 | Promoter | 7 | 90130232 | 90209447 | 79216 | 1 | ENSMUST00 | 0 | Picalm | ENSMUSG00000039361.9 |
| chr7 | 34912063 | 34913223 | 34.36702 | 111.64683 | 107.76208 | Promoter | 7 | 34912379 | 35044710 | 132332 | 1 | ENSMUST00 | 0 | Pepd | ENSMUSG00000063931.9 |
| chr8 | 79398146 | 79399848 | 34.36702 | 111.64683 | 107.76208 | Promoter | 8 | 79338395 | 79399518 | 61124 | 2 | ENSMUST00 | 0 | Smad1 | ENSMUSG00000031681.9 |
| chr9 | 56041506 | 56042817 | 34.36702 | 111.64683 | 107.76208 | Promoter | 9 | 56041845 | 56061882 | 20038 | 1 | ENSMUST00 | 0 | Rcn2 | ENSMUSG00000032320.7 |
| chr6 | 108212049 | 108214484 | 34.95504 | 111.6348 | 107.75649 | Promoter | 6 | 108213118 | 108551109 | 337992 | 1 | ENSMUST00 | 0 | Itpr1 | ENSMUSG00000030102.8 |
| chr17 | 31564766 | 31566088 | 33.17231 | 111.61063 | 107.73259 | Promoter | 17 | 31564801 | 31607680 | 42880 | 1 | ENSMUST00 | 0 | Pknox1 | ENSMUSG00000006705.7 |
| chr14 | 34200921 | 34201713 | 35.5046 | 111.54979 | 107.67184 | Promoter | 14 | 34194441 | 34201633 | 7193 | 2 | ENSMUST00 | 0 | Gprin2 | ENSMUSG00000071531.2 |
| chr9 | 100643537 | 100645131 | 36.08632 | 111.51659 | 107.63924 | Promoter | 9 | 100643600 | 100739597 | 95998 | 1 | ENSMUST00 | 0 | Stag1 | ENSMUSG00000037286.10 |
| chr13 | 63562847 | 63568718 | 33.69019 | 111.49343 | 107.61708 | Promoter | 13 | 63511533 | 63565520 | 53988 | 2 | ENSMUST00 | 0 | Ptch1 | ENSMUSG00000021466.6 |
| chr2 | 91053659 | 91055010 | 33.69019 | 111.49343 | 107.61708 | Promoter | 2 | 91054009 | 91059432 | 5424 | 1 | ENSMUST00 | 0 | Psmc3 | ENSMUSG00000002102.10 |
| chr2 | 148672076 | 148673503 | 33.69019 | 111.49343 | 107.61708 | Promoter | 2 | 148672601 | 148676027 | 3427 | 1 | ENSMUST00 | 0 | Nxt1 | ENSMUSG00000036992.5 |
| chr2 | 157367116 | 157368359 | 33.69019 | 111.49343 | 107.61708 | Promoter | 2 | 157367594 | 157396763 | 29170 | 1 | ENSMUST00 | 0 | Manbal | ENSMUSG00000063019.5 |
| chr3 | 138142532 | 138145013 | 33.69019 | 111.49343 | 107.61708 | Promoter | 3 | 138143448 | 138159821 | 16374 | 1 | ENSMUST00 | 0 | Trmt10a | ENSMUSG00000004127.7 |
| chr3 | 122617382 | 122620944 | 33.69019 | 111.49343 | 107.61708 | Promoter | 3 | 122538719 | 122619667 | 80949 | 2 | ENSMUST00 | 0 | Fnbp1l | ENSMUSG00000039735.10 |
| chr15 | 86212867 | 86215206 | 31.94275 | 111.4613 | 107.58682 | Promoter | 15 | 86214459 | 86498503 | 284045 | 1 | ENSMUST00 | 0 | Tbc1d22a | ENSMUSG00000051864.9 |
| chr16 | 56028147 | 56030547 | 31.94275 | 111.4613 | 107.58682 | Promoter | 16 | 56015506 | 56029696 | 14191 | 2 | ENSMUST00 | 0 | Pcnp | ENSMUSG00000071533.6 |
| chr18 | 23751642 | 23754155 | 31.94275 | 111.4613 | 107.58682 | Promoter | 18 | 23752333 | 23774684 | 22352 | 1 | ENSMUST00 | 0 | Mapre2 | ENSMUSG00000024277.9 |
| chr2 | 26139348 | 26141046 | 31.94275 | 111.4613 | 107.58682 | Promoter | 2 | 26136814 | 26140521 | 3708 | 2 | ENSMUST00 | 0 | C330006A16I | ENSMUSG00000087679.2 |
| chr1 | 78488119 | 78489734 | 30.26787 | 111.42836 | 107.5545 | Promoter | 1 | 78417975 | 78488859 | 70885 | 2 | ENSMUST00 | 0 | Farsb | ENSMUSG00000026245.11 |
| chr1 | 187215072 | 187216193 | 30.26787 | 111.42836 | 107.5545 | Promoter | 1 | 187044648 | 187215465 | 170818 | 2 | ENSMUST00 | 0 | Spata17 | ENSMUSG00000026611.12 |
| chr10 | 86300140 | 86302829 | 30.26787 | 111.42836 | 107.5545 | Promoter | 10 | 86300372 | 86349506 | 49135 | 1 | ENSMUST00 | 0 | Timp3 | ENSMUSG00000020044.8 |
| chr10 | 120226667 | 120228099 | 30.26787 | 111.42836 | 107.5545 | Promoter | 10 | 120227070 | 120232071 | 5002 | 1 | ENSMUST00 | 0 | Llph | ENSMUSG00000020224.9 |
| chr11 | 103951021 | 103954455 | 30.26787 | 111.42836 | 107.5545 | Promoter | 11 | 103821782 | 103954056 | 132275 | 2 | ENSMUST00 | 0 | Nsf | ENSMUSG00000034187.13 |
| chr11 | 64079007 | 64080159 | 30.26787 | 111.42836 | 107.5545 | Promoter | 11 | 63962627 | 64079466 | 116840 | 2 | ENSMUST00 | 0 | Cox10 | ENSMUSG00000042148.8 |
| chr11 | 75453976 | 75455090 | 35.43268 | 111.42518 | 107.5545 | Promoter | 11 | 75440944 | 75454717 | 13774 | 2 | ENSMUST00 | 0 | Wdr81 | ENSMUSG00000045374.13 |
| chr12 | 69892027 | 69893734 | 30.26787 | 111.42836 | 107.5545 | Promoter | 12 | 69893105 | 69964085 | 70981 | 1 | ENSMUST00 | 0 | Atl1 | ENSMUSG00000021066.8 |
| chr13 | 73603140 | 73605160 | 30.26787 | 111.42836 | 107.5545 | Promoter | 13 | 73604002 | 73620639 | 16638 | 1 | ENSMUST00 | 0 | Clptm1l | ENSMUSG00000021610.7 |
| chr14 | 75283583 | 75285322 | 30.26787 | 111.42836 | 107.5545 | Promoter | 14 | 75284373 | 75344426 | 60054 | 1 | ENSMUST00 | 0 | Zc3h13 | ENSMUSG00000022000.10 |
| chr2 | 34753418 | 34755238 | 30.26787 | 111.42836 | 107.5545 | Promoter | 2 | 34676184 | 34755232 | 79049 | 2 | ENSMUST00 | 0 | Gapvd1 | ENSMUSG00000026867.13 |
| chr3 | 60500555 | 60503503 | 30.26787 | 111.42836 | 107.5545 | Promoter | 3 | 60501252 | 60629748 | 128497 | 1 | ENSMUST00 | 0 | Mbnl1 | ENSMUSG00000027763.8 |
| chr3 | 138527203 | 138528587 | 30.26787 | 111.42836 | 107.5545 | Promoter | 3 | 138526415 | 138526494 | 80 | 1 | ENSMUST00 | 789 | Mir1956 | ENSMUSG00000088373.2 |
| chr4 | 107922052 | 107923649 | 30.26787 | 111.42836 | 107.5545 | Promoter | 4 | 107903981 | 107923610 | 19630 | 2 | ENSMUST00 | 0 | Cpt2 | ENSMUSG00000028607.11 |
| chr4 | 134128046 | 134129392 | 30.26787 | 111.42836 | 107.5545 | Promoter | 4 | 134127406 | 134128789 | 1384 | 2 | ENSMUST00 | 0 | Sh3bgrl3 | ENSMUSG00000028843.8 |
| chr4 | 132638569 | 132639890 | 30.26787 | 111.42836 | 107.5545 | Promoter | 4 | 132638987 | 132694883 | 55897 | 1 | ENSMUST00 | 0 | Eya3 | ENSMUSG00000028886.10 |
| chr4 | 3677777 | 3679180 | 30.26787 | 111.42836 | 107.5545 | Promoter | 4 | 3678115 | 3791613 | 113499 | 1 | ENSMUST00 | 0 | Lyn | ENSMUSG00000042228.9 |
| chr4 | 143411893 | 143413428 | 30.26787 | 111.42836 | 107.5545 | Promoter | 4 | 143412426 | 143421091 | 8666 | 1 | ENSMUST00 | 0 | Pramef8 | ENSMUSG00000046862.9 |
| chr5 | 145231137 | 145232649 | 30.26787 | 111.42836 | 107.5545 | Promoter | 5 | 145231723 | 145247302 | 15580 | 1 | ENSMUST00 | 0 | Zfp655 | ENSMUSG00000007812.11 |
| chr6 | 37870693 | 37872083 | 30.26787 | 111.42836 | 107.5545 | Promoter | 6 | 37870811 | 37921505 | 50695 | 1 | ENSMUST00 | 0 | Trim24 | ENSMUSG00000029833.12 |
| chr8 | 69995345 | 69996366 | 30.26787 | 111.42836 | 107.5545 | Promoter | 8 | 69907069 | 69996379 | 89311 | 2 | ENSMUST00 | 13 | Gatad2a | ENSMUSG00000036180.9 |
| chr8 | 105899942 | 105901307 | 30.26787 | 111.42836 | 107.5545 | Promoter | 8 | 105900441 | 105931778 | 31338 | 1 | ENSMUST00 | 0 | Pskh1 | ENSMUSG00000048310.8 |
| chr9 | 56994400 | 56996703 | 30.26787 | 111.42836 | 107.5545 | Promoter | 9 | 56994968 | 57062805 | 67838 | 1 | ENSMUST00 | 0 | Ptpn9 | ENSMUSG00000032290.6 |
| chr10 | 128337679 | 128338932 | 34.81759 | 111.39526 | 107.52535 | Promoter | 10 | 128337832 | 128362479 | 24648 | 1 | ENSMUST00 | 0 | Cs | ENSMUSG00000005683.7 |
| chr11 | 76178676 | 76180354 | 34.81759 | 111.39526 | 107.52535 | Promoter | 11 | 76046226 | 76179647 | 133422 | 2 | ENSMUST00 | 0 | Vps53 | ENSMUSG00000017288.10 |
| chr11 | 60698688 | 60700396 | 34.81759 | 111.39526 | 107.52535 | Promoter | 11 | 60699723 | 60714186 | 14464 | 1 | ENSMUST00 | 0 | Llgl1 | ENSMUSG00000020536.7 |
| chr12 | 102876356 | 102879146 | 34.81759 | 111.39526 | 107.52535 | Promoter | 12 | 102784656 | 102878406 | 93751 | 2 | ENSMUST00 | 0 | Btbd7 | ENSMUSG00000041702.6 |
| chr17 | 46327328 | 46328690 | 34.81759 | 111.39526 | 107.52535 | Promoter | 17 | 46303221 | 46328023 | 24803 | 2 | ENSMUST00 | 0 | Abcc10 | ENSMUSG00000032842.8 |
| chr5 | 140701639 | 140702698 | 34.81759 | 111.39526 | 107.52535 | Promoter | 5 | 140661827 | 140702378 | 40552 | 2 | ENSMUST00 | 0 | Iqce | ENSMUSG00000036555.9 |
| chr6 | 28480251 | 28481228 | 34.81759 | 111.39526 | 107.52535 | Promoter | 6 | 28480348 | 28877723 | 397376 | 1 | ENSMUST00 | 0 | Snd1 | ENSMUSG00000001424.9 |
| chr7 | 16313234 | 16314485 | 34.81759 | 111.39526 | 107.52535 | Promoter | 7 | 16313517 | 16317697 | 4181 | 1 | ENSMUST00 | 0 | Bbc3 | ENSMUSG00000002083.7 |
| chr15 | 38077360 | 38079609 | 36.01109 | 111.38685 | 107.51837 | Promoter | 15 | 37967328 | 38078853 | 111526 | 2 | ENSMUST00 | 0 | Ubr5 | ENSMUSG00000037487.6 |
| chr11 | 120466761 | 120468595 | 27.18312 | 111.36749 | 107.49936 | Promoter | 11 | 120467635 | 120475319 | 7685 | 1 | ENSMUST00 | 0 | Hgs | ENSMUSG00000025793.10 |
| chr11 | 106036199 | 106037906 | 27.18312 | 111.36749 | 107.49936 | Promoter | 11 | 106036872 | 106059324 | 22453 | 1 | ENSMUST00 | 0 | Dcaf7 | ENSMUSG00000049354.8 |
| chr11 | 120631883 | 120634076 | 27.18312 | 111.36749 | 107.49936 | Promoter | 11 | 120625117 | 120633600 | 8484 | 2 | ENSMUST00 | 0 | Mafg | ENSMUSG00000051510.8 |
| chr12 | 59060367 | 59061713 | 27.18312 | 111.36749 | 107.49936 | Promoter | 12 | 59043093 | 59061461 | 18369 | 2 | ENSMUST00 | 0 | Trappc6b | ENSMUSG00000020993.8 |
| chr13 | 24936852 | 24938257 | 27.18312 | 111.36749 | 107.49936 | Promoter | 13 | 24907579 | 24937661 | 30083 | 2 | ENSMUST00 | 0 | Aldh5a1 | ENSMUSG00000035936.6 |
| chr15 | 36495983 | 36497122 | 27.18312 | 111.36749 | 107.49936 | Promoter | 15 | 36477668 | 36496791 | 19124 | 2 | ENSMUST00 | 0 | Ankrd46 | ENSMUSG00000048307.7 |
| chr17 | 25239511 | 25240902 | 27.18312 | 111.36749 | 107.49936 | Promoter | 17 | 25240170 | 25242798 | 2629 | 1 | ENSMUST00 | 0 | Tsr3 | ENSMUSG00000015126.9 |
| chr17 | 84511480 | 84513349 | 27.18312 | 111.36749 | 107.49936 | Promoter | 17 | 84511895 | 84622142 | 110248 | 1 | ENSMUST00 | 0 | Plekhh2 | ENSMUSG00000040852.5 |
| chr2 | 31571910 | 31573549 | 27.18312 | 111.36749 | 107.49936 | Promoter | 2 | 31572651 | 31617526 | 44876 | 1 | ENSMUST00 | 0 | Fubp3 | ENSMUSG00000026843.10 |
| chr3 | 145924240 | 145926076 | 27.18312 | 111.36749 | 107.49936 | Promoter | 3 | 145924378 | 145934283 | 9906 | 1 | ENSMUST00 | 0 | Bcl10 | ENSMUSG00000028191.10 |
| chr6 | 40434915 | 40436523 | 27.18312 | 111.36749 | 107.49936 | Promoter | 6 | 40401375 | 40436135 | 34761 | 2 | ENSMUST00 | 0 | E330009J07F | ENSMUSG00000037172.9 |
| chr6 | 35252601 | 35253613 | 27.18312 | 111.36749 | 107.49936 | Promoter | 6 | 35252654 | 35263496 | 10843 | 1 | ENSMUST00 | 0 | 1810058I24R | ENSMUSG00000073155.7 |
| chr8 | 107292711 | 107294946 | 27.18312 | 111.36749 | 107.49936 | Promoter | 8 | 107293470 | 107329307 | 35838 | 1 | ENSMUST00 | 0 | Nfat5 | ENSMUSG00000003847.11 |
| chr8 | 88696931 | 88697911 | 27.18312 | 111.36749 | 107.49936 | Promoter | 8 | 88697028 | 88751945 | 54918 | 1 | ENSMUST00 | 0 | Cyld | ENSMUSG00000036712.9 |
| chr10 | 123075268 | 123077129 | 34.74927 | 111.27614 | 107.40929 | Promoter | 10 | 122992061 | 123076505 | 84445 | 2 | ENSMUST00 | 0 | Mon2 | ENSMUSG00000034602.8 |
| chr1 | 52726241 | 52727775 | 28.59262 | 111.22587 | 107.35971 | Promoter | 1 | 52656286 | 52727462 | 71177 | 2 | ENSMUST00 | 0 | Mfsd6 | ENSMUSG00000041439.10 |
| chr11 | 116622477 | 116624455 | 28.59262 | 111.22587 | 107.35971 | Promoter | 11 | 116598165 | 116624200 | 26036 | 2 | ENSMUST00 | 0 | Rhbdf2 | ENSMUSG00000020806.10 |
| chr13 | 81656629 | 81659028 | 28.59262 | 111.22587 | 107.35971 | Promoter | 13 | 81657806 | 81671888 | 14083 | 1 | ENSMUST00 | 0 | Lysmd3 | ENSMUSG00000035840.6 |
| chr15 | 76349778 | 76352229 | 28.59262 | 111.22587 | 107.35971 | Promoter | 15 | 76347040 | 76351110 | 4071 | 2 | ENSMUST00 | 0 | Sharpin | ENSMUSG00000022552.10 |
| chr16 | 91372405 | 91373495 | 28.59262 | 111.22587 | 107.35971 | Promoter | 16 | 91372783 | 91405589 | 32807 | 1 | ENSMUST00 | 0 | Ifnar2 | ENSMUSG00000022971.13 |
| chr17 | 24469635 | 24471499 | 28.59262 | 111.22587 | 107.35971 | Promoter | 17 | 24470473 | 24471591 | 1119 | 1 | ENSMUST00 | 0 | Pgp | ENSMUSG00000043445.6 |
| chr18 | 38296359 | 38297841 | 28.59262 | 111.22587 | 107.35971 | Promoter | 18 | 38296635 | 38317847 | 21213 | 1 | ENSMUST00 | 0 | Rnf14 | ENSMUSG00000060450.8 |
| chr2 | 109917200 | 109921050 | 28.59262 | 111.22587 | 107.35971 | Promoter | 2 | 109917647 | 110014257 | 96611 | 1 | ENSMUST00 | 0 | Lgr4 | ENSMUSG00000050199.8 |
| chr3 | 30968253 | 30969738 | 28.59262 | 111.22587 | 107.35971 | Promoter | 3 | 30899295 | 30969415 | 70121 | 2 | ENSMUST00 | 0 | Phc3 | ENSMUSG00000037652.10 |
| chr4 | 86611335 | 86612624 | 28.59262 | 111.22587 | 107.35971 | Promoter | 4 | 86578855 | 86612055 | 33201 | 2 | ENSMUST00 | 0 | Haus6 | ENSMUSG00000038047.13 |
| chr9 | 109875378 | 109876629 | 28.59262 | 111.22587 | 107.35971 | Promoter | 9 | 109875579 | 109893888 | 18310 | 1 | ENSMUST00 | 0 | Cdc25a | ENSMUSG00000032477.9 |
| chr2 | 174463205 | 174464469 | 31.25232 | 111.22263 | 107.35811 | Promoter | 2 | 174461072 | 174464105 | 3034 | 2 | ENSMUST00 | 0 | Atp5e | ENSMUSG00000016252.7 |
| chr4 | 45407590 | 45409701 | 31.25232 | 111.22263 | 107.35811 | Promoter | 4 | 45395923 | 45408761 | 12839 | 2 | ENSMUST00 | 0 | Slc25a51 | ENSMUSG00000045973.13 |
| chr10 | 61647907 | 61649648 | 35.2897 | 111.17733 | 107.31322 | Promoter | 10 | 61648552 | 61674168 | 25617 | 1 | ENSMUST00 | 0 | Ppa1 | ENSMUSG00000020089.8 |
| chr11 | 58937697 | 58938949 | 34.68122 | 111.15741 | 107.2936 | Promoter | 11 | 58932288 | 58938916 | 6629 | 2 | ENSMUST00 | 0 | Rnf187 | ENSMUSG00000020496.8 |
| chr1 | 92906695 | 92907971 | 32.87541 | 111.07999 | 107.21675 | Promoter | 1 | 92906981 | 92908435 | 1455 | 1 | ENSMUST00 | 0 | Dusp28 | ENSMUSG00000047067.7 |
| chr1 | 64690907 | 64692184 | 32.87541 | 111.07999 | 107.21675 | Promoter | 1 | 64691183 | 64711453 | 20271 | 1 | ENSMUST00 | 0 | Ccnyl1 | ENSMUSG00000070871.5 |
| chr11 | 97987688 | 97989380 | 32.87541 | 111.07999 | 107.21675 | Downstream | 11 | 97923238 | 97986444 | 63207 | 2 | ENSMUST00 | -1245 | Plxdc1 | ENSMUSG00000017417.9 |
| chr2 | 103483845 | 103485827 | 32.87541 | 111.07999 | 107.21675 | Promoter | 2 | 103453849 | 103485160 | 31312 | 2 | ENSMUST00 | 0 | Cat | ENSMUSG00000027187.5 |
| chr2 | 37422026 | 37423429 | 32.87541 | 111.07999 | 107.21675 | Promoter | 2 | 37370069 | 37422903 | 52835 | 2 | ENSMUST00 | 0 | Rc3h2 | ENSMUSG00000075376.5 |
| chr18 | 4375235 | 4376514 | 35.21864 | 111.05409 | 107.192 | Promoter | 18 | 4375592 | 4397330 | 21739 | 1 | ENSMUST00 | 0 | Mtpap | ENSMUSG00000024234.6 |
| chr11 | 120186648 | 120190389 | 30.60583 | 111.04385 | 107.18215 | Promoter | 11 | 120185400 | 120188751 | 3352 | 2 | ENSMUST00 | 0 | 2810410L24F | ENSMUSG00000075389.3 |
| chr17 | 65950415 | 65951253 | 30.60583 | 111.04385 | 107.18215 | Promoter | 17 | 65923066 | 65951187 | 28122 | 2 | ENSMUST00 | 0 | Twsg1 | ENSMUSG00000024098.5 |
| chr2 | 72285363 | 72286768 | 34.61343 | 111.03911 | 107.17758 | Promoter | 2 | 72285637 | 72442610 | 156974 | 1 | ENSMUST00 | 0 | Zak | ENSMUSG00000004085.9 |
| chr18 | 77064771 | 77066818 | 35.78724 | 111.00061 | 107.13931 | Promoter | 18 | 77065208 | 77155708 | 90501 | 1 | ENSMUST00 | 0 | Pias2 | ENSMUSG00000025423.10 |
| chr3 | 130059441 | 130061742 | 35.78724 | 111.00061 | 107.13931 | Promoter | 3 | 129982759 | 130060907 | 78149 | 2 | ENSMUST00 | 0 | Sec24b | ENSMUSG00000001052.10 |
| chr6 | 54971060 | 54973296 | 35.78724 | 111.00061 | 107.13931 | Promoter | 6 | 54923942 | 54971661 | 47720 | 2 | ENSMUST00 | 0 | Nod1 | ENSMUSG00000038058.9 |

| | | | | | | | | | | | | | | | |
|---|---|---|---|---|---|---|---|---|---|---|---|---|---|---|---|
| chr13 | 25019457 | 25020800 | 35.14787 | 110.9313 | 107.0704 | Promoter | 13 | 24992483 | 25020379 | 27897 | 2 | ENSMUST00 | 0 | Mrs2 | ENSMUSG00000021339.3 |
| chr18 | 3336441 | 3338545 | 35.14787 | 110.9313 | 107.0704 | Promoter | 18 | 3266354 | 3337679 | 71326 | 2 | ENSMUST00 | 0 | Crem | ENSMUSG00000063889.11 |
| chr2 | 120562969 | 120563923 | 24.69612 | 110.86208 | 107.00209 | Promoter | 2 | 120506820 | 120563807 | 56988 | 2 | ENSMUST00 | 0 | Zfp106 | ENSMUSG00000027288.11 |
| chr7 | 114253808 | 114255033 | 24.69612 | 110.86208 | 107.00209 | Promoter | 7 | 114215559 | 114254680 | 39122 | 2 | ENSMUST00 | 0 | Copb1 | ENSMUSG00000030754.8 |
| chr1 | 131137640 | 131139080 | 33.91391 | 110.8507 | 106.99079 | Promoter | 1 | 131127455 | 131138340 | 10886 | 2 | ENSMUST00 | 0 | Dyrk3 | ENSMUSG00000016526.8 |
| chr11 | 54100609 | 54101866 | 33.91391 | 110.8507 | 106.99079 | Promoter | 11 | 54100924 | 54131652 | 30729 | 1 | ENSMUST00 | 0 | P4ha2 | ENSMUSG00000018906.9 |
| chr11 | 30771296 | 30773413 | 33.91391 | 110.8507 | 106.99079 | Promoter | 11 | 30771726 | 30880361 | 108636 | 1 | ENSMUST00 | 0 | Psme4 | ENSMUSG00000040850.12 |
| chr19 | 3999790 | 4001275 | 33.91391 | 110.8507 | 106.99079 | Promoter | 19 | 4000580 | 4002103 | 1524 | 1 | ENSMUST00 | 0 | Nudt8 | ENSMUSG00000024869.4 |
| chr4 | 133574113 | 133575391 | 33.91391 | 110.8507 | 106.99079 | Promoter | 4 | 133574745 | 133584243 | 9499 | 1 | ENSMUST00 | 0 | Gpatch3 | ENSMUSG00000028850.2 |
| chr6 | 97204495 | 97205669 | 33.91391 | 110.8507 | 106.99079 | Promoter | 6 | 97183670 | 97205560 | 21891 | 2 | ENSMUST00 | 0 | Uba3 | ENSMUSG00000030061.11 |
| chr7 | 139388995 | 139389982 | 33.91391 | 110.8507 | 106.99079 | Promoter | 7 | 139389109 | 139579652 | 190544 | 1 | ENSMUST00 | 0 | Inpp5a | ENSMUSG00000025477.8 |
| chr9 | 124422515 | 124426377 | 33.91391 | 110.8507 | 106.99079 | Promoter | 9 | 124422622 | 124424154 | 1533 | 2 | ENSMUST00 | 0 | Ppp2r3d | ENSMUSG00000093803.2 |
| chr1 | 91458226 | 91460023 | 35.07738 | 110.80895 | 106.9512 | Promoter | 1 | 91415982 | 91459324 | 43343 | 2 | ENSMUST00 | 0 | Per2 | ENSMUSG00000055866.9 |
| chr10 | 12963461 | 12964767 | 35.07738 | 110.80895 | 106.9512 | Promoter | 10 | 12939983 | 12964259 | 24277 | 2 | ENSMUST00 | 0 | Stx11 | ENSMUSG00000039232.6 |
| chr4 | 116876655 | 116878267 | 35.07738 | 110.80895 | 106.9512 | Promoter | 4 | 116877376 | 116989264 | 111889 | 1 | ENSMUST00 | 0 | Zswim5 | ENSMUSG00000033948.3 |
| chr7 | 13036853 | 13039040 | 35.07738 | 110.80895 | 106.9512 | Promoter | 7 | 13035120 | 13038275 | 3156 | 2 | ENSMUST00 | 0 | Ube2m | ENSMUSG00000005575.10 |
| chr4 | 134913993 | 134915235 | 32.11694 | 110.747 | 106.89115 | Promoter | 4 | 134897849 | 134915024 | 17176 | 2 | ENSMUST00 | 0 | Tmem50a | ENSMUSG00000028822.6 |
| chr5 | 122353779 | 122355233 | 32.11694 | 110.747 | 106.89115 | Promoter | 5 | 122354369 | 122364984 | 10616 | 1 | ENSMUST00 | 0 | Vps29 | ENSMUSG00000029462.13 |
| chr5 | 96160231 | 96163458 | 32.11694 | 110.747 | 106.89115 | Promoter | 5 | 96070333 | 96161990 | 91658 | 2 | ENSMUST00 | 0 | Cnot6l | ENSMUSG00000034724.12 |
| chr2 | 30952810 | 30953880 | 35.63955 | 110.7456 | 106.89037 | Promoter | 2 | 30952959 | 30959015 | 6057 | 1 | ENSMUST00 | 0 | Tor1b | ENSMUSG00000026848.9 |
| chr2 | 119593731 | 119595705 | 33.84988 | 110.73796 | 106.88306 | Promoter | 2 | 119594297 | 119600746 | 6450 | 1 | ENSMUST00 | 0 | 1700020I14R | ENSMUSG00000085438.1 |
| chr6 | 70790221 | 70792994 | 33.84988 | 110.73796 | 106.88306 | Promoter | 6 | 70765720 | 70792232 | 26513 | 2 | ENSMUST00 | 0 | Rpia | ENSMUSG00000053604.5 |
| chr15 | 58822098 | 58824291 | 33.25798 | 110.72714 | 106.87233 | Promoter | 15 | 58782269 | 58823530 | 41262 | 2 | ENSMUST00 | 0 | Tmem65 | ENSMUSG00000062373.7 |
| chr9 | 64810764 | 64812508 | 33.78609 | 110.6256 | 106.77135 | Promoter | 9 | 64811340 | 64919667 | 108328 | 1 | ENSMUST00 | 0 | Dennd4a | ENSMUSG00000053641.8 |
| chr12 | 44268371 | 44270172 | 34.93725 | 110.5656 | 106.71194 | Promoter | 12 | 44269186 | 44312244 | 43059 | 1 | ENSMUST00 | 0 | Pnpla8 | ENSMUSG00000036257.9 |
| chr10 | 95414179 | 95418016 | 35.49308 | 110.49252 | 106.63927 | Promoter | 10 | 95411490 | 95415484 | 3995 | 2 | ENSMUST00 | 0 | Socs2 | ENSMUSG00000020027.13 |
| chr16 | 91484460 | 91486279 | 31.40916 | 110.4849 | 106.632 | Promoter | 16 | 91485238 | 91507441 | 22204 | 1 | ENSMUST00 | 0 | Ifnar1 | ENSMUSG00000022967.7 |
| chr2 | 103760437 | 103761988 | 31.40916 | 110.4849 | 106.632 | Promoter | 2 | 103721256 | 103761270 | 40015 | 2 | ENSMUST00 | 0 | Nat10 | ENSMUSG00000027185.9 |
| chr4 | 45528278 | 45531712 | 31.40916 | 110.4849 | 106.632 | Promoter | 4 | 45423278 | 45530330 | 107053 | 2 | ENSMUST00 | 0 | Shb | ENSMUSG00000044813.10 |
| chr3 | 69125615 | 69127992 | 34.86761 | 110.44456 | 106.5922 | Promoter | 3 | 69067149 | 69127113 | 59965 | 2 | ENSMUST00 | 0 | Kpna4 | ENSMUSG00000027782.5 |
| chr1 | 82724492 | 82725933 | 33.07612 | 110.40385 | 106.55179 | Promoter | 1 | 82724890 | 82752390 | 27501 | 1 | ENSMUST00 | 0 | Mff | ENSMUSG00000026150.9 |
| chr11 | 115812381 | 115815436 | 33.07612 | 110.40385 | 106.55179 | Promoter | 11 | 115814724 | 115823094 | 8371 | 1 | ENSMUST00 | 0 | Tsen54 | ENSMUSG00000020781.9 |
| chr11 | 53350131 | 53352443 | 33.07612 | 110.40385 | 106.55179 | Promoter | 11 | 53350833 | 53421830 | 70998 | 1 | ENSMUST00 | 0 | Aff4 | ENSMUSG00000049470.8 |
| chr12 | 111814116 | 111814909 | 33.07612 | 110.40385 | 106.55179 | Promoter | 12 | 111814170 | 111828388 | 14219 | 1 | ENSMUST00 | 0 | Zfyve21 | ENSMUSG00000021286.7 |
| chr15 | 26894129 | 26896386 | 33.07612 | 110.40385 | 106.55179 | Promoter | 15 | 26543084 | 26895049 | 351966 | 2 | ENSMUST00 | 0 | Fbxl7 | ENSMUSG00000043556.9 |
| chr16 | 18425518 | 18427105 | 33.07612 | 110.40385 | 106.55179 | Promoter | 16 | 18407346 | 18426852 | 19507 | 2 | ENSMUST00 | 0 | Comt | ENSMUSG00000000326.7 |
| chr17 | 27133321 | 27134253 | 33.07612 | 110.40385 | 106.55179 | Promoter | 17 | 27122663 | 27133916 | 11254 | 2 | ENSMUST00 | 0 | Uqcc2 | ENSMUSG00000024208.10 |
| chr18 | 35118136 | 35120892 | 33.07612 | 110.40385 | 106.55179 | Promoter | 18 | 35118888 | 35254773 | 135886 | 1 | ENSMUST00 | 0 | Ctnna1 | ENSMUSG00000037815.6 |
| chr4 | 126262294 | 126262967 | 33.07612 | 110.40385 | 106.55179 | Promoter | 4 | 126262325 | 126275883 | 13559 | 1 | ENSMUST00 | 0 | Trappc3 | ENSMUSG00000028847.8 |
| chr5 | 43780884 | 43782943 | 33.07612 | 110.40385 | 106.55179 | Promoter | 5 | 43744616 | 43782118 | 37503 | 2 | ENSMUST00 | 0 | Fbxl5 | ENSMUSG00000039753.11 |
| chr7 | 44891585 | 44893164 | 33.07612 | 110.40385 | 106.55179 | Promoter | 7 | 44879391 | 44892397 | 13007 | 2 | ENSMUST00 | 0 | Med25 | ENSMUSG00000002968.7 |
| chr1 | 132315711 | 132317998 | 31.91637 | 110.38313 | 106.53261 | Promoter | 1 | 132316126 | 132333293 | 17168 | 1 | ENSMUST00 | 0 | Nuak2 | ENSMUSG00000009772.10 |
| chr1 | 75210303 | 75211459 | 31.91637 | 110.38313 | 106.53261 | Promoter | 1 | 75198243 | 75210778 | 12536 | 2 | ENSMUST00 | 0 | Glb1l | ENSMUSG00000026200.8 |
| chr1 | 166000437 | 166002688 | 31.91637 | 110.38313 | 106.53261 | Promoter | 1 | 165875457 | 166002609 | 127153 | 2 | ENSMUST00 | 0 | Pou2f1 | ENSMUSG00000026565.13 |
| chr1 | 163779596 | 163780335 | 31.91637 | 110.38313 | 106.53261 | Promoter | 1 | 163779583 | 163917109 | 137527 | 1 | ENSMUST00 | 14 | Kifap3 | ENSMUSG00000026585.8 |
| chr1 | 5082795 | 5083891 | 31.91637 | 110.38313 | 106.53261 | Promoter | 1 | 5083173 | 5162549 | 79377 | 1 | ENSMUST00 | 0 | Atp6v1h | ENSMUSG00000033793.7 |
| chr10 | 58497512 | 58498602 | 31.91637 | 110.38313 | 106.53261 | Promoter | 10 | 58497970 | 58576242 | 78273 | 1 | ENSMUST00 | 0 | Ccdc138 | ENSMUSG00000038010.8 |
| chr10 | 19933409 | 19936698 | 31.91637 | 110.38313 | 106.53261 | Promoter | 10 | 19934472 | 20142753 | 208282 | 1 | ENSMUST00 | 0 | Map3k5 | ENSMUSG00000071369.5 |
| chr11 | 77606424 | 77608552 | 31.91637 | 110.38313 | 106.53261 | Promoter | 11 | 77529162 | 77607815 | 78654 | 2 | ENSMUST00 | 0 | Taok1 | ENSMUSG00000017291.8 |
| chr11 | 4894677 | 4896003 | 31.91637 | 110.38313 | 106.53261 | Promoter | 11 | 4895320 | 4928867 | 33548 | 1 | ENSMUST00 | 0 | Thoc5 | ENSMUSG00000034274.6 |
| chr11 | 73137620 | 73138653 | 31.91637 | 110.38313 | 106.53261 | Promoter | 11 | 73135485 | 73138294 | 2810 | 2 | ENSMUST00 | 0 | Gsg2 | ENSMUSG00000050107.2 |
| chr13 | 92844494 | 92845356 | 31.91637 | 110.38313 | 106.53261 | Promoter | 13 | 92844787 | 92858230 | 13444 | 1 | ENSMUST00 | 0 | Mtx3 | ENSMUSG00000021704.7 |
| chr15 | 76671572 | 76672462 | 31.91637 | 110.38313 | 106.53261 | Promoter | 15 | 76671615 | 76694919 | 23305 | 1 | ENSMUST00 | 0 | Ppp1r16a | ENSMUSG00000033819.10 |
| chr15 | 78405455 | 78407074 | 31.91637 | 110.38313 | 106.53261 | Promoter | 15 | 78399556 | 78405859 | 6304 | 2 | ENSMUST00 | 0 | Tst | ENSMUSG00000044986.8 |
| chr16 | 4419243 | 4421071 | 31.91637 | 110.38313 | 106.53261 | Promoter | 16 | 4287529 | 4419808 | 132280 | 2 | ENSMUST00 | 0 | Adcy9 | ENSMUSG00000005580.6 |
| chr16 | 44173160 | 44174315 | 31.91637 | 110.38313 | 106.53261 | Promoter | 16 | 44173246 | 44227465 | 54220 | 1 | ENSMUST00 | 0 | Gm608 | ENSMUSG00000068284.9 |
| chr18 | 6239314 | 6242188 | 31.91637 | 110.38313 | 106.53261 | Promoter | 18 | 6201002 | 6241523 | 40522 | 2 | ENSMUST00 | 0 | Kif5b | ENSMUSG00000006740.8 |
| chr18 | 38249506 | 38251021 | 31.91637 | 110.38313 | 106.53261 | Promoter | 18 | 38250249 | 38262629 | 12381 | 1 | ENSMUST00 | 0 | 0610009O20 | ENSMUSG00000024442.5 |
| chr18 | 35598305 | 35599706 | 31.91637 | 110.38313 | 106.53261 | Promoter | 18 | 35598617 | 35617186 | 18570 | 1 | ENSMUST00 | 0 | Paip2 | ENSMUSG00000037058.9 |
| chr18 | 38337936 | 38339622 | 31.91637 | 110.38313 | 106.53261 | Promoter | 18 | 38327558 | 38339003 | 11446 | 2 | ENSMUST00 | 0 | Gnpda1 | ENSMUSG00000052102.9 |
| chr19 | 5911416 | 5913321 | 31.91637 | 110.38313 | 106.53261 | Promoter | 19 | 5896516 | 5912871 | 16356 | 2 | ENSMUST00 | 0 | Dpf2 | ENSMUSG00000024826.9 |
| chr19 | 4148113 | 4149802 | 31.91637 | 110.38313 | 106.53261 | Promoter | 19 | 4148619 | 4150504 | 1886 | 1 | ENSMUST00 | 0 | Coro1b | ENSMUSG00000024835.9 |
| chr19 | 38930241 | 38931722 | 31.91637 | 110.38313 | 106.53261 | Promoter | 19 | 38930915 | 38971051 | 40137 | 1 | ENSMUST00 | 0 | Hells | ENSMUSG00000025001.6 |
| chr19 | 41846750 | 41848862 | 31.91637 | 110.38313 | 106.53261 | Promoter | 19 | 41845978 | 41848132 | 2155 | 2 | ENSMUST00 | 0 | Frat2 | ENSMUSG00000047604.2 |
| chr2 | 71786545 | 71788373 | 31.91637 | 110.38313 | 106.53261 | Promoter | 2 | 71786929 | 71820943 | 34015 | 1 | ENSMUST00 | 0 | Itga6 | ENSMUSG00000027111.10 |
| chr2 | 131210000 | 131211274 | 31.91637 | 110.38313 | 106.53261 | Promoter | 2 | 131210370 | 131213002 | 2633 | 1 | ENSMUST00 | 0 | Ap5s1 | ENSMUSG00000068264.6 |
| chr3 | 14578291 | 14580268 | 31.91637 | 110.38313 | 106.53261 | Promoter | 3 | 14578671 | 14606308 | 27638 | 1 | ENSMUST00 | 0 | E2f5 | ENSMUSG00000027552.9 |
| chr4 | 133369618 | 133371516 | 31.91637 | 110.38313 | 106.53261 | Promoter | 4 | 133369706 | 133423702 | 53997 | 1 | ENSMUST00 | 0 | Slc9a1 | ENSMUSG00000028854.9 |
| chr5 | 134312869 | 134316109 | 31.91637 | 110.38313 | 106.53261 | Promoter | 5 | 134237834 | 134314760 | 76927 | 2 | ENSMUST00 | 0 | Gtf2i | ENSMUSG00000060261.10 |
| chr5 | 14514361 | 14515748 | 31.91637 | 110.38313 | 106.53261 | Promoter | 5 | 14514918 | 14795993 | 281076 | 1 | ENSMUST00 | 0 | Pclo | ENSMUSG00000061601.10 |
| chr6 | 120463001 | 120464153 | 31.91637 | 110.38313 | 106.53261 | Promoter | 6 | 120463197 | 120483727 | 20531 | 1 | ENSMUST00 | 0 | Il17ra | ENSMUSG00000002897.3 |
| chr6 | 124964714 | 124966415 | 31.91637 | 110.38313 | 106.53261 | Promoter | 6 | 124958413 | 124965538 | 7126 | 2 | ENSMUST00 | 0 | Cops7a | ENSMUSG00000030127.10 |
| chr6 | 115852833 | 115854067 | 31.91637 | 110.38313 | 106.53261 | Promoter | 6 | 115840697 | 115853371 | 12675 | 2 | ENSMUST00 | 0 | Mbd4 | ENSMUSG00000030322.7 |
| chr6 | 54680902 | 54682590 | 31.91637 | 110.38313 | 106.53261 | Promoter | 6 | 54681624 | 54703851 | 22228 | 1 | ENSMUST00 | 0 | Mturn | ENSMUSG00000038065.8 |
| chr8 | 47532529 | 47534431 | 31.91637 | 110.38313 | 106.53261 | Promoter | 8 | 47533664 | 47551279 | 17616 | 1 | ENSMUST00 | 0 | Rwdd4a | ENSMUSG00000031568.11 |
| chr8 | 106209902 | 106211779 | 31.91637 | 110.38313 | 106.53261 | Promoter | 8 | 106200438 | 106210933 | 10496 | 2 | ENSMUST00 | 0 | Slc7a6os | ENSMUSG00000033106.5 |
| chr8 | 3676017 | 3677566 | 31.91637 | 110.38313 | 106.53261 | Promoter | 8 | 3676299 | 3681255 | 4957 | 1 | ENSMUST00 | 0 | Trappc5 | ENSMUSG00000040236.3 |
| chr9 | 53383794 | 53385192 | 31.91637 | 110.38313 | 106.53261 | Promoter | 9 | 53384023 | 53401867 | 17845 | 1 | ENSMUST00 | 0 | Kdelc2 | ENSMUSG00000034487.3 |
| chr10 | 17947067 | 17948455 | 34.79824 | 110.32398 | 106.47989 | Promoter | 10 | 17900466 | 17948067 | 47602 | 2 | ENSMUST00 | 0 | Heca | ENSMUSG00000039879.8 |
| chr11 | 31671232 | 31672519 | 34.79824 | 110.32398 | 106.47989 | Promoter | 11 | 31665181 | 31671885 | 6705 | 2 | ENSMUST00 | 0 | Bod1 | ENSMUSG00000044502.11 |
| chr19 | 4510390 | 4511178 | 34.79824 | 110.32398 | 106.47989 | Promoter | 19 | 4510472 | 4621752 | 111281 | 1 | ENSMUST00 | 0 | Pcx | ENSMUSG00000024892.10 |
| chr10 | 120208618 | 120209998 | 35.34781 | 110.24136 | 106.39787 | Promoter | 10 | 120208750 | 120224917 | 16168 | 1 | ENSMUST00 | 0 | Tmbim4 | ENSMUSG00000020225.5 |
| chr11 | 117653596 | 117656293 | 35.34781 | 110.24136 | 106.39787 | Promoter | 11 | 117654289 | 117763437 | 109149 | 1 | ENSMUST00 | 0 | Tnrc6c | ENSMUSG00000025571.8 |
| chr11 | 87357598 | 87359414 | 35.34781 | 110.24136 | 106.39787 | Promoter | 11 | 87226906 | 87359023 | 132118 | 2 | ENSMUST00 | 0 | Ppm1e | ENSMUSG00000046442.4 |
| chr14 | 75786215 | 75787868 | 35.34781 | 110.24136 | 106.39787 | Promoter | 14 | 75761999 | 75787037 | 25039 | 2 | ENSMUST00 | 0 | Slc25a30 | ENSMUSG00000022003.6 |
| chr2 | 30133098 | 30134976 | 35.34781 | 110.24136 | 106.39787 | Promoter | 2 | 30133746 | 30137296 | 3551 | 1 | ENSMUST00 | 0 | Tbc1d13 | ENSMUSG00000039678.7 |
| chr4 | 115827538 | 115828949 | 25.71568 | 110.22335 | 106.38094 | Promoter | 4 | 115828092 | 115836185 | 8094 | 1 | ENSMUST00 | 0 | Mob3c | ENSMUSG00000028709.3 |
| chr7 | 35646560 | 35647846 | 25.71568 | 110.22335 | 106.38094 | Promoter | 7 | 35641982 | 35647521 | 5540 | 2 | ENSMUST00 | 0 | Pdcd5 | ENSMUSG00000030417.10 |
| chr9 | 15305543 | 15307003 | 25.71568 | 110.22335 | 106.38094 | Promoter | 9 | 15283337 | 15306167 | 22831 | 2 | ENSMUST00 | 0 | 4931406C07I | ENSMUSG00000031938.9 |
| chr11 | 75172287 | 75174081 | 34.14626 | 110.2223 | 106.38018 | Promoter | 11 | 75173665 | 75173765 | 101 | 1 | ENSMUST00 | 0 | Mir132 | ENSMUSG00000065537.1 |
| chr12 | 75817574 | 75819634 | 34.14626 | 110.2223 | 106.38018 | Promoter | 12 | 75818134 | 75880506 | 62373 | 1 | ENSMUST00 | 0 | Syne2 | ENSMUSG00000063450.9 |
| chr15 | 98662100 | 98663226 | 34.14626 | 110.2223 | 106.38018 | Promoter | 15 | 98645134 | 98662889 | 17756 | 2 | ENSMUST00 | 0 | Ddx23 | ENSMUSG00000003360.9 |
| chr16 | 18088404 | 18089910 | 34.14626 | 110.2223 | 106.38018 | Promoter | 16 | 18060357 | 18089163 | 28807 | 2 | ENSMUST00 | 0 | Prodh | ENSMUSG00000003526.6 |
| chr19 | 4257116 | 4257744 | 34.14626 | 110.2223 | 106.38018 | Distal Interge | 19 | 4263818 | 4263878 | 61 | 2 | ENSMUST00 | 6134 | Mir6985 | ENSMUSG00000099290.1 |
| chr5 | 33651424 | 33652689 | 34.14626 | 110.2223 | 106.38018 | Promoter | 5 | 33634952 | 33652345 | 17394 | 2 | ENSMUST00 | 0 | Slbp | ENSMUSG00000004642.8 |
| chr5 | 76950025 | 76952599 | 34.14626 | 110.2223 | 106.38018 | Promoter | 5 | 76913249 | 76951572 | 38324 | 2 | ENSMUST00 | 0 | Ppat | ENSMUSG00000029246.9 |
| chr5 | 123343837 | 123346514 | 34.14626 | 110.2223 | 106.38018 | Promoter | 5 | 123344448 | 123374082 | 29635 | 1 | ENSMUST00 | 0 | Bcl7a | ENSMUSG00000029438.4 |
| chr7 | 143069312 | 143070519 | 34.14626 | 110.2223 | 106.38018 | Promoter | 7 | 143069341 | 143070546 | 1206 | 1 | ENSMUST00 | 0 | Tssc4 | ENSMUSG00000045752.7 |
| chr4 | 100095760 | 100097247 | 34.72914 | 110.20382 | 106.3632 | Promoter | 4 | 100095791 | 100444765 | 348975 | 1 | ENSMUST00 | 0 | Ror1 | ENSMUSG00000035305.5 |
| chr1 | 9545377 | 9546776 | 23.57607 | 110.17163 | 106.33151 | Promoter | 1 | 9545408 | 9547455 | 2048 | 1 | ENSMUST00 | 0 | Rrs1 | ENSMUSG00000061024.8 |
| chr9 | 114932730 | 114934401 | 30.12659 | 110.14252 | 106.30244 | Promoter | 9 | 114899341 | 114933987 | 34647 | 2 | ENSMUST00 | 0 | Gpd1l | ENSMUSG00000050627.8 |
| chr10 | 110918934 | 110921076 | 35.27562 | 110.11647 | 106.27654 | Promoter | 10 | 110920177 | 110939599 | 19423 | 1 | ENSMUST00 | 0 | Csrp2 | ENSMUSG00000020186.6 |
| chr18 | 24205060 | 24207222 | 35.27562 | 110.11647 | 106.27654 | Promoter | 18 | 24205344 | 24286818 | 81475 | 1 | ENSMUST00 | 0 | Galnt1 | ENSMUSG00000000420.10 |
| chr11 | 118353231 | 118355882 | 34.66032 | 110.08408 | 106.24455 | Promoter | 11 | 118301069 | 118355506 | 54438 | 2 | ENSMUST00 | 0 | Timp2 | ENSMUSG00000017466.4 |
| chr1 | 9797518 | 9799601 | 35.84351 | 110.06287 | 106.22365 | Promoter | 1 | 9798107 | 9900843 | 102737 | 1 | ENSMUST00 | 0 | Sgk3 | ENSMUSG00000025915.9 |
| chr1 | 172026431 | 172028021 | 35.84351 | 110.06287 | 106.22365 | Promoter | 1 | 172000960 | 172027295 | 26336 | 2 | ENSMUST00 | 0 | Vangl2 | ENSMUSG00000026556.10 |
| chr10 | 25359196 | 25361334 | 35.84351 | 110.06287 | 106.22365 | Promoter | 10 | 25359798 | 25523519 | 163722 | 1 | ENSMUST00 | 0 | Epb4.1l2 | ENSMUSG00000019978.10 |
| chr10 | 34417473 | 34419144 | 35.84351 | 110.06287 | 106.22365 | Promoter | 10 | 34303609 | 34418539 | 114931 | 2 | ENSMUST00 | 0 | Nt5dc1 | ENSMUSG00000039480.8 |
| chr11 | 53430099 | 53431593 | 35.84351 | 110.06287 | 106.22365 | Promoter | 11 | 53431023 | 53436902 | 5880 | 1 | ENSMUST00 | 0 | Gdf9 | ENSMUSG00000018238.6 |
| chr14 | 70598881 | 70600254 | 35.84351 | 110.06287 | 106.22365 | Promoter | 14 | 70583296 | 70599835 | 16540 | 2 | ENSMUST00 | 0 | Fam160b2 | ENSMUSG00000022095.9 |
| chr16 | 14361174 | 14362412 | 35.84351 | 110.06287 | 106.22365 | Promoter | 16 | 14361558 | 14473902 | 112345 | 1 | ENSMUST00 | 0 | Abcc1 | ENSMUSG00000023088.11 |
| chr5 | 112276637 | 112277521 | 35.84351 | 110.06287 | 106.22365 | Promoter | 5 | 112276691 | 112315361 | 38671 | 1 | ENSMUST00 | 0 | Tpst2 | ENSMUSG00000029344.9 |
| chr5 | 143817080 | 143818885 | 35.84351 | 110.06287 | 106.22365 | Promoter | 5 | 143817788 | 143904256 | 86469 | 1 | ENSMUST00 | 0 | Eif2ak1 | ENSMUSG00000029613.10 |
| chr6 | 51270156 | 51271460 | 35.84351 | 110.06287 | 106.22365 | Promoter | 6 | 51269812 | 51269910 | 99 | 2 | ENSMUST00 | -247 | Mir148a | ENSMUSG00000065505.1 |
| chr8 | 70872784 | 70874067 | 35.84351 | 110.06287 | 106.22365 | Promoter | 8 | 70868227 | 70873490 | 5264 | 2 | ENSMUST00 | 0 | Ccdc124 | ENSMUSG00000007721.5 |
| chr9 | 8002688 | 8005179 | 35.84351 | 110.06287 | 106.22365 | Promoter | 9 | 7931999 | 8004588 | 72590 | 2 | ENSMUST00 | 0 | Yap1 | ENSMUSG00000053110.8 |
| chr14 | 55590291 | 55592805 | 29.54372 | 110.04967 | 106.21311 | Promoter | 14 | 55591708 | 55603693 | 11986 | 1 | ENSMUST00 | 0 | Rnf31 | ENSMUSG00000047098.12 |
| chr11 | 120436715 | 120438083 | 32.29726 | 110.04213 | 106.20562 | Promoter | 11 | 120380370 | 120437708 | 57339 | 2 | ENSMUST00 | 0 | Nploc4 | ENSMUSG00000039703.10 |
| chr13 | 55099316 | 55100664 | 32.29726 | 110.04213 | 106.20562 | Promoter | 13 | 55027880 | 55100295 | 72416 | 2 | ENSMUST00 | 0 | Uimc1 | ENSMUSG00000025878.11 |
| chr8 | 8658981 | 8662087 | 32.29726 | 110.04213 | 106.20562 | Promoter | 8 | 8617434 | 8660773 | 43340 | 2 | ENSMUST00 | 0 | Efnb2 | ENSMUSG00000001300.10 |
| chr9 | 37347727 | 37349150 | 34.01509 | 109.99249 | 106.15683 | Promoter | 9 | 37275835 | 37348392 | 72558 | 2 | ENSMUST00 | 0 | Ccdc15 | ENSMUSG00000034303.6 |
| chr7 | 25218622 | 25222086 | 35.20372 | 109.99207 | 106.15649 | Promoter | 7 | 25221451 | 25227507 | 6057 | 1 | ENSMUST00 | 0 | Zfp526 | ENSMUSG00000046541.8 |
| chr15 | 79686797 | 79688562 | 34.59177 | 109.96477 | 106.12952 | Promoter | 15 | 79674250 | 79687872 | 13623 | 2 | ENSMUST00 | 0 | Josd1 | ENSMUSG00000022426.5 |
| chr9 | 66591521 | 66593918 | 35.77904 | 109.9518 | 106.11687 | Promoter | 9 | 66514637 | 66593142 | 78506 | 2 | ENSMUST00 | 0 | Usp3 | ENSMUSG00000032376.7 |
| chr5 | 146220447 | 146221984 | 33.94988 | 109.87815 | 106.04343 | Promoter | 5 | 146209192 | 146221555 | 12364 | 2 | ENSMUST00 | 0 | Rnf6 | ENSMUSG00000029634.10 |
| chr1 | 136624228 | 136626150 | 33.94791 | 109.87469 | 106.04007 | Promoter | 1 | 136624901 | 136630053 | 5153 | 1 | ENSMUST00 | 0 | Zfp281 | ENSMUSG00000041483.9 |
| chr11 | 54437572 | 54439590 | 33.94791 | 109.87469 | 106.04007 | Promoter | 11 | 54438199 | 54518235 | 80037 | 1 | ENSMUST00 | 0 | Fnip1 | ENSMUSG00000035992.10 |
| chr11 | 94989305 | 94994756 | 33.94791 | 109.87469 | 106.04007 | Promoter | 11 | 94991035 | 95006899 | 15865 | 1 | ENSMUST00 | 0 | Ppp1r9b | ENSMUSG00000038976.7 |

| | | | | | | | | | | | | | | | |
|---|---|---|---|---|---|---|---|---|---|---|---|---|---|---|---|
| chr11 | 5761289 | 5762854 | 33.94791 | 109.87469 | 106.04007 | Promoter | 11 | 5713417 | 5762376 | 48960 | 2 | ENSMUST00 | 0 | Urgcp | ENSMUSG00000049680.12 |
| chr12 | 117843263 | 117844345 | 33.94791 | 109.87469 | 106.04007 | Promoter | 12 | 117843515 | 117878706 | 35192 | 1 | ENSMUST00 | 0 | Cdca7l | ENSMUSG00000021175.10 |
| chr12 | 110849707 | 110851180 | 33.94791 | 109.87469 | 106.04007 | Promoter | 12 | 110850279 | 110869998 | 19720 | 1 | ENSMUST00 | 0 | Zfp839 | ENSMUSG00000021271.4 |
| chr13 | 63239341 | 63244158 | 33.94791 | 109.87469 | 106.04007 | Exon (ENSMU | 13 | 63282142 | 63326096 | 43955 | 1 | ENSMUST00 | -37984 | 2010111I01R | ENSMUSG00000021458.11 |
| chr13 | 54302347 | 54304259 | 33.94791 | 109.87469 | 106.04007 | Distal Interge | 13 | 54371349 | 54383917 | 12569 | 1 | ENSMUST00 | -67090 | Cplx2 | ENSMUSG00000025867.8 |
| chr13 | 107887775 | 107890104 | 33.94791 | 109.87469 | 106.04007 | Promoter | 13 | 107724618 | 107890064 | 165447 | 2 | ENSMUST00 | 0 | Zswim6 | ENSMUSG00000032846.9 |
| chr13 | 67525573 | 67526669 | 33.94791 | 109.87469 | 106.04007 | Promoter | 13 | 67515781 | 67526177 | 10397 | 2 | ENSMUST00 | 0 | Zfp87 | ENSMUSG00000097333.2 |
| chr15 | 43477125 | 43478158 | 33.94791 | 109.87469 | 106.04007 | Promoter | 15 | 43477229 | 43527777 | 50549 | 1 | ENSMUST00 | 0 | Emc2 | ENSMUSG00000022337.6 |
| chr15 | 35370761 | 35372510 | 33.94791 | 109.87469 | 106.04007 | Promoter | 15 | 35371546 | 35931228 | 559683 | 1 | ENSMUST00 | 0 | Vps13b | ENSMUSG00000037646.9 |
| chr16 | 32277025 | 32278537 | 33.94791 | 109.87469 | 106.04007 | Promoter | 16 | 32277459 | 32289194 | 11736 | 1 | ENSMUST00 | 0 | Rnf168 | ENSMUSG00000014074.12 |
| chr16 | 22160989 | 22164041 | 33.94791 | 109.87469 | 106.04007 | Promoter | 16 | 22059009 | 22163299 | 104291 | 2 | ENSMUST00 | 0 | Igf2bp2 | ENSMUSG00000033581.11 |
| chr17 | 23828084 | 23829590 | 33.94791 | 109.87469 | 106.04007 | Promoter | 17 | 23824740 | 23829109 | 4370 | 2 | ENSMUST00 | 0 | Tceb2 | ENSMUSG00000055839.5 |
| chr18 | 36793391 | 36794618 | 33.94791 | 109.87469 | 106.04007 | Promoter | 18 | 36793923 | 36799660 | 5738 | 1 | ENSMUST00 | 0 | Zmat2 | ENSMUSG00000001383.8 |
| chr19 | 5294471 | 5295473 | 33.94791 | 109.87469 | 106.04007 | Promoter | 19 | 5273923 | 5295455 | 21533 | 2 | ENSMUST00 | 0 | Sf3b2 | ENSMUSG00000024853.9 |
| chr19 | 5728681 | 5729823 | 33.94791 | 109.87469 | 106.04007 | Promoter | 19 | 5728087 | 5729633 | 1547 | 2 | ENSMUST00 | 0 | Fam89b | ENSMUSG00000024939.6 |
| chr19 | 53329381 | 53330780 | 33.94791 | 109.87469 | 106.04007 | Promoter | 19 | 53329413 | 53375810 | 46398 | 1 | ENSMUST00 | 0 | Mxi1 | ENSMUSG00000025025.8 |
| chr19 | 45363722 | 45365113 | 33.94791 | 109.87469 | 106.04007 | Promoter | 19 | 45363734 | 45533341 | 169608 | 1 | ENSMUST00 | 0 | Btrc | ENSMUSG00000025217.9 |
| chr2 | 120153103 | 120155266 | 33.94791 | 109.87469 | 106.04007 | Promoter | 2 | 120089175 | 120154606 | 65432 | 2 | ENSMUST00 | 0 | Ehd4 | ENSMUSG00000027293.8 |
| chr4 | 119108327 | 119109801 | 33.94791 | 109.87469 | 106.04007 | Promoter | 4 | 119108711 | 119137330 | 28620 | 1 | ENSMUST00 | 0 | Slc2a1 | ENSMUSG00000028645.5 |
| chr4 | 155066244 | 155067763 | 33.94791 | 109.87469 | 106.04007 | Promoter | 4 | 155067016 | 155072433 | 5418 | 1 | ENSMUST00 | 0 | Pex10 | ENSMUSG00000029047.8 |
| chr4 | 118178793 | 118180839 | 33.94791 | 109.87469 | 106.04007 | Promoter | 4 | 118136957 | 118180043 | 43087 | 2 | ENSMUST00 | 0 | Kdm4a | ENSMUSG00000033326.10 |
| chr6 | 116628227 | 116629310 | 33.94791 | 109.87469 | 106.04007 | Promoter | 6 | 116624016 | 116628999 | 4984 | 2 | ENSMUST00 | 0 | Zfp422 | ENSMUSG00000059878.7 |
| chr7 | 67371358 | 67373918 | 33.94791 | 109.87469 | 106.04007 | Promoter | 7 | 67231163 | 67372858 | 141696 | 2 | ENSMUST00 | 0 | Mef2a | ENSMUSG00000030557.11 |
| chr7 | 105632662 | 105634117 | 33.94791 | 109.87469 | 106.04007 | Promoter | 7 | 105604463 | 105633508 | 29046 | 2 | ENSMUST00 | 0 | Trim3 | ENSMUSG00000036989.10 |
| chr7 | 5037968 | 5039081 | 33.94791 | 109.87469 | 106.04007 | Promoter | 7 | 5034439 | 5038446 | 4008 | 2 | ENSMUST00 | 0 | Zfp784 | ENSMUSG00000043290.6 |
| chr7 | 105399362 | 105400078 | 33.94791 | 109.87469 | 106.04007 | Promoter | 7 | 105371211 | 105400054 | 28844 | 2 | ENSMUST00 | 0 | Fam160a2 | ENSMUSG00000044465.13 |
| chr8 | 46162924 | 46164395 | 33.94791 | 109.87469 | 106.04007 | Promoter | 8 | 46163651 | 46195590 | 31940 | 1 | ENSMUST00 | 0 | 4933411K20I | ENSMUSG00000031631.10 |
| chr12 | 112644742 | 112646905 | 35.13211 | 109.86813 | 106.03947 | Promoter | 12 | 112644828 | 112649152 | 4325 | 1 | ENSMUST00 | 0 | Siva1 | ENSMUSG00000064326.7 |
| chr13 | 9763025 | 9765435 | 35.13211 | 109.86813 | 106.03947 | Promoter | 13 | 9684836 | 9764423 | 79588 | 2 | ENSMUST00 | 0 | Zmynd11 | ENSMUSG00000021156.11 |
| chr3 | 96219035 | 96222442 | 35.13211 | 109.86813 | 106.03947 | Promoter | 3 | 96196863 | 96219709 | 22847 | 2 | ENSMUST00 | 0 | Bola1 | ENSMUSG00000015943.6 |
| chr2 | 20966651 | 20970031 | 34.52349 | 109.84589 | 106.01766 | Promoter | 2 | 20847919 | 20967721 | 119803 | 2 | ENSMUST00 | 0 | Arhgap21 | ENSMUSG00000036591.10 |
| chr9 | 50616356 | 50618031 | 34.52349 | 109.84589 | 106.01766 | Promoter | 9 | 50617321 | 50624997 | 7677 | 1 | ENSMUST00 | 0 | Pih1d2 | ENSMUSG00000000167.9 |
| chr1 | 61638253 | 61641326 | 26.89699 | 109.78668 | 105.95901 | Promoter | 1 | 61638824 | 62078259 | 439436 | 1 | ENSMUST00 | 0 | Pard3b | ENSMUSG00000052062.9 |
| chr10 | 12866850 | 12869855 | 26.89699 | 109.78668 | 105.95901 | Distal Interge | 10 | 12382188 | 12861735 | 479548 | 2 | ENSMUST00 | -5116 | Utrn | ENSMUSG00000019820.9 |
| chr13 | 81710251 | 81712428 | 26.89699 | 109.78668 | 105.95901 | Promoter | 13 | 81673843 | 81711013 | 37171 | 2 | ENSMUST00 | 0 | Polr3g | ENSMUSG00000035834.6 |
| chr17 | 35879709 | 35880552 | 26.89699 | 109.78668 | 105.95901 | Promoter | 17 | 35879819 | 35892670 | 12852 | 1 | ENSMUST00 | 0 | Dhx16 | ENSMUSG00000024422.9 |
| chr3 | 100161592 | 100163238 | 26.89699 | 109.78668 | 105.95901 | Promoter | 3 | 100162381 | 100206981 | 44601 | 1 | ENSMUST00 | 0 | Gdap2 | ENSMUSG00000027865.5 |
| chr6 | 100670213 | 100671482 | 26.89699 | 109.78668 | 105.95901 | Promoter | 6 | 100573081 | 100671157 | 98077 | 2 | ENSMUST00 | 0 | Shq1 | ENSMUSG00000035378.12 |
| chr1 | 13371872 | 13374991 | 29.92778 | 109.76675 | 105.9399 | Promoter | 1 | 13139105 | 13372439 | 233335 | 2 | ENSMUST00 | 0 | Ncoa2 | ENSMUSG00000005886.9 |
| chr1 | 85793126 | 85795106 | 29.92778 | 109.76675 | 105.9399 | Promoter | 1 | 85793441 | 85851576 | 58136 | 1 | ENSMUST00 | 0 | Cab39 | ENSMUSG00000036707.11 |
| chr1 | 92641003 | 92642384 | 29.92778 | 109.76675 | 105.9399 | Promoter | 1 | 92637145 | 92641923 | 4779 | 2 | ENSMUST00 | 0 | Myeov2 | ENSMUSG00000073616.5 |
| chr10 | 76441739 | 76443274 | 29.92778 | 109.76675 | 105.9399 | Promoter | 10 | 76351254 | 76442912 | 91659 | 2 | ENSMUST00 | 0 | Pcnt | ENSMUSG00000001151.9 |
| chr11 | 4848399 | 4849540 | 29.92778 | 109.76675 | 105.9399 | Promoter | 11 | 4765845 | 4849536 | 83692 | 2 | ENSMUST00 | 0 | Nf2 | ENSMUSG00000009073.11 |
| chr11 | 107188394 | 107189750 | 29.92778 | 109.76675 | 105.9399 | Promoter | 11 | 107166663 | 107189381 | 22719 | 2 | ENSMUST00 | 0 | Nol11 | ENSMUSG00000018433.9 |
| chr11 | 67454716 | 67457354 | 29.92778 | 109.76675 | 105.9399 | Promoter | 11 | 67455437 | 67683460 | 228024 | 1 | ENSMUST00 | 0 | Gas7 | ENSMUSG00000033066.10 |
| chr11 | 65787211 | 65788689 | 29.92778 | 109.76675 | 105.9399 | Promoter | 11 | 65688243 | 65788297 | 100055 | 2 | ENSMUST00 | 0 | Map2k4 | ENSMUSG00000033352.6 |
| chr11 | 69795520 | 69796207 | 29.92778 | 109.76675 | 105.9399 | Promoter | 11 | 69784036 | 69795943 | 11908 | 2 | ENSMUST00 | 0 | Chrnb1 | ENSMUSG00000041189.9 |
| chr16 | 4886161 | 4886997 | 29.92778 | 109.76675 | 105.9399 | Promoter | 16 | 4886100 | 4938296 | 52197 | 1 | ENSMUST00 | 62 | Mgrn1 | ENSMUSG00000022517.11 |
| chr16 | 90220255 | 90221966 | 29.92778 | 109.76675 | 105.9399 | Promoter | 16 | 90220742 | 90226322 | 5581 | 1 | ENSMUST00 | 0 | Sod1 | ENSMUSG00000022982.9 |
| chr2 | 127246948 | 127249138 | 29.92778 | 109.76675 | 105.9399 | Promoter | 2 | 127240938 | 127247816 | 6879 | 2 | ENSMUST00 | 0 | Ciao1 | ENSMUSG00000003662.5 |
| chr2 | 135658344 | 135660234 | 29.92778 | 109.76675 | 105.9399 | Promoter | 2 | 135659011 | 135742751 | 83741 | 1 | ENSMUST00 | 0 | Plcb4 | ENSMUSG00000039943.11 |
| chr3 | 87525082 | 87526619 | 29.92778 | 109.76675 | 105.9399 | Promoter | 3 | 87525407 | 87540156 | 14750 | 1 | ENSMUST00 | 0 | Etv3 | ENSMUSG00000003382.13 |
| chr3 | 84038826 | 84040973 | 29.92778 | 109.76675 | 105.9399 | Promoter | 3 | 83898287 | 84040161 | 141875 | 2 | ENSMUST00 | 0 | D930015E06I | ENSMUSG00000033767.9 |
| chr3 | 79284979 | 79287963 | 29.92778 | 109.76675 | 105.9399 | Distal Interge | 3 | 79345376 | 79464129 | 118754 | 1 | ENSMUST00 | -57413 | Gm17359 | ENSMUSG00000091685.1 |
| chr4 | 82858685 | 82859999 | 29.92778 | 109.76675 | 105.9399 | Promoter | 4 | 82798738 | 82859673 | 60936 | 2 | ENSMUST00 | 0 | Zdhhc21 | ENSMUSG00000028403.10 |
| chr4 | 132328653 | 132330129 | 29.92778 | 109.76675 | 105.9399 | Promoter | 4 | 132311763 | 132329489 | 17727 | 2 | ENSMUST00 | 0 | Trnau1ap | ENSMUSG00000028898.10 |
| chr4 | 46506749 | 46508135 | 29.92778 | 109.76675 | 105.9399 | 3' UTR | 4 | 46495155 | 46495285 | 131 | 1 | ENSMUST00 | 11595 | Gm22247 | ENSMUSG00000064410.1 |
| chr5 | 124628313 | 124629960 | 29.92778 | 109.76675 | 105.9399 | Promoter | 5 | 124629067 | 124722144 | 93078 | 1 | ENSMUST00 | 0 | Atp6v0a2 | ENSMUSG00000038023.8 |
| chr6 | 116207319 | 116208881 | 29.92778 | 109.76675 | 105.9399 | Promoter | 6 | 116208033 | 116262668 | 54636 | 1 | ENSMUST00 | 0 | Fam21 | ENSMUSG00000024104.9 |
| chr6 | 136803563 | 136805037 | 29.92778 | 109.76675 | 105.9399 | Promoter | 6 | 136801553 | 136804431 | 2879 | 2 | ENSMUST00 | 0 | Hist4h4 | ENSMUSG00000096010.1 |
| chr7 | 111028422 | 111029808 | 29.92778 | 109.76675 | 105.9399 | Promoter | 7 | 111028951 | 111056377 | 27427 | 1 | ENSMUST00 | 0 | Ctr9 | ENSMUSG00000005609.10 |
| chr7 | 66059897 | 66061516 | 29.92778 | 109.76675 | 105.9399 | Promoter | 7 | 66060299 | 66069526 | 9228 | 1 | ENSMUST00 | 0 | Snrpa1 | ENSMUSG00000030512.8 |
| chr9 | 108490636 | 108491934 | 29.92778 | 109.76675 | 105.9399 | Promoter | 9 | 108490676 | 108501697 | 11022 | 1 | ENSMUST00 | 0 | Usp19 | ENSMUSG00000006676.11 |
| chr13 | 100014961 | 100016231 | 31.57133 | 109.7552 | 105.93209 | Promoter | 13 | 99948533 | 100015639 | 67107 | 2 | ENSMUST00 | 0 | Mccc2 | ENSMUSG00000021646.8 |
| chr15 | 83148053 | 83149978 | 31.57133 | 109.7552 | 105.93209 | Promoter | 15 | 83125978 | 83149331 | 23354 | 2 | ENSMUST00 | 0 | Poldip3 | ENSMUSG00000041815.8 |
| chr18 | 36782287 | 36783970 | 31.57133 | 109.7552 | 105.93209 | Promoter | 18 | 36766530 | 36783205 | 16676 | 2 | ENSMUST00 | 0 | Hars | ENSMUSG00000001380.6 |
| chr5 | 65696596 | 65698919 | 35.06079 | 109.74464 | 105.9221 | Promoter | 5 | 65615260 | 65697856 | 82597 | 2 | ENSMUST00 | 0 | Pds5a | ENSMUSG00000029202.9 |
| chr10 | 128091204 | 128094095 | 33.28429 | 109.73968 | 105.91746 | Promoter | 10 | 128092783 | 128129303 | 36521 | 1 | ENSMUST00 | 0 | Baz2a | ENSMUSG00000040054.10 |
| chr14 | 45351108 | 45352008 | 33.28429 | 109.73968 | 105.91746 | Promoter | 14 | 45351473 | 45373585 | 22113 | 1 | ENSMUST00 | 0 | Styx | ENSMUSG00000053205.8 |
| chr5 | 22549343 | 22550862 | 33.28429 | 109.73968 | 105.91746 | Promoter | 5 | 22486485 | 22550304 | 63820 | 2 | ENSMUST00 | 0 | Orc5 | ENSMUSG00000029012.6 |
| chr6 | 87837982 | 87839537 | 33.28429 | 109.73968 | 105.91746 | Promoter | 6 | 87818447 | 87838759 | 20313 | 2 | ENSMUST00 | 0 | Isy1 | ENSMUSG00000030056.6 |
| chr16 | 17232910 | 17234712 | 35.62953 | 109.69414 | 105.87395 | Promoter | 16 | 17233572 | 17263430 | 29859 | 1 | ENSMUST00 | 0 | Hic2 | ENSMUSG00000050240.9 |
| chr6 | 148441860 | 148445082 | 35.62953 | 109.69414 | 105.87395 | Promoter | 6 | 148232430 | 148444389 | 211960 | 2 | ENSMUST00 | 0 | Tmtc1 | ENSMUSG00000030306.9 |
| chr10 | 80674492 | 80676385 | 34.38775 | 109.60938 | 105.79044 | Distal Interge | 10 | 80665332 | 80671976 | 6645 | 2 | ENSMUST00 | -2517 | Mknk2 | ENSMUSG00000020190.9 |
| chr3 | 69043976 | 69045799 | 34.38775 | 109.60938 | 105.79044 | Promoter | 3 | 69035288 | 69044755 | 9468 | 2 | ENSMUST00 | 0 | Trim59 | ENSMUSG00000034317.9 |
| chr11 | 116138092 | 116138868 | 28.28183 | 109.60786 | 105.79031 | Promoter | 11 | 116131817 | 116138868 | 7052 | 2 | ENSMUST00 | 0 | Mrpl38 | ENSMUSG00000020775.8 |
| chr13 | 22043145 | 22043866 | 28.28183 | 109.60786 | 105.79031 | Promoter | 13 | 22043214 | 22043676 | 463 | 1 | ENSMUST00 | 0 | Hist1h2bj | ENSMUSG00000069300.3 |
| chr13 | 41605781 | 41607035 | 28.28183 | 109.60786 | 105.79031 | Promoter | 13 | 41606216 | 41641347 | 35132 | 1 | ENSMUST00 | 0 | Tmem170b | ENSMUSG00000087370.2 |
| chr15 | 75940688 | 75941885 | 28.28183 | 109.60786 | 105.79031 | Promoter | 15 | 75940952 | 75949377 | 8426 | 1 | ENSMUST00 | 0 | Zfp623 | ENSMUSG00000050846.8 |
| chr16 | 20516331 | 20518870 | 28.28183 | 109.60786 | 105.79031 | Promoter | 16 | 20516982 | 20531650 | 14669 | 1 | ENSMUST00 | 0 | Dvl3 | ENSMUSG00000003233.6 |
| chr17 | 34116580 | 34119365 | 28.28183 | 109.60786 | 105.79031 | Promoter | 17 | 34112363 | 34118489 | 6127 | 2 | ENSMUST00 | 0 | Brd2 | ENSMUSG00000024335.14 |
| chr17 | 33759353 | 33760455 | 28.28183 | 109.60786 | 105.79031 | Promoter | 17 | 33746750 | 33760306 | 13557 | 2 | ENSMUST00 | 0 | Rab11b | ENSMUSG00000077450.7 |
| chr18 | 36559278 | 36560278 | 28.28183 | 109.60786 | 105.79031 | Promoter | 18 | 36559987 | 36583681 | 23695 | 1 | ENSMUST00 | 0 | Ankhd1 | ENSMUSG00000024483.13 |
| chr2 | 160730735 | 160732558 | 28.28183 | 109.60786 | 105.79031 | Promoter | 2 | 160731300 | 160762502 | 31203 | 1 | ENSMUST00 | 0 | Plcg1 | ENSMUSG00000016933.12 |
| chr3 | 133309986 | 133311194 | 28.28183 | 109.60786 | 105.79031 | Promoter | 3 | 133310110 | 133378078 | 67969 | 1 | ENSMUST00 | 0 | Ppa2 | ENSMUSG00000028013.11 |
| chr5 | 135369244 | 135370772 | 28.28183 | 109.60786 | 105.79031 | Promoter | 5 | 135369953 | 135376798 | 6846 | 1 | ENSMUST00 | 0 | Nsun5 | ENSMUSG00000000916.10 |
| chr5 | 36830014 | 36830930 | 28.28183 | 109.60786 | 105.79031 | Promoter | 5 | 36806921 | 36830653 | 23733 | 2 | ENSMUST00 | 0 | Man2b2 | ENSMUSG00000029119.4 |
| chr5 | 36465079 | 36466142 | 28.28183 | 109.60786 | 105.79031 | Promoter | 5 | 36464998 | 36474080 | 9083 | 1 | ENSMUST00 | 82 | Grpel1 | ENSMUSG00000029198.3 |
| chr6 | 40470723 | 40472224 | 28.28183 | 109.60786 | 105.79031 | Promoter | 6 | 40471352 | 40481823 | 10472 | 1 | ENSMUST00 | 0 | Ssbp1 | ENSMUSG00000029911.9 |
| chr18 | 67640413 | 67642182 | 35.55525 | 109.56605 | 105.75019 | Promoter | 18 | 67641599 | 67654162 | 12564 | 1 | ENSMUST00 | 0 | Psmg2 | ENSMUSG00000024537.4 |
| chr9 | 114639214 | 114640185 | 30.8931 | 109.53436 | 105.71891 | Promoter | 9 | 114585878 | 114640200 | 54323 | 2 | ENSMUST00 | 15 | Cnot10 | ENSMUSG00000056167.6 |
| chr12 | 112721427 | 112723136 | 34.91903 | 109.49904 | 105.68399 | Promoter | 12 | 112722174 | 112746591 | 24418 | 1 | ENSMUST00 | 0 | Cep170b | ENSMUSG00000072825.5 |
| chr7 | 102441375 | 102442573 | 34.91903 | 109.49904 | 105.68399 | Promoter | 7 | 102441695 | 102469771 | 28077 | 1 | ENSMUST00 | 0 | Rrm1 | ENSMUSG00000030978.9 |
| chr7 | 35554712 | 35556253 | 34.91903 | 109.49904 | 105.68399 | Promoter | 7 | 35547521 | 35555194 | 7674 | 2 | ENSMUST00 | 0 | Nudt19 | ENSMUSG00000034875.5 |
| chr11 | 59162784 | 59164239 | 34.32027 | 109.49174 | 105.67705 | Promoter | 11 | 59155369 | 59163739 | 8371 | 2 | ENSMUST00 | 0 | Iba57 | ENSMUSG00000049287.5 |
| chr6 | 92213776 | 92215455 | 34.32027 | 109.49174 | 105.67705 | Promoter | 6 | 92186712 | 92214925 | 28214 | 2 | ENSMUST00 | 0 | Zfyve20 | ENSMUSG00000014550.9 |
| chr18 | 31609020 | 31610283 | 34.84858 | 109.37691 | 105.56304 | Promoter | 18 | 31580168 | 31609902 | 29735 | 2 | ENSMUST00 | 0 | Slc25a46 | ENSMUSG00000024259.8 |
| chr7 | 122020495 | 122021521 | 34.84858 | 109.37691 | 105.56304 | Promoter | 7 | 121986722 | 122021167 | 34446 | 2 | ENSMUST00 | 0 | Gga2 | ENSMUSG00000030872.9 |
| chr7 | 132851815 | 132852798 | 34.84858 | 109.37691 | 105.56304 | Promoter | 7 | 132827457 | 132852673 | 25217 | 2 | ENSMUST00 | 0 | Mettl10 | ENSMUSG00000030960.11 |
| chr4 | 134467498 | 134469579 | 34.25306 | 109.37451 | 105.56091 | Promoter | 4 | 134468320 | 134473843 | 5524 | 1 | ENSMUST00 | 0 | Stmn1 | ENSMUSG00000028832.6 |
| chr8 | 22859496 | 22861135 | 34.25306 | 109.37451 | 105.56091 | Promoter | 8 | 22859535 | 22943259 | 83725 | 1 | ENSMUST00 | 0 | Kat6a | ENSMUSG00000031540.8 |
| chr2 | 30364179 | 30365016 | 30.25803 | 109.37234 | 105.55905 | Promoter | 2 | 30364233 | 30385521 | 21289 | 1 | ENSMUST00 | 0 | Fam73b | ENSMUSG00000026858.13 |
| chr7 | 127875160 | 127877572 | 30.25803 | 109.37234 | 105.55905 | Promoter | 7 | 127865170 | 127876828 | 11659 | 2 | ENSMUST00 | 0 | Zfp668 | ENSMUSG00000049728.9 |
| chr9 | 13827687 | 13828673 | 30.25803 | 109.37234 | 105.55905 | Promoter | 9 | 13827716 | 13846541 | 18826 | 1 | ENSMUST00 | 0 | Fam76b | ENSMUSG00000037808.7 |
| chr10 | 94514294 | 94516659 | 32.48404 | 109.34737 | 105.53431 | Promoter | 10 | 94514857 | 94590956 | 76100 | 1 | ENSMUST00 | 0 | Tmcc3 | ENSMUSG00000020023.12 |
| chr13 | 29855005 | 29855892 | 32.48404 | 109.34737 | 105.53431 | Promoter | 13 | 29191746 | 29855647 | 663902 | 2 | ENSMUST00 | 0 | Cdkal1 | ENSMUSG00000006191.12 |
| chr13 | 81782643 | 81784311 | 32.48404 | 109.34737 | 105.53431 | Promoter | 13 | 81783292 | 81797143 | 13852 | 1 | ENSMUST00 | 0 | Cetn3 | ENSMUSG00000021537.8 |
| chr13 | 73330672 | 73331957 | 32.48404 | 109.34737 | 105.53431 | Promoter | 13 | 73331009 | 73332175 | 1167 | 1 | ENSMUST00 | 0 | Mrpl36 | ENSMUSG00000021607.8 |
| chr18 | 67464224 | 67465553 | 32.48404 | 109.34737 | 105.53431 | Promoter | 18 | 67464849 | 67480581 | 15733 | 1 | ENSMUST00 | 0 | Slmo1 | ENSMUSG00000024530.7 |
| chr2 | 164822569 | 164823635 | 32.48404 | 109.34737 | 105.53431 | Promoter | 2 | 164822686 | 164826871 | 4186 | 1 | ENSMUST00 | 0 | Zswim1 | ENSMUSG00000017764.2 |
| chr7 | 38270329 | 38272115 | 32.48404 | 109.34737 | 105.53431 | Promoter | 7 | 38262820 | 38271348 | 8529 | 2 | ENSMUST00 | 0 | Pop4 | ENSMUSG00000030423.6 |
| chr8 | 105825929 | 105828096 | 32.48404 | 109.34737 | 105.53431 | Promoter | 8 | 105827744 | 105844676 | 16933 | 1 | ENSMUST00 | 0 | Tsnaxip1 | ENSMUSG00000031893.6 |
| chr9 | 119401529 | 119403875 | 32.48404 | 109.34737 | 105.53431 | Promoter | 9 | 119402444 | 119433501 | 31058 | 1 | ENSMUST00 | 0 | Acvr2b | ENSMUSG00000061393.8 |
| chr11 | 26386933 | 26387962 | 24.44916 | 109.34614 | 105.53412 | Promoter | 11 | 26387202 | 26422460 | 35259 | 1 | ENSMUST00 | 0 | Fancl | ENSMUSG00000004018.4 |
| chr19 | 53310158 | 53314217 | 35.4076 | 109.31139 | 105.49952 | Promoter | 19 | 53310506 | 53375810 | 65305 | 1 | ENSMUST00 | 0 | Mxi1 | ENSMUSG00000025025.8 |
| chr9 | 108783389 | 108785213 | 35.4076 | 109.31139 | 105.49952 | Distal Interge | 9 | 108795242 | 108795388 | 147 | 2 | ENSMUST00 | 10175 | Gm23034 | ENSMUSG00000088626.2 |
| chr10 | 93588768 | 93590059 | 34.77841 | 109.25525 | 105.44401 | Promoter | 10 | 93583029 | 93589706 | 6678 | 2 | ENSMUST00 | 0 | Snrpf | ENSMUSG00000020018.6 |
| chr8 | 116943321 | 116944464 | 29.10187 | 109.2007 | 105.38981 | Promoter | 8 | 116943393 | 116960445 | 17053 | 1 | ENSMUST00 | 0 | Atmin | ENSMUSG00000047388.10 |
| chr18 | 6489051 | 6491620 | 35.33424 | 109.18479 | 105.37405 | Promoter | 18 | 6435953 | 6490856 | 54904 | 2 | ENSMUST00 | 0 | Epc1 | ENSMUSG00000024240.8 |
| chr7 | 4811383 | 4812694 | 35.33424 | 109.18479 | 105.37405 | Promoter | 7 | 4808155 | 4812351 | 4197 | 2 | ENSMUST00 | 0 | Ube2s | ENSMUSG00000060860.7 |
| chr8 | 54529007 | 54530608 | 35.33424 | 109.18479 | 105.37405 | Promoter | 8 | 54520433 | 54529998 | 9566 | 2 | ENSMUST00 | 0 | Spcs3 | ENSMUSG00000054408.9 |
| chr3 | 122245528 | 122246629 | 34.11942 | 109.14127 | 105.33086 | Promoter | 3 | 122245557 | 122269984 | 24428 | 1 | ENSMUST00 | 0 | Gclm | ENSMUSG00000028124.10 |
| chr3 | 55112014 | 55113386 | 34.70852 | 109.13401 | 105.32389 | Promoter | 3 | 55112108 | 55137322 | 25215 | 1 | ENSMUST00 | 0 | Spg20 | ENSMUSG00000036580.10 |
| chr10 | 128231529 | 128232831 | 33.50033 | 109.08844 | 105.27855 | Promoter | 10 | 128232065 | 128252941 | 20877 | 1 | ENSMUST00 | 0 | Timeless | ENSMUSG00000039994.10 |
| chr13 | 60601515 | 60602641 | 33.50033 | 109.08844 | 105.27855 | Promoter | 13 | 60601947 | 60763185 | 161239 | 1 | ENSMUST00 | 0 | Dapk1 | ENSMUSG00000021559.8 |
| chr15 | 36281706 | 36283694 | 33.50033 | 109.08844 | 105.27855 | Promoter | 15 | 36239934 | 36283147 | 43214 | 2 | ENSMUST00 | 0 | Rnf19a | ENSMUSG00000022280.8 |
| chr4 | 89136849 | 89138351 | 33.50033 | 109.08844 | 105.27855 | Promoter | 4 | 89137122 | 89181081 | 43960 | 1 | ENSMUST00 | 0 | Mtap | ENSMUSG00000062937.5 |
| chr17 | 6654460 | 6656315 | 20.93824 | 109.0712 | 105.26324 | Promoter | 17 | 6648941 | 6655939 | 6999 | 2 | ENSMUST00 | 0 | Dynlt1f | ENSMUSG00000095677.1 |

| | | | | | | | | | | | | | | | |
|---|---|---|---|---|---|---|---|---|---|---|---|---|---|---|---|
| chr3 | 32736102 | 32736844 | 31.7391 | 109.03407 | 105.22643 | Promoter | 3 | 32736990 | 32751566 | 14577 | 1 | ENSMUST00 | -146 | Ndufb5 | ENSMUSG00000027673.7 |
| chr4 | 59545854 | 59549379 | 31.7391 | 109.03407 | 105.22643 | Promoter | 4 | 59471868 | 59549364 | 77497 | 2 | ENSMUST00 | 0 | Ptbp3 | ENSMUSG00000028382.10 |
| chr7 | 4792221 | 4794069 | 31.7391 | 109.03407 | 105.22643 | Promoter | 7 | 4792874 | 4794562 | 1689 | 1 | ENSMUST00 | 0 | Rpl28 | ENSMUSG00000030432.7 |
| chr11 | 60726256 | 60729003 | 34.63892 | 109.01322 | 105.20635 | Promoter | 11 | 60714143 | 60727263 | 13121 | 2 | ENSMUST00 | 0 | Flii | ENSMUSG00000002812.4 |
| chr12 | 91778477 | 91779493 | 34.63892 | 109.01322 | 105.20635 | Promoter | 12 | 91714082 | 91779137 | 65056 | 2 | ENSMUST00 | 0 | Ston2 | ENSMUSG00000020961.9 |
| chr14 | 26638056 | 26640031 | 34.63892 | 109.01322 | 105.20635 | Promoter | 14 | 26638074 | 26657258 | 19185 | 1 | ENSMUST00 | 0 | Arf4 | ENSMUSG00000021877.6 |
| chr2 | 105398455 | 105399983 | 34.63892 | 109.01322 | 105.20635 | Promoter | 2 | 105386291 | 105399319 | 13029 | 2 | ENSMUST00 | 0 | Rcn1 | ENSMUSG00000005973.6 |
| chr4 | 135493806 | 135495302 | 34.63892 | 109.01322 | 105.20635 | Promoter | 4 | 135494790 | 135519349 | 24560 | 1 | ENSMUST00 | 0 | Stpg1 | ENSMUSG00000028801.9 |
| chr4 | 155219895 | 155222771 | 34.63892 | 109.01322 | 105.20635 | Promoter | 4 | 155154075 | 155222535 | 68461 | 2 | ENSMUST00 | 0 | Ski | ENSMUSG00000029050.10 |
| chr6 | 39591947 | 39593585 | 34.63892 | 109.01322 | 105.20635 | Promoter | 6 | 39592574 | 39603382 | 10809 | 1 | ENSMUST00 | 0 | Ndufb2 | ENSMUSG00000002416.8 |
| chr11 | 51635019 | 51636571 | 33.43708 | 108.9771 | 105.17175 | Promoter | 11 | 51623671 | 51635647 | 11977 | 2 | ENSMUST00 | 0 | Rmnd5b | ENSMUSG00000001054.4 |
| chr11 | 79338660 | 79340697 | 35.18842 | 108.93304 | 105.12779 | Promoter | 11 | 79339693 | 79345186 | 5494 | 1 | ENSMUST00 | 0 | Nf1 | ENSMUSG00000020716.10 |
| chr10 | 116949413 | 116951139 | 34.56959 | 108.89287 | 105.0881 | Promoter | 10 | 116905784 | 116950380 | 44597 | 2 | ENSMUST00 | 0 | Rab3ip | ENSMUSG00000064181.5 |
| chr15 | 91572223 | 91573629 | 34.56959 | 108.89287 | 105.0881 | Promoter | 15 | 91267696 | 91573261 | 305566 | 2 | ENSMUST00 | 0 | Slc2a13 | ENSMUSG00000036298.10 |
| chr13 | 23368333 | 23369895 | 33.37407 | 108.86614 | 105.06176 | Promoter | 13 | 23353103 | 23369208 | 16106 | 2 | ENSMUST00 | 0 | Zfp322a | ENSMUSG00000046351.5 |
| chr13 | 18947976 | 18949900 | 35.11596 | 108.8079 | 105.00359 | Promoter | 13 | 18948371 | 19150913 | 202543 | 1 | ENSMUST00 | 0 | Amph | ENSMUSG00000021314.7 |
| chr8 | 107435790 | 107437401 | 35.11596 | 108.8079 | 105.00359 | Promoter | 8 | 107436398 | 107558594 | 122197 | 1 | ENSMUST00 | 0 | Wwp2 | ENSMUSG00000031930.10 |
| chr3 | 88141285 | 88144231 | 33.92091 | 108.79443 | 104.99046 | Promoter | 3 | 88142372 | 88172086 | 29715 | 1 | ENSMUST00 | 0 | Mef2d | ENSMUSG00000001419.12 |
| chr15 | 89591232 | 89592401 | 31.04394 | 108.79036 | 104.98655 | Promoter | 15 | 89582533 | 89591923 | 9391 | 2 | ENSMUST00 | 0 | Rabl2 | ENSMUSG00000022621.11 |
| chr4 | 123016294 | 123017787 | 31.04394 | 108.79036 | 104.98655 | Promoter | 4 | 123016597 | 123054949 | 38353 | 1 | ENSMUST00 | 0 | Trit1 | ENSMUSG00000028653.11 |
| chr7 | 141454899 | 141456677 | 31.04394 | 108.79036 | 104.98655 | Promoter | 7 | 141455198 | 141460743 | 5546 | 1 | ENSMUST00 | 0 | Pnpla2 | ENSMUSG00000025509.10 |
| chr9 | 20641954 | 20645680 | 31.04394 | 108.79036 | 104.98655 | Promoter | 9 | 20618118 | 20642313 | 24196 | 2 | ENSMUST00 | 0 | Fbxl12 | ENSMUSG00000066892.8 |
| chr4 | 106315477 | 106318241 | 33.31129 | 108.75554 | 104.95251 | Promoter | 4 | 106316213 | 106441317 | 125105 | 1 | ENSMUST00 | 0 | Usp24 | ENSMUSG00000028514.10 |
| chr14 | 62837574 | 62838781 | 23.34493 | 108.68487 | 104.88192 | Promoter | 14 | 62837690 | 62956886 | 119197 | 1 | ENSMUST00 | 0 | Wdfy2 | ENSMUSG00000014547.9 |
| chr1 | 189728162 | 189732470 | 25.45057 | 108.67924 | 104.87682 | Promoter | 1 | 189728268 | 189876689 | 148422 | 1 | ENSMUST00 | 0 | Ptpn14 | ENSMUSG00000026604.12 |
| chr11 | 119298749 | 119300721 | 25.45057 | 108.67924 | 104.87682 | Promoter | 11 | 119288363 | 119300089 | 11727 | 2 | ENSMUST00 | 0 | Eif4a3 | ENSMUSG00000025580.10 |
| chr12 | 8640412 | 8641670 | 25.45057 | 108.67924 | 104.87682 | Distal Interge | 12 | 8674134 | 8751454 | 77321 | 1 | ENSMUST00 | -32464 | Pum2 | ENSMUSG00000020594.9 |
| chr19 | 41800952 | 41802458 | 25.45057 | 108.67924 | 104.87682 | Promoter | 19 | 41766588 | 41802047 | 35460 | 2 | ENSMUST00 | 0 | Arhgap19 | ENSMUSG00000025154.9 |
| chr2 | 180582646 | 180583426 | 25.45057 | 108.67924 | 104.87682 | Promoter | 2 | 180582712 | 180586287 | 3576 | 1 | ENSMUST00 | 0 | Mrgbp | ENSMUSG00000027569.10 |
| chr1 | 36555717 | 36559135 | 31.54088 | 108.67455 | 104.87239 | Promoter | 1 | 36548642 | 36558381 | 9740 | 2 | ENSMUST00 | 0 | Sema4c | ENSMUSG00000026121.8 |
| chr1 | 132006980 | 132007372 | 31.54088 | 108.67455 | 104.87239 | Promoter | 1 | 132007607 | 132026448 | 18842 | 1 | ENSMUST00 | 0 | Elk4 | ENSMUSG00000026436.10 |
| chr1 | 20819645 | 20820373 | 31.54088 | 108.67455 | 104.87239 | Promoter | 1 | 20802968 | 20820312 | 17345 | 2 | ENSMUST00 | 0 | Mcm3 | ENSMUSG00000041859.10 |
| chr10 | 85871484 | 85872772 | 31.54088 | 108.67455 | 104.87239 | Promoter | 10 | 85871831 | 85889096 | 17266 | 1 | ENSMUST00 | 0 | Pwp1 | ENSMUSG00000001785.6 |
| chr10 | 127040927 | 127042546 | 31.54088 | 108.67455 | 104.87239 | Promoter | 10 | 127041414 | 127043182 | 1769 | 1 | ENSMUST00 | 0 | Mettl1 | ENSMUSG00000006732.9 |
| chr11 | 119943036 | 119944833 | 31.54088 | 108.67455 | 104.87239 | Promoter | 11 | 119943116 | 119960532 | 17417 | 1 | ENSMUST00 | 0 | Baiap2 | ENSMUSG00000025372.11 |
| chr11 | 120491501 | 120492705 | 31.54088 | 108.67455 | 104.87239 | Promoter | 11 | 120491840 | 120499187 | 7348 | 1 | ENSMUST00 | 0 | Slc25a10 | ENSMUSG00000025792.4 |
| chr12 | 76709398 | 76711013 | 31.54088 | 108.67455 | 104.87239 | Promoter | 12 | 76580488 | 76710547 | 130060 | 2 | ENSMUST00 | 0 | Sptb | ENSMUSG00000021061.10 |
| chr13 | 68581239 | 68582933 | 31.54088 | 108.67455 | 104.87239 | Promoter | 13 | 68582248 | 68592279 | 10032 | 1 | ENSMUST00 | 0 | Fastkd3 | ENSMUSG00000021532.10 |
| chr13 | 104177288 | 104179575 | 31.54088 | 108.67455 | 104.87239 | Promoter | 13 | 104142149 | 104178469 | 36321 | 2 | ENSMUST00 | 0 | Trappc13 | ENSMUSG00000021711.11 |
| chr13 | 117023675 | 117025980 | 31.54088 | 108.67455 | 104.87239 | Promoter | 13 | 116854824 | 117025516 | 170693 | 2 | ENSMUST00 | 0 | Parp8 | ENSMUSG00000021725.8 |
| chr14 | 25457181 | 25461653 | 31.54088 | 108.67455 | 104.87239 | Promoter | 14 | 25459185 | 25538492 | 79308 | 1 | ENSMUST00 | 0 | Zmiz1 | ENSMUSG00000007817.9 |
| chr14 | 118136391 | 118138211 | 31.54088 | 108.67455 | 104.87239 | Promoter | 14 | 118137127 | 118164232 | 27106 | 1 | ENSMUST00 | 0 | Gpr180 | ENSMUSG00000022131.2 |
| chr14 | 47276211 | 47277770 | 31.54088 | 108.67455 | 104.87239 | Promoter | 14 | 47240944 | 47276790 | 35847 | 2 | ENSMUST00 | 0 | Wdhd1 | ENSMUSG00000037572.11 |
| chr17 | 28176587 | 28177973 | 31.54088 | 108.67455 | 104.87239 | Promoter | 17 | 28177207 | 28205886 | 28680 | 1 | ENSMUST00 | 0 | Zfp523 | ENSMUSG00000024220.8 |
| chr17 | 66446496 | 66450714 | 31.54088 | 108.67455 | 104.87239 | Promoter | 17 | 66336982 | 66449750 | 112769 | 2 | ENSMUST00 | 0 | Mtcl1 | ENSMUSG00000052105.11 |
| chr19 | 44106519 | 44108483 | 31.54088 | 108.67455 | 104.87239 | Promoter | 19 | 44073335 | 44107480 | 34146 | 2 | ENSMUST00 | 0 | Chuk | ENSMUSG00000025199.11 |
| chr19 | 40894179 | 40895684 | 31.54088 | 108.67455 | 104.87239 | Promoter | 19 | 40894705 | 40917947 | 23243 | 1 | ENSMUST00 | 0 | Zfp518a | ENSMUSG00000049164.6 |
| chr2 | 90903727 | 90905438 | 31.54088 | 108.67455 | 104.87239 | Promoter | 2 | 90904740 | 90910561 | 5822 | 1 | ENSMUST00 | 0 | Kbtbd4 | ENSMUSG00000005505.7 |
| chr2 | 32757165 | 32757998 | 31.54088 | 108.67455 | 104.87239 | Promoter | 2 | 32757234 | 32762244 | 5011 | 1 | ENSMUST00 | 0 | Tor2a | ENSMUSG00000009563.11 |
| chr3 | 36150590 | 36151824 | 31.54088 | 108.67455 | 104.87239 | Distal Interge | 3 | 36159522 | 36170335 | 10814 | 1 | ENSMUST00 | -7698 | D3Ertd254e | ENSMUSG00000033883.7 |
| chr4 | 145193812 | 145195132 | 31.54088 | 108.67455 | 104.87239 | Promoter | 4 | 144972622 | 145195005 | 222384 | 2 | ENSMUST00 | 0 | Vps13d | ENSMUSG00000020220.11 |
| chr4 | 154295799 | 154300705 | 31.54088 | 108.67455 | 104.87239 | Promoter | 4 | 154278486 | 154299892 | 21407 | 2 | ENSMUST00 | 0 | Arhgef16 | ENSMUSG00000029032.7 |
| chr5 | 135777646 | 135778401 | 31.54088 | 108.67455 | 104.87239 | Promoter | 5 | 135747220 | 135778238 | 31019 | 2 | ENSMUST00 | 0 | Styxl1 | ENSMUSG00000019178.12 |
| chr5 | 52189080 | 52191292 | 31.54088 | 108.67455 | 104.87239 | Promoter | 5 | 52150209 | 52190519 | 40311 | 2 | ENSMUST00 | 0 | Dhx15 | ENSMUSG00000029169.7 |
| chr5 | 69556482 | 69557799 | 31.54088 | 108.67455 | 104.87239 | Promoter | 5 | 69556924 | 69573104 | 16181 | 1 | ENSMUST00 | 0 | Guf1 | ENSMUSG00000029208.11 |
| chr5 | 136198124 | 136199653 | 31.54088 | 108.67455 | 104.87239 | Promoter | 5 | 136178127 | 136198963 | 20837 | 2 | ENSMUST00 | 0 | Prkrip1 | ENSMUSG00000039737.6 |
| chr7 | 19507045 | 19509433 | 31.54088 | 108.67455 | 104.87239 | Promoter | 7 | 19508670 | 19516138 | 7469 | 1 | ENSMUST00 | 0 | Trappc6a | ENSMUSG00000002043.11 |
| chr7 | 38106694 | 38107599 | 31.54088 | 108.67455 | 104.87239 | Promoter | 7 | 38097984 | 38107534 | 9551 | 2 | ENSMUST00 | 0 | Ccne1 | ENSMUSG00000002068.11 |
| chr7 | 19714708 | 19715498 | 31.54088 | 108.67455 | 104.87239 | Promoter | 7 | 19701313 | 19715438 | 14126 | 2 | ENSMUST00 | 0 | Tomm40 | ENSMUSG00000002984.11 |
| chr7 | 65861608 | 65863627 | 31.54088 | 108.67455 | 104.87239 | Promoter | 7 | 65861734 | 65952984 | 91251 | 1 | ENSMUST00 | 0 | Pcsk6 | ENSMUSG00000030513.9 |
| chr7 | 126565521 | 126567070 | 31.54088 | 108.67455 | 104.87239 | Promoter | 7 | 126546455 | 126566411 | 19957 | 2 | ENSMUST00 | 0 | Eif3c | ENSMUSG00000030738.10 |
| chr8 | 33599519 | 33601159 | 31.54088 | 108.67455 | 104.87239 | Promoter | 8 | 33599621 | 33619794 | 20174 | 1 | ENSMUST00 | 0 | Ppp2cb | ENSMUSG00000009630.9 |
| chr8 | 111875830 | 111876802 | 31.54088 | 108.67455 | 104.87239 | Promoter | 8 | 111864898 | 111876675 | 11778 | 2 | ENSMUST00 | 0 | Tmem170 | ENSMUSG00000031953.1 |
| chr9 | 7183574 | 7185575 | 31.54088 | 108.67455 | 104.87239 | Promoter | 9 | 7184566 | 7207031 | 22466 | 1 | ENSMUST00 | 0 | Dcun1d5 | ENSMUSG00000032002.9 |
| chr9 | 102352850 | 102354848 | 31.54088 | 108.67455 | 104.87239 | Promoter | 9 | 101922128 | 102354693 | 432566 | 2 | ENSMUST00 | 0 | Ephb1 | ENSMUSG00000032537.9 |
| chr9 | 106821829 | 106823113 | 31.54088 | 108.67455 | 104.87239 | Promoter | 9 | 106821874 | 106880443 | 58570 | 1 | ENSMUST00 | 0 | Vprbp | ENSMUSG00000040325.11 |
| chr1 | 34010872 | 34012641 | 32.67761 | 108.66347 | 104.86716 | Promoter | 1 | 34011822 | 34308661 | 296840 | 1 | ENSMUST00 | 0 | Dst | ENSMUSG00000026131.12 |
| chr10 | 88826000 | 88827293 | 32.67761 | 108.66347 | 104.86716 | Promoter | 10 | 88746607 | 88826814 | 80208 | 2 | ENSMUST00 | 0 | Utp20 | ENSMUSG00000004356.7 |
| chr11 | 83302172 | 83303781 | 32.67761 | 108.66347 | 104.86716 | Promoter | 11 | 83298030 | 83302586 | 4557 | 2 | ENSMUST00 | 0 | Pex12 | ENSMUSG00000018733.5 |
| chr14 | 25841710 | 25843211 | 32.67761 | 108.66347 | 104.86716 | Promoter | 14 | 25842156 | 25886804 | 44649 | 1 | ENSMUST00 | 0 | Anxa11 | ENSMUSG00000021866.10 |
| chr19 | 8735275 | 8736504 | 32.67761 | 108.66347 | 104.86716 | Promoter | 19 | 8735839 | 8740623 | 4785 | 1 | ENSMUST00 | 0 | Wdr74 | ENSMUSG00000042729.9 |
| chr3 | 63928887 | 63929594 | 32.67761 | 108.66347 | 104.86716 | Promoter | 3 | 63914684 | 63929472 | 14789 | 2 | ENSMUST00 | 0 | E130311K13I | ENSMUSG00000048581.7 |
| chr7 | 114115691 | 114118263 | 32.67761 | 108.66347 | 104.86716 | Promoter | 7 | 114046785 | 114117781 | 70997 | 2 | ENSMUST00 | 0 | Rras2 | ENSMUSG00000055723.5 |
| chr14 | 70288628 | 70289902 | 34.43176 | 108.65345 | 104.85863 | Promoter | 14 | 70217898 | 70289449 | 71552 | 2 | ENSMUST00 | 0 | Ppp3cc | ENSMUSG00000022092.10 |
| chr1 | 151343938 | 151345916 | 30.39373 | 108.60838 | 104.8139 | Promoter | 1 | 151344477 | 151364422 | 19946 | 1 | ENSMUST00 | 0 | Ivns1abp | ENSMUSG00000023150.9 |
| chr1 | 86358246 | 86360433 | 30.39373 | 108.60838 | 104.8139 | Promoter | 1 | 86344719 | 86359400 | 14682 | 2 | ENSMUST00 | 0 | Ncl | ENSMUSG00000026234.7 |
| chr12 | 20920326 | 20921715 | 30.39373 | 108.60838 | 104.8139 | Promoter | 12 | 20875640 | 20921262 | 45623 | 2 | ENSMUST00 | 0 | 3110053B16I | ENSMUSG00000085642.2 |
| chr14 | 87415190 | 87418026 | 30.39373 | 108.60838 | 104.8139 | Promoter | 14 | 87416639 | 87514766 | 98128 | 1 | ENSMUST00 | 0 | Tdrd3 | ENSMUSG00000022019.10 |
| chr3 | 81932356 | 81933286 | 30.39373 | 108.60838 | 104.8139 | Promoter | 3 | 81932601 | 81956725 | 24125 | 1 | ENSMUST00 | 0 | Ctso | ENSMUSG00000028015.2 |
| chr15 | 83555016 | 83556174 | 34.97194 | 108.55902 | 104.76495 | Promoter | 15 | 83546797 | 83555711 | 8915 | 2 | ENSMUST00 | 0 | Mcat | ENSMUSG00000048755.8 |
| chr11 | 29525995 | 29527141 | 33.18644 | 108.53539 | 104.74159 | Promoter | 11 | 29526408 | 29533397 | 6990 | 1 | ENSMUST00 | 0 | Mtif2 | ENSMUSG00000020459.9 |
| chr11 | 116412272 | 116413687 | 34.36326 | 108.53439 | 104.74065 | Promoter | 11 | 116336353 | 116413004 | 76652 | 2 | ENSMUST00 | 0 | Rnf157 | ENSMUSG00000052949.9 |
| chr18 | 60848151 | 60849055 | 34.36326 | 108.53439 | 104.74065 | Promoter | 18 | 60813755 | 60848929 | 35175 | 2 | ENSMUST00 | 0 | Tcof1 | ENSMUSG00000024613.11 |
| chr7 | 122100946 | 122102271 | 34.36326 | 108.53439 | 104.74065 | Promoter | 7 | 122085403 | 122101886 | 16484 | 2 | ENSMUST00 | 0 | Ndufab1 | ENSMUSG00000030869.6 |
| chr2 | 118745142 | 118747076 | 33.7247 | 108.45113 | 104.65779 | Promoter | 2 | 118745758 | 118748810 | 3053 | 1 | ENSMUST00 | 0 | 5430417L22F | ENSMUSG00000074918.4 |
| chr4 | 52438330 | 52439908 | 33.7247 | 108.45113 | 104.65779 | Promoter | 4 | 52439243 | 52488260 | 49018 | 1 | ENSMUST00 | 0 | Smc2 | ENSMUSG00000028312.14 |
| chr4 | 116650295 | 116652083 | 33.7247 | 108.45113 | 104.65779 | Promoter | 4 | 116636510 | 116651680 | 15171 | 2 | ENSMUST00 | 0 | Akr1a1 | ENSMUSG00000028692.9 |
| chr9 | 52047271 | 52050025 | 33.7247 | 108.45113 | 104.65779 | Promoter | 9 | 52047941 | 52075177 | 27237 | 1 | ENSMUST00 | 0 | Rdx | ENSMUSG00000032050.11 |
| chr9 | 78587317 | 78588583 | 33.7247 | 108.45113 | 104.65779 | Promoter | 9 | 78536488 | 78588041 | 51554 | 2 | ENSMUST00 | 0 | Slc17a5 | ENSMUSG00000049624.8 |
| chr11 | 116827118 | 116828605 | 34.90036 | 108.43528 | 104.64344 | Promoter | 11 | 116803062 | 116828046 | 24985 | 2 | ENSMUST00 | 0 | Mxra7 | ENSMUSG00000020814.8 |
| chr15 | 27629823 | 27631338 | 34.90036 | 108.43528 | 104.64344 | Promoter | 15 | 27605919 | 27630693 | 24775 | 2 | ENSMUST00 | 0 | Otulin | ENSMUSG00000046034.7 |
| chr19 | 6241598 | 6242537 | 34.90036 | 108.43528 | 104.64344 | Promoter | 19 | 6241668 | 6262335 | 20668 | 1 | ENSMUST00 | 0 | Atg2a | ENSMUSG00000024773.9 |
| chr2 | 60123837 | 60126567 | 34.90036 | 108.43528 | 104.64344 | Promoter | 2 | 59899363 | 60125697 | 226335 | 2 | ENSMUST00 | 0 | Baz2b | ENSMUSG00000026987.11 |
| chr7 | 138908947 | 138910265 | 34.90036 | 108.43528 | 104.64344 | Promoter | 7 | 138890836 | 138909506 | 18671 | 2 | ENSMUST00 | 0 | Bnip3 | ENSMUSG00000078566.3 |
| chr11 | 82763563 | 82765645 | 34.29503 | 108.41577 | 104.62504 | Promoter | 11 | 82719250 | 82764306 | 45057 | 2 | ENSMUST00 | 0 | Cct6b | ENSMUSG00000020698.6 |
| chr1 | 93803311 | 93804724 | 31.91277 | 108.3221 | 104.53201 | Promoter | 1 | 93803965 | 93822095 | 18131 | 1 | ENSMUST00 | 0 | Ing5 | ENSMUSG00000026283.8 |
| chr12 | 8973605 | 8974550 | 31.91277 | 108.3221 | 104.53201 | Promoter | 12 | 8973892 | 9028847 | 54956 | 1 | ENSMUST00 | 0 | Wdr35 | ENSMUSG00000066643.7 |
| chr19 | 41932612 | 41934160 | 31.91277 | 108.3221 | 104.53201 | Promoter | 19 | 41933472 | 41944095 | 10624 | 1 | ENSMUST00 | 0 | Zdhhc16 | ENSMUSG00000025157.7 |
| chr7 | 24971656 | 24972749 | 31.91277 | 108.3221 | 104.53201 | Promoter | 7 | 24969750 | 24972728 | 2979 | 2 | ENSMUST00 | 0 | Rabac1 | ENSMUSG00000003380.10 |
| chr10 | 57486338 | 57487441 | 34.82909 | 108.31201 | 104.52283 | Promoter | 10 | 57486385 | 57513133 | 26749 | 1 | ENSMUST00 | 0 | Hsf2 | ENSMUSG00000019878.7 |
| chr3 | 69721465 | 69723120 | 34.22706 | 108.29755 | 104.50865 | Promoter | 3 | 69721985 | 69724461 | 2477 | 1 | ENSMUST00 | 0 | Nmd3 | ENSMUSG00000027787.9 |
| chr5 | 129724482 | 129726168 | 34.22706 | 108.29755 | 104.50865 | Promoter | 5 | 129725063 | 129758327 | 33265 | 1 | ENSMUST00 | 0 | Gbas | ENSMUSG00000029432.7 |
| chr6 | 43665308 | 43666768 | 34.22706 | 108.29755 | 104.50865 | Promoter | 6 | 43345001 | 43666278 | 321278 | 2 | ENSMUST00 | 0 | Tpk1 | ENSMUSG00000029735.12 |
| chr18 | 73814766 | 73816183 | 32.44109 | 108.24224 | 104.45363 | Promoter | 18 | 73770040 | 73815392 | 45353 | 2 | ENSMUST00 | 0 | Me2 | ENSMUSG00000024556.3 |
| chr10 | 82622357 | 82623644 | 35.38397 | 108.24114 | 104.45254 | Promoter | 10 | 82619851 | 82622926 | 3076 | 2 | ENSMUST00 | 0 | 1190007I07R | ENSMUSG00000063320.6 |
| chr11 | 76434132 | 76437894 | 35.38397 | 108.24114 | 104.45254 | Promoter | 11 | 76423048 | 76437220 | 14173 | 2 | ENSMUST00 | 0 | Abr | ENSMUSG00000017631.13 |
| chr12 | 111165749 | 111167378 | 35.38397 | 108.24114 | 104.45254 | Promoter | 12 | 111166370 | 111267153 | 100784 | 1 | ENSMUST00 | 0 | Traf3 | ENSMUSG00000021277.11 |
| chr13 | 48869692 | 48872023 | 35.38397 | 108.24114 | 104.45254 | Promoter | 13 | 48801750 | 48870885 | 69136 | 2 | ENSMUST00 | 0 | Phf2 | ENSMUSG00000038025.7 |
| chr13 | 55592693 | 55594174 | 35.38397 | 108.24114 | 104.45254 | Promoter | 13 | 55593135 | 55597663 | 4529 | 1 | ENSMUST00 | 0 | Tmed9 | ENSMUSG00000058569.7 |
| chr14 | 70350670 | 70351850 | 35.38397 | 108.24114 | 104.45254 | Promoter | 14 | 70303469 | 70351425 | 47957 | 2 | ENSMUST00 | 0 | Slc39a14 | ENSMUSG00000022094.10 |
| chr19 | 32485186 | 32486937 | 35.38397 | 108.24114 | 104.45254 | Promoter | 19 | 32485769 | 32515370 | 29602 | 1 | ENSMUST00 | 0 | Minpp1 | ENSMUSG00000024896.8 |
| chr2 | 164496980 | 164498263 | 35.38397 | 108.24114 | 104.45254 | Promoter | 2 | 164497520 | 164508301 | 10782 | 1 | ENSMUST00 | 0 | Pigt | ENSMUSG00000017721.10 |
| chr2 | 30205248 | 30206666 | 35.38397 | 108.24114 | 104.45254 | Promoter | 2 | 30185124 | 30205847 | 20724 | 2 | ENSMUST00 | 0 | Ccbl1 | ENSMUSG00000039648.9 |
| chr4 | 132768278 | 132769206 | 35.38397 | 108.24114 | 104.45254 | Promoter | 4 | 132768332 | 132776710 | 8379 | 1 | ENSMUST00 | 0 | Rpa2 | ENSMUSG00000028884.9 |
| chr5 | 108693790 | 108695271 | 35.38397 | 108.24114 | 104.45254 | Promoter | 5 | 108694229 | 108706950 | 12722 | 1 | ENSMUST00 | 0 | Fgfrl1 | ENSMUSG00000008090.9 |
| chr5 | 134098943 | 134100390 | 35.38397 | 108.24114 | 104.45254 | Promoter | 5 | 134099711 | 134144343 | 44633 | 1 | ENSMUST00 | 0 | Gatsl2 | ENSMUSG00000015944.8 |
| chr5 | 142813148 | 142817670 | 35.38397 | 108.24114 | 104.45254 | Promoter | 5 | 142754789 | 142817662 | 62874 | 2 | ENSMUST00 | 0 | Tnrc18 | ENSMUSG00000039477.11 |
| chr5 | 142463474 | 142464783 | 35.38397 | 108.24114 | 104.45254 | Promoter | 5 | 142463931 | 142478715 | 14785 | 1 | ENSMUST00 | 0 | Ap5z1 | ENSMUSG00000039623.9 |
| chr5 | 38683969 | 38685238 | 35.38397 | 108.24114 | 104.45254 | Promoter | 5 | 38668484 | 38684750 | 16267 | 2 | ENSMUST00 | 0 | Zfp518b | ENSMUSG00000046572.5 |
| chr7 | 25247802 | 25251702 | 35.38397 | 108.24114 | 104.45254 | Promoter | 7 | 25242561 | 25250761 | 8201 | 2 | ENSMUST00 | 0 | Erf | ENSMUSG00000040857.10 |
| chr8 | 83651941 | 83653788 | 35.38397 | 108.24114 | 104.45254 | Promoter | 8 | 83652678 | 83664788 | 12111 | 1 | ENSMUST00 | 0 | Gipc1 | ENSMUSG00000019433.8 |
| chr11 | 61761317 | 61762420 | 33.59515 | 108.2242 | 104.43813 | Promoter | 11 | 61729650 | 61762088 | 32439 | 2 | ENSMUST00 | 0 | Prpsap2 | ENSMUSG00000020528.9 |
| chr1 | 179545202 | 179547773 | 26.61085 | 108.21047 | 104.42454 | Promoter | 1 | 179528055 | 179546267 | 18213 | 2 | ENSMUST00 | 0 | Tfb2m | ENSMUSG00000026492.6 |
| chr10 | 127676990 | 127677796 | 26.61085 | 108.21047 | 104.42454 | Promoter | 10 | 127677064 | 127695168 | 18105 | 1 | ENSMUST00 | 0 | Tmem194 | ENSMUSG00000040195.6 |
| chr11 | 20543047 | 20544428 | 26.61085 | 108.21047 | 104.42454 | Promoter | 11 | 20543253 | 20597020 | 53768 | 1 | ENSMUST00 | 0 | Sertad2 | ENSMUSG00000049800.8 |
| chr11 | 50236154 | 50237650 | 26.61085 | 108.21047 | 104.42454 | Promoter | 11 | 50237009 | 50288839 | 51831 | 1 | ENSMUST00 | 0 | Gm26542 | ENSMUSG00000097887.1 |
| chr15 | 73183476 | 73185498 | 26.61085 | 108.21047 | 104.42454 | Promoter | 15 | 73095844 | 73184840 | 88997 | 2 | ENSMUST00 | 0 | Ago2 | ENSMUSG00000036698.10 |
| chr16 | 92057556 | 92059912 | 26.61085 | 108.21047 | 104.42454 | Promoter | 16 | 92058270 | 92112227 | 53958 | 1 | ENSMUST00 | 0 | Mrps6 | ENSMUSG00000039680.9 |

| | | | | | | | | | | | | | | | |
|---|---|---|---|---|---|---|---|---|---|---|---|---|---|---|---|
| chr16 | 21793830 | 21795312 | 26.61085 | 108.21047 | 104.42454 | Promoter | 16 | 21794347 | 21806123 | 11777 | 1 | ENSMUST00 | 0 | 1300002E11I | ENSMUSG00000097290.2 |
| chr17 | 25941413 | 25942368 | 26.61085 | 108.21047 | 104.42454 | Promoter | 17 | 25926428 | 25941883 | 15456 | 2 | ENSMUST00 | 0 | Pigq | ENSMUSG00000025728.9 |
| chr19 | 53944799 | 53946110 | 26.61085 | 108.21047 | 104.42454 | Promoter | 19 | 53944876 | 54033268 | 88393 | 1 | ENSMUST00 | 0 | Shoc2 | ENSMUSG00000024976.8 |
| chr4 | 53825699 | 53826939 | 26.61085 | 108.21047 | 104.42454 | Promoter | 4 | 53826045 | 53862019 | 35975 | 1 | ENSMUST00 | 0 | Tmem38b | ENSMUSG00000028420.8 |
| chr5 | 34549141 | 34550254 | 26.61085 | 108.21047 | 104.42454 | Promoter | 5 | 34549576 | 34563625 | 14050 | 1 | ENSMUST00 | 0 | Sh3bp2 | ENSMUSG00000054520.10 |
| chr9 | 71484364 | 71486018 | 26.61085 | 108.21047 | 104.42454 | Promoter | 9 | 71478437 | 71485935 | 7499 | 2 | ENSMUST00 | 0 | Polr2m | ENSMUSG00000032199.7 |
| chr11 | 101094903 | 101095918 | 34.15937 | 108.17976 | 104.39504 | Promoter | 11 | 101091823 | 101095436 | 3614 | 2 | ENSMUST00 | 0 | Psmc3ip | ENSMUSG00000019303.9 |
| chr2 | 35978342 | 35980211 | 34.15937 | 108.17976 | 104.39504 | Promoter | 2 | 35751241 | 35979624 | 228384 | 2 | ENSMUST00 | 0 | Ttll11 | ENSMUSG00000026885.8 |
| chr5 | 99976691 | 99979009 | 35.32033 | 108.13151 | 104.34708 | Promoter | 5 | 99958426 | 99978938 | 20513 | 2 | ENSMUST00 | 0 | Hnrnpd | ENSMUSG00000000568.10 |
| chr1 | 64121784 | 64122677 | 33.5288 | 108.10788 | 104.32592 | Promoter | 1 | 64079103 | 64122282 | 43180 | 2 | ENSMUST00 | 0 | Klf7 | ENSMUSG00000025959.8 |
| chr1 | 63175611 | 63177736 | 29.58769 | 108.11006 | 104.32592 | Promoter | 1 | 63176825 | 63180486 | 3662 | 1 | ENSMUST00 | 0 | Eef1b2 | ENSMUSG00000025967.11 |
| chr1 | 4784947 | 4786386 | 29.58769 | 108.11006 | 104.32592 | Promoter | 1 | 4773206 | 4785710 | 12505 | 2 | ENSMUST00 | 0 | Mrpl15 | ENSMUSG00000033845.8 |
| chr10 | 79988286 | 79990465 | 33.5288 | 108.10788 | 104.32592 | Promoter | 10 | 79988584 | 79996062 | 7479 | 1 | ENSMUST00 | 0 | Cnn2 | ENSMUSG00000004665.5 |
| chr11 | 101169063 | 101171430 | 33.5288 | 108.10788 | 104.32592 | Promoter | 11 | 101170523 | 101178316 | 7794 | 1 | ENSMUST00 | 0 | Cntnap1 | ENSMUSG00000017167.6 |
| chr11 | 69632880 | 69633976 | 29.58769 | 108.11006 | 104.32592 | Promoter | 11 | 69632990 | 69653297 | 20308 | 1 | ENSMUST00 | 0 | Fxr2 | ENSMUSG00000018765.5 |
| chr11 | 29130145 | 29131524 | 33.5288 | 108.10788 | 104.32592 | Promoter | 11 | 29130744 | 29161680 | 30937 | 1 | ENSMUST00 | 0 | Pnpt1 | ENSMUSG00000020464.10 |
| chr11 | 65162059 | 65163092 | 29.58769 | 108.11006 | 104.32592 | Promoter | 11 | 65002039 | 65162961 | 160923 | 2 | ENSMUST00 | 0 | Arhgap44 | ENSMUSG00000033389.11 |
| chr12 | 101911905 | 101913759 | 29.58769 | 108.11006 | 104.32592 | Promoter | 12 | 101834043 | 101913267 | 79225 | 2 | ENSMUST00 | 0 | Trip11 | ENSMUSG00000021188.9 |
| chr12 | 102756811 | 102759183 | 29.58769 | 108.11006 | 104.32592 | Promoter | 12 | 102757975 | 102777701 | 19727 | 1 | ENSMUST00 | 0 | Ubr7 | ENSMUSG00000041712.6 |
| chr13 | 62964530 | 62965530 | 33.5288 | 108.10788 | 104.32592 | Distal Interge | 13 | 63014934 | 63299656 | 284723 | 1 | ENSMUST00 | -49404 | 2010111I01R | ENSMUSG00000021458.11 |
| chr14 | 57825764 | 57827010 | 29.58769 | 108.11006 | 104.32592 | Promoter | 14 | 57806561 | 57826163 | 19603 | 2 | ENSMUST00 | 0 | Ska3 | ENSMUSG00000021965.2 |
| chr14 | 103098163 | 103100145 | 33.5288 | 108.10788 | 104.32592 | Promoter | 14 | 103080239 | 103099554 | 19316 | 2 | ENSMUST00 | 0 | Fbxl3 | ENSMUSG00000022124.9 |
| chr14 | 54617254 | 54618266 | 33.5288 | 108.10788 | 104.32592 | Promoter | 14 | 54614120 | 54617995 | 3876 | 2 | ENSMUST00 | 0 | Psmb5 | ENSMUSG00000022193.6 |
| chr15 | 66968167 | 66970335 | 29.58769 | 108.11006 | 104.32592 | Promoter | 15 | 66929321 | 66969640 | 40320 | 2 | ENSMUST00 | 0 | Ndrg1 | ENSMUSG00000005125.7 |
| chr15 | 82345970 | 82347455 | 33.5288 | 108.10788 | 104.32592 | Promoter | 15 | 82346014 | 82349091 | 3078 | 1 | ENSMUST00 | 0 | Smdt1 | ENSMUSG00000022452.9 |
| chr15 | 76606796 | 76607621 | 29.58769 | 108.11006 | 104.32592 | Promoter | 15 | 76595809 | 76607591 | 11783 | 2 | ENSMUST00 | 0 | Cpsf1 | ENSMUSG00000034022.7 |
| chr15 | 74708602 | 74709818 | 29.58769 | 108.11006 | 104.32592 | Promoter | 15 | 74709161 | 74710372 | 1212 | 1 | ENSMUST00 | 0 | 4933427E11I | ENSMUSG00000044276.2 |
| chr15 | 27680994 | 27682108 | 29.58769 | 108.11006 | 104.32592 | Promoter | 15 | 27655071 | 27681542 | 26472 | 2 | ENSMUST00 | 0 | Fam105a | ENSMUSG00000056069.8 |
| chr17 | 46680020 | 46681746 | 29.58769 | 108.11006 | 104.32592 | Promoter | 17 | 46681137 | 46683105 | 1969 | 1 | ENSMUST00 | 0 | Mea1 | ENSMUSG00000002768.7 |
| chr17 | 66518531 | 66520403 | 29.58769 | 108.11006 | 104.32592 | Promoter | 17 | 66494512 | 66519670 | 25159 | 2 | ENSMUST00 | 0 | Rab12 | ENSMUSG00000023460.7 |
| chr17 | 26067642 | 26070245 | 33.5288 | 108.10788 | 104.32592 | Promoter | 17 | 25989036 | 26069177 | 80142 | 2 | ENSMUST00 | 0 | Rab11fip3 | ENSMUSG00000037098.12 |
| chr17 | 25832014 | 25834183 | 33.5288 | 108.10788 | 104.32592 | Promoter | 17 | 25830636 | 25833361 | 2726 | 2 | ENSMUST00 | 0 | Stub1 | ENSMUSG00000039615.8 |
| chr18 | 11656858 | 11658394 | 29.58769 | 108.11006 | 104.32592 | Promoter | 18 | 11657349 | 11743207 | 85859 | 1 | ENSMUST00 | 0 | Rbbp8 | ENSMUSG00000041238.10 |
| chr19 | 40830383 | 40832269 | 33.5288 | 108.10788 | 104.32592 | Promoter | 19 | 40831279 | 40848572 | 17294 | 1 | ENSMUST00 | 0 | Ccnj | ENSMUSG00000025010.8 |
| chr19 | 47730652 | 47732360 | 33.5288 | 108.10788 | 104.32592 | Promoter | 19 | 47731756 | 47735588 | 3833 | 1 | ENSMUST00 | 0 | Sfr1 | ENSMUSG00000025066.9 |
| chr2 | 118388524 | 118389863 | 29.58769 | 108.11006 | 104.32592 | Promoter | 2 | 118388618 | 118428213 | 39596 | 1 | ENSMUST00 | 0 | Eif2ak4 | ENSMUSG00000005102.8 |
| chr2 | 90579154 | 90581189 | 29.58769 | 108.11006 | 104.32592 | Promoter | 2 | 90432889 | 90580647 | 147759 | 2 | ENSMUST00 | 0 | Ptprj | ENSMUSG00000025314.11 |
| chr2 | 73213550 | 73215039 | 29.58769 | 108.11006 | 104.32592 | Promoter | 2 | 73092801 | 73214447 | 121647 | 2 | ENSMUST00 | 0 | Ola1 | ENSMUSG00000027108.10 |
| chr2 | 129065326 | 129066998 | 33.5288 | 108.10788 | 104.32592 | Promoter | 2 | 129065942 | 129096283 | 30342 | 1 | ENSMUST00 | 0 | Ttl | ENSMUSG00000027394.8 |
| chr2 | 174075923 | 174077510 | 33.5288 | 108.10788 | 104.32592 | Promoter | 2 | 174076308 | 174092433 | 16126 | 1 | ENSMUST00 | 0 | Stx16 | ENSMUSG00000027522.10 |
| chr2 | 28446214 | 28447644 | 33.5288 | 108.10788 | 104.32592 | Promoter | 2 | 28446800 | 28450506 | 3707 | 1 | ENSMUST00 | 0 | Ppp1r26 | ENSMUSG00000035829.8 |
| chr2 | 166793414 | 166794519 | 33.5288 | 108.10788 | 104.32592 | Promoter | 2 | 166793767 | 166799505 | 5739 | 1 | ENSMUST00 | 0 | Trp53rk | ENSMUSG00000042854.9 |
| chr2 | 49450752 | 49453348 | 29.58769 | 108.11006 | 104.32592 | Promoter | 2 | 49451486 | 49551948 | 100463 | 1 | ENSMUST00 | 0 | Epc2 | ENSMUSG00000069495.7 |
| chr3 | 158035595 | 158037426 | 29.58769 | 108.11006 | 104.32592 | Promoter | 3 | 158010473 | 158036639 | 26167 | 2 | ENSMUST00 | 0 | Srsf11 | ENSMUSG00000055436.13 |
| chr4 | 148558045 | 148559195 | 29.58769 | 108.11006 | 104.32592 | Promoter | 4 | 148558429 | 148582401 | 23973 | 1 | ENSMUST00 | 0 | Exosc10 | ENSMUSG00000017264.11 |
| chr4 | 106678141 | 106679264 | 29.58769 | 108.11006 | 104.32592 | Promoter | 4 | 106662256 | 106678944 | 16689 | 2 | ENSMUST00 | 0 | Ttc4 | ENSMUSG00000025413.8 |
| chr4 | 107879222 | 107880603 | 29.58769 | 108.11006 | 104.32592 | Promoter | 4 | 107879755 | 107887424 | 7670 | 1 | ENSMUST00 | 0 | Magoh | ENSMUSG00000028609.6 |
| chr4 | 137047733 | 137049094 | 29.58769 | 108.11006 | 104.32592 | Promoter | 4 | 136979732 | 137048695 | 68964 | 2 | ENSMUST00 | 0 | Zbtb40 | ENSMUSG00000060862.9 |
| chr5 | 64802420 | 64806226 | 29.58769 | 108.11006 | 104.32592 | Promoter | 5 | 64803523 | 64830129 | 26607 | 1 | ENSMUST00 | 0 | Klf3 | ENSMUSG00000029178.9 |
| chr6 | 54594686 | 54596059 | 33.5288 | 108.10788 | 104.32592 | Promoter | 6 | 54595111 | 54645839 | 50729 | 1 | ENSMUST00 | 0 | Plekha8 | ENSMUSG00000005225.10 |
| chr7 | 19768283 | 19771284 | 29.58769 | 108.11006 | 104.32592 | Promoter | 7 | 19756131 | 19770547 | 14417 | 2 | ENSMUST00 | 0 | Bcam | ENSMUSG00000002980.9 |
| chr7 | 27591673 | 27593253 | 29.58769 | 108.11006 | 104.32592 | Promoter | 7 | 27591746 | 27639450 | 47705 | 1 | ENSMUST00 | 0 | Akt2 | ENSMUSG00000004056.10 |
| chr7 | 126928356 | 126929847 | 29.58769 | 108.11006 | 104.32592 | Promoter | 7 | 126928879 | 126945631 | 16753 | 1 | ENSMUST00 | 0 | Kctd13 | ENSMUSG00000030685.5 |
| chr7 | 143738218 | 143740816 | 29.58769 | 108.11006 | 104.32592 | Promoter | 7 | 143688762 | 143740341 | 51580 | 2 | ENSMUST00 | 0 | Osbpl5 | ENSMUSG00000037606.12 |
| chr7 | 5014072 | 5016177 | 33.5288 | 108.10788 | 104.32592 | Promoter | 7 | 5015501 | 5018486 | 2986 | 1 | ENSMUST00 | 0 | Zfp524 | ENSMUSG00000051184.6 |
| chr8 | 60992445 | 60994609 | 33.5288 | 108.10788 | 104.32592 | Promoter | 8 | 60993195 | 61131339 | 138145 | 1 | ENSMUST00 | 0 | Nek1 | ENSMUSG00000031644.13 |
| chr8 | 88520943 | 88522438 | 33.5288 | 108.10788 | 104.32592 | Promoter | 8 | 88521344 | 88594887 | 73544 | 1 | ENSMUST00 | 0 | Nkd1 | ENSMUSG00000031661.6 |
| chr8 | 60889868 | 60891265 | 33.5288 | 108.10788 | 104.32592 | Promoter | 8 | 60890418 | 60907575 | 17158 | 1 | ENSMUST00 | 0 | 2700029M09 | ENSMUSG00000038005.9 |
| chr9 | 73112935 | 73114437 | 29.58769 | 108.11006 | 104.32592 | Promoter | 9 | 73113426 | 73122712 | 9287 | 1 | ENSMUST00 | 0 | Rsl24d1 | ENSMUSG00000032215.10 |
| chr9 | 72661484 | 72665268 | 29.58769 | 108.11006 | 104.32592 | Promoter | 9 | 72662346 | 72749852 | 87507 | 1 | ENSMUST00 | 0 | Nedd4 | ENSMUSG00000032216.9 |
| chr9 | 95857164 | 95858685 | 29.58769 | 108.11006 | 104.32592 | Promoter | 9 | 95857597 | 95868312 | 10716 | 1 | ENSMUST00 | 0 | Atr | ENSMUSG00000032409.9 |
| chr9 | 22070511 | 22071901 | 33.5288 | 108.10788 | 104.32592 | Promoter | 9 | 22055460 | 22071345 | 15886 | 2 | ENSMUST00 | 0 | Zfp653 | ENSMUSG00000038895.10 |
| chr9 | 45828055 | 45829186 | 33.5288 | 108.10788 | 104.32592 | Promoter | 9 | 45766946 | 45828691 | 61746 | 2 | ENSMUST00 | 0 | Cep164 | ENSMUSG00000043987.12 |
| chr12 | 110590657 | 110592416 | 31.1999 | 108.05416 | 104.27962 | Distal Interge | 12 | 110601452 | 110666945 | 65494 | 1 | ENSMUST00 | -9036 | Dync1h1 | ENSMUSG00000018707.8 |
| chr12 | 100723877 | 100725193 | 31.1999 | 108.05416 | 104.27962 | Promoter | 12 | 100549778 | 100725028 | 175251 | 2 | ENSMUST00 | 0 | Rps6ka5 | ENSMUSG00000021180.8 |
| chr13 | 18716811 | 18717905 | 31.1999 | 108.05416 | 104.27962 | Promoter | 13 | 18717292 | 18866809 | 149518 | 1 | ENSMUST00 | 0 | Vps41 | ENSMUSG00000041236.7 |
| chr14 | 121377820 | 121380208 | 31.1999 | 108.05416 | 104.27962 | Promoter | 14 | 121286343 | 121379334 | 92992 | 2 | ENSMUST00 | 0 | Stk24 | ENSMUSG00000063410.7 |
| chr8 | 106935129 | 106937196 | 31.1999 | 108.05416 | 104.27962 | Promoter | 8 | 106935750 | 107014192 | 78443 | 1 | ENSMUST00 | 0 | Sntb2 | ENSMUSG00000041308.5 |
| chr8 | 68276198 | 68280543 | 31.1999 | 108.05416 | 104.27962 | Promoter | 8 | 68276567 | 68347694 | 71128 | 1 | ENSMUST00 | 0 | Sh2d4a | ENSMUSG00000053886.3 |
| chr13 | 3538044 | 3539605 | 33.46659 | 107.99879 | 104.22513 | Promoter | 13 | 3538075 | 3566261 | 28187 | 1 | ENSMUST00 | 0 | Gdi2 | ENSMUSG00000021218.9 |
| chr17 | 56997256 | 56998063 | 33.46659 | 107.99879 | 104.22513 | Promoter | 17 | 56997339 | 56999311 | 1973 | 1 | ENSMUST00 | 0 | Alkbh7 | ENSMUSG00000002661.9 |
| chr1 | 143776236 | 143778292 | 27.97104 | 107.99458 | 104.2211 | Promoter | 1 | 143777272 | 143807305 | 30034 | 1 | ENSMUST00 | 0 | Uchl5 | ENSMUSG00000018189.7 |
| chr11 | 109610494 | 109611486 | 27.97104 | 107.99458 | 104.2211 | Promoter | 11 | 109573331 | 109611432 | 38102 | 2 | ENSMUST00 | 0 | Wipi1 | ENSMUSG00000041895.10 |
| chr11 | 100471596 | 100473123 | 27.97104 | 107.99458 | 104.2211 | Promoter | 11 | 100462614 | 100472741 | 10128 | 2 | ENSMUST00 | 0 | Klhl11 | ENSMUSG00000048732.5 |
| chr17 | 24850465 | 24851465 | 27.97104 | 107.99458 | 104.2211 | Promoter | 17 | 24850490 | 24864450 | 13961 | 1 | ENSMUST00 | 0 | Hagh | ENSMUSG00000024158.12 |
| chr19 | 9982044 | 9983828 | 27.97104 | 107.99458 | 104.2211 | Promoter | 19 | 9982703 | 9985092 | 2390 | 1 | ENSMUST00 | 0 | Fth1 | ENSMUSG00000024661.6 |
| chr2 | 180119314 | 180120317 | 27.97104 | 107.99458 | 104.2211 | Promoter | 2 | 180119335 | 180137085 | 17751 | 1 | ENSMUST00 | 0 | Osbpl2 | ENSMUSG00000039050.8 |
| chr4 | 34613891 | 34615909 | 27.97104 | 107.99458 | 104.2211 | Promoter | 4 | 34614957 | 34660167 | 45211 | 1 | ENSMUST00 | 0 | Rars2 | ENSMUSG00000028292.9 |
| chr5 | 112342400 | 112343885 | 27.97104 | 107.99458 | 104.2211 | Promoter | 5 | 112337392 | 112343020 | 5629 | 2 | ENSMUST00 | 0 | Srrd | ENSMUSG00000029346.7 |
| chr9 | 44879791 | 44881351 | 27.97104 | 107.99458 | 104.2211 | Promoter | 9 | 44803355 | 44881274 | 77920 | 2 | ENSMUST00 | 0 | Kmt2a | ENSMUSG00000002028.7 |
| chr9 | 123112407 | 123114045 | 27.97104 | 107.99458 | 104.2211 | Promoter | 9 | 123113215 | 123136129 | 22915 | 1 | ENSMUST00 | 0 | Exosc7 | ENSMUSG00000025785.4 |
| chr9 | 106367929 | 106370134 | 27.97104 | 107.99458 | 104.2211 | Promoter | 9 | 106368632 | 106375724 | 7093 | 1 | ENSMUST00 | 0 | Dusp7 | ENSMUSG00000053716.8 |
| chr13 | 119789745 | 119791716 | 32.87838 | 107.99121 | 104.21945 | Promoter | 13 | 119765187 | 119790805 | 25619 | 2 | ENSMUST00 | 0 | Zfp131 | ENSMUSG00000094870.2 |
| chr15 | 98092443 | 98094045 | 32.87838 | 107.99121 | 104.21945 | Promoter | 15 | 98092589 | 98132447 | 39859 | 1 | ENSMUST00 | 0 | Pfkm | ENSMUSG00000033065.8 |
| chr19 | 4011854 | 4012763 | 32.87838 | 107.99121 | 104.21945 | Promoter | 19 | 4007497 | 4012666 | 5170 | 2 | ENSMUST00 | 0 | Ndufv1 | ENSMUSG00000037916.8 |
| chr3 | 145575887 | 145577178 | 32.87838 | 107.99121 | 104.21945 | Promoter | 3 | 145576208 | 145605245 | 29038 | 1 | ENSMUST00 | 0 | Znhit6 | ENSMUSG00000074182.4 |
| chr9 | 95510814 | 95512717 | 32.87838 | 107.99121 | 104.21945 | Promoter | 9 | 95456898 | 95511894 | 54997 | 2 | ENSMUST00 | 0 | U2surp | ENSMUSG00000032407.8 |
| chr13 | 113793817 | 113795802 | 34.02479 | 107.94543 | 104.17527 | Promoter | 13 | 113794505 | 114157461 | 362957 | 1 | ENSMUST00 | 0 | Arl15 | ENSMUSG00000042348.9 |
| chr14 | 54912185 | 54913452 | 34.02479 | 107.94543 | 104.17527 | Promoter | 14 | 54906727 | 54913132 | 6406 | 2 | ENSMUST00 | 0 | Slc22a17 | ENSMUSG00000022199.10 |
| chr16 | 87439758 | 87441306 | 34.02479 | 107.94543 | 104.17527 | Promoter | 16 | 87433407 | 87440573 | 7167 | 2 | ENSMUST00 | 0 | Rwdd2b | ENSMUSG00000041079.6 |
| chr11 | 51059117 | 51059984 | 35.17274 | 107.8772 | 104.10775 | Promoter | 11 | 51059257 | 51071362 | 12106 | 1 | ENSMUST00 | 0 | Zfp354a | ENSMUSG00000020364.9 |
| chr5 | 122899282 | 122901119 | 35.17274 | 107.8772 | 104.10775 | Promoter | 5 | 122870668 | 122900355 | 29688 | 2 | ENSMUST00 | 0 | Kdm2b | ENSMUSG00000029475.12 |
| chr11 | 117809434 | 117811136 | 30.53388 | 107.8511 | 104.08218 | Promoter | 11 | 117809668 | 117814283 | 4616 | 1 | ENSMUST00 | 0 | Syngr2 | ENSMUSG00000048277.10 |
| chr4 | 118132585 | 118135492 | 24.2022 | 107.83455 | 104.06592 | Promoter | 4 | 117932154 | 118134890 | 202737 | 2 | ENSMUST00 | 0 | St3gal3 | ENSMUSG00000028538.7 |
| chr12 | 79028669 | 79030046 | 33.9579 | 107.82887 | 104.06033 | Promoter | 12 | 79029163 | 79081648 | 52486 | 1 | ENSMUST00 | 0 | Plekhh1 | ENSMUSG00000060716.6 |
| chr2 | 132144191 | 132145924 | 33.9579 | 107.82887 | 104.06033 | Promoter | 2 | 132052496 | 132145108 | 92613 | 2 | ENSMUST00 | 0 | Slc23a2 | ENSMUSG00000027340.10 |
| chr7 | 80024655 | 80028372 | 33.9579 | 107.82887 | 104.06033 | Promoter | 7 | 80024814 | 80092751 | 67938 | 1 | ENSMUST00 | 0 | Zfp710 | ENSMUSG00000048897.10 |
| chr8 | 86623979 | 86625540 | 33.9579 | 107.82887 | 104.06033 | Promoter | 8 | 86624043 | 86716631 | 92589 | 1 | ENSMUST00 | 0 | Lonp2 | ENSMUSG00000047866.13 |
| chr14 | 20640227 | 20641954 | 34.54686 | 107.82351 | 104.0565 | Promoter | 14 | 20641009 | 20644048 | 3040 | 1 | ENSMUST00 | 0 | RP24-155O1 | ENSMUSG00000099966.1 |
| chr1 | 86154131 | 86155428 | 32.75675 | 107.77596 | 104.00922 | Promoter | 1 | 86154780 | 86246326 | 91547 | 1 | ENSMUST00 | 0 | Armc9 | ENSMUSG00000062590.8 |
| chr19 | 6986900 | 6988151 | 33.89127 | 107.71272 | 103.94643 | Promoter | 19 | 6982473 | 6987629 | 5157 | 2 | ENSMUST00 | 0 | Vegfb | ENSMUSG00000024962.8 |
| chr4 | 136469495 | 136471700 | 34.47702 | 107.70252 | 103.93674 | Promoter | 4 | 136469761 | 136549315 | 79555 | 1 | ENSMUST00 | 0 | Luzp1 | ENSMUSG00000001089.9 |
| chr5 | 41842950 | 41845617 | 34.47702 | 107.70252 | 103.93674 | Promoter | 5 | 41787542 | 41844168 | 56627 | 2 | ENSMUST00 | 0 | Bod1l | ENSMUSG00000061755.7 |
| chr7 | 127707789 | 127709220 | 34.47702 | 107.70252 | 103.93674 | Promoter | 7 | 127663353 | 127708785 | 45433 | 2 | ENSMUST00 | 0 | Bcl7c | ENSMUSG00000030814.12 |
| chr10 | 118141340 | 118142516 | 32.09266 | 107.61995 | 103.85494 | Promoter | 10 | 118141787 | 118168997 | 27211 | 1 | ENSMUST00 | 0 | Mdm1 | ENSMUSG00000020212.8 |
| chr11 | 84179012 | 84180621 | 32.09266 | 107.61995 | 103.85494 | Promoter | 11 | 84179831 | 84214195 | 34365 | 1 | ENSMUST00 | 0 | Acaca | ENSMUSG00000020532.13 |
| chr11 | 103696863 | 103698327 | 32.09266 | 107.61995 | 103.85494 | Promoter | 11 | 103676849 | 103697898 | 21050 | 2 | ENSMUST00 | 0 | Gosr2 | ENSMUSG00000020946.8 |
| chr13 | 96670116 | 96671529 | 32.09266 | 107.61995 | 103.85494 | Promoter | 13 | 96648967 | 96670936 | 21970 | 2 | ENSMUST00 | 0 | Hmgcr | ENSMUSG00000021670.9 |
| chr5 | 117286664 | 117288029 | 32.09266 | 107.61995 | 103.85494 | Promoter | 5 | 117282651 | 117287625 | 4975 | 2 | ENSMUST00 | 0 | Pebp1 | ENSMUSG00000032959.7 |
| chr9 | 54698960 | 54700445 | 32.09266 | 107.61995 | 103.85494 | Promoter | 9 | 54699514 | 54709336 | 9823 | 1 | ENSMUST00 | 0 | Dnaja4 | ENSMUSG00000032285.10 |
| chr12 | 4916822 | 4918831 | 29.32506 | 107.61204 | 103.84809 | Promoter | 12 | 4917404 | 5044047 | 126644 | 1 | ENSMUST00 | 0 | Atad2b | ENSMUSG00000052812.4 |
| chr15 | 98952176 | 98954298 | 33.82489 | 107.59698 | 103.83312 | Promoter | 15 | 98949841 | 98953551 | 3711 | 2 | ENSMUST00 | 0 | Tuba1a | ENSMUSG00000072235.5 |
| chr7 | 12818207 | 12819221 | 33.82489 | 107.59698 | 103.83312 | Promoter | 7 | 12804977 | 12818854 | 13878 | 2 | ENSMUST00 | 0 | Zfp329 | ENSMUSG00000057894.10 |
| chr8 | 109691717 | 109693283 | 27.7677 | 107.59161 | 103.82806 | Promoter | 8 | 109671322 | 109693294 | 21973 | 2 | ENSMUST00 | 11 | Ist1 | ENSMUSG00000031729.5 |
| chr8 | 40861959 | 40863338 | 34.40746 | 107.58196 | 103.81843 | Promoter | 8 | 40862396 | 40918056 | 55661 | 1 | ENSMUST00 | 0 | Slc7a2 | ENSMUSG00000031596.10 |
| chr2 | 127008343 | 127009585 | 34.95366 | 107.49944 | 103.73631 | Promoter | 2 | 127008717 | 127037344 | 28628 | 1 | ENSMUST00 | 0 | Ap4e1 | ENSMUSG00000001998.10 |
| chr17 | 10839745 | 10841170 | 33.75878 | 107.48165 | 103.71877 | Promoter | 17 | 10840384 | 11557234 | 716851 | 1 | ENSMUST00 | 0 | Park2 | ENSMUSG00000023826.9 |
| chr18 | 49978679 | 49980374 | 33.75878 | 107.48165 | 103.71877 | Promoter | 18 | 49979427 | 50091001 | 111575 | 1 | ENSMUST00 | 0 | Tnfaip8 | ENSMUSG00000062210.8 |
| chr8 | 33927502 | 33930859 | 33.75878 | 107.48165 | 103.71877 | Promoter | 8 | 33782643 | 33929863 | 147221 | 2 | ENSMUST00 | 0 | Rbpms | ENSMUSG00000031586.11 |
| chr4 | 132352621 | 132354151 | 34.33818 | 107.46186 | 103.69923 | Promoter | 4 | 132337823 | 132353605 | 15783 | 2 | ENSMUST00 | 0 | Rcc1 | ENSMUSG00000028896.8 |
| chr2 | 126673657 | 126676639 | 33.69293 | 107.36673 | 103.60477 | Promoter | 2 | 126627442 | 126675568 | 48127 | 2 | ENSMUST00 | 0 | Gabpb1 | ENSMUSG00000027361.10 |
| chr5 | 34369227 | 34371027 | 33.69293 | 107.36673 | 103.60477 | Promoter | 5 | 34369933 | 34486456 | 116524 | 1 | ENSMUST00 | 0 | Fam193a | ENSMUSG00000037210.11 |
| chr17 | 31657432 | 31659338 | 34.26917 | 107.34218 | 103.58047 | Promoter | 17 | 31647082 | 31658754 | 11673 | 2 | ENSMUST00 | 0 | U2af1 | ENSMUSG00000061613.7 |
| chr5 | 113993025 | 113994545 | 34.26917 | 107.34218 | 103.58047 | Promoter | 5 | 113937094 | 113993894 | 56801 | 2 | ENSMUST00 | 0 | Ssh1 | ENSMUSG00000042121.11 |
| chr1 | 153548984 | 153550082 | 33.08674 | 107.33151 | 103.57004 | Promoter | 1 | 153503015 | 153550045 | 47031 | 2 | ENSMUST00 | 0 | Npl | ENSMUSG00000042684.8 |
| chr1 | 54925084 | 54926821 | 33.08674 | 107.33151 | 103.57004 | Promoter | 1 | 54645340 | 54926387 | 281048 | 2 | ENSMUST00 | 0 | Ankrd44 | ENSMUSG00000052331.9 |
| chr17 | 55891367 | 55892903 | 33.08674 | 107.33151 | 103.57004 | Promoter | 17 | 55892120 | 55898928 | 6809 | 1 | ENSMUST00 | 0 | Zfp959 | ENSMUSG00000003198.9 |
| chr19 | 37550051 | 37551353 | 33.08674 | 107.33151 | 103.57004 | Promoter | 19 | 37550418 | 37683245 | 132828 | 1 | ENSMUST00 | 0 | Exoc6 | ENSMUSG00000053799.6 |

| | | | | | | | | | | | | | | | |
|---|---|---|---|---|---|---|---|---|---|---|---|---|---|---|---|
| chr2 | 119741625 | 119743529 | 33.08674 | 107.33151 | 103.57004 | Promoter | 2 | 119742337 | 119751263 | 8927 | 1 | ENSMUST00 | 0 | Itpka | ENSMUSG00000027296.7 |
| chr3 | 122728814 | 122731114 | 33.08674 | 107.33151 | 103.57004 | Promoter | 3 | 122729158 | 122859374 | 130217 | 1 | ENSMUST00 | 0 | Pde5a | ENSMUSG00000053965.6 |
| chr4 | 105156812 | 105160389 | 33.08674 | 107.33151 | 103.57004 | Promoter | 4 | 105157347 | 105232764 | 75418 | 1 | ENSMUST00 | 0 | Ppap2b | ENSMUSG00000028517.8 |
| chr5 | 32459049 | 32460692 | 33.08674 | 107.33151 | 103.57004 | Promoter | 5 | 32459152 | 32517433 | 58282 | 1 | ENSMUST00 | 0 | Ppp1cb | ENSMUSG00000014956.10 |
| chr5 | 106695254 | 106697207 | 33.08674 | 107.33151 | 103.57004 | Promoter | 5 | 106616739 | 106696830 | 80092 | 2 | ENSMUST00 | 0 | Zfp644 | ENSMUSG00000049606.11 |
| chr7 | 47007232 | 47009188 | 33.08674 | 107.33151 | 103.57004 | Promoter | 7 | 46990396 | 47008411 | 18016 | 2 | ENSMUST00 | 0 | Spty2d1 | ENSMUSG00000049516.7 |
| chr8 | 22192598 | 22193978 | 33.08674 | 107.33151 | 103.57004 | Promoter | 8 | 22192809 | 22220843 | 28035 | 1 | ENSMUST00 | 0 | Vps36 | ENSMUSG00000031479.8 |
| chr13 | 21753345 | 21754894 | 31.36125 | 107.32627 | 103.56666 | Promoter | 13 | 21753435 | 21753827 | 393 | 2 | ENSMUST00 | 0 | Hist1h2ak | ENSMUSG00000063021.3 |
| chr5 | 135632628 | 135633357 | 31.36125 | 107.32627 | 103.56666 | Promoter | 5 | 135632654 | 135646448 | 13795 | 1 | ENSMUST00 | 0 | Rhbdd2 | ENSMUSG00000039917.6 |
| chr9 | 82829415 | 82830730 | 31.36125 | 107.32627 | 103.56666 | Promoter | 9 | 82829540 | 82847687 | 18148 | 1 | ENSMUST00 | 0 | Irak1bp1 | ENSMUSG00000032251.7 |
| chr15 | 10469446 | 10471180 | 33.62733 | 107.25218 | 103.49312 | Promoter | 15 | 10446756 | 10470516 | 23761 | 2 | ENSMUST00 | 0 | Dnajc21 | ENSMUSG00000044224.11 |
| chr9 | 85326142 | 85328587 | 33.62733 | 107.25218 | 103.49312 | Promoter | 9 | 85320439 | 85327124 | 6686 | 2 | ENSMUST00 | 0 | Fam46a | ENSMUSG00000032265.9 |
| chr5 | 100156800 | 100160103 | 34.80912 | 107.25002 | 103.49115 | Promoter | 5 | 100077872 | 100159808 | 81937 | 2 | ENSMUST00 | 0 | Tmem150c | ENSMUSG00000050640.11 |
| chr1 | 74504943 | 74507160 | 34.20045 | 107.22294 | 103.46436 | Promoter | 1 | 74506058 | 74530842 | 24785 | 1 | ENSMUST00 | 0 | Rqcd1 | ENSMUSG00000026174.9 |
| chr1 | 93235486 | 93236791 | 34.20045 | 107.22294 | 103.46436 | Promoter | 1 | 93235841 | 93301065 | 65225 | 1 | ENSMUST00 | 0 | Sned1 | ENSMUSG00000047793.8 |
| chr10 | 63336957 | 63339812 | 34.20045 | 107.22294 | 103.46436 | Promoter | 10 | 63319007 | 63336992 | 17986 | 2 | ENSMUST00 | 0 | Sirt1 | ENSMUSG00000020063.11 |
| chr19 | 8929002 | 8930617 | 34.20045 | 107.22294 | 103.46436 | Promoter | 19 | 8929694 | 8941563 | 11870 | 1 | ENSMUST00 | 0 | Eml3 | ENSMUSG00000071647.4 |
| chr2 | 38643089 | 38644443 | 34.20045 | 107.22294 | 103.46436 | Promoter | 2 | 38588036 | 38644087 | 56052 | 2 | ENSMUST00 | 0 | Psmb7 | ENSMUSG00000026750.5 |
| chr4 | 43560786 | 43563861 | 34.20045 | 107.22294 | 103.46436 | Promoter | 4 | 43531519 | 43562422 | 30904 | 2 | ENSMUST00 | 0 | Tln1 | ENSMUSG00000028465.11 |
| chr10 | 85183451 | 85186083 | 25.18546 | 107.13963 | 103.3826 | Promoter | 10 | 85131702 | 85185054 | 53353 | 2 | ENSMUST00 | 0 | Cry1 | ENSMUSG00000020038.9 |
| chr11 | 55204247 | 55205368 | 25.18546 | 107.13963 | 103.3826 | Promoter | 11 | 55204350 | 55236330 | 31981 | 1 | ENSMUST00 | 0 | Slc36a1 | ENSMUSG00000020261.10 |
| chr11 | 86356114 | 86358387 | 25.18546 | 107.13963 | 103.3826 | Promoter | 11 | 86267033 | 86357602 | 90570 | 2 | ENSMUST00 | 0 | Med13 | ENSMUSG00000034297.9 |
| chr17 | 71000626 | 71002523 | 25.18546 | 107.13963 | 103.3826 | Promoter | 17 | 70994294 | 71002165 | 7872 | 2 | ENSMUST00 | 0 | Myl12a | ENSMUSG00000024048.9 |
| chr7 | 141474160 | 141476599 | 25.18546 | 107.13963 | 103.3826 | Promoter | 7 | 141475240 | 141493427 | 18188 | 1 | ENSMUST00 | 0 | Tspan4 | ENSMUSG00000025511.9 |
| chr6 | 118418720 | 118420152 | 33.56199 | 107.13804 | 103.38137 | Promoter | 6 | 118383383 | 118419417 | 36035 | 2 | ENSMUST00 | 0 | Bms1 | ENSMUSG00000030138.6 |
| chr15 | 96283762 | 96289319 | 34.73729 | 107.12603 | 103.36953 | Promoter | 15 | 96287518 | 96352038 | 64521 | 1 | ENSMUST00 | 0 | Arid2 | ENSMUSG00000033237.13 |
| chr5 | 140388513 | 140389686 | 34.73729 | 107.12603 | 103.36953 | Promoter | 5 | 140340304 | 140389251 | 48948 | 2 | ENSMUST00 | 0 | Snx8 | ENSMUSG00000029560.8 |
| chr6 | 122338950 | 122340676 | 34.73729 | 107.12603 | 103.36953 | Promoter | 6 | 122317738 | 122339658 | 21921 | 2 | ENSMUST00 | 0 | Phc1 | ENSMUSG00000040669.9 |
| chr11 | 74830423 | 74832317 | 30.67871 | 107.10089 | 103.34502 | Promoter | 11 | 74830920 | 74845725 | 14806 | 1 | ENSMUST00 | 0 | Mnt | ENSMUSG00000000282.7 |
| chr15 | 67173622 | 67176826 | 30.67871 | 107.10089 | 103.34502 | Distal Interge | 15 | 67226769 | 67377094 | 150326 | 1 | ENSMUST00 | -49943 | RP23-163N4. | ENSMUSG00000102069.1 |
| chr5 | 135933350 | 135935455 | 30.67871 | 107.10089 | 103.34502 | Promoter | 5 | 135908409 | 135934599 | 26191 | 2 | ENSMUST00 | 0 | Ywhag | ENSMUSG00000051391.8 |
| chr17 | 46704061 | 46705063 | 33.4969 | 107.02429 | 103.26894 | Promoter | 17 | 46682991 | 46705002 | 22012 | 2 | ENSMUST00 | 0 | Ppp2r5d | ENSMUSG00000059409.8 |
| chr7 | 105580532 | 105581844 | 33.4969 | 107.02429 | 103.26894 | Promoter | 7 | 105558483 | 105581653 | 23171 | 2 | ENSMUST00 | 0 | Apbb1 | ENSMUSG00000037032.10 |
| chr5 | 121835543 | 121838063 | 34.66576 | 107.00249 | 103.24738 | Promoter | 5 | 121815488 | 121836825 | 21338 | 2 | ENSMUST00 | 0 | Sh2b3 | ENSMUSG00000042594.11 |
| chr1 | 163929283 | 163930774 | 31.16539 | 106.97112 | 103.21663 | Promoter | 1 | 163929765 | 163955126 | 25362 | 1 | ENSMUST00 | 0 | Scyl3 | ENSMUSG00000026584.9 |
| chr10 | 67095916 | 67097940 | 31.16539 | 106.97112 | 103.21663 | Promoter | 10 | 67009189 | 67096988 | 87800 | 2 | ENSMUST00 | 0 | Reep3 | ENSMUSG00000019873.7 |
| chr10 | 21295455 | 21297112 | 31.16539 | 106.97112 | 103.21663 | Promoter | 10 | 21295979 | 21311198 | 15220 | 1 | ENSMUST00 | 0 | Hbs1l | ENSMUSG00000019977.10 |
| chr13 | 119428264 | 119430336 | 31.16539 | 106.97112 | 103.21663 | Promoter | 13 | 119428601 | 119457391 | 28791 | 1 | ENSMUST00 | 0 | Paip1 | ENSMUSG00000025451.10 |
| chr13 | 112660256 | 112661388 | 31.16539 | 106.97112 | 103.21663 | Promoter | 13 | 112660766 | 112738743 | 77978 | 1 | ENSMUST00 | 0 | Slc38a9 | ENSMUSG00000047789.4 |
| chr13 | 100102898 | 100104640 | 31.16539 | 106.97112 | 103.21663 | Promoter | 13 | 100017994 | 100104070 | 86077 | 2 | ENSMUST00 | 0 | Bdp1 | ENSMUSG00000049658.8 |
| chr14 | 14702478 | 14704027 | 31.16539 | 106.97112 | 103.21663 | Promoter | 14 | 14703025 | 14799943 | 96919 | 1 | ENSMUST00 | 0 | Slc4a7 | ENSMUSG00000021733.9 |
| chr14 | 32159418 | 32161018 | 31.16539 | 106.97112 | 103.21663 | Promoter | 14 | 32159865 | 32179855 | 19991 | 1 | ENSMUST00 | 0 | Ncoa4 | ENSMUSG00000056234.9 |
| chr16 | 91646065 | 91649433 | 31.16539 | 106.97112 | 103.21663 | Promoter | 16 | 91647506 | 91679207 | 31702 | 1 | ENSMUST00 | 0 | Son | ENSMUSG00000022961.12 |
| chr17 | 33358117 | 33359422 | 31.16539 | 106.97112 | 103.21663 | Promoter | 17 | 33329342 | 33358878 | 29537 | 2 | ENSMUST00 | 0 | Zfp81 | ENSMUSG00000003929.10 |
| chr17 | 24140408 | 24142347 | 31.16539 | 106.97112 | 103.21663 | Promoter | 17 | 24073680 | 24141595 | 67916 | 2 | ENSMUST00 | 0 | Pdpk1 | ENSMUSG00000024122.11 |
| chr17 | 35120282 | 35122490 | 31.16539 | 106.97112 | 103.21663 | Promoter | 17 | 35116196 | 35121429 | 5234 | 2 | ENSMUST00 | 0 | Csnk2b | ENSMUSG00000024387.8 |
| chr19 | 5566994 | 5568966 | 31.16539 | 106.97112 | 103.21663 | Promoter | 19 | 5568074 | 5571261 | 3188 | 1 | ENSMUST00 | 0 | Ap5b1 | ENSMUSG00000049562.5 |
| chr2 | 34770899 | 34771870 | 31.16539 | 106.97112 | 103.21663 | Promoter | 2 | 34771970 | 34772710 | 741 | 1 | ENSMUST00 | -100 | Hspa5 | ENSMUSG00000026864.8 |
| chr2 | 92374485 | 92376159 | 31.16539 | 106.97112 | 103.21663 | Promoter | 2 | 92374676 | 92381216 | 6541 | 1 | ENSMUST00 | 0 | Pex16 | ENSMUSG00000027222.9 |
| chr2 | 104095745 | 104097982 | 31.16539 | 106.97112 | 103.21663 | Promoter | 2 | 104095801 | 104115349 | 19549 | 1 | ENSMUST00 | 0 | Cd59a | ENSMUSG00000032679.7 |
| chr3 | 103789748 | 103791963 | 31.16539 | 106.97112 | 103.21663 | Promoter | 3 | 103739815 | 103791563 | 51749 | 2 | ENSMUST00 | 0 | Hipk1 | ENSMUSG00000008730.12 |
| chr3 | 89912092 | 89913983 | 31.16539 | 106.97112 | 103.21663 | Promoter | 3 | 89869324 | 89913162 | 43839 | 2 | ENSMUST00 | 0 | Il6ra | ENSMUSG00000027947.7 |
| chr3 | 104218748 | 104221302 | 31.16539 | 106.97112 | 103.21663 | Promoter | 3 | 104013259 | 104220374 | 207116 | 2 | ENSMUST00 | 0 | Magi3 | ENSMUSG00000052539.10 |
| chr4 | 155623921 | 155625589 | 31.16539 | 106.97112 | 103.21663 | Promoter | 4 | 155624854 | 155630340 | 5487 | 1 | ENSMUST00 | 0 | Cdk11b | ENSMUSG00000029062.9 |
| chr4 | 139129787 | 139131399 | 31.16539 | 106.97112 | 103.21663 | Promoter | 4 | 139101814 | 139131113 | 29300 | 2 | ENSMUST00 | 0 | Minos1 | ENSMUSG00000050608.6 |
| chr4 | 98395386 | 98396772 | 31.16539 | 106.97112 | 103.21663 | Promoter | 4 | 98395785 | 98405677 | 9893 | 1 | ENSMUST00 | 0 | Inadl | ENSMUSG00000061859.11 |
| chr5 | 130028838 | 130030023 | 31.16539 | 106.97112 | 103.21663 | Promoter | 5 | 130029290 | 130060789 | 31500 | 1 | ENSMUST00 | 0 | Crcp | ENSMUSG00000025532.10 |
| chr5 | 129942250 | 129942973 | 31.16539 | 106.97112 | 103.21663 | Promoter | 5 | 129942113 | 129982891 | 40779 | 1 | ENSMUST00 | 138 | Vkorc1l1 | ENSMUSG00000066735.5 |
| chr6 | 31398336 | 31399951 | 31.16539 | 106.97112 | 103.21663 | Promoter | 6 | 31398735 | 31516811 | 118077 | 1 | ENSMUST00 | 0 | Mkln1 | ENSMUSG00000025609.10 |
| chr6 | 89595015 | 89595706 | 31.16539 | 106.97112 | 103.21663 | Promoter | 6 | 89383146 | 89595652 | 212507 | 2 | ENSMUST00 | 0 | Chchd6 | ENSMUSG00000030086.11 |
| chr6 | 145249108 | 145251136 | 31.16539 | 106.97112 | 103.21663 | Promoter | 6 | 145216699 | 145250239 | 33541 | 2 | ENSMUST00 | 0 | Kras | ENSMUSG00000030265.9 |
| chr6 | 128842150 | 128844024 | 31.16539 | 106.97112 | 103.21663 | Promoter | 6 | 128843786 | 128846959 | 3174 | 1 | ENSMUST00 | 0 | Gm26656 | ENSMUSG00000096931.1 |
| chr7 | 112427013 | 112429707 | 31.16539 | 106.97112 | 103.21663 | Promoter | 7 | 112427505 | 112589629 | 162125 | 1 | ENSMUST00 | 0 | Parva | ENSMUSG00000030770.9 |
| chr7 | 80370176 | 80371869 | 31.16539 | 106.97112 | 103.21663 | Promoter | 7 | 80349098 | 80371375 | 22278 | 2 | ENSMUST00 | 0 | Man2a2 | ENSMUSG00000038886.9 |
| chr7 | 79273144 | 79274150 | 31.16539 | 106.97112 | 103.21663 | Promoter | 7 | 79273201 | 79365508 | 92308 | 1 | ENSMUST00 | 0 | Abhd2 | ENSMUSG00000039202.7 |
| chr7 | 34387343 | 34390805 | 31.16539 | 106.97112 | 103.21663 | Promoter | 7 | 34344646 | 34389540 | 44895 | 2 | ENSMUST00 | 0 | Lsm14a | ENSMUSG00000066568.7 |
| chr8 | 70301887 | 70303713 | 31.16539 | 106.97112 | 103.21663 | Promoter | 8 | 70302518 | 70312990 | 10473 | 1 | ENSMUST00 | 0 | Cope | ENSMUSG00000055681.9 |
| chr9 | 21149019 | 21150250 | 31.16539 | 106.97112 | 103.21663 | Promoter | 9 | 21138541 | 21149906 | 11366 | 2 | ENSMUST00 | 0 | Cdc37 | ENSMUSG00000019471.9 |
| chr9 | 120570891 | 120572455 | 31.16539 | 106.97112 | 103.21663 | Promoter | 9 | 120571517 | 120574653 | 3137 | 1 | ENSMUST00 | 0 | Rpl14 | ENSMUSG00000025794.8 |
| chr9 | 122293466 | 122295123 | 31.16539 | 106.97112 | 103.21663 | Promoter | 9 | 122175879 | 122294374 | 118496 | 2 | ENSMUST00 | 0 | Ano10 | ENSMUSG00000037949.7 |
| chr9 | 21988524 | 21990166 | 31.16539 | 106.97112 | 103.21663 | Promoter | 9 | 21971527 | 21989453 | 17927 | 2 | ENSMUST00 | 0 | Rgl3 | ENSMUSG00000040146.8 |
| chr9 | 108078441 | 108079690 | 31.16539 | 106.97112 | 103.21663 | Promoter | 9 | 108051672 | 108079301 | 27630 | 2 | ENSMUST00 | 0 | Rnf123 | ENSMUSG00000041528.10 |
| chr9 | 21072811 | 21074046 | 31.16539 | 106.97112 | 103.21663 | Promoter | 9 | 21067520 | 21073514 | 5995 | 2 | ENSMUST00 | 0 | Fdx1l | ENSMUSG00000079677.1 |
| chr16 | 35982507 | 35984385 | 31.14594 | 106.93581 | 103.18726 | Promoter | 16 | 35983315 | 36037131 | 53817 | 1 | ENSMUST00 | 0 | Kpna1 | ENSMUSG00000022905.7 |
| chr10 | 36973845 | 36975536 | 32.27911 | 106.92835 | 103.17982 | Promoter | 10 | 36974544 | 37001889 | 27346 | 1 | ENSMUST00 | 0 | Hdac2 | ENSMUSG00000019777.11 |
| chr11 | 4637155 | 4638771 | 32.27911 | 106.92835 | 103.17982 | Promoter | 11 | 4637747 | 4683385 | 45639 | 1 | ENSMUST00 | 0 | Ascc2 | ENSMUSG00000020412.11 |
| chr18 | 61210759 | 61212026 | 32.27911 | 106.92835 | 103.17982 | Promoter | 18 | 61192919 | 61211612 | 18694 | 2 | ENSMUST00 | 0 | Slc26a2 | ENSMUSG00000034320.9 |
| chr7 | 116442665 | 116444451 | 32.27911 | 106.92835 | 103.17982 | Promoter | 7 | 116337276 | 116443458 | 106183 | 2 | ENSMUST00 | 0 | Pik3c2a | ENSMUSG00000030660.8 |
| chr8 | 105880700 | 105881719 | 32.27911 | 106.92835 | 103.17982 | Promoter | 8 | 105880881 | 105893225 | 12345 | 1 | ENSMUST00 | 0 | Edc4 | ENSMUSG00000036270.11 |
| chr12 | 83520016 | 83521210 | 33.43206 | 106.91093 | 103.1637 | Promoter | 12 | 83520466 | 83541988 | 21523 | 1 | ENSMUST00 | 0 | Dcaf4 | ENSMUSG00000021222.8 |
| chr14 | 70152889 | 70154166 | 32.83827 | 106.89391 | 103.14682 | Promoter | 14 | 70138177 | 70153811 | 15635 | 2 | ENSMUST00 | 0 | Ccar2 | ENSMUSG00000033712.5 |
| chr18 | 9707115 | 9709019 | 34.59452 | 106.87945 | 103.13243 | Promoter | 18 | 9707648 | 9877995 | 170348 | 1 | ENSMUST00 | 0 | Colec12 | ENSMUSG00000036103.8 |
| chr16 | 20651158 | 20652659 | 33.99591 | 106.86781 | 103.1211 | Promoter | 16 | 20651652 | 20663414 | 11763 | 1 | ENSMUST00 | 0 | Psmd2 | ENSMUSG00000006998.10 |
| chr18 | 69345746 | 69346752 | 33.99591 | 106.86781 | 103.1211 | Promoter | 18 | 69345771 | 69487850 | 142080 | 1 | ENSMUST00 | 0 | Tcf4 | ENSMUSG00000053477.10 |
| chr6 | 83506331 | 83507511 | 32.22038 | 106.82387 | 103.07761 | Promoter | 6 | 83480217 | 83506969 | 26753 | 2 | ENSMUST00 | 0 | Dguok | ENSMUSG00000014554.8 |
| chr15 | 63997162 | 63998791 | 34.52358 | 106.75686 | 103.01088 | Promoter | 15 | 63930944 | 63997969 | 67026 | 2 | ENSMUST00 | 0 | Fam49b | ENSMUSG00000022378.8 |
| chr6 | 42691997 | 42693318 | 34.52358 | 106.75686 | 103.01088 | Promoter | 6 | 42669942 | 42693087 | 23146 | 2 | ENSMUST00 | 0 | Fam115a | ENSMUSG00000036667.9 |
| chr6 | 128361968 | 128363462 | 22.15647 | 106.74507 | 102.9996 | Promoter | 6 | 128362967 | 128368962 | 5996 | 1 | ENSMUST00 | 0 | Foxm1 | ENSMUSG00000001517.9 |
| chr18 | 11838767 | 11842359 | 28.35087 | 106.74432 | 102.99886 | Promoter | 18 | 11839220 | 11945621 | 106402 | 1 | ENSMUST00 | 0 | Cables1 | ENSMUSG00000040957.9 |
| chr10 | 60830021 | 60831584 | 33.30314 | 106.68535 | 102.93993 | Promoter | 10 | 60762610 | 60831581 | 68972 | 2 | ENSMUST00 | 0 | Unc5b | ENSMUSG00000020099.7 |
| chr12 | 56344852 | 56346314 | 33.30314 | 106.68535 | 102.93993 | Promoter | 12 | 56328306 | 56345894 | 17589 | 2 | ENSMUST00 | 0 | Mbip | ENSMUSG00000021028.7 |
| chr18 | 6764641 | 6766162 | 33.30314 | 106.68535 | 102.93993 | Promoter | 18 | 6765205 | 6790231 | 25027 | 1 | ENSMUST00 | 0 | Rab18 | ENSMUSG00000073639.5 |
| chr2 | 25364801 | 25365954 | 33.30314 | 106.68535 | 102.93993 | Promoter | 2 | 25359889 | 25365682 | 5794 | 2 | ENSMUST00 | 0 | Uap1l1 | ENSMUSG00000026956.10 |
| chr3 | 30855641 | 30856601 | 33.30314 | 106.68535 | 102.93993 | Promoter | 3 | 30855950 | 30879461 | 23512 | 1 | ENSMUST00 | 0 | Gpr160 | ENSMUSG00000037661.9 |
| chr4 | 32799634 | 32800936 | 33.30314 | 106.68535 | 102.93993 | Promoter | 4 | 32800253 | 32801383 | 1131 | 1 | ENSMUST00 | 0 | Lyrm2 | ENSMUSG00000045854.4 |
| chr5 | 135778559 | 135779650 | 33.30314 | 106.68535 | 102.93993 | Promoter | 5 | 135778662 | 135790398 | 11737 | 1 | ENSMUST00 | 0 | Mdh2 | ENSMUSG00000019179.6 |
| chr5 | 36747653 | 36748934 | 33.30314 | 106.68535 | 102.93993 | Promoter | 5 | 36747374 | 36748679 | 1306 | 2 | ENSMUST00 | 0 | Bloc1s4 | ENSMUSG00000060708.3 |
| chr9 | 71590867 | 71592639 | 33.30314 | 106.68535 | 102.93993 | Promoter | 9 | 71504347 | 71592360 | 88014 | 2 | ENSMUST00 | 0 | Myzap | ENSMUSG00000041361.8 |
| chr1 | 191223989 | 191225334 | 26.32471 | 106.6389 | 102.89503 | Promoter | 1 | 191224474 | 191225686 | 1213 | 1 | ENSMUST00 | 0 | D730003I15F | ENSMUSG00000073478.5 |
| chr11 | 62455956 | 62459233 | 26.32471 | 106.6389 | 102.89503 | Promoter | 11 | 62458460 | 62514417 | 55958 | 1 | ENSMUST00 | 0 | Pigl | ENSMUSG00000014245.8 |
| chr11 | 22512039 | 22513396 | 26.32471 | 106.6389 | 102.89503 | Promoter | 11 | 22512088 | 22519234 | 7147 | 1 | ENSMUST00 | 0 | Tmem17 | ENSMUSG00000049904.7 |
| chr12 | 13268801 | 13270170 | 26.32471 | 106.6389 | 102.89503 | Promoter | 12 | 13269127 | 13583810 | 314684 | 1 | ENSMUST00 | 0 | Nbas | ENSMUSG00000020576.9 |
| chr17 | 25133328 | 25134022 | 26.32471 | 106.6389 | 102.89503 | Promoter | 17 | 25133391 | 25162104 | 28714 | 1 | ENSMUST00 | 0 | Clcn7 | ENSMUSG00000036636.4 |
| chr2 | 74577860 | 74579489 | 26.32471 | 106.6389 | 102.89503 | Promoter | 2 | 74520291 | 74578948 | 58658 | 2 | ENSMUST00 | 0 | Lnp | ENSMUSG00000009207.10 |
| chr2 | 30981257 | 30983331 | 26.32471 | 106.6389 | 102.89503 | Promoter | 2 | 30972833 | 30981941 | 9109 | 2 | ENSMUST00 | 0 | BC005624 | ENSMUSG00000026851.8 |
| chr1 | 143701477 | 143703505 | 34.45293 | 106.63475 | 102.89172 | Promoter | 1 | 143598800 | 143702893 | 104094 | 2 | ENSMUST00 | 0 | Cdc73 | ENSMUSG00000026361.8 |
| chr3 | 57573515 | 57576914 | 34.45293 | 106.63475 | 102.89172 | Promoter | 3 | 57455649 | 57575828 | 120180 | 2 | ENSMUST00 | 0 | Wwtr1 | ENSMUSG00000027803.9 |
| chr8 | 123858622 | 123860243 | 34.45293 | 106.63475 | 102.89172 | Promoter | 8 | 123840831 | 123859530 | 18700 | 2 | ENSMUST00 | 0 | Ccsap | ENSMUSG00000031971.10 |
| chr9 | 118925107 | 118929928 | 34.45293 | 106.63475 | 102.89172 | Promoter | 9 | 118926453 | 119043998 | 117546 | 1 | ENSMUST00 | 0 | Ctdspl | ENSMUSG00000047409.8 |
| chr4 | 101354509 | 101357858 | 33.86091 | 106.63318 | 102.89114 | Promoter | 4 | 101346531 | 101356248 | 9718 | 2 | ENSMUST00 | 0 | E130102H24I | ENSMUSG00000086782.2 |
| chr13 | 97137178 | 97138877 | 31.52828 | 106.60726 | 102.86559 | Promoter | 13 | 97137937 | 97155477 | 17541 | 1 | ENSMUST00 | 0 | Gfm2 | ENSMUSG00000021666.11 |
| chr5 | 134183660 | 134184868 | 31.52828 | 106.60726 | 102.86559 | Promoter | 5 | 134184019 | 134224355 | 40337 | 1 | ENSMUST00 | 0 | Gtf2ird2 | ENSMUSG00000015942.4 |
| chr5 | 76904825 | 76906192 | 31.52828 | 106.60726 | 102.86559 | Promoter | 5 | 76873659 | 76905427 | 31769 | 2 | ENSMUST00 | 0 | Aasdh | ENSMUSG00000055923.11 |
| chr9 | 21592227 | 21593964 | 31.52828 | 106.60726 | 102.86559 | Promoter | 9 | 21588709 | 21592831 | 4123 | 2 | ENSMUST00 | 0 | Yipf2 | ENSMUSG00000032182.9 |
| chr2 | 61711261 | 61712790 | 33.23905 | 106.57314 | 102.83228 | Promoter | 2 | 61711694 | 61800376 | 88683 | 1 | ENSMUST00 | 0 | Psmd14 | ENSMUSG00000026914.10 |
| chr1 | 63112781 | 63116118 | 29.2476 | 106.45832 | 102.71833 | Promoter | 1 | 63047801 | 63114267 | 66467 | 2 | ENSMUST00 | 0 | Ino80d | ENSMUSG00000040865.10 |
| chr11 | 75587816 | 75589321 | 29.2476 | 106.45832 | 102.71833 | Promoter | 11 | 75588097 | 75628804 | 40708 | 1 | ENSMUST00 | 0 | Pitpna | ENSMUSG00000017781.12 |
| chr11 | 115823561 | 115824840 | 29.2476 | 106.45832 | 102.71833 | Promoter | 11 | 115824049 | 115855780 | 31732 | 1 | ENSMUST00 | 0 | Llgl2 | ENSMUSG00000020782.13 |
| chr11 | 94676447 | 94677907 | 29.2476 | 106.45832 | 102.71833 | Promoter | 11 | 94663851 | 94677515 | 13665 | 2 | ENSMUST00 | 0 | Xylt2 | ENSMUSG00000020868.10 |
| chr13 | 66904045 | 66905759 | 29.2476 | 106.45832 | 102.71833 | Promoter | 13 | 66900617 | 66905378 | 4762 | 2 | ENSMUST00 | 0 | Uqcrb | ENSMUSG00000021520.4 |
| chr13 | 54574538 | 54576156 | 29.2476 | 106.45832 | 102.71833 | Promoter | 13 | 54575015 | 54581128 | 6114 | 1 | ENSMUST00 | 0 | Arl10 | ENSMUSG00000025870.5 |
| chr13 | 45001036 | 45002726 | 29.2476 | 106.45832 | 102.71833 | Promoter | 13 | 44922079 | 45002096 | 80018 | 2 | ENSMUST00 | 0 | Dtnbp1 | ENSMUSG00000057531.8 |
| chr14 | 74731818 | 74733133 | 29.2476 | 106.45832 | 102.71833 | Promoter | 14 | 74732297 | 74750765 | 18469 | 1 | ENSMUST00 | 0 | Esd | ENSMUSG00000021996.11 |
| chr15 | 67176959 | 67179707 | 29.2476 | 106.45832 | 102.71833 | Distal Interge | 15 | 67226769 | 67377094 | 150326 | 1 | ENSMUST00 | -47062 | RP23-163N4. | ENSMUSG00000102069.1 |
| chr16 | 16982800 | 16984633 | 29.2476 | 106.45832 | 102.71833 | Promoter | 16 | 16983382 | 17044749 | 61368 | 1 | ENSMUST00 | 0 | Mapk1 | ENSMUSG00000063358.10 |
| chr17 | 64600157 | 64602660 | 29.2476 | 106.45832 | 102.71833 | Promoter | 17 | 64600736 | 64755110 | 154375 | 1 | ENSMUST00 | 0 | Man2a1 | ENSMUSG00000024085.8 |
| chr18 | 46958425 | 46959506 | 29.2476 | 106.45832 | 102.71833 | Promoter | 18 | 46958862 | 47087992 | 129131 | 1 | ENSMUST00 | 0 | Commd10 | ENSMUSG00000042705.8 |
| chr2 | 150786247 | 150787742 | 29.2476 | 106.45832 | 102.71833 | Promoter | 2 | 150786735 | 150831758 | 45024 | 1 | ENSMUST00 | 0 | Pygb | ENSMUSG00000033059.7 |

| | | | | | | | | | | | | | | | |
|---|---|---|---|---|---|---|---|---|---|---|---|---|---|---|---|
| chr2 | 26963536 | 26965076 | 29.2476 | 106.45832 | 102.71833 | Promoter | 2 | 26953563 | 26964386 | 10824 | 2 | ENSMUST00 | 0 | Rexo4 | ENSMUSG00000052406.9 |
| chr2 | 33430325 | 33431829 | 29.2476 | 106.45832 | 102.71833 | Promoter | 2 | 33406108 | 33431324 | 25217 | 2 | ENSMUST00 | 0 | Zbtb34 | ENSMUSG00000068966.5 |
| chr3 | 144570490 | 144571614 | 29.2476 | 106.45832 | 102.71833 | Promoter | 3 | 144570693 | 144597661 | 26969 | 1 | ENSMUST00 | 0 | Sep-15 | ENSMUSG00000037072.10 |
| chr3 | 97657492 | 97659356 | 29.2476 | 106.45832 | 102.71833 | Promoter | 3 | 97658193 | 97670799 | 12607 | 1 | ENSMUST00 | 0 | Prkab2 | ENSMUSG00000038205.7 |
| chr3 | 88774813 | 88775822 | 29.2476 | 106.45832 | 102.71833 | Promoter | 3 | 88744700 | 88775164 | 30465 | 2 | ENSMUST00 | 0 | Syt11 | ENSMUSG00000068923.9 |
| chr4 | 55349975 | 55351158 | 29.2476 | 106.45832 | 102.71833 | Promoter | 4 | 55350043 | 55392237 | 42195 | 1 | ENSMUST00 | 0 | Rad23b | ENSMUSG00000028426.10 |
| chr4 | 43874975 | 43876536 | 29.2476 | 106.45832 | 102.71833 | Promoter | 4 | 43875530 | 43944745 | 69216 | 1 | ENSMUST00 | 0 | Reck | ENSMUSG00000028476.8 |
| chr4 | 150909156 | 150910322 | 29.2476 | 106.45832 | 102.71833 | Promoter | 4 | 150897133 | 150909926 | 12794 | 2 | ENSMUST00 | 0 | Park7 | ENSMUSG00000028964.9 |
| chr4 | 86873870 | 86875346 | 29.2476 | 106.45832 | 102.71833 | Promoter | 4 | 86874396 | 86920875 | 46480 | 1 | ENSMUST00 | 0 | Acer2 | ENSMUSG00000038007.9 |
| chr4 | 141722638 | 141723984 | 29.2476 | 106.45832 | 102.71833 | Promoter | 4 | 141677549 | 141723419 | 45871 | 2 | ENSMUST00 | 0 | Ddi2 | ENSMUSG00000078515.4 |
| chr5 | 120586812 | 120588712 | 29.2476 | 106.45832 | 102.71833 | Promoter | 5 | 120534160 | 120588613 | 54454 | 2 | ENSMUST00 | 0 | Tpcn1 | ENSMUSG00000032741.8 |
| chr5 | 135393587 | 135395145 | 29.2476 | 106.45832 | 102.71833 | Promoter | 5 | 135376141 | 135394510 | 18370 | 2 | ENSMUST00 | 0 | Pom121 | ENSMUSG00000053293.6 |
| chr5 | 129886167 | 129887389 | 29.2476 | 106.45832 | 102.71833 | Promoter | 5 | 129881159 | 129887273 | 6115 | 2 | ENSMUST00 | 0 | Chchd2 | ENSMUSG00000070493.3 |
| chr6 | 91877489 | 91878947 | 29.2476 | 106.45832 | 102.71833 | Promoter | 6 | 91878053 | 91899843 | 21791 | 1 | ENSMUST00 | 0 | Ccdc174 | ENSMUSG00000034083.11 |
| chr6 | 127453204 | 127454489 | 29.2476 | 106.45832 | 102.71833 | Promoter | 6 | 127453624 | 127472028 | 18405 | 1 | ENSMUST00 | 0 | Parp11 | ENSMUSG00000037997.7 |
| chr7 | 126797639 | 126799355 | 29.2476 | 106.45832 | 102.71833 | Promoter | 7 | 126795234 | 126799254 | 4021 | 2 | ENSMUST00 | 0 | Aldoa | ENSMUSG00000030695.8 |
| chr8 | 94036106 | 94038234 | 29.2476 | 106.45832 | 102.71833 | Promoter | 8 | 94017770 | 94037021 | 19252 | 2 | ENSMUST00 | 0 | Nudt21 | ENSMUSG00000031754.9 |
| chr8 | 111642915 | 111644635 | 29.2476 | 106.45832 | 102.71833 | Promoter | 8 | 111643443 | 111671008 | 27566 | 1 | ENSMUST00 | 0 | Zfp1 | ENSMUSG00000055835.9 |
| chr9 | 44406531 | 44408619 | 29.2476 | 106.45832 | 102.71833 | Promoter | 9 | 44407714 | 44410405 | 2692 | 1 | ENSMUST00 | 0 | Rps25 | ENSMUSG00000009927.8 |
| chr9 | 22116245 | 22117724 | 29.2476 | 106.45832 | 102.71833 | Promoter | 9 | 22112990 | 22117148 | 4159 | 2 | ENSMUST00 | 0 | Elof1 | ENSMUSG00000013822.6 |
| chr9 | 100544788 | 100547004 | 29.2476 | 106.45832 | 102.71833 | Promoter | 9 | 100492293 | 100546134 | 53842 | 2 | ENSMUST00 | 0 | Nck1 | ENSMUSG00000032475.10 |
| chr9 | 108566205 | 108568464 | 29.2476 | 106.45832 | 102.71833 | Promoter | 9 | 108565866 | 108567342 | 1477 | 2 | ENSMUST00 | 0 | Ndufaf3 | ENSMUSG00000070283.3 |
| chr11 | 3288485 | 3291480 | 34.92444 | 106.42502 | 102.68874 | Promoter | 11 | 3288874 | 3290192 | 1319 | 1 | ENSMUST00 | 0 | Patz1 | ENSMUSG00000020453.12 |
| chr11 | 119084733 | 119086327 | 34.92444 | 106.42502 | 102.68874 | Promoter | 11 | 119077573 | 119086221 | 8649 | 2 | ENSMUST00 | 0 | Cbx4 | ENSMUSG00000039989.7 |
| chr12 | 76764979 | 76766448 | 34.92444 | 106.42502 | 102.68874 | Promoter | 12 | 76765589 | 76920659 | 155071 | 1 | ENSMUST00 | 0 | Fntb | ENSMUSG00000033373.11 |
| chr16 | 33684190 | 33685739 | 34.92444 | 106.42502 | 102.68874 | Promoter | 16 | 33684466 | 33722913 | 38448 | 1 | ENSMUST00 | 0 | Heg1 | ENSMUSG00000075254.6 |
| chr17 | 83350894 | 83352559 | 34.92444 | 106.42502 | 102.68874 | Promoter | 17 | 83350931 | 83480361 | 129431 | 1 | ENSMUST00 | 0 | Eml4 | ENSMUSG00000032624.10 |
| chr2 | 164832882 | 164834605 | 34.92444 | 106.42502 | 102.68874 | Promoter | 2 | 164832887 | 164833759 | 873 | 1 | ENSMUST00 | 0 | Ctsa | ENSMUSG00000017760.10 |
| chr2 | 32981149 | 32982972 | 34.92444 | 106.42502 | 102.68874 | Promoter | 2 | 32972990 | 32982082 | 9093 | 2 | ENSMUST00 | 0 | Slc2a8 | ENSMUSG00000026791.9 |
| chr5 | 110135737 | 110136728 | 34.92444 | 106.42502 | 102.68874 | Promoter | 5 | 110135840 | 110171972 | 36133 | 1 | ENSMUST00 | 0 | Chfr | ENSMUSG00000014668.10 |
| chr6 | 86792202 | 86794077 | 34.92444 | 106.42502 | 102.68874 | Promoter | 6 | 86736840 | 86793584 | 56745 | 2 | ENSMUST00 | 0 | Anxa4 | ENSMUSG00000029994.12 |
| chr6 | 83101111 | 83102693 | 34.92444 | 106.42502 | 102.68874 | Promoter | 6 | 83101617 | 83108377 | 6761 | 1 | ENSMUST00 | 0 | Ccdc142 | ENSMUSG00000079511.4 |
| chr8 | 120486986 | 120490241 | 34.92444 | 106.42502 | 102.68874 | Promoter | 8 | 120488447 | 120581390 | 92944 | 1 | ENSMUST00 | 0 | Gse1 | ENSMUSG00000031822.13 |
| chr8 | 84235808 | 84237146 | 34.92444 | 106.42502 | 102.68874 | Promoter | 8 | 84210942 | 84237042 | 26101 | 2 | ENSMUST00 | 0 | Zswim4 | ENSMUSG00000035671.5 |
| chr9 | 37144737 | 37147348 | 34.92444 | 106.42502 | 102.68874 | Promoter | 9 | 36890982 | 37147289 | 256308 | 2 | ENSMUST00 | 0 | Pknox2 | ENSMUSG00000035934.10 |
| chr9 | 33728137 | 33728494 | 34.92444 | 106.42502 | 102.68874 | Promoter | 9 | 33728247 | 33728850 | 604 | 1 | ENSMUST00 | 0 | Gm10698 | ENSMUSG00000079884.6 |
| chr18 | 24019506 | 24021414 | 31.98759 | 106.40915 | 102.67528 | Promoter | 18 | 24009702 | 24020810 | 11109 | 2 | ENSMUST00 | 0 | Zfp191 | ENSMUSG00000051469.9 |
| chr11 | 121228401 | 121229619 | 34.31248 | 106.39187 | 102.65826 | Promoter | 11 | 121222588 | 121229305 | 6718 | 2 | ENSMUST00 | 0 | BC017643 | ENSMUSG00000039294.9 |
| chr10 | 59220670 | 59223467 | 27.66025 | 106.38611 | 102.65278 | Promoter | 10 | 59164113 | 59221847 | 57735 | 2 | ENSMUST00 | 0 | Sep-10 | ENSMUSG00000019917.9 |
| chr10 | 82698621 | 82700018 | 27.66025 | 106.38611 | 102.65278 | Promoter | 10 | 82699007 | 82742428 | 43422 | 1 | ENSMUST00 | 0 | Hcfc2 | ENSMUSG00000020246.8 |
| chr11 | 78037515 | 78038718 | 27.66025 | 106.38611 | 102.65278 | Promoter | 11 | 78037931 | 78060434 | 22504 | 1 | ENSMUST00 | 0 | Flot2 | ENSMUSG00000061981.9 |
| chr12 | 35534046 | 35535469 | 27.66025 | 106.38611 | 102.65278 | Promoter | 12 | 35497974 | 35535038 | 37065 | 2 | ENSMUST00 | 0 | Ahr | ENSMUSG00000019256.12 |
| chr12 | 55124043 | 55125539 | 27.66025 | 106.38611 | 102.65278 | Promoter | 12 | 55124528 | 55142078 | 17551 | 1 | ENSMUST00 | 0 | Fam177a | ENSMUSG00000095595.1 |
| chr15 | 79690814 | 79691865 | 27.66025 | 106.38611 | 102.65278 | Promoter | 15 | 79690895 | 79721478 | 30584 | 1 | ENSMUST00 | 0 | Gtpbp1 | ENSMUSG00000042535.8 |
| chr17 | 56626194 | 56627894 | 27.66025 | 106.38611 | 102.65278 | Promoter | 17 | 56614301 | 56626903 | 12603 | 2 | ENSMUST00 | 0 | Lonp1 | ENSMUSG00000041168.9 |
| chr19 | 45748670 | 45749806 | 27.66025 | 106.38611 | 102.65278 | Promoter | 19 | 45747734 | 45749591 | 1858 | 2 | ENSMUST00 | 0 | Npm3 | ENSMUSG00000056209.7 |
| chr2 | 14073137 | 14075226 | 27.66025 | 106.38611 | 102.65278 | Promoter | 2 | 14074100 | 14148338 | 74239 | 1 | ENSMUST00 | 0 | Stam | ENSMUSG00000026718.12 |
| chr2 | 130423439 | 130425305 | 27.66025 | 106.38611 | 102.65278 | Promoter | 2 | 130424339 | 130438453 | 14115 | 1 | ENSMUST00 | 0 | Vps16 | ENSMUSG00000027411.12 |
| chr3 | 107630637 | 107632300 | 27.66025 | 106.38611 | 102.65278 | Promoter | 3 | 107612532 | 107631710 | 19179 | 2 | ENSMUST00 | 0 | Strip1 | ENSMUSG00000014601.9 |
| chr3 | 94581275 | 94582913 | 27.66025 | 106.38611 | 102.65278 | Promoter | 3 | 94497544 | 94582716 | 85173 | 2 | ENSMUST00 | 0 | Snx27 | ENSMUSG00000028136.10 |
| chr4 | 148087183 | 148088332 | 27.66025 | 106.38611 | 102.65278 | Promoter | 4 | 148077061 | 148088031 | 10971 | 2 | ENSMUST00 | 0 | Agtrap | ENSMUSG00000029007.8 |
| chr4 | 135271193 | 135272820 | 27.66025 | 106.38611 | 102.65278 | Promoter | 4 | 135213969 | 135272814 | 58846 | 2 | ENSMUST00 | 0 | Clic4 | ENSMUSG00000037242.8 |
| chr5 | 107287431 | 107289736 | 27.66025 | 106.38611 | 102.65278 | Promoter | 5 | 107106570 | 107289629 | 183060 | 2 | ENSMUST00 | 0 | Tgfbr3 | ENSMUSG00000029287.9 |
| chr6 | 122874095 | 122875405 | 27.66025 | 106.38611 | 102.65278 | Promoter | 6 | 122874557 | 122888934 | 14378 | 1 | ENSMUST00 | 0 | Necap1 | ENSMUSG00000030327.6 |
| chr7 | 4136986 | 4138227 | 27.66025 | 106.38611 | 102.65278 | Promoter | 7 | 4137039 | 4142183 | 5145 | 1 | ENSMUST00 | 0 | Leng8 | ENSMUSG00000035545.8 |
| chr8 | 109868032 | 109869607 | 27.66025 | 106.38611 | 102.65278 | Promoter | 8 | 109868542 | 109904971 | 36430 | 1 | ENSMUST00 | 0 | Phlpp2 | ENSMUSG00000031732.7 |
| chr9 | 106684656 | 106686403 | 27.66025 | 106.38611 | 102.65278 | Promoter | 9 | 106658746 | 106685682 | 26937 | 2 | ENSMUST00 | 0 | Tex264 | ENSMUSG00000040813.10 |
| chr13 | 54621256 | 54622835 | 30.82847 | 106.35825 | 102.62659 | Promoter | 13 | 54621784 | 54664068 | 42285 | 1 | ENSMUST00 | 0 | Faf2 | ENSMUSG00000025873.10 |
| chr9 | 55148832 | 55150580 | 30.82847 | 106.35825 | 102.62659 | Promoter | 9 | 55148854 | 55207529 | 58676 | 1 | ENSMUST00 | 0 | Ube2q2 | ENSMUSG00000032307.11 |
| chr5 | 139150127 | 139151032 | 32.53288 | 106.35484 | 102.62381 | Promoter | 5 | 139150223 | 139186510 | 36288 | 1 | ENSMUST00 | 0 | Heatr2 | ENSMUSG00000025857.6 |
| chr2 | 157203730 | 157205350 | 33.11161 | 106.34985 | 102.61888 | Promoter | 2 | 157145893 | 157204512 | 58620 | 2 | ENSMUST00 | 0 | Rbl1 | ENSMUSG00000027641.8 |
| chr10 | 31313314 | 31314055 | 33.10969 | 106.34647 | 102.61562 | Promoter | 10 | 31313382 | 31328204 | 14823 | 1 | ENSMUST00 | 0 | Hddc2 | ENSMUSG00000000295.7 |
| chr11 | 45955059 | 45955668 | 33.10969 | 106.34647 | 102.61562 | Promoter | 11 | 45946843 | 45955188 | 8346 | 2 | ENSMUST00 | 0 | Thg1l | ENSMUSG00000011254.11 |
| chr11 | 100440528 | 100441216 | 33.10969 | 106.34647 | 102.61562 | Promoter | 11 | 100422321 | 100441089 | 18769 | 2 | ENSMUST00 | 0 | Nt5c3b | ENSMUSG00000017176.12 |
| chr11 | 96318131 | 96320429 | 33.10969 | 106.34647 | 102.61562 | Promoter | 11 | 96318267 | 96321638 | 3372 | 1 | ENSMUST00 | 0 | Hoxb4 | ENSMUSG00000038692.8 |
| chr14 | 24490363 | 24491710 | 33.10969 | 106.34647 | 102.61562 | Promoter | 14 | 24490681 | 24495432 | 4752 | 1 | ENSMUST00 | 0 | Rps24 | ENSMUSG00000025290.11 |
| chr14 | 52019298 | 52021224 | 33.10969 | 106.34647 | 102.61562 | Promoter | 14 | 52006077 | 52020733 | 14657 | 2 | ENSMUST00 | 0 | Zfp219 | ENSMUSG00000049295.11 |
| chr15 | 85036337 | 85037731 | 33.10969 | 106.34647 | 102.61562 | Promoter | 15 | 85037062 | 85062828 | 25767 | 1 | ENSMUST00 | 0 | Fam118a | ENSMUSG00000022434.7 |
| chr15 | 78842114 | 78845803 | 33.10969 | 106.34647 | 102.61562 | Promoter | 15 | 78842647 | 78850902 | 8256 | 1 | ENSMUST00 | 0 | Cdc42ep1 | ENSMUSG00000049521.1 |
| chr15 | 99125784 | 99127374 | 33.10969 | 106.34647 | 102.61562 | Promoter | 15 | 99126578 | 99213206 | 86629 | 1 | ENSMUST00 | 0 | Spats2 | ENSMUSG00000051934.4 |
| chr16 | 38562109 | 38563688 | 33.10969 | 106.34647 | 102.61562 | Promoter | 16 | 38562812 | 38592162 | 29351 | 1 | ENSMUST00 | 0 | Tmem39a | ENSMUSG00000002845.9 |
| chr16 | 17508212 | 17509862 | 33.10969 | 106.34647 | 102.61562 | Promoter | 16 | 17508688 | 17526333 | 17646 | 1 | ENSMUST00 | 0 | Lztr1 | ENSMUSG00000022761.5 |
| chr16 | 21332479 | 21334089 | 33.10969 | 106.34647 | 102.61562 | Distal Interge | 16 | 21247518 | 21296872 | 49355 | 2 | ENSMUST00 | -35608 | Gm16863 | ENSMUSG00000096918.1 |
| chr17 | 25784914 | 25785870 | 33.10969 | 106.34647 | 102.61562 | Promoter | 17 | 25779843 | 25785540 | 5698 | 2 | ENSMUST00 | 0 | Haghl | ENSMUSG00000061046.4 |
| chr2 | 31949970 | 31953731 | 33.10969 | 106.34647 | 102.61562 | Promoter | 2 | 31950260 | 31973442 | 23183 | 1 | ENSMUST00 | 0 | Aif1l | ENSMUSG00000001864.8 |
| chr2 | 164804497 | 164805846 | 33.10969 | 106.34647 | 102.61562 | Promoter | 2 | 164792768 | 164804882 | 12115 | 2 | ENSMUST00 | 0 | Acot8 | ENSMUSG00000017307.10 |
| chr2 | 156143420 | 156145158 | 33.10969 | 106.34647 | 102.61562 | Promoter | 2 | 156123639 | 156144186 | 20548 | 2 | ENSMUST00 | 0 | Nfs1 | ENSMUSG00000027618.12 |
| chr2 | 129592138 | 129595051 | 33.10969 | 106.34647 | 102.61562 | Promoter | 2 | 129592835 | 129608992 | 16158 | 1 | ENSMUST00 | 0 | Sirpa | ENSMUSG00000037902.13 |
| chr2 | 93187257 | 93189185 | 33.10969 | 106.34647 | 102.61562 | Promoter | 2 | 93187548 | 93198919 | 11372 | 1 | ENSMUST00 | 0 | Trp53i11 | ENSMUSG00000068735.9 |
| chr3 | 37418934 | 37419636 | 33.10969 | 106.34647 | 102.61562 | Promoter | 3 | 37404910 | 37419580 | 14671 | 2 | ENSMUST00 | 0 | Nudt6 | ENSMUSG00000050174.10 |
| chr4 | 101419025 | 101420579 | 33.10969 | 106.34647 | 102.61562 | Promoter | 4 | 101419277 | 101466995 | 47719 | 1 | ENSMUST00 | 0 | Ak4 | ENSMUSG00000028527.13 |
| chr4 | 150854624 | 150856217 | 33.10969 | 106.34647 | 102.61562 | Promoter | 4 | 150855073 | 150855814 | 742 | 1 | ENSMUST00 | 0 | Errfi1 | ENSMUSG00000028967.5 |
| chr4 | 148443954 | 148445078 | 33.10969 | 106.34647 | 102.61562 | Promoter | 4 | 148434495 | 148444771 | 10277 | 2 | ENSMUST00 | 0 | Ubiad1 | ENSMUSG00000047719.2 |
| chr5 | 138263274 | 138264832 | 33.10969 | 106.34647 | 102.61562 | Promoter | 5 | 138259656 | 138263994 | 4339 | 2 | ENSMUST00 | 0 | BC037034 | ENSMUSG00000036948.12 |
| chr7 | 34652958 | 34655928 | 33.10969 | 106.34647 | 102.61562 | Promoter | 7 | 34639014 | 34654398 | 15385 | 2 | ENSMUST00 | 0 | Kctd15 | ENSMUSG00000030499.4 |
| chr7 | 27165284 | 27167620 | 33.10969 | 106.34647 | 102.61562 | Promoter | 7 | 27158658 | 27166802 | 8145 | 2 | ENSMUST00 | 0 | Egln2 | ENSMUSG00000058709.6 |
| chr8 | 8688529 | 8690897 | 33.10969 | 106.34647 | 102.61562 | Promoter | 8 | 8666577 | 8690537 | 23961 | 2 | ENSMUST00 | 0 | Arglu1 | ENSMUSG00000040459.10 |
| chr8 | 121116146 | 121123525 | 33.10969 | 106.34647 | 102.61562 | Promoter | 8 | 121116171 | 121118895 | 2725 | 1 | ENSMUST00 | 0 | Foxc2 | ENSMUSG00000046714.7 |
| chr8 | 104394767 | 104396288 | 33.10969 | 106.34647 | 102.61562 | Promoter | 8 | 104348191 | 104395807 | 47617 | 2 | ENSMUST00 | 0 | Cmtm4 | ENSMUSG00000096188.1 |
| chr9 | 114688118 | 114690077 | 33.10969 | 106.34647 | 102.61562 | Promoter | 9 | 114688790 | 114724302 | 35513 | 1 | ENSMUST00 | 0 | Dync1li1 | ENSMUSG00000032435.8 |
| chr9 | 121995462 | 121996156 | 33.10969 | 106.34647 | 102.61562 | Promoter | 9 | 121981606 | 121996026 | 14421 | 2 | ENSMUST00 | 0 | Pomgnt2 | ENSMUSG00000066235.6 |
| chr17 | 29006909 | 29008516 | 23.95524 | 106.32738 | 102.60156 | Promoter | 17 | 28970885 | 29007937 | 37053 | 2 | ENSMUST00 | 0 | Stk38 | ENSMUSG00000024006.11 |
| chr13 | 98262196 | 98264201 | 33.6604 | 106.28434 | 102.55876 | Promoter | 13 | 98263158 | 98273853 | 10696 | 1 | ENSMUST00 | 0 | Ankra2 | ENSMUSG00000021661.10 |
| chr14 | 20673707 | 20675377 | 33.6604 | 106.28434 | 102.55876 | Promoter | 14 | 20674321 | 20694850 | 20530 | 1 | ENSMUST00 | 0 | Sec24c | ENSMUSG00000039367.9 |
| chr17 | 26123419 | 26124426 | 34.24269 | 106.27111 | 102.54579 | Promoter | 17 | 26123520 | 26126613 | 3094 | 1 | ENSMUST00 | 0 | Mrpl28 | ENSMUSG00000024181.8 |
| chr16 | 70313499 | 70315103 | 32.47248 | 106.24805 | 102.52307 | Promoter | 16 | 70313949 | 70569716 | 255768 | 1 | ENSMUST00 | 0 | Gbe1 | ENSMUSG00000022707.11 |
| chr17 | 26676095 | 26677386 | 32.47248 | 106.24805 | 102.52307 | Promoter | 17 | 26676396 | 26699644 | 23249 | 1 | ENSMUST00 | 0 | Atp6v0e | ENSMUSG00000015575.9 |
| chr3 | 108084894 | 108087049 | 32.47248 | 106.24805 | 102.52307 | Promoter | 3 | 108074062 | 108086627 | 12566 | 2 | ENSMUST00 | 0 | Ampd2 | ENSMUSG00000027889.12 |
| chr3 | 40540442 | 40541329 | 32.47248 | 106.24805 | 102.52307 | Promoter | 3 | 40540767 | 40704774 | 164008 | 1 | ENSMUST00 | 0 | Intu | ENSMUSG00000060798.4 |
| chr7 | 144552932 | 144554372 | 32.47248 | 106.24805 | 102.52307 | Promoter | 7 | 144476758 | 144553729 | 76972 | 2 | ENSMUST00 | 0 | Ppfia1 | ENSMUSG00000037519.11 |
| chr7 | 24884130 | 24885761 | 32.47248 | 106.24805 | 102.52307 | Promoter | 7 | 24884371 | 24889806 | 5436 | 1 | ENSMUST00 | 0 | Rps19 | ENSMUSG00000040952.11 |
| chr7 | 67222099 | 67223466 | 32.47248 | 106.24805 | 102.52307 | Promoter | 7 | 67222544 | 67228468 | 5925 | 1 | ENSMUST00 | 0 | Lysmd4 | ENSMUSG00000043831.6 |
| chr1 | 136344371 | 136346807 | 33.04826 | 106.23876 | 102.51532 | Promoter | 1 | 136268123 | 136346104 | 77982 | 2 | ENSMUST00 | 0 | Camsap2 | ENSMUSG00000041570.9 |
| chr5 | 19223861 | 19232405 | 34.78864 | 106.1911 | 102.46782 | Promoter | 5 | 19227046 | 20702903 | 1475858 | 1 | ENSMUST00 | 0 | Magi2 | ENSMUSG00000040003.12 |
| chr9 | 67046899 | 67049946 | 34.78864 | 106.1911 | 102.46782 | Promoter | 9 | 67022590 | 67049267 | 26678 | 2 | ENSMUST00 | 0 | Tpm1 | ENSMUSG00000032366.10 |
| chr11 | 70763721 | 70765043 | 30.17466 | 106.17292 | 102.44986 | Promoter | 11 | 70764209 | 70772928 | 8720 | 1 | ENSMUST00 | 0 | Zfp3 | ENSMUSG00000043602.5 |
| chr12 | 112976024 | 112977315 | 34.17318 | 106.15081 | 102.42828 | Promoter | 12 | 112976482 | 112978930 | 2449 | 1 | ENSMUST00 | 0 | Btbd6 | ENSMUSG00000002803.8 |
| chr12 | 99391404 | 99393876 | 34.17318 | 106.15081 | 102.42828 | Promoter | 12 | 99194980 | 99393797 | 198818 | 2 | ENSMUST00 | 0 | Foxn3 | ENSMUSG00000033713.6 |
| chr5 | 45449460 | 45450241 | 34.17318 | 106.15081 | 102.42828 | Promoter | 5 | 45434032 | 45450235 | 16204 | 2 | ENSMUST00 | 0 | Qdpr | ENSMUSG00000015806.7 |
| chr3 | 102995208 | 102996541 | 32.98514 | 106.12805 | 102.40588 | Promoter | 3 | 102995728 | 103003914 | 8187 | 1 | ENSMUST00 | 0 | Sike1 | ENSMUSG00000027854.7 |
| chr4 | 101647700 | 101649031 | 32.98514 | 106.12805 | 102.40588 | Promoter | 4 | 101647718 | 101659364 | 11647 | 1 | ENSMUST00 | 0 | Leprot | ENSMUSG00000035212.9 |
| chr17 | 34592255 | 34596534 | 34.71595 | 106.06588 | 102.34386 | Promoter | 17 | 34592263 | 34597400 | 5138 | 1 | ENSMUST00 | 0 | Pbx2 | ENSMUSG00000034673.9 |
| chr2 | 106692866 | 106695403 | 34.71595 | 106.06588 | 102.34386 | Promoter | 2 | 106693269 | 106868356 | 175088 | 1 | ENSMUST00 | 0 | Mpped2 | ENSMUSG00000016386.10 |
| chr4 | 132463229 | 132464465 | 34.71595 | 106.06588 | 102.34386 | Promoter | 4 | 132458731 | 132463921 | 5191 | 2 | ENSMUST00 | 0 | Med18 | ENSMUSG00000066042.4 |
| chr1 | 194975913 | 194977642 | 33.52805 | 106.05383 | 102.33234 | Promoter | 1 | 194976375 | 195021493 | 45119 | 1 | ENSMUST00 | 0 | A330023F24 | ENSMUSG00000096929.2 |
| chr10 | 128058154 | 128060485 | 33.52805 | 106.05383 | 102.33234 | Promoter | 10 | 128058954 | 128077272 | 18319 | 1 | ENSMUST00 | 0 | Ptges3 | ENSMUSG00000071072.7 |
| chr13 | 41249177 | 41250862 | 33.52805 | 106.05383 | 102.33234 | Promoter | 13 | 41249860 | 41276574 | 26715 | 1 | ENSMUST00 | 0 | Smim13 | ENSMUSG00000091264.2 |
| chr2 | 105122726 | 105134202 | 33.52805 | 106.05383 | 102.33234 | Promoter | 2 | 105126529 | 105173612 | 47084 | 1 | ENSMUST00 | 0 | Wt1 | ENSMUSG00000016458.8 |
| chr2 | 121169593 | 121171927 | 33.52805 | 106.05383 | 102.33234 | Promoter | 2 | 121170654 | 121184430 | 13777 | 1 | ENSMUST00 | 0 | Tubgcp4 | ENSMUSG00000027263.7 |
| chr6 | 49073330 | 49074698 | 33.52805 | 106.05383 | 102.33234 | Promoter | 6 | 49073795 | 49084717 | 10923 | 1 | ENSMUST00 | 0 | Malsu1 | ENSMUSG00000029815.8 |
| chr7 | 126474394 | 126476021 | 33.52805 | 106.05383 | 102.33234 | Promoter | 7 | 126466994 | 126475130 | 8137 | 2 | ENSMUST00 | 0 | Sh2b1 | ENSMUSG00000030733.9 |
| chr8 | 84659950 | 84663752 | 33.52805 | 106.05383 | 102.33234 | Promoter | 8 | 84661331 | 84662852 | 1522 | 2 | ENSMUST00 | 0 | Ier2 | ENSMUSG00000053560.4 |
| chr19 | 46043949 | 46046177 | 34.10395 | 106.03094 | 102.3112 | Promoter | 19 | 46032593 | 46045214 | 12622 | 2 | ENSMUST00 | 0 | Ldb1 | ENSMUSG00000025223.9 |
| chr8 | 95487773 | 95489217 | 34.10395 | 106.03094 | 102.3112 | Promoter | 8 | 95446096 | 95488820 | 42725 | 2 | ENSMUST00 | 0 | Csnk2a2 | ENSMUSG00000046707.8 |
| chr16 | 56075165 | 56076480 | 34.64357 | 105.94113 | 102.22192 | Promoter | 16 | 56075409 | 56126544 | 51136 | 1 | ENSMUST00 | 0 | Senp7 | ENSMUSG00000052917.9 |
| chr9 | 44325844 | 44327884 | 34.64357 | 105.94113 | 102.22192 | Promoter | 9 | 44326845 | 44333600 | 6756 | 1 | ENSMUST00 | 0 | Dpagt1 | ENSMUSG00000032123.4 |
| chr5 | 121847865 | 121849996 | 34.03501 | 105.91151 | 102.19291 | Promoter | 5 | 121849028 | 121854599 | 5572 | 1 | ENSMUST00 | 0 | Fam109a | ENSMUSG00000044134.5 |
| chr10 | 30802117 | 30803889 | 31.70129 | 105.8978 | 102.17969 | Promoter | 10 | 30645584 | 30803107 | 157524 | 2 | ENSMUST00 | 0 | Ncoa7 | ENSMUSG00000039697.10 |

| | | | | | | | | | | | | | | | |
|---|---|---|---|---|---|---|---|---|---|---|---|---|---|---|---|
| chr10 | 86777964 | 86780307 | 31.70129 | 105.8978 | 102.17969 | Promoter | 10 | 86779005 | 86838389 | 59385 | 1 | ENSMUST00 | 0 | Nt5dc3 | ENSMUSG00000054027.7 |
| chr1 | 191025038 | 191026751 | 33.96634 | 105.79253 | 102.076 | Promoter | 1 | 191005832 | 191026190 | 20359 | 2 | ENSMUST00 | 0 | Mfsd7b | ENSMUSG00000066595.4 |
| chr11 | 115475189 | 115476342 | 33.96634 | 105.79253 | 102.076 | Promoter | 11 | 115475667 | 115490467 | 14801 | 1 | ENSMUST00 | 0 | Armc7 | ENSMUSG00000057219.6 |
| chr15 | 85734614 | 85736405 | 22.88266 | 105.72421 | 102.00799 | Promoter | 15 | 85734983 | 85777632 | 42650 | 1 | ENSMUST00 | 0 | Ppara | ENSMUSG00000022383.8 |
| chr1 | 60409123 | 60410773 | 34.49972 | 105.69308 | 101.97717 | Promoter | 1 | 60409619 | 60481158 | 71540 | 1 | ENSMUST00 | 0 | Abi2 | ENSMUSG00000026782.10 |
| chr4 | 108327397 | 108328719 | 34.49972 | 105.69308 | 101.97717 | Promoter | 4 | 108328140 | 108341544 | 13405 | 1 | ENSMUST00 | 0 | Coa7 | ENSMUSG00000048351.9 |
| chr19 | 8740880 | 8742746 | 33.89795 | 105.67399 | 101.95845 | Promoter | 19 | 8741413 | 8755586 | 14174 | 1 | ENSMUST00 | 0 | Stx5a | ENSMUSG00000010110.12 |
| chr19 | 21271332 | 21275517 | 33.89795 | 105.67399 | 101.95845 | Promoter | 19 | 21272278 | 21282289 | 10012 | 1 | ENSMUST00 | 0 | Zfand5 | ENSMUSG00000024750.5 |
| chr2 | 69821915 | 69823594 | 33.89795 | 105.67399 | 101.95845 | Promoter | 2 | 69821944 | 69824323 | 2380 | 1 | ENSMUST00 | 0 | Klhl23 | ENSMUSG00000042155.3 |
| chr7 | 134670248 | 134672039 | 33.89795 | 105.67399 | 101.95845 | Promoter | 7 | 134670687 | 135173639 | 502953 | 1 | ENSMUST00 | 0 | Dock1 | ENSMUSG00000058325.5 |
| chr11 | 119267921 | 119268857 | 30.9834 | 105.62366 | 101.90845 | Promoter | 11 | 119268026 | 119285454 | 17429 | 1 | ENSMUST00 | 0 | Gaa | ENSMUSG00000025579.9 |
| chr17 | 34186475 | 34188545 | 30.9834 | 105.62366 | 101.90845 | Promoter | 17 | 34187553 | 34197225 | 9673 | 1 | ENSMUST00 | 0 | Tap1 | ENSMUSG00000037321.12 |
| chr4 | 152007699 | 152011153 | 30.9834 | 105.62366 | 101.90845 | Promoter | 4 | 152008803 | 152017680 | 8878 | 1 | ENSMUST00 | 0 | Klhl21 | ENSMUSG00000073700.3 |
| chr15 | 25983685 | 25986055 | 24.92035 | 105.60457 | 101.88994 | Promoter | 15 | 25984366 | 25998482 | 14117 | 1 | ENSMUST00 | 0 | Zfp622 | ENSMUSG00000052253.5 |
| chr7 | 59227911 | 59230196 | 24.92035 | 105.60457 | 101.88994 | Promoter | 7 | 59228752 | 59306727 | 77976 | 1 | ENSMUST00 | 0 | Ube3a | ENSMUSG00000025326.8 |
| chr1 | 46852212 | 46853473 | 33.26644 | 105.59763 | 101.88331 | Promoter | 1 | 46807544 | 46854046 | 46503 | 2 | ENSMUST00 | 573 | Slc39a10 | ENSMUSG00000025986.6 |
| chr1 | 23256193 | 23259443 | 33.26644 | 105.59763 | 101.88331 | Distal Interge | 1 | 23245211 | 23245331 | 121 | 1 | ENSMUST00 | 10983 | Gm24176 | ENSMUSG00000089553.1 |
| chr6 | 97210699 | 97212326 | 33.26644 | 105.59763 | 101.88331 | Promoter | 6 | 97210689 | 97233315 | 22627 | 1 | ENSMUST00 | 11 | Arl6ip5 | ENSMUSG00000035199.6 |
| chr13 | 113614178 | 113618647 | 32.67316 | 105.57997 | 101.8659 | Promoter | 13 | 113592180 | 113618564 | 26385 | 2 | ENSMUST00 | 0 | Snx18 | ENSMUSG00000042364.10 |
| chr15 | 98830661 | 98831981 | 32.67316 | 105.57997 | 101.8659 | Promoter | 15 | 98812798 | 98831508 | 18711 | 2 | ENSMUST00 | 0 | Prkag1 | ENSMUSG00000067713.6 |
| chr16 | 13818427 | 13820142 | 32.67316 | 105.57997 | 101.8659 | Promoter | 16 | 13819251 | 13835451 | 16201 | 1 | ENSMUST00 | 0 | Ntan1 | ENSMUSG00000022681.9 |
| chr17 | 79894559 | 79896570 | 32.67316 | 105.57997 | 101.8659 | Promoter | 17 | 79848392 | 79896028 | 47637 | 2 | ENSMUST00 | 0 | Atl2 | ENSMUSG00000059811.7 |
| chr3 | 88553099 | 88554546 | 32.67316 | 105.57997 | 101.8659 | Promoter | 3 | 88553716 | 88569725 | 16010 | 1 | ENSMUST00 | 0 | Ubqln4 | ENSMUSG00000008604.6 |
| chr3 | 132682978 | 132685271 | 32.67316 | 105.57997 | 101.8659 | Promoter | 3 | 132660498 | 132683879 | 23382 | 2 | ENSMUST00 | 0 | Aimp1 | ENSMUSG00000028029.6 |
| chr6 | 29768392 | 29769584 | 32.67316 | 105.57997 | 101.8659 | Promoter | 6 | 29768443 | 29867419 | 98977 | 1 | ENSMUST00 | 0 | Ahcyl2 | ENSMUSG00000029772.12 |
| chr9 | 94536192 | 94539228 | 32.67316 | 105.57997 | 101.8659 | Promoter | 9 | 94517864 | 94538081 | 20218 | 2 | ENSMUST00 | 0 | 1190002N15 | ENSMUSG00000045414.7 |
| chr3 | 100488212 | 100489978 | 34.42823 | 105.56979 | 101.85753 | Promoter | 3 | 100451628 | 100489198 | 37571 | 2 | ENSMUST00 | 0 | Fam46c | ENSMUSG00000044468.9 |
| chr9 | 107707503 | 107710482 | 34.42823 | 105.56979 | 101.85753 | Promoter | 9 | 107681502 | 107710475 | 28974 | 2 | ENSMUST00 | 0 | Sema3f | ENSMUSG00000034684.7 |
| chr17 | 24719189 | 24719973 | 30.92995 | 105.52683 | 101.81509 | Promoter | 17 | 24719676 | 24719803 | 128 | 2 | ENSMUST00 | 0 | Snora78 | ENSMUSG00000089255.1 |
| chr19 | 10576378 | 10578104 | 32.05589 | 105.50996 | 101.79823 | Promoter | 19 | 10570478 | 10577362 | 6885 | 2 | ENSMUST00 | 0 | Tmem138 | ENSMUSG00000024666.5 |
| chr11 | 102603096 | 102607889 | 33.76198 | 105.43819 | 101.727 | Promoter | 11 | 102604396 | 102608058 | 3663 | 1 | ENSMUST00 | 0 | Fzd2 | ENSMUSG00000050288.6 |
| chr13 | 75706984 | 75708992 | 33.76198 | 105.43819 | 101.727 | Promoter | 13 | 75707484 | 75772356 | 64873 | 1 | ENSMUST00 | 0 | Ell2 | ENSMUSG00000001542.6 |
| chr15 | 58033599 | 58034490 | 33.76198 | 105.43819 | 101.727 | Promoter | 15 | 58022259 | 58034315 | 12057 | 2 | ENSMUST00 | 0 | 9130401M01 | ENSMUSG00000101892.1 |
| chr17 | 47610467 | 47612451 | 33.76198 | 105.43819 | 101.727 | Promoter | 17 | 47611589 | 47650289 | 38701 | 1 | ENSMUST00 | 0 | Gm20517 | ENSMUSG00000023984.12 |
| chr3 | 135422276 | 135424307 | 33.76198 | 105.43819 | 101.727 | Promoter | 3 | 135406412 | 135423925 | 17514 | 2 | ENSMUST00 | 0 | Cisd2 | ENSMUSG00000028165.8 |
| chr3 | 88579332 | 88580391 | 33.76198 | 105.43819 | 101.727 | Promoter | 3 | 88579671 | 88588397 | 8727 | 1 | ENSMUST00 | 0 | Ssr2 | ENSMUSG00000041355.8 |
| chr6 | 72233454 | 72235881 | 33.76198 | 105.43819 | 101.727 | Promoter | 6 | 72206177 | 72235577 | 29401 | 2 | ENSMUST00 | 0 | Atoh8 | ENSMUSG00000037621.7 |
| chr7 | 80900501 | 80901947 | 33.76198 | 105.43819 | 101.727 | Promoter | 7 | 80890723 | 80901269 | 10547 | 2 | ENSMUST00 | 0 | Wdr73 | ENSMUSG00000025722.11 |
| chr4 | 134244758 | 134245936 | 30.31349 | 105.41657 | 101.70665 | Promoter | 4 | 134243289 | 134245592 | 2304 | 2 | ENSMUST00 | 0 | Zfp593 | ENSMUSG00000028840.10 |
| chr11 | 32221136 | 32222863 | 33.13715 | 105.37189 | 101.66248 | Promoter | 11 | 32209585 | 32222300 | 12716 | 2 | ENSMUST00 | 0 | Rhbdf1 | ENSMUSG00000020282.13 |
| chr6 | 57580791 | 57581804 | 33.13715 | 105.37189 | 101.66248 | Promoter | 6 | 57580992 | 57665136 | 84145 | 1 | ENSMUST00 | 0 | Herc6 | ENSMUSG00000029798.9 |
| chr5 | 3341034 | 3344969 | 32.55001 | 105.36325 | 101.65399 | Promoter | 5 | 3343893 | 3523218 | 179326 | 1 | ENSMUST00 | 0 | Cdk6 | ENSMUSG00000040274.6 |
| chr12 | 78225441 | 78228015 | 33.69441 | 105.32094 | 101.61211 | Promoter | 12 | 78226655 | 78684767 | 458113 | 1 | ENSMUST00 | 0 | Gphn | ENSMUSG00000047454.6 |
| chr8 | 123211850 | 123213553 | 21.93925 | 105.29307 | 101.58453 | Promoter | 8 | 123204264 | 123212760 | 8497 | 2 | ENSMUST00 | 0 | Chmp1a | ENSMUSG00000000743.8 |
| chr1 | 75168055 | 75169352 | 30.78991 | 105.27288 | 101.56436 | Promoter | 1 | 75168646 | 75170752 | 2107 | 1 | ENSMUST00 | 0 | Zfand2b | ENSMUSG00000026197.7 |
| chr1 | 84837968 | 84840939 | 30.78991 | 105.27288 | 101.56436 | Promoter | 1 | 84721189 | 84839304 | 118116 | 2 | ENSMUST00 | 0 | Trip12 | ENSMUSG00000026219.9 |
| chr10 | 70095817 | 70098606 | 30.78991 | 105.27288 | 101.56436 | Promoter | 10 | 70097121 | 70193200 | 96080 | 1 | ENSMUST00 | 0 | Ccdc6 | ENSMUSG00000048701.8 |
| chr11 | 20201342 | 20203232 | 30.78991 | 105.27288 | 101.56436 | Promoter | 11 | 20201432 | 20226856 | 25425 | 1 | ENSMUST00 | 0 | Rab1 | ENSMUSG00000020149.12 |
| chr11 | 106611514 | 106613072 | 29.6869 | 105.26892 | 101.56436 | Promoter | 11 | 106502147 | 106612930 | 110784 | 2 | ENSMUST00 | 0 | Tex2 | ENSMUSG00000040548.10 |
| chr12 | 108179724 | 108180617 | 29.6869 | 105.26892 | 101.56436 | Promoter | 12 | 108179738 | 108203359 | 23622 | 1 | ENSMUST00 | 0 | Ccnk | ENSMUSG00000021258.9 |
| chr13 | 74061625 | 74062582 | 30.78991 | 105.27288 | 101.56436 | Promoter | 13 | 74036495 | 74062285 | 25791 | 2 | ENSMUST00 | 0 | Cep72 | ENSMUSG00000021572.8 |
| chr13 | 23581534 | 23586947 | 30.78991 | 105.27288 | 101.56436 | Promoter | 13 | 23581598 | 23581990 | 393 | 1 | ENSMUST00 | 0 | Hist1h4d | ENSMUSG00000061482.6 |
| chr14 | 57798183 | 57800517 | 30.78991 | 105.27288 | 101.56436 | Promoter | 14 | 57798218 | 57804969 | 6752 | 1 | ENSMUST00 | 0 | Sap18 | ENSMUSG00000021963.11 |
| chr14 | 67007547 | 67009614 | 30.78991 | 105.27288 | 101.56436 | Promoter | 14 | 66985239 | 67008856 | 23618 | 2 | ENSMUST00 | 0 | Bnip3l | ENSMUSG00000022051.10 |
| chr14 | 121797106 | 121798657 | 30.78991 | 105.27288 | 101.56436 | Promoter | 14 | 121542039 | 121797734 | 255696 | 2 | ENSMUST00 | 0 | Dock9 | ENSMUSG00000025558.10 |
| chr15 | 41788058 | 41790113 | 30.78991 | 105.27288 | 101.56436 | Promoter | 15 | 41789034 | 41861045 | 72012 | 1 | ENSMUST00 | 0 | Oxr1 | ENSMUSG00000022307.10 |
| chr15 | 98608565 | 98610228 | 30.78991 | 105.27288 | 101.56436 | Promoter | 15 | 98589994 | 98607633 | 17640 | 2 | ENSMUST00 | -933 | Adcy6 | ENSMUSG00000022994.8 |
| chr17 | 79019683 | 79021424 | 30.78991 | 105.27288 | 101.56436 | Promoter | 17 | 78949405 | 79020816 | 71412 | 2 | ENSMUST00 | 0 | Prkd3 | ENSMUSG00000024070.10 |
| chr17 | 33032825 | 33033649 | 30.78991 | 105.27288 | 101.56436 | Promoter | 17 | 33016863 | 33033402 | 16540 | 2 | ENSMUST00 | 0 | Zfp763 | ENSMUSG00000067430.7 |
| chr18 | 67289111 | 67290354 | 30.78991 | 105.27288 | 101.56436 | Promoter | 18 | 67289253 | 67319146 | 29894 | 1 | ENSMUST00 | 0 | Impa2 | ENSMUSG00000024525.6 |
| chr18 | 12120780 | 12121829 | 30.78991 | 105.27288 | 101.56436 | Promoter | 18 | 11981303 | 12121480 | 140178 | 2 | ENSMUST00 | 0 | Tmem241 | ENSMUSG00000049411.8 |
| chr19 | 4125868 | 4127236 | 30.78991 | 105.27288 | 101.56436 | Promoter | 19 | 4125934 | 4132285 | 6352 | 1 | ENSMUST00 | 0 | Tmem134 | ENSMUSG00000024845.12 |
| chr19 | 23140239 | 23143387 | 30.78991 | 105.27288 | 101.56436 | Promoter | 19 | 23141226 | 23166911 | 25686 | 1 | ENSMUST00 | 0 | Klf9 | ENSMUSG00000033863.1 |
| chr4 | 130662855 | 130664831 | 30.78991 | 105.27288 | 101.56436 | Promoter | 4 | 130663321 | 130781564 | 118244 | 1 | ENSMUST00 | 0 | Pum1 | ENSMUSG00000028580.10 |
| chr4 | 126752441 | 126754692 | 30.78991 | 105.27288 | 101.56436 | Promoter | 4 | 126753544 | 126791894 | 38351 | 1 | ENSMUST00 | 0 | AU040320 | ENSMUSG00000028830.9 |
| chr4 | 131836342 | 131838375 | 30.78991 | 105.27288 | 101.56436 | Promoter | 4 | 131768457 | 131838288 | 69832 | 2 | ENSMUST00 | 0 | Ptpru | ENSMUSG00000028909.12 |
| chr4 | 155802847 | 155804319 | 30.78991 | 105.27288 | 101.56436 | Promoter | 4 | 155802878 | 155803642 | 765 | 1 | ENSMUST00 | 0 | Mrpl20 | ENSMUSG00000029066.7 |
| chr5 | 34573033 | 34575112 | 30.78991 | 105.27288 | 101.56436 | Promoter | 5 | 34573664 | 34632308 | 58645 | 1 | ENSMUST00 | 0 | Add1 | ENSMUSG00000029106.9 |
| chr5 | 21645773 | 21646765 | 30.78991 | 105.27288 | 101.56436 | Promoter | 5 | 21645813 | 21661616 | 15804 | 1 | ENSMUST00 | 0 | Armc10 | ENSMUSG00000038525.11 |
| chr5 | 45492745 | 45494309 | 30.78991 | 105.27288 | 101.56436 | Promoter | 5 | 45493374 | 45512674 | 19301 | 1 | ENSMUST00 | 0 | Lap3 | ENSMUSG00000039682.7 |
| chr5 | 146832555 | 146834377 | 30.78991 | 105.27288 | 101.56436 | Promoter | 5 | 146832890 | 146837032 | 4143 | 1 | ENSMUST00 | 0 | Rpl21 | ENSMUSG00000041453.7 |
| chr6 | 87009352 | 87010440 | 30.78991 | 105.27288 | 101.56436 | Promoter | 6 | 87009836 | 87028454 | 18619 | 1 | ENSMUST00 | 0 | Nfu1 | ENSMUSG00000029993.11 |
| chr6 | 129532540 | 129534683 | 30.78991 | 105.27288 | 101.56436 | Promoter | 6 | 129533200 | 129542331 | 9132 | 1 | ENSMUST00 | 0 | Gabarapl1 | ENSMUSG00000030161.6 |
| chr6 | 73247964 | 73249716 | 30.78991 | 105.27288 | 101.56436 | Promoter | 6 | 73248382 | 73276911 | 28530 | 1 | ENSMUST00 | 0 | Suclg1 | ENSMUSG00000052738.9 |
| chr7 | 99182220 | 99183208 | 30.78991 | 105.27288 | 101.56436 | Promoter | 7 | 99153663 | 99182713 | 29051 | 2 | ENSMUST00 | 0 | Dgat2 | ENSMUSG00000030747.4 |
| chr7 | 28370828 | 28372772 | 30.78991 | 105.27288 | 101.56436 | Promoter | 7 | 28359604 | 28372233 | 12630 | 2 | ENSMUST00 | 0 | Plekhg2 | ENSMUSG00000037552.12 |
| chr7 | 118447355 | 118448957 | 30.78991 | 105.27288 | 101.56436 | Distal Interge | 7 | 118381856 | 118443552 | 61697 | 2 | ENSMUST00 | -3804 | Syt17 | ENSMUSG00000058420.5 |
| chr7 | 73555379 | 73558486 | 30.78991 | 105.27288 | 101.56436 | Promoter | 7 | 73539801 | 73558395 | 18595 | 2 | ENSMUST00 | 0 | 1810026B05 | ENSMUSG00000098496.2 |
| chr9 | 57589080 | 57590210 | 30.78991 | 105.27288 | 101.56436 | Promoter | 9 | 57589452 | 57596233 | 6782 | 1 | ENSMUST00 | 0 | Ulk3 | ENSMUSG00000032308.8 |
| chr16 | 33055841 | 33057426 | 27.07707 | 105.22646 | 101.52403 | Promoter | 16 | 33015385 | 33056190 | 40806 | 2 | ENSMUST00 | 0 | Iqcg | ENSMUSG00000035578.10 |
| chr1 | 74302782 | 74304983 | 33.62711 | 105.2041 | 101.50168 | Promoter | 1 | 74288247 | 74304405 | 16159 | 2 | ENSMUST00 | 0 | Tmbim1 | ENSMUSG00000006301.12 |
| chr11 | 70451574 | 70452744 | 31.8806 | 105.19856 | 101.49664 | Promoter | 11 | 70451919 | 70453240 | 1322 | 1 | ENSMUST00 | 0 | Med11 | ENSMUSG00000018923.8 |
| chr11 | 100319937 | 100321297 | 31.8806 | 105.19856 | 101.49664 | Promoter | 11 | 100320209 | 100320989 | 781 | 1 | ENSMUST00 | 0 | Eif1 | ENSMUSG00000035530.8 |
| chr14 | 75844263 | 75845074 | 31.8806 | 105.19856 | 101.49664 | Promoter | 14 | 75843379 | 75844964 | 1586 | 2 | ENSMUST00 | 0 | Gm4285 | ENSMUSG00000085829.1 |
| chr19 | 45559956 | 45561138 | 31.8806 | 105.19856 | 101.49664 | Promoter | 19 | 45552275 | 45560531 | 8257 | 2 | ENSMUST00 | 0 | Poll | ENSMUSG00000025218.6 |
| chr2 | 23571194 | 23572480 | 31.8806 | 105.19856 | 101.49664 | Promoter | 2 | 23506220 | 23572104 | 65885 | 2 | ENSMUST00 | 0 | Spopl | ENSMUSG00000026771.9 |
| chr9 | 35210385 | 35212079 | 31.8806 | 105.19856 | 101.49664 | Promoter | 9 | 35211155 | 35217014 | 5860 | 1 | ENSMUST00 | 0 | Srpr | ENSMUSG00000032042.6 |
| chr8 | 110617975 | 110619522 | 29.09955 | 105.17428 | 101.47362 | Promoter | 8 | 110618638 | 110720396 | 101759 | 1 | ENSMUST00 | 0 | Vac14 | ENSMUSG00000010936.10 |
| chr8 | 78507901 | 78510468 | 28.0287 | 105.12276 | 101.42252 | Promoter | 8 | 78509335 | 78734005 | 224671 | 1 | ENSMUST00 | 0 | Slc10a7 | ENSMUSG00000031684.10 |
| chr4 | 62285924 | 62287996 | 33.56007 | 105.08768 | 101.38747 | Promoter | 4 | 62286449 | 62298411 | 11963 | 1 | ENSMUST00 | 0 | Slc31a2 | ENSMUSG00000066152.6 |
| chr3 | 103809519 | 103810080 | 34.14524 | 105.08128 | 101.38137 | Promoter | 3 | 103809520 | 103822023 | 12504 | 1 | ENSMUST00 | 0 | Ap4b1 | ENSMUSG00000032952.6 |
| chr10 | 9899546 | 9902121 | 26.03857 | 105.07202 | 101.37238 | Promoter | 10 | 9755547 | 9900683 | 145137 | 2 | ENSMUST00 | 0 | Stxbp5 | ENSMUSG00000019790.11 |
| chr10 | 80899740 | 80900929 | 26.03857 | 105.07202 | 101.37238 | Promoter | 10 | 80899450 | 80900969 | 1520 | 2 | ENSMUST00 | 40 | Timm13 | ENSMUSG00000020219.6 |
| chr10 | 18318175 | 18319701 | 26.03857 | 105.07202 | 101.37238 | Distal Interge | 10 | 18390322 | 18524289 | 133968 | 1 | ENSMUST00 | -70621 | Nhsl1 | ENSMUSG00000039835.10 |
| chr11 | 90249309 | 90250643 | 26.03857 | 105.07202 | 101.37238 | Promoter | 11 | 90249456 | 90268465 | 19010 | 1 | ENSMUST00 | 0 | Mmd | ENSMUSG00000003948.11 |
| chr11 | 86806985 | 86808025 | 26.03857 | 105.07202 | 101.37238 | Promoter | 11 | 86768846 | 86807746 | 38901 | 2 | ENSMUST00 | 0 | Dhx40 | ENSMUSG00000018425.13 |
| chr2 | 4918699 | 4919626 | 26.03857 | 105.07202 | 101.37238 | Promoter | 2 | 4919019 | 4938730 | 19712 | 1 | ENSMUST00 | 0 | Phyh | ENSMUSG00000026664.7 |
| chr3 | 65527589 | 65530267 | 26.03857 | 105.07202 | 101.37238 | Promoter | 3 | 65528410 | 65555145 | 26736 | 1 | ENSMUST00 | 0 | Tiparp | ENSMUSG00000034640.4 |
| chr4 | 43577870 | 43579625 | 26.03857 | 105.07202 | 101.37238 | Promoter | 4 | 43566928 | 43578089 | 11162 | 2 | ENSMUST00 | 0 | Gba2 | ENSMUSG00000028467.10 |
| chr4 | 12906702 | 12907574 | 26.03857 | 105.07202 | 101.37238 | Promoter | 4 | 12906838 | 12980873 | 74036 | 1 | ENSMUST00 | 0 | Triqk | ENSMUSG00000055963.7 |
| chr5 | 129007637 | 129009002 | 26.03857 | 105.07202 | 101.37238 | Promoter | 5 | 128984557 | 129008574 | 24018 | 2 | ENSMUST00 | 0 | Stx2 | ENSMUSG00000029428.8 |
| chr6 | 37439782 | 37442252 | 26.03857 | 105.07202 | 101.37238 | Promoter | 6 | 37327255 | 37442146 | 114892 | 2 | ENSMUST00 | 0 | Creb3l2 | ENSMUSG00000038648.5 |
| chr8 | 27023076 | 27024558 | 26.03857 | 105.07202 | 101.37238 | Promoter | 8 | 27023799 | 27039433 | 15635 | 1 | ENSMUST00 | 0 | Erlin2 | ENSMUSG00000031483.7 |
| chr8 | 27042133 | 27043546 | 26.03857 | 105.07202 | 101.37238 | Promoter | 8 | 27042555 | 27050240 | 7686 | 1 | ENSMUST00 | 0 | Prosc | ENSMUSG00000031485.9 |
| chr11 | 60536603 | 60538667 | 32.94511 | 105.03617 | 101.33749 | Promoter | 11 | 60537978 | 60558512 | 20535 | 1 | ENSMUST00 | 0 | Alkbh5 | ENSMUSG00000042650.3 |
| chr1 | 171080070 | 171081267 | 14.06175 | 105.00215 | 101.30357 | Promoter | 1 | 171079840 | 171079901 | 62 | 2 | ENSMUST00 | -170 | Gm27844 | ENSMUSG00000098965.1 |
| chr6 | 141249145 | 141251922 | 34.07522 | 104.96031 | 101.26198 | Promoter | 6 | 141249269 | 141505940 | 256672 | 1 | ENSMUST00 | 0 | Pde3a | ENSMUSG00000041741.9 |
| chr8 | 70509389 | 70511406 | 32.30648 | 104.93403 | 101.23596 | Promoter | 8 | 70508263 | 70510350 | 2088 | 2 | ENSMUST00 | 0 | Uba52 | ENSMUSG00000090137.3 |
| chr10 | 84916703 | 84918437 | 32.88158 | 104.92503 | 101.22703 | Promoter | 10 | 84917616 | 85016446 | 98831 | 1 | ENSMUST00 | 0 | Ric8b | ENSMUSG00000035620.9 |
| chr13 | 21710012 | 21711411 | 32.88158 | 104.92503 | 101.22703 | Distal Interge | 13 | 21715763 | 21716143 | 381 | 2 | ENSMUST00 | 4732 | Hist1h2bl | ENSMUSG00000094338.1 |
| chr16 | 43979193 | 43980269 | 32.88158 | 104.92503 | 101.22703 | Promoter | 16 | 43969078 | 43979093 | 10016 | 2 | ENSMUST00 | -101 | Zdhhc23 | ENSMUSG00000036304.8 |
| chr17 | 56934576 | 56935985 | 32.88158 | 104.92503 | 101.22703 | Promoter | 17 | 56892612 | 56935388 | 42777 | 2 | ENSMUST00 | 0 | Mllt1 | ENSMUSG00000024212.8 |
| chr17 | 6960123 | 6961910 | 32.88158 | 104.92503 | 101.22703 | Promoter | 17 | 6954965 | 6961156 | 6192 | 2 | ENSMUST00 | 0 | Tagap1 | ENSMUSG00000052031.7 |
| chr4 | 118620213 | 118621499 | 32.88158 | 104.92503 | 101.22703 | Promoter | 4 | 118620799 | 118627776 | 6978 | 1 | ENSMUST00 | 0 | Ebna1bp2 | ENSMUSG00000028729.10 |
| chr5 | 32784971 | 32786245 | 32.88158 | 104.92503 | 101.22703 | Promoter | 5 | 32736301 | 32785585 | 49285 | 2 | ENSMUST00 | 0 | Pisd | ENSMUSG00000023452.13 |
| chr6 | 106768594 | 106770191 | 32.88158 | 104.92503 | 101.22703 | Promoter | 6 | 106769120 | 106777681 | 8562 | 1 | ENSMUST00 | 0 | Trnt1 | ENSMUSG00000013736.11 |
| chr6 | 35132576 | 35134663 | 32.88158 | 104.92503 | 101.22703 | Promoter | 6 | 35022066 | 35133737 | 111672 | 2 | ENSMUST00 | 0 | Cnot4 | ENSMUSG00000038784.8 |
| chr17 | 27819348 | 27820669 | 31.14379 | 104.89771 | 101.20105 | Promoter | 17 | 27751232 | 27820558 | 69327 | 2 | ENSMUST00 | 0 | D17Wsu92e | ENSMUSG00000056692.7 |
| chr18 | 53743447 | 53745668 | 31.14379 | 104.89771 | 101.20105 | Promoter | 18 | 53681724 | 53744547 | 62824 | 2 | ENSMUST00 | 0 | Cep120 | ENSMUSG00000048799.7 |
| chr19 | 6057225 | 6058915 | 31.14379 | 104.89771 | 101.20105 | Promoter | 19 | 6053630 | 6057751 | 4122 | 2 | ENSMUST00 | 0 | Mrpl49 | ENSMUSG00000007338.9 |
| chr2 | 140169580 | 140171663 | 31.14379 | 104.89771 | 101.20105 | Promoter | 2 | 140170649 | 140203689 | 33041 | 1 | ENSMUST00 | 0 | Ndufaf5 | ENSMUSG00000027384.6 |
| chr4 | 83323194 | 83324767 | 31.14379 | 104.89771 | 101.20105 | Promoter | 4 | 83220300 | 83324255 | 103956 | 2 | ENSMUST00 | 0 | Ttc39b | ENSMUSG00000038172.9 |
| chr5 | 114941716 | 114942920 | 31.14379 | 104.89771 | 101.20105 | Promoter | 5 | 114942158 | 114949783 | 7626 | 1 | ENSMUST00 | 0 | 2210016L21 | ENSMUSG00000029559.10 |
| chr8 | 105297176 | 105298591 | 31.14379 | 104.89771 | 101.20105 | Promoter | 8 | 105297663 | 105305369 | 7707 | 1 | ENSMUST00 | 0 | E2f4 | ENSMUSG00000014859.8 |
| chr13 | 108213922 | 108215375 | 34.00549 | 104.83981 | 101.14426 | Promoter | 13 | 108214404 | 108287105 | 72702 | 1 | ENSMUST00 | 0 | Elovl7 | ENSMUSG00000021696.8 |
| chr17 | 59012479 | 59013604 | 34.00549 | 104.83981 | 101.14426 | Promoter | 17 | 58999618 | 59013372 | 13755 | 2 | ENSMUST00 | 0 | Nudt12 | ENSMUSG00000024228.7 |
| chr17 | 35666938 | 35668137 | 34.00549 | 104.83981 | 101.14426 | Promoter | 17 | 35655634 | 35667592 | 11959 | 2 | ENSMUST00 | 0 | Vars2 | ENSMUSG00000038838.10 |
| chr2 | 112454162 | 112455814 | 34.00549 | 104.83981 | 101.14426 | Promoter | 2 | 112454997 | 112472162 | 17166 | 1 | ENSMUST00 | 0 | Emc7 | ENSMUSG00000055943.5 |

| | | | | | | | | | | | | | | | |
|---|---|---|---|---|---|---|---|---|---|---|---|---|---|---|---|
| chr3 | 90476084 | 90477052 | 34.00549 | 104.83981 | 101.14426 | Promoter | 3 | 90476201 | 90488379 | 12179 | 1 | ENSMUST00 | 0 | Ilf2 | ENSMUSG00000001016.8 |
| chr5 | 115326237 | 115328553 | 34.00549 | 104.83981 | 101.14426 | Promoter | 5 | 115327177 | 115333080 | 5904 | 1 | ENSMUST00 | 0 | Srsf9 | ENSMUSG00000029538.8 |
| chr1 | 185203116 | 185205017 | 23.70827 | 104.82465 | 101.13002 | Promoter | 1 | 185204168 | 185286743 | 82576 | 1 | ENSMUST00 | 0 | Rab3gap2 | ENSMUSG00000039318.7 |
| chr13 | 55512304 | 55513824 | 23.70827 | 104.82465 | 101.13002 | Promoter | 13 | 55495795 | 55513438 | 17644 | 2 | ENSMUST00 | 0 | Pdlim7 | ENSMUSG00000021493.10 |
| chr3 | 65956813 | 65959201 | 23.70827 | 104.82465 | 101.13002 | Promoter | 3 | 65946151 | 65958162 | 12012 | 2 | ENSMUST00 | 0 | Ccnl1 | ENSMUSG00000027829.10 |
| chr3 | 57651085 | 57652152 | 23.70827 | 104.82465 | 101.13002 | Promoter | 3 | 57642904 | 57651693 | 8790 | 2 | ENSMUST00 | 0 | Commd2 | ENSMUSG00000036513.10 |
| chr8 | 70698975 | 70701534 | 23.70827 | 104.82465 | 101.13002 | Promoter | 8 | 70699538 | 70700080 | 543 | 2 | ENSMUST00 | 0 | Gm11175 | ENSMUSG00000080058.1 |
| chr1 | 37719170 | 37720553 | 28.90751 | 104.81161 | 101.11721 | Promoter | 1 | 37611677 | 37720085 | 108409 | 2 | ENSMUST00 | 0 | 2010300C02I | ENSMUSG00000026090.11 |
| chr1 | 180568491 | 180569887 | 28.90751 | 104.81161 | 101.11721 | Promoter | 1 | 180568955 | 180601253 | 32299 | 1 | ENSMUST00 | 0 | Parp1 | ENSMUSG00000026496.6 |
| chr1 | 133024460 | 133026209 | 28.90751 | 104.81161 | 101.11721 | Promoter | 1 | 132959484 | 133025338 | 65855 | 2 | ENSMUST00 | 0 | Mdm4 | ENSMUSG00000054387.8 |
| chr10 | 22272927 | 22274323 | 28.90751 | 104.81161 | 101.11721 | Promoter | 10 | 22273346 | 22288849 | 15504 | 1 | ENSMUST00 | 0 | H60b | ENSMUSG00000075297.5 |
| chr11 | 53769560 | 53771265 | 28.90751 | 104.81161 | 101.11721 | Promoter | 11 | 53770014 | 53777323 | 7310 | 1 | ENSMUST00 | 0 | Irf1 | ENSMUSG00000018899.11 |
| chr11 | 119913059 | 119914872 | 28.90751 | 104.81161 | 101.11721 | Promoter | 11 | 119913441 | 119917008 | 3568 | 1 | ENSMUST00 | 0 | Chmp6 | ENSMUSG00000025371.7 |
| chr12 | 84100637 | 84101532 | 28.90751 | 104.81161 | 101.11721 | Promoter | 12 | 84100654 | 84111349 | 10696 | 1 | ENSMUST00 | 0 | Acot6 | ENSMUSG00000043487.3 |
| chr13 | 66932235 | 66934137 | 28.90751 | 104.81161 | 101.11721 | Promoter | 13 | 66932830 | 66998401 | 65572 | 1 | ENSMUST00 | 0 | Ptdss1 | ENSMUSG00000021518.3 |
| chr16 | 90809703 | 90810502 | 28.90751 | 104.81161 | 101.11721 | Promoter | 16 | 90751527 | 90810392 | 58866 | 2 | ENSMUST00 | 0 | Urb1 | ENSMUSG00000039929.9 |
| chr17 | 34586640 | 34592164 | 28.90751 | 104.81161 | 101.11721 | Promoter | 17 | 34586945 | 34587501 | 557 | 1 | ENSMUST00 | 0 | Notch4 | ENSMUSG00000015468.9 |
| chr18 | 32067200 | 32068691 | 28.90751 | 104.81161 | 101.11721 | Promoter | 18 | 32067734 | 32104331 | 36598 | 1 | ENSMUST00 | 0 | Iws1 | ENSMUSG00000024384.3 |
| chr18 | 80933292 | 80934494 | 28.90751 | 104.81161 | 101.11721 | Promoter | 18 | 80734143 | 80934058 | 199916 | 2 | ENSMUST00 | 0 | Atp9b | ENSMUSG00000024566.8 |
| chr2 | 80638530 | 80639693 | 28.90751 | 104.81161 | 101.11721 | Promoter | 2 | 80638816 | 80658902 | 20087 | 1 | ENSMUST00 | 0 | Nup35 | ENSMUSG00000026999.9 |
| chr2 | 146511465 | 146512506 | 28.90751 | 104.81161 | 101.11721 | Promoter | 2 | 146239879 | 146512004 | 272126 | 2 | ENSMUST00 | 0 | Ralgapa2 | ENSMUSG00000037110.14 |
| chr3 | 94953471 | 94954644 | 28.90751 | 104.81161 | 101.11721 | Promoter | 3 | 94954075 | 94959496 | 5422 | 1 | ENSMUST00 | 0 | Rfx5 | ENSMUSG00000005774.7 |
| chr4 | 133751327 | 133754985 | 28.90751 | 104.81161 | 101.11721 | Promoter | 4 | 133679008 | 133753611 | 74604 | 2 | ENSMUST00 | 0 | Arid1a | ENSMUSG00000007880.11 |
| chr5 | 122820560 | 122821729 | 28.90751 | 104.81161 | 101.11721 | Promoter | 5 | 122787469 | 122821339 | 33871 | 2 | ENSMUST00 | 0 | Anapc5 | ENSMUSG00000029472.9 |
| chr5 | 144100241 | 144101315 | 28.90751 | 104.81161 | 101.11721 | Promoter | 5 | 144100436 | 144188204 | 87769 | 1 | ENSMUST00 | 0 | Lmtk2 | ENSMUSG00000038970.7 |
| chr6 | 34176058 | 34177139 | 28.90751 | 104.81161 | 101.11721 | Promoter | 6 | 34153380 | 34177111 | 23732 | 2 | ENSMUST00 | 0 | Slc35b4 | ENSMUSG00000018999.9 |
| chr6 | 42348851 | 42351089 | 28.90751 | 104.81161 | 101.11721 | Promoter | 6 | 42349828 | 42358392 | 8565 | 1 | ENSMUST00 | 0 | Zyx | ENSMUSG00000029860.11 |
| chr6 | 83108467 | 83109941 | 28.90751 | 104.81161 | 101.11721 | Promoter | 6 | 83109095 | 83109939 | 845 | 1 | ENSMUST00 | 0 | Mrpl53 | ENSMUSG00000030037.8 |
| chr7 | 45123273 | 45124400 | 28.90751 | 104.81161 | 101.11721 | Promoter | 7 | 45122388 | 45124389 | 2002 | 2 | ENSMUST00 | 0 | Rps11 | ENSMUSG00000003429.10 |
| chr7 | 25297151 | 25298419 | 28.90751 | 104.81161 | 101.11721 | Promoter | 7 | 25295049 | 25297312 | 2264 | 2 | ENSMUST00 | 0 | Pafah1b3 | ENSMUSG00000005447.7 |
| chr7 | 101905830 | 101906676 | 28.90751 | 104.81161 | 101.11721 | Promoter | 7 | 101905837 | 101911901 | 6065 | 1 | ENSMUST00 | 0 | Lamtor1 | ENSMUSG00000030842.6 |
| chr8 | 64732771 | 64734174 | 28.90751 | 104.81161 | 101.11721 | Promoter | 8 | 64718139 | 64733792 | 15654 | 2 | ENSMUST00 | 0 | Msmo1 | ENSMUSG00000031604.5 |
| chr9 | 57261442 | 57262990 | 28.90751 | 104.81161 | 101.11721 | Promoter | 9 | 57252322 | 57262599 | 10278 | 2 | ENSMUST00 | 0 | 1700017B05I | ENSMUSG00000032300.6 |
| chr1 | 150392105 | 150394544 | 27.34946 | 104.78247 | 101.09169 | Promoter | 1 | 150392838 | 150449815 | 56978 | 1 | ENSMUST00 | 0 | Tpr | ENSMUSG00000006005.12 |
| chr10 | 40256686 | 40258314 | 27.34946 | 104.78247 | 101.09169 | Promoter | 10 | 40249203 | 40257665 | 8463 | 2 | ENSMUST00 | 0 | Gtf3c6 | ENSMUSG00000019837.7 |
| chr11 | 23894200 | 23895678 | 27.34946 | 104.78247 | 101.09169 | Promoter | 11 | 23862646 | 23895253 | 32608 | 2 | ENSMUST00 | 0 | Papolg | ENSMUSG00000020273.8 |
| chr11 | 76201511 | 76202751 | 27.34946 | 104.78247 | 101.09169 | Promoter | 11 | 76202015 | 76209416 | 7402 | 1 | ENSMUST00 | 0 | Fam57a | ENSMUSG00000069808.8 |
| chr13 | 75942785 | 75944428 | 27.34946 | 104.78247 | 101.09169 | Promoter | 13 | 75869537 | 75943867 | 74331 | 2 | ENSMUST00 | 0 | Rhobtb3 | ENSMUSG00000021589.7 |
| chr17 | 79614607 | 79615964 | 27.34946 | 104.78247 | 101.09169 | Promoter | 17 | 79614900 | 79682152 | 67253 | 1 | ENSMUST00 | 0 | Rmdn2 | ENSMUSG00000036368.7 |
| chr3 | 31093380 | 31096252 | 27.34946 | 104.78247 | 101.09169 | Promoter | 3 | 31095058 | 31122577 | 27520 | 1 | ENSMUST00 | 0 | Skil | ENSMUSG00000027660.11 |
| chr3 | 88949487 | 88952169 | 27.34946 | 104.78247 | 101.09169 | Promoter | 3 | 88950622 | 89079375 | 128754 | 1 | ENSMUST00 | 0 | Ash1l | ENSMUSG00000028053.8 |
| chr3 | 36862581 | 36863954 | 27.34946 | 104.78247 | 101.09169 | Promoter | 3 | 36863106 | 37053033 | 189928 | 1 | ENSMUST00 | 0 | 4932438A13I | ENSMUSG00000037270.12 |
| chr4 | 125126469 | 125128407 | 27.34946 | 104.78247 | 101.09169 | Promoter | 4 | 125118423 | 125127840 | 9418 | 2 | ENSMUST00 | 0 | Zc3h12a | ENSMUSG00000042677.7 |
| chr6 | 134034454 | 134036769 | 27.34946 | 104.78247 | 101.09169 | Promoter | 6 | 134035700 | 134270158 | 234459 | 1 | ENSMUST00 | 0 | Etv6 | ENSMUSG00000030199.11 |
| chr7 | 81827970 | 81829981 | 27.34946 | 104.78247 | 101.09169 | Promoter | 7 | 81792075 | 81829431 | 37357 | 2 | ENSMUST00 | 0 | Btbd1 | ENSMUSG00000025103.7 |
| chr9 | 20887702 | 20889004 | 27.34946 | 104.78247 | 101.09169 | Promoter | 9 | 20888175 | 20892178 | 4004 | 1 | ENSMUST00 | 0 | Ppan | ENSMUSG00000004100.6 |
| chr9 | 57076296 | 57077594 | 27.34946 | 104.78247 | 101.09169 | Promoter | 9 | 57076376 | 57128364 | 51989 | 1 | ENSMUST00 | 0 | Sin3a | ENSMUSG00000042557.9 |
| chr9 | 75408649 | 75410600 | 27.34946 | 104.78247 | 101.09169 | Promoter | 9 | 75386902 | 75409359 | 22458 | 2 | ENSMUST00 | 0 | Mapk6 | ENSMUSG00000042688.11 |
| chr17 | 66075855 | 66077884 | 33.36056 | 104.74094 | 101.05196 | Promoter | 17 | 65967501 | 66077046 | 109546 | 2 | ENSMUST00 | 0 | Ankrd12 | ENSMUSG00000034647.9 |
| chr11 | 55469582 | 55470757 | 32.18608 | 104.72149 | 101.03278 | Promoter | 11 | 55469685 | 55504838 | 35154 | 1 | ENSMUST00 | 0 | G3bp1 | ENSMUSG00000018583.5 |
| chr1 | 87852728 | 87854508 | 30.45704 | 104.66754 | 100.97924 | Promoter | 1 | 87853287 | 87944490 | 91204 | 1 | ENSMUST00 | 0 | Dgkd | ENSMUSG00000070738.5 |
| chr11 | 62076979 | 62077897 | 30.45704 | 104.66754 | 100.97924 | Promoter | 11 | 62077024 | 62223005 | 145982 | 1 | ENSMUST00 | 0 | Specc1 | ENSMUSG00000042331.8 |
| chr12 | 88360163 | 88361325 | 30.45704 | 104.66754 | 100.97924 | Promoter | 12 | 88360554 | 88461719 | 101166 | 1 | ENSMUST00 | 0 | Adck1 | ENSMUSG00000021044.9 |
| chr14 | 65097178 | 65098671 | 30.45704 | 104.66754 | 100.97924 | Promoter | 14 | 65052060 | 65098106 | 46047 | 2 | ENSMUST00 | 0 | Extl3 | ENSMUSG00000021978.8 |
| chr2 | 132815143 | 132816695 | 30.45704 | 104.66754 | 100.97924 | Promoter | 2 | 132816141 | 132843895 | 27755 | 1 | ENSMUST00 | 0 | Mcm8 | ENSMUSG00000027353.9 |
| chr4 | 99714998 | 99716279 | 30.45704 | 104.66754 | 100.97924 | Promoter | 4 | 99715664 | 99730223 | 14560 | 1 | ENSMUST00 | 0 | Alg6 | ENSMUSG00000073792.6 |
| chr15 | 99456565 | 99458437 | 33.29458 | 104.62617 | 100.93848 | Promoter | 15 | 99422035 | 99457748 | 35714 | 2 | ENSMUST00 | 0 | Nckap5l | ENSMUSG00000023009.9 |
| chr10 | 81425223 | 81427495 | 34.46491 | 104.61457 | 100.92722 | Promoter | 10 | 81396186 | 81427187 | 31002 | 2 | ENSMUST00 | 0 | Nfic | ENSMUSG00000055053.12 |
| chr13 | 56702432 | 56704287 | 34.46491 | 104.61457 | 100.92722 | Promoter | 13 | 56703010 | 56742377 | 39368 | 1 | ENSMUST00 | 0 | Smad5 | ENSMUSG00000021540.11 |
| chr17 | 57274696 | 57277105 | 34.46491 | 104.61457 | 100.92722 | Distal Interge | 17 | 57279100 | 57327950 | 48851 | 1 | ENSMUST00 | -1995 | Vav1 | ENSMUSG00000034116.12 |
| chr18 | 34623373 | 34625328 | 34.46491 | 104.61457 | 100.92722 | Promoter | 18 | 34598615 | 34624601 | 25987 | 2 | ENSMUST00 | 0 | Brd8 | ENSMUSG00000003778.9 |
| chr18 | 35829308 | 35833126 | 34.46491 | 104.61457 | 100.92722 | Promoter | 18 | 35829818 | 35861687 | 31870 | 1 | ENSMUST00 | 0 | Cxxc5 | ENSMUSG00000046668.8 |
| chr19 | 5741479 | 5742777 | 34.46491 | 104.61457 | 100.92722 | Promoter | 19 | 5740904 | 5758532 | 17629 | 1 | ENSMUST00 | 576 | Ltbp3 | ENSMUSG00000024940.10 |
| chr2 | 154372080 | 154373167 | 34.46491 | 104.61457 | 100.92722 | Promoter | 2 | 154335380 | 154373010 | 37631 | 2 | ENSMUST00 | 0 | Cdk5rap1 | ENSMUSG00000027487.6 |
| chr3 | 89714676 | 89715826 | 34.46491 | 104.61457 | 100.92722 | Promoter | 3 | 89715022 | 89753446 | 38425 | 1 | ENSMUST00 | 0 | Adar | ENSMUSG00000027951.11 |
| chr4 | 116220461 | 116221244 | 34.46491 | 104.61457 | 100.92722 | Promoter | 4 | 116221618 | 116259576 | 37959 | 1 | ENSMUST00 | -374 | Pik3r3 | ENSMUSG00000028698.8 |
| chr5 | 146230632 | 146232287 | 34.46491 | 104.61457 | 100.92722 | Promoter | 5 | 146231230 | 146302874 | 71645 | 1 | ENSMUST00 | 0 | Cdk8 | ENSMUSG00000029635.10 |
| chr7 | 80946663 | 80948309 | 34.46491 | 104.61457 | 100.92722 | Promoter | 7 | 80904889 | 80947545 | 42657 | 2 | ENSMUST00 | 0 | Sec11a | ENSMUSG00000025724.7 |
| chr8 | 41373761 | 41375419 | 34.46491 | 104.61457 | 100.92722 | Promoter | 8 | 41340197 | 41374773 | 34577 | 2 | ENSMUST00 | 0 | Asah1 | ENSMUSG00000031591.9 |
| chr9 | 63376955 | 63378575 | 34.46491 | 104.61457 | 100.92722 | Promoter | 9 | 63163769 | 63377902 | 214134 | 2 | ENSMUST00 | 0 | Map2k5 | ENSMUSG00000058444.7 |
| chr8 | 70905581 | 70906912 | 33.86687 | 104.60014 | 100.91516 | Promoter | 8 | 70905974 | 70917526 | 11553 | 1 | ENSMUST00 | 0 | Map1s | ENSMUSG00000019261.2 |
| chr9 | 22225080 | 22226309 | 33.86687 | 104.60014 | 100.91516 | Promoter | 9 | 22225714 | 22243197 | 17484 | 1 | ENSMUST00 | 0 | Zfp809 | ENSMUSG00000057982.7 |
| chr1 | 132476575 | 132478169 | 32.69058 | 104.59052 | 100.90593 | Promoter | 1 | 132477365 | 132504490 | 27126 | 1 | ENSMUST00 | 0 | Rbbp5 | ENSMUSG00000026439.9 |
| chr1 | 134405521 | 134406555 | 32.69058 | 104.59052 | 100.90593 | Promoter | 1 | 134405559 | 134407990 | 2432 | 1 | ENSMUST00 | 0 | Cyb5r1 | ENSMUSG00000026456.13 |
| chr1 | 155812138 | 155813540 | 32.69058 | 104.59052 | 100.90593 | Promoter | 1 | 155778155 | 155812883 | 34729 | 2 | ENSMUST00 | 0 | Qsox1 | ENSMUSG00000033684.9 |
| chr1 | 165193163 | 165195083 | 32.69058 | 104.59052 | 100.90593 | Promoter | 1 | 165174341 | 165194433 | 20093 | 2 | ENSMUST00 | 0 | Sft2d2 | ENSMUSG00000040848.6 |
| chr10 | 79682099 | 79683765 | 32.69058 | 104.59052 | 100.90593 | Promoter | 10 | 79682195 | 79688394 | 6200 | 1 | ENSMUST00 | 0 | Cdc34 | ENSMUSG00000020307.9 |
| chr10 | 40246194 | 40247719 | 32.69058 | 104.59052 | 100.90593 | Promoter | 10 | 40223246 | 40247002 | 23757 | 2 | ENSMUST00 | 0 | Rpf2 | ENSMUSG00000038510.7 |
| chr13 | 98813742 | 98815937 | 32.69058 | 104.59052 | 100.90593 | Promoter | 13 | 98723407 | 98815449 | 92043 | 2 | ENSMUST00 | 0 | Fcho2 | ENSMUSG00000041685.10 |
| chr15 | 58415024 | 58416723 | 32.69058 | 104.59052 | 100.90593 | Promoter | 15 | 58415468 | 58457801 | 42334 | 1 | ENSMUST00 | 0 | D15Ertd621e | ENSMUSG00000037119.3 |
| chr15 | 89122321 | 89123657 | 32.69058 | 104.59052 | 100.90593 | Promoter | 15 | 89098357 | 89123087 | 24731 | 2 | ENSMUST00 | 0 | Tubgcp6 | ENSMUSG00000051786.9 |
| chr17 | 47628770 | 47630664 | 32.69058 | 104.59052 | 100.90593 | Promoter | 17 | 47630690 | 47686738 | 56049 | 1 | ENSMUST00 | -26 | Usp49 | ENSMUSG00000090115.2 |
| chr18 | 46279728 | 46281220 | 32.69058 | 104.59052 | 100.90593 | Promoter | 18 | 46239949 | 46280850 | 40902 | 2 | ENSMUST00 | 0 | Pggt1b | ENSMUSG00000024477.3 |
| chr19 | 15984196 | 15985563 | 32.69058 | 104.59052 | 100.90593 | Promoter | 19 | 15955774 | 15984989 | 29216 | 2 | ENSMUST00 | 0 | Cep78 | ENSMUSG00000041491.7 |
| chr2 | 30391708 | 30393784 | 32.69058 | 104.59052 | 100.90593 | Promoter | 2 | 30392254 | 30400529 | 8276 | 1 | ENSMUST00 | 0 | Dolpp1 | ENSMUSG00000026856.9 |
| chr2 | 104590447 | 104591512 | 32.69058 | 104.59052 | 100.90593 | Promoter | 2 | 104590523 | 104665429 | 74907 | 1 | ENSMUST00 | 0 | Cstf3 | ENSMUSG00000027176.8 |
| chr2 | 144368284 | 144369979 | 32.69058 | 104.59052 | 100.90593 | Promoter | 2 | 144368983 | 144407676 | 38694 | 1 | ENSMUST00 | 0 | Csrp2bp | ENSMUSG00000027425.13 |
| chr4 | 155490668 | 155493066 | 32.69058 | 104.59052 | 100.90593 | Promoter | 4 | 155491361 | 155559269 | 67909 | 1 | ENSMUST00 | 0 | Gnb1 | ENSMUSG00000029064.10 |
| chr5 | 30906784 | 30908449 | 32.69058 | 104.59052 | 100.90593 | Promoter | 5 | 30905885 | 30907788 | 1904 | 2 | ENSMUST00 | 0 | Ost4 | ENSMUSG00000038803.5 |
| chr6 | 83571740 | 83573137 | 32.69058 | 104.59052 | 100.90593 | Promoter | 6 | 83543206 | 83572504 | 29299 | 2 | ENSMUST00 | 0 | Stambp | ENSMUSG00000006906.9 |
| chr6 | 87913750 | 87914946 | 32.69058 | 104.59052 | 100.90593 | Promoter | 6 | 87913935 | 87936629 | 22695 | 1 | ENSMUST00 | 0 | Hmces | ENSMUSG00000030060.9 |
| chr6 | 39116813 | 39119136 | 32.69058 | 104.59052 | 100.90593 | Promoter | 6 | 39086410 | 39118349 | 31940 | 2 | ENSMUST00 | 0 | Parp12 | ENSMUSG00000038507.6 |
| chr8 | 64848977 | 64850628 | 32.69058 | 104.59052 | 100.90593 | Promoter | 8 | 64739863 | 64849818 | 109956 | 2 | ENSMUST00 | 0 | Klhl2 | ENSMUSG00000031605.7 |
| chr9 | 44360770 | 44362079 | 32.69058 | 104.59052 | 100.90593 | Promoter | 9 | 44348106 | 44361670 | 13565 | 2 | ENSMUST00 | 0 | Vps11 | ENSMUSG00000032127.8 |
| chr9 | 121718570 | 121720727 | 32.69058 | 104.59052 | 100.90593 | Promoter | 9 | 121719169 | 121742718 | 23550 | 1 | ENSMUST00 | 0 | Nktr | ENSMUSG00000032525.9 |
| chr9 | 75070770 | 75072219 | 32.69058 | 104.59052 | 100.90593 | Promoter | 9 | 75071015 | 75117308 | 46294 | 1 | ENSMUST00 | 0 | Myo5a | ENSMUSG00000034593.11 |
| chr1 | 127204578 | 127207504 | 33.22886 | 104.51182 | 100.83223 | Promoter | 1 | 127205015 | 127388212 | 183198 | 1 | ENSMUST00 | 0 | Mgat5 | ENSMUSG00000036155.8 |
| chr11 | 72134751 | 72136239 | 33.22886 | 104.51182 | 100.83223 | Promoter | 11 | 72047528 | 72135721 | 88194 | 2 | ENSMUST00 | 0 | Pitpnm3 | ENSMUSG00000040543.11 |
| chr16 | 11202379 | 11204032 | 33.22886 | 104.51182 | 100.83223 | Promoter | 16 | 11192970 | 11203331 | 10362 | 2 | ENSMUST00 | 0 | Rsl1d1 | ENSMUSG00000005846.11 |
| chr10 | 30199924 | 30200973 | 32.06657 | 104.51031 | 100.83098 | Promoter | 10 | 30196011 | 30200540 | 4530 | 2 | ENSMUST00 | 0 | Cenpw | ENSMUSG00000075266.4 |
| chr12 | 76254776 | 76256315 | 32.06657 | 104.51031 | 100.83098 | Promoter | 12 | 76255232 | 76319820 | 64589 | 1 | ENSMUST00 | 0 | Mthfd1 | ENSMUSG00000021048.6 |
| chr16 | 90385388 | 90387479 | 32.06657 | 104.51031 | 100.83098 | Promoter | 16 | 90386397 | 90499553 | 113157 | 1 | ENSMUST00 | 0 | Hunk | ENSMUSG00000053414.7 |
| chr3 | 95992078 | 95997273 | 32.06657 | 104.51031 | 100.83098 | Promoter | 3 | 95988429 | 95996001 | 7573 | 2 | ENSMUST00 | 0 | Plekho1 | ENSMUSG00000015745.4 |
| chr9 | 7763992 | 7764964 | 32.06657 | 104.51031 | 100.83098 | Promoter | 9 | 7764041 | 7794333 | 30293 | 1 | ENSMUST00 | 0 | Tmem123 | ENSMUSG00000050912.10 |
| chr12 | 113097661 | 113099119 | 34.40292 | 104.50784 | 100.82993 | Promoter | 12 | 113098278 | 113137205 | 38928 | 1 | ENSMUST00 | 0 | Mta1 | ENSMUSG00000021144.9 |
| chr14 | 32201043 | 32202975 | 29.81544 | 104.49987 | 100.82211 | Promoter | 14 | 32180162 | 32201898 | 21737 | 2 | ENSMUST00 | 0 | Timm23 | ENSMUSG00000013701.7 |
| chr10 | 95501140 | 95502039 | 32.62992 | 104.48419 | 100.80671 | Promoter | 10 | 95480806 | 95501927 | 21122 | 2 | ENSMUST00 | 0 | Mrpl42 | ENSMUSG00000062981.4 |
| chr19 | 43689367 | 43690591 | 33.79798 | 104.48097 | 100.80365 | Promoter | 19 | 43689672 | 43733697 | 44026 | 1 | ENSMUST00 | 0 | Entpd7 | ENSMUSG00000025192.8 |
| chr1 | 74587078 | 74588189 | 33.1634 | 104.39787 | 100.72088 | Promoter | 1 | 74565127 | 74588028 | 22902 | 2 | ENSMUST00 | 0 | Zfp142 | ENSMUSG00000026135.12 |
| chr2 | 26408352 | 26409246 | 29.21465 | 104.38775 | 100.71102 | Promoter | 2 | 26396249 | 26409200 | 12952 | 2 | ENSMUST00 | 0 | Inpp5e | ENSMUSG00000026925.11 |
| chr19 | 6045560 | 6048074 | 33.72937 | 104.36225 | 100.686 | Promoter | 19 | 6046576 | 6053712 | 7137 | 1 | ENSMUST00 | 0 | Syvn1 | ENSMUSG00000024807.12 |
| chr2 | 16355582 | 16359803 | 33.72937 | 104.36225 | 100.686 | Promoter | 2 | 16356304 | 16755839 | 399536 | 1 | ENSMUST00 | 0 | Plxdc2 | ENSMUSG00000026748.8 |
| chr13 | 67305784 | 67306929 | 28.65092 | 104.32529 | 100.64934 | Promoter | 13 | 67292450 | 67306412 | 13963 | 2 | ENSMUST00 | 0 | Zfp457 | ENSMUSG00000055341.8 |
| chr8 | 84839859 | 84840734 | 28.65092 | 104.32529 | 100.64934 | Promoter | 8 | 84834652 | 84840641 | 5990 | 2 | ENSMUST00 | 0 | Rad23a | ENSMUSG00000003813.10 |
| chr12 | 98746611 | 98748039 | 33.09821 | 104.28432 | 100.60858 | Promoter | 12 | 98746968 | 98787774 | 40807 | 1 | ENSMUST00 | 0 | Zc3h14 | ENSMUSG00000021012.10 |
| chr15 | 85668507 | 85673353 | 33.09821 | 104.28432 | 100.60858 | Distal Interge | 15 | 85639747 | 85645934 | 6188 | 1 | ENSMUST00 | 28761 | RP23-64J7.2 | ENSMUSG00000101828.1 |
| chr16 | 11322315 | 11324140 | 33.09821 | 104.28432 | 100.60858 | Promoter | 16 | 11322908 | 11755472 | 432565 | 1 | ENSMUST00 | 0 | Snx29 | ENSMUSG00000071669.9 |
| chr2 | 121270666 | 121272040 | 33.0982 | 104.28432 | 100.60858 | Promoter | 2 | 121194838 | 121271341 | 76504 | 2 | ENSMUST00 | 0 | Trp53bp1 | ENSMUSG00000043909.11 |
| chr5 | 110879341 | 110880511 | 33.09821 | 104.28432 | 100.60858 | Promoter | 5 | 110879803 | 111289779 | 409977 | 1 | ENSMUST00 | 0 | Ttc28 | ENSMUSG00000033209.12 |
| chr6 | 38550233 | 38551299 | 33.09821 | 104.28432 | 100.60858 | Promoter | 6 | 38551334 | 38605040 | 53707 | 1 | ENSMUST00 | -35 | Luc7l2 | ENSMUSG00000029823.11 |
| chr8 | 72474318 | 72476077 | 33.09821 | 104.28432 | 100.60858 | Promoter | 8 | 72460489 | 72475233 | 14745 | 2 | ENSMUST00 | 0 | Cherp | ENSMUSG00000052488.6 |
| chr7 | 24081395 | 24082851 | 22.65152 | 104.2504 | 100.57662 | Promoter | 7 | 24081924 | 24107698 | 25775 | 1 | ENSMUST00 | 0 | Zfp180 | ENSMUSG00000057101.6 |
| chr18 | 62179460 | 62180731 | 33.66105 | 104.24397 | 100.57021 | Promoter | 18 | 62177817 | 62179959 | 2143 | 2 | ENSMUST00 | 0 | Adrb2 | ENSMUSG00000045730.3 |
| chr6 | 28831812 | 28833824 | 33.66105 | 104.24397 | 100.57021 | Promoter | 6 | 28828133 | 28831747 | 3615 | 2 | ENSMUST00 | -66 | Lrrc4 | ENSMUSG00000049939.6 |
| chr13 | 52979288 | 52982918 | 34.18774 | 104.13715 | 100.46495 | Promoter | 13 | 52967209 | 52981073 | 13865 | 2 | ENSMUST00 | 0 | Nfil3 | ENSMUSG00000056749.7 |
| chr9 | 106477225 | 106478088 | 34.18774 | 104.13715 | 100.46495 | Promoter | 9 | 106477269 | 106485424 | 8156 | 1 | ENSMUST00 | 0 | Rrp9 | ENSMUSG00000041506.9 |
| chr12 | 73286052 | 73287550 | 24.65524 | 104.07409 | 100.40247 | Promoter | 12 | 73280410 | 73286711 | 6302 | 2 | ENSMUST00 | 0 | Trmt5 | ENSMUSG00000034442.10 |
| chr14 | 20702046 | 20703649 | 24.65524 | 104.07409 | 100.40247 | Promoter | 14 | 20703027 | 20704417 | 1391 | 1 | ENSMUST00 | 0 | Chchd1 | ENSMUSG00000063787.3 |

| | | | | | | | | | | | | | | | |
|---|---|---|---|---|---|---|---|---|---|---|---|---|---|---|---|
| chr16 | 95990153 | 95991577 | 24.65524 | 104.07409 | 100.40247 | Promoter | 16 | 95979933 | 95990960 | 11028 | 2 | ENSMUST00 | 0 | Psmg1 | ENSMUSG00000022913.9 |
| chr2 | 91182039 | 91183580 | 24.65524 | 104.07409 | 100.40247 | Promoter | 2 | 91137360 | 91182848 | 45489 | 2 | ENSMUST00 | 0 | Madd | ENSMUSG00000040687.11 |
| chr5 | 123394150 | 123396353 | 24.65524 | 104.07409 | 100.40247 | Promoter | 5 | 123394798 | 123446896 | 52099 | 1 | ENSMUST00 | 0 | Mlxip | ENSMUSG00000038342.10 |
| chr6 | 57691329 | 57692533 | 24.65524 | 104.07409 | 100.40247 | Promoter | 6 | 57686576 | 57692078 | 5503 | 2 | ENSMUST00 | 0 | Pyurf | ENSMUSG00000043162.7 |
| chr8 | 80738046 | 80740342 | 32.96857 | 104.0584 | 100.38712 | Promoter | 8 | 80698507 | 80739497 | 40991 | 2 | ENSMUST00 | 0 | Smarca5 | ENSMUSG00000031715.9 |
| chr4 | 53631039 | 53632373 | 32.38209 | 104.04922 | 100.37821 | Promoter | 4 | 53631471 | 53707009 | 75539 | 1 | ENSMUST00 | 0 | Fsd1l | ENSMUSG00000054752.11 |
| chr15 | 99223127 | 99225653 | 34.11661 | 104.01456 | 100.34361 | Promoter | 15 | 99224976 | 99242817 | 17842 | 1 | ENSMUST00 | 0 | Kcnh3 | ENSMUSG00000037579.6 |
| chr4 | 41568168 | 41569606 | 34.11661 | 104.01456 | 100.34361 | Promoter | 4 | 41517691 | 41569509 | 51819 | 2 | ENSMUST00 | 0 | Fam219a | ENSMUSG00000028439.9 |
| chr9 | 20491789 | 20493539 | 34.11661 | 104.01456 | 100.34361 | Promoter | 9 | 20468549 | 20492746 | 24198 | 2 | ENSMUST00 | 0 | Zfp426 | ENSMUSG00000059475.8 |
| chr1 | 52816600 | 52818107 | 32.90413 | 103.94604 | 100.27567 | Promoter | 1 | 52785427 | 52817688 | 32262 | 2 | ENSMUST00 | 0 | Inpp1 | ENSMUSG00000026102.4 |
| chr6 | 65042284 | 65043527 | 32.90413 | 103.94604 | 100.27567 | Promoter | 6 | 65042667 | 65116050 | 73384 | 1 | ENSMUST00 | 0 | Smarcad1 | ENSMUSG00000029920.7 |
| chr19 | 41829166 | 41831508 | 30.60556 | 103.92632 | 100.25623 | Promoter | 19 | 41830168 | 41830809 | 642 | 1 | ENSMUST00 | 0 | Frat1 | ENSMUSG00000067199.3 |
| chr5 | 122500881 | 122503112 | 30.60556 | 103.92632 | 100.25623 | Promoter | 5 | 122442702 | 122502225 | 59524 | 2 | ENSMUST00 | 0 | Atp2a2 | ENSMUSG00000029467.10 |
| chr11 | 78094238 | 78095478 | 34.04577 | 103.89243 | 100.2229 | Promoter | 11 | 78094660 | 78156700 | 62041 | 1 | ENSMUST00 | 0 | Fam222b | ENSMUSG00000037750.11 |
| chr8 | 85298510 | 85300590 | 34.04577 | 103.89243 | 100.2229 | Promoter | 8 | 85260397 | 85299513 | 39117 | 2 | ENSMUST00 | 0 | Vps35 | ENSMUSG00000031696.8 |
| chr11 | 67921383 | 67922888 | 33.45771 | 103.89173 | 100.22243 | Promoter | 11 | 67854523 | 67922153 | 67631 | 2 | ENSMUST00 | 0 | Usp43 | ENSMUSG00000020905.6 |
| chr8 | 46738444 | 46742094 | 33.45771 | 103.89173 | 100.22243 | Promoter | 8 | 46739745 | 46847458 | 107714 | 1 | ENSMUST00 | 0 | Irf2 | ENSMUSG00000031627.7 |
| chr10 | 53749886 | 53751730 | 32.25957 | 103.83389 | 100.16508 | Promoter | 10 | 53633145 | 53750880 | 117736 | 2 | ENSMUST00 | 0 | Fam184a | ENSMUSG00000019856.8 |
| chr12 | 113014500 | 113015637 | 32.25957 | 103.83389 | 100.16508 | Promoter | 12 | 113014508 | 113074401 | 59894 | 1 | ENSMUST00 | 0 | Pacs2 | ENSMUSG00000021143.9 |
| chr16 | 36070577 | 36072663 | 32.25957 | 103.83389 | 100.16508 | Promoter | 16 | 36043844 | 36071515 | 27672 | 2 | ENSMUST00 | 0 | Fam162a | ENSMUSG00000003955.7 |
| chr3 | 36571453 | 36572046 | 32.25957 | 103.83389 | 100.16508 | Promoter | 3 | 36565978 | 36571927 | 5950 | 2 | ENSMUST00 | 0 | Ccna2 | ENSMUSG00000027715.4 |
| chr5 | 73646992 | 73648042 | 32.25957 | 103.83389 | 100.16508 | Promoter | 5 | 73632749 | 73647713 | 14965 | 2 | ENSMUST00 | 0 | Sgcb | ENSMUSG00000029156.10 |
| chr6 | 52639334 | 52640742 | 32.25957 | 103.83389 | 100.16508 | Promoter | 6 | 52546228 | 52640389 | 94162 | 2 | ENSMUST00 | 0 | Hibadh | ENSMUSG00000029776.10 |
| chr6 | 4086193 | 4087389 | 32.25957 | 103.83389 | 100.16508 | Promoter | 6 | 4076899 | 4086972 | 10074 | 2 | ENSMUST00 | 0 | Bet1 | ENSMUSG00000032757.4 |
| chr6 | 54816612 | 54818738 | 32.83994 | 103.83406 | 100.16508 | Promoter | 6 | 54816916 | 54893500 | 76585 | 1 | ENSMUST00 | 0 | Znrf2 | ENSMUSG00000058446.9 |
| chr8 | 70211568 | 70212393 | 32.25957 | 103.83389 | 100.16508 | Promoter | 8 | 70184340 | 70212256 | 27917 | 2 | ENSMUST00 | 0 | Slc25a42 | ENSMUSG00000002346.11 |
| chr8 | 122550054 | 122551803 | 32.25957 | 103.83389 | 100.16508 | Promoter | 8 | 122481698 | 122551329 | 69632 | 2 | ENSMUST00 | 0 | Piezo1 | ENSMUSG00000014444.11 |
| chr15 | 80012006 | 80015223 | 33.97523 | 103.77077 | 100.10409 | Promoter | 15 | 79995900 | 80014808 | 18909 | 2 | ENSMUST00 | 0 | Pdgfb | ENSMUSG00000000489.6 |
| chr7 | 16875289 | 16875877 | 33.97523 | 103.77077 | 100.10409 | Promoter | 7 | 16875317 | 16887301 | 11985 | 1 | ENSMUST00 | 0 | Dact3 | ENSMUSG00000078794.3 |
| chr4 | 141138228 | 141139871 | 29.94827 | 103.73744 | 100.07098 | Promoter | 4 | 141113001 | 141139727 | 26727 | 2 | ENSMUST00 | 0 | Szrd1 | ENSMUSG00000040842.12 |
| chr6 | 85914902 | 85916630 | 29.94827 | 103.73744 | 100.07098 | Promoter | 6 | 85915738 | 85927608 | 11871 | 1 | ENSMUST00 | 0 | Tprkb | ENSMUSG00000054226.8 |
| chr11 | 16507148 | 16508821 | 32.77601 | 103.72247 | 100.0565 | Promoter | 11 | 16500530 | 16508484 | 7955 | 2 | ENSMUST00 | 0 | Sec61g | ENSMUSG00000078974.5 |
| chr1 | 182018483 | 182020835 | 33.32351 | 103.65904 | 99.99326 | Promoter | 1 | 181904445 | 182019980 | 115536 | 2 | ENSMUST00 | 0 | Enah | ENSMUSG00000022995.11 |
| chr1 | 161033938 | 161036613 | 33.32351 | 103.65904 | 99.99326 | Promoter | 1 | 161008794 | 161034849 | 26056 | 2 | ENSMUST00 | 0 | Zbtb37 | ENSMUSG00000043467.14 |
| chr10 | 93888741 | 93890501 | 33.32351 | 103.65904 | 99.99326 | Promoter | 10 | 93858489 | 93889814 | 31326 | 2 | ENSMUST00 | 0 | Metap2 | ENSMUSG00000036112.9 |
| chr2 | 104815665 | 104818377 | 33.32351 | 103.65904 | 99.99326 | Promoter | 2 | 104754795 | 104816696 | 61902 | 2 | ENSMUST00 | 0 | Qser1 | ENSMUSG00000074994.4 |
| chr5 | 52668987 | 52670046 | 33.32351 | 103.65904 | 99.99326 | Promoter | 5 | 52640087 | 52669729 | 29643 | 2 | ENSMUST00 | 0 | Sepsecs | ENSMUSG00000029173.7 |
| chr5 | 108312006 | 108313586 | 33.32351 | 103.65904 | 99.99326 | Promoter | 5 | 108312609 | 108348233 | 35625 | 1 | ENSMUST00 | 0 | Pigg | ENSMUSG00000029263.10 |
| chr4 | 136052141 | 136053418 | 29.33348 | 103.60687 | 99.9429 | Promoter | 4 | 136028265 | 136052685 | 24421 | 2 | ENSMUST00 | 0 | Rpl11 | ENSMUSG00000059291.10 |
| chr6 | 99520242 | 99523905 | 29.33348 | 103.60687 | 99.9429 | Promoter | 6 | 98929189 | 99520997 | 591809 | 2 | ENSMUST00 | 0 | Foxp1 | ENSMUSG00000030067.12 |
| chr7 | 141446958 | 141448497 | 29.33348 | 103.60687 | 99.9429 | Promoter | 7 | 141447370 | 141451585 | 4216 | 1 | ENSMUST00 | 0 | Rplp2 | ENSMUSG00000025508.8 |
| chr1 | 181143562 | 181145514 | 30.41442 | 103.5799 | 99.9162 | Promoter | 1 | 181093423 | 181144157 | 50735 | 2 | ENSMUST00 | 0 | Nvl | ENSMUSG00000026516.7 |
| chr1 | 58391838 | 58394196 | 30.41442 | 103.5799 | 99.9162 | Promoter | 1 | 58392898 | 58407353 | 14456 | 1 | ENSMUST00 | 0 | Bzw1 | ENSMUSG00000051223.9 |
| chr10 | 10557459 | 10558341 | 30.41442 | 103.5799 | 99.9162 | Promoter | 10 | 10545011 | 10558233 | 13223 | 2 | ENSMUST00 | 0 | Rab32 | ENSMUSG00000019832.3 |
| chr10 | 79096648 | 79097890 | 30.41442 | 103.5799 | 99.9162 | Promoter | 10 | 79064374 | 79097600 | 33227 | 2 | ENSMUST00 | 0 | 2610008E11I | ENSMUSG00000060301.5 |
| chr11 | 104468564 | 104469932 | 30.41442 | 103.5799 | 99.9162 | Promoter | 11 | 104333501 | 104468861 | 135361 | 2 | ENSMUST00 | 0 | Kansl1 | ENSMUSG00000018412.11 |
| chr11 | 106255111 | 106257244 | 30.41442 | 103.5799 | 99.9162 | Promoter | 11 | 106249142 | 106256079 | 6938 | 2 | ENSMUST00 | 0 | Ftsj3 | ENSMUSG00000020706.8 |
| chr11 | 22858157 | 22860897 | 30.41442 | 103.5799 | 99.9162 | Promoter | 11 | 22834747 | 22860101 | 25355 | 2 | ENSMUST00 | 0 | B3gnt2 | ENSMUSG00000051650.10 |
| chr12 | 50647570 | 50649686 | 30.41442 | 103.5799 | 99.9162 | Promoter | 12 | 50341231 | 50649223 | 307993 | 2 | ENSMUST00 | 0 | Prkd1 | ENSMUSG00000002688.7 |
| chr12 | 99883014 | 99885191 | 30.41442 | 103.5799 | 99.9162 | Promoter | 12 | 99717531 | 99883442 | 165912 | 2 | ENSMUST00 | 0 | Efcab11 | ENSMUSG00000021176.5 |
| chr13 | 70636576 | 70638601 | 30.41442 | 103.5799 | 99.9162 | Promoter | 13 | 70588689 | 70637634 | 48946 | 2 | ENSMUST00 | 0 | BC018507 | ENSMUSG00000034525.6 |
| chr15 | 73421126 | 73424166 | 30.41442 | 103.5799 | 99.9162 | Promoter | 15 | 73205105 | 73423191 | 218087 | 2 | ENSMUST00 | 0 | Ptk2 | ENSMUSG00000022607.9 |
| chr15 | 99725578 | 99726497 | 30.41442 | 103.5799 | 99.9162 | Promoter | 15 | 99725618 | 99728136 | 2519 | 1 | ENSMUST00 | 0 | Cox14 | ENSMUSG00000023020.2 |
| chr15 | 102980785 | 102982803 | 30.41442 | 103.5799 | 99.9162 | 5' UTR | 15 | 102977032 | 102985099 | 8068 | 1 | ENSMUST00 | 3754 | Hoxc9 | ENSMUSG00000036139.6 |
| chr16 | 5012992 | 5013607 | 30.41442 | 103.5799 | 99.9162 | Promoter | 16 | 5008730 | 5013517 | 4788 | 2 | ENSMUST00 | 0 | Rogdi | ENSMUSG00000022540.11 |
| chr19 | 28989833 | 28991359 | 30.41442 | 103.5799 | 99.9162 | Promoter | 19 | 28990494 | 29017569 | 27076 | 1 | ENSMUST00 | 0 | Cdc37l1 | ENSMUSG00000024780.6 |
| chr19 | 10880728 | 10882347 | 30.41442 | 103.5799 | 99.9162 | Promoter | 19 | 10870660 | 10882001 | 11342 | 2 | ENSMUST00 | 0 | Tmem109 | ENSMUSG00000034659.9 |
| chr2 | 121866208 | 121868007 | 30.41442 | 103.5799 | 99.9162 | Promoter | 2 | 121866970 | 121936205 | 69236 | 1 | ENSMUST00 | 0 | Casc4 | ENSMUSG00000060227.10 |
| chr2 | 156720047 | 156723142 | 30.41442 | 103.5799 | 99.9162 | Promoter | 2 | 156720572 | 156749091 | 28520 | 1 | ENSMUST00 | 0 | Dlgap4 | ENSMUSG00000061689.10 |
| chr2 | 29826439 | 29828177 | 30.41442 | 103.5799 | 99.9162 | Promoter | 2 | 29827342 | 29843374 | 16033 | 1 | ENSMUST00 | 0 | Urm1 | ENSMUSG00000069020.4 |
| chr3 | 100680989 | 100686556 | 30.41442 | 103.5799 | 99.9162 | Promoter | 3 | 100562208 | 100685503 | 123296 | 2 | ENSMUST00 | 0 | Man1a2 | ENSMUSG00000008763.11 |
| chr3 | 135689869 | 135692324 | 30.41442 | 103.5799 | 99.9162 | Promoter | 3 | 135584655 | 135691546 | 106892 | 2 | ENSMUST00 | 0 | Nfkb1 | ENSMUSG00000028163.12 |
| chr4 | 21775904 | 21777133 | 30.41442 | 103.5799 | 99.9162 | Promoter | 4 | 21776261 | 21784840 | 8580 | 1 | ENSMUST00 | 0 | Usp45 | ENSMUSG00000040455.12 |
| chr5 | 120613053 | 120614224 | 30.41442 | 103.5799 | 99.9162 | Promoter | 5 | 120613130 | 120628590 | 15461 | 1 | ENSMUST00 | 0 | Ddx54 | ENSMUSG00000029599.10 |
| chr5 | 139483738 | 139484539 | 30.41442 | 103.5799 | 99.9162 | Promoter | 5 | 139471216 | 139484517 | 13302 | 2 | ENSMUST00 | 0 | Zfand2a | ENSMUSG00000053581.8 |
| chr6 | 142756393 | 142757636 | 30.41442 | 103.5799 | 99.9162 | Promoter | 6 | 142756686 | 142775714 | 19029 | 1 | ENSMUST00 | 0 | Cmas | ENSMUSG00000030282.9 |
| chr7 | 6382653 | 6383974 | 30.41442 | 103.5799 | 99.9162 | Promoter | 7 | 6383318 | 6396633 | 13316 | 1 | ENSMUST00 | 0 | Zfp28 | ENSMUSG00000062861.7 |
| chr7 | 25680168 | 25681143 | 30.41442 | 103.5799 | 99.9162 | Promoter | 7 | 25681158 | 25686558 | 5401 | 1 | ENSMUST00 | -15 | B9d2 | ENSMUSG00000063439.6 |
| chr7 | 28277660 | 28278372 | 30.41442 | 103.5799 | 99.9162 | Promoter | 7 | 28277706 | 28280129 | 2424 | 1 | ENSMUST00 | 0 | Eid2b | ENSMUSG00000070705.2 |
| chr8 | 128356451 | 128361175 | 30.41442 | 103.5799 | 99.9162 | Promoter | 8 | 128359073 | 128505462 | 146390 | 1 | ENSMUST00 | 0 | Nrp1 | ENSMUSG00000025810.8 |
| chr8 | 95081119 | 95081904 | 30.41442 | 103.5799 | 99.9162 | Promoter | 8 | 95081201 | 95099874 | 18674 | 1 | ENSMUST00 | 0 | Katnb1 | ENSMUSG00000031787.7 |
| chr8 | 123040842 | 123042311 | 30.41442 | 103.5799 | 99.9162 | Promoter | 8 | 123042466 | 123044602 | 2137 | 1 | ENSMUST00 | -155 | RP23-460F21 | ENSMUSG00000099881.1 |
| chr9 | 55208507 | 55209941 | 30.41442 | 103.5799 | 99.9162 | Promoter | 9 | 55208925 | 55224433 | 15509 | 1 | ENSMUST00 | 0 | Fbxo22 | ENSMUSG00000032309.10 |
| chr9 | 73039068 | 73040618 | 30.41442 | 103.5799 | 99.9162 | Promoter | 9 | 73007419 | 73040378 | 32960 | 2 | ENSMUST00 | 0 | Pigb | ENSMUSG00000079469.5 |
| chr16 | 76370364 | 76374292 | 33.25682 | 103.54333 | 99.8849 | Promoter | 16 | 76287400 | 76373827 | 86428 | 2 | ENSMUST00 | 0 | Nrip1 | ENSMUSG00000048490.8 |
| chr14 | 66123977 | 66124544 | 33.83502 | 103.52887 | 99.8707 | Promoter | 14 | 66084377 | 66124500 | 40124 | 2 | ENSMUST00 | 0 | Ephx2 | ENSMUSG00000022040.7 |
| chr1 | 24099193 | 24100931 | 25.75243 | 103.50989 | 99.85224 | Promoter | 1 | 24011245 | 24100194 | 88950 | 2 | ENSMUST00 | 0 | Fam135a | ENSMUSG00000026153.10 |
| chr10 | 59950517 | 59952770 | 25.75243 | 103.50989 | 99.85224 | Promoter | 10 | 59949675 | 59951770 | 2096 | 2 | ENSMUST00 | 0 | Ddit4 | ENSMUSG00000020108.3 |
| chr11 | 54865974 | 54867430 | 25.75243 | 103.50989 | 99.85224 | Promoter | 11 | 54866383 | 54870501 | 4119 | 1 | ENSMUST00 | 0 | Hint1 | ENSMUSG00000020267.6 |
| chr16 | 78375631 | 78377323 | 25.75243 | 103.50989 | 99.85224 | Promoter | 16 | 78359860 | 78376810 | 16951 | 2 | ENSMUST00 | 0 | Btg3 | ENSMUSG00000022863.9 |
| chr3 | 40845976 | 40847610 | 25.75243 | 103.50989 | 99.85224 | Promoter | 3 | 40818070 | 40846885 | 28816 | 2 | ENSMUST00 | 0 | Mfsd8 | ENSMUSG00000025759.7 |
| chr6 | 71543403 | 71545051 | 25.75243 | 103.50989 | 99.85224 | Promoter | 6 | 71543797 | 71581579 | 37783 | 1 | ENSMUST00 | 0 | Chmp3 | ENSMUSG00000053119.6 |
| chr8 | 111258097 | 111259935 | 25.75243 | 103.50989 | 99.85224 | Promoter | 8 | 111154421 | 111259216 | 104796 | 2 | ENSMUST00 | 0 | Glg1 | ENSMUSG00000003316.9 |
| chr8 | 22059673 | 22061551 | 25.75243 | 103.50989 | 99.85224 | Promoter | 8 | 21992785 | 22060305 | 67521 | 2 | ENSMUST00 | 0 | Atp7b | ENSMUSG00000006567.6 |
| chr8 | 56550132 | 56552391 | 25.75243 | 103.50989 | 99.85224 | Promoter | 8 | 56551090 | 56571125 | 20036 | 1 | ENSMUST00 | 0 | Fbxo8 | ENSMUSG00000038206.7 |
| chr9 | 25149881 | 25152473 | 25.75243 | 103.50989 | 99.85224 | Promoter | 9 | 25108130 | 25151781 | 43652 | 2 | ENSMUST00 | 0 | Herpud2 | ENSMUSG00000008429.7 |
| chr9 | 57536855 | 57538456 | 25.75243 | 103.50989 | 99.85224 | Promoter | 9 | 57537528 | 57543187 | 5660 | 1 | ENSMUST00 | 0 | Fam219b | ENSMUSG00000032305.9 |
| chr16 | 4639898 | 4641726 | 32.64888 | 103.50044 | 99.84376 | Promoter | 16 | 4639989 | 4687292 | 47304 | 1 | ENSMUST00 | 0 | Dnaja3 | ENSMUSG00000004069.11 |
| chr1 | 171101881 | 171102987 | 31.48209 | 103.47416 | 99.8177 | Distal Interge | 1 | 171113918 | 171126829 | 12912 | 1 | ENSMUST00 | -10931 | 1700009P17I | ENSMUSG00000026649.9 |
| chr1 | 164248386 | 164249924 | 31.48209 | 103.47416 | 99.8177 | Promoter | 1 | 164249046 | 164265385 | 16340 | 1 | ENSMUST00 | 0 | Slc19a2 | ENSMUSG00000040918.7 |
| chr11 | 75347755 | 75349013 | 31.48209 | 103.47416 | 99.8177 | Promoter | 11 | 75298166 | 75348324 | 50159 | 2 | ENSMUST00 | 0 | Rpa1 | ENSMUSG00000000751.8 |
| chr12 | 51347377 | 51348896 | 31.48209 | 103.47416 | 99.8177 | Promoter | 12 | 51348061 | 51376986 | 28926 | 1 | ENSMUST00 | 0 | G2e3 | ENSMUSG00000035293.8 |
| chr16 | 38346628 | 38347635 | 31.48209 | 103.47416 | 99.8177 | Promoter | 16 | 38346991 | 38352763 | 5773 | 1 | ENSMUST00 | 0 | Cox17 | ENSMUSG00000046516.10 |
| chr17 | 72917390 | 72919689 | 31.48209 | 103.47416 | 99.8177 | Promoter | 17 | 72918305 | 72941942 | 23638 | 1 | ENSMUST00 | 0 | Lbh | ENSMUSG00000024063.7 |
| chr19 | 45659211 | 45660899 | 31.48209 | 103.47416 | 99.8177 | Promoter | 19 | 45578254 | 45660312 | 82059 | 2 | ENSMUST00 | 0 | Fbxw4 | ENSMUSG00000040913.5 |
| chr2 | 30358513 | 30359650 | 31.48209 | 103.47416 | 99.8177 | Promoter | 2 | 30344809 | 30359240 | 14432 | 2 | ENSMUST00 | 0 | Sh3glb2 | ENSMUSG00000026860.11 |
| chr6 | 3498030 | 3499113 | 31.48209 | 103.47416 | 99.8177 | Promoter | 6 | 3498382 | 3603315 | 104934 | 1 | ENSMUST00 | 0 | Ccdc132 | ENSMUSG00000001376.12 |
| chr7 | 123377961 | 123378620 | 31.48209 | 103.47416 | 99.8177 | Promoter | 7 | 123377982 | 123430358 | 52377 | 1 | ENSMUST00 | 0 | Lcmt1 | ENSMUSG00000030763.6 |
| chr9 | 122950202 | 122951544 | 31.48209 | 103.47416 | 99.8177 | Promoter | 9 | 122945089 | 122951000 | 5912 | 2 | ENSMUST00 | 0 | 1110059G10 | ENSMUSG00000032551.6 |
| chr3 | 90293103 | 90294459 | 33.19039 | 103.42804 | 99.77289 | Distal Interge | 3 | 90278008 | 90278670 | 663 | 1 | ENSMUST00 | 15096 | Dennd4b | ENSMUSG00000042404.11 |
| chr3 | 144203799 | 144206219 | 33.76535 | 103.4086 | 99.75368 | Promoter | 3 | 144188530 | 144205220 | 16691 | 2 | ENSMUST00 | 0 | Lmo4 | ENSMUSG00000028266.12 |
| chr11 | 115707334 | 115709306 | 23.46131 | 103.32642 | 99.67202 | Promoter | 11 | 115644045 | 115708597 | 64553 | 2 | ENSMUST00 | 0 | Grb2 | ENSMUSG00000059923.8 |
| chr3 | 96239518 | 96240302 | 23.46131 | 103.32642 | 99.67202 | Promoter | 3 | 96239749 | 96240276 | 528 | 2 | ENSMUST00 | 0 | Hist2h2aa1 | ENSMUSG00000063954.7 |
| chr9 | 21264611 | 21266112 | 23.46131 | 103.32642 | 99.67202 | Promoter | 9 | 21265299 | 21274368 | 9070 | 1 | ENSMUST00 | 0 | Atg4d | ENSMUSG00000002820.5 |
| chr17 | 15526508 | 15528256 | 33.12423 | 103.31316 | 99.65884 | Promoter | 17 | 15519208 | 15527281 | 8074 | 2 | ENSMUST00 | 0 | Pdcd2 | ENSMUSG00000014771.7 |
| chr10 | 7792084 | 7793742 | 33.69596 | 103.2888 | 99.6348 | Promoter | 10 | 7792891 | 7823135 | 30245 | 1 | ENSMUST00 | 0 | Ppil4 | ENSMUSG00000015757.5 |
| chr8 | 105635762 | 105638088 | 33.69596 | 103.2888 | 99.6348 | Promoter | 8 | 105636568 | 105681647 | 45080 | 1 | ENSMUST00 | 0 | Ctcf | ENSMUSG00000005698.10 |
| chr13 | 67637112 | 67638371 | 32.52273 | 103.27992 | 99.62622 | Promoter | 13 | 67617001 | 67637791 | 20791 | 2 | ENSMUST00 | 0 | A530054K11I | ENSMUSG00000021510.10 |
| chr11 | 79253142 | 79254985 | 30.75929 | 103.19345 | 99.54015 | Promoter | 11 | 79239372 | 79254671 | 15300 | 2 | ENSMUST00 | 0 | Wsb1 | ENSMUSG00000017677.6 |
| chr16 | 94568436 | 94571673 | 30.75929 | 103.19345 | 99.54015 | Promoter | 16 | 94570010 | 94695067 | 125058 | 1 | ENSMUST00 | 0 | Dyrk1a | ENSMUSG00000022897.9 |
| chr19 | 5106004 | 5107122 | 30.75929 | 103.19345 | 99.54015 | Promoter | 19 | 5099207 | 5106996 | 7790 | 2 | ENSMUST00 | 0 | Rab1b | ENSMUSG00000024870.5 |
| chr6 | 29608825 | 29610408 | 30.75929 | 103.19345 | 99.54015 | Promoter | 6 | 29540827 | 29609607 | 68781 | 2 | ENSMUST00 | 0 | Tnpo3 | ENSMUSG00000012535.9 |
| chr11 | 73198447 | 73200105 | 27.03867 | 103.18376 | 99.53125 | Promoter | 11 | 73183596 | 73199042 | 15447 | 2 | ENSMUST00 | 0 | Ctns | ENSMUSG00000005949.4 |
| chr11 | 76216820 | 76217929 | 27.03867 | 103.18376 | 99.53125 | Promoter | 11 | 76217608 | 76218666 | 1059 | 1 | ENSMUST00 | 0 | Dbil5 | ENSMUSG00000038057.4 |
| chr13 | 55569705 | 55571873 | 27.03867 | 103.18376 | 99.53125 | Promoter | 13 | 55539319 | 55571120 | 31802 | 2 | ENSMUST00 | 0 | Fam193b | ENSMUSG00000021495.8 |
| chr13 | 112800784 | 112802503 | 27.03867 | 103.18376 | 99.53125 | Promoter | 13 | 112800894 | 112867881 | 66988 | 1 | ENSMUST00 | 0 | Ppap2a | ENSMUSG00000021759.8 |
| chr13 | 12565502 | 12566770 | 27.03867 | 103.18376 | 99.53125 | Promoter | 13 | 12565883 | 12609526 | 43644 | 1 | ENSMUST00 | 0 | Ero1lb | ENSMUSG00000057069.6 |
| chr15 | 58932941 | 58934443 | 27.03867 | 103.18376 | 99.53125 | Promoter | 15 | 58933810 | 58939489 | 5680 | 1 | ENSMUST00 | 0 | Ndufb9 | ENSMUSG00000022354.3 |
| chr18 | 34372240 | 34373485 | 27.03867 | 103.18376 | 99.53125 | Promoter | 18 | 34344889 | 34373415 | 28527 | 2 | ENSMUST00 | 0 | Reep5 | ENSMUSG00000005873.5 |
| chr2 | 145934018 | 145935221 | 27.03867 | 103.18376 | 99.53125 | Promoter | 2 | 145917479 | 145935014 | 17536 | 2 | ENSMUST00 | 0 | Crnkl1 | ENSMUSG00000001767.5 |
| chr2 | 77279943 | 77281168 | 27.03867 | 103.18376 | 99.53125 | Promoter | 2 | 77186555 | 77280592 | 94038 | 2 | ENSMUST00 | 0 | Sestd1 | ENSMUSG00000042272.12 |
| chr3 | 61362950 | 61366087 | 27.03867 | 103.18376 | 99.53125 | Promoter | 3 | 61364507 | 61368703 | 4197 | 1 | ENSMUST00 | 0 | Rap2b | ENSMUSG00000036894.2 |
| chr5 | 74067293 | 74069360 | 27.03867 | 103.18376 | 99.53125 | Promoter | 5 | 73998463 | 74068420 | 69958 | 2 | ENSMUST00 | 0 | Usp46 | ENSMUSG00000054814.9 |
| chr6 | 146724505 | 146725882 | 27.03867 | 103.18376 | 99.53125 | Promoter | 6 | 146724995 | 146778812 | 53818 | 1 | ENSMUST00 | 0 | Stk38l | ENSMUSG00000001630.8 |
| chr6 | 59208815 | 59209796 | 27.03867 | 103.18376 | 99.53125 | Promoter | 6 | 59208870 | 59212033 | 3164 | 1 | ENSMUST00 | 0 | Tigd2 | ENSMUSG00000049232.3 |
| chr7 | 127206978 | 127208714 | 27.03867 | 103.18376 | 99.53125 | Promoter | 7 | 127197465 | 127208468 | 11004 | 2 | ENSMUST00 | 0 | Tbc1d10b | ENSMUSG00000042492.7 |
| chr8 | 119838819 | 119841220 | 27.03867 | 103.18376 | 99.53125 | Promoter | 8 | 119809225 | 119840522 | 31298 | 2 | ENSMUST00 | 0 | Cotl1 | ENSMUSG00000031827.7 |
| chr9 | 13826148 | 13827508 | 27.03867 | 103.18376 | 99.53125 | Promoter | 9 | 13807792 | 13826991 | 19200 | 2 | ENSMUST00 | 0 | Cep57 | ENSMUSG00000031922.7 |

| | | | | | | | | | | | | | | | |
|---|---|---|---|---|---|---|---|---|---|---|---|---|---|---|---|
| chr1 | 186748495 | 186749995 | 28.56742 | 103.16996 | 99.51957 | Promoter | 1 | 186720978 | 186749358 | 28381 | 2 | ENSMUST00 | 0 | Rrp15 | ENSMUSG00000001305.5 |
| chr1 | 38128784 | 38130448 | 28.56742 | 103.16996 | 99.51957 | Promoter | 1 | 38052787 | 38129662 | 76876 | 2 | ENSMUST00 | 0 | Rev1 | ENSMUSG00000026082.6 |
| chr1 | 82315985 | 82317296 | 28.56742 | 103.16996 | 99.51957 | Promoter | 1 | 82316452 | 82445366 | 128915 | 1 | ENSMUST00 | 0 | Rhbdd1 | ENSMUSG00000026142.10 |
| chr10 | 42761507 | 42763100 | 28.56742 | 103.16996 | 99.51957 | Promoter | 10 | 42761615 | 42830392 | 68778 | 1 | ENSMUST00 | 0 | Sec63 | ENSMUSG00000019802.8 |
| chr10 | 28073969 | 28076790 | 32.46003 | 103.17023 | 99.51957 | Promoter | 10 | 28074820 | 28597397 | 522578 | 1 | ENSMUST00 | 0 | Ptprk | ENSMUSG00000019889.9 |
| chr10 | 20148389 | 20150329 | 28.56742 | 103.16996 | 99.51957 | Promoter | 10 | 20148471 | 20278699 | 130229 | 1 | ENSMUST00 | 0 | Map7 | ENSMUSG00000019996.11 |
| chr10 | 61782647 | 61784301 | 28.56742 | 103.16996 | 99.51957 | Promoter | 10 | 61738665 | 61784147 | 45483 | 2 | ENSMUST00 | 0 | H2afy2 | ENSMUSG00000020086.6 |
| chr10 | 81036845 | 81037439 | 32.46003 | 103.17023 | 99.51957 | Promoter | 10 | 81028538 | 81037426 | 8889 | 2 | ENSMUST00 | 0 | Slc39a3 | ENSMUSG00000046822.7 |
| chr11 | 102229308 | 102230763 | 28.56742 | 103.16996 | 99.51957 | Promoter | 11 | 102194432 | 102230166 | 35735 | 2 | ENSMUST00 | 0 | Hdac5 | ENSMUSG00000008855.12 |
| chr11 | 76242987 | 76244430 | 28.56742 | 103.16996 | 99.51957 | Promoter | 11 | 76120987 | 76243675 | 122689 | 2 | ENSMUST00 | 0 | Glod4 | ENSMUSG00000017286.10 |
| chr11 | 51583944 | 51585416 | 28.56742 | 103.16996 | 99.51957 | Promoter | 11 | 51584757 | 51603254 | 18498 | 1 | ENSMUST00 | 0 | Phykpl | ENSMUSG00000020359.8 |
| chr11 | 58903451 | 58904664 | 28.56742 | 103.16996 | 99.51957 | Promoter | 11 | 58888153 | 58904225 | 16073 | 2 | ENSMUST00 | 0 | Zfp39 | ENSMUSG00000037001.10 |
| chr11 | 95858153 | 95860265 | 28.56742 | 103.16996 | 99.51957 | Promoter | 11 | 95858162 | 95859174 | 1013 | 2 | ENSMUST00 | 0 | Gm11527 | ENSMUSG00000085578.1 |
| chr12 | 83949954 | 83951946 | 28.56742 | 103.16996 | 99.51957 | Promoter | 12 | 83950608 | 83952953 | 2346 | 1 | ENSMUST00 | 0 | 2410016O06 | ENSMUSG00000046791.7 |
| chr14 | 47648782 | 47650473 | 28.56742 | 103.16996 | 99.51957 | Promoter | 14 | 47649311 | 47736557 | 87247 | 1 | ENSMUST00 | 0 | Ktn1 | ENSMUSG00000021843.11 |
| chr16 | 32002157 | 32003778 | 28.56742 | 103.16996 | 99.51957 | Promoter | 16 | 31962506 | 32003287 | 40782 | 2 | ENSMUST00 | 0 | Senp5 | ENSMUSG00000022772.6 |
| chr17 | 34971678 | 34972891 | 28.56742 | 103.16996 | 99.51957 | Promoter | 17 | 34972703 | 34979285 | 6583 | 1 | ENSMUST00 | 0 | Hspa1l | ENSMUSG00000007033.4 |
| chr17 | 46890621 | 46892494 | 28.56742 | 103.16996 | 99.51957 | Promoter | 17 | 46890621 | 46892463 | 1843 | 1 | ENSMUST00 | 1 | Tbcc | ENSMUSG00000036430.7 |
| chr19 | 10182386 | 10184244 | 28.56742 | 103.16996 | 99.51957 | Promoter | 19 | 10182888 | 10196870 | 13983 | 1 | ENSMUST00 | 0 | Fads1 | ENSMUSG00000010663.4 |
| chr19 | 3851080 | 3852947 | 28.56742 | 103.16996 | 99.51957 | Promoter | 19 | 3851773 | 3894365 | 42593 | 1 | ENSMUST00 | 0 | Chka | ENSMUSG00000024843.10 |
| chr2 | 25557062 | 25558826 | 28.56742 | 103.16996 | 99.51957 | Promoter | 2 | 25557847 | 25562082 | 4236 | 1 | ENSMUST00 | 0 | Edf1 | ENSMUSG00000015092.4 |
| chr2 | 51972334 | 51973672 | 32.46003 | 103.17023 | 99.51957 | Promoter | 2 | 51948487 | 51973224 | 24738 | 2 | ENSMUST00 | 0 | Nmi | ENSMUSG00000026946.4 |
| chr2 | 68860798 | 68863593 | 28.56742 | 103.16996 | 99.51957 | Promoter | 2 | 68861441 | 69114282 | 252842 | 1 | ENSMUST00 | 0 | Cers6 | ENSMUSG00000027035.5 |
| chr2 | 74697475 | 74699399 | 28.56742 | 103.16996 | 99.51957 | Promoter | 2 | 74697727 | 74700208 | 2482 | 1 | ENSMUST00 | 0 | Hoxd9 | ENSMUSG00000043342.9 |
| chr3 | 103967695 | 103969065 | 28.56742 | 103.16996 | 99.51957 | Promoter | 3 | 103968110 | 104007488 | 39379 | 1 | ENSMUST00 | 0 | Phtf1 | ENSMUSG00000058388.9 |
| chr3 | 103019827 | 103021910 | 28.56742 | 103.16996 | 99.51957 | Promoter | 3 | 103020546 | 103058189 | 37644 | 1 | ENSMUST00 | 0 | Csde1 | ENSMUSG00000068823.6 |
| chr5 | 124352090 | 124355494 | 28.56742 | 103.16996 | 99.51957 | Promoter | 5 | 124345417 | 124354671 | 9255 | 2 | ENSMUST00 | 0 | Cdk2ap1 | ENSMUSG00000029394.6 |
| chr5 | 122850121 | 122850888 | 28.56742 | 103.16996 | 99.51957 | Promoter | 5 | 122850188 | 122868945 | 18758 | 1 | ENSMUST00 | 0 | Rnf34 | ENSMUSG00000029474.6 |
| chr6 | 134641015 | 134641911 | 28.56742 | 103.16996 | 99.51957 | Promoter | 6 | 134640940 | 134711184 | 70245 | 1 | ENSMUST00 | 76 | Loh12cr1 | ENSMUSG00000042992.10 |
| chr7 | 12977339 | 12978767 | 32.46003 | 103.17023 | 99.51957 | Promoter | 7 | 12977848 | 12983396 | 5549 | 1 | ENSMUST00 | 0 | Zfp446 | ENSMUSG00000033961.10 |
| chr7 | 45525112 | 45526849 | 28.56742 | 103.16996 | 99.51957 | Promoter | 7 | 45526330 | 45549569 | 23240 | 1 | ENSMUST00 | 0 | Plekha4 | ENSMUSG00000040428.12 |
| chr8 | 122475096 | 122476986 | 28.56742 | 103.16996 | 99.51957 | Promoter | 8 | 122466147 | 122476064 | 9918 | 2 | ENSMUST00 | 0 | Rnf166 | ENSMUSG00000014470.4 |
| chr9 | 110100259 | 110101122 | 28.56742 | 103.16996 | 99.51957 | Promoter | 9 | 110084320 | 110101087 | 16768 | 2 | ENSMUST00 | 0 | Dhx30 | ENSMUSG00000032480.12 |
| chr2 | 181364391 | 181366453 | 31.83799 | 103.0912 | 99.44543 | Promoter | 2 | 181364928 | 181380794 | 15867 | 1 | ENSMUST00 | 0 | Zgpat | ENSMUSG00000027582.11 |
| chr17 | 26780865 | 26781966 | 32.99268 | 103.08465 | 99.4389 | Promoter | 17 | 26781079 | 26792521 | 11443 | 1 | ENSMUST00 | 0 | Bnip1 | ENSMUSG00000024191.9 |
| chr11 | 52282532 | 52283722 | 33.55804 | 103.05054 | 99.40515 | Promoter | 11 | 52252371 | 52282571 | 30201 | 2 | ENSMUST00 | 0 | Tcf7 | ENSMUSG00000000782.10 |
| chr11 | 51855275 | 51859109 | 33.55804 | 103.05054 | 99.40515 | Promoter | 11 | 51813455 | 51857653 | 44199 | 2 | ENSMUST00 | 0 | Jade2 | ENSMUSG00000020387.10 |
| chr14 | 61599012 | 61600029 | 33.55804 | 103.05054 | 99.40515 | Promoter | 14 | 61599512 | 61605946 | 6435 | 1 | ENSMUST00 | 0 | Trim13 | ENSMUSG00000035235.7 |
| chr16 | 93603346 | 93605110 | 33.55804 | 103.05054 | 99.40515 | Promoter | 16 | 93583457 | 93603791 | 20335 | 2 | ENSMUST00 | 0 | Setd4 | ENSMUSG00000022948.9 |
| chr3 | 145648777 | 145650997 | 33.55804 | 103.05054 | 99.40515 | Promoter | 3 | 145646971 | 145649985 | 3015 | 2 | ENSMUST00 | 0 | Cyr61 | ENSMUSG00000028195.3 |
| chr9 | 14276184 | 14277840 | 33.55804 | 103.05054 | 99.40515 | Promoter | 9 | 14276301 | 14326138 | 49838 | 1 | ENSMUST00 | 0 | Sesn3 | ENSMUSG00000032009.7 |
| chr9 | 50527769 | 50528826 | 33.55804 | 103.05054 | 99.40515 | Promoter | 9 | 50521618 | 50528641 | 7024 | 2 | ENSMUST00 | 0 | Pts | ENSMUSG00000032067.5 |
| chr1 | 52118891 | 52120761 | 30.08562 | 102.9821 | 99.33765 | Promoter | 1 | 52119440 | 52155412 | 35973 | 1 | ENSMUST00 | 0 | Stat1 | ENSMUSG00000026104.9 |
| chr11 | 54521936 | 54523905 | 30.08562 | 102.9821 | 99.33765 | Promoter | 11 | 54522847 | 54699285 | 176439 | 1 | ENSMUST00 | 0 | Rapgef6 | ENSMUSG00000037533.12 |
| chr14 | 30714921 | 30717340 | 30.08562 | 102.9821 | 99.33765 | Promoter | 14 | 30715167 | 30815084 | 99918 | 1 | ENSMUST00 | 0 | Sfmbt1 | ENSMUSG00000006527.9 |
| chr15 | 38299118 | 38302130 | 30.08562 | 102.9821 | 99.33765 | Promoter | 15 | 38294413 | 38300707 | 6295 | 2 | ENSMUST00 | 0 | Klf10 | ENSMUSG00000037465.9 |
| chr8 | 44936734 | 44939539 | 30.08562 | 102.9821 | 99.33765 | Intron (ENSM | 8 | 44935447 | 45052257 | 116811 | 1 | ENSMUST00 | 1288 | Fat1 | ENSMUSG00000070047.7 |
| chr8 | 11757637 | 11759504 | 32.33533 | 102.95196 | 99.30833 | Promoter | 8 | 11758354 | 11835219 | 76866 | 1 | ENSMUST00 | 0 | Arhgef7 | ENSMUSG00000031511.9 |
| chr4 | 119421416 | 119423230 | 33.48951 | 102.93208 | 99.28856 | Promoter | 4 | 119418530 | 119422358 | 3829 | 2 | ENSMUST00 | 0 | Ppcs | ENSMUSG00000028636.9 |
| chr7 | 101933239 | 101934720 | 33.48951 | 102.93208 | 99.28856 | Promoter | 7 | 101912990 | 101933806 | 20817 | 2 | ENSMUST00 | 0 | Lrrc51 | ENSMUSG00000064307.7 |
| chr11 | 17257066 | 17258492 | 31.71955 | 102.88206 | 99.23879 | Promoter | 11 | 17257579 | 17262910 | 5332 | 1 | ENSMUST00 | 0 | C1d | ENSMUSG00000000581.3 |
| chr19 | 5687926 | 5692029 | 32.27334 | 102.84337 | 99.20039 | Promoter | 19 | 5689131 | 5702862 | 13732 | 1 | ENSMUST00 | 0 | Map3k11 | ENSMUSG00000004054.8 |
| chr1 | 182124638 | 182125968 | 32.27147 | 102.84008 | 99.19724 | Promoter | 1 | 182124737 | 182131919 | 7183 | 1 | ENSMUST00 | 0 | Srp9 | ENSMUSG00000026511.6 |
| chr12 | 87299220 | 87300746 | 32.27147 | 102.84008 | 99.19724 | Promoter | 12 | 87278638 | 87299705 | 21068 | 2 | ENSMUST00 | 0 | Ism2 | ENSMUSG00000050671.7 |
| chr14 | 13960176 | 13962110 | 32.27147 | 102.84008 | 99.19724 | Promoter | 14 | 13949014 | 13961225 | 12212 | 2 | ENSMUST00 | 0 | Thoc7 | ENSMUSG00000053453.8 |
| chr15 | 99771336 | 99772882 | 32.27147 | 102.84008 | 99.19724 | Promoter | 15 | 99734881 | 99772462 | 37582 | 2 | ENSMUST00 | 0 | Cers5 | ENSMUSG00000023021.10 |
| chr17 | 51177960 | 51179858 | 32.27147 | 102.84008 | 99.19724 | Promoter | 17 | 50733127 | 51179349 | 446223 | 2 | ENSMUST00 | 0 | Tbc1d5 | ENSMUSG00000023923.9 |
| chr17 | 36031269 | 36032981 | 32.27147 | 102.84008 | 99.19724 | Promoter | 17 | 36029773 | 36032541 | 2769 | 2 | ENSMUST00 | 0 | H2-T23 | ENSMUSG00000067212.8 |
| chr18 | 36678392 | 36680341 | 32.27147 | 102.84008 | 99.19724 | Promoter | 18 | 36671160 | 36679366 | 8207 | 2 | ENSMUST00 | 0 | Sra1 | ENSMUSG00000006050.6 |
| chr18 | 57877616 | 57879893 | 32.27147 | 102.84008 | 99.19724 | Promoter | 18 | 57878678 | 57946821 | 68144 | 1 | ENSMUST00 | 0 | Slc12a2 | ENSMUSG00000024597.10 |
| chr18 | 14782562 | 14784378 | 32.27147 | 102.84008 | 99.19724 | Promoter | 18 | 14783245 | 14900359 | 117115 | 1 | ENSMUST00 | 0 | Taf4b | ENSMUSG00000054321.6 |
| chr19 | 47050052 | 47051230 | 32.27147 | 102.84008 | 99.19724 | Promoter | 19 | 47033619 | 47050845 | 17227 | 2 | ENSMUST00 | 0 | Pcgf6 | ENSMUSG00000025050.7 |
| chr19 | 4213990 | 4215760 | 32.27147 | 102.84008 | 99.19724 | Promoter | 19 | 4214238 | 4223490 | 9253 | 1 | ENSMUST00 | 0 | Clcf1 | ENSMUSG00000040663.8 |
| chr2 | 71872647 | 71874146 | 32.27147 | 102.84008 | 99.19724 | Promoter | 2 | 71873224 | 71903858 | 30635 | 1 | ENSMUST00 | 0 | Pdk1 | ENSMUSG00000006494.6 |
| chr2 | 26315294 | 26316499 | 32.27147 | 102.84008 | 99.19724 | Promoter | 2 | 26315515 | 26348237 | 32723 | 1 | ENSMUST00 | 0 | Gpsm1 | ENSMUSG00000026930.10 |
| chr5 | 123014659 | 123016346 | 32.27147 | 102.84008 | 99.19724 | Promoter | 5 | 123007579 | 123015080 | 7502 | 2 | ENSMUST00 | 0 | Kdm2b | ENSMUSG00000029475.12 |
| chr5 | 73490170 | 73492849 | 32.27147 | 102.84008 | 99.19724 | Promoter | 5 | 73491026 | 73560794 | 69769 | 1 | ENSMUST00 | 0 | Dcun1d4 | ENSMUSG00000051674.9 |
| chr5 | 36795370 | 36797310 | 32.27147 | 102.84008 | 99.19724 | Promoter | 5 | 36794867 | 36796772 | 1906 | 2 | ENSMUST00 | 0 | Mrfap1 | ENSMUSG00000055302.5 |
| chr6 | 87810377 | 87812633 | 32.27147 | 102.84008 | 99.19724 | Promoter | 6 | 87788853 | 87811779 | 22927 | 2 | ENSMUST00 | 0 | Rab43 | ENSMUSG00000030055.11 |
| chr6 | 100287608 | 100288345 | 32.27147 | 102.84008 | 99.19724 | Promoter | 6 | 100228566 | 100287441 | 58876 | 2 | ENSMUST00 | -168 | Rybp | ENSMUSG00000072872.2 |
| chr7 | 28436601 | 28437531 | 32.27147 | 102.84008 | 99.19724 | Promoter | 7 | 28437447 | 28446546 | 9100 | 1 | ENSMUST00 | 0 | Gmfg | ENSMUSG00000060791.10 |
| chr8 | 27127935 | 27129062 | 32.27147 | 102.84008 | 99.19724 | Promoter | 8 | 27123832 | 27128632 | 4801 | 2 | ENSMUST00 | 0 | Brf2 | ENSMUSG00000031487.4 |
| chr8 | 48842559 | 48844996 | 32.27147 | 102.84008 | 99.19724 | Promoter | 8 | 48227682 | 48844026 | 616345 | 2 | ENSMUST00 | 0 | Tenm3 | ENSMUSG00000031561.10 |
| chr9 | 75625127 | 75626647 | 32.27147 | 102.84008 | 99.19724 | Promoter | 9 | 75625732 | 75637773 | 12042 | 1 | ENSMUST00 | 0 | Lysmd2 | ENSMUSG00000032184.4 |
| chr9 | 108691764 | 108693954 | 32.27147 | 102.84008 | 99.19724 | Promoter | 9 | 108692143 | 108749512 | 57370 | 1 | ENSMUST00 | 0 | Prkar2a | ENSMUSG00000032601.8 |
| chr9 | 95749469 | 95750806 | 32.27147 | 102.84008 | 99.19724 | Promoter | 9 | 95705082 | 95750358 | 45277 | 2 | ENSMUST00 | 0 | Trpc1 | ENSMUSG00000032839.8 |
| chrX | 53669506 | 53670749 | 32.27147 | 102.84008 | 99.19724 | Promoter | 20 | 53669177 | 53670408 | 1232 | 2 | ENSMUST00 | 0 | Cxx1b | ENSMUSG00000067924.4 |
| chr3 | 89164269 | 89166387 | 29.45621 | 102.83202 | 99.19408 | Promoter | 3 | 89164795 | 89176921 | 12127 | 1 | ENSMUST00 | 0 | Clk2 | ENSMUSG00000068917.7 |
| chr7 | 46845314 | 46847259 | 29.45621 | 102.83202 | 99.19408 | Promoter | 7 | 46845839 | 46855627 | 9789 | 1 | ENSMUST00 | 0 | Ldha | ENSMUSG00000063229.9 |
| chr2 | 18676027 | 18678781 | 33.42125 | 102.81408 | 99.17643 | Promoter | 2 | 18677018 | 18686629 | 9612 | 1 | ENSMUST00 | 0 | Bmi1 | ENSMUSG00000026739.8 |
| chr4 | 151107451 | 151109285 | 33.42125 | 102.81408 | 99.17643 | Promoter | 4 | 151059814 | 151108576 | 48763 | 2 | ENSMUST00 | 0 | Camta1 | ENSMUSG00000014592.14 |
| chr7 | 5060226 | 5063489 | 33.42125 | 102.81408 | 99.17643 | Promoter | 7 | 5062143 | 5079945 | 17803 | 1 | ENSMUST00 | 0 | U2af2 | ENSMUSG00000030435.10 |
| chr10 | 83647819 | 83649464 | 34.00537 | 102.80987 | 99.17244 | Promoter | 10 | 83600033 | 83648738 | 48706 | 2 | ENSMUST00 | 0 | Appl2 | ENSMUSG00000020263.9 |
| chr10 | 62725516 | 62726603 | 34.00537 | 102.80987 | 99.17244 | Promoter | 10 | 62659043 | 62725801 | 66759 | 2 | ENSMUST00 | 0 | Stox1 | ENSMUSG00000036923.7 |
| chr11 | 51649934 | 51651437 | 34.00537 | 102.80987 | 99.17244 | Promoter | 11 | 51643063 | 51650842 | 7780 | 2 | ENSMUST00 | 0 | N4bp3 | ENSMUSG00000001053.10 |
| chr11 | 121203972 | 121205344 | 34.00537 | 102.80987 | 99.17244 | Promoter | 11 | 121177591 | 121204711 | 27121 | 2 | ENSMUST00 | 0 | Ogfod3 | ENSMUSG00000025169.6 |
| chr12 | 24679777 | 24682165 | 34.00537 | 102.80987 | 99.17244 | Promoter | 12 | 24665838 | 24681813 | 15976 | 2 | ENSMUST00 | 0 | Cys1 | ENSMUSG00000062563.9 |
| chr16 | 24392713 | 24394688 | 34.00537 | 102.80987 | 99.17244 | Promoter | 16 | 24393350 | 24992576 | 599227 | 1 | ENSMUST00 | 0 | Lpp | ENSMUSG00000033306.9 |
| chr17 | 34030328 | 34032845 | 34.00537 | 102.80987 | 99.17244 | Promoter | 17 | 34028267 | 34031690 | 3424 | 2 | ENSMUST00 | 0 | Slc39a7 | ENSMUSG00000024327.10 |
| chr3 | 32364866 | 32366223 | 34.00537 | 102.80987 | 99.17244 | Promoter | 3 | 32334796 | 32365620 | 30825 | 2 | ENSMUST00 | 0 | Zmat3 | ENSMUSG00000027663.7 |
| chr3 | 53016343 | 53017515 | 34.00537 | 102.80987 | 99.17244 | Promoter | 3 | 52982123 | 53017223 | 35101 | 2 | ENSMUST00 | 0 | Cog6 | ENSMUSG00000027742.8 |
| chr3 | 90051276 | 90052549 | 34.00537 | 102.80987 | 99.17244 | Promoter | 3 | 90051636 | 90062018 | 10383 | 1 | ENSMUST00 | 0 | 4933434E20I | ENSMUSG00000027942.11 |
| chr6 | 28420719 | 28422295 | 34.00537 | 102.80987 | 99.17244 | Promoter | 6 | 28416091 | 28421724 | 5634 | 2 | ENSMUST00 | 0 | Gcc1 | ENSMUSG00000029708.9 |
| chr6 | 117167809 | 117170507 | 34.00537 | 102.80987 | 99.17244 | Promoter | 6 | 117168535 | 117175069 | 6535 | 1 | ENSMUST00 | 0 | Cxcl12 | ENSMUSG00000061353.6 |
| chr7 | 141466983 | 141469576 | 34.00537 | 102.80987 | 99.17244 | Promoter | 7 | 141467362 | 141471479 | 4118 | 1 | ENSMUST00 | 0 | Cd151 | ENSMUSG00000025510.8 |
| chr7 | 118583975 | 118585062 | 34.00537 | 102.80987 | 99.17244 | Promoter | 7 | 118535843 | 118584736 | 48894 | 2 | ENSMUST00 | 0 | Tmc7 | ENSMUSG00000042246.4 |
| chr8 | 70527096 | 70528644 | 34.00537 | 102.80987 | 99.17244 | Promoter | 8 | 70527724 | 70535328 | 7605 | 1 | ENSMUST00 | 0 | Fkbp8 | ENSMUSG00000019428.11 |
| chr8 | 106682794 | 106683671 | 34.00537 | 102.80987 | 99.17244 | Promoter | 8 | 106683068 | 106851439 | 168372 | 1 | ENSMUST00 | 0 | Tango6 | ENSMUSG00000041949.5 |
| chr8 | 115705637 | 115708698 | 34.00537 | 102.80987 | 99.17244 | Promoter | 8 | 115682942 | 115707096 | 24155 | 2 | ENSMUST00 | 0 | Maf | ENSMUSG00000055435.6 |
| chr10 | 13192290 | 13193446 | 22.42038 | 102.78101 | 99.1459 | Promoter | 10 | 13178640 | 13193137 | 14498 | 2 | ENSMUST00 | 0 | Ltv1 | ENSMUSG00000019814.4 |
| chr10 | 128565011 | 128566823 | 31.66066 | 102.77801 | 99.14294 | Promoter | 10 | 128557766 | 128565934 | 8169 | 2 | ENSMUST00 | 0 | Pa2g4 | ENSMUSG00000025364.7 |
| chr19 | 36925411 | 36928022 | 31.66066 | 102.77801 | 99.14294 | Promoter | 19 | 36926079 | 37012752 | 86674 | 1 | ENSMUST00 | 0 | Btaf1 | ENSMUSG00000040565.7 |
| chr2 | 12418253 | 12419764 | 31.66066 | 102.77801 | 99.14294 | Promoter | 2 | 12347263 | 12419444 | 72182 | 2 | ENSMUST00 | 0 | Fam188a | ENSMUSG00000026767.7 |
| chr5 | 123136690 | 123143966 | 31.66066 | 102.77801 | 99.14294 | Promoter | 5 | 123142193 | 123149981 | 7789 | 1 | ENSMUST00 | 0 | Setd1b | ENSMUSG00000038384.11 |
| chr7 | 46033517 | 46034659 | 31.66066 | 102.77801 | 99.14294 | Promoter | 7 | 46033698 | 46084212 | 50515 | 1 | ENSMUST00 | 0 | Nomo1 | ENSMUSG00000030835.6 |
| chr9 | 21290440 | 21291318 | 31.66066 | 102.77801 | 99.14294 | Promoter | 9 | 21288463 | 21291209 | 2747 | 2 | ENSMUST00 | 0 | Cdkn2d | ENSMUSG00000096472.1 |
| chr15 | 79773737 | 79775005 | 32.79732 | 102.7449 | 99.11129 | Promoter | 15 | 79761453 | 79774383 | 12931 | 2 | ENSMUST00 | 0 | Dnal4 | ENSMUSG00000022420.10 |
| chr8 | 25993540 | 25994439 | 32.79732 | 102.7449 | 99.11129 | Promoter | 8 | 25980604 | 25994121 | 13518 | 2 | ENSMUST00 | 0 | Pomk | ENSMUSG00000037251.4 |
| chr2 | 114200861 | 114201793 | 28.86686 | 102.73642 | 99.10302 | Promoter | 2 | 114193461 | 114201469 | 8009 | 2 | ENSMUST00 | 0 | Zfp770 | ENSMUSG00000040321.3 |
| chr10 | 30599860 | 30601247 | 33.94421 | 102.70458 | 99.07153 | Promoter | 10 | 30534225 | 30600749 | 66525 | 2 | ENSMUST00 | 0 | Trmt11 | ENSMUSG00000019792.7 |
| chr19 | 45074346 | 45076160 | 33.35327 | 102.69651 | 99.06359 | Promoter | 19 | 45075241 | 45079289 | 4049 | 1 | ENSMUST00 | 0 | Kazald1 | ENSMUSG00000025213.6 |
| chr7 | 142386717 | 142388441 | 33.35327 | 102.69651 | 99.06359 | Promoter | 7 | 142375910 | 142388038 | 12129 | 2 | ENSMUST00 | 0 | Ctsd | ENSMUSG00000007891.10 |
| chr17 | 74293736 | 74296377 | 31.60199 | 102.67427 | 99.04176 | Promoter | 17 | 74200706 | 74294883 | 94178 | 2 | ENSMUST00 | 0 | Memo1 | ENSMUSG00000058704.7 |
| chr12 | 105031242 | 105032421 | 32.73271 | 102.63245 | 98.99996 | Promoter | 12 | 105031075 | 105031349 | 275 | 2 | ENSMUST00 | 0 | Scarna13 | ENSMUSG00000088789.1 |
| chr8 | 83165020 | 83166443 | 33.87315 | 102.58221 | 98.95009 | Promoter | 8 | 83165352 | 83272422 | 107071 | 1 | ENSMUST00 | 0 | Tbc1d9 | ENSMUSG00000031709.10 |
| chr1 | 36367800 | 36369502 | 24.39013 | 102.54827 | 98.91666 | Promoter | 1 | 36335730 | 36369157 | 33428 | 2 | ENSMUST00 | 0 | Kansl3 | ENSMUSG00000010453.7 |
| chr1 | 36691042 | 36692379 | 24.39013 | 102.54827 | 98.91666 | Promoter | 1 | 36691487 | 36693385 | 1899 | 1 | ENSMUST00 | 0 | Cox5b | ENSMUSG00000061518.5 |
| chr17 | 27906960 | 27910842 | 24.39013 | 102.54827 | 98.91666 | Promoter | 17 | 27901122 | 27909242 | 8121 | 2 | ENSMUST00 | 0 | Taf11 | ENSMUSG00000024218.7 |
| chr18 | 44827764 | 44829696 | 24.39013 | 102.54827 | 98.91666 | Promoter | 18 | 44828665 | 44889720 | 61056 | 1 | ENSMUST00 | 0 | Ythdc2 | ENSMUSG00000034653.8 |
| chr4 | 40947210 | 40949211 | 24.39013 | 102.54827 | 98.91666 | Promoter | 4 | 40936398 | 40948281 | 11884 | 2 | ENSMUST00 | 0 | Bag1 | ENSMUSG00000028416.6 |
| chr5 | 75043434 | 75045043 | 24.39013 | 102.54827 | 98.91666 | Promoter | 5 | 75001026 | 75044774 | 43749 | 2 | ENSMUST00 | 0 | Chic2 | ENSMUSG00000029229.7 |
| chr7 | 132858964 | 132860046 | 24.39013 | 102.54827 | 98.91666 | Promoter | 7 | 132859225 | 132885111 | 25887 | 1 | ENSMUST00 | 0 | Fam175b | ENSMUSG00000030965.13 |
| chr8 | 84256296 | 84257689 | 24.39013 | 102.54827 | 98.91666 | Promoter | 8 | 84249906 | 84257324 | 7419 | 2 | ENSMUST00 | 0 | Mri1 | ENSMUSG00000004996.4 |
| chr11 | 84128853 | 84130280 | 32.66836 | 102.52041 | 98.88917 | Promoter | 11 | 84078920 | 84129600 | 50681 | 2 | ENSMUST00 | 0 | Tada2a | ENSMUSG00000018651.9 |
| chr16 | 59638561 | 59639962 | 32.66836 | 102.52041 | 98.88917 | Promoter | 16 | 59612949 | 59639167 | 26219 | 2 | ENSMUST00 | 0 | Arl6 | ENSMUSG00000022722.13 |
| chr5 | 147075302 | 147078299 | 32.66836 | 102.52041 | 98.88917 | Promoter | 5 | 147016655 | 147076586 | 59932 | 2 | ENSMUST00 | 0 | Lnx2 | ENSMUSG00000016520.7 |
| chr6 | 115992161 | 115995454 | 32.66836 | 102.52041 | 98.88917 | Promoter | 6 | 115954811 | 115995005 | 40195 | 2 | ENSMUST00 | 0 | Plxnd1 | ENSMUSG00000030123.10 |
| chr10 | 95939944 | 95941817 | 30.91854 | 102.46954 | 98.8397 | Promoter | 10 | 95940663 | 96045518 | 104856 | 1 | ENSMUST00 | 0 | Eea1 | ENSMUSG00000036499.8 |

| | | | | | | | | | | | | | | | |
|---|---|---|---|---|---|---|---|---|---|---|---|---|---|---|---|
| chr11 | 43528218 | 43529735 | 30.91854 | 102.46954 | 98.8397 | Promoter | 11 | 43528784 | 43586997 | 58214 | 1 | ENSMUST00 | 0 | Ccnjl | ENSMUSG00000044707.7 |
| chr11 | 78182563 | 78183922 | 30.91854 | 102.46954 | 98.8397 | Promoter | 11 | 78180932 | 78183077 | 2146 | 2 | ENSMUST00 | 0 | Rpl23a | ENSMUSG00000058546.8 |
| chr13 | 104227893 | 104229418 | 30.91854 | 102.46954 | 98.8397 | Promoter | 13 | 104205121 | 104228844 | 23724 | 2 | ENSMUST00 | 0 | Ppwd1 | ENSMUSG00000021713.8 |
| chr2 | 128818031 | 128819335 | 30.91854 | 102.46954 | 98.8397 | Promoter | 2 | 128818118 | 128854261 | 36144 | 1 | ENSMUST00 | 0 | Tmem87b | ENSMUSG00000014353.9 |
| chr2 | 153345733 | 153347755 | 30.91854 | 102.46954 | 98.8397 | Promoter | 2 | 153345845 | 153404007 | 58163 | 1 | ENSMUST00 | 0 | Asxl1 | ENSMUSG00000042548.8 |
| chr3 | 95014242 | 95016985 | 30.91854 | 102.46954 | 98.8397 | Promoter | 3 | 95006702 | 95015238 | 8537 | 2 | ENSMUST00 | 0 | Zfp687 | ENSMUSG00000019338.8 |
| chr8 | 107424203 | 107425061 | 30.91854 | 102.46954 | 98.8397 | Promoter | 8 | 107412486 | 107425041 | 12556 | 2 | ENSMUST00 | 0 | Nob1 | ENSMUSG00000003848.9 |
| chr17 | 87442999 | 87447868 | 33.21814 | 102.46268 | 98.83372 | Promoter | 17 | 87433412 | 87446935 | 13524 | 2 | ENSMUST00 | 0 | Calm2 | ENSMUSG00000036438.7 |
| chr8 | 68061707 | 68062981 | 32.02775 | 102.41263 | 98.78432 | Promoter | 8 | 67790470 | 68062286 | 271817 | 2 | ENSMUST00 | 0 | Psd3 | ENSMUSG00000030465.13 |
| chr12 | 71014262 | 71015695 | 32.60426 | 102.40874 | 98.78054 | Promoter | 12 | 70991615 | 71015596 | 23982 | 2 | ENSMUST00 | 0 | 3110056K07I | ENSMUSG00000085622.3 |
| chr5 | 149050983 | 149053595 | 30.86329 | 102.37074 | 98.74274 | Promoter | 5 | 149046702 | 149053040 | 6339 | 2 | ENSMUST00 | 0 | Hmgb1 | ENSMUSG00000066551.7 |
| chr15 | 99874555 | 99875959 | 33.15098 | 102.34641 | 98.71844 | Promoter | 15 | 99778470 | 99875428 | 96959 | 2 | ENSMUST00 | 0 | Lima1 | ENSMUSG00000023022.8 |
| chr2 | 27677305 | 27678495 | 33.15098 | 102.34641 | 98.71844 | Promoter | 2 | 27677262 | 27761417 | 84156 | 1 | ENSMUST00 | 44 | Rxra | ENSMUSG00000015846.9 |
| chr4 | 132260423 | 132262171 | 33.15098 | 102.34641 | 98.71844 | Promoter | 4 | 132222946 | 132261521 | 38576 | 2 | ENSMUST00 | 0 | Gmeb1 | ENSMUSG00000028901.8 |
| chr5 | 137509828 | 137511544 | 33.15098 | 102.34641 | 98.71844 | Intron (ENSM | 5 | 137501439 | 137502518 | 1080 | 2 | ENSMUST00 | -7311 | Pop7 | ENSMUSG00000029715.5 |
| chr17 | 6947623 | 6949113 | 25.6494 | 102.3303 | 98.70287 | Promoter | 17 | 6904716 | 6948356 | 43641 | 2 | ENSMUST00 | 0 | Rsph3b | ENSMUSG00000023806.9 |
| chr4 | 126147596 | 126150575 | 31.96693 | 102.30585 | 98.67843 | Promoter | 4 | 126147744 | 126149874 | 2131 | 1 | ENSMUST00 | 0 | Eva1b | ENSMUSG00000050212.4 |
| chr11 | 69117442 | 69119156 | 30.22771 | 102.23431 | 98.60725 | Downstream | 11 | 69120404 | 69124055 | 3652 | 1 | ENSMUST00 | -1248 | Hes7 | ENSMUSG00000023781.2 |
| chr3 | 32737005 | 32737938 | 30.22771 | 102.23431 | 98.60725 | Promoter | 3 | 32737071 | 32748900 | 11830 | 1 | ENSMUST00 | 0 | Ndufb5 | ENSMUSG00000027673.7 |
| chr16 | 16599022 | 16600812 | 33.0841 | 102.23056 | 98.60399 | Promoter | 16 | 16422563 | 16599978 | 177416 | 2 | ENSMUST00 | 0 | Fgd4 | ENSMUSG00000022788.11 |
| chr7 | 130518559 | 130520450 | 33.0841 | 102.23056 | 98.60399 | Promoter | 7 | 130391526 | 130519530 | 128005 | 2 | ENSMUST00 | 0 | Ate1 | ENSMUSG00000030850.10 |
| chr13 | 23423328 | 23424988 | 32.4768 | 102.1866 | 98.56064 | Promoter | 13 | 23421539 | 23423866 | 2328 | 2 | ENSMUST00 | 0 | Abt1 | ENSMUSG00000036376.2 |
| chr9 | 20606949 | 20608085 | 32.4768 | 102.1866 | 98.56064 | Promoter | 9 | 20607481 | 20617271 | 9791 | 1 | ENSMUST00 | 0 | Fbxl12os | ENSMUSG00000062470.9 |
| chr15 | 3582852 | 3583587 | 33.01748 | 102.11515 | 98.48943 | Promoter | 15 | 3317760 | 3583230 | 265471 | 2 | ENSMUST00 | 0 | Ghr | ENSMUSG00000055737.7 |
| chr2 | 19656994 | 19659619 | 33.01748 | 102.11515 | 98.48943 | Promoter | 2 | 19657752 | 19660583 | 2832 | 1 | ENSMUST00 | 0 | Otud1 | ENSMUSG00000043415.5 |
| chr1 | 155243730 | 155245165 | 31.84599 | 102.09332 | 98.46812 | Promoter | 1 | 155212471 | 155244444 | 31974 | 2 | ENSMUST00 | 0 | BC034090 | ENSMUSG00000033722.4 |
| chr1 | 156673054 | 156674892 | 31.84599 | 102.09332 | 98.46812 | Promoter | 1 | 156653617 | 156674325 | 20709 | 2 | ENSMUST00 | 0 | Tor3a | ENSMUSG00000060519.6 |
| chr11 | 72488626 | 72490519 | 31.84599 | 102.09332 | 98.46812 | Promoter | 11 | 72451638 | 72489904 | 38267 | 2 | ENSMUST00 | 0 | Spns2 | ENSMUSG00000040447.10 |
| chr17 | 24803885 | 24804973 | 31.84599 | 102.09332 | 98.46812 | Promoter | 17 | 24804382 | 24839787 | 35406 | 1 | ENSMUST00 | 0 | Meiob | ENSMUSG00000024155.7 |
| chr18 | 4352435 | 4354157 | 31.84599 | 102.09332 | 98.46812 | Promoter | 18 | 4331327 | 4352953 | 21627 | 2 | ENSMUST00 | 0 | Map3k8 | ENSMUSG00000024235.5 |
| chr2 | 158409269 | 158410905 | 31.84599 | 102.09332 | 98.46812 | Promoter | 2 | 158409848 | 158499253 | 89406 | 1 | ENSMUST00 | 0 | Ralgapb | ENSMUSG00000027652.10 |
| chr2 | 151541564 | 151543754 | 31.84599 | 102.09332 | 98.46812 | Promoter | 2 | 151542483 | 151561692 | 19210 | 1 | ENSMUST00 | 0 | Fkbp1a | ENSMUSG00000032966.9 |
| chr3 | 57735249 | 57737120 | 31.84599 | 102.09332 | 98.46812 | Promoter | 3 | 57736066 | 57835233 | 99168 | 1 | ENSMUST00 | 0 | Rnf13 | ENSMUSG00000036503.9 |
| chr3 | 158030930 | 158032283 | 31.84599 | 102.09332 | 98.46812 | Promoter | 3 | 158010919 | 158031726 | 20808 | 2 | ENSMUST00 | 0 | Srsf11 | ENSMUSG00000055436.13 |
| chr4 | 137387272 | 137389203 | 31.84599 | 102.09332 | 98.46812 | Promoter | 4 | 137387266 | 137388382 | 1117 | 2 | ENSMUST00 | 0 | RP23-246F18 | ENSMUSG00000101493.1 |
| chr5 | 98030106 | 98031628 | 31.84599 | 102.09332 | 98.46812 | Promoter | 5 | 97882783 | 98030733 | 147951 | 2 | ENSMUST00 | 0 | Antxr2 | ENSMUSG00000029338.9 |
| chr6 | 115134572 | 115135637 | 31.84599 | 102.09332 | 98.46812 | Promoter | 6 | 115134902 | 115276559 | 141658 | 1 | ENSMUST00 | 0 | Syn2 | ENSMUSG00000009394.8 |
| chr6 | 31219869 | 31221288 | 31.84599 | 102.09332 | 98.46812 | Promoter | 6 | 31220351 | 31337247 | 116897 | 1 | ENSMUST00 | 0 | 2210408F21I | ENSMUSG00000087380.2 |
| chr12 | 64917326 | 64918820 | 32.41345 | 102.0761 | 98.45256 | Promoter | 12 | 64917911 | 64924591 | 6681 | 1 | ENSMUST00 | 0 | Gm527 | ENSMUSG00000047227.4 |
| chr11 | 23664733 | 23667134 | 29.58305 | 102.06358 | 98.44028 | Promoter | 11 | 23665674 | 23732876 | 67203 | 1 | ENSMUST00 | 0 | Pus10 | ENSMUSG00000020280.7 |
| chr2 | 136890535 | 136892781 | 29.58305 | 102.06358 | 98.44028 | Promoter | 2 | 136873780 | 136891382 | 17603 | 2 | ENSMUST00 | 0 | Mkks | ENSMUSG00000027274.11 |
| chr6 | 47763624 | 47764161 | 8.30651 | 102.0048 | 98.38202 | Distal Interge | 6 | 47773329 | 47773429 | 101 | 1 | ENSMUST00 | -9168 | Gm24975 | ENSMUSG00000094807.1 |
| chr11 | 86543896 | 86545818 | 32.35035 | 101.96598 | 98.34379 | Promoter | 11 | 86544991 | 86567359 | 22369 | 1 | ENSMUST00 | 0 | Tubd1 | ENSMUSG00000020513.10 |
| chr10 | 23786112 | 23788131 | 25.4663 | 101.95255 | 98.33054 | Promoter | 10 | 23785211 | 23787195 | 1985 | 2 | ENSMUST00 | 0 | Rps12 | ENSMUSG00000061983.6 |
| chr11 | 4135153 | 4136015 | 25.4663 | 101.95255 | 98.33054 | Promoter | 11 | 4135202 | 4141172 | 5971 | 1 | ENSMUST00 | 0 | Rnf215 | ENSMUSG00000003581.9 |
| chr12 | 51377022 | 51378274 | 25.4663 | 101.95255 | 98.33054 | Promoter | 12 | 51377580 | 51450096 | 72517 | 1 | ENSMUST00 | 0 | Scfd1 | ENSMUSG00000020952.9 |
| chr13 | 32337977 | 32339540 | 25.4663 | 101.95255 | 98.33054 | Promoter | 13 | 31819579 | 32338712 | 519134 | 2 | ENSMUST00 | 0 | Gmds | ENSMUSG00000038372.8 |
| chr14 | 61172490 | 61173715 | 25.4663 | 101.95255 | 98.33054 | Promoter | 14 | 61172990 | 61195663 | 22674 | 1 | ENSMUST00 | 0 | Sacs | ENSMUSG00000048279.12 |
| chr17 | 36257111 | 36258175 | 25.4663 | 101.95255 | 98.33054 | Promoter | 17 | 36255645 | 36257856 | 2212 | 2 | ENSMUST00 | 0 | Rpp21 | ENSMUSG00000024446.8 |
| chr17 | 12916373 | 12917402 | 25.4663 | 101.95255 | 98.33054 | Promoter | 17 | 12916382 | 12925067 | 8686 | 1 | ENSMUST00 | 0 | Tcp1 | ENSMUSG00000068039.7 |
| chr19 | 3574095 | 3576469 | 25.4663 | 101.95255 | 98.33054 | Promoter | 19 | 3454933 | 3575749 | 120817 | 2 | ENSMUST00 | 0 | Ppp6r3 | ENSMUSG00000024908.9 |
| chr2 | 94436427 | 94438271 | 25.4663 | 101.95255 | 98.33054 | Promoter | 2 | 94411682 | 94438136 | 26455 | 2 | ENSMUST00 | 0 | Api5 | ENSMUSG00000027193.7 |
| chr6 | 67266963 | 67268876 | 25.4663 | 101.95255 | 98.33054 | Promoter | 6 | 67266979 | 67289302 | 22324 | 1 | ENSMUST00 | 0 | Serbp1 | ENSMUSG00000036371.4 |
| chr7 | 28337743 | 28339287 | 25.4663 | 101.95255 | 98.33054 | Promoter | 7 | 28314898 | 28338719 | 23822 | 2 | ENSMUST00 | 0 | Supt5 | ENSMUSG00000003435.8 |
| chr6 | 125215490 | 125216295 | 28.98007 | 101.95001 | 98.32897 | Promoter | 6 | 125215551 | 125222306 | 6756 | 1 | ENSMUST00 | 0 | Vamp1 | ENSMUSG00000030337.11 |
| chr1 | 191574722 | 191576556 | 30.03893 | 101.89226 | 98.2715 | Promoter | 1 | 191575763 | 191622672 | 46910 | 1 | ENSMUST00 | 0 | Ints7 | ENSMUSG00000037461.5 |
| chr1 | 54556793 | 54558256 | 30.03893 | 101.89226 | 98.2715 | Promoter | 1 | 54472994 | 54557682 | 84689 | 2 | ENSMUST00 | 0 | Pgap1 | ENSMUSG00000073678.4 |
| chr1 | 119525551 | 119526807 | 30.03893 | 101.89226 | 98.2715 | Promoter | 1 | 119520090 | 119526359 | 6270 | 2 | ENSMUST00 | 0 | Gm27184 | ENSMUSG00000098447.2 |
| chr10 | 20951930 | 20953388 | 30.03893 | 101.89226 | 98.2715 | Promoter | 10 | 20952547 | 21080428 | 127882 | 1 | ENSMUST00 | 0 | Ahi1 | ENSMUSG00000019986.11 |
| chr10 | 81384364 | 81385334 | 30.03893 | 101.89226 | 98.2715 | Promoter | 10 | 81384428 | 81388352 | 3925 | 1 | ENSMUST00 | 0 | Dohh | ENSMUSG00000078440.4 |
| chr11 | 72214454 | 72215992 | 30.03893 | 101.89226 | 98.2715 | Promoter | 11 | 72211724 | 72215592 | 3869 | 2 | ENSMUST00 | 0 | Med31 | ENSMUSG00000020801.8 |
| chr12 | 4082253 | 4083211 | 30.03893 | 101.89226 | 98.2715 | Promoter | 12 | 4082574 | 4110612 | 28039 | 1 | ENSMUST00 | 0 | Dnajc27 | ENSMUSG00000020657.10 |
| chr12 | 78861768 | 78863108 | 30.03893 | 101.89226 | 98.2715 | Promoter | 12 | 78861819 | 78887010 | 25192 | 1 | ENSMUST00 | 0 | Eif2s1 | ENSMUSG00000021116.9 |
| chr15 | 37005733 | 37008177 | 30.03893 | 101.89226 | 98.2715 | Promoter | 15 | 36997031 | 37007402 | 10372 | 2 | ENSMUST00 | 0 | Zfp706 | ENSMUSG00000062397.7 |
| chr16 | 33061959 | 33063463 | 30.03893 | 101.89226 | 98.2715 | Promoter | 16 | 33062521 | 33127695 | 65175 | 1 | ENSMUST00 | 0 | Lmln | ENSMUSG00000022802.2 |
| chr18 | 23803418 | 23805095 | 30.03893 | 101.89226 | 98.2715 | Promoter | 18 | 23803984 | 23893857 | 89874 | 1 | ENSMUST00 | 0 | Mapre2 | ENSMUSG00000024277.9 |
| chr18 | 80361986 | 80364088 | 30.03893 | 101.89226 | 98.2715 | Promoter | 18 | 80322679 | 80363336 | 40658 | 2 | ENSMUST00 | 0 | Kcng2 | ENSMUSG00000059852.7 |
| chr19 | 46960757 | 46963137 | 30.03893 | 101.89226 | 98.2715 | Promoter | 19 | 46886831 | 46962351 | 75521 | 2 | ENSMUST00 | 0 | Nt5c2 | ENSMUSG00000025041.11 |
| chr2 | 93461321 | 93462514 | 30.03893 | 101.89226 | 98.2715 | Promoter | 2 | 93419111 | 93462411 | 43301 | 2 | ENSMUST00 | 0 | Cd82 | ENSMUSG00000027215.8 |
| chr2 | 126874723 | 126876927 | 30.03893 | 101.89226 | 98.2715 | Promoter | 2 | 126791565 | 126876230 | 84666 | 2 | ENSMUST00 | 0 | Trpm7 | ENSMUSG00000027365.9 |
| chr2 | 3118032 | 3119629 | 30.03893 | 101.89226 | 98.2715 | Promoter | 2 | 3118382 | 3227806 | 109425 | 1 | ENSMUST00 | 0 | Fam171a1 | ENSMUSG00000050530.9 |
| chr2 | 170130125 | 170131972 | 30.03893 | 101.89226 | 98.2715 | Promoter | 2 | 170108643 | 170131220 | 22578 | 2 | ENSMUST00 | 0 | Zfp217 | ENSMUSG00000052056.9 |
| chr4 | 129334195 | 129335430 | 30.03893 | 101.89226 | 98.2715 | Promoter | 4 | 129307100 | 129335370 | 28271 | 2 | ENSMUST00 | 0 | Rbbp4 | ENSMUSG00000057236.10 |
| chr5 | 25705056 | 25706319 | 30.03893 | 101.89226 | 98.2715 | Promoter | 5 | 25689812 | 25705825 | 16014 | 2 | ENSMUST00 | 0 | Xrcc2 | ENSMUSG00000028933.7 |
| chr5 | 121451584 | 121453273 | 30.03893 | 101.89226 | 98.2715 | Promoter | 5 | 121451893 | 121524183 | 72291 | 1 | ENSMUST00 | 0 | Tmem116 | ENSMUSG00000029452.13 |
| chr6 | 6577340 | 6579136 | 30.03893 | 101.89226 | 98.2715 | Promoter | 6 | 6557294 | 6578658 | 21365 | 2 | ENSMUST00 | 0 | Shfm1 | ENSMUSG00000042541.9 |
| chr7 | 110221194 | 110222833 | 30.03893 | 101.89226 | 98.2715 | Promoter | 7 | 110221703 | 110283503 | 61801 | 1 | ENSMUST00 | 0 | Swap70 | ENSMUSG00000031015.7 |
| chr7 | 30072094 | 30073231 | 30.03893 | 101.89226 | 98.2715 | Promoter | 7 | 30056037 | 30072900 | 16864 | 2 | ENSMUST00 | 0 | Zfp82 | ENSMUSG00000098022.2 |
| chr8 | 123064785 | 123066376 | 30.03893 | 101.89226 | 98.2715 | Promoter | 8 | 123065179 | 123097760 | 32582 | 1 | ENSMUST00 | 0 | Spg7 | ENSMUSG00000000738.12 |
| chr8 | 22805201 | 22809095 | 30.03893 | 101.89226 | 98.2715 | Promoter | 8 | 22787354 | 22805654 | 18301 | 2 | ENSMUST00 | 0 | Ap3m2 | ENSMUSG00000031539.5 |
| chr9 | 57708441 | 57709366 | 30.03893 | 101.89226 | 98.2715 | Promoter | 9 | 57708540 | 57752499 | 43960 | 1 | ENSMUST00 | 0 | Edc3 | ENSMUSG00000038957.8 |
| chr9 | 65826248 | 65828549 | 30.03893 | 101.89226 | 98.2715 | Promoter | 9 | 65692391 | 65827564 | 135174 | 2 | ENSMUST00 | 0 | Zfp609 | ENSMUSG00000040524.9 |
| chr9 | 96477435 | 96478907 | 30.03893 | 101.89226 | 98.2715 | Promoter | 9 | 96470937 | 96478675 | 7739 | 2 | ENSMUST00 | 0 | Rnf7 | ENSMUSG00000051234.6 |
| chr14 | 50923896 | 50925701 | 32.88505 | 101.88557 | 98.27007 | Promoter | 14 | 50906478 | 50924834 | 18357 | 2 | ENSMUST00 | 0 | Osgep | ENSMUSG00000006289.9 |
| chr13 | 33934956 | 33936478 | 31.72596 | 101.88219 | 98.2679 | Promoter | 13 | 33917918 | 33936083 | 18166 | 2 | ENSMUST00 | 0 | Serpinb6a | ENSMUSG00000060147.10 |
| chr2 | 155235845 | 155237665 | 27.88396 | 101.87045 | 98.25623 | Promoter | 2 | 155236533 | 155250272 | 13740 | 1 | ENSMUST00 | 0 | Dynlrb1 | ENSMUSG00000047459.9 |
| chr12 | 76532849 | 76534984 | 33.45292 | 101.85796 | 98.2438 | Promoter | 12 | 76533560 | 76579039 | 45480 | 1 | ENSMUST00 | 0 | Plekhg3 | ENSMUSG00000052609.8 |
| chr14 | 20793027 | 20794619 | 33.45292 | 101.85796 | 98.2438 | Promoter | 14 | 20734875 | 20794088 | 59214 | 2 | ENSMUST00 | 0 | Camk2g | ENSMUSG00000021820.11 |
| chr4 | 46388736 | 46390523 | 32.28748 | 101.85625 | 98.2423 | Promoter | 4 | 46376505 | 46389418 | 12914 | 2 | ENSMUST00 | 0 | 5830415F09I | ENSMUSG00000028331.7 |
| chr18 | 46740400 | 46743200 | 23.21435 | 101.83275 | 98.21896 | Promoter | 18 | 46741917 | 46790826 | 48910 | 1 | ENSMUST00 | 0 | Ap3s1 | ENSMUSG00000024480.7 |
| chr8 | 107587386 | 107588794 | 23.21435 | 101.83275 | 98.21896 | Promoter | 8 | 107580381 | 107588482 | 8102 | 2 | ENSMUST00 | 0 | Psmd7 | ENSMUSG00000039067.4 |
| chr12 | 16894002 | 16896666 | 32.81923 | 101.77142 | 98.15781 | Promoter | 12 | 16894978 | 16988274 | 93297 | 1 | ENSMUST00 | 0 | Rock2 | ENSMUSG00000020580.9 |
| chr7 | 27195262 | 27196830 | 32.81923 | 101.77142 | 98.15781 | Promoter | 7 | 27187007 | 27195760 | 8754 | 2 | ENSMUST00 | 0 | Snrpa | ENSMUSG00000061479.10 |
| chr8 | 119394376 | 119396080 | 32.81923 | 101.77142 | 98.15781 | Promoter | 8 | 119394878 | 119411102 | 16225 | 1 | ENSMUST00 | 0 | Mlycd | ENSMUSG00000074064.6 |
| chr1 | 191182292 | 191186167 | 31.08358 | 101.75523 | 98.14187 | Promoter | 1 | 191170296 | 191183340 | 13045 | 2 | ENSMUST00 | 0 | Atf3 | ENSMUSG00000026628.8 |
| chr10 | 81059550 | 81060575 | 31.08358 | 101.75523 | 98.14187 | Promoter | 10 | 81044075 | 81060178 | 16104 | 2 | ENSMUST00 | 0 | Sgta | ENSMUSG00000004937.5 |
| chr12 | 83596150 | 83597284 | 31.08358 | 101.75523 | 98.14187 | Promoter | 12 | 83546941 | 83597147 | 50207 | 2 | ENSMUST00 | 0 | Zfyve1 | ENSMUSG00000042628.7 |
| chr14 | 56776691 | 56779122 | 31.08358 | 101.75523 | 98.14187 | Promoter | 14 | 56722441 | 56778316 | 55876 | 2 | ENSMUST00 | 0 | Pspc1 | ENSMUSG00000021938.6 |
| chr2 | 165286909 | 165287951 | 31.08358 | 101.75523 | 98.14187 | Promoter | 2 | 165276554 | 165287755 | 11202 | 2 | ENSMUST00 | 0 | Slc35c2 | ENSMUSG00000017664.11 |
| chr2 | 114174415 | 114175414 | 33.38388 | 101.73887 | 98.12704 | Promoter | 2 | 114101170 | 114175339 | 74170 | 2 | ENSMUST00 | 0 | Aqr | ENSMUSG00000040383.11 |
| chr9 | 107871754 | 107873634 | 32.75367 | 101.65766 | 98.0462 | Promoter | 9 | 107773559 | 107872724 | 99166 | 2 | ENSMUST00 | 0 | Rbm6 | ENSMUSG00000032582.9 |
| chr11 | 109362828 | 109365045 | 32.16249 | 101.63789 | 98.02663 | Promoter | 11 | 109362831 | 109380475 | 17645 | 1 | ENSMUST00 | 0 | Gna13 | ENSMUSG00000020611.9 |
| chr1 | 190927800 | 190929765 | 33.31514 | 101.62023 | 98.00909 | Promoter | 1 | 190928492 | 190938435 | 9944 | 1 | ENSMUST00 | 0 | Angel2 | ENSMUSG00000026634.11 |
| chr9 | 70140571 | 70142290 | 33.31514 | 101.62023 | 98.00909 | Promoter | 9 | 70089310 | 70141578 | 52269 | 2 | ENSMUST00 | 0 | Fam81a | ENSMUSG00000032224.9 |
| chr10 | 127311072 | 127312491 | 26.72788 | 101.58999 | 97.97913 | Promoter | 10 | 127296221 | 127311786 | 15566 | 2 | ENSMUST00 | 0 | Mars | ENSMUSG00000040354.9 |
| chr11 | 80089011 | 80090131 | 26.72788 | 101.58999 | 97.97913 | Promoter | 11 | 80089400 | 80135791 | 46392 | 1 | ENSMUST00 | 0 | Atad5 | ENSMUSG00000017550.9 |
| chr12 | 54984607 | 54986836 | 26.72788 | 101.58999 | 97.97913 | Promoter | 12 | 54892989 | 54986336 | 93348 | 2 | ENSMUST00 | 0 | Baz1a | ENSMUSG00000035021.8 |
| chr14 | 67930568 | 67934296 | 26.72788 | 101.58999 | 97.97913 | Promoter | 14 | 67751928 | 67933572 | 181645 | 2 | ENSMUST00 | 0 | Dock5 | ENSMUSG00000044447.4 |
| chr15 | 79741496 | 79743083 | 26.72788 | 101.58999 | 97.97913 | Promoter | 15 | 79724070 | 79742456 | 18387 | 2 | ENSMUST00 | 0 | Sun2 | ENSMUSG00000042524.8 |
| chr16 | 23224518 | 23227486 | 26.72788 | 101.58999 | 97.97913 | Promoter | 16 | 23224740 | 23360348 | 135609 | 1 | ENSMUST00 | 0 | St6gal1 | ENSMUSG00000022885.10 |
| chr17 | 35000585 | 35002024 | 26.72788 | 101.58999 | 97.97913 | Promoter | 17 | 35000987 | 35016322 | 15336 | 1 | ENSMUST00 | 0 | Vars | ENSMUSG00000007029.11 |
| chr17 | 29660044 | 29661644 | 26.72788 | 101.58999 | 97.97913 | Promoter | 17 | 29660595 | 29705979 | 45385 | 1 | ENSMUST00 | 0 | Cmtr1 | ENSMUSG00000024019.12 |
| chr19 | 23757870 | 23759680 | 26.72788 | 101.58999 | 97.97913 | Promoter | 19 | 23758876 | 23949597 | 190722 | 1 | ENSMUST00 | 0 | Apba1 | ENSMUSG00000024897.8 |
| chr19 | 44068955 | 44070157 | 26.72788 | 101.58999 | 97.97913 | Promoter | 19 | 44034944 | 44069526 | 34583 | 2 | ENSMUST00 | 0 | Erlin1 | ENSMUSG00000025198.11 |
| chr4 | 21727046 | 21728585 | 26.72788 | 101.58999 | 97.97913 | Promoter | 4 | 21727701 | 21750546 | 22846 | 1 | ENSMUST00 | 0 | Ccnc | ENSMUSG00000028252.15 |
| chr4 | 86856343 | 86858339 | 26.72788 | 101.58999 | 97.97913 | Promoter | 4 | 86854660 | 86857412 | 2753 | 2 | ENSMUST00 | 0 | Rps6 | ENSMUSG00000028495.9 |
| chr4 | 129058051 | 129059547 | 26.72788 | 101.58999 | 97.97913 | Promoter | 4 | 129058271 | 129084526 | 26256 | 1 | ENSMUST00 | 0 | Rnf19b | ENSMUSG00000028793.10 |
| chr5 | 123907208 | 123908092 | 26.72788 | 101.58999 | 97.97913 | Promoter | 5 | 123907269 | 123908508 | 1240 | 1 | ENSMUST00 | 0 | Denr | ENSMUSG00000023106.10 |
| chr5 | 24586142 | 24588039 | 26.72788 | 101.58999 | 97.97913 | Promoter | 5 | 24586741 | 24592487 | 5747 | 1 | ENSMUST00 | 0 | Chpf2 | ENSMUSG00000038181.11 |
| chr6 | 87042081 | 87043758 | 26.72788 | 101.58999 | 97.97913 | Promoter | 6 | 87042846 | 87055424 | 12579 | 1 | ENSMUST00 | 0 | Gfpt1 | ENSMUSG00000029992.9 |
| chr6 | 118454724 | 118456063 | 26.72788 | 101.58999 | 97.97913 | Promoter | 6 | 118427319 | 118455528 | 28210 | 2 | ENSMUST00 | 0 | Zfp248 | ENSMUSG00000030145.9 |
| chr7 | 44495874 | 44497219 | 26.72788 | 101.58999 | 97.97913 | Promoter | 7 | 44489937 | 44496529 | 6593 | 2 | ENSMUST00 | 0 | Emc10 | ENSMUSG00000008140.12 |
| chr8 | 3492673 | 3494001 | 26.72788 | 101.58999 | 97.97913 | Promoter | 8 | 3493138 | 3497208 | 4071 | 1 | ENSMUST00 | 0 | Zfp358 | ENSMUSG00000047264.7 |
| chr9 | 69289292 | 69291949 | 26.72788 | 101.58999 | 97.97913 | Promoter | 9 | 69289682 | 69379809 | 90128 | 1 | ENSMUST00 | 0 | Rora | ENSMUSG00000032238.12 |
| chr1 | 181150788 | 181151670 | 28.22734 | 101.53345 | 97.92473 | Promoter | 1 | 181150931 | 181168994 | 18064 | 1 | ENSMUST00 | 0 | Cnih4 | ENSMUSG00000062169.8 |
| chr10 | 50591981 | 50593604 | 28.22734 | 101.53345 | 97.92473 | Promoter | 10 | 50592669 | 50851202 | 258534 | 1 | ENSMUST00 | 0 | Ascc3 | ENSMUSG00000038774.8 |
| chr11 | 84512655 | 84514216 | 28.22734 | 101.53345 | 97.92473 | Promoter | 11 | 84422855 | 84513522 | 90668 | 2 | ENSMUST00 | 0 | Aatf | ENSMUSG00000018697.9 |
| chr11 | 106778703 | 106779939 | 28.22734 | 101.53345 | 97.92473 | Promoter | 11 | 106768253 | 106779515 | 11263 | 2 | ENSMUST00 | 0 | Polg2 | ENSMUSG00000020718.7 |

| | | | | | | | | | | | | | | | |
|---|---|---|---|---|---|---|---|---|---|---|---|---|---|---|---|
| chr11 | 115534905 | 115536788 | 28.22734 | 101.53345 | 97.92473 | Promoter | 11 | 115523102 | 115535817 | 12716 | 2 | ENSMUST00 | 0 | Sumo2 | ENSMUSG00000020738.11 |
| chr11 | 103266683 | 103268189 | 28.22734 | 101.53345 | 97.92473 | Promoter | 11 | 103219762 | 103267472 | 47711 | 2 | ENSMUST00 | 0 | Map3k14 | ENSMUSG00000020941.7 |
| chr11 | 120573364 | 120574265 | 28.22734 | 101.53345 | 97.92473 | Promoter | 11 | 120560298 | 120573253 | 12956 | 2 | ENSMUST00 | -112 | P4hb | ENSMUSG00000025130.6 |
| chr12 | 112828574 | 112829497 | 28.22734 | 101.53345 | 97.92473 | Promoter | 12 | 112820237 | 112829389 | 9153 | 2 | ENSMUST00 | 0 | Cdca4 | ENSMUSG00000047832.8 |
| chr14 | 45219632 | 45221251 | 28.22734 | 101.53345 | 97.92473 | Promoter | 14 | 45134907 | 45220328 | 85422 | 2 | ENSMUST00 | 0 | Txndc16 | ENSMUSG00000021830.8 |
| chr14 | 27507768 | 27509136 | 28.22734 | 101.53345 | 97.92473 | Promoter | 14 | 27482412 | 27508460 | 26049 | 2 | ENSMUST00 | 0 | Ccdc66 | ENSMUSG00000046753.9 |
| chr18 | 40218507 | 40219431 | 28.22734 | 101.53345 | 97.92473 | Promoter | 18 | 40204865 | 40219399 | 14535 | 2 | ENSMUST00 | 0 | Yipf5 | ENSMUSG00000024487.4 |
| chr19 | 5117175 | 5118826 | 28.22734 | 101.53345 | 97.92473 | Promoter | 19 | 5107746 | 5118297 | 10552 | 2 | ENSMUST00 | 0 | Klc2 | ENSMUSG00000024862.11 |
| chr19 | 47917779 | 47920263 | 28.22734 | 101.53345 | 97.92473 | Promoter | 19 | 47786580 | 47919287 | 132708 | 2 | ENSMUST00 | 0 | Wdr96 | ENSMUSG00000044948.11 |
| chr2 | 126151549 | 126152953 | 28.22734 | 101.53345 | 97.92473 | Promoter | 2 | 126152141 | 126165276 | 13136 | 1 | ENSMUST00 | 0 | Dtwd1 | ENSMUSG00000023330.7 |
| chr2 | 5844120 | 5846008 | 28.22734 | 101.53345 | 97.92473 | Promoter | 2 | 5845019 | 5871895 | 26877 | 1 | ENSMUST00 | 0 | Nudt5 | ENSMUSG00000025817.7 |
| chr2 | 177464698 | 177465628 | 28.22734 | 101.53345 | 97.92473 | Promoter | 2 | 177464733 | 177478129 | 13397 | 1 | ENSMUST00 | 0 | Gm14420 | ENSMUSG00000078866.5 |
| chr4 | 135855184 | 135857243 | 28.22734 | 101.53345 | 97.92473 | Promoter | 4 | 135855747 | 135865815 | 10069 | 1 | ENSMUST00 | 0 | Srsf10 | ENSMUSG00000028676.12 |
| chr4 | 19708046 | 19709648 | 28.22734 | 101.53345 | 97.92473 | Promoter | 4 | 19608303 | 19708993 | 100691 | 2 | ENSMUST00 | 0 | Wwp1 | ENSMUSG00000041058.10 |
| chr5 | 30886978 | 30889835 | 28.22734 | 101.53345 | 97.92473 | Promoter | 5 | 30888694 | 30891207 | 2514 | 1 | ENSMUST00 | 0 | Agbl5 | ENSMUSG00000029165.11 |
| chr5 | 5513956 | 5515239 | 28.22734 | 101.53345 | 97.92473 | Promoter | 5 | 5505015 | 5514789 | 9775 | 2 | ENSMUST00 | 0 | Cldn12 | ENSMUSG00000046798.9 |
| chr6 | 70844451 | 70845559 | 28.22734 | 101.53345 | 97.92473 | Promoter | 6 | 70844515 | 70905245 | 60731 | 1 | ENSMUST00 | 0 | Eif2ak3 | ENSMUSG00000031668.9 |
| chr6 | 113075230 | 113076760 | 28.22734 | 101.53345 | 97.92473 | Promoter | 6 | 113067659 | 113077177 | 9519 | 2 | ENSMUST00 | 417 | Gt(ROSA)26 | ENSMUSG00000086429.4 |
| chr7 | 118242001 | 118243629 | 28.22734 | 101.53345 | 97.92473 | Promoter | 7 | 118200696 | 118243627 | 42932 | 2 | ENSMUST00 | 0 | Smg1 | ENSMUSG00000030655.9 |
| chr7 | 102267172 | 102268649 | 28.22734 | 101.53345 | 97.92473 | Promoter | 7 | 102267824 | 102436848 | 169025 | 1 | ENSMUST00 | 0 | Stim1 | ENSMUSG00000030987.4 |
| chr7 | 127344643 | 127346973 | 28.22734 | 101.53345 | 97.92473 | Promoter | 7 | 127342797 | 127345314 | 2518 | 2 | ENSMUST00 | 0 | Zfp768 | ENSMUSG00000047371.6 |
| chr8 | 105757781 | 105759401 | 28.22734 | 101.53345 | 97.92473 | Promoter | 8 | 105713854 | 105758664 | 44811 | 2 | ENSMUST00 | 0 | Gfod2 | ENSMUSG00000013150.10 |
| chr8 | 57651425 | 57653684 | 28.22734 | 101.53345 | 97.92473 | Promoter | 8 | 57523828 | 57653031 | 129204 | 2 | ENSMUST00 | 0 | Galnt7 | ENSMUSG00000031608.8 |
| chr8 | 105936542 | 105938522 | 28.22734 | 101.53345 | 97.92473 | Promoter | 8 | 105935728 | 105938392 | 2665 | 2 | ENSMUST00 | 0 | Psmb10 | ENSMUSG00000031897.8 |
| chr8 | 79710841 | 79712537 | 28.22734 | 101.53345 | 97.92473 | Promoter | 8 | 79711820 | 79777321 | 65502 | 1 | ENSMUST00 | 0 | Anapc10 | ENSMUSG00000036977.7 |
| chr3 | 108571311 | 108572720 | 33.24668 | 101.50206 | 97.89681 | Promoter | 3 | 108571698 | 108581370 | 9673 | 1 | ENSMUST00 | 0 | Taf13 | ENSMUSG00000048100.8 |
| chr10 | 7589305 | 7590761 | 30.3748 | 101.49462 | 97.88963 | Promoter | 10 | 7589800 | 7608567 | 18768 | 1 | ENSMUST00 | 0 | Lrp11 | ENSMUSG00000019796.7 |
| chr4 | 45083577 | 45085029 | 30.3748 | 101.49462 | 97.88963 | Promoter | 4 | 45034247 | 45084604 | 50358 | 2 | ENSMUST00 | 0 | Fbxo10 | ENSMUSG00000048232.7 |
| chr4 | 147124721 | 147125521 | 14.86491 | 101.45053 | 97.84645 | Promoter | 4 | 147125256 | 147125816 | 561 | 1 | ENSMUST00 | 0 | Gm13162 | ENSMUSG00000082779.1 |
| chr15 | 78934764 | 78935809 | 32.62334 | 101.43139 | 97.82731 | Promoter | 15 | 78934933 | 78943634 | 8702 | 1 | ENSMUST00 | 0 | Nol12 | ENSMUSG00000033099.4 |
| chr1 | 34841471 | 34843649 | 32.03847 | 101.42104 | 97.81718 | Promoter | 1 | 34813219 | 34843050 | 29832 | 2 | ENSMUST00 | 0 | Fam168b | ENSMUSG00000037503.7 |
| chr1 | 43091387 | 43093290 | 32.03847 | 101.42104 | 97.81718 | Promoter | 1 | 43092588 | 43093835 | 1248 | 1 | ENSMUST00 | 0 | 8430432A02 | ENSMUSG00000041878.3 |
| chr19 | 5087724 | 5089304 | 32.03847 | 101.42104 | 97.81718 | Promoter | 19 | 5088538 | 5092879 | 4342 | 1 | ENSMUST00 | 0 | Yif1a | ENSMUSG00000024875.4 |
| chr3 | 113629253 | 113630972 | 32.03847 | 101.42104 | 97.81718 | Promoter | 3 | 113605067 | 113629900 | 24834 | 2 | ENSMUST00 | 0 | Rnpc3 | ENSMUSG00000027981.9 |
| chr5 | 143548668 | 143549562 | 32.03847 | 101.42104 | 97.81718 | Promoter | 5 | 143548706 | 143564525 | 15820 | 1 | ENSMUST00 | 0 | Fam220a | ENSMUSG00000083012.3 |
| chr11 | 102295679 | 102297953 | 33.1785 | 101.38432 | 97.78168 | Promoter | 11 | 102289300 | 102296631 | 7332 | 2 | ENSMUST00 | 0 | Atxn7l3 | ENSMUSG00000059995.7 |
| chr17 | 24752281 | 24755341 | 33.1785 | 101.38432 | 97.78168 | Promoter | 17 | 24753003 | 24758683 | 5681 | 1 | ENSMUST00 | 0 | Hs3st6 | ENSMUSG00000039628.8 |
| chr6 | 53067479 | 53069129 | 33.1785 | 101.38432 | 97.78168 | Promoter | 6 | 52768797 | 53068631 | 299835 | 2 | ENSMUST00 | 0 | Jazf1 | ENSMUSG00000063568.6 |
| chr9 | 15493121 | 15494847 | 33.1785 | 101.38432 | 97.78168 | Distal Interge | 9 | 15505487 | 15545260 | 39774 | 1 | ENSMUST00 | -10640 | Smco4 | ENSMUSG00000058173.6 |
| chr1 | 92933726 | 92935362 | 32.55857 | 101.31886 | 97.7165 | Promoter | 1 | 92934376 | 92947941 | 13566 | 1 | ENSMUST00 | 0 | Capn10 | ENSMUSG00000026270.6 |
| chr17 | 84957997 | 84959482 | 32.55857 | 101.31886 | 97.7165 | Promoter | 17 | 84958001 | 85014776 | 56776 | 1 | ENSMUST00 | 0 | Ppm1b | ENSMUSG00000061130.7 |
| chr17 | 24426215 | 24427651 | 31.97681 | 101.31316 | 97.71113 | Promoter | 17 | 24426683 | 24439316 | 12634 | 1 | ENSMUST00 | 0 | Eci1 | ENSMUSG00000024132.5 |
| chr18 | 73702684 | 73704403 | 31.97681 | 101.31316 | 97.71113 | Promoter | 18 | 73639009 | 73703780 | 64772 | 2 | ENSMUST00 | 0 | Smad4 | ENSMUSG00000024515.8 |
| chr10 | 86703997 | 86705619 | 29.71419 | 101.30199 | 97.70005 | Promoter | 10 | 86690840 | 86705509 | 14670 | 2 | ENSMUST00 | 0 | Hsp90b1 | ENSMUSG00000020048.8 |
| chr10 | 80917475 | 80918510 | 29.71419 | 101.30199 | 97.70005 | Promoter | 10 | 80901203 | 80918245 | 17043 | 2 | ENSMUST00 | 0 | Lmnb2 | ENSMUSG00000062075.8 |
| chr11 | 115403196 | 115404587 | 29.71419 | 101.30199 | 97.70005 | Promoter | 11 | 115403752 | 115412674 | 8923 | 1 | ENSMUST00 | 0 | Ict1 | ENSMUSG00000018858.10 |
| chr3 | 88057578 | 88058938 | 29.71419 | 101.30199 | 97.70005 | Promoter | 3 | 88056523 | 88058495 | 1973 | 2 | ENSMUST00 | 0 | Apoa1bp | ENSMUSG00000028070.6 |
| chr3 | 89429914 | 89431194 | 29.71419 | 101.30199 | 97.70005 | Promoter | 3 | 89430214 | 89432743 | 2530 | 1 | ENSMUST00 | 0 | Pygo2 | ENSMUSG00000047824.8 |
| chr4 | 132273801 | 132275297 | 29.71419 | 101.30199 | 97.70005 | Promoter | 4 | 132274375 | 132295766 | 21392 | 1 | ENSMUST00 | 0 | Taf12 | ENSMUSG00000028899.6 |
| chr5 | 32134342 | 32139172 | 29.71419 | 101.30199 | 97.70005 | Promoter | 5 | 32136472 | 32157831 | 21360 | 1 | ENSMUST00 | 0 | Fosl2 | ENSMUSG00000029135.9 |
| chr1 | 64531819 | 64534191 | 33.1106 | 101.26704 | 97.66567 | Promoter | 1 | 64532645 | 64597695 | 65051 | 1 | ENSMUST00 | 0 | Creb1 | ENSMUSG00000025958.9 |
| chr16 | 36827704 | 36829155 | 33.1106 | 101.26704 | 97.66567 | Promoter | 16 | 36828385 | 36831565 | 3181 | 1 | ENSMUST00 | 0 | Iqcb1 | ENSMUSG00000022837.8 |
| chr18 | 24528898 | 24530722 | 33.1106 | 101.26704 | 97.66567 | Promoter | 18 | 24484962 | 24530204 | 45243 | 2 | ENSMUST00 | 0 | Rprd1a | ENSMUSG00000040446.3 |
| chr2 | 71388694 | 71389921 | 33.1106 | 101.26704 | 97.66567 | Promoter | 2 | 71388958 | 71441622 | 52665 | 1 | ENSMUST00 | 0 | Hat1 | ENSMUSG00000027018.6 |
| chr3 | 84580512 | 84583491 | 33.1106 | 101.26704 | 97.66567 | Promoter | 3 | 84496093 | 84582534 | 86442 | 2 | ENSMUST00 | 0 | Arfip1 | ENSMUSG00000074513.4 |
| chr4 | 149953980 | 149955073 | 33.1106 | 101.26704 | 97.66567 | Promoter | 4 | 149896283 | 149955043 | 58761 | 2 | ENSMUST00 | 0 | Spsb1 | ENSMUSG00000039911.8 |
| chr6 | 146887899 | 146889364 | 33.1106 | 101.26704 | 97.66567 | Promoter | 6 | 146888487 | 147032025 | 143539 | 1 | ENSMUST00 | 0 | Ppfibp1 | ENSMUSG00000016487.10 |
| chr2 | 151008552 | 151009746 | 32.49405 | 101.20673 | 97.60635 | Promoter | 2 | 150934519 | 151009398 | 74880 | 2 | ENSMUST00 | 0 | Ninl | ENSMUSG00000068115.8 |
| chr9 | 47529352 | 47532275 | 32.49405 | 101.20673 | 97.60635 | Promoter | 9 | 47530173 | 47853385 | 323213 | 1 | ENSMUST00 | 0 | Cadm1 | ENSMUSG00000032076.13 |
| chr2 | 25054278 | 25055297 | 29.09699 | 101.16944 | 97.56942 | Promoter | 2 | 25054355 | 25062881 | 8527 | 1 | ENSMUST00 | 0 | Nsmf | ENSMUSG00000006476.14 |
| chr3 | 37312066 | 37313316 | 29.09699 | 101.16944 | 97.56942 | Promoter | 3 | 37312519 | 37404866 | 92348 | 1 | ENSMUST00 | 0 | Fgf2 | ENSMUSG00000037225.8 |
| chr8 | 48553777 | 48556246 | 31.85421 | 101.09849 | 97.49914 | Promoter | 8 | 48334753 | 48555693 | 220941 | 2 | ENSMUST00 | 0 | Tenm3 | ENSMUSG00000031561.10 |
| chr1 | 13588480 | 13590440 | 31.85236 | 101.09525 | 97.496 | Promoter | 1 | 13564698 | 13589910 | 25213 | 2 | ENSMUST00 | 0 | Tram1 | ENSMUSG00000025935.5 |
| chr1 | 91051607 | 91058713 | 31.85236 | 101.09525 | 97.496 | Promoter | 1 | 91053444 | 91128944 | 75501 | 1 | ENSMUST00 | 0 | Lrrfip1 | ENSMUSG00000026305.10 |
| chr10 | 127739030 | 127740414 | 31.85236 | 101.09525 | 97.496 | Promoter | 10 | 127739538 | 127747335 | 7798 | 1 | ENSMUST00 | 0 | Zbtb39 | ENSMUSG00000044617.7 |
| chr11 | 69579501 | 69582098 | 31.85236 | 101.09525 | 97.496 | Promoter | 11 | 69577708 | 69580255 | 2548 | 2 | ENSMUST00 | 0 | Wrap53 | ENSMUSG00000041346.6 |
| chr11 | 58199041 | 58200075 | 31.85236 | 101.09525 | 97.496 | Promoter | 11 | 58199618 | 58222771 | 23154 | 1 | ENSMUST00 | 0 | Irgm2 | ENSMUSG00000069874.6 |
| chr12 | 80109381 | 80113014 | 31.85236 | 101.09525 | 97.496 | Promoter | 12 | 80107760 | 80113013 | 5254 | 2 | ENSMUST00 | 0 | Zfp36l1 | ENSMUSG00000021127.6 |
| chr12 | 83485304 | 83489226 | 31.85236 | 101.09525 | 97.496 | Promoter | 12 | 83213751 | 83487716 | 273966 | 2 | ENSMUST00 | 0 | Dpf3 | ENSMUSG00000021221.10 |
| chr12 | 111758148 | 111760750 | 28.51907 | 101.09029 | 97.496 | Promoter | 12 | 111758849 | 111806775 | 47927 | 1 | ENSMUST00 | 0 | Klc1 | ENSMUSG00000021288.13 |
| chr12 | 112130010 | 112131148 | 31.85236 | 101.09525 | 97.496 | Promoter | 12 | 112130880 | 112130955 | 76 | 1 | ENSMUST00 | 0 | Mir203 | ENSMUSG00000065574.1 |
| chr12 | 81483636 | 81485527 | 31.85236 | 101.09525 | 97.496 | Promoter | 12 | 81359026 | 81485110 | 126085 | 2 | ENSMUST00 | 0 | Cox16 | ENSMUSG00000091803.2 |
| chr13 | 67359944 | 67361004 | 31.85236 | 101.09525 | 97.496 | Promoter | 13 | 67360671 | 67365679 | 5009 | 1 | ENSMUST00 | 0 | Gm17039 | ENSMUSG00000092103.2 |
| chr16 | 75592585 | 75594022 | 31.85236 | 101.09525 | 97.496 | Promoter | 16 | 75592844 | 75601695 | 8852 | 1 | ENSMUST00 | 0 | Rbm11 | ENSMUSG00000032940.11 |
| chr17 | 53478468 | 53481041 | 31.85236 | 101.09525 | 97.496 | Promoter | 17 | 53479234 | 53507671 | 28438 | 1 | ENSMUST00 | 0 | Rab5a | ENSMUSG00000017831.7 |
| chr19 | 6968628 | 6970505 | 31.85236 | 101.09525 | 97.496 | Promoter | 19 | 6953714 | 6969759 | 16046 | 2 | ENSMUST00 | 0 | Plcb3 | ENSMUSG00000024960.8 |
| chr2 | 76646657 | 76648968 | 31.85236 | 101.09525 | 97.496 | Promoter | 2 | 76629898 | 76648015 | 18118 | 2 | ENSMUST00 | 0 | Prkra | ENSMUSG00000002731.6 |
| chr2 | 25210115 | 25211701 | 31.85236 | 101.09525 | 97.496 | Promoter | 2 | 25199712 | 25211489 | 11778 | 2 | ENSMUST00 | 0 | Nelfb | ENSMUSG00000013465.13 |
| chr2 | 143914317 | 143916573 | 31.85236 | 101.09525 | 97.496 | Promoter | 2 | 143915320 | 143943324 | 28005 | 1 | ENSMUST00 | 0 | Dstn | ENSMUSG00000015932.8 |
| chr2 | 163602069 | 163603114 | 31.85236 | 101.09525 | 97.496 | Promoter | 2 | 163602314 | 163619012 | 16699 | 1 | ENSMUST00 | 0 | Ttpal | ENSMUSG00000017679.11 |
| chr2 | 31759357 | 31761932 | 31.85236 | 101.09525 | 97.496 | Promoter | 2 | 31759943 | 31804227 | 44285 | 1 | ENSMUST00 | 0 | Abl1 | ENSMUSG00000026842.11 |
| chr2 | 36135650 | 36137375 | 31.85236 | 101.09525 | 97.496 | Promoter | 2 | 36136389 | 36190647 | 54259 | 1 | ENSMUST00 | 0 | Mrrf | ENSMUSG00000026887.4 |
| chr2 | 59160059 | 59162590 | 28.51907 | 101.09029 | 97.496 | Promoter | 2 | 59160850 | 59355204 | 194355 | 1 | ENSMUST00 | 0 | Pkp4 | ENSMUSG00000026991.15 |
| chr2 | 148731521 | 148732711 | 31.85236 | 101.09525 | 97.496 | Promoter | 2 | 148693985 | 148732413 | 38429 | 2 | ENSMUST00 | 0 | Napb | ENSMUSG00000027438.9 |
| chr2 | 120976796 | 120978057 | 31.85236 | 101.09525 | 97.496 | Promoter | 2 | 120977017 | 121007852 | 30836 | 1 | ENSMUST00 | 0 | Tmem62 | ENSMUSG00000054484.9 |
| chr3 | 41563120 | 41566467 | 31.85236 | 101.09525 | 97.496 | Promoter | 3 | 41563383 | 41616862 | 53480 | 1 | ENSMUST00 | 0 | Jade1 | ENSMUSG00000025764.9 |
| chr3 | 146404092 | 146405945 | 31.85236 | 101.09525 | 97.496 | Promoter | 3 | 146404642 | 146440139 | 35498 | 1 | ENSMUST00 | 0 | Ssx2ip | ENSMUSG00000036825.7 |
| chr4 | 59580508 | 59582421 | 31.85236 | 101.09525 | 97.496 | Promoter | 4 | 59581563 | 59618689 | 37127 | 1 | ENSMUST00 | 0 | Hsdl2 | ENSMUSG00000028383.12 |
| chr4 | 46304404 | 46305687 | 31.85236 | 101.09525 | 97.496 | Intron (ENSM | 4 | 46327296 | 46327765 | 470 | 1 | ENSMUST00 | -21609 | Gm12447 | ENSMUSG00000080816.1 |
| chr7 | 97842956 | 97846713 | 31.85236 | 101.09525 | 97.496 | Promoter | 7 | 97842935 | 97912381 | 69447 | 1 | ENSMUST00 | 22 | Pak1 | ENSMUSG00000030774.8 |
| chr9 | 66834217 | 66834807 | 31.85236 | 101.09525 | 97.496 | Promoter | 9 | 66814994 | 66834726 | 19733 | 2 | ENSMUST00 | 0 | Aph1c | ENSMUSG00000053040.8 |
| chr19 | 5838337 | 5845488 | 27.97678 | 101.05884 | 97.46488 | Promoter | 19 | 5824708 | 5845478 | 20771 | 2 | ENSMUST00 | 0 | Neat1 | ENSMUSG00000092274.2 |
| chr3 | 95658222 | 95661738 | 27.97678 | 101.05884 | 97.46488 | Promoter | 3 | 95658721 | 95663176 | 4456 | 1 | ENSMUST00 | 0 | Mcl1 | ENSMUSG00000038612.11 |
| chr14 | 47472460 | 47473449 | 31.25476 | 101.05123 | 97.45738 | Promoter | 14 | 47472561 | 47531962 | 59402 | 1 | ENSMUST00 | 0 | Fbxo34 | ENSMUSG00000037536.8 |
| chr15 | 102099897 | 102107616 | 31.25476 | 101.05123 | 97.45738 | Promoter | 15 | 102102988 | 102116391 | 13404 | 1 | ENSMUST00 | 0 | Tenc1 | ENSMUSG00000037003.10 |
| chr2 | 132575235 | 132578510 | 31.25476 | 101.05123 | 97.45738 | Promoter | 2 | 132529082 | 132578178 | 49097 | 2 | ENSMUST00 | 0 | Gpcpd1 | ENSMUSG00000027346.10 |
| chr2 | 127586902 | 127588170 | 31.25476 | 101.05123 | 97.45738 | Promoter | 2 | 127587222 | 127606829 | 19608 | 1 | ENSMUST00 | 0 | Mrps5 | ENSMUSG00000027374.7 |
| chr3 | 97763573 | 97768572 | 31.25476 | 101.05123 | 97.45738 | Promoter | 3 | 97738854 | 97768009 | 29156 | 2 | ENSMUST00 | 0 | Pde4dip | ENSMUSG00000038170.10 |
| chr5 | 30231760 | 30233605 | 31.25476 | 101.05123 | 97.45738 | Promoter | 5 | 30232581 | 30272012 | 39432 | 1 | ENSMUST00 | 0 | Ept1 | ENSMUSG00000075703.10 |
| chr6 | 30509099 | 30510160 | 31.25476 | 101.05123 | 97.45738 | Promoter | 6 | 30479053 | 30509704 | 30652 | 2 | ENSMUST00 | 0 | Tmem209 | ENSMUSG00000029782.13 |
| chr7 | 35055378 | 35057248 | 31.25476 | 101.05123 | 97.45738 | Promoter | 7 | 35046422 | 35056503 | 10082 | 2 | ENSMUST00 | 0 | Cebpg | ENSMUSG00000056216.8 |
| chr11 | 106192974 | 106193624 | 32.97563 | 101.03377 | 97.44133 | Promoter | 11 | 106163351 | 106193536 | 30186 | 2 | ENSMUST00 | 0 | Strada | ENSMUSG00000069631.9 |
| chr11 | 75189684 | 75190592 | 32.97563 | 101.03377 | 97.44133 | Promoter | 11 | 75177643 | 75190519 | 12877 | 2 | ENSMUST00 | 0 | Dph1 | ENSMUSG00000078789.4 |
| chr18 | 63708304 | 63709486 | 32.97563 | 101.03377 | 97.44133 | Promoter | 18 | 63708695 | 63989760 | 281066 | 1 | ENSMUST00 | 0 | Wdr7 | ENSMUSG00000040560.9 |
| chr6 | 137251894 | 137256757 | 32.97563 | 101.03377 | 97.44133 | Promoter | 6 | 137252299 | 137464629 | 212331 | 1 | ENSMUST00 | 0 | Ptpro | ENSMUSG00000030223.9 |
| chr11 | 73047761 | 73048865 | 24.12502 | 101.02714 | 97.43493 | Promoter | 11 | 73047783 | 73080217 | 32435 | 1 | ENSMUST00 | 0 | 1200014J11 | ENSMUSG00000020783.9 |
| chr14 | 52196202 | 52197962 | 24.12502 | 101.02714 | 97.43493 | Promoter | 14 | 52160419 | 52197239 | 36821 | 2 | ENSMUST00 | 0 | Supt16 | ENSMUSG00000035726.7 |
| chr18 | 68299186 | 68301290 | 24.12502 | 101.02714 | 97.43493 | Promoter | 18 | 68300355 | 68324852 | 24498 | 1 | ENSMUST00 | 0 | Rnmt | ENSMUSG00000009535.8 |
| chr1 | 184998999 | 185000324 | 33.54585 | 101.01101 | 97.41918 | Promoter | 1 | 184896443 | 184999549 | 103107 | 2 | ENSMUST00 | 0 | Mark1 | ENSMUSG00000026620.6 |
| chr10 | 127380110 | 127381752 | 33.54585 | 101.01101 | 97.41918 | Promoter | 10 | 127380327 | 127444391 | 64065 | 1 | ENSMUST00 | 0 | R3hdm2 | ENSMUSG00000025404.10 |
| chr10 | 127195052 | 127196188 | 33.54585 | 101.01101 | 97.41918 | Promoter | 10 | 127190378 | 127195709 | 5332 | 2 | ENSMUST00 | 0 | Dtx3 | ENSMUSG00000040415.10 |
| chr10 | 76031985 | 76033514 | 33.54585 | 101.01101 | 97.41918 | Promoter | 10 | 76032654 | 76043234 | 10581 | 1 | ENSMUST00 | 0 | Zfp280b | ENSMUSG00000049764.7 |
| chr11 | 74770368 | 74771645 | 33.54585 | 101.01101 | 97.41918 | Promoter | 11 | 74770830 | 74817894 | 47065 | 1 | ENSMUST00 | 0 | Mettl16 | ENSMUSG00000010554.9 |
| chr11 | 69416361 | 69418915 | 33.54585 | 101.01101 | 97.41918 | Downstream | 11 | 69398508 | 69413675 | 15168 | 2 | ENSMUST00 | -2687 | Kdm6b | ENSMUSG00000018476.7 |
| chr11 | 102025982 | 102027728 | 33.54585 | 101.01101 | 97.41918 | Promoter | 11 | 101999652 | 102026094 | 26443 | 2 | ENSMUST00 | 0 | Mpp3 | ENSMUSG00000052373.9 |
| chr11 | 88715830 | 88719254 | 33.54585 | 101.01101 | 97.41918 | Promoter | 11 | 88339382 | 88718229 | 378848 | 2 | ENSMUST00 | 0 | Msi2 | ENSMUSG00000069769.8 |
| chr14 | 87140405 | 87141744 | 33.54585 | 101.01101 | 97.41918 | Promoter | 14 | 86655367 | 87141224 | 485858 | 2 | ENSMUST00 | 0 | Diap3 | ENSMUSG00000022021.8 |
| chr16 | 87698632 | 87700430 | 33.54585 | 101.01101 | 97.41918 | Promoter | 16 | 87698945 | 87733346 | 34402 | 1 | ENSMUST00 | 0 | Bach1 | ENSMUSG00000025612.5 |
| chr17 | 27017685 | 27018575 | 33.54585 | 101.01101 | 97.41918 | Promoter | 17 | 27018013 | 27026693 | 8681 | 1 | ENSMUST00 | 0 | Ggnbp1 | ENSMUSG00000048731.9 |
| chr19 | 46337945 | 46338771 | 33.54585 | 101.01101 | 97.41918 | Promoter | 19 | 46329812 | 46338660 | 8849 | 2 | ENSMUST00 | 0 | Cuedc2 | ENSMUSG00000036748.10 |
| chr3 | 89959985 | 89960959 | 33.54585 | 101.01101 | 97.41918 | Promoter | 3 | 89939481 | 89960629 | 21149 | 2 | ENSMUST00 | 0 | Atp8b2 | ENSMUSG00000060671.7 |
| chr4 | 148158990 | 148161122 | 33.54585 | 101.01101 | 97.41918 | Promoter | 4 | 148152800 | 148159595 | 6796 | 2 | ENSMUST00 | 0 | Fbxo44 | ENSMUSG00000029001.10 |
| chr5 | 125177014 | 125179760 | 33.54585 | 101.01101 | 97.41918 | Promoter | 5 | 125017153 | 125179214 | 162062 | 2 | ENSMUST00 | 0 | Ncor2 | ENSMUSG00000029478.11 |
| chr1 | 183296405 | 183298299 | 19.5713 | 100.97023 | 97.3811 | Promoter | 1 | 183297060 | 183324501 | 27442 | 1 | ENSMUST00 | 0 | Aida | ENSMUSG00000042901.5 |
| chr14 | 79426350 | 79427272 | 21.28758 | 100.96285 | 97.37373 | Promoter | 14 | 79426511 | 79431039 | 4529 | 1 | ENSMUST00 | 0 | Kbtbd7 | ENSMUSG00000043881.8 |
| chr3 | 88834896 | 88835948 | 30.05765 | 100.92319 | 97.33411 | Promoter | 3 | 88835231 | 88910103 | 74873 | 1 | ENSMUST00 | 0 | Gon4l | ENSMUSG00000054199.11 |
| chr3 | 90247626 | 90249164 | 31.13914 | 100.84679 | 97.25846 | Promoter | 3 | 90248172 | 90253612 | 5441 | 1 | ENSMUST00 | 0 | Slc39a1 | ENSMUSG00000052310.9 |
| chr2 | 91526441 | 91527615 | 31.67206 | 100.77918 | 97.19132 | Promoter | 2 | 91526762 | 91620664 | 93903 | 1 | ENSMUST00 | 0 | Ckap5 | ENSMUSG00000040549.11 |

| | | | | | | | | | | | | | | | |
|---|---|---|---|---|---|---|---|---|---|---|---|---|---|---|---|
| chr12 | 116047153 | 116049091 | 30.52716 | 100.76356 | 97.17578 | Promoter | 12 | 116047724 | 116060592 | 12869 | 1 | ENSMUST00 | 0 | Zfp386 | ENSMUSG00000042063.11 |
| chr13 | 48261011 | 48263120 | 30.52716 | 100.76356 | 97.17578 | Promoter | 13 | 48261427 | 48264036 | 2610 | 1 | ENSMUST00 | 0 | Id4 | ENSMUSG00000021379.1 |
| chr3 | 19627648 | 19629625 | 30.52716 | 100.76356 | 97.17578 | Promoter | 3 | 19610368 | 19628677 | 18310 | 2 | ENSMUST00 | 0 | 1700064H15I | ENSMUSG00000082766.1 |
| chr8 | 83593648 | 83595091 | 30.52716 | 100.76356 | 97.17578 | Promoter | 8 | 83571700 | 83594429 | 22730 | 2 | ENSMUST00 | 0 | Tecr | ENSMUSG00000031708.10 |
| chr8 | 105263838 | 105265434 | 30.52716 | 100.76356 | 97.17578 | Promoter | 8 | 105258286 | 105264609 | 6324 | 2 | ENSMUST00 | 0 | Tradd | ENSMUSG00000031887.8 |
| chr9 | 36767956 | 36768690 | 30.52716 | 100.76356 | 97.17578 | Promoter | 9 | 36767910 | 36782675 | 14766 | 1 | ENSMUST00 | 47 | Gm26787 | ENSMUSG00000096993.2 |
| chr15 | 101077078 | 101079386 | 32.23851 | 100.76218 | 97.17527 | Distal Interge | 15 | 101050194 | 101054399 | 4206 | 2 | ENSMUST00 | -22680 | Fignl2 | ENSMUSG00000095440.1 |
| chr17 | 84465500 | 84466677 | 32.23851 | 100.76218 | 97.17527 | Promoter | 17 | 84190056 | 84466196 | 276141 | 2 | ENSMUST00 | 0 | Thada | ENSMUSG00000024251.9 |
| chr17 | 24885498 | 24886412 | 32.23851 | 100.76218 | 97.17527 | Promoter | 17 | 24882611 | 24886350 | 3740 | 2 | ENSMUST00 | 0 | Nubp2 | ENSMUSG00000039183.5 |
| chr2 | 48813188 | 48815735 | 32.23851 | 100.76218 | 97.17527 | Promoter | 2 | 48814109 | 48903269 | 89161 | 1 | ENSMUST00 | 0 | Acvr2a | ENSMUSG00000052155.5 |
| chr2 | 78868289 | 78870641 | 32.77523 | 100.68713 | 97.10149 | Promoter | 2 | 78869047 | 78921293 | 52247 | 1 | ENSMUST00 | 0 | Ube2e3 | ENSMUSG00000027011.9 |
| chr1 | 167000262 | 167003097 | 33.34559 | 100.66625 | 97.08096 | Promoter | 1 | 167001417 | 167091302 | 89886 | 1 | ENSMUST00 | 0 | Fam78b | ENSMUSG00000060568.9 |
| chr14 | 55015477 | 55016249 | 33.34559 | 100.66625 | 97.08096 | Promoter | 14 | 55015454 | 55024135 | 8682 | 1 | ENSMUST00 | 24 | Ngdn | ENSMUSG00000022204.8 |
| chr5 | 129669228 | 129670541 | 33.34559 | 100.66625 | 97.08096 | Promoter | 5 | 129654593 | 129670099 | 15507 | 2 | ENSMUST00 | 0 | Zfp11 | ENSMUSG00000051034.7 |
| chr4 | 44232600 | 44234015 | 32.70897 | 100.57243 | 96.98774 | Promoter | 4 | 44152434 | 44233789 | 81356 | 2 | ENSMUST00 | 0 | Rnf38 | ENSMUSG00000035696.10 |
| chr11 | 116130136 | 116131506 | 29.84986 | 100.54769 | 96.96338 | Promoter | 11 | 116121846 | 116131124 | 9279 | 2 | ENSMUST00 | 0 | Trim65 | ENSMUSG00000054517.4 |
| chr9 | 59290797 | 59292489 | 29.84986 | 100.54769 | 96.96338 | Promoter | 9 | 59291572 | 59316199 | 24628 | 1 | ENSMUST00 | 0 | Adpgk | ENSMUSG00000025236.10 |
| chr10 | 82762672 | 82764547 | 32.64297 | 100.45817 | 96.87489 | Promoter | 10 | 82748701 | 82764144 | 15444 | 2 | ENSMUST00 | 0 | Nfyb | ENSMUSG00000020248.13 |
| chr3 | 103278682 | 103281247 | 32.64297 | 100.45817 | 96.87489 | Promoter | 3 | 103279293 | 103358768 | 79476 | 1 | ENSMUST00 | 0 | Trim33 | ENSMUSG00000033014.7 |
| chr1 | 192136182 | 192138954 | 25.18016 | 100.40004 | 96.81754 | Promoter | 1 | 192098656 | 192138010 | 39355 | 2 | ENSMUST00 | 0 | Rcor3 | ENSMUSG00000037395.10 |
| chr17 | 33683865 | 33685869 | 25.18016 | 100.40004 | 96.81754 | Promoter | 17 | 33646235 | 33685392 | 39158 | 2 | ENSMUST00 | 0 | Hnrnpm | ENSMUSG00000059208.9 |
| chr19 | 4282703 | 4283507 | 25.18016 | 100.40004 | 96.81754 | Promoter | 19 | 4270180 | 4283135 | 12956 | 2 | ENSMUST00 | 0 | Ankrd13d | ENSMUSG00000005986.10 |
| chr6 | 86668165 | 86669251 | 25.18016 | 100.40004 | 96.81754 | Promoter | 6 | 86647044 | 86669159 | 22116 | 2 | ENSMUST00 | 0 | Mxd1 | ENSMUSG00000001156.7 |
| chr6 | 29916461 | 29917797 | 25.18016 | 100.40004 | 96.81754 | Promoter | 6 | 29917013 | 29945554 | 28542 | 1 | ENSMUST00 | 0 | Strip2 | ENSMUSG00000039629.9 |
| chr7 | 100925652 | 100932907 | 25.18016 | 100.40004 | 96.81754 | Promoter | 7 | 100869746 | 100932161 | 62416 | 2 | ENSMUST00 | 0 | Arhgef17 | ENSMUSG00000032875.7 |
| chr7 | 105743540 | 105744790 | 25.18016 | 100.40004 | 96.81754 | Promoter | 7 | 105739393 | 105744361 | 4969 | 2 | ENSMUST00 | 0 | Taf10 | ENSMUSG00000043866.5 |
| chr9 | 122351245 | 122352433 | 25.18016 | 100.40004 | 96.81754 | Promoter | 9 | 122351608 | 122381524 | 29917 | 1 | ENSMUST00 | 0 | Abhd5 | ENSMUSG00000032540.10 |
| chr15 | 82243721 | 82245060 | 29.21782 | 100.39505 | 96.81345 | Promoter | 15 | 82233779 | 82244747 | 10969 | 2 | ENSMUST00 | 0 | Cenpm | ENSMUSG00000068101.5 |
| chr7 | 127375200 | 127376114 | 29.21782 | 100.39505 | 96.81345 | Promoter | 7 | 127372564 | 127376050 | 3487 | 2 | ENSMUST00 | 0 | Zfp747 | ENSMUSG00000054381.4 |
| chr8 | 86744704 | 86746872 | 29.21782 | 100.39505 | 96.81345 | Promoter | 8 | 86723940 | 86746006 | 22067 | 2 | ENSMUST00 | 0 | Siah1a | ENSMUSG00000036840.4 |
| chr11 | 6266806 | 6268347 | 31.43241 | 100.35837 | 96.77719 | Promoter | 11 | 6258919 | 6267772 | 8854 | 2 | ENSMUST00 | 0 | Ddx56 | ENSMUSG00000004393.5 |
| chr11 | 120579222 | 120582581 | 31.43241 | 100.35837 | 96.77719 | Promoter | 11 | 120578104 | 120581624 | 3521 | 2 | ENSMUST00 | 0 | Arhgdia | ENSMUSG00000025132.8 |
| chr11 | 75509802 | 75510755 | 31.43241 | 100.35837 | 96.77719 | Promoter | 11 | 75510094 | 75513168 | 3075 | 1 | ENSMUST00 | 0 | Rilp | ENSMUSG00000038195.6 |
| chr13 | 119408593 | 119409890 | 31.43241 | 100.35837 | 96.77719 | Promoter | 13 | 119335448 | 119408997 | 73550 | 2 | ENSMUST00 | 0 | Nnt | ENSMUSG00000025453.11 |
| chr14 | 76413767 | 76416715 | 31.43241 | 100.35837 | 96.77719 | Promoter | 14 | 76414961 | 76507762 | 92802 | 1 | ENSMUST00 | 0 | Tsc22d1 | ENSMUSG00000022010.14 |
| chr19 | 24998983 | 25000509 | 31.43241 | 100.35837 | 96.77719 | Promoter | 19 | 24999529 | 25202432 | 202904 | 1 | ENSMUST00 | 0 | Dock8 | ENSMUSG00000052085.6 |
| chr2 | 164460919 | 164462087 | 31.43241 | 100.35837 | 96.77719 | Promoter | 2 | 164460971 | 164465508 | 4538 | 1 | ENSMUST00 | 0 | Sys1 | ENSMUSG00000045503.11 |
| chr8 | 110902051 | 110902699 | 31.43241 | 100.35837 | 96.77719 | Promoter | 8 | 110882463 | 110902459 | 19997 | 2 | ENSMUST00 | 0 | Fuk | ENSMUSG00000033703.8 |
| chr1 | 161727049 | 161727715 | 22.96739 | 100.34364 | 96.76433 | Promoter | 1 | 161726242 | 161727489 | 1248 | 2 | ENSMUST00 | 0 | Gm15429 | ENSMUSG00000040323.13 |
| chr11 | 75732399 | 75734879 | 22.96739 | 100.34364 | 96.76433 | Promoter | 11 | 75732869 | 75765845 | 32977 | 1 | ENSMUST00 | 0 | Ywhae | ENSMUSG00000020849.8 |
| chr8 | 123224194 | 123225624 | 28.62665 | 100.29842 | 96.71969 | Promoter | 8 | 123224841 | 123232250 | 7410 | 1 | ENSMUST00 | 0 | Cdk10 | ENSMUSG00000033862.6 |
| chr7 | 98814693 | 98816760 | 27.06219 | 100.28806 | 96.70959 | Distal Interge | 7 | 98835112 | 98846587 | 11476 | 1 | ENSMUST00 | -18352 | Wnt11 | ENSMUSG00000015957.6 |
| chr8 | 95351992 | 95353528 | 27.06219 | 100.28806 | 96.70959 | Promoter | 8 | 95352268 | 95374293 | 22026 | 1 | ENSMUST00 | 0 | Mmp15 | ENSMUSG00000031790.7 |
| chr19 | 6117454 | 6119032 | 27.55201 | 100.24977 | 96.67152 | Promoter | 19 | 6116004 | 6118586 | 2583 | 2 | ENSMUST00 | 0 | Sac3d1 | ENSMUSG00000024790.7 |
| chr15 | 76710078 | 76711301 | 31.92468 | 100.21517 | 96.63724 | Promoter | 15 | 76710623 | 76713570 | 2948 | 1 | ENSMUST00 | 0 | Lrrc14 | ENSMUSG00000033728.9 |
| chr1 | 44101623 | 44102830 | 29.66345 | 100.21 | 96.63225 | Promoter | 1 | 44086613 | 44102391 | 15779 | 2 | ENSMUST00 | 0 | Tex30 | ENSMUSG00000026049.6 |
| chr1 | 118387964 | 118390405 | 29.66345 | 100.21 | 96.63225 | Promoter | 1 | 118389058 | 118609432 | 220375 | 1 | ENSMUST00 | 0 | Clasp1 | ENSMUSG00000064302.8 |
| chr10 | 127662238 | 127666785 | 29.66345 | 100.21 | 96.63225 | Promoter | 10 | 127660918 | 127666710 | 5793 | 2 | ENSMUST00 | 0 | Nab2 | ENSMUSG00000025402.7 |
| chr10 | 122096005 | 122097350 | 29.66345 | 100.21 | 96.63225 | Promoter | 10 | 122081260 | 122097156 | 15897 | 2 | ENSMUST00 | 0 | Tmem5 | ENSMUSG00000034620.12 |
| chr10 | 127120354 | 127121235 | 29.66345 | 100.21 | 96.63225 | Promoter | 10 | 127094259 | 127121160 | 26902 | 2 | ENSMUST00 | 0 | Os9 | ENSMUSG00000040462.7 |
| chr11 | 100574589 | 100575730 | 29.66345 | 100.21 | 96.63225 | Promoter | 11 | 100574904 | 100581728 | 6825 | 1 | ENSMUST00 | 0 | Cnp | ENSMUSG00000006782.11 |
| chr11 | 62280815 | 62282196 | 29.66345 | 100.21 | 96.63225 | Promoter | 11 | 62267224 | 62281375 | 14152 | 2 | ENSMUST00 | 0 | Zswim7 | ENSMUSG00000014243.5 |
| chr11 | 3408169 | 3409742 | 29.66345 | 100.21 | 96.63225 | Promoter | 11 | 3344256 | 3409174 | 64919 | 2 | ENSMUST00 | 0 | Limk2 | ENSMUSG00000020451.12 |
| chr13 | 54789169 | 54790514 | 29.66345 | 100.21 | 96.63225 | Promoter | 13 | 54789377 | 54796776 | 7400 | 1 | ENSMUST00 | 0 | Tspan17 | ENSMUSG00000025875.9 |
| chr13 | 58157315 | 58158795 | 29.66345 | 100.21 | 96.63225 | Promoter | 13 | 58157649 | 58164693 | 7045 | 1 | ENSMUST00 | 0 | Idnk | ENSMUSG00000050002.8 |
| chr14 | 73509938 | 73510748 | 29.66345 | 100.21 | 96.63225 | Promoter | 14 | 73510049 | 73518545 | 8497 | 1 | ENSMUST00 | 0 | Med4 | ENSMUSG00000022109.5 |
| chr15 | 98803793 | 98809569 | 29.66345 | 100.21 | 96.63225 | Promoter | 15 | 98803782 | 98807925 | 4144 | 2 | ENSMUST00 | 0 | Ddn | ENSMUSG00000059213.6 |
| chr16 | 20732505 | 20734317 | 29.66345 | 100.21 | 96.63225 | Promoter | 16 | 20733127 | 20738152 | 5026 | 1 | ENSMUST00 | 0 | Chrd | ENSMUSG00000006958.8 |
| chr16 | 27388323 | 27390008 | 29.66345 | 100.21 | 96.63225 | Promoter | 16 | 27388869 | 27452218 | 63350 | 1 | ENSMUST00 | 0 | Ccdc50 | ENSMUSG00000038127.9 |
| chr17 | 46628274 | 46630084 | 29.66345 | 100.21 | 96.63225 | Promoter | 17 | 46564451 | 46629504 | 65054 | 2 | ENSMUST00 | 0 | Ptk7 | ENSMUSG00000023972.9 |
| chr19 | 41262676 | 41264809 | 29.66345 | 100.21 | 96.63225 | Promoter | 19 | 41210842 | 41264004 | 53163 | 2 | ENSMUST00 | 0 | Tm9sf3 | ENSMUSG00000025016.10 |
| chr19 | 12595622 | 12597597 | 29.66345 | 100.21 | 96.63225 | Promoter | 19 | 12585868 | 12596566 | 10699 | 2 | ENSMUST00 | 0 | A330040F15I | ENSMUSG00000086213.1 |
| chr2 | 35336294 | 35337626 | 29.66345 | 100.21 | 96.63225 | Promoter | 2 | 35313986 | 35336976 | 22991 | 2 | ENSMUST00 | 0 | Stom | ENSMUSG00000026880.10 |
| chr2 | 36048205 | 36049802 | 29.66345 | 100.21 | 96.63225 | Promoter | 2 | 36049458 | 36079718 | 30261 | 1 | ENSMUST00 | 0 | Morn5 | ENSMUSG00000026894.4 |
| chr2 | 77814761 | 77817865 | 29.66345 | 100.21 | 96.63225 | Promoter | 2 | 77411100 | 77816816 | 405717 | 2 | ENSMUST00 | 0 | Zfp385b | ENSMUSG00000027016.12 |
| chr2 | 120352207 | 120353328 | 29.66345 | 100.21 | 96.63225 | Promoter | 2 | 120316461 | 120353125 | 36665 | 2 | ENSMUST00 | 0 | Vps39 | ENSMUSG00000027291.10 |
| chr2 | 153648932 | 153650679 | 29.66345 | 100.21 | 96.63225 | Promoter | 2 | 153649450 | 153687730 | 38281 | 1 | ENSMUST00 | 0 | Dnmt3b | ENSMUSG00000027478.10 |
| chr2 | 165033630 | 165035229 | 29.66345 | 100.21 | 96.63225 | Promoter | 2 | 165000357 | 165034867 | 34511 | 2 | ENSMUST00 | 0 | Ncoa5 | ENSMUSG00000039804.10 |
| chr3 | 90052818 | 90053990 | 29.66345 | 100.21 | 96.63225 | Promoter | 3 | 90052829 | 90058651 | 5823 | 1 | ENSMUST00 | 0 | 4933434E20I | ENSMUSG00000027942.11 |
| chr4 | 132509245 | 132510546 | 29.66345 | 100.21 | 96.63225 | Promoter | 4 | 132492032 | 132510501 | 18470 | 2 | ENSMUST00 | 0 | Sesn2 | ENSMUSG00000028893.8 |
| chr4 | 151988077 | 151989205 | 29.66345 | 100.21 | 96.63225 | Promoter | 4 | 151982639 | 151988997 | 6359 | 2 | ENSMUST00 | 0 | Thap3 | ENSMUSG00000039759.5 |
| chr4 | 117251477 | 117252620 | 29.66345 | 100.21 | 96.63225 | Promoter | 4 | 117251951 | 117268580 | 16630 | 1 | ENSMUST00 | 0 | Tmem53 | ENSMUSG00000048772.10 |
| chr5 | 103691558 | 103694062 | 29.66345 | 100.21 | 96.63225 | Promoter | 5 | 103692374 | 103816428 | 124055 | 1 | ENSMUST00 | 0 | Aff1 | ENSMUSG00000029313.13 |
| chr5 | 117318562 | 117320661 | 29.66345 | 100.21 | 96.63225 | Promoter | 5 | 117319266 | 117355005 | 35740 | 1 | ENSMUST00 | 0 | Vsig10 | ENSMUSG00000066894.9 |
| chr6 | 35177153 | 35178417 | 29.66345 | 100.21 | 96.63225 | Promoter | 6 | 35177430 | 35247449 | 70020 | 1 | ENSMUST00 | 0 | Nup205 | ENSMUSG00000038759.10 |
| chr6 | 71907767 | 71909828 | 29.66345 | 100.21 | 96.63225 | Promoter | 6 | 71909053 | 71979356 | 70304 | 1 | ENSMUST00 | 0 | Polr1a | ENSMUSG00000049553.9 |
| chr7 | 16387340 | 16387863 | 29.66345 | 100.21 | 96.63225 | Promoter | 7 | 16327053 | 16387797 | 60745 | 2 | ENSMUST00 | 0 | Sae1 | ENSMUSG00000052833.8 |
| chr8 | 46617390 | 46619181 | 29.66345 | 100.21 | 96.63225 | Promoter | 8 | 46617449 | 46638706 | 21258 | 1 | ENSMUST00 | 0 | Casp3 | ENSMUSG00000031628.8 |
| chr9 | 59484660 | 59487428 | 29.66345 | 100.21 | 96.63225 | Promoter | 9 | 59388258 | 59486614 | 98357 | 2 | ENSMUST00 | 0 | Arih1 | ENSMUSG00000025234.6 |
| chr9 | 4308674 | 4310563 | 29.66345 | 100.21 | 96.63225 | Promoter | 9 | 4309743 | 4331721 | 21979 | 1 | ENSMUST00 | 0 | Kbtbd3 | ENSMUSG00000025893.7 |
| chr9 | 44733443 | 44735510 | 29.66345 | 100.21 | 96.63225 | Promoter | 9 | 44686308 | 44735198 | 48891 | 2 | ENSMUST00 | 0 | Phldb1 | ENSMUSG00000048537.10 |
| chr7 | 100226921 | 100228355 | 31.31394 | 100.15002 | 96.57744 | Promoter | 7 | 100227394 | 100275358 | 47965 | 1 | ENSMUST00 | 0 | Pgm2l1 | ENSMUSG00000030729.11 |
| chr10 | 128747459 | 128749169 | 32.44658 | 100.11784 | 96.54536 | Promoter | 10 | 128747879 | 128766568 | 18690 | 1 | ENSMUST00 | 0 | Wibg | ENSMUSG00000064030.9 |
| chr10 | 82353887 | 82354857 | 32.44658 | 100.11784 | 96.54536 | Distal Interge | 10 | 82378593 | 82381237 | 2645 | 1 | ENSMUST00 | -23736 | Gm4924 | ENSMUSG00000073427.3 |
| chr11 | 69681078 | 69682590 | 32.44658 | 100.11784 | 96.54536 | Promoter | 11 | 69673115 | 69681846 | 8732 | 2 | ENSMUST00 | 0 | Senp3 | ENSMUSG00000005204.7 |
| chr3 | 88531122 | 88532849 | 32.44658 | 100.11784 | 96.54536 | Promoter | 3 | 88532395 | 88541396 | 9002 | 1 | ENSMUST00 | 0 | Mex3a | ENSMUSG00000074480.4 |
| chr4 | 41712735 | 41714205 | 32.44658 | 100.11784 | 96.54536 | Promoter | 4 | 41712033 | 41713534 | 1502 | 2 | ENSMUST00 | 0 | Rpp25l | ENSMUSG00000036114.3 |
| chr5 | 35104751 | 35106285 | 32.44658 | 100.11784 | 96.54536 | Promoter | 5 | 35091490 | 35105766 | 14277 | 2 | ENSMUST00 | 0 | Lrpap1 | ENSMUSG00000029103.11 |
| chr7 | 25316780 | 25318001 | 32.44658 | 100.11784 | 96.54536 | Promoter | 7 | 25317164 | 25365917 | 48754 | 1 | ENSMUST00 | 0 | Megf8 | ENSMUSG00000045039.9 |
| chr8 | 77548938 | 77550394 | 32.44658 | 100.11784 | 96.54536 | Promoter | 8 | 77549397 | 77581338 | 31942 | 1 | ENSMUST00 | 0 | Prmt10 | ENSMUSG00000037134.11 |
| chr15 | 102202622 | 102204939 | 30.68508 | 100.04181 | 96.47092 | Promoter | 15 | 102203648 | 102215605 | 11958 | 1 | ENSMUST00 | 0 | Zfp740 | ENSMUSG00000046897.11 |
| chr8 | 85079913 | 85081646 | 30.68508 | 100.04181 | 96.47092 | Promoter | 8 | 85075035 | 85080632 | 5598 | 2 | ENSMUST00 | 0 | Wdr83 | ENSMUSG00000005150.11 |
| chr8 | 13889527 | 13890626 | 30.68508 | 100.04181 | 96.47092 | Promoter | 8 | 13884788 | 13890271 | 5484 | 2 | ENSMUST00 | 0 | Coprs | ENSMUSG00000031458.7 |
| chr3 | 75952519 | 75957408 | 32.38164 | 100.00523 | 96.43553 | Promoter | 3 | 75876183 | 75956949 | 80767 | 2 | ENSMUST00 | 0 | Golim4 | ENSMUSG00000034109.10 |
| chr10 | 128909451 | 128910702 | 26.41709 | 100.00125 | 96.43182 | Promoter | 10 | 128909866 | 128912816 | 2951 | 1 | ENSMUST00 | 0 | Cd63 | ENSMUSG00000025351.8 |
| chr12 | 91587910 | 91591057 | 26.41709 | 100.00125 | 96.43182 | Promoter | 12 | 91555916 | 91590009 | 34094 | 2 | ENSMUST00 | 0 | Gtf2a1 | ENSMUSG00000020962.8 |
| chr13 | 59768265 | 59770791 | 26.41709 | 100.00125 | 96.43182 | Promoter | 13 | 59755409 | 59769810 | 14402 | 2 | ENSMUST00 | 0 | Isca1 | ENSMUSG00000044792.6 |
| chr2 | 158624587 | 158625889 | 26.41709 | 100.00125 | 96.43182 | Promoter | 2 | 158624888 | 158629157 | 4270 | 1 | ENSMUST00 | 0 | Actr5 | ENSMUSG00000037761.11 |
| chr3 | 142764529 | 142766473 | 26.41709 | 100.00125 | 96.43182 | Promoter | 3 | 142765226 | 142783603 | 18378 | 1 | ENSMUST00 | 0 | Gtf2b | ENSMUSG00000028271.8 |
| chr5 | 118245193 | 118246258 | 26.41709 | 100.00125 | 96.43182 | Promoter | 5 | 118245227 | 118266093 | 20867 | 1 | ENSMUST00 | 0 | 2410131K14I | ENSMUSG00000032840.7 |
| chr5 | 53997850 | 54000404 | 26.41709 | 100.00125 | 96.43182 | Promoter | 5 | 53998417 | 54121057 | 122641 | 1 | ENSMUST00 | 0 | Stim2 | ENSMUSG00000039156.14 |
| chr8 | 25840092 | 25841976 | 26.41709 | 100.00125 | 96.43182 | Promoter | 8 | 25816191 | 25840779 | 24589 | 2 | ENSMUST00 | 0 | Ash2l | ENSMUSG00000031575.12 |
| chr8 | 95887452 | 95888719 | 26.41709 | 100.00125 | 96.43182 | Promoter | 8 | 95864134 | 95888547 | 24414 | 2 | ENSMUST00 | 0 | Got2 | ENSMUSG00000031672.7 |
| chr9 | 45983749 | 45985474 | 26.41709 | 100.00125 | 96.43182 | Promoter | 9 | 45965311 | 45984871 | 19561 | 2 | ENSMUST00 | 0 | Pafah1b2 | ENSMUSG00000003131.6 |
| chr9 | 40801224 | 40802473 | 26.41709 | 100.00125 | 96.43182 | Promoter | 9 | 40801235 | 40805200 | 3966 | 1 | ENSMUST00 | 0 | Hspa8 | ENSMUSG00000015656.11 |
| chr2 | 32287238 | 32289480 | 31.80086 | 99.99902 | 96.43139 | Promoter | 2 | 32287384 | 32297776 | 10393 | 1 | ENSMUST00 | 0 | Golga2 | ENSMUSG00000002546.11 |
| chr15 | 76870579 | 76871809 | 32.93275 | 99.9547 | 96.38725 | Promoter | 15 | 76852148 | 76871435 | 19288 | 2 | ENSMUST00 | 0 | Zfp251 | ENSMUSG00000022526.7 |
| chr16 | 13436798 | 13438361 | 32.93275 | 99.9547 | 96.38725 | Distal Interge | 16 | 13449523 | 13449601 | 79 | 1 | ENSMUST00 | -11162 | Mir193b | ENSMUSG00000076972.2 |
| chr1 | 75478421 | 75480081 | 27.88725 | 99.90214 | 96.33501 | Promoter | 1 | 75474569 | 75479271 | 4703 | 2 | ENSMUST00 | 0 | Chpf | ENSMUSG00000032997.11 |
| chr1 | 86703168 | 86704679 | 27.88725 | 99.90214 | 96.33501 | Promoter | 1 | 86703808 | 86748331 | 44524 | 1 | ENSMUST00 | 0 | Dis3l2 | ENSMUSG00000053333.10 |
| chr10 | 80622599 | 80624420 | 27.88725 | 99.90214 | 96.33501 | Promoter | 10 | 80622780 | 80640771 | 17992 | 1 | ENSMUST00 | 0 | Csnk1g2 | ENSMUSG00000003345.10 |
| chr10 | 127030208 | 127030924 | 27.88725 | 99.90214 | 96.33501 | Promoter | 10 | 127011572 | 127030803 | 19232 | 2 | ENSMUST00 | 0 | Tsfm | ENSMUSG00000040521.6 |
| chr11 | 69408142 | 69412281 | 27.88725 | 99.90214 | 96.33501 | 5' UTR | 11 | 69398508 | 69413675 | 15168 | 2 | ENSMUST00 | 1394 | Kdm6b | ENSMUSG00000018476.7 |
| chr14 | 14011969 | 14013852 | 27.88725 | 99.90214 | 96.33501 | Promoter | 14 | 14012491 | 14107296 | 94806 | 1 | ENSMUST00 | 0 | Atxn7 | ENSMUSG00000021738.4 |
| chr14 | 45656729 | 45658743 | 27.88725 | 99.90214 | 96.33501 | Promoter | 14 | 45593174 | 45658143 | 64970 | 2 | ENSMUST00 | 0 | Ddhd1 | ENSMUSG00000037697.13 |
| chr17 | 53566162 | 53568773 | 27.88725 | 99.90214 | 96.33501 | Promoter | 17 | 53566861 | 53624491 | 57631 | 1 | ENSMUST00 | 0 | Kat2b | ENSMUSG00000000708.9 |
| chr17 | 33718047 | 33718789 | 27.88725 | 99.90214 | 96.33501 | Promoter | 17 | 33685692 | 33718670 | 32979 | 2 | ENSMUST00 | 0 | Mar-02 | ENSMUSG00000079557.5 |
| chr18 | 21001005 | 21001722 | 27.88725 | 99.90214 | 96.33501 | Promoter | 18 | 21001341 | 21028223 | 26883 | 1 | ENSMUST00 | 0 | Rnf138 | ENSMUSG00000024317.9 |
| chr18 | 37643084 | 37644787 | 27.88725 | 99.90214 | 96.33501 | Promoter | 18 | 37640902 | 37644204 | 3303 | 2 | ENSMUST00 | 0 | Taf7 | ENSMUSG00000051316.7 |
| chr18 | 50052968 | 50055720 | 27.88725 | 99.90214 | 96.33501 | Promoter | 18 | 50053282 | 50091782 | 38501 | 1 | ENSMUST00 | 0 | Tnfaip8 | ENSMUSG00000062210.8 |
| chr19 | 17835376 | 17838194 | 27.88725 | 99.90214 | 96.33501 | Promoter | 19 | 17432832 | 17837632 | 404801 | 2 | ENSMUST00 | 0 | Pcsk5 | ENSMUSG00000024713.10 |
| chr2 | 75659232 | 75661067 | 27.88725 | 99.90214 | 96.33501 | Promoter | 2 | 75659261 | 75665839 | 6579 | 1 | ENSMUST00 | 0 | Hnrnpa3 | ENSMUSG00000059005.8 |
| chr2 | 69860961 | 69862760 | 27.88725 | 99.90214 | 96.33501 | Promoter | 2 | 69861562 | 69871786 | 10225 | 1 | ENSMUST00 | 0 | Ssb | ENSMUSG00000068882.8 |
| chr4 | 45341634 | 45343307 | 27.88725 | 99.90214 | 96.33501 | Promoter | 4 | 45316954 | 45342732 | 25779 | 2 | ENSMUST00 | 0 | Exosc3 | ENSMUSG00000028322.6 |
| chr4 | 151043578 | 151044724 | 27.88725 | 99.90214 | 96.33501 | Promoter | 4 | 151003652 | 151044665 | 41014 | 2 | ENSMUST00 | 0 | Per3 | ENSMUSG00000028957.7 |
| chr4 | 155600964 | 155602448 | 27.88725 | 99.90214 | 96.33501 | Promoter | 4 | 155601416 | 155623340 | 21925 | 1 | ENSMUST00 | 0 | Slc35e2 | ENSMUSG00000042202.10 |
| chr5 | 122158261 | 122159841 | 27.88725 | 99.90214 | 96.33501 | Promoter | 5 | 122158278 | 122175273 | 16996 | 1 | ENSMUST00 | 0 | Ppp1cc | ENSMUSG00000004455.11 |
| chr5 | 91961707 | 91963799 | 27.88725 | 99.90214 | 96.33501 | Promoter | 5 | 91948904 | 91962556 | 13653 | 2 | ENSMUST00 | 0 | Rchy1 | ENSMUSG00000029397.10 |

| | | | | | | | | | | | | | | | |
|---|---|---|---|---|---|---|---|---|---|---|---|---|---|---|---|
| chr5 | 125475256 | 125476543 | 27.88725 | 99.90214 | 96.33501 | Promoter | 5 | 125475814 | 125517412 | 41599 | 1 | ENSMUST00 | 0 | Aacs | ENSMUSG00000029482.3 |
| chr5 | 144254325 | 144259060 | 27.88725 | 99.90214 | 96.33501 | Promoter | 5 | 144255225 | 144446757 | 191533 | 1 | ENSMUST00 | 0 | Bri3 | ENSMUSG00000047843.11 |
| chr6 | 88724035 | 88725421 | 27.88725 | 99.90214 | 96.33501 | Promoter | 6 | 88724412 | 88828360 | 103949 | 1 | ENSMUST00 | 0 | Mgll | ENSMUSG00000033174.12 |
| chr6 | 48395103 | 48396285 | 27.88725 | 99.90214 | 96.33501 | Promoter | 6 | 48395586 | 48419545 | 23960 | 1 | ENSMUST00 | 0 | Krba1 | ENSMUSG00000042810.9 |
| chr7 | 35753775 | 35754912 | 27.88725 | 99.90214 | 96.33501 | Promoter | 7 | 35685165 | 35754454 | 69290 | 2 | ENSMUST00 | 0 | Dpy19l3 | ENSMUSG00000043671.9 |
| chr7 | 144941140 | 144943434 | 27.88725 | 99.90214 | 96.33501 | Distal Interge | 7 | 144929931 | 144939869 | 9939 | 2 | ENSMUST00 | -1272 | Ccnd1 | ENSMUSG00000070348.4 |
| chr8 | 33731482 | 33733039 | 32.31696 | 99.89304 | 96.32935 | Promoter | 8 | 33731833 | 33777164 | 45332 | 1 | ENSMUST00 | 0 | Gtf2e2 | ENSMUSG00000031585.8 |
| chr6 | 87887849 | 87888619 | 31.7393 | 99.89149 | 96.32799 | Promoter | 6 | 87887928 | 87890759 | 2832 | 1 | ENSMUST00 | 0 | Copg1 | ENSMUSG00000030058.12 |
| chr6 | 15720519 | 15722168 | 31.7393 | 99.89149 | 96.32799 | Promoter | 6 | 15720661 | 15802165 | 81505 | 1 | ENSMUST00 | 0 | Mdfic | ENSMUSG00000041390.13 |
| chr13 | 52928873 | 52930167 | 21.95811 | 99.8556 | 96.2922 | Promoter | 13 | 52835119 | 52929661 | 94543 | 2 | ENSMUST00 | 0 | Auh | ENSMUSG00000021460.11 |
| chr19 | 18670839 | 18672337 | 21.95811 | 99.8556 | 96.2922 | Promoter | 19 | 18670780 | 18704788 | 34009 | 1 | ENSMUST00 | 60 | 2410127L17I | ENSMUSG00000024726.9 |
| chr15 | 75968756 | 75969982 | 31.1379 | 99.84005 | 96.2767 | Promoter | 15 | 75969123 | 75975749 | 6627 | 1 | ENSMUST00 | 0 | Zfp707 | ENSMUSG00000034429.9 |
| chr17 | 70989464 | 70991264 | 29.99031 | 99.80121 | 96.2382 | Promoter | 17 | 70973920 | 70990787 | 16868 | 2 | ENSMUST00 | 0 | Myl12b | ENSMUSG00000034868.8 |
| chr8 | 18845670 | 18847232 | 29.99031 | 99.80121 | 96.2382 | Promoter | 8 | 18846277 | 18891361 | 45085 | 1 | ENSMUST00 | 0 | Agpat5 | ENSMUSG00000031467.5 |
| chr12 | 86082275 | 86083475 | 32.25254 | 99.78123 | 96.21911 | Promoter | 12 | 86082561 | 86162458 | 79898 | 1 | ENSMUST00 | 0 | Ift43 | ENSMUSG00000007867.8 |
| chr5 | 64229858 | 64231190 | 32.25254 | 99.78123 | 96.21911 | Promoter | 5 | 64230299 | 64351482 | 121184 | 1 | ENSMUST00 | 0 | Tbc1d1 | ENSMUSG00000029174.13 |
| chr2 | 27514311 | 27516190 | 31.61691 | 99.67752 | 96.11594 | Promoter | 2 | 27515157 | 27520653 | 5497 | 1 | ENSMUST00 | 0 | Wdr5 | ENSMUSG00000026917.10 |
| chr3 | 8922912 | 8924319 | 31.61691 | 99.67752 | 96.11594 | Promoter | 3 | 8802146 | 8923918 | 121773 | 2 | ENSMUST00 | 0 | Mrps28 | ENSMUSG00000040269.5 |
| chr5 | 135688309 | 135690083 | 31.61691 | 99.67752 | 96.11594 | Promoter | 5 | 135689036 | 135735326 | 46291 | 1 | ENSMUST00 | 0 | Por | ENSMUSG00000005514.9 |
| chr8 | 105325610 | 105327102 | 31.61691 | 99.67752 | 96.11594 | Promoter | 8 | 105326354 | 105329057 | 2704 | 1 | ENSMUST00 | 0 | Tmem208 | ENSMUSG00000014856.7 |
| chr9 | 121704441 | 121705820 | 31.61691 | 99.67752 | 96.11594 | Promoter | 9 | 121680047 | 121705029 | 24983 | 2 | ENSMUST00 | 0 | Sec22c | ENSMUSG00000061536.7 |
| chr11 | 93954825 | 93956484 | 32.18837 | 99.66983 | 96.10942 | Promoter | 11 | 93949814 | 93956259 | 6446 | 2 | ENSMUST00 | 0 | Nme2 | ENSMUSG00000020857.6 |
| chr17 | 22314221 | 22315478 | 32.18837 | 99.66983 | 96.10942 | Distal Interge | 17 | 22337989 | 22361400 | 23412 | 2 | ENSMUST00 | 45922 | Zfp944 | ENSMUSG00000033972.9 |
| chr11 | 30024227 | 30026706 | 29.34276 | 99.62727 | 96.06711 | Promoter | 11 | 29743048 | 30026033 | 282986 | 2 | ENSMUST00 | 0 | Eml6 | ENSMUSG00000044072.9 |
| chr12 | 111537199 | 111537960 | 29.34276 | 99.62727 | 96.06711 | Promoter | 12 | 111538101 | 111546753 | 8653 | 1 | ENSMUST00 | -141 | Eif5 | ENSMUSG00000021282.11 |
| chr5 | 143111903 | 143113458 | 29.34276 | 99.62727 | 96.06711 | Promoter | 5 | 142990893 | 143113020 | 122128 | 2 | ENSMUST00 | 0 | Rnf216 | ENSMUSG00000045078.8 |
| chr9 | 121759121 | 121761008 | 32.73014 | 99.60502 | 96.04542 | Promoter | 9 | 121760033 | 121771742 | 11710 | 1 | ENSMUST00 | 0 | Zfp651 | ENSMUSG00000013419.6 |
| chr11 | 84915665 | 84916848 | 31.55606 | 99.5711 | 96.01173 | Promoter | 11 | 84910950 | 84916366 | 5417 | 2 | ENSMUST00 | 0 | Znhit3 | ENSMUSG00000020526.7 |
| chr4 | 153974348 | 153976041 | 28.73777 | 99.51218 | 95.95325 | Promoter | 4 | 153964449 | 153975126 | 10678 | 2 | ENSMUST00 | 0 | Dffb | ENSMUSG00000029027.3 |
| chr14 | 56886554 | 56888224 | 23.85991 | 99.51073 | 95.95212 | Promoter | 14 | 56887795 | 56962579 | 74785 | 1 | ENSMUST00 | 0 | Zmym2 | ENSMUSG00000021945.8 |
| chr17 | 56584951 | 56586211 | 23.85991 | 99.51073 | 95.95212 | Promoter | 17 | 56584974 | 56606219 | 21246 | 1 | ENSMUST00 | 0 | Safb | ENSMUSG00000071054.7 |
| chr2 | 131491475 | 131492673 | 23.85991 | 99.51073 | 95.95212 | Promoter | 2 | 131491496 | 131520243 | 28748 | 1 | ENSMUST00 | 0 | Smox | ENSMUSG00000027333.13 |
| chr3 | 37639368 | 37642407 | 23.85991 | 99.51073 | 95.95212 | Promoter | 3 | 37639947 | 37644598 | 4652 | 1 | ENSMUST00 | 0 | Spry1 | ENSMUSG00000037211.7 |
| chr7 | 34195961 | 34197031 | 23.85991 | 99.51073 | 95.95212 | Promoter | 7 | 34184499 | 34196661 | 12163 | 2 | ENSMUST00 | 0 | Pdcd2l | ENSMUSG00000002635.6 |
| chr7 | 89979712 | 89981930 | 23.85991 | 99.51073 | 95.95212 | Promoter | 7 | 89954656 | 89980983 | 26328 | 2 | ENSMUST00 | 0 | Eed | ENSMUSG00000030619.8 |
| chr10 | 127059084 | 127060802 | 32.66316 | 99.48935 | 95.93111 | Promoter | 10 | 127055011 | 127060184 | 5174 | 2 | ENSMUST00 | 0 | Mar-09 | ENSMUSG00000040502.5 |
| chr13 | 24280200 | 24281526 | 32.66316 | 99.48935 | 95.93111 | Promoter | 13 | 24012659 | 24280795 | 268137 | 2 | ENSMUST00 | 0 | Lrrc16a | ENSMUSG00000021338.12 |
| chr18 | 37805761 | 37807729 | 32.66316 | 99.48935 | 95.93111 | Promoter | 18 | 37806410 | 37841873 | 35464 | 1 | ENSMUST00 | 0 | Pcdhga12 | ENSMUSG00000023036.9 |
| chr2 | 25500351 | 25502049 | 32.66316 | 99.48935 | 95.93111 | Promoter | 2 | 25498651 | 25500613 | 1963 | 2 | ENSMUST00 | 0 | C8g | ENSMUSG00000015083.6 |
| chr6 | 83124497 | 83125481 | 32.66316 | 99.48935 | 95.93111 | Promoter | 6 | 83121765 | 83125431 | 3667 | 2 | ENSMUST00 | 0 | Ino80b | ENSMUSG00000030034.6 |
| chr7 | 39588355 | 39589139 | 32.66316 | 99.48935 | 95.93111 | Promoter | 7 | 39588931 | 39589717 | 787 | 1 | ENSMUST00 | 0 | Gm2058 | ENSMUSG00000074903.5 |
| chr1 | 121567130 | 121568339 | 31.49545 | 99.46503 | 95.90771 | Promoter | 1 | 121553835 | 121567989 | 14155 | 2 | ENSMUST00 | 0 | Ddx18 | ENSMUSG00000001674.6 |
| chr15 | 6707265 | 6709859 | 31.49545 | 99.46503 | 95.90771 | Promoter | 15 | 6708381 | 6800398 | 92018 | 1 | ENSMUST00 | 0 | Rictor | ENSMUSG00000050310.8 |
| chr1 | 184033118 | 184034192 | 27.13995 | 99.45934 | 95.90212 | Promoter | 1 | 184033030 | 184034301 | 1272 | 2 | ENSMUST00 | 109 | 1700056E22I | ENSMUSG00000044854.6 |
| chr15 | 76068809 | 76069842 | 27.13995 | 99.45934 | 95.90212 | Promoter | 15 | 76047158 | 76069754 | 22597 | 2 | ENSMUST00 | 0 | Scrib | ENSMUSG00000022568.11 |
| chr10 | 81642746 | 81644258 | 32.06079 | 99.44822 | 95.89121 | Promoter | 10 | 81643803 | 81647474 | 3672 | 1 | ENSMUST00 | 0 | Ankrd24 | ENSMUSG00000054708.12 |
| chr2 | 11500699 | 11503640 | 31.43507 | 99.35931 | 95.8029 | Promoter | 2 | 11471430 | 11502025 | 30596 | 2 | ENSMUST00 | 0 | Pfkfb3 | ENSMUSG00000026773.14 |
| chr1 | 93477516 | 93480357 | 31.43325 | 99.35611 | 95.79983 | Promoter | 1 | 93478964 | 93510260 | 31297 | 1 | ENSMUST00 | 0 | Sep-02 | ENSMUSG00000026276.12 |
| chr1 | 156938566 | 156940198 | 31.43325 | 99.35611 | 95.79983 | Promoter | 1 | 156804166 | 156939602 | 135437 | 2 | ENSMUST00 | 0 | Ralgps2 | ENSMUSG00000026594.9 |
| chr10 | 88378807 | 88380508 | 31.43325 | 99.35611 | 95.79983 | Promoter | 10 | 88340563 | 88379080 | 38518 | 2 | ENSMUST00 | 0 | Dram1 | ENSMUSG00000020057.2 |
| chr10 | 128803556 | 128806228 | 31.43325 | 99.35611 | 95.79983 | Promoter | 10 | 128805676 | 128819446 | 13771 | 1 | ENSMUST00 | 0 | Dnajc14 | ENSMUSG00000025354.4 |
| chr10 | 117843919 | 117846716 | 31.43325 | 99.35611 | 95.79983 | Promoter | 10 | 117814471 | 117845935 | 31465 | 2 | ENSMUST00 | 0 | Rap1b | ENSMUSG00000052681.7 |
| chr11 | 80208996 | 80210259 | 31.43325 | 99.35611 | 95.79983 | Promoter | 11 | 80209019 | 80266812 | 57794 | 1 | ENSMUST00 | 0 | Rhot1 | ENSMUSG00000017686.11 |
| chr11 | 86200505 | 86202460 | 31.43325 | 99.35611 | 95.79983 | Promoter | 11 | 86058138 | 86201193 | 143056 | 2 | ENSMUST00 | 0 | Brip1 | ENSMUSG00000034329.11 |
| chr11 | 115366826 | 115368055 | 31.43325 | 99.35611 | 95.79983 | Promoter | 11 | 115347707 | 115367719 | 20013 | 2 | ENSMUST00 | 0 | Hid1 | ENSMUSG00000034586.9 |
| chr14 | 122912508 | 122913604 | 31.43325 | 99.35611 | 95.79983 | Promoter | 14 | 122816223 | 122913120 | 96898 | 2 | ENSMUST00 | 0 | Ggact | ENSMUSG00000041625.9 |
| chr16 | 8636510 | 8638437 | 31.43325 | 99.35611 | 95.79983 | Promoter | 16 | 8637674 | 8657531 | 19858 | 1 | ENSMUST00 | 0 | Pmm2 | ENSMUSG00000022711.10 |
| chr17 | 46889042 | 46890426 | 31.43325 | 99.35611 | 95.79983 | Promoter | 17 | 46889216 | 46890405 | 1190 | 2 | ENSMUST00 | 0 | A330017A19 | ENSMUSG00000073388.2 |
| chr19 | 4962075 | 4962761 | 31.43325 | 99.35611 | 95.79983 | Promoter | 19 | 4962306 | 4966995 | 4690 | 1 | ENSMUST00 | 0 | Mrpl11 | ENSMUSG00000024902.4 |
| chr3 | 7366038 | 7368264 | 31.43325 | 99.35611 | 95.79983 | Promoter | 3 | 7366604 | 7445366 | 78763 | 1 | ENSMUST00 | 0 | Pkia | ENSMUSG00000027499.7 |
| chr3 | 58690452 | 58692965 | 31.43325 | 99.35611 | 95.79983 | Promoter | 3 | 58674938 | 58692400 | 17463 | 2 | ENSMUST00 | 0 | Siah2 | ENSMUSG00000036432.8 |
| chr7 | 56018956 | 56020119 | 31.43325 | 99.35611 | 95.79983 | Promoter | 7 | 55977567 | 56019954 | 42388 | 2 | ENSMUST00 | 0 | Nipa1 | ENSMUSG00000047037.5 |
| chr7 | 80260738 | 80262097 | 31.43325 | 99.35611 | 95.79983 | Promoter | 7 | 80261215 | 80265374 | 4160 | 1 | ENSMUST00 | 0 | Ngrn | ENSMUSG00000047084.8 |
| chr8 | 66485738 | 66487009 | 31.43325 | 99.35611 | 95.79983 | Promoter | 8 | 66474330 | 66486497 | 12168 | 2 | ENSMUST00 | 0 | Tma16 | ENSMUSG00000025591.5 |
| chr9 | 59539139 | 59540326 | 31.43325 | 99.35611 | 95.79983 | Promoter | 9 | 59539667 | 59565105 | 25439 | 1 | ENSMUST00 | 0 | Hexa | ENSMUSG00000025232.7 |
| chr9 | 86464292 | 86465201 | 31.43325 | 99.35611 | 95.79983 | Promoter | 9 | 86307306 | 86464950 | 157645 | 2 | ENSMUST00 | 0 | Ube2cbp | ENSMUSG00000032415.9 |
| chr9 | 72806838 | 72808468 | 31.43325 | 99.35611 | 95.79983 | Promoter | 9 | 72806874 | 72917291 | 110418 | 1 | ENSMUST00 | 0 | Prtg | ENSMUSG00000036030.9 |
| chr1 | 189921639 | 189922858 | 30.84885 | 99.33006 | 95.77879 | Promoter | 1 | 189880492 | 189922363 | 41872 | 2 | ENSMUST00 | 0 | Smyd2 | ENSMUSG00000026603.8 |
| chr10 | 80319474 | 80320632 | 30.84885 | 99.33006 | 95.77879 | Promoter | 10 | 80318254 | 80320537 | 2284 | 2 | ENSMUST00 | 0 | 2310011J03I | ENSMUSG00000020133.8 |
| chr11 | 62878821 | 62880274 | 30.84885 | 99.33006 | 95.77879 | Promoter | 11 | 62879455 | 62895185 | 15731 | 1 | ENSMUST00 | 0 | Tvp23b | ENSMUSG00000014177.5 |
| chr18 | 60525701 | 60527274 | 30.84885 | 99.33006 | 95.77879 | Promoter | 18 | 60526221 | 60558762 | 32542 | 1 | ENSMUST00 | 0 | Dctn4 | ENSMUSG00000024603.7 |
| chr7 | 45872722 | 45874235 | 30.84885 | 99.33006 | 95.77879 | Promoter | 7 | 45872772 | 45883731 | 10960 | 1 | ENSMUST00 | 0 | Kdelr1 | ENSMUSG00000002778.7 |
| chr9 | 108936628 | 108937631 | 30.84885 | 99.33006 | 95.77879 | Promoter | 9 | 108936648 | 108949640 | 12993 | 1 | ENSMUST00 | 0 | Uqcrc1 | ENSMUSG00000025651.9 |
| chr15 | 101223942 | 101225305 | 32.53002 | 99.25928 | 95.7094 | Promoter | 15 | 101224207 | 101232755 | 8549 | 1 | ENSMUST00 | 0 | Grasp | ENSMUSG00000000531.4 |
| chr5 | 100673438 | 100675140 | 32.53002 | 99.25928 | 95.7094 | Promoter | 5 | 100654723 | 100674284 | 19562 | 2 | ENSMUST00 | 0 | Coq2 | ENSMUSG00000029319.7 |
| chr11 | 59449770 | 59451008 | 31.93423 | 99.22817 | 95.67865 | Promoter | 11 | 59407134 | 59449999 | 42866 | 2 | ENSMUST00 | 0 | Snap47 | ENSMUSG00000009894.10 |
| chr19 | 4927464 | 4928856 | 31.93423 | 99.22817 | 95.67865 | Promoter | 19 | 4907229 | 4928287 | 21059 | 2 | ENSMUST00 | 0 | Dpp3 | ENSMUSG00000063904.3 |
| chr2 | 132247043 | 132248310 | 31.93423 | 99.22817 | 95.67865 | Promoter | 2 | 132239492 | 132247654 | 8163 | 2 | ENSMUST00 | 0 | Tmem230 | ENSMUSG00000027341.5 |
| chr7 | 105787188 | 105788106 | 31.93423 | 99.22817 | 95.67865 | Promoter | 7 | 105752989 | 105787550 | 34562 | 2 | ENSMUST00 | 0 | Dchs1 | ENSMUSG00000036862.9 |
| chr1 | 88007807 | 88008854 | 33.08631 | 99.21803 | 95.66874 | Promoter | 1 | 87945121 | 88008551 | 63431 | 2 | ENSMUST00 | 0 | Usp40 | ENSMUSG00000005501.9 |
| chr11 | 74837494 | 74839155 | 33.08631 | 99.21803 | 95.66874 | Promoter | 11 | 74837468 | 74843775 | 6308 | 1 | ENSMUST00 | 27 | Mnt | ENSMUSG00000000282.7 |
| chr11 | 85138661 | 85140048 | 33.08631 | 99.21803 | 95.66874 | Promoter | 11 | 84984442 | 85140038 | 155597 | 2 | ENSMUST00 | 0 | Usp32 | ENSMUSG00000000804.9 |
| chr14 | 25767764 | 25769441 | 33.08631 | 99.21803 | 95.66874 | Promoter | 14 | 25711642 | 25769039 | 57398 | 2 | ENSMUST00 | 0 | Zcchc24 | ENSMUSG00000055538.7 |
| chr14 | 115040106 | 115042806 | 33.08631 | 99.21803 | 95.66874 | Promoter | 14 | 115042879 | 115046727 | 3849 | 1 | ENSMUST00 | -73 | Mir17hg | ENSMUSG00000089726.2 |
| chr4 | 43668905 | 43670089 | 33.08631 | 99.21803 | 95.66874 | Promoter | 4 | 43659622 | 43669141 | 9520 | 2 | ENSMUST00 | 0 | Fam221b | ENSMUSG00000043633.6 |
| chr8 | 12756624 | 12758223 | 33.08631 | 99.21803 | 95.66874 | Promoter | 8 | 12757014 | 12868728 | 111715 | 1 | ENSMUST00 | 0 | Atp11a | ENSMUSG00000031441.10 |
| chr6 | 148895238 | 148896628 | 31.87132 | 99.11874 | 95.57203 | Promoter | 6 | 148842492 | 148896237 | 53746 | 2 | ENSMUST00 | 0 | Caprin2 | ENSMUSG00000030309.11 |
| chr13 | 69738105 | 69740890 | 33.0268 | 99.11563 | 95.56912 | Promoter | 13 | 69702835 | 69739897 | 37063 | 2 | ENSMUST00 | 0 | Ube2ql1 | ENSMUSG00000052981.5 |
| chr9 | 64281205 | 64282333 | 33.0268 | 99.11563 | 95.56912 | Promoter | 9 | 64281607 | 64304792 | 23186 | 1 | ENSMUST00 | 0 | Tipin | ENSMUSG00000032397.6 |
| chr10 | 67547675 | 67550046 | 30.13579 | 99.06311 | 95.51676 | Promoter | 10 | 67544512 | 67548955 | 4444 | 2 | ENSMUST00 | 0 | Ado | ENSMUSG00000057134.4 |
| chr2 | 168954334 | 168956001 | 30.13579 | 99.06311 | 95.51676 | Promoter | 2 | 168893331 | 168955587 | 62257 | 2 | ENSMUST00 | 0 | Zfp64 | ENSMUSG00000027551.10 |
| chr2 | 92431256 | 92434116 | 30.13579 | 99.06311 | 95.51676 | Promoter | 2 | 92403646 | 92434043 | 30398 | 2 | ENSMUST00 | 0 | Cry2 | ENSMUSG00000068742.6 |
| chr2 | 34799175 | 34800304 | 30.13579 | 99.06311 | 95.51676 | Promoter | 2 | 34777556 | 34799912 | 22357 | 2 | ENSMUST00 | 0 | Rabepk | ENSMUSG00000070953.8 |
| chr4 | 45184329 | 45185591 | 30.13579 | 99.06311 | 95.51676 | Promoter | 4 | 45184875 | 45261900 | 77026 | 1 | ENSMUST00 | 0 | Frmpd1 | ENSMUSG00000035615.7 |
| chr8 | 108714360 | 108715770 | 30.13579 | 99.06311 | 95.51676 | Promoter | 8 | 108714644 | 108961630 | 246987 | 1 | ENSMUST00 | 0 | Zfhx3 | ENSMUSG00000038872.9 |
| chr7 | 140855492 | 140858230 | 32.39795 | 99.03095 | 95.4856 | Promoter | 7 | 140853384 | 140856383 | 3000 | 2 | ENSMUST00 | 0 | Bet1l | ENSMUSG00000025484.8 |
| chr10 | 79915955 | 79917517 | 31.80867 | 99.00969 | 95.46461 | Promoter | 10 | 79916971 | 79922273 | 5303 | 1 | ENSMUST00 | 0 | Kiss1r | ENSMUSG00000035773.5 |
| chr11 | 85352326 | 85353737 | 31.80867 | 99.00969 | 95.46461 | Promoter | 11 | 85353167 | 85510991 | 157825 | 1 | ENSMUST00 | 0 | Bcas3 | ENSMUSG00000059439.10 |
| chr17 | 51811751 | 51813952 | 31.80866 | 99.00968 | 95.46461 | Promoter | 17 | 51739476 | 51812680 | 73205 | 2 | ENSMUST00 | 0 | Satb1 | ENSMUSG00000023927.10 |
| chr3 | 41083359 | 41084082 | 31.80867 | 99.00969 | 95.46461 | Promoter | 3 | 41066326 | 41083046 | 16721 | 2 | ENSMUST00 | -314 | Pgrmc2 | ENSMUSG00000049940.7 |
| chr3 | 123507313 | 123509354 | 31.80867 | 99.00969 | 95.46461 | Promoter | 3 | 123507937 | 123508066 | 130 | 2 | ENSMUST00 | 0 | Snora24 | ENSMUSG00000092730.1 |
| chr8 | 70522446 | 70523697 | 31.80867 | 99.00969 | 95.46461 | Promoter | 8 | 70508272 | 70523163 | 14892 | 2 | ENSMUST00 | 0 | Kxd1 | ENSMUSG00000055553.11 |
| chr4 | 80910516 | 80912537 | 32.95766 | 98.99662 | 95.4528 | Promoter | 4 | 80910646 | 80955628 | 44983 | 1 | ENSMUST00 | 0 | Lurap1l | ENSMUSG00000048706.3 |
| chr2 | 26020809 | 26022200 | 31.19586 | 98.93996 | 95.39634 | Promoter | 2 | 25998543 | 26021747 | 23205 | 2 | ENSMUST00 | 0 | Ubac1 | ENSMUSG00000036352.11 |
| chr11 | 59847557 | 59848814 | 32.33232 | 98.9174 | 95.3739 | Promoter | 11 | 59848073 | 59876533 | 28461 | 1 | ENSMUST00 | 0 | Nt5m | ENSMUSG00000032615.9 |
| chr17 | 27056273 | 27059567 | 32.33232 | 98.9174 | 95.3739 | Promoter | 17 | 27057304 | 27122223 | 64920 | 1 | ENSMUST00 | 0 | Itpr3 | ENSMUSG00000042644.8 |
| chr5 | 124477289 | 124478449 | 32.33232 | 98.9174 | 95.3739 | Promoter | 5 | 124463265 | 124478366 | 15102 | 2 | ENSMUST00 | 0 | Rilpl2 | ENSMUSG00000029401.7 |
| chr11 | 113750511 | 113752445 | 32.8888 | 98.87808 | 95.33504 | Promoter | 11 | 113726850 | 113751318 | 24469 | 2 | ENSMUST00 | 0 | Cdc42ep4 | ENSMUSG00000041598.7 |
| chr13 | 63430637 | 63432256 | 32.8888 | 98.87808 | 95.33504 | Promoter | 13 | 63304709 | 63431745 | 127037 | 2 | ENSMUST00 | 0 | Fancc | ENSMUSG00000021461.11 |
| chr16 | 30586693 | 30588498 | 32.8888 | 98.87808 | 95.33504 | Promoter | 16 | 30560494 | 30587592 | 27099 | 2 | ENSMUST00 | 0 | Lsg1 | ENSMUSG00000022538.14 |
| chr19 | 29521423 | 29523832 | 32.8888 | 98.87808 | 95.33504 | Promoter | 19 | 29522282 | 29606829 | 84548 | 1 | ENSMUST00 | 0 | C030046E11I | ENSMUSG00000038658.10 |
| chr3 | 135485185 | 135486284 | 32.8888 | 98.87808 | 95.33504 | Promoter | 3 | 135485611 | 135555566 | 69956 | 1 | ENSMUST00 | 0 | Manba | ENSMUSG00000028164.10 |
| chr1 | 39576052 | 39577945 | 29.47202 | 98.86652 | 95.32386 | Promoter | 1 | 39551296 | 39577405 | 26110 | 2 | ENSMUST00 | 0 | Rnf149 | ENSMUSG00000048234.8 |
| chr10 | 80292108 | 80293604 | 29.47202 | 98.86652 | 95.32386 | Promoter | 10 | 80292453 | 80294114 | 1662 | 1 | ENSMUST00 | 0 | Rps15 | ENSMUSG00000063457.9 |
| chr11 | 97703384 | 97704134 | 29.47202 | 98.86652 | 95.32386 | Promoter | 11 | 97703399 | 97713500 | 10102 | 1 | ENSMUST00 | 0 | Psmb3 | ENSMUSG00000069744.7 |
| chr19 | 6306058 | 6307381 | 29.47202 | 98.86652 | 95.32386 | Promoter | 19 | 6306456 | 6325652 | 19197 | 1 | ENSMUST00 | 0 | Cdc42bpg | ENSMUSG00000024769.7 |
| chr2 | 91962693 | 91963743 | 29.47202 | 98.86652 | 95.32386 | Promoter | 2 | 91932826 | 91963657 | 30832 | 2 | ENSMUST00 | 0 | Dgkz | ENSMUSG00000040479.6 |
| chr6 | 134897406 | 134899080 | 29.47202 | 98.86652 | 95.32386 | Promoter | 6 | 134869093 | 134897815 | 28723 | 2 | ENSMUST00 | 0 | Gpr19 | ENSMUSG00000032641.12 |
| chr11 | 64434865 | 64438804 | 22.72043 | 98.85921 | 95.31703 | Promoter | 11 | 64435332 | 64522841 | 87510 | 1 | ENSMUST00 | 0 | Hs3st3a1 | ENSMUSG00000047759.6 |
| chr8 | 26157472 | 26159346 | 22.72043 | 98.85921 | 95.31703 | Promoter | 8 | 26158212 | 26162940 | 4729 | 1 | ENSMUST00 | 0 | Thap1 | ENSMUSG00000037214.6 |
| chr1 | 15805135 | 15806856 | 24.89402 | 98.85246 | 95.31046 | Promoter | 1 | 15805646 | 15844052 | 38407 | 1 | ENSMUST00 | 0 | Terf1 | ENSMUSG00000025925.9 |
| chr1 | 60180506 | 60181850 | 24.89402 | 98.85246 | 95.31046 | Promoter | 1 | 60180599 | 60338328 | 157730 | 1 | ENSMUST00 | 0 | Nbeal1 | ENSMUSG00000073664.6 |
| chr10 | 52380966 | 52382996 | 24.89402 | 98.85246 | 95.31046 | Promoter | 10 | 52337024 | 52382124 | 45101 | 2 | ENSMUST00 | 0 | Gopc | ENSMUSG00000019861.9 |
| chr10 | 82127649 | 82128617 | 24.89402 | 98.85246 | 95.31046 | Distal Interge | 10 | 82138253 | 82152011 | 13759 | 1 | ENSMUST00 | -9636 | AU041133 | ENSMUSG00000078435.2 |
| chr12 | 85823896 | 85825603 | 24.89402 | 98.85246 | 95.31046 | Promoter | 12 | 85824659 | 85857405 | 32747 | 1 | ENSMUST00 | 0 | Ttll5 | ENSMUSG00000012609.13 |
| chr16 | 49854841 | 49857032 | 24.89402 | 98.85246 | 95.31046 | Promoter | 16 | 49855618 | 49915010 | 59393 | 1 | ENSMUST00 | 0 | Cd47 | ENSMUSG00000055447.13 |
| chr4 | 126967024 | 126968587 | 24.89402 | 98.85246 | 95.31046 | Promoter | 4 | 126861939 | 126968135 | 106197 | 2 | ENSMUST00 | 0 | Zmym4 | ENSMUSG00000042446.11 |
| chr6 | 30896016 | 30897439 | 24.89402 | 98.85246 | 95.31046 | Promoter | 6 | 30747554 | 30896739 | 149186 | 2 | ENSMUST00 | 0 | Copg2 | ENSMUSG00000025607.10 |

| | | | | | | | | | | | | | | | |
|---|---|---|---|---|---|---|---|---|---|---|---|---|---|---|---|
| chr8 | 106010514 | 106011979 | 24.89402 | 98.85246 | 95.31046 | Promoter | 8 | 106011507 | 106053819 | 42313 | 1 | ENSMUST00 | 0 | Dus2 | ENSMUSG00000031901.7 |
| chr8 | 47287267 | 47289869 | 24.89402 | 98.85246 | 95.31046 | Promoter | 8 | 47180048 | 47289402 | 109355 | 2 | ENSMUST00 | 0 | Stox2 | ENSMUSG00000038143.9 |
| chr1 | 57969340 | 57970392 | 32.26695 | 98.80429 | 95.26329 | Promoter | 1 | 57955101 | 57970084 | 14984 | 2 | ENSMUST00 | 0 | Kctd18 | ENSMUSG00000054770.11 |
| chr2 | 57114096 | 57117611 | 32.26695 | 98.80429 | 95.26329 | Promoter | 2 | 57106830 | 57115038 | 8209 | 2 | ENSMUST00 | 0 | Nr4a2 | ENSMUSG00000026826.8 |
| chr4 | 65604538 | 65605674 | 31.68408 | 98.79272 | 95.25197 | Promoter | 4 | 65604986 | 65616238 | 11253 | 1 | ENSMUST00 | 0 | Trim32 | ENSMUSG00000051675.8 |
| chr6 | 56923096 | 56924656 | 31.68408 | 98.79272 | 95.25197 | Promoter | 6 | 56882400 | 56923932 | 41533 | 2 | ENSMUST00 | 0 | Nt5c3 | ENSMUSG00000029780.9 |
| chr11 | 94472650 | 94474728 | 32.82023 | 98.76002 | 95.21949 | Promoter | 11 | 94408391 | 94474100 | 65710 | 2 | ENSMUST00 | 0 | Cacna1g | ENSMUSG00000020866.12 |
| chr8 | 48109196 | 48110812 | 31.07761 | 98.73238 | 95.19199 | Promoter | 8 | 48109958 | 48141665 | 31708 | 1 | ENSMUST00 | 0 | Dctd | ENSMUSG00000031562.10 |
| chr14 | 73142491 | 73144603 | 28.8526 | 98.7319 | 95.19158 | Promoter | 14 | 73142536 | 73164719 | 22184 | 1 | ENSMUST00 | 0 | Rcbtb2 | ENSMUSG00000022106.9 |
| chr17 | 24935707 | 24937390 | 28.8526 | 98.7319 | 95.19158 | Promoter | 17 | 24892153 | 24936938 | 44786 | 2 | ENSMUST00 | 0 | Mapk8ip3 | ENSMUSG00000024163.12 |
| chr2 | 127133421 | 127134430 | 28.8526 | 98.7319 | 95.19158 | Promoter | 2 | 127103809 | 127133954 | 30146 | 2 | ENSMUST00 | 0 | Ncaph | ENSMUSG00000034906.10 |
| chr5 | 102068756 | 102070662 | 28.8526 | 98.7319 | 95.19158 | Promoter | 5 | 101832956 | 102069921 | 236966 | 2 | ENSMUST00 | 0 | Wdfy3 | ENSMUSG00000043940.9 |
| chr9 | 83924637 | 83925761 | 28.8526 | 98.7319 | 95.19158 | Promoter | 9 | 83925145 | 84124239 | 199095 | 1 | ENSMUST00 | 0 | Bckdhb | ENSMUSG00000032263.9 |
| chr9 | 89704222 | 89705572 | 28.8526 | 98.7319 | 95.19158 | Promoter | 9 | 89699205 | 89705068 | 5864 | 2 | ENSMUST00 | 0 | Tmed3 | ENSMUSG00000032353.8 |
| chr5 | 23712012 | 23713165 | 32.20185 | 98.69158 | 95.15165 | Promoter | 5 | 23710991 | 23711061 | 71 | 2 | ENSMUST00 | -952 | Mir3096 | ENSMUSG00000096832.1 |
| chr2 | 155818434 | 155819678 | 31.62215 | 98.68481 | 95.14514 | Promoter | 2 | 155817730 | 155819426 | 1697 | 2 | ENSMUST00 | 0 | BC029722 | ENSMUSG00000074649.6 |
| chr9 | 44964733 | 44965620 | 28.27324 | 98.65216 | 95.1127 | Promoter | 9 | 44923131 | 44965560 | 42430 | 2 | ENSMUST00 | 0 | Ube4a | ENSMUSG00000059890.10 |
| chr13 | 54071575 | 54072789 | 32.75195 | 98.6424 | 95.10313 | Promoter | 13 | 54071845 | 54108346 | 36502 | 1 | ENSMUST00 | 0 | Sfxn1 | ENSMUSG00000021474.8 |
| chr17 | 86166332 | 86169593 | 32.75195 | 98.6424 | 95.10313 | Promoter | 17 | 86167785 | 86657919 | 490135 | 1 | ENSMUST00 | 0 | Prkce | ENSMUSG00000045038.9 |
| chr5 | 108131611 | 108133259 | 27.22006 | 98.6343 | 95.09524 | Promoter | 5 | 108132914 | 108232947 | 100034 | 1 | ENSMUST00 | 0 | Ccdc18 | ENSMUSG00000056531.7 |
| chr10 | 81377526 | 81378535 | 31.01882 | 98.6291 | 95.09013 | Promoter | 10 | 81366222 | 81378516 | 12295 | 2 | ENSMUST00 | 0 | Fzr1 | ENSMUSG00000020235.11 |
| chr16 | 49698661 | 49699981 | 31.01882 | 98.6291 | 95.09013 | Promoter | 16 | 49699233 | 49765126 | 65894 | 1 | ENSMUST00 | 0 | Ift57 | ENSMUSG00000032965.5 |
| chr17 | 83705691 | 83707110 | 31.01882 | 98.6291 | 95.09013 | Promoter | 17 | 83706163 | 83792873 | 86711 | 1 | ENSMUST00 | 0 | Mta3 | ENSMUSG00000055817.12 |
| chr5 | 136961829 | 136962820 | 31.01882 | 98.6291 | 95.09013 | Promoter | 5 | 136962077 | 136966234 | 4158 | 1 | ENSMUST00 | 0 | Fis1 | ENSMUSG00000019054.8 |
| chr9 | 75037326 | 75038664 | 31.01882 | 98.6291 | 95.09013 | Promoter | 9 | 75037614 | 75060294 | 22681 | 1 | ENSMUST00 | 0 | Arpp19 | ENSMUSG00000007656.8 |
| chr15 | 37791004 | 37792729 | 32.13701 | 98.5793 | 95.04197 | Promoter | 15 | 37366175 | 37792000 | 425826 | 2 | ENSMUST00 | 0 | Ncald | ENSMUSG00000051359.9 |
| chr5 | 50057827 | 50059574 | 31.56046 | 98.57728 | 95.04015 | Promoter | 5 | 49959951 | 50058996 | 99046 | 2 | ENSMUST00 | 0 | Gpr125 | ENSMUSG00000029090.8 |
| chr9 | 23222397 | 23223906 | 31.56046 | 98.57728 | 95.04015 | Promoter | 9 | 23223076 | 23485202 | 262127 | 1 | ENSMUST00 | 0 | Bmper | ENSMUSG00000031963.7 |
| chr1 | 181841117 | 181843213 | 29.28796 | 98.53321 | 94.99628 | Promoter | 1 | 181815316 | 181842401 | 27086 | 2 | ENSMUST00 | 0 | Lbr | ENSMUSG00000004880.6 |
| chr1 | 34849606 | 34850820 | 29.28796 | 98.53321 | 94.99628 | Promoter | 1 | 34849976 | 34873956 | 23981 | 1 | ENSMUST00 | 0 | Plekhb2 | ENSMUSG00000026123.6 |
| chr1 | 34438762 | 34440576 | 29.28796 | 98.53321 | 94.99628 | Promoter | 1 | 34439851 | 34449356 | 9506 | 1 | ENSMUST00 | 0 | Imp4 | ENSMUSG00000026127.8 |
| chr1 | 156558296 | 156559937 | 29.28796 | 98.53321 | 94.99628 | Promoter | 1 | 156558786 | 156625376 | 66591 | 1 | ENSMUST00 | 0 | Abl2 | ENSMUSG00000026596.8 |
| chr10 | 81166899 | 81167857 | 29.28796 | 98.53321 | 94.99628 | Promoter | 10 | 81155672 | 81167721 | 12050 | 2 | ENSMUST00 | 0 | Pias4 | ENSMUSG00000004934.8 |
| chr10 | 125965700 | 125967874 | 29.28796 | 98.53321 | 94.99628 | Promoter | 10 | 125966219 | 126015359 | 49141 | 1 | ENSMUST00 | 0 | Lrig3 | ENSMUSG00000020105.8 |
| chr10 | 111472714 | 111474509 | 29.28796 | 98.53321 | 94.99628 | Promoter | 10 | 111473192 | 111496886 | 23695 | 1 | ENSMUST00 | 0 | Nap1l1 | ENSMUSG00000058799.9 |
| chr11 | 100618769 | 100620432 | 29.28796 | 98.53321 | 94.99628 | Promoter | 11 | 100582818 | 100620168 | 37351 | 2 | ENSMUST00 | 0 | Dnajc7 | ENSMUSG00000014195.10 |
| chr11 | 85233236 | 85235725 | 29.28796 | 98.53321 | 94.99628 | Promoter | 11 | 85187262 | 85235130 | 47869 | 2 | ENSMUST00 | 0 | Appbp2 | ENSMUSG00000018481.9 |
| chr12 | 84969862 | 84971593 | 29.28796 | 98.53321 | 94.99628 | Promoter | 12 | 84970897 | 84983303 | 12407 | 1 | ENSMUST00 | 0 | Fcf1 | ENSMUSG00000021243.9 |
| chr13 | 90088873 | 90090502 | 29.28796 | 98.53321 | 94.99628 | Promoter | 13 | 90089123 | 90112257 | 23135 | 1 | ENSMUST00 | 0 | Tmem167 | ENSMUSG00000012422.9 |
| chr13 | 21199268 | 21200908 | 29.28796 | 98.53321 | 94.99628 | Distal Interge | 13 | 21190092 | 21192475 | 2384 | 1 | ENSMUST00 | 9177 | Trim27 | ENSMUSG00000021326.6 |
| chr13 | 73937258 | 73938823 | 29.28796 | 98.53321 | 94.99628 | Promoter | 13 | 73912462 | 73937767 | 25306 | 2 | ENSMUST00 | 0 | Trip13 | ENSMUSG00000021569.8 |
| chr13 | 73327793 | 73329104 | 29.28796 | 98.53321 | 94.99628 | Promoter | 13 | 73319874 | 73328542 | 8669 | 2 | ENSMUST00 | 0 | Ndufs6 | ENSMUSG00000021606.7 |
| chr13 | 23534750 | 23535801 | 29.28796 | 98.53321 | 94.99628 | Promoter | 13 | 23535434 | 23535860 | 427 | 1 | ENSMUST00 | 0 | Hist1h3g | ENSMUSG00000099517.1 |
| chr15 | 97963406 | 97965116 | 29.28796 | 98.53321 | 94.99628 | Promoter | 15 | 97964229 | 97970286 | 6058 | 1 | ENSMUST00 | 0 | Tmem106c | ENSMUSG00000052369.7 |
| chr15 | 76947906 | 76949370 | 29.28796 | 98.53321 | 94.99628 | Promoter | 15 | 76948539 | 76950729 | 2191 | 1 | ENSMUST00 | 0 | 1110038F14I | ENSMUSG00000063236.5 |
| chr2 | 25428578 | 25430241 | 29.28796 | 98.53321 | 94.99628 | Promoter | 2 | 25428703 | 25448540 | 19838 | 1 | ENSMUST00 | 0 | Abca2 | ENSMUSG00000026944.13 |
| chr2 | 129197808 | 129199982 | 29.28796 | 98.53321 | 94.99628 | Promoter | 2 | 129198764 | 129211616 | 12853 | 1 | ENSMUST00 | 0 | Slc20a1 | ENSMUSG00000027397.9 |
| chr2 | 71452849 | 71453973 | 29.28796 | 98.53321 | 94.99628 | Promoter | 2 | 71453276 | 71525191 | 71916 | 1 | ENSMUST00 | 0 | Metap1d | ENSMUSG00000041921.11 |
| chr3 | 87884369 | 87885651 | 29.28796 | 98.53321 | 94.99628 | Promoter | 3 | 87858900 | 87885608 | 26709 | 2 | ENSMUST00 | 0 | Prcc | ENSMUSG00000004895.8 |
| chr3 | 98427166 | 98428732 | 29.28796 | 98.53321 | 94.99628 | 3' UTR | 3 | 98447198 | 98448976 | 1779 | 2 | ENSMUST00 | 20244 | Gm4450 | ENSMUSG00000090817.2 |
| chr4 | 53158422 | 53160317 | 29.28796 | 98.53321 | 94.99628 | Promoter | 4 | 53030787 | 53159895 | 129109 | 2 | ENSMUST00 | 0 | Abca1 | ENSMUSG00000015243.4 |
| chr4 | 137992884 | 137995121 | 29.28796 | 98.53321 | 94.99628 | Promoter | 4 | 137993022 | 138105419 | 112398 | 1 | ENSMUST00 | 0 | Eif4g3 | ENSMUSG00000028760.11 |
| chr5 | 3927741 | 3929337 | 29.28796 | 98.53321 | 94.99628 | Promoter | 5 | 3928054 | 3954765 | 26712 | 1 | ENSMUST00 | 0 | Akap9 | ENSMUSG00000040407.12 |
| chr5 | 138622453 | 138623785 | 29.28796 | 98.53321 | 94.99628 | Promoter | 5 | 138622859 | 138640823 | 17965 | 1 | ENSMUST00 | 0 | A430033K04I | ENSMUSG00000056014.10 |
| chr6 | 88189480 | 88190234 | 29.28796 | 98.53321 | 94.99628 | Distal Interge | 6 | 88193891 | 88207030 | 13140 | 1 | ENSMUST00 | -3657 | Gata2 | ENSMUSG00000015053.9 |
| chr6 | 88841141 | 88842056 | 29.28796 | 98.53321 | 94.99628 | Promoter | 6 | 88835915 | 88841935 | 6021 | 2 | ENSMUST00 | 0 | Abtb1 | ENSMUSG00000030083.9 |
| chr7 | 48788641 | 48790126 | 29.28796 | 98.53321 | 94.99628 | Promoter | 7 | 48789003 | 48827440 | 38438 | 1 | ENSMUST00 | 0 | Zdhhc13 | ENSMUSG00000030471.12 |
| chr7 | 126883515 | 126885205 | 29.28796 | 98.53321 | 94.99628 | Promoter | 7 | 126865678 | 126884703 | 19026 | 2 | ENSMUST00 | 0 | Taok2 | ENSMUSG00000059981.6 |
| chr8 | 70282389 | 70283850 | 29.28796 | 98.53321 | 94.99628 | Promoter | 8 | 70282827 | 70286530 | 3704 | 1 | ENSMUST00 | 0 | Homer3 | ENSMUSG00000003573.10 |
| chr8 | 22410907 | 22412288 | 29.28796 | 98.53321 | 94.99628 | Promoter | 8 | 22411340 | 22429665 | 18326 | 1 | ENSMUST00 | 0 | Mrps31 | ENSMUSG00000031533.3 |
| chr8 | 74993050 | 74994752 | 29.28796 | 98.53321 | 94.99628 | Promoter | 8 | 74993356 | 75031972 | 38617 | 1 | ENSMUST00 | 0 | Hmgxb4 | ENSMUSG00000034518.8 |
| chr8 | 71922378 | 71923838 | 29.28796 | 98.53321 | 94.99628 | Promoter | 8 | 71922852 | 71934629 | 11778 | 1 | ENSMUST00 | 0 | Zfp617 | ENSMUSG00000066880.10 |
| chr9 | 99575435 | 99576541 | 29.28796 | 98.53321 | 94.99628 | Promoter | 9 | 99575799 | 99584501 | 8703 | 1 | ENSMUST00 | 0 | Dbr1 | ENSMUSG00000032469.7 |
| chr9 | 20975370 | 20977071 | 29.28796 | 98.53321 | 94.99628 | Promoter | 9 | 20965952 | 20976793 | 10842 | 2 | ENSMUST00 | 0 | S1pr2 | ENSMUSG00000043895.5 |
| chr14 | 65261170 | 65263071 | 32.68394 | 98.52523 | 94.99333 | Promoter | 14 | 65201026 | 65262463 | 61438 | 2 | ENSMUST00 | 0 | Fzd3 | ENSMUSG00000007989.6 |
| chr7 | 120676997 | 120678514 | 31.49902 | 98.4701 | 94.93838 | Promoter | 7 | 120677620 | 120734854 | 57235 | 1 | ENSMUST00 | 0 | BC030336 | ENSMUSG00000046096.6 |
| chr1 | 138962926 | 138964541 | 32.07243 | 98.46742 | 94.9359 | Promoter | 1 | 138963436 | 139063719 | 100284 | 1 | ENSMUST00 | 0 | Dennd1b | ENSMUSG00000056268.10 |
| chr3 | 106721230 | 106723023 | 32.07243 | 98.46742 | 94.9359 | Promoter | 3 | 106721682 | 106736574 | 14893 | 1 | ENSMUST00 | 0 | Lrif1 | ENSMUSG00000056260.10 |
| chr19 | 4755824 | 4757634 | 30.90191 | 98.42355 | 94.89227 | Promoter | 19 | 4756543 | 4762876 | 6334 | 1 | ENSMUST00 | 0 | Rbm4b | ENSMUSG00000033760.11 |
| chr7 | 80687626 | 80689456 | 30.90191 | 98.42355 | 94.89227 | Promoter | 7 | 80586627 | 80688877 | 102251 | 2 | ENSMUST00 | 0 | Crtc3 | ENSMUSG00000030527.10 |
| chr1 | 182281371 | 182283460 | 26.1063 | 98.41757 | 94.88632 | Promoter | 1 | 182275772 | 182282804 | 7033 | 2 | ENSMUST00 | 0 | Degs1 | ENSMUSG00000038633.5 |
| chr10 | 78351249 | 78352952 | 26.1063 | 98.41757 | 94.88632 | Promoter | 10 | 78269178 | 78352442 | 83265 | 2 | ENSMUST00 | 0 | Agpat3 | ENSMUSG00000001211.9 |
| chr11 | 70646200 | 70647062 | 26.1063 | 98.41757 | 94.88632 | Promoter | 11 | 70644196 | 70647058 | 2863 | 2 | ENSMUST00 | 0 | Slc25a11 | ENSMUSG00000014606.9 |
| chr11 | 5064372 | 5065809 | 26.1063 | 98.41757 | 94.88632 | Promoter | 11 | 5054132 | 5065294 | 11163 | 2 | ENSMUST00 | 0 | Gas2l1 | ENSMUSG00000034201.8 |
| chr14 | 62331739 | 62332866 | 26.1063 | 98.41757 | 94.88632 | Promoter | 14 | 62332068 | 62372992 | 40925 | 1 | ENSMUST00 | 0 | Rnaseh2b | ENSMUSG00000021932.8 |
| chr14 | 70173369 | 70177377 | 26.1063 | 98.41757 | 94.88632 | Promoter | 14 | 70164231 | 70176819 | 12589 | 2 | ENSMUST00 | 0 | Pdlim2 | ENSMUSG00000022090.5 |
| chr19 | 58859113 | 58861512 | 26.1063 | 98.41757 | 94.88632 | Promoter | 19 | 58795751 | 58860984 | 65234 | 2 | ENSMUST00 | 0 | Hspa12a | ENSMUSG00000025092.6 |
| chr2 | 32259334 | 32260614 | 26.1063 | 98.41757 | 94.88632 | Promoter | 2 | 32255002 | 32260159 | 5158 | 2 | ENSMUST00 | 0 | Uck1 | ENSMUSG00000002550.11 |
| chr2 | 121139280 | 121141380 | 26.1063 | 98.41757 | 94.88632 | Promoter | 2 | 121140428 | 121145514 | 5087 | 1 | ENSMUST00 | 0 | Adal | ENSMUSG00000027259.10 |
| chr2 | 126706747 | 126707944 | 26.1063 | 98.41757 | 94.88632 | Promoter | 2 | 126707328 | 126759248 | 51921 | 1 | ENSMUST00 | 0 | Usp8 | ENSMUSG00000027363.10 |
| chr4 | 117835153 | 117836940 | 26.1063 | 98.41757 | 94.88632 | Promoter | 4 | 117835333 | 117875196 | 39864 | 1 | ENSMUST00 | 0 | Slc6a9 | ENSMUSG00000028542.12 |
| chr4 | 57299328 | 57302225 | 26.1063 | 98.41757 | 94.88632 | Promoter | 4 | 57190841 | 57301837 | 110997 | 2 | ENSMUST00 | 0 | Ptpn3 | ENSMUSG00000038764.9 |
| chr9 | 113931013 | 113932730 | 26.1063 | 98.41757 | 94.88632 | Promoter | 9 | 113931564 | 113976817 | 45254 | 1 | ENSMUST00 | 0 | Ubp1 | ENSMUSG00000009741.9 |
| chr9 | 105642577 | 105643629 | 26.1063 | 98.41757 | 94.88632 | Promoter | 9 | 105642978 | 105679320 | 36343 | 1 | ENSMUST00 | 0 | Pik3r4 | ENSMUSG00000032571.8 |
| chr12 | 111677482 | 111678811 | 32.61622 | 98.40852 | 94.879 | Promoter | 12 | 111678105 | 111683902 | 5798 | 1 | ENSMUST00 | 0 | Trmt61a | ENSMUSG00000060950.6 |
| chr19 | 6276749 | 6279190 | 21.72697 | 98.39969 | 94.87038 | Promoter | 19 | 6276925 | 6281392 | 4468 | 1 | ENSMUST00 | 0 | Ehd1 | ENSMUSG00000024772.9 |
| chr19 | 45005529 | 45007479 | 21.72697 | 98.39969 | 94.87038 | Promoter | 19 | 45005014 | 45006442 | 1429 | 2 | ENSMUST00 | 0 | Mrpl43 | ENSMUSG00000025208.8 |
| chr5 | 138085011 | 138085914 | 21.72697 | 98.39969 | 94.87038 | Promoter | 5 | 138085084 | 138107822 | 22739 | 1 | ENSMUST00 | 0 | Zkscan1 | ENSMUSG00000029729.7 |
| chr10 | 18021720 | 18023486 | 32.00811 | 98.35596 | 94.82688 | Promoter | 10 | 18011263 | 18023252 | 11990 | 2 | ENSMUST00 | 0 | Abracl | ENSMUSG00000078453.2 |
| chr11 | 76026815 | 76028460 | 32.00811 | 98.35596 | 94.82688 | Promoter | 11 | 76019194 | 76027782 | 8589 | 2 | ENSMUST00 | 0 | Fam101b | ENSMUSG00000020846.6 |
| chr11 | 120232940 | 120238211 | 32.00811 | 98.35596 | 94.82688 | Promoter | 11 | 120232947 | 120292282 | 59336 | 1 | ENSMUST00 | 0 | Bahcc1 | ENSMUSG00000039741.10 |
| chr4 | 127060456 | 127061634 | 32.00811 | 98.35596 | 94.82688 | Promoter | 4 | 127047094 | 127061152 | 14059 | 2 | ENSMUST00 | 0 | Zmym1 | ENSMUSG00000043872.9 |
| chr7 | 133708247 | 133710181 | 32.00811 | 98.35596 | 94.82688 | Promoter | 7 | 133686243 | 133709094 | 22852 | 2 | ENSMUST00 | 0 | Uros | ENSMUSG00000030979.7 |
| chr9 | 56937002 | 56938607 | 32.00811 | 98.35596 | 94.82688 | Promoter | 9 | 56937475 | 56938398 | 924 | 1 | ENSMUST00 | 0 | Imp3 | ENSMUSG00000032288.9 |
| chr10 | 75666959 | 75668360 | 30.28657 | 98.33401 | 94.80609 | Intron (ENSM | 10 | 75636619 | 75644008 | 7390 | 2 | ENSMUST00 | -22952 | Susd2 | ENSMUSG00000006342.9 |
| chr11 | 103016419 | 103017187 | 30.28657 | 98.33401 | 94.80609 | Promoter | 11 | 102994047 | 103017168 | 23122 | 2 | ENSMUST00 | 0 | Dcakd | ENSMUSG00000020935.3 |
| chr14 | 69706783 | 69707701 | 30.28657 | 98.33401 | 94.80609 | Promoter | 14 | 69697307 | 69707584 | 10278 | 2 | ENSMUST00 | 0 | R3hcc1 | ENSMUSG00000034194.11 |
| chr3 | 51558835 | 51561958 | 30.28657 | 98.33401 | 94.80609 | Promoter | 3 | 51515319 | 51560878 | 45560 | 2 | ENSMUST00 | 0 | Setd7 | ENSMUSG00000037111.8 |
| chr5 | 115010549 | 115012400 | 30.28657 | 98.33401 | 94.80609 | Promoter | 5 | 115011137 | 115098790 | 87654 | 1 | ENSMUST00 | 0 | Sppl3 | ENSMUSG00000029550.8 |
| chr5 | 135962842 | 135963512 | 30.28657 | 98.33401 | 94.80609 | Promoter | 5 | 135960224 | 135962327 | 2104 | 2 | ENSMUST00 | -516 | Srcrb4d | ENSMUSG00000029699.8 |
| chr5 | 115559447 | 115560373 | 30.28657 | 98.33401 | 94.80609 | Promoter | 5 | 115559467 | 115563727 | 4261 | 1 | ENSMUST00 | 0 | Rplp0 | ENSMUSG00000067274.5 |
| chr4 | 48044233 | 48048412 | 30.84378 | 98.32127 | 94.79446 | Promoter | 4 | 48045153 | 48086445 | 41293 | 1 | ENSMUST00 | 0 | Nr4a3 | ENSMUSG00000028341.4 |
| chr1 | 57405719 | 57407662 | 27.54716 | 98.2761 | 94.74957 | Promoter | 1 | 57406328 | 57417953 | 11626 | 1 | ENSMUST00 | 0 | 9430016H08I | ENSMUSG00000025971.6 |
| chr10 | 82177916 | 82179221 | 27.54716 | 98.2761 | 94.74957 | Promoter | 10 | 82178235 | 82198385 | 20151 | 1 | ENSMUST00 | 0 | Gm15608 | ENSMUSG00000087481.2 |
| chr11 | 8663227 | 8664945 | 27.54716 | 98.2761 | 94.74957 | Promoter | 11 | 8431652 | 8664535 | 232884 | 2 | ENSMUST00 | 0 | Tns3 | ENSMUSG00000020422.8 |
| chr11 | 103100966 | 103102757 | 27.54716 | 98.2761 | 94.74957 | Promoter | 11 | 103070304 | 103101658 | 31355 | 2 | ENSMUST00 | 0 | Plcd3 | ENSMUSG00000020937.9 |
| chr11 | 120457377 | 120459007 | 27.54716 | 98.2761 | 94.74957 | Promoter | 11 | 120456606 | 120458068 | 1463 | 2 | ENSMUST00 | 0 | Oxld1 | ENSMUSG00000039670.2 |
| chr12 | 72084484 | 72085510 | 27.54716 | 98.2761 | 94.74957 | Promoter | 12 | 72073428 | 72085439 | 12012 | 2 | ENSMUST00 | 0 | L3hypdh | ENSMUSG00000019718.3 |
| chr12 | 4592382 | 4594262 | 27.54716 | 98.2761 | 94.74957 | Promoter | 12 | 4593008 | 4713950 | 120943 | 1 | ENSMUST00 | 0 | Itsn2 | ENSMUSG00000020640.9 |
| chr12 | 26414174 | 26415373 | 27.54716 | 98.2761 | 94.74957 | Promoter | 12 | 26306797 | 26415256 | 108460 | 2 | ENSMUST00 | 0 | Rnf144a | ENSMUSG00000020642.7 |
| chr12 | 17881272 | 17883012 | 27.54716 | 98.2761 | 94.74957 | Distal Interge | 12 | 17924294 | 17925668 | 1375 | 1 | ENSMUST00 | -41282 | B430203G13 | ENSMUSG00000067356.5 |
| chr15 | 97704990 | 97706312 | 27.54716 | 98.2761 | 94.74957 | Promoter | 15 | 97675105 | 97705822 | 30718 | 2 | ENSMUST00 | 0 | Rpap3 | ENSMUSG00000022466.5 |
| chr16 | 4624053 | 4625167 | 27.54716 | 98.2761 | 94.74957 | Promoter | 16 | 4616464 | 4624936 | 8473 | 2 | ENSMUST00 | 0 | Pam16 | ENSMUSG00000014301.7 |
| chr16 | 96145333 | 96146075 | 27.54716 | 98.2761 | 94.74957 | Promoter | 16 | 96145407 | 96146764 | 1358 | 1 | ENSMUST00 | 0 | Wrb | ENSMUSG00000023147.10 |
| chr17 | 46383068 | 46384821 | 27.54716 | 98.2761 | 94.74957 | Promoter | 17 | 46383731 | 46414814 | 31084 | 1 | ENSMUST00 | 0 | Zfp318 | ENSMUSG00000015597.11 |
| chr19 | 24223944 | 24225914 | 27.54716 | 98.2761 | 94.74957 | Promoter | 19 | 24094523 | 24225026 | 130504 | 2 | ENSMUST00 | 0 | Tjp2 | ENSMUSG00000024812.9 |
| chr2 | 154602815 | 154603790 | 27.54716 | 98.2761 | 94.74957 | Promoter | 2 | 154585758 | 154603708 | 17951 | 2 | ENSMUST00 | 0 | Pxmp4 | ENSMUSG00000000876.6 |
| chr2 | 143062580 | 143064002 | 27.54716 | 98.2761 | 94.74957 | Promoter | 2 | 143063039 | 143072853 | 9815 | 1 | ENSMUST00 | 0 | Snrpb2 | ENSMUSG00000008333.7 |
| chr2 | 105013606 | 105017444 | 27.54716 | 98.2761 | 94.74957 | Promoter | 2 | 104999656 | 105017080 | 17425 | 2 | ENSMUST00 | 0 | Eif3m | ENSMUSG00000027170.7 |
| chr2 | 103566306 | 103569145 | 27.54716 | 98.2761 | 94.74957 | Promoter | 2 | 103566310 | 103718423 | 152114 | 1 | ENSMUST00 | 0 | Abtb2 | ENSMUSG00000032724.5 |
| chr2 | 152104864 | 152106303 | 27.54716 | 98.2761 | 94.74957 | Promoter | 2 | 152105516 | 152109211 | 3696 | 1 | ENSMUST00 | 0 | Srxn1 | ENSMUSG00000032802.8 |
| chr2 | 34370530 | 34373633 | 27.54716 | 98.2761 | 94.74957 | Promoter | 2 | 34171457 | 34372038 | 200582 | 2 | ENSMUST00 | 0 | Pbx3 | ENSMUSG00000038718.10 |
| chr3 | 63975856 | 63977361 | 27.54716 | 98.2761 | 94.74957 | Promoter | 3 | 63976106 | 64022579 | 46474 | 1 | ENSMUST00 | 0 | Gmps | ENSMUSG00000027823.8 |
| chr3 | 153723569 | 153725538 | 27.54716 | 98.2761 | 94.74957 | Promoter | 3 | 153205405 | 153725062 | 519658 | 2 | ENSMUST00 | 0 | St6galnac3 | ENSMUSG00000052544.5 |
| chr5 | 107986196 | 107988211 | 27.54716 | 98.2761 | 94.74957 | Promoter | 5 | 107908053 | 107987085 | 79033 | 2 | ENSMUST00 | 0 | Fam69a | ENSMUSG00000029270.5 |
| chr5 | 103752781 | 103755575 | 27.54716 | 98.2761 | 94.74957 | Promoter | 5 | 103754162 | 103855322 | 101161 | 1 | ENSMUST00 | 0 | Aff1 | ENSMUSG00000029313.13 |
| chr5 | 150673763 | 150674767 | 27.54716 | 98.2761 | 94.74957 | Promoter | 5 | 150673827 | 150810667 | 136841 | 1 | ENSMUST00 | 0 | Pds5b | ENSMUSG00000034021.10 |
| chr6 | 29163882 | 29165103 | 27.54716 | 98.2761 | 94.74957 | Promoter | 6 | 29123573 | 29165006 | 41434 | 2 | ENSMUST00 | 0 | Rbm28 | ENSMUSG00000029701.10 |

| | | | | | | | | | | | | | | | |
|---|---|---|---|---|---|---|---|---|---|---|---|---|---|---|---|
| chr6 | 134791180 | 134792650 | 27.54716 | 98.2761 | 94.74957 | Promoter | 6 | 134715473 | 134792625 | 77153 | 2 | ENSMUST00 | 0 | Dusp16 | ENSMUSG00000030203.12 |
| chr7 | 45465934 | 45467296 | 27.54716 | 98.2761 | 94.74957 | Promoter | 7 | 45461695 | 45466898 | 5204 | 2 | ENSMUST00 | 0 | Bax | ENSMUSG00000003873.10 |
| chr7 | 82865501 | 82869284 | 27.54716 | 98.2761 | 94.74957 | Promoter | 7 | 82867333 | 82871563 | 4231 | 1 | ENSMUST00 | 0 | Mex3b | ENSMUSG00000057706.6 |
| chr7 | 45703819 | 45709044 | 27.54716 | 98.2761 | 94.74957 | Promoter | 7 | 45705088 | 45710155 | 5068 | 1 | ENSMUST00 | 0 | Dbp | ENSMUSG00000059824.5 |
| chr7 | 127935036 | 127936269 | 27.54716 | 98.2761 | 94.74957 | Promoter | 7 | 127932638 | 127935479 | 2842 | 2 | ENSMUST00 | 0 | Prss36 | ENSMUSG00000070371.6 |
| chr8 | 105565120 | 105567512 | 27.54716 | 98.2761 | 94.74957 | Promoter | 8 | 105524465 | 105566047 | 41583 | 2 | ENSMUST00 | 0 | Atp6v0d1 | ENSMUSG00000013160.7 |
| chr8 | 111730354 | 111732664 | 27.54716 | 98.2761 | 94.74957 | Intron (ENSM | 8 | 111710485 | 111743849 | 33365 | 2 | ENSMUST00 | 11185 | Bcar1 | ENSMUSG00000031955.9 |
| chr8 | 79028069 | 79031248 | 27.54716 | 98.2761 | 94.74957 | Promoter | 8 | 79028437 | 79071837 | 43401 | 1 | ENSMUST00 | 0 | Zfp827 | ENSMUSG00000071064.8 |
| chr9 | 110407056 | 110408833 | 27.54716 | 98.2761 | 94.74957 | Promoter | 9 | 110385089 | 110408210 | 23122 | 2 | ENSMUST00 | 0 | Ptpn23 | ENSMUSG00000036057.7 |
| chr9 | 106885477 | 106887821 | 27.54716 | 98.2761 | 94.74957 | Promoter | 9 | 106880918 | 106887428 | 6511 | 2 | ENSMUST00 | 0 | Rbm15b | ENSMUSG00000074102.3 |
| chr6 | 33249030 | 33249796 | 31.37684 | 98.25688 | 94.7338 | Promoter | 6 | 33249085 | 33973979 | 724895 | 1 | ENSMUST00 | 0 | Exoc4 | ENSMUSG00000029763.12 |
| chr6 | 47771783 | 47772516 | 8.30211 | 98.20082 | 94.67821 | Promoter | 6 | 47773329 | 47773429 | 101 | 1 | ENSMUST00 | -813 | Gm24975 | ENSMUSG00000094807.1 |
| chr16 | 91464576 | 91465705 | 32.48162 | 98.17645 | 94.65383 | Promoter | 16 | 91465104 | 91470121 | 5018 | 1 | ENSMUST00 | 0 | A930006K02I | ENSMUSG00000084866.1 |
| chr19 | 24476142 | 24478155 | 31.88024 | 98.13424 | 94.612 | Promoter | 19 | 24476502 | 24477356 | 855 | 2 | ENSMUST00 | 0 | Fam122a | ENSMUSG00000074922.1 |
| chr18 | 50127855 | 50128857 | 29.60582 | 98.11335 | 94.5914 | Promoter | 18 | 50128201 | 50196269 | 68069 | 1 | ENSMUST00 | 0 | Hsd17b4 | ENSMUSG00000024507.6 |
| chr6 | 114917117 | 114921811 | 29.60582 | 98.11335 | 94.5914 | Promoter | 6 | 114861424 | 114921821 | 60398 | 2 | ENSMUST00 | 10 | Vgll4 | ENSMUSG00000030315.10 |
| chr7 | 128523225 | 128525180 | 29.60582 | 98.11335 | 94.5914 | Promoter | 7 | 128523583 | 128546977 | 23395 | 1 | ENSMUST00 | 0 | Bag3 | ENSMUSG00000030847.7 |
| chr7 | 45717180 | 45719165 | 29.60582 | 98.11335 | 94.5914 | Promoter | 7 | 45709467 | 45718002 | 8536 | 2 | ENSMUST00 | 0 | Sphk2 | ENSMUSG00000057342.8 |
| chr7 | 25268350 | 25269389 | 32.41473 | 98.06107 | 94.53979 | Promoter | 7 | 25268387 | 25271272 | 2886 | 1 | ENSMUST00 | 0 | Cic | ENSMUSG00000005442.8 |
| chr2 | 180700691 | 180702845 | 31.25561 | 98.0451 | 94.52408 | Promoter | 2 | 180657964 | 180702350 | 44387 | 2 | ENSMUST00 | 0 | Dido1 | ENSMUSG00000038914.10 |
| chr2 | 5894821 | 5895828 | 31.81669 | 98.02396 | 94.50304 | Promoter | 2 | 5870987 | 5895432 | 24446 | 2 | ENSMUST00 | 0 | Sec61a2 | ENSMUSG00000025816.10 |
| chr2 | 13572823 | 13576831 | 31.81669 | 98.02396 | 94.50304 | Promoter | 2 | 13573927 | 13574738 | 812 | 1 | ENSMUST00 | 0 | Vim | ENSMUSG00000026728.4 |
| chr12 | 17544060 | 17544701 | 30.67071 | 98.01645 | 94.49575 | Promoter | 12 | 17544873 | 17551502 | 6630 | 1 | ENSMUST00 | -172 | Odc1 | ENSMUSG00000011179.7 |
| chr1 | 152088689 | 152091255 | 23.5948 | 97.99914 | 94.47849 | Promoter | 1 | 151852403 | 152089970 | 237568 | 2 | ENSMUST00 | 0 | 1700025G04 | ENSMUSG00000032666.11 |
| chr10 | 71344330 | 71345439 | 23.5948 | 97.99914 | 94.47849 | Promoter | 10 | 71330480 | 71344954 | 14475 | 2 | ENSMUST00 | 0 | Cisd1 | ENSMUSG00000037710.8 |
| chr11 | 59226686 | 59228835 | 23.5948 | 97.99914 | 94.47849 | Promoter | 11 | 59211412 | 59228162 | 16751 | 2 | ENSMUST00 | 0 | Arf1 | ENSMUSG00000048076.9 |
| chr18 | 37934343 | 37935812 | 23.5948 | 97.99914 | 94.47849 | Promoter | 18 | 37843601 | 37935476 | 91876 | 2 | ENSMUST00 | 0 | Diap1 | ENSMUSG00000024456.11 |
| chr10 | 80854482 | 80856036 | 28.97134 | 97.95801 | 94.43773 | Promoter | 10 | 80855275 | 80868705 | 13431 | 1 | ENSMUST00 | 0 | Sppl2b | ENSMUSG00000035206.9 |
| chr5 | 124095050 | 124095851 | 28.97134 | 97.95801 | 94.43773 | Promoter | 5 | 124061530 | 124095798 | 34269 | 2 | ENSMUST00 | 0 | Abcb9 | ENSMUSG00000029408.8 |
| chr7 | 135715635 | 135716557 | 32.34813 | 97.94613 | 94.42644 | Promoter | 7 | 135689788 | 135716379 | 26592 | 2 | ENSMUST00 | 0 | Mki67 | ENSMUSG00000031004.7 |
| chr10 | 29312648 | 29314111 | 31.19535 | 97.93977 | 94.4203 | Promoter | 10 | 29313166 | 29345560 | 32395 | 1 | ENSMUST00 | 0 | Echdc1 | ENSMUSG00000019883.5 |
| chr15 | 85336032 | 85337422 | 31.19535 | 97.93977 | 94.4203 | Promoter | 15 | 85336381 | 85463836 | 127456 | 1 | ENSMUST00 | 0 | Atxn10 | ENSMUSG00000016541.9 |
| chr2 | 144555321 | 144557008 | 31.19535 | 97.93977 | 94.4203 | Promoter | 2 | 144556229 | 144568027 | 11799 | 1 | ENSMUST00 | 0 | Sec23b | ENSMUSG00000027429.11 |
| chr3 | 95754160 | 95754927 | 31.19535 | 97.93977 | 94.4203 | Promoter | 3 | 95739976 | 95754977 | 15002 | 2 | ENSMUST00 | 50 | Tars2 | ENSMUSG00000028107.9 |
| chr14 | 5811977 | 5813258 | 20.07858 | 97.92787 | 94.40955 | Distal Interge | 14 | 5607620 | 5807982 | 200363 | 2 | ENSMUST00 | -3996 | Gm3383 | ENSMUSG00000096629.2 |
| chr11 | 30953354 | 30955080 | 31.75339 | 97.91409 | 94.3958 | Promoter | 11 | 30954395 | 31102704 | 148310 | 1 | ENSMUST00 | 0 | Asb3 | ENSMUSG00000020305.8 |
| chr3 | 86141359 | 86142620 | 28.37855 | 97.86033 | 94.34225 | Promoter | 3 | 86137940 | 86142702 | 4763 | 2 | ENSMUST00 | 82 | Rps3a1 | ENSMUSG00000028081.6 |
| chr7 | 125443975 | 125445550 | 28.37855 | 97.86033 | 94.34225 | Promoter | 7 | 125444676 | 125462265 | 17590 | 1 | ENSMUST00 | 0 | Kdm8 | ENSMUSG00000030752.3 |
| chr3 | 157533711 | 157535409 | 26.81296 | 97.85361 | 94.33582 | Promoter | 3 | 157534160 | 157548390 | 14231 | 1 | ENSMUST00 | 0 | Zranb2 | ENSMUSG00000028180.13 |
| chr11 | 102429983 | 102431352 | 32.28179 | 97.83163 | 94.31397 | Promoter | 11 | 102430315 | 102436809 | 6495 | 1 | ENSMUST00 | 0 | Grn | ENSMUSG00000034708.6 |
| chr12 | 28749799 | 28750775 | 27.82348 | 97.81387 | 94.29646 | Promoter | 12 | 28690629 | 28750394 | 59766 | 2 | ENSMUST00 | 0 | Trappc12 | ENSMUSG00000020628.10 |
| chr13 | 73466857 | 73468175 | 31.69034 | 97.80461 | 94.28741 | Promoter | 13 | 73467197 | 73516422 | 49226 | 1 | ENSMUST00 | 0 | Lpcat1 | ENSMUSG00000021608.9 |
| chr2 | 180966622 | 180968168 | 31.69034 | 97.80461 | 94.28741 | Promoter | 2 | 180967225 | 180982458 | 15234 | 1 | ENSMUST00 | 0 | Arfgap1 | ENSMUSG00000027575.14 |
| chr10 | 89731414 | 89733039 | 31.07551 | 97.73014 | 94.2132 | Promoter | 10 | 89711973 | 89732295 | 20323 | 2 | ENSMUST00 | 0 | Actr6 | ENSMUSG00000019948.3 |
| chr12 | 76403128 | 76406376 | 31.07551 | 97.73014 | 94.2132 | Promoter | 12 | 76404176 | 76406934 | 2759 | 1 | ENSMUST00 | 0 | Hspa2 | ENSMUSG00000059970.6 |
| chr14 | 55577811 | 55578944 | 32.21572 | 97.71757 | 94.20073 | Promoter | 14 | 55578123 | 55581346 | 3224 | 1 | ENSMUST00 | 0 | Psme1 | ENSMUSG00000022216.11 |
| chr7 | 28049526 | 28050131 | 32.21572 | 97.71757 | 94.20073 | Promoter | 7 | 28041707 | 28050101 | 8395 | 2 | ENSMUST00 | 0 | Psmc4 | ENSMUSG00000030603.10 |
| chr16 | 94369721 | 94372158 | 31.62754 | 97.69553 | 94.17948 | Promoter | 16 | 94358763 | 94370289 | 11527 | 2 | ENSMUST00 | 0 | Pigp | ENSMUSG00000022940.11 |
| chr1 | 172311315 | 172313237 | 31.01414 | 97.6227 | 94.10706 | Promoter | 1 | 172311771 | 172319841 | 8071 | 1 | ENSMUST00 | 0 | Igsf8 | ENSMUSG00000038034.10 |
| chr10 | 68275857 | 68280673 | 31.01414 | 97.6227 | 94.10706 | Promoter | 10 | 68095593 | 68278726 | 183134 | 2 | ENSMUST00 | 0 | Arid5b | ENSMUSG00000019947.9 |
| chr11 | 97661507 | 97664564 | 31.01414 | 97.6227 | 94.10706 | Promoter | 11 | 97663414 | 97685463 | 22050 | 1 | ENSMUST00 | 0 | Mllt6 | ENSMUSG00000038437.6 |
| chr12 | 103531957 | 103533229 | 31.01414 | 97.6227 | 94.10706 | Promoter | 12 | 103532283 | 103613831 | 81549 | 1 | ENSMUST00 | 0 | Ppp4r4 | ENSMUSG00000021209.7 |
| chr12 | 9029631 | 9031447 | 31.01414 | 97.6227 | 94.10706 | Promoter | 12 | 9029997 | 9036394 | 6398 | 1 | ENSMUST00 | 0 | Ttc32 | ENSMUSG00000066637.1 |
| chr13 | 30659404 | 30660874 | 31.01414 | 97.6227 | 94.10706 | Promoter | 13 | 30659999 | 30711231 | 51233 | 1 | ENSMUST00 | 0 | Dusp22 | ENSMUSG00000069255.7 |
| chr14 | 21993562 | 21996021 | 31.01414 | 97.6227 | 94.10706 | Distal Interge | 14 | 21983962 | 21989601 | 5640 | 2 | ENSMUST00 | -3962 | Zfp503 | ENSMUSG00000039081.9 |
| chr14 | 102980932 | 102983668 | 31.01414 | 97.6227 | 94.10706 | Promoter | 14 | 102982549 | 102982632 | 84 | 2 | ENSMUST00 | 0 | Mir5130 | ENSMUSG00000092901.1 |
| chr15 | 37960142 | 37961118 | 31.01414 | 97.6227 | 94.10706 | Promoter | 15 | 37923952 | 37961103 | 37152 | 2 | ENSMUST00 | 0 | Rrm2b | ENSMUSG00000022292.10 |
| chr18 | 67723607 | 67725123 | 31.01414 | 97.6227 | 94.10706 | Promoter | 18 | 67665511 | 67724573 | 59063 | 2 | ENSMUST00 | 0 | Ptpn2 | ENSMUSG00000024539.12 |
| chr18 | 67799533 | 67800892 | 31.01414 | 97.6227 | 94.10706 | Promoter | 18 | 67800107 | 67885168 | 85062 | 1 | ENSMUST00 | 0 | Cep192 | ENSMUSG00000024542.8 |
| chr19 | 56396578 | 56397904 | 31.01414 | 97.6227 | 94.10706 | Promoter | 19 | 56397129 | 56442344 | 45216 | 1 | ENSMUST00 | 0 | Casp7 | ENSMUSG00000025076.9 |
| chr2 | 76674806 | 76676198 | 31.01414 | 97.6227 | 94.10706 | Promoter | 2 | 76675281 | 76696828 | 21548 | 1 | ENSMUST00 | 0 | Plekha3 | ENSMUSG00000002733.8 |
| chr2 | 180171482 | 180172920 | 31.01414 | 97.6227 | 94.10706 | Promoter | 2 | 180171485 | 180176286 | 4802 | 1 | ENSMUST00 | 0 | Adrm1 | ENSMUSG00000039041.10 |
| chr3 | 94442821 | 94444075 | 31.01414 | 97.6227 | 94.10706 | Promoter | 3 | 94443318 | 94446841 | 3524 | 1 | ENSMUST00 | 0 | Mrpl9 | ENSMUSG00000028140.8 |
| chr3 | 38485055 | 38485573 | 31.01414 | 97.6227 | 94.10706 | Promoter | 3 | 38449261 | 38484844 | 35584 | 2 | ENSMUST00 | -212 | Ankrd50 | ENSMUSG00000044864.11 |
| chr4 | 156202698 | 156204105 | 31.01414 | 97.6227 | 94.10706 | Promoter | 4 | 156203304 | 156205151 | 1848 | 1 | ENSMUST00 | 0 | AW011738 | ENSMUSG00000078349.2 |
| chr6 | 38353230 | 38355708 | 31.01414 | 97.6227 | 94.10706 | Promoter | 6 | 38305286 | 38354273 | 48988 | 2 | ENSMUST00 | 0 | Zc3hav1 | ENSMUSG00000029826.9 |
| chr6 | 85431671 | 85432599 | 31.01414 | 97.6227 | 94.10706 | Promoter | 6 | 85431989 | 85446435 | 14447 | 1 | ENSMUST00 | 0 | Smyd5 | ENSMUSG00000033706.8 |
| chr8 | 127063243 | 127065529 | 31.01414 | 97.6227 | 94.10706 | Promoter | 8 | 127063893 | 127611970 | 548078 | 1 | ENSMUST00 | 0 | Pard3 | ENSMUSG00000025812.12 |
| chr8 | 116978265 | 116979486 | 31.01414 | 97.6227 | 94.10706 | Promoter | 8 | 116969599 | 116978943 | 9345 | 2 | ENSMUST00 | 0 | 1700030J22F | ENSMUSG00000031847.7 |
| chr9 | 119982428 | 119983802 | 31.01414 | 97.6227 | 94.10706 | Intron (ENSM | 9 | 119971166 | 119977250 | 6085 | 2 | ENSMUST00 | -5179 | Csrnp1 | ENSMUSG00000032515.7 |
| chr9 | 57071455 | 57074688 | 31.01414 | 97.6227 | 94.10706 | Promoter | 9 | 57072040 | 57089496 | 17457 | 1 | ENSMUST00 | 0 | Sin3a | ENSMUSG00000042557.9 |
| chr15 | 34441922 | 34444253 | 30.44295 | 97.61457 | 94.10351 | Promoter | 15 | 34440505 | 34442908 | 2404 | 2 | ENSMUST00 | 0 | Rpl30 | ENSMUSG00000058600.8 |
| chr16 | 93883326 | 93884864 | 30.44295 | 97.61457 | 94.10351 | Promoter | 16 | 93883901 | 93906115 | 22215 | 1 | ENSMUST00 | 0 | Chaf1b | ENSMUSG00000022945.6 |
| chr3 | 88716584 | 88718376 | 30.44295 | 97.61457 | 94.10351 | Promoter | 3 | 88716838 | 88729727 | 12890 | 1 | ENSMUST00 | 0 | Rit1 | ENSMUSG00000028057.9 |
| chr17 | 27555151 | 27557949 | 32.14993 | 97.60393 | 94.09416 | Promoter | 17 | 27556620 | 27563674 | 7055 | 1 | ENSMUST00 | 0 | Hmga1 | ENSMUSG00000046711.10 |
| chr4 | 123686316 | 123687833 | 32.14993 | 97.60393 | 94.09416 | Distal Interge | 4 | 123349720 | 123684360 | 334641 | 2 | ENSMUST00 | -1957 | Macf1 | ENSMUSG00000028649.12 |
| chr4 | 150086980 | 150088393 | 32.14993 | 97.60393 | 94.09416 | Promoter | 4 | 150087365 | 150105927 | 18563 | 1 | ENSMUST00 | 0 | Gpr157 | ENSMUSG00000047875.6 |
| chr5 | 72167426 | 72168293 | 32.14993 | 97.60393 | 94.09416 | Promoter | 5 | 72157000 | 72168183 | 11184 | 2 | ENSMUST00 | 0 | Commd8 | ENSMUSG00000029213.7 |
| chr7 | 123368151 | 123370934 | 32.14993 | 97.60393 | 94.09416 | Promoter | 7 | 123279150 | 123369915 | 90766 | 2 | ENSMUST00 | 0 | Arhgap17 | ENSMUSG00000030766.10 |
| chr11 | 101118885 | 101119919 | 32.62678 | 97.43105 | 93.92233 | Promoter | 11 | 101096322 | 101119843 | 23522 | 2 | ENSMUST00 | 0 | Fam134c | ENSMUSG00000017802.9 |
| chr11 | 59191203 | 59192601 | 32.62678 | 97.43105 | 93.92233 | Promoter | 11 | 59183875 | 59191952 | 8078 | 2 | ENSMUST00 | 0 | Guk1 | ENSMUSG00000020444.14 |
| chr16 | 10313103 | 10314401 | 32.62678 | 97.43105 | 93.92233 | Promoter | 16 | 10281749 | 10313968 | 32220 | 2 | ENSMUST00 | 0 | Emp2 | ENSMUSG00000022505.8 |
| chr17 | 56079589 | 56080712 | 32.62678 | 97.43105 | 93.92233 | Promoter | 17 | 56079657 | 56100596 | 20940 | 1 | ENSMUST00 | 0 | Hdgfrp2 | ENSMUSG00000002833.10 |
| chr19 | 5601647 | 5602621 | 32.62678 | 97.43105 | 93.92233 | Promoter | 19 | 5601873 | 5603439 | 1567 | 1 | ENSMUST00 | 0 | Rnaseh2c | ENSMUSG00000024925.5 |
| chr2 | 166805014 | 166806509 | 32.62678 | 97.43105 | 93.92233 | Promoter | 2 | 166805588 | 166898052 | 92465 | 1 | ENSMUST00 | 0 | Arfgef2 | ENSMUSG00000074582.5 |
| chr3 | 94757534 | 94758837 | 32.62678 | 97.43105 | 93.92233 | Promoter | 3 | 94720023 | 94758116 | 38094 | 2 | ENSMUST00 | 0 | Gm15264 | ENSMUSG00000081355.1 |
| chr4 | 59437774 | 59438840 | 32.62678 | 97.43105 | 93.92233 | Promoter | 4 | 59314683 | 59438633 | 123951 | 2 | ENSMUST00 | 0 | Susd1 | ENSMUSG00000038578.10 |
| chr4 | 121097396 | 121098624 | 32.62678 | 97.43105 | 93.92233 | Promoter | 4 | 121059237 | 121098241 | 39005 | 2 | ENSMUST00 | 0 | Zmpste24 | ENSMUSG00000043207.5 |
| chr5 | 132541111 | 132544655 | 32.62678 | 97.43105 | 93.92233 | Promoter | 5 | 131437333 | 132543344 | 1106012 | 2 | ENSMUST00 | 0 | Auts2 | ENSMUSG00000029673.12 |
| chr11 | 96064220 | 96065900 | 22.47347 | 97.37944 | 93.87287 | Promoter | 11 | 96047432 | 96065388 | 17957 | 2 | ENSMUST00 | 0 | Ube2z | ENSMUSG00000014349.7 |
| chr13 | 5861507 | 5863975 | 32.01914 | 97.37792 | 93.87154 | Promoter | 13 | 5861489 | 5870393 | 8905 | 1 | ENSMUST00 | 19 | Klf6 | ENSMUSG00000000078.6 |
| chr14 | 57056396 | 57058454 | 32.01914 | 97.37792 | 93.87154 | Promoter | 14 | 57035608 | 57058030 | 22423 | 2 | ENSMUST00 | 0 | Gja3 | ENSMUSG00000048582.6 |
| chr6 | 122811026 | 122812125 | 32.01914 | 97.37792 | 93.87154 | Downstream | 6 | 122819914 | 122845366 | 25453 | 1 | ENSMUST00 | -7789 | Foxj2 | ENSMUSG00000003154.10 |
| chr11 | 101086676 | 101087863 | 31.44063 | 97.37058 | 93.86444 | Promoter | 11 | 101087277 | 101092207 | 4931 | 1 | ENSMUST00 | 0 | Mlx | ENSMUSG00000017801.10 |
| chr12 | 99627219 | 99628409 | 29.74442 | 97.36826 | 93.86232 | Promoter | 12 | 99626053 | 99627974 | 1922 | 2 | ENSMUST00 | 0 | RP24-342G5. | ENSMUSG00000100211.1 |
| chr13 | 107021328 | 107023109 | 29.74442 | 97.36826 | 93.86232 | Promoter | 13 | 106958996 | 107022118 | 63123 | 2 | ENSMUST00 | 0 | Kif2a | ENSMUSG00000021693.14 |
| chr4 | 137276713 | 137278585 | 32.5681 | 97.3301 | 93.82507 | Promoter | 4 | 137277489 | 137299726 | 22238 | 1 | ENSMUST00 | 0 | Wnt4 | ENSMUSG00000036856.4 |
| chr10 | 80797167 | 80799728 | 24.60788 | 97.30984 | 93.80507 | Promoter | 10 | 80798198 | 80804924 | 6727 | 1 | ENSMUST00 | 0 | Sf3a2 | ENSMUSG00000020211.10 |
| chr11 | 104440325 | 104443479 | 24.60788 | 97.30984 | 93.80507 | Promoter | 11 | 104333229 | 104442291 | 109063 | 2 | ENSMUST00 | 0 | Kansl1 | ENSMUSG00000018412.11 |
| chr13 | 3917543 | 3918683 | 24.60788 | 97.30984 | 93.80507 | Promoter | 13 | 3882565 | 3918220 | 35656 | 2 | ENSMUST00 | 0 | Net1 | ENSMUSG00000021215.9 |
| chr16 | 35768281 | 35771151 | 24.60788 | 97.30984 | 93.80507 | Promoter | 16 | 35694062 | 35769356 | 75295 | 2 | ENSMUST00 | 0 | Dirc2 | ENSMUSG00000022848.8 |
| chr17 | 74394874 | 74396426 | 24.60788 | 97.30984 | 93.80507 | Promoter | 17 | 74395608 | 74424229 | 28622 | 1 | ENSMUST00 | 0 | Slc30a6 | ENSMUSG00000024069.7 |
| chr4 | 32615322 | 32616144 | 24.60788 | 97.30984 | 93.80507 | Promoter | 4 | 32615451 | 32631810 | 16360 | 1 | ENSMUST00 | 0 | Casp8ap2 | ENSMUSG00000028282.7 |
| chr4 | 107252934 | 107254526 | 24.60788 | 97.30984 | 93.80507 | Promoter | 4 | 107233514 | 107253532 | 20019 | 2 | ENSMUST00 | 0 | Lrrc42 | ENSMUSG00000028617.5 |
| chr5 | 110268875 | 110270012 | 24.60788 | 97.30984 | 93.80507 | Promoter | 5 | 110259130 | 110269913 | 10784 | 2 | ENSMUST00 | 0 | Pgam5 | ENSMUSG00000029500.11 |
| chr6 | 149100192 | 149102240 | 24.60788 | 97.30984 | 93.80507 | Promoter | 6 | 148988071 | 149101680 | 113610 | 2 | ENSMUST00 | 0 | Dennd5b | ENSMUSG00000030313.10 |
| chr7 | 142094161 | 142096036 | 24.60788 | 97.30984 | 93.80507 | Promoter | 7 | 142079490 | 142095272 | 15783 | 2 | ENSMUST00 | 0 | Dusp8 | ENSMUSG00000037887.6 |
| chr7 | 127614375 | 127616118 | 24.60788 | 97.30984 | 93.80507 | Promoter | 7 | 127607031 | 127614433 | 7403 | 2 | ENSMUST00 | 0 | Zfp629 | ENSMUSG00000045639.9 |
| chr9 | 96630303 | 96631963 | 24.60788 | 97.30984 | 93.80507 | Promoter | 9 | 96539300 | 96631617 | 92318 | 2 | ENSMUST00 | 0 | Rasa2 | ENSMUSG00000032413.7 |
| chr13 | 111685596 | 111687859 | 31.95414 | 97.26555 | 93.76166 | Promoter | 13 | 111686178 | 111718596 | 32419 | 1 | ENSMUST00 | 0 | Mier3 | ENSMUSG00000032727.8 |
| chr8 | 3278295 | 3280203 | 31.95414 | 97.26555 | 93.76166 | Promoter | 8 | 3150922 | 3279617 | 128696 | 2 | ENSMUST00 | 0 | Insr | ENSMUSG00000005534.9 |
| chr13 | 23453461 | 23454353 | 31.37882 | 97.26304 | 93.75943 | Downstream | 13 | 23456992 | 23465875 | 8884 | 2 | ENSMUST00 | 11522 | Btn1a1 | ENSMUSG00000000706.10 |
| chr14 | 55559308 | 55561525 | 31.37882 | 97.26304 | 93.75943 | Promoter | 14 | 55560006 | 55570065 | 10060 | 1 | ENSMUST00 | 0 | Dcaf11 | ENSMUSG00000022214.9 |
| chr16 | 20693151 | 20693986 | 31.37882 | 97.26304 | 93.75943 | Promoter | 16 | 20693282 | 20699283 | 6002 | 1 | ENSMUST00 | 0 | Fam131a | ENSMUSG00000050821.8 |
| chr19 | 8770661 | 8771878 | 31.37882 | 97.26304 | 93.75943 | Promoter | 19 | 8770996 | 8772475 | 1480 | 1 | ENSMUST00 | 0 | Nxf1 | ENSMUSG00000010097.8 |
| chr6 | 83349072 | 83350186 | 31.88941 | 97.15359 | 93.65204 | Promoter | 6 | 83349147 | 83358396 | 9250 | 1 | ENSMUST00 | 0 | Bola3 | ENSMUSG00000045160.9 |
| chr1 | 119420266 | 119422956 | 32.43201 | 97.0959 | 93.5947 | Promoter | 1 | 119415465 | 119422248 | 6784 | 2 | ENSMUST00 | 0 | Inhbb | ENSMUSG00000037035.5 |
| chr10 | 40682986 | 40683970 | 32.43201 | 97.0959 | 93.5947 | Promoter | 10 | 40683282 | 40811083 | 127802 | 1 | ENSMUST00 | 0 | Mettl24 | ENSMUSG00000045555.3 |
| chr17 | 33628988 | 33630087 | 32.43201 | 97.0959 | 93.5947 | Promoter | 17 | 33629090 | 33631716 | 2627 | 1 | ENSMUST00 | 0 | Zfp414 | ENSMUSG00000073423.4 |
| chr15 | 102325753 | 102327357 | 28.48738 | 97.07422 | 93.57343 | Promoter | 15 | 102326116 | 102331873 | 5758 | 1 | ENSMUST00 | 0 | Pfdn5 | ENSMUSG00000001289.7 |
| chr4 | 128883298 | 128884769 | 28.48738 | 97.07422 | 93.57343 | Promoter | 4 | 128883580 | 128903565 | 19986 | 1 | ENSMUST00 | 0 | Trim62 | ENSMUSG00000041000.8 |
| chr8 | 13907172 | 13908404 | 28.48738 | 97.07422 | 93.57343 | Promoter | 8 | 13907806 | 13940521 | 32716 | 1 | ENSMUST00 | 0 | Fbxo25 | ENSMUSG00000038365.7 |
| chr9 | 119444581 | 119445858 | 28.48738 | 97.07422 | 93.57343 | Promoter | 9 | 119444923 | 119465517 | 20595 | 1 | ENSMUST00 | 0 | Exog | ENSMUSG00000042787.8 |
| chr7 | 66688676 | 66690647 | 31.82494 | 97.04205 | 93.54184 | Promoter | 7 | 66644565 | 66688884 | 44320 | 2 | ENSMUST00 | 0 | Asb7 | ENSMUSG00000030509.11 |
| chr9 | 114779700 | 114782404 | 31.82494 | 97.04205 | 93.54184 | Promoter | 9 | 114756837 | 114781993 | 25157 | 2 | ENSMUST00 | 0 | Cmtm7 | ENSMUSG00000032436.11 |
| chr4 | 119319846 | 119321257 | 26.88811 | 97.02409 | 93.52411 | Promoter | 4 | 119300010 | 119320484 | 20475 | 2 | ENSMUST00 | 0 | Ppih | ENSMUSG00000060288.8 |
| chr10 | 93833509 | 93834612 | 27.91982 | 97.01129 | 93.51138 | Intron (ENSM | 10 | 93831555 | 93848524 | 16970 | 1 | ENSMUST00 | 1955 | Usp44 | ENSMUSG00000020020.7 |
| chr11 | 22981480 | 22982775 | 27.91982 | 97.01129 | 93.51138 | Promoter | 11 | 22896136 | 22982382 | 86247 | 2 | ENSMUST00 | 0 | Commd1 | ENSMUSG00000051355.13 |

| | | | | | | | | | | | | | | | |
|---|---|---|---|---|---|---|---|---|---|---|---|---|---|---|---|
| chr17 | 13007841 | 13009063 | 27.91982 | 97.01129 | 93.51138 | Promoter | 17 | 13007839 | 13018119 | 10281 | 1 | ENSMUST00 | 3 | Sod2 | ENSMUSG00000006818.4 |
| chr7 | 116092636 | 116094195 | 27.91982 | 97.01129 | 93.51138 | Promoter | 7 | 116093301 | 116105210 | 11910 | 1 | ENSMUST00 | 0 | 1110004F10I | ENSMUSG00000030663.7 |
| chr10 | 61475289 | 61476610 | 27.38781 | 96.99624 | 93.49658 | Promoter | 10 | 61475833 | 61582226 | 106394 | 1 | ENSMUST00 | 0 | Lrrc20 | ENSMUSG00000037151.7 |
| chr10 | 56228199 | 56229435 | 31.19482 | 96.94263 | 93.44326 | Promoter | 10 | 56014298 | 56228869 | 214392 | 2 | ENSMUST00 | 0 | Tbc1d32 | ENSMUSG00000038122.8 |
| chr11 | 121420875 | 121422162 | 30.60524 | 96.90557 | 93.40668 | Promoter | 11 | 121421401 | 121431288 | 9888 | 1 | ENSMUST00 | 0 | Fn3krp | ENSMUSG00000039253.7 |
| chr13 | 63572249 | 63573850 | 30.60524 | 96.90557 | 93.40668 | Distal Interge | 13 | 63511533 | 63565520 | 53988 | 2 | ENSMUST00 | -6730 | Ptch1 | ENSMUSG00000021466.6 |
| chr17 | 35125969 | 35127369 | 30.60524 | 96.90557 | 93.40668 | Promoter | 17 | 35126373 | 35126910 | 538 | 1 | ENSMUST00 | 0 | D17H6S53E | ENSMUSG00000043311.7 |
| chr3 | 142529802 | 142531213 | 30.60524 | 96.90557 | 93.40668 | Promoter | 3 | 142530338 | 142550083 | 19746 | 1 | ENSMUST00 | 0 | Gbp7 | ENSMUSG00000040253.10 |
| chr4 | 154024913 | 154026694 | 30.60524 | 96.90557 | 93.40668 | Promoter | 4 | 154020470 | 154025616 | 5147 | 2 | ENSMUST00 | 0 | Smim1 | ENSMUSG00000078350.6 |
| chr9 | 106170267 | 106173420 | 30.60524 | 96.90557 | 93.40668 | Promoter | 9 | 106170928 | 106191136 | 20209 | 1 | ENSMUST00 | 0 | Wdr82 | ENSMUSG00000020257.7 |
| chr1 | 171064455 | 171065442 | 11.07378 | 96.90156 | 93.40425 | Promoter | 1 | 171057828 | 171064935 | 7108 | 2 | ENSMUST00 | 0 | Fcgr3 | ENSMUSG00000059498.8 |
| chr1 | 58711052 | 58714555 | 28.91247 | 96.86193 | 93.36482 | Promoter | 1 | 58711508 | 58713886 | 2379 | 1 | ENSMUST00 | 0 | Cflar | ENSMUSG00000026031.10 |
| chr1 | 192769875 | 192771502 | 28.91247 | 96.86193 | 93.36482 | Promoter | 1 | 192512828 | 192771213 | 258386 | 2 | ENSMUST00 | 0 | Hhat | ENSMUSG00000037375.11 |
| chr1 | 105355638 | 105357120 | 28.91247 | 96.86193 | 93.36482 | Promoter | 1 | 105276914 | 105356670 | 79757 | 2 | ENSMUST00 | 0 | Rnf152 | ENSMUSG00000047496.6 |
| chr1 | 135313070 | 135314254 | 28.91247 | 96.86193 | 93.36482 | Promoter | 1 | 135295213 | 135313778 | 18566 | 2 | ENSMUST00 | 0 | Timm17a | ENSMUSG00000062580.4 |
| chr10 | 128376389 | 128377970 | 28.91247 | 96.86193 | 93.36482 | Promoter | 10 | 128377115 | 128385496 | 8382 | 1 | ENSMUST00 | 0 | Ankrd52 | ENSMUSG00000014498.4 |
| chr10 | 75859421 | 75860642 | 28.91247 | 96.86193 | 93.36482 | Promoter | 10 | 75859353 | 75860250 | 898 | 2 | ENSMUST00 | 0 | Mif | ENSMUSG00000033307.6 |
| chr10 | 41886943 | 41888211 | 28.91247 | 96.86193 | 93.36482 | Promoter | 10 | 41887439 | 41908424 | 20986 | 1 | ENSMUST00 | 0 | Sesn1 | ENSMUSG00000038332.7 |
| chr11 | 70970151 | 70971198 | 28.91247 | 96.86193 | 93.36482 | Promoter | 11 | 70970213 | 70977833 | 7621 | 1 | ENSMUST00 | 0 | Rpain | ENSMUSG00000018449.7 |
| chr11 | 80778230 | 80780044 | 28.91247 | 96.86193 | 93.36482 | Promoter | 11 | 80482126 | 80780025 | 297900 | 2 | ENSMUST00 | 0 | Myo1d | ENSMUSG00000035441.9 |
| chr11 | 79703487 | 79704556 | 28.91247 | 96.86193 | 93.36482 | Promoter | 11 | 79704630 | 79705420 | 791 | 1 | ENSMUST00 | -74 | Gm11205 | ENSMUSG00000084969.1 |
| chr12 | 78748881 | 78751313 | 28.91247 | 96.86193 | 93.36482 | Promoter | 12 | 78748947 | 78840711 | 91765 | 1 | ENSMUST00 | 0 | Mpp5 | ENSMUSG00000021112.8 |
| chr12 | 102742911 | 102744352 | 28.91247 | 96.86193 | 93.36482 | Promoter | 12 | 102743760 | 102747561 | 3802 | 1 | ENSMUST00 | 0 | Tmem251 | ENSMUSG00000046675.6 |
| chr13 | 12257055 | 12258247 | 28.91247 | 96.86193 | 93.36482 | Promoter | 13 | 12186542 | 12258113 | 71572 | 2 | ENSMUST00 | 0 | Mtr | ENSMUSG00000021311.8 |
| chr14 | 103069501 | 103071221 | 28.91247 | 96.86193 | 93.36482 | Promoter | 14 | 103070216 | 103077579 | 7364 | 1 | ENSMUST00 | 0 | Cln5 | ENSMUSG00000022125.6 |
| chr14 | 8080276 | 8081204 | 28.91247 | 96.86193 | 93.36482 | Promoter | 14 | 8080367 | 8091834 | 11468 | 1 | ENSMUST00 | 0 | Rpp14 | ENSMUSG00000023156.3 |
| chr14 | 119006416 | 119008128 | 28.91247 | 96.86193 | 93.36482 | Promoter | 14 | 118985039 | 119007465 | 22427 | 2 | ENSMUST00 | 0 | Uggt2 | ENSMUSG00000042104.13 |
| chr17 | 24695460 | 24696926 | 28.91247 | 96.86193 | 93.36482 | Promoter | 17 | 24696234 | 24700529 | 4296 | 1 | ENSMUST00 | 0 | Noxo1 | ENSMUSG00000019320.8 |
| chr17 | 32319923 | 32322060 | 28.91247 | 96.86193 | 93.36482 | Promoter | 17 | 32303676 | 32321153 | 17478 | 2 | ENSMUST00 | 0 | Akap8 | ENSMUSG00000024045.5 |
| chr17 | 69416699 | 69418386 | 28.91247 | 96.86193 | 93.36482 | Promoter | 17 | 69417402 | 69421632 | 4231 | 1 | ENSMUST00 | 0 | C030034I22R | ENSMUSG00000073374.3 |
| chr19 | 9898690 | 9900139 | 28.91247 | 96.86193 | 93.36482 | Promoter | 19 | 9872297 | 9899533 | 27237 | 2 | ENSMUST00 | 0 | Incenp | ENSMUSG00000024660.8 |
| chr2 | 122147170 | 122149548 | 28.91247 | 96.86193 | 93.36482 | Promoter | 2 | 122147686 | 122153083 | 5398 | 1 | ENSMUST00 | 0 | B2m | ENSMUSG00000060802.8 |
| chr2 | 84669973 | 84671112 | 28.91247 | 96.86193 | 93.36482 | Promoter | 2 | 84669215 | 84670770 | 1556 | 2 | ENSMUST00 | 0 | 2700094K13I | ENSMUSG00000076437.5 |
| chr3 | 95433809 | 95435481 | 28.91247 | 96.86193 | 93.36482 | Promoter | 3 | 95434388 | 95491622 | 57235 | 1 | ENSMUST00 | 0 | Arnt | ENSMUSG00000015522.13 |
| chr3 | 121722702 | 121724803 | 28.91247 | 96.86193 | 93.36482 | Promoter | 3 | 121723537 | 121735052 | 11516 | 1 | ENSMUST00 | 0 | F3 | ENSMUSG00000028128.9 |
| chr3 | 40949721 | 40951569 | 28.91247 | 96.86193 | 93.36482 | Promoter | 3 | 40950631 | 40977793 | 27163 | 1 | ENSMUST00 | 0 | Larp1b | ENSMUSG00000037814.9 |
| chr4 | 34882101 | 34883867 | 28.91247 | 96.86193 | 93.36482 | Promoter | 4 | 34803113 | 34882930 | 79818 | 2 | ENSMUST00 | 0 | Zfp292 | ENSMUSG00000039967.9 |
| chr5 | 115564886 | 115565983 | 28.91247 | 96.86193 | 93.36482 | Promoter | 5 | 115565254 | 115622656 | 57403 | 1 | ENSMUST00 | 0 | Gcn1l1 | ENSMUSG00000041638.13 |
| chr5 | 143269182 | 143270604 | 28.91247 | 96.86193 | 93.36482 | Promoter | 5 | 143249697 | 143270022 | 20326 | 2 | ENSMUST00 | 0 | Zfp316 | ENSMUSG00000046658.11 |
| chr5 | 88564232 | 88566691 | 28.91247 | 96.86193 | 93.36482 | Distal Interge | 5 | 88554483 | 88556083 | 1601 | 1 | ENSMUST00 | 9750 | Utp3 | ENSMUSG00000070697.3 |
| chr6 | 95718032 | 95719391 | 28.91247 | 96.86193 | 93.36482 | Promoter | 6 | 95474134 | 95718837 | 244704 | 2 | ENSMUST00 | 0 | Suclg2 | ENSMUSG00000061838.5 |
| chr7 | 142059991 | 142061607 | 28.91247 | 96.86193 | 93.36482 | Promoter | 7 | 142008553 | 142061034 | 52482 | 2 | ENSMUST00 | 0 | Mob2 | ENSMUSG00000025147.6 |
| chr7 | 130546352 | 130547429 | 28.91247 | 96.86193 | 93.36482 | Promoter | 7 | 130532523 | 130547381 | 14859 | 2 | ENSMUST00 | 0 | Nsmce4a | ENSMUSG00000040331.10 |
| chr7 | 126271603 | 126273389 | 28.91247 | 96.86193 | 93.36482 | Promoter | 7 | 126272619 | 126294999 | 22381 | 1 | ENSMUST00 | 0 | Sbk1 | ENSMUSG00000042978.9 |
| chr7 | 12897732 | 12898483 | 28.91247 | 96.86193 | 93.36482 | Promoter | 7 | 12897815 | 12908239 | 10425 | 1 | ENSMUST00 | 0 | Zscan22 | ENSMUSG00000054715.5 |
| chr8 | 85492134 | 85493453 | 28.91247 | 96.86193 | 93.36482 | Promoter | 8 | 85492576 | 85527560 | 34985 | 1 | ENSMUST00 | 0 | Gpt2 | ENSMUSG00000031700.6 |
| chr8 | 71950413 | 71951789 | 28.91247 | 96.86193 | 93.36482 | Promoter | 8 | 71951038 | 71970333 | 19296 | 1 | ENSMUST00 | 0 | Zfp961 | ENSMUSG00000052446.11 |
| chr9 | 21367262 | 21369079 | 28.91247 | 96.86193 | 93.36482 | Promoter | 9 | 21368019 | 21401803 | 33785 | 1 | ENSMUST00 | 0 | Ilf3 | ENSMUSG00000032178.8 |
| chr9 | 14380027 | 14381955 | 28.91247 | 96.86193 | 93.36482 | Promoter | 9 | 14353990 | 14381242 | 27253 | 2 | ENSMUST00 | 0 | Endod1 | ENSMUSG00000037419.8 |
| chr9 | 8042356 | 8043289 | 28.91247 | 96.86193 | 93.36482 | Promoter | 9 | 8021672 | 8042823 | 21152 | 2 | ENSMUST00 | 0 | 9230110C19I | ENSMUSG00000053070.4 |
| chr1 | 106758646 | 106760410 | 25.79552 | 96.83904 | 93.34669 | Promoter | 1 | 106720459 | 106759727 | 39269 | 2 | ENSMUST00 | 0 | Kdsr | ENSMUSG00000009905.5 |
| chr1 | 82839079 | 82840598 | 25.79552 | 96.83904 | 93.34669 | Promoter | 1 | 82839483 | 82901182 | 61700 | 1 | ENSMUST00 | 0 | Agfg1 | ENSMUSG00000026159.8 |
| chr1 | 182516139 | 182517998 | 25.79552 | 96.83904 | 93.34669 | Promoter | 1 | 182467256 | 182517495 | 50240 | 2 | ENSMUST00 | 0 | Capn2 | ENSMUSG00000026509.10 |
| chr10 | 128269913 | 128271255 | 25.79552 | 96.83904 | 93.34669 | Promoter | 10 | 128270576 | 128292849 | 22274 | 1 | ENSMUST00 | 0 | Stat2 | ENSMUSG00000040033.10 |
| chr11 | 80153779 | 80154721 | 25.79552 | 96.83904 | 93.34669 | Promoter | 11 | 80154105 | 80178958 | 24854 | 1 | ENSMUST00 | 0 | Adap2 | ENSMUSG00000020709.10 |
| chr11 | 119389663 | 119391370 | 25.79552 | 96.83904 | 93.34669 | Promoter | 11 | 119390472 | 119390561 | 90 | 1 | ENSMUST00 | 0 | Mir1932 | ENSMUSG00000088015.1 |
| chr14 | 70544958 | 70545891 | 25.79552 | 96.83904 | 93.34669 | Promoter | 14 | 70545251 | 70548933 | 3683 | 1 | ENSMUST00 | 0 | Reep4 | ENSMUSG00000033589.3 |
| chr16 | 21946729 | 21947835 | 25.79552 | 96.83904 | 93.34669 | Promoter | 16 | 21934327 | 21947552 | 13226 | 2 | ENSMUST00 | 0 | Tmem41a | ENSMUSG00000022856.8 |
| chr19 | 5636775 | 5638669 | 25.79552 | 96.83904 | 93.34669 | Promoter | 19 | 5637483 | 5648130 | 10648 | 1 | ENSMUST00 | 0 | Rela | ENSMUSG00000024927.7 |
| chr3 | 131271139 | 131272691 | 25.79552 | 96.83904 | 93.34669 | Promoter | 3 | 131233419 | 131272101 | 38683 | 2 | ENSMUST00 | 0 | Hadh | ENSMUSG00000027984.8 |
| chr4 | 147904068 | 147905402 | 25.79552 | 96.83904 | 93.34669 | Promoter | 4 | 147873599 | 147904687 | 31089 | 2 | ENSMUST00 | 0 | Mfn2 | ENSMUSG00000029020.8 |
| chr5 | 145139074 | 145141215 | 25.79552 | 96.83904 | 93.34669 | Promoter | 5 | 145128769 | 145140238 | 11470 | 2 | ENSMUST00 | 0 | Pdap1 | ENSMUSG00000029623.9 |
| chr5 | 121220002 | 121221164 | 25.79552 | 96.83904 | 93.34669 | Promoter | 5 | 121220219 | 121368573 | 148355 | 1 | ENSMUST00 | 0 | Gm15800 | ENSMUSG00000042744.11 |
| chr6 | 71439112 | 71440634 | 25.79552 | 96.83904 | 93.34669 | Promoter | 6 | 71388634 | 71440637 | 52004 | 2 | ENSMUST00 | 3 | Rmnd5a | ENSMUSG00000002222.9 |
| chr6 | 48444505 | 48446173 | 25.79552 | 96.83904 | 93.34669 | Promoter | 6 | 48427697 | 48445464 | 17768 | 2 | ENSMUST00 | 0 | Zfp467 | ENSMUSG00000068551.7 |
| chr7 | 28741106 | 28742576 | 25.79552 | 96.83904 | 93.34669 | Promoter | 7 | 28739641 | 28741770 | 2130 | 2 | ENSMUST00 | 0 | Mrps12 | ENSMUSG00000045948.8 |
| chr7 | 127511504 | 127513494 | 25.79552 | 96.83904 | 93.34669 | Promoter | 7 | 127511983 | 127527040 | 15058 | 1 | ENSMUST00 | 0 | Srcap | ENSMUSG00000053877.7 |
| chr9 | 54950323 | 54951673 | 25.79552 | 96.83904 | 93.34669 | Promoter | 9 | 54950790 | 54958030 | 7241 | 1 | ENSMUST00 | 0 | Psma4 | ENSMUSG00000032301.8 |
| chr11 | 60219782 | 60220851 | 31.63308 | 96.70988 | 93.22008 | Promoter | 11 | 60199089 | 60220604 | 21516 | 2 | ENSMUST00 | 0 | Srebf1 | ENSMUSG00000020538.10 |
| chr14 | 78731087 | 78733615 | 20.63592 | 96.67229 | 93.18274 | Distal Interge | 14 | 78569609 | 78725089 | 155481 | 2 | ENSMUST00 | -5999 | Dgkh | ENSMUSG00000034731.9 |
| chr1 | 93511315 | 93512892 | 27.20707 | 96.65539 | 93.16586 | Promoter | 1 | 93512079 | 93621976 | 109898 | 1 | ENSMUST00 | 0 | Farp2 | ENSMUSG00000034066.8 |
| chr1 | 184731227 | 184733192 | 27.20707 | 96.65539 | 93.16586 | Promoter | 1 | 184727140 | 184732619 | 5480 | 2 | ENSMUST00 | 0 | Hlx | ENSMUSG00000039377.6 |
| chr10 | 78463827 | 78465439 | 27.20707 | 96.65539 | 93.16586 | Promoter | 10 | 78436744 | 78464975 | 28232 | 2 | ENSMUST00 | 0 | Pdxk | ENSMUSG00000032788.9 |
| chr11 | 120377964 | 120379202 | 27.20707 | 96.65539 | 93.16586 | Promoter | 11 | 120369055 | 120378735 | 9681 | 2 | ENSMUST00 | 0 | 2310003H01I | ENSMUSG00000025384.10 |
| chr12 | 78860566 | 78861640 | 27.20707 | 96.65539 | 93.16586 | Promoter | 12 | 78842989 | 78861638 | 18650 | 2 | ENSMUST00 | 0 | Atp6v1d | ENSMUSG00000021114.8 |
| chr12 | 84616765 | 84618108 | 27.20707 | 96.65539 | 93.16586 | Promoter | 12 | 84602531 | 84617466 | 14936 | 2 | ENSMUST00 | 0 | Abcd4 | ENSMUSG00000021240.5 |
| chr17 | 24478308 | 24479534 | 27.20707 | 96.65539 | 93.16586 | Promoter | 17 | 24473550 | 24479078 | 5529 | 2 | ENSMUST00 | 0 | Mlst8 | ENSMUSG00000024142.9 |
| chr18 | 13969413 | 13973642 | 27.20707 | 96.65539 | 93.16586 | Promoter | 18 | 13687013 | 13972733 | 285721 | 2 | ENSMUST00 | 0 | Zfp521 | ENSMUSG00000024420.8 |
| chr2 | 26603398 | 26604791 | 27.20707 | 96.65539 | 93.16586 | Promoter | 2 | 26593057 | 26604211 | 11155 | 2 | ENSMUST00 | 0 | Agpat2 | ENSMUSG00000026922.8 |
| chr2 | 154891616 | 154893464 | 27.20707 | 96.65539 | 93.16586 | Promoter | 2 | 154871410 | 154892935 | 21526 | 2 | ENSMUST00 | 0 | Eif2s2 | ENSMUSG00000074656.7 |
| chr3 | 59005156 | 59007686 | 27.20707 | 96.65539 | 93.16586 | Promoter | 3 | 59006978 | 59093788 | 86811 | 1 | ENSMUST00 | 0 | Med12l | ENSMUSG00000056476.8 |
| chr6 | 71493411 | 71495597 | 27.20707 | 96.65539 | 93.16586 | Promoter | 6 | 71493894 | 71510881 | 16988 | 1 | ENSMUST00 | 0 | Rnf103 | ENSMUSG00000052656.7 |
| chr8 | 31111227 | 31112953 | 27.20707 | 96.65539 | 93.16586 | Promoter | 8 | 31111820 | 31131482 | 19663 | 1 | ENSMUST00 | 0 | Rnf122 | ENSMUSG00000039328.9 |
| chr8 | 25200274 | 25202264 | 27.20707 | 96.65539 | 93.16586 | Promoter | 8 | 25154552 | 25201449 | 46898 | 2 | ENSMUST00 | 0 | Tacc1 | ENSMUSG00000065954.5 |
| chr8 | 84200501 | 84204316 | 27.20707 | 96.65539 | 93.16586 | Distal Interge | 8 | 84192774 | 84197667 | 4894 | 2 | ENSMUST00 | -2835 | Gm26887 | ENSMUSG00000097120.1 |
| chr9 | 48479750 | 48481477 | 27.20707 | 96.65539 | 93.16586 | Promoter | 9 | 48468514 | 48480611 | 12098 | 2 | ENSMUST00 | 0 | Rexo2 | ENSMUSG00000032026.6 |
| chr17 | 40933813 | 40935462 | 32.16322 | 96.63297 | 93.14677 | Promoter | 17 | 40923051 | 40935047 | 11997 | 2 | ENSMUST00 | 0 | Cenpq | ENSMUSG00000023919.7 |
| chr4 | 138434165 | 138435568 | 32.16322 | 96.63297 | 93.14677 | Promoter | 4 | 138434671 | 138442265 | 7595 | 1 | ENSMUST00 | 0 | Mul1 | ENSMUSG00000041241.7 |
| chr8 | 70658804 | 70660437 | 32.16322 | 96.63297 | 93.14677 | Promoter | 8 | 70646436 | 70659738 | 13303 | 2 | ENSMUST00 | 0 | Pgpep1 | ENSMUSG00000056204.7 |
| chr10 | 67535269 | 67539267 | 29.88806 | 96.63187 | 93.14587 | Promoter | 10 | 67535475 | 67540246 | 4772 | 1 | ENSMUST00 | 0 | Egr2 | ENSMUSG00000037868.10 |
| chr13 | 50417527 | 50418576 | 29.88806 | 96.63187 | 93.14587 | Promoter | 13 | 50417864 | 50433769 | 15906 | 1 | ENSMUST00 | 0 | Fbxw17 | ENSMUSG00000037816.5 |
| chr2 | 57237553 | 57239193 | 29.88806 | 96.63187 | 93.14587 | Promoter | 2 | 57237635 | 57367853 | 130219 | 1 | ENSMUST00 | 0 | Gpd2 | ENSMUSG00000026827.7 |
| chr7 | 118128541 | 118130153 | 29.88806 | 96.63187 | 93.14587 | Promoter | 7 | 118118891 | 118129625 | 10735 | 2 | ENSMUST00 | 0 | Arl6ip1 | ENSMUSG00000030654.7 |
| chr11 | 83963960 | 83965228 | 31.56964 | 96.59996 | 93.11521 | Promoter | 11 | 83964428 | 84040954 | 76527 | 1 | ENSMUST00 | 0 | Synrg | ENSMUSG00000034940.10 |
| chr2 | 30471582 | 30474710 | 31.56964 | 96.59996 | 93.11521 | Promoter | 2 | 30471537 | 30474219 | 2683 | 1 | ENSMUST00 | 0 | Ier5l | ENSMUSG00000089762.3 |
| chr4 | 109000638 | 109001894 | 31.56964 | 96.59996 | 93.11521 | Promoter | 4 | 109000655 | 109061777 | 61123 | 1 | ENSMUST00 | 0 | Nrd1 | ENSMUSG00000053510.7 |
| chr7 | 111081145 | 111083078 | 31.56964 | 96.59996 | 93.11521 | Promoter | 7 | 111070568 | 111083030 | 12463 | 2 | ENSMUST00 | 0 | Eif4g2 | ENSMUSG00000005610.11 |
| chr9 | 105131129 | 105132266 | 31.56964 | 96.59996 | 93.11521 | Promoter | 9 | 105128903 | 105131312 | 2410 | 2 | ENSMUST00 | 0 | Nudt16 | ENSMUSG00000032565.8 |
| chr2 | 181592050 | 181593379 | 32.09672 | 96.51835 | 93.03485 | Promoter | 2 | 181582103 | 181592803 | 10701 | 2 | ENSMUST00 | 0 | Znf512b | ENSMUSG00000000823.11 |
| chr5 | 5748835 | 5750108 | 32.09672 | 96.51835 | 93.03485 | Promoter | 5 | 5736317 | 5749326 | 13010 | 2 | ENSMUST00 | 0 | Steap1 | ENSMUSG00000015652.4 |
| chr17 | 75551107 | 75551900 | 23.32969 | 96.49238 | 93.00913 | Promoter | 17 | 75544467 | 75551796 | 7330 | 2 | ENSMUST00 | 0 | Fam98a | ENSMUSG00000002017.9 |
| chr2 | 146855421 | 146856407 | 23.32969 | 96.49238 | 93.00913 | Promoter | 2 | 146855864 | 146970097 | 114234 | 1 | ENSMUST00 | 0 | Kiz | ENSMUSG00000074749.5 |
| chr2 | 177205881 | 177206528 | 23.32969 | 96.49238 | 93.00913 | Promoter | 2 | 177194849 | 177206421 | 11573 | 2 | ENSMUST00 | 0 | Gm14410 | ENSMUSG00000078870.4 |
| chr4 | 42998907 | 43000474 | 23.32969 | 96.49238 | 93.00913 | Promoter | 4 | 42993712 | 43000433 | 6722 | 2 | ENSMUST00 | 0 | Vcp | ENSMUSG00000028452.7 |
| chr5 | 3543740 | 3544744 | 23.32969 | 96.49238 | 93.00913 | Promoter | 5 | 3543844 | 3569810 | 25967 | 1 | ENSMUST00 | 0 | Fam133b | ENSMUSG00000058503.7 |
| chr9 | 45953182 | 45955922 | 23.32969 | 96.49238 | 93.00913 | Promoter | 9 | 45937857 | 45955247 | 17391 | 2 | ENSMUST00 | 0 | Sidt2 | ENSMUSG00000034908.10 |
| chr5 | 123973277 | 123975236 | 31.50646 | 96.49045 | 93.00759 | Promoter | 5 | 123973628 | 124003211 | 29584 | 1 | ENSMUST00 | 0 | Hip1r | ENSMUSG00000000915.10 |
| chr10 | 83359694 | 83360997 | 29.22133 | 96.43108 | 92.94852 | Promoter | 10 | 83360221 | 83368835 | 8615 | 1 | ENSMUST00 | 0 | D10Wsu102e | ENSMUSG00000020255.7 |
| chr11 | 59306344 | 59308012 | 29.22133 | 96.43108 | 92.94852 | Promoter | 11 | 59306928 | 59333552 | 26625 | 1 | ENSMUST00 | 0 | Wnt9a | ENSMUSG00000000126.6 |
| chr12 | 108791687 | 108792746 | 29.22133 | 96.43108 | 92.94852 | Promoter | 12 | 108792973 | 108816632 | 23660 | 1 | ENSMUST00 | -227 | Yy1 | ENSMUSG00000021264.11 |
| chr5 | 143908877 | 143910578 | 29.22133 | 96.43108 | 92.94852 | Promoter | 5 | 143902704 | 143909839 | 7136 | 2 | ENSMUST00 | 0 | Aimp2 | ENSMUSG00000029610.8 |
| chr6 | 83165140 | 83166832 | 29.22133 | 96.43108 | 92.94852 | Promoter | 6 | 83165920 | 83200116 | 34197 | 1 | ENSMUST00 | 0 | Dctn1 | ENSMUSG00000031865.10 |
| chr16 | 56716231 | 56717497 | 30.89292 | 96.41597 | 92.93404 | Promoter | 16 | 56690332 | 56717450 | 27119 | 2 | ENSMUST00 | 0 | Tfg | ENSMUSG00000022757.13 |
| chr11 | 107130199 | 107132795 | 32.03048 | 96.40417 | 92.92239 | Promoter | 11 | 107033081 | 107131922 | 98842 | 2 | ENSMUST00 | 0 | Bptf | ENSMUSG00000040481.11 |
| chr8 | 15026934 | 15029068 | 32.03048 | 96.40417 | 92.92239 | Promoter | 8 | 15027290 | 15033333 | 6044 | 1 | ENSMUST00 | 0 | Kbtbd11 | ENSMUSG00000055675.6 |
| chr2 | 34825403 | 34826864 | 30.83324 | 96.31171 | 92.83044 | Promoter | 2 | 34805178 | 34826073 | 20896 | 2 | ENSMUST00 | 0 | Fbxw2 | ENSMUSG00000035949.10 |
| chr11 | 78244811 | 78246806 | 28.59991 | 96.29427 | 92.81311 | Promoter | 11 | 78206746 | 78245987 | 39242 | 2 | ENSMUST00 | 0 | Supt6 | ENSMUSG00000002052.6 |
| chr2 | 93333540 | 93334966 | 28.59991 | 96.29427 | 92.81311 | Promoter | 2 | 93201760 | 93334487 | 132728 | 2 | ENSMUST00 | 0 | Tspan18 | ENSMUSG00000027217.8 |
| chr7 | 98177725 | 98178598 | 28.59991 | 96.29427 | 92.81311 | Promoter | 7 | 98121559 | 98178274 | 56716 | 2 | ENSMUST00 | 0 | Capn5 | ENSMUSG00000035547.9 |
| chr15 | 79931972 | 79933094 | 31.96453 | 96.29042 | 92.80983 | Promoter | 15 | 79915807 | 79932652 | 16846 | 2 | ENSMUST00 | 0 | Cbx7 | ENSMUSG00000053411.11 |
| chr4 | 154963727 | 154965130 | 31.96453 | 96.29042 | 92.80983 | Promoter | 4 | 154964123 | 154980871 | 16749 | 1 | ENSMUST00 | 0 | Pank4 | ENSMUSG00000029056.8 |
| chr4 | 148150673 | 148152220 | 31.96453 | 96.29042 | 92.80983 | Promoter | 4 | 148145716 | 148151908 | 6193 | 2 | ENSMUST00 | 0 | Fbxo6 | ENSMUSG00000055401.9 |
| chr7 | 44849167 | 44850909 | 31.38084 | 96.2726 | 92.79221 | Promoter | 7 | 44849224 | 44855421 | 6198 | 1 | ENSMUST00 | 0 | Akt1s1 | ENSMUSG00000011096.11 |
| chr1 | 172081970 | 172083760 | 30.77379 | 96.20782 | 92.72798 | Promoter | 1 | 172066013 | 172082795 | 16783 | 2 | ENSMUST00 | 0 | Ncstn | ENSMUSG00000003458.7 |
| chr2 | 112492346 | 112494140 | 30.77379 | 96.20782 | 92.72798 | Promoter | 2 | 112492964 | 112634573 | 141610 | 1 | ENSMUST00 | 0 | Aven | ENSMUSG00000003604.9 |
| chr4 | 84883849 | 84885458 | 30.77379 | 96.20782 | 92.72798 | Promoter | 4 | 84884309 | 85131921 | 247613 | 1 | ENSMUST00 | 0 | Cntln | ENSMUSG00000038070.10 |
| chr7 | 45830103 | 45831240 | 30.77379 | 96.20782 | 92.72798 | Promoter | 7 | 45825227 | 45830789 | 5563 | 2 | ENSMUST00 | 0 | Grwd1 | ENSMUSG00000053801.11 |
| chr9 | 103201179 | 103202396 | 30.77379 | 96.20782 | 92.72798 | Promoter | 9 | 103188032 | 103202137 | 14106 | 2 | ENSMUST00 | 0 | Srprb | ENSMUSG00000032553.9 |

| | | | | | | | | | | | | | | | |
|---|---|---|---|---|---|---|---|---|---|---|---|---|---|---|---|
| chr17 | 33954580 | 33955679 | 26.96557 | 96.19833 | 92.71962 | Promoter | 17 | 33951999 | 33955212 | 3214 | 2 | ENSMUST00 | 0 | Rps18 | ENSMUSG00000008668.9 |
| chr14 | 75845256 | 75846538 | 27.47569 | 96.18372 | 92.70509 | Promoter | 14 | 75845302 | 75848525 | 3224 | 1 | ENSMUST00 | 0 | Tpt1 | ENSMUSG00000060126.9 |
| chr2 | 120403160 | 120405558 | 27.47569 | 96.18372 | 92.70509 | Promoter | 2 | 120355312 | 120404069 | 48758 | 2 | ENSMUST00 | 0 | Tmem87a | ENSMUSG00000033808.11 |
| chr3 | 65658315 | 65659888 | 31.89884 | 96.17711 | 92.69861 | Promoter | 3 | 65659288 | 65659426 | 139 | 1 | ENSMUST00 | 0 | Mir8120 | ENSMUSG00000098944.1 |
| chr5 | 114099830 | 114101172 | 31.89884 | 96.17711 | 92.69861 | Promoter | 5 | 114100333 | 114123506 | 23174 | 1 | ENSMUST00 | 0 | Usp30 | ENSMUSG00000029592.7 |
| chr18 | 36734573 | 36735835 | 31.31841 | 96.16425 | 92.68594 | Promoter | 18 | 36735068 | 36742391 | 7324 | 1 | ENSMUST00 | 0 | Tmco6 | ENSMUSG00000006850.7 |
| chr7 | 126486726 | 126488354 | 31.31841 | 96.16425 | 92.68594 | Promoter | 7 | 126487355 | 126490730 | 3376 | 1 | ENSMUST00 | 0 | Tufm | ENSMUSG00000073838.5 |
| chr10 | 80589113 | 80590391 | 31.83343 | 96.06423 | 92.58627 | Promoter | 10 | 80583654 | 80590341 | 6688 | 2 | ENSMUST00 | 0 | Abhd17a | ENSMUSG00000003346.9 |
| chr10 | 72654498 | 72655923 | 31.83343 | 96.06423 | 92.58627 | Promoter | 10 | 72654845 | 72674964 | 20120 | 1 | ENSMUST00 | 0 | Zwint | ENSMUSG00000019923.8 |
| chr19 | 8892616 | 8893726 | 31.83343 | 96.06423 | 92.58627 | Promoter | 19 | 8892987 | 8897890 | 4904 | 1 | ENSMUST00 | 0 | Ints5 | ENSMUSG00000071652.6 |
| chr5 | 122284135 | 122285620 | 31.83343 | 96.06423 | 92.58627 | Promoter | 5 | 122284365 | 122324281 | 39917 | 1 | ENSMUST00 | 0 | Pptc7 | ENSMUSG00000038582.10 |
| chr4 | 128727024 | 128730437 | 31.25623 | 96.05631 | 92.57859 | Promoter | 4 | 128727581 | 128752881 | 25301 | 1 | ENSMUST00 | 0 | Phc2 | ENSMUSG00000028796.12 |
| chr3 | 32395974 | 32398506 | 30.09291 | 96.00345 | 92.52599 | Promoter | 3 | 32397671 | 32466107 | 68437 | 1 | ENSMUST00 | 0 | Pik3ca | ENSMUSG00000027665.8 |
| chr11 | 58008900 | 58010805 | 30.65557 | 96.00109 | 92.52367 | Promoter | 11 | 58009064 | 58062031 | 52968 | 1 | ENSMUST00 | 0 | Larp1 | ENSMUSG00000037331.10 |
| chr17 | 57030151 | 57031593 | 31.76828 | 95.95177 | 92.47445 | Promoter | 17 | 57021058 | 57031507 | 10450 | 2 | ENSMUST00 | 0 | Khsrp | ENSMUSG00000007670.9 |
| chr1 | 171410882 | 171412889 | 22.22651 | 95.90443 | 92.42858 | Promoter | 1 | 171411313 | 171416977 | 5665 | 1 | ENSMUST00 | 0 | Usf1 | ENSMUSG00000026641.8 |
| chr1 | 128102166 | 128104140 | 22.22651 | 95.90443 | 92.42858 | Promoter | 1 | 127954184 | 128102462 | 148279 | 2 | ENSMUST00 | 0 | Zranb3 | ENSMUSG00000036086.11 |
| chr11 | 62731116 | 62732190 | 30.03704 | 95.90482 | 92.42858 | Promoter | 11 | 62700356 | 62731885 | 31530 | 2 | ENSMUST00 | 0 | Zfp287 | ENSMUSG00000005267.8 |
| chr3 | 84155278 | 84155788 | 30.03704 | 95.90482 | 92.42858 | Promoter | 3 | 84087934 | 84155786 | 67853 | 2 | ENSMUST00 | 0 | Mnd1 | ENSMUSG00000033752.6 |
| chr5 | 144013952 | 144014954 | 30.03704 | 95.90482 | 92.42858 | Promoter | 5 | 143987916 | 144014853 | 26938 | 2 | ENSMUST00 | 0 | Ccz1 | ENSMUSG00000029617.9 |
| chr6 | 128837556 | 128839860 | 30.03704 | 95.90482 | 92.42858 | Distal Interge | 6 | 128843786 | 128846959 | 3174 | 1 | ENSMUST00 | -3926 | Gm26656 | ENSMUSG00000096931.1 |
| chr7 | 132575447 | 132576749 | 30.03704 | 95.90482 | 92.42858 | Promoter | 7 | 132557475 | 132576398 | 18924 | 2 | ENSMUST00 | 0 | Oat | ENSMUSG00000030934.8 |
| chr7 | 45714137 | 45715334 | 22.22651 | 95.90443 | 92.42858 | Promoter | 7 | 45710285 | 45715207 | 4923 | 2 | ENSMUST00 | 0 | Sphk2 | ENSMUSG00000057342.8 |
| chr7 | 39543998 | 39545210 | 30.03704 | 95.90482 | 92.42858 | Distal Interge | 7 | 39552496 | 39556305 | 3810 | 1 | ENSMUST00 | -7286 | RP23-402K24 | ENSMUSG00000099697.1 |
| chr8 | 72219388 | 72220648 | 30.03704 | 95.90482 | 92.42858 | Promoter | 8 | 72219730 | 72223764 | 4035 | 1 | ENSMUST00 | 0 | Fam32a | ENSMUSG00000003039.8 |
| chr9 | 99435672 | 99437648 | 30.03704 | 95.90482 | 92.42858 | Promoter | 9 | 99385420 | 99437381 | 51962 | 2 | ENSMUST00 | 0 | Mras | ENSMUSG00000032470.11 |
| chr9 | 108808289 | 108809045 | 30.03704 | 95.90482 | 92.42858 | Promoter | 9 | 108808380 | 108818366 | 9987 | 1 | ENSMUST00 | 0 | Nckipsd | ENSMUSG00000032598.7 |
| chr12 | 111907130 | 111908809 | 30.59681 | 95.89824 | 92.42339 | Promoter | 12 | 111828458 | 111908055 | 79598 | 2 | ENSMUST00 | 0 | Ppp1r13b | ENSMUSG00000021285.9 |
| chr8 | 46151196 | 46153054 | 30.59681 | 95.89824 | 92.42339 | Promoter | 8 | 46151771 | 46170191 | 18421 | 1 | ENSMUST00 | 0 | 4933411K20I | ENSMUSG00000031631.10 |
| chr1 | 72236158 | 72237397 | 30.59503 | 95.89513 | 92.42037 | Promoter | 1 | 72236946 | 72237136 | 191 | 1 | ENSMUST00 | 0 | Gm23472 | ENSMUSG00000094050.1 |
| chr11 | 100859990 | 100861109 | 30.59503 | 95.89513 | 92.42037 | Promoter | 11 | 100860345 | 100863220 | 2876 | 1 | ENSMUST00 | 0 | Stat5a | ENSMUSG00000004043.9 |
| chr11 | 98836489 | 98837525 | 30.59503 | 95.89513 | 92.42037 | Promoter | 11 | 98836785 | 98853697 | 16913 | 1 | ENSMUST00 | 0 | Rapgefl1 | ENSMUSG00000038020.5 |
| chr14 | 56810418 | 56811481 | 30.59503 | 95.89513 | 92.42037 | Promoter | 14 | 56792929 | 56811464 | 18536 | 2 | ENSMUST00 | 0 | Zmym5 | ENSMUSG00000040123.11 |
| chr15 | 8108704 | 8110145 | 30.59503 | 95.89513 | 92.42037 | Promoter | 15 | 8109313 | 8159857 | 50545 | 1 | ENSMUST00 | 0 | Nup155 | ENSMUSG00000022142.7 |
| chr17 | 56764186 | 56765623 | 30.59503 | 95.89513 | 92.42037 | Promoter | 17 | 56764751 | 56770092 | 5342 | 1 | ENSMUST00 | 0 | Dus3l | ENSMUSG00000007603.8 |
| chr17 | 47501313 | 47502596 | 30.59503 | 95.89513 | 92.42037 | Promoter | 17 | 47488050 | 47502287 | 14238 | 2 | ENSMUST00 | 0 | Taf8 | ENSMUSG00000023980.6 |
| chr17 | 22360747 | 22362108 | 30.59503 | 95.89513 | 92.42037 | Promoter | 17 | 22337989 | 22361400 | 23412 | 2 | ENSMUST00 | 0 | Zfp944 | ENSMUSG00000033972.9 |
| chr18 | 53245095 | 53246206 | 30.59503 | 95.89513 | 92.42037 | Promoter | 18 | 53245662 | 53390823 | 145162 | 1 | ENSMUST00 | 0 | Snx24 | ENSMUSG00000024535.10 |
| chr19 | 3685204 | 3686997 | 30.59503 | 95.89513 | 92.42037 | Promoter | 19 | 3584828 | 3686556 | 101729 | 2 | ENSMUST00 | 0 | Lrp5 | ENSMUSG00000024913.11 |
| chr19 | 44554460 | 44555878 | 30.59503 | 95.89513 | 92.42037 | Promoter | 19 | 44548572 | 44555426 | 6855 | 2 | ENSMUST00 | 0 | Ndufb8 | ENSMUSG00000025204.5 |
| chr2 | 125672595 | 125674020 | 30.59503 | 95.89513 | 92.42037 | Promoter | 2 | 125627447 | 125673632 | 46186 | 2 | ENSMUST00 | 0 | Shc4 | ENSMUSG00000035109.9 |
| chr2 | 153527885 | 153530666 | 30.59503 | 95.89513 | 92.42037 | Promoter | 2 | 153407462 | 153529971 | 122510 | 2 | ENSMUST00 | 0 | Nol4l | ENSMUSG00000061411.7 |
| chr2 | 84742264 | 84744578 | 30.59503 | 95.89513 | 92.42037 | Promoter | 2 | 84742169 | 84743655 | 1487 | 2 | ENSMUST00 | 0 | Gm19426 | ENSMUSG00000097187.1 |
| chr3 | 116507276 | 116508509 | 30.59503 | 95.89513 | 92.42037 | Promoter | 3 | 116488963 | 116508208 | 19246 | 2 | ENSMUST00 | 0 | Rtca | ENSMUSG00000000339.9 |
| chr3 | 152395164 | 152397484 | 30.59503 | 95.89513 | 92.42037 | Promoter | 3 | 152395473 | 152459474 | 64002 | 1 | ENSMUST00 | 0 | Zzz3 | ENSMUSG00000039068.11 |
| chr5 | 114690140 | 114691487 | 30.59503 | 95.89513 | 92.42037 | Promoter | 5 | 114669403 | 114690984 | 21582 | 2 | ENSMUST00 | 0 | Gltp | ENSMUSG00000011884.8 |
| chr5 | 31697125 | 31698833 | 30.59503 | 95.89513 | 92.42037 | Promoter | 5 | 31624439 | 31697627 | 73189 | 2 | ENSMUST00 | 0 | Rbks | ENSMUSG00000029136.7 |
| chr5 | 145113817 | 145115268 | 30.59503 | 95.89513 | 92.42037 | Promoter | 5 | 145114215 | 145129699 | 15485 | 1 | ENSMUST00 | 0 | Arpc1b | ENSMUSG00000029622.11 |
| chr6 | 113604325 | 113605680 | 30.59503 | 95.89513 | 92.42037 | Promoter | 6 | 113604772 | 113616948 | 12177 | 1 | ENSMUST00 | 0 | Brk1 | ENSMUSG00000033940.7 |
| chr6 | 59425318 | 59426818 | 30.59503 | 95.89513 | 92.42037 | Promoter | 6 | 59352461 | 59426290 | 73830 | 2 | ENSMUST00 | 0 | Gprin3 | ENSMUSG00000045441.4 |
| chr7 | 68748543 | 68749370 | 30.59503 | 95.89513 | 92.42037 | Promoter | 7 | 68737045 | 68749241 | 12197 | 2 | ENSMUST00 | 0 | Arrdc4 | ENSMUSG00000042659.9 |
| chr7 | 30050815 | 30051657 | 30.59503 | 95.89513 | 92.42037 | Downstream | 7 | 30038053 | 30044734 | 6682 | 2 | ENSMUST00 | -6082 | Zfp14 | ENSMUSG00000053985.9 |
| chr8 | 11549980 | 11551097 | 30.59503 | 95.89513 | 92.42037 | Promoter | 8 | 11514021 | 11550771 | 36751 | 2 | ENSMUST00 | 0 | Cars2 | ENSMUSG00000056228.9 |
| chr9 | 108853637 | 108854924 | 30.59503 | 95.89513 | 92.42037 | Promoter | 9 | 108854037 | 108912558 | 58522 | 1 | ENSMUST00 | 0 | Slc26a6 | ENSMUSG00000023259.11 |
| chr2 | 139677037 | 139679067 | 31.13259 | 95.84155 | 92.37124 | Promoter | 2 | 139678178 | 139758580 | 80403 | 1 | ENSMUST00 | 0 | Ism1 | ENSMUSG00000074766.5 |
| chr16 | 31080144 | 31081702 | 31.7034 | 95.83972 | 92.36961 | Promoter | 16 | 30955627 | 31081432 | 125806 | 2 | ENSMUST00 | 0 | Xxylt1 | ENSMUSG00000047434.8 |
| chr8 | 92855065 | 92855993 | 31.7034 | 95.83972 | 92.36961 | Promoter | 8 | 92855350 | 92919279 | 63930 | 1 | ENSMUST00 | 0 | Lpcat2 | ENSMUSG00000033192.4 |
| chr1 | 179667616 | 179668998 | 24.32174 | 95.77222 | 92.30251 | Promoter | 1 | 179668210 | 179687189 | 18980 | 1 | ENSMUST00 | 0 | Sccpdh | ENSMUSG00000038936.8 |
| chr17 | 46554523 | 46556263 | 24.32174 | 95.77222 | 92.30251 | Promoter | 17 | 46546839 | 46556162 | 9324 | 2 | ENSMUST00 | 0 | Srf | ENSMUSG00000015605.5 |
| chr2 | 122374944 | 122376729 | 24.32174 | 95.77222 | 92.30251 | Distal Interge | 2 | 122349128 | 122369162 | 20035 | 2 | ENSMUST00 | -5783 | Shf | ENSMUSG00000033256.9 |
| chr2 | 153239613 | 153242107 | 24.32174 | 95.77222 | 92.30251 | Promoter | 2 | 153241533 | 153270247 | 28715 | 1 | ENSMUST00 | 0 | Pofut1 | ENSMUSG00000046020.8 |
| chr3 | 56182369 | 56184233 | 24.32174 | 95.77222 | 92.30251 | Promoter | 3 | 55625205 | 56183701 | 558497 | 2 | ENSMUST00 | 0 | Nbea | ENSMUSG00000027799.8 |
| chr4 | 123282579 | 123284220 | 24.32174 | 95.77222 | 92.30251 | Promoter | 4 | 123282825 | 123298925 | 16101 | 1 | ENSMUST00 | 0 | Pabpc4 | ENSMUSG00000011257.14 |
| chr6 | 91472668 | 91474413 | 24.32174 | 95.77222 | 92.30251 | Promoter | 6 | 91473703 | 91488459 | 14757 | 1 | ENSMUST00 | 0 | Tmem43 | ENSMUSG00000030095.5 |
| chr7 | 80268932 | 80270499 | 24.32174 | 95.77222 | 92.30251 | Promoter | 7 | 80269649 | 80291579 | 21931 | 1 | ENSMUST00 | 0 | Vps33b | ENSMUSG00000030534.8 |
| chr9 | 79758867 | 79760573 | 31.07114 | 95.73475 | 92.26586 | Promoter | 9 | 79755241 | 79759853 | 4613 | 2 | ENSMUST00 | 0 | Cox7a2 | ENSMUSG00000032330.6 |
| chr10 | 13982367 | 13983535 | 31.63878 | 95.72812 | 92.25946 | Intron (ENSN | 10 | 13967024 | 14079675 | 112652 | 1 | ENSMUST00 | 15344 | Hivep2 | ENSMUSG00000015501.5 |
| chr12 | 46817868 | 46820117 | 31.63878 | 95.72812 | 92.25946 | Promoter | 12 | 46694517 | 46818775 | 124259 | 2 | ENSMUST00 | 0 | Nova1 | ENSMUSG00000021047.6 |
| chr19 | 29647505 | 29648672 | 31.63878 | 95.72812 | 92.25946 | Promoter | 19 | 29608214 | 29648415 | 40202 | 2 | ENSMUST00 | 0 | Ermp1 | ENSMUSG00000046324.7 |
| chr9 | 45838320 | 45839724 | 31.63878 | 95.72812 | 92.25946 | Promoter | 9 | 45838580 | 45862484 | 23905 | 1 | ENSMUST00 | 0 | Bace1 | ENSMUSG00000032086.7 |
| chr1 | 120263582 | 120265737 | 29.35304 | 95.67908 | 92.21078 | Promoter | 1 | 120190757 | 120265255 | 74499 | 2 | ENSMUST00 | 0 | Steap3 | ENSMUSG00000026389.11 |
| chr1 | 7088071 | 7088834 | 29.35304 | 95.67908 | 92.21078 | Promoter | 1 | 7088920 | 7173628 | 84709 | 1 | ENSMUST00 | -86 | Pcmtd1 | ENSMUSG00000051285.12 |
| chr12 | 13248352 | 13249753 | 29.35304 | 95.67908 | 92.21078 | Promoter | 12 | 13219308 | 13249213 | 29906 | 2 | ENSMUST00 | 0 | Ddx1 | ENSMUSG00000037149.9 |
| chr4 | 66402455 | 66404785 | 29.35304 | 95.67908 | 92.21078 | Promoter | 4 | 65380803 | 66404483 | 1023681 | 2 | ENSMUST00 | 0 | Astn2 | ENSMUSG00000028373.11 |
| chr4 | 116685139 | 116686728 | 29.35304 | 95.67908 | 92.21078 | Promoter | 4 | 116685544 | 116700822 | 15279 | 1 | ENSMUST00 | 0 | Prdx1 | ENSMUSG00000028691.7 |
| chr6 | 13676723 | 13678457 | 29.35304 | 95.67908 | 92.21078 | Promoter | 6 | 13625677 | 13677966 | 52290 | 2 | ENSMUST00 | 0 | B630005N14 | ENSMUSG00000042742.5 |
| chr7 | 30121250 | 30122569 | 29.35304 | 95.67908 | 92.21078 | Promoter | 7 | 30121942 | 30134665 | 12724 | 1 | ENSMUST00 | 0 | Zfp382 | ENSMUSG00000074220.4 |
| chr8 | 4237910 | 4239927 | 29.35304 | 95.67908 | 92.21078 | Promoter | 8 | 4238740 | 4247896 | 9157 | 1 | ENSMUST00 | 0 | Map2k7 | ENSMUSG00000002948.13 |
| chr9 | 110738126 | 110744546 | 29.35304 | 95.67908 | 92.21078 | Promoter | 9 | 110722088 | 110742610 | 20523 | 2 | ENSMUST00 | 0 | Pth1r | ENSMUSG00000032492.9 |
| chr11 | 11685209 | 11686868 | 32.16725 | 95.65015 | 92.18271 | Promoter | 11 | 11685925 | 11772112 | 86188 | 1 | ENSMUST00 | 0 | Ikzf1 | ENSMUSG00000018654.12 |
| chr11 | 119040284 | 119040989 | 32.16725 | 95.65015 | 92.18271 | Promoter | 11 | 119036305 | 119040969 | 4665 | 2 | ENSMUST00 | 0 | Cbx8 | ENSMUSG00000025578.9 |
| chr12 | 83632162 | 83633479 | 32.16725 | 95.65015 | 92.18271 | Promoter | 12 | 83632252 | 83683123 | 50872 | 1 | ENSMUST00 | 0 | Rbm25 | ENSMUSG00000010608.10 |
| chr14 | 46760078 | 46761094 | 32.16725 | 95.65015 | 92.18271 | Promoter | 14 | 46760541 | 46771525 | 10985 | 1 | ENSMUST00 | 0 | Cdkn3 | ENSMUSG00000037628.9 |
| chr14 | 14345517 | 14346893 | 32.16725 | 95.65015 | 92.18271 | Promoter | 14 | 14346495 | 14355490 | 8996 | 1 | ENSMUST00 | 0 | Il3ra | ENSMUSG00000068758.7 |
| chr16 | 13754154 | 13755338 | 32.16725 | 95.65015 | 92.18271 | Distal Interge | 16 | 13715059 | 13730983 | 15925 | 2 | ENSMUST00 | -23172 | Pla2g10 | ENSMUSG00000022683.8 |
| chr17 | 28800335 | 28802507 | 32.16725 | 95.65015 | 92.18271 | Promoter | 17 | 28801090 | 28838546 | 37457 | 1 | ENSMUST00 | 0 | Brpf3 | ENSMUSG00000063952.10 |
| chr19 | 53528084 | 53530077 | 32.16725 | 95.65015 | 92.18271 | Promoter | 19 | 53529109 | 53542431 | 13323 | 1 | ENSMUST00 | 0 | Dusp5 | ENSMUSG00000034765.6 |
| chr2 | 176798712 | 176799522 | 32.16725 | 95.65015 | 92.18271 | Promoter | 2 | 176798612 | 176808618 | 10007 | 1 | ENSMUST00 | 101 | Gm14295 | ENSMUSG00000078877.6 |
| chr4 | 147847823 | 147848975 | 32.16725 | 95.65015 | 92.18271 | Promoter | 4 | 147822986 | 147848366 | 25381 | 2 | ENSMUST00 | 0 | Zfp933 | ENSMUSG00000059423.8 |
| chr5 | 34512942 | 34514474 | 32.16725 | 95.65015 | 92.18271 | Promoter | 5 | 34496087 | 34513963 | 17877 | 2 | ENSMUST00 | 0 | Tnip2 | ENSMUSG00000059866.8 |
| chr8 | 111337324 | 111338558 | 32.16725 | 95.65015 | 92.18271 | Promoter | 8 | 111311797 | 111338177 | 26381 | 2 | ENSMUST00 | 0 | Mlkl | ENSMUSG00000012519.9 |
| chrX | 6047176 | 6048520 | 32.16725 | 95.65015 | 92.18271 | Promoter | 20 | 6047453 | 6055110 | 7658 | 1 | ENSMUST00 | 0 | Nudt11 | ENSMUSG00000073295.4 |
| chr11 | 95258496 | 95260049 | 31.00994 | 95.62833 | 92.16279 | Distal Interge | 11 | 95261529 | 95269265 | 7737 | 1 | ENSMUST00 | -1480 | Tac4 | ENSMUSG00000020872.4 |
| chr17 | 6106150 | 6108092 | 31.00994 | 95.62833 | 92.16279 | Promoter | 17 | 6106437 | 6138792 | 32356 | 1 | ENSMUST00 | 0 | Tulp4 | ENSMUSG00000034377.5 |
| chr15 | 57975179 | 57976978 | 31.57443 | 95.61693 | 92.15154 | Distal Interge | 15 | 57985419 | 58011009 | 25591 | 1 | ENSMUST00 | -8441 | Fam83a | ENSMUSG00000051225.4 |
| chr2 | 90678746 | 90680070 | 31.57443 | 95.61693 | 92.15154 | 5' UTR | 2 | 90677215 | 90736328 | 59114 | 1 | ENSMUST00 | 1532 | Nup160 | ENSMUSG00000051329.8 |
| chr3 | 152339838 | 152340784 | 31.57443 | 95.61693 | 92.15154 | Promoter | 3 | 152273459 | 152340407 | 66949 | 2 | ENSMUST00 | 0 | Fam73a | ENSMUSG00000054942.8 |
| chr5 | 92277532 | 92278322 | 31.57443 | 95.61693 | 92.15154 | Promoter | 5 | 92257659 | 92278170 | 20512 | 2 | ENSMUST00 | 0 | Naaa | ENSMUSG00000029413.9 |
| chr5 | 139371444 | 139372564 | 31.57443 | 95.61693 | 92.15154 | Intron (ENSN | 5 | 139369650 | 139369745 | 96 | 2 | ENSMUST00 | -1700 | Mir339 | ENSMUSG00000065593.1 |
| chr3 | 98381893 | 98383460 | 30.42184 | 95.59177 | 92.12663 | Promoter | 3 | 98382481 | 98431948 | 49468 | 1 | ENSMUST00 | 0 | Zfp697 | ENSMUSG00000050064.9 |
| chr19 | 6140301 | 6141783 | 30.94897 | 95.52228 | 92.05737 | Promoter | 19 | 6134374 | 6141140 | 6767 | 2 | ENSMUST00 | 0 | Arl2 | ENSMUSG00000024944.5 |
| chr11 | 119942167 | 119942815 | 28.71633 | 95.52087 | 92.05702 | Promoter | 11 | 119942763 | 120002621 | 59859 | 1 | ENSMUST00 | 0 | Baiap2 | ENSMUSG00000025372.11 |
| chr14 | 78848611 | 78849970 | 28.71633 | 95.52087 | 92.05702 | Promoter | 14 | 78849178 | 79202310 | 353133 | 1 | ENSMUST00 | 0 | Vwa8 | ENSMUSG00000058997.7 |
| chr19 | 59942619 | 59944211 | 28.71633 | 95.52087 | 92.05702 | Promoter | 19 | 59902884 | 59943654 | 40771 | 2 | ENSMUST00 | 0 | Rab11fip2 | ENSMUSG00000040022.9 |
| chr3 | 83055444 | 83056344 | 28.71633 | 95.52087 | 92.05702 | Promoter | 3 | 83055522 | 83072355 | 16834 | 1 | ENSMUST00 | 0 | Plrg1 | ENSMUSG00000027998.6 |
| chr5 | 5693212 | 5694665 | 28.71633 | 95.52087 | 92.05702 | Promoter | 5 | 5665954 | 5694060 | 28107 | 2 | ENSMUST00 | 0 | Steap2 | ENSMUSG00000015653.8 |
| chr9 | 123020357 | 123022072 | 28.71633 | 95.52087 | 92.05702 | Promoter | 9 | 123021326 | 123023490 | 2165 | 1 | ENSMUST00 | 0 | Tmem42 | ENSMUSG00000066233.5 |
| chr15 | 59314322 | 59315989 | 31.51034 | 95.50614 | 92.04281 | Promoter | 15 | 59315077 | 59328958 | 13882 | 1 | ENSMUST00 | 0 | Sqle | ENSMUSG00000022351.9 |
| chr5 | 110284644 | 110286789 | 31.51034 | 95.50614 | 92.04281 | Promoter | 5 | 110286306 | 110337474 | 51169 | 1 | ENSMUST00 | 0 | Pole | ENSMUSG00000007080.9 |
| chr2 | 104711578 | 104712863 | 21.26469 | 95.5017 | 92.03861 | Promoter | 2 | 104678017 | 104712156 | 34140 | 2 | ENSMUST00 | 0 | Tcp11l1 | ENSMUSG00000027175.5 |
| chr5 | 96996701 | 96998913 | 21.26469 | 95.5017 | 92.03861 | Promoter | 5 | 96997689 | 97069649 | 71961 | 1 | ENSMUST00 | 0 | Bmp2k | ENSMUSG00000034663.8 |
| chr7 | 4148241 | 4151710 | 21.26469 | 95.5017 | 92.03861 | Promoter | 7 | 4147915 | 4150374 | 2460 | 2 | ENSMUST00 | 0 | Leng9 | ENSMUSG00000043432.7 |
| chr11 | 76397725 | 76399811 | 28.12215 | 95.42207 | 91.9593 | Promoter | 11 | 76257198 | 76399140 | 141943 | 2 | ENSMUST00 | 0 | Nxn | ENSMUSG00000020844.6 |
| chr17 | 56217823 | 56219558 | 28.12215 | 95.42207 | 91.9593 | Promoter | 17 | 56186682 | 56218889 | 32208 | 2 | ENSMUST00 | 0 | Dpp9 | ENSMUSG00000001229.7 |
| chr3 | 90212135 | 90214248 | 28.12215 | 95.42207 | 91.9593 | Promoter | 3 | 90213695 | 90226385 | 12691 | 1 | ENSMUST00 | 0 | Rab13 | ENSMUSG00000027935.9 |
| chr7 | 4788688 | 4789872 | 31.4465 | 95.39577 | 91.93361 | Promoter | 7 | 4784595 | 4789656 | 5062 | 2 | ENSMUST00 | 0 | Tmem238 | ENSMUSG00000030431.7 |
| chr19 | 5366022 | 5367764 | 27.5664 | 95.37588 | 91.91415 | Promoter | 19 | 5366813 | 5371511 | 4699 | 1 | ENSMUST00 | 0 | Eif1ad | ENSMUSG00000024841.7 |
| chr15 | 102256411 | 102258508 | 31.97522 | 95.31979 | 91.85828 | Promoter | 15 | 102234938 | 102257517 | 22580 | 2 | ENSMUST00 | 0 | Rarg | ENSMUSG00000001288.9 |
| chr15 | 85810839 | 85812186 | 30.24887 | 95.28834 | 91.82743 | Promoter | 15 | 85806972 | 85811696 | 4725 | 2 | ENSMUST00 | 0 | Cdpf1 | ENSMUSG00000064284.7 |
| chr16 | 59471585 | 59472836 | 30.24887 | 95.28834 | 91.82743 | Promoter | 16 | 59471775 | 59492461 | 20687 | 1 | ENSMUST00 | 0 | Mina | ENSMUSG00000022724.10 |
| chr4 | 41465045 | 41465680 | 31.38293 | 95.2858 | 91.82495 | Promoter | 4 | 41465151 | 41480926 | 15776 | 1 | ENSMUST00 | 0 | Nudt2 | ENSMUSG00000028443.6 |
| chr9 | 43239020 | 43240233 | 31.38293 | 95.2858 | 91.82495 | Promoter | 9 | 43221278 | 43239816 | 18539 | 2 | ENSMUST00 | 0 | Oaf | ENSMUSG00000032014.5 |
| chr1 | 59762382 | 59765962 | 25.48472 | 95.2657 | 91.8051 | Promoter | 1 | 59763400 | 59879014 | 115615 | 1 | ENSMUST00 | 0 | Bmpr2 | ENSMUSG00000067336.6 |
| chr10 | 13552480 | 13554033 | 25.48472 | 95.2657 | 91.8051 | Promoter | 10 | 13552894 | 13563376 | 10483 | 1 | ENSMUST00 | 0 | Adat2 | ENSMUSG00000019808.8 |
| chr10 | 71347157 | 71349107 | 25.48472 | 95.2657 | 91.8051 | Promoter | 10 | 71347763 | 71385885 | 38123 | 1 | ENSMUST00 | 0 | Ipmk | ENSMUSG00000060733.8 |
| chr11 | 116108781 | 116110664 | 25.48472 | 95.2657 | 91.8051 | Promoter | 11 | 116105752 | 116110237 | 4486 | 2 | ENSMUST00 | 0 | Trim47 | ENSMUSG00000020773.6 |
| chr11 | 116580541 | 116581616 | 25.48472 | 95.2657 | 91.8051 | Promoter | 11 | 116537740 | 116581447 | 43708 | 2 | ENSMUST00 | 0 | Ube2o | ENSMUSG00000020802.8 |
| chr13 | 55714413 | 55715531 | 25.48472 | 95.2657 | 91.8051 | Promoter | 13 | 55714650 | 55726221 | 11572 | 1 | ENSMUST00 | 0 | Txndc15 | ENSMUSG00000021497.9 |

| | | | | | | | | | | | | | | | |
|---|---|---|---|---|---|---|---|---|---|---|---|---|---|---|---|
| chr15 | 101283352 | 101285088 | 25.48472 | 95.2657 | 91.8051 | Promoter | 15 | 101284301 | 101290934 | 6634 | 1 | ENSMUST00 | 0 | Atg101 | ENSMUSG00000037204.6 |
| chr16 | 20498071 | 20499669 | 25.48472 | 95.2657 | 91.8051 | Promoter | 16 | 20498817 | 20509955 | 11139 | 1 | ENSMUST00 | 0 | Eif2b5 | ENSMUSG00000003235.7 |
| chr16 | 23127320 | 23128515 | 25.48472 | 95.2657 | 91.8051 | Promoter | 16 | 23113943 | 23127737 | 13795 | 2 | ENSMUST00 | 0 | Rfc4 | ENSMUSG00000022881.9 |
| chr16 | 90933837 | 90935564 | 25.48472 | 95.2657 | 91.8051 | Promoter | 16 | 90925809 | 90934827 | 9019 | 2 | ENSMUST00 | 0 | 1110004E09I | ENSMUSG00000022972.4 |
| chr18 | 34220262 | 34222439 | 25.48472 | 95.2657 | 91.8051 | Promoter | 18 | 34220924 | 34313506 | 92583 | 1 | ENSMUST00 | 0 | Apc | ENSMUSG00000005871.9 |
| chr19 | 23273105 | 23274085 | 25.48472 | 95.2657 | 91.8051 | Promoter | 19 | 23206441 | 23273897 | 67457 | 2 | ENSMUST00 | 0 | Smc5 | ENSMUSG00000024943.8 |
| chr2 | 118478601 | 118480175 | 25.48472 | 95.2657 | 91.8051 | Promoter | 2 | 118475850 | 118479711 | 3862 | 2 | ENSMUST00 | 0 | Srp14 | ENSMUSG00000009549.9 |
| chr2 | 119046523 | 119047735 | 25.48472 | 95.2657 | 91.8051 | Promoter | 2 | 119047119 | 119069271 | 22153 | 1 | ENSMUST00 | 0 | Casc5 | ENSMUSG00000027326.8 |
| chr3 | 8666061 | 8668353 | 25.48472 | 95.2657 | 91.8051 | Promoter | 3 | 8663362 | 8667039 | 3678 | 2 | ENSMUST00 | 0 | Hey1 | ENSMUSG00000040289.3 |
| chr5 | 53809403 | 53810342 | 25.48472 | 95.2657 | 91.8051 | Promoter | 5 | 53809627 | 53904380 | 94754 | 1 | ENSMUST00 | 0 | Tbc1d19 | ENSMUSG00000039178.6 |
| chr6 | 50565606 | 50567370 | 25.48472 | 95.2657 | 91.8051 | Promoter | 6 | 50562563 | 50566538 | 3976 | 2 | ENSMUST00 | 0 | Cycs | ENSMUSG00000063694.5 |
| chr8 | 23256419 | 23257782 | 25.48472 | 95.2657 | 91.8051 | Promoter | 8 | 23241353 | 23257021 | 15669 | 2 | ENSMUST00 | 0 | Golga7 | ENSMUSG00000015341.11 |
| chr8 | 27085211 | 27087724 | 25.48472 | 95.2657 | 91.8051 | Promoter | 8 | 27085583 | 27123435 | 37853 | 1 | ENSMUST00 | 0 | Gpr124 | ENSMUSG00000031486.9 |
| chr12 | 81593966 | 81595618 | 31.90856 | 95.20506 | 91.74643 | Promoter | 12 | 81573557 | 81595006 | 21450 | 2 | ENSMUST00 | 0 | Med6 | ENSMUSG00000002679.9 |
| chr2 | 49619087 | 49620154 | 31.90856 | 95.20506 | 91.74643 | Promoter | 2 | 49619298 | 49774778 | 155481 | 1 | ENSMUST00 | 0 | Kif5c | ENSMUSG00000026764.10 |
| chr6 | 113442252 | 113443389 | 31.90856 | 95.20506 | 91.74643 | Promoter | 6 | 113442569 | 113448229 | 5661 | 1 | ENSMUST00 | 0 | Jagn1 | ENSMUSG00000051256.9 |
| chr10 | 128410784 | 128412444 | 28.53699 | 95.19626 | 91.7378 | Promoter | 10 | 128411616 | 128441438 | 29823 | 1 | ENSMUST00 | 0 | Rnf41 | ENSMUSG00000025373.8 |
| chr10 | 128367888 | 128369602 | 28.53699 | 95.19626 | 91.7378 | Promoter | 10 | 128363097 | 128370032 | 6936 | 2 | ENSMUST00 | 430 | Coq10a | ENSMUSG00000039914.5 |
| chr12 | 110978291 | 110979566 | 28.53699 | 95.19626 | 91.7378 | Promoter | 12 | 110975353 | 110979040 | 3688 | 2 | ENSMUST00 | 0 | Ankrd9 | ENSMUSG00000037904.9 |
| chr13 | 55321107 | 55323065 | 28.53699 | 95.19626 | 91.7378 | Promoter | 13 | 55322055 | 55325271 | 3217 | 1 | ENSMUST00 | 0 | Prelid1 | ENSMUSG00000021486.6 |
| chr14 | 18269737 | 18271010 | 28.53699 | 95.19626 | 91.7378 | Promoter | 14 | 18268946 | 18270626 | 1681 | 2 | ENSMUST00 | 0 | Rpl15 | ENSMUSG00000012405.10 |
| chr14 | 16248824 | 16250539 | 28.53699 | 95.19626 | 91.7378 | Promoter | 14 | 16249280 | 16311926 | 62647 | 1 | ENSMUST00 | 0 | Ngly1 | ENSMUSG00000021785.7 |
| chr14 | 122982323 | 122984015 | 28.53699 | 95.19626 | 91.7378 | Promoter | 14 | 122918971 | 122983283 | 64313 | 2 | ENSMUST00 | 0 | Tmtc4 | ENSMUSG00000041594.12 |
| chr15 | 79440865 | 79442525 | 28.53699 | 95.19626 | 91.7378 | Promoter | 15 | 79417856 | 79441999 | 24144 | 2 | ENSMUST00 | 0 | Csnk1e | ENSMUSG00000022433.13 |
| chr15 | 88750934 | 88752603 | 28.53699 | 95.19626 | 91.7378 | Promoter | 15 | 88751660 | 88784516 | 32857 | 1 | ENSMUST00 | 0 | Zbed4 | ENSMUSG00000034333.10 |
| chr16 | 17276273 | 17277170 | 28.53699 | 95.19626 | 91.7378 | Promoter | 16 | 17276300 | 17278660 | 2361 | 1 | ENSMUST00 | 0 | Tmem191c | ENSMUSG00000055692.13 |
| chr17 | 23725979 | 23727245 | 28.53699 | 95.19626 | 91.7378 | Promoter | 17 | 23726336 | 23736729 | 10394 | 1 | ENSMUST00 | 0 | Pkmyt1 | ENSMUSG00000023908.7 |
| chr17 | 24413610 | 24415692 | 28.53699 | 95.19626 | 91.7378 | Promoter | 17 | 24414675 | 24425895 | 11221 | 1 | ENSMUST00 | 0 | Rnps1 | ENSMUSG00000034681.11 |
| chr18 | 54985294 | 54988807 | 28.53699 | 95.19626 | 91.7378 | 5' UTR | 18 | 54888045 | 54990180 | 102136 | 2 | ENSMUST00 | 1373 | Zfp608 | ENSMUSG00000052713.8 |
| chr2 | 120969913 | 120971260 | 28.53699 | 95.19626 | 91.7378 | Promoter | 2 | 120860269 | 120970715 | 110447 | 2 | ENSMUST00 | 0 | Ubr1 | ENSMUSG00000027272.5 |
| chr2 | 119799062 | 119801370 | 28.53699 | 95.19626 | 91.7378 | Promoter | 2 | 119799239 | 119805650 | 6412 | 1 | ENSMUST00 | 0 | Tyro3 | ENSMUSG00000027298.12 |
| chr2 | 158768008 | 158768690 | 28.53699 | 95.19626 | 91.7378 | Promoter | 2 | 158768093 | 158786637 | 18545 | 1 | ENSMUST00 | 0 | Fam83d | ENSMUSG00000027654.2 |
| chr2 | 34406362 | 34407501 | 28.53699 | 95.19626 | 91.7378 | Promoter | 2 | 34406771 | 34444408 | 37638 | 1 | ENSMUST00 | 0 | Mapkap1 | ENSMUSG00000038696.9 |
| chr2 | 166712878 | 166713884 | 28.53699 | 95.19626 | 91.7378 | Promoter | 2 | 166566342 | 166713832 | 147491 | 2 | ENSMUST00 | 0 | Prex1 | ENSMUSG00000039621.8 |
| chr5 | 114707703 | 114708580 | 28.53699 | 95.19626 | 91.7378 | Promoter | 5 | 114707760 | 114722327 | 14568 | 1 | ENSMUST00 | 0 | Tchp | ENSMUSG00000002486.10 |
| chr5 | 21786262 | 21787620 | 28.53699 | 95.19626 | 91.7378 | Promoter | 5 | 21785283 | 21803787 | 18505 | 1 | ENSMUST00 | 980 | Psmc2 | ENSMUSG00000028932.7 |
| chr5 | 65434910 | 65436584 | 28.53699 | 95.19626 | 91.7378 | Promoter | 5 | 65413202 | 65435731 | 22530 | 2 | ENSMUST00 | 0 | Ugdh | ENSMUSG00000029201.9 |
| chr6 | 47877134 | 47878849 | 28.53699 | 95.19626 | 91.7378 | Promoter | 6 | 47877204 | 47908485 | 31282 | 1 | ENSMUST00 | 0 | Zfp282 | ENSMUSG00000025821.9 |
| chr6 | 65671465 | 65673440 | 28.53699 | 95.19626 | 91.7378 | Promoter | 6 | 65671590 | 65706930 | 35341 | 1 | ENSMUST00 | 0 | Ndnf | ENSMUSG00000049001.4 |
| chr7 | 4684288 | 4685518 | 28.53699 | 95.19626 | 91.7378 | Promoter | 7 | 4660521 | 4684963 | 24443 | 2 | ENSMUST00 | 0 | Hspbp1 | ENSMUSG00000063802.4 |
| chr8 | 106415187 | 106415996 | 28.53699 | 95.19626 | 91.7378 | Promoter | 8 | 106415339 | 106425889 | 10551 | 1 | ENSMUST00 | 0 | Zfp90 | ENSMUSG00000031907.8 |
| chr9 | 65214597 | 65216055 | 28.53699 | 95.19626 | 91.7378 | Promoter | 9 | 65214690 | 65239219 | 24530 | 1 | ENSMUST00 | 0 | Parp16 | ENSMUSG00000032392.10 |
| chr9 | 51764398 | 51766274 | 28.53699 | 95.19626 | 91.7378 | Promoter | 9 | 51765337 | 51790249 | 24913 | 1 | ENSMUST00 | 0 | Arhgap20 | ENSMUSG00000053199.8 |
| chr1 | 36470838 | 36472511 | 30.19165 | 95.18787 | 91.73401 | Promoter | 1 | 36471620 | 36508764 | 37145 | 1 | ENSMUST00 | 0 | Cnnm4 | ENSMUSG00000037408.9 |
| chr10 | 80038985 | 80039798 | 30.19165 | 95.18787 | 91.73401 | Promoter | 10 | 80035949 | 80039710 | 3762 | 2 | ENSMUST00 | 0 | Polr2e | ENSMUSG00000004667.13 |
| chr10 | 94147520 | 94148719 | 30.19165 | 95.18787 | 91.73401 | Promoter | 10 | 94148023 | 94197211 | 49189 | 1 | ENSMUST00 | 0 | Nr2c1 | ENSMUSG00000005897.9 |
| chr10 | 127290751 | 127292540 | 30.19165 | 95.18787 | 91.73401 | Promoter | 10 | 127290774 | 127296288 | 5515 | 1 | ENSMUST00 | 0 | Ddit3 | ENSMUSG00000025408.10 |
| chr11 | 120597382 | 120599283 | 30.19165 | 95.18787 | 91.73401 | Promoter | 11 | 120592121 | 120598365 | 6245 | 2 | ENSMUST00 | 0 | Alyref | ENSMUSG00000025134.2 |
| chr19 | 61140231 | 61141320 | 30.19165 | 95.18787 | 91.73401 | Promoter | 19 | 61053840 | 61140795 | 86956 | 2 | ENSMUST00 | 0 | 5830428H23I | ENSMUSG00000074733.9 |
| chr7 | 128460002 | 128461770 | 30.19165 | 95.18787 | 91.73401 | Promoter | 7 | 128439777 | 128461672 | 21896 | 2 | ENSMUST00 | 0 | Tial1 | ENSMUSG00000030846.10 |
| chr9 | 44112814 | 44113565 | 30.19165 | 95.18787 | 91.73401 | Promoter | 9 | 44066327 | 44113500 | 47174 | 2 | ENSMUST00 | 0 | Rnf26 | ENSMUSG00000053128.9 |
| chr3 | 94974132 | 94975930 | 30.70748 | 95.10176 | 91.64964 | Promoter | 3 | 94974731 | 95006843 | 32113 | 1 | ENSMUST00 | 0 | Pi4kb | ENSMUSG00000038861.9 |
| chr12 | 84038265 | 84039142 | 31.84217 | 95.09076 | 91.6388 | Promoter | 12 | 84038379 | 84044723 | 6345 | 1 | ENSMUST00 | 0 | Acot4 | ENSMUSG00000052392.4 |
| chr9 | 55326284 | 55327538 | 31.25655 | 95.06709 | 91.61534 | Promoter | 9 | 55221301 | 55326749 | 105449 | 2 | ENSMUST00 | 0 | Nrg4 | ENSMUSG00000032311.12 |
| chr1 | 21077190 | 21079961 | 26.86698 | 95.04005 | 91.5885 | Promoter | 1 | 20996299 | 21079229 | 82931 | 2 | ENSMUST00 | 0 | Tram2 | ENSMUSG00000041779.5 |
| chr1 | 40790126 | 40791201 | 26.86698 | 95.04005 | 91.5885 | Promoter | 1 | 40772038 | 40790663 | 18626 | 2 | ENSMUST00 | 0 | Mfsd9 | ENSMUSG00000041945.7 |
| chr1 | 180813696 | 180814720 | 26.86698 | 95.04005 | 91.5885 | Promoter | 1 | 180803024 | 180813943 | 10920 | 2 | ENSMUST00 | 0 | H3f3a | ENSMUSG00000060743.7 |
| chr10 | 69211966 | 69213907 | 26.86698 | 95.04005 | 91.5885 | Promoter | 10 | 69212634 | 69291791 | 79158 | 1 | ENSMUST00 | 0 | Rhobtb1 | ENSMUSG00000019944.9 |
| chr10 | 41302583 | 41304291 | 26.86698 | 95.04005 | 91.5885 | Promoter | 10 | 41188172 | 41303241 | 115070 | 2 | ENSMUST00 | 0 | Fig4 | ENSMUSG00000038417.3 |
| chr11 | 61266257 | 61267900 | 26.86698 | 95.04005 | 91.5885 | Promoter | 11 | 61223417 | 61267185 | 43769 | 2 | ENSMUST00 | 0 | Aldh3a2 | ENSMUSG00000010025.13 |
| chr11 | 106918841 | 106920785 | 26.86698 | 95.04005 | 91.5885 | Promoter | 11 | 106820066 | 106920715 | 100650 | 2 | ENSMUST00 | 0 | Smurf2 | ENSMUSG00000018363.11 |
| chr11 | 57800608 | 57802887 | 26.86698 | 95.04005 | 91.5885 | Promoter | 11 | 57801637 | 57810217 | 8581 | 1 | ENSMUST00 | 0 | Sap30l | ENSMUSG00000020519.5 |
| chr11 | 113682857 | 113685142 | 26.86698 | 95.04005 | 91.5885 | Promoter | 11 | 113684412 | 113694647 | 10236 | 1 | ENSMUST00 | 0 | D11Wsu47e | ENSMUSG00000041623.12 |
| chr11 | 120149746 | 120151354 | 26.86698 | 95.04005 | 91.5885 | Promoter | 11 | 120103960 | 120151346 | 47387 | 2 | ENSMUST00 | 0 | Slc38a10 | ENSMUSG00000061306.10 |
| chr12 | 4840928 | 4842094 | 26.86698 | 95.04005 | 91.5885 | Promoter | 12 | 4833175 | 4841595 | 8421 | 2 | ENSMUST00 | 0 | Fkbp1b | ENSMUSG00000020635.7 |
| chr14 | 101639794 | 101640933 | 26.86698 | 95.04005 | 91.5885 | Promoter | 14 | 101633766 | 101640686 | 6921 | 2 | ENSMUST00 | 0 | Commd6 | ENSMUSG00000075486.4 |
| chr14 | 79451301 | 79452111 | 26.86698 | 95.04005 | 91.5885 | Promoter | 14 | 79451835 | 79454816 | 2982 | 1 | ENSMUST00 | 0 | Zbtbd6 | ENSMUSG00000075502.1 |
| chr16 | 18065862 | 18067235 | 26.86698 | 95.04005 | 91.5885 | Promoter | 16 | 18066403 | 18071363 | 4961 | 1 | ENSMUST00 | 0 | Dgcr6 | ENSMUSG00000003531.9 |
| chr16 | 44557656 | 44559430 | 26.86698 | 95.04005 | 91.5885 | Promoter | 16 | 44485045 | 44558870 | 73826 | 2 | ENSMUST00 | 0 | Boc | ENSMUSG00000022687.6 |
| chr16 | 37776641 | 37777898 | 26.86698 | 95.04005 | 91.5885 | Promoter | 16 | 37776873 | 37836514 | 59642 | 1 | ENSMUST00 | 0 | Fstl1 | ENSMUSG00000022816.6 |
| chr16 | 10541909 | 10543316 | 26.86698 | 95.04005 | 91.5885 | Promoter | 16 | 10502531 | 10543054 | 40524 | 2 | ENSMUST00 | 0 | Dexi | ENSMUSG00000038055.7 |
| chr18 | 10028791 | 10030110 | 26.86698 | 95.04005 | 91.5885 | Promoter | 18 | 9995594 | 10030067 | 34474 | 2 | ENSMUST00 | 0 | Usp14 | ENSMUSG00000047879.11 |
| chr2 | 155517941 | 155518867 | 26.86698 | 95.04005 | 91.5885 | Promoter | 2 | 155504773 | 155518237 | 13465 | 2 | ENSMUST00 | 0 | Ggt7 | ENSMUSG00000027603.10 |
| chr2 | 25261996 | 25263624 | 26.86698 | 95.04005 | 91.5885 | Promoter | 2 | 25262618 | 25269885 | 7268 | 1 | ENSMUST00 | 0 | Tprn | ENSMUSG00000048707.9 |
| chr2 | 125624336 | 125625729 | 26.86698 | 95.04005 | 91.5885 | Promoter | 2 | 125563088 | 125625113 | 62026 | 2 | ENSMUST00 | 0 | Cep152 | ENSMUSG00000068394.4 |
| chr4 | 12171134 | 12172245 | 26.86698 | 95.04005 | 91.5885 | Promoter | 4 | 12153409 | 12172015 | 18607 | 2 | ENSMUST00 | 0 | Fam92a | ENSMUSG00000028218.14 |
| chr4 | 43442217 | 43444119 | 26.86698 | 95.04005 | 91.5885 | Promoter | 4 | 43443321 | 43447794 | 4474 | 1 | ENSMUST00 | 0 | Tesk1 | ENSMUSG00000028458.7 |
| chr4 | 83416858 | 83418806 | 26.86698 | 95.04005 | 91.5885 | Promoter | 4 | 83417724 | 83453340 | 35617 | 1 | ENSMUST00 | 0 | Snapc3 | ENSMUSG00000028483.8 |
| chr4 | 152297280 | 152298074 | 26.86698 | 95.04005 | 91.5885 | Promoter | 4 | 152297304 | 152307120 | 9817 | 1 | ENSMUST00 | 0 | Icmt | ENSMUSG00000039662.11 |
| chr4 | 138972203 | 138973837 | 26.86698 | 95.04005 | 91.5885 | Promoter | 4 | 138972888 | 138990720 | 17833 | 1 | ENSMUST00 | 0 | Tmco4 | ENSMUSG00000041143.11 |
| chr6 | 124931319 | 124932573 | 26.86698 | 95.04005 | 91.5885 | Promoter | 6 | 124931388 | 124936138 | 4751 | 1 | ENSMUST00 | 0 | Mlf2 | ENSMUSG00000030120.9 |
| chr6 | 83077577 | 83079311 | 26.86698 | 95.04005 | 91.5885 | Promoter | 6 | 83077869 | 83078995 | 1127 | 1 | ENSMUST00 | 0 | Pcgf1 | ENSMUSG00000069678.5 |
| chr7 | 19024333 | 19025407 | 26.86698 | 95.04005 | 91.5885 | Promoter | 7 | 19024377 | 19054617 | 30241 | 1 | ENSMUST00 | 0 | Sympk | ENSMUSG00000023118.10 |
| chr7 | 126428298 | 126429917 | 26.86698 | 95.04005 | 91.5885 | Promoter | 7 | 126428759 | 126445901 | 17143 | 1 | ENSMUST00 | 0 | Rabep2 | ENSMUSG00000030727.7 |
| chr7 | 63939161 | 63940138 | 26.86698 | 95.04005 | 91.5885 | Promoter | 7 | 63886347 | 63938915 | 52569 | 2 | ENSMUST00 | -247 | Klf13 | ENSMUSG00000052040.9 |
| chr7 | 24593435 | 24594385 | 26.86698 | 95.04005 | 91.5885 | 5' UTR | 7 | 24583838 | 24587641 | 3804 | 2 | ENSMUST00 | -5795 | Zfp575 | ENSMUSG00000066721.3 |
| chr9 | 108305026 | 108306057 | 26.86698 | 95.04005 | 91.5885 | Promoter | 9 | 108302959 | 108305951 | 2993 | 2 | ENSMUST00 | 0 | Tcta | ENSMUSG00000039461.7 |
| chr10 | 34296759 | 34298214 | 30.6477 | 94.99754 | 91.54929 | Promoter | 10 | 34297442 | 34301320 | 3879 | 1 | ENSMUST00 | 0 | Tspyl4 | ENSMUSG00000039485.7 |
| chr5 | 21185878 | 21186941 | 30.6477 | 94.99754 | 91.54929 | Promoter | 5 | 21186267 | 21291701 | 105435 | 1 | ENSMUST00 | 0 | Gsap | ENSMUSG00000039934.8 |
| chr10 | 60277281 | 60278511 | 23.06458 | 94.99054 | 91.54246 | Promoter | 10 | 60277627 | 60302594 | 24968 | 1 | ENSMUST00 | 0 | Psap | ENSMUSG00000004207.9 |
| chr18 | 12235572 | 12237162 | 23.06458 | 94.99054 | 91.54246 | Promoter | 18 | 12189693 | 12236400 | 46708 | 2 | ENSMUST00 | 0 | Npc1 | ENSMUSG00000024413.8 |
| chr7 | 30230749 | 30233116 | 31.77605 | 94.9769 | 91.52924 | Promoter | 7 | 30224131 | 30232272 | 8142 | 2 | ENSMUST00 | 0 | Tbcb | ENSMUSG00000006095.7 |
| chr11 | 43681948 | 43683166 | 31.19374 | 94.95832 | 91.51084 | Promoter | 11 | 43681998 | 43707191 | 25194 | 1 | ENSMUST00 | 0 | Pwwp2a | ENSMUSG00000044950.7 |
| chr15 | 36609230 | 36609985 | 31.19374 | 94.95832 | 91.51084 | Promoter | 15 | 36603496 | 36608973 | 5478 | 2 | ENSMUST00 | -258 | Pabpc1 | ENSMUSG00000022283.9 |
| chr11 | 76945142 | 76946667 | 29.48955 | 94.93548 | 91.48817 | Promoter | 11 | 76945504 | 76987379 | 41876 | 1 | ENSMUST00 | 0 | Blmh | ENSMUSG00000020840.5 |
| chr13 | 67268581 | 67269639 | 29.48955 | 94.93548 | 91.48817 | Promoter | 13 | 67254918 | 67269068 | 14151 | 2 | ENSMUST00 | 0 | Zfp458 | ENSMUSG00000055480.6 |
| chr16 | 3743206 | 3744968 | 29.48955 | 94.93548 | 91.48817 | Promoter | 16 | 3744093 | 3750781 | 6689 | 1 | ENSMUST00 | 0 | Zfp263 | ENSMUSG00000022529.6 |
| chr5 | 124444651 | 124446796 | 30.58814 | 94.89368 | 91.44746 | Promoter | 5 | 124445493 | 124460883 | 15391 | 1 | ENSMUST00 | 0 | Setd8 | ENSMUSG00000049327.12 |
| chr11 | 114850502 | 114853302 | 31.71022 | 94.86349 | 91.41748 | Promoter | 11 | 114851152 | 114863733 | 12582 | 1 | ENSMUST00 | 0 | Gprc5c | ENSMUSG00000051043.11 |
| chr13 | 55471214 | 55473074 | 31.13118 | 94.84997 | 91.40416 | 5' UTR | 13 | 55477681 | 55477776 | 96 | 2 | ENSMUST00 | 4702 | Mir6944 | ENSMUSG00000098341.1 |
| chr16 | 18811243 | 18813326 | 31.13118 | 94.84997 | 91.40416 | Promoter | 16 | 18780447 | 18811972 | 31526 | 2 | ENSMUST00 | 0 | Cdc45 | ENSMUSG00000000028.9 |
| chr19 | 36083207 | 36084627 | 31.13118 | 94.84997 | 91.40416 | Promoter | 19 | 36083716 | 36104772 | 21057 | 1 | ENSMUST00 | 0 | Rpp30 | ENSMUSG00000024800.7 |
| chr5 | 134582676 | 134583394 | 31.13118 | 94.84997 | 91.40416 | Promoter | 5 | 134582690 | 134598328 | 15639 | 1 | ENSMUST00 | 0 | Rfc2 | ENSMUSG00000023104.6 |
| chr5 | 145190630 | 145191996 | 31.13118 | 94.84997 | 91.40416 | Promoter | 5 | 145183706 | 145191592 | 7887 | 2 | ENSMUST00 | 0 | Atp5j2 | ENSMUSG00000038690.10 |
| chr11 | 102155638 | 102156471 | 28.83684 | 94.75451 | 91.30991 | Promoter | 11 | 102152546 | 102156404 | 3859 | 2 | ENSMUST00 | 0 | Tmem101 | ENSMUSG00000020921.6 |
| chr19 | 40270513 | 40271964 | 28.83684 | 94.75451 | 91.30991 | Promoter | 19 | 40221173 | 40271616 | 50444 | 2 | ENSMUST00 | 0 | Pdlim1 | ENSMUSG00000055044.7 |
| chr4 | 100776119 | 100778615 | 28.83684 | 94.75451 | 91.30991 | Promoter | 4 | 100776675 | 101004544 | 227870 | 1 | ENSMUST00 | 0 | Cachd1 | ENSMUSG00000028532.9 |
| chr9 | 96718877 | 96720320 | 28.83684 | 94.75451 | 91.30991 | Promoter | 9 | 96686258 | 96719850 | 33593 | 2 | ENSMUST00 | 0 | Zbtb38 | ENSMUSG00000040433.11 |
| chr10 | 61382020 | 61383994 | 31.64465 | 94.75051 | 91.30649 | Promoter | 10 | 61319657 | 61383523 | 63867 | 2 | ENSMUST00 | 0 | Pald1 | ENSMUSG00000020092.8 |
| chr4 | 136462373 | 136463335 | 31.64465 | 94.75051 | 91.30649 | Promoter | 4 | 136462250 | 136463977 | 1728 | 1 | ENSMUST00 | 124 | 6030445D17I | ENSMUSG00000066178.2 |
| chr7 | 127194919 | 127196530 | 31.06887 | 94.74199 | 91.29819 | Promoter | 7 | 127191660 | 127196000 | 4341 | 2 | ENSMUST00 | 0 | Cd2bp2 | ENSMUSG00000042502.5 |
| chr11 | 20830006 | 20832131 | 28.22848 | 94.63612 | 91.19299 | Promoter | 11 | 20823576 | 20831056 | 7481 | 2 | ENSMUST00 | 0 | Lgalsl | ENSMUSG00000042363.9 |
| chr18 | 53861654 | 53863702 | 28.22848 | 94.63612 | 91.19299 | Promoter | 18 | 53862113 | 53955684 | 93572 | 1 | ENSMUST00 | 0 | Csnk1g3 | ENSMUSG00000073563.2 |
| chr3 | 95105324 | 95107428 | 28.22848 | 94.63612 | 91.19299 | Promoter | 3 | 95058530 | 95106875 | 48346 | 2 | ENSMUST00 | 0 | Pip5k1a | ENSMUSG00000028126.11 |
| chr9 | 78175862 | 78177230 | 28.22848 | 94.63612 | 91.19299 | Promoter | 9 | 78175914 | 78178879 | 2966 | 1 | ENSMUST00 | 0 | C920006O11 | ENSMUSG00000097574.1 |
| chr15 | 88861608 | 88864034 | 31.00681 | 94.63441 | 91.19181 | Promoter | 15 | 88862186 | 88865718 | 3533 | 1 | ENSMUST00 | 0 | Pim3 | ENSMUSG00000035828.10 |
| chr13 | 55535668 | 55537037 | 29.85283 | 94.59187 | 91.14945 | Promoter | 13 | 55530410 | 55536658 | 6249 | 2 | ENSMUST00 | 0 | Ddx41 | ENSMUSG00000021494.8 |
| chr5 | 31613825 | 31614695 | 29.85283 | 94.59187 | 91.14945 | Promoter | 5 | 31613939 | 31622644 | 8706 | 1 | ENSMUST00 | 0 | Mrpl33 | ENSMUSG00000029142.9 |
| chr11 | 100737545 | 100738217 | 30.41087 | 94.58424 | 91.14193 | Promoter | 11 | 100715009 | 100738215 | 23207 | 2 | ENSMUST00 | 0 | Rab5c | ENSMUSG00000019173.6 |
| chr9 | 67043055 | 67043963 | 27.6601 | 94.57302 | 91.13082 | Promoter | 9 | 67022590 | 67043907 | 21318 | 2 | ENSMUST00 | 0 | Tpm1 | ENSMUSG00000032366.10 |
| chr5 | 34760835 | 34762760 | 27.12789 | 94.55903 | 91.11707 | Promoter | 5 | 34761740 | 34820754 | 59015 | 1 | ENSMUST00 | 0 | Htt | ENSMUSG00000029104.10 |
| chr7 | 81788685 | 81789981 | 27.12789 | 94.55903 | 91.11707 | Promoter | 7 | 81782182 | 81789478 | 7297 | 2 | ENSMUST00 | 0 | 3110040N11 | ENSMUSG00000025102.9 |
| chr7 | 79148383 | 79149564 | 30.945 | 94.52721 | 91.08539 | Promoter | 7 | 79133769 | 79149060 | 15292 | 2 | ENSMUST00 | 0 | Mfge8 | ENSMUSG00000030605.10 |
| chr10 | 122044483 | 122048243 | 31.51432 | 94.52584 | 91.08421 | Promoter | 10 | 121780991 | 122047308 | 266318 | 2 | ENSMUST00 | 0 | Srgap1 | ENSMUSG00000020121.10 |
| chr10 | 126900716 | 126901734 | 31.51432 | 94.52584 | 91.08421 | Promoter | 10 | 126868430 | 126901371 | 32942 | 2 | ENSMUST00 | 0 | Xrcc6bp1 | ENSMUSG00000025436.6 |
| chr1 | 38987135 | 38989246 | 30.35223 | 94.4818 | 91.04041 | Promoter | 1 | 38987814 | 38997236 | 9423 | 1 | ENSMUST00 | 0 | Pdcl3 | ENSMUSG00000026078.5 |
| chr12 | 98736301 | 98738314 | 30.35223 | 94.4818 | 91.04041 | Promoter | 12 | 98676741 | 98737405 | 60665 | 2 | ENSMUST00 | 0 | Ptpn21 | ENSMUSG00000021009.9 |
| chr14 | 54258802 | 54260140 | 30.35223 | 94.4818 | 91.04041 | Promoter | 14 | 54259232 | 54269169 | 9938 | 1 | ENSMUST00 | 0 | Abhd4 | ENSMUSG00000040997.8 |

| | | | | | | | | | | | | | | | |
|---|---|---|---|---|---|---|---|---|---|---|---|---|---|---|---|
| chr2 | 85047255 | 85048940 | 30.35223 | 94.4818 | 91.04041 | Promoter | 2 | 85048022 | 85052134 | 4113 | 1 | ENSMUST00 | 0 | Tnks1bp1 | ENSMUSG00000033955.7 |
| chr4 | 139351755 | 139353339 | 30.35223 | 94.4818 | 91.04041 | Promoter | 4 | 139347435 | 139352576 | 5142 | 2 | ENSMUST00 | 0 | Mrto4 | ENSMUSG00000028741.8 |
| chr5 | 27790221 | 27792376 | 30.35223 | 94.4818 | 91.04041 | Promoter | 5 | 27740667 | 27791339 | 50673 | 2 | ENSMUST00 | 0 | Paxip1 | ENSMUSG00000002221.7 |
| chr6 | 56796286 | 56798543 | 30.35223 | 94.4818 | 91.04041 | Promoter | 6 | 56777524 | 56797736 | 20213 | 2 | ENSMUST00 | 0 | Kbtbd2 | ENSMUSG00000059486.6 |
| chr8 | 124862295 | 124863810 | 21.97955 | 94.4342 | 90.99384 | Promoter | 8 | 124839355 | 124863029 | 23675 | 2 | ENSMUST00 | 0 | 2810004N23 | ENSMUSG00000031984.7 |
| chr13 | 54589495 | 54591533 | 31.44956 | 94.41415 | 90.97417 | Promoter | 13 | 54590207 | 54591158 | 952 | 1 | ENSMUST00 | 0 | Higd2a | ENSMUSG00000025868.6 |
| chr7 | 24133602 | 24134815 | 30.29382 | 94.37968 | 90.93996 | Promoter | 7 | 24134174 | 24142266 | 8093 | 1 | ENSMUST00 | 0 | Zfp235 | ENSMUSG00000047603.8 |
| chr8 | 3515071 | 3517492 | 30.82211 | 94.31399 | 90.87436 | Promoter | 8 | 3515384 | 3544266 | 28883 | 1 | ENSMUST00 | 0 | Pnpla6 | ENSMUSG00000004565.9 |
| chr10 | 117375634 | 117377721 | 31.38507 | 94.30289 | 90.86349 | Promoter | 10 | 117344673 | 117376970 | 32298 | 2 | ENSMUST00 | 0 | Cpsf6 | ENSMUSG00000055531.7 |
| chr12 | 108893365 | 108894771 | 29.68625 | 94.29817 | 90.859 | Promoter | 12 | 108860030 | 108894137 | 34108 | 2 | ENSMUST00 | 0 | Wars | ENSMUSG00000021266.11 |
| chr7 | 81453319 | 81455625 | 30.23563 | 94.27794 | 90.8388 | Promoter | 7 | 81347026 | 81454758 | 107733 | 2 | ENSMUST00 | 0 | Cpeb1 | ENSMUSG00000025586.11 |
| chr19 | 5423832 | 5426468 | 29.11804 | 94.27461 | 90.83557 | Promoter | 19 | 5422805 | 5423923 | 1119 | 2 | ENSMUST00 | 0 | Drap1 | ENSMUSG00000024914.11 |
| chr1 | 95313212 | 95314559 | 24.0356 | 94.23969 | 90.80066 | Promoter | 1 | 95313623 | 95335284 | 21662 | 1 | ENSMUST00 | 0 | Fam174a | ENSMUSG00000051185.9 |
| chr1 | 133421916 | 133424490 | 24.0356 | 94.23969 | 90.80066 | Promoter | 1 | 133382303 | 133424377 | 42075 | 2 | ENSMUST00 | 0 | Sox13 | ENSMUSG00000070643.6 |
| chr10 | 93452710 | 93454355 | 24.0356 | 94.23969 | 90.80066 | Promoter | 10 | 93453396 | 93484896 | 31501 | 1 | ENSMUST00 | 0 | Lta4h | ENSMUSG00000015889.7 |
| chr10 | 62919971 | 62921526 | 24.0356 | 94.23969 | 90.80066 | Promoter | 10 | 62920633 | 62946498 | 25866 | 1 | ENSMUST00 | 0 | Slc25a16 | ENSMUSG00000071253.3 |
| chr13 | 108157847 | 108159463 | 24.0356 | 94.23969 | 90.80066 | Promoter | 13 | 108158731 | 108195364 | 36634 | 1 | ENSMUST00 | 0 | Ercc8 | ENSMUSG00000021694.13 |
| chr16 | 57606175 | 57607420 | 24.0356 | 94.23969 | 90.80066 | Promoter | 16 | 57302199 | 57606825 | 304627 | 2 | ENSMUST00 | 0 | Cmss1 | ENSMUSG00000022748.7 |
| chr16 | 93832171 | 93833436 | 24.0356 | 94.23969 | 90.80066 | Promoter | 16 | 93832121 | 93876072 | 43952 | 1 | ENSMUST00 | 51 | Morc3 | ENSMUSG00000039456.6 |
| chr2 | 93821553 | 93822638 | 24.0356 | 94.23969 | 90.80066 | Promoter | 2 | 93661028 | 93822549 | 161522 | 2 | ENSMUST00 | 0 | Ext2 | ENSMUSG00000027198.11 |
| chr3 | 79628914 | 79629682 | 24.0356 | 94.23969 | 90.80066 | Promoter | 3 | 79629079 | 79632817 | 3739 | 1 | ENSMUST00 | 0 | 4930579G24 | ENSMUSG00000027811.8 |
| chr3 | 146498729 | 146501117 | 24.0356 | 94.23969 | 90.80066 | Promoter | 3 | 146457203 | 146499753 | 42551 | 2 | ENSMUST00 | 0 | Spata1 | ENSMUSG00000028188.8 |
| chr3 | 10350735 | 10351367 | 24.0356 | 94.23969 | 90.80066 | Promoter | 3 | 10339953 | 10351317 | 11365 | 2 | ENSMUST00 | 0 | Zfand1 | ENSMUSG00000039795.9 |
| chr4 | 117886355 | 117887755 | 24.0356 | 94.23969 | 90.80066 | Promoter | 4 | 117884326 | 117887329 | 3004 | 2 | ENSMUST00 | 0 | Atp6v0b | ENSMUSG00000033379.8 |
| chr4 | 140700984 | 140703222 | 24.0356 | 94.23969 | 90.80066 | Promoter | 4 | 140701473 | 140710470 | 8998 | 1 | ENSMUST00 | 0 | Rcc2 | ENSMUSG00000040945.8 |
| chr5 | 29569158 | 29570344 | 24.0356 | 94.23969 | 90.80066 | Promoter | 5 | 29569242 | 29676077 | 106836 | 1 | ENSMUST00 | 0 | Ube3c | ENSMUSG00000039000.4 |
| chr7 | 31150227 | 31151795 | 24.0356 | 94.23969 | 90.80066 | Promoter | 7 | 31130127 | 31151012 | 20886 | 2 | ENSMUST00 | 0 | Gramd1a | ENSMUSG00000001248.9 |
| chr9 | 114730348 | 114732085 | 24.0356 | 94.23969 | 90.80066 | Promoter | 9 | 114731203 | 114749343 | 18141 | 1 | ENSMUST00 | 0 | Cmtm6 | ENSMUSG00000032434.8 |
| chr6 | 30302352 | 30304619 | 30.76103 | 94.20794 | 90.76972 | Promoter | 6 | 30211289 | 30304539 | 93251 | 2 | ENSMUST00 | 0 | Ube2h | ENSMUSG00000039159.11 |
| chr1 | 33757109 | 33758128 | 29.63113 | 94.2009 | 90.76292 | Promoter | 1 | 33745484 | 33757795 | 12312 | 2 | ENSMUST00 | 0 | Bag2 | ENSMUSG00000042215.8 |
| chr14 | 30950835 | 30952082 | 29.63113 | 94.2009 | 90.76292 | Promoter | 14 | 30951486 | 30987439 | 35954 | 1 | ENSMUST00 | 0 | Nek4 | ENSMUSG00000021918.9 |
| chr17 | 57247969 | 57250927 | 29.63113 | 94.2009 | 90.76292 | Promoter | 17 | 57249451 | 57263697 | 14247 | 1 | ENSMUST00 | 0 | Trip10 | ENSMUSG00000019487.10 |
| chr2 | 150748998 | 150749763 | 29.63113 | 94.2009 | 90.76292 | Promoter | 2 | 150749042 | 150771675 | 22634 | 1 | ENSMUST00 | 0 | Entpd6 | ENSMUSG00000033068.11 |
| chr6 | 86437720 | 86439429 | 29.63113 | 94.2009 | 90.76292 | Promoter | 6 | 86438374 | 86470388 | 32015 | 1 | ENSMUST00 | 0 | C87436 | ENSMUSG00000046679.11 |
| chr8 | 3499872 | 3501326 | 29.63113 | 94.2009 | 90.76292 | Promoter | 8 | 3500457 | 3515232 | 14776 | 1 | ENSMUST00 | 0 | Mcoln1 | ENSMUSG00000004567.10 |
| chr8 | 69900972 | 69903639 | 29.63113 | 94.2009 | 90.76292 | Promoter | 8 | 69894188 | 69902558 | 8371 | 2 | ENSMUST00 | 0 | Ndufa13 | ENSMUSG00000036199.8 |
| chr13 | 104285383 | 104288917 | 31.32084 | 94.19205 | 90.75528 | Promoter | 13 | 104287873 | 104494763 | 206891 | 1 | ENSMUST00 | 0 | Adamts6 | ENSMUSG00000046169.9 |
| chr4 | 46535290 | 46536540 | 31.32084 | 94.19205 | 90.75528 | Promoter | 4 | 46493781 | 46536122 | 42342 | 2 | ENSMUST00 | 0 | Trim14 | ENSMUSG00000039853.13 |
| chr11 | 101081815 | 101083384 | 30.17592 | 94.17345 | 90.73757 | Promoter | 11 | 101082565 | 101086619 | 4055 | 1 | ENSMUST00 | 0 | Coasy | ENSMUSG00000001755.7 |
| chr11 | 55032702 | 55033970 | 30.17592 | 94.17345 | 90.73757 | Promoter | 11 | 54979108 | 55033445 | 54338 | 2 | ENSMUST00 | 0 | Anxa6 | ENSMUSG00000018340.8 |
| chr11 | 120617109 | 120618261 | 30.17592 | 94.17345 | 90.73757 | Promoter | 11 | 120610087 | 120617936 | 7850 | 2 | ENSMUST00 | 0 | Pcyt2 | ENSMUSG00000025137.10 |
| chr11 | 77077526 | 77079024 | 30.17592 | 94.17345 | 90.73757 | Promoter | 11 | 77044292 | 77078435 | 34144 | 2 | ENSMUST00 | 0 | Ccdc55 | ENSMUSG00000037958.8 |
| chr12 | 30910612 | 30912594 | 30.17592 | 94.17345 | 90.73757 | Promoter | 12 | 30911668 | 30960018 | 48351 | 1 | ENSMUST00 | 0 | Sh3yl1 | ENSMUSG00000020669.10 |
| chr12 | 87265582 | 87267562 | 30.17592 | 94.17345 | 90.73757 | Promoter | 12 | 87266746 | 87273997 | 7252 | 1 | ENSMUST00 | 0 | Ahsa1 | ENSMUSG00000021037.6 |
| chr13 | 69610732 | 69611538 | 30.17592 | 94.17345 | 90.73757 | Promoter | 13 | 69573449 | 69611442 | 37994 | 2 | ENSMUST00 | 0 | Srd5a1 | ENSMUSG00000021594.10 |
| chr13 | 14038732 | 14040316 | 30.17592 | 94.17345 | 90.73757 | Promoter | 13 | 13997949 | 14039579 | 41631 | 2 | ENSMUST00 | 0 | Tbce | ENSMUSG00000039233.7 |
| chr13 | 23561789 | 23562486 | 30.17592 | 94.17345 | 90.73757 | Promoter | 13 | 23561534 | 23562369 | 836 | 2 | ENSMUST00 | 0 | Hist1h3e | ENSMUSG00000069273.2 |
| chr13 | 17992320 | 17993900 | 30.17592 | 94.17345 | 90.73757 | Promoter | 13 | 17986640 | 17993351 | 6712 | 2 | ENSMUST00 | 0 | Yae1d1 | ENSMUSG00000075054.4 |
| chr15 | 102244836 | 102248267 | 30.17592 | 94.17345 | 90.73757 | Promoter | 15 | 102234940 | 102246500 | 11561 | 2 | ENSMUST00 | 0 | Rarg | ENSMUSG00000001288.9 |
| chr16 | 37539029 | 37540718 | 30.17592 | 94.17345 | 90.73757 | Promoter | 16 | 37539885 | 37572385 | 32501 | 1 | ENSMUST00 | 0 | Rabl3 | ENSMUSG00000022827.9 |
| chr16 | 34572257 | 34574303 | 30.17592 | 94.17345 | 90.73757 | Promoter | 16 | 34049689 | 34573532 | 523844 | 2 | ENSMUST00 | 0 | Kalrn | ENSMUSG00000061751.10 |
| chr17 | 35134562 | 35136490 | 30.17592 | 94.17345 | 90.73757 | Promoter | 17 | 35135178 | 35147305 | 12128 | 1 | ENSMUST00 | 0 | Bag6 | ENSMUSG00000024392.12 |
| chr18 | 43392549 | 43393845 | 30.17592 | 94.17345 | 90.73757 | Promoter | 18 | 43324288 | 43393377 | 69090 | 2 | ENSMUST00 | 0 | Dpysl3 | ENSMUSG00000024501.14 |
| chr19 | 42090349 | 42091418 | 30.17592 | 94.17345 | 90.73757 | Promoter | 19 | 42090435 | 42122218 | 31784 | 1 | ENSMUST00 | 0 | Pi4k2a | ENSMUSG00000025178.8 |
| chr19 | 36918249 | 36919478 | 30.17592 | 94.17345 | 90.73757 | Promoter | 19 | 36917550 | 36919615 | 2066 | 2 | ENSMUST00 | 137 | Fgfbp3 | ENSMUSG00000047632.10 |
| chr2 | 119971973 | 119973511 | 30.17592 | 94.17345 | 90.73757 | Promoter | 2 | 119972699 | 120027408 | 54710 | 1 | ENSMUST00 | 0 | Mapkbp1 | ENSMUSG00000033902.11 |
| chr2 | 32362481 | 32363897 | 30.17592 | 94.17345 | 90.73757 | Promoter | 2 | 32363010 | 32367407 | 4398 | 1 | ENSMUST00 | 0 | Ciz1 | ENSMUSG00000039205.11 |
| chr2 | 29802524 | 29803400 | 30.17592 | 94.17345 | 90.73757 | Promoter | 2 | 29802634 | 29817522 | 14889 | 1 | ENSMUST00 | 0 | Slc27a4 | ENSMUSG00000059316.2 |
| chr2 | 119872141 | 119873656 | 30.17592 | 94.17345 | 90.73757 | Distal Interg | 2 | 119878069 | 119878403 | 335 | 2 | ENSMUST00 | 4747 | Gm13998 | ENSMUSG00000081056.1 |
| chr3 | 87171121 | 87175434 | 30.17592 | 94.17345 | 90.73757 | Promoter | 3 | 87078593 | 87174700 | 96108 | 2 | ENSMUST00 | 0 | Kirrel | ENSMUSG00000041734.10 |
| chr4 | 120746669 | 120747856 | 30.17592 | 94.17345 | 90.73757 | Promoter | 4 | 120696138 | 120747248 | 51111 | 2 | ENSMUST00 | 0 | Kcnq4 | ENSMUSG00000028631.7 |
| chr4 | 5643981 | 5645206 | 30.17592 | 94.17345 | 90.73757 | Promoter | 4 | 5644090 | 5798382 | 154293 | 1 | ENSMUST00 | 0 | Fam110b | ENSMUSG00000049119.9 |
| chr4 | 120286553 | 120288405 | 30.17592 | 94.17345 | 90.73757 | Promoter | 4 | 120267079 | 120287349 | 20271 | 2 | ENSMUST00 | 0 | Foxo6 | ENSMUSG00000052135.8 |
| chr5 | 38038465 | 38040150 | 30.17592 | 94.17345 | 90.73757 | Promoter | 5 | 38038742 | 38137765 | 99024 | 1 | ENSMUST00 | 0 | Stx18 | ENSMUSG00000029125.9 |
| chr5 | 92602620 | 92603582 | 30.17592 | 94.17345 | 90.73757 | Promoter | 5 | 92603051 | 92606579 | 3529 | 1 | ENSMUST00 | 0 | Stbd1 | ENSMUSG00000047963.6 |
| chr6 | 4902738 | 4904537 | 30.17592 | 94.17345 | 90.73757 | Promoter | 6 | 4902917 | 5159590 | 256674 | 1 | ENSMUST00 | 0 | Ppp1r9a | ENSMUSG00000032827.10 |
| chr7 | 105639767 | 105641351 | 30.17592 | 94.17345 | 90.73757 | Promoter | 7 | 105634203 | 105640416 | 6214 | 2 | ENSMUST00 | 0 | Arfip2 | ENSMUSG00000030881.9 |
| chr7 | 41632488 | 41634349 | 30.17592 | 94.17345 | 90.73757 | Promoter | 7 | 41633203 | 41647548 | 14346 | 1 | ENSMUST00 | 0 | Zfp788 | ENSMUSG00000074165.5 |
| chr7 | 4914514 | 4916194 | 30.17592 | 94.17345 | 90.73757 | Promoter | 7 | 4915217 | 4922003 | 6787 | 1 | ENSMUST00 | 0 | Zfp628 | ENSMUSG00000074406.4 |
| chr8 | 116920849 | 116922423 | 30.17592 | 94.17345 | 90.73757 | Promoter | 8 | 116888685 | 116921455 | 32771 | 2 | ENSMUST00 | 0 | Cmc2 | ENSMUSG00000014633.10 |
| chr8 | 33653035 | 33654537 | 30.17592 | 94.17345 | 90.73757 | Promoter | 8 | 33653238 | 33698163 | 44926 | 1 | ENSMUST00 | 0 | Gsr | ENSMUSG00000031584.11 |
| chr8 | 116732022 | 116733738 | 30.17592 | 94.17345 | 90.73757 | Promoter | 8 | 116568724 | 116732991 | 164268 | 2 | ENSMUST00 | 0 | Cdyl2 | ENSMUSG00000031758.9 |
| chr8 | 22185261 | 22186153 | 30.17592 | 94.17345 | 90.73757 | Promoter | 8 | 22168152 | 22185819 | 17668 | 2 | ENSMUST00 | 0 | Ckap2 | ENSMUSG00000037725.7 |
| chr9 | 35569470 | 35570915 | 30.17592 | 94.17345 | 90.73757 | Promoter | 9 | 35560820 | 35570398 | 9579 | 2 | ENSMUST00 | 0 | Hyls1 | ENSMUSG00000050555.9 |
| chr11 | 6443550 | 6444540 | 31.25687 | 94.08162 | 90.65025 | Promoter | 11 | 6427229 | 6444443 | 17215 | 2 | ENSMUST00 | 0 | H2afv | ENSMUSG00000041126.11 |
| chr2 | 175141256 | 175142162 | 31.25687 | 94.08162 | 90.65025 | Promoter | 2 | 175129493 | 175141999 | 12507 | 2 | ENSMUST00 | 0 | Gm14399 | ENSMUSG00000090093.3 |
| chr10 | 105572740 | 105575603 | 30.6396 | 93.99698 | 90.56607 | Promoter | 10 | 105187663 | 105574435 | 386773 | 2 | ENSMUST00 | 0 | Tmtc2 | ENSMUSG00000036019.8 |
| chr1 | 131244552 | 131246593 | 28.96167 | 93.99567 | 90.56499 | Promoter | 1 | 131176410 | 131245258 | 68849 | 2 | ENSMUST00 | 0 | Rassf5 | ENSMUSG00000026430.10 |
| chr10 | 80701379 | 80702000 | 28.96167 | 93.99567 | 90.56499 | Promoter | 10 | 80685256 | 80701820 | 16565 | 2 | ENSMUST00 | 0 | Mob3a | ENSMUSG00000003348.9 |
| chr13 | 74349381 | 74350958 | 28.96167 | 93.99567 | 90.56499 | Promoter | 13 | 74322254 | 74350280 | 28027 | 2 | ENSMUST00 | 0 | Sdha | ENSMUSG00000021577.8 |
| chr18 | 75342214 | 75343517 | 28.96167 | 93.99567 | 90.56499 | Distal Interg | 18 | 75365332 | 75367088 | 1757 | 2 | ENSMUST00 | 23571 | Gm20544 | ENSMUSG00000092569.1 |
| chr7 | 110613377 | 110615752 | 31.19317 | 93.97161 | 90.54182 | Promoter | 7 | 110308013 | 110614922 | 306910 | 2 | ENSMUST00 | 0 | Sbf2 | ENSMUSG00000038371.9 |
| chr11 | 83440161 | 83441032 | 31.70772 | 93.8754 | 90.44607 | Promoter | 11 | 83440131 | 83440828 | 698 | 1 | ENSMUST00 | 31 | 1700020L24 | ENSMUSG00000035085.5 |
| chr12 | 112941531 | 112942503 | 31.70772 | 93.8754 | 90.44607 | Promoter | 12 | 112934733 | 112942118 | 7386 | 2 | ENSMUST00 | 0 | Nudt14 | ENSMUSG00000002804.3 |
| chr13 | 101693541 | 101695297 | 31.70772 | 93.8754 | 90.44607 | Promoter | 13 | 101689040 | 101694600 | 5561 | 2 | ENSMUST00 | 0 | Pik3r1 | ENSMUSG00000041417.10 |
| chr14 | 57570910 | 57572174 | 31.70772 | 93.8754 | 90.44607 | Promoter | 14 | 57549598 | 57571569 | 21972 | 2 | ENSMUST00 | 0 | N6amt2 | ENSMUSG00000021951.5 |
| chr16 | 4212030 | 4214851 | 31.70772 | 93.8754 | 90.44607 | Promoter | 16 | 4081328 | 4213407 | 132080 | 2 | ENSMUST00 | 0 | Crebbp | ENSMUSG00000022521.9 |
| chr19 | 6334640 | 6336004 | 30.00511 | 93.87433 | 90.44607 | Promoter | 19 | 6334979 | 6340880 | 5902 | 1 | ENSMUST00 | 0 | Men1 | ENSMUSG00000024947.11 |
| chr2 | 154406634 | 154408534 | 31.70772 | 93.8754 | 90.44607 | Promoter | 2 | 154376313 | 154408081 | 31769 | 2 | ENSMUST00 | 0 | Snta1 | ENSMUSG00000027488.7 |
| chr3 | 123385144 | 123386655 | 31.70772 | 93.8754 | 90.44607 | Promoter | 3 | 123364645 | 123385948 | 21304 | 2 | ENSMUST00 | 0 | Mettl14 | ENSMUSG00000028114.11 |
| chr4 | 88031234 | 88034901 | 31.70772 | 93.8754 | 90.44607 | Promoter | 4 | 87769925 | 88033364 | 263440 | 2 | ENSMUST00 | 0 | Mllt3 | ENSMUSG00000028496.12 |
| chr5 | 135488346 | 135489702 | 31.70772 | 93.8754 | 90.44607 | Intron (ENSM | 5 | 135406523 | 135545122 | 138600 | 2 | ENSMUST00 | 55420 | Hip1 | ENSMUSG00000039959.9 |
| chr6 | 114642620 | 114644013 | 31.70772 | 93.8754 | 90.44607 | Promoter | 6 | 114643097 | 114860614 | 217518 | 1 | ENSMUST00 | 0 | Atg7 | ENSMUSG00000030314.11 |
| chr7 | 66079616 | 66080501 | 31.70772 | 93.8754 | 90.44607 | Promoter | 7 | 66079649 | 66089412 | 9764 | 1 | ENSMUST00 | 0 | Vimp | ENSMUSG00000075701.8 |
| chr8 | 71272013 | 71273842 | 31.70772 | 93.8754 | 90.44607 | Promoter | 8 | 71272714 | 71360712 | 87999 | 1 | ENSMUST00 | 0 | Myo9b | ENSMUSG00000004677.10 |
| chr9 | 114977880 | 114979450 | 31.70772 | 93.8754 | 90.44607 | Promoter | 9 | 114978569 | 115232223 | 253655 | 1 | ENSMUST00 | 0 | Osbpl10 | ENSMUSG00000040875.7 |
| chr1 | 180903735 | 180905066 | 28.33848 | 93.85651 | 90.42933 | Promoter | 1 | 180904274 | 180908087 | 3814 | 1 | ENSMUST00 | 0 | Pycr2 | ENSMUSG00000026520.7 |
| chr12 | 59011130 | 59012052 | 28.33848 | 93.85651 | 90.42933 | Promoter | 12 | 58958383 | 59012017 | 53635 | 2 | ENSMUST00 | 0 | Sec23a | ENSMUSG00000020986.8 |
| chr3 | 145758375 | 145759815 | 28.33848 | 93.85651 | 90.42933 | Promoter | 3 | 145758675 | 145894277 | 135603 | 1 | ENSMUST00 | 0 | Ddah1 | ENSMUSG00000028194.10 |
| chr7 | 3692185 | 3695281 | 28.33848 | 93.85651 | 90.42933 | Promoter | 7 | 3693375 | 3701023 | 7649 | 1 | ENSMUST00 | 0 | Tsen34 | ENSMUSG00000035585.11 |
| chr10 | 79766064 | 79768018 | 26.2242 | 93.81972 | 90.39303 | Promoter | 10 | 79758515 | 79766952 | 8438 | 2 | ENSMUST00 | 0 | Rnf126 | ENSMUSG00000035890.8 |
| chr10 | 80248693 | 80250377 | 30.51912 | 93.78751 | 90.36088 | Promoter | 10 | 80249121 | 80256793 | 7673 | 1 | ENSMUST00 | 0 | Ndufs7 | ENSMUSG00000020153.9 |
| chr12 | 8313281 | 8314141 | 30.51912 | 93.78751 | 90.36088 | Promoter | 12 | 8313433 | 8343824 | 30392 | 1 | ENSMUST00 | 0 | Hs1bp3 | ENSMUSG00000020605.8 |
| chr13 | 51651070 | 51652438 | 30.51912 | 93.78751 | 90.36088 | Promoter | 13 | 51651697 | 51684044 | 32348 | 1 | ENSMUST00 | 0 | Secisbp2 | ENSMUSG00000035139.9 |
| chr2 | 144550282 | 144550984 | 30.51912 | 93.78751 | 90.36088 | Promoter | 2 | 144542265 | 144550869 | 8605 | 2 | ENSMUST00 | 0 | Rbbp9 | ENSMUSG00000027428.9 |
| chr5 | 30666340 | 30667506 | 30.51912 | 93.78751 | 90.36088 | Promoter | 5 | 30666777 | 30674378 | 7602 | 1 | ENSMUST00 | 0 | Cenpa | ENSMUSG00000029177.4 |
| chr5 | 122778520 | 122779989 | 30.51912 | 93.78751 | 90.36088 | Promoter | 5 | 122731173 | 122779410 | 48238 | 2 | ENSMUST00 | 0 | Camkk2 | ENSMUSG00000029471.9 |
| chr9 | 107296194 | 107297688 | 30.51912 | 93.78751 | 90.36088 | Promoter | 9 | 107296710 | 107302784 | 6075 | 1 | ENSMUST00 | 0 | Cish | ENSMUSG00000032578.7 |
| chr9 | 64960469 | 64961684 | 30.51912 | 93.78751 | 90.36088 | Promoter | 9 | 64960832 | 64986974 | 26143 | 1 | ENSMUST00 | 0 | Vwa9 | ENSMUSG00000034263.7 |
| chr10 | 24149036 | 24150224 | 27.75693 | 93.7755 | 90.3501 | Promoter | 10 | 24149317 | 24188959 | 39643 | 1 | ENSMUST00 | 0 | Stx7 | ENSMUSG00000019998.5 |
| chr11 | 101665012 | 101668610 | 27.75693 | 93.7755 | 90.3501 | Promoter | 11 | 101665541 | 101667832 | 2292 | 1 | ENSMUST00 | 0 | Arl4d | ENSMUSG00000034936.2 |
| chr19 | 41895457 | 41896740 | 27.75693 | 93.7755 | 90.3501 | Promoter | 19 | 41862852 | 41896153 | 33302 | 2 | ENSMUST00 | 0 | Rrp12 | ENSMUSG00000035049.3 |
| chr6 | 47655632 | 47655927 | 10.19355 | 93.77309 | 90.34816 | Distal Interg | 6 | 47637143 | 47637219 | 77 | 1 | ENSMUST00 | 18490 | Gm25283 | ENSMUSG00000087897.1 |
| chr7 | 133775865 | 133777162 | 26.70311 | 93.76232 | 90.33739 | Promoter | 7 | 133720942 | 133776803 | 55862 | 2 | ENSMUST00 | 0 | Dhx32 | ENSMUSG00000030986.7 |
| chr1 | 23382281 | 23383443 | 25.17393 | 93.69764 | 90.27328 | Promoter | 1 | 23366424 | 23383175 | 16752 | 2 | ENSMUST00 | 0 | Ogfrl1 | ENSMUSG00000026158.6 |
| chr11 | 75651407 | 75657466 | 25.17393 | 93.69764 | 90.27328 | Promoter | 11 | 75651510 | 75673910 | 22401 | 1 | ENSMUST00 | 0 | Myo1c | ENSMUSG00000017774.14 |
| chr11 | 120712903 | 120714574 | 25.17393 | 93.69764 | 90.27328 | Promoter | 11 | 120710942 | 120713732 | 2791 | 2 | ENSMUST00 | 0 | Stra13 | ENSMUSG00000025144.12 |
| chr13 | 21362385 | 21363516 | 25.17393 | 93.69764 | 90.27328 | Promoter | 13 | 21362827 | 21369824 | 6998 | 1 | ENSMUST00 | 0 | Zscan12 | ENSMUSG00000036721.8 |
| chr14 | 57663945 | 57666077 | 25.17393 | 93.69764 | 90.27328 | Promoter | 14 | 57581002 | 57665430 | 84429 | 2 | ENSMUST00 | 0 | Xpo4 | ENSMUSG00000021952.10 |
| chr15 | 60822622 | 60825350 | 25.17393 | 93.69764 | 90.27328 | Promoter | 15 | 60818994 | 60825080 | 6087 | 2 | ENSMUST00 | 0 | Fam84b | ENSMUSG00000072568.4 |
| chr16 | 91596520 | 91597737 | 25.17393 | 93.69764 | 90.27328 | Promoter | 16 | 91575758 | 91597664 | 21907 | 2 | ENSMUST00 | 0 | Tmem50b | ENSMUSG00000022964.9 |
| chr17 | 47737000 | 47739098 | 25.17393 | 93.69764 | 90.27328 | Promoter | 17 | 47737030 | 47792419 | 55390 | 1 | ENSMUST00 | 0 | Tfeb | ENSMUSG00000023990.13 |
| chr18 | 74999858 | 75001288 | 25.17393 | 93.69764 | 90.27328 | Promoter | 18 | 75000477 | 75003381 | 2905 | 1 | ENSMUST00 | 0 | Rpl17 | ENSMUSG00000062328.7 |
| chr2 | 155955858 | 155957255 | 25.17393 | 93.69764 | 90.27328 | Promoter | 2 | 155956458 | 155986613 | 30156 | 1 | ENSMUST00 | 0 | Cep250 | ENSMUSG00000038241.11 |
| chr2 | 17729605 | 17732624 | 25.17393 | 93.69764 | 90.27328 | Promoter | 2 | 17343909 | 17731043 | 387135 | 2 | ENSMUST00 | 0 | Nebl | ENSMUSG00000053702.11 |
| chr4 | 56945931 | 56947685 | 25.17393 | 93.69764 | 90.27328 | Promoter | 4 | 56866923 | 56947437 | 80515 | 2 | ENSMUST00 | 0 | Tmem245 | ENSMUSG00000055296.9 |
| chr6 | 94698891 | 94701598 | 25.17393 | 93.69764 | 90.27328 | Promoter | 6 | 94604529 | 94700158 | 95630 | 2 | ENSMUST00 | 0 | Lrig1 | ENSMUSG00000030029.9 |
| chr7 | 144914444 | 144915918 | 25.17393 | 93.69764 | 90.27328 | Promoter | 7 | 144915100 | 144930748 | 15649 | 1 | ENSMUST00 | 0 | Oraov1 | ENSMUSG00000031072.8 |

| | | | | | | | | | | | | | | | |
|---|---|---|---|---|---|---|---|---|---|---|---|---|---|---|---|
| chr7 | 109985350 | 109987042 | 25.17393 | 93.69764 | 90.27328 | Promoter | 7 | 109972187 | 109986451 | 14265 | 2 | ENSMUST00 | 0 | Tmem41b | ENSMUSG00000047554.8 |
| chr18 | 32376592 | 32378381 | 31.58442 | 93.66337 | 90.24092 | Promoter | 18 | 32377217 | 32435736 | 58520 | 1 | ENSMUST00 | 0 | Bin1 | ENSMUSG00000024381.10 |
| chr12 | 112673011 | 112674859 | 30.39959 | 93.57949 | 90.15746 | Promoter | 12 | 112653821 | 112674276 | 20456 | 2 | ENSMUST00 | 0 | Akt1 | ENSMUSG00000001729.9 |
| chr11 | 98399904 | 98400956 | 31.51843 | 93.54985 | 90.12806 | Promoter | 11 | 98388677 | 98400490 | 11814 | 2 | ENSMUST00 | 0 | Pgap3 | ENSMUSG00000038208.8 |
| chr2 | 30093188 | 30094333 | 31.51843 | 93.54985 | 90.12806 | Promoter | 2 | 30090944 | 30093648 | 2705 | 2 | ENSMUST00 | 0 | Zdhhc12 | ENSMUSG00000015335.11 |
| chr6 | 72413818 | 72415019 | 31.51843 | 93.54985 | 90.12806 | Promoter | 6 | 72414308 | 72430712 | 16405 | 1 | ENSMUST00 | 0 | Ggcx | ENSMUSG00000053460.7 |
| chr10 | 127063552 | 127064638 | 28.1615 | 93.53628 | 90.11485 | Promoter | 10 | 127063603 | 127067002 | 3400 | 1 | ENSMUST00 | 0 | Cdk4 | ENSMUSG00000006728.7 |
| chr10 | 81770370 | 81771566 | 28.1615 | 93.53628 | 90.11485 | Distal Interge | 10 | 81745983 | 81747503 | 1521 | 2 | ENSMUST00 | -22868 | Gm3055 | ENSMUSG00000094622.1 |
| chr11 | 58953638 | 58956401 | 28.1615 | 93.53628 | 90.11485 | Promoter | 11 | 58954685 | 58973098 | 18414 | 1 | ENSMUST00 | 0 | Trim17 | ENSMUSG00000036964.9 |
| chr12 | 28649202 | 28650442 | 28.1615 | 93.53628 | 90.11485 | Promoter | 12 | 28649602 | 28659591 | 9990 | 1 | ENSMUST00 | 0 | Rnaseh1 | ENSMUSG00000020630.8 |
| chr14 | 64948455 | 64950969 | 28.1615 | 93.53628 | 90.11485 | Promoter | 14 | 64811600 | 64949850 | 138251 | 2 | ENSMUST00 | 0 | Hmbox1 | ENSMUSG00000021972.9 |
| chr15 | 76368511 | 76369676 | 28.1615 | 93.53628 | 90.11485 | Promoter | 15 | 76368898 | 76371403 | 2506 | 1 | ENSMUST00 | 0 | Fam203a | ENSMUSG00000022554.6 |
| chr15 | 59647395 | 59650262 | 28.1615 | 93.53628 | 90.11485 | Promoter | 15 | 59648350 | 59656550 | 8201 | 1 | ENSMUST00 | 0 | Trib1 | ENSMUSG00000032501.8 |
| chr16 | 22438273 | 22439688 | 28.1615 | 93.53628 | 90.11485 | Promoter | 16 | 22401864 | 22439616 | 37753 | 2 | ENSMUST00 | 0 | Etv5 | ENSMUSG00000013089.10 |
| chr17 | 47008952 | 47011261 | 28.1615 | 93.53628 | 90.11485 | Promoter | 17 | 46928292 | 47010532 | 82241 | 2 | ENSMUST00 | 0 | Ubr2 | ENSMUSG00000023977.9 |
| chr17 | 47368416 | 47369691 | 28.1615 | 93.53628 | 90.11485 | Promoter | 17 | 47368887 | 47378679 | 9793 | 1 | ENSMUST00 | 0 | Mrps10 | ENSMUSG00000034729.11 |
| chr18 | 34650773 | 34652232 | 30.94091 | 93.53562 | 90.11485 | Promoter | 18 | 34630951 | 34651735 | 20785 | 2 | ENSMUST00 | 0 | Cdc23 | ENSMUSG00000024370.11 |
| chr18 | 15062362 | 15065257 | 28.1615 | 93.53628 | 90.11485 | Promoter | 18 | 14968685 | 15063582 | 94898 | 2 | ENSMUST00 | 0 | Kctd1 | ENSMUSG00000036225.9 |
| chr18 | 73753844 | 73755108 | 28.1615 | 93.53628 | 90.11485 | Promoter | 18 | 73735038 | 73754479 | 19442 | 2 | ENSMUST00 | 0 | Elac1 | ENSMUSG00000036941.2 |
| chr19 | 5874539 | 5875650 | 28.1615 | 93.53628 | 90.11485 | Promoter | 19 | 5849702 | 5875274 | 25573 | 2 | ENSMUST00 | 0 | Frmd8 | ENSMUSG00000024816.7 |
| chr19 | 12500787 | 12502023 | 28.1615 | 93.53628 | 90.11485 | Promoter | 19 | 12466341 | 12501996 | 35656 | 2 | ENSMUST00 | 0 | Dtx4 | ENSMUSG00000039982.7 |
| chr19 | 18712461 | 18714024 | 28.1615 | 93.53628 | 90.11485 | Promoter | 19 | 18713193 | 18718428 | 5236 | 1 | ENSMUST00 | 0 | D030056L22I | ENSMUSG00000047044.7 |
| chr19 | 33391413 | 33392646 | 28.1615 | 93.53628 | 90.11485 | Promoter | 19 | 33137747 | 33392265 | 254519 | 2 | ENSMUST00 | 0 | Rnls | ENSMUSG00000071573.8 |
| chr19 | 60755532 | 60757412 | 28.1615 | 93.53628 | 90.11485 | Promoter | 19 | 60755987 | 60759913 | 3927 | 1 | ENSMUST00 | 0 | Nanos1 | ENSMUSG00000072437.3 |
| chr2 | 164745339 | 164746885 | 28.1615 | 93.53628 | 90.11485 | Promoter | 2 | 164745983 | 164768607 | 22625 | 1 | ENSMUST00 | 0 | Dnttip1 | ENSMUSG00000017299.8 |
| chr3 | 35931692 | 35933728 | 28.1615 | 93.53628 | 90.11485 | Promoter | 3 | 35892105 | 35932334 | 40230 | 2 | ENSMUST00 | 0 | Dcun1d1 | ENSMUSG00000027708.9 |
| chr3 | 9608405 | 9610425 | 28.1615 | 93.53628 | 90.11485 | Promoter | 3 | 9427011 | 9610085 | 183075 | 2 | ENSMUST00 | 0 | Zfp704 | ENSMUSG00000040209.7 |
| chr4 | 129286769 | 129288550 | 28.1615 | 93.53628 | 90.11485 | Promoter | 4 | 129287617 | 129308559 | 20943 | 1 | ENSMUST00 | 0 | Sync | ENSMUSG00000001333.9 |
| chr4 | 33030273 | 33032302 | 28.1615 | 93.53628 | 90.11485 | Promoter | 4 | 33031416 | 33052363 | 20948 | 1 | ENSMUST00 | 0 | Ube2j1 | ENSMUSG00000028277.8 |
| chr4 | 94613865 | 94615126 | 28.1615 | 93.53628 | 90.11485 | Promoter | 4 | 94614491 | 94664698 | 50208 | 1 | ENSMUST00 | 0 | Ift74 | ENSMUSG00000028576.7 |
| chr4 | 125084460 | 125085865 | 28.1615 | 93.53628 | 90.11485 | Promoter | 4 | 125085134 | 125113238 | 28105 | 1 | ENSMUST00 | 0 | Meaf6 | ENSMUSG00000028863.8 |
| chr4 | 149103856 | 149105144 | 28.1615 | 93.53628 | 90.11485 | Promoter | 4 | 149104146 | 149118870 | 14725 | 1 | ENSMUST00 | 0 | Dffa | ENSMUSG00000028974.8 |
| chr4 | 44981043 | 44982002 | 28.1615 | 93.53628 | 90.11485 | Promoter | 4 | 44981395 | 44990734 | 9340 | 1 | ENSMUST00 | 0 | Grhpr | ENSMUSG00000035637.9 |
| chr4 | 126201231 | 126202738 | 28.1615 | 93.53628 | 90.11485 | Promoter | 4 | 126172843 | 126202589 | 29747 | 2 | ENSMUST00 | 0 | Thrap3 | ENSMUSG00000043962.11 |
| chr5 | 34948690 | 34949989 | 28.1615 | 93.53628 | 90.11485 | Promoter | 5 | 34949445 | 35033595 | 84151 | 1 | ENSMUST00 | 0 | Rgs12 | ENSMUSG00000029101.9 |
| chr5 | 36483232 | 36484174 | 28.1615 | 93.53628 | 90.11485 | Promoter | 5 | 36475742 | 36484112 | 8371 | 2 | ENSMUST00 | 0 | Tada2b | ENSMUSG00000029196.9 |
| chr5 | 150118265 | 150119526 | 30.94091 | 93.53562 | 90.11485 | Distal Interge | 5 | 150080622 | 150082184 | 1563 | 1 | ENSMUST00 | 37644 | Rxfp2 | ENSMUSG00000053368.8 |
| chr7 | 80401665 | 80403431 | 28.1615 | 93.53628 | 90.11485 | Promoter | 7 | 80389194 | 80402762 | 13569 | 2 | ENSMUST00 | 0 | Furin | ENSMUSG00000030530.10 |
| chr7 | 30321546 | 30322797 | 28.1615 | 93.53628 | 90.11485 | Promoter | 7 | 30321740 | 30323472 | 1733 | 1 | ENSMUST00 | 0 | E130208F15I | ENSMUSG00000074210.3 |
| chr8 | 9976849 | 9978849 | 28.1615 | 93.53628 | 90.11485 | Promoter | 8 | 9977707 | 9992155 | 14449 | 1 | ENSMUST00 | 0 | Abhd13 | ENSMUSG00000040396.7 |
| chr8 | 3587339 | 3588506 | 28.1615 | 93.53628 | 90.11485 | Promoter | 8 | 3587450 | 3609074 | 21625 | 1 | ENSMUST00 | 0 | Camsap3 | ENSMUSG00000044433.10 |
| chr9 | 86695180 | 86696353 | 28.1615 | 93.53628 | 90.11485 | Promoter | 9 | 86581371 | 86695953 | 114583 | 2 | ENSMUST00 | 0 | Me1 | ENSMUSG00000032418.10 |
| chr9 | 57939633 | 57941400 | 28.1615 | 93.53628 | 90.11485 | Promoter | 9 | 57940113 | 57962864 | 22752 | 1 | ENSMUST00 | 0 | Sema7a | ENSMUSG00000038264.7 |
| chr10 | 39730769 | 39733802 | 22.79947 | 93.49365 | 90.07696 | Promoter | 10 | 39732118 | 39875211 | 143094 | 1 | ENSMUST00 | 0 | Rev3l | ENSMUSG00000019841.10 |
| chr10 | 59878813 | 59880788 | 22.79947 | 93.49365 | 90.07696 | Promoter | 10 | 59879556 | 59899302 | 19747 | 1 | ENSMUST00 | 0 | Dnajb12 | ENSMUSG00000020109.8 |
| chr13 | 90922498 | 90923934 | 22.79947 | 93.49365 | 90.07696 | Promoter | 13 | 90923131 | 90924942 | 1812 | 1 | ENSMUST00 | 0 | Rps23 | ENSMUSG00000049517.7 |
| chr9 | 123716958 | 123717687 | 22.79947 | 93.49365 | 90.07696 | Promoter | 9 | 123694416 | 123717625 | 23210 | 2 | ENSMUST00 | 0 | Lztfl1 | ENSMUSG00000025245.8 |
| chr12 | 21372741 | 21374479 | 29.77807 | 93.47606 | 90.05975 | Promoter | 12 | 21323509 | 21373593 | 50085 | 2 | ENSMUST00 | 0 | Adam17 | ENSMUSG00000052593.10 |
| chr17 | 12768928 | 12770076 | 29.77807 | 93.47606 | 90.05975 | Promoter | 17 | 12682406 | 12769664 | 87259 | 2 | ENSMUST00 | 0 | Igf2r | ENSMUSG00000023830.8 |
| chr17 | 50508569 | 50510296 | 29.77807 | 93.47606 | 90.05975 | Promoter | 17 | 50509547 | 50688493 | 178947 | 1 | ENSMUST00 | 0 | Plcl2 | ENSMUSG00000038910.6 |
| chr5 | 45668827 | 45670551 | 29.77807 | 93.47606 | 90.05975 | Promoter | 5 | 45669922 | 45700544 | 30623 | 1 | ENSMUST00 | 0 | Ncapg | ENSMUSG00000015880.9 |
| chr9 | 110656072 | 110657490 | 29.77807 | 93.47606 | 90.05975 | Promoter | 9 | 110656503 | 110711590 | 55088 | 1 | ENSMUST00 | 0 | Ccdc12 | ENSMUSG00000019659.4 |
| chr1 | 165641628 | 165643253 | 26.52689 | 93.43019 | 90.01563 | Distal Interge | 1 | 165592181 | 165634541 | 42361 | 2 | ENSMUST00 | -7088 | Mpzl1 | ENSMUSG00000026566.10 |
| chr1 | 135429257 | 135430977 | 26.52689 | 93.43019 | 90.01563 | Promoter | 1 | 135382312 | 135430499 | 48188 | 2 | ENSMUST00 | 0 | Ipo9 | ENSMUSG00000041879.8 |
| chr10 | 83543505 | 83544857 | 26.52689 | 93.43019 | 90.01563 | Promoter | 10 | 83543941 | 83596469 | 52529 | 1 | ENSMUST00 | 0 | A230046K03I | ENSMUSG00000034560.5 |
| chr11 | 83298225 | 83299602 | 30.87849 | 93.42764 | 90.01563 | Promoter | 11 | 83294642 | 83298977 | 4336 | 2 | ENSMUST00 | 0 | Pex12 | ENSMUSG00000018733.5 |
| chr11 | 32266754 | 32267959 | 26.52689 | 93.43019 | 90.01563 | Promoter | 11 | 32225628 | 32267689 | 42062 | 2 | ENSMUST00 | 0 | Nprl3 | ENSMUSG00000020289.10 |
| chr11 | 31870855 | 31874873 | 26.52689 | 93.43019 | 90.01563 | Promoter | 11 | 31872211 | 31931577 | 59367 | 1 | ENSMUST00 | 0 | Cpeb4 | ENSMUSG00000020300.9 |
| chr11 | 118417946 | 118419122 | 26.52689 | 93.43019 | 90.01563 | Promoter | 11 | 118406289 | 118419052 | 12764 | 2 | ENSMUST00 | 0 | Cant1 | ENSMUSG00000025575.9 |
| chr11 | 101301558 | 101302372 | 26.52689 | 93.43019 | 90.01563 | Promoter | 11 | 101285952 | 101302286 | 16335 | 2 | ENSMUST00 | 0 | Becn1 | ENSMUSG00000035086.8 |
| chr15 | 79228973 | 79229946 | 26.52689 | 93.43019 | 90.01563 | Promoter | 15 | 79229173 | 79249484 | 20312 | 1 | ENSMUST00 | 0 | Pick1 | ENSMUSG00000068206.7 |
| chr18 | 64488271 | 64489481 | 26.52689 | 93.43019 | 90.01563 | Promoter | 18 | 64456550 | 64489066 | 32517 | 2 | ENSMUST00 | 0 | Fech | ENSMUSG00000024588.8 |
| chr19 | 8722564 | 8723338 | 26.52689 | 93.43019 | 90.01563 | Promoter | 19 | 8706882 | 8723369 | 16488 | 2 | ENSMUST00 | 31 | Slc3a2 | ENSMUSG00000010095.8 |
| chr19 | 10895211 | 10896661 | 26.52689 | 93.43019 | 90.01563 | Promoter | 19 | 10895231 | 10905532 | 10302 | 1 | ENSMUST00 | 0 | Prpf19 | ENSMUSG00000024735.8 |
| chr19 | 4162710 | 4163785 | 26.52689 | 93.43019 | 90.01563 | Promoter | 19 | 4156713 | 4163354 | 6642 | 2 | ENSMUST00 | 0 | Rps6kb2 | ENSMUSG00000024830.11 |
| chr19 | 31663317 | 31665214 | 26.52689 | 93.43019 | 90.01563 | Promoter | 19 | 30568547 | 31664370 | 1095824 | 2 | ENSMUST00 | 0 | Prkg1 | ENSMUSG00000052920.9 |
| chr19 | 4397356 | 4398674 | 26.52689 | 93.43019 | 90.01563 | Promoter | 19 | 4314419 | 4398285 | 83867 | 2 | ENSMUST00 | 0 | Kdm2a | ENSMUSG00000054611.11 |
| chr2 | 25982006 | 25983386 | 26.52689 | 93.43019 | 90.01563 | Promoter | 2 | 25926838 | 25983282 | 56445 | 2 | ENSMUST00 | 0 | Camsap1 | ENSMUSG00000026933.12 |
| chr2 | 121806207 | 121807707 | 26.52689 | 93.43019 | 90.01563 | Promoter | 2 | 121545529 | 121807087 | 261559 | 2 | ENSMUST00 | 0 | Frmd5 | ENSMUSG00000027238.11 |
| chr2 | 145674554 | 145676544 | 26.52689 | 93.43019 | 90.01563 | Promoter | 2 | 145675367 | 145714989 | 39623 | 1 | ENSMUST00 | 0 | BC039771 | ENSMUSG00000087594.1 |
| chr3 | 34080600 | 34082064 | 26.52689 | 93.43019 | 90.01563 | Promoter | 3 | 34056020 | 34081312 | 25293 | 2 | ENSMUST00 | 0 | Dnajc19 | ENSMUSG00000027679.8 |
| chr4 | 72198600 | 72202177 | 26.52689 | 93.43019 | 90.01563 | Promoter | 4 | 72117142 | 72200919 | 83778 | 2 | ENSMUST00 | 0 | Tle1 | ENSMUSG00000008305.13 |
| chr4 | 109664951 | 109667518 | 26.52689 | 93.43019 | 90.01563 | Promoter | 4 | 109660876 | 109665372 | 4497 | 2 | ENSMUST00 | 0 | Cdkn2c | ENSMUSG00000028551.9 |
| chr4 | 108578239 | 108580007 | 26.52689 | 93.43019 | 90.01563 | Promoter | 4 | 108579423 | 108595439 | 16017 | 1 | ENSMUST00 | 0 | Orc1 | ENSMUSG00000028587.13 |
| chr5 | 135994738 | 135995738 | 26.52689 | 93.43019 | 90.01563 | Promoter | 5 | 135994800 | 136032872 | 38073 | 1 | ENSMUST00 | 0 | Dtx2 | ENSMUSG00000004947.10 |
| chr5 | 24029506 | 24031016 | 26.52689 | 93.43019 | 90.01563 | Promoter | 5 | 23915277 | 24030312 | 115036 | 2 | ENSMUST00 | 0 | Fam126a | ENSMUSG00000028995.9 |
| chr5 | 107597049 | 107598312 | 26.52689 | 93.43019 | 90.01563 | Promoter | 5 | 107548967 | 107597639 | 48673 | 2 | ENSMUST00 | 0 | Glmn | ENSMUSG00000029276.8 |
| chr5 | 92042169 | 92043742 | 26.52689 | 93.43019 | 90.01563 | Promoter | 5 | 92006074 | 92043041 | 36968 | 2 | ENSMUST00 | 0 | Cdkl2 | ENSMUSG00000029403.9 |
| chr6 | 92183232 | 92184448 | 26.52689 | 93.43019 | 90.01563 | Promoter | 6 | 92169525 | 92184033 | 14509 | 2 | ENSMUST00 | 0 | Mrps25 | ENSMUSG00000014551.8 |
| chr6 | 17693660 | 17695108 | 30.87849 | 93.42764 | 90.01563 | Promoter | 6 | 17693947 | 17943022 | 249076 | 1 | ENSMUST00 | 0 | St7 | ENSMUSG00000029534.12 |
| chr6 | 42264469 | 42266101 | 26.52689 | 93.43019 | 90.01563 | Promoter | 6 | 42264985 | 42282508 | 17524 | 1 | ENSMUST00 | 0 | Casp2 | ENSMUSG00000029863.8 |
| chr7 | 121706329 | 121708050 | 26.52689 | 93.43019 | 90.01563 | Promoter | 7 | 121707075 | 121708013 | 939 | 1 | ENSMUST00 | 0 | 1700069B07I | ENSMUSG00000053081.3 |
| chr8 | 72239485 | 72241092 | 26.52689 | 93.43019 | 90.01563 | Promoter | 8 | 72240018 | 72257385 | 17368 | 1 | ENSMUST00 | 0 | Ap1m1 | ENSMUSG00000003033.9 |
| chr8 | 104533603 | 104535957 | 26.52689 | 93.43019 | 90.01563 | Promoter | 8 | 104511028 | 104534636 | 23609 | 2 | ENSMUST00 | 0 | Nae1 | ENSMUSG00000031878.12 |
| chr9 | 107554606 | 107556283 | 26.52689 | 93.43019 | 90.01563 | Promoter | 9 | 107554656 | 107562263 | 7608 | 1 | ENSMUST00 | 0 | Rassf1 | ENSMUSG00000010067.8 |
| chr9 | 101250702 | 101251977 | 26.52689 | 93.43019 | 90.01563 | Promoter | 9 | 101105084 | 101251795 | 146712 | 2 | ENSMUST00 | 0 | Ppp2r3a | ENSMUSG00000043154.10 |
| chr9 | 20520577 | 20522228 | 26.52689 | 93.43019 | 90.01563 | Promoter | 9 | 20495068 | 20521417 | 26350 | 2 | ENSMUST00 | 0 | Zfp266 | ENSMUSG00000060510.8 |
| chr5 | 140829591 | 140830475 | 30.22203 | 93.27019 | 89.85963 | Promoter | 5 | 140758408 | 140830431 | 72024 | 2 | ENSMUST00 | 0 | Gna12 | ENSMUSG00000000149.6 |
| chr10 | 128625655 | 128626818 | 29.09105 | 93.24493 | 89.8345 | Promoter | 10 | 128624530 | 128626506 | 1977 | 2 | ENSMUST00 | 0 | Rps26 | ENSMUSG00000025362.5 |
| chr14 | 55681200 | 55682448 | 29.09105 | 93.24493 | 89.8345 | Promoter | 14 | 55679083 | 55681817 | 2735 | 2 | ENSMUST00 | 0 | Tinf2 | ENSMUSG00000007589.8 |
| chr17 | 35088938 | 35090122 | 29.09105 | 93.24493 | 89.8345 | Promoter | 17 | 35089263 | 35091826 | 2564 | 1 | ENSMUST00 | 0 | Abhd16a | ENSMUSG00000007036.10 |
| chr7 | 30562463 | 30564201 | 29.09105 | 93.24493 | 89.8345 | Promoter | 7 | 30561868 | 30563184 | 1317 | 2 | ENSMUST00 | 0 | Psenen | ENSMUSG00000036835.6 |
| chr7 | 44467585 | 44468728 | 29.09105 | 93.24493 | 89.8345 | Promoter | 7 | 44467980 | 44471652 | 3673 | 1 | ENSMUST00 | 0 | Josd2 | ENSMUSG00000038695.7 |
| chr8 | 75033611 | 75034936 | 29.09105 | 93.24493 | 89.8345 | Promoter | 8 | 75033686 | 75070121 | 36436 | 1 | ENSMUST00 | 0 | Tom1 | ENSMUSG00000042870.9 |
| chr3 | 88552107 | 88553013 | 30.75439 | 93.21284 | 89.80341 | Promoter | 3 | 88549820 | 88552859 | 3040 | 2 | ENSMUST00 | 0 | Lamtor2 | ENSMUSG00000028062.10 |
| chr5 | 124326967 | 124327933 | 30.75439 | 93.21284 | 89.80341 | Promoter | 5 | 124252677 | 124327849 | 75173 | 2 | ENSMUST00 | 0 | Mphosph9 | ENSMUSG00000038126.12 |
| chr8 | 83996357 | 83998054 | 31.32211 | 93.21193 | 89.80268 | Promoter | 8 | 83997672 | 84000386 | 2715 | 1 | ENSMUST00 | 0 | Samd1 | ENSMUSG00000079003.1 |
| chr11 | 115932592 | 115934597 | 30.16331 | 93.1678 | 89.7588 | Promoter | 11 | 115892595 | 115933477 | 40883 | 2 | ENSMUST00 | 0 | Recql5 | ENSMUSG00000020752.6 |
| chr1 | 86487725 | 86488470 | 30.69271 | 93.10602 | 89.69717 | Distal Interge | 1 | 86473659 | 86474006 | 348 | 1 | ENSMUST00 | 14067 | RP23-403I3.6 | ENSMUSG00000101328.1 |
| chr10 | 81232790 | 81233661 | 30.69271 | 93.10602 | 89.69717 | Promoter | 10 | 81233155 | 81237553 | 4399 | 1 | ENSMUST00 | 0 | Zfr2 | ENSMUSG00000034949.13 |
| chr13 | 23750936 | 23751547 | 30.69271 | 93.10602 | 89.69717 | Promoter | 13 | 23751126 | 23751598 | 473 | 1 | ENSMUST00 | 0 | Hist1h2ab | ENSMUSG00000061615.2 |
| chr14 | 63605924 | 63607510 | 30.69271 | 93.10602 | 89.69717 | Promoter | 14 | 63606503 | 63820809 | 214307 | 1 | ENSMUST00 | 0 | Xkr6 | ENSMUSG00000035067.9 |
| chr14 | 47188655 | 47189862 | 30.69271 | 93.10602 | 89.69717 | Promoter | 14 | 47153895 | 47189413 | 35519 | 2 | ENSMUST00 | 0 | Gch1 | ENSMUSG00000037580.9 |
| chr5 | 33936454 | 33937405 | 30.69271 | 93.10602 | 89.69717 | Promoter | 5 | 33896936 | 33936413 | 39478 | 2 | ENSMUST00 | -42 | Nelfa | ENSMUSG00000029111.6 |
| chr6 | 29215492 | 29216667 | 30.69271 | 93.10602 | 89.69717 | Promoter | 6 | 29200434 | 29216331 | 15898 | 2 | ENSMUST00 | 0 | Impdh1 | ENSMUSG00000003500.8 |
| chr6 | 83455308 | 83457331 | 30.69271 | 93.10602 | 89.69717 | Promoter | 6 | 83368042 | 83456392 | 88351 | 2 | ENSMUST00 | 0 | Tet3 | ENSMUSG00000034832.10 |
| chr7 | 30625383 | 30626226 | 30.69271 | 93.10602 | 89.69717 | Promoter | 7 | 30616974 | 30626151 | 9178 | 2 | ENSMUST00 | 0 | Cox6b1 | ENSMUSG00000036751.6 |
| chr12 | 84450126 | 84452383 | 31.25721 | 93.10017 | 89.69231 | Promoter | 12 | 84430723 | 84450950 | 20228 | 2 | ENSMUST00 | 0 | Aldh6a1 | ENSMUSG00000021238.10 |
| chr2 | 25706352 | 25707563 | 31.25721 | 93.10017 | 89.69231 | Promoter | 2 | 25706739 | 25707721 | 983 | 1 | ENSMUST00 | 0 | Bmyc | ENSMUSG00000049086.8 |
| chr13 | 38527177 | 38528831 | 28.45235 | 93.08366 | 89.67602 | Promoter | 13 | 38500273 | 38527914 | 27642 | 2 | ENSMUST00 | 0 | Txndc5 | ENSMUSG00000038991.11 |
| chr17 | 28951668 | 28953760 | 28.45235 | 93.08366 | 89.67602 | Promoter | 17 | 28951950 | 28964121 | 12172 | 1 | ENSMUST00 | 0 | Kctd20 | ENSMUSG00000005936.13 |
| chr5 | 20881099 | 20882123 | 28.45235 | 93.08366 | 89.67602 | Promoter | 5 | 20788747 | 20882077 | 93331 | 2 | ENSMUST00 | 0 | Phtf2 | ENSMUSG00000039987.10 |
| chr6 | 22355441 | 22356405 | 28.45235 | 93.08366 | 89.67602 | Promoter | 6 | 22306520 | 22356166 | 49647 | 2 | ENSMUST00 | 0 | Fam3c | ENSMUSG00000029672.11 |
| chr7 | 35334087 | 35335639 | 28.45235 | 93.08366 | 89.67602 | Promoter | 7 | 35334170 | 35392289 | 58120 | 1 | ENSMUST00 | 0 | Rhpn2 | ENSMUSG00000030494.8 |
| chr9 | 54916841 | 54917915 | 28.45235 | 93.08366 | 89.67602 | Promoter | 9 | 54917283 | 54949924 | 32642 | 1 | ENSMUST00 | 0 | Hykk | ENSMUSG00000035878.8 |
| chr6 | 47812239 | 47814216 | 31.19258 | 92.98883 | 89.58222 | Promoter | 6 | 47796142 | 47813512 | 17371 | 2 | ENSMUST00 | 0 | Pdia4 | ENSMUSG00000025823.8 |
| chr11 | 84818196 | 84819934 | 27.85705 | 92.98368 | 89.57738 | Promoter | 11 | 84813061 | 84819515 | 6455 | 2 | ENSMUST00 | 0 | Mrm1 | ENSMUSG00000018405.7 |
| chr16 | 32877139 | 32879191 | 27.85705 | 92.98368 | 89.57738 | Promoter | 16 | 32877500 | 32891409 | 13910 | 1 | ENSMUST00 | 0 | Fyttd1 | ENSMUSG00000022800.8 |
| chr17 | 21733276 | 21734361 | 27.85705 | 92.98368 | 89.57738 | Promoter | 17 | 21733680 | 21736081 | 2402 | 1 | ENSMUST00 | 0 | Zfp229 | ENSMUSG00000061544.8 |
| chr8 | 69774229 | 69775194 | 27.85705 | 92.98368 | 89.57738 | Promoter | 8 | 69763843 | 69774911 | 11069 | 2 | ENSMUST00 | 0 | Zfp866 | ENSMUSG00000043090.13 |
| chr15 | 32243266 | 32247730 | 21.73259 | 92.96881 | 89.56307 | Promoter | 15 | 32244813 | 32696341 | 451529 | 1 | ENSMUST00 | 0 | Sema5a | ENSMUSG00000022231.9 |
| chr17 | 45432557 | 45434542 | 21.73259 | 92.96881 | 89.56307 | Promoter | 17 | 45391892 | 45433707 | 41816 | 2 | ENSMUST00 | 0 | Cdc5l | ENSMUSG00000023932.8 |
| chr15 | 102670144 | 102671188 | 26.7801 | 92.93985 | 89.5344 | Promoter | 15 | 102662864 | 102671127 | 8264 | 2 | ENSMUST00 | 0 | Atp5g2 | ENSMUSG00000062683.6 |
| chr3 | 30595061 | 30596020 | 26.7801 | 92.93985 | 89.5344 | Promoter | 3 | 30595346 | 30595757 | 412 | 2 | ENSMUST00 | 0 | Gm24413 | ENSMUSG00000065361.1 |
| chr3 | 96246722 | 96247388 | 27.30088 | 92.93774 | 89.53244 | Promoter | 3 | 96246685 | 96248504 | 1820 | 1 | ENSMUST00 | 38 | Hist2h3c1 | ENSMUSG00000093769.3 |
| chr8 | 14888012 | 14890072 | 30.5701 | 92.89357 | 89.48846 | Promoter | 8 | 14888045 | 14895059 | 7015 | 1 | ENSMUST00 | 0 | Cln8 | ENSMUSG00000026317.7 |
| chr14 | 103345093 | 103347425 | 30.50916 | 92.78792 | 89.38336 | Promoter | 14 | 103113411 | 103346418 | 233008 | 2 | ENSMUST00 | 0 | Mycbp2 | ENSMUSG00000033004.9 |
| chr2 | 91069362 | 91071119 | 31.06412 | 92.76744 | 89.36313 | Promoter | 2 | 91061791 | 91070229 | 8439 | 2 | ENSMUST00 | 0 | Slc39a13 | ENSMUSG00000002105.10 |

| | | | | | | | | | | | | | | | |
|---|---|---|---|---|---|---|---|---|---|---|---|---|---|---|---|
| chr11 | 87109172 | 87110316 | 29.93067 | 92.76174 | 89.35763 | Promoter | 11 | 87109234 | 87124635 | 15402 | 1 | ENSMUST00 | 0 | Ska2 | ENSMUSG00000020492.6 |
| chr17 | 56461389 | 56463239 | 29.93067 | 92.76174 | 89.35763 | Promoter | 17 | 56412431 | 56462216 | 49786 | 2 | ENSMUST00 | 0 | Ptprs | ENSMUSG00000013236.11 |
| chr8 | 31167764 | 31169441 | 29.93067 | 92.76174 | 89.35763 | Promoter | 8 | 31159468 | 31168724 | 9257 | 2 | ENSMUST00 | 0 | Mak16 | ENSMUSG00000031578.4 |
| chr1 | 88698405 | 88702774 | 23.74947 | 92.7123 | 89.30923 | Promoter | 1 | 88673125 | 88701876 | 28752 | 2 | ENSMUST00 | 0 | Arl4c | ENSMUSG00000049866.7 |
| chr11 | 94629366 | 94630841 | 23.74947 | 92.7123 | 89.30923 | Promoter | 11 | 94629767 | 94645216 | 15450 | 1 | ENSMUST00 | 0 | Lrrc59 | ENSMUSG00000020869.8 |
| chr12 | 52005712 | 52006850 | 23.74947 | 92.7123 | 89.30923 | Promoter | 12 | 51988312 | 52006501 | 18190 | 2 | ENSMUST00 | 0 | Dtd2 | ENSMUSG00000020956.9 |
| chr13 | 29983304 | 29986029 | 23.74947 | 92.7123 | 89.30923 | Promoter | 13 | 29906575 | 29985668 | 79094 | 2 | ENSMUST00 | 0 | E2f3 | ENSMUSG00000016477.12 |
| chr13 | 74807412 | 74808982 | 23.74947 | 92.7123 | 89.30923 | Promoter | 13 | 74694286 | 74807921 | 113636 | 2 | ENSMUST00 | 0 | Cast | ENSMUSG00000021585.8 |
| chr19 | 4810388 | 4812472 | 23.74947 | 92.7123 | 89.30923 | Promoter | 19 | 4800569 | 4811634 | 11066 | 2 | ENSMUST00 | 0 | Rbm14 | ENSMUSG00000006456.10 |
| chr2 | 32693401 | 32694844 | 23.74947 | 92.7123 | 89.30923 | Promoter | 2 | 32682609 | 32694185 | 11577 | 2 | ENSMUST00 | 0 | Fpgs | ENSMUSG00000009566.7 |
| chr2 | 174472236 | 174473056 | 23.74947 | 92.7123 | 89.30923 | Promoter | 2 | 174465585 | 174472898 | 7314 | 2 | ENSMUST00 | 0 | Slmo2 | ENSMUSG00000016257.10 |
| chr3 | 22075480 | 22078230 | 23.74947 | 92.7123 | 89.30923 | Promoter | 3 | 22076652 | 22216594 | 139943 | 1 | ENSMUST00 | 0 | Tbl1xr1 | ENSMUSG00000027630.8 |
| chr5 | 30647389 | 30648960 | 23.74947 | 92.7123 | 89.30923 | Promoter | 5 | 30647933 | 30659731 | 11799 | 1 | ENSMUST00 | 0 | Slc35f6 | ENSMUSG00000029175.12 |
| chr1 | 51477533 | 51478508 | 30.44846 | 92.68264 | 89.28044 | Promoter | 1 | 51465862 | 51478399 | 12538 | 2 | ENSMUST00 | 0 | Nabp1 | ENSMUSG00000026107.6 |
| chr5 | 31047525 | 31049639 | 30.44846 | 92.68264 | 89.28044 | Promoter | 5 | 31036036 | 31048002 | 11967 | 2 | ENSMUST00 | 0 | Slc5a6 | ENSMUSG00000006641.7 |
| chr11 | 104549416 | 104551043 | 20.80241 | 92.62246 | 89.22079 | Promoter | 11 | 104502745 | 104550483 | 47739 | 2 | ENSMUST00 | 0 | Cdc27 | ENSMUSG00000020687.8 |
| chr2 | 154436395 | 154437605 | 20.80241 | 92.62246 | 89.22079 | Promoter | 2 | 154436481 | 154539356 | 102876 | 1 | ENSMUST00 | 0 | Cbfa2t2 | ENSMUSG00000038533.10 |
| chr15 | 10981214 | 10982460 | 30.388 | 92.57774 | 89.17617 | Promoter | 15 | 10981756 | 10996624 | 14869 | 1 | ENSMUST00 | 0 | Amacr | ENSMUSG00000022244.6 |
| chr15 | 89088657 | 89090103 | 30.93671 | 92.54772 | 89.14647 | Promoter | 15 | 89089084 | 89098180 | 9097 | 1 | ENSMUST00 | 0 | Selo | ENSMUSG00000035757.11 |
| chr1 | 52498999 | 52501178 | 29.22523 | 92.5029 | 89.10189 | Promoter | 1 | 52457294 | 52500300 | 43007 | 2 | ENSMUST00 | 0 | Nab1 | ENSMUSG00000002881.9 |
| chr10 | 43172654 | 43175535 | 29.22523 | 92.5029 | 89.10189 | Promoter | 10 | 43002500 | 43174530 | 172031 | 2 | ENSMUST00 | 0 | Sobp | ENSMUSG00000038248.8 |
| chr15 | 102624179 | 102626066 | 29.22523 | 92.5029 | 89.10189 | Promoter | 15 | 102523843 | 102625067 | 101225 | 2 | ENSMUST00 | 0 | Gm28047 | ENSMUSG00000052414.9 |
| chr17 | 23739513 | 23741047 | 29.22523 | 92.5029 | 89.10189 | Promoter | 17 | 23736186 | 23740330 | 4145 | 2 | ENSMUST00 | 0 | Paqr4 | ENSMUSG00000023909.3 |
| chr17 | 23546333 | 23547640 | 29.22523 | 92.5029 | 89.10189 | Distal Interge | 17 | 23551073 | 23553488 | 2416 | 2 | ENSMUST00 | 5848 | Casp16 | ENSMUSG00000067882.8 |
| chr5 | 130078896 | 130080200 | 29.22523 | 92.5029 | 89.10189 | Promoter | 5 | 130079370 | 130135729 | 56360 | 1 | ENSMUST00 | 0 | Tpst1 | ENSMUSG00000034118.10 |
| chr14 | 54253171 | 54254753 | 29.75854 | 92.4608 | 89.06126 | Promoter | 14 | 54235479 | 54254104 | 18626 | 2 | ENSMUST00 | 0 | Dad1 | ENSMUSG00000022174.8 |
| chr1 | 156473664 | 156474600 | 29.75681 | 92.45776 | 89.05831 | Promoter | 1 | 156424525 | 156474331 | 49807 | 2 | ENSMUST00 | 0 | Soat1 | ENSMUSG00000026600.7 |
| chr10 | 128193936 | 128195391 | 29.75681 | 92.45776 | 89.05831 | Promoter | 10 | 128194457 | 128209939 | 15483 | 1 | ENSMUST00 | 0 | Gls2 | ENSMUSG00000044005.8 |
| chr11 | 52004208 | 52005092 | 29.75681 | 92.45776 | 89.05831 | Promoter | 11 | 52004221 | 52033802 | 29582 | 1 | ENSMUST00 | 0 | Cdkl3 | ENSMUSG00000020389.13 |
| chr11 | 33512805 | 33514033 | 29.75681 | 92.45776 | 89.05831 | Promoter | 11 | 33211795 | 33513746 | 301952 | 2 | ENSMUST00 | 0 | Ranbp17 | ENSMUSG00000040594.13 |
| chr11 | 70969181 | 70970072 | 29.75681 | 92.45776 | 89.05831 | Promoter | 11 | 70943058 | 70969959 | 26902 | 2 | ENSMUST00 | 0 | Nup88 | ENSMUSG00000040667.6 |
| chr12 | 105783997 | 105786120 | 29.75681 | 92.45776 | 89.05831 | Promoter | 12 | 105784694 | 105838807 | 54114 | 1 | ENSMUST00 | 0 | Papola | ENSMUSG00000021111.10 |
| chr12 | 17347679 | 17349106 | 29.75681 | 92.45776 | 89.05831 | Promoter | 12 | 17348458 | 17430094 | 81637 | 1 | ENSMUST00 | 0 | Nol10 | ENSMUSG00000061458.8 |
| chr12 | 55079554 | 55081389 | 29.75681 | 92.45776 | 89.05831 | Promoter | 12 | 55045661 | 55080110 | 34450 | 2 | ENSMUST00 | 0 | 2700097O09 | ENSMUSG00000062198.4 |
| chr13 | 44730600 | 44734844 | 29.75681 | 92.45776 | 89.05831 | Promoter | 13 | 44730774 | 44920108 | 189335 | 1 | ENSMUST00 | 0 | Jarid2 | ENSMUSG00000038518.10 |
| chr16 | 30599184 | 30602189 | 29.75681 | 92.45776 | 89.05831 | Promoter | 16 | 30599723 | 30602797 | 3075 | 1 | ENSMUST00 | 0 | Fam43a | ENSMUSG00000046546.3 |
| chr17 | 35916918 | 35919053 | 29.75681 | 92.45776 | 89.05831 | Promoter | 17 | 35916998 | 35924295 | 7298 | 1 | ENSMUST00 | 0 | Ppp1r10 | ENSMUSG00000039220.11 |
| chr17 | 17825989 | 17829531 | 29.75681 | 92.45776 | 89.05831 | Promoter | 17 | 17830188 | 17830257 | 70 | 1 | ENSMUST00 | -657 | Mir99b | ENSMUSG00000065396.1 |
| chr2 | 113847972 | 113849522 | 29.75681 | 92.45776 | 89.05831 | Promoter | 2 | 113831492 | 113848661 | 17170 | 2 | ENSMUST00 | 0 | Arhgap11a | ENSMUSG00000041219.9 |
| chr4 | 117681524 | 117682659 | 29.75681 | 92.45776 | 89.05831 | Promoter | 4 | 117674681 | 117682253 | 7573 | 2 | ENSMUST00 | 0 | Dmap1 | ENSMUSG00000009640.6 |
| chr4 | 109475549 | 109477184 | 29.75681 | 92.45776 | 89.05831 | Promoter | 4 | 109451098 | 109476675 | 25578 | 2 | ENSMUST00 | 0 | Rnf11 | ENSMUSG00000028557.5 |
| chr4 | 116149764 | 116151273 | 29.75681 | 92.45776 | 89.05831 | Promoter | 4 | 116150518 | 116159844 | 9327 | 1 | ENSMUST00 | 0 | Pomgnt1 | ENSMUSG00000028700.9 |
| chr5 | 5558827 | 5560157 | 29.75681 | 92.45776 | 89.05831 | Promoter | 5 | 5537454 | 5559538 | 22085 | 2 | ENSMUST00 | 0 | Gtpbp10 | ENSMUSG00000040464.10 |
| chr5 | 25529773 | 25530757 | 29.75681 | 92.45776 | 89.05831 | Promoter | 5 | 25530013 | 25531466 | 1454 | 1 | ENSMUST00 | 0 | RP23-405O1 | ENSMUSG00000101856.1 |
| chr7 | 79742393 | 79743430 | 29.75681 | 92.45776 | 89.05831 | Promoter | 7 | 79735957 | 79743080 | 7124 | 2 | ENSMUST00 | 0 | Pex11a | ENSMUSG00000030545.8 |
| chr7 | 70357618 | 70362933 | 29.75681 | 92.45776 | 89.05831 | Promoter | 7 | 70351946 | 70360593 | 8648 | 2 | ENSMUST00 | 0 | Nr2f2 | ENSMUSG00000030551.8 |
| chr7 | 19920197 | 19921509 | 29.75681 | 92.45776 | 89.05831 | Promoter | 7 | 19903578 | 19921160 | 17583 | 2 | ENSMUST00 | 0 | Pvr | ENSMUSG00000040511.8 |
| chr9 | 119936825 | 119937927 | 29.75681 | 92.45776 | 89.05831 | Promoter | 9 | 119925673 | 119937558 | 11886 | 2 | ENSMUST00 | 0 | Gorasp1 | ENSMUSG00000032513.7 |
| chr10 | 119453220 | 119455580 | 30.8734 | 92.43849 | 89.0432 | Promoter | 10 | 119454034 | 120087260 | 633227 | 1 | ENSMUST00 | 0 | Grip1 | ENSMUSG00000034813.11 |
| chr13 | 28949862 | 28954154 | 30.8734 | 92.43849 | 89.0432 | Promoter | 13 | 28948919 | 28953713 | 4795 | 2 | ENSMUST00 | 0 | Sox4 | ENSMUSG00000076431.4 |
| chr2 | 177497900 | 177499216 | 30.8734 | 92.43849 | 89.0432 | Promoter | 2 | 177498215 | 177508869 | 10655 | 1 | ENSMUST00 | 0 | Gm14403 | ENSMUSG00000094786.1 |
| chr7 | 29907874 | 29909004 | 30.8734 | 92.43849 | 89.0432 | Promoter | 7 | 29908223 | 29916813 | 8591 | 1 | ENSMUST00 | 0 | ZNF383 | ENSMUSG00000099689.1 |
| chr1 | 153331293 | 153333837 | 19.98426 | 92.42262 | 89.02813 | Promoter | 1 | 153218922 | 153332786 | 113865 | 2 | ENSMUST00 | 0 | Lamc1 | ENSMUSG00000026478.9 |
| chr3 | 101109427 | 101110666 | 30.2678 | 92.36906 | 88.97462 | Promoter | 3 | 101040232 | 101110278 | 70047 | 2 | ENSMUST00 | 0 | Ptgfrn | ENSMUSG00000027864.8 |
| chr14 | 99298002 | 99299904 | 29.7016 | 92.36115 | 88.96692 | Promoter | 14 | 99298691 | 99313405 | 14715 | 1 | ENSMUST00 | 0 | Klf5 | ENSMUSG00000005148.7 |
| chr10 | 79983154 | 79984959 | 30.81035 | 92.32966 | 88.93555 | Promoter | 10 | 79977118 | 79984330 | 7213 | 2 | ENSMUST00 | 0 | Tmem259 | ENSMUSG00000013858.9 |
| chr14 | 55634877 | 55636107 | 30.81035 | 92.32966 | 88.93555 | Promoter | 14 | 55625400 | 55635684 | 10285 | 2 | ENSMUST00 | 0 | Ipo4 | ENSMUSG00000002319.10 |
| chr1 | 91297809 | 91299047 | 28.5703 | 92.31807 | 88.92419 | Promoter | 1 | 91298338 | 91321075 | 22738 | 1 | ENSMUST00 | 0 | Scly | ENSMUSG00000026307.7 |
| chr15 | 5115562 | 5117636 | 28.5703 | 92.31807 | 88.92419 | Promoter | 15 | 5116613 | 5119140 | 2528 | 1 | ENSMUST00 | 0 | Rpl37 | ENSMUSG00000041841.7 |
| chr18 | 35702362 | 35703608 | 28.5703 | 92.31807 | 88.92419 | Promoter | 18 | 35671103 | 35703144 | 32042 | 2 | ENSMUST00 | 0 | Dnajc18 | ENSMUSG00000024350.5 |
| chr6 | 48601477 | 48602808 | 28.5703 | 92.31807 | 88.92419 | Distal Interge | 6 | 48594319 | 48596623 | 2305 | 1 | ENSMUST00 | 7159 | Repin1 | ENSMUSG00000052751.11 |
| chr7 | 27472589 | 27474624 | 28.5703 | 92.31807 | 88.92419 | Promoter | 7 | 27473768 | 27477359 | 3592 | 1 | ENSMUST00 | 0 | Sertad3 | ENSMUSG00000055200.6 |
| chr9 | 72132652 | 72133982 | 28.5703 | 92.31807 | 88.92419 | Distal Interge | 9 | 72128587 | 72129391 | 805 | 1 | ENSMUST00 | 4066 | Gm7863 | ENSMUSG00000099253.1 |
| chr19 | 46760836 | 46761355 | 30.20806 | 92.26528 | 88.87222 | Promoter | 19 | 46761596 | 46878795 | 117200 | 1 | ENSMUST00 | -241 | Cnnm2 | ENSMUSG00000064105.7 |
| chr11 | 100711979 | 100712568 | 29.64488 | 92.26185 | 88.869 | Promoter | 11 | 100704746 | 100712173 | 7428 | 2 | ENSMUST00 | 0 | Kat2a | ENSMUSG00000020918.9 |
| chr9 | 77909795 | 77911783 | 25.89225 | 92.22567 | 88.8329 | Distal Interge | 9 | 77917364 | 77984519 | 67156 | 1 | ENSMUST00 | -5581 | Elovl5 | ENSMUSG00000032349.8 |
| chr7 | 16452759 | 16453648 | 30.74755 | 92.22124 | 88.82853 | Promoter | 7 | 16452779 | 16455489 | 2711 | 1 | ENSMUST00 | 0 | Tmem160 | ENSMUSG00000019158.8 |
| chr9 | 30942383 | 30943953 | 30.74755 | 92.22124 | 88.82853 | Promoter | 9 | 30942562 | 30962856 | 20295 | 1 | ENSMUST00 | 0 | Adamts8 | ENSMUSG00000031994.8 |
| chr16 | 5254553 | 5256620 | 29.06336 | 92.21697 | 88.82447 | Promoter | 16 | 5244152 | 5255983 | 11832 | 2 | ENSMUST00 | 0 | Fam86 | ENSMUSG00000022544.8 |
| chr1 | 152399483 | 152401550 | 27.96063 | 92.19791 | 88.80544 | Promoter | 1 | 152399830 | 152509818 | 109989 | 1 | ENSMUST00 | 0 | Colgalt2 | ENSMUSG00000032649.9 |
| chr4 | 122835637 | 122836931 | 27.96063 | 92.19791 | 88.80544 | Promoter | 4 | 122836242 | 122859078 | 22837 | 1 | ENSMUST00 | 0 | Ppt1 | ENSMUSG00000028657.9 |
| chr1 | 135282967 | 135284430 | 24.86315 | 92.1349 | 88.74321 | Promoter | 1 | 135262712 | 135284084 | 21373 | 2 | ENSMUST00 | 0 | Rnpep | ENSMUSG00000041926.10 |
| chr1 | 134494021 | 134495459 | 24.86315 | 92.1349 | 88.74321 | Promoter | 1 | 134494648 | 134508774 | 14127 | 1 | ENSMUST00 | 0 | Rabif | ENSMUSG00000042229.9 |
| chr11 | 80382405 | 80385417 | 24.86315 | 92.1349 | 88.74321 | Promoter | 11 | 80383279 | 80396245 | 12967 | 1 | ENSMUST00 | 0 | Zfp207 | ENSMUSG00000017421.13 |
| chr11 | 9010305 | 9011731 | 27.39171 | 92.13462 | 88.74321 | Promoter | 11 | 8993137 | 9011160 | 18024 | 2 | ENSMUST00 | 0 | Hus1 | ENSMUSG00000020413.6 |
| chr12 | 78747954 | 78748798 | 24.86315 | 92.1349 | 88.74321 | Promoter | 12 | 78748947 | 78840711 | 91765 | 1 | ENSMUST00 | -149 | Mpp5 | ENSMUSG00000021112.8 |
| chr15 | 10485017 | 10486692 | 24.86315 | 92.1349 | 88.74321 | Promoter | 15 | 10474779 | 10485947 | 11169 | 2 | ENSMUST00 | 0 | Brix1 | ENSMUSG00000022247.7 |
| chr18 | 60603340 | 60613557 | 27.39171 | 92.13462 | 88.74321 | Promoter | 18 | 60600207 | 60610105 | 9899 | 2 | ENSMUST00 | 0 | Synpo | ENSMUSG00000043079.12 |
| chr2 | 26351234 | 26352873 | 24.86315 | 92.1349 | 88.74321 | Promoter | 2 | 26352187 | 26352775 | 589 | 2 | ENSMUST00 | 0 | Card9 | ENSMUSG00000026928.9 |
| chr2 | 167420983 | 167422465 | 24.86315 | 92.1349 | 88.74321 | Promoter | 2 | 167421712 | 167477000 | 55289 | 1 | ENSMUST00 | 0 | Slc9a8 | ENSMUSG00000039463.9 |
| chr4 | 109200480 | 109202667 | 24.86315 | 92.1349 | 88.74321 | Promoter | 4 | 109061655 | 109202217 | 140563 | 2 | ENSMUST00 | 0 | Osbpl9 | ENSMUSG00000028559.12 |
| chr4 | 138453728 | 138456452 | 24.86315 | 92.1349 | 88.74321 | Promoter | 4 | 138454314 | 138460123 | 5810 | 1 | ENSMUST00 | 0 | Camk2n1 | ENSMUSG00000046447.3 |
| chr5 | 73292784 | 73293753 | 24.86315 | 92.1349 | 88.74321 | Promoter | 5 | 73292794 | 73309356 | 16563 | 1 | ENSMUST00 | 0 | Ociad1 | ENSMUSG00000029152.8 |
| chr5 | 139199935 | 139201685 | 24.86315 | 92.1349 | 88.74321 | Promoter | 5 | 139200637 | 139249840 | 49204 | 1 | ENSMUST00 | 0 | Sun1 | ENSMUSG00000036817.9 |
| chr5 | 28316314 | 28318412 | 27.39171 | 92.13462 | 88.74321 | Promoter | 5 | 28317121 | 28352868 | 35748 | 1 | ENSMUST00 | 0 | Rbm33 | ENSMUSG00000048271.9 |
| chr6 | 94282364 | 94284879 | 24.86315 | 92.1349 | 88.74321 | Promoter | 6 | 93675455 | 94283322 | 607868 | 2 | ENSMUST00 | 0 | Magi1 | ENSMUSG00000045095.11 |
| chr8 | 72420659 | 72422185 | 24.86315 | 92.1349 | 88.74321 | Promoter | 8 | 72341001 | 72421474 | 80474 | 2 | ENSMUST00 | 0 | Eps15l1 | ENSMUSG00000006276.9 |
| chr8 | 121577810 | 121578884 | 24.86315 | 92.1349 | 88.74321 | Promoter | 8 | 121549440 | 121578806 | 29367 | 2 | ENSMUST00 | 0 | Fbxo31 | ENSMUSG00000052934.8 |
| chr9 | 123478684 | 123479821 | 24.86315 | 92.1349 | 88.74321 | Promoter | 9 | 123478701 | 123521548 | 42848 | 1 | ENSMUST00 | 0 | Limd1 | ENSMUSG00000025239.2 |
| chr9 | 43102050 | 43107425 | 27.39171 | 92.13462 | 88.74321 | Promoter | 9 | 42963842 | 43105718 | 141877 | 2 | ENSMUST00 | 0 | Arhgef12 | ENSMUSG00000059495.8 |
| chr1 | 74283791 | 74284772 | 31.24818 | 92.1069 | 88.71768 | Promoter | 1 | 74279840 | 74284738 | 4899 | 2 | ENSMUST00 | 0 | Aamp | ENSMUSG00000006299.8 |
| chr1 | 16518543 | 16520190 | 31.24818 | 92.1069 | 88.71768 | Promoter | 1 | 16228674 | 16519306 | 290633 | 2 | ENSMUST00 | 0 | Stau2 | ENSMUSG00000025920.14 |
| chr11 | 100413912 | 100415193 | 31.24818 | 92.1069 | 88.71768 | Promoter | 11 | 100408456 | 100414836 | 6381 | 2 | ENSMUST00 | 0 | Leprel4 | ENSMUSG00000006931.10 |
| chr11 | 51688104 | 51689576 | 31.24818 | 92.1069 | 88.71768 | Promoter | 11 | 51685386 | 51688653 | 3268 | 2 | ENSMUST00 | 0 | 0610009B22 | ENSMUSG00000007777.4 |
| chr12 | 53252318 | 53254087 | 31.24818 | 92.1069 | 88.71768 | Intron (ENSM | 12 | 53248677 | 54072175 | 823499 | 1 | ENSMUST00 | 3642 | Npas3 | ENSMUSG00000021010.7 |
| chr15 | 102990385 | 102993111 | 31.24818 | 92.1069 | 88.71768 | Promoter | 15 | 102990607 | 102993821 | 3215 | 1 | ENSMUST00 | 0 | Hoxc8 | ENSMUSG00000001657.7 |
| chr4 | 136247308 | 136248988 | 31.24818 | 92.1069 | 88.71768 | Promoter | 4 | 136247729 | 136271304 | 23576 | 1 | ENSMUST00 | 0 | Tcea3 | ENSMUSG00000001604.9 |
| chr4 | 43039333 | 43040348 | 31.24818 | 92.1069 | 88.71768 | Promoter | 4 | 43032414 | 43040288 | 7875 | 2 | ENSMUST00 | 0 | Fam214b | ENSMUSG00000036002.7 |
| chr14 | 118131916 | 118133153 | 30.08928 | 92.05882 | 88.67145 | Promoter | 14 | 118111911 | 118132765 | 20855 | 2 | ENSMUST00 | 0 | Tgds | ENSMUSG00000022130.9 |
| chr16 | 4678825 | 4680120 | 30.08928 | 92.05882 | 88.67145 | Promoter | 16 | 4626133 | 4679777 | 53645 | 2 | ENSMUST00 | 0 | Coro7 | ENSMUSG00000039637.9 |
| chr6 | 5725457 | 5726859 | 30.08928 | 92.05882 | 88.67145 | Promoter | 6 | 5725639 | 6028039 | 302401 | 1 | ENSMUST00 | 0 | Dync1i1 | ENSMUSG00000029757.11 |
| chr4 | 146610889 | 146611537 | 17.87554 | 92.05333 | 88.66702 | Promoter | 4 | 146610961 | 146658160 | 47200 | 1 | ENSMUST00 | 0 | Gm13248 | ENSMUSG00000063245.6 |
| chr17 | 27634180 | 27635882 | 30.62272 | 92.00563 | 88.61945 | Promoter | 17 | 27630422 | 27635276 | 4855 | 2 | ENSMUST00 | 0 | Rps10 | ENSMUSG00000052146.10 |
| chr6 | 50109410 | 50111194 | 30.62272 | 92.00563 | 88.61945 | Promoter | 6 | 50110241 | 50198598 | 88358 | 1 | ENSMUST00 | 0 | Mpp6 | ENSMUSG00000038388.9 |
| chr1 | 165235872 | 165237874 | 22.53436 | 92.00179 | 88.61587 | Promoter | 1 | 165214922 | 165236943 | 22022 | 2 | ENSMUST00 | 0 | Tiprl | ENSMUSG00000040843.5 |
| chr15 | 34494573 | 34496010 | 22.53436 | 92.00179 | 88.61587 | Promoter | 15 | 34484021 | 34495255 | 11235 | 2 | ENSMUST00 | 0 | Hrsp12 | ENSMUSG00000022323.6 |
| chr17 | 55878000 | 55879719 | 22.53436 | 92.00179 | 88.61587 | Promoter | 17 | 55864892 | 55878930 | 14039 | 2 | ENSMUST00 | 0 | Zfp119a | ENSMUSG00000057835.7 |
| chr10 | 77605823 | 77607318 | 29.476 | 91.96591 | 88.58034 | Promoter | 10 | 77606097 | 77618331 | 12235 | 1 | ENSMUST00 | 0 | Sumo3 | ENSMUSG00000020265.10 |
| chr6 | 47668287 | 47668554 | 16.43574 | 91.94095 | 88.5556 | Distal Interge | 6 | 47637143 | 47637219 | 77 | 1 | ENSMUST00 | 31145 | Gm25283 | ENSMUSG00000087897.1 |
| chr7 | 101310088 | 101310911 | 30.56069 | 91.89841 | 88.51308 | Distal Interge | 7 | 101317086 | 101346626 | 29541 | 1 | ENSMUST00 | -6175 | Stard10 | ENSMUSG00000030688.9 |
| chr9 | 124281002 | 124282804 | 27.25512 | 91.88436 | 88.49937 | Distal Interge | 9 | 124293831 | 124311899 | 18069 | 2 | ENSMUST00 | 29095 | 2010315B03 | ENSMUSG00000074829.5 |
| chr10 | 7725607 | 7726872 | 27.78601 | 91.88203 | 88.49705 | Promoter | 10 | 7726000 | 7763149 | 37150 | 1 | ENSMUST00 | 0 | Katna1 | ENSMUSG00000019794.8 |
| chr10 | 93640662 | 93642462 | 27.78601 | 91.88203 | 88.49705 | Promoter | 10 | 93640681 | 93644957 | 4277 | 1 | ENSMUST00 | 0 | Ntn4 | ENSMUSG00000020019.4 |
| chr10 | 111505983 | 111508558 | 27.78601 | 91.88203 | 88.49705 | Promoter | 10 | 111506286 | 111508645 | 2360 | 1 | ENSMUST00 | 0 | Phlda1 | ENSMUSG00000020205.8 |
| chr11 | 74637560 | 74638919 | 27.78601 | 91.88203 | 88.49705 | 3' UTR | 11 | 74649495 | 74656551 | 7057 | 1 | ENSMUST00 | -10576 | Cluh | ENSMUSG00000020741.12 |
| chr12 | 84217669 | 84219521 | 27.78601 | 91.88203 | 88.49705 | Promoter | 12 | 84149176 | 84218881 | 69706 | 2 | ENSMUST00 | 0 | Elmsan1 | ENSMUSG00000042507.9 |
| chr13 | 58384406 | 58385635 | 27.78601 | 91.88203 | 88.49705 | Promoter | 13 | 58380046 | 58385225 | 5180 | 2 | ENSMUST00 | 0 | 2210016F16 | ENSMUSG00000021550.5 |
| chr13 | 49681387 | 49683105 | 27.78601 | 91.88203 | 88.49705 | Promoter | 13 | 49682100 | 49734267 | 52168 | 1 | ENSMUST00 | 0 | Iars | ENSMUSG00000037851.8 |
| chr13 | 98353706 | 98359675 | 27.78601 | 91.88203 | 88.49705 | Promoter | 13 | 98354242 | 98356703 | 2462 | 1 | ENSMUST00 | 0 | Foxd1 | ENSMUSG00000078302.4 |
| chr13 | 12649806 | 12651375 | 27.78601 | 91.88203 | 88.49705 | Promoter | 13 | 12619084 | 12650297 | 31214 | 2 | ENSMUST00 | 0 | Gpr137b-ps | ENSMUSG00000097715.1 |
| chr14 | 57397972 | 57399204 | 27.78601 | 91.88203 | 88.49705 | Promoter | 14 | 57275034 | 57398483 | 123450 | 2 | ENSMUST00 | 0 | Cryl1 | ENSMUSG00000021947.10 |
| chr14 | 61680629 | 61683530 | 27.78601 | 91.88203 | 88.49705 | Promoter | 14 | 61602839 | 61682373 | 79535 | 2 | ENSMUST00 | 0 | Dleu2 | ENSMUSG00000097589.4 |
| chr15 | 80947928 | 80949299 | 27.78601 | 91.88203 | 88.49705 | Promoter | 15 | 80948490 | 80970947 | 22458 | 1 | ENSMUST00 | 0 | Adsl | ENSMUSG00000022407.3 |
| chr15 | 91048925 | 91050373 | 27.78601 | 91.88203 | 88.49705 | Promoter | 15 | 90933276 | 91049948 | 116673 | 2 | ENSMUST00 | 0 | Kif21a | ENSMUSG00000022629.11 |
| chr15 | 78876271 | 78878467 | 27.78601 | 91.88203 | 88.49705 | Promoter | 15 | 78877190 | 78894585 | 17396 | 1 | ENSMUST00 | 0 | Gga1 | ENSMUSG00000033128.8 |
| chr16 | 17759171 | 17760419 | 27.78601 | 91.88203 | 88.49705 | Promoter | 16 | 17759618 | 17771827 | 12210 | 1 | ENSMUST00 | 0 | Klhl22 | ENSMUSG00000022750.12 |
| chr16 | 96158967 | 96159866 | 27.78601 | 91.88203 | 88.49705 | 3' UTR | 16 | 96145476 | 96147584 | 2109 | 2 | ENSMUST00 | -11384 | Gm15317 | ENSMUSG00000087282.1 |
| chr17 | 57010552 | 57012337 | 27.78601 | 91.88203 | 88.49705 | Promoter | 17 | 57003405 | 57011288 | 7884 | 2 | ENSMUST00 | 0 | Gtf2f1 | ENSMUSG00000002658.9 |

| | | | | | | | | | | | | | | | |
|---|---|---|---|---|---|---|---|---|---|---|---|---|---|---|---|
| chr17 | 43666540 | 43667878 | 27.78601 | 91.88203 | 88.49705 | Promoter | 17 | 43641900 | 43667015 | 25116 | 2 | ENSMUST00 | 0 | Slc25a27 | ENSMUSG00000023912.9 |
| chr17 | 30575193 | 30577241 | 27.78601 | 91.88203 | 88.49705 | Promoter | 17 | 30215524 | 30576287 | 360764 | 2 | ENSMUST00 | 0 | Btbd9 | ENSMUSG00000062202.8 |
| chr18 | 9210613 | 9214860 | 27.78601 | 91.88203 | 88.49705 | Promoter | 18 | 9212856 | 9216201 | 3346 | 1 | ENSMUST00 | 0 | Fzd8 | ENSMUSG00000036904.5 |
| chr19 | 7240402 | 7241621 | 27.78601 | 91.88203 | 88.49705 | Promoter | 19 | 7225668 | 7241222 | 15555 | 2 | ENSMUST00 | 0 | Naa40 | ENSMUSG00000024764.9 |
| chr19 | 40611503 | 40612421 | 27.78601 | 91.88203 | 88.49705 | Promoter | 19 | 40596446 | 40612233 | 15788 | 2 | ENSMUST00 | 0 | Tctn3 | ENSMUSG00000025008.10 |
| chr2 | 94178038 | 94179142 | 27.78601 | 91.88203 | 88.49705 | Distal Interge | 2 | 94184027 | 94184130 | 104 | 2 | ENSMUST00 | 4988 | Gm23630 | ENSMUSG00000093033.1 |
| chr3 | 103808753 | 103809440 | 27.78601 | 91.88203 | 88.49705 | Promoter | 3 | 103800605 | 103809378 | 8774 | 2 | ENSMUST00 | 0 | Dclre1b | ENSMUSG00000027845.10 |
| chr3 | 157946510 | 157948743 | 27.78601 | 91.88203 | 88.49705 | Promoter | 3 | 157947414 | 158005910 | 58497 | 1 | ENSMUST00 | 0 | Ankrd13c | ENSMUSG00000039988.9 |
| chr3 | 14610429 | 14611323 | 27.78601 | 91.88203 | 88.49705 | Promoter | 3 | 14606284 | 14611285 | 5002 | 2 | ENSMUST00 | 0 | 1810022K09I | ENSMUSG00000078784.3 |
| chr3 | 108554052 | 108554913 | 27.78601 | 91.88203 | 88.49705 | Promoter | 3 | 108554338 | 108554751 | 414 | 2 | ENSMUST00 | 0 | Scarna2 | ENSMUSG00000088185.1 |
| chr4 | 109675990 | 109677632 | 27.78601 | 91.88203 | 88.49705 | Promoter | 4 | 109676588 | 109963960 | 287373 | 1 | ENSMUST00 | 0 | Faf1 | ENSMUSG00000010517.7 |
| chr4 | 53011420 | 53012767 | 27.78601 | 91.88203 | 88.49705 | Promoter | 4 | 53011880 | 53022060 | 10181 | 1 | ENSMUST00 | 0 | Nipsnap3b | ENSMUSG00000015247.5 |
| chr4 | 117891481 | 117892086 | 27.78601 | 91.88203 | 88.49705 | Promoter | 4 | 117888643 | 117892032 | 3390 | 2 | ENSMUST00 | 0 | Dph2 | ENSMUSG00000028540.6 |
| chr4 | 70409774 | 70411085 | 27.78601 | 91.88203 | 88.49705 | Promoter | 4 | 70216856 | 70410443 | 193588 | 2 | ENSMUST00 | 0 | Cdk5rap2 | ENSMUSG00000039298.11 |
| chr6 | 97806110 | 97807782 | 27.78601 | 91.88203 | 88.49705 | Promoter | 6 | 97807058 | 97942179 | 135122 | 1 | ENSMUST00 | 0 | Mitf | ENSMUSG00000035158.10 |
| chr7 | 127470082 | 127472628 | 27.78601 | 91.88203 | 88.49705 | Promoter | 7 | 127470371 | 127476759 | 6389 | 1 | ENSMUST00 | 0 | Prr14 | ENSMUSG00000030822.10 |
| chr7 | 25618809 | 25620007 | 27.78601 | 91.88203 | 88.49705 | Promoter | 7 | 25619414 | 25625551 | 6138 | 1 | ENSMUST00 | 0 | Atp5sl | ENSMUSG00000057229.6 |
| chr7 | 73616872 | 73618375 | 27.78601 | 91.88203 | 88.49705 | Distal Interge | 7 | 73645985 | 73646089 | 105 | 2 | ENSMUST00 | 27714 | Gm26176 | ENSMUSG00000065729.1 |
| chr8 | 71908250 | 71909533 | 27.78601 | 91.88203 | 88.49705 | Promoter | 8 | 71908635 | 71932495 | 23861 | 1 | ENSMUST00 | 0 | Zfp617 | ENSMUSG00000066880.10 |
| chr9 | 35266797 | 35268694 | 27.78601 | 91.88203 | 88.49705 | Promoter | 9 | 35267865 | 35275965 | 8101 | 1 | ENSMUST00 | 0 | Rpusd4 | ENSMUSG00000032044.4 |
| chr9 | 64339957 | 64341585 | 27.78601 | 91.88203 | 88.49705 | Promoter | 9 | 64306756 | 64341257 | 34502 | 2 | ENSMUST00 | 0 | Dis3l | ENSMUSG00000032396.12 |
| chr9 | 114389854 | 114391090 | 27.78601 | 91.88203 | 88.49705 | Promoter | 9 | 114375134 | 114390675 | 15542 | 2 | ENSMUST00 | 0 | Crtap | ENSMUSG00000032431.6 |
| chr1 | 160195160 | 160195982 | 26.18681 | 91.82585 | 88.44535 | Promoter | 1 | 160195215 | 160199820 | 4606 | 1 | ENSMUST00 | 0 | Mrps14 | ENSMUSG00000058267.8 |
| chr1 | 58652252 | 58653336 | 26.18681 | 91.82585 | 88.44535 | Promoter | 1 | 58652641 | 58655831 | 3191 | 1 | ENSMUST00 | 0 | RP24-285E3. | ENSMUSG00000102059.1 |
| chr11 | 3451200 | 3453231 | 26.18681 | 91.82585 | 88.44535 | Promoter | 11 | 3415982 | 3452363 | 36382 | 2 | ENSMUST00 | 0 | Rnf185 | ENSMUSG00000020448.11 |
| chr11 | 103116270 | 103119164 | 26.18681 | 91.82585 | 88.44535 | Promoter | 11 | 103116231 | 103119725 | 3495 | 1 | ENSMUST00 | 40 | Hexim1 | ENSMUSG00000048878.6 |
| chr12 | 5373404 | 5376248 | 26.18681 | 91.82585 | 88.44535 | Promoter | 12 | 5077472 | 5375682 | 298211 | 2 | ENSMUST00 | 0 | Klhl29 | ENSMUSG00000020627.9 |
| chr12 | 52097424 | 52098312 | 26.18681 | 91.82585 | 88.44535 | Promoter | 12 | 52097737 | 52312744 | 215008 | 1 | ENSMUST00 | 0 | Nubpl | ENSMUSG00000035142.13 |
| chr12 | 4816870 | 4818608 | 26.18681 | 91.82585 | 88.44535 | Promoter | 12 | 4817608 | 4827658 | 10051 | 1 | ENSMUST00 | 0 | Sf3b6 | ENSMUSG00000037361.7 |
| chr13 | 55622271 | 55623942 | 26.18681 | 91.82585 | 88.44535 | Promoter | 13 | 55623005 | 55632411 | 9407 | 1 | ENSMUST00 | 0 | Caml | ENSMUSG00000021501.4 |
| chr13 | 103918287 | 103922058 | 26.18681 | 91.82585 | 88.44535 | Promoter | 13 | 103818787 | 103920514 | 101728 | 2 | ENSMUST00 | 0 | Erbb2ip | ENSMUSG00000021709.9 |
| chr15 | 55111809 | 55113638 | 26.18681 | 91.82585 | 88.44535 | Promoter | 15 | 55112317 | 55259271 | 146955 | 1 | ENSMUST00 | 0 | Deptor | ENSMUSG00000022419.10 |
| chr16 | 38452104 | 38453454 | 26.18681 | 91.82585 | 88.44535 | Promoter | 16 | 38445399 | 38452689 | 7291 | 2 | ENSMUST00 | 0 | Adprh | ENSMUSG00000002844.8 |
| chr16 | 91405640 | 91407210 | 26.18681 | 91.82585 | 88.44535 | Promoter | 16 | 91406164 | 91425822 | 19659 | 1 | ENSMUST00 | 0 | Il10rb | ENSMUSG00000022969.8 |
| chr17 | 33919171 | 33920968 | 26.18681 | 91.82585 | 88.44535 | Promoter | 17 | 33919332 | 33929288 | 9957 | 1 | ENSMUST00 | 0 | Tapbp | ENSMUSG00000024308.9 |
| chr17 | 24162821 | 24163851 | 26.18681 | 91.82585 | 88.44535 | Promoter | 17 | 24155833 | 24163730 | 7898 | 2 | ENSMUST00 | 0 | Amdhd2 | ENSMUSG00000036820.6 |
| chr17 | 28328018 | 28329379 | 26.18681 | 91.82585 | 88.44535 | Promoter | 17 | 28328471 | 28331032 | 2562 | 1 | ENSMUST00 | 0 | Rpl10a | ENSMUSG00000037805.9 |
| chr17 | 6978383 | 6979605 | 26.18681 | 91.82585 | 88.44535 | Promoter | 17 | 6978860 | 6998193 | 19334 | 1 | ENSMUST00 | 0 | Rnaset2b | ENSMUSG00000094724.2 |
| chr2 | 120608534 | 120610256 | 26.18681 | 91.82585 | 88.44535 | Promoter | 2 | 120609383 | 120619906 | 10524 | 1 | ENSMUST00 | 0 | Haus2 | ENSMUSG00000027285.8 |
| chr2 | 33637941 | 33643219 | 26.18681 | 91.82585 | 88.44535 | Promoter | 2 | 33560965 | 33640511 | 79547 | 2 | ENSMUST00 | 0 | Lmx1b | ENSMUSG00000038765.8 |
| chr3 | 96196650 | 96197954 | 26.18681 | 91.82585 | 88.44535 | Promoter | 3 | 96196588 | 96197586 | 999 | 2 | ENSMUST00 | 0 | Bola1 | ENSMUSG00000015943.6 |
| chr4 | 62501735 | 62502741 | 26.18681 | 91.82585 | 88.44535 | Promoter | 4 | 62499008 | 62502276 | 3269 | 2 | ENSMUST00 | 0 | Hdhd3 | ENSMUSG00000038422.2 |
| chr4 | 155693984 | 155695391 | 26.18681 | 91.82585 | 88.44535 | Promoter | 4 | 155694342 | 155696481 | 2140 | 1 | ENSMUST00 | 0 | B930041F14I | ENSMUSG00000074738.2 |
| chr6 | 21850969 | 21852979 | 26.18681 | 91.82585 | 88.44535 | Promoter | 6 | 21771395 | 21852088 | 80694 | 2 | ENSMUST00 | 0 | Tspan12 | ENSMUSG00000029669.9 |
| chr6 | 118138293 | 118139593 | 26.18681 | 91.82585 | 88.44535 | Promoter | 6 | 118107452 | 118139140 | 31689 | 2 | ENSMUST00 | 0 | Csgalnact2 | ENSMUSG00000042042.9 |
| chr7 | 114414652 | 114417189 | 26.18681 | 91.82585 | 88.44535 | Promoter | 7 | 114415281 | 114539251 | 123971 | 1 | ENSMUST00 | 0 | Pde3b | ENSMUSG00000030671.8 |
| chr7 | 107370116 | 107371837 | 26.18681 | 91.82585 | 88.44535 | Promoter | 7 | 107370728 | 107436336 | 65609 | 1 | ENSMUST00 | 0 | Syt9 | ENSMUSG00000062542.6 |
| chr8 | 124946790 | 124950147 | 26.18681 | 91.82585 | 88.44535 | Promoter | 8 | 124908596 | 124949254 | 40659 | 2 | ENSMUST00 | 0 | Egln1 | ENSMUSG00000031987.5 |
| chr9 | 69397656 | 69398853 | 26.18681 | 91.82585 | 88.44535 | Promoter | 9 | 69397906 | 69433122 | 35217 | 1 | ENSMUST00 | 0 | Narg2 | ENSMUSG00000032235.10 |
| chr9 | 8133662 | 8134918 | 26.18681 | 91.82585 | 88.44535 | Promoter | 9 | 8076633 | 8134294 | 57662 | 2 | ENSMUST00 | 0 | AK129341 | ENSMUSG00000040729.6 |
| chr12 | 33146768 | 33148505 | 30.4989 | 91.7916 | 88.41422 | Promoter | 12 | 33147693 | 33251146 | 103454 | 1 | ENSMUST00 | 0 | Atxn7l1 | ENSMUSG00000020564.11 |
| chr3 | 135211828 | 135213077 | 30.4989 | 91.7916 | 88.41422 | Promoter | 3 | 135212563 | 135273540 | 60978 | 1 | ENSMUST00 | 0 | Cenpe | ENSMUSG00000045328.7 |
| chr4 | 141793602 | 141794545 | 30.4989 | 91.7916 | 88.41422 | Promoter | 4 | 141793612 | 141798086 | 4475 | 1 | ENSMUST00 | 0 | Casp9 | ENSMUSG00000028914.8 |
| chr1 | 90998212 | 90999695 | 31.06164 | 91.78615 | 88.409 | Promoter | 1 | 90998737 | 91108640 | 109904 | 1 | ENSMUST00 | 0 | Lrrfip1 | ENSMUSG00000026305.10 |
| chr10 | 128673422 | 128674818 | 31.06164 | 91.78615 | 88.409 | Promoter | 10 | 128669887 | 128673918 | 4032 | 2 | ENSMUST00 | 0 | Suox | ENSMUSG00000049858.7 |
| chr17 | 26917006 | 26917751 | 31.06164 | 91.78615 | 88.409 | Promoter | 17 | 26917091 | 26932579 | 15489 | 1 | ENSMUST00 | 0 | Kifc5b | ENSMUSG00000024301.9 |
| chr8 | 11558328 | 11559201 | 31.06164 | 91.78615 | 88.409 | Intron (ENSM | 8 | 11556066 | 11563250 | 7185 | 1 | ENSMUST00 | 2263 | Ing1 | ENSMUSG00000045969.7 |
| chr1 | 37539932 | 37541579 | 29.36448 | 91.77025 | 88.3935 | Promoter | 1 | 37439340 | 37541016 | 101677 | 2 | ENSMUST00 | 0 | Mgat4a | ENSMUSG00000026110.10 |
| chr1 | 128359124 | 128360407 | 29.36448 | 91.77025 | 88.3935 | Promoter | 1 | 128331590 | 128359664 | 28075 | 2 | ENSMUST00 | 0 | Mcm6 | ENSMUSG00000026355.6 |
| chr11 | 23769609 | 23771991 | 29.36448 | 91.77025 | 88.3935 | Promoter | 11 | 23741514 | 23770970 | 29457 | 2 | ENSMUST00 | 0 | Rel | ENSMUSG00000020275.8 |
| chr11 | 102437370 | 102438734 | 29.36448 | 91.77025 | 88.3935 | 3' UTR | 11 | 102438748 | 102439777 | 1030 | 2 | ENSMUST00 | 1043 | Fam171a2 | ENSMUSG00000034685.4 |
| chr12 | 81531833 | 81533139 | 29.36448 | 91.77025 | 88.3935 | Promoter | 12 | 81358860 | 81532905 | 174046 | 2 | ENSMUST00 | 0 | Gm20498 | ENSMUSG00000021139.12 |
| chr16 | 45842591 | 45845351 | 29.36448 | 91.77025 | 88.3935 | Promoter | 16 | 45746243 | 45844378 | 98136 | 2 | ENSMUST00 | 0 | Phldb2 | ENSMUSG00000033149.11 |
| chr2 | 49787410 | 49788169 | 29.36448 | 91.77025 | 88.3935 | Promoter | 2 | 49787688 | 49948849 | 161162 | 1 | ENSMUST00 | 0 | Lypd6b | ENSMUSG00000026765.7 |
| chr4 | 107066242 | 107067705 | 29.36448 | 91.77025 | 88.3935 | Promoter | 4 | 107066988 | 107078247 | 11260 | 1 | ENSMUST00 | 0 | Cyb5rl | ENSMUSG00000028621.12 |
| chr4 | 16013032 | 16014530 | 29.36448 | 91.77025 | 88.3935 | Promoter | 4 | 15997121 | 16013888 | 16768 | 2 | ENSMUST00 | 0 | Osgin2 | ENSMUSG00000041153.4 |
| chr8 | 4275117 | 4276486 | 29.36448 | 91.77025 | 88.3935 | Promoter | 8 | 4259731 | 4275885 | 16155 | 2 | ENSMUST00 | 0 | Timm44 | ENSMUSG00000002949.9 |
| chr14 | 8213589 | 8215726 | 30.43737 | 91.68518 | 88.31002 | Promoter | 14 | 8214148 | 8223567 | 9420 | 1 | ENSMUST00 | 0 | Kctd6 | ENSMUSG00000021752.7 |
| chr1 | 62702681 | 62704220 | 30.99688 | 91.67474 | 88.2998 | Promoter | 1 | 62703285 | 62739932 | 36648 | 1 | ENSMUST00 | 0 | Nrp2 | ENSMUSG00000025969.10 |
| chr5 | 77405039 | 77408352 | 25.08176 | 91.61557 | 88.24099 | Promoter | 5 | 77349240 | 77408045 | 58806 | 2 | ENSMUST00 | 0 | Igfbp7 | ENSMUSG00000036256.8 |
| chr11 | 77493384 | 77494525 | 30.37608 | 91.57915 | 88.20459 | Promoter | 11 | 77493562 | 77507786 | 14225 | 1 | ENSMUST00 | 0 | Git1 | ENSMUSG00000011877.8 |
| chr11 | 69695084 | 69696451 | 30.37608 | 91.57915 | 88.20459 | Promoter | 11 | 69682577 | 69696098 | 13522 | 2 | ENSMUST00 | 0 | Tnfsf12Tnfsf | ENSMUSG00000018752.14 |
| chr7 | 25282142 | 25283253 | 30.37608 | 91.57915 | 88.20459 | Promoter | 7 | 25282179 | 25294145 | 11967 | 1 | ENSMUST00 | 0 | Cic | ENSMUSG00000005442.8 |
| chr17 | 33939725 | 33941436 | 28.69254 | 91.56027 | 88.18607 | Promoter | 17 | 33938821 | 33940343 | 1523 | 2 | ENSMUST00 | 0 | Pfdn6 | ENSMUSG00000024309.9 |
| chr17 | 67352761 | 67355220 | 28.69254 | 91.56027 | 88.18607 | Promoter | 17 | 66666848 | 67354459 | 687612 | 2 | ENSMUST00 | 0 | Ptprm | ENSMUSG00000033278.9 |
| chr2 | 26902239 | 26903850 | 28.69254 | 91.56027 | 88.18607 | Promoter | 2 | 26890435 | 26902879 | 12445 | 2 | ENSMUST00 | 0 | Surf6 | ENSMUSG00000036160.8 |
| chr2 | 11602268 | 11603483 | 28.69254 | 91.56027 | 88.18607 | Promoter | 2 | 11585439 | 11603235 | 17797 | 2 | ENSMUST00 | 0 | Rbm17 | ENSMUSG00000037197.6 |
| chr3 | 19187442 | 19189116 | 28.69254 | 91.56027 | 88.18607 | Promoter | 3 | 19188110 | 19220817 | 32708 | 1 | ENSMUST00 | 0 | Mtfr1 | ENSMUSG00000027601.8 |
| chr3 | 107278562 | 107280094 | 28.69254 | 91.56027 | 88.18607 | Promoter | 3 | 107278858 | 107284079 | 5222 | 1 | ENSMUST00 | 0 | Lamtor5 | ENSMUSG00000087260.1 |
| chr6 | 87671311 | 87672842 | 28.69254 | 91.56027 | 88.18607 | Promoter | 6 | 87628429 | 87672168 | 43740 | 2 | ENSMUST00 | 0 | Aplf | ENSMUSG00000030051.5 |
| chr8 | 84965470 | 84966188 | 28.69254 | 91.56027 | 88.18607 | Promoter | 8 | 84956610 | 84966011 | 9402 | 2 | ENSMUST00 | 0 | Rnaseh2a | ENSMUSG00000052926.10 |
| chr12 | 100198919 | 100201212 | 21.48562 | 91.50835 | 88.13527 | Promoter | 12 | 100199435 | 100209806 | 10372 | 1 | ENSMUST00 | 0 | Calm1 | ENSMUSG00000001175.8 |
| chr18 | 57354231 | 57355888 | 21.48562 | 91.50835 | 88.13527 | Promoter | 18 | 57354733 | 57392719 | 37987 | 1 | ENSMUST00 | 0 | Prrc1 | ENSMUSG00000024594.8 |
| chr6 | 83325284 | 83326888 | 25.50501 | 91.48401 | 88.11109 | Promoter | 6 | 83326016 | 83343776 | 17761 | 1 | ENSMUST00 | 0 | Mob1a | ENSMUSG00000043131.9 |
| chr7 | 25236886 | 25239305 | 30.31504 | 91.47351 | 88.10067 | Promoter | 7 | 25228258 | 25237851 | 9594 | 2 | ENSMUST00 | 0 | Gsk3a | ENSMUSG00000057177.7 |
| chr9 | 90269919 | 90271221 | 30.86817 | 91.45322 | 88.08056 | Promoter | 9 | 90202027 | 90270769 | 68743 | 2 | ENSMUST00 | 0 | Tbc1d2b | ENSMUSG00000037410.8 |
| chr9 | 48834371 | 48836926 | 30.86817 | 91.45322 | 88.08056 | Promoter | 9 | 48654311 | 48835945 | 181635 | 2 | ENSMUST00 | 0 | Zbtb16 | ENSMUSG00000066687.4 |
| chr11 | 102696829 | 102698408 | 28.06786 | 91.41865 | 88.0463 | Promoter | 11 | 102684686 | 102697782 | 13097 | 2 | ENSMUST00 | 0 | Ccdc43 | ENSMUSG00000020925.11 |
| chr13 | 64161342 | 64163393 | 28.06786 | 91.41865 | 88.0463 | Promoter | 13 | 64161866 | 64186536 | 24671 | 1 | ENSMUST00 | 0 | Habp4 | ENSMUSG00000021476.8 |
| chr16 | 20610201 | 20612182 | 28.06786 | 91.41865 | 88.0463 | Promoter | 16 | 20611593 | 20619007 | 7415 | 1 | ENSMUST00 | 0 | Ece2 | ENSMUSG00000022842.11 |
| chr16 | 94327749 | 94329658 | 28.06786 | 91.41865 | 88.0463 | Promoter | 16 | 94328420 | 94336935 | 8516 | 1 | ENSMUST00 | 0 | Ripply3 | ENSMUSG00000022941.8 |
| chr2 | 91648776 | 91651859 | 28.06786 | 91.41865 | 88.0463 | Promoter | 2 | 91649860 | 91672326 | 22467 | 1 | ENSMUST00 | 0 | Arhgap1 | ENSMUSG00000027247.11 |
| chr6 | 34780162 | 34781417 | 28.06786 | 91.41865 | 88.0463 | Promoter | 6 | 34780432 | 34857932 | 77501 | 1 | ENSMUST00 | 0 | Agbl3 | ENSMUSG00000038836.10 |
| chr7 | 67645361 | 67646264 | 28.06786 | 91.41865 | 88.0463 | Promoter | 7 | 67618062 | 67645268 | 27207 | 2 | ENSMUST00 | -94 | Lrrc28 | ENSMUSG00000030556.7 |
| chr5 | 114444225 | 114444941 | 25.95436 | 91.38634 | 88.01462 | Promoter | 5 | 114444269 | 114453149 | 8881 | 1 | ENSMUST00 | 0 | Mvk | ENSMUSG00000041939.9 |
| chr10 | 12922325 | 12923501 | 29.14396 | 91.38276 | 88.01111 | Promoter | 10 | 12916648 | 12923100 | 6453 | 2 | ENSMUST00 | 0 | B230208H11 | ENSMUSG00000097378.2 |
| chr12 | 111442259 | 111444418 | 30.25424 | 91.36826 | 87.99667 | Promoter | 12 | 111442469 | 111455018 | 12550 | 1 | ENSMUST00 | 0 | Tnfaip2 | ENSMUSG00000021281.10 |
| chr7 | 41498525 | 41500525 | 30.25424 | 91.36826 | 87.99667 | Promoter | 7 | 41478880 | 41499890 | 21011 | 2 | ENSMUST00 | 0 | AW146154 | ENSMUSG00000074166.6 |
| chr15 | 76625129 | 76626788 | 27.48563 | 91.33697 | 87.96664 | Promoter | 15 | 76622088 | 76626027 | 3940 | 2 | ENSMUST00 | 0 | Tonsl | ENSMUSG00000059323.8 |
| chr4 | 123662851 | 123665633 | 27.48563 | 91.33697 | 87.96664 | Promoter | 4 | 123456349 | 123664752 | 208404 | 2 | ENSMUST00 | 0 | Macf1 | ENSMUSG00000028649.12 |
| chr5 | 38259797 | 38261236 | 27.48563 | 91.33697 | 87.96664 | Promoter | 5 | 38260185 | 38269622 | 9438 | 1 | ENSMUST00 | 0 | Tmem128 | ENSMUSG00000067365.5 |
| chr8 | 13869309 | 13870531 | 26.43231 | 91.32629 | 87.95647 | Promoter | 8 | 13869641 | 13881639 | 11999 | 1 | ENSMUST00 | 0 | Champ1 | ENSMUSG00000047710.7 |
| chr10 | 79554260 | 79555448 | 26.94166 | 91.30816 | 87.93849 | Promoter | 10 | 79540245 | 79555091 | 14847 | 2 | ENSMUST00 | 0 | Mier2 | ENSMUSG00000042570.9 |
| chr13 | 100491648 | 100492949 | 26.94166 | 91.30816 | 87.93849 | Promoter | 13 | 100460218 | 100492338 | 32121 | 2 | ENSMUST00 | 0 | Gtf2h2 | ENSMUSG00000021639.11 |
| chr1 | 161875523 | 161876835 | 23.46333 | 91.19009 | 87.82124 | Promoter | 1 | 161816112 | 161876661 | 60550 | 2 | ENSMUST00 | 0 | Suco | ENSMUSG00000040297.7 |
| chr13 | 38959640 | 38961007 | 23.46333 | 91.19009 | 87.82124 | Promoter | 13 | 38932140 | 38960537 | 28398 | 2 | ENSMUST00 | 0 | Slc35b3 | ENSMUSG00000021432.9 |
| chr18 | 9957507 | 9958955 | 23.46333 | 91.19009 | 87.82124 | Promoter | 18 | 9958180 | 9995484 | 37305 | 1 | ENSMUST00 | 0 | Thoc1 | ENSMUSG00000024287.7 |
| chr4 | 117018933 | 117020146 | 23.46333 | 91.19009 | 87.82124 | Promoter | 4 | 117019402 | 117086852 | 67451 | 1 | ENSMUST00 | 0 | Eif2b3 | ENSMUSG00000028683.9 |
| chr6 | 113306728 | 113308496 | 20.57128 | 91.19001 | 87.82124 | Promoter | 6 | 113307181 | 113324857 | 17677 | 1 | ENSMUST00 | 0 | Brpf1 | ENSMUSG00000001632.11 |
| chr8 | 121107519 | 121108793 | 23.46333 | 91.19009 | 87.82124 | Promoter | 8 | 121097555 | 121108295 | 10741 | 2 | ENSMUST00 | 0 | Mthfsd | ENSMUSG00000031816.9 |
| chr9 | 66059514 | 66061149 | 23.46333 | 91.19009 | 87.82124 | Promoter | 9 | 66060169 | 66066629 | 6461 | 1 | ENSMUST00 | 0 | Ppib | ENSMUSG00000032383.6 |
| chr8 | 70838618 | 70839627 | 30.13338 | 91.15889 | 87.79092 | Promoter | 8 | 70835155 | 70839720 | 4566 | 2 | ENSMUST00 | 93 | Arrdc2 | ENSMUSG00000002910.10 |
| chr2 | 18025808 | 18029357 | 29.56612 | 91.1473 | 87.77953 | Promoter | 2 | 18027926 | 18028760 | 835 | 2 | ENSMUST00 | 0 | A930004D18 | ENSMUSG00000054057.2 |
| chr7 | 92873936 | 92875664 | 29.56612 | 91.1473 | 87.77953 | Promoter | 7 | 92857529 | 92874213 | 16685 | 2 | ENSMUST00 | 0 | 4632434I11R | ENSMUSG00000030641.9 |
| chr11 | 69095226 | 69097021 | 30.07331 | 91.05476 | 87.68732 | Promoter | 11 | 69095217 | 69106458 | 11242 | 1 | ENSMUST00 | 10 | Per1 | ENSMUSG00000020893.12 |
| chr14 | 77036071 | 77037177 | 30.07331 | 91.05476 | 87.68732 | Promoter | 14 | 77036772 | 77112204 | 75433 | 1 | ENSMUST00 | 0 | Ccdc122 | ENSMUSG00000034795.9 |
| chr7 | 45569998 | 45571389 | 30.07331 | 91.05476 | 87.68732 | Promoter | 7 | 45570153 | 45589710 | 19558 | 1 | ENSMUST00 | 0 | Bcat2 | ENSMUSG00000030826.12 |
| chr7 | 81344058 | 81345731 | 30.07331 | 91.05476 | 87.68732 | Promoter | 7 | 81342732 | 81344531 | 1800 | 2 | ENSMUST00 | 0 | Rps17 | ENSMUSG00000061787.9 |
| chr2 | 145902873 | 145904471 | 29.50911 | 91.04776 | 87.68056 | Promoter | 2 | 145903241 | 145916425 | 13185 | 1 | ENSMUST00 | 0 | Naa20 | ENSMUSG00000002728.9 |
| chr3 | 95110639 | 95111926 | 29.50911 | 91.04776 | 87.68056 | Promoter | 3 | 95111022 | 95123051 | 12030 | 1 | ENSMUST00 | 0 | Vps72 | ENSMUSG00000008958.9 |
| chr3 | 89962488 | 89964514 | 29.50911 | 91.04776 | 87.68056 | Promoter | 3 | 89939481 | 89963508 | 24028 | 2 | ENSMUST00 | 0 | Atp8b2 | ENSMUSG00000060671.7 |
| chr4 | 151862125 | 151863367 | 29.50911 | 91.04776 | 87.68056 | Promoter | 4 | 151059525 | 151861876 | 802352 | 2 | ENSMUST00 | -250 | Camta1 | ENSMUSG00000014592.14 |
| chr5 | 129908219 | 129909160 | 29.50911 | 91.04776 | 87.68056 | Promoter | 5 | 129908568 | 129910396 | 1829 | 1 | ENSMUST00 | 0 | Nupr1l | ENSMUSG00000095789.1 |
| chr7 | 29179346 | 29180586 | 29.50911 | 91.04776 | 87.68056 | Promoter | 7 | 29174188 | 29180552 | 6365 | 2 | ENSMUST00 | 0 | Psmd8 | ENSMUSG00000030591.11 |
| chr7 | 114275235 | 114276795 | 29.50911 | 91.04776 | 87.68056 | Promoter | 7 | 114264606 | 114276118 | 11513 | 2 | ENSMUST00 | 0 | Psma1 | ENSMUSG00000030751.12 |
| chr7 | 84688896 | 84690552 | 29.50911 | 91.04776 | 87.68056 | Promoter | 7 | 84689640 | 84776549 | 86910 | 1 | ENSMUST00 | 0 | 2610206C17I | ENSMUSG00000085236.2 |
| chr8 | 113634558 | 113636457 | 29.50911 | 91.04776 | 87.68056 | Promoter | 8 | 113635586 | 113645178 | 9593 | 1 | ENSMUST00 | 0 | Mon1b | ENSMUSG00000078908.4 |
| chr9 | 44378636 | 44380622 | 19.76704 | 91.01538 | 87.64919 | Promoter | 9 | 44379490 | 44392369 | 12880 | 1 | ENSMUST00 | 0 | Hyou1 | ENSMUSG00000032115.8 |

| | | | | | | | | | | | | | | | |
|---|---|---|---|---|---|---|---|---|---|---|---|---|---|---|---|
| chr9 | 62534315 | 62537345 | 30.01348 | 90.95103 | 87.5851 | Promoter | 9 | 62419492 | 62537044 | 117553 | 2 | ENSMUST00 | 0 | Coro2b | ENSMUSG00000041729.10 |
| chr3 | 144718529 | 144720795 | 29.95388 | 90.84766 | 87.48237 | Promoter | 3 | 144688844 | 144720323 | 31480 | 2 | ENSMUST00 | 0 | Sh3glb1 | ENSMUSG00000037062.9 |
| chr1 | 156309730 | 156319004 | 28.81932 | 90.81086 | 87.44578 | Promoter | 1 | 156310727 | 156328035 | 17309 | 1 | ENSMUST00 | 0 | Nphs2 | ENSMUSG00000026602.4 |
| chr13 | 114817558 | 114819439 | 28.81932 | 90.81086 | 87.44578 | Promoter | 13 | 114818236 | 114828723 | 10488 | 1 | ENSMUST00 | 0 | Mocs2 | ENSMUSG00000015536.9 |
| chr13 | 67779361 | 67780383 | 28.81932 | 90.81086 | 87.44578 | Promoter | 13 | 67779754 | 67789078 | 9325 | 1 | ENSMUST00 | 0 | Zfp493 | ENSMUSG00000090659.3 |
| chr16 | 35938251 | 35939914 | 28.81932 | 90.81086 | 87.44578 | Promoter | 16 | 35938470 | 35972605 | 34136 | 1 | ENSMUST00 | 0 | Parp9 | ENSMUSG00000022906.10 |
| chr17 | 29263357 | 29264677 | 28.81932 | 90.81086 | 87.44578 | Promoter | 17 | 29250803 | 29264158 | 13356 | 2 | ENSMUST00 | 0 | Ppil1 | ENSMUSG00000024007.9 |
| chr2 | 155073743 | 155074933 | 28.81932 | 90.81086 | 87.44578 | Promoter | 2 | 155059310 | 155074497 | 15188 | 2 | ENSMUST00 | 0 | Ahcy | ENSMUSG00000027597.10 |
| chr2 | 29869074 | 29870047 | 28.81932 | 90.81086 | 87.44578 | Promoter | 2 | 29869164 | 29871355 | 2192 | 1 | ENSMUST00 | 0 | Cercam | ENSMUSG00000039787.8 |
| chr5 | 88764418 | 88765711 | 30.48835 | 90.7988 | 87.43492 | Promoter | 5 | 88764996 | 88783281 | 18286 | 1 | ENSMUST00 | 0 | Dck | ENSMUSG00000029366.9 |
| chr10 | 61679667 | 61681197 | 29.3377 | 90.74815 | 87.38457 | Promoter | 10 | 61680310 | 61693295 | 12986 | 1 | ENSMUST00 | 0 | Sar1a | ENSMUSG00000020088.9 |
| chr10 | 121350567 | 121351903 | 29.3377 | 90.74815 | 87.38457 | Distal Interge | 10 | 121365090 | 121397245 | 32156 | 1 | ENSMUST00 | -13187 | Gns | ENSMUSG00000034707.6 |
| chr11 | 76762717 | 76763684 | 29.3377 | 90.74815 | 87.38457 | Promoter | 11 | 76726602 | 76763579 | 36978 | 2 | ENSMUST00 | 0 | Gosr1 | ENSMUSG00000010392.8 |
| chr11 | 55417325 | 55420160 | 29.3377 | 90.74815 | 87.38457 | Promoter | 11 | 55394500 | 55419898 | 25399 | 2 | ENSMUST00 | 0 | Sparc | ENSMUSG00000018593.7 |
| chr11 | 120672681 | 120674029 | 29.3377 | 90.74815 | 87.38457 | Promoter | 11 | 120672973 | 120674346 | 1374 | 1 | ENSMUST00 | 0 | Aspscr1 | ENSMUSG00000025142.12 |
| chr15 | 53343924 | 53346154 | 29.3377 | 90.74815 | 87.38457 | Promoter | 15 | 53068664 | 53345659 | 276996 | 2 | ENSMUST00 | 0 | Ext1 | ENSMUSG00000061731.4 |
| chr18 | 33462104 | 33464772 | 29.3377 | 90.74815 | 87.38457 | Promoter | 18 | 33437019 | 33463634 | 26616 | 2 | ENSMUST00 | 0 | Nrep | ENSMUSG00000042834.9 |
| chr2 | 37358172 | 37359760 | 29.3377 | 90.74815 | 87.38457 | Promoter | 2 | 37350074 | 37359332 | 9259 | 2 | ENSMUST00 | 0 | Pdcl | ENSMUSG00000009030.9 |
| chr2 | 116060041 | 116064346 | 29.3377 | 90.74815 | 87.38457 | Promoter | 2 | 115996056 | 116062360 | 66305 | 2 | ENSMUST00 | 0 | Meis2 | ENSMUSG00000027210.15 |
| chr2 | 11552855 | 11554333 | 29.3377 | 90.74815 | 87.38457 | Distal Interge | 2 | 11585439 | 11585990 | 552 | 2 | ENSMUST00 | 31657 | Rbm17 | ENSMUSG00000037197.6 |
| chr2 | 29124443 | 29126022 | 29.3377 | 90.74815 | 87.38457 | Promoter | 2 | 29124992 | 29182471 | 57480 | 1 | ENSMUST00 | 0 | Setx | ENSMUSG00000043535.8 |
| chr3 | 145987290 | 145989994 | 29.3377 | 90.74815 | 87.38457 | Promoter | 3 | 145987870 | 146021720 | 33851 | 1 | ENSMUST00 | 0 | Syde2 | ENSMUSG00000036863.7 |
| chr3 | 96600614 | 96602004 | 29.3377 | 90.74815 | 87.38457 | Promoter | 3 | 96601133 | 96625352 | 24220 | 1 | ENSMUST00 | 0 | Lix1l | ENSMUSG00000049288.3 |
| chr5 | 4103644 | 4105223 | 29.3377 | 90.74815 | 87.38457 | Promoter | 5 | 4081145 | 4104746 | 23602 | 2 | ENSMUST00 | 0 | Cyp51 | ENSMUSG00000001467.4 |
| chr5 | 109557542 | 109559389 | 29.3377 | 90.74815 | 87.38457 | Promoter | 5 | 109554711 | 109558993 | 4283 | 2 | ENSMUST00 | 0 | Crlf2 | ENSMUSG00000033467.8 |
| chr8 | 105346518 | 105349557 | 29.3377 | 90.74815 | 87.38457 | Promoter | 8 | 105329163 | 105347909 | 18747 | 2 | ENSMUST00 | 0 | Fhod1 | ENSMUSG00000014778.5 |
| chr9 | 77655043 | 77656067 | 29.3377 | 90.74815 | 87.38457 | 3' UTR | 9 | 77636500 | 77660127 | 23628 | 1 | ENSMUST00 | 18544 | Klhl31 | ENSMUSG00000044938.8 |
| chr1 | 175624830 | 175626156 | 28.22945 | 90.73652 | 87.3771 | Promoter | 1 | 175600374 | 175625635 | 25262 | 2 | ENSMUST00 | 0 | Fh1 | ENSMUSG00000026526.9 |
| chr10 | 24711471 | 24712723 | 30.42596 | 90.69117 | 87.33178 | Promoter | 10 | 24637914 | 24712158 | 74245 | 2 | ENSMUST00 | 0 | Enpp1 | ENSMUSG00000037370.8 |
| chr13 | 95616110 | 95618852 | 30.42596 | 90.69117 | 87.33178 | Promoter | 13 | 95601804 | 95618459 | 16656 | 2 | ENSMUST00 | 0 | F2r | ENSMUSG00000048376.5 |
| chr17 | 25843814 | 25844907 | 30.42596 | 90.69117 | 87.33178 | Promoter | 17 | 25838457 | 25844851 | 6395 | 2 | ENSMUST00 | 0 | Rhot2 | ENSMUSG00000025733.12 |
| chr4 | 125065400 | 125067636 | 30.42596 | 90.69117 | 87.33178 | Promoter | 4 | 125055338 | 125065703 | 10366 | 2 | ENSMUST00 | 0 | Dnali1 | ENSMUSG00000042707.6 |
| chr5 | 100845079 | 100847111 | 30.42596 | 90.69117 | 87.33178 | Promoter | 5 | 100845713 | 100846810 | 1098 | 1 | ENSMUST00 | 0 | Agpat9 | ENSMUSG00000029314.9 |
| chr9 | 102834367 | 102836715 | 30.42596 | 90.69117 | 87.33178 | Promoter | 9 | 102834917 | 102907542 | 72626 | 1 | ENSMUST00 | 0 | Ryk | ENSMUSG00000032547.7 |
| chr12 | 110485064 | 110486904 | 28.17892 | 90.64638 | 87.2878 | Promoter | 12 | 110485739 | 110583061 | 97323 | 1 | ENSMUST00 | 0 | Ppp2r5c | ENSMUSG00000017843.8 |
| chr14 | 76503316 | 76505898 | 28.17892 | 90.64638 | 87.2878 | Promoter | 14 | 76504214 | 76507763 | 3550 | 1 | ENSMUST00 | 0 | Tsc22d1 | ENSMUSG00000022010.14 |
| chr7 | 30728799 | 30730420 | 28.17892 | 90.64638 | 87.2878 | Promoter | 7 | 30727701 | 30729534 | 1834 | 2 | ENSMUST00 | 0 | Tmem147 | ENSMUSG00000006315.8 |
| chr8 | 89040662 | 89045860 | 28.17892 | 90.64638 | 87.2878 | Promoter | 8 | 89027248 | 89044162 | 16915 | 2 | ENSMUST00 | 0 | Sall1 | ENSMUSG00000031665.6 |
| chr9 | 21852095 | 21853205 | 28.17892 | 90.64638 | 87.2878 | Promoter | 9 | 21800184 | 21852635 | 52452 | 2 | ENSMUST00 | 0 | Dock6 | ENSMUSG00000032198.8 |
| chr2 | 101796152 | 101798205 | 29.8354 | 90.64201 | 87.2842 | Promoter | 2 | 101714285 | 101797707 | 83423 | 2 | ENSMUST00 | 0 | Prr5l | ENSMUSG00000032841.10 |
| chr2 | 121955227 | 121957775 | 29.8354 | 90.64201 | 87.2842 | Promoter | 2 | 121956001 | 122013642 | 57642 | 1 | ENSMUST00 | 0 | Ctdspl2 | ENSMUSG00000033411.11 |
| chr2 | 153631784 | 153633126 | 29.8354 | 90.64201 | 87.2842 | Promoter | 2 | 153616933 | 153632781 | 15849 | 2 | ENSMUST00 | 0 | Commd7 | ENSMUSG00000056941.12 |
| chr17 | 35241818 | 35242927 | 28.71271 | 90.62266 | 87.26511 | Promoter | 17 | 35241870 | 35253680 | 11811 | 1 | ENSMUST00 | 0 | Ddx39b | ENSMUSG00000019432.10 |
| chr1 | 58585293 | 58587521 | 24.55236 | 90.57755 | 87.22021 | Promoter | 1 | 58586384 | 58595964 | 9581 | 1 | ENSMUST00 | 0 | Ndufb3 | ENSMUSG00000026032.8 |
| chr1 | 75505204 | 75506553 | 24.55236 | 90.57755 | 87.22021 | Promoter | 1 | 75485825 | 75506361 | 20537 | 2 | ENSMUST00 | 0 | Obsl1 | ENSMUSG00000026211.12 |
| chr11 | 53706462 | 53707792 | 24.55236 | 90.57755 | 87.22021 | Promoter | 11 | 53649519 | 53707319 | 57801 | 2 | ENSMUST00 | 0 | Rad50 | ENSMUSG00000020380.11 |
| chr14 | 64440031 | 64441524 | 24.55236 | 90.57755 | 87.22021 | 5' UTR | 14 | 64122623 | 64455903 | 333281 | 2 | ENSMUST00 | 14379 | Msra | ENSMUSG00000054733.8 |
| chr14 | 31128013 | 31129010 | 24.55236 | 90.57755 | 87.22021 | Promoter | 14 | 31088869 | 31128930 | 40062 | 2 | ENSMUST00 | 0 | Smim4 | ENSMUSG00000058351.6 |
| chr15 | 98634150 | 98635460 | 24.55236 | 90.57755 | 87.22021 | Promoter | 15 | 98634764 | 98644530 | 9767 | 1 | ENSMUST00 | 0 | Cacnb3 | ENSMUSG00000003352.8 |
| chr15 | 83122305 | 83122853 | 24.55236 | 90.57755 | 87.22021 | Promoter | 15 | 83113433 | 83122801 | 9369 | 2 | ENSMUST00 | 0 | Rrp7a | ENSMUSG00000018040.4 |
| chr15 | 81960281 | 81961421 | 24.55236 | 90.57755 | 87.22021 | Promoter | 15 | 81951108 | 81960869 | 9762 | 2 | ENSMUST00 | 0 | Pmm1 | ENSMUSG00000022474.9 |
| chr15 | 84987053 | 84988586 | 24.55236 | 90.57755 | 87.22021 | Promoter | 15 | 84943936 | 84988000 | 44065 | 2 | ENSMUST00 | 0 | 5031439G07 | ENSMUSG00000036046.9 |
| chr2 | 105903837 | 105905507 | 24.55236 | 90.57755 | 87.22021 | Promoter | 2 | 105701027 | 105904564 | 203538 | 2 | ENSMUST00 | 0 | Elp4 | ENSMUSG00000027167.6 |
| chr2 | 25331751 | 25333538 | 24.55236 | 90.57755 | 87.22021 | Promoter | 2 | 25332338 | 25343280 | 10943 | 1 | ENSMUST00 | 0 | Man1b1 | ENSMUSG00000036646.8 |
| chr4 | 22357091 | 22358524 | 24.55236 | 90.57755 | 87.22021 | Promoter | 4 | 22357543 | 22376725 | 19183 | 1 | ENSMUST00 | 0 | Fbxl4 | ENSMUSG00000040410.9 |
| chr4 | 129695887 | 129697149 | 24.55236 | 90.57755 | 87.22021 | Promoter | 4 | 129676355 | 129696796 | 20442 | 2 | ENSMUST00 | 0 | Tmem39b | ENSMUSG00000053730.10 |
| chr5 | 3595303 | 3596981 | 24.55236 | 90.57755 | 87.22021 | Promoter | 5 | 3596066 | 3637230 | 41165 | 1 | ENSMUST00 | 0 | Pex1 | ENSMUSG00000005907.9 |
| chr5 | 139380120 | 139383385 | 24.55236 | 90.57755 | 87.22021 | Promoter | 5 | 139380581 | 139396415 | 15835 | 1 | ENSMUST00 | 0 | Gpr146 | ENSMUSG00000044197.8 |
| chr6 | 147090907 | 147092605 | 24.55236 | 90.57755 | 87.22021 | Promoter | 6 | 147091379 | 147112778 | 21400 | 1 | ENSMUST00 | 0 | Klhl42 | ENSMUSG00000040102.9 |
| chr6 | 115544234 | 115545177 | 24.55236 | 90.57755 | 87.22021 | Promoter | 6 | 115544664 | 115554272 | 9609 | 1 | ENSMUST00 | 0 | Tsen2 | ENSMUSG00000042389.8 |
| chr7 | 128237092 | 128238963 | 24.55236 | 90.57755 | 87.22021 | Promoter | 7 | 128237357 | 128245100 | 7744 | 1 | ENSMUST00 | 0 | Armc5 | ENSMUSG00000042178.5 |
| chr7 | 141413270 | 141414145 | 24.55236 | 90.57755 | 87.22021 | Promoter | 7 | 141408135 | 141414090 | 5956 | 2 | ENSMUST00 | 0 | Pddc1 | ENSMUSG00000051007.9 |
| chr8 | 84902697 | 84903908 | 24.55236 | 90.57755 | 87.22021 | Promoter | 8 | 84901928 | 84905295 | 3368 | 1 | ENSMUST00 | 770 | Klf1 | ENSMUSG00000054191.7 |
| chr3 | 108185509 | 108187025 | 29.22734 | 90.55506 | 87.19949 | Promoter | 3 | 108186335 | 108191776 | 5442 | 1 | ENSMUST00 | 0 | Amigo1 | ENSMUSG00000050947.8 |
| chr3 | 153911873 | 153913720 | 26.01842 | 90.55051 | 87.195 | Promoter | 3 | 153907289 | 153912203 | 4915 | 2 | ENSMUST00 | 0 | Rabggtb | ENSMUSG00000038975.9 |
| chr11 | 76493190 | 76495482 | 27.58279 | 90.54513 | 87.1897 | Promoter | 11 | 76419104 | 76495456 | 76353 | 2 | ENSMUST00 | 0 | Abr | ENSMUSG00000017631.13 |
| chr2 | 32381188 | 32382428 | 27.58279 | 90.54513 | 87.1897 | Promoter | 2 | 32377097 | 32381862 | 4766 | 2 | ENSMUST00 | 0 | 1110008P14 | ENSMUSG00000039195.4 |
| chr10 | 59702528 | 59703890 | 22.26925 | 90.51499 | 87.16024 | Promoter | 10 | 59702563 | 59744896 | 42334 | 1 | ENSMUST00 | 0 | Micu1 | ENSMUSG00000020111.10 |
| chr15 | 80263587 | 80264806 | 22.26925 | 90.51499 | 87.16024 | Promoter | 15 | 80260614 | 80264306 | 3693 | 2 | ENSMUST00 | 0 | Rps19bp1 | ENSMUSG00000051518.7 |
| chr17 | 74338195 | 74340082 | 22.26925 | 90.51499 | 87.16024 | Promoter | 17 | 74338987 | 74391113 | 52127 | 1 | ENSMUST00 | 0 | Spast | ENSMUSG00000024068.6 |
| chr3 | 68492555 | 68495979 | 22.26925 | 90.51499 | 87.16024 | Promoter | 3 | 68493491 | 68626231 | 132741 | 1 | ENSMUST00 | 0 | Schip1 | ENSMUSG00000027777.10 |
| chr15 | 76511007 | 76512009 | 26.50616 | 90.50309 | 87.1487 | Promoter | 15 | 76502015 | 76511951 | 9937 | 2 | ENSMUST00 | 0 | Dgat1 | ENSMUSG00000022555.6 |
| chr2 | 127444054 | 127445059 | 26.50616 | 90.50309 | 87.1487 | Promoter | 2 | 127436215 | 127444550 | 8336 | 2 | ENSMUST00 | 0 | Fahd2a | ENSMUSG00000027371.5 |
| chr4 | 134102506 | 134102969 | 26.50616 | 90.50309 | 87.1487 | Promoter | 4 | 134102570 | 134125165 | 22596 | 1 | ENSMUST00 | 0 | Ubxn11 | ENSMUSG00000012126.11 |
| chr1 | 131962458 | 131963909 | 30.30193 | 90.47713 | 87.1233 | Promoter | 1 | 131962967 | 131982969 | 20003 | 1 | ENSMUST00 | 0 | Slc45a3 | ENSMUSG00000026435.10 |
| chr8 | 119557720 | 119559274 | 30.30193 | 90.47713 | 87.1233 | Promoter | 8 | 119508156 | 119558761 | 50606 | 2 | ENSMUST00 | 0 | Mbtps1 | ENSMUSG00000031835.10 |
| chr12 | 112692915 | 112694138 | 29.71786 | 90.43782 | 87.08427 | Distal Interge | 12 | 112678878 | 112679970 | 1093 | 1 | ENSMUST00 | 14038 | Zbtb42 | ENSMUSG00000037638.5 |
| chr18 | 4920198 | 4922438 | 29.71786 | 90.43782 | 87.08427 | Promoter | 18 | 4920540 | 5119291 | 198752 | 1 | ENSMUST00 | 0 | Svil | ENSMUSG00000024236.12 |
| chr5 | 31179015 | 31180907 | 29.71786 | 90.43782 | 87.08427 | Promoter | 5 | 31156005 | 31180132 | 24128 | 2 | ENSMUST00 | 0 | Gtf3c2 | ENSMUSG00000029144.11 |
| chr11 | 109472077 | 109474269 | 30.2403 | 90.37072 | 87.01736 | Promoter | 11 | 109473374 | 109489999 | 16626 | 1 | ENSMUST00 | 0 | Arsg | ENSMUSG00000020604.8 |
| chr5 | 114130275 | 114131743 | 30.2403 | 90.37072 | 87.01736 | Promoter | 5 | 114130386 | 114139323 | 8938 | 1 | ENSMUST00 | 0 | Ung | ENSMUSG00000029591.9 |
| chr8 | 46210151 | 46211994 | 30.2403 | 90.37072 | 87.01736 | Promoter | 8 | 46206797 | 46211284 | 4488 | 2 | ENSMUST00 | 0 | Slc25a4 | ENSMUSG00000031633.4 |
| chr9 | 103007566 | 103008928 | 30.2403 | 90.37072 | 87.01736 | Promoter | 9 | 103008002 | 103090451 | 82450 | 1 | ENSMUST00 | 0 | Slco2a1 | ENSMUSG00000032548.9 |
| chr1 | 74284949 | 74285941 | 30.78865 | 90.34473 | 86.99165 | Promoter | 1 | 74285045 | 74353684 | 68640 | 1 | ENSMUST00 | 0 | Pnkd | ENSMUSG00000026179.9 |
| chr10 | 40882172 | 40885028 | 30.78865 | 90.34473 | 86.99165 | Promoter | 10 | 40883475 | 40938569 | 55095 | 1 | ENSMUST00 | 0 | Wasf1 | ENSMUSG00000019831.9 |
| chr17 | 26414273 | 26415574 | 30.78865 | 90.34473 | 86.99165 | Promoter | 17 | 26414829 | 26441795 | 26967 | 1 | ENSMUST00 | 0 | Neurl1b | ENSMUSG00000034413.9 |
| chr2 | 172938608 | 172940993 | 30.78865 | 90.34473 | 86.99165 | Promoter | 2 | 172868012 | 172940321 | 72310 | 2 | ENSMUST00 | 0 | Bmp7 | ENSMUSG00000008999.7 |
| chr2 | 166786014 | 166787197 | 30.78865 | 90.34473 | 86.99165 | Distal Interge | 2 | 166792531 | 166794388 | 1858 | 1 | ENSMUST00 | -5334 | Trp53rk | ENSMUSG00000042854.9 |
| chr4 | 133812841 | 133815309 | 30.78865 | 90.34473 | 86.99165 | Distal Interge | 4 | 133816420 | 133817614 | 1195 | 1 | ENSMUST00 | -1111 | Gm12974 | ENSMUSG00000084789.1 |
| chr5 | 8055753 | 8057709 | 30.78865 | 90.34473 | 86.99165 | Promoter | 5 | 8056527 | 8069151 | 12625 | 1 | ENSMUST00 | 0 | Sri | ENSMUSG00000003161.10 |
| chr5 | 131536784 | 131539135 | 30.78865 | 90.34473 | 86.99165 | Promoter | 5 | 131439298 | 131538687 | 99390 | 2 | ENSMUST00 | 0 | Auts2 | ENSMUSG00000029673.12 |
| chr5 | 137293142 | 137295178 | 30.78865 | 90.34473 | 86.99165 | Promoter | 5 | 137294629 | 137295665 | 1037 | 1 | ENSMUST00 | 0 | Ufsp1 | ENSMUSG00000051502.6 |
| chr7 | 16286309 | 16287057 | 30.78865 | 90.34473 | 86.99165 | Promoter | 7 | 16274045 | 16286790 | 12746 | 2 | ENSMUST00 | 0 | Ccdc9 | ENSMUSG00000041375.13 |
| chr9 | 31385609 | 31386970 | 30.78865 | 90.34473 | 86.99165 | Promoter | 9 | 31386192 | 31421333 | 35142 | 1 | ENSMUST00 | 0 | Nfrkb | ENSMUSG00000042185.8 |
| chr11 | 96285812 | 96287443 | 29.65943 | 90.33628 | 86.98486 | Promoter | 11 | 96286623 | 96290162 | 3540 | 1 | ENSMUST00 | 0 | Hoxb7 | ENSMUSG00000038721.8 |
| chr12 | 80434574 | 80437085 | 29.65943 | 90.33628 | 86.98486 | Promoter | 12 | 80335847 | 80436601 | 100755 | 2 | ENSMUST00 | 0 | Dcaf5 | ENSMUSG00000049106.6 |
| chr14 | 16242820 | 16244027 | 29.65943 | 90.33628 | 86.98486 | Promoter | 14 | 16239915 | 16243331 | 3417 | 2 | ENSMUST00 | 0 | Oxsm | ENSMUSG00000021786.7 |
| chr16 | 30549927 | 30550858 | 30.17891 | 90.26469 | 86.91433 | Promoter | 16 | 30511855 | 30549946 | 38092 | 2 | ENSMUST00 | 0 | Tmem44 | ENSMUSG00000022537.12 |
| chr2 | 119662219 | 119663430 | 30.17891 | 90.26469 | 86.91433 | Promoter | 2 | 119655446 | 119662820 | 7375 | 2 | ENSMUST00 | 0 | Ndufaf1 | ENSMUSG00000027305.4 |
| chr8 | 106149855 | 106151141 | 30.17891 | 90.26469 | 86.91433 | Promoter | 8 | 106150399 | 106164713 | 14315 | 1 | ENSMUST00 | 0 | Pla2g15 | ENSMUSG00000031903.6 |
| chr15 | 81810073 | 81812169 | 29.06085 | 90.26336 | 86.91321 | Promoter | 15 | 81811046 | 81821256 | 10211 | 1 | ENSMUST00 | 0 | Tef | ENSMUSG00000022389.9 |
| chr11 | 58171317 | 58172395 | 30.73327 | 90.24956 | 86.89949 | Promoter | 11 | 58171661 | 58179565 | 7905 | 1 | ENSMUST00 | 0 | Mrpl22 | ENSMUSG00000020514.3 |
| chr6 | 83032551 | 83034569 | 30.73327 | 90.24956 | 86.89949 | Promoter | 6 | 83034173 | 83037555 | 3383 | 1 | ENSMUST00 | 0 | Loxl3 | ENSMUSG00000000693.5 |
| chr18 | 36759747 | 36760823 | 29.60123 | 90.23508 | 86.88511 | Promoter | 18 | 36760239 | 36763708 | 3470 | 1 | ENSMUST00 | 0 | Wdr55 | ENSMUSG00000042660.7 |
| chr5 | 24100027 | 24101627 | 29.60123 | 90.23508 | 86.88511 | Promoter | 5 | 24100590 | 24161223 | 60634 | 1 | ENSMUST00 | 0 | Klhl7 | ENSMUSG00000028986.7 |
| chr1 | 20890081 | 20891582 | 27.41053 | 90.23362 | 86.88379 | Promoter | 1 | 20890606 | 20939650 | 49045 | 1 | ENSMUST00 | 0 | Paqr8 | ENSMUSG00000025931.10 |
| chr1 | 37889628 | 37891186 | 27.41053 | 90.23362 | 86.88379 | Promoter | 1 | 37890477 | 37898535 | 8059 | 1 | ENSMUST00 | 0 | Mrpl30 | ENSMUSG00000026087.6 |
| chr11 | 88998994 | 89000981 | 27.41053 | 90.23362 | 86.88379 | Promoter | 11 | 88999376 | 89013702 | 14327 | 1 | ENSMUST00 | 0 | Trim25 | ENSMUSG00000000275.11 |
| chr12 | 55819950 | 55821461 | 27.41053 | 90.23362 | 86.88379 | Promoter | 12 | 55602929 | 55821167 | 218239 | 2 | ENSMUST00 | 0 | Ralgapa1 | ENSMUSG00000021027.10 |
| chr17 | 33842822 | 33843840 | 27.41053 | 90.23362 | 86.88379 | Promoter | 17 | 33843091 | 33849774 | 6684 | 1 | ENSMUST00 | 0 | Cd320 | ENSMUSG00000002308.10 |
| chr18 | 79108575 | 79110521 | 27.41053 | 90.23362 | 86.88379 | Promoter | 18 | 78750380 | 79109391 | 359012 | 2 | ENSMUST00 | 0 | Setbp1 | ENSMUSG00000024548.10 |
| chr19 | 41481135 | 41484006 | 27.41053 | 90.23362 | 86.88379 | Promoter | 19 | 41482645 | 41562246 | 79602 | 1 | ENSMUST00 | 0 | Lcor | ENSMUSG00000025019.10 |
| chr19 | 46394581 | 46396186 | 27.41053 | 90.23362 | 86.88379 | Promoter | 19 | 46376814 | 46395735 | 18922 | 2 | ENSMUST00 | 0 | Actr1a | ENSMUSG00000025228.4 |
| chr19 | 53676291 | 53677707 | 27.41053 | 90.23362 | 86.88379 | Promoter | 19 | 53677306 | 53843763 | 166458 | 1 | ENSMUST00 | 0 | Rbm20 | ENSMUSG00000043639.9 |
| chr2 | 92458898 | 92460809 | 27.41053 | 90.23362 | 86.88379 | Promoter | 2 | 92452764 | 92459794 | 7031 | 2 | ENSMUST00 | 0 | Slc35c1 | ENSMUSG00000049922.8 |
| chr2 | 157077375 | 157079761 | 27.41053 | 90.23362 | 86.88379 | Promoter | 2 | 157015799 | 157079254 | 63456 | 2 | ENSMUST00 | 0 | Soga1 | ENSMUSG00000055485.6 |
| chr2 | 90916805 | 90918306 | 27.41053 | 90.23362 | 86.88379 | Promoter | 2 | 90908716 | 90917857 | 9142 | 2 | ENSMUST00 | 0 | Ptpmt1 | ENSMUSG00000063235.11 |
| chr2 | 18765448 | 18767736 | 27.41053 | 90.23362 | 86.88379 | Distal Interge | 2 | 18742715 | 18760862 | 18148 | 2 | ENSMUST00 | -4587 | Gm20539 | ENSMUSG00000092353.1 |
| chr3 | 131564762 | 131565597 | 27.41053 | 90.23362 | 86.88379 | Promoter | 3 | 131564768 | 131643670 | 78903 | 1 | ENSMUST00 | 0 | Papss1 | ENSMUSG00000028032.9 |
| chr3 | 144198116 | 144199381 | 27.41053 | 90.23362 | 86.88379 | Intron (ENSM | 3 | 144201623 | 144202300 | 678 | 2 | ENSMUST00 | 2919 | Lmo4 | ENSMUSG00000028266.12 |
| chr6 | 148830615 | 148831891 | 27.41053 | 90.23362 | 86.88379 | Promoter | 6 | 148770683 | 148831467 | 60785 | 2 | ENSMUST00 | 0 | Ipo8 | ENSMUSG00000040029.9 |
| chr7 | 43579211 | 43580522 | 27.41053 | 90.23362 | 86.88379 | Promoter | 7 | 43579610 | 43593235 | 13626 | 1 | ENSMUST00 | 0 | Zfp719 | ENSMUSG00000030469.9 |
| chr7 | 116503862 | 116505056 | 27.41053 | 90.23362 | 86.88379 | Promoter | 7 | 116504369 | 116540554 | 36186 | 1 | ENSMUST00 | 0 | Nucb2 | ENSMUSG00000030659.9 |
| chr8 | 70864890 | 70866158 | 27.41053 | 90.23362 | 86.88379 | 3' UTR | 8 | 70863127 | 70865428 | 2302 | 1 | ENSMUST00 | 1764 | A230052G05 | ENSMUSG00000020887.3 |
| chr8 | 105851349 | 105852120 | 27.41053 | 90.23362 | 86.88379 | Promoter | 8 | 105844678 | 105852008 | 7331 | 2 | ENSMUST00 | 0 | Cenpt | ENSMUSG00000036672.4 |
| chr8 | 26976009 | 26979489 | 27.41053 | 90.23362 | 86.88379 | Promoter | 8 | 26977336 | 26981461 | 4126 | 1 | ENSMUST00 | 0 | Zfp703 | ENSMUSG00000085795.2 |
| chr9 | 105394719 | 105396315 | 27.41053 | 90.23362 | 86.88379 | Promoter | 9 | 105395392 | 105405758 | 10367 | 1 | ENSMUST00 | 0 | Aste1 | ENSMUSG00000032567.9 |
| chr9 | 53535946 | 53537950 | 27.41053 | 90.23362 | 86.88379 | Promoter | 9 | 53537047 | 53574342 | 37296 | 1 | ENSMUST00 | 0 | Npat | ENSMUSG00000033054.7 |

| | | | | | | | | | | | | | | | |
|---|---|---|---|---|---|---|---|---|---|---|---|---|---|---|---|
| chr9 | 108049175 | 108050073 | 27.41053 | 90.23362 | 86.88379 | Promoter | 9 | 108049269 | 108049581 | 313 | 1 | ENSMUST00 | 0 | Gmppb | ENSMUSG00000070284.5 |
| chr1 | 9699653 | 9701408 | 25.84672 | 90.2271 | 86.88143 | Promoter | 1 | 9667415 | 9700209 | 32795 | 2 | ENSMUST00 | 0 | Mybl1 | ENSMUSG00000025912.11 |
| chr1 | 167349624 | 167350909 | 25.84672 | 90.2271 | 86.88143 | Promoter | 1 | 167349991 | 167368530 | 18540 | 1 | ENSMUST00 | 0 | Aldh9a1 | ENSMUSG00000026687.9 |
| chr10 | 80225979 | 80227278 | 25.84672 | 90.2271 | 86.88143 | Promoter | 10 | 80226434 | 80243903 | 17470 | 1 | ENSMUST00 | 0 | Mum1 | ENSMUSG00000020156.11 |
| chr11 | 51619706 | 51620498 | 25.84672 | 90.2271 | 86.88143 | Promoter | 11 | 51619735 | 51623714 | 3980 | 1 | ENSMUST00 | 0 | Nhp2 | ENSMUSG00000001056.3 |
| chr11 | 72206093 | 72208424 | 25.84672 | 90.2271 | 86.88143 | Promoter | 11 | 72207547 | 72210015 | 2469 | 1 | ENSMUST00 | 0 | Txndc17 | ENSMUSG00000020803.3 |
| chr11 | 121451320 | 121452786 | 25.84672 | 90.2271 | 86.88143 | Promoter | 11 | 121451949 | 121617164 | 165216 | 1 | ENSMUST00 | 0 | Tbcd | ENSMUSG00000039230.9 |
| chr12 | 103388611 | 103389464 | 25.84672 | 90.2271 | 86.88143 | Promoter | 12 | 103388682 | 103406339 | 17658 | 1 | ENSMUST00 | 0 | Otub2 | ENSMUSG00000021203.10 |
| chr13 | 21811753 | 21812466 | 25.84672 | 90.2271 | 86.88143 | Promoter | 13 | 21811746 | 21812147 | 402 | 1 | ENSMUST00 | 8 | Hist1h4m | ENSMUSG00000069306.5 |
| chr14 | 70074380 | 70079100 | 25.84672 | 90.2271 | 86.88143 | Promoter | 14 | 70077445 | 70080157 | 2713 | 1 | ENSMUST00 | 0 | Egr3 | ENSMUSG00000033730.3 |
| chr15 | 31600631 | 31602918 | 25.84672 | 90.2271 | 86.88143 | Promoter | 15 | 31590800 | 31601847 | 11048 | 2 | ENSMUST00 | 0 | Cct5 | ENSMUSG00000022234.9 |
| chr15 | 90223598 | 90225155 | 25.84672 | 90.2271 | 86.88143 | Promoter | 15 | 90224311 | 90230554 | 6244 | 1 | ENSMUST00 | 0 | Alg10b | ENSMUSG00000075470.1 |
| chr16 | 13109230 | 13110710 | 25.84672 | 90.2271 | 86.88143 | Promoter | 16 | 13109684 | 13148734 | 39051 | 1 | ENSMUST00 | 0 | Ercc4 | ENSMUSG00000022545.8 |
| chr16 | 58727491 | 58729008 | 25.84672 | 90.2271 | 86.88143 | Promoter | 16 | 58727910 | 58734251 | 6342 | 1 | ENSMUST00 | 0 | Cldn25 | ENSMUSG00000022744.7 |
| chr16 | 91729283 | 91730791 | 25.84672 | 90.2271 | 86.88143 | Promoter | 16 | 91729353 | 91871651 | 142299 | 1 | ENSMUST00 | 0 | Itsn1 | ENSMUSG00000022957.14 |
| chr17 | 28349620 | 28351838 | 25.84672 | 90.2271 | 86.88143 | Promoter | 17 | 28331671 | 28350346 | 18676 | 2 | ENSMUST00 | 0 | Tead3 | ENSMUSG00000002249.12 |
| chr17 | 29716438 | 29717471 | 25.84672 | 90.2271 | 86.88143 | Promoter | 17 | 29689378 | 29717009 | 27632 | 2 | ENSMUST00 | 0 | Ccdc167 | ENSMUSG00000024018.12 |
| chr17 | 83861359 | 83863132 | 25.84672 | 90.2271 | 86.88143 | Distal Interge | 17 | 83851675 | 83854393 | 2719 | 2 | ENSMUST00 | -6967 | C430042M11 | ENSMUSG00000097254.1 |
| chr18 | 76160877 | 76161480 | 25.84672 | 90.2271 | 86.88143 | Distal Interge | 18 | 76170552 | 76170652 | 101 | 1 | ENSMUST00 | -9072 | Mir6358 | ENSMUSG00000098276.1 |
| chr19 | 6974159 | 6976424 | 25.84672 | 90.2271 | 86.88143 | Promoter | 19 | 6975048 | 6977324 | 2277 | 1 | ENSMUST00 | 0 | Ppp1r14b | ENSMUSG00000056612.6 |
| chr2 | 15054450 | 15056235 | 25.84672 | 90.2271 | 86.88143 | Promoter | 2 | 15055282 | 15082456 | 27175 | 1 | ENSMUST00 | 0 | Arl5b | ENSMUSG00000017418.8 |
| chr3 | 88913350 | 88914844 | 25.84672 | 90.2271 | 86.88143 | Promoter | 3 | 88905107 | 88913931 | 8825 | 2 | ENSMUST00 | 0 | Msto1 | ENSMUSG00000068922.9 |
| chr4 | 135685906 | 135687177 | 25.84672 | 90.2271 | 86.88143 | Promoter | 4 | 135686287 | 135708181 | 21895 | 1 | ENSMUST00 | 0 | Ifnlr1 | ENSMUSG00000062157.6 |
| chr7 | 81932850 | 81934959 | 25.84672 | 90.2271 | 86.88143 | Promoter | 7 | 81881251 | 81934473 | 53223 | 2 | ENSMUST00 | 0 | Hdgfrp3 | ENSMUSG00000025104.8 |
| chr7 | 45458297 | 45460393 | 25.84672 | 90.2271 | 86.88143 | Promoter | 7 | 45457944 | 45459884 | 1941 | 2 | ENSMUST00 | 0 | Ftl1 | ENSMUSG00000050708.9 |
| chr8 | 13286832 | 13289049 | 25.84672 | 90.2271 | 86.88143 | Promoter | 8 | 13288013 | 13322924 | 34912 | 1 | ENSMUST00 | 0 | Tmco3 | ENSMUSG00000038497.6 |
| chr8 | 40423157 | 40424706 | 25.84672 | 90.2271 | 86.88143 | Promoter | 8 | 40423815 | 40485725 | 61911 | 1 | ENSMUST00 | 0 | Zdhhc2 | ENSMUSG00000039470.10 |
| chr9 | 107533968 | 107536510 | 25.84672 | 90.2271 | 86.88143 | Promoter | 9 | 107533945 | 107538656 | 4712 | 1 | ENSMUST00 | 24 | Tmem115 | ENSMUSG00000010045.1 |
| chr9 | 53609834 | 53610732 | 25.84672 | 90.2271 | 86.88143 | Promoter | 9 | 53580522 | 53610382 | 29861 | 2 | ENSMUST00 | 0 | Acat1 | ENSMUSG00000032047.5 |
| chr9 | 77750140 | 77751518 | 25.84672 | 90.2271 | 86.88143 | Distal Interge | 9 | 77754535 | 77794485 | 39951 | 1 | ENSMUST00 | -3017 | Gclc | ENSMUSG00000032350.8 |
| chr15 | 103239056 | 103241486 | 29.00577 | 90.16677 | 86.82411 | Promoter | 15 | 103191544 | 103239816 | 48273 | 2 | ENSMUST00 | 0 | Cbx5 | ENSMUSG00000009575.9 |
| chr4 | 99120070 | 99121403 | 30.11778 | 90.15907 | 86.81646 | Promoter | 4 | 98936665 | 99120915 | 184251 | 2 | ENSMUST00 | 0 | Dock7 | ENSMUSG00000028556.10 |
| chr5 | 148398764 | 148400373 | 30.66893 | 90.13895 | 86.79655 | Promoter | 5 | 148327410 | 148399904 | 72495 | 2 | ENSMUST00 | 0 | Slc7a1 | ENSMUSG00000041313.9 |
| chr3 | 104788434 | 104790298 | 29.54326 | 90.13423 | 86.79193 | Promoter | 3 | 104789011 | 104794459 | 5449 | 1 | ENSMUST00 | 0 | Rhoc | ENSMUSG00000002233.8 |
| chr7 | 131560290 | 131561157 | 29.54326 | 90.13423 | 86.79193 | Promoter | 7 | 131560391 | 131571898 | 11508 | 1 | ENSMUST00 | 0 | Bub3 | ENSMUSG00000066979.5 |
| chr1 | 181015848 | 181017585 | 28.9509 | 90.07051 | 86.72839 | Promoter | 1 | 180989554 | 181017569 | 28016 | 2 | ENSMUST00 | 0 | Ephx1 | ENSMUSG00000038776.8 |
| chr10 | 39898529 | 39899580 | 28.9509 | 90.07051 | 86.72839 | Promoter | 10 | 39890018 | 39899238 | 9221 | 2 | ENSMUST00 | 0 | AA474331 | ENSMUSG00000096687.2 |
| chr11 | 54860049 | 54861355 | 28.9509 | 90.07051 | 86.72839 | Promoter | 11 | 54826866 | 54860916 | 34051 | 2 | ENSMUST00 | 0 | Lyrm7 | ENSMUSG00000020268.8 |
| chr17 | 56989494 | 56991165 | 28.9509 | 90.07051 | 86.72839 | Promoter | 17 | 56990264 | 56996371 | 6108 | 1 | ENSMUST00 | 0 | Clpp | ENSMUSG00000002660.7 |
| chr3 | 105799997 | 105801764 | 28.9509 | 90.07051 | 86.72839 | Promoter | 3 | 105727267 | 105801330 | 74064 | 2 | ENSMUST00 | 0 | Rap1a | ENSMUSG00000068798.6 |
| chr4 | 34686600 | 34687906 | 28.9509 | 90.07051 | 86.72839 | Promoter | 4 | 34663257 | 34687438 | 24182 | 2 | ENSMUST00 | 0 | Slc35a1 | ENSMUSG00000028293.9 |
| chr6 | 125144855 | 125146037 | 28.9509 | 90.07051 | 86.72839 | Promoter | 6 | 125145241 | 125161782 | 16542 | 1 | ENSMUST00 | 0 | Iffo1 | ENSMUSG00000038271.12 |
| chr8 | 105700255 | 105702068 | 28.9509 | 90.07051 | 86.72839 | Promoter | 8 | 105701154 | 105703493 | 2340 | 1 | ENSMUST00 | 0 | Pard6a | ENSMUSG00000005699.10 |
| chr9 | 90162466 | 90164113 | 28.9509 | 90.07051 | 86.72839 | Promoter | 9 | 90163069 | 90200100 | 37032 | 1 | ENSMUST00 | 0 | Adamts7 | ENSMUSG00000032363.10 |
| chr1 | 133045387 | 133047073 | 30.05689 | 90.05383 | 86.71302 | Promoter | 1 | 133045667 | 133067229 | 21563 | 1 | ENSMUST00 | 0 | Pik3c2b | ENSMUSG00000026447.11 |
| chr15 | 76090740 | 76092009 | 30.05689 | 90.05383 | 86.71302 | Promoter | 15 | 76085595 | 76090013 | 4419 | 2 | ENSMUST00 | -728 | Nrbp2 | ENSMUSG00000075590.2 |
| chr6 | 72361642 | 72362476 | 30.05689 | 90.05383 | 86.71302 | Promoter | 6 | 72359714 | 72362393 | 2680 | 2 | ENSMUST00 | 0 | Rnf181 | ENSMUSG00000055850.7 |
| chr9 | 21759876 | 21761053 | 30.05689 | 90.05383 | 86.71302 | Promoter | 9 | 21755442 | 21760286 | 4845 | 2 | ENSMUST00 | 0 | Spc24 | ENSMUSG00000074476.5 |
| chr1 | 165633299 | 165635197 | 21.23866 | 90.05283 | 86.71227 | Promoter | 1 | 165592181 | 165634541 | 42361 | 2 | ENSMUST00 | 0 | Mpzl1 | ENSMUSG00000026566.10 |
| chr14 | 21075275 | 21076773 | 30.60485 | 90.02879 | 86.68851 | Intron (ENSM | 14 | 21052574 | 21448568 | 395995 | 1 | ENSMUST00 | 22702 | Adk | ENSMUSG00000039197.9 |
| chr17 | 69156516 | 69157768 | 30.60485 | 90.02879 | 86.68851 | Promoter | 17 | 69156810 | 69289985 | 133176 | 1 | ENSMUST00 | 0 | Epb4.1l3 | ENSMUSG00000024044.10 |
| chr19 | 23686825 | 23688717 | 25.17802 | 89.90526 | 86.56598 | Promoter | 19 | 23687400 | 23721129 | 33730 | 1 | ENSMUST00 | 0 | Ptar1 | ENSMUSG00000074925.3 |
| chr10 | 78425215 | 78426300 | 28.29403 | 89.8816 | 86.54237 | Promoter | 10 | 78425670 | 78427619 | 1950 | 1 | ENSMUST00 | 0 | Cstb | ENSMUSG00000005054.6 |
| chr15 | 89178489 | 89181434 | 28.29403 | 89.8816 | 86.54237 | Downstream | 15 | 89185465 | 89185537 | 73 | 2 | ENSMUST00 | 4103 | Mir6958 | ENSMUSG00000098663.1 |
| chr16 | 38741965 | 38742942 | 28.29403 | 89.8816 | 86.54237 | Promoter | 16 | 38742264 | 38769049 | 26786 | 1 | ENSMUST00 | 0 | B4galt4 | ENSMUSG00000022793.11 |
| chr7 | 143599137 | 143600712 | 28.29403 | 89.8816 | 86.54237 | Promoter | 7 | 143557230 | 143600090 | 42861 | 2 | ENSMUST00 | 0 | Cars | ENSMUSG00000010755.11 |
| chr8 | 91799641 | 91802556 | 28.29403 | 89.8816 | 86.54237 | Promoter | 8 | 91798525 | 91801915 | 3391 | 2 | ENSMUST00 | 0 | Irx3 | ENSMUSG00000031734.12 |
| chr8 | 85430895 | 85433408 | 28.29403 | 89.8816 | 86.54237 | Promoter | 8 | 85408759 | 85432841 | 24083 | 2 | ENSMUST00 | 0 | 4921524J17R | ENSMUSG00000036934.8 |
| chr11 | 52167719 | 52168683 | 29.93585 | 89.84451 | 86.50629 | Intron (ENSM | 11 | 52155378 | 52156283 | 906 | 2 | ENSMUST00 | -11437 | Olfr1372-ps1 | ENSMUSG00000084141.1 |
| chr11 | 94327395 | 94329063 | 29.37071 | 89.8338 | 86.49576 | Promoter | 11 | 94328001 | 94341841 | 13841 | 1 | ENSMUST00 | 0 | Ankrd40 | ENSMUSG00000020864.8 |
| chr2 | 125122790 | 125123998 | 29.37071 | 89.8338 | 86.49576 | Promoter | 2 | 125086271 | 125123595 | 37325 | 2 | ENSMUST00 | 0 | Myef2 | ENSMUSG00000027201.11 |
| chr5 | 23843490 | 23844969 | 30.4775 | 89.80972 | 86.47182 | Promoter | 5 | 23838944 | 23844161 | 5218 | 2 | ENSMUST00 | 0 | Tomm7 | ENSMUSG00000028998.6 |
| chr1 | 133609373 | 133610411 | 25.61729 | 89.79346 | 86.4557 | Promoter | 1 | 133603871 | 133610291 | 6421 | 2 | ENSMUST00 | 0 | Snrpe | ENSMUSG00000090553.3 |
| chr2 | 104122441 | 104123372 | 25.61729 | 89.79346 | 86.4557 | Promoter | 2 | 104122769 | 104124749 | 1981 | 1 | ENSMUST00 | 0 | A930018P22I | ENSMUSG00000032671.5 |
| chr5 | 104571294 | 104572374 | 28.78753 | 89.78359 | 86.4459 | Distal Interge | 5 | 104540571 | 104552074 | 11504 | 1 | ENSMUST00 | 30724 | D930016D06 | ENSMUSG00000097392.1 |
| chr17 | 25187913 | 25189669 | 20.34014 | 89.76241 | 86.42476 | Promoter | 17 | 25188397 | 25234443 | 46047 | 1 | ENSMUST00 | 0 | Unkl | ENSMUSG00000015127.9 |
| chr4 | 150007535 | 150009402 | 27.68336 | 89.75954 | 86.42194 | Promoter | 4 | 149979475 | 150009023 | 29549 | 2 | ENSMUST00 | 0 | H6pd | ENSMUSG00000028980.9 |
| chr3 | 152346111 | 152347383 | 29.31364 | 89.73435 | 86.39749 | Promoter | 3 | 152346478 | 152393612 | 47135 | 1 | ENSMUST00 | 0 | Usp33 | ENSMUSG00000025437.10 |
| chr5 | 24576525 | 24578236 | 30.41422 | 89.70082 | 86.36422 | Promoter | 5 | 24565341 | 24577467 | 12127 | 2 | ENSMUST00 | 0 | Abcf2 | ENSMUSG00000028953.9 |
| chr18 | 36319794 | 36320996 | 27.1142 | 89.69608 | 86.35963 | Distal Interge | 18 | 36345619 | 36366720 | 21102 | 1 | ENSMUST00 | -24623 | Cystm1 | ENSMUSG00000046727.7 |
| chr2 | 3713081 | 3714311 | 27.1142 | 89.69608 | 86.35963 | Promoter | 2 | 3713458 | 3782142 | 68685 | 1 | ENSMUST00 | 0 | Fam107b | ENSMUSG00000026655.9 |
| chr2 | 23067595 | 23069970 | 27.1142 | 89.69608 | 86.35963 | Promoter | 2 | 23068168 | 23114510 | 46343 | 1 | ENSMUST00 | 0 | Acbd5 | ENSMUSG00000026781.10 |
| chr6 | 91515061 | 91516824 | 27.1142 | 89.69608 | 86.35963 | Promoter | 6 | 91489305 | 91515884 | 26580 | 2 | ENSMUST00 | 0 | Xpc | ENSMUSG00000030094.7 |
| chr1 | 162739099 | 162741067 | 23.17719 | 89.67316 | 86.33752 | Promoter | 1 | 162670725 | 162740495 | 69771 | 2 | ENSMUST00 | 0 | Prrc2c | ENSMUSG00000040225.10 |
| chr1 | 79760642 | 79762273 | 23.17719 | 89.67316 | 86.33752 | Promoter | 1 | 79702262 | 79761769 | 59508 | 2 | ENSMUST00 | 0 | Wdfy1 | ENSMUSG00000073643.6 |
| chr10 | 81495430 | 81497133 | 23.17719 | 89.67316 | 86.33752 | Promoter | 10 | 81486249 | 81496392 | 10144 | 2 | ENSMUST00 | 0 | Ncln | ENSMUSG00000020238.9 |
| chr11 | 96034340 | 96035728 | 23.17719 | 89.67316 | 86.33752 | Promoter | 11 | 96034885 | 96047430 | 12546 | 1 | ENSMUST00 | 0 | Snf8 | ENSMUSG00000006058.5 |
| chr11 | 115381437 | 115382858 | 23.17719 | 89.67316 | 86.33752 | Promoter | 11 | 115381916 | 115396132 | 14217 | 1 | ENSMUST00 | 0 | Cdr2l | ENSMUSG00000050910.5 |
| chr11 | 101253109 | 101254459 | 23.17719 | 89.67316 | 86.33752 | Promoter | 11 | 101253707 | 101258049 | 4343 | 1 | ENSMUST00 | 0 | Vps25 | ENSMUSG00000078656.6 |
| chr12 | 51828430 | 51830566 | 23.17719 | 89.67316 | 86.33752 | Promoter | 12 | 51743722 | 51829536 | 85815 | 2 | ENSMUST00 | 0 | Hectd1 | ENSMUSG00000035247.10 |
| chr16 | 8736868 | 8739534 | 23.17719 | 89.67316 | 86.33752 | Promoter | 16 | 8689600 | 8738420 | 48821 | 2 | ENSMUST00 | 0 | Usp7 | ENSMUSG00000022710.12 |
| chr2 | 172439724 | 172441491 | 23.17719 | 89.67316 | 86.33752 | Promoter | 2 | 172440556 | 172469908 | 29353 | 1 | ENSMUST00 | 0 | Rtfdc1 | ENSMUSG00000027502.6 |
| chr4 | 126532569 | 126534044 | 23.17719 | 89.67316 | 86.33752 | Promoter | 4 | 126489541 | 126533472 | 43932 | 2 | ENSMUST00 | 0 | Ago4 | ENSMUSG00000042500.11 |
| chr4 | 108847424 | 108849115 | 23.17719 | 89.67316 | 86.33752 | Promoter | 4 | 108847785 | 108849413 | 1629 | 1 | ENSMUST00 | 0 | Kti12 | ENSMUSG00000073775.4 |
| chr5 | 130256958 | 130257794 | 23.17719 | 89.67316 | 86.33752 | Promoter | 5 | 130257035 | 130277749 | 20715 | 1 | ENSMUST00 | 0 | Tyw1 | ENSMUSG00000056310.9 |
| chr6 | 11906440 | 11908484 | 23.17719 | 89.67316 | 86.33752 | Promoter | 6 | 11907809 | 11933410 | 25602 | 1 | ENSMUST00 | 0 | Phf14 | ENSMUSG00000029629.11 |
| chr9 | 44417144 | 44418636 | 23.17719 | 89.67316 | 86.33752 | Promoter | 9 | 44410160 | 44418003 | 7844 | 2 | ENSMUST00 | 0 | Ccdc84 | ENSMUSG00000043923.7 |
| chr12 | 3806158 | 3809012 | 29.81577 | 89.63673 | 86.30187 | Promoter | 12 | 3806498 | 3867484 | 60987 | 1 | ENSMUST00 | 0 | Dnmt3a | ENSMUSG00000020661.10 |
| chr12 | 20614560 | 20615896 | 29.81577 | 89.63673 | 86.30187 | Distal Interge | 12 | 20804562 | 20804708 | 147 | 2 | ENSMUST00 | 188812 | RP23-152O2. | ENSMUSG00000101813.1 |
| chr13 | 51845444 | 51849264 | 29.81577 | 89.63673 | 86.30187 | Promoter | 13 | 51846678 | 51848468 | 1791 | 1 | ENSMUST00 | 0 | Gadd45g | ENSMUSG00000021453.2 |
| chr4 | 41274452 | 41275941 | 29.81577 | 89.63673 | 86.30187 | Promoter | 4 | 41194313 | 41275144 | 80832 | 2 | ENSMUST00 | 0 | Ubap2 | ENSMUSG00000028433.9 |
| chr5 | 93091910 | 93098273 | 29.81577 | 89.63673 | 86.30187 | Promoter | 5 | 93093457 | 93174958 | 81502 | 1 | ENSMUST00 | 0 | Sep-11 | ENSMUSG00000058013.8 |
| chr8 | 4492808 | 4494348 | 29.81577 | 89.63673 | 86.30187 | Promoter | 8 | 4493026 | 4518664 | 25639 | 1 | ENSMUST00 | 0 | Cers4 | ENSMUSG00000008206.10 |
| chr8 | 81015123 | 81015937 | 29.81577 | 89.63673 | 86.30187 | Promoter | 8 | 80980734 | 81014906 | 34173 | 2 | ENSMUST00 | -218 | Usp38 | ENSMUSG00000038250.8 |
| chr9 | 22002914 | 22003927 | 29.81577 | 89.63673 | 86.30187 | Promoter | 9 | 22003035 | 22014219 | 11185 | 1 | ENSMUST00 | 0 | Prkcsh | ENSMUSG00000003402.8 |
| chr9 | 43115803 | 43116749 | 29.81577 | 89.63673 | 86.30187 | Promoter | 9 | 43108653 | 43116570 | 7918 | 2 | ENSMUST00 | 0 | Tmem136 | ENSMUSG00000048503.6 |
| chr15 | 50888324 | 50890636 | 29.25679 | 89.63524 | 86.30125 | Promoter | 15 | 50654752 | 50889691 | 234940 | 2 | ENSMUST00 | 0 | Trps1 | ENSMUSG00000038679.11 |
| chr2 | 174109904 | 174111766 | 29.25679 | 89.63524 | 86.30125 | Promoter | 2 | 174110349 | 174123070 | 12722 | 1 | ENSMUST00 | 0 | Npepl1 | ENSMUSG00000039263.11 |
| chr2 | 23155399 | 23157490 | 28.67964 | 89.59386 | 86.26003 | Promoter | 2 | 23156369 | 23199260 | 42892 | 1 | ENSMUST00 | 0 | Yme1l1 | ENSMUSG00000026775.4 |
| chr5 | 146948130 | 146949643 | 30.35121 | 89.59235 | 86.25857 | Promoter | 5 | 146948657 | 146955614 | 6958 | 1 | ENSMUST00 | 0 | Gtf3a | ENSMUSG00000016503.11 |
| chr19 | 24555259 | 24557043 | 29.7561 | 89.53341 | 86.19995 | Promoter | 19 | 24294794 | 24555842 | 261049 | 2 | ENSMUST00 | 0 | Pip5k1b | ENSMUSG00000024867.9 |
| chr19 | 24030106 | 24031612 | 30.28845 | 89.48431 | 86.15106 | Promoter | 19 | 23972751 | 24031019 | 58269 | 2 | ENSMUST00 | 0 | Fam189a2 | ENSMUSG00000071604.4 |
| chr19 | 36409130 | 36410665 | 29.69666 | 89.43046 | 86.09753 | Promoter | 19 | 36409748 | 36456192 | 46445 | 1 | ENSMUST00 | 0 | Pcgf5 | ENSMUSG00000024805.10 |
| chr9 | 96362708 | 96364402 | 29.69666 | 89.43046 | 86.09753 | Promoter | 9 | 96332655 | 96364442 | 31788 | 2 | ENSMUST00 | 40 | Atp1b3 | ENSMUSG00000032412.8 |
| chr4 | 133887125 | 133888391 | 30.22595 | 89.37666 | 86.04396 | Promoter | 4 | 133847290 | 133887797 | 40508 | 2 | ENSMUST00 | 0 | Rps6ka1 | ENSMUSG00000003644.12 |
| chr10 | 85386715 | 85388887 | 29.08755 | 89.33995 | 86.00743 | Promoter | 10 | 85386814 | 85660292 | 273479 | 1 | ENSMUST00 | 0 | Btbd11 | ENSMUSG00000020042.10 |
| chr11 | 88862882 | 88865272 | 29.08755 | 89.33995 | 86.00743 | Promoter | 11 | 88830792 | 88864586 | 33795 | 2 | ENSMUST00 | 0 | Akap1 | ENSMUSG00000018428.10 |
| chr11 | 119041153 | 119041789 | 29.08755 | 89.33995 | 86.00743 | Promoter | 11 | 119036305 | 119040969 | 4665 | 2 | ENSMUST00 | -185 | Cbx8 | ENSMUSG00000025578.9 |
| chr17 | 48410026 | 48411454 | 29.08755 | 89.33995 | 86.00743 | Promoter | 17 | 48410080 | 48417266 | 7187 | 1 | ENSMUST00 | 0 | Oard1 | ENSMUSG00000040771.8 |
| chr18 | 37969012 | 37970288 | 29.08755 | 89.33995 | 86.00743 | Promoter | 18 | 37957434 | 37969774 | 12341 | 2 | ENSMUST00 | 0 | Fchsd1 | ENSMUSG00000038524.8 |
| chr2 | 165492333 | 165493875 | 29.08755 | 89.33995 | 86.00743 | Promoter | 2 | 165490110 | 165493324 | 3215 | 2 | ENSMUST00 | 0 | 2810408M09 | ENSMUSG00000039725.6 |
| chr5 | 137501484 | 137503362 | 29.08755 | 89.33995 | 86.00743 | Promoter | 5 | 137501438 | 137502425 | 988 | 2 | ENSMUST00 | 0 | Pop7 | ENSMUSG00000029715.5 |
| chr5 | 121618332 | 121619845 | 29.08755 | 89.33995 | 86.00743 | Promoter | 5 | 121598284 | 121618938 | 20655 | 2 | ENSMUST00 | 0 | Acad12 | ENSMUSG00000042647.8 |
| chr3 | 10366530 | 10367720 | 30.16371 | 89.26943 | 85.93817 | Promoter | 3 | 10366907 | 10391005 | 24099 | 1 | ENSMUST00 | 0 | Chmp4c | ENSMUSG00000027536.5 |
| chr8 | 124019985 | 124022062 | 29.03158 | 89.24218 | 85.91109 | Promoter | 8 | 124021508 | 124048504 | 26997 | 1 | ENSMUST00 | 0 | Urb2 | ENSMUSG00000031976.9 |
| chr3 | 129969532 | 129971010 | 29.5785 | 89.22566 | 85.89466 | Promoter | 3 | 129914960 | 129970206 | 55247 | 2 | ENSMUST00 | 0 | Ccdc109b | ENSMUSG00000027994.9 |
| chr7 | 35396032 | 35397680 | 29.5785 | 89.22566 | 85.89466 | Promoter | 7 | 35397035 | 35438687 | 41653 | 1 | ENSMUST00 | 0 | Cep89 | ENSMUSG00000023072.9 |
| chr7 | 126897697 | 126898685 | 29.5785 | 89.22566 | 85.89466 | Promoter | 7 | 126886220 | 126898253 | 12034 | 2 | ENSMUST00 | 0 | Tmem219 | ENSMUSG00000060538.9 |
| chr13 | 108315740 | 108316882 | 24.80701 | 89.18987 | 85.85909 | Promoter | 13 | 108316332 | 108389585 | 73254 | 1 | ENSMUST00 | 0 | Depdc1b | ENSMUSG00000021697.7 |
| chr4 | 41464243 | 41464954 | 30.10173 | 89.16261 | 85.83184 | Promoter | 4 | 41390745 | 41464840 | 74096 | 2 | ENSMUST00 | 0 | Kif24 | ENSMUSG00000028438.11 |
| chr16 | 36040667 | 36042231 | 28.41342 | 89.1249 | 85.7944 | Promoter | 16 | 36041190 | 36042973 | 1784 | 1 | ENSMUST00 | 0 | Wdr5b | ENSMUSG00000034379.8 |
| chr17 | 35673214 | 35673783 | 28.41342 | 89.1249 | 85.7944 | Promoter | 17 | 35667730 | 35673739 | 6010 | 2 | ENSMUST00 | 0 | Gtf2h4 | ENSMUSG00000001524.9 |
| chr3 | 108721002 | 108722934 | 28.41342 | 89.1249 | 85.7944 | Promoter | 3 | 108678638 | 108722309 | 43672 | 2 | ENSMUST00 | 0 | Gpsm2 | ENSMUSG00000027883.10 |
| chr3 | 88620710 | 88623767 | 28.41342 | 89.1249 | 85.7944 | Promoter | 3 | 88621106 | 88648052 | 26947 | 1 | ENSMUST00 | 0 | Arhgef2 | ENSMUSG00000028059.9 |
| chr4 | 129541578 | 129543395 | 28.41342 | 89.1249 | 85.7944 | Promoter | 4 | 129516104 | 129542713 | 26610 | 2 | ENSMUST00 | 0 | Hdac1 | ENSMUSG00000028800.10 |
| chr6 | 112695923 | 112697100 | 28.41342 | 89.1249 | 85.7944 | Promoter | 6 | 112619850 | 112696686 | 76837 | 2 | ENSMUST00 | 0 | Rad18 | ENSMUSG00000030254.11 |
| chr8 | 122376081 | 122377570 | 28.41342 | 89.1249 | 85.7944 | Promoter | 8 | 122376609 | 122417359 | 40751 | 1 | ENSMUST00 | 0 | Zc3h18 | ENSMUSG00000017478.10 |
| chr4 | 154960482 | 154961833 | 30.04 | 89.0562 | 85.72701 | Promoter | 4 | 154960923 | 154962371 | 1449 | 1 | ENSMUST00 | 0 | Hes5 | ENSMUSG00000048001.7 |

| | | | | | | | | | | | | | | | |
|---|---|---|---|---|---|---|---|---|---|---|---|---|---|---|---|
| chr10 | 82047825 | 82048909 | 28.92027 | 89.04764 | 85.7187 | Distal Interge | 10 | 82058417 | 82061329 | 2913 | 1 | ENSMUST00 | -9508 | Zfp873 | ENSMUSG00000061371.6 |
| chr1 | 65177109 | 65180019 | 28.91859 | 89.0447 | 85.71584 | Promoter | 1 | 65158616 | 65179181 | 20566 | 2 | ENSMUST00 | 0 | Idh1 | ENSMUSG00000025950.11 |
| chr1 | 169968711 | 169969974 | 28.91859 | 89.0447 | 85.71584 | Promoter | 1 | 169949535 | 169969241 | 19707 | 2 | ENSMUST00 | 0 | Hsd17b7 | ENSMUSG00000026675.7 |
| chr1 | 36546526 | 36547277 | 28.91859 | 89.0447 | 85.71584 | Promoter | 1 | 36538173 | 36547201 | 9029 | 2 | ENSMUST00 | 0 | Ankrd39 | ENSMUSG00000079610.3 |
| chr11 | 94495773 | 94500513 | 28.91859 | 89.0447 | 85.71584 | Promoter | 11 | 94489599 | 94499974 | 10376 | 2 | ENSMUST00 | 0 | Epn3 | ENSMUSG00000010080.10 |
| chr11 | 115627558 | 115628197 | 28.91859 | 89.0447 | 85.71584 | Promoter | 11 | 115614178 | 115628145 | 13968 | 2 | ENSMUST00 | 0 | Slc25a19 | ENSMUSG00000020744.8 |
| chr12 | 55154489 | 55156508 | 28.91859 | 89.0447 | 85.71584 | Promoter | 12 | 55155104 | 55189213 | 34110 | 1 | ENSMUST00 | 0 | Srp54b | ENSMUSG00000079108.5 |
| chr13 | 41015784 | 41016985 | 28.91859 | 89.0447 | 85.71584 | Promoter | 13 | 41016250 | 41022575 | 6326 | 1 | ENSMUST00 | 0 | Tmem14c | ENSMUSG00000021361.6 |
| chr17 | 56074251 | 56075442 | 28.91859 | 89.0447 | 85.71584 | Promoter | 17 | 56067045 | 56075028 | 7984 | 2 | ENSMUST00 | 0 | Ubxn6 | ENSMUSG00000019578.6 |
| chr17 | 88529505 | 88530968 | 28.91859 | 89.0447 | 85.71584 | Promoter | 17 | 88530118 | 88588367 | 58250 | 1 | ENSMUST00 | 0 | Ppp1r21 | ENSMUSG00000034709.8 |
| chr18 | 16806869 | 16809934 | 28.91859 | 89.0447 | 85.71584 | Promoter | 18 | 16588877 | 16809246 | 220370 | 2 | ENSMUST00 | 0 | Cdh2 | ENSMUSG00000024304.9 |
| chr2 | 130449526 | 130451893 | 28.91859 | 89.0447 | 85.71584 | Promoter | 2 | 130450278 | 130554026 | 103749 | 1 | ENSMUST00 | 0 | Ptpra | ENSMUSG00000027303.12 |
| chr2 | 181287155 | 181288638 | 28.91859 | 89.0447 | 85.71584 | Promoter | 2 | 181251449 | 181288035 | 36587 | 2 | ENSMUST00 | 0 | Gmeb2 | ENSMUSG00000038705.8 |
| chr2 | 64096333 | 64099107 | 28.91859 | 89.0447 | 85.71584 | Promoter | 2 | 63971507 | 64098012 | 126506 | 2 | ENSMUST00 | 0 | Fign | ENSMUSG00000075324.8 |
| chr4 | 116806883 | 116808307 | 28.91859 | 89.0447 | 85.71584 | Promoter | 4 | 116807723 | 116819431 | 11709 | 1 | ENSMUST00 | 0 | Mutyh | ENSMUSG00000028687.12 |
| chr4 | 119173076 | 119174292 | 28.91859 | 89.0447 | 85.71584 | Promoter | 4 | 119169513 | 119173856 | 4344 | 2 | ENSMUST00 | 0 | Zfp691 | ENSMUSG00000045268.8 |
| chr5 | 30949346 | 30950582 | 28.91859 | 89.0447 | 85.71584 | Promoter | 5 | 30950066 | 30955091 | 5026 | 1 | ENSMUST00 | 0 | Abhd1 | ENSMUSG00000006638.9 |
| chr5 | 104507986 | 104509227 | 28.91859 | 89.0447 | 85.71584 | Promoter | 5 | 104508352 | 104522383 | 14032 | 1 | ENSMUST00 | 0 | BC005561 | ENSMUSG00000079065.2 |
| chr5 | 99906235 | 99907018 | 28.91859 | 89.0447 | 85.71584 | Distal Interge | 5 | 99928283 | 99928381 | 99 | 2 | ENSMUST00 | 21363 | Gm22838 | ENSMUSG00000089062.1 |
| chr7 | 16119437 | 16120524 | 28.91859 | 89.0447 | 85.71584 | Promoter | 7 | 16119895 | 16121524 | 1630 | 1 | ENSMUST00 | 0 | Kptn | ENSMUSG00000006021.5 |
| chr7 | 83891929 | 83893016 | 28.91859 | 89.0447 | 85.71584 | Promoter | 7 | 83892000 | 83901532 | 9533 | 1 | ENSMUST00 | 0 | Mesdc2 | ENSMUSG00000038503.9 |
| chr8 | 70139326 | 70140529 | 28.91859 | 89.0447 | 85.71584 | Promoter | 8 | 70139499 | 70147256 | 7758 | 1 | ENSMUST00 | 0 | 2310045N01 | ENSMUSG00000002345.12 |
| chr9 | 59655687 | 59657702 | 28.91859 | 89.0447 | 85.71584 | Promoter | 9 | 59656368 | 59679375 | 23008 | 1 | ENSMUST00 | 0 | Pkm | ENSMUSG00000032294.11 |
| chr9 | 110333216 | 110334287 | 28.91859 | 89.0447 | 85.71584 | Promoter | 9 | 110333288 | 110373081 | 39794 | 1 | ENSMUST00 | 0 | Scap | ENSMUSG00000032485.9 |
| chr4 | 11253039 | 11254812 | 22.00414 | 89.03333 | 85.70822 | Promoter | 4 | 11199158 | 11254258 | 55101 | 2 | ENSMUST00 | 0 | Ints8 | ENSMUSG00000040738.5 |
| chr6 | 135011599 | 135012354 | 22.00414 | 89.03333 | 85.70822 | Promoter | 6 | 135011612 | 135023776 | 12165 | 1 | ENSMUST00 | 0 | Ddx47 | ENSMUSG00000030204.5 |
| chr1 | 186116654 | 186117652 | 24.24157 | 89.02564 | 85.70093 | Promoter | 1 | 186087731 | 186117310 | 29580 | 2 | ENSMUST00 | 0 | Lyplal1 | ENSMUSG00000039246.7 |
| chr10 | 115313463 | 115316288 | 24.24157 | 89.02564 | 85.70093 | Promoter | 10 | 115289863 | 115315591 | 25729 | 2 | ENSMUST00 | 0 | Rab21 | ENSMUSG00000020132.9 |
| chr11 | 97041499 | 97042462 | 24.24157 | 89.02564 | 85.70093 | Promoter | 11 | 97041560 | 97049213 | 7654 | 1 | ENSMUST00 | 0 | Mrpl10 | ENSMUSG00000001445.5 |
| chr11 | 117872882 | 117873723 | 24.24157 | 89.02564 | 85.70093 | Promoter | 11 | 117867952 | 117873459 | 5508 | 2 | ENSMUST00 | 0 | Tha1 | ENSMUSG00000017713.9 |
| chr11 | 78327871 | 78329155 | 24.24157 | 89.02564 | 85.70093 | Promoter | 11 | 78328415 | 78342782 | 14368 | 1 | ENSMUST00 | 0 | Pigs | ENSMUSG00000041958.10 |
| chr12 | 4768645 | 4769603 | 24.24157 | 89.02564 | 85.70093 | Promoter | 12 | 4769295 | 4778266 | 8972 | 1 | ENSMUST00 | 0 | Pfn4 | ENSMUSG00000020639.8 |
| chr13 | 23571319 | 23572523 | 24.24157 | 89.02564 | 85.70093 | Promoter | 13 | 23571408 | 23571884 | 477 | 1 | ENSMUST00 | 0 | Hist1h2bg | ENSMUSG00000058385.7 |
| chr13 | 46668898 | 46670479 | 24.24157 | 89.02564 | 85.70093 | Promoter | 13 | 46669522 | 46678056 | 8535 | 1 | ENSMUST00 | 0 | C78339 | ENSMUSG00000069237.8 |
| chr15 | 98566754 | 98568959 | 24.24157 | 89.02564 | 85.70093 | Promoter | 15 | 98543211 | 98567636 | 24426 | 2 | ENSMUST00 | 0 | Ccnt1 | ENSMUSG00000011960.7 |
| chr16 | 90726770 | 90728024 | 24.24157 | 89.02564 | 85.70093 | Promoter | 16 | 90719312 | 90727404 | 8093 | 2 | ENSMUST00 | 0 | Mis18a | ENSMUSG00000022978.11 |
| chr17 | 5941320 | 5942472 | 24.24157 | 89.02564 | 85.70093 | Promoter | 17 | 5941378 | 6039390 | 98013 | 1 | ENSMUST00 | 0 | Synj2 | ENSMUSG00000023805.11 |
| chr17 | 86962102 | 86964034 | 24.24157 | 89.02564 | 85.70093 | Promoter | 17 | 86963082 | 87000069 | 36988 | 1 | ENSMUST00 | 0 | Rhoq | ENSMUSG00000024143.9 |
| chr17 | 25984935 | 25986517 | 24.24157 | 89.02564 | 85.70093 | Promoter | 17 | 25959556 | 25985680 | 26125 | 2 | ENSMUST00 | 0 | Capn15 | ENSMUSG00000037326.9 |
| chr19 | 40512533 | 40514324 | 24.24157 | 89.02564 | 85.70093 | Distal Interge | 19 | 40553427 | 40554741 | 1315 | 2 | ENSMUST00 | 40417 | Aldh18a1 | ENSMUSG00000025007.8 |
| chr19 | 44145674 | 44146784 | 24.24157 | 89.02564 | 85.70093 | Promoter | 19 | 44139266 | 44146446 | 7181 | 2 | ENSMUST00 | 0 | Bloc1s2 | ENSMUSG00000057506.4 |
| chr2 | 101678047 | 101679533 | 24.24157 | 89.02564 | 85.70093 | Promoter | 2 | 101678429 | 101686235 | 7807 | 1 | ENSMUST00 | 0 | Traf6 | ENSMUSG00000027164.8 |
| chr2 | 164784984 | 164786753 | 24.24157 | 89.02564 | 85.70093 | Promoter | 2 | 164785823 | 164792487 | 6665 | 1 | ENSMUST00 | 0 | Snx21 | ENSMUSG00000050373.8 |
| chr3 | 135437447 | 135438173 | 24.24157 | 89.02564 | 85.70093 | Promoter | 3 | 135436226 | 135437759 | 1534 | 2 | ENSMUST00 | 0 | 4930539J05R | ENSMUSG00000097032.2 |
| chr4 | 31963245 | 31965103 | 24.24157 | 89.02564 | 85.70093 | Promoter | 4 | 31964097 | 32023467 | 59371 | 1 | ENSMUST00 | 0 | Map3k7 | ENSMUSG00000028284.8 |
| chr5 | 124115848 | 124116954 | 24.24157 | 89.02564 | 85.70093 | Promoter | 5 | 124116089 | 124117042 | 954 | 1 | ENSMUST00 | 0 | Arl6ip4 | ENSMUSG00000029404.10 |
| chr6 | 18512890 | 18515216 | 24.24157 | 89.02564 | 85.70093 | Promoter | 6 | 18366547 | 18514869 | 148323 | 2 | ENSMUST00 | 0 | Cttnbp2 | ENSMUSG00000000416.11 |
| chr6 | 113530947 | 113532074 | 24.24157 | 89.02564 | 85.70093 | Promoter | 6 | 113514874 | 113531638 | 16765 | 2 | ENSMUST00 | 0 | Emc3 | ENSMUSG00000030286.5 |
| chr6 | 85137245 | 85138120 | 24.24157 | 89.02564 | 85.70093 | Promoter | 6 | 85133678 | 85137766 | 4089 | 2 | ENSMUST00 | 0 | Spr | ENSMUSG00000033735.7 |
| chr9 | 61370729 | 61374567 | 24.24157 | 89.02564 | 85.70093 | Promoter | 9 | 61372366 | 61418497 | 46132 | 1 | ENSMUST00 | 0 | Tle3 | ENSMUSG00000032280.11 |
| chr13 | 103333218 | 103335263 | 27.78755 | 88.98067 | 85.65775 | Promoter | 13 | 102732486 | 103334192 | 601707 | 2 | ENSMUST00 | 0 | Mast4 | ENSMUSG00000034751.10 |
| chr13 | 47014163 | 47015244 | 27.78755 | 88.98067 | 85.65775 | Promoter | 13 | 47012558 | 47014850 | 2293 | 2 | ENSMUST00 | 0 | Nhlrc1 | ENSMUSG00000044231.2 |
| chr15 | 88818558 | 88819435 | 27.78755 | 88.98067 | 85.65775 | Promoter | 15 | 88805244 | 88819318 | 14075 | 2 | ENSMUST00 | 0 | Alg12 | ENSMUSG00000035845.12 |
| chr17 | 56039798 | 56040882 | 27.78755 | 88.98067 | 85.65775 | Promoter | 17 | 56040416 | 56068026 | 27611 | 1 | ENSMUST00 | 0 | Chaf1a | ENSMUSG00000002835.8 |
| chr2 | 69789115 | 69790449 | 27.78755 | 88.98067 | 85.65775 | Promoter | 2 | 69789623 | 69795653 | 6031 | 1 | ENSMUST00 | 0 | Phospho2 | ENSMUSG00000027088.5 |
| chr4 | 40267762 | 40270252 | 27.78755 | 88.98067 | 85.65775 | Promoter | 4 | 40269569 | 40270940 | 1372 | 1 | ENSMUST00 | 0 | Toporsos | ENSMUSG00000028407.3 |
| chr5 | 96209312 | 96210945 | 27.78755 | 88.98067 | 85.65775 | Promoter | 5 | 96209493 | 96266727 | 57235 | 1 | ENSMUST00 | 0 | Mrpl1 | ENSMUSG00000029486.8 |
| chr7 | 140115662 | 140117006 | 27.78755 | 88.98067 | 85.65775 | Promoter | 7 | 140105710 | 140116476 | 10767 | 2 | ENSMUST00 | 0 | Echs1 | ENSMUSG00000025465.7 |
| chr7 | 137410007 | 137411686 | 27.78755 | 88.98067 | 85.65775 | Promoter | 7 | 137375574 | 137410756 | 35183 | 2 | ENSMUST00 | 0 | 9430038I01R | ENSMUSG00000040139.9 |
| chr7 | 19003795 | 19005127 | 27.78755 | 88.98067 | 85.65775 | Promoter | 7 | 19004065 | 19006761 | 2697 | 1 | ENSMUST00 | 0 | Irf2bp1 | ENSMUSG00000044030.3 |
| chr14 | 11160851 | 11162642 | 29.97852 | 88.95019 | 85.62817 | Promoter | 14 | 9550100 | 11162032 | 1611933 | 2 | ENSMUST00 | 0 | Fhit | ENSMUSG00000060579.7 |
| chr14 | 56571341 | 56572315 | 29.97852 | 88.95019 | 85.62817 | Promoter | 14 | 56526761 | 56571846 | 45086 | 2 | ENSMUST00 | 0 | Cenpj | ENSMUSG00000064128.6 |
| chr3 | 93520371 | 93523048 | 29.97852 | 88.95019 | 85.62817 | Promoter | 3 | 93520488 | 93526287 | 5800 | 1 | ENSMUST00 | 0 | S100a11 | ENSMUSG00000027907.4 |
| chr7 | 25266992 | 25267991 | 29.97852 | 88.95019 | 85.62817 | Promoter | 7 | 25267704 | 25294159 | 26456 | 1 | ENSMUST00 | 0 | Cic | ENSMUSG00000005442.8 |
| chr9 | 32539982 | 32542054 | 29.97852 | 88.95019 | 85.62817 | Promoter | 9 | 32422204 | 32541595 | 119392 | 2 | ENSMUST00 | 0 | Fli1 | ENSMUSG00000016087.8 |
| chr2 | 167061714 | 167063859 | 29.403 | 88.9212 | 85.59992 | Promoter | 2 | 167035793 | 167063015 | 27223 | 2 | ENSMUST00 | 0 | Znfx1 | ENSMUSG00000039501.9 |
| chr11 | 97765791 | 97767137 | 27.20494 | 88.89825 | 85.57716 | Promoter | 11 | 97745615 | 97766556 | 20942 | 2 | ENSMUST00 | 0 | Cwc25 | ENSMUSG00000018541.5 |
| chr15 | 76537923 | 76539844 | 27.20494 | 88.89825 | 85.57716 | Promoter | 15 | 76535730 | 76538746 | 3017 | 2 | ENSMUST00 | 0 | Fbxl6 | ENSMUSG00000022559.7 |
| chr5 | 149635405 | 149637058 | 27.20494 | 88.89825 | 85.57716 | Promoter | 5 | 149614287 | 149636171 | 21885 | 2 | ENSMUST00 | 0 | Hsph1 | ENSMUSG00000029657.10 |
| chr18 | 75005427 | 75006690 | 26.15274 | 88.89023 | 85.56958 | Promoter | 18 | 75005900 | 75009933 | 4034 | 1 | ENSMUST00 | 0 | BC031181 | ENSMUSG00000036299.4 |
| chr1 | 74712846 | 74714683 | 26.66126 | 88.87019 | 85.5497 | Promoter | 1 | 74713574 | 74737892 | 24319 | 1 | ENSMUST00 | 0 | Cyp27a1 | ENSMUSG00000026170.6 |
| chr2 | 161109244 | 161110158 | 26.66126 | 88.87019 | 85.5497 | Promoter | 2 | 161008034 | 161109075 | 101042 | 2 | ENSMUST00 | -170 | Chd6 | ENSMUSG00000057133.9 |
| chr8 | 104250540 | 104251779 | 26.66126 | 88.87019 | 85.5497 | Promoter | 8 | 104250903 | 104263443 | 12541 | 1 | ENSMUST00 | 0 | Cklf | ENSMUSG00000054400.10 |
| chr9 | 4376477 | 4377172 | 26.66126 | 88.87019 | 85.5497 | Distal Interge | 9 | 4371269 | 4371335 | 67 | 2 | ENSMUST00 | -5143 | Gm23992 | ENSMUSG00000094086.1 |
| chr6 | 24526536 | 24528331 | 28.80981 | 88.85442 | 85.53428 | Promoter | 6 | 24518666 | 24527570 | 8905 | 2 | ENSMUST00 | 0 | Ndufa5 | ENSMUSG00000023089.7 |
| chr6 | 118065871 | 118067211 | 28.80981 | 88.85442 | 85.53428 | Promoter | 6 | 118066385 | 118091546 | 25162 | 1 | ENSMUST00 | 0 | Rasgef1a | ENSMUSG00000030134.9 |
| chr9 | 15709114 | 15710996 | 29.34497 | 88.82044 | 85.50058 | Promoter | 9 | 15709738 | 15738579 | 28842 | 1 | ENSMUST00 | 0 | Slc36a4 | ENSMUSG00000043885.8 |
| chr2 | 25271712 | 25273274 | 28.20397 | 88.75508 | 85.43546 | Promoter | 2 | 25272478 | 25285915 | 13438 | 1 | ENSMUST00 | 0 | Anapc2 | ENSMUSG00000026965.7 |
| chr4 | 43030580 | 43031848 | 29.85632 | 88.73936 | 85.41974 | Promoter | 4 | 43027690 | 43031710 | 4021 | 2 | ENSMUST00 | 0 | Stoml2 | ENSMUSG00000028455.10 |
| chr9 | 49796389 | 49799150 | 29.85632 | 88.73936 | 85.41974 | Promoter | 9 | 49504361 | 49798729 | 294369 | 2 | ENSMUST00 | 0 | Ncam1 | ENSMUSG00000039542.10 |
| chr1 | 17096332 | 17098208 | 29.28716 | 88.72003 | 85.40064 | Promoter | 1 | 16994938 | 17097889 | 102952 | 2 | ENSMUST00 | 0 | Jph1 | ENSMUSG00000042686.5 |
| chr5 | 30921470 | 30922282 | 29.79559 | 88.63454 | 85.31541 | Promoter | 5 | 30921556 | 30931246 | 9691 | 1 | ENSMUST00 | 0 | Khk | ENSMUSG00000029162.10 |
| chr7 | 27178237 | 27179485 | 29.79559 | 88.63454 | 85.31541 | Promoter | 7 | 27168424 | 27178896 | 10473 | 2 | ENSMUST00 | 0 | Rab4b | ENSMUSG00000053291.10 |
| chr1 | 82289450 | 82292285 | 25.50663 | 88.63404 | 85.31516 | Promoter | 1 | 82233101 | 82291416 | 58316 | 2 | ENSMUST00 | 0 | Irs1 | ENSMUSG00000055980.2 |
| chr10 | 128589190 | 128590713 | 25.50663 | 88.63404 | 85.31516 | Promoter | 10 | 128567523 | 128589652 | 22130 | 2 | ENSMUST00 | 0 | Erbb3 | ENSMUSG00000018166.8 |
| chr10 | 43220403 | 43222284 | 25.50663 | 88.63404 | 85.31516 | Promoter | 10 | 43221486 | 43405395 | 183910 | 1 | ENSMUST00 | 0 | Pdss2 | ENSMUSG00000038240.8 |
| chr10 | 29360820 | 29362611 | 25.50663 | 88.63404 | 85.31516 | Promoter | 10 | 29344178 | 29362442 | 18265 | 2 | ENSMUST00 | 0 | Rnf146 | ENSMUSG00000038876.6 |
| chr10 | 22148088 | 22150989 | 25.50663 | 88.63404 | 85.31516 | Promoter | 10 | 22113050 | 22148819 | 35770 | 2 | ENSMUST00 | 0 | E030030I06R | ENSMUSG00000097327.2 |
| chr11 | 50429896 | 50431640 | 25.50663 | 88.63404 | 85.31516 | Promoter | 11 | 50389286 | 50431125 | 41840 | 2 | ENSMUST00 | 0 | Rufy1 | ENSMUSG00000020375.4 |
| chr11 | 68691089 | 68693279 | 25.50663 | 88.63404 | 85.31516 | Promoter | 11 | 68691559 | 68699259 | 7701 | 1 | ENSMUST00 | 0 | Myh10 | ENSMUSG00000020900.10 |
| chr11 | 120086093 | 120086892 | 25.50663 | 88.63404 | 85.31516 | Promoter | 11 | 120064430 | 120086827 | 22398 | 2 | ENSMUST00 | 0 | Cep131 | ENSMUSG00000039781.9 |
| chr12 | 116404853 | 116405992 | 25.50663 | 88.63404 | 85.31516 | Promoter | 12 | 116405402 | 116463531 | 58130 | 1 | ENSMUST00 | 0 | Ncapg2 | ENSMUSG00000042029.6 |
| chr13 | 100107426 | 100108677 | 25.50663 | 88.63404 | 85.31516 | Promoter | 13 | 100107997 | 100114233 | 6237 | 1 | ENSMUST00 | 0 | Serf1 | ENSMUSG00000021643.9 |
| chr14 | 29978251 | 29979281 | 25.50663 | 88.63404 | 85.31516 | Promoter | 14 | 29978337 | 29993221 | 14885 | 1 | ENSMUST00 | 0 | Actr8 | ENSMUSG00000015971.4 |
| chr14 | 66866432 | 66869441 | 25.50663 | 88.63404 | 85.31516 | Promoter | 14 | 66802864 | 66868688 | 65825 | 2 | ENSMUST00 | 0 | Dpysl2 | ENSMUSG00000022048.8 |
| chr14 | 24486268 | 24487147 | 25.50663 | 88.63404 | 85.31516 | Promoter | 14 | 24448694 | 24487046 | 38353 | 2 | ENSMUST00 | 0 | Polr3a | ENSMUSG00000025280.7 |
| chr16 | 3908245 | 3909661 | 25.50663 | 88.63404 | 85.31516 | Promoter | 16 | 3895179 | 3908653 | 13475 | 2 | ENSMUST00 | 0 | 1700037C18I | ENSMUSG00000005983.10 |
| chr17 | 5840733 | 5842592 | 25.50663 | 88.63404 | 85.31516 | Promoter | 17 | 5841328 | 5931033 | 89706 | 1 | ENSMUST00 | 0 | Snx9 | ENSMUSG00000002365.9 |
| chr19 | 42128190 | 42129561 | 25.50663 | 88.63404 | 85.31516 | Promoter | 19 | 42123273 | 42129059 | 5787 | 2 | ENSMUST00 | 0 | Avpi1 | ENSMUSG00000018821.3 |
| chr19 | 57452694 | 57453540 | 25.50663 | 88.63404 | 85.31516 | Promoter | 19 | 57452906 | 57488910 | 36005 | 1 | ENSMUST00 | 0 | Trub1 | ENSMUSG00000025086.7 |
| chr2 | 26121900 | 26123466 | 25.50663 | 88.63404 | 85.31516 | Promoter | 2 | 26055535 | 26122818 | 67284 | 2 | ENSMUST00 | 0 | Nacc2 | ENSMUSG00000026932.9 |
| chr2 | 35108876 | 35110289 | 25.50663 | 88.63404 | 85.31516 | Promoter | 2 | 35109492 | 35130713 | 21222 | 1 | ENSMUST00 | 0 | Cntrl | ENSMUSG00000057110.10 |
| chr5 | 146844279 | 146848368 | 25.50663 | 88.63404 | 85.31516 | Promoter | 5 | 146845071 | 146847726 | 2656 | 1 | ENSMUST00 | 0 | Rasl11a | ENSMUSG00000029641.7 |
| chr6 | 134919565 | 134922632 | 25.50663 | 88.63404 | 85.31516 | Promoter | 6 | 134920401 | 134925513 | 5113 | 1 | ENSMUST00 | 0 | Cdkn1b | ENSMUSG00000003031.9 |
| chr6 | 99027636 | 99029292 | 25.50663 | 88.63404 | 85.31516 | Promoter | 6 | 98929466 | 99028258 | 98793 | 2 | ENSMUST00 | 0 | Foxp1 | ENSMUSG00000030067.12 |
| chr6 | 128200042 | 128201537 | 25.50663 | 88.63404 | 85.31516 | Promoter | 6 | 128187846 | 128200623 | 12778 | 2 | ENSMUST00 | 0 | Gm10010 | ENSMUSG00000056771.6 |
| chr7 | 101663367 | 101664391 | 25.50663 | 88.63404 | 85.31516 | Promoter | 7 | 101663633 | 101722922 | 59290 | 1 | ENSMUST00 | 0 | Clpb | ENSMUSG00000001829.11 |
| chr7 | 45332526 | 45333989 | 25.50663 | 88.63404 | 85.31516 | Promoter | 7 | 45303155 | 45333780 | 30626 | 2 | ENSMUST00 | 0 | Trpm4 | ENSMUSG00000038260.9 |
| chr9 | 87254716 | 87256055 | 25.50663 | 88.63404 | 85.31516 | Promoter | 9 | 87189577 | 87255536 | 65960 | 2 | ENSMUST00 | 0 | Cep162 | ENSMUSG00000056919.9 |
| chr1 | 164306496 | 164308372 | 29.22959 | 88.61999 | 85.30408 | Promoter | 1 | 164289800 | 164307484 | 17685 | 2 | ENSMUST00 | 0 | Blzf1 | ENSMUSG00000026577.8 |
| chr11 | 119490697 | 119492351 | 29.22959 | 88.61999 | 85.30408 | Promoter | 11 | 119491347 | 119511423 | 20077 | 1 | ENSMUST00 | 0 | Endov | ENSMUSG00000039850.11 |
| chr14 | 67714976 | 67715747 | 29.22958 | 88.61999 | 85.30408 | Promoter | 14 | 67676355 | 67715610 | 39256 | 2 | ENSMUST00 | 0 | Cdca2 | ENSMUSG00000048922.13 |
| chr16 | 35489892 | 35491182 | 29.22959 | 88.61999 | 85.30408 | Promoter | 16 | 35397313 | 35490873 | 93561 | 2 | ENSMUST00 | 0 | Pdia5 | ENSMUSG00000022844.7 |
| chr17 | 80944259 | 80945584 | 29.22958 | 88.61999 | 85.30408 | Promoter | 17 | 80944632 | 81001816 | 57185 | 1 | ENSMUST00 | 0 | Tmem178 | ENSMUSG00000024245.4 |
| chr19 | 8888290 | 8889526 | 29.22959 | 88.61999 | 85.30408 | Promoter | 19 | 8888538 | 8888685 | 148 | 2 | ENSMUST00 | 0 | Gm25822 | ENSMUSG00000070167.1 |
| chr3 | 127552453 | 127553971 | 29.22959 | 88.61999 | 85.30408 | Promoter | 3 | 127536954 | 127553348 | 16395 | 2 | ENSMUST00 | 0 | Larp7 | ENSMUSG00000027968.7 |
| chr4 | 62407602 | 62409705 | 29.22959 | 88.61999 | 85.30408 | Promoter | 4 | 62408797 | 62426990 | 18194 | 1 | ENSMUST00 | 0 | Prpf4 | ENSMUSG00000066148.3 |
| chr6 | 32586391 | 32588544 | 29.22959 | 88.61999 | 85.30408 | Promoter | 6 | 32144268 | 32588192 | 443925 | 2 | ENSMUST00 | 0 | Plxna4 | ENSMUSG00000029765.11 |
| chr7 | 30308171 | 30309572 | 29.22959 | 88.61999 | 85.30408 | Promoter | 7 | 30308717 | 30314304 | 5588 | 1 | ENSMUST00 | 0 | Alkbh6 | ENSMUSG00000042831.8 |
| chr7 | 143829592 | 143830480 | 29.22959 | 88.61999 | 85.30408 | Promoter | 7 | 143829626 | 143848410 | 18785 | 1 | ENSMUST00 | 0 | Dhcr7 | ENSMUSG00000058454.9 |
| chr5 | 135063491 | 135065117 | 20.9917 | 88.60235 | 85.28741 | Promoter | 5 | 135052957 | 135064666 | 11710 | 2 | ENSMUST00 | 0 | Wbscr22 | ENSMUSG00000005378.12 |
| chr5 | 105875939 | 105878057 | 20.9917 | 88.60235 | 85.28741 | Promoter | 5 | 105876565 | 105914780 | 38216 | 1 | ENSMUST00 | 0 | Zfp326 | ENSMUSG00000029290.7 |
| chr1 | 80338787 | 80341787 | 27.03504 | 88.5911 | 85.27629 | Promoter | 1 | 80264923 | 80340480 | 75558 | 2 | ENSMUST00 | 0 | Cul3 | ENSMUSG00000004364.9 |
| chr1 | 92180566 | 92181428 | 27.03504 | 88.5911 | 85.27629 | Promoter | 1 | 91928779 | 92180341 | 251563 | 2 | ENSMUST00 | -226 | Hdac4 | ENSMUSG00000026313.10 |
| chr10 | 121585897 | 121587605 | 27.03504 | 88.5911 | 85.27629 | Promoter | 10 | 121546455 | 121586794 | 40340 | 2 | ENSMUST00 | 0 | Tbk1 | ENSMUSG00000020115.2 |
| chr10 | 4432512 | 4433545 | 27.03504 | 88.5911 | 85.27629 | Promoter | 10 | 4432513 | 4455140 | 22628 | 1 | ENSMUST00 | 0 | 1700052N19 | ENSMUSG00000061759.10 |
| chr11 | 35769023 | 35770839 | 27.03504 | 88.5911 | 85.27629 | Promoter | 11 | 35769484 | 35791285 | 21802 | 1 | ENSMUST00 | 0 | Pank3 | ENSMUSG00000018846.8 |
| chr11 | 80141468 | 80142824 | 27.03504 | 88.5911 | 85.27629 | Promoter | 11 | 80136678 | 80142175 | 5498 | 2 | ENSMUST00 | 0 | Tefm | ENSMUSG00000046909.9 |

| | | | | | | | | | | | | | | | |
|---|---|---|---|---|---|---|---|---|---|---|---|---|---|---|---|
| chr12 | 8770910 | 8773220 | 27.03504 | 88.5911 | 85.27629 | Promoter | 12 | 8771323 | 8793715 | 22393 | 1 | ENSMUST00 | 0 | Sdc1 | ENSMUSG00000020592.9 |
| chr13 | 35905489 | 35906429 | 27.03504 | 88.5911 | 85.27629 | Promoter | 13 | 35893472 | 35906359 | 12888 | 2 | ENSMUST00 | 0 | Rpp40 | ENSMUSG00000021418.10 |
| chr14 | 52013893 | 52015600 | 30.32912 | 88.589 | 85.27629 | 5' UTR | 14 | 52016865 | 52019787 | 2923 | 1 | ENSMUST00 | -1265 | Tmem253 | ENSMUSG00000072571.2 |
| chr14 | 100284160 | 100286947 | 30.32912 | 88.589 | 85.27629 | Distal Interge | 14 | 100219486 | 100219789 | 304 | 1 | ENSMUST00 | 64675 | Gm16260 | ENSMUSG00000084991.1 |
| chr15 | 44747875 | 44748912 | 30.32912 | 88.589 | 85.27629 | 5' UTR | 15 | 44671859 | 44752458 | 80600 | 2 | ENSMUST00 | 3546 | Sybu | ENSMUSG00000022340.9 |
| chr15 | 73644244 | 73646291 | 30.32912 | 88.589 | 85.27629 | Promoter | 15 | 73605593 | 73645762 | 40170 | 2 | ENSMUST00 | 0 | Slc45a4 | ENSMUSG00000079020.4 |
| chr15 | 79027416 | 79030756 | 27.03504 | 88.5911 | 85.27629 | Promoter | 15 | 79028212 | 79030498 | 2287 | 1 | ENSMUST00 | 0 | H1f0 | ENSMUSG00000096210.1 |
| chr17 | 46545596 | 46546716 | 27.03504 | 88.5911 | 85.27629 | Promoter | 17 | 46500605 | 46546388 | 45784 | 2 | ENSMUST00 | 0 | Cul9 | ENSMUSG00000040327.11 |
| chr19 | 5609269 | 5610577 | 27.03504 | 88.5911 | 85.27629 | Promoter | 19 | 5603017 | 5610094 | 7078 | 2 | ENSMUST00 | 0 | Kat5 | ENSMUSG00000024926.9 |
| chr19 | 4792775 | 4793921 | 27.03504 | 88.5911 | 85.27629 | Promoter | 19 | 4784293 | 4793901 | 9609 | 2 | ENSMUST00 | 0 | Rbm4 | ENSMUSG00000094936.2 |
| chr2 | 106002542 | 106003829 | 27.03504 | 88.5911 | 85.27629 | Promoter | 2 | 105966709 | 106003549 | 36841 | 2 | ENSMUST00 | 0 | Dnajc24 | ENSMUSG00000027166.9 |
| chr2 | 152029391 | 152030080 | 27.03504 | 88.5911 | 85.27629 | Distal Interge | 2 | 151999897 | 152009246 | 9350 | 1 | ENSMUST00 | 29495 | Slc52a3 | ENSMUSG00000027463.9 |
| chr2 | 125781288 | 125783313 | 27.03504 | 88.5911 | 85.27629 | Promoter | 2 | 125736986 | 125782870 | 45885 | 2 | ENSMUST00 | 0 | Secisbp2l | ENSMUSG00000035093.11 |
| chr2 | 163658124 | 163659535 | 30.32912 | 88.589 | 85.27629 | Promoter | 2 | 163658386 | 163723758 | 65373 | 1 | ENSMUST00 | 0 | Pkig | ENSMUSG00000035268.9 |
| chr2 | 18055597 | 18058275 | 30.32912 | 88.589 | 85.27629 | Promoter | 2 | 18054193 | 18056613 | 2421 | 2 | ENSMUST00 | 0 | 1810059C17 | ENSMUSG00000086340.1 |
| chr3 | 104967665 | 104968520 | 27.03504 | 88.5911 | 85.27629 | Distal Interge | 3 | 104944805 | 104961709 | 16905 | 2 | ENSMUST00 | -5957 | Wnt2b | ENSMUSG00000027840.4 |
| chr3 | 89391380 | 89396724 | 30.32912 | 88.589 | 85.27629 | Promoter | 3 | 89377644 | 89392069 | 14426 | 2 | ENSMUST00 | 0 | Zbtb7b | ENSMUSG00000028042.10 |
| chr3 | 97226662 | 97228545 | 27.03504 | 88.5911 | 85.27629 | Promoter | 3 | 97203662 | 97227364 | 23703 | 2 | ENSMUST00 | 0 | Bcl9 | ENSMUSG00000038256.10 |
| chr4 | 6452287 | 6454754 | 30.32912 | 88.589 | 85.27629 | Promoter | 4 | 6396207 | 6454271 | 58065 | 2 | ENSMUST00 | 0 | Nsmaf | ENSMUSG00000028245.10 |
| chr4 | 123717632 | 123718719 | 27.03504 | 88.5911 | 85.27629 | Promoter | 4 | 123712710 | 123718186 | 5477 | 2 | ENSMUST00 | 0 | Ndufs5 | ENSMUSG00000028648.8 |
| chr4 | 155086550 | 155087285 | 27.03504 | 88.5911 | 85.27629 | Promoter | 4 | 155086577 | 155112181 | 25605 | 1 | ENSMUST00 | 0 | Morn1 | ENSMUSG00000029049.9 |
| chr5 | 137530397 | 137533556 | 27.03504 | 88.5911 | 85.27629 | Promoter | 5 | 137483020 | 137533242 | 50223 | 2 | ENSMUST00 | 0 | Epo | ENSMUSG00000029711.8 |
| chr5 | 138154912 | 138155829 | 27.03504 | 88.5911 | 85.27629 | Promoter | 5 | 138139702 | 138155744 | 16043 | 2 | ENSMUST00 | 0 | Zfp113 | ENSMUSG00000037007.12 |
| chr5 | 144964984 | 144966586 | 30.32912 | 88.589 | 85.27629 | Promoter | 5 | 144876495 | 144965839 | 89345 | 2 | ENSMUST00 | 0 | Smurf1 | ENSMUSG00000038780.9 |
| chr5 | 21700358 | 21702125 | 27.03504 | 88.5911 | 85.27629 | Promoter | 5 | 21662901 | 21701394 | 38494 | 2 | ENSMUST00 | 0 | Napepld | ENSMUSG00000044968.11 |
| chr5 | 143730802 | 143733135 | 30.32912 | 88.589 | 85.27629 | Promoter | 5 | 143710325 | 143732280 | 21956 | 2 | ENSMUST00 | 0 | Usp42 | ENSMUSG00000051306.7 |
| chr6 | 34877381 | 34878524 | 27.03504 | 88.5911 | 85.27629 | Promoter | 6 | 34871771 | 34878021 | 6251 | 2 | ENSMUST00 | 0 | 3110062M04 | ENSMUSG00000046806.8 |
| chr7 | 81212149 | 81214474 | 27.03504 | 88.5911 | 85.27629 | Promoter | 7 | 81213596 | 81334533 | 120938 | 1 | ENSMUST00 | 0 | Pde8a | ENSMUSG00000025584.12 |
| chr7 | 92740196 | 92743123 | 27.03504 | 88.5911 | 85.27629 | Promoter | 7 | 92741603 | 92837117 | 95515 | 1 | ENSMUST00 | 0 | Rab30 | ENSMUSG00000030643.8 |
| chr7 | 97399686 | 97400928 | 27.03504 | 88.5911 | 85.27629 | Promoter | 7 | 97400003 | 97407802 | 7800 | 1 | ENSMUST00 | 0 | Ndufc2 | ENSMUSG00000030647.8 |
| chr7 | 119793387 | 119794738 | 30.32912 | 88.589 | 85.27629 | Promoter | 7 | 119794006 | 119848943 | 54938 | 1 | ENSMUST00 | 0 | 2610020H08 | ENSMUSG00000030924.10 |
| chr7 | 7171264 | 7172088 | 27.03504 | 88.5911 | 85.27629 | Promoter | 7 | 7171330 | 7183562 | 12233 | 1 | ENSMUST00 | 0 | Zfp418 | ENSMUSG00000034538.7 |
| chr7 | 109959166 | 109960911 | 27.03504 | 88.5911 | 85.27629 | Promoter | 7 | 109893780 | 109960400 | 66621 | 2 | ENSMUST00 | 0 | Dennd5a | ENSMUSG00000035901.9 |
| chr7 | 30534310 | 30535717 | 30.32912 | 88.589 | 85.27629 | Promoter | 7 | 30522226 | 30534180 | 11955 | 2 | ENSMUST00 | -131 | Arhgap33 | ENSMUSG00000036882.6 |
| chr7 | 19572424 | 19574065 | 30.32912 | 88.589 | 85.27629 | Promoter | 7 | 19564946 | 19573823 | 8878 | 2 | ENSMUST00 | 0 | Gemin7 | ENSMUSG00000044709.5 |
| chr7 | 4740091 | 4741544 | 30.32912 | 88.589 | 85.27629 | Promoter | 7 | 4740115 | 4747514 | 7400 | 1 | ENSMUST00 | 0 | Suv420h2 | ENSMUSG00000059851.10 |
| chr8 | 105254023 | 105255259 | 27.03504 | 88.5911 | 85.27629 | Promoter | 8 | 105252638 | 105255153 | 2516 | 2 | ENSMUST00 | 0 | B3gnt9 | ENSMUSG00000069920.5 |
| chr9 | 21546884 | 21547906 | 27.03504 | 88.5911 | 85.27629 | Promoter | 9 | 21546894 | 21589467 | 42574 | 1 | ENSMUST00 | 0 | Carm1 | ENSMUSG00000032185.9 |
| chr9 | 57552087 | 57552876 | 30.32912 | 88.589 | 85.27629 | Promoter | 9 | 57544256 | 57552763 | 8508 | 2 | ENSMUST00 | 0 | Mpi | ENSMUSG00000032306.9 |
| chr9 | 107984961 | 107986547 | 27.03504 | 88.5911 | 85.27629 | Promoter | 9 | 107984227 | 107985879 | 1653 | 2 | ENSMUST00 | 0 | Fam212a | ENSMUSG00000042106.4 |
| chr11 | 86483912 | 86485744 | 28.64569 | 88.56696 | 85.25784 | Promoter | 11 | 86484657 | 86499025 | 14369 | 1 | ENSMUST00 | 0 | Rnft1 | ENSMUSG00000020521.7 |
| chr11 | 96788747 | 96790143 | 29.73511 | 88.5301 | 85.22105 | Promoter | 11 | 96789127 | 96803969 | 14843 | 1 | ENSMUST00 | 0 | Cbx1 | ENSMUSG00000018666.8 |
| chr17 | 47694413 | 47695873 | 29.73511 | 88.5301 | 85.22105 | Promoter | 17 | 47695198 | 47704286 | 9089 | 1 | ENSMUST00 | 0 | Frs3 | ENSMUSG00000023266.6 |
| chr11 | 106065447 | 106066855 | 22.1708 | 88.48181 | 85.17323 | Promoter | 11 | 106066061 | 106073612 | 7552 | 1 | ENSMUST00 | 0 | Taco1 | ENSMUSG00000001983.7 |
| chr3 | 20034336 | 20035567 | 30.21119 | 88.38634 | 85.07815 | Promoter | 3 | 19995945 | 20035315 | 39371 | 2 | ENSMUST00 | 0 | Hps3 | ENSMUSG00000027615.9 |
| chr14 | 55478576 | 55479508 | 28.53732 | 88.37692 | 85.06884 | Promoter | 14 | 55478758 | 55490338 | 11581 | 1 | ENSMUST00 | 0 | Dhrs4 | ENSMUSG00000022210.6 |
| chr4 | 116052857 | 116054293 | 28.53732 | 88.37692 | 85.06884 | Promoter | 4 | 116032842 | 116053876 | 21035 | 2 | ENSMUST00 | 0 | Nsun4 | ENSMUSG00000028706.9 |
| chr5 | 76139804 | 76141310 | 28.53732 | 88.37692 | 85.06884 | Promoter | 5 | 76140271 | 76151515 | 11245 | 1 | ENSMUST00 | 0 | Srd5a3 | ENSMUSG00000029233.10 |
| chr7 | 41598721 | 41599941 | 28.53732 | 88.37692 | 85.06884 | Promoter | 7 | 41599261 | 41628533 | 29273 | 1 | ENSMUST00 | 0 | 2610021A01 | ENSMUSG00000091474.1 |
| chr9 | 108596863 | 108598133 | 28.53732 | 88.37692 | 85.06884 | Promoter | 9 | 108578826 | 108597600 | 18775 | 2 | ENSMUST00 | 0 | P4htm | ENSMUSG00000006675.5 |
| chr1 | 66862165 | 66864406 | 20.109 | 88.33971 | 85.03291 | Promoter | 1 | 66830839 | 66863277 | 32439 | 2 | ENSMUST00 | 0 | Acadl | ENSMUSG00000026003.5 |
| chr3 | 121953085 | 121955451 | 20.109 | 88.33971 | 85.03291 | Promoter | 3 | 121953229 | 122016677 | 63449 | 1 | ENSMUST00 | 0 | Arhgap29 | ENSMUSG00000039831.11 |
| chr4 | 155085258 | 155086435 | 20.109 | 88.33971 | 85.03291 | Promoter | 4 | 155074110 | 155086371 | 12262 | 2 | ENSMUST00 | 0 | Rer1 | ENSMUSG00000029048.3 |
| chr9 | 103363917 | 103365715 | 20.109 | 88.33971 | 85.03291 | Promoter | 9 | 103353104 | 103364897 | 11794 | 2 | ENSMUST00 | 0 | Cdv3 | ENSMUSG00000032803.9 |
| chr1 | 106712428 | 106714820 | 29.05819 | 88.32191 | 85.01541 | Promoter | 1 | 106538178 | 106714274 | 176097 | 2 | ENSMUST00 | 0 | Bcl2 | ENSMUSG00000057329.7 |
| chr11 | 120098903 | 120099754 | 29.05819 | 88.32191 | 85.01541 | Promoter | 11 | 120098928 | 120100705 | 1778 | 1 | ENSMUST00 | 0 | 1810043H04 | ENSMUSG00000078572.3 |
| chr14 | 120275080 | 120277905 | 29.55512 | 88.21911 | 84.9134 | Promoter | 14 | 120275669 | 120431697 | 156029 | 1 | ENSMUST00 | 0 | Mbnl2 | ENSMUSG00000022139.11 |
| chr1 | 171044018 | 171045451 | 27.89552 | 88.209 | 84.90351 | Distal Interge | 1 | 171051174 | 171058054 | 6881 | 2 | ENSMUST00 | 12603 | Fcgr3 | ENSMUSG00000059498.8 |
| chr10 | 86685002 | 86686127 | 27.89552 | 88.209 | 84.90351 | Promoter | 10 | 86685601 | 86694806 | 9206 | 1 | ENSMUST00 | 0 | 1810014B01 | ENSMUSG00000097412.1 |
| chr11 | 67965353 | 67967226 | 27.89552 | 88.209 | 84.90351 | Promoter | 11 | 67966193 | 68207148 | 240956 | 1 | ENSMUST00 | 0 | Stx8 | ENSMUSG00000020903.8 |
| chr16 | 18498196 | 18499519 | 27.89552 | 88.209 | 84.90351 | Promoter | 16 | 18498768 | 18566679 | 67912 | 1 | ENSMUST00 | 0 | Gnb1l | ENSMUSG00000000884.11 |
| chr17 | 34897900 | 34900008 | 27.89552 | 88.209 | 84.90351 | Promoter | 17 | 34898469 | 34914051 | 15583 | 1 | ENSMUST00 | 0 | Ehmt2 | ENSMUSG00000013787.10 |
| chr19 | 8785821 | 8786418 | 27.89552 | 88.209 | 84.90351 | Promoter | 19 | 8774354 | 8786316 | 11963 | 2 | ENSMUST00 | 0 | Taf6l | ENSMUSG00000003680.10 |
| chr3 | 95159474 | 95162067 | 27.89552 | 88.209 | 84.90351 | Promoter | 3 | 95160457 | 95164542 | 4086 | 1 | ENSMUST00 | 0 | Sema6c | ENSMUSG00000038777.14 |
| chr5 | 110779699 | 110781025 | 27.89552 | 88.209 | 84.90351 | Promoter | 5 | 110773667 | 110779967 | 6301 | 2 | ENSMUST00 | 0 | Pus1 | ENSMUSG00000029507.11 |
| chr11 | 5837050 | 5838383 | 25.28022 | 88.20625 | 84.90166 | Promoter | 11 | 5827860 | 5837789 | 9930 | 2 | ENSMUST00 | 0 | Polm | ENSMUSG00000020474.6 |
| chr1 | 175691698 | 175692850 | 22.89105 | 88.16154 | 84.85722 | Promoter | 1 | 175662421 | 175692776 | 30356 | 2 | ENSMUST00 | 0 | Opn3 | ENSMUSG00000026525.9 |
| chr13 | 55329368 | 55330650 | 22.89105 | 88.16154 | 84.85722 | Promoter | 13 | 55325168 | 55329758 | 4591 | 2 | ENSMUST00 | 0 | Mxd3 | ENSMUSG00000021485.8 |
| chr13 | 67608978 | 67610109 | 22.89105 | 88.16154 | 84.85722 | Promoter | 13 | 67589443 | 67609707 | 20265 | 2 | ENSMUST00 | 0 | AA987161 | ENSMUSG00000058093.7 |
| chr17 | 17063537 | 17064886 | 22.89105 | 88.16154 | 84.85722 | Promoter | 17 | 17064113 | 17089628 | 25516 | 1 | ENSMUST00 | 0 | Zfp960 | ENSMUSG00000096696.1 |
| chr18 | 9448898 | 9450908 | 22.89105 | 88.16154 | 84.85722 | Promoter | 18 | 9314044 | 9450150 | 136107 | 2 | ENSMUST00 | 0 | Ccny | ENSMUSG00000024286.7 |
| chr18 | 84684779 | 84686215 | 22.89105 | 88.16154 | 84.85722 | Promoter | 18 | 84667470 | 84685633 | 18164 | 2 | ENSMUST00 | 0 | Cndp2 | ENSMUSG00000024644.11 |
| chr2 | 119034515 | 119035578 | 22.89105 | 88.16154 | 84.85722 | Promoter | 2 | 119034790 | 119039769 | 4980 | 1 | ENSMUST00 | 0 | Rpusd2 | ENSMUSG00000027324.1 |
| chr7 | 114562339 | 114563407 | 22.89105 | 88.16154 | 84.85722 | Promoter | 7 | 114550166 | 114562972 | 12807 | 2 | ENSMUST00 | 0 | Cyp2r1 | ENSMUSG00000030670.10 |
| chr7 | 127232200 | 127233944 | 22.89105 | 88.16154 | 84.85722 | Promoter | 7 | 127233061 | 127238179 | 5119 | 1 | ENSMUST00 | 0 | Zfp553 | ENSMUSG00000045598.9 |
| chr7 | 139247202 | 139249854 | 22.89105 | 88.16154 | 84.85722 | Promoter | 7 | 139248482 | 139267253 | 18772 | 1 | ENSMUST00 | 0 | Pwwp2b | ENSMUSG00000060260.7 |
| chr14 | 20478679 | 20480317 | 25.73672 | 88.11626 | 84.8129 | Promoter | 14 | 20455261 | 20480133 | 24873 | 2 | ENSMUST00 | 0 | Anxa7 | ENSMUSG00000021814.11 |
| chr6 | 72615470 | 72617444 | 25.73672 | 88.11626 | 84.8129 | Promoter | 6 | 72608432 | 72617000 | 8569 | 2 | ENSMUST00 | 0 | Tgoln1 | ENSMUSG00000056429.4 |
| chr10 | 117745534 | 117746867 | 27.29887 | 88.10642 | 84.80337 | Promoter | 10 | 117733679 | 117746358 | 12680 | 2 | ENSMUST00 | 0 | Slc35e3 | ENSMUSG00000060181.5 |
| chr7 | 80294374 | 80294938 | 27.29887 | 88.10642 | 84.80337 | Promoter | 7 | 80294450 | 80316259 | 21810 | 1 | ENSMUST00 | 0 | Prc1 | ENSMUSG00000038943.11 |
| chr8 | 92355053 | 92358743 | 27.29887 | 88.10642 | 84.80337 | Promoter | 8 | 92326038 | 92355831 | 29794 | 2 | ENSMUST00 | 0 | Crnde | ENSMUSG00000031736.5 |
| chr11 | 96943379 | 96944651 | 30.02262 | 88.06211 | 84.75982 | Promoter | 11 | 96937825 | 96943779 | 5955 | 2 | ENSMUST00 | 0 | Pnpo | ENSMUSG00000018659.7 |
| chr16 | 23986923 | 23989929 | 30.02262 | 88.06211 | 84.75982 | Promoter | 16 | 23965052 | 23988852 | 23801 | 2 | ENSMUST00 | 0 | Bcl6 | ENSMUSG00000022508.5 |
| chr2 | 119288276 | 119289498 | 30.02262 | 88.06211 | 84.75982 | Promoter | 2 | 119288740 | 119295487 | 6748 | 1 | ENSMUST00 | 0 | Vps18 | ENSMUSG00000034216.7 |
| chr12 | 21416080 | 21418715 | 26.74277 | 88.06109 | 84.75893 | Promoter | 12 | 21390071 | 21417637 | 27567 | 2 | ENSMUST00 | 0 | Ywhaq | ENSMUSG00000076432.7 |
| chr2 | 140394864 | 140396429 | 26.74277 | 88.06109 | 84.75893 | Promoter | 2 | 140395309 | 142392966 | 1997658 | 1 | ENSMUST00 | 0 | Macrod2 | ENSMUSG00000068205.9 |
| chr3 | 40893351 | 40895033 | 26.74277 | 88.06109 | 84.75893 | Promoter | 3 | 40894145 | 40938138 | 43994 | 1 | ENSMUST00 | 0 | 3110057O12 | ENSMUSG00000037818.11 |
| chr17 | 37269977 | 37271191 | 29.43634 | 88.01369 | 84.71214 | Promoter | 17 | 37270220 | 37274484 | 4265 | 1 | ENSMUST00 | 0 | H2-M3 | ENSMUSG00000016206.6 |
| chr1 | 195241487 | 195242029 | 6.08306 | 87.96767 | 84.66632 | Promoter | 1 | 195240910 | 195241007 | 98 | 2 | ENSMUST00 | -481 | Gm27940 | ENSMUSG00000099208.1 |
| chr8 | 79293953 | 79295699 | 28.83278 | 87.92931 | 84.62811 | Promoter | 8 | 79266424 | 79294956 | 28533 | 2 | ENSMUST00 | 0 | Mmaa | ENSMUSG00000037022.2 |
| chr4 | 134702765 | 134704907 | 29.37731 | 87.91154 | 84.61048 | Promoter | 4 | 134561690 | 134704290 | 142601 | 2 | ENSMUST00 | 0 | Man1c1 | ENSMUSG00000037306.8 |
| chr19 | 43675078 | 43676446 | 28.77697 | 87.83199 | 84.53201 | Promoter | 19 | 43663822 | 43674881 | 11060 | 2 | ENSMUST00 | -198 | Slc25a28 | ENSMUSG00000040414.7 |
| chr8 | 71487578 | 71489429 | 28.77697 | 87.83199 | 84.53201 | Promoter | 8 | 71488103 | 71493400 | 5298 | 1 | ENSMUST00 | 0 | Gtpbp3 | ENSMUSG00000007610.10 |
| chr14 | 56667534 | 56669410 | 29.31851 | 87.80975 | 84.5099 | Promoter | 14 | 56668248 | 56697430 | 29183 | 1 | ENSMUST00 | 0 | Mphosph8 | ENSMUSG00000079184.5 |
| chr11 | 34832903 | 34834594 | 29.83638 | 87.74163 | 84.44196 | Promoter | 11 | 34809190 | 34833641 | 24452 | 2 | ENSMUST00 | 0 | Spdl1 | ENSMUSG00000069910.2 |
| chr11 | 62538296 | 62539629 | 29.25995 | 87.70835 | 84.40896 | Promoter | 11 | 62524946 | 62539261 | 14316 | 2 | ENSMUST00 | 0 | Cenpv | ENSMUSG00000018509.3 |
| chr6 | 124755579 | 124757546 | 29.25995 | 87.70835 | 84.40896 | Promoter | 6 | 124739189 | 124756021 | 16833 | 2 | ENSMUST00 | 0 | Atn1 | ENSMUSG00000004263.10 |
| chr1 | 33669291 | 33670761 | 28.666 | 87.63837 | 84.3392 | Promoter | 1 | 33453810 | 33669795 | 215986 | 2 | ENSMUST00 | 0 | Prim2 | ENSMUSG00000026134.11 |
| chr13 | 34344051 | 34345646 | 28.666 | 87.63837 | 84.3392 | Promoter | 13 | 34179158 | 34345182 | 166025 | 2 | ENSMUST00 | 0 | Slc22a23 | ENSMUSG00000038267.9 |
| chr2 | 3283218 | 3283980 | 28.666 | 87.63837 | 84.3392 | Promoter | 2 | 3284212 | 3328877 | 44666 | 1 | ENSMUST00 | -232 | Nmt2 | ENSMUSG00000026643.11 |
| chr5 | 138255010 | 138256431 | 28.666 | 87.63837 | 84.3392 | Promoter | 5 | 138255608 | 138259395 | 3788 | 1 | ENSMUST00 | 0 | Lamtor4 | ENSMUSG00000050552.7 |
| chr5 | 110978341 | 110980520 | 28.666 | 87.63837 | 84.3392 | Intron (ENSM | 5 | 111004144 | 111004252 | 109 | 1 | ENSMUST00 | -23624 | Mir701 | ENSMUSG00000076163.1 |
| chr7 | 141391636 | 141393012 | 28.666 | 87.63837 | 84.3392 | Promoter | 7 | 141392199 | 141402968 | 10770 | 1 | ENSMUST00 | 0 | Taldo1 | ENSMUSG00000025503.4 |
| chr7 | 38227135 | 38228402 | 28.666 | 87.63837 | 84.3392 | Promoter | 7 | 38220654 | 38227994 | 7341 | 2 | ENSMUST00 | 0 | Plekhf1 | ENSMUSG00000074170.4 |
| chr8 | 70775837 | 70778039 | 28.666 | 87.63837 | 84.3392 | Promoter | 8 | 70768176 | 70776713 | 8538 | 2 | ENSMUST00 | 0 | Pik3r2 | ENSMUSG00000031834.10 |
| chr11 | 98863105 | 98864606 | 29.20162 | 87.6073 | 84.30928 | Promoter | 11 | 98863638 | 98892299 | 28662 | 1 | ENSMUST00 | 0 | Wipf2 | ENSMUSG00000038013.9 |
| chr13 | 23683198 | 23683993 | 29.20162 | 87.6073 | 84.30928 | Promoter | 13 | 23683449 | 23683924 | 476 | 1 | ENSMUST00 | 0 | Hist1h2ac | ENSMUSG00000069270.5 |
| chr5 | 8621993 | 8623333 | 29.20162 | 87.6073 | 84.30928 | Promoter | 5 | 8490334 | 8622952 | 132619 | 2 | ENSMUST00 | 0 | Rundc3b | ENSMUSG00000040570.9 |
| chr1 | 139421471 | 139423147 | 21.73903 | 87.55687 | 84.25908 | Promoter | 1 | 139422383 | 139453005 | 30623 | 1 | ENSMUST00 | 0 | Zbtb41 | ENSMUSG00000033964.7 |
| chr14 | 30479073 | 30480785 | 21.73903 | 87.55687 | 84.25908 | Promoter | 14 | 30479565 | 30527056 | 47492 | 1 | ENSMUST00 | 0 | Dcp1a | ENSMUSG00000021962.8 |
| chr18 | 75818942 | 75821711 | 21.73903 | 87.55687 | 84.25908 | Promoter | 18 | 75820178 | 76148561 | 328384 | 1 | ENSMUST00 | 0 | Zbtb7c | ENSMUSG00000044646.9 |
| chr2 | 75831720 | 75833180 | 21.73903 | 87.55687 | 84.25908 | Promoter | 2 | 75832177 | 75931350 | 99174 | 1 | ENSMUST00 | 0 | Agps | ENSMUSG00000042410.10 |
| chr5 | 77308798 | 77311242 | 21.73903 | 87.55687 | 84.25908 | Promoter | 5 | 77310147 | 77349324 | 39178 | 1 | ENSMUST00 | 0 | Polr2b | ENSMUSG00000029250.5 |
| chr15 | 58213813 | 58215815 | 29.14352 | 87.50663 | 84.2095 | Promoter | 15 | 58175879 | 58214892 | 39014 | 2 | ENSMUST00 | 0 | Fbxo32 | ENSMUSG00000022358.6 |
| chr9 | 63756312 | 63758698 | 29.14352 | 87.50663 | 84.2095 | Promoter | 9 | 63646767 | 63757994 | 111228 | 2 | ENSMUST00 | 0 | Smad3 | ENSMUSG00000032402.7 |
| chr1 | 24004866 | 24005978 | 23.93078 | 87.47927 | 84.18233 | Promoter | 1 | 23995939 | 24005656 | 9718 | 2 | ENSMUST00 | 0 | 1110058L19 | ENSMUSG00000026154.4 |
| chr10 | 128743100 | 128744571 | 23.93078 | 87.47927 | 84.18233 | Promoter | 10 | 128720141 | 128744056 | 23916 | 2 | ENSMUST00 | 0 | Dgka | ENSMUSG00000025357.7 |
| chr11 | 60351504 | 60353003 | 23.93078 | 87.47927 | 84.18233 | Promoter | 11 | 60226714 | 60352905 | 126192 | 2 | ENSMUST00 | 0 | Tom1l2 | ENSMUSG00000000538.13 |
| chr13 | 91460982 | 91462809 | 23.93078 | 87.47927 | 84.18233 | Promoter | 13 | 91461050 | 91703429 | 242380 | 1 | ENSMUST00 | 0 | Ssbp2 | ENSMUSG00000003992.9 |
| chr13 | 54503289 | 54504756 | 23.93078 | 87.47927 | 84.18233 | Promoter | 13 | 54503779 | 54551290 | 47512 | 1 | ENSMUST00 | 0 | Simc1 | ENSMUSG00000043183.13 |
| chr14 | 63859568 | 63860954 | 23.93078 | 87.47927 | 84.18233 | Promoter | 14 | 63860312 | 63919859 | 59548 | 1 | ENSMUST00 | 0 | Pinx1 | ENSMUSG00000021958.4 |
| chr16 | 3846647 | 3847711 | 23.93078 | 87.47927 | 84.18233 | Promoter | 16 | 3847268 | 3858880 | 11613 | 1 | ENSMUST00 | 0 | Zfp174 | ENSMUSG00000054939.6 |
| chr18 | 31788480 | 31790095 | 23.93078 | 87.47927 | 84.18233 | Promoter | 18 | 31789159 | 31796642 | 7484 | 1 | ENSMUST00 | 0 | Polr2d | ENSMUSG00000024258.3 |
| chr18 | 24653102 | 24654660 | 23.93078 | 87.47927 | 84.18233 | Promoter | 18 | 24653691 | 24701556 | 47866 | 1 | ENSMUST00 | 0 | Mocos | ENSMUSG00000039616.9 |
| chr2 | 130273567 | 130275245 | 23.93078 | 87.47927 | 84.18233 | Promoter | 2 | 130274430 | 130279313 | 4884 | 1 | ENSMUST00 | 0 | Nop56 | ENSMUSG00000027405.11 |
| chr2 | 129129012 | 129130508 | 23.93078 | 87.47927 | 84.18233 | Promoter | 2 | 129129700 | 129134134 | 4435 | 1 | ENSMUST00 | 0 | Chchd5 | ENSMUSG00000037938.4 |
| chr4 | 41754961 | 41756541 | 23.93078 | 87.47927 | 84.18233 | Promoter | 4 | 41755228 | 41758695 | 3468 | 1 | ENSMUST00 | 0 | Galt | ENSMUSG00000036073.12 |
| chr7 | 36695587 | 36698932 | 23.93078 | 87.47927 | 84.18233 | Promoter | 7 | 36698118 | 36773553 | 75436 | 1 | ENSMUST00 | 0 | Tshz3 | ENSMUSG00000021217.7 |

| | | | | | | | | | | | | | | | |
|---|---|---|---|---|---|---|---|---|---|---|---|---|---|---|---|
| chr7 | 13009256 | 13010609 | 23.93078 | 87.47927 | 84.18233 | Promoter | 7 | 13004936 | 13009822 | 4887 | 2 | ENSMUST00 | 0 | Zbtb45 | ENSMUSG00000049600.7 |
| chr8 | 123805532 | 123806981 | 23.93078 | 87.47927 | 84.18233 | Promoter | 8 | 123805985 | 123835287 | 29303 | 1 | ENSMUST00 | 0 | Rab4a | ENSMUSG00000019478.10 |
| chr9 | 61913515 | 61915287 | 23.93078 | 87.47927 | 84.18233 | Promoter | 9 | 61913284 | 61914538 | 1255 | 2 | ENSMUST00 | 0 | Rplp1 | ENSMUSG00000007892.7 |
| chr9 | 119321480 | 119323666 | 23.93078 | 87.47927 | 84.18233 | Promoter | 9 | 119238432 | 119322427 | 83996 | 2 | ENSMUST00 | 0 | Oxsr1 | ENSMUSG00000036737.7 |
| chr9 | 110653365 | 110654625 | 23.93078 | 87.47927 | 84.18233 | Promoter | 9 | 110624789 | 110654161 | 29373 | 2 | ENSMUST00 | 0 | Nbeal2 | ENSMUSG00000056724.9 |
| chr16 | 72662877 | 72665011 | 28.00751 | 87.44511 | 84.14996 | Promoter | 16 | 72663149 | 73046095 | 382947 | 1 | ENSMUST00 | 0 | Robo1 | ENSMUSG00000022883.10 |
| chr2 | 122027707 | 122028413 | 28.00751 | 87.44511 | 84.14996 | Promoter | 2 | 122028546 | 122056598 | 28053 | 1 | ENSMUST00 | -133 | Eif3j1 | ENSMUSG00000027236.8 |
| chr3 | 123446595 | 123447803 | 28.00751 | 87.44511 | 84.14996 | Promoter | 3 | 123446913 | 123506602 | 59690 | 1 | ENSMUST00 | 0 | Prss12 | ENSMUSG00000027978.8 |
| chr6 | 126939092 | 126940812 | 28.00751 | 87.44511 | 84.14996 | Promoter | 6 | 126923050 | 126939587 | 16538 | 2 | ENSMUST00 | 0 | Rad51ap1 | ENSMUSG00000030346.11 |
| chr8 | 121652509 | 121653931 | 28.00751 | 87.44511 | 84.14996 | Promoter | 8 | 121598703 | 121652901 | 54199 | 2 | ENSMUST00 | 0 | Zcchc14 | ENSMUSG00000061410.6 |
| chr11 | 4873681 | 4874761 | 29.08566 | 87.40631 | 84.11242 | Promoter | 11 | 4873951 | 4894200 | 20250 | 1 | ENSMUST00 | 0 | Nipsnap1 | ENSMUSG00000034285.10 |
| chr17 | 33955795 | 33956779 | 25.33372 | 87.3614 | 84.0677 | Promoter | 17 | 33955519 | 33956001 | 483 | 2 | ENSMUST00 | 0 | Rps18 | ENSMUSG00000008668.9 |
| chr3 | 95216676 | 95218334 | 28.50113 | 87.35039 | 84.05681 | Promoter | 3 | 95181766 | 95217709 | 35944 | 2 | ENSMUST00 | 0 | Gabpb2 | ENSMUSG00000038766.11 |
| chr1 | 85894103 | 85895855 | 28.49948 | 87.34749 | 84.05397 | Promoter | 1 | 85894281 | 85908675 | 14395 | 1 | ENSMUST00 | 0 | Itm2c | ENSMUSG00000026223.10 |
| chr11 | 96074697 | 96075738 | 28.49948 | 87.34749 | 84.05397 | Promoter | 11 | 96068852 | 96075670 | 6819 | 2 | ENSMUST00 | 0 | Atp5g1 | ENSMUSG00000006057.10 |
| chr12 | 69168116 | 69169221 | 28.49948 | 87.34749 | 84.05397 | Promoter | 12 | 69168814 | 69179010 | 10197 | 1 | ENSMUST00 | 0 | Lrr1 | ENSMUSG00000034883.8 |
| chr14 | 63122106 | 63124349 | 28.49948 | 87.34749 | 84.05397 | Promoter | 14 | 63122462 | 63145919 | 23458 | 1 | ENSMUST00 | 0 | Ctsb | ENSMUSG00000021939.7 |
| chr17 | 23192260 | 23193947 | 28.49948 | 87.34749 | 84.05397 | Promoter | 17 | 23173869 | 23193252 | 19384 | 2 | ENSMUST00 | 0 | Zfp40 | ENSMUSG00000002617.9 |
| chr17 | 10313109 | 10319334 | 28.49948 | 87.34749 | 84.05397 | Promoter | 17 | 10206471 | 10319361 | 112891 | 2 | ENSMUST00 | 27 | Qk | ENSMUSG00000062078.9 |
| chr19 | 3906455 | 3907711 | 28.49948 | 87.34749 | 84.05397 | Promoter | 19 | 3896050 | 3906991 | 10942 | 2 | ENSMUST00 | 0 | Tcirg1 | ENSMUSG00000001750.10 |
| chr19 | 9134608 | 9136070 | 28.49948 | 87.34749 | 84.05397 | Promoter | 19 | 9109868 | 9135636 | 25769 | 2 | ENSMUST00 | 0 | Asrgl1 | ENSMUSG00000024654.8 |
| chr19 | 11965745 | 11967003 | 28.49948 | 87.34749 | 84.05397 | Promoter | 19 | 11965844 | 11994105 | 28262 | 1 | ENSMUST00 | 0 | Osbp | ENSMUSG00000024687.10 |
| chr19 | 46689613 | 46690621 | 28.49948 | 87.34749 | 84.05397 | Promoter | 19 | 46689906 | 46703382 | 13477 | 1 | ENSMUST00 | 0 | 2010012O05 | ENSMUSG00000062376.7 |
| chr2 | 73385746 | 73386551 | 28.49948 | 87.34749 | 84.05397 | Promoter | 2 | 73341958 | 73386412 | 44455 | 2 | ENSMUST00 | 0 | Gpr155 | ENSMUSG00000041762.11 |
| chr4 | 46488882 | 46490309 | 28.49948 | 87.34749 | 84.05397 | Promoter | 4 | 46489248 | 46503632 | 14385 | 1 | ENSMUST00 | 0 | Nans | ENSMUSG00000028334.5 |
| chr5 | 111416272 | 111419191 | 28.49948 | 87.34749 | 84.05397 | Promoter | 5 | 111417362 | 111418073 | 712 | 1 | ENSMUST00 | 0 | Mn1 | ENSMUSG00000070576.4 |
| chr6 | 72597573 | 72598983 | 28.49948 | 87.34749 | 84.05397 | Promoter | 6 | 72598475 | 72608425 | 9951 | 1 | ENSMUST00 | 0 | Retsat | ENSMUSG00000056666.8 |
| chr7 | 58657824 | 58659125 | 28.49948 | 87.34749 | 84.05397 | Promoter | 7 | 58658202 | 58829426 | 171225 | 1 | ENSMUST00 | 0 | Atp10a | ENSMUSG00000025324.7 |
| chr8 | 111027202 | 111028251 | 28.49948 | 87.34749 | 84.05397 | Promoter | 8 | 111003188 | 111027764 | 24577 | 2 | ENSMUST00 | 0 | Ddx19b | ENSMUSG00000033658.11 |
| chr9 | 109073136 | 109074690 | 28.49948 | 87.34749 | 84.05397 | Promoter | 9 | 109057933 | 109074071 | 16139 | 2 | ENSMUST00 | 0 | Atrip | ENSMUSG00000025646.3 |
| chr9 | 51076314 | 51077616 | 28.49948 | 87.34749 | 84.05397 | Promoter | 9 | 51056780 | 51077094 | 20315 | 2 | ENSMUST00 | 0 | Layn | ENSMUSG00000060594.5 |
| chr9 | 101073177 | 101073780 | 28.49948 | 87.34749 | 84.05397 | Promoter | 9 | 101074101 | 101101785 | 27685 | 1 | ENSMUST00 | -321 | Msl2 | ENSMUSG00000066415.4 |
| chr14 | 65592319 | 65593487 | 27.39617 | 87.32104 | 84.03119 | Promoter | 14 | 65530446 | 65593112 | 62667 | 2 | ENSMUST00 | 0 | Elp3 | ENSMUSG00000022031.5 |
| chr15 | 53346264 | 53347038 | 27.39617 | 87.32104 | 84.03119 | Promoter | 15 | 53064038 | 53346159 | 282122 | 2 | ENSMUST00 | -106 | Ext1 | ENSMUSG00000061731.4 |
| chr17 | 3326502 | 3327735 | 27.39617 | 87.32104 | 84.03119 | Promoter | 17 | 3326573 | 3519397 | 192825 | 1 | ENSMUST00 | 0 | Tiam2 | ENSMUSG00000023800.9 |
| chr18 | 74778092 | 74780533 | 27.39617 | 87.32104 | 84.03119 | Promoter | 18 | 74779197 | 74806207 | 27011 | 1 | ENSMUST00 | 0 | Acaa2 | ENSMUSG00000036880.9 |
| chr4 | 15944626 | 15945984 | 27.39617 | 87.32104 | 84.03119 | Promoter | 4 | 15917240 | 15945507 | 28268 | 2 | ENSMUST00 | 0 | Decr1 | ENSMUSG00000028223.8 |
| chr5 | 35757434 | 35758701 | 27.39617 | 87.32104 | 84.03119 | Promoter | 5 | 35757880 | 35884466 | 126587 | 1 | ENSMUST00 | 0 | Ablim2 | ENSMUSG00000029095.12 |
| chr7 | 118712468 | 118713097 | 27.39617 | 87.32104 | 84.03119 | Promoter | 7 | 118712552 | 118737024 | 24473 | 1 | ENSMUST00 | 0 | Ccp110 | ENSMUSG00000033904.10 |
| chr6 | 52190914 | 52191800 | 29.59164 | 87.32004 | 84.03093 | Promoter | 6 | 52189671 | 52191753 | 2083 | 2 | ENSMUST00 | 0 | Hoxa4 | ENSMUSG00000000942.10 |
| chr7 | 27486916 | 27490677 | 29.02802 | 87.30634 | 84.01743 | Promoter | 7 | 27487375 | 27489507 | 2133 | 1 | ENSMUST00 | 0 | Sertad1 | ENSMUSG00000008384.8 |
| chr19 | 34526383 | 34527533 | 25.79933 | 87.28319 | 83.99445 | Promoter | 19 | 34492318 | 34527474 | 35157 | 2 | ENSMUST00 | 0 | Lipa | ENSMUSG00000024781.10 |
| chr6 | 119387786 | 119388742 | 25.79933 | 87.28319 | 83.99445 | Promoter | 6 | 119353765 | 119388686 | 34922 | 2 | ENSMUST00 | 0 | Adipor2 | ENSMUSG00000030168.8 |
| chr14 | 27335535 | 27337019 | 26.8271 | 87.25735 | 83.9688 | Intron (ENSM | 14 | 27380732 | 27380850 | 119 | 2 | ENSMUST00 | 43831 | Gm23633 | ENSMUSG00000093034.1 |
| chr19 | 47865149 | 47866289 | 26.8271 | 87.25735 | 83.9688 | Promoter | 19 | 47865545 | 47886324 | 20780 | 1 | ENSMUST00 | 0 | Gsto2 | ENSMUSG00000025069.9 |
| chr1 | 191820986 | 191822282 | 26.29604 | 87.24678 | 83.95876 | Promoter | 1 | 191821444 | 191833050 | 11607 | 1 | ENSMUST00 | 0 | Nek2 | ENSMUSG00000026622.10 |
| chr3 | 20154289 | 20155423 | 26.29604 | 87.24678 | 83.95876 | Promoter | 3 | 20122084 | 20155317 | 33234 | 2 | ENSMUST00 | 0 | Gyg | ENSMUSG00000019528.13 |
| chr8 | 75108793 | 75110118 | 26.29604 | 87.24678 | 83.95876 | Promoter | 8 | 75109528 | 75128439 | 18912 | 1 | ENSMUST00 | 0 | Mcm5 | ENSMUSG00000005410.5 |
| chr11 | 70647243 | 70648576 | 29.53108 | 87.21563 | 83.92796 | Promoter | 11 | 70645231 | 70647479 | 2249 | 2 | ENSMUST00 | 0 | Slc25a11 | ENSMUSG00000014606.9 |
| chr11 | 69325980 | 69326924 | 29.53108 | 87.21563 | 83.92796 | Promoter | 11 | 69326258 | 69333042 | 6785 | 1 | ENSMUST00 | 0 | Kcnab3 | ENSMUSG00000018470.3 |
| chr18 | 60747065 | 60749034 | 29.53108 | 87.21563 | 83.92796 | Distal Interge | 18 | 60774510 | 60778546 | 4037 | 1 | ENSMUST00 | -25476 | Rps14 | ENSMUSG00000024608.5 |
| chr18 | 82909434 | 82915656 | 29.53108 | 87.21563 | 83.92796 | Promoter | 18 | 82910879 | 83005314 | 94436 | 1 | ENSMUST00 | 0 | Zfp516 | ENSMUSG00000058881.7 |
| chr4 | 9769848 | 9771306 | 29.53108 | 87.21563 | 83.92796 | Promoter | 4 | 9770585 | 9821951 | 51367 | 1 | ENSMUST00 | 0 | 4930412C18I | ENSMUSG00000085558.2 |
| chr8 | 71370848 | 71371955 | 28.97061 | 87.20673 | 83.91971 | Promoter | 8 | 71371298 | 71374382 | 3085 | 1 | ENSMUST00 | 0 | Ocel1 | ENSMUSG00000002396.8 |
| chr16 | 56037006 | 56038363 | 20.74474 | 87.15693 | 83.87018 | Promoter | 16 | 56032609 | 56037819 | 5211 | 2 | ENSMUST00 | 0 | Trmt10c | ENSMUSG00000044763.8 |
| chr11 | 78549980 | 78551211 | 29.47077 | 87.11163 | 83.82504 | Promoter | 11 | 78541817 | 78550777 | 8961 | 2 | ENSMUST00 | 0 | Tmem97 | ENSMUSG00000037278.9 |
| chr16 | 55838318 | 55839457 | 29.47077 | 87.11163 | 83.82504 | Promoter | 16 | 55811375 | 55838899 | 27525 | 2 | ENSMUST00 | 0 | Nfkbiz | ENSMUSG00000035356.11 |
| chr1 | 161130561 | 161131835 | 25.16654 | 87.04672 | 83.76057 | Promoter | 1 | 161088375 | 161131511 | 43137 | 2 | ENSMUST00 | 0 | Klhl20 | ENSMUSG00000026705.11 |
| chr10 | 20347291 | 20348106 | 25.16654 | 87.04672 | 83.76057 | Promoter | 10 | 20347819 | 20361669 | 13851 | 1 | ENSMUST00 | 0 | Mtfr2 | ENSMUSG00000019992.7 |
| chr10 | 24594963 | 24597717 | 25.16654 | 87.04672 | 83.76057 | Promoter | 10 | 24595442 | 24598678 | 3237 | 1 | ENSMUST00 | 0 | Ctgf | ENSMUSG00000019997.6 |
| chr11 | 101467402 | 101468763 | 25.16654 | 87.04672 | 83.76057 | Promoter | 11 | 101468175 | 101471853 | 3679 | 1 | ENSMUST00 | 0 | Rnd2 | ENSMUSG00000001313.7 |
| chr11 | 101627645 | 101628686 | 25.16654 | 87.04672 | 83.76057 | Promoter | 11 | 101627942 | 101636100 | 8159 | 1 | ENSMUST00 | 0 | Rdm1 | ENSMUSG00000010362.4 |
| chr12 | 85598428 | 85600604 | 25.16654 | 87.04672 | 83.76057 | Promoter | 12 | 85599105 | 85639878 | 40774 | 1 | ENSMUST00 | 0 | Jdp2 | ENSMUSG00000034271.10 |
| chr13 | 111724511 | 111726223 | 25.16654 | 87.04672 | 83.76057 | Promoter | 13 | 111725517 | 111735124 | 9608 | 1 | ENSMUST00 | 0 | Gm15287 | ENSMUSG00000086480.1 |
| chr13 | 67551988 | 67553547 | 25.16654 | 87.04672 | 83.76057 | Promoter | 13 | 67538641 | 67553138 | 14498 | 2 | ENSMUST00 | 0 | Zfp748 | ENSMUSG00000095432.3 |
| chr15 | 85878650 | 85880068 | 25.16654 | 87.04672 | 83.76057 | Promoter | 15 | 85879312 | 85892598 | 13287 | 1 | ENSMUST00 | 0 | Trmu | ENSMUSG00000022386.7 |
| chr15 | 81663541 | 81665072 | 25.16654 | 87.04672 | 83.76057 | Promoter | 15 | 81663889 | 81688315 | 24427 | 1 | ENSMUST00 | 0 | L3mbtl2 | ENSMUSG00000022394.9 |
| chr15 | 93397435 | 93399514 | 25.16654 | 87.04672 | 83.76057 | Promoter | 15 | 93386097 | 93398301 | 12205 | 2 | ENSMUST00 | 0 | Zcrb1 | ENSMUSG00000022635.4 |
| chr15 | 90678118 | 90680514 | 25.16654 | 87.04672 | 83.76057 | Promoter | 15 | 90487482 | 90679371 | 191890 | 2 | ENSMUST00 | 0 | Cpne8 | ENSMUSG00000052560.10 |
| chr15 | 76924945 | 76926154 | 25.16654 | 87.04672 | 83.76057 | Distal Interge | 15 | 76910851 | 76918010 | 7160 | 2 | ENSMUST00 | -6936 | Zfp647 | ENSMUSG00000054967.6 |
| chr17 | 71597789 | 71599268 | 25.16654 | 87.04672 | 83.76057 | Promoter | 17 | 71557027 | 71598755 | 41729 | 2 | ENSMUST00 | 0 | Trmt61b | ENSMUSG00000085492.2 |
| chr17 | 21531730 | 21533285 | 25.16654 | 87.04672 | 83.76057 | Intron (ENSM | 17 | 21528194 | 21535878 | 7685 | 2 | ENSMUST00 | 2593 | 9330136K24I | ENSMUSG00000097781.1 |
| chr18 | 31930574 | 31933785 | 25.16654 | 87.04672 | 83.76057 | Promoter | 18 | 31931507 | 31958619 | 27113 | 1 | ENSMUST00 | 0 | Lims2 | ENSMUSG00000024395.7 |
| chr2 | 121008044 | 121009281 | 25.16654 | 87.04672 | 83.76057 | Promoter | 2 | 121008403 | 121016904 | 8502 | 1 | ENSMUST00 | 0 | Ccndbp1 | ENSMUSG00000023572.11 |
| chr4 | 41683408 | 41684199 | 25.16654 | 87.04672 | 83.76057 | Intron (ENSM | 4 | 41657498 | 41695442 | 37945 | 2 | ENSMUST00 | 11243 | Cntfr | ENSMUSG00000028444.12 |
| chr4 | 126467729 | 126469026 | 25.16654 | 87.04672 | 83.76057 | Promoter | 4 | 126435012 | 126468583 | 33572 | 2 | ENSMUST00 | 0 | Ago1 | ENSMUSG00000041530.11 |
| chr4 | 124656736 | 124658371 | 25.16654 | 87.04672 | 83.76057 | Promoter | 4 | 124656807 | 124660655 | 3849 | 1 | ENSMUST00 | 0 | Pou3f1 | ENSMUSG00000090125.3 |
| chr6 | 30390483 | 30391506 | 25.16654 | 87.04672 | 83.76057 | Promoter | 6 | 30366380 | 30391023 | 24644 | 2 | ENSMUST00 | 0 | Zc3hc1 | ENSMUSG00000039130.13 |
| chr7 | 46639184 | 46640418 | 25.16654 | 87.04672 | 83.76057 | Promoter | 7 | 46443168 | 46639807 | 196640 | 2 | ENSMUST00 | 0 | Sergef | ENSMUSG00000030839.5 |
| chr7 | 27258187 | 27259515 | 25.16654 | 87.04672 | 83.76057 | Promoter | 7 | 27258433 | 27266801 | 8369 | 1 | ENSMUST00 | 0 | Numbl | ENSMUSG00000063160.7 |
| chr7 | 101969394 | 101971896 | 25.16654 | 87.04672 | 83.76057 | Promoter | 7 | 101969843 | 102014959 | 45117 | 1 | ENSMUST00 | 0 | Numa1 | ENSMUSG00000066306.7 |
| chr8 | 117801395 | 117802861 | 25.16654 | 87.04672 | 83.76057 | Promoter | 8 | 117791645 | 117801929 | 10285 | 2 | ENSMUST00 | 0 | Mphosph6 | ENSMUSG00000031843.1 |
| chr8 | 110079316 | 110080321 | 25.16654 | 87.04672 | 83.76057 | Promoter | 8 | 110079764 | 110090181 | 10418 | 1 | ENSMUST00 | 0 | Zfp612 | ENSMUSG00000044676.9 |
| chr13 | 104816275 | 104818088 | 28.85647 | 87.00858 | 83.7254 | Promoter | 13 | 104631140 | 104816953 | 185814 | 2 | ENSMUST00 | 0 | Cwc27 | ENSMUSG00000021715.7 |
| chr2 | 32235727 | 32237616 | 28.85647 | 87.00858 | 83.7254 | Promoter | 2 | 32236590 | 32255005 | 18416 | 1 | ENSMUST00 | 0 | Pomt1 | ENSMUSG00000039254.11 |
| chr2 | 27676111 | 27677129 | 29.4107 | 87.008 | 83.72497 | Promoter | 2 | 27676440 | 27741898 | 65459 | 1 | ENSMUST00 | 0 | Rxra | ENSMUSG00000015846.9 |
| chr4 | 58943316 | 58944136 | 29.4107 | 87.008 | 83.72497 | Promoter | 4 | 58943628 | 58958355 | 14728 | 1 | ENSMUST00 | 0 | Zkscan16 | ENSMUSG00000038630.8 |
| chr1 | 16656086 | 16657782 | 26.65955 | 86.95457 | 83.67183 | Promoter | 1 | 16641725 | 16656884 | 15160 | 2 | ENSMUST00 | 0 | Tceb1 | ENSMUSG00000079658.4 |
| chr10 | 71158323 | 71160162 | 26.65955 | 86.95457 | 83.67183 | Promoter | 10 | 70922832 | 71159700 | 236869 | 2 | ENSMUST00 | 0 | Bicc1 | ENSMUSG00000014329.9 |
| chr10 | 88356579 | 88357610 | 26.65955 | 86.95457 | 83.67183 | Promoter | 10 | 88322804 | 88356996 | 34193 | 2 | ENSMUST00 | 0 | Dram1 | ENSMUSG00000020057.2 |
| chr11 | 29546901 | 29548934 | 26.65955 | 86.95457 | 83.67183 | Promoter | 11 | 29545846 | 29547860 | 2015 | 2 | ENSMUST00 | 0 | Rps27a | ENSMUSG00000020460.10 |
| chr11 | 98959620 | 98961343 | 26.65955 | 86.95457 | 83.67183 | Promoter | 11 | 98960412 | 98974942 | 14531 | 1 | ENSMUST00 | 0 | Rara | ENSMUSG00000037992.11 |
| chr11 | 48815920 | 48818558 | 26.65955 | 86.95457 | 83.67183 | Promoter | 11 | 48806404 | 48817353 | 10950 | 2 | ENSMUST00 | 0 | Trim41 | ENSMUSG00000040365.9 |
| chr12 | 84192322 | 84194307 | 26.65955 | 86.95457 | 83.67183 | Promoter | 12 | 84149278 | 84194015 | 44738 | 2 | ENSMUST00 | 0 | Elmsan1 | ENSMUSG00000042507.9 |
| chr12 | 70967738 | 70970109 | 26.65955 | 86.95457 | 83.67183 | Distal Interge | 12 | 70974621 | 70996347 | 21727 | 1 | ENSMUST00 | -4512 | Psma3 | ENSMUSG00000060073.4 |
| chr13 | 36116667 | 36118632 | 26.65955 | 86.95457 | 83.67183 | Promoter | 13 | 36117411 | 36537592 | 420182 | 1 | ENSMUST00 | 0 | Fars2 | ENSMUSG00000021420.7 |
| chr14 | 19750569 | 19752607 | 26.65955 | 86.95457 | 83.67183 | Promoter | 14 | 19751257 | 19811787 | 60531 | 1 | ENSMUST00 | 0 | Nid2 | ENSMUSG00000021806.3 |
| chr15 | 102510182 | 102511139 | 26.65955 | 86.95457 | 83.67183 | Promoter | 15 | 102498776 | 102510700 | 11925 | 2 | ENSMUST00 | 0 | Map3k12 | ENSMUSG00000023050.3 |
| chr19 | 46599005 | 46600299 | 26.65955 | 86.95457 | 83.67183 | Promoter | 19 | 46599084 | 46657389 | 58306 | 1 | ENSMUST00 | 0 | Wbp1l | ENSMUSG00000047731.12 |
| chr2 | 24962288 | 24963116 | 26.65955 | 86.95457 | 83.67183 | Promoter | 2 | 24962400 | 24972163 | 9764 | 1 | ENSMUST00 | 0 | Dph7 | ENSMUSG00000026975.5 |
| chr2 | 169632988 | 169635980 | 26.65955 | 86.95457 | 83.67183 | Promoter | 2 | 169633013 | 169913093 | 280081 | 1 | ENSMUST00 | 0 | Tshz2 | ENSMUSG00000047907.11 |
| chr3 | 90231187 | 90232526 | 26.65955 | 86.95457 | 83.67183 | Promoter | 3 | 90231597 | 90235838 | 4242 | 1 | ENSMUST00 | 0 | Jtb | ENSMUSG00000027937.10 |
| chr3 | 96629299 | 96630871 | 26.65955 | 86.95457 | 83.67183 | Promoter | 3 | 96629928 | 96633779 | 3852 | 1 | ENSMUST00 | 0 | Rbm8a | ENSMUSG00000038374.6 |
| chr3 | 67514339 | 67515861 | 26.65955 | 86.95457 | 83.67183 | Promoter | 3 | 67478886 | 67515523 | 36638 | 2 | ENSMUST00 | 0 | Rarres1 | ENSMUSG00000049404.6 |
| chr4 | 118408792 | 118410205 | 26.65955 | 86.95457 | 83.67183 | Promoter | 4 | 118409337 | 118415111 | 5775 | 1 | ENSMUST00 | 0 | Med8 | ENSMUSG00000006392.11 |
| chr4 | 95966720 | 95968092 | 26.65955 | 86.95457 | 83.67183 | Promoter | 4 | 95967240 | 96025413 | 58174 | 1 | ENSMUST00 | 0 | Hook1 | ENSMUSG00000028572.8 |
| chr4 | 129819570 | 129821757 | 26.65955 | 86.95457 | 83.67183 | Promoter | 4 | 129820221 | 129849978 | 29758 | 1 | ENSMUST00 | 0 | Ptp4a2 | ENSMUSG00000028788.9 |
| chr4 | 152177486 | 152178864 | 26.65955 | 86.95457 | 83.67183 | Promoter | 4 | 152178134 | 152271852 | 93719 | 1 | ENSMUST00 | 0 | Acot7 | ENSMUSG00000028937.9 |
| chr5 | 120116125 | 120117223 | 26.65955 | 86.95457 | 83.67183 | Promoter | 5 | 120116510 | 120197979 | 81470 | 1 | ENSMUST00 | 0 | Rbm19 | ENSMUSG00000029594.9 |
| chr5 | 138193791 | 138195196 | 26.65955 | 86.95457 | 83.67183 | Promoter | 5 | 138194314 | 138195621 | 1308 | 1 | ENSMUST00 | 0 | Mblac1 | ENSMUSG00000049285.4 |
| chr6 | 114967501 | 114970246 | 26.65955 | 86.95457 | 83.67183 | Promoter | 6 | 114862321 | 114969180 | 106860 | 2 | ENSMUST00 | 0 | Vgll4 | ENSMUSG00000030315.10 |
| chr6 | 125008334 | 125010316 | 26.65955 | 86.95457 | 83.67183 | Promoter | 6 | 125009145 | 125036180 | 27036 | 1 | ENSMUST00 | 0 | Zfp384 | ENSMUSG00000038346.13 |
| chr7 | 45865939 | 45866990 | 26.65955 | 86.95457 | 83.67183 | Promoter | 7 | 45831883 | 45867129 | 35247 | 2 | ENSMUST00 | 139 | Grin2d | ENSMUSG00000002771.10 |
| chr7 | 3617096 | 3618088 | 26.65955 | 86.95457 | 83.67183 | Promoter | 7 | 3617373 | 3620161 | 2789 | 1 | ENSMUST00 | 0 | Ndufa3 | ENSMUSG00000035674.8 |
| chr7 | 49758797 | 49759670 | 26.65955 | 86.95457 | 83.67183 | Promoter | 7 | 49759106 | 49773999 | 14894 | 1 | ENSMUST00 | 0 | Htatip2 | ENSMUSG00000039745.7 |
| chr7 | 24902634 | 24904205 | 26.65955 | 86.95457 | 83.67183 | Promoter | 7 | 24902912 | 24926592 | 23681 | 1 | ENSMUST00 | 0 | Arhgef1 | ENSMUSG00000040940.13 |
| chr7 | 15921023 | 15923182 | 26.65955 | 86.95457 | 83.67183 | Promoter | 7 | 15917210 | 15922371 | 5162 | 2 | ENSMUST00 | 0 | Sepw1 | ENSMUSG00000041571.8 |
| chr7 | 25657902 | 25658993 | 26.65955 | 86.95457 | 83.67183 | Promoter | 7 | 25629852 | 25658761 | 28910 | 2 | ENSMUST00 | 0 | Bckdha | ENSMUSG00000060376.6 |
| chr8 | 121950115 | 121951651 | 26.65955 | 86.95457 | 83.67183 | Promoter | 8 | 121950492 | 122026177 | 75686 | 1 | ENSMUST00 | 0 | Banp | ENSMUSG00000025316.11 |
| chr8 | 61487618 | 61488425 | 26.65955 | 86.95457 | 83.67183 | Promoter | 8 | 61487734 | 61503482 | 15749 | 1 | ENSMUST00 | 0 | Cbr4 | ENSMUSG00000031641.8 |
| chr9 | 66125734 | 66127424 | 26.65955 | 86.95457 | 83.67183 | Promoter | 9 | 66126611 | 66138955 | 12345 | 1 | ENSMUST00 | 0 | Fam96a | ENSMUSG00000032381.5 |
| chr1 | 93801314 | 93802529 | 28.79973 | 86.91003 | 83.63136 | Promoter | 1 | 93792576 | 93801934 | 9359 | 2 | ENSMUST00 | 0 | Dtymk | ENSMUSG00000026281.10 |
| chr11 | 20331759 | 20333631 | 28.79974 | 86.91003 | 83.63136 | Promoter | 11 | 20302180 | 20332713 | 30534 | 2 | ENSMUST00 | 0 | Slc1a4 | ENSMUSG00000020142.7 |
| chr16 | 13447301 | 13448787 | 28.79974 | 86.91003 | 83.63136 | Promoter | 16 | 13449523 | 13449601 | 79 | 1 | ENSMUST00 | -736 | Mir193b | ENSMUSG00000076972.2 |
| chr3 | 95817553 | 95819386 | 28.79974 | 86.91003 | 83.63136 | Promoter | 3 | 95759873 | 95818953 | 59081 | 2 | ENSMUST00 | 0 | Rprd2 | ENSMUSG00000028106.9 |
| chr7 | 129591302 | 129592736 | 28.79974 | 86.91003 | 83.63136 | Promoter | 7 | 129591863 | 129635738 | 43876 | 1 | ENSMUST00 | 0 | Wdr11 | ENSMUSG00000042055.8 |
| chr1 | 171246458 | 171247900 | 29.86959 | 86.83979 | 83.56236 | Promoter | 1 | 171234853 | 171247122 | 12270 | 2 | ENSMUST00 | 0 | Ndufs2 | ENSMUSG00000013593.7 |
| chr10 | 94943292 | 94945671 | 29.86959 | 86.83979 | 83.56236 | Promoter | 10 | 94790866 | 94944578 | 153713 | 2 | ENSMUST00 | 0 | Plxnc1 | ENSMUSG00000074785.4 |

| | | | | | | | | | | | | | | | |
|---|---|---|---|---|---|---|---|---|---|---|---|---|---|---|---|
| chr2 | 26379956 | 26380890 | 29.86959 | 86.83979 | 83.56236 | Promoter | 2 | 26362765 | 26380649 | 17885 | 2 | ENSMUST00 | 0 | Snapc4 | ENSMUSG00000036281.8 |
| chr4 | 45297042 | 45297754 | 29.86959 | 86.83979 | 83.56236 | Promoter | 4 | 45297127 | 45316131 | 19005 | 1 | ENSMUST00 | 0 | Trmt10b | ENSMUSG00000035601.4 |
| chr4 | 120282409 | 120283440 | 29.86959 | 86.83979 | 83.56236 | Intron (ENSM | 4 | 120267079 | 120287349 | 20271 | 2 | ENSMUST00 | 3909 | Foxo6 | ENSMUSG00000052135.8 |
| chr5 | 86064854 | 86065631 | 29.86959 | 86.83979 | 83.56236 | Promoter | 5 | 86012043 | 86065583 | 53541 | 2 | ENSMUST00 | 0 | Cenpc1 | ENSMUSG00000029253.8 |
| chr6 | 118478769 | 118479810 | 29.86959 | 86.83979 | 83.56236 | Promoter | 6 | 118461950 | 118479320 | 17371 | 2 | ENSMUST00 | 0 | Zfp9 | ENSMUSG00000072623.5 |
| chr7 | 16309242 | 16310716 | 29.86959 | 86.83979 | 83.56236 | Promoter | 7 | 16309616 | 16318205 | 8590 | 1 | ENSMUST00 | 0 | Bbc3 | ENSMUSG00000002083.7 |
| chr7 | 89632371 | 89633483 | 29.86959 | 86.83979 | 83.56236 | Promoter | 7 | 89632392 | 89810197 | 177806 | 1 | ENSMUST00 | 0 | Me3 | ENSMUSG00000030621.12 |
| chr17 | 3082964 | 3085057 | 19.11308 | 86.81667 | 83.5409 | Promoter | 17 | 3076578 | 3084082 | 7505 | 2 | ENSMUST00 | 0 | Pisd-ps2 | ENSMUSG00000023795.11 |
| chr3 | 65665753 | 65666870 | 28.74323 | 86.81182 | 83.53605 | Promoter | 3 | 65666228 | 65686179 | 19952 | 1 | ENSMUST00 | 0 | Lekr1 | ENSMUSG00000074579.8 |
| chr4 | 45971834 | 45973478 | 28.74323 | 86.81182 | 83.53605 | Promoter | 4 | 45972233 | 46034761 | 62529 | 1 | ENSMUST00 | 0 | Tdrd7 | ENSMUSG00000035517.12 |
| chr19 | 44544905 | 44546097 | 29.2913 | 86.80193 | 83.52631 | Promoter | 19 | 44516957 | 44545864 | 28908 | 2 | ENSMUST00 | 0 | Sec31b | ENSMUSG00000051984.8 |
| chr9 | 27298739 | 27300804 | 29.81586 | 86.74751 | 83.47216 | Promoter | 9 | 27299228 | 27330135 | 30908 | 1 | ENSMUST00 | 0 | Igsf9b | ENSMUSG00000034275.12 |
| chr2 | 127207783 | 127209230 | 28.68694 | 86.71395 | 83.43872 | Promoter | 2 | 127208386 | 127222717 | 14332 | 1 | ENSMUST00 | 0 | Snrnp200 | ENSMUSG00000003660.10 |
| chr10 | 34019816 | 34020609 | 29.23196 | 86.69946 | 83.42436 | Promoter | 10 | 33996555 | 34019624 | 23070 | 2 | ENSMUST00 | -193 | Rwdd1 | ENSMUSG00000019782.9 |
| chr3 | 109122707 | 109123960 | 29.23196 | 86.69946 | 83.42436 | Promoter | 3 | 109123149 | 109168457 | 45309 | 1 | ENSMUST00 | 0 | Slc25a24 | ENSMUSG00000040322.6 |
| chr10 | 77416895 | 77418831 | 28.12373 | 86.68959 | 83.41467 | Promoter | 10 | 77290726 | 77418265 | 127540 | 2 | ENSMUST00 | 0 | Adarb1 | ENSMUSG00000020262.10 |
| chr10 | 128821844 | 128822726 | 28.12373 | 86.68959 | 83.41467 | Promoter | 10 | 128821771 | 128877638 | 55868 | 1 | ENSMUST00 | 74 | Sarnp | ENSMUSG00000078427.2 |
| chr11 | 5861239 | 5863555 | 28.12373 | 86.68959 | 83.41467 | Promoter | 11 | 5861947 | 5872088 | 10142 | 1 | ENSMUST00 | 0 | Aebp1 | ENSMUSG00000020473.8 |
| chr13 | 65277836 | 65279571 | 28.12373 | 86.68959 | 83.41467 | Promoter | 13 | 65278814 | 65297795 | 18982 | 1 | ENSMUST00 | 0 | Zfp369 | ENSMUSG00000021514.12 |
| chr18 | 38599522 | 38602400 | 28.12373 | 86.68959 | 83.41467 | Promoter | 18 | 38586268 | 38601268 | 15001 | 2 | ENSMUST00 | 0 | Spry4 | ENSMUSG00000024427.6 |
| chr2 | 115581105 | 115582329 | 28.12373 | 86.68959 | 83.41467 | Promoter | 2 | 115581716 | 115778768 | 197053 | 1 | ENSMUST00 | 0 | BC052040 | ENSMUSG00000040282.8 |
| chr7 | 13033890 | 13035078 | 28.12373 | 86.68959 | 83.41467 | Promoter | 7 | 13032006 | 13034777 | 2772 | 2 | ENSMUST00 | 0 | Chmp2a | ENSMUSG00000033916.4 |
| chr11 | 50223936 | 50226450 | 22.60491 | 86.65532 | 83.38163 | Promoter | 11 | 50225335 | 50235103 | 9769 | 1 | ENSMUST00 | 0 | Mgat4b | ENSMUSG00000036620.9 |
| chr11 | 30648808 | 30649813 | 22.60491 | 86.65532 | 83.38163 | Promoter | 11 | 30505991 | 30649587 | 143597 | 2 | ENSMUST00 | 0 | Acyp2 | ENSMUSG00000060923.5 |
| chr12 | 95691821 | 95698644 | 22.60491 | 86.65532 | 83.38163 | Promoter | 12 | 95692226 | 95785215 | 92990 | 1 | ENSMUST00 | 0 | Flrt2 | ENSMUSG00000047414.6 |
| chr15 | 3995190 | 3995797 | 22.60491 | 86.65532 | 83.38163 | Promoter | 15 | 3982036 | 3995752 | 13717 | 2 | ENSMUST00 | 0 | AW549877 | ENSMUSG00000041935.9 |
| chr4 | 120569090 | 120570929 | 22.60491 | 86.65532 | 83.38163 | Promoter | 4 | 120539868 | 120570276 | 30409 | 2 | ENSMUST00 | 0 | Ctps | ENSMUSG00000028633.7 |
| chr5 | 124578504 | 124579886 | 22.60491 | 86.65532 | 83.38163 | Promoter | 5 | 124579140 | 124597680 | 18541 | 1 | ENSMUST00 | 0 | Gtf2h3 | ENSMUSG00000029387.5 |
| chr7 | 120842456 | 120844000 | 22.60491 | 86.65532 | 83.38163 | Promoter | 7 | 120842831 | 120907219 | 64389 | 1 | ENSMUST00 | 0 | Eef2k | ENSMUSG00000035064.11 |
| chr11 | 119354793 | 119356014 | 28.63087 | 86.61642 | 83.34377 | Promoter | 11 | 119343425 | 119355502 | 12078 | 2 | ENSMUST00 | 0 | Sgsh | ENSMUSG00000005043.8 |
| chr4 | 126162443 | 126163467 | 29.17286 | 86.59738 | 83.32484 | Promoter | 4 | 126150602 | 126162626 | 12025 | 2 | ENSMUST00 | 0 | Sh3d21 | ENSMUSG00000073758.5 |
| chr5 | 135009113 | 135010551 | 29.17286 | 86.59738 | 83.32484 | Promoter | 5 | 135009152 | 135012021 | 2870 | 1 | ENSMUST00 | 0 | Abhd11 | ENSMUSG00000040532.9 |
| chr1 | 165461194 | 165462209 | 27.49702 | 86.54257 | 83.27029 | Promoter | 1 | 165461224 | 165477694 | 16471 | 1 | ENSMUST00 | 0 | Mpc2 | ENSMUSG00000026568.5 |
| chr10 | 8885006 | 8886830 | 27.49702 | 86.54257 | 83.27029 | Promoter | 10 | 8722219 | 8886070 | 163852 | 2 | ENSMUST00 | 0 | Sash1 | ENSMUSG00000015305.5 |
| chr5 | 36988297 | 36989768 | 27.49702 | 86.54257 | 83.27029 | Promoter | 5 | 36966104 | 36988982 | 22879 | 2 | ENSMUST00 | 0 | Wfs1 | ENSMUSG00000039474.8 |
| chr7 | 66108901 | 66110654 | 27.49702 | 86.54257 | 83.27029 | Promoter | 7 | 66109515 | 66173789 | 64275 | 1 | ENSMUST00 | 0 | Chsy1 | ENSMUSG00000032640.9 |
| chr9 | 75440979 | 75442254 | 27.49702 | 86.54257 | 83.27029 | Promoter | 9 | 75441524 | 75466432 | 24909 | 1 | ENSMUST00 | 0 | Leo1 | ENSMUSG00000042487.5 |
| chr16 | 11065844 | 11067095 | 29.69128 | 86.53342 | 83.26198 | Promoter | 16 | 10963343 | 11066157 | 102815 | 2 | ENSMUST00 | 0 | Litaf | ENSMUSG00000022500.9 |
| chr16 | 20620717 | 20622730 | 29.69128 | 86.53342 | 83.26198 | Promoter | 16 | 20619215 | 20621287 | 2073 | 2 | ENSMUST00 | 0 | Camk2n2 | ENSMUSG00000051146.2 |
| chr5 | 25099478 | 25101663 | 29.69128 | 86.53342 | 83.26198 | Promoter | 5 | 24862744 | 25100642 | 237899 | 2 | ENSMUST00 | 0 | Prkag2 | ENSMUSG00000028944.9 |
| chr5 | 142817908 | 142824082 | 29.69128 | 86.53342 | 83.26198 | Promoter | 5 | 142754789 | 142817662 | 62874 | 2 | ENSMUST00 | -247 | Tnrc18 | ENSMUSG00000039477.11 |
| chr5 | 124031065 | 124032535 | 29.69128 | 86.53342 | 83.26198 | Promoter | 5 | 124004641 | 124032260 | 27620 | 2 | ENSMUST00 | 0 | Vps37b | ENSMUSG00000066278.5 |
| chr6 | 125349048 | 125350835 | 29.69128 | 86.53342 | 83.26198 | Promoter | 6 | 125349362 | 125350035 | 674 | 1 | ENSMUST00 | 0 | Tnfrsf1a | ENSMUSG00000030341.12 |
| chr4 | 99193514 | 99195055 | 25.38893 | 86.51994 | 83.2489 | Promoter | 4 | 99193934 | 99259787 | 65854 | 1 | ENSMUST00 | 0 | Atg4c | ENSMUSG00000028550.10 |
| chr1 | 159231413 | 159233676 | 28.01773 | 86.50358 | 83.23278 | Promoter | 1 | 159232326 | 159347580 | 115255 | 1 | ENSMUST00 | 0 | Rfwd2 | ENSMUSG00000040782.7 |
| chr19 | 6908992 | 6910951 | 29.114 | 86.49567 | 83.2249 | Promoter | 19 | 6906697 | 6909978 | 3282 | 2 | ENSMUST00 | 0 | Prdx5 | ENSMUSG00000024953.11 |
| chr13 | 102903474 | 102906807 | 26.91438 | 86.45937 | 83.18881 | Promoter | 13 | 102735002 | 102906106 | 171105 | 2 | ENSMUST00 | 0 | Mast4 | ENSMUSG00000034751.10 |
| chr16 | 38902161 | 38902911 | 26.91438 | 86.45937 | 83.18881 | Promoter | 16 | 38902316 | 39027159 | 124844 | 1 | ENSMUST00 | 0 | Igsf11 | ENSMUSG00000022790.8 |
| chr19 | 12795661 | 12796166 | 26.91438 | 86.45937 | 83.18881 | Promoter | 19 | 12763660 | 12796126 | 32467 | 2 | ENSMUST00 | 0 | Zfp91 | ENSMUSG00000024695.10 |
| chr19 | 6950499 | 6952904 | 26.91438 | 86.45937 | 83.18881 | Promoter | 19 | 6950767 | 6951874 | 1108 | 1 | ENSMUST00 | 0 | Bad | ENSMUSG00000024959.8 |
| chr3 | 89997952 | 89999307 | 26.91438 | 86.45937 | 83.18881 | Promoter | 3 | 89995457 | 89998686 | 3230 | 2 | ENSMUST00 | 0 | Hax1 | ENSMUSG00000027944.9 |
| chr8 | 69624639 | 69625923 | 26.91438 | 86.45937 | 83.18881 | Promoter | 8 | 69610857 | 69625548 | 14692 | 2 | ENSMUST00 | 0 | Zfp868 | ENSMUSG00000060427.10 |
| chr8 | 126473965 | 126475285 | 26.91438 | 86.45937 | 83.18881 | Promoter | 8 | 126426652 | 126475065 | 48414 | 2 | ENSMUST00 | 0 | Gm17296 | ENSMUSG00000090290.1 |
| chr16 | 20548026 | 20548513 | 25.86399 | 86.45414 | 83.18411 | Promoter | 16 | 20548577 | 20561379 | 12803 | 1 | ENSMUST00 | -64 | Abcf3 | ENSMUSG00000003234.10 |
| chr7 | 67644584 | 67645257 | 25.86399 | 86.45414 | 83.18411 | Promoter | 7 | 67513411 | 67645231 | 131821 | 2 | ENSMUST00 | 0 | Lrrc28 | ENSMUSG00000030556.7 |
| chr11 | 121237187 | 121238048 | 26.37134 | 86.4321 | 83.16229 | Promoter | 11 | 121237253 | 121255856 | 18604 | 1 | ENSMUST00 | 0 | Narf | ENSMUSG00000000056.7 |
| chr11 | 97702515 | 97703210 | 26.37134 | 86.4321 | 83.16229 | Promoter | 11 | 97703399 | 97713500 | 10102 | 1 | ENSMUST00 | -189 | Psmb3 | ENSMUSG00000069744.7 |
| chr18 | 67244310 | 67246272 | 26.37134 | 86.4321 | 83.16229 | Promoter | 18 | 67225530 | 67245830 | 20301 | 2 | ENSMUST00 | 0 | Mppe1 | ENSMUSG00000062526.3 |
| chr7 | 55793661 | 55794907 | 26.37134 | 86.4321 | 83.16229 | Promoter | 7 | 55794148 | 55831447 | 37300 | 1 | ENSMUST00 | 0 | Tubgcp5 | ENSMUSG00000033790.10 |
| chr15 | 57911762 | 57913026 | 28.51938 | 86.42239 | 83.15316 | Promoter | 15 | 57912199 | 57970061 | 57863 | 1 | ENSMUST00 | 0 | Tbc1d31 | ENSMUSG00000022364.9 |
| chr6 | 128142694 | 128144365 | 28.51938 | 86.42239 | 83.15316 | Promoter | 6 | 127961396 | 128143563 | 182168 | 2 | ENSMUST00 | 0 | Tspan9 | ENSMUSG00000030352.9 |
| chr17 | 46673639 | 46674675 | 29.05538 | 86.39434 | 83.12526 | Promoter | 17 | 46667458 | 46674255 | 6798 | 2 | ENSMUST00 | 0 | Rrp36 | ENSMUSG00000023971.8 |
| chr3 | 144201264 | 144203047 | 28.46397 | 86.32589 | 83.05701 | Promoter | 3 | 144188530 | 144202317 | 13788 | 2 | ENSMUST00 | 0 | Lmo4 | ENSMUSG00000028266.12 |
| chr11 | 12464163 | 12465333 | 29.56773 | 86.32103 | 83.05225 | Promoter | 11 | 12236608 | 12464804 | 228197 | 2 | ENSMUST00 | 0 | Cobl | ENSMUSG00000020173.12 |
| chr7 | 127421244 | 127422700 | 27.91253 | 86.3188 | 83.05016 | Promoter | 7 | 127418967 | 127422027 | 3061 | 2 | ENSMUST00 | 0 | Zfp688 | ENSMUSG00000045251.8 |
| chr17 | 28223534 | 28224238 | 28.99699 | 86.29337 | 83.02477 | Exon (ENSMU | 17 | 28232754 | 28301469 | 68716 | 1 | ENSMUST00 | -8516 | Ppard | ENSMUSG00000002250.10 |
| chr2 | 131261792 | 131263701 | 29.50634 | 86.21544 | 82.94717 | Promoter | 2 | 131262495 | 131283437 | 20943 | 1 | ENSMUST00 | 0 | Pank2 | ENSMUSG00000037514.12 |
| chr11 | 103028472 | 103029571 | 28.93884 | 86.19278 | 82.92467 | Promoter | 11 | 103028585 | 103057484 | 28900 | 1 | ENSMUST00 | 0 | Nmt1 | ENSMUSG00000020936.8 |
| chr12 | 16999550 | 17000515 | 28.93884 | 86.19278 | 82.92467 | Promoter | 12 | 16988648 | 17000032 | 11385 | 2 | ENSMUST00 | 0 | Pqlc3 | ENSMUSG00000045679.8 |
| chr12 | 17174111 | 17177123 | 28.93884 | 86.19278 | 82.92467 | Promoter | 12 | 17172100 | 17176888 | 4789 | 2 | ENSMUST00 | 0 | Kcnf1 | ENSMUSG00000051726.6 |
| chr17 | 24849449 | 24850386 | 28.93884 | 86.19278 | 82.92467 | Promoter | 17 | 24848896 | 24850302 | 1407 | 2 | ENSMUST00 | 0 | Fahd1 | ENSMUSG00000045316.4 |
| chr17 | 32890799 | 32892159 | 28.93884 | 86.19278 | 82.92467 | Distal Interge | 17 | 32879228 | 32886083 | 6856 | 2 | ENSMUST00 | -4717 | Zfp870 | ENSMUSG00000095325.1 |
| chr19 | 42779292 | 42780311 | 28.93884 | 86.19278 | 82.92467 | Promoter | 19 | 42755105 | 42779978 | 24874 | 2 | ENSMUST00 | 0 | Hps1 | ENSMUSG00000025188.10 |
| chr2 | 45115710 | 45117667 | 28.93884 | 86.19278 | 82.92467 | Promoter | 2 | 44983699 | 45117395 | 133697 | 2 | ENSMUST00 | 0 | Zeb2 | ENSMUSG00000026872.11 |
| chr2 | 91255775 | 91257804 | 28.93884 | 86.19278 | 82.92467 | Promoter | 2 | 91255954 | 91261569 | 5616 | 1 | ENSMUST00 | 0 | Pacsin3 | ENSMUSG00000027257.8 |
| chr5 | 139343962 | 139345242 | 28.93884 | 86.19278 | 82.92467 | Promoter | 5 | 139336189 | 139345233 | 9045 | 2 | ENSMUST00 | 0 | Cox19 | ENSMUSG00000045438.7 |
| chr1 | 135765813 | 135767149 | 27.80811 | 86.13519 | 82.86789 | Promoter | 1 | 135766119 | 135769136 | 3018 | 1 | ENSMUST00 | 0 | Phlda3 | ENSMUSG00000041801.5 |
| chr7 | 127840892 | 127842835 | 28.35378 | 86.13386 | 82.86658 | Promoter | 7 | 127841764 | 127849019 | 7256 | 1 | ENSMUST00 | 0 | Stx4a | ENSMUSG00000030805.9 |
| chr17 | 86752839 | 86756072 | 28.88092 | 86.09254 | 82.82554 | Promoter | 17 | 86753907 | 86833410 | 79504 | 1 | ENSMUST00 | 0 | Epas1 | ENSMUSG00000024140.9 |
| chr1 | 172562102 | 172563770 | 21.47392 | 86.08567 | 82.81886 | 3' UTR | 1 | 172559906 | 172563716 | 3811 | 1 | ENSMUST00 | 2197 | Vsig8 | ENSMUSG00000049598.4 |
| chr14 | 26969342 | 26971279 | 21.47392 | 86.08567 | 82.81886 | Promoter | 14 | 26918988 | 26971232 | 52245 | 2 | ENSMUST00 | 0 | Appl1 | ENSMUSG00000040760.5 |
| chr16 | 17921338 | 17922831 | 21.47392 | 86.08567 | 82.81886 | Downstream | 16 | 17925682 | 17927343 | 1662 | 2 | ENSMUST00 | 4512 | Slc25a1 | ENSMUSG00000003528.9 |
| chr16 | 17404953 | 17406958 | 21.47392 | 86.08567 | 82.81886 | Promoter | 16 | 17406003 | 17430826 | 24824 | 1 | ENSMUST00 | 0 | Snap29 | ENSMUSG00000022765.8 |
| chr16 | 94312366 | 94314005 | 21.47392 | 86.08567 | 82.81886 | Promoter | 16 | 94129195 | 94313563 | 184369 | 2 | ENSMUST00 | 0 | Hlcs | ENSMUSG00000040820.10 |
| chr17 | 44187799 | 44192338 | 21.47392 | 86.08567 | 82.81886 | Promoter | 17 | 44188572 | 44280168 | 91597 | 1 | ENSMUST00 | 0 | Clic5 | ENSMUSG00000023959.8 |
| chr3 | 99140544 | 99141906 | 27.7562 | 86.04385 | 82.77744 | Promoter | 3 | 99141068 | 99220203 | 79136 | 1 | ENSMUST00 | 0 | Wars2 | ENSMUSG00000004233.9 |
| chr5 | 100518379 | 100519423 | 28.299 | 86.03832 | 82.77194 | Promoter | 5 | 100518443 | 100532095 | 13653 | 1 | ENSMUST00 | 0 | Cops4 | ENSMUSG00000035297.9 |
| chr12 | 24498062 | 24499146 | 29.38432 | 86.00549 | 82.73921 | Promoter | 12 | 24498581 | 24558571 | 59991 | 1 | ENSMUST00 | 0 | Taf1b | ENSMUSG00000059669.7 |
| chr10 | 128321898 | 128323638 | 28.24444 | 85.94312 | 82.67722 | Promoter | 10 | 128322459 | 128327187 | 4729 | 1 | ENSMUST00 | 0 | Cnpy2 | ENSMUSG00000025381.2 |
| chr12 | 108757506 | 108758117 | 28.24444 | 85.94312 | 82.67722 | Distal Interge | 12 | 108792973 | 108816632 | 23660 | 1 | ENSMUST00 | -34856 | Yy1 | ENSMUSG00000021264.11 |
| chr15 | 102997422 | 103001867 | 28.24444 | 85.94312 | 82.67722 | Promoter | 15 | 102998257 | 103010768 | 12512 | 1 | ENSMUST00 | 0 | Hoxc6 | ENSMUSG00000001661.5 |
| chr15 | 35295490 | 35297547 | 28.24444 | 85.94312 | 82.67722 | Promoter | 15 | 35296098 | 35303304 | 7207 | 1 | ENSMUST00 | 0 | Osr2 | ENSMUSG00000022330.4 |
| chr16 | 57548939 | 57549947 | 28.24444 | 85.94312 | 82.67722 | Promoter | 16 | 57549242 | 57572804 | 23563 | 1 | ENSMUST00 | 0 | Filip1l | ENSMUSG00000043336.9 |
| chr19 | 3912160 | 3912841 | 28.24444 | 85.94312 | 82.67722 | Promoter | 19 | 3908870 | 3912717 | 3848 | 2 | ENSMUST00 | 0 | Ndufs8 | ENSMUSG00000059734.6 |
| chr2 | 68470868 | 68472728 | 28.24444 | 85.94312 | 82.67722 | Promoter | 2 | 68210445 | 68472268 | 261824 | 2 | ENSMUST00 | 0 | Stk39 | ENSMUSG00000027030.10 |
| chr5 | 33721432 | 33723845 | 28.24444 | 85.94312 | 82.67722 | Promoter | 5 | 33721724 | 33737067 | 15344 | 1 | ENSMUST00 | 0 | Fgfr3 | ENSMUSG00000054252.12 |
| chr7 | 141539056 | 141540467 | 28.24444 | 85.94312 | 82.67722 | Promoter | 7 | 141493136 | 141539816 | 46681 | 2 | ENSMUST00 | 0 | Chid1 | ENSMUSG00000025512.9 |
| chr10 | 84439467 | 84441707 | 23.61999 | 85.9385 | 82.67356 | Promoter | 10 | 84370905 | 84440597 | 69693 | 2 | ENSMUST00 | 0 | Nuak1 | ENSMUSG00000020032.9 |
| chr11 | 83472283 | 83474074 | 23.61999 | 85.9385 | 82.67356 | Promoter | 11 | 83473086 | 83489213 | 16128 | 1 | ENSMUST00 | 0 | Taf15 | ENSMUSG00000020680.10 |
| chr12 | 108334256 | 108334858 | 23.61999 | 85.9385 | 82.67356 | Promoter | 12 | 108334377 | 108362231 | 27855 | 1 | ENSMUST00 | 0 | Cyp46a1 | ENSMUSG00000021259.4 |
| chr14 | 118088592 | 118090263 | 23.61999 | 85.9385 | 82.67356 | Distal Interge | 14 | 118107831 | 118117100 | 9270 | 1 | ENSMUST00 | -17568 | Gm26773 | ENSMUSG00000097358.1 |
| chr16 | 17208423 | 17210710 | 23.61999 | 85.9385 | 82.67356 | Promoter | 16 | 17208135 | 17213921 | 5787 | 1 | ENSMUST00 | 289 | Rimbp3 | ENSMUSG00000071636.6 |
| chr17 | 26138184 | 26139766 | 23.61999 | 85.9385 | 82.67356 | Promoter | 17 | 26138688 | 26195811 | 57124 | 1 | ENSMUST00 | 0 | Axin1 | ENSMUSG00000024182.11 |
| chr19 | 38223425 | 38224996 | 23.61999 | 85.9385 | 82.67356 | Promoter | 19 | 38188481 | 38224138 | 35658 | 2 | ENSMUST00 | 0 | Fra10ac1 | ENSMUSG00000054237.5 |
| chr2 | 152669171 | 152670751 | 23.61999 | 85.9385 | 82.67356 | Promoter | 2 | 152669461 | 152708670 | 39210 | 1 | ENSMUST00 | 0 | H13 | ENSMUSG00000019188.11 |
| chr3 | 88324806 | 88325916 | 23.61999 | 85.9385 | 82.67356 | Promoter | 3 | 88325023 | 88331313 | 6291 | 1 | ENSMUST00 | 0 | 0610031J06F | ENSMUSG00000001418.8 |
| chr3 | 28780885 | 28782289 | 23.61999 | 85.9385 | 82.67356 | Promoter | 3 | 28781311 | 28798846 | 17536 | 1 | ENSMUST00 | 0 | Eif5a2 | ENSMUSG00000050192.7 |
| chr3 | 96499421 | 96503414 | 23.61999 | 85.9385 | 82.67356 | Distal Interge | 3 | 96488729 | 96488893 | 165 | 1 | ENSMUST00 | 10693 | Gm22614 | ENSMUSG00000094812.1 |
| chr5 | 24444466 | 24445612 | 23.61999 | 85.9385 | 82.67356 | Promoter | 5 | 24438440 | 24445194 | 6755 | 2 | ENSMUST00 | 0 | Fastk | ENSMUSG00000028959.9 |
| chr6 | 6216289 | 6217432 | 23.61999 | 85.9385 | 82.67356 | Promoter | 6 | 6041218 | 6217118 | 175901 | 2 | ENSMUST00 | 0 | Slc25a13 | ENSMUSG00000015112.10 |
| chr8 | 22397785 | 22399225 | 23.61999 | 85.9385 | 82.67356 | Promoter | 8 | 22375554 | 22398621 | 23068 | 2 | ENSMUST00 | 0 | Slc25a15 | ENSMUSG00000031482.8 |
| chr9 | 110244859 | 110246631 | 23.61999 | 85.9385 | 82.67356 | 5' UTR | 9 | 110243783 | 110262575 | 18793 | 1 | ENSMUST00 | 1077 | Cspg5 | ENSMUSG00000032482.5 |
| chr11 | 99385348 | 99386841 | 28.76577 | 85.89317 | 82.63004 | Promoter | 11 | 99387340 | 99387897 | 558 | 1 | ENSMUST00 | -499 | Gm11940 | ENSMUSG00000087692.1 |
| chr13 | 100729796 | 100731298 | 29.2633 | 85.79715 | 82.53433 | Promoter | 13 | 100697027 | 100730942 | 33916 | 2 | ENSMUST00 | 0 | Cdk7 | ENSMUSG00000069089.7 |
| chr14 | 18271132 | 18271928 | 28.70854 | 85.79402 | 82.53136 | Promoter | 14 | 18269304 | 18271391 | 2088 | 2 | ENSMUST00 | 0 | Rpl15 | ENSMUSG00000012405.10 |
| chr19 | 8879994 | 8881359 | 28.70854 | 85.79402 | 82.53136 | Promoter | 19 | 8880016 | 8880933 | 918 | 2 | ENSMUST00 | 0 | AI462493 | ENSMUSG00000071654.5 |
| chr5 | 138186452 | 138187933 | 24.99138 | 85.77554 | 82.51305 | Promoter | 5 | 138187485 | 138193918 | 6434 | 1 | ENSMUST00 | 0 | Cnpy4 | ENSMUSG00000036968.10 |
| chr16 | 55894677 | 55895890 | 27.6016 | 85.77155 | 82.50918 | Promoter | 16 | 55839953 | 55895279 | 55327 | 2 | ENSMUST00 | 0 | Nxpe3 | ENSMUSG00000075033.3 |
| chr17 | 47139299 | 47142406 | 27.6016 | 85.77155 | 82.50918 | Promoter | 17 | 47140875 | 47329536 | 188662 | 1 | ENSMUST00 | 0 | Trerf1 | ENSMUSG00000064043.8 |
| chr2 | 163419689 | 163420757 | 27.6016 | 85.77155 | 82.50918 | Promoter | 2 | 163419795 | 163420361 | 567 | 1 | ENSMUST00 | 0 | RP23-117O1 | ENSMUSG00000099632.1 |
| chr3 | 27182540 | 27184235 | 27.6016 | 85.77155 | 82.50918 | Promoter | 3 | 27182965 | 27244958 | 61994 | 1 | ENSMUST00 | 0 | Nceh1 | ENSMUSG00000027698.9 |
| chr5 | 130002290 | 130003250 | 27.6016 | 85.77155 | 82.50918 | Promoter | 5 | 129989011 | 130003049 | 14039 | 2 | ENSMUST00 | 0 | Gusb | ENSMUSG00000025534.12 |
| chr7 | 99857968 | 99859477 | 27.6016 | 85.77155 | 82.50918 | Promoter | 7 | 99859218 | 99917824 | 58607 | 1 | ENSMUST00 | 0 | Xrra1 | ENSMUSG00000035211.8 |
| chr9 | 75313818 | 75314989 | 27.6016 | 85.77155 | 82.50918 | Exon (ENSMU | 9 | 75311395 | 75344964 | 33570 | 1 | ENSMUST00 | 2424 | Gnb5 | ENSMUSG00000032192.8 |
| chr9 | 97017209 | 97019396 | 27.6016 | 85.77155 | 82.50918 | Promoter | 9 | 96943482 | 97018843 | 75362 | 2 | ENSMUST00 | 0 | Spsb4 | ENSMUSG00000046997.5 |
| chr12 | 3307840 | 3310000 | 20.49778 | 85.71664 | 82.45538 | Promoter | 12 | 3247430 | 3309969 | 62540 | 2 | ENSMUST00 | 0 | Rab10 | ENSMUSG00000020671.8 |
| chr14 | 32513234 | 32514216 | 20.49778 | 85.71664 | 82.45538 | Promoter | 14 | 32513521 | 32580989 | 67469 | 1 | ENSMUST00 | 0 | Ercc6 | ENSMUSG00000054051.6 |
| chr6 | 116192457 | 116194100 | 20.49778 | 85.71664 | 82.45538 | Promoter | 6 | 116018618 | 116193486 | 174869 | 2 | ENSMUST00 | 0 | Tmcc1 | ENSMUSG00000030126.12 |

| | | | | | | | | | | | | | | | |
|---|---|---|---|---|---|---|---|---|---|---|---|---|---|---|---|
| chr1 | 36067902 | 36069968 | 29.20317 | 85.69358 | 82.43262 | Promoter | 1 | 36068400 | 36106446 | 38047 | 1 | ENSMUST00 | 0 | Hs6st1 | ENSMUSG00000045216.6 |
| chr11 | 40733557 | 40734781 | 27.0048 | 85.66756 | 82.40694 | Promoter | 11 | 40733667 | 40740046 | 6380 | 1 | ENSMUST00 | 0 | Nudcd2 | ENSMUSG00000020328.5 |
| chr12 | 83987286 | 83988883 | 27.0048 | 85.66756 | 82.40694 | Promoter | 12 | 83987861 | 83993873 | 6013 | 1 | ENSMUST00 | 0 | Acot2 | ENSMUSG00000021226.7 |
| chr7 | 29815327 | 29816905 | 27.0048 | 85.66756 | 82.40694 | Promoter | 7 | 29816072 | 29832042 | 15971 | 1 | ENSMUST00 | 0 | Zfp790 | ENSMUSG00000011427.10 |
| chr11 | 24078881 | 24082282 | 28.08037 | 85.65662 | 82.39679 | Promoter | 11 | 24080670 | 24173558 | 92889 | 1 | ENSMUST00 | 0 | Bcl11a | ENSMUSG00000000861.10 |
| chr11 | 5443230 | 5445570 | 28.08037 | 85.65662 | 82.39679 | Promoter | 11 | 5276324 | 5444847 | 168524 | 2 | ENSMUST00 | 0 | Znrf3 | ENSMUSG00000041961.9 |
| chr15 | 81360196 | 81361269 | 28.08037 | 85.65662 | 82.39679 | Promoter | 15 | 81318921 | 81360765 | 41845 | 2 | ENSMUST00 | 0 | Slc25a17 | ENSMUSG00000022404.7 |
| chr16 | 94525785 | 94527653 | 28.08037 | 85.65662 | 82.39679 | Promoter | 16 | 94497783 | 94526830 | 29048 | 2 | ENSMUST00 | 0 | Dscr3 | ENSMUSG00000022898.7 |
| chr16 | 62853976 | 62855300 | 28.08037 | 85.65662 | 82.39679 | Promoter | 16 | 62854307 | 62929346 | 75040 | 1 | ENSMUST00 | 0 | Pros1 | ENSMUSG00000022912.8 |
| chr17 | 21008449 | 21009772 | 28.08037 | 85.65662 | 82.39679 | Promoter | 17 | 21008941 | 21028852 | 19912 | 1 | ENSMUST00 | 0 | Zfp160 | ENSMUSG00000067942.5 |
| chr18 | 53417224 | 53418581 | 28.08037 | 85.65662 | 82.39679 | Promoter | 18 | 53406341 | 53418007 | 11667 | 2 | ENSMUST00 | 0 | Ppic | ENSMUSG00000024538.7 |
| chr19 | 7056278 | 7057898 | 28.08037 | 85.65662 | 82.39679 | Promoter | 19 | 7056768 | 7198061 | 141294 | 1 | ENSMUST00 | 0 | Macrod1 | ENSMUSG00000036278.6 |
| chr2 | 129296675 | 129298001 | 28.08037 | 85.65662 | 82.39679 | Promoter | 2 | 129268210 | 129297212 | 29003 | 2 | ENSMUST00 | 0 | Ckap2l | ENSMUSG00000048327.6 |
| chr3 | 96261973 | 96263403 | 28.08037 | 85.65662 | 82.39679 | Promoter | 3 | 96261682 | 96263311 | 1630 | 2 | ENSMUST00 | 0 | Hist2h4 | ENSMUSG00000091405.2 |
| chr3 | 36474815 | 36475713 | 28.08037 | 85.65662 | 82.39679 | Promoter | 3 | 36475461 | 36475524 | 64 | 2 | ENSMUST00 | 0 | Mir7009 | ENSMUSG00000098808.1 |
| chr5 | 36529746 | 36530852 | 28.08037 | 85.65662 | 82.39679 | Promoter | 5 | 36490605 | 36530623 | 40019 | 2 | ENSMUST00 | 0 | Tbc1d14 | ENSMUSG00000029192.12 |
| chr5 | 123747425 | 123749472 | 28.08037 | 85.65662 | 82.39679 | Promoter | 5 | 123728426 | 123749389 | 20964 | 2 | ENSMUST00 | 0 | Rsrc2 | ENSMUSG00000029422.9 |
| chr6 | 142345281 | 142346714 | 28.08037 | 85.65662 | 82.39679 | Promoter | 6 | 142345654 | 142363257 | 17604 | 1 | ENSMUST00 | 0 | Pyroxd1 | ENSMUSG00000041671.8 |
| chr7 | 25394610 | 25396560 | 28.08037 | 85.65662 | 82.39679 | Promoter | 7 | 25379527 | 25395987 | 16461 | 2 | ENSMUST00 | 0 | Lipe | ENSMUSG00000003123.10 |
| chr7 | 3628894 | 3630671 | 28.08037 | 85.65662 | 82.39679 | Promoter | 7 | 3620324 | 3629884 | 9561 | 2 | ENSMUST00 | 0 | Tfpt | ENSMUSG00000006335.11 |
| chr7 | 19181038 | 19182936 | 28.08037 | 85.65662 | 82.39679 | Promoter | 7 | 19181159 | 19206482 | 25324 | 1 | ENSMUST00 | 0 | Eml2 | ENSMUSG00000040811.10 |
| chr7 | 29853057 | 29853986 | 28.08037 | 85.65662 | 82.39679 | Promoter | 7 | 29833620 | 29853669 | 20050 | 2 | ENSMUST00 | 0 | Zfp940 | ENSMUSG00000050855.11 |
| chr8 | 68880055 | 68883509 | 28.08037 | 85.65662 | 82.39679 | Promoter | 8 | 68880491 | 68906934 | 26444 | 1 | ENSMUST00 | 0 | Lpl | ENSMUSG00000015568.10 |
| chr17 | 68001890 | 68004635 | 25.93082 | 85.62936 | 82.37304 | Promoter | 17 | 67842713 | 68004120 | 161408 | 2 | ENSMUST00 | 0 | Arhgap28 | ENSMUSG00000024043.8 |
| chr1 | 134962060 | 134963089 | 26.44925 | 85.62254 | 82.36654 | Promoter | 1 | 134962565 | 134974162 | 11598 | 1 | ENSMUST00 | 0 | Ube2t | ENSMUSG00000026429.8 |
| chr4 | 21879202 | 21880393 | 26.44925 | 85.62254 | 82.36654 | Promoter | 4 | 21879673 | 21912162 | 32490 | 1 | ENSMUST00 | 0 | Coq3 | ENSMUSG00000028247.2 |
| chr9 | 51278067 | 51279570 | 26.44925 | 85.62254 | 82.36654 | Promoter | 9 | 51278513 | 51284210 | 5698 | 1 | ENSMUST00 | 0 | 2010007H06 | ENSMUSG00000044694.2 |
| chr10 | 62338804 | 62341149 | 28.59476 | 85.59679 | 82.34128 | Promoter | 10 | 62268855 | 62340542 | 71688 | 2 | ENSMUST00 | 0 | Hk1 | ENSMUSG00000037012.13 |
| chr6 | 49367191 | 49368630 | 28.59476 | 85.59679 | 82.34128 | Promoter | 6 | 49367739 | 49389905 | 22167 | 1 | ENSMUST00 | 0 | Fam221a | ENSMUSG00000047115.10 |
| chr4 | 136205955 | 136207125 | 29.14328 | 85.59041 | 82.33509 | Promoter | 4 | 136206365 | 136245216 | 38852 | 1 | ENSMUST00 | 0 | Asap3 | ENSMUSG00000036995.7 |
| chr11 | 53858798 | 53859561 | 29.08364 | 85.48761 | 82.2328 | Promoter | 11 | 53784291 | 53859144 | 74854 | 2 | ENSMUST00 | 0 | Gm12216 | ENSMUSG00000081769.3 |
| chr17 | 25860890 | 25862012 | 29.08364 | 85.48761 | 82.2328 | Promoter | 17 | 25844771 | 25861383 | 16613 | 2 | ENSMUST00 | 0 | Wdr90 | ENSMUSG00000073434.6 |
| chr2 | 178413634 | 178415302 | 29.08364 | 85.48761 | 82.2328 | Promoter | 2 | 178411206 | 178414472 | 3267 | 2 | ENSMUST00 | 0 | Ppp1r3d | ENSMUSG00000049999.4 |
| chr4 | 124751028 | 124752650 | 29.08364 | 85.48761 | 82.2328 | Promoter | 4 | 124735730 | 124751354 | 15625 | 2 | ENSMUST00 | 0 | Gm12915 | ENSMUSG00000086589.1 |
| chr6 | 117878860 | 117880769 | 29.08364 | 85.48761 | 82.2328 | Distal Interge | 6 | 117863084 | 117870734 | 7651 | 1 | ENSMUST00 | 15777 | Zfp239 | ENSMUSG00000042097.12 |
| chr5 | 31494271 | 31495602 | 27.97474 | 85.47195 | 82.2178 | Promoter | 5 | 31494741 | 31512903 | 18163 | 1 | ENSMUST00 | 0 | Gpn1 | ENSMUSG00000064037.8 |
| chr1 | 172632236 | 172633275 | 24.82645 | 85.46523 | 82.21117 | Promoter | 1 | 172630769 | 172632974 | 2206 | 2 | ENSMUST00 | 0 | Dusp23 | ENSMUSG00000026544.5 |
| chr1 | 180777278 | 180778774 | 24.82645 | 85.46523 | 82.21117 | Distal Interge | 1 | 180802568 | 180809483 | 6916 | 1 | ENSMUST00 | -23794 | Gm17275 | ENSMUSG00000097347.1 |
| chr11 | 119055633 | 119057493 | 24.82645 | 85.46523 | 82.21117 | Distal Interge | 11 | 119036305 | 119040969 | 4665 | 2 | ENSMUST00 | -14665 | Cbx8 | ENSMUSG00000025578.9 |
| chr11 | 116842041 | 116843458 | 24.82645 | 85.46523 | 82.21117 | Promoter | 11 | 116837432 | 116843281 | 5850 | 2 | ENSMUST00 | 0 | Jmjd6 | ENSMUSG00000056962.6 |
| chr13 | 67499778 | 67500818 | 24.82645 | 85.46523 | 82.21117 | Promoter | 13 | 67490167 | 67500503 | 10337 | 2 | ENSMUST00 | 0 | Zfp58 | ENSMUSG00000071291.5 |
| chr15 | 103272831 | 103273692 | 24.82645 | 85.46523 | 82.21117 | Promoter | 15 | 103272911 | 103299862 | 26952 | 1 | ENSMUST00 | 0 | Copz1 | ENSMUSG00000060992.8 |
| chr18 | 12643179 | 12644524 | 24.82645 | 85.46523 | 82.21117 | Promoter | 18 | 12643344 | 12689445 | 46102 | 1 | ENSMUST00 | 0 | Ttc39c | ENSMUSG00000024424.9 |
| chr19 | 8712136 | 8714441 | 24.82645 | 85.46523 | 82.21117 | Promoter | 19 | 8706882 | 8713897 | 7016 | 2 | ENSMUST00 | 0 | Slc3a2 | ENSMUSG00000010095.8 |
| chr19 | 14598489 | 14599256 | 24.82645 | 85.46523 | 82.21117 | Promoter | 19 | 14448150 | 14598051 | 149902 | 2 | ENSMUST00 | -439 | Tle4 | ENSMUSG00000024642.11 |
| chr19 | 24173413 | 24175288 | 24.82645 | 85.46523 | 82.21117 | Intron (ENSM | 19 | 24094523 | 24225026 | 130504 | 2 | ENSMUST00 | 49738 | Tjp2 | ENSMUSG00000024812.9 |
| chr2 | 122028561 | 122029777 | 24.82645 | 85.46523 | 82.21117 | Promoter | 2 | 122028652 | 122029328 | 677 | 1 | ENSMUST00 | 0 | Eif3j1 | ENSMUSG00000027236.8 |
| chr2 | 155825684 | 155827241 | 24.82645 | 85.46523 | 82.21117 | Promoter | 2 | 155819832 | 155826925 | 7094 | 2 | ENSMUST00 | 0 | Eif6 | ENSMUSG00000027613.10 |
| chr2 | 150607893 | 150609197 | 24.82645 | 85.46523 | 82.21117 | Promoter | 2 | 150583080 | 150608567 | 25488 | 2 | ENSMUST00 | 0 | Apmap | ENSMUSG00000033096.7 |
| chr2 | 122348388 | 122349553 | 24.82645 | 85.46523 | 82.21117 | 3' UTR | 2 | 122349504 | 122354131 | 4628 | 2 | ENSMUST00 | 4578 | Shf | ENSMUSG00000033256.9 |
| chr3 | 86919574 | 86921174 | 24.82645 | 85.46523 | 82.21117 | Promoter | 3 | 86786150 | 86920884 | 134735 | 2 | ENSMUST00 | 0 | Dclk2 | ENSMUSG00000028078.9 |
| chr3 | 85574092 | 85574815 | 24.82645 | 85.46523 | 82.21117 | Promoter | 3 | 85574119 | 85646579 | 72461 | 1 | ENSMUST00 | 0 | Pet112 | ENSMUSG00000028085.7 |
| chr4 | 143160943 | 143162268 | 24.82645 | 85.46523 | 82.21117 | Exon (ENSMU | 4 | 143166603 | 143167396 | 794 | 1 | ENSMUST00 | -4335 | Gm13039 | ENSMUSG00000083826.1 |
| chr5 | 148550262 | 148553141 | 24.82645 | 85.46523 | 82.21117 | Promoter | 5 | 148504631 | 148552788 | 48158 | 2 | ENSMUST00 | 0 | Ubl3 | ENSMUSG00000001687.10 |
| chr5 | 129715087 | 129716184 | 24.82645 | 85.46523 | 82.21117 | Promoter | 5 | 129715497 | 129718694 | 3198 | 1 | ENSMUST00 | 0 | Mrps17 | ENSMUSG00000034211.9 |
| chr6 | 90715816 | 90716965 | 24.82645 | 85.46523 | 82.21117 | Promoter | 6 | 90659598 | 90716529 | 56932 | 2 | ENSMUST00 | 0 | Iqsec1 | ENSMUSG00000034312.8 |
| chr8 | 104508986 | 104510025 | 24.82645 | 85.46523 | 82.21117 | Promoter | 8 | 104446719 | 104509887 | 63169 | 2 | ENSMUST00 | 0 | Ccdc79 | ENSMUSG00000052616.5 |
| chr8 | 69653962 | 69655366 | 24.82645 | 85.46523 | 82.21117 | Distal Interge | 8 | 69659263 | 69664170 | 4908 | 1 | ENSMUST00 | -3897 | Zfp964 | ENSMUSG00000091764.1 |
| chr9 | 64016675 | 64022981 | 24.82645 | 85.46523 | 82.21117 | Promoter | 9 | 63953076 | 64022059 | 68984 | 2 | ENSMUST00 | 0 | Smad6 | ENSMUSG00000036867.7 |
| chr10 | 84533060 | 84534572 | 29.02424 | 85.38521 | 82.13421 | Promoter | 10 | 84526305 | 84533888 | 7584 | 2 | ENSMUST00 | 0 | Ckap4 | ENSMUSG00000046841.4 |
| chr1 | 178317405 | 178320432 | 27.92142 | 85.37866 | 82.12785 | Promoter | 1 | 178319130 | 178322335 | 3206 | 1 | ENSMUST00 | 0 | Cox20 | ENSMUSG00000026500.6 |
| chr1 | 38626226 | 38628260 | 26.28407 | 85.32412 | 82.07336 | Promoter | 1 | 38177326 | 38627185 | 449860 | 2 | ENSMUST00 | 0 | Aff3 | ENSMUSG00000037138.11 |
| chr10 | 94198282 | 94199942 | 26.28407 | 85.32412 | 82.07336 | Promoter | 10 | 94198955 | 94221443 | 22489 | 1 | ENSMUST00 | 0 | Ndufa12 | ENSMUSG00000020022.11 |
| chr11 | 115186754 | 115187994 | 26.28407 | 85.32412 | 82.07336 | Promoter | 11 | 115182832 | 115187332 | 4501 | 2 | ENSMUST00 | 0 | Nat9 | ENSMUSG00000015542.12 |
| chr11 | 69366750 | 69369724 | 26.28407 | 85.32412 | 82.07336 | Promoter | 11 | 69343273 | 69369406 | 26134 | 2 | ENSMUST00 | 0 | Chd3 | ENSMUSG00000018474.12 |
| chr11 | 29246191 | 29247874 | 26.28407 | 85.32412 | 82.07336 | Promoter | 11 | 29221532 | 29247409 | 25878 | 2 | ENSMUST00 | 0 | Ccdc104 | ENSMUSG00000020462.9 |
| chr11 | 59808878 | 59810986 | 26.28407 | 85.32412 | 82.07336 | Promoter | 11 | 59791408 | 59810016 | 18609 | 2 | ENSMUST00 | 0 | Flcn | ENSMUSG00000032633.7 |
| chr11 | 58847077 | 58848128 | 26.28407 | 85.32412 | 82.07336 | Promoter | 11 | 58847158 | 58861956 | 14799 | 1 | ENSMUST00 | 0 | Gm12258 | ENSMUSG00000072915.3 |
| chr12 | 111484468 | 111487284 | 26.28407 | 85.32412 | 82.07336 | Promoter | 12 | 111484611 | 111485823 | 1213 | 2 | ENSMUST00 | 0 | Gm266 | ENSMUSG00000010529.5 |
| chr12 | 24572192 | 24573771 | 26.28407 | 85.32412 | 82.07336 | Promoter | 12 | 24572287 | 24617387 | 45101 | 1 | ENSMUST00 | 0 | Grhl1 | ENSMUSG00000020656.10 |
| chr12 | 108554275 | 108555764 | 26.28407 | 85.32412 | 82.07336 | Promoter | 12 | 108554720 | 108688513 | 133794 | 1 | ENSMUST00 | 0 | Evl | ENSMUSG00000021262.9 |
| chr15 | 89354959 | 89356871 | 26.28407 | 85.32412 | 82.07336 | Promoter | 15 | 89355719 | 89372826 | 17108 | 1 | ENSMUST00 | 0 | Ncaph2 | ENSMUSG00000008690.10 |
| chr16 | 16146226 | 16147296 | 26.28407 | 85.32412 | 82.07336 | Promoter | 16 | 15889227 | 16146851 | 257625 | 2 | ENSMUST00 | 0 | Spidr | ENSMUSG00000041974.9 |
| chr17 | 23676112 | 23677653 | 26.28407 | 85.32412 | 82.07336 | Promoter | 17 | 23675447 | 23677449 | 2003 | 2 | ENSMUST00 | 0 | Tnfrsf12a | ENSMUSG00000023905.9 |
| chr2 | 150667485 | 150668773 | 26.28407 | 85.32412 | 82.07336 | Promoter | 2 | 150618105 | 150668258 | 50154 | 2 | ENSMUST00 | 0 | Acss1 | ENSMUSG00000027452.6 |
| chr3 | 32509946 | 32511591 | 26.28407 | 85.32412 | 82.07336 | Distal Interge | 3 | 32515295 | 32520685 | 5391 | 1 | ENSMUST00 | -3704 | Zfp639 | ENSMUSG00000027667.8 |
| chr3 | 94837352 | 94839024 | 26.28407 | 85.32412 | 82.07336 | Promoter | 3 | 94837567 | 94882326 | 44760 | 1 | ENSMUST00 | 0 | Pogz | ENSMUSG00000038902.9 |
| chr4 | 83485372 | 83487207 | 26.28407 | 85.32412 | 82.07336 | Promoter | 4 | 83455680 | 83486459 | 30780 | 2 | ENSMUST00 | 0 | Psip1 | ENSMUSG00000028484.11 |
| chr4 | 135986291 | 135987789 | 26.28407 | 85.32412 | 82.07336 | Promoter | 4 | 135975460 | 135987264 | 11805 | 2 | ENSMUST00 | 0 | Pithd1 | ENSMUSG00000028669.10 |
| chr4 | 130107031 | 130108407 | 26.28407 | 85.32412 | 82.07336 | Promoter | 4 | 130107556 | 130128134 | 20579 | 1 | ENSMUST00 | 0 | Pef1 | ENSMUSG00000028779.11 |
| chr4 | 133276832 | 133278394 | 26.28407 | 85.32412 | 82.07336 | Promoter | 4 | 133266045 | 133277792 | 11748 | 2 | ENSMUST00 | 0 | Tmem222 | ENSMUSG00000028857.11 |
| chr4 | 133240506 | 133242188 | 26.28407 | 85.32412 | 82.07336 | Promoter | 4 | 133240818 | 133252929 | 12112 | 1 | ENSMUST00 | 0 | Map3k6 | ENSMUSG00000028862.6 |
| chr4 | 154141650 | 154143092 | 26.28407 | 85.32412 | 82.07336 | Promoter | 4 | 154142372 | 154156816 | 14445 | 1 | ENSMUST00 | 0 | Wrap73 | ENSMUSG00000029029.9 |
| chr4 | 107434005 | 107435717 | 26.28407 | 85.32412 | 82.07336 | Promoter | 4 | 107434591 | 107436036 | 1446 | 1 | ENSMUST00 | 0 | Glis1 | ENSMUSG00000034762.4 |
| chr5 | 137745433 | 137747808 | 26.28407 | 85.32412 | 82.07336 | Promoter | 5 | 137745730 | 137768450 | 22721 | 1 | ENSMUST00 | 0 | Tsc22d4 | ENSMUSG00000029723.11 |
| chr6 | 72354790 | 72356034 | 26.28407 | 85.32412 | 82.07336 | Promoter | 6 | 72355447 | 72359762 | 4316 | 1 | ENSMUST00 | 0 | Tmem150a | ENSMUSG00000055912.3 |
| chr7 | 110626970 | 110629632 | 26.28407 | 85.32412 | 82.07336 | Promoter | 7 | 110627661 | 110629820 | 2160 | 1 | ENSMUST00 | 0 | Adm | ENSMUSG00000030790.9 |
| chr8 | 110919321 | 110921240 | 26.28407 | 85.32412 | 82.07336 | Promoter | 8 | 110919922 | 110969823 | 49902 | 1 | ENSMUST00 | 0 | St3gal2 | ENSMUSG00000031749.7 |
| chr9 | 66511174 | 66512787 | 26.28407 | 85.32412 | 82.07336 | 3' UTR | 9 | 66508459 | 66514593 | 6135 | 2 | ENSMUST00 | 1806 | Fbxl22 | ENSMUSG00000050503.8 |
| chr5 | 118559542 | 118561980 | 28.96508 | 85.28318 | 82.03651 | Promoter | 5 | 118560719 | 118765435 | 204717 | 1 | ENSMUST00 | 0 | Med13l | ENSMUSG00000018076.9 |
| chr6 | 47768143 | 47768503 | 9.28662 | 85.25 | 82.00354 | Distal Interge | 6 | 47773329 | 47773429 | 101 | 1 | ENSMUST00 | -4826 | Gm24975 | ENSMUSG00000094807.1 |
| chr11 | 6273729 | 6275465 | 28.36989 | 85.2065 | 81.96005 | Promoter | 11 | 6270369 | 6274870 | 4502 | 2 | ENSMUST00 | 0 | Tmed4 | ENSMUSG00000004394.7 |
| chr13 | 21505925 | 21506836 | 28.36989 | 85.2065 | 81.96005 | Promoter | 13 | 21504382 | 21505056 | 675 | 2 | ENSMUST00 | -870 | Zfp389 | ENSMUSG00000082396.3 |
| chr14 | 122533621 | 122534902 | 28.36989 | 85.2065 | 81.96005 | Promoter | 14 | 122534324 | 122582977 | 48654 | 1 | ENSMUST00 | 0 | Pcca | ENSMUSG00000041650.10 |
| chr4 | 102568208 | 102575841 | 28.36989 | 85.2065 | 81.96005 | Promoter | 4 | 102570095 | 102605847 | 35753 | 1 | ENSMUST00 | 0 | Pde4b | ENSMUSG00000028525.11 |
| chr6 | 118561639 | 118562832 | 28.36989 | 85.2065 | 81.96005 | Promoter | 6 | 118501308 | 118562226 | 60919 | 2 | ENSMUST00 | 0 | Ankrd26 | ENSMUSG00000007827.10 |
| chr6 | 83440905 | 83443443 | 28.36989 | 85.2065 | 81.96005 | Promoter | 6 | 83362373 | 83441675 | 79303 | 2 | ENSMUST00 | 0 | Tet3 | ENSMUSG00000034832.10 |
| chr7 | 45017877 | 45019667 | 28.36989 | 85.2065 | 81.96005 | Promoter | 7 | 45017978 | 45021646 | 3669 | 1 | ENSMUST00 | 0 | Rras | ENSMUSG00000038387.7 |
| chr9 | 107399489 | 107400750 | 28.36989 | 85.2065 | 81.96005 | Promoter | 9 | 107399612 | 107529346 | 129735 | 1 | ENSMUST00 | 0 | Cacna2d2 | ENSMUSG00000010066.10 |
| chr1 | 161070832 | 161071893 | 27.81538 | 85.193 | 81.94753 | Promoter | 1 | 161070767 | 161086724 | 15958 | 1 | ENSMUST00 | 66 | Cenpl | ENSMUSG00000026708.11 |
| chr5 | 64159212 | 64161446 | 28.90617 | 85.18155 | 81.93615 | Promoter | 5 | 64159451 | 64351480 | 192030 | 1 | ENSMUST00 | 0 | Tbc1d1 | ENSMUSG00000029174.13 |
| chr7 | 126759577 | 126760602 | 28.90617 | 85.18155 | 81.93615 | Promoter | 7 | 126759601 | 126764845 | 5245 | 1 | ENSMUST00 | 0 | Mapk3 | ENSMUSG00000063065.8 |
| chr9 | 25481120 | 25482139 | 28.90617 | 85.18155 | 81.93615 | Promoter | 9 | 25481547 | 25604110 | 122564 | 1 | ENSMUST00 | 0 | Eepd1 | ENSMUSG00000036611.5 |
| chr1 | 59119469 | 59120710 | 22.31878 | 85.15456 | 81.90936 | Promoter | 1 | 59100590 | 59120094 | 19505 | 2 | ENSMUST00 | 0 | Tmem237 | ENSMUSG00000038079.9 |
| chr10 | 59615385 | 59617197 | 22.31878 | 85.15456 | 81.90936 | Promoter | 10 | 59446984 | 59616692 | 169709 | 2 | ENSMUST00 | 0 | Mcu | ENSMUSG00000009647.8 |
| chr10 | 81175938 | 81177970 | 22.31878 | 85.15456 | 81.90936 | Promoter | 10 | 81176631 | 81182507 | 5877 | 1 | ENSMUST00 | 0 | Eef2 | ENSMUSG00000034994.9 |
| chr11 | 97186741 | 97188025 | 22.31878 | 85.15456 | 81.90936 | Promoter | 11 | 97159714 | 97187881 | 28168 | 2 | ENSMUST00 | 0 | Kpnb1 | ENSMUSG00000001440.5 |
| chr11 | 84067823 | 84068916 | 22.31878 | 85.15456 | 81.90936 | Promoter | 11 | 84048041 | 84068357 | 20317 | 2 | ENSMUST00 | 0 | Dusp14 | ENSMUSG00000018648.10 |
| chr11 | 71003453 | 71005291 | 22.31878 | 85.15456 | 81.90936 | Promoter | 11 | 70984091 | 71004437 | 20347 | 2 | ENSMUST00 | 0 | Dhx33 | ENSMUSG00000040620.10 |
| chr12 | 111966196 | 111967098 | 22.31878 | 85.15456 | 81.90936 | Promoter | 12 | 111961376 | 111966977 | 5602 | 2 | ENSMUST00 | 0 | 2010107E04 | ENSMUSG00000021290.5 |
| chr14 | 105894686 | 105899120 | 22.31878 | 85.15456 | 81.90936 | Promoter | 14 | 105891949 | 105896819 | 4871 | 2 | ENSMUST00 | 0 | Spry2 | ENSMUSG00000022114.4 |
| chr19 | 10303885 | 10305229 | 22.31878 | 85.15456 | 81.90936 | Promoter | 19 | 10245265 | 10304877 | 59613 | 2 | ENSMUST00 | 0 | Dagla | ENSMUSG00000035735.5 |
| chr2 | 92599547 | 92603771 | 22.31878 | 85.15456 | 81.90936 | Promoter | 2 | 92599707 | 92615250 | 15544 | 1 | ENSMUST00 | 0 | Chst1 | ENSMUSG00000027221.5 |
| chr8 | 23207317 | 23209189 | 22.31878 | 85.15456 | 81.90936 | Promoter | 8 | 23172946 | 23208453 | 35508 | 2 | ENSMUST00 | 0 | Agpat6 | ENSMUSG00000031545.5 |
| chr8 | 71881569 | 71882618 | 22.31878 | 85.15456 | 81.90936 | Promoter | 8 | 71882068 | 71892565 | 10498 | 1 | ENSMUST00 | 0 | Zfp709 | ENSMUSG00000056019.7 |
| chr9 | 44881494 | 44883305 | 22.31878 | 85.15456 | 81.90936 | Promoter | 9 | 44841563 | 44881296 | 39734 | 2 | ENSMUST00 | -199 | Kmt2a | ENSMUSG00000002028.7 |
| chr9 | 58822227 | 58824265 | 22.31878 | 85.15456 | 81.90936 | Promoter | 9 | 58823512 | 58860955 | 37444 | 1 | ENSMUST00 | 0 | Hcn4 | ENSMUSG00000032338.8 |
| chr1 | 166409016 | 166410272 | 28.31422 | 85.10979 | 81.86543 | Promoter | 1 | 166384622 | 166409782 | 25161 | 2 | ENSMUST00 | 0 | Pogk | ENSMUSG00000040596.10 |
| chr2 | 66256085 | 66257392 | 28.31422 | 85.10979 | 81.86543 | Promoter | 2 | 66184327 | 66256617 | 72291 | 2 | ENSMUST00 | 0 | Ttc21b | ENSMUSG00000034848.12 |
| chr8 | 125569988 | 125570633 | 28.31422 | 85.10979 | 81.86543 | Distal Interge | 8 | 125418063 | 125492710 | 74648 | 2 | ENSMUST00 | -77279 | Sipa1l2 | ENSMUSG00000001995.8 |
| chr11 | 87737445 | 87738342 | 27.76267 | 85.10065 | 81.85645 | Promoter | 11 | 87737552 | 87743617 | 6066 | 1 | ENSMUST00 | 0 | Supt4a | ENSMUSG00000020485.9 |
| chr11 | 69121276 | 69124643 | 27.76267 | 85.10065 | 81.85645 | Promoter | 11 | 69120054 | 69122589 | 2536 | 2 | ENSMUST00 | 0 | 9130213A22 | ENSMUSG00000097386.1 |
| chr1 | 7398087 | 7398800 | 29.41006 | 85.09721 | 81.85307 | Promoter | 1 | 7349406 | 7397801 | 48396 | 2 | ENSMUST00 | -287 | Gm26901 | ENSMUSG00000097797.1 |
| chr15 | 64381829 | 64383394 | 29.41006 | 85.09721 | 81.85307 | Promoter | 15 | 64088539 | 64382919 | 294381 | 2 | ENSMUST00 | 0 | Asap1 | ENSMUSG00000022377.11 |
| chr16 | 32643583 | 32644599 | 29.41006 | 85.09721 | 81.85307 | Promoter | 16 | 32643874 | 32668279 | 24406 | 1 | ENSMUST00 | 0 | Tnk2 | ENSMUSG00000022791.11 |
| chr16 | 18629329 | 18630303 | 29.41006 | 85.09721 | 81.85307 | Promoter | 16 | 18621811 | 18629938 | 8128 | 2 | ENSMUST00 | 0 | Sep-05 | ENSMUSG00000072214.6 |
| chr17 | 79051976 | 79052822 | 29.41006 | 85.09721 | 81.85307 | Promoter | 17 | 79051906 | 79090243 | 38338 | 1 | ENSMUST00 | 71 | Qpct | ENSMUSG00000024084.7 |
| chr18 | 24470500 | 24471384 | 29.41006 | 85.09721 | 81.85307 | Promoter | 18 | 24470861 | 24477765 | 6905 | 1 | ENSMUST00 | 0 | 2700062C07 | ENSMUSG00000024273.7 |

| | | | | | | | | | | | | | | | |
|---|---|---|---|---|---|---|---|---|---|---|---|---|---|---|---|
| chr5 | 138992846 | 138996442 | 29.41006 | 85.09721 | 81.85307 | Promoter | 5 | 138976014 | 138995037 | 19024 | 2 | ENSMUST00 | 0 | Pdgfa | ENSMUSG00000025856.10 |
| chr5 | 137349196 | 137351472 | 29.41006 | 85.09721 | 81.85307 | Promoter | 5 | 137350109 | 137374528 | 24420 | 1 | ENSMUST00 | 0 | Ephb4 | ENSMUSG00000029710.10 |
| chr6 | 29693777 | 29694874 | 29.41006 | 85.09721 | 81.85307 | Promoter | 6 | 29694222 | 29718559 | 24338 | 1 | ENSMUST00 | 0 | Tspan33 | ENSMUSG00000001763.9 |
| chr6 | 132463619 | 132463896 | 29.41006 | 85.09721 | 81.85307 | Distal Interge | 6 | 132361041 | 132364134 | 3094 | 2 | ENSMUST00 | -99486 | Gm8882 | ENSMUSG00000030143.7 |
| chr7 | 34131933 | 34133828 | 29.41006 | 85.09721 | 81.85307 | Promoter | 7 | 34109543 | 34133268 | 23726 | 2 | ENSMUST00 | 0 | Wtip | ENSMUSG00000036459.10 |
| chr10 | 75935440 | 75936535 | 28.84749 | 85.08029 | 81.83772 | Promoter | 10 | 75935573 | 75937725 | 2153 | 1 | ENSMUST00 | 0 | Chchd10 | ENSMUSG00000049422.6 |
| chr11 | 105943387 | 105944660 | 28.84749 | 85.08029 | 81.83772 | Promoter | 11 | 105933702 | 105944412 | 10711 | 2 | ENSMUST00 | 0 | Cyb561 | ENSMUSG00000019590.11 |
| chr4 | 141663133 | 141664825 | 28.25877 | 85.01341 | 81.77112 | Promoter | 4 | 141625734 | 141664118 | 38385 | 2 | ENSMUST00 | 0 | Plekhm2 | ENSMUSG00000028917.9 |
| chr4 | 129513471 | 129514770 | 28.25877 | 85.01341 | 81.77112 | Promoter | 4 | 129513581 | 129515985 | 2405 | 1 | ENSMUST00 | 0 | Marcksl1 | ENSMUSG00000047945.6 |
| chr10 | 33950095 | 33951710 | 27.71015 | 85.00858 | 81.76643 | Promoter | 10 | 33926936 | 33951212 | 24277 | 2 | ENSMUST00 | 0 | Zufsp | ENSMUSG00000039531.7 |
| chr11 | 95823751 | 95825261 | 27.71015 | 85.00858 | 81.76643 | Promoter | 11 | 95824499 | 95830701 | 6203 | 1 | ENSMUST00 | 0 | Phospho1 | ENSMUSG00000050860.6 |
| chr13 | 45507198 | 45508834 | 27.71015 | 85.00858 | 81.76643 | Promoter | 13 | 45507444 | 45546382 | 38939 | 1 | ENSMUST00 | 0 | Gmpr | ENSMUSG00000000253.7 |
| chr15 | 99029288 | 99031062 | 27.71015 | 85.00858 | 81.76643 | Promoter | 15 | 99029891 | 99038105 | 8215 | 1 | ENSMUST00 | 0 | Tuba1c | ENSMUSG00000043091.8 |
| chr7 | 127041959 | 127042865 | 27.71015 | 85.00858 | 81.76643 | Promoter | 7 | 127027732 | 127042420 | 14689 | 2 | ENSMUST00 | 0 | Kif22 | ENSMUSG00000030677.7 |
| chr7 | 101347632 | 101348783 | 27.71015 | 85.00858 | 81.76643 | Promoter | 7 | 101348067 | 101411894 | 63828 | 1 | ENSMUST00 | 0 | Arap1 | ENSMUSG00000032812.11 |
| chr8 | 36993213 | 36994518 | 27.71015 | 85.00858 | 81.76643 | Promoter | 8 | 36993575 | 36995531 | 1957 | 1 | ENSMUST00 | 0 | AI429214 | ENSMUSG00000074384.3 |
| chr9 | 107632708 | 107635411 | 27.71015 | 85.00858 | 81.76643 | Promoter | 9 | 107614138 | 107635342 | 21205 | 2 | ENSMUST00 | 0 | Gnai2 | ENSMUSG00000032562.8 |
| chr11 | 55460352 | 55461216 | 25.04113 | 84.92952 | 81.6889 | Promoter | 11 | 55446644 | 55461197 | 14554 | 2 | ENSMUST00 | 0 | Atox1 | ENSMUSG00000018585.9 |
| chr11 | 11461573 | 11462733 | 25.04113 | 84.92952 | 81.6889 | Promoter | 11 | 11462094 | 11488537 | 26444 | 1 | ENSMUST00 | 0 | 4930415F15I | ENSMUSG00000020191.6 |
| chr7 | 7297989 | 7299897 | 25.04113 | 84.92952 | 81.6889 | Promoter | 7 | 7282316 | 7299511 | 17196 | 2 | ENSMUST00 | 0 | Clcn4-2 | ENSMUSG00000000605.8 |
| chr7 | 100492519 | 100494857 | 28.20354 | 84.91737 | 81.67689 | Promoter | 7 | 100493337 | 100502020 | 8684 | 1 | ENSMUST00 | 0 | Ucp2 | ENSMUSG00000033685.7 |
| chr14 | 51062791 | 51066193 | 27.65783 | 84.91682 | 81.67644 | Distal Interge | 14 | 51070311 | 51071442 | 1132 | 2 | ENSMUST00 | 5249 | Olfr750 | ENSMUSG00000035932.5 |
| chr15 | 76330669 | 76332093 | 27.09851 | 84.88239 | 81.64215 | Promoter | 15 | 76331231 | 76334907 | 3677 | 1 | ENSMUST00 | 0 | Gpaa1 | ENSMUSG00000022561.7 |
| chr4 | 106839087 | 106840723 | 27.09851 | 84.88239 | 81.64215 | Downstream | 4 | 106841761 | 106847920 | 6160 | 2 | ENSMUST00 | 7197 | Gm12746 | ENSMUSG00000086940.1 |
| chr4 | 152026890 | 152028328 | 27.60571 | 84.82537 | 81.58639 | Promoter | 4 | 152019774 | 152027661 | 7888 | 2 | ENSMUST00 | 0 | Zbtb48 | ENSMUSG00000028952.7 |
| chr4 | 117914129 | 117915849 | 26.52989 | 84.81848 | 81.57964 | Promoter | 4 | 117894486 | 117914999 | 20514 | 2 | ENSMUST00 | 0 | Ipo13 | ENSMUSG00000033365.9 |
| chr5 | 4756381 | 4759090 | 26.52989 | 84.81848 | 81.57964 | Promoter | 5 | 4753873 | 4758035 | 4163 | 2 | ENSMUST00 | 0 | Fzd1 | ENSMUSG00000044674.5 |
| chr7 | 19381893 | 19382966 | 26.52989 | 84.81848 | 81.57964 | Promoter | 7 | 19382010 | 19395694 | 13685 | 1 | ENSMUST00 | 0 | Ercc2 | ENSMUSG00000030400.12 |
| chr1 | 131523158 | 131527435 | 25.99992 | 84.80917 | 81.57088 | Promoter | 1 | 131285251 | 131527347 | 242097 | 2 | ENSMUST00 | 0 | Srgap2 | ENSMUSG00000026425.10 |
| chr1 | 163994206 | 163995669 | 25.99992 | 84.80917 | 81.57088 | Promoter | 1 | 163994889 | 163997243 | 2355 | 1 | ENSMUST00 | 0 | Mettl18 | ENSMUSG00000041396.8 |
| chr14 | 33319411 | 33320665 | 25.99992 | 84.80917 | 81.57088 | Promoter | 14 | 33319703 | 33369934 | 50232 | 1 | ENSMUST00 | 0 | Arhgap22 | ENSMUSG00000063506.8 |
| chr4 | 148139571 | 148141195 | 25.99992 | 84.80917 | 81.57088 | Promoter | 4 | 148140182 | 148145121 | 4940 | 1 | ENSMUST00 | 0 | Mad2l2 | ENSMUSG00000029003.6 |
| chr7 | 99535128 | 99537920 | 25.99992 | 84.80917 | 81.57088 | Promoter | 7 | 99535466 | 99606771 | 71306 | 1 | ENSMUST00 | 0 | Arrb1 | ENSMUSG00000018909.10 |
| chr10 | 89442963 | 89444403 | 29.23449 | 84.79565 | 81.55782 | Promoter | 10 | 89408823 | 89443967 | 35145 | 2 | ENSMUST00 | 0 | Gas2l3 | ENSMUSG00000074802.5 |
| chr11 | 78465380 | 78466347 | 29.23449 | 84.79565 | 81.55782 | Promoter | 11 | 78465697 | 78471945 | 6249 | 1 | ENSMUST00 | 0 | Slc46a1 | ENSMUSG00000020829.9 |
| chr8 | 38659941 | 38661317 | 28.67288 | 84.77877 | 81.54134 | Promoter | 8 | 37522298 | 38661508 | 1139211 | 2 | ENSMUST00 | 191 | Sgcz | ENSMUSG00000039539.8 |
| chr4 | 103113428 | 103115228 | 28.09372 | 84.72628 | 81.48907 | Promoter | 4 | 103114390 | 103115904 | 1515 | 1 | ENSMUST00 | 0 | Mier1 | ENSMUSG00000028522.11 |
| chr1 | 183220772 | 183221590 | 23.47096 | 84.72127 | 81.48416 | Promoter | 1 | 183086264 | 183221529 | 135266 | 2 | ENSMUST00 | 0 | Disp1 | ENSMUSG00000030768.7 |
| chr8 | 35587571 | 35589216 | 23.47096 | 84.72127 | 81.48416 | Promoter | 8 | 35587798 | 35679449 | 91652 | 1 | ENSMUST00 | 0 | Mfhas1 | ENSMUSG00000070056.5 |
| chr10 | 86021772 | 86022997 | 21.20881 | 84.61981 | 81.38317 | Promoter | 10 | 86021972 | 86051873 | 29902 | 1 | ENSMUST00 | 0 | Fbxo7 | ENSMUSG00000001786.9 |
| chr11 | 6291088 | 6292643 | 21.20881 | 84.61981 | 81.38317 | Promoter | 11 | 6291633 | 6356642 | 65010 | 1 | ENSMUST00 | 0 | Ogdh | ENSMUSG00000020456.12 |
| chr17 | 24631895 | 24633295 | 21.20881 | 84.61981 | 81.38317 | Promoter | 17 | 24595937 | 24632629 | 36693 | 2 | ENSMUST00 | 0 | Tsc2 | ENSMUSG00000002496.9 |
| chr7 | 30193627 | 30195415 | 21.20881 | 84.61981 | 81.38317 | Promoter | 7 | 30186942 | 30195164 | 8223 | 2 | ENSMUST00 | 0 | Capns1 | ENSMUSG00000001794.7 |
| chr7 | 16399095 | 16403914 | 21.20881 | 84.61981 | 81.38317 | Promoter | 7 | 16401196 | 16437694 | 36499 | 1 | ENSMUST00 | 0 | Zc3h4 | ENSMUSG00000059273.7 |
| chr9 | 63398366 | 63399901 | 21.20881 | 84.61981 | 81.38317 | Promoter | 9 | 63394447 | 63398827 | 4381 | 2 | ENSMUST00 | 0 | 2300009A05I | ENSMUSG00000032403.8 |
| chr11 | 115412283 | 115413838 | 27.98475 | 84.53653 | 81.30061 | 3' UTR | 11 | 115410160 | 115410908 | 749 | 1 | ENSMUST00 | 2124 | Ict1 | ENSMUSG00000018858.10 |
| chr13 | 21832784 | 21833857 | 16.92661 | 84.48347 | 81.24769 | Promoter | 13 | 21833743 | 21836665 | 2923 | 1 | ENSMUST00 | 0 | Hist1h2br | ENSMUSG00000069303.5 |
| chr16 | 98081618 | 98082956 | 29.05239 | 84.48264 | 81.24685 | Distal Interge | 16 | 98008629 | 98009008 | 380 | 1 | ENSMUST00 | 72990 | B230307C23I | ENSMUSG00000080717.4 |
| chr1 | 171328255 | 171330422 | 28.50037 | 84.48055 | 81.24493 | Promoter | 1 | 171329145 | 171342331 | 13187 | 1 | ENSMUST00 | 0 | Dedd | ENSMUSG00000013973.7 |
| chr15 | 75920962 | 75921570 | 28.50037 | 84.48055 | 81.24493 | Promoter | 15 | 75916463 | 75921560 | 5098 | 2 | ENSMUST00 | 0 | Pycrl | ENSMUSG00000022571.8 |
| chr3 | 88424399 | 88425681 | 28.50037 | 84.48055 | 81.24493 | Promoter | 3 | 88410498 | 88425139 | 14642 | 2 | ENSMUST00 | 0 | Slc25a44 | ENSMUSG00000050144.8 |
| chr4 | 149518122 | 149519059 | 28.50037 | 84.48055 | 81.24493 | Promoter | 4 | 149518236 | 149566437 | 48202 | 1 | ENSMUST00 | 0 | Ctnnbip1 | ENSMUSG00000028988.8 |
| chr5 | 34168859 | 34169522 | 28.50037 | 84.48055 | 81.24493 | Promoter | 5 | 34007198 | 34169448 | 162251 | 2 | ENSMUST00 | 0 | Poln | ENSMUSG00000045102.8 |
| chr9 | 122805110 | 122806474 | 27.39917 | 84.46252 | 81.22775 | Promoter | 9 | 122805539 | 122836334 | 30796 | 1 | ENSMUST00 | 0 | Tcaim | ENSMUSG00000046603.10 |
| chr1 | 72710755 | 72712518 | 23.3092 | 84.4034 | 81.16875 | Promoter | 1 | 72711260 | 72713809 | 2550 | 1 | ENSMUST00 | 0 | Rpl37a | ENSMUSG00000046330.9 |
| chr1 | 133363252 | 133364156 | 23.3092 | 84.4034 | 81.16875 | Promoter | 1 | 133363572 | 133380336 | 16765 | 1 | ENSMUST00 | 0 | Etnk2 | ENSMUSG00000070644.8 |
| chr10 | 88146453 | 88147529 | 23.3092 | 84.4034 | 81.16875 | Promoter | 10 | 88146992 | 88178388 | 31397 | 1 | ENSMUST00 | 0 | Nup37 | ENSMUSG00000035351.10 |
| chr11 | 7063589 | 7065749 | 23.3092 | 84.4034 | 81.16875 | Promoter | 11 | 7063489 | 7178506 | 115018 | 1 | ENSMUST00 | 101 | Adcy1 | ENSMUSG00000020431.5 |
| chr12 | 113139865 | 113141336 | 23.3092 | 84.4034 | 81.16875 | Promoter | 12 | 113140236 | 113145504 | 5269 | 1 | ENSMUST00 | 0 | Crip2 | ENSMUSG00000006356.6 |
| chr13 | 3892601 | 3893908 | 23.3092 | 84.4034 | 81.16875 | Promoter | 13 | 3882565 | 3893578 | 11014 | 2 | ENSMUST00 | 0 | Net1 | ENSMUSG00000021215.9 |
| chr14 | 8097842 | 8099268 | 23.3092 | 84.4034 | 81.16875 | Promoter | 14 | 8098213 | 8165111 | 66899 | 1 | ENSMUST00 | 0 | Pxk | ENSMUSG00000033885.9 |
| chr15 | 89373196 | 89374492 | 23.3092 | 84.4034 | 81.16875 | Promoter | 15 | 89371639 | 89373818 | 2180 | 2 | ENSMUST00 | 0 | Sco2 | ENSMUSG00000091780.2 |
| chr17 | 8147318 | 8148171 | 23.3092 | 84.4034 | 81.16875 | Promoter | 17 | 8128610 | 8148097 | 19488 | 2 | ENSMUST00 | 0 | Rnaset2a | ENSMUSG00000095687.1 |
| chr18 | 75366329 | 75373500 | 23.3092 | 84.4034 | 81.16875 | Promoter | 18 | 75367529 | 75395935 | 28407 | 1 | ENSMUST00 | 0 | Smad7 | ENSMUSG00000025880.6 |
| chr2 | 69205583 | 69206754 | 23.3092 | 84.4034 | 81.16875 | Promoter | 2 | 69193895 | 69206135 | 12241 | 2 | ENSMUST00 | 0 | Spc25 | ENSMUSG00000005233.11 |
| chr3 | 54755279 | 54756891 | 23.3092 | 84.4034 | 81.16875 | Promoter | 3 | 54755582 | 54801257 | 45676 | 1 | ENSMUST00 | 0 | Smad9 | ENSMUSG00000027796.2 |
| chr3 | 116006495 | 116008826 | 23.3092 | 84.4034 | 81.16875 | Promoter | 3 | 116007462 | 116029016 | 21555 | 1 | ENSMUST00 | 0 | Extl2 | ENSMUSG00000027963.9 |
| chr4 | 122995308 | 122997677 | 23.3092 | 84.4034 | 81.16875 | Promoter | 4 | 122995652 | 122996739 | 1088 | 1 | ENSMUST00 | 0 | Mycl | ENSMUSG00000028654.8 |
| chr5 | 144222689 | 144224417 | 23.3092 | 84.4034 | 81.16875 | Promoter | 5 | 144194442 | 144223615 | 29174 | 2 | ENSMUST00 | 0 | Tecpr1 | ENSMUSG00000066621.7 |
| chr8 | 126943948 | 126946384 | 23.3092 | 84.4034 | 81.16875 | Promoter | 8 | 126944847 | 126944980 | 134 | 2 | ENSMUST00 | 0 | Gm26397 | ENSMUSG00000065637.1 |
| chr9 | 62121399 | 62123449 | 23.3092 | 84.4034 | 81.16875 | Promoter | 9 | 62057248 | 62122610 | 65363 | 2 | ENSMUST00 | 0 | Glce | ENSMUSG00000032252.9 |
| chr9 | 82972816 | 82976631 | 23.3092 | 84.4034 | 81.16875 | Promoter | 9 | 82866159 | 82975516 | 109358 | 2 | ENSMUST00 | 0 | Phip | ENSMUSG00000032253.8 |
| chr1 | 192091878 | 192093127 | 28.44333 | 84.38186 | 81.14881 | Promoter | 1 | 191997430 | 192092559 | 95130 | 2 | ENSMUST00 | 0 | Traf5 | ENSMUSG00000026637.8 |
| chr5 | 121592937 | 121594485 | 28.44333 | 84.38186 | 81.14881 | Promoter | 5 | 121566027 | 121593824 | 27798 | 2 | ENSMUST00 | 0 | Aldh2 | ENSMUSG00000029455.9 |
| chr18 | 74063646 | 74065684 | 28.9922 | 84.37911 | 81.14628 | Promoter | 18 | 73928486 | 74064392 | 135907 | 2 | ENSMUST00 | 0 | Mapk4 | ENSMUSG00000024558.7 |
| chr13 | 21721642 | 21722491 | 27.34802 | 84.37254 | 81.13987 | Promoter | 13 | 21721457 | 21721808 | 352 | 2 | ENSMUST00 | 0 | Hist1h2aj | ENSMUSG00000080076.4 |
| chr12 | 8921298 | 8924062 | 20.25082 | 84.28156 | 81.04926 | Promoter | 12 | 8921664 | 8938742 | 17079 | 1 | ENSMUST00 | 0 | Laptm4a | ENSMUSG00000020585.4 |
| chr7 | 132315520 | 132317720 | 20.25082 | 84.28156 | 81.04926 | Promoter | 7 | 132235780 | 132316714 | 80935 | 2 | ENSMUST00 | 0 | Chst15 | ENSMUSG00000030930.8 |
| chr8 | 106168195 | 106170002 | 20.25082 | 84.28156 | 81.04926 | Promoter | 8 | 106168875 | 106198704 | 29830 | 1 | ENSMUST00 | 0 | Slc7a6 | ENSMUSG00000031904.4 |
| chr11 | 101552780 | 101554398 | 28.93225 | 84.27598 | 81.04385 | Promoter | 11 | 101552858 | 101581951 | 29094 | 1 | ENSMUST00 | 0 | Nbr1 | ENSMUSG00000017119.14 |
| chr14 | 20392850 | 20394150 | 28.93225 | 84.27598 | 81.04385 | Promoter | 14 | 20391231 | 20393555 | 2325 | 2 | ENSMUST00 | 0 | Mrps16 | ENSMUSG00000049960.3 |
| chr2 | 74704033 | 74706369 | 28.93225 | 84.27598 | 81.04385 | Promoter | 2 | 74704615 | 74706812 | 2198 | 1 | ENSMUST00 | 0 | Hoxd8 | ENSMUSG00000027102.4 |
| chr8 | 93809849 | 93811435 | 28.93225 | 84.27598 | 81.04385 | Promoter | 8 | 93809966 | 93950403 | 140438 | 1 | ENSMUST00 | 0 | Gnao1 | ENSMUSG00000031748.10 |
| chr12 | 73584164 | 73586065 | 27.82288 | 84.25431 | 81.02234 | Promoter | 12 | 73584797 | 73778185 | 193389 | 1 | ENSMUST00 | 0 | Prkch | ENSMUSG00000021108.12 |
| chr14 | 51984317 | 51986017 | 27.82288 | 84.25431 | 81.02234 | Promoter | 14 | 51984719 | 51987467 | 2749 | 1 | ENSMUST00 | 0 | Arhgef40 | ENSMUSG00000004562.11 |
| chr19 | 6920899 | 6922749 | 27.82288 | 84.25431 | 81.02234 | Promoter | 19 | 6910977 | 6921808 | 10832 | 2 | ENSMUST00 | 0 | Esrra | ENSMUSG00000024955.8 |
| chr2 | 158360817 | 158361635 | 27.82288 | 84.25431 | 81.02234 | Promoter | 2 | 158355307 | 158361510 | 6204 | 2 | ENSMUST00 | 0 | 9430008C03I | ENSMUSG00000085385.2 |
| chr6 | 38298370 | 38299589 | 27.82288 | 84.25431 | 81.02234 | Promoter | 6 | 38287396 | 38299259 | 11864 | 2 | ENSMUST00 | 0 | Zc3hav1l | ENSMUSG00000047749.2 |
| chr6 | 86364623 | 86366430 | 27.82288 | 84.25431 | 81.02234 | Promoter | 6 | 86365683 | 86370057 | 4375 | 1 | ENSMUST00 | 0 | Fam136a | ENSMUSG00000057497.7 |
| chr6 | 134983348 | 134984394 | 27.82288 | 84.25431 | 81.02234 | 5' UTR | 6 | 134982001 | 134986836 | 4836 | 1 | ENSMUST00 | 1348 | Apold1 | ENSMUSG00000090698.1 |
| chr7 | 16046337 | 16049003 | 27.82288 | 84.25431 | 81.02234 | Distal Interge | 7 | 16071942 | 16096328 | 24387 | 1 | ENSMUST00 | -22939 | Zfp541 | ENSMUSG00000078796.3 |
| chr1 | 180851100 | 180851884 | 24.64903 | 84.19559 | 80.96455 | Promoter | 1 | 180851151 | 180868097 | 16947 | 1 | ENSMUST00 | 0 | Sde2 | ENSMUSG00000038806.6 |
| chr10 | 120201051 | 120202300 | 24.64903 | 84.19559 | 80.96455 | Promoter | 10 | 120201590 | 120203215 | 1626 | 1 | ENSMUST00 | 0 | Tmbim4 | ENSMUSG00000020225.5 |
| chr7 | 7120751 | 7121715 | 28.32993 | 84.18556 | 80.95464 | Promoter | 7 | 7114690 | 7121445 | 6756 | 2 | ENSMUST00 | 0 | Zfp954 | ENSMUSG00000062116.8 |
| chr17 | 56303188 | 56304605 | 27.1957 | 84.10433 | 80.87383 | Promoter | 17 | 56303321 | 56323486 | 20166 | 1 | ENSMUST00 | 0 | Uhrf1 | ENSMUSG00000001228.9 |
| chr5 | 140704407 | 140704830 | 27.1957 | 84.10433 | 80.87383 | Promoter | 5 | 140705011 | 140719379 | 14369 | 1 | ENSMUST00 | -181 | Brat1 | ENSMUSG00000000148.12 |
| chr5 | 122707005 | 122708548 | 27.1957 | 84.10433 | 80.87383 | Promoter | 5 | 122707584 | 122729044 | 21461 | 1 | ENSMUST00 | 0 | P2rx4 | ENSMUSG00000029470.10 |
| chr6 | 124662145 | 124663822 | 27.1957 | 84.10433 | 80.87383 | Promoter | 6 | 124663027 | 124704370 | 41344 | 1 | ENSMUST00 | 0 | Lpcat3 | ENSMUSG00000004270.8 |
| chr8 | 121585003 | 121585533 | 27.1957 | 84.10433 | 80.87383 | Intron (ENSM | 8 | 121581765 | 121582191 | 427 | 1 | ENSMUST00 | 3239 | Gm17786 | ENSMUSG00000097987.1 |
| chr14 | 21847509 | 21848985 | 27.716 | 84.06778 | 80.83876 | Promoter | 14 | 21845861 | 21848977 | 3117 | 2 | ENSMUST00 | 0 | Comtd1 | ENSMUSG00000021773.5 |
| chr13 | 43558962 | 43561028 | 18.68094 | 84.05138 | 80.82243 | Promoter | 13 | 43538412 | 43560191 | 21780 | 2 | ENSMUST00 | 0 | Mcur1 | ENSMUSG00000021371.4 |
| chr10 | 81626885 | 81628214 | 26.61343 | 84.02032 | 80.79139 | Promoter | 10 | 81621785 | 81627320 | 5536 | 2 | ENSMUST00 | 0 | Sirt6 | ENSMUSG00000034748.11 |
| chr11 | 19919639 | 19920788 | 26.61343 | 84.02032 | 80.79139 | Distal Interge | 11 | 19924375 | 20024026 | 99652 | 1 | ENSMUST00 | -3587 | Spred2 | ENSMUSG00000045671.12 |
| chr15 | 58872196 | 58873319 | 26.61343 | 84.02032 | 80.79139 | Promoter | 15 | 58872649 | 58876781 | 4133 | 1 | ENSMUST00 | 0 | Trmt12 | ENSMUSG00000037085.7 |
| chr16 | 50731759 | 50733214 | 26.61343 | 84.02032 | 80.79139 | Promoter | 16 | 50719294 | 50732773 | 13480 | 2 | ENSMUST00 | 0 | 5330426P16I | ENSMUSG00000022639.9 |
| chr19 | 34921589 | 34922788 | 26.61343 | 84.02032 | 80.79139 | Promoter | 19 | 34922358 | 34975731 | 53374 | 1 | ENSMUST00 | 0 | Kif20b | ENSMUSG00000024795.10 |
| chr8 | 83441742 | 83443146 | 26.61343 | 84.02032 | 80.79139 | Promoter | 8 | 83434493 | 83442648 | 8156 | 2 | ENSMUST00 | 0 | Scoc | ENSMUSG00000063253.5 |
| chr14 | 79586786 | 79588351 | 25.5656 | 84.01802 | 80.7898 | Promoter | 14 | 79587691 | 79629755 | 42065 | 1 | ENSMUST00 | 0 | Sugt1 | ENSMUSG00000022024.9 |
| chr15 | 79658185 | 79659768 | 25.5656 | 84.01802 | 80.7898 | Promoter | 15 | 79659227 | 79667660 | 8434 | 1 | ENSMUST00 | 0 | Cby1 | ENSMUSG00000022428.7 |
| chr1 | 63730061 | 63731245 | 26.0714 | 83.99387 | 80.766 | Promoter | 1 | 63730614 | 63754655 | 24042 | 1 | ENSMUST00 | 0 | Fastkd2 | ENSMUSG00000025962.10 |
| chr3 | 96557994 | 96562539 | 26.0714 | 83.99387 | 80.766 | Promoter | 3 | 96559498 | 96560013 | 516 | 1 | ENSMUST00 | 0 | Txnip | ENSMUSG00000038393.9 |
| chr19 | 7416554 | 7418702 | 27.66286 | 83.97498 | 80.7477 | Promoter | 19 | 7417625 | 7425904 | 8280 | 1 | ENSMUST00 | 0 | 2700081O15 | ENSMUSG00000053080.10 |
| chr1 | 171269902 | 171271264 | 27.66126 | 83.97217 | 80.74496 | Promoter | 1 | 171270328 | 171276895 | 6568 | 1 | ENSMUST00 | 0 | B4galt3 | ENSMUSG00000052423.9 |
| chr10 | 126927626 | 126929244 | 27.66126 | 83.97217 | 80.74496 | Distal Interge | 10 | 126887249 | 126901371 | 14123 | 2 | ENSMUST00 | -26256 | Xrcc6bp1 | ENSMUSG00000025436.6 |
| chr10 | 81137634 | 81138894 | 27.66126 | 83.97217 | 80.74496 | Promoter | 10 | 81137953 | 81148458 | 10506 | 1 | ENSMUST00 | 0 | Zbtb7a | ENSMUSG00000035011.10 |
| chr11 | 46311482 | 46313023 | 27.66126 | 83.97217 | 80.74496 | Promoter | 11 | 46193850 | 46312358 | 118509 | 2 | ENSMUST00 | 0 | Cyfip2 | ENSMUSG00000020340.11 |
| chr15 | 83463171 | 83465159 | 27.66126 | 83.97217 | 80.74496 | Promoter | 15 | 83375607 | 83464606 | 89000 | 2 | ENSMUST00 | 0 | Pacsin2 | ENSMUSG00000016664.9 |
| chr15 | 89172791 | 89174246 | 27.66126 | 83.97217 | 80.74496 | Distal Interge | 15 | 89155549 | 89171120 | 15572 | 2 | ENSMUST00 | -1672 | Plxnb2 | ENSMUSG00000036606.11 |
| chr16 | 90829810 | 90831614 | 27.66126 | 83.97217 | 80.74496 | Promoter | 16 | 90830808 | 90905109 | 74302 | 1 | ENSMUST00 | 0 | Eva1c | ENSMUSG00000039903.11 |
| chr17 | 56325969 | 56327059 | 27.66126 | 83.97217 | 80.74496 | Promoter | 17 | 56326062 | 56356056 | 29995 | 1 | ENSMUST00 | 0 | Kdm4b | ENSMUSG00000024201.7 |
| chr17 | 65771973 | 65773014 | 27.66126 | 83.97217 | 80.74496 | Promoter | 17 | 65651726 | 65772752 | 121027 | 2 | ENSMUST00 | 0 | Rab31 | ENSMUSG00000056515.8 |
| chr18 | 36669542 | 36670664 | 27.66126 | 83.97217 | 80.74496 | Promoter | 18 | 36667246 | 36670286 | 3041 | 2 | ENSMUST00 | 0 | Sra1 | ENSMUSG00000006050.6 |
| chr2 | 174438027 | 174439283 | 27.66126 | 83.97217 | 80.74496 | Promoter | 2 | 174427493 | 174439039 | 11547 | 2 | ENSMUST00 | 0 | Ctsz | ENSMUSG00000016256.10 |
| chr3 | 142168586 | 142169710 | 27.66126 | 83.97217 | 80.74496 | Promoter | 3 | 141840367 | 142169213 | 328847 | 2 | ENSMUST00 | 0 | Bmpr1b | ENSMUSG00000052430.10 |
| chr4 | 147867549 | 147869781 | 27.66126 | 83.97217 | 80.74496 | Promoter | 4 | 147860778 | 147868719 | 7942 | 2 | ENSMUST00 | 0 | Miip | ENSMUSG00000029022.13 |
| chr6 | 149408692 | 149410015 | 27.66126 | 83.97217 | 80.74496 | Promoter | 6 | 149408984 | 149563329 | 154346 | 1 | ENSMUST00 | 0 | Bicd1 | ENSMUSG00000003452.10 |
| chr6 | 90987314 | 90989227 | 27.66126 | 83.97217 | 80.74496 | Distal Interge | 6 | 91013317 | 91017429 | 4113 | 2 | ENSMUST00 | 28202 | Nup210 | ENSMUSG00000030091.12 |
| chr6 | 43265083 | 43266670 | 27.66126 | 83.97217 | 80.74496 | Promoter | 6 | 43265582 | 43289320 | 23739 | 1 | ENSMUST00 | 0 | Arhgef5 | ENSMUSG00000033542.8 |

| | | | | | | | | | | | | | | | |
|---|---|---|---|---|---|---|---|---|---|---|---|---|---|---|---|
| chr7 | 49778215 | 49778931 | 27.66126 | 83.97217 | 80.74496 | Promoter | 7 | 49778346 | 49782505 | 4160 | 1 | ENSMUST00 | 0 | Prmt3 | ENSMUSG00000030505.11 |
| chr7 | 102222429 | 102223527 | 27.66126 | 83.97217 | 80.74496 | Promoter | 7 | 102222440 | 102236167 | 13728 | 1 | ENSMUST00 | 0 | Pgap2 | ENSMUSG00000030990.12 |
| chr7 | 140953561 | 140956714 | 27.66126 | 83.97217 | 80.74496 | Promoter | 7 | 140954839 | 140955961 | 1123 | 2 | ENSMUST00 | 0 | Ifitm2 | ENSMUSG00000060591.8 |
| chr8 | 71464912 | 71465606 | 27.66126 | 83.97217 | 80.74496 | Promoter | 8 | 71464926 | 71465752 | 827 | 1 | ENSMUST00 | 0 | Mrpl34 | ENSMUSG00000034880.6 |
| chr15 | 81744733 | 81746324 | 28.16151 | 83.89376 | 80.67007 | Promoter | 15 | 81744848 | 81796269 | 51422 | 1 | ENSMUST00 | 0 | Zc3h7b | ENSMUSG00000022390.10 |
| chr1 | 131152837 | 131153921 | 24.48636 | 83.88964 | 80.66612 | Promoter | 1 | 131153181 | 131158902 | 5722 | 1 | ENSMUST00 | 0 | Eif2d | ENSMUSG00000026427.11 |
| chr10 | 9674014 | 9676226 | 24.48636 | 83.88964 | 80.66612 | Promoter | 10 | 9627259 | 9675208 | 47950 | 2 | ENSMUST00 | 0 | Samd5 | ENSMUSG00000060487.5 |
| chr11 | 65806297 | 65808006 | 24.48636 | 83.88964 | 80.66612 | Promoter | 11 | 65807175 | 65829238 | 22064 | 1 | ENSMUST00 | 0 | Zkscan6 | ENSMUSG00000018347.11 |
| chr11 | 115563642 | 115565044 | 24.48636 | 83.88964 | 80.66612 | Promoter | 11 | 115564434 | 115583985 | 19552 | 1 | ENSMUST00 | 0 | Nup85 | ENSMUSG00000020739.6 |
| chr12 | 84772154 | 84773058 | 24.48636 | 83.88964 | 80.66612 | Promoter | 12 | 84754560 | 84773112 | 18553 | 2 | ENSMUST00 | 54 | Npc2 | ENSMUSG00000021242.8 |
| chr13 | 97240304 | 97242323 | 24.48636 | 83.88964 | 80.66612 | Promoter | 13 | 97241105 | 97253039 | 11935 | 1 | ENSMUST00 | 0 | Enc1 | ENSMUSG00000041773.7 |
| chr13 | 62466106 | 62467284 | 24.48636 | 83.88964 | 80.66612 | Promoter | 13 | 62454323 | 62466709 | 12387 | 2 | ENSMUST00 | 0 | Zfp935 | ENSMUSG00000055228.7 |
| chr14 | 104521848 | 104523065 | 24.48636 | 83.88964 | 80.66612 | Promoter | 14 | 104477534 | 104522666 | 45133 | 2 | ENSMUST00 | 0 | Rnf219 | ENSMUSG00000022120.3 |
| chr17 | 44734011 | 44736771 | 24.48636 | 83.88964 | 80.66612 | Promoter | 17 | 44495987 | 44736176 | 240190 | 2 | ENSMUST00 | 0 | Runx2 | ENSMUSG00000039153.11 |
| chr3 | 88022473 | 88024031 | 24.48636 | 83.88964 | 80.66612 | Promoter | 3 | 88022638 | 88023673 | 1036 | 2 | ENSMUST00 | 0 | Hapln2 | ENSMUSG00000004894.5 |
| chr4 | 149773400 | 149774273 | 24.48636 | 83.88964 | 80.66612 | Promoter | 4 | 149744036 | 149774277 | 30242 | 2 | ENSMUST00 | 4 | Slc25a33 | ENSMUSG00000028982.9 |
| chr6 | 86848422 | 86851621 | 24.48636 | 83.88964 | 80.66612 | Promoter | 6 | 86849517 | 86991864 | 142348 | 1 | ENSMUST00 | 0 | Aak1 | ENSMUSG00000057230.8 |
| chr6 | 38636221 | 38637766 | 24.48636 | 83.88964 | 80.66612 | Promoter | 6 | 38626660 | 38637239 | 10580 | 2 | ENSMUST00 | 0 | Klrg2 | ENSMUSG00000071537.3 |
| chr7 | 109703217 | 109704397 | 24.48636 | 83.88964 | 80.66612 | Promoter | 7 | 109703690 | 109712189 | 8500 | 1 | ENSMUST00 | 0 | Akip1 | ENSMUSG00000031023.5 |
| chr9 | 21917580 | 21918741 | 24.48636 | 83.88964 | 80.66612 | Promoter | 9 | 21907491 | 21918192 | 10702 | 2 | ENSMUST00 | 0 | Rab3d | ENSMUSG00000019066.8 |
| chr15 | 81190967 | 81191490 | 28.6362 | 83.76622 | 80.54601 | Promoter | 15 | 81190853 | 81192756 | 1904 | 1 | ENSMUST00 | 115 | 4930483J18R | ENSMUSG00000047938.7 |
| chr19 | 4614686 | 4616300 | 28.6362 | 83.76622 | 80.54601 | Promoter | 19 | 4611785 | 4615667 | 3883 | 2 | ENSMUST00 | 0 | Lrfn4 | ENSMUSG00000045045.7 |
| chr3 | 20057338 | 20058558 | 28.6362 | 83.76622 | 80.54601 | Promoter | 3 | 20057811 | 20117281 | 59471 | 1 | ENSMUST00 | 0 | Hltf | ENSMUSG00000002428.7 |
| chr4 | 129247693 | 129249005 | 28.6362 | 83.76622 | 80.54601 | Promoter | 4 | 129219578 | 129248443 | 28866 | 2 | ENSMUST00 | 0 | C77080 | ENSMUSG00000050390.7 |
| chr8 | 119604419 | 119605303 | 28.05034 | 83.70095 | 80.48139 | Promoter | 8 | 119597975 | 119605222 | 7248 | 2 | ENSMUST00 | 0 | Taf1c | ENSMUSG00000031832.9 |
| chr1 | 111864110 | 111865465 | 25.90858 | 83.69981 | 80.48042 | Promoter | 1 | 111858702 | 111864768 | 6067 | 2 | ENSMUST00 | 0 | Dsel | ENSMUSG00000038702.6 |
| chr1 | 168429977 | 168432206 | 25.90858 | 83.69981 | 80.48042 | Promoter | 1 | 168153542 | 168431897 | 278356 | 2 | ENSMUST00 | 0 | Pbx1 | ENSMUSG00000052534.10 |
| chr10 | 80655644 | 80656595 | 25.90858 | 83.69981 | 80.48042 | Promoter | 10 | 80642617 | 80656494 | 13878 | 2 | ENSMUST00 | 0 | Btbd2 | ENSMUSG00000003344.9 |
| chr10 | 70440157 | 70441617 | 25.90858 | 83.69981 | 80.48042 | Promoter | 10 | 70440668 | 70558729 | 118062 | 1 | ENSMUST00 | 0 | Fam13c | ENSMUSG00000043259.9 |
| chr11 | 6546029 | 6548154 | 25.90858 | 83.69981 | 80.48042 | Promoter | 11 | 6546887 | 6596744 | 49858 | 1 | ENSMUST00 | 0 | Ccm2 | ENSMUSG00000000378.10 |
| chr11 | 81151425 | 81153765 | 25.90858 | 83.69981 | 80.48042 | Promoter | 11 | 80880169 | 81153321 | 273153 | 2 | ENSMUST00 | 0 | Asic2 | ENSMUSG00000020704.8 |
| chr12 | 18609655 | 18611204 | 25.90858 | 83.69981 | 80.48042 | Distal Interge | 12 | 18514738 | 18534191 | 19454 | 1 | ENSMUST00 | 94918 | 5730507C01R | ENSMUSG00000073197.3 |
| chr13 | 17694330 | 17694872 | 25.90858 | 83.69981 | 80.48042 | Promoter | 13 | 16857855 | 17694732 | 836878 | 2 | ENSMUST00 | 0 | Sugct | ENSMUSG00000055137.6 |
| chr14 | 122181104 | 122182249 | 25.90858 | 83.69981 | 80.48042 | Promoter | 14 | 122181694 | 122402232 | 220539 | 1 | ENSMUST00 | 0 | Clybl | ENSMUSG00000025545.9 |
| chr17 | 33908374 | 33910645 | 25.90858 | 83.69981 | 80.48042 | Promoter | 17 | 33909414 | 33911423 | 2010 | 1 | ENSMUST00 | 0 | Daxx | ENSMUSG00000002307.10 |
| chr2 | 122701481 | 122703325 | 25.90858 | 83.69981 | 80.48042 | Promoter | 2 | 122681233 | 122702639 | 21407 | 2 | ENSMUST00 | 0 | Slc30a4 | ENSMUSG00000005802.7 |
| chr3 | 88042631 | 88043693 | 25.90858 | 83.69981 | 80.48042 | Promoter | 3 | 88043108 | 88055990 | 12883 | 1 | ENSMUST00 | 0 | Gpatch4 | ENSMUSG00000028069.11 |
| chr3 | 97867118 | 97868651 | 25.90858 | 83.69981 | 80.48042 | Promoter | 3 | 97746693 | 97868248 | 121556 | 2 | ENSMUST00 | 0 | Pde4dip | ENSMUSG00000038170.10 |
| chr4 | 150371430 | 150372679 | 27.50468 | 83.69844 | 80.48042 | Intron (ENSM | 4 | 150406394 | 150470381 | 63988 | 1 | ENSMUST00 | -33715 | Rere | ENSMUSG00000039852.11 |
| chr4 | 155734052 | 155735926 | 25.90858 | 83.69981 | 80.48042 | Promoter | 4 | 155734800 | 155740142 | 5343 | 1 | ENSMUST00 | 0 | Tmem240 | ENSMUSG00000084845.4 |
| chr5 | 114379830 | 114381806 | 25.90858 | 83.69981 | 80.48042 | Promoter | 5 | 114363567 | 114380515 | 16949 | 2 | ENSMUST00 | 0 | Kctd10 | ENSMUSG00000001098.10 |
| chr6 | 92939881 | 92944204 | 25.90858 | 83.69981 | 80.48042 | Promoter | 6 | 92772699 | 92943492 | 170794 | 2 | ENSMUST00 | 0 | Adamts9 | ENSMUSG00000030022.9 |
| chr7 | 44856729 | 44858016 | 25.90858 | 83.69981 | 80.48042 | Promoter | 7 | 44857139 | 44862650 | 5512 | 1 | ENSMUST00 | 0 | Pnkp | ENSMUSG00000002963.11 |
| chr7 | 98308665 | 98310071 | 25.90858 | 83.69981 | 80.48042 | Promoter | 7 | 98213658 | 98309528 | 95871 | 2 | ENSMUST00 | 0 | Acer3 | ENSMUSG00000030760.9 |
| chr7 | 19358771 | 19362302 | 25.90858 | 83.69981 | 80.48042 | Promoter | 7 | 19359749 | 19361316 | 1568 | 1 | ENSMUST00 | 0 | Ppp1r13l | ENSMUSG00000040734.9 |
| chr7 | 5050922 | 5052722 | 25.90858 | 83.69981 | 80.48042 | Promoter | 7 | 5051532 | 5053722 | 2191 | 1 | ENSMUST00 | 0 | Zfp580 | ENSMUSG00000055633.5 |
| chr8 | 54076910 | 54078170 | 25.90858 | 83.69981 | 80.48042 | Promoter | 8 | 54077532 | 54186454 | 108923 | 1 | ENSMUST00 | 0 | Vegfc | ENSMUSG00000031520.5 |
| chr9 | 71215273 | 71217229 | 25.90858 | 83.69981 | 80.48042 | Promoter | 9 | 71215789 | 71296243 | 80455 | 1 | ENSMUST00 | 0 | Aldh1a2 | ENSMUSG00000013584.5 |
| chr10 | 52232734 | 52234679 | 22.03264 | 83.65932 | 80.44393 | Promoter | 10 | 52233619 | 52321376 | 87758 | 1 | ENSMUST00 | 0 | Dcbld1 | ENSMUSG00000019891.8 |
| chr10 | 81128971 | 81129919 | 22.03264 | 83.65932 | 80.44393 | Distal Interge | 10 | 81135220 | 81144496 | 9277 | 1 | ENSMUST00 | -5301 | Zbtb7a | ENSMUSG00000035011.10 |
| chr12 | 70452544 | 70453891 | 22.03264 | 83.65932 | 80.44393 | Promoter | 12 | 70453095 | 70468040 | 14946 | 1 | ENSMUST00 | 0 | Tmx1 | ENSMUSG00000021072.7 |
| chr13 | 112926380 | 112928598 | 22.03264 | 83.65932 | 80.44393 | Promoter | 13 | 112867780 | 112927380 | 59601 | 2 | ENSMUST00 | 0 | Skiv2l2 | ENSMUSG00000016018.3 |
| chr15 | 99294285 | 99296025 | 22.03264 | 83.65932 | 80.44393 | Promoter | 15 | 99295087 | 99305965 | 10879 | 1 | ENSMUST00 | 0 | Prpf40b | ENSMUSG00000023007.10 |
| chr2 | 157133813 | 157135626 | 22.03264 | 83.65932 | 80.44393 | Promoter | 2 | 157097533 | 157135265 | 37733 | 2 | ENSMUST00 | 0 | Samhd1 | ENSMUSG00000027639.11 |
| chr2 | 151740435 | 151741794 | 22.03264 | 83.65932 | 80.44393 | Promoter | 2 | 151715812 | 151741310 | 25499 | 2 | ENSMUST00 | 0 | Psmf1 | ENSMUSG00000032869.10 |
| chr3 | 95855120 | 95856342 | 22.03264 | 83.65932 | 80.44393 | Promoter | 3 | 95830124 | 95855885 | 25762 | 2 | ENSMUST00 | 0 | Prpf3 | ENSMUSG00000015748.8 |
| chr6 | 122819193 | 122821504 | 22.03264 | 83.65932 | 80.44393 | Promoter | 6 | 122819914 | 122845366 | 25453 | 1 | ENSMUST00 | 0 | Foxj2 | ENSMUSG00000003154.10 |
| chr7 | 112953937 | 112957582 | 22.03264 | 83.65932 | 80.44393 | Promoter | 7 | 112953962 | 112957457 | 3496 | 1 | ENSMUST00 | 0 | Rassf10 | ENSMUSG00000098132.1 |
| chr10 | 127287777 | 127290398 | 27.94004 | 83.5095 | 80.29546 | Promoter | 10 | 127281956 | 127288890 | 6935 | 2 | ENSMUST00 | 0 | Mbd6 | ENSMUSG00000025409.9 |
| chr11 | 69015548 | 69016760 | 27.94004 | 83.5095 | 80.29546 | Promoter | 11 | 69015911 | 69036473 | 20563 | 1 | ENSMUST00 | 0 | Ctc1 | ENSMUSG00000020898.13 |
| chr13 | 100785849 | 100786933 | 27.94004 | 83.5095 | 80.29546 | Promoter | 13 | 100778650 | 100786570 | 7921 | 2 | ENSMUST00 | 0 | Ccnb1 | ENSMUSG00000041431.11 |
| chr16 | 92696643 | 92698812 | 27.94004 | 83.5095 | 80.29546 | Promoter | 16 | 92603656 | 92697328 | 93673 | 2 | ENSMUST00 | 0 | Runx1 | ENSMUSG00000022952.11 |
| chr2 | 11705089 | 11706956 | 27.94004 | 83.5095 | 80.29546 | Promoter | 2 | 11705287 | 11733977 | 28691 | 1 | ENSMUST00 | 0 | Il15ra | ENSMUSG00000023206.11 |
| chr2 | 28582020 | 28583594 | 27.94004 | 83.5095 | 80.29546 | Promoter | 2 | 28566311 | 28583189 | 16879 | 2 | ENSMUST00 | 0 | Gtf3c5 | ENSMUSG00000026816.9 |
| chr4 | 155360805 | 155362252 | 27.94004 | 83.5095 | 80.29546 | Promoter | 4 | 155261721 | 155361361 | 99641 | 2 | ENSMUST00 | 0 | Prkcz | ENSMUSG00000029053.11 |
| chr5 | 137115516 | 137116832 | 27.94004 | 83.5095 | 80.29546 | Promoter | 5 | 137105644 | 137116207 | 10564 | 2 | ENSMUST00 | 0 | Trim56 | ENSMUSG00000043279.9 |
| chr18 | 46311332 | 46312310 | 27.88522 | 83.41428 | 80.20149 | Promoter | 18 | 46282151 | 46311928 | 29778 | 2 | ENSMUST00 | 0 | Ccdc112 | ENSMUSG00000071855.4 |
| chr10 | 78590821 | 78592452 | 28.40368 | 83.36528 | 80.15265 | Promoter | 10 | 78584503 | 78591964 | 7462 | 2 | ENSMUST00 | 0 | Syde1 | ENSMUSG00000032714.6 |
| chr1 | 54193963 | 54195437 | 28.95052 | 83.36136 | 80.14891 | Promoter | 1 | 53806876 | 54194048 | 387173 | 2 | ENSMUST00 | 0 | Hecw2 | ENSMUSG00000042807.10 |
| chr11 | 60045661 | 60046559 | 28.95052 | 83.36136 | 80.14891 | Promoter | 11 | 59970614 | 60046477 | 75864 | 2 | ENSMUST00 | 0 | Pemt | ENSMUSG00000000301.11 |
| chr14 | 52315306 | 52316762 | 28.95052 | 83.36136 | 80.14891 | Promoter | 14 | 52311172 | 52316323 | 5152 | 2 | ENSMUST00 | 0 | Sall2 | ENSMUSG00000049532.8 |
| chr16 | 38711971 | 38713697 | 28.95052 | 83.36136 | 80.14891 | Promoter | 16 | 38598340 | 38713274 | 114935 | 2 | ENSMUST00 | 0 | Arhgap31 | ENSMUSG00000022799.4 |
| chr19 | 44562423 | 44563784 | 28.95052 | 83.36136 | 80.14891 | Promoter | 19 | 44562850 | 44563952 | 1103 | 1 | ENSMUST00 | 0 | Hif1an | ENSMUSG00000036450.4 |
| chrX | 163909011 | 163910239 | 28.95052 | 83.36136 | 80.14891 | Promoter | 20 | 163909017 | 163933666 | 24650 | 1 | ENSMUST00 | 0 | Ap1s2 | ENSMUSG00000031367.10 |
| chr4 | 62699665 | 62702487 | 24.69334 | 83.34509 | 80.13418 | Promoter | 4 | 62699680 | 62701153 | 1474 | 1 | ENSMUST00 | 0 | Rgs3 | ENSMUSG00000059810.13 |
| chr16 | 30387757 | 30389792 | 27.29656 | 83.33401 | 80.12321 | Promoter | 16 | 30312423 | 30388530 | 76108 | 2 | ENSMUST00 | 0 | Atp13a3 | ENSMUSG00000022533.8 |
| chr17 | 3556940 | 3557897 | 27.29656 | 83.33401 | 80.12321 | Promoter | 17 | 3519263 | 3557713 | 38451 | 2 | ENSMUST00 | 0 | Tfb1m | ENSMUSG00000036983.6 |
| chr5 | 143464014 | 143465328 | 27.29656 | 83.33401 | 80.12321 | Promoter | 5 | 143464493 | 143504414 | 39922 | 1 | ENSMUST00 | 0 | Daglb | ENSMUSG00000039206.11 |
| chr6 | 66535340 | 66536027 | 27.29656 | 83.33401 | 80.12321 | Promoter | 6 | 66535390 | 66541033 | 5644 | 1 | ENSMUST00 | 0 | Mad2l1 | ENSMUSG00000029910.9 |
| chr7 | 128417382 | 128418878 | 27.29656 | 83.33401 | 80.12321 | Promoter | 7 | 128373621 | 128418173 | 44553 | 2 | ENSMUST00 | 0 | Rgs10 | ENSMUSG00000030844.6 |
| chr8 | 36094331 | 36095970 | 27.29656 | 83.33401 | 80.12321 | Promoter | 8 | 36094828 | 36147787 | 52960 | 1 | ENSMUST00 | 0 | D8Ertd82e | ENSMUSG00000050271.7 |
| chr8 | 90886723 | 90888073 | 27.29656 | 83.33401 | 80.12321 | Intron (ENSM | 8 | 90907939 | 90908415 | 477 | 2 | ENSMUST00 | 20342 | Gm6658 | ENSMUSG00000074171.1 |
| chr9 | 108452250 | 108452840 | 27.29656 | 83.33401 | 80.12321 | Promoter | 9 | 108447658 | 108452377 | 4720 | 2 | ENSMUST00 | 0 | Klhdc8b | ENSMUSG00000032609.7 |
| chr6 | 117916314 | 117918503 | 25.14555 | 83.24793 | 80.0387 | Promoter | 6 | 117916981 | 117924382 | 7402 | 1 | ENSMUST00 | 0 | Hnrnpf | ENSMUSG00000042079.8 |
| chr11 | 97415033 | 97416334 | 26.7 | 83.22852 | 80.01951 | Promoter | 11 | 97415533 | 97502402 | 86870 | 1 | ENSMUST00 | 0 | Arhgap23 | ENSMUSG00000049807.11 |
| chr13 | 85289533 | 85290252 | 26.7 | 83.22852 | 80.01951 | Promoter | 13 | 85214780 | 85289029 | 74250 | 2 | ENSMUST00 | -505 | Rasa1 | ENSMUSG00000021549.8 |
| chr16 | 84702735 | 84703691 | 26.7 | 83.22852 | 80.01951 | Promoter | 16 | 84703167 | 84715245 | 12079 | 1 | ENSMUST00 | 0 | Gm26667 | ENSMUSG00000097418.2 |
| chr6 | 121183324 | 121184834 | 26.7 | 83.22852 | 80.01951 | Promoter | 6 | 121183667 | 121196185 | 12519 | 1 | ENSMUST00 | 0 | Pex26 | ENSMUSG00000067825.6 |
| chr17 | 34204429 | 34205851 | 25.62849 | 83.19241 | 79.98436 | Promoter | 17 | 34204567 | 34216318 | 11752 | 1 | ENSMUST00 | 0 | Tap2 | ENSMUSG00000024339.7 |
| chr3 | 139205213 | 139206353 | 25.62849 | 83.19241 | 79.98436 | Promoter | 3 | 139205694 | 139709610 | 503917 | 1 | ENSMUST00 | 0 | Stpg2 | ENSMUSG00000047940.8 |
| chr5 | 92309245 | 92310576 | 25.62849 | 83.19241 | 79.98436 | Promoter | 5 | 92284010 | 92310024 | 26015 | 2 | ENSMUST00 | 0 | Sdad1 | ENSMUSG00000029415.1 |
| chr6 | 134830125 | 134831005 | 25.62849 | 83.19241 | 79.98436 | Promoter | 6 | 134830154 | 134858931 | 28778 | 1 | ENSMUST00 | 0 | Crebl2 | ENSMUSG00000032652.8 |
| chr10 | 25296481 | 25297459 | 26.1454 | 83.18384 | 79.97624 | Promoter | 10 | 25283923 | 25297073 | 13151 | 2 | ENSMUST00 | 0 | Akap7 | ENSMUSG00000039166.10 |
| chr11 | 21570978 | 21572906 | 26.1454 | 83.18384 | 79.97624 | Promoter | 11 | 21572235 | 21664385 | 92151 | 1 | ENSMUST00 | 0 | Wdpcp | ENSMUSG00000020319.4 |
| chr11 | 68900974 | 68902659 | 26.1454 | 83.18384 | 79.97624 | Promoter | 11 | 68901583 | 68904534 | 2952 | 1 | ENSMUST00 | 0 | Rpl26 | ENSMUSG00000060938.9 |
| chr17 | 25578689 | 25579887 | 26.1454 | 83.18384 | 79.97624 | Promoter | 17 | 25579174 | 25662826 | 83653 | 1 | ENSMUST00 | 0 | Lmf1 | ENSMUSG00000002279.13 |
| chr4 | 135962895 | 135964814 | 26.1454 | 83.18384 | 79.97624 | Promoter | 4 | 135963727 | 135968178 | 4452 | 1 | ENSMUST00 | 0 | Gale | ENSMUSG00000028671.11 |
| chr6 | 128798583 | 128800204 | 26.1454 | 83.18384 | 79.97624 | Distal Interge | 6 | 128784764 | 128789215 | 4452 | 2 | ENSMUST00 | -9369 | Klrb1c | ENSMUSG00000030325.11 |
| chr11 | 97659512 | 97661428 | 28.83795 | 83.16807 | 79.96098 | Promoter | 11 | 97644184 | 97660367 | 16184 | 2 | ENSMUST00 | 0 | Gm11613 | ENSMUSG00000085586.1 |
| chr12 | 113156348 | 113156806 | 28.83795 | 83.16807 | 79.96098 | Promoter | 12 | 113156421 | 113165456 | 9036 | 1 | ENSMUST00 | 0 | 4930427A07R | ENSMUSG00000037466.9 |
| chr5 | 110057391 | 110058634 | 28.83795 | 83.16807 | 79.96098 | Distal Interge | 5 | 110046232 | 110046486 | 255 | 2 | ENSMUST00 | -10906 | Gm17655 | ENSMUSG00000090963.1 |
| chr2 | 129100894 | 129101797 | 28.28884 | 83.16705 | 79.96007 | Promoter | 2 | 129100995 | 129126594 | 25600 | 1 | ENSMUST00 | 0 | Polr1b | ENSMUSG00000027395.10 |
| chr12 | 10390672 | 10391830 | 20.9437 | 83.15932 | 79.95254 | Promoter | 12 | 10390780 | 10395562 | 4783 | 1 | ENSMUST00 | 0 | Rdh14 | ENSMUSG00000020621.5 |
| chr19 | 7164595 | 7166503 | 20.9437 | 83.15932 | 79.95254 | Intron (ENSM | 19 | 7199540 | 7200464 | 925 | 2 | ENSMUST00 | 33961 | Otub1 | ENSMUSG00000024767.6 |
| chr8 | 36610903 | 36614533 | 20.9437 | 83.15932 | 79.95254 | Promoter | 8 | 36567751 | 36613943 | 46193 | 2 | ENSMUST00 | 0 | Dlc1 | ENSMUSG00000031523.11 |
| chr9 | 21002313 | 21003705 | 20.9437 | 83.15932 | 79.95254 | Promoter | 9 | 21002737 | 21008835 | 6099 | 1 | ENSMUST00 | 0 | Mrpl4 | ENSMUSG00000003299.9 |
| chr11 | 116843598 | 116844460 | 27.72203 | 83.13062 | 79.92423 | Promoter | 11 | 116843673 | 116849629 | 5957 | 1 | ENSMUST00 | 0 | Mettl23 | ENSMUSG00000090266.5 |
| chr3 | 115713724 | 115715586 | 27.72203 | 83.13062 | 79.92423 | Promoter | 3 | 115710433 | 115715055 | 4623 | 2 | ENSMUST00 | 0 | S1pr1 | ENSMUSG00000045092.7 |
| chr8 | 9975661 | 9976429 | 27.72203 | 83.13062 | 79.92423 | Promoter | 8 | 9970020 | 9976323 | 6304 | 2 | ENSMUST00 | 0 | Lig4 | ENSMUSG00000049717.8 |
| chr1 | 172499800 | 172502632 | 28.23176 | 83.06847 | 79.86218 | Promoter | 1 | 172500047 | 172505934 | 5888 | 1 | ENSMUST00 | 0 | Tagln2 | ENSMUSG00000026547.10 |
| chr19 | 8837082 | 8838490 | 28.23176 | 83.06847 | 79.86218 | Promoter | 19 | 8837467 | 8848683 | 11217 | 1 | ENSMUST00 | 0 | Bscl2 | ENSMUSG00000071657.7 |
| chr6 | 137754427 | 137755439 | 28.23176 | 83.06847 | 79.86218 | Promoter | 6 | 137754542 | 137837891 | 83350 | 1 | ENSMUST00 | 0 | Dera | ENSMUSG00000030225.9 |
| chr1 | 40680316 | 40682232 | 28.7777 | 83.06458 | 79.8585 | Promoter | 1 | 40681712 | 40768885 | 87174 | 1 | ENSMUST00 | 0 | Slc9a2 | ENSMUSG00000026062.7 |
| chr17 | 22865878 | 22867248 | 28.7777 | 83.06458 | 79.8585 | Promoter | 17 | 22846697 | 22867134 | 20438 | 2 | ENSMUST00 | 0 | Zfp945 | ENSMUSG00000059142.10 |
| chr19 | 8251712 | 8252895 | 28.7777 | 83.06458 | 79.8585 | Distal Interge | 19 | 8160304 | 8218900 | 58597 | 2 | ENSMUST00 | -32813 | Slc22a29 | ENSMUSG00000075044.6 |
| chr7 | 127260210 | 127260775 | 28.7777 | 83.06458 | 79.8585 | Promoter | 7 | 127256959 | 127260709 | 3751 | 2 | ENSMUST00 | 0 | Dctpp1 | ENSMUSG00000042462.4 |
| chr8 | 105600584 | 105607833 | 28.7777 | 83.06458 | 79.8585 | Promoter | 8 | 105605229 | 105622194 | 16966 | 1 | ENSMUST00 | 0 | Fam65a | ENSMUSG00000038604.8 |
| chr10 | 81570656 | 81572272 | 28.7177 | 82.96152 | 79.75617 | Downstream | 10 | 81574561 | 81590466 | 15906 | 1 | ENSMUST00 | -2289 | Tle2 | ENSMUSG00000034771.10 |
| chr2 | 154568805 | 154570548 | 28.7177 | 82.96152 | 79.75617 | Promoter | 2 | 154559407 | 154569892 | 10486 | 2 | ENSMUST00 | 0 | E2f1 | ENSMUSG00000027490.12 |
| chr1 | 89579858 | 89582737 | 22.99841 | 82.87405 | 79.66897 | Promoter | 1 | 89580236 | 89895277 | 315042 | 1 | ENSMUST00 | 0 | Agap1 | ENSMUSG00000055013.9 |
| chr11 | 58306725 | 58308128 | 22.99841 | 82.87405 | 79.66897 | Promoter | 11 | 58307069 | 58314613 | 7545 | 1 | ENSMUST00 | 0 | Zfp692 | ENSMUSG00000037243.12 |
| chr13 | 74208061 | 74209360 | 22.99841 | 82.87405 | 79.66897 | Promoter | 13 | 74169805 | 74208700 | 38896 | 2 | ENSMUST00 | 0 | Exoc3 | ENSMUSG00000034152.5 |
| chr16 | 17910680 | 17911425 | 22.99841 | 82.87405 | 79.66897 | Promoter | 16 | 17900711 | 17911348 | 10638 | 2 | ENSMUST00 | 0 | Dgcr14 | ENSMUSG00000003527.8 |
| chr16 | 33380162 | 33382282 | 22.99841 | 82.87405 | 79.66897 | Promoter | 16 | 33380775 | 33503903 | 123129 | 1 | ENSMUST00 | 0 | Zfp148 | ENSMUSG00000022811.11 |
| chr19 | 37375447 | 37376930 | 22.99841 | 82.87405 | 79.66897 | Promoter | 19 | 37376403 | 37421859 | 45457 | 1 | ENSMUST00 | 0 | Kif11 | ENSMUSG00000012443.3 |
| chr3 | 63295209 | 63297475 | 22.99841 | 82.87405 | 79.66897 | Promoter | 3 | 63295872 | 63382231 | 86360 | 1 | ENSMUST00 | 0 | Mme | ENSMUSG00000027820.7 |
| chr5 | 122371498 | 122373147 | 22.99841 | 82.87405 | 79.66897 | Promoter | 5 | 122364580 | 122372364 | 7785 | 2 | ENSMUST00 | 0 | Fam216a | ENSMUSG00000029463.5 |
| chr6 | 113623555 | 113624970 | 22.99841 | 82.87405 | 79.66897 | Promoter | 6 | 113623959 | 113631633 | 7675 | 1 | ENSMUST00 | 0 | Vhl | ENSMUSG00000033933.8 |

| | | | | | | | | | | | | | | | |
|---|---|---|---|---|---|---|---|---|---|---|---|---|---|---|---|
| chr6 | 49319068 | 49320144 | 22.99841 | 82.87405 | 79.66897 | Promoter | 6 | 49319274 | 49341582 | 22309 | 1 | ENSMUST00 | 0 | Ccdc126 | ENSMUSG00000050786.8 |
| chr7 | 127776169 | 127778338 | 22.99841 | 82.87405 | 79.66897 | Promoter | 7 | 127776670 | 127800122 | 23453 | 1 | ENSMUST00 | 0 | Setd1a | ENSMUSG00000042308.9 |
| chr9 | 15357105 | 15358278 | 22.99841 | 82.87405 | 79.66897 | Promoter | 9 | 15316915 | 15357788 | 40874 | 2 | ENSMUST00 | 0 | 5830418K08I | ENSMUSG00000046111.11 |
| chr6 | 128354529 | 128356336 | 28.1183 | 82.87242 | 79.66893 | Promoter | 6 | 128321140 | 128355851 | 34712 | 2 | ENSMUST00 | 0 | Tulp3 | ENSMUSG00000001521.7 |
| chr7 | 25718404 | 25719600 | 28.65795 | 82.85884 | 79.65552 | Promoter | 7 | 25711106 | 25719088 | 7983 | 2 | ENSMUST00 | 0 | Ccdc97 | ENSMUSG00000002608.3 |
| chr13 | 14629710 | 14630866 | 28.06191 | 82.77493 | 79.57206 | Promoter | 13 | 14630245 | 14638202 | 7958 | 1 | ENSMUST00 | 0 | AW209491 | ENSMUSG00000039182.5 |
| chr15 | 36173499 | 36174698 | 28.06191 | 82.77493 | 79.57206 | Promoter | 15 | 36174010 | 36177007 | 2998 | 1 | ENSMUST00 | 0 | Polr2k | ENSMUSG00000045996.7 |
| chr17 | 24220123 | 24221335 | 28.06191 | 82.77493 | 79.57206 | Promoter | 17 | 24215054 | 24220790 | 5737 | 2 | ENSMUST00 | 0 | 1600002H07I | ENSMUSG00000024118.8 |
| chr5 | 138754087 | 138756196 | 28.06191 | 82.77493 | 79.57206 | Promoter | 5 | 138754514 | 138810077 | 55564 | 1 | ENSMUST00 | 0 | Fam20c | ENSMUSG00000025854.10 |
| chr7 | 16944385 | 16945377 | 28.06191 | 82.77493 | 79.57206 | Promoter | 7 | 16944682 | 16948828 | 4147 | 1 | ENSMUST00 | 0 | Pnmal2 | ENSMUSG00000070802.5 |
| chr9 | 14613601 | 14615334 | 28.06191 | 82.77493 | 79.57206 | Promoter | 9 | 14541966 | 14615000 | 73035 | 2 | ENSMUST00 | 0 | Amotl1 | ENSMUSG00000013076.10 |
| chr11 | 69036969 | 69037934 | 27.50739 | 82.75696 | 79.55487 | Distal Interge | 11 | 69034879 | 69036180 | 1302 | 1 | ENSMUST00 | 2091 | Ctc1 | ENSMUSG00000020898.13 |
| chr14 | 16572513 | 16576178 | 19.18445 | 82.69909 | 79.49731 | Promoter | 14 | 16430842 | 16575472 | 144631 | 2 | ENSMUST00 | 0 | Rarb | ENSMUSG00000017491.8 |
| chr2 | 30123739 | 30125160 | 19.18445 | 82.69909 | 79.49731 | Promoter | 2 | 30097283 | 30124454 | 27172 | 2 | ENSMUST00 | 0 | Zer1 | ENSMUSG00000039686.9 |
| chr15 | 77152314 | 77153833 | 28.00574 | 82.6778 | 79.47607 | Promoter | 15 | 77078991 | 77153812 | 74822 | 2 | ENSMUST00 | 0 | Rbfox2 | ENSMUSG00000033565.11 |
| chr2 | 75937799 | 75939059 | 18.46372 | 82.67341 | 79.47188 | Promoter | 2 | 75935849 | 75938585 | 2737 | 2 | ENSMUST00 | 0 | Ttc30b | ENSMUSG00000075273.4 |
| chr14 | 69804386 | 69806027 | 28.53919 | 82.65468 | 79.45325 | Promoter | 14 | 69784990 | 69805530 | 20541 | 2 | ENSMUST00 | 0 | Rhobtb2 | ENSMUSG00000022075.6 |
| chr13 | 67080753 | 67082022 | 27.9498 | 82.58102 | 79.37986 | Promoter | 13 | 67069398 | 67081512 | 12115 | 2 | ENSMUST00 | 0 | Zfp708 | ENSMUSG00000058883.9 |
| chr3 | 133542885 | 133545353 | 27.9498 | 82.58102 | 79.37986 | Promoter | 3 | 133463679 | 133544390 | 80712 | 2 | ENSMUST00 | 0 | Tet2 | ENSMUSG00000040943.8 |
| chr11 | 57962276 | 57963391 | 27.40132 | 82.57204 | 79.37107 | Distal Interge | 11 | 57969342 | 57969612 | 271 | 1 | ENSMUST00 | -5951 | Gm26025 | ENSMUSG00000089280.1 |
| chr2 | 18671836 | 18672324 | 27.40132 | 82.57204 | 79.37107 | Promoter | 2 | 18672384 | 18676230 | 3847 | 1 | ENSMUST00 | -60 | Commd3 | ENSMUSG00000051154.6 |
| chr15 | 99651111 | 99652018 | 28.48018 | 82.55319 | 79.35308 | Promoter | 15 | 99620496 | 99651643 | 31148 | 2 | ENSMUST00 | 0 | Racgap1 | ENSMUSG00000023015.8 |
| chr16 | 18127073 | 18128250 | 28.48018 | 82.55319 | 79.35308 | Promoter | 16 | 18127642 | 18152408 | 24767 | 1 | ENSMUST00 | 0 | Rtn4r | ENSMUSG00000043811.5 |
| chr19 | 46356699 | 46358297 | 28.48018 | 82.55319 | 79.35308 | Promoter | 19 | 46356880 | 46375252 | 18373 | 1 | ENSMUST00 | 0 | Tmem180 | ENSMUSG00000025227.9 |
| chr10 | 95323306 | 95324150 | 24.73908 | 82.49783 | 79.2979 | Promoter | 10 | 95174746 | 95324097 | 149352 | 2 | ENSMUST00 | 0 | Cradd | ENSMUSG00000045867.9 |
| chr16 | 8470484 | 8471552 | 24.73908 | 82.49783 | 79.2979 | Promoter | 16 | 8470788 | 8490684 | 19897 | 1 | ENSMUST00 | 0 | Mettl22 | ENSMUSG00000039345.10 |
| chr13 | 67723525 | 67724645 | 27.89408 | 82.48458 | 79.28481 | Promoter | 13 | 67705306 | 67724272 | 18967 | 2 | ENSMUST00 | 0 | Zfp65 | ENSMUSG00000071281.5 |
| chr17 | 24488162 | 24489370 | 27.89408 | 82.48458 | 79.28481 | Promoter | 17 | 24488783 | 24508905 | 20123 | 1 | ENSMUST00 | 0 | Caskin1 | ENSMUSG00000033597.8 |
| chr11 | 115491314 | 115492539 | 26.78979 | 82.44355 | 79.24416 | Promoter | 11 | 115490420 | 115491815 | 1396 | 2 | ENSMUST00 | 0 | Nt5c | ENSMUSG00000020736.7 |
| chr12 | 112808491 | 112809807 | 26.78979 | 82.44355 | 79.24416 | Promoter | 12 | 112808975 | 112815710 | 6736 | 1 | ENSMUST00 | 0 | BC022687 | ENSMUSG00000037594.5 |
| chr15 | 81925571 | 81926298 | 26.78979 | 82.44355 | 79.24416 | Promoter | 15 | 81915030 | 81926213 | 11184 | 2 | ENSMUST00 | 0 | Polr3h | ENSMUSG00000022476.7 |
| chr4 | 59549506 | 59550194 | 26.78979 | 82.44355 | 79.24416 | Promoter | 4 | 59471868 | 59549364 | 77497 | 2 | ENSMUST00 | -143 | Ptbp3 | ENSMUSG00000028382.10 |
| chr8 | 84147141 | 84148466 | 26.78979 | 82.44355 | 79.24416 | Promoter | 8 | 84148025 | 84162347 | 14323 | 1 | ENSMUST00 | 0 | 4930432K21I | ENSMUSG00000008129.9 |
| chr8 | 90347017 | 90349010 | 26.78979 | 82.44355 | 79.24416 | Promoter | 8 | 90247040 | 90348343 | 101304 | 2 | ENSMUST00 | 0 | Tox3 | ENSMUSG00000043668.8 |
| chr1 | 59684376 | 59686148 | 25.20037 | 82.41283 | 79.21442 | Promoter | 1 | 59684971 | 59692666 | 7696 | 1 | ENSMUST00 | 0 | Nop58 | ENSMUSG00000026020.4 |
| chr17 | 88255832 | 88257141 | 25.20037 | 82.41283 | 79.21442 | Distal Interge | 17 | 88342388 | 88342482 | 95 | 2 | ENSMUST00 | 85341 | Gm22542 | ENSMUSG00000088791.1 |
| chr2 | 37430089 | 37431854 | 25.20037 | 82.41283 | 79.21442 | Promoter | 2 | 37425500 | 37430919 | 5420 | 2 | ENSMUST00 | 0 | Zbtb6 | ENSMUSG00000066798.3 |
| chr3 | 54692163 | 54693652 | 25.20037 | 82.41283 | 79.21442 | Promoter | 3 | 54693105 | 54716837 | 23733 | 1 | ENSMUST00 | 0 | Supt20 | ENSMUSG00000027751.9 |
| chr7 | 98702178 | 98704054 | 25.20037 | 82.41283 | 79.21442 | Promoter | 7 | 98703103 | 98715367 | 12265 | 1 | ENSMUST00 | 0 | Prkrir | ENSMUSG00000030753.10 |
| chr19 | 46304097 | 46306268 | 27.29606 | 82.38838 | 79.1904 | Promoter | 19 | 46304737 | 46312090 | 7354 | 1 | ENSMUST00 | 0 | Nfkb2 | ENSMUSG00000025225.9 |
| chr9 | 120933357 | 120935851 | 27.83859 | 82.38849 | 79.1904 | Promoter | 9 | 120933400 | 120960507 | 27108 | 1 | ENSMUST00 | 0 | Ctnnb1 | ENSMUSG00000006932.11 |
| chr10 | 80602807 | 80603731 | 26.22205 | 82.37943 | 79.18155 | Promoter | 10 | 80602880 | 80607654 | 4775 | 1 | ENSMUST00 | 0 | Adat3 | ENSMUSG00000035370.7 |
| chr15 | 99251025 | 99253132 | 26.22205 | 82.37943 | 79.18155 | Promoter | 15 | 99242818 | 99251961 | 9144 | 2 | ENSMUST00 | 0 | Mcrs1 | ENSMUSG00000037570.10 |
| chr2 | 25402830 | 25403648 | 26.22205 | 82.37943 | 79.18155 | Promoter | 2 | 25403071 | 25409494 | 6424 | 1 | ENSMUST00 | 0 | Npdc1 | ENSMUSG00000015094.11 |
| chr3 | 137867258 | 137868612 | 26.22205 | 82.37943 | 79.18155 | Promoter | 3 | 137867675 | 137916557 | 48883 | 1 | ENSMUST00 | 0 | Dnajb14 | ENSMUSG00000074212.3 |
| chr2 | 33215963 | 33217985 | 25.69356 | 82.37146 | 79.1743 | Promoter | 2 | 33216069 | 33247717 | 31649 | 1 | ENSMUST00 | 0 | Angptl2 | ENSMUSG00000004105.6 |
| chr2 | 98666177 | 98667372 | 3.57929 | 82.36507 | 79.1683 | Promoter | 2 | 98666547 | 98667301 | 755 | 2 | ENSMUST00 | 0 | Gm10800 | ENSMUSG00000075014.1 |
| chr2 | 98662174 | 98663034 | 3.57929 | 82.36507 | 79.1683 | Promoter | 2 | 98662237 | 98664083 | 1847 | 1 | ENSMUST00 | 0 | Gm10801 | ENSMUSG00000075015.3 |
| chr9 | 2999956 | 3001822 | 3.57929 | 82.36507 | 79.1683 | Promoter | 9 | 3000922 | 3002330 | 1409 | 1 | ENSMUST00 | 0 | Gm10722 | ENSMUSG00000091028.1 |
| chr1 | 34801511 | 34802534 | 24.14628 | 82.32003 | 79.12344 | Promoter | 1 | 34801704 | 34813309 | 11606 | 1 | ENSMUST00 | 0 | Arhgef4 | ENSMUSG00000037509.13 |
| chr11 | 98438192 | 98439084 | 24.14628 | 82.32003 | 79.12344 | Promoter | 11 | 98437708 | 98438991 | 1284 | 2 | ENSMUST00 | 0 | Mien1 | ENSMUSG00000002580.7 |
| chr11 | 75178193 | 75179315 | 24.14628 | 82.32003 | 79.12344 | Promoter | 11 | 75175942 | 75178835 | 2894 | 2 | ENSMUST00 | 0 | Ovca2 | ENSMUSG00000038268.4 |
| chr13 | 23554700 | 23556159 | 24.14628 | 82.32003 | 79.12344 | Promoter | 13 | 23555087 | 23555830 | 744 | 1 | ENSMUST00 | 0 | Hist1h1d | ENSMUSG00000052565.6 |
| chr14 | 55671126 | 55672885 | 24.14628 | 82.32003 | 79.12344 | Promoter | 14 | 55672235 | 55678750 | 6516 | 1 | ENSMUST00 | 0 | Gmpr2 | ENSMUSG00000002326.5 |
| chr16 | 15862257 | 15863860 | 24.14628 | 82.32003 | 79.12344 | Promoter | 16 | 15848441 | 15863347 | 14907 | 2 | ENSMUST00 | 0 | Mzt2 | ENSMUSG00000022671.7 |
| chr16 | 38087776 | 38088856 | 24.14628 | 82.32003 | 79.12344 | Promoter | 16 | 38089001 | 38246084 | 157084 | 1 | ENSMUST00 | -145 | Gsk3b | ENSMUSG00000022812.8 |
| chr16 | 65814792 | 65816194 | 24.14628 | 82.32003 | 79.12344 | Promoter | 16 | 65815633 | 65863057 | 47425 | 1 | ENSMUST00 | 0 | Vgll3 | ENSMUSG00000091243.1 |
| chr17 | 64332312 | 64332961 | 24.14628 | 82.32003 | 79.12344 | Promoter | 17 | 64281005 | 64331916 | 50912 | 2 | ENSMUST00 | -397 | Pja2 | ENSMUSG00000024083.10 |
| chr17 | 84184922 | 84188829 | 24.14628 | 82.32003 | 79.12344 | Promoter | 17 | 84183931 | 84187947 | 4017 | 2 | ENSMUST00 | 0 | Zfp36l2 | ENSMUSG00000045817.8 |
| chr18 | 56572380 | 56574181 | 24.14628 | 82.32003 | 79.12344 | Promoter | 18 | 56572822 | 56587712 | 14891 | 1 | ENSMUST00 | 0 | Phax | ENSMUSG00000008301.6 |
| chr18 | 35661566 | 35662435 | 24.14628 | 82.32003 | 79.12344 | Promoter | 18 | 35656690 | 35662186 | 5497 | 2 | ENSMUST00 | 0 | Spata24 | ENSMUSG00000024352.6 |
| chr19 | 60811097 | 60812456 | 24.14628 | 82.32003 | 79.12344 | Promoter | 19 | 60811585 | 60830864 | 19280 | 1 | ENSMUST00 | 0 | Fam45a | ENSMUSG00000024993.10 |
| chr19 | 6995397 | 6996691 | 24.14628 | 82.32003 | 79.12344 | Promoter | 19 | 6993018 | 6996037 | 3020 | 2 | ENSMUST00 | 0 | Nudt22 | ENSMUSG00000037349.4 |
| chr19 | 37173708 | 37176086 | 24.14628 | 82.32003 | 79.12344 | Promoter | 19 | 37021297 | 37174095 | 152799 | 2 | ENSMUST00 | 0 | Cpeb3 | ENSMUSG00000039652.11 |
| chr3 | 123266862 | 123268531 | 24.14628 | 82.32003 | 79.12344 | Promoter | 3 | 123267496 | 123365635 | 98140 | 1 | ENSMUST00 | 0 | Sec24d | ENSMUSG00000039234.7 |
| chr3 | 127895360 | 127896376 | 24.14628 | 82.32003 | 79.12344 | Promoter | 3 | 127869688 | 127896323 | 26636 | 2 | ENSMUST00 | 0 | 5730508B09I | ENSMUSG00000050549.7 |
| chr7 | 28168874 | 28170713 | 24.14628 | 82.32003 | 79.12344 | Promoter | 7 | 28169748 | 28179269 | 9522 | 1 | ENSMUST00 | 0 | Fbl | ENSMUSG00000046865.6 |
| chr7 | 66839626 | 66840776 | 24.14628 | 82.32003 | 79.12344 | Promoter | 7 | 66839735 | 66969555 | 129821 | 1 | ENSMUST00 | 0 | Adamts17 | ENSMUSG00000058145.10 |
| chr7 | 52005354 | 52006907 | 24.14628 | 82.32003 | 79.12344 | Promoter | 7 | 51997165 | 52006018 | 8854 | 2 | ENSMUST00 | 0 | Svip | ENSMUSG00000074093.3 |
| chr8 | 111933176 | 111934090 | 24.14628 | 82.32003 | 79.12344 | Promoter | 8 | 111912018 | 111933791 | 21774 | 2 | ENSMUST00 | 0 | Tmem231 | ENSMUSG00000031951.7 |
| chr5 | 32713275 | 32714410 | 23.17682 | 82.30755 | 79.11359 | Distal Interge | 5 | 32724352 | 32736194 | 11843 | 1 | ENSMUST00 | -9942 | Gm2420 | ENSMUSG00000082305.1 |
| chr18 | 76240774 | 76242782 | 27.24373 | 82.29701 | 79.10307 | Promoter | 18 | 76241580 | 76305731 | 64152 | 1 | ENSMUST00 | 0 | Smad2 | ENSMUSG00000024563.10 |
| chr11 | 96849555 | 96850735 | 27.24215 | 82.29425 | 79.10038 | Promoter | 11 | 96849870 | 96851590 | 1721 | 1 | ENSMUST00 | 0 | Copz2 | ENSMUSG00000018672.10 |
| chr12 | 105032567 | 105033749 | 27.24215 | 82.29425 | 79.10038 | Promoter | 12 | 105032689 | 105040910 | 8222 | 1 | ENSMUST00 | 0 | Glrx5 | ENSMUSG00000021102.3 |
| chr12 | 85485608 | 85487883 | 27.24215 | 82.29425 | 79.10038 | Distal Interge | 12 | 85474911 | 85476063 | 1153 | 1 | ENSMUST00 | 10698 | Fos | ENSMUSG00000021250.8 |
| chr13 | 98889912 | 98891282 | 27.24215 | 82.29425 | 79.10038 | Promoter | 13 | 98839019 | 98891040 | 52022 | 2 | ENSMUST00 | 0 | Tnpo1 | ENSMUSG00000009470.11 |
| chr13 | 24638238 | 24639466 | 27.78331 | 82.29276 | 79.10038 | Promoter | 13 | 24638648 | 24733816 | 95169 | 1 | ENSMUST00 | 0 | Fam65b | ENSMUSG00000036006.14 |
| chr13 | 47122064 | 47123515 | 27.24215 | 82.29425 | 79.10038 | Promoter | 13 | 47122720 | 47247991 | 125272 | 1 | ENSMUST00 | 0 | Rnf144b | ENSMUSG00000038068.9 |
| chr16 | 52450004 | 52454370 | 27.24215 | 82.29425 | 79.10038 | Promoter | 16 | 52248996 | 52454074 | 205079 | 2 | ENSMUST00 | 0 | Alcam | ENSMUSG00000022636.8 |
| chr17 | 35049563 | 35050923 | 27.78331 | 82.29276 | 79.10038 | Promoter | 17 | 35049966 | 35058749 | 8784 | 1 | ENSMUST00 | 0 | Clic1 | ENSMUSG00000007041.9 |
| chr17 | 31057350 | 31059147 | 27.24215 | 82.29425 | 79.10038 | Promoter | 17 | 31057698 | 31115777 | 58080 | 1 | ENSMUST00 | 0 | Abcg1 | ENSMUSG00000024030.6 |
| chr2 | 32966378 | 32967701 | 27.24215 | 82.29425 | 79.10038 | Distal Interge | 2 | 32963389 | 32964035 | 647 | 1 | ENSMUST00 | 2990 | Rpl12 | ENSMUSG00000038900.12 |
| chr2 | 4651042 | 4652227 | 27.24215 | 82.29425 | 79.10038 | Promoter | 2 | 4622058 | 4652086 | 30029 | 2 | ENSMUST00 | 0 | Prpf18 | ENSMUSG00000039449.9 |
| chr4 | 130359115 | 130360023 | 27.24215 | 82.29425 | 79.10038 | Promoter | 4 | 130315383 | 130359943 | 44561 | 2 | ENSMUST00 | 0 | Zcchc17 | ENSMUSG00000028772.14 |
| chr4 | 40238727 | 40240422 | 27.24215 | 82.29425 | 79.10038 | Promoter | 4 | 40204849 | 40239828 | 34980 | 2 | ENSMUST00 | 0 | Ddx58 | ENSMUSG00000040296.10 |
| chr5 | 138170353 | 138172591 | 27.24215 | 82.29425 | 79.10038 | Promoter | 5 | 138172002 | 138178708 | 6707 | 1 | ENSMUST00 | 0 | Ap4m1 | ENSMUSG00000019518.5 |
| chr5 | 135167981 | 135169578 | 27.24215 | 82.29425 | 79.10038 | Promoter | 5 | 135168283 | 135181855 | 13573 | 1 | ENSMUST00 | 0 | Bcl7b | ENSMUSG00000029681.7 |
| chr5 | 122263710 | 122264482 | 27.24215 | 82.29425 | 79.10038 | Promoter | 5 | 122241287 | 122264460 | 23174 | 2 | ENSMUST00 | 0 | Tctn1 | ENSMUSG00000038593.13 |
| chr7 | 84409363 | 84410707 | 27.24215 | 82.29425 | 79.10038 | Promoter | 7 | 84246281 | 84409959 | 163679 | 2 | ENSMUST00 | 0 | Arnt2 | ENSMUSG00000015709.8 |
| chr7 | 120834657 | 120836487 | 27.24215 | 82.29425 | 79.10038 | Distal Interge | 7 | 120842831 | 120907219 | 64389 | 1 | ENSMUST00 | -6344 | Eef2k | ENSMUSG00000035064.11 |
| chr8 | 11476845 | 11479251 | 27.24215 | 82.29425 | 79.10038 | Promoter | 8 | 11453392 | 11478638 | 25247 | 2 | ENSMUST00 | 0 | Rab20 | ENSMUSG00000031504.5 |
| chr9 | 119357292 | 119358064 | 27.24215 | 82.29425 | 79.10038 | Promoter | 9 | 119357381 | 119393797 | 36417 | 1 | ENSMUST00 | 0 | Xylb | ENSMUSG00000035769.8 |
| chr3 | 107759461 | 107760773 | 27.1916 | 82.20596 | 79.01564 | Promoter | 3 | 107741048 | 107760236 | 19189 | 2 | ENSMUST00 | 0 | Csf1 | ENSMUSG00000014599.5 |
| chr11 | 62549794 | 62551074 | 21.7465 | 82.16968 | 78.97962 | Promoter | 11 | 62550047 | 62551041 | 995 | 2 | ENSMUST00 | 0 | Gm12279 | ENSMUSG00000087120.1 |
| chr14 | 18330705 | 18332441 | 21.7465 | 82.16968 | 78.97962 | Promoter | 14 | 18282722 | 18331859 | 49138 | 2 | ENSMUST00 | 0 | Ube2e1 | ENSMUSG00000021774.7 |
| chr14 | 31493999 | 31495974 | 21.7465 | 82.16968 | 78.97962 | Promoter | 14 | 31495079 | 31509858 | 14780 | 1 | ENSMUST00 | 0 | Eaf1 | ENSMUSG00000021890.5 |
| chr2 | 19371327 | 19372096 | 21.7465 | 82.16968 | 78.97962 | Promoter | 2 | 19371440 | 19383284 | 11845 | 1 | ENSMUST00 | 0 | Msrb2 | ENSMUSG00000023094.9 |
| chr2 | 180069990 | 180071542 | 21.7465 | 82.16968 | 78.97962 | Promoter | 2 | 180070588 | 180085895 | 15308 | 1 | ENSMUST00 | 0 | Mtg2 | ENSMUSG00000039069.14 |
| chr3 | 32529073 | 32530325 | 21.7465 | 82.16968 | 78.97962 | Promoter | 3 | 32529465 | 32579239 | 49775 | 1 | ENSMUST00 | 0 | Mfn1 | ENSMUSG00000027668.8 |
| chr5 | 143179842 | 143181208 | 21.7465 | 82.16968 | 78.97962 | Promoter | 5 | 143181017 | 143205075 | 24059 | 1 | ENSMUST00 | 0 | 4933411G11 | ENSMUSG00000029586.4 |
| chr5 | 121204229 | 121205346 | 21.7465 | 82.16968 | 78.97962 | Promoter | 5 | 121204481 | 121209241 | 4761 | 1 | ENSMUST00 | 0 | Rpl6 | ENSMUSG00000029614.9 |
| chr5 | 109751073 | 109752140 | 21.7465 | 82.16968 | 78.97962 | Promoter | 5 | 109735990 | 109751597 | 15608 | 2 | ENSMUST00 | 0 | 4930522L14I | ENSMUSG00000072762.4 |
| chr1 | 72211410 | 72212622 | 25.53309 | 82.08177 | 78.89293 | Promoter | 1 | 72159442 | 72212307 | 52866 | 2 | ENSMUST00 | 0 | Mreg | ENSMUSG00000039395.8 |
| chr1 | 160044086 | 160045572 | 25.53309 | 82.08177 | 78.89293 | Promoter | 1 | 160044384 | 160071536 | 27153 | 1 | ENSMUST00 | 0 | 4930523C07I | ENSMUSG00000090394.3 |
| chr10 | 18743271 | 18744336 | 25.53309 | 82.08177 | 78.89293 | Promoter | 10 | 18588011 | 18743758 | 155748 | 2 | ENSMUST00 | 0 | D10Bwg1379 | ENSMUSG00000019852.6 |
| chr11 | 50930543 | 50932040 | 25.53309 | 82.08177 | 78.89293 | Promoter | 11 | 50921823 | 50931633 | 9811 | 2 | ENSMUST00 | 0 | Zfp354b | ENSMUSG00000020335.8 |
| chr12 | 73123438 | 73124676 | 25.53309 | 82.08177 | 78.89293 | Promoter | 12 | 73123717 | 73233548 | 109832 | 1 | ENSMUST00 | 0 | Mnat1 | ENSMUSG00000021103.7 |
| chr12 | 85270031 | 85270927 | 25.53309 | 82.08177 | 78.89293 | Promoter | 12 | 85234529 | 85270564 | 36036 | 2 | ENSMUST00 | 0 | Mlh3 | ENSMUSG00000021245.9 |
| chr14 | 76009828 | 76011321 | 25.53309 | 82.08177 | 78.89293 | Promoter | 14 | 75896937 | 76010865 | 113929 | 2 | ENSMUST00 | 0 | Gtf2f2 | ENSMUSG00000067995.4 |
| chr18 | 12168096 | 12169590 | 25.53309 | 82.08177 | 78.89293 | Promoter | 18 | 12168717 | 12181116 | 12400 | 1 | ENSMUST00 | 0 | 3110002H16I | ENSMUSG00000024410.9 |
| chr19 | 46572370 | 46575355 | 25.53309 | 82.08177 | 78.89293 | Promoter | 19 | 46531109 | 46573085 | 41977 | 2 | ENSMUST00 | 0 | Arl3 | ENSMUSG00000025035.8 |
| chr2 | 167348132 | 167349979 | 25.53309 | 82.08177 | 78.89293 | Promoter | 2 | 167298444 | 167349183 | 50740 | 2 | ENSMUST00 | 0 | B4galt5 | ENSMUSG00000017929.8 |
| chr2 | 181679788 | 181680978 | 25.53309 | 82.08177 | 78.89293 | Promoter | 2 | 181680310 | 181687347 | 7038 | 1 | ENSMUST00 | 0 | Tcea2 | ENSMUSG00000059540.10 |
| chr4 | 155868656 | 155870211 | 25.53309 | 82.08177 | 78.89293 | Promoter | 4 | 155869546 | 155888685 | 19140 | 1 | ENSMUST00 | 0 | Cpsf3l | ENSMUSG00000029034.4 |
| chr5 | 136083512 | 136084617 | 25.53309 | 82.08177 | 78.89293 | Promoter | 5 | 136083916 | 136111860 | 27945 | 1 | ENSMUST00 | 0 | Rasa4 | ENSMUSG00000004952.8 |
| chr5 | 124598741 | 124599359 | 25.53309 | 82.08177 | 78.89293 | Promoter | 5 | 124598749 | 124602080 | 3332 | 1 | ENSMUST00 | 0 | Tctn2 | ENSMUSG00000029386.10 |
| chr6 | 135197714 | 135198661 | 25.53309 | 82.08177 | 78.89293 | Promoter | 6 | 135197977 | 135236242 | 38266 | 1 | ENSMUST00 | 0 | 8430419L09I | ENSMUSG00000030207.10 |
| chr6 | 61179826 | 61181514 | 25.53309 | 82.08177 | 78.89293 | Promoter | 6 | 61180324 | 62382865 | 1202542 | 1 | ENSMUST00 | 0 | Ccser1 | ENSMUSG00000039578.11 |
| chr7 | 25278343 | 25279876 | 25.53309 | 82.08177 | 78.89293 | Promoter | 7 | 25278422 | 25284969 | 6548 | 1 | ENSMUST00 | 0 | Cic | ENSMUSG00000005442.8 |
| chr7 | 130645344 | 130646821 | 25.53309 | 82.08177 | 78.89293 | Intron (ENSM | 7 | 130692556 | 130764784 | 72229 | 1 | ENSMUST00 | -45735 | Tacc2 | ENSMUSG00000030852.10 |
| chr7 | 42504984 | 42506214 | 25.53309 | 82.08177 | 78.89293 | Promoter | 7 | 42473386 | 42505726 | 32341 | 2 | ENSMUST00 | 0 | Zfp141 | ENSMUSG00000092416.1 |
| chr9 | 7872437 | 7873684 | 25.53309 | 82.08177 | 78.89293 | Promoter | 9 | 7848699 | 7873024 | 24326 | 2 | ENSMUST00 | 0 | Birc3 | ENSMUSG00000032000.12 |
| chr9 | 44084344 | 44085603 | 25.53309 | 82.08177 | 78.89293 | Promoter | 9 | 44084950 | 44093926 | 8977 | 1 | ENSMUST00 | 0 | Usp2 | ENSMUSG00000032010.9 |
| chr3 | 134239268 | 134240836 | 28.18876 | 82.05154 | 78.86629 | Promoter | 3 | 134240505 | 134240583 | 79 | 2 | ENSMUST00 | 0 | Mir1895 | ENSMUSG00000084564.1 |
| chr12 | 86946143 | 86947252 | 27.08794 | 82.02477 | 78.8401 | Promoter | 12 | 86947043 | 86962158 | 15116 | 1 | ENSMUST00 | 0 | Cipc | ENSMUSG00000034157.7 |
| chr13 | 49652992 | 49653956 | 27.08794 | 82.02477 | 78.8401 | Promoter | 13 | 49653350 | 49679015 | 25666 | 1 | ENSMUST00 | 0 | Nol8 | ENSMUSG00000021392.7 |
| chr3 | 27153244 | 27153909 | 28.07385 | 81.85352 | 78.66954 | Promoter | 3 | 27097222 | 27153861 | 56640 | 2 | ENSMUST00 | 0 | Ect2 | ENSMUSG00000027699.14 |
| chr7 | 24529736 | 24531913 | 28.07385 | 81.85352 | 78.66954 | Promoter | 7 | 24530648 | 24538599 | 7952 | 1 | ENSMUST00 | 0 | Irgq | ENSMUSG00000041037.7 |

| | | | | | | | | | | | | | | | |
|---|---|---|---|---|---|---|---|---|---|---|---|---|---|---|---|
| chr11 | 16951305 | 16951975 | 27.5102 | 81.81912 | 78.63541 | Promoter | 11 | 16951375 | 16954447 | 3073 | 1 | ENSMUST00 | 0 | Fbxo48 | ENSMUSG00000044966.4 |
| chr14 | 54553529 | 54554959 | 27.5102 | 81.81912 | 78.63541 | Promoter | 14 | 54541785 | 54554361 | 12577 | 2 | ENSMUST00 | 0 | Haus4 | ENSMUSG00000022177.8 |
| chr5 | 37337843 | 37338935 | 27.5102 | 81.81912 | 78.63541 | Promoter | 5 | 37338499 | 37425055 | 86557 | 1 | ENSMUST00 | 0 | Evc2 | ENSMUSG00000050248.11 |
| chr11 | 32347309 | 32348974 | 24.34554 | 81.76674 | 78.58393 | Promoter | 11 | 32347820 | 32428173 | 80354 | 1 | ENSMUST00 | 0 | Sh3pxd2b | ENSMUSG00000040711.7 |
| chr15 | 84922584 | 84924126 | 20.67859 | 81.70428 | 78.522 | Promoter | 15 | 84923428 | 84942955 | 19528 | 1 | ENSMUST00 | 0 | Nup50 | ENSMUSG00000016619.9 |
| chr3 | 95041610 | 95043054 | 20.67859 | 81.70428 | 78.522 | Promoter | 3 | 95032694 | 95042614 | 9921 | 2 | ENSMUST00 | 0 | Psmd4 | ENSMUSG00000005625.10 |
| chr8 | 111033744 | 111034867 | 20.67859 | 81.70428 | 78.522 | Promoter | 8 | 111033879 | 111045124 | 11246 | 1 | ENSMUST00 | 0 | Aars | ENSMUSG00000031960.9 |
| chr11 | 4703541 | 4705462 | 26.88298 | 81.66597 | 78.48403 | Promoter | 11 | 4704678 | 4737669 | 32992 | 1 | ENSMUST00 | 0 | Zmat5 | ENSMUSG00000009076.5 |
| chr11 | 116087178 | 116087997 | 26.88298 | 81.66597 | 78.48403 | Promoter | 11 | 116081829 | 116086995 | 5167 | 2 | ENSMUST00 | -184 | Wbp2 | ENSMUSG00000034341.12 |
| chr2 | 120027013 | 120028358 | 26.88298 | 81.66597 | 78.48403 | Promoter | 2 | 120027493 | 120043033 | 15541 | 1 | ENSMUST00 | 0 | Gm28042 | ENSMUSG00000033852.11 |
| chr5 | 151368629 | 151369885 | 26.88298 | 81.66597 | 78.48403 | Promoter | 5 | 151368864 | 151369172 | 309 | 2 | ENSMUST00 | 0 | D730045B01 | ENSMUSG00000054622.4 |
| chr9 | 106247236 | 106248391 | 26.88298 | 81.66597 | 78.48403 | Promoter | 9 | 106233455 | 106247730 | 14276 | 2 | ENSMUST00 | 0 | Alas1 | ENSMUSG00000032786.9 |
| chr5 | 143328488 | 143330078 | 24.78633 | 81.65401 | 78.47337 | Promoter | 5 | 143316489 | 143329256 | 12768 | 2 | ENSMUST00 | 0 | Zdhhc4 | ENSMUSG00000001844.5 |
| chr1 | 135687608 | 135688623 | 28.49099 | 81.63235 | 78.4519 | Promoter | 1 | 135606447 | 135688105 | 81659 | 2 | ENSMUST00 | 0 | Nav1 | ENSMUSG00000090399.1 |
| chr14 | 69767260 | 69767922 | 28.49099 | 81.63235 | 78.4519 | Promoter | 14 | 69767472 | 69784403 | 16932 | 1 | ENSMUST00 | 0 | Tnfrsf10b | ENSMUSG00000022074.5 |
| chr14 | 54925688 | 54927212 | 28.49099 | 81.63235 | 78.4519 | Promoter | 14 | 54916543 | 54926788 | 10246 | 2 | ENSMUST00 | 0 | Efs | ENSMUSG00000022203.6 |
| chr15 | 76246734 | 76247770 | 28.49099 | 81.63235 | 78.4519 | Promoter | 15 | 76246807 | 76249904 | 3098 | 1 | ENSMUST00 | 0 | Grina | ENSMUSG00000022564.6 |
| chr17 | 45594883 | 45595981 | 28.49099 | 81.63235 | 78.4519 | Promoter | 17 | 45585203 | 45595570 | 10368 | 2 | ENSMUST00 | 0 | Slc29a1 | ENSMUSG00000023942.10 |
| chr17 | 37045570 | 37046946 | 28.49099 | 81.63235 | 78.4519 | Promoter | 17 | 37045966 | 37075067 | 29102 | 1 | ENSMUST00 | 0 | Gabbr1 | ENSMUSG00000024462.12 |
| chr18 | 46526120 | 46526817 | 28.49099 | 81.63235 | 78.4519 | Promoter | 18 | 46501746 | 46525971 | 24226 | 2 | ENSMUST00 | -150 | Fem1c | ENSMUSG00000033319.5 |
| chr19 | 4439010 | 4440014 | 28.49099 | 81.63235 | 78.4519 | Promoter | 19 | 4425459 | 4439432 | 13974 | 2 | ENSMUST00 | 0 | Rhod | ENSMUSG00000041845.5 |
| chr19 | 8920094 | 8921667 | 28.49099 | 81.63235 | 78.4519 | Promoter | 19 | 8920374 | 8927234 | 6861 | 1 | ENSMUST00 | 0 | B3gat3 | ENSMUSG00000071649.5 |
| chr2 | 21204889 | 21206438 | 27.40245 | 81.632 | 78.4519 | Promoter | 2 | 21180731 | 21205365 | 24635 | 2 | ENSMUST00 | 0 | Enkur | ENSMUSG00000026679.3 |
| chr5 | 96792343 | 96794781 | 28.49099 | 81.63235 | 78.4519 | Promoter | 5 | 96793385 | 96845968 | 52584 | 1 | ENSMUST00 | 0 | Anxa3 | ENSMUSG00000029484.8 |
| chr5 | 37027988 | 37029567 | 28.49099 | 81.63235 | 78.4519 | Promoter | 5 | 37028335 | 37091447 | 63113 | 1 | ENSMUST00 | 0 | Jakmip1 | ENSMUSG00000063646.8 |
| chr7 | 89940455 | 89941624 | 28.49099 | 81.63235 | 78.4519 | Promoter | 7 | 89917529 | 89941185 | 23657 | 2 | ENSMUST00 | 0 | l7Rn6 | ENSMUSG00000062797.6 |
| chr8 | 71462915 | 71464611 | 27.40245 | 81.632 | 78.4519 | Promoter | 8 | 71456700 | 71463657 | 6958 | 2 | ENSMUST00 | 0 | Abhd8 | ENSMUSG00000007950.8 |
| chr8 | 3620446 | 3621394 | 28.49099 | 81.63235 | 78.4519 | Promoter | 8 | 3608421 | 3621305 | 12885 | 2 | ENSMUST00 | 0 | Xab2 | ENSMUSG00000019470.7 |
| chr8 | 125733809 | 125734983 | 28.49099 | 81.63235 | 78.4519 | Promoter | 8 | 125734203 | 125748235 | 14033 | 1 | ENSMUST00 | 0 | Ntpcr | ENSMUSG00000031851.9 |
| chrX | 7318483 | 7319382 | 28.49099 | 81.63235 | 78.4519 | Promoter | 20 | 7159469 | 7319233 | 159765 | 2 | ENSMUST00 | 0 | Clcn5 | ENSMUSG00000004317.9 |
| chrX | 152232960 | 152234571 | 28.49099 | 81.63235 | 78.4519 | Promoter | 20 | 152233020 | 152273768 | 40749 | 1 | ENSMUST00 | 0 | Kdm5c | ENSMUSG00000025332.9 |
| chr16 | 15636693 | 15638435 | 25.25706 | 81.58189 | 78.40311 | Promoter | 16 | 15637866 | 15684749 | 46884 | 1 | ENSMUST00 | 0 | Prkdc | ENSMUSG00000022672.8 |
| chr8 | 107047636 | 107049189 | 25.25706 | 81.58189 | 78.40311 | Promoter | 8 | 107046520 | 107048614 | 2095 | 2 | ENSMUST00 | 0 | Pdf | ENSMUSG00000078931.4 |
| chr16 | 26581293 | 26582646 | 26.30149 | 81.58109 | 78.4027 | Promoter | 16 | 26581704 | 26725264 | 143561 | 1 | ENSMUST00 | 0 | Il1rap | ENSMUSG00000022514.9 |
| chr17 | 26939737 | 26940231 | 26.30149 | 81.58109 | 78.4027 | Promoter | 17 | 26937973 | 26939478 | 1506 | 2 | ENSMUST00 | -260 | Cuta | ENSMUSG00000024194.10 |
| chr3 | 101376757 | 101378196 | 26.30149 | 81.58109 | 78.4027 | Promoter | 3 | 101377125 | 101463060 | 85936 | 1 | ENSMUST00 | 0 | Igsf3 | ENSMUSG00000042035.6 |
| chr5 | 149183302 | 149184274 | 26.30149 | 81.58109 | 78.4027 | Promoter | 5 | 149119565 | 149184063 | 64499 | 2 | ENSMUST00 | 0 | 5730422E09I | ENSMUSG00000097191.1 |
| chr7 | 127386192 | 127387231 | 26.30149 | 81.58109 | 78.4027 | Promoter | 7 | 127382260 | 127387166 | 4907 | 2 | ENSMUST00 | 0 | 9130019O22 | ENSMUSG00000030823.6 |
| chr18 | 76943673 | 76945062 | 26.83222 | 81.57701 | 78.3995 | Promoter | 18 | 76944115 | 76962053 | 17939 | 1 | ENSMUST00 | 0 | Hdhd2 | ENSMUSG00000025421.9 |
| chr14 | 63177488 | 63177972 | 27.90323 | 81.55928 | 78.38181 | Promoter | 14 | 63145153 | 63177793 | 32641 | 2 | ENSMUST00 | 0 | Fdft1 | ENSMUSG00000021273.9 |
| chr7 | 84605357 | 84606814 | 27.90323 | 81.55928 | 78.38181 | Promoter | 7 | 84585159 | 84605827 | 20669 | 2 | ENSMUST00 | 0 | Fah | ENSMUSG00000030630.10 |
| chr10 | 7390330 | 7393128 | 25.76091 | 81.55552 | 78.37826 | Distal Interge | 10 | 7445391 | 7473361 | 27971 | 2 | ENSMUST00 | 80233 | Ulbp1 | ENSMUSG00000079685.4 |
| chr5 | 114127243 | 114128311 | 25.76091 | 81.55552 | 78.37826 | Promoter | 5 | 114123926 | 114128176 | 4251 | 2 | ENSMUST00 | 0 | Alkbh2 | ENSMUSG00000044339.7 |
| chr7 | 45153704 | 45155178 | 25.76091 | 81.55552 | 78.37826 | Promoter | 7 | 45154303 | 45160069 | 5767 | 1 | ENSMUST00 | 0 | Pih1d1 | ENSMUSG00000003423.9 |
| chr7 | 73618591 | 73619476 | 25.76091 | 81.55552 | 78.37826 | Distal Interge | 7 | 73645985 | 73646089 | 105 | 2 | ENSMUST00 | 26613 | Gm26176 | ENSMUSG00000065729.1 |
| chr16 | 42875791 | 42876932 | 27.34889 | 81.53893 | 78.36224 | Promoter | 16 | 42875881 | 43463175 | 587295 | 1 | ENSMUST00 | 0 | Zbtb20 | ENSMUSG00000022708.11 |
| chr14 | 67071434 | 67072796 | 27.29554 | 81.44618 | 78.26986 | Promoter | 14 | 67014057 | 67072471 | 58415 | 2 | ENSMUST00 | 0 | Ppp2r2a | ENSMUSG00000022052.8 |
| chr7 | 28375275 | 28379742 | 28.38021 | 81.44218 | 78.26595 | Promoter | 7 | 28376784 | 28379255 | 2472 | 2 | ENSMUST00 | 0 | Zfp36 | ENSMUSG00000044786.5 |
| chr8 | 4252660 | 4253872 | 19.7569 | 81.42724 | 78.25113 | Promoter | 8 | 4253102 | 4256219 | 3118 | 1 | ENSMUST00 | 0 | Snapc2 | ENSMUSG00000011837.3 |
| chr13 | 64311532 | 64312714 | 26.73128 | 81.39996 | 78.22401 | Promoter | 13 | 64291836 | 64312710 | 20875 | 2 | ENSMUST00 | 0 | Aaed1 | ENSMUSG00000021482.9 |
| chr17 | 46297255 | 46298762 | 27.79064 | 81.36491 | 78.18899 | Promoter | 17 | 46297421 | 46301005 | 3585 | 1 | ENSMUST00 | 0 | Dlk2 | ENSMUSG00000047428.9 |
| chr13 | 25054929 | 25057285 | 27.2424 | 81.35376 | 78.17802 | Promoter | 13 | 25056004 | 25121523 | 65520 | 1 | ENSMUST00 | 0 | Dcdc2a | ENSMUSG00000035910.10 |
| chr1 | 131827249 | 131828792 | 22.68762 | 81.35053 | 78.1749 | Promoter | 1 | 131827493 | 131830276 | 2784 | 1 | ENSMUST00 | 0 | Slc41a1 | ENSMUSG00000013275.9 |
| chr1 | 30948406 | 30950686 | 22.68762 | 81.35053 | 78.1749 | Promoter | 1 | 30940302 | 30949770 | 9469 | 2 | ENSMUST00 | 0 | Ptp4a1 | ENSMUSG00000026064.10 |
| chr11 | 101416477 | 101417754 | 22.68762 | 81.35053 | 78.1749 | Promoter | 11 | 101406839 | 101417615 | 10777 | 2 | ENSMUST00 | 0 | Aarsd1 | ENSMUSG00000075528.8 |
| chr13 | 100615532 | 100617288 | 22.68762 | 81.35053 | 78.1749 | Promoter | 13 | 100595957 | 100616944 | 20988 | 2 | ENSMUST00 | 0 | Marveld2 | ENSMUSG00000021636.9 |
| chr13 | 8885151 | 8886558 | 22.68762 | 81.35053 | 78.1749 | Promoter | 13 | 8885501 | 8892451 | 6951 | 1 | ENSMUST00 | 0 | Idi1 | ENSMUSG00000058258.9 |
| chr14 | 54640971 | 54641938 | 22.68762 | 81.35053 | 78.1749 | Promoter | 14 | 54631992 | 54641364 | 9373 | 2 | ENSMUST00 | 0 | Cdh24 | ENSMUSG00000059674.6 |
| chr15 | 76079693 | 76081549 | 22.68762 | 81.35053 | 78.1749 | Promoter | 15 | 76070184 | 76080946 | 10763 | 2 | ENSMUST00 | 0 | Puf60 | ENSMUSG00000002524.10 |
| chr18 | 73572684 | 73574834 | 22.68762 | 81.35053 | 78.1749 | Promoter | 18 | 73573044 | 73592575 | 19532 | 1 | ENSMUST00 | 0 | Mex3c | ENSMUSG00000037253.8 |
| chr2 | 91264604 | 91266283 | 22.68762 | 81.35053 | 78.1749 | Promoter | 2 | 91264974 | 91276931 | 11958 | 1 | ENSMUST00 | 0 | Arfgap2 | ENSMUSG00000027255.9 |
| chr2 | 18672417 | 18673195 | 22.68762 | 81.35053 | 78.1749 | Promoter | 2 | 18672423 | 18676231 | 3809 | 1 | ENSMUST00 | 0 | Commd3 | ENSMUSG00000051154.6 |
| chr4 | 118359487 | 118360710 | 22.68762 | 81.35053 | 78.1749 | Promoter | 4 | 118359999 | 118361294 | 1296 | 1 | ENSMUST00 | 0 | Hyi | ENSMUSG00000006395.11 |
| chr4 | 14863572 | 14865346 | 22.68762 | 81.35053 | 78.1749 | Promoter | 4 | 14864076 | 14915176 | 51101 | 1 | ENSMUST00 | 0 | Tmem55a | ENSMUSG00000028221.3 |
| chr6 | 54429012 | 54432013 | 22.68762 | 81.35053 | 78.1749 | Promoter | 6 | 54269681 | 54430221 | 160541 | 2 | ENSMUST00 | 0 | 9130019P16I | ENSMUSG00000073067.5 |
| chr8 | 82773478 | 82774728 | 22.68762 | 81.35053 | 78.1749 | Promoter | 8 | 82763621 | 82774160 | 10540 | 2 | ENSMUST00 | 0 | Zfp330 | ENSMUSG00000031711.6 |
| chr8 | 15010824 | 15012332 | 22.68762 | 81.35053 | 78.1749 | Promoter | 8 | 15011025 | 15033333 | 22309 | 1 | ENSMUST00 | 0 | Kbtbd11 | ENSMUSG00000055675.6 |
| chr19 | 46348294 | 46349707 | 28.32091 | 81.34037 | 78.16626 | Promoter | 19 | 46348201 | 46350302 | 2102 | 2 | ENSMUST00 | 595 | RP23-62I23.3 | ENSMUSG00000099655.1 |
| chr3 | 88536138 | 88537358 | 18.95331 | 81.30177 | 78.12804 | Promoter | 3 | 88536301 | 88536382 | 82 | 2 | ENSMUST00 | 0 | Mir1905 | ENSMUSG00000084728.1 |
| chr5 | 20882357 | 20883362 | 18.95331 | 81.30177 | 78.12804 | Promoter | 5 | 20882191 | 20886870 | 4680 | 1 | ENSMUST00 | 167 | Tmem60 | ENSMUSG00000045435.8 |
| chr1 | 164457085 | 164459055 | 27.73468 | 81.26827 | 78.09463 | Promoter | 1 | 164437267 | 164458355 | 21089 | 2 | ENSMUST00 | 0 | Atp1b1 | ENSMUSG00000026576.7 |
| chr19 | 6398550 | 6400677 | 28.26186 | 81.23897 | 78.06556 | Promoter | 19 | 6399340 | 6415210 | 15871 | 1 | ENSMUST00 | 0 | Rasgrp2 | ENSMUSG00000032946.11 |
| chr4 | 41348146 | 41349973 | 26.63109 | 81.22407 | 78.05081 | Promoter | 4 | 41348996 | 41389766 | 40771 | 1 | ENSMUST00 | 0 | Ubap1 | ENSMUSG00000028437.9 |
| chr1 | 93305067 | 93306324 | 28.20306 | 81.13795 | 77.96497 | Promoter | 1 | 93299211 | 93305863 | 6653 | 2 | ENSMUST00 | 0 | Mterfd2 | ENSMUSG00000026273.8 |
| chr11 | 116198099 | 116199882 | 27.08421 | 81.07839 | 77.90557 | Promoter | 11 | 116171888 | 116198801 | 26914 | 2 | ENSMUST00 | 0 | Acox1 | ENSMUSG00000020777.11 |
| chr6 | 52215657 | 52218495 | 27.08421 | 81.07839 | 77.90557 | Promoter | 6 | 52216440 | 52217452 | 1013 | 2 | ENSMUST00 | 0 | Hoxa7 | ENSMUSG00000038236.7 |
| chr1 | 166253860 | 166256724 | 27.62344 | 81.07606 | 77.90333 | Promoter | 1 | 166254139 | 166316823 | 62685 | 1 | ENSMUST00 | 0 | Ildr2 | ENSMUSG00000040612.8 |
| chr4 | 116221357 | 116223082 | 27.62344 | 81.07606 | 77.90333 | Promoter | 4 | 116221618 | 116259576 | 37959 | 1 | ENSMUST00 | 0 | Pik3r3 | ENSMUSG00000028698.8 |
| chr5 | 97110863 | 97111913 | 27.56815 | 80.98046 | 77.80884 | Promoter | 5 | 97082329 | 97111596 | 29268 | 2 | ENSMUST00 | 0 | Paqr3 | ENSMUSG00000055725.6 |
| chr2 | 122764897 | 122766045 | 28.08619 | 80.9371 | 77.76572 | Promoter | 2 | 122765237 | 122809569 | 44333 | 1 | ENSMUST00 | 0 | Sqrdl | ENSMUSG00000005803.9 |
| chr11 | 106083606 | 106084248 | 24.38567 | 80.91491 | 77.74369 | Promoter | 11 | 106084613 | 106155446 | 70834 | 1 | ENSMUST00 | -365 | Map3k3 | ENSMUSG00000020700.11 |
| chr15 | 75908350 | 75909515 | 24.38567 | 80.91491 | 77.74369 | Promoter | 15 | 75894796 | 75909355 | 14560 | 2 | ENSMUST00 | 0 | Eef1d | ENSMUSG00000055762.10 |
| chr17 | 15563607 | 15565165 | 24.38567 | 80.91491 | 77.74369 | Promoter | 17 | 15553224 | 15564340 | 11117 | 2 | ENSMUST00 | 0 | Prdm9 | ENSMUSG00000051977.9 |
| chr1 | 143738928 | 143740484 | 26.97976 | 80.89639 | 77.72532 | Promoter | 1 | 143739029 | 143746776 | 7748 | 1 | ENSMUST00 | 0 | Glrx2 | ENSMUSG00000018196.13 |
| chr10 | 75211632 | 75212747 | 26.97976 | 80.89639 | 77.72532 | Promoter | 10 | 75212073 | 75312399 | 100327 | 1 | ENSMUST00 | 0 | Specc1l | ENSMUSG00000033444.8 |
| chr2 | 57181368 | 57182501 | 26.97976 | 80.89639 | 77.72532 | Promoter | 2 | 57127479 | 57181754 | 54276 | 2 | ENSMUST00 | 0 | BB557941 | ENSMUSG00000084938.1 |
| chr7 | 73739902 | 73741430 | 26.97976 | 80.89639 | 77.72532 | Promoter | 7 | 73740307 | 73776919 | 36613 | 1 | ENSMUST00 | 0 | Fam174b | ENSMUSG00000078670.2 |
| chr8 | 105595140 | 105600368 | 26.97976 | 80.89639 | 77.72532 | Distal Interge | 8 | 105605229 | 105622194 | 16966 | 1 | ENSMUST00 | -4861 | Fam65a | ENSMUSG00000038604.8 |
| chr9 | 42471420 | 42472586 | 24.83515 | 80.81389 | 77.64394 | Promoter | 9 | 42412316 | 42472258 | 59943 | 2 | ENSMUST00 | 0 | Tbcel | ENSMUSG00000037287.9 |
| chr8 | 121651240 | 121652363 | 27.45824 | 80.79031 | 77.62074 | Promoter | 8 | 121598703 | 121652901 | 54199 | 2 | ENSMUST00 | 538 | Zcchc14 | ENSMUSG00000061410.6 |
| chr7 | 19228284 | 19228710 | 26.38389 | 80.78929 | 77.61987 | Promoter | 7 | 19228334 | 19247868 | 19535 | 1 | ENSMUST00 | 0 | Opa3 | ENSMUSG00000052214.8 |
| chr1 | 82583771 | 82590897 | 23.80619 | 80.75649 | 77.58801 | Promoter | 1 | 82448423 | 82586849 | 138427 | 2 | ENSMUST00 | 0 | Col4a4 | ENSMUSG00000067158.9 |
| chr10 | 30841853 | 30844572 | 23.80619 | 80.75649 | 77.58801 | Promoter | 10 | 30832359 | 30842801 | 10443 | 2 | ENSMUST00 | 0 | Hey2 | ENSMUSG00000019789.8 |
| chr11 | 69991006 | 69992775 | 23.80619 | 80.75649 | 77.58801 | Promoter | 11 | 69991143 | 69994951 | 3809 | 1 | ENSMUST00 | 0 | Gabarap | ENSMUSG00000018567.9 |
| chr11 | 116335078 | 116335905 | 25.31571 | 80.75538 | 77.58801 | Promoter | 11 | 116330704 | 116335399 | 4696 | 2 | ENSMUST00 | 0 | Foxj1 | ENSMUSG00000034227.7 |
| chr11 | 77512623 | 77514094 | 23.80619 | 80.75649 | 77.58801 | Promoter | 11 | 77508099 | 77513370 | 5272 | 2 | ENSMUST00 | 0 | Trp53i13 | ENSMUSG00000044328.8 |
| chr12 | 69724145 | 69725595 | 23.80619 | 80.75649 | 77.58801 | Promoter | 12 | 69690436 | 69724874 | 34439 | 2 | ENSMUST00 | 0 | L2hgdh | ENSMUSG00000020988.8 |
| chr13 | 23445146 | 23447042 | 23.80619 | 80.75649 | 77.58801 | Distal Interge | 13 | 23427975 | 23431022 | 3048 | 2 | ENSMUST00 | -14125 | C230035I16R | ENSMUSG00000085024.1 |
| chr17 | 23673248 | 23675090 | 23.80619 | 80.75649 | 77.58801 | Promoter | 17 | 23673596 | 23675227 | 1632 | 1 | ENSMUST00 | 0 | Hcfc1r1 | ENSMUSG00000023904.5 |
| chr17 | 24723619 | 24724481 | 25.31571 | 80.75538 | 77.58801 | Promoter | 17 | 24722060 | 24724420 | 2361 | 2 | ENSMUST00 | 0 | Ndufb10 | ENSMUSG00000040048.9 |
| chr18 | 74267664 | 74269096 | 23.80619 | 80.75649 | 77.58801 | Promoter | 18 | 74268272 | 74282685 | 14414 | 1 | ENSMUST00 | 0 | Mbd1 | ENSMUSG00000024561.8 |
| chr19 | 26603929 | 26608671 | 23.80619 | 80.75649 | 77.58801 | Promoter | 19 | 26605115 | 26778321 | 173207 | 1 | ENSMUST00 | 0 | Smarca2 | ENSMUSG00000024921.11 |
| chr19 | 6340456 | 6341900 | 23.80619 | 80.75649 | 77.58801 | Promoter | 19 | 6341135 | 6355612 | 14478 | 1 | ENSMUST00 | 0 | Map4k2 | ENSMUSG00000024948.9 |
| chr3 | 36065828 | 36066842 | 23.80619 | 80.75649 | 77.58801 | Promoter | 3 | 36065979 | 36094944 | 28966 | 1 | ENSMUST00 | 0 | Acad9 | ENSMUSG00000027710.9 |
| chr4 | 104366910 | 104369011 | 23.80619 | 80.75649 | 77.58801 | Promoter | 4 | 104367113 | 104679496 | 312384 | 1 | ENSMUST00 | 0 | Dab1 | ENSMUSG00000028519.11 |
| chr5 | 71699782 | 71701650 | 23.80619 | 80.75649 | 77.58801 | Promoter | 5 | 71699932 | 72138389 | 438458 | 1 | ENSMUST00 | 0 | Gabrb1 | ENSMUSG00000029212.7 |
| chr5 | 99979164 | 99979864 | 23.80619 | 80.75649 | 77.58801 | Promoter | 5 | 99979061 | 99979663 | 603 | 1 | ENSMUST00 | 104 | 4930524J08R | ENSMUSG00000035364.1 |
| chr6 | 86683657 | 86684907 | 23.80619 | 80.75649 | 77.58801 | Promoter | 6 | 86675221 | 86684522 | 9302 | 2 | ENSMUST00 | 0 | Snrnp27 | ENSMUSG00000001158.8 |
| chr6 | 113482373 | 113484218 | 25.31571 | 80.75538 | 77.58801 | Promoter | 6 | 113483297 | 113493338 | 10042 | 1 | ENSMUST00 | 0 | Creld1 | ENSMUSG00000030284.6 |
| chr7 | 19307857 | 19310719 | 23.80619 | 80.75649 | 77.58801 | Promoter | 7 | 19302721 | 19310045 | 7325 | 2 | ENSMUST00 | 0 | Fosb | ENSMUSG00000003545.2 |
| chr7 | 45433582 | 45434632 | 23.80619 | 80.75649 | 77.58801 | Promoter | 7 | 45421898 | 45434623 | 12726 | 2 | ENSMUST00 | 0 | Ruvbl2 | ENSMUSG00000003868.7 |
| chr7 | 144469753 | 144471284 | 23.80619 | 80.75649 | 77.58801 | Promoter | 7 | 144435733 | 144470745 | 35013 | 2 | ENSMUST00 | 0 | Cttn | ENSMUSG00000031078.10 |
| chr7 | 27978497 | 27979759 | 23.80619 | 80.75649 | 77.58801 | Distal Interge | 7 | 27962708 | 27975368 | 12661 | 2 | ENSMUST00 | -3130 | Zfp780b | ENSMUSG00000063047.7 |
| chr8 | 25719928 | 25720990 | 23.80619 | 80.75649 | 77.58801 | Promoter | 8 | 25720061 | 25724887 | 4827 | 1 | ENSMUST00 | 0 | Ppapdc1b | ENSMUSG00000031570.11 |
| chr8 | 69996593 | 69997305 | 25.31571 | 80.75538 | 77.58801 | Promoter | 8 | 69907069 | 69996379 | 89311 | 2 | ENSMUST00 | -215 | Gatad2a | ENSMUSG00000036180.9 |
| chr8 | 25753663 | 25755164 | 23.80619 | 80.75649 | 77.58801 | Promoter | 8 | 25725324 | 25754280 | 28957 | 2 | ENSMUST00 | 0 | Ddhd2 | ENSMUSG00000061313.7 |
| chr9 | 110475529 | 110477398 | 23.80619 | 80.75649 | 77.58801 | Promoter | 9 | 110476994 | 110522131 | 45138 | 1 | ENSMUST00 | 0 | Kif9 | ENSMUSG00000032489.12 |
| chr9 | 72958252 | 72959286 | 25.31571 | 80.75538 | 77.58801 | Promoter | 9 | 72958852 | 73016319 | 57468 | 1 | ENSMUST00 | 0 | Dyx1c1Ccpg1 | ENSMUSG00000089865.2 |
| chr1 | 171079216 | 171079981 | 11.23412 | 80.75291 | 77.58743 | Promoter | 1 | 171079840 | 171079901 | 62 | 2 | ENSMUST00 | 0 | Gm27844 | ENSMUSG00000098965.1 |
| chr3 | 129327289 | 129333393 | 25.83068 | 80.74496 | 77.57947 | Promoter | 3 | 129269175 | 129332720 | 63546 | 2 | ENSMUST00 | 0 | Enpep | ENSMUSG00000028024.9 |
| chr5 | 138116298 | 138117872 | 25.83068 | 80.74496 | 77.57947 | Promoter | 5 | 138116903 | 138134213 | 17311 | 1 | ENSMUST00 | 0 | Zscan21 | ENSMUSG00000037017.10 |
| chr8 | 69399027 | 69399988 | 25.83068 | 80.74496 | 77.57947 | Promoter | 8 | 69399725 | 69400020 | 296 | 1 | ENSMUST00 | 0 | Gm25661 | ENSMUSG00000088329.1 |
| chr13 | 99514671 | 99517115 | 21.46036 | 80.68572 | 77.52129 | Promoter | 13 | 99421464 | 99516602 | 95139 | 2 | ENSMUST00 | 0 | Map1b | ENSMUSG00000052727.5 |
| chr15 | 88819536 | 88820434 | 21.46036 | 80.68572 | 77.52129 | Promoter | 15 | 88819646 | 88826681 | 7036 | 1 | ENSMUST00 | 0 | Creld2 | ENSMUSG00000023272.3 |
| chr16 | 25058670 | 25059871 | 21.46036 | 80.68572 | 77.52129 | Promoter | 16 | 25059639 | 25069058 | 9420 | 1 | ENSMUST00 | 0 | A230028O05 | ENSMUSG00000096960.1 |
| chr5 | 88553392 | 88554295 | 21.46036 | 80.68572 | 77.52129 | Promoter | 5 | 88554483 | 88556083 | 1601 | 1 | ENSMUST00 | -188 | Utp3 | ENSMUSG00000070697.3 |
| chr6 | 86194626 | 86196458 | 21.46036 | 80.68572 | 77.52129 | Promoter | 6 | 86195251 | 86275449 | 80199 | 1 | ENSMUST00 | 0 | Tgfa | ENSMUSG00000029999.9 |
| chr7 | 13123515 | 13124268 | 21.46036 | 80.68572 | 77.52129 | Distal Interge | 7 | 13131471 | 13132358 | 888 | 2 | ENSMUST00 | 8090 | Vmn1r87 | ENSMUSG00000070815.1 |

| | | | | | | | | | | | | | | | |
|---|---|---|---|---|---|---|---|---|---|---|---|---|---|---|---|
| chr9 | 37656139 | 37657688 | 21.46036 | 80.68572 | 77.52129 | Promoter | 9 | 37649182 | 37657213 | 8032 | 2 | ENSMUST00 | 0 | Tbrg1 | ENSMUSG00000011114.12 |
| chr10 | 45485336 | 45486940 | 26.82304 | 80.62296 | 77.45952 | Distal Interge | 10 | 45376620 | 45470201 | 93582 | 2 | ENSMUST00 | -15136 | Lin28b | ENSMUSG00000063804.7 |
| chr11 | 58866823 | 58867853 | 26.82304 | 80.62296 | 77.45952 | Promoter | 11 | 58867216 | 58883288 | 16073 | 1 | ENSMUST00 | 0 | 2810021J22R | ENSMUSG00000020491.10 |
| chr11 | 120097608 | 120098790 | 26.82304 | 80.62296 | 77.45952 | Promoter | 11 | 120090531 | 120098731 | 8201 | 2 | ENSMUST00 | 0 | Enthd2 | ENSMUSG00000025377.9 |
| chr15 | 38473271 | 38473881 | 26.82304 | 80.62296 | 77.45952 | Distal Interge | 15 | 38490473 | 38490547 | 75 | 2 | ENSMUST00 | 16666 | Mir6951 | ENSMUSG00000099110.1 |
| chr17 | 25791720 | 25793145 | 26.82304 | 80.62296 | 77.45952 | Promoter | 17 | 25790508 | 25792284 | 1777 | 2 | ENSMUST00 | 0 | Fam173a | ENSMUSG00000057411.8 |
| chr19 | 5923998 | 5925355 | 26.82304 | 80.62296 | 77.45952 | Promoter | 19 | 5917556 | 5924816 | 7261 | 2 | ENSMUST00 | 0 | Cdc42ep2 | ENSMUSG00000045664.4 |
| chr3 | 106482181 | 106483132 | 26.82304 | 80.62296 | 77.45952 | Promoter | 3 | 106482405 | 106487974 | 5570 | 1 | ENSMUST00 | 0 | Dennd2d | ENSMUSG00000027901.7 |
| chr3 | 89458727 | 89459745 | 26.82304 | 80.62296 | 77.45952 | Promoter | 3 | 89459118 | 89469013 | 9896 | 1 | ENSMUST00 | 0 | Pmvk | ENSMUSG00000027952.11 |
| chr5 | 118244557 | 118245070 | 26.82304 | 80.62296 | 77.45952 | Promoter | 5 | 118190736 | 118244861 | 54126 | 2 | ENSMUST00 | 0 | Rnft2 | ENSMUSG00000032850.11 |
| chr7 | 64373364 | 64374517 | 26.82304 | 80.62296 | 77.45952 | Promoter | 7 | 64346758 | 64373503 | 26746 | 2 | ENSMUST00 | 0 | Fan1 | ENSMUSG00000033458.12 |
| chr7 | 98360068 | 98361628 | 26.82304 | 80.62296 | 77.45952 | Promoter | 7 | 98350668 | 98361288 | 10621 | 2 | ENSMUST00 | 0 | Tsku | ENSMUSG00000049580.7 |
| chr7 | 28013457 | 28014834 | 26.82304 | 80.62296 | 77.45952 | Promoter | 7 | 27984854 | 28014115 | 29262 | 2 | ENSMUST00 | 0 | Zfp850 | ENSMUSG00000096916.2 |
| chr8 | 45974837 | 45976220 | 26.82304 | 80.62296 | 77.45952 | Promoter | 8 | 45975528 | 45996958 | 21431 | 1 | ENSMUST00 | 0 | Ufsp2 | ENSMUSG00000031634.7 |
| chr9 | 30426784 | 30428168 | 26.82304 | 80.62296 | 77.45952 | Promoter | 9 | 30427329 | 30466726 | 39398 | 1 | ENSMUST00 | 0 | Snx19 | ENSMUSG00000031993.6 |
| chr4 | 97757030 | 97758446 | 27.3492 | 80.60155 | 77.44116 | Promoter | 4 | 97754168 | 97757389 | 3222 | 2 | ENSMUST00 | 0 | Gm12676 | ENSMUSG00000085710.1 |
| chr5 | 31201290 | 31203128 | 27.3492 | 80.60155 | 77.44116 | Promoter | 5 | 31198981 | 31202303 | 3323 | 2 | ENSMUST00 | 0 | Zfp513 | ENSMUSG00000043059.11 |
| chr6 | 90462123 | 90463911 | 27.3492 | 80.60155 | 77.44116 | Promoter | 6 | 90462576 | 90475238 | 12663 | 1 | ENSMUST00 | 0 | Klf15 | ENSMUSG00000030087.6 |
| chr8 | 70233576 | 70235234 | 23.2161 | 80.55299 | 77.39275 | Promoter | 8 | 70220172 | 70234466 | 14295 | 2 | ENSMUST00 | 0 | Armc6 | ENSMUSG00000002343.6 |
| chr12 | 79295504 | 79297884 | 27.85533 | 80.53999 | 77.37978 | Promoter | 12 | 79297351 | 79327463 | 30113 | 1 | ENSMUST00 | 0 | Rad51b | ENSMUSG00000059060.8 |
| chr1 | 33719716 | 33720760 | 25.15761 | 80.47005 | 77.31023 | Promoter | 1 | 33719887 | 33726891 | 7005 | 1 | ENSMUST00 | 0 | Rab23 | ENSMUSG00000004768.9 |
| chr1 | 74816511 | 74817596 | 25.15761 | 80.47005 | 77.31023 | Distal Interge | 1 | 74793363 | 74803495 | 10133 | 1 | ENSMUST00 | 23149 | Wnt10a | ENSMUSG00000026167.9 |
| chr1 | 131867165 | 131868198 | 25.15761 | 80.47005 | 77.31023 | Promoter | 1 | 131867224 | 131872739 | 5516 | 1 | ENSMUST00 | 0 | Rab29 | ENSMUSG00000026433.8 |
| chr10 | 120474296 | 120477856 | 25.15761 | 80.47005 | 77.31023 | Promoter | 10 | 120361275 | 120476469 | 115195 | 2 | ENSMUST00 | 0 | Hmga2 | ENSMUSG00000056758.9 |
| chr12 | 81780327 | 81781786 | 25.15761 | 80.47005 | 77.31023 | Promoter | 12 | 81714950 | 81781170 | 66221 | 2 | ENSMUST00 | 0 | Map3k9 | ENSMUSG00000042724.7 |
| chr13 | 21401899 | 21403173 | 25.15761 | 80.47005 | 77.31023 | Promoter | 13 | 21387013 | 21402713 | 15701 | 2 | ENSMUST00 | 0 | Zkscan3 | ENSMUSG00000021327.13 |
| chr15 | 68927803 | 68930333 | 25.15761 | 80.47005 | 77.31023 | Promoter | 15 | 68928420 | 69093512 | 165093 | 1 | ENSMUST00 | 0 | Khdrbs3 | ENSMUSG00000022332.7 |
| chr16 | 4593688 | 4596406 | 25.15761 | 80.47005 | 77.31023 | Promoter | 16 | 4594713 | 4615957 | 21245 | 1 | ENSMUST00 | 0 | Glis2 | ENSMUSG00000014303.8 |
| chr16 | 46009644 | 46011113 | 25.15761 | 80.47005 | 77.31023 | Promoter | 16 | 45959263 | 46010218 | 50956 | 2 | ENSMUST00 | 0 | Plcxd2 | ENSMUSG00000087141.1 |
| chr17 | 55712007 | 55712707 | 25.15761 | 80.47005 | 77.31023 | Promoter | 17 | 55652025 | 55712628 | 60604 | 2 | ENSMUST00 | 0 | Pot1b | ENSMUSG00000024174.8 |
| chr18 | 33213183 | 33214422 | 25.15761 | 80.47005 | 77.31023 | Promoter | 18 | 33199355 | 33213862 | 14508 | 2 | ENSMUST00 | 0 | Stard4 | ENSMUSG00000024378.4 |
| chr18 | 80046481 | 80048014 | 25.15761 | 80.47005 | 77.31023 | Promoter | 18 | 80046895 | 80119640 | 72746 | 1 | ENSMUST00 | 0 | Pard6g | ENSMUSG00000056214.8 |
| chr19 | 58942770 | 58944181 | 25.15761 | 80.47005 | 77.31023 | Promoter | 19 | 58943425 | 58971419 | 27995 | 1 | ENSMUST00 | 0 | Eno4 | ENSMUSG00000048029.7 |
| chr19 | 40872916 | 40873873 | 25.15761 | 80.47005 | 77.31023 | Exon (ENSMU | 19 | 40868386 | 40874071 | 5686 | 1 | ENSMUST00 | 4531 | E030044B06I | ENSMUSG00000097740.1 |
| chr2 | 179441800 | 179443894 | 25.15761 | 80.47005 | 77.31023 | Promoter | 2 | 179442431 | 179899373 | 456943 | 1 | ENSMUST00 | 0 | Cdh4 | ENSMUSG00000000305.7 |
| chr2 | 168741242 | 168742724 | 25.15761 | 80.47005 | 77.31023 | Promoter | 2 | 168634438 | 168741800 | 107363 | 2 | ENSMUST00 | 0 | Atp9a | ENSMUSG00000027546.10 |
| chr2 | 156196153 | 156197761 | 25.15761 | 80.47005 | 77.31023 | Promoter | 2 | 156196466 | 156254310 | 57845 | 1 | ENSMUST00 | 0 | Phf20 | ENSMUSG00000038116.11 |
| chr2 | 154612811 | 154614277 | 25.15761 | 80.47005 | 77.31023 | Promoter | 2 | 154613297 | 154646821 | 33525 | 1 | ENSMUST00 | 0 | Zfp341 | ENSMUSG00000059842.7 |
| chr2 | 150135779 | 150137150 | 25.15761 | 80.47005 | 77.31023 | Promoter | 2 | 150114406 | 150136678 | 22273 | 2 | ENSMUST00 | 0 | Zfp120 | ENSMUSG00000068134.8 |
| chr3 | 35999793 | 36001137 | 25.15761 | 80.47005 | 77.31023 | Promoter | 3 | 35959312 | 36000609 | 41298 | 2 | ENSMUST00 | 0 | Mccc1 | ENSMUSG00000027709.4 |
| chr4 | 3573746 | 3575814 | 25.15761 | 80.47005 | 77.31023 | Promoter | 4 | 3549041 | 3574809 | 25769 | 2 | ENSMUST00 | 0 | Tmem68 | ENSMUSG00000028232.8 |
| chr4 | 134186247 | 134187154 | 25.15761 | 80.47005 | 77.31023 | Promoter | 4 | 134129858 | 134187112 | 57255 | 2 | ENSMUST00 | 0 | Cep85 | ENSMUSG00000037443.8 |
| chr5 | 30959152 | 30960708 | 25.15761 | 80.47005 | 77.31023 | Promoter | 5 | 30950853 | 30959234 | 8382 | 2 | ENSMUST00 | 0 | Preb | ENSMUSG00000045302.8 |
| chr5 | 123930269 | 123931336 | 25.15761 | 80.47005 | 77.31023 | Promoter | 5 | 123930689 | 123936645 | 5957 | 1 | ENSMUST00 | 0 | Ccdc62 | ENSMUSG00000061882.7 |
| chr7 | 44373719 | 44375025 | 25.15761 | 80.47005 | 77.31023 | Promoter | 7 | 44360099 | 44374514 | 14416 | 2 | ENSMUST00 | 0 | 1700008O03 | ENSMUSG00000008028.10 |
| chr7 | 35333177 | 35333959 | 25.15761 | 80.47005 | 77.31023 | Promoter | 7 | 35334170 | 35392289 | 58120 | 1 | ENSMUST00 | -211 | Rhpn2 | ENSMUSG00000030494.8 |
| chr7 | 141215258 | 141216658 | 25.15761 | 80.47005 | 77.31023 | Promoter | 7 | 141215860 | 141218658 | 2799 | 1 | ENSMUST00 | 0 | Rassf7 | ENSMUSG00000038618.6 |
| chr7 | 73374798 | 73376759 | 25.15761 | 80.47005 | 77.31023 | Promoter | 7 | 73375520 | 73419899 | 44380 | 1 | ENSMUST00 | 0 | Rgma | ENSMUSG00000070509.10 |
| chr7 | 12779972 | 12780813 | 25.15761 | 80.47005 | 77.31023 | Distal Interge | 7 | 12768092 | 12775658 | 7567 | 2 | ENSMUST00 | -4315 | Zscan18 | ENSMUSG00000070822.4 |
| chr8 | 75212723 | 75214161 | 25.15761 | 80.47005 | 77.31023 | Promoter | 8 | 75213944 | 75224113 | 10170 | 1 | ENSMUST00 | 0 | Rasd2 | ENSMUSG00000034472.8 |
| chr8 | 61927541 | 61928613 | 25.15761 | 80.47005 | 77.31023 | Promoter | 8 | 61928087 | 61963742 | 35656 | 1 | ENSMUST00 | 0 | Ddx60 | ENSMUSG00000037921.9 |
| chr9 | 66794589 | 66796275 | 25.15761 | 80.47005 | 77.31023 | Promoter | 9 | 66775202 | 66795490 | 20289 | 2 | ENSMUST00 | 0 | Aph1b | ENSMUSG00000032375.10 |
| chr9 | 19622050 | 19622942 | 25.15761 | 80.47005 | 77.31023 | Distal Interge | 9 | 19641224 | 19648048 | 6825 | 1 | ENSMUST00 | -18282 | Zfp317 | ENSMUSG00000057551.8 |
| chr9 | 22084690 | 22086122 | 25.15761 | 80.47005 | 77.31023 | Promoter | 9 | 22072246 | 22085402 | 13157 | 2 | ENSMUST00 | 0 | Ecsit | ENSMUSG00000066839.6 |
| chr12 | 111813362 | 111813984 | 27.79821 | 80.44167 | 77.28537 | Promoter | 12 | 111803193 | 111813841 | 10649 | 2 | ENSMUST00 | 0 | Xrcc3 | ENSMUSG00000021287.7 |
| chr4 | 28812513 | 28815985 | 27.79821 | 80.44167 | 77.28537 | Promoter | 4 | 28813131 | 28947047 | 133917 | 1 | ENSMUST00 | 0 | Epha7 | ENSMUSG00000028289.7 |
| chr12 | 71830218 | 71830892 | 27.24102 | 80.41412 | 77.25797 | Promoter | 12 | 71831078 | 71992367 | 161290 | 1 | ENSMUST00 | -186 | Daam1 | ENSMUSG00000034574.9 |
| chr19 | 10018186 | 10019690 | 27.24102 | 80.41412 | 77.25797 | Promoter | 19 | 10018193 | 10035456 | 17264 | 1 | ENSMUST00 | 0 | Rab3il1 | ENSMUSG00000024663.12 |
| chr17 | 81669148 | 81670762 | 27.74132 | 80.34372 | 77.18787 | Distal Interge | 17 | 81388691 | 81649607 | 260917 | 2 | ENSMUST00 | -19542 | Slc8a1 | ENSMUSG00000054640.9 |
| chr19 | 46075397 | 46076916 | 27.74132 | 80.34372 | 77.18787 | Promoter | 19 | 46075863 | 46085528 | 9666 | 1 | ENSMUST00 | 0 | Nolc1 | ENSMUSG00000015176.9 |
| chr5 | 33983108 | 33984220 | 27.74132 | 80.34372 | 77.18787 | Promoter | 5 | 33983437 | 33985009 | 1573 | 1 | ENSMUST00 | 0 | Gm1673 | ENSMUSG00000070858.6 |
| chr6 | 39557112 | 39558181 | 27.74132 | 80.34372 | 77.18787 | Promoter | 6 | 39462378 | 39557867 | 95490 | 2 | ENSMUST00 | 0 | Dennd2a | ENSMUSG00000038456.4 |
| chr2 | 75981255 | 75982222 | 27.18725 | 80.32091 | 77.16563 | Promoter | 2 | 75978247 | 75981967 | 3721 | 2 | ENSMUST00 | 0 | Ttc30a1 | ENSMUSG00000075271.4 |
| chr10 | 99261081 | 99265157 | 20.41348 | 80.25478 | 77.09972 | Promoter | 10 | 99263231 | 99267488 | 4258 | 1 | ENSMUST00 | 0 | Dusp6 | ENSMUSG00000019960.7 |
| chr11 | 100526782 | 100528524 | 20.41348 | 80.25478 | 77.09972 | Promoter | 11 | 100476353 | 100528000 | 51648 | 2 | ENSMUST00 | 0 | Acly | ENSMUSG00000020917.12 |
| chr14 | 120476887 | 120479540 | 20.41348 | 80.25478 | 77.09972 | Promoter | 14 | 120478461 | 120507194 | 28734 | 1 | ENSMUST00 | 0 | Rap2a | ENSMUSG00000051615.8 |
| chr2 | 148680257 | 148681868 | 27.13369 | 80.22803 | 77.07352 | Promoter | 2 | 148681023 | 148692949 | 11927 | 1 | ENSMUST00 | 0 | Gzf1 | ENSMUSG00000027439.9 |
| chr11 | 75530546 | 75532710 | 23.99775 | 80.19453 | 77.04016 | Promoter | 11 | 75531694 | 75543323 | 11630 | 1 | ENSMUST00 | 0 | Slc43a2 | ENSMUSG00000038178.11 |
| chr12 | 112619682 | 112620467 | 26.56992 | 80.18054 | 77.02628 | Promoter | 12 | 112620045 | 112641339 | 21295 | 1 | ENSMUST00 | 0 | Adssl1 | ENSMUSG00000011148.8 |
| chr2 | 160367241 | 160369416 | 27.62823 | 80.1489 | 76.99468 | Promoter | 2 | 160363703 | 160367065 | 3363 | 2 | ENSMUST00 | -177 | Mafb | ENSMUSG00000074622.4 |
| chr6 | 125164309 | 125166835 | 24.4673 | 80.14021 | 76.9862 | Promoter | 6 | 125161853 | 125166467 | 4615 | 2 | ENSMUST00 | 0 | Gapdh | ENSMUSG00000057666.13 |
| chr10 | 91180987 | 91183079 | 27.08035 | 80.13548 | 76.98148 | Distal Interge | 10 | 91160417 | 91171582 | 11166 | 2 | ENSMUST00 | -9406 | Tmpo | ENSMUSG00000019961.11 |
| chr11 | 6625269 | 6626927 | 27.08035 | 80.13548 | 76.98148 | Promoter | 11 | 6615598 | 6626043 | 10446 | 2 | ENSMUST00 | 0 | Tbrg4 | ENSMUSG00000000384.10 |
| chr11 | 69235064 | 69236519 | 27.08035 | 80.13548 | 76.98148 | Promoter | 11 | 69218117 | 69237022 | 18906 | 2 | ENSMUST00 | 503 | Gucy2e | ENSMUSG00000020890.6 |
| chr11 | 119022563 | 119023907 | 27.08035 | 80.13548 | 76.98148 | Promoter | 11 | 119022962 | 119031270 | 8309 | 1 | ENSMUST00 | 0 | Cbx2 | ENSMUSG00000025577.7 |
| chr2 | 110017452 | 110018514 | 27.08035 | 80.13548 | 76.98148 | Promoter | 2 | 110017817 | 110045316 | 27500 | 1 | ENSMUST00 | 0 | Ccdc34 | ENSMUSG00000027160.11 |
| chr3 | 122923822 | 122925193 | 27.08035 | 80.13548 | 76.98148 | Promoter | 3 | 122924242 | 122926194 | 1953 | 1 | ENSMUST00 | 0 | 1810037I17R | ENSMUSG00000054091.8 |
| chr5 | 89882037 | 89883791 | 27.08035 | 80.13548 | 76.98148 | Promoter | 5 | 89673841 | 89883334 | 209494 | 2 | ENSMUST00 | 0 | Adamts3 | ENSMUSG00000043635.7 |
| chr1 | 92910170 | 92912267 | 24.4271 | 80.06635 | 76.91331 | Promoter | 1 | 92910783 | 92920841 | 10059 | 1 | ENSMUST00 | 0 | Rnpepl1 | ENSMUSG00000026269.9 |
| chr6 | 116263822 | 116264669 | 24.4271 | 80.06635 | 76.91331 | Promoter | 6 | 116264222 | 116330298 | 66077 | 1 | ENSMUST00 | 0 | Zfand4 | ENSMUSG00000042213.11 |
| chr14 | 119137389 | 119139165 | 27.57204 | 80.05205 | 76.89922 | Promoter | 14 | 119138415 | 119869594 | 731180 | 1 | ENSMUST00 | 0 | Hs6st3 | ENSMUSG00000053465.5 |
| chr8 | 70791832 | 70793049 | 27.02721 | 80.04324 | 76.89059 | Promoter | 8 | 70778117 | 70792433 | 14317 | 2 | ENSMUST00 | 0 | Mast3 | ENSMUSG00000031833.9 |
| chr1 | 37429187 | 37431192 | 19.50994 | 80.00812 | 76.85559 | Promoter | 1 | 37430170 | 37439124 | 8955 | 1 | ENSMUST00 | 0 | Unc50 | ENSMUSG00000026111.10 |
| chr5 | 24907736 | 24909252 | 19.50994 | 80.00812 | 76.85559 | Promoter | 5 | 24862748 | 24908469 | 45722 | 2 | ENSMUST00 | 0 | Prkag2 | ENSMUSG00000028944.9 |
| chr1 | 34459177 | 34460393 | 26.46939 | 80.00455 | 76.85218 | Promoter | 1 | 34459762 | 34461056 | 1295 | 1 | ENSMUST00 | 0 | Ptpn18 | ENSMUSG00000026126.10 |
| chr10 | 78068651 | 78070259 | 26.46939 | 80.00455 | 76.85218 | Promoter | 10 | 78069360 | 78079525 | 10166 | 1 | ENSMUST00 | 0 | Icosl | ENSMUSG00000000732.8 |
| chr18 | 12861677 | 12863060 | 26.46939 | 80.00455 | 76.85218 | Promoter | 18 | 12755314 | 12862673 | 107360 | 2 | ENSMUST00 | 0 | Osbpl1a | ENSMUSG00000044252.12 |
| chr2 | 121289045 | 121290238 | 26.46939 | 80.00455 | 76.85218 | Promoter | 2 | 121289600 | 121310832 | 21233 | 1 | ENSMUST00 | 0 | Map1a | ENSMUSG00000027254.8 |
| chr18 | 66290977 | 66292649 | 24.88564 | 79.9777 | 76.82633 | Promoter | 18 | 66045302 | 66291446 | 246145 | 2 | ENSMUST00 | 0 | Ccbe1 | ENSMUSG00000046318.10 |
| chr3 | 89417931 | 89420270 | 24.88564 | 79.9777 | 76.82633 | Promoter | 3 | 89415472 | 89418383 | 2912 | 2 | ENSMUST00 | 0 | Cks1b | ENSMUSG00000028044.6 |
| chr10 | 120112319 | 120113648 | 27.51607 | 79.95555 | 76.80458 | Promoter | 10 | 120083608 | 120112987 | 29380 | 2 | ENSMUST00 | 0 | Helb | ENSMUSG00000020228.6 |
| chr10 | 81574805 | 81575907 | 27.51607 | 79.95555 | 76.80458 | Promoter | 10 | 81575260 | 81590470 | 15211 | 1 | ENSMUST00 | 0 | Tle2 | ENSMUSG00000034771.10 |
| chr11 | 77489086 | 77490311 | 27.51607 | 79.95555 | 76.80458 | Promoter | 11 | 77470485 | 77489666 | 19182 | 2 | ENSMUST00 | 0 | Ankrd13b | ENSMUSG00000037907.11 |
| chr13 | 114930908 | 114932478 | 27.51607 | 79.95555 | 76.80458 | Promoter | 13 | 114835916 | 114932041 | 96126 | 2 | ENSMUST00 | 0 | Itga2 | ENSMUSG00000015533.8 |
| chr11 | 68973019 | 68974189 | 26.97428 | 79.95133 | 76.80052 | Promoter | 11 | 68972484 | 68973789 | 1306 | 2 | ENSMUST00 | 0 | Rangrf | ENSMUSG00000032892.9 |
| chr11 | 59471460 | 59472616 | 25.90299 | 79.94019 | 76.78951 | Promoter | 11 | 59461197 | 59472474 | 11278 | 2 | ENSMUST00 | 0 | Zfp867 | ENSMUSG00000054519.8 |
| chr13 | 67060203 | 67061680 | 25.90299 | 79.94019 | 76.78951 | Promoter | 13 | 67038594 | 67061170 | 22577 | 2 | ENSMUST00 | 0 | Zfp712 | ENSMUSG00000090641.1 |
| chr17 | 34931249 | 34932414 | 25.90299 | 79.94019 | 76.78951 | Promoter | 17 | 34931253 | 34935921 | 4669 | 1 | ENSMUST00 | 0 | Neu1 | ENSMUSG00000007038.5 |
| chr19 | 38818370 | 38819734 | 25.37642 | 79.93365 | 76.78373 | Promoter | 19 | 38788128 | 38819237 | 31110 | 2 | ENSMUST00 | 0 | Noc3l | ENSMUSG00000024999.7 |
| chr5 | 8996006 | 8997859 | 25.37642 | 79.93365 | 76.78373 | Promoter | 5 | 8966033 | 8997324 | 31292 | 2 | ENSMUST00 | 0 | Crot | ENSMUSG00000003623.4 |
| chr5 | 23782853 | 23784133 | 25.37642 | 79.93365 | 76.78373 | Promoter | 5 | 23740648 | 23783711 | 43064 | 2 | ENSMUST00 | 0 | Pus7 | ENSMUSG00000057541.9 |
| chr8 | 123477139 | 123479106 | 18.02928 | 79.93274 | 76.78327 | Promoter | 8 | 123477903 | 123503916 | 26014 | 1 | ENSMUST00 | 0 | Afg3l1 | ENSMUSG00000031967.10 |
| chr11 | 3537868 | 3539726 | 28.03146 | 79.91027 | 76.76085 | Promoter | 11 | 3517523 | 3539239 | 21717 | 2 | ENSMUST00 | 0 | Smtn | ENSMUSG00000020439.12 |
| chr15 | 89139588 | 89141123 | 28.03146 | 79.91027 | 76.76085 | Promoter | 15 | 89130585 | 89140703 | 10119 | 2 | ENSMUST00 | 0 | Mapk12 | ENSMUSG00000022610.9 |
| chr15 | 98933157 | 98935385 | 28.03146 | 79.91027 | 76.76085 | Promoter | 15 | 98931425 | 98934565 | 3141 | 2 | ENSMUST00 | 0 | Tuba1b | ENSMUSG00000023004.8 |
| chr2 | 34913146 | 34914714 | 28.03146 | 79.91027 | 76.76085 | Promoter | 2 | 34893757 | 34913976 | 20220 | 2 | ENSMUST00 | 0 | Phf19 | ENSMUSG00000026873.4 |
| chr2 | 116064436 | 116065867 | 28.03146 | 79.91027 | 76.76085 | Promoter | 2 | 115863064 | 116065047 | 201984 | 2 | ENSMUST00 | 0 | Meis2 | ENSMUSG00000027210.15 |
| chr2 | 92182752 | 92186025 | 28.03146 | 79.91027 | 76.76085 | Promoter | 2 | 92184106 | 92361829 | 177724 | 1 | ENSMUST00 | 0 | Phf21a | ENSMUSG00000058318.10 |
| chr4 | 128805149 | 128807225 | 28.03146 | 79.91027 | 76.76085 | Promoter | 4 | 128773088 | 128806045 | 32958 | 2 | ENSMUST00 | 0 | Zfp362 | ENSMUSG00000028799.10 |
| chr5 | 99728110 | 99729245 | 28.03146 | 79.91027 | 76.76085 | Promoter | 5 | 99297244 | 99729045 | 431802 | 2 | ENSMUST00 | 0 | A930011G23 | ENSMUSG00000089809.3 |
| chr6 | 8777263 | 8778576 | 28.03146 | 79.91027 | 76.76085 | Promoter | 6 | 8630527 | 8778154 | 147628 | 2 | ENSMUST00 | 0 | Ica1 | ENSMUSG00000062995.7 |
| chr6 | 120492232 | 120494265 | 28.03146 | 79.91027 | 76.76085 | Promoter | 6 | 120488939 | 120493807 | 4869 | 2 | ENSMUST00 | 0 | Cecr6 | ENSMUSG00000094626.1 |
| chr7 | 112225499 | 112226787 | 28.03146 | 79.91027 | 76.76085 | Promoter | 7 | 112225856 | 112355194 | 129339 | 1 | ENSMUST00 | 0 | Mical2 | ENSMUSG00000038244.8 |
| chr7 | 16174832 | 16175809 | 28.03146 | 79.91027 | 76.76085 | Promoter | 7 | 16175090 | 16178793 | 3704 | 1 | ENSMUST00 | 0 | Meis3 | ENSMUSG00000041420.13 |
| chr9 | 32031756 | 32033342 | 28.03146 | 79.91027 | 76.76085 | Distal Interge | 9 | 32116136 | 32268446 | 152311 | 1 | ENSMUST00 | -82794 | Arhgap32 | ENSMUSG00000041444.9 |
| chr9 | 58487725 | 58488315 | 28.03146 | 79.91027 | 76.76085 | Promoter | 9 | 58488603 | 58499742 | 11140 | 1 | ENSMUST00 | 0 | 6030419C18I | ENSMUSG00000066607.4 |
| chrX | 13280005 | 13282849 | 28.03146 | 79.91027 | 76.76085 | Promoter | 20 | 13280970 | 13294052 | 13083 | 1 | ENSMUST00 | 0 | Ddx3x | ENSMUSG00000000787.7 |
| chrX | 74312999 | 74313879 | 28.03146 | 79.91027 | 76.76085 | Promoter | 20 | 74313033 | 74320149 | 7117 | 1 | ENSMUST00 | 0 | Fam50a | ENSMUSG00000001962.8 |
| chrX | 8175164 | 8176464 | 28.03146 | 79.91027 | 76.76085 | Promoter | 20 | 8154472 | 8176181 | 21710 | 2 | ENSMUST00 | 0 | Tbc1d25 | ENSMUSG00000039201.4 |
| chr4 | 97751131 | 97752475 | 27.46033 | 79.85941 | 76.71169 | Intron (ENSM | 4 | 97754168 | 97757389 | 3222 | 2 | ENSMUST00 | 4914 | Gm12676 | ENSMUSG00000085710.1 |
| chr1 | 119503795 | 119504823 | 22.37683 | 79.83291 | 76.68538 | Promoter | 1 | 119470305 | 119504794 | 34490 | 2 | ENSMUST00 | 0 | Ralb | ENSMUSG00000004451.9 |
| chr10 | 58323527 | 58324339 | 22.37683 | 79.83291 | 76.68538 | Promoter | 10 | 58323466 | 58424691 | 101226 | 1 | ENSMUST00 | 62 | Lims1 | ENSMUSG00000019920.12 |
| chr11 | 70668946 | 70670273 | 22.37683 | 79.83291 | 76.68538 | Promoter | 11 | 70663771 | 70669416 | 5646 | 2 | ENSMUST00 | 0 | Spag7 | ENSMUSG00000018287.7 |
| chr11 | 32532209 | 32534043 | 22.37683 | 79.83291 | 76.68538 | Promoter | 11 | 32533305 | 32624587 | 91283 | 1 | ENSMUST00 | 0 | Stk10 | ENSMUSG00000020272.8 |
| chr13 | 67483325 | 67484694 | 22.37683 | 79.83291 | 76.68538 | Promoter | 13 | 67471514 | 67484225 | 12712 | 2 | ENSMUST00 | 0 | Zfp874b | ENSMUSG00000059839.8 |
| chr14 | 31217560 | 31220214 | 22.37683 | 79.83291 | 76.68538 | Promoter | 14 | 31217860 | 31230350 | 12491 | 1 | ENSMUST00 | 0 | Sema3g | ENSMUSG00000021904.5 |
| chr19 | 7038988 | 7039985 | 22.37683 | 79.83291 | 76.68538 | Promoter | 19 | 7020696 | 7040026 | 19331 | 2 | ENSMUST00 | 41 | Stip1 | ENSMUSG00000024966.8 |
| chr2 | 119547092 | 119548523 | 22.37683 | 79.83291 | 76.68538 | Promoter | 2 | 119547697 | 119587027 | 39331 | 1 | ENSMUST00 | 0 | Chp1 | ENSMUSG00000014077.8 |

| | | | | | | | | | | | | | | | |
|---|---|---|---|---|---|---|---|---|---|---|---|---|---|---|---|
| chr2 | 157737355 | 157738113 | 22.37683 | 79.83291 | 76.68538 | Promoter | 2 | 157737401 | 157891614 | 154214 | 1 | ENSMUST00 | 0 | Ctnnbl1 | ENSMUSG00000027649.10 |
| chr3 | 103101985 | 103103380 | 22.37683 | 79.83291 | 76.68538 | Promoter | 3 | 103102604 | 103169769 | 67166 | 1 | ENSMUST00 | 0 | Dennd2c | ENSMUSG00000007379.10 |
| chr3 | 89349013 | 89350604 | 22.37683 | 79.83291 | 76.68538 | Promoter | 3 | 89338542 | 89349982 | 11441 | 2 | ENSMUST00 | 0 | Adam15 | ENSMUSG00000028041.12 |
| chr6 | 88627137 | 88627981 | 22.37683 | 79.83291 | 76.68538 | Promoter | 6 | 88547340 | 88627442 | 80103 | 2 | ENSMUST00 | 0 | Kbtbd12 | ENSMUSG00000033182.7 |
| chr6 | 34475683 | 34477021 | 22.37683 | 79.83291 | 76.68538 | Promoter | 6 | 34476207 | 34505613 | 29407 | 1 | ENSMUST00 | 0 | Bpgm | ENSMUSG00000038871.5 |
| chr8 | 123101676 | 123103515 | 22.37683 | 79.83291 | 76.68538 | Promoter | 8 | 123102350 | 123105244 | 2895 | 1 | ENSMUST00 | 0 | Rpl13 | ENSMUSG00000000740.6 |
| chr9 | 120576838 | 120577979 | 22.37683 | 79.83291 | 76.68538 | Promoter | 9 | 120577346 | 120578072 | 727 | 1 | ENSMUST00 | 0 | 5830454E08I | ENSMUSG00000052658.4 |
| chr9 | 100569791 | 100571278 | 22.37683 | 79.83291 | 76.68538 | Promoter | 9 | 100552188 | 100571090 | 18903 | 2 | ENSMUST00 | 0 | Slc35g2 | ENSMUSG00000070287.3 |
| chr11 | 60139224 | 60140901 | 27.98104 | 79.82376 | 76.67776 | Promoter | 11 | 60140082 | 60199195 | 59114 | 1 | ENSMUST00 | 0 | Rai1 | ENSMUSG00000062115.10 |
| chr12 | 51592925 | 51594931 | 27.98104 | 79.82376 | 76.67776 | Promoter | 12 | 51593341 | 51605773 | 12433 | 1 | ENSMUST00 | 0 | Coch | ENSMUSG00000020953.11 |
| chr4 | 150913674 | 150915272 | 27.40481 | 79.76363 | 76.61784 | Promoter | 4 | 150897133 | 150914437 | 17305 | 2 | ENSMUST00 | 0 | Park7 | ENSMUSG00000028964.9 |
| chr5 | 24451349 | 24453343 | 26.32003 | 79.74272 | 76.59713 | Promoter | 5 | 24452177 | 24502047 | 49871 | 1 | ENSMUST00 | 0 | Agap3 | ENSMUSG00000023353.9 |
| chr4 | 20008062 | 20009027 | 27.92246 | 79.72324 | 76.57767 | Promoter | 4 | 20008357 | 20030785 | 22429 | 1 | ENSMUST00 | 0 | Ttpa | ENSMUSG00000073988.8 |
| chr15 | 76293950 | 76295402 | 27.86412 | 79.62311 | 76.47798 | Promoter | 15 | 76294434 | 76296896 | 2463 | 1 | ENSMUST00 | 0 | Smpd5 | ENSMUSG00000071724.4 |
| chr17 | 25773713 | 25774689 | 27.86412 | 79.62311 | 76.47798 | Promoter | 17 | 25773776 | 25782791 | 9016 | 1 | ENSMUST00 | 0 | Narfl | ENSMUSG00000002280.5 |
| chr2 | 33886553 | 33887927 | 27.29445 | 79.57311 | 76.42841 | Promoter | 2 | 33825048 | 33886845 | 61798 | 2 | ENSMUST00 | 0 | Mvb12b | ENSMUSG00000038740.4 |
| chr2 | 156064974 | 156066119 | 23.62199 | 79.492 | 76.34769 | Promoter | 2 | 156065175 | 156069501 | 4327 | 1 | ENSMUST00 | 0 | Spag4 | ENSMUSG00000038180.6 |
| chr7 | 131371083 | 131371957 | 23.62199 | 79.492 | 76.34769 | Promoter | 7 | 131371141 | 131387839 | 16699 | 1 | ENSMUST00 | 0 | Pstk | ENSMUSG00000063179.7 |
| chr7 | 65644835 | 65645593 | 27.2396 | 79.47839 | 76.33418 | Promoter | 7 | 65644898 | 65683814 | 38917 | 1 | ENSMUST00 | 0 | Tarsl2 | ENSMUSG00000030515.8 |
| chr1 | 172126209 | 172127445 | 27.18497 | 79.384 | 76.2402 | Promoter | 1 | 172126755 | 172135283 | 8529 | 1 | ENSMUST00 | 0 | Pex19 | ENSMUSG00000003464.8 |
| chr14 | 52304058 | 52305438 | 27.18497 | 79.384 | 76.2402 | Promoter | 14 | 52294668 | 52305083 | 10416 | 2 | ENSMUST00 | 0 | Mettl3 | ENSMUSG00000022160.12 |
| chr4 | 42949218 | 42950597 | 27.18497 | 79.384 | 76.2402 | Promoter | 4 | 42949814 | 42959425 | 9612 | 1 | ENSMUST00 | 0 | Dnajb5 | ENSMUSG00000036052.9 |
| chr5 | 149184565 | 149185728 | 27.18497 | 79.384 | 76.2402 | Promoter | 5 | 149184663 | 149215430 | 30768 | 1 | ENSMUST00 | 0 | Uspl1 | ENSMUSG00000041264.11 |
| chr15 | 89148739 | 89150078 | 27.69056 | 79.32504 | 76.18215 | Promoter | 15 | 89142486 | 89149606 | 7121 | 2 | ENSMUST00 | 0 | Mapk11 | ENSMUSG00000053137.6 |
| chr7 | 81858985 | 81859871 | 27.69056 | 79.32504 | 76.18215 | Promoter | 7 | 81859001 | 81884071 | 25071 | 1 | ENSMUST00 | 0 | Tm6sf1 | ENSMUSG00000038623.4 |
| chr1 | 165706914 | 165708815 | 26.60951 | 79.31685 | 76.1741 | Promoter | 1 | 165648945 | 165708094 | 59150 | 2 | ENSMUST00 | 0 | Rcsd1 | ENSMUSG00000040723.9 |
| chr13 | 98205680 | 98206849 | 26.60951 | 79.31685 | 76.1741 | Promoter | 13 | 97898595 | 98206165 | 307571 | 2 | ENSMUST00 | 0 | Arhgef28 | ENSMUSG00000021662.9 |
| chr2 | 27885967 | 27887278 | 26.60951 | 79.31685 | 76.1741 | Promoter | 2 | 27886425 | 28039514 | 153090 | 1 | ENSMUST00 | 0 | Col5a1 | ENSMUSG00000026837.10 |
| chr18 | 56974768 | 56975654 | 26.0748 | 79.312 | 76.16934 | Promoter | 18 | 56955833 | 56975368 | 19536 | 2 | ENSMUST00 | 0 | C330018D20I | ENSMUSG00000024592.10 |
| chr19 | 8967138 | 8968191 | 26.0748 | 79.312 | 76.16934 | Promoter | 19 | 8967570 | 8972584 | 5015 | 1 | ENSMUST00 | 0 | Eef1g | ENSMUSG00000071644.5 |
| chr7 | 44548189 | 44548860 | 26.0748 | 79.312 | 76.16934 | Promoter | 7 | 44532746 | 44548847 | 16102 | 2 | ENSMUST00 | 0 | Pold1 | ENSMUSG00000038644.9 |
| chr11 | 82889573 | 82890631 | 26.5582 | 79.22746 | 76.08501 | Promoter | 11 | 82876434 | 82890589 | 14156 | 2 | ENSMUST00 | 0 | Rad51d | ENSMUSG00000018841.12 |
| chr13 | 68997906 | 68999860 | 26.5582 | 79.22746 | 76.08501 | Promoter | 13 | 68620043 | 68999541 | 379499 | 2 | ENSMUST00 | 0 | Adcy2 | ENSMUSG00000021536.7 |
| chr13 | 67127885 | 67128892 | 26.5582 | 79.22746 | 76.08501 | Promoter | 13 | 67128228 | 67141787 | 13560 | 1 | ENSMUST00 | 0 | Zfp759 | ENSMUSG00000057396.6 |
| chr5 | 69591559 | 69592538 | 26.5582 | 79.22746 | 76.08501 | Promoter | 5 | 69575001 | 69592337 | 17337 | 2 | ENSMUST00 | 0 | Gnpda2 | ENSMUSG00000029209.10 |
| chr9 | 108578036 | 108579239 | 26.5582 | 79.22746 | 76.08501 | Promoter | 9 | 108572313 | 108578670 | 6358 | 2 | ENSMUST00 | 0 | Wdr6 | ENSMUSG00000066357.5 |
| chr13 | 28957933 | 28960010 | 27.63319 | 79.22645 | 76.08477 | Distal Interge | 13 | 28948919 | 28953713 | 4795 | 2 | ENSMUST00 | -4221 | Sox4 | ENSMUSG00000076431.4 |
| chr4 | 155150178 | 155151714 | 24.46993 | 79.22128 | 76.07978 | Downstream | 4 | 155154075 | 155188296 | 34222 | 2 | ENSMUST00 | 36582 | Ski | ENSMUSG00000029050.10 |
| chr11 | 69845891 | 69847275 | 21.17422 | 79.20749 | 76.06625 | Promoter | 11 | 69846376 | 69851036 | 4661 | 1 | ENSMUST00 | 0 | Plscr3 | ENSMUSG00000019461.7 |
| chr12 | 30583902 | 30585668 | 21.17422 | 79.20749 | 76.06625 | Promoter | 12 | 30584443 | 30591213 | 6771 | 1 | ENSMUST00 | 0 | Tmem18 | ENSMUSG00000043061.9 |
| chr13 | 94975953 | 94977107 | 21.17422 | 79.20749 | 76.06625 | Promoter | 13 | 94976344 | 95014531 | 38188 | 1 | ENSMUST00 | 0 | Wdr41 | ENSMUSG00000042015.11 |
| chr14 | 70158320 | 70159598 | 21.17422 | 79.20749 | 76.06625 | Promoter | 14 | 70154405 | 70159502 | 5098 | 2 | ENSMUST00 | 0 | 9930012K11I | ENSMUSG00000044551.8 |
| chr16 | 4789130 | 4790637 | 21.17422 | 79.20749 | 76.06625 | Promoter | 16 | 4765461 | 4789990 | 24530 | 2 | ENSMUST00 | 0 | Cdip1 | ENSMUSG00000004071.7 |
| chr2 | 6871587 | 6873131 | 21.17422 | 79.20749 | 76.06625 | Promoter | 2 | 6546859 | 6872597 | 325739 | 2 | ENSMUST00 | 0 | Celf2 | ENSMUSG00000002107.13 |
| chr3 | 131302547 | 131303599 | 21.17422 | 79.20749 | 76.06625 | Promoter | 3 | 131290491 | 131303227 | 12737 | 2 | ENSMUST00 | 0 | Cyp2u1 | ENSMUSG00000027983.9 |
| chr4 | 132532706 | 132534067 | 21.17422 | 79.20749 | 76.06625 | Promoter | 4 | 132530555 | 132533488 | 2934 | 2 | ENSMUST00 | 0 | Atpif1 | ENSMUSG00000054428.7 |
| chr5 | 140606439 | 140608441 | 21.17422 | 79.20749 | 76.06625 | Promoter | 5 | 140607341 | 140615543 | 8203 | 1 | ENSMUST00 | 0 | Lfng | ENSMUSG00000029570.1 |
| chr8 | 76898535 | 76903248 | 21.17422 | 79.20749 | 76.06625 | Promoter | 8 | 76899442 | 76908413 | 8972 | 1 | ENSMUST00 | 0 | Nr3c2 | ENSMUSG00000031618.8 |
| chr1 | 39367167 | 39368903 | 23.4661 | 79.19912 | 76.05865 | Promoter | 1 | 39367851 | 39371908 | 4058 | 1 | ENSMUST00 | 0 | Rpl31 | ENSMUSG00000073702.6 |
| chr10 | 5823125 | 5825026 | 23.4661 | 79.19912 | 76.05865 | Promoter | 10 | 5811887 | 5823910 | 12024 | 2 | ENSMUST00 | 0 | Mtrf1l | ENSMUSG00000019774.8 |
| chr10 | 118867715 | 118870312 | 23.4661 | 79.19912 | 76.05865 | Promoter | 10 | 118859349 | 118868903 | 9555 | 2 | ENSMUST00 | 0 | Dyrk2 | ENSMUSG00000028630.8 |
| chr10 | 7667173 | 7668076 | 23.4661 | 79.19912 | 76.05865 | Promoter | 10 | 7667503 | 7678881 | 11379 | 1 | ENSMUST00 | 0 | Nup43 | ENSMUSG00000040034.10 |
| chr11 | 69900342 | 69902635 | 23.4661 | 79.19912 | 76.05865 | Promoter | 11 | 69897352 | 69900894 | 3543 | 2 | ENSMUST00 | 0 | 2810408A11I | ENSMUSG00000018570.12 |
| chr12 | 100186599 | 100188479 | 23.4661 | 79.19912 | 76.05865 | Promoter | 12 | 100187968 | 100193540 | 5573 | 1 | ENSMUST00 | 0 | Gm10433 | ENSMUSG00000072884.5 |
| chr17 | 15374183 | 15377041 | 23.4661 | 79.19912 | 76.05865 | Promoter | 17 | 15367354 | 15375976 | 8623 | 2 | ENSMUST00 | 0 | Dll1 | ENSMUSG00000014773.8 |
| chr17 | 45555335 | 45556614 | 23.4661 | 79.19912 | 76.05865 | Promoter | 17 | 45555716 | 45563169 | 7454 | 1 | ENSMUST00 | 0 | Nfkbie | ENSMUSG00000023947.7 |
| chr18 | 64515716 | 64517784 | 23.4661 | 79.19912 | 76.05865 | Promoter | 18 | 64499665 | 64516557 | 16893 | 2 | ENSMUST00 | 0 | Nars | ENSMUSG00000024587.9 |
| chr3 | 129532063 | 129534638 | 23.4661 | 79.19912 | 76.05865 | Promoter | 3 | 129532386 | 129638495 | 106110 | 1 | ENSMUST00 | 0 | Elovl6 | ENSMUSG00000041220.6 |
| chr4 | 24966127 | 24967507 | 23.4661 | 79.19912 | 76.05865 | Promoter | 4 | 24966407 | 25007300 | 40894 | 1 | ENSMUST00 | 0 | Gpr63 | ENSMUSG00000040372.2 |
| chr6 | 119543362 | 119544750 | 23.4661 | 79.19912 | 76.05865 | Promoter | 6 | 119432531 | 119544347 | 111817 | 2 | ENSMUST00 | 0 | Wnt5b | ENSMUSG00000030170.9 |
| chr7 | 19148410 | 19149307 | 23.4661 | 79.19912 | 76.05865 | Promoter | 7 | 19140217 | 19149196 | 8980 | 2 | ENSMUST00 | 0 | Qpctl | ENSMUSG00000030407.1 |
| chr8 | 13785055 | 13786498 | 23.4661 | 79.19912 | 76.05865 | Promoter | 8 | 13785615 | 13798538 | 12924 | 1 | ENSMUST00 | 0 | Upf3a | ENSMUSG00000038398.7 |
| chr9 | 56634833 | 56636173 | 23.4661 | 79.19912 | 76.05865 | Promoter | 9 | 56619088 | 56635628 | 16541 | 2 | ENSMUST00 | 0 | Lingo1 | ENSMUSG00000049556.4 |
| chr9 | 67632276 | 67634639 | 23.4661 | 79.19912 | 76.05865 | Distal Interge | 9 | 67217087 | 67559703 | 342617 | 2 | ENSMUST00 | -72574 | Tln2 | ENSMUSG00000052698.9 |
| chr10 | 128409073 | 128409854 | 24.93786 | 79.14572 | 76.00787 | Promoter | 10 | 128401395 | 128409728 | 8334 | 2 | ENSMUST00 | 0 | Nabp2 | ENSMUSG00000025374.8 |
| chr14 | 55105638 | 55106777 | 25.97798 | 79.14165 | 76.00417 | Promoter | 14 | 55098578 | 55106534 | 7957 | 2 | ENSMUST00 | 0 | Ap1g2 | ENSMUSG00000040701.10 |
| chr4 | 155784003 | 155784774 | 25.97798 | 79.14165 | 76.00417 | 3' UTR | 4 | 155781591 | 155785874 | 4284 | 2 | ENSMUST00 | 1100 | Tmem88b | ENSMUSG00000073680.2 |
| chr5 | 129846830 | 129847747 | 25.97798 | 79.14165 | 76.00417 | Promoter | 5 | 129846990 | 129863951 | 16962 | 1 | ENSMUST00 | 0 | Sumf2 | ENSMUSG00000025538.10 |
| chr9 | 75610419 | 75611479 | 25.97798 | 79.14165 | 76.00417 | Promoter | 9 | 75565621 | 75611325 | 45705 | 2 | ENSMUST00 | 0 | Tmod2 | ENSMUSG00000032186.9 |
| chr4 | 132344787 | 132345841 | 25.4393 | 79.11702 | 75.98065 | Promoter | 4 | 132331919 | 132345747 | 13829 | 2 | ENSMUST00 | 0 | Rcc1 | ENSMUSG00000028896.8 |
| chr8 | 47412486 | 47414325 | 25.4393 | 79.11702 | 75.98065 | Promoter | 8 | 47413129 | 47415359 | 2231 | 1 | ENSMUST00 | 0 | Gm10083 | ENSMUSG00000060559.5 |
| chr9 | 72925014 | 72926508 | 27.02239 | 79.1029 | 75.96719 | Promoter | 9 | 72925645 | 72952181 | 26537 | 1 | ENSMUST00 | 0 | Pygo1 | ENSMUSG00000034910.5 |
| chr14 | 45218754 | 45219478 | 26.45618 | 79.04958 | 75.91403 | Promoter | 14 | 45134448 | 45219394 | 84947 | 2 | ENSMUST00 | 0 | Txndc16 | ENSMUSG00000021830.8 |
| chr16 | 14159566 | 14159981 | 26.96863 | 79.00988 | 75.87456 | Promoter | 16 | 14159626 | 14159692 | 67 | 1 | ENSMUST00 | 0 | Mir484 | ENSMUSG00000070074.1 |
| chr1 | 127868509 | 127869519 | 26.40393 | 78.95839 | 75.82329 | Promoter | 1 | 127868773 | 127943868 | 75096 | 1 | ENSMUST00 | 0 | Rab3gap1 | ENSMUSG00000036104.6 |
| chr11 | 114675270 | 114676405 | 26.40393 | 78.95839 | 75.82329 | Promoter | 11 | 114675431 | 114720977 | 45547 | 1 | ENSMUST00 | 0 | Ttyh2 | ENSMUSG00000034714.9 |
| chr14 | 54516726 | 54517467 | 26.40393 | 78.95839 | 75.82329 | Promoter | 14 | 54511206 | 54517435 | 6230 | 2 | ENSMUST00 | 0 | Prmt5 | ENSMUSG00000023110.7 |
| chr15 | 75703357 | 75704139 | 26.40393 | 78.95839 | 75.82329 | Promoter | 15 | 75704280 | 75714408 | 10129 | 1 | ENSMUST00 | -141 | Rhpn1 | ENSMUSG00000022580.7 |
| chr17 | 40879593 | 40881031 | 26.40393 | 78.95839 | 75.82329 | Promoter | 17 | 40875482 | 40880558 | 5077 | 2 | ENSMUST00 | 0 | 9130008F23I | ENSMUSG00000054951.2 |
| chr19 | 38054962 | 38055492 | 26.40393 | 78.95839 | 75.82329 | Promoter | 19 | 38055025 | 38074425 | 19401 | 1 | ENSMUST00 | 0 | Cep55 | ENSMUSG00000024989.9 |
| chr19 | 46135475 | 46137449 | 26.40393 | 78.95839 | 75.82329 | Promoter | 19 | 46136495 | 46137289 | 795 | 2 | ENSMUST00 | 0 | Pitx3 | ENSMUSG00000025229.10 |
| chr2 | 164769302 | 164770309 | 26.40393 | 78.95839 | 75.82329 | Promoter | 2 | 164769898 | 164772907 | 3010 | 1 | ENSMUST00 | 0 | Ube2c | ENSMUSG00000001403.8 |
| chr2 | 29787072 | 29788791 | 26.40393 | 78.95839 | 75.82329 | Promoter | 2 | 29787493 | 29797935 | 10443 | 1 | ENSMUST00 | 0 | Coq4 | ENSMUSG00000026798.5 |
| chr2 | 156546933 | 156548474 | 26.40393 | 78.95839 | 75.82329 | Promoter | 2 | 156547584 | 156568156 | 20573 | 1 | ENSMUST00 | 0 | Aar2 | ENSMUSG00000027628.5 |
| chr6 | 91440430 | 91441757 | 26.40393 | 78.95839 | 75.82329 | Promoter | 6 | 91430460 | 91440854 | 10395 | 2 | ENSMUST00 | 0 | 4930471M09 | ENSMUSG00000097693.1 |
| chr7 | 27928557 | 27929955 | 26.40393 | 78.95839 | 75.82329 | Promoter | 7 | 27907392 | 27929430 | 22039 | 2 | ENSMUST00 | 0 | 1700049G17 | ENSMUSG00000070709.7 |
| chr8 | 70130974 | 70131966 | 26.40393 | 78.95839 | 75.82329 | Promoter | 8 | 70131333 | 70133739 | 2407 | 1 | ENSMUST00 | 0 | Nr2c2ap | ENSMUSG00000071078.5 |
| chrX | 163957631 | 163959170 | 26.40393 | 78.95839 | 75.82329 | Promoter | 20 | 163935443 | 163958589 | 23147 | 2 | ENSMUST00 | 0 | Zrsr2 | ENSMUSG00000031370.7 |
| chr12 | 12937456 | 12942953 | 27.46248 | 78.93296 | 75.80077 | Promoter | 12 | 12936096 | 12940616 | 4521 | 2 | ENSMUST00 | 0 | Mycn | ENSMUSG00000037169.9 |
| chr6 | 124813168 | 124814519 | 26.91508 | 78.9172 | 75.78516 | Promoter | 6 | 124810586 | 124814296 | 3711 | 2 | ENSMUST00 | 0 | Tpi1 | ENSMUSG00000023456.9 |
| chr1 | 161070062 | 161070740 | 25.83409 | 78.88815 | 75.75626 | Promoter | 1 | 161040601 | 161070658 | 30058 | 2 | ENSMUST00 | 0 | Dars2 | ENSMUSG00000026709.5 |
| chr18 | 88893282 | 88894599 | 26.35494 | 78.87288 | 75.741 | Promoter | 18 | 88668695 | 88894339 | 225645 | 2 | ENSMUST00 | 0 | Socs6 | ENSMUSG00000056153.9 |
| chr1 | 60042521 | 60043601 | 24.78212 | 78.86476 | 75.73294 | Promoter | 1 | 59989068 | 60043087 | 54020 | 2 | ENSMUST00 | 0 | Ica1l | ENSMUSG00000026018.7 |
| chr1 | 130461445 | 130463349 | 24.78212 | 78.86476 | 75.73294 | Promoter | 1 | 130439027 | 130462744 | 23718 | 2 | ENSMUST00 | 0 | Cd55 | ENSMUSG00000026399.7 |
| chr1 | 162477270 | 162478324 | 24.78212 | 78.86476 | 75.73294 | Promoter | 1 | 161987302 | 162478034 | 490733 | 2 | ENSMUST00 | 0 | Dnm3 | ENSMUSG00000040265.11 |
| chr1 | 89454009 | 89456011 | 24.78212 | 78.86476 | 75.73294 | Promoter | 1 | 89454806 | 89897617 | 442812 | 1 | ENSMUST00 | 0 | Agap1 | ENSMUSG00000055013.9 |
| chr10 | 127165095 | 127166396 | 24.78212 | 78.86476 | 75.73294 | Promoter | 10 | 127165156 | 127172340 | 7185 | 1 | ENSMUST00 | 0 | B4galnt1 | ENSMUSG00000006731.9 |
| chr11 | 4986471 | 4987778 | 24.78212 | 78.86476 | 75.73294 | Promoter | 11 | 4986824 | 5042729 | 55906 | 1 | ENSMUST00 | 0 | Ap1b1 | ENSMUSG00000009090.12 |
| chr11 | 117848353 | 117849591 | 24.78212 | 78.86476 | 75.73294 | Promoter | 11 | 117849251 | 117855732 | 6482 | 1 | ENSMUST00 | 0 | Birc5 | ENSMUSG00000017716.10 |
| chr13 | 62558158 | 62559082 | 24.78212 | 78.86476 | 75.73294 | Distal Interge | 13 | 62543434 | 62543503 | 70 | 2 | ENSMUST00 | -14656 | Gm22615 | ENSMUSG00000094815.1 |
| chr15 | 80132695 | 80134176 | 24.78212 | 78.86476 | 75.73294 | Promoter | 15 | 80133154 | 80161701 | 28548 | 1 | ENSMUST00 | 0 | Tab1 | ENSMUSG00000022414.7 |
| chr17 | 56507130 | 56507891 | 24.78212 | 78.86476 | 75.73294 | Distal Interge | 17 | 56511248 | 56512483 | 1236 | 2 | ENSMUST00 | 4592 | Znrf4 | ENSMUSG00000044526.2 |
| chr17 | 83631115 | 83633458 | 24.78212 | 78.86476 | 75.73294 | Promoter | 17 | 83585957 | 83631895 | 45939 | 2 | ENSMUST00 | 0 | Kcng3 | ENSMUSG00000045053.1 |
| chr18 | 32239351 | 32241232 | 24.78212 | 78.86476 | 75.73294 | Promoter | 18 | 32240300 | 32270151 | 29852 | 1 | ENSMUST00 | 0 | Ercc3 | ENSMUSG00000024382.8 |
| chr19 | 44134936 | 44135949 | 24.78212 | 78.86476 | 75.73294 | Promoter | 19 | 44108644 | 44135876 | 27233 | 2 | ENSMUST00 | 0 | Cwf19l1 | ENSMUSG00000025200.6 |
| chr2 | 12300984 | 12302218 | 24.78212 | 78.86476 | 75.73294 | Promoter | 2 | 12106632 | 12301920 | 195289 | 2 | ENSMUST00 | 0 | Itga8 | ENSMUSG00000026768.5 |
| chr2 | 102185400 | 102186830 | 24.78212 | 78.86476 | 75.73294 | Promoter | 2 | 101950203 | 102186358 | 236156 | 2 | ENSMUST00 | 0 | Ldlrad3 | ENSMUSG00000048058.12 |
| chr2 | 31258576 | 31259600 | 24.78212 | 78.86476 | 75.73294 | Intron (ENSM | 2 | 31246166 | 31292319 | 46154 | 1 | ENSMUST00 | 12411 | Ncs1 | ENSMUSG00000062661.6 |
| chr2 | 131909450 | 131910584 | 24.78212 | 78.86476 | 75.73294 | Promoter | 2 | 131909928 | 131938429 | 28502 | 1 | ENSMUST00 | 0 | Prnp | ENSMUSG00000079037.4 |
| chr3 | 97158037 | 97159684 | 24.78212 | 78.86476 | 75.73294 | Promoter | 3 | 97158777 | 97176574 | 17798 | 1 | ENSMUST00 | 0 | Acp6 | ENSMUSG00000028093.10 |
| chr4 | 152477051 | 152478573 | 24.78212 | 78.86476 | 75.73294 | Promoter | 4 | 152390744 | 152477871 | 87128 | 2 | ENSMUST00 | 0 | Kcnab2 | ENSMUSG00000028931.6 |
| chr5 | 110839100 | 110840690 | 24.78212 | 78.86476 | 75.73294 | Promoter | 5 | 110840017 | 110874133 | 34117 | 1 | ENSMUST00 | 0 | Chek2 | ENSMUSG00000029521.3 |
| chr5 | 62765401 | 62766855 | 24.78212 | 78.86476 | 75.73294 | Promoter | 5 | 62602445 | 62766198 | 163754 | 2 | ENSMUST00 | 0 | Arap2 | ENSMUSG00000037999.10 |
| chr6 | 55203062 | 55203968 | 24.78212 | 78.86476 | 75.73294 | Promoter | 6 | 55203383 | 55320222 | 116840 | 1 | ENSMUST00 | 0 | Fam188b | ENSMUSG00000038022.9 |
| chr7 | 125551556 | 125552977 | 24.78212 | 78.86476 | 75.73294 | Promoter | 7 | 125552282 | 125579472 | 27191 | 1 | ENSMUST00 | 0 | Il4ra | ENSMUSG00000030748.8 |
| chr8 | 102783585 | 102786947 | 24.78212 | 78.86476 | 75.73294 | Promoter | 8 | 102632995 | 102785111 | 152117 | 2 | ENSMUST00 | 0 | Cdh11 | ENSMUSG00000031673.5 |
| chr9 | 100642044 | 100643414 | 24.78212 | 78.86476 | 75.73294 | Promoter | 9 | 100643448 | 100796791 | 153344 | 1 | ENSMUST00 | -34 | Stag1 | ENSMUSG00000037286.10 |
| chr8 | 45332530 | 45333879 | 26.86175 | 78.82483 | 75.69657 | Promoter | 8 | 45304946 | 45333216 | 28271 | 2 | ENSMUST00 | 0 | Cyp4v3 | ENSMUSG00000079057.3 |
| chr9 | 88522164 | 88523627 | 26.86175 | 78.82483 | 75.69657 | Promoter | 9 | 88521050 | 88522929 | 1880 | 2 | ENSMUST00 | 0 | Snhg5 | ENSMUSG00000097195.3 |
| chr11 | 70432049 | 70433420 | 20.14837 | 78.81087 | 75.68276 | Promoter | 11 | 70432635 | 70435996 | 3362 | 1 | ENSMUST00 | 0 | Arrb2 | ENSMUSG00000060216.10 |
| chr13 | 60173585 | 60177569 | 20.14837 | 78.81087 | 75.68276 | Promoter | 13 | 60174405 | 60177365 | 2961 | 2 | ENSMUST00 | 0 | Gas1 | ENSMUSG00000052957.7 |
| chr15 | 98532844 | 98534818 | 20.14837 | 78.81087 | 75.68276 | Promoter | 15 | 98518821 | 98534244 | 15424 | 2 | ENSMUST00 | 0 | Kansl2 | ENSMUSG00000022992.11 |
| chr2 | 32845892 | 32847353 | 20.14837 | 78.81087 | 75.68276 | Promoter | 2 | 32787602 | 32847239 | 59638 | 2 | ENSMUST00 | 0 | Stxbp1 | ENSMUSG00000026797.10 |
| chr2 | 156839609 | 156842066 | 20.14837 | 78.81087 | 75.68276 | Promoter | 2 | 156840007 | 156855570 | 15564 | 1 | ENSMUST00 | 0 | Tgif2 | ENSMUSG00000062175.8 |
| chr3 | 95227430 | 95229926 | 20.14837 | 78.81087 | 75.68276 | Promoter | 3 | 95228732 | 95236409 | 7678 | 1 | ENSMUST00 | 0 | Cdc42se1 | ENSMUSG00000046722.9 |
| chr8 | 111853408 | 111854783 | 20.14837 | 78.81087 | 75.68276 | Promoter | 8 | 111768473 | 111854310 | 85838 | 2 | ENSMUST00 | 0 | Cfdp1 | ENSMUSG00000031954.5 |
| chr1 | 183388545 | 183389796 | 27.34985 | 78.73917 | 75.61149 | Promoter | 1 | 183388981 | 183409353 | 20373 | 1 | ENSMUST00 | 0 | Taf1a | ENSMUSG00000072258.5 |
| chr11 | 23067253 | 23068481 | 27.34985 | 78.73917 | 75.61149 | Promoter | 11 | 23067998 | 23075574 | 7577 | 1 | ENSMUST00 | 0 | Gm12059 | ENSMUSG00000085665.1 |

| | | | | | | | | | | | | | | | |
|---|---|---|---|---|---|---|---|---|---|---|---|---|---|---|---|
| chr10 | 80405880 | 80407455 | 26.80862 | 78.7328 | 75.60527 | Promoter | 10 | 80402997 | 80406766 | 3770 | 2 | ENSMUST00 | 0 | Uqcr11 | ENSMUSG00000020163.7 |
| chr15 | 96709302 | 96710655 | 26.25447 | 78.69737 | 75.57 | Distal Interge | 15 | 96687394 | 96699698 | 12305 | 2 | ENSMUST00 | -9605 | Slc38a2 | ENSMUSG00000022462.6 |
| chr11 | 102284865 | 102285829 | 27.29387 | 78.64282 | 75.51551 | Promoter | 11 | 102284931 | 102289237 | 4307 | 1 | ENSMUST00 | 0 | Tmub2 | ENSMUSG00000034757.10 |
| chr13 | 111490250 | 111491172 | 27.29387 | 78.64282 | 75.51551 | Promoter | 13 | 111425680 | 111490111 | 64432 | 2 | ENSMUST00 | -140 | Gpbp1 | ENSMUSG00000032745.12 |
| chr14 | 50061249 | 50062970 | 27.29387 | 78.64282 | 75.51551 | Distal Interge | 14 | 50083714 | 50084679 | 966 | 2 | ENSMUST00 | 21709 | Olfr726 | ENSMUSG00000060523.2 |
| chr16 | 17833274 | 17834518 | 27.29387 | 78.64282 | 75.51551 | Downstream | 16 | 17835276 | 17838364 | 3089 | 2 | ENSMUST00 | 3846 | Car15 | ENSMUSG00000090236.2 |
| chr4 | 151860740 | 151861826 | 27.29387 | 78.64282 | 75.51551 | Promoter | 4 | 150917322 | 151861669 | 944348 | 2 | ENSMUST00 | 0 | Camta1 | ENSMUSG00000014592.14 |
| chr4 | 89294261 | 89295701 | 26.75571 | 78.6411 | 75.51434 | Promoter | 4 | 89274541 | 89294653 | 20113 | 2 | ENSMUST00 | 0 | Cdkn2a | ENSMUSG00000044303.6 |
| chr9 | 95405075 | 95408037 | 26.703 | 78.54972 | 75.42343 | Promoter | 9 | 95399292 | 95406722 | 7431 | 2 | ENSMUST00 | 0 | Chst2 | ENSMUSG00000033350.7 |
| chr10 | 61715082 | 61715770 | 27.23813 | 78.54684 | 75.42068 | Promoter | 10 | 61715263 | 61738653 | 23391 | 1 | ENSMUST00 | 0 | Aifm2 | ENSMUSG00000020085.10 |
| chr8 | 85059691 | 85061009 | 18.49104 | 78.52299 | 75.39706 | Promoter | 8 | 85060055 | 85067121 | 7067 | 1 | ENSMUST00 | 0 | Fbxw9 | ENSMUSG00000008167.9 |
| chr9 | 98296463 | 98297381 | 24.06788 | 78.48502 | 75.35915 | Promoter | 9 | 98296583 | 98411434 | 114852 | 1 | ENSMUST00 | 0 | Nmnat3 | ENSMUSG00000032456.8 |
| chr9 | 119102059 | 119103232 | 24.06788 | 78.48502 | 75.35915 | Promoter | 9 | 119102478 | 119125295 | 22818 | 1 | ENSMUST00 | 0 | Dlec1 | ENSMUSG00000038060.10 |
| chr1 | 37864293 | 37864898 | 26.6505 | 78.45867 | 75.33303 | Promoter | 1 | 37832406 | 37864836 | 32431 | 2 | ENSMUST00 | 0 | Tsga10 | ENSMUSG00000060771.9 |
| chr1 | 191989623 | 191990539 | 26.6505 | 78.45867 | 75.33303 | Promoter | 1 | 191986832 | 191990138 | 3307 | 2 | ENSMUST00 | 0 | Gm26670 | ENSMUSG00000097416.1 |
| chr11 | 90687001 | 90688119 | 26.6505 | 78.45867 | 75.33303 | Promoter | 11 | 90643465 | 90687607 | 44143 | 2 | ENSMUST00 | 0 | Tom1l1 | ENSMUSG00000020541.7 |
| chr16 | 92300602 | 92302068 | 26.6505 | 78.45867 | 75.33303 | Promoter | 16 | 92301286 | 92313041 | 11756 | 1 | ENSMUST00 | 0 | Smim11 | ENSMUSG00000051989.6 |
| chr2 | 106973606 | 106974761 | 26.6505 | 78.45867 | 75.33303 | Promoter | 2 | 106962527 | 106974424 | 11898 | 2 | ENSMUST00 | 0 | Arl14ep | ENSMUSG00000027122.10 |
| chr7 | 30094381 | 30095835 | 26.6505 | 78.45867 | 75.33303 | Promoter | 7 | 30094777 | 30104291 | 9515 | 1 | ENSMUST00 | 0 | Zfp260 | ENSMUSG00000049421.8 |
| chr7 | 99351184 | 99353540 | 26.6505 | 78.45867 | 75.33303 | Promoter | 7 | 99345375 | 99353230 | 7856 | 2 | ENSMUST00 | 0 | Serpinh1 | ENSMUSG00000070436.6 |
| chr7 | 45413339 | 45414538 | 26.6505 | 78.45867 | 75.33303 | Promoter | 7 | 45413695 | 45417178 | 3484 | 1 | ENSMUST00 | 0 | Ntf5 | ENSMUSG00000074121.2 |
| chr7 | 142283685 | 142285408 | 26.6505 | 78.45867 | 75.33303 | Distal Interge | 7 | 142280268 | 142280939 | 672 | 2 | ENSMUST00 | -2747 | Gm10013 | ENSMUSG00000089731.3 |
| chr8 | 70673266 | 70673970 | 26.6505 | 78.45867 | 75.33303 | Promoter | 8 | 70673432 | 70678048 | 4617 | 1 | ENSMUST00 | 0 | Lsm4 | ENSMUSG00000031848.9 |
| chr15 | 78899411 | 78900583 | 27.18262 | 78.45122 | 75.32643 | Promoter | 15 | 78899667 | 78919032 | 19366 | 1 | ENSMUST00 | 0 | Sh3bp1 | ENSMUSG00000022436.10 |
| chr7 | 31208263 | 31209130 | 27.18262 | 78.45122 | 75.32643 | Distal Interge | 7 | 31182037 | 31182823 | 787 | 1 | ENSMUST00 | 26227 | RP23-412I17 | ENSMUSG00000099536.1 |
| chr11 | 3912589 | 3914725 | 24.51421 | 78.37994 | 75.25539 | Promoter | 11 | 3907021 | 3914664 | 7644 | 2 | ENSMUST00 | 0 | Slc35e4 | ENSMUSG00000048807.2 |
| chr15 | 80082426 | 80083401 | 24.51421 | 78.37994 | 75.25539 | Promoter | 15 | 80077794 | 80082583 | 4790 | 2 | ENSMUST00 | 0 | Rpl3 | ENSMUSG00000060036.8 |
| chr19 | 4838649 | 4839994 | 24.51421 | 78.37994 | 75.25539 | Promoter | 19 | 4825366 | 4839322 | 13957 | 2 | ENSMUST00 | 0 | Ccs | ENSMUSG00000034108.5 |
| chr4 | 133756056 | 133757903 | 24.51421 | 78.37994 | 75.25539 | Promoter | 4 | 133679941 | 133756769 | 76829 | 2 | ENSMUST00 | 0 | Arid1a | ENSMUSG00000007880.11 |
| chr7 | 45062040 | 45063248 | 24.51421 | 78.37994 | 75.25539 | Promoter | 7 | 45062429 | 45077416 | 14988 | 1 | ENSMUST00 | 0 | Nosip | ENSMUSG00000003421.7 |
| chr8 | 83972078 | 83972808 | 24.51421 | 78.37994 | 75.25539 | Promoter | 8 | 83972978 | 83996445 | 23468 | 1 | ENSMUST00 | -170 | Prkaca | ENSMUSG00000005469.9 |
| chr19 | 4099427 | 4101190 | 26.59821 | 78.36794 | 75.24377 | Promoter | 19 | 4099998 | 4113965 | 13968 | 1 | ENSMUST00 | 0 | Pitpnm1 | ENSMUSG00000024851.8 |
| chr10 | 110744846 | 110746074 | 26.05581 | 78.34985 | 75.22605 | Promoter | 10 | 110745439 | 110748146 | 2708 | 1 | ENSMUST00 | 0 | E2f7 | ENSMUSG00000020185.11 |
| chr16 | 85800671 | 85803715 | 26.05581 | 78.34985 | 75.22605 | Promoter | 16 | 85793827 | 85803113 | 9287 | 2 | ENSMUST00 | 0 | Adamts1 | ENSMUSG00000022893.9 |
| chr4 | 99828492 | 99830254 | 26.05581 | 78.34985 | 75.22605 | Promoter | 4 | 99765402 | 99829186 | 63785 | 2 | ENSMUST00 | 0 | Itgb3bp | ENSMUSG00000028549.12 |
| chr5 | 51551775 | 51553975 | 26.05581 | 78.34985 | 75.22605 | Promoter | 5 | 51454249 | 51553921 | 99673 | 2 | ENSMUST00 | 0 | Ppargc1a | ENSMUSG00000029167.8 |
| chr6 | 34028850 | 34030028 | 26.05581 | 78.34985 | 75.22605 | Promoter | 6 | 34029448 | 34134034 | 104587 | 1 | ENSMUST00 | 0 | Lrguk | ENSMUSG00000056215.7 |
| chr7 | 45135798 | 45136549 | 26.05581 | 78.34985 | 75.22605 | Promoter | 7 | 45125922 | 45136231 | 10310 | 2 | ENSMUST00 | 0 | Rpl13a | ENSMUSG00000074129.7 |
| chr1 | 54438271 | 54439277 | 22.06604 | 78.32127 | 75.1984 | Promoter | 1 | 54396004 | 54438971 | 42968 | 2 | ENSMUST00 | 0 | Gtf3c3 | ENSMUSG00000041303.7 |
| chr10 | 62601290 | 62602995 | 22.06604 | 78.32127 | 75.1984 | Promoter | 10 | 62580248 | 62602298 | 22051 | 2 | ENSMUST00 | 0 | Ddx21 | ENSMUSG00000020075.7 |
| chr11 | 19018184 | 19021149 | 22.06604 | 78.32127 | 75.1984 | Promoter | 11 | 18879817 | 19018985 | 139169 | 2 | ENSMUST00 | 0 | Meis1 | ENSMUSG00000020160.13 |
| chr12 | 32378503 | 32381087 | 22.06604 | 78.32127 | 75.1984 | Promoter | 12 | 32378704 | 32382943 | 4240 | 1 | ENSMUST00 | 0 | Ccdc71l | ENSMUSG00000090946.3 |
| chr14 | 73524637 | 73525512 | 22.06604 | 78.32127 | 75.1984 | Promoter | 14 | 73525319 | 73560543 | 35225 | 1 | ENSMUST00 | 0 | Sucla2 | ENSMUSG00000022110.8 |
| chr14 | 55584681 | 55585616 | 22.06604 | 78.32127 | 75.1984 | Promoter | 14 | 55581516 | 55584946 | 3431 | 2 | ENSMUST00 | 0 | Emc9 | ENSMUSG00000022217.9 |
| chr17 | 27565197 | 27566231 | 22.06604 | 78.32127 | 75.1984 | Promoter | 17 | 27563575 | 27565732 | 2158 | 2 | ENSMUST00 | 0 | AI413582 | ENSMUSG00000062753.8 |
| chr4 | 154635029 | 154637946 | 22.06604 | 78.32127 | 75.1984 | Promoter | 4 | 154316125 | 154636866 | 320742 | 2 | ENSMUST00 | 0 | Prdm16 | ENSMUSG00000039410.11 |
| chr5 | 117388392 | 117389363 | 22.06604 | 78.32127 | 75.1984 | Promoter | 5 | 117378103 | 117389047 | 10945 | 2 | ENSMUST00 | 0 | Rfc5 | ENSMUSG00000029363.9 |
| chr5 | 150593462 | 150594948 | 22.06604 | 78.32127 | 75.1984 | Promoter | 5 | 150571643 | 150594525 | 22883 | 2 | ENSMUST00 | 0 | N4bp2l1 | ENSMUSG00000041132.8 |
| chr7 | 141192831 | 141195274 | 22.06604 | 78.32127 | 75.1984 | Promoter | 7 | 141189105 | 141193960 | 4856 | 2 | ENSMUST00 | 0 | Hras | ENSMUSG00000025499.12 |
| chr7 | 90475283 | 90477161 | 22.06604 | 78.32127 | 75.1984 | Promoter | 7 | 90468194 | 90476001 | 7808 | 2 | ENSMUST00 | 0 | Tmem126b | ENSMUSG00000030614.7 |
| chr7 | 27673368 | 27675137 | 22.06604 | 78.32127 | 75.1984 | Promoter | 7 | 27656375 | 27674594 | 18220 | 2 | ENSMUST00 | 0 | Map3k10 | ENSMUSG00000040390.8 |
| chr8 | 71396597 | 71397592 | 22.06604 | 78.32127 | 75.1984 | Promoter | 8 | 71396855 | 71404772 | 7918 | 1 | ENSMUST00 | 0 | Babam1 | ENSMUSG00000031820.8 |
| chr8 | 41133006 | 41134407 | 22.06604 | 78.32127 | 75.1984 | Promoter | 8 | 40990914 | 41133726 | 142813 | 2 | ENSMUST00 | 0 | Mtus1 | ENSMUSG00000045636.11 |
| chr9 | 107887603 | 107888866 | 22.06604 | 78.32127 | 75.1984 | Promoter | 9 | 107888129 | 107903125 | 14997 | 1 | ENSMUST00 | 0 | Mon1a | ENSMUSG00000032583.2 |
| chr9 | 58554690 | 58555860 | 22.06604 | 78.32127 | 75.1984 | Promoter | 9 | 58524302 | 58555132 | 30831 | 2 | ENSMUST00 | 0 | Cd276 | ENSMUSG00000035914.5 |
| chr7 | 30168946 | 30170898 | 24.99193 | 78.31828 | 75.19689 | Promoter | 7 | 30161274 | 30169727 | 8454 | 2 | ENSMUST00 | 0 | Zfp146 | ENSMUSG00000037029.7 |
| chr11 | 117778803 | 117781710 | 25.50448 | 78.30592 | 75.18496 | Promoter | 11 | 117765988 | 117779648 | 13661 | 2 | ENSMUST00 | 0 | Tmc6 | ENSMUSG00000025572.12 |
| chr11 | 97021341 | 97022635 | 25.50448 | 78.30592 | 75.18496 | 5' UTR | 11 | 97018733 | 97024731 | 5999 | 1 | ENSMUST00 | 2609 | Sp6 | ENSMUSG00000038560.7 |
| chr2 | 153444761 | 153446242 | 25.50448 | 78.30592 | 75.18496 | Promoter | 2 | 153417826 | 153445581 | 27756 | 2 | ENSMUST00 | 0 | Nol4l | ENSMUSG00000061411.7 |
| chr5 | 145200795 | 145201950 | 25.50448 | 78.30592 | 75.18496 | Promoter | 5 | 145194946 | 145201868 | 6923 | 2 | ENSMUST00 | 0 | Zkscan14 | ENSMUSG00000029627.11 |
| chr7 | 79715217 | 79716305 | 25.50448 | 78.30592 | 75.18496 | Promoter | 7 | 79698098 | 79715720 | 17623 | 2 | ENSMUST00 | 0 | Kif7 | ENSMUSG00000050382.9 |
| chr8 | 106135923 | 106138736 | 25.50448 | 78.30592 | 75.18496 | Promoter | 8 | 106130319 | 106136974 | 6656 | 2 | ENSMUST00 | 0 | Esrp2 | ENSMUSG00000084128.5 |
| chr9 | 49485457 | 49486535 | 26.54612 | 78.27752 | 75.1573 | Promoter | 9 | 49436963 | 49486225 | 49263 | 2 | ENSMUST00 | 0 | Ttc12 | ENSMUSG00000040219.4 |
| chr11 | 62248327 | 62249985 | 27.07226 | 78.26105 | 75.141 | Promoter | 11 | 62248984 | 62266453 | 17470 | 1 | ENSMUST00 | 0 | Adora2b | ENSMUSG00000018500.2 |
| chr11 | 98457270 | 98458375 | 27.57193 | 78.19525 | 75.07539 | Distal Interge | 11 | 98453789 | 98455125 | 1337 | 1 | ENSMUST00 | 3482 | Grb7 | ENSMUSG00000019312.5 |
| chr17 | 5439388 | 5441013 | 27.57193 | 78.19525 | 75.07539 | Promoter | 17 | 5410870 | 5440243 | 29374 | 2 | ENSMUST00 | 0 | Tmem242 | ENSMUSG00000004945.10 |
| chr17 | 56425075 | 56426001 | 27.57193 | 78.19525 | 75.07539 | Exon (ENSMU | 17 | 56418846 | 56418909 | 64 | 2 | ENSMUST00 | -6167 | Mir6977 | ENSMUSG00000098339.1 |
| chr2 | 126932250 | 126933908 | 27.57193 | 78.19525 | 75.07539 | Promoter | 2 | 126890391 | 126933235 | 42845 | 2 | ENSMUST00 | 0 | Sppl2a | ENSMUSG00000027366.7 |
| chr2 | 152847921 | 152848580 | 27.57193 | 78.19525 | 75.07539 | Promoter | 2 | 152847964 | 152895321 | 47358 | 1 | ENSMUST00 | 0 | Tpx2 | ENSMUSG00000027469.11 |
| chr2 | 155472648 | 155474429 | 27.57193 | 78.19525 | 75.07539 | Promoter | 2 | 155390665 | 155473806 | 83142 | 2 | ENSMUST00 | 0 | Ncoa6 | ENSMUSG00000038369.9 |
| chr4 | 62965204 | 62966197 | 27.57193 | 78.19525 | 75.07539 | Promoter | 4 | 62965573 | 63115748 | 150176 | 1 | ENSMUST00 | 0 | Zfp618 | ENSMUSG00000028358.10 |
| chr8 | 85082980 | 85083798 | 27.57193 | 78.19525 | 75.07539 | Promoter | 8 | 85083269 | 85098739 | 15471 | 1 | ENSMUST00 | 0 | Man2b1 | ENSMUSG00000005142.9 |
| chr9 | 106453266 | 106454683 | 27.57193 | 78.19525 | 75.07539 | Promoter | 9 | 106453291 | 106461094 | 7804 | 1 | ENSMUST00 | 0 | Pcbp4 | ENSMUSG00000023495.7 |
| chr9 | 103111207 | 103113371 | 27.57193 | 78.19525 | 75.07539 | Promoter | 9 | 103111787 | 103162657 | 50871 | 1 | ENSMUST00 | 0 | Rab6b | ENSMUSG00000032549.7 |
| chr9 | 18472999 | 18474072 | 27.57193 | 78.19525 | 75.07539 | Promoter | 9 | 18456192 | 18473559 | 17368 | 2 | ENSMUST00 | 0 | Zfp558 | ENSMUSG00000074500.6 |
| chrX | 85573920 | 85574890 | 27.57193 | 78.19525 | 75.07539 | Promoter | 20 | 85574022 | 85634469 | 60448 | 1 | ENSMUST00 | 0 | Tab3 | ENSMUSG00000035476.4 |
| chrX | 103012691 | 103014310 | 27.57193 | 78.19525 | 75.07539 | Promoter | 20 | 103013698 | 103014387 | 690 | 1 | ENSMUST00 | 0 | Pabpc1l2b-ps | ENSMUSG00000084842.3 |
| chr16 | 18343048 | 18344368 | 26.49423 | 78.18742 | 75.06905 | Promoter | 16 | 18300827 | 18343927 | 43101 | 2 | ENSMUST00 | 0 | Tango2 | ENSMUSG00000013539.8 |
| chr11 | 119090174 | 119091704 | 27.01742 | 78.16648 | 75.04827 | Distal Interge | 11 | 119081373 | 119086221 | 4849 | 2 | ENSMUST00 | -3954 | Cbx4 | ENSMUSG00000039989.7 |
| chr9 | 44267348 | 44268373 | 27.52234 | 78.1102 | 74.99217 | Promoter | 9 | 44252713 | 44268187 | 15475 | 2 | ENSMUST00 | 0 | Nlrx1 | ENSMUSG00000032109.9 |
| chr14 | 61359907 | 61360988 | 26.39107 | 78.00815 | 74.89062 | Promoter | 14 | 61339763 | 61360445 | 20683 | 2 | ENSMUST00 | 0 | Ebpl | ENSMUSG00000021928.8 |
| chr2 | 6129615 | 6130668 | 26.39107 | 78.00815 | 74.89062 | Promoter | 2 | 6100025 | 6130146 | 30122 | 2 | ENSMUST00 | 0 | Proser2 | ENSMUSG00000045319.8 |
| chr4 | 133632405 | 133633763 | 23.27964 | 77.93649 | 74.81927 | Promoter | 4 | 133605299 | 133633469 | 28171 | 2 | ENSMUST00 | 0 | Zdhhc18 | ENSMUSG00000037553.9 |
| chr5 | 66619941 | 66620884 | 26.33979 | 77.91899 | 74.80188 | Distal Interge | 5 | 66298861 | 66618828 | 319968 | 2 | ENSMUST00 | -1114 | Apbb2 | ENSMUSG00000029207.11 |
| chr11 | 113200994 | 113202047 | 27.40733 | 77.91289 | 74.79585 | Promoter | 11 | 113043895 | 113201822 | 157928 | 2 | ENSMUST00 | 0 | 2610035D17 | ENSMUSG00000087259.2 |
| chr16 | 8513058 | 8514438 | 27.40733 | 77.91289 | 74.79585 | Promoter | 16 | 8513429 | 8621568 | 108140 | 1 | ENSMUST00 | 0 | Abat | ENSMUSG00000057880.7 |
| chr2 | 180103287 | 180104905 | 27.40733 | 77.91289 | 74.79585 | Promoter | 2 | 180099465 | 180104488 | 5024 | 2 | ENSMUST00 | 0 | Hrh3 | ENSMUSG00000039059.6 |
| chr3 | 146450143 | 146451147 | 27.40733 | 77.91289 | 74.79585 | Promoter | 3 | 146450470 | 146465849 | 15380 | 1 | ENSMUST00 | 0 | Ctbs | ENSMUSG00000028189.7 |
| chr4 | 116824403 | 116825465 | 27.40733 | 77.91289 | 74.79585 | Promoter | 4 | 116824903 | 116830603 | 5701 | 1 | ENSMUST00 | 0 | Gm12996 | ENSMUSG00000086417.1 |
| chr10 | 26772092 | 26774866 | 26.85422 | 77.8849 | 74.7682 | Promoter | 10 | 26772477 | 26918648 | 146172 | 1 | ENSMUST00 | 0 | Arhgap18 | ENSMUSG00000039031.10 |
| chr7 | 111121782 | 111123426 | 26.85422 | 77.8849 | 74.7682 | Distal Interge | 7 | 111070568 | 111083030 | 12463 | 2 | ENSMUST00 | -38753 | Eif4g2 | ENSMUSG00000005610.11 |
| chr11 | 69836901 | 69838304 | 26.80025 | 77.79173 | 74.67541 | Promoter | 11 | 69823122 | 69837784 | 14663 | 2 | ENSMUST00 | 0 | Nlgn2 | ENSMUSG00000051790.10 |
| chr9 | 21411431 | 21412584 | 26.80025 | 77.79173 | 74.67541 | Promoter | 9 | 21411837 | 21420277 | 8441 | 1 | ENSMUST00 | 0 | Qtrt1 | ENSMUSG00000002825.6 |
| chr1 | 75521023 | 75522382 | 20.88808 | 77.73511 | 74.61916 | Promoter | 1 | 75521529 | 75537335 | 15807 | 1 | ENSMUST00 | 0 | Stk11ip | ENSMUSG00000026213.10 |
| chr11 | 22990063 | 22991717 | 20.88808 | 77.73511 | 74.61916 | Promoter | 11 | 22990519 | 23003780 | 13262 | 1 | ENSMUST00 | 0 | Cct4 | ENSMUSG00000007739.5 |
| chr17 | 56613301 | 56614568 | 20.88808 | 77.73511 | 74.61916 | Promoter | 17 | 56613395 | 56614243 | 849 | 1 | ENSMUST00 | 0 | Rpl36 | ENSMUSG00000057863.5 |
| chr2 | 167014823 | 167016046 | 20.88808 | 77.73511 | 74.61916 | Promoter | 2 | 167015193 | 167034947 | 19755 | 1 | ENSMUST00 | 0 | Ddx27 | ENSMUSG00000017999.11 |
| chr7 | 74553987 | 74555444 | 20.88808 | 77.73511 | 74.61916 | Promoter | 7 | 74275419 | 74554633 | 279215 | 2 | ENSMUST00 | 0 | Slco3a1 | ENSMUSG00000025790.9 |
| chr7 | 109671273 | 109672671 | 20.88808 | 77.73511 | 74.61916 | Distal Interge | 7 | 109703690 | 109712189 | 8500 | 1 | ENSMUST00 | -31019 | Akip1 | ENSMUSG00000031023.5 |
| chr9 | 83833982 | 83835194 | 20.88808 | 77.73511 | 74.61916 | Promoter | 9 | 83834689 | 83869493 | 34805 | 1 | ENSMUST00 | 0 | Ttk | ENSMUSG00000038379.10 |
| chr8 | 84689204 | 84690070 | 26.7465 | 77.69891 | 74.58387 | Promoter | 8 | 84689247 | 84699787 | 10541 | 1 | ENSMUST00 | 0 | Trmt1 | ENSMUSG00000001909.10 |
| chr19 | 29361260 | 29362431 | 26.18714 | 77.65332 | 74.53902 | Promoter | 19 | 29348599 | 29361970 | 13372 | 2 | ENSMUST00 | 0 | Plgrkt | ENSMUSG00000016495.7 |
| chr10 | 100015316 | 100016774 | 23.12601 | 77.64799 | 74.53377 | Promoter | 10 | 100015630 | 100100413 | 84784 | 1 | ENSMUST00 | 0 | Kitl | ENSMUSG00000019966.12 |
| chr10 | 81559124 | 81561616 | 23.12601 | 77.64799 | 74.53377 | Promoter | 10 | 81559494 | 81566362 | 6869 | 1 | ENSMUST00 | 0 | Aes | ENSMUSG00000054452.9 |
| chr11 | 94548415 | 94549345 | 23.12601 | 77.64799 | 74.53377 | Promoter | 11 | 94539798 | 94549199 | 9402 | 2 | ENSMUST00 | 0 | Rsad1 | ENSMUSG00000039096.5 |
| chr13 | 55726885 | 55728189 | 23.12601 | 77.64799 | 74.53377 | Promoter | 13 | 55727368 | 55776828 | 49461 | 1 | ENSMUST00 | 0 | Pcbd2 | ENSMUSG00000021496.7 |
| chr13 | 109902335 | 109904319 | 23.12601 | 77.64799 | 74.53377 | Promoter | 13 | 109903102 | 109951387 | 48286 | 1 | ENSMUST00 | 0 | Pde4d | ENSMUSG00000021699.12 |
| chr13 | 97070974 | 97072321 | 23.12601 | 77.64799 | 74.53377 | Promoter | 13 | 97071643 | 97129521 | 57879 | 1 | ENSMUST00 | 0 | Fam169a | ENSMUSG00000041817.8 |
| chr15 | 102721189 | 102722153 | 23.12601 | 77.64799 | 74.53377 | Promoter | 15 | 102707585 | 102722131 | 14547 | 2 | ENSMUST00 | 0 | Calcoco1 | ENSMUSG00000023055.4 |
| chr17 | 23699230 | 23700464 | 23.12601 | 77.64799 | 74.53377 | Distal Interge | 17 | 23705058 | 23706339 | 1282 | 2 | ENSMUST00 | 5875 | 1520401A03 | ENSMUSG00000043747.7 |
| chr19 | 25236314 | 25239245 | 23.12601 | 77.64799 | 74.53377 | Promoter | 19 | 25236975 | 25434496 | 197522 | 1 | ENSMUST00 | 0 | Kank1 | ENSMUSG00000032702.11 |
| chr2 | 102399957 | 102401748 | 23.12601 | 77.64799 | 74.53377 | Promoter | 2 | 102300119 | 102400900 | 100782 | 2 | ENSMUST00 | 0 | Trim44 | ENSMUSG00000027189.11 |
| chr2 | 155356572 | 155358047 | 23.12601 | 77.64799 | 74.53377 | Promoter | 2 | 155278243 | 155357430 | 79188 | 2 | ENSMUST00 | 0 | Pigu | ENSMUSG00000038383.11 |
| chr4 | 48042390 | 48044094 | 23.12601 | 77.64799 | 74.53377 | Distal Interge | 4 | 48045153 | 48086445 | 41293 | 1 | ENSMUST00 | -1059 | Nr4a3 | ENSMUSG00000028341.4 |
| chr4 | 124700324 | 124701319 | 23.12601 | 77.64799 | 74.53377 | Promoter | 4 | 124700701 | 124708094 | 7394 | 1 | ENSMUST00 | 0 | Fhl3 | ENSMUSG00000032643.7 |
| chr4 | 136284651 | 136286770 | 23.12601 | 77.64799 | 74.53377 | Promoter | 4 | 136284683 | 136290327 | 5645 | 1 | ENSMUST00 | 0 | Zfp46 | ENSMUSG00000051351.9 |
| chr5 | 33656992 | 33659141 | 23.12601 | 77.64799 | 74.53377 | Promoter | 5 | 33653216 | 33657977 | 4762 | 2 | ENSMUST00 | 0 | Tmem129 | ENSMUSG00000019295.7 |
| chr5 | 36694931 | 36696494 | 23.12601 | 77.64799 | 74.53377 | Promoter | 5 | 36600504 | 36695969 | 95466 | 2 | ENSMUST00 | 0 | D5Ertd579e | ENSMUSG00000029190.14 |
| chr5 | 36582041 | 36582816 | 23.12601 | 77.64799 | 74.53377 | Promoter | 5 | 36490604 | 36582397 | 91794 | 2 | ENSMUST00 | 0 | Tbc1d14 | ENSMUSG00000029192.12 |
| chr6 | 47953064 | 47954174 | 23.12601 | 77.64799 | 74.53377 | Promoter | 6 | 47953390 | 47965300 | 11911 | 1 | ENSMUST00 | 0 | Zfp956 | ENSMUSG00000045466.13 |
| chr7 | 82647978 | 82649359 | 23.12601 | 77.64799 | 74.53377 | Promoter | 7 | 82648614 | 82674555 | 25942 | 1 | ENSMUST00 | 0 | Eftud1 | ENSMUSG00000038563.8 |
| chr7 | 29280794 | 29282413 | 23.12601 | 77.64799 | 74.53377 | Promoter | 7 | 29256331 | 29281977 | 25647 | 2 | ENSMUST00 | 0 | Spint2 | ENSMUSG00000074227.6 |
| chr7 | 39552060 | 39553247 | 23.12601 | 77.64799 | 74.53377 | Promoter | 7 | 39552496 | 39556305 | 3810 | 1 | ENSMUST00 | 0 | RP23-402K24 | ENSMUSG00000099697.1 |
| chr8 | 4676905 | 4678922 | 23.12601 | 77.64799 | 74.53377 | Promoter | 8 | 4677539 | 4678365 | 827 | 2 | ENSMUST00 | 0 | Gm7461 | ENSMUSG00000048484.4 |
| chr1 | 162547847 | 162549019 | 24.1047 | 77.63509 | 74.52327 | Promoter | 1 | 162532127 | 162548551 | 16425 | 2 | ENSMUST00 | 0 | Mettl13 | ENSMUSG00000026694.14 |
| chr16 | 30063120 | 30067742 | 24.1047 | 77.63509 | 74.52327 | Promoter | 16 | 30064384 | 30065569 | 1186 | 1 | ENSMUST00 | 0 | Hes1 | ENSMUSG00000022528.7 |
| chr19 | 8723448 | 8724334 | 24.1047 | 77.63509 | 74.52327 | Promoter | 19 | 8723993 | 8724064 | 72 | 1 | ENSMUST00 | 0 | Gm25855 | ENSMUSG00000064721.4 |
| chr4 | 124714395 | 124715568 | 26.69297 | 77.60641 | 74.495 | Promoter | 4 | 124714776 | 124732460 | 17685 | 1 | ENSMUST00 | 0 | Sf3a3 | ENSMUSG00000028902.4 |

| | | | | | | | | | | | | | | | |
|---|---|---|---|---|---|---|---|---|---|---|---|---|---|---|---|
| chr6 | 29211215 | 29212626 | 26.69297 | 77.60641 | 74.495 | Promoter | 6 | 29200437 | 29212249 | 11813 | 2 | ENSMUST00 | 0 | Impdh1 | ENSMUSG00000003500.8 |
| chr17 | 29346218 | 29348377 | 26.13664 | 77.56537 | 74.45414 | Promoter | 17 | 29332072 | 29347904 | 15833 | 2 | ENSMUST00 | 0 | Mtch1 | ENSMUSG00000024012.12 |
| chr2 | 73386735 | 73387181 | 26.13664 | 77.56537 | 74.45414 | Promoter | 2 | 73386970 | 73420498 | 33529 | 1 | ENSMUST00 | 0 | D230022J07F | ENSMUSG00000087176.1 |
| chr6 | 41604462 | 41605908 | 26.13664 | 77.56537 | 74.45414 | Promoter | 6 | 41605482 | 41620509 | 15028 | 1 | ENSMUST00 | 0 | Ephb6 | ENSMUSG00000029869.7 |
| chr6 | 145864843 | 145866819 | 26.13664 | 77.56537 | 74.45414 | Promoter | 6 | 145858243 | 145865558 | 7316 | 2 | ENSMUST00 | 0 | Bhlhe41 | ENSMUSG00000030256.6 |
| chr7 | 100536530 | 100537475 | 26.13664 | 77.56537 | 74.45414 | Promoter | 7 | 100537100 | 100540369 | 3270 | 1 | ENSMUST00 | 0 | Coa4 | ENSMUSG00000044881.6 |
| chr12 | 86078492 | 86079616 | 24.56002 | 77.54259 | 74.4321 | Promoter | 12 | 86056744 | 86079041 | 22298 | 2 | ENSMUST00 | 0 | Tgfb3 | ENSMUSG00000021253.6 |
| chr2 | 25455822 | 25461073 | 24.56002 | 77.54259 | 74.4321 | Promoter | 2 | 25456838 | 25458676 | 1839 | 1 | ENSMUST00 | 0 | Clic3 | ENSMUSG00000015093.10 |
| chr12 | 59012234 | 59013989 | 27.18019 | 77.52287 | 74.41271 | Promoter | 12 | 59013393 | 59028470 | 15078 | 1 | ENSMUST00 | 0 | Gemin2 | ENSMUSG00000060121.10 |
| chr2 | 92388689 | 92389914 | 27.18019 | 77.52287 | 74.41271 | Exon (ENSMU | 2 | 92387384 | 92387446 | 63 | 2 | ENSMUST00 | -1244 | Mir7000 | ENSMUSG00000099093.1 |
| chr7 | 28267611 | 28268724 | 27.18019 | 77.52287 | 74.41271 | Promoter | 7 | 28267881 | 28269272 | 1392 | 1 | ENSMUST00 | 0 | Eid2 | ENSMUSG00000046058.7 |
| chr11 | 75630258 | 75632213 | 25.57207 | 77.50074 | 74.3909 | Promoter | 11 | 75630988 | 75648871 | 17884 | 1 | ENSMUST00 | 0 | Inpp5k | ENSMUSG00000006127.4 |
| chr12 | 38778143 | 38781472 | 25.57207 | 77.50074 | 74.3909 | Promoter | 12 | 38779380 | 38866249 | 86870 | 1 | ENSMUST00 | 0 | Etv1 | ENSMUSG00000004151.11 |
| chr14 | 31018221 | 31020204 | 25.57207 | 77.50074 | 74.3909 | Promoter | 14 | 31019138 | 31118901 | 99764 | 1 | ENSMUST00 | 0 | Pbrm1 | ENSMUSG00000042323.11 |
| chr15 | 68257894 | 68259920 | 25.57207 | 77.50074 | 74.3909 | Promoter | 15 | 68083766 | 68258856 | 175091 | 2 | ENSMUST00 | 0 | Zfat | ENSMUSG00000022335.11 |
| chr16 | 43889115 | 43890424 | 25.57207 | 77.50074 | 74.3909 | Promoter | 16 | 43889800 | 43964314 | 74515 | 1 | ENSMUST00 | 0 | 2610015P09I | ENSMUSG00000022701.12 |
| chr2 | 181186784 | 181188108 | 25.57207 | 77.50074 | 74.3909 | Promoter | 2 | 181187247 | 181188771 | 1525 | 1 | ENSMUST00 | 0 | Ppdpf | ENSMUSG00000016344.9 |
| chr2 | 31245344 | 31246825 | 25.57207 | 77.50074 | 74.3909 | Promoter | 2 | 31245823 | 31295989 | 50167 | 1 | ENSMUST00 | 0 | Ncs1 | ENSMUSG00000062661.6 |
| chr8 | 84989803 | 84991156 | 25.57207 | 77.50074 | 74.3909 | Promoter | 8 | 84990595 | 85003364 | 12770 | 1 | ENSMUST00 | 0 | Hook2 | ENSMUSG00000052566.7 |
| chr11 | 116671178 | 116672541 | 25.04793 | 77.4957 | 74.38671 | Promoter | 11 | 116671660 | 116675798 | 4139 | 1 | ENSMUST00 | 0 | 1810032O08 | ENSMUSG00000020812.12 |
| chr15 | 76457167 | 76458476 | 25.04793 | 77.4957 | 74.38671 | Promoter | 15 | 76457438 | 76459468 | 2031 | 1 | ENSMUST00 | 0 | Scx | ENSMUSG00000034161.7 |
| chr4 | 24850287 | 24851504 | 25.04793 | 77.4957 | 74.38671 | Promoter | 4 | 24612554 | 24851124 | 238571 | 2 | ENSMUST00 | 0 | Klhl32 | ENSMUSG00000040387.11 |
| chr8 | 3630558 | 3631845 | 25.04793 | 77.4957 | 74.38671 | Promoter | 8 | 3630955 | 3643644 | 12690 | 1 | ENSMUST00 | 0 | Stxbp2 | ENSMUSG00000004626.9 |
| chr11 | 87592061 | 87593337 | 26.08635 | 77.47771 | 74.36935 | Promoter | 11 | 87592162 | 87616302 | 24141 | 1 | ENSMUST00 | 0 | Mtmr4 | ENSMUSG00000018401.12 |
| chr7 | 30279798 | 30281238 | 17.00533 | 77.38029 | 74.27235 | Promoter | 7 | 30280094 | 30290237 | 10144 | 1 | ENSMUST00 | 0 | Thap8 | ENSMUSG00000013928.7 |
| chr10 | 76237092 | 76238425 | 19.88326 | 77.37266 | 74.26472 | Promoter | 10 | 76207222 | 76237865 | 30644 | 2 | ENSMUST00 | 0 | Prmt2 | ENSMUSG00000020230.9 |
| chr11 | 6474409 | 6476644 | 19.88326 | 77.37266 | 74.26472 | Promoter | 11 | 6475425 | 6490918 | 15494 | 1 | ENSMUST00 | 0 | Gm11973 | ENSMUSG00000086839.2 |
| chr2 | 39004267 | 39005987 | 19.88326 | 77.37266 | 74.26472 | Promoter | 2 | 39001580 | 39005624 | 4045 | 2 | ENSMUST00 | 0 | Rpl35 | ENSMUSG00000062997.6 |
| chr4 | 155830691 | 155832625 | 19.88326 | 77.37266 | 74.26472 | Promoter | 4 | 155831272 | 155832739 | 1468 | 1 | ENSMUST00 | 0 | Aurkaip1 | ENSMUSG00000065990.7 |
| chr5 | 137599660 | 137602084 | 19.88326 | 77.37266 | 74.26472 | Promoter | 5 | 137596645 | 137600708 | 4064 | 2 | ENSMUST00 | 0 | Mospd3 | ENSMUSG00000037221.8 |
| chr6 | 71440869 | 71441545 | 19.88326 | 77.37266 | 74.26472 | Promoter | 6 | 71388634 | 71440637 | 52004 | 2 | ENSMUST00 | -233 | Rmnd5a | ENSMUSG00000002222.9 |
| chr6 | 54039287 | 54040535 | 26.53366 | 77.33094 | 74.22335 | Promoter | 6 | 54039554 | 54067798 | 28245 | 1 | ENSMUST00 | 0 | Chn2 | ENSMUSG00000004633.12 |
| chr10 | 127069397 | 127071246 | 25.98482 | 77.30067 | 74.19341 | Promoter | 10 | 127067317 | 127070261 | 2945 | 2 | ENSMUST00 | 0 | Tspan31 | ENSMUSG00000006736.8 |
| chr11 | 50601623 | 50603606 | 25.98482 | 77.30067 | 74.19341 | Promoter | 11 | 50602084 | 50807573 | 205490 | 1 | ENSMUST00 | 0 | Adamts2 | ENSMUSG00000036545.8 |
| chr14 | 29721447 | 29723107 | 25.98482 | 77.30067 | 74.19341 | Promoter | 14 | 28904943 | 29721864 | 816922 | 2 | ENSMUST00 | 0 | Cacna2d3 | ENSMUSG00000021991.8 |
| chr15 | 31452958 | 31454212 | 25.98482 | 77.30067 | 74.19341 | Promoter | 15 | 31441210 | 31453689 | 12480 | 2 | ENSMUST00 | 0 | Ropn1l | ENSMUSG00000022236.9 |
| chr15 | 96461063 | 96461897 | 25.98482 | 77.30067 | 74.19341 | Promoter | 15 | 96411698 | 96460843 | 49146 | 2 | ENSMUST00 | -221 | Scaf11 | ENSMUSG00000033228.7 |
| chr17 | 17373492 | 17375488 | 25.98482 | 77.30067 | 74.19341 | Promoter | 17 | 17374332 | 17394898 | 20567 | 1 | ENSMUST00 | 0 | Chd1 | ENSMUSG00000023852.8 |
| chr18 | 36744012 | 36745688 | 25.98482 | 77.30067 | 74.19341 | Promoter | 18 | 36742332 | 36744557 | 2226 | 2 | ENSMUST00 | 0 | Ndufa2 | ENSMUSG00000014294.4 |
| chr18 | 80712341 | 80713735 | 25.98482 | 77.30067 | 74.19341 | Promoter | 18 | 80606205 | 80713071 | 106867 | 2 | ENSMUST00 | 0 | Nfatc1 | ENSMUSG00000033016.10 |
| chr2 | 31670641 | 31671452 | 25.98482 | 77.30067 | 74.19341 | Promoter | 2 | 31670715 | 31681350 | 10636 | 1 | ENSMUST00 | 0 | Exosc2 | ENSMUSG00000039356.10 |
| chr4 | 129500396 | 129501680 | 25.98482 | 77.30067 | 74.19341 | Intron (ENSM | 4 | 129491969 | 129509746 | 17778 | 2 | ENSMUST00 | 8066 | Gm12979 | ENSMUSG00000086336.1 |
| chr6 | 5297228 | 5298667 | 25.98482 | 77.30067 | 74.19341 | Promoter | 6 | 5264147 | 5298455 | 34309 | 2 | ENSMUST00 | 0 | Pon2 | ENSMUSG00000032667.12 |
| chr7 | 35214583 | 35215853 | 25.98482 | 77.30067 | 74.19341 | Promoter | 7 | 35200878 | 35215370 | 14493 | 2 | ENSMUST00 | 0 | Lrp3 | ENSMUSG00000001802.11 |
| chr7 | 80186223 | 80187680 | 25.98482 | 77.30067 | 74.19341 | Promoter | 7 | 80186841 | 80221832 | 34992 | 1 | ENSMUST00 | 0 | Sema4b | ENSMUSG00000030539.8 |
| chr7 | 81523006 | 81524204 | 25.98482 | 77.30067 | 74.19341 | Promoter | 7 | 81523550 | 81529554 | 6005 | 1 | ENSMUST00 | 0 | 2900076A07I | ENSMUSG00000097277.2 |
| chr8 | 71366516 | 71368084 | 25.98482 | 77.30067 | 74.19341 | Promoter | 8 | 71366848 | 71369729 | 2882 | 1 | ENSMUST00 | 0 | Use1 | ENSMUSG00000002395.9 |
| chr8 | 104540605 | 104541754 | 25.98482 | 77.30067 | 74.19341 | Promoter | 8 | 104540807 | 104550343 | 9537 | 1 | ENSMUST00 | 0 | Car7 | ENSMUSG00000031883.7 |
| chr9 | 44772876 | 44773729 | 25.98482 | 77.30067 | 74.19341 | Promoter | 9 | 44772935 | 44792714 | 19780 | 1 | ENSMUST00 | 0 | Ift46 | ENSMUSG00000002031.6 |
| chr1 | 195130337 | 195132022 | 24.40663 | 77.266 | 74.1616 | Promoter | 1 | 195103789 | 195131570 | 27782 | 2 | ENSMUST00 | 0 | Cr1l | ENSMUSG00000016481.10 |
| chr1 | 60342766 | 60344158 | 24.40663 | 77.266 | 74.1616 | Promoter | 1 | 60343323 | 60372707 | 29385 | 1 | ENSMUST00 | 0 | Cyp20a1 | ENSMUSG00000049439.8 |
| chr1 | 90431359 | 90438623 | 24.40663 | 77.266 | 74.1616 | Distal Interge | 1 | 90453394 | 90454501 | 1108 | 2 | ENSMUST00 | 15878 | RP24-154L11 | ENSMUSG00000101997.1 |
| chr10 | 75772714 | 75773424 | 24.40663 | 77.266 | 74.1616 | Promoter | 10 | 75771230 | 75773034 | 1805 | 2 | ENSMUST00 | 0 | Ddt | ENSMUSG00000001666.8 |
| chr11 | 78343121 | 78344003 | 24.40663 | 77.266 | 74.1616 | Promoter | 11 | 78343482 | 78349164 | 5683 | 1 | ENSMUST00 | 0 | Unc119 | ENSMUSG00000002058.8 |
| chr13 | 74008885 | 74010801 | 24.40663 | 77.266 | 74.1616 | Promoter | 13 | 74009407 | 74031371 | 21965 | 1 | ENSMUST00 | 0 | Tppp | ENSMUSG00000021573.9 |
| chr14 | 60176802 | 60177825 | 24.40663 | 77.266 | 74.1616 | Promoter | 14 | 60086582 | 60177539 | 90958 | 2 | ENSMUST00 | 0 | Atp8a2 | ENSMUSG00000021983.10 |
| chr17 | 36167367 | 36168675 | 24.40663 | 77.266 | 74.1616 | Promoter | 17 | 36164443 | 36168523 | 4081 | 2 | ENSMUST00 | 0 | Gm8909 | ENSMUSG00000073402.6 |
| chr18 | 70529710 | 70531341 | 24.40663 | 77.266 | 74.1616 | Promoter | 18 | 70508680 | 70530138 | 21459 | 2 | ENSMUST00 | 0 | Poli | ENSMUSG00000038425.12 |
| chr2 | 91236355 | 91237807 | 24.40663 | 77.266 | 74.1616 | Promoter | 2 | 91211588 | 91236907 | 25320 | 2 | ENSMUST00 | 0 | Ddb2 | ENSMUSG00000002109.9 |
| chr4 | 116596656 | 116598461 | 24.40663 | 77.266 | 74.1616 | Promoter | 4 | 116596730 | 116600266 | 3537 | 1 | ENSMUST00 | 0 | Ccdc17 | ENSMUSG00000034035.13 |
| chr5 | 35582450 | 35583832 | 24.40663 | 77.266 | 74.1616 | Promoter | 5 | 35583040 | 35613801 | 30762 | 1 | ENSMUST00 | 0 | Acox3 | ENSMUSG00000029098.11 |
| chr5 | 36581265 | 36581929 | 24.40663 | 77.266 | 74.1616 | Promoter | 5 | 36490604 | 36581803 | 91200 | 2 | ENSMUST00 | 0 | Tbc1d14 | ENSMUSG00000029192.12 |
| chr5 | 120611934 | 120612873 | 24.40663 | 77.266 | 74.1616 | Promoter | 5 | 120594305 | 120612339 | 18035 | 2 | ENSMUST00 | 0 | Rita1 | ENSMUSG00000029600.5 |
| chr5 | 67426148 | 67428221 | 24.40663 | 77.266 | 74.1616 | Promoter | 5 | 67392147 | 67427799 | 35653 | 2 | ENSMUST00 | 0 | Bend4 | ENSMUSG00000092060.1 |
| chr6 | 17463213 | 17464736 | 24.40663 | 77.266 | 74.1616 | Promoter | 6 | 17463800 | 17573980 | 110181 | 1 | ENSMUST00 | 0 | Met | ENSMUSG00000009376.10 |
| chr7 | 116033322 | 116034862 | 24.40663 | 77.266 | 74.1616 | Intron (ENSM | 7 | 116031979 | 116032550 | 572 | 1 | ENSMUST00 | 1344 | Sox6os | ENSMUSG00000030664.3 |
| chr7 | 126861005 | 126863038 | 24.40663 | 77.266 | 74.1616 | Promoter | 7 | 126851580 | 126861463 | 9884 | 2 | ENSMUST00 | 0 | Ino80e | ENSMUSG00000030689.10 |
| chr7 | 127406173 | 127406855 | 24.40663 | 77.266 | 74.1616 | Promoter | 7 | 127403668 | 127406822 | 3155 | 2 | ENSMUST00 | 0 | Zfp764 | ENSMUSG00000045757.6 |
| chr7 | 7209419 | 7210038 | 24.40663 | 77.266 | 74.1616 | Promoter | 7 | 7202122 | 7209998 | 7877 | 2 | ENSMUST00 | 0 | Zfp772 | ENSMUSG00000066838.6 |
| chr8 | 84066764 | 84067958 | 24.40663 | 77.266 | 74.1616 | Promoter | 8 | 84066836 | 84096992 | 30157 | 1 | ENSMUST00 | 0 | Rfx1 | ENSMUSG00000031706.6 |
| chr8 | 14910723 | 14912537 | 24.40663 | 77.266 | 74.1616 | Promoter | 8 | 14911663 | 15001085 | 89423 | 1 | ENSMUST00 | 0 | Arhgef10 | ENSMUSG00000071176.5 |
| chr3 | 95588802 | 95589687 | 27.01228 | 77.23428 | 74.13344 | Promoter | 3 | 95588934 | 95619242 | 30309 | 1 | ENSMUST00 | 0 | Golph3l | ENSMUSG00000046519.10 |
| chr11 | 48837053 | 48838781 | 17.59484 | 77.21284 | 74.11221 | Promoter | 11 | 48837465 | 48850555 | 13091 | 1 | ENSMUST00 | 0 | Trim7 | ENSMUSG00000040350.10 |
| chr13 | 105053662 | 105055291 | 26.42849 | 77.14894 | 74.04846 | Promoter | 13 | 104947153 | 105054930 | 107778 | 2 | ENSMUST00 | 0 | Rgs7bp | ENSMUSG00000021719.8 |
| chr16 | 17489151 | 17490431 | 26.42849 | 77.14894 | 74.04846 | Promoter | 16 | 17489611 | 17507485 | 17875 | 1 | ENSMUST00 | 0 | Aifm3 | ENSMUSG00000022763.11 |
| chr11 | 100622779 | 100623594 | 18.2599 | 77.14165 | 74.04132 | Promoter | 11 | 100622876 | 100627607 | 4732 | 1 | ENSMUST00 | 0 | Nkiras2 | ENSMUSG00000017837.7 |
| chr4 | 129376396 | 129378534 | 18.2599 | 77.14165 | 74.04132 | Promoter | 4 | 129353628 | 129378116 | 24489 | 2 | ENSMUST00 | 0 | Zbtb8a | ENSMUSG00000028807.2 |
| chr4 | 88785084 | 88785605 | 18.2599 | 77.14165 | 74.04132 | Promoter | 4 | 88785709 | 88786257 | 549 | 1 | ENSMUST00 | -104 | Gm13276 | ENSMUSG00000099545.1 |
| chr19 | 31762329 | 31765338 | 26.95677 | 77.13882 | 74.03856 | Promoter | 19 | 30567551 | 31765033 | 1197483 | 2 | ENSMUST00 | 0 | Prkg1 | ENSMUSG00000052920.9 |
| chr4 | 49520206 | 49521914 | 26.95677 | 77.13882 | 74.03856 | Promoter | 4 | 49521176 | 49531517 | 10342 | 1 | ENSMUST00 | 0 | Zfp189 | ENSMUSG00000039634.7 |
| chr2 | 102450101 | 102452417 | 25.88708 | 77.13007 | 74.02996 | Promoter | 2 | 102449366 | 102452499 | 3134 | 2 | ENSMUST00 | 82 | Fjx1 | ENSMUSG00000075012.4 |
| chr17 | 24720090 | 24720644 | 23.30216 | 77.06895 | 73.96892 | Promoter | 17 | 24720110 | 24721929 | 1820 | 1 | ENSMUST00 | 0 | Rps2 | ENSMUSG00000044533.10 |
| chr12 | 75668930 | 75669583 | 26.90149 | 77.04372 | 73.94399 | Promoter | 12 | 75630596 | 75669537 | 38942 | 2 | ENSMUST00 | 0 | Wdr89 | ENSMUSG00000045690.8 |
| chr7 | 51861279 | 51862822 | 25.78858 | 76.95798 | 73.85856 | Promoter | 7 | 51862015 | 51994975 | 132961 | 1 | ENSMUST00 | 0 | Gas2 | ENSMUSG00000030498.9 |
| chr11 | 43425573 | 43426753 | 26.84644 | 76.94898 | 73.84959 | Promoter | 11 | 43420250 | 43426251 | 6002 | 2 | ENSMUST00 | 0 | Pttg1 | ENSMUSG00000020415.11 |
| chr11 | 62647832 | 62649284 | 26.84644 | 76.94898 | 73.84959 | Promoter | 11 | 62604884 | 62648523 | 43640 | 2 | ENSMUST00 | 0 | Lrrc75a | ENSMUSG00000046417.9 |
| chr19 | 46038359 | 46040333 | 26.84644 | 76.94898 | 73.84959 | Promoter | 19 | 46032594 | 46039621 | 7028 | 2 | ENSMUST00 | 0 | Ldb1 | ENSMUSG00000025223.9 |
| chr5 | 114657776 | 114658683 | 26.84644 | 76.94898 | 73.84959 | Promoter | 5 | 114622152 | 114658421 | 36270 | 2 | ENSMUST00 | 0 | Trpv4 | ENSMUSG00000014158.6 |
| chr2 | 145860076 | 145863605 | 23.70866 | 76.9101 | 73.81126 | Promoter | 2 | 145862644 | 145878974 | 16331 | 1 | ENSMUST00 | 0 | Rin2 | ENSMUSG00000001768.10 |
| chr9 | 121276862 | 121278187 | 23.70866 | 76.9101 | 73.81126 | Promoter | 9 | 120964454 | 121277197 | 312744 | 2 | ENSMUST00 | 0 | Ulk4 | ENSMUSG00000040936.9 |
| chr2 | 62408166 | 62412568 | 26.27231 | 76.87836 | 73.77971 | Promoter | 2 | 62330073 | 62412231 | 82159 | 2 | ENSMUST00 | 0 | Dpp4 | ENSMUSG00000035000.8 |
| chr2 | 15048699 | 15049735 | 26.7916 | 76.85461 | 73.75612 | Promoter | 2 | 15049395 | 15078371 | 28977 | 1 | ENSMUST00 | 0 | Arl5b | ENSMUSG00000017418.8 |
| chr7 | 127744610 | 127747241 | 26.7916 | 76.85461 | 73.75612 | Promoter | 7 | 127745201 | 127768927 | 23727 | 1 | ENSMUST00 | 0 | Fbxl19 | ENSMUSG00000030811.9 |
| chr11 | 49202015 | 49204776 | 21.75525 | 76.8157 | 73.7174 | Promoter | 11 | 49203292 | 49218553 | 15262 | 1 | ENSMUST00 | 0 | Zfp62 | ENSMUSG00000046311.8 |
| chr13 | 15462764 | 15465151 | 21.75525 | 76.8157 | 73.7174 | Promoter | 13 | 15463235 | 15715091 | 251857 | 1 | ENSMUST00 | 0 | Gli3 | ENSMUSG00000021318.10 |
| chr13 | 92529903 | 92531684 | 21.75525 | 76.8157 | 73.7174 | Promoter | 13 | 92487108 | 92530868 | 43761 | 2 | ENSMUST00 | 0 | Zfyve16 | ENSMUSG00000021706.9 |
| chr16 | 44346842 | 44348178 | 21.75525 | 76.8157 | 73.7174 | Promoter | 16 | 44347121 | 44388497 | 41377 | 1 | ENSMUST00 | 0 | Spice1 | ENSMUSG00000043065.7 |
| chr17 | 26561044 | 26562480 | 21.75525 | 76.8157 | 73.7174 | Promoter | 17 | 26561489 | 26656931 | 95443 | 1 | ENSMUST00 | 0 | Ergic1 | ENSMUSG00000001576.9 |
| chr17 | 35699302 | 35701573 | 21.75525 | 76.8157 | 73.7174 | Promoter | 17 | 35681567 | 35700610 | 19044 | 2 | ENSMUST00 | 0 | Ddr1 | ENSMUSG00000003534.12 |
| chr17 | 34951628 | 34953049 | 21.75525 | 76.8157 | 73.7174 | Promoter | 17 | 34950241 | 34952448 | 2208 | 2 | ENSMUST00 | 0 | 1110038B12I | ENSMUSG00000092203.2 |
| chr2 | 181693179 | 181694305 | 21.75525 | 76.8157 | 73.7174 | Promoter | 2 | 181688422 | 181693920 | 5499 | 2 | ENSMUST00 | 0 | Rgs19 | ENSMUSG00000002458.8 |
| chr8 | 111392792 | 111394218 | 21.75525 | 76.8157 | 73.7174 | Promoter | 8 | 111345141 | 111393824 | 48684 | 2 | ENSMUST00 | 0 | Fa2h | ENSMUSG00000033579.11 |
| chr9 | 26998819 | 27000333 | 21.75525 | 76.8157 | 73.7174 | Promoter | 9 | 26974135 | 26999566 | 25432 | 2 | ENSMUST00 | 0 | Acad8 | ENSMUSG00000031969.10 |
| chr13 | 23710162 | 23710971 | 26.22066 | 76.7888 | 73.6919 | Promoter | 13 | 23703840 | 23710854 | 7015 | 2 | ENSMUST00 | 0 | Hfe | ENSMUSG00000006611.9 |
| chr13 | 43365238 | 43366264 | 26.22066 | 76.7888 | 73.6919 | Distal Interge | 13 | 43370710 | 43395202 | 24493 | 1 | ENSMUST00 | -4446 | Sirt5 | ENSMUSG00000054021.5 |
| chr14 | 31130528 | 31131738 | 26.22066 | 76.7888 | 73.6919 | Distal Interge | 14 | 31124802 | 31128930 | 4129 | 2 | ENSMUST00 | -1599 | Smim4 | ENSMUSG00000058351.6 |
| chr15 | 41446789 | 41448543 | 26.22065 | 76.7888 | 73.6919 | Promoter | 15 | 41447482 | 41861045 | 413564 | 1 | ENSMUST00 | 0 | Oxr1 | ENSMUSG00000022307.10 |
| chr17 | 78916000 | 78917238 | 26.22066 | 76.7888 | 73.6919 | Promoter | 17 | 78916499 | 78921037 | 4539 | 1 | ENSMUST00 | 0 | Cebpzos | ENSMUSG00000062691.5 |
| chr19 | 8840256 | 8841299 | 26.22066 | 76.7888 | 73.6919 | Promoter | 19 | 8840519 | 8848666 | 8148 | 1 | ENSMUST00 | 0 | Bscl2 | ENSMUSG00000071657.7 |
| chr2 | 25546078 | 25547023 | 26.22066 | 76.7888 | 73.6919 | Promoter | 2 | 25517982 | 25546940 | 28959 | 2 | ENSMUST00 | 0 | Traf2 | ENSMUSG00000026942.8 |
| chr5 | 31054270 | 31055389 | 24.14278 | 76.78871 | 73.6919 | Promoter | 5 | 31054780 | 31078479 | 23700 | 1 | ENSMUST00 | 0 | Cad | ENSMUSG00000013629.11 |
| chr5 | 53266685 | 53267919 | 24.14278 | 76.78871 | 73.6919 | Promoter | 5 | 53267106 | 53278487 | 11382 | 1 | ENSMUST00 | 0 | Smim20 | ENSMUSG00000061461.7 |
| chr8 | 71569284 | 71571112 | 26.22066 | 76.7888 | 73.6919 | Promoter | 8 | 71568927 | 71586708 | 17782 | 1 | ENSMUST00 | 358 | Slc27a1 | ENSMUSG00000031808.6 |
| chr13 | 21782848 | 21783999 | 24.60744 | 76.70956 | 73.61404 | Promoter | 13 | 21782960 | 21783370 | 411 | 2 | ENSMUST00 | 0 | Hist1h3i | ENSMUSG00000101972.1 |
| chr2 | 38511327 | 38512271 | 24.60744 | 76.70956 | 73.61404 | Promoter | 2 | 38511643 | 38588041 | 76399 | 1 | ENSMUST00 | 0 | Nek6 | ENSMUSG00000026749.6 |
| chr7 | 118740282 | 118741378 | 25.64223 | 76.702 | 73.60689 | Promoter | 7 | 118740732 | 118784055 | 43324 | 1 | ENSMUST00 | 0 | 9030624J02F | ENSMUSG00000030982.12 |
| chr2 | 50295404 | 50297372 | 26.1692 | 76.69956 | 73.60538 | Promoter | 2 | 50279881 | 50296695 | 16815 | 2 | ENSMUST00 | 0 | Mmadhc | ENSMUSG00000026766.11 |
| chr1 | 132880058 | 132881648 | 25.10597 | 76.67836 | 73.5843 | Promoter | 1 | 132880273 | 132940005 | 59733 | 1 | ENSMUST00 | 0 | Lrrn2 | ENSMUSG00000026443.3 |
| chr10 | 79746074 | 79746659 | 25.10597 | 76.67836 | 73.5843 | Promoter | 10 | 79736123 | 79746573 | 10451 | 2 | ENSMUST00 | 0 | Polrmt | ENSMUSG00000020329.7 |
| chr11 | 26592904 | 26594047 | 25.10597 | 76.67836 | 73.5843 | Promoter | 11 | 26471404 | 26593920 | 122517 | 2 | ENSMUST00 | 0 | Vrk2 | ENSMUSG00000064090.9 |
| chr13 | 49146617 | 49148299 | 25.10597 | 76.67836 | 73.5843 | Promoter | 13 | 49036303 | 49148012 | 111710 | 2 | ENSMUST00 | 0 | Wnk2 | ENSMUSG00000037989.10 |
| chr2 | 103969199 | 103971275 | 25.10597 | 76.67836 | 73.5843 | Promoter | 2 | 103969528 | 103981092 | 11565 | 1 | ENSMUST00 | 0 | Lmo2 | ENSMUSG00000032698.10 |
| chr3 | 75557062 | 75557823 | 25.10597 | 76.67836 | 73.5843 | Promoter | 3 | 75557547 | 75616709 | 59163 | 1 | ENSMUST00 | 0 | Serpini1 | ENSMUSG00000027834.10 |
| chr5 | 130254286 | 130255526 | 25.10597 | 76.67836 | 73.5843 | Promoter | 5 | 130246405 | 130255405 | 9001 | 2 | ENSMUST00 | 0 | Sbds | ENSMUSG00000025337.5 |
| chr5 | 100428474 | 100430552 | 25.10597 | 76.67836 | 73.5843 | Promoter | 5 | 100420842 | 100429513 | 8672 | 2 | ENSMUST00 | 0 | 5430416N02 | ENSMUSG00000097772.2 |
| chr9 | 108338609 | 108340230 | 25.10597 | 76.67836 | 73.5843 | Promoter | 9 | 108339080 | 108340338 | 1259 | 1 | ENSMUST00 | 0 | Gpx1 | ENSMUSG00000063856.6 |
| chr11 | 116273124 | 116274837 | 26.68262 | 76.66692 | 73.57355 | Promoter | 11 | 116245166 | 116274135 | 28970 | 2 | ENSMUST00 | 0 | Srp68 | ENSMUSG00000020780.10 |
| chr15 | 55556621 | 55558207 | 25.59381 | 76.61724 | 73.52408 | Promoter | 15 | 55557408 | 55626423 | 69016 | 1 | ENSMUST00 | 0 | Mtbp | ENSMUSG00000022369.8 |

| | | | | | | | | | | | | | | | |
|---|---|---|---|---|---|---|---|---|---|---|---|---|---|---|---|
| chr4 | 116507049 | 116508324 | 26.11796 | 76.61063 | 73.51752 | Promoter | 4 | 116507549 | 116538235 | 30687 | 1 | ENSMUST00 | 0 | Ipp | ENSMUSG00000028696.7 |
| chr11 | 97150144 | 97152028 | 26.62845 | 76.57359 | 73.48064 | Promoter | 11 | 97137099 | 97151036 | 13938 | 2 | ENSMUST00 | 0 | Tbkbp1 | ENSMUSG00000038517.10 |
| chr9 | 59352804 | 59353875 | 26.06691 | 76.52203 | 73.42928 | Promoter | 9 | 59321966 | 59353508 | 31543 | 2 | ENSMUST00 | 0 | Bbs4 | ENSMUSG00000025235.7 |
| chr4 | 147131779 | 147132874 | 13.10216 | 76.5132 | 73.42055 | Promoter | 4 | 147132038 | 147180579 | 48542 | 1 | ENSMUST00 | 0 | Gm13139 | ENSMUSG00000067916.4 |
| chr1 | 172205664 | 172207142 | 27.11239 | 76.48743 | 73.39479 | Promoter | 1 | 172196728 | 172206804 | 10077 | 2 | ENSMUST00 | 0 | Pea15a | ENSMUSG00000013698.7 |
| chr11 | 76622106 | 76624259 | 27.11239 | 76.48743 | 73.39479 | Promoter | 11 | 76479057 | 76622314 | 143258 | 2 | ENSMUST00 | 0 | Abr | ENSMUSG00000017631.13 |
| chr12 | 4132822 | 4134093 | 27.11239 | 76.48743 | 73.39479 | Promoter | 12 | 4133103 | 4213525 | 80423 | 1 | ENSMUST00 | 0 | Adcy3 | ENSMUSG00000020654.10 |
| chr13 | 112987057 | 112988572 | 27.11239 | 76.48743 | 73.39479 | Promoter | 13 | 112987802 | 112990778 | 2977 | 1 | ENSMUST00 | 0 | Ccno | ENSMUSG00000042417.4 |
| chr15 | 97830319 | 97832239 | 27.11239 | 76.48743 | 73.39479 | Promoter | 15 | 97792664 | 97831494 | 38831 | 2 | ENSMUST00 | 0 | Hdac7 | ENSMUSG00000022475.13 |
| chr7 | 25659104 | 25659885 | 27.11239 | 76.48743 | 73.39479 | Promoter | 7 | 25659153 | 25668031 | 8879 | 1 | ENSMUST00 | 0 | Exosc5 | ENSMUSG00000061286.6 |
| chr8 | 84663906 | 84665749 | 27.11239 | 76.48743 | 73.39479 | Promoter | 8 | 84665128 | 84669406 | 4279 | 1 | ENSMUST00 | 0 | Gm26664 | ENSMUSG00000097193.1 |
| chr9 | 106202779 | 106203827 | 27.11239 | 76.48743 | 73.39479 | Promoter | 9 | 106203108 | 106215389 | 12282 | 1 | ENSMUST00 | 0 | Twf2 | ENSMUSG00000023277.6 |
| chr9 | 20868419 | 20869407 | 27.11239 | 76.48743 | 73.39479 | Promoter | 9 | 20868642 | 20874307 | 5666 | 1 | ENSMUST00 | 0 | A230050P20 | ENSMUSG00000038884.9 |
| chr13 | 63271624 | 63282991 | 26.57451 | 76.48061 | 73.3894 | Promoter | 13 | 63282142 | 63326096 | 43955 | 1 | ENSMUST00 | 0 | 2010111I01R | ENSMUSG00000021458.11 |
| chr15 | 102331366 | 102332565 | 26.57451 | 76.48061 | 73.3894 | Promoter | 15 | 102331709 | 102338139 | 6431 | 1 | ENSMUST00 | 0 | Myg1 | ENSMUSG00000001285.7 |
| chr16 | 3871717 | 3873084 | 26.57451 | 76.48061 | 73.3894 | Promoter | 16 | 3872375 | 3900811 | 28437 | 1 | ENSMUST00 | 0 | Naa60 | ENSMUSG00000005982.8 |
| chr17 | 25016032 | 25017159 | 27.06363 | 76.40382 | 73.31287 | Promoter | 17 | 25016091 | 25092801 | 76711 | 1 | ENSMUST00 | 0 | Ift140 | ENSMUSG00000024169.10 |
| chr4 | 21930870 | 21932043 | 27.06363 | 76.40382 | 73.31287 | Promoter | 4 | 21931329 | 21994146 | 62818 | 1 | ENSMUST00 | 0 | Faxc | ENSMUSG00000028246.8 |
| chr10 | 125388733 | 125389820 | 22.93729 | 76.38734 | 73.29665 | Distal Interge | 10 | 125226464 | 125328963 | 102500 | 2 | ENSMUST00 | -59771 | Slc16a7 | ENSMUSG00000020102.9 |
| chr6 | 47650747 | 47651674 | 21.50605 | 76.3248 | 73.2343 | Distal Interge | 6 | 47637143 | 47637219 | 77 | 1 | ENSMUST00 | 13605 | Gm25283 | ENSMUSG00000087897.1 |
| chr2 | 84648190 | 84651309 | 27.00697 | 76.30663 | 73.21615 | Promoter | 2 | 84600071 | 84650745 | 50675 | 2 | ENSMUST00 | 0 | Ctnnd1 | ENSMUSG00000034101.9 |
| chr10 | 85927711 | 85929091 | 20.60195 | 76.26862 | 73.17842 | Promoter | 10 | 85928774 | 85946206 | 17433 | 1 | ENSMUST00 | 0 | Ascl4 | ENSMUSG00000085111.3 |
| chr11 | 116029765 | 116031490 | 20.60195 | 76.26862 | 73.17842 | Promoter | 11 | 116030322 | 116061214 | 30893 | 1 | ENSMUST00 | 0 | Unk | ENSMUSG00000020770.8 |
| chr12 | 104864253 | 104865506 | 20.60195 | 76.26862 | 73.17842 | Promoter | 12 | 104763114 | 104865076 | 101963 | 2 | ENSMUST00 | 0 | Clmn | ENSMUSG00000021097.9 |
| chr13 | 55210377 | 55213952 | 20.60195 | 76.26862 | 73.17842 | Promoter | 13 | 55209782 | 55318325 | 108544 | 1 | ENSMUST00 | 596 | Nsd1 | ENSMUSG00000021488.7 |
| chr13 | 51644323 | 51646300 | 20.60195 | 76.26862 | 73.17842 | Promoter | 13 | 51645232 | 51650660 | 5429 | 1 | ENSMUST00 | 0 | Cks2 | ENSMUSG00000062248.5 |
| chr16 | 14162717 | 14164701 | 20.60195 | 76.26862 | 73.17842 | Promoter | 16 | 14163275 | 14192928 | 29654 | 1 | ENSMUST00 | 0 | Nde1 | ENSMUSG00000022678.11 |
| chr19 | 10203466 | 10204864 | 20.60195 | 76.26862 | 73.17842 | Promoter | 19 | 10199132 | 10204169 | 5038 | 2 | ENSMUST00 | 0 | Fen1 | ENSMUSG00000024742.10 |
| chr19 | 7340227 | 7342485 | 20.60195 | 76.26862 | 73.17842 | Promoter | 19 | 7275396 | 7341819 | 66424 | 2 | ENSMUST00 | 0 | Mark2 | ENSMUSG00000024969.10 |
| chr7 | 127769280 | 127770956 | 20.60195 | 76.26862 | 73.17842 | Promoter | 7 | 127769815 | 127774692 | 4878 | 1 | ENSMUST00 | 0 | Orai3 | ENSMUSG00000043964.9 |
| chr8 | 84237227 | 84237958 | 20.60195 | 76.26862 | 73.17842 | Promoter | 8 | 84210942 | 84237042 | 26101 | 2 | ENSMUST00 | -186 | Zswim4 | ENSMUSG00000035671.5 |
| chr11 | 3963302 | 3964820 | 26.95054 | 76.20985 | 73.12051 | Promoter | 11 | 3963975 | 3980004 | 16030 | 1 | ENSMUST00 | 0 | Pes1 | ENSMUSG00000020430.7 |
| chr2 | 18046379 | 18049120 | 26.95054 | 76.20985 | 73.12051 | Promoter | 2 | 18044083 | 18049051 | 4969 | 2 | ENSMUST00 | 0 | Skida1 | ENSMUSG00000054074.9 |
| chr16 | 34920992 | 34925857 | 23.32542 | 76.20397 | 73.11496 | Exon (ENSMU | 16 | 34935854 | 34939329 | 3476 | 2 | ENSMUST00 | 13472 | RP23-80O10. | ENSMUSG00000100593.1 |
| chr17 | 8800574 | 8803185 | 23.32542 | 76.20397 | 73.11496 | Promoter | 17 | 8801693 | 8899189 | 97497 | 1 | ENSMUST00 | 0 | Pde10a | ENSMUSG00000023868.10 |
| chr10 | 3365843 | 3367564 | 25.8647 | 76.17062 | 73.08194 | Promoter | 10 | 3366150 | 3464975 | 98826 | 1 | ENSMUST00 | 0 | Ppp1r14c | ENSMUSG00000040653.5 |
| chr7 | 16221325 | 16222244 | 25.8647 | 76.17062 | 73.08194 | Promoter | 7 | 16197147 | 16222037 | 24891 | 2 | ENSMUST00 | 0 | Dhx34 | ENSMUSG00000006019.10 |
| chr6 | 83831033 | 83831815 | 26.3609 | 76.11214 | 73.02366 | Promoter | 6 | 83805401 | 83831741 | 26341 | 2 | ENSMUST00 | 0 | Paip2b | ENSMUSG00000045896.9 |
| chr1 | 191062243 | 191063529 | 22.78592 | 76.10319 | 73.01487 | Promoter | 1 | 191045826 | 191062932 | 17107 | 2 | ENSMUST00 | 0 | Tatdn3 | ENSMUSG00000026632.12 |
| chr1 | 39649894 | 39651348 | 22.78592 | 76.10319 | 73.01487 | Promoter | 1 | 39618406 | 39651182 | 32777 | 2 | ENSMUST00 | 0 | Creg2 | ENSMUSG00000050967.4 |
| chr11 | 121672549 | 121673918 | 22.78592 | 76.10319 | 73.01487 | Promoter | 11 | 121616197 | 121673153 | 56957 | 2 | ENSMUST00 | 0 | B3gntl1 | ENSMUSG00000046605.9 |
| chr11 | 80476339 | 80478051 | 22.78592 | 76.10319 | 73.01487 | Promoter | 11 | 80477023 | 80481184 | 4162 | 1 | ENSMUST00 | 0 | Cdk5r1 | ENSMUSG00000048895.13 |
| chr12 | 25095179 | 25096936 | 22.78592 | 76.10319 | 73.01487 | Promoter | 12 | 25093801 | 25096092 | 2292 | 2 | ENSMUST00 | 0 | Id2 | ENSMUSG00000020644.8 |
| chr12 | 72069853 | 72071359 | 22.78592 | 76.10319 | 73.01487 | Promoter | 12 | 72069618 | 72070991 | 1374 | 2 | ENSMUST00 | 0 | Gpr135 | ENSMUSG00000043398.4 |
| chr15 | 102518102 | 102519426 | 22.78592 | 76.10319 | 73.01487 | Promoter | 15 | 102518192 | 102523674 | 5483 | 1 | ENSMUST00 | 0 | Tarbp2 | ENSMUSG00000023051.5 |
| chr15 | 102470489 | 102471870 | 22.78592 | 76.10319 | 73.01487 | Promoter | 15 | 102470632 | 102500059 | 29428 | 1 | ENSMUST00 | 0 | Pcbp2 | ENSMUSG00000056851.8 |
| chr17 | 26938724 | 26939594 | 22.78592 | 76.10319 | 73.01487 | Promoter | 17 | 26937973 | 26939468 | 1496 | 2 | ENSMUST00 | 0 | Cuta | ENSMUSG00000024194.10 |
| chr17 | 35915629 | 35916728 | 22.78592 | 76.10319 | 73.01487 | Promoter | 17 | 35910379 | 35916389 | 6011 | 2 | ENSMUST00 | 0 | Mrps18b | ENSMUSG00000024436.11 |
| chr18 | 84851321 | 84852574 | 22.78592 | 76.10319 | 73.01487 | Promoter | 18 | 84851338 | 84880401 | 29064 | 1 | ENSMUST00 | 0 | Cyb5a | ENSMUSG00000024646.8 |
| chr19 | 53943307 | 53944593 | 22.78592 | 76.10319 | 73.01487 | Promoter | 19 | 53944306 | 54033268 | 88963 | 1 | ENSMUST00 | 0 | Shoc2 | ENSMUSG00000024976.8 |
| chr2 | 181863775 | 181865150 | 22.78592 | 76.10319 | 73.01487 | Promoter | 2 | 181864337 | 181870830 | 6494 | 1 | ENSMUST00 | 0 | Polr3k | ENSMUSG00000038628.8 |
| chr2 | 175206326 | 175207000 | 22.78592 | 76.10319 | 73.01487 | Promoter | 2 | 175191065 | 175206903 | 15839 | 2 | ENSMUST00 | 0 | Gm14391 | ENSMUSG00000078903.5 |
| chr3 | 127789334 | 127791477 | 22.78592 | 76.10319 | 73.01487 | Promoter | 3 | 127789872 | 127798392 | 8521 | 1 | ENSMUST00 | 0 | Tifa | ENSMUSG00000046688.9 |
| chr3 | 14611402 | 14611874 | 22.78592 | 76.10319 | 73.01487 | Promoter | 3 | 14606284 | 14611285 | 5002 | 2 | ENSMUST00 | -118 | 1810022K09 | ENSMUSG00000078784.3 |
| chr5 | 24423040 | 24423662 | 22.78592 | 76.10319 | 73.01487 | Promoter | 5 | 24418250 | 24423530 | 5281 | 2 | ENSMUST00 | 0 | Cdk5 | ENSMUSG00000028969.6 |
| chr6 | 131387180 | 131389119 | 22.78592 | 76.10319 | 73.01487 | Promoter | 6 | 131364858 | 131388450 | 23593 | 2 | ENSMUST00 | 0 | Ybx3 | ENSMUSG00000030189.10 |
| chr6 | 71822785 | 71824077 | 22.78592 | 76.10319 | 73.01487 | Promoter | 6 | 71813003 | 71823810 | 10808 | 2 | ENSMUST00 | 0 | Mrpl35 | ENSMUSG00000052962.2 |
| chr7 | 79934570 | 79936261 | 22.78592 | 76.10319 | 73.01487 | Promoter | 7 | 79925361 | 79935359 | 9999 | 2 | ENSMUST00 | 0 | 2610034B18 | ENSMUSG00000039043.5 |
| chr7 | 28435601 | 28436397 | 22.78592 | 76.10319 | 73.01487 | Intron (ENSM | 7 | 28437447 | 28446546 | 9100 | 1 | ENSMUST00 | -1050 | Gmfg | ENSMUSG00000060791.10 |
| chr8 | 122610865 | 122612393 | 22.78592 | 76.10319 | 73.01487 | Promoter | 8 | 122611626 | 122615588 | 3963 | 1 | ENSMUST00 | 0 | Trappc2l | ENSMUSG00000015013.8 |
| chr8 | 70593499 | 70595422 | 22.78592 | 76.10319 | 73.01487 | Promoter | 8 | 70594481 | 70597288 | 2808 | 1 | ENSMUST00 | 0 | Isyna1 | ENSMUSG00000019139.9 |
| chr9 | 107541192 | 107542682 | 22.78592 | 76.10319 | 73.01487 | Promoter | 9 | 107542209 | 107545704 | 3496 | 1 | ENSMUST00 | 0 | Nprl2 | ENSMUSG00000010057.3 |
| chr11 | 59963913 | 59965603 | 25.81463 | 76.08353 | 72.99751 | Promoter | 11 | 59963181 | 59964944 | 1764 | 2 | ENSMUST00 | 0 | Rasd1 | ENSMUSG00000049892.7 |
| chr11 | 119227871 | 119229195 | 23.73948 | 76.05544 | 72.96951 | Promoter | 11 | 119228572 | 119265212 | 36641 | 1 | ENSMUST00 | 0 | Ccdc40 | ENSMUSG00000039963.13 |
| chr4 | 134000237 | 134001349 | 23.73948 | 76.05544 | 72.96951 | Promoter | 4 | 133969028 | 134000890 | 31863 | 2 | ENSMUST00 | 0 | Dhdds | ENSMUSG00000012117.8 |
| chr7 | 49245452 | 49248461 | 23.73948 | 76.05544 | 72.96951 | Promoter | 7 | 49246189 | 49409643 | 163455 | 1 | ENSMUST00 | 0 | Nav2 | ENSMUSG00000052512.10 |
| chr1 | 34472564 | 34473417 | 26.30804 | 76.02086 | 72.93523 | Promoter | 1 | 34472736 | 34475733 | 2998 | 1 | ENSMUST00 | 0 | Ptpn18 | ENSMUSG00000026126.10 |
| chr11 | 53324333 | 53325373 | 26.30804 | 76.02086 | 72.93523 | Promoter | 11 | 53324679 | 53333301 | 8623 | 1 | ENSMUST00 | 0 | Zcchc10 | ENSMUSG00000018239.4 |
| chr13 | 62382716 | 62383700 | 26.30804 | 76.02086 | 72.93523 | Promoter | 13 | 62368328 | 62383166 | 14839 | 2 | ENSMUST00 | 0 | Gm3604 | ENSMUSG00000094942.2 |
| chr3 | 87919203 | 87920400 | 26.30804 | 76.02086 | 72.93523 | Promoter | 3 | 87919506 | 87923662 | 4157 | 1 | ENSMUST00 | 0 | Mrpl24 | ENSMUSG00000019710.8 |
| chr6 | 124829087 | 124830563 | 26.30804 | 76.02086 | 72.93523 | Promoter | 6 | 124829547 | 124832664 | 3118 | 1 | ENSMUST00 | 0 | Cdca3 | ENSMUSG00000023505.8 |
| chr16 | 22439906 | 22440859 | 24.18217 | 75.9461 | 72.86143 | Promoter | 16 | 22381309 | 22439719 | 58411 | 2 | ENSMUST00 | -188 | Etv5 | ENSMUSG00000013089.10 |
| chr8 | 53511396 | 53512355 | 24.18217 | 75.9461 | 72.86143 | Promoter | 8 | 53511702 | 53523421 | 11720 | 1 | ENSMUST00 | 0 | Aga | ENSMUSG00000031521.4 |
| chr11 | 61021904 | 61023783 | 19.61815 | 75.94017 | 72.85583 | Promoter | 11 | 61022564 | 61071131 | 48568 | 1 | ENSMUST00 | 0 | Kcnj12 | ENSMUSG00000042529.9 |
| chr14 | 70191374 | 70192726 | 19.61815 | 75.94017 | 72.85583 | Exon (ENSMU | 14 | 70205324 | 70217659 | 12336 | 1 | ENSMUST00 | -12598 | Gm26908 | ENSMUSG00000097243.1 |
| chr15 | 83349550 | 83350723 | 19.61815 | 75.94017 | 72.85583 | Promoter | 15 | 83299740 | 83350247 | 50508 | 2 | ENSMUST00 | 0 | Arfgap3 | ENSMUSG00000054277.7 |
| chr2 | 24918542 | 24920152 | 19.61815 | 75.94017 | 72.85583 | Promoter | 2 | 24790769 | 24919609 | 128841 | 2 | ENSMUST00 | 0 | Ehmt1 | ENSMUSG00000036893.11 |
| chr7 | 35317509 | 35318641 | 19.61815 | 75.94017 | 72.85583 | Promoter | 7 | 35276536 | 35318440 | 41905 | 2 | ENSMUST00 | 0 | Gpatch1 | ENSMUSG00000063808.9 |
| chr9 | 21091000 | 21092579 | 19.61815 | 75.94017 | 72.85583 | Promoter | 9 | 21074168 | 21091988 | 17821 | 2 | ENSMUST00 | 0 | Raver1 | ENSMUSG00000010205.10 |
| chr17 | 46253492 | 46254618 | 26.78267 | 75.92174 | 72.83789 | Promoter | 17 | 46254165 | 46257316 | 3152 | 1 | ENSMUST00 | 0 | Lrrc73 | ENSMUSG00000071073.4 |
| chr9 | 40455021 | 40456468 | 26.78267 | 75.92174 | 72.83789 | Promoter | 9 | 40293245 | 40455670 | 162426 | 2 | ENSMUST00 | 0 | Gramd1b | ENSMUSG00000040111.9 |
| chr1 | 37865298 | 37866022 | 25.71508 | 75.91022 | 72.8265 | Promoter | 1 | 37865299 | 37871388 | 6090 | 1 | ENSMUST00 | 0 | Gm26805 | ENSMUSG00000097268.1 |
| chr4 | 41740373 | 41741683 | 25.71508 | 75.91022 | 72.8265 | Promoter | 4 | 41738493 | 41741359 | 2867 | 2 | ENSMUST00 | 0 | Sigmar1 | ENSMUSG00000036078.11 |
| chr5 | 140417875 | 140419085 | 25.71508 | 75.91022 | 72.8265 | Promoter | 5 | 140419305 | 140443358 | 24054 | 1 | ENSMUST00 | -220 | Eif3b | ENSMUSG00000056076.9 |
| chr5 | 106574021 | 106575115 | 25.71508 | 75.91022 | 72.8265 | Promoter | 5 | 106500894 | 106574715 | 73822 | 2 | ENSMUST00 | 0 | Gm28050 | ENSMUSG00000099061.1 |
| chr10 | 78574095 | 78575626 | 24.65656 | 75.88111 | 72.79811 | Promoter | 10 | 78574500 | 78584498 | 9999 | 1 | ENSMUST00 | 0 | Ilvbl | ENSMUSG00000032763.10 |
| chr2 | 109280275 | 109281543 | 24.65656 | 75.88111 | 72.79811 | Promoter | 2 | 109280738 | 109341747 | 61010 | 1 | ENSMUST00 | 0 | Kif18a | ENSMUSG00000027115.9 |
| chr6 | 113326034 | 113326727 | 24.65656 | 75.88111 | 72.79811 | Promoter | 6 | 113308666 | 113326747 | 18082 | 2 | ENSMUST00 | 20 | Gm15492 | ENSMUSG00000085012.1 |
| chr10 | 81266206 | 81267114 | 25.16617 | 75.86667 | 72.78426 | Promoter | 10 | 81264713 | 81266934 | 2222 | 2 | ENSMUST00 | 0 | Mrpl54 | ENSMUSG00000034932.4 |
| chr15 | 75929091 | 75930247 | 25.16617 | 75.86667 | 72.78426 | Promoter | 15 | 75924684 | 75929730 | 5047 | 2 | ENSMUST00 | 0 | Tsta3 | ENSMUSG00000022570.5 |
| chr3 | 95624921 | 95625958 | 25.16617 | 75.86667 | 72.78426 | Promoter | 3 | 95624980 | 95630493 | 5514 | 1 | ENSMUST00 | 0 | Ensa | ENSMUSG00000038619.7 |
| chr4 | 134496595 | 134497573 | 25.16617 | 75.86667 | 72.78426 | Promoter | 4 | 134496697 | 134508813 | 12117 | 1 | ENSMUST00 | 0 | Paqr7 | ENSMUSG00000037348.10 |
| chr5 | 64092528 | 64093954 | 25.16617 | 75.86667 | 72.78426 | Promoter | 5 | 64092936 | 64128355 | 35420 | 1 | ENSMUST00 | 0 | Pgm1 | ENSMUSG00000029171.8 |
| chr11 | 33161949 | 33163764 | 18.76905 | 75.7838 | 72.70259 | Promoter | 11 | 33152287 | 33163206 | 10920 | 2 | ENSMUST00 | 0 | Npm1 | ENSMUSG00000057113.8 |
| chr17 | 26243366 | 26244679 | 18.76905 | 75.7838 | 72.70259 | Promoter | 17 | 26211822 | 26244223 | 32402 | 2 | ENSMUST00 | 0 | Itfg3 | ENSMUSG00000024187.9 |
| chr17 | 46646171 | 46646812 | 18.02876 | 75.76575 | 72.68469 | Promoter | 17 | 46646229 | 46650132 | 3904 | 1 | ENSMUST00 | 0 | Mrpl2 | ENSMUSG00000002767.8 |
| chr5 | 5663481 | 5664453 | 26.1507 | 75.74902 | 72.66802 | Promoter | 5 | 5579278 | 5664239 | 84962 | 2 | ENSMUST00 | 0 | A330021E22 | ENSMUSG00000040473.10 |
| chr5 | 108433548 | 108434787 | 26.1507 | 75.74902 | 72.66802 | Promoter | 5 | 108433244 | 108434448 | 1205 | 2 | ENSMUST00 | 0 | Atp5k | ENSMUSG00000050856.11 |
| chr1 | 152623682 | 152625635 | 24.03115 | 75.67385 | 72.59326 | Promoter | 1 | 152516951 | 152625217 | 108267 | 2 | ENSMUST00 | 0 | Rgl1 | ENSMUSG00000026482.8 |
| chr11 | 87403994 | 87405030 | 24.03115 | 75.67385 | 72.59326 | Promoter | 11 | 87376645 | 87404523 | 27879 | 2 | ENSMUST00 | 0 | Rad51c | ENSMUSG00000007646.8 |
| chr11 | 51998950 | 52000765 | 24.03115 | 75.67385 | 72.59326 | Promoter | 11 | 51985497 | 52000484 | 14988 | 2 | ENSMUST00 | 0 | Ube2b | ENSMUSG00000020390.7 |
| chr11 | 96777059 | 96777646 | 24.03115 | 75.67385 | 72.59326 | Promoter | 11 | 96767555 | 96777531 | 9977 | 2 | ENSMUST00 | 0 | Snx11 | ENSMUSG00000020876.9 |
| chr11 | 79961481 | 79963264 | 24.03115 | 75.67385 | 72.59326 | Promoter | 11 | 79932321 | 79962379 | 30059 | 2 | ENSMUST00 | 0 | Utp6 | ENSMUSG00000035575.6 |
| chr12 | 98627633 | 98628884 | 24.03115 | 75.67385 | 72.59326 | Promoter | 12 | 98628157 | 98669815 | 41659 | 1 | ENSMUST00 | 0 | Spata7 | ENSMUSG00000021007.8 |
| chr12 | 85472966 | 85475150 | 24.03115 | 75.67385 | 72.59326 | Promoter | 12 | 85473890 | 85477273 | 3384 | 1 | ENSMUST00 | 0 | Fos | ENSMUSG00000021250.8 |
| chr13 | 67682782 | 67684085 | 24.03115 | 75.67385 | 72.59326 | Promoter | 13 | 67667437 | 67683467 | 16031 | 2 | ENSMUST00 | 0 | Zfp738 | ENSMUSG00000048280.12 |
| chr14 | 57103551 | 57105089 | 24.03115 | 75.67385 | 72.59326 | Promoter | 14 | 57098600 | 57104702 | 6103 | 2 | ENSMUST00 | 0 | Gjb2 | ENSMUSG00000046352.7 |
| chr15 | 12738458 | 12740528 | 24.03115 | 75.67385 | 72.59326 | Promoter | 15 | 12443331 | 12739924 | 296594 | 2 | ENSMUST00 | 0 | Pdzd2 | ENSMUSG00000022197.10 |
| chr15 | 93275405 | 93276823 | 24.03115 | 75.67385 | 72.59326 | Promoter | 15 | 93239742 | 93275161 | 35420 | 2 | ENSMUST00 | -245 | Gxylt1 | ENSMUSG00000036197.10 |
| chr15 | 75745907 | 75748880 | 24.03115 | 75.67385 | 72.59326 | Promoter | 15 | 75746843 | 75747922 | 1080 | 2 | ENSMUST00 | 0 | Mafa | ENSMUSG00000047591.4 |
| chr16 | 37653045 | 37654797 | 24.03115 | 75.67385 | 72.59326 | Promoter | 16 | 37647170 | 37654453 | 7284 | 2 | ENSMUST00 | 0 | Ndufb4 | ENSMUSG00000022820.3 |
| chr18 | 70471561 | 70473046 | 24.03115 | 75.67385 | 72.59326 | Promoter | 18 | 70453140 | 70472480 | 19341 | 2 | ENSMUST00 | 0 | 4930503L19 | ENSMUSG00000044906.4 |
| chr18 | 35873085 | 35874233 | 24.03115 | 75.67385 | 72.59326 | Distal Interge | 18 | 35830389 | 35831093 | 705 | 2 | ENSMUST00 | -41993 | RP24-326K1 | ENSMUSG00000100320.1 |
| chr2 | 131178711 | 131180611 | 24.03115 | 75.67385 | 72.59326 | Promoter | 2 | 131175182 | 131180067 | 4886 | 2 | ENSMUST00 | 0 | Cenpb | ENSMUSG00000068267.5 |
| chr2 | 130663878 | 130665005 | 24.03115 | 75.67385 | 72.59326 | Promoter | 2 | 130653960 | 130664659 | 10700 | 2 | ENSMUST00 | 0 | Ddrgk1 | ENSMUSG00000068290.6 |
| chr2 | 26639227 | 26640357 | 24.03115 | 75.67385 | 72.59326 | Promoter | 2 | 26639189 | 26639326 | 138 | 2 | ENSMUST00 | 0 | Snora17 | ENSMUSG00000077192.2 |
| chr3 | 104780720 | 104782171 | 24.03115 | 75.67385 | 72.59326 | Promoter | 3 | 104781056 | 104786018 | 4963 | 1 | ENSMUST00 | 0 | Ppm1j | ENSMUSG00000002228.6 |
| chr3 | 117868381 | 117869632 | 24.03115 | 75.67385 | 72.59326 | Promoter | 3 | 117781497 | 117868936 | 87440 | 2 | ENSMUST00 | 0 | Snx7 | ENSMUSG00000028007.8 |
| chr4 | 48584659 | 48586353 | 24.03115 | 75.67385 | 72.59326 | Promoter | 4 | 48585174 | 48663131 | 77958 | 1 | ENSMUST00 | 0 | Tmeff1 | ENSMUSG00000028347.9 |
| chr4 | 133038350 | 133039857 | 24.03115 | 75.67385 | 72.59326 | Promoter | 4 | 133039493 | 133077242 | 37750 | 1 | ENSMUST00 | 0 | Ahdc1 | ENSMUSG00000037692.9 |
| chr4 | 8646552 | 8648739 | 24.03115 | 75.67385 | 72.59326 | Distal Interge | 4 | 8672633 | 8673582 | 950 | 2 | ENSMUST00 | 24843 | Gm11809 | ENSMUSG00000082122.1 |
| chr5 | 18358905 | 18360944 | 24.03115 | 75.67385 | 72.59326 | Promoter | 5 | 18265135 | 18360413 | 95279 | 2 | ENSMUST00 | 0 | Gnai1 | ENSMUSG00000057614.5 |
| chr6 | 100833184 | 100834843 | 24.03115 | 75.67385 | 72.59326 | Promoter | 6 | 100833638 | 100868717 | 35080 | 1 | ENSMUST00 | 0 | Ppp4r2 | ENSMUSG00000052144.6 |
| chr8 | 90876446 | 90877999 | 24.03115 | 75.67385 | 72.59326 | Intron (ENSM | 8 | 90907939 | 90908415 | 477 | 2 | ENSMUST00 | 30416 | Gm6658 | ENSMUSG00000074171.1 |
| chr9 | 32695428 | 32698295 | 24.03115 | 75.67385 | 72.59326 | Promoter | 9 | 32696022 | 32757820 | 61799 | 1 | ENSMUST00 | 0 | Ets1 | ENSMUSG00000032035.10 |
| chr9 | 21509504 | 21511024 | 24.03115 | 75.67385 | 72.59326 | Promoter | 9 | 21507380 | 21510186 | 2807 | 2 | ENSMUST00 | 0 | Tmed1 | ENSMUSG00000032180.10 |
| chr11 | 20542079 | 20542945 | 25.56719 | 75.65247 | 72.57522 | Promoter | 11 | 20543253 | 20597020 | 53768 | 1 | ENSMUST00 | -308 | Sertad2 | ENSMUSG00000049800.8 |
| chr1 | 92831029 | 92833403 | 25.56571 | 75.64988 | 72.57267 | Promoter | 1 | 92831645 | 92860779 | 29135 | 1 | ENSMUST00 | 0 | Gpc1 | ENSMUSG00000034220.7 |

| | | | | | | | | | | | | | | | |
|---|---|---|---|---|---|---|---|---|---|---|---|---|---|---|---|
| chr1 | 79670988 | 79672518 | 25.56571 | 75.64988 | 72.57267 | Promoter | 1 | 79606876 | 79671972 | 65097 | 2 | ENSMUST00 | 0 | Ap1s3 | ENSMUSG00000054702.9 |
| chr10 | 127621282 | 127622474 | 25.56571 | 75.64988 | 72.57267 | Promoter | 10 | 127587627 | 127621148 | 33522 | 2 | ENSMUST00 | -135 | Lrp1 | ENSMUSG00000040249.10 |
| chr11 | 70014694 | 70015713 | 25.56571 | 75.64988 | 72.57267 | Promoter | 11 | 70010183 | 70015408 | 5226 | 2 | ENSMUST00 | 0 | Acadvl | ENSMUSG00000018574.9 |
| chr11 | 51289361 | 51290454 | 25.56571 | 75.64988 | 72.57267 | Promoter | 11 | 51289920 | 51583918 | 293999 | 1 | ENSMUST00 | 0 | Col23a1 | ENSMUSG00000063564.8 |
| chr11 | 48854958 | 48857221 | 25.56571 | 75.64988 | 72.57267 | Distal Interge | 11 | 48843648 | 48853493 | 9846 | 2 | ENSMUST00 | -1466 | Gm16170 | ENSMUSG00000086245.1 |
| chr12 | 24974466 | 24975655 | 25.56571 | 75.64988 | 72.57267 | Promoter | 12 | 24974932 | 25059697 | 84766 | 1 | ENSMUST00 | 0 | Kidins220 | ENSMUSG00000036333.10 |
| chr13 | 3537313 | 3537923 | 25.56571 | 75.64988 | 72.57267 | Promoter | 13 | 3538075 | 3566261 | 28187 | 1 | ENSMUST00 | -152 | Gdi2 | ENSMUSG00000021218.9 |
| chr13 | 89738844 | 89741663 | 25.56571 | 75.64988 | 72.57267 | Promoter | 13 | 89725434 | 89740397 | 14964 | 2 | ENSMUST00 | 0 | Vcan | ENSMUSG00000021614.11 |
| chr16 | 17576106 | 17576965 | 25.56571 | 75.64988 | 72.57267 | Promoter | 16 | 17572018 | 17576721 | 4704 | 2 | ENSMUST00 | 0 | Slc7a4 | ENSMUSG00000022756.10 |
| chr17 | 31295863 | 31297198 | 25.56571 | 75.64988 | 72.57267 | Promoter | 17 | 31296219 | 31350696 | 54478 | 1 | ENSMUST00 | 0 | Slc37a1 | ENSMUSG00000024036.10 |
| chr18 | 67609568 | 67611350 | 25.56571 | 75.64988 | 72.57267 | Distal Interge | 18 | 67617397 | 67641336 | 23940 | 2 | ENSMUST00 | 29986 | Cep76 | ENSMUSG00000073542.2 |
| chr19 | 6076399 | 6077750 | 25.56571 | 75.64988 | 72.57267 | Promoter | 19 | 6067994 | 6077197 | 9204 | 2 | ENSMUST00 | 0 | Vps51 | ENSMUSG00000024797.7 |
| chr2 | 28533088 | 28533934 | 25.56571 | 75.64988 | 72.57267 | Promoter | 2 | 28533355 | 28553066 | 19712 | 1 | ENSMUST00 | 0 | Ralgds | ENSMUSG00000026821.10 |
| chr2 | 30807859 | 30808651 | 25.56571 | 75.64988 | 72.57267 | Promoter | 2 | 30807941 | 30823017 | 15077 | 1 | ENSMUST00 | 0 | Ntmt1 | ENSMUSG00000026857.4 |
| chr2 | 10372318 | 10375667 | 25.56571 | 75.64988 | 72.57267 | Promoter | 2 | 10372436 | 10439137 | 66702 | 1 | ENSMUST00 | 0 | Sfmbt2 | ENSMUSG00000061186.10 |
| chr3 | 152191653 | 152193826 | 25.56571 | 75.64988 | 72.57267 | Promoter | 3 | 152183955 | 152193803 | 9849 | 2 | ENSMUST00 | 0 | Dnajb4 | ENSMUSG00000028035.8 |
| chr3 | 89177416 | 89178289 | 25.56571 | 75.64988 | 72.57267 | Promoter | 3 | 89177473 | 89182765 | 5293 | 1 | ENSMUST00 | 0 | Scamp3 | ENSMUSG00000028049.10 |
| chr4 | 62524414 | 62525862 | 25.56571 | 75.64988 | 72.57267 | Promoter | 4 | 62522649 | 62525068 | 2420 | 2 | ENSMUST00 | 0 | Pole3 | ENSMUSG00000028394.4 |
| chr4 | 81442015 | 81443104 | 25.56571 | 75.64988 | 72.57267 | Promoter | 4 | 81278500 | 81442805 | 164306 | 2 | ENSMUST00 | 0 | Mpdz | ENSMUSG00000028402.12 |
| chr4 | 118290285 | 118291695 | 25.56571 | 75.64988 | 72.57267 | Promoter | 4 | 118208213 | 118291405 | 83193 | 2 | ENSMUST00 | 0 | Ptprf | ENSMUSG00000033295.8 |
| chr4 | 47056745 | 47057908 | 25.56571 | 75.64988 | 72.57267 | Promoter | 4 | 47015669 | 47057306 | 41638 | 2 | ENSMUST00 | 0 | Anks6 | ENSMUSG00000066191.6 |
| chr5 | 24525602 | 24527538 | 25.56571 | 75.64988 | 72.57267 | Promoter | 5 | 24502757 | 24527202 | 24446 | 2 | ENSMUST00 | 0 | Gbx1 | ENSMUSG00000067724.5 |
| chr6 | 31564130 | 31565675 | 25.56571 | 75.64988 | 72.57267 | Promoter | 6 | 31519488 | 31563981 | 44494 | 2 | ENSMUST00 | -150 | Podxl | ENSMUSG00000025608.9 |
| chr7 | 141122679 | 141123729 | 25.56571 | 75.64988 | 72.57267 | Promoter | 7 | 141122382 | 141154456 | 32075 | 1 | ENSMUST00 | 298 | Ptdss2 | ENSMUSG00000025495.9 |
| chr7 | 56049558 | 56051192 | 25.56571 | 75.64988 | 72.57267 | Promoter | 7 | 56050155 | 56231794 | 181640 | 1 | ENSMUST00 | 0 | Herc2 | ENSMUSG00000030451.10 |
| chr9 | 60712746 | 60713581 | 25.56571 | 75.64988 | 72.57267 | Promoter | 9 | 60712989 | 60738801 | 25813 | 1 | ENSMUST00 | 0 | Larp6 | ENSMUSG00000034839.5 |
| chr19 | 6014421 | 6016110 | 26.04686 | 75.56942 | 72.49508 | Promoter | 19 | 5988546 | 6015247 | 26702 | 2 | ENSMUST00 | 0 | Capn1 | ENSMUSG00000024942.11 |
| chr2 | 101628448 | 101629492 | 25.42099 | 75.39734 | 72.3238 | Promoter | 2 | 101560781 | 101628990 | 68210 | 2 | ENSMUST00 | 0 | B230118H07 | ENSMUSG00000027165.11 |
| chr11 | 80183321 | 80184415 | 23.34944 | 75.34173 | 72.26852 | Promoter | 11 | 80183851 | 80199757 | 15907 | 1 | ENSMUST00 | 0 | Rnf135 | ENSMUSG00000020707.6 |
| chr5 | 65199640 | 65200386 | 23.34944 | 75.34173 | 72.26852 | Promoter | 5 | 65199696 | 65260415 | 60720 | 1 | ENSMUST00 | 0 | Wdr19 | ENSMUSG00000037890.9 |
| chr6 | 145934002 | 145934968 | 23.34944 | 75.34173 | 72.26852 | Promoter | 6 | 145934122 | 145965223 | 31102 | 1 | ENSMUST00 | 0 | Sspn | ENSMUSG00000030255.8 |
| chr7 | 27447900 | 27448417 | 23.34944 | 75.34173 | 72.26852 | Promoter | 7 | 27447978 | 27466144 | 18167 | 1 | ENSMUST00 | 0 | Blvrb | ENSMUSG00000040466.10 |
| chr11 | 87447966 | 87448754 | 21.90749 | 75.32819 | 72.25517 | Promoter | 11 | 87448627 | 87448842 | 216 | 1 | ENSMUST00 | 0 | Rnu3b3 | ENSMUSG00000099291.1 |
| chr9 | 106428781 | 106430242 | 21.90749 | 75.32819 | 72.25517 | Promoter | 9 | 106429454 | 106431568 | 2115 | 1 | ENSMUST00 | 0 | Rpl29 | ENSMUSG00000048758.8 |
| chr1 | 194618995 | 194621050 | 21.44446 | 75.31629 | 72.24329 | Promoter | 1 | 194619829 | 194816869 | 197041 | 1 | ENSMUST00 | 0 | Plxna2 | ENSMUSG00000026640.7 |
| chr11 | 82907659 | 82908659 | 21.44446 | 75.31629 | 72.24329 | Promoter | 11 | 82900768 | 82908411 | 7644 | 2 | ENSMUST00 | 0 | Nle1 | ENSMUSG00000020692.9 |
| chr12 | 4233182 | 4234847 | 21.44446 | 75.31629 | 72.24329 | Promoter | 12 | 4197822 | 4233965 | 36144 | 2 | ENSMUST00 | 0 | Cenpo | ENSMUSG00000020652.12 |
| chr13 | 100775287 | 100776483 | 21.44446 | 75.31629 | 72.24329 | Promoter | 13 | 100759674 | 100775899 | 16226 | 2 | ENSMUST00 | 0 | Cenph | ENSMUSG00000045273.8 |
| chr14 | 11551784 | 11556756 | 21.44446 | 75.31629 | 72.24329 | Promoter | 14 | 11553532 | 12242041 | 688510 | 1 | ENSMUST00 | 0 | Ptprg | ENSMUSG00000021745.8 |
| chr15 | 39196825 | 39199510 | 21.44446 | 75.31629 | 72.24329 | Promoter | 15 | 39198332 | 39681940 | 483609 | 1 | ENSMUST00 | 0 | Rims2 | ENSMUSG00000037386.10 |
| chr17 | 55931660 | 55932584 | 21.44446 | 75.31629 | 72.24329 | Distal Interge | 17 | 55938381 | 55945259 | 6879 | 2 | ENSMUST00 | 12675 | Zfp119b | ENSMUSG00000062101.7 |
| chr18 | 77794248 | 77795438 | 21.44446 | 75.31629 | 72.24329 | Promoter | 18 | 77794545 | 77882007 | 87463 | 1 | ENSMUST00 | 0 | Pstpip2 | ENSMUSG00000025429.8 |
| chr2 | 24974134 | 24976414 | 21.44446 | 75.31629 | 72.24329 | Promoter | 2 | 24976033 | 25054057 | 78025 | 1 | ENSMUST00 | 0 | Pnpla7 | ENSMUSG00000036833.11 |
| chr3 | 58576362 | 58577992 | 21.44446 | 75.31629 | 72.24329 | Promoter | 3 | 58576636 | 58593133 | 16498 | 1 | ENSMUST00 | 0 | Selt | ENSMUSG00000075700.9 |
| chr5 | 137333094 | 137334240 | 21.44446 | 75.31629 | 72.24329 | Promoter | 5 | 137314558 | 137333597 | 19040 | 2 | ENSMUST00 | 0 | Slc12a9 | ENSMUSG00000037344.8 |
| chr11 | 60454154 | 60455260 | 26.39899 | 75.26234 | 72.19086 | Promoter | 11 | 60454591 | 60461853 | 7263 | 1 | ENSMUST00 | 0 | Drg2 | ENSMUSG00000020537.4 |
| chr11 | 106159606 | 106161040 | 26.39899 | 75.26234 | 72.19086 | Promoter | 11 | 106156256 | 106160142 | 3887 | 2 | ENSMUST00 | 0 | Limd2 | ENSMUSG00000040699.8 |
| chr13 | 3633655 | 3634709 | 26.39899 | 75.26234 | 72.19086 | Promoter | 13 | 3634032 | 3651774 | 17743 | 1 | ENSMUST00 | 0 | Asb13 | ENSMUSG00000033781.6 |
| chr19 | 28677815 | 28680438 | 26.39899 | 75.26234 | 72.19086 | Promoter | 19 | 28258851 | 28680077 | 421227 | 2 | ENSMUST00 | 0 | Glis3 | ENSMUSG00000052942.8 |
| chr2 | 130642161 | 130643208 | 26.39899 | 75.26234 | 72.19086 | Promoter | 2 | 130632842 | 130642803 | 9962 | 2 | ENSMUST00 | 0 | Lzts3 | ENSMUSG00000037703.9 |
| chr4 | 152107604 | 152109436 | 25.32445 | 75.22865 | 72.15769 | Promoter | 4 | 152107636 | 152115394 | 7759 | 1 | ENSMUST00 | 0 | Plekhg5 | ENSMUSG00000039713.11 |
| chr16 | 34690361 | 34691460 | 25.84161 | 75.21405 | 72.14314 | Promoter | 16 | 34690616 | 34725194 | 34579 | 1 | ENSMUST00 | 0 | Ccdc14 | ENSMUSG00000022833.6 |
| chr10 | 77110823 | 77114286 | 25.79081 | 75.126 | 72.05579 | Promoter | 10 | 77052181 | 77113676 | 61496 | 2 | ENSMUST00 | 0 | Col18a1 | ENSMUSG00000001435.9 |
| chr3 | 88409469 | 88410900 | 25.79081 | 75.126 | 72.05579 | Promoter | 3 | 88394143 | 88410316 | 16174 | 2 | ENSMUST00 | 0 | Pmf1 | ENSMUSG00000028066.10 |
| chr10 | 62110512 | 62111216 | 24.22294 | 75.10754 | 72.03808 | Promoter | 10 | 62107655 | 62111050 | 3396 | 2 | ENSMUST00 | 0 | 2010107G23 | ENSMUSG00000020083.8 |
| chr3 | 27938231 | 27939763 | 24.22294 | 75.10754 | 72.03808 | Promoter | 3 | 27938695 | 28133239 | 194545 | 1 | ENSMUST00 | 0 | Pld1 | ENSMUSG00000027695.11 |
| chr7 | 132811861 | 132814347 | 24.22294 | 75.10754 | 72.03808 | Promoter | 7 | 132712082 | 132813528 | 101447 | 2 | ENSMUST00 | 0 | Fam53b | ENSMUSG00000030956.10 |
| chr10 | 127642238 | 127644543 | 26.29138 | 75.07713 | 72.00807 | Promoter | 10 | 127642986 | 127660957 | 17972 | 1 | ENSMUST00 | 0 | Stat6 | ENSMUSG00000002147.13 |
| chr13 | 43170882 | 43172192 | 25.22864 | 75.06109 | 71.99223 | Promoter | 13 | 43151744 | 43171501 | 19758 | 2 | ENSMUST00 | 0 | Tbc1d7 | ENSMUSG00000021368.9 |
| chr7 | 28466889 | 28467949 | 25.22864 | 75.06109 | 71.99223 | 3' UTR | 7 | 28458658 | 28468242 | 9585 | 1 | ENSMUST00 | 8232 | Lrfn1 | ENSMUSG00000030600.10 |
| chr17 | 33089253 | 33089945 | 24.70746 | 75.05766 | 71.98973 | Promoter | 17 | 33089310 | 33106203 | 16894 | 1 | ENSMUST00 | 0 | Zfp563 | ENSMUSG00000067424.7 |
| chr4 | 40702286 | 40703968 | 24.70746 | 75.05766 | 71.98973 | Promoter | 4 | 40682382 | 40702914 | 20533 | 2 | ENSMUST00 | 0 | Aptx | ENSMUSG00000028411.10 |
| chr5 | 137045243 | 137046377 | 24.70746 | 75.05766 | 71.98973 | Promoter | 5 | 137034993 | 137046135 | 11143 | 2 | ENSMUST00 | 0 | Ap1s1 | ENSMUSG00000004849.9 |
| chr8 | 105707549 | 105708330 | 24.70746 | 75.05766 | 71.98973 | Promoter | 8 | 105703655 | 105707933 | 4279 | 2 | ENSMUST00 | 0 | Enkd1 | ENSMUSG00000013155.9 |
| chr9 | 79793550 | 79794200 | 24.70746 | 75.05766 | 71.98973 | Promoter | 9 | 79793641 | 79803235 | 9595 | 1 | ENSMUST00 | 0 | 4930429F24I | ENSMUSG00000086918.1 |
| chr10 | 53596026 | 53596728 | 25.7402 | 75.03825 | 71.97101 | Promoter | 10 | 53596961 | 53609213 | 12253 | 1 | ENSMUST00 | -233 | Asf1a | ENSMUSG00000019857.6 |
| chr10 | 88200722 | 88201573 | 26.18464 | 74.89332 | 71.82663 | Promoter | 10 | 88201093 | 88246158 | 45066 | 1 | ENSMUST00 | 0 | Ccdc53 | ENSMUSG00000020056.11 |
| chr19 | 29366664 | 29367831 | 25.63958 | 74.86369 | 71.79715 | Promoter | 19 | 29358376 | 29367308 | 8933 | 2 | ENSMUST00 | 0 | Plgrkt | ENSMUSG00000016495.7 |
| chr11 | 87736508 | 87737347 | 25.08628 | 74.81181 | 71.74545 | Promoter | 11 | 87737552 | 87743617 | 6066 | 1 | ENSMUST00 | -205 | Supt4a | ENSMUSG00000020485.9 |
| chr1 | 167392967 | 167394793 | 20.31581 | 74.8081 | 71.74177 | Promoter | 1 | 167371966 | 167393841 | 21876 | 2 | ENSMUST00 | 0 | Mgst3 | ENSMUSG00000026688.5 |
| chr11 | 57517580 | 57519099 | 20.31581 | 74.8081 | 71.74177 | Promoter | 11 | 57518664 | 57533815 | 15152 | 1 | ENSMUST00 | 0 | Mfap3 | ENSMUSG00000020522.8 |
| chr11 | 95512463 | 95515642 | 20.31581 | 74.8081 | 71.74177 | Promoter | 11 | 95509845 | 95514570 | 4726 | 2 | ENSMUST00 | 0 | Nxph3 | ENSMUSG00000046719.7 |
| chr12 | 65172151 | 65173446 | 20.31581 | 74.8081 | 71.74177 | Promoter | 12 | 65132734 | 65172580 | 39847 | 2 | ENSMUST00 | 0 | Mis18bp1 | ENSMUSG00000047534.11 |
| chr17 | 33823736 | 33825347 | 20.31581 | 74.8081 | 71.74177 | Promoter | 17 | 33824591 | 33838316 | 13726 | 1 | ENSMUST00 | 0 | Ndufa7 | ENSMUSG00000041881.7 |
| chr18 | 67204723 | 67207000 | 20.31581 | 74.8081 | 71.74177 | Intron (ENSM | 18 | 67174885 | 67174991 | 107 | 2 | ENSMUST00 | -29733 | Gm23512 | ENSMUSG00000095972.1 |
| chr2 | 73528721 | 73530153 | 20.31581 | 74.8081 | 71.74177 | Promoter | 2 | 73429610 | 73529734 | 100125 | 2 | ENSMUST00 | 0 | Wipf1 | ENSMUSG00000075284.5 |
| chr5 | 137683658 | 137684817 | 20.31581 | 74.8081 | 71.74177 | Promoter | 5 | 137650483 | 137684726 | 34244 | 2 | ENSMUST00 | 0 | Agfg2 | ENSMUSG00000029722.10 |
| chr7 | 29953107 | 29954850 | 20.31581 | 74.8081 | 71.74177 | Promoter | 7 | 29931711 | 29954087 | 22377 | 2 | ENSMUST00 | 0 | Zfp74 | ENSMUSG00000059975.9 |
| chr4 | 75277417 | 75278561 | 26.1316 | 74.80193 | 71.73632 | Promoter | 4 | 75277354 | 75278305 | 952 | 2 | ENSMUST00 | 0 | Tmem261 | ENSMUSG00000028398.8 |
| chr11 | 115276364 | 115277051 | 26.65286 | 74.78688 | 71.72144 | Promoter | 11 | 115268024 | 115276914 | 8891 | 2 | ENSMUST00 | 0 | Fdxr | ENSMUSG00000018861.3 |
| chr11 | 101008912 | 101010834 | 26.65286 | 74.78688 | 71.72144 | Promoter | 11 | 101009452 | 101021410 | 11959 | 1 | ENSMUST00 | 0 | Atp6v0a1 | ENSMUSG00000019302.11 |
| chr12 | 101818214 | 101819644 | 26.65286 | 74.78688 | 71.72144 | Promoter | 12 | 101746565 | 101819055 | 72491 | 2 | ENSMUST00 | 0 | Fbln5 | ENSMUSG00000021186.8 |
| chr17 | 26948318 | 26949056 | 26.65286 | 74.78688 | 71.72144 | Promoter | 17 | 26948056 | 26948119 | 64 | 2 | ENSMUST00 | -200 | Mir3083 | ENSMUSG00000093118.1 |
| chr3 | 68572019 | 68576372 | 26.65286 | 74.78688 | 71.72144 | Promoter | 3 | 68572245 | 68626464 | 54220 | 1 | ENSMUST00 | 0 | Schip1 | ENSMUSG00000027777.10 |
| chr5 | 117413423 | 117414465 | 26.65286 | 74.78688 | 71.72144 | Promoter | 5 | 117414000 | 117555942 | 141943 | 1 | ENSMUST00 | 0 | Ksr2 | ENSMUSG00000061578.8 |
| chr6 | 135065110 | 135066008 | 26.65286 | 74.78688 | 71.72144 | Promoter | 6 | 135065651 | 135084709 | 19059 | 1 | ENSMUST00 | 0 | Gprc5a | ENSMUSG00000046733.7 |
| chr7 | 4776455 | 4777620 | 26.65286 | 74.78688 | 71.72144 | Promoter | 7 | 4773055 | 4778141 | 5087 | 2 | ENSMUST00 | 521 | Il11 | ENSMUSG00000004371.9 |
| chr7 | 37770206 | 37772620 | 26.65286 | 74.78688 | 71.72144 | Promoter | 7 | 37479122 | 37770980 | 291859 | 2 | ENSMUST00 | 0 | Zfp536 | ENSMUSG00000043456.11 |
| chr9 | 37612835 | 37614176 | 26.65286 | 74.78688 | 71.72144 | Promoter | 9 | 37613847 | 37648318 | 34472 | 1 | ENSMUST00 | 0 | Siae | ENSMUSG00000001942.7 |
| chrX | 47912348 | 47913327 | 26.65286 | 74.78688 | 71.72144 | Promoter | 20 | 47912387 | 47965868 | 53482 | 1 | ENSMUST00 | 0 | Ocrl | ENSMUSG00000001173.10 |
| chrX | 160136967 | 160138880 | 26.65286 | 74.78688 | 71.72144 | Promoter | 20 | 160122209 | 160138413 | 16205 | 2 | ENSMUST00 | 0 | Pdha1 | ENSMUSG00000031299.10 |
| chrX | 74084207 | 74085691 | 26.65286 | 74.78688 | 71.72144 | Promoter | 20 | 74026592 | 74085638 | 59047 | 2 | ENSMUST00 | 0 | Mecp2 | ENSMUSG00000031393.11 |
| chrX | 144317476 | 144319227 | 26.65286 | 74.78688 | 71.72144 | Promoter | 20 | 144317812 | 144325197 | 7386 | 1 | ENSMUST00 | 0 | Alg13 | ENSMUSG00000041718.10 |
| chr8 | 19979636 | 19980779 | 21.16908 | 74.77386 | 71.70985 | Distal Interge | 8 | 20122365 | 20126624 | 4260 | 1 | ENSMUST00 | -141586 | Gm21769 | ENSMUSG00000094614.1 |
| chr2 | 69712946 | 69713838 | 16.59556 | 74.72475 | 71.66077 | Promoter | 2 | 69705085 | 69713498 | 8414 | 2 | ENSMUST00 | 0 | Fastkd1 | ENSMUSG00000027086.11 |
| chr11 | 26209894 | 26211068 | 26.07877 | 74.71087 | 71.64691 | Promoter | 11 | 25617668 | 26210569 | 592902 | 2 | ENSMUST00 | 0 | 5730522E02I | ENSMUSG00000032985.10 |
| chr3 | 95687298 | 95688332 | 26.60493 | 74.70471 | 71.64091 | Promoter | 3 | 95676201 | 95687917 | 11717 | 2 | ENSMUST00 | 0 | Adamtsl4 | ENSMUSG00000015850.6 |
| chr4 | 155651243 | 155655658 | 26.60493 | 74.70471 | 71.64091 | Promoter | 4 | 155650655 | 155653384 | 2730 | 2 | ENSMUST00 | 0 | Mmp23 | ENSMUSG00000029061.2 |
| chr7 | 29859911 | 29860761 | 26.60493 | 74.70471 | 71.64091 | Promoter | 7 | 29859979 | 29876505 | 16527 | 1 | ENSMUST00 | 0 | Zfp420 | ENSMUSG00000058402.8 |
| chr13 | 119462051 | 119463603 | 22.972 | 74.64658 | 71.58298 | Promoter | 13 | 119462759 | 119486117 | 23359 | 1 | ENSMUST00 | 0 | 4833420G17 | ENSMUSG00000062822.7 |
| chr12 | 31633878 | 31635103 | 26.02615 | 74.62016 | 71.5567 | Promoter | 12 | 31595354 | 31634597 | 39244 | 2 | ENSMUST00 | 0 | Bcap29 | ENSMUSG00000020650.9 |
| chr17 | 46281444 | 46283018 | 26.02615 | 74.62016 | 71.5567 | Promoter | 17 | 46257851 | 46283026 | 25176 | 2 | ENSMUST00 | 8 | Tjap1 | ENSMUSG00000012296.10 |
| chr11 | 78188241 | 78189701 | 26.54922 | 74.60921 | 71.54591 | Promoter | 11 | 78188430 | 78192188 | 3759 | 1 | ENSMUST00 | 0 | Rab34 | ENSMUSG00000002059.12 |
| chr12 | 100158813 | 100160040 | 26.54922 | 74.60921 | 71.54591 | Promoter | 12 | 100125452 | 100159653 | 34202 | 2 | ENSMUST00 | 0 | Nrde2 | ENSMUSG00000021179.7 |
| chr3 | 138526131 | 138527087 | 25.49012 | 74.6041 | 71.54089 | Promoter | 3 | 138526191 | 138557599 | 31409 | 1 | ENSMUST00 | 0 | Eif4e | ENSMUSG00000028156.8 |
| chr3 | 67462977 | 67464335 | 25.49012 | 74.6041 | 71.54089 | Promoter | 3 | 67457998 | 67463926 | 5929 | 2 | ENSMUST00 | 0 | Lxn | ENSMUSG00000047557.2 |
| chr1 | 136130929 | 136131999 | 22.44583 | 74.56482 | 71.50171 | Promoter | 1 | 136131389 | 136177998 | 46610 | 1 | ENSMUST00 | 0 | Kif21b | ENSMUSG00000041642.13 |
| chr10 | 56114711 | 56116412 | 22.44583 | 74.56482 | 71.50171 | 5' UTR | 10 | 56106917 | 56116880 | 9964 | 1 | ENSMUST00 | 7795 | Msl3l2 | ENSMUSG00000047669.7 |
| chr11 | 43747194 | 43748425 | 22.44583 | 74.56482 | 71.50171 | Promoter | 11 | 43729512 | 43748008 | 18497 | 2 | ENSMUST00 | 0 | Ttc1 | ENSMUSG00000041278.5 |
| chr13 | 21944656 | 21945794 | 22.44583 | 74.56482 | 71.50171 | Promoter | 13 | 21945094 | 21960485 | 15392 | 1 | ENSMUST00 | 0 | Zfp184 | ENSMUSG00000006720.11 |
| chr13 | 49652256 | 49652833 | 22.44583 | 74.56482 | 71.50171 | Promoter | 13 | 49464027 | 49652746 | 188720 | 2 | ENSMUST00 | 0 | Cenpp | ENSMUSG00000021391.9 |
| chr14 | 55094506 | 55095451 | 22.44583 | 74.56482 | 71.50171 | Promoter | 14 | 55094784 | 55098986 | 4203 | 1 | ENSMUST00 | 0 | Thtpa | ENSMUSG00000045691.8 |
| chr2 | 103072739 | 103074465 | 22.44583 | 74.56482 | 71.50171 | Promoter | 2 | 103073675 | 103092644 | 18970 | 1 | ENSMUST00 | 0 | Apip | ENSMUSG00000010911.7 |
| chr4 | 117120054 | 117121362 | 22.44583 | 74.56482 | 71.50171 | Promoter | 4 | 117120706 | 117121265 | 560 | 2 | ENSMUST00 | 0 | Btbd19 | ENSMUSG00000073771.6 |
| chr6 | 29272332 | 29273265 | 22.44583 | 74.56482 | 71.50171 | Promoter | 6 | 29272488 | 29275186 | 2699 | 1 | ENSMUST00 | 0 | Hilpda | ENSMUSG00000043421.8 |
| chr7 | 30459332 | 30463950 | 22.44583 | 74.56482 | 71.50171 | Promoter | 7 | 30459725 | 30487223 | 27499 | 1 | ENSMUST00 | 0 | Nphs1 | ENSMUSG00000006649.11 |
| chr8 | 122775339 | 122776260 | 22.44583 | 74.56482 | 71.50171 | Promoter | 8 | 122775505 | 122817880 | 42376 | 1 | ENSMUST00 | 0 | Acsf3 | ENSMUSG00000015016.7 |
| chr8 | 85035567 | 85038616 | 22.44583 | 74.56482 | 71.50171 | Promoter | 8 | 85036915 | 85057583 | 20669 | 1 | ENSMUST00 | 0 | Tnpo2 | ENSMUSG00000031691.8 |
| chr8 | 71485346 | 71486066 | 22.44583 | 74.56482 | 71.50171 | Promoter | 8 | 71476023 | 71486067 | 10045 | 2 | ENSMUST00 | 1 | Ano8 | ENSMUSG00000034863.7 |
| chr9 | 65908312 | 65910257 | 22.44583 | 74.56482 | 71.50171 | Promoter | 9 | 65908974 | 66004168 | 95195 | 1 | ENSMUST00 | 0 | Csnk1g1 | ENSMUSG00000032384.10 |
| chr9 | 85841971 | 85844734 | 22.44583 | 74.56482 | 71.50171 | Promoter | 9 | 85842380 | 85847040 | 4661 | 1 | ENSMUST00 | 0 | Tpbg | ENSMUSG00000035274.8 |
| chr11 | 98550566 | 98551756 | 25.97375 | 74.52978 | 71.46892 | Promoter | 11 | 98551097 | 98558665 | 7569 | 1 | ENSMUST00 | 0 | Zpbp2 | ENSMUSG00000017195.10 |
| chr10 | 40141250 | 40142645 | 19.35304 | 74.51352 | 71.45322 | Promoter | 10 | 40033535 | 40142254 | 108720 | 2 | ENSMUST00 | 0 | Slc16a10 | ENSMUSG00000019838.10 |
| chr10 | 96616250 | 96616891 | 19.35304 | 74.51352 | 71.45322 | Promoter | 10 | 96617001 | 96622813 | 5813 | 1 | ENSMUST00 | -110 | Btg1 | ENSMUSG00000036478.7 |
| chr11 | 98192483 | 98193831 | 19.35304 | 74.51352 | 71.45322 | Promoter | 11 | 98152154 | 98193260 | 41107 | 2 | ENSMUST00 | 0 | Med1 | ENSMUSG00000018160.10 |
| chr13 | 107890303 | 107892333 | 19.35304 | 74.51352 | 71.45322 | Promoter | 13 | 107724618 | 107890064 | 165447 | 2 | ENSMUST00 | -240 | Zswim6 | ENSMUSG00000032846.9 |
| chr17 | 29268447 | 29269771 | 19.35304 | 74.51352 | 71.45322 | Promoter | 17 | 29268788 | 29302881 | 34094 | 1 | ENSMUST00 | 0 | BC004004 | ENSMUSG00000052712.11 |
| chr7 | 64391514 | 64393386 | 19.35304 | 74.51352 | 71.45322 | Promoter | 7 | 64376576 | 64392268 | 15693 | 2 | ENSMUST00 | 0 | Mphosph10 | ENSMUSG00000030521.9 |

| | | | | | | | | | | | | | | | |
|---|---|---|---|---|---|---|---|---|---|---|---|---|---|---|---|
| chr7 | 130263579 | 130266780 | 19.35304 | 74.51352 | 71.45322 | Promoter | 7 | 130162512 | 130266245 | 103734 | 2 | ENSMUST00 | 0 | Fgfr2 | ENSMUSG00000030849.12 |
| chr11 | 53979317 | 53980607 | 23.37426 | 74.48241 | 71.42245 | Promoter | 11 | 53949965 | 53980332 | 30368 | 2 | ENSMUST00 | 0 | Slc22a21 | ENSMUSG00000063652.5 |
| chr12 | 32060375 | 32061738 | 23.37426 | 74.48241 | 71.42245 | Promoter | 12 | 31958476 | 32061235 | 102760 | 2 | ENSMUST00 | 0 | Prkar2b | ENSMUSG00000002997.10 |
| chr13 | 23542865 | 23545672 | 25.39144 | 74.43253 | 71.37302 | Promoter | 13 | 23542970 | 23543357 | 388 | 2 | ENSMUST00 | 0 | Hist1h2bh | ENSMUSG00000064168.4 |
| chr5 | 146962650 | 146964279 | 26.43852 | 74.41935 | 71.35993 | Promoter | 5 | 146951573 | 146963785 | 12213 | 2 | ENSMUST00 | 0 | Mtif3 | ENSMUSG00000016510.6 |
| chr18 | 77767203 | 77767861 | 17.79762 | 74.39539 | 71.33608 | Promoter | 18 | 77757567 | 77767780 | 10214 | 2 | ENSMUST00 | 0 | Haus1 | ENSMUSG00000041840.7 |
| chr18 | 46197970 | 46199750 | 18.52209 | 74.38695 | 71.32771 | Promoter | 18 | 46165302 | 46198818 | 33517 | 2 | ENSMUST00 | 0 | Trim36 | ENSMUSG00000033949.6 |
| chr11 | 78031905 | 78032988 | 23.80432 | 74.35631 | 71.29721 | Promoter | 11 | 78032280 | 78037866 | 5587 | 1 | ENSMUST00 | 0 | Dhrs13 | ENSMUSG00000020834.12 |
| chr17 | 33816999 | 33818499 | 25.86957 | 74.34999 | 71.29122 | 5' UTR | 17 | 33821830 | 33822275 | 446 | 1 | ENSMUST00 | -3331 | Kank3 | ENSMUSG00000042099.10 |
| chr4 | 95052498 | 95053357 | 25.86957 | 74.34999 | 71.29122 | Promoter | 4 | 95052951 | 95060658 | 7708 | 1 | ENSMUST00 | 0 | Junos | ENSMUSG00000087366.2 |
| chr9 | 65140715 | 65142580 | 25.86957 | 74.34999 | 71.29122 | Promoter | 9 | 65141154 | 65185857 | 44704 | 1 | ENSMUST00 | 0 | Igdcc3 | ENSMUSG00000032394.5 |
| chr5 | 5379620 | 5381052 | 25.34239 | 74.34718 | 71.28911 | Promoter | 5 | 4803391 | 5380197 | 576807 | 2 | ENSMUST00 | 0 | Cdk14 | ENSMUSG00000028926.10 |
| chr5 | 24604312 | 24605347 | 25.34239 | 74.34718 | 71.28911 | Distal Interge | 5 | 24592622 | 24602012 | 9391 | 2 | ENSMUST00 | -2301 | Smarcd3 | ENSMUSG00000028949.8 |
| chr3 | 36475860 | 36477050 | 26.38351 | 74.32498 | 71.26698 | Promoter | 3 | 36475937 | 36482299 | 6363 | 1 | ENSMUST00 | 0 | 1810062G17 | ENSMUSG00000027713.6 |
| chr1 | 135374356 | 135375499 | 25.29353 | 74.26213 | 71.20484 | Promoter | 1 | 135371057 | 135375237 | 4181 | 2 | ENSMUST00 | 0 | Shisa4 | ENSMUSG00000041889.7 |
| chr1 | 186703443 | 186705953 | 25.29353 | 74.26213 | 71.20484 | Promoter | 1 | 186705695 | 186707033 | 1339 | 1 | ENSMUST00 | 0 | RP24-276N2 | ENSMUSG00000100393.1 |
| chr12 | 28674926 | 28675952 | 25.29353 | 74.26213 | 71.20484 | Promoter | 12 | 28675231 | 28682175 | 6945 | 1 | ENSMUST00 | 0 | Adi1 | ENSMUSG00000020629.7 |
| chr13 | 13392538 | 13394560 | 25.29353 | 74.26213 | 71.20484 | Promoter | 13 | 13357623 | 13393624 | 36002 | 2 | ENSMUST00 | 0 | Gpr137b | ENSMUSG00000021306.10 |
| chr15 | 83563248 | 83564704 | 25.29353 | 74.26213 | 71.20484 | Promoter | 15 | 83563573 | 83574201 | 10629 | 1 | ENSMUST00 | 0 | Tspo | ENSMUSG00000041736.6 |
| chr3 | 57290579 | 57295802 | 25.29353 | 74.26213 | 71.20484 | Promoter | 3 | 57285611 | 57294889 | 9279 | 2 | ENSMUST00 | 0 | Tm4sf1 | ENSMUSG00000027800.9 |
| chr7 | 144937673 | 144940744 | 25.29353 | 74.26213 | 71.20484 | Promoter | 7 | 144929931 | 144939869 | 9939 | 2 | ENSMUST00 | 0 | Ccnd1 | ENSMUSG00000070348.4 |
| chr10 | 128821080 | 128821713 | 24.76026 | 74.23955 | 71.18308 | Promoter | 10 | 128817457 | 128821631 | 4175 | 2 | ENSMUST00 | 0 | Ormdl2 | ENSMUSG00000025353.4 |
| chr13 | 21974257 | 21975102 | 24.76026 | 74.23955 | 71.18308 | Distal Interge | 13 | 21981194 | 21988734 | 7541 | 1 | ENSMUST00 | -6092 | Pom121l2 | ENSMUSG00000016982.6 |
| chr17 | 12118585 | 12120526 | 24.76026 | 74.23955 | 71.18308 | Promoter | 17 | 12118704 | 12210710 | 92007 | 1 | ENSMUST00 | 0 | Agpat4 | ENSMUSG00000023827.3 |
| chr17 | 36836199 | 36837810 | 24.76026 | 74.23955 | 71.18308 | Promoter | 17 | 36837134 | 36837855 | 722 | 1 | ENSMUST00 | 0 | Trim26 | ENSMUSG00000024457.11 |
| chr19 | 12796321 | 12797320 | 24.76026 | 74.23955 | 71.18308 | Promoter | 19 | 12767020 | 12796126 | 29107 | 2 | ENSMUST00 | -196 | Zfp91 | ENSMUSG00000024695.10 |
| chr19 | 12004808 | 12006757 | 24.76026 | 74.23955 | 71.18308 | Distal Interge | 19 | 12035808 | 12036740 | 933 | 2 | ENSMUST00 | 29983 | Olfr1423 | ENSMUSG00000067529.1 |
| chr3 | 10087877 | 10088366 | 24.76026 | 74.23955 | 71.18308 | Promoter | 3 | 10088173 | 10092564 | 4392 | 1 | ENSMUST00 | 0 | Gm9833 | ENSMUSG00000049230.7 |
| chr4 | 132394676 | 132398208 | 24.76026 | 74.23955 | 71.18308 | Promoter | 4 | 132377461 | 132398199 | 20739 | 2 | ENSMUST00 | 0 | Phactr4 | ENSMUSG00000066043.8 |
| chr8 | 83607299 | 83608079 | 24.76026 | 74.23955 | 71.18308 | Promoter | 8 | 83608175 | 83611902 | 3728 | 1 | ENSMUST00 | -96 | Dnajb1 | ENSMUSG00000005483.9 |
| chr9 | 3335055 | 3336304 | 25.76622 | 74.17153 | 71.11615 | Promoter | 9 | 3335478 | 3387050 | 51573 | 1 | ENSMUST00 | 0 | Alkbh8 | ENSMUSG00000025899.8 |
| chr3 | 159839398 | 159841336 | 24.71438 | 74.15884 | 71.1036 | Promoter | 3 | 159839737 | 159935833 | 96097 | 1 | ENSMUST00 | 0 | Wls | ENSMUSG00000028173.6 |
| chr2 | 13543589 | 13545024 | 26.27418 | 74.13734 | 71.08211 | Promoter | 2 | 13509014 | 13544664 | 35651 | 2 | ENSMUST00 | 0 | Trdmt1 | ENSMUSG00000026723.5 |
| chr1 | 75381728 | 75383103 | 23.65566 | 74.08845 | 71.03342 | Promoter | 1 | 75382114 | 75388345 | 6232 | 1 | ENSMUST00 | 0 | Speg | ENSMUSG00000026207.11 |
| chr1 | 69105028 | 69108834 | 23.65566 | 74.08845 | 71.03342 | Promoter | 1 | 68032186 | 69107756 | 1075571 | 2 | ENSMUST00 | 0 | Erbb4 | ENSMUSG00000062209.10 |
| chr10 | 43900909 | 43902472 | 23.65566 | 74.08845 | 71.03342 | Promoter | 10 | 43874188 | 43901745 | 27558 | 2 | ENSMUST00 | 0 | Qrsl1 | ENSMUSG00000019863.6 |
| chr10 | 60752055 | 60753394 | 23.65566 | 74.08845 | 71.03342 | Promoter | 10 | 60712072 | 60752782 | 40711 | 2 | ENSMUST00 | 0 | Slc29a3 | ENSMUSG00000020100.10 |
| chr10 | 40024670 | 40025645 | 23.65566 | 74.08845 | 71.03342 | Promoter | 10 | 40001575 | 40025268 | 23694 | 2 | ENSMUST00 | 0 | AI317395 | ENSMUSG00000038522.6 |
| chr11 | 108424572 | 108425870 | 23.65566 | 74.08845 | 71.03342 | Promoter | 11 | 108425192 | 108860615 | 435424 | 1 | ENSMUST00 | 0 | Cep112 | ENSMUSG00000020728.12 |
| chr12 | 84589871 | 84590777 | 23.65566 | 74.08845 | 71.03342 | Intron (ENSM | 12 | 84569840 | 84595457 | 25618 | 1 | ENSMUST00 | 20032 | Vsx2 | ENSMUSG00000021239.7 |
| chr13 | 3358153 | 3361680 | 23.65566 | 74.08845 | 71.03342 | Promoter | 13 | 3361029 | 3361533 | 505 | 1 | ENSMUST00 | 0 | Gm16505 | ENSMUSG00000096107.1 |
| chr14 | 51883870 | 51885654 | 23.65566 | 74.08845 | 71.03342 | Promoter | 14 | 51884842 | 51891868 | 7027 | 1 | ENSMUST00 | 0 | Mettl17 | ENSMUSG00000004561.9 |
| chr16 | 4557998 | 4560889 | 23.65566 | 74.08845 | 71.03342 | Promoter | 16 | 4544661 | 4559720 | 15060 | 2 | ENSMUST00 | 0 | Tfap4 | ENSMUSG00000005718.7 |
| chr16 | 91728054 | 91729199 | 23.65566 | 74.08845 | 71.03342 | Promoter | 16 | 91689322 | 91728975 | 39654 | 2 | ENSMUST00 | 0 | Cryzl1 | ENSMUSG00000058240.8 |
| chr17 | 25808581 | 25809868 | 23.65566 | 74.08845 | 71.03342 | Promoter | 17 | 25809085 | 25821244 | 12160 | 1 | ENSMUST00 | 0 | Fbxl16 | ENSMUSG00000025738.7 |
| chr18 | 10705994 | 10707368 | 23.65566 | 74.08845 | 71.03342 | Promoter | 18 | 10644411 | 10706771 | 62361 | 2 | ENSMUST00 | 0 | Abhd3 | ENSMUSG00000002475.10 |
| chr18 | 75695237 | 75698016 | 23.65566 | 74.08845 | 71.03342 | Promoter | 18 | 75431224 | 75697696 | 266473 | 2 | ENSMUST00 | 0 | Ctif | ENSMUSG00000052928.8 |
| chr4 | 155846801 | 155848451 | 23.65566 | 74.08845 | 71.03342 | Promoter | 4 | 155847402 | 155859303 | 11902 | 1 | ENSMUST00 | 0 | Dvl1 | ENSMUSG00000029071.11 |
| chr5 | 24685889 | 24687425 | 23.65566 | 74.08845 | 71.03342 | Promoter | 5 | 24685815 | 24710555 | 24741 | 1 | ENSMUST00 | 75 | Nub1 | ENSMUSG00000028954.7 |
| chr5 | 140331173 | 140332313 | 23.65566 | 74.08845 | 71.03342 | Promoter | 5 | 140327674 | 140331898 | 4225 | 2 | ENSMUST00 | 0 | Ftsj2 | ENSMUSG00000029557.7 |
| chr5 | 72913155 | 72914239 | 23.65566 | 74.08845 | 71.03342 | Promoter | 5 | 72914304 | 72978829 | 64526 | 1 | ENSMUST00 | -65 | Slain2 | ENSMUSG00000036087.13 |
| chr5 | 135077676 | 135078736 | 23.65566 | 74.08845 | 71.03342 | Promoter | 5 | 135072900 | 135078266 | 5367 | 2 | ENSMUST00 | 0 | Vps37d | ENSMUSG00000043614.8 |
| chr6 | 120916286 | 120917807 | 23.65566 | 74.08845 | 71.03342 | Promoter | 6 | 120891930 | 120916836 | 24907 | 2 | ENSMUST00 | 0 | Bid | ENSMUSG00000004446.7 |
| chr6 | 57533792 | 57535516 | 23.65566 | 74.08845 | 71.03342 | Promoter | 6 | 57506502 | 57535426 | 28925 | 2 | ENSMUST00 | 0 | Ppm1k | ENSMUSG00000037826.3 |
| chr7 | 45628098 | 45629263 | 23.65566 | 74.08845 | 71.03342 | Promoter | 7 | 45627488 | 45639093 | 11606 | 1 | ENSMUST00 | 611 | Rasip1 | ENSMUSG00000044562.10 |
| chr8 | 83899124 | 83901392 | 23.65566 | 74.08845 | 71.03342 | Promoter | 8 | 83900105 | 83941954 | 41850 | 1 | ENSMUST00 | 0 | Lphn1 | ENSMUSG00000013033.11 |
| chr9 | 90054333 | 90055498 | 23.65566 | 74.08845 | 71.03342 | Promoter | 9 | 90054506 | 90076087 | 21582 | 1 | ENSMUST00 | 0 | Ctsh | ENSMUSG00000032359.9 |
| chr9 | 109832255 | 109833458 | 23.65566 | 74.08845 | 71.03342 | Promoter | 9 | 109832749 | 109842529 | 9781 | 1 | ENSMUST00 | 0 | Nme6 | ENSMUSG00000032478.9 |
| chr19 | 43939281 | 43940742 | 26.21985 | 74.04404 | 70.99222 | Distal Interge | 19 | 43932316 | 43932432 | 117 | 1 | ENSMUST00 | 6966 | n-R5s21 | ENSMUSG00000084641.1 |
| chr8 | 84004168 | 84005290 | 25.14806 | 74.00871 | 70.95702 | Promoter | 8 | 84005628 | 84005764 | 137 | 1 | ENSMUST00 | -338 | Mir8111 | ENSMUSG00000098823.1 |
| chr10 | 14544762 | 14545897 | 25.1466 | 74.00616 | 70.95453 | Promoter | 10 | 14402585 | 14545036 | 142452 | 2 | ENSMUST00 | 0 | Gpr126 | ENSMUSG00000039116.4 |
| chr11 | 97985916 | 97986757 | 25.1466 | 74.00616 | 70.95453 | Promoter | 11 | 97923238 | 97986444 | 63207 | 2 | ENSMUST00 | 0 | Plxdc1 | ENSMUSG00000017417.9 |
| chr13 | 41605028 | 41605689 | 25.1466 | 74.00616 | 70.95453 | Promoter | 13 | 41606216 | 41641347 | 35132 | 1 | ENSMUST00 | -527 | Tmem170b | ENSMUSG00000087370.2 |
| chr14 | 61648941 | 61649715 | 25.1466 | 74.00616 | 70.95453 | Promoter | 14 | 61648950 | 61667312 | 18363 | 1 | ENSMUST00 | 0 | Gm27017 | ENSMUSG00000097997.1 |
| chr17 | 16950870 | 16952365 | 25.1466 | 74.00616 | 70.95453 | Promoter | 17 | 16951539 | 16974157 | 22619 | 1 | ENSMUST00 | 0 | BC002059 | ENSMUSG00000060149.6 |
| chr17 | 47410244 | 47410827 | 25.1466 | 74.00616 | 70.95453 | Promoter | 17 | 47410363 | 47410740 | 378 | 1 | ENSMUST00 | 0 | Gm5814 | ENSMUSG00000096361.1 |
| chr19 | 11769912 | 11771101 | 25.1466 | 74.00616 | 70.95453 | Promoter | 19 | 11770415 | 11774944 | 4530 | 1 | ENSMUST00 | 0 | Mrpl16 | ENSMUSG00000024683.5 |
| chr19 | 24279685 | 24281055 | 25.1466 | 74.00616 | 70.95453 | Promoter | 19 | 24261453 | 24280605 | 19153 | 2 | ENSMUST00 | 0 | Fxn | ENSMUSG00000059363.5 |
| chr2 | 127584152 | 127585102 | 25.1466 | 74.00616 | 70.95453 | Promoter | 2 | 127575533 | 127584669 | 9137 | 2 | ENSMUST00 | 0 | Zfp661 | ENSMUSG00000034800.10 |
| chr2 | 92058378 | 92060209 | 25.1466 | 74.00616 | 70.95453 | Promoter | 2 | 92054989 | 92059369 | 4381 | 2 | ENSMUST00 | 0 | 2900072N19 | ENSMUSG00000087018.1 |
| chr4 | 54944740 | 54946604 | 25.1466 | 74.00616 | 70.95453 | Promoter | 4 | 54945048 | 55008314 | 63267 | 1 | ENSMUST00 | 0 | Zfp462 | ENSMUSG00000060206.6 |
| chr5 | 134687933 | 134689442 | 25.1466 | 74.00616 | 70.95453 | Promoter | 5 | 134656039 | 134688590 | 32552 | 2 | ENSMUST00 | 0 | Limk1 | ENSMUSG00000029674.8 |
| chr5 | 120510942 | 120512134 | 25.1466 | 74.00616 | 70.95453 | Promoter | 5 | 120511168 | 120534024 | 22857 | 1 | ENSMUST00 | 0 | Slc8b1 | ENSMUSG00000032754.9 |
| chr5 | 81019533 | 81023055 | 25.1466 | 74.00616 | 70.95453 | Promoter | 5 | 81020138 | 81388784 | 368647 | 1 | ENSMUST00 | 0 | Lphn3 | ENSMUSG00000037605.11 |
| chr6 | 91116178 | 91117315 | 25.1466 | 74.00616 | 70.95453 | Promoter | 6 | 91013068 | 91116829 | 103762 | 2 | ENSMUST00 | 0 | Nup210 | ENSMUSG00000030091.12 |
| chr7 | 143648986 | 143649725 | 25.1466 | 74.00616 | 70.95453 | Promoter | 7 | 143634808 | 143649661 | 14854 | 2 | ENSMUST00 | 0 | Tnfrsf22 | ENSMUSG00000010751.9 |
| chr7 | 105553586 | 105555108 | 25.1466 | 74.00616 | 70.95453 | Promoter | 7 | 105554360 | 105558388 | 4029 | 1 | ENSMUST00 | 0 | Smpd1 | ENSMUSG00000037049.8 |
| chr11 | 5877581 | 5878300 | 26.11187 | 73.85856 | 70.81015 | Promoter | 11 | 5872180 | 5878292 | 6113 | 2 | ENSMUST00 | 0 | Pold2 | ENSMUSG00000020471.6 |
| chr15 | 76715314 | 76716153 | 25.05201 | 73.84118 | 70.7929 | 3' UTR | 15 | 76716207 | 76718416 | 2210 | 2 | ENSMUST00 | 2263 | Lrrc24 | ENSMUSG00000033707.8 |
| chr6 | 117862878 | 117863582 | 25.05201 | 73.84118 | 70.7929 | Promoter | 6 | 117863077 | 117870739 | 7663 | 1 | ENSMUST00 | 0 | Zfp239 | ENSMUSG00000042097.12 |
| chr15 | 3978820 | 3979702 | 24.53253 | 73.83858 | 70.79035 | Promoter | 15 | 3965445 | 3979573 | 14129 | 2 | ENSMUST00 | 0 | Fbxo4 | ENSMUSG00000022184.8 |
| chr16 | 14316617 | 14317905 | 21.13367 | 73.82313 | 70.77491 | Promoter | 16 | 14299244 | 14317347 | 18104 | 2 | ENSMUST00 | 0 | Fopnl | ENSMUSG00000022677.9 |
| chr2 | 152949781 | 152952321 | 21.13367 | 73.82313 | 70.77491 | Promoter | 2 | 152940997 | 152951456 | 10460 | 2 | ENSMUST00 | 0 | Dusp15 | ENSMUSG00000042662.11 |
| chr2 | 52741060 | 52743122 | 21.13367 | 73.82313 | 70.77491 | Promoter | 2 | 52691664 | 52742200 | 50537 | 2 | ENSMUST00 | 0 | Stam2 | ENSMUSG00000055371.12 |
| chr2 | 153160857 | 153162449 | 21.13367 | 73.82313 | 70.77491 | Promoter | 2 | 153161303 | 153204862 | 43560 | 1 | ENSMUST00 | 0 | Tm9sf4 | ENSMUSG00000068040.5 |
| chr2 | 13271534 | 13272156 | 21.13367 | 73.82313 | 70.77491 | Promoter | 2 | 13271602 | 13324791 | 53190 | 1 | ENSMUST00 | 0 | Gm13270 | ENSMUSG00000086358.1 |
| chr5 | 124540331 | 124541863 | 21.13367 | 73.82313 | 70.77491 | Promoter | 5 | 124540695 | 124550506 | 9812 | 1 | ENSMUST00 | 0 | Tmed2 | ENSMUSG00000029390.8 |
| chr8 | 125669323 | 125670600 | 21.13367 | 73.82313 | 70.77491 | Promoter | 8 | 125669818 | 125673359 | 3542 | 1 | ENSMUST00 | 0 | Map10 | ENSMUSG00000050930.5 |
| chr9 | 71769462 | 71771959 | 21.13367 | 73.82313 | 70.77491 | Promoter | 9 | 71626509 | 71771602 | 145094 | 2 | ENSMUST00 | 0 | Cgnl1 | ENSMUSG00000032232.9 |
| chr9 | 57910323 | 57911706 | 21.13367 | 73.82313 | 70.77491 | Promoter | 9 | 57910986 | 57929968 | 18983 | 1 | ENSMUST00 | 0 | Ubl7 | ENSMUSG00000055720.9 |
| chr19 | 27216675 | 27219273 | 25.56199 | 73.81844 | 70.77154 | Promoter | 19 | 27217020 | 27234856 | 17837 | 1 | ENSMUST00 | 0 | Vldlr | ENSMUSG00000024924.9 |
| chr17 | 21679450 | 21680613 | 22.99022 | 73.77998 | 70.73319 | Distal Interge | 17 | 21690766 | 21691024 | 259 | 1 | ENSMUST00 | -10153 | Gm10509 | ENSMUSG00000073448.1 |
| chr9 | 64178892 | 64180171 | 24.48749 | 73.75915 | 70.7127 | Promoter | 9 | 64179297 | 64182684 | 3388 | 1 | ENSMUST00 | 0 | Snapc5 | ENSMUSG00000032398.6 |
| chr14 | 30825422 | 30826479 | 25.00426 | 73.75784 | 70.7114 | Promoter | 14 | 30825590 | 30876986 | 51397 | 1 | ENSMUST00 | 0 | Tmem110 | ENSMUSG00000066526.6 |
| chr17 | 24250691 | 24252009 | 25.00426 | 73.75784 | 70.7114 | Promoter | 17 | 24223232 | 24251409 | 28178 | 2 | ENSMUST00 | 0 | Ccnf | ENSMUSG00000072082.6 |
| chr7 | 29231970 | 29232978 | 25.51143 | 73.73096 | 70.68457 | Promoter | 7 | 29221928 | 29232522 | 10595 | 2 | ENSMUST00 | 0 | Kcnk6 | ENSMUSG00000046410.9 |
| chr8 | 127439260 | 127441222 | 25.51143 | 73.73096 | 70.68457 | Intron (ENSM | 8 | 127447746 | 127474513 | 26768 | 1 | ENSMUST00 | -6524 | Pard3 | ENSMUSG00000025812.12 |
| chr11 | 12035826 | 12038123 | 26.00477 | 73.67448 | 70.62826 | Promoter | 11 | 11930508 | 12037401 | 106894 | 2 | ENSMUST00 | 0 | Grb10 | ENSMUSG00000020176.12 |
| chr7 | 126622952 | 126625762 | 23.39993 | 73.62621 | 70.58024 | Promoter | 7 | 126623249 | 126625676 | 2428 | 2 | ENSMUST00 | 0 | Nupr1 | ENSMUSG00000030717.4 |
| chr8 | 104590930 | 104592113 | 23.39993 | 73.62621 | 70.58024 | Promoter | 8 | 104591468 | 104596836 | 5369 | 1 | ENSMUST00 | 0 | Pdp2 | ENSMUSG00000048371.7 |
| chr1 | 72283836 | 72284524 | 25.95155 | 73.58297 | 70.53738 | Promoter | 1 | 72284369 | 72303104 | 18736 | 1 | ENSMUST00 | 0 | Tmem169 | ENSMUSG00000026188.6 |
| chr18 | 65580239 | 65581655 | 25.95155 | 73.58297 | 70.53738 | Promoter | 18 | 65580382 | 65623519 | 43138 | 1 | ENSMUST00 | 0 | Zfp532 | ENSMUSG00000042439.7 |
| chr19 | 36903975 | 36904907 | 25.95155 | 73.58297 | 70.53738 | Distal Interge | 19 | 36917550 | 36919615 | 2066 | 2 | ENSMUST00 | 14708 | Fgfbp3 | ENSMUSG00000047632.10 |
| chr10 | 81292865 | 81293889 | 23.83845 | 73.51238 | 70.46748 | Promoter | 10 | 81292963 | 81319972 | 27010 | 1 | ENSMUST00 | 0 | Pip5k1c | ENSMUSG00000034902.12 |
| chr7 | 141326953 | 141329130 | 23.83845 | 73.51238 | 70.46748 | Promoter | 7 | 141327739 | 141333286 | 5548 | 1 | ENSMUST00 | 0 | Tmem80 | ENSMUSG00000025505.11 |
| chr7 | 127015838 | 127017400 | 23.83845 | 73.51238 | 70.46748 | Promoter | 7 | 127001480 | 127016484 | 15005 | 2 | ENSMUST00 | 0 | Pagr1b | ENSMUSG00000092534.3 |
| chr4 | 155249526 | 155251162 | 24.86209 | 73.50948 | 70.465 | Promoter | 4 | 155249802 | 155256419 | 6618 | 1 | ENSMUST00 | 0 | 2610002J02 | ENSMUSG00000073684.8 |
| chr10 | 75316699 | 75317813 | 25.89855 | 73.4918 | 70.44736 | Promoter | 10 | 75316943 | 75334788 | 17846 | 1 | ENSMUST00 | 0 | Adora2a | ENSMUSG00000020178.5 |
| chr2 | 181837469 | 181838742 | 25.36096 | 73.47037 | 70.42609 | Promoter | 2 | 181837854 | 181857461 | 19608 | 1 | ENSMUST00 | 0 | Pcmtd2 | ENSMUSG00000027589.9 |
| chr2 | 177759218 | 177759991 | 25.36096 | 73.47037 | 70.42609 | Promoter | 2 | 177759288 | 177769962 | 10675 | 1 | ENSMUST00 | 0 | Gm14322 | ENSMUSG00000078864.5 |
| chr3 | 145653019 | 145653876 | 25.36096 | 73.47037 | 70.42609 | Distal Interge | 3 | 145650312 | 145652093 | 1782 | 1 | ENSMUST00 | 2708 | Gm17501 | ENSMUSG00000097183.1 |
| chr6 | 113638389 | 113638982 | 25.36096 | 73.47037 | 70.42609 | Promoter | 6 | 113638467 | 113695009 | 56543 | 1 | ENSMUST00 | 0 | Irak2 | ENSMUSG00000060477.9 |
| chr6 | 30459513 | 30460254 | 25.36096 | 73.47037 | 70.42609 | Promoter | 6 | 30460128 | 30463261 | 3134 | 1 | ENSMUST00 | 0 | 1700012C14 | ENSMUSG00000085470.1 |
| chr9 | 50727707 | 50728565 | 25.36096 | 73.47037 | 70.42609 | Promoter | 9 | 50662752 | 50727975 | 65224 | 2 | ENSMUST00 | 0 | Dixdc1 | ENSMUSG00000032064.12 |
| chr1 | 133163562 | 133164662 | 24.30896 | 73.44389 | 70.40032 | Distal Interge | 1 | 133181321 | 133301219 | 119899 | 1 | ENSMUST00 | -16659 | Plekha6 | ENSMUSG00000041757.10 |
| chr13 | 5860614 | 5861246 | 24.30896 | 73.44389 | 70.40032 | Promoter | 13 | 5861489 | 5870393 | 8905 | 1 | ENSMUST00 | -243 | Klf6 | ENSMUSG00000000078.6 |
| chr16 | 33966228 | 33967555 | 24.30896 | 73.44389 | 70.40032 | Promoter | 16 | 33954782 | 33967038 | 12257 | 2 | ENSMUST00 | 0 | Umps | ENSMUSG00000022814.6 |
| chr7 | 80323423 | 80324792 | 24.30896 | 73.44389 | 70.40032 | Promoter | 7 | 80316596 | 80324444 | 7849 | 2 | ENSMUST00 | 0 | Rccd1 | ENSMUSG00000038930.6 |
| chr3 | 41741443 | 41743295 | 24.81506 | 73.42724 | 70.38435 | Promoter | 3 | 41742611 | 41759389 | 16779 | 1 | ENSMUST00 | 0 | D3Ertd751e | ENSMUSG00000025766.9 |
| chr3 | 95427567 | 95428438 | 24.81506 | 73.42724 | 70.38435 | Promoter | 3 | 95427671 | 95431087 | 3417 | 1 | ENSMUST00 | 0 | Gm4349 | ENSMUSG00000097532.1 |
| chr4 | 148038133 | 148039704 | 24.81506 | 73.42724 | 70.38435 | Promoter | 4 | 148039077 | 148056206 | 17130 | 1 | ENSMUST00 | 0 | Mthfr | ENSMUSG00000029009.12 |
| chr9 | 21320297 | 21321765 | 24.81506 | 73.42724 | 70.38435 | Distal Interge | 9 | 21295457 | 21312333 | 16877 | 2 | ENSMUST00 | -7965 | Ap1m2 | ENSMUSG00000003309.8 |
| chr4 | 129048436 | 129049847 | 25.84577 | 73.40098 | 70.35904 | Downstream | 4 | 129050097 | 129057983 | 7887 | 2 | ENSMUST00 | 8136 | 1700086P04 | ENSMUSG00000085085.2 |
| chr10 | 39133430 | 39134478 | 20.02967 | 73.35366 | 70.31202 | Promoter | 10 | 39134023 | 39151058 | 17036 | 1 | ENSMUST00 | 0 | Tube1 | ENSMUSG00000019845.7 |
| chr11 | 107027494 | 107028820 | 20.02967 | 73.35366 | 70.31202 | Promoter | 11 | 107028118 | 107030438 | 2321 | 1 | ENSMUST00 | 0 | 1810010H24 | ENSMUSG00000078607.3 |
| chr2 | 164438320 | 164443697 | 20.02967 | 73.35366 | 70.31202 | Promoter | 2 | 164424247 | 164443186 | 18940 | 2 | ENSMUST00 | 0 | Sdc4 | ENSMUSG00000017009.3 |
| chr3 | 122418997 | 122422602 | 20.02967 | 73.35366 | 70.31202 | Promoter | 3 | 122419780 | 122530180 | 110401 | 1 | ENSMUST00 | 0 | Bcar3 | ENSMUSG00000028121.4 |
| chr7 | 110061710 | 110062924 | 20.02967 | 73.35366 | 70.31202 | Promoter | 7 | 110061710 | 110095387 | 33678 | 1 | ENSMUST00 | 1 | Zfp143 | ENSMUSG00000061079.8 |

| | | | | | | | | | | | | | | | |
|---|---|---|---|---|---|---|---|---|---|---|---|---|---|---|---|
| chr11 | 51967584 | 51968496 | 22.2526 | 73.30839 | 70.26766 | Promoter | 11 | 51967661 | 51977334 | 9674 | 1 | ENSMUST00 | 0 | Cdkn2aipnl | ENSMUSG00000020392.8 |
| chr16 | 84734500 | 84735983 | 25.26163 | 73.29816 | 70.2575 | Promoter | 16 | 84717576 | 84735742 | 18167 | 2 | ENSMUST00 | 0 | Mrpl39 | ENSMUSG00000022889.7 |
| chr4 | 63215161 | 63216250 | 25.26163 | 73.29816 | 70.2575 | Promoter | 4 | 63215412 | 63275093 | 59682 | 1 | ENSMUST00 | 0 | Col27a1 | ENSMUSG00000045672.10 |
| chr11 | 96298899 | 96299642 | 25.74084 | 73.22035 | 70.17986 | Promoter | 11 | 96299171 | 96301568 | 2398 | 1 | ENSMUST00 | 0 | Hoxb6 | ENSMUSG00000000690.5 |
| chr3 | 121967379 | 121969883 | 25.74084 | 73.22035 | 70.17986 | Promoter | 3 | 121967830 | 122009914 | 42085 | 1 | ENSMUST00 | 0 | Arhgap29 | ENSMUSG00000039831.11 |
| chr17 | 65782365 | 65784357 | 16.94318 | 73.17311 | 70.13294 | Promoter | 17 | 65782573 | 65803912 | 21340 | 1 | ENSMUST00 | 0 | Ppp4r1 | ENSMUSG00000061950.12 |
| chr11 | 83291080 | 83292571 | 25.68869 | 73.13053 | 70.09039 | Promoter | 11 | 83291699 | 83294631 | 2933 | 1 | ENSMUST00 | 0 | AA465934 | ENSMUSG00000093483.1 |
| chr17 | 24707039 | 24708553 | 25.68869 | 73.13053 | 70.09039 | Promoter | 17 | 24697949 | 24707660 | 9712 | 2 | ENSMUST00 | 0 | Tbl3 | ENSMUSG00000040688.11 |
| chr2 | 62645474 | 62646806 | 22.61859 | 73.09582 | 70.05593 | Promoter | 2 | 62595798 | 62646211 | 50414 | 2 | ENSMUST00 | 0 | Ifih1 | ENSMUSG00000026896.9 |
| chr5 | 142366791 | 142368168 | 22.61859 | 73.09582 | 70.05593 | Distal Interge | 5 | 142401497 | 142462014 | 60518 | 1 | ENSMUST00 | -33329 | Foxk1 | ENSMUSG00000056493.8 |
| chr11 | 61683598 | 61684846 | 26.19333 | 73.09374 | 70.054 | Promoter | 11 | 61684091 | 61686858 | 2768 | 1 | ENSMUST00 | 0 | Fam83g | ENSMUSG00000042377.8 |
| chr12 | 72663805 | 72665580 | 26.19333 | 73.09374 | 70.054 | Promoter | 12 | 72650356 | 72664828 | 14473 | 2 | ENSMUST00 | 0 | Dhrs7 | ENSMUSG00000021094.9 |
| chr12 | 105008329 | 105010500 | 19.08793 | 73.09275 | 70.054 | Promoter | 12 | 104929949 | 105009809 | 79861 | 2 | ENSMUST00 | 0 | Syne3 | ENSMUSG00000054150.6 |
| chr13 | 64431923 | 64433090 | 26.19333 | 73.09374 | 70.054 | Promoter | 13 | 64432553 | 64439720 | 7168 | 1 | ENSMUST00 | 0 | Cdk20 | ENSMUSG00000021483.7 |
| chr14 | 54431085 | 54433050 | 26.19333 | 73.09374 | 70.054 | Promoter | 14 | 54431604 | 54441258 | 9655 | 1 | ENSMUST00 | 0 | Mmp14 | ENSMUSG00000000957.8 |
| chr14 | 51088104 | 51091567 | 19.08793 | 73.09275 | 70.054 | Promoter | 14 | 51091077 | 51106151 | 15075 | 1 | ENSMUST00 | 0 | Rnase4 | ENSMUSG00000021876.9 |
| chr14 | 46386588 | 46388392 | 26.19333 | 73.09374 | 70.054 | Promoter | 14 | 46387519 | 46389282 | 1764 | 1 | ENSMUST00 | 0 | Gm15222 | ENSMUSG00000086746.1 |
| chr15 | 6386203 | 6388628 | 26.19333 | 73.09374 | 70.054 | Promoter | 15 | 6386598 | 6440705 | 54108 | 1 | ENSMUST00 | 0 | Dab2 | ENSMUSG00000022150.11 |
| chr17 | 56829772 | 56831129 | 26.19333 | 73.09374 | 70.054 | Promoter | 17 | 56775897 | 56831008 | 55112 | 2 | ENSMUST00 | 0 | Rfx2 | ENSMUSG00000024206.9 |
| chr17 | 33810173 | 33811324 | 26.19333 | 73.09374 | 70.054 | Promoter | 17 | 33810520 | 33822918 | 12399 | 1 | ENSMUST00 | 0 | Kank3 | ENSMUSG00000042099.10 |
| chr7 | 141265471 | 141266718 | 26.19333 | 73.09374 | 70.054 | Promoter | 7 | 141262706 | 141266421 | 3716 | 2 | ENSMUST00 | 0 | Irf7 | ENSMUSG00000025498.9 |
| chr7 | 24269820 | 24270770 | 26.19333 | 73.09374 | 70.054 | Promoter | 7 | 24270420 | 24276817 | 6398 | 1 | ENSMUST00 | 0 | Zfp93 | ENSMUSG00000055305.10 |
| chr7 | 24610391 | 24611848 | 19.08793 | 73.09275 | 70.054 | Promoter | 7 | 24611328 | 24629288 | 17961 | 1 | ENSMUST00 | 0 | Phldb3 | ENSMUSG00000074277.3 |
| chr8 | 22475915 | 22477695 | 19.08793 | 73.09275 | 70.054 | Promoter | 8 | 22462615 | 22476879 | 14265 | 2 | ENSMUST00 | 0 | Smim19 | ENSMUSG00000031534.9 |
| chr8 | 44934665 | 44936566 | 26.19333 | 73.09374 | 70.054 | Promoter | 8 | 44935447 | 45052257 | 116811 | 1 | ENSMUST00 | 0 | Fat1 | ENSMUSG00000070047.7 |
| chr9 | 117249955 | 117253836 | 26.19333 | 73.09374 | 70.054 | Promoter | 9 | 116572746 | 117252463 | 679718 | 2 | ENSMUST00 | 0 | Rbms3 | ENSMUSG00000039607.10 |
| chr9 | 3531610 | 3532694 | 26.19333 | 73.09374 | 70.054 | Promoter | 9 | 3532354 | 3897342 | 364989 | 1 | ENSMUST00 | 0 | Gucy1a2 | ENSMUSG00000041624.9 |
| chrX | 93831268 | 93832563 | 26.19333 | 73.09374 | 70.054 | Promoter | 20 | 93764607 | 93832201 | 67595 | 2 | ENSMUST00 | 0 | Pdk3 | ENSMUSG00000035232.8 |
| chrX | 12672910 | 12674275 | 26.19333 | 73.09374 | 70.054 | Promoter | 20 | 12654879 | 12673668 | 18790 | 2 | ENSMUST00 | 0 | 1810030O07 | ENSMUSG00000044148.6 |
| chr1 | 75218367 | 75220073 | 22.10575 | 73.03297 | 69.99519 | Promoter | 1 | 75214228 | 75219294 | 5067 | 2 | ENSMUST00 | 0 | Tuba4a | ENSMUSG00000026202.8 |
| chr1 | 184882283 | 184883885 | 22.10575 | 73.03297 | 69.99519 | Promoter | 1 | 184871926 | 184883218 | 11293 | 2 | ENSMUST00 | 0 | C130074G19 | ENSMUSG00000039349.5 |
| chr1 | 74124171 | 74125317 | 22.10575 | 73.03297 | 69.99519 | Promoter | 1 | 73910231 | 74124447 | 214217 | 2 | ENSMUST00 | 0 | Tns1 | ENSMUSG00000055322.9 |
| chr1 | 171281295 | 171281917 | 22.10575 | 73.03297 | 69.99519 | Promoter | 1 | 171280109 | 171281186 | 1078 | 2 | ENSMUST00 | -110 | Ppox | ENSMUSG00000062729.6 |
| chr11 | 68384591 | 68387483 | 22.10575 | 73.03297 | 69.99519 | Promoter | 11 | 68209364 | 68386826 | 177463 | 2 | ENSMUST00 | 0 | Ntn1 | ENSMUSG00000020902.7 |
| chr11 | 119546666 | 119548665 | 22.10575 | 73.03297 | 69.99519 | Promoter | 11 | 119538719 | 119547753 | 9035 | 2 | ENSMUST00 | 0 | Nptx1 | ENSMUSG00000025582.4 |
| chr12 | 100519792 | 100520779 | 22.10575 | 73.03297 | 69.99519 | Promoter | 12 | 100300771 | 100520822 | 220052 | 2 | ENSMUST00 | 43 | Ttc7b | ENSMUSG00000033530.7 |
| chr13 | 21787499 | 21787923 | 22.10575 | 73.03297 | 69.99519 | Promoter | 13 | 21787508 | 21787976 | 469 | 1 | ENSMUST00 | 0 | Hist1h2bp | ENSMUSG00000069308.7 |
| chr14 | 73237671 | 73239739 | 22.10575 | 73.03297 | 69.99519 | Promoter | 14 | 73237895 | 73243294 | 5400 | 1 | ENSMUST00 | 0 | Lpar6 | ENSMUSG00000033446.7 |
| chr15 | 84557188 | 84558145 | 22.10575 | 73.03297 | 69.99519 | Promoter | 15 | 84553398 | 84557823 | 4426 | 2 | ENSMUST00 | 0 | Ldoc1l | ENSMUSG00000055745.4 |
| chr16 | 20425246 | 20426842 | 22.10575 | 73.03297 | 69.99519 | Promoter | 16 | 20331303 | 20426394 | 95092 | 2 | ENSMUST00 | 0 | Abcc5 | ENSMUSG00000022822.10 |
| chr16 | 93607589 | 93608539 | 22.10575 | 73.03297 | 69.99519 | Promoter | 16 | 93607837 | 93610505 | 2669 | 1 | ENSMUST00 | 0 | Cbr1 | ENSMUSG00000051483.8 |
| chr4 | 34777600 | 34778460 | 22.10575 | 73.03297 | 69.99519 | Promoter | 4 | 34768664 | 34778270 | 9607 | 2 | ENSMUST00 | 0 | Smim8 | ENSMUSG00000028295.9 |
| chr4 | 133212078 | 133213226 | 22.10575 | 73.03297 | 69.99519 | Promoter | 4 | 133212131 | 133221770 | 9640 | 1 | ENSMUST00 | 0 | Cd164l2 | ENSMUSG00000028865.8 |
| chr5 | 30072847 | 30073989 | 22.10575 | 73.03297 | 69.99519 | Promoter | 5 | 30058202 | 30073581 | 15380 | 2 | ENSMUST00 | 0 | Tyms | ENSMUSG00000025747.7 |
| chr6 | 131292577 | 131293822 | 22.10575 | 73.03297 | 69.99519 | Promoter | 6 | 131284388 | 131293244 | 8857 | 2 | ENSMUST00 | 0 | Magohb | ENSMUSG00000030188.7 |
| chr9 | 103304624 | 103305792 | 22.10575 | 73.03297 | 69.99519 | Promoter | 9 | 103305215 | 103350428 | 45214 | 1 | ENSMUST00 | 0 | Topbp1 | ENSMUSG00000032555.4 |
| chr5 | 76588487 | 76589753 | 17.56648 | 73.03062 | 69.99512 | Promoter | 5 | 76588698 | 76644908 | 56211 | 1 | ENSMUST00 | 0 | Cep135 | ENSMUSG00000036403.10 |
| chr7 | 79659408 | 79661045 | 17.56648 | 73.03062 | 69.99512 | Promoter | 7 | 79660196 | 79698134 | 37939 | 1 | ENSMUST00 | 0 | Ticrr | ENSMUSG00000046591.9 |
| chr18 | 36017700 | 36019090 | 18.27513 | 72.99583 | 69.9605 | 3' UTR | 18 | 35987019 | 36006084 | 19066 | 1 | ENSMUST00 | 30682 | Psd2 | ENSMUSG00000024347.11 |
| chr5 | 146793690 | 146794900 | 18.27513 | 72.99583 | 69.9605 | Promoter | 5 | 146734809 | 146795006 | 60198 | 2 | ENSMUST00 | 106 | Usp12 | ENSMUSG00000029640.12 |
| chr8 | 110997039 | 110998471 | 18.27513 | 72.99583 | 69.9605 | Promoter | 8 | 110974996 | 110997800 | 22805 | 2 | ENSMUST00 | 0 | Ddx19a | ENSMUSG00000015023.7 |
| chr17 | 36950746 | 36952126 | 25.58503 | 72.95191 | 69.91682 | Promoter | 17 | 36948356 | 36951713 | 3358 | 2 | ENSMUST00 | 0 | Ppp1r11 | ENSMUSG00000036398.8 |
| chr12 | 119236888 | 119239435 | 23.00904 | 72.91611 | 69.8813 | Promoter | 12 | 119158022 | 119238802 | 80781 | 2 | ENSMUST00 | 0 | Itgb8 | ENSMUSG00000025321.9 |
| chr13 | 84221590 | 84223059 | 25.53352 | 72.8631 | 69.82863 | Promoter | 13 | 84222314 | 84295870 | 73557 | 1 | ENSMUST00 | 0 | Tmem161b | ENSMUSG00000035762.10 |
| chr1 | 189687748 | 189688831 | 26.03697 | 72.82576 | 69.79147 | Promoter | 1 | 189640599 | 189688086 | 47488 | 2 | ENSMUST00 | 0 | Cenpf | ENSMUSG00000026605.9 |
| chr11 | 69621632 | 69622631 | 26.03697 | 72.82576 | 69.79147 | Promoter | 11 | 69622024 | 69623870 | 1847 | 1 | ENSMUST00 | 0 | Sat2 | ENSMUSG00000069835.5 |
| chr15 | 98108058 | 98109151 | 26.03697 | 72.82576 | 69.79147 | Promoter | 15 | 98108471 | 98132447 | 23977 | 1 | ENSMUST00 | 0 | Pfkm | ENSMUSG00000033065.8 |
| chr2 | 32352625 | 32353735 | 26.03697 | 72.82576 | 69.79147 | Promoter | 2 | 32308471 | 32353247 | 44777 | 2 | ENSMUST00 | 0 | Dnm1 | ENSMUSG00000026825.12 |
| chr13 | 74389957 | 74391013 | 23.42648 | 72.77338 | 69.73964 | Distal Interge | 13 | 74371426 | 74376566 | 5141 | 2 | ENSMUST00 | -13392 | Zfp72 | ENSMUSG00000069184.2 |
| chr19 | 4906067 | 4906702 | 23.42648 | 72.77338 | 69.73964 | Promoter | 19 | 4886898 | 4906627 | 19730 | 2 | ENSMUST00 | 0 | Bbs1 | ENSMUSG00000006464.8 |
| chr6 | 72379564 | 72380905 | 23.42648 | 72.77338 | 69.73964 | Promoter | 6 | 72368049 | 72380468 | 12420 | 2 | ENSMUST00 | 0 | Vamp5 | ENSMUSG00000073002.4 |
| chr8 | 107095512 | 107097189 | 23.42648 | 72.77338 | 69.73964 | Promoter | 8 | 107069400 | 107096545 | 27146 | 2 | ENSMUST00 | 0 | Terf2 | ENSMUSG00000031921.12 |
| chr11 | 103360131 | 103361046 | 25.4311 | 72.68642 | 69.65319 | Promoter | 11 | 103331497 | 103360900 | 29404 | 2 | ENSMUST00 | 0 | Arhgap27 | ENSMUSG00000034255.12 |
| chr12 | 112829598 | 112830264 | 25.4311 | 72.68642 | 69.65319 | Promoter | 12 | 112820237 | 112829389 | 9153 | 2 | ENSMUST00 | -210 | Cdca4 | ENSMUSG00000047832.8 |
| chr7 | 139212604 | 139213790 | 25.4311 | 72.68642 | 69.65319 | Promoter | 7 | 139212988 | 139242979 | 29992 | 1 | ENSMUST00 | 0 | Lrrc27 | ENSMUSG00000015980.9 |
| chr7 | 18909717 | 18910468 | 25.4311 | 72.68642 | 69.65319 | Promoter | 7 | 18890883 | 18910415 | 19533 | 2 | ENSMUST00 | 0 | Ccdc61 | ENSMUSG00000074358.4 |
| chr8 | 84799139 | 84800864 | 25.4311 | 72.68642 | 69.65319 | Promoter | 8 | 84708614 | 84800140 | 91527 | 2 | ENSMUST00 | 0 | Nfix | ENSMUSG00000001911.11 |
| chr15 | 78173350 | 78174161 | 23.87381 | 72.67255 | 69.64001 | Promoter | 15 | 78159465 | 78174108 | 14644 | 2 | ENSMUST00 | 0 | Ift27 | ENSMUSG00000016637.6 |
| chr6 | 90604548 | 90605461 | 23.87381 | 72.67255 | 69.64001 | Promoter | 6 | 90604725 | 90646412 | 41688 | 1 | ENSMUST00 | 0 | Slc41a3 | ENSMUSG00000030089.10 |
| chr7 | 101320580 | 101321978 | 23.87381 | 72.67255 | 69.64001 | Promoter | 7 | 101321113 | 101346308 | 25196 | 1 | ENSMUST00 | 0 | Stard10 | ENSMUSG00000030688.9 |
| chr11 | 30985860 | 30987022 | 25.92862 | 72.63998 | 69.60807 | Promoter | 11 | 30976707 | 30986350 | 9644 | 2 | ENSMUST00 | 0 | Chac2 | ENSMUSG00000020309.6 |
| chr13 | 37778006 | 37779602 | 24.87197 | 72.62122 | 69.58942 | Promoter | 13 | 37778400 | 37893849 | 115450 | 1 | ENSMUST00 | 0 | Rreb1 | ENSMUSG00000039087.11 |
| chr14 | 31205406 | 31207302 | 24.87197 | 72.62122 | 69.58942 | Promoter | 14 | 31170930 | 31206801 | 35872 | 2 | ENSMUST00 | 0 | Nisch | ENSMUSG00000021910.10 |
| chr10 | 21158821 | 21161478 | 24.35436 | 72.61947 | 69.58835 | Promoter | 10 | 21124936 | 21160905 | 35970 | 2 | ENSMUST00 | 0 | Myb | ENSMUSG00000019982.9 |
| chr14 | 79397530 | 79398542 | 24.35436 | 72.61947 | 69.58835 | Promoter | 14 | 79397772 | 79423650 | 25879 | 1 | ENSMUST00 | 0 | Mtrf1 | ENSMUSG00000022022.2 |
| chr15 | 76576110 | 76577014 | 24.35436 | 72.61947 | 69.58835 | Promoter | 15 | 76576358 | 76595816 | 19459 | 1 | ENSMUST00 | 0 | Adck5 | ENSMUSG00000022550.12 |
| chr17 | 21449880 | 21450957 | 24.35436 | 72.61947 | 69.58835 | Promoter | 17 | 21450352 | 21465591 | 15240 | 1 | ENSMUST00 | 0 | Zfp51 | ENSMUSG00000023892.7 |
| chr18 | 20961215 | 20962973 | 24.35436 | 72.61947 | 69.58835 | Intron (ENSN | 18 | 20944625 | 20983848 | 39224 | 1 | ENSMUST00 | 16591 | Rnf125 | ENSMUSG00000033107.5 |
| chr18 | 34840320 | 34841334 | 24.35436 | 72.61947 | 69.58835 | Promoter | 18 | 34840589 | 34847463 | 6875 | 1 | ENSMUST00 | 0 | Reep2 | ENSMUSG00000038555.7 |
| chr3 | 155092535 | 155094049 | 24.35436 | 72.61947 | 69.58835 | Promoter | 3 | 155093434 | 155194278 | 100845 | 1 | ENSMUST00 | 0 | Lrriq3 | ENSMUSG00000028182.9 |
| chr3 | 94786091 | 94787429 | 24.35436 | 72.61947 | 69.58835 | Promoter | 3 | 94760069 | 94786463 | 26395 | 2 | ENSMUST00 | 0 | Cgn | ENSMUSG00000068876.9 |
| chr4 | 57845075 | 57848262 | 24.35436 | 72.61947 | 69.58835 | Promoter | 4 | 57845247 | 57896982 | 51736 | 1 | ENSMUST00 | 0 | Akap2 | ENSMUSG00000038729.15 |
| chr7 | 43442913 | 43444736 | 24.35436 | 72.61947 | 69.58835 | Promoter | 7 | 43444108 | 43457800 | 13693 | 1 | ENSMUST00 | 0 | Etfb | ENSMUSG00000004610.3 |
| chr1 | 127306652 | 127307946 | 25.3802 | 72.59858 | 69.5683 | Promoter | 1 | 127306762 | 127483704 | 176943 | 1 | ENSMUST00 | 0 | Mgat5 | ENSMUSG00000036155.8 |
| chr10 | 62885627 | 62887639 | 24.8241 | 72.5379 | 69.50792 | Intron (ENSN | 10 | 62804570 | 62880014 | 75445 | 2 | ENSMUST00 | -5614 | Tet1 | ENSMUSG00000047146.10 |
| chr1 | 72306763 | 72307946 | 23.28017 | 72.50986 | 69.48009 | Promoter | 1 | 72307421 | 72394952 | 87532 | 1 | ENSMUST00 | 0 | Xrcc5 | ENSMUSG00000026187.8 |
| chr1 | 87509293 | 87510649 | 23.28017 | 72.50986 | 69.48009 | Promoter | 1 | 87476834 | 87510362 | 33529 | 2 | ENSMUST00 | 0 | Ngef | ENSMUSG00000026259.9 |
| chr10 | 93309491 | 93311606 | 23.28017 | 72.50986 | 69.48009 | Promoter | 10 | 93247414 | 93311135 | 63722 | 2 | ENSMUST00 | 0 | Elk3 | ENSMUSG00000008398.9 |
| chr11 | 49250056 | 49251442 | 23.28017 | 72.50986 | 69.48009 | Promoter | 11 | 49250512 | 49263030 | 12519 | 1 | ENSMUST00 | 0 | Mgat1 | ENSMUSG00000020346.11 |
| chr11 | 73018603 | 73019752 | 23.28017 | 72.50986 | 69.48009 | Promoter | 11 | 73019008 | 73042062 | 23055 | 1 | ENSMUST00 | 0 | Camkk1 | ENSMUSG00000020785.12 |
| chr11 | 58214493 | 58215673 | 23.28017 | 72.50986 | 69.48009 | Promoter | 11 | 58215028 | 58222779 | 7752 | 1 | ENSMUST00 | 0 | Irgm2 | ENSMUSG00000069874.6 |
| chr19 | 43752357 | 43753913 | 23.28017 | 72.50986 | 69.48009 | Promoter | 19 | 43752996 | 43768631 | 15636 | 1 | ENSMUST00 | 0 | Cutc | ENSMUSG00000025193.9 |
| chr2 | 166153906 | 166155964 | 23.28017 | 72.50986 | 69.48009 | Promoter | 2 | 166073089 | 166155663 | 82575 | 2 | ENSMUST00 | 0 | Sulf2 | ENSMUSG00000006800.8 |
| chr3 | 105450974 | 105453152 | 23.28017 | 72.50986 | 69.48009 | Promoter | 3 | 105452330 | 105674002 | 221673 | 1 | ENSMUST00 | 0 | Kcnd3 | ENSMUSG00000040896.11 |
| chr4 | 138912971 | 138914013 | 23.28017 | 72.50986 | 69.48009 | Promoter | 4 | 138895379 | 138913945 | 18567 | 2 | ENSMUST00 | 0 | Otud3 | ENSMUSG00000041161.8 |
| chr5 | 109923113 | 109924529 | 23.28017 | 72.50986 | 69.48009 | Promoter | 5 | 109923400 | 109943936 | 20537 | 1 | ENSMUST00 | 0 | Gm15446 | ENSMUSG00000090015.3 |
| chr6 | 48047316 | 48048137 | 23.28017 | 72.50986 | 69.48009 | Promoter | 6 | 48024188 | 48048114 | 23927 | 2 | ENSMUST00 | 0 | Zfp777 | ENSMUSG00000071477.5 |
| chr7 | 39517280 | 39518457 | 23.28017 | 72.50986 | 69.48009 | Promoter | 7 | 39517766 | 39540415 | 22650 | 1 | ENSMUST00 | 0 | Zfp619 | ENSMUSG00000068959.5 |
| chr8 | 123442872 | 123443754 | 23.28017 | 72.50986 | 69.48009 | Promoter | 8 | 123442986 | 123460515 | 17530 | 1 | ENSMUST00 | 0 | Def8 | ENSMUSG00000001482.9 |
| chr8 | 70760335 | 70760970 | 23.28017 | 72.50986 | 69.48009 | Promoter | 8 | 70758649 | 70760921 | 2273 | 2 | ENSMUST00 | 0 | Mpv17l2 | ENSMUSG00000035559.8 |
| chr4 | 120924201 | 120925562 | 25.82118 | 72.45564 | 69.42885 | Promoter | 4 | 120921196 | 120925005 | 3810 | 2 | ENSMUST00 | 0 | Exo5 | ENSMUSG00000028629.4 |
| chr2 | 18688735 | 18690594 | 24.72893 | 72.3721 | 69.34575 | Distal Interge | 2 | 18694032 | 18698884 | 4853 | 1 | ENSMUST00 | -3438 | Gm13334 | ENSMUSG00000087182.2 |
| chr11 | 59263814 | 59264639 | 24.72749 | 72.3696 | 69.3433 | Intron (ENSN | 11 | 59248033 | 59290752 | 42720 | 2 | ENSMUST00 | 26113 | Wnt3a | ENSMUSG00000009900.7 |
| chr15 | 103013754 | 103015861 | 24.72749 | 72.3696 | 69.3433 | Promoter | 15 | 103013815 | 103017429 | 3615 | 1 | ENSMUST00 | 0 | Hoxc5 | ENSMUSG00000022485.3 |
| chr15 | 98880737 | 98881405 | 24.72749 | 72.3696 | 69.3433 | Promoter | 15 | 98877778 | 98881404 | 3627 | 2 | ENSMUST00 | 0 | Rhebl1 | ENSMUSG00000023755.9 |
| chr2 | 80616660 | 80618108 | 24.72749 | 72.3696 | 69.3433 | Promoter | 2 | 80617236 | 80658906 | 41671 | 1 | ENSMUST00 | 0 | Nup35 | ENSMUSG00000026999.9 |
| chr3 | 84258819 | 84260075 | 24.72749 | 72.3696 | 69.3433 | Promoter | 3 | 84160440 | 84259812 | 99373 | 2 | ENSMUST00 | 0 | Trim2 | ENSMUSG00000027993.11 |
| chr4 | 54946727 | 54950127 | 24.72749 | 72.3696 | 69.3433 | Promoter | 4 | 54947976 | 55083563 | 135588 | 1 | ENSMUST00 | 0 | Zfp462 | ENSMUSG00000060206.6 |
| chr6 | 29467299 | 29468653 | 24.72749 | 72.3696 | 69.3433 | Promoter | 6 | 29467718 | 29470512 | 2795 | 1 | ENSMUST00 | 0 | Atp6v1f | ENSMUSG00000004285.7 |
| chr6 | 82041283 | 82043111 | 24.72749 | 72.3696 | 69.3433 | Promoter | 6 | 82041623 | 82093099 | 51477 | 1 | ENSMUST00 | 0 | Eva1a | ENSMUSG00000035104.9 |
| chr7 | 65155442 | 65156992 | 24.72749 | 72.3696 | 69.3433 | Promoter | 7 | 64756113 | 65156570 | 400458 | 2 | ENSMUST00 | 0 | Fam189a1 | ENSMUSG00000030518.12 |
| chr7 | 3414812 | 3415922 | 24.72749 | 72.3696 | 69.3433 | Promoter | 7 | 3415066 | 3415156 | 91 | 1 | ENSMUST00 | 0 | Gm23450 | ENSMUSG00000094210.1 |
| chr8 | 84893193 | 84893862 | 24.72749 | 72.3696 | 69.3433 | Promoter | 8 | 84890738 | 84893285 | 2548 | 2 | ENSMUST00 | 0 | Gcdh | ENSMUSG00000003809.9 |
| chr8 | 45409716 | 45410824 | 24.72749 | 72.3696 | 69.3433 | Promoter | 8 | 45395665 | 45410539 | 14875 | 2 | ENSMUST00 | 0 | Tlr3 | ENSMUSG00000031639.6 |
| chr8 | 117498027 | 117499046 | 24.72749 | 72.3696 | 69.3433 | Promoter | 8 | 117498291 | 117635140 | 136850 | 1 | ENSMUST00 | 0 | Plcg2 | ENSMUSG00000034330.9 |
| chr8 | 91313466 | 91314567 | 24.72749 | 72.3696 | 69.3433 | Promoter | 8 | 91313532 | 91668439 | 354908 | 1 | ENSMUST00 | 0 | Fto | ENSMUSG00000055932.9 |
| chr10 | 60001946 | 60003837 | 20.82289 | 72.3363 | 69.31286 | Promoter | 10 | 59987409 | 60003135 | 15727 | 2 | ENSMUST00 | 0 | Anapc16 | ENSMUSG00000020107.5 |
| chr10 | 130412221 | 130412640 | 20.82289 | 72.3363 | 69.31286 | Promoter | 10 | 130412499 | 130413312 | 814 | 1 | ENSMUST00 | 0 | Gm9770 | ENSMUSG00000040540.5 |
| chr10 | 13323002 | 13324565 | 20.82289 | 72.3363 | 69.31286 | Promoter | 10 | 13207717 | 13324160 | 116444 | 2 | ENSMUST00 | 0 | Phactr2 | ENSMUSG00000062866.10 |
| chr11 | 121236329 | 121237017 | 20.82289 | 72.3363 | 69.31286 | Promoter | 11 | 121237253 | 121255856 | 18604 | 1 | ENSMUST00 | -236 | Narf | ENSMUSG00000000056.7 |
| chr11 | 94653480 | 94654642 | 20.82289 | 72.3363 | 69.31286 | Promoter | 11 | 94653767 | 94660089 | 6323 | 1 | ENSMUST00 | 0 | Mrpl27 | ENSMUSG00000024414.8 |
| chr12 | 116262451 | 116263413 | 20.82289 | 72.3363 | 69.31286 | Promoter | 12 | 116207050 | 116263025 | 55976 | 2 | ENSMUST00 | 0 | Wdr60 | ENSMUSG00000042050.7 |
| chr16 | 85899317 | 85901820 | 20.82289 | 72.3363 | 69.31286 | Promoter | 16 | 85858170 | 85901125 | 42956 | 2 | ENSMUST00 | 0 | Adamts5 | ENSMUSG00000022894.5 |
| chr19 | 41911360 | 41913199 | 20.82289 | 72.3363 | 69.31286 | Promoter | 19 | 41911871 | 41918664 | 6794 | 1 | ENSMUST00 | 0 | Pgam1 | ENSMUSG00000011752.6 |
| chr2 | 26445563 | 26446532 | 20.82289 | 72.3363 | 69.31286 | Promoter | 2 | 26445696 | 26459379 | 13684 | 1 | ENSMUST00 | 0 | 0610009E02I | ENSMUSG00000086714.1 |
| chr3 | 145118077 | 145119514 | 20.82289 | 72.3363 | 69.31286 | Promoter | 3 | 145118589 | 145153915 | 35327 | 1 | ENSMUST00 | 0 | Odf2l | ENSMUSG00000028256.11 |
| chr6 | 124808299 | 124810012 | 20.82289 | 72.3363 | 69.31286 | Promoter | 6 | 124808661 | 124810619 | 1959 | 1 | ENSMUST00 | 0 | Spsb2 | ENSMUSG00000038451.8 |
| chr4 | 20041510 | 20042682 | 24.68161 | 72.28964 | 69.2675 | Promoter | 4 | 20042052 | 20066750 | 24699 | 1 | ENSMUST00 | 0 | Ggh | ENSMUSG00000073987.4 |

| | | | | | | | | | | | | | | | |
|---|---|---|---|---|---|---|---|---|---|---|---|---|---|---|---|
| chr9 | 65101038 | 65103015 | 25.71461 | 72.27271 | 69.25061 | Promoter | 9 | 65101486 | 65137940 | 36455 | 1 | ENSMUST00 | 0 | Igdcc4 | ENSMUSG00000032816.9 |
| chr3 | 108443858 | 108445673 | 22.63099 | 72.22495 | 69.20312 | Promoter | 3 | 108424865 | 108445198 | 20334 | 2 | ENSMUST00 | 0 | Sars | ENSMUSG00000068739.8 |
| chr8 | 46551979 | 46552654 | 22.63099 | 72.22495 | 69.20312 | Promoter | 8 | 46552055 | 46580007 | 27953 | 1 | ENSMUST00 | 0 | Cenpu | ENSMUSG00000031629.8 |
| chr8 | 120228411 | 120230255 | 25.66166 | 72.18176 | 69.16018 | Promoter | 8 | 120204434 | 120228230 | 23797 | 2 | ENSMUST00 | -182 | A330074K22I | ENSMUSG00000097960.1 |
| chr7 | 16842573 | 16844283 | 25.12873 | 72.16408 | 69.14261 | Promoter | 7 | 16842902 | 16870461 | 27560 | 1 | ENSMUST00 | 0 | Prkd2 | ENSMUSG00000041187.11 |
| chr17 | 46110628 | 46111724 | 24.58752 | 72.12553 | 69.10418 | Promoter | 17 | 46110986 | 46128910 | 17925 | 1 | ENSMUST00 | 0 | Mrps18a | ENSMUSG00000023967.4 |
| chr1 | 160213072 | 160213802 | 25.07903 | 72.07811 | 69.05697 | Promoter | 1 | 160202367 | 160212892 | 10526 | 2 | ENSMUST00 | -181 | Cacybp | ENSMUSG00000014226.5 |
| chr11 | 101983072 | 101985233 | 23.0285 | 72.05518 | 69.03415 | Promoter | 11 | 101971143 | 101983503 | 12361 | 2 | ENSMUST00 | 0 | Dusp3 | ENSMUSG00000003518.8 |
| chr11 | 69070136 | 69071381 | 23.0285 | 72.05518 | 69.03415 | Promoter | 11 | 69070806 | 69073030 | 2225 | 1 | ENSMUST00 | 0 | Tmem107 | ENSMUSG00000020895.9 |
| chr2 | 144164562 | 144166242 | 23.0285 | 72.05518 | 69.03415 | Distal Interge | 2 | 144174315 | 144179086 | 4772 | 2 | ENSMUST00 | 12844 | Gm5535 | ENSMUSG00000074758.4 |
| chr3 | 61002193 | 61003795 | 23.0285 | 72.05518 | 69.03415 | Promoter | 3 | 61002795 | 61008979 | 6185 | 1 | ENSMUST00 | 0 | P2ry1 | ENSMUSG00000027765.1 |
| chr17 | 33092120 | 33093457 | 20.67151 | 72.03884 | 69.0182 | Promoter | 17 | 33092458 | 33105058 | 12601 | 1 | ENSMUST00 | 0 | Zfp563 | ENSMUSG00000067424.7 |
| chr7 | 97480918 | 97481811 | 21.57688 | 72.03428 | 69.01365 | Promoter | 7 | 97480956 | 97541397 | 60442 | 1 | ENSMUST00 | 0 | Ints4 | ENSMUSG00000025133.8 |
| chr19 | 30174568 | 30175616 | 25.55642 | 72.00093 | 68.98035 | Promoter | 19 | 30098449 | 30175418 | 76970 | 2 | ENSMUST00 | 0 | Gldc | ENSMUSG00000024827.9 |
| chr8 | 13338544 | 13339623 | 25.55642 | 72.00093 | 68.98035 | Promoter | 8 | 13339674 | 13377702 | 38029 | 1 | ENSMUST00 | -51 | Tfdp1 | ENSMUSG00000038482.10 |
| chr15 | 82338111 | 82339414 | 23.45396 | 71.92415 | 68.90385 | Promoter | 15 | 82338959 | 82349059 | 10101 | 1 | ENSMUST00 | 0 | Smdt1 | ENSMUSG00000022452.9 |
| chr15 | 33686873 | 33688146 | 23.45396 | 71.92415 | 68.90385 | Promoter | 15 | 33683876 | 33687883 | 4008 | 2 | ENSMUST00 | 0 | Tspyl5 | ENSMUSG00000038984.3 |
| chr17 | 70848566 | 70853866 | 23.45396 | 71.92415 | 68.90385 | Promoter | 17 | 70844205 | 70849790 | 5586 | 2 | ENSMUST00 | 0 | Tgif1 | ENSMUSG00000047407.12 |
| chr7 | 38182798 | 38184390 | 23.45396 | 71.92415 | 68.90385 | Promoter | 7 | 38183217 | 38197563 | 14347 | 1 | ENSMUST00 | 0 | 1600014C10I | ENSMUSG00000054676.10 |
| chr3 | 59343873 | 59344690 | 25.50411 | 71.91103 | 68.89117 | Promoter | 3 | 59316735 | 59344256 | 27522 | 2 | ENSMUST00 | 0 | Igsf10 | ENSMUSG00000036334.7 |
| chr11 | 69063131 | 69064397 | 24.98023 | 71.90708 | 68.88776 | Distal Interge | 11 | 69057718 | 69060483 | 2766 | 2 | ENSMUST00 | -2649 | 9330160F10I | ENSMUSG00000072809.2 |
| chr1 | 132298350 | 132299550 | 19.74353 | 71.90538 | 68.88615 | Promoter | 1 | 132298626 | 132307357 | 8732 | 1 | ENSMUST00 | 0 | Klhdc8a | ENSMUSG00000042115.4 |
| chr1 | 179516989 | 179518360 | 19.74353 | 71.90538 | 68.88615 | Promoter | 1 | 178951960 | 179518034 | 566075 | 2 | ENSMUST00 | 0 | Smyd3 | ENSMUSG00000055067.10 |
| chr11 | 101442399 | 101443270 | 19.74353 | 71.90538 | 68.88615 | Promoter | 11 | 101442438 | 101443934 | 1497 | 1 | ENSMUST00 | 0 | Rpl27 | ENSMUSG00000063316.8 |
| chr13 | 103773260 | 103775065 | 19.74353 | 71.90538 | 68.88615 | Distal Interge | 13 | 103741615 | 103764504 | 22890 | 2 | ENSMUST00 | -8757 | Srek1 | ENSMUSG00000032621.7 |
| chr15 | 77969917 | 77971584 | 19.74353 | 71.90538 | 68.88615 | Promoter | 15 | 77958998 | 77970824 | 11827 | 2 | ENSMUST00 | 0 | Eif3d | ENSMUSG00000016554.8 |
| chr17 | 32992675 | 32994158 | 19.74353 | 71.90538 | 68.88615 | Promoter | 17 | 32993129 | 32997582 | 4454 | 1 | ENSMUST00 | 0 | Zfp952 | ENSMUSG00000053390.10 |
| chr6 | 126848088 | 126849679 | 19.74353 | 71.90538 | 68.88615 | Promoter | 6 | 126821863 | 126849144 | 27282 | 2 | ENSMUST00 | 0 | Ndufa9 | ENSMUSG00000000399.7 |
| chr7 | 117380196 | 117382139 | 19.74353 | 71.90538 | 68.88615 | Promoter | 7 | 117380979 | 117635185 | 254207 | 1 | ENSMUST00 | 0 | Xylt1 | ENSMUSG00000030657.6 |
| chr7 | 34229243 | 34230723 | 19.74353 | 71.90538 | 68.88615 | Promoter | 7 | 34202122 | 34230281 | 28160 | 2 | ENSMUST00 | 0 | Gpi1 | ENSMUSG00000036427.4 |
| chr11 | 29514307 | 29515477 | 23.91045 | 71.83717 | 68.8187 | Promoter | 11 | 29511757 | 29515031 | 3275 | 2 | ENSMUST00 | 0 | Prorsd1 | ENSMUSG00000032673.5 |
| chr17 | 25920218 | 25921077 | 23.91045 | 71.83717 | 68.8187 | Promoter | 17 | 25883639 | 25919727 | 36089 | 2 | ENSMUST00 | -492 | Rab40c | ENSMUSG00000025730.9 |
| chr12 | 102282779 | 102284358 | 24.93112 | 71.82204 | 68.80418 | Promoter | 12 | 102283048 | 102390855 | 107808 | 1 | ENSMUST00 | 0 | Rin3 | ENSMUSG00000044456.11 |
| chr12 | 69363809 | 69364743 | 24.93112 | 71.82204 | 68.80418 | Distal Interge | 12 | 69361182 | 69361481 | 300 | 2 | ENSMUST00 | -2329 | Gm26191 | ENSMUSG00000099250.1 |
| chr13 | 74563363 | 74564601 | 24.93112 | 71.82204 | 68.80418 | Distal Interge | 13 | 74639872 | 74691875 | 52004 | 1 | ENSMUST00 | -75271 | Erap1 | ENSMUSG00000021583.6 |
| chr14 | 62555064 | 62556501 | 25.45202 | 71.82146 | 68.80418 | Promoter | 14 | 62555737 | 62588523 | 32787 | 1 | ENSMUST00 | 0 | Fam124a | ENSMUSG00000035184.10 |
| chr15 | 79803764 | 79805110 | 24.93112 | 71.82204 | 68.80418 | Promoter | 15 | 79786352 | 79804709 | 18358 | 2 | ENSMUST00 | 0 | Nptxr | ENSMUSG00000022421.13 |
| chr17 | 42315825 | 42317062 | 24.93112 | 71.82204 | 68.80418 | Promoter | 17 | 42315947 | 42507741 | 191795 | 1 | ENSMUST00 | 0 | Ptchd4 | ENSMUSG00000042256.4 |
| chr19 | 5040687 | 5042027 | 25.45202 | 71.82146 | 68.80418 | Promoter | 19 | 5041404 | 5049917 | 8514 | 1 | ENSMUST00 | 0 | Brms1 | ENSMUSG00000080268.3 |
| chr6 | 72538340 | 72538858 | 24.93112 | 71.82204 | 68.80418 | Distal Interge | 6 | 72544391 | 72562983 | 18593 | 1 | ENSMUST00 | -5533 | Capg | ENSMUSG00000056737.9 |
| chr7 | 24481368 | 24483109 | 24.93112 | 71.82204 | 68.80418 | Promoter | 7 | 24482023 | 24504539 | 22517 | 1 | ENSMUST00 | 0 | Cadm4 | ENSMUSG00000054793.7 |
| chr7 | 27355311 | 27357563 | 24.93112 | 71.82204 | 68.80418 | Promoter | 7 | 27342133 | 27356019 | 13887 | 2 | ENSMUST00 | 0 | Shkbp1 | ENSMUSG00000089832.3 |
| chr8 | 70629543 | 70630276 | 24.93112 | 71.82204 | 68.80418 | Exon (ENSMU | 8 | 70629393 | 70632095 | 2703 | 2 | ENSMUST00 | 1819 | Gdf15 | ENSMUSG00000038508.7 |
| chr9 | 60836859 | 60842659 | 24.93112 | 71.82204 | 68.80418 | Promoter | 9 | 60821855 | 60838200 | 16346 | 2 | ENSMUST00 | 0 | Gm9869 | ENSMUSG00000052143.7 |
| chr13 | 62129321 | 62130457 | 24.40147 | 71.80055 | 68.78354 | Promoter | 13 | 62129916 | 62173419 | 43504 | 1 | ENSMUST00 | 0 | Zfp808 | ENSMUSG00000074867.3 |
| chr7 | 67758534 | 67759924 | 25.40015 | 71.73223 | 68.71624 | Promoter | 7 | 67730161 | 67759742 | 29582 | 2 | ENSMUST00 | 0 | Synm | ENSMUSG00000030554.10 |
| chr16 | 13667412 | 13668408 | 18.82282 | 71.67798 | 68.66222 | Promoter | 16 | 13537964 | 13668170 | 130207 | 2 | ENSMUST00 | 0 | Parn | ENSMUSG00000022685.8 |
| chr4 | 126609373 | 126610249 | 18.82282 | 71.67798 | 68.66222 | Promoter | 4 | 126609818 | 126614371 | 4554 | 1 | ENSMUST00 | 0 | 5730409E04I | ENSMUSG00000073755.4 |
| chr7 | 100371160 | 100372007 | 18.82282 | 71.67798 | 68.66222 | Promoter | 7 | 100326737 | 100371896 | 45160 | 2 | ENSMUST00 | 0 | Ppme1 | ENSMUSG00000030718.8 |
| chr8 | 123235367 | 123236593 | 18.82282 | 71.67798 | 68.66222 | Promoter | 8 | 123232255 | 123236206 | 3952 | 2 | ENSMUST00 | 0 | Spata2l | ENSMUSG00000033594.10 |
| chr9 | 108569535 | 108570677 | 18.82282 | 71.67798 | 68.66222 | Promoter | 9 | 108569892 | 108572771 | 2880 | 1 | ENSMUST00 | 0 | Dalrd3 | ENSMUSG00000019039.8 |
| chr13 | 45078716 | 45079143 | 17.33535 | 71.67154 | 68.65611 | Promoter | 13 | 45078498 | 45079127 | 630 | 2 | ENSMUST00 | 0 | Hsp25-ps1 | ENSMUSG00000078915.1 |
| chr14 | 20660725 | 20662033 | 24.83347 | 71.65281 | 68.63744 | Exon (ENSMU | 14 | 20658948 | 20664574 | 5627 | 2 | ENSMUST00 | 2541 | Synpo2l | ENSMUSG00000039376.8 |
| chr2 | 32570777 | 32571537 | 24.83347 | 71.65281 | 68.63744 | Promoter | 2 | 32570858 | 32573563 | 2706 | 1 | ENSMUST00 | 0 | Dpm2 | ENSMUSG00000026810.7 |
| chr19 | 26845437 | 26845941 | 25.34849 | 71.64335 | 68.62808 | Distal Interge | 19 | 26768801 | 26823907 | 55107 | 2 | ENSMUST00 | -21531 | 4931403E22I | ENSMUSG00000093772.1 |
| chr13 | 76017591 | 76019304 | 24.3095 | 71.63969 | 68.62457 | Promoter | 13 | 76001535 | 76018562 | 17028 | 2 | ENSMUST00 | 0 | Rfesd | ENSMUSG00000043190.8 |
| chr10 | 77978010 | 77979172 | 18.02817 | 71.61055 | 68.59548 | Promoter | 10 | 77978524 | 77985434 | 6911 | 1 | ENSMUST00 | 0 | 1810043G02 | ENSMUSG00000020284.11 |
| chr13 | 94057297 | 94058880 | 18.02817 | 71.61055 | 68.59548 | Promoter | 13 | 94057796 | 94195409 | 137614 | 1 | ENSMUST00 | 0 | Lhfpl2 | ENSMUSG00000045312.6 |
| chr18 | 67390202 | 67391868 | 18.02817 | 71.61055 | 68.59548 | Promoter | 18 | 67390717 | 67402748 | 12032 | 1 | ENSMUST00 | 0 | Tubb6 | ENSMUSG00000001473.6 |
| chr4 | 128654558 | 128655651 | 18.02817 | 71.61055 | 68.59548 | Promoter | 4 | 128654702 | 128752881 | 98180 | 1 | ENSMUST00 | 0 | Phc2 | ENSMUSG00000028796.12 |
| chr4 | 120666281 | 120667632 | 18.02817 | 71.61055 | 68.59548 | Promoter | 4 | 120666572 | 120667820 | 1249 | 1 | ENSMUST00 | 0 | Cited4 | ENSMUSG00000070803.6 |
| chr11 | 101552053 | 101552666 | 25.29703 | 71.55479 | 68.54001 | Promoter | 11 | 101552149 | 101581951 | 29803 | 1 | ENSMUST00 | 0 | Nbr1 | ENSMUSG00000017119.14 |
| chr7 | 18990641 | 18992635 | 25.29703 | 71.55479 | 68.54001 | Promoter | 7 | 18991245 | 18992496 | 1252 | 1 | ENSMUST00 | 0 | Mypop | ENSMUSG00000048481.10 |
| chr11 | 86256732 | 86258058 | 21.76566 | 71.50774 | 68.49327 | Promoter | 11 | 86210681 | 86257550 | 46870 | 2 | ENSMUST00 | 0 | Ints2 | ENSMUSG00000018068.8 |
| chr11 | 114065630 | 114067369 | 21.76566 | 71.50774 | 68.49327 | Promoter | 11 | 113776374 | 114066218 | 289845 | 2 | ENSMUST00 | 0 | Sdk2 | ENSMUSG00000041592.11 |
| chr15 | 27465975 | 27467367 | 21.76566 | 71.50774 | 68.49327 | Promoter | 15 | 27466677 | 27594909 | 128233 | 1 | ENSMUST00 | 0 | Ank | ENSMUSG00000022265.6 |
| chr17 | 24527234 | 24527975 | 21.76566 | 71.50774 | 68.49327 | Promoter | 17 | 24508562 | 24527938 | 19377 | 2 | ENSMUST00 | 0 | Traf7 | ENSMUSG00000052752.10 |
| chr18 | 30272433 | 30273733 | 21.76566 | 71.50774 | 68.49327 | Promoter | 18 | 30272747 | 30348126 | 75380 | 1 | ENSMUST00 | 0 | Pik3c3 | ENSMUSG00000033628.10 |
| chr2 | 122610739 | 122611707 | 21.76566 | 71.50774 | 68.49327 | Promoter | 2 | 122594467 | 122611303 | 16837 | 2 | ENSMUST00 | 0 | Gatm | ENSMUSG00000027199.9 |
| chr2 | 136678336 | 136680011 | 21.76566 | 71.50774 | 68.49327 | Distal Interge | 2 | 136713453 | 136782428 | 68976 | 1 | ENSMUST00 | -33442 | Snap25 | ENSMUSG00000027273.8 |
| chr2 | 150909203 | 150910351 | 21.76566 | 71.50774 | 68.49327 | Promoter | 2 | 150909594 | 150931280 | 21687 | 1 | ENSMUST00 | 0 | Gins1 | ENSMUSG00000027454.5 |
| chr3 | 116807428 | 116808817 | 21.76566 | 71.50774 | 68.49327 | Promoter | 3 | 116739999 | 116808166 | 68168 | 2 | ENSMUST00 | 0 | Agl | ENSMUSG00000033400.9 |
| chr3 | 110141967 | 110143972 | 21.76566 | 71.50774 | 68.49327 | Promoter | 3 | 109780040 | 110143004 | 362965 | 2 | ENSMUST00 | 0 | Ntng1 | ENSMUSG00000059857.10 |
| chr4 | 133599275 | 133600441 | 21.76566 | 71.50774 | 68.49327 | Downstream | 4 | 133600556 | 133602168 | 1613 | 2 | ENSMUST00 | 1727 | Sfn | ENSMUSG00000047281.3 |
| chr7 | 43561859 | 43563041 | 21.76566 | 71.50774 | 68.49327 | Promoter | 7 | 43562256 | 43575461 | 13206 | 1 | ENSMUST00 | 0 | Zfp658 | ENSMUSG00000056592.9 |
| chr7 | 12834261 | 12835688 | 21.76566 | 71.50774 | 68.49327 | Promoter | 7 | 12834761 | 12850577 | 15817 | 1 | ENSMUST00 | 0 | Zfp110 | ENSMUSG00000058638.8 |
| chr8 | 69749288 | 69750483 | 21.76566 | 71.50774 | 68.49327 | Promoter | 8 | 69741639 | 69749962 | 8324 | 2 | ENSMUST00 | 0 | Zfp963 | ENSMUSG00000092260.2 |
| chr8 | 83874821 | 83876133 | 21.76566 | 71.50774 | 68.49327 | Distal Interge | 8 | 83887079 | 83899501 | 12423 | 2 | ENSMUST00 | 23368 | Gm26721 | ENSMUSG00000096943.1 |
| chr10 | 80141816 | 80143104 | 25.7338 | 71.40817 | 68.39619 | Promoter | 10 | 80142315 | 80145818 | 3504 | 1 | ENSMUST00 | 0 | Atp5d | ENSMUSG00000003072.10 |
| chr10 | 81267921 | 81269195 | 25.7338 | 71.40817 | 68.39619 | Promoter | 10 | 81268172 | 81273246 | 5075 | 1 | ENSMUST00 | 0 | Apba3 | ENSMUSG00000004931.10 |
| chr10 | 127188230 | 127190051 | 25.7338 | 71.40817 | 68.39619 | Promoter | 10 | 127182521 | 127189823 | 7303 | 2 | ENSMUST00 | 0 | Arhgef25 | ENSMUSG00000019467.8 |
| chr11 | 102761199 | 102762462 | 25.7338 | 71.40817 | 68.39619 | Promoter | 11 | 102761439 | 102780262 | 18824 | 1 | ENSMUST00 | 0 | Adam11 | ENSMUSG00000020926.11 |
| chr11 | 102446437 | 102447794 | 25.7338 | 71.40817 | 68.39619 | Promoter | 11 | 102436981 | 102447682 | 10702 | 2 | ENSMUST00 | 0 | Fam171a2 | ENSMUSG00000034685.4 |
| chr12 | 54738455 | 54739299 | 25.7338 | 71.40817 | 68.39619 | Promoter | 12 | 54738459 | 54738622 | 164 | 2 | ENSMUST00 | 0 | Gm22317 | ENSMUSG00000096206.1 |
| chr14 | 69502012 | 69503357 | 25.7338 | 71.40817 | 68.39619 | Promoter | 14 | 69363504 | 69503320 | 139817 | 2 | ENSMUST00 | 0 | Gm28049 | ENSMUSG00000093954.3 |
| chr15 | 83526301 | 83527847 | 25.7338 | 71.40817 | 68.39619 | Promoter | 15 | 83526862 | 83544635 | 17774 | 1 | ENSMUST00 | 0 | Bik | ENSMUSG00000016758.3 |
| chr17 | 35516164 | 35517735 | 25.7338 | 71.40817 | 68.39619 | Promoter | 17 | 35517100 | 35530996 | 13897 | 1 | ENSMUST00 | 0 | Cchcr1 | ENSMUSG00000040312.9 |
| chr2 | 86019949 | 86020925 | 25.7338 | 71.40817 | 68.39619 | Distal Interge | 2 | 86007778 | 86008710 | 933 | 1 | ENSMUST00 | 12172 | Olfr1032 | ENSMUSG00000042796.1 |
| chr2 | 151701955 | 151703200 | 25.7338 | 71.40817 | 68.39619 | Promoter | 2 | 151702008 | 151707310 | 5303 | 1 | ENSMUST00 | 0 | Tmem74b | ENSMUSG00000044364.5 |
| chr2 | 150362205 | 150363193 | 25.7338 | 71.40817 | 68.39619 | Promoter | 2 | 150310935 | 150362765 | 51831 | 2 | ENSMUST00 | 0 | 3300002I08R | ENSMUSG00000063364.5 |
| chr4 | 134370510 | 134372917 | 25.7338 | 71.40817 | 68.39619 | Promoter | 4 | 134356372 | 134372558 | 16187 | 2 | ENSMUST00 | 0 | Extl1 | ENSMUSG00000028838.6 |
| chr5 | 52470936 | 52471906 | 25.7338 | 71.40817 | 68.39619 | Promoter | 5 | 52374651 | 52471543 | 96893 | 2 | ENSMUST00 | 0 | Ccdc149 | ENSMUSG00000045790.5 |
| chr5 | 37822966 | 37824987 | 25.7338 | 71.40817 | 68.39619 | Promoter | 5 | 37820485 | 37824583 | 4099 | 2 | ENSMUST00 | 0 | Msx1 | ENSMUSG00000048450.10 |
| chr7 | 132610198 | 132611085 | 25.7338 | 71.40817 | 68.39619 | Promoter | 7 | 132610638 | 132630626 | 19989 | 1 | ENSMUST00 | 0 | Lhpp | ENSMUSG00000030946.7 |
| chr7 | 12421962 | 12422969 | 25.7338 | 71.40817 | 68.39619 | Promoter | 7 | 12415153 | 12422751 | 7599 | 2 | ENSMUST00 | 0 | Zfp551 | ENSMUSG00000034071.9 |
| chrX | 71555539 | 71557518 | 25.7338 | 71.40817 | 68.39619 | Promoter | 20 | 71555918 | 71560676 | 4759 | 1 | ENSMUST00 | 0 | Hmgb3 | ENSMUSG00000015217.6 |
| chrX | 6399688 | 6401191 | 25.7338 | 71.40817 | 68.39619 | Promoter | 20 | 6399854 | 6633837 | 233984 | 1 | ENSMUST00 | 0 | Shroom4 | ENSMUSG00000068270.10 |
| chr1 | 135728677 | 135729930 | 25.19474 | 71.37865 | 68.36819 | Promoter | 1 | 135729147 | 135752232 | 23086 | 1 | ENSMUST00 | 0 | Csrp1 | ENSMUSG00000026421.9 |
| chr1 | 193172724 | 193174122 | 22.64381 | 71.35664 | 68.34631 | Promoter | 1 | 193173469 | 193176298 | 2830 | 1 | ENSMUST00 | 0 | A130010J15I | ENSMUSG00000079144.6 |
| chr19 | 47578934 | 47580860 | 22.64381 | 71.35664 | 68.34631 | Promoter | 19 | 47579678 | 47645246 | 65569 | 1 | ENSMUST00 | 0 | Slk | ENSMUSG00000025060.9 |
| chr3 | 121171056 | 121172163 | 22.64381 | 71.35664 | 68.34631 | Promoter | 3 | 121155400 | 121171695 | 16296 | 2 | ENSMUST00 | 0 | Rwdd3 | ENSMUSG00000028133.12 |
| chr4 | 57433813 | 57434810 | 22.64381 | 71.35664 | 68.34631 | Promoter | 4 | 57434247 | 57712016 | 277770 | 1 | ENSMUST00 | 0 | Palm2 | ENSMUSG00000090053.4 |
| chr8 | 84871771 | 84873009 | 25.63373 | 71.23676 | 68.22707 | Promoter | 8 | 84872111 | 84887446 | 15336 | 1 | ENSMUST00 | 0 | Syce2 | ENSMUSG00000003824.7 |
| chr7 | 137313067 | 137314953 | 25.09328 | 71.20383 | 68.1943 | Promoter | 7 | 137193673 | 137314445 | 120773 | 2 | ENSMUST00 | 0 | Ebf3 | ENSMUSG00000010476.8 |
| chr10 | 11342945 | 11344338 | 23.04863 | 71.19738 | 68.188 | Promoter | 10 | 11343404 | 11459644 | 116241 | 1 | ENSMUST00 | 0 | Epm2a | ENSMUSG00000055493.4 |
| chr17 | 56756832 | 56757982 | 23.04863 | 71.19738 | 68.188 | Promoter | 17 | 56751325 | 56757530 | 6206 | 2 | ENSMUST00 | 0 | Nrtn | ENSMUSG00000039481.5 |
| chr19 | 5963624 | 5964739 | 23.04863 | 71.19738 | 68.188 | Promoter | 19 | 5940543 | 5964206 | 23664 | 2 | ENSMUST00 | 0 | Pola2 | ENSMUSG00000024833.9 |
| chr3 | 88297064 | 88298281 | 23.04863 | 71.19738 | 68.188 | Promoter | 3 | 88297116 | 88321767 | 24652 | 1 | ENSMUST00 | 0 | Cct3 | ENSMUSG00000001416.8 |
| chr2 | 153741215 | 153742321 | 24.0377 | 71.16332 | 68.15427 | Promoter | 2 | 153741274 | 153773310 | 32037 | 1 | ENSMUST00 | 0 | Mapre1 | ENSMUSG00000027479.9 |
| chr8 | 33516961 | 33517709 | 25.58018 | 71.14499 | 68.13602 | Promoter | 8 | 33517324 | 33573558 | 56235 | 1 | ENSMUST00 | 0 | Tex15 | ENSMUSG00000009628.9 |
| chr1 | 91250241 | 91251404 | 15.50981 | 71.12668 | 68.11806 | Promoter | 1 | 91250304 | 91275479 | 25176 | 1 | ENSMUST00 | 0 | Ube2f | ENSMUSG00000034343.9 |
| chr12 | 69361531 | 69362179 | 25.04286 | 71.11688 | 68.10826 | Promoter | 12 | 69361182 | 69361481 | 300 | 2 | ENSMUST00 | -51 | Gm26191 | ENSMUSG00000099250.1 |
| chr13 | 40730137 | 40731718 | 25.04286 | 71.11688 | 68.10826 | Promoter | 13 | 40715675 | 40730418 | 14744 | 2 | ENSMUST00 | 0 | Tfap2a | ENSMUSG00000021359.10 |
| chr14 | 118179999 | 118181108 | 25.04286 | 71.11688 | 68.10826 | Distal Interge | 14 | 118137127 | 118164232 | 27106 | 1 | ENSMUST00 | 42873 | Gpr180 | ENSMUSG00000022131.2 |
| chr18 | 54989542 | 54993168 | 25.04286 | 71.11688 | 68.10826 | Promoter | 18 | 54888045 | 54990180 | 102136 | 2 | ENSMUST00 | 0 | Zfp608 | ENSMUSG00000052713.8 |
| chr3 | 69598180 | 69599335 | 23.48243 | 71.0788 | 68.07035 | Promoter | 3 | 69574186 | 69598861 | 24676 | 2 | ENSMUST00 | 0 | B3galnt1 | ENSMUSG00000043300.1 |
| chr7 | 19149437 | 19150428 | 23.48243 | 71.0788 | 68.07035 | Promoter | 7 | 19149722 | 19152735 | 3014 | 1 | ENSMUST00 | 0 | Snrpd2 | ENSMUSG00000040824.2 |
| chr9 | 51103329 | 51104096 | 23.48243 | 71.0788 | 68.07035 | Promoter | 9 | 51103562 | 51103645 | 84 | 2 | ENSMUST00 | 0 | Mir34b | ENSMUSG00000065492.1 |
| chr6 | 119107305 | 119109398 | 25.52684 | 71.05358 | 68.04573 | Promoter | 6 | 118592464 | 119108189 | 515726 | 2 | ENSMUST00 | 0 | Cacna1c | ENSMUSG00000051331.9 |
| chr10 | 74966689 | 74967900 | 24.99264 | 71.03027 | 68.02251 | Promoter | 10 | 74967177 | 75016907 | 49731 | 1 | ENSMUST00 | 0 | Gnaz | ENSMUSG00000040009.5 |
| chr10 | 13473618 | 13475037 | 24.99264 | 71.03027 | 68.02251 | Promoter | 10 | 13213395 | 13474396 | 261002 | 2 | ENSMUST00 | 0 | Phactr2 | ENSMUSG00000062866.10 |
| chr17 | 21383356 | 21384429 | 24.99264 | 71.03027 | 68.02251 | Promoter | 17 | 21383748 | 21399262 | 15515 | 1 | ENSMUST00 | 0 | Zfp677 | ENSMUSG00000062743.4 |
| chr4 | 141544698 | 141545900 | 24.99264 | 71.03027 | 68.02251 | Promoter | 4 | 141546189 | 141547892 | 1704 | 1 | ENSMUST00 | -289 | B330016D10 | ENSMUSG00000048406.5 |
| chr4 | 130519734 | 130520891 | 24.99264 | 71.03027 | 68.02251 | Distal Interge | 4 | 130539063 | 130557988 | 18926 | 2 | ENSMUST00 | 37097 | Nkain1 | ENSMUSG00000078532.4 |
| chr5 | 102860005 | 102862831 | 24.99264 | 71.03027 | 68.02251 | Exon (ENSMU | 5 | 102845007 | 102897829 | 52823 | 1 | ENSMUST00 | 14999 | Arhgap24 | ENSMUSG00000057315.9 |
| chr6 | 56831538 | 56832862 | 24.99264 | 71.03027 | 68.02251 | Promoter | 6 | 56832059 | 56879358 | 47300 | 1 | ENSMUST00 | 0 | Fkbp9 | ENSMUSG00000029781.7 |
| chr8 | 15059401 | 15062365 | 24.99264 | 71.03027 | 68.02251 | Promoter | 8 | 15060190 | 15073840 | 13651 | 1 | ENSMUST00 | 0 | Myom2 | ENSMUSG00000031461.4 |
| chr8 | 90828109 | 90829753 | 24.99264 | 71.03027 | 68.02251 | Promoter | 8 | 90828835 | 91054508 | 225674 | 1 | ENSMUST00 | 0 | Chd9 | ENSMUSG00000056608.8 |

| | | | | | | | | | | | | | | | |
|---|---|---|---|---|---|---|---|---|---|---|---|---|---|---|---|
| chr1 | 60179757 | 60180364 | 23.94845 | 71.00658 | 67.99948 | Promoter | 1 | 60180599 | 60338328 | 157730 | 1 | ENSMUST00 | -235 | Nbeal1 | ENSMUSG00000073664.6 |
| chr11 | 69045318 | 69046399 | 23.94845 | 71.00658 | 67.99948 | Promoter | 11 | 69045647 | 69051664 | 6018 | 1 | ENSMUST00 | 0 | Aurkb | ENSMUSG00000020897.7 |
| chr15 | 83724317 | 83725388 | 23.94845 | 71.00658 | 67.99948 | Promoter | 15 | 83602583 | 83725021 | 122439 | 2 | ENSMUST00 | 0 | Scube1 | ENSMUSG00000016763.9 |
| chr17 | 66123165 | 66124386 | 23.94845 | 71.00658 | 67.99948 | Promoter | 17 | 66123520 | 66152167 | 28648 | 1 | ENSMUST00 | 0 | Ddx11 | ENSMUSG00000035842.9 |
| chr17 | 33393847 | 33395161 | 23.94845 | 71.00658 | 67.99948 | Promoter | 17 | 33360507 | 33394378 | 33872 | 2 | ENSMUST00 | 0 | Zfp101 | ENSMUSG00000055240.11 |
| chr3 | 84479213 | 84481047 | 23.94845 | 71.00658 | 67.99948 | Promoter | 3 | 84444035 | 84479491 | 35457 | 2 | ENSMUST00 | 0 | Fhdc1 | ENSMUSG00000041842.10 |
| chr1 | 127676929 | 127678248 | 24.45041 | 70.98759 | 67.98131 | Promoter | 1 | 127486546 | 127678096 | 191551 | 2 | ENSMUST00 | 0 | Tmem163 | ENSMUSG00000026347.8 |
| chr11 | 23254787 | 23255543 | 24.45041 | 70.98759 | 67.98131 | Promoter | 11 | 23256041 | 23297597 | 41557 | 1 | ENSMUST00 | -498 | Xpo1 | ENSMUSG00000020290.9 |
| chr16 | 64872848 | 64874373 | 24.45041 | 70.98759 | 67.98131 | Distal Interge | 16 | 64852702 | 64855986 | 3285 | 1 | ENSMUST00 | 20147 | Cggbp1 | ENSMUSG00000054604.7 |
| chr4 | 144891998 | 144895332 | 25.47373 | 70.96252 | 67.95692 | Promoter | 4 | 144892827 | 144927642 | 34816 | 1 | ENSMUST00 | 0 | Dhrs3 | ENSMUSG00000066026.9 |
| chr9 | 96888740 | 96890149 | 24.94261 | 70.94398 | 67.93848 | Promoter | 9 | 96823336 | 96889413 | 66078 | 2 | ENSMUST00 | 0 | Pxylp1 | ENSMUSG00000043587.10 |
| chr1 | 134075527 | 134079138 | 22.90469 | 70.93823 | 67.93287 | Promoter | 1 | 134075170 | 134079120 | 3951 | 2 | ENSMUST00 | 0 | Btg2 | ENSMUSG00000020423.6 |
| chr10 | 82858525 | 82860999 | 22.90469 | 70.93823 | 67.93287 | Promoter | 10 | 82859206 | 82897724 | 38519 | 1 | ENSMUST00 | 0 | Txnrd1 | ENSMUSG00000020250.9 |
| chr11 | 69604857 | 69606538 | 22.90469 | 70.93823 | 67.93287 | Promoter | 11 | 69599736 | 69605829 | 6094 | 2 | ENSMUST00 | 0 | Atp1b2 | ENSMUSG00000041329.8 |
| chr15 | 36938198 | 36939286 | 22.90469 | 70.93823 | 67.93287 | Distal Interge | 15 | 36870501 | 36879816 | 9316 | 2 | ENSMUST00 | -58383 | Gm10384 | ENSMUSG00000072618.2 |
| chr16 | 35870459 | 35872171 | 22.90469 | 70.93823 | 67.93287 | Promoter | 16 | 35832874 | 35871544 | 38671 | 2 | ENSMUST00 | 0 | Parp14 | ENSMUSG00000034422.9 |
| chr16 | 14158558 | 14159239 | 22.90469 | 70.93823 | 67.93287 | Promoter | 16 | 14109166 | 14159274 | 50109 | 2 | ENSMUST00 | 35 | Marf1 | ENSMUSG00000060657.7 |
| chr17 | 7945193 | 7946321 | 22.90469 | 70.93823 | 67.93287 | Promoter | 17 | 7945653 | 7979824 | 34172 | 1 | ENSMUST00 | 0 | Rsph3a | ENSMUSG00000073471.2 |
| chr18 | 42261214 | 42262776 | 22.90469 | 70.93823 | 67.93287 | Promoter | 18 | 42202350 | 42262122 | 59773 | 2 | ENSMUST00 | 0 | Lars | ENSMUSG00000024493.8 |
| chr2 | 117342204 | 117343178 | 22.90469 | 70.93823 | 67.93287 | Promoter | 2 | 117279993 | 117343001 | 63009 | 2 | ENSMUST00 | 0 | Rasgrp1 | ENSMUSG00000027347.13 |
| chr2 | 144594009 | 144594865 | 22.90469 | 70.93823 | 67.93287 | Promoter | 2 | 144594054 | 144595366 | 1313 | 1 | ENSMUST00 | 0 | Gm561 | ENSMUSG00000074754.4 |
| chr2 | 175337669 | 175338313 | 22.90469 | 70.93823 | 67.93287 | Promoter | 2 | 175321955 | 175338197 | 16243 | 2 | ENSMUST00 | 0 | Gm4631 | ENSMUSG00000078899.4 |
| chr2 | 116975328 | 116976313 | 22.90469 | 70.93823 | 67.93287 | Promoter | 2 | 116972575 | 116976428 | 3854 | 2 | ENSMUST00 | 115 | RP23-326F22 | ENSMUSG00000099470.1 |
| chr3 | 7612514 | 7613917 | 22.90469 | 70.93823 | 67.93287 | Promoter | 3 | 7572077 | 7613427 | 41351 | 2 | ENSMUST00 | 0 | Il7 | ENSMUSG00000040329.9 |
| chr5 | 96373764 | 96375990 | 22.90469 | 70.93823 | 67.93287 | Promoter | 5 | 96373955 | 96784728 | 410774 | 1 | ENSMUST00 | 0 | Fras1 | ENSMUSG00000034687.4 |
| chr5 | 31295128 | 31296346 | 22.90469 | 70.93823 | 67.93287 | Promoter | 5 | 31292255 | 31295877 | 3623 | 2 | ENSMUST00 | 0 | Fndc4 | ENSMUSG00000038552.9 |
| chr7 | 3703579 | 3704989 | 22.90469 | 70.93823 | 67.93287 | Promoter | 7 | 3703993 | 3706901 | 2909 | 1 | ENSMUST00 | 0 | Rps9 | ENSMUSG00000006333.9 |
| chr7 | 118115197 | 118116797 | 22.90469 | 70.93823 | 67.93287 | Promoter | 7 | 118104378 | 118115256 | 10879 | 2 | ENSMUST00 | 0 | Rps15a | ENSMUSG00000008683.11 |
| chr7 | 128246452 | 128247624 | 22.90469 | 70.93823 | 67.93287 | Promoter | 7 | 128246812 | 128253711 | 6900 | 1 | ENSMUST00 | 0 | Tgfb1i1 | ENSMUSG00000030782.10 |
| chr7 | 6171255 | 6173219 | 22.90469 | 70.93823 | 67.93287 | Promoter | 7 | 6172226 | 6183951 | 11726 | 1 | ENSMUST00 | 0 | Zfp444 | ENSMUSG00000044876.9 |
| chr7 | 19119792 | 19120605 | 22.90469 | 70.93823 | 67.93287 | Promoter | 7 | 19119859 | 19138261 | 18403 | 1 | ENSMUST00 | 0 | Fbxo46 | ENSMUSG00000050428.6 |
| chr9 | 21130643 | 21131837 | 22.90469 | 70.93823 | 67.93287 | Distal Interge | 9 | 21104069 | 21127346 | 23278 | 2 | ENSMUST00 | -3298 | Tyk2 | ENSMUSG00000032175.10 |
| chrX | 36873986 | 36875275 | 22.90469 | 70.93823 | 67.93287 | Promoter | 20 | 36848550 | 36874111 | 25562 | 2 | ENSMUST00 | 0 | C330007P06I | ENSMUSG00000006423.10 |
| chr1 | 88205233 | 88206133 | 24.40335 | 70.90572 | 67.90328 | Promoter | 1 | 88204732 | 88205782 | 1051 | 2 | ENSMUST00 | 0 | Dnajb3 | ENSMUSG00000081984.3 |
| chr8 | 85699887 | 85701561 | 24.89279 | 70.85799 | 67.85581 | Promoter | 8 | 85700905 | 85701021 | 117 | 1 | ENSMUST00 | 0 | Mir8109 | ENSMUSG00000098388.1 |
| chr1 | 86433982 | 86435081 | 20.5121 | 70.85588 | 67.85384 | Distal Interge | 1 | 86426329 | 86428052 | 1724 | 1 | ENSMUST00 | 7654 | 1700019O17 | ENSMUSG00000036574.5 |
| chr10 | 67979029 | 67980169 | 20.5121 | 70.85588 | 67.85384 | Promoter | 10 | 67979570 | 68043868 | 64299 | 1 | ENSMUST00 | 0 | Rtkn2 | ENSMUSG00000037846.13 |
| chr11 | 60175495 | 60177680 | 20.5121 | 70.85588 | 67.85384 | Promoter | 11 | 60175879 | 60199197 | 23319 | 1 | ENSMUST00 | 0 | Rai1 | ENSMUSG00000062115.10 |
| chr12 | 71309332 | 71310789 | 20.5121 | 70.85588 | 67.85384 | Promoter | 12 | 71309884 | 71320107 | 10224 | 1 | ENSMUST00 | 0 | Dact1 | ENSMUSG00000044548.6 |
| chr16 | 94289487 | 94290497 | 20.5121 | 70.85588 | 67.85384 | Intron (ENSN | 16 | 94129306 | 94287856 | 158551 | 2 | ENSMUST00 | -1632 | Hlcs | ENSMUSG00000040820.10 |
| chr17 | 46751480 | 46752624 | 20.5121 | 70.85588 | 67.85384 | Promoter | 17 | 46735705 | 46752214 | 16510 | 2 | ENSMUST00 | 0 | Cnpy3 | ENSMUSG00000023973.8 |
| chr3 | 108200038 | 108201466 | 20.5121 | 70.85588 | 67.85384 | Promoter | 3 | 108195687 | 108200834 | 5148 | 2 | ENSMUST00 | 0 | Cyb561d1 | ENSMUSG00000048796.7 |
| chr6 | 83866959 | 83868257 | 20.5121 | 70.85588 | 67.85384 | Distal Interge | 6 | 83874379 | 83875380 | 1002 | 2 | ENSMUST00 | 7123 | Gm5138 | ENSMUSG00000079501.3 |
| chr7 | 4500798 | 4502864 | 20.5121 | 70.85588 | 67.85384 | Promoter | 7 | 4481520 | 4501361 | 19842 | 2 | ENSMUST00 | 0 | Ppp1r12c | ENSMUSG00000019254.11 |
| chr11 | 93967612 | 93968449 | 24.84317 | 70.77231 | 67.77174 | Promoter | 11 | 93960668 | 93968272 | 7605 | 2 | ENSMUST00 | 0 | Nme1 | ENSMUSG00000037601.6 |
| chr6 | 28479479 | 28480083 | 24.84317 | 70.77231 | 67.77174 | Promoter | 6 | 28480348 | 28877723 | 397376 | 1 | ENSMUST00 | -265 | Snd1 | ENSMUSG00000001424.9 |
| chr7 | 102210370 | 102211123 | 24.84317 | 70.77231 | 67.77174 | Promoter | 7 | 102210372 | 102226445 | 16074 | 1 | ENSMUST00 | 0 | Pgap2 | ENSMUSG00000030990.12 |
| chr11 | 90002186 | 90003289 | 24.30838 | 70.74035 | 67.73997 | Promoter | 11 | 89982665 | 90002894 | 20230 | 2 | ENSMUST00 | 0 | Pctp | ENSMUSG00000020553.8 |
| chr13 | 55152536 | 55153874 | 24.30838 | 70.74035 | 67.73997 | Promoter | 13 | 55152640 | 55168759 | 16120 | 1 | ENSMUST00 | 0 | Fgfr4 | ENSMUSG00000005320.9 |
| chr18 | 41950572 | 41951583 | 24.30838 | 70.74035 | 67.73997 | Promoter | 18 | 41875696 | 41951194 | 75499 | 2 | ENSMUST00 | 0 | Prelid2 | ENSMUSG00000056671.6 |
| chr18 | 12955151 | 12956564 | 24.30838 | 70.74035 | 67.73997 | Distal Interge | 18 | 12961499 | 12961805 | 307 | 1 | ENSMUST00 | -4935 | Gm26407 | ENSMUSG00000089129.1 |
| chr2 | 181241481 | 181242514 | 24.30838 | 70.74035 | 67.73997 | Promoter | 2 | 181227615 | 181242027 | 14413 | 2 | ENSMUST00 | 0 | Helz2 | ENSMUSG00000027580.12 |
| chr2 | 84886283 | 84888418 | 24.30838 | 70.74035 | 67.73997 | Promoter | 2 | 84871924 | 84886710 | 14787 | 2 | ENSMUST00 | 0 | Rtn4rl2 | ENSMUSG00000050896.7 |
| chr3 | 28262261 | 28263926 | 24.30838 | 70.74035 | 67.73997 | Promoter | 3 | 28263214 | 28670585 | 407372 | 1 | ENSMUST00 | 0 | Tnik | ENSMUSG00000027692.11 |
| chr4 | 155961847 | 155963148 | 24.30838 | 70.74035 | 67.73997 | Promoter | 4 | 155962318 | 155966629 | 4312 | 1 | ENSMUST00 | 0 | Fam132a | ENSMUSG00000023571.4 |
| chr4 | 102759017 | 102763519 | 24.30838 | 70.74035 | 67.73997 | Promoter | 4 | 102759222 | 102883347 | 124126 | 1 | ENSMUST00 | 0 | Sgip1 | ENSMUSG00000028524.16 |
| chr4 | 108031243 | 108032486 | 24.30838 | 70.74035 | 67.73997 | Promoter | 4 | 108014791 | 108032090 | 17300 | 2 | ENSMUST00 | 0 | Podn | ENSMUSG00000028600.10 |
| chr4 | 154139476 | 154141532 | 24.30838 | 70.74035 | 67.73997 | Promoter | 4 | 154056253 | 154140208 | 83956 | 2 | ENSMUST00 | 0 | Trp73 | ENSMUSG00000029026.11 |
| chr4 | 44234096 | 44234951 | 24.30838 | 70.74035 | 67.73997 | Promoter | 4 | 44152434 | 44233789 | 81356 | 2 | ENSMUST00 | -308 | Rnf38 | ENSMUSG00000035696.10 |
| chr5 | 137312954 | 137314692 | 24.30838 | 70.74035 | 67.73997 | Promoter | 5 | 137309899 | 137314241 | 4343 | 2 | ENSMUST00 | 0 | Trip6 | ENSMUSG00000023348.10 |
| chr5 | 142550294 | 142551407 | 24.30838 | 70.74035 | 67.73997 | Promoter | 5 | 142484839 | 142551098 | 66260 | 2 | ENSMUST00 | 0 | Radil | ENSMUSG00000029576.12 |
| chr5 | 33995688 | 33997721 | 24.30838 | 70.74035 | 67.73997 | Promoter | 5 | 33995984 | 34005916 | 9933 | 1 | ENSMUST00 | 0 | Nat8l | ENSMUSG00000048142.8 |
| chrX | 152768035 | 152770247 | 24.30838 | 70.74035 | 67.73997 | Promoter | 20 | 152609509 | 152769461 | 159953 | 2 | ENSMUST00 | 0 | Shroom2 | ENSMUSG00000045180.8 |
| chrX | 136270049 | 136271023 | 24.30838 | 70.74035 | 67.73997 | Promoter | 20 | 136270253 | 136271978 | 1726 | 1 | ENSMUST00 | 0 | Ngfrap1 | ENSMUSG00000046432.7 |
| chr4 | 149585970 | 149588285 | 25.31572 | 70.69148 | 67.6937 | Promoter | 4 | 149586468 | 149648899 | 62432 | 1 | ENSMUST00 | 0 | Clstn1 | ENSMUSG00000039953.8 |
| chr4 | 126096411 | 126097218 | 25.31572 | 70.69148 | 67.6937 | Promoter | 4 | 126096623 | 126098584 | 1962 | 1 | ENSMUST00 | 0 | Lsm10 | ENSMUSG00000050188.8 |
| chr13 | 42916148 | 42917655 | 22.27177 | 70.67676 | 67.6792 | Intron (ENSN | 13 | 42868338 | 42869226 | 889 | 2 | ENSMUST00 | -46923 | Gm15812 | ENSMUSG00000085276.2 |
| chr2 | 35661012 | 35662284 | 24.74451 | 70.60188 | 67.60455 | Promoter | 2 | 35661350 | 35708847 | 47498 | 1 | ENSMUST00 | 0 | Dab2ip | ENSMUSG00000026883.12 |
| chr1 | 74600681 | 74601850 | 24.21694 | 70.581 | 67.58389 | Promoter | 1 | 74593748 | 74601372 | 7625 | 2 | ENSMUST00 | 0 | Rnf25 | ENSMUSG00000026171.7 |
| chr9 | 66834886 | 66835777 | 25.21146 | 70.51252 | 67.51562 | Promoter | 9 | 66814994 | 66834726 | 19733 | 2 | ENSMUST00 | -161 | Aph1c | ENSMUSG00000053040.8 |
| chr3 | 132948145 | 132950666 | 16.50874 | 70.50747 | 67.5107 | Promoter | 3 | 132881745 | 132950291 | 68547 | 2 | ENSMUST00 | 0 | Npnt | ENSMUSG00000040998.13 |
| chr15 | 81522003 | 81524194 | 22.65707 | 70.49107 | 67.49436 | Distal Interge | 15 | 81499084 | 81499548 | 465 | 2 | ENSMUST00 | -22456 | Gm5218 | ENSMUSG00000091198.1 |
| chr9 | 83145513 | 83146848 | 22.65707 | 70.49107 | 67.49436 | Promoter | 9 | 83109948 | 83146470 | 36523 | 2 | ENSMUST00 | 0 | Hmgn3 | ENSMUSG00000066456.9 |
| chr10 | 3972798 | 3974211 | 19.45739 | 70.46334 | 67.46697 | Promoter | 10 | 3973118 | 4156111 | 182994 | 1 | ENSMUST00 | 0 | Mthfd1l | ENSMUSG00000040675.12 |
| chr15 | 88732641 | 88734627 | 19.45739 | 70.46334 | 67.46697 | Promoter | 15 | 88687300 | 88734233 | 46934 | 2 | ENSMUST00 | 0 | Brd1 | ENSMUSG00000022387.11 |
| chr6 | 25808383 | 25809801 | 19.45739 | 70.46334 | 67.46697 | Promoter | 6 | 25743737 | 25809226 | 65490 | 2 | ENSMUST00 | 0 | Pot1a | ENSMUSG00000029676.10 |
| chr6 | 85587283 | 85588437 | 19.45739 | 70.46334 | 67.46697 | Promoter | 6 | 85587531 | 85702753 | 115223 | 1 | ENSMUST00 | 0 | Alms1 | ENSMUSG00000063810.5 |
| chr7 | 102686722 | 102687967 | 19.45739 | 70.46334 | 67.46697 | Promoter | 7 | 102677580 | 102687326 | 9747 | 2 | ENSMUST00 | 0 | Trim68 | ENSMUSG00000073968.3 |
| chr8 | 122567163 | 122568548 | 19.45739 | 70.46334 | 67.46697 | Promoter | 8 | 122568015 | 122573128 | 5114 | 1 | ENSMUST00 | 0 | Cdt1 | ENSMUSG00000006585.2 |
| chr8 | 68734332 | 68735824 | 19.45739 | 70.46334 | 67.46697 | Promoter | 8 | 68356781 | 68735146 | 378366 | 2 | ENSMUST00 | 0 | Csgalnact1 | ENSMUSG00000036356.9 |
| chr9 | 22388550 | 22389727 | 23.06947 | 70.34298 | 67.34766 | Promoter | 9 | 22331214 | 22389206 | 57993 | 2 | ENSMUST00 | 0 | Anln | ENSMUSG00000036777.7 |
| chr17 | 35045923 | 35046624 | 24.07899 | 70.34031 | 67.34542 | Promoter | 17 | 35028605 | 35046622 | 18018 | 2 | ENSMUST00 | 0 | Msh5 | ENSMUSG00000007035.10 |
| chr10 | 81285314 | 81286310 | 25.10806 | 70.33492 | 67.34009 | Intron (ENSN | 10 | 81273207 | 81291267 | 18061 | 2 | ENSMUST00 | 4957 | Tjp3 | ENSMUSG00000034917.7 |
| chr9 | 50752870 | 50754879 | 25.10806 | 70.33492 | 67.34009 | Promoter | 9 | 50752758 | 50756633 | 3876 | 1 | ENSMUST00 | 113 | Cryab | ENSMUSG00000032060.9 |
| chr1 | 192854608 | 192856186 | 17.10421 | 70.31819 | 67.3235 | Promoter | 1 | 192844431 | 192855752 | 11322 | 2 | ENSMUST00 | 0 | Sertad4 | ENSMUSG00000016262.9 |
| chr11 | 97781302 | 97783180 | 17.10421 | 70.31819 | 67.3235 | Promoter | 11 | 97781639 | 97781775 | 137 | 2 | ENSMUST00 | 0 | Snora21 | ENSMUSG00000064901.1 |
| chr9 | 44396250 | 44397700 | 17.10421 | 70.31819 | 67.3235 | Promoter | 9 | 44395797 | 44395915 | 119 | 2 | ENSMUST00 | -336 | Gm26306 | ENSMUSG00000088398.1 |
| chr17 | 12959996 | 12961192 | 18.55771 | 70.26926 | 67.27464 | Promoter | 17 | 12942890 | 12960747 | 17858 | 2 | ENSMUST00 | 0 | Acat2 | ENSMUSG00000023832.8 |
| chr5 | 135023204 | 135024594 | 18.55771 | 70.26926 | 67.27464 | Promoter | 5 | 135023482 | 135051100 | 27619 | 1 | ENSMUST00 | 0 | Stx1a | ENSMUSG00000007207.5 |
| chr5 | 23417333 | 23419520 | 18.55771 | 70.26926 | 67.27464 | Intron (ENSN | 5 | 23422404 | 23433309 | 10906 | 2 | ENSMUST00 | 13789 | 5031425E22I | ENSMUSG00000073147.3 |
| chr6 | 122801209 | 122802532 | 18.55771 | 70.26926 | 67.27464 | Promoter | 6 | 122737085 | 122801640 | 64556 | 2 | ENSMUST00 | 0 | Slc2a3 | ENSMUSG00000003153.5 |
| chr9 | 22156224 | 22157709 | 18.55771 | 70.26926 | 67.27464 | Promoter | 9 | 22156846 | 22158358 | 1513 | 1 | ENSMUST00 | 0 | Pigyl | ENSMUSG00000010607.8 |
| chr9 | 44334630 | 44336041 | 18.55771 | 70.26926 | 67.27464 | Promoter | 9 | 44334694 | 44336077 | 1384 | 1 | ENSMUST00 | 0 | H2afx | ENSMUSG00000049932.3 |
| chr9 | 56506171 | 56507750 | 24.03335 | 70.26061 | 67.26643 | Distal Interge | 9 | 56418644 | 56496884 | 78241 | 1 | ENSMUST00 | 87528 | Hmg20a | ENSMUSG00000032329.3 |
| chr12 | 86947335 | 86948361 | 21.5679 | 70.25594 | 67.2618 | Promoter | 12 | 86947343 | 86965362 | 18020 | 1 | ENSMUST00 | 0 | Cipc | ENSMUSG00000034157.7 |
| chr11 | 119391851 | 119394080 | 25.05667 | 70.24663 | 67.25255 | Promoter | 11 | 119393100 | 119487415 | 94316 | 1 | ENSMUST00 | 0 | Rnf213 | ENSMUSG00000070327.8 |
| chr13 | 21478635 | 21479819 | 25.05667 | 70.24663 | 67.25255 | Promoter | 13 | 21478907 | 21485507 | 6601 | 1 | ENSMUST00 | 0 | Zkscan4 | ENSMUSG00000054931.6 |
| chr13 | 78197737 | 78199944 | 25.05667 | 70.24663 | 67.25255 | Promoter | 13 | 78188973 | 78199757 | 10785 | 2 | ENSMUST00 | 0 | Nr2f1 | ENSMUSG00000069171.8 |
| chr18 | 38211156 | 38212817 | 25.05667 | 70.24663 | 67.25255 | Promoter | 18 | 38185915 | 38212053 | 26139 | 2 | ENSMUST00 | 0 | Pcdh1 | ENSMUSG00000051375.10 |
| chr7 | 45339213 | 45340222 | 25.05667 | 70.24663 | 67.25255 | Promoter | 7 | 45339281 | 45340762 | 1482 | 2 | ENSMUST00 | 540 | Ppfia3 | ENSMUSG00000003863.11 |
| chr19 | 55253032 | 55254583 | 23.51194 | 70.23763 | 67.24405 | Promoter | 19 | 55253369 | 55296627 | 43259 | 1 | ENSMUST00 | 0 | Acsl5 | ENSMUSG00000024981.5 |
| chr2 | 112261831 | 112262761 | 23.51194 | 70.23763 | 67.24405 | Promoter | 2 | 112261926 | 112263269 | 1344 | 1 | ENSMUST00 | 0 | Nop10 | ENSMUSG00000027133.3 |
| chr6 | 52011818 | 52012598 | 23.51194 | 70.23763 | 67.24405 | Promoter | 6 | 51859166 | 52012549 | 153384 | 2 | ENSMUST00 | 0 | Skap2 | ENSMUSG00000059182.5 |
| chr7 | 6363007 | 6363881 | 23.51194 | 70.23763 | 67.24405 | Distal Interge | 7 | 6371364 | 6379491 | 8128 | 1 | ENSMUST00 | -7483 | Zfp78 | ENSMUSG00000055150.9 |
| chr11 | 83941557 | 83942838 | 17.78121 | 70.23115 | 67.23819 | Promoter | 11 | 83942062 | 83963088 | 21027 | 1 | ENSMUST00 | 0 | Ddx52 | ENSMUSG00000020677.7 |
| chr11 | 69039288 | 69041871 | 17.78121 | 70.23115 | 67.23819 | Distal Interge | 11 | 69045647 | 69051664 | 6018 | 1 | ENSMUST00 | -3776 | Aurkb | ENSMUSG00000020897.7 |
| chr2 | 104849125 | 104850173 | 17.78121 | 70.23115 | 67.23819 | Promoter | 2 | 104830741 | 104849876 | 19136 | 2 | ENSMUST00 | 0 | Prrg4 | ENSMUSG00000027171.5 |
| chr1 | 75545925 | 75546995 | 23.98789 | 70.18118 | 67.18839 | Promoter | 1 | 75546266 | 75562172 | 15907 | 1 | ENSMUST00 | 0 | Slc4a3 | ENSMUSG00000006576.11 |
| chr1 | 79857873 | 79859155 | 24.50127 | 70.18111 | 67.18839 | Promoter | 1 | 79794197 | 79858696 | 64500 | 2 | ENSMUST00 | 0 | Serpine2 | ENSMUSG00000026249.5 |
| chr1 | 86486943 | 86487503 | 24.50127 | 70.18111 | 67.18839 | Distal Interge | 1 | 86473659 | 86474006 | 348 | 1 | ENSMUST00 | 13285 | RP23-403I3.6 | ENSMUSG00000101328.1 |
| chr10 | 13500729 | 13501497 | 24.50127 | 70.18111 | 67.18839 | Promoter | 10 | 13501001 | 13515543 | 14543 | 1 | ENSMUST00 | 0 | Fuca2 | ENSMUSG00000019810.10 |
| chr10 | 81606332 | 81606813 | 23.98789 | 70.18118 | 67.18839 | Promoter | 10 | 81606303 | 81609836 | 3534 | 1 | ENSMUST00 | 30 | BC025920 | ENSMUSG00000074862.6 |
| chr11 | 78301219 | 78302279 | 23.98789 | 70.18118 | 67.18839 | Promoter | 11 | 78301529 | 78322457 | 20929 | 1 | ENSMUST00 | 0 | Spag5 | ENSMUSG00000002055.4 |
| chr11 | 70020904 | 70021553 | 24.50127 | 70.18111 | 67.18839 | Intron (ENSN | 11 | 70018864 | 70031149 | 12286 | 1 | ENSMUST00 | 2041 | Dlg4 | ENSMUSG00000020886.11 |
| chr13 | 58112415 | 58113896 | 24.50127 | 70.18111 | 67.18839 | Promoter | 13 | 58004221 | 58113592 | 109372 | 2 | ENSMUST00 | 0 | Klhl3 | ENSMUSG00000014164.9 |
| chr14 | 8002583 | 8003712 | 23.98789 | 70.18118 | 67.18839 | Promoter | 14 | 8002902 | 8056555 | 53654 | 1 | ENSMUST00 | 0 | Abhd6 | ENSMUSG00000025277.8 |
| chr14 | 116925098 | 116927440 | 24.50127 | 70.18111 | 67.18839 | Promoter | 14 | 116925297 | 117979529 | 1054233 | 1 | ENSMUST00 | 0 | Gpc6 | ENSMUSG00000058571.5 |
| chr15 | 79504784 | 79506214 | 24.50127 | 70.18111 | 67.18839 | Promoter | 15 | 79483718 | 79505241 | 21524 | 2 | ENSMUST00 | 0 | Kcnj4 | ENSMUSG00000044216.6 |
| chr17 | 27820756 | 27821417 | 23.98789 | 70.18118 | 67.18839 | Promoter | 17 | 27751232 | 27820558 | 69327 | 2 | ENSMUST00 | -199 | D17Wsu92e | ENSMUSG00000056692.7 |
| chr2 | 118548996 | 118550448 | 23.98789 | 70.18118 | 67.18839 | Promoter | 2 | 118528757 | 118549687 | 20931 | 2 | ENSMUST00 | 0 | Bmf | ENSMUSG00000040093.10 |
| chr3 | 107895364 | 107896351 | 23.98789 | 70.18118 | 67.18839 | Promoter | 3 | 107895821 | 107898686 | 2866 | 1 | ENSMUST00 | 0 | Gstm5 | ENSMUSG00000004032.5 |
| chr9 | 22164585 | 22165544 | 24.50127 | 70.18111 | 67.18839 | Distal Interge | 9 | 22156946 | 22158325 | 1380 | 1 | ENSMUST00 | 7640 | Pigyl | ENSMUSG00000010607.8 |
| chr9 | 44720798 | 44721735 | 23.98789 | 70.18118 | 67.18839 | Promoter | 9 | 44716124 | 44721424 | 5301 | 2 | ENSMUST00 | 0 | Phldb1 | ENSMUSG00000048537.10 |
| chr19 | 4509642 | 4510212 | 25.0055 | 70.15867 | 67.16741 | Promoter | 19 | 4510472 | 4621752 | 111281 | 1 | ENSMUST00 | -260 | Pcx | ENSMUSG00000024892.10 |
| chr6 | 100286055 | 100287492 | 25.0055 | 70.15867 | 67.16741 | Promoter | 6 | 100228566 | 100287441 | 58876 | 2 | ENSMUST00 | 0 | Rybp | ENSMUSG00000072872.2 |
| chr1 | 64119053 | 64121494 | 21.42557 | 69.98924 | 66.99868 | Promoter | 1 | 64029447 | 64121491 | 92045 | 2 | ENSMUST00 | 0 | Klf7 | ENSMUSG00000025959.8 |
| chr10 | 82984757 | 82986471 | 21.42557 | 69.98924 | 66.99868 | Promoter | 10 | 82985498 | 83195900 | 210403 | 1 | ENSMUST00 | 0 | Chst11 | ENSMUSG00000034612.7 |
| chr11 | 60830214 | 60831240 | 21.42557 | 69.98924 | 66.99868 | Promoter | 11 | 60830631 | 60858423 | 27793 | 1 | ENSMUST00 | 0 | Dhrs7b | ENSMUSG00000042569.8 |

| | | | | | | | | | | | | | | | |
|---|---|---|---|---|---|---|---|---|---|---|---|---|---|---|---|
| chr14 | 69390377 | 69391553 | 21.42557 | 69.98924 | 66.99868 | Promoter | 14 | 69390854 | 69480287 | 89434 | 1 | ENSMUST00 | 0 | Gm21451 | ENSMUSG00000094526.1 |
| chr15 | 81466302 | 81467458 | 21.42557 | 69.98924 | 66.99868 | Promoter | 15 | 81466316 | 81476369 | 10054 | 1 | ENSMUST00 | 0 | Rbx1 | ENSMUSG00000022400.8 |
| chr16 | 17069675 | 17070569 | 21.42557 | 69.98924 | 66.99868 | Promoter | 16 | 17069696 | 17086736 | 17041 | 1 | ENSMUST00 | 0 | Ypel1 | ENSMUSG00000022773.11 |
| chr2 | 84839354 | 84841115 | 21.42557 | 69.98924 | 66.99868 | Promoter | 2 | 84839395 | 84850142 | 10748 | 1 | ENSMUST00 | 0 | Slc43a1 | ENSMUSG00000027075.11 |
| chr3 | 96238498 | 96239143 | 21.42557 | 69.98924 | 66.99868 | Promoter | 3 | 96238108 | 96239127 | 1020 | 2 | ENSMUST00 | 0 | Hist2h3c2 | ENSMUSG00000081058.5 |
| chr3 | 94820977 | 94822042 | 21.42557 | 69.98924 | 66.99868 | Distal Interge | 3 | 94832977 | 94833805 | 829 | 1 | ENSMUST00 | -10935 | Gm15263 | ENSMUSG00000084058.1 |
| chr4 | 107367443 | 107368458 | 21.42557 | 69.98924 | 66.99868 | Promoter | 4 | 107367784 | 107416346 | 48563 | 1 | ENSMUST00 | 0 | Ndc1 | ENSMUSG00000028614.9 |
| chr5 | 92808748 | 92813129 | 21.42557 | 69.98924 | 66.99868 | Promoter | 5 | 92809378 | 92965318 | 155941 | 1 | ENSMUST00 | 0 | Shroom3 | ENSMUSG00000029381.10 |
| chr7 | 46709938 | 46710759 | 21.42557 | 69.98924 | 66.99868 | Promoter | 7 | 46686108 | 46710680 | 24573 | 2 | ENSMUST00 | 0 | Saal1 | ENSMUSG00000006763.7 |
| chr7 | 18957376 | 18958691 | 21.42557 | 69.98924 | 66.99868 | 3' UTR | 7 | 18962415 | 18965173 | 2759 | 2 | ENSMUST00 | 6482 | Gm26821 | ENSMUSG00000097038.1 |
| chr8 | 46616644 | 46617278 | 21.42557 | 69.98924 | 66.99868 | Promoter | 8 | 46575579 | 46617200 | 41622 | 2 | ENSMUST00 | 0 | Primpol | ENSMUSG00000038225.9 |
| chr8 | 36457200 | 36458776 | 21.42557 | 69.98924 | 66.99868 | Promoter | 8 | 36457548 | 36512956 | 55409 | 1 | ENSMUST00 | 0 | 6430573F11 | ENSMUSG00000039620.11 |
| chr8 | 94917111 | 94918353 | 21.42557 | 69.98924 | 66.99868 | Promoter | 8 | 94902869 | 94918098 | 15230 | 2 | ENSMUST00 | 0 | Ccdc102a | ENSMUSG00000063605.5 |
| chr1 | 16687810 | 16689283 | 23.85253 | 69.94444 | 66.95615 | Promoter | 1 | 16688051 | 16691947 | 3897 | 1 | ENSMUST00 | 0 | Ly96 | ENSMUSG00000025779.5 |
| chr1 | 184034360 | 184035976 | 24.31009 | 69.8498 | 66.86187 | Promoter | 1 | 184034381 | 184075636 | 41256 | 1 | ENSMUST00 | 0 | Dusp10 | ENSMUSG00000039384.8 |
| chr14 | 50944087 | 50945775 | 24.31009 | 69.8498 | 66.86187 | Promoter | 14 | 50944302 | 50953412 | 9111 | 1 | ENSMUST00 | 0 | Pnp | ENSMUSG00000021871.12 |
| chr2 | 26502084 | 26504362 | 24.31009 | 69.8498 | 66.86187 | Promoter | 2 | 26457903 | 26503822 | 45920 | 2 | ENSMUST00 | 0 | Notch1 | ENSMUSG00000026923.10 |
| chr4 | 14825500 | 14827022 | 24.31009 | 69.8498 | 66.86187 | Promoter | 4 | 14809498 | 14826587 | 17090 | 2 | ENSMUST00 | 0 | Otud6b | ENSMUSG00000040550.11 |
| chr10 | 81342803 | 81343699 | 24.80288 | 69.81013 | 66.8223 | Promoter | 10 | 81337762 | 81343266 | 5505 | 2 | ENSMUST00 | 0 | Gipc3 | ENSMUSG00000034872.5 |
| chr13 | 48512957 | 48513786 | 22.27859 | 69.80411 | 66.81643 | Promoter | 13 | 48487647 | 48513436 | 25790 | 2 | ENSMUST00 | 0 | Zfp169 | ENSMUSG00000050954.10 |
| chr13 | 62760193 | 62760995 | 25.27427 | 69.73028 | 66.74294 | Promoter | 13 | 62760739 | 62760846 | 108 | 1 | ENSMUST00 | 0 | Mir713 | ENSMUSG00000096098.1 |
| chr14 | 69336773 | 69337747 | 25.27427 | 69.73028 | 66.74294 | Promoter | 14 | 69337174 | 69367270 | 30097 | 1 | ENSMUST00 | 0 | Entpd4 | ENSMUSG00000095463.3 |
| chr19 | 6933513 | 6934451 | 25.27427 | 69.73028 | 66.74294 | Promoter | 19 | 6925710 | 6934515 | 8806 | 2 | ENSMUST00 | 64 | Kcnk4 | ENSMUSG00000024957.8 |
| chr2 | 25180327 | 25182244 | 25.27427 | 69.73028 | 66.74294 | Promoter | 2 | 25180758 | 25183339 | 2582 | 1 | ENSMUST00 | 0 | Nrarp | ENSMUSG00000078202.3 |
| chr2 | 73892851 | 73893412 | 25.27427 | 69.73028 | 66.74294 | Promoter | 2 | 73892996 | 73900850 | 7855 | 1 | ENSMUST00 | 0 | Gm10822 | ENSMUSG00000097377.1 |
| chr5 | 115429202 | 115430988 | 25.27427 | 69.73028 | 66.74294 | Promoter | 5 | 115429599 | 115455698 | 26100 | 1 | ENSMUST00 | 0 | Msi1 | ENSMUSG00000054256.6 |
| chr6 | 6955350 | 6956778 | 25.27427 | 69.73028 | 66.74294 | Promoter | 6 | 6956004 | 7042979 | 86976 | 1 | ENSMUST00 | 0 | Acn9 | ENSMUSG00000042505.6 |
| chr7 | 27552707 | 27553827 | 25.27427 | 69.73028 | 66.74294 | Promoter | 7 | 27532000 | 27553212 | 21213 | 2 | ENSMUST00 | 0 | Pld3 | ENSMUSG00000003363.10 |
| chr7 | 96209775 | 96212128 | 25.27427 | 69.73028 | 66.74294 | Promoter | 7 | 96210146 | 96438966 | 228821 | 1 | ENSMUST00 | 0 | Tenm4 | ENSMUSG00000048078.11 |
| chr9 | 107587668 | 107588446 | 25.27427 | 69.73028 | 66.74294 | Promoter | 9 | 107587718 | 107593038 | 5321 | 1 | ENSMUST00 | 0 | Ifrd2 | ENSMUSG00000010048.5 |
| chr9 | 83805510 | 83806659 | 25.27427 | 69.73028 | 66.74294 | Promoter | 9 | 83778692 | 83806252 | 27561 | 2 | ENSMUST00 | 0 | Elovl4 | ENSMUSG00000032262.8 |
| chr9 | 88326968 | 88328103 | 25.27427 | 69.73028 | 66.74294 | Promoter | 9 | 88327197 | 88372092 | 44896 | 1 | ENSMUST00 | 0 | Nt5e | ENSMUSG00000032420.7 |
| chr9 | 108661808 | 108663010 | 25.27427 | 69.73028 | 66.74294 | Promoter | 9 | 108662098 | 108684641 | 22544 | 1 | ENSMUST00 | 0 | Slc25a20 | ENSMUSG00000032602.5 |
| chr9 | 124311612 | 124312922 | 25.27427 | 69.73028 | 66.74294 | Promoter | 9 | 124291804 | 124312696 | 20893 | 2 | ENSMUST00 | 0 | 2010315B03 | ENSMUSG00000074829.5 |
| chrX | 38455467 | 38456479 | 25.27427 | 69.73028 | 66.74294 | Promoter | 20 | 38401357 | 38456454 | 55098 | 2 | ENSMUST00 | 0 | Lamp2 | ENSMUSG00000016534.10 |
| chrX | 36111740 | 36113074 | 25.27427 | 69.73028 | 66.74294 | Promoter | 20 | 36112110 | 36171259 | 59150 | 1 | ENSMUST00 | 0 | Il13ra1 | ENSMUSG00000017057.9 |
| chrX | 52911776 | 52913925 | 25.27427 | 69.73028 | 66.74294 | Promoter | 20 | 52912266 | 52956943 | 44678 | 1 | ENSMUST00 | 0 | Phf6 | ENSMUSG00000025626.11 |
| chrX | 13070252 | 13073605 | 25.27427 | 69.73028 | 66.74294 | Promoter | 20 | 13071498 | 13123449 | 51952 | 1 | ENSMUST00 | 0 | Usp9x | ENSMUSG00000031010.12 |
| chrX | 56454847 | 56456286 | 25.27427 | 69.73028 | 66.74294 | Promoter | 20 | 56454857 | 56507840 | 52984 | 1 | ENSMUST00 | 0 | Ddx26b | ENSMUSG00000035967.10 |
| chrX | 60403062 | 60404044 | 25.27427 | 69.73028 | 66.74294 | Promoter | 20 | 60223293 | 60403981 | 180689 | 2 | ENSMUST00 | 0 | Atp11c | ENSMUSG00000062949.8 |
| chrX | 71815952 | 71817658 | 25.27427 | 69.73028 | 66.74294 | Promoter | 20 | 71816758 | 71824706 | 7949 | 1 | ENSMUST00 | 0 | Vma21 | ENSMUSG00000073131.7 |
| chr4 | 152138374 | 152139201 | 24.70279 | 69.63779 | 66.65218 | Exon (ENSMU | 4 | 152121121 | 152136325 | 15205 | 2 | ENSMUST00 | -2050 | Espn | ENSMUSG00000028943.12 |
| chr2 | 119236856 | 119238257 | 22.67078 | 69.62844 | 66.64306 | Promoter | 2 | 119237362 | 119249514 | 12153 | 1 | ENSMUST00 | 0 | Spint1 | ENSMUSG00000027315.8 |
| chr7 | 125491064 | 125492014 | 22.67078 | 69.62844 | 66.64306 | Promoter | 7 | 125467640 | 125491596 | 23957 | 2 | ENSMUST00 | 0 | Nsmce1 | ENSMUSG00000030750.8 |
| chr8 | 70438562 | 70440049 | 22.67078 | 69.62844 | 66.64306 | Promoter | 8 | 70382355 | 70439579 | 57225 | 2 | ENSMUST00 | 0 | Crtc1 | ENSMUSG00000003575.8 |
| chr13 | 98390311 | 98391925 | 25.17599 | 69.56199 | 66.57701 | Distal Interge | 13 | 98402095 | 98403083 | 989 | 2 | ENSMUST00 | 11158 | Gm2445 | ENSMUSG00000098113.1 |
| chr2 | 125866000 | 125867056 | 25.17599 | 69.56199 | 66.57701 | Promoter | 2 | 125866107 | 125929570 | 63464 | 1 | ENSMUST00 | 0 | Galk2 | ENSMUSG00000027207.10 |
| chr4 | 127235892 | 127236965 | 24.65305 | 69.55211 | 66.56724 | 3' UTR | 4 | 127243784 | 127247809 | 4026 | 1 | ENSMUST00 | -6819 | Smim12 | ENSMUSG00000042380.8 |
| chr13 | 14062233 | 14063503 | 23.09106 | 69.49222 | 66.50756 | Promoter | 13 | 14052449 | 14063401 | 10953 | 2 | ENSMUST00 | 0 | Ggps1 | ENSMUSG00000021302.9 |
| chr18 | 69343931 | 69345637 | 23.09106 | 69.49222 | 66.50756 | Promoter | 18 | 69344146 | 69684343 | 340198 | 1 | ENSMUST00 | 0 | Tcf4 | ENSMUSG00000053477.10 |
| chr2 | 114654182 | 114655559 | 23.09106 | 69.49222 | 66.50756 | Promoter | 2 | 114516416 | 114654953 | 138538 | 2 | ENSMUST00 | 0 | Dph6 | ENSMUSG00000057147.8 |
| chr5 | 93043671 | 93045528 | 23.09106 | 69.49222 | 66.50756 | Promoter | 5 | 93041123 | 93045022 | 3900 | 2 | ENSMUST00 | 0 | Sowahb | ENSMUSG00000045314.5 |
| chr5 | 31153661 | 31154902 | 23.09106 | 69.49222 | 66.50756 | Promoter | 5 | 31140660 | 31154251 | 13592 | 2 | ENSMUST00 | 0 | Mpv17 | ENSMUSG00000090262.2 |
| chr12 | 108305971 | 108306910 | 25.12339 | 69.47189 | 66.48775 | Promoter | 12 | 108306270 | 108328300 | 22031 | 1 | ENSMUST00 | 0 | Hhipl1 | ENSMUSG00000021260.7 |
| chr12 | 84148042 | 84148694 | 25.12339 | 69.47189 | 66.48775 | Promoter | 12 | 84146131 | 84148489 | 2359 | 2 | ENSMUST00 | 0 | Pnma1 | ENSMUSG00000054383.2 |
| chr17 | 46152634 | 46153928 | 25.12339 | 69.47189 | 66.48775 | Promoter | 17 | 46147390 | 46153553 | 6164 | 2 | ENSMUST00 | 0 | Mad2l1bp | ENSMUSG00000034509.8 |
| chr8 | 125909739 | 125911484 | 25.12339 | 69.47189 | 66.48775 | Promoter | 8 | 125910450 | 125947439 | 36990 | 1 | ENSMUST00 | 0 | BC021891 | ENSMUSG00000031853.4 |
| chr13 | 23684160 | 23685889 | 24.60351 | 69.46672 | 66.48286 | Promoter | 13 | 23684199 | 23692488 | 8290 | 1 | ENSMUST00 | 0 | Hist1h2bc | ENSMUSG00000018102.4 |
| chr4 | 32614791 | 32615243 | 24.60351 | 69.46672 | 66.48286 | Promoter | 4 | 32615451 | 32631810 | 16360 | 1 | ENSMUST00 | -208 | Casp8ap2 | ENSMUSG00000028282.7 |
| chr1 | 137898312 | 137901031 | 23.54255 | 69.40097 | 66.41734 | Promoter | 1 | 137898595 | 137899729 | 1135 | 1 | ENSMUST00 | 0 | Gm4258 | ENSMUSG00000098243.3 |
| chr10 | 63381460 | 63382823 | 23.54255 | 69.40097 | 66.41734 | Promoter | 10 | 63319005 | 63381704 | 62700 | 2 | ENSMUST00 | 0 | Sirt1 | ENSMUSG00000020063.11 |
| chr11 | 22971710 | 22973143 | 23.54255 | 69.40097 | 66.41734 | Promoter | 11 | 22900023 | 22972859 | 72837 | 2 | ENSMUST00 | 0 | Commd1 | ENSMUSG00000051355.13 |
| chr11 | 87073213 | 87074855 | 23.54255 | 69.40097 | 66.41734 | Promoter | 11 | 87033867 | 87074062 | 40196 | 2 | ENSMUST00 | 0 | Gdpd1 | ENSMUSG00000061666.6 |
| chr18 | 82474741 | 82476320 | 23.54255 | 69.40097 | 66.41734 | Promoter | 18 | 82475146 | 82558703 | 83558 | 1 | ENSMUST00 | 0 | Mbp | ENSMUSG00000041607.11 |
| chr4 | 127076728 | 127077872 | 23.54255 | 69.40097 | 66.41734 | Promoter | 4 | 127077383 | 127124370 | 46988 | 1 | ENSMUST00 | 0 | Zmym6 | ENSMUSG00000042408.8 |
| chr5 | 33377779 | 33379570 | 23.54255 | 69.40097 | 66.41734 | Promoter | 5 | 33378549 | 33410519 | 31971 | 1 | ENSMUST00 | 0 | Uvssa | ENSMUSG00000037355.9 |
| chr6 | 83114925 | 83116453 | 23.54255 | 69.40097 | 66.41734 | Promoter | 6 | 83115496 | 83118898 | 3403 | 1 | ENSMUST00 | 0 | Mogs | ENSMUSG00000030036.7 |
| chr7 | 3664496 | 3666141 | 23.54255 | 69.40097 | 66.41734 | Promoter | 7 | 3660098 | 3665840 | 5743 | 2 | ENSMUST00 | 0 | Leng1 | ENSMUSG00000078813.3 |
| chr8 | 13200086 | 13201411 | 23.54255 | 69.40097 | 66.41734 | Promoter | 8 | 13176869 | 13200624 | 23756 | 2 | ENSMUST00 | 0 | Grtp1 | ENSMUSG00000038515.9 |
| chr1 | 175880304 | 175881272 | 20.20131 | 69.382 | 66.39919 | Promoter | 1 | 175880778 | 175911396 | 30619 | 1 | ENSMUST00 | 0 | Exo1 | ENSMUSG00000039748.6 |
| chr11 | 96915837 | 96916595 | 24.55417 | 69.38167 | 66.39919 | Promoter | 11 | 96907426 | 96916496 | 9071 | 2 | ENSMUST00 | 0 | Cdk5rap3 | ENSMUSG00000018669.9 |
| chr11 | 96830148 | 96830883 | 24.55417 | 69.38167 | 66.39919 | Promoter | 11 | 96817414 | 96829968 | 12555 | 2 | ENSMUST00 | -181 | Nfe2l1 | ENSMUSG00000038615.12 |
| chr13 | 73763388 | 73764824 | 24.55417 | 69.38167 | 66.39919 | Promoter | 13 | 73763697 | 73816743 | 53047 | 1 | ENSMUST00 | 0 | Slc12a7 | ENSMUSG00000017756.8 |
| chr13 | 64248123 | 64249296 | 20.20131 | 69.382 | 66.39919 | Promoter | 13 | 64192545 | 64248526 | 55982 | 2 | ENSMUST00 | 0 | Cdc14b | ENSMUSG00000033102.9 |
| chr13 | 34129135 | 34131581 | 24.55417 | 69.38167 | 66.39919 | Promoter | 13 | 34127008 | 34130354 | 3347 | 2 | ENSMUST00 | 0 | Tubb2b | ENSMUSG00000045136.5 |
| chr13 | 78163045 | 78164161 | 20.20131 | 69.382 | 66.39919 | 3' UTR | 13 | 78189872 | 78190443 | 572 | 2 | ENSMUST00 | 26282 | Nr2f1 | ENSMUSG00000069171.8 |
| chr14 | 101729087 | 101730561 | 20.20131 | 69.382 | 66.39919 | Promoter | 14 | 101729957 | 101934710 | 204754 | 1 | ENSMUST00 | 0 | Lmo7 | ENSMUSG00000033060.10 |
| chr15 | 85652927 | 85654833 | 20.20131 | 69.382 | 66.39919 | Distal Interge | 15 | 85639747 | 85645934 | 6188 | 1 | ENSMUST00 | 13181 | RP23-64J7.2 | ENSMUSG00000101828.1 |
| chr19 | 24960896 | 24961679 | 20.20131 | 69.382 | 66.39919 | Promoter | 19 | 24919916 | 24961616 | 41701 | 2 | ENSMUST00 | 0 | Cbwd1 | ENSMUSG00000024878.8 |
| chr2 | 26909810 | 26912049 | 20.20131 | 69.382 | 66.39919 | Promoter | 2 | 26905262 | 26910677 | 5416 | 2 | ENSMUST00 | 0 | Med22 | ENSMUSG00000015776.7 |
| chr4 | 62207811 | 62209358 | 20.20131 | 69.382 | 66.39919 | Promoter | 4 | 62189540 | 62208446 | 18907 | 2 | ENSMUST00 | 0 | Zfp37 | ENSMUSG00000028389.5 |
| chr5 | 139251817 | 139253823 | 20.20131 | 69.382 | 66.39919 | Promoter | 5 | 139252324 | 139270051 | 17728 | 1 | ENSMUST00 | 0 | Get4 | ENSMUSG00000025858.7 |
| chr7 | 97737410 | 97738709 | 20.20131 | 69.382 | 66.39919 | Promoter | 7 | 97726379 | 97738247 | 11869 | 2 | ENSMUST00 | 0 | Aqp11 | ENSMUSG00000042797.8 |
| chr8 | 119910112 | 119911760 | 20.20131 | 69.382 | 66.39919 | Promoter | 8 | 119910360 | 119957555 | 47196 | 1 | ENSMUST00 | 0 | Usp10 | ENSMUSG00000031826.14 |
| chr1 | 77514048 | 77516422 | 22.5292 | 69.37365 | 66.39268 | Promoter | 1 | 77367185 | 77515088 | 147904 | 2 | ENSMUST00 | 0 | Epha4 | ENSMUSG00000026235.9 |
| chr1 | 125433585 | 125435601 | 22.5292 | 69.37365 | 66.39268 | Promoter | 1 | 125392906 | 125435547 | 42642 | 2 | ENSMUST00 | 0 | Actr3 | ENSMUSG00000026341.11 |
| chr10 | 121032940 | 121034794 | 22.5292 | 69.37365 | 66.39268 | Promoter | 10 | 121033960 | 121100650 | 66691 | 1 | ENSMUST00 | 0 | Wif1 | ENSMUSG00000020218.6 |
| chr11 | 95414187 | 95415400 | 22.5292 | 69.37365 | 66.39268 | Promoter | 11 | 95414243 | 95474335 | 60093 | 1 | ENSMUST00 | 0 | Spop | ENSMUSG00000057522.10 |
| chr14 | 58073800 | 58076799 | 22.5292 | 69.37365 | 66.39268 | Promoter | 14 | 58075115 | 58110174 | 35060 | 1 | ENSMUST00 | 0 | Fgf9 | ENSMUSG00000021974.9 |
| chr14 | 104336851 | 104337137 | 22.5292 | 69.37365 | 66.39268 | Distal Interge | 14 | 104463973 | 104465365 | 1393 | 2 | ENSMUST00 | 128228 | Pou4f1 | ENSMUSG00000048349.9 |
| chr17 | 25114989 | 25116341 | 22.5292 | 69.37365 | 66.39268 | Promoter | 17 | 25099570 | 25115907 | 16338 | 2 | ENSMUST00 | 0 | Telo2 | ENSMUSG00000024170.10 |
| chr17 | 29886537 | 29888883 | 22.5292 | 69.37365 | 66.39268 | Promoter | 17 | 29827956 | 29888629 | 60674 | 2 | ENSMUST00 | 0 | Mdga1 | ENSMUSG00000043557.10 |
| chr18 | 36514845 | 36516217 | 22.5292 | 69.37365 | 66.39268 | Promoter | 18 | 36504929 | 36515805 | 10877 | 2 | ENSMUST00 | 0 | Hbegf | ENSMUSG00000024486.5 |
| chr19 | 4096503 | 4097199 | 22.5292 | 69.37365 | 66.39268 | Promoter | 19 | 4097182 | 4099017 | 1836 | 1 | ENSMUST00 | 0 | Cdk2ap2 | ENSMUSG00000024856.5 |
| chr2 | 91201938 | 91203471 | 22.5292 | 69.37365 | 66.39268 | Promoter | 2 | 91202885 | 91208313 | 5429 | 1 | ENSMUST00 | 0 | Acp2 | ENSMUSG00000002103.10 |
| chr2 | 30048435 | 30049494 | 22.5292 | 69.37365 | 66.39268 | Promoter | 2 | 30031546 | 30048881 | 17336 | 2 | ENSMUST00 | 0 | Wdr34 | ENSMUSG00000039715.8 |
| chr3 | 108839254 | 108840617 | 22.5292 | 69.37365 | 66.39268 | Promoter | 3 | 108793176 | 108840511 | 47336 | 2 | ENSMUST00 | 0 | Stxbp3a | ENSMUSG00000027882.13 |
| chr3 | 131110142 | 131112789 | 22.5292 | 69.37365 | 66.39268 | Promoter | 3 | 131110471 | 131224356 | 113886 | 1 | ENSMUST00 | 0 | Lef1 | ENSMUSG00000027985.9 |
| chr4 | 139622845 | 139623738 | 22.5292 | 69.37365 | 66.39268 | Promoter | 4 | 139622866 | 139649690 | 26825 | 1 | ENSMUST00 | 0 | Aldh4a1 | ENSMUSG00000028737.10 |
| chr4 | 142083653 | 142084938 | 22.5292 | 69.37365 | 66.39268 | Promoter | 4 | 142030992 | 142084304 | 53313 | 2 | ENSMUST00 | 0 | Tmem51 | ENSMUSG00000040616.3 |
| chr4 | 12139930 | 12140530 | 22.5292 | 69.37365 | 66.39268 | Promoter | 4 | 12140264 | 12144688 | 4425 | 1 | ENSMUST00 | 0 | Rbm12b1 | ENSMUSG00000046667.9 |
| chr4 | 89310286 | 89311501 | 22.5292 | 69.37365 | 66.39268 | Promoter | 4 | 89306289 | 89311032 | 4744 | 2 | ENSMUST00 | 0 | Cdkn2b | ENSMUSG00000073802.4 |
| chr4 | 35155675 | 35157544 | 22.5292 | 69.37365 | 66.39268 | Promoter | 4 | 34949074 | 35157484 | 208411 | 2 | ENSMUST00 | 0 | Mob3b | ENSMUSG00000073910.5 |
| chr5 | 90930648 | 90932111 | 22.5292 | 69.37365 | 66.39268 | Promoter | 5 | 90931196 | 91021360 | 90165 | 1 | ENSMUST00 | 0 | Mthfd2l | ENSMUSG00000029376.7 |
| chr6 | 145862672 | 145864243 | 22.5292 | 69.37365 | 66.39268 | 3' UTR | 6 | 145863505 | 145865302 | 1798 | 2 | ENSMUST00 | 1059 | Bhlhe41 | ENSMUSG00000030256.6 |
| chr6 | 117905652 | 117906561 | 22.5292 | 69.37365 | 66.39268 | Promoter | 6 | 117906790 | 117923798 | 17009 | 1 | ENSMUST00 | -229 | Hnrnpf | ENSMUSG00000042079.8 |
| chr7 | 97371343 | 97372245 | 22.5292 | 69.37365 | 66.39268 | Promoter | 7 | 97371606 | 97392153 | 20548 | 1 | ENSMUST00 | 0 | Alg8 | ENSMUSG00000035704.12 |
| chr7 | 100372111 | 100372991 | 22.5292 | 69.37365 | 66.39268 | Promoter | 7 | 100372233 | 100469461 | 97229 | 1 | ENSMUST00 | 0 | C2cd3 | ENSMUSG00000047248.14 |
| chr8 | 22165798 | 22166749 | 22.5292 | 69.37365 | 66.39268 | Promoter | 8 | 22128283 | 22166412 | 38130 | 2 | ENSMUST00 | 0 | Nek3 | ENSMUSG00000031478.11 |
| chr8 | 31186857 | 31187993 | 22.5292 | 69.37365 | 66.39268 | Promoter | 8 | 31187331 | 31261483 | 74153 | 1 | ENSMUST00 | 0 | Fut10 | ENSMUSG00000046152.11 |
| chr9 | 122864928 | 122866263 | 22.5292 | 69.37365 | 66.39268 | Distal Interge | 9 | 122851914 | 122862128 | 10215 | 2 | ENSMUST00 | -2801 | Zfp445 | ENSMUSG00000047036.7 |
| chr13 | 67754365 | 67755559 | 24.02885 | 69.3614 | 66.38335 | Promoter | 13 | 67747800 | 67755134 | 7335 | 2 | ENSMUST00 | 0 | Zfp85 | ENSMUSG00000058331.9 |
| chr18 | 35887544 | 35890163 | 24.02885 | 69.3614 | 66.38335 | Distal Interge | 18 | 35830389 | 35831093 | 705 | 2 | ENSMUST00 | -56452 | RP24-326K1 | ENSMUSG00000100320.1 |
| chr6 | 40324965 | 40326179 | 24.02885 | 69.3614 | 66.38335 | Promoter | 6 | 40325478 | 40396762 | 71285 | 1 | ENSMUST00 | 0 | Agk | ENSMUSG00000029916.8 |
| chr8 | 84847106 | 84847731 | 24.02885 | 69.3614 | 66.38335 | Promoter | 8 | 84847188 | 84848415 | 1228 | 1 | ENSMUST00 | 0 | 1700122E12 | ENSMUSG00000087685.1 |
| chr19 | 61225202 | 61227892 | 23.49892 | 69.32433 | 66.34698 | Promoter | 19 | 61224402 | 61228418 | 4017 | 2 | ENSMUST00 | 526 | Csf2ra | ENSMUSG00000059326.6 |
| chr15 | 12548931 | 12550332 | 24.50502 | 69.29691 | 66.31957 | Promoter | 15 | 12407282 | 12549278 | 141997 | 2 | ENSMUST00 | 0 | Pdzd2 | ENSMUSG00000022197.10 |
| chr19 | 5893650 | 5894481 | 24.50502 | 69.29691 | 66.31957 | Promoter | 19 | 5891139 | 5894107 | 2969 | 2 | ENSMUST00 | 0 | Tigd3 | ENSMUSG00000044390.4 |
| chr6 | 71707487 | 71708627 | 24.50502 | 69.29691 | 66.31957 | Promoter | 6 | 71707561 | 71810710 | 103150 | 1 | ENSMUST00 | 0 | Reep1 | ENSMUSG00000052852.7 |
| chr11 | 109361906 | 109362668 | 24.45608 | 69.21248 | 66.23546 | Promoter | 11 | 109362831 | 109380475 | 17645 | 1 | ENSMUST00 | -163 | Gna13 | ENSMUSG00000020611.9 |
| chr5 | 124328089 | 124328649 | 21.91255 | 69.13551 | 66.15882 | Promoter | 5 | 124328098 | 124341844 | 13747 | 1 | ENSMUST00 | 0 | 2810006K23 | ENSMUSG00000047635.6 |
| chr1 | 46853640 | 46854715 | 23.88927 | 69.11851 | 66.14216 | Promoter | 1 | 46807544 | 46854046 | 46503 | 2 | ENSMUST00 | 0 | Slc39a10 | ENSMUSG00000025986.6 |
| chr1 | 72225567 | 72226163 | 23.88927 | 69.11851 | 66.14216 | Promoter | 1 | 72226240 | 72226430 | 191 | 1 | ENSMUST00 | -77 | Gm25360 | ENSMUSG00000094655.1 |
| chr10 | 80475045 | 80475836 | 23.88927 | 69.11851 | 66.14216 | Promoter | 10 | 80476352 | 80476455 | 104 | 1 | ENSMUST00 | -516 | Gm25044 | ENSMUSG00000089228.1 |
| chr11 | 97029795 | 97030755 | 23.88927 | 69.11851 | 66.14216 | Promoter | 11 | 97029938 | 97033958 | 4021 | 1 | ENSMUST00 | 0 | Scrn2 | ENSMUSG00000020877.6 |
| chr13 | 34076532 | 34078549 | 23.88927 | 69.11851 | 66.14216 | Promoter | 13 | 34074301 | 34078008 | 3708 | 2 | ENSMUST00 | 0 | Tubb2a | ENSMUSG00000058672.6 |
| chr14 | 115091719 | 115092797 | 23.88927 | 69.11851 | 66.14216 | Promoter | 14 | 115092215 | 115134210 | 41996 | 1 | ENSMUST00 | 0 | Gpc5 | ENSMUSG00000022112.9 |
| chr14 | 55114573 | 55115695 | 23.88927 | 69.11851 | 66.14216 | Promoter | 14 | 55107277 | 55115552 | 8276 | 2 | ENSMUST00 | 0 | Jph4 | ENSMUSG00000022208.6 |

| | | | | | | | | | | | | | | | |
|---|---|---|---|---|---|---|---|---|---|---|---|---|---|---|---|
| chr15 | 89127842 | 89129292 | 23.88927 | 69.11851 | 66.14216 | Promoter | 15 | 89123307 | 89128653 | 5347 | 2 | ENSMUST00 | 0 | Hdac10 | ENSMUSG00000062906.7 |
| chr16 | 36874271 | 36874938 | 23.88927 | 69.11851 | 66.14216 | Promoter | 16 | 36792884 | 36874826 | 81943 | 2 | ENSMUST00 | 0 | Eaf2 | ENSMUSG00000022838.8 |
| chr18 | 82691477 | 82693572 | 23.88927 | 69.11851 | 66.14216 | Promoter | 18 | 82593593 | 82692883 | 99291 | 2 | ENSMUST00 | 0 | Zfp236 | ENSMUSG00000041258.11 |
| chr19 | 5406783 | 5407807 | 23.88927 | 69.11851 | 66.14216 | Promoter | 19 | 5406815 | 5417683 | 10869 | 1 | ENSMUST00 | 0 | 4930481A15 | ENSMUSG00000086938.3 |
| chr4 | 120032346 | 120033678 | 23.88927 | 69.11851 | 66.14216 | Intron (ENSM | 4 | 120094489 | 120134557 | 40069 | 1 | ENSMUST00 | -60811 | Hivep3 | ENSMUSG00000028634.11 |
| chr4 | 137468321 | 137469378 | 23.88927 | 69.11851 | 66.14216 | Promoter | 4 | 137468769 | 137570630 | 101862 | 1 | ENSMUST00 | 0 | Hspg2 | ENSMUSG00000028763.12 |
| chr5 | 142960079 | 142961488 | 23.88927 | 69.11851 | 66.14216 | Promoter | 5 | 142960343 | 142973185 | 12843 | 1 | ENSMUST00 | 0 | Fscn1 | ENSMUSG00000029581.9 |
| chr5 | 114175390 | 114176597 | 23.88927 | 69.11851 | 66.14216 | Promoter | 5 | 114175906 | 114190338 | 14433 | 1 | ENSMUST00 | 0 | Acacb | ENSMUSG00000042010.11 |
| chr6 | 108064806 | 108065649 | 23.88927 | 69.11851 | 66.14216 | Promoter | 6 | 108065045 | 108077122 | 12078 | 1 | ENSMUST00 | 0 | Setmar | ENSMUSG00000034639.6 |
| chr6 | 96116095 | 96117294 | 23.88927 | 69.11851 | 66.14216 | Promoter | 6 | 96115311 | 96656467 | 541157 | 1 | ENSMUST00 | 785 | Fam19a1 | ENSMUSG00000059187.7 |
| chr8 | 105518200 | 105519332 | 23.88927 | 69.11851 | 66.14216 | Promoter | 8 | 105518746 | 105523984 | 5239 | 1 | ENSMUST00 | 0 | Hsd11b2 | ENSMUSG00000031891.5 |
| chr8 | 122282038 | 122285122 | 23.88927 | 69.11851 | 66.14216 | Promoter | 8 | 122282141 | 122337251 | 55111 | 1 | ENSMUST00 | 0 | Zfpm1 | ENSMUSG00000049577.9 |
| chr9 | 107580290 | 107581431 | 23.88927 | 69.11851 | 66.14216 | Promoter | 9 | 107580818 | 107586832 | 6015 | 1 | ENSMUST00 | 0 | Hyal3 | ENSMUSG00000036091.3 |
| chr18 | 88971425 | 88972587 | 24.91517 | 69.11501 | 66.14118 | Promoter | 18 | 88971790 | 89131013 | 159224 | 1 | ENSMUST00 | 0 | Rttn | ENSMUSG00000023066.9 |
| chr2 | 25574161 | 25575623 | 24.91517 | 69.11501 | 66.14118 | Promoter | 2 | 25570174 | 25574845 | 4672 | 2 | ENSMUST00 | 0 | Mamdc4 | ENSMUSG00000026941.11 |
| chr8 | 104340179 | 104341479 | 24.91517 | 69.11501 | 66.14118 | Promoter | 8 | 104340594 | 104347672 | 7079 | 1 | ENSMUST00 | 0 | Cmtm3 | ENSMUSG00000031875.6 |
| chr7 | 35234171 | 35235263 | 24.35876 | 69.04449 | 66.07082 | Distal Interge | 7 | 35217235 | 35217360 | 126 | 1 | ENSMUST00 | 16937 | Gm24505 | ENSMUSG00000088307.1 |
| chr1 | 58795215 | 58796359 | 19.17126 | 69.02763 | 66.05415 | Promoter | 1 | 58795374 | 58847503 | 52130 | 1 | ENSMUST00 | 0 | Casp8 | ENSMUSG00000026029.9 |
| chr1 | 71556723 | 71557836 | 19.17126 | 69.02763 | 66.05415 | Promoter | 1 | 71557150 | 71579631 | 22482 | 1 | ENSMUST00 | 0 | Atic | ENSMUSG00000026192.8 |
| chr1 | 75236435 | 75237423 | 19.17126 | 69.02763 | 66.05415 | Promoter | 1 | 75236465 | 75244804 | 8340 | 1 | ENSMUST00 | 0 | Dnajb2 | ENSMUSG00000026203.11 |
| chr11 | 8499137 | 8504359 | 19.17126 | 69.02763 | 66.05415 | Intron (ENSM | 11 | 8451070 | 8468675 | 17606 | 2 | ENSMUST00 | -30463 | Tns3 | ENSMUSG00000020422.8 |
| chr12 | 36381280 | 36382483 | 19.17126 | 69.02763 | 66.05415 | Promoter | 12 | 36381519 | 36689444 | 307926 | 1 | ENSMUST00 | 0 | Ispd | ENSMUSG00000043153.5 |
| chr14 | 61037400 | 61038207 | 19.17126 | 69.02763 | 66.05415 | Promoter | 14 | 60963834 | 61037987 | 74154 | 2 | ENSMUST00 | 0 | Tnfrsf19 | ENSMUSG00000060548.7 |
| chr17 | 32946535 | 32947924 | 19.17126 | 69.02763 | 66.05415 | Promoter | 17 | 32924688 | 32947402 | 22715 | 2 | ENSMUST00 | 0 | Cyp4f13 | ENSMUSG00000024055.9 |
| chr17 | 36957679 | 36959243 | 19.17126 | 69.02763 | 66.05415 | Promoter | 17 | 36958592 | 36965622 | 7031 | 1 | ENSMUST00 | 0 | Znrd1as | ENSMUSG00000036214.8 |
| chr4 | 133095363 | 133097697 | 19.17126 | 69.02763 | 66.05415 | Distal Interge | 4 | 133085474 | 133089146 | 3673 | 1 | ENSMUST00 | 9890 | Gm26615 | ENSMUSG00000097088.1 |
| chr7 | 118532564 | 118533836 | 19.17126 | 69.02763 | 66.05415 | Promoter | 7 | 118509659 | 118533301 | 23643 | 2 | ENSMUST00 | 0 | Coq7 | ENSMUSG00000030652.5 |
| chr6 | 125231134 | 125232314 | 24.81235 | 68.93864 | 65.96623 | Promoter | 6 | 125231276 | 125239934 | 8659 | 1 | ENSMUST00 | 0 | Vamp1 | ENSMUSG00000030337.11 |
| chr13 | 67450997 | 67452249 | 22.28564 | 68.93403 | 65.96172 | Promoter | 13 | 67424549 | 67451585 | 27037 | 2 | ENSMUST00 | 0 | Zfp874a | ENSMUSG00000069206.8 |
| chr3 | 89101174 | 89101985 | 22.28564 | 68.93403 | 65.96172 | Promoter | 3 | 89093592 | 89101967 | 8376 | 2 | ENSMUST00 | 0 | Fdps | ENSMUSG00000059743.8 |
| chr4 | 116017072 | 116018149 | 22.28564 | 68.93403 | 65.96172 | Promoter | 4 | 115967145 | 116017675 | 50531 | 2 | ENSMUST00 | 0 | Faah | ENSMUSG00000034171.8 |
| chr10 | 44003320 | 44004521 | 18.2926 | 68.86671 | 65.89487 | Promoter | 10 | 44004154 | 44004701 | 548 | 1 | ENSMUST00 | 0 | Gm10812 | ENSMUSG00000075162.3 |
| chr11 | 32226473 | 32227086 | 18.2926 | 68.86671 | 65.89487 | Promoter | 11 | 32226505 | 32232022 | 5518 | 1 | ENSMUST00 | 0 | Mpg | ENSMUSG00000020287.10 |
| chr12 | 73111613 | 73113805 | 18.2926 | 68.86671 | 65.89487 | Promoter | 12 | 73099609 | 73113425 | 13817 | 2 | ENSMUST00 | 0 | Six4 | ENSMUSG00000034460.9 |
| chr12 | 84416817 | 84417928 | 18.2926 | 68.86671 | 65.89487 | Promoter | 12 | 84417196 | 84418578 | 1383 | 1 | ENSMUST00 | 0 | Rnf113a2 | ENSMUSG00000098134.1 |
| chr6 | 34910113 | 34911221 | 18.2926 | 68.86671 | 65.89487 | Promoter | 6 | 34880426 | 34910876 | 30451 | 2 | ENSMUST00 | 0 | Wdr91 | ENSMUSG00000058486.6 |
| chr7 | 132930467 | 132932593 | 18.2926 | 68.86671 | 65.89487 | Promoter | 7 | 132931142 | 132986391 | 55250 | 1 | ENSMUST00 | 0 | Zranb1 | ENSMUSG00000030967.8 |
| chr15 | 57891485 | 57893195 | 17.53424 | 68.85774 | 65.88623 | Promoter | 15 | 57869502 | 57892418 | 22917 | 2 | ENSMUST00 | 0 | Derl1 | ENSMUSG00000022365.5 |
| chr16 | 20715653 | 20718713 | 17.53424 | 68.85774 | 65.88623 | Promoter | 16 | 20717665 | 20722267 | 4603 | 1 | ENSMUST00 | 0 | Polr2h | ENSMUSG00000021018.7 |
| chr2 | 180257337 | 180258149 | 17.53424 | 68.85774 | 65.88623 | Promoter | 2 | 180257377 | 180258445 | 1069 | 1 | ENSMUST00 | 0 | Rps21 | ENSMUSG00000039001.7 |
| chr7 | 43671486 | 43673049 | 17.53424 | 68.85774 | 65.88623 | Promoter | 7 | 43672016 | 43678298 | 6283 | 1 | ENSMUST00 | 0 | Ctu1 | ENSMUSG00000038888.8 |
| chr9 | 13246922 | 13247810 | 17.53424 | 68.85774 | 65.88623 | Promoter | 9 | 13246982 | 13289409 | 42428 | 1 | ENSMUST00 | 0 | Ccdc82 | ENSMUSG00000079084.4 |
| chr9 | 22475377 | 22476602 | 24.76126 | 68.85096 | 65.8796 | Promoter | 9 | 22475715 | 22888280 | 412566 | 1 | ENSMUST00 | 0 | Bbs9 | ENSMUSG00000035919.11 |
| chr11 | 102774091 | 102775253 | 22.68498 | 68.76898 | 65.7979 | Promoter | 11 | 102774117 | 102776401 | 2285 | 1 | ENSMUST00 | 0 | Adam11 | ENSMUSG00000020926.11 |
| chr2 | 118598079 | 118598685 | 22.68498 | 68.76898 | 65.7979 | Promoter | 2 | 118598211 | 118641591 | 43381 | 1 | ENSMUST00 | 0 | Bub1b | ENSMUSG00000040084.9 |
| chr4 | 135872022 | 135874455 | 22.68498 | 68.76898 | 65.7979 | Promoter | 4 | 135870918 | 135873850 | 2933 | 2 | ENSMUST00 | 0 | Pnrc2 | ENSMUSG00000028675.7 |
| chr9 | 123366391 | 123367540 | 22.68498 | 68.76898 | 65.7979 | Promoter | 9 | 123366940 | 123462664 | 95725 | 1 | ENSMUST00 | 0 | Lars2 | ENSMUSG00000035202.7 |
| chr2 | 33370398 | 33371796 | 24.71037 | 68.76362 | 65.79295 | Promoter | 2 | 33133421 | 33371481 | 238061 | 2 | ENSMUST00 | 0 | Ralgps1 | ENSMUSG00000038831.11 |
| chr11 | 116305985 | 116307207 | 21.22555 | 68.73992 | 65.76938 | Promoter | 11 | 116288001 | 116306734 | 18734 | 2 | ENSMUST00 | 0 | Exoc7 | ENSMUSG00000020792.10 |
| chr9 | 57560513 | 57561714 | 24.16644 | 68.71214 | 65.74169 | Promoter | 9 | 57560944 | 57588797 | 27854 | 1 | ENSMUST00 | 0 | Scamp2 | ENSMUSG00000040188.9 |
| chr2 | 101885880 | 101886828 | 23.11343 | 68.64539 | 65.67522 | Promoter | 2 | 101886247 | 101896940 | 10694 | 1 | ENSMUST00 | 0 | Commd9 | ENSMUSG00000027163.8 |
| chr9 | 108991894 | 108992377 | 23.11343 | 68.64539 | 65.67522 | Promoter | 9 | 108991902 | 109010572 | 18671 | 1 | ENSMUST00 | 0 | Pfkfb4 | ENSMUSG00000025648.12 |
| chr8 | 122432190 | 122433189 | 24.11884 | 68.62979 | 65.66022 | Promoter | 8 | 122424772 | 122432940 | 8169 | 2 | ENSMUST00 | 0 | Cyba | ENSMUSG00000006519.10 |
| chr10 | 81070028 | 81070753 | 24.60923 | 68.58992 | 65.62046 | Promoter | 10 | 81070035 | 81082559 | 12525 | 1 | ENSMUST00 | 0 | Thop1 | ENSMUSG00000004929.7 |
| chr1 | 191312579 | 191313204 | 23.5743 | 68.56921 | 65.60024 | Intron (ENSM | 1 | 191306789 | 191318194 | 11406 | 2 | ENSMUST00 | 4990 | Nenf | ENSMUSG00000037499.9 |
| chr18 | 10051557 | 10053003 | 23.5743 | 68.56921 | 65.60024 | Distal Interge | 18 | 10021704 | 10045119 | 23416 | 2 | ENSMUST00 | -6439 | Usp14 | ENSMUSG00000047879.11 |
| chr2 | 133551874 | 133553509 | 23.5743 | 68.56921 | 65.60024 | Promoter | 2 | 133552159 | 133562885 | 10727 | 1 | ENSMUST00 | 0 | Bmp2 | ENSMUSG00000027358.6 |
| chr5 | 33018136 | 33018705 | 23.5743 | 68.56921 | 65.60024 | Promoter | 5 | 33018816 | 33027966 | 9151 | 1 | ENSMUST00 | -111 | Ywhah | ENSMUSG00000018965.10 |
| chr8 | 116992467 | 116994099 | 23.5743 | 68.56921 | 65.60024 | Promoter | 8 | 116981810 | 116993537 | 11728 | 2 | ENSMUST00 | 0 | Gcsh | ENSMUSG00000034424.5 |
| chr10 | 23348112 | 23350902 | 24.07142 | 68.54774 | 65.57966 | Promoter | 10 | 23104168 | 23349895 | 245728 | 2 | ENSMUST00 | 0 | Eya4 | ENSMUSG00000010461.9 |
| chr11 | 69394552 | 69396853 | 24.07142 | 68.54774 | 65.57966 | Promoter | 11 | 69391925 | 69395631 | 3707 | 2 | ENSMUST00 | 0 | Cyb5d1 | ENSMUSG00000044795.7 |
| chr14 | 61045367 | 61046913 | 24.07142 | 68.54774 | 65.57966 | Promoter | 14 | 60963834 | 61046855 | 83022 | 2 | ENSMUST00 | 0 | Tnfrsf19 | ENSMUSG00000060548.7 |
| chr2 | 127070198 | 127071325 | 24.07142 | 68.54774 | 65.57966 | Promoter | 2 | 127070665 | 127087870 | 17206 | 1 | ENSMUST00 | 0 | Blvra | ENSMUSG00000001999.10 |
| chr2 | 163054198 | 163055226 | 24.07142 | 68.54774 | 65.57966 | Promoter | 2 | 163054687 | 163084688 | 30002 | 1 | ENSMUST00 | 0 | Mybl2 | ENSMUSG00000017861.6 |
| chr2 | 32311860 | 32312859 | 24.07142 | 68.54774 | 65.57966 | Promoter | 2 | 32309485 | 32312265 | 2781 | 2 | ENSMUST00 | 0 | Dnm1 | ENSMUSG00000026825.12 |
| chr2 | 152343299 | 152344669 | 24.07142 | 68.54774 | 65.57966 | Promoter | 2 | 152337422 | 152344032 | 6611 | 2 | ENSMUST00 | 0 | Trib3 | ENSMUSG00000032715.9 |
| chr2 | 121354397 | 121355435 | 24.07142 | 68.54774 | 65.57966 | Promoter | 2 | 121310561 | 121355330 | 44770 | 2 | ENSMUST00 | 0 | Ppip5k1 | ENSMUSG00000033526.11 |
| chr10 | 121641006 | 121642571 | 24.55897 | 68.50356 | 65.53616 | Promoter | 10 | 121641588 | 121677301 | 35714 | 1 | ENSMUST00 | 0 | D930020B18 | ENSMUSG00000047642.9 |
| chr14 | 7817745 | 7819052 | 24.55897 | 68.50356 | 65.53616 | Promoter | 14 | 7817957 | 7951588 | 133632 | 1 | ENSMUST00 | 0 | Flnb | ENSMUSG00000025278.8 |
| chr2 | 131233487 | 131234939 | 21.55834 | 68.48431 | 65.51707 | Promoter | 2 | 131234063 | 131248022 | 13960 | 1 | ENSMUST00 | 0 | Mavs | ENSMUSG00000037523.8 |
| chr1 | 171437150 | 171440146 | 21.08548 | 68.47759 | 65.51048 | Promoter | 1 | 171437535 | 171464603 | 27069 | 1 | ENSMUST00 | 0 | F11r | ENSMUSG00000038235.4 |
| chr10 | 67911967 | 67912778 | 21.08548 | 68.47759 | 65.51048 | Promoter | 10 | 67886103 | 67912662 | 26560 | 2 | ENSMUST00 | 0 | Zfp365 | ENSMUSG00000037855.10 |
| chr16 | 18347868 | 18349368 | 21.08548 | 68.47759 | 65.51048 | Promoter | 16 | 18348182 | 18407076 | 58895 | 1 | ENSMUST00 | 0 | Arvcf | ENSMUSG00000000325.10 |
| chr19 | 4942387 | 4943623 | 21.08548 | 68.47759 | 65.51048 | Promoter | 19 | 4930651 | 4943092 | 12442 | 2 | ENSMUST00 | 0 | Peli3 | ENSMUSG00000024901.9 |
| chr2 | 6212609 | 6213761 | 21.08548 | 68.47759 | 65.51048 | Promoter | 2 | 6188465 | 6213033 | 24569 | 2 | ENSMUST00 | 0 | Echdc3 | ENSMUSG00000039063.4 |
| chr3 | 96057350 | 96058848 | 21.08548 | 68.47759 | 65.51048 | Promoter | 3 | 95999832 | 96058466 | 58635 | 2 | ENSMUST00 | 0 | Vps45 | ENSMUSG00000015747.5 |
| chr3 | 79628001 | 79628815 | 21.08548 | 68.47759 | 65.51048 | Promoter | 3 | 79603803 | 79628697 | 24895 | 2 | ENSMUST00 | 0 | Etfdh | ENSMUSG00000027809.9 |
| chr4 | 124936304 | 124937952 | 21.08548 | 68.47759 | 65.51048 | Promoter | 4 | 124918465 | 124936917 | 18453 | 2 | ENSMUST00 | 0 | Cdca8 | ENSMUSG00000028873.11 |
| chr7 | 27731140 | 27732175 | 21.08548 | 68.47759 | 65.51048 | Promoter | 7 | 27731389 | 27753702 | 22314 | 1 | ENSMUST00 | 0 | Zfp60 | ENSMUSG00000037640.10 |
| chr5 | 109690220 | 109691416 | 24.45906 | 68.33184 | 65.3672 | Promoter | 5 | 109674545 | 109691041 | 16497 | 2 | ENSMUST00 | 0 | 5430403G16 | ENSMUSG00000072763.2 |
| chr9 | 89211168 | 89211897 | 24.45906 | 68.33184 | 65.3672 | Promoter | 9 | 89211383 | 89240115 | 28733 | 1 | ENSMUST00 | 0 | Mthfs | ENSMUSG00000066442.11 |
| chr12 | 80113160 | 80116456 | 21.91337 | 68.25864 | 65.29428 | Promoter | 12 | 80107760 | 80113013 | 5254 | 2 | ENSMUST00 | -148 | Zfp36l1 | ENSMUSG00000021127.6 |
| chr6 | 65778665 | 65779629 | 21.91337 | 68.25864 | 65.29428 | Promoter | 6 | 65778988 | 65936373 | 157386 | 1 | ENSMUST00 | 0 | Prdm5 | ENSMUSG00000029913.9 |
| chr5 | 22343696 | 22345699 | 24.40942 | 68.24645 | 65.28231 | Promoter | 5 | 21884454 | 22344702 | 460249 | 2 | ENSMUST00 | 0 | Reln | ENSMUSG00000042453.9 |
| chr7 | 102564866 | 102565822 | 24.40942 | 68.24645 | 65.28231 | Promoter | 7 | 102557922 | 102565469 | 7548 | 2 | ENSMUST00 | 0 | Trim21 | ENSMUSG00000030966.7 |
| chr5 | 88674798 | 88676601 | 15.57114 | 68.18018 | 65.21632 | Promoter | 5 | 88659448 | 88676171 | 16724 | 2 | ENSMUST00 | 0 | Grsf1 | ENSMUSG00000044221.9 |
| chr19 | 61199567 | 61200137 | 4.59443 | 68.17046 | 65.20663 | Distal Interge | 19 | 61183890 | 61183955 | 66 | 1 | ENSMUST00 | 15678 | Gm6020 | ENSMUSG00000069475.1 |
| chr15 | 36727085 | 36728327 | 24.35997 | 68.16138 | 65.19754 | Distal Interge | 15 | 36790712 | 36792414 | 1703 | 2 | ENSMUST00 | 64087 | Ywhaz | ENSMUSG00000022285.11 |
| chr9 | 65675887 | 65677358 | 24.35997 | 68.16138 | 65.19754 | Promoter | 9 | 65676548 | 65690300 | 13753 | 1 | ENSMUST00 | 0 | Oaz2 | ENSMUSG00000040652.10 |
| chr10 | 84621752 | 84623247 | 22.29294 | 68.06667 | 65.1033 | Promoter | 10 | 84622437 | 84727177 | 104741 | 1 | ENSMUST00 | 0 | Polr3b | ENSMUSG00000034453.7 |
| chr5 | 21372292 | 21373711 | 22.29294 | 68.06667 | 65.1033 | Promoter | 5 | 21372642 | 21378374 | 5733 | 1 | ENSMUST00 | 0 | Fgl2 | ENSMUSG00000039899.5 |
| chr8 | 69394845 | 69396049 | 22.29294 | 68.06667 | 65.1033 | Distal Interge | 8 | 69399725 | 69400020 | 296 | 1 | ENSMUST00 | -3676 | Gm25661 | ENSMUSG00000088329.1 |
| chr11 | 98338360 | 98339625 | 24.81473 | 68.0602 | 65.09722 | Distal Interge | 11 | 98325415 | 98329648 | 4234 | 2 | ENSMUST00 | -8713 | Neurod2 | ENSMUSG00000038255.6 |
| chr15 | 87623999 | 87625775 | 24.81473 | 68.0602 | 65.09722 | Promoter | 15 | 87625230 | 87759336 | 134107 | 1 | ENSMUST00 | 0 | Fam19a5 | ENSMUSG00000054863.8 |
| chr2 | 84798207 | 84799862 | 24.81473 | 68.0602 | 65.09722 | Promoter | 2 | 84798828 | 84810176 | 11349 | 1 | ENSMUST00 | 0 | Ube2l6 | ENSMUSG00000027078.9 |
| chr2 | 118813873 | 118814746 | 24.81473 | 68.0602 | 65.09722 | Promoter | 2 | 118814003 | 118837696 | 23694 | 1 | ENSMUST00 | 0 | Knstrn | ENSMUSG00000027331.10 |
| chr4 | 136171617 | 136173503 | 24.81473 | 68.0602 | 65.09722 | Promoter | 4 | 136172394 | 136196057 | 23664 | 1 | ENSMUST00 | 0 | E2f2 | ENSMUSG00000018983.9 |
| chr5 | 34525486 | 34526710 | 24.81473 | 68.0602 | 65.09722 | Promoter | 5 | 34525838 | 34551700 | 25863 | 1 | ENSMUST00 | 0 | Sh3bp2 | ENSMUSG00000054520.10 |
| chr6 | 29526293 | 29527809 | 24.81473 | 68.0602 | 65.09722 | Promoter | 6 | 29526625 | 29537319 | 10695 | 1 | ENSMUST00 | 0 | Irf5 | ENSMUSG00000029771.7 |
| chrX | 12587201 | 12588431 | 24.81473 | 68.0602 | 65.09722 | Promoter | 20 | 12587801 | 12596581 | 8781 | 1 | ENSMUST00 | 0 | Atp6ap2 | ENSMUSG00000031007.2 |
| chrX | 73965220 | 73968256 | 24.81473 | 68.0602 | 65.09722 | Promoter | 20 | 73942792 | 73966357 | 23566 | 2 | ENSMUST00 | 0 | Hcfc1 | ENSMUSG00000031386.9 |
| chrX | 106219016 | 106220227 | 24.81473 | 68.0602 | 65.09722 | Promoter | 20 | 106206883 | 106219901 | 13019 | 2 | ENSMUST00 | 0 | Taf9b | ENSMUSG00000047242.9 |
| chrX | 136033350 | 136034096 | 24.81473 | 68.0602 | 65.09722 | Promoter | 20 | 136033367 | 136034946 | 1580 | 1 | ENSMUST00 | 0 | Arxes1 | ENSMUSG00000048355.2 |
| chr4 | 62479669 | 62480443 | 24.26167 | 67.9922 | 65.03062 | Promoter | 4 | 62480053 | 62497298 | 17246 | 1 | ENSMUST00 | 0 | Bspry | ENSMUSG00000028392.10 |
| chr1 | 134555703 | 134557642 | 19.89052 | 67.91473 | 64.9535 | Distal Interge | 1 | 134560171 | 134635285 | 75115 | 1 | ENSMUST00 | -2529 | Kdm5b | ENSMUSG00000042207.12 |
| chr10 | 31445275 | 31446108 | 19.89052 | 67.91473 | 64.9535 | Promoter | 10 | 31332380 | 31445921 | 113542 | 2 | ENSMUST00 | 0 | Tpd52l1 | ENSMUSG00000000296.7 |
| chr10 | 100487513 | 100488167 | 19.89052 | 67.91473 | 64.9535 | Promoter | 10 | 100488289 | 100573655 | 85367 | 1 | ENSMUST00 | -122 | Cep290 | ENSMUSG00000019971.9 |
| chr10 | 100588659 | 100589507 | 19.89052 | 67.91473 | 64.9535 | Promoter | 10 | 100572274 | 100589259 | 16986 | 2 | ENSMUST00 | 0 | 4930430F08 | ENSMUSG00000046567.9 |
| chr12 | 12261863 | 12263016 | 19.89052 | 67.91473 | 64.9535 | Promoter | 12 | 12262139 | 12376359 | 114221 | 1 | ENSMUST00 | 0 | Fam49a | ENSMUSG00000020589.11 |
| chr3 | 121262251 | 121263451 | 19.89052 | 67.91473 | 64.9535 | Promoter | 3 | 121202008 | 121263159 | 61152 | 2 | ENSMUST00 | 0 | Tmem56 | ENSMUSG00000028132.10 |
| chr5 | 125404310 | 125407021 | 19.89052 | 67.91473 | 64.9535 | Distal Interge | 5 | 125385977 | 125390202 | 4226 | 2 | ENSMUST00 | -14109 | Ubc | ENSMUSG00000008348.9 |
| chr5 | 53212386 | 53214037 | 19.89052 | 67.91473 | 64.9535 | Promoter | 5 | 53107084 | 53213452 | 106369 | 2 | ENSMUST00 | 0 | Sel1l3 | ENSMUSG00000029189.6 |
| chr5 | 114443548 | 114444071 | 19.89052 | 67.91473 | 64.9535 | Promoter | 5 | 114431034 | 114444059 | 13026 | 2 | ENSMUST00 | 0 | Mmab | ENSMUSG00000029575.12 |
| chr7 | 13277932 | 13279594 | 19.89052 | 67.91473 | 64.9535 | Promoter | 7 | 13278826 | 13311425 | 32600 | 1 | ENSMUST00 | 0 | Lig1 | ENSMUSG00000056394.12 |
| chr8 | 71628786 | 71629956 | 19.89052 | 67.91473 | 64.9535 | 3' UTR | 8 | 71631047 | 71631106 | 60 | 2 | ENSMUST00 | 1150 | Mir6769b | ENSMUSG00000098976.1 |
| chr11 | 30754685 | 30755562 | 22.69969 | 67.91293 | 64.95291 | Promoter | 11 | 30754572 | 30754961 | 390 | 2 | ENSMUST00 | 0 | Gm12101 | ENSMUSG00000083990.1 |
| chr2 | 74762574 | 74763758 | 22.69969 | 67.91293 | 64.95291 | Promoter | 2 | 74762980 | 74765142 | 2163 | 1 | ENSMUST00 | 0 | Hoxd1 | ENSMUSG00000042448.5 |
| chr3 | 88138472 | 88139892 | 22.69969 | 67.91293 | 64.95291 | Distal Interge | 3 | 88142372 | 88172086 | 29715 | 1 | ENSMUST00 | -2480 | Mef2d | ENSMUSG00000001419.12 |
| chr7 | 126799476 | 126800481 | 23.69868 | 67.90152 | 64.94212 | Promoter | 7 | 126795235 | 126800447 | 5213 | 2 | ENSMUST00 | 0 | Aldoa | ENSMUSG00000030695.8 |
| chr17 | 34264372 | 34265289 | 24.71824 | 67.89503 | 64.9357 | Exon (ENSMU | 17 | 34263209 | 34269418 | 6210 | 1 | ENSMUST00 | 1164 | H2-Ab1 | ENSMUSG00000073421.5 |
| chr14 | 120910190 | 120912364 | 16.0743 | 67.86449 | 64.90525 | Promoter | 14 | 120911194 | 120948042 | 36849 | 1 | ENSMUST00 | 0 | Ipo5 | ENSMUSG00000030662.9 |
| chr10 | 79532838 | 79534077 | 21.2154 | 67.84969 | 64.89048 | Promoter | 10 | 79526430 | 79533742 | 7313 | 2 | ENSMUST00 | 0 | Ppap2c | ENSMUSG00000052151.6 |
| chr11 | 75966189 | 75967304 | 21.2154 | 67.84969 | 64.89048 | Distal Interge | 11 | 75906572 | 75938429 | 31858 | 2 | ENSMUST00 | -27761 | Rph3al | ENSMUSG00000020847.10 |
| chr11 | 58097331 | 58098325 | 24.16416 | 67.82425 | 64.86514 | Promoter | 11 | 58097104 | 58097773 | 670 | 2 | ENSMUST00 | 0 | Gm12247 | ENSMUSG00000081223.1 |
| chr14 | 62761387 | 62762328 | 23.6529 | 67.82199 | 64.86302 | Promoter | 14 | 62676330 | 62761112 | 84783 | 2 | ENSMUST00 | -276 | Ints6 | ENSMUSG00000035161.6 |
| chr1 | 75179724 | 75180728 | 22.15371 | 67.81625 | 64.85734 | Promoter | 1 | 75171717 | 75180392 | 8676 | 2 | ENSMUST00 | 0 | Abcb6 | ENSMUSG00000026198.10 |
| chr10 | 13868363 | 13869834 | 22.15371 | 67.81625 | 64.85734 | Promoter | 10 | 13647054 | 13868980 | 221927 | 2 | ENSMUST00 | 0 | Aig1 | ENSMUSG00000019806.8 |
| chr11 | 61504771 | 61505604 | 22.15371 | 67.81625 | 64.85734 | Promoter | 11 | 61505144 | 61512931 | 7788 | 1 | ENSMUST00 | 0 | B9d1 | ENSMUSG00000001039.7 |

| | | | | | | | | | | | | | | | |
|---|---|---|---|---|---|---|---|---|---|---|---|---|---|---|---|
| chr12 | 73053350 | 73054533 | 22.15371 | 67.81625 | 64.85734 | Promoter | 12 | 73043531 | 73053887 | 10357 | 2 | ENSMUST00 | 0 | Six1 | ENSMUSG00000051367.8 |
| chr13 | 41355780 | 41359729 | 22.15371 | 67.81625 | 64.85734 | Promoter | 13 | 41309916 | 41359001 | 49086 | 2 | ENSMUST00 | 0 | Nedd9 | ENSMUSG00000021365.9 |
| chr13 | 34992948 | 34994858 | 22.15371 | 67.81625 | 64.85734 | Promoter | 13 | 34977748 | 34994144 | 16397 | 2 | ENSMUST00 | 0 | Eci2 | ENSMUSG00000021417.9 |
| chr14 | 58070134 | 58071635 | 22.15371 | 67.81625 | 64.85734 | Promoter | 14 | 58070547 | 58112337 | 41791 | 1 | ENSMUST00 | 0 | Fgf9 | ENSMUSG00000021974.9 |
| chr15 | 103168825 | 103169601 | 22.15371 | 67.81625 | 64.85734 | Distal Interge | 15 | 103153307 | 103158002 | 4696 | 2 | ENSMUST00 | -10824 | Smug1 | ENSMUSG00000036061.9 |
| chr17 | 67697029 | 67698070 | 22.15371 | 67.81625 | 64.85734 | Promoter | 17 | 67697265 | 67822645 | 125381 | 1 | ENSMUST00 | 0 | Lama1 | ENSMUSG00000032796.7 |
| chr17 | 36205206 | 36206489 | 22.15371 | 67.81625 | 64.85734 | Promoter | 17 | 36201996 | 36206339 | 4344 | 2 | ENSMUST00 | 0 | Gm8835 | ENSMUSG00000092457.3 |
| chr19 | 54044572 | 54047741 | 22.15371 | 67.81625 | 64.85734 | Promoter | 19 | 54045182 | 54048982 | 3801 | 1 | ENSMUST00 | 0 | Adra2a | ENSMUSG00000033717.5 |
| chr2 | 71366842 | 71368486 | 22.15371 | 67.81625 | 64.85734 | Promoter | 2 | 71271063 | 71367749 | 96687 | 2 | ENSMUST00 | 0 | Slc25a12 | ENSMUSG00000027010.11 |
| chr2 | 155591686 | 155593225 | 22.15371 | 67.81625 | 64.85734 | Promoter | 2 | 155563181 | 155592810 | 29630 | 2 | ENSMUST00 | 0 | Gss | ENSMUSG00000027610.12 |
| chr2 | 116066009 | 116067477 | 22.15371 | 67.81625 | 64.85734 | Promoter | 2 | 116067225 | 116085911 | 18687 | 1 | ENSMUST00 | 0 | G630016G05 | ENSMUSG00000086308.1 |
| chr3 | 89266395 | 89267119 | 22.15371 | 67.81625 | 64.85734 | Promoter | 3 | 89266552 | 89267077 | 526 | 1 | ENSMUST00 | 0 | Dpm3 | ENSMUSG00000042737.4 |
| chr5 | 34188018 | 34188670 | 22.15371 | 67.81625 | 64.85734 | Promoter | 5 | 34176579 | 34187720 | 11142 | 2 | ENSMUST00 | -299 | Mxd4 | ENSMUSG00000037235.8 |
| chr6 | 52238979 | 52240843 | 22.15371 | 67.81625 | 64.85734 | Promoter | 6 | 52232082 | 52240795 | 8714 | 2 | ENSMUST00 | 0 | Hoxa10 | ENSMUSG00000000938.11 |
| chr6 | 48085517 | 48086971 | 22.15371 | 67.81625 | 64.85734 | Promoter | 6 | 48062395 | 48086593 | 24199 | 2 | ENSMUST00 | 0 | Zfp746 | ENSMUSG00000057691.6 |
| chr6 | 47663853 | 47664155 | 22.15371 | 67.81625 | 64.85734 | Distal Interge | 6 | 47637143 | 47637219 | 77 | 1 | ENSMUST00 | 26711 | Gm25283 | ENSMUSG00000087897.1 |
| chr7 | 29288969 | 29289636 | 22.15371 | 67.81625 | 64.85734 | Promoter | 7 | 29289320 | 29293390 | 4071 | 1 | ENSMUST00 | 0 | Ppp1r14a | ENSMUSG00000037166.4 |
| chr7 | 27507246 | 27508158 | 22.15371 | 67.81625 | 64.85734 | Promoter | 7 | 27507801 | 27520034 | 12234 | 1 | ENSMUST00 | 0 | Prx | ENSMUSG00000053198.8 |
| chr8 | 83740029 | 83742098 | 22.15371 | 67.81625 | 64.85734 | Promoter | 8 | 83723251 | 83741326 | 18076 | 2 | ENSMUST00 | 0 | Cd97 | ENSMUSG00000002885.9 |
| chr8 | 122576232 | 122577300 | 22.15371 | 67.81625 | 64.85734 | Promoter | 8 | 122574637 | 122576907 | 2271 | 2 | ENSMUST00 | 0 | Aprt | ENSMUSG00000006589.7 |
| chr8 | 72570135 | 72572933 | 22.15371 | 67.81625 | 64.85734 | Promoter | 8 | 72572103 | 72587282 | 15180 | 1 | ENSMUST00 | 0 | Tmem38a | ENSMUSG00000031791.7 |
| chr8 | 120535456 | 120538941 | 22.15371 | 67.81625 | 64.85734 | Promoter | 8 | 120535857 | 120579480 | 43624 | 1 | ENSMUST00 | 0 | Gse1 | ENSMUSG00000031822.13 |
| chr8 | 35825655 | 35827155 | 22.15371 | 67.81625 | 64.85734 | Promoter | 8 | 35824709 | 35826559 | 1851 | 2 | ENSMUST00 | 0 | Cldn23 | ENSMUSG00000055976.5 |
| chr2 | 93955911 | 93957490 | 24.6666 | 67.80661 | 64.85043 | Promoter | 2 | 93955810 | 93957049 | 1240 | 2 | ENSMUST00 | 0 | Gm13889 | ENSMUSG00000087006.3 |
| chr4 | 55530022 | 55533908 | 24.6666 | 67.80661 | 64.85043 | Promoter | 4 | 55527143 | 55532466 | 5324 | 2 | ENSMUST00 | 0 | Klf4 | ENSMUSG00000003032.8 |
| chr1 | 93685234 | 93686792 | 23.13664 | 67.80279 | 64.84689 | Promoter | 1 | 93685660 | 93695764 | 10105 | 1 | ENSMUST00 | 0 | Bok | ENSMUSG00000026278.9 |
| chr11 | 101245876 | 101246906 | 23.13664 | 67.80279 | 64.84689 | Promoter | 11 | 101246028 | 101248250 | 2223 | 1 | ENSMUST00 | 0 | Ramp2 | ENSMUSG00000001240.8 |
| chr3 | 49756025 | 49757785 | 23.13664 | 67.80279 | 64.84689 | Promoter | 3 | 49743296 | 49757316 | 14021 | 2 | ENSMUST00 | 0 | Pcdh18 | ENSMUSG00000037892.8 |
| chr17 | 35195427 | 35196374 | 23.60729 | 67.74274 | 64.78755 | Promoter | 17 | 35194972 | 35195049 | 78 | 1 | ENSMUST00 | 456 | Mir6973a | ENSMUSG00000098291.1 |
| chr5 | 15933989 | 15935703 | 23.60729 | 67.74274 | 64.78755 | Promoter | 5 | 15934691 | 16374511 | 439821 | 1 | ENSMUST00 | 0 | Cacna2d1 | ENSMUSG00000040118.10 |
| chr6 | 99691965 | 99693676 | 23.60729 | 67.74274 | 64.78755 | Promoter | 6 | 99692679 | 99693818 | 1140 | 1 | ENSMUST00 | 0 | Gpr27 | ENSMUSG00000072875.1 |
| chr8 | 94532544 | 94533595 | 23.60729 | 67.74274 | 64.78755 | Promoter | 8 | 94532990 | 94570529 | 37540 | 1 | ENSMUST00 | 0 | Cpne2 | ENSMUSG00000034361.4 |
| chr10 | 110455341 | 110456443 | 24.1157 | 67.74074 | 64.78616 | Promoter | 10 | 109899103 | 110456204 | 557102 | 2 | ENSMUST00 | 0 | Nav3 | ENSMUSG00000020181.10 |
| chr15 | 79833220 | 79834719 | 24.1157 | 67.74074 | 64.78616 | Promoter | 15 | 79823896 | 79834688 | 10793 | 2 | ENSMUST00 | 0 | Cbx6 | ENSMUSG00000089715.6 |
| chr17 | 25796309 | 25797389 | 24.1157 | 67.74074 | 64.78616 | Promoter | 17 | 25794571 | 25797045 | 2475 | 2 | ENSMUST00 | 0 | Metrn | ENSMUSG00000002274.7 |
| chr2 | 32775247 | 32776504 | 24.1157 | 67.74074 | 64.78616 | Promoter | 2 | 32757026 | 32775383 | 18358 | 2 | ENSMUST00 | 0 | Ttc16 | ENSMUSG00000039021.10 |
| chr10 | 80385919 | 80387636 | 24.61517 | 67.71854 | 64.76461 | Promoter | 10 | 80380357 | 80387651 | 7295 | 2 | ENSMUST00 | 15 | Mex3d | ENSMUSG00000048696.10 |
| chr2 | 61655178 | 61657128 | 24.56396 | 67.6308 | 64.67719 | Distal Interge | 2 | 61643831 | 61649977 | 6147 | 1 | ENSMUST00 | 11348 | Tank | ENSMUSG00000064289.10 |
| chr11 | 77762905 | 77764356 | 16.64193 | 67.62904 | 64.67553 | Promoter | 11 | 77763246 | 77865980 | 102735 | 1 | ENSMUST00 | 0 | Myo18a | ENSMUSG00000000631.14 |
| chr13 | 67194280 | 67195093 | 16.64193 | 67.62904 | 64.67553 | Promoter | 13 | 67194506 | 67209298 | 14793 | 1 | ENSMUST00 | 0 | Zfp455 | ENSMUSG00000051037.10 |
| chr13 | 62785268 | 62786273 | 21.55333 | 67.6013 | 64.64786 | Distal Interge | 13 | 62791507 | 62791613 | 107 | 1 | ENSMUST00 | -5234 | Gm25448 | ENSMUSG00000095825.1 |
| chr11 | 68967300 | 68968611 | 18.88512 | 67.59834 | 64.64505 | Promoter | 11 | 68968131 | 68972516 | 4386 | 1 | ENSMUST00 | 0 | Slc25a35 | ENSMUSG00000018740.9 |
| chr11 | 70539548 | 70540732 | 18.88512 | 67.59834 | 64.64505 | Promoter | 11 | 70540064 | 70541183 | 1120 | 1 | ENSMUST00 | 0 | Pld2 | ENSMUSG00000020828.8 |
| chr11 | 72440680 | 72442086 | 18.88512 | 67.59834 | 64.64505 | Promoter | 11 | 72441355 | 72451768 | 10414 | 1 | ENSMUST00 | 0 | Mybbp1a | ENSMUSG00000040463.11 |
| chr13 | 97197795 | 97198703 | 18.88512 | 67.59834 | 64.64505 | Promoter | 13 | 97176332 | 97198357 | 22026 | 2 | ENSMUST00 | 0 | Hexb | ENSMUSG00000021665.7 |
| chr2 | 30658018 | 30659016 | 18.88512 | 67.59834 | 64.64505 | Exon (ENSMU | 2 | 30658468 | 30669908 | 11441 | 2 | ENSMUST00 | 10892 | Gm14486 | ENSMUSG00000087268.1 |
| chr6 | 57535719 | 57536412 | 18.88512 | 67.59834 | 64.64505 | Promoter | 6 | 57506502 | 57535426 | 28925 | 2 | ENSMUST00 | -294 | Ppm1k | ENSMUSG00000037826.3 |
| chr7 | 102064411 | 102066178 | 18.88512 | 67.59834 | 64.64505 | Promoter | 7 | 102065713 | 102096416 | 30704 | 1 | ENSMUST00 | 0 | Trpc2 | ENSMUSG00000070425.7 |
| chrX | 103165034 | 103165655 | 18.88512 | 67.59834 | 64.64505 | Promoter | 20 | 103163110 | 103165407 | 2298 | 2 | ENSMUST00 | 0 | Gm6206 | ENSMUSG00000048949.8 |
| chr7 | 27804196 | 27805515 | 24.01936 | 67.57464 | 64.62208 | Distal Interge | 7 | 27810813 | 27818669 | 7857 | 1 | ENSMUST00 | -5298 | Zfp626 | ENSMUSG00000030604.8 |
| chr3 | 10012125 | 10013647 | 24.51296 | 67.54341 | 64.59099 | Promoter | 3 | 10012548 | 10016607 | 4060 | 1 | ENSMUST00 | 0 | Fabp5 | ENSMUSG00000027533.10 |
| chr5 | 147893748 | 147895374 | 20.58102 | 67.52638 | 64.57406 | Promoter | 5 | 147878437 | 147894815 | 16379 | 2 | ENSMUST00 | 0 | Slc46a3 | ENSMUSG00000029650.5 |
| chr1 | 132390559 | 132392052 | 23.47016 | 67.50422 | 64.55198 | Promoter | 1 | 132356315 | 132391281 | 34967 | 2 | ENSMUST00 | 0 | Tmcc2 | ENSMUSG00000042066.10 |
| chr11 | 77514569 | 77516040 | 23.47016 | 67.50422 | 64.55198 | Promoter | 11 | 77515121 | 77520045 | 4925 | 1 | ENSMUST00 | 0 | Abhd15 | ENSMUSG00000000686.11 |
| chr11 | 113565227 | 113566061 | 23.47016 | 67.50422 | 64.55198 | Promoter | 11 | 113244853 | 113565756 | 320904 | 2 | ENSMUST00 | 0 | Slc39a11 | ENSMUSG00000041654.10 |
| chr14 | 51912893 | 51914272 | 23.47016 | 67.50422 | 64.55198 | Promoter | 14 | 51905271 | 51913488 | 8218 | 2 | ENSMUST00 | 0 | Ndrg2 | ENSMUSG00000004558.9 |
| chr15 | 34306458 | 34307261 | 23.47016 | 67.50422 | 64.55198 | Promoter | 15 | 34306680 | 34436238 | 129559 | 1 | ENSMUST00 | 0 | Matn2 | ENSMUSG00000022324.9 |
| chr17 | 44078541 | 44079337 | 23.47016 | 67.50422 | 64.55198 | Promoter | 17 | 44078813 | 44086567 | 7755 | 1 | ENSMUST00 | 0 | Enpp5 | ENSMUSG00000023960.8 |
| chr18 | 77185315 | 77186278 | 23.47016 | 67.50422 | 64.55198 | Promoter | 18 | 77185853 | 77255450 | 69598 | 1 | ENSMUST00 | 0 | St8sia5 | ENSMUSG00000025425.12 |
| chr19 | 7267778 | 7269372 | 23.47016 | 67.50422 | 64.55198 | Promoter | 19 | 7268301 | 7275225 | 6925 | 1 | ENSMUST00 | 0 | Rcor2 | ENSMUSG00000024968.8 |
| chr2 | 127335645 | 127336933 | 23.47016 | 67.50422 | 64.55198 | Promoter | 2 | 127336159 | 127338376 | 2218 | 1 | ENSMUST00 | 0 | Dusp2 | ENSMUSG00000027368.6 |
| chr2 | 127142816 | 127143808 | 23.47016 | 67.50422 | 64.55198 | Promoter | 2 | 127138772 | 127143442 | 4671 | 2 | ENSMUST00 | 0 | Itpripl1 | ENSMUSG00000074825.4 |
| chr5 | 136907666 | 136908800 | 23.47016 | 67.50422 | 64.55198 | Promoter | 5 | 136908150 | 136913244 | 5095 | 1 | ENSMUST00 | 0 | Ift22 | ENSMUSG00000007987.8 |
| chr7 | 45366411 | 45367908 | 23.47016 | 67.50422 | 64.55198 | Promoter | 7 | 45339126 | 45366714 | 27589 | 2 | ENSMUST00 | 0 | Ppfia3 | ENSMUSG00000003863.11 |
| chr7 | 111083205 | 111084051 | 23.47016 | 67.50422 | 64.55198 | Promoter | 7 | 111070568 | 111083030 | 12463 | 2 | ENSMUST00 | -176 | Eif4g2 | ENSMUSG00000005610.11 |
| chr8 | 104170268 | 104171332 | 23.47016 | 67.50422 | 64.55198 | Promoter | 8 | 104170513 | 104219092 | 48580 | 1 | ENSMUST00 | 0 | Bean1 | ENSMUSG00000031872.8 |
| chr8 | 123653446 | 123654610 | 23.47016 | 67.50422 | 64.55198 | Promoter | 8 | 123653929 | 123655985 | 2057 | 1 | ENSMUST00 | 0 | Rhou | ENSMUSG00000039960.5 |
| chr1 | 136414780 | 136415964 | 17.28728 | 67.4904 | 64.54075 | Promoter | 1 | 136415271 | 136417792 | 2522 | 1 | ENSMUST00 | 0 | Ddx59 | ENSMUSG00000026404.7 |
| chr16 | 21787345 | 21788245 | 17.28728 | 67.4904 | 64.54075 | Promoter | 16 | 21761287 | 21787844 | 26558 | 2 | ENSMUST00 | 0 | Ehhadh | ENSMUSG00000022853.6 |
| chr19 | 47854432 | 47855663 | 17.28728 | 67.4904 | 64.54075 | Promoter | 19 | 47854970 | 47864790 | 9821 | 1 | ENSMUST00 | 0 | Gsto1 | ENSMUSG00000025068.7 |
| chr7 | 30551563 | 30553162 | 17.28728 | 67.4904 | 64.54075 | Promoter | 7 | 30552178 | 30555440 | 3263 | 1 | ENSMUST00 | 0 | Hspb6 | ENSMUSG00000036854.9 |
| chr11 | 118289150 | 118290206 | 18.02749 | 67.47041 | 64.5209 | Promoter | 11 | 118262388 | 118290180 | 27793 | 2 | ENSMUST00 | 0 | Usp36 | ENSMUSG00000033909.12 |
| chr11 | 48852866 | 48853692 | 18.02749 | 67.47041 | 64.5209 | Promoter | 11 | 48843648 | 48853493 | 9846 | 2 | ENSMUST00 | 0 | Gm16170 | ENSMUSG00000086245.1 |
| chr16 | 45742277 | 45743436 | 18.02749 | 67.47041 | 64.5209 | Promoter | 16 | 45729725 | 45742955 | 13231 | 2 | ENSMUST00 | 0 | Abhd10 | ENSMUSG00000033157.11 |
| chr18 | 5333642 | 5334839 | 18.02749 | 67.47041 | 64.5209 | Promoter | 18 | 5210031 | 5334439 | 124409 | 2 | ENSMUST00 | 0 | Zfp438 | ENSMUSG00000050945.7 |
| chr7 | 44304267 | 44305249 | 18.02749 | 67.47041 | 64.5209 | 3' UTR | 7 | 44303770 | 44306959 | 3190 | 2 | ENSMUST00 | 1710 | Clec11a | ENSMUSG00000004473.9 |
| chr7 | 127253646 | 127254844 | 18.02749 | 67.47041 | 64.5209 | 3' UTR | 7 | 127256959 | 127260709 | 3751 | 2 | ENSMUST00 | 5865 | Dctpp1 | ENSMUSG00000042462.4 |
| chr5 | 72203003 | 72204225 | 21.91422 | 67.38415 | 64.43526 | Promoter | 5 | 72203329 | 72230475 | 27147 | 1 | ENSMUST00 | 0 | Atp10d | ENSMUSG00000046808.12 |
| chr12 | 44328558 | 44330404 | 24.41159 | 67.36964 | 64.42106 | Promoter | 12 | 44328885 | 44601846 | 272962 | 1 | ENSMUST00 | 0 | Nrcam | ENSMUSG00000020598.10 |
| chr7 | 126648973 | 126650146 | 23.82897 | 67.246 | 64.29786 | Promoter | 7 | 126649309 | 126664004 | 14696 | 1 | ENSMUST00 | 0 | Ccdc101 | ENSMUSG00000030714.9 |
| chr5 | 105731574 | 105732860 | 22.30049 | 67.20228 | 64.25432 | Promoter | 5 | 105731784 | 105811848 | 80065 | 1 | ENSMUST00 | 0 | Lrrc8d | ENSMUSG00000046079.11 |
| chr10 | 70598571 | 70600290 | 24.26109 | 67.11149 | 64.1642 | Promoter | 10 | 70557682 | 70599291 | 41610 | 2 | ENSMUST00 | 0 | Phyhipl | ENSMUSG00000037747.4 |
| chr11 | 94210365 | 94211075 | 22.71493 | 67.06055 | 64.11349 | Promoter | 11 | 94211454 | 94215495 | 4042 | 1 | ENSMUST00 | -379 | Tob1 | ENSMUSG00000037573.5 |
| chr17 | 22423609 | 22425123 | 22.71493 | 67.06055 | 64.11349 | Promoter | 17 | 22424222 | 22447416 | 23195 | 1 | ENSMUST00 | 0 | Zfp946 | ENSMUSG00000071266.8 |
| chr18 | 80150219 | 80151770 | 22.71493 | 67.06055 | 64.11349 | Promoter | 18 | 80126311 | 80151482 | 25172 | 2 | ENSMUST00 | 0 | Adnp2 | ENSMUSG00000053950.10 |
| chr2 | 131159441 | 131161142 | 22.71493 | 67.06055 | 64.11349 | Promoter | 2 | 131146324 | 131160034 | 13711 | 2 | ENSMUST00 | 0 | 1700037H04 | ENSMUSG00000027327.11 |
| chr9 | 73103164 | 73104069 | 22.71493 | 67.06055 | 64.11349 | Promoter | 9 | 73102510 | 73104192 | 1683 | 1 | ENSMUST00 | 655 | Khdc3 | ENSMUSG00000092622.3 |
| chr11 | 102818144 | 102819718 | 24.21134 | 67.02609 | 64.07967 | Promoter | 11 | 102799579 | 102819179 | 19601 | 2 | ENSMUST00 | 0 | Gjc1 | ENSMUSG00000034520.9 |
| chr13 | 42049007 | 42050458 | 23.68814 | 67.0026 | 64.0563 | Promoter | 13 | 42049334 | 42050287 | 954 | 1 | ENSMUST00 | 0 | RP24-87A4.1 | ENSMUSG00000100071.1 |
| chr2 | 3423757 | 3424851 | 23.68814 | 67.0026 | 64.0563 | Promoter | 2 | 3424131 | 3461111 | 36981 | 1 | ENSMUST00 | 0 | Dclre1c | ENSMUSG00000026648.13 |
| chr5 | 124482408 | 124484009 | 23.68814 | 67.0026 | 64.0563 | Promoter | 5 | 124483134 | 124491124 | 7991 | 1 | ENSMUST00 | 0 | Snrnp35 | ENSMUSG00000029402.8 |
| chr10 | 82811958 | 82812501 | 20.74539 | 66.97287 | 64.02668 | Promoter | 10 | 82812123 | 82824242 | 12120 | 1 | ENSMUST00 | 0 | 1700028I16R | ENSMUSG00000056821.8 |
| chr10 | 82240618 | 82241707 | 20.74539 | 66.97287 | 64.02668 | Promoter | 10 | 82224850 | 82241273 | 16424 | 2 | ENSMUST00 | 0 | Zfp938 | ENSMUSG00000062931.10 |
| chr11 | 63130660 | 63133452 | 20.74539 | 66.97287 | 64.02668 | Promoter | 11 | 63131512 | 63159547 | 28036 | 1 | ENSMUST00 | 0 | Pmp22 | ENSMUSG00000018217.7 |
| chr11 | 78751097 | 78752111 | 20.74539 | 66.97287 | 64.02668 | Promoter | 11 | 78750506 | 78751729 | 1224 | 2 | ENSMUST00 | 0 | Fam58b | ENSMUSG00000049489.5 |
| chr14 | 46084150 | 46084802 | 20.74539 | 66.97287 | 64.02668 | Promoter | 14 | 46084126 | 46084815 | 690 | 1 | ENSMUST00 | 25 | Gm1821 | ENSMUSG00000044285.7 |
| chr14 | 54882829 | 54884007 | 20.74539 | 66.97287 | 64.02668 | Promoter | 14 | 54883377 | 54888234 | 4858 | 1 | ENSMUST00 | 0 | Bcl2l2 | ENSMUSG00000089682.4 |
| chr16 | 18234256 | 18236098 | 20.74539 | 66.97287 | 64.02668 | Promoter | 16 | 18220754 | 18235136 | 14383 | 2 | ENSMUST00 | 0 | Zdhhc8 | ENSMUSG00000060166.5 |
| chr17 | 43801261 | 43802494 | 20.74539 | 66.97287 | 64.02668 | Promoter | 17 | 43801851 | 44039516 | 237666 | 1 | ENSMUST00 | 0 | Rcan2 | ENSMUSG00000039601.10 |
| chr3 | 66973388 | 66973920 | 20.74539 | 66.97287 | 64.02668 | 3' UTR | 3 | 66972780 | 66978447 | 5668 | 2 | ENSMUST00 | 4527 | Shox2 | ENSMUSG00000027833.11 |
| chr3 | 127498329 | 127499295 | 20.74539 | 66.97287 | 64.02668 | Promoter | 3 | 126923957 | 127499115 | 575159 | 2 | ENSMUST00 | 0 | Ank2 | ENSMUSG00000032826.11 |
| chr4 | 11703933 | 11705523 | 20.74539 | 66.97287 | 64.02668 | Promoter | 4 | 11704457 | 11714752 | 10296 | 1 | ENSMUST00 | 0 | Gem | ENSMUSG00000028214.8 |
| chr4 | 34886470 | 34887745 | 20.74539 | 66.97287 | 64.02668 | Distal Interge | 4 | 34803113 | 34882960 | 79848 | 2 | ENSMUST00 | -3511 | Zfp292 | ENSMUSG00000039967.9 |
| chr5 | 71656804 | 71658689 | 20.74539 | 66.97287 | 64.02668 | Promoter | 5 | 71569734 | 71658308 | 88575 | 2 | ENSMUST00 | 0 | Gabra4 | ENSMUSG00000029211.7 |
| chr5 | 103655363 | 103656018 | 20.74539 | 66.97287 | 64.02668 | Promoter | 5 | 103648039 | 103656010 | 7972 | 2 | ENSMUST00 | 0 | 1700016H13 | ENSMUSG00000029320.6 |
| chr5 | 36484541 | 36485546 | 20.74539 | 66.97287 | 64.02668 | Promoter | 5 | 36484588 | 36488172 | 3585 | 1 | ENSMUST00 | 0 | Ccdc96 | ENSMUSG00000050677.2 |
| chr6 | 50596277 | 50597083 | 20.74539 | 66.97287 | 64.02668 | Promoter | 6 | 50573302 | 50596632 | 23331 | 2 | ENSMUST00 | 0 | 4921507P07 | ENSMUSG00000029828.4 |
| chr6 | 137169061 | 137169767 | 20.74539 | 66.97287 | 64.02668 | Promoter | 6 | 137054831 | 137169732 | 114902 | 2 | ENSMUST00 | 0 | Rerg | ENSMUSG00000030222.8 |
| chr8 | 84975875 | 84979344 | 20.74539 | 66.97287 | 64.02668 | Promoter | 8 | 84976909 | 84978748 | 1840 | 2 | ENSMUST00 | 0 | Junb | ENSMUSG00000052837.5 |
| chr8 | 32008396 | 32009876 | 20.74539 | 66.97287 | 64.02668 | Promoter | 8 | 32008504 | 32015759 | 7256 | 1 | ENSMUST00 | 0 | Gm26578 | ENSMUSG00000097928.1 |
| chr9 | 109037832 | 109039215 | 20.74539 | 66.97287 | 64.02668 | Promoter | 9 | 109038565 | 109057719 | 19155 | 1 | ENSMUST00 | 0 | Shisa5 | ENSMUSG00000025647.11 |
| chr9 | 72438074 | 72439029 | 20.74539 | 66.97287 | 64.02668 | Promoter | 9 | 72438526 | 72452742 | 14217 | 1 | ENSMUST00 | 0 | Mns1 | ENSMUSG00000032221.9 |
| chr9 | 22411101 | 22411967 | 20.74539 | 66.97287 | 64.02668 | Promoter | 9 | 22411577 | 22444677 | 33101 | 1 | ENSMUST00 | 0 | 9530077C05 | ENSMUSG00000036411.9 |
| chr9 | 24502352 | 24504092 | 20.74539 | 66.97287 | 64.02668 | Promoter | 9 | 24411778 | 24503140 | 91363 | 2 | ENSMUST00 | 0 | Dpy19l1 | ENSMUSG00000043067.10 |
| chr10 | 69352037 | 69353348 | 23.16072 | 66.96479 | 64.02055 | Promoter | 10 | 69336635 | 69352938 | 16304 | 2 | ENSMUST00 | 0 | Cdk1 | ENSMUSG00000019942.7 |
| chr10 | 81648679 | 81649789 | 23.16072 | 66.96479 | 64.02055 | Promoter | 10 | 81649069 | 81667361 | 18293 | 1 | ENSMUST00 | 0 | Gm10778 | ENSMUSG00000096856.2 |
| chr15 | 76904068 | 76905202 | 23.16072 | 66.96479 | 64.02055 | Promoter | 15 | 76904071 | 76906318 | 2248 | 1 | ENSMUST00 | 0 | Rpl8 | ENSMUSG00000003970.8 |
| chr17 | 26089549 | 26090527 | 23.16072 | 66.96479 | 64.02055 | Promoter | 17 | 26081211 | 26090300 | 9090 | 2 | ENSMUST00 | 0 | Decr2 | ENSMUSG00000036775.7 |
| chr2 | 83812364 | 83813184 | 23.16072 | 66.96479 | 64.02055 | Promoter | 2 | 83812636 | 83883486 | 70851 | 1 | ENSMUST00 | 0 | Fam171b | ENSMUSG00000048388.3 |
| chr5 | 130023644 | 130024663 | 23.16072 | 66.96479 | 64.02055 | Promoter | 5 | 130011258 | 130024381 | 13124 | 2 | ENSMUST00 | 0 | Asl | ENSMUSG00000025533.10 |
| chr6 | 53817748 | 53821510 | 23.16072 | 66.96479 | 64.02055 | Promoter | 6 | 53815466 | 53820830 | 5365 | 2 | ENSMUST00 | 0 | Tril | ENSMUSG00000043496.6 |
| chr16 | 21310720 | 21311470 | 24.16179 | 66.94102 | 63.99775 | Distal Interge | 16 | 21247518 | 21296872 | 49355 | 2 | ENSMUST00 | -13849 | Gm16863 | ENSMUSG00000096918.1 |
| chr7 | 24399214 | 24400413 | 24.16179 | 66.94102 | 63.99775 | Promoter | 7 | 24399619 | 24408273 | 8655 | 1 | ENSMUST00 | 0 | Smg9 | ENSMUSG00000002210.6 |
| chr9 | 110879851 | 110880693 | 20.27321 | 66.94082 | 63.99775 | Promoter | 9 | 110879870 | 110882722 | 2853 | 1 | ENSMUST00 | 0 | Als2cl | ENSMUSG00000044037.10 |
| chr3 | 108383400 | 108384170 | 23.64158 | 66.92205 | 63.97926 | Promoter | 3 | 108383839 | 108388179 | 4341 | 1 | ENSMUST00 | 0 | Psrc1 | ENSMUSG00000068744.7 |
| chr1 | 75384399 | 75387322 | 24.11245 | 66.85627 | 63.91418 | Promoter | 1 | 75384700 | 75387948 | 3249 | 1 | ENSMUST00 | 0 | Speg | ENSMUSG00000026207.11 |
| chr2 | 156420374 | 156422132 | 24.11245 | 66.85627 | 63.91418 | Promoter | 2 | 156420909 | 156543214 | 122306 | 1 | ENSMUST00 | 0 | Epb4.1l1 | ENSMUSG00000027624.12 |
| chr17 | 28522419 | 28523541 | 24.0633 | 66.77184 | 63.83004 | Promoter | 17 | 28523257 | 28525347 | 2091 | 1 | ENSMUST00 | 0 | E230001N04 | ENSMUSG00000066170.6 |

| | | | | | | | | | | | | | | | |
|---|---|---|---|---|---|---|---|---|---|---|---|---|---|---|---|
| chr7 | 44853779 | 44855457 | 23.54898 | 66.76177 | 63.82013 | Promoter | 7 | 44852920 | 44854644 | 1725 | 1 | ENSMUST00 | 860 | Akt1s1 | ENSMUSG00000011096.11 |
| chr11 | 71019547 | 71020536 | 21.54815 | 66.72034 | 63.77881 | Promoter | 11 | 71009933 | 71019841 | 9909 | 2 | ENSMUST00 | 0 | Derl2 | ENSMUSG00000018442.8 |
| chr5 | 115235208 | 115236090 | 21.54815 | 66.72034 | 63.77881 | Promoter | 5 | 115235836 | 115240991 | 5156 | 1 | ENSMUST00 | 0 | Pop5 | ENSMUSG00000060152.9 |
| chr11 | 69323956 | 69324634 | 23.96561 | 66.60394 | 63.66297 | Promoter | 11 | 69323980 | 69325793 | 1814 | 1 | ENSMUST00 | 0 | Trappc1 | ENSMUSG00000049299.9 |
| chr17 | 21488599 | 21489847 | 23.96561 | 66.60394 | 63.66297 | Promoter | 17 | 21485177 | 21488875 | 3699 | 2 | ENSMUST00 | 0 | Gm26753 | ENSMUSG00000097759.1 |
| chr3 | 36551911 | 36553152 | 23.96561 | 66.60394 | 63.66297 | Promoter | 3 | 36552606 | 36563271 | 10666 | 1 | ENSMUST00 | 0 | Exosc9 | ENSMUSG00000027714.6 |
| chr2 | 84726571 | 84727937 | 22.94371 | 66.5854 | 63.64468 | Promoter | 2 | 84722104 | 84727242 | 5139 | 2 | ENSMUST00 | 0 | Clp1 | ENSMUSG00000027079.10 |
| chr13 | 33964344 | 33965896 | 23.91706 | 66.52044 | 63.57986 | Promoter | 13 | 33964659 | 33988465 | 23807 | 1 | ENSMUST00 | 0 | Nqo2 | ENSMUSG00000046949.10 |
| chr3 | 107333310 | 107334618 | 21.91509 | 66.5122 | 63.57177 | Promoter | 3 | 107326110 | 107333289 | 7180 | 2 | ENSMUST00 | -22 | Rbm15 | ENSMUSG00000048109.9 |
| chr11 | 100355127 | 100356777 | 19.57973 | 66.4542 | 63.51413 | Promoter | 11 | 100347327 | 100356128 | 8802 | 2 | ENSMUST00 | 0 | Hap1 | ENSMUSG00000006930.10 |
| chr13 | 64128692 | 64129858 | 19.57973 | 66.4542 | 63.51413 | Promoter | 13 | 64096310 | 64129330 | 33021 | 2 | ENSMUST00 | 0 | Slc35d2 | ENSMUSG00000033114.10 |
| chr15 | 103162876 | 103163733 | 19.57973 | 66.4542 | 63.51413 | Distal Interg | 15 | 103153307 | 103158002 | 4696 | 2 | ENSMUST00 | -4875 | Smug1 | ENSMUSG00000036061.9 |
| chr17 | 62879599 | 62884335 | 19.57973 | 66.4542 | 63.51413 | Promoter | 17 | 62604184 | 62881317 | 277134 | 2 | ENSMUST00 | 0 | Efna5 | ENSMUSG00000048915.7 |
| chr18 | 56431280 | 56432909 | 19.57973 | 66.4542 | 63.51413 | Promoter | 18 | 56432132 | 56503792 | 71661 | 1 | ENSMUST00 | 0 | Gramd3 | ENSMUSG00000001700.9 |
| chr6 | 144780525 | 144782962 | 19.57973 | 66.4542 | 63.51413 | Promoter | 6 | 143843739 | 144781977 | 938239 | 2 | ENSMUST00 | 0 | Sox5 | ENSMUSG00000041540.11 |
| chr8 | 85024492 | 85025924 | 19.57973 | 66.4542 | 63.51413 | Promoter | 8 | 85017934 | 85025278 | 7345 | 2 | ENSMUST00 | 0 | Asna1 | ENSMUSG00000052456.8 |
| chr17 | 23548772 | 23551443 | 23.86871 | 66.43727 | 63.49846 | 3' UTR | 17 | 23551073 | 23553488 | 2416 | 2 | ENSMUST00 | 2045 | Casp16 | ENSMUSG00000067882.8 |
| chr10 | 79874058 | 79874856 | 24.3552 | 66.39809 | 63.45943 | Promoter | 10 | 79860475 | 79874570 | 14096 | 2 | ENSMUST00 | 0 | BC005764 | ENSMUSG00000035835.9 |
| chr10 | 81192017 | 81193242 | 24.3552 | 66.39809 | 63.45943 | 3' UTR | 10 | 81194609 | 81198439 | 3831 | 1 | ENSMUST00 | -1367 | Atcayos | ENSMUSG00000085779.1 |
| chr11 | 100770104 | 100771089 | 24.3552 | 66.39809 | 63.45943 | Promoter | 11 | 100766032 | 100770957 | 4926 | 2 | ENSMUST00 | 0 | Ghdc | ENSMUSG00000017747.8 |
| chr11 | 69578523 | 69579388 | 24.3552 | 66.39809 | 63.45943 | Promoter | 11 | 69561759 | 69579324 | 17566 | 2 | ENSMUST00 | 0 | Wrap53 | ENSMUSG00000041346.6 |
| chr15 | 102355503 | 102356091 | 24.3552 | 66.39809 | 63.45943 | Downstream | 15 | 102338247 | 102350759 | 12513 | 2 | ENSMUST00 | -4745 | Aaas | ENSMUSG00000036678.7 |
| chr15 | 103304505 | 103305880 | 24.3552 | 66.39809 | 63.45943 | Downstream | 15 | 103308236 | 103310438 | 2203 | 2 | ENSMUST00 | 4558 | Gpr84 | ENSMUSG00000063234.4 |
| chr17 | 34893816 | 34895421 | 24.3552 | 66.39809 | 63.45943 | Promoter | 17 | 34894559 | 34896867 | 2309 | 1 | ENSMUST00 | 0 | Zbtb12 | ENSMUSG00000049823.9 |
| chr4 | 63494549 | 63496504 | 24.3552 | 66.39809 | 63.45943 | Promoter | 4 | 63414910 | 63495951 | 81042 | 2 | ENSMUST00 | 0 | Whrn | ENSMUSG00000039137.13 |
| chr6 | 88874011 | 88875352 | 24.3552 | 66.39809 | 63.45943 | Promoter | 6 | 88842565 | 88874045 | 31481 | 2 | ENSMUST00 | 0 | Podxl2 | ENSMUSG00000033152.8 |
| chr7 | 141414302 | 141414730 | 24.3552 | 66.39809 | 63.45943 | Promoter | 7 | 141408184 | 141414136 | 5953 | 2 | ENSMUST00 | -167 | Pddc1 | ENSMUSG00000051007.9 |
| chr7 | 6285947 | 6287260 | 24.3552 | 66.39809 | 63.45943 | Promoter | 7 | 6286579 | 6307883 | 21305 | 1 | ENSMUST00 | 0 | Zfp667 | ENSMUSG00000054893.10 |
| chr8 | 111521456 | 111522207 | 24.3552 | 66.39809 | 63.45943 | Promoter | 8 | 111448784 | 111522107 | 73324 | 2 | ENSMUST00 | 0 | Wdr59 | ENSMUSG00000031959.9 |
| chrX | 20617730 | 20618826 | 24.3552 | 66.39809 | 63.45943 | Promoter | 20 | 20617783 | 20650901 | 33119 | 1 | ENSMUST00 | 0 | Rbm10 | ENSMUSG00000031060.11 |
| chrX | 160501750 | 160503740 | 24.3552 | 66.39809 | 63.45943 | Promoter | 20 | 160502166 | 160591141 | 88976 | 1 | ENSMUST00 | 0 | Phka2 | ENSMUSG00000031295.8 |
| chrX | 162760381 | 162761708 | 24.3552 | 66.39809 | 63.45943 | Promoter | 20 | 162760402 | 162779092 | 18691 | 1 | ENSMUST00 | 0 | Rbbp7 | ENSMUSG00000031353.8 |
| chrX | 99469695 | 99470948 | 24.3552 | 66.39809 | 63.45943 | Promoter | 20 | 99465738 | 99470687 | 4950 | 2 | ENSMUST00 | 0 | Pja1 | ENSMUSG00000034403.11 |
| chrX | 20364120 | 20365207 | 24.3552 | 66.39809 | 63.45943 | Promoter | 20 | 20364481 | 20400853 | 36373 | 1 | ENSMUST00 | 0 | Rp2h | ENSMUSG00000060090.11 |
| chr11 | 97404289 | 97405289 | 23.82055 | 66.35441 | 63.41718 | Distal Interg | 11 | 97415533 | 97502402 | 86870 | 1 | ENSMUST00 | -10244 | Arhgap23 | ENSMUSG00000049807.11 |
| chr10 | 117628811 | 117630031 | 22.30831 | 66.34106 | 63.40397 | Promoter | 10 | 117629500 | 117635373 | 5874 | 1 | ENSMUST00 | 0 | Cpm | ENSMUSG00000020183.6 |
| chr6 | 38253205 | 38254651 | 22.30831 | 66.34106 | 63.40397 | Promoter | 6 | 38123176 | 38254009 | 130834 | 2 | ENSMUST00 | 0 | D630045J12I | ENSMUSG00000063455.11 |
| chr5 | 113220424 | 113221566 | 24.3114 | 66.32314 | 63.38662 | Distal Interg | 5 | 113226909 | 113239263 | 12355 | 1 | ENSMUST00 | -5343 | Tmem211 | ENSMUSG00000066964.2 |
| chr1 | 9847993 | 9849006 | 21.77823 | 66.26614 | 63.32996 | Promoter | 1 | 9848272 | 9900843 | 52572 | 1 | ENSMUST00 | 0 | Sgk3 | ENSMUSG00000025915.9 |
| chr1 | 161968857 | 161970072 | 21.77823 | 66.26614 | 63.32996 | Promoter | 1 | 161969186 | 161973435 | 4250 | 1 | ENSMUST00 | 0 | Pigc | ENSMUSG00000026698.7 |
| chr11 | 116918492 | 116920145 | 21.77823 | 66.26614 | 63.32996 | Promoter | 11 | 116918863 | 116986948 | 68086 | 1 | ENSMUST00 | 0 | Mgat5b | ENSMUSG00000043857.11 |
| chr15 | 74720791 | 74722000 | 21.77823 | 66.26614 | 63.32996 | Promoter | 15 | 74721204 | 74724639 | 3436 | 1 | ENSMUST00 | 0 | Them6 | ENSMUSG00000056665.2 |
| chr17 | 30611469 | 30612679 | 21.77823 | 66.26614 | 63.32996 | Promoter | 17 | 30592866 | 30612659 | 19794 | 2 | ENSMUST00 | 0 | Glo1 | ENSMUSG00000024026.7 |
| chr17 | 33977452 | 33979158 | 21.77823 | 66.26614 | 63.32996 | Promoter | 17 | 33974831 | 33978790 | 3960 | 2 | ENSMUST00 | 0 | H2-K2 | ENSMUSG00000067203.7 |
| chr19 | 45017892 | 45019345 | 21.77823 | 66.26614 | 63.32996 | Promoter | 19 | 45018135 | 45027104 | 8970 | 1 | ENSMUST00 | 0 | Lzts2 | ENSMUSG00000035342.8 |
| chr2 | 153492067 | 153493444 | 21.77823 | 66.26614 | 63.32996 | Promoter | 2 | 153459746 | 153492517 | 32772 | 2 | ENSMUST00 | 0 | Nol4l | ENSMUSG00000061411.7 |
| chr4 | 143297953 | 143299748 | 21.77823 | 66.26614 | 63.32996 | Promoter | 4 | 143267431 | 143299564 | 32134 | 2 | ENSMUST00 | 0 | Pdpn | ENSMUSG00000028583.9 |
| chr5 | 138160735 | 138161817 | 21.77823 | 66.26614 | 63.32996 | Promoter | 5 | 138161071 | 138164646 | 3576 | 1 | ENSMUST00 | 0 | Cops6 | ENSMUSG00000019494.9 |
| chr5 | 141240751 | 141242376 | 21.77823 | 66.26614 | 63.32996 | Promoter | 5 | 141241490 | 142215586 | 974097 | 1 | ENSMUST00 | 0 | Sdk1 | ENSMUSG00000039683.11 |
| chr5 | 139907631 | 139909529 | 21.77823 | 66.26614 | 63.32996 | Promoter | 5 | 139907943 | 139974711 | 66769 | 1 | ENSMUST00 | 0 | Elfn1 | ENSMUSG00000048988.7 |
| chr5 | 75977571 | 75979173 | 21.77823 | 66.26614 | 63.32996 | Promoter | 5 | 75932827 | 75978458 | 45632 | 2 | ENSMUST00 | 0 | Kdr | ENSMUSG00000062960.7 |
| chr6 | 48048221 | 48049301 | 21.77823 | 66.26614 | 63.32996 | Promoter | 6 | 48024205 | 48048911 | 24707 | 2 | ENSMUST00 | 0 | Zfp777 | ENSMUSG00000071477.5 |
| chr7 | 101450511 | 101451804 | 21.77823 | 66.26614 | 63.32996 | Exon (ENSMU | 7 | 101475376 | 101475443 | 68 | 1 | ENSMUST00 | -23572 | Mir139 | ENSMUSG00000065446.1 |
| chr8 | 70506017 | 70506799 | 21.77823 | 66.26614 | 63.32996 | Promoter | 8 | 70504292 | 70506721 | 2430 | 2 | ENSMUST00 | 0 | 2810428I15R | ENSMUSG00000058833.6 |
| chr9 | 42263561 | 42264865 | 21.77823 | 66.26614 | 63.32996 | Promoter | 9 | 42254177 | 42264300 | 10124 | 2 | ENSMUST00 | 0 | Sc5d | ENSMUSG00000032018.8 |
| chr9 | 99629371 | 99630536 | 21.77823 | 66.26614 | 63.32996 | Promoter | 9 | 99629496 | 99637796 | 8301 | 1 | ENSMUST00 | 0 | Dzip1l | ENSMUSG00000037784.9 |
| chr10 | 5805615 | 5806682 | 22.73073 | 66.21214 | 63.27877 | Promoter | 10 | 5799160 | 5805600 | 6441 | 2 | ENSMUST00 | -16 | Fbxo5 | ENSMUSG00000019773.7 |
| chr3 | 89246323 | 89247126 | 22.73073 | 66.21214 | 63.27877 | Promoter | 3 | 89246434 | 89249709 | 3276 | 1 | ENSMUST00 | 0 | Krtcap2 | ENSMUSG00000042747.7 |
| chr5 | 122157659 | 122158175 | 22.73073 | 66.21214 | 63.27877 | Promoter | 5 | 122158278 | 122175273 | 16996 | 1 | ENSMUST00 | -103 | Ppp1cc | ENSMUSG00000004455.11 |
| chr1 | 120120081 | 120122069 | 18.59898 | 66.17558 | 63.2431 | Promoter | 1 | 120113280 | 120121078 | 7799 | 2 | ENSMUST00 | 0 | Dbi | ENSMUSG00000026385.11 |
| chr11 | 102948140 | 102949075 | 18.59898 | 66.17558 | 63.2431 | Intron (ENSM | 11 | 102939264 | 102946688 | 7425 | 2 | ENSMUST00 | -1453 | C1ql1 | ENSMUSG00000045532.5 |
| chr16 | 11908415 | 11909852 | 18.59898 | 66.17558 | 63.2431 | Promoter | 16 | 11803721 | 11909423 | 105703 | 2 | ENSMUST00 | 0 | Cpped1 | ENSMUSG00000065979.7 |
| chr17 | 46645075 | 46646076 | 18.59898 | 66.17558 | 63.2431 | Promoter | 17 | 46630631 | 46645144 | 14514 | 2 | ENSMUST00 | 0 | Klc4 | ENSMUSG00000003546.9 |
| chr17 | 25568768 | 25571582 | 18.59898 | 66.17558 | 63.2431 | Promoter | 17 | 25565892 | 25570686 | 4795 | 2 | ENSMUST00 | 0 | Sox8 | ENSMUSG00000024176.5 |
| chr17 | 25823394 | 25824341 | 18.59898 | 66.17558 | 63.2431 | Promoter | 17 | 25823627 | 25828730 | 5104 | 1 | ENSMUST00 | 0 | Wdr24 | ENSMUSG00000025737.5 |
| chr18 | 37954158 | 37956169 | 18.59898 | 66.17558 | 63.2431 | Promoter | 18 | 37936972 | 37954988 | 18017 | 2 | ENSMUST00 | 0 | Hdac3 | ENSMUSG00000024454.10 |
| chr19 | 53140463 | 53142585 | 18.59898 | 66.17558 | 63.2431 | Promoter | 19 | 53140445 | 53247399 | 106955 | 1 | ENSMUST00 | 19 | Add3 | ENSMUSG00000025026.9 |
| chr6 | 128423781 | 128425420 | 18.59898 | 66.17558 | 63.2431 | Promoter | 6 | 128409444 | 128424931 | 15488 | 2 | ENSMUST00 | 0 | Itfg2 | ENSMUSG00000001518.10 |
| chr9 | 122116993 | 122118488 | 18.59898 | 66.17558 | 63.2431 | Promoter | 9 | 122117283 | 122137176 | 19894 | 1 | ENSMUST00 | 0 | Snrk | ENSMUSG00000038145.11 |
| chr2 | 30771231 | 30772294 | 22.68861 | 66.13821 | 63.20662 | Distal Interg | 2 | 30795468 | 30796664 | 1197 | 2 | ENSMUST00 | 24370 | 1700001O22 | ENSMUSG00000044320.9 |
| chr9 | 108189782 | 108190729 | 23.18573 | 66.13177 | 63.2002 | Promoter | 9 | 108096022 | 108190384 | 94363 | 2 | ENSMUST00 | 0 | Bsn | ENSMUSG00000032589.9 |
| chr4 | 156233883 | 156235233 | 17.04032 | 66.12922 | 63.19824 | Promoter | 4 | 156229044 | 156234857 | 5814 | 2 | ENSMUST00 | 0 | Klhl17 | ENSMUSG00000078484.3 |
| chr8 | 111991651 | 111992564 | 17.04032 | 66.12922 | 63.19824 | Promoter | 8 | 111966908 | 111992302 | 25395 | 2 | ENSMUST00 | 0 | Adat1 | ENSMUSG00000031949.12 |
| chr9 | 61945954 | 61947259 | 17.04032 | 66.12922 | 63.19824 | Promoter | 9 | 61917280 | 61946799 | 29520 | 2 | ENSMUST00 | 0 | Kif23 | ENSMUSG00000032254.9 |
| chr18 | 62921844 | 62923342 | 23.67723 | 66.10762 | 63.1768 | Promoter | 18 | 62922327 | 62953195 | 30869 | 1 | ENSMUST00 | 0 | Apcdd1 | ENSMUSG00000071847.7 |
| chr2 | 5137432 | 5138467 | 23.67723 | 66.10762 | 63.1768 | Promoter | 2 | 5137776 | 5230878 | 93103 | 1 | ENSMUST00 | 0 | Ccdc3 | ENSMUSG00000026676.7 |
| chr2 | 122467324 | 122467999 | 23.67723 | 66.10762 | 63.1768 | Promoter | 2 | 122466935 | 122467526 | 592 | 2 | ENSMUST00 | 0 | Bambi-ps1 | ENSMUSG00000081219.1 |
| chr9 | 34488374 | 34489689 | 23.67723 | 66.10762 | 63.1768 | Promoter | 9 | 34488733 | 35036716 | 547984 | 1 | ENSMUST00 | 0 | Kirrel3 | ENSMUSG00000032036.10 |
| chr1 | 180194971 | 180196465 | 17.76238 | 66.08042 | 63.15025 | Promoter | 1 | 180165238 | 180196020 | 30783 | 2 | ENSMUST00 | 0 | Adck3 | ENSMUSG00000026489.8 |
| chr11 | 75168275 | 75169318 | 17.76238 | 66.08042 | 63.15025 | Promoter | 11 | 75167153 | 75168146 | 994 | 2 | ENSMUST00 | -130 | Hic1 | ENSMUSG00000043099.4 |
| chr13 | 30136106 | 30136896 | 21.19411 | 66.07429 | 63.14441 | Promoter | 13 | 30136498 | 30246717 | 110220 | 1 | ENSMUST00 | 0 | Mboat1 | ENSMUSG00000038732.9 |
| chr4 | 129618457 | 129619553 | 21.19411 | 66.07429 | 63.14441 | Promoter | 4 | 129615126 | 129619099 | 3974 | 2 | ENSMUST00 | 0 | Iqcc | ENSMUSG00000040795.7 |
| chr7 | 84779299 | 84779781 | 23.14111 | 66.05422 | 63.12461 | Distal Interg | 7 | 84775703 | 84776020 | 318 | 1 | ENSMUST00 | 3597 | 2610206C17I | ENSMUSG00000085236.2 |
| chr1 | 59481280 | 59481937 | 23.62984 | 66.02597 | 63.09645 | Promoter | 1 | 59482424 | 59486955 | 4532 | 1 | ENSMUST00 | -487 | Fzd7 | ENSMUSG00000041075.8 |
| chr11 | 83270713 | 83271739 | 24.10907 | 65.97681 | 63.0475 | Promoter | 11 | 83271498 | 83271688 | 191 | 1 | ENSMUST00 | 0 | Gm23444 | ENSMUSG00000064856.1 |
| chr17 | 37002318 | 37003101 | 24.10907 | 65.97681 | 63.0475 | Promoter | 17 | 37002482 | 37009442 | 6961 | 1 | ENSMUST00 | 0 | Zfp57 | ENSMUSG00000036036.10 |
| chr17 | 22817155 | 22818376 | 24.10907 | 65.97681 | 63.0475 | Promoter | 17 | 22784798 | 22817741 | 32944 | 2 | ENSMUST00 | 0 | Gm16386 | ENSMUSG00000096975.1 |
| chr11 | 3123203 | 3124473 | 13.87731 | 65.92068 | 62.99162 | Distal Interg | 11 | 3125904 | 3129396 | 3493 | 1 | ENSMUST00 | -1431 | Pisd-ps1 | ENSMUSG00000082286.5 |
| chr10 | 85828913 | 85829995 | 23.05105 | 65.89761 | 62.96862 | Distal Interg | 10 | 85871831 | 85889096 | 17266 | 1 | ENSMUST00 | -41836 | Pwp1 | ENSMUSG00000001785.6 |
| chr10 | 78184366 | 78185716 | 23.05105 | 65.89761 | 62.96862 | Promoter | 10 | 78170910 | 78185149 | 14240 | 2 | ENSMUST00 | 0 | Pwp2 | ENSMUSG00000032834.10 |
| chr11 | 97315793 | 97316314 | 23.05105 | 65.89761 | 62.96862 | Promoter | 11 | 97315908 | 97321725 | 5818 | 1 | ENSMUST00 | 0 | Mrpl45 | ENSMUSG00000018882.7 |
| chr12 | 3959782 | 3960655 | 23.05105 | 65.89761 | 62.96862 | 3' UTR | 12 | 3954951 | 3960618 | 5668 | 1 | ENSMUST00 | 4832 | Pomc | ENSMUSG00000020660.5 |
| chr13 | 35967945 | 35969796 | 23.05105 | 65.89761 | 62.96862 | Promoter | 13 | 35967853 | 35970388 | 2536 | 1 | ENSMUST00 | 93 | Ppp1r3g | ENSMUSG00000050423.10 |
| chr14 | 32165334 | 32166047 | 23.05105 | 65.89761 | 62.96862 | Promoter | 14 | 32165589 | 32179855 | 14267 | 1 | ENSMUST00 | 0 | Ncoa4 | ENSMUSG00000056234.9 |
| chr15 | 95526988 | 95528909 | 23.05105 | 65.89761 | 62.96862 | Promoter | 15 | 95219451 | 95528706 | 309256 | 2 | ENSMUST00 | 0 | Nell2 | ENSMUSG00000022454.11 |
| chr15 | 102350148 | 102351433 | 23.05105 | 65.89761 | 62.96862 | Promoter | 15 | 102338247 | 102350759 | 12513 | 2 | ENSMUST00 | 0 | Aaas | ENSMUSG00000036678.7 |
| chr16 | 8828855 | 8829800 | 23.05105 | 65.89761 | 62.96862 | Promoter | 16 | 8830100 | 8858916 | 28817 | 1 | ENSMUST00 | -300 | 1810013L24I | ENSMUSG00000022507.5 |
| chr19 | 46147538 | 46149252 | 23.05105 | 65.89761 | 62.96862 | Promoter | 19 | 46135685 | 46148326 | 12642 | 2 | ENSMUST00 | 0 | Pitx3 | ENSMUSG00000025229.10 |
| chr19 | 8797711 | 8799075 | 23.05105 | 65.89761 | 62.96862 | Promoter | 19 | 8793129 | 8798557 | 5429 | 2 | ENSMUST00 | 0 | Polr2g | ENSMUSG00000071662.3 |
| chr2 | 28699401 | 28700587 | 23.05105 | 65.89761 | 62.96862 | Promoter | 2 | 28700164 | 28813165 | 113002 | 1 | ENSMUST00 | 0 | Ak8 | ENSMUSG00000026807.8 |
| chr2 | 181598474 | 181599566 | 23.05105 | 65.89761 | 62.96862 | Promoter | 2 | 181595218 | 181599212 | 3995 | 2 | ENSMUST00 | 0 | Samd10 | ENSMUSG00000038605.11 |
| chr4 | 110351079 | 110352094 | 23.05105 | 65.89761 | 62.96862 | Promoter | 4 | 110203737 | 110351909 | 148173 | 2 | ENSMUST00 | 0 | Elavl4 | ENSMUSG00000028546.12 |
| chr5 | 125340179 | 125341485 | 23.05105 | 65.89761 | 62.96862 | Promoter | 5 | 125277087 | 125341094 | 64008 | 2 | ENSMUST00 | 0 | Scarb1 | ENSMUSG00000037936.10 |
| chr6 | 5495847 | 5497159 | 23.05105 | 65.89761 | 62.96862 | Promoter | 6 | 5483351 | 5496309 | 12959 | 2 | ENSMUST00 | 0 | Pdk4 | ENSMUSG00000019577.4 |
| chr7 | 45896985 | 45897906 | 23.05105 | 65.89761 | 62.96862 | Promoter | 7 | 45897081 | 45917413 | 20333 | 1 | ENSMUST00 | 0 | Tmem143 | ENSMUSG00000002781.12 |
| chr8 | 84985294 | 84986495 | 23.05105 | 65.89761 | 62.96862 | Distal Interg | 8 | 84990595 | 85003364 | 12770 | 1 | ENSMUST00 | -4100 | Hook2 | ENSMUSG00000052566.7 |
| chr9 | 44478008 | 44480080 | 23.05105 | 65.89761 | 62.96862 | Promoter | 9 | 44478778 | 44482180 | 3403 | 1 | ENSMUST00 | 0 | C030014I23R | ENSMUSG00000060380.2 |
| chr9 | 63123642 | 63124322 | 23.05105 | 65.89761 | 62.96862 | Distal Interg | 9 | 63133283 | 63133389 | 107 | 1 | ENSMUST00 | -8961 | Gm25064 | ENSMUSG00000094201.1 |
| chr11 | 78114982 | 78116222 | 23.53563 | 65.86355 | 62.93705 | Promoter | 11 | 78115606 | 78156700 | 41095 | 1 | ENSMUST00 | 0 | Fam222b | ENSMUSG00000037750.11 |
| chr14 | 77156243 | 77157902 | 21.54279 | 65.84157 | 62.9152 | Promoter | 14 | 77156763 | 77721760 | 564998 | 1 | ENSMUST00 | 0 | Enox1 | ENSMUSG00000022012.8 |
| chr17 | 21707291 | 21708374 | 21.54279 | 65.84157 | 62.9152 | Promoter | 17 | 21707741 | 21725636 | 17896 | 1 | ENSMUST00 | 0 | Zfp760 | ENSMUSG00000067928.6 |
| chr3 | 131489555 | 131490997 | 21.54279 | 65.84157 | 62.9152 | Distal Interg | 3 | 131564768 | 131643670 | 78903 | 1 | ENSMUST00 | -73771 | Papss1 | ENSMUSG00000028032.9 |
| chr15 | 89498433 | 89500609 | 24.00916 | 65.80567 | 62.87967 | Promoter | 15 | 89499623 | 89560261 | 60639 | 1 | ENSMUST00 | 0 | Shank3 | ENSMUSG00000022623.10 |
| chr5 | 43515384 | 43518831 | 23.48881 | 65.78277 | 62.85688 | Promoter | 5 | 43515569 | 43616585 | 101017 | 1 | ENSMUST00 | 0 | C1qtnf7 | ENSMUSG00000061535.9 |
| chr10 | 86705710 | 86706864 | 21.916 | 65.643 | 62.71761 | Promoter | 10 | 86705811 | 86724584 | 18774 | 1 | ENSMUST00 | 0 | BC030307 | ENSMUSG00000044937.9 |
| chr19 | 34191999 | 34193663 | 21.916 | 65.643 | 62.71761 | Promoter | 19 | 34192229 | 34240333 | 48105 | 1 | ENSMUST00 | 0 | Stambpl1 | ENSMUSG00000024776.12 |
| chr4 | 84545134 | 84546146 | 21.916 | 65.643 | 62.71761 | Promoter | 4 | 84411154 | 84546290 | 135137 | 2 | ENSMUST00 | 144 | Bnc2 | ENSMUSG00000028487.13 |
| chr5 | 108668563 | 108669885 | 21.916 | 65.643 | 62.71761 | Promoter | 5 | 108646693 | 108669672 | 22980 | 2 | ENSMUST00 | 0 | Dgkq | ENSMUSG00000004815.7 |
| chr7 | 139187550 | 139189051 | 21.916 | 65.643 | 62.71761 | Promoter | 7 | 139103638 | 139188517 | 84880 | 2 | ENSMUST00 | 0 | Stk32c | ENSMUSG00000015981.7 |
| chr13 | 93143859 | 93144810 | 23.91008 | 65.63585 | 62.71086 | Promoter | 13 | 93040713 | 93144724 | 104012 | 2 | ENSMUST00 | 0 | Cmya5 | ENSMUSG00000047419.5 |
| chr14 | 118854492 | 118855622 | 23.91008 | 65.63585 | 62.71086 | Promoter | 14 | 118854732 | 118875489 | 20758 | 1 | ENSMUST00 | 0 | Cldn10 | ENSMUSG00000022132.10 |
| chr19 | 44388072 | 44389078 | 23.91008 | 65.63585 | 62.71086 | Distal Interg | 19 | 44394451 | 44407709 | 13259 | 2 | ENSMUST00 | 18631 | Scd1 | ENSMUSG00000037071.2 |
| chr8 | 95141640 | 95142860 | 23.91008 | 65.63585 | 62.71086 | Promoter | 8 | 95099832 | 95142540 | 42709 | 2 | ENSMUST00 | 0 | Kifc3 | ENSMUSG00000031788.8 |
| chr1 | 151571107 | 151571892 | 22.31642 | 65.48328 | 62.55884 | Promoter | 1 | 151571186 | 151644898 | 73713 | 1 | ENSMUST00 | 0 | Fam129a | ENSMUSG00000026483.8 |
| chr14 | 55014352 | 55015181 | 22.31642 | 65.48328 | 62.55884 | Promoter | 14 | 55015454 | 55024135 | 8682 | 1 | ENSMUST00 | -273 | Ngdn | ENSMUSG00000022204.8 |
| chr10 | 52690237 | 52691512 | 20.4053 | 65.4752 | 62.55133 | Promoter | 10 | 52690533 | 53111622 | 421090 | 1 | ENSMUST00 | 0 | Slc35f1 | ENSMUSG00000038602.7 |
| chr11 | 101278209 | 101279587 | 20.4053 | 65.4752 | 62.55133 | Promoter | 11 | 101277968 | 101279114 | 1147 | 2 | ENSMUST00 | 0 | Coa3 | ENSMUSG00000017188.3 |
| chr11 | 72041629 | 72042829 | 20.4053 | 65.4752 | 62.55133 | Promoter | 11 | 72042032 | 72043282 | 1251 | 1 | ENSMUST00 | 0 | Fam64a | ENSMUSG00000020808.3 |

| | | | | | | | | | | | | | | |
|---|---|---|---|---|---|---|---|---|---|---|---|---|---|---|
| chr11 | 97883479 | 97884137 | 20.4053 | 65.4752 | 62.55133 | Promoter | 11 | 97853826 | 97884154 | 30329 | 2 | ENSMUST00 | 17 | Fbxo47 | ENSMUSG00000070336.3 |
| chr13 | 91460078 | 91460804 | 20.4053 | 65.4752 | 62.55133 | Promoter | 13 | 91460283 | 91688866 | 228584 | 1 | ENSMUST00 | 0 | Ssbp2 | ENSMUSG00000003992.9 |
| chr13 | 56521973 | 56523166 | 20.4053 | 65.4752 | 62.55133 | Promoter | 13 | 56522472 | 56527467 | 4996 | 1 | ENSMUST00 | 0 | Fbxl21 | ENSMUSG00000035509.12 |
| chr15 | 96640997 | 96642347 | 20.4053 | 65.4752 | 62.55133 | Promoter | 15 | 96571418 | 96641962 | 70545 | 2 | ENSMUST00 | 0 | Slc38a1 | ENSMUSG00000023169.9 |
| chr15 | 102295828 | 102296669 | 20.4053 | 65.4752 | 62.55133 | Promoter | 15 | 102296293 | 102324356 | 28064 | 1 | ENSMUST00 | 0 | Espl1 | ENSMUSG00000058290.3 |
| chr15 | 85625428 | 85626331 | 20.4053 | 65.4752 | 62.55133 | Distal Interge | 15 | 85639747 | 85645934 | 6188 | 1 | ENSMUST00 | -13416 | RP23-64J7.2 | ENSMUSG00000101828.1 |
| chr17 | 24204929 | 24206345 | 20.4053 | 65.4752 | 62.55133 | Promoter | 17 | 24175431 | 24205562 | 30132 | 2 | ENSMUST00 | 0 | Tbc1d24 | ENSMUSG00000036473.10 |
| chr17 | 13759895 | 13761892 | 20.4053 | 65.4752 | 62.55133 | Promoter | 17 | 13489273 | 13761378 | 272106 | 2 | ENSMUST00 | 0 | Tcte2 | ENSMUSG00000038347.9 |
| chr19 | 6978978 | 6980047 | 20.4053 | 65.4752 | 62.55133 | Promoter | 19 | 6977741 | 6980440 | 2700 | 2 | ENSMUST00 | 393 | Fkbp2 | ENSMUSG00000056629.10 |
| chr2 | 44926157 | 44927009 | 20.4053 | 65.4752 | 62.55133 | Promoter | 2 | 44791193 | 44927149 | 135957 | 2 | ENSMUST00 | 140 | Gtdc1 | ENSMUSG00000036890.8 |
| chr3 | 89730311 | 89731567 | 20.4053 | 65.4752 | 62.55133 | Promoter | 3 | 89730748 | 89751406 | 20659 | 1 | ENSMUST00 | 0 | Adar | ENSMUSG00000027951.11 |
| chr3 | 81036384 | 81037850 | 20.4053 | 65.4752 | 62.55133 | Promoter | 3 | 81036416 | 81214040 | 177625 | 1 | ENSMUST00 | 0 | Pdgfc | ENSMUSG00000028019.4 |
| chr3 | 152264840 | 152266333 | 20.4053 | 65.4752 | 62.55133 | Promoter | 3 | 152236986 | 152266320 | 29335 | 2 | ENSMUST00 | 0 | Nexn | ENSMUSG00000039103.8 |
| chr3 | 96269603 | 96270370 | 20.4053 | 65.4752 | 62.55133 | Promoter | 3 | 96269721 | 96279001 | 9281 | 1 | ENSMUST00 | 0 | Hist2h2bb | ENSMUSG00000050936.4 |
| chr6 | 83136694 | 83138101 | 20.4053 | 65.4752 | 62.55133 | Promoter | 6 | 83137089 | 83152579 | 15491 | 1 | ENSMUST00 | 0 | Rtkn | ENSMUSG00000034930.10 |
| chr7 | 113765642 | 113767217 | 20.4053 | 65.4752 | 62.55133 | Promoter | 7 | 113765998 | 114043370 | 277373 | 1 | ENSMUST00 | 0 | Spon1 | ENSMUSG00000038156.10 |
| chr7 | 24111845 | 24113268 | 20.4053 | 65.4752 | 62.55133 | Promoter | 7 | 24112314 | 24127952 | 15639 | 1 | ENSMUST00 | 0 | Zfp112 | ENSMUSG00000052675.5 |
| chr8 | 87472162 | 87473695 | 20.4053 | 65.4752 | 62.55133 | Promoter | 8 | 87468854 | 87472592 | 3739 | 2 | ENSMUST00 | 0 | Cbln1 | ENSMUSG00000031654.10 |
| chr9 | 123259525 | 123261205 | 20.4053 | 65.4752 | 62.55133 | Promoter | 9 | 123259058 | 123260789 | 1732 | 2 | ENSMUST00 | 0 | Tmem158 | ENSMUSG00000054871.4 |
| chr10 | 116472943 | 116474490 | 23.81181 | 65.46733 | 62.54538 | Promoter | 10 | 116417861 | 116473878 | 56018 | 2 | ENSMUST00 | 0 | Kcnmb4 | ENSMUSG00000054934.5 |
| chr6 | 121245119 | 121246418 | 23.81181 | 65.46733 | 62.54538 | Promoter | 6 | 121245906 | 121270916 | 25011 | 1 | ENSMUST00 | 0 | Usp18 | ENSMUSG00000030107.9 |
| chr6 | 125039772 | 125040409 | 23.30336 | 65.46256 | 62.54073 | Promoter | 6 | 125039836 | 125046185 | 6350 | 1 | ENSMUST00 | 0 | Ing4 | ENSMUSG00000030330.10 |
| chr15 | 67039226 | 67040096 | 23.76298 | 65.38354 | 62.46201 | Distal Interge | 15 | 66946537 | 66969640 | 23104 | 2 | ENSMUST00 | -69587 | Ndrg1 | ENSMUSG00000005125.7 |
| chr1 | 143640102 | 143644670 | 22.74714 | 65.36804 | 62.44672 | Promoter | 1 | 143606161 | 143640581 | 34421 | 2 | ENSMUST00 | 0 | Cdc73 | ENSMUSG00000026361.8 |
| chr11 | 118427431 | 118430359 | 22.74714 | 65.36804 | 62.44672 | Promoter | 11 | 118428203 | 118443843 | 15641 | 1 | ENSMUST00 | 0 | C1qtnf1 | ENSMUSG00000017446.9 |
| chr13 | 74493446 | 74494689 | 22.74714 | 65.36804 | 62.44672 | Distal Interge | 13 | 74480517 | 74482943 | 2427 | 2 | ENSMUST00 | -10504 | Zfp825 | ENSMUSG00000069208.5 |
| chr2 | 28192692 | 28193602 | 22.74714 | 65.36804 | 62.44672 | Promoter | 2 | 28192992 | 28214425 | 21434 | 1 | ENSMUST00 | 0 | Olfm1 | ENSMUSG00000026833.13 |
| chr4 | 127168318 | 127169773 | 22.74714 | 65.36804 | 62.44672 | Promoter | 4 | 127169204 | 127237022 | 67819 | 1 | ENSMUST00 | 0 | Dlgap3 | ENSMUSG00000042388.9 |
| chr5 | 115193220 | 115194674 | 22.74714 | 65.36804 | 62.44672 | Promoter | 5 | 115168689 | 115194381 | 25693 | 2 | ENSMUST00 | 0 | Cabp1 | ENSMUSG00000029544.11 |
| chr6 | 38532769 | 38533838 | 22.74714 | 65.36804 | 62.44672 | Promoter | 6 | 38533502 | 38539448 | 5947 | 1 | ENSMUST00 | 0 | 1110001J03R | ENSMUSG00000019689.4 |
| chr7 | 30559014 | 30559681 | 22.74714 | 65.36804 | 62.44672 | Promoter | 7 | 30555441 | 30559646 | 4206 | 2 | ENSMUST00 | 0 | Lin37 | ENSMUSG00000036845.7 |
| chr9 | 3403443 | 3404852 | 22.74714 | 65.36804 | 62.44672 | Promoter | 9 | 3404085 | 3479236 | 75152 | 1 | ENSMUST00 | 0 | Cwf19l2 | ENSMUSG00000025898.5 |
| chr11 | 72960879 | 72961822 | 23.21173 | 65.30416 | 62.38367 | Promoter | 11 | 72961169 | 72993043 | 31875 | 1 | ENSMUST00 | 0 | Atp2a3 | ENSMUSG00000020788.10 |
| chr12 | 11149236 | 11150975 | 23.21173 | 65.30416 | 62.38367 | Promoter | 12 | 11090202 | 11150373 | 60172 | 2 | ENSMUST00 | 0 | Kcns3 | ENSMUSG00000043673.5 |
| chr13 | 105293720 | 105294627 | 23.21173 | 65.30416 | 62.38367 | Distal Interge | 13 | 105149352 | 105271039 | 121688 | 2 | ENSMUST00 | -22682 | Rnf180 | ENSMUSG00000021720.10 |
| chr4 | 136835062 | 136836866 | 23.21173 | 65.30416 | 62.38367 | Promoter | 4 | 136647539 | 136835988 | 188450 | 2 | ENSMUST00 | 0 | Ephb2 | ENSMUSG00000028664.9 |
| chr7 | 13053993 | 13055125 | 23.21173 | 65.30416 | 62.38367 | Promoter | 7 | 13042303 | 13054764 | 12462 | 2 | ENSMUST00 | 0 | Mzf1 | ENSMUSG00000030380.11 |
| chr11 | 5057743 | 5058645 | 23.71435 | 65.30009 | 62.38026 | Promoter | 11 | 5058128 | 5060385 | 2258 | 1 | ENSMUST00 | 0 | Rasl10a | ENSMUSG00000034209.4 |
| chr2 | 121456715 | 121457893 | 23.71435 | 65.30009 | 62.38026 | Promoter | 2 | 121456963 | 121458672 | 1710 | 1 | ENSMUST00 | 0 | Hypk | ENSMUSG00000027245.6 |
| chr15 | 102516392 | 102517958 | 23.16618 | 65.22538 | 62.30607 | Promoter | 15 | 102497649 | 102516836 | 19188 | 2 | ENSMUST00 | 0 | Map3k12 | ENSMUSG00000023050.3 |
| chr2 | 121442062 | 121443385 | 23.66592 | 65.21694 | 62.29776 | Promoter | 2 | 121439027 | 121442601 | 3575 | 2 | ENSMUST00 | 0 | Ell3 | ENSMUSG00000027246.7 |
| chr1 | 171502954 | 171504470 | 21.18292 | 65.18933 | 62.27029 | Promoter | 1 | 171503478 | 171504750 | 1273 | 1 | ENSMUST00 | 0 | Alyref2 | ENSMUSG00000060244.1 |
| chr5 | 108903150 | 108903971 | 21.18292 | 65.18933 | 62.27029 | Intron (ENSM | 5 | 108872655 | 108892380 | 19726 | 2 | ENSMUST00 | -10771 | Vmn2r-ps21 | ENSMUSG00000093510.1 |
| chr9 | 20726444 | 20728001 | 23.12081 | 65.14688 | 62.22808 | Promoter | 9 | 20667986 | 20728219 | 60234 | 2 | ENSMUST00 | 218 | Olfm2 | ENSMUSG00000032172.7 |
| chr5 | 108366449 | 108368339 | 23.61768 | 65.13411 | 62.21548 | Distal Interge | 5 | 108369858 | 108378239 | 8382 | 1 | ENSMUST00 | -1519 | Gm10419 | ENSMUSG00000072769.2 |
| chr1 | 40266288 | 40267027 | 23.56964 | 65.0516 | 62.13324 | Promoter | 1 | 40266586 | 40316198 | 49613 | 1 | ENSMUST00 | 0 | Il1r1 | ENSMUSG00000026072.7 |
| chr1 | 44118127 | 44120328 | 19.26894 | 65.0005 | 62.08229 | Promoter | 1 | 44106546 | 44118808 | 12263 | 2 | ENSMUST00 | 0 | Kdelc1 | ENSMUSG00000026047.8 |
| chr7 | 116031412 | 116032817 | 19.26894 | 65.0005 | 62.08229 | Promoter | 7 | 116031979 | 116032550 | 572 | 1 | ENSMUST00 | 0 | Sox6os | ENSMUSG00000030664.3 |
| chr6 | 51432306 | 51433522 | 23.5218 | 64.96938 | 62.05244 | Promoter | 6 | 51432670 | 51458768 | 26099 | 1 | ENSMUST00 | 0 | Nfe2l3 | ENSMUSG00000029832.11 |
| chr11 | 22287381 | 22288365 | 21.53724 | 64.96514 | 62.04835 | Promoter | 11 | 22005828 | 22286795 | 280968 | 2 | ENSMUST00 | -587 | Ehbp1 | ENSMUSG00000042302.9 |
| chr15 | 83405838 | 83406219 | 16.17966 | 64.96381 | 62.04733 | Promoter | 15 | 83405612 | 83406105 | 494 | 2 | ENSMUST00 | 0 | Gm5417 | ENSMUSG00000049414.6 |
| chr7 | 122158538 | 122160077 | 16.17966 | 64.96381 | 62.04733 | Promoter | 7 | 122159437 | 122169875 | 10439 | 1 | ENSMUST00 | 0 | Plk1 | ENSMUSG00000030867.6 |
| chr5 | 34636252 | 34637483 | 23.47415 | 64.88747 | 61.97117 | Promoter | 5 | 34633642 | 34637189 | 3548 | 2 | ENSMUST00 | 0 | Mfsd10 | ENSMUSG00000001082.7 |
| chr11 | 48799136 | 48801644 | 21.91694 | 64.77673 | 61.86085 | Promoter | 11 | 48800332 | 48804974 | 4643 | 1 | ENSMUST00 | 0 | Gnb2l1 | ENSMUSG00000020372.10 |
| chr17 | 35841571 | 35842377 | 21.91694 | 64.77673 | 61.86085 | Promoter | 17 | 35841676 | 35846031 | 4356 | 1 | ENSMUST00 | 0 | Mdc1 | ENSMUSG00000061607.9 |
| chr2 | 11171884 | 11172604 | 21.91694 | 64.77673 | 61.86085 | Promoter | 2 | 11172113 | 11263085 | 90973 | 1 | ENSMUST00 | 0 | Prkcq | ENSMUSG00000026778.8 |
| chr2 | 38997863 | 38998600 | 21.91694 | 64.77673 | 61.86085 | Promoter | 2 | 38998319 | 39001582 | 3264 | 1 | ENSMUST00 | 0 | Wdr38 | ENSMUSG00000035295.5 |
| chr4 | 145745099 | 145745560 | 14.54045 | 64.75945 | 61.8442 | Promoter | 4 | 145745746 | 145747401 | 1656 | 1 | ENSMUST00 | -186 | Gm13244 | ENSMUSG00000082334.1 |
| chr1 | 9966951 | 9968548 | 18.31284 | 64.75944 | 61.84419 | Promoter | 1 | 9960163 | 9967932 | 7770 | 2 | ENSMUST00 | 0 | Tcf24 | ENSMUSG00000099032.2 |
| chr11 | 88203626 | 88205228 | 18.31284 | 64.75944 | 61.84419 | Promoter | 11 | 88204388 | 88211507 | 7120 | 1 | ENSMUST00 | 0 | Mrps23 | ENSMUSG00000023723.12 |
| chr12 | 80757398 | 80759652 | 18.31284 | 64.75944 | 61.84419 | Promoter | 12 | 80758406 | 80761267 | 2862 | 1 | ENSMUST00 | 0 | Gm26796 | ENSMUSG00000097575.1 |
| chr13 | 54694268 | 54694933 | 18.31284 | 64.75944 | 61.84419 | Promoter | 13 | 54679410 | 54693907 | 14498 | 2 | ENSMUST00 | -362 | Rnf44 | ENSMUSG00000034928.11 |
| chr15 | 76701128 | 76702212 | 18.31284 | 64.75944 | 61.84419 | Promoter | 15 | 76701542 | 76704239 | 2698 | 1 | ENSMUST00 | 0 | Mfsd3 | ENSMUSG00000019080.7 |
| chr3 | 94309394 | 94310950 | 18.31284 | 64.75944 | 61.84419 | Promoter | 3 | 94310089 | 94332540 | 22452 | 1 | ENSMUST00 | 0 | Them4 | ENSMUSG00000028145.7 |
| chr9 | 44798958 | 44799935 | 18.31284 | 64.75944 | 61.84419 | Promoter | 9 | 44793776 | 44799242 | 5467 | 2 | ENSMUST00 | 0 | Tmem25 | ENSMUSG00000002032.11 |
| chr9 | 20581018 | 20582070 | 18.31284 | 64.75944 | 61.84419 | Promoter | 9 | 20581325 | 20595192 | 13868 | 1 | ENSMUST00 | 0 | Zfp846 | ENSMUSG00000058192.10 |
| chr10 | 80190052 | 80191250 | 23.89567 | 64.74409 | 61.82953 | Distal Interge | 10 | 80179482 | 80190010 | 10529 | 1 | ENSMUST00 | 10571 | Efna2 | ENSMUSG00000003070.6 |
| chr10 | 128668703 | 128669550 | 23.89567 | 64.74409 | 61.82953 | Downstream | 10 | 128669887 | 128673918 | 4032 | 2 | ENSMUST00 | 4368 | Suox | ENSMUSG00000049858.7 |
| chr16 | 31428479 | 31430227 | 23.89567 | 64.74409 | 61.82953 | Promoter | 16 | 31428748 | 31458901 | 30154 | 1 | ENSMUST00 | 0 | Bdh1 | ENSMUSG00000046598.9 |
| chr17 | 78881781 | 78882948 | 23.89567 | 64.74409 | 61.82953 | Promoter | 17 | 78852564 | 78882573 | 30010 | 2 | ENSMUST00 | 0 | Eif2ak2 | ENSMUSG00000024079.4 |
| chr2 | 74690898 | 74693568 | 23.89567 | 64.74409 | 61.82953 | Promoter | 2 | 74691924 | 74695105 | 3182 | 1 | ENSMUST00 | 0 | Hoxd10 | ENSMUSG00000050368.4 |
| chr2 | 175275080 | 175275809 | 23.89567 | 64.74409 | 61.82953 | Promoter | 2 | 175275125 | 175286805 | 11681 | 1 | ENSMUST00 | 0 | Gm14440 | ENSMUSG00000078901.3 |
| chr2 | 38823156 | 38825507 | 23.89567 | 64.74409 | 61.82953 | Promoter | 2 | 38823257 | 38838600 | 15344 | 1 | ENSMUST00 | 0 | Nr6a1os | ENSMUSG00000085201.1 |
| chrX | 139870916 | 139871958 | 23.89567 | 64.74409 | 61.82953 | Promoter | 20 | 139821632 | 139871654 | 50023 | 2 | ENSMUST00 | 0 | Morc4 | ENSMUSG00000031434.10 |
| chrX | 18161640 | 18162378 | 23.89567 | 64.74409 | 61.82953 | Promoter | 20 | 18162575 | 18279936 | 117362 | 1 | ENSMUST00 | -197 | Kdm6a | ENSMUSG00000037369.11 |
| chrX | 41910403 | 41912427 | 23.89567 | 64.74409 | 61.82953 | Promoter | 20 | 41794991 | 41911901 | 116911 | 2 | ENSMUST00 | 0 | Thoc2 | ENSMUSG00000037475.10 |
| chrX | 36644727 | 36645512 | 23.89567 | 64.74409 | 61.82953 | Promoter | 20 | 36608314 | 36645395 | 37082 | 2 | ENSMUST00 | 0 | Akap17b | ENSMUSG00000059708.7 |
| chrX | 60404142 | 60405175 | 23.89567 | 64.74409 | 61.82953 | Promoter | 20 | 60223293 | 60403981 | 180689 | 2 | ENSMUST00 | -162 | Atp11c | ENSMUSG00000062949.8 |
| chr10 | 4431665 | 4432359 | 21.40274 | 64.72344 | 61.81032 | Promoter | 10 | 4426094 | 4432344 | 6251 | 2 | ENSMUST00 | 0 | Rmnd1 | ENSMUSG00000019763.6 |
| chr10 | 18844766 | 18845515 | 21.40274 | 64.72344 | 61.81032 | Promoter | 10 | 18845071 | 18857073 | 12003 | 1 | ENSMUST00 | 0 | Perp | ENSMUSG00000019851.7 |
| chr10 | 39959584 | 39960542 | 21.40274 | 64.72344 | 61.81032 | Promoter | 10 | 39946907 | 39960153 | 13247 | 2 | ENSMUST00 | 0 | G630090E17 | ENSMUSG00000071335.4 |
| chr11 | 97188127 | 97189029 | 21.40274 | 64.72344 | 61.81032 | Promoter | 11 | 97159714 | 97187881 | 28168 | 2 | ENSMUST00 | -247 | Kpnb1 | ENSMUSG00000001440.5 |
| chr11 | 87590719 | 87591887 | 21.40274 | 64.72344 | 61.81032 | Promoter | 11 | 87592162 | 87616302 | 24141 | 1 | ENSMUST00 | -275 | Mtmr4 | ENSMUSG00000018401.12 |
| chr11 | 111064881 | 111066677 | 21.40274 | 64.72344 | 61.81032 | Promoter | 11 | 111066164 | 111076821 | 10658 | 1 | ENSMUST00 | 0 | Kcnj2 | ENSMUSG00000041695.2 |
| chr12 | 30372789 | 30373491 | 21.40274 | 64.72344 | 61.81032 | Promoter | 12 | 30174482 | 30373328 | 198847 | 2 | ENSMUST00 | 0 | Sntg2 | ENSMUSG00000020672.8 |
| chr13 | 109440167 | 109441959 | 21.40274 | 64.72344 | 61.81032 | Promoter | 13 | 109442183 | 109951364 | 509182 | 1 | ENSMUST00 | -224 | Pde4d | ENSMUSG00000021699.12 |
| chr15 | 53901151 | 53903072 | 21.40274 | 64.72344 | 61.81032 | Promoter | 15 | 53461621 | 53902537 | 440917 | 2 | ENSMUST00 | 0 | Samd12 | ENSMUSG00000058656.7 |
| chr17 | 33523880 | 33524638 | 21.40274 | 64.72344 | 61.81032 | Promoter | 17 | 33524204 | 33553768 | 29565 | 1 | ENSMUST00 | 0 | Adamts10 | ENSMUSG00000024299.11 |
| chr17 | 88597214 | 88598641 | 21.40274 | 64.72344 | 61.81032 | Promoter | 17 | 88597684 | 88635829 | 38146 | 1 | ENSMUST00 | 0 | Ston1 | ENSMUSG00000033855.10 |
| chr19 | 10555165 | 10556626 | 21.40274 | 64.72344 | 61.81032 | Promoter | 19 | 10533865 | 10555736 | 21872 | 2 | ENSMUST00 | 0 | Tmem216 | ENSMUSG00000024667.6 |
| chr2 | 176708214 | 176708823 | 21.40274 | 64.72344 | 61.81032 | Promoter | 2 | 176708328 | 176721813 | 13486 | 1 | ENSMUST00 | 0 | Gm14305 | ENSMUSG00000078878.4 |
| chr4 | 129136559 | 129137531 | 21.40274 | 64.72344 | 61.81032 | Promoter | 4 | 129136999 | 129144593 | 7595 | 1 | ENSMUST00 | 0 | Fndc5 | ENSMUSG00000001334.9 |
| chr4 | 106560261 | 106561867 | 21.40274 | 64.72344 | 61.81032 | Promoter | 4 | 106561038 | 106589113 | 28076 | 1 | ENSMUST00 | 0 | Dhcr24 | ENSMUSG00000034926.3 |
| chr6 | 134631905 | 134632850 | 21.40274 | 64.72344 | 61.81032 | Promoter | 6 | 134609210 | 134632488 | 23279 | 2 | ENSMUST00 | 0 | Mansc1 | ENSMUSG00000032718.3 |
| chr7 | 107594323 | 107595630 | 21.40274 | 64.72344 | 61.81032 | Promoter | 7 | 107595051 | 107744894 | 149844 | 1 | ENSMUST00 | 0 | Ppfibp2 | ENSMUSG00000036528.9 |
| chr7 | 143458479 | 143460972 | 21.40274 | 64.72344 | 61.81032 | Promoter | 7 | 143458350 | 143461050 | 2701 | 2 | ENSMUST00 | 78 | Cdkn1c | ENSMUSG00000037664.7 |
| chr7 | 19094120 | 19096125 | 21.40274 | 64.72344 | 61.81032 | Promoter | 7 | 19094594 | 19098549 | 3956 | 1 | ENSMUST00 | 0 | Six5 | ENSMUSG00000040841.5 |
| chr7 | 28981153 | 28981891 | 21.40274 | 64.72344 | 61.81032 | Promoter | 7 | 28971373 | 28981814 | 10442 | 2 | ENSMUST00 | 0 | Eif3k | ENSMUSG00000053565.9 |
| chr7 | 65526935 | 65528127 | 21.40274 | 64.72344 | 61.81032 | Distal Interge | 7 | 65582031 | 65618869 | 36839 | 1 | ENSMUST00 | -53904 | BC046251 | ENSMUSG00000086662.2 |
| chr1 | 86586777 | 86587797 | 17.49727 | 64.69686 | 61.78638 | Promoter | 1 | 86587100 | 86606500 | 19401 | 1 | ENSMUST00 | 0 | Cops7b | ENSMUSG00000026240.7 |
| chr17 | 57059228 | 57061163 | 17.49727 | 64.69686 | 61.78638 | Promoter | 17 | 57043711 | 57059863 | 16153 | 2 | ENSMUST00 | 0 | Slc25a23 | ENSMUSG00000046329.9 |
| chr4 | 103313729 | 103314546 | 17.49727 | 64.69686 | 61.78638 | Promoter | 4 | 103313812 | 103366443 | 52632 | 1 | ENSMUST00 | 0 | Oma1 | ENSMUSG00000035069.3 |
| chr9 | 7836210 | 7838078 | 17.49727 | 64.69686 | 61.78638 | Promoter | 9 | 7818227 | 7837057 | 18831 | 2 | ENSMUST00 | 0 | Birc2 | ENSMUSG00000057367.7 |
| chr9 | 4375774 | 4376361 | 17.49727 | 64.69686 | 61.78638 | Distal Interge | 9 | 4371269 | 4371335 | 67 | 2 | ENSMUST00 | -4440 | Gm23992 | ENSMUSG00000094086.1 |
| chr7 | 46957725 | 46959184 | 22.8523 | 64.68157 | 61.77136 | Promoter | 7 | 46923539 | 46958527 | 34989 | 2 | ENSMUST00 | 0 | Uevld | ENSMUSG00000043262.9 |
| chr13 | 113292857 | 113293708 | 23.85269 | 64.67058 | 61.76043 | Distal Interge | 13 | 113317084 | 113370515 | 53432 | 1 | ENSMUST00 | -23376 | BC067074 | ENSMUSG00000021763.11 |
| chr11 | 106193709 | 106194230 | 23.33235 | 64.64359 | 61.73353 | Promoter | 11 | 106183902 | 106193572 | 9671 | 2 | ENSMUST00 | -138 | Strada | ENSMUSG00000069631.9 |
| chr1 | 74386524 | 74387500 | 22.32483 | 64.62918 | 61.71925 | Distal Interge | 1 | 74383600 | 74385197 | 1598 | 1 | ENSMUST00 | 2925 | Slc11a1 | ENSMUSG00000026177.6 |
| chr10 | 128302721 | 128304467 | 22.32483 | 64.62918 | 61.71925 | Promoter | 10 | 128303335 | 128321358 | 18024 | 1 | ENSMUST00 | 0 | Pan2 | ENSMUSG00000005682.8 |
| chr12 | 98733722 | 98735478 | 22.32483 | 64.62918 | 61.71925 | Promoter | 12 | 98676741 | 98734645 | 57905 | 2 | ENSMUST00 | 0 | Ptpn21 | ENSMUSG00000021009.9 |
| chr5 | 14025087 | 14028582 | 22.32483 | 64.62918 | 61.71925 | Promoter | 5 | 14025276 | 14254314 | 229039 | 1 | ENSMUST00 | 0 | Sema3e | ENSMUSG00000063531.6 |
| chr7 | 45798003 | 45798772 | 22.32483 | 64.62918 | 61.71925 | Exon (ENSMU | 7 | 45787904 | 45792586 | 4683 | 1 | ENSMUST00 | 10100 | Lmtk3 | ENSMUSG00000062044.8 |
| chr9 | 58311829 | 58313301 | 22.32483 | 64.62918 | 61.71925 | Promoter | 9 | 58287723 | 58313212 | 25490 | 2 | ENSMUST00 | 0 | Loxl1 | ENSMUSG00000032334.9 |
| chr11 | 101784049 | 101785755 | 23.80275 | 64.58516 | 61.676 | Promoter | 11 | 101769742 | 101785310 | 15569 | 2 | ENSMUST00 | 0 | Etv4 | ENSMUSG00000017724.9 |
| chr17 | 73804453 | 73805776 | 23.28547 | 64.56288 | 61.65382 | Promoter | 17 | 73804841 | 73832093 | 27253 | 1 | ENSMUST00 | 0 | Ehd3 | ENSMUSG00000024065.7 |
| chr13 | 62652596 | 62653642 | 20.83488 | 64.55455 | 61.64568 | Distal Interge | 13 | 62624194 | 62624301 | 108 | 1 | ENSMUST00 | 28403 | Gm24020 | ENSMUSG00000095020.1 |
| chr8 | 119777613 | 119778471 | 20.83488 | 64.55455 | 61.64568 | Promoter | 8 | 119760587 | 119778416 | 17830 | 2 | ENSMUST00 | 0 | Tldc1 | ENSMUSG00000034105.8 |
| chr18 | 20745431 | 20746612 | 22.76417 | 64.5286 | 61.61989 | Promoter | 18 | 20684599 | 20746404 | 61806 | 2 | ENSMUST00 | 0 | B4galt6 | ENSMUSG00000056124.5 |
| chr4 | 85205049 | 85206196 | 22.76417 | 64.5286 | 61.61989 | Promoter | 4 | 85205126 | 85399529 | 194404 | 1 | ENSMUST00 | 0 | Sh3gl2 | ENSMUSG00000028488.10 |
| chr4 | 129105090 | 129107018 | 22.76417 | 64.5286 | 61.61989 | Promoter | 4 | 129105548 | 129111626 | 6079 | 1 | ENSMUST00 | 0 | Tmem54 | ENSMUSG00000028786.10 |
| chr2 | 4558843 | 4560357 | 23.75302 | 64.50008 | 61.59195 | Promoter | 2 | 4559754 | 4597989 | 38236 | 1 | ENSMUST00 | 0 | Frmd4a | ENSMUSG00000026657.11 |
| chr16 | 4880656 | 4881356 | 23.23877 | 64.48248 | 61.57465 | Promoter | 16 | 4874778 | 4880315 | 5538 | 2 | ENSMUST00 | -342 | Ubald1 | ENSMUSG00000039568.6 |
| chr17 | 35058835 | 35060326 | 23.23877 | 64.48248 | 61.57465 | Promoter | 17 | 35059035 | 35062091 | 3057 | 1 | ENSMUST00 | 0 | Ddah2 | ENSMUSG00000007039.7 |
| chr2 | 90889316 | 90890526 | 23.23877 | 64.48248 | 61.57465 | 5' UTR | 2 | 90885860 | 90890525 | 4666 | 1 | ENSMUST00 | 3457 | C1qtnf4 | ENSMUSG00000040794.5 |
| chr4 | 98321436 | 98322450 | 23.23877 | 64.48248 | 61.57465 | Promoter | 4 | 98321448 | 98331791 | 10344 | 1 | ENSMUST00 | 0 | 0610025J13R | ENSMUSG00000046683.11 |
| chr4 | 150030018 | 150031436 | 23.23877 | 64.48248 | 61.57465 | Distal Interge | 4 | 150049013 | 150056976 | 7964 | 1 | ENSMUST00 | -17577 | Gm13067 | ENSMUSG00000086665.1 |
| chr5 | 148994297 | 148995572 | 23.23877 | 64.48248 | 61.57465 | Promoter | 5 | 148893775 | 148995147 | 101373 | 2 | ENSMUST00 | 0 | Katnal1 | ENSMUSG00000041298.10 |
| chr5 | 39643780 | 39645490 | 23.23877 | 64.48248 | 61.57465 | Promoter | 5 | 39613939 | 39644634 | 30696 | 2 | ENSMUST00 | 0 | Hs3st1 | ENSMUSG00000051022.7 |

| | | | | | | | | | | | | | | | |
|---|---|---|---|---|---|---|---|---|---|---|---|---|---|---|---|
| chr3 | 95893407 | 95895298 | 23.7035 | 64.41534 | 61.50821 | Promoter | 3 | 95893969 | 95896758 | 2790 | 1 | ENSMUST00 | 0 | Aph1a | ENSMUSG00000015750.10 |
| chr4 | 132308628 | 132309796 | 23.7035 | 64.41534 | 61.50821 | Promoter | 4 | 132309465 | 132309600 | 136 | 1 | ENSMUST00 | 0 | Snora16a | ENSMUSG00000065097.1 |
| chr15 | 80797921 | 80799937 | 23.19226 | 64.40237 | 61.49533 | Intron (ENSM | 15 | 80711319 | 80941086 | 229768 | 1 | ENSMUST00 | 86603 | Tnrc6b | ENSMUSG00000047888.9 |
| chr3 | 137623372 | 137624485 | 23.14593 | 64.32256 | 61.41582 | Promoter | 3 | 137623672 | 137628333 | 4662 | 1 | ENSMUST00 | 0 | Ddit4l | ENSMUSG00000046818.7 |
| chr4 | 119492004 | 119493020 | 23.14593 | 64.32256 | 61.41582 | Promoter | 4 | 119465283 | 119492628 | 27346 | 2 | ENSMUST00 | 0 | Rimkla | ENSMUSG00000048899.8 |
| chr5 | 25529018 | 25529675 | 23.14593 | 64.32256 | 61.41582 | Promoter | 5 | 25530013 | 25531466 | 1454 | 1 | ENSMUST00 | -338 | RP23-405O1 | ENSMUSG00000101856.1 |
| chr1 | 45925183 | 45926367 | 21.17136 | 64.3064 | 61.3998 | Promoter | 1 | 45908068 | 45925582 | 17515 | 2 | ENSMUST00 | 0 | Slc40a1 | ENSMUSG00000025993.5 |
| chr18 | 66458212 | 66459004 | 21.17136 | 64.3064 | 61.3998 | Promoter | 18 | 66458604 | 66465550 | 6947 | 1 | ENSMUST00 | 0 | Pmaip1 | ENSMUSG00000024521.7 |
| chr9 | 60521536 | 60522296 | 21.17136 | 64.3064 | 61.3998 | Promoter | 9 | 60428226 | 60522046 | 93821 | 2 | ENSMUST00 | 0 | Thsd4 | ENSMUSG00000032289.10 |
| chr1 | 170255176 | 170255999 | 22.63194 | 64.29881 | 61.39256 | Promoter | 1 | 170254049 | 170255739 | 1691 | 2 | ENSMUST00 | 0 | Gm6345 | ENSMUSG00000096964.1 |
| chr11 | 77462140 | 77463101 | 22.63194 | 64.29881 | 61.39256 | Promoter | 11 | 77462411 | 77470484 | 8074 | 1 | ENSMUST00 | 0 | Coro6 | ENSMUSG00000020836.10 |
| chr11 | 40694060 | 40695488 | 22.63194 | 64.29881 | 61.39256 | Promoter | 11 | 40679314 | 40695203 | 15890 | 2 | ENSMUST00 | 0 | Mat2b | ENSMUSG00000042032.8 |
| chr11 | 103114825 | 103116009 | 22.63194 | 64.29881 | 61.39256 | Promoter | 11 | 103116231 | 103119725 | 3495 | 1 | ENSMUST00 | -222 | Hexim1 | ENSMUSG00000048878.6 |
| chr2 | 25983537 | 25984187 | 22.63194 | 64.29881 | 61.39256 | Promoter | 2 | 25929726 | 25983282 | 53557 | 2 | ENSMUST00 | -256 | Camsap1 | ENSMUSG00000026933.12 |
| chr2 | 166713988 | 166714722 | 22.63194 | 64.29881 | 61.39256 | Promoter | 2 | 166566342 | 166713832 | 147491 | 2 | ENSMUST00 | -157 | Prex1 | ENSMUSG00000039621.8 |
| chr2 | 30707595 | 30708517 | 22.63194 | 64.29881 | 61.39256 | Distal Interge | 2 | 30713150 | 30717043 | 3894 | 2 | ENSMUST00 | 8526 | Gm14488 | ENSMUSG00000075416.5 |
| chr4 | 139192025 | 139192789 | 22.63194 | 64.29881 | 61.39256 | Promoter | 4 | 139192899 | 139291818 | 98920 | 1 | ENSMUST00 | -110 | Capzb | ENSMUSG00000028745.13 |
| chr5 | 136952722 | 136954222 | 22.63194 | 64.29881 | 61.39256 | Promoter | 5 | 136953275 | 136966232 | 12958 | 1 | ENSMUST00 | 0 | Fis1 | ENSMUSG00000019054.8 |
| chr6 | 97486822 | 97488123 | 22.63194 | 64.29881 | 61.39256 | Promoter | 6 | 97288360 | 97487805 | 199446 | 2 | ENSMUST00 | 0 | Frmd4b | ENSMUSG00000030064.11 |
| chr7 | 81706209 | 81707397 | 22.63194 | 64.29881 | 61.39256 | Promoter | 7 | 81600481 | 81706925 | 106445 | 2 | ENSMUST00 | 0 | Homer2 | ENSMUSG00000025813.8 |
| chr7 | 104244137 | 104245216 | 22.63194 | 64.29881 | 61.39256 | Promoter | 7 | 104244457 | 104262234 | 17778 | 1 | ENSMUST00 | 0 | Trim34a | ENSMUSG00000056144.8 |
| chr4 | 101292040 | 101293747 | 23.60507 | 64.24684 | 61.34294 | Exon (ENSMU | 4 | 101284221 | 101285203 | 983 | 1 | ENSMUST00 | 7820 | Gm12801 | ENSMUSG00000085863.1 |
| chr6 | 122485575 | 122487005 | 23.60507 | 64.24684 | 61.34294 | Promoter | 6 | 122453101 | 122486313 | 33213 | 2 | ENSMUST00 | 0 | Rimklb | ENSMUSG00000040649.10 |
| chr16 | 84773321 | 84775332 | 23.09979 | 64.24302 | 61.33921 | Promoter | 16 | 84774123 | 84825928 | 51806 | 1 | ENSMUST00 | 0 | Jam2 | ENSMUSG00000053062.10 |
| chr1 | 167340282 | 167340802 | 16.5442 | 64.21555 | 61.31196 | Promoter | 1 | 167340369 | 167340424 | 56 | 2 | ENSMUST00 | 0 | Gm25911 | ENSMUSG00000088246.1 |
| chr16 | 17806821 | 17807977 | 21.53151 | 64.09126 | 61.18795 | 3' UTR | 16 | 17797282 | 17808287 | 11006 | 1 | ENSMUST00 | 9540 | Scarf2 | ENSMUSG00000012017.3 |
| chr2 | 130667327 | 130668368 | 21.53151 | 64.09126 | 61.18795 | Promoter | 2 | 130667610 | 130681614 | 14005 | 1 | ENSMUST00 | 0 | Itpa | ENSMUSG00000074797.6 |
| chr7 | 27332275 | 27333899 | 23.50745 | 64.07966 | 61.17685 | Promoter | 7 | 27305140 | 27333663 | 28524 | 2 | ENSMUST00 | 0 | Ltbp4 | ENSMUSG00000040488.11 |
| chr11 | 116694015 | 116695007 | 22.96246 | 64.00615 | 61.10351 | Promoter | 11 | 116675303 | 116694868 | 19566 | 2 | ENSMUST00 | 0 | St6galnac2 | ENSMUSG00000057286.5 |
| chr7 | 19403628 | 19404355 | 22.96246 | 64.00615 | 61.10351 | Promoter | 7 | 19394437 | 19404104 | 9668 | 2 | ENSMUST00 | 0 | Klc3 | ENSMUSG00000040714.8 |
| chr10 | 76961345 | 76962056 | 20.06521 | 63.98471 | 61.08233 | Promoter | 10 | 76761857 | 76961788 | 199932 | 2 | ENSMUST00 | 0 | Pcbp3 | ENSMUSG00000001120.10 |
| chr11 | 55097930 | 55098556 | 20.06521 | 63.98471 | 61.08233 | Promoter | 11 | 55098115 | 55113029 | 14915 | 1 | ENSMUST00 | 0 | Gm2a | ENSMUSG00000000594.7 |
| chr11 | 4266382 | 4267454 | 20.06521 | 63.98471 | 61.08233 | Promoter | 11 | 4266796 | 4268249 | 1454 | 1 | ENSMUST00 | 0 | Lif | ENSMUSG00000034394.9 |
| chr11 | 95310432 | 95311102 | 20.06521 | 63.98471 | 61.08233 | Promoter | 11 | 95310429 | 95324496 | 14068 | 1 | ENSMUST00 | 4 | Gm11520 | ENSMUSG00000085143.1 |
| chr13 | 115101042 | 115102480 | 20.06521 | 63.98471 | 61.08233 | Promoter | 13 | 114958079 | 115101964 | 143886 | 2 | ENSMUST00 | 0 | Itga1 | ENSMUSG00000042284.9 |
| chr2 | 69884960 | 69886184 | 20.06521 | 63.98471 | 61.08233 | Promoter | 2 | 69871198 | 69885615 | 14418 | 2 | ENSMUST00 | 0 | Mettl5 | ENSMUSG00000051730.10 |
| chr3 | 90432562 | 90433609 | 20.06521 | 63.98471 | 61.08233 | Promoter | 3 | 90391380 | 90433636 | 42257 | 2 | ENSMUST00 | 27 | Ints3 | ENSMUSG00000027933.6 |
| chr3 | 37348216 | 37349254 | 20.06521 | 63.98471 | 61.08233 | Promoter | 3 | 37348346 | 37405377 | 57032 | 1 | ENSMUST00 | 0 | Fgf2 | ENSMUSG00000037225.8 |
| chr4 | 48539714 | 48541193 | 20.06521 | 63.98471 | 61.08233 | Promoter | 4 | 48539935 | 48561918 | 21984 | 1 | ENSMUST00 | 0 | Msantd3 | ENSMUSG00000039693.6 |
| chr5 | 129895201 | 129896182 | 20.06521 | 63.98471 | 61.08233 | Promoter | 5 | 129895723 | 129903623 | 7901 | 1 | ENSMUST00 | 0 | Zbed5 | ENSMUSG00000034173.8 |
| chr6 | 54326642 | 54328336 | 20.06521 | 63.98471 | 61.08233 | Promoter | 6 | 54327012 | 54330200 | 3189 | 1 | ENSMUST00 | 0 | Prr15 | ENSMUSG00000045725.4 |
| chr7 | 29905698 | 29906978 | 20.06521 | 63.98471 | 61.08233 | Promoter | 7 | 29893333 | 29906572 | 13240 | 2 | ENSMUST00 | 0 | Zfp27 | ENSMUSG00000062040.9 |
| chr13 | 21717439 | 21718124 | 13.18946 | 63.94324 | 61.04279 | Promoter | 13 | 21717659 | 21718069 | 411 | 1 | ENSMUST00 | 0 | Hist1h3h | ENSMUSG00000101355.1 |
| chr11 | 69323258 | 69323819 | 23.41064 | 63.91375 | 61.01354 | Promoter | 11 | 69299487 | 69323775 | 24289 | 2 | ENSMUST00 | 0 | Cntrob | ENSMUSG00000032782.14 |
| chr11 | 67025097 | 67025690 | 21.91791 | 63.91367 | 61.01354 | Promoter | 11 | 67025154 | 67034790 | 9637 | 1 | ENSMUST00 | 0 | Tmem220 | ENSMUSG00000050270.7 |
| chr15 | 8568508 | 8570040 | 21.91791 | 63.91367 | 61.01354 | Distal Interge | 15 | 8529135 | 8529434 | 300 | 2 | ENSMUST00 | -39075 | Gm2348 | ENSMUSG00000078850.1 |
| chr18 | 12740951 | 12741782 | 21.91791 | 63.91367 | 61.01354 | Promoter | 18 | 12741324 | 12755146 | 13823 | 1 | ENSMUST00 | 0 | Cabyr | ENSMUSG00000024430.8 |
| chr18 | 23986802 | 23987690 | 21.91791 | 63.91367 | 61.01354 | Distal Interge | 18 | 23989632 | 24005376 | 15745 | 1 | ENSMUST00 | -1942 | Zfp35 | ENSMUSG00000063281.9 |
| chr19 | 23447071 | 23448521 | 23.41064 | 63.91375 | 61.01354 | Promoter | 19 | 23302609 | 23448322 | 145714 | 2 | ENSMUST00 | 0 | Mamdc2 | ENSMUSG00000033207.6 |
| chr10 | 80138590 | 80139651 | 22.33355 | 63.77912 | 60.87984 | Promoter | 10 | 80138632 | 80142400 | 3769 | 1 | ENSMUST00 | 0 | Atp5d | ENSMUSG00000003072.10 |
| chr15 | 82127285 | 82128669 | 22.33355 | 63.77912 | 60.87984 | Promoter | 15 | 82127922 | 82142200 | 14279 | 1 | ENSMUST00 | 0 | Ccdc134 | ENSMUSG00000068114.6 |
| chr2 | 25621448 | 25622233 | 22.33355 | 63.77912 | 60.87984 | Promoter | 2 | 25620067 | 25621985 | 1919 | 2 | ENSMUST00 | 0 | Tmem141 | ENSMUSG00000026939.7 |
| chr2 | 127481915 | 127483499 | 22.33355 | 63.77912 | 60.87984 | Promoter | 2 | 127458130 | 127482499 | 24370 | 2 | ENSMUST00 | 0 | Kcnip3 | ENSMUSG00000079056.7 |
| chr4 | 110397436 | 110398121 | 22.33355 | 63.77912 | 60.87984 | Promoter | 4 | 110397661 | 110955596 | 557936 | 1 | ENSMUST00 | 0 | Agbl4 | ENSMUSG00000061298.9 |
| chr16 | 42874637 | 42875395 | 22.82676 | 63.77181 | 60.87331 | Promoter | 16 | 42725702 | 42875587 | 149886 | 2 | ENSMUST00 | 192 | 4932412D23I | ENSMUSG00000075070.6 |
| chr12 | 113013711 | 113014344 | 22.29138 | 63.70553 | 60.80725 | Promoter | 12 | 113014508 | 113074401 | 59894 | 1 | ENSMUST00 | -164 | Pacs2 | ENSMUSG00000021143.9 |
| chr1 | 180331432 | 180333654 | 22.78188 | 63.69425 | 60.796 | Promoter | 1 | 180326970 | 180330550 | 3581 | 2 | ENSMUST00 | -883 | Gm5069 | ENSMUSG00000055676.3 |
| chr13 | 51170779 | 51171816 | 22.78188 | 63.69425 | 60.796 | Promoter | 13 | 51171025 | 51175187 | 4163 | 1 | ENSMUST00 | 0 | Nxnl2 | ENSMUSG00000021396.4 |
| chr14 | 55659840 | 55660682 | 22.78188 | 63.69425 | 60.796 | Promoter | 14 | 55657879 | 55660508 | 2630 | 2 | ENSMUST00 | 0 | Mdp1 | ENSMUSG00000002329.6 |
| chr19 | 8802444 | 8803097 | 22.78188 | 63.69425 | 60.796 | Promoter | 19 | 8802530 | 8804854 | 2325 | 1 | ENSMUST00 | 0 | Zbtb3 | ENSMUSG00000071661.6 |
| chr3 | 96244657 | 96245389 | 22.78188 | 63.69425 | 60.796 | Promoter | 3 | 96245497 | 96246082 | 586 | 1 | ENSMUST00 | -108 | Hist2h2aa2 | ENSMUSG00000064220.6 |
| chr5 | 135148518 | 135149494 | 22.78188 | 63.69425 | 60.796 | Promoter | 5 | 135149657 | 135163873 | 14217 | 1 | ENSMUST00 | -163 | Tbl2 | ENSMUSG00000005374.8 |
| chr7 | 90302017 | 90302856 | 22.78188 | 63.69425 | 60.796 | Distal Interge | 7 | 90293734 | 90293846 | 113 | 1 | ENSMUST00 | 8284 | n-R5s155 | ENSMUSG00000084480.1 |
| chr8 | 45358163 | 45359051 | 22.78188 | 63.69425 | 60.796 | Promoter | 8 | 45336717 | 45358865 | 22149 | 2 | ENSMUST00 | 0 | Fam149a | ENSMUSG00000070044.10 |
| chr1 | 66174814 | 66175822 | 23.26691 | 63.66726 | 60.76967 | Promoter | 1 | 66175273 | 66380617 | 205345 | 1 | ENSMUST00 | 0 | Map2 | ENSMUSG00000015222.12 |
| chr10 | 13090292 | 13091986 | 23.26691 | 63.66726 | 60.76967 | Promoter | 10 | 13090691 | 13124860 | 34170 | 1 | ENSMUST00 | 0 | Plagl1 | ENSMUSG00000019817.13 |
| chr17 | 3314035 | 3314882 | 23.26691 | 63.66726 | 60.76967 | Distal Interge | 17 | 3326573 | 3519397 | 192825 | 1 | ENSMUST00 | -11691 | Tiam2 | ENSMUSG00000023800.9 |
| chr19 | 38395575 | 38396391 | 23.26691 | 63.66726 | 60.76967 | Promoter | 19 | 38395980 | 38405607 | 9628 | 1 | ENSMUST00 | 0 | Slc35g1 | ENSMUSG00000044026.2 |
| chr7 | 79391611 | 79392924 | 23.26691 | 63.66726 | 60.76967 | Promoter | 7 | 79391929 | 79401886 | 9958 | 1 | ENSMUST00 | 0 | Fanci | ENSMUSG00000039187.11 |
| chr7 | 4836060 | 4837007 | 23.26691 | 63.66726 | 60.76967 | Promoter | 7 | 4825552 | 4836174 | 10623 | 2 | ENSMUST00 | 0 | Shisa7 | ENSMUSG00000053550.8 |
| chr9 | 45054781 | 45055886 | 23.26691 | 63.66726 | 60.76967 | Promoter | 9 | 45055186 | 45077436 | 22251 | 1 | ENSMUST00 | 0 | Mpzl3 | ENSMUSG00000070305.5 |
| chr14 | 76556033 | 76557095 | 20.8177 | 63.66574 | 60.7686 | Promoter | 14 | 76532814 | 76556687 | 23874 | 2 | ENSMUST00 | 0 | Serp2 | ENSMUSG00000052584.9 |
| chr7 | 142021191 | 142026097 | 20.8177 | 63.66574 | 60.7686 | Intron (ENSM | 7 | 141998661 | 142001441 | 2781 | 1 | ENSMUST00 | 22531 | Brsk2 | ENSMUSG00000053046.9 |
| chr5 | 105518730 | 105520178 | 15.94852 | 63.64039 | 60.74356 | Promoter | 5 | 105519388 | 105613018 | 93631 | 1 | ENSMUST00 | 0 | Lrrc8c | ENSMUSG00000054720.7 |
| chr15 | 103322998 | 103324152 | 22.24938 | 63.63219 | 60.73542 | Intron (ENSM | 15 | 103308236 | 103310438 | 2203 | 2 | ENSMUST00 | -12561 | Gpr84 | ENSMUSG00000063234.4 |
| chr11 | 4159944 | 4161381 | 18.95815 | 63.55374 | 60.65727 | Promoter | 11 | 4160350 | 4182541 | 22192 | 1 | ENSMUST00 | 0 | Sf3a1 | ENSMUSG00000002129.6 |
| chr11 | 22000785 | 22002392 | 18.95815 | 63.55374 | 60.65727 | Promoter | 11 | 21994764 | 22001615 | 6852 | 2 | ENSMUST00 | 0 | Otx1 | ENSMUSG00000005917.10 |
| chr17 | 32964939 | 32966677 | 18.95815 | 63.55374 | 60.65727 | Promoter | 17 | 32965814 | 32979233 | 13420 | 1 | ENSMUST00 | 0 | Zfp472 | ENSMUSG00000053600.7 |
| chr19 | 40587950 | 40589676 | 18.95815 | 63.55374 | 60.65727 | Promoter | 19 | 40550257 | 40588369 | 38113 | 2 | ENSMUST00 | 0 | Aldh18a1 | ENSMUSG00000025007.8 |
| chr19 | 8965979 | 8966930 | 18.95815 | 63.55374 | 60.65727 | Promoter | 19 | 8967041 | 8978479 | 11439 | 1 | ENSMUST00 | -111 | Eef1g | ENSMUSG00000071644.5 |
| chr2 | 91457134 | 91459385 | 18.95815 | 63.55374 | 60.65727 | Promoter | 2 | 91457511 | 91479287 | 21777 | 1 | ENSMUST00 | 0 | Lrp4 | ENSMUSG00000027253.10 |
| chr6 | 52191910 | 52192593 | 18.95815 | 63.55374 | 60.65727 | Promoter | 6 | 52189671 | 52191753 | 2083 | 2 | ENSMUST00 | -158 | Hoxa4 | ENSMUSG00000000942.10 |
| chr7 | 38107709 | 38108320 | 18.95815 | 63.55374 | 60.65727 | Promoter | 7 | 38097984 | 38107534 | 9551 | 2 | ENSMUST00 | -176 | Ccne1 | ENSMUSG00000002068.11 |
| chr7 | 41392551 | 41394049 | 18.95815 | 63.55374 | 60.65727 | Promoter | 7 | 41372923 | 41393260 | 20338 | 2 | ENSMUST00 | 0 | AI987944 | ENSMUSG00000056383.9 |
| chr9 | 54733800 | 54735023 | 18.95815 | 63.55374 | 60.65727 | Promoter | 9 | 54714735 | 54734482 | 19748 | 2 | ENSMUST00 | 0 | Wdr61 | ENSMUSG00000061559.10 |
| chr15 | 80090658 | 80091946 | 23.17207 | 63.50449 | 60.60925 | Promoter | 15 | 80091334 | 80113441 | 22108 | 1 | ENSMUST00 | 0 | Syngr1 | ENSMUSG00000022415.7 |
| chr13 | 119965374 | 119966467 | 14.34657 | 63.50294 | 60.60785 | Distal Interge | 13 | 119962244 | 119963227 | 984 | 1 | ENSMUST00 | 3131 | Gm20784 | ENSMUSG00000095367.1 |
| chr6 | 47757752 | 47758418 | 14.34657 | 63.50294 | 60.60785 | Distal Interge | 6 | 47773329 | 47773429 | 101 | 1 | ENSMUST00 | -14911 | Gm24975 | ENSMUSG00000094807.1 |
| chr11 | 117598673 | 117600062 | 22.16583 | 63.48624 | 60.59115 | Distal Interge | 11 | 117611247 | 117613857 | 2611 | 1 | ENSMUST00 | -11185 | 2900041M22 | ENSMUSG00000054418.6 |
| chr12 | 55301909 | 55303744 | 16.5464 | 63.42566 | 60.53069 | Promoter | 12 | 55280814 | 55303000 | 22187 | 2 | ENSMUST00 | 0 | Ppp2r3c | ENSMUSG00000021022.8 |
| chr7 | 101011613 | 101012522 | 21.1594 | 63.42564 | 60.53069 | Promoter | 7 | 100996572 | 101011991 | 15420 | 2 | ENSMUST00 | 0 | P2ry2 | ENSMUSG00000032860.4 |
| chr8 | 45999323 | 46000786 | 21.1594 | 63.42564 | 60.53069 | Promoter | 8 | 45996902 | 45999867 | 2966 | 2 | ENSMUST00 | 0 | Ankrd37 | ENSMUSG00000050914.10 |
| chr2 | 4400894 | 4401782 | 22.60412 | 63.38673 | 60.49239 | Promoter | 2 | 4400975 | 4614043 | 213069 | 1 | ENSMUST00 | 0 | Frmd4a | ENSMUSG00000026657.11 |
| chr11 | 23007291 | 23008863 | 18.0267 | 63.35003 | 60.45576 | Promoter | 11 | 23007531 | 23020236 | 12706 | 1 | ENSMUST00 | 0 | Fam161a | ENSMUSG00000049811.10 |
| chr12 | 98919977 | 98921028 | 18.0267 | 63.35003 | 60.45576 | Promoter | 12 | 98920574 | 98983232 | 62659 | 1 | ENSMUST00 | 0 | Ttc8 | ENSMUSG00000021013.10 |
| chr13 | 100669114 | 100670221 | 18.0267 | 63.35003 | 60.45576 | Promoter | 13 | 100669481 | 100697240 | 27760 | 1 | ENSMUST00 | 0 | Ccdc125 | ENSMUSG00000048924.8 |
| chr16 | 90142378 | 90144645 | 18.0267 | 63.35003 | 60.45576 | Distal Interge | 16 | 90220742 | 90226322 | 5581 | 1 | ENSMUST00 | -76097 | Sod1 | ENSMUSG00000022982.9 |
| chr17 | 22080846 | 22081566 | 18.0267 | 63.35003 | 60.45576 | Distal Interge | 17 | 22039028 | 22040539 | 1512 | 2 | ENSMUST00 | -40308 | Gm7052 | ENSMUSG00000092147.1 |
| chr2 | 32137420 | 32138013 | 18.0267 | 63.35003 | 60.45576 | Distal Interge | 2 | 32151082 | 32234537 | 83456 | 1 | ENSMUST00 | -13069 | Prrc2b | ENSMUSG00000039262.11 |
| chr9 | 108507349 | 108508757 | 18.0267 | 63.35003 | 60.45576 | Promoter | 9 | 108508005 | 108515941 | 7937 | 1 | ENSMUST00 | 0 | Qars | ENSMUSG00000032604.9 |
| chr11 | 116434028 | 116435508 | 17.23216 | 63.31984 | 60.4264 | Promoter | 11 | 116434094 | 116439077 | 4984 | 1 | ENSMUST00 | 0 | Ubald2 | ENSMUSG00000050628.6 |
| chr7 | 100862774 | 100863626 | 17.23216 | 63.31984 | 60.4264 | Promoter | 7 | 100845848 | 100863446 | 17599 | 2 | ENSMUST00 | 0 | Relt | ENSMUSG00000008318.4 |
| chr9 | 18291639 | 18293431 | 17.23216 | 63.31984 | 60.4264 | Promoter | 9 | 18292267 | 18314000 | 21734 | 1 | ENSMUST00 | 0 | Chordc1 | ENSMUSG00000001774.7 |
| chr3 | 100969039 | 100970106 | 22.51627 | 63.23458 | 60.34166 | Promoter | 3 | 100938860 | 100969663 | 30804 | 2 | ENSMUST00 | 0 | Ttf2 | ENSMUSG00000033222.11 |
| chr10 | 85127497 | 85128711 | 21.52556 | 63.2201 | 60.32722 | Promoter | 10 | 85119435 | 85128027 | 8593 | 2 | ENSMUST00 | 0 | Mterfd3 | ENSMUSG00000049038.2 |
| chr12 | 84773183 | 84774111 | 21.52556 | 63.2201 | 60.32722 | Promoter | 12 | 84773270 | 84775089 | 1820 | 1 | ENSMUST00 | 0 | Isca2 | ENSMUSG00000021241.6 |
| chr17 | 33981090 | 33981826 | 21.52556 | 63.2201 | 60.32722 | Promoter | 17 | 33980486 | 33980744 | 259 | 2 | ENSMUST00 | -347 | Gm4152 | ENSMUSG00000092512.1 |
| chr4 | 17852871 | 17854099 | 21.52556 | 63.2201 | 60.32722 | Promoter | 4 | 17852893 | 18117457 | 264565 | 1 | ENSMUST00 | 0 | Mmp16 | ENSMUSG00000028226.10 |
| chr10 | 44528998 | 44530261 | 21.02725 | 63.18829 | 60.29588 | Promoter | 10 | 44446717 | 44528501 | 81785 | 2 | ENSMUST00 | -498 | Prdm1 | ENSMUSG00000038151.7 |
| chr11 | 96978006 | 96978625 | 21.02725 | 63.18829 | 60.29588 | Promoter | 11 | 96953341 | 96977711 | 24371 | 2 | ENSMUST00 | -296 | Sp2 | ENSMUSG00000018678.7 |
| chr11 | 97628183 | 97630377 | 21.02725 | 63.18829 | 60.29588 | Promoter | 11 | 97627389 | 97629702 | 2314 | 2 | ENSMUST00 | 0 | E130012A19I | ENSMUSG00000043439.5 |
| chr12 | 112928440 | 112930203 | 21.02725 | 63.18829 | 60.29588 | Promoter | 12 | 112908590 | 112929495 | 20906 | 2 | ENSMUST00 | 0 | Jag2 | ENSMUSG00000002799.5 |
| chr13 | 24799815 | 24800731 | 21.02725 | 63.18829 | 60.29588 | Promoter | 13 | 24801657 | 24814013 | 12357 | 1 | ENSMUST00 | -926 | BC005537 | ENSMUSG00000019132.5 |
| chr13 | 31557253 | 31560191 | 21.02725 | 63.18829 | 60.29588 | Promoter | 13 | 31558324 | 31560976 | 2653 | 1 | ENSMUST00 | 0 | Foxq1 | ENSMUSG00000038415.9 |
| chr14 | 58071755 | 58073331 | 21.02725 | 63.18829 | 60.29588 | Promoter | 14 | 58072686 | 58110174 | 37489 | 1 | ENSMUST00 | 0 | Fgf9 | ENSMUSG00000021974.9 |
| chr14 | 55853819 | 55855746 | 21.02725 | 63.18829 | 60.29588 | Promoter | 14 | 55854036 | 55874291 | 20256 | 1 | ENSMUST00 | 0 | Nynrin | ENSMUSG00000075592.4 |
| chr14 | 43925131 | 43925901 | 21.02725 | 63.18829 | 60.29588 | Promoter | 14 | 43925390 | 43933411 | 8022 | 1 | ENSMUST00 | 0 | Gm8113 | ENSMUSG00000089901.1 |
| chr15 | 102965979 | 102968245 | 21.02725 | 63.18829 | 60.29588 | Promoter | 15 | 102966796 | 102971893 | 5098 | 1 | ENSMUST00 | 0 | Hoxc10 | ENSMUSG00000022484.7 |
| chr15 | 76232001 | 76233096 | 21.02725 | 63.18829 | 60.29588 | Promoter | 15 | 76188464 | 76232045 | 43582 | 2 | ENSMUST00 | 0 | Plec | ENSMUSG00000022565.10 |
| chr15 | 80173046 | 80174497 | 21.02725 | 63.18829 | 60.29588 | Promoter | 15 | 80173721 | 80215519 | 41799 | 1 | ENSMUST00 | 0 | Mgat3 | ENSMUSG00000042428.6 |
| chr2 | 132028950 | 132030583 | 21.02725 | 63.18829 | 60.29588 | Promoter | 2 | 131989415 | 132030001 | 40587 | 2 | ENSMUST00 | 0 | Rassf2 | ENSMUSG00000027339.10 |
| chr2 | 84677366 | 84678132 | 21.02725 | 63.18829 | 60.29588 | Promoter | 2 | 84671316 | 84678118 | 6803 | 2 | ENSMUST00 | 0 | Tmx2 | ENSMUSG00000050043.11 |
| chr2 | 94273044 | 94273945 | 21.02725 | 63.18829 | 60.29588 | Distal Interge | 2 | 94260966 | 94264758 | 3793 | 2 | ENSMUST00 | -8287 | E530001K10I | ENSMUSG00000075020.5 |
| chr3 | 90513691 | 90515996 | 21.02725 | 63.18829 | 60.29588 | Promoter | 3 | 90514435 | 90524581 | 10147 | 1 | ENSMUST00 | 0 | S100a13 | ENSMUSG00000042312.9 |
| chr3 | 92080275 | 92082186 | 21.02725 | 63.18829 | 60.29588 | Promoter | 3 | 92080271 | 92083142 | 2872 | 2 | ENSMUST00 | 956 | Lor | ENSMUSG00000043165.7 |
| chr4 | 41123800 | 41124400 | 21.02725 | 63.18829 | 60.29588 | Promoter | 4 | 41114427 | 41124455 | 10029 | 2 | ENSMUST00 | 55 | Nol6 | ENSMUSG00000028430.9 |
| chr4 | 117483869 | 117485136 | 21.02725 | 63.18829 | 60.29588 | Intron (ENSM | 4 | 117490235 | 117495011 | 4777 | 2 | ENSMUST00 | 9875 | Rnf220 | ENSMUSG00000028677.13 |
| chr4 | 135433323 | 135434728 | 21.02725 | 63.18829 | 60.29588 | Promoter | 4 | 135412308 | 135433853 | 21546 | 2 | ENSMUST00 | 0 | Rcan3 | ENSMUSG00000059713.7 |

| | | | | | | | | | | | | | | | |
|---|---|---|---|---|---|---|---|---|---|---|---|---|---|---|---|
| chr5 | 28165273 | 28167260 | 21.02725 | 63.18829 | 60.29588 | Promoter | 5 | 28165694 | 28172166 | 6473 | 1 | ENSMUST00 | 0 | En2 | ENSMUSG00000039095.8 |
| chr5 | 98185118 | 98186704 | 21.02725 | 63.18829 | 60.29588 | Promoter | 5 | 98185296 | 98185984 | 689 | 2 | ENSMUST00 | 0 | A730035I17F | ENSMUSG00000045238.7 |
| chr7 | 19075406 | 19077361 | 21.02725 | 63.18829 | 60.29588 | Promoter | 7 | 19076249 | 19082776 | 6528 | 1 | ENSMUST00 | 0 | Dmwd | ENSMUSG00000030410.11 |
| chr7 | 114635653 | 114636558 | 21.02725 | 63.18829 | 60.29588 | Promoter | 7 | 114631480 | 114636357 | 4878 | 2 | ENSMUST00 | 0 | Calca | ENSMUSG00000030669.8 |
| chr7 | 119894701 | 119896756 | 21.02725 | 63.18829 | 60.29588 | Promoter | 7 | 119895861 | 119916754 | 20894 | 1 | ENSMUST00 | 0 | Lyrm1 | ENSMUSG00000030922.6 |
| chr7 | 35117750 | 35120763 | 21.02725 | 63.18829 | 60.29588 | Promoter | 7 | 35119293 | 35121926 | 2634 | 1 | ENSMUST00 | 0 | Cebpa | ENSMUSG00000034957.9 |
| chr7 | 89517043 | 89518500 | 21.02725 | 63.18829 | 60.29588 | Promoter | 7 | 89507785 | 89517586 | 9802 | 2 | ENSMUST00 | 0 | Prss23 | ENSMUSG00000039405.6 |
| chr7 | 12880632 | 12881773 | 21.02725 | 63.18829 | 60.29588 | Promoter | 7 | 12881177 | 12891841 | 10665 | 1 | ENSMUST00 | 0 | Zfp128 | ENSMUSG00000060397.6 |
| chr9 | 35175171 | 35176500 | 21.02725 | 63.18829 | 60.29588 | Promoter | 9 | 35124408 | 35176061 | 51654 | 2 | ENSMUST00 | 0 | Dcps | ENSMUSG00000032040.10 |
| chr9 | 101033842 | 101035401 | 21.02725 | 63.18829 | 60.29588 | Promoter | 9 | 100982032 | 101034877 | 52846 | 2 | ENSMUST00 | 0 | Pccb | ENSMUSG00000032527.8 |
| chr11 | 103027534 | 103028369 | 23.43614 | 63.09834 | 60.20888 | Promoter | 11 | 102999644 | 103028304 | 28661 | 2 | ENSMUST00 | 0 | Dcakd | ENSMUSG00000020935.3 |
| chr13 | 54948747 | 54950005 | 23.43614 | 63.09834 | 60.20888 | Promoter | 13 | 54949411 | 55006018 | 56608 | 1 | ENSMUST00 | 0 | Unc5a | ENSMUSG00000025876.10 |
| chr14 | 69283729 | 69285101 | 23.43614 | 63.09834 | 60.20888 | Promoter | 14 | 69244176 | 69284967 | 40792 | 2 | ENSMUST00 | 0 | Slc25a37 | ENSMUSG00000034248.7 |
| chr15 | 100761312 | 100762307 | 23.43614 | 63.09834 | 60.20888 | Promoter | 15 | 100761747 | 100823968 | 62222 | 1 | ENSMUST00 | 0 | Slc4a8 | ENSMUSG00000023032.7 |
| chr2 | 119337545 | 119338441 | 23.43614 | 63.09834 | 60.20888 | Distal Interge | 2 | 119328538 | 119332153 | 3616 | 1 | ENSMUST00 | 9008 | Dll4 | ENSMUSG00000027314.6 |
| chr2 | 146221101 | 146223600 | 23.43614 | 63.09834 | 60.20888 | Promoter | 2 | 146221921 | 146225020 | 3100 | 1 | ENSMUST00 | 0 | Insm1 | ENSMUSG00000068154.5 |
| chr2 | 93014233 | 93015383 | 23.43614 | 63.09834 | 60.20888 | Promoter | 2 | 93009973 | 93013910 | 3938 | 2 | ENSMUST00 | -324 | Prdm11 | ENSMUSG00000075028.6 |
| chr4 | 137913305 | 137914277 | 23.43614 | 63.09834 | 60.20888 | Promoter | 4 | 137913471 | 137935387 | 21917 | 1 | ENSMUST00 | 0 | Ece1 | ENSMUSG00000057530.9 |
| chr5 | 30711168 | 30713618 | 23.43614 | 63.09834 | 60.20888 | Promoter | 5 | 30711564 | 30799375 | 87812 | 1 | ENSMUST00 | 0 | Dpysl5 | ENSMUSG00000029168.9 |
| chr6 | 29396281 | 29397152 | 23.43614 | 63.09834 | 60.20888 | Promoter | 6 | 29396309 | 29399449 | 3141 | 1 | ENSMUST00 | 0 | Ccdc136 | ENSMUSG00000029769.11 |
| chr6 | 128034264 | 128035425 | 23.43614 | 63.09834 | 60.20888 | Promoter | 6 | 127965747 | 128034592 | 68846 | 2 | ENSMUST00 | 0 | Tspan9 | ENSMUSG00000030352.9 |
| chr7 | 74012486 | 74014302 | 23.43614 | 63.09834 | 60.20888 | Promoter | 7 | 73939119 | 74013682 | 74564 | 2 | ENSMUST00 | 0 | St8sia2 | ENSMUSG00000025789.4 |
| chr7 | 80339335 | 80340983 | 23.43614 | 63.09834 | 60.20888 | Promoter | 7 | 80325295 | 80340219 | 14925 | 2 | ENSMUST00 | 0 | Unc45a | ENSMUSG00000030533.10 |
| chrX | 73785927 | 73787885 | 23.43614 | 63.09834 | 60.20888 | Promoter | 20 | 73778963 | 73786897 | 7935 | 2 | ENSMUST00 | 0 | Idh3g | ENSMUSG00000002010.12 |
| chrX | 77510423 | 77512224 | 23.43614 | 63.09834 | 60.20888 | Promoter | 20 | 77511013 | 77662983 | 151971 | 1 | ENSMUST00 | 0 | Tbl1x | ENSMUSG00000025246.8 |
| chrX | 151343760 | 151345169 | 23.43614 | 63.09834 | 60.20888 | Promoter | 20 | 151344175 | 151471842 | 127668 | 1 | ENSMUST00 | 0 | Fam120c | ENSMUSG00000025262.8 |
| chrX | 35888509 | 35889983 | 23.43614 | 63.09834 | 60.20888 | Promoter | 20 | 35888832 | 36076562 | 187731 | 1 | ENSMUST00 | 0 | Dock11 | ENSMUSG00000031093.9 |
| chrX | 7841753 | 7843408 | 23.43614 | 63.09834 | 60.20888 | Promoter | 20 | 7841831 | 7876626 | 34796 | 1 | ENSMUST00 | 0 | Otud5 | ENSMUSG00000031154.10 |
| chrX | 99136135 | 99141180 | 23.43614 | 63.09834 | 60.20888 | Promoter | 20 | 99138131 | 99147230 | 9100 | 1 | ENSMUST00 | 0 | Efnb1 | ENSMUSG00000031217.8 |
| chrX | 106011313 | 106011816 | 23.43614 | 63.09834 | 60.20888 | Promoter | 20 | 105970478 | 106011786 | 41309 | 2 | ENSMUST00 | 0 | Magt1 | ENSMUSG00000031232.11 |
| chrX | 53268993 | 53269848 | 23.43614 | 63.09834 | 60.20888 | Promoter | 20 | 53243415 | 53269805 | 26391 | 2 | ENSMUST00 | 0 | Fam122b | ENSMUSG00000036022.10 |
| chrX | 18162564 | 18165181 | 23.43614 | 63.09834 | 60.20888 | Promoter | 20 | 18162575 | 18279936 | 117362 | 1 | ENSMUST00 | 0 | Kdm6a | ENSMUSG00000037369.11 |
| chrX | 103979894 | 103981279 | 23.43614 | 63.09834 | 60.20888 | Promoter | 20 | 103957163 | 103981284 | 24122 | 2 | ENSMUST00 | 5 | Rlim | ENSMUSG00000056537.6 |
| chrX | 159372169 | 159373328 | 23.43614 | 63.09834 | 60.20888 | Promoter | 20 | 159372178 | 159376558 | 4381 | 1 | ENSMUST00 | 0 | Eif1ax | ENSMUSG00000067194.6 |
| chrX | 73227742 | 73229483 | 23.43614 | 63.09834 | 60.20888 | Promoter | 20 | 73228291 | 73230795 | 2505 | 1 | ENSMUST00 | 0 | F8a | ENSMUSG00000078317.6 |
| chr14 | 24003674 | 24005097 | 21.91892 | 63.05404 | 60.16603 | Promoter | 14 | 23299381 | 24004205 | 704825 | 2 | ENSMUST00 | 0 | Kcnma1 | ENSMUSG00000063142.8 |
| chr15 | 6873356 | 6875149 | 21.91892 | 63.05404 | 60.16603 | Promoter | 15 | 6813577 | 6874969 | 61393 | 2 | ENSMUST00 | 0 | Osmr | ENSMUSG00000022146.7 |
| chr19 | 46707078 | 46708255 | 21.91892 | 63.05404 | 60.16603 | Promoter | 19 | 46707443 | 46741095 | 33653 | 1 | ENSMUST00 | 0 | As3mt | ENSMUSG00000003559.7 |
| chr19 | 3912973 | 3913685 | 21.91892 | 63.05404 | 60.16603 | Promoter | 19 | 3908870 | 3912717 | 3848 | 2 | ENSMUST00 | -257 | Ndufs8 | ENSMUSG00000059734.6 |
| chr2 | 5942205 | 5943119 | 21.91892 | 63.05404 | 60.16603 | Promoter | 2 | 5896115 | 5942792 | 46678 | 2 | ENSMUST00 | 0 | Dhtkd1 | ENSMUSG00000025815.8 |
| chr4 | 129335566 | 129336753 | 21.91892 | 63.05404 | 60.16603 | Promoter | 4 | 129335695 | 129349871 | 14177 | 1 | ENSMUST00 | 0 | Zbtb8os | ENSMUSG00000057572.10 |
| chr5 | 76856384 | 76858338 | 21.91892 | 63.05404 | 60.16603 | Promoter | 5 | 76858842 | 76866491 | 7650 | 1 | ENSMUST00 | -504 | C530008M17 | ENSMUSG00000036377.13 |
| chr7 | 143822072 | 143822884 | 21.91892 | 63.05404 | 60.16603 | Promoter | 7 | 143795584 | 143822841 | 27258 | 2 | ENSMUST00 | 0 | Nadsyn1 | ENSMUSG00000031090.9 |
| chr9 | 27154754 | 27155702 | 21.91892 | 63.05404 | 60.16603 | Promoter | 9 | 27097385 | 27155421 | 58037 | 2 | ENSMUST00 | 0 | Jam3 | ENSMUSG00000031990.9 |
| chr1 | 171293910 | 171295312 | 20.47566 | 63.04224 | 60.15488 | Promoter | 1 | 171288563 | 171295024 | 6462 | 2 | ENSMUST00 | 0 | Ufc1 | ENSMUSG00000062963.5 |
| chr11 | 72410970 | 72411994 | 20.47566 | 63.04224 | 60.15488 | Promoter | 11 | 72389164 | 72411711 | 22548 | 2 | ENSMUST00 | 0 | Smtnl2 | ENSMUSG00000045667.9 |
| chr13 | 23516819 | 23519594 | 20.47566 | 63.04224 | 60.15488 | Distal Interge | 13 | 23527011 | 23530943 | 3933 | 2 | ENSMUST00 | 11349 | Gm11335 | ENSMUSG00000085016.1 |
| chr10 | 57794384 | 57795660 | 23.39399 | 63.02628 | 60.13908 | Promoter | 10 | 57794568 | 57807470 | 12903 | 1 | ENSMUST00 | 0 | Smpdl3a | ENSMUSG00000019872.8 |
| chr4 | 11387144 | 11388540 | 22.89212 | 63.02343 | 60.13633 | Promoter | 4 | 11332038 | 11386783 | 54746 | 2 | ENSMUST00 | -362 | Esrp1 | ENSMUSG00000040728.10 |
| chr9 | 44707832 | 44708957 | 22.38578 | 63.0083 | 60.12133 | Promoter | 9 | 44703573 | 44708427 | 4855 | 2 | ENSMUST00 | 0 | Phldb1 | ENSMUSG00000048537.10 |
| chr15 | 35293585 | 35294811 | 23.34501 | 62.94254 | 60.0558 | Distal Interge | 15 | 35296098 | 35303304 | 7207 | 1 | ENSMUST00 | -1287 | Osr2 | ENSMUSG00000022330.4 |
| chr19 | 7040331 | 7041000 | 23.34501 | 62.94254 | 60.0558 | Promoter | 19 | 7020696 | 7040026 | 19331 | 2 | ENSMUST00 | -306 | Stip1 | ENSMUSG00000024966.8 |
| chr18 | 38238767 | 38239758 | 22.34262 | 62.9334 | 60.04675 | Exon (ENSMU | 18 | 38244828 | 38245382 | 555 | 1 | ENSMUST00 | -5070 | Gm10268 | ENSMUSG00000066475.6 |
| chr5 | 151367775 | 151368540 | 22.34262 | 62.9334 | 60.04675 | Promoter | 5 | 151368675 | 151380912 | 12238 | 1 | ENSMUST00 | -135 | 1700028E10I | ENSMUSG00000097321.2 |
| chr12 | 118198504 | 118199248 | 22.8003 | 62.86544 | 59.97937 | Promoter | 12 | 117877982 | 118199043 | 321062 | 2 | ENSMUST00 | 0 | Dnah11 | ENSMUSG00000018581.10 |
| chr4 | 116993601 | 116994502 | 22.8003 | 62.86544 | 59.97937 | Promoter | 4 | 116989965 | 116994413 | 4449 | 2 | ENSMUST00 | 0 | Urod | ENSMUSG00000028684.9 |
| chr5 | 134455170 | 134457151 | 22.8003 | 62.86544 | 59.97937 | Promoter | 5 | 134357669 | 134456262 | 98594 | 2 | ENSMUST00 | 0 | Gtf2ird1 | ENSMUSG00000023079.9 |
| chr7 | 45537977 | 45538826 | 22.8003 | 62.86544 | 59.97937 | Exon (ENSMU | 7 | 45530571 | 45554032 | 23462 | 1 | ENSMUST00 | 7407 | Plekha4 | ENSMUSG00000040428.12 |
| chr9 | 118606450 | 118607555 | 22.8003 | 62.86544 | 59.97937 | Promoter | 9 | 118606690 | 118901003 | 294314 | 1 | ENSMUST00 | 0 | Itga9 | ENSMUSG00000039115.8 |
| chr9 | 95844679 | 95845549 | 22.8003 | 62.86544 | 59.97937 | Promoter | 9 | 95752642 | 95845286 | 92645 | 2 | ENSMUST00 | 0 | Pls1 | ENSMUSG00000049493.8 |
| chr17 | 46781856 | 46783237 | 23.29623 | 62.85914 | 59.97368 | Promoter | 17 | 46773907 | 46782656 | 8750 | 2 | ENSMUST00 | 0 | Rpl7l1 | ENSMUSG00000063888.6 |
| chr8 | 94178656 | 94180513 | 23.29623 | 62.85914 | 59.97368 | Promoter | 8 | 94179089 | 94180325 | 1237 | 1 | ENSMUST00 | 0 | Mt1 | ENSMUSG00000031765.7 |
| chr1 | 178527358 | 178530919 | 20.79993 | 62.77878 | 59.89386 | Promoter | 1 | 178529125 | 178932730 | 403606 | 1 | ENSMUST00 | 0 | Kif26b | ENSMUSG00000026494.7 |
| chr11 | 69331855 | 69332661 | 23.24766 | 62.77607 | 59.89146 | Promoter | 11 | 69329825 | 69332122 | 2298 | 2 | ENSMUST00 | 0 | Kcnab3os | ENSMUSG00000086900.1 |
| chr7 | 67647357 | 67648016 | 22.21412 | 62.71022 | 59.82578 | Promoter | 7 | 67647410 | 67672564 | 25155 | 1 | ENSMUST00 | 0 | Ttc23 | ENSMUSG00000030555.10 |
| chr8 | 71469105 | 71470205 | 22.70921 | 62.70861 | 59.8242 | Promoter | 8 | 71469199 | 71472246 | 3048 | 1 | ENSMUST00 | 0 | Dda1 | ENSMUSG00000074247.5 |
| chr1 | 72254269 | 72255424 | 22.21283 | 62.70797 | 59.8237 | Promoter | 1 | 72255008 | 72255198 | 191 | 1 | ENSMUST00 | 0 | Gm25939 | ENSMUSG00000093843.1 |
| chr11 | 85140193 | 85141092 | 22.21283 | 62.70797 | 59.8237 | Promoter | 11 | 85081176 | 85140161 | 58986 | 2 | ENSMUST00 | -33 | Usp32 | ENSMUSG00000000804.9 |
| chr11 | 115277266 | 115277988 | 22.21283 | 62.70797 | 59.8237 | Promoter | 11 | 115268024 | 115277050 | 9027 | 2 | ENSMUST00 | -217 | Fdxr | ENSMUSG00000018861.3 |
| chr12 | 112949963 | 112951035 | 22.21283 | 62.70797 | 59.8237 | Promoter | 12 | 112946363 | 112950512 | 4150 | 2 | ENSMUST00 | 0 | Gm26583 | ENSMUSG00000097476.1 |
| chr13 | 98365666 | 98367402 | 22.21283 | 62.70797 | 59.8237 | Distal Interge | 13 | 98354242 | 98356703 | 2462 | 1 | ENSMUST00 | 11425 | Foxd1 | ENSMUSG00000078302.4 |
| chr14 | 55822836 | 55824209 | 22.21283 | 62.70797 | 59.8237 | Promoter | 14 | 55824795 | 55833943 | 9149 | 1 | ENSMUST00 | -586 | Nfatc4 | ENSMUSG00000023411.6 |
| chr14 | 21988023 | 21988467 | 22.21283 | 62.70797 | 59.8237 | 5' UTR | 14 | 21983962 | 21989601 | 5640 | 2 | ENSMUST00 | 1134 | Zfp503 | ENSMUSG00000039081.9 |
| chr2 | 178077492 | 178078612 | 22.21283 | 62.70797 | 59.8237 | Promoter | 2 | 178067695 | 178078425 | 10731 | 2 | ENSMUST00 | 0 | Zfp931 | ENSMUSG00000078861.3 |
| chr3 | 5218270 | 5219919 | 22.21283 | 62.70797 | 59.8237 | Promoter | 3 | 5218526 | 5413165 | 194640 | 1 | ENSMUST00 | 0 | Zfhx4 | ENSMUSG00000025255.13 |
| chr4 | 58552134 | 58553904 | 22.21283 | 62.70797 | 59.8237 | Promoter | 4 | 58435255 | 58553311 | 118057 | 2 | ENSMUST00 | 0 | Lpar1 | ENSMUSG00000038668.9 |
| chr6 | 84587347 | 84589663 | 22.21283 | 62.70797 | 59.8237 | 5' UTR | 6 | 84571414 | 84593810 | 22397 | 2 | ENSMUST00 | 4147 | Cyp26b1 | ENSMUSG00000063415.7 |
| chr7 | 68302694 | 68303491 | 22.21283 | 62.70797 | 59.8237 | Intron (ENSM | 7 | 68300113 | 68363087 | 62975 | 1 | ENSMUST00 | 2582 | Fam169b | ENSMUSG00000074071.4 |
| chr8 | 22974162 | 22975667 | 22.21283 | 62.70797 | 59.8237 | Promoter | 8 | 22974844 | 23149933 | 175090 | 1 | ENSMUST00 | 0 | Ank1 | ENSMUSG00000031543.13 |
| chr8 | 94875489 | 94876384 | 22.21283 | 62.70797 | 59.8237 | Promoter | 8 | 94863828 | 94876312 | 12485 | 2 | ENSMUST00 | 0 | Dok4 | ENSMUSG00000040631.7 |
| chr8 | 86623413 | 86623857 | 22.21283 | 62.70797 | 59.8237 | Promoter | 8 | 86624043 | 86716631 | 92589 | 1 | ENSMUST00 | -186 | Lonp2 | ENSMUSG00000047866.13 |
| chr9 | 42943675 | 42945076 | 22.21283 | 62.70797 | 59.8237 | Promoter | 9 | 42518135 | 42944493 | 426359 | 2 | ENSMUST00 | 0 | Grik4 | ENSMUSG00000032017.9 |
| chr9 | 107950489 | 107951499 | 22.21283 | 62.70797 | 59.8237 | Promoter | 9 | 107950963 | 107972268 | 21306 | 1 | ENSMUST00 | 0 | Traip | ENSMUSG00000032586.4 |
| chr12 | 98574228 | 98575364 | 22.66394 | 62.63062 | 59.74885 | Intron (ENSM | 12 | 98433994 | 98577940 | 143947 | 2 | ENSMUST00 | 2576 | Kcnk10 | ENSMUSG00000033854.9 |
| chr17 | 28079753 | 28080398 | 22.66394 | 62.63062 | 59.74885 | Promoter | 17 | 28066751 | 28080567 | 13817 | 2 | ENSMUST00 | 169 | Tcp11 | ENSMUSG00000062859.9 |
| chr16 | 93710993 | 93711494 | 23.15112 | 62.61088 | 59.72924 | Promoter | 16 | 93711907 | 93810585 | 98679 | 1 | ENSMUST00 | -413 | Dopey2 | ENSMUSG00000022946.8 |
| chr12 | 3235019 | 3236318 | 22.61885 | 62.5529 | 59.67142 | Promoter | 12 | 3235504 | 3241410 | 5907 | 1 | ENSMUST00 | 0 | Rab10os | ENSMUSG00000079179.4 |
| chr4 | 115962618 | 115963438 | 21.14701 | 62.5472 | 59.66584 | Distal Interge | 4 | 115915120 | 115939926 | 24807 | 2 | ENSMUST00 | -22693 | Dmbx1 | ENSMUSG00000028707.10 |
| chr9 | 20459072 | 20460621 | 21.14701 | 62.5472 | 59.66584 | Promoter | 9 | 20432972 | 20460160 | 27189 | 2 | ENSMUST00 | 0 | Zfp26 | ENSMUSG00000063108.9 |
| chr1 | 177442303 | 177443552 | 19.72513 | 62.50151 | 59.62064 | Promoter | 1 | 177429047 | 177443270 | 14224 | 2 | ENSMUST00 | 0 | Gm26801 | ENSMUSG00000097892.1 |
| chr10 | 94311546 | 94312854 | 19.72513 | 62.50151 | 59.62064 | Promoter | 10 | 94311949 | 94430897 | 118949 | 1 | ENSMUST00 | 0 | Tmcc3 | ENSMUSG00000020023.12 |
| chr11 | 50174812 | 50175473 | 19.72513 | 62.50151 | 59.62064 | Promoter | 11 | 50174849 | 50200115 | 25267 | 1 | ENSMUST00 | 0 | 3010026O09 | ENSMUSG00000020381.5 |
| chr11 | 78079786 | 78081012 | 19.72513 | 62.50151 | 59.62064 | Promoter | 11 | 78073376 | 78080383 | 7008 | 2 | ENSMUST00 | 0 | Eral1 | ENSMUSG00000020832.9 |
| chr14 | 27622263 | 27623083 | 19.72513 | 62.50151 | 59.62064 | Promoter | 14 | 27622442 | 28478537 | 856096 | 1 | ENSMUST00 | 0 | Erc2 | ENSMUSG00000040640.9 |
| chr15 | 98676338 | 98677490 | 19.72513 | 62.50151 | 59.62064 | Promoter | 15 | 98663421 | 98677460 | 14040 | 2 | ENSMUST00 | 0 | Rnd1 | ENSMUSG00000054855.8 |
| chr16 | 17530073 | 17531784 | 19.72513 | 62.50151 | 59.62064 | Promoter | 16 | 17527982 | 17531136 | 3155 | 2 | ENSMUST00 | 0 | Thap7 | ENSMUSG00000022760.10 |
| chr17 | 35865184 | 35867409 | 19.72513 | 62.50151 | 59.62064 | Promoter | 17 | 35865593 | 35875591 | 9999 | 1 | ENSMUST00 | 0 | Ppp1r18 | ENSMUSG00000034595.12 |
| chr17 | 36144789 | 36146054 | 19.72513 | 62.50151 | 59.62064 | Promoter | 17 | 36145018 | 36162374 | 17357 | 1 | ENSMUST00 | 0 | 2410017I17R | ENSMUSG00000038311.8 |
| chr17 | 80372881 | 80374126 | 19.72513 | 62.50151 | 59.62064 | Promoter | 17 | 80373218 | 80373541 | 324 | 2 | ENSMUST00 | 0 | Gm10190 | ENSMUSG00000066938.1 |
| chr18 | 24466329 | 24467929 | 19.72513 | 62.50151 | 59.62064 | Distal Interge | 18 | 24470861 | 24477765 | 6905 | 1 | ENSMUST00 | -2932 | 2700062C07I | ENSMUSG00000024273.7 |
| chr19 | 11120933 | 11121746 | 19.72513 | 62.50151 | 59.62064 | Promoter | 19 | 11121510 | 11122425 | 916 | 1 | ENSMUST00 | 0 | RP24-512H1 | ENSMUSG00000099998.1 |
| chr2 | 32960956 | 32962491 | 19.72513 | 62.50151 | 59.62064 | Promoter | 2 | 32925216 | 32961255 | 36040 | 2 | ENSMUST00 | 0 | Lrsam1 | ENSMUSG00000026792.11 |
| chr2 | 127182479 | 127183437 | 19.72513 | 62.50151 | 59.62064 | Downstream | 2 | 127202567 | 127208254 | 5688 | 2 | ENSMUST00 | 24817 | 1810024B03I | ENSMUSG00000044145.2 |
| chr3 | 95950530 | 95951708 | 19.72513 | 62.50151 | 59.62064 | Distal Interge | 3 | 95944204 | 95945523 | 1320 | 1 | ENSMUST00 | 6327 | Anp32e | ENSMUSG00000015749.7 |
| chr3 | 94412473 | 94414007 | 19.72513 | 62.50151 | 59.62064 | Promoter | 3 | 94413296 | 94432633 | 19338 | 1 | ENSMUST00 | 0 | Tdrkh | ENSMUSG00000041912.7 |
| chr6 | 124744930 | 124746565 | 19.72513 | 62.50151 | 59.62064 | Promoter | 6 | 124745172 | 124745625 | 454 | 2 | ENSMUST00 | 0 | Atn1 | ENSMUSG00000004263.10 |
| chr7 | 105480927 | 105482479 | 19.72513 | 62.50151 | 59.62064 | Promoter | 7 | 105480607 | 105482197 | 1591 | 2 | ENSMUST00 | 0 | Prkcdbp | ENSMUSG00000037060.2 |
| chr7 | 48131735 | 48132282 | 19.72513 | 62.50151 | 59.62064 | Distal Interge | 7 | 48116772 | 48117441 | 670 | 2 | ENSMUST00 | -14295 | RP23-62M19 | ENSMUSG00000101352.1 |
| chr15 | 102376529 | 102377484 | 22.08709 | 62.48929 | 59.61029 | Distal Interge | 15 | 102373880 | 102373983 | 104 | 1 | ENSMUST00 | 2650 | Gm23590 | ENSMUSG00000088286.1 |
| chr2 | 157423815 | 157424803 | 22.08709 | 62.48929 | 59.61029 | Promoter | 2 | 157424116 | 157455707 | 31592 | 1 | ENSMUST00 | 0 | Src | ENSMUSG00000027646.10 |
| chr15 | 98727315 | 98728505 | 22.57394 | 62.47547 | 59.59652 | Promoter | 15 | 98724373 | 98728198 | 3826 | 2 | ENSMUST00 | 0 | Fkbp11 | ENSMUSG00000003355.6 |
| chr4 | 122886181 | 122887223 | 22.57394 | 62.47547 | 59.59652 | Promoter | 4 | 122859047 | 122886056 | 27010 | 2 | ENSMUST00 | -126 | Cap1 | ENSMUSG00000028656.9 |
| chr4 | 43957286 | 43958457 | 23.05539 | 62.44698 | 59.56813 | Promoter | 4 | 43957401 | 43979118 | 21718 | 1 | ENSMUST00 | 0 | Glipr2 | ENSMUSG00000028480.9 |
| chr19 | 5473603 | 5475286 | 20.14468 | 62.43501 | 59.55626 | Promoter | 19 | 5473973 | 5477578 | 3606 | 1 | ENSMUST00 | 0 | Efemp2 | ENSMUSG00000024909.9 |
| chr11 | 55468395 | 55469249 | 21.5587 | 62.4211 | 59.54246 | Promoter | 11 | 55469685 | 55504838 | 35154 | 1 | ENSMUST00 | -436 | G3bp1 | ENSMUSG00000018583.5 |
| chr18 | 36453843 | 36454966 | 22.52921 | 62.39832 | 59.51973 | Promoter | 18 | 36403679 | 36454495 | 50817 | 2 | ENSMUST00 | 0 | Pfdn1 | ENSMUSG00000024346.5 |
| chr13 | 62684450 | 62685389 | 21.5194 | 62.35189 | 59.47348 | Distal Interge | 13 | 62700108 | 62701046 | 939 | 2 | ENSMUST00 | 15657 | Gm5665 | ENSMUSG00000074863.4 |
| chr5 | 9265761 | 9266601 | 21.5194 | 62.35189 | 59.47348 | Promoter | 5 | 9266118 | 9479736 | 213619 | 1 | ENSMUST00 | 0 | 9330182L06 | ENSMUSG00000056004.11 |
| chr9 | 37208253 | 37208859 | 21.5194 | 62.35189 | 59.47348 | Promoter | 9 | 37208333 | 37223210 | 14878 | 1 | ENSMUST00 | 0 | Tmem218 | ENSMUSG00000032121.8 |
| chr7 | 16272560 | 16274343 | 21.96151 | 62.27057 | 59.39301 | Promoter | 7 | 16272014 | 16273692 | 1679 | 2 | ENSMUST00 | 0 | Prr24 | ENSMUSG00000091811.1 |
| chr9 | 40799626 | 40800806 | 21.96151 | 62.27057 | 59.39301 | Promoter | 9 | 40800984 | 40805200 | 4217 | 1 | ENSMUST00 | -178 | Hspa8 | ENSMUSG00000015656.11 |
| chr10 | 22644821 | 22645805 | 22.44027 | 62.24483 | 59.36732 | Promoter | 10 | 22645011 | 22704285 | 59275 | 1 | ENSMUST00 | 0 | Slc2a12 | ENSMUSG00000037490.5 |
| chr14 | 20928550 | 20929186 | 22.91325 | 62.20349 | 59.32608 | Promoter | 14 | 20929433 | 21033673 | 104241 | 1 | ENSMUST00 | -247 | Vcl | ENSMUSG00000021823.9 |
| chr11 | 3535468 | 3536634 | 21.91997 | 62.19815 | 59.32085 | Promoter | 11 | 3517523 | 3537398 | 19876 | 2 | ENSMUST00 | 764 | Smtn | ENSMUSG00000020439.12 |
| chr17 | 26095049 | 26095767 | 21.91997 | 62.19815 | 59.32085 | Promoter | 17 | 26091734 | 26095508 | 3775 | 2 | ENSMUST00 | 0 | Nme4 | ENSMUSG00000024177.8 |
| chr18 | 60500849 | 60501922 | 21.91997 | 62.19815 | 59.32085 | Promoter | 18 | 60474193 | 60501983 | 27791 | 2 | ENSMUST00 | 61 | Smim3 | ENSMUSG00000038059.6 |
| chr18 | 6136234 | 6136978 | 21.91997 | 62.19815 | 59.32085 | Promoter | 18 | 6025217 | 6136098 | 110882 | 2 | ENSMUST00 | -137 | Arhgap12 | ENSMUSG00000041225.11 |
| chr19 | 57007763 | 57008670 | 21.91997 | 62.19815 | 59.32085 | Promoter | 19 | 56912361 | 57008198 | 95838 | 2 | ENSMUST00 | 0 | Afap1l2 | ENSMUSG00000025083.12 |

| | | | | | | | | | | | | | | | |
|---|---|---|---|---|---|---|---|---|---|---|---|---|---|---|---|
| chr3 | 96593397 | 96594507 | 21.91997 | 62.19815 | 59.32085 | Promoter | 3 | 96577872 | 96594181 | 16310 | 2 | ENSMUST00 | 0 | Polr3gl | ENSMUSG00000028104.9 |
| chr4 | 156109168 | 156109790 | 21.91997 | 62.19815 | 59.32085 | Promoter | 4 | 156109982 | 156127265 | 17284 | 1 | ENSMUST00 | -192 | 9430015G10 | ENSMUSG00000059939.8 |
| chr11 | 70525085 | 70526193 | 22.39606 | 62.16851 | 59.29202 | Promoter | 11 | 70525367 | 70526424 | 1058 | 1 | ENSMUST00 | 0 | Psmb6 | ENSMUSG00000018286.6 |
| chr8 | 13493638 | 13494541 | 22.39606 | 62.16851 | 59.29202 | Promoter | 8 | 13465374 | 13494535 | 29162 | 2 | ENSMUST00 | 0 | Gas6 | ENSMUSG00000031451.5 |
| chr10 | 75836789 | 75837882 | 18.64736 | 62.11404 | 59.23804 | Distal Interge | 10 | 75831845 | 75834881 | 3037 | 2 | ENSMUST00 | -1909 | Gstt2 | ENSMUSG00000033318.6 |
| chr12 | 21542953 | 21544077 | 18.64736 | 62.11404 | 59.23804 | Distal Interge | 12 | 21417911 | 21419803 | 1893 | 1 | ENSMUST00 | 125043 | Gm4419 | ENSMUSG00000097055.1 |
| chr13 | 67172593 | 67173883 | 18.64736 | 62.11404 | 59.23804 | Promoter | 13 | 67173207 | 67183126 | 9920 | 1 | ENSMUST00 | 0 | Rsl1 | ENSMUSG00000058900.4 |
| chr18 | 15149737 | 15151710 | 18.64736 | 62.11404 | 59.23804 | Promoter | 18 | 14968685 | 15151446 | 182762 | 2 | ENSMUST00 | 0 | Kctd1 | ENSMUSG00000036225.9 |
| chr19 | 5731148 | 5732118 | 18.64736 | 62.11404 | 59.23804 | Promoter | 19 | 5728088 | 5731713 | 3626 | 2 | ENSMUST00 | 0 | Sssca1 | ENSMUSG00000079478.3 |
| chr2 | 128943333 | 128944466 | 18.64736 | 62.11404 | 59.23804 | Promoter | 2 | 128926268 | 128944077 | 17810 | 2 | ENSMUST00 | 0 | Zc3h8 | ENSMUSG00000027387.11 |
| chr6 | 124740221 | 124741831 | 18.64736 | 62.11404 | 59.23804 | Promoter | 6 | 124739183 | 124740695 | 1513 | 2 | ENSMUST00 | 0 | Grcc10 | ENSMUSG00000072772.3 |
| chr10 | 69398308 | 69400147 | 22.35204 | 62.09245 | 59.21756 | Distal Interge | 10 | 69336635 | 69352938 | 16304 | 2 | ENSMUST00 | -45371 | Cdk1 | ENSMUSG00000019942.7 |
| chr5 | 121384670 | 121386247 | 22.35204 | 62.09245 | 59.21756 | Promoter | 5 | 121371725 | 121385632 | 13908 | 2 | ENSMUST00 | 0 | Trafd1 | ENSMUSG00000042726.9 |
| chr8 | 110217852 | 110218480 | 22.35203 | 62.09245 | 59.21756 | Promoter | 8 | 110217989 | 110224486 | 6498 | 1 | ENSMUST00 | 0 | Cmtr2 | ENSMUSG00000046441.5 |
| chr5 | 136135180 | 136137068 | 16.29944 | 62.08345 | 59.20918 | Promoter | 5 | 136122772 | 136136074 | 13303 | 2 | ENSMUST00 | 0 | Lrwd1 | ENSMUSG00000029703.5 |
| chr6 | 100704452 | 100706031 | 22.81947 | 62.04271 | 59.16861 | Promoter | 6 | 100704734 | 100805081 | 100348 | 1 | ENSMUST00 | 0 | Gxylt2 | ENSMUSG00000030074.8 |
| chr9 | 40530678 | 40531652 | 22.81947 | 62.04271 | 59.16861 | Promoter | 9 | 40293239 | 40531383 | 238145 | 2 | ENSMUST00 | 0 | Gramd1b | ENSMUSG00000040111.9 |
| chr13 | 115088957 | 115091242 | 22.30817 | 62.01665 | 59.14301 | Promoter | 13 | 115088357 | 115090188 | 1832 | 2 | ENSMUST00 | 0 | Pelo | ENSMUSG00000042275.8 |
| chr2 | 132686461 | 132687606 | 22.30817 | 62.01665 | 59.14301 | Promoter | 2 | 132686931 | 132751055 | 64125 | 1 | ENSMUST00 | 0 | 1110034G24 | ENSMUSG00000044991.5 |
| chr1 | 59482026 | 59483998 | 16.96705 | 61.94942 | 59.07605 | Promoter | 1 | 59482424 | 59486955 | 4532 | 1 | ENSMUST00 | 0 | Fzd7 | ENSMUSG00000041075.8 |
| chr11 | 120932301 | 120933163 | 16.96705 | 61.94942 | 59.07605 | Promoter | 11 | 120826529 | 120932872 | 106344 | 2 | ENSMUST00 | 0 | Ccdc57 | ENSMUSG00000048445.6 |
| chr17 | 21566408 | 21567855 | 16.96705 | 61.94942 | 59.07605 | Promoter | 17 | 21567046 | 21588697 | 21652 | 1 | ENSMUST00 | 0 | Zfp948 | ENSMUSG00000067931.5 |
| chr17 | 34958204 | 34960170 | 16.96705 | 61.94942 | 59.07605 | Promoter | 17 | 34956436 | 34959238 | 2803 | 2 | ENSMUST00 | 0 | Hspa1b | ENSMUSG00000090877.3 |
| chr8 | 72318583 | 72321527 | 16.96705 | 61.94942 | 59.07605 | Promoter | 8 | 72319033 | 72321656 | 2624 | 1 | ENSMUST00 | 0 | Klf2 | ENSMUSG00000055148.7 |
| chr14 | 41013064 | 41014319 | 17.74057 | 61.94745 | 59.07436 | Promoter | 14 | 40993740 | 41013775 | 20036 | 2 | ENSMUST00 | 0 | Fam213a | ENSMUSG00000021792.10 |
| chr17 | 26508607 | 26509136 | 17.74057 | 61.94745 | 59.07436 | Promoter | 17 | 26505590 | 26508519 | 2930 | 2 | ENSMUST00 | -89 | Dusp1 | ENSMUSG00000024190.6 |
| chr5 | 116421455 | 116423156 | 17.74057 | 61.94745 | 59.07436 | Promoter | 5 | 116408491 | 116422864 | 14374 | 2 | ENSMUST00 | 0 | Hspb8 | ENSMUSG00000041548.4 |
| chr9 | 53771247 | 53772322 | 20.78155 | 61.8938 | 59.02139 | Promoter | 9 | 53771535 | 53818161 | 46627 | 1 | ENSMUST00 | 0 | Slc35f2 | ENSMUSG00000042195.8 |
| chr5 | 107687856 | 107688720 | 22.72645 | 61.88314 | 59.01102 | Promoter | 5 | 107659097 | 107688015 | 28919 | 2 | ENSMUST00 | 0 | Gm26692 | ENSMUSG00000097035.1 |
| chr10 | 59969839 | 59970559 | 22.17762 | 61.79087 | 58.9192 | Distal Interge | 10 | 59949675 | 59951770 | 2096 | 2 | ENSMUST00 | -18070 | Ddit4 | ENSMUSG00000020108.3 |
| chr3 | 95870869 | 95872043 | 21.13419 | 61.67127 | 58.79994 | Promoter | 3 | 95862634 | 95871519 | 8886 | 2 | ENSMUST00 | 0 | Mrps21 | ENSMUSG00000054312.5 |
| chr5 | 24394123 | 24394939 | 21.13419 | 61.67127 | 58.79994 | Promoter | 5 | 24394155 | 24409945 | 15791 | 1 | ENSMUST00 | 0 | Abcb8 | ENSMUSG00000028973.13 |
| chr1 | 185331829 | 185332940 | 20.65177 | 61.66082 | 58.78991 | Promoter | 1 | 185332149 | 185357777 | 25629 | 1 | ENSMUST00 | 0 | Bpnt1 | ENSMUSG00000026617.10 |
| chr1 | 191097816 | 191098806 | 20.65177 | 61.66082 | 58.78991 | Promoter | 1 | 191098414 | 191108943 | 10530 | 1 | ENSMUST00 | 0 | Batf3 | ENSMUSG00000026630.4 |
| chr1 | 165281262 | 165282706 | 20.65177 | 61.66082 | 58.78991 | Distal Interge | 1 | 165301863 | 165322930 | 21068 | 1 | ENSMUST00 | -19157 | Gpr161 | ENSMUSG00000040836.10 |
| chr1 | 93824772 | 93826075 | 20.65177 | 61.66082 | 58.78991 | Promoter | 1 | 93824909 | 93826458 | 1550 | 1 | ENSMUST00 | 0 | D2hgdh | ENSMUSG00000073609.4 |
| chr13 | 6647037 | 6649458 | 20.65177 | 61.66082 | 58.78991 | Promoter | 13 | 6579768 | 6648777 | 69010 | 2 | ENSMUST00 | 0 | Pfkp | ENSMUSG00000021196.8 |
| chr13 | 109685162 | 109687571 | 20.65177 | 61.66082 | 58.78991 | Promoter | 13 | 109686156 | 109951474 | 265319 | 1 | ENSMUST00 | 0 | Pde4d | ENSMUSG00000021699.12 |
| chr17 | 87636111 | 87637290 | 20.65177 | 61.66082 | 58.78991 | Promoter | 17 | 87635979 | 87651106 | 15128 | 1 | ENSMUST00 | 133 | Epcam | ENSMUSG00000045394.8 |
| chr2 | 13270663 | 13271398 | 20.65177 | 61.66082 | 58.78991 | Promoter | 2 | 13076967 | 13271315 | 194349 | 2 | ENSMUST00 | 0 | Rsu1 | ENSMUSG00000026727.5 |
| chr2 | 74738622 | 74741151 | 20.65177 | 61.66082 | 58.78991 | Promoter | 2 | 74739619 | 74747900 | 8282 | 1 | ENSMUST00 | 0 | Hoxd3 | ENSMUSG00000079277.4 |
| chr3 | 65391460 | 65392682 | 20.65177 | 61.66082 | 58.78991 | Promoter | 3 | 65379655 | 65392623 | 12969 | 2 | ENSMUST00 | 0 | Ssr3 | ENSMUSG00000027828.7 |
| chr3 | 96708069 | 96708622 | 20.65177 | 61.66082 | 58.78991 | Promoter | 3 | 96705892 | 96708560 | 2669 | 2 | ENSMUST00 | 0 | Nudt17 | ENSMUSG00000028100.6 |
| chr3 | 107035703 | 107037207 | 20.65177 | 61.66082 | 58.78991 | Promoter | 3 | 107036162 | 107038129 | 1968 | 1 | ENSMUST00 | 0 | Kcna3 | ENSMUSG00000047959.3 |
| chr4 | 138261857 | 138263010 | 20.65177 | 61.66082 | 58.78991 | Promoter | 4 | 138262232 | 138301492 | 39261 | 1 | ENSMUST00 | 0 | Kif17 | ENSMUSG00000028758.7 |
| chr5 | 72867770 | 72869007 | 20.65177 | 61.66082 | 58.78991 | Promoter | 5 | 72755718 | 72868452 | 112735 | 2 | ENSMUST00 | 0 | Tec | ENSMUSG00000029217.11 |
| chr8 | 84969075 | 84970575 | 20.65177 | 61.66082 | 58.78991 | Promoter | 8 | 84969587 | 84974834 | 5248 | 1 | ENSMUST00 | 0 | Prdx2 | ENSMUSG00000005161.10 |
| chr8 | 67947543 | 67950750 | 20.65177 | 61.66082 | 58.78991 | Intron (ENSM | 8 | 67903648 | 67951879 | 48232 | 2 | ENSMUST00 | 1129 | Psd3 | ENSMUSG00000030465.13 |
| chr8 | 84010752 | 84012547 | 20.65177 | 61.66082 | 58.78991 | Promoter | 8 | 84010222 | 84012158 | 1937 | 2 | ENSMUST00 | 0 | 2210011C24I | ENSMUSG00000074217.3 |
| chr9 | 106198701 | 106199614 | 20.65177 | 61.66082 | 58.78991 | Promoter | 9 | 106194172 | 106198942 | 4771 | 2 | ENSMUST00 | 0 | Ppm1m | ENSMUSG00000020253.9 |
| chr9 | 83440355 | 83441682 | 20.65177 | 61.66082 | 58.78991 | Promoter | 9 | 83391364 | 83441127 | 49764 | 2 | ENSMUST00 | 0 | Lca5 | ENSMUSG00000032258.10 |
| chr9 | 96364501 | 96365263 | 20.65177 | 61.66082 | 58.78991 | Promoter | 9 | 96332655 | 96364442 | 31788 | 2 | ENSMUST00 | -60 | Atp1b3 | ENSMUSG00000032412.8 |
| chr9 | 63132093 | 63133795 | 20.65177 | 61.66082 | 58.78991 | Promoter | 9 | 63133283 | 63133389 | 107 | 1 | ENSMUST00 | 0 | Gm25064 | ENSMUSG00000094201.1 |
| chrX | 155262446 | 155263332 | 20.65177 | 61.66082 | 58.78991 | Promoter | 20 | 155262505 | 155297646 | 35142 | 1 | ENSMUST00 | 0 | Acot9 | ENSMUSG00000025287.10 |
| chr6 | 119388835 | 119389615 | 19.25321 | 61.61364 | 58.74545 | Promoter | 6 | 119353765 | 119388686 | 34922 | 2 | ENSMUST00 | -150 | Adipor2 | ENSMUSG00000030168.8 |
| chr4 | 147442923 | 147443929 | 12.85555 | 61.54892 | 58.68096 | Promoter | 4 | 147443471 | 147444029 | 559 | 1 | ENSMUST00 | 0 | Gm13161 | ENSMUSG00000082248.1 |
| chr11 | 84523693 | 84526470 | 21.51301 | 61.48686 | 58.61925 | Promoter | 11 | 84518284 | 84525535 | 7252 | 2 | ENSMUST00 | 0 | Lhx1 | ENSMUSG00000018698.10 |
| chr2 | 50065979 | 50067249 | 21.51301 | 61.48686 | 58.61925 | Promoter | 2 | 50066429 | 50193569 | 127141 | 1 | ENSMUST00 | 0 | Lypd6 | ENSMUSG00000050447.10 |
| chr6 | 92705404 | 92706488 | 21.51301 | 61.48686 | 58.61925 | Promoter | 6 | 92370892 | 92706145 | 335254 | 2 | ENSMUST00 | 0 | Prickle2 | ENSMUSG00000030020.8 |
| chr11 | 109312006 | 109312746 | 22.97661 | 61.46103 | 58.5941 | Distal Interge | 11 | 109225355 | 109298129 | 72775 | 2 | ENSMUST00 | -13878 | Rgs9 | ENSMUSG00000020599.8 |
| chr11 | 76997321 | 76998998 | 22.97661 | 61.46103 | 58.5941 | Promoter | 11 | 76998603 | 77032339 | 33737 | 1 | ENSMUST00 | 0 | Slc6a4 | ENSMUSG00000020838.7 |
| chr11 | 99023633 | 99024847 | 22.97661 | 61.46103 | 58.5941 | Promoter | 11 | 98992943 | 99024189 | 31247 | 2 | ENSMUST00 | 0 | Top2a | ENSMUSG00000020914.12 |
| chr2 | 181038826 | 181039612 | 22.97661 | 61.46103 | 58.5941 | Promoter | 2 | 181018380 | 181039187 | 20808 | 2 | ENSMUST00 | 0 | Chrna4 | ENSMUSG00000027577.9 |
| chr2 | 167188384 | 167189902 | 22.97661 | 61.46103 | 58.5941 | Promoter | 2 | 167103378 | 167188799 | 85422 | 2 | ENSMUST00 | 0 | Kcnb1 | ENSMUSG00000050556.8 |
| chr3 | 152477894 | 152479035 | 22.97661 | 61.46103 | 58.5941 | Exon (ENSMU | 3 | 152457417 | 152458535 | 1119 | 1 | ENSMUST00 | 20478 | Zzz3 | ENSMUSG00000039068.11 |
| chr7 | 25072030 | 25073077 | 22.97661 | 61.46103 | 58.5941 | Promoter | 7 | 25009853 | 25072369 | 62517 | 2 | ENSMUST00 | 0 | Grik5 | ENSMUSG00000003378.8 |
| chr7 | 70350687 | 70352400 | 22.97661 | 61.46103 | 58.5941 | 3' UTR | 7 | 70351946 | 70360593 | 8648 | 2 | ENSMUST00 | 8193 | Nr2f2 | ENSMUSG00000030551.8 |
| chr8 | 95852893 | 95853824 | 22.97661 | 61.46103 | 58.5941 | Promoter | 8 | 95835920 | 95853539 | 17620 | 2 | ENSMUST00 | 0 | Slc38a7 | ENSMUSG00000036534.5 |
| chr8 | 31089441 | 31090243 | 22.97661 | 61.46103 | 58.5941 | Promoter | 8 | 31089471 | 31097047 | 7577 | 1 | ENSMUST00 | 0 | Dusp26 | ENSMUSG00000039661.9 |
| chrX | 74367926 | 74368772 | 22.97661 | 61.46103 | 58.5941 | Promoter | 20 | 74365718 | 74368673 | 2956 | 2 | ENSMUST00 | 0 | Ubl4 | ENSMUSG00000015290.10 |
| chrX | 142388849 | 142389115 | 22.97661 | 61.46103 | 58.5941 | Promoter | 20 | 142317993 | 142390344 | 72352 | 2 | ENSMUST00 | 0 | Acsl4 | ENSMUSG00000031278.7 |
| chrX | 101532695 | 101533585 | 22.97661 | 61.46103 | 58.5941 | Promoter | 20 | 101532734 | 101601034 | 68301 | 1 | ENSMUST00 | 0 | Taf1 | ENSMUSG00000031314.12 |
| chr10 | 80179157 | 80180165 | 22.93528 | 61.39042 | 58.52499 | Promoter | 10 | 80179482 | 80190010 | 10529 | 1 | ENSMUST00 | 0 | Efna2 | ENSMUSG00000003070.6 |
| chr12 | 51844288 | 51845215 | 21.43343 | 61.34734 | 58.48203 | Distal Interge | 12 | 51743722 | 51829536 | 85815 | 2 | ENSMUST00 | -14753 | Hectd1 | ENSMUSG00000035247.10 |
| chr14 | 103650039 | 103651136 | 21.92106 | 61.34632 | 58.48102 | Promoter | 14 | 103650228 | 103704907 | 54680 | 1 | ENSMUST00 | 0 | Slain1 | ENSMUSG00000055717.8 |
| chr19 | 44835788 | 44836990 | 21.92106 | 61.34632 | 58.48102 | Promoter | 19 | 44828493 | 44835938 | 7446 | 2 | ENSMUST00 | 0 | 1700039E22I | ENSMUSG00000092369.1 |
| chr5 | 124248582 | 124250472 | 21.92106 | 61.34632 | 58.48102 | Promoter | 5 | 124118690 | 124249760 | 131071 | 2 | ENSMUST00 | 0 | Pitpnm2 | ENSMUSG00000029406.10 |
| chr12 | 84113944 | 84114922 | 19.51381 | 61.26624 | 58.40183 | Promoter | 12 | 84114366 | 84139838 | 25473 | 1 | ENSMUST00 | 0 | Dnal1 | ENSMUSG00000042523.7 |
| chr1 | 72243554 | 72244622 | 20.42851 | 61.25884 | 58.39448 | Promoter | 1 | 72244213 | 72244403 | 191 | 1 | ENSMUST00 | 0 | Gm24497 | ENSMUSG00000093956.1 |
| chr11 | 83462482 | 83463312 | 20.42851 | 61.25884 | 58.39448 | Promoter | 11 | 83440768 | 83463022 | 22255 | 2 | ENSMUST00 | 0 | Mmp28 | ENSMUSG00000020682.12 |
| chr9 | 38717767 | 38719795 | 20.42851 | 61.25884 | 58.39448 | Promoter | 9 | 38718268 | 38743337 | 25070 | 1 | ENSMUST00 | 0 | Vwa5a | ENSMUSG00000023186.9 |
| chr2 | 152626525 | 152627552 | 22.36183 | 61.25668 | 58.3926 | Promoter | 2 | 152626951 | 152635194 | 8244 | 1 | ENSMUST00 | 0 | Rem1 | ENSMUSG00000000359.3 |
| chr9 | 42123512 | 42124548 | 22.36183 | 61.25668 | 58.3926 | Promoter | 9 | 41964720 | 42124297 | 159578 | 2 | ENSMUST00 | 0 | Sorl1 | ENSMUSG00000049313.8 |
| chr1 | 87186472 | 87187264 | 22.83944 | 61.22662 | 58.36318 | Exon (ENSMU | 1 | 87183313 | 87188405 | 5093 | 1 | ENSMUST00 | 3160 | Prss56 | ENSMUSG00000036480.9 |
| chr13 | 67420946 | 67421927 | 22.83944 | 61.22662 | 58.36318 | Promoter | 13 | 67405722 | 67421418 | 15697 | 2 | ENSMUST00 | 0 | Zfp459 | ENSMUSG00000055560.7 |
| chr15 | 76899239 | 76900771 | 22.83944 | 61.22662 | 58.36318 | Promoter | 15 | 76899910 | 76901305 | 1396 | 1 | ENSMUST00 | 0 | Commd5 | ENSMUSG00000055041.7 |
| chr19 | 10100545 | 10102386 | 22.83944 | 61.22662 | 58.36318 | Promoter | 19 | 10064164 | 10101503 | 37340 | 2 | ENSMUST00 | 0 | Fads2 | ENSMUSG00000024665.7 |
| chr1 | 92178778 | 92180363 | 21.79372 | 61.12524 | 58.26245 | Promoter | 1 | 91928779 | 92180341 | 251563 | 2 | ENSMUST00 | 0 | Hdac4 | ENSMUSG00000026313.10 |
| chr1 | 163402860 | 163404270 | 21.79372 | 61.12524 | 58.26245 | Promoter | 1 | 163384908 | 163403669 | 18762 | 2 | ENSMUST00 | 0 | Gorab | ENSMUSG00000040124.5 |
| chr10 | 121309954 | 121311322 | 21.79372 | 61.12524 | 58.26245 | Promoter | 10 | 121263824 | 121311189 | 47366 | 2 | ENSMUST00 | 0 | Tbc1d30 | ENSMUSG00000052302.5 |
| chr12 | 17992390 | 17993459 | 21.79372 | 61.12524 | 58.26245 | Distal Interge | 12 | 17949458 | 17949548 | 91 | 2 | ENSMUST00 | -42843 | Gm23913 | ENSMUSG00000077876.2 |
| chr14 | 118705611 | 118706713 | 21.79372 | 61.12524 | 58.26245 | Promoter | 14 | 118482692 | 118706219 | 223528 | 2 | ENSMUST00 | 0 | Abcc4 | ENSMUSG00000032849.8 |
| chr15 | 96642436 | 96643385 | 21.79372 | 61.12524 | 58.26245 | Promoter | 15 | 96571418 | 96642913 | 71496 | 2 | ENSMUST00 | 0 | Slc38a1 | ENSMUSG00000023169.9 |
| chr17 | 35909382 | 35910365 | 21.79372 | 61.12524 | 58.26245 | Promoter | 17 | 35897595 | 35910075 | 12481 | 2 | ENSMUST00 | 0 | Atat1 | ENSMUSG00000024426.12 |
| chr19 | 42085737 | 42086612 | 21.79372 | 61.12524 | 58.26245 | Promoter | 19 | 42074939 | 42086370 | 11432 | 2 | ENSMUST00 | 0 | Morn4 | ENSMUSG00000049670.8 |
| chr2 | 26627864 | 26628866 | 21.79372 | 61.12524 | 58.26245 | Promoter | 2 | 26628457 | 26636497 | 8041 | 1 | ENSMUST00 | 0 | Fam69b | ENSMUSG00000036186.5 |
| chr3 | 97296446 | 97298749 | 21.79372 | 61.12524 | 58.26245 | Promoter | 3 | 97284908 | 97297778 | 12871 | 2 | ENSMUST00 | 0 | Gm17608 | ENSMUSG00000097143.1 |
| chr4 | 97771503 | 97773573 | 21.79372 | 61.12524 | 58.26245 | Promoter | 4 | 97772734 | 98000466 | 227733 | 1 | ENSMUST00 | 0 | Nfia | ENSMUSG00000028565.13 |
| chr4 | 131920283 | 131922489 | 21.79372 | 61.12524 | 58.26245 | Promoter | 4 | 131921771 | 131923140 | 1370 | 1 | ENSMUST00 | 0 | Tmem200b | ENSMUSG00000070720.3 |
| chr5 | 135250447 | 135251586 | 21.79372 | 61.12524 | 58.26245 | Promoter | 5 | 135248938 | 135251230 | 2293 | 2 | ENSMUST00 | 0 | Fzd9 | ENSMUSG00000049551.2 |
| chr6 | 87980374 | 87982160 | 21.79372 | 61.12524 | 58.26245 | Promoter | 6 | 87980421 | 87981637 | 1217 | 2 | ENSMUST00 | 0 | H1fx | ENSMUSG00000044927.6 |
| chr7 | 73390852 | 73392583 | 21.79372 | 61.12524 | 58.26245 | Promoter | 7 | 73391160 | 73409451 | 18292 | 1 | ENSMUST00 | 0 | Rgma | ENSMUSG00000070509.10 |
| chr9 | 73044245 | 73045292 | 21.79372 | 61.12524 | 58.26245 | Promoter | 9 | 73044854 | 73097629 | 52776 | 1 | ENSMUST00 | 0 | Rab27a | ENSMUSG00000032202.6 |
| chr9 | 86879244 | 86880928 | 21.79372 | 61.12524 | 58.26245 | Promoter | 9 | 86765923 | 86880439 | 114517 | 2 | ENSMUST00 | 0 | Snap91 | ENSMUSG00000033419.10 |
| chrX | 48453190 | 48454762 | 21.79372 | 61.12524 | 58.26245 | Promoter | 20 | 48411046 | 48454169 | 43124 | 2 | ENSMUST00 | 0 | Elf4 | ENSMUSG00000031103.7 |
| chr15 | 36164446 | 36165253 | 22.2725 | 61.10292 | 58.24237 | Promoter | 15 | 36150060 | 36164884 | 14825 | 2 | ENSMUST00 | 0 | Fbxo43 | ENSMUSG00000048230.5 |
| chr17 | 78199554 | 78200068 | 22.2725 | 61.10292 | 58.24237 | Promoter | 17 | 78200248 | 78376592 | 176345 | 1 | ENSMUST00 | -180 | Crim1 | ENSMUSG00000024074.7 |
| chr1 | 10993391 | 10994521 | 22.74441 | 61.06413 | 58.20372 | Promoter | 1 | 10993465 | 11303681 | 310217 | 1 | ENSMUST00 | 0 | Prex2 | ENSMUSG00000048960.8 |
| chr7 | 29211282 | 29212104 | 22.74441 | 61.06413 | 58.20372 | Promoter | 7 | 29196750 | 29211495 | 14746 | 2 | ENSMUST00 | 0 | Catsperg1 | ENSMUSG00000049676.9 |
| chr5 | 73650501 | 73651960 | 21.75328 | 61.05499 | 58.1947 | Promoter | 5 | 73651380 | 73679096 | 27717 | 1 | ENSMUST00 | 0 | Spata18 | ENSMUSG00000029155.11 |
| chr11 | 45980011 | 45981033 | 22.2281 | 61.02647 | 58.16623 | Promoter | 11 | 45980310 | 46017994 | 37685 | 1 | ENSMUST00 | 0 | Sox30 | ENSMUSG00000040489.5 |
| chr1 | 14308804 | 14310699 | 19.38504 | 61.02572 | 58.1656 | Promoter | 1 | 14168954 | 14310200 | 141247 | 2 | ENSMUST00 | 0 | Eya1 | ENSMUSG00000025932.9 |
| chr1 | 118458684 | 118460044 | 19.38504 | 61.02572 | 58.1656 | Promoter | 1 | 118459141 | 118459265 | 125 | 2 | ENSMUST00 | 0 | Gm22710 | ENSMUSG00000080542.1 |
| chr10 | 127521935 | 127522640 | 19.38504 | 61.02572 | 58.1656 | Promoter | 10 | 127517123 | 127522444 | 5322 | 2 | ENSMUST00 | 0 | Shmt2 | ENSMUSG00000025403.4 |
| chr11 | 69098805 | 69100459 | 19.38504 | 61.02572 | 58.1656 | Promoter | 11 | 69098948 | 69109960 | 11013 | 1 | ENSMUST00 | 0 | Per1 | ENSMUSG00000020893.12 |
| chr13 | 59674664 | 59676081 | 19.38504 | 61.02572 | 58.1656 | Promoter | 13 | 59634996 | 59675784 | 40789 | 2 | ENSMUST00 | 0 | Golm1 | ENSMUSG00000021556.6 |
| chr14 | 57524652 | 57526082 | 19.38504 | 61.02572 | 58.1656 | Promoter | 14 | 57524829 | 57543165 | 18337 | 1 | ENSMUST00 | 0 | Il17d | ENSMUSG00000050222.10 |
| chr16 | 91618422 | 91619161 | 19.38504 | 61.02572 | 58.1656 | Promoter | 16 | 91614257 | 91618999 | 4743 | 2 | ENSMUST00 | 0 | Dnajc28 | ENSMUSG00000039763.9 |
| chr17 | 8525349 | 8526722 | 19.38504 | 61.02572 | 58.1656 | Promoter | 17 | 8525372 | 8981671 | 456300 | 1 | ENSMUST00 | 0 | Pde10a | ENSMUSG00000023868.10 |
| chr17 | 12915287 | 12916199 | 19.38504 | 61.02572 | 58.1656 | Promoter | 17 | 12913846 | 12915624 | 1779 | 2 | ENSMUST00 | 0 | Mrpl18 | ENSMUSG00000057388.4 |
| chr2 | 164856602 | 164858134 | 19.38504 | 61.02572 | 58.1656 | Promoter | 2 | 164839518 | 164857542 | 18025 | 2 | ENSMUST00 | 0 | Pltp | ENSMUSG00000017754.8 |
| chr2 | 127788438 | 127789597 | 19.38504 | 61.02572 | 58.1656 | Promoter | 2 | 127740732 | 127788854 | 48123 | 2 | ENSMUST00 | 0 | Nphp1 | ENSMUSG00000027378.11 |
| chr2 | 119786821 | 119788084 | 19.38504 | 61.02572 | 58.1656 | Promoter | 2 | 119763304 | 119787504 | 24201 | 2 | ENSMUST00 | 0 | Rpap1 | ENSMUSG00000034032.10 |
| chr2 | 119167027 | 119168187 | 19.38504 | 61.02572 | 58.1656 | Promoter | 2 | 119167773 | 119172390 | 4618 | 1 | ENSMUST00 | 0 | Gchfr | ENSMUSG00000046814.3 |
| chr3 | 109026385 | 109028112 | 19.38504 | 61.02572 | 58.1656 | Promoter | 3 | 108970997 | 109027273 | 56277 | 2 | ENSMUST00 | 0 | Fam102b | ENSMUSG00000040339.5 |
| chr4 | 99929156 | 99930731 | 19.38504 | 61.02572 | 58.1656 | Promoter | 4 | 99929414 | 99987294 | 57881 | 1 | ENSMUST00 | 0 | Pgm2 | ENSMUSG00000025791.13 |
| chr4 | 128961951 | 128962962 | 19.38504 | 61.02572 | 58.1656 | Promoter | 4 | 128930233 | 128962420 | 32188 | 2 | ENSMUST00 | 0 | Adc | ENSMUSG00000028789.11 |
| chr4 | 14930141 | 14931139 | 19.38504 | 61.02572 | 58.1656 | Promoter | 4 | 14930661 | 14953030 | 22370 | 1 | ENSMUST00 | 0 | Gm11837 | ENSMUSG00000086587.2 |

| | | | | | | | | | | | | | | | |
|---|---|---|---|---|---|---|---|---|---|---|---|---|---|---|---|
| chr5 | 31451928 | 31453552 | 19.38504 | 61.02572 | 58.1656 | Promoter | 5 | 31452436 | 31481749 | 29314 | 1 | ENSMUST00 | 0 | Zfp512 | ENSMUSG00000062761.4 |
| chr7 | 4995549 | 4996856 | 19.38504 | 61.02572 | 58.1656 | Promoter | 7 | 4983483 | 4996055 | 12573 | 2 | ENSMUST00 | 0 | Zfp579 | ENSMUSG00000051550.10 |
| chr8 | 31149119 | 31151505 | 19.38504 | 61.02572 | 58.1656 | Promoter | 8 | 31150316 | 31164702 | 14387 | 1 | ENSMUST00 | 0 | Tti2 | ENSMUSG00000031577.9 |
| chr8 | 94097748 | 94099004 | 19.38504 | 61.02572 | 58.1656 | Promoter | 8 | 94067954 | 94098811 | 30858 | 2 | ENSMUST00 | 0 | Bbs2 | ENSMUSG00000031755.5 |
| chr8 | 70607750 | 70610214 | 19.38504 | 61.02572 | 58.1656 | Promoter | 8 | 70597491 | 70608314 | 10824 | 2 | ENSMUST00 | 0 | Ssbp4 | ENSMUSG00000070003.6 |
| chr9 | 113792584 | 113794246 | 19.38504 | 61.02572 | 58.1656 | Distal Interge | 9 | 113776182 | 113776298 | 117 | 2 | ENSMUST00 | -16287 | Gm25059 | ENSMUSG00000087992.1 |
| chr19 | 5796007 | 5804263 | 15.48624 | 61.01235 | 58.15405 | Promoter | 19 | 5797562 | 5798549 | 988 | 1 | ENSMUST00 | 0 | Gm20417 | ENSMUSG00000092267.1 |
| chr16 | 93682573 | 93684551 | 20.76252 | 61.01094 | 58.15269 | Promoter | 16 | 93683215 | 93690990 | 7776 | 1 | ENSMUST00 | 0 | Cbr3 | ENSMUSG00000022947.7 |
| chr4 | 134922877 | 134923436 | 21.71174 | 60.98278 | 58.12498 | Promoter | 4 | 134923592 | 134927671 | 4080 | 1 | ENSMUST00 | -156 | Rsrp1 | ENSMUSG00000037266.13 |
| chr2 | 37775966 | 37778479 | 22.18388 | 60.95028 | 58.09254 | Promoter | 2 | 37776249 | 37799103 | 22855 | 1 | ENSMUST00 | 0 | Crb2 | ENSMUSG00000035403.8 |
| chr5 | 31290315 | 31291682 | 22.18388 | 60.95028 | 58.09254 | Promoter | 5 | 31253280 | 31291114 | 37835 | 2 | ENSMUST00 | 0 | Ift172 | ENSMUSG00000038564.7 |
| chr5 | 77459962 | 77460461 | 22.65015 | 60.90288 | 58.04541 | Distal Interge | 5 | 77511178 | 77511564 | 387 | 2 | ENSMUST00 | 51103 | Pea15b | ENSMUSG00000047652.4 |
| chr16 | 26371091 | 26371876 | 22.13983 | 60.87438 | 58.01701 | Promoter | 16 | 26356642 | 26371841 | 15200 | 2 | ENSMUST00 | 0 | Cldn1 | ENSMUSG00000022512.2 |
| chr17 | 35836718 | 35838317 | 21.62913 | 60.83906 | 57.9818 | Promoter | 17 | 35833921 | 35838306 | 4386 | 2 | ENSMUST00 | 0 | Tubb5 | ENSMUSG00000001525.10 |
| chr17 | 24736523 | 24737434 | 22.09595 | 60.79875 | 57.94164 | Promoter | 17 | 24736642 | 24742778 | 6137 | 1 | ENSMUST00 | 0 | Msrb1 | ENSMUSG00000075705.7 |
| chr7 | 30664482 | 30665631 | 21.1209 | 60.79805 | 57.94103 | Promoter | 7 | 30653711 | 30664933 | 11223 | 2 | ENSMUST00 | 0 | Haus5 | ENSMUSG00000078762.5 |
| chr9 | 57467317 | 57468435 | 21.1209 | 60.79805 | 57.94103 | Promoter | 9 | 57441328 | 57468024 | 26697 | 2 | ENSMUST00 | 0 | Scamp5 | ENSMUSG00000040722.6 |
| chr7 | 100606850 | 100607386 | 21.58806 | 60.76757 | 57.91108 | Promoter | 7 | 100607410 | 100641268 | 33859 | 1 | ENSMUST00 | -24 | Rab6a | ENSMUSG00000030704.9 |
| chr8 | 109607763 | 109609294 | 21.58806 | 60.76757 | 57.91108 | Promoter | 8 | 109593248 | 109608673 | 15426 | 2 | ENSMUST00 | 0 | Dhodh | ENSMUSG00000031730.12 |
| chr8 | 129233519 | 129235104 | 21.54714 | 60.69631 | 57.84021 | Promoter | 8 | 129216354 | 129234046 | 17693 | 2 | ENSMUST00 | 0 | 2610044O15 | ENSMUSG00000071302.5 |
| chr11 | 6006195 | 6007325 | 18.33657 | 60.6815 | 57.82544 | Intron (ENSM | 11 | 5990395 | 6000448 | 10054 | 2 | ENSMUST00 | -5748 | Camk2b | ENSMUSG00000057897.9 |
| chr12 | 76837212 | 76838449 | 18.33657 | 60.6815 | 57.82544 | Promoter | 12 | 76837414 | 76921412 | 83999 | 1 | ENSMUST00 | 0 | Fntb | ENSMUSG00000033373.11 |
| chr13 | 55487090 | 55488351 | 18.33657 | 60.6815 | 57.82544 | Promoter | 13 | 55473429 | 55488111 | 14683 | 2 | ENSMUST00 | 0 | Dbn1 | ENSMUSG00000034675.12 |
| chr13 | 12702430 | 12703024 | 18.33657 | 60.6815 | 57.82544 | Promoter | 13 | 12702362 | 12702589 | 228 | 1 | ENSMUST00 | 69 | Gm2399 | ENSMUSG00000078141.2 |
| chr14 | 59439678 | 59441779 | 18.33657 | 60.6815 | 57.82544 | Promoter | 14 | 59440981 | 59548903 | 107923 | 1 | ENSMUST00 | 0 | Cab39l | ENSMUSG00000021981.9 |
| chr14 | 27114386 | 27115616 | 18.33657 | 60.6815 | 57.82544 | Distal Interge | 14 | 27039001 | 27107286 | 68286 | 1 | ENSMUST00 | 75386 | Il17rd | ENSMUSG00000040717.5 |
| chr16 | 4003174 | 4004370 | 18.33657 | 60.6815 | 57.82544 | Promoter | 16 | 4001003 | 4003770 | 2768 | 2 | ENSMUST00 | 0 | Slx4 | ENSMUSG00000039738.11 |
| chr3 | 95133404 | 95134075 | 18.33657 | 60.6815 | 57.82544 | Promoter | 3 | 95129536 | 95134011 | 4476 | 2 | ENSMUST00 | 0 | Scnm1 | ENSMUSG00000092607.4 |
| chr4 | 84674350 | 84676199 | 18.33657 | 60.6815 | 57.82544 | Promoter | 4 | 84275095 | 84675034 | 399940 | 2 | ENSMUST00 | 0 | Bnc2 | ENSMUSG00000028487.13 |
| chr6 | 58640162 | 58641165 | 18.33657 | 60.6815 | 57.82544 | Promoter | 6 | 58640546 | 58664783 | 24238 | 1 | ENSMUST00 | 0 | Abcg2 | ENSMUSG00000029802.8 |
| chr7 | 44951540 | 44952944 | 18.33657 | 60.6815 | 57.82544 | Exon (ENSMU | 7 | 44943261 | 44958032 | 14772 | 1 | ENSMUST00 | 8280 | Tsks | ENSMUSG00000059891.8 |
| chr10 | 130541711 | 130542753 | 22.51023 | 60.66335 | 57.80838 | Distal Interge | 10 | 130557348 | 130557464 | 117 | 2 | ENSMUST00 | 14711 | Gm23793 | ENSMUSG00000096766.1 |
| chr4 | 44300809 | 44301535 | 22.51023 | 60.66335 | 57.80838 | Promoter | 4 | 44300876 | 44364675 | 63800 | 1 | ENSMUST00 | 0 | Melk | ENSMUSG00000035683.8 |
| chr3 | 30011921 | 30013391 | 20.08725 | 60.64133 | 57.78656 | Promoter | 3 | 29951299 | 30013156 | 61858 | 2 | ENSMUST00 | 0 | Mecom | ENSMUSG00000027684.11 |
| chr1 | 171032991 | 171034770 | 21.50638 | 60.6253 | 57.77077 | Distal Interge | 1 | 171027965 | 171028129 | 165 | 1 | ENSMUST00 | 5027 | Gm26110 | ENSMUSG00000065905.1 |
| chr11 | 78826527 | 78827336 | 21.50638 | 60.6253 | 57.77077 | Promoter | 11 | 78826584 | 78846870 | 20287 | 1 | ENSMUST00 | 0 | Lyrm9 | ENSMUSG00000072640.6 |
| chr6 | 118593253 | 118594248 | 21.50638 | 60.6253 | 57.77077 | 3' UTR | 6 | 118592304 | 118595426 | 3123 | 2 | ENSMUST00 | 1178 | Cacna1c | ENSMUSG00000051331.9 |
| chr8 | 95080175 | 95080787 | 21.50638 | 60.6253 | 57.77077 | Promoter | 8 | 95081201 | 95099874 | 18674 | 1 | ENSMUST00 | -414 | Katnb1 | ENSMUSG00000031787.7 |
| chr10 | 5804803 | 5805508 | 16.70194 | 60.58573 | 57.73199 | Promoter | 10 | 5799160 | 5805600 | 6441 | 2 | ENSMUST00 | 92 | Fbxo5 | ENSMUSG00000019773.7 |
| chr10 | 37135907 | 37137976 | 16.70194 | 60.58573 | 57.73199 | Promoter | 10 | 37133375 | 37138920 | 5546 | 2 | ENSMUST00 | 944 | Marcks | ENSMUSG00000069662.5 |
| chr11 | 77347879 | 77349056 | 16.70194 | 60.58573 | 57.73199 | Promoter | 11 | 77348280 | 77413635 | 65356 | 1 | ENSMUST00 | 0 | Ssh2 | ENSMUSG00000037926.10 |
| chr12 | 101028372 | 101029660 | 16.70194 | 60.58573 | 57.73199 | Promoter | 12 | 100912700 | 101028983 | 116284 | 2 | ENSMUST00 | 0 | Ccdc88c | ENSMUSG00000021182.9 |
| chr19 | 3935126 | 3936326 | 16.70194 | 60.58573 | 57.73199 | Promoter | 19 | 3935186 | 3949339 | 14154 | 1 | ENSMUST00 | 0 | Unc93b1 | ENSMUSG00000036908.11 |
| chr2 | 5713426 | 5715268 | 16.70194 | 60.58573 | 57.73199 | Promoter | 2 | 5293457 | 5714515 | 421059 | 2 | ENSMUST00 | 0 | Camk1d | ENSMUSG00000039145.11 |
| chr6 | 97615795 | 97617988 | 16.70194 | 60.58573 | 57.73199 | Promoter | 6 | 97286867 | 97617541 | 330675 | 2 | ENSMUST00 | 0 | Frmd4b | ENSMUSG00000030064.11 |
| chr9 | 107567738 | 107570259 | 16.70194 | 60.58573 | 57.73199 | Promoter | 9 | 107569117 | 107572776 | 3660 | 1 | ENSMUST00 | 0 | Hyal2 | ENSMUSG00000010047.7 |
| chr14 | 21500795 | 21501734 | 17.45443 | 60.55182 | 57.69857 | Promoter | 14 | 21500853 | 21621401 | 120549 | 1 | ENSMUST00 | 0 | Kat6b | ENSMUSG00000021767.11 |
| chr15 | 84852984 | 84856941 | 17.45443 | 60.55182 | 57.69857 | Promoter | 15 | 84785381 | 84855110 | 69730 | 2 | ENSMUST00 | 0 | Phf21b | ENSMUSG00000016624.10 |
| chr4 | 116143868 | 116144863 | 17.45443 | 60.55182 | 57.69857 | Promoter | 4 | 116132406 | 116144609 | 12204 | 2 | ENSMUST00 | 0 | Lurap1 | ENSMUSG00000028701.4 |
| chr4 | 40278654 | 40280055 | 17.45443 | 60.55182 | 57.69857 | Promoter | 4 | 40270591 | 40279421 | 8831 | 2 | ENSMUST00 | 0 | Ndufb6 | ENSMUSG00000071014.5 |
| chr9 | 36726073 | 36727366 | 17.45443 | 60.55182 | 57.69857 | Promoter | 9 | 36708482 | 36726374 | 17893 | 2 | ENSMUST00 | 0 | Chek1 | ENSMUSG00000032113.10 |
| chr19 | 47463319 | 47465034 | 22.41791 | 60.5052 | 57.65255 | Promoter | 19 | 47260174 | 47464411 | 204238 | 2 | ENSMUST00 | 0 | Sh3pxd2a | ENSMUSG00000053617.6 |
| chr4 | 82504132 | 82507030 | 22.41791 | 60.5052 | 57.65255 | Promoter | 4 | 82290173 | 82505308 | 215136 | 2 | ENSMUST00 | 0 | Nfib | ENSMUSG00000008575.12 |
| chr5 | 117119390 | 117119927 | 22.41791 | 60.5052 | 57.65255 | Promoter | 5 | 117120129 | 117275217 | 155089 | 1 | ENSMUST00 | -202 | Taok3 | ENSMUSG00000061288.8 |
| chr11 | 102189364 | 102190440 | 21.92219 | 60.49891 | 57.64639 | Promoter | 11 | 102189620 | 102194081 | 4462 | 1 | ENSMUST00 | 0 | G6pc3 | ENSMUSG00000034793.10 |
| chr13 | 52582909 | 52583810 | 21.92219 | 60.49891 | 57.64639 | Promoter | 13 | 52583173 | 52648792 | 65620 | 1 | ENSMUST00 | 0 | Syk | ENSMUSG00000021457.9 |
| chr15 | 102406202 | 102407269 | 21.92219 | 60.49891 | 57.64639 | Promoter | 15 | 102406328 | 102432679 | 26352 | 1 | ENSMUST00 | 0 | Sp1 | ENSMUSG00000001280.7 |
| chr2 | 35201180 | 35201824 | 21.92219 | 60.49891 | 57.64639 | Promoter | 2 | 35180205 | 35201120 | 20916 | 2 | ENSMUST00 | -61 | Rab14 | ENSMUSG00000026878.11 |
| chr6 | 120293805 | 120295125 | 21.92219 | 60.49891 | 57.64639 | Promoter | 6 | 120203094 | 120294559 | 91466 | 2 | ENSMUST00 | 0 | B4galnt3 | ENSMUSG00000041372.9 |
| chr7 | 88429918 | 88431011 | 21.92219 | 60.49891 | 57.64639 | Promoter | 7 | 88430273 | 88491466 | 61194 | 1 | ENSMUST00 | 0 | Rab38 | ENSMUSG00000030559.7 |
| chr8 | 121906697 | 121908197 | 21.92219 | 60.49891 | 57.64639 | Promoter | 8 | 121881150 | 121907694 | 26545 | 2 | ENSMUST00 | 0 | Slc7a5 | ENSMUSG00000040010.10 |
| chr8 | 69208655 | 69209811 | 21.92219 | 60.49891 | 57.64639 | Distal Interge | 8 | 69226343 | 69228980 | 2638 | 1 | ENSMUST00 | -16532 | Zfp930 | ENSMUSG00000059897.4 |
| chr2 | 120628545 | 120629650 | 22.37203 | 60.42657 | 57.57468 | Promoter | 2 | 120629126 | 120682782 | 53657 | 1 | ENSMUST00 | 0 | Stard9 | ENSMUSG00000033705.11 |
| chr17 | 17909720 | 17910766 | 21.87917 | 60.42461 | 57.57317 | Intron (ENSM | 17 | 17911039 | 17911947 | 909 | 2 | ENSMUST00 | 1181 | Gm7535 | ENSMUSG00000090957.1 |
| chr2 | 20519278 | 20521509 | 21.87917 | 60.42461 | 57.57317 | Promoter | 2 | 20519791 | 20809885 | 290095 | 1 | ENSMUST00 | 0 | Etl4 | ENSMUSG00000036617.11 |
| chr9 | 108478517 | 108480814 | 19.47644 | 60.36021 | 57.50917 | Promoter | 9 | 108479862 | 108490530 | 10669 | 1 | ENSMUST00 | 0 | Lamb2 | ENSMUSG00000052911.4 |
| chr9 | 70678383 | 70678768 | 21.75113 | 60.20329 | 57.35284 | Promoter | 9 | 70678997 | 70778382 | 99386 | 1 | ENSMUST00 | -229 | Adam10 | ENSMUSG00000054693.9 |
| chr11 | 120304949 | 120305940 | 22.23551 | 60.19247 | 57.34209 | Distal Interge | 11 | 120289793 | 120292167 | 2375 | 1 | ENSMUST00 | 15157 | Bahcc1 | ENSMUSG00000039741.10 |
| chr11 | 62575384 | 62576384 | 20.27628 | 60.14116 | 57.29097 | Promoter | 11 | 62575547 | 62581304 | 5758 | 1 | ENSMUST00 | 0 | Trpv2 | ENSMUSG00000018507.11 |
| chr11 | 80809976 | 80811357 | 20.27628 | 60.14116 | 57.29097 | Promoter | 11 | 80810175 | 80822033 | 11859 | 1 | ENSMUST00 | 0 | Tmem98 | ENSMUSG00000035413.8 |
| chr14 | 34309864 | 34310546 | 20.27628 | 60.14116 | 57.29097 | Promoter | 14 | 34237041 | 34310503 | 73463 | 2 | ENSMUST00 | 0 | Fam35a | ENSMUSG00000041471.8 |
| chr15 | 85645614 | 85646702 | 20.27628 | 60.14116 | 57.29097 | Exon (ENSMU | 15 | 85639747 | 85645934 | 6188 | 1 | ENSMUST00 | 5868 | RP23-64J7.2 | ENSMUSG00000101828.1 |
| chr16 | 97996552 | 97997922 | 20.27628 | 60.14116 | 57.29097 | Promoter | 16 | 97997323 | 98010000 | 12678 | 1 | ENSMUST00 | 0 | B230307C23I | ENSMUSG00000080717.4 |
| chr17 | 17004585 | 17006014 | 20.27628 | 60.14116 | 57.29097 | Distal Interge | 17 | 16972910 | 16973770 | 861 | 1 | ENSMUST00 | 31676 | BC002059 | ENSMUSG00000060149.6 |
| chr2 | 167661681 | 167662263 | 20.27628 | 60.14116 | 57.29097 | Promoter | 2 | 167662109 | 167667697 | 5589 | 1 | ENSMUST00 | 0 | Gm14320 | ENSMUSG00000085914.1 |
| chr3 | 121291051 | 121292667 | 20.27628 | 60.14116 | 57.29097 | Promoter | 3 | 121291817 | 121362011 | 70195 | 1 | ENSMUST00 | 0 | Alg14 | ENSMUSG00000039887.7 |
| chr4 | 13742710 | 13744561 | 20.27628 | 60.14116 | 57.29097 | Promoter | 4 | 13743436 | 13893649 | 150214 | 1 | ENSMUST00 | 0 | Runx1t1 | ENSMUSG00000006586.10 |
| chr5 | 33695092 | 33696139 | 20.27628 | 60.14116 | 57.29097 | Distal Interge | 5 | 33669569 | 33672201 | 2633 | 1 | ENSMUST00 | 25524 | Tacc3 | ENSMUSG00000037313.11 |
| chr5 | 136170095 | 136171831 | 20.27628 | 60.14116 | 57.29097 | Promoter | 5 | 136147464 | 136170656 | 23193 | 2 | ENSMUST00 | 0 | Orai2 | ENSMUSG00000039747.7 |
| chr5 | 109866235 | 109867588 | 20.27628 | 60.14116 | 57.29097 | Distal Interge | 5 | 109886231 | 109891137 | 4907 | 1 | ENSMUST00 | -18643 | Gm26779 | ENSMUSG00000097140.1 |
| chr5 | 136918742 | 136919902 | 20.27628 | 60.14116 | 57.29097 | Promoter | 5 | 136919146 | 136937106 | 17961 | 1 | ENSMUST00 | 0 | 4933404O12 | ENSMUSG00000097908.2 |
| chr7 | 38061000 | 38061862 | 20.27628 | 60.14116 | 57.29097 | Distal Interge | 7 | 38098811 | 38099369 | 559 | 2 | ENSMUST00 | 37507 | Ccne1 | ENSMUSG00000002068.11 |
| chr7 | 27859993 | 27861362 | 20.27628 | 60.14116 | 57.29097 | Distal Interge | 7 | 27862557 | 27879567 | 17011 | 1 | ENSMUST00 | -1195 | Zfp607 | ENSMUSG00000020420.7 |
| chr8 | 123514693 | 123515506 | 20.27628 | 60.14116 | 57.29097 | Promoter | 8 | 123504718 | 123515455 | 10738 | 2 | ENSMUST00 | 0 | Dbndd1 | ENSMUSG00000031970.11 |
| chr8 | 84414241 | 84415968 | 20.27628 | 60.14116 | 57.29097 | Promoter | 8 | 84415364 | 84640246 | 224883 | 1 | ENSMUST00 | 0 | Cacna1a | ENSMUSG00000034656.11 |
| chr8 | 72160295 | 72160929 | 20.27628 | 60.14116 | 57.29097 | Promoter | 8 | 72147601 | 72160855 | 13255 | 2 | ENSMUST00 | 0 | Gm16091 | ENSMUSG00000087502.1 |
| chr9 | 35558231 | 35560124 | 20.27628 | 60.14116 | 57.29097 | Promoter | 9 | 35541847 | 35558528 | 16682 | 2 | ENSMUST00 | 0 | Ddx25 | ENSMUSG00000032101.5 |
| chr9 | 7176497 | 7177355 | 20.27628 | 60.14116 | 57.29097 | Promoter | 9 | 6928503 | 7177046 | 248544 | 2 | ENSMUST00 | 0 | Dync2h1 | ENSMUSG00000047193.10 |
| chr9 | 14751392 | 14752460 | 20.27628 | 60.14116 | 57.29097 | Promoter | 9 | 14748459 | 14752122 | 3664 | 2 | ENSMUST00 | 0 | Fut4 | ENSMUSG00000049307.5 |
| chr9 | 57501755 | 57504739 | 20.27628 | 60.14116 | 57.29097 | Promoter | 9 | 57504102 | 57505447 | 1346 | 1 | ENSMUST00 | 0 | Rpp25 | ENSMUSG00000062309.7 |
| chr11 | 3370747 | 3371857 | 20.74282 | 60.13039 | 57.28273 | Promoter | 11 | 3344256 | 3371376 | 27121 | 2 | ENSMUST00 | 0 | Limk2 | ENSMUSG00000020451.12 |
| chr11 | 69340229 | 69341261 | 21.70878 | 60.13001 | 57.28273 | Promoter | 11 | 69340461 | 69342641 | 2181 | 1 | ENSMUST00 | 0 | Chd3os | ENSMUSG00000043419.10 |
| chr14 | 63178123 | 63178718 | 20.74282 | 60.13039 | 57.28273 | Promoter | 14 | 63145153 | 63177793 | 32641 | 2 | ENSMUST00 | -331 | Fdft1 | ENSMUSG00000021273.9 |
| chr17 | 34309237 | 34310272 | 20.74282 | 60.13039 | 57.28273 | Exon (ENSMU | 17 | 34305877 | 34316199 | 10323 | 1 | ENSMUST00 | 3361 | H2-Eb1 | ENSMUSG00000060586.10 |
| chr3 | 89278368 | 89280098 | 20.74282 | 60.13039 | 57.28273 | Promoter | 3 | 89271733 | 89279645 | 7913 | 2 | ENSMUST00 | 0 | Efna1 | ENSMUSG00000027954.4 |
| chr4 | 132842194 | 132843679 | 20.74282 | 60.13039 | 57.28273 | Promoter | 4 | 132826929 | 132843169 | 16241 | 2 | ENSMUST00 | 0 | Ppp1r8 | ENSMUSG00000028882.8 |
| chr6 | 54565710 | 54566766 | 19.75722 | 60.04038 | 57.19346 | Promoter | 6 | 54505890 | 54566489 | 60600 | 2 | ENSMUST00 | 0 | Scrn1 | ENSMUSG00000019124.5 |
| chr7 | 42692241 | 42693493 | 19.75722 | 60.04038 | 57.19346 | Promoter | 7 | 42660105 | 42692718 | 32614 | 2 | ENSMUST00 | 0 | Gm5595 | ENSMUSG00000069727.4 |
| chr11 | 101732893 | 101733889 | 22.10065 | 59.96099 | 57.11441 | Promoter | 11 | 101732919 | 101735696 | 2778 | 1 | ENSMUST00 | 0 | Dhx8 | ENSMUSG00000034931.10 |
| chr11 | 117265778 | 117267655 | 22.10065 | 59.96099 | 57.11441 | Promoter | 11 | 117266246 | 117362313 | 96068 | 1 | ENSMUST00 | 0 | Sep-09 | ENSMUSG00000059248.8 |
| chr12 | 82616645 | 82618014 | 22.05606 | 59.8844 | 57.03867 | Promoter | 12 | 82616925 | 83092731 | 475807 | 1 | ENSMUST00 | 0 | Rgs6 | ENSMUSG00000021219.11 |
| chr4 | 114821453 | 114822346 | 22.05606 | 59.8844 | 57.03867 | Promoter | 4 | 114821722 | 114856166 | 34445 | 1 | ENSMUST00 | 0 | Gm12830 | ENSMUSG00000055198.3 |
| chr7 | 143293497 | 143296592 | 22.05606 | 59.8844 | 57.03867 | Promoter | 7 | 143293932 | 143294155 | 224 | 1 | ENSMUST00 | 0 | Gm27539 | ENSMUSG00000099240.1 |
| chr1 | 12692880 | 12693967 | 22.51707 | 59.83231 | 56.98677 | Promoter | 1 | 12692532 | 12852489 | 159958 | 1 | ENSMUST00 | 349 | Sulf1 | ENSMUSG00000016918.10 |
| chr1 | 164898967 | 164899567 | 22.51707 | 59.83231 | 56.98677 | Distal Interge | 1 | 164931644 | 164935527 | 3884 | 2 | ENSMUST00 | 35960 | Xcl1 | ENSMUSG00000026573.7 |
| chr12 | 44221584 | 44222391 | 22.51707 | 59.83231 | 56.98677 | Promoter | 12 | 44221370 | 44312092 | 90723 | 1 | ENSMUST00 | 215 | Pnpla8 | ENSMUSG00000036257.9 |
| chr16 | 67619517 | 67621341 | 22.51707 | 59.83231 | 56.98677 | Promoter | 16 | 66655421 | 67620908 | 965488 | 2 | ENSMUST00 | 0 | Cadm2 | ENSMUSG00000064115.8 |
| chr17 | 34626596 | 34628611 | 22.51707 | 59.83231 | 56.98677 | Promoter | 17 | 34616662 | 34627435 | 10774 | 2 | ENSMUST00 | 0 | Ppt2 | ENSMUSG00000015474.11 |
| chr17 | 23598594 | 23599797 | 22.51707 | 59.83231 | 56.98677 | Promoter | 17 | 23575852 | 23599487 | 23636 | 2 | ENSMUST00 | 0 | Zfp13 | ENSMUSG00000062012.8 |
| chr19 | 34290290 | 34291652 | 22.51707 | 59.83231 | 56.98677 | Promoter | 19 | 34290659 | 34309350 | 18692 | 1 | ENSMUST00 | 0 | Fas | ENSMUSG00000024778.7 |
| chr3 | 79841921 | 79843054 | 22.51707 | 59.83231 | 56.98677 | Promoter | 3 | 79813153 | 79842662 | 29510 | 2 | ENSMUST00 | 0 | Tmem144 | ENSMUSG00000027956.6 |
| chr4 | 148130346 | 148131392 | 22.51707 | 59.83231 | 56.98677 | Promoter | 4 | 148130384 | 148140791 | 10408 | 1 | ENSMUST00 | 0 | Mad2l2 | ENSMUSG00000029003.6 |
| chr5 | 35278708 | 35281037 | 22.51707 | 59.83231 | 56.98677 | Promoter | 5 | 35278566 | 35281761 | 3196 | 1 | ENSMUST00 | 143 | Adra2c | ENSMUSG00000045318.5 |
| chr8 | 71988349 | 71989103 | 22.51707 | 59.83231 | 56.98677 | 3' UTR | 8 | 71988483 | 71992915 | 4433 | 2 | ENSMUST00 | 3812 | Cyp4f18 | ENSMUSG00000003484.4 |
| chr8 | 3467229 | 3467945 | 22.51707 | 59.83231 | 56.98677 | Promoter | 8 | 3457105 | 3467680 | 10576 | 2 | ENSMUST00 | 0 | Pex11g | ENSMUSG00000069633.7 |
| chr8 | 105612839 | 105614792 | 22.51707 | 59.83231 | 56.98677 | Promoter | 8 | 105615466 | 105615573 | 108 | 1 | ENSMUST00 | -674 | Mir1966 | ENSMUSG00000089432.1 |
| chrX | 7762650 | 7764496 | 22.51707 | 59.83231 | 56.98677 | Promoter | 20 | 7762658 | 7775202 | 12545 | 1 | ENSMUST00 | 0 | Tfe3 | ENSMUSG00000000134.12 |
| chrX | 36597823 | 36599108 | 22.51707 | 59.83231 | 56.98677 | Promoter | 20 | 36598206 | 36606079 | 7874 | 1 | ENSMUST00 | 0 | Pgrmc1 | ENSMUSG00000006373.10 |
| chrX | 9272313 | 9273362 | 22.51707 | 59.83231 | 56.98677 | Promoter | 20 | 9272756 | 9313250 | 40495 | 1 | ENSMUST00 | 0 | Xk | ENSMUSG00000015342.6 |
| chrX | 36795379 | 36796755 | 22.51707 | 59.83231 | 56.98677 | Promoter | 20 | 36795651 | 36798807 | 3157 | 1 | ENSMUST00 | 0 | Slc25a5 | ENSMUSG00000016319.3 |
| chrX | 37126563 | 37127864 | 22.51707 | 59.83231 | 56.98677 | Promoter | 20 | 37126795 | 37150746 | 23952 | 1 | ENSMUST00 | 0 | Nkap | ENSMUSG00000016409.4 |
| chrX | 20616575 | 20617600 | 22.51707 | 59.83231 | 56.98677 | Promoter | 20 | 20617503 | 20650888 | 33386 | 1 | ENSMUST00 | 0 | Rbm10 | ENSMUSG00000031060.11 |
| chrX | 101429680 | 101430429 | 22.51707 | 59.83231 | 56.98677 | Promoter | 20 | 101429735 | 101442431 | 12697 | 1 | ENSMUST00 | 0 | Nono | ENSMUSG00000031311.12 |
| chrX | 162887628 | 162888483 | 22.51707 | 59.83231 | 56.98677 | Promoter | 20 | 162857057 | 162888447 | 31391 | 2 | ENSMUST00 | 0 | Syap1 | ENSMUSG00000031357.3 |
| chrX | 77794773 | 77796146 | 22.51707 | 59.83231 | 56.98677 | Promoter | 20 | 77771455 | 77795972 | 24518 | 2 | ENSMUST00 | 0 | Prkx | ENSMUSG00000035725.8 |
| chrX | 48343740 | 48345229 | 22.51707 | 59.83231 | 56.98677 | Promoter | 20 | 48343770 | 48408049 | 64280 | 1 | ENSMUST00 | 0 | Bcorl1 | ENSMUSG00000036959.10 |
| chrX | 20424963 | 20426636 | 22.51707 | 59.83231 | 56.98677 | Promoter | 20 | 20425688 | 20519939 | 94252 | 1 | ENSMUST00 | 0 | Jade3 | ENSMUSG00000037315.9 |
| chrX | 145504092 | 145506026 | 22.51707 | 59.83231 | 56.98677 | Promoter | 20 | 145446425 | 145505144 | 58720 | 2 | ENSMUST00 | 0 | Amot | ENSMUSG00000041688.11 |

| | | | | | | | | | | | | | | | |
|---|---|---|---|---|---|---|---|---|---|---|---|---|---|---|---|
| chr9 | 104336789 | 104338232 | 22.01165 | 59.80809 | 56.96389 | Promoter | 9 | 104288316 | 104337728 | 49413 | 2 | ENSMUST00 | 0 | Acpp | ENSMUSG00000032561.9 |
| chr11 | 103217688 | 103218579 | 21.4995 | 59.76749 | 56.92346 | Promoter | 11 | 103208127 | 103218432 | 10306 | 2 | ENSMUST00 | 0 | Spata32 | ENSMUSG00000044787.7 |
| chr15 | 98295453 | 98296554 | 21.4995 | 59.76749 | 56.92346 | Promoter | 15 | 98285588 | 98296130 | 10543 | 2 | ENSMUST00 | 0 | Zfp641 | ENSMUSG00000022987.6 |
| chr15 | 76973870 | 76974491 | 21.4995 | 59.76749 | 56.92346 | Distal Interge | 15 | 76948539 | 76950729 | 2191 | 1 | ENSMUST00 | 25332 | 1110038F14I | ENSMUSG00000063236.5 |
| chr2 | 30061791 | 30063403 | 21.4995 | 59.76749 | 56.92346 | Promoter | 2 | 30061853 | 30072577 | 10725 | 1 | ENSMUST00 | 0 | Set | ENSMUSG00000054766.8 |
| chr6 | 112946109 | 112947557 | 21.4995 | 59.76749 | 56.92346 | Promoter | 6 | 112717971 | 112947266 | 229296 | 2 | ENSMUST00 | 0 | Srgap3 | ENSMUSG00000030257.11 |
| chr17 | 22165296 | 22166172 | 20.05708 | 59.74671 | 56.90333 | Promoter | 17 | 22144359 | 22166190 | 21832 | 2 | ENSMUST00 | 18 | Zfp947 | ENSMUSG00000063383.5 |
| chr7 | 19228805 | 19229686 | 20.05708 | 59.74671 | 56.90333 | Promoter | 7 | 19228383 | 19256543 | 28161 | 1 | ENSMUST00 | 423 | Opa3 | ENSMUSG00000052214.8 |
| chr4 | 88756840 | 88757747 | 21.96742 | 59.73206 | 56.88897 | Promoter | 4 | 88757844 | 88758380 | 537 | 1 | ENSMUST00 | -97 | Gm13286 | ENSMUSG00000081501.5 |
| chr2 | 62663853 | 62664702 | 15.2551 | 59.70792 | 56.86497 | Promoter | 2 | 62664285 | 62694109 | 29825 | 1 | ENSMUST00 | 0 | Gca | ENSMUSG00000026893.4 |
| chr3 | 14533323 | 14534756 | 15.2551 | 59.70792 | 56.86497 | Promoter | 3 | 14533788 | 14572658 | 38871 | 1 | ENSMUST00 | 0 | Lrrcc1 | ENSMUSG00000027550.9 |
| chr4 | 45012721 | 45013528 | 21.45812 | 59.69573 | 56.85284 | Promoter | 4 | 45012830 | 45014399 | 1570 | 1 | ENSMUST00 | 0 | 1700055D18I | ENSMUSG00000070997.4 |
| chr1 | 158355443 | 158356657 | 21.92336 | 59.65632 | 56.81357 | Promoter | 1 | 158245271 | 158356260 | 110990 | 2 | ENSMUST00 | 0 | Brinp2 | ENSMUSG00000004031.8 |
| chr1 | 136259841 | 136261055 | 21.92336 | 59.65632 | 56.81357 | Promoter | 1 | 136258914 | 136260873 | 1960 | 2 | ENSMUST00 | 0 | Gpr25 | ENSMUSG00000052759.6 |
| chr17 | 57061663 | 57062839 | 21.92336 | 59.65632 | 56.81357 | Promoter | 17 | 57062231 | 57065918 | 3688 | 1 | ENSMUST00 | 0 | Crb3 | ENSMUSG00000044279.10 |
| chr3 | 10342299 | 10343121 | 21.92336 | 59.65632 | 56.81357 | Exon (ENSMU | 3 | 10339955 | 10344727 | 4773 | 2 | ENSMUST00 | 1606 | Zfand1 | ENSMUSG00000039795.9 |
| chr4 | 57142712 | 57143905 | 21.92336 | 59.65632 | 56.81357 | Promoter | 4 | 56991972 | 57143437 | 151466 | 2 | ENSMUST00 | 0 | Epb4.1l4b | ENSMUSG00000028434.7 |
| chr9 | 65579586 | 65581133 | 21.92336 | 59.65632 | 56.81357 | Promoter | 9 | 65554386 | 65580040 | 25655 | 2 | ENSMUST00 | 0 | Plekho2 | ENSMUSG00000050721.8 |
| chr1 | 38897569 | 38898453 | 22.38265 | 59.60271 | 56.76062 | Promoter | 1 | 38863877 | 38898160 | 34284 | 2 | ENSMUST00 | 0 | Chst10 | ENSMUSG00000026080.8 |
| chr13 | 43784668 | 43785693 | 22.38265 | 59.60271 | 56.76062 | Promoter | 13 | 43785107 | 43803130 | 18024 | 1 | ENSMUST00 | 0 | Cd83 | ENSMUSG00000015396.3 |
| chr1 | 178275750 | 178276846 | 21.87949 | 59.58086 | 56.73908 | Promoter | 1 | 178276093 | 178276639 | 547 | 2 | ENSMUST00 | 0 | 4930527J03R | ENSMUSG00000070489.4 |
| chr1 | 86441607 | 86443281 | 19.04495 | 59.55748 | 56.71581 | Distal Interge | 1 | 86426329 | 86428052 | 1724 | 1 | ENSMUST00 | 15279 | 1700019O17 | ENSMUSG00000036574.5 |
| chr10 | 80621560 | 80622431 | 19.04495 | 59.55748 | 56.71581 | Promoter | 10 | 80622780 | 80640771 | 17992 | 1 | ENSMUST00 | -349 | Csnk1g2 | ENSMUSG00000003345.10 |
| chr10 | 70244489 | 70246103 | 19.04495 | 59.55748 | 56.71581 | Promoter | 10 | 70245100 | 70285968 | 40869 | 1 | ENSMUST00 | 0 | Slc16a9 | ENSMUSG00000037762.6 |
| chr10 | 5192240 | 5195052 | 19.04495 | 59.55748 | 56.71581 | Distal Interge | 10 | 5069192 | 5071975 | 2784 | 1 | ENSMUST00 | 123049 | Gm10097 | ENSMUSG00000096553.1 |
| chr11 | 87461732 | 87462557 | 19.04495 | 59.55748 | 56.71581 | Promoter | 11 | 87462286 | 87462500 | 215 | 1 | ENSMUST00 | 0 | Rnu3b4 | ENSMUSG00000098641.1 |
| chr12 | 72535810 | 72536947 | 19.04495 | 59.55748 | 56.71581 | Promoter | 12 | 72536357 | 72580213 | 43857 | 1 | ENSMUST00 | 0 | Pcnxl4 | ENSMUSG00000034501.6 |
| chr12 | 4843291 | 4844254 | 19.04495 | 59.55748 | 56.71581 | Promoter | 12 | 4843303 | 4852913 | 9611 | 1 | ENSMUST00 | 0 | BC068281 | ENSMUSG00000051721.4 |
| chr16 | 74408228 | 74412370 | 19.04495 | 59.55748 | 56.71581 | Promoter | 16 | 73892306 | 74410989 | 518684 | 2 | ENSMUST00 | 0 | Robo2 | ENSMUSG00000052516.12 |
| chr17 | 63849148 | 63849911 | 19.04495 | 59.55748 | 56.71581 | Distal Interge | 17 | 63863300 | 63863791 | 492 | 2 | ENSMUST00 | 13880 | A930002H24 | ENSMUSG00000045506.3 |
| chr19 | 54114760 | 54115385 | 19.04495 | 59.55748 | 56.71581 | Distal Interge | 19 | 54045182 | 54048982 | 3801 | 1 | ENSMUST00 | 69579 | Adra2a | ENSMUSG00000033717.5 |
| chr2 | 119325079 | 119327754 | 19.04495 | 59.55748 | 56.71581 | Promoter | 2 | 119325784 | 119335962 | 10179 | 1 | ENSMUST00 | 0 | Dll4 | ENSMUSG00000027314.6 |
| chr3 | 116512667 | 116513742 | 19.04495 | 59.55748 | 56.71581 | Promoter | 3 | 116513070 | 116549976 | 36907 | 1 | ENSMUST00 | 0 | Dbt | ENSMUSG00000000340.6 |
| chr4 | 107801651 | 107804211 | 19.04495 | 59.55748 | 56.71581 | Promoter | 4 | 107802261 | 107874167 | 71907 | 1 | ENSMUST00 | 0 | Lrp8 | ENSMUSG00000028613.9 |
| chr5 | 77094535 | 77095508 | 19.04495 | 59.55748 | 56.71581 | Promoter | 5 | 77086988 | 77095253 | 8266 | 2 | ENSMUST00 | 0 | Hopx | ENSMUSG00000059325.9 |
| chr7 | 34811244 | 34813377 | 19.04495 | 59.55748 | 56.71581 | Promoter | 7 | 34674468 | 34812686 | 138219 | 2 | ENSMUST00 | 0 | Chst8 | ENSMUSG00000060402.6 |
| chr9 | 21545661 | 21546699 | 19.04495 | 59.55748 | 56.71581 | Promoter | 9 | 21546894 | 21589467 | 42574 | 1 | ENSMUST00 | -195 | Carm1 | ENSMUSG00000032185.9 |
| chr9 | 50767428 | 50769019 | 19.04495 | 59.55748 | 56.71581 | Promoter | 9 | 50768236 | 50773348 | 5113 | 1 | ENSMUST00 | 0 | Fdxacb1 | ENSMUSG00000037845.9 |
| chr9 | 20384397 | 20385770 | 19.04495 | 59.55748 | 56.71581 | Promoter | 9 | 20345136 | 20385177 | 40042 | 2 | ENSMUST00 | 0 | Zfp560 | ENSMUSG00000045519.9 |
| chr10 | 103367489 | 103369582 | 21.37461 | 59.55079 | 56.71094 | Promoter | 10 | 103367808 | 103419378 | 51571 | 1 | ENSMUST00 | 0 | Slc6a15 | ENSMUSG00000019894.8 |
| chr10 | 76530871 | 76532111 | 21.37461 | 59.55079 | 56.71094 | Promoter | 10 | 76531588 | 76535568 | 3981 | 1 | ENSMUST00 | 0 | Lss | ENSMUSG00000033105.7 |
| chr11 | 116012188 | 116013235 | 21.37461 | 59.55079 | 56.71094 | Promoter | 11 | 116008457 | 116012719 | 4263 | 2 | ENSMUST00 | 0 | Galk1 | ENSMUSG00000020766.4 |
| chr11 | 119036853 | 119037823 | 21.37461 | 59.55079 | 56.71094 | 3' UTR | 11 | 119039053 | 119040831 | 1779 | 2 | ENSMUST00 | 3008 | Cbx8 | ENSMUSG00000025578.9 |
| chr13 | 67399111 | 67400344 | 21.37461 | 59.55079 | 56.71094 | Promoter | 13 | 67389309 | 67399750 | 10442 | 2 | ENSMUST00 | 0 | Zfp429 | ENSMUSG00000078994.3 |
| chr15 | 97907246 | 97908866 | 21.37461 | 59.55079 | 56.71094 | Promoter | 15 | 97854425 | 97908310 | 53886 | 2 | ENSMUST00 | 0 | Vdr | ENSMUSG00000022479.10 |
| chr16 | 17146647 | 17147957 | 21.37461 | 59.55079 | 56.71094 | Promoter | 16 | 17139064 | 17147229 | 8166 | 2 | ENSMUST00 | 0 | Ccdc116 | ENSMUSG00000022768.10 |
| chr17 | 36314848 | 36315780 | 21.37461 | 59.55079 | 56.71094 | Distal Interge | 17 | 36323201 | 36326145 | 2945 | 2 | ENSMUST00 | 10365 | H2-M10.1 | ENSMUSG00000024448.9 |
| chr19 | 34746696 | 34747457 | 21.37461 | 59.55079 | 56.71094 | Promoter | 19 | 34668403 | 34747289 | 78887 | 2 | ENSMUST00 | 0 | Slc16a12 | ENSMUSG00000009378.3 |
| chr2 | 28205247 | 28206399 | 21.37461 | 59.55079 | 56.71094 | Promoter | 2 | 28205659 | 28214425 | 8767 | 1 | ENSMUST00 | 0 | Olfm1 | ENSMUSG00000026833.13 |
| chr2 | 94158236 | 94158718 | 21.37461 | 59.55079 | 56.71094 | Promoter | 2 | 94032689 | 94157964 | 125276 | 2 | ENSMUST00 | -273 | Hsd17b12 | ENSMUSG00000027195.5 |
| chr3 | 14886008 | 14887153 | 21.37461 | 59.55079 | 56.71094 | Promoter | 3 | 14886426 | 14900769 | 14344 | 1 | ENSMUST00 | 0 | Car2 | ENSMUSG00000027562.7 |
| chr3 | 96597281 | 96598839 | 21.37461 | 59.55079 | 56.71094 | Promoter | 3 | 96596636 | 96599775 | 3140 | 1 | ENSMUST00 | 646 | Ankrd34a | ENSMUSG00000049097.9 |
| chr4 | 130792040 | 130793287 | 21.37461 | 59.55079 | 56.71094 | Promoter | 4 | 130792537 | 130826319 | 33783 | 1 | ENSMUST00 | 0 | Sdc3 | ENSMUSG00000025743.9 |
| chr5 | 136881536 | 136883244 | 21.37461 | 59.55079 | 56.71094 | Promoter | 5 | 136741759 | 136883121 | 141363 | 2 | ENSMUST00 | 0 | Col26a1 | ENSMUSG00000004415.10 |
| chr6 | 30958391 | 30959512 | 21.37461 | 59.55079 | 56.71094 | Promoter | 6 | 30956023 | 30959078 | 3056 | 2 | ENSMUST00 | 0 | Klf14 | ENSMUSG00000073209.4 |
| chr7 | 88277640 | 88278814 | 21.37461 | 59.55079 | 56.71094 | Promoter | 7 | 88278085 | 88310888 | 32804 | 1 | ENSMUST00 | 0 | Ctsc | ENSMUSG00000030560.11 |
| chr7 | 44384080 | 44385000 | 21.37461 | 59.55079 | 56.71094 | Promoter | 7 | 44384102 | 44400029 | 15928 | 1 | ENSMUST00 | 0 | Syt3 | ENSMUSG00000030731.8 |
| chr9 | 120582202 | 120582810 | 21.37461 | 59.55079 | 56.71094 | Distal Interge | 9 | 120577346 | 120578072 | 727 | 1 | ENSMUST00 | 4857 | 5830454E08I | ENSMUSG00000052658.4 |
| chrX | 68678124 | 68679977 | 21.37461 | 59.55079 | 56.71094 | Promoter | 20 | 68678541 | 68717963 | 39423 | 1 | ENSMUST00 | 0 | Fmr1 | ENSMUSG00000000838.12 |
| chr4 | 130573364 | 130574393 | 22.33599 | 59.52296 | 56.68532 | Promoter | 4 | 130531602 | 130573966 | 42365 | 2 | ENSMUST00 | 0 | Nkain1 | ENSMUSG00000078532.4 |
| chr7 | 66427081 | 66428260 | 22.33599 | 59.52296 | 56.68532 | Promoter | 7 | 66390892 | 66427517 | 36626 | 2 | ENSMUST00 | 0 | Aldh1a3 | ENSMUSG00000015134.9 |
| chr4 | 58205390 | 58206908 | 20.37803 | 59.48265 | 56.64523 | Promoter | 4 | 58042442 | 58206596 | 164155 | 2 | ENSMUST00 | 0 | Svep1 | ENSMUSG00000028369.10 |
| chr3 | 135824972 | 135826805 | 21.33495 | 59.48191 | 56.64483 | Promoter | 3 | 135825279 | 135888386 | 63108 | 1 | ENSMUST00 | 0 | Slc39a8 | ENSMUSG00000053897.10 |
| chr4 | 146603758 | 146604404 | 14.83195 | 59.47061 | 56.63358 | Promoter | 4 | 146604212 | 146604776 | 565 | 1 | ENSMUST00 | 0 | Gm13240 | ENSMUSG00000096366.1 |
| chr6 | 54992306 | 54993434 | 19.43785 | 59.45512 | 56.6181 | Promoter | 6 | 54982580 | 54992950 | 10371 | 2 | ENSMUST00 | 0 | Ggct | ENSMUSG00000002797.7 |
| chr12 | 33302526 | 33303732 | 21.79225 | 59.43074 | 56.59393 | Promoter | 12 | 33302515 | 33368273 | 65759 | 1 | ENSMUST00 | 12 | Atxn7l1 | ENSMUSG00000020564.11 |
| chr11 | 25099986 | 25100286 | 15.80552 | 59.41874 | 56.58204 | Distal Interge | 11 | 25232626 | 25367133 | 134508 | 1 | ENSMUST00 | -132340 | 4933427E13I | ENSMUSG00000033015.11 |
| chr17 | 94749275 | 94750618 | 15.80552 | 59.41874 | 56.58204 | Promoter | 17 | 94727080 | 94749892 | 22813 | 2 | ENSMUST00 | 0 | Mettl4 | ENSMUSG00000055660.8 |
| chr15 | 103182393 | 103182735 | 21.29421 | 59.41112 | 56.57455 | Distal Interge | 15 | 103153307 | 103158002 | 4696 | 2 | ENSMUST00 | -24392 | Smug1 | ENSMUSG00000036061.9 |
| chr11 | 116167477 | 116168650 | 22.24324 | 59.3644 | 56.5279 | Promoter | 11 | 116142285 | 116167780 | 25496 | 2 | ENSMUST00 | 0 | Fbf1 | ENSMUSG00000020776.13 |
| chr2 | 118861546 | 118862430 | 22.24324 | 59.3644 | 56.5279 | Promoter | 2 | 118861954 | 118882909 | 20956 | 1 | ENSMUST00 | 0 | Ivd | ENSMUSG00000027332.6 |
| chr6 | 13838329 | 13839204 | 21.7489 | 59.35609 | 56.51971 | Promoter | 6 | 13835068 | 13838454 | 3387 | 2 | ENSMUST00 | 0 | Gpr85 | ENSMUSG00000048216.8 |
| chr6 | 47740203 | 47740550 | 14.45332 | 59.30346 | 56.46724 | Distal Interge | 6 | 47773329 | 47773429 | 101 | 1 | ENSMUST00 | -32779 | Gm24975 | ENSMUSG00000094807.1 |
| chr5 | 125055981 | 125058881 | 22.19715 | 59.28558 | 56.44936 | Promoter | 5 | 125030800 | 125058400 | 27601 | 2 | ENSMUST00 | 0 | Ncor2 | ENSMUSG00000029478.11 |
| chr5 | 145203613 | 145205310 | 22.19715 | 59.28558 | 56.44936 | Promoter | 5 | 145204562 | 145221750 | 17189 | 1 | ENSMUST00 | 0 | Zkscan5 | ENSMUSG00000055991.10 |
| chr1 | 51914973 | 51916278 | 18.02578 | 59.25625 | 56.42029 | Promoter | 1 | 51749765 | 51916071 | 166307 | 2 | ENSMUST00 | 0 | Myo1b | ENSMUSG00000018417.9 |
| chr10 | 83336897 | 83338171 | 18.02578 | 59.25625 | 56.42029 | Promoter | 10 | 83230848 | 83337882 | 107035 | 2 | ENSMUST00 | 0 | Slc41a2 | ENSMUSG00000034591.5 |
| chr10 | 82629283 | 82630833 | 18.02578 | 59.25625 | 56.42029 | Promoter | 10 | 82629828 | 82650324 | 20497 | 1 | ENSMUST00 | 0 | Tdg | ENSMUSG00000034674.13 |
| chr11 | 98728850 | 98729459 | 18.02578 | 59.25625 | 56.42029 | Promoter | 11 | 98704591 | 98729435 | 24845 | 2 | ENSMUST00 | 0 | Med24 | ENSMUSG00000017210.9 |
| chr15 | 100614147 | 100615159 | 18.02578 | 59.25625 | 56.42029 | Promoter | 15 | 100614140 | 100615110 | 971 | 2 | ENSMUST00 | 0 | C330013E15I | ENSMUSG00000097093.1 |
| chr16 | 55965761 | 55967413 | 18.02578 | 59.25625 | 56.42029 | Promoter | 16 | 55966275 | 55971435 | 5161 | 1 | ENSMUST00 | 0 | Rpl24 | ENSMUSG00000098274.2 |
| chr18 | 3506950 | 3507774 | 18.02578 | 59.25625 | 56.42029 | Promoter | 18 | 3507957 | 3516404 | 8448 | 1 | ENSMUST00 | -183 | Bambi | ENSMUSG00000024232.1 |
| chr18 | 73571379 | 73572465 | 18.02578 | 59.25625 | 56.42029 | Promoter | 18 | 73573044 | 73592575 | 19532 | 1 | ENSMUST00 | -579 | Mex3c | ENSMUSG00000037253.8 |
| chr19 | 7557359 | 7558288 | 18.02578 | 59.25625 | 56.42029 | Promoter | 19 | 7557459 | 7581728 | 24270 | 1 | ENSMUST00 | 0 | Pla2g16 | ENSMUSG00000060675.8 |
| chr6 | 39810232 | 39811219 | 18.02578 | 59.25625 | 56.42029 | Promoter | 6 | 39801804 | 39810988 | 9185 | 2 | ENSMUST00 | 0 | Mrps33 | ENSMUSG00000029918.8 |
| chr7 | 68128511 | 68130116 | 18.02578 | 59.25625 | 56.42029 | Intron (ENSM | 7 | 68236608 | 68264233 | 27626 | 2 | ENSMUST00 | 134117 | Pgpep1l | ENSMUSG00000030553.2 |
| chr7 | 63754712 | 63755800 | 18.02578 | 59.25625 | 56.42029 | Promoter | 7 | 63754712 | 63757925 | 3214 | 1 | ENSMUST00 | 1 | Otud7a | ENSMUSG00000033510.9 |
| chr11 | 105967580 | 105968446 | 20.72239 | 59.25234 | 56.41743 | Promoter | 11 | 105967945 | 105989964 | 22020 | 1 | ENSMUST00 | 0 | Ace | ENSMUSG00000020681.9 |
| chr13 | 21167008 | 21170327 | 20.72239 | 59.25234 | 56.41743 | Distal Interge | 13 | 21179445 | 21180416 | 972 | 1 | ENSMUST00 | -9118 | Trim27 | ENSMUSG00000021326.6 |
| chr2 | 165387626 | 165388570 | 20.72239 | 59.25234 | 56.41743 | Promoter | 2 | 165374264 | 165388259 | 13996 | 2 | ENSMUST00 | 0 | Zfp334 | ENSMUSG00000017667.9 |
| chr5 | 93266136 | 93267130 | 20.72239 | 59.25234 | 56.41743 | Promoter | 5 | 93267257 | 93276231 | 8975 | 1 | ENSMUST00 | -127 | Ccng2 | ENSMUSG00000029385.9 |
| chr4 | 46038637 | 46039904 | 18.8759 | 59.24087 | 56.40649 | Promoter | 4 | 46038935 | 46116032 | 77098 | 1 | ENSMUST00 | 0 | Tmod1 | ENSMUSG00000028328.8 |
| chr4 | 59188732 | 59190495 | 16.43683 | 59.22888 | 56.39454 | Promoter | 4 | 59189257 | 59222833 | 33577 | 1 | ENSMUST00 | 0 | Ugcg | ENSMUSG00000028381.8 |
| chr7 | 48137234 | 48138685 | 16.43683 | 59.22888 | 56.39454 | Distal Interge | 7 | 48116772 | 48117441 | 670 | 2 | ENSMUST00 | -19794 | RP23-62M19 | ENSMUSG00000101352.1 |
| chr13 | 74462197 | 74463605 | 22.15125 | 59.20708 | 56.37309 | Distal Interge | 13 | 74480517 | 74482943 | 2427 | 2 | ENSMUST00 | 19338 | Zfp825 | ENSMUSG00000069208.5 |
| chr19 | 45300973 | 45302769 | 22.15125 | 59.20708 | 56.37309 | Promoter | 19 | 45300463 | 45303034 | 2572 | 2 | ENSMUST00 | 265 | RP23-188J8.1 | ENSMUSG00000100618.1 |
| chr6 | 28215131 | 28215985 | 22.15125 | 59.20708 | 56.37309 | Promoter | 6 | 28215555 | 28260862 | 45308 | 1 | ENSMUST00 | 0 | 6530409C15I | ENSMUSG00000043340.3 |
| chr11 | 109425910 | 109426774 | 17.16829 | 59.16322 | 56.32936 | Promoter | 11 | 109425946 | 109438148 | 12203 | 1 | ENSMUST00 | 0 | Amz2 | ENSMUSG00000020610.11 |
| chr12 | 69790220 | 69791544 | 17.16829 | 59.16322 | 56.32936 | Promoter | 12 | 69746848 | 69790707 | 43860 | 2 | ENSMUST00 | 0 | Cdkl1 | ENSMUSG00000020990.4 |
| chr12 | 104472324 | 104473489 | 17.16829 | 59.16322 | 56.32936 | Promoter | 12 | 104471209 | 104473330 | 2122 | 2 | ENSMUST00 | 0 | Gsc | ENSMUSG00000021095.5 |
| chr15 | 76197865 | 76200595 | 17.16829 | 59.16322 | 56.32936 | Promoter | 15 | 76170974 | 76199835 | 28862 | 2 | ENSMUST00 | 0 | Plec | ENSMUSG00000022565.10 |
| chr19 | 5084979 | 5085826 | 17.16829 | 59.16322 | 56.32936 | Promoter | 19 | 5071035 | 5085487 | 14453 | 2 | ENSMUST00 | 0 | Tmem151a | ENSMUSG00000061451.8 |
| chr3 | 10330668 | 10331623 | 17.16829 | 59.16322 | 56.32936 | Promoter | 3 | 10313540 | 10331439 | 17900 | 2 | ENSMUST00 | 0 | Impa1 | ENSMUSG00000027531.11 |
| chr3 | 14641270 | 14642919 | 17.16829 | 59.16322 | 56.32936 | Promoter | 3 | 14641727 | 14663002 | 21276 | 1 | ENSMUST00 | 0 | Car13 | ENSMUSG00000027555.8 |
| chr8 | 36731870 | 36731965 | 17.16829 | 59.16322 | 56.32936 | Promoter | 8 | 36567751 | 36733054 | 165304 | 2 | ENSMUST00 | 0 | Dlc1 | ENSMUSG00000031523.11 |
| chr16 | 44172251 | 44172968 | 19.72203 | 59.14077 | 56.30748 | Promoter | 16 | 44173246 | 44227465 | 54220 | 1 | ENSMUST00 | -278 | Gm608 | ENSMUSG00000068284.9 |
| chr4 | 123342314 | 123343684 | 19.72203 | 59.14077 | 56.30748 | Promoter | 4 | 123312645 | 123342803 | 30159 | 2 | ENSMUST00 | 0 | Bmp8a | ENSMUSG00000032726.7 |
| chr2 | 32360860 | 32362104 | 21.61986 | 59.13376 | 56.30069 | Promoter | 2 | 32363010 | 32367407 | 4398 | 1 | ENSMUST00 | -906 | Ciz1 | ENSMUSG00000039205.11 |
| chr5 | 122951325 | 122952765 | 22.10555 | 59.12887 | 56.29591 | Promoter | 5 | 122932575 | 122952506 | 19932 | 2 | ENSMUST00 | 0 | Kdm2b | ENSMUSG00000029475.12 |
| chr8 | 70138496 | 70139218 | 22.10555 | 59.12887 | 56.29591 | Promoter | 8 | 70130812 | 70139197 | 8386 | 2 | ENSMUST00 | 0 | Rfxank | ENSMUSG00000036120.9 |
| chr1 | 64690203 | 64690718 | 21.57718 | 59.06017 | 56.22787 | Promoter | 1 | 64690455 | 64725644 | 35190 | 1 | ENSMUST00 | 0 | Ccnyl1 | ENSMUSG00000070871.5 |
| chr1 | 4571432 | 4572261 | 21.0928 | 59.0607 | 56.22787 | Distal Interge | 1 | 4529017 | 4529123 | 107 | 1 | ENSMUST00 | 42416 | Gm22307 | ENSMUSG00000096126.1 |
| chr15 | 98985117 | 98986470 | 21.0928 | 59.0607 | 56.22787 | Promoter | 15 | 98985965 | 98989281 | 3317 | 1 | ENSMUST00 | 0 | 4930578M01 | ENSMUSG00000097587.1 |
| chr6 | 85936553 | 85937879 | 21.0928 | 59.0607 | 56.22787 | Distal Interge | 6 | 85932681 | 85933379 | 699 | 2 | ENSMUST00 | -3175 | Nat8b | ENSMUSG00000089634.1 |
| chr8 | 123318125 | 123319092 | 21.0928 | 59.0607 | 56.22787 | Promoter | 8 | 123268300 | 123318576 | 50277 | 2 | ENSMUST00 | 0 | Fanca | ENSMUSG00000032815.10 |
| chr11 | 48825828 | 48827167 | 20.5724 | 58.98804 | 56.15619 | Promoter | 11 | 48826140 | 48851187 | 25048 | 1 | ENSMUST00 | 0 | Trim7 | ENSMUSG00000040350.10 |
| chr11 | 62551230 | 62554112 | 22.0147 | 58.97336 | 56.14162 | Promoter | 11 | 62551676 | 62552853 | 1178 | 1 | ENSMUST00 | 0 | Ubb | ENSMUSG00000019505.7 |
| chr18 | 46727599 | 46728609 | 21.0133 | 58.92216 | 56.09053 | Promoter | 18 | 46713205 | 46728342 | 15138 | 2 | ENSMUST00 | 0 | Cdo1 | ENSMUSG00000033022.7 |
| chr10 | 79996429 | 79997945 | 21.49234 | 58.91379 | 56.0822 | Promoter | 10 | 79996494 | 80015572 | 19079 | 1 | ENSMUST00 | 0 | Abca7 | ENSMUSG00000035722.10 |
| chr16 | 14704873 | 14707329 | 21.49234 | 58.91379 | 56.0822 | Promoter | 16 | 14705852 | 14709395 | 3544 | 1 | ENSMUST00 | 0 | Snai2 | ENSMUSG00000022676.6 |
| chr7 | 16294945 | 16295821 | 21.49234 | 58.91379 | 56.0822 | Distal Interge | 7 | 16282755 | 16286795 | 4041 | 2 | ENSMUST00 | -8151 | Ccdc9 | ENSMUSG00000041375.13 |
| chr9 | 21031044 | 21033818 | 21.49234 | 58.91379 | 56.0822 | Promoter | 9 | 21032038 | 21039036 | 6999 | 1 | ENSMUST00 | 0 | Icam5 | ENSMUSG00000032174.4 |
| chr4 | 120730797 | 120731965 | 21.96955 | 58.89605 | 56.06502 | Intron (ENSM | 4 | 120696138 | 120747248 | 51111 | 2 | ENSMUST00 | 15283 | Kcnq4 | ENSMUSG00000028631.7 |
| chr4 | 108406051 | 108407135 | 20.02586 | 58.85371 | 56.02288 | Promoter | 4 | 108400390 | 108406961 | 6572 | 2 | ENSMUST00 | 0 | Gpx7 | ENSMUSG00000028597.11 |
| chr7 | 24316135 | 24317143 | 20.02586 | 58.85371 | 56.02288 | Promoter | 7 | 24301704 | 24316605 | 14902 | 2 | ENSMUST00 | 0 | Zfp94 | ENSMUSG00000074282.5 |
| chr8 | 88453839 | 88454852 | 21.45017 | 58.84098 | 56.01045 | Distal Interge | 8 | 88521344 | 88594887 | 73544 | 1 | ENSMUST00 | -66492 | Nkd1 | ENSMUSG00000031661.6 |
| chr7 | 27232642 | 27233642 | 21.92459 | 58.81904 | 55.9886 | Promoter | 7 | 27233023 | 27257950 | 24928 | 1 | ENSMUST00 | 0 | Adck4 | ENSMUSG00000003762.8 |
| chr6 | 42644695 | 42645610 | 21.40816 | 58.76844 | 55.9385 | Promoter | 6 | 42623019 | 42645254 | 22236 | 2 | ENSMUST00 | 0 | Fam115c | ENSMUSG00000029851.4 |

| | | | | | | | | | | | | | | | |
|---|---|---|---|---|---|---|---|---|---|---|---|---|---|---|---|
| chr4 | 106651039 | 106651672 | 21.87981 | 58.7423 | 55.91244 | Promoter | 4 | 106651069 | 106653539 | 2471 | 1 | ENSMUST00 | 0 | Pars2 | ENSMUSG00000043572.7 |
| chr12 | 117533667 | 117534635 | 21.36632 | 58.69615 | 55.86647 | Intron (ENSM | 12 | 117516479 | 117756978 | 240500 | 1 | ENSMUST00 | 17189 | Rapgef5 | ENSMUSG00000041992.8 |
| chr17 | 26662990 | 26663863 | 21.36632 | 58.69615 | 55.86647 | Distal Interge | 17 | 26676396 | 26699644 | 23249 | 1 | ENSMUST00 | -12533 | Atp6v0e | ENSMUSG00000015575.9 |
| chr18 | 37143450 | 37144693 | 21.83522 | 58.66587 | 55.83627 | Promoter | 18 | 37143969 | 37187657 | 43689 | 1 | ENSMUST00 | 0 | Pcdha10 | ENSMUSG00000007440.9 |
| chr1 | 58444707 | 58446248 | 19.90079 | 58.62946 | 55.80003 | Promoter | 1 | 58430994 | 58445429 | 14436 | 2 | ENSMUST00 | 0 | Ppil3 | ENSMUSG00000026035.10 |
| chr1 | 71652644 | 71654070 | 19.90079 | 58.62946 | 55.80003 | Promoter | 1 | 71585520 | 71653172 | 67653 | 2 | ENSMUST00 | 0 | Fn1 | ENSMUSG00000026193.10 |
| chr10 | 128015096 | 128015692 | 19.90079 | 58.62946 | 55.80003 | Promoter | 10 | 128015168 | 128029983 | 14816 | 1 | ENSMUST00 | 0 | Prim1 | ENSMUSG00000025395.9 |
| chr10 | 79613614 | 79614290 | 19.90079 | 58.62946 | 55.80003 | Promoter | 10 | 79606857 | 79614018 | 7162 | 2 | ENSMUST00 | 0 | C2cd4c | ENSMUSG00000045912.8 |
| chr12 | 76962441 | 76963203 | 19.90079 | 58.62946 | 55.80003 | Promoter | 12 | 76937269 | 76962248 | 24980 | 2 | ENSMUST00 | -194 | Max | ENSMUSG00000059436.7 |
| chr14 | 79486710 | 79487981 | 19.90079 | 58.62946 | 55.80003 | Intron (ENSM | 14 | 79459937 | 79481268 | 21332 | 2 | ENSMUST00 | -5443 | Wbp4 | ENSMUSG00000022023.5 |
| chr14 | 20082326 | 20083648 | 19.90079 | 58.62946 | 55.80003 | Promoter | 14 | 20075646 | 20083172 | 7527 | 2 | ENSMUST00 | 0 | Saysd1 | ENSMUSG00000045107.4 |
| chr17 | 17234300 | 17235950 | 19.90079 | 58.62946 | 55.80003 | Distal Interge | 17 | 17294443 | 17318104 | 23662 | 1 | ENSMUST00 | -58493 | Gm6712 | ENSMUSG00000072761.5 |
| chr2 | 112284259 | 112285182 | 19.90079 | 58.62946 | 55.80003 | Promoter | 2 | 112284577 | 112363156 | 78580 | 1 | ENSMUST00 | 0 | Slc12a6 | ENSMUSG00000027130.10 |
| chr4 | 44168264 | 44168962 | 19.90079 | 58.62946 | 55.80003 | Promoter | 4 | 44126215 | 44168339 | 42125 | 2 | ENSMUST00 | 0 | Rnf38 | ENSMUSG00000035696.10 |
| chr5 | 8798003 | 8799073 | 19.90079 | 58.62946 | 55.80003 | Promoter | 5 | 8798147 | 8866314 | 68168 | 1 | ENSMUST00 | 0 | Abcb1b | ENSMUSG00000028970.5 |
| chr5 | 122849418 | 122849958 | 19.90079 | 58.62946 | 55.80003 | Promoter | 5 | 122850188 | 122868945 | 18758 | 1 | ENSMUST00 | -230 | Rnf34 | ENSMUSG00000029474.6 |
| chr5 | 151650455 | 151651684 | 19.90079 | 58.62946 | 55.80003 | Promoter | 5 | 151642756 | 151651242 | 8487 | 2 | ENSMUST00 | 0 | Rfc3 | ENSMUSG00000033970.10 |
| chr5 | 110386393 | 110387736 | 19.90079 | 58.62946 | 55.80003 | Promoter | 5 | 110361754 | 110387127 | 25374 | 2 | ENSMUST00 | 0 | Fbrsl1 | ENSMUSG00000043323.11 |
| chr6 | 71143314 | 71144570 | 19.90079 | 58.62946 | 55.80003 | Promoter | 6 | 71128166 | 71144346 | 16181 | 2 | ENSMUST00 | 0 | Thnsl2 | ENSMUSG00000054474.10 |
| chr7 | 44895753 | 44896806 | 19.90079 | 58.62946 | 55.80003 | Promoter | 7 | 44896079 | 44900623 | 4545 | 1 | ENSMUST00 | 0 | Fuz | ENSMUSG00000011658.9 |
| chr7 | 100893076 | 100894901 | 19.90079 | 58.62946 | 55.80003 | Intron (ENSM | 7 | 100882306 | 100882409 | 104 | 2 | ENSMUST00 | -10668 | Mir3102 | ENSMUSG00000093296.1 |
| chr8 | 14089716 | 14091085 | 19.90079 | 58.62946 | 55.80003 | Promoter | 8 | 14027565 | 14090327 | 62763 | 2 | ENSMUST00 | 0 | Erich1 | ENSMUSG00000051978.8 |
| chr2 | 35581533 | 35583251 | 20.35144 | 58.59763 | 55.77069 | Promoter | 2 | 35582840 | 35730992 | 148153 | 1 | ENSMUST00 | 0 | Dab2ip | ENSMUSG00000026883.12 |
| chr9 | 106159717 | 106160690 | 20.35144 | 58.59763 | 55.77069 | Exon (ENSMU | 9 | 106152869 | 106158138 | 5270 | 2 | ENSMUST00 | -1580 | Glyctk | ENSMUSG00000020258.10 |
| chr11 | 78176183 | 78179336 | 13.57108 | 58.53591 | 55.70979 | Promoter | 11 | 78166106 | 78176611 | 10506 | 2 | ENSMUST00 | 0 | Nek8 | ENSMUSG00000017405.9 |
| chr2 | 11534837 | 11536076 | 21.70252 | 58.43828 | 55.61241 | Distal Interge | 2 | 11477967 | 11502101 | 24135 | 2 | ENSMUST00 | -32737 | Pfkfb3 | ENSMUSG00000026773.14 |
| chr15 | 85858919 | 85860146 | 15.02397 | 58.41 | 55.58427 | Promoter | 15 | 85859707 | 85876572 | 16866 | 1 | ENSMUST00 | 0 | Gtse1 | ENSMUSG00000022385.10 |
| chr1 | 74332151 | 74333246 | 21.20057 | 58.40949 | 55.58382 | Promoter | 1 | 74332610 | 74353686 | 21077 | 1 | ENSMUST00 | 0 | Pnkd | ENSMUSG00000026179.9 |
| chr1 | 171280294 | 171281198 | 20.7012 | 58.37697 | 55.55136 | Promoter | 1 | 171275990 | 171281186 | 5197 | 2 | ENSMUST00 | 0 | Ppox | ENSMUSG00000062729.6 |
| chr13 | 74616744 | 74618124 | 20.7012 | 58.37697 | 55.55136 | Distal Interge | 13 | 74639872 | 74691875 | 52004 | 1 | ENSMUST00 | -21748 | Erap1 | ENSMUSG00000021583.6 |
| chr17 | 25432876 | 25434527 | 20.7012 | 58.37697 | 55.55136 | Promoter | 17 | 25374285 | 25433783 | 59499 | 2 | ENSMUST00 | 0 | Cacna1h | ENSMUSG00000024112.11 |
| chr4 | 114924951 | 114926133 | 20.7012 | 58.37697 | 55.55136 | Promoter | 4 | 114925147 | 114926013 | 867 | 2 | ENSMUST00 | 0 | Foxe3 | ENSMUSG00000044518.3 |
| chr8 | 84639597 | 84640208 | 20.7012 | 58.37697 | 55.55136 | Promoter | 8 | 84638991 | 84639434 | 444 | 1 | ENSMUST00 | 607 | Cacna1a | ENSMUSG00000034656.11 |
| chr11 | 88973699 | 88974671 | 16.78291 | 58.37249 | 55.54753 | Promoter | 11 | 88973935 | 88991613 | 17679 | 1 | ENSMUST00 | 0 | Coil | ENSMUSG00000033983.8 |
| chr17 | 56199916 | 56200182 | 21.65864 | 58.36299 | 55.53803 | Intron (ENSM | 17 | 56175744 | 56183920 | 8177 | 2 | ENSMUST00 | -15997 | D17Wsu104e | ENSMUSG00000019579.8 |
| chr15 | 98180775 | 98181775 | 21.11866 | 58.26765 | 55.44304 | Distal Interge | 15 | 98167806 | 98170134 | 2329 | 1 | ENSMUST00 | 12970 | Ccdc184 | ENSMUSG00000029875.4 |
| chr11 | 24075913 | 24078659 | 19.68565 | 58.24254 | 55.41799 | Promoter | 11 | 24078056 | 24173194 | 95139 | 1 | ENSMUST00 | 0 | Bcl11a | ENSMUSG00000000861.10 |
| chr7 | 80246319 | 80246837 | 19.68565 | 58.24254 | 55.41799 | Promoter | 7 | 80246376 | 80260821 | 14446 | 1 | ENSMUST00 | 0 | Ttll13 | ENSMUSG00000045467.8 |
| chr12 | 113151461 | 113152599 | 21.57142 | 58.21323 | 55.38894 | Promoter | 12 | 113152012 | 113153877 | 1866 | 1 | ENSMUST00 | 0 | Crip1 | ENSMUSG00000006360.7 |
| chr7 | 44748081 | 44749367 | 21.57142 | 58.21323 | 55.38894 | Promoter | 7 | 44748413 | 44777515 | 29103 | 1 | ENSMUST00 | 0 | Vrk3 | ENSMUSG00000002205.10 |
| chr9 | 59577636 | 59578658 | 21.57142 | 58.21323 | 55.38894 | Promoter | 9 | 59577917 | 59607292 | 29376 | 1 | ENSMUST00 | 0 | Celf6 | ENSMUSG00000032297.6 |
| chr12 | 76822081 | 76822978 | 22.05754 | 58.21236 | 55.38818 | Promoter | 12 | 76797960 | 76822553 | 24594 | 2 | ENSMUST00 | 0 | Rab15 | ENSMUSG00000021062.10 |
| chr12 | 86421349 | 86422647 | 22.05754 | 58.21236 | 55.38818 | Promoter | 12 | 86421644 | 86515522 | 93879 | 1 | ENSMUST00 | 0 | Esrrb | ENSMUSG00000021255.12 |
| chr17 | 27511994 | 27513684 | 22.05754 | 58.21236 | 55.38818 | Promoter | 17 | 27425904 | 27513341 | 87438 | 2 | ENSMUST00 | 0 | Grm4 | ENSMUSG00000063239.11 |
| chr19 | 45014893 | 45017754 | 22.05754 | 58.21236 | 55.38818 | Promoter | 19 | 45015176 | 45027104 | 11929 | 1 | ENSMUST00 | 0 | Lzts2 | ENSMUSG00000035342.8 |
| chr2 | 35461025 | 35462807 | 22.05754 | 58.21236 | 55.38818 | Promoter | 2 | 35400179 | 35461266 | 61088 | 2 | ENSMUST00 | 0 | Ggta1 | ENSMUSG00000035778.12 |
| chr2 | 127791824 | 127792813 | 22.05754 | 58.21236 | 55.38818 | Promoter | 2 | 127791388 | 127792488 | 1101 | 2 | ENSMUST00 | 0 | 1500011K16I | ENSMUSG00000051319.6 |
| chr3 | 89159692 | 89160297 | 22.05754 | 58.21236 | 55.38818 | Promoter | 3 | 89146074 | 89160196 | 14123 | 2 | ENSMUST00 | 0 | Hcn3 | ENSMUSG00000028051.5 |
| chr6 | 127561115 | 127561758 | 22.05754 | 58.21236 | 55.38818 | Distal Interge | 6 | 127577975 | 127629938 | 51964 | 1 | ENSMUST00 | -16217 | Efcab4b | ENSMUSG00000061414.4 |
| chr7 | 110768357 | 110769407 | 22.05754 | 58.21236 | 55.38818 | Promoter | 7 | 110768343 | 110791269 | 22927 | 1 | ENSMUST00 | 15 | Ampd3 | ENSMUSG00000005686.11 |
| chr7 | 19576859 | 19578264 | 22.05754 | 58.21236 | 55.38818 | Promoter | 7 | 19577287 | 19580656 | 3370 | 1 | ENSMUST00 | 0 | Zfp296 | ENSMUSG00000011267.7 |
| chr8 | 94171990 | 94173502 | 22.05754 | 58.21236 | 55.38818 | Promoter | 8 | 94172618 | 94173567 | 950 | 1 | ENSMUST00 | 0 | Mt2 | ENSMUSG00000031762.6 |
| chrX | 20687911 | 20689733 | 22.05754 | 58.21236 | 55.38818 | Promoter | 20 | 20687954 | 20699880 | 11927 | 1 | ENSMUST00 | 0 | Cdk16 | ENSMUSG00000031065.9 |
| chrX | 107815505 | 107816688 | 22.05754 | 58.21236 | 55.38818 | Promoter | 20 | 107782751 | 107816334 | 33584 | 2 | ENSMUST00 | 0 | 2610002M06 | ENSMUSG00000031242.7 |
| chrX | 94212066 | 94213274 | 22.05754 | 58.21236 | 55.38818 | Promoter | 20 | 94188707 | 94212709 | 24003 | 2 | ENSMUST00 | 0 | Eif2s3x | ENSMUSG00000035150.10 |
| chrX | 8073771 | 8074610 | 22.05754 | 58.21236 | 55.38818 | Promoter | 20 | 8061435 | 8074258 | 12824 | 2 | ENSMUST00 | 0 | Suv39h1 | ENSMUSG00000039231.13 |
| chrX | 94234167 | 94235882 | 22.05754 | 58.21236 | 55.38818 | Promoter | 20 | 94234599 | 94238157 | 3559 | 1 | ENSMUST00 | 0 | Klhl15 | ENSMUSG00000043929.11 |
| chrX | 95956174 | 95956935 | 22.05754 | 58.21236 | 55.38818 | Promoter | 20 | 95935335 | 95956913 | 21579 | 2 | ENSMUST00 | 0 | Las1l | ENSMUSG00000057421.7 |
| chrX | 56597362 | 56598025 | 22.05754 | 58.21236 | 55.38818 | Promoter | 20 | 56585512 | 56598069 | 12558 | 2 | ENSMUST00 | 44 | Mmgt1 | ENSMUSG00000061273.3 |
| chr13 | 67107831 | 67109205 | 21.07794 | 58.1971 | 55.37426 | Promoter | 13 | 67069399 | 67108549 | 39151 | 2 | ENSMUST00 | 0 | RSLCAN18 | ENSMUSG00000100235.1 |
| chr7 | 28596642 | 28598238 | 21.07794 | 58.1971 | 55.37426 | Promoter | 7 | 28399549 | 28598144 | 198596 | 2 | ENSMUST00 | 0 | Pak4 | ENSMUSG00000030602.7 |
| chr7 | 145300780 | 145302012 | 21.07794 | 58.1971 | 55.37426 | Promoter | 7 | 145300828 | 145309557 | 8730 | 1 | ENSMUST00 | 0 | Mrgprf | ENSMUSG00000031070.10 |
| chr9 | 56437433 | 56437718 | 21.52807 | 58.13876 | 55.3166 | Intron (ENSM | 9 | 56418644 | 56496884 | 78241 | 1 | ENSMUST00 | 18790 | Hmg20a | ENSMUSG00000032329.3 |
| chr11 | 32204944 | 32205875 | 18.70486 | 58.09692 | 55.27494 | Promoter | 11 | 32205415 | 32208984 | 3570 | 1 | ENSMUST00 | 0 | Snrnp25 | ENSMUSG00000040767.5 |
| chr12 | 69404298 | 69406539 | 18.70486 | 58.09692 | 55.27494 | Distal Interge | 12 | 69371835 | 69372458 | 624 | 2 | ENSMUST00 | -31841 | Gm9887 | ENSMUSG00000052673.1 |
| chr13 | 6547649 | 6548804 | 18.70486 | 58.09692 | 55.27494 | Promoter | 13 | 6548157 | 6580139 | 31983 | 1 | ENSMUST00 | 0 | Pitrm1 | ENSMUSG00000021193.9 |
| chr15 | 37135021 | 37136216 | 18.70486 | 58.09692 | 55.27494 | Distal Interge | 15 | 37233036 | 37333647 | 100612 | 1 | ENSMUST00 | -96820 | Grhl2 | ENSMUSG00000022286.10 |
| chr18 | 80985485 | 80987981 | 18.70486 | 58.09692 | 55.27494 | Promoter | 18 | 80966376 | 80986578 | 20203 | 2 | ENSMUST00 | 0 | Sall3 | ENSMUSG00000024565.8 |
| chr18 | 61707110 | 61707860 | 18.70486 | 58.09692 | 55.27494 | Promoter | 18 | 61696837 | 61707635 | 10799 | 2 | ENSMUST00 | 0 | Pcyox1l | ENSMUSG00000024579.2 |
| chr18 | 24120871 | 24122263 | 18.70486 | 58.09692 | 55.27494 | Promoter | 18 | 24104757 | 24121953 | 17197 | 2 | ENSMUST00 | 0 | Ino80c | ENSMUSG00000047989.6 |
| chr2 | 144330377 | 144332324 | 18.70486 | 58.09692 | 55.27494 | Promoter | 2 | 144305175 | 144331695 | 26521 | 2 | ENSMUST00 | 0 | Ovol2 | ENSMUSG00000037279.8 |
| chr2 | 29054051 | 29055377 | 18.70486 | 58.09692 | 55.27494 | Promoter | 2 | 28926949 | 29055024 | 128076 | 2 | ENSMUST00 | 0 | 1700101E01I | ENSMUSG00000079502.3 |
| chr4 | 119414935 | 119415914 | 18.70486 | 58.09692 | 55.27494 | Promoter | 4 | 119323841 | 119415521 | 91681 | 2 | ENSMUST00 | 0 | Ccdc30 | ENSMUSG00000028637.10 |
| chr4 | 141059952 | 141061398 | 15.55856 | 58.09642 | 55.27494 | Promoter | 4 | 141016637 | 141060550 | 43914 | 2 | ENSMUST00 | 0 | Crocc | ENSMUSG00000040860.11 |
| chr4 | 47437444 | 47438190 | 18.70486 | 58.09692 | 55.27494 | Distal Interge | 4 | 47453632 | 47454953 | 1322 | 1 | ENSMUST00 | -15442 | Gm12430 | ENSMUSG00000081152.1 |
| chr5 | 98329057 | 98329625 | 18.70486 | 58.09692 | 55.27494 | Promoter | 5 | 98329354 | 98801910 | 472557 | 1 | ENSMUST00 | 0 | 1700007G11 | ENSMUSG00000057816.3 |
| chr7 | 6330784 | 6331810 | 18.70486 | 58.09692 | 55.27494 | Promoter | 7 | 6315660 | 6331285 | 15626 | 2 | ENSMUST00 | 0 | Zfp583 | ENSMUSG00000030443.11 |
| chr7 | 45509504 | 45510885 | 18.70486 | 58.09692 | 55.27494 | Promoter | 7 | 45492874 | 45510406 | 17533 | 2 | ENSMUST00 | 0 | Nucb1 | ENSMUSG00000030824.10 |
| chr7 | 58398508 | 58399911 | 18.70486 | 58.09692 | 55.27494 | Distal Interge | 7 | 58148096 | 58148227 | 132 | 2 | ENSMUST00 | -250282 | Gm26288 | ENSMUSG00000077356.1 |
| chr8 | 124751238 | 124752087 | 18.70486 | 58.09692 | 55.27494 | Promoter | 8 | 124740257 | 124751809 | 11553 | 2 | ENSMUST00 | 0 | Fam89a | ENSMUSG00000043068.5 |
| chr8 | 71476070 | 71477063 | 18.70486 | 58.09692 | 55.27494 | 3' UTR | 8 | 71470411 | 71476098 | 5688 | 1 | ENSMUST00 | 5660 | Dda1 | ENSMUSG00000074247.5 |
| chr8 | 129262985 | 129264150 | 18.70486 | 58.09692 | 55.27494 | Distal Interge | 8 | 129252998 | 129253597 | 600 | 1 | ENSMUST00 | 9988 | Gm21399 | ENSMUSG00000094530.1 |
| chr9 | 65629075 | 65631311 | 18.70486 | 58.09692 | 55.27494 | Promoter | 9 | 65629648 | 65660528 | 30881 | 1 | ENSMUST00 | 0 | Rbpms2 | ENSMUSG00000032387.9 |
| chr9 | 54981246 | 54982209 | 18.70486 | 58.09692 | 55.27494 | Promoter | 9 | 54980880 | 55007779 | 26900 | 1 | ENSMUST00 | 367 | Chrna5 | ENSMUSG00000035594.9 |
| chr9 | 78481906 | 78482475 | 18.70486 | 58.09692 | 55.27494 | Promoter | 9 | 78478449 | 78481724 | 3276 | 2 | ENSMUST00 | -183 | Eef1a1 | ENSMUSG00000037742.9 |
| chr3 | 88771231 | 88772027 | 21.97177 | 58.06593 | 55.24583 | Promoter | 3 | 88744701 | 88772599 | 27899 | 2 | ENSMUST00 | 572 | Syt11 | ENSMUSG00000068923.9 |
| chr2 | 118110556 | 118112456 | 21.4849 | 58.06458 | 55.24457 | Promoter | 2 | 118111876 | 118127133 | 15258 | 1 | ENSMUST00 | 0 | Thbs1 | ENSMUSG00000040152.8 |
| chr9 | 50745778 | 50746844 | 21.4849 | 58.06458 | 55.24457 | Promoter | 9 | 50739691 | 50746521 | 6831 | 2 | ENSMUST00 | 0 | 2310030G06 | ENSMUSG00000032062.1 |
| chr4 | 143389309 | 143389825 | 21.92587 | 57.98755 | 55.16829 | Promoter | 4 | 143389659 | 143390416 | 758 | 1 | ENSMUST00 | 0 | Anp32b-ps1 | ENSMUSG00000081792.4 |
| chr7 | 27837846 | 27839317 | 21.92587 | 57.98755 | 55.16829 | Distal Interge | 7 | 27842855 | 27855086 | 12232 | 1 | ENSMUST00 | -3538 | Zfp59 | ENSMUSG00000078779.3 |
| chr1 | 195017069 | 195018317 | 20.9555 | 57.98476 | 55.16582 | Promoter | 1 | 195017399 | 195037908 | 20510 | 1 | ENSMUST00 | 0 | A330023F24I | ENSMUSG00000096929.2 |
| chr10 | 63386235 | 63387236 | 20.9555 | 57.98476 | 55.16582 | Promoter | 10 | 63386563 | 63408840 | 22278 | 1 | ENSMUST00 | 0 | Dnajc12 | ENSMUSG00000036764.6 |
| chr11 | 49609025 | 49610617 | 20.9555 | 57.98476 | 55.16582 | Promoter | 11 | 49609263 | 49628941 | 19679 | 1 | ENSMUST00 | 0 | Flt4 | ENSMUSG00000020357.3 |
| chr11 | 105362936 | 105363544 | 20.9555 | 57.98476 | 55.16582 | Promoter | 11 | 105363252 | 105390642 | 27391 | 1 | ENSMUST00 | 0 | Gm11638 | ENSMUSG00000086487.1 |
| chr13 | 63249311 | 63254013 | 20.9555 | 57.98476 | 55.16582 | Intron (ENSM | 13 | 63282142 | 63326096 | 43955 | 1 | ENSMUST00 | -28129 | 2010111I01R | ENSMUSG00000021458.11 |
| chr13 | 98503874 | 98505195 | 20.9555 | 57.98476 | 55.16582 | Distal Interge | 13 | 98489348 | 98492001 | 2654 | 2 | ENSMUST00 | -11874 | Gm10320 | ENSMUSG00000092116.1 |
| chr19 | 47178464 | 47179677 | 20.9555 | 57.98476 | 55.16582 | Promoter | 19 | 47178820 | 47259440 | 80621 | 1 | ENSMUST00 | 0 | Neurl1a | ENSMUSG00000006435.10 |
| chr2 | 13010014 | 13011697 | 20.9555 | 57.98476 | 55.16582 | Promoter | 2 | 13003457 | 13011747 | 8291 | 2 | ENSMUST00 | 50 | C1ql3 | ENSMUSG00000049630.5 |
| chr2 | 21141410 | 21142657 | 20.9555 | 57.98476 | 55.16582 | Promoter | 2 | 21044808 | 21141954 | 97147 | 2 | ENSMUST00 | 0 | Gm13377 | ENSMUSG00000086821.3 |
| chr3 | 129724731 | 129725669 | 20.9555 | 57.98476 | 55.16582 | Exon (ENSMU | 3 | 129677575 | 129755322 | 77748 | 2 | ENSMUST00 | 29653 | Egf | ENSMUSG00000028017.3 |
| chr3 | 116861301 | 116862041 | 20.9555 | 57.98476 | 55.16582 | Distal Interge | 3 | 116878227 | 116903750 | 25524 | 1 | ENSMUST00 | -16186 | Frrs1 | ENSMUSG00000033386.6 |
| chr4 | 62470501 | 62471308 | 20.9555 | 57.98476 | 55.16582 | Promoter | 4 | 62452632 | 62470887 | 18256 | 2 | ENSMUST00 | 0 | Wdr31 | ENSMUSG00000028391.11 |
| chr4 | 116820878 | 116821929 | 20.9555 | 57.98476 | 55.16582 | Promoter | 4 | 116819904 | 116821707 | 1804 | 2 | ENSMUST00 | 0 | Hpdl | ENSMUSG00000043155.4 |
| chr6 | 134916606 | 134917371 | 20.9555 | 57.98476 | 55.16582 | Distal Interge | 6 | 134920401 | 134925513 | 5113 | 1 | ENSMUST00 | -3030 | Cdkn1b | ENSMUSG00000003031.9 |
| chr6 | 137648440 | 137649503 | 20.9555 | 57.98476 | 55.16582 | Promoter | 6 | 137477245 | 137649285 | 172041 | 2 | ENSMUST00 | 0 | Eps8 | ENSMUSG00000015766.9 |
| chr7 | 107664446 | 107667691 | 20.9555 | 57.98476 | 55.16582 | Intron (ENSM | 7 | 107623971 | 107625161 | 1191 | 2 | ENSMUST00 | -39286 | 5330417H12I | ENSMUSG00000049740.2 |
| chr9 | 42896787 | 42898140 | 20.9555 | 57.98476 | 55.16582 | Intron (ENSM | 9 | 42862051 | 42862129 | 79 | 2 | ENSMUST00 | -34659 | Gm27509 | ENSMUSG00000076355.2 |
| chrX | 23693026 | 23693954 | 20.9555 | 57.98476 | 55.16582 | Promoter | 20 | 23693051 | 23805990 | 112940 | 1 | ENSMUST00 | 0 | Wdr44 | ENSMUSG00000036769.9 |
| chr17 | 17120848 | 17122006 | 19.99354 | 57.96247 | 55.14572 | Promoter | 17 | 17121415 | 17146831 | 25417 | 1 | ENSMUST00 | 0 | Zfp97 | ENSMUSG00000095990.1 |
| chr2 | 119028580 | 119029472 | 19.99354 | 57.96247 | 55.14572 | Promoter | 2 | 119017779 | 119029393 | 11615 | 2 | ENSMUST00 | 0 | Ccdc32 | ENSMUSG00000039983.8 |
| chr1 | 72824443 | 72825771 | 21.88015 | 57.90947 | 55.09324 | Promoter | 1 | 72824503 | 72852474 | 27972 | 1 | ENSMUST00 | 0 | Igfbp2 | ENSMUSG00000039323.13 |
| chr14 | 54602842 | 54603818 | 16.17172 | 57.87897 | 55.06284 | Intron (ENSM | 14 | 54583663 | 54605908 | 22246 | 2 | ENSMUST00 | 2090 | 4931414P19I | ENSMUSG00000022179.4 |
| chr10 | 80930057 | 80931209 | 17.71499 | 57.83842 | 55.02271 | Promoter | 10 | 80930091 | 80932204 | 2114 | 1 | ENSMUST00 | 0 | Gadd45b | ENSMUSG00000015312.8 |
| chr13 | 23436216 | 23438696 | 17.71499 | 57.83842 | 55.02271 | Distal Interge | 13 | 23427975 | 23431022 | 3048 | 2 | ENSMUST00 | -5195 | C230035I16R | ENSMUSG00000085024.1 |
| chr15 | 101253655 | 101254885 | 17.71499 | 57.83842 | 55.02271 | Distal Interge | 15 | 101266846 | 101274792 | 7947 | 1 | ENSMUST00 | -11961 | Nr4a1 | ENSMUSG00000023034.6 |
| chr15 | 75615707 | 75617294 | 17.71499 | 57.83842 | 55.02271 | Promoter | 15 | 75616679 | 75625300 | 8622 | 1 | ENSMUST00 | 0 | Zfp41 | ENSMUSG00000047003.9 |
| chr19 | 57314342 | 57315314 | 17.71499 | 57.83842 | 55.02271 | Promoter | 19 | 57037251 | 57314919 | 277669 | 2 | ENSMUST00 | 0 | Ablim1 | ENSMUSG00000025085.11 |
| chr2 | 165233419 | 165235246 | 17.71499 | 57.83842 | 55.02271 | Promoter | 2 | 165111507 | 165234853 | 123347 | 2 | ENSMUST00 | 0 | Cdh22 | ENSMUSG00000053166.9 |
| chr6 | 83156002 | 83156727 | 17.71499 | 57.83842 | 55.02271 | Promoter | 6 | 83156404 | 83162975 | 6572 | 1 | ENSMUST00 | 0 | 1700003E16I | ENSMUSG00000030030.4 |
| chr9 | 21722916 | 21724671 | 17.71499 | 57.83842 | 55.02271 | Promoter | 9 | 21723576 | 21749916 | 26341 | 1 | ENSMUST00 | 0 | Ldlr | ENSMUSG00000032193.8 |
| chr10 | 81182941 | 81183458 | 16.88215 | 57.7818 | 54.96725 | Promoter | 10 | 81183007 | 81193197 | 10191 | 1 | ENSMUST00 | 0 | Dapk3 | ENSMUSG00000034974.7 |
| chr11 | 50841244 | 50841845 | 16.88215 | 57.7818 | 54.96725 | Promoter | 11 | 50832031 | 50841552 | 9522 | 2 | ENSMUST00 | 0 | Zfp879 | ENSMUSG00000044296.5 |
| chr13 | 13438186 | 13439077 | 16.88215 | 57.7818 | 54.96725 | Promoter | 13 | 13437602 | 13512275 | 74674 | 1 | ENSMUST00 | 585 | Nid1 | ENSMUSG00000005397.7 |
| chr13 | 100551427 | 100553367 | 16.88215 | 57.7818 | 54.96725 | Promoter | 13 | 100496507 | 100552498 | 55992 | 2 | ENSMUST00 | 0 | Ocln | ENSMUSG00000021638.7 |
| chr2 | 134786019 | 134787187 | 16.88215 | 57.7818 | 54.96725 | Promoter | 2 | 134786159 | 135475258 | 689100 | 1 | ENSMUST00 | 0 | Plcb1 | ENSMUSG00000051177.11 |
| chr5 | 24601323 | 24603193 | 16.88215 | 57.7818 | 54.96725 | Promoter | 5 | 24590818 | 24601976 | 11159 | 2 | ENSMUST00 | 0 | Smarcd3 | ENSMUSG00000028949.8 |
| chr7 | 141171961 | 141173303 | 16.88215 | 57.7818 | 54.96725 | Promoter | 7 | 141160328 | 141172851 | 12524 | 2 | ENSMUST00 | 0 | Rnh1 | ENSMUSG00000038650.9 |
| chr7 | 27689403 | 27689873 | 16.88215 | 57.7818 | 54.96725 | Promoter | 7 | 27689340 | 27706484 | 17145 | 1 | ENSMUST00 | 64 | C030039L03I | ENSMUSG00000057093.9 |

| | | | | | | | | | | | | | | | |
|---|---|---|---|---|---|---|---|---|---|---|---|---|---|---|---|
| chr7 | 139388179 | 139388840 | 21.31392 | 57.7705 | 54.95658 | Promoter | 7 | 139389109 | 139579652 | 190544 | 1 | ENSMUST00 | -269 | Inpp5a | ENSMUSG00000025477.8 |
| chr8 | 70796627 | 70798163 | 21.31392 | 57.7705 | 54.95658 | Promoter | 8 | 70798135 | 70798192 | 58 | 1 | ENSMUST00 | 0 | Mir7240 | ENSMUSG00000099031.1 |
| chr14 | 55091378 | 55092748 | 20.32388 | 57.71489 | 54.90119 | Promoter | 14 | 55060262 | 55092324 | 32063 | 2 | ENSMUST00 | 0 | Zfhx2 | ENSMUSG00000040721.9 |
| chr15 | 32920480 | 32923199 | 20.32388 | 57.71489 | 54.90119 | Promoter | 15 | 32920723 | 33034730 | 114008 | 1 | ENSMUST00 | 0 | Sdc2 | ENSMUSG00000022261.5 |
| chr4 | 5797750 | 5798789 | 20.32388 | 57.71489 | 54.90119 | Promoter | 4 | 5798084 | 5798531 | 448 | 1 | ENSMUST00 | 0 | Fam110b | ENSMUSG00000049119.9 |
| chr5 | 32863170 | 32864374 | 20.32388 | 57.71489 | 54.90119 | Promoter | 5 | 32863701 | 32887048 | 23348 | 1 | ENSMUST00 | 0 | Depdc5 | ENSMUSG00000037426.12 |
| chr7 | 70330931 | 70332393 | 20.32388 | 57.71489 | 54.90119 | Distal Interge | 7 | 70351946 | 70360593 | 8648 | 2 | ENSMUST00 | 28200 | Nr2f2 | ENSMUSG00000030551.8 |
| chr8 | 23149750 | 23150286 | 20.32388 | 57.71489 | 54.90119 | Promoter | 8 | 23138794 | 23149585 | 10792 | 2 | ENSMUST00 | -166 | Gm15816 | ENSMUSG00000086741.1 |
| chr12 | 31950570 | 31951169 | 21.2716 | 57.69764 | 54.88451 | Promoter | 12 | 31926254 | 31950535 | 24282 | 2 | ENSMUST00 | -36 | Hbp1 | ENSMUSG00000002996.12 |
| chr10 | 90828821 | 90830153 | 21.74415 | 57.67707 | 54.86403 | Promoter | 10 | 90829111 | 90941465 | 112355 | 1 | ENSMUST00 | 0 | Anks1b | ENSMUSG00000058589.9 |
| chr2 | 35621712 | 35622983 | 21.74415 | 57.67707 | 54.86403 | Promoter | 2 | 35621981 | 35730992 | 109012 | 1 | ENSMUST00 | 0 | Dab2ip | ENSMUSG00000026883.12 |
| chr9 | 59749942 | 59750763 | 21.74415 | 57.67707 | 54.86403 | Promoter | 9 | 59734259 | 59750649 | 16391 | 2 | ENSMUST00 | 0 | Senp8 | ENSMUSG00000051705.7 |
| chr14 | 32084642 | 32086564 | 19.35681 | 57.64819 | 54.83527 | Promoter | 14 | 32080566 | 32085609 | 5044 | 2 | ENSMUST00 | 0 | Dph3 | ENSMUSG00000021905.8 |
| chr19 | 60860902 | 60861931 | 21.22944 | 57.62505 | 54.81237 | Promoter | 19 | 60837277 | 60861427 | 24151 | 2 | ENSMUST00 | 0 | Sfxn4 | ENSMUSG00000063698.4 |
| chr19 | 5559894 | 5561298 | 21.69919 | 57.6002 | 54.78762 | Promoter | 19 | 5549137 | 5560575 | 11439 | 2 | ENSMUST00 | 0 | Ovol1 | ENSMUSG00000024922.1 |
| chr7 | 63211691 | 63212782 | 21.65441 | 57.52362 | 54.71128 | Promoter | 7 | 63098692 | 63212526 | 113835 | 2 | ENSMUST00 | 0 | Chrna7 | ENSMUSG00000030525.7 |
| chr14 | 54652953 | 54653965 | 20.67921 | 57.50454 | 54.69232 | Promoter | 14 | 54642161 | 54653616 | 11456 | 2 | ENSMUST00 | 0 | Acin1 | ENSMUSG00000022185.13 |
| chr19 | 4200842 | 4202377 | 20.67921 | 57.50454 | 54.69232 | Promoter | 19 | 4195198 | 4201603 | 6406 | 2 | ENSMUST00 | 0 | Rad9a | ENSMUSG00000024824.6 |
| chr2 | 170153300 | 170154119 | 20.67921 | 57.50454 | 54.69232 | Distal Interge | 2 | 170120719 | 170148103 | 27385 | 2 | ENSMUST00 | -5198 | Zfp217 | ENSMUSG00000052056.9 |
| chr6 | 34383947 | 34385289 | 20.67921 | 57.50454 | 54.69232 | Promoter | 6 | 34384218 | 34396950 | 12733 | 1 | ENSMUST00 | 0 | Akr1b10 | ENSMUSG00000061758.8 |
| chr8 | 117673228 | 117673898 | 20.67921 | 57.50454 | 54.69232 | Promoter | 8 | 117661402 | 117673689 | 12288 | 2 | ENSMUST00 | 0 | Sdr42e1 | ENSMUSG00000034308.8 |
| chr11 | 120643188 | 120643821 | 21.60982 | 57.44734 | 54.63594 | Promoter | 11 | 120635712 | 120643670 | 7959 | 2 | ENSMUST00 | 0 | Pycr1 | ENSMUSG00000025140.10 |
| chr3 | 40948945 | 40949626 | 21.60982 | 57.44734 | 54.63594 | Distal Interge | 3 | 40950631 | 40977793 | 27163 | 1 | ENSMUST00 | -1005 | Larp1b | ENSMUSG00000037814.9 |
| chr18 | 5162128 | 5163848 | 21.56542 | 57.37136 | 54.56027 | Exon (ENSMU | 18 | 5162880 | 5165731 | 2852 | 2 | ENSMUST00 | 1883 | Gm26682 | ENSMUSG00000097888.1 |
| chr3 | 156560459 | 156561242 | 21.56542 | 57.37136 | 54.56027 | Promoter | 3 | 156546761 | 156561582 | 14822 | 2 | ENSMUST00 | 340 | 4930570G19 | ENSMUSG00000085084.2 |
| chr1 | 72582695 | 72583963 | 19.64801 | 57.34576 | 54.5348 | Promoter | 1 | 72583251 | 72585991 | 2741 | 1 | ENSMUST00 | 0 | Smarcal1 | ENSMUSG00000039354.11 |
| chr11 | 94601426 | 94602381 | 19.64801 | 57.34576 | 54.5348 | Promoter | 11 | 94557102 | 94601871 | 44770 | 2 | ENSMUST00 | 0 | Acsf2 | ENSMUSG00000076435.3 |
| chr10 | 29535013 | 29536190 | 21.06249 | 57.33728 | 54.52658 | Promoter | 10 | 29453109 | 29535867 | 82759 | 2 | ENSMUST00 | 0 | Rspo3 | ENSMUSG00000019880.9 |
| chr18 | 35751662 | 35752528 | 21.06249 | 57.33728 | 54.52658 | Distal Interge | 18 | 35733678 | 35740554 | 6877 | 2 | ENSMUST00 | -11109 | Tmem173 | ENSMUSG00000024349.9 |
| chr19 | 5510725 | 5511536 | 21.06249 | 57.33728 | 54.52658 | Promoter | 19 | 5495278 | 5510489 | 15212 | 2 | ENSMUST00 | -237 | Snx32 | ENSMUSG00000056185.4 |
| chr2 | 93849274 | 93850375 | 21.06249 | 57.33728 | 54.52658 | Promoter | 2 | 93833467 | 93849941 | 16475 | 2 | ENSMUST00 | 0 | Accs | ENSMUSG00000040272.9 |
| chr8 | 114133594 | 114134346 | 21.06249 | 57.33728 | 54.52658 | Promoter | 8 | 114133618 | 114152312 | 18695 | 1 | ENSMUST00 | 0 | Nudt7 | ENSMUSG00000031767.8 |
| chr9 | 122310500 | 122311253 | 21.06249 | 57.33728 | 54.52658 | Promoter | 9 | 122306130 | 122310928 | 4799 | 2 | ENSMUST00 | 0 | Gm26797 | ENSMUSG00000097572.1 |
| chr11 | 103170531 | 103171521 | 21.52119 | 57.29567 | 54.48558 | Promoter | 11 | 103171107 | 103198901 | 27795 | 1 | ENSMUST00 | 0 | Fmnl1 | ENSMUSG00000055805.9 |
| chr7 | 45215589 | 45216644 | 21.52119 | 57.29567 | 54.48558 | Promoter | 7 | 45215753 | 45233619 | 17867 | 1 | ENSMUST00 | 0 | Tead2 | ENSMUSG00000030796.10 |
| chr1 | 171110763 | 171112407 | 21.47715 | 57.22026 | 54.41045 | Distal Interge | 1 | 171113918 | 171126829 | 12912 | 1 | ENSMUST00 | -1511 | 1700009P17I | ENSMUSG00000026649.9 |
| chr15 | 86032848 | 86035051 | 21.47715 | 57.22026 | 54.41045 | Promoter | 15 | 85898758 | 86033777 | 135020 | 2 | ENSMUST00 | 0 | Celsr1 | ENSMUSG00000016028.9 |
| chr17 | 35993807 | 35995359 | 21.47715 | 57.22026 | 54.41045 | Promoter | 17 | 35994453 | 36038435 | 43983 | 1 | ENSMUST00 | 0 | A930015D03 | ENSMUSG00000092368.1 |
| chr8 | 95677448 | 95678620 | 21.47715 | 57.22026 | 54.41045 | Promoter | 8 | 95678085 | 95714956 | 36872 | 1 | ENSMUST00 | 0 | Ndrg4 | ENSMUSG00000036564.11 |
| chr7 | 99141486 | 99141993 | 21.43328 | 57.14514 | 54.33587 | Promoter | 7 | 98886743 | 99141144 | 254402 | 2 | ENSMUST00 | -343 | Uvrag | ENSMUSG00000035354.8 |
| chr8 | 88696151 | 88696778 | 21.43328 | 57.14514 | 54.33587 | Promoter | 8 | 88697028 | 88751945 | 54918 | 1 | ENSMUST00 | -250 | Cyld | ENSMUSG00000036712.9 |
| chr1 | 128590520 | 128593468 | 19.52531 | 57.12588 | 54.31675 | Promoter | 1 | 128588199 | 128592290 | 4092 | 2 | ENSMUST00 | 0 | Cxcr4 | ENSMUSG00000045382.6 |
| chr10 | 75589036 | 75591503 | 19.52531 | 57.12588 | 54.31675 | Promoter | 10 | 75589340 | 75617200 | 27861 | 1 | ENSMUST00 | 0 | Ggt5 | ENSMUSG00000006344.11 |
| chr10 | 128546563 | 128547789 | 19.52531 | 57.12588 | 54.31675 | Promoter | 10 | 128541963 | 128547774 | 5812 | 2 | ENSMUST00 | 0 | Zc3h10 | ENSMUSG00000039810.6 |
| chr10 | 86498271 | 86499152 | 19.52531 | 57.12588 | 54.31675 | Promoter | 10 | 86055125 | 86498856 | 443732 | 2 | ENSMUST00 | 0 | Syn3 | ENSMUSG00000059602.9 |
| chr11 | 48833520 | 48834497 | 19.52531 | 57.12588 | 54.31675 | Intron (ENSM | 11 | 48837465 | 48850555 | 13091 | 1 | ENSMUST00 | -2968 | Trim7 | ENSMUSG00000040350.10 |
| chr14 | 21988610 | 21990214 | 19.52531 | 57.12588 | 54.31675 | Promoter | 14 | 21983962 | 21989601 | 5640 | 2 | ENSMUST00 | 0 | Zfp503 | ENSMUSG00000039081.9 |
| chr14 | 52118544 | 52119654 | 19.52531 | 57.12588 | 54.31675 | Promoter | 14 | 52118977 | 52119558 | 582 | 1 | ENSMUST00 | 0 | Rpgrip1 | ENSMUSG00000057132.10 |
| chr16 | 30510208 | 30511499 | 19.52531 | 57.12588 | 54.31675 | Downstream | 16 | 30513970 | 30540909 | 26940 | 2 | ENSMUST00 | 29410 | Tmem44 | ENSMUSG00000022537.12 |
| chr17 | 87745446 | 87746878 | 19.52531 | 57.12588 | 54.31675 | 3' UTR | 17 | 87696541 | 87717678 | 21138 | 1 | ENSMUST00 | 48906 | Msh2 | ENSMUSG00000024151.8 |
| chr2 | 150450591 | 150451911 | 19.52531 | 57.12588 | 54.31675 | Promoter | 2 | 150407141 | 150451486 | 44346 | 2 | ENSMUST00 | 0 | Zfp442 | ENSMUSG00000068130.6 |
| chr2 | 118676806 | 118677874 | 19.52531 | 57.12588 | 54.31675 | Promoter | 2 | 118677229 | 118698018 | 20790 | 1 | ENSMUST00 | 0 | Pak6 | ENSMUSG00000074923.5 |
| chr3 | 108209908 | 108211309 | 19.52531 | 57.12588 | 54.31675 | Promoter | 3 | 108202222 | 108210486 | 8265 | 2 | ENSMUST00 | 0 | Atxn7l2 | ENSMUSG00000048997.12 |
| chr4 | 129639863 | 129641496 | 19.52531 | 57.12588 | 54.31675 | Promoter | 4 | 129626078 | 129641065 | 14988 | 2 | ENSMUST00 | 0 | Txlna | ENSMUSG00000053841.10 |
| chr4 | 57567668 | 57568619 | 19.52531 | 57.12588 | 54.31675 | Promoter | 4 | 57568179 | 57896982 | 328804 | 1 | ENSMUST00 | 0 | Palm2Akap2 | ENSMUSG00000089945.5 |
| chr5 | 115731087 | 115732982 | 19.52531 | 57.12588 | 54.31675 | Promoter | 5 | 115648175 | 115731621 | 83447 | 2 | ENSMUST00 | 0 | Ccdc64 | ENSMUSG00000041609.11 |
| chr6 | 124828247 | 124828989 | 19.52531 | 57.12588 | 54.31675 | Promoter | 6 | 124824136 | 124828895 | 4760 | 2 | ENSMUST00 | 0 | Usp5 | ENSMUSG00000038429.5 |
| chr7 | 143501704 | 143503089 | 19.52531 | 57.12588 | 54.31675 | Promoter | 7 | 143501545 | 143502541 | 997 | 2 | ENSMUST00 | 0 | Phlda2 | ENSMUSG00000010760.4 |
| chr7 | 109780974 | 109781998 | 19.52531 | 57.12588 | 54.31675 | Promoter | 7 | 109758059 | 109781545 | 23487 | 2 | ENSMUST00 | 0 | Nrip3 | ENSMUSG00000034825.8 |
| chr8 | 34806492 | 34810291 | 19.52531 | 57.12588 | 54.31675 | Promoter | 8 | 34807297 | 34819894 | 12598 | 1 | ENSMUST00 | 0 | Dusp4 | ENSMUSG00000031530.6 |
| chr8 | 23058112 | 23059056 | 19.52531 | 57.12588 | 54.31675 | Promoter | 8 | 23058259 | 23150497 | 92239 | 1 | ENSMUST00 | 0 | Ank1 | ENSMUSG00000031543.13 |
| chr9 | 110304494 | 110305925 | 19.52531 | 57.12588 | 54.31675 | Promoter | 9 | 110305192 | 110322092 | 16901 | 1 | ENSMUST00 | 0 | Elp6 | ENSMUSG00000054836.8 |
| chr2 | 122233894 | 122235351 | 14.79283 | 57.11871 | 54.31201 | Promoter | 2 | 122234749 | 122265340 | 30592 | 1 | ENSMUST00 | 0 | Sord | ENSMUSG00000027227.7 |
| chr9 | 9238203 | 9239549 | 14.79283 | 57.11871 | 54.31201 | Promoter | 9 | 8994329 | 9239101 | 244773 | 2 | ENSMUST00 | 0 | Arhgap42 | ENSMUSG00000050730.11 |
| chr3 | 129752368 | 129755481 | 20.44714 | 57.09995 | 54.29332 | Promoter | 3 | 129677575 | 129755322 | 77748 | 2 | ENSMUST00 | 0 | Egf | ENSMUSG00000028017.3 |
| chr3 | 88254293 | 88254954 | 19.96007 | 57.07313 | 54.26652 | Promoter | 3 | 88242874 | 88254709 | 11836 | 2 | ENSMUST00 | 0 | Rhbg | ENSMUSG00000001417.5 |
| chr4 | 43009833 | 43010972 | 19.96007 | 57.07313 | 54.26652 | Promoter | 4 | 43002343 | 43010355 | 8013 | 2 | ENSMUST00 | 0 | Fancg | ENSMUSG00000028453.5 |
| chr7 | 70369313 | 70370722 | 21.3896 | 57.0703 | 54.26409 | Intron (ENSM | 7 | 70354439 | 70366735 | 12297 | 2 | ENSMUST00 | -2579 | Nr2f2 | ENSMUSG00000030551.8 |
| chr2 | 154557669 | 154559308 | 20.89815 | 57.05354 | 54.24761 | Promoter | 2 | 154544399 | 154558851 | 14453 | 2 | ENSMUST00 | 0 | Necab3 | ENSMUSG00000027489.10 |
| chr4 | 123139142 | 123140214 | 21.34609 | 56.99575 | 54.18992 | Promoter | 4 | 123127115 | 123139951 | 12837 | 2 | ENSMUST00 | 0 | Ppie | ENSMUSG00000028651.7 |
| chr11 | 98774164 | 98775366 | 20.85746 | 56.98322 | 54.17754 | Promoter | 11 | 98767932 | 98775333 | 7402 | 2 | ENSMUST00 | 0 | Nr1d1 | ENSMUSG00000020889.11 |
| chr3 | 146149816 | 146150506 | 20.81693 | 56.91315 | 54.10769 | Promoter | 3 | 146149833 | 146195513 | 45681 | 1 | ENSMUST00 | 0 | Mcoln2 | ENSMUSG00000011008.8 |
| chr9 | 106447118 | 106449172 | 20.77656 | 56.84333 | 54.03812 | Promoter | 9 | 106447788 | 106451609 | 3822 | 1 | ENSMUST00 | 0 | Abhd14b | ENSMUSG00000042073.9 |
| chr3 | 53041138 | 53042152 | 20.2953 | 56.83459 | 54.02946 | Promoter | 3 | 53041528 | 53261679 | 220152 | 1 | ENSMUST00 | 0 | Lhfp | ENSMUSG00000048332.8 |
| chr7 | 99827718 | 99828727 | 20.2953 | 56.83459 | 54.02946 | Promoter | 7 | 99811439 | 99828417 | 16979 | 2 | ENSMUST00 | 0 | Neu3 | ENSMUSG00000035239.6 |
| chr10 | 23796517 | 23797521 | 15.31159 | 56.78092 | 53.97643 | Promoter | 10 | 23796986 | 23827968 | 30983 | 1 | ENSMUST00 | 0 | Slc18b1 | ENSMUSG00000037455.11 |
| chr10 | 58812228 | 58813758 | 21.21663 | 56.77375 | 53.96939 | Promoter | 10 | 58813359 | 59050020 | 236662 | 1 | ENSMUST00 | 0 | Sh3rf3 | ENSMUSG00000037990.13 |
| chr19 | 43523562 | 43525376 | 13.72719 | 56.76396 | 53.95979 | Promoter | 19 | 43499752 | 43524605 | 24854 | 2 | ENSMUST00 | 0 | Got1 | ENSMUSG00000025190.8 |
| chr16 | 18581216 | 18582925 | 19.31422 | 56.74648 | 53.94233 | 3' UTR | 16 | 18581713 | 18586969 | 5257 | 2 | ENSMUST00 | 4044 | Tbx1 | ENSMUSG00000009097.9 |
| chr3 | 89182855 | 89183747 | 19.31422 | 56.74648 | 53.94233 | Promoter | 3 | 89183143 | 89184679 | 1537 | 1 | ENSMUST00 | 0 | Fam189b | ENSMUSG00000032657.10 |
| chr4 | 117288453 | 117289927 | 20.69629 | 56.7044 | 53.90051 | Promoter | 4 | 117271464 | 117288772 | 17309 | 2 | ENSMUST00 | 0 | Rnf220 | ENSMUSG00000028677.13 |
| chr11 | 4094614 | 4095769 | 18.36477 | 56.64417 | 53.84045 | Promoter | 11 | 4091480 | 4095445 | 3966 | 2 | ENSMUST00 | 0 | Mtfp1 | ENSMUSG00000004748.5 |
| chr11 | 48869670 | 48872632 | 18.36477 | 56.64417 | 53.84045 | Promoter | 11 | 48861968 | 48871683 | 9716 | 2 | ENSMUST00 | 0 | Irgm1 | ENSMUSG00000046879.6 |
| chr15 | 102692259 | 102693634 | 18.36477 | 56.64417 | 53.84045 | Distal Interge | 15 | 102706972 | 102709902 | 2931 | 2 | ENSMUST00 | 16268 | Calcoco1 | ENSMUSG00000023055.4 |
| chr15 | 81693459 | 81695071 | 18.36477 | 56.64417 | 53.84045 | Promoter | 15 | 81678296 | 81694028 | 15733 | 2 | ENSMUST00 | 0 | Chadl | ENSMUSG00000063765.6 |
| chr18 | 24708503 | 24710423 | 18.36477 | 56.64417 | 53.84045 | Promoter | 18 | 24709445 | 25133500 | 424056 | 1 | ENSMUST00 | 0 | Fhod3 | ENSMUSG00000034295.9 |
| chr19 | 6991854 | 6993065 | 18.36477 | 56.64417 | 53.84045 | Promoter | 19 | 6987911 | 6992272 | 4362 | 2 | ENSMUST00 | 0 | Dnajc4 | ENSMUSG00000024963.6 |
| chr4 | 45018316 | 45019389 | 18.36477 | 56.64417 | 53.84045 | Promoter | 4 | 45018583 | 45033692 | 15110 | 1 | ENSMUST00 | 0 | Polr1e | ENSMUSG00000028318.9 |
| chr4 | 21688125 | 21689306 | 18.36477 | 56.64417 | 53.84045 | Distal Interge | 4 | 21677481 | 21685963 | 8483 | 2 | ENSMUST00 | -2163 | Prdm13 | ENSMUSG00000040478.9 |
| chr4 | 107314316 | 107315256 | 18.36477 | 56.64417 | 53.84045 | Promoter | 4 | 107314363 | 107359822 | 45460 | 1 | ENSMUST00 | 0 | Yipf1 | ENSMUSG00000057375.8 |
| chr4 | 127125797 | 127126519 | 18.36477 | 56.64417 | 53.84045 | Promoter | 4 | 127126046 | 127129644 | 3599 | 1 | ENSMUST00 | 0 | Gm12942 | ENSMUSG00000070737.5 |
| chr6 | 7692466 | 7693443 | 18.36477 | 56.64417 | 53.84045 | Promoter | 6 | 7675169 | 7692871 | 17703 | 2 | ENSMUST00 | 0 | Asns | ENSMUSG00000029752.7 |
| chr6 | 108658527 | 108661954 | 18.36477 | 56.64417 | 53.84045 | Promoter | 6 | 108660629 | 108666925 | 6297 | 1 | ENSMUST00 | 0 | Bhlhe40 | ENSMUSG00000030103.6 |
| chr7 | 145283332 | 145284140 | 18.36477 | 56.64417 | 53.84045 | Promoter | 7 | 145253923 | 145283927 | 30005 | 2 | ENSMUST00 | 0 | Tpcn2 | ENSMUSG00000048677.8 |
| chr9 | 123717772 | 123718283 | 18.36477 | 56.64417 | 53.84045 | Promoter | 9 | 123697012 | 123717625 | 20614 | 2 | ENSMUST00 | -148 | Lztfl1 | ENSMUSG00000025245.8 |
| chr9 | 40414422 | 40415228 | 18.36477 | 56.64417 | 53.84045 | Intron (ENSM | 9 | 40317511 | 40392450 | 74940 | 2 | ENSMUST00 | -21973 | Gramd1b | ENSMUSG00000040111.9 |
| chr5 | 124439909 | 124440688 | 19.71305 | 56.63538 | 53.8334 | Promoter | 5 | 124439930 | 124462308 | 22379 | 1 | ENSMUST00 | 0 | Setd8 | ENSMUSG00000049327.12 |
| chr10 | 80928632 | 80929826 | 20.65638 | 56.63529 | 53.83331 | Promoter | 10 | 80930091 | 80932204 | 2114 | 1 | ENSMUST00 | -265 | Gadd45b | ENSMUSG00000015312.8 |
| chr3 | 90292388 | 90292926 | 20.65638 | 56.63529 | 53.83331 | Distal Interge | 3 | 90278008 | 90278670 | 663 | 1 | ENSMUST00 | 14381 | Dennd4b | ENSMUSG00000042404.11 |
| chr3 | 96263509 | 96264194 | 20.65638 | 56.63529 | 53.83331 | Promoter | 3 | 96261682 | 96263311 | 1630 | 2 | ENSMUST00 | -199 | Hist2h4 | ENSMUSG00000091405.2 |
| chr10 | 34483224 | 34485079 | 21.59801 | 56.60135 | 53.80005 | Promoter | 10 | 34483400 | 34611226 | 127827 | 1 | ENSMUST00 | 0 | Frk | ENSMUSG00000019779.9 |
| chr11 | 102087123 | 102089604 | 21.59801 | 56.60135 | 53.80005 | Promoter | 11 | 102057015 | 102088515 | 31501 | 2 | ENSMUST00 | 0 | Mpp2 | ENSMUSG00000017314.7 |
| chr11 | 116653464 | 116654389 | 21.59801 | 56.60135 | 53.80005 | Promoter | 11 | 116645595 | 116654313 | 8719 | 2 | ENSMUST00 | 0 | Cygb | ENSMUSG00000020810.5 |
| chr12 | 81630687 | 81632079 | 21.59801 | 56.60135 | 53.80005 | Promoter | 12 | 81631369 | 81664941 | 33573 | 1 | ENSMUST00 | 0 | Ttc9 | ENSMUSG00000042734.5 |
| chr12 | 54713478 | 54714886 | 21.59801 | 56.60135 | 53.80005 | Promoter | 12 | 54714218 | 54714381 | 164 | 1 | ENSMUST00 | 0 | Gm23804 | ENSMUSG00000094826.1 |
| chr13 | 31628983 | 31630033 | 21.59801 | 56.60135 | 53.80005 | Intron (ENSM | 13 | 31626446 | 31626543 | 98 | 2 | ENSMUST00 | -2441 | Gm27516 | ENSMUSG00000098266.1 |
| chr14 | 26206365 | 26207119 | 21.59801 | 56.60135 | 53.80005 | Promoter | 14 | 26203369 | 26207041 | 3673 | 2 | ENSMUST00 | 0 | Tmem254b | ENSMUSG00000021867.11 |
| chr14 | 60506281 | 60507719 | 21.59801 | 56.60135 | 53.80005 | Exon (ENSMU | 14 | 60485208 | 60599031 | 113824 | 1 | ENSMUST00 | 21074 | RP24-127N1 | ENSMUSG00000101797.1 |
| chr17 | 21909047 | 21910121 | 21.59801 | 56.60135 | 53.80005 | Promoter | 17 | 21879570 | 21909926 | 30357 | 2 | ENSMUST00 | 0 | 2210404O09 | ENSMUSG00000078546.3 |
| chr19 | 44511286 | 44512337 | 21.59801 | 56.60135 | 53.80005 | 3' UTR | 19 | 44517644 | 44520593 | 2950 | 2 | ENSMUST00 | 8256 | Sec31b | ENSMUSG00000051984.8 |
| chr2 | 30077582 | 30078298 | 21.59801 | 56.60135 | 53.80005 | Promoter | 2 | 30077684 | 30081125 | 3442 | 1 | ENSMUST00 | 0 | Pkn3 | ENSMUSG00000026785.9 |
| chr2 | 155018305 | 155020039 | 21.59801 | 56.60135 | 53.80005 | Promoter | 2 | 155018266 | 155027104 | 8839 | 1 | ENSMUST00 | 40 | Gm14226 | ENSMUSG00000084897.1 |
| chr4 | 129987553 | 129988458 | 21.59801 | 56.60135 | 53.80005 | Intron (ENSM | 4 | 129985268 | 130022633 | 37366 | 1 | ENSMUST00 | 2286 | Bai2 | ENSMUSG00000028782.9 |
| chr4 | 149675474 | 149676671 | 21.59801 | 56.60135 | 53.80005 | Promoter | 4 | 149649171 | 149676044 | 26874 | 2 | ENSMUST00 | 0 | Pik3cd | ENSMUSG00000039936.13 |
| chr6 | 115774489 | 115775073 | 21.59801 | 56.60135 | 53.80005 | Promoter | 6 | 115774538 | 115804893 | 30356 | 1 | ENSMUST00 | 0 | Cand2 | ENSMUSG00000030319.7 |
| chr7 | 19822220 | 19823046 | 21.59801 | 56.60135 | 53.80005 | Promoter | 7 | 19808462 | 19822770 | 14309 | 2 | ENSMUST00 | 0 | Bcl3 | ENSMUSG00000053175.11 |
| chr9 | 114562275 | 114564649 | 21.59801 | 56.60135 | 53.80005 | Promoter | 9 | 114507133 | 114564369 | 57237 | 2 | ENSMUST00 | 0 | Trim71 | ENSMUSG00000079259.2 |
| chrX | 151800846 | 151801654 | 21.59801 | 56.60135 | 53.80005 | Promoter | 20 | 151800883 | 151804511 | 3629 | 1 | ENSMUST00 | 0 | Huwe1 | ENSMUSG00000025261.12 |
| chrX | 151046631 | 151048116 | 21.59801 | 56.60135 | 53.80005 | Promoter | 20 | 151047170 | 151089521 | 42352 | 1 | ENSMUST00 | 0 | Fgd1 | ENSMUSG00000025265.9 |
| chrX | 52987728 | 52989091 | 21.59801 | 56.60135 | 53.80005 | Promoter | 20 | 52988137 | 53021659 | 33523 | 1 | ENSMUST00 | 0 | Hprt | ENSMUSG00000025630.8 |
| chrX | 47891758 | 47892570 | 21.59801 | 56.60135 | 53.80005 | Promoter | 20 | 47809368 | 47892519 | 83152 | 2 | ENSMUST00 | 0 | Smarca1 | ENSMUSG00000031099.11 |
| chrX | 57391747 | 57393024 | 21.59801 | 56.60135 | 53.80005 | Promoter | 20 | 57383530 | 57392981 | 9452 | 2 | ENSMUST00 | 0 | Rbmx | ENSMUSG00000031134.11 |
| chrX | 8144353 | 8146212 | 21.59801 | 56.60135 | 53.80005 | Promoter | 20 | 8138978 | 8145697 | 6720 | 2 | ENSMUST00 | 0 | Rbm3 | ENSMUSG00000031167.11 |
| chrX | 8251803 | 8253291 | 21.59801 | 56.60135 | 53.80005 | Promoter | 20 | 8238668 | 8252406 | 13739 | 2 | ENSMUST00 | 0 | Ftsj1 | ENSMUSG00000031171.6 |
| chrX | 169319768 | 169320406 | 21.59801 | 56.60135 | 53.80005 | Promoter | 20 | 169250193 | 169320372 | 70180 | 2 | ENSMUST00 | 0 | Hccs | ENSMUSG00000031352.5 |
| chrX | 168672052 | 168674729 | 21.59801 | 56.60135 | 53.80005 | Promoter | 20 | 168654117 | 168673898 | 19782 | 2 | ENSMUST00 | 0 | Msl3 | ENSMUSG00000031358.12 |
| chrX | 102643741 | 102645047 | 21.59801 | 56.60135 | 53.80005 | Promoter | 20 | 102513975 | 102644246 | 130272 | 2 | ENSMUST00 | 0 | Phka1 | ENSMUSG00000034055.11 |
| chrX | 129749389 | 129750968 | 21.59801 | 56.60135 | 53.80005 | Promoter | 20 | 129749742 | 130180471 | 430730 | 1 | ENSMUST00 | 0 | Diap2 | ENSMUSG00000034480.13 |

| | | | | | | | | | | | | | | | |
|---|---|---|---|---|---|---|---|---|---|---|---|---|---|---|---|
| chrX | 20291251 | 20292098 | 21.59801 | 56.60135 | 53.80005 | Promoter | 20 | 20105755 | 20291807 | 186053 | 2 | ENSMUST00 | 0 | Slc9a7 | ENSMUSG00000037341.8 |
| chrX | 159532608 | 159533488 | 21.59801 | 56.60135 | 53.80005 | Promoter | 20 | 159532668 | 159593081 | 60414 | 1 | ENSMUST00 | 0 | A830080D01 | ENSMUSG00000044150.7 |
| chrX | 38189347 | 38190847 | 21.59801 | 56.60135 | 53.80005 | Promoter | 20 | 38189793 | 38197046 | 7254 | 1 | ENSMUST00 | 0 | Zbtb33 | ENSMUSG00000048047.3 |
| chrX | 166509715 | 166510472 | 21.59801 | 56.60135 | 53.80005 | Promoter | 20 | 166499812 | 166510478 | 10667 | 2 | ENSMUST00 | 6 | Tceanc | ENSMUSG00000051224.8 |
| chrX | 104482463 | 104485053 | 21.59801 | 56.60135 | 53.80005 | Promoter | 20 | 104482818 | 104507490 | 24673 | 1 | ENSMUST00 | 0 | Uprt | ENSMUSG00000073016.3 |
| chr14 | 65833453 | 65833985 | 20.61663 | 56.56642 | 53.76645 | Promoter | 14 | 65819038 | 65833994 | 14957 | 2 | ENSMUST00 | 9 | Esco2 | ENSMUSG00000022034.9 |
| chr16 | 17646147 | 17646967 | 20.61663 | 56.56642 | 53.76645 | Promoter | 16 | 17646470 | 17649415 | 2946 | 1 | ENSMUST00 | 0 | Ccdc74a | ENSMUSG00000041617.5 |
| chr17 | 51831671 | 51833605 | 15.90661 | 56.5361 | 53.73628 | Promoter | 17 | 51736187 | 51832672 | 96486 | 2 | ENSMUST00 | 0 | Satb1 | ENSMUSG00000023927.10 |
| chr2 | 125246816 | 125248232 | 15.90661 | 56.5361 | 53.73628 | Promoter | 2 | 125247190 | 125257748 | 10559 | 1 | ENSMUST00 | 0 | Dut | ENSMUSG00000027203.10 |
| chr3 | 127779785 | 127780770 | 15.90661 | 56.5361 | 53.73628 | Promoter | 3 | 127670399 | 127780527 | 110129 | 2 | ENSMUST00 | 0 | Alpk1 | ENSMUSG00000028028.6 |
| chr9 | 107231033 | 107232285 | 15.90661 | 56.5361 | 53.73628 | Promoter | 9 | 106892834 | 107231909 | 339076 | 2 | ENSMUST00 | 0 | Dock3 | ENSMUSG00000039716.7 |
| chr11 | 89100705 | 89101521 | 21.55916 | 56.53507 | 53.73548 | Distal Interge | 11 | 89087048 | 89087151 | 104 | 2 | ENSMUST00 | -13555 | Gm22534 | ENSMUSG00000088798.1 |
| chr6 | 50455174 | 50456558 | 18.73102 | 56.5065 | 53.70699 | Promoter | 6 | 50296799 | 50456170 | 159372 | 2 | ENSMUST00 | 0 | Osbpl3 | ENSMUSG00000029822.10 |
| chr18 | 23989012 | 23990303 | 21.04643 | 56.48158 | 53.68221 | Promoter | 18 | 23989632 | 24005376 | 15745 | 1 | ENSMUST00 | 0 | Zfp35 | ENSMUSG00000063281.9 |
| chr3 | 96277788 | 96278601 | 21.04643 | 56.48158 | 53.68221 | 3' UTR | 3 | 96267075 | 96270289 | 3215 | 2 | ENSMUST00 | -7500 | Gm20628 | ENSMUSG00000093656.1 |
| chr5 | 38158912 | 38160110 | 21.04643 | 56.48158 | 53.68221 | Promoter | 5 | 38137193 | 38159467 | 22275 | 2 | ENSMUST00 | 0 | Nsg1 | ENSMUSG00000029126.4 |
| chr7 | 45016468 | 45017564 | 21.04643 | 56.48158 | 53.68221 | Promoter | 7 | 45002948 | 45016363 | 13416 | 2 | ENSMUST00 | -106 | Scaf1 | ENSMUSG00000038406.10 |
| chr7 | 83631425 | 83632921 | 21.04643 | 56.48158 | 53.68221 | Promoter | 7 | 83631959 | 83642318 | 10360 | 1 | ENSMUST00 | 0 | Stard5 | ENSMUSG00000046027.11 |
| chr12 | 57575849 | 57576569 | 19.60905 | 56.45056 | 53.6518 | Promoter | 12 | 57575817 | 57737928 | 162112 | 1 | ENSMUST00 | 33 | Ttc6 | ENSMUSG00000046782.9 |
| chr2 | 66123899 | 66125366 | 19.60905 | 56.45056 | 53.6518 | Promoter | 2 | 66082766 | 66124994 | 42229 | 2 | ENSMUST00 | 0 | Galnt3 | ENSMUSG00000026994.4 |
| chr2 | 157456913 | 157457681 | 19.60905 | 56.45056 | 53.6518 | Promoter | 2 | 157457085 | 157470355 | 13271 | 1 | ENSMUST00 | 0 | Src | ENSMUSG00000027646.10 |
| chr7 | 44970351 | 44970794 | 19.60905 | 56.45056 | 53.6518 | Exon (ENSMU | 7 | 44959373 | 44974851 | 15479 | 2 | ENSMUST00 | 4057 | Cpt1c | ENSMUSG00000007783.9 |
| chr1 | 138856365 | 138857236 | 17.4042 | 56.42813 | 53.62976 | Promoter | 1 | 138851979 | 138856854 | 4876 | 2 | ENSMUST00 | 0 | 2310009B15I | ENSMUSG00000079283.1 |
| chr11 | 77492500 | 77493299 | 20.53639 | 56.4273 | 53.62976 | Promoter | 11 | 77493562 | 77507786 | 14225 | 1 | ENSMUST00 | -263 | Git1 | ENSMUSG00000011877.8 |
| chr11 | 20200618 | 20201079 | 20.53639 | 56.4273 | 53.62976 | Promoter | 11 | 20201432 | 20226856 | 25425 | 1 | ENSMUST00 | -353 | Rab1 | ENSMUSG00000020149.12 |
| chr11 | 30197643 | 30198889 | 20.53639 | 56.4273 | 53.62976 | Promoter | 11 | 30106785 | 30198257 | 91473 | 2 | ENSMUST00 | 0 | Sptbn1 | ENSMUSG00000020315.13 |
| chr11 | 96270998 | 96272761 | 20.53639 | 56.4273 | 53.62976 | Promoter | 11 | 96271457 | 96276595 | 5139 | 1 | ENSMUST00 | 0 | Hoxb9 | ENSMUSG00000020875.9 |
| chr12 | 91383813 | 91384959 | 17.4042 | 56.42813 | 53.62976 | Promoter | 12 | 90998492 | 91384409 | 385918 | 2 | ENSMUST00 | 0 | Cep128 | ENSMUSG00000061533.10 |
| chr16 | 44062938 | 44063625 | 20.53639 | 56.4273 | 53.62976 | Distal Interge | 16 | 43980355 | 44027945 | 47591 | 2 | ENSMUST00 | -34994 | Gramd1c | ENSMUSG00000036292.8 |
| chr17 | 34602775 | 34603756 | 20.53639 | 56.4273 | 53.62976 | Promoter | 17 | 34601091 | 34603690 | 2600 | 2 | ENSMUST00 | 0 | Rnf5 | ENSMUSG00000015478.8 |
| chr17 | 34197335 | 34198797 | 20.53639 | 56.4273 | 53.62976 | Promoter | 17 | 34197721 | 34200918 | 3198 | 1 | ENSMUST00 | 0 | Psmb8 | ENSMUSG00000024338.9 |
| chr17 | 94833969 | 94835327 | 20.53639 | 56.4273 | 53.62976 | Promoter | 17 | 94759944 | 94834799 | 74856 | 2 | ENSMUST00 | 0 | Gm1976 | ENSMUSG00000066057.6 |
| chr19 | 60874075 | 60874892 | 17.4042 | 56.42813 | 53.62976 | Promoter | 19 | 60864051 | 60874556 | 10506 | 2 | ENSMUST00 | 0 | Prdx3 | ENSMUSG00000024997.7 |
| chr2 | 104742362 | 104743204 | 17.4042 | 56.42813 | 53.62976 | Promoter | 2 | 104721784 | 104742878 | 21095 | 2 | ENSMUST00 | 0 | Depdc7 | ENSMUSG00000027173.7 |
| chr3 | 65973958 | 65975550 | 17.4042 | 56.42813 | 53.62976 | Distal Interge | 3 | 65946151 | 65958249 | 12099 | 2 | ENSMUST00 | -15710 | Ccnl1 | ENSMUSG00000027829.10 |
| chr4 | 63409582 | 63410994 | 20.53639 | 56.4273 | 53.62976 | Distal Interge | 4 | 63367125 | 63403354 | 36230 | 2 | ENSMUST00 | -6229 | Akna | ENSMUSG00000039158.11 |
| chr4 | 149136915 | 149137713 | 20.53639 | 56.4273 | 53.62976 | Promoter | 4 | 149127121 | 149137548 | 10428 | 2 | ENSMUST00 | 0 | Apitd1 | ENSMUSG00000073705.5 |
| chr5 | 66675778 | 66677132 | 20.53639 | 56.4273 | 53.62976 | Promoter | 5 | 66676098 | 66687231 | 11134 | 1 | ENSMUST00 | 0 | Uchl1 | ENSMUSG00000029223.9 |
| chr5 | 140608552 | 140610115 | 20.53639 | 56.4273 | 53.62976 | Intron (ENSM | 5 | 140607341 | 140615543 | 8203 | 1 | ENSMUST00 | 1212 | Lfng | ENSMUSG00000029570.1 |
| chr5 | 139324944 | 139325865 | 17.4042 | 56.42813 | 53.62976 | Promoter | 5 | 139271876 | 139325622 | 53747 | 2 | ENSMUST00 | 0 | Adap1 | ENSMUSG00000056413.11 |
| chr6 | 38380761 | 38381881 | 20.53639 | 56.4273 | 53.62976 | Promoter | 6 | 38381469 | 38427647 | 46179 | 1 | ENSMUST00 | 0 | Ttc26 | ENSMUSG00000056832.9 |
| chr8 | 75093119 | 75094099 | 20.53639 | 56.4273 | 53.62976 | Promoter | 8 | 75093591 | 75100596 | 7006 | 1 | ENSMUST00 | 0 | Hmox1 | ENSMUSG00000005413.7 |
| chr8 | 22227115 | 22227798 | 17.4042 | 56.42813 | 53.62976 | Promoter | 8 | 22227313 | 22261334 | 34022 | 1 | ENSMUST00 | 0 | Thsd1 | ENSMUSG00000031480.10 |
| chr9 | 119092493 | 119093965 | 17.4042 | 56.42813 | 53.62976 | Promoter | 9 | 119071527 | 119093496 | 21970 | 2 | ENSMUST00 | 0 | Plcd1 | ENSMUSG00000010660.2 |
| chr13 | 23430426 | 23432630 | 21.00431 | 56.40921 | 53.61382 | Promoter | 13 | 23427975 | 23431022 | 3048 | 2 | ENSMUST00 | 0 | C230035I16R | ENSMUSG00000085024.1 |
| chr1 | 6382520 | 6383355 | 16.59601 | 56.40767 | 53.61239 | Intron (ENSM | 1 | 6359331 | 6394731 | 35401 | 1 | ENSMUST00 | 23190 | Fam150a | ENSMUSG00000087247.2 |
| chr10 | 111297960 | 111299142 | 16.59601 | 56.40767 | 53.61239 | Promoter | 10 | 111298679 | 111301736 | 3058 | 1 | ENSMUST00 | 0 | Bbs10 | ENSMUSG00000035759.3 |
| chr12 | 4037994 | 4039133 | 16.59601 | 56.40767 | 53.61239 | Promoter | 12 | 3962554 | 4038915 | 76362 | 2 | ENSMUST00 | 0 | Efr3b | ENSMUSG00000020658.9 |
| chr4 | 43025093 | 43025900 | 16.59601 | 56.40767 | 53.61239 | Promoter | 4 | 43017635 | 43025756 | 8122 | 2 | ENSMUST00 | 0 | Pigo | ENSMUSG00000028454.11 |
| chr7 | 19344647 | 19345778 | 16.59601 | 56.40767 | 53.61239 | Promoter | 7 | 19344778 | 19356524 | 11747 | 1 | ENSMUST00 | 0 | Ercc1 | ENSMUSG00000003549.4 |
| chr7 | 19282455 | 19283197 | 16.59601 | 56.40767 | 53.61239 | Promoter | 7 | 19282662 | 19296164 | 13503 | 1 | ENSMUST00 | 0 | Rtn2 | ENSMUSG00000030401.10 |
| chr8 | 71205473 | 71206176 | 16.59601 | 56.40767 | 53.61239 | Distal Interge | 8 | 71256038 | 71256653 | 616 | 2 | ENSMUST00 | 50477 | Haus8 | ENSMUSG00000035439.8 |
| chr14 | 52243947 | 52245093 | 20.96235 | 56.3371 | 53.54275 | Distal Interge | 14 | 52198451 | 52237791 | 39341 | 2 | ENSMUST00 | -6157 | Chd8 | ENSMUSG00000053754.8 |
| chr4 | 145774244 | 145774822 | 14.11932 | 56.26894 | 53.47484 | Promoter | 4 | 145774719 | 145775283 | 565 | 1 | ENSMUST00 | 0 | Gm13243 | ENSMUSG00000094417.1 |
| chr11 | 97050565 | 97051738 | 21.37974 | 56.22882 | 53.43486 | Promoter | 11 | 97050628 | 97068904 | 18277 | 1 | ENSMUST00 | 0 | Osbpl7 | ENSMUSG00000038534.12 |
| chr17 | 68840805 | 68842224 | 21.37974 | 56.22882 | 53.43486 | Exon (ENSMU | 17 | 68837136 | 68843138 | 6003 | 1 | ENSMUST00 | 3670 | Tmem200c | ENSMUSG00000095407.1 |
| chr17 | 21845023 | 21846086 | 20.42014 | 56.22556 | 53.43172 | Promoter | 17 | 21816876 | 21845759 | 28884 | 2 | ENSMUST00 | 0 | Zfp820 | ENSMUSG00000069743.4 |
| chr11 | 87578063 | 87578884 | 19.92537 | 56.18586 | 53.39219 | Promoter | 11 | 87578401 | 87590539 | 12139 | 1 | ENSMUST00 | 0 | Sep-04 | ENSMUSG00000020486.13 |
| chr12 | 72407776 | 72408968 | 19.92537 | 56.18586 | 53.39219 | Promoter | 12 | 72211752 | 72409054 | 197303 | 2 | ENSMUST00 | 86 | Rtn1 | ENSMUSG00000021087.12 |
| chr2 | 180256442 | 180257196 | 19.92537 | 56.18586 | 53.39219 | Promoter | 2 | 180257377 | 180258445 | 1069 | 1 | ENSMUST00 | -181 | Rps21 | ENSMUSG00000039001.7 |
| chr3 | 88214883 | 88216290 | 19.92537 | 56.18586 | 53.39219 | Promoter | 3 | 88215589 | 88215696 | 108 | 2 | ENSMUST00 | 0 | Gm25641 | ENSMUSG00000092876.1 |
| chr4 | 34729724 | 34730516 | 19.92537 | 56.18586 | 53.39219 | Promoter | 4 | 34688559 | 34730206 | 41648 | 2 | ENSMUST00 | 0 | 1700003M02 | ENSMUSG00000028294.10 |
| chr11 | 79660266 | 79661139 | 18.99158 | 56.16371 | 53.37056 | Promoter | 11 | 79660562 | 79683498 | 22937 | 1 | ENSMUST00 | 0 | Rab11fip4 | ENSMUSG00000017639.8 |
| chr15 | 75981175 | 75982500 | 18.99158 | 56.16371 | 53.37056 | Promoter | 15 | 75979872 | 75982455 | 2584 | 2 | ENSMUST00 | 0 | Ccdc166 | ENSMUSG00000098176.1 |
| chr9 | 44133925 | 44135926 | 18.99158 | 56.16371 | 53.37056 | Promoter | 9 | 44134469 | 44138934 | 4466 | 1 | ENSMUST00 | 0 | Mcam | ENSMUSG00000032135.9 |
| chr5 | 24423745 | 24426392 | 21.33535 | 56.153 | 53.36009 | Promoter | 5 | 24423837 | 24427612 | 3776 | 1 | ENSMUST00 | 0 | Slc4a2 | ENSMUSG00000028962.9 |
| chr7 | 127572645 | 127574176 | 21.29115 | 56.07748 | 53.28482 | Promoter | 7 | 127573340 | 127578621 | 5282 | 1 | ENSMUST00 | 0 | Phkg2 | ENSMUSG00000030815.6 |
| chr13 | 40916900 | 40918236 | 21.24712 | 56.00225 | 53.20983 | Promoter | 13 | 40917634 | 40960891 | 43258 | 1 | ENSMUST00 | 0 | Gcnt2 | ENSMUSG00000021360.10 |
| chr11 | 76570744 | 76571776 | 18.43306 | 55.95763 | 53.16539 | Promoter | 11 | 76416734 | 76571535 | 154802 | 2 | ENSMUST00 | 0 | Abr | ENSMUSG00000017631.13 |
| chr13 | 21440418 | 21441471 | 20.26563 | 55.95697 | 53.16479 | Promoter | 13 | 21421275 | 21440901 | 19627 | 2 | ENSMUST00 | 0 | Pgbd1 | ENSMUSG00000055313.9 |
| chr14 | 65343139 | 65343787 | 20.26563 | 55.95697 | 53.16479 | Distal Interge | 14 | 65358676 | 65398930 | 40255 | 1 | ENSMUST00 | -14889 | Zfp395 | ENSMUSG00000034522.9 |
| chr14 | 50807147 | 50808551 | 20.26563 | 55.95697 | 53.16479 | Promoter | 14 | 50807946 | 50821300 | 13355 | 1 | ENSMUST00 | 0 | Parp2 | ENSMUSG00000036023.5 |
| chr18 | 67088074 | 67089317 | 20.26563 | 55.95697 | 53.16479 | Promoter | 18 | 67088336 | 67226792 | 138457 | 1 | ENSMUST00 | 0 | Gnal | ENSMUSG00000024524.11 |
| chr2 | 30469735 | 30471289 | 20.26563 | 55.95697 | 53.16479 | Downstream | 2 | 30471537 | 30474219 | 2683 | 2 | ENSMUST00 | 2930 | Ier5l | ENSMUSG00000089762.3 |
| chr3 | 89202916 | 89203496 | 20.26563 | 55.95697 | 53.16479 | Promoter | 3 | 89202928 | 89208679 | 5752 | 1 | ENSMUST00 | 0 | Gba | ENSMUSG00000028048.6 |
| chr12 | 14151778 | 14152799 | 21.20328 | 55.92731 | 53.13585 | Promoter | 12 | 14147599 | 14152038 | 4440 | 2 | ENSMUST00 | 0 | Fam84a | ENSMUSG00000020607.6 |
| chr17 | 56474360 | 56476713 | 21.20328 | 55.92731 | 53.13585 | Promoter | 17 | 56412433 | 56476483 | 64051 | 2 | ENSMUST00 | 0 | Ptprs | ENSMUSG00000013236.11 |
| chr9 | 59680104 | 59680672 | 20.22737 | 55.89037 | 53.0991 | Promoter | 9 | 59680144 | 59713923 | 33780 | 1 | ENSMUST00 | 0 | Gramd2 | ENSMUSG00000074259.5 |
| chr11 | 95009394 | 95010240 | 21.15962 | 55.85266 | 53.06144 | Promoter | 11 | 95009879 | 95026087 | 16209 | 1 | ENSMUST00 | 0 | Samd14 | ENSMUSG00000047181.7 |
| chr9 | 50698317 | 50699375 | 21.15962 | 55.85266 | 53.06144 | Promoter | 9 | 50662756 | 50699362 | 36607 | 2 | ENSMUST00 | 0 | Dixdc1 | ENSMUSG00000032064.12 |
| chr4 | 136142962 | 136145178 | 19.27017 | 55.84607 | 53.05495 | Promoter | 4 | 136143497 | 136145392 | 1896 | 1 | ENSMUST00 | 0 | Id3 | ENSMUSG00000007872.3 |
| chr7 | 19420947 | 19421717 | 19.27017 | 55.84607 | 53.05495 | Promoter | 7 | 19418812 | 19421417 | 2606 | 2 | ENSMUST00 | 0 | A930016O22 | ENSMUSG00000040705.1 |
| chr15 | 98893717 | 98895109 | 20.67329 | 55.83958 | 53.04871 | 3' UTR | 15 | 98893031 | 98898540 | 5510 | 2 | ENSMUST00 | 3431 | Dhh | ENSMUSG00000023000.4 |
| chr4 | 63558298 | 63559257 | 14.56169 | 55.83414 | 53.04337 | Promoter | 4 | 63558781 | 63586357 | 27577 | 1 | ENSMUST00 | 0 | 6330416G13 | ENSMUSG00000045917.12 |
| chr6 | 47780786 | 47781347 | 19.70982 | 55.80626 | 53.0156 | Promoter | 6 | 47781624 | 47781725 | 102 | 1 | ENSMUST00 | -277 | Rny3 | ENSMUSG00000064945.1 |
| chr10 | 22158277 | 22159656 | 20.63265 | 55.76951 | 52.97896 | Promoter | 10 | 22158569 | 22183917 | 25349 | 1 | ENSMUST00 | 0 | Raet1e | ENSMUSG00000053219.10 |
| chr11 | 116086763 | 116087033 | 20.63265 | 55.76951 | 52.97896 | Promoter | 11 | 116078573 | 116086995 | 8423 | 2 | ENSMUST00 | 0 | Wbp2 | ENSMUSG00000034341.12 |
| chr3 | 31056300 | 31057454 | 20.63265 | 55.76951 | 52.97896 | Distal Interge | 3 | 31038966 | 31039028 | 63 | 1 | ENSMUST00 | 17335 | Mir7008 | ENSMUSG00000098890.1 |
| chr4 | 114999349 | 115000515 | 20.63265 | 55.76951 | 52.97896 | Promoter | 4 | 115000159 | 115043196 | 43038 | 1 | ENSMUST00 | 0 | Stil | ENSMUSG00000028718.11 |
| chr5 | 137627921 | 137628752 | 20.63265 | 55.76951 | 52.97896 | Promoter | 5 | 137612506 | 137628956 | 16451 | 2 | ENSMUST00 | 204 | Fbxo24 | ENSMUSG00000089984.2 |
| chr11 | 23498177 | 23498540 | 21.07283 | 55.70422 | 52.91429 | Promoter | 11 | 23488547 | 23498030 | 9484 | 2 | ENSMUST00 | -148 | Ahsa2 | ENSMUSG00000020288.8 |
| chr2 | 151979576 | 151980566 | 21.07283 | 55.70422 | 52.91429 | Promoter | 2 | 151969398 | 151980219 | 10822 | 2 | ENSMUST00 | 0 | Fam110a | ENSMUSG00000027459.11 |
| chr3 | 95881776 | 95883244 | 21.07283 | 55.70422 | 52.91429 | Promoter | 3 | 95878503 | 95882205 | 3703 | 2 | ENSMUST00 | 0 | Ciart | ENSMUSG00000038550.5 |
| chr1 | 21218527 | 21219653 | 21.02971 | 55.63043 | 52.84086 | Promoter | 1 | 21218575 | 21230167 | 11593 | 1 | ENSMUST00 | 0 | Tmem14a | ENSMUSG00000025933.10 |
| chr1 | 23761809 | 23763464 | 19.14982 | 55.63055 | 52.84086 | Promoter | 1 | 23761926 | 23849158 | 87233 | 1 | ENSMUST00 | 0 | B3gat2 | ENSMUSG00000026156.8 |
| chr10 | 18545620 | 18546277 | 19.14982 | 55.63055 | 52.84086 | Promoter | 10 | 18540123 | 18546076 | 5954 | 2 | ENSMUST00 | 0 | Hebp2 | ENSMUSG00000019853.5 |
| chr10 | 91187503 | 91188452 | 19.14982 | 55.63055 | 52.84086 | Distal Interge | 10 | 91160417 | 91171582 | 11166 | 2 | ENSMUST00 | -15922 | Tmpo | ENSMUSG00000019961.11 |
| chr11 | 98357889 | 98359357 | 19.14982 | 55.63055 | 52.84086 | Promoter | 11 | 98358368 | 98381099 | 22732 | 1 | ENSMUST00 | 0 | Stard3 | ENSMUSG00000018167.5 |
| chr11 | 119524618 | 119525063 | 19.14982 | 55.63055 | 52.84086 | Distal Interge | 11 | 119501978 | 119507389 | 5412 | 1 | ENSMUST00 | 22641 | Endov | ENSMUSG00000039850.11 |
| chr11 | 75170188 | 75170675 | 19.14982 | 55.63055 | 52.84086 | Promoter | 11 | 75164565 | 75169519 | 4955 | 2 | ENSMUST00 | -670 | Hic1 | ENSMUSG00000043099.4 |
| chr13 | 13783834 | 13785333 | 19.14982 | 55.63055 | 52.84086 | Promoter | 13 | 13784059 | 13806113 | 22055 | 1 | ENSMUST00 | 0 | Gng4 | ENSMUSG00000021303.8 |
| chr13 | 45871618 | 45872678 | 19.14982 | 55.63055 | 52.84086 | Intron (ENSM | 13 | 45549758 | 45964991 | 415234 | 2 | ENSMUST00 | 92313 | Atxn1 | ENSMUSG00000046876.10 |
| chr13 | 62015417 | 62016461 | 19.14982 | 55.63055 | 52.84086 | Distal Interge | 13 | 62013035 | 62013144 | 110 | 2 | ENSMUST00 | -2274 | Gm25649 | ENSMUSG00000094096.1 |
| chr15 | 79327467 | 79328736 | 19.14982 | 55.63055 | 52.84086 | Promoter | 15 | 79286228 | 79327964 | 41737 | 2 | ENSMUST00 | 0 | Pla2g6 | ENSMUSG00000042632.11 |
| chr16 | 38476131 | 38477433 | 19.14982 | 55.63055 | 52.84086 | Intron (ENSM | 16 | 38458933 | 38486933 | 28001 | 1 | ENSMUST00 | 17199 | Cd80 | ENSMUSG00000075122.4 |
| chr17 | 35342245 | 35343652 | 19.14982 | 55.63055 | 52.84086 | Promoter | 17 | 35342242 | 35345794 | 3553 | 1 | ENSMUST00 | 4 | H2-Q2 | ENSMUSG00000091705.3 |
| chr19 | 5071646 | 5072829 | 19.14982 | 55.63055 | 52.84086 | Promoter | 19 | 5070859 | 5072132 | 1274 | 2 | ENSMUST00 | 0 | Tmem151a | ENSMUSG00000061451.8 |
| chr19 | 12009975 | 12013473 | 19.14982 | 55.63055 | 52.84086 | Distal Interge | 19 | 12035808 | 12036740 | 933 | 2 | ENSMUST00 | 23267 | Olfr1423 | ENSMUSG00000067529.1 |
| chr2 | 24473135 | 24475336 | 19.14982 | 55.63055 | 52.84086 | Promoter | 2 | 24421520 | 24475097 | 53578 | 2 | ENSMUST00 | 0 | Pax8 | ENSMUSG00000026976.10 |
| chr4 | 43045142 | 43046346 | 19.14982 | 55.63055 | 52.84086 | Promoter | 4 | 43032414 | 43045797 | 13384 | 2 | ENSMUST00 | 0 | Fam214b | ENSMUSG00000036002.7 |
| chr5 | 109844619 | 109845276 | 19.14982 | 55.63055 | 52.84086 | Distal Interge | 5 | 109886231 | 109891137 | 4907 | 1 | ENSMUST00 | -40955 | Gm26779 | ENSMUSG00000097140.1 |
| chr6 | 91156460 | 91157255 | 19.14982 | 55.63055 | 52.84086 | Promoter | 6 | 91156665 | 91174692 | 18028 | 1 | ENSMUST00 | 0 | Hdac11 | ENSMUSG00000034245.5 |
| chr6 | 52012707 | 52013310 | 19.14982 | 55.63055 | 52.84086 | Promoter | 6 | 51859166 | 52012549 | 153384 | 2 | ENSMUST00 | -159 | Skap2 | ENSMUSG00000059182.5 |
| chr7 | 100590651 | 100591651 | 19.14982 | 55.63055 | 52.84086 | Promoter | 7 | 100550583 | 100591426 | 40844 | 2 | ENSMUST00 | 0 | Mrpl48 | ENSMUSG00000030706.11 |
| chr7 | 43465030 | 43466623 | 19.14982 | 55.63055 | 52.84086 | Exon (ENSMU | 7 | 43463151 | 43471913 | 8763 | 1 | ENSMUST00 | 1880 | Vsig10l | ENSMUSG00000070604.3 |
| chr9 | 21161156 | 21161755 | 19.14982 | 55.63055 | 52.84086 | Distal Interge | 9 | 21165714 | 21213248 | 47535 | 1 | ENSMUST00 | -3959 | Pde4a | ENSMUSG00000032177.11 |
| chr9 | 13296261 | 13298513 | 19.14982 | 55.63055 | 52.84086 | Distal Interge | 9 | 13275516 | 13275622 | 107 | 1 | ENSMUST00 | 20746 | Gm26189 | ENSMUSG00000095068.1 |
| chr3 | 83789476 | 83790166 | 18.67955 | 55.59657 | 52.80968 | Promoter | 3 | 83773853 | 83789956 | 16104 | 2 | ENSMUST00 | 0 | Gm26771 | ENSMUSG00000097250.1 |
| chr1 | 118627523 | 118628593 | 19.5687 | 55.55706 | 52.77042 | Promoter | 1 | 118627945 | 118637766 | 9822 | 1 | ENSMUST00 | 0 | Tfcp2l1 | ENSMUSG00000026380.10 |
| chr1 | 119912773 | 119913697 | 19.5687 | 55.55706 | 52.77042 | Promoter | 1 | 119907890 | 119913208 | 5319 | 2 | ENSMUST00 | 0 | Tmem177 | ENSMUSG00000036975.5 |
| chr14 | 55761825 | 55762933 | 20.98676 | 55.55692 | 52.77042 | Promoter | 14 | 55761428 | 55763229 | 1802 | 1 | ENSMUST00 | 398 | Ltb4r2 | ENSMUSG00000040432.4 |
| chr19 | 15924003 | 15925843 | 19.5687 | 55.55706 | 52.77042 | Promoter | 19 | 15904678 | 15925059 | 20382 | 2 | ENSMUST00 | 0 | Psat1 | ENSMUSG00000024640.4 |
| chr10 | 128082089 | 128083044 | 20.47165 | 55.49172 | 52.70563 | Promoter | 10 | 128083273 | 128090391 | 7119 | 1 | ENSMUST00 | -229 | Atp5b | ENSMUSG00000025393.7 |
| chr19 | 4037316 | 4038461 | 15.06463 | 55.47236 | 52.68647 | Promoter | 19 | 4035407 | 4037921 | 2515 | 2 | ENSMUST00 | 0 | Gstp1 | ENSMUSG00000060803.5 |
| chr17 | 37028426 | 37029167 | 20.90138 | 55.41071 | 52.6251 | Distal Interge | 17 | 37010743 | 37023398 | 12656 | 2 | ENSMUST00 | -5029 | Mog | ENSMUSG00000076439.7 |
| chr1 | 181255919 | 181257413 | 19.88939 | 55.30079 | 52.51555 | Distal Interge | 1 | 181241988 | 181242320 | 333 | 2 | ENSMUST00 | -13600 | Rpl35a-ps2 | ENSMUSG00000081214.3 |
| chr11 | 95666922 | 95667800 | 19.88939 | 55.30079 | 52.51555 | Promoter | 11 | 95666957 | 95680773 | 13817 | 1 | ENSMUST00 | 0 | Phb | ENSMUSG00000038846.6 |
| chr16 | 44331889 | 44333411 | 19.88939 | 55.30079 | 52.51555 | Promoter | 16 | 44240180 | 44332991 | 92812 | 2 | ENSMUST00 | 0 | Sidt1 | ENSMUSG00000022696.12 |

| | | | | | | | | | | | | | | | |
|---|---|---|---|---|---|---|---|---|---|---|---|---|---|---|---|
| chr2 | 38339114 | 38340086 | 19.88939 | 55.30079 | 52.51555 | Promoter | 2 | 38339281 | 38369199 | 29919 | 1 | ENSMUST00 | 0 | Lhx2 | ENSMUSG00000000247.6 |
| chr2 | 168049306 | 168050962 | 19.88939 | 55.30079 | 52.51555 | Distal Interge | 2 | 168064820 | 168064897 | 78 | 2 | ENSMUST00 | 13935 | Gm24327 | ENSMUSG00000092849.1 |
| chr9 | 106464773 | 106465852 | 19.88939 | 55.30079 | 52.51555 | Promoter | 9 | 106463962 | 106465938 | 1977 | 2 | ENSMUST00 | 86 | Gpr62 | ENSMUSG00000091735.2 |
| chr10 | 43578187 | 43581161 | 20.8167 | 55.26561 | 52.48107 | Promoter | 10 | 43579169 | 43584262 | 5094 | 1 | ENSMUST00 | 0 | Cd24a | ENSMUSG00000047139.8 |
| chr3 | 129901175 | 129902288 | 18.9428 | 55.25868 | 52.47425 | Promoter | 3 | 129901425 | 129914103 | 12679 | 1 | ENSMUST00 | 0 | Casp6 | ENSMUSG00000027997.8 |
| chr5 | 88886438 | 88887943 | 18.9428 | 55.25868 | 52.47425 | Promoter | 5 | 88886818 | 89239653 | 352836 | 1 | ENSMUST00 | 0 | Slc4a4 | ENSMUSG00000060961.9 |
| chr13 | 21786741 | 21787408 | 15.6415 | 55.20039 | 52.41629 | Promoter | 13 | 21786826 | 21787218 | 393 | 2 | ENSMUST00 | 0 | Hist1h2an | ENSMUSG00000069309.2 |
| chr2 | 122377525 | 122378653 | 15.6415 | 55.20039 | 52.41629 | Distal Interge | 2 | 122385489 | 122385640 | 152 | 2 | ENSMUST00 | 6987 | Gm24409 | ENSMUSG00000088506.1 |
| chr4 | 98816819 | 98818200 | 15.6415 | 55.20039 | 52.41629 | Promoter | 4 | 98754898 | 98817537 | 62640 | 2 | ENSMUST00 | 0 | Kank4 | ENSMUSG00000035407.8 |
| chr5 | 110094118 | 110095082 | 15.6415 | 55.20039 | 52.41629 | Distal Interge | 5 | 110099969 | 110103815 | 3847 | 1 | ENSMUST00 | -4887 | Plcxd1 | ENSMUSG00000064247.9 |
| chr1 | 156040804 | 156041431 | 18.02468 | 55.19937 | 52.41551 | Promoter | 1 | 156040890 | 156051669 | 10780 | 1 | ENSMUST00 | 0 | Tor1aip2 | ENSMUSG00000050565.11 |
| chr12 | 84284837 | 84286013 | 18.02468 | 55.19937 | 52.41551 | Promoter | 12 | 84285232 | 84315832 | 30601 | 1 | ENSMUST00 | 0 | Ptgr2 | ENSMUSG00000072946.7 |
| chr19 | 4268382 | 4269562 | 18.02468 | 55.19937 | 52.41551 | Promoter | 19 | 4261668 | 4269172 | 7505 | 2 | ENSMUST00 | 0 | Ssh3 | ENSMUSG00000034616.11 |
| chr19 | 8818123 | 8819393 | 18.02468 | 55.19937 | 52.41551 | Promoter | 19 | 8809075 | 8819333 | 10259 | 2 | ENSMUST00 | 0 | Ttc9c | ENSMUSG00000071660.5 |
| chr3 | 83841021 | 83841872 | 18.02468 | 55.19937 | 52.41551 | Promoter | 3 | 83836273 | 83841767 | 5495 | 2 | ENSMUST00 | 0 | Tlr2 | ENSMUSG00000027995.9 |
| chr3 | 89411195 | 89412567 | 18.02468 | 55.19937 | 52.41551 | Promoter | 3 | 89401004 | 89411870 | 10867 | 2 | ENSMUST00 | 0 | Flad1 | ENSMUSG00000042642.8 |
| chr4 | 43631185 | 43632004 | 18.02468 | 55.19937 | 52.41551 | Promoter | 4 | 43631935 | 43651244 | 19310 | 1 | ENSMUST00 | 0 | Npr2 | ENSMUSG00000028469.10 |
| chr5 | 24729777 | 24731419 | 18.02468 | 55.19937 | 52.41551 | Promoter | 5 | 24711738 | 24730727 | 18990 | 2 | ENSMUST00 | 0 | Wdr86 | ENSMUSG00000055235.6 |
| chr7 | 24298640 | 24300207 | 18.02468 | 55.19937 | 52.41551 | Promoter | 7 | 24291039 | 24299675 | 8637 | 2 | ENSMUST00 | 0 | Zfp61 | ENSMUSG00000050605.12 |
| chr1 | 171379651 | 171381313 | 20.77461 | 55.19346 | 52.4113 | Exon (ENSMU | 1 | 171388954 | 171410240 | 21287 | 1 | ENSMUST00 | -7641 | Arhgap30 | ENSMUSG00000048865.11 |
| chr17 | 80223875 | 80225115 | 20.7327 | 55.12159 | 52.33961 | Promoter | 17 | 80224441 | 80226818 | 2378 | 1 | ENSMUST00 | 0 | Gemin6 | ENSMUSG00000055760.7 |
| chr14 | 55603783 | 55604920 | 20.23482 | 55.08224 | 52.30038 | Promoter | 14 | 55603985 | 55610030 | 6046 | 1 | ENSMUST00 | 0 | Irf9 | ENSMUSG00000002325.9 |
| chr8 | 70803851 | 70805345 | 20.23482 | 55.08224 | 52.30038 | Distal Interge | 8 | 70808449 | 70821424 | 12976 | 1 | ENSMUST00 | -3104 | Il12rb1 | ENSMUSG00000000791.8 |
| chr9 | 109081350 | 109083201 | 20.23482 | 55.08224 | 52.30038 | Promoter | 9 | 109082496 | 109093363 | 10868 | 1 | ENSMUST00 | 0 | Ccdc51 | ENSMUSG00000025645.7 |
| chr2 | 34249435 | 34250326 | 20.69095 | 55.04997 | 52.26868 | Intron (ENSM | 2 | 34287771 | 34304440 | 16670 | 1 | ENSMUST00 | -37445 | C79798 | ENSMUSG00000084908.2 |
| chr13 | 21452567 | 21453886 | 16.30988 | 55.04095 | 52.25988 | Promoter | 13 | 21442175 | 21453727 | 11553 | 2 | ENSMUST00 | 0 | Zscan26 | ENSMUSG00000022228.8 |
| chr13 | 43615542 | 43616454 | 16.30988 | 55.04095 | 52.25988 | Promoter | 13 | 43615710 | 43670945 | 55236 | 1 | ENSMUST00 | 0 | Rnf182 | ENSMUSG00000044164.2 |
| chr14 | 3049118 | 3049979 | 16.30988 | 55.04095 | 52.25988 | Promoter | 14 | 3049285 | 3076840 | 27556 | 1 | ENSMUST00 | 0 | Gm2897 | ENSMUSG00000079410.4 |
| chr6 | 13870378 | 13872359 | 16.30988 | 55.04095 | 52.25988 | Promoter | 6 | 13867735 | 13871518 | 3784 | 2 | ENSMUST00 | 0 | 2610001J05R | ENSMUSG00000052419.7 |
| chr11 | 102880366 | 102881751 | 17.09341 | 55.02552 | 52.24499 | Promoter | 11 | 102838473 | 102880967 | 42495 | 2 | ENSMUST00 | 0 | Eftud2 | ENSMUSG00000020929.9 |
| chr12 | 105962580 | 105963742 | 17.09341 | 55.02552 | 52.24499 | Distal Interge | 12 | 106010263 | 106077410 | 67148 | 1 | ENSMUST00 | -46521 | Vrk1 | ENSMUSG00000021115.9 |
| chr15 | 76655745 | 76657556 | 17.09341 | 55.02552 | 52.24499 | Promoter | 15 | 76646927 | 76656920 | 9994 | 2 | ENSMUST00 | 0 | Cyhr1 | ENSMUSG00000053929.11 |
| chr19 | 32595343 | 32596330 | 17.09341 | 55.02552 | 52.24499 | Distal Interge | 19 | 32620005 | 32667187 | 47183 | 1 | ENSMUST00 | -23675 | Papss2 | ENSMUSG00000024899.6 |
| chr2 | 113326973 | 113328143 | 17.09341 | 55.02552 | 52.24499 | Promoter | 2 | 113327736 | 113716767 | 389032 | 1 | ENSMUST00 | 0 | Fmn1 | ENSMUSG00000044042.13 |
| chr2 | 144458685 | 144459566 | 17.09341 | 55.02552 | 52.24499 | Promoter | 2 | 144459280 | 144467794 | 8515 | 1 | ENSMUST00 | 0 | Zfp133-ps | ENSMUSG00000083674.3 |
| chr4 | 120854676 | 120855486 | 17.09341 | 55.02552 | 52.24499 | Promoter | 4 | 120854816 | 120884444 | 29629 | 1 | ENSMUST00 | 0 | Rims3 | ENSMUSG00000032890.12 |
| chr5 | 122987524 | 122989984 | 17.09341 | 55.02552 | 52.24499 | Promoter | 5 | 122870668 | 122989099 | 118432 | 2 | ENSMUST00 | 0 | Kdm2b | ENSMUSG00000029475.12 |
| chr6 | 52210986 | 52214025 | 17.09341 | 55.02552 | 52.24499 | Promoter | 6 | 52169062 | 52213115 | 44054 | 2 | ENSMUST00 | 0 | Hoxa3 | ENSMUSG00000079560.8 |
| chr7 | 122288850 | 122289900 | 17.09341 | 55.02552 | 52.24499 | Promoter | 7 | 122289297 | 122628655 | 339359 | 1 | ENSMUST00 | 0 | Prkcb | ENSMUSG00000052889.6 |
| chr8 | 94771750 | 94772906 | 17.09341 | 55.02552 | 52.24499 | Promoter | 8 | 94772009 | 94778945 | 6937 | 1 | ENSMUST00 | 0 | Cx3cl1 | ENSMUSG00000031778.7 |
| chr9 | 35116129 | 35117433 | 17.09341 | 55.02552 | 52.24499 | Promoter | 9 | 35046579 | 35116810 | 70232 | 2 | ENSMUST00 | 0 | St3gal4 | ENSMUSG00000032038.6 |
| chr9 | 104001869 | 104002467 | 17.09341 | 55.02552 | 52.24499 | Promoter | 9 | 104002544 | 104043811 | 41268 | 1 | ENSMUST00 | -77 | Nphp3 | ENSMUSG00000032558.13 |
| chr9 | 107340165 | 107341059 | 17.09341 | 55.02552 | 52.24499 | Promoter | 9 | 107340602 | 107349683 | 9082 | 1 | ENSMUST00 | 0 | 6430571L13R | ENSMUSG00000037977.5 |
| chr9 | 56950068 | 56951690 | 17.09341 | 55.02552 | 52.24499 | Promoter | 9 | 56950878 | 56983206 | 32329 | 1 | ENSMUST00 | 0 | Snupn | ENSMUSG00000055334.4 |
| chr9 | 72491490 | 72492403 | 17.09341 | 55.02552 | 52.24499 | Promoter | 9 | 72450394 | 72491956 | 41563 | 2 | ENSMUST00 | 0 | Tex9 | ENSMUSG00000090626.4 |
| chr1 | 42952733 | 42953876 | 21.13848 | 54.99947 | 52.22001 | Promoter | 1 | 42952872 | 42955155 | 2284 | 1 | ENSMUST00 | 0 | Gpr45 | ENSMUSG00000041907.9 |
| chr11 | 19026439 | 19027727 | 21.13848 | 54.99947 | 52.22001 | Distal Interge | 11 | 18879817 | 19018985 | 139169 | 2 | ENSMUST00 | -7455 | Meis1 | ENSMUSG00000020160.13 |
| chr11 | 101259782 | 101261361 | 21.13848 | 54.99947 | 52.22001 | Promoter | 11 | 101260567 | 101277409 | 16843 | 1 | ENSMUST00 | 0 | Wnk4 | ENSMUSG00000035112.12 |
| chr11 | 113618587 | 113619979 | 21.13848 | 54.99947 | 52.22001 | Promoter | 11 | 113619342 | 113626028 | 6687 | 1 | ENSMUST00 | 0 | Sstr2 | ENSMUSG00000047904.6 |
| chr13 | 21810029 | 21811180 | 21.13848 | 54.99947 | 52.22001 | Promoter | 13 | 21806412 | 21810199 | 3788 | 2 | ENSMUST00 | 0 | Hist1h2bq | ENSMUSG00000069307.5 |
| chr14 | 111674793 | 111676534 | 21.13848 | 54.99947 | 52.22001 | Promoter | 14 | 111675115 | 111683134 | 8020 | 1 | ENSMUST00 | 0 | Slitrk5 | ENSMUSG00000033214.9 |
| chr15 | 84719815 | 84721020 | 21.13848 | 54.99947 | 52.22001 | Promoter | 15 | 84720052 | 84772207 | 52156 | 1 | ENSMUST00 | 0 | Arhgap8 | ENSMUSG00000078954.4 |
| chr17 | 69075440 | 69076694 | 21.13848 | 54.99947 | 52.22001 | Distal Interge | 17 | 69156810 | 69289985 | 133176 | 1 | ENSMUST00 | -80116 | Epb4.1l3 | ENSMUSG00000024044.10 |
| chr17 | 80482568 | 80483679 | 21.13848 | 54.99947 | 52.22001 | Distal Interge | 17 | 80393752 | 80480453 | 86702 | 2 | ENSMUST00 | -2116 | Sos1 | ENSMUSG00000024241.6 |
| chr18 | 44811896 | 44812505 | 21.13848 | 54.99947 | 52.22001 | Promoter | 18 | 44425061 | 44812182 | 387122 | 2 | ENSMUST00 | 0 | Mcc | ENSMUSG00000071856.5 |
| chr2 | 45111990 | 45113601 | 21.13848 | 54.99947 | 52.22001 | Promoter | 2 | 44983634 | 45113279 | 129646 | 2 | ENSMUST00 | 0 | Zeb2 | ENSMUSG00000026872.11 |
| chr2 | 180892819 | 180894892 | 21.13848 | 54.99947 | 52.22001 | Promoter | 2 | 180894031 | 180894117 | 87 | 1 | ENSMUST00 | 0 | Mir124a-3 | ENSMUSG00000065454.1 |
| chr8 | 84103873 | 84104844 | 21.13848 | 54.99947 | 52.22001 | Promoter | 8 | 84097075 | 84104773 | 7699 | 2 | ENSMUST00 | 0 | Dcaf15 | ENSMUSG00000037103.7 |
| chr8 | 71536900 | 71538161 | 21.13848 | 54.99947 | 52.22001 | Promoter | 8 | 71534264 | 71537437 | 3174 | 2 | ENSMUST00 | 0 | Bst2 | ENSMUSG00000046718.7 |
| chr8 | 3621512 | 3621933 | 21.13848 | 54.99947 | 52.22001 | Promoter | 8 | 3621551 | 3624231 | 2681 | 1 | ENSMUST00 | 0 | Pet100 | ENSMUSG00000087687.2 |
| chr9 | 122572099 | 122572964 | 21.13848 | 54.99947 | 52.22001 | Promoter | 9 | 122572497 | 122678965 | 106469 | 1 | ENSMUST00 | 0 | 9530059O14 | ENSMUSG00000097736.2 |
| chrX | 16617908 | 16620088 | 21.13848 | 54.99947 | 52.22001 | Promoter | 20 | 16619698 | 16687818 | 68121 | 1 | ENSMUST00 | 0 | Maoa | ENSMUSG00000025037.6 |
| chrX | 113184768 | 113185956 | 21.13848 | 54.99947 | 52.22001 | Promoter | 20 | 113040593 | 113185517 | 144925 | 2 | ENSMUST00 | 0 | Chm | ENSMUSG00000025531.9 |
| chrX | 48623115 | 48624262 | 21.13848 | 54.99947 | 52.22001 | Promoter | 20 | 48623413 | 48631036 | 7624 | 1 | ENSMUST00 | 0 | Slc25a14 | ENSMUSG00000031105.11 |
| chrX | 160993660 | 160995693 | 21.13848 | 54.99947 | 52.22001 | Promoter | 20 | 160784308 | 160994702 | 210395 | 2 | ENSMUST00 | 0 | Cdkl5 | ENSMUSG00000031292.9 |
| chrX | 104412652 | 104414068 | 21.13848 | 54.99947 | 52.22001 | Promoter | 20 | 104280657 | 104413846 | 133190 | 2 | ENSMUST00 | 0 | Abcb7 | ENSMUSG00000031333.7 |
| chrX | 72986865 | 72988251 | 21.13848 | 54.99947 | 52.22001 | Promoter | 20 | 72987339 | 73031543 | 44205 | 1 | ENSMUST00 | 0 | Zfp185 | ENSMUSG00000031351.10 |
| chrX | 73921079 | 73922554 | 21.13848 | 54.99947 | 52.22001 | Promoter | 20 | 73916873 | 73921944 | 5072 | 2 | ENSMUST00 | 0 | Naa10 | ENSMUSG00000031388.9 |
| chrX | 101640055 | 101642007 | 21.13848 | 54.99947 | 52.22001 | Promoter | 20 | 101640060 | 101682940 | 42881 | 1 | ENSMUST00 | 0 | Ogt | ENSMUSG00000034160.8 |
| chrX | 71491872 | 71493555 | 21.13848 | 54.99947 | 52.22001 | Promoter | 20 | 71420060 | 71492645 | 72586 | 2 | ENSMUST00 | 0 | Cd99l2 | ENSMUSG00000035776.9 |
| chrX | 12076787 | 12081284 | 21.13848 | 54.99947 | 52.22001 | Promoter | 20 | 12036740 | 12080553 | 43814 | 2 | ENSMUST00 | 0 | Bcor | ENSMUSG00000040363.9 |
| chrX | 152143983 | 152145711 | 21.13848 | 54.99947 | 52.22001 | Promoter | 20 | 152144268 | 152225235 | 80968 | 1 | ENSMUST00 | 0 | Iqsec2 | ENSMUSG00000041115.11 |
| chrX | 137049118 | 137050290 | 21.13848 | 54.99947 | 52.22001 | Promoter | 20 | 137049594 | 137082503 | 32910 | 1 | ENSMUST00 | 0 | Fam199x | ENSMUSG00000042595.7 |
| chr3 | 105704253 | 105705864 | 19.22456 | 54.94707 | 52.16912 | Promoter | 3 | 105704599 | 105720842 | 16244 | 1 | ENSMUST00 | 0 | Fam212b | ENSMUSG00000048458.8 |
| chr9 | 56926890 | 56929065 | 19.22456 | 54.94707 | 52.16912 | Promoter | 9 | 56917200 | 56928371 | 11172 | 2 | ENSMUST00 | 0 | Snx33 | ENSMUSG00000032733.5 |
| chr8 | 83955649 | 83956131 | 19.67028 | 54.91714 | 52.13961 | Promoter | 8 | 83955714 | 83967847 | 12134 | 1 | ENSMUST00 | 0 | Asf1b | ENSMUSG00000005470.8 |
| chr1 | 75124518 | 75125711 | 20.60796 | 54.90754 | 52.13 | Promoter | 1 | 75125136 | 75125200 | 65 | 2 | ENSMUST00 | 0 | Nhej1 | ENSMUSG00000026162.7 |
| chr10 | 56377095 | 56379177 | 20.60796 | 54.90754 | 52.13 | Promoter | 10 | 56377300 | 56390419 | 13120 | 1 | ENSMUST00 | 0 | Gja1 | ENSMUSG00000050953.9 |
| chr13 | 52977991 | 52979106 | 20.60796 | 54.90754 | 52.13 | Intron (ENSM | 13 | 52967209 | 52981073 | 13865 | 2 | ENSMUST00 | 1967 | Nfil3 | ENSMUSG00000056749.7 |
| chr17 | 85687187 | 85689367 | 20.60796 | 54.90754 | 52.13 | Promoter | 17 | 85684277 | 85688274 | 3998 | 2 | ENSMUST00 | 0 | Six2 | ENSMUSG00000024134.11 |
| chr17 | 47909208 | 47910013 | 20.60796 | 54.90754 | 52.13 | Intron (ENSM | 17 | 47922378 | 47923429 | 1052 | 1 | ENSMUST00 | -12365 | Gm15556 | ENSMUSG00000085932.1 |
| chr1 | 87263912 | 87265128 | 20.11728 | 54.87862 | 52.10172 | Promoter | 1 | 87264363 | 87310836 | 46474 | 1 | ENSMUST00 | 0 | Efhd1 | ENSMUSG00000026255.10 |
| chr10 | 77032569 | 77033943 | 20.11728 | 54.87862 | 52.10172 | Promoter | 10 | 77032684 | 77050432 | 17749 | 1 | ENSMUST00 | 0 | Slc19a1 | ENSMUSG00000001436.10 |
| chr10 | 117792951 | 117793526 | 20.11728 | 54.87862 | 52.10172 | Promoter | 10 | 117750643 | 117792705 | 42063 | 2 | ENSMUST00 | -247 | Nup107 | ENSMUSG00000052798.7 |
| chr10 | 61977942 | 61979412 | 20.11728 | 54.87862 | 52.10172 | Promoter | 10 | 61838236 | 61979073 | 140838 | 2 | ENSMUST00 | 0 | Col13a1 | ENSMUSG00000058806.9 |
| chr13 | 92483403 | 92484400 | 20.11728 | 54.87862 | 52.10172 | Promoter | 13 | 92449622 | 92484031 | 34410 | 2 | ENSMUST00 | 0 | Fam151b | ENSMUSG00000034334.7 |
| chr15 | 79030840 | 79031298 | 20.11728 | 54.87862 | 52.10172 | Promoter | 15 | 79030874 | 79038348 | 7475 | 1 | ENSMUST00 | 0 | Gcat | ENSMUSG00000006378.8 |
| chr15 | 34678309 | 34679063 | 20.11728 | 54.87862 | 52.10172 | Promoter | 15 | 34572799 | 34678706 | 105908 | 2 | ENSMUST00 | 0 | Nipal2 | ENSMUSG00000038879.7 |
| chr15 | 103034081 | 103036049 | 20.11728 | 54.87862 | 52.10172 | Promoter | 15 | 103034395 | 103036852 | 2458 | 1 | ENSMUST00 | 0 | Hoxc4 | ENSMUSG00000075394.4 |
| chr17 | 35165145 | 35166016 | 20.11728 | 54.87862 | 52.10172 | Promoter | 17 | 35165119 | 35167186 | 2068 | 1 | ENSMUST00 | 27 | Gm17705 | ENSMUSG00000090936.2 |
| chr17 | 32113436 | 32114667 | 20.11728 | 54.87862 | 52.10172 | Promoter | 17 | 32113429 | 32141839 | 28411 | 1 | ENSMUST00 | 8 | Gm17276 | ENSMUSG00000097305.1 |
| chr3 | 89269869 | 89270833 | 20.11728 | 54.87862 | 52.10172 | Promoter | 3 | 89268246 | 89270479 | 2234 | 2 | ENSMUST00 | 0 | Slc50a1 | ENSMUSG00000027953.8 |
| chr3 | 156561377 | 156562528 | 20.11728 | 54.87862 | 52.10172 | Promoter | 3 | 156561794 | 157316445 | 754652 | 1 | ENSMUST00 | 0 | Negr1 | ENSMUSG00000040037.8 |
| chr5 | 136243696 | 136245529 | 20.11728 | 54.87862 | 52.10172 | Promoter | 5 | 136218149 | 136244903 | 26755 | 2 | ENSMUST00 | 0 | Sh2b2 | ENSMUSG00000005057.9 |
| chr5 | 66745342 | 66747077 | 20.11728 | 54.87862 | 52.10172 | Promoter | 5 | 66745835 | 67056073 | 310239 | 1 | ENSMUST00 | 0 | Limch1 | ENSMUSG00000037736.13 |
| chr5 | 23615898 | 23617353 | 20.11728 | 54.87862 | 52.10172 | Promoter | 5 | 23503356 | 23616571 | 113216 | 2 | ENSMUST00 | 0 | Srpk2 | ENSMUSG00000062604.7 |
| chr9 | 114025933 | 114026928 | 20.11728 | 54.87862 | 52.10172 | Promoter | 9 | 113963637 | 114026729 | 63093 | 2 | ENSMUST00 | 0 | Fbxl2 | ENSMUSG00000032507.10 |
| chr17 | 34644726 | 34645479 | 20.56672 | 54.83671 | 52.06188 | Promoter | 17 | 34644764 | 34646324 | 1561 | 1 | ENSMUST00 | 0 | Fkbpl | ENSMUSG00000033739.8 |
| chr17 | 21535333 | 21536358 | 20.56672 | 54.83671 | 52.06188 | Promoter | 17 | 21528194 | 21535878 | 7685 | 2 | ENSMUST00 | 0 | 9330136K24R | ENSMUSG00000097781.1 |
| chr19 | 4231890 | 4232260 | 20.56672 | 54.83671 | 52.06188 | Promoter | 19 | 4231899 | 4233631 | 1733 | 1 | ENSMUST00 | 0 | Pold4 | ENSMUSG00000024854.3 |
| chr14 | 19584540 | 19585580 | 21.01229 | 54.78426 | 52.00962 | Promoter | 14 | 19563801 | 19585135 | 21335 | 2 | ENSMUST00 | 0 | Gm2237 | ENSMUSG00000093979.2 |
| chr7 | 104352353 | 104353745 | 20.52563 | 54.76614 | 51.99178 | Promoter | 7 | 104338754 | 104353362 | 14609 | 2 | ENSMUST00 | 0 | Trim12c | ENSMUSG00000057143.9 |
| chr8 | 47714237 | 47714899 | 20.04161 | 54.7474 | 51.97314 | Promoter | 8 | 47709344 | 47713931 | 4588 | 2 | ENSMUST00 | -307 | Cdkn2aip | ENSMUSG00000038069.5 |
| chr5 | 32132837 | 32134178 | 20.96848 | 54.70951 | 51.93532 | Distal Interge | 5 | 32136472 | 32157831 | 21360 | 1 | ENSMUST00 | -2294 | Fosl2 | ENSMUSG00000029135.9 |
| chr13 | 76055260 | 76057031 | 18.62636 | 54.68748 | 51.91348 | Promoter | 13 | 76054851 | 76056996 | 2146 | 2 | ENSMUST00 | 0 | Gpr150 | ENSMUSG00000045509.5 |
| chr13 | 48878227 | 48879165 | 19.52687 | 54.66541 | 51.89166 | Downstream | 13 | 48871112 | 48875460 | 4349 | 1 | ENSMUST00 | 7116 | Phf2os1 | ENSMUSG00000086889.1 |
| chr15 | 3270402 | 3272276 | 19.52687 | 54.66541 | 51.89166 | Promoter | 15 | 3270767 | 3280508 | 9742 | 1 | ENSMUST00 | 0 | Sepp1 | ENSMUSG00000064373.6 |
| chr17 | 71526937 | 71527595 | 19.52687 | 54.66541 | 51.89166 | Promoter | 17 | 71526896 | 71556543 | 29648 | 1 | ENSMUST00 | 42 | Mettl4-ps1 | ENSMUSG00000079356.4 |
| chr7 | 24236857 | 24237741 | 19.52687 | 54.66541 | 51.89166 | Promoter | 7 | 24228048 | 24236632 | 8585 | 2 | ENSMUST00 | -226 | Zfp109 | ENSMUSG00000074283.4 |
| chr11 | 89301432 | 89303232 | 20.92485 | 54.63506 | 51.8618 | Promoter | 11 | 89300638 | 89302332 | 1695 | 2 | ENSMUST00 | 0 | Nog | ENSMUSG00000048616.4 |
| chr14 | 75834423 | 75835535 | 20.92485 | 54.63506 | 51.8618 | Distal Interge | 14 | 75843379 | 75844964 | 1586 | 2 | ENSMUST00 | 9429 | Gm4285 | ENSMUSG00000085829.1 |
| chr2 | 165326644 | 165327257 | 20.44396 | 54.62578 | 51.85264 | Promoter | 2 | 165315672 | 165326479 | 10808 | 2 | ENSMUST00 | -166 | Elmo2 | ENSMUSG00000017670.11 |
| chr16 | 4586213 | 4586723 | 20.88141 | 54.5609 | 51.78793 | Distal Interge | 16 | 4594713 | 4615957 | 21245 | 1 | ENSMUST00 | -7990 | Glis2 | ENSMUSG00000014303.8 |
| chr18 | 34887042 | 34887995 | 20.40337 | 54.55598 | 51.78317 | Distal Interge | 18 | 34861207 | 34864956 | 3750 | 1 | ENSMUST00 | 25836 | Egr1 | ENSMUSG00000038418.7 |
| chr6 | 38662591 | 38663553 | 20.83814 | 54.48703 | 51.7144 | Promoter | 6 | 38663069 | 38680864 | 17796 | 1 | ENSMUST00 | 0 | Clec2l | ENSMUSG00000079598.3 |
| chr13 | 98572501 | 98573514 | 19.85205 | 54.41815 | 51.64575 | Distal Interge | 13 | 98560354 | 98560461 | 108 | 1 | ENSMUST00 | 12148 | Gm22698 | ENSMUSG00000093116.1 |
| chr18 | 36801339 | 36803017 | 19.85205 | 54.41815 | 51.64575 | Promoter | 18 | 36801866 | 36802008 | 143 | 1 | ENSMUST00 | 0 | Vaultrc5 | ENSMUSG00000065145.1 |
| chr2 | 158794168 | 158794648 | 19.85205 | 54.41815 | 51.64575 | Promoter | 2 | 158794807 | 158858214 | 63408 | 1 | ENSMUST00 | -159 | Dhx35 | ENSMUSG00000027655.9 |
| chr2 | 167777365 | 167778095 | 19.85205 | 54.41815 | 51.64575 | Promoter | 2 | 167777467 | 167782569 | 5103 | 1 | ENSMUST00 | 0 | Gm14321 | ENSMUSG00000084797.1 |
| chr3 | 130180278 | 130181636 | 19.85205 | 54.41815 | 51.64575 | Promoter | 3 | 130180891 | 130551024 | 370134 | 1 | ENSMUST00 | 0 | Col25a1 | ENSMUSG00000058897.12 |
| chr7 | 68275563 | 68276607 | 19.85205 | 54.41815 | 51.64575 | Promoter | 7 | 68275995 | 68353646 | 77652 | 1 | ENSMUST00 | 0 | Fam169b | ENSMUSG00000074071.4 |
| chr7 | 4876914 | 4877704 | 19.85205 | 54.41815 | 51.64575 | Promoter | 7 | 4877053 | 4895717 | 18665 | 1 | ENSMUST00 | 0 | Isoc2a | ENSMUSG00000086784.1 |
| chr8 | 121127717 | 121129604 | 19.85205 | 54.41815 | 51.64575 | Promoter | 8 | 121127940 | 121130644 | 2705 | 1 | ENSMUST00 | 0 | Foxl1 | ENSMUSG00000097084.1 |
| chr2 | 179177140 | 179177482 | 20.32266 | 54.41713 | 51.64543 | Promoter | 2 | 179177425 | 179177880 | 456 | 1 | ENSMUST00 | 0 | Gm14292 | ENSMUSG00000061897.5 |
| chr7 | 19332600 | 19333531 | 20.79506 | 54.41344 | 51.64182 | Distal Interge | 7 | 19344778 | 19356524 | 11747 | 1 | ENSMUST00 | -11247 | Ercc1 | ENSMUSG00000003549.4 |
| chr9 | 92275235 | 92278970 | 18.89232 | 54.3548 | 51.58332 | Promoter | 9 | 92275602 | 92297731 | 22130 | 1 | ENSMUST00 | 0 | Plscr2 | ENSMUSG00000032372.9 |
| chr14 | 54687028 | 54687670 | 20.28255 | 54.34807 | 51.57687 | Promoter | 14 | 54642728 | 54686931 | 44204 | 2 | ENSMUST00 | -98 | Acin1 | ENSMUSG00000022185.13 |
| chr15 | 37961208 | 37961613 | 20.70941 | 54.26712 | 51.49624 | Promoter | 15 | 37946752 | 37961318 | 14567 | 2 | ENSMUST00 | 0 | Rrm2b | ENSMUSG00000022292.10 |

| | | | | | | | | | | | | | | | |
|---|---|---|---|---|---|---|---|---|---|---|---|---|---|---|---|
| chr4 | 116123290 | 116124365 | 20.2028 | 54.2107 | 51.43995 | Promoter | 4 | 116123840 | 116138850 | 15011 | 1 | ENSMUST00 | 0 | Pomgnt1 | ENSMUSG00000028700.9 |
| chr5 | 140704987 | 140705733 | 20.2028 | 54.2107 | 51.43995 | Promoter | 5 | 140705011 | 140719379 | 14369 | 1 | ENSMUST00 | 0 | Brat1 | ENSMUSG00000000148.12 |
| chr5 | 129778788 | 129779623 | 20.2028 | 54.2107 | 51.43995 | 5' UTR | 5 | 129765558 | 129771615 | 6058 | 2 | ENSMUST00 | -7174 | Psph | ENSMUSG00000029446.9 |
| chr5 | 52114733 | 52116391 | 20.2028 | 54.2107 | 51.43995 | Distal Interge | 5 | 52135902 | 52136017 | 116 | 2 | ENSMUST00 | 19626 | Mir6417 | ENSMUSG00000098726.1 |
| chr10 | 90969441 | 90970508 | 20.66686 | 54.19438 | 51.42418 | 3' UTR | 10 | 90989313 | 90991590 | 2278 | 2 | ENSMUST00 | 21082 | Apaf1 | ENSMUSG00000019979.7 |
| chr15 | 62037741 | 62038388 | 20.66686 | 54.19438 | 51.42418 | Promoter | 15 | 62037986 | 62249528 | 211543 | 1 | ENSMUST00 | 0 | Pvt1 | ENSMUSG00000097039.3 |
| chr6 | 115360762 | 115362314 | 14.81767 | 54.17085 | 51.40078 | Promoter | 6 | 115361221 | 115490401 | 129181 | 1 | ENSMUST00 | 0 | Pparg | ENSMUSG00000000440.7 |
| chr1 | 75487272 | 75488169 | 18.77433 | 54.14365 | 51.37372 | Promoter | 1 | 75486851 | 75487802 | 952 | 2 | ENSMUST00 | 0 | Obsl1 | ENSMUSG00000026211.12 |
| chr1 | 132138961 | 132139885 | 18.77433 | 54.14365 | 51.37372 | Promoter | 1 | 132112237 | 132139684 | 27448 | 2 | ENSMUST00 | 0 | Cdk18 | ENSMUSG00000026437.6 |
| chr1 | 175692980 | 175693562 | 18.77433 | 54.14365 | 51.37372 | Promoter | 1 | 175662421 | 175692776 | 30356 | 2 | ENSMUST00 | -205 | Opn3 | ENSMUSG00000026525.9 |
| chr1 | 182763639 | 182764659 | 18.77433 | 54.14365 | 51.37372 | Promoter | 1 | 182763860 | 182854040 | 90181 | 1 | ENSMUST00 | 0 | Susd4 | ENSMUSG00000038576.10 |
| chr1 | 52844968 | 52845904 | 18.77433 | 54.14365 | 51.37372 | Promoter | 1 | 52845013 | 52920981 | 75969 | 1 | ENSMUST00 | 0 | Hibch | ENSMUSG00000041426.7 |
| chr10 | 80172765 | 80173576 | 18.77433 | 54.14365 | 51.37372 | Promoter | 10 | 80172944 | 80175146 | 2203 | 1 | ENSMUST00 | 0 | 1600002K03I | ENSMUSG00000035595.6 |
| chr10 | 79793233 | 79794127 | 18.77433 | 54.14365 | 51.37372 | Promoter | 10 | 79793572 | 79820896 | 27325 | 1 | ENSMUST00 | 0 | Palm | ENSMUSG00000035863.8 |
| chr11 | 58948124 | 58949431 | 18.77433 | 54.14365 | 51.37372 | Promoter | 11 | 58948920 | 58949533 | 614 | 1 | ENSMUST00 | 0 | Hist3h2ba | ENSMUSG00000056895.4 |
| chr11 | 117199219 | 117200136 | 18.77433 | 54.14365 | 51.37372 | Promoter | 11 | 117199661 | 117362325 | 162665 | 1 | ENSMUST00 | 0 | Sep-09 | ENSMUSG00000059248.8 |
| chr14 | 63435957 | 63436764 | 18.77433 | 54.14365 | 51.37372 | Promoter | 14 | 63436394 | 63465498 | 29105 | 1 | ENSMUST00 | 0 | Fam167a | ENSMUSG00000035095.11 |
| chr14 | 61686797 | 61687458 | 18.77433 | 54.14365 | 51.37372 | Distal Interge | 14 | 61602839 | 61682373 | 79535 | 2 | ENSMUST00 | -4425 | Dleu2 | ENSMUSG00000097589.4 |
| chr15 | 72805816 | 72811358 | 18.77433 | 54.14365 | 51.37372 | Intron (ENSM | 15 | 72721321 | 72721387 | 67 | 2 | ENSMUST00 | -84430 | Gm28020 | ENSMUSG00000088364.2 |
| chr16 | 75766135 | 75767980 | 18.77433 | 54.14365 | 51.37372 | Promoter | 16 | 75755190 | 75766821 | 11632 | 2 | ENSMUST00 | 0 | Hspa13 | ENSMUSG00000032932.9 |
| chr19 | 44725985 | 44726780 | 18.77433 | 54.14365 | 51.37372 | Distal Interge | 19 | 44756045 | 44837851 | 81807 | 1 | ENSMUST00 | -29265 | Pax2 | ENSMUSG00000004231.10 |
| chr2 | 67565176 | 67566948 | 18.77433 | 54.14365 | 51.37372 | Promoter | 2 | 67565871 | 67810894 | 245024 | 1 | ENSMUST00 | 0 | B3galt1 | ENSMUSG00000034780.6 |
| chr2 | 180118670 | 180119157 | 18.77433 | 54.14365 | 51.37372 | Promoter | 2 | 180119306 | 180162680 | 43375 | 1 | ENSMUST00 | -149 | Osbpl2 | ENSMUSG00000039050.8 |
| chr2 | 109277879 | 109278600 | 18.77433 | 54.14365 | 51.37372 | Promoter | 2 | 109092297 | 109278290 | 185994 | 2 | ENSMUST00 | 0 | Mettl15 | ENSMUSG00000057234.4 |
| chr3 | 96556721 | 96557800 | 18.77433 | 54.14365 | 51.37372 | Promoter | 3 | 96555765 | 96557815 | 2051 | 2 | ENSMUST00 | 15 | Gm15441 | ENSMUSG00000074398.5 |
| chr4 | 133910275 | 133911377 | 18.77433 | 54.14365 | 51.37372 | Distal Interge | 4 | 133847290 | 133887797 | 40508 | 2 | ENSMUST00 | -22479 | Rps6ka1 | ENSMUSG00000003644.12 |
| chr6 | 121272080 | 121273002 | 18.77433 | 54.14365 | 51.37372 | Distal Interge | 6 | 121245906 | 121270916 | 25011 | 1 | ENSMUST00 | 26175 | Usp18 | ENSMUSG00000030107.9 |
| chr6 | 47788220 | 47789028 | 18.77433 | 54.14365 | 51.37372 | Promoter | 6 | 47788069 | 47788180 | 112 | 2 | ENSMUST00 | -41 | Rny1 | ENSMUSG00000065701.1 |
| chr7 | 142967306 | 142969434 | 18.77433 | 54.14365 | 51.37372 | Promoter | 7 | 142966822 | 142969264 | 2443 | 2 | ENSMUST00 | 0 | Ascl2 | ENSMUSG00000009248.5 |
| chr7 | 45395996 | 45396388 | 18.77433 | 54.14365 | 51.37372 | Promoter | 7 | 45376462 | 45395692 | 19231 | 2 | ENSMUST00 | -305 | Snrnp70 | ENSMUSG00000063511.8 |
| chr8 | 94702746 | 94704001 | 18.77433 | 54.14365 | 51.37372 | Distal Interge | 8 | 94698016 | 94698285 | 270 | 2 | ENSMUST00 | -4462 | Gm22846 | ENSMUSG00000088331.1 |
| chr8 | 46686640 | 46687317 | 18.77433 | 54.14365 | 51.37372 | Distal Interge | 8 | 46730972 | 46739515 | 8544 | 2 | ENSMUST00 | 52198 | Gm16675 | ENSMUSG00000097534.1 |
| chr16 | 97356333 | 97357944 | 20.12367 | 54.0743 | 51.30701 | Promoter | 16 | 97356728 | 97439012 | 82285 | 1 | ENSMUST00 | 0 | Bace2 | ENSMUSG00000040605.6 |
| chr1 | 23271151 | 23276131 | 19.17732 | 54.04958 | 51.28253 | Promoter | 1 | 23272264 | 23272345 | 82 | 1 | ENSMUST00 | 0 | Mir30a | ENSMUSG00000065405.2 |
| chr13 | 62623693 | 62624626 | 19.17732 | 54.04958 | 51.28253 | Promoter | 13 | 62624194 | 62624301 | 108 | 1 | ENSMUST00 | 0 | Gm24020 | ENSMUSG00000095020.1 |
| chr18 | 67133165 | 67134477 | 20.58227 | 54.04973 | 51.28253 | Promoter | 18 | 67133826 | 67226792 | 92967 | 1 | ENSMUST00 | 0 | Gnal | ENSMUSG00000024524.11 |
| chr6 | 56568606 | 56569441 | 19.17732 | 54.04958 | 51.28253 | Distal Interge | 6 | 56583715 | 56583846 | 132 | 2 | ENSMUST00 | 14405 | Gm24709 | ENSMUSG00000096087.1 |
| chr7 | 141437009 | 141437898 | 19.17732 | 54.04958 | 51.28253 | Promoter | 7 | 141429744 | 141437892 | 8149 | 2 | ENSMUST00 | 0 | Slc25a22 | ENSMUSG00000019082.12 |
| chr7 | 62463429 | 62464681 | 20.58227 | 54.04973 | 51.28253 | Promoter | 7 | 62461871 | 62464510 | 2640 | 2 | ENSMUST00 | 0 | Peg12 | ENSMUSG00000070526.2 |
| chr8 | 106870004 | 106871560 | 19.17732 | 54.04958 | 51.28253 | Promoter | 8 | 106870242 | 106882902 | 12661 | 1 | ENSMUST00 | 0 | Has3 | ENSMUSG00000031910.9 |
| chr12 | 108333194 | 108334007 | 20.08434 | 54.00646 | 51.24007 | Promoter | 12 | 108334377 | 108362231 | 27855 | 1 | ENSMUST00 | -370 | Cyp46a1 | ENSMUSG00000021259.4 |
| chr11 | 20455141 | 20456649 | 15.37639 | 53.87199 | 51.10612 | Distal Interge | 11 | 20446412 | 20446810 | 399 | 1 | ENSMUST00 | 8730 | Gm12034 | ENSMUSG00000083878.3 |
| chr4 | 126057932 | 126059069 | 20.00613 | 53.87149 | 51.10585 | Promoter | 4 | 126058565 | 126092195 | 33631 | 1 | ENSMUST00 | 0 | Oscp1 | ENSMUSG00000042616.7 |
| chr17 | 67952886 | 67956202 | 20.45667 | 53.83482 | 51.06929 | Intron (ENSM | 17 | 67845110 | 67950940 | 105831 | 2 | ENSMUST00 | -1947 | Arhgap28 | ENSMUSG00000024043.8 |
| chr5 | 150195615 | 150197097 | 20.45667 | 53.83482 | 51.06929 | Distal Interge | 5 | 150259930 | 150497752 | 237823 | 1 | ENSMUST00 | -62833 | Fry | ENSMUSG00000056602.8 |
| chr4 | 147808915 | 147810080 | 12.07892 | 53.79575 | 51.03043 | Promoter | 4 | 147753974 | 147809788 | 55815 | 2 | ENSMUST00 | 0 | Gm13157 | ENSMUSG00000078495.5 |
| chr3 | 100922357 | 100922987 | 18.57137 | 53.77931 | 51.01399 | Promoter | 3 | 100922475 | 100934011 | 11537 | 1 | ENSMUST00 | 0 | Trim45 | ENSMUSG00000033233.12 |
| chr7 | 139683849 | 139684776 | 18.57137 | 53.77931 | 51.01399 | Promoter | 7 | 139678270 | 139683817 | 5548 | 2 | ENSMUST00 | -33 | 9330101J02R | ENSMUSG00000049571.11 |
| chr2 | 38926655 | 38927318 | 20.41515 | 53.76371 | 50.99926 | Promoter | 2 | 38723370 | 38927688 | 204319 | 2 | ENSMUST00 | 370 | Nr6a1 | ENSMUSG00000063972.8 |
| chr1 | 54250657 | 54251334 | 17.6846 | 53.76268 | 50.99834 | Promoter | 1 | 54250683 | 54278961 | 28279 | 1 | ENSMUST00 | 0 | Ccdc150 | ENSMUSG00000025983.6 |
| chr1 | 75435729 | 75436668 | 17.6846 | 53.76268 | 50.99834 | Promoter | 1 | 75435930 | 75443174 | 7245 | 1 | ENSMUST00 | 0 | Gmppa | ENSMUSG00000033021.11 |
| chr12 | 45073476 | 45075217 | 17.6846 | 53.76268 | 50.99834 | Promoter | 12 | 44852488 | 45074112 | 221625 | 2 | ENSMUST00 | 0 | Stxbp6 | ENSMUSG00000046314.10 |
| chr12 | 69502212 | 69503540 | 17.6846 | 53.76268 | 50.99834 | Distal Interge | 12 | 69561229 | 69561337 | 109 | 2 | ENSMUST00 | 57797 | Gm23018 | ENSMUSG00000095776.1 |
| chr13 | 38344943 | 38347015 | 17.6846 | 53.76268 | 50.99834 | Promoter | 13 | 38345716 | 38499728 | 154013 | 1 | ENSMUST00 | 0 | Bmp6 | ENSMUSG00000039004.5 |
| chr13 | 10359615 | 10361337 | 17.6846 | 53.76268 | 50.99834 | Promoter | 13 | 9875486 | 10360844 | 485359 | 2 | ENSMUST00 | 0 | Chrm3 | ENSMUSG00000046159.10 |
| chr15 | 32778424 | 32779830 | 17.6846 | 53.76268 | 50.99834 | Distal Interge | 15 | 32920723 | 33034730 | 114008 | 1 | ENSMUST00 | -140893 | Sdc2 | ENSMUSG00000022261.5 |
| chr16 | 17815300 | 17816035 | 17.6846 | 53.76268 | 50.99834 | Distal Interge | 16 | 17797282 | 17808287 | 11006 | 1 | ENSMUST00 | 18019 | Scarf2 | ENSMUSG00000012017.3 |
| chr16 | 20240581 | 20241822 | 17.6846 | 53.76268 | 50.99834 | Promoter | 16 | 20233310 | 20241358 | 8049 | 2 | ENSMUST00 | 0 | Map6d1 | ENSMUSG00000041205.7 |
| chr18 | 61554545 | 61555087 | 17.6846 | 53.76268 | 50.99834 | Promoter | 18 | 61555274 | 61590061 | 34788 | 1 | ENSMUST00 | -187 | Csnk1a1 | ENSMUSG00000024576.9 |
| chr2 | 69585169 | 69586565 | 17.6846 | 53.76268 | 50.99834 | Promoter | 2 | 69424340 | 69586065 | 161726 | 2 | ENSMUST00 | 0 | Lrp2 | ENSMUSG00000027070.9 |
| chr2 | 120566919 | 120568583 | 17.6846 | 53.76268 | 50.99834 | Promoter | 2 | 120567671 | 120601255 | 33585 | 1 | ENSMUST00 | 0 | Snap23 | ENSMUSG00000027287.9 |
| chr4 | 119813872 | 119815377 | 17.6846 | 53.76268 | 50.99834 | Promoter | 4 | 119814331 | 119998432 | 184102 | 1 | ENSMUST00 | 0 | Hivep3 | ENSMUSG00000028634.11 |
| chr4 | 129622917 | 129625414 | 17.6846 | 53.76268 | 50.99834 | Promoter | 4 | 129619274 | 129623947 | 4674 | 2 | ENSMUST00 | 0 | Ccdc28b | ENSMUSG00000028795.10 |
| chr6 | 149141230 | 149142150 | 17.6846 | 53.76268 | 50.99834 | Promoter | 6 | 149141513 | 149151166 | 9654 | 1 | ENSMUST00 | 0 | Mettl20 | ENSMUSG00000039958.12 |
| chr7 | 79721202 | 79721934 | 17.6846 | 53.76268 | 50.99834 | 3' UTR | 7 | 79698098 | 79715720 | 17623 | 2 | ENSMUST00 | -5483 | Kif7 | ENSMUSG00000050382.9 |
| chr8 | 3220969 | 3221812 | 17.6846 | 53.76268 | 50.99834 | Intron (ENSM | 8 | 3259704 | 3260641 | 938 | 1 | ENSMUST00 | -37892 | Gm16180 | ENSMUSG00000086065.1 |
| chr1 | 21960440 | 21962504 | 16.02374 | 53.68177 | 50.91932 | Promoter | 1 | 21398403 | 21961581 | 563179 | 2 | ENSMUST00 | 0 | Kcnq5 | ENSMUSG00000028033.11 |
| chr10 | 128211095 | 128212197 | 16.02374 | 53.68177 | 50.91932 | Promoter | 10 | 128209908 | 128211794 | 1887 | 2 | ENSMUST00 | 0 | Spryd4 | ENSMUSG00000051346.8 |
| chr12 | 84069359 | 84070056 | 16.02374 | 53.68177 | 50.91932 | Promoter | 12 | 84069325 | 84076020 | 6696 | 1 | ENSMUST00 | 35 | Acot5 | ENSMUSG00000042540.7 |
| chr5 | 36397524 | 36398268 | 16.02374 | 53.68177 | 50.91932 | Promoter | 5 | 36017180 | 36398124 | 380945 | 2 | ENSMUST00 | 0 | Sorcs2 | ENSMUSG00000029093.9 |
| chr14 | 75015949 | 75016694 | 19.88996 | 53.67079 | 50.90882 | Promoter | 14 | 75016027 | 75052532 | 36506 | 1 | ENSMUST00 | 0 | 5031414D18I | ENSMUSG00000034959.7 |
| chr7 | 45922817 | 45924562 | 19.41141 | 53.64976 | 50.88786 | Promoter | 7 | 45924072 | 45931607 | 7536 | 1 | ENSMUST00 | 0 | Ccdc114 | ENSMUSG00000040189.10 |
| chr11 | 59661712 | 59664872 | 16.78262 | 53.63071 | 50.86883 | Promoter | 11 | 59662495 | 59780860 | 118366 | 1 | ENSMUST00 | 0 | Mprip | ENSMUSG00000005417.12 |
| chr11 | 104230662 | 104231268 | 16.78262 | 53.63071 | 50.86883 | Promoter | 11 | 104231390 | 104332090 | 100701 | 1 | ENSMUST00 | -122 | Mapt | ENSMUSG00000018411.12 |
| chr11 | 69879287 | 69881992 | 16.78262 | 53.63071 | 50.86883 | Promoter | 11 | 69878261 | 69881406 | 3146 | 2 | ENSMUST00 | 0 | Kctd11 | ENSMUSG00000046731.4 |
| chr2 | 177897039 | 177897634 | 16.78262 | 53.63071 | 50.86883 | Promoter | 2 | 177897096 | 177905784 | 8689 | 1 | ENSMUST00 | 0 | Gm14327 | ENSMUSG00000074521.4 |
| chr5 | 137787023 | 137787654 | 16.78262 | 53.63071 | 50.86883 | Promoter | 5 | 137787798 | 137822621 | 34824 | 1 | ENSMUST00 | -144 | Zcwpw1 | ENSMUSG00000037108.8 |
| chr6 | 29767601 | 29768298 | 16.78262 | 53.63071 | 50.86883 | Promoter | 6 | 29768011 | 29768727 | 717 | 1 | ENSMUST00 | 0 | Ahcyl2 | ENSMUSG00000029772.12 |
| chr8 | 33641088 | 33642607 | 16.78262 | 53.63071 | 50.86883 | Promoter | 8 | 33619586 | 33641976 | 22391 | 2 | ENSMUST00 | 0 | Ubxn8 | ENSMUSG00000052906.6 |
| chr9 | 57147428 | 57149059 | 16.78262 | 53.63071 | 50.86883 | Promoter | 9 | 57142800 | 57148127 | 5328 | 2 | ENSMUST00 | 0 | Neil1 | ENSMUSG00000032298.8 |
| chr11 | 75460689 | 75461242 | 20.29158 | 53.552 | 50.79138 | Promoter | 11 | 75461539 | 75466676 | 5138 | 1 | ENSMUST00 | -297 | Mir22hg | ENSMUSG00000085148.1 |
| chr2 | 5950319 | 5951076 | 20.29158 | 53.552 | 50.79138 | Promoter | 2 | 5951469 | 6056703 | 105235 | 1 | ENSMUST00 | -393 | Upf2 | ENSMUSG00000043241.9 |
| chr11 | 95309015 | 95309668 | 19.81326 | 53.53814 | 50.77763 | Promoter | 11 | 95274259 | 95309602 | 35344 | 2 | ENSMUST00 | 0 | Kat7 | ENSMUSG00000038909.11 |
| chr14 | 51082670 | 51083781 | 19.81326 | 53.53814 | 50.77763 | Distal Interge | 14 | 51091077 | 51106151 | 15075 | 1 | ENSMUST00 | -7296 | Rnase4 | ENSMUSG00000021876.9 |
| chr17 | 29237464 | 29238786 | 19.81326 | 53.53814 | 50.77763 | Promoter | 17 | 29156550 | 29237797 | 81248 | 2 | ENSMUST00 | 0 | Cpne5 | ENSMUSG00000024008.10 |
| chr4 | 135353427 | 135354334 | 19.81326 | 53.53814 | 50.77763 | Promoter | 4 | 135320484 | 135353321 | 32838 | 2 | ENSMUST00 | -107 | Srrm1 | ENSMUSG00000028809.11 |
| chr9 | 6168375 | 6169084 | 19.81326 | 53.53814 | 50.77763 | Promoter | 9 | 6168612 | 6377519 | 208908 | 1 | ENSMUST00 | 0 | Pdgfd | ENSMUSG00000032006.9 |
| chrX | 170009224 | 170009853 | 6.40876 | 53.53633 | 50.77636 | Promoter | 20 | 170009659 | 170019281 | 9623 | 1 | ENSMUST00 | 0 | Gm21887 | ENSMUSG00000095562.2 |
| chr10 | 62651390 | 62652194 | 20.25072 | 53.48195 | 50.722 | Promoter | 10 | 62616023 | 62651218 | 35196 | 2 | ENSMUST00 | -173 | Ddx50 | ENSMUSG00000020076.6 |
| chr14 | 60377812 | 60379347 | 18.84007 | 53.45216 | 50.69242 | Promoter | 14 | 60378286 | 60381003 | 2718 | 1 | ENSMUST00 | 0 | Amer2 | ENSMUSG00000021986.7 |
| chr3 | 90220275 | 90221220 | 18.84007 | 53.45216 | 50.69242 | Promoter | 3 | 90220787 | 90225969 | 5183 | 1 | ENSMUST00 | 0 | Rab13 | ENSMUSG00000027935.9 |
| chr6 | 113471067 | 113472016 | 18.84007 | 53.45216 | 50.69242 | Promoter | 6 | 113471427 | 113483140 | 11714 | 1 | ENSMUST00 | 0 | Il17rc | ENSMUSG00000030281.11 |
| chr13 | 19618978 | 19619910 | 20.21003 | 53.41216 | 50.65275 | Promoter | 13 | 19591709 | 19619830 | 28122 | 2 | ENSMUST00 | 0 | Epdr1 | ENSMUSG00000002808.6 |
| chr14 | 84442979 | 84444732 | 20.67895 | 53.40692 | 50.64761 | Promoter | 14 | 84443563 | 84537060 | 93498 | 1 | ENSMUST00 | 0 | Pcdh17 | ENSMUSG00000035566.6 |
| chr15 | 98918338 | 98918836 | 20.67895 | 53.40692 | 50.64761 | Promoter | 15 | 98903921 | 98918231 | 14311 | 2 | ENSMUST00 | -108 | Lmbr1l | ENSMUSG00000022999.9 |
| chr17 | 57105261 | 57105947 | 20.67895 | 53.40692 | 50.64761 | Promoter | 17 | 57105385 | 57107757 | 2373 | 1 | ENSMUST00 | 0 | Tnfsf9 | ENSMUSG00000035678.7 |
| chr18 | 23310009 | 23311168 | 20.67895 | 53.40692 | 50.64761 | Distal Interge | 18 | 23415415 | 23604230 | 188816 | 1 | ENSMUST00 | -104247 | Dtna | ENSMUSG00000024302.10 |
| chr3 | 80913147 | 80913922 | 20.67895 | 53.40692 | 50.64761 | Promoter | 3 | 80843599 | 80913614 | 70016 | 2 | ENSMUST00 | 0 | Glrb | ENSMUSG00000028020.11 |
| chr4 | 55650856 | 55651769 | 20.67895 | 53.40692 | 50.64761 | Distal Interge | 4 | 55605149 | 55636901 | 31753 | 1 | ENSMUST00 | 45708 | Gm12506 | ENSMUSG00000085375.1 |
| chr5 | 29467722 | 29468746 | 20.67895 | 53.40692 | 50.64761 | Distal Interge | 5 | 29473034 | 29478470 | 5437 | 2 | ENSMUST00 | 9724 | Mnx1 | ENSMUSG00000001566.9 |
| chr6 | 52175412 | 52176523 | 20.67895 | 53.40692 | 50.64761 | Promoter | 6 | 52175235 | 52215076 | 39842 | 1 | ENSMUST00 | 178 | Hoxaas3 | ENSMUSG00000085696.3 |
| chr7 | 25315191 | 25316112 | 20.67895 | 53.40692 | 50.64761 | 3' UTR | 7 | 25317164 | 25365917 | 48754 | 1 | ENSMUST00 | -1052 | Megf8 | ENSMUSG00000045039.9 |
| chr8 | 71676145 | 71676980 | 20.67895 | 53.40692 | 50.64761 | Promoter | 8 | 71676296 | 71688556 | 12261 | 1 | ENSMUST00 | 0 | Jak3 | ENSMUSG00000031805.12 |
| chrX | 112600460 | 112602098 | 20.67895 | 53.40692 | 50.64761 | Promoter | 20 | 112600526 | 112626073 | 25548 | 1 | ENSMUST00 | 0 | Zfp711 | ENSMUSG00000025529.9 |
| chrX | 81070309 | 81071644 | 20.67895 | 53.40692 | 50.64761 | Promoter | 20 | 81070698 | 81094776 | 24079 | 1 | ENSMUST00 | 0 | Tmem47 | ENSMUSG00000025666.11 |
| chrX | 96095680 | 96097279 | 20.67895 | 53.40692 | 50.64761 | Promoter | 20 | 96096042 | 96168552 | 72511 | 1 | ENSMUST00 | 0 | Msn | ENSMUSG00000031207.11 |
| chrX | 140540370 | 140543204 | 20.67895 | 53.40692 | 50.64761 | Promoter | 20 | 140539528 | 140542668 | 3141 | 2 | ENSMUST00 | 0 | Tsc22d3 | ENSMUSG00000031431.8 |
| chrX | 70364527 | 70365416 | 20.67895 | 53.40692 | 50.64761 | Promoter | 20 | 70343069 | 70365084 | 22016 | 2 | ENSMUST00 | 0 | Ids | ENSMUSG00000035847.10 |
| chrX | 48512817 | 48514323 | 20.67895 | 53.40692 | 50.64761 | Promoter | 20 | 48474944 | 48513426 | 38483 | 2 | ENSMUST00 | 0 | Aifm1 | ENSMUSG00000036932.9 |
| chrX | 162643170 | 162643946 | 20.67895 | 53.40692 | 50.64761 | Promoter | 20 | 162411954 | 162643601 | 231648 | 2 | ENSMUST00 | 0 | Reps2 | ENSMUSG00000040855.10 |
| chrX | 159626654 | 159628266 | 20.67895 | 53.40692 | 50.64761 | Promoter | 20 | 159627272 | 159978069 | 350798 | 1 | ENSMUST00 | 0 | Sh3kbp1 | ENSMUSG00000040990.12 |
| chr11 | 116532365 | 116534374 | 20.16949 | 53.34262 | 50.58483 | Promoter | 11 | 116532444 | 116536674 | 4231 | 1 | ENSMUST00 | 0 | Sphk1 | ENSMUSG00000061878.10 |
| chr2 | 14603812 | 14605255 | 20.16949 | 53.34262 | 50.58483 | Promoter | 2 | 14604306 | 14987908 | 383603 | 1 | ENSMUST00 | 0 | Cacnb2 | ENSMUSG00000057914.10 |
| chr7 | 72672768 | 72673082 | 20.16949 | 53.34262 | 50.58483 | Promoter | 7 | 72672818 | 72674278 | 1461 | 1 | ENSMUST00 | 0 | Kansl2-ps | ENSMUSG00000097530.1 |
| chr1 | 134646242 | 134647283 | 19.69817 | 53.33889 | 50.58166 | Promoter | 1 | 134646677 | 134741829 | 95153 | 1 | ENSMUST00 | 0 | Syt2 | ENSMUSG00000026452.10 |
| chr10 | 75780513 | 75781480 | 19.69817 | 53.33889 | 50.58166 | Promoter | 10 | 75774115 | 75781414 | 7300 | 2 | ENSMUST00 | 0 | Gstt3 | ENSMUSG00000001665.5 |
| chr10 | 58394347 | 58395666 | 19.69817 | 53.33889 | 50.58166 | Promoter | 10 | 58394373 | 58421783 | 27411 | 1 | ENSMUST00 | 0 | Lims1 | ENSMUSG00000019920.12 |
| chr10 | 60826421 | 60827506 | 19.69817 | 53.33889 | 50.58166 | Intron (ENSM | 10 | 60762610 | 60831581 | 68972 | 2 | ENSMUST00 | 4075 | Unc5b | ENSMUSG00000020099.7 |
| chr11 | 28852855 | 28854039 | 19.69817 | 53.33889 | 50.58166 | Promoter | 11 | 28853204 | 28926743 | 73540 | 1 | ENSMUST00 | 0 | Efemp1 | ENSMUSG00000020467.10 |
| chr11 | 87108240 | 87108850 | 19.69817 | 53.33889 | 50.58166 | Promoter | 11 | 87089153 | 87108708 | 19556 | 2 | ENSMUST00 | 0 | Prr11 | ENSMUSG00000020493.8 |
| chr13 | 23697582 | 23698486 | 19.69817 | 53.33889 | 50.58166 | Promoter | 13 | 23698060 | 23698454 | 395 | 2 | ENSMUST00 | 0 | Hist1h4c | ENSMUSG00000060678.4 |
| chr14 | 59625056 | 59626481 | 19.69817 | 53.33889 | 50.58166 | Promoter | 14 | 59625281 | 59631658 | 6378 | 1 | ENSMUST00 | 0 | Shisa2 | ENSMUSG00000044461.5 |
| chr14 | 118234864 | 118237398 | 19.69817 | 53.33889 | 50.58166 | Promoter | 14 | 118233232 | 118237030 | 3799 | 2 | ENSMUST00 | 0 | Sox21 | ENSMUSG00000061517.8 |
| chr15 | 76722485 | 76725119 | 19.69817 | 53.33889 | 50.58166 | Promoter | 15 | 76721466 | 76723845 | 2380 | 2 | ENSMUST00 | 0 | Lrrc24 | ENSMUSG00000033707.8 |
| chr15 | 100303932 | 100305487 | 19.69817 | 53.33889 | 50.58166 | Promoter | 15 | 100304817 | 100314348 | 9532 | 1 | ENSMUST00 | 0 | Mettl7a1 | ENSMUSG00000054619.6 |
| chr17 | 36110876 | 36111934 | 19.69817 | 53.33889 | 50.58166 | Promoter | 17 | 36109234 | 36111658 | 2425 | 2 | ENSMUST00 | 0 | C920025E04I | ENSMUSG00000073405.6 |
| chr19 | 5295609 | 5296117 | 19.69817 | 53.33889 | 50.58166 | Promoter | 19 | 5273923 | 5295455 | 21533 | 2 | ENSMUST00 | -155 | Sf3b2 | ENSMUSG00000024853.9 |
| chr19 | 10456412 | 10458156 | 19.69817 | 53.33889 | 50.58166 | Promoter | 19 | 10455371 | 10457447 | 2077 | 2 | ENSMUST00 | 0 | Lrrc10b | ENSMUSG00000090291.2 |

| | | | | | | | | | | | | | | | |
|---|---|---|---|---|---|---|---|---|---|---|---|---|---|---|---|
| chr3 | 146851908 | 146852954 | 19.69817 | 53.33889 | 50.58166 | Promoter | 3 | 146852367 | 146940256 | 87890 | 1 | ENSMUST00 | 0 | Ttll7 | ENSMUSG00000036745.10 |
| chr4 | 86669182 | 86670516 | 19.69817 | 53.33889 | 50.58166 | Promoter | 4 | 86656565 | 86670060 | 13496 | 2 | ENSMUST00 | 0 | Plin2 | ENSMUSG00000028494.7 |
| chr5 | 52476179 | 52476916 | 19.69817 | 53.33889 | 50.58166 | Distal Interge | 5 | 52374651 | 52471543 | 96893 | 2 | ENSMUST00 | -4637 | Ccdc149 | ENSMUSG00000045790.5 |
| chr7 | 7300374 | 7300976 | 19.23353 | 53.33828 | 50.58166 | Promoter | 7 | 7282316 | 7299511 | 17196 | 2 | ENSMUST00 | -864 | Clcn4-2 | ENSMUSG00000000605.8 |
| chr7 | 144895312 | 144897307 | 19.69817 | 53.33889 | 50.58166 | Promoter | 7 | 144896532 | 144900953 | 4422 | 1 | ENSMUST00 | 0 | Fgf15 | ENSMUSG00000031073.5 |
| chr7 | 29169161 | 29171818 | 19.69817 | 53.33889 | 50.58166 | Promoter | 7 | 29170220 | 29173933 | 3714 | 1 | ENSMUST00 | 0 | Ggn | ENSMUSG00000031493.9 |
| chr8 | 122443655 | 122444733 | 19.69817 | 53.33889 | 50.58166 | Promoter | 8 | 122433596 | 122443422 | 9827 | 2 | ENSMUST00 | -234 | Mvd | ENSMUSG00000006517.4 |
| chr8 | 120735776 | 120736926 | 19.69817 | 53.33889 | 50.58166 | Promoter | 8 | 120736358 | 120756694 | 20337 | 1 | ENSMUST00 | 0 | Irf8 | ENSMUSG00000041515.4 |
| chr11 | 70668185 | 70668815 | 20.12913 | 53.27335 | 50.5182 | Promoter | 11 | 70664003 | 70669218 | 5216 | 2 | ENSMUST00 | 403 | Spag7 | ENSMUSG00000018287.7 |
| chr4 | 11558399 | 11559409 | 20.59854 | 53.26988 | 50.51485 | Promoter | 4 | 11558922 | 11593653 | 34732 | 1 | ENSMUST00 | 0 | Rad54b | ENSMUSG00000078773.6 |
| chr15 | 85634793 | 85635329 | 18.26114 | 53.21968 | 50.46473 | Distal Interge | 15 | 85639747 | 85645934 | 6188 | 1 | ENSMUST00 | -4418 | RP23-64J7.2 | ENSMUSG00000101828.1 |
| chr19 | 58453143 | 58456144 | 18.26114 | 53.21968 | 50.46473 | Promoter | 19 | 58235604 | 58455398 | 219795 | 2 | ENSMUST00 | 0 | Gfra1 | ENSMUSG00000025089.10 |
| chr5 | 123507981 | 123509104 | 18.26114 | 53.21968 | 50.46473 | Promoter | 5 | 123508395 | 123514011 | 5617 | 1 | ENSMUST00 | 0 | Gm15751 | ENSMUSG00000086753.1 |
| chr14 | 54780833 | 54782193 | 20.08892 | 53.20432 | 50.4496 | Promoter | 14 | 54722216 | 54781886 | 59671 | 2 | ENSMUST00 | 0 | Slc7a8 | ENSMUSG00000022180.6 |
| chr13 | 103096647 | 103097446 | 20.5555 | 53.19651 | 50.4419 | Promoter | 13 | 102794128 | 103096828 | 302701 | 2 | ENSMUST00 | 0 | Mast4 | ENSMUSG00000034751.10 |
| chr3 | 45376055 | 45379083 | 20.5555 | 53.19651 | 50.4419 | Promoter | 3 | 45378398 | 45399594 | 21197 | 1 | ENSMUST00 | 0 | Pcdh10 | ENSMUSG00000049100.10 |
| chr5 | 41767831 | 41768602 | 20.5555 | 53.19651 | 50.4419 | Distal Interge | 5 | 41761483 | 41764501 | 3019 | 2 | ENSMUST00 | -3331 | Nkx3-2 | ENSMUSG00000049691.8 |
| chr6 | 72389183 | 72391497 | 20.5555 | 53.19651 | 50.4419 | Promoter | 6 | 72385223 | 72390703 | 5481 | 2 | ENSMUST00 | 0 | Vamp8 | ENSMUSG00000050732.9 |
| chr2 | 156474878 | 156476371 | 19.12836 | 53.15374 | 50.39941 | Promoter | 2 | 156475550 | 156499674 | 24125 | 1 | ENSMUST00 | 0 | Epb4.1l1 | ENSMUSG00000027624.12 |
| chr2 | 118372616 | 118374429 | 19.12836 | 53.15374 | 50.39941 | Promoter | 2 | 118277110 | 118373419 | 96310 | 2 | ENSMUST00 | 0 | Gpr176 | ENSMUSG00000040133.2 |
| chr4 | 137861800 | 137862868 | 19.12836 | 53.15374 | 50.39941 | Promoter | 4 | 137862237 | 137965229 | 102993 | 1 | ENSMUST00 | 0 | Ece1 | ENSMUSG00000057530.9 |
| chr4 | 114901057 | 114903863 | 19.12836 | 53.15374 | 50.39941 | Promoter | 4 | 114817161 | 114902859 | 85699 | 2 | ENSMUST00 | 0 | 9130410C08I | ENSMUSG00000085323.1 |
| chr7 | 16737852 | 16739254 | 20.04887 | 53.13555 | 50.38177 | Promoter | 7 | 16738410 | 16749290 | 10881 | 1 | ENSMUST00 | 0 | Ap2s1 | ENSMUSG00000008036.10 |
| chr11 | 69935210 | 69936511 | 20.51264 | 53.12344 | 50.36976 | Promoter | 11 | 69935796 | 69941605 | 5810 | 1 | ENSMUST00 | 0 | Ybx2 | ENSMUSG00000018554.8 |
| chr2 | 181319293 | 181320346 | 17.73747 | 53.06287 | 50.30949 | Promoter | 2 | 181319739 | 181356616 | 36878 | 1 | ENSMUST00 | 0 | Rtel1 | ENSMUSG00000038685.13 |
| chr5 | 24447457 | 24448224 | 17.73747 | 53.06287 | 50.30949 | Promoter | 5 | 24445458 | 24447854 | 2397 | 2 | ENSMUST00 | 0 | Tmub1 | ENSMUSG00000028958.9 |
| chr7 | 12961280 | 12962340 | 19.96926 | 52.99875 | 50.24557 | Promoter | 7 | 12948592 | 12961768 | 13177 | 2 | ENSMUST00 | 0 | 1810019N24 | ENSMUSG00000097115.2 |
| chr5 | 90890807 | 90891600 | 20.42746 | 52.97813 | 50.22504 | Promoter | 5 | 90891241 | 90893115 | 1875 | 1 | ENSMUST00 | 0 | Cxcl1 | ENSMUSG00000029380.10 |
| chr12 | 111670564 | 111672617 | 17.2615 | 52.96976 | 50.21678 | Promoter | 12 | 111669355 | 111672338 | 2984 | 2 | ENSMUST00 | 0 | Ckb | ENSMUSG00000001270.8 |
| chr18 | 38185112 | 38186528 | 20.38514 | 52.9059 | 50.15308 | 3' UTR | 18 | 38188888 | 38203163 | 14276 | 2 | ENSMUST00 | 16635 | Pcdh1 | ENSMUSG00000051375.10 |
| chr17 | 4907803 | 4909035 | 19.43846 | 52.88828 | 50.13554 | Distal Interge | 17 | 4994332 | 5097733 | 103402 | 1 | ENSMUST00 | -85297 | Arid1b | ENSMUSG00000069729.8 |
| chr17 | 36082963 | 36084288 | 19.43846 | 52.88828 | 50.13554 | Promoter | 17 | 36080189 | 36084223 | 4035 | 2 | ENSMUST00 | 0 | H2-Bl | ENSMUSG00000073406.5 |
| chr3 | 8766691 | 8767982 | 19.43846 | 52.88828 | 50.13554 | Distal Interge | 3 | 8679359 | 8679477 | 119 | 2 | ENSMUST00 | -87215 | Gm23670 | ENSMUSG00000088094.1 |
| chr4 | 41639574 | 41640212 | 19.43846 | 52.88828 | 50.13554 | Promoter | 4 | 41638144 | 41640324 | 2181 | 2 | ENSMUST00 | 112 | Enho | ENSMUSG00000028445.7 |
| chr5 | 47982701 | 47985505 | 19.43846 | 52.88828 | 50.13554 | Promoter | 5 | 47983155 | 47990332 | 7178 | 1 | ENSMUST00 | 0 | Slit2 | ENSMUSG00000031558.10 |
| chr5 | 149031105 | 149031748 | 19.43846 | 52.88828 | 50.13554 | Distal Interge | 5 | 148995651 | 149015813 | 20163 | 2 | ENSMUST00 | -15293 | Gm15411 | ENSMUSG00000085971.1 |
| chr6 | 146795267 | 146796559 | 19.43846 | 52.88828 | 50.13554 | Promoter | 6 | 146796055 | 146805578 | 9524 | 1 | ENSMUST00 | 0 | Arntl2 | ENSMUSG00000040187.10 |
| chr1 | 185454750 | 185455722 | 18.51447 | 52.87213 | 50.12019 | Promoter | 1 | 185454848 | 185468762 | 13915 | 1 | ENSMUST00 | 0 | Slc30a10 | ENSMUSG00000026614.6 |
| chr11 | 98051992 | 98053609 | 18.51447 | 52.87213 | 50.12019 | Promoter | 11 | 98036623 | 98053462 | 16840 | 2 | ENSMUST00 | 0 | Stac2 | ENSMUSG00000017400.5 |
| chr18 | 35552973 | 35553908 | 18.51447 | 52.87213 | 50.12019 | Promoter | 18 | 35553371 | 35558316 | 4946 | 1 | ENSMUST00 | 0 | Snhg4 | ENSMUSG00000097769.2 |
| chr4 | 33188741 | 33189861 | 18.51447 | 52.87213 | 50.12019 | Promoter | 4 | 33174230 | 33189433 | 15204 | 2 | ENSMUST00 | 0 | Pm20d2 | ENSMUSG00000054659.8 |
| chr8 | 46551401 | 46551820 | 18.51447 | 52.87213 | 50.12019 | Promoter | 8 | 46552055 | 46580007 | 27953 | 1 | ENSMUST00 | -235 | Cenpu | ENSMUSG00000031629.8 |
| chr3 | 84228184 | 84229304 | 19.89027 | 52.86293 | 50.11123 | Intron (ENSM | 3 | 84160441 | 84220889 | 60449 | 2 | ENSMUST00 | -7296 | Trim2 | ENSMUSG00000027993.11 |
| chr19 | 50677906 | 50679093 | 20.34299 | 52.83396 | 50.08234 | Promoter | 19 | 50143301 | 50678646 | 535346 | 2 | ENSMUST00 | 0 | Sorcs1 | ENSMUSG00000043531.10 |
| chr3 | 25143455 | 25144537 | 20.34299 | 52.83396 | 50.08234 | Distal Interge | 3 | 25211016 | 25212605 | 1590 | 2 | ENSMUST00 | 68068 | Gm10259 | ENSMUSG00000069083.4 |
| chr4 | 146263185 | 146263997 | 15.22315 | 52.78796 | 50.03657 | Promoter | 4 | 146263559 | 146264190 | 632 | 1 | ENSMUST00 | 0 | Gm13168 | ENSMUSG00000081460.1 |
| chr11 | 87639012 | 87639621 | 19.8119 | 52.72808 | 49.97682 | Intron (ENSM | 11 | 87617164 | 87659542 | 42379 | 1 | ENSMUST00 | 21849 | Hsf5 | ENSMUSG00000070345.3 |
| chr15 | 81802642 | 81804673 | 20.25921 | 52.6909 | 49.93975 | Promoter | 15 | 81802707 | 81806049 | 3343 | 1 | ENSMUST00 | 0 | Tef | ENSMUSG00000022389.9 |
| chr4 | 108070413 | 108071809 | 17.96111 | 52.67509 | 49.92405 | Promoter | 4 | 108044378 | 108071363 | 26986 | 2 | ENSMUST00 | 0 | Scp2 | ENSMUSG00000028603.10 |
| chr1 | 171221519 | 171222502 | 18.39885 | 52.66533 | 49.91441 | Promoter | 1 | 171217802 | 171222184 | 4383 | 2 | ENSMUST00 | 0 | Tomm40l | ENSMUSG00000005674.4 |
| chr1 | 195091802 | 195092875 | 18.39885 | 52.66533 | 49.91441 | Promoter | 1 | 195041285 | 195092248 | 50964 | 2 | ENSMUST00 | 0 | Cd46 | ENSMUSG00000016493.8 |
| chr1 | 155740637 | 155743345 | 18.39885 | 52.66533 | 49.91441 | 5' UTR | 1 | 155698031 | 155751710 | 53680 | 2 | ENSMUST00 | 8365 | Lhx4 | ENSMUSG00000026468.9 |
| chr1 | 21166285 | 21167113 | 18.39885 | 52.66533 | 49.91441 | Distal Interge | 1 | 21178988 | 21187427 | 8440 | 1 | ENSMUST00 | -11875 | RP23-80N7.1 | ENSMUSG00000100819.1 |
| chr10 | 19712162 | 19712955 | 18.39885 | 52.66533 | 49.91441 | Promoter | 10 | 19712587 | 19760053 | 47467 | 1 | ENSMUST00 | 0 | Il20ra | ENSMUSG00000020007.3 |
| chr10 | 81811954 | 81812659 | 18.39885 | 52.66533 | 49.91441 | Promoter | 10 | 81812186 | 81826490 | 14305 | 1 | ENSMUST00 | 0 | Gm4767 | ENSMUSG00000094076.2 |
| chr11 | 4946942 | 4948152 | 18.39885 | 52.66533 | 49.91441 | Promoter | 11 | 4938754 | 4948064 | 9311 | 2 | ENSMUST00 | 0 | Nefh | ENSMUSG00000020396.8 |
| chr13 | 42050686 | 42051679 | 18.39885 | 52.66533 | 49.91441 | Promoter | 13 | 42052021 | 42185026 | 133006 | 1 | ENSMUST00 | -342 | Hivep1 | ENSMUSG00000021366.7 |
| chr13 | 67830660 | 67831434 | 18.39885 | 52.66533 | 49.91441 | Promoter | 13 | 67796594 | 67830985 | 34392 | 2 | ENSMUST00 | 0 | 4930525G20 | ENSMUSG00000097276.1 |
| chr15 | 61984706 | 61987357 | 18.39885 | 52.66533 | 49.91441 | Promoter | 15 | 61985341 | 61990352 | 5012 | 1 | ENSMUST00 | 0 | Myc | ENSMUSG00000022346.9 |
| chr17 | 35469918 | 35471169 | 18.39885 | 52.66533 | 49.91441 | Promoter | 17 | 35470089 | 35474563 | 4475 | 1 | ENSMUST00 | 0 | H2-Q10 | ENSMUSG00000067235.9 |
| chr19 | 4332597 | 4334428 | 18.39885 | 52.66533 | 49.91441 | Promoter | 19 | 4324906 | 4334001 | 9096 | 2 | ENSMUST00 | 0 | Kdm2a | ENSMUSG00000054611.11 |
| chr2 | 18821353 | 18822362 | 18.39885 | 52.66533 | 49.91441 | Distal Interge | 2 | 18799245 | 18801818 | 2574 | 2 | ENSMUST00 | -19536 | Carlr | ENSMUSG00000097638.1 |
| chr2 | 74727178 | 74727798 | 18.39885 | 52.66533 | 49.91441 | Promoter | 2 | 74727074 | 74729123 | 2050 | 1 | ENSMUST00 | 105 | Hoxd4 | ENSMUSG00000101174.1 |
| chr3 | 108225864 | 108227336 | 18.39885 | 52.66533 | 49.91441 | Promoter | 3 | 108211472 | 108226648 | 15177 | 2 | ENSMUST00 | 0 | Sypl2 | ENSMUSG00000027887.9 |
| chr5 | 151189646 | 151190537 | 18.39885 | 52.66533 | 49.91441 | Promoter | 5 | 151037628 | 151190194 | 152567 | 2 | ENSMUST00 | 0 | Stard13 | ENSMUSG00000016128.9 |
| chr5 | 34287792 | 34288812 | 18.39885 | 52.66533 | 49.91441 | Promoter | 5 | 34194874 | 34288326 | 93453 | 2 | ENSMUST00 | 0 | Zfyve28 | ENSMUSG00000037224.10 |
| chr5 | 24750232 | 24752112 | 18.39885 | 52.66533 | 49.91441 | Promoter | 5 | 24751003 | 24751324 | 322 | 2 | ENSMUST00 | 0 | Crygn | ENSMUSG00000038135.4 |
| chr7 | 6343408 | 6344193 | 18.39885 | 52.66533 | 49.91441 | Distal Interge | 7 | 6315660 | 6331285 | 15626 | 2 | ENSMUST00 | -12124 | Zfp583 | ENSMUSG00000030443.11 |
| chr7 | 29766855 | 29767723 | 18.39885 | 52.66533 | 49.91441 | Promoter | 7 | 29768552 | 29779821 | 11270 | 1 | ENSMUST00 | -829 | Zfp84 | ENSMUSG00000046185.10 |
| chr7 | 93079670 | 93081372 | 18.39885 | 52.66533 | 49.91441 | Promoter | 7 | 93079879 | 93081715 | 1837 | 1 | ENSMUST00 | 0 | Fam181b | ENSMUSG00000051515.8 |
| chr7 | 42642109 | 42643251 | 18.39885 | 52.66533 | 49.91441 | Promoter | 7 | 42609526 | 42642588 | 33063 | 2 | ENSMUST00 | 0 | 9830147E19I | ENSMUSG00000074158.4 |
| chr7 | 73538075 | 73541788 | 18.39885 | 52.66533 | 49.91441 | Promoter | 7 | 73426691 | 73541746 | 115056 | 2 | ENSMUST00 | 0 | Chd2 | ENSMUSG00000078671.5 |
| chr8 | 122442663 | 122443547 | 18.39885 | 52.66533 | 49.91441 | Promoter | 8 | 122433596 | 122443422 | 9827 | 2 | ENSMUST00 | 0 | Mvd | ENSMUSG00000006517.4 |
| chr9 | 119983901 | 119984798 | 18.39885 | 52.66533 | 49.91441 | Intron (ENSM | 9 | 119971166 | 119977250 | 6085 | 2 | ENSMUST00 | -6652 | Csrnp1 | ENSMUSG00000032515.7 |
| chr9 | 107337899 | 107338921 | 18.39885 | 52.66533 | 49.91441 | Promoter | 9 | 107327692 | 107338317 | 10626 | 2 | ENSMUST00 | 0 | Hemk1 | ENSMUSG00000032579.9 |
| chr9 | 45369887 | 45371230 | 18.39885 | 52.66533 | 49.91441 | Promoter | 9 | 45370185 | 45396158 | 25974 | 1 | ENSMUST00 | 0 | Fxyd6 | ENSMUSG00000066705.6 |
| chr9 | 22327380 | 22328388 | 18.39885 | 52.66533 | 49.91441 | Downstream | 9 | 22320314 | 22321410 | 1097 | 2 | ENSMUST00 | -5971 | Gm10182 | ENSMUSG00000094627.1 |
| chr10 | 128547879 | 128549100 | 19.77295 | 52.66102 | 49.91236 | Promoter | 10 | 128548120 | 128549102 | 983 | 2 | ENSMUST00 | 2 | Rpl41 | ENSMUSG00000093674.2 |
| chr11 | 8623718 | 8624944 | 19.77295 | 52.66102 | 49.91236 | Intron (ENSM | 11 | 8431652 | 8664535 | 232884 | 2 | ENSMUST00 | 39591 | Tns3 | ENSMUSG00000020422.8 |
| chr11 | 6528014 | 6528785 | 19.77295 | 52.66102 | 49.91236 | Promoter | 11 | 6528033 | 6528165 | 133 | 2 | ENSMUST00 | 0 | Gm22457 | ENSMUSG00000064513.1 |
| chr11 | 58959775 | 58960689 | 19.77295 | 52.66102 | 49.91236 | Intron (ENSM | 11 | 58961812 | 58962222 | 411 | 1 | ENSMUST00 | -1123 | Gm12260 | ENSMUSG00000080152.4 |
| chr12 | 25099209 | 25101142 | 19.77295 | 52.66102 | 49.91236 | Distal Interge | 12 | 25093801 | 25096092 | 2292 | 2 | ENSMUST00 | -3118 | Id2 | ENSMUSG00000020644.8 |
| chr12 | 27338585 | 27339824 | 19.77295 | 52.66101 | 49.91236 | 3' UTR | 12 | 27334264 | 27342574 | 8311 | 2 | ENSMUST00 | 2750 | Sox11 | ENSMUSG00000063632.6 |
| chr14 | 50931125 | 50931992 | 19.77295 | 52.66101 | 49.91236 | Promoter | 14 | 50927215 | 50930856 | 3642 | 2 | ENSMUST00 | -270 | Tmem55b | ENSMUSG00000035953.8 |
| chr17 | 33950616 | 33951977 | 19.77295 | 52.66101 | 49.91236 | Promoter | 17 | 33949909 | 33951477 | 1569 | 2 | ENSMUST00 | 0 | B3galt4 | ENSMUSG00000067370.3 |
| chr7 | 110163997 | 110164938 | 19.77295 | 52.66102 | 49.91236 | Distal Interge | 7 | 109998412 | 110151202 | 152791 | 2 | ENSMUST00 | -12796 | RP24-272D1! | ENSMUSG00000101585.1 |
| chr8 | 45627304 | 45628303 | 19.77295 | 52.66102 | 49.91236 | Promoter | 8 | 45627745 | 45745807 | 118063 | 1 | ENSMUST00 | 0 | Sorbs2 | ENSMUSG00000031626.11 |
| chr3 | 86998763 | 86999954 | 20.21758 | 52.61978 | 49.87169 | Promoter | 3 | 86995834 | 86999309 | 3476 | 2 | ENSMUST00 | 0 | Cd1d1 | ENSMUSG00000028076.7 |
| chr2 | 127831364 | 127832306 | 19.73415 | 52.59419 | 49.84622 | Promoter | 2 | 127801122 | 127831865 | 30744 | 2 | ENSMUST00 | 0 | Bub1 | ENSMUSG00000027379.8 |
| chr13 | 113044785 | 113046798 | 15.11128 | 52.55099 | 49.80313 | Promoter | 13 | 113042763 | 113046388 | 3626 | 2 | ENSMUST00 | 0 | Gpx8 | ENSMUSG00000021760.3 |
| chr14 | 54893354 | 54893892 | 18.78595 | 52.55086 | 49.80313 | Promoter | 14 | 54894148 | 54898156 | 4009 | 1 | ENSMUST00 | -256 | Pabpn1 | ENSMUSG00000022194.10 |
| chr16 | 37011163 | 37012278 | 15.11128 | 52.55099 | 49.80313 | Promoter | 16 | 37011786 | 37094992 | 83207 | 1 | ENSMUST00 | 0 | Polq | ENSMUSG00000034206.10 |
| chr2 | 119112165 | 119113292 | 15.11128 | 52.55099 | 49.80313 | Promoter | 2 | 119112793 | 119123846 | 11054 | 1 | ENSMUST00 | 0 | Rad51 | ENSMUSG00000027323.5 |
| chr5 | 34659465 | 34660171 | 18.78595 | 52.55086 | 49.80313 | Promoter | 5 | 34638532 | 34660148 | 21617 | 2 | ENSMUST00 | 0 | Nop14 | ENSMUSG00000036693.9 |
| chr6 | 113418553 | 113419900 | 18.78595 | 52.55086 | 49.80313 | Promoter | 6 | 113415319 | 113419348 | 4030 | 2 | ENSMUST00 | 0 | Rpusd3 | ENSMUSG00000051169.9 |
| chr1 | 164275320 | 164276174 | 13.46766 | 52.51953 | 49.7725 | Promoter | 1 | 164275585 | 164287847 | 12263 | 1 | ENSMUST00 | 0 | Ccdc181 | ENSMUSG00000026578.6 |
| chr11 | 49901520 | 49902492 | 20.13483 | 52.47836 | 49.7314 | Promoter | 11 | 49901835 | 49980222 | 78388 | 1 | ENSMUST00 | 0 | Rasgef1c | ENSMUSG00000020374.11 |
| chr11 | 66984558 | 66985689 | 20.13483 | 52.47836 | 49.7314 | Distal Interge | 11 | 67002086 | 67011625 | 9540 | 1 | ENSMUST00 | -16397 | Gm12299 | ENSMUSG00000085926.1 |
| chr14 | 70658934 | 70660134 | 20.13483 | 52.47836 | 49.7314 | 3' UTR | 14 | 70647302 | 70653084 | 5783 | 2 | ENSMUST00 | -5851 | Npm2 | ENSMUSG00000047911.5 |
| chr5 | 35056563 | 35057538 | 20.13483 | 52.47836 | 49.7314 | Promoter | 5 | 35056813 | 35080964 | 24152 | 1 | ENSMUST00 | 0 | Dok7 | ENSMUSG00000044716.7 |
| chr6 | 124903738 | 124904839 | 20.13483 | 52.47836 | 49.7314 | 3' UTR | 6 | 124908599 | 124910051 | 1453 | 2 | ENSMUST00 | 5212 | Lag3 | ENSMUSG00000030124.2 |
| chr7 | 44986832 | 44987564 | 20.13483 | 52.47836 | 49.7314 | Promoter | 7 | 44986900 | 44991916 | 5017 | 1 | ENSMUST00 | 0 | Gm15545 | ENSMUSG00000087138.1 |
| chr12 | 54728991 | 54729929 | 12.60172 | 52.46623 | 49.71969 | Promoter | 12 | 54729602 | 54729765 | 164 | 1 | ENSMUST00 | 0 | Gm26444 | ENSMUSG00000093815.1 |
| chr8 | 122335652 | 122336940 | 20.05275 | 52.33802 | 49.5918 | Exon (ENSMU | 8 | 122333786 | 122335761 | 1976 | 1 | ENSMUST00 | 1867 | Zfpm1 | ENSMUSG00000049577.9 |
| chr1 | 189342645 | 189344089 | 17.34451 | 52.33425 | 49.58816 | Promoter | 1 | 189207931 | 189343353 | 135423 | 2 | ENSMUST00 | 0 | Kcnk2 | ENSMUSG00000037624.10 |
| chr10 | 44458183 | 44459278 | 17.34451 | 52.33425 | 49.58816 | Promoter | 10 | 44437177 | 44458748 | 21572 | 2 | ENSMUST00 | 0 | Prdm1 | ENSMUSG00000038151.7 |
| chr10 | 22058265 | 22059231 | 17.34451 | 52.33425 | 49.58816 | Distal Interge | 10 | 22066307 | 22068079 | 1773 | 2 | ENSMUST00 | 8848 | 4930444G20 | ENSMUSG00000069712.5 |
| chr12 | 69227716 | 69228488 | 17.34451 | 52.33425 | 49.58816 | Promoter | 12 | 69201779 | 69228190 | 26412 | 2 | ENSMUST00 | 0 | Pole2 | ENSMUSG00000020974.5 |
| chr14 | 76506045 | 76507699 | 17.34451 | 52.33425 | 49.58816 | Promoter | 14 | 76505098 | 76507002 | 1905 | 1 | ENSMUST00 | 948 | Tsc22d1 | ENSMUSG00000022010.14 |
| chr16 | 34745215 | 34745786 | 17.34451 | 52.33425 | 49.58816 | Promoter | 16 | 34745210 | 34877836 | 132627 | 1 | ENSMUST00 | 6 | Mylk | ENSMUSG00000022836.10 |
| chr17 | 29489687 | 29490498 | 17.34451 | 52.33425 | 49.58816 | Promoter | 17 | 29490812 | 29496111 | 5300 | 1 | ENSMUST00 | -314 | Pim1 | ENSMUSG00000024014.7 |
| chr17 | 56609150 | 56610552 | 17.34451 | 52.33425 | 49.58816 | Promoter | 17 | 56607452 | 56609771 | 2320 | 2 | ENSMUST00 | 0 | 2410015M20 | ENSMUSG00000049760.5 |
| chr17 | 23924482 | 23925483 | 17.34451 | 52.33425 | 49.58816 | Distal Interge | 17 | 23917458 | 23919441 | 1984 | 2 | ENSMUST00 | 7025 | Dcpp3 | ENSMUSG00000057417.6 |
| chr17 | 22738421 | 22740255 | 17.34451 | 52.33425 | 49.58816 | Promoter | 17 | 22739944 | 22751509 | 11566 | 1 | ENSMUST00 | 0 | Gm5493 | ENSMUSG00000098164.1 |
| chr4 | 154898399 | 154899526 | 17.34451 | 52.33425 | 49.58816 | Promoter | 4 | 154895504 | 154899038 | 3535 | 2 | ENSMUST00 | 0 | Fam213b | ENSMUSG00000029059.4 |
| chr4 | 33923493 | 33925292 | 17.34451 | 52.33425 | 49.58816 | Promoter | 4 | 33924593 | 33935316 | 10724 | 1 | ENSMUST00 | 0 | Cnr1 | ENSMUSG00000044288.6 |
| chr5 | 137740937 | 137742017 | 17.34451 | 52.33425 | 49.58816 | Promoter | 5 | 137730913 | 137741607 | 10695 | 2 | ENSMUST00 | 0 | Nyap1 | ENSMUSG00000045348.10 |
| chr9 | 65884237 | 65885594 | 17.34451 | 52.33425 | 49.58816 | Promoter | 9 | 65831136 | 65885055 | 53920 | 2 | ENSMUST00 | 0 | Trip4 | ENSMUSG00000032386.10 |
| chr9 | 57439339 | 57440186 | 17.34451 | 52.33425 | 49.58816 | Promoter | 9 | 57412668 | 57440114 | 27447 | 2 | ENSMUST00 | 0 | Ppcdc | ENSMUSG00000063849.5 |
| chr9 | 63108809 | 63109798 | 17.34451 | 52.33425 | 49.58816 | Distal Interge | 9 | 63133283 | 63133389 | 107 | 1 | ENSMUST00 | -23485 | Gm25064 | ENSMUSG00000094201.1 |
| chr12 | 23009739 | 23010495 | 15.7376 | 52.33027 | 49.58585 | Promoter | 12 | 22953997 | 23010243 | 56247 | 2 | ENSMUST00 | 0 | 2410018L13I | ENSMUSG00000073164.6 |
| chr14 | 70618113 | 70618914 | 15.7376 | 52.33027 | 49.58585 | 5' UTR | 14 | 70602185 | 70627065 | 24881 | 2 | ENSMUST00 | 8151 | Dmtn | ENSMUSG00000022099.11 |
| chr14 | 78724500 | 78725859 | 15.7376 | 52.33027 | 49.58585 | Promoter | 14 | 78569609 | 78725089 | 155481 | 2 | ENSMUST00 | 0 | Dgkh | ENSMUSG00000034731.9 |
| chr14 | 32856657 | 32857567 | 15.7376 | 52.33027 | 49.58585 | Promoter | 14 | 32856756 | 32939487 | 82732 | 1 | ENSMUST00 | 0 | Vstm4 | ENSMUSG00000050666.8 |
| chr2 | 32082901 | 32084991 | 15.7376 | 52.33027 | 49.58585 | Promoter | 2 | 32050055 | 32084669 | 34615 | 2 | ENSMUST00 | 0 | Fam78a | ENSMUSG00000050592.8 |
| chr7 | 127013650 | 127014751 | 15.7376 | 52.33027 | 49.58585 | Promoter | 7 | 126986860 | 127014621 | 27762 | 2 | ENSMUST00 | 0 | Mvp | ENSMUSG00000030681.13 |
| chr9 | 110624023 | 110625607 | 15.7376 | 52.33027 | 49.58585 | Promoter | 9 | 110621137 | 110624393 | 3257 | 2 | ENSMUST00 | 0 | Nradd | ENSMUSG00000032491.10 |
| chr11 | 103102978 | 103103575 | 19.58047 | 52.32923 | 49.58531 | Promoter | 11 | 103103021 | 103105466 | 2446 | 1 | ENSMUST00 | 0 | Acbd4 | ENSMUSG00000056938.11 |

| | | | | | | | | | | | | | | | |
|---|---|---|---|---|---|---|---|---|---|---|---|---|---|---|---|
| chr10 | 62865033 | 62866107 | 18.19994 | 52.30853 | 49.5647 | Promoter | 10 | 62839097 | 62865304 | 26208 | 2 | ENSMUST00 | 0 | Tet1 | ENSMUSG00000047146.10 |
| chr2 | 116075411 | 116076396 | 18.19994 | 52.30853 | 49.5647 | Intron (ENSM | 2 | 116067968 | 116085910 | 17943 | 1 | ENSMUST00 | 7444 | G630016G05 | ENSMUSG00000086308.1 |
| chr10 | 93081228 | 93081728 | 20.01196 | 52.26824 | 49.52467 | Promoter | 10 | 92775619 | 93081618 | 306000 | 2 | ENSMUST00 | 0 | 4930485B16 | ENSMUSG00000020014.10 |
| chr9 | 20544090 | 20545340 | 20.01196 | 52.26824 | 49.52467 | Intron (ENSM | 9 | 20495068 | 20521417 | 26350 | 2 | ENSMUST00 | -22674 | Zfp266 | ENSMUSG00000060510.8 |
| chr7 | 73558783 | 73559920 | 19.54243 | 52.26357 | 49.52013 | Promoter | 7 | 73539801 | 73558395 | 18595 | 2 | ENSMUST00 | -389 | 1810026B05 | ENSMUSG00000098496.2 |
| chr10 | 60106252 | 60107797 | 19.07758 | 52.25967 | 49.51631 | Promoter | 10 | 60106468 | 60133908 | 27441 | 1 | ENSMUST00 | 0 | Spock2 | ENSMUSG00000058297.11 |
| chr11 | 69008066 | 69008797 | 19.07758 | 52.25967 | 49.51631 | Promoter | 11 | 68985697 | 69008460 | 22764 | 2 | ENSMUST00 | 0 | Pfas | ENSMUSG00000020899.10 |
| chr6 | 14900632 | 14902079 | 19.07758 | 52.25967 | 49.51631 | Promoter | 6 | 14901349 | 15441977 | 540629 | 1 | ENSMUST00 | 0 | Foxp2 | ENSMUSG00000029563.11 |
| chr6 | 86028552 | 86029359 | 19.07758 | 52.25967 | 49.51631 | Distal Interge | 6 | 86017191 | 86020996 | 3806 | 1 | ENSMUST00 | 11362 | Figla | ENSMUSG00000030001.3 |
| chr8 | 68793287 | 68793815 | 19.07758 | 52.25967 | 49.51631 | Promoter | 8 | 68793929 | 68829410 | 35482 | 1 | ENSMUST00 | -114 | Ints10 | ENSMUSG00000031864.10 |
| chr10 | 60218504 | 60219508 | 16.47183 | 52.24387 | 49.50113 | Promoter | 10 | 60181532 | 60219260 | 37729 | 2 | ENSMUST00 | 0 | Chst3 | ENSMUSG00000057337.8 |
| chr14 | 55490480 | 55491847 | 16.47183 | 52.24387 | 49.50113 | Promoter | 14 | 55491093 | 55508246 | 17154 | 1 | ENSMUST00 | 0 | Lrrc16b | ENSMUSG00000022211.8 |
| chr4 | 20777270 | 20778731 | 16.47183 | 52.24387 | 49.50113 | Promoter | 4 | 20118874 | 20778567 | 659694 | 2 | ENSMUST00 | 0 | Nkain3 | ENSMUSG00000055761.9 |
| chr4 | 88760058 | 88760619 | 16.47183 | 52.24387 | 49.50113 | Promoter | 4 | 88760119 | 88761581 | 1463 | 1 | ENSMUST00 | 0 | Gm13283 | ENSMUSG00000100505.1 |
| chr7 | 98834790 | 98835529 | 16.47183 | 52.24387 | 49.50113 | Promoter | 7 | 98835112 | 98846587 | 11476 | 1 | ENSMUST00 | 0 | Wnt11 | ENSMUSG00000015957.6 |
| chr8 | 84065929 | 84066555 | 16.47183 | 52.24387 | 49.50113 | Promoter | 8 | 84066836 | 84096992 | 30157 | 1 | ENSMUST00 | -281 | Rfx1 | ENSMUSG00000031706.6 |
| chr15 | 64920781 | 64923006 | 19.97134 | 52.19873 | 49.45697 | Promoter | 15 | 64699042 | 64922296 | 223255 | 2 | ENSMUST00 | 0 | Adcy8 | ENSMUSG00000022376.7 |
| chr11 | 69921657 | 69922644 | 19.93088 | 52.12947 | 49.38807 | Promoter | 11 | 69916720 | 69921958 | 5239 | 2 | ENSMUST00 | 0 | Eif5a | ENSMUSG00000078812.5 |
| chr2 | 52675430 | 52676764 | 19.93088 | 52.12947 | 49.38807 | Promoter | 2 | 52428320 | 52676582 | 248263 | 2 | ENSMUST00 | 0 | Cacnb4 | ENSMUSG00000017412.10 |
| chr7 | 30089768 | 30091197 | 19.89059 | 52.06047 | 49.31924 | Promoter | 7 | 30056034 | 30090410 | 34377 | 2 | ENSMUST00 | 0 | Gm26920 | ENSMUSG00000058447.8 |
| chr13 | 67332011 | 67333072 | 19.3917 | 52.00318 | 49.26217 | Promoter | 13 | 67312998 | 67332547 | 19550 | 2 | ENSMUST00 | 0 | Zfp595 | ENSMUSG00000057842.8 |
| chr13 | 48544202 | 48545798 | 19.3917 | 52.00318 | 49.26217 | Distal Interge | 13 | 48538171 | 48538270 | 100 | 2 | ENSMUST00 | -5933 | Mirlet7a-1 | ENSMUSG00000065421.2 |
| chr7 | 100582519 | 100583374 | 19.3917 | 52.00318 | 49.26217 | Promoter | 7 | 100545751 | 100583102 | 37352 | 2 | ENSMUST00 | 0 | Mrpl48 | ENSMUSG00000030706.11 |
| chr9 | 64385390 | 64386659 | 19.3917 | 52.00318 | 49.26217 | Promoter | 9 | 64385626 | 64700975 | 315350 | 1 | ENSMUST00 | 0 | Megf11 | ENSMUSG00000036466.12 |
| chr10 | 81929884 | 81930928 | 19.85046 | 51.99174 | 49.25125 | Distal Interge | 10 | 81905073 | 81910899 | 5827 | 2 | ENSMUST00 | -18986 | Zfp781 | ENSMUSG00000096718.1 |
| chr4 | 132302863 | 132303566 | 19.85046 | 51.99174 | 49.25125 | Promoter | 4 | 132302197 | 132303385 | 1189 | 2 | ENSMUST00 | 0 | Rab42 | ENSMUSG00000089687.2 |
| chr13 | 21833990 | 21834707 | 19.81049 | 51.92325 | 49.18326 | Promoter | 13 | 21833758 | 21837530 | 3773 | 1 | ENSMUST00 | 233 | Hist1h2br | ENSMUSG00000069303.5 |
| chr1 | 187608590 | 187609891 | 20.21941 | 51.82391 | 49.08421 | Promoter | 1 | 187608791 | 188043502 | 434712 | 1 | ENSMUST00 | 0 | Esrrg | ENSMUSG00000026610.8 |
| chr11 | 69546420 | 69547501 | 20.21941 | 51.82391 | 49.08421 | Intron (ENSM | 11 | 69420809 | 69549108 | 128300 | 2 | ENSMUST00 | 1607 | Dnah2 | ENSMUSG00000005237.8 |
| chr11 | 69415361 | 69416249 | 20.21941 | 51.82391 | 49.08421 | Distal Interge | 11 | 69398508 | 69413675 | 15168 | 2 | ENSMUST00 | -1687 | Kdm6b | ENSMUSG00000018476.7 |
| chr11 | 54303906 | 54304505 | 20.21941 | 51.82391 | 49.08421 | Promoter | 11 | 54304005 | 54361535 | 57531 | 1 | ENSMUST00 | 0 | Acsl6 | ENSMUSG00000020333.12 |
| chr13 | 114458240 | 114460083 | 20.21941 | 51.82391 | 49.08421 | Promoter | 13 | 114452262 | 114458730 | 6469 | 2 | ENSMUST00 | 0 | Fst | ENSMUSG00000021765.7 |
| chr13 | 9093226 | 9093624 | 20.21941 | 51.82391 | 49.08421 | Promoter | 13 | 9093881 | 9171104 | 77224 | 1 | ENSMUST00 | -257 | Larp4b | ENSMUSG00000033499.8 |
| chr14 | 7773884 | 7774597 | 20.21941 | 51.82391 | 49.08421 | Promoter | 14 | 7764818 | 7774069 | 9252 | 2 | ENSMUST00 | 0 | Gm10044 | ENSMUSG00000072723.5 |
| chr15 | 89059206 | 89060479 | 20.21941 | 51.82391 | 49.08421 | Promoter | 15 | 89059734 | 89069878 | 10145 | 1 | ENSMUST00 | 0 | Panx2 | ENSMUSG00000058441.7 |
| chr4 | 8238607 | 8239312 | 20.21941 | 51.82391 | 49.08421 | Promoter | 4 | 8143362 | 8239041 | 95680 | 2 | ENSMUST00 | 0 | Car8 | ENSMUSG00000041261.9 |
| chr5 | 147724596 | 147726689 | 20.21941 | 51.82391 | 49.08421 | Promoter | 5 | 147561604 | 147726011 | 164408 | 2 | ENSMUST00 | 0 | Flt1 | ENSMUSG00000029648.8 |
| chr7 | 15966389 | 15967632 | 20.21941 | 51.82391 | 49.08421 | Promoter | 7 | 15946977 | 15967567 | 20591 | 2 | ENSMUST00 | 0 | Ehd2 | ENSMUSG00000074364.5 |
| chrX | 53369514 | 53370565 | 20.21941 | 51.82391 | 49.08421 | Promoter | 20 | 53344598 | 53370502 | 25905 | 2 | ENSMUST00 | 0 | Mospd1 | ENSMUSG00000023074.6 |
| chrX | 158042028 | 158044073 | 20.21941 | 51.82391 | 49.08421 | Promoter | 20 | 157821436 | 158043294 | 221859 | 2 | ENSMUST00 | 0 | Cnksr2 | ENSMUSG00000025658.11 |
| chrX | 50840775 | 50842647 | 20.21941 | 51.82391 | 49.08421 | Promoter | 20 | 50841047 | 50893097 | 52051 | 1 | ENSMUST00 | 0 | 2610018G03 | ENSMUSG00000031112.5 |
| chrX | 7670173 | 7671711 | 20.21941 | 51.82391 | 49.08421 | Promoter | 20 | 7667941 | 7671390 | 3450 | 2 | ENSMUST00 | 0 | Plp2 | ENSMUSG00000031146.6 |
| chrX | 101419086 | 101420481 | 20.21941 | 51.82391 | 49.08421 | Promoter | 20 | 101404384 | 101419798 | 15415 | 2 | ENSMUST00 | 0 | Zmym3 | ENSMUSG00000031310.11 |
| chrX | 142965604 | 142967368 | 20.21941 | 51.82391 | 49.08421 | Promoter | 20 | 142851146 | 142966728 | 115583 | 2 | ENSMUST00 | 0 | Ammecr1 | ENSMUSG00000042225.3 |
| chrX | 104199573 | 104201696 | 20.21941 | 51.82391 | 49.08421 | Promoter | 20 | 104077434 | 104201185 | 123752 | 2 | ENSMUST00 | 0 | C77370 | ENSMUSG00000046449.10 |
| chrX | 122396661 | 122397969 | 20.21941 | 51.82391 | 49.08421 | Promoter | 20 | 122394565 | 122397401 | 2837 | 2 | ENSMUST00 | 0 | Nap1l3 | ENSMUSG00000055733.6 |
| chrX | 12760788 | 12762958 | 20.21941 | 51.82391 | 49.08421 | Promoter | 20 | 12675369 | 12762073 | 86705 | 2 | ENSMUST00 | 0 | Med14 | ENSMUSG00000064127.8 |
| chr10 | 80164488 | 80165117 | 19.27906 | 51.80829 | 49.07 | Promoter | 10 | 80165985 | 80170316 | 4332 | 1 | ENSMUST00 | -868 | Cirbp | ENSMUSG00000045193.8 |
| chr10 | 37138147 | 37141657 | 19.27906 | 51.80829 | 49.07 | Promoter | 10 | 37139402 | 37143366 | 3965 | 1 | ENSMUST00 | 0 | 5930403N24 | ENSMUSG00000046463.4 |
| chr11 | 82386895 | 82389905 | 19.27906 | 51.80829 | 49.07 | Promoter | 11 | 82388900 | 82446327 | 57428 | 1 | ENSMUST00 | 0 | Tmem132e | ENSMUSG00000020701.7 |
| chr11 | 75193308 | 75195378 | 19.27906 | 51.80829 | 49.07 | Promoter | 11 | 75193783 | 75267769 | 73987 | 1 | ENSMUST00 | 0 | Rtn4rl1 | ENSMUSG00000045287.6 |
| chr11 | 94148737 | 94149674 | 19.27906 | 51.80829 | 49.07 | Intron (ENSM | 11 | 94135660 | 94138845 | 3186 | 1 | ENSMUST00 | 13078 | B230206L02 | ENSMUSG00000086003.2 |
| chr11 | 98294205 | 98295077 | 19.27906 | 51.80829 | 49.07 | Distal Interge | 11 | 98285382 | 98287444 | 2063 | 1 | ENSMUST00 | 8824 | Gm20644 | ENSMUSG00000093673.1 |
| chr13 | 54955846 | 54956750 | 19.27906 | 51.80829 | 49.07 | Intron (ENSM | 13 | 54949411 | 55006018 | 56608 | 1 | ENSMUST00 | 6436 | Unc5a | ENSMUSG00000025876.10 |
| chr14 | 40967191 | 40967920 | 19.27906 | 51.80829 | 49.07 | Promoter | 14 | 40906445 | 40966807 | 60363 | 2 | ENSMUST00 | -385 | Tspan14 | ENSMUSG00000037824.5 |
| chr18 | 32938690 | 32939857 | 19.27906 | 51.80829 | 49.07 | Promoter | 18 | 32939041 | 33195767 | 256727 | 1 | ENSMUST00 | 0 | Camk4 | ENSMUSG00000038128.6 |
| chr18 | 35653884 | 35655647 | 19.27906 | 51.80829 | 49.07 | Promoter | 18 | 35650351 | 35655240 | 4890 | 2 | ENSMUST00 | 0 | Prob1 | ENSMUSG00000073600.2 |
| chr18 | 25486570 | 25487611 | 19.27906 | 51.80829 | 49.07 | 3' UTR | 18 | 25391801 | 25392332 | 532 | 1 | ENSMUST00 | 94770 | Gm16558 | ENSMUSG00000089908.1 |
| chr18 | 65934764 | 65935655 | 19.27906 | 51.80829 | 49.07 | 3' UTR | 18 | 65937538 | 65940447 | 2910 | 1 | ENSMUST00 | -1883 | Raxos1 | ENSMUSG00000097907.1 |
| chr19 | 4854679 | 4855850 | 19.27906 | 51.80829 | 49.07 | Promoter | 19 | 4855129 | 4860912 | 5784 | 1 | ENSMUST00 | 0 | Ctsf | ENSMUSG00000083282.2 |
| chr2 | 156613214 | 156614522 | 19.27906 | 51.80829 | 49.07 | Promoter | 2 | 156613705 | 156764361 | 150657 | 1 | ENSMUST00 | 0 | Dlgap4 | ENSMUSG00000061689.10 |
| chr2 | 104409725 | 104410963 | 19.27906 | 51.80829 | 49.07 | Promoter | 2 | 104143073 | 104410334 | 267262 | 2 | ENSMUST00 | 0 | D430041D05 | ENSMUSG00000068373.9 |
| chr3 | 98338685 | 98340237 | 19.27906 | 51.80829 | 49.07 | Promoter | 3 | 98313170 | 98339990 | 26821 | 2 | ENSMUST00 | 0 | Phgdh | ENSMUSG00000053398.6 |
| chr4 | 156077382 | 156078257 | 19.27906 | 51.80829 | 49.07 | Distal Interge | 4 | 156052715 | 156059802 | 7088 | 2 | ENSMUST00 | -17581 | Gm13648 | ENSMUSG00000086549.1 |
| chr5 | 45639019 | 45639955 | 19.27906 | 51.80829 | 49.07 | Promoter | 5 | 45529706 | 45639501 | 109796 | 2 | ENSMUST00 | 0 | Fam184b | ENSMUSG00000015879.7 |
| chr5 | 134932181 | 134932859 | 19.27906 | 51.80829 | 49.07 | Promoter | 5 | 134932368 | 134942637 | 10270 | 1 | ENSMUST00 | 0 | Wbscr27 | ENSMUSG00000040557.9 |
| chr5 | 139459923 | 139460949 | 19.27906 | 51.80829 | 49.07 | Promoter | 5 | 139359739 | 139460502 | 100764 | 2 | ENSMUST00 | 0 | 3110082I17R | ENSMUSG00000053553.7 |
| chr5 | 32857720 | 32858391 | 19.27906 | 51.80829 | 49.07 | Distal Interge | 5 | 32789218 | 32854256 | 65039 | 2 | ENSMUST00 | -3465 | Prr14l | ENSMUSG00000054280.9 |
| chr7 | 80342997 | 80343659 | 19.27906 | 51.80829 | 49.07 | Promoter | 7 | 80343137 | 80346097 | 2961 | 1 | ENSMUST00 | 0 | Hddc3 | ENSMUSG00000030532.5 |
| chr7 | 44359562 | 44360672 | 19.27906 | 51.80829 | 49.07 | 3' UTR | 7 | 44350762 | 44352057 | 1296 | 1 | ENSMUST00 | 8801 | Shank1 | ENSMUSG00000038738.10 |
| chr8 | 67494611 | 67495537 | 19.27906 | 51.80829 | 49.07 | Promoter | 8 | 67494867 | 67502576 | 7710 | 1 | ENSMUST00 | 0 | Nat2 | ENSMUSG00000051147.9 |
| chr9 | 21183946 | 21184567 | 19.27906 | 51.80829 | 49.07 | Promoter | 9 | 21184101 | 21193498 | 9398 | 1 | ENSMUST00 | 0 | Pde4a | ENSMUSG00000032177.11 |
| chr9 | 70421251 | 70422027 | 19.27906 | 51.80829 | 49.07 | Promoter | 9 | 70407689 | 70421554 | 13866 | 2 | ENSMUST00 | 0 | Ccnb2 | ENSMUSG00000032218.6 |
| chr9 | 106821212 | 106821746 | 19.27906 | 51.80829 | 49.07 | Promoter | 9 | 106821874 | 106880443 | 58570 | 1 | ENSMUST00 | -128 | Vprbp | ENSMUSG00000040325.11 |
| chr10 | 121476470 | 121476958 | 19.73103 | 51.78704 | 49.0507 | Promoter | 10 | 121410350 | 121476250 | 65901 | 2 | ENSMUST00 | -221 | Rassf3 | ENSMUSG00000025795.7 |
| chr18 | 84091473 | 84092668 | 19.73103 | 51.78704 | 49.0507 | Intron (ENSM | 18 | 84088158 | 84097514 | 9357 | 1 | ENSMUST00 | 3316 | Zadh2 | ENSMUSG00000049090.2 |
| chr7 | 62419235 | 62420498 | 19.73103 | 51.78704 | 49.0507 | Promoter | 7 | 62417593 | 62420139 | 2547 | 2 | ENSMUST00 | 0 | Mkrn3 | ENSMUSG00000070527.3 |
| chr8 | 70158317 | 70158910 | 19.73103 | 51.78704 | 49.0507 | Intron (ENSM | 8 | 70152781 | 70167487 | 14707 | 1 | ENSMUST00 | 5537 | Mef2b | ENSMUSG00000079033.4 |
| chr6 | 83120740 | 83122376 | 17.89592 | 51.76047 | 49.0247 | Promoter | 6 | 83119044 | 83121559 | 2516 | 2 | ENSMUST00 | 0 | Wbp1 | ENSMUSG00000030035.9 |
| chr8 | 72492870 | 72493635 | 17.89592 | 51.76047 | 49.0247 | Promoter | 8 | 72492934 | 72505499 | 12566 | 1 | ENSMUST00 | 0 | Gm17435 | ENSMUSG00000097000.1 |
| chr18 | 35918794 | 35920648 | 19.69154 | 51.71931 | 48.98378 | Distal Interge | 18 | 35952824 | 35952907 | 84 | 1 | ENSMUST00 | -32176 | Gm23882 | ENSMUSG00000080529.1 |
| chr5 | 91139205 | 91139991 | 20.14079 | 51.68999 | 48.95457 | Promoter | 5 | 91139615 | 91148432 | 8818 | 1 | ENSMUST00 | 0 | Areg | ENSMUSG00000029378.4 |
| chr8 | 84183303 | 84184333 | 19.6522 | 51.65182 | 48.91653 | Distal Interge | 8 | 84178863 | 84178968 | 106 | 2 | ENSMUST00 | -4336 | Mir181c | ENSMUSG00000065483.1 |
| chr11 | 74521980 | 74523361 | 18.72985 | 51.651 | 48.91581 | Intron (ENSM | 11 | 74437043 | 74590124 | 153082 | 2 | ENSMUST00 | 66763 | Rap1gap2 | ENSMUSG00000038807.12 |
| chr14 | 60634125 | 60635526 | 18.72985 | 51.651 | 48.91581 | Promoter | 14 | 60634243 | 60764549 | 130307 | 1 | ENSMUST00 | 0 | Spata13 | ENSMUSG00000021990.10 |
| chr15 | 55905198 | 55907229 | 18.72985 | 51.651 | 48.91581 | Promoter | 15 | 55638843 | 55906949 | 268107 | 2 | ENSMUST00 | 0 | Sntb1 | ENSMUSG00000060429.6 |
| chr19 | 6127537 | 6128305 | 18.72985 | 51.651 | 48.91581 | Promoter | 19 | 6119399 | 6128304 | 8906 | 2 | ENSMUST00 | 0 | Snx15 | ENSMUSG00000024787.8 |
| chr1 | 93635894 | 93636374 | 17.38968 | 51.62804 | 48.89336 | Promoter | 1 | 93619751 | 93635722 | 15972 | 2 | ENSMUST00 | -173 | Stk25 | ENSMUSG00000026277.8 |
| chr10 | 128890662 | 128892018 | 20.09871 | 51.61829 | 48.88371 | Promoter | 10 | 128882295 | 128891718 | 9424 | 2 | ENSMUST00 | 0 | Gdf11 | ENSMUSG00000025352.6 |
| chr9 | 54559677 | 54560474 | 20.09871 | 51.61829 | 48.88371 | Promoter | 9 | 54544794 | 54560218 | 15425 | 2 | ENSMUST00 | 0 | Cib2 | ENSMUSG00000037493.6 |
| chr14 | 52278606 | 52279500 | 14.32375 | 51.58946 | 48.85514 | Promoter | 14 | 52279146 | 52296391 | 17246 | 1 | ENSMUST00 | 0 | Tox4 | ENSMUSG00000016831.11 |
| chr12 | 113074415 | 113075319 | 16.12455 | 51.56509 | 48.83098 | Promoter | 12 | 113074502 | 113088914 | 14413 | 1 | ENSMUST00 | 0 | Tex22 | ENSMUSG00000012211.8 |
| chr9 | 40685726 | 40686705 | 20.05681 | 51.54688 | 48.81284 | Promoter | 9 | 40685962 | 40761060 | 75099 | 1 | ENSMUST00 | 0 | Clmp | ENSMUSG00000032024.10 |
| chr3 | 116251414 | 116252762 | 20.01508 | 51.47575 | 48.74197 | Promoter | 3 | 116249654 | 116253484 | 3831 | 2 | ENSMUST00 | 722 | Gpr88 | ENSMUSG00000068696.5 |
| chr4 | 116708496 | 116709212 | 19.97352 | 51.4049 | 48.67135 | Promoter | 4 | 116708509 | 116715099 | 6591 | 1 | ENSMUST00 | 0 | Ccdc163 | ENSMUSG00000028689.9 |
| chr13 | 23763107 | 23764270 | 18.13663 | 51.39822 | 48.66477 | Promoter | 13 | 23763717 | 23764358 | 642 | 1 | ENSMUST00 | 0 | Hist1h1a | ENSMUSG00000049539.2 |
| chr16 | 22656675 | 22657628 | 19.49643 | 51.38433 | 48.65112 | Promoter | 16 | 22468461 | 22657173 | 188713 | 2 | ENSMUST00 | 0 | Dgkg | ENSMUSG00000022861.11 |
| chr11 | 87595219 | 87596051 | 19.02488 | 51.36755 | 48.63442 | Promoter | 11 | 87595652 | 87601077 | 5426 | 1 | ENSMUST00 | 0 | Mtmr4 | ENSMUSG00000018401.12 |
| chr14 | 70205110 | 70207222 | 19.02488 | 51.36755 | 48.63442 | Promoter | 14 | 70205324 | 70217659 | 12336 | 1 | ENSMUST00 | 0 | Gm26908 | ENSMUSG00000097243.1 |
| chr17 | 49562125 | 49564376 | 19.02488 | 51.36755 | 48.63442 | Promoter | 17 | 49456022 | 49564337 | 108316 | 2 | ENSMUST00 | 0 | Daam2 | ENSMUSG00000040260.7 |
| chr2 | 37788131 | 37791502 | 19.02488 | 51.36755 | 48.63442 | Promoter | 2 | 37791171 | 37793523 | 2353 | 1 | ENSMUST00 | 0 | Crb2 | ENSMUSG00000035403.8 |
| chr15 | 76697607 | 76698111 | 19.41946 | 51.25203 | 48.51994 | Promoter | 15 | 76696764 | 76699674 | 2911 | 1 | ENSMUST00 | 844 | Gpt | ENSMUSG00000022546.4 |
| chr15 | 50869392 | 50870337 | 14.84617 | 51.23753 | 48.50555 | Intron (ENSM | 15 | 50660435 | 50882806 | 222372 | 2 | ENSMUST00 | 12469 | Trps1 | ENSMUSG00000038679.11 |
| chr3 | 142700303 | 142702025 | 14.84617 | 51.23753 | 48.50555 | Promoter | 3 | 142701051 | 142744527 | 43477 | 1 | ENSMUST00 | 0 | Ccbl2 | ENSMUSG00000040213.8 |
| chr7 | 99482588 | 99483693 | 14.84617 | 51.23753 | 48.50555 | Promoter | 7 | 99477904 | 99483651 | 5748 | 2 | ENSMUST00 | 0 | Rps3 | ENSMUSG00000030744.7 |
| chr10 | 80806831 | 80807728 | 18.02336 | 51.19578 | 48.46415 | Exon (ENSMU | 10 | 80805248 | 80807648 | 2401 | 1 | ENSMUST00 | 1584 | Amh | ENSMUSG00000035262.4 |
| chr11 | 90388817 | 90391517 | 18.02336 | 51.19578 | 48.46415 | Promoter | 11 | 90336536 | 90390895 | 54360 | 2 | ENSMUST00 | 0 | Hlf | ENSMUSG00000003949.11 |
| chr11 | 120720726 | 120722245 | 18.02336 | 51.19578 | 48.46415 | Promoter | 11 | 120721470 | 120723969 | 2500 | 1 | ENSMUST00 | 0 | Rac3 | ENSMUSG00000018012.6 |
| chr11 | 120310720 | 120311507 | 18.02336 | 51.19578 | 48.46415 | Distal Interge | 11 | 120289793 | 120292167 | 2375 | 1 | ENSMUST00 | 20928 | Bahcc1 | ENSMUSG00000039741.10 |
| chr12 | 75669735 | 75670464 | 18.02336 | 51.19578 | 48.46415 | Promoter | 12 | 75630596 | 75669537 | 38942 | 2 | ENSMUST00 | -199 | Wdr89 | ENSMUSG00000045690.8 |
| chr12 | 84009221 | 84010045 | 18.02336 | 51.19578 | 48.46415 | Promoter | 12 | 84009502 | 84017670 | 8169 | 1 | ENSMUST00 | 0 | Acot1 | ENSMUSG00000072949.5 |
| chr14 | 34363326 | 34364236 | 18.02336 | 51.19578 | 48.46415 | Distal Interge | 14 | 34351881 | 34355433 | 3553 | 2 | ENSMUST00 | -7894 | Fam25c | ENSMUSG00000043681.5 |
| chr15 | 79891619 | 79892918 | 18.02336 | 51.19578 | 48.46415 | Promoter | 15 | 79891659 | 79905547 | 13889 | 1 | ENSMUST00 | 0 | Apobec3 | ENSMUSG00000009585.12 |
| chr15 | 75861839 | 75862989 | 18.02336 | 51.19578 | 48.46415 | Promoter | 15 | 75862339 | 75867404 | 5066 | 1 | ENSMUST00 | 0 | Gsdmd | ENSMUSG00000022575.4 |
| chr18 | 56561767 | 56563208 | 18.02336 | 51.19578 | 48.46415 | Promoter | 18 | 56562443 | 56587189 | 24747 | 1 | ENSMUST00 | 0 | Phax | ENSMUSG00000008301.6 |
| chr19 | 44755477 | 44757601 | 18.02336 | 51.19578 | 48.46415 | Promoter | 19 | 44756045 | 44837851 | 81807 | 1 | ENSMUST00 | 0 | Pax2 | ENSMUSG00000004231.10 |
| chr19 | 6276149 | 6276600 | 18.02336 | 51.19578 | 48.46415 | Promoter | 19 | 6268795 | 6276605 | 7811 | 2 | ENSMUST00 | 5 | Gm14963 | ENSMUSG00000085196.1 |
| chr2 | 36104558 | 36105718 | 18.02336 | 51.19578 | 48.46415 | Promoter | 2 | 36081953 | 36105271 | 23319 | 2 | ENSMUST00 | 0 | Lhx6 | ENSMUSG00000026890.14 |
| chr4 | 21766290 | 21767267 | 18.02336 | 51.19578 | 48.46415 | Promoter | 4 | 21757382 | 21767212 | 9831 | 2 | ENSMUST00 | 0 | Tstd3 | ENSMUSG00000028251.5 |
| chr4 | 149698048 | 149698982 | 18.02336 | 51.19578 | 48.46415 | Promoter | 4 | 149649168 | 149698698 | 49531 | 2 | ENSMUST00 | 0 | Pik3cd | ENSMUSG00000039936.13 |
| chr4 | 88782300 | 88782678 | 18.02336 | 51.19578 | 48.46415 | Promoter | 4 | 88782131 | 88783592 | 1462 | 1 | ENSMUST00 | 170 | Ifnz | ENSMUSG00000096854.1 |
| chr6 | 54592713 | 54593676 | 18.02336 | 51.19578 | 48.46415 | Promoter | 6 | 54577604 | 54593177 | 15574 | 2 | ENSMUST00 | 0 | Fkbp14 | ENSMUSG00000038074.11 |
| chr6 | 54698757 | 54699893 | 18.02336 | 51.19578 | 48.46415 | 3' UTR | 6 | 54680930 | 54681917 | 988 | 2 | ENSMUST00 | -16841 | RP23-13A15. | ENSMUSG00000101651.1 |
| chr7 | 45434803 | 45435328 | 18.02336 | 51.19578 | 48.46415 | Promoter | 7 | 45434844 | 45456619 | 21776 | 1 | ENSMUST00 | 0 | Gys1 | ENSMUSG00000003865.9 |
| chr7 | 98838134 | 98839457 | 18.02336 | 51.19578 | 48.46415 | Promoter | 7 | 98838845 | 98853751 | 14907 | 1 | ENSMUST00 | 0 | Wnt11 | ENSMUSG00000015957.6 |
| chr7 | 57590489 | 57592145 | 19.84988 | 51.19398 | 48.46415 | Promoter | 7 | 57590518 | 57828802 | 238285 | 1 | ENSMUST00 | 0 | Gabrb3 | ENSMUSG00000033676.8 |
| chr7 | 4812907 | 4813792 | 18.02336 | 51.19578 | 48.46415 | Promoter | 7 | 4808155 | 4812351 | 4197 | 2 | ENSMUST00 | -557 | Ube2s | ENSMUSG00000060860.7 |
| chr8 | 111629932 | 111630657 | 18.02336 | 51.19578 | 48.46415 | Promoter | 8 | 111625503 | 111630335 | 4833 | 2 | ENSMUST00 | 0 | Ldhd | ENSMUSG00000031958.11 |

| | | | | | | | | | | | | | | | |
|---|---|---|---|---|---|---|---|---|---|---|---|---|---|---|---|
| chr9 | 31130567 | 31132119 | 18.02336 | 51.19578 | 48.46415 | Promoter | 9 | 31089402 | 31131853 | 42452 | 2 | ENSMUST00 | 0 | St14 | ENSMUSG00000031995.3 |
| chrX | 161716902 | 161718697 | 18.02336 | 51.19578 | 48.46415 | Promoter | 20 | 161717069 | 161779494 | 62426 | 1 | ENSMUST00 | 0 | Rai2 | ENSMUSG00000043518.7 |
| chr1 | 86045759 | 86046745 | 18.39457 | 51.06108 | 48.33263 | Promoter | 1 | 86045863 | 86053353 | 7491 | 1 | ENSMUST00 | 0 | 2810459M11 | ENSMUSG00000026227.6 |
| chr2 | 128863594 | 128864410 | 18.39457 | 51.06108 | 48.33263 | Promoter | 2 | 128863932 | 128897030 | 33099 | 1 | ENSMUST00 | 0 | Fbln7 | ENSMUSG00000027386.4 |
| chr5 | 115301092 | 115301866 | 18.39457 | 51.06108 | 48.33263 | Promoter | 5 | 115301120 | 115319208 | 18089 | 1 | ENSMUST00 | 0 | Gm13830 | ENSMUSG00000086368.1 |
| chr4 | 12087306 | 12088431 | 19.30515 | 51.05533 | 48.32729 | Promoter | 4 | 12039355 | 12087957 | 48603 | 2 | ENSMUST00 | 0 | Tmem67 | ENSMUSG00000049488.9 |
| chr10 | 80702221 | 80703270 | 18.81137 | 50.99623 | 48.2684 | Promoter | 10 | 80702693 | 80705382 | 2690 | 1 | ENSMUST00 | 0 | Izumo4 | ENSMUSG00000055862.6 |
| chr1 | 43163168 | 43164278 | 15.45146 | 50.98658 | 48.25883 | Promoter | 1 | 43123074 | 43163961 | 40888 | 2 | ENSMUST00 | 0 | Fhl2 | ENSMUSG00000008136.9 |
| chr2 | 130294730 | 130296281 | 15.45146 | 50.98658 | 48.25883 | Promoter | 2 | 130295169 | 130370481 | 75313 | 1 | ENSMUST00 | 0 | Ebf4 | ENSMUSG00000053552.9 |
| chr5 | 25222552 | 25223940 | 15.45146 | 50.98658 | 48.25883 | Promoter | 5 | 25222882 | 25265918 | 43037 | 1 | ENSMUST00 | 0 | Galnt11 | ENSMUSG00000038072.9 |
| chr5 | 137189213 | 137189789 | 15.45146 | 50.98658 | 48.25883 | Distal Interge | 5 | 137208813 | 137212389 | 3577 | 2 | ENSMUST00 | 22600 | A630081J09R | ENSMUSG00000094840.1 |
| chr7 | 19227340 | 19228152 | 15.45146 | 50.98658 | 48.25883 | Promoter | 7 | 19228334 | 19247868 | 19535 | 1 | ENSMUST00 | -182 | Opa3 | ENSMUSG00000052214.8 |
| chr7 | 78576814 | 78579420 | 15.45146 | 50.98658 | 48.25883 | Promoter | 7 | 78578830 | 78581034 | 2205 | 1 | ENSMUST00 | 0 | Gm9885 | ENSMUSG00000052629.5 |
| chr7 | 3290519 | 3291885 | 15.45146 | 50.98658 | 48.25883 | Promoter | 7 | 3290556 | 3299345 | 8790 | 1 | ENSMUST00 | 0 | Myadm | ENSMUSG00000068566.7 |
| chr8 | 41040522 | 41042270 | 15.45146 | 50.98658 | 48.25883 | Promoter | 8 | 40991052 | 41041866 | 50815 | 2 | ENSMUST00 | 0 | Mtus1 | ENSMUSG00000045636.11 |
| chr8 | 84912839 | 84913660 | 19.22969 | 50.92535 | 48.19825 | Promoter | 8 | 84913391 | 84922915 | 9525 | 1 | ENSMUST00 | 0 | Dnase2a | ENSMUSG00000003812.8 |
| chr11 | 75999686 | 76000307 | 19.68739 | 50.91653 | 48.1895 | Promoter | 11 | 75999912 | 76003569 | 3658 | 1 | ENSMUST00 | 0 | 1700016K19R | ENSMUSG00000053783.5 |
| chr15 | 74518940 | 74520094 | 19.68739 | 50.91653 | 48.1895 | Intron (ENSM | 15 | 74516831 | 74589464 | 72634 | 1 | ENSMUST00 | 2110 | Bai1 | ENSMUSG00000034730.11 |
| chr11 | 120726821 | 120727387 | 17.00442 | 50.91425 | 48.18763 | Promoter | 11 | 120725399 | 120727243 | 1845 | 2 | ENSMUST00 | 0 | Dcxr | ENSMUSG00000039450.6 |
| chr12 | 74316771 | 74317292 | 17.00442 | 50.91425 | 48.18763 | Promoter | 12 | 74315110 | 74316394 | 1285 | 2 | ENSMUST00 | -378 | Gm11042 | ENSMUSG00000079061.3 |
| chr13 | 23760650 | 23762774 | 17.00442 | 50.91425 | 48.18763 | Promoter | 13 | 23760692 | 23761230 | 539 | 2 | ENSMUST00 | 0 | Hist1h4a | ENSMUSG00000060093.6 |
| chr14 | 63943003 | 63944658 | 17.00442 | 50.91425 | 48.18763 | Promoter | 14 | 63943674 | 63950732 | 7059 | 1 | ENSMUST00 | 0 | Sox7 | ENSMUSG00000063060.5 |
| chr18 | 47366217 | 47370796 | 17.00442 | 50.91425 | 48.18763 | Promoter | 18 | 47245254 | 47368830 | 123577 | 2 | ENSMUST00 | 0 | Sema6a | ENSMUSG00000019647.11 |
| chr19 | 30539372 | 30540014 | 17.00442 | 50.91425 | 48.18763 | Promoter | 19 | 30538928 | 30539554 | 627 | 2 | ENSMUST00 | 0 | Ppp1r2-ps3 | ENSMUSG00000058816.4 |
| chr2 | 155050417 | 155051491 | 17.00442 | 50.91425 | 48.18763 | 3' UTR | 2 | 155047615 | 155051011 | 3397 | 1 | ENSMUST00 | 2803 | a | ENSMUSG00000027596.5 |
| chr4 | 108165083 | 108166200 | 17.00442 | 50.91425 | 48.18763 | Promoter | 4 | 108165437 | 108179308 | 13872 | 1 | ENSMUST00 | 0 | Echdc2 | ENSMUSG00000028601.13 |
| chr4 | 124741377 | 124742388 | 17.00442 | 50.91425 | 48.18763 | Promoter | 4 | 124741850 | 124801511 | 59662 | 1 | ENSMUST00 | 0 | Inpp5b | ENSMUSG00000028894.13 |
| chr4 | 120814986 | 120816353 | 17.00442 | 50.91425 | 48.18763 | Promoter | 4 | 120757438 | 120815712 | 58275 | 2 | ENSMUST00 | 0 | Nfyc | ENSMUSG00000032897.12 |
| chr4 | 126556620 | 126557683 | 17.00442 | 50.91425 | 48.18763 | Promoter | 4 | 126556935 | 126593903 | 36969 | 1 | ENSMUST00 | 0 | Clspn | ENSMUSG00000042489.10 |
| chr5 | 106925543 | 106926395 | 17.00442 | 50.91425 | 48.18763 | Promoter | 5 | 106840194 | 106926321 | 86128 | 2 | ENSMUST00 | 0 | Hfm1 | ENSMUSG00000043410.11 |
| chr7 | 101896043 | 101897252 | 17.00442 | 50.91425 | 48.18763 | Promoter | 7 | 101896340 | 101899540 | 3201 | 1 | ENSMUST00 | 0 | Anapc15 | ENSMUSG00000030649.12 |
| chr7 | 126582714 | 126583897 | 17.00442 | 50.91425 | 48.18763 | Promoter | 7 | 126571400 | 126583247 | 11848 | 2 | ENSMUST00 | 0 | Cln3 | ENSMUSG00000030720.11 |
| chr9 | 106367148 | 106367760 | 17.00442 | 50.91425 | 48.18763 | Promoter | 9 | 106368632 | 106375724 | 7093 | 1 | ENSMUST00 | -872 | Dusp7 | ENSMUSG00000053716.8 |
| chr11 | 69124925 | 69126298 | 16.16105 | 50.86513 | 48.14017 | Promoter | 11 | 69125896 | 69149115 | 23220 | 1 | ENSMUST00 | 0 | Aloxe3 | ENSMUSG00000020892.5 |
| chr12 | 110843843 | 110846184 | 16.16105 | 50.86513 | 48.14017 | Promoter | 12 | 110842276 | 110845216 | 2941 | 2 | ENSMUST00 | 0 | Gm26912 | ENSMUSG00000097832.1 |
| chr13 | 21778789 | 21779358 | 16.16105 | 50.86513 | 48.14017 | Downstream | 13 | 21779883 | 21780625 | 743 | 2 | ENSMUST00 | 1267 | Hist1h1b | ENSMUSG00000058773.2 |
| chr15 | 79516024 | 79516982 | 16.16105 | 50.86513 | 48.14017 | Promoter | 15 | 79516408 | 79527739 | 11332 | 1 | ENSMUST00 | 0 | Kdelr3 | ENSMUSG00000010830.7 |
| chr17 | 15830077 | 15831606 | 16.16105 | 50.86513 | 48.14017 | Distal Interge | 17 | 15806253 | 15826586 | 20334 | 2 | ENSMUST00 | -3492 | Rgmb | ENSMUSG00000048027.7 |
| chr17 | 21516073 | 21517433 | 16.16105 | 50.86513 | 48.14017 | Distal Interge | 17 | 21528194 | 21535878 | 7685 | 2 | ENSMUST00 | 18445 | 9330136K24R | ENSMUSG00000097781.1 |
| chr18 | 34408838 | 34409878 | 16.16105 | 50.86513 | 48.14017 | Promoter | 18 | 34409423 | 34441794 | 32372 | 1 | ENSMUST00 | 0 | Pkd2l2 | ENSMUSG00000014503.9 |
| chr19 | 45234747 | 45236371 | 16.16105 | 50.86513 | 48.14017 | Promoter | 19 | 45232684 | 45235812 | 3129 | 2 | ENSMUST00 | 0 | Lbx1 | ENSMUSG00000025216.9 |
| chr4 | 150047890 | 150049456 | 16.16105 | 50.86513 | 48.14017 | Promoter | 4 | 150049013 | 150056976 | 7964 | 1 | ENSMUST00 | 0 | Gm13067 | ENSMUSG00000086665.1 |
| chr5 | 135008374 | 135008934 | 16.16105 | 50.86513 | 48.14017 | Promoter | 5 | 135009152 | 135012021 | 2870 | 1 | ENSMUST00 | -218 | Abhd11 | ENSMUSG00000040532.9 |
| chr7 | 30972858 | 30974247 | 16.16105 | 50.86513 | 48.14017 | Promoter | 7 | 30957775 | 30973464 | 15690 | 2 | ENSMUST00 | 0 | Lsr | ENSMUSG00000001247.11 |
| chr7 | 12949121 | 12950310 | 16.16105 | 50.86513 | 48.14017 | Promoter | 7 | 12949690 | 12950066 | 377 | 1 | ENSMUST00 | 0 | 1810019N24 | ENSMUSG00000097115.2 |
| chr8 | 69183634 | 69184817 | 16.16105 | 50.86513 | 48.14017 | Promoter | 8 | 69132669 | 69184225 | 51557 | 2 | ENSMUST00 | 0 | Lzts1 | ENSMUSG00000036306.6 |
| chr9 | 40638266 | 40638896 | 16.16105 | 50.86513 | 48.14017 | Distal Interge | 9 | 40670613 | 40671014 | 402 | 1 | ENSMUST00 | -31717 | Gm21915 | ENSMUSG00000094204.1 |
| chr3 | 90537113 | 90538338 | 19.64718 | 50.84783 | 48.12388 | Promoter | 3 | 90537254 | 90543148 | 5895 | 1 | ENSMUST00 | 0 | S100a16 | ENSMUSG00000074457.5 |
| chr11 | 102924633 | 102925610 | 17.82851 | 50.8465 | 48.12268 | Promoter | 11 | 102905529 | 102925124 | 19596 | 2 | ENSMUST00 | 0 | Kif18b | ENSMUSG00000051378.10 |
| chr11 | 9048273 | 9049093 | 19.60713 | 50.77938 | 48.05589 | Promoter | 11 | 9048594 | 9069356 | 20763 | 1 | ENSMUST00 | 0 | Gm11992 | ENSMUSG00000040978.4 |
| chr2 | 33543919 | 33544917 | 19.60713 | 50.77938 | 48.05589 | Distal Interge | 2 | 33489782 | 33503632 | 13851 | 1 | ENSMUST00 | 54138 | Gm13530 | ENSMUSG00000086939.1 |
| chr17 | 17178711 | 17180059 | 13.63714 | 50.76526 | 48.04188 | Distal Interge | 17 | 17121415 | 17146831 | 25417 | 1 | ENSMUST00 | 57297 | Zfp97 | ENSMUSG00000095990.1 |
| chr1 | 183438856 | 183439355 | 18.67167 | 50.75271 | 48.02938 | Distal Interge | 1 | 183418503 | 183437053 | 18551 | 1 | ENSMUST00 | 20354 | Hhipl2 | ENSMUSG00000053461.4 |
| chr14 | 65149167 | 65150135 | 18.67167 | 50.75271 | 48.02938 | Distal Interge | 14 | 65052060 | 65098106 | 46047 | 2 | ENSMUST00 | -51062 | Extl3 | ENSMUSG00000021978.8 |
| chr15 | 79546856 | 79547519 | 18.67167 | 50.75271 | 48.02938 | Promoter | 15 | 79527707 | 79546741 | 19035 | 2 | ENSMUST00 | -116 | Ddx17 | ENSMUSG00000055065.6 |
| chr13 | 41079347 | 41079863 | 19.56725 | 50.7112 | 47.98855 | Promoter | 13 | 41025120 | 41079706 | 54587 | 2 | ENSMUST00 | 0 | Mak | ENSMUSG00000021363.8 |
| chr15 | 102746412 | 102747635 | 19.52753 | 50.64327 | 47.92091 | Distal Interge | 15 | 102706779 | 102722178 | 15400 | 2 | ENSMUST00 | -24235 | Calcoco1 | ENSMUSG00000023055.4 |
| chr9 | 89210602 | 89210952 | 19.52753 | 50.64327 | 47.92091 | Promoter | 9 | 89210676 | 89239981 | 29306 | 1 | ENSMUST00 | 0 | Mthfs | ENSMUSG00000066442.11 |
| chr6 | 47660199 | 47660483 | 6.79673 | 50.61199 | 47.88982 | Distal Interge | 6 | 47637143 | 47637219 | 77 | 1 | ENSMUST00 | 23057 | Gm25283 | ENSMUSG00000087897.1 |
| chr9 | 114367864 | 114368718 | 19.04358 | 50.60438 | 47.88221 | Promoter | 9 | 114368303 | 114377229 | 8927 | 1 | ENSMUST00 | 0 | 4930520O04 | ENSMUSG00000074039.5 |
| chr14 | 72602247 | 72603533 | 18.07108 | 50.48883 | 47.76702 | Promoter | 14 | 72539530 | 72602962 | 63433 | 2 | ENSMUST00 | 0 | Fndc3a | ENSMUSG00000033487.9 |
| chr18 | 5989344 | 5990027 | 18.07108 | 50.48883 | 47.76702 | Distal Interge | 18 | 6032960 | 6034707 | 1748 | 2 | ENSMUST00 | 44680 | Arhgap12 | ENSMUSG00000041225.11 |
| chr7 | 24254329 | 24255279 | 18.07108 | 50.48883 | 47.76702 | Distal Interge | 7 | 24257644 | 24261914 | 4271 | 1 | ENSMUST00 | -2365 | Zfp108 | ENSMUSG00000030486.8 |
| chr4 | 144586176 | 144586487 | 18.97014 | 50.47753 | 47.75603 | Promoter | 4 | 144587244 | 144588632 | 1389 | 1 | ENSMUST00 | -757 | Gm13127 | ENSMUSG00000070610.4 |
| chr10 | 80260517 | 80262231 | 19.37025 | 50.37409 | 47.65327 | Promoter | 10 | 80258151 | 80261012 | 2862 | 2 | ENSMUST00 | 0 | Gamt | ENSMUSG00000020150.8 |
| chr9 | 58248714 | 58250214 | 18.89727 | 50.35157 | 47.63088 | Promoter | 9 | 58218076 | 58249786 | 31711 | 2 | ENSMUST00 | 0 | Pml | ENSMUSG00000036986.11 |
| chr3 | 93443512 | 93448142 | 14.07679 | 50.30981 | 47.58934 | 3' UTR | 3 | 93442330 | 93449077 | 6748 | 1 | ENSMUST00 | 1183 | Tchh | ENSMUSG00000052415.5 |
| chr1 | 37871724 | 37872945 | 18.85995 | 50.28701 | 47.56678 | Promoter | 1 | 37871738 | 37875343 | 3606 | 1 | ENSMUST00 | 0 | Lipt1 | ENSMUSG00000037216.4 |
| chr1 | 120072898 | 120074442 | 18.85995 | 50.28701 | 47.56678 | Promoter | 1 | 120067377 | 120074074 | 6698 | 2 | ENSMUST00 | 0 | Tmem37 | ENSMUSG00000050777.7 |
| chr11 | 102572951 | 102573533 | 18.85995 | 50.28701 | 47.56678 | Downstream | 11 | 102576491 | 102579119 | 2629 | 2 | ENSMUST00 | 5586 | Gm11627 | ENSMUSG00000078640.4 |
| chr2 | 145167300 | 145168685 | 18.85995 | 50.28701 | 47.56678 | Promoter | 2 | 145167754 | 145642166 | 474413 | 1 | ENSMUST00 | 0 | Slc24a3 | ENSMUSG00000063873.5 |
| chr3 | 151836798 | 151837872 | 18.85995 | 50.28701 | 47.56678 | Promoter | 3 | 151798918 | 151837630 | 38713 | 2 | ENSMUST00 | 0 | Ptgfr | ENSMUSG00000028036.5 |
| chr4 | 129984360 | 129985631 | 18.85995 | 50.28701 | 47.56678 | Promoter | 4 | 129984870 | 130022633 | 37764 | 1 | ENSMUST00 | 0 | Bai2 | ENSMUSG00000028782.9 |
| chr5 | 100499667 | 100501061 | 18.85995 | 50.28701 | 47.56678 | Promoter | 5 | 100441918 | 100500592 | 58675 | 2 | ENSMUST00 | 0 | Lin54 | ENSMUSG00000035310.10 |
| chr6 | 92480119 | 92481672 | 18.85995 | 50.28701 | 47.56678 | Promoter | 6 | 92375384 | 92481392 | 106009 | 2 | ENSMUST00 | 0 | Prickle2 | ENSMUSG00000030020.8 |
| chr7 | 141902590 | 141903005 | 18.85995 | 50.28701 | 47.56678 | Promoter | 7 | 141881583 | 141902487 | 20905 | 2 | ENSMUST00 | -104 | Tollip | ENSMUSG00000025139.8 |
| chr7 | 30421476 | 30423111 | 18.85995 | 50.28701 | 47.56678 | Promoter | 7 | 30421732 | 30428746 | 7015 | 1 | ENSMUST00 | 0 | Nfkbid | ENSMUSG00000036931.9 |
| chr1 | 187651427 | 187652308 | 19.75988 | 50.25065 | 47.53236 | Intron (ENSM | 1 | 187609696 | 187793742 | 184047 | 1 | ENSMUST00 | 41732 | Esrrg | ENSMUSG00000026610.8 |
| chr10 | 39612829 | 39613850 | 19.75988 | 50.25065 | 47.53236 | Promoter | 10 | 39612934 | 39655307 | 42374 | 1 | ENSMUST00 | 0 | Traf3ip2 | ENSMUSG00000019842.6 |
| chr10 | 80348006 | 80348705 | 19.75988 | 50.25065 | 47.53236 | Promoter | 10 | 80339793 | 80348412 | 8620 | 2 | ENSMUST00 | 0 | Adamtsl5 | ENSMUSG00000043822.13 |
| chr11 | 107011809 | 107012825 | 19.75988 | 50.25065 | 47.53236 | Distal Interge | 11 | 107003909 | 107004740 | 832 | 2 | ENSMUST00 | -7070 | Gm11708 | ENSMUSG00000083356.1 |
| chr13 | 101801359 | 101802093 | 19.75988 | 50.25065 | 47.53236 | Distal Interge | 13 | 101680563 | 101768217 | 87655 | 2 | ENSMUST00 | -33143 | Pik3r1 | ENSMUSG00000041417.10 |
| chr15 | 25412086 | 25413982 | 19.75988 | 50.25065 | 47.53236 | Promoter | 15 | 25363285 | 25413764 | 50480 | 2 | ENSMUST00 | 0 | Basp1 | ENSMUSG00000045763.7 |
| chr18 | 58209256 | 58210616 | 19.75988 | 50.25065 | 47.53236 | Promoter | 18 | 58008623 | 58209926 | 201304 | 2 | ENSMUST00 | 0 | Fbn2 | ENSMUSG00000024598.8 |
| chr2 | 162660168 | 162661408 | 19.75988 | 50.25065 | 47.53236 | Promoter | 2 | 161521990 | 162661147 | 1139158 | 2 | ENSMUST00 | 0 | Ptprt | ENSMUSG00000053141.11 |
| chr2 | 31653889 | 31654842 | 19.75988 | 50.25065 | 47.53236 | 3' UTR | 2 | 31660470 | 31661427 | 958 | 1 | ENSMUST00 | -5628 | Gm13424 | ENSMUSG00000083143.1 |
| chr4 | 65123536 | 65127080 | 19.75988 | 50.25065 | 47.53236 | Promoter | 4 | 65124174 | 65357509 | 233336 | 1 | ENSMUST00 | 0 | Pappa | ENSMUSG00000028370.7 |
| chr4 | 129120896 | 129121613 | 19.75988 | 50.25065 | 47.53236 | Promoter | 4 | 129111576 | 129121291 | 9716 | 2 | ENSMUST00 | 0 | Hpca | ENSMUSG00000028785.7 |
| chr5 | 112275912 | 112276452 | 19.75988 | 50.25065 | 47.53236 | Promoter | 5 | 112276691 | 112315361 | 38671 | 1 | ENSMUST00 | -239 | Tpst2 | ENSMUSG00000029344.9 |
| chr5 | 142701860 | 142702619 | 19.75988 | 50.25065 | 47.53236 | Promoter | 5 | 142702101 | 142722490 | 20390 | 1 | ENSMUST00 | 0 | Slc29a4 | ENSMUSG00000050822.7 |
| chr6 | 72552561 | 72553439 | 19.75988 | 50.25065 | 47.53236 | Promoter | 6 | 72552938 | 72558257 | 5320 | 1 | ENSMUST00 | 0 | Capg | ENSMUSG00000056737.9 |
| chr7 | 25787969 | 25789058 | 19.75988 | 50.25065 | 47.53236 | Promoter | 7 | 25757273 | 25788687 | 31415 | 2 | ENSMUST00 | 0 | Axl | ENSMUSG00000002602.11 |
| chr7 | 105751515 | 105752458 | 19.75988 | 50.25065 | 47.53236 | Promoter | 7 | 105744847 | 105752207 | 7361 | 2 | ENSMUST00 | 0 | Tpp1 | ENSMUSG00000030894.4 |
| chr8 | 120588562 | 120589160 | 19.75988 | 50.25065 | 47.53236 | Promoter | 8 | 120578633 | 120588980 | 10348 | 2 | ENSMUST00 | 0 | Gins2 | ENSMUSG00000031821.6 |
| chr8 | 89006293 | 89007087 | 19.75988 | 50.25065 | 47.53236 | Distal Interge | 8 | 89024735 | 89024831 | 97 | 1 | ENSMUST00 | -17648 | Mir8110 | ENSMUSG00000099078.1 |
| chrX | 100776682 | 100777626 | 19.75988 | 50.25065 | 47.53236 | Promoter | 20 | 100777285 | 100818408 | 41124 | 1 | ENSMUST00 | 0 | Dlg3 | ENSMUSG00000000881.7 |
| chrX | 71364259 | 71365317 | 19.75988 | 50.25065 | 47.53236 | Promoter | 20 | 71364760 | 71388756 | 23997 | 1 | ENSMUST00 | 0 | Mtmr1 | ENSMUSG00000015214.9 |
| chrX | 73672693 | 73674323 | 19.75988 | 50.25065 | 47.53236 | Promoter | 20 | 73673150 | 73682502 | 9353 | 1 | ENSMUST00 | 0 | Slc6a8 | ENSMUSG00000019558.9 |
| chrX | 151803265 | 151804859 | 19.75988 | 50.25065 | 47.53236 | Promoter | 20 | 151803327 | 151853201 | 49875 | 1 | ENSMUST00 | 0 | Huwe1 | ENSMUSG00000025261.12 |
| chrX | 105120156 | 105121800 | 19.75988 | 50.25065 | 47.53236 | Promoter | 20 | 105120378 | 105123927 | 3550 | 1 | ENSMUST00 | 0 | Magee1 | ENSMUSG00000031227.8 |
| chrX | 139610210 | 139611250 | 19.75988 | 50.25065 | 47.53236 | Promoter | 20 | 139610620 | 139673145 | 62526 | 1 | ENSMUST00 | 0 | Rnf128 | ENSMUSG00000031438.6 |
| chrX | 51204522 | 51206287 | 19.75988 | 50.25065 | 47.53236 | Promoter | 20 | 51117269 | 51205832 | 88564 | 2 | ENSMUST00 | 0 | Mbnl3 | ENSMUSG00000036109.12 |
| chrX | 48208001 | 48209281 | 19.75988 | 50.25065 | 47.53236 | Promoter | 20 | 48171969 | 48208687 | 36719 | 2 | ENSMUST00 | 0 | Zdhhc9 | ENSMUSG00000036985.7 |
| chrX | 166439936 | 166441336 | 19.75988 | 50.25065 | 47.53236 | Promoter | 20 | 166390033 | 166440704 | 50672 | 2 | ENSMUST00 | 0 | Ofd1 | ENSMUSG00000040586.8 |
| chrX | 135742045 | 135743544 | 19.75988 | 50.25065 | 47.53236 | Promoter | 20 | 135742733 | 135761990 | 19258 | 1 | ENSMUST00 | 0 | Gprasp1 | ENSMUSG00000043384.10 |
| chrX | 153036735 | 153038188 | 19.75988 | 50.25065 | 47.53236 | Promoter | 20 | 153036166 | 153037575 | 1410 | 2 | ENSMUST00 | 0 | Mageh1 | ENSMUSG00000047238.4 |
| chrX | 94635772 | 94636746 | 19.75988 | 50.25065 | 47.53236 | Promoter | 20 | 94636069 | 94638559 | 2491 | 1 | ENSMUST00 | 0 | Gspt2 | ENSMUSG00000071723.7 |
| chrX | 101849005 | 101850558 | 19.75988 | 50.25065 | 47.53236 | Promoter | 20 | 101849385 | 101949635 | 100251 | 1 | ENSMUST00 | 0 | Nhsl2 | ENSMUSG00000079481.6 |
| chrX | 94121286 | 94123843 | 19.75988 | 50.25065 | 47.53236 | Promoter | 20 | 94074631 | 94123407 | 48777 | 2 | ENSMUST00 | 0 | Zfx | ENSMUSG00000079509.5 |
| chr11 | 54902545 | 54903266 | 19.29256 | 50.24099 | 47.52407 | Promoter | 11 | 54902772 | 54910377 | 7606 | 1 | ENSMUST00 | 0 | Gpx3 | ENSMUSG00000018339.6 |
| chr11 | 5151758 | 5152539 | 19.29256 | 50.24099 | 47.52407 | Promoter | 11 | 5106265 | 5152257 | 45993 | 2 | ENSMUST00 | 0 | Emid1 | ENSMUSG00000034164.12 |
| chr9 | 13245325 | 13245891 | 18.82496 | 50.22645 | 47.51003 | Promoter | 9 | 13242790 | 13245741 | 2952 | 2 | ENSMUST00 | 0 | Jrkl | ENSMUSG00000079083.2 |
| chr17 | 31563690 | 31564428 | 19.72434 | 50.19016 | 47.47384 | Promoter | 17 | 31564749 | 31605481 | 40733 | 1 | ENSMUST00 | -321 | Pknox1 | ENSMUSG00000006705.7 |
| chr2 | 163749577 | 163750631 | 19.72434 | 50.19016 | 47.47384 | Promoter | 2 | 163726584 | 163750239 | 23656 | 2 | ENSMUST00 | 0 | Ada | ENSMUSG00000017697.3 |
| chr14 | 93890254 | 93891752 | 18.33134 | 50.1574 | 47.44131 | Distal Interge | 14 | 93015512 | 93888732 | 873221 | 2 | ENSMUST00 | -1523 | Pcdh9 | ENSMUSG00000055421.3 |
| chr15 | 83099604 | 83100808 | 18.33134 | 50.1574 | 47.44131 | Promoter | 15 | 83100204 | 83116674 | 16471 | 1 | ENSMUST00 | 0 | Serhl | ENSMUSG00000058586.6 |
| chr4 | 145463595 | 145464468 | 17.44328 | 50.15193 | 47.43638 | Promoter | 4 | 145464204 | 145465487 | 1284 | 1 | ENSMUST00 | 0 | Smarca5-ps | ENSMUSG00000052912.7 |
| chr7 | 80534444 | 80535755 | 13.03321 | 50.05443 | 47.33913 | Promoter | 7 | 80454993 | 80534998 | 80006 | 2 | ENSMUST00 | 0 | Blm | ENSMUSG00000030528.7 |
| chr7 | 31126362 | 31127506 | 19.64192 | 50.04983 | 47.33455 | Promoter | 7 | 31116524 | 31126844 | 10321 | 2 | ENSMUST00 | 0 | Scn1b | ENSMUSG00000019194.9 |
| chr7 | 27508301 | 27508964 | 19.64192 | 50.04983 | 47.33455 | Promoter | 7 | 27502718 | 27508122 | 5405 | 2 | ENSMUST00 | -180 | Gm15541 | ENSMUSG00000087371.1 |
| chr13 | 109260086 | 109261287 | 19.17718 | 50.04316 | 47.32813 | Promoter | 13 | 109260654 | 109951364 | 690711 | 1 | ENSMUST00 | 0 | Pde4d | ENSMUSG00000021699.12 |
| chr3 | 96268974 | 96269493 | 19.60097 | 49.98008 | 47.26519 | Promoter | 3 | 96269721 | 96279001 | 9281 | 1 | ENSMUST00 | -228 | Hist2h2bb | ENSMUSG00000050936.4 |
| chr17 | 21422784 | 21423775 | 19.13903 | 49.9777 | 47.26295 | Promoter | 17 | 21423227 | 21435640 | 12414 | 1 | ENSMUST00 | 0 | Zfp54 | ENSMUSG00000023882.10 |
| chr11 | 58301401 | 58303425 | 14.58106 | 49.93175 | 47.21736 | Distal Interge | 11 | 58307069 | 58314613 | 7545 | 1 | ENSMUST00 | -3644 | Zfp692 | ENSMUSG00000037243.12 |
| chr13 | 13437424 | 13437964 | 14.58106 | 49.93175 | 47.21736 | Promoter | 13 | 13437602 | 13512275 | 74674 | 1 | ENSMUST00 | 0 | Nid1 | ENSMUSG00000005397.7 |
| chr4 | 132731964 | 132732720 | 14.58106 | 49.93175 | 47.21736 | Promoter | 4 | 132724897 | 132732546 | 7650 | 2 | ENSMUST00 | 0 | Xkr8 | ENSMUSG00000037752.4 |
| chr11 | 16751845 | 16753270 | 19.56019 | 49.91061 | 47.19658 | Promoter | 11 | 16752203 | 16887923 | 135721 | 1 | ENSMUST00 | 0 | Egfr | ENSMUSG00000020122.11 |
| chr1 | 171195742 | 171196335 | 18.61129 | 49.85614 | 47.14222 | Promoter | 1 | 171173262 | 171196268 | 23007 | 2 | ENSMUST00 | 0 | Pcp4l1 | ENSMUSG00000038370.6 |

| | | | | | | | | | | | | | | | |
|---|---|---|---|---|---|---|---|---|---|---|---|---|---|---|---|
| chr13 | 16011699 | 16012757 | 18.61129 | 49.85614 | 47.14222 | Promoter | 13 | 16011851 | 16027188 | 15338 | 1 | ENSMUST00 | 0 | Inhba | ENSMUSG00000041324.8 |
| chr15 | 102975608 | 102978084 | 18.61129 | 49.85614 | 47.14222 | Promoter | 15 | 102976439 | 102981669 | 5231 | 1 | ENSMUST00 | 0 | Hoxc9 | ENSMUSG00000036139.6 |
| chr19 | 37434061 | 37436006 | 18.61129 | 49.85614 | 47.14222 | Promoter | 19 | 37434810 | 37440731 | 5922 | 1 | ENSMUST00 | 0 | Hhex | ENSMUSG00000024986.6 |
| chr3 | 94363293 | 94364443 | 18.61129 | 49.85614 | 47.14222 | Promoter | 3 | 94362444 | 94364567 | 2124 | 1 | ENSMUST00 | 850 | C2cd4d | ENSMUSG00000091648.1 |
| chr8 | 3671978 | 3672606 | 18.61129 | 49.85614 | 47.14222 | Promoter | 8 | 3671188 | 3672336 | 1149 | 2 | ENSMUST00 | 0 | Fcor | ENSMUSG00000089665.1 |
| chr4 | 155774045 | 155775002 | 19.06318 | 49.84748 | 47.13424 | Promoter | 4 | 155768149 | 155774698 | 6550 | 2 | ENSMUST00 | 0 | Vwa1 | ENSMUSG00000042116.3 |
| chr1 | 151754475 | 151755108 | 18.57615 | 49.79513 | 47.08208 | Promoter | 1 | 151748949 | 151755018 | 6070 | 2 | ENSMUST00 | 0 | RP23-370J16 | ENSMUSG00000100594.1 |
| chr2 | 32586689 | 32587897 | 18.57615 | 49.79513 | 47.08208 | Promoter | 2 | 32587095 | 32599698 | 12604 | 1 | ENSMUST00 | 0 | St6galnac4 | ENSMUSG00000079442.7 |
| chr11 | 101448329 | 101449398 | 17.24266 | 49.78797 | 47.07495 | Promoter | 11 | 101448407 | 101458698 | 10292 | 1 | ENSMUST00 | 0 | Ifi35 | ENSMUSG00000010358.8 |
| chr5 | 108448531 | 108449829 | 17.24266 | 49.78797 | 47.07495 | Promoter | 5 | 108441054 | 108448891 | 7838 | 2 | ENSMUST00 | 0 | Mfsd7a | ENSMUSG00000029490.3 |
| chr12 | 93928499 | 93929325 | 14.51174 | 49.78466 | 47.07175 | Promoter | 12 | 93927004 | 93929102 | 2099 | 2 | ENSMUST00 | 0 | Gm9726 | ENSMUSG00000094935.1 |
| chr7 | 118993772 | 118995615 | 19.47913 | 49.77247 | 47.05964 | Promoter | 7 | 118972040 | 118995211 | 23172 | 2 | ENSMUST00 | 0 | Gprc5b | ENSMUSG00000008734.8 |
| chr1 | 74588268 | 74588770 | 17.64787 | 49.73517 | 47.02245 | Promoter | 1 | 74588289 | 74592124 | 3836 | 1 | ENSMUST00 | 0 | Bcs1l | ENSMUSG00000026172.7 |
| chr1 | 144003517 | 144005075 | 17.64787 | 49.73517 | 47.02245 | Promoter | 1 | 143999338 | 144004161 | 4824 | 2 | ENSMUST00 | 0 | Rgs2 | ENSMUSG00000026360.4 |
| chr1 | 74783909 | 74784660 | 17.64787 | 49.73517 | 47.02245 | 3' UTR | 1 | 74782439 | 74784602 | 2164 | 1 | ENSMUST00 | 1471 | Wnt6 | ENSMUSG00000033227.7 |
| chr1 | 7089010 | 7090932 | 17.64787 | 49.73517 | 47.02245 | Promoter | 1 | 7089104 | 7160902 | 71799 | 1 | ENSMUST00 | 0 | Pcmtd1 | ENSMUSG00000051285.12 |
| chr10 | 81167982 | 81169233 | 17.64787 | 49.73517 | 47.02245 | Promoter | 10 | 81153266 | 81167923 | 14658 | 2 | ENSMUST00 | -60 | Pias4 | ENSMUSG00000004934.8 |
| chr11 | 69419049 | 69420647 | 17.64787 | 49.73517 | 47.02245 | Downstream | 11 | 69398508 | 69413675 | 15168 | 2 | ENSMUST00 | -5375 | Kdm6b | ENSMUSG00000018476.7 |
| chr11 | 43681095 | 43681749 | 17.64787 | 49.73517 | 47.02245 | Promoter | 11 | 43681998 | 43707191 | 25194 | 1 | ENSMUST00 | -249 | Pwwp2a | ENSMUSG00000044950.7 |
| chr13 | 59876075 | 59878826 | 17.64787 | 49.73517 | 47.02245 | Distal Interge | 13 | 59771879 | 59823147 | 51269 | 2 | ENSMUST00 | -52929 | Zcchc6 | ENSMUSG00000035248.8 |
| chr15 | 76229079 | 76230123 | 17.64787 | 49.73517 | 47.02245 | Promoter | 15 | 76170975 | 76229497 | 58523 | 2 | ENSMUST00 | 0 | Plec | ENSMUSG00000022565.10 |
| chr17 | 46143764 | 46144614 | 17.64787 | 49.73517 | 47.02245 | Promoter | 17 | 46129271 | 46144216 | 14946 | 2 | ENSMUST00 | 0 | Rsph9 | ENSMUSG00000023966.3 |
| chr17 | 32924464 | 32925285 | 18.54115 | 49.73434 | 47.02245 | Promoter | 17 | 32925332 | 32925885 | 554 | 2 | ENSMUST00 | 600 | Cyp4f13 | ENSMUSG00000024055.9 |
| chr17 | 34263175 | 34263688 | 17.64787 | 49.73517 | 47.02245 | Promoter | 17 | 34263209 | 34269418 | 6210 | 1 | ENSMUST00 | 0 | H2-Ab1 | ENSMUSG00000073421.5 |
| chr19 | 4794065 | 4794570 | 17.64787 | 49.73517 | 47.02245 | Promoter | 19 | 4784293 | 4793901 | 9609 | 2 | ENSMUST00 | -165 | Rbm4 | ENSMUSG00000094936.2 |
| chr2 | 91115261 | 91115681 | 17.64787 | 49.73517 | 47.02245 | 3' UTR | 2 | 91118144 | 91136510 | 18367 | 1 | ENSMUST00 | -2463 | Mybpc3 | ENSMUSG00000002100.10 |
| chr2 | 80446457 | 80448477 | 17.64787 | 49.73517 | 47.02245 | Promoter | 2 | 80411970 | 80447625 | 35656 | 2 | ENSMUST00 | 0 | Frzb | ENSMUSG00000027004.3 |
| chr3 | 96414291 | 96414965 | 18.54115 | 49.73434 | 47.02245 | Promoter | 3 | 96414437 | 96414859 | 423 | 2 | ENSMUST00 | 0 | Terc | ENSMUSG00000064796.1 |
| chr3 | 46447441 | 46448314 | 17.64787 | 49.73517 | 47.02245 | Promoter | 3 | 46442197 | 46447941 | 5745 | 2 | ENSMUST00 | 0 | Pabpc4l | ENSMUSG00000090919.1 |
| chr6 | 48504258 | 48505082 | 17.64787 | 49.73517 | 47.02245 | Distal Interge | 6 | 48493780 | 48496206 | 2427 | 1 | ENSMUST00 | 10479 | Sspo | ENSMUSG00000029797.7 |
| chr6 | 51612270 | 51613066 | 17.64787 | 49.73517 | 47.02245 | Distal Interge | 6 | 51621632 | 51621708 | 77 | 1 | ENSMUST00 | -8566 | Gm22914 | ENSMUSG00000094528.1 |
| chr8 | 109693365 | 109693957 | 17.64787 | 49.73517 | 47.02245 | Promoter | 8 | 109671322 | 109693294 | 21973 | 2 | ENSMUST00 | -72 | Ist1 | ENSMUSG00000031729.5 |
| chr8 | 94433779 | 94434684 | 17.64787 | 49.73517 | 47.02245 | Promoter | 8 | 94422899 | 94434157 | 11259 | 2 | ENSMUST00 | 0 | 9330175E14I | ENSMUSG00000097194.1 |
| chr9 | 113930277 | 113930872 | 17.64787 | 49.73517 | 47.02245 | Promoter | 9 | 113930934 | 113977199 | 46266 | 1 | ENSMUST00 | -62 | Ubp1 | ENSMUSG00000009741.9 |
| chr9 | 119156432 | 119157076 | 17.64787 | 49.73517 | 47.02245 | Promoter | 9 | 119148023 | 119157093 | 9071 | 2 | ENSMUST00 | 17 | Acaa1b | ENSMUSG00000010651.4 |
| chr9 | 41950494 | 41951589 | 17.64787 | 49.73517 | 47.02245 | Distal Interge | 9 | 41969739 | 41974587 | 4849 | 2 | ENSMUST00 | 22998 | Sorl1 | ENSMUSG00000049313.8 |
| chr9 | 56324531 | 56325132 | 17.64787 | 49.73517 | 47.02245 | Intron (ENSM | 9 | 56257890 | 56267780 | 9891 | 1 | ENSMUST00 | 66642 | Peak1os | ENSMUSG00000097111.1 |
| chrX | 8095931 | 8096571 | 17.64787 | 49.73517 | 47.02245 | Promoter | 20 | 8097087 | 8106701 | 9615 | 1 | ENSMUST00 | -516 | Gm6787 | ENSMUSG00000086404.1 |
| chr16 | 81200246 | 81201154 | 18.98792 | 49.7182 | 47.00772 | Promoter | 16 | 81200697 | 81597805 | 397109 | 1 | ENSMUST00 | 0 | Ncam2 | ENSMUSG00000022762.12 |
| chr17 | 83276028 | 83277166 | 16.77503 | 49.70845 | 46.99806 | Distal Interge | 17 | 83269971 | 83270105 | 135 | 1 | ENSMUST00 | 6058 | Gm24240 | ENSMUSG00000088410.1 |
| chr11 | 116085758 | 116086662 | 19.43886 | 49.70381 | 46.99345 | Promoter | 11 | 116083096 | 116086972 | 3877 | 2 | ENSMUST00 | 310 | Wbp2 | ENSMUSG00000034341.12 |
| chr14 | 25606620 | 25608499 | 15.16532 | 49.65084 | 46.94072 | Promoter | 14 | 25607357 | 25643657 | 36301 | 1 | ENSMUST00 | 0 | Zmiz1 | ENSMUSG00000007817.9 |
| chr14 | 65805752 | 65806297 | 15.16532 | 49.65084 | 46.94072 | Promoter | 14 | 65805837 | 65817822 | 11986 | 1 | ENSMUST00 | 0 | Pbk | ENSMUSG00000022033.4 |
| chr2 | 177397378 | 177398585 | 15.16532 | 49.65084 | 46.94072 | Promoter | 2 | 177387901 | 177398320 | 10420 | 2 | ENSMUST00 | 0 | Gm14418 | ENSMUSG00000078867.4 |
| chr4 | 88640755 | 88641212 | 15.16532 | 49.65084 | 46.94072 | Downstream | 4 | 88643641 | 88644459 | 819 | 2 | ENSMUST00 | 3247 | Ifna13 | ENSMUSG00000063376.5 |
| chr5 | 109795773 | 109796769 | 15.16532 | 49.65084 | 46.94072 | Distal Interge | 5 | 109757522 | 109757931 | 410 | 2 | ENSMUST00 | -37843 | Gm8508 | ENSMUSG00000049477.4 |
| chr18 | 52825014 | 52828158 | 19.39874 | 49.63541 | 46.92579 | Intron (ENSM | 18 | 52837964 | 52915586 | 77623 | 1 | ENSMUST00 | -9806 | Sncaip | ENSMUSG00000024534.10 |
| chr6 | 124856897 | 124857761 | 19.39874 | 49.63541 | 46.92579 | Promoter | 6 | 124841089 | 124857752 | 16664 | 2 | ENSMUST00 | 0 | Leprel2 | ENSMUSG00000023191.4 |
| chr11 | 101963509 | 101964404 | 18.91326 | 49.58984 | 46.88035 | 3' UTR | 11 | 101962458 | 101967015 | 4558 | 2 | ENSMUST00 | 2611 | Sost | ENSMUSG00000001494.6 |
| chr11 | 3331738 | 3332542 | 18.91326 | 49.58984 | 46.88035 | Promoter | 11 | 3332072 | 3333679 | 1608 | 1 | ENSMUST00 | 0 | Pik3ip1 | ENSMUSG00000034614.9 |
| chr18 | 70052833 | 70053619 | 18.91326 | 49.58984 | 46.88035 | Promoter | 18 | 69983656 | 70053588 | 69933 | 2 | ENSMUST00 | 0 | Rab27b | ENSMUSG00000024511.10 |
| chr2 | 75658153 | 75658982 | 17.13008 | 49.58303 | 46.87408 | Promoter | 2 | 75659244 | 75659317 | 74 | 1 | ENSMUST00 | -262 | Gm24574 | ENSMUSG00000093242.1 |
| chr1 | 136465900 | 136466795 | 19.3588 | 49.56728 | 46.8587 | Promoter | 1 | 136466343 | 136495916 | 29574 | 1 | ENSMUST00 | 0 | Kif14 | ENSMUSG00000041498.8 |
| chr6 | 47942754 | 47943601 | 18.87615 | 49.526 | 46.81754 | Promoter | 6 | 47943171 | 47954549 | 11379 | 1 | ENSMUST00 | 0 | Zfp783 | ENSMUSG00000072653.7 |
| chr1 | 161876914 | 161877795 | 13.406 | 49.51596 | 46.80759 | Promoter | 1 | 161816112 | 161876661 | 60550 | 2 | ENSMUST00 | -254 | Suco | ENSMUSG00000040297.7 |
| chr1 | 171070490 | 171070896 | 13.406 | 49.51596 | 46.80759 | Downstream | 1 | 171072226 | 171072287 | 62 | 2 | ENSMUST00 | 1391 | Gm27552 | ENSMUSG00000098356.1 |
| chr1 | 190168694 | 190171152 | 16.66433 | 49.50283 | 46.79451 | Promoter | 1 | 190118035 | 190170714 | 52680 | 2 | ENSMUST00 | 0 | Prox1 | ENSMUSG00000010175.8 |
| chr13 | 37857152 | 37858663 | 16.66433 | 49.50283 | 46.79451 | Promoter | 13 | 37857934 | 37893914 | 35981 | 1 | ENSMUST00 | 0 | Rreb1 | ENSMUSG00000039087.11 |
| chr3 | 36688981 | 36690356 | 16.66433 | 49.50283 | 46.79451 | Promoter | 3 | 36620482 | 36690167 | 69686 | 2 | ENSMUST00 | 0 | Trpc3 | ENSMUSG00000027716.8 |
| chr4 | 156031080 | 156032314 | 16.66433 | 49.50283 | 46.79451 | Promoter | 4 | 156023727 | 156031643 | 7917 | 2 | ENSMUST00 | 0 | Gm16008 | ENSMUSG00000087381.3 |
| chr5 | 57715045 | 57720059 | 16.66433 | 49.50283 | 46.79451 | Promoter | 5 | 57718021 | 57729465 | 11445 | 1 | ENSMUST00 | 0 | Pcdh7 | ENSMUSG00000029108.9 |
| chr7 | 4477917 | 4479213 | 16.66433 | 49.50283 | 46.79451 | Promoter | 7 | 4477666 | 4479244 | 1579 | 1 | ENSMUST00 | 252 | Eps8l1 | ENSMUSG00000006154.8 |
| chr9 | 88858425 | 88859397 | 16.66433 | 49.50283 | 46.79451 | Promoter | 9 | 88840236 | 88858866 | 18631 | 2 | ENSMUST00 | 0 | 9330159M07 | ENSMUSG00000097177.2 |
| chr7 | 49974868 | 49975780 | 19.31902 | 49.4994 | 46.79264 | Promoter | 7 | 49975069 | 50280036 | 304968 | 1 | ENSMUST00 | 0 | Nell1 | ENSMUSG00000055409.8 |
| chr1 | 181352275 | 181353393 | 15.85026 | 49.49464 | 46.78799 | Promoter | 1 | 181352628 | 181459909 | 107282 | 1 | ENSMUST00 | 0 | Cnih3 | ENSMUSG00000026514.8 |
| chr10 | 39171283 | 39172006 | 15.85026 | 49.49464 | 46.78799 | Distal Interge | 10 | 39150971 | 39163794 | 12824 | 2 | ENSMUST00 | -7490 | Wisp3 | ENSMUSG00000062074.6 |
| chr11 | 78535381 | 78536261 | 15.85026 | 49.49464 | 46.78799 | Promoter | 11 | 78522850 | 78536271 | 13422 | 2 | ENSMUST00 | 10 | Tnfaip1 | ENSMUSG00000017615.7 |
| chr11 | 109439719 | 109440803 | 15.85026 | 49.49464 | 46.78799 | Promoter | 11 | 109440156 | 109452813 | 12658 | 1 | ENSMUST00 | 0 | Amz2 | ENSMUSG00000020610.11 |
| chr13 | 46501688 | 46502622 | 15.85026 | 49.49464 | 46.78799 | Promoter | 13 | 46501848 | 46649642 | 147795 | 1 | ENSMUST00 | 0 | Cap2 | ENSMUSG00000021373.10 |
| chr18 | 7003755 | 7005881 | 15.85026 | 49.49464 | 46.78799 | Promoter | 18 | 6934966 | 7004780 | 69815 | 2 | ENSMUST00 | 0 | Mkx | ENSMUSG00000061013.5 |
| chr7 | 16311865 | 16313153 | 15.85026 | 49.49464 | 46.78799 | Promoter | 7 | 16313517 | 16317697 | 4181 | 1 | ENSMUST00 | -364 | Bbc3 | ENSMUSG00000002083.7 |
| chr7 | 110629711 | 110631062 | 15.85026 | 49.49464 | 46.78799 | 3' UTR | 7 | 110628028 | 110628551 | 524 | 1 | ENSMUST00 | 1684 | Adm | ENSMUSG00000030790.9 |
| chr8 | 4258335 | 4259772 | 15.85026 | 49.49464 | 46.78799 | Promoter | 8 | 4257648 | 4259274 | 1627 | 2 | ENSMUST00 | 0 | Ctxn1 | ENSMUSG00000048644.7 |
| chr11 | 102455661 | 102456449 | 18.83918 | 49.46239 | 46.75671 | Promoter | 11 | 102453304 | 102456190 | 2887 | 2 | ENSMUST00 | 0 | Itga2b | ENSMUSG00000034664.8 |
| chr13 | 92425718 | 92426998 | 19.2794 | 49.4318 | 46.72622 | Promoter | 13 | 92425896 | 92441658 | 15763 | 1 | ENSMUST00 | 0 | Ankrd34b | ENSMUSG00000045034.7 |
| chr13 | 12519482 | 12520657 | 19.2794 | 49.4318 | 46.72622 | Promoter | 13 | 12471209 | 12520450 | 49242 | 2 | ENSMUST00 | 0 | Edaradd | ENSMUSG00000095105.1 |
| chr5 | 72647230 | 72648317 | 19.2794 | 49.4318 | 46.72622 | Promoter | 5 | 72647796 | 72671078 | 23283 | 1 | ENSMUST00 | 0 | Nipal1 | ENSMUSG00000067219.5 |
| chr14 | 63164409 | 63165221 | 18.80236 | 49.39901 | 46.69355 | Exon (ENSMU | 14 | 63145153 | 63177793 | 32641 | 2 | ENSMUST00 | 12572 | Fdft1 | ENSMUSG00000021273.9 |
| chr13 | 40722313 | 40723573 | 17.45709 | 49.39342 | 46.68804 | Exon (ENSMU | 13 | 40727808 | 40735162 | 7355 | 1 | ENSMUST00 | -4235 | Gm26688 | ENSMUSG00000097752.2 |
| chr19 | 34550380 | 34551364 | 17.45709 | 49.39342 | 46.68804 | Promoter | 19 | 34550694 | 34573192 | 22499 | 1 | ENSMUST00 | 0 | Ifit2 | ENSMUSG00000045932.7 |
| chr2 | 130696844 | 130698000 | 17.45709 | 49.39342 | 46.68804 | Promoter | 2 | 130684113 | 130697519 | 13407 | 2 | ENSMUST00 | 0 | Slc4a11 | ENSMUSG00000074796.5 |
| chr7 | 6282967 | 6284008 | 17.45709 | 49.39342 | 46.68804 | Distal Interge | 7 | 6286579 | 6307883 | 21305 | 1 | ENSMUST00 | -2571 | Zfp667 | ENSMUSG00000054893.10 |
| chr9 | 107288792 | 107290348 | 17.45709 | 49.39342 | 46.68804 | Promoter | 9 | 107254927 | 107289877 | 34951 | 2 | ENSMUST00 | 0 | Mapkapk3 | ENSMUSG00000032577.11 |
| chr10 | 128499054 | 128499748 | 19.23994 | 49.36444 | 46.65932 | Promoter | 10 | 128499364 | 128504242 | 4879 | 1 | ENSMUST00 | 0 | A430046D13 | ENSMUSG00000097353.1 |
| chr4 | 134396341 | 134396847 | 19.23994 | 49.36444 | 46.65932 | Promoter | 4 | 134396370 | 134418072 | 21703 | 1 | ENSMUST00 | 0 | Pafah2 | ENSMUSG00000037366.9 |
| chr5 | 121842506 | 121843411 | 19.23994 | 49.36444 | 46.65932 | Distal Interge | 5 | 121849028 | 121854599 | 5572 | 1 | ENSMUST00 | -5617 | Fam109a | ENSMUSG00000044134.5 |
| chr7 | 141481690 | 141482574 | 19.23994 | 49.36444 | 46.65932 | 5' UTR | 7 | 141476400 | 141493421 | 17022 | 1 | ENSMUST00 | 5291 | Tspan4 | ENSMUSG00000025511.9 |
| chr9 | 98424229 | 98425440 | 18.72915 | 49.27289 | 46.56846 | 5' UTR | 9 | 98422961 | 98446575 | 23615 | 1 | ENSMUST00 | 1269 | Rbp1 | ENSMUSG00000046402.10 |
| chr7 | 16780879 | 16782693 | 18.26577 | 49.25508 | 46.55071 | Promoter | 7 | 16781346 | 16798257 | 16912 | 1 | ENSMUST00 | 0 | Slc1a5 | ENSMUSG00000001918.12 |
| chr19 | 38053901 | 38054867 | 18.6565 | 49.14766 | 46.44425 | Promoter | 19 | 38051774 | 38054528 | 2755 | 2 | ENSMUST00 | 0 | I830134H01F | ENSMUSG00000069554.4 |
| chr14 | 54605589 | 54606514 | 19.08372 | 49.09757 | 46.39423 | Promoter | 14 | 54583663 | 54605908 | 22246 | 2 | ENSMUST00 | 0 | 4931414P19F | ENSMUSG00000022179.4 |
| chr5 | 139735730 | 139736407 | 18.62039 | 49.08537 | 46.38214 | Promoter | 5 | 139706693 | 139736336 | 29644 | 2 | ENSMUST00 | 0 | Micall2 | ENSMUSG00000036718.12 |
| chr14 | 50869917 | 50871102 | 13.82983 | 49.0377 | 46.33454 | Promoter | 14 | 50824062 | 50870560 | 46499 | 2 | ENSMUST00 | 0 | Tep1 | ENSMUSG00000006281.7 |
| chr19 | 4476542 | 4477536 | 13.82983 | 49.0377 | 46.33454 | Promoter | 19 | 4445908 | 4477447 | 31540 | 2 | ENSMUST00 | 0 | Syt12 | ENSMUSG00000049303.5 |
| chr8 | 70754657 | 70755419 | 13.82983 | 49.0377 | 46.33454 | Promoter | 8 | 70754679 | 70758677 | 3999 | 1 | ENSMUST00 | 0 | Rab3a | ENSMUSG00000031840.7 |
| chr7 | 113206430 | 113207171 | 19.04506 | 49.03148 | 46.32841 | Promoter | 7 | 113207465 | 113314122 | 106658 | 1 | ENSMUST00 | -294 | Arntl | ENSMUSG00000055116.7 |
| chr9 | 123528545 | 123529543 | 18.58442 | 49.02329 | 46.32032 | Promoter | 9 | 123529882 | 123592598 | 62717 | 1 | ENSMUST00 | -339 | Sacm1l | ENSMUSG00000025240.8 |
| chr7 | 6729668 | 6730727 | 17.68659 | 49.02076 | 46.31784 | Promoter | 7 | 6703892 | 6730421 | 26530 | 2 | ENSMUST00 | 0 | Peg3 | ENSMUSG00000002265.10 |
| chr18 | 20557803 | 20558701 | 19.00656 | 48.96563 | 46.26303 | Promoter | 18 | 20558074 | 20604521 | 46448 | 1 | ENSMUST00 | 0 | Dsg2 | ENSMUSG00000044393.10 |
| chr11 | 74829633 | 74830333 | 18.54859 | 48.96143 | 46.25894 | Promoter | 11 | 74830920 | 74845725 | 14806 | 1 | ENSMUST00 | -587 | Mnt | ENSMUSG00000000282.7 |
| chr2 | 30845081 | 30845822 | 18.54859 | 48.96143 | 46.25894 | Promoter | 2 | 30845444 | 30878657 | 33214 | 1 | ENSMUST00 | 0 | Prrx2 | ENSMUSG00000039476.8 |
| chr11 | 97450104 | 97452307 | 17.16547 | 48.86851 | 46.16669 | Promoter | 11 | 97450160 | 97502400 | 52241 | 1 | ENSMUST00 | 0 | Arhgap23 | ENSMUSG00000049807.11 |
| chr17 | 25726986 | 25727459 | 17.16547 | 48.86851 | 46.16669 | Promoter | 17 | 25718926 | 25727371 | 8446 | 2 | ENSMUST00 | 0 | Chtf18 | ENSMUSG00000019214.8 |
| chr8 | 20557501 | 20558087 | 10.02373 | 48.85992 | 46.15827 | Promoter | 8 | 20556695 | 20557333 | 639 | 2 | ENSMUST00 | -169 | Gm21182 | ENSMUSG00000095766.1 |
| chr8 | 22858604 | 22859344 | 18.47733 | 48.83834 | 46.1367 | Promoter | 8 | 22859535 | 22943259 | 83725 | 1 | ENSMUST00 | -191 | Kat6a | ENSMUSG00000031540.8 |
| chr2 | 91444186 | 91444787 | 15.87453 | 48.7791 | 46.0777 | Promoter | 2 | 91275068 | 91444622 | 169555 | 2 | ENSMUST00 | 0 | 1110051M20 | ENSMUSG00000040591.13 |
| chr10 | 83533257 | 83534381 | 18.44084 | 48.77526 | 46.0739 | Promoter | 10 | 83487450 | 83534140 | 46691 | 2 | ENSMUST00 | 0 | Aldh1l2 | ENSMUSG00000020256.9 |
| chr11 | 3863478 | 3864305 | 18.44084 | 48.77526 | 46.0739 | Promoter | 11 | 3703731 | 3863903 | 160173 | 2 | ENSMUST00 | 0 | Osbp2 | ENSMUSG00000020435.12 |
| chr11 | 33146998 | 33148481 | 18.44084 | 48.77526 | 46.0739 | Promoter | 11 | 33117430 | 33147400 | 29971 | 2 | ENSMUST00 | 0 | Fgf18 | ENSMUSG00000057967.7 |
| chr12 | 70226918 | 70228127 | 18.44084 | 48.77526 | 46.0739 | Promoter | 12 | 70190811 | 70227683 | 36873 | 2 | ENSMUST00 | 0 | Pygl | ENSMUSG00000021069.11 |
| chr12 | 19387558 | 19388842 | 18.44084 | 48.77526 | 46.0739 | Promoter | 12 | 19387809 | 19390286 | 2478 | 1 | ENSMUST00 | 0 | Gm5784 | ENSMUSG00000093908.1 |
| chr13 | 94702294 | 94703317 | 18.44084 | 48.77526 | 46.0739 | Distal Interge | 13 | 94788943 | 94842899 | 53957 | 1 | ENSMUST00 | -85626 | Tbca | ENSMUSG00000042043.5 |
| chr13 | 58358554 | 58359564 | 18.44084 | 48.77526 | 46.0739 | Distal Interge | 13 | 58287516 | 58354862 | 67347 | 2 | ENSMUST00 | -3693 | Kif27 | ENSMUSG00000060176.3 |
| chr14 | 63193342 | 63194028 | 18.44084 | 48.77526 | 46.0739 | Promoter | 14 | 63182443 | 63193541 | 11099 | 2 | ENSMUST00 | 0 | Neil2 | ENSMUSG00000035121.5 |
| chr19 | 4120833 | 4121578 | 18.44084 | 48.77526 | 46.0739 | Promoter | 19 | 4114464 | 4121575 | 7112 | 2 | ENSMUST00 | 0 | Aip | ENSMUSG00000024847.9 |
| chr19 | 44988855 | 44989791 | 18.44084 | 48.77526 | 46.0739 | Promoter | 19 | 44989101 | 45003397 | 14297 | 1 | ENSMUST00 | 0 | Sema4g | ENSMUSG00000025207.10 |
| chr2 | 33631177 | 33632414 | 18.44084 | 48.77526 | 46.0739 | Promoter | 2 | 33567156 | 33632283 | 65128 | 2 | ENSMUST00 | 0 | Lmx1b | ENSMUSG00000038765.8 |
| chr4 | 153481128 | 153483155 | 18.44084 | 48.77526 | 46.0739 | Promoter | 4 | 153373221 | 153482811 | 109591 | 2 | ENSMUST00 | 0 | Ajap1 | ENSMUSG00000039546.9 |
| chr4 | 153118908 | 153120529 | 18.44084 | 48.77526 | 46.0739 | Distal Interge | 4 | 153145590 | 153146943 | 1354 | 2 | ENSMUST00 | 26414 | Gm13173 | ENSMUSG00000085648.1 |
| chr5 | 120812022 | 120813285 | 18.44084 | 48.77526 | 46.0739 | Promoter | 5 | 120800194 | 120812514 | 12321 | 2 | ENSMUST00 | 0 | Oas1c | ENSMUSG00000001166.12 |
| chr5 | 107359351 | 107360408 | 18.44084 | 48.77526 | 46.0739 | Promoter | 5 | 107359837 | 107386361 | 26525 | 1 | ENSMUST00 | 0 | Brdt | ENSMUSG00000029279.10 |
| chr6 | 42324147 | 42325306 | 18.44084 | 48.77526 | 46.0739 | Promoter | 6 | 42315312 | 42324640 | 9329 | 2 | ENSMUST00 | 0 | Fam131b | ENSMUSG00000029861.12 |
| chr6 | 121210440 | 121211763 | 18.44084 | 48.77526 | 46.0739 | Promoter | 6 | 121210734 | 121226854 | 16121 | 1 | ENSMUST00 | 0 | Tuba8 | ENSMUSG00000030137.7 |
| chr6 | 29178916 | 29179800 | 18.44084 | 48.77526 | 46.0739 | Promoter | 6 | 29169232 | 29179584 | 10353 | 2 | ENSMUST00 | 0 | Prrt4 | ENSMUSG00000079654.3 |
| chrX | 85776910 | 85777661 | 18.44084 | 48.77526 | 46.0739 | Promoter | 20 | 85701937 | 85776819 | 74883 | 2 | ENSMUST00 | -92 | Gyk | ENSMUSG00000025059.11 |
| chr6 | 48554354 | 48555344 | 18.89198 | 48.76956 | 46.07009 | Promoter | 6 | 48554796 | 48570722 | 15927 | 1 | ENSMUST00 | 0 | Lrrc61 | ENSMUSG00000073096.6 |
| chr6 | 47739814 | 47740114 | 6.03039 | 48.75505 | 46.05571 | Distal Interge | 6 | 47773329 | 47773429 | 101 | 1 | ENSMUST00 | -33215 | Gm24975 | ENSMUSG00000094807.1 |
| chr9 | 7763441 | 7763867 | 18.40662 | 48.71608 | 46.01675 | Promoter | 9 | 7764041 | 7794333 | 30293 | 1 | ENSMUST00 | -174 | Tmem123 | ENSMUSG00000050912.10 |

| | | | | | | | | | | | | | | | |
|---|---|---|---|---|---|---|---|---|---|---|---|---|---|---|---|
| chr12 | 54655411 | 54657033 | 18.85409 | 48.70469 | 46.00539 | Promoter | 12 | 54645374 | 54656572 | 11199 | 2 | ENSMUST00 | 0 | Sptssa | ENSMUSG00000044408.6 |
| chr13 | 53472200 | 53473099 | 18.85409 | 48.70469 | 46.00539 | Promoter | 13 | 53466884 | 53473074 | 6191 | 2 | ENSMUST00 | 0 | Msx2 | ENSMUSG00000021469.9 |
| chr13 | 28884070 | 28885838 | 18.85409 | 48.70469 | 46.00539 | Promoter | 13 | 28460778 | 28885620 | 424843 | 2 | ENSMUST00 | 0 | 2610307P16I | ENSMUSG00000085936.2 |
| chr15 | 101410897 | 101413261 | 18.85409 | 48.70469 | 46.00539 | Promoter | 15 | 101411043 | 101411983 | 941 | 1 | ENSMUST00 | 0 | Krt7 | ENSMUSG00000023039.11 |
| chr7 | 14623140 | 14623611 | 18.85409 | 48.70469 | 46.00539 | Promoter | 7 | 14530652 | 14622479 | 91828 | 2 | ENSMUST00 | -662 | Nlrp5-ps | ENSMUSG00000041596.7 |
| chr8 | 82401878 | 82403319 | 18.85409 | 48.70469 | 46.00539 | Promoter | 8 | 82331637 | 82403222 | 71586 | 2 | ENSMUST00 | 0 | Il15 | ENSMUSG00000031712.8 |
| chr1 | 83037782 | 83038696 | 19.30035 | 48.68736 | 45.98855 | Promoter | 1 | 83012523 | 83038448 | 25926 | 2 | ENSMUST00 | 0 | Slc19a3 | ENSMUSG00000038496.11 |
| chr10 | 29146646 | 29147804 | 19.30035 | 48.68736 | 45.98855 | 5' UTR | 10 | 29143282 | 29144194 | 913 | 2 | ENSMUST00 | -2453 | Gm9996 | ENSMUSG00000056316.2 |
| chr11 | 69966010 | 69967136 | 19.30035 | 48.68736 | 45.98855 | Promoter | 11 | 69966918 | 69967652 | 735 | 1 | ENSMUST00 | 0 | Cldn7 | ENSMUSG00000018569.7 |
| chr13 | 120096753 | 120097057 | 19.30035 | 48.68736 | 45.98855 | Downstream | 13 | 120099349 | 120100332 | 984 | 2 | ENSMUST00 | 3275 | Gm21378 | ENSMUSG00000095709.1 |
| chr18 | 77564720 | 77566297 | 19.30035 | 48.68736 | 45.98855 | Promoter | 18 | 77456110 | 77565136 | 109027 | 2 | ENSMUST00 | 0 | Rnf165 | ENSMUSG00000025427.9 |
| chr18 | 74960528 | 74961975 | 19.30035 | 48.68736 | 45.98855 | Promoter | 18 | 74939322 | 74961263 | 21942 | 2 | ENSMUST00 | 0 | Lipg | ENSMUSG00000053846.4 |
| chr19 | 5725465 | 5726721 | 19.30035 | 48.68736 | 45.98855 | Promoter | 19 | 5707376 | 5726317 | 18942 | 2 | ENSMUST00 | 0 | Ehbp1l1 | ENSMUSG00000024937.9 |
| chr2 | 58356830 | 58358149 | 19.30035 | 48.68736 | 45.98855 | Promoter | 2 | 58267453 | 58357895 | 90443 | 2 | ENSMUST00 | 0 | Acvr1c | ENSMUSG00000026834.8 |
| chr4 | 40854197 | 40854723 | 19.30035 | 48.68736 | 45.98855 | Promoter | 4 | 40809154 | 40854005 | 44852 | 2 | ENSMUST00 | -193 | B4galt1 | ENSMUSG00000028413.8 |
| chr4 | 141538723 | 141539706 | 19.30035 | 48.68736 | 45.98855 | Promoter | 4 | 141467890 | 141538597 | 70708 | 2 | ENSMUST00 | -127 | Spen | ENSMUSG00000040761.11 |
| chr5 | 147957175 | 147958331 | 19.30035 | 48.68736 | 45.98855 | Promoter | 5 | 147957320 | 148316065 | 358746 | 1 | ENSMUST00 | 0 | Mtus2 | ENSMUSG00000029651.11 |
| chr7 | 25076277 | 25077009 | 19.30035 | 48.68736 | 45.98855 | Promoter | 7 | 25077205 | 25083492 | 6288 | 1 | ENSMUST00 | -196 | Zfp574 | ENSMUSG00000045252.10 |
| chr8 | 84025841 | 84026307 | 19.30035 | 48.68736 | 45.98855 | Promoter | 8 | 84026075 | 84027049 | 975 | 1 | ENSMUST00 | 0 | Palm3 | ENSMUSG00000047986.7 |
| chr9 | 91364331 | 91365365 | 19.30035 | 48.68736 | 45.98855 | Promoter | 9 | 91358058 | 91365768 | 7711 | 2 | ENSMUST00 | 403 | Zic1 | ENSMUSG00000032368.9 |
| chr9 | 91361995 | 91362631 | 19.30035 | 48.68736 | 45.98855 | Promoter | 9 | 91362413 | 91378915 | 16503 | 1 | ENSMUST00 | 0 | Zic4 | ENSMUSG00000036972.9 |
| chrX | 74428042 | 74430275 | 19.30035 | 48.68736 | 45.98855 | Promoter | 20 | 74429281 | 74443357 | 14077 | 1 | ENSMUST00 | 0 | Ikbkg | ENSMUSG00000004221.11 |
| chrX | 17571391 | 17572275 | 19.30035 | 48.68736 | 45.98855 | Promoter | 20 | 17557524 | 17571641 | 14118 | 2 | ENSMUST00 | 0 | Fundc1 | ENSMUSG00000025040.8 |
| chrX | 111462500 | 111463737 | 19.30035 | 48.68736 | 45.98855 | Promoter | 20 | 111388192 | 111463043 | 74852 | 2 | ENSMUST00 | 0 | Rps6ka6 | ENSMUSG00000025665.11 |
| chrX | 38563569 | 38565117 | 19.30035 | 48.68736 | 45.98855 | Promoter | 20 | 38533274 | 38564525 | 31252 | 2 | ENSMUST00 | 0 | Cul4b | ENSMUSG00000031095.10 |
| chrX | 7697123 | 7698072 | 19.30035 | 48.68736 | 45.98855 | Promoter | 20 | 7697131 | 7710262 | 13132 | 1 | ENSMUST00 | 0 | Gpkow | ENSMUSG00000031148.9 |
| chrX | 151520665 | 151522628 | 19.30035 | 48.68736 | 45.98855 | Promoter | 20 | 151520672 | 151570807 | 50136 | 1 | ENSMUST00 | 0 | Phf8 | ENSMUSG00000041229.10 |
| chrX | 7374969 | 7376714 | 19.30035 | 48.68736 | 45.98855 | Promoter | 20 | 7371276 | 7375826 | 4551 | 2 | ENSMUST00 | 0 | Usp27x | ENSMUSG00000046269.11 |
| chrX | 98937406 | 98939211 | 19.30035 | 48.68736 | 45.98855 | Promoter | 20 | 98937761 | 98944906 | 7146 | 1 | ENSMUST00 | 0 | Yipf6 | ENSMUSG00000047694.7 |
| chrX | 134756578 | 134757470 | 19.30035 | 48.68736 | 45.98855 | Promoter | 20 | 134756595 | 134761425 | 4831 | 1 | ENSMUST00 | 0 | Armcx3 | ENSMUSG00000049047.6 |
| chrX | 56609297 | 56610422 | 19.30035 | 48.68736 | 45.98855 | Promoter | 20 | 56609757 | 56633579 | 23823 | 1 | ENSMUST00 | 0 | Slc9a6 | ENSMUSG00000060681.9 |
| chrX | 135885422 | 135886593 | 19.30035 | 48.68736 | 45.98855 | Promoter | 20 | 135885851 | 135891078 | 5228 | 1 | ENSMUST00 | 0 | Bhlhb9 | ENSMUSG00000072964.8 |
| chrX | 166478509 | 166480235 | 19.30035 | 48.68736 | 45.98855 | Promoter | 20 | 166457253 | 166479867 | 22615 | 2 | ENSMUST00 | 0 | Rab9 | ENSMUSG00000079316.5 |
| chrX | 123102881 | 123104946 | 19.30035 | 48.68736 | 45.98855 | Promoter | 20 | 123103495 | 123145651 | 42157 | 1 | ENSMUST00 | 0 | 3110007F17I | ENSMUSG00000079450.6 |
| chr13 | 23573311 | 23574630 | 17.93284 | 48.67312 | 45.97572 | Promoter | 13 | 23573736 | 23574196 | 461 | 2 | ENSMUST00 | 0 | Hist1h2bf | ENSMUSG00000069268.4 |
| chr15 | 99093529 | 99094203 | 17.93284 | 48.67312 | 45.97572 | Promoter | 15 | 99093170 | 99100917 | 7748 | 1 | ENSMUST00 | 360 | Dnajc22 | ENSMUSG00000038009.7 |
| chr18 | 37879850 | 37880387 | 17.93284 | 48.67312 | 45.97572 | Intron (ENSM | 18 | 37896157 | 37897942 | 1786 | 2 | ENSMUST00 | 17555 | Diap1 | ENSMUSG00000024456.11 |
| chr19 | 23134219 | 23136818 | 17.93284 | 48.67312 | 45.97572 | Promoter | 19 | 23134900 | 23135717 | 818 | 1 | ENSMUST00 | 0 | 2410080I02R | ENSMUSG00000044387.5 |
| chr2 | 155929228 | 155930716 | 17.93284 | 48.67312 | 45.97572 | Promoter | 2 | 155846894 | 155930060 | 83167 | 2 | ENSMUST00 | 0 | Uqcc1 | ENSMUSG00000005882.13 |
| chr7 | 45488439 | 45489284 | 17.93284 | 48.67312 | 45.97572 | Promoter | 7 | 45472956 | 45488805 | 15850 | 2 | ENSMUST00 | 0 | Dhdh | ENSMUSG00000011382.6 |
| chr8 | 69584487 | 69585158 | 13.98842 | 48.66103 | 45.96413 | Distal Interge | 8 | 69612420 | 69614063 | 1644 | 2 | ENSMUST00 | 28905 | Zfp868 | ENSMUSG00000060427.10 |
| chr18 | 10324866 | 10325953 | 14.31595 | 48.63377 | 45.93701 | Promoter | 18 | 10325177 | 10502412 | 177236 | 1 | ENSMUST00 | 0 | Greb1l | ENSMUSG00000042942.10 |
| chr19 | 38041811 | 38043957 | 14.31595 | 48.63377 | 45.93701 | Promoter | 19 | 37899036 | 38043577 | 144542 | 2 | ENSMUST00 | 0 | Myof | ENSMUSG00000048612.10 |
| chr4 | 12136061 | 12137120 | 14.31595 | 48.63377 | 45.93701 | Distal Interge | 4 | 12140264 | 12144688 | 4425 | 1 | ENSMUST00 | -3144 | Rbm12b1 | ENSMUSG00000046667.9 |
| chr6 | 29170054 | 29171542 | 14.31595 | 48.63377 | 45.93701 | Exon (ENSMU | 6 | 29123573 | 29165006 | 41434 | 2 | ENSMUST00 | -5049 | Rbm28 | ENSMUSG00000029701.10 |
| chr8 | 85119162 | 85120048 | 14.31595 | 48.63377 | 45.93701 | Promoter | 8 | 85109647 | 85119637 | 9991 | 2 | ENSMUST00 | 0 | Zfp791 | ENSMUSG00000074194.3 |
| chr15 | 7397715 | 7398736 | 19.26563 | 48.62831 | 45.9318 | Promoter | 15 | 7206120 | 7398304 | 192185 | 2 | ENSMUST00 | 0 | Egflam | ENSMUSG00000042961.8 |
| chr4 | 108387514 | 108388356 | 19.26563 | 48.62831 | 45.9318 | Distal Interge | 4 | 108367777 | 108383349 | 15573 | 2 | ENSMUST00 | -4166 | Fam159a | ENSMUSG00000059816.5 |
| chr10 | 19849128 | 19851849 | 18.33645 | 48.59467 | 45.89824 | Promoter | 10 | 19847917 | 19851459 | 3543 | 2 | ENSMUST00 | 0 | Slc35d3 | ENSMUSG00000050473.4 |
| chr17 | 56276023 | 56277505 | 18.77877 | 48.57565 | 45.87926 | Promoter | 17 | 56269319 | 56276786 | 7468 | 2 | ENSMUST00 | 0 | Ticam1 | ENSMUSG00000047123.8 |
| chr9 | 120010981 | 120011832 | 18.77877 | 48.57565 | 45.87926 | Intron (ENSM | 9 | 120013755 | 120023598 | 9844 | 2 | ENSMUST00 | 11766 | Xirp1 | ENSMUSG00000079243.2 |
| chr2 | 26091165 | 26092272 | 18.70405 | 48.44756 | 45.75203 | Promoter | 2 | 26055535 | 26092177 | 36643 | 2 | ENSMUST00 | 0 | Nacc2 | ENSMUSG00000026932.9 |
| chr1 | 105971539 | 105972408 | 18.66691 | 48.38385 | 45.68857 | Distal Interge | 1 | 105955775 | 105956310 | 536 | 2 | ENSMUST00 | -15230 | RP24-79F8.2 | ENSMUSG00000099448.1 |
| chr6 | 17280878 | 17282201 | 18.66691 | 48.38385 | 45.68857 | Promoter | 6 | 17281185 | 17289115 | 7931 | 1 | ENSMUST00 | 0 | Cav2 | ENSMUSG00000000058.6 |
| chr6 | 113469297 | 113470104 | 18.66691 | 48.38385 | 45.68857 | Exon (ENSMU | 6 | 113471427 | 113483140 | 11714 | 1 | ENSMUST00 | -1323 | Il17rc | ENSMUSG00000030281.11 |
| chr17 | 79712319 | 79716277 | 16.88344 | 48.35773 | 45.66254 | Promoter | 17 | 79706953 | 79715041 | 8089 | 2 | ENSMUST00 | 0 | Cyp1b1 | ENSMUSG00000024087.3 |
| chr15 | 74487214 | 74488131 | 19.1053 | 48.35543 | 45.66036 | Distal Interge | 15 | 74516195 | 74589464 | 73270 | 1 | ENSMUST00 | -28064 | Bai1 | ENSMUSG00000034730.11 |
| chr13 | 55464059 | 55464918 | 18.19771 | 48.35426 | 45.65929 | Promoter | 13 | 55464267 | 55473155 | 8889 | 1 | ENSMUST00 | 0 | Prr7 | ENSMUSG00000034686.7 |
| chr2 | 84871851 | 84872692 | 18.19771 | 48.35426 | 45.65929 | 3' UTR | 2 | 84880523 | 84886476 | 5954 | 2 | ENSMUST00 | 13784 | Rtn4rl2 | ENSMUSG00000050896.7 |
| chr7 | 24207504 | 24208541 | 18.19771 | 48.35426 | 45.65929 | Promoter | 7 | 24193947 | 24208149 | 14203 | 2 | ENSMUST00 | 0 | Zfp111 | ENSMUSG00000087598.4 |
| chr9 | 67830708 | 67831879 | 18.19771 | 48.35426 | 45.65929 | Promoter | 9 | 67830532 | 67832330 | 1799 | 2 | ENSMUST00 | 451 | C2cd4a | ENSMUSG00000047990.5 |
| chr15 | 33082802 | 33084204 | 14.87918 | 48.32321 | 45.62892 | Promoter | 15 | 33083129 | 33594552 | 511424 | 1 | ENSMUST00 | 0 | Cpq | ENSMUSG00000039007.9 |
| chr16 | 22844179 | 22845152 | 14.87918 | 48.32321 | 45.62892 | Intron (ENSM | 16 | 22813215 | 22857569 | 44355 | 2 | ENSMUST00 | 12417 | Tbccd1 | ENSMUSG00000004462.6 |
| chr6 | 88465098 | 88466262 | 14.87918 | 48.32321 | 45.62892 | Promoter | 6 | 88465409 | 88497572 | 32164 | 1 | ENSMUST00 | 0 | Ruvbl1 | ENSMUSG00000030079.10 |
| chr7 | 81493138 | 81494081 | 14.87918 | 48.32321 | 45.62892 | Promoter | 7 | 81460401 | 81493925 | 33525 | 2 | ENSMUST00 | 0 | Ap3b2 | ENSMUSG00000062444.9 |
| chr4 | 118436854 | 118437912 | 18.62992 | 48.32038 | 45.6266 | Promoter | 4 | 118432901 | 118437352 | 4452 | 2 | ENSMUST00 | 0 | Cdc20 | ENSMUSG00000006398.10 |
| chr1 | 133033393 | 133034371 | 19.06563 | 48.28789 | 45.59428 | Distal Interge | 1 | 133009151 | 133030561 | 21411 | 2 | ENSMUST00 | -2833 | Mdm4 | ENSMUSG00000054387.8 |
| chr12 | 85287825 | 85289333 | 19.06563 | 48.28789 | 45.59428 | Promoter | 12 | 85272547 | 85288438 | 15892 | 2 | ENSMUST00 | 0 | Acyp1 | ENSMUSG00000008822.10 |
| chr1 | 62706055 | 62707835 | 17.27239 | 48.28369 | 45.59018 | Intron (ENSM | 1 | 62703702 | 62818695 | 114994 | 1 | ENSMUST00 | 2354 | Nrp2 | ENSMUSG00000025969.10 |
| chr11 | 77981332 | 77982525 | 17.27239 | 48.28369 | 45.59018 | Promoter | 11 | 77982754 | 78026214 | 43461 | 1 | ENSMUST00 | -229 | Phf12 | ENSMUSG00000037791.10 |
| chr12 | 67221348 | 67222975 | 17.27239 | 48.28369 | 45.59018 | Promoter | 12 | 66469568 | 67222549 | 752982 | 2 | ENSMUST00 | 0 | Mdga2 | ENSMUSG00000034912.11 |
| chr12 | 78802591 | 78805573 | 17.27239 | 48.28369 | 45.59018 | Intron (ENSM | 12 | 78752687 | 78752788 | 102 | 2 | ENSMUST00 | -49804 | Gm25225 | ENSMUSG00000088822.1 |
| chr14 | 104467129 | 104468396 | 17.27239 | 48.28369 | 45.59018 | Promoter | 14 | 104461676 | 104467999 | 6324 | 2 | ENSMUST00 | 0 | Pou4f1 | ENSMUSG00000048349.9 |
| chr14 | 67188201 | 67189254 | 17.27239 | 48.28369 | 45.59018 | Distal Interge | 14 | 67232437 | 67233129 | 693 | 2 | ENSMUST00 | 43875 | Gm20675 | ENSMUSG00000093744.1 |
| chr16 | 42189822 | 42190876 | 17.27239 | 48.28369 | 45.59018 | Distal Interge | 16 | 42144025 | 42174283 | 30259 | 1 | ENSMUST00 | 45798 | Lsamp | ENSMUSG00000061080.6 |
| chr16 | 21246024 | 21246848 | 17.27239 | 48.28369 | 45.59018 | Downstream | 16 | 21256013 | 21256066 | 54 | 2 | ENSMUST00 | 9218 | Mir7680 | ENSMUSG00000098400.1 |
| chr17 | 35234794 | 35236541 | 17.27239 | 48.28369 | 45.59018 | Promoter | 17 | 35235088 | 35237805 | 2718 | 1 | ENSMUST00 | 0 | Atp6v1g2 | ENSMUSG00000024403.11 |
| chr18 | 67343268 | 67344146 | 17.27239 | 48.28369 | 45.59018 | Promoter | 18 | 67343564 | 67367794 | 24231 | 1 | ENSMUST00 | 0 | Cidea | ENSMUSG00000024526.8 |
| chr4 | 115062666 | 115063696 | 17.27239 | 48.28369 | 45.59018 | 5' UTR | 4 | 115059508 | 115070469 | 10962 | 1 | ENSMUST00 | 3159 | Tal1 | ENSMUSG00000028717.7 |
| chr5 | 24348080 | 24348971 | 17.27239 | 48.28369 | 45.59018 | Exon (ENSMU | 5 | 24319589 | 24351604 | 32016 | 2 | ENSMUST00 | 2633 | Kcnh2 | ENSMUSG00000038319.9 |
| chr5 | 114822963 | 114823742 | 17.27239 | 48.28369 | 45.59018 | Promoter | 5 | 114809036 | 114823460 | 14425 | 2 | ENSMUST00 | 0 | Gm20499 | ENSMUSG00000092252.1 |
| chr6 | 21985113 | 21987485 | 17.27239 | 48.28369 | 45.59018 | Promoter | 6 | 21985916 | 22240003 | 254088 | 1 | ENSMUST00 | 0 | Cped1 | ENSMUSG00000062980.10 |
| chr7 | 126808782 | 126809590 | 17.27239 | 48.28369 | 45.59018 | Promoter | 7 | 126800790 | 126809108 | 8319 | 2 | ENSMUST00 | 0 | Gm15676 | ENSMUSG00000085583.1 |
| chr8 | 25256163 | 25256631 | 17.27239 | 48.28369 | 45.59018 | Distal Interge | 8 | 25154552 | 25201449 | 46898 | 2 | ENSMUST00 | -54715 | Tacc1 | ENSMUSG00000065954.5 |
| chr9 | 44287470 | 44288577 | 17.27239 | 48.28369 | 45.59018 | Promoter | 9 | 44273188 | 44287544 | 14357 | 2 | ENSMUST00 | 0 | Abcg4 | ENSMUSG00000032131.10 |
| chr9 | 20878312 | 20878946 | 17.27239 | 48.28369 | 45.59018 | Promoter | 9 | 20873809 | 20879727 | 5919 | 2 | ENSMUST00 | 781 | Angptl6 | ENSMUSG00000038742.6 |
| chr16 | 11143973 | 11144240 | 3.97618 | 48.25122 | 45.56001 | Promoter | 16 | 11144125 | 11144181 | 57 | 1 | ENSMUST00 | 0 | Gm23935 | ENSMUSG00000076258.1 |
| chr11 | 103809516 | 103810661 | 15.53947 | 48.13258 | 45.44172 | Intron (ENSM | 11 | 103822934 | 103824100 | 1167 | 2 | ENSMUST00 | 13439 | Nsf | ENSMUSG00000034187.13 |
| chr13 | 41219968 | 41220867 | 15.53947 | 48.13258 | 45.44172 | Promoter | 13 | 41182495 | 41220405 | 37911 | 2 | ENSMUST00 | 0 | Elovl2 | ENSMUSG00000021364.10 |
| chr13 | 75839868 | 75840968 | 15.53947 | 48.13258 | 45.44172 | Promoter | 13 | 75839868 | 75850151 | 10284 | 1 | ENSMUST00 | 1 | Glrx | ENSMUSG00000021591.7 |
| chr17 | 29093483 | 29095648 | 15.53947 | 48.13258 | 45.44172 | Promoter | 17 | 29093772 | 29100722 | 6951 | 1 | ENSMUST00 | 0 | Cdkn1a | ENSMUSG00000023067.8 |
| chr17 | 21650177 | 21651177 | 15.53947 | 48.13258 | 45.44172 | Distal Interge | 17 | 21657582 | 21662540 | 4959 | 1 | ENSMUST00 | -6405 | 3110052M02 | ENSMUSG00000035868.8 |
| chr17 | 27027519 | 27028691 | 15.53947 | 48.13258 | 45.44172 | Promoter | 17 | 27019812 | 27028578 | 8767 | 2 | ENSMUST00 | 0 | Bak1 | ENSMUSG00000057789.8 |
| chr19 | 43440264 | 43441765 | 15.53947 | 48.13258 | 45.44172 | Promoter | 19 | 43440436 | 43497212 | 56777 | 1 | ENSMUST00 | 0 | Cnnm1 | ENSMUSG00000025189.8 |
| chr5 | 32745569 | 32747337 | 15.53947 | 48.13258 | 45.44172 | Promoter | 5 | 32736424 | 32746312 | 9889 | 2 | ENSMUST00 | 0 | Pisd | ENSMUSG00000023452.13 |
| chr6 | 136518390 | 136520007 | 15.53947 | 48.13258 | 45.44172 | Promoter | 6 | 136518802 | 136611038 | 92237 | 1 | ENSMUST00 | 0 | Atf7ip | ENSMUSG00000030213.7 |
| chr6 | 67266025 | 67266877 | 15.53947 | 48.13258 | 45.44172 | Promoter | 6 | 67266979 | 67289302 | 22324 | 1 | ENSMUST00 | -102 | Serbp1 | ENSMUSG00000036371.4 |
| chr9 | 98031988 | 98033615 | 15.53947 | 48.13258 | 45.44172 | Promoter | 9 | 97444395 | 98033181 | 588787 | 2 | ENSMUST00 | 0 | Clstn2 | ENSMUSG00000032452.7 |
| chr9 | 108795574 | 108798004 | 15.53947 | 48.13258 | 45.44172 | Promoter | 9 | 108795994 | 108806333 | 10340 | 1 | ENSMUST00 | 0 | Ip6k2 | ENSMUSG00000032599.7 |
| chr15 | 91672736 | 91674386 | 17.61183 | 48.10911 | 45.41927 | Promoter | 15 | 91673175 | 91816120 | 142946 | 1 | ENSMUST00 | 0 | Lrrk2 | ENSMUSG00000036273.10 |
| chr1 | 156420479 | 156421401 | 16.32424 | 48.10018 | 45.4107 | Promoter | 1 | 156323509 | 156421159 | 97651 | 2 | ENSMUST00 | 0 | Axdnd1 | ENSMUSG00000026601.8 |
| chr1 | 74854639 | 74856159 | 16.32424 | 48.10018 | 45.4107 | Promoter | 1 | 74854934 | 74857732 | 2799 | 1 | ENSMUST00 | 0 | Cdk5r2 | ENSMUSG00000090071.4 |
| chr10 | 63099833 | 63100820 | 16.32424 | 48.10018 | 45.4107 | Promoter | 10 | 63100156 | 63100605 | 450 | 1 | ENSMUST00 | 0 | Atoh7 | ENSMUSG00000036816.3 |
| chr11 | 69758740 | 69759758 | 16.32424 | 48.10018 | 45.4107 | Promoter | 11 | 69733997 | 69758637 | 24641 | 2 | ENSMUST00 | -104 | Polr2a | ENSMUSG00000005198.10 |
| chr12 | 80168115 | 80168828 | 16.32424 | 48.10018 | 45.4107 | 3' UTR | 12 | 80120548 | 80132844 | 12297 | 2 | ENSMUST00 | -35272 | 2310015A10I | ENSMUSG00000097729.1 |
| chr3 | 95356009 | 95357528 | 16.32424 | 48.10018 | 45.4107 | Promoter | 3 | 95323525 | 95357155 | 33631 | 2 | ENSMUST00 | 0 | Setdb1 | ENSMUSG00000015697.9 |
| chr5 | 69340534 | 69342037 | 16.32424 | 48.10018 | 45.4107 | Promoter | 5 | 69109285 | 69341709 | 232425 | 2 | ENSMUST00 | 0 | Kctd8 | ENSMUSG00000037653.6 |
| chr7 | 139949976 | 139951693 | 16.32424 | 48.10018 | 45.4107 | Promoter | 7 | 139950396 | 139963327 | 12932 | 1 | ENSMUST00 | 0 | Gm16201 | ENSMUSG00000086742.2 |
| chr9 | 108831838 | 108833065 | 16.32424 | 48.10018 | 45.4107 | Exon (ENSMU | 9 | 108826320 | 108852963 | 26644 | 1 | ENSMUST00 | 5519 | Celsr3 | ENSMUSG00000023473.6 |
| chr9 | 14275598 | 14275998 | 18.94761 | 48.08681 | 45.3989 | Promoter | 9 | 14276301 | 14326138 | 49838 | 1 | ENSMUST00 | -303 | Sesn3 | ENSMUSG00000032009.7 |
| chr11 | 53478643 | 53480402 | 18.48341 | 48.06877 | 45.38095 | Promoter | 11 | 53476573 | 53480274 | 3702 | 2 | ENSMUST00 | 0 | Sowaha | ENSMUSG00000044352.6 |
| chr18 | 36725264 | 36726884 | 18.48341 | 48.06877 | 45.38095 | Promoter | 18 | 36725074 | 36726736 | 1663 | 2 | ENSMUST00 | 0 | Cd14 | ENSMUSG00000051439.6 |
| chr4 | 116707754 | 116708410 | 18.48341 | 48.06877 | 45.38095 | Promoter | 4 | 116702279 | 116708406 | 6128 | 2 | ENSMUST00 | 0 | Mmachc | ENSMUSG00000028690.4 |
| chr19 | 47313175 | 47315281 | 18.90859 | 48.02031 | 45.33304 | Exon (ENSMU | 19 | 47273519 | 47273589 | 71 | 2 | ENSMUST00 | -39587 | Mir6995 | ENSMUSG00000098786.1 |
| chr6 | 49822027 | 49823019 | 18.90859 | 48.02031 | 45.33304 | Promoter | 6 | 49822710 | 49829507 | 6798 | 1 | ENSMUST00 | 0 | Npy | ENSMUSG00000029819.6 |
| chr8 | 70172359 | 70172788 | 18.90859 | 48.02031 | 45.33304 | Promoter | 8 | 70172405 | 70183667 | 11263 | 1 | ENSMUST00 | 0 | Tmem161a | ENSMUSG00000002342.12 |
| chr11 | 66167781 | 66168605 | 18.86974 | 47.95406 | 45.26699 | Promoter | 11 | 65831282 | 66168533 | 337252 | 2 | ENSMUST00 | 0 | Dnah9 | ENSMUSG00000056752.11 |
| chr7 | 63444294 | 63445444 | 18.86974 | 47.95406 | 45.26699 | Promoter | 7 | 63444751 | 63759028 | 314278 | 1 | ENSMUST00 | 0 | Otud7a | ENSMUSG00000033510.9 |
| chr14 | 20180394 | 20181858 | 17.08566 | 47.94947 | 45.2625 | Promoter | 14 | 20140058 | 20181782 | 41725 | 2 | ENSMUST00 | 0 | Kcnk5 | ENSMUSG00000023243.8 |
| chr18 | 90579141 | 90580293 | 17.08566 | 47.94947 | 45.2625 | Distal Interge | 18 | 90510154 | 90543267 | 33114 | 1 | ENSMUST00 | 68988 | Tmx3 | ENSMUSG00000024614.6 |
| chr13 | 21715096 | 21716609 | 17.95989 | 47.94115 | 45.25451 | Promoter | 13 | 21716422 | 21716814 | 393 | 1 | ENSMUST00 | 0 | Hist1h2ai | ENSMUSG00000071516.2 |
| chr11 | 121434257 | 121435431 | 18.79251 | 47.82232 | 45.13606 | Promoter | 11 | 121434913 | 121450491 | 15579 | 1 | ENSMUST00 | 0 | Fn3k | ENSMUSG00000025175.7 |
| chr5 | 118380404 | 118381632 | 18.79251 | 47.82232 | 45.13606 | Distal Interge | 5 | 118418727 | 118418818 | 92 | 2 | ENSMUST00 | 37186 | Gm26455 | ENSMUSG00000093041.1 |
| chr6 | 82772928 | 82774934 | 18.79251 | 47.82232 | 45.13606 | Promoter | 6 | 82725025 | 82774454 | 49430 | 2 | ENSMUST00 | 0 | Hk2 | ENSMUSG00000000628.5 |
| chr7 | 143685214 | 143685949 | 18.79251 | 47.82232 | 45.13606 | Promoter | 7 | 143665809 | 143685872 | 20064 | 2 | ENSMUST00 | 0 | Tnfrsf23 | ENSMUSG00000037613.10 |
| chr7 | 24904348 | 24904948 | 18.79251 | 47.82232 | 45.13606 | Promoter | 7 | 24904358 | 24926589 | 22232 | 1 | ENSMUST00 | 0 | Arhgef1 | ENSMUSG00000040940.13 |
| chr8 | 125053893 | 125054769 | 18.79251 | 47.82232 | 45.13606 | Promoter | 8 | 125054195 | 125261151 | 206957 | 1 | ENSMUST00 | 0 | Disc1 | ENSMUSG00000043051.12 |
| chr11 | 96076096 | 96076468 | 17.85986 | 47.76698 | 45.0813 | Promoter | 11 | 96068852 | 96075670 | 6819 | 2 | ENSMUST00 | -427 | Atp5g1 | ENSMUSG00000006057.10 |

| | | | | | | | | | | | | | | | |
|---|---|---|---|---|---|---|---|---|---|---|---|---|---|---|---|
| chr12 | 57196551 | 57197296 | 17.85986 | 47.76698 | 45.0813 | Promoter | 12 | 56712634 | 57197472 | 484839 | 2 | ENSMUST00 | 176 | Slc25a21 | ENSMUSG00000035472.8 |
| chr17 | 80553359 | 80554123 | 17.85986 | 47.76698 | 45.0813 | Intron (ENSM | 17 | 80523550 | 80563834 | 40285 | 2 | ENSMUST00 | 9711 | Cdkl4 | ENSMUSG00000033966.7 |
| chr5 | 146385010 | 146385701 | 17.85986 | 47.76698 | 45.0813 | Promoter | 5 | 146384985 | 146472411 | 87427 | 1 | ENSMUST00 | 26 | Wasf3 | ENSMUSG00000029636.8 |
| chr6 | 114397453 | 114398494 | 17.85986 | 47.76698 | 45.0813 | Promoter | 6 | 114397936 | 114483296 | 85361 | 1 | ENSMUST00 | 0 | Hrh1 | ENSMUSG00000053004.8 |
| chr7 | 126780668 | 126781314 | 17.85986 | 47.76698 | 45.0813 | Promoter | 7 | 126781483 | 126785359 | 3877 | 1 | ENSMUST00 | -169 | Tbx6 | ENSMUSG00000030699.11 |
| chr7 | 104314494 | 104315994 | 17.85986 | 47.76698 | 45.0813 | Promoter | 7 | 104299897 | 104315455 | 15559 | 2 | ENSMUST00 | 0 | Trim12a | ENSMUSG00000066258.8 |
| chr14 | 34661195 | 34662596 | 18.7159 | 47.69159 | 45.00676 | Distal Interge | 14 | 34673928 | 34747983 | 74056 | 1 | ENSMUST00 | -11332 | Wapal | ENSMUSG00000041408.11 |
| chr1 | 64737999 | 64739348 | 18.1972 | 47.5761 | 44.89159 | Promoter | 1 | 64730558 | 64737751 | 7194 | 2 | ENSMUST00 | -249 | Fzd5 | ENSMUSG00000045005.9 |
| chr10 | 123264442 | 123265808 | 18.63992 | 47.56184 | 44.87756 | Promoter | 10 | 123264076 | 123741204 | 477129 | 1 | ENSMUST00 | 367 | Fam19a2 | ENSMUSG00000044071.7 |
| chr13 | 22040633 | 22041421 | 17.3021 | 47.56202 | 44.87756 | Promoter | 13 | 22040636 | 22041362 | 727 | 2 | ENSMUST00 | 0 | Hist1h4i | ENSMUSG00000060639.5 |
| chr18 | 46573984 | 46574695 | 17.3021 | 47.56202 | 44.87756 | Promoter | 18 | 46559155 | 46574533 | 15379 | 2 | ENSMUST00 | 0 | Ticam2 | ENSMUSG00000056130.4 |
| chr8 | 4779168 | 4779598 | 17.3021 | 47.56202 | 44.87756 | Promoter | 8 | 4735980 | 4779534 | 43555 | 2 | ENSMUST00 | 0 | Shcbp1 | ENSMUSG00000022322.7 |
| chr4 | 151044842 | 151045319 | 18.16205 | 47.51548 | 44.83143 | Promoter | 4 | 151024190 | 151044665 | 20476 | 2 | ENSMUST00 | -178 | Per3 | ENSMUSG00000028957.7 |
| chr5 | 36464270 | 36464888 | 18.16205 | 47.51548 | 44.83143 | Promoter | 5 | 36447611 | 36464821 | 17211 | 2 | ENSMUST00 | 0 | 2210406O10 | ENSMUSG00000039563.6 |
| chr9 | 64084430 | 64085342 | 18.16205 | 47.51548 | 44.83143 | Intron (ENSM | 9 | 64080761 | 64086961 | 6201 | 1 | ENSMUST00 | 3670 | Scarletltr | ENSMUSG00000093759.1 |
| chr1 | 91179575 | 91180319 | 18.60216 | 47.49733 | 44.81332 | Promoter | 1 | 91179822 | 91206790 | 26969 | 1 | ENSMUST00 | 0 | Ramp1 | ENSMUSG00000034353.9 |
| chr17 | 34135566 | 34135930 | 18.12703 | 47.45507 | 44.77117 | Promoter | 17 | 34135182 | 34139101 | 3920 | 1 | ENSMUST00 | 385 | H2-DMa | ENSMUSG00000037649.9 |
| chr5 | 76657346 | 76658471 | 18.12703 | 47.45507 | 44.77117 | Promoter | 5 | 76657683 | 76840881 | 183199 | 1 | ENSMUST00 | 0 | C530008M17 | ENSMUSG00000036377.13 |
| chr1 | 132190047 | 132191155 | 16.80025 | 47.43613 | 44.75273 | Promoter | 1 | 132191431 | 132259227 | 67797 | 1 | ENSMUST00 | -276 | Lemd1 | ENSMUSG00000079330.3 |
| chr19 | 5487532 | 5489063 | 16.80025 | 47.43613 | 44.75273 | Promoter | 19 | 5482345 | 5488016 | 5672 | 2 | ENSMUST00 | 0 | Mus81 | ENSMUSG00000024906.4 |
| chr7 | 28825273 | 28826429 | 16.80025 | 47.43613 | 44.75273 | Promoter | 7 | 28825349 | 28826656 | 1308 | 1 | ENSMUST00 | 0 | Ech1 | ENSMUSG00000053898.7 |
| chr7 | 34233690 | 34234987 | 16.80025 | 47.43613 | 44.75273 | Promoter | 7 | 34234377 | 34240207 | 5831 | 1 | ENSMUST00 | 0 | Gm12758 | ENSMUSG00000085105.1 |
| chr10 | 80387745 | 80388393 | 18.56455 | 47.43306 | 44.74982 | Promoter | 10 | 80380355 | 80387659 | 7305 | 2 | ENSMUST00 | -87 | Mex3d | ENSMUSG00000048696.10 |
| chr13 | 54192010 | 54192739 | 18.56455 | 47.43306 | 44.74982 | Promoter | 13 | 54192129 | 54222432 | 30304 | 1 | ENSMUST00 | 0 | Hrh2 | ENSMUSG00000034987.3 |
| chr11 | 120315894 | 120317217 | 16.34629 | 47.3758 | 44.69273 | Distal Interge | 11 | 120289793 | 120292167 | 2375 | 1 | ENSMUST00 | 26102 | Bahcc1 | ENSMUSG00000039741.10 |
| chr15 | 89475744 | 89476685 | 14.05084 | 47.34375 | 44.66087 | Promoter | 15 | 89472528 | 89476113 | 3586 | 2 | ENSMUST00 | 0 | Arsa | ENSMUSG00000022620.9 |
| chr18 | 77046531 | 77047777 | 14.05084 | 47.34375 | 44.66087 | Promoter | 18 | 76977148 | 77047308 | 70161 | 2 | ENSMUST00 | 0 | Katnal2 | ENSMUSG00000025420.8 |
| chr1 | 165844405 | 165845742 | 18.02173 | 47.27326 | 44.5908 | Intron (ENSM | 1 | 165858216 | 165861555 | 3340 | 1 | ENSMUST00 | -12474 | Cd247 | ENSMUSG00000005763.10 |
| chr1 | 133417877 | 133418807 | 18.02173 | 47.27326 | 44.5908 | Intron (ENSM | 1 | 133382303 | 133424377 | 42075 | 2 | ENSMUST00 | 5570 | Sox13 | ENSMUSG00000070643.6 |
| chr11 | 109425224 | 109425788 | 18.02173 | 47.27326 | 44.5908 | Promoter | 11 | 109425946 | 109438148 | 12203 | 1 | ENSMUST00 | -158 | Amz2 | ENSMUSG00000020610.11 |
| chr12 | 76324449 | 76325281 | 18.02173 | 47.27326 | 44.5908 | Promoter | 12 | 76324891 | 76334151 | 9261 | 1 | ENSMUST00 | 0 | Akap5 | ENSMUSG00000021057.10 |
| chr13 | 55181899 | 55182864 | 18.02173 | 47.27326 | 44.5908 | Distal Interge | 13 | 55156117 | 55156904 | 788 | 1 | ENSMUST00 | 25783 | Fgfr4 | ENSMUSG00000005320.9 |
| chr13 | 59557574 | 59557965 | 18.02173 | 47.27326 | 44.5908 | Promoter | 13 | 59531182 | 59557347 | 26166 | 2 | ENSMUST00 | -228 | Agtpbp1 | ENSMUSG00000021557.9 |
| chr13 | 83730584 | 83731490 | 18.02173 | 47.27326 | 44.5908 | Intron (ENSM | 13 | 83723442 | 83729582 | 6141 | 2 | ENSMUST00 | -1003 | Gm26803 | ENSMUSG00000097265.1 |
| chr14 | 18892782 | 18894436 | 18.02173 | 47.27326 | 44.5908 | Promoter | 14 | 18573577 | 18893776 | 320200 | 2 | ENSMUST00 | 0 | Ube2e2 | ENSMUSG00000058317.6 |
| chr18 | 23414765 | 23415951 | 18.02173 | 47.27326 | 44.5908 | Promoter | 18 | 23415415 | 23604230 | 188816 | 1 | ENSMUST00 | 0 | Dtna | ENSMUSG00000024302.10 |
| chr19 | 10868950 | 10870186 | 18.02173 | 47.27326 | 44.5908 | Promoter | 19 | 10857822 | 10869940 | 12119 | 2 | ENSMUST00 | 0 | Tmem132a | ENSMUSG00000024736.9 |
| chr19 | 23892822 | 23893629 | 18.02173 | 47.27326 | 44.5908 | Exon (ENSMU | 19 | 23758876 | 23949597 | 190722 | 1 | ENSMUST00 | 133947 | Apba1 | ENSMUSG00000024897.8 |
| chr2 | 156392750 | 156393038 | 18.02173 | 47.27326 | 44.5908 | Distal Interge | 2 | 156386741 | 156387222 | 482 | 1 | ENSMUST00 | 6010 | Rpl21-ps9 | ENSMUSG00000084017.1 |
| chr4 | 55349442 | 55349872 | 18.02173 | 47.27326 | 44.5908 | Promoter | 4 | 55350043 | 55392237 | 42195 | 1 | ENSMUST00 | -171 | Rad23b | ENSMUSG00000028426.10 |
| chr4 | 103563401 | 103564179 | 18.02173 | 47.27326 | 44.5908 | Promoter | 4 | 103563402 | 103564529 | 1128 | 1 | ENSMUST00 | 0 | Gm12715 | ENSMUSG00000059195.10 |
| chr6 | 88898170 | 88898753 | 18.02173 | 47.27326 | 44.5908 | Promoter | 6 | 88883475 | 88898780 | 15306 | 2 | ENSMUST00 | 27 | Mcm2 | ENSMUSG00000002870.7 |
| chr6 | 21948939 | 21949473 | 18.02173 | 47.27326 | 44.5908 | Promoter | 6 | 21949571 | 21976038 | 26468 | 1 | ENSMUST00 | -98 | Ing3 | ENSMUSG00000029670.7 |
| chr6 | 39381020 | 39381852 | 18.02173 | 47.27326 | 44.5908 | Promoter | 6 | 39381175 | 39390380 | 9206 | 1 | ENSMUST00 | 0 | Rab19 | ENSMUSG00000029923.4 |
| chr6 | 64728558 | 64730281 | 18.02173 | 47.27326 | 44.5908 | Promoter | 6 | 64729146 | 64731233 | 2088 | 1 | ENSMUST00 | 0 | Atoh1 | ENSMUSG00000073043.4 |
| chr6 | 53286463 | 53288445 | 18.02173 | 47.27326 | 44.5908 | Distal Interge | 6 | 53433974 | 53434137 | 164 | 1 | ENSMUST00 | -145529 | Gm25818 | ENSMUSG00000096147.1 |
| chr7 | 4163577 | 4164847 | 18.02173 | 47.27326 | 44.5908 | Promoter | 7 | 4151551 | 4164539 | 12989 | 2 | ENSMUST00 | 0 | Cdc42ep5 | ENSMUSG00000063838.6 |
| chr9 | 111311397 | 111313081 | 18.02173 | 47.27326 | 44.5908 | Promoter | 9 | 111311739 | 111395774 | 84036 | 1 | ENSMUST00 | 0 | Trank1 | ENSMUSG00000062296.4 |
| chrX | 20662871 | 20664098 | 18.02173 | 47.27326 | 44.5908 | Promoter | 20 | 20662898 | 20683179 | 20282 | 1 | ENSMUST00 | 0 | Uba1 | ENSMUSG00000001924.10 |
| chrX | 48593603 | 48594826 | 18.02173 | 47.27326 | 44.5908 | Promoter | 20 | 48541625 | 48594373 | 52749 | 2 | ENSMUST00 | 0 | Zfp280c | ENSMUSG00000036916.8 |
| chr11 | 51615182 | 51616076 | 17.53433 | 47.19838 | 44.51795 | Distal Interge | 11 | 51619735 | 51623714 | 3980 | 1 | ENSMUST00 | -3659 | Nhp2 | ENSMUSG00000001056.3 |
| chr19 | 33589465 | 33590527 | 17.53433 | 47.19838 | 44.51795 | Promoter | 19 | 33517740 | 33590154 | 72415 | 2 | ENSMUST00 | 0 | Lipo1 | ENSMUSG00000024766.9 |
| chr10 | 56106696 | 56107390 | 18.41563 | 47.17836 | 44.49838 | Promoter | 10 | 56106917 | 56116880 | 9964 | 1 | ENSMUST00 | 0 | Msl3l2 | ENSMUSG00000047669.7 |
| chr11 | 68939432 | 68940863 | 18.41563 | 47.17836 | 44.49838 | Downstream | 11 | 68944527 | 68947380 | 2854 | 2 | ENSMUST00 | 6517 | Arhgef15 | ENSMUSG00000052921.8 |
| chr18 | 76378707 | 76379499 | 18.41563 | 47.17836 | 44.49838 | Distal Interge | 18 | 76262492 | 76302642 | 40151 | 1 | ENSMUST00 | 116216 | Smad2 | ENSMUSG00000024563.10 |
| chr4 | 109656755 | 109658163 | 18.41563 | 47.17836 | 44.49838 | Promoter | 4 | 109651529 | 109657827 | 6299 | 2 | ENSMUST00 | 0 | 9630013D21 | ENSMUSG00000059027.9 |
| chr5 | 98253723 | 98254866 | 18.41563 | 47.17836 | 44.49838 | Promoter | 5 | 98254184 | 98277030 | 22847 | 1 | ENSMUST00 | 0 | Fgf5 | ENSMUSG00000029337.1 |
| chr8 | 123518425 | 123519534 | 18.41563 | 47.17836 | 44.49838 | Promoter | 8 | 123518835 | 123536649 | 17815 | 1 | ENSMUST00 | 0 | Gas8 | ENSMUSG00000040220.9 |
| chr1 | 78203332 | 78204277 | 18.84082 | 47.13427 | 44.45482 | Distal Interge | 1 | 78101270 | 78197134 | 95865 | 2 | ENSMUST00 | -6199 | Pax3 | ENSMUSG00000004872.10 |
| chr10 | 79780286 | 79781351 | 18.84082 | 47.13427 | 44.45482 | Promoter | 10 | 79780581 | 79782607 | 2027 | 1 | ENSMUST00 | 0 | Fstl3 | ENSMUSG00000020325.10 |
| chr14 | 26066783 | 26067511 | 18.84082 | 47.13427 | 44.45482 | Promoter | 14 | 26063919 | 26067355 | 3437 | 2 | ENSMUST00 | 0 | Tmem254c | ENSMUSG00000072680.6 |
| chr15 | 28158610 | 28159545 | 18.84082 | 47.13427 | 44.45482 | Distal Interge | 15 | 28203752 | 28472045 | 268294 | 1 | ENSMUST00 | -44207 | Dnah5 | ENSMUSG00000022262.7 |
| chr17 | 42891932 | 42892257 | 18.84082 | 47.13427 | 44.45482 | Distal Interge | 17 | 42792951 | 42876424 | 83474 | 2 | ENSMUST00 | -15509 | Cd2ap | ENSMUSG00000061665.6 |
| chr19 | 53325354 | 53326134 | 18.84082 | 47.13427 | 44.45482 | Intron (ENSM | 19 | 53329413 | 53375810 | 46398 | 1 | ENSMUST00 | -3279 | Mxi1 | ENSMUSG00000025025.8 |
| chr2 | 13792857 | 13794193 | 18.84082 | 47.13427 | 44.45482 | Promoter | 2 | 13651020 | 13793808 | 142789 | 2 | ENSMUST00 | 0 | St8sia6 | ENSMUSG00000003418.10 |
| chr2 | 92007631 | 92008416 | 18.84082 | 47.13427 | 44.45482 | Intron (ENSM | 2 | 91982328 | 92024502 | 42175 | 2 | ENSMUST00 | 16086 | Creb3l1 | ENSMUSG00000027230.9 |
| chr3 | 94372718 | 94373954 | 18.84082 | 47.13427 | 44.45482 | Promoter | 3 | 94372794 | 94397840 | 25047 | 1 | ENSMUST00 | 0 | Rorc | ENSMUSG00000028150.9 |
| chr3 | 127257496 | 127258209 | 18.84082 | 47.13427 | 44.45482 | Intron (ENSM | 3 | 126923957 | 127225932 | 301976 | 2 | ENSMUST00 | -31565 | Ank2 | ENSMUSG00000032826.11 |
| chr4 | 134551500 | 134552505 | 18.84082 | 47.13427 | 44.45482 | Promoter | 4 | 134537892 | 134552166 | 14275 | 2 | ENSMUST00 | 0 | Sepn1 | ENSMUSG00000050989.9 |
| chr5 | 137338125 | 137339159 | 18.84082 | 47.13427 | 44.45482 | Distal Interge | 5 | 137329768 | 137333597 | 3830 | 2 | ENSMUST00 | -4529 | Slc12a9 | ENSMUSG00000037344.8 |
| chr5 | 24351205 | 24352105 | 18.84082 | 47.13427 | 44.45482 | Promoter | 5 | 24319589 | 24351604 | 32016 | 2 | ENSMUST00 | 0 | Kcnh2 | ENSMUSG00000038319.9 |
| chr7 | 96212520 | 96213657 | 18.84082 | 47.13427 | 44.45482 | Promoter | 7 | 96211671 | 96911093 | 699423 | 1 | ENSMUST00 | 850 | Tenm4 | ENSMUSG00000048078.11 |
| chrX | 10717005 | 10719476 | 18.84082 | 47.13427 | 44.45482 | Promoter | 20 | 10717390 | 10719690 | 2301 | 1 | ENSMUST00 | 0 | Mid1ip1 | ENSMUSG00000008035.7 |
| chrX | 74281767 | 74283502 | 18.84082 | 47.13427 | 44.45482 | Promoter | 20 | 74281912 | 74282865 | 954 | 1 | ENSMUST00 | 0 | Taz | ENSMUSG00000009995.12 |
| chrX | 36327630 | 36330397 | 18.84082 | 47.13427 | 44.45482 | Promoter | 20 | 36328353 | 36342611 | 14259 | 1 | ENSMUST00 | 0 | Lonrf3 | ENSMUSG00000016239.6 |
| chrX | 37190506 | 37192515 | 18.84082 | 47.13427 | 44.45482 | Promoter | 20 | 37187588 | 37191163 | 3576 | 2 | ENSMUST00 | 0 | Ndufa1 | ENSMUSG00000016427.7 |
| chrX | 155215198 | 155216468 | 18.84082 | 47.13427 | 44.45482 | Promoter | 20 | 155213132 | 155216449 | 3318 | 2 | ENSMUST00 | 0 | Sat1 | ENSMUSG00000025283.10 |
| chrX | 167381725 | 167383017 | 18.84082 | 47.13427 | 44.45482 | Promoter | 20 | 167346322 | 167382749 | 36428 | 2 | ENSMUST00 | 0 | Prps2 | ENSMUSG00000025742.5 |
| chrX | 42149209 | 42150570 | 18.84082 | 47.13427 | 44.45482 | Promoter | 20 | 42149317 | 42277185 | 127869 | 1 | ENSMUST00 | 0 | Stag2 | ENSMUSG00000025862.9 |
| chrX | 13845424 | 13847667 | 18.84082 | 47.13427 | 44.45482 | Promoter | 20 | 13517080 | 13846541 | 329462 | 2 | ENSMUST00 | 0 | Cask | ENSMUSG00000031012.12 |
| chrX | 48694475 | 48695457 | 18.84082 | 47.13427 | 44.45482 | Promoter | 20 | 48695004 | 48710723 | 15720 | 1 | ENSMUST00 | 0 | Rbmx2 | ENSMUSG00000031107.6 |
| chrX | 10215717 | 10216848 | 18.84082 | 47.13427 | 44.45482 | Promoter | 20 | 10158216 | 10216453 | 58238 | 2 | ENSMUST00 | 0 | Rpgr | ENSMUSG00000031174.12 |
| chrX | 134058855 | 134059973 | 18.84082 | 47.13427 | 44.45482 | Promoter | 20 | 134059187 | 134086046 | 26860 | 1 | ENSMUST00 | 0 | Cstf2 | ENSMUSG00000031256.6 |
| chrX | 159255753 | 159257406 | 18.84082 | 47.13427 | 44.45482 | Promoter | 20 | 159255782 | 159311525 | 55744 | 1 | ENSMUST00 | 0 | Rps6ka3 | ENSMUSG00000031309.10 |
| chrX | 133686703 | 133689068 | 18.84082 | 47.13427 | 44.45482 | Promoter | 20 | 133582863 | 133688993 | 106131 | 2 | ENSMUST00 | 0 | Pcdh19 | ENSMUSG00000051323.11 |
| chrX | 135008769 | 135009718 | 18.84082 | 47.13427 | 44.45482 | Promoter | 20 | 134971372 | 135009185 | 37814 | 2 | ENSMUST00 | 0 | Zmat1 | ENSMUSG00000052676.11 |
| chrX | 99974986 | 99976711 | 18.84082 | 47.13427 | 44.45482 | Promoter | 20 | 99975606 | 100311299 | 335694 | 1 | ENSMUST00 | 0 | Eda | ENSMUSG00000059327.4 |
| chr15 | 8279160 | 8280317 | 18.37877 | 47.11527 | 44.43725 | Distal Interge | 15 | 8247103 | 8252976 | 5874 | 1 | ENSMUST00 | 32058 | 2410089E03I | ENSMUSG00000039801.6 |
| chr6 | 120038862 | 120040121 | 12.94372 | 47.03971 | 44.36203 | Promoter | 6 | 119923969 | 120038672 | 114704 | 2 | ENSMUST00 | -191 | Wnk1 | ENSMUSG00000045962.11 |
| chr7 | 29818873 | 29819672 | 17.88563 | 47.03791 | 44.36029 | Intron (ENSM | 7 | 29816072 | 29832042 | 15971 | 1 | ENSMUST00 | 2802 | Zfp790 | ENSMUSG00000011427.10 |
| chr15 | 101046148 | 101047343 | 17.00309 | 47.03089 | 44.35329 | Downstream | 15 | 101050194 | 101054399 | 4206 | 2 | ENSMUST00 | 7056 | Fignl2 | ENSMUSG00000095440.1 |
| chr19 | 6067004 | 6068333 | 17.00309 | 47.03089 | 44.35329 | Promoter | 19 | 6062821 | 6067803 | 4983 | 2 | ENSMUST00 | 0 | Tm7sf2 | ENSMUSG00000024799.9 |
| chr19 | 16029003 | 16029584 | 17.00309 | 47.03089 | 44.35329 | Distal Interge | 19 | 15985090 | 16010912 | 25823 | 1 | ENSMUST00 | 43914 | C130060C02I | ENSMUSG00000097557.1 |
| chr7 | 13043453 | 13044508 | 17.00309 | 47.03089 | 44.35329 | 3' UTR | 7 | 13035120 | 13038275 | 3156 | 2 | ENSMUST00 | -5179 | Ube2m | ENSMUSG00000005575.10 |
| chr2 | 152942785 | 152943788 | 18.76756 | 47.00972 | 44.33237 | Promoter | 2 | 152941022 | 152944556 | 3535 | 2 | ENSMUST00 | 768 | Dusp15 | ENSMUSG00000042662.11 |
| chr7 | 14622078 | 14623016 | 18.30549 | 46.98978 | 44.31301 | Promoter | 7 | 14530652 | 14622479 | 91828 | 2 | ENSMUST00 | 0 | Nlrp5-ps | ENSMUSG00000041596.7 |
| chr14 | 21990294 | 21992314 | 18.72834 | 46.94304 | 44.26641 | Promoter | 14 | 21983962 | 21989601 | 5640 | 2 | ENSMUST00 | -694 | Zfp503 | ENSMUSG00000039081.9 |
| chr17 | 26202712 | 26203863 | 18.72834 | 46.94304 | 44.26641 | Promoter | 17 | 26202951 | 26211315 | 8365 | 1 | ENSMUST00 | 0 | Rgs11 | ENSMUSG00000024186.10 |
| chr5 | 110342434 | 110343458 | 18.26907 | 46.92738 | 44.25112 | Promoter | 5 | 110339812 | 110343035 | 3224 | 2 | ENSMUST00 | 0 | P2rx2 | ENSMUSG00000029503.11 |
| chr7 | 7277551 | 7278399 | 8.37265 | 46.86385 | 44.18798 | Distal Interge | 7 | 7265498 | 7269335 | 3838 | 2 | ENSMUST00 | -8217 | Gm18190 | ENSMUSG00000098900.1 |
| chr17 | 94864402 | 94865307 | 17.78412 | 46.86213 | 44.18626 | Distal Interge | 17 | 94869873 | 94869988 | 116 | 2 | ENSMUST00 | 4681 | Gm23102 | ENSMUSG00000065658.1 |
| chr3 | 83765798 | 83766977 | 17.78412 | 46.86213 | 44.18626 | Promoter | 3 | 83766321 | 83774316 | 7996 | 1 | ENSMUST00 | 0 | Sfrp2 | ENSMUSG00000027996.8 |
| chr7 | 44929975 | 44930461 | 17.78412 | 46.86213 | 44.18626 | Promoter | 7 | 44900373 | 44929496 | 29124 | 2 | ENSMUST00 | -480 | Ap2a1 | ENSMUSG00000060279.8 |
| chr9 | 37384307 | 37385045 | 17.78412 | 46.86213 | 44.18626 | 3' UTR | 9 | 37401897 | 37414019 | 12123 | 1 | ENSMUST00 | -16852 | Robo4 | ENSMUSG00000032125.14 |
| chr1 | 34750119 | 34751103 | 16.8969 | 46.84153 | 44.16631 | Promoter | 1 | 34750881 | 34807144 | 56264 | 1 | ENSMUST00 | 0 | Arhgef4 | ENSMUSG00000037509.13 |
| chr1 | 82282881 | 82284308 | 16.8969 | 46.84153 | 44.16631 | Intron (ENSM | 1 | 82233101 | 82291416 | 58316 | 2 | ENSMUST00 | 7108 | Irs1 | ENSMUSG00000055980.2 |
| chr10 | 20903904 | 20905847 | 16.8969 | 46.84153 | 44.16631 | Distal Interge | 10 | 20952547 | 21080428 | 127882 | 1 | ENSMUST00 | -46700 | Ahi1 | ENSMUSG00000019986.11 |
| chr10 | 4266071 | 4267207 | 16.8969 | 46.84153 | 44.16631 | Promoter | 10 | 4266329 | 4359468 | 93140 | 1 | ENSMUST00 | 0 | Akap12 | ENSMUSG00000038587.8 |
| chr10 | 115361706 | 115362980 | 16.8969 | 46.84153 | 44.16631 | Promoter | 10 | 115340739 | 115362262 | 21524 | 2 | ENSMUST00 | 0 | Tmem19 | ENSMUSG00000069520.5 |
| chr11 | 72266173 | 72267481 | 16.8969 | 46.84153 | 44.16631 | Promoter | 11 | 72241994 | 72266602 | 24609 | 2 | ENSMUST00 | 0 | Slc13a5 | ENSMUSG00000020805.8 |
| chr11 | 82990865 | 82992051 | 16.8969 | 46.84153 | 44.16631 | Promoter | 11 | 82979629 | 82991830 | 12202 | 2 | ENSMUST00 | 0 | Slfn9 | ENSMUSG00000069793.6 |
| chr13 | 117220483 | 117221539 | 16.8969 | 46.84153 | 44.16631 | Promoter | 13 | 117220625 | 117274098 | 53474 | 1 | ENSMUST00 | 0 | Emb | ENSMUSG00000021728.7 |
| chr13 | 4608510 | 4609539 | 16.8969 | 46.84153 | 44.16631 | Promoter | 13 | 4590757 | 4609174 | 18418 | 2 | ENSMUST00 | 0 | Akr1e1 | ENSMUSG00000045410.12 |
| chr15 | 78447838 | 78448446 | 16.8969 | 46.84153 | 44.16631 | Intron (ENSM | 15 | 78433982 | 78437579 | 3598 | 1 | ENSMUST00 | 13857 | Kctd17 | ENSMUSG00000033287.10 |
| chr15 | 103255040 | 103255575 | 16.8969 | 46.84153 | 44.16631 | Promoter | 15 | 103248212 | 103255439 | 7228 | 2 | ENSMUST00 | 0 | Nfe2 | ENSMUSG00000058794.7 |
| chr16 | 31932898 | 31934158 | 16.8969 | 46.84153 | 44.16631 | Promoter | 16 | 31933851 | 31946046 | 12196 | 1 | ENSMUST00 | 0 | Pigz | ENSMUSG00000045625.4 |
| chr16 | 32200234 | 32201333 | 16.8969 | 46.84153 | 44.16631 | Distal Interge | 16 | 32186186 | 32186949 | 764 | 1 | ENSMUST00 | 14049 | Bex6 | ENSMUSG00000075269.4 |
| chr17 | 71782421 | 71783433 | 16.8969 | 46.84153 | 44.16631 | Promoter | 17 | 71781947 | 71858351 | 76405 | 1 | ENSMUST00 | 475 | Clip4 | ENSMUSG00000024059.9 |
| chr2 | 119270826 | 119271469 | 16.8969 | 46.84153 | 44.16631 | Promoter | 2 | 119269201 | 119271272 | 2072 | 2 | ENSMUST00 | 0 | Rhov | ENSMUSG00000034226.7 |
| chr2 | 61811598 | 61812742 | 16.8969 | 46.84153 | 44.16631 | 3' UTR | 2 | 61806273 | 61811801 | 5529 | 1 | ENSMUST00 | 5326 | Tbr1 | ENSMUSG00000035033.10 |
| chr2 | 60283735 | 60284626 | 16.8969 | 46.84153 | 44.16631 | Promoter | 2 | 60251993 | 60284418 | 32426 | 2 | ENSMUST00 | 0 | Cd302 | ENSMUSG00000060703.7 |
| chr2 | 93470078 | 93470742 | 16.8969 | 46.84153 | 44.16631 | Distal Interge | 2 | 93468439 | 93468675 | 237 | 1 | ENSMUST00 | 1640 | Gm10804 | ENSMUSG00000075025.5 |
| chr3 | 99253147 | 99254326 | 16.8969 | 46.84153 | 44.16631 | Promoter | 3 | 99253760 | 99354259 | 100500 | 1 | ENSMUST00 | 0 | Tbx15 | ENSMUSG00000027868.6 |
| chr3 | 109339973 | 109341428 | 16.8969 | 46.84153 | 44.16631 | Promoter | 3 | 109340653 | 109685698 | 345046 | 1 | ENSMUST00 | 0 | Vav3 | ENSMUSG00000033721.11 |
| chr5 | 151232958 | 151234226 | 16.8969 | 46.84153 | 44.16631 | Promoter | 5 | 151063546 | 151233774 | 170229 | 2 | ENSMUST00 | 0 | Stard13 | ENSMUSG00000016128.9 |
| chr5 | 77015696 | 77016579 | 16.8969 | 46.84153 | 44.16631 | Promoter | 5 | 77016023 | 77061522 | 45500 | 1 | ENSMUST00 | 0 | 1700023E05I | ENSMUSG00000029248.9 |
| chr5 | 92682736 | 92683875 | 16.8969 | 46.84153 | 44.16631 | Promoter | 5 | 92683435 | 92965314 | 281880 | 1 | ENSMUST00 | 0 | Shroom3 | ENSMUSG00000029381.10 |
| chr5 | 28070416 | 28071178 | 16.8969 | 46.84153 | 44.16631 | Promoter | 5 | 28019465 | 28071105 | 51641 | 2 | ENSMUST00 | 0 | Gm26608 | ENSMUSG00000097673.1 |

| | | | | | | | | | | | | | | | |
|---|---|---|---|---|---|---|---|---|---|---|---|---|---|---|---|
| chr6 | 51523540 | 51524635 | 16.8969 | 46.84153 | 44.16631 | Promoter | 6 | 51523901 | 51590679 | 66779 | 1 | ENSMUST00 | 0 | Snx10 | ENSMUSG00000038301.10 |
| chr8 | 121114739 | 121115965 | 16.8969 | 46.84153 | 44.16631 | Promoter | 8 | 121116171 | 121118895 | 2725 | 1 | ENSMUST00 | -206 | Foxc2 | ENSMUSG00000046714.7 |
| chr2 | 65022103 | 65023166 | 18.65041 | 46.81047 | 44.13735 | Promoter | 2 | 64912476 | 65022782 | 110307 | 2 | ENSMUST00 | 0 | Grb14 | ENSMUSG00000026888.9 |
| chr5 | 134571532 | 134572669 | 18.19667 | 46.80325 | 44.13025 | Distal Interge | 5 | 134582690 | 134598328 | 15639 | 1 | ENSMUST00 | -10021 | Rfc2 | ENSMUSG00000023104.6 |
| chr1 | 51478593 | 51479239 | 15.22868 | 46.77911 | 44.10619 | Promoter | 1 | 51471129 | 51478425 | 7297 | 2 | ENSMUST00 | -169 | Nabp1 | ENSMUSG00000026107.6 |
| chr13 | 46735464 | 46736953 | 15.22868 | 46.77911 | 44.10619 | Distal Interge | 13 | 46679905 | 46727940 | 48036 | 2 | ENSMUST00 | -7525 | Nup153 | ENSMUSG00000021374.9 |
| chr14 | 22019289 | 22020203 | 15.22868 | 46.77911 | 44.10619 | Promoter | 14 | 22019712 | 23056085 | 1036374 | 1 | ENSMUST00 | 0 | 1700112E06I | ENSMUSG00000063458.7 |
| chr19 | 3743929 | 3744538 | 15.22868 | 46.77911 | 44.10619 | Distal Interge | 19 | 3767421 | 3808829 | 41409 | 1 | ENSMUST00 | -22883 | Suv420h1 | ENSMUSG00000045098.13 |
| chr2 | 60382702 | 60383580 | 15.22868 | 46.77911 | 44.10619 | Promoter | 2 | 60292103 | 60383303 | 91201 | 2 | ENSMUST00 | 0 | Ly75 | ENSMUSG00000026980.10 |
| chr7 | 98489601 | 98490689 | 15.22868 | 46.77911 | 44.10619 | Distal Interge | 7 | 98494222 | 98501831 | 7610 | 1 | ENSMUST00 | -3533 | Lrrc32 | ENSMUSG00000090958.1 |
| chr13 | 91370313 | 91371539 | 17.7171 | 46.74591 | 44.0739 | Exon (ENSMU | 13 | 91373842 | 91374294 | 453 | 2 | ENSMUST00 | 2755 | RP23-241P22 | ENSMUSG00000099340.1 |
| chr15 | 52039937 | 52040738 | 18.61169 | 46.74458 | 44.0726 | Promoter | 15 | 52040107 | 52045722 | 5616 | 1 | ENSMUST00 | 0 | Aard | ENSMUSG00000068522.4 |
| chr16 | 37384268 | 37385223 | 18.61169 | 46.74458 | 44.0726 | Promoter | 16 | 37114942 | 37384923 | 269982 | 2 | ENSMUST00 | 0 | Stxbp5l | ENSMUSG00000022829.9 |
| chr4 | 141367848 | 141368683 | 18.61169 | 46.74458 | 44.0726 | Promoter | 4 | 141368220 | 141384175 | 15956 | 1 | ENSMUST00 | 0 | Fam131c | ENSMUSG00000006218.4 |
| chr1 | 190978128 | 190979621 | 15.98415 | 46.70646 | 44.03467 | Promoter | 1 | 190947648 | 190978998 | 31351 | 2 | ENSMUST00 | 0 | Vash2 | ENSMUSG00000037568.7 |
| chr11 | 69661810 | 69663290 | 15.98415 | 46.70646 | 44.03467 | Promoter | 11 | 69656697 | 69662642 | 5946 | 2 | ENSMUST00 | 0 | Mpdu1 | ENSMUSG00000018761.9 |
| chr11 | 70458977 | 70460075 | 15.98415 | 46.70646 | 44.03467 | Promoter | 11 | 70453983 | 70459984 | 6002 | 2 | ENSMUST00 | 0 | Cxcl16 | ENSMUSG00000018920.6 |
| chr11 | 55077234 | 55078353 | 15.98415 | 46.70646 | 44.03467 | Promoter | 11 | 55049731 | 55078107 | 28377 | 2 | ENSMUST00 | 0 | Ccdc69 | ENSMUSG00000049588.8 |
| chr13 | 21326587 | 21327420 | 15.98415 | 46.70646 | 44.03467 | Distal Interge | 13 | 21317258 | 21319624 | 2367 | 1 | ENSMUST00 | 9330 | Gpx6 | ENSMUSG00000004341.7 |
| chr13 | 29986304 | 29987095 | 15.98415 | 46.70646 | 44.03467 | Promoter | 13 | 29906575 | 29985668 | 79094 | 2 | ENSMUST00 | -637 | E2f3 | ENSMUSG00000016477.12 |
| chr13 | 21734298 | 21735906 | 15.98415 | 46.70646 | 44.03467 | Promoter | 13 | 21735064 | 21735837 | 774 | 1 | ENSMUST00 | 0 | Hist1h4j | ENSMUSG00000067455.5 |
| chr17 | 23585465 | 23586805 | 15.98415 | 46.70646 | 44.03467 | Promoter | 17 | 23575845 | 23586232 | 10388 | 2 | ENSMUST00 | 0 | Zfp13 | ENSMUSG00000062012.8 |
| chr2 | 91921594 | 91922691 | 15.98415 | 46.70646 | 44.03467 | Promoter | 2 | 91920528 | 91920928 | 401 | 2 | ENSMUST00 | -667 | Gm10805 | ENSMUSG00000075036.1 |
| chr3 | 53656106 | 53658302 | 15.98415 | 46.70646 | 44.03467 | Promoter | 3 | 53513939 | 53657355 | 143417 | 2 | ENSMUST00 | 0 | Frem2 | ENSMUSG00000037016.7 |
| chr4 | 55514837 | 55515383 | 15.98415 | 46.70646 | 44.03467 | Distal Interge | 4 | 55508294 | 55508708 | 415 | 1 | ENSMUST00 | 6544 | Gm12507 | ENSMUSG00000082952.1 |
| chr4 | 133659625 | 133660921 | 15.98415 | 46.70646 | 44.03467 | 3' UTR | 4 | 133654002 | 133654257 | 256 | 1 | ENSMUST00 | 5624 | Gm22662 | ENSMUSG00000093031.2 |
| chr5 | 110109876 | 110110828 | 15.98415 | 46.70646 | 44.03467 | Promoter | 5 | 110110092 | 110129794 | 19703 | 1 | ENSMUST00 | 0 | Zfp605 | ENSMUSG00000023284.10 |
| chr5 | 102845002 | 102848980 | 15.98415 | 46.70646 | 44.03467 | Promoter | 5 | 102845007 | 102897829 | 52823 | 1 | ENSMUST00 | 0 | Arhgap24 | ENSMUSG00000057315.9 |
| chr5 | 149228600 | 149229612 | 15.98415 | 46.70646 | 44.03467 | Distal Interge | 5 | 149231332 | 149237984 | 6653 | 1 | ENSMUST00 | -1720 | RP23-408A1. | ENSMUSG00000100783.1 |
| chr6 | 34862367 | 34863819 | 15.98415 | 46.70646 | 44.03467 | Promoter | 6 | 34863146 | 34871587 | 8442 | 1 | ENSMUST00 | 0 | Tmem140 | ENSMUSG00000057137.4 |
| chr7 | 44847892 | 44849023 | 15.98415 | 46.70646 | 44.03467 | Promoter | 7 | 44848991 | 44855408 | 6418 | 1 | ENSMUST00 | 0 | Akt1s1 | ENSMUSG00000011096.11 |
| chr9 | 78615445 | 78616389 | 15.98415 | 46.70646 | 44.03467 | Promoter | 9 | 78615546 | 78716253 | 100708 | 1 | ENSMUST00 | 0 | Cd109 | ENSMUSG00000046186.8 |
| chr11 | 105293126 | 105294450 | 18.57313 | 46.67894 | 44.00877 | Promoter | 11 | 105292668 | 105343871 | 51204 | 1 | ENSMUST00 | 459 | Mrc2 | ENSMUSG00000020695.9 |
| chr16 | 77643859 | 77644877 | 18.57313 | 46.67894 | 44.00877 | Intron (ENSM | 16 | 77646273 | 77646343 | 71 | 1 | ENSMUST00 | -1396 | Mir125b-2 | ENSMUSG00000065472.1 |
| chr2 | 103564927 | 103566225 | 17.22045 | 46.64732 | 43.97726 | Promoter | 2 | 103566310 | 103718423 | 152114 | 1 | ENSMUST00 | -85 | Abtb2 | ENSMUSG00000032724.5 |
| chr1 | 119504905 | 119505393 | 18.05357 | 46.55767 | 43.88821 | Promoter | 1 | 119481133 | 119504794 | 23662 | 2 | ENSMUST00 | -112 | Ralb | ENSMUSG00000004451.9 |
| chr1 | 52952372 | 52953382 | 18.05357 | 46.55767 | 43.88821 | Promoter | 1 | 52922324 | 52952844 | 30521 | 2 | ENSMUST00 | 0 | 1700019D03I | ENSMUSG00000043629.7 |
| chr19 | 5662563 | 5664184 | 18.05357 | 46.55767 | 43.88821 | Promoter | 19 | 5651185 | 5663707 | 12523 | 2 | ENSMUST00 | 0 | Sipa1 | ENSMUSG00000056917.7 |
| chr16 | 28445015 | 28447058 | 18.49648 | 46.54843 | 43.8795 | Promoter | 16 | 28160120 | 28445227 | 285108 | 2 | ENSMUST00 | 0 | Fgf12 | ENSMUSG00000022523.9 |
| chr12 | 113155178 | 113156098 | 16.71423 | 46.51484 | 43.84613 | Promoter | 12 | 113156421 | 113165456 | 9036 | 1 | ENSMUST00 | -323 | 4930427A07I | ENSMUSG00000037466.9 |
| chr6 | 34598171 | 34599963 | 16.71423 | 46.51484 | 43.84613 | Promoter | 6 | 34598500 | 34775454 | 176955 | 1 | ENSMUST00 | 0 | Cald1 | ENSMUSG00000029761.11 |
| chr2 | 153224712 | 153225555 | 18.01814 | 46.49681 | 43.82834 | Promoter | 2 | 153222370 | 153225441 | 3072 | 2 | ENSMUST00 | 0 | Tspyl3 | ENSMUSG00000074671.6 |
| chr3 | 27984106 | 27985779 | 18.01814 | 46.49681 | 43.82834 | Promoter | 3 | 27984173 | 28132050 | 147878 | 1 | ENSMUST00 | 0 | Pld1 | ENSMUSG00000027695.11 |
| chr11 | 5485478 | 5486080 | 18.45839 | 46.48355 | 43.81516 | Distal Interge | 11 | 5495192 | 5495446 | 255 | 2 | ENSMUST00 | 9366 | Gm11962 | ENSMUSG00000085232.3 |
| chr8 | 83998252 | 83998820 | 18.45839 | 46.48355 | 43.81516 | Promoter | 8 | 83997672 | 84000386 | 2715 | 1 | ENSMUST00 | 581 | Samd1 | ENSMUSG00000079003.1 |
| chr7 | 10494871 | 10495551 | 17.98285 | 46.43617 | 43.76802 | Promoter | 7 | 10487229 | 10495394 | 8166 | 2 | ENSMUST00 | 0 | Zik1 | ENSMUSG00000030393.8 |
| chr18 | 25754076 | 25754530 | 18.42046 | 46.41892 | 43.75084 | Promoter | 18 | 25477632 | 25753983 | 276352 | 2 | ENSMUST00 | -94 | Celf4 | ENSMUSG00000024268.10 |
| chr5 | 121838190 | 121839252 | 18.42046 | 46.41892 | 43.75084 | Distal Interge | 5 | 121815648 | 121836859 | 21212 | 2 | ENSMUST00 | -1332 | Sh2b3 | ENSMUSG00000042594.11 |
| chr2 | 18587001 | 18588257 | 18.34506 | 46.29042 | 43.62275 | Distal Interge | 2 | 18568126 | 18568198 | 73 | 1 | ENSMUST00 | 18876 | Gm13353 | ENSMUSG00000081181.1 |
| chr4 | 138304060 | 138304627 | 18.34506 | 46.29042 | 43.62275 | Promoter | 4 | 138304730 | 138312628 | 7899 | 1 | ENSMUST00 | -103 | Ddost | ENSMUSG00000028757.4 |
| chr5 | 44100105 | 44101861 | 18.34506 | 46.29042 | 43.62275 | Promoter | 5 | 43993625 | 44101736 | 108112 | 2 | ENSMUST00 | 0 | Prom1 | ENSMUSG00000029086.10 |
| chr8 | 124433676 | 124434584 | 18.34506 | 46.29042 | 43.62275 | Promoter | 8 | 124369049 | 124433936 | 64888 | 2 | ENSMUST00 | 0 | Pgbd5 | ENSMUSG00000050751.9 |
| chr2 | 30711573 | 30712506 | 17.45395 | 46.28863 | 43.62135 | Downstream | 2 | 30713150 | 30717043 | 3894 | 2 | ENSMUST00 | 4537 | Gm14488 | ENSMUSG00000075416.5 |
| chr4 | 141559134 | 141560104 | 17.45395 | 46.28863 | 43.62135 | Distal Interge | 4 | 141546189 | 141547892 | 1704 | 1 | ENSMUST00 | 12946 | B330016D10 | ENSMUSG00000048406.5 |
| chr6 | 86669549 | 86670136 | 17.45395 | 46.28863 | 43.62135 | Promoter | 6 | 86647044 | 86669159 | 22116 | 2 | ENSMUST00 | -391 | Mxd1 | ENSMUSG00000001156.7 |
| chr17 | 49614528 | 49615540 | 18.3076 | 46.22653 | 43.55991 | Promoter | 17 | 49615136 | 49755388 | 140253 | 1 | ENSMUST00 | 0 | Kif6 | ENSMUSG00000023999.9 |
| chr7 | 109865264 | 109866076 | 18.3076 | 46.22653 | 43.55991 | Promoter | 7 | 109798676 | 109865586 | 66911 | 2 | ENSMUST00 | 0 | Scube2 | ENSMUSG00000007279.9 |
| chr1 | 157458183 | 157459060 | 18.27028 | 46.16288 | 43.49645 | Promoter | 1 | 157458580 | 157489060 | 30481 | 1 | ENSMUST00 | 0 | BC026585 | ENSMUSG00000033488.6 |
| chr4 | 8911065 | 8912002 | 18.27028 | 46.16288 | 43.49645 | Distal Interge | 4 | 8941259 | 8941656 | 398 | 1 | ENSMUST00 | -29257 | Rps18-ps2 | ENSMUSG00000083786.1 |
| chr3 | 108044452 | 108045354 | 16.91761 | 46.11281 | 43.44656 | Promoter | 3 | 108040408 | 108044894 | 4487 | 2 | ENSMUST00 | 0 | Gstm4 | ENSMUSG00000027890.12 |
| chr4 | 148966100 | 148968885 | 16.91761 | 46.11281 | 43.44656 | Exon (ENSMU | 4 | 148960360 | 148966998 | 6639 | 1 | ENSMUST00 | 5741 | Gm17029 | ENSMUSG00000086740.1 |
| chr7 | 29238327 | 29238999 | 16.91761 | 46.11281 | 43.44656 | Promoter | 7 | 29238329 | 29247529 | 9201 | 1 | ENSMUST00 | 0 | Yif1b | ENSMUSG00000030588.6 |
| chr8 | 70610927 | 70612123 | 16.91761 | 46.11281 | 43.44656 | Distal Interge | 8 | 70597491 | 70608314 | 10824 | 2 | ENSMUST00 | -2614 | Ssbp4 | ENSMUSG00000070003.6 |
| chr7 | 25815840 | 25817023 | 18.23312 | 46.09947 | 43.43353 | Promoter | 7 | 25802475 | 25816653 | 14179 | 2 | ENSMUST00 | 0 | Cyp2s1 | ENSMUSG00000040703.6 |
| chr6 | 49213612 | 49215549 | 13.78573 | 46.06182 | 43.39605 | Promoter | 6 | 49085221 | 49214954 | 129734 | 2 | ENSMUST00 | 0 | Igf2bp3 | ENSMUSG00000029814.8 |
| chr8 | 69832324 | 69833060 | 13.78573 | 46.06182 | 43.39605 | Promoter | 8 | 69832491 | 69872292 | 39802 | 1 | ENSMUST00 | 0 | Pbx4 | ENSMUSG00000031860.12 |
| chr9 | 78442429 | 78443926 | 15.9985 | 45.97639 | 43.31121 | Promoter | 9 | 78430526 | 78443204 | 12679 | 2 | ENSMUST00 | 0 | Mb21d1 | ENSMUSG00000032344.12 |
| chr16 | 17914174 | 17915238 | 18.15925 | 45.97338 | 43.30825 | Promoter | 16 | 17913107 | 17915059 | 1953 | 2 | ENSMUST00 | 0 | Gsc2 | ENSMUSG00000022738.6 |
| chr17 | 32505773 | 32507065 | 18.15925 | 45.97338 | 43.30825 | Promoter | 17 | 32506462 | 32528894 | 22433 | 1 | ENSMUST00 | 0 | Cyp4f17 | ENSMUSG00000091586.1 |
| chr10 | 81633258 | 81634137 | 17.70547 | 45.95868 | 43.29366 | Promoter | 10 | 81633699 | 81643023 | 9325 | 1 | ENSMUST00 | 0 | Ankrd24 | ENSMUSG00000054708.12 |
| chr10 | 119691612 | 119692930 | 17.70547 | 45.95868 | 43.29366 | Intron (ENSM | 10 | 119746063 | 119761699 | 15637 | 2 | ENSMUST00 | 68769 | Grip1os2 | ENSMUSG00000084764.1 |
| chr6 | 116107389 | 116108281 | 17.70547 | 45.95868 | 43.29366 | Promoter | 6 | 116043768 | 116107567 | 63800 | 2 | ENSMUST00 | 0 | Tmcc1 | ENSMUSG00000030126.12 |
| chr9 | 37207707 | 37208136 | 17.70547 | 45.95868 | 43.29366 | Promoter | 9 | 37208333 | 37223210 | 14878 | 1 | ENSMUST00 | -197 | Tmem218 | ENSMUSG00000032121.8 |
| chr8 | 29218998 | 29219998 | 18.12254 | 45.91068 | 43.24614 | Promoter | 8 | 28646717 | 29219338 | 572622 | 2 | ENSMUST00 | 0 | Unc5d | ENSMUSG00000063626.5 |
| chr4 | 97745647 | 97746407 | 17.6714 | 45.89991 | 43.23549 | Intron (ENSM | 4 | 97754168 | 97757389 | 3222 | 2 | ENSMUST00 | 10982 | Gm12676 | ENSMUSG00000085710.1 |
| chr9 | 72449206 | 72450162 | 18.08597 | 45.84822 | 43.18387 | Exon (ENSMU | 9 | 72441079 | 72441779 | 701 | 2 | ENSMUST00 | -7428 | Gm27231 | ENSMUSG00000098778.1 |
| chr15 | 79927444 | 79928362 | 12.71259 | 45.81302 | 43.1488 | Intron (ENSM | 15 | 79918921 | 79932516 | 13596 | 2 | ENSMUST00 | 4154 | Cbx7 | ENSMUSG00000053411.11 |
| chr1 | 34434002 | 34435403 | 17.60262 | 45.78123 | 43.11721 | Promoter | 1 | 34433121 | 34433199 | 79 | 2 | ENSMUST00 | -804 | Mir5103 | ENSMUSG00000092852.1 |
| chr11 | 31870012 | 31870741 | 17.60262 | 45.78123 | 43.11721 | Distal Interge | 11 | 31872211 | 31931577 | 59367 | 1 | ENSMUST00 | -1470 | Cpeb4 | ENSMUSG00000020300.9 |
| chr11 | 97574739 | 97575836 | 17.60262 | 45.78123 | 43.11721 | Promoter | 11 | 97509340 | 97575215 | 65876 | 2 | ENSMUST00 | 0 | Srcin1 | ENSMUSG00000038453.10 |
| chr11 | 101645951 | 101646655 | 17.60262 | 45.78123 | 43.11721 | Promoter | 11 | 101645740 | 101645930 | 191 | 2 | ENSMUST00 | -22 | Gm23971 | ENSMUSG00000065251.1 |
| chr14 | 69555053 | 69556025 | 17.60262 | 45.78123 | 43.11721 | Promoter | 14 | 69555424 | 69585520 | 30097 | 1 | ENSMUST00 | 0 | Gm21685 | ENSMUSG00000022066.11 |
| chr14 | 13933256 | 13934208 | 17.60262 | 45.78123 | 43.11721 | Promoter | 14 | 13932894 | 13949558 | 16665 | 1 | ENSMUST00 | 363 | Gm11100 | ENSMUSG00000079357.3 |
| chr15 | 77857605 | 77858242 | 17.60262 | 45.78123 | 43.11721 | Distal Interge | 15 | 77760587 | 77842175 | 81589 | 2 | ENSMUST00 | -15431 | Myh9 | ENSMUSG00000022443.11 |
| chr15 | 82365056 | 82365351 | 17.60262 | 45.78123 | 43.11721 | Distal Interge | 15 | 82350139 | 82354291 | 4153 | 2 | ENSMUST00 | -10766 | Ndufa6 | ENSMUSG00000022450.5 |
| chr15 | 101266077 | 101268132 | 17.60262 | 45.78123 | 43.11721 | Promoter | 15 | 101266846 | 101274792 | 7947 | 1 | ENSMUST00 | 0 | Nr4a1 | ENSMUSG00000023034.6 |
| chr15 | 84167520 | 84168405 | 17.60262 | 45.78123 | 43.11721 | Promoter | 15 | 84167816 | 84189512 | 21697 | 1 | ENSMUST00 | 0 | Pnpla3 | ENSMUSG00000041653.4 |
| chr15 | 76242804 | 76243599 | 17.60262 | 45.78123 | 43.11721 | Promoter | 15 | 76233003 | 76243420 | 10418 | 2 | ENSMUST00 | 0 | Parp10 | ENSMUSG00000063268.7 |
| chr16 | 10192591 | 10193435 | 17.60262 | 45.78123 | 43.11721 | Promoter | 16 | 10192712 | 10243051 | 50340 | 1 | ENSMUST00 | 0 | Atf7ip2 | ENSMUSG00000039200.10 |
| chr17 | 74527497 | 74528154 | 17.60262 | 45.78123 | 43.11721 | Promoter | 17 | 74528295 | 74703356 | 175062 | 1 | ENSMUST00 | -141 | Birc6 | ENSMUSG00000024073.9 |
| chr19 | 28834599 | 28835525 | 17.60262 | 45.78123 | 43.11721 | Promoter | 19 | 28835049 | 28879301 | 44253 | 1 | ENSMUST00 | 0 | Slc1a1 | ENSMUSG00000024935.5 |
| chr2 | 5012387 | 5012758 | 17.60262 | 45.78123 | 43.11721 | Promoter | 2 | 5006511 | 5012753 | 6243 | 2 | ENSMUST00 | 0 | Mcm10 | ENSMUSG00000026669.9 |
| chr2 | 120850593 | 120851016 | 17.60262 | 45.78123 | 43.11721 | Promoter | 2 | 120850705 | 120866650 | 15946 | 1 | ENSMUST00 | 0 | AV039307 | ENSMUSG00000087519.3 |
| chr3 | 127632339 | 127634596 | 17.60262 | 45.78123 | 43.11721 | Promoter | 3 | 127633136 | 127635631 | 2496 | 1 | ENSMUST00 | 0 | Neurog2 | ENSMUSG00000027967.7 |
| chr4 | 36949976 | 36952371 | 17.60262 | 45.78123 | 43.11721 | Promoter | 4 | 35707406 | 36951223 | 1243818 | 2 | ENSMUST00 | 0 | Lingo2 | ENSMUSG00000045083.9 |
| chr4 | 53196158 | 53196726 | 17.60262 | 45.78123 | 43.11721 | Distal Interge | 4 | 53205066 | 53217561 | 12496 | 2 | ENSMUST00 | 20835 | 4930412L05I | ENSMUSG00000087192.2 |
| chr4 | 22487447 | 22488873 | 17.60262 | 45.78123 | 43.11721 | Promoter | 4 | 22482780 | 22488366 | 5587 | 2 | ENSMUST00 | 0 | Pou3f2 | ENSMUSG00000095139.2 |
| chr5 | 43661972 | 43663748 | 17.60262 | 45.78123 | 43.11721 | Promoter | 5 | 43662379 | 43740972 | 78594 | 1 | ENSMUST00 | 0 | Cc2d2a | ENSMUSG00000039765.10 |
| chr5 | 128599806 | 128601582 | 17.60262 | 45.78123 | 43.11721 | Promoter | 5 | 128579106 | 128600687 | 21582 | 2 | ENSMUST00 | 0 | 5930412G12 | ENSMUSG00000072591.5 |
| chr6 | 88196796 | 88199942 | 17.60262 | 45.78123 | 43.11721 | Promoter | 6 | 88198664 | 88207030 | 8367 | 1 | ENSMUST00 | 0 | Gata2 | ENSMUSG00000015053.9 |
| chr6 | 87335081 | 87335924 | 17.60262 | 45.78123 | 43.11721 | Promoter | 6 | 87133854 | 87335775 | 201922 | 2 | ENSMUST00 | 0 | Antxr1 | ENSMUSG00000033420.10 |
| chr6 | 71270850 | 71271480 | 17.60262 | 45.78123 | 43.11721 | Promoter | 6 | 71271671 | 71271801 | 131 | 1 | ENSMUST00 | -191 | Mir8112 | ENSMUSG00000098623.1 |
| chr7 | 81991499 | 81993036 | 17.60262 | 45.78123 | 43.11721 | Promoter | 7 | 81966672 | 81992299 | 25628 | 2 | ENSMUST00 | 0 | Bnc1 | ENSMUSG00000025105.8 |
| chr7 | 12803015 | 12804280 | 17.60262 | 45.78123 | 43.11721 | Downstream | 7 | 12806073 | 12808075 | 2003 | 2 | ENSMUST00 | 3795 | Zfp329 | ENSMUSG00000057894.10 |
| chr7 | 143444764 | 143446666 | 17.60262 | 45.78123 | 43.11721 | Promoter | 7 | 143447605 | 143447665 | 61 | 1 | ENSMUST00 | -939 | Gm27901 | ENSMUSG00000099103.1 |
| chr8 | 81341690 | 81342919 | 17.60262 | 45.78123 | 43.11721 | Promoter | 8 | 81342556 | 81745890 | 403335 | 1 | ENSMUST00 | 0 | Inpp4b | ENSMUSG00000037940.10 |
| chr9 | 21615400 | 21615987 | 17.60262 | 45.78123 | 43.11721 | Promoter | 9 | 21616169 | 21704217 | 88049 | 1 | ENSMUST00 | -182 | Smarca4 | ENSMUSG00000032187.11 |
| chr17 | 6316560 | 6317784 | 17.13582 | 45.73338 | 43.07122 | Promoter | 17 | 6310547 | 6317474 | 6928 | 2 | ENSMUST00 | 0 | Dynlt1a | ENSMUSG00000092074.1 |
| chr6 | 125039050 | 125039686 | 17.13582 | 45.73338 | 43.07122 | Promoter | 6 | 125039760 | 125049002 | 9243 | 1 | ENSMUST00 | -74 | Ing4 | ENSMUSG00000030330.10 |
| chr7 | 135537761 | 135538603 | 17.13582 | 45.73338 | 43.07122 | Promoter | 7 | 135537824 | 135686294 | 148471 | 1 | ENSMUST00 | 0 | Ptpre | ENSMUSG00000041836.9 |
| chr15 | 100983554 | 100984459 | 18.01328 | 45.724 | 43.06234 | Exon (ENSMU | 15 | 100955398 | 101045929 | 90532 | 1 | ENSMUST00 | 28157 | Scn8a | ENSMUSG00000023033.9 |
| chr4 | 139622375 | 139622748 | 18.01328 | 45.724 | 43.06234 | Promoter | 4 | 139622866 | 139649690 | 26825 | 1 | ENSMUST00 | -118 | Aldh4a1 | ENSMUSG00000028737.10 |
| chr12 | 100725410 | 100726035 | 14.30691 | 45.69287 | 43.03132 | Promoter | 12 | 100549778 | 100725028 | 175251 | 2 | ENSMUST00 | -383 | Rps6ka5 | ENSMUSG00000021180.8 |
| chr14 | 65331642 | 65332471 | 14.30691 | 45.69287 | 43.03132 | Distal Interge | 14 | 65358676 | 65398930 | 40255 | 1 | ENSMUST00 | -26205 | Zfp395 | ENSMUSG00000034522.9 |
| chr14 | 25377781 | 25378838 | 14.30691 | 45.69287 | 43.03132 | Promoter | 14 | 25355801 | 25378635 | 22835 | 2 | ENSMUST00 | 0 | Gm26660 | ENSMUSG00000097601.1 |
| chr17 | 31386037 | 31386819 | 14.30691 | 45.69287 | 43.03132 | Promoter | 17 | 31386234 | 31476310 | 90077 | 1 | ENSMUST00 | 0 | Pde9a | ENSMUSG00000041119.6 |
| chr3 | 97628394 | 97629222 | 14.30691 | 45.69287 | 43.03132 | Promoter | 3 | 97628804 | 97655282 | 26479 | 1 | ENSMUST00 | 0 | Fmo5 | ENSMUSG00000028088.9 |
| chr11 | 104607595 | 104608566 | 17.97716 | 45.66223 | 43.00122 | Promoter | 11 | 104608000 | 104670476 | 62477 | 1 | ENSMUST00 | 0 | Itgb3 | ENSMUSG00000020689.4 |
| chr14 | 118910196 | 118910780 | 17.97716 | 45.66223 | 43.00122 | Intron (ENSM | 14 | 118875520 | 118923167 | 47648 | 2 | ENSMUST00 | 12387 | Dzip1 | ENSMUSG00000042156.10 |
| chr5 | 98854756 | 98855542 | 17.97716 | 45.66223 | 43.00122 | Promoter | 5 | 98854439 | 98880960 | 26522 | 1 | ENSMUST00 | 318 | Bmp3 | ENSMUSG00000029335.4 |
| chr9 | 98954684 | 98957106 | 17.97716 | 45.66223 | 43.00122 | Promoter | 9 | 98955607 | 98956847 | 1241 | 1 | ENSMUST00 | 0 | Foxl2 | ENSMUSG00000050397.3 |
| chr1 | 135818557 | 135819086 | 16.62524 | 45.5939 | 42.93356 | Promoter | 1 | 135818598 | 135833342 | 14745 | 1 | ENSMUST00 | 0 | Lad1 | ENSMUSG00000041782.9 |
| chr17 | 28141701 | 28143481 | 16.62524 | 45.5939 | 42.93356 | Promoter | 17 | 28142316 | 28174852 | 32537 | 1 | ENSMUST00 | 0 | Scube3 | ENSMUSG00000038677.8 |
| chr1 | 167688371 | 167689424 | 18.38128 | 45.59164 | 42.93153 | Promoter | 1 | 167689237 | 167848741 | 159505 | 1 | ENSMUST00 | 0 | Lmx1a | ENSMUSG00000026686.9 |
| chr14 | 28504424 | 28507723 | 18.38128 | 45.59164 | 42.93153 | Promoter | 14 | 28504750 | 28527447 | 22698 | 1 | ENSMUST00 | 0 | Wnt5a | ENSMUSG00000021994.9 |
| chr14 | 7173751 | 7174920 | 18.38128 | 45.59164 | 42.93153 | Promoter | 14 | 7153511 | 7174533 | 21023 | 2 | ENSMUST00 | 0 | Gm3512 | ENSMUSG00000090363.3 |
| chr15 | 8709698 | 8711004 | 18.38128 | 45.59164 | 42.93153 | Promoter | 15 | 8634124 | 8710764 | 76641 | 2 | ENSMUST00 | 0 | Slc1a3 | ENSMUSG00000005360.9 |
| chr15 | 82211767 | 82213072 | 18.38128 | 45.59164 | 42.93153 | Promoter | 15 | 82206952 | 82212815 | 5864 | 2 | ENSMUST00 | 0 | AI848285 | ENSMUSG00000096883.1 |

| | | | | | | | | | | | | | | | |
|---|---|---|---|---|---|---|---|---|---|---|---|---|---|---|---|
| chr15 | 103146571 | 103147229 | 18.38128 | 45.59164 | 42.93153 | Promoter | 15 | 103123070 | 103146828 | 23759 | 2 | ENSMUST00 | 0 | D930007P13I | ENSMUSG00000097003.1 |
| chr17 | 70834385 | 70835379 | 18.38128 | 45.59164 | 42.93153 | Promoter | 17 | 70834664 | 70836044 | 1381 | 1 | ENSMUST00 | 0 | Gm20703 | ENSMUSG00000093622.1 |
| chr2 | 168280606 | 168281519 | 18.38128 | 45.59164 | 42.93153 | Promoter | 2 | 168260117 | 168281132 | 21016 | 2 | ENSMUST00 | 0 | Kcng1 | ENSMUSG00000074575.4 |
| chr2 | 175869225 | 175869797 | 18.38128 | 45.59164 | 42.93153 | Promoter | 2 | 175869245 | 175879891 | 10647 | 1 | ENSMUST00 | 0 | Gm6710 | ENSMUSG00000078887.5 |
| chr4 | 118260493 | 118261379 | 18.38128 | 45.59164 | 42.93153 | Intron (ENSM | 4 | 118209005 | 118236445 | 27441 | 2 | ENSMUST00 | -24049 | Ptprf | ENSMUSG00000033295.8 |
| chr4 | 152296589 | 152297152 | 18.38128 | 45.59164 | 42.93153 | Promoter | 4 | 152297227 | 152307121 | 9895 | 1 | ENSMUST00 | -75 | Icmt | ENSMUSG00000039662.11 |
| chr4 | 126286438 | 126287270 | 18.38128 | 45.59164 | 42.93153 | Promoter | 4 | 126286793 | 126314330 | 27538 | 1 | ENSMUST00 | 0 | Col8a2 | ENSMUSG00000056174.8 |
| chr5 | 98167011 | 98167873 | 18.38128 | 45.59164 | 42.93153 | Distal Interge | 5 | 98180869 | 98187448 | 6580 | 1 | ENSMUST00 | -12996 | Prdm8 | ENSMUSG00000035456.6 |
| chr6 | 127108932 | 127110032 | 18.38128 | 45.59164 | 42.93153 | Promoter | 6 | 127085116 | 127109550 | 24435 | 2 | ENSMUST00 | 0 | 9630033F20I | ENSMUSG00000038028.8 |
| chr7 | 101820682 | 101822603 | 18.38128 | 45.59164 | 42.93153 | 3' UTR | 7 | 101818313 | 101822726 | 4414 | 1 | ENSMUST00 | 2370 | Phox2a | ENSMUSG00000007946.10 |
| chr7 | 18925887 | 18926597 | 18.38128 | 45.59164 | 42.93153 | Promoter | 7 | 18925888 | 18962057 | 36170 | 1 | ENSMUST00 | 0 | Nova2 | ENSMUSG00000030411.10 |
| chr8 | 88627037 | 88628051 | 18.38128 | 45.59164 | 42.93153 | 3' UTR | 8 | 88626828 | 88636128 | 9301 | 2 | ENSMUST00 | 8077 | Snx20 | ENSMUSG00000031662.7 |
| chr9 | 30921524 | 30922879 | 18.38128 | 45.59164 | 42.93153 | Promoter | 9 | 30899155 | 30922452 | 23298 | 2 | ENSMUST00 | 0 | Adamts15 | ENSMUSG00000033453.7 |
| chrX | 140664601 | 140665872 | 18.38128 | 45.59164 | 42.93153 | Promoter | 20 | 140664956 | 140767715 | 102760 | 1 | ENSMUST00 | 0 | Mid2 | ENSMUSG00000000266.6 |
| chrX | 74254683 | 74255724 | 18.38128 | 45.59164 | 42.93153 | Promoter | 20 | 74254687 | 74257887 | 3201 | 1 | ENSMUST00 | 0 | Emd | ENSMUSG00000001964.9 |
| chrX | 94540721 | 94542785 | 18.38128 | 45.59164 | 42.93153 | Promoter | 20 | 94535474 | 94542143 | 6670 | 2 | ENSMUST00 | 0 | Maged1 | ENSMUSG00000025151.11 |
| chrX | 152015863 | 152017508 | 18.38128 | 45.59164 | 42.93153 | Promoter | 20 | 152004582 | 152016295 | 11714 | 2 | ENSMUST00 | 0 | Ribc1 | ENSMUSG00000025257.5 |
| chrX | 112311378 | 112312070 | 18.38128 | 45.59164 | 42.93153 | Promoter | 20 | 112311366 | 112372430 | 61065 | 1 | ENSMUST00 | 13 | Apool | ENSMUSG00000025525.7 |
| chrX | 105928697 | 105929771 | 18.38128 | 45.59164 | 42.93153 | Promoter | 20 | 105797615 | 105929397 | 131783 | 2 | ENSMUST00 | 0 | Atrx | ENSMUSG00000031229.11 |
| chrX | 141475093 | 141476973 | 18.38128 | 45.59164 | 42.93153 | Promoter | 20 | 141475385 | 141689234 | 213850 | 1 | ENSMUST00 | 0 | Col4a5 | ENSMUSG00000031274.11 |
| chrX | 103356107 | 103357299 | 18.38128 | 45.59164 | 42.93153 | Promoter | 20 | 103356476 | 103396092 | 39617 | 1 | ENSMUST00 | 0 | Chic1 | ENSMUSG00000031327.10 |
| chrX | 71214989 | 71215841 | 18.38128 | 45.59164 | 42.93153 | Promoter | 20 | 71215006 | 71260810 | 45805 | 1 | ENSMUST00 | 0 | Mtm1 | ENSMUSG00000031337.11 |
| chrX | 7573333 | 7574848 | 18.38128 | 45.59164 | 42.93153 | Promoter | 20 | 7573600 | 7581015 | 7416 | 1 | ENSMUST00 | 0 | Ppp1r3fos | ENSMUSG00000039545.10 |
| chrX | 135796047 | 135796987 | 18.38128 | 45.59164 | 42.93153 | Promoter | 20 | 135797588 | 135803466 | 5879 | 1 | ENSMUST00 | -601 | Gprasp1 | ENSMUSG00000043384.10 |
| chrX | 134540801 | 134542148 | 18.38128 | 45.59164 | 42.93153 | Promoter | 20 | 134537256 | 134541865 | 4610 | 2 | ENSMUST00 | 0 | Timm8a1 | ENSMUSG00000048007.4 |
| chrX | 167206266 | 167210665 | 18.38128 | 45.59164 | 42.93153 | Promoter | 20 | 167207093 | 167209315 | 2223 | 2 | ENSMUST00 | 0 | Tmsb4x | ENSMUSG00000049775.11 |
| chrX | 134686134 | 134687343 | 18.38128 | 45.59164 | 42.93153 | Promoter | 20 | 134686519 | 134696757 | 10239 | 1 | ENSMUST00 | 0 | Armcx4 | ENSMUSG00000049804.9 |
| chrX | 102250067 | 102252696 | 18.38128 | 45.59164 | 42.93153 | Promoter | 20 | 102247381 | 102252181 | 4801 | 2 | ENSMUST00 | 0 | Cited1 | ENSMUSG00000051159.11 |
| chrX | 8892266 | 8893915 | 18.38128 | 45.59164 | 42.93153 | Promoter | 20 | 8892382 | 8894964 | 2583 | 1 | ENSMUST00 | 0 | B630019K06 | ENSMUSG00000052364.4 |
| chrX | 48256092 | 48257602 | 18.38128 | 45.59164 | 42.93153 | Promoter | 20 | 48256862 | 48282453 | 25592 | 1 | ENSMUST00 | 0 | Utp14a | ENSMUSG00000063785.7 |
| chrX | 70476151 | 70477520 | 18.38128 | 45.59164 | 42.93153 | Promoter | 20 | 70459753 | 70477180 | 17428 | 2 | ENSMUST00 | 0 | BC023829 | ENSMUSG00000073139.4 |
| chrX | 151169495 | 151169948 | 18.38128 | 45.59164 | 42.93153 | Promoter | 20 | 151169700 | 151171764 | 2065 | 1 | ENSMUST00 | 0 | Gm15138 | ENSMUSG00000086286.1 |
| chr7 | 72306035 | 72306916 | 17.04263 | 45.57019 | 42.91153 | Promoter | 7 | 72077830 | 72306595 | 228766 | 2 | ENSMUST00 | 0 | Mctp2 | ENSMUSG00000032776.8 |
| chr10 | 68134465 | 68136910 | 16.165 | 45.52484 | 42.86639 | Exon (ENSMU | 10 | 68043968 | 68051046 | 7079 | 1 | ENSMUST00 | 90498 | Rtkn2 | ENSMUSG00000037846.13 |
| chr18 | 73622810 | 73623454 | 16.165 | 45.52484 | 42.86639 | Downstream | 18 | 73624928 | 73625037 | 110 | 1 | ENSMUST00 | -1474 | Gm22065 | ENSMUSG00000076941.1 |
| chr9 | 44965716 | 44966312 | 16.165 | 45.52484 | 42.86639 | Promoter | 9 | 44925848 | 44965600 | 39753 | 2 | ENSMUST00 | -117 | Ube4a | ENSMUSG00000059890.10 |
| chr10 | 80399731 | 80400305 | 14.91789 | 45.43439 | 42.77634 | Promoter | 10 | 80392539 | 80399550 | 7012 | 2 | ENSMUST00 | -182 | Mbd3 | ENSMUSG00000035478.9 |
| chr13 | 61803887 | 61804707 | 14.91789 | 45.43439 | 42.77634 | Promoter | 13 | 61804141 | 61804248 | 108 | 2 | ENSMUST00 | 0 | Gm23739 | ENSMUSG00000096212.1 |
| chr14 | 55539974 | 55541250 | 14.91789 | 45.43439 | 42.77634 | Promoter | 14 | 55540266 | 55550017 | 9752 | 1 | ENSMUST00 | 0 | Pck2 | ENSMUSG00000040618.6 |
| chr15 | 75677956 | 75678746 | 14.91789 | 45.43439 | 42.77634 | Promoter | 15 | 75657035 | 75678800 | 21766 | 2 | ENSMUST00 | 0 | Top1mt | ENSMUSG00000000934.8 |
| chr2 | 152735847 | 152738277 | 14.91789 | 45.43439 | 42.77634 | Promoter | 2 | 152736251 | 152737410 | 1160 | 1 | ENSMUST00 | 0 | Id1 | ENSMUSG00000042745.9 |
| chr3 | 95928665 | 95929089 | 14.91789 | 45.43439 | 42.77634 | Promoter | 3 | 95929246 | 95947390 | 18145 | 1 | ENSMUST00 | -157 | Anp32e | ENSMUSG00000015749.7 |
| chr4 | 107889275 | 107890655 | 14.91789 | 45.43439 | 42.77634 | Promoter | 4 | 107889813 | 107899384 | 9572 | 1 | ENSMUST00 | 0 | 0610037L13F | ENSMUSG00000028608.10 |
| chr6 | 134792843 | 134793179 | 14.91789 | 45.43439 | 42.77634 | Promoter | 6 | 134715473 | 134792625 | 77153 | 2 | ENSMUST00 | -219 | Dusp16 | ENSMUSG00000030203.12 |
| chrX | 144275894 | 144276672 | 14.91789 | 45.43439 | 42.77634 | Distal Interge | 20 | 144268049 | 144268204 | 156 | 2 | ENSMUST00 | -7691 | Gm15067 | ENSMUSG00000081897.1 |
| chr10 | 75797350 | 75798652 | 16.52141 | 45.4089 | 42.75185 | Promoter | 10 | 75783813 | 75798584 | 14772 | 2 | ENSMUST00 | 0 | Gstt1 | ENSMUSG00000001663.5 |
| chr12 | 87388574 | 87389074 | 16.52141 | 45.4089 | 42.75185 | Promoter | 12 | 87305058 | 87388355 | 83298 | 2 | ENSMUST00 | -220 | Sptlc2 | ENSMUSG00000021036.5 |
| chr13 | 74522137 | 74523114 | 16.52141 | 45.4089 | 42.75185 | Distal Interge | 13 | 74480517 | 74482943 | 2427 | 2 | ENSMUST00 | -39195 | Zfp825 | ENSMUSG00000069208.5 |
| chr15 | 79042600 | 79043543 | 16.52141 | 45.4089 | 42.75185 | Promoter | 15 | 79041885 | 79043558 | 1674 | 1 | ENSMUST00 | 716 | Gcat | ENSMUSG00000006378.8 |
| chr15 | 74671925 | 74672932 | 16.52141 | 45.4089 | 42.75185 | Promoter | 15 | 74669083 | 74672568 | 3486 | 2 | ENSMUST00 | 0 | Arc | ENSMUSG00000022602.9 |
| chr16 | 16896510 | 16897476 | 16.52141 | 45.4089 | 42.75185 | Promoter | 16 | 16896574 | 16915194 | 18621 | 1 | ENSMUST00 | 0 | Ppm1f | ENSMUSG00000026181.11 |
| chr17 | 45736363 | 45737272 | 16.52141 | 45.4089 | 42.75185 | Intron (ENSM | 17 | 45734506 | 45781251 | 46746 | 1 | ENSMUST00 | 1858 | F630040K05I | ENSMUSG00000097727.2 |
| chr19 | 4169619 | 4170188 | 16.52141 | 45.4089 | 42.75185 | Promoter | 19 | 4169566 | 4174284 | 4719 | 1 | ENSMUST00 | 54 | Gm17552 | ENSMUSG00000097233.1 |
| chr2 | 179974884 | 179975870 | 16.52141 | 45.4089 | 42.75185 | Promoter | 2 | 179912152 | 179975559 | 63408 | 2 | ENSMUST00 | 0 | Taf4a | ENSMUSG00000039117.11 |
| chr2 | 152931092 | 152933192 | 16.52141 | 45.4089 | 42.75185 | Promoter | 2 | 152931898 | 152933208 | 1311 | 2 | ENSMUST00 | 16 | Foxs1 | ENSMUSG00000074676.2 |
| chr2 | 76338684 | 76339272 | 16.52141 | 45.4089 | 42.75185 | Promoter | 2 | 75989912 | 76338774 | 348863 | 2 | ENSMUST00 | 0 | Pde11a | ENSMUSG00000075270.6 |
| chr2 | 118975797 | 118976914 | 16.52141 | 45.4089 | 42.75185 | Distal Interge | 2 | 118997068 | 119000684 | 3617 | 1 | ENSMUST00 | -20154 | Gm14089 | ENSMUSG00000084802.1 |
| chr5 | 8893300 | 8894172 | 16.52141 | 45.4089 | 42.75185 | Promoter | 5 | 8893721 | 8959225 | 65505 | 1 | ENSMUST00 | 0 | Abcb4 | ENSMUSG00000042476.8 |
| chr6 | 70843649 | 70844180 | 16.52141 | 45.4089 | 42.75185 | Promoter | 6 | 70844515 | 70905245 | 60731 | 1 | ENSMUST00 | -335 | Eif2ak3 | ENSMUSG00000031668.9 |
| chr6 | 29984813 | 29985641 | 16.52141 | 45.4089 | 42.75185 | Promoter | 6 | 29985329 | 29993530 | 8202 | 1 | ENSMUST00 | 0 | 1700023L04F | ENSMUSG00000045709.10 |
| chr9 | 76566554 | 76567582 | 16.52141 | 45.4089 | 42.75185 | Promoter | 9 | 76490054 | 76567116 | 77063 | 2 | ENSMUST00 | 0 | Fam83b | ENSMUSG00000032358.7 |
| chr9 | 32223834 | 32227622 | 16.52141 | 45.4089 | 42.75185 | Promoter | 9 | 32224249 | 32264245 | 39997 | 1 | ENSMUST00 | 0 | Arhgap32 | ENSMUSG00000041444.9 |
| chr1 | 168432526 | 168432982 | 17.37054 | 45.37996 | 42.72523 | Promoter | 1 | 168119364 | 168432270 | 312907 | 2 | ENSMUST00 | -257 | Pbx1 | ENSMUSG00000052534.10 |
| chr11 | 59786975 | 59787807 | 17.37054 | 45.37996 | 42.72523 | Promoter | 11 | 59783897 | 59787645 | 3749 | 2 | ENSMUST00 | 0 | Pld6 | ENSMUSG00000043648.7 |
| chr2 | 31845645 | 31846888 | 17.37054 | 45.37996 | 42.72523 | Promoter | 2 | 31813290 | 31846005 | 32716 | 2 | ENSMUST00 | 0 | Fibcd1 | ENSMUSG00000026841.7 |
| chr2 | 34961236 | 34961902 | 17.37054 | 45.37996 | 42.72523 | Promoter | 2 | 34943256 | 34961585 | 18330 | 2 | ENSMUST00 | 0 | Traf1 | ENSMUSG00000026875.9 |
| chr3 | 87910387 | 87911255 | 17.37054 | 45.37996 | 42.72523 | Promoter | 3 | 87910034 | 87914182 | 4149 | 1 | ENSMUST00 | 354 | Hdgf | ENSMUSG00000004897.11 |
| chr10 | 29533295 | 29534895 | 18.23346 | 45.34046 | 42.68647 | Promoter | 10 | 29453109 | 29535867 | 82759 | 2 | ENSMUST00 | 972 | Rspo3 | ENSMUSG00000019880.9 |
| chr11 | 87852781 | 87853881 | 15.64407 | 45.32187 | 42.668 | Promoter | 11 | 87853215 | 87863803 | 10589 | 1 | ENSMUST00 | 0 | Mks1 | ENSMUSG00000034121.7 |
| chr11 | 94520983 | 94521703 | 15.64407 | 45.32187 | 42.668 | Promoter | 11 | 94501347 | 94521742 | 20396 | 2 | ENSMUST00 | 39 | Mycbpap | ENSMUSG00000039110.9 |
| chr13 | 119631546 | 119632308 | 15.64407 | 45.32187 | 42.668 | Intron (ENSM | 13 | 119623819 | 119654354 | 30536 | 1 | ENSMUST00 | 7728 | Ccl28 | ENSMUSG00000074715.2 |
| chr15 | 36210217 | 36210922 | 15.64407 | 45.32187 | 42.668 | Exon (ENSMU | 15 | 36205260 | 36205322 | 63 | 1 | ENSMUST00 | 4958 | Gm23754 | ENSMUSG00000088201.1 |
| chr16 | 19760030 | 19761621 | 15.64407 | 45.32187 | 42.668 | Promoter | 16 | 19760208 | 19772233 | 12026 | 1 | ENSMUST00 | 0 | B3gnt5 | ENSMUSG00000022686.9 |
| chr16 | 18626018 | 18626977 | 15.64407 | 45.32187 | 42.668 | Intron (ENSM | 16 | 18621811 | 18629938 | 8128 | 2 | ENSMUST00 | 2961 | Sep-05 | ENSMUSG00000072214.6 |
| chr17 | 36946810 | 36947789 | 15.64407 | 45.32187 | 42.668 | 3' UTR | 17 | 36945078 | 36947286 | 2209 | 1 | ENSMUST00 | 1733 | Rnf39 | ENSMUSG00000036492.7 |
| chr2 | 131422949 | 131423445 | 15.64407 | 45.32187 | 42.668 | Distal Interge | 2 | 131414921 | 131415245 | 325 | 2 | ENSMUST00 | -7705 | Gm14283 | ENSMUSG00000081652.1 |
| chr4 | 109406379 | 109407463 | 15.64407 | 45.32187 | 42.668 | Promoter | 4 | 109406623 | 109428134 | 21512 | 1 | ENSMUST00 | 0 | Ttc39a | ENSMUSG00000028555.10 |
| chr4 | 108999883 | 109000456 | 15.64407 | 45.32187 | 42.668 | Promoter | 4 | 109000655 | 109061777 | 61123 | 1 | ENSMUST00 | -199 | Nrd1 | ENSMUSG00000053510.7 |
| chr5 | 30622200 | 30623232 | 15.64407 | 45.32187 | 42.668 | 3' UTR | 5 | 30647933 | 30659731 | 11799 | 1 | ENSMUST00 | -24701 | Slc35f6 | ENSMUSG00000029175.12 |
| chr5 | 114979186 | 114980466 | 15.64407 | 45.32187 | 42.668 | Exon (ENSMU | 5 | 114968705 | 114971094 | 2390 | 2 | ENSMUST00 | -8093 | Hnf1a | ENSMUSG00000029556.7 |
| chr6 | 122810365 | 122810935 | 15.64407 | 45.32187 | 42.668 | Downstream | 6 | 122737085 | 122801640 | 64556 | 2 | ENSMUST00 | -8726 | Slc2a3 | ENSMUSG00000003153.5 |
| chr6 | 118196606 | 118197761 | 15.64407 | 45.32187 | 42.668 | Promoter | 6 | 118151748 | 118197744 | 45997 | 2 | ENSMUST00 | 0 | Ret | ENSMUSG00000030110.8 |
| chr6 | 67533927 | 67536284 | 15.64407 | 45.32187 | 42.668 | Promoter | 6 | 67534059 | 67535822 | 1764 | 2 | ENSMUST00 | 0 | Tacstd2 | ENSMUSG00000051397.4 |
| chr1 | 5018761 | 5020599 | 18.15775 | 45.21172 | 42.55974 | Promoter | 1 | 4909576 | 5019539 | 109964 | 2 | ENSMUST00 | 0 | Rgs20 | ENSMUSG00000002459.12 |
| chr19 | 5067956 | 5068600 | 18.15775 | 45.21172 | 42.55974 | Promoter | 19 | 5068078 | 5070637 | 2560 | 1 | ENSMUST00 | 0 | Cd248 | ENSMUSG00000056481.6 |
| chr12 | 33957327 | 33958679 | 12.16433 | 45.21117 | 42.5593 | Promoter | 12 | 33957671 | 33959817 | 2147 | 1 | ENSMUST00 | 0 | Twist1 | ENSMUSG00000035799.5 |
| chr8 | 84908134 | 84909194 | 12.16433 | 45.21117 | 42.5593 | Promoter | 8 | 84908560 | 84911469 | 2910 | 1 | ENSMUST00 | 0 | Dnase2a | ENSMUSG00000003812.8 |
| chr11 | 79590789 | 79591724 | 17.69329 | 45.17611 | 42.52464 | Promoter | 11 | 79591212 | 79698023 | 106812 | 1 | ENSMUST00 | 0 | Rab11fip4 | ENSMUSG00000017639.8 |
| chr6 | 42709334 | 42710127 | 18.12012 | 45.14773 | 42.49636 | Promoter | 6 | 42668002 | 42710067 | 42066 | 2 | ENSMUST00 | 0 | Fam115a | ENSMUSG00000036667.9 |
| chr1 | 153751482 | 153752879 | 17.65844 | 45.11633 | 42.46507 | Promoter | 1 | 153751952 | 153760898 | 8947 | 1 | ENSMUST00 | 0 | Rnasel | ENSMUSG00000066800.6 |
| chr7 | 92670075 | 92670717 | 17.65844 | 45.11633 | 42.46507 | Promoter | 7 | 92643712 | 92669933 | 26222 | 2 | ENSMUST00 | -143 | Pcf11 | ENSMUSG00000041328.10 |
| chr2 | 30177878 | 30178893 | 16.34281 | 45.08974 | 42.43857 | Promoter | 2 | 30173453 | 30178429 | 4977 | 2 | ENSMUST00 | 0 | D2Wsu81e | ENSMUSG00000039660.11 |
| chr15 | 79261165 | 79262134 | 17.62372 | 45.05676 | 42.40596 | Promoter | 15 | 79259716 | 79261297 | 1582 | 2 | ENSMUST00 | 0 | Baiap2l2 | ENSMUSG00000018126.10 |
| chr16 | 35540419 | 35542272 | 17.62372 | 45.05676 | 42.40596 | Promoter | 16 | 35541145 | 35648140 | 106996 | 1 | ENSMUST00 | 0 | Sema5b | ENSMUSG00000052133.11 |
| chr17 | 24687012 | 24687865 | 17.62372 | 45.05676 | 42.40596 | Promoter | 17 | 24686633 | 24688199 | 1567 | 2 | ENSMUST00 | 334 | Syngr3 | ENSMUSG00000007021.4 |
| chr18 | 84850493 | 84851186 | 17.62372 | 45.05676 | 42.40596 | Promoter | 18 | 84851338 | 84880401 | 29064 | 1 | ENSMUST00 | -152 | Cyb5a | ENSMUSG00000024646.8 |
| chr2 | 180954236 | 180955116 | 17.62372 | 45.05676 | 42.40596 | Promoter | 2 | 180934772 | 180954699 | 19928 | 2 | ENSMUST00 | 0 | Nkain4 | ENSMUSG00000027574.10 |
| chr4 | 139074592 | 139075826 | 17.62372 | 45.05676 | 42.40596 | Promoter | 4 | 139061108 | 139075570 | 14463 | 2 | ENSMUST00 | 0 | Htr6 | ENSMUSG00000028747.5 |
| chr7 | 46100418 | 46101074 | 18.04534 | 45.02049 | 42.3703 | Promoter | 7 | 46098725 | 46099897 | 1173 | 2 | ENSMUST00 | -522 | Kcnj11 | ENSMUSG00000096146.1 |
| chr6 | 127887518 | 127888088 | 18.00818 | 44.95724 | 42.30725 | Promoter | 6 | 127887589 | 127953031 | 65443 | 1 | ENSMUST00 | 0 | Tspan11 | ENSMUSG00000030351.4 |
| chr10 | 43021009 | 43022292 | 17.55469 | 44.93826 | 42.28838 | Exon (ENSMU | 10 | 42939881 | 42958721 | 18841 | 1 | ENSMUST00 | 81129 | Scml4 | ENSMUSG00000044770.9 |
| chr11 | 7062914 | 7063460 | 17.97118 | 44.89424 | 42.24445 | Promoter | 11 | 7063489 | 7178506 | 115018 | 1 | ENSMUST00 | -29 | Adcy1 | ENSMUSG00000020431.5 |
| chr11 | 96340714 | 96341283 | 17.97118 | 44.89424 | 42.24445 | Promoter | 11 | 96340945 | 96341774 | 830 | 1 | ENSMUST00 | 0 | Hoxb3 | ENSMUSG00000048763.6 |
| chr1 | 38835598 | 38836828 | 17.4862 | 44.82059 | 42.1711 | Promoter | 1 | 38793669 | 38836090 | 42422 | 2 | ENSMUST00 | 0 | Lonrf2 | ENSMUSG00000048814.5 |
| chr1 | 132067516 | 132068285 | 17.04805 | 44.82024 | 42.17082 | Promoter | 1 | 132022806 | 132067911 | 45106 | 2 | ENSMUST00 | 0 | Mfsd4 | ENSMUSG00000059149.12 |
| chr11 | 3761662 | 3762496 | 17.04805 | 44.82024 | 42.17082 | Intron (ENSM | 11 | 3717852 | 3773789 | 55938 | 2 | ENSMUST00 | 11293 | Osbp2 | ENSMUSG00000020435.12 |
| chr19 | 6084044 | 6085628 | 17.04805 | 44.82024 | 42.17082 | Promoter | 19 | 6084983 | 6091777 | 6795 | 1 | ENSMUST00 | 0 | Cdca5 | ENSMUSG00000024791.5 |
| chr5 | 24474087 | 24475301 | 17.04805 | 44.82024 | 42.17082 | Intron (ENSM | 5 | 24452177 | 24502047 | 49871 | 1 | ENSMUST00 | 21911 | Agap3 | ENSMUSG00000023353.9 |
| chr5 | 117362867 | 117364306 | 17.04805 | 44.82024 | 42.17082 | Promoter | 5 | 117363522 | 117377807 | 14286 | 1 | ENSMUST00 | 0 | Wsb2 | ENSMUSG00000029364.10 |
| chrX | 142630711 | 142631903 | 17.04805 | 44.82024 | 42.17082 | Exon (ENSMU | 20 | 142631483 | 142634015 | 2533 | 2 | ENSMUST00 | 2112 | Gm5763 | ENSMUSG00000048600.7 |
| chr1 | 163311933 | 163313000 | 13.52062 | 44.78815 | 42.13932 | Promoter | 1 | 163252866 | 163313668 | 60803 | 2 | ENSMUST00 | 668 | Prrx1 | ENSMUSG00000026586.11 |
| chr17 | 36041824 | 36043783 | 13.52062 | 44.78815 | 42.13932 | Promoter | 17 | 36037128 | 36042500 | 5373 | 2 | ENSMUST00 | 0 | H2-T22 | ENSMUSG00000056116.13 |
| chr3 | 89337430 | 89338226 | 13.52062 | 44.78815 | 42.13932 | Promoter | 3 | 89333390 | 89338028 | 4639 | 2 | ENSMUST00 | 0 | Efna4 | ENSMUSG00000028040.7 |
| chr8 | 110721059 | 110721907 | 13.52062 | 44.78815 | 42.13932 | Promoter | 8 | 110721476 | 110741400 | 19925 | 1 | ENSMUST00 | 0 | Mtss1l | ENSMUSG00000033763.9 |
| chr1 | 55225946 | 55227077 | 17.89762 | 44.76895 | 42.12037 | Promoter | 1 | 55170159 | 55226782 | 56624 | 2 | ENSMUST00 | 0 | Rftn2 | ENSMUSG00000025978.9 |
| chr1 | 169531574 | 169532226 | 17.89762 | 44.76895 | 42.12037 | Promoter | 1 | 169497934 | 169531444 | 33531 | 2 | ENSMUST00 | -111 | Nuf2 | ENSMUSG00000026683.9 |
| chr13 | 38150957 | 38153283 | 17.89762 | 44.76895 | 42.12037 | Promoter | 13 | 38151328 | 38198577 | 47250 | 1 | ENSMUST00 | 0 | Dsp | ENSMUSG00000054889.4 |
| chr9 | 119087813 | 119089048 | 17.89762 | 44.76895 | 42.12037 | Intron (ENSM | 9 | 119071527 | 119093496 | 21970 | 2 | ENSMUST00 | 4448 | Plcd1 | ENSMUSG00000010660.2 |
| chr9 | 67759714 | 67760836 | 17.89762 | 44.76895 | 42.12037 | Promoter | 9 | 67758712 | 67760208 | 1497 | 2 | ENSMUST00 | 0 | M5C1000I18 | ENSMUSG00000055125.6 |
| chr16 | 17804205 | 17805003 | 16.98498 | 44.71047 | 42.06234 | Exon (ENSMU | 16 | 17797282 | 17808287 | 11006 | 1 | ENSMUST00 | 6924 | Scarf2 | ENSMUSG00000012017.3 |
| chr4 | 137795885 | 137796518 | 17.41825 | 44.70375 | 42.05574 | Promoter | 4 | 137741733 | 137796384 | 54652 | 2 | ENSMUST00 | 0 | Alpl | ENSMUSG00000028766.5 |
| chr15 | 103085884 | 103086945 | 16.53312 | 44.67334 | 42.02541 | Distal Interge | 15 | 103096151 | 103102111 | 5961 | 2 | ENSMUST00 | 15166 | RP24-89G21 | ENSMUSG00000101356.1 |
| chr9 | 21181427 | 21181966 | 16.53312 | 44.67334 | 42.02541 | Promoter | 9 | 21181616 | 21211627 | 30012 | 1 | ENSMUST00 | 0 | Pde4a | ENSMUSG00000032177.11 |
| chr9 | 122902886 | 122903688 | 16.53312 | 44.67334 | 42.02541 | Promoter | 9 | 122885080 | 122903184 | 18105 | 2 | ENSMUST00 | 0 | A530083I20F | ENSMUSG00000097322.2 |
| chr2 | 118702445 | 118704619 | 15.26979 | 44.6239 | 41.97645 | Promoter | 2 | 118699103 | 118703963 | 4861 | 2 | ENSMUST00 | 0 | Ankrd63 | ENSMUSG00000078137.1 |
| chr7 | 29307559 | 29308299 | 16.0698 | 44.59954 | 41.95213 | Promoter | 7 | 29307924 | 29316896 | 8973 | 1 | ENSMUST00 | 0 | Dpf1 | ENSMUSG00000030584.8 |
| chr6 | 21214081 | 21215743 | 17.78841 | 44.58281 | 41.93571 | Promoter | 6 | 21215503 | 21729805 | 514303 | 1 | ENSMUST00 | 0 | Kcnd2 | ENSMUSG00000060882.5 |
| chr8 | 77723660 | 77724600 | 17.78841 | 44.58281 | 41.93571 | Promoter | 8 | 77663031 | 77724436 | 61406 | 2 | ENSMUST00 | 0 | Ednra | ENSMUSG00000031616.7 |

| | | | | | | | | | | | | | | | |
|---|---|---|---|---|---|---|---|---|---|---|---|---|---|---|---|
| chr11 | 52003475 | 52004117 | 17.28391 | 44.47248 | 41.82566 | Promoter | 11 | 52004221 | 52033802 | 29582 | 1 | ENSMUST00 | -104 | Cdkl3 | ENSMUSG00000020389.13 |
| chr19 | 10388504 | 10389864 | 17.28391 | 44.47248 | 41.82566 | Promoter | 19 | 10389090 | 10453181 | 64092 | 1 | ENSMUST00 | 0 | Syt7 | ENSMUSG00000024743.9 |
| chr6 | 124805331 | 124806580 | 17.28391 | 44.47248 | 41.82566 | Distal Interge | 6 | 124808661 | 124810619 | 1959 | 1 | ENSMUST00 | -2081 | Spsb2 | ENSMUSG00000038451.8 |
| chr6 | 29432990 | 29433840 | 17.28391 | 44.47248 | 41.82566 | Promoter | 6 | 29433153 | 29461888 | 28736 | 1 | ENSMUST00 | 0 | Flnc | ENSMUSG00000068699.7 |
| chr7 | 19473851 | 19474684 | 17.28391 | 44.47248 | 41.82566 | Distal Interge | 7 | 19460797 | 19460935 | 139 | 2 | ENSMUST00 | -12917 | Gm23099 | ENSMUSG00000065388.1 |
| chr9 | 40730639 | 40731944 | 17.28391 | 44.47248 | 41.82566 | Intron (ENSM | 9 | 40718906 | 40719986 | 1081 | 2 | ENSMUST00 | -10654 | Gm16095 | ENSMUSG00000086014.1 |
| chr11 | 3202687 | 3203561 | 8.307 | 44.44816 | 41.80192 | Promoter | 11 | 3202689 | 3213434 | 10746 | 1 | ENSMUST00 | 0 | Eif4enif1 | ENSMUSG00000020454.12 |
| chr1 | 34005763 | 34006858 | 17.68052 | 44.39876 | 41.75258 | Promoter | 1 | 34005880 | 34174795 | 168916 | 1 | ENSMUST00 | 0 | Dst | ENSMUSG00000026131.12 |
| chr10 | 128174944 | 128177059 | 14.02077 | 44.3905 | 41.74443 | Promoter | 10 | 128129470 | 128176568 | 47099 | 2 | ENSMUST00 | 0 | Rbms2 | ENSMUSG00000040043.10 |
| chr17 | 30901288 | 30902252 | 14.02077 | 44.3905 | 41.74443 | Promoter | 17 | 30901867 | 30936510 | 34644 | 1 | ENSMUST00 | 0 | Glp1r | ENSMUSG00000024027.7 |
| chr17 | 6827297 | 6828817 | 14.02077 | 44.3905 | 41.74443 | Promoter | 17 | 6798390 | 6827990 | 29601 | 2 | ENSMUST00 | 0 | Gm2885 | ENSMUSG00000097103.1 |
| chr19 | 43879292 | 43880119 | 14.02077 | 44.3905 | 41.74443 | Intron (ENSM | 19 | 43857245 | 43857351 | 107 | 2 | ENSMUST00 | -21942 | Gm24400 | ENSMUSG00000088490.1 |
| chr6 | 113326915 | 113327680 | 14.02077 | 44.3905 | 41.74443 | Promoter | 6 | 113326972 | 113334188 | 7217 | 1 | ENSMUST00 | 0 | Ogg1 | ENSMUSG00000030271.9 |
| chr11 | 101120042 | 101120889 | 17.18351 | 44.2994 | 41.65401 | Promoter | 11 | 101119938 | 101126419 | 6482 | 1 | ENSMUST00 | 105 | Tubg1 | ENSMUSG00000035198.9 |
| chr11 | 69838434 | 69839193 | 17.18351 | 44.2994 | 41.65401 | Promoter | 11 | 69838514 | 69839613 | 1100 | 1 | ENSMUST00 | 0 | Tmem256 | ENSMUSG00000070394.5 |
| chr14 | 118924900 | 118925765 | 17.18351 | 44.2994 | 41.65401 | Promoter | 14 | 118876108 | 118925314 | 49207 | 2 | ENSMUST00 | 0 | Dzip1 | ENSMUSG00000042156.10 |
| chr15 | 25364079 | 25364894 | 17.18351 | 44.2994 | 41.65401 | 3' UTR | 15 | 25363285 | 25413764 | 50480 | 2 | ENSMUST00 | 48870 | Basp1 | ENSMUSG00000045763.7 |
| chr15 | 103101488 | 103102633 | 17.18351 | 44.2994 | 41.65401 | Promoter | 15 | 103102237 | 103105069 | 2833 | 1 | ENSMUST00 | 0 | RP24-89G21. | ENSMUSG00000099758.1 |
| chr17 | 15380084 | 15381280 | 17.18351 | 44.2994 | 41.65401 | Distal Interge | 17 | 15374794 | 15376872 | 2079 | 2 | ENSMUST00 | -3213 | Dll1 | ENSMUSG00000014773.8 |
| chr2 | 28512794 | 28514028 | 17.18351 | 44.2994 | 41.65401 | Promoter | 2 | 28513125 | 28542950 | 29826 | 1 | ENSMUST00 | 0 | Ralgds | ENSMUSG00000026821.10 |
| chr2 | 70508762 | 70509795 | 17.18351 | 44.2994 | 41.65401 | Promoter | 2 | 70508819 | 70540884 | 32066 | 1 | ENSMUST00 | 0 | Erich2 | ENSMUSG00000075302.5 |
| chr2 | 150310975 | 150312075 | 17.18351 | 44.2994 | 41.65401 | 3' UTR | 2 | 150305761 | 150306516 | 756 | 1 | ENSMUST00 | 5215 | Gm14125 | ENSMUSG00000084320.2 |
| chr3 | 87885932 | 87886332 | 17.18351 | 44.2994 | 41.65401 | Promoter | 3 | 87858900 | 87885608 | 26709 | 2 | ENSMUST00 | -325 | Prcc | ENSMUSG00000004895.8 |
| chr4 | 58986362 | 58987364 | 17.18351 | 44.2994 | 41.65401 | Promoter | 4 | 58965439 | 58987119 | 21681 | 2 | ENSMUST00 | 0 | Ptgr1 | ENSMUSG00000028378.6 |
| chr5 | 101663398 | 101665950 | 17.18351 | 44.2994 | 41.65401 | Promoter | 5 | 101659177 | 101664996 | 5820 | 2 | ENSMUST00 | 0 | Nkx6-1 | ENSMUSG00000035187.9 |
| chr6 | 113494140 | 113495221 | 17.18351 | 44.2994 | 41.65401 | 3' UTR | 6 | 113492019 | 113493335 | 1317 | 1 | ENSMUST00 | 2122 | Creld1 | ENSMUSG00000030284.6 |
| chr6 | 119329143 | 119331143 | 17.18351 | 44.2994 | 41.65401 | Promoter | 6 | 119330769 | 119344042 | 13274 | 1 | ENSMUST00 | 0 | Cacna2d4 | ENSMUSG00000041460.9 |
| chr6 | 125081526 | 125081992 | 17.18351 | 44.2994 | 41.65401 | Exon (ENSMU | 6 | 125071277 | 125082472 | 11196 | 1 | ENSMUST00 | 10250 | Lpar5 | ENSMUSG00000067714.8 |
| chr7 | 119739897 | 119740178 | 17.18351 | 44.2994 | 41.65401 | Promoter | 7 | 119739926 | 119742226 | 2301 | 1 | ENSMUST00 | 0 | Gm5601 | ENSMUSG00000044341.7 |
| chr7 | 141949193 | 141950689 | 17.18351 | 44.2994 | 41.65401 | Promoter | 7 | 141949751 | 141999005 | 49255 | 1 | ENSMUST00 | 0 | Brsk2 | ENSMUSG00000053046.9 |
| chr9 | 54068054 | 54069515 | 17.18351 | 44.2994 | 41.65401 | Promoter | 9 | 54025606 | 54068411 | 42806 | 2 | ENSMUST00 | 0 | Tnfaip8l3 | ENSMUSG00000074345.3 |
| chr1 | 83159554 | 83160069 | 16.73731 | 44.27835 | 41.63479 | Promoter | 1 | 83159752 | 83179824 | 20073 | 1 | ENSMUST00 | 0 | Daw1 | ENSMUSG00000053161.11 |
| chr2 | 60036737 | 60037756 | 16.73731 | 44.27835 | 41.63479 | Promoter | 2 | 60037141 | 60038837 | 1697 | 1 | ENSMUST00 | 0 | Gm13572 | ENSMUSG00000085975.1 |
| chr8 | 70492678 | 70493568 | 16.73731 | 44.27835 | 41.63479 | Promoter | 8 | 70493158 | 70504081 | 10924 | 1 | ENSMUST00 | 0 | Crlf1 | ENSMUSG00000007888.10 |
| chr10 | 127759754 | 127761591 | 17.60932 | 44.27721 | 41.6341 | Promoter | 10 | 127759787 | 127792694 | 32908 | 1 | ENSMUST00 | 0 | Gm28046 | ENSMUSG00000099009.1 |
| chr19 | 47536467 | 47537802 | 17.60932 | 44.27721 | 41.6341 | Promoter | 19 | 47501033 | 47537119 | 36087 | 2 | ENSMUST00 | 0 | Obfc1 | ENSMUSG00000042694.12 |
| chr12 | 11455575 | 11456518 | 17.57394 | 44.21677 | 41.57382 | Promoter | 12 | 11456079 | 11462928 | 6850 | 1 | ENSMUST00 | 0 | Rad51ap2 | ENSMUSG00000086022.1 |
| chr17 | 80127395 | 80128061 | 16.24739 | 44.16666 | 41.52389 | Promoter | 17 | 80127471 | 80185101 | 57631 | 1 | ENSMUST00 | 0 | Galm | ENSMUSG00000035473.5 |
| chr4 | 129238793 | 129239660 | 16.24739 | 44.16666 | 41.52389 | Promoter | 4 | 129219578 | 129239174 | 19597 | 2 | ENSMUST00 | 0 | C77080 | ENSMUSG00000050390.7 |
| chr1 | 14755034 | 14756154 | 17.53869 | 44.15654 | 41.51399 | Promoter | 1 | 14753346 | 14755990 | 2645 | 2 | ENSMUST00 | 0 | Msc | ENSMUSG00000025930.6 |
| chr1 | 192834556 | 192835495 | 17.53869 | 44.15654 | 41.51399 | Promoter | 1 | 192833169 | 192834719 | 1551 | 2 | ENSMUST00 | 0 | A730013G03 | ENSMUSG00000047384.2 |
| chr11 | 53456881 | 53457657 | 17.53869 | 44.15654 | 41.51399 | Promoter | 11 | 53457205 | 53467766 | 10562 | 1 | ENSMUST00 | 0 | Shroom1 | ENSMUSG00000018387.7 |
| chr11 | 43766966 | 43767392 | 17.53869 | 44.15654 | 41.51399 | Distal Interge | 11 | 43729512 | 43748008 | 18497 | 2 | ENSMUST00 | -18959 | Ttc1 | ENSMUSG00000041278.5 |
| chr12 | 31264923 | 31266287 | 17.53869 | 44.15654 | 41.51399 | Promoter | 12 | 31265234 | 31329493 | 64260 | 1 | ENSMUST00 | 0 | Lamb1 | ENSMUSG00000002900.10 |
| chr16 | 77329083 | 77331387 | 17.53869 | 44.15654 | 41.51399 | Promoter | 16 | 77329180 | 77680187 | 351008 | 1 | ENSMUST00 | 0 | 2810055G20 | ENSMUSG00000090386.4 |
| chr2 | 160911747 | 160912648 | 15.40564 | 44.13477 | 41.49272 | Promoter | 2 | 160906437 | 160912317 | 5881 | 2 | ENSMUST00 | 0 | Emilin3 | ENSMUSG00000050700.6 |
| chr1 | 12990500 | 12992953 | 14.6071 | 44.0986 | 41.45673 | Promoter | 1 | 12866549 | 12992650 | 126102 | 2 | ENSMUST00 | 0 | Slco5a1 | ENSMUSG00000025938.11 |
| chr10 | 75559770 | 75560540 | 14.6071 | 44.0986 | 41.45673 | Promoter | 10 | 75550125 | 75560330 | 10206 | 2 | ENSMUST00 | 0 | Lrrc75b | ENSMUSG00000046807.9 |
| chr11 | 79704650 | 79705267 | 14.6071 | 44.0986 | 41.45673 | Promoter | 11 | 79704630 | 79705420 | 791 | 1 | ENSMUST00 | 21 | Gm11205 | ENSMUSG00000084969.1 |
| chr13 | 51792785 | 51793794 | 14.6071 | 44.0986 | 41.45673 | Promoter | 13 | 51701246 | 51793673 | 92428 | 2 | ENSMUST00 | 0 | Sema4d | ENSMUSG00000021451.10 |
| chr16 | 30308354 | 30309386 | 14.6071 | 44.0986 | 41.45673 | Exon (ENSMU | 16 | 30307686 | 30310779 | 3094 | 2 | ENSMUST00 | 1393 | Gp5 | ENSMUSG00000047953.6 |
| chr17 | 84956541 | 84957558 | 14.6071 | 44.0986 | 41.45673 | Promoter | 17 | 84956741 | 85023991 | 67251 | 1 | ENSMUST00 | 0 | Ppm1b | ENSMUSG00000061130.7 |
| chr7 | 27330451 | 27331846 | 14.6071 | 44.0986 | 41.45673 | Intron (ENSM | 7 | 27305998 | 27333613 | 27616 | 2 | ENSMUST00 | 1767 | Ltbp4 | ENSMUSG00000040488.11 |
| chr7 | 16052049 | 16053426 | 14.6071 | 44.0986 | 41.45673 | Distal Interge | 7 | 16071942 | 16096328 | 24387 | 1 | ENSMUST00 | -18516 | Zfp541 | ENSMUSG00000078796.3 |
| chr1 | 12691712 | 12692719 | 17.92175 | 44.05972 | 41.41886 | Promoter | 1 | 12692277 | 12861192 | 168916 | 1 | ENSMUST00 | 0 | Sulf1 | ENSMUSG00000016918.10 |
| chr1 | 177322626 | 177323317 | 17.92175 | 44.05972 | 41.41886 | Distal Interge | 1 | 177022116 | 177258203 | 236088 | 2 | ENSMUST00 | -64424 | Akt3 | ENSMUSG00000019699.10 |
| chr1 | 55363020 | 55363973 | 17.92175 | 44.05972 | 41.41886 | Promoter | 1 | 55300069 | 55363319 | 63251 | 2 | ENSMUST00 | 0 | Boll | ENSMUSG00000025977.10 |
| chr11 | 105125994 | 105126324 | 17.92175 | 44.05972 | 41.41886 | Promoter | 11 | 105126425 | 105140394 | 13970 | 1 | ENSMUST00 | -101 | Mettl2 | ENSMUSG00000020691.8 |
| chr11 | 101171610 | 101172765 | 17.92175 | 44.05972 | 41.41886 | Promoter | 11 | 101162679 | 101171351 | 8673 | 2 | ENSMUST00 | -260 | Plekhh3 | ENSMUSG00000035172.10 |
| chr11 | 119086431 | 119086956 | 17.92175 | 44.05972 | 41.41886 | Promoter | 11 | 119081373 | 119086221 | 4849 | 2 | ENSMUST00 | -211 | Cbx4 | ENSMUSG00000039989.7 |
| chr12 | 102948695 | 102950272 | 17.92175 | 44.05972 | 41.41886 | Promoter | 12 | 102948859 | 103184001 | 235143 | 1 | ENSMUST00 | 0 | Unc79 | ENSMUSG00000021198.11 |
| chr12 | 108765419 | 108766724 | 17.92175 | 44.05972 | 41.41886 | Distal Interge | 12 | 108792973 | 108816632 | 23660 | 1 | ENSMUST00 | -26249 | Yy1 | ENSMUSG00000021264.11 |
| chr15 | 88837190 | 88838108 | 17.92175 | 44.05972 | 41.41886 | Distal Interge | 15 | 88835300 | 88835406 | 107 | 1 | ENSMUST00 | 1891 | Gm23144 | ENSMUSG00000093831.1 |
| chr17 | 83350276 | 83350807 | 17.92175 | 44.05972 | 41.41886 | Promoter | 17 | 83350931 | 83480361 | 129431 | 1 | ENSMUST00 | -124 | Eml4 | ENSMUSG00000032624.10 |
| chr17 | 32162155 | 32162976 | 17.92175 | 44.05972 | 41.41886 | Intron (ENSM | 17 | 32120820 | 32166880 | 46061 | 2 | ENSMUST00 | 3904 | Notch3 | ENSMUSG00000038146.7 |
| chr17 | 69475459 | 69475891 | 17.92175 | 44.05972 | 41.41886 | Intron (ENSM | 17 | 69439326 | 69489233 | 49908 | 1 | ENSMUST00 | 36134 | A330050F15 | ENSMUSG00000091636.1 |
| chr2 | 30807144 | 30807622 | 17.92175 | 44.05972 | 41.41886 | Promoter | 2 | 30807826 | 30823033 | 15208 | 1 | ENSMUST00 | -204 | Ntmt1 | ENSMUSG00000026857.4 |
| chr2 | 70474235 | 70475430 | 17.92175 | 44.05972 | 41.41886 | Promoter | 2 | 70474923 | 70477729 | 2807 | 1 | ENSMUST00 | 0 | Sp5 | ENSMUSG00000075304.2 |
| chr4 | 122960305 | 122961349 | 17.92175 | 44.05972 | 41.41886 | Promoter | 4 | 122946850 | 122961152 | 14303 | 2 | ENSMUST00 | 0 | Mfsd2a | ENSMUSG00000028655.6 |
| chr4 | 46664438 | 46664986 | 17.92175 | 44.05972 | 41.41886 | 3' UTR | 4 | 46632580 | 46650209 | 17630 | 2 | ENSMUST00 | -14230 | Tbc1d2 | ENSMUSG00000039813.9 |
| chr5 | 99036519 | 99037492 | 17.92175 | 44.05972 | 41.41886 | Promoter | 5 | 98929773 | 99037351 | 107579 | 2 | ENSMUST00 | 0 | Prkg2 | ENSMUSG00000029334.9 |
| chr6 | 87590248 | 87591207 | 17.92175 | 44.05972 | 41.41886 | Promoter | 6 | 87578593 | 87590701 | 12109 | 2 | ENSMUST00 | 0 | Prokr1 | ENSMUSG00000049409.7 |
| chr8 | 71725301 | 71726001 | 17.92175 | 44.05972 | 41.41886 | Promoter | 8 | 71708394 | 71725716 | 17323 | 2 | ENSMUST00 | 0 | Fcho1 | ENSMUSG00000070000.8 |
| chr9 | 107606557 | 107607592 | 17.92175 | 44.05972 | 41.41886 | Promoter | 9 | 107598354 | 107606656 | 8303 | 2 | ENSMUST00 | 0 | Sema3b | ENSMUSG00000057969.10 |
| chrX | 20848525 | 20849511 | 17.92175 | 44.05972 | 41.41886 | Promoter | 20 | 20848543 | 20860519 | 11977 | 1 | ENSMUST00 | 0 | Araf | ENSMUSG00000001127.7 |
| chrX | 20950224 | 20951147 | 17.92175 | 44.05972 | 41.41886 | Promoter | 20 | 20933395 | 20950608 | 17214 | 2 | ENSMUST00 | 0 | Elk1 | ENSMUSG00000009406.8 |
| chrX | 75873307 | 75875446 | 17.92175 | 44.05972 | 41.41886 | Promoter | 20 | 75785654 | 75875182 | 89529 | 2 | ENSMUST00 | 0 | Pls3 | ENSMUSG00000016382.10 |
| chrX | 152001783 | 152002585 | 17.92175 | 44.05972 | 41.41886 | Promoter | 20 | 152001845 | 152004442 | 2598 | 1 | ENSMUST00 | 0 | Hsd17b10 | ENSMUSG00000025260.5 |
| chrX | 151016646 | 151018076 | 17.92175 | 44.05972 | 41.41886 | Promoter | 20 | 150983141 | 151017322 | 34182 | 2 | ENSMUST00 | 0 | Gnl3l | ENSMUSG00000025266.6 |
| chrX | 42150683 | 42153221 | 17.92175 | 44.05972 | 41.41886 | Promoter | 20 | 42150701 | 42277185 | 126485 | 1 | ENSMUST00 | 0 | Stag2 | ENSMUSG00000025862.9 |
| chrX | 57325797 | 57326841 | 17.92175 | 44.05972 | 41.41886 | Promoter | 20 | 57248987 | 57326599 | 77613 | 2 | ENSMUST00 | 0 | Arhgef6 | ENSMUSG00000031133.7 |
| chrX | 7750460 | 7751176 | 17.92175 | 44.05972 | 41.41886 | Promoter | 20 | 7738460 | 7750905 | 12446 | 2 | ENSMUST00 | 0 | Ccdc120 | ENSMUSG00000031150.7 |
| chrX | 7877866 | 7878914 | 17.92175 | 44.05972 | 41.41886 | Promoter | 20 | 7878262 | 7880322 | 2061 | 1 | ENSMUST00 | 0 | Pim2 | ENSMUSG00000031155.9 |
| chrX | 8206074 | 8207019 | 17.92175 | 44.05972 | 41.41886 | Promoter | 20 | 8193848 | 8206525 | 12678 | 2 | ENSMUST00 | 0 | Porcn | ENSMUSG00000031169.8 |
| chrX | 99042356 | 99043269 | 17.92175 | 44.05972 | 41.41886 | Promoter | 20 | 99042581 | 99074728 | 32148 | 1 | ENSMUST00 | 0 | Stard8 | ENSMUSG00000031216.8 |
| chrX | 140956024 | 140956742 | 17.92175 | 44.05972 | 41.41886 | Promoter | 20 | 140948429 | 140956689 | 8261 | 2 | ENSMUST00 | 0 | Psmd10 | ENSMUSG00000031429.9 |
| chrX | 94177439 | 94178702 | 17.92175 | 44.05972 | 41.41886 | Distal Interge | 20 | 94199641 | 94212651 | 13011 | 2 | ENSMUST00 | 33949 | Eif2s3x | ENSMUSG00000035150.10 |
| chrX | 20920167 | 20921068 | 17.92175 | 44.05972 | 41.41886 | Promoter | 20 | 20860511 | 20921004 | 60494 | 2 | ENSMUST00 | 0 | Syn1 | ENSMUSG00000037217.10 |
| chrX | 12091578 | 12092219 | 17.92175 | 44.05972 | 41.41886 | Intron (ENSM | 20 | 12036740 | 12080553 | 43814 | 2 | ENSMUST00 | -11026 | Bcor | ENSMUSG00000040363.9 |
| chrX | 12127816 | 12129066 | 17.92175 | 44.05972 | 41.41886 | Promoter | 20 | 12058981 | 12128400 | 69420 | 2 | ENSMUST00 | 0 | Bcor | ENSMUSG00000040363.9 |
| chrX | 137037403 | 137038705 | 17.92175 | 44.05972 | 41.41886 | Promoter | 20 | 136981116 | 137038300 | 57185 | 2 | ENSMUST00 | 0 | Slc25a53 | ENSMUSG00000044348.9 |
| chrX | 101221672 | 101222606 | 17.92175 | 44.05972 | 41.41886 | Promoter | 20 | 101097786 | 101222553 | 124768 | 2 | ENSMUST00 | 0 | Snx12 | ENSMUSG00000046032.11 |
| chrX | 153006655 | 153007849 | 17.92175 | 44.05972 | 41.41886 | Promoter | 20 | 153006469 | 153009416 | 2948 | 1 | ENSMUST00 | 187 | Usp51 | ENSMUSG00000067215.5 |
| chrX | 95025694 | 95027004 | 17.92175 | 44.05972 | 41.41886 | Promoter | 20 | 95022510 | 95026682 | 4173 | 2 | ENSMUST00 | 0 | Spin4 | ENSMUSG00000071722.4 |
| chrX | 103493504 | 103494554 | 17.92175 | 44.05972 | 41.41886 | Promoter | 20 | 103493558 | 103506425 | 12868 | 1 | ENSMUST00 | 0 | Jpx | ENSMUSG00000097571.3 |
| chr11 | 103362467 | 103363175 | 17.46863 | 44.03677 | 41.39737 | Promoter | 11 | 103331498 | 103363692 | 32195 | 2 | ENSMUST00 | 0 | Arhgap27 | ENSMUSG00000034255.12 |
| chr10 | 127533907 | 127535067 | 16.14593 | 43.98602 | 41.34693 | Promoter | 10 | 127525473 | 127534559 | 9087 | 2 | ENSMUST00 | 0 | Nxph4 | ENSMUSG00000040258.6 |
| chr10 | 127326022 | 127327272 | 16.14593 | 43.98602 | 41.34693 | Exon (ENSMU | 10 | 127323727 | 127329943 | 6217 | 1 | ENSMUST00 | 2296 | Arhgap9 | ENSMUSG00000040345.9 |
| chr10 | 124288068 | 124289218 | 16.14593 | 43.98602 | 41.34693 | Distal Interge | 10 | 124455059 | 124455317 | 259 | 1 | ENSMUST00 | -165841 | Gm23777 | ENSMUSG00000089619.1 |
| chr10 | 5549973 | 5550939 | 16.14593 | 43.98602 | 41.34693 | Promoter | 10 | 5336949 | 5550692 | 213744 | 2 | ENSMUST00 | 0 | Syne1 | ENSMUSG00000096054.2 |
| chr11 | 77800559 | 77802076 | 16.14593 | 43.98602 | 41.34693 | Promoter | 11 | 77801298 | 77865837 | 64540 | 1 | ENSMUST00 | 0 | Myo18a | ENSMUSG00000000631.14 |
| chr13 | 63244274 | 63245622 | 16.14593 | 43.98602 | 41.34693 | Intron (ENSM | 13 | 63282142 | 63326096 | 43955 | 1 | ENSMUST00 | -36520 | 2010111I01R | ENSMUSG00000021458.11 |
| chr15 | 7813062 | 7814102 | 16.14593 | 43.98602 | 41.34693 | Intron (ENSM | 15 | 7811011 | 7837575 | 26565 | 1 | ENSMUST00 | 2052 | Gdnf | ENSMUSG00000022144.3 |
| chr15 | 85580363 | 85582124 | 16.14593 | 43.98602 | 41.34693 | Promoter | 15 | 85535439 | 85580729 | 45291 | 2 | ENSMUST00 | 0 | Wnt7b | ENSMUSG00000022382.9 |
| chr17 | 47832749 | 47834530 | 16.14593 | 43.98602 | 41.34693 | Promoter | 17 | 47815328 | 47833241 | 17914 | 2 | ENSMUST00 | 0 | Mdfi | ENSMUSG00000032717.9 |
| chr17 | 26946217 | 26947265 | 16.14593 | 43.98602 | 41.34693 | Promoter | 17 | 26948056 | 26948119 | 64 | 2 | ENSMUST00 | 854 | Mir3083 | ENSMUSG00000093118.1 |
| chr18 | 56755851 | 56756882 | 16.14593 | 43.98602 | 41.34693 | Distal Interge | 18 | 56769717 | 56801688 | 31972 | 1 | ENSMUST00 | -12835 | Gm15345 | ENSMUSG00000087597.1 |
| chr2 | 3418559 | 3419503 | 16.14593 | 43.98602 | 41.34693 | Promoter | 2 | 3409043 | 3419019 | 9977 | 2 | ENSMUST00 | 0 | Meig1 | ENSMUSG00000026650.10 |
| chr2 | 93642426 | 93643610 | 16.14593 | 43.98602 | 41.34693 | Promoter | 2 | 93642388 | 93681339 | 38952 | 1 | ENSMUST00 | 39 | Alx4 | ENSMUSG00000040310.7 |
| chr2 | 113440160 | 113441633 | 16.14593 | 43.98602 | 41.34693 | Promoter | 2 | 113441064 | 113709855 | 268792 | 1 | ENSMUST00 | 0 | Fmn1 | ENSMUSG00000044042.13 |
| chr2 | 30801081 | 30802017 | 16.14593 | 43.98602 | 41.34693 | Promoter | 2 | 30795301 | 30801707 | 6407 | 2 | ENSMUST00 | 0 | 1700001O22 | ENSMUSG00000044320.9 |
| chr3 | 124320828 | 124321483 | 16.14593 | 43.98602 | 41.34693 | Promoter | 3 | 124321037 | 124323260 | 2224 | 1 | ENSMUST00 | 0 | Tram1l1 | ENSMUSG00000044528.4 |
| chr4 | 9843696 | 9845446 | 16.14593 | 43.98602 | 41.34693 | Promoter | 4 | 9844372 | 9862345 | 17974 | 1 | ENSMUST00 | 0 | Gdf6 | ENSMUSG00000051279.2 |
| chr4 | 88867944 | 88868447 | 16.14593 | 43.98602 | 41.34693 | Promoter | 4 | 88867882 | 88868379 | 498 | 2 | ENSMUST00 | 0 | 4930553M12 | ENSMUSG00000054351.1 |
| chr4 | 140813959 | 140814941 | 16.14593 | 43.98602 | 41.34693 | 3' UTR | 4 | 140810993 | 140817511 | 6519 | 1 | ENSMUST00 | 2967 | Gm13032 | ENSMUSG00000085918.1 |
| chr5 | 30104850 | 30105636 | 16.14593 | 43.98602 | 41.34693 | Promoter | 5 | 30105161 | 30118378 | 13218 | 1 | ENSMUST00 | 0 | Gareml | ENSMUSG00000044576.6 |
| chr6 | 101377007 | 101378628 | 16.14593 | 43.98602 | 41.34693 | Promoter | 6 | 101149609 | 101377897 | 228289 | 2 | ENSMUST00 | 0 | Pdzrn3 | ENSMUSG00000035357.11 |
| chr7 | 118243892 | 118244454 | 16.14593 | 43.98602 | 41.34693 | Promoter | 7 | 118243865 | 118245551 | 1687 | 1 | ENSMUST00 | 28 | 4930583K01I | ENSMUSG00000055159.7 |
| chr7 | 30650347 | 30650947 | 16.14593 | 43.98602 | 41.34693 | Promoter | 7 | 30650385 | 30652711 | 2327 | 1 | ENSMUST00 | 0 | Gm26610 | ENSMUSG00000097674.1 |
| chr8 | 71670669 | 71672081 | 16.14593 | 43.98602 | 41.34693 | Promoter | 8 | 71624417 | 71671757 | 47341 | 2 | ENSMUST00 | 0 | Unc13a | ENSMUSG00000034799.11 |
| chr8 | 64692485 | 64693243 | 16.14593 | 43.98602 | 41.34693 | Promoter | 8 | 64592558 | 64693040 | 100483 | 2 | ENSMUST00 | 0 | Cpe | ENSMUSG00000037852.7 |
| chrX | 101253735 | 101255612 | 16.14593 | 43.98602 | 41.34693 | Promoter | 20 | 101254528 | 101260873 | 6346 | 1 | ENSMUST00 | 0 | Foxo4 | ENSMUSG00000042903.7 |
| chrX | 101273700 | 101274973 | 16.14593 | 43.98602 | 41.34693 | Promoter | 20 | 101274030 | 101297465 | 23436 | 1 | ENSMUST00 | 0 | Med12 | ENSMUSG00000079487.6 |
| chr5 | 29195452 | 29196494 | 17.4338 | 43.97722 | 41.34015 | Promoter | 5 | 29195992 | 29203135 | 7144 | 1 | ENSMUST00 | 0 | Rnf32 | ENSMUSG00000029130.7 |
| chr7 | 18839929 | 18840777 | 17.4338 | 43.97722 | 41.34015 | Promoter | 7 | 18839966 | 18858653 | 18688 | 1 | ENSMUST00 | 0 | Mill2 | ENSMUSG00000040987.9 |
| chr1 | 33907302 | 33908701 | 15.30398 | 43.94659 | 41.30965 | Promoter | 1 | 33908225 | 34183904 | 275680 | 1 | ENSMUST00 | 0 | Dst | ENSMUSG00000026131.12 |
| chr11 | 18877713 | 18878078 | 15.30398 | 43.94659 | 41.30965 | Downstream | 11 | 18854043 | 18874309 | 20267 | 2 | ENSMUST00 | -3405 | 8430419K02I | ENSMUSG00000085743.1 |
| chr12 | 80131939 | 80132873 | 15.30398 | 43.94659 | 41.30965 | Promoter | 12 | 80120548 | 80132844 | 12297 | 2 | ENSMUST00 | 0 | 2310015A10I | ENSMUSG00000097729.1 |

| | | | | | | | | | | | | | | | |
|---|---|---|---|---|---|---|---|---|---|---|---|---|---|---|---|
| chr15 | 76878873 | 76879841 | 15.30398 | 43.94659 | 41.30965 | Promoter | 15 | 76879259 | 76892392 | 13134 | 1 | ENSMUST00 | 0 | Zfp7 | ENSMUSG00000033669.7 |
| chr15 | 80254431 | 80255089 | 15.30398 | 43.94659 | 41.30965 | Promoter | 15 | 80255184 | 80257540 | 2357 | 1 | ENSMUST00 | -95 | Atf4 | ENSMUSG00000042406.7 |
| chr15 | 76639563 | 76639988 | 15.30398 | 43.94659 | 41.30965 | Promoter | 15 | 76626238 | 76639958 | 13721 | 2 | ENSMUST00 | 0 | Tonsl | ENSMUSG00000059323.8 |
| chr17 | 87530230 | 87532496 | 15.30398 | 43.94659 | 41.30965 | Distal Interge | 17 | 87490748 | 87502616 | 11869 | 1 | ENSMUST00 | 39483 | RP23-243O1 | ENSMUSG00000100847.1 |
| chr18 | 69592339 | 69594600 | 15.30398 | 43.94659 | 41.30965 | Promoter | 18 | 69593417 | 69683252 | 89836 | 1 | ENSMUST00 | 0 | Tcf4 | ENSMUSG00000053477.10 |
| chr4 | 88763063 | 88763667 | 15.30398 | 43.94659 | 41.30965 | Promoter | 4 | 88763709 | 88771606 | 7898 | 1 | ENSMUST00 | -42 | Gm13284 | ENSMUSG00000082789.3 |
| chr6 | 88499874 | 88500453 | 15.30398 | 43.94659 | 41.30965 | Distal Interge | 6 | 88504801 | 88506371 | 1571 | 2 | ENSMUST00 | 5918 | Sec61a1 | ENSMUSG00000030082.9 |
| chr6 | 24955049 | 24956514 | 15.30398 | 43.94659 | 41.30965 | Promoter | 6 | 24951141 | 24956297 | 5157 | 2 | ENSMUST00 | 0 | Tmem229a | ENSMUSG00000048022.11 |
| chr8 | 121125330 | 121126765 | 15.30398 | 43.94659 | 41.30965 | Intron (ENSM | 8 | 121127940 | 121130644 | 2705 | 1 | ENSMUST00 | -1175 | Foxl1 | ENSMUSG00000097084.1 |
| chr10 | 79852880 | 79854274 | 16.95696 | 43.908 | 41.27272 | Promoter | 10 | 79854427 | 79864432 | 10006 | 1 | ENSMUST00 | -153 | Ptbp1 | ENSMUSG00000006498.11 |
| chr10 | 63243089 | 63243701 | 16.95696 | 43.908 | 41.27272 | Promoter | 10 | 63243797 | 63317881 | 74085 | 1 | ENSMUST00 | -96 | Herc4 | ENSMUSG00000020064.8 |
| chr11 | 31368365 | 31370319 | 16.95696 | 43.908 | 41.27272 | Promoter | 11 | 31357307 | 31370074 | 12768 | 2 | ENSMUST00 | 0 | Stc2 | ENSMUSG00000020303.2 |
| chr11 | 107025535 | 107026285 | 16.95696 | 43.908 | 41.27272 | Promoter | 11 | 107025681 | 107025907 | 227 | 1 | ENSMUST00 | 0 | Gm11705 | ENSMUSG00000086772.1 |
| chr18 | 7692569 | 7693676 | 16.95696 | 43.908 | 41.27272 | Distal Interge | 18 | 7347962 | 7626863 | 278902 | 2 | ENSMUST00 | -65707 | Mpp7 | ENSMUSG00000057440.7 |
| chr2 | 25291872 | 25292980 | 16.95696 | 43.908 | 41.27272 | Promoter | 2 | 25291220 | 25298925 | 7706 | 1 | ENSMUST00 | 653 | Grin1os | ENSMUSG00000085830.1 |
| chr4 | 117882974 | 117883799 | 16.95696 | 43.908 | 41.27272 | Promoter | 4 | 117869260 | 117883155 | 13896 | 2 | ENSMUST00 | 0 | B4galt2 | ENSMUSG00000028541.9 |
| chr4 | 126202918 | 126203353 | 16.95696 | 43.908 | 41.27272 | Promoter | 4 | 126164082 | 126202760 | 38679 | 2 | ENSMUST00 | -159 | Thrap3 | ENSMUSG00000043962.11 |
| chr14 | 79300605 | 79301938 | 17.81477 | 43.87807 | 41.24343 | Promoter | 14 | 79288756 | 79301645 | 12890 | 2 | ENSMUST00 | 0 | Rgcc | ENSMUSG00000022018.7 |
| chr19 | 29501028 | 29501592 | 17.81477 | 43.87807 | 41.24343 | Intron (ENSM | 19 | 29503663 | 29521989 | 18327 | 2 | ENSMUST00 | 20397 | A930007I19R | ENSMUSG00000097855.2 |
| chr2 | 152143070 | 152144393 | 17.81477 | 43.87807 | 41.24343 | Promoter | 2 | 152143561 | 152149097 | 5537 | 1 | ENSMUST00 | 0 | Tcf15 | ENSMUSG00000068079.5 |
| chr7 | 140048638 | 140050028 | 17.36457 | 43.85876 | 41.22438 | Promoter | 7 | 140046157 | 140049089 | 2933 | 2 | ENSMUST00 | 0 | Msx3 | ENSMUSG00000025469.10 |
| chr8 | 69822000 | 69823147 | 17.77762 | 43.81499 | 41.18071 | Promoter | 8 | 69822429 | 69831101 | 8673 | 1 | ENSMUST00 | 0 | Lpar2 | ENSMUSG00000031861.10 |
| chr8 | 61759431 | 61760301 | 17.29589 | 43.74115 | 41.10756 | Promoter | 8 | 61513252 | 61760089 | 246838 | 2 | ENSMUST00 | 0 | Palld | ENSMUSG00000058056.10 |
| chr1 | 166378713 | 166379179 | 16.82961 | 43.68747 | 41.0541 | Promoter | 1 | 166379097 | 166393621 | 14525 | 1 | ENSMUST00 | 0 | Tada1 | ENSMUSG00000026563.8 |
| chr16 | 92399134 | 92400236 | 15.97138 | 43.67439 | 41.04113 | Promoter | 16 | 92391953 | 92400077 | 8125 | 2 | ENSMUST00 | 0 | Rcan1 | ENSMUSG00000022951.10 |
| chr5 | 43807866 | 43808565 | 17.66712 | 43.62722 | 40.9943 | Intron (ENSM | 5 | 43818886 | 43825404 | 6519 | 1 | ENSMUST00 | -10321 | Bst1 | ENSMUSG00000029082.12 |
| chr8 | 41038344 | 41040438 | 17.22774 | 43.62438 | 40.99156 | Promoter | 8 | 41014376 | 41039587 | 25212 | 2 | ENSMUST00 | 0 | Mtus1 | ENSMUSG00000045636.11 |
| chr1 | 78310039 | 78310931 | 17.19387 | 43.56631 | 40.93357 | Promoter | 1 | 78310345 | 78420289 | 109945 | 1 | ENSMUST00 | 0 | Sgpp2 | ENSMUSG00000032908.9 |
| chr11 | 102623539 | 102624414 | 17.19387 | 43.56631 | 40.93357 | Promoter | 11 | 102621318 | 102624366 | 3049 | 2 | ENSMUST00 | 0 | 2810433D01 | ENSMUSG00000086283.2 |
| chr19 | 57197103 | 57197837 | 17.19387 | 43.56631 | 40.93357 | Promoter | 19 | 57033274 | 57197631 | 164358 | 2 | ENSMUST00 | 0 | Ablim1 | ENSMUSG00000025085.11 |
| chr6 | 145340930 | 145341971 | 17.19387 | 43.56631 | 40.93357 | Distal Interge | 6 | 145315613 | 145316608 | 996 | 2 | ENSMUST00 | -24323 | Gm15707 | ENSMUSG00000083962.1 |
| chr7 | 45369930 | 45371040 | 17.19387 | 43.56631 | 40.93357 | Promoter | 7 | 45367891 | 45370583 | 2693 | 2 | ENSMUST00 | 0 | Lin7b | ENSMUSG00000003872.7 |
| chr1 | 182959377 | 182960327 | 17.63059 | 43.56512 | 40.9329 | Intron (ENSM | 1 | 182954788 | 182976044 | 21257 | 1 | ENSMUST00 | 4590 | Tlr5 | ENSMUSG00000079164.3 |
| chr5 | 135106555 | 135107454 | 13.25551 | 43.5229 | 40.89079 | Promoter | 5 | 135106891 | 135123042 | 16152 | 1 | ENSMUST00 | 0 | Mlxipl | ENSMUSG00000005373.8 |
| chr16 | 23982418 | 23983788 | 17.59421 | 43.50326 | 40.87146 | Intron (ENSM | 16 | 23965052 | 23988852 | 23801 | 2 | ENSMUST00 | 5064 | Bcl6 | ENSMUSG00000022508.5 |
| chr11 | 30884899 | 30885770 | 17.55798 | 43.44164 | 40.81004 | Promoter | 11 | 30885358 | 30893729 | 8372 | 1 | ENSMUST00 | 0 | Gpr75 | ENSMUSG00000043999.6 |
| chr5 | 24684829 | 24685667 | 17.55798 | 43.44164 | 40.81004 | Promoter | 5 | 24685815 | 24710555 | 24741 | 1 | ENSMUST00 | -148 | Nub1 | ENSMUSG00000028954.7 |
| chr9 | 64049678 | 64050557 | 16.64214 | 43.36206 | 40.73079 | Promoter | 9 | 64049829 | 64054100 | 4272 | 1 | ENSMUST00 | 0 | 1110036E04 | ENSMUSG00000097036.1 |
| chr10 | 79936883 | 79937621 | 17.45018 | 43.2582 | 40.62776 | Intron (ENSM | 10 | 79930424 | 79931218 | 795 | 1 | ENSMUST00 | 6460 | Arid3a | ENSMUSG00000019564.7 |
| chr4 | 62702958 | 62704727 | 17.45018 | 43.2582 | 40.62776 | 3' UTR | 4 | 62705981 | 62725737 | 19757 | 1 | ENSMUST00 | -1254 | Gm11210 | ENSMUSG00000085223.1 |
| chr6 | 89362742 | 89363328 | 17.45018 | 43.2582 | 40.62776 | Promoter | 6 | 89316316 | 89362613 | 46298 | 2 | ENSMUST00 | -130 | Plxna1 | ENSMUSG00000030084.6 |
| chr1 | 63272916 | 63274416 | 16.14863 | 43.24384 | 40.61377 | Promoter | 1 | 63273265 | 63314576 | 41312 | 1 | ENSMUST00 | 0 | Zdbf2 | ENSMUSG00000027520.10 |
| chr15 | 36374035 | 36375538 | 16.14863 | 43.24384 | 40.61377 | Distal Interge | 15 | 36308144 | 36308956 | 813 | 2 | ENSMUST00 | -65080 | Rpl7a-ps3 | ENSMUSG00000068579.4 |
| chr6 | 39871178 | 39873476 | 15.30291 | 43.20574 | 40.57601 | Promoter | 6 | 39873271 | 40255075 | 381805 | 1 | ENSMUST00 | 0 | Tmem178b | ENSMUSG00000057716.6 |
| chr17 | 5189499 | 5190676 | 15.70458 | 43.19576 | 40.56621 | Intron (ENSM | 17 | 5110781 | 5112119 | 1339 | 2 | ENSMUST00 | -77381 | Gm15599 | ENSMUSG00000081783.1 |
| chr2 | 150217757 | 150218424 | 15.70458 | 43.19576 | 40.56621 | Promoter | 2 | 150218073 | 150240725 | 22653 | 1 | ENSMUST00 | 0 | Zfp937 | ENSMUSG00000060336.6 |
| chr3 | 118561956 | 118563094 | 15.70458 | 43.19576 | 40.56621 | Promoter | 3 | 118562129 | 119432918 | 870790 | 1 | ENSMUST00 | 0 | Dpyd | ENSMUSG00000033308.11 |
| chr3 | 90433743 | 90434232 | 17.37905 | 43.13707 | 40.50774 | Promoter | 3 | 90391380 | 90433636 | 42257 | 2 | ENSMUST00 | -108 | Ints3 | ENSMUSG00000027933.6 |
| chr12 | 103241883 | 103243036 | 13.73463 | 43.09688 | 40.46774 | Promoter | 12 | 103196908 | 103242146 | 45239 | 2 | ENSMUST00 | 0 | Prima1 | ENSMUSG00000041669.10 |
| chr12 | 99964487 | 99966311 | 13.73463 | 43.09688 | 40.46774 | Promoter | 12 | 99964499 | 100062682 | 98184 | 1 | ENSMUST00 | 0 | Kcnk13 | ENSMUSG00000045404.10 |
| chr18 | 67288324 | 67289010 | 13.73463 | 43.09688 | 40.46774 | Promoter | 18 | 67289253 | 67319146 | 29894 | 1 | ENSMUST00 | -243 | Impa2 | ENSMUSG00000024525.6 |
| chr2 | 180455867 | 180456781 | 13.73463 | 43.09688 | 40.46774 | Promoter | 2 | 180456245 | 180474867 | 18623 | 1 | ENSMUST00 | 0 | Slco4a1 | ENSMUSG00000038963.10 |
| chr3 | 37204100 | 37204763 | 13.73463 | 43.09688 | 40.46774 | Distal Interge | 3 | 37199979 | 37200979 | 1001 | 2 | ENSMUST00 | -3122 | Gm13621 | ENSMUSG00000081634.1 |
| chr4 | 46601221 | 46602300 | 13.73463 | 43.09688 | 40.46774 | Promoter | 4 | 46536943 | 46602202 | 65260 | 2 | ENSMUST00 | 0 | Coro2a | ENSMUSG00000028337.9 |
| chr4 | 47091669 | 47092843 | 13.73463 | 43.09688 | 40.46774 | Promoter | 4 | 47091909 | 47123070 | 31162 | 1 | ENSMUST00 | 0 | Galnt12 | ENSMUSG00000039774.7 |
| chr3 | 60527647 | 60528528 | 17.3437 | 43.07685 | 40.44819 | Intron (ENSM | 3 | 60501252 | 60629748 | 128497 | 1 | ENSMUST00 | 26396 | Mbnl1 | ENSMUSG00000027763.8 |
| chr6 | 83710552 | 83711584 | 17.30849 | 43.01685 | 40.38835 | Promoter | 6 | 83711264 | 83738313 | 27050 | 1 | ENSMUST00 | 0 | Vax2 | ENSMUSG00000034777.2 |
| chr11 | 97018410 | 97019205 | 16.86235 | 42.99675 | 40.36834 | Promoter | 11 | 97018733 | 97024731 | 5999 | 1 | ENSMUST00 | 0 | Sp6 | ENSMUSG00000038560.7 |
| chr13 | 21779454 | 21780876 | 16.86235 | 42.99675 | 40.36834 | Promoter | 13 | 21779883 | 21780625 | 743 | 2 | ENSMUST00 | 0 | Hist1h1b | ENSMUSG00000058773.2 |
| chr13 | 84347485 | 84349633 | 17.27343 | 42.95708 | 40.32916 | Distal Interge | 13 | 84339149 | 84340113 | 965 | 2 | ENSMUST00 | -7373 | Gm26927 | ENSMUSG00000098245.1 |
| chr14 | 18326962 | 18327382 | 17.23851 | 42.89754 | 40.26978 | Promoter | 14 | 18285213 | 18327291 | 42079 | 2 | ENSMUST00 | 0 | Ube2e1 | ENSMUSG00000021774.7 |
| chr15 | 61157707 | 61158533 | 17.23851 | 42.89754 | 40.26978 | Distal Interge | 15 | 61357871 | 61358004 | 134 | 2 | ENSMUST00 | 199471 | Gm24696 | ENSMUSG00000065812.1 |
| chr3 | 78966754 | 78967418 | 17.20373 | 42.83822 | 40.21063 | Promoter | 3 | 78966356 | 78966799 | 444 | 2 | ENSMUST00 | 0 | Gm9762 | ENSMUSG00000037096.4 |
| chr10 | 68541390 | 68542024 | 16.33881 | 42.83351 | 40.20603 | Promoter | 10 | 68430953 | 68541896 | 110944 | 2 | ENSMUST00 | 0 | 1700040L02 | ENSMUSG00000019945.4 |
| chr3 | 95819497 | 95820267 | 16.33881 | 42.83351 | 40.20603 | Promoter | 3 | 95759873 | 95818953 | 59081 | 2 | ENSMUST00 | -545 | Rprd2 | ENSMUSG00000028106.9 |
| chr1 | 77513322 | 77513852 | 16.7644 | 42.82802 | 40.20103 | Intron (ENSM | 1 | 77367189 | 77515079 | 147891 | 2 | ENSMUST00 | 1227 | Epha4 | ENSMUSG00000026235.9 |
| chr1 | 155082926 | 155084017 | 16.7644 | 42.82802 | 40.20103 | Promoter | 1 | 155063447 | 155083266 | 19820 | 2 | ENSMUST00 | 0 | Gm9530 | ENSMUSG00000096950.2 |
| chr10 | 78779759 | 78780765 | 16.7644 | 42.82802 | 40.20103 | Promoter | 10 | 78780496 | 78814825 | 34330 | 1 | ENSMUST00 | 0 | Slc1a6 | ENSMUSG00000005357.8 |
| chr11 | 121847508 | 121848408 | 16.7644 | 42.82802 | 40.20103 | Distal Interge | 11 | 121856773 | 121861227 | 4455 | 1 | ENSMUST00 | -8365 | Gm12586 | ENSMUSG00000084401.1 |
| chr12 | 85705233 | 85706182 | 16.7644 | 42.82802 | 40.20103 | Intron (ENSM | 12 | 85718477 | 85718595 | 119 | 2 | ENSMUST00 | 12413 | n-R5s64 | ENSMUSG00000075973.1 |
| chr17 | 56831298 | 56831710 | 16.7644 | 42.82802 | 40.20103 | Promoter | 17 | 56775897 | 56831008 | 55112 | 2 | ENSMUST00 | -291 | Rfx2 | ENSMUSG00000024206.9 |
| chr17 | 66017176 | 66017649 | 16.7644 | 42.82802 | 40.20103 | Intron (ENSM | 17 | 66049786 | 66050640 | 855 | 2 | ENSMUST00 | 32991 | Ankrd12 | ENSMUSG00000034647.9 |
| chr17 | 24206452 | 24207031 | 16.7644 | 42.82802 | 40.20103 | Promoter | 17 | 24175431 | 24205562 | 30132 | 2 | ENSMUST00 | -891 | Tbc1d24 | ENSMUSG00000036473.10 |
| chr17 | 24528210 | 24528903 | 16.7644 | 42.82802 | 40.20103 | Promoter | 17 | 24528476 | 24528553 | 78 | 1 | ENSMUST00 | 0 | Snord60 | ENSMUSG00000064768.1 |
| chr17 | 35185787 | 35186488 | 16.7644 | 42.82802 | 40.20103 | Intron (ENSM | 17 | 35185095 | 35188439 | 3345 | 2 | ENSMUST00 | 1951 | Lst1 | ENSMUSG00000073412.6 |
| chr3 | 29082356 | 29083287 | 16.7644 | 42.82802 | 40.20103 | Promoter | 3 | 29082577 | 29671627 | 589051 | 1 | ENSMUST00 | 0 | Egfem1 | ENSMUSG00000063600.9 |
| chr4 | 148041068 | 148042129 | 16.7644 | 42.82802 | 40.20103 | Promoter | 4 | 148041191 | 148059551 | 18361 | 1 | ENSMUST00 | 0 | Mthfr | ENSMUSG00000029009.12 |
| chr4 | 86930659 | 86931435 | 16.7644 | 42.82802 | 40.20103 | Promoter | 4 | 86930691 | 86934822 | 4132 | 1 | ENSMUST00 | 0 | Acer2 | ENSMUSG00000038007.9 |
| chr5 | 37336499 | 37337317 | 16.7644 | 42.82802 | 40.20103 | Promoter | 5 | 37299171 | 37336894 | 37724 | 2 | ENSMUST00 | 0 | Evc | ENSMUSG00000029122.6 |
| chr5 | 24383298 | 24384025 | 16.7644 | 42.82802 | 40.20103 | Promoter | 5 | 24366192 | 24383582 | 17391 | 2 | ENSMUST00 | 0 | Gm15587 | ENSMUSG00000086769.2 |
| chr6 | 5255570 | 5256501 | 16.7644 | 42.82802 | 40.20103 | Promoter | 6 | 5220852 | 5256233 | 35382 | 2 | ENSMUST00 | 0 | Pon3 | ENSMUSG00000029759.4 |
| chr6 | 59024080 | 59024888 | 16.7644 | 42.82802 | 40.20103 | Promoter | 6 | 58932090 | 59024549 | 92460 | 2 | ENSMUST00 | 0 | Fam13a | ENSMUSG00000037709.8 |
| chr9 | 74975429 | 74976867 | 16.7644 | 42.82802 | 40.20103 | Promoter | 9 | 74976111 | 75032468 | 56358 | 1 | ENSMUST00 | 0 | Fam214a | ENSMUSG00000034858.10 |
| chrX | 73824175 | 73825239 | 16.7644 | 42.82802 | 40.20103 | Promoter | 20 | 73793359 | 73824969 | 31611 | 2 | ENSMUST00 | 0 | Pdzd4 | ENSMUSG00000002006.7 |
| chrX | 74245610 | 74247227 | 16.7644 | 42.82802 | 40.20103 | Promoter | 20 | 74223461 | 74246398 | 22938 | 2 | ENSMUST00 | 0 | Flna | ENSMUSG00000031328.10 |
| chrX | 162828425 | 162829462 | 16.7644 | 42.82802 | 40.20103 | Promoter | 20 | 162778919 | 162829454 | 50536 | 2 | ENSMUST00 | 0 | Txlng | ENSMUSG00000038344.9 |
| chrX | 142681290 | 142683291 | 16.7644 | 42.82802 | 40.20103 | Promoter | 20 | 142681406 | 142843494 | 162089 | 1 | ENSMUST00 | 0 | Tmem164 | ENSMUSG00000047045.12 |
| chr2 | 25371869 | 25372995 | 12.59502 | 42.79506 | 40.1699 | Promoter | 2 | 25372321 | 25378213 | 5893 | 1 | ENSMUST00 | 0 | Sapcd2 | ENSMUSG00000026955.8 |
| chr1 | 125911335 | 125912478 | 14.29631 | 42.77194 | 40.14703 | Promoter | 1 | 125867622 | 125912214 | 44593 | 2 | ENSMUST00 | 0 | Lypd1 | ENSMUSG00000026344.4 |
| chr11 | 4217797 | 4218874 | 14.29631 | 42.77194 | 40.14703 | Promoter | 11 | 4218225 | 4222409 | 4185 | 1 | ENSMUST00 | 0 | Gatsl3 | ENSMUSG00000020424.3 |
| chr11 | 120346386 | 120347030 | 14.29631 | 42.77194 | 40.14703 | Promoter | 11 | 120346382 | 120346977 | 596 | 2 | ENSMUST00 | 0 | Actg1 | ENSMUSG00000062825.10 |
| chr11 | 50358918 | 50359947 | 14.29631 | 42.77194 | 40.14703 | 5' UTR | 11 | 50357785 | 50357879 | 95 | 2 | ENSMUST00 | -1040 | Mir804 | ENSMUSG00000076450.1 |
| chr16 | 50736275 | 50737752 | 14.29631 | 42.77194 | 40.14703 | Distal Interge | 16 | 50719294 | 50732773 | 13480 | 2 | ENSMUST00 | -3503 | 5330426P16 | ENSMUSG00000022639.9 |
| chr16 | 42907033 | 42911476 | 14.29631 | 42.77194 | 40.14703 | Promoter | 16 | 42907392 | 43072468 | 165077 | 1 | ENSMUST00 | 0 | Zbtb20 | ENSMUSG00000022708.11 |
| chr17 | 28517555 | 28518704 | 14.29631 | 42.77194 | 40.14703 | Promoter | 17 | 28399105 | 28517524 | 118420 | 2 | ENSMUST00 | -32 | Fkbp5 | ENSMUSG00000024222.11 |
| chr17 | 3557985 | 3558477 | 14.29631 | 42.77194 | 40.14703 | Promoter | 17 | 3519263 | 3557713 | 38451 | 2 | ENSMUST00 | -273 | Tfb1m | ENSMUSG00000036983.6 |
| chr17 | 25727689 | 25728337 | 14.29631 | 42.77194 | 40.14703 | Promoter | 17 | 25727751 | 25731456 | 3706 | 1 | ENSMUST00 | 0 | Rpusd1 | ENSMUSG00000041199.2 |
| chr6 | 134829342 | 134829976 | 14.29631 | 42.77194 | 40.14703 | Promoter | 6 | 134830154 | 134858931 | 28778 | 1 | ENSMUST00 | -178 | Crebl2 | ENSMUSG00000032652.8 |
| chr7 | 127894949 | 127895958 | 14.29631 | 42.77194 | 40.14703 | Promoter | 7 | 127886227 | 127895454 | 9228 | 2 | ENSMUST00 | 0 | Gm21974 | ENSMUSG00000030804.8 |
| chr1 | 134987538 | 134988202 | 15.0633 | 42.76468 | 40.14062 | Exon (ENSMU | 1 | 134971956 | 134972857 | 902 | 1 | ENSMUST00 | 15583 | Ube2t | ENSMUSG00000026429.8 |
| chr10 | 24223147 | 24224066 | 15.86955 | 42.74938 | 40.12535 | Promoter | 10 | 24223517 | 24302790 | 79274 | 1 | ENSMUST00 | 0 | Moxd1 | ENSMUSG00000020000.7 |
| chr10 | 80676724 | 80677626 | 15.86955 | 42.74938 | 40.12535 | Distal Interge | 10 | 80681838 | 80687176 | 5339 | 1 | ENSMUST00 | -4212 | Gm26541 | ENSMUSG00000097889.1 |
| chr19 | 61151417 | 61152466 | 15.86955 | 42.74938 | 40.12535 | Distal Interge | 19 | 61114984 | 61140840 | 25857 | 2 | ENSMUST00 | -10578 | 5830428H23 | ENSMUSG00000074733.9 |
| chr3 | 55054968 | 55055786 | 15.44655 | 42.73007 | 40.10627 | Promoter | 3 | 55045469 | 55055055 | 9587 | 2 | ENSMUST00 | 0 | Ccna1 | ENSMUSG00000027793.2 |
| chr17 | 46556370 | 46556965 | 17.13459 | 42.72025 | 40.09655 | Promoter | 17 | 46546839 | 46556162 | 9324 | 2 | ENSMUST00 | -209 | Srf | ENSMUSG00000015605.5 |
| chr15 | 101845766 | 101846382 | 17.10022 | 42.66158 | 40.03807 | 3' UTR | 15 | 101845428 | 101850786 | 5359 | 2 | ENSMUST00 | 4404 | Krt1 | ENSMUSG00000046834.6 |
| chr3 | 66980702 | 66981759 | 17.10022 | 42.66158 | 40.03807 | Promoter | 3 | 66971727 | 66981318 | 9592 | 2 | ENSMUST00 | 0 | Shox2 | ENSMUSG00000027833.11 |
| chr4 | 132514550 | 132515571 | 17.10022 | 42.66158 | 40.03807 | Distal Interge | 4 | 132492032 | 132510501 | 18470 | 2 | ENSMUST00 | -4050 | Sesn2 | ENSMUSG00000028893.8 |
| chr7 | 143437720 | 143438769 | 17.10022 | 42.66158 | 40.03807 | Distal Interge | 7 | 143447605 | 143447665 | 61 | 1 | ENSMUST00 | -8836 | Gm27901 | ENSMUSG00000099103.1 |
| chr9 | 22050478 | 22051328 | 17.10022 | 42.66158 | 40.03807 | Promoter | 9 | 22015005 | 22052023 | 37019 | 2 | ENSMUST00 | 695 | Elavl3 | ENSMUSG00000003410.7 |
| chr1 | 136007302 | 136008656 | 14.96389 | 42.58086 | 39.95799 | Promoter | 1 | 136008219 | 136035030 | 26812 | 1 | ENSMUST00 | 0 | Tmem9 | ENSMUSG00000026411.12 |
| chr1 | 180850115 | 180850818 | 14.96389 | 42.58086 | 39.95799 | Promoter | 1 | 180851151 | 180868097 | 16947 | 1 | ENSMUST00 | -333 | Sde2 | ENSMUSG00000038806.6 |
| chr11 | 78697648 | 78698322 | 14.96389 | 42.58086 | 39.95799 | Promoter | 11 | 78567168 | 78697373 | 130206 | 2 | ENSMUST00 | -276 | Nlk | ENSMUSG00000017376.10 |
| chr13 | 67728531 | 67729741 | 14.96389 | 42.58086 | 39.95799 | Promoter | 13 | 67729262 | 67755714 | 26453 | 1 | ENSMUST00 | 0 | Zfp85os | ENSMUSG00000044081.7 |
| chr19 | 56721342 | 56724468 | 14.96389 | 42.58086 | 39.95799 | Promoter | 19 | 56722372 | 56724862 | 2491 | 1 | ENSMUST00 | 0 | Adrb1 | ENSMUSG00000035283.4 |
| chr2 | 76672383 | 76673586 | 14.96389 | 42.58086 | 39.95799 | Promoter | 2 | 76663044 | 76673070 | 10027 | 2 | ENSMUST00 | 0 | Fkbp7 | ENSMUSG00000002732.9 |
| chr2 | 26586251 | 26588302 | 14.96389 | 42.58086 | 39.95799 | Promoter | 2 | 26586630 | 26592681 | 6052 | 1 | ENSMUST00 | 0 | Egfl7 | ENSMUSG00000026921.13 |
| chr3 | 89436276 | 89437465 | 14.96389 | 42.58086 | 39.95799 | Promoter | 3 | 89436706 | 89450952 | 14247 | 1 | ENSMUST00 | 0 | Pbxip1 | ENSMUSG00000042613.4 |
| chr3 | 68869128 | 68870633 | 14.96389 | 42.58086 | 39.95799 | Promoter | 3 | 68869586 | 68872163 | 2578 | 1 | ENSMUST00 | 0 | 1110032F04 | ENSMUSG00000046999.2 |
| chr4 | 41502473 | 41503143 | 14.96389 | 42.58086 | 39.95799 | Promoter | 4 | 41495604 | 41503076 | 7473 | 2 | ENSMUST00 | 0 | AI464131 | ENSMUSG00000046312.4 |
| chr6 | 28134897 | 28136135 | 14.96389 | 42.58086 | 39.95799 | Promoter | 6 | 27934377 | 28135178 | 200802 | 2 | ENSMUST00 | 0 | Grm8 | ENSMUSG00000024211.10 |
| chr6 | 125493775 | 125495138 | 14.96389 | 42.58086 | 39.95799 | Promoter | 6 | 125460266 | 125494791 | 34526 | 2 | ENSMUST00 | 0 | Cd9 | ENSMUSG00000030342.8 |
| chr6 | 145339893 | 145340769 | 14.96389 | 42.58086 | 39.95799 | Distal Interge | 6 | 145315613 | 145316608 | 996 | 2 | ENSMUST00 | -23286 | Gm15707 | ENSMUSG00000083962.1 |
| chr9 | 106280457 | 106281720 | 14.96389 | 42.58086 | 39.95799 | Promoter | 9 | 106281061 | 106349879 | 68819 | 1 | ENSMUST00 | 0 | Poc1a | ENSMUSG00000023345.10 |
| chr9 | 96536896 | 96537429 | 14.96389 | 42.58086 | 39.95799 | Promoter | 9 | 96537161 | 96537854 | 694 | 1 | ENSMUST00 | 0 | RP24-91A16. | ENSMUSG00000101615.1 |
| chr1 | 134513225 | 134514120 | 15.77044 | 42.5731 | 39.95169 | Distal Interge | 1 | 134518081 | 134518563 | 483 | 1 | ENSMUST00 | -3961 | Gm15454 | ENSMUSG00000096712.1 |
| chr10 | 68282568 | 68283301 | 15.77044 | 42.5731 | 39.95169 | Distal Interge | 10 | 68095593 | 68278726 | 183134 | 2 | ENSMUST00 | -3843 | Arid5b | ENSMUSG00000019947.9 |
| chr11 | 95990773 | 95991742 | 15.77044 | 42.5731 | 39.95169 | Intron (ENSM | 11 | 95973979 | 95982611 | 8633 | 2 | ENSMUST00 | -8163 | Igf2bp1 | ENSMUSG00000013415.4 |

| | | | | | | | | | | | | | | | |
|---|---|---|---|---|---|---|---|---|---|---|---|---|---|---|---|
| chr12 | 78906694 | 78907242 | 15.77044 | 42.5731 | 39.95169 | Promoter | 12 | 78888691 | 78906964 | 18274 | 2 | ENSMUST00 | 0 | Plek2 | ENSMUSG00000021118.7 |
| chr13 | 51568717 | 51569764 | 15.77044 | 42.5731 | 39.95169 | Distal Interge | 13 | 51431041 | 51567084 | 136044 | 2 | ENSMUST00 | -1634 | Shc3 | ENSMUSG00000021448.6 |
| chr14 | 122463410 | 122465727 | 15.77044 | 42.5731 | 39.95169 | Promoter | 14 | 122456795 | 122465677 | 8883 | 2 | ENSMUST00 | 0 | Zic5 | ENSMUSG00000041703.7 |
| chr15 | 41830850 | 41832770 | 15.77044 | 42.5731 | 39.95169 | Promoter | 15 | 41830958 | 41861045 | 30088 | 1 | ENSMUST00 | 0 | Oxr1 | ENSMUSG00000022307.10 |
| chr17 | 67947219 | 67951695 | 15.77044 | 42.5731 | 39.95169 | Promoter | 17 | 67845110 | 67950940 | 105831 | 2 | ENSMUST00 | 0 | Arhgap28 | ENSMUSG00000024043.8 |
| chr17 | 24548890 | 24549850 | 15.77044 | 42.5731 | 39.95169 | Promoter | 17 | 24549950 | 24596514 | 46565 | 1 | ENSMUST00 | -100 | Pkd1 | ENSMUSG00000032855.5 |
| chr17 | 34604317 | 34605145 | 15.77044 | 42.5731 | 39.95169 | Promoter | 17 | 34604844 | 34611396 | 6553 | 1 | ENSMUST00 | 0 | Agpat1 | ENSMUSG00000034254.12 |
| chr19 | 7143059 | 7144600 | 15.77044 | 42.5731 | 39.95169 | Intron (ENSM | 19 | 7092014 | 7105729 | 13716 | 2 | ENSMUST00 | -37331 | Flrt1 | ENSMUSG00000047787.7 |
| chr2 | 77169910 | 77170720 | 15.77044 | 42.5731 | 39.95169 | Promoter | 2 | 77009902 | 77170635 | 160734 | 2 | ENSMUST00 | 0 | Ccdc141 | ENSMUSG00000044033.11 |
| chr3 | 18054071 | 18055883 | 15.77044 | 42.5731 | 39.95169 | Promoter | 3 | 18054174 | 18057517 | 3344 | 1 | ENSMUST00 | 0 | Bhlhe22 | ENSMUSG00000025128.7 |
| chr3 | 159495045 | 159495850 | 15.77044 | 42.5731 | 39.95169 | Promoter | 3 | 159495433 | 159529955 | 34523 | 1 | ENSMUST00 | 0 | Depdc1a | ENSMUSG00000028175.10 |
| chr3 | 97824080 | 97824657 | 15.77044 | 42.5731 | 39.95169 | Exon (ENSMU | 3 | 97820982 | 97868197 | 47216 | 2 | ENSMUST00 | 43540 | Pde4dip | ENSMUSG00000038170.10 |
| chr4 | 129555572 | 129557250 | 15.77044 | 42.5731 | 39.95169 | Promoter | 4 | 129548895 | 129555848 | 6954 | 2 | ENSMUST00 | 0 | Lck | ENSMUSG00000000409.9 |
| chr4 | 134159897 | 134160486 | 15.77044 | 42.5731 | 39.95169 | Intron (ENSM | 4 | 134155901 | 134156921 | 1021 | 2 | ENSMUST00 | -2977 | Cep85 | ENSMUSG00000037443.8 |
| chr8 | 120633713 | 120634858 | 15.77044 | 42.5731 | 39.95169 | Promoter | 8 | 120608602 | 120634492 | 25891 | 2 | ENSMUST00 | 0 | 1190005I06R | ENSMUSG00000043687.10 |
| chr9 | 8899649 | 8902155 | 15.77044 | 42.5731 | 39.95169 | Promoter | 9 | 8899833 | 8968611 | 68779 | 1 | ENSMUST00 | 0 | Pgr | ENSMUSG00000031870.11 |
| chr13 | 22042760 | 22043062 | 17.03191 | 42.54489 | 39.9255 | Promoter | 13 | 22042460 | 22042944 | 485 | 2 | ENSMUST00 | 0 | Hist1h2ag | ENSMUSG00000069301.5 |
| chr1 | 182258180 | 182258461 | 17.46222 | 42.53877 | 39.91948 | Distal Interge | 1 | 182279004 | 182282224 | 3221 | 2 | ENSMUST00 | 23763 | Degs1 | ENSMUSG00000038633.5 |
| chr10 | 80494631 | 80495787 | 17.46222 | 42.53877 | 39.91948 | Promoter | 10 | 80494835 | 80517276 | 22442 | 1 | ENSMUST00 | 0 | Onecut3 | ENSMUSG00000045518.8 |
| chr13 | 95222477 | 95223520 | 17.46222 | 42.53877 | 39.91948 | Promoter | 13 | 95024505 | 95223066 | 198562 | 2 | ENSMUST00 | 0 | Pde8b | ENSMUSG00000021684.11 |
| chr17 | 33916115 | 33917028 | 17.46222 | 42.53877 | 39.91948 | Promoter | 17 | 33916171 | 33919324 | 3154 | 1 | ENSMUST00 | 0 | Zbtb22 | ENSMUSG00000051390.9 |
| chr17 | 35257950 | 35258582 | 17.46222 | 42.53877 | 39.91948 | Distal Interge | 17 | 35262730 | 35267499 | 4770 | 1 | ENSMUST00 | -4148 | H2-D1 | ENSMUSG00000073411.6 |
| chr19 | 4064062 | 4064821 | 17.46222 | 42.53877 | 39.91948 | Distal Interge | 19 | 4057477 | 4059446 | 1970 | 2 | ENSMUST00 | -4617 | BC021614 | ENSMUSG00000058216.7 |
| chr2 | 92400838 | 92401651 | 17.46222 | 42.53877 | 39.91948 | Promoter | 2 | 92383680 | 92401263 | 17584 | 2 | ENSMUST00 | 0 | Mapk8ip1 | ENSMUSG00000027223.10 |
| chr3 | 52101179 | 52102153 | 17.46222 | 42.53877 | 39.91948 | Intron (ENSM | 3 | 51687611 | 52104891 | 417281 | 2 | ENSMUST00 | 2738 | Maml3 | ENSMUSG00000061143.10 |
| chr4 | 42916794 | 42917719 | 17.46222 | 42.53877 | 39.91948 | Promoter | 4 | 42917251 | 42944752 | 27502 | 1 | ENSMUST00 | 0 | N28178 | ENSMUSG00000036062.8 |
| chr5 | 116445494 | 116447174 | 17.46222 | 42.53877 | 39.91948 | Promoter | 5 | 116442432 | 116446844 | 4413 | 2 | ENSMUST00 | 0 | Srrm4 | ENSMUSG00000063919.7 |
| chr7 | 28451692 | 28452490 | 17.46222 | 42.53877 | 39.91948 | Promoter | 7 | 28451980 | 28459285 | 7306 | 1 | ENSMUST00 | 0 | Lrfn1 | ENSMUSG00000030600.10 |
| chr7 | 144678442 | 144679142 | 17.46222 | 42.53877 | 39.91948 | Promoter | 7 | 144655601 | 144678885 | 23285 | 2 | ENSMUST00 | 0 | Ano1 | ENSMUSG00000031075.12 |
| chr7 | 7188036 | 7188561 | 17.46222 | 42.53877 | 39.91948 | Distal Interge | 7 | 7171330 | 7183562 | 12233 | 1 | ENSMUST00 | 16707 | Zfp418 | ENSMUSG00000034538.7 |
| chr8 | 4216872 | 4217552 | 17.46222 | 42.53877 | 39.91948 | Promoter | 8 | 4209543 | 4217312 | 7770 | 2 | ENSMUST00 | 0 | BC068157 | ENSMUSG00000064125.9 |
| chr9 | 91381036 | 91382009 | 17.46222 | 42.53877 | 39.91948 | Promoter | 9 | 91382410 | 91388044 | 5635 | 1 | ENSMUST00 | -401 | Zic4 | ENSMUSG00000036972.9 |
| chrX | 49287553 | 49288332 | 17.46222 | 42.53877 | 39.91948 | Promoter | 20 | 49009707 | 49288212 | 278506 | 2 | ENSMUST00 | 0 | Enox2 | ENSMUSG00000031109.11 |
| chrX | 7898746 | 7899988 | 17.46222 | 42.53877 | 39.91948 | Promoter | 20 | 7894519 | 7899269 | 4751 | 2 | ENSMUST00 | 0 | Pqbp1 | ENSMUSG00000031157.5 |
| chrX | 69360177 | 69360994 | 17.46222 | 42.53877 | 39.91948 | Promoter | 20 | 69360294 | 69868037 | 507744 | 1 | ENSMUST00 | 0 | Aff2 | ENSMUSG00000031189.7 |
| chrX | 162901567 | 162902756 | 17.46222 | 42.53877 | 39.91948 | Promoter | 20 | 162901605 | 163032325 | 130721 | 1 | ENSMUST00 | 0 | Ctps2 | ENSMUSG00000031360.9 |
| chrX | 164419753 | 164420387 | 17.46222 | 42.53877 | 39.91948 | Promoter | 20 | 164419793 | 164433911 | 14119 | 1 | ENSMUST00 | 0 | Piga | ENSMUSG00000031381.10 |
| chrX | 74023137 | 74024274 | 17.46222 | 42.53877 | 39.91948 | Promoter | 20 | 74013914 | 74023918 | 10005 | 2 | ENSMUST00 | 0 | Irak1 | ENSMUSG00000031392.13 |
| chrX | 140598132 | 140600658 | 17.46222 | 42.53877 | 39.91948 | Promoter | 20 | 140540262 | 140600659 | 60398 | 2 | ENSMUST00 | 1 | Tsc22d3 | ENSMUSG00000031431.8 |
| chrX | 139684244 | 139686102 | 17.46222 | 42.53877 | 39.91948 | Promoter | 20 | 139684996 | 139753405 | 68410 | 1 | ENSMUST00 | 0 | Tbc1d8b | ENSMUSG00000042473.8 |
| chrX | 85047907 | 85049126 | 17.46222 | 42.53877 | 39.91948 | Promoter | 20 | 85048295 | 85055597 | 7303 | 1 | ENSMUST00 | 0 | Dmd | ENSMUSG00000045103.12 |
| chrX | 157597665 | 157599272 | 17.46222 | 42.53877 | 39.91948 | Promoter | 20 | 157547824 | 157598715 | 50892 | 2 | ENSMUST00 | 0 | Mbtps2 | ENSMUSG00000046873.13 |
| chrX | 153497465 | 153499227 | 17.46222 | 42.53877 | 39.91948 | Promoter | 20 | 153498227 | 153501570 | 3344 | 1 | ENSMUST00 | 0 | Ubqln2 | ENSMUSG00000050148.9 |
| chrX | 95443457 | 95445327 | 17.46222 | 42.53877 | 39.91948 | Promoter | 20 | 95420318 | 95444872 | 24555 | 2 | ENSMUST00 | 0 | Amer1 | ENSMUSG00000050332.7 |
| chrX | 134750555 | 134751664 | 17.46222 | 42.53877 | 39.91948 | Promoter | 20 | 134748461 | 134751331 | 2871 | 2 | ENSMUST00 | 0 | Armcx6 | ENSMUSG00000050394.9 |
| chrX | 21061380 | 21062544 | 17.46222 | 42.53877 | 39.91948 | Promoter | 20 | 21028939 | 21062003 | 33065 | 2 | ENSMUST00 | 0 | Zfp182 | ENSMUSG00000054737.6 |
| chrX | 102504302 | 102505471 | 17.46222 | 42.53877 | 39.91948 | Promoter | 20 | 102284639 | 102505359 | 220721 | 2 | ENSMUST00 | 0 | Hdac8 | ENSMUSG00000067567.7 |
| chrX | 57052914 | 57054288 | 17.46222 | 42.53877 | 39.91948 | Promoter | 20 | 57053583 | 57057206 | 3624 | 1 | ENSMUST00 | 0 | Htatsf1 | ENSMUSG00000067873.6 |
| chrX | 56346063 | 56347209 | 17.46222 | 42.53877 | 39.91948 | Promoter | 20 | 56346400 | 56365674 | 19275 | 1 | ENSMUST00 | 0 | Zfp449 | ENSMUSG00000073176.4 |
| chrX | 102119445 | 102120058 | 17.46222 | 42.53877 | 39.91948 | Promoter | 20 | 102119447 | 102127669 | 8223 | 1 | ENSMUST00 | 0 | Pin4 | ENSMUSG00000079480.3 |
| chr3 | 8664465 | 8665320 | 16.99796 | 42.48687 | 39.86913 | 3' UTR | 3 | 8663362 | 8667039 | 3678 | 2 | ENSMUST00 | 1719 | Hey1 | ENSMUSG00000040289.3 |
| chr9 | 81632454 | 81633933 | 16.99796 | 42.48687 | 39.86913 | Promoter | 9 | 81628291 | 81633828 | 5538 | 2 | ENSMUST00 | 0 | Htr1b | ENSMUSG00000049511.5 |
| chr10 | 79985067 | 79986304 | 16.54337 | 42.44649 | 39.82893 | Promoter | 10 | 79985506 | 79985612 | 107 | 1 | ENSMUST00 | 0 | Gm26243 | ENSMUSG00000095615.1 |
| chr15 | 81465557 | 81466212 | 16.54337 | 42.44649 | 39.82893 | Promoter | 15 | 81466316 | 81476369 | 10054 | 1 | ENSMUST00 | -104 | Rbx1 | ENSMUSG00000022400.8 |
| chr8 | 23410911 | 23412471 | 16.54337 | 42.44649 | 39.82893 | Promoter | 8 | 23411502 | 23449632 | 38131 | 1 | ENSMUST00 | 0 | Sfrp1 | ENSMUSG00000031548.7 |
| chr17 | 7169205 | 7169866 | 16.96414 | 42.42906 | 39.81211 | Promoter | 17 | 7170115 | 7303313 | 133199 | 1 | ENSMUST00 | -249 | Rps6ka2 | ENSMUSG00000023809.9 |
| chr3 | 66219954 | 66221741 | 17.39432 | 42.42359 | 39.80675 | Promoter | 3 | 66220245 | 66221430 | 1186 | 1 | ENSMUST00 | 0 | Ptx3 | ENSMUSG00000027832.5 |
| chr4 | 145585109 | 145585732 | 13.37499 | 42.36604 | 39.7494 | Promoter | 4 | 145585166 | 145625345 | 40180 | 1 | ENSMUST00 | 0 | Gm13212 | ENSMUSG00000078502.5 |
| chr11 | 96284031 | 96284998 | 17.35798 | 42.36193 | 39.74529 | 3' UTR | 11 | 96282725 | 96284693 | 1969 | 1 | ENSMUST00 | 1307 | Hoxb8 | ENSMUSG00000056648.5 |
| chr16 | 37684332 | 37685347 | 17.35798 | 42.36193 | 39.74529 | Distal Interge | 16 | 37647170 | 37654453 | 7284 | 2 | ENSMUST00 | -29880 | Ndufb4 | ENSMUSG00000022820.3 |
| chr5 | 139129375 | 139129992 | 17.35798 | 42.36193 | 39.74529 | Promoter | 5 | 139017306 | 139129759 | 112454 | 2 | ENSMUST00 | 0 | Prkar1b | ENSMUSG00000025855.8 |
| chr6 | 122297169 | 122297803 | 17.35798 | 42.36193 | 39.74529 | Distal Interge | 6 | 122305544 | 122305798 | 255 | 2 | ENSMUST00 | 7995 | Gm27826 | ENSMUSG00000093023.2 |
| chr9 | 90125920 | 90127035 | 17.35798 | 42.36193 | 39.74529 | Distal Interge | 9 | 90118937 | 90119807 | 871 | 1 | ENSMUST00 | 6984 | Gm6223 | ENSMUSG00000096647.2 |
| chr17 | 26714994 | 26715492 | 16.04633 | 42.32132 | 39.70501 | Promoter | 17 | 26715650 | 26776635 | 60986 | 1 | ENSMUST00 | -158 | Crebrf | ENSMUSG00000048249.9 |
| chr13 | 23526188 | 23526970 | 16.89691 | 42.31408 | 39.69809 | Downstream | 13 | 23527011 | 23530943 | 3933 | 2 | ENSMUST00 | 3973 | Gm11335 | ENSMUSG00000085016.1 |
| chr4 | 88937867 | 88938878 | 17.32179 | 42.30051 | 39.68461 | Distal Interge | 4 | 88910557 | 88910662 | 106 | 2 | ENSMUST00 | -27206 | Mir31 | ENSMUSG00000065408.1 |
| chr7 | 4865656 | 4867011 | 15.19686 | 42.27666 | 39.66088 | Promoter | 7 | 4844959 | 4866193 | 21235 | 2 | ENSMUST00 | 0 | Isoc2b | ENSMUSG00000052605.6 |
| chr18 | 82811792 | 82812590 | 15.59995 | 42.26899 | 39.65329 | Distal Interge | 18 | 82855894 | 82870604 | 14711 | 1 | ENSMUST00 | -43304 | 4930445N18 | ENSMUSG00000097738.1 |
| chr2 | 27442386 | 27442963 | 15.59995 | 42.26899 | 39.65329 | Downstream | 2 | 27450549 | 27450630 | 82 | 2 | ENSMUST00 | 7667 | Mir7578 | ENSMUSG00000098812.1 |
| chr3 | 88296175 | 88296929 | 15.59995 | 42.26899 | 39.65329 | Promoter | 3 | 88282760 | 88296838 | 14079 | 2 | ENSMUST00 | 0 | Tsacc | ENSMUSG00000010538.9 |
| chr12 | 83631427 | 83632065 | 12.9904 | 42.26624 | 39.65078 | Promoter | 12 | 83631339 | 83646232 | 14894 | 1 | ENSMUST00 | 89 | Rbm25 | ENSMUSG00000010608.10 |
| chr9 | 102689705 | 102690990 | 12.9904 | 42.26624 | 39.65078 | Distal Interge | 9 | 102692885 | 102693834 | 950 | 1 | ENSMUST00 | -1895 | RP24-221A14 | ENSMUSG00000099491.1 |
| chr9 | 118478380 | 118479448 | 17.28575 | 42.23933 | 39.6242 | Promoter | 9 | 118478861 | 118482757 | 3897 | 1 | ENSMUST00 | 0 | Eomes | ENSMUSG00000032446.9 |
| chr10 | 5921715 | 5922605 | 16.8302 | 42.19991 | 39.5849 | Promoter | 10 | 5825663 | 5922386 | 96724 | 2 | ENSMUST00 | 0 | Rgs17 | ENSMUSG00000019775.12 |
| chr11 | 4096791 | 4097403 | 16.8302 | 42.19991 | 39.5849 | 3' UTR | 11 | 4091480 | 4095445 | 3966 | 2 | ENSMUST00 | -1347 | Mtfp1 | ENSMUSG00000004748.5 |
| chr12 | 79190953 | 79192085 | 16.76403 | 42.08657 | 39.472 | Promoter | 12 | 79175551 | 79190961 | 15411 | 2 | ENSMUST00 | 0 | Rdh11 | ENSMUSG00000066441.9 |
| chr17 | 55445333 | 55446292 | 16.76403 | 42.08657 | 39.472 | Promoter | 17 | 55445382 | 55483032 | 37651 | 1 | ENSMUST00 | 0 | St6gal2 | ENSMUSG00000024172.5 |
| chr7 | 82174596 | 82175560 | 16.76403 | 42.08657 | 39.472 | Promoter | 7 | 82174796 | 82307419 | 132624 | 1 | ENSMUST00 | 0 | Sh3gl3 | ENSMUSG00000030638.8 |
| chr1 | 133800021 | 133801596 | 17.10778 | 41.93702 | 39.32339 | Promoter | 1 | 133699457 | 133801041 | 101585 | 2 | ENSMUST00 | 0 | Atp2b4 | ENSMUSG00000026463.12 |
| chr10 | 35711374 | 35711869 | 17.10778 | 41.93702 | 39.32339 | Promoter | 10 | 35708691 | 35711891 | 3201 | 2 | ENSMUST00 | 22 | Amd2 | ENSMUSG00000063953.2 |
| chr13 | 22028821 | 22029159 | 17.10778 | 41.93702 | 39.32339 | Intron (ENSM | 13 | 22026699 | 22031610 | 4912 | 1 | ENSMUST00 | 2123 | Gm11292 | ENSMUSG00000043145.6 |
| chr9 | 96436926 | 96438702 | 17.10778 | 41.93702 | 39.32339 | Promoter | 9 | 96435783 | 96438138 | 2356 | 2 | ENSMUST00 | 0 | BC043934 | ENSMUSG00000056418.3 |
| chr1 | 44951123 | 44951386 | 16.66573 | 41.91806 | 39.30453 | Distal Interge | 1 | 44924149 | 44924893 | 745 | 2 | ENSMUST00 | -26231 | RP23-231B19 | ENSMUSG00000101251.1 |
| chr16 | 4756741 | 4757383 | 16.66573 | 41.91806 | 39.30453 | Promoter | 16 | 4756845 | 4764684 | 7840 | 1 | ENSMUST00 | 0 | Hmox2 | ENSMUSG00000004070.10 |
| chr3 | 88492514 | 88493067 | 16.66573 | 41.91806 | 39.30453 | Promoter | 3 | 88483206 | 88493312 | 10107 | 2 | ENSMUST00 | 245 | Lmna | ENSMUSG00000028063.10 |
| chr1 | 136233900 | 136234692 | 16.23624 | 41.91433 | 39.30089 | Promoter | 1 | 136213531 | 136234264 | 20734 | 2 | ENSMUST00 | 0 | 5730559C18R | ENSMUSG00000041605.11 |
| chr12 | 109052707 | 109054022 | 16.23624 | 41.91433 | 39.30089 | Promoter | 12 | 109037652 | 109053581 | 15930 | 2 | ENSMUST00 | 0 | Begain | ENSMUSG00000040867.10 |
| chr13 | 32965180 | 32966767 | 16.23624 | 41.91433 | 39.30089 | Promoter | 13 | 32965209 | 32979067 | 13859 | 1 | ENSMUST00 | 0 | Serpinb6b | ENSMUSG00000042842.5 |
| chr18 | 12973384 | 12973859 | 16.23624 | 41.91433 | 39.30089 | Intron (ENSM | 18 | 12972252 | 12992948 | 20697 | 1 | ENSMUST00 | 1133 | Impact | ENSMUSG00000024423.5 |
| chr2 | 71580951 | 71581554 | 16.23624 | 41.91433 | 39.30089 | Distal Interge | 2 | 71543408 | 71546754 | 3347 | 2 | ENSMUST00 | -34198 | Dlx2 | ENSMUSG00000023391.8 |
| chr2 | 97466311 | 97466883 | 16.23624 | 41.91433 | 39.30089 | Promoter | 2 | 97467657 | 97631664 | 164008 | 1 | ENSMUST00 | -774 | Lrrc4c | ENSMUSG00000050587.9 |
| chr5 | 73256876 | 73257621 | 16.23624 | 41.91433 | 39.30089 | Promoter | 5 | 73193828 | 73256618 | 62791 | 2 | ENSMUST00 | -259 | Fryl | ENSMUSG00000070733.8 |
| chr7 | 28982014 | 28982456 | 16.23624 | 41.91433 | 39.30089 | Promoter | 7 | 28971373 | 28981814 | 10442 | 2 | ENSMUST00 | -201 | Eif3k | ENSMUSG00000053565.9 |
| chr11 | 78194473 | 78195324 | 17.07262 | 41.87726 | 39.26436 | Promoter | 11 | 78194704 | 78205467 | 10764 | 1 | ENSMUST00 | 0 | Proca1 | ENSMUSG00000044122.9 |
| chr1 | 22533192 | 22534045 | 16.20615 | 41.86206 | 39.24932 | Exon (ENSMU | 1 | 22288422 | 22432843 | 144422 | 2 | ENSMUST00 | -100350 | Rims1 | ENSMUSG00000041670.10 |
| chr13 | 13232607 | 13232896 | 14.95512 | 41.83493 | 39.22221 | Promoter | 13 | 13232158 | 13232895 | 738 | 2 | ENSMUST00 | 0 | Gm2423 | ENSMUSG00000061724.4 |
| chr1 | 180768151 | 180769015 | 15.76414 | 41.82454 | 39.21188 | Distal Interge | 1 | 180802568 | 180809483 | 6916 | 1 | ENSMUST00 | -33553 | Gm17275 | ENSMUSG00000097347.1 |
| chr19 | 3388772 | 3389396 | 15.76414 | 41.82454 | 39.21188 | Promoter | 19 | 3388857 | 3407823 | 18967 | 1 | ENSMUST00 | 0 | Mtl5 | ENSMUSG00000024905.10 |
| chr16 | 30069736 | 30070434 | 13.44849 | 41.81219 | 39.19989 | Distal Interge | 16 | 30066338 | 30067319 | 982 | 1 | ENSMUST00 | 3399 | Hes1 | ENSMUSG00000022528.7 |
| chr6 | 120356593 | 120357803 | 13.44849 | 41.81219 | 39.19989 | Promoter | 6 | 120324323 | 120357429 | 33107 | 2 | ENSMUST00 | 0 | Ccdc77 | ENSMUSG00000030177.9 |
| chr2 | 118256643 | 118257177 | 15.33936 | 41.80194 | 39.19019 | Promoter | 2 | 118204888 | 118256966 | 52079 | 2 | ENSMUST00 | 0 | Fsip1 | ENSMUSG00000027344.8 |
| chr1 | 172309908 | 172310236 | 17.00274 | 41.75843 | 39.14683 | Intron (ENSM | 1 | 172311771 | 172319841 | 8071 | 1 | ENSMUST00 | -1535 | Igsf8 | ENSMUSG00000038034.10 |
| chr12 | 85745947 | 85746842 | 17.00274 | 41.75843 | 39.14683 | Promoter | 12 | 85746539 | 85813585 | 67047 | 1 | ENSMUST00 | 0 | Mfsd7c | ENSMUSG00000034258.3 |
| chr18 | 45558299 | 45560518 | 17.00274 | 41.75843 | 39.14683 | Promoter | 18 | 45559469 | 45671249 | 111781 | 1 | ENSMUST00 | 0 | Kcnn2 | ENSMUSG00000054477.9 |
| chr4 | 133480007 | 133480599 | 17.00274 | 41.75843 | 39.14683 | Promoter | 4 | 133480132 | 133487938 | 7807 | 1 | ENSMUST00 | 0 | Fam46b | ENSMUSG00000046694.6 |
| chr14 | 17172359 | 17173495 | 16.11655 | 41.70626 | 39.09511 | Distal Interge | 14 | 17660960 | 18036559 | 375600 | 1 | ENSMUST00 | -487465 | Thrb | ENSMUSG00000021779.11 |
| chr13 | 73846859 | 73847642 | 16.93343 | 41.6405 | 39.02962 | Promoter | 13 | 73818534 | 73847631 | 29098 | 2 | ENSMUST00 | 0 | Nkd2 | ENSMUSG00000021567.9 |
| chr7 | 70345833 | 70346633 | 16.93343 | 41.6405 | 39.02962 | Distal Interge | 7 | 70351946 | 70360593 | 8648 | 2 | ENSMUST00 | 13960 | Nr2f2 | ENSMUSG00000030551.8 |
| chr10 | 78752348 | 78753058 | 16.89899 | 41.58187 | 38.97116 | Promoter | 10 | 78746924 | 78753067 | 6144 | 2 | ENSMUST00 | 9 | Ccdc105 | ENSMUSG00000078442.2 |
| chr15 | 75893843 | 75894709 | 16.44079 | 41.53171 | 38.92122 | Promoter | 15 | 75890964 | 75894481 | 3518 | 2 | ENSMUST00 | 0 | Naprt1 | ENSMUSG00000022574.6 |
| chr5 | 107403099 | 107404121 | 16.44079 | 41.53171 | 38.92122 | Promoter | 5 | 107403496 | 107430035 | 26540 | 1 | ENSMUST00 | 0 | Ephx4 | ENSMUSG00000033805.7 |
| chr4 | 117494120 | 117495657 | 16.83052 | 41.46527 | 38.85544 | Promoter | 4 | 117490235 | 117495011 | 4777 | 2 | ENSMUST00 | 0 | Rnf220 | ENSMUSG00000028677.13 |
| chr1 | 127308107 | 127310766 | 13.98552 | 41.45461 | 38.84487 | Intron (ENSM | 1 | 127306762 | 127483704 | 176943 | 1 | ENSMUST00 | 1346 | Mgat5 | ENSMUSG00000036155.8 |
| chr10 | 81316916 | 81317538 | 13.98552 | 41.45461 | 38.84487 | Promoter | 10 | 81316842 | 81318013 | 1172 | 1 | ENSMUST00 | 75 | Pip5k1c | ENSMUSG00000034902.12 |
| chr12 | 17265979 | 17266372 | 13.98552 | 41.45461 | 38.84487 | Promoter | 12 | 17266545 | 17281810 | 15266 | 1 | ENSMUST00 | -173 | Pdia6 | ENSMUSG00000020571.7 |
| chr12 | 112802017 | 112802797 | 13.98552 | 41.45461 | 38.84487 | Promoter | 12 | 112781010 | 112802657 | 21648 | 2 | ENSMUST00 | 0 | Ahnak2 | ENSMUSG00000072812.4 |
| chr17 | 71310315 | 71311529 | 13.98552 | 41.45461 | 38.84487 | Promoter | 17 | 71252176 | 71310965 | 58790 | 2 | ENSMUST00 | 0 | Emilin2 | ENSMUSG00000024053.10 |
| chr2 | 119350772 | 119351629 | 13.98552 | 41.45461 | 38.84487 | Promoter | 2 | 119351229 | 119354381 | 3153 | 1 | ENSMUST00 | 0 | Chac1 | ENSMUSG00000027313.3 |
| chr3 | 100921334 | 100922086 | 13.98552 | 41.45461 | 38.84487 | Promoter | 3 | 100922202 | 100931702 | 9501 | 1 | ENSMUST00 | -116 | Trim45 | ENSMUSG00000033233.12 |
| chr4 | 109153150 | 109157434 | 13.98552 | 41.45461 | 38.84487 | Promoter | 4 | 109062087 | 109156610 | 94524 | 2 | ENSMUST00 | 0 | Osbpl9 | ENSMUSG00000028559.12 |
| chr4 | 16164084 | 16164594 | 13.98552 | 41.45461 | 38.84487 | Promoter | 4 | 16164110 | 16266225 | 102116 | 1 | ENSMUST00 | 0 | A530072M11 | ENSMUSG00000085112.1 |
| chr9 | 72274530 | 72276395 | 13.98552 | 41.45461 | 38.84487 | Promoter | 9 | 72274860 | 72362891 | 88032 | 1 | ENSMUST00 | 0 | Zfp280d | ENSMUSG00000038535.12 |
| chr7 | 18892344 | 18892773 | 15.96941 | 41.44991 | 38.84101 | Promoter | 7 | 18890884 | 18892546 | 1663 | 2 | ENSMUST00 | 0 | Ccdc61 | ENSMUSG00000074358.4 |
| chr1 | 20951200 | 20952243 | 15.9403 | 41.39913 | 38.79041 | Promoter | 1 | 20951626 | 20955527 | 3902 | 1 | ENSMUST00 | 0 | Efhc1 | ENSMUSG00000041809.5 |
| chr10 | 42017978 | 42018807 | 15.9403 | 41.39913 | 38.79041 | Promoter | 10 | 41914990 | 42018436 | 103447 | 2 | ENSMUST00 | 0 | Armc2 | ENSMUSG00000071324.6 |
| chr2 | 151575569 | 151576775 | 15.9403 | 41.39913 | 38.79041 | Intron (ENSM | 2 | 151572622 | 151590005 | 17384 | 1 | ENSMUST00 | 2948 | Sdcbp2 | ENSMUSG00000027456.8 |
| chr7 | 4136298 | 4136888 | 15.9403 | 41.39913 | 38.79041 | Promoter | 7 | 4126340 | 4136926 | 10587 | 2 | ENSMUST00 | 38 | D030047H15 | ENSMUSG00000085886.1 |
| chr1 | 15892401 | 15893023 | 16.34529 | 41.36735 | 38.7591 | Promoter | 1 | 15853862 | 15892722 | 38861 | 2 | ENSMUST00 | 0 | Sbspon | ENSMUSG00000032719.4 |

| | | | | | | | | | | | | | | | |
|---|---|---|---|---|---|---|---|---|---|---|---|---|---|---|---|
| chr1 | 91366248 | 91366936 | 16.34529 | 41.36735 | 38.7591 | Promoter | 1 | 91366430 | 91374217 | 7788 | 1 | ENSMUST00 | 0 | Fam132b | ENSMUSG00000047443.9 |
| chr10 | 81482245 | 81482889 | 16.34529 | 41.36735 | 38.7591 | Promoter | 10 | 81459227 | 81482709 | 23483 | 2 | ENSMUST00 | 0 | Celf5 | ENSMUSG00000034818.11 |
| chr12 | 27341240 | 27343696 | 16.34529 | 41.36735 | 38.7591 | Promoter | 12 | 27334264 | 27342574 | 8311 | 2 | ENSMUST00 | 0 | Sox11 | ENSMUSG00000063632.6 |
| chr12 | 54734000 | 54735323 | 16.34529 | 41.36735 | 38.7591 | Promoter | 12 | 54734555 | 54734718 | 164 | 2 | ENSMUST00 | 0 | Gm24305 | ENSMUSG00000095590.1 |
| chr13 | 3478013 | 3478760 | 16.34529 | 41.36735 | 38.7591 | Promoter | 13 | 3478230 | 3486177 | 7948 | 1 | ENSMUST00 | 0 | 2810429I04R | ENSMUSG00000097709.2 |
| chr19 | 36056974 | 36058001 | 16.34529 | 41.36735 | 38.7591 | Promoter | 19 | 35958734 | 36057340 | 98607 | 2 | ENSMUST00 | 0 | Htr7 | ENSMUSG00000024798.10 |
| chr2 | 25395810 | 25396416 | 16.34529 | 41.36735 | 38.7591 | Promoter | 2 | 25395874 | 25401321 | 5448 | 1 | ENSMUST00 | 0 | Entpd2 | ENSMUSG00000015085.8 |
| chr2 | 158146116 | 158146456 | 16.34529 | 41.36735 | 38.7591 | Promoter | 2 | 158116402 | 158146436 | 30035 | 2 | ENSMUST00 | 0 | Tgm2 | ENSMUSG00000037820.10 |
| chr2 | 161177254 | 161178197 | 16.34529 | 41.36735 | 38.7591 | Promoter | 2 | 161179064 | 161216577 | 37514 | 1 | ENSMUST00 | -867 | Gm14243 | ENSMUSG00000067745.4 |
| chr4 | 148806885 | 148807657 | 16.34529 | 41.36735 | 38.7591 | Intron (ENSM | 4 | 148804465 | 148839746 | 35282 | 1 | ENSMUST00 | 2421 | Casz1 | ENSMUSG00000028977.11 |
| chr5 | 66259636 | 66260355 | 16.34529 | 41.36735 | 38.7591 | Promoter | 5 | 66259912 | 66297062 | 37151 | 1 | ENSMUST00 | 0 | Nsun7 | ENSMUSG00000029206.8 |
| chr6 | 127149169 | 127151101 | 16.34529 | 41.36735 | 38.7591 | Promoter | 6 | 127125779 | 127151048 | 25270 | 2 | ENSMUST00 | 0 | Ccnd2 | ENSMUSG00000000184.9 |
| chr7 | 28732424 | 28733381 | 16.34529 | 41.36735 | 38.7591 | Promoter | 7 | 28732500 | 28737482 | 4983 | 1 | ENSMUST00 | 0 | Fbxo17 | ENSMUSG00000030598.10 |
| chr7 | 45521661 | 45522268 | 16.34529 | 41.36735 | 38.7591 | Exon (ENSMU | 7 | 45522916 | 45523769 | 854 | 2 | ENSMUST00 | 1501 | Ppp1r15a | ENSMUSG00000040435.10 |
| chr7 | 141060643 | 141061515 | 16.34529 | 41.36735 | 38.7591 | Promoter | 7 | 141061274 | 141072119 | 10846 | 1 | ENSMUST00 | 0 | B4galnt4 | ENSMUSG00000055629.4 |
| chr7 | 75781439 | 75782469 | 16.34529 | 41.36735 | 38.7591 | Promoter | 7 | 75769762 | 75782099 | 12338 | 2 | ENSMUST00 | 0 | AU020206 | ENSMUSG00000097415.1 |
| chr8 | 24438048 | 24439492 | 16.34529 | 41.36735 | 38.7591 | Promoter | 8 | 24437180 | 24438984 | 1805 | 2 | ENSMUST00 | 0 | 1810011O10 | ENSMUSG00000056313.5 |
| chr9 | 107551259 | 107552030 | 16.34529 | 41.36735 | 38.7591 | Promoter | 9 | 107551555 | 107562263 | 10709 | 1 | ENSMUST00 | 0 | Rassf1 | ENSMUSG00000010067.8 |
| chr9 | 66069010 | 66069834 | 16.34529 | 41.36735 | 38.7591 | Promoter | 9 | 66065176 | 66069731 | 4556 | 2 | ENSMUST00 | 0 | Snx22 | ENSMUSG00000039452.5 |
| chr9 | 108264086 | 108265067 | 16.34529 | 41.36735 | 38.7591 | Promoter | 9 | 108205958 | 108263924 | 57967 | 2 | ENSMUST00 | -163 | Dag1 | ENSMUSG00000039952.10 |
| chrX | 71049698 | 71052480 | 16.34529 | 41.36735 | 38.7591 | Promoter | 20 | 71050256 | 71154717 | 104462 | 1 | ENSMUST00 | 0 | Mamld1 | ENSMUSG00000059401.8 |
| chr5 | 123749665 | 123750060 | 16.76261 | 41.34956 | 38.74308 | Promoter | 5 | 123749726 | 123821593 | 71868 | 1 | ENSMUST00 | 0 | Kntc1 | ENSMUSG00000029414.7 |
| chr2 | 25224949 | 25225717 | 16.72886 | 41.29202 | 38.68602 | Promoter | 2 | 25222160 | 25224702 | 2543 | 2 | ENSMUST00 | -248 | Tubb4b | ENSMUSG00000036752.4 |
| chr2 | 143034759 | 143035696 | 16.69524 | 41.2347 | 38.62885 | Distal Interge | 2 | 143063039 | 143072853 | 9815 | 1 | ENSMUST00 | -27343 | Snrpb2 | ENSMUSG00000008333.7 |
| chr10 | 62946603 | 62947482 | 14.6238 | 41.22485 | 38.6191 | Promoter | 10 | 62947026 | 62974185 | 27160 | 1 | ENSMUST00 | 0 | Dna2 | ENSMUSG00000036875.10 |
| chr12 | 83922093 | 83922794 | 14.6238 | 41.22485 | 38.6191 | Promoter | 12 | 83794034 | 83921934 | 127901 | 2 | ENSMUST00 | -160 | Numb | ENSMUSG00000021224.9 |
| chr16 | 94084811 | 94086164 | 14.6238 | 41.22485 | 38.6191 | Promoter | 16 | 94085260 | 94127032 | 41773 | 1 | ENSMUST00 | 0 | Sim2 | ENSMUSG00000062713.7 |
| chr17 | 6601259 | 6601680 | 14.6238 | 41.22485 | 38.6191 | Promoter | 17 | 6601671 | 6609679 | 8009 | 1 | ENSMUST00 | 0 | Dynlt1c | ENSMUSG00000000579.9 |
| chr19 | 5447378 | 5448950 | 14.6238 | 41.22485 | 38.6191 | Promoter | 19 | 5447703 | 5455945 | 8243 | 1 | ENSMUST00 | 0 | Fosl1 | ENSMUSG00000024912.5 |
| chr3 | 145923480 | 145924025 | 14.6238 | 41.22485 | 38.6191 | Promoter | 3 | 145924378 | 145934283 | 9906 | 1 | ENSMUST00 | -353 | Bcl10 | ENSMUSG00000028191.10 |
| chr4 | 138249948 | 138251008 | 14.6238 | 41.22485 | 38.6191 | Promoter | 4 | 138250435 | 138301967 | 51533 | 1 | ENSMUST00 | 0 | Kif17 | ENSMUSG00000028758.7 |
| chr6 | 24543105 | 24545291 | 14.6238 | 41.22485 | 38.6191 | Intron (ENSM | 6 | 24556072 | 24573164 | 17093 | 1 | ENSMUST00 | -10781 | Asb15 | ENSMUSG00000029685.10 |
| chr6 | 39256659 | 39257527 | 14.6238 | 41.22485 | 38.6191 | Distal Interge | 6 | 39265844 | 39265942 | 99 | 2 | ENSMUST00 | 8415 | Gm26008 | ENSMUSG00000089422.1 |
| chr6 | 129237778 | 129238619 | 14.6238 | 41.22485 | 38.6191 | Promoter | 6 | 129208412 | 129237948 | 29537 | 2 | ENSMUST00 | 0 | 2310001H17I | ENSMUSG00000097354.2 |
| chr7 | 27446331 | 27446890 | 14.6238 | 41.22485 | 38.6191 | Promoter | 7 | 27356383 | 27446599 | 90217 | 2 | ENSMUST00 | 0 | Sptbn4 | ENSMUSG00000011751.11 |
| chr7 | 99625504 | 99627205 | 14.6238 | 41.22485 | 38.6191 | Promoter | 7 | 99624082 | 99627103 | 3022 | 2 | ENSMUST00 | 0 | Gm4980 | ENSMUSG00000096606.2 |
| chr8 | 12873270 | 12874325 | 14.6238 | 41.22485 | 38.6191 | Promoter | 8 | 12873806 | 12880146 | 6341 | 1 | ENSMUST00 | 0 | Mcf2l | ENSMUSG00000031442.15 |
| chr9 | 21165061 | 21166447 | 14.6238 | 41.22485 | 38.6191 | Promoter | 9 | 21165714 | 21213248 | 47535 | 1 | ENSMUST00 | 0 | Pde4a | ENSMUSG00000032177.11 |
| chr10 | 43002283 | 43003461 | 16.66176 | 41.17759 | 38.57329 | 3' UTR | 10 | 42939881 | 42958721 | 18841 | 1 | ENSMUST00 | 62403 | Scml4 | ENSMUSG00000044770.9 |
| chr3 | 8963869 | 8964497 | 16.66176 | 41.17759 | 38.57329 | Promoter | 3 | 8929598 | 8964046 | 34449 | 2 | ENSMUST00 | 0 | Tpd52 | ENSMUSG00000027506.10 |
| chr6 | 88193507 | 88194515 | 16.66176 | 41.17759 | 38.57329 | Promoter | 6 | 88193891 | 88207030 | 13140 | 1 | ENSMUST00 | 0 | Gata2 | ENSMUSG00000015053.9 |
| chr7 | 45699729 | 45701036 | 16.66176 | 41.17759 | 38.57329 | Promoter | 7 | 45699843 | 45704682 | 4840 | 1 | ENSMUST00 | 0 | Car11 | ENSMUSG00000003273.8 |
| chr9 | 88599150 | 88600133 | 16.66176 | 41.17759 | 38.57329 | Promoter | 9 | 88595325 | 88599243 | 3919 | 2 | ENSMUST00 | 0 | 9430037G07 | ENSMUSG00000097073.2 |
| chr1 | 105780405 | 105781169 | 15.39495 | 41.17039 | 38.56658 | Promoter | 1 | 105780718 | 105847981 | 67264 | 1 | ENSMUST00 | 0 | Tnfrsf11a | ENSMUSG00000026321.7 |
| chr1 | 172340612 | 172341400 | 15.39495 | 41.17039 | 38.56658 | Promoter | 1 | 172341210 | 172374085 | 32876 | 1 | ENSMUST00 | 0 | Kcnj10 | ENSMUSG00000044708.5 |
| chr1 | 176274640 | 176275801 | 15.39495 | 41.17039 | 38.56658 | Promoter | 1 | 175962306 | 176275298 | 312993 | 2 | ENSMUST00 | 0 | Pld5 | ENSMUSG00000055214.10 |
| chr10 | 7181972 | 7183538 | 15.39495 | 41.17039 | 38.56658 | Intron (ENSM | 10 | 7119063 | 7212237 | 93175 | 2 | ENSMUST00 | 28699 | Cnksr3 | ENSMUSG00000015202.8 |
| chr10 | 70094788 | 70095709 | 15.39495 | 41.17039 | 38.56658 | Distal Interge | 10 | 70097121 | 70193200 | 96080 | 1 | ENSMUST00 | -1412 | Ccdc6 | ENSMUSG00000048701.8 |
| chr11 | 68853234 | 68853893 | 15.39495 | 41.17039 | 38.56658 | Promoter | 11 | 68821434 | 68853135 | 31702 | 2 | ENSMUST00 | -100 | Ndel1 | ENSMUSG00000018736.9 |
| chr11 | 66525341 | 66526900 | 15.39495 | 41.17039 | 38.56658 | Promoter | 11 | 66211725 | 66525795 | 314071 | 2 | ENSMUST00 | 0 | Shisa6 | ENSMUSG00000053930.8 |
| chr12 | 31558921 | 31559896 | 15.39495 | 41.17039 | 38.56658 | Promoter | 12 | 31519827 | 31559969 | 40143 | 2 | ENSMUST00 | 73 | Slc26a4 | ENSMUSG00000020651.7 |
| chr12 | 73044328 | 73047056 | 15.39495 | 41.17039 | 38.56658 | Promoter | 12 | 73043774 | 73045291 | 1518 | 2 | ENSMUST00 | 0 | Six1 | ENSMUSG00000051367.8 |
| chr13 | 73536148 | 73537118 | 15.39495 | 41.17039 | 38.56658 | Promoter | 13 | 73536747 | 73578672 | 41926 | 1 | ENSMUST00 | 0 | Slc6a3 | ENSMUSG00000021609.6 |
| chr13 | 42866208 | 42867521 | 15.39495 | 41.17039 | 38.56658 | Promoter | 13 | 42866247 | 43135304 | 269058 | 1 | ENSMUST00 | 0 | Phactr1 | ENSMUSG00000054728.11 |
| chr13 | 23570413 | 23571218 | 15.39495 | 41.17039 | 38.56658 | Promoter | 13 | 23570662 | 23571121 | 460 | 2 | ENSMUST00 | 0 | Hist1h2ae | ENSMUSG00000069272.5 |
| chr14 | 7104799 | 7105748 | 15.39495 | 41.17039 | 38.56658 | Distal Interge | 14 | 7083208 | 7100621 | 17414 | 2 | ENSMUST00 | -4179 | Gm3696 | ENSMUSG00000092167.3 |
| chr15 | 37458319 | 37460172 | 15.39495 | 41.17039 | 38.56658 | Promoter | 15 | 37366218 | 37459360 | 93143 | 2 | ENSMUST00 | 0 | Ncald | ENSMUSG00000051359.9 |
| chr16 | 34095427 | 34096307 | 15.39495 | 41.17039 | 38.56658 | Promoter | 16 | 33974258 | 34095987 | 121730 | 2 | ENSMUST00 | 0 | Kalrn | ENSMUSG00000061751.10 |
| chr17 | 69968953 | 69970217 | 15.39495 | 41.17039 | 38.56658 | Promoter | 17 | 69969073 | 70821413 | 852341 | 1 | ENSMUST00 | 0 | Dlgap1 | ENSMUSG00000003279.11 |
| chr18 | 46464920 | 46465941 | 15.39495 | 41.17039 | 38.56658 | Promoter | 18 | 46465215 | 46465790 | 576 | 2 | ENSMUST00 | 0 | Mospd4 | ENSMUSG00000046487.6 |
| chr19 | 20388965 | 20390989 | 15.39495 | 41.17039 | 38.56658 | Promoter | 19 | 20373428 | 20390944 | 17517 | 2 | ENSMUST00 | 0 | Anxa1 | ENSMUSG00000024659.9 |
| chr2 | 168117297 | 168117736 | 15.39495 | 41.17039 | 38.56658 | Distal Interge | 2 | 168148017 | 168153533 | 5517 | 2 | ENSMUST00 | 35797 | Gm14235 | ENSMUSG00000085267.1 |
| chr3 | 100825118 | 100827041 | 15.39495 | 41.17039 | 38.56658 | Promoter | 3 | 100825459 | 100895679 | 70221 | 1 | ENSMUST00 | 0 | Vtcn1 | ENSMUSG00000051076.7 |
| chr4 | 41760142 | 41761120 | 15.39495 | 41.17039 | 38.56658 | Promoter | 4 | 41760196 | 41769474 | 9279 | 1 | ENSMUST00 | 0 | Il11ra1 | ENSMUSG00000073889.5 |
| chr5 | 138365410 | 138365674 | 15.39495 | 41.17039 | 38.56658 | Promoter | 5 | 138364260 | 138365552 | 1293 | 2 | ENSMUST00 | 0 | Got2-ps1 | ENSMUSG00000080935.3 |
| chr7 | 140941081 | 140942637 | 15.39495 | 41.17039 | 38.56658 | Promoter | 7 | 140941391 | 140947656 | 6266 | 1 | ENSMUST00 | 0 | Athl1 | ENSMUSG00000062031.7 |
| chr8 | 86585276 | 86585854 | 15.39495 | 41.17039 | 38.56658 | Distal Interge | 8 | 86533961 | 86580686 | 46726 | 2 | ENSMUST00 | -4591 | Abcc12 | ENSMUSG00000036872.11 |
| chr9 | 78191361 | 78192508 | 15.39495 | 41.17039 | 38.56658 | Promoter | 9 | 78191966 | 78209348 | 17383 | 1 | ENSMUST00 | 0 | Gsta4 | ENSMUSG00000032348.5 |
| chr9 | 108289989 | 108290874 | 15.39495 | 41.17039 | 38.56658 | Promoter | 9 | 108290448 | 108296498 | 6051 | 1 | ENSMUST00 | 0 | Nicn1 | ENSMUSG00000032606.6 |
| chr9 | 74848371 | 74849583 | 15.39495 | 41.17039 | 38.56658 | Promoter | 9 | 74848437 | 74860134 | 11698 | 1 | ENSMUST00 | 0 | Gm16551 | ENSMUSG00000066477.8 |
| chrX | 153359318 | 153360577 | 15.39495 | 41.17039 | 38.56658 | Promoter | 20 | 153359613 | 153395797 | 36185 | 1 | ENSMUST00 | 0 | Klf8 | ENSMUSG00000041649.8 |
| chrX | 106187110 | 106188039 | 15.39495 | 41.17039 | 38.56658 | Promoter | 20 | 106187163 | 106203661 | 16499 | 1 | ENSMUST00 | 0 | Pgk1 | ENSMUSG00000062070.7 |
| chr16 | 17802510 | 17803686 | 15.73948 | 41.04805 | 38.4465 | Exon (ENSMU | 16 | 17797282 | 17808287 | 11006 | 1 | ENSMUST00 | 5229 | Scarf2 | ENSMUSG00000012017.3 |
| chr10 | 93539629 | 93540696 | 17.00269 | 41.02908 | 38.42754 | Promoter | 10 | 93523338 | 93540033 | 16696 | 2 | ENSMUST00 | 0 | Amdhd1 | ENSMUSG00000015890.2 |
| chr11 | 100545213 | 100546009 | 17.00269 | 41.02908 | 38.42754 | Promoter | 11 | 100545607 | 100571963 | 26357 | 1 | ENSMUST00 | 0 | Ttc25 | ENSMUSG00000006784.9 |
| chr11 | 103311925 | 103312530 | 17.00269 | 41.02908 | 38.42754 | Distal Interge | 11 | 103333229 | 103333395 | 167 | 2 | ENSMUST00 | 20865 | Arhgap27 | ENSMUSG00000034255.12 |
| chr11 | 6064895 | 6066080 | 17.00269 | 41.02908 | 38.42754 | Promoter | 11 | 5969644 | 6065748 | 96105 | 2 | ENSMUST00 | 0 | Camk2b | ENSMUSG00000057897.9 |
| chr14 | 7567919 | 7568783 | 17.00269 | 41.02908 | 38.42754 | Promoter | 14 | 7545151 | 7568566 | 23416 | 2 | ENSMUST00 | 0 | Gm3558 | ENSMUSG00000079364.3 |
| chr15 | 84445823 | 84446678 | 17.00269 | 41.02908 | 38.42754 | Promoter | 15 | 84380047 | 84446545 | 66499 | 2 | ENSMUST00 | 0 | 1810041L15F | ENSMUSG00000062760.5 |
| chr15 | 100443628 | 100444535 | 17.00269 | 41.02908 | 38.42754 | Distal Interge | 15 | 100449684 | 100449793 | 110 | 1 | ENSMUST00 | -5149 | n-R5s43 | ENSMUSG00000084688.1 |
| chr18 | 12305183 | 12306014 | 17.00269 | 41.02908 | 38.42754 | Promoter | 18 | 12252362 | 12305711 | 53350 | 2 | ENSMUST00 | 0 | Ankrd29 | ENSMUSG00000057766.8 |
| chr2 | 155775484 | 155776154 | 17.00269 | 41.02908 | 38.42754 | Promoter | 2 | 155775342 | 155818366 | 43025 | 1 | ENSMUST00 | 143 | Mmp24 | ENSMUSG00000027612.7 |
| chr3 | 58636252 | 58637357 | 17.00269 | 41.02908 | 38.42754 | Promoter | 3 | 58616300 | 58637207 | 20908 | 2 | ENSMUST00 | 0 | Erich6 | ENSMUSG00000070471.3 |
| chr5 | 135888320 | 135889132 | 17.00269 | 41.02908 | 38.42754 | Promoter | 5 | 135888018 | 135889561 | 1544 | 1 | ENSMUST00 | 303 | Hspb1 | ENSMUSG00000004951.10 |
| chr7 | 25178834 | 25180107 | 17.00269 | 41.02908 | 38.42754 | Distal Interge | 7 | 25120618 | 25159922 | 39305 | 2 | ENSMUST00 | -18913 | Pou2f2 | ENSMUSG00000008496.13 |
| chr7 | 47132080 | 47132998 | 17.00269 | 41.02908 | 38.42754 | Promoter | 7 | 47077798 | 47132326 | 54529 | 2 | ENSMUST00 | 0 | Ptpn5 | ENSMUSG00000030854.10 |
| chr7 | 142658304 | 142659721 | 17.00269 | 41.02908 | 38.42754 | Promoter | 7 | 142650769 | 142659489 | 8721 | 2 | ENSMUST00 | 0 | Igf2 | ENSMUSG00000048583.11 |
| chr9 | 120303424 | 120304710 | 17.00269 | 41.02908 | 38.42754 | Promoter | 9 | 120303936 | 120474841 | 170906 | 1 | ENSMUST00 | 0 | Myrip | ENSMUSG00000041794.7 |
| chr9 | 95559137 | 95560707 | 17.00269 | 41.02908 | 38.42754 | Promoter | 9 | 95559657 | 95568023 | 8367 | 1 | ENSMUST00 | 0 | Paqr9 | ENSMUSG00000064225.6 |
| chrX | 52163494 | 52165503 | 17.00269 | 41.02908 | 38.42754 | Promoter | 20 | 52053021 | 52165252 | 112232 | 2 | ENSMUST00 | 0 | Gpc4 | ENSMUSG00000031119.4 |
| chrX | 7884032 | 7884970 | 17.00269 | 41.02908 | 38.42754 | Promoter | 20 | 7884041 | 7894492 | 10452 | 1 | ENSMUST00 | 0 | Slc35a2 | ENSMUSG00000031156.12 |
| chrX | 8192147 | 8193775 | 17.00269 | 41.02908 | 38.42754 | Promoter | 20 | 8185329 | 8193512 | 8184 | 2 | ENSMUST00 | 0 | Ebp | ENSMUSG00000031168.9 |
| chrX | 75415868 | 75417305 | 17.00269 | 41.02908 | 38.42754 | Promoter | 20 | 75410444 | 75416543 | 6100 | 2 | ENSMUST00 | 0 | Mtcp1 | ENSMUSG00000031200.11 |
| chrX | 109011548 | 109014121 | 17.00269 | 41.02908 | 38.42754 | Promoter | 20 | 109004534 | 109013404 | 8871 | 2 | ENSMUST00 | 0 | Hmgn5 | ENSMUSG00000031245.8 |
| chrX | 106027255 | 106028197 | 17.00269 | 41.02908 | 38.42754 | Promoter | 20 | 106027276 | 106124917 | 97642 | 1 | ENSMUST00 | 0 | Atp7a | ENSMUSG00000033792.7 |
| chrX | 7604989 | 7605424 | 17.00269 | 41.02908 | 38.42754 | Promoter | 20 | 7558602 | 7605381 | 46780 | 2 | ENSMUST00 | 0 | Ppp1r3f | ENSMUSG00000039556.11 |
| chrX | 134601177 | 134602157 | 17.00269 | 41.02908 | 38.42754 | Promoter | 20 | 134601179 | 134607054 | 5876 | 1 | ENSMUST00 | 0 | Hnrnph2 | ENSMUSG00000045427.8 |
| chrX | 108832866 | 108834360 | 17.00269 | 41.02908 | 38.42754 | Promoter | 20 | 108737016 | 108834348 | 97333 | 2 | ENSMUST00 | 0 | Brwd3 | ENSMUSG00000063663.6 |
| chrX | 133897922 | 133898454 | 17.00269 | 41.02908 | 38.42754 | Promoter | 20 | 133891068 | 133898429 | 7362 | 2 | ENSMUST00 | 0 | Tspan6 | ENSMUSG00000067377.7 |
| chrX | 150477935 | 150478691 | 17.00269 | 41.02908 | 38.42754 | Distal Interge | 20 | 150463726 | 150466156 | 2431 | 1 | ENSMUST00 | 14210 | 3110067C02F | ENSMUSG00000081187.1 |
| chrX | 75764116 | 75765210 | 17.00269 | 41.02908 | 38.42754 | Promoter | 20 | 75706932 | 75764699 | 57768 | 2 | ENSMUST00 | 0 | 4933407K13I | ENSMUSG00000087396.2 |
| chrY | 897595 | 898726 | 17.00269 | 41.02908 | 38.42754 | Promoter | 21 | 897788 | 917605 | 19818 | 1 | ENSMUST00 | 0 | Kdm5d | ENSMUSG00000056673.9 |
| chr11 | 87421949 | 87422861 | 12.72529 | 41.01831 | 38.41836 | Promoter | 11 | 87422867 | 87423030 | 164 | 1 | ENSMUST00 | -6 | Rnu1a1 | ENSMUSG00000095969.1 |
| chr6 | 137272934 | 137275402 | 12.72529 | 41.01831 | 38.41836 | Intron (ENSM | 6 | 137252299 | 137464629 | 212331 | 1 | ENSMUST00 | 20636 | Ptpro | ENSMUSG00000030223.9 |
| chr7 | 141461557 | 141462588 | 12.72529 | 41.01831 | 38.41836 | Promoter | 7 | 141461895 | 141465029 | 3135 | 1 | ENSMUST00 | 0 | Efcab4a | ENSMUSG00000048200.6 |
| chr15 | 103058507 | 103060089 | 16.56211 | 41.00751 | 38.40777 | Distal Interge | 15 | 103034395 | 103036852 | 2458 | 1 | ENSMUST00 | 24113 | Hoxc4 | ENSMUSG00000075394.4 |
| chr10 | 75950875 | 75951997 | 16.12979 | 40.99568 | 38.39605 | Distal Interge | 10 | 75954514 | 76009591 | 55078 | 1 | ENSMUST00 | -2517 | Gm5134 | ENSMUSG00000033255.8 |
| chr17 | 23785858 | 23786812 | 16.12979 | 40.99568 | 38.39605 | Promoter | 17 | 23776916 | 23786081 | 9166 | 2 | ENSMUST00 | 0 | Flywch2 | ENSMUSG00000023911.7 |
| chr18 | 32559358 | 32560222 | 16.12979 | 40.99568 | 38.39605 | Promoter | 18 | 32528322 | 32560034 | 31713 | 2 | ENSMUST00 | 0 | Gypc | ENSMUSG00000090523.2 |
| chr5 | 135545622 | 135546032 | 16.12979 | 40.99568 | 38.39605 | Promoter | 5 | 135406523 | 135545122 | 138600 | 2 | ENSMUST00 | -501 | Hip1 | ENSMUSG00000039959.9 |
| chr7 | 30143506 | 30144064 | 16.12979 | 40.99568 | 38.39605 | Distal Interge | 7 | 30125615 | 30133545 | 7931 | 1 | ENSMUST00 | 17892 | Zfp382 | ENSMUSG00000074220.4 |
| chr13 | 74770224 | 74771108 | 16.97211 | 40.97725 | 38.37832 | Intron (ENSM | 13 | 74694286 | 74807921 | 113636 | 2 | ENSMUST00 | 36813 | Cast | ENSMUSG00000021585.8 |
| chr4 | 43499322 | 43500136 | 16.93657 | 40.91702 | 38.31829 | Promoter | 4 | 43496142 | 43499695 | 3554 | 2 | ENSMUST00 | 0 | Arhgef39 | ENSMUSG00000051517.9 |
| chr13 | 95523996 | 95525452 | 15.65496 | 40.89989 | 38.30132 | Promoter | 13 | 95511732 | 95525240 | 13509 | 2 | ENSMUST00 | 0 | F2rl1 | ENSMUSG00000021678.8 |
| chr5 | 137644938 | 137646119 | 15.65496 | 40.89989 | 38.30132 | Promoter | 5 | 137643032 | 137645714 | 2683 | 2 | ENSMUST00 | 0 | Irs3 | ENSMUSG00000047182.5 |
| chr11 | 106613192 | 106613790 | 15.22852 | 40.8738 | 38.27569 | Promoter | 11 | 106568098 | 106613423 | 45326 | 2 | ENSMUST00 | 0 | Tex2 | ENSMUSG00000040548.10 |
| chr3 | 96497223 | 96497868 | 15.22852 | 40.8738 | 38.27569 | Distal Interge | 3 | 96488729 | 96488893 | 165 | 1 | ENSMUST00 | 8495 | Gm22614 | ENSMUSG00000094812.1 |
| chr15 | 82340950 | 82341601 | 16.90119 | 40.85703 | 38.25915 | Promoter | 15 | 82340986 | 82349007 | 8022 | 1 | ENSMUST00 | 0 | Smdt1 | ENSMUSG00000022452.9 |
| chr16 | 97610374 | 97611477 | 16.90119 | 40.85703 | 38.25915 | Promoter | 16 | 97564684 | 97611195 | 46512 | 2 | ENSMUST00 | 0 | Tmprss2 | ENSMUSG00000000385.7 |
| chr18 | 15759782 | 15760426 | 16.90119 | 40.85703 | 38.25915 | Promoter | 18 | 15451924 | 15760157 | 308234 | 2 | ENSMUST00 | 0 | Chst9 | ENSMUSG00000047161.7 |
| chr8 | 12394922 | 12395972 | 16.90119 | 40.85703 | 38.25915 | Promoter | 8 | 12395295 | 12400126 | 4832 | 1 | ENSMUST00 | 0 | Sox1 | ENSMUSG00000096014.1 |
| chr7 | 27485942 | 27486823 | 16.43109 | 40.78362 | 38.18623 | Promoter | 7 | 27486910 | 27490316 | 3407 | 1 | ENSMUST00 | -87 | Sertad1 | ENSMUSG00000008384.8 |
| chr11 | 79973309 | 79974662 | 16.83086 | 40.73776 | 38.14046 | Distal Interge | 11 | 79933953 | 79962390 | 28438 | 2 | ENSMUST00 | -10920 | Utp6 | ENSMUSG00000035575.6 |
| chr1 | 193152749 | 193153978 | 16.79592 | 40.67848 | 38.08141 | Promoter | 1 | 193153112 | 193166868 | 13757 | 1 | ENSMUST00 | 0 | Irf6 | ENSMUSG00000026638.10 |
| chr2 | 170287072 | 170287903 | 16.76111 | 40.61943 | 38.02256 | Distal Interge | 2 | 170285035 | 170285880 | 846 | 1 | ENSMUST00 | 2038 | Gm14270 | ENSMUSG00000084013.1 |
| chr12 | 21783440 | 21784809 | 16.33418 | 40.6178 | 38.021 | Distal Interge | 12 | 21776159 | 21776707 | 549 | 2 | ENSMUST00 | -6734 | Gm5633 | ENSMUSG00000091105.1 |
| chr10 | 80656757 | 80657091 | 16.72646 | 40.56061 | 37.96445 | Promoter | 10 | 80642617 | 80657071 | 14455 | 2 | ENSMUST00 | 0 | Btbd2 | ENSMUSG00000003344.9 |
| chr10 | 77901982 | 77903681 | 13.16235 | 40.53663 | 37.94056 | Promoter | 10 | 77897575 | 77902536 | 4962 | 2 | ENSMUST00 | 0 | Lrrc3 | ENSMUSG00000051652.4 |

| | | | | | | | | | | | | | | | |
|---|---|---|---|---|---|---|---|---|---|---|---|---|---|---|---|
| chr11 | 35979365 | 35981000 | 13.16235 | 40.53663 | 37.94056 | Promoter | 11 | 35838400 | 35980527 | 142128 | 2 | ENSMUST00 | 0 | Wwc1 | ENSMUSG00000018849.6 |
| chr13 | 119755239 | 119756642 | 13.16235 | 40.53663 | 37.94056 | Promoter | 13 | 119710094 | 119755882 | 45789 | 2 | ENSMUST00 | 0 | Nim1k | ENSMUSG00000095930.1 |
| chr14 | 20294589 | 20295313 | 13.16235 | 40.53663 | 37.94056 | Promoter | 14 | 20294690 | 20317575 | 22886 | 1 | ENSMUST00 | 0 | Nudt13 | ENSMUSG00000021809.9 |
| chr17 | 73709348 | 73710868 | 13.16235 | 40.53663 | 37.94056 | Promoter | 17 | 73493228 | 73710451 | 217224 | 2 | ENSMUST00 | 0 | Galnt14 | ENSMUSG00000024064.8 |
| chr18 | 63386057 | 63387399 | 13.16235 | 40.53663 | 37.94056 | Promoter | 18 | 63010213 | 63387183 | 376971 | 2 | ENSMUST00 | 0 | Piezo2 | ENSMUSG00000041482.11 |
| chr2 | 163202708 | 163203711 | 13.16235 | 40.53663 | 37.94056 | Promoter | 2 | 163203125 | 163324170 | 121046 | 1 | ENSMUST00 | 0 | Tox2 | ENSMUSG00000074607.6 |
| chr15 | 48789549 | 48790198 | 16.69194 | 40.50202 | 37.90651 | Intron (ENSM | 15 | 47580637 | 48791933 | 1211297 | 2 | ENSMUST00 | 1735 | Csmd3 | ENSMUSG00000022311.10 |
| chr1 | 95667025 | 95668366 | 15.83033 | 40.47729 | 37.8819 | Promoter | 1 | 95587682 | 95667571 | 79890 | 2 | ENSMUST00 | 0 | St8sia4 | ENSMUSG00000040710.5 |
| chr11 | 4118213 | 4118877 | 15.83033 | 40.47729 | 37.8819 | Promoter | 11 | 4097039 | 4118831 | 21793 | 2 | ENSMUST00 | 0 | Sec14l2 | ENSMUSG00000003585.8 |
| chr11 | 50174279 | 50174664 | 15.83033 | 40.47729 | 37.8819 | Promoter | 11 | 50174444 | 50178348 | 3905 | 1 | ENSMUST00 | 0 | 3010026O09 | ENSMUSG00000020381.5 |
| chr6 | 91978407 | 91979681 | 15.83033 | 40.47729 | 37.8819 | Promoter | 6 | 91978878 | 92076004 | 97127 | 1 | ENSMUST00 | 0 | Fgd5 | ENSMUSG00000034037.9 |
| chr13 | 119689658 | 119690178 | 16.2384 | 40.45376 | 37.85896 | Promoter | 13 | 119690462 | 119708086 | 17625 | 1 | ENSMUST00 | -284 | Hmgcs1 | ENSMUSG00000093930.1 |
| chr17 | 32492227 | 32492815 | 16.2384 | 40.45376 | 37.85896 | Exon (ENSMU | 17 | 32506462 | 32528894 | 22433 | 1 | ENSMUST00 | -13647 | Cyp4f17 | ENSMUSG00000091586.1 |
| chr2 | 168589884 | 168590651 | 16.65757 | 40.44366 | 37.84893 | Promoter | 2 | 168476410 | 168590260 | 113851 | 2 | ENSMUST00 | 0 | Nfatc2 | ENSMUSG00000027544.11 |
| chr4 | 109116279 | 109118694 | 14.97413 | 40.41832 | 37.82371 | Promoter | 4 | 109061145 | 109118022 | 56878 | 2 | ENSMUST00 | 0 | Osbpl9 | ENSMUSG00000028559.12 |
| chr5 | 24842964 | 24843392 | 14.97413 | 40.41832 | 37.82371 | Promoter | 5 | 24802823 | 24842624 | 39802 | 2 | ENSMUST00 | -341 | Rheb | ENSMUSG00000028945.8 |
| chr19 | 45046467 | 45048099 | 15.37965 | 40.41568 | 37.82122 | Promoter | 19 | 45047576 | 45056383 | 8808 | 1 | ENSMUST00 | 0 | Sfxn3 | ENSMUSG00000025212.11 |
| chr3 | 129393223 | 129393932 | 15.37965 | 40.41568 | 37.82122 | Distal Interge | 3 | 129298594 | 129332720 | 34127 | 2 | ENSMUST00 | -60504 | Enpep | ENSMUSG00000028024.9 |
| chr5 | 116419828 | 116421247 | 15.37965 | 40.41568 | 37.82122 | Intron (ENSM | 5 | 116408491 | 116422864 | 14374 | 2 | ENSMUST00 | 1617 | Hspb8 | ENSMUSG00000041548.4 |
| chr9 | 14274996 | 14275511 | 16.62334 | 40.38552 | 37.79139 | Promoter | 9 | 14276301 | 14326138 | 49838 | 1 | ENSMUST00 | -790 | Sesn3 | ENSMUSG00000032009.7 |
| chr15 | 79357079 | 79358282 | 12.1011 | 40.35582 | 37.7618 | 3' UTR | 15 | 79348061 | 79359072 | 11012 | 1 | ENSMUST00 | 9019 | Maff | ENSMUSG00000042622.8 |
| chr1 | 42713057 | 42714553 | 16.58925 | 40.3276 | 37.73373 | Promoter | 1 | 42707061 | 42713454 | 6394 | 2 | ENSMUST00 | 0 | Pou3f3os2 | ENSMUSG00000097063.2 |
| chr19 | 44284311 | 44285094 | 16.58925 | 40.3276 | 37.73373 | Distal Interge | 19 | 44293676 | 44306864 | 13189 | 1 | ENSMUST00 | -8582 | Scd2 | ENSMUSG00000025203.5 |
| chr15 | 73724395 | 73725364 | 16.5553 | 40.26991 | 37.6762 | Promoter | 15 | 73724930 | 73758766 | 33837 | 1 | ENSMUST00 | 0 | Ptp4a3 | ENSMUSG00000059895.7 |
| chr2 | 158667020 | 158668469 | 16.5553 | 40.26991 | 37.6762 | Promoter | 2 | 158667119 | 158766334 | 99216 | 1 | ENSMUST00 | 0 | Ppp1r16b | ENSMUSG00000037754.8 |
| chr7 | 114020375 | 114020965 | 16.5553 | 40.26991 | 37.6762 | Intron (ENSM | 7 | 114046785 | 114117781 | 70997 | 2 | ENSMUST00 | 96816 | Rras2 | ENSMUSG00000055723.5 |
| chr7 | 71705310 | 71705898 | 16.11243 | 40.23772 | 37.64439 | Distal Interge | 7 | 71370333 | 71379813 | 9481 | 1 | ENSMUST00 | 334978 | RP24-173C11 | ENSMUSG00000099853.1 |
| chr1 | 118481424 | 118482973 | 16.48782 | 40.15519 | 37.56212 | Intron (ENSM | 1 | 118478604 | 118479591 | 988 | 2 | ENSMUST00 | -1834 | RP24-553C6. | ENSMUSG00000100770.1 |
| chr11 | 120184503 | 120186230 | 13.67473 | 40.1468 | 37.55384 | Exon (ENSMU | 11 | 120185400 | 120188751 | 3352 | 2 | ENSMUST00 | 2521 | 2810410L24 | ENSMUSG00000075389.3 |
| chr15 | 7809876 | 7811553 | 13.67473 | 40.1468 | 37.55384 | Promoter | 15 | 7811011 | 7837575 | 26565 | 1 | ENSMUST00 | 0 | Gdnf | ENSMUSG00000022144.3 |
| chr15 | 85831894 | 85832784 | 13.67473 | 40.1468 | 37.55384 | Promoter | 15 | 85832306 | 85858822 | 26517 | 1 | ENSMUST00 | 0 | Ttc38 | ENSMUSG00000035944.12 |
| chr17 | 47672176 | 47673092 | 13.67473 | 40.1468 | 37.55384 | Promoter | 17 | 47672720 | 47681056 | 8337 | 1 | ENSMUST00 | 0 | Usp49 | ENSMUSG00000090115.2 |
| chr18 | 37814265 | 37816104 | 13.67473 | 40.1468 | 37.55384 | Promoter | 18 | 37815079 | 37841870 | 26792 | 1 | ENSMUST00 | 0 | Pcdhga12 | ENSMUSG00000023036.9 |
| chr4 | 155839481 | 155841295 | 13.67473 | 40.1468 | 37.55384 | Promoter | 4 | 155839680 | 155842130 | 2451 | 1 | ENSMUST00 | 0 | Mxra8 | ENSMUSG00000029070.4 |
| chr5 | 104020933 | 104022665 | 13.67473 | 40.1468 | 37.55384 | Promoter | 5 | 103989762 | 104021919 | 32158 | 2 | ENSMUST00 | 0 | Hsd17b11 | ENSMUSG00000029311.11 |
| chr6 | 127330017 | 127330893 | 13.67473 | 40.1468 | 37.55384 | Distal Interge | 6 | 127282063 | 127282509 | 447 | 2 | ENSMUST00 | -47509 | Gm7308 | ENSMUSG00000081113.1 |
| chr7 | 30373735 | 30374093 | 13.67473 | 40.1468 | 37.55384 | Distal Interge | 7 | 30355514 | 30362772 | 7259 | 2 | ENSMUST00 | -10964 | Lrfn3 | ENSMUSG00000036957.7 |
| chr8 | 105288985 | 105290387 | 13.67473 | 40.1468 | 37.55384 | Promoter | 8 | 105280410 | 105289528 | 9119 | 2 | ENSMUST00 | 0 | 4931428F04 | ENSMUSG00000014837.9 |
| chr4 | 95556964 | 95558007 | 16.05018 | 40.13083 | 37.5387 | Promoter | 4 | 95557507 | 95926939 | 369433 | 1 | ENSMUST00 | 0 | Fggy | ENSMUSG00000028573.13 |
| chr10 | 89872994 | 89874358 | 16.01923 | 40.07766 | 37.48569 | Promoter | 10 | 89873509 | 90970219 | 1096711 | 1 | ENSMUST00 | 0 | Anks1b | ENSMUSG00000058589.9 |
| chr10 | 8761269 | 8762607 | 16.01923 | 40.07766 | 37.48569 | Intron (ENSM | 10 | 8824011 | 8824147 | 137 | 2 | ENSMUST00 | 61540 | Gm25410 | ENSMUSG00000065692.1 |
| chr4 | 53217574 | 53218523 | 16.01923 | 40.07766 | 37.48569 | Promoter | 4 | 53214557 | 53217857 | 3301 | 2 | ENSMUST00 | 0 | 4930412L05 | ENSMUSG00000087192.2 |
| chr8 | 69328900 | 69329440 | 16.01923 | 40.07766 | 37.48569 | Distal Interge | 8 | 69300776 | 69303162 | 2387 | 1 | ENSMUST00 | 28125 | D130040H23 | ENSMUSG00000079038.2 |
| chr9 | 107553604 | 107554526 | 16.01923 | 40.07766 | 37.48569 | Promoter | 9 | 107554656 | 107562263 | 7608 | 1 | ENSMUST00 | -130 | Rassf1 | ENSMUSG00000010067.8 |
| chr5 | 23386552 | 23387148 | 16.42088 | 40.04133 | 37.44989 | Exon (ENSMU | 5 | 23422404 | 23433309 | 10906 | 2 | ENSMUST00 | 46161 | 5031425E22 | ENSMUSG00000073147.3 |
| chr2 | 72296487 | 72298952 | 15.54179 | 39.97545 | 37.38426 | Promoter | 2 | 72297901 | 72355622 | 57722 | 1 | ENSMUST00 | 0 | Zak | ENSMUSG00000004085.9 |
| chr5 | 115483894 | 115484982 | 15.54179 | 39.97545 | 37.38426 | Promoter | 5 | 115478010 | 115484297 | 6288 | 2 | ENSMUST00 | 0 | Sirt4 | ENSMUSG00000029524.11 |
| chr9 | 57074897 | 57076055 | 14.72811 | 39.97517 | 37.38426 | Promoter | 9 | 57075338 | 57128366 | 53029 | 1 | ENSMUST00 | 0 | Sin3a | ENSMUSG00000042557.9 |
| chr4 | 155769708 | 155770765 | 15.11386 | 39.94567 | 37.35505 | Promoter | 4 | 155769969 | 155776161 | 6193 | 1 | ENSMUST00 | 0 | Gm26840 | ENSMUSG00000097728.1 |
| chr9 | 48266870 | 48267537 | 15.11386 | 39.94567 | 37.35505 | Distal Interge | 9 | 48319144 | 48340834 | 21691 | 2 | ENSMUST00 | 73297 | Nxpe2 | ENSMUSG00000032028.10 |
| chr1 | 74770804 | 74772444 | 15.92618 | 39.91766 | 37.3274 | Promoter | 1 | 74771892 | 74785322 | 13431 | 1 | ENSMUST00 | 0 | Wnt6 | ENSMUSG00000033227.7 |
| chr1 | 90203977 | 90206221 | 15.92618 | 39.91766 | 37.3274 | Promoter | 1 | 90203980 | 90216751 | 12772 | 1 | ENSMUST00 | 0 | Ackr3 | ENSMUSG00000044337.5 |
| chr10 | 21144680 | 21145568 | 15.92618 | 39.91766 | 37.3274 | Promoter | 10 | 21145094 | 21154374 | 9281 | 1 | ENSMUST00 | 0 | Gm26577 | ENSMUSG00000097923.1 |
| chr11 | 120948429 | 120949719 | 15.92618 | 39.91766 | 37.3274 | Promoter | 11 | 120948480 | 120958775 | 10296 | 1 | ENSMUST00 | 0 | Slc16a3 | ENSMUSG00000025161.11 |
| chr12 | 103433887 | 103435079 | 15.92618 | 39.91766 | 37.3274 | Promoter | 12 | 103434211 | 103440207 | 5997 | 1 | ENSMUST00 | 0 | Ifi27 | ENSMUSG00000064215.8 |
| chr12 | 24096368 | 24097680 | 15.92618 | 39.91766 | 37.3274 | Promoter | 12 | 24044701 | 24096968 | 52268 | 2 | ENSMUST00 | 0 | 9030624G23 | ENSMUSG00000073158.3 |
| chr14 | 33923135 | 33924145 | 15.92618 | 39.91766 | 37.3274 | Promoter | 14 | 33923587 | 33935280 | 11694 | 1 | ENSMUST00 | 0 | Gdf10 | ENSMUSG00000021943.6 |
| chr15 | 97053965 | 97056019 | 15.92618 | 39.91766 | 37.3274 | Promoter | 15 | 96994823 | 97055956 | 61134 | 2 | ENSMUST00 | 0 | Slc38a4 | ENSMUSG00000022464.8 |
| chr17 | 23592399 | 23593246 | 15.92618 | 39.91766 | 37.3274 | Intron (ENSM | 17 | 23575845 | 23586232 | 10388 | 2 | ENSMUST00 | -6168 | Zfp13 | ENSMUSG00000062012.8 |
| chr17 | 47973525 | 47974624 | 15.92618 | 39.91766 | 37.3274 | Distal Interge | 17 | 48003379 | 48003572 | 194 | 2 | ENSMUST00 | 28948 | Gm14871 | ENSMUSG00000082492.1 |
| chr18 | 57132860 | 57133844 | 15.92618 | 39.91766 | 37.3274 | Promoter | 18 | 57133090 | 57297467 | 164378 | 1 | ENSMUST00 | 0 | Megf10 | ENSMUSG00000024593.10 |
| chr19 | 59075104 | 59076053 | 15.92618 | 39.91766 | 37.3274 | Promoter | 19 | 58973358 | 59076069 | 102712 | 2 | ENSMUST00 | 16 | 4930506M07 | ENSMUSG00000041362.11 |
| chr7 | 109169466 | 109170643 | 15.92618 | 39.91766 | 37.3274 | Promoter | 7 | 109138572 | 109170308 | 31737 | 2 | ENSMUST00 | 0 | Lmo1 | ENSMUSG00000036111.7 |
| chr7 | 45204316 | 45204907 | 15.92618 | 39.91766 | 37.3274 | Promoter | 7 | 45183624 | 45204892 | 21269 | 2 | ENSMUST00 | 0 | Ccdc155 | ENSMUSG00000038292.12 |
| chr8 | 126584020 | 126584892 | 15.92618 | 39.91766 | 37.3274 | Intron (ENSM | 8 | 126588296 | 126593986 | 5691 | 2 | ENSMUST00 | 9094 | Irf2bp2 | ENSMUSG00000051495.7 |
| chr9 | 41207407 | 41207993 | 15.92618 | 39.91766 | 37.3274 | Intron (ENSM | 9 | 41206137 | 41217564 | 11428 | 1 | ENSMUST00 | 1271 | RP24-531B2 | ENSMUSG00000100837.1 |
| chrX | 73342224 | 73343318 | 15.92618 | 39.91766 | 37.3274 | Promoter | 20 | 73342621 | 73359080 | 16460 | 1 | ENSMUST00 | 0 | Zfp275 | ENSMUSG00000031365.8 |
| chr1 | 155145995 | 155146842 | 14.28371 | 39.87881 | 37.29034 | Promoter | 1 | 155127878 | 155146783 | 18906 | 2 | ENSMUST00 | 0 | Mr1 | ENSMUSG00000026471.9 |
| chr11 | 107793136 | 107795006 | 14.28371 | 39.87881 | 37.29034 | Promoter | 11 | 107732357 | 107794557 | 62201 | 2 | ENSMUST00 | 0 | Cacng4 | ENSMUSG00000020723.3 |
| chr11 | 68399515 | 68400667 | 14.28371 | 39.87881 | 37.29034 | Promoter | 11 | 68209364 | 68400823 | 191460 | 2 | ENSMUST00 | 156 | Ntn1 | ENSMUSG00000020902.7 |
| chr11 | 80681163 | 80682790 | 14.28371 | 39.87881 | 37.29034 | Exon (ENSMU | 11 | 80673667 | 80780003 | 106337 | 2 | ENSMUST00 | 97213 | Myo1d | ENSMUSG00000035441.9 |
| chr12 | 9570047 | 9570942 | 14.28371 | 39.87881 | 37.29034 | Promoter | 12 | 9565236 | 9570585 | 5350 | 2 | ENSMUST00 | 0 | RP23-228C1 | ENSMUSG00000099360.1 |
| chr15 | 99074174 | 99075319 | 14.28371 | 39.87881 | 37.29034 | Promoter | 15 | 99074973 | 99083407 | 8435 | 1 | ENSMUST00 | 0 | Troap | ENSMUSG00000032783.8 |
| chr17 | 25828527 | 25829660 | 14.28371 | 39.87881 | 37.29034 | Promoter | 17 | 25828867 | 25831840 | 2974 | 1 | ENSMUST00 | 0 | Jmjd8 | ENSMUSG00000025736.9 |
| chr17 | 23790181 | 23791199 | 14.28371 | 39.87881 | 37.29034 | Promoter | 17 | 23790662 | 23808518 | 17857 | 1 | ENSMUST00 | 0 | Srrm2 | ENSMUSG00000039218.11 |
| chr3 | 36521111 | 36521695 | 14.28371 | 39.87881 | 37.29034 | Promoter | 3 | 36515057 | 36521813 | 6757 | 2 | ENSMUST00 | 118 | Gm11549 | ENSMUSG00000085007.1 |
| chr4 | 152130570 | 152131958 | 14.28371 | 39.87881 | 37.29034 | Promoter | 4 | 152120870 | 152131865 | 10996 | 2 | ENSMUST00 | 0 | Espn | ENSMUSG00000028943.12 |
| chr5 | 108368775 | 108369662 | 14.28371 | 39.87881 | 37.29034 | Promoter | 5 | 108369858 | 108378239 | 8382 | 1 | ENSMUST00 | -196 | Gm10419 | ENSMUSG00000072769.2 |
| chr7 | 73536788 | 73537796 | 14.28371 | 39.87881 | 37.29034 | Promoter | 7 | 73519523 | 73537621 | 18099 | 2 | ENSMUST00 | 0 | Chd2 | ENSMUSG00000078671.5 |
| chr8 | 91312844 | 91313348 | 14.28371 | 39.87881 | 37.29034 | Promoter | 8 | 91217030 | 91313262 | 96233 | 2 | ENSMUST00 | 0 | Rpgrip1l | ENSMUSG00000033282.8 |
| chr8 | 71558435 | 71559376 | 14.28371 | 39.87881 | 37.29034 | Promoter | 8 | 71554303 | 71558871 | 4569 | 2 | ENSMUST00 | 0 | Tmem221 | ENSMUSG00000043664.6 |
| chr9 | 111056021 | 111057753 | 13.85327 | 39.78301 | 37.19628 | Promoter | 9 | 111054835 | 111057270 | 2436 | 2 | ENSMUST00 | 0 | Ccrl2 | ENSMUSG00000043953.8 |
| chr8 | 64204720 | 64207047 | 12.46018 | 39.77933 | 37.19262 | Promoter | 8 | 64014770 | 64205993 | 191224 | 2 | ENSMUST00 | 0 | Tll1 | ENSMUSG00000053626.4 |
| chr1 | 46066472 | 46067289 | 15.01947 | 39.77812 | 37.19162 | Promoter | 1 | 46066738 | 46373546 | 306809 | 1 | ENSMUST00 | 0 | Dnah7b | ENSMUSG00000041144.10 |
| chr10 | 127341518 | 127342336 | 15.01947 | 39.77812 | 37.19162 | Promoter | 10 | 127329889 | 127341589 | 11701 | 2 | ENSMUST00 | 0 | Gli1 | ENSMUSG00000025407.6 |
| chr10 | 81605436 | 81606146 | 15.01947 | 39.77812 | 37.19162 | Promoter | 10 | 81606270 | 81620703 | 14434 | 1 | ENSMUST00 | -124 | BC025920 | ENSMUSG00000074862.6 |
| chr12 | 70824542 | 70825277 | 15.01947 | 39.77812 | 37.19162 | Promoter | 12 | 70825514 | 70902232 | 76719 | 1 | ENSMUST00 | -237 | Frmd6 | ENSMUSG00000048285.8 |
| chr12 | 54709610 | 54710327 | 15.01947 | 39.77812 | 37.19162 | Promoter | 12 | 54710308 | 54710471 | 164 | 1 | ENSMUST00 | 0 | Gm22634 | ENSMUSG00000096214.1 |
| chr13 | 46418090 | 46419504 | 15.01947 | 39.77812 | 37.19162 | Promoter | 13 | 46418300 | 46431090 | 12791 | 1 | ENSMUST00 | 0 | Rbm24 | ENSMUSG00000038132.5 |
| chr14 | 36918174 | 36919751 | 15.01947 | 39.77812 | 37.19162 | Promoter | 14 | 36879143 | 36919333 | 40191 | 2 | ENSMUST00 | 0 | Ccser2 | ENSMUSG00000058690.9 |
| chr16 | 5131703 | 5132833 | 15.01947 | 39.77812 | 37.19162 | Promoter | 16 | 5086291 | 5132481 | 46191 | 2 | ENSMUST00 | 0 | Ppl | ENSMUSG00000039457.3 |
| chr17 | 25081688 | 25083565 | 15.01947 | 39.77812 | 37.19162 | Promoter | 17 | 25057702 | 25081181 | 23480 | 2 | ENSMUST00 | -508 | Tmem204 | ENSMUSG00000024168.7 |
| chr18 | 53463703 | 53465379 | 15.01947 | 39.77812 | 37.19162 | Promoter | 18 | 53464546 | 53575857 | 111312 | 1 | ENSMUST00 | 0 | Prdm6 | ENSMUSG00000069378.8 |
| chr18 | 82855363 | 82856644 | 15.01947 | 39.77812 | 37.19162 | Promoter | 18 | 82855894 | 82870604 | 14711 | 1 | ENSMUST00 | 0 | 4930445N18 | ENSMUSG00000097738.1 |
| chr3 | 107458539 | 107460440 | 15.01947 | 39.77812 | 37.19162 | Promoter | 3 | 107438303 | 107458898 | 20596 | 2 | ENSMUST00 | 0 | Kcnc4 | ENSMUSG00000027895.8 |
| chr4 | 136180058 | 136180616 | 15.01947 | 39.77812 | 37.19162 | Promoter | 4 | 136180783 | 136186807 | 6025 | 1 | ENSMUST00 | -167 | E2f2 | ENSMUSG00000018983.9 |
| chr4 | 73790394 | 73791263 | 15.01947 | 39.77812 | 37.19162 | Promoter | 4 | 73714579 | 73790602 | 76024 | 2 | ENSMUST00 | 0 | Rasef | ENSMUSG00000043003.8 |
| chr4 | 63744609 | 63745629 | 15.01947 | 39.77812 | 37.19162 | Promoter | 4 | 63727084 | 63745113 | 18030 | 2 | ENSMUST00 | 0 | Tnfsf15 | ENSMUSG00000050395.8 |
| chr4 | 88550314 | 88550967 | 15.01947 | 39.77812 | 37.19162 | Distal Interge | 4 | 88557673 | 88558245 | 573 | 2 | ENSMUST00 | 7278 | Ifna15 | ENSMUSG00000096011.1 |
| chr9 | 106437894 | 106438939 | 15.01947 | 39.77812 | 37.19162 | Promoter | 9 | 106432981 | 106438319 | 5339 | 2 | ENSMUST00 | 0 | Acy1 | ENSMUSG00000023262.7 |
| chr9 | 78488672 | 78489472 | 15.01947 | 39.77812 | 37.19162 | Promoter | 9 | 78479649 | 78489151 | 9503 | 2 | ENSMUST00 | 0 | Eef1a1 | ENSMUSG00000037742.9 |
| chr9 | 109058794 | 109059890 | 15.01947 | 39.77812 | 37.19162 | Promoter | 9 | 109057933 | 109059734 | 1802 | 2 | ENSMUST00 | 0 | Trex1 | ENSMUSG00000049734.8 |
| chr9 | 110780486 | 110782085 | 15.01947 | 39.77812 | 37.19162 | Distal Interge | 9 | 110769001 | 110769794 | 794 | 1 | ENSMUST00 | 11486 | Myl3 | ENSMUSG00000059741.8 |
| chr4 | 129810738 | 129811390 | 16.22329 | 39.70485 | 37.12044 | Promoter | 4 | 129811219 | 129839232 | 28014 | 1 | ENSMUST00 | 0 | Ptp4a2 | ENSMUSG00000028788.9 |
| chr6 | 48443941 | 48444382 | 16.22329 | 39.70485 | 37.12044 | Promoter | 6 | 48427697 | 48444036 | 16340 | 2 | ENSMUST00 | 0 | Zfp467 | ENSMUSG00000068551.7 |
| chr17 | 53688796 | 53690412 | 14.15692 | 39.64408 | 37.06027 | Promoter | 17 | 53674786 | 53689333 | 14548 | 2 | ENSMUST00 | 0 | Sgol1 | ENSMUSG00000023940.9 |
| chr8 | 83954804 | 83955365 | 15.74599 | 39.60725 | 37.02347 | Promoter | 8 | 83932329 | 83955205 | 22877 | 2 | ENSMUST00 | 0 | Gm10644 | ENSMUSG00000074219.2 |
| chr19 | 8134178 | 8135169 | 16.15848 | 39.59435 | 37.01059 | Distal Interge | 19 | 8062209 | 8131982 | 69774 | 2 | ENSMUST00 | -2197 | Slc22a28 | ENSMUSG00000063590.3 |
| chr16 | 97762788 | 97764160 | 15.7162 | 39.55585 | 36.97221 | Promoter | 16 | 97741933 | 97763491 | 21559 | 2 | ENSMUST00 | 0 | Ripk4 | ENSMUSG00000005251.9 |
| chr4 | 129584635 | 129585487 | 15.7162 | 39.55585 | 36.97221 | Distal Interge | 4 | 129576815 | 129578549 | 1735 | 2 | ENSMUST00 | -6087 | Fam167b | ENSMUSG00000050493.8 |
| chr6 | 52203683 | 52204833 | 15.7162 | 39.55585 | 36.97221 | Promoter | 6 | 52201754 | 52204587 | 2834 | 2 | ENSMUST00 | 0 | Hoxa5 | ENSMUSG00000038253.6 |
| chr1 | 172550086 | 172550983 | 16.54316 | 39.53096 | 36.94807 | Promoter | 1 | 172538133 | 172550669 | 12537 | 2 | ENSMUST00 | 0 | 4933439K11 | ENSMUSG00000085184.4 |
| chr11 | 4849669 | 4850249 | 16.54316 | 39.53096 | 36.94807 | Promoter | 11 | 4765845 | 4849536 | 83692 | 2 | ENSMUST00 | -134 | Nf2 | ENSMUSG00000009073.11 |
| chr11 | 17051585 | 17052745 | 16.54316 | 39.53096 | 36.94807 | Promoter | 11 | 16971464 | 17052381 | 80918 | 2 | ENSMUST00 | 0 | Plek | ENSMUSG00000020120.10 |
| chr13 | 56752856 | 56753975 | 16.54316 | 39.53096 | 36.94807 | Distal Interge | 13 | 56733017 | 56738993 | 5977 | 1 | ENSMUST00 | 19840 | Smad5 | ENSMUSG00000021540.11 |
| chr13 | 78202990 | 78203865 | 16.54316 | 39.53096 | 36.94807 | Promoter | 13 | 78202787 | 78279391 | 76605 | 1 | ENSMUST00 | 204 | A830082K12 | ENSMUSG00000087143.4 |
| chr15 | 94403292 | 94404783 | 16.54316 | 39.53096 | 36.94807 | Promoter | 15 | 94270163 | 94404258 | 134096 | 2 | ENSMUST00 | 0 | Adamts20 | ENSMUSG00000022449.9 |
| chr17 | 56668497 | 56669004 | 16.54316 | 39.53096 | 36.94807 | Distal Interge | 17 | 56673225 | 56711769 | 38545 | 1 | ENSMUST00 | -4221 | Ranbp3 | ENSMUSG00000002372.8 |
| chr17 | 24208747 | 24209588 | 16.54316 | 39.53096 | 36.94807 | Promoter | 17 | 24203848 | 24209387 | 5540 | 2 | ENSMUST00 | 0 | Ntn3 | ENSMUSG00000079662.1 |
| chr17 | 70817900 | 70818433 | 16.54316 | 39.53096 | 36.94807 | 3' UTR | 17 | 70834664 | 70836044 | 1381 | 1 | ENSMUST00 | -16231 | Gm20703 | ENSMUSG00000093622.1 |
| chr19 | 12517969 | 12518526 | 16.54316 | 39.53096 | 36.94807 | Distal Interge | 19 | 12466341 | 12501996 | 35656 | 2 | ENSMUST00 | -15974 | Dtx4 | ENSMUSG00000039982.7 |
| chr2 | 19553633 | 19554407 | 16.54316 | 39.53096 | 36.94807 | Promoter | 2 | 19462837 | 19553914 | 91078 | 2 | ENSMUST00 | 0 | 4921504E06 | ENSMUSG00000026734.9 |
| chr2 | 176226466 | 176226940 | 16.54316 | 39.53096 | 36.94807 | Promoter | 2 | 176226569 | 176240328 | 13760 | 1 | ENSMUST00 | 0 | 2210418O10 | ENSMUSG00000078894.4 |
| chr4 | 91371227 | 91372351 | 16.54316 | 39.53096 | 36.94807 | Promoter | 4 | 91250763 | 91372034 | 121272 | 2 | ENSMUST00 | 0 | Elavl2 | ENSMUSG00000008489.13 |
| chr4 | 141213596 | 141214549 | 16.54316 | 39.53096 | 36.94807 | Promoter | 4 | 141213956 | 141220148 | 6193 | 1 | ENSMUST00 | 0 | Rsg1 | ENSMUSG00000073733.4 |
| chr5 | 139484636 | 139485130 | 16.54316 | 39.53096 | 36.94807 | Promoter | 5 | 139471211 | 139484549 | 13339 | 2 | ENSMUST00 | -88 | Zfand2a | ENSMUSG00000053581.8 |
| chr8 | 105275937 | 105276737 | 16.54316 | 39.53096 | 36.94807 | Promoter | 8 | 105276447 | 105281939 | 5493 | 1 | ENSMUST00 | 0 | Nol3 | ENSMUSG00000014776.6 |
| chr8 | 15062447 | 15063102 | 16.54316 | 39.53096 | 36.94807 | Intron (ENSM | 8 | 15060190 | 15073840 | 13651 | 1 | ENSMUST00 | 2258 | Myom2 | ENSMUSG00000031461.4 |

| | | | | | | | | | | | | | | | |
|---|---|---|---|---|---|---|---|---|---|---|---|---|---|---|---|
| chrX | 161162553 | 161163380 | 16.54316 | 39.53096 | 36.94807 | Promoter | 20 | 161162750 | 161256952 | 94203 | 1 | ENSMUST00 | 0 | Scml2 | ENSMUSG00000000037.11 |
| chrX | 93631600 | 93632724 | 16.54316 | 39.53096 | 36.94807 | Promoter | 20 | 93304767 | 93632155 | 327389 | 2 | ENSMUST00 | 0 | Pola1 | ENSMUSG00000006678.6 |
| chrX | 150587545 | 150588831 | 16.54316 | 39.53096 | 36.94807 | Promoter | 20 | 150520352 | 150588082 | 67731 | 2 | ENSMUST00 | 0 | Apex2 | ENSMUSG00000025269.11 |
| chrX | 7789795 | 7790688 | 16.54316 | 39.53096 | 36.94807 | Promoter | 20 | 7789994 | 7812483 | 22490 | 1 | ENSMUST00 | 0 | Gripap1 | ENSMUSG00000031153.11 |
| chrX | 7841360 | 7841643 | 16.54316 | 39.53096 | 36.94807 | Promoter | 20 | 7841364 | 7842008 | 645 | 1 | ENSMUST00 | 0 | Otud5 | ENSMUSG00000031154.10 |
| chrX | 8131828 | 8133166 | 16.54316 | 39.53096 | 36.94807 | Promoter | 20 | 8123301 | 8132808 | 9508 | 2 | ENSMUST00 | 0 | Wdr13 | ENSMUSG00000031166.8 |
| chrX | 100494296 | 100495101 | 16.54316 | 39.53096 | 36.94807 | Promoter | 20 | 100494355 | 100499397 | 5043 | 1 | ENSMUST00 | 0 | Igbp1 | ENSMUSG00000031221.7 |
| chrX | 140456594 | 140457493 | 16.54316 | 39.53096 | 36.94807 | Promoter | 20 | 140456613 | 140465442 | 8830 | 1 | ENSMUST00 | 0 | Prps1 | ENSMUSG00000031432.3 |
| chrX | 36902646 | 36903906 | 16.54316 | 39.53096 | 36.94807 | Promoter | 20 | 36887540 | 36902899 | 15360 | 2 | ENSMUST00 | 0 | Nkrf | ENSMUSG00000044149.6 |
| chrX | 37047520 | 37049681 | 16.54316 | 39.53096 | 36.94807 | Promoter | 20 | 37048845 | 37050418 | 1574 | 1 | ENSMUST00 | 0 | Sowahd | ENSMUSG00000044400.1 |
| chrX | 140956889 | 140957534 | 16.54316 | 39.53096 | 36.94807 | Promoter | 20 | 140956907 | 141164270 | 207364 | 1 | ENSMUST00 | 0 | Atg4a | ENSMUSG00000079418.3 |
| chrX | 60547102 | 60548340 | 16.54316 | 39.53096 | 36.94807 | Promoter | 20 | 60548019 | 60548849 | 831 | 1 | ENSMUST00 | 0 | Gm715 | ENSMUSG00000082226.3 |
| chrX | 162735193 | 162736464 | 16.54316 | 39.53096 | 36.94807 | Promoter | 20 | 162735078 | 162736613 | 1536 | 2 | ENSMUST00 | 149 | Gm15201 | ENSMUSG00000086034.1 |
| chrY | 1285124 | 1287078 | 16.54316 | 39.53096 | 36.94807 | Promoter | 21 | 1260771 | 1286629 | 25859 | 2 | ENSMUST00 | 0 | Ddx3y | ENSMUSG00000069045.6 |
| chr1 | 75387422 | 75389391 | 15.26358 | 39.48916 | 36.90792 | 3' UTR | 1 | 75385610 | 75432304 | 46695 | 1 | ENSMUST00 | 1813 | Speg | ENSMUSG00000026207.11 |
| chr11 | 98703830 | 98705879 | 15.26358 | 39.48916 | 36.90792 | Promoter | 11 | 98704843 | 98706480 | 1638 | 2 | ENSMUST00 | 601 | Med24 | ENSMUSG00000017210.9 |
| chr15 | 83510461 | 83511252 | 15.26358 | 39.48916 | 36.90792 | Promoter | 15 | 83483772 | 83510893 | 27122 | 2 | ENSMUST00 | 0 | Ttll1 | ENSMUSG00000022442.10 |
| chr17 | 50189417 | 50190751 | 14.8571 | 39.48906 | 36.90792 | Promoter | 17 | 49992257 | 50190674 | 198418 | 2 | ENSMUST00 | 0 | Rftn1 | ENSMUSG00000039316.9 |
| chr2 | 168912105 | 168912980 | 15.26358 | 39.48916 | 36.90792 | Promoter | 2 | 168906842 | 168912633 | 5792 | 2 | ENSMUST00 | 0 | Zfp64 | ENSMUSG00000027551.10 |
| chr3 | 152165408 | 152166214 | 15.26358 | 39.48916 | 36.90792 | Promoter | 3 | 152130434 | 152165750 | 35317 | 2 | ENSMUST00 | 0 | Gipc2 | ENSMUSG00000039131.10 |
| chr9 | 121839157 | 121839725 | 15.26358 | 39.48916 | 36.90792 | Promoter | 9 | 121797627 | 121839461 | 41835 | 2 | ENSMUST00 | 0 | Ccdc13 | ENSMUSG00000079235.5 |
| chr11 | 9117790 | 9119443 | 16.09418 | 39.48465 | 36.90396 | Promoter | 11 | 9118103 | 9136164 | 18062 | 1 | ENSMUST00 | 0 | Upp1 | ENSMUSG00000020407.8 |
| chr13 | 56609323 | 56609874 | 16.5134 | 39.48057 | 36.89997 | Promoter | 13 | 56609603 | 56639339 | 29737 | 1 | ENSMUST00 | 0 | Tgfbi | ENSMUSG00000035493.9 |
| chr8 | 115159244 | 115160420 | 16.5134 | 39.48057 | 36.89997 | Intron (ENSM | 8 | 115052814 | 115052913 | 100 | 2 | ENSMUST00 | -106332 | Gm22556 | ENSMUSG00000088428.1 |
| chr10 | 4333687 | 4335629 | 16.06223 | 39.4301 | 36.8496 | Intron (ENSM | 10 | 4367475 | 4387462 | 19988 | 2 | ENSMUST00 | 51833 | Zbtb2 | ENSMUSG00000075327.4 |
| chr17 | 9764667 | 9765360 | 16.06223 | 39.4301 | 36.8496 | Distal Interge | 17 | 9666497 | 9669704 | 3208 | 2 | ENSMUST00 | -94964 | Pabpc6 | ENSMUSG00000046173.2 |
| chr7 | 19002606 | 19003412 | 16.06223 | 39.4301 | 36.8496 | Promoter | 7 | 19004065 | 19006761 | 2697 | 1 | ENSMUST00 | -653 | Irf2bp1 | ENSMUSG00000044030.3 |
| chr19 | 5653889 | 5654944 | 16.47883 | 39.42202 | 36.84161 | Exon (ENSMU | 19 | 5651185 | 5663707 | 12523 | 2 | ENSMUST00 | 8763 | Sipa1 | ENSMUSG00000056917.7 |
| chr3 | 96669883 | 96670763 | 16.47883 | 39.42202 | 36.84161 | Promoter | 3 | 96670131 | 96690468 | 20338 | 1 | ENSMUST00 | 0 | Ankrd35 | ENSMUSG00000038354.8 |
| chr13 | 119487894 | 119489135 | 10.11917 | 39.41373 | 36.83339 | Promoter | 13 | 119487941 | 119495505 | 7565 | 1 | ENSMUST00 | 0 | Gm7120 | ENSMUSG00000074634.6 |
| chr19 | 4698106 | 4698714 | 16.0304 | 39.37575 | 36.79544 | Promoter | 19 | 4625841 | 4698668 | 72828 | 2 | ENSMUST00 | 0 | Gm960 | ENSMUSG00000071691.5 |
| chr17 | 56124208 | 56125071 | 16.4444 | 39.3637 | 36.78348 | Exon (ENSMU | 17 | 56119680 | 56121946 | 2267 | 2 | ENSMUST00 | -2263 | Lrg1 | ENSMUSG00000037095.7 |
| chr19 | 4097293 | 4098050 | 15.59818 | 39.35201 | 36.77204 | Promoter | 19 | 4097401 | 4098465 | 1065 | 1 | ENSMUST00 | 0 | Cdk2ap2 | ENSMUSG00000024856.5 |
| chr11 | 69802553 | 69804725 | 12.87622 | 39.27038 | 36.69063 | Promoter | 11 | 69796073 | 69801857 | 5785 | 2 | ENSMUST00 | -697 | Fgf11 | ENSMUSG00000042826.8 |
| chr11 | 115698853 | 115699962 | 12.87622 | 39.27038 | 36.69063 | Promoter | 11 | 115644045 | 115699307 | 55263 | 2 | ENSMUST00 | 0 | Grb2 | ENSMUSG00000059923.8 |
| chr12 | 118836477 | 118837548 | 12.87622 | 39.27038 | 36.69063 | Distal Interge | 12 | 118846329 | 118852578 | 6250 | 1 | ENSMUST00 | -8781 | Sp8 | ENSMUSG00000048562.6 |
| chr9 | 37460097 | 37461136 | 16.37597 | 39.24776 | 36.66856 | Distal Interge | 9 | 37416045 | 37433163 | 17119 | 2 | ENSMUST00 | -26935 | Robo3 | ENSMUSG00000032128.9 |
| chr11 | 54027815 | 54028717 | 13.94198 | 39.24413 | 36.66504 | Promoter | 11 | 53983123 | 54028090 | 44968 | 2 | ENSMUST00 | 0 | Slc22a4 | ENSMUSG00000020334.6 |
| chr16 | 96200156 | 96201054 | 15.90433 | 39.1603 | 36.58143 | Promoter | 16 | 96200470 | 96228933 | 28464 | 1 | ENSMUST00 | 0 | Sh3bgr | ENSMUSG00000040666.12 |
| chr2 | 44556641 | 44557550 | 15.90433 | 39.16029 | 36.58143 | Distal Interge | 2 | 44565133 | 44571347 | 6215 | 2 | ENSMUST00 | 13797 | Gtdc1 | ENSMUSG00000036890.8 |
| chr2 | 167537609 | 167538780 | 16.30811 | 39.13273 | 36.55445 | Promoter | 2 | 167538195 | 167542814 | 4620 | 1 | ENSMUST00 | 0 | Snai1 | ENSMUSG00000042821.7 |
| chr17 | 35320266 | 35321338 | 15.84204 | 39.05372 | 36.47581 | Promoter | 17 | 35320405 | 35325099 | 4695 | 1 | ENSMUST00 | 0 | H2-Q1 | ENSMUSG00000079507.5 |
| chr10 | 108443290 | 108444613 | 15.42443 | 39.05124 | 36.47339 | Distal Interge | 10 | 108428067 | 108428160 | 94 | 1 | ENSMUST00 | 15224 | Gm25591 | ENSMUSG00000093762.1 |
| chr12 | 85697928 | 85698747 | 15.42443 | 39.05124 | 36.47339 | Intron (ENSM | 12 | 85686669 | 85709087 | 22419 | 1 | ENSMUST00 | 11260 | Batf | ENSMUSG00000034266.5 |
| chr13 | 40732890 | 40734280 | 15.42443 | 39.05124 | 36.47339 | Promoter | 13 | 40715675 | 40733823 | 18149 | 2 | ENSMUST00 | 0 | Tfap2a | ENSMUSG00000021359.10 |
| chr4 | 43405677 | 43408085 | 15.42443 | 39.05124 | 36.47339 | Promoter | 4 | 43406435 | 43427088 | 20654 | 1 | ENSMUST00 | 0 | Rusc2 | ENSMUSG00000035969.10 |
| chr17 | 8340401 | 8341644 | 14.99516 | 39.01754 | 36.44046 | Promoter | 17 | 8340406 | 8344112 | 3707 | 1 | ENSMUST00 | 0 | Prr18 | ENSMUSG00000055945.7 |
| chr3 | 146684970 | 146685850 | 14.99516 | 39.01754 | 36.44046 | Promoter | 3 | 146645195 | 146685592 | 40398 | 2 | ENSMUST00 | 0 | Samd13 | ENSMUSG00000048652.7 |
| chr4 | 120267912 | 120268986 | 14.99516 | 39.01754 | 36.44046 | 3' UTR | 4 | 120267079 | 120287349 | 20271 | 2 | ENSMUST00 | 18363 | Foxo6 | ENSMUSG00000052135.8 |
| chr9 | 60687282 | 60688502 | 15.39584 | 39.00168 | 36.42487 | Promoter | 9 | 60568859 | 60688124 | 119266 | 2 | ENSMUST00 | 0 | Lrrc49 | ENSMUSG00000047766.10 |
| chr2 | 19275969 | 19276688 | 15.81108 | 39.00072 | 36.42392 | Intron (ENSM | 2 | 19344317 | 19349202 | 4886 | 1 | ENSMUST00 | -67629 | 4930447M23 | ENSMUSG00000087674.2 |
| chr17 | 26884764 | 26885840 | 15.36737 | 38.95228 | 36.37565 | Promoter | 17 | 26860287 | 26886175 | 25889 | 2 | ENSMUST00 | 335 | Gm17382 | ENSMUSG00000078838.2 |
| chr3 | 96486156 | 96487789 | 10.65394 | 38.93504 | 36.35848 | Promoter | 3 | 96486555 | 96486719 | 165 | 2 | ENSMUST00 | 0 | Gm25890 | ENSMUSG00000095260.1 |
| chr1 | 52113378 | 52113806 | 13.73347 | 38.8536 | 36.27722 | Intron (ENSM | 1 | 52119440 | 52155412 | 35973 | 1 | ENSMUST00 | -5634 | Stat1 | ENSMUSG00000026104.9 |
| chr6 | 124863267 | 124864173 | 16.14089 | 38.84904 | 36.27268 | Promoter | 6 | 124858444 | 124863917 | 5474 | 2 | ENSMUST00 | 0 | Gpr162 | ENSMUSG00000038390.9 |
| chr1 | 22805483 | 22806478 | 13.36394 | 38.84871 | 36.27245 | Promoter | 1 | 22286251 | 22805724 | 519474 | 2 | ENSMUST00 | 0 | Rims1 | ENSMUSG00000041670.10 |
| chr1 | 161170568 | 161171086 | 13.36394 | 38.84871 | 36.27245 | Distal Interge | 1 | 161156843 | 161159314 | 2472 | 2 | ENSMUST00 | -11255 | 4930469G21 | ENSMUSG00000049160.10 |
| chr11 | 95830246 | 95831655 | 13.36394 | 38.84871 | 36.27245 | Promoter | 11 | 95830763 | 95831592 | 830 | 1 | ENSMUST00 | 0 | Phospho1 | ENSMUSG00000050860.6 |
| chr13 | 76383790 | 76385194 | 13.36394 | 38.84871 | 36.27245 | Promoter | 13 | 76384535 | 77031810 | 647276 | 1 | ENSMUST00 | 0 | Mctp1 | ENSMUSG00000021596.11 |
| chr18 | 74207271 | 74208128 | 13.36394 | 38.84871 | 36.27245 | Promoter | 18 | 74195299 | 74207818 | 12520 | 2 | ENSMUST00 | 0 | Ska1 | ENSMUSG00000036223.10 |
| chr2 | 170132897 | 170133813 | 13.36394 | 38.84871 | 36.27245 | Promoter | 2 | 170120619 | 170133945 | 13327 | 2 | ENSMUST00 | 132 | Zfp217 | ENSMUSG00000052056.9 |
| chr2 | 18031167 | 18032419 | 13.36394 | 38.84871 | 36.27245 | Promoter | 2 | 18032533 | 18039633 | 7101 | 1 | ENSMUST00 | -114 | Gm13330 | ENSMUSG00000085319.1 |
| chr3 | 157924517 | 157925556 | 13.36394 | 38.84871 | 36.27245 | Promoter | 3 | 157925269 | 157927147 | 1879 | 1 | ENSMUST00 | 0 | 1810013D15I | ENSMUSG00000028177.9 |
| chr3 | 88177499 | 88178632 | 13.36394 | 38.84871 | 36.27245 | Promoter | 3 | 88171560 | 88177785 | 6226 | 2 | ENSMUST00 | 0 | 1700113A16I | ENSMUSG00000096935.1 |
| chr5 | 24387463 | 24388401 | 13.36394 | 38.84871 | 36.27245 | Promoter | 5 | 24387105 | 24387966 | 862 | 2 | ENSMUST00 | 0 | Atg9b | ENSMUSG00000038295.9 |
| chr6 | 135167677 | 135168588 | 13.36394 | 38.84871 | 36.27245 | Promoter | 6 | 135137519 | 135168215 | 30697 | 2 | ENSMUST00 | 0 | Hebp1 | ENSMUSG00000042770.7 |
| chr12 | 116485636 | 116486601 | 16.10786 | 38.79296 | 36.21762 | Promoter | 12 | 116485720 | 116773330 | 287611 | 1 | ENSMUST00 | 0 | Ptprn2 | ENSMUSG00000056553.9 |
| chr2 | 152962141 | 152962790 | 16.10786 | 38.79296 | 36.21762 | Promoter | 2 | 152962485 | 153007340 | 44856 | 1 | ENSMUST00 | 0 | Ttll9 | ENSMUSG00000074673.10 |
| chr4 | 44703225 | 44704814 | 16.10786 | 38.79296 | 36.21762 | Promoter | 4 | 44679440 | 44704006 | 24567 | 2 | ENSMUST00 | 0 | Pax5 | ENSMUSG00000014030.10 |
| chr11 | 49243444 | 49244480 | 15.68842 | 38.79056 | 36.21557 | Promoter | 11 | 49244191 | 49263022 | 18832 | 1 | ENSMUST00 | 0 | Mgat1 | ENSMUSG00000020346.11 |
| chr11 | 85843964 | 85844877 | 15.59767 | 38.63485 | 36.0603 | Downstream | 11 | 85846790 | 85850333 | 3544 | 2 | ENSMUST00 | 5456 | Gm11444 | ENSMUSG00000069785.4 |
| chr16 | 56717583 | 56718127 | 11.32576 | 38.62201 | 36.04802 | Promoter | 16 | 56690332 | 56717450 | 27119 | 2 | ENSMUST00 | -134 | Tfg | ENSMUSG00000022757.13 |
| chr14 | 6037588 | 6038673 | 14.36889 | 38.61326 | 36.0393 | Promoter | 14 | 6016759 | 6038209 | 21451 | 2 | ENSMUST00 | 0 | Gm8206 | ENSMUSG00000091568.2 |
| chr1 | 133181174 | 133181969 | 15.56766 | 38.58331 | 36.00948 | Promoter | 1 | 133181321 | 133301219 | 119899 | 1 | ENSMUST00 | 0 | Plekha6 | ENSMUSG00000041757.10 |
| chr10 | 94545392 | 94546191 | 15.14328 | 38.56274 | 35.98906 | Promoter | 10 | 94545431 | 94578922 | 33492 | 1 | ENSMUST00 | 0 | Tmcc3 | ENSMUSG00000020023.12 |
| chr2 | 74746552 | 74747523 | 15.14328 | 38.56274 | 35.98906 | Promoter | 2 | 74745722 | 74748442 | 2721 | 1 | ENSMUST00 | 831 | Hoxd3 | ENSMUSG00000079277.4 |
| chr4 | 134251001 | 134252508 | 15.14328 | 38.56274 | 35.98906 | Promoter | 4 | 134251706 | 134253023 | 1318 | 2 | ENSMUST00 | 515 | Grrp1 | ENSMUSG00000050105.5 |
| chr4 | 133497531 | 133498909 | 15.14328 | 38.56274 | 35.98906 | Promoter | 4 | 133491100 | 133498550 | 7451 | 2 | ENSMUST00 | 0 | Trnp1 | ENSMUSG00000056596.8 |
| chr11 | 96861404 | 96862432 | 14.73601 | 38.55974 | 35.98649 | Downstream | 11 | 96854131 | 96861203 | 7073 | 1 | ENSMUST00 | 7274 | Copz2 | ENSMUSG00000018672.10 |
| chr6 | 142507058 | 142508304 | 14.73601 | 38.55974 | 35.98649 | Promoter | 6 | 142490249 | 142507957 | 17709 | 2 | ENSMUST00 | 0 | Ldhb | ENSMUSG00000030246.6 |
| chr15 | 83250797 | 83251939 | 12.19507 | 38.54947 | 35.97643 | Promoter | 15 | 83226726 | 83251731 | 25006 | 2 | ENSMUST00 | 0 | A4galt | ENSMUSG00000047878.7 |
| chr5 | 63811992 | 63812814 | 12.19507 | 38.54947 | 35.97643 | Promoter | 5 | 63812495 | 63899619 | 87125 | 1 | ENSMUST00 | 0 | 0610040J01R | ENSMUSG00000060512.3 |
| chr1 | 104993815 | 104994565 | 13.94362 | 38.54297 | 35.97016 | 3' UTR | 1 | 104911431 | 104912333 | 903 | 2 | ENSMUST00 | -81483 | Gm15699 | ENSMUSG00000089866.1 |
| chr11 | 64978317 | 64978864 | 13.94362 | 38.54297 | 35.97016 | Promoter | 11 | 64979038 | 65002069 | 23032 | 1 | ENSMUST00 | -174 | Elac2 | ENSMUSG00000020549.9 |
| chr11 | 3514172 | 3515438 | 13.94362 | 38.54297 | 35.97016 | Promoter | 11 | 3514684 | 3517348 | 2665 | 1 | ENSMUST00 | 0 | Selm | ENSMUSG00000075702.9 |
| chr12 | 32819564 | 32820103 | 13.94362 | 38.54297 | 35.97016 | Promoter | 12 | 32820335 | 32853369 | 33035 | 1 | ENSMUST00 | -232 | Nampt | ENSMUSG00000020572.7 |
| chr15 | 76659480 | 76661269 | 13.94362 | 38.54297 | 35.97016 | Promoter | 15 | 76660641 | 76668196 | 7556 | 1 | ENSMUST00 | 0 | Kifc2 | ENSMUSG00000004187.10 |
| chr16 | 91597899 | 91598244 | 13.94362 | 38.54297 | 35.97016 | Promoter | 16 | 91580202 | 91597800 | 17599 | 2 | ENSMUST00 | -100 | Tmem50b | ENSMUSG00000022964.9 |
| chr4 | 152338334 | 152339533 | 13.94362 | 38.54297 | 35.97016 | Promoter | 4 | 152338651 | 152390194 | 51544 | 1 | ENSMUST00 | 0 | Chd5 | ENSMUSG00000005045.11 |
| chr5 | 66069504 | 66070243 | 13.94362 | 38.54297 | 35.97016 | Intron (ENSM | 5 | 66027220 | 66080984 | 53765 | 2 | ENSMUST00 | 10741 | Rbm47 | ENSMUSG00000070780.6 |
| chr7 | 34629812 | 34631162 | 13.94362 | 38.54297 | 35.97016 | Distal Interge | 7 | 34639015 | 34652837 | 13823 | 2 | ENSMUST00 | 21675 | Kctd15 | ENSMUSG00000030499.4 |
| chr13 | 73626646 | 73627384 | 15.94471 | 38.51573 | 35.94438 | Promoter | 13 | 73627001 | 73649041 | 22041 | 1 | ENSMUST00 | 0 | Tert | ENSMUSG00000021611.8 |
| chr19 | 55643218 | 55644045 | 15.94471 | 38.51573 | 35.94438 | Exon (ENSMU | 19 | 55640589 | 55702204 | 61616 | 2 | ENSMUST00 | 58159 | 4930552P12I | ENSMUSG00000097632.1 |
| chr1 | 72873479 | 72874846 | 15.50707 | 38.47921 | 35.908 | Promoter | 1 | 72858065 | 72874275 | 16211 | 2 | ENSMUST00 | 0 | Igfbp5 | ENSMUSG00000026185.8 |
| chr11 | 76612455 | 76613341 | 15.50707 | 38.47921 | 35.908 | Promoter | 11 | 76479113 | 76613395 | 134283 | 2 | ENSMUST00 | 54 | Abr | ENSMUSG00000017631.13 |
| chr11 | 98981927 | 98983052 | 15.50707 | 38.47921 | 35.908 | Promoter | 11 | 98982180 | 98983016 | 837 | 2 | ENSMUST00 | 0 | Gjd3 | ENSMUSG00000047197.1 |
| chr11 | 97546418 | 97547365 | 15.50707 | 38.47921 | 35.908 | Intron (ENSM | 11 | 97566711 | 97576186 | 9476 | 1 | ENSMUST00 | -19346 | Gm11612 | ENSMUSG00000086394.1 |
| chr12 | 102128688 | 102129752 | 15.50707 | 38.47921 | 35.908 | Promoter | 12 | 102128733 | 102267091 | 138359 | 1 | ENSMUST00 | 0 | Slc24a4 | ENSMUSG00000041771.8 |
| chr17 | 8524255 | 8525207 | 15.50707 | 38.47921 | 35.908 | Promoter | 17 | 8525372 | 8981671 | 456300 | 1 | ENSMUST00 | -165 | Pde10a | ENSMUSG00000023868.10 |
| chr18 | 69198079 | 69198930 | 15.50707 | 38.47921 | 35.908 | Distal Interge | 18 | 69173987 | 69174120 | 134 | 2 | ENSMUST00 | -23960 | Gm23536 | ENSMUSG00000088166.1 |
| chr19 | 37696983 | 37699244 | 15.50707 | 38.47921 | 35.908 | Promoter | 19 | 37697808 | 37701528 | 3721 | 1 | ENSMUST00 | 0 | Cyp26a1 | ENSMUSG00000024987.5 |
| chr19 | 6530956 | 6532503 | 15.50707 | 38.47921 | 35.908 | Promoter | 19 | 6530711 | 6532011 | 1301 | 2 | ENSMUST00 | 0 | Gm14964 | ENSMUSG00000052188.6 |
| chr2 | 126552529 | 126553545 | 15.50707 | 38.47921 | 35.908 | Promoter | 2 | 126553590 | 126564685 | 11096 | 1 | ENSMUST00 | -45 | Slc27a2 | ENSMUSG00000027359.11 |
| chr2 | 140670634 | 140671600 | 15.50707 | 38.47921 | 35.908 | Promoter | 2 | 140650914 | 140671450 | 20537 | 2 | ENSMUST00 | 0 | Flrt3 | ENSMUSG00000051379.7 |
| chr3 | 134235553 | 134236537 | 15.50707 | 38.47921 | 35.908 | Promoter | 3 | 134236491 | 134262090 | 25600 | 1 | ENSMUST00 | 0 | Cxxc4 | ENSMUSG00000044365.11 |
| chr4 | 134081796 | 134082364 | 15.50707 | 38.47921 | 35.908 | Promoter | 4 | 134082312 | 134091096 | 8785 | 1 | ENSMUST00 | 0 | Aim1l | ENSMUSG00000012123.10 |
| chr4 | 152273869 | 152274536 | 15.50707 | 38.47921 | 35.908 | Promoter | 4 | 152274232 | 152285337 | 11106 | 1 | ENSMUST00 | 0 | Gpr153 | ENSMUSG00000042804.8 |
| chr5 | 137031245 | 137033291 | 15.50707 | 38.47921 | 35.908 | Promoter | 5 | 137030701 | 137031407 | 707 | 1 | ENSMUST00 | 545 | Vgf | ENSMUSG00000037428.9 |
| chr5 | 143757857 | 143758937 | 15.50707 | 38.47921 | 35.908 | Promoter | 5 | 143758325 | 143766974 | 8650 | 1 | ENSMUST00 | 0 | D130017N08 | ENSMUSG00000097287.1 |
| chr6 | 36809394 | 36810394 | 15.50707 | 38.47921 | 35.908 | Promoter | 6 | 36715663 | 36811361 | 95699 | 2 | ENSMUST00 | 967 | Ptn | ENSMUSG00000029838.8 |
| chr6 | 124996608 | 124997571 | 15.50707 | 38.47921 | 35.908 | Promoter | 6 | 124996694 | 125003096 | 6403 | 1 | ENSMUST00 | 0 | Pianp | ENSMUSG00000030329.6 |
| chr7 | 109174576 | 109175924 | 15.50707 | 38.47921 | 35.908 | Distal Interge | 7 | 109138572 | 109170308 | 31737 | 2 | ENSMUST00 | -4269 | Lmo1 | ENSMUSG00000036111.7 |
| chr7 | 25626852 | 25628244 | 15.50707 | 38.47921 | 35.908 | Promoter | 7 | 25627624 | 25629490 | 1867 | 1 | ENSMUST00 | 0 | B3gnt8 | ENSMUSG00000059479.6 |
| chr9 | 79717796 | 79719888 | 15.50707 | 38.47921 | 35.908 | Promoter | 9 | 79598991 | 79718518 | 119528 | 2 | ENSMUST00 | 0 | Col12a1 | ENSMUSG00000032332.12 |
| chr9 | 100454200 | 100454877 | 15.50707 | 38.47921 | 35.908 | Downstream | 9 | 100450322 | 100451912 | 1591 | 1 | ENSMUST00 | 3879 | RP23-368K22 | ENSMUSG00000101287.1 |
| chrX | 95711373 | 95712338 | 15.50707 | 38.47921 | 35.908 | Promoter | 20 | 95711678 | 95924758 | 213081 | 1 | ENSMUST00 | 0 | Zc3h12b | ENSMUSG00000035045.12 |
| chr18 | 38631193 | 38632129 | 15.88037 | 38.4063 | 35.83701 | Intron (ENSM | 18 | 38627287 | 38642930 | 15644 | 1 | ENSMUST00 | 3907 | Arhgap26 | ENSMUSG00000036452.12 |
| chr1 | 86669597 | 86670931 | 14.64398 | 38.39655 | 35.82735 | Promoter | 1 | 86666291 | 86670571 | 4281 | 2 | ENSMUST00 | 0 | Nppc | ENSMUSG00000026241.5 |
| chr1 | 153185628 | 153186411 | 14.64398 | 38.39655 | 35.82735 | Promoter | 1 | 153122756 | 153186047 | 63292 | 2 | ENSMUST00 | 0 | Lamc2 | ENSMUSG00000026479.8 |
| chr10 | 127262805 | 127263474 | 14.64398 | 38.39655 | 35.82735 | Promoter | 10 | 127225696 | 127263363 | 37668 | 2 | ENSMUST00 | 0 | Kif5a | ENSMUSG00000074657.4 |
| chr11 | 46165919 | 46166762 | 14.64398 | 38.39655 | 35.82735 | Promoter | 11 | 46148155 | 46166508 | 18354 | 2 | ENSMUST00 | 0 | Nipal4 | ENSMUSG00000020411.2 |
| chr11 | 100319280 | 100319746 | 14.64398 | 38.39655 | 35.82735 | Promoter | 11 | 100319885 | 100322096 | 2212 | 1 | ENSMUST00 | -139 | Eif1 | ENSMUSG00000035530.8 |
| chr11 | 3330937 | 3331508 | 14.64398 | 38.39655 | 35.82735 | Promoter | 11 | 3305426 | 3331178 | 25753 | 2 | ENSMUST00 | 0 | Gm12592 | ENSMUSG00000053263.3 |
| chr11 | 96004422 | 96005307 | 14.64398 | 38.39655 | 35.82735 | Promoter | 11 | 96005889 | 96008055 | 2167 | 1 | ENSMUST00 | -582 | RP23-235J5. | ENSMUSG00000099928.1 |
| chr12 | 85455512 | 85456149 | 14.64398 | 38.39655 | 35.82735 | Distal Interge | 12 | 85473890 | 85477273 | 3384 | 1 | ENSMUST00 | -17741 | Fos | ENSMUSG00000021250.8 |
| chr13 | 23463520 | 23465180 | 14.64398 | 38.39655 | 35.82735 | Promoter | 13 | 23456992 | 23465875 | 8884 | 2 | ENSMUST00 | 695 | Btn1a1 | ENSMUSG00000000706.10 |
| chr13 | 59584388 | 59585130 | 14.64398 | 38.39655 | 35.82735 | Promoter | 13 | 59532043 | 59585211 | 53169 | 2 | ENSMUST00 | 81 | Agtpbp1 | ENSMUSG00000021557.9 |
| chr14 | 54479291 | 54479981 | 14.64398 | 38.39655 | 35.82735 | 3' UTR | 14 | 54477471 | 54479117 | 1647 | 1 | ENSMUST00 | 1821 | Rem2 | ENSMUSG00000022176.6 |

| | | | | | | | | | | | | | | | |
|---|---|---|---|---|---|---|---|---|---|---|---|---|---|---|---|
| chr14 | 66635187 | 66635868 | 14.64398 | 38.39655 | 35.82735 | Promoter | 14 | 66635251 | 66733462 | 98212 | 1 | ENSMUST00 | 0 | Adra1a | ENSMUSG00000045875.7 |
| chr14 | 6108068 | 6108956 | 14.64398 | 38.39655 | 35.82735 | Promoter | 14 | 6087295 | 6108665 | 21371 | 2 | ENSMUST00 | 0 | Gm3468 | ENSMUSG00000094634.2 |
| chr17 | 46496571 | 46497214 | 14.64398 | 38.39655 | 35.82735 | Promoter | 17 | 46496789 | 46499618 | 2830 | 1 | ENSMUST00 | 0 | Dnph1 | ENSMUSG00000040658.6 |
| chr19 | 26747465 | 26748613 | 14.64398 | 38.39655 | 35.82735 | Promoter | 19 | 26748268 | 26777476 | 29209 | 1 | ENSMUST00 | 0 | Smarca2 | ENSMUSG00000024921.11 |
| chr2 | 92008547 | 92009082 | 14.64398 | 38.39655 | 35.82735 | Intron (ENSM | 2 | 91982328 | 92024502 | 42175 | 2 | ENSMUST00 | 15420 | Creb3l1 | ENSMUSG00000027230.9 |
| chr4 | 43523271 | 43524667 | 14.64398 | 38.39655 | 35.82735 | Promoter | 4 | 43514711 | 43523438 | 8728 | 2 | ENSMUST00 | 0 | Tpm2 | ENSMUSG00000028464.11 |
| chr4 | 136021894 | 136022300 | 14.64398 | 38.39655 | 35.82735 | Promoter | 4 | 136003368 | 136021763 | 18396 | 2 | ENSMUST00 | -132 | Tceb3 | ENSMUSG00000028668.5 |
| chr4 | 155223453 | 155223772 | 14.64398 | 38.39655 | 35.82735 | Promoter | 4 | 155154075 | 155222592 | 68518 | 2 | ENSMUST00 | -862 | Ski | ENSMUSG00000029050.10 |
| chr5 | 53995121 | 53995884 | 14.64398 | 38.39655 | 35.82735 | Distal Interge | 5 | 53998417 | 54121057 | 122641 | 1 | ENSMUST00 | -2533 | Stim2 | ENSMUSG00000039156.14 |
| chr7 | 127636649 | 127637757 | 14.64398 | 38.39655 | 35.82735 | Distal Interge | 7 | 127608756 | 127615797 | 7042 | 2 | ENSMUST00 | -20853 | Zfp629 | ENSMUSG00000045639.9 |
| chr7 | 116039235 | 116040440 | 14.64398 | 38.39655 | 35.82735 | Promoter | 7 | 115471344 | 116038744 | 567401 | 2 | ENSMUST00 | -492 | Sox6 | ENSMUSG00000051910.8 |
| chr11 | 97884236 | 97884966 | 15.84839 | 38.35189 | 35.78466 | Promoter | 11 | 97884594 | 97893811 | 9218 | 1 | ENSMUST00 | 0 | Gm11630 | ENSMUSG00000086917.1 |
| chr15 | 12204201 | 12204883 | 15.84839 | 38.35189 | 35.78466 | Promoter | 15 | 12205028 | 12272240 | 67213 | 1 | ENSMUST00 | -145 | Mtmr12 | ENSMUSG00000039458.9 |
| chr4 | 74013134 | 74014153 | 15.84839 | 38.35189 | 35.78466 | Promoter | 4 | 74013442 | 74202214 | 188773 | 1 | ENSMUST00 | 0 | Frmd3 | ENSMUSG00000049122.12 |
| chr11 | 7213223 | 7214316 | 15.30262 | 38.12728 | 35.5612 | Promoter | 11 | 7206086 | 7213897 | 7812 | 2 | ENSMUST00 | 0 | Igfbp3 | ENSMUSG00000020427.6 |
| chr19 | 19110177 | 19111709 | 15.30262 | 38.12728 | 35.5612 | Promoter | 19 | 18930605 | 19111196 | 180592 | 2 | ENSMUST00 | 0 | Rorb | ENSMUSG00000036192.10 |
| chr5 | 115652443 | 115653059 | 15.30262 | 38.12728 | 35.5612 | Promoter | 5 | 115648177 | 115652986 | 4810 | 2 | ENSMUST00 | 0 | Ccdc64 | ENSMUSG00000041609.11 |
| chr6 | 112489092 | 112490309 | 15.30262 | 38.12728 | 35.5612 | Promoter | 6 | 112473684 | 112489808 | 16125 | 2 | ENSMUST00 | 0 | Oxtr | ENSMUSG00000049112.8 |
| chr8 | 123117222 | 123117932 | 15.30262 | 38.12728 | 35.5612 | Promoter | 8 | 123117374 | 123135180 | 17807 | 1 | ENSMUST00 | 0 | Cpne7 | ENSMUSG00000034796.9 |
| chr10 | 117905555 | 117906960 | 14.48567 | 38.11502 | 35.5495 | Distal Interge | 10 | 117814471 | 117845935 | 31465 | 2 | ENSMUST00 | -59621 | Rap1b | ENSMUSG00000052681.7 |
| chr17 | 27826256 | 27826938 | 15.69042 | 38.08285 | 35.51791 | Distal Interge | 17 | 27751232 | 27820558 | 69327 | 2 | ENSMUST00 | -5699 | D17Wsu92e | ENSMUSG00000056692.7 |
| chr11 | 78924790 | 78925459 | 16.08362 | 38.04472 | 35.47989 | Intron (ENSM | 11 | 78920787 | 78960254 | 39468 | 1 | ENSMUST00 | 4004 | Nos2 | ENSMUSG00000020826.6 |
| chr11 | 52764400 | 52765208 | 16.08362 | 38.04472 | 35.47989 | Promoter | 11 | 52764634 | 53188538 | 423905 | 1 | ENSMUST00 | 0 | Fstl4 | ENSMUSG00000036264.9 |
| chr12 | 53248191 | 53248818 | 16.08362 | 38.04472 | 35.47989 | Promoter | 12 | 53248677 | 54072175 | 823499 | 1 | ENSMUST00 | 0 | Npas3 | ENSMUSG00000021010.7 |
| chr12 | 20815010 | 20816139 | 16.08362 | 38.04472 | 35.47989 | Promoter | 12 | 20814257 | 20815727 | 1471 | 2 | ENSMUST00 | 0 | 1700030C10I | ENSMUSG00000091071.2 |
| chr13 | 22005809 | 22007213 | 16.08362 | 38.04472 | 35.47989 | Promoter | 13 | 22002926 | 22006415 | 3490 | 2 | ENSMUST00 | 0 | Prss16 | ENSMUSG00000006179.3 |
| chr13 | 94882509 | 94883870 | 16.08362 | 38.04472 | 35.47989 | Promoter | 13 | 94882600 | 94883552 | 953 | 1 | ENSMUST00 | 0 | Otp | ENSMUSG00000021685.11 |
| chr13 | 116309356 | 116310235 | 16.08362 | 38.04472 | 35.47989 | Promoter | 13 | 116298281 | 116309689 | 11409 | 2 | ENSMUST00 | 0 | Isl1 | ENSMUSG00000042258.8 |
| chr14 | 6889379 | 6890249 | 16.08362 | 38.04472 | 35.47989 | Promoter | 14 | 6868502 | 6889962 | 21461 | 2 | ENSMUST00 | 0 | Gm3667 | ENSMUSG00000090691.3 |
| chr15 | 44618498 | 44619033 | 16.08362 | 38.04472 | 35.47989 | Promoter | 15 | 44619641 | 44641024 | 21384 | 1 | ENSMUST00 | -608 | Ebag9 | ENSMUSG00000022339.8 |
| chr15 | 84231266 | 84232526 | 16.08362 | 38.04472 | 35.47989 | Promoter | 15 | 84232043 | 84315688 | 83646 | 1 | ENSMUST00 | 0 | Parvb | ENSMUSG00000022438.6 |
| chr15 | 99817939 | 99820181 | 16.08362 | 38.04472 | 35.47989 | Promoter | 15 | 99778470 | 99820143 | 41674 | 2 | ENSMUST00 | 0 | Lima1 | ENSMUSG00000023022.8 |
| chr15 | 82379548 | 82380266 | 16.08362 | 38.04472 | 35.47989 | Promoter | 15 | 82370528 | 82380260 | 9733 | 2 | ENSMUST00 | 0 | Cyp2d22 | ENSMUSG00000061740.7 |
| chr17 | 83545672 | 83546607 | 16.08362 | 38.04472 | 35.47989 | Distal Interge | 17 | 83501919 | 83514333 | 12415 | 2 | ENSMUST00 | -31340 | Cox7a2l | ENSMUSG00000024248.8 |
| chr17 | 48931575 | 48932116 | 16.08362 | 38.04472 | 35.47989 | Promoter | 17 | 48932379 | 49097588 | 165210 | 1 | ENSMUST00 | -263 | Lrfn2 | ENSMUSG00000040490.4 |
| chr2 | 71117719 | 71118666 | 16.08362 | 38.04472 | 35.47989 | Promoter | 2 | 71117923 | 71142926 | 25004 | 1 | ENSMUST00 | 0 | Cybrd1 | ENSMUSG00000027015.4 |
| chr2 | 169735950 | 169736561 | 16.08362 | 38.04472 | 35.47989 | Intron (ENSM | 2 | 169633676 | 169888504 | 254829 | 1 | ENSMUST00 | 102275 | Tshz2 | ENSMUSG00000047907.11 |
| chr2 | 155027109 | 155027562 | 16.08362 | 38.04472 | 35.47989 | Intron (ENSM | 2 | 155018266 | 155027104 | 8839 | 1 | ENSMUST00 | 8844 | Gm14226 | ENSMUSG00000084897.1 |
| chr3 | 21935258 | 21936108 | 16.08362 | 38.04472 | 35.47989 | Distal Interge | 3 | 22076652 | 22216594 | 139943 | 1 | ENSMUST00 | -140544 | Tbl1xr1 | ENSMUSG00000027630.8 |
| chr3 | 117359674 | 117360803 | 16.08362 | 38.04472 | 35.47989 | Promoter | 3 | 117319145 | 117360876 | 41732 | 2 | ENSMUST00 | 73 | D3Bwg0562e | ENSMUSG00000044667.8 |
| chr4 | 82513102 | 82513677 | 16.08362 | 38.04472 | 35.47989 | Intron (ENSM | 4 | 82508016 | 82547292 | 39277 | 1 | ENSMUST00 | 5087 | Gm11266 | ENSMUSG00000087413.1 |
| chr5 | 86065747 | 86067028 | 16.08362 | 38.04472 | 35.47989 | Promoter | 5 | 86012043 | 86065583 | 53541 | 2 | ENSMUST00 | -165 | Cenpc1 | ENSMUSG00000029253.8 |
| chr5 | 91517463 | 91518423 | 16.08362 | 38.04472 | 35.47989 | Promoter | 5 | 91517615 | 91626894 | 109280 | 1 | ENSMUST00 | 0 | Parm1 | ENSMUSG00000034981.9 |
| chr5 | 135867751 | 135868589 | 16.08362 | 38.04472 | 35.47989 | 3' UTR | 5 | 135866617 | 135868209 | 1593 | 1 | ENSMUST00 | 1135 | Srrm3 | ENSMUSG00000039860.14 |
| chr7 | 120597631 | 120598215 | 16.08362 | 38.04472 | 35.47989 | Promoter | 7 | 120597625 | 120598475 | 851 | 1 | ENSMUST00 | 7 | E130201H02I | ENSMUSG00000044141.8 |
| chr9 | 78527333 | 78528473 | 16.08362 | 38.04472 | 35.47989 | Distal Interge | 9 | 78479649 | 78489151 | 9503 | 2 | ENSMUST00 | -38183 | Eef1a1 | ENSMUSG00000037742.9 |
| chrX | 151095928 | 151096899 | 16.08362 | 38.04472 | 35.47989 | Promoter | 20 | 151087094 | 151096543 | 9450 | 2 | ENSMUST00 | 0 | Tsr2 | ENSMUSG00000025264.7 |
| chrX | 21088608 | 21089799 | 16.08362 | 38.04472 | 35.47989 | Promoter | 20 | 21079174 | 21089229 | 10056 | 2 | ENSMUST00 | 0 | Zfp300 | ENSMUSG00000031079.7 |
| chrX | 75514246 | 75515231 | 16.08362 | 38.04472 | 35.47989 | Promoter | 20 | 75514299 | 75524046 | 9748 | 1 | ENSMUST00 | 0 | Vbp1 | ENSMUSG00000031197.6 |
| chrX | 98890055 | 98891324 | 16.08362 | 38.04472 | 35.47989 | Promoter | 20 | 98554277 | 98890985 | 336709 | 2 | ENSMUST00 | 0 | Ophn1 | ENSMUSG00000031214.8 |
| chrX | 109094875 | 109096274 | 16.08362 | 38.04472 | 35.47989 | Promoter | 20 | 109095365 | 109162471 | 67107 | 1 | ENSMUST00 | 0 | Sh3bgrl | ENSMUSG00000031246.9 |
| chrX | 159988066 | 159988852 | 16.08362 | 38.04472 | 35.47989 | Promoter | 20 | 159988433 | 160123351 | 134919 | 1 | ENSMUST00 | 0 | Map3k15 | ENSMUSG00000031303.8 |
| chrX | 166238777 | 166239575 | 16.08362 | 38.04472 | 35.47989 | Promoter | 20 | 166238911 | 166388988 | 150078 | 1 | ENSMUST00 | 0 | Gpm6b | ENSMUSG00000031342.12 |
| chrX | 163908206 | 163908912 | 16.08362 | 38.04472 | 35.47989 | Promoter | 20 | 163909017 | 163933666 | 24650 | 1 | ENSMUST00 | -105 | Ap1s2 | ENSMUSG00000031367.10 |
| chrX | 75129972 | 75131008 | 16.08362 | 38.04472 | 35.47989 | Promoter | 20 | 75110648 | 75130844 | 20197 | 2 | ENSMUST00 | 0 | Mpp1 | ENSMUSG00000031402.7 |
| chrX | 75095132 | 75096641 | 16.08362 | 38.04472 | 35.47989 | Promoter | 20 | 75095854 | 75109777 | 13924 | 1 | ENSMUST00 | 0 | Dkc1 | ENSMUSG00000031403.9 |
| chrX | 134717563 | 134718523 | 16.08362 | 38.04472 | 35.47989 | Promoter | 20 | 134717963 | 134721908 | 3946 | 1 | ENSMUST00 | 0 | Armcx1 | ENSMUSG00000033460.9 |
| chrX | 48340813 | 48342607 | 16.08362 | 38.04472 | 35.47989 | Promoter | 20 | 48341358 | 48408049 | 66692 | 1 | ENSMUST00 | 0 | Bcorl1 | ENSMUSG00000036959.10 |
| chrX | 164075842 | 164076545 | 16.08362 | 38.04472 | 35.47989 | Promoter | 20 | 164070705 | 164076121 | 5417 | 2 | ENSMUST00 | 0 | Siah1b | ENSMUSG00000040749.6 |
| chrX | 159414375 | 159415569 | 16.08362 | 38.04472 | 35.47989 | Promoter | 20 | 159414572 | 159498757 | 84186 | 1 | ENSMUST00 | 0 | Map7d2 | ENSMUSG00000041020.9 |
| chrX | 136244748 | 136245928 | 16.08362 | 38.04472 | 35.47989 | Promoter | 20 | 136245079 | 136247139 | 2061 | 1 | ENSMUST00 | 0 | Wbp5 | ENSMUSG00000042712.5 |
| chrX | 51017427 | 51019512 | 16.08362 | 38.04472 | 35.47989 | Promoter | 20 | 51003912 | 51018018 | 14107 | 2 | ENSMUST00 | 0 | Rap2c | ENSMUSG00000050029.7 |
| chrX | 52612296 | 52614474 | 16.08362 | 38.04472 | 35.47989 | Promoter | 20 | 52272426 | 52613921 | 341496 | 2 | ENSMUST00 | 0 | Gpc3 | ENSMUSG00000055653.8 |
| chrX | 75155536 | 75156011 | 16.08362 | 38.04472 | 35.47989 | Promoter | 20 | 75155819 | 75156273 | 455 | 1 | ENSMUST00 | 0 | Gm8503 | ENSMUSG00000082244.1 |
| chrX | 152326990 | 152327564 | 16.08362 | 38.04472 | 35.47989 | Promoter | 20 | 152298500 | 152327495 | 28996 | 2 | ENSMUST00 | 0 | Kantr | ENSMUSG00000087403.3 |
| chrX | 48284800 | 48285343 | 16.08362 | 38.04472 | 35.47989 | Promoter | 20 | 48276792 | 48285426 | 8635 | 2 | ENSMUST00 | 83 | 9530027J09F | ENSMUSG00000097197.1 |
| chr1 | 178368575 | 178369724 | 12.59008 | 38.01365 | 35.45054 | Distal Interge | 1 | 178363401 | 178363521 | 121 | 2 | ENSMUST00 | -5055 | Gm27458 | ENSMUSG00000099303.1 |
| chr10 | 4712098 | 4712991 | 12.59008 | 38.01365 | 35.45054 | Promoter | 10 | 4712651 | 4784824 | 72174 | 1 | ENSMUST00 | 0 | Esr1 | ENSMUSG00000019768.11 |
| chr15 | 103502456 | 103503556 | 12.59008 | 38.01365 | 35.45054 | Promoter | 15 | 103503298 | 103530052 | 26755 | 1 | ENSMUST00 | 0 | Pde1b | ENSMUSG00000022489.5 |
| chr18 | 52527968 | 52530011 | 12.59008 | 38.01365 | 35.45054 | Promoter | 18 | 52516069 | 52529867 | 13799 | 2 | ENSMUST00 | 0 | Lox | ENSMUSG00000024529.8 |
| chr5 | 52113384 | 52114289 | 12.59008 | 38.01365 | 35.45054 | Distal Interge | 5 | 52135902 | 52136017 | 116 | 2 | ENSMUST00 | 21728 | Mir6417 | ENSMUSG00000098726.1 |
| chr6 | 52244858 | 52245631 | 12.59008 | 38.01365 | 35.45054 | Promoter | 6 | 52245243 | 52249769 | 4527 | 1 | ENSMUST00 | 0 | Hoxa11os | ENSMUSG00000086427.3 |
| chr2 | 121448585 | 121449040 | 11.60718 | 37.9516 | 35.3891 | Promoter | 2 | 121449195 | 121453420 | 4226 | 1 | ENSMUST00 | -155 | Serf2 | ENSMUSG00000074884.7 |
| chr7 | 19749861 | 19750714 | 11.60718 | 37.9516 | 35.3891 | Promoter | 7 | 19724165 | 19749573 | 25409 | 2 | ENSMUST00 | -289 | Pvrl2 | ENSMUSG00000062300.8 |
| chr10 | 80355854 | 80357192 | 15.98761 | 37.88226 | 35.32005 | Promoter | 10 | 80356459 | 80365489 | 9031 | 1 | ENSMUST00 | 0 | Plk5 | ENSMUSG00000035486.9 |
| chr15 | 94628841 | 94629705 | 15.98761 | 37.88226 | 35.32005 | Promoter | 15 | 94629185 | 95096097 | 466913 | 1 | ENSMUST00 | 0 | Tmem117 | ENSMUSG00000063296.4 |
| chr2 | 143545567 | 143546690 | 15.95428 | 37.82584 | 35.264 | Promoter | 2 | 143546156 | 143816285 | 270130 | 1 | ENSMUST00 | 0 | Pcsk2 | ENSMUSG00000027419.9 |
| chr7 | 6361971 | 6362862 | 15.95428 | 37.82584 | 35.264 | Distal Interge | 7 | 6371364 | 6379491 | 8128 | 1 | ENSMUST00 | -8502 | Zfp78 | ENSMUSG00000055150.9 |
| chr19 | 7612064 | 7612784 | 15.53557 | 37.81869 | 35.25693 | Promoter | 19 | 7612541 | 7614745 | 2205 | 1 | ENSMUST00 | 0 | Hrasls5 | ENSMUSG00000024973.11 |
| chr1 | 57436276 | 57437484 | 15.47449 | 37.71435 | 35.15295 | Distal Interge | 1 | 57422244 | 57422373 | 130 | 2 | ENSMUST00 | -13904 | Gm22371 | ENSMUSG00000077776.1 |
| chr13 | 80962659 | 80963447 | 15.47449 | 37.71435 | 35.15295 | Promoter | 13 | 80962888 | 81126424 | 163537 | 1 | ENSMUST00 | 0 | 9330111N05 | ENSMUSG00000097098.2 |
| chr15 | 102399060 | 102400347 | 15.47449 | 37.71435 | 35.15295 | Distal Interge | 15 | 102406143 | 102408235 | 2093 | 1 | ENSMUST00 | -5796 | Sp1 | ENSMUSG00000001280.7 |
| chr8 | 91805255 | 91805890 | 15.47449 | 37.71435 | 35.15295 | Exon (ENSMU | 8 | 91802175 | 91805215 | 3041 | 1 | ENSMUST00 | 3081 | Irx3os | ENSMUSG00000046413.7 |
| chr9 | 41228890 | 41229328 | 15.47448 | 37.71435 | 35.15295 | Distal Interge | 9 | 41216948 | 41217566 | 619 | 1 | ENSMUST00 | 11943 | RP24-531B2 | ENSMUSG00000100837.1 |
| chr10 | 61272966 | 61273639 | 14.24369 | 37.68267 | 35.12186 | Promoter | 10 | 61197112 | 61273438 | 76327 | 2 | ENSMUST00 | 0 | Adamts14 | ENSMUSG00000059901.7 |
| chr19 | 5489205 | 5489624 | 14.24369 | 37.68267 | 35.12186 | Promoter | 19 | 5482832 | 5488402 | 5571 | 2 | ENSMUST00 | -804 | Mus81 | ENSMUSG00000024906.4 |
| chr11 | 101626762 | 101627326 | 15.85511 | 37.65791 | 35.09732 | Promoter | 11 | 101627195 | 101630900 | 3706 | 1 | ENSMUST00 | 0 | Rdm1 | ENSMUSG00000010362.4 |
| chr8 | 95702701 | 95704003 | 15.85511 | 37.65791 | 35.09732 | Promoter | 8 | 95703037 | 95715119 | 12083 | 1 | ENSMUST00 | 0 | Ndrg4 | ENSMUSG00000036564.11 |
| chr6 | 63255838 | 63257886 | 15.01852 | 37.63644 | 35.07596 | Promoter | 6 | 63255876 | 63314071 | 58196 | 1 | ENSMUST00 | 0 | Grid2 | ENSMUSG00000071424.4 |
| chr7 | 25848162 | 25848720 | 15.01852 | 37.63644 | 35.07596 | Distal Interge | 7 | 25828587 | 25828877 | 291 | 2 | ENSMUST00 | -19286 | Gm4613 | ENSMUSG00000090467.2 |
| chr11 | 115859158 | 115859905 | 14.61067 | 37.63035 | 35.07038 | Distal Interge | 11 | 115861612 | 115892592 | 30981 | 1 | ENSMUST00 | -1707 | Myo15b | ENSMUSG00000034427.11 |
| chr11 | 106920883 | 106921302 | 14.61067 | 37.63035 | 35.07038 | Promoter | 11 | 106920658 | 106920761 | 104 | 2 | ENSMUST00 | -123 | Gm27322 | ENSMUSG00000099125.1 |
| chr18 | 74443774 | 74444853 | 14.61067 | 37.63035 | 35.07038 | Intron (ENSM | 18 | 74442607 | 74770805 | 328199 | 1 | ENSMUST00 | 1168 | Myo5b | ENSMUSG00000025885.13 |
| chr19 | 41384765 | 41385378 | 14.61067 | 37.63035 | 35.07038 | Promoter | 19 | 41274218 | 41385070 | 110853 | 2 | ENSMUST00 | 0 | Pik3ap1 | ENSMUSG00000025017.9 |
| chr6 | 6005787 | 6006432 | 14.61067 | 37.63035 | 35.07038 | Exon (ENSMU | 6 | 5963909 | 5977393 | 13485 | 2 | ENSMUST00 | -28395 | 1700019G24 | ENSMUSG00000085416.1 |
| chr19 | 46308963 | 46309253 | 15.82232 | 37.60237 | 35.04273 | Exon (ENSMU | 19 | 46305695 | 46312090 | 6396 | 1 | ENSMUST00 | 3269 | Nfkb2 | ENSMUSG00000025225.9 |
| chr1 | 86503793 | 86505070 | 13.05315 | 37.5606 | 35.00106 | Distal Interge | 1 | 86526726 | 86530712 | 3987 | 1 | ENSMUST00 | -21656 | Ptma | ENSMUSG00000026238.9 |
| chr13 | 23551125 | 23552074 | 13.05315 | 37.5606 | 35.00106 | Promoter | 13 | 23551258 | 23551648 | 391 | 2 | ENSMUST00 | 0 | Hist1h4f | ENSMUSG00000069274.3 |
| chr15 | 3581766 | 3582717 | 13.05315 | 37.5606 | 35.00106 | Promoter | 15 | 3319009 | 3581842 | 262834 | 2 | ENSMUST00 | 0 | Ghr | ENSMUSG00000055737.7 |
| chr17 | 56761821 | 56762532 | 13.05315 | 37.5606 | 35.00106 | Distal Interge | 17 | 56764751 | 56770092 | 5342 | 1 | ENSMUST00 | -2219 | Dus3l | ENSMUSG00000007603.8 |
| chr4 | 117182271 | 117183179 | 13.05315 | 37.5606 | 35.00106 | Promoter | 4 | 117159639 | 117182639 | 23001 | 2 | ENSMUST00 | 0 | Kif2c | ENSMUSG00000028678.8 |
| chr4 | 130307794 | 130309178 | 13.05315 | 37.5606 | 35.00106 | Promoter | 4 | 130308595 | 130315463 | 6869 | 1 | ENSMUST00 | 0 | Fabp3 | ENSMUSG00000028773.8 |
| chr8 | 84638530 | 84639416 | 15.78968 | 37.54704 | 34.98843 | Promoter | 8 | 84638991 | 84639434 | 444 | 1 | ENSMUST00 | 0 | Cacna1a | ENSMUSG00000034656.11 |
| chr8 | 36532711 | 36533637 | 15.75716 | 37.49194 | 34.93351 | Distal Interge | 8 | 36505489 | 36512593 | 7105 | 1 | ENSMUST00 | 27223 | 6430573F11I | ENSMUSG00000039620.11 |
| chr7 | 27542374 | 27543302 | 15.32385 | 37.45674 | 34.89842 | Promoter | 7 | 27541740 | 27542745 | 1006 | 2 | ENSMUST00 | 0 | Pld3 | ENSMUSG00000003363.10 |
| chr11 | 44617917 | 44618811 | 15.72478 | 37.43704 | 34.87882 | Promoter | 11 | 44618116 | 44646820 | 28705 | 1 | ENSMUST00 | 0 | Ebf1 | ENSMUSG00000057098.9 |
| chr6 | 97311581 | 97312764 | 15.72478 | 37.43704 | 34.87882 | Promoter | 6 | 97287082 | 97312380 | 25299 | 2 | ENSMUST00 | 0 | Frmd4b | ENSMUSG00000030064.11 |
| chr8 | 69887294 | 69889519 | 15.72478 | 37.43704 | 34.87882 | Promoter | 8 | 69880369 | 69887687 | 7319 | 2 | ENSMUST00 | 0 | Cilp2 | ENSMUSG00000044006.8 |
| chr2 | 61453969 | 61454647 | 13.6939 | 37.38395 | 34.82589 | Distal Interge | 2 | 61492766 | 61492894 | 129 | 1 | ENSMUST00 | -38119 | Gm22338 | ENSMUSG00000088340.1 |
| chr1 | 97976586 | 97977623 | 11.92996 | 37.32891 | 34.77106 | Promoter | 1 | 97821100 | 97977255 | 156156 | 2 | ENSMUST00 | 0 | Pam | ENSMUSG00000026335.11 |
| chr1 | 155973396 | 155974279 | 11.92996 | 37.32891 | 34.77106 | Promoter | 1 | 155953442 | 155972890 | 19449 | 2 | ENSMUST00 | -507 | Cep350 | ENSMUSG00000033671.12 |
| chr11 | 69011917 | 69012538 | 11.92996 | 37.32891 | 34.77106 | Distal Interge | 11 | 69015911 | 69036473 | 20563 | 1 | ENSMUST00 | -3373 | Ctc1 | ENSMUSG00000020898.13 |
| chr10 | 7867155 | 7867812 | 15.66042 | 37.32788 | 34.77026 | Exon (ENSMU | 10 | 7848361 | 7848481 | 121 | 2 | ENSMUST00 | -18675 | Gm24726 | ENSMUSG00000088857.1 |
| chr13 | 22050126 | 22051551 | 15.66042 | 37.32788 | 34.77026 | Distal Interge | 13 | 22057346 | 22057862 | 517 | 2 | ENSMUST00 | 6311 | Vmn1r-ps92 | ENSMUSG00000083873.1 |
| chr19 | 17356049 | 17356821 | 15.66042 | 37.32788 | 34.77026 | Promoter | 19 | 17326141 | 17356667 | 30527 | 2 | ENSMUST00 | 0 | Gcnt1 | ENSMUSG00000038843.12 |
| chr4 | 124985979 | 124987322 | 15.66042 | 37.32788 | 34.77026 | Promoter | 4 | 124986430 | 125009099 | 22670 | 1 | ENSMUST00 | 0 | Rspo1 | ENSMUSG00000028871.7 |
| chr7 | 100178425 | 100179373 | 15.66042 | 37.32788 | 34.77026 | Promoter | 7 | 100178683 | 100184869 | 6187 | 1 | ENSMUST00 | 0 | Kcne3 | ENSMUSG00000035165.8 |
| chr11 | 85832779 | 85833988 | 15.23487 | 37.30433 | 34.74706 | Promoter | 11 | 85832551 | 85841948 | 9398 | 1 | ENSMUST00 | 229 | Tbx2 | ENSMUSG00000000093.6 |
| chr16 | 44015748 | 44016791 | 15.23487 | 37.30433 | 34.74706 | Promoter | 16 | 43980355 | 44016436 | 36082 | 2 | ENSMUST00 | 0 | Gramd1c | ENSMUSG00000036292.8 |
| chr14 | 75130450 | 75131599 | 15.62844 | 37.27362 | 34.7164 | Promoter | 14 | 75084303 | 75130881 | 46579 | 2 | ENSMUST00 | 0 | Lrrc63 | ENSMUSG00000021997.4 |
| chr15 | 98992450 | 98993902 | 15.62844 | 37.27362 | 34.7164 | Distal Interge | 15 | 98985965 | 98989281 | 3317 | 1 | ENSMUST00 | 6486 | 4930578M01 | ENSMUSG00000097587.1 |
| chr17 | 31856014 | 31856714 | 15.62844 | 37.27362 | 34.7164 | Promoter | 17 | 31844248 | 31855792 | 11545 | 2 | ENSMUST00 | -223 | Sik1 | ENSMUSG00000024042.6 |
| chr11 | 60114432 | 60115777 | 14.00966 | 37.26205 | 34.70494 | Promoter | 11 | 60112219 | 60115230 | 3012 | 2 | ENSMUST00 | 0 | 4930412M03 | ENSMUSG00000051008.5 |
| chr18 | 61535608 | 61536934 | 14.00966 | 37.26205 | 34.70494 | Promoter | 18 | 61493794 | 61536536 | 42743 | 2 | ENSMUST00 | 0 | Arhgef37 | ENSMUSG00000045094.7 |
| chr15 | 97663845 | 97664335 | 15.59658 | 37.21957 | 34.66261 | Distal Interge | 15 | 97675105 | 97705822 | 30718 | 2 | ENSMUST00 | 41487 | Rpap3 | ENSMUSG00000022466.5 |
| chr1 | 169531126 | 169531482 | 13.60354 | 37.21757 | 34.66072 | Promoter | 1 | 169497934 | 169531464 | 33531 | 2 | ENSMUST00 | 0 | Nuf2 | ENSMUSG00000026683.9 |
| chr10 | 112270109 | 112271212 | 13.60354 | 37.21757 | 34.66072 | Promoter | 10 | 112271123 | 112466304 | 195182 | 1 | ENSMUST00 | 0 | Kcnc2 | ENSMUSG00000035681.6 |
| chr11 | 20531889 | 20533096 | 13.60354 | 37.21757 | 34.66072 | Distal Interge | 11 | 20543253 | 20597020 | 53768 | 1 | ENSMUST00 | -10157 | Sertad2 | ENSMUSG00000049800.8 |

| | | | | | | | | | | | | | | | |
|---|---|---|---|---|---|---|---|---|---|---|---|---|---|---|---|
| chr12 | 14149609 | 14150936 | 13.60354 | 37.21757 | 34.66072 | 5' UTR | 12 | 14147599 | 14152038 | 4440 | 2 | ENSMUST00 | 1102 | Fam84a | ENSMUSG00000020607.6 |
| chr12 | 17011214 | 17011843 | 13.60354 | 37.21757 | 34.66072 | Promoter | 12 | 17003319 | 17011727 | 8409 | 2 | ENSMUST00 | 0 | 2410004P03I | ENSMUSG00000071398.3 |
| chr13 | 47193392 | 47194928 | 13.60354 | 37.21757 | 34.66072 | Promoter | 13 | 47194003 | 47247991 | 53989 | 1 | ENSMUST00 | 0 | Rnf144b | ENSMUSG00000038068.9 |
| chr16 | 13940421 | 13941256 | 13.60354 | 37.21757 | 34.66072 | Promoter | 16 | 13940630 | 13948723 | 8094 | 1 | ENSMUST00 | 0 | Mpv17l | ENSMUSG00000022679.8 |
| chr17 | 88626424 | 88629225 | 13.60354 | 37.21757 | 34.66072 | Promoter | 17 | 88626555 | 88645724 | 19170 | 1 | ENSMUST00 | 0 | Ston1 | ENSMUSG00000033855.10 |
| chr3 | 36572140 | 36572836 | 13.60354 | 37.21757 | 34.66072 | Promoter | 3 | 36564866 | 36572150 | 7285 | 2 | ENSMUST00 | 0 | Ccna2 | ENSMUSG00000027715.4 |
| chr4 | 101496262 | 101497630 | 13.60354 | 37.21757 | 34.66072 | Promoter | 4 | 101496648 | 101642799 | 146152 | 1 | ENSMUST00 | 0 | Dnajc6 | ENSMUSG00000028528.11 |
| chr8 | 84197247 | 84198456 | 13.60354 | 37.21757 | 34.66072 | Promoter | 8 | 84192774 | 84197667 | 4894 | 2 | ENSMUST00 | 0 | Gm26887 | ENSMUSG00000097120.1 |
| chr1 | 183146159 | 183147549 | 15.17612 | 37.20362 | 34.64814 | Promoter | 1 | 183086270 | 183147461 | 61192 | 2 | ENSMUST00 | 0 | Disp1 | ENSMUSG00000030768.7 |
| chr14 | 55814847 | 55815738 | 15.17612 | 37.20362 | 34.64814 | Distal Interge | 14 | 55824795 | 55833943 | 9149 | 1 | ENSMUST00 | -9057 | Nfatc4 | ENSMUSG00000023411.6 |
| chr17 | 29889004 | 29889859 | 15.17612 | 37.20362 | 34.64814 | Promoter | 17 | 29850562 | 29889950 | 39389 | 2 | ENSMUST00 | 91 | Mdga1 | ENSMUSG00000043557.10 |
| chr5 | 118027361 | 118028840 | 14.35816 | 37.18478 | 34.62976 | Promoter | 5 | 118027743 | 118061878 | 34136 | 1 | ENSMUST00 | 0 | Tesc | ENSMUSG00000029359.8 |
| chr5 | 131615984 | 131616853 | 15.56486 | 37.16572 | 34.61091 | Promoter | 5 | 131437788 | 131616605 | 178818 | 2 | ENSMUST00 | 0 | Auts2 | ENSMUSG00000029673.12 |
| chr10 | 80825982 | 80826298 | 14.74478 | 37.16129 | 34.60659 | Promoter | 10 | 80826656 | 80829290 | 2635 | 1 | ENSMUST00 | -358 | Oaz1 | ENSMUSG00000035242.10 |
| chr16 | 50363548 | 50364971 | 14.74478 | 37.16129 | 34.60659 | Intron (ENSM | 16 | 50197159 | 50330993 | 133835 | 2 | ENSMUST00 | -32556 | Bbx | ENSMUSG00000022641.10 |
| chr4 | 83315154 | 83316017 | 14.74478 | 37.16129 | 34.60659 | Intron (ENSM | 4 | 83259198 | 83324189 | 64992 | 2 | ENSMUST00 | 8172 | Ttc39b | ENSMUSG00000038172.9 |
| chr6 | 30737734 | 30738494 | 14.74478 | 37.16129 | 34.60659 | Promoter | 6 | 30738050 | 30744394 | 6345 | 1 | ENSMUST00 | 0 | Mest | ENSMUSG00000051855.10 |
| chr12 | 113000904 | 113001494 | 15.5018 | 37.05862 | 34.50454 | Promoter | 12 | 112959862 | 113000621 | 40760 | 2 | ENSMUST00 | -284 | Brf1 | ENSMUSG00000011158.7 |
| chr14 | 66324882 | 66325663 | 15.5018 | 37.05862 | 34.50454 | Distal Interge | 14 | 66344296 | 66356756 | 12461 | 1 | ENSMUST00 | -18633 | Stmn4 | ENSMUSG00000022044.9 |
| chr8 | 107402824 | 107403825 | 15.5018 | 37.05862 | 34.50454 | Promoter | 8 | 107388225 | 107403206 | 14982 | 2 | ENSMUST00 | 0 | Nqo1 | ENSMUSG00000003849.8 |
| chr1 | 74397527 | 74398277 | 15.08796 | 37.05233 | 34.49837 | Distal Interge | 1 | 74394310 | 74394394 | 85 | 1 | ENSMUST00 | 3218 | Mir26b | ENSMUSG00000065468.1 |
| chr10 | 41476033 | 41477908 | 15.08796 | 37.05233 | 34.49837 | Promoter | 10 | 41476314 | 41487030 | 10717 | 1 | ENSMUST00 | 0 | Mical1 | ENSMUSG00000019823.12 |
| chr11 | 54068682 | 54069337 | 15.08796 | 37.05233 | 34.49837 | Promoter | 11 | 54054928 | 54069014 | 14087 | 2 | ENSMUST00 | 0 | Pdlim4 | ENSMUSG00000020388.7 |
| chr11 | 3931448 | 3932506 | 15.08796 | 37.05233 | 34.49837 | Promoter | 11 | 3917192 | 3931789 | 14598 | 2 | ENSMUST00 | 0 | Tcn2 | ENSMUSG00000020432.7 |
| chr11 | 72301385 | 72302721 | 15.08796 | 37.05233 | 34.49837 | Promoter | 11 | 72301629 | 72309061 | 7433 | 1 | ENSMUST00 | 0 | Xaf1 | ENSMUSG00000040483.10 |
| chr14 | 34768320 | 34770010 | 15.08796 | 37.05233 | 34.49837 | Exon (ENSMU | 14 | 34760227 | 34770553 | 10327 | 1 | ENSMUST00 | 8094 | A930038B10 | ENSMUSG00000097310.1 |
| chr16 | 18586258 | 18587332 | 15.08796 | 37.05233 | 34.49837 | Promoter | 16 | 18581713 | 18586969 | 5257 | 2 | ENSMUST00 | 0 | Tbx1 | ENSMUSG00000009097.9 |
| chr17 | 55958575 | 55959968 | 15.08796 | 37.05233 | 34.49837 | Promoter | 17 | 55959187 | 55967949 | 8763 | 1 | ENSMUST00 | 0 | Ccdc94 | ENSMUSG00000003208.8 |
| chr17 | 20944553 | 20945245 | 15.08796 | 37.05233 | 34.49837 | Promoter | 17 | 20945311 | 20965916 | 20606 | 1 | ENSMUST00 | -66 | Ppp2r1a | ENSMUSG00000007564.9 |
| chr18 | 32147411 | 32147934 | 15.08796 | 37.05233 | 34.49837 | Distal Interge | 18 | 32153476 | 32160427 | 6952 | 1 | ENSMUST00 | -5542 | Gm26717 | ENSMUSG00000097363.1 |
| chr2 | 127655573 | 127656684 | 15.08796 | 37.05233 | 34.49837 | Promoter | 2 | 127634874 | 127656219 | 21346 | 2 | ENSMUST00 | 0 | Mal | ENSMUSG00000027375.9 |
| chr2 | 148406833 | 148408646 | 15.08796 | 37.05233 | 34.49837 | Promoter | 2 | 148404466 | 148408188 | 3723 | 2 | ENSMUST00 | 0 | Thbd | ENSMUSG00000074743.4 |
| chr2 | 73584727 | 73585698 | 15.08796 | 37.05233 | 34.49837 | Promoter | 2 | 73584850 | 73589187 | 4338 | 1 | ENSMUST00 | 0 | Chrna1os | ENSMUSG00000086382.1 |
| chr4 | 136424289 | 136425176 | 15.08796 | 37.05233 | 34.49837 | Promoter | 4 | 136424612 | 136441970 | 17359 | 1 | ENSMUST00 | 0 | Htr1d | ENSMUSG00000070687.6 |
| chr6 | 72362563 | 72363086 | 15.08796 | 37.05233 | 34.49837 | Promoter | 6 | 72360245 | 72362721 | 2477 | 2 | ENSMUST00 | 0 | Rnf181 | ENSMUSG00000055850.7 |
| chr6 | 120579532 | 120580461 | 15.08796 | 37.05233 | 34.49837 | Distal Interge | 6 | 120509494 | 120531319 | 21826 | 2 | ENSMUST00 | -48214 | Cecr5 | ENSMUSG00000058979.6 |
| chr7 | 73399209 | 73400163 | 15.08796 | 37.05233 | 34.49837 | Intron (ENSM | 7 | 73391227 | 73419899 | 28673 | 1 | ENSMUST00 | 7983 | Rgma | ENSMUSG00000070509.10 |
| chr8 | 104570572 | 104571264 | 15.08796 | 37.05233 | 34.49837 | Distal Interge | 8 | 104591468 | 104596836 | 5369 | 1 | ENSMUST00 | -20204 | Pdp2 | ENSMUSG00000048371.7 |
| chr8 | 70697986 | 70698758 | 15.08796 | 37.05233 | 34.49837 | Promoter | 8 | 70697739 | 70700616 | 2878 | 1 | ENSMUST00 | 248 | Jund | ENSMUSG00000071076.5 |
| chr9 | 53340339 | 53341883 | 15.08796 | 37.05233 | 34.49837 | Promoter | 9 | 53341084 | 53368955 | 27872 | 1 | ENSMUST00 | 0 | Exph5 | ENSMUSG00000034584.3 |
| chr9 | 65391003 | 65391718 | 15.08796 | 37.05233 | 34.49837 | Promoter | 9 | 65388684 | 65391652 | 2969 | 2 | ENSMUST00 | 0 | Kbtbd13 | ENSMUSG00000054978.3 |
| chr9 | 32940340 | 32940784 | 15.08796 | 37.05233 | 34.49837 | Distal Interge | 9 | 32904937 | 32928966 | 24030 | 2 | ENSMUST00 | -11375 | Gm27162 | ENSMUSG00000099025.1 |
| chr4 | 146239373 | 146240167 | 10.64379 | 37.03648 | 34.48422 | Promoter | 4 | 146239954 | 146241633 | 1680 | 1 | ENSMUST00 | 0 | Gm13169 | ENSMUSG00000084127.1 |
| chr1 | 134298720 | 134299437 | 14.26849 | 37.02594 | 34.4737 | 3' UTR | 1 | 134296789 | 134304808 | 8020 | 2 | ENSMUST00 | 5371 | Ppfia4 | ENSMUSG00000026458.8 |
| chr1 | 187997337 | 187998108 | 14.26849 | 37.02594 | 34.4737 | Promoter | 1 | 187997827 | 188214885 | 217059 | 1 | ENSMUST00 | 0 | Esrrg | ENSMUSG00000026610.8 |
| chr12 | 81332367 | 81333403 | 14.26849 | 37.02594 | 34.4737 | Promoter | 12 | 81197915 | 81333164 | 135250 | 2 | ENSMUST00 | 0 | Slc8a3 | ENSMUSG00000079055.5 |
| chr13 | 67892587 | 67892876 | 14.26849 | 37.02594 | 34.4737 | Distal Interge | 13 | 67863326 | 67863923 | 598 | 1 | ENSMUST00 | 29262 | BC048507 | ENSMUSG00000064063.5 |
| chr17 | 43568010 | 43569102 | 14.26849 | 37.02594 | 34.4737 | Promoter | 17 | 43568098 | 43600591 | 32494 | 1 | ENSMUST00 | 0 | Pla2g7 | ENSMUSG00000023913.12 |
| chr18 | 65430226 | 65430779 | 14.26849 | 37.02594 | 34.4737 | Promoter | 18 | 65430963 | 65478823 | 47861 | 1 | ENSMUST00 | -184 | Malt1 | ENSMUSG00000032688.7 |
| chr19 | 46317440 | 46318465 | 14.26849 | 37.02594 | 34.4737 | Intron (ENSM | 19 | 46312087 | 46327156 | 15070 | 2 | ENSMUST00 | 8691 | Psd | ENSMUSG00000037126.10 |
| chr2 | 164960770 | 164961185 | 14.26849 | 37.02594 | 34.4737 | Promoter | 2 | 164960816 | 164974540 | 13725 | 1 | ENSMUST00 | 0 | Slc12a5 | ENSMUSG00000017740.11 |
| chr3 | 50906023 | 50906810 | 14.26849 | 37.02594 | 34.4737 | Distal Interge | 3 | 50838581 | 50838715 | 135 | 2 | ENSMUST00 | -67309 | Gm24503 | ENSMUSG00000089079.1 |
| chr4 | 128811604 | 128812168 | 14.26849 | 37.02594 | 34.4737 | Distal Interge | 4 | 128777084 | 128806045 | 28962 | 2 | ENSMUST00 | -5560 | Zfp362 | ENSMUSG00000028799.10 |
| chr4 | 33209041 | 33209844 | 14.26849 | 37.02594 | 34.4737 | Promoter | 4 | 33209263 | 33231293 | 22031 | 1 | ENSMUST00 | 0 | Srsf12 | ENSMUSG00000054679.3 |
| chr4 | 123200919 | 123201913 | 14.26849 | 37.02594 | 34.4737 | Promoter | 4 | 123201504 | 123216275 | 14772 | 1 | ENSMUST00 | 0 | Nt5c1a | ENSMUSG00000054958.6 |
| chr4 | 151386347 | 151387508 | 14.26849 | 37.02594 | 34.4737 | Intron (ENSM | 4 | 151376470 | 151379582 | 3113 | 1 | ENSMUST00 | 9878 | 4930589P08I | ENSMUSG00000085593.1 |
| chr5 | 115488835 | 115490242 | 14.26849 | 37.02594 | 34.4737 | Promoter | 5 | 115489459 | 115489599 | 141 | 1 | ENSMUST00 | 0 | Gm24265 | ENSMUSG00000096243.1 |
| chr6 | 51379282 | 51380226 | 14.26849 | 37.02594 | 34.4737 | Exon (ENSMU | 6 | 51379710 | 51392777 | 13068 | 2 | ENSMUST00 | 12551 | Gm26677 | ENSMUSG00000097598.1 |
| chr7 | 140102025 | 140102732 | 14.26849 | 37.02594 | 34.4737 | Promoter | 7 | 140096770 | 140102403 | 5634 | 2 | ENSMUST00 | 0 | Fuom | ENSMUSG00000025466.13 |
| chr7 | 34570293 | 34571186 | 14.26849 | 37.02594 | 34.4737 | Distal Interge | 7 | 34547206 | 34556204 | 8999 | 1 | ENSMUST00 | 23088 | Gm12784 | ENSMUSG00000086631.1 |
| chr5 | 114922968 | 114923653 | 14.25959 | 37.01015 | 34.45982 | Promoter | 5 | 114923240 | 114937912 | 14673 | 1 | ENSMUST00 | 0 | Oasl1 | ENSMUSG00000041827.10 |
| chr8 | 60954034 | 60955093 | 15.05997 | 37.00426 | 34.45401 | Promoter | 8 | 60911896 | 60954748 | 42853 | 2 | ENSMUST00 | 0 | Clcn3 | ENSMUSG00000004319.10 |
| chr3 | 88831165 | 88831978 | 13.48279 | 36.99454 | 34.44434 | Promoter | 3 | 88800688 | 88832368 | 31681 | 2 | ENSMUST00 | 390 | 1500004A13I | ENSMUSG00000098912.2 |
| chr7 | 111779164 | 111780393 | 13.48279 | 36.99454 | 34.44434 | Promoter | 7 | 111471661 | 111779977 | 308317 | 2 | ENSMUST00 | 0 | Galnt18 | ENSMUSG00000038296.9 |
| chr5 | 135961931 | 135962393 | 15.43924 | 36.95233 | 34.40221 | Promoter | 5 | 135960224 | 135962327 | 2104 | 2 | ENSMUST00 | 0 | Srcrb4d | ENSMUSG00000029699.8 |
| chr7 | 141168086 | 141169179 | 15.43924 | 36.95233 | 34.40221 | Promoter | 7 | 141160328 | 141168936 | 8609 | 2 | ENSMUST00 | 0 | Rnh1 | ENSMUSG00000038650.9 |
| chr18 | 12949163 | 12949453 | 4.49855 | 36.92913 | 34.37911 | Distal Interge | 18 | 12870218 | 12941841 | 71624 | 2 | ENSMUST00 | -7323 | Osbpl1a | ENSMUSG00000044252.12 |
| chr10 | 45335418 | 45336262 | 15.40816 | 36.89948 | 34.3495 | Promoter | 10 | 45335762 | 45369708 | 33947 | 1 | ENSMUST00 | 0 | Bves | ENSMUSG00000071317.3 |
| chr11 | 61342082 | 61343105 | 13.7832 | 36.85258 | 34.30273 | Promoter | 11 | 61301631 | 61342860 | 41230 | 2 | ENSMUST00 | 0 | Slc47a2 | ENSMUSG00000069855.4 |
| chr12 | 24592715 | 24593894 | 15.37719 | 36.84684 | 34.29707 | Intron (ENSM | 12 | 24580749 | 24617391 | 36643 | 1 | ENSMUST00 | 11967 | Grhl1 | ENSMUSG00000020656.10 |
| chr2 | 60552535 | 60553715 | 15.37719 | 36.84684 | 34.29707 | Promoter | 2 | 60417543 | 60553308 | 135766 | 2 | ENSMUST00 | 0 | Pla2r1 | ENSMUSG00000054580.9 |
| chr3 | 88137961 | 88138340 | 15.37719 | 36.84684 | 34.29707 | Distal Interge | 3 | 88142372 | 88172086 | 29715 | 1 | ENSMUST00 | -4032 | Mef2d | ENSMUSG00000001419.12 |
| chr10 | 87492594 | 87493800 | 15.34636 | 36.79438 | 34.24475 | Promoter | 10 | 87490819 | 87493660 | 2842 | 2 | ENSMUST00 | 0 | Ascl1 | ENSMUSG00000020052.7 |
| chr11 | 61956776 | 61957458 | 15.34636 | 36.79438 | 34.24475 | Promoter | 11 | 61956779 | 62141726 | 184948 | 1 | ENSMUST00 | 0 | Specc1 | ENSMUSG00000042331.8 |
| chr13 | 23449301 | 23450597 | 15.34636 | 36.79438 | 34.24475 | Distal Interge | 13 | 23456992 | 23465875 | 8884 | 2 | ENSMUST00 | 15278 | Btn1a1 | ENSMUSG00000000706.10 |
| chr14 | 47417921 | 47418619 | 15.34636 | 36.79438 | 34.24475 | Promoter | 14 | 47387779 | 47418407 | 30629 | 2 | ENSMUST00 | 0 | Dlgap5 | ENSMUSG00000037544.8 |
| chr17 | 87796906 | 87798947 | 15.34636 | 36.79438 | 34.24475 | Promoter | 17 | 87745837 | 87797994 | 52158 | 2 | ENSMUST00 | 0 | Kcnk12 | ENSMUSG00000050138.7 |
| chr18 | 69570451 | 69571235 | 15.34636 | 36.79438 | 34.24475 | Intron (ENSM | 18 | 69563335 | 69636964 | 73630 | 1 | ENSMUST00 | 7117 | Tcf4 | ENSMUSG00000053477.10 |
| chr10 | 78481695 | 78482652 | 12.30394 | 36.76666 | 34.21749 | Downstream | 10 | 78484570 | 78487842 | 3273 | 2 | ENSMUST00 | 5190 | Gm9978 | ENSMUSG00000055704.4 |
| chr10 | 83722344 | 83723468 | 12.30394 | 36.76666 | 34.21749 | Promoter | 10 | 83722865 | 83762761 | 39897 | 1 | ENSMUST00 | 0 | 1500009L16I | ENSMUSG00000087651.2 |
| chr11 | 96351914 | 96353821 | 12.30394 | 36.76666 | 34.21749 | Promoter | 11 | 96351631 | 96354012 | 2382 | 1 | ENSMUST00 | 284 | Hoxb2 | ENSMUSG00000075588.6 |
| chr12 | 91683606 | 91686349 | 12.30394 | 36.76666 | 34.21749 | Promoter | 12 | 91684665 | 91693590 | 8926 | 1 | ENSMUST00 | 0 | Gm16876 | ENSMUSG00000086689.1 |
| chr15 | 75641982 | 75642949 | 12.30394 | 36.76666 | 34.21749 | Promoter | 15 | 75642129 | 75647437 | 5309 | 1 | ENSMUST00 | 0 | 2810039B14I | ENSMUSG00000099034.2 |
| chr17 | 34398729 | 34399253 | 12.30394 | 36.76666 | 34.21749 | Promoter | 17 | 34398820 | 34460734 | 61915 | 1 | ENSMUST00 | 0 | BC051142 | ENSMUSG00000057246.9 |
| chr2 | 102658165 | 102659605 | 12.30394 | 36.76666 | 34.21749 | Promoter | 2 | 102658659 | 102790784 | 132126 | 1 | ENSMUST00 | 0 | Slc1a2 | ENSMUSG00000005089.10 |
| chr5 | 108001128 | 108001665 | 12.30394 | 36.76666 | 34.21749 | Promoter | 5 | 108002260 | 108002486 | 227 | 1 | ENSMUST00 | -595 | Gm9850 | ENSMUSG00000084855.2 |
| chr8 | 107114369 | 107115495 | 12.30394 | 36.76666 | 34.21749 | Distal Interge | 8 | 107075517 | 107096547 | 21031 | 2 | ENSMUST00 | -17823 | Terf2 | ENSMUSG00000031921.12 |
| chr7 | 70364822 | 70366760 | 11.36022 | 36.76307 | 34.21434 | Promoter | 7 | 70354439 | 70366735 | 12297 | 2 | ENSMUST00 | 0 | Nr2f2 | ENSMUSG00000030551.8 |
| chr8 | 25038210 | 25038958 | 14.11424 | 36.75194 | 34.20333 | Promoter | 8 | 25025451 | 25038962 | 13512 | 2 | ENSMUST00 | 4 | Htra4 | ENSMUSG00000037406.6 |
| chr7 | 105650475 | 105651400 | 14.88904 | 36.71027 | 34.16194 | Promoter | 7 | 105650827 | 105721799 | 70973 | 1 | ENSMUST00 | 0 | Dnhd1 | ENSMUSG00000030882.12 |
| chr19 | 59169142 | 59170482 | 14.48083 | 36.70091 | 34.15315 | Promoter | 19 | 59166187 | 59170056 | 3870 | 2 | ENSMUST00 | 0 | Vax1 | ENSMUSG00000006270.7 |
| chr19 | 33761502 | 33762078 | 14.48083 | 36.70091 | 34.15315 | Promoter | 19 | 33555167 | 33761951 | 206785 | 2 | ENSMUST00 | 0 | Lipo1 | ENSMUSG00000024766.9 |
| chr2 | 125505290 | 125506524 | 14.48083 | 36.70091 | 34.15315 | Promoter | 2 | 125300594 | 125506385 | 205792 | 2 | ENSMUST00 | 0 | Fbn1 | ENSMUSG00000027204.8 |
| chr2 | 35132175 | 35132958 | 14.48083 | 36.70091 | 34.15315 | Promoter | 2 | 35132234 | 35178822 | 46589 | 1 | ENSMUST00 | 0 | Cntrl | ENSMUSG00000057110.10 |
| chr3 | 51396333 | 51397054 | 14.48083 | 36.70091 | 34.15315 | Promoter | 3 | 51388415 | 51396502 | 8088 | 2 | ENSMUST00 | 0 | Mgarp | ENSMUSG00000037161.9 |
| chr11 | 117832202 | 117832625 | 15.28505 | 36.69006 | 34.14264 | Promoter | 11 | 117832230 | 117837231 | 5002 | 1 | ENSMUST00 | 0 | Afmid | ENSMUSG00000017718.10 |
| chr17 | 17837498 | 17839072 | 14.45496 | 36.65565 | 34.10837 | Promoter | 17 | 17837504 | 17839071 | 1568 | 1 | ENSMUST00 | 0 | Spaca6 | ENSMUSG00000080316.5 |
| chr16 | 4718567 | 4719580 | 15.22423 | 36.58649 | 34.03935 | Promoter | 16 | 4710059 | 4719109 | 9051 | 2 | ENSMUST00 | 0 | Nmral1 | ENSMUSG00000063445.9 |
| chr10 | 75892874 | 75893789 | 15.62409 | 36.57069 | 34.02363 | Promoter | 10 | 75893398 | 75895941 | 2544 | 1 | ENSMUST00 | 0 | Derl3 | ENSMUSG00000009092.4 |
| chr11 | 102393046 | 102393977 | 15.62409 | 36.57069 | 34.02363 | Promoter | 11 | 102393403 | 102401754 | 8352 | 1 | ENSMUST00 | 0 | Rundc3a | ENSMUSG00000006575.9 |
| chr11 | 110166679 | 110168650 | 15.62409 | 36.57069 | 34.02363 | Promoter | 11 | 110100749 | 110168196 | 67448 | 2 | ENSMUST00 | 0 | Abca9 | ENSMUSG00000041797.7 |
| chr12 | 57545240 | 57546360 | 15.62409 | 36.57069 | 34.02363 | Promoter | 12 | 57540631 | 57546121 | 5491 | 2 | ENSMUST00 | 0 | Foxa1 | ENSMUSG00000035451.6 |
| chr12 | 18514300 | 18515437 | 15.62409 | 36.57069 | 34.02363 | Promoter | 12 | 18514738 | 18534191 | 19454 | 1 | ENSMUST00 | 0 | 5730507C01I | ENSMUSG00000073197.3 |
| chr13 | 98015595 | 98016881 | 15.62409 | 36.57069 | 34.02363 | Intron (ENSM | 13 | 97898595 | 98206165 | 307571 | 2 | ENSMUST00 | 189284 | Arhgef28 | ENSMUSG00000021662.9 |
| chr13 | 114456886 | 114457881 | 15.62409 | 36.57069 | 34.02363 | Promoter | 13 | 114452262 | 114458730 | 6469 | 2 | ENSMUST00 | 849 | Fst | ENSMUSG00000021765.7 |
| chr14 | 122474804 | 122476455 | 15.62409 | 36.57069 | 34.02363 | Promoter | 14 | 122475435 | 122479852 | 4418 | 1 | ENSMUST00 | 0 | Zic2 | ENSMUSG00000061524.8 |
| chr15 | 75566414 | 75567244 | 15.62409 | 36.57069 | 34.02363 | Promoter | 15 | 75551268 | 75566648 | 15381 | 2 | ENSMUST00 | 0 | Ly6h | ENSMUSG00000022577.11 |
| chr16 | 37916082 | 37916811 | 15.62409 | 36.57069 | 34.02363 | Promoter | 16 | 37916496 | 38007527 | 91032 | 1 | ENSMUST00 | 0 | Gpr156 | ENSMUSG00000046961.7 |
| chr17 | 56005183 | 56006139 | 15.62409 | 36.57069 | 34.02363 | Promoter | 17 | 56005711 | 56011924 | 6214 | 1 | ENSMUST00 | 0 | Mpnd | ENSMUSG00000003199.11 |
| chr17 | 26200703 | 26201702 | 15.62409 | 36.57069 | 34.02363 | Promoter | 17 | 26196226 | 26201143 | 4918 | 2 | ENSMUST00 | 0 | Arhgdig | ENSMUSG00000073433.5 |
| chr17 | 9167217 | 9168590 | 15.62409 | 36.57069 | 34.02363 | Promoter | 17 | 9147719 | 9167836 | 20118 | 2 | ENSMUST00 | 0 | 6530411M01 | ENSMUSG00000097049.1 |
| chr2 | 164432671 | 164438188 | 15.62409 | 36.57069 | 34.02363 | Intron (ENSM | 2 | 164424247 | 164443186 | 18940 | 2 | ENSMUST00 | 4998 | Sdc4 | ENSMUSG00000017009.3 |
| chr2 | 130634758 | 130635618 | 15.62409 | 36.57069 | 34.02363 | 3' UTR | 2 | 130632839 | 130638095 | 5257 | 2 | ENSMUST00 | 2477 | Lzts3 | ENSMUSG00000037703.9 |
| chr2 | 18607971 | 18609779 | 15.62409 | 36.57069 | 34.02363 | Distal Interge | 2 | 18641786 | 18642102 | 317 | 2 | ENSMUST00 | 32323 | Gm13355 | ENSMUSG00000082943.1 |
| chr3 | 9334652 | 9335076 | 15.62409 | 36.57069 | 34.02363 | Distal Interge | 3 | 9402175 | 9402485 | 311 | 1 | ENSMUST00 | -67099 | Gm22604 | ENSMUSG00000088229.1 |
| chr5 | 135887419 | 135888188 | 15.62409 | 36.57069 | 34.02363 | Promoter | 5 | 135887919 | 135889563 | 1645 | 1 | ENSMUST00 | 0 | Hspb1 | ENSMUSG00000004951.10 |
| chr5 | 136622974 | 136623489 | 15.62409 | 36.57069 | 34.02363 | Distal Interge | 5 | 136613702 | 136615328 | 1627 | 2 | ENSMUST00 | -7647 | 4731417B20I | ENSMUSG00000046548.2 |
| chr7 | 16285599 | 16286129 | 15.62409 | 36.57069 | 34.02363 | Promoter | 7 | 16282824 | 16286025 | 3202 | 2 | ENSMUST00 | 0 | Ccdc9 | ENSMUSG00000041375.13 |
| chrX | 100625403 | 100626653 | 15.62409 | 36.57069 | 34.02363 | Promoter | 20 | 100622883 | 100625938 | 3056 | 2 | ENSMUST00 | 0 | Pdzd11 | ENSMUSG00000015668.8 |
| chrX | 150812483 | 150813955 | 15.62409 | 36.57069 | 34.02363 | Promoter | 20 | 150806421 | 150812968 | 6548 | 2 | ENSMUST00 | 0 | Maged2 | ENSMUSG00000025268.10 |
| chrX | 7657113 | 7658374 | 15.62409 | 36.57069 | 34.02363 | Promoter | 20 | 7657260 | 7667958 | 10699 | 1 | ENSMUST00 | 0 | Prickle3 | ENSMUSG00000031145.10 |
| chrX | 106015463 | 106016543 | 15.62409 | 36.57069 | 34.02363 | Promoter | 20 | 106015700 | 106022450 | 6751 | 1 | ENSMUST00 | 0 | Cox7b | ENSMUSG00000031231.4 |
| chrX | 72917746 | 72918482 | 15.62409 | 36.57069 | 34.02363 | Promoter | 20 | 72913532 | 72918411 | 4880 | 2 | ENSMUST00 | 0 | Cetn2 | ENSMUSG00000031347.7 |
| chrX | 85573408 | 85573813 | 15.62409 | 36.57069 | 34.02363 | Promoter | 20 | 85574022 | 85634469 | 60448 | 1 | ENSMUST00 | -209 | Tab3 | ENSMUSG00000035476.4 |
| chrX | 152341792 | 152343312 | 15.62409 | 36.57069 | 34.02363 | Promoter | 20 | 152336852 | 152342425 | 5574 | 2 | ENSMUST00 | 0 | Tspyl2 | ENSMUSG00000041096.8 |
| chrX | 153237509 | 153238606 | 15.62409 | 36.57069 | 34.02363 | Promoter | 20 | 153237748 | 153396132 | 158385 | 1 | ENSMUST00 | 0 | Klf8 | ENSMUSG00000041649.8 |
| chrX | 70385091 | 70386514 | 15.62409 | 36.57069 | 34.02363 | Promoter | 20 | 70385877 | 70389417 | 3541 | 1 | ENSMUST00 | 0 | 1110012L19I | ENSMUSG00000045237.5 |
| chrX | 53054948 | 53059241 | 15.62409 | 36.57069 | 34.02363 | Promoter | 20 | 53055046 | 53057190 | 2145 | 2 | ENSMUST00 | 0 | C430049B03I | ENSMUSG00000087365.2 |

| | | | | | | | | | | | | | | | |
|---|---|---|---|---|---|---|---|---|---|---|---|---|---|---|---|
| chr12 | 85471798 | 85472829 | 14.80502 | 36.56548 | 34.02003 | Distal Interge | 12 | 85473890 | 85477273 | 3384 | 1 | ENSMUST00 | -1061 | Fos | ENSMUSG00000021250.8 |
| chr15 | 79690017 | 79690458 | 15.194 | 36.53499 | 33.98956 | Promoter | 15 | 79690895 | 79721478 | 30584 | 1 | ENSMUST00 | -437 | Gtpbp1 | ENSMUSG00000042535.8 |
| chr17 | 70754976 | 70755855 | 15.194 | 36.53499 | 33.98956 | Promoter | 17 | 70745329 | 70755582 | 10254 | 2 | ENSMUST00 | 0 | 5031415H12I | ENSMUSG00000097282.1 |
| chr6 | 31549758 | 31552254 | 15.59599 | 36.52319 | 33.97784 | Intron (ENSM | 6 | 31519488 | 31563981 | 44494 | 2 | ENSMUST00 | 11727 | Podxl | ENSMUSG00000025608.9 |
| chr7 | 16946319 | 16947742 | 13.56395 | 36.45369 | 33.90862 | 3' UTR | 7 | 16944682 | 16948828 | 4147 | 1 | ENSMUST00 | 1638 | Pnmal2 | ENSMUSG00000070802.5 |
| chr9 | 20877677 | 20878122 | 15.53082 | 36.41302 | 33.86811 | Exon (ENSMU | 9 | 20873809 | 20879727 | 5919 | 2 | ENSMUST00 | 1605 | Angptl6 | ENSMUSG00000038742.6 |
| chr11 | 63923140 | 63923793 | 15.10403 | 36.38161 | 33.83695 | Promoter | 11 | 63885792 | 63922290 | 36499 | 2 | ENSMUST00 | -851 | Hs3st3b1 | ENSMUSG00000070407.5 |
| chr4 | 155562706 | 155563361 | 15.10403 | 36.38161 | 33.83695 | Promoter | 4 | 155562378 | 155590997 | 28620 | 1 | ENSMUST00 | 329 | Nadk | ENSMUSG00000029063.11 |
| chr1 | 171077733 | 171078475 | 5.98627 | 36.36336 | 33.81879 | Downstream | 1 | 171079840 | 171079901 | 62 | 2 | ENSMUST00 | 1426 | Gm27844 | ENSMUSG00000098965.1 |
| chr11 | 120659687 | 120661470 | 13.87847 | 36.33116 | 33.78668 | Promoter | 11 | 120653788 | 120660606 | 6819 | 2 | ENSMUST00 | 0 | Notum | ENSMUSG00000042988.5 |
| chr15 | 38470059 | 38471389 | 13.87847 | 36.33116 | 33.78668 | Distal Interge | 15 | 38490473 | 38490547 | 75 | 2 | ENSMUST00 | 19158 | Mir6951 | ENSMUSG00000099110.1 |
| chr17 | 32099818 | 32100619 | 13.87847 | 36.33116 | 33.78668 | Promoter | 17 | 32076549 | 32100286 | 23738 | 2 | ENSMUST00 | 0 | Pdxk-ps | ENSMUSG00000002791.11 |
| chr2 | 153345044 | 153345640 | 15.07427 | 36.33085 | 33.78652 | Promoter | 2 | 153341157 | 153345810 | 4654 | 2 | ENSMUST00 | 170 | 2500004C02I | ENSMUSG00000073236.4 |
| chr5 | 20701623 | 20702915 | 15.07427 | 36.33085 | 33.78652 | Promoter | 5 | 20702277 | 20706049 | 3773 | 1 | ENSMUST00 | 0 | Magi2 | ENSMUSG00000073174.5 |
| chr2 | 181669993 | 181671838 | 15.4662 | 36.30373 | 33.75949 | Promoter | 2 | 181669836 | 181671640 | 1805 | 2 | ENSMUST00 | 0 | Sox18 | ENSMUSG00000046470.5 |
| chr1 | 180330478 | 180331292 | 12.74236 | 36.28267 | 33.73857 | Promoter | 1 | 180330485 | 180424802 | 94318 | 1 | ENSMUST00 | 0 | Itpkb | ENSMUSG00000038855.9 |
| chr11 | 120046527 | 120047209 | 12.74236 | 36.28267 | 33.73857 | Promoter | 11 | 120007313 | 120047070 | 39758 | 2 | ENSMUST00 | 0 | Aatk | ENSMUSG00000025375.10 |
| chr17 | 14202801 | 14204122 | 12.74236 | 36.28267 | 33.73857 | Promoter | 17 | 14195231 | 14203831 | 8601 | 2 | ENSMUST00 | 0 | Dact2 | ENSMUSG00000048826.6 |
| chr19 | 45057619 | 45059254 | 12.74236 | 36.28267 | 33.73857 | Distal Interge | 19 | 45047576 | 45056383 | 8808 | 1 | ENSMUST00 | 10044 | Sfxn3 | ENSMUSG00000025212.11 |
| chr2 | 36230166 | 36231210 | 12.74236 | 36.28267 | 33.73857 | Promoter | 2 | 36230426 | 36252272 | 21847 | 1 | ENSMUST00 | 0 | Ptgs1 | ENSMUSG00000047250.8 |
| chr3 | 122483254 | 122485690 | 12.74236 | 36.28267 | 33.73857 | Intron (ENSM | 3 | 122464727 | 122464882 | 156 | 1 | ENSMUST00 | 18528 | Gm25153 | ENSMUSG00000087743.1 |
| chr5 | 108692046 | 108692570 | 12.74236 | 36.28267 | 33.73857 | Distal Interge | 5 | 108694229 | 108706950 | 12722 | 1 | ENSMUST00 | -1659 | Fgfrl1 | ENSMUSG00000008090.9 |
| chr5 | 12383040 | 12384295 | 12.74236 | 36.28267 | 33.73857 | Promoter | 5 | 12383166 | 12588943 | 205778 | 1 | ENSMUST00 | 0 | Sema3d | ENSMUSG00000040254.7 |
| chr7 | 19715878 | 19716497 | 12.74236 | 36.28267 | 33.73857 | Promoter | 7 | 19701313 | 19715438 | 14126 | 2 | ENSMUST00 | -441 | Tomm40 | ENSMUSG00000002984.11 |
| chr4 | 134249595 | 134250166 | 15.04464 | 36.28028 | 33.73701 | Downstream | 4 | 134251706 | 134253023 | 1318 | 2 | ENSMUST00 | 2857 | Grrp1 | ENSMUSG00000050105.5 |
| chr6 | 125380150 | 125380811 | 15.04464 | 36.28028 | 33.73701 | Promoter | 6 | 125362660 | 125380793 | 18134 | 2 | ENSMUST00 | 0 | Plekhg6 | ENSMUSG00000038167.6 |
| chr6 | 83268078 | 83268644 | 15.04464 | 36.28028 | 33.73701 | Exon (ENSMU | 6 | 83277393 | 83288451 | 11059 | 1 | ENSMUST00 | -8749 | Slc4a5 | ENSMUSG00000068323.7 |
| chr1 | 74098188 | 74099467 | 14.22617 | 36.25443 | 33.71164 | Intron (ENSM | 1 | 73997055 | 74096868 | 99814 | 2 | ENSMUST00 | -1321 | Tns1 | ENSMUSG00000055322.9 |
| chr9 | 7595003 | 7595454 | 14.22617 | 36.25443 | 33.71164 | Distal Interge | 9 | 7571463 | 7581885 | 10423 | 1 | ENSMUST00 | 23541 | Mmp27 | ENSMUSG00000070323.6 |
| chr13 | 92130874 | 92132084 | 14.61261 | 36.2332 | 33.69079 | Promoter | 13 | 91884432 | 92131494 | 247063 | 2 | ENSMUST00 | 0 | Rasgrf2 | ENSMUSG00000021708.11 |
| chr13 | 23745698 | 23747402 | 14.61261 | 36.2332 | 33.69079 | Promoter | 13 | 23746790 | 23747241 | 452 | 1 | ENSMUST00 | 0 | Hist1h2bb | ENSMUSG00000075031.3 |
| chr15 | 61987445 | 61988323 | 14.61261 | 36.2332 | 33.69079 | Promoter | 15 | 61987422 | 61990253 | 2832 | 1 | ENSMUST00 | 24 | Myc | ENSMUSG00000022346.9 |
| chr17 | 35077460 | 35078270 | 14.61261 | 36.2332 | 33.69079 | Promoter | 17 | 35071477 | 35078065 | 6589 | 2 | ENSMUST00 | 0 | Ly6g6d | ENSMUSG00000073413.4 |
| chr18 | 65703856 | 65704519 | 14.61261 | 36.2332 | 33.69079 | Intron (ENSM | 18 | 65698270 | 65751601 | 53332 | 1 | ENSMUST00 | 5587 | Oacyl | ENSMUSG00000046610.9 |
| chr3 | 85573429 | 85573952 | 14.61261 | 36.2332 | 33.69079 | Promoter | 3 | 85556046 | 85573945 | 17900 | 2 | ENSMUST00 | 0 | Gm15535 | ENSMUSG00000085527.1 |
| chr4 | 101646671 | 101647201 | 14.61261 | 36.2332 | 33.69079 | Promoter | 4 | 101647718 | 101659364 | 11647 | 1 | ENSMUST00 | -517 | Leprot | ENSMUSG00000035212.9 |
| chr4 | 140610139 | 140611652 | 14.61261 | 36.2332 | 33.69079 | Exon (ENSMU | 4 | 140593169 | 140617065 | 23897 | 2 | ENSMUST00 | 5413 | Arhgef10l | ENSMUSG00000040964.11 |
| chr7 | 30525831 | 30526848 | 14.61261 | 36.2332 | 33.69079 | Exon (ENSMU | 7 | 30522226 | 30534180 | 11955 | 2 | ENSMUST00 | 7332 | Arhgap33 | ENSMUSG00000036882.6 |
| chr2 | 152707371 | 152708201 | 9.56722 | 36.15532 | 33.61367 | 3' UTR | 2 | 152694733 | 152696692 | 1960 | 2 | ENSMUST00 | -10680 | Gm26841 | ENSMUSG00000097722.1 |
| chr6 | 102563890 | 102564740 | 11.66485 | 36.11788 | 33.57639 | Distal Interge | 6 | 102163306 | 102464667 | 301362 | 2 | ENSMUST00 | -99224 | Cntn3 | ENSMUSG00000030075.8 |
| chr12 | 73949495 | 73950067 | 13.35156 | 36.06488 | 33.52377 | Promoter | 12 | 73926481 | 73949785 | 23305 | 2 | ENSMUST00 | 0 | Gm15283 | ENSMUSG00000087700.1 |
| chr11 | 88187255 | 88188228 | 15.2755 | 35.98095 | 33.44006 | Promoter | 11 | 88187821 | 88193004 | 5184 | 1 | ENSMUST00 | 0 | Cuedc1 | ENSMUSG00000018378.8 |
| chr16 | 4541442 | 4541913 | 15.2755 | 35.98095 | 33.44006 | Promoter | 16 | 4487579 | 4541816 | 54238 | 2 | ENSMUST00 | 0 | Srl | ENSMUSG00000022519.8 |
| chr12 | 73545466 | 73546636 | 13.65044 | 35.92182 | 33.38125 | Promoter | 12 | 73543114 | 73546395 | 3282 | 2 | ENSMUST00 | 0 | Tmem30b | ENSMUSG00000034435.5 |
| chr10 | 77165554 | 77166644 | 13.26345 | 35.90286 | 33.3624 | Promoter | 10 | 77052178 | 77166548 | 114371 | 2 | ENSMUST00 | 0 | Col18a1 | ENSMUSG00000001435.9 |
| chr11 | 78497041 | 78498266 | 13.26345 | 35.90286 | 33.3624 | Promoter | 11 | 78472330 | 78497458 | 25129 | 2 | ENSMUST00 | 0 | Sarm1 | ENSMUSG00000050132.8 |
| chr13 | 47042695 | 47043214 | 13.26345 | 35.90286 | 33.3624 | Promoter | 13 | 47025170 | 47043210 | 18041 | 2 | ENSMUST00 | 0 | Tpmt | ENSMUSG00000021376.5 |
| chr15 | 101478972 | 101479805 | 13.26345 | 35.90286 | 33.3624 | 3' UTR | 15 | 101473478 | 101479983 | 6506 | 1 | ENSMUST00 | 5495 | Krt86 | ENSMUSG00000067614.4 |
| chr16 | 4474527 | 4475390 | 13.26345 | 35.90286 | 33.3624 | Distal Interge | 16 | 4493603 | 4523063 | 29461 | 2 | ENSMUST00 | 47673 | Srl | ENSMUSG00000022519.8 |
| chr18 | 34751743 | 34752186 | 13.26345 | 35.90286 | 33.3624 | Promoter | 18 | 34751809 | 34756662 | 4854 | 1 | ENSMUST00 | 0 | 2010110K18I | ENSMUSG00000097755.1 |
| chr2 | 118761847 | 118763098 | 13.26345 | 35.90286 | 33.3624 | Promoter | 2 | 118754158 | 118762661 | 8504 | 2 | ENSMUST00 | 0 | A430105I19F | ENSMUSG00000045838.8 |
| chr2 | 131545623 | 131546454 | 13.26345 | 35.90286 | 33.3624 | 3' UTR | 2 | 131561614 | 131563179 | 1566 | 1 | ENSMUST00 | -15160 | Gm14285 | ENSMUSG00000087524.1 |
| chr4 | 135273078 | 135273685 | 13.26345 | 35.90286 | 33.3624 | Promoter | 4 | 135223464 | 135272814 | 49351 | 2 | ENSMUST00 | -265 | Clic4 | ENSMUSG00000037242.8 |
| chr4 | 6989721 | 6991384 | 13.26345 | 35.90286 | 33.3624 | Promoter | 4 | 6686353 | 6990799 | 304447 | 2 | ENSMUST00 | 0 | Tox | ENSMUSG00000041272.6 |
| chr5 | 64505373 | 64506387 | 13.26345 | 35.90286 | 33.3624 | Distal Interge | 5 | 64456328 | 64467946 | 11619 | 2 | ENSMUST00 | -37428 | RP23-195B3. | ENSMUSG00000100911.1 |
| chr6 | 122602035 | 122603053 | 13.26345 | 35.90286 | 33.3624 | Promoter | 6 | 122577792 | 122602444 | 24653 | 2 | ENSMUST00 | 0 | Apobec1 | ENSMUSG00000040613.9 |
| chr7 | 38084593 | 38084906 | 13.26345 | 35.90286 | 33.3624 | Distal Interge | 7 | 38098811 | 38099369 | 559 | 2 | ENSMUST00 | 14463 | Ccne1 | ENSMUSG00000002068.11 |
| chr8 | 84186616 | 84187375 | 13.26345 | 35.90286 | 33.3624 | Distal Interge | 8 | 84178863 | 84178968 | 106 | 2 | ENSMUST00 | -7649 | Mir181c | ENSMUSG00000065483.1 |
| chr9 | 20651171 | 20651879 | 13.26345 | 35.90286 | 33.3624 | Promoter | 9 | 20652130 | 20666584 | 14455 | 1 | ENSMUST00 | -251 | Pin1 | ENSMUSG00000032171.6 |
| chrX | 37109504 | 37110376 | 13.26345 | 35.90286 | 33.3624 | Promoter | 20 | 37091678 | 37110322 | 18645 | 2 | ENSMUST00 | 0 | Upf3b | ENSMUSG00000036572.11 |
| chr2 | 143862373 | 143863177 | 14.81168 | 35.88207 | 33.34299 | Promoter | 2 | 143826528 | 143862943 | 36416 | 2 | ENSMUST00 | 0 | Bfsp1 | ENSMUSG00000027420.8 |
| chr14 | 65968470 | 65969026 | 15.21298 | 35.87502 | 33.336 | Promoter | 14 | 65968483 | 65981548 | 13066 | 1 | ENSMUST00 | 0 | Clu | ENSMUSG00000022037.9 |
| chr5 | 35391544 | 35392592 | 15.21298 | 35.87502 | 33.336 | 3' UTR | 5 | 35389117 | 35392861 | 3745 | 1 | ENSMUST00 | 2428 | Hmx1 | ENSMUSG00000067438.4 |
| chr9 | 16499580 | 16502412 | 15.21298 | 35.87502 | 33.336 | Distal Interge | 9 | 15910205 | 16378231 | 468027 | 2 | ENSMUST00 | -121350 | Fat3 | ENSMUSG00000074505.4 |
| chr16 | 17796819 | 17797871 | 14.78306 | 35.83308 | 33.29445 | Promoter | 16 | 17797282 | 17808287 | 11006 | 1 | ENSMUST00 | 0 | Scarf2 | ENSMUSG00000012017.3 |
| chr15 | 98167575 | 98168613 | 15.18191 | 35.82236 | 33.28377 | Promoter | 15 | 98167806 | 98170134 | 2329 | 1 | ENSMUST00 | 0 | Ccdc184 | ENSMUSG00000029875.4 |
| chr11 | 109297763 | 109298271 | 13.98032 | 35.82108 | 33.28258 | Promoter | 11 | 109225355 | 109298129 | 72775 | 2 | ENSMUST00 | 0 | Rgs9 | ENSMUSG00000020599.8 |
| chr2 | 75137348 | 75138099 | 14.75456 | 35.78426 | 33.24598 | Promoter | 2 | 75124871 | 75137798 | 12928 | 2 | ENSMUST00 | 0 | Gm13652 | ENSMUSG00000087333.1 |
| chr9 | 21926813 | 21927406 | 14.75456 | 35.78426 | 33.24598 | Promoter | 9 | 21927471 | 21935872 | 8402 | 1 | ENSMUST00 | -65 | Ccdc159 | ENSMUSG00000006241.10 |
| chr9 | 21243977 | 21245284 | 14.75456 | 35.78426 | 33.24598 | 5' UTR | 9 | 21242912 | 21248443 | 5532 | 2 | ENSMUST00 | 3159 | S1pr5 | ENSMUSG00000045087.8 |
| chr11 | 115297255 | 115297860 | 14.34627 | 35.7714 | 33.23358 | Promoter | 11 | 115279622 | 115297514 | 17893 | 2 | ENSMUST00 | 0 | Fads6 | ENSMUSG00000044788.10 |
| chr19 | 5509975 | 5510643 | 14.34627 | 35.7714 | 33.23358 | Promoter | 19 | 5495278 | 5510489 | 15212 | 2 | ENSMUST00 | 0 | Snx32 | ENSMUSG00000056185.4 |
| chr2 | 156180424 | 156181016 | 14.34627 | 35.7714 | 33.23358 | Promoter | 2 | 156174774 | 156180238 | 5465 | 2 | ENSMUST00 | -187 | Rbm39 | ENSMUSG00000027620.11 |
| chr2 | 84424494 | 84425582 | 14.34627 | 35.7714 | 33.23358 | Promoter | 2 | 84330626 | 84425411 | 94786 | 2 | ENSMUST00 | 0 | Calcrl | ENSMUSG00000059588.8 |
| chr4 | 123233410 | 123234037 | 14.34627 | 35.7714 | 33.23358 | Promoter | 4 | 123233556 | 123249875 | 16320 | 1 | ENSMUST00 | 0 | Heyl | ENSMUSG00000032744.4 |
| chr4 | 140957484 | 140958054 | 14.34627 | 35.7714 | 33.23358 | Promoter | 4 | 140947666 | 140957902 | 10237 | 2 | ENSMUST00 | 0 | Gm13031 | ENSMUSG00000087698.1 |
| chr10 | 97313184 | 97314105 | 13.89301 | 35.6666 | 33.12951 | Distal Interge | 10 | 97479500 | 97518162 | 38663 | 1 | ENSMUST00 | -165395 | Dcn | ENSMUSG00000019929.10 |
| chr10 | 75833576 | 75835210 | 13.89301 | 35.6666 | 33.12951 | Promoter | 10 | 75831845 | 75834881 | 3037 | 2 | ENSMUST00 | 0 | Gstt2 | ENSMUSG00000033318.6 |
| chr10 | 42632603 | 42633925 | 13.89301 | 35.6666 | 33.12951 | Intron (ENSM | 10 | 42678916 | 42702459 | 23544 | 1 | ENSMUST00 | -44991 | Ostm1 | ENSMUSG00000038280.7 |
| chr11 | 62820166 | 62821348 | 15.08946 | 35.66558 | 33.12951 | Promoter | 11 | 62820231 | 62842936 | 22706 | 1 | ENSMUST00 | 0 | Trim16 | ENSMUSG00000047821.11 |
| chr12 | 80518308 | 80519778 | 13.89301 | 35.6666 | 33.12951 | Promoter | 12 | 80518990 | 80603896 | 84907 | 1 | ENSMUST00 | 0 | Galnt16 | ENSMUSG00000021130.7 |
| chr12 | 5119273 | 5119865 | 13.89301 | 35.6666 | 33.12951 | Intron (ENSM | 12 | 4999266 | 4999368 | 103 | 2 | ENSMUST00 | -119906 | Gm23167 | ENSMUSG00000087780.1 |
| chr13 | 74166838 | 74167319 | 13.89301 | 35.6666 | 33.12951 | Downstream | 13 | 74169805 | 74208700 | 38896 | 2 | ENSMUST00 | 41381 | Exoc3 | ENSMUSG00000034152.5 |
| chr13 | 112100438 | 112101481 | 13.89301 | 35.6666 | 33.12951 | Distal Interge | 13 | 112005502 | 112017332 | 11831 | 1 | ENSMUST00 | 94937 | Gm15323 | ENSMUSG00000086092.1 |
| chr14 | 55899691 | 55900883 | 13.89301 | 35.6666 | 33.12951 | Promoter | 14 | 55897286 | 55900232 | 2947 | 2 | ENSMUST00 | 0 | Sdr39u1 | ENSMUSG00000022223.8 |
| chr14 | 34819124 | 34820803 | 13.89301 | 35.6666 | 33.12951 | Promoter | 14 | 34819825 | 35583379 | 763555 | 1 | ENSMUST00 | 0 | Grid1 | ENSMUSG00000041078.6 |
| chr18 | 32162290 | 32162808 | 13.89301 | 35.6666 | 33.12951 | Promoter | 18 | 32163089 | 32236751 | 73663 | 1 | ENSMUST00 | -281 | Map3k2 | ENSMUSG00000024383.8 |
| chr19 | 44247327 | 44248258 | 13.89301 | 35.6666 | 33.12951 | Distal Interge | 19 | 44203288 | 44244016 | 40729 | 1 | ENSMUST00 | 44040 | Scd3 | ENSMUSG00000025202.7 |
| chr2 | 131174549 | 131174883 | 13.89301 | 35.6666 | 33.12951 | Promoter | 2 | 131171611 | 131174653 | 3043 | 2 | ENSMUST00 | 0 | Spef1 | ENSMUSG00000027329.4 |
| chr3 | 70104096 | 70104899 | 13.89301 | 35.6666 | 33.12951 | Distal Interge | 3 | 70007613 | 70028708 | 21096 | 1 | ENSMUST00 | 96484 | Otol1 | ENSMUSG00000027788.6 |
| chr3 | 40531184 | 40532253 | 13.89301 | 35.6666 | 33.12951 | Promoter | 3 | 40531286 | 40654617 | 123332 | 1 | ENSMUST00 | 0 | Intu | ENSMUSG00000060798.4 |
| chr4 | 95160997 | 95162146 | 13.89301 | 35.6666 | 33.12951 | Exon (ENSMU | 4 | 95151740 | 95157503 | 5764 | 1 | ENSMUST00 | 9258 | RP23-77I17.4 | ENSMUSG00000100189.1 |
| chr6 | 29484845 | 29485283 | 13.89301 | 35.6666 | 33.12951 | Promoter | 6 | 29482036 | 29484860 | 2825 | 2 | ENSMUST00 | 0 | Kcp | ENSMUSG00000059022.9 |
| chr7 | 45128991 | 45129861 | 13.89301 | 35.6666 | 33.12951 | Promoter | 7 | 45125558 | 45128748 | 3191 | 2 | ENSMUST00 | -244 | Rpl13a | ENSMUSG00000074129.7 |
| chr9 | 123678146 | 123679098 | 13.89301 | 35.6666 | 33.12951 | Promoter | 9 | 123678439 | 123781965 | 103527 | 1 | ENSMUST00 | 0 | Ccr9 | ENSMUSG00000029530.10 |
| chr17 | 12741181 | 12742643 | 14.6697 | 35.63879 | 33.10369 | Promoter | 17 | 12741311 | 12860122 | 118812 | 1 | ENSMUST00 | 0 | Airn | ENSMUSG00000078247.3 |
| chr1 | 171222739 | 171223191 | 14.66885 | 35.63733 | 33.10225 | Promoter | 1 | 171217802 | 171222514 | 4713 | 2 | ENSMUST00 | -226 | Tomm40l | ENSMUSG00000005674.4 |
| chr1 | 134093180 | 134093692 | 14.66885 | 35.63733 | 33.10225 | Distal Interge | 1 | 134075170 | 134079120 | 3951 | 2 | ENSMUST00 | -14061 | Btg2 | ENSMUSG00000020423.6 |
| chr11 | 101551536 | 101551888 | 14.66885 | 35.63733 | 33.10225 | Promoter | 11 | 101488764 | 101551879 | 63116 | 2 | ENSMUST00 | 0 | Brca1 | ENSMUSG00000017146.7 |
| chr12 | 87101752 | 87102702 | 14.66885 | 35.63733 | 33.10225 | Promoter | 12 | 87097531 | 87102522 | 4992 | 2 | ENSMUST00 | 0 | Ngb | ENSMUSG00000021032.8 |
| chr12 | 76176972 | 76177994 | 14.66885 | 35.63733 | 33.10225 | Promoter | 12 | 76120419 | 76177259 | 56841 | 2 | ENSMUST00 | 0 | Esr2 | ENSMUSG00000021055.8 |
| chr14 | 103349292 | 103350308 | 14.66885 | 35.63733 | 33.10225 | Distal Interge | 14 | 103311695 | 103346814 | 35120 | 2 | ENSMUST00 | -2479 | Mycbp2 | ENSMUSG00000033004.9 |
| chr15 | 55306497 | 55307313 | 14.66885 | 35.63733 | 33.10225 | Promoter | 15 | 55307750 | 55520802 | 213053 | 1 | ENSMUST00 | -437 | Col14a1 | ENSMUSG00000022371.10 |
| chr15 | 99670262 | 99671023 | 14.66885 | 35.63733 | 33.10225 | Promoter | 15 | 99670718 | 99701127 | 30410 | 1 | ENSMUST00 | 0 | Asic1 | ENSMUSG00000023017.9 |
| chr15 | 81584642 | 81585068 | 14.66885 | 35.63733 | 33.10225 | Promoter | 15 | 81585351 | 81588344 | 2994 | 1 | ENSMUST00 | -283 | Ep300 | ENSMUSG00000055024.7 |
| chr17 | 45599350 | 45599978 | 14.66885 | 35.63733 | 33.10225 | Promoter | 17 | 45585200 | 45599603 | 14404 | 2 | ENSMUST00 | 0 | Slc29a1 | ENSMUSG00000023942.10 |
| chr17 | 68837024 | 68837736 | 14.66885 | 35.63733 | 33.10225 | Promoter | 17 | 68837136 | 68843138 | 6003 | 1 | ENSMUST00 | 0 | Tmem200c | ENSMUSG00000095407.1 |
| chr18 | 35939606 | 35939925 | 14.66885 | 35.63733 | 33.10225 | Distal Interge | 18 | 35952824 | 35952907 | 84 | 1 | ENSMUST00 | -12899 | Gm23882 | ENSMUSG00000080529.1 |
| chr2 | 90782922 | 90783482 | 14.66885 | 35.63733 | 33.10225 | Promoter | 2 | 90783238 | 90814269 | 31032 | 1 | ENSMUST00 | 0 | Agbl2 | ENSMUSG00000040812.9 |
| chr2 | 118663250 | 118663945 | 14.66885 | 35.63733 | 33.10225 | Promoter | 2 | 118663303 | 118698020 | 34718 | 1 | ENSMUST00 | 0 | Pak6 | ENSMUSG00000074923.5 |
| chr3 | 30507549 | 30509587 | 14.66885 | 35.63733 | 33.10225 | Promoter | 3 | 29963074 | 30509514 | 546441 | 2 | ENSMUST00 | 0 | Mecom | ENSMUSG00000027684.11 |
| chr3 | 83127156 | 83129397 | 14.66885 | 35.63733 | 33.10225 | Promoter | 3 | 83125276 | 83129246 | 3971 | 2 | ENSMUST00 | 0 | Gm10710 | ENSMUSG00000074517.2 |
| chr4 | 152312974 | 152314051 | 14.66885 | 35.63733 | 33.10225 | Promoter | 4 | 152311557 | 152313403 | 1847 | 2 | ENSMUST00 | 0 | Rnf207 | ENSMUSG00000058498.7 |
| chr5 | 107960707 | 107961330 | 14.66885 | 35.63733 | 33.10225 | Intron (ENSM | 5 | 107914398 | 107972998 | 58601 | 2 | ENSMUST00 | 11668 | Fam69a | ENSMUSG00000029270.5 |
| chr5 | 92897654 | 92898726 | 14.66885 | 35.63733 | 33.10225 | Promoter | 5 | 92897993 | 92965308 | 67316 | 1 | ENSMUST00 | 0 | Shroom3 | ENSMUSG00000029381.10 |
| chr5 | 127241440 | 127242164 | 14.66885 | 35.63733 | 33.10225 | Promoter | 5 | 127241808 | 127565793 | 323986 | 1 | ENSMUST00 | 0 | Tmem132c | ENSMUSG00000034324.11 |
| chr5 | 140649160 | 140649698 | 14.66885 | 35.63733 | 33.10225 | Promoter | 5 | 140620586 | 140649031 | 28446 | 2 | ENSMUST00 | -130 | Ttyh3 | ENSMUSG00000036565.3 |
| chr5 | 30588173 | 30589030 | 14.66885 | 35.63733 | 33.10225 | Promoter | 5 | 30588170 | 30625270 | 37101 | 1 | ENSMUST00 | 4 | Kcnk3 | ENSMUSG00000049265.6 |
| chr6 | 88074669 | 88075628 | 14.66885 | 35.63733 | 33.10225 | Distal Interge | 6 | 88084473 | 88105304 | 20832 | 1 | ENSMUST00 | -8845 | Rpn1 | ENSMUSG00000030062.6 |
| chr6 | 84584701 | 84586936 | 14.66885 | 35.63733 | 33.10225 | Intron (ENSM | 6 | 84571414 | 84593810 | 22397 | 2 | ENSMUST00 | 6874 | Cyp26b1 | ENSMUSG00000063415.7 |
| chr7 | 28294336 | 28295031 | 14.66885 | 35.63733 | 33.10225 | Promoter | 7 | 28293599 | 28294481 | 883 | 2 | ENSMUST00 | 0 | Dll3 | ENSMUSG00000003436.11 |
| chr7 | 99380483 | 99381351 | 14.66885 | 35.63733 | 33.10225 | Promoter | 7 | 99381549 | 99460983 | 79435 | 1 | ENSMUST00 | -198 | Gdpd5 | ENSMUSG00000035314.8 |
| chr9 | 21291496 | 21291922 | 14.66885 | 35.63733 | 33.10225 | Promoter | 9 | 21288463 | 21291209 | 2747 | 2 | ENSMUST00 | -288 | Cdkn2d | ENSMUSG00000096472.1 |
| chrX | 38634493 | 38635118 | 14.66885 | 35.63733 | 33.10225 | Promoter | 20 | 38630785 | 38635087 | 4303 | 2 | ENSMUST00 | 0 | C1galt1c1 | ENSMUSG00000048970.9 |
| chrX | 157490958 | 157492445 | 14.66885 | 35.63733 | 33.10225 | Promoter | 20 | 157443855 | 157492287 | 48433 | 2 | ENSMUST00 | 0 | Sms | ENSMUSG00000071708.6 |
| chr10 | 81319785 | 81320699 | 15.05889 | 35.61372 | 33.08038 | Promoter | 10 | 81321103 | 81326251 | 5149 | 1 | ENSMUST00 | -404 | Cactin | ENSMUSG00000034889.7 |
| chr17 | 23749055 | 23749686 | 15.05889 | 35.61372 | 33.08038 | Promoter | 17 | 23749236 | 23754065 | 4830 | 1 | ENSMUST00 | 0 | 9530082P21I | ENSMUSG00000097006.2 |
| chr2 | 156362087 | 156363149 | 10.03929 | 35.57722 | 33.04412 | Intron (ENSM | 2 | 156347202 | 156347840 | 639 | 2 | ENSMUST00 | -14248 | Gm14253 | ENSMUSG00000082321.3 |

| | | | | | | | | | | | | | | | |
|---|---|---|---|---|---|---|---|---|---|---|---|---|---|---|---|
| chr10 | 75932124 | 75932953 | 12.0178 | 35.52963 | 32.99667 | Promoter | 10 | 75923222 | 75932475 | 9254 | 2 | ENSMUST00 | 0 | Mmp11 | ENSMUSG00000000901.10 |
| chr11 | 114764944 | 114765658 | 12.0178 | 35.52963 | 32.99667 | Promoter | 11 | 114765388 | 114786179 | 20792 | 1 | ENSMUST00 | 0 | Kif19a | ENSMUSG00000010021.8 |
| chr16 | 29813613 | 29814424 | 12.0178 | 35.52963 | 32.99667 | Intron (ENSM | 16 | 29811058 | 29837814 | 26757 | 1 | ENSMUST00 | 2556 | Gm26834 | ENSMUSG00000097307.1 |
| chr9 | 65586713 | 65588441 | 13.42979 | 35.52332 | 32.99081 | Promoter | 9 | 65587160 | 65595964 | 8805 | 1 | ENSMUST00 | 0 | Pif1 | ENSMUSG00000041064.8 |
| chr11 | 102466964 | 102467901 | 14.99812 | 35.51057 | 32.97813 | Promoter | 11 | 102456472 | 102467199 | 10728 | 2 | ENSMUST00 | 0 | Itga2b | ENSMUSG00000034664.8 |
| chr4 | 115056388 | 115058097 | 14.58582 | 35.49482 | 32.96247 | Promoter | 4 | 115056426 | 115071755 | 15330 | 1 | ENSMUST00 | 0 | Tal1 | ENSMUSG00000028717.7 |
| chr18 | 37824941 | 37825891 | 14.96792 | 35.45929 | 32.92697 | Exon (ENSMU | 18 | 37819546 | 37841873 | 22328 | 1 | ENSMUST00 | 5396 | Pcdhga12 | ENSMUSG00000023036.9 |
| chr4 | 147406873 | 147407593 | 9.09097 | 35.43891 | 32.90672 | Intron (ENSM | 4 | 147378929 | 147379574 | 646 | 1 | ENSMUST00 | 27945 | Gm8178 | ENSMUSG00000096354.1 |
| chr15 | 78428587 | 78429115 | 14.93785 | 35.4082 | 32.87602 | Promoter | 15 | 78428628 | 78438909 | 10282 | 1 | ENSMUST00 | 0 | Kctd17 | ENSMUSG00000033287.10 |
| chr4 | 129406148 | 129406871 | 13.74281 | 35.40012 | 32.86803 | Distal Interge | 4 | 129426269 | 129432529 | 6261 | 2 | ENSMUST00 | 25658 | Zbtb8b | ENSMUSG00000048485.7 |
| chr7 | 102512845 | 102513281 | 13.74281 | 35.40012 | 32.86803 | Distal Interge | 7 | 102493823 | 102494773 | 951 | 2 | ENSMUST00 | -18073 | Olfr545 | ENSMUSG00000044824.5 |
| chr1 | 172490262 | 172490572 | 14.90789 | 35.35729 | 32.82543 | Promoter | 1 | 172491087 | 172491980 | 894 | 1 | ENSMUST00 | -515 | Igsf9 | ENSMUSG00000037995.10 |
| chr1 | 161195682 | 161196521 | 14.90789 | 35.35729 | 32.82543 | Distal Interge | 1 | 161156843 | 161159314 | 2472 | 2 | ENSMUST00 | -36369 | 4930469G21 | ENSMUSG00000049160.10 |
| chr15 | 102458717 | 102460824 | 14.90789 | 35.35729 | 32.82543 | Promoter | 15 | 102459028 | 102460588 | 1561 | 1 | ENSMUST00 | 0 | Prr13 | ENSMUSG00000023048.8 |
| chr3 | 51700857 | 51701793 | 14.90789 | 35.35729 | 32.82543 | Intron (ENSM | 3 | 51693771 | 51695090 | 1320 | 1 | ENSMUST00 | 7087 | Gm10729 | ENSMUSG00000074603.2 |
| chr11 | 112782058 | 112783156 | 14.08946 | 35.32397 | 32.79257 | Promoter | 11 | 112782224 | 112787760 | 5537 | 1 | ENSMUST00 | 0 | Sox9 | ENSMUSG00000000567.5 |
| chr14 | 30008620 | 30009339 | 14.08946 | 35.32397 | 32.79257 | Promoter | 14 | 29996141 | 30008813 | 12673 | 2 | ENSMUST00 | 0 | Il17rb | ENSMUSG00000015966.11 |
| chr18 | 41860669 | 41861840 | 14.08946 | 35.32397 | 32.79257 | Distal Interge | 18 | 41875696 | 41951194 | 75499 | 2 | ENSMUST00 | 89354 | Prelid2 | ENSMUSG00000056671.6 |
| chr6 | 72655030 | 72655884 | 14.08946 | 35.32397 | 32.79257 | Intron (ENSM | 6 | 72648815 | 72671404 | 22590 | 2 | ENSMUST00 | 15520 | Tcf7l1 | ENSMUSG00000055799.8 |
| chr7 | 139978424 | 139979252 | 14.08946 | 35.32397 | 32.79257 | Promoter | 7 | 139972303 | 139978723 | 6421 | 2 | ENSMUST00 | 0 | 6430531B16 | ENSMUSG00000073795.5 |
| chr11 | 104132332 | 104133810 | 14.87805 | 35.30658 | 32.77555 | Promoter | 11 | 104132855 | 104175523 | 42669 | 1 | ENSMUST00 | 0 | Crhr1 | ENSMUSG00000018634.10 |
| chr15 | 56695535 | 56695970 | 14.47545 | 35.30513 | 32.77419 | Promoter | 15 | 56665627 | 56694539 | 28913 | 2 | ENSMUST00 | -997 | Has2 | ENSMUSG00000022367.7 |
| chr5 | 36712796 | 36713593 | 14.47545 | 35.30513 | 32.77419 | Distal Interge | 5 | 36671399 | 36696024 | 24626 | 2 | ENSMUST00 | -16773 | D5Ertd579e | ENSMUSG00000029190.14 |
| chr7 | 126949503 | 126950104 | 14.47545 | 35.30513 | 32.77419 | Promoter | 7 | 126945989 | 126949582 | 3594 | 2 | ENSMUST00 | 0 | Asphd1 | ENSMUSG00000046378.6 |
| chr8 | 90954670 | 90956951 | 14.47545 | 35.30513 | 32.77419 | Exon (ENSMU | 8 | 90907939 | 90908415 | 477 | 2 | ENSMUST00 | -46256 | Gm6658 | ENSMUSG00000074171.1 |
| chr9 | 99357613 | 99358669 | 14.47545 | 35.30513 | 32.77419 | Promoter | 9 | 99309963 | 99358518 | 48556 | 2 | ENSMUST00 | 0 | Esyt3 | ENSMUSG00000037681.8 |
| chr4 | 118543399 | 118544596 | 10.42562 | 35.28478 | 32.75441 | Promoter | 4 | 118540941 | 118543728 | 2788 | 2 | ENSMUST00 | 0 | Tmem125 | ENSMUSG00000050854.4 |
| chr10 | 34127320 | 34127887 | 14.81873 | 35.20571 | 32.67548 | Promoter | 10 | 34126069 | 34127972 | 1904 | 2 | ENSMUST00 | 85 | Fam26f | ENSMUSG00000046031.6 |
| chr17 | 35051064 | 35051895 | 14.81873 | 35.20571 | 32.67548 | Promoter | 17 | 35052450 | 35055741 | 3292 | 1 | ENSMUST00 | -555 | Clic1 | ENSMUSG00000007041.9 |
| chr9 | 123924504 | 123925876 | 6.5408 | 35.1478 | 32.61777 | Distal Interge | 9 | 123962126 | 123968692 | 6567 | 2 | ENSMUST00 | 42816 | Ccr1 | ENSMUSG00000025804.4 |
| chr10 | 60346690 | 60347418 | 15.16456 | 35.10921 | 32.57929 | Promoter | 10 | 60346851 | 60372684 | 25834 | 1 | ENSMUST00 | 0 | 4632428N05 | ENSMUSG00000020101.9 |
| chr11 | 83409404 | 83410117 | 15.16456 | 35.10921 | 32.57929 | Promoter | 11 | 83409688 | 83421037 | 11350 | 1 | ENSMUST00 | 0 | Rasl10b | ENSMUSG00000020684.9 |
| chr13 | 75797721 | 75798042 | 15.16456 | 35.10921 | 32.57929 | Distal Interge | 13 | 75768068 | 75768197 | 130 | 1 | ENSMUST00 | 29654 | Gm23127 | ENSMUSG00000077449.1 |
| chr14 | 68124142 | 68125178 | 15.16456 | 35.10921 | 32.57929 | Promoter | 14 | 68082590 | 68124846 | 42257 | 2 | ENSMUST00 | 0 | Nefm | ENSMUSG00000022054.6 |
| chr15 | 99055261 | 99056206 | 15.16456 | 35.10921 | 32.57929 | Promoter | 15 | 99055174 | 99058978 | 3805 | 1 | ENSMUST00 | 88 | Prph | ENSMUSG00000023484.8 |
| chr16 | 29839272 | 29839659 | 15.16456 | 35.10921 | 32.57929 | Distal Interge | 16 | 29811058 | 29837814 | 26757 | 1 | ENSMUST00 | 28215 | Gm26834 | ENSMUSG00000097307.1 |
| chr17 | 65540686 | 65541201 | 15.16456 | 35.10921 | 32.57929 | Promoter | 17 | 65256005 | 65540782 | 284778 | 2 | ENSMUST00 | 0 | Tmem232 | ENSMUSG00000045036.9 |
| chr18 | 76543667 | 76544947 | 15.16456 | 35.10921 | 32.57929 | Distal Interge | 18 | 76262492 | 76302642 | 40151 | 1 | ENSMUST00 | 281176 | Smad2 | ENSMUSG00000024563.10 |
| chr19 | 53314417 | 53314840 | 15.16456 | 35.10921 | 32.57929 | Intron (ENSM | 19 | 53310506 | 53375810 | 65305 | 1 | ENSMUST00 | 3912 | Mxi1 | ENSMUSG00000025025.8 |
| chr2 | 176927493 | 176927925 | 15.16456 | 35.10921 | 32.57929 | Promoter | 2 | 176892509 | 176927794 | 35286 | 2 | ENSMUST00 | 0 | Gm14296 | ENSMUSG00000074527.6 |
| chr2 | 175800466 | 175801058 | 15.16456 | 35.10921 | 32.57929 | Promoter | 2 | 175791243 | 175800961 | 9719 | 2 | ENSMUST00 | 0 | Gm14288 | ENSMUSG00000078889.6 |
| chr2 | 175372440 | 175372857 | 15.16456 | 35.10921 | 32.57929 | Promoter | 2 | 175372462 | 175384201 | 11740 | 1 | ENSMUST00 | 0 | Gm4723 | ENSMUSG00000078898.4 |
| chr2 | 176152766 | 176153477 | 15.16456 | 35.10921 | 32.57929 | Distal Interge | 2 | 176144300 | 176149825 | 5526 | 2 | ENSMUST00 | -2942 | Gm2004 | ENSMUSG00000095648.2 |
| chr4 | 46991172 | 46992205 | 15.16456 | 35.10921 | 32.57929 | Promoter | 4 | 46662318 | 46991873 | 329556 | 2 | ENSMUST00 | 0 | Gabbr2 | ENSMUSG00000039809.9 |
| chr4 | 124575138 | 124576197 | 15.16456 | 35.10921 | 32.57929 | Intron (ENSM | 4 | 124569190 | 124574645 | 5456 | 1 | ENSMUST00 | 5949 | 4933407E24 | ENSMUSG00000073759.6 |
| chr6 | 107529484 | 107530616 | 15.16456 | 35.10921 | 32.57929 | Promoter | 6 | 107529726 | 107570228 | 40503 | 1 | ENSMUST00 | 0 | Lrrn1 | ENSMUSG00000034648.8 |
| chr8 | 70212762 | 70213258 | 15.16456 | 35.10921 | 32.57929 | Promoter | 8 | 70192369 | 70212305 | 19937 | 2 | ENSMUST00 | -458 | Slc25a42 | ENSMUSG00000002346.11 |
| chr8 | 125995148 | 125996018 | 15.16456 | 35.10921 | 32.57929 | Promoter | 8 | 125995102 | 126030683 | 35582 | 1 | ENSMUST00 | 47 | Kcnk1 | ENSMUSG00000033998.8 |
| chrX | 74304772 | 74305988 | 15.16456 | 35.10921 | 32.57929 | Promoter | 20 | 74304998 | 74311862 | 6865 | 1 | ENSMUST00 | 0 | Gdi1 | ENSMUSG00000015291.5 |
| chrX | 56730978 | 56731656 | 15.16456 | 35.10921 | 32.57929 | Promoter | 20 | 56731787 | 56793346 | 61560 | 1 | ENSMUST00 | -131 | Fhl1 | ENSMUSG00000023092.11 |
| chrX | 105079225 | 105080343 | 15.16456 | 35.10921 | 32.57929 | Promoter | 20 | 105079756 | 105085186 | 5431 | 1 | ENSMUST00 | 0 | Pbdc1 | ENSMUSG00000031226.8 |
| chrX | 102188452 | 102189481 | 15.16456 | 35.10921 | 32.57929 | Promoter | 20 | 102184941 | 102189394 | 4454 | 2 | ENSMUST00 | 0 | Rps4x | ENSMUSG00000031320.9 |
| chrX | 78582957 | 78583978 | 15.16456 | 35.10921 | 32.57929 | Promoter | 20 | 78449613 | 78583782 | 134170 | 2 | ENSMUST00 | 0 | Prrg1 | ENSMUSG00000047996.11 |
| chrX | 164979680 | 164980321 | 15.16456 | 35.10921 | 32.57929 | Promoter | 20 | 164936318 | 164980297 | 43980 | 2 | ENSMUST00 | 0 | Mospd2 | ENSMUSG00000061778.5 |
| chrX | 75232790 | 75233129 | 15.16456 | 35.10921 | 32.57929 | Promoter | 20 | 75233063 | 75234312 | 1250 | 1 | ENSMUST00 | 0 | Gm6039 | ENSMUSG00000083736.1 |
| chr12 | 3791498 | 3792459 | 14.75988 | 35.10558 | 32.57729 | Distal Interge | 12 | 3779548 | 3781398 | 1851 | 1 | ENSMUST00 | 11951 | Dtnb | ENSMUSG00000071454.8 |
| chr9 | 121285581 | 121286394 | 14.75988 | 35.10558 | 32.57729 | Distal Interge | 9 | 121102544 | 121277197 | 174654 | 2 | ENSMUST00 | -8385 | Ulk4 | ENSMUSG00000040936.9 |
| chr2 | 91078578 | 91079094 | 15.13729 | 35.06316 | 32.53499 | Distal Interge | 2 | 91082390 | 91114366 | 31977 | 1 | ENSMUST00 | -3296 | Spi1 | ENSMUSG00000002111.8 |
| chr1 | 180386917 | 180387574 | 12.43157 | 35.01519 | 32.48718 | Intron (ENSM | 1 | 180326970 | 180330550 | 3581 | 2 | ENSMUST00 | -56368 | Gm5069 | ENSMUSG00000055676.3 |
| chr12 | 75308014 | 75309297 | 12.43157 | 35.01519 | 32.48718 | Promoter | 12 | 75308322 | 75395817 | 87496 | 1 | ENSMUST00 | 0 | Rhoj | ENSMUSG00000046768.8 |
| chr2 | 180333637 | 180335240 | 12.43157 | 35.01519 | 32.48718 | Promoter | 2 | 180325133 | 180334699 | 9567 | 2 | ENSMUST00 | 0 | Gata5 | ENSMUSG00000015627.5 |
| chr5 | 66150356 | 66151714 | 12.43157 | 35.01519 | 32.48718 | Promoter | 5 | 66018602 | 66150898 | 132297 | 2 | ENSMUST00 | 0 | Rbm47 | ENSMUSG00000070780.6 |
| chr4 | 129460901 | 129461396 | 15.10559 | 35.00965 | 32.48246 | Promoter | 4 | 129461581 | 129488498 | 26918 | 1 | ENSMUST00 | -185 | Bsdc1 | ENSMUSG00000040859.9 |
| chr5 | 66337298 | 66337890 | 15.10559 | 35.00965 | 32.48246 | Promoter | 5 | 66301872 | 66337663 | 35792 | 2 | ENSMUST00 | 0 | Apbb2 | ENSMUSG00000029207.11 |
| chr17 | 25951469 | 25952377 | 13.51324 | 34.99091 | 32.46387 | Promoter | 17 | 25951472 | 25963610 | 12139 | 1 | ENSMUST00 | 0 | D630044L22 | ENSMUSG00000097405.1 |
| chr2 | 119896566 | 119898614 | 13.51324 | 34.99091 | 32.46387 | Promoter | 2 | 119897228 | 119967749 | 70522 | 1 | ENSMUST00 | 0 | Mga | ENSMUSG00000033943.10 |
| chr4 | 31823781 | 31824864 | 15.07403 | 34.95635 | 32.42956 | Distal Interge | 4 | 31964097 | 32023467 | 59371 | 1 | ENSMUST00 | -139233 | Map3k7 | ENSMUSG00000028284.8 |
| chr10 | 67185348 | 67187978 | 11.39974 | 34.91658 | 32.39005 | Promoter | 10 | 67185749 | 67216171 | 30423 | 1 | ENSMUST00 | 0 | Jmjd1c | ENSMUSG00000037876.11 |
| chr4 | 147101652 | 147102593 | 11.39974 | 34.91658 | 32.39005 | Promoter | 4 | 147102327 | 147103981 | 1655 | 1 | ENSMUST00 | 0 | Gm13142 | ENSMUSG00000081405.1 |
| chr6 | 125330708 | 125331390 | 11.39974 | 34.91658 | 32.39005 | Exon (ENSMU | 6 | 125321521 | 125344612 | 23092 | 1 | ENSMUST00 | 9188 | Scnn1a | ENSMUSG00000030340.11 |
| chr9 | 32409074 | 32409671 | 11.39974 | 34.91658 | 32.39005 | Distal Interge | 9 | 32393985 | 32399192 | 5208 | 1 | ENSMUST00 | 15090 | Kcnj1 | ENSMUSG00000041248.8 |
| chr12 | 8298015 | 8298941 | 15.0426 | 34.90326 | 32.37702 | Exon (ENSMU | 12 | 8297934 | 8301954 | 4021 | 2 | ENSMUST00 | 3013 | Gdf7 | ENSMUSG00000037660.4 |
| chr6 | 70824167 | 70825168 | 15.0426 | 34.90326 | 32.37702 | Distal Interge | 6 | 70844515 | 70905245 | 60731 | 1 | ENSMUST00 | -19347 | Eif2ak3 | ENSMUSG00000031668.9 |
| chr5 | 142928889 | 142929624 | 13.84168 | 34.89006 | 32.36391 | Distal Interge | 5 | 142903115 | 142906754 | 3640 | 2 | ENSMUST00 | -22136 | Actb | ENSMUSG00000029580.9 |
| chr5 | 72580086 | 72581105 | 13.84168 | 34.89006 | 32.36391 | Promoter | 5 | 72577114 | 72581084 | 3971 | 2 | ENSMUST00 | 0 | Zar1 | ENSMUSG00000063935.2 |
| chr17 | 32165972 | 32167122 | 14.61479 | 34.85841 | 32.33254 | Promoter | 17 | 32120820 | 32166880 | 46061 | 2 | ENSMUST00 | 0 | Notch3 | ENSMUSG00000038146.7 |
| chr3 | 89196151 | 89196673 | 14.61479 | 34.85841 | 32.33254 | Distal Interge | 3 | 89201167 | 89201273 | 107 | 2 | ENSMUST00 | 4600 | Gm23269 | ENSMUSG00000094489.1 |
| chr1 | 157086256 | 157086927 | 15.01131 | 34.85038 | 32.32499 | Distal Interge | 1 | 157099313 | 157108388 | 9076 | 2 | ENSMUST00 | 21461 | Tex35 | ENSMUSG00000026592.8 |
| chr10 | 128933540 | 128934382 | 15.01131 | 34.85038 | 32.32499 | Promoter | 10 | 128933813 | 128958277 | 24465 | 1 | ENSMUST00 | 0 | Itga7 | ENSMUSG00000025348.8 |
| chr18 | 64339875 | 64340797 | 15.01131 | 34.85038 | 32.32499 | Promoter | 18 | 64340364 | 64398486 | 58123 | 1 | ENSMUST00 | 0 | Onecut2 | ENSMUSG00000045991.13 |
| chr2 | 27540322 | 27540901 | 15.01131 | 34.85038 | 32.32499 | Promoter | 2 | 27540426 | 27543362 | 2937 | 1 | ENSMUST00 | 0 | Gm13421 | ENSMUSG00000085929.1 |
| chr10 | 60831673 | 60832323 | 14.20671 | 34.84183 | 32.31654 | Promoter | 10 | 60762610 | 60831581 | 68972 | 2 | ENSMUST00 | -93 | Unc5b | ENSMUSG00000020099.7 |
| chr11 | 94044197 | 94044760 | 14.20671 | 34.84183 | 32.31654 | Promoter | 11 | 94044205 | 94126080 | 81876 | 1 | ENSMUST00 | 0 | Spag9 | ENSMUSG00000020859.11 |
| chr18 | 35652386 | 35653533 | 14.20671 | 34.84183 | 32.31654 | Promoter | 18 | 35650355 | 35652006 | 1652 | 2 | ENSMUST00 | -381 | Prob1 | ENSMUSG00000073600.2 |
| chr8 | 47242647 | 47244577 | 14.20671 | 34.84183 | 32.31654 | Intron (ENSM | 8 | 47180048 | 47289402 | 109355 | 2 | ENSMUST00 | 44825 | Stox2 | ENSMUSG00000038143.9 |
| chr11 | 115373770 | 115374713 | 14.55755 | 34.76078 | 32.23622 | Distal Interge | 11 | 115347707 | 115367756 | 20050 | 2 | ENSMUST00 | -6015 | Hid1 | ENSMUSG00000034586.9 |
| chr1 | 131909903 | 131910322 | 14.9491 | 34.74524 | 32.2208 | Promoter | 1 | 131910534 | 131932679 | 22146 | 1 | ENSMUST00 | -212 | Nucks1 | ENSMUSG00000026434.7 |
| chr11 | 60222448 | 60223187 | 14.9491 | 34.74524 | 32.2208 | Promoter | 11 | 60206950 | 60222581 | 15632 | 2 | ENSMUST00 | 0 | Srebf1 | ENSMUSG00000020538.10 |
| chr15 | 36179267 | 36179812 | 14.91819 | 34.69299 | 32.16871 | Promoter | 15 | 36179368 | 36235610 | 56243 | 1 | ENSMUST00 | 0 | Spag1 | ENSMUSG00000037617.6 |
| chr14 | 62836870 | 62837274 | 14.50075 | 34.66385 | 32.13967 | Promoter | 14 | 62837690 | 62956886 | 119197 | 1 | ENSMUST00 | -416 | Wdfy2 | ENSMUSG00000014547.9 |
| chr3 | 96180864 | 96181436 | 14.88741 | 34.64093 | 32.11682 | Promoter | 3 | 96181151 | 96195521 | 14371 | 1 | ENSMUST00 | 0 | Sv2a | ENSMUSG00000038486.8 |
| chr1 | 134060670 | 134061705 | 14.47252 | 34.61563 | 32.09162 | Distal Interge | 1 | 134075170 | 134079120 | 3951 | 2 | ENSMUST00 | 17415 | Btg2 | ENSMUSG00000020423.6 |
| chr1 | 135584578 | 135585850 | 12.92336 | 34.59913 | 32.07519 | Promoter | 1 | 135434580 | 135585355 | 150776 | 2 | ENSMUST00 | 0 | Nav1 | ENSMUSG00000009418.10 |
| chr10 | 80114764 | 80115501 | 12.92336 | 34.59913 | 32.07519 | Promoter | 10 | 80115803 | 80130469 | 14667 | 1 | ENSMUST00 | -302 | Stk11 | ENSMUSG00000003068.10 |
| chr11 | 97531873 | 97532303 | 12.92336 | 34.59913 | 32.07519 | Exon (ENSMU | 11 | 97519612 | 97521895 | 2284 | 2 | ENSMUST00 | -9979 | Gm11611 | ENSMUSG00000087042.1 |
| chr14 | 55520359 | 55521186 | 12.92336 | 34.59913 | 32.07519 | 3' UTR | 14 | 55518978 | 55524191 | 5214 | 2 | ENSMUST00 | 3005 | Nrl | ENSMUSG00000040632.10 |
| chr15 | 103214892 | 103216001 | 12.92336 | 34.59913 | 32.07519 | Promoter | 15 | 103196687 | 103215352 | 18666 | 2 | ENSMUST00 | 0 | Cbx5 | ENSMUSG00000009575.9 |
| chr15 | 78174426 | 78175163 | 12.92336 | 34.59913 | 32.07519 | Promoter | 15 | 78159465 | 78174108 | 14644 | 2 | ENSMUST00 | -319 | Ift27 | ENSMUSG00000016637.6 |
| chr17 | 46552796 | 46553732 | 12.92336 | 34.59913 | 32.07519 | Exon (ENSMU | 17 | 46546839 | 46556162 | 9324 | 2 | ENSMUST00 | 2430 | Srf | ENSMUSG00000015605.5 |
| chr18 | 6666397 | 6667067 | 12.92336 | 34.59913 | 32.07519 | Distal Interge | 18 | 6603633 | 6638966 | 35334 | 1 | ENSMUST00 | 62765 | 4921524L21 | ENSMUSG00000039540.8 |
| chr19 | 34474298 | 34475845 | 12.92336 | 34.59913 | 32.07519 | Promoter | 19 | 34473786 | 34475135 | 1350 | 2 | ENSMUST00 | 0 | Ch25h | ENSMUSG00000050370.4 |
| chr19 | 48204986 | 48205750 | 12.92336 | 34.59913 | 32.07519 | Promoter | 19 | 48206025 | 48805505 | 599481 | 1 | ENSMUST00 | -275 | Sorcs3 | ENSMUSG00000063434.5 |
| chr2 | 74732605 | 74733925 | 12.92336 | 34.59913 | 32.07519 | Promoter | 2 | 74732913 | 74744214 | 11302 | 1 | ENSMUST00 | 0 | Hoxd3 | ENSMUSG00000079277.4 |
| chr3 | 88685028 | 88685594 | 12.92336 | 34.59913 | 32.07519 | Promoter | 3 | 88685794 | 88712933 | 27140 | 1 | ENSMUST00 | -200 | 2810403A07 | ENSMUSG00000028060.10 |
| chr4 | 46527324 | 46528139 | 12.92336 | 34.59913 | 32.07519 | Intron (ENSM | 4 | 46494819 | 46534414 | 39596 | 2 | ENSMUST00 | 6275 | Trim14 | ENSMUSG00000039853.13 |
| chr4 | 35099818 | 35100578 | 12.92336 | 34.59913 | 32.07519 | Intron (ENSM | 4 | 35152056 | 35154005 | 1950 | 1 | ENSMUST00 | -51478 | Ifnk | ENSMUSG00000042993.6 |
| chr5 | 114570268 | 114572884 | 12.92336 | 34.59913 | 32.07519 | Intron (ENSM | 5 | 114568017 | 114594678 | 26662 | 1 | ENSMUST00 | 2252 | Fam222a | ENSMUSG00000041930.7 |
| chr8 | 40926107 | 40926838 | 12.92336 | 34.59913 | 32.07519 | Promoter | 8 | 40926212 | 40939944 | 13733 | 1 | ENSMUST00 | 0 | Pdgfrl | ENSMUSG00000031595.9 |
| chr5 | 91402334 | 91403210 | 14.85675 | 34.58908 | 32.06662 | Promoter | 5 | 91357261 | 91402994 | 45734 | 2 | ENSMUST00 | 0 | Btc | ENSMUSG00000082361.3 |
| chr7 | 18893607 | 18894213 | 14.0502 | 34.57104 | 32.04866 | 3' UTR | 7 | 18890884 | 18892546 | 1663 | 2 | ENSMUST00 | -1062 | Ccdc61 | ENSMUSG00000074358.4 |
| chr18 | 80714491 | 80715784 | 14.82622 | 34.53743 | 32.01512 | Distal Interge | 18 | 80606205 | 80713071 | 106867 | 2 | ENSMUST00 | -1421 | Nfatc1 | ENSMUSG00000033016.10 |
| chr2 | 133554247 | 133554993 | 14.41639 | 34.51971 | 31.99752 | 5' UTR | 2 | 133552159 | 133562885 | 10727 | 1 | ENSMUST00 | 2089 | Bmp2 | ENSMUSG00000027358.6 |
| chr5 | 100719181 | 100720119 | 14.41639 | 34.51971 | 31.99752 | Promoter | 5 | 100679484 | 100719716 | 40233 | 2 | ENSMUST00 | 0 | Hpse | ENSMUSG00000035273.9 |
| chr11 | 16257340 | 16258042 | 14.76554 | 34.43472 | 31.91294 | Promoter | 11 | 16257724 | 16284186 | 26463 | 1 | ENSMUST00 | 0 | Vstm2a | ENSMUSG00000048834.10 |
| chr13 | 91741312 | 91741987 | 14.76554 | 34.43472 | 31.91294 | Promoter | 13 | 91741512 | 91786148 | 44637 | 1 | ENSMUST00 | 0 | Acot12 | ENSMUSG00000021620.4 |
| chr8 | 41217625 | 41218366 | 14.76554 | 34.43472 | 31.91294 | Promoter | 8 | 41218139 | 41220894 | 2756 | 1 | ENSMUST00 | 0 | Gm16348 | ENSMUSG00000085445.1 |
| chr17 | 56139628 | 56140521 | 13.94776 | 34.39338 | 31.87212 | Promoter | 17 | 56123085 | 56140343 | 17259 | 2 | ENSMUST00 | 0 | Sema6b | ENSMUSG00000001227.6 |
| chr6 | 122491329 | 122492171 | 13.94776 | 34.39338 | 31.87212 | Distal Interge | 6 | 122453101 | 122486313 | 33213 | 2 | ENSMUST00 | -5017 | Rimklb | ENSMUSG00000040649.10 |
| chr7 | 45406099 | 45407198 | 13.94776 | 34.39338 | 31.87212 | Promoter | 7 | 45405960 | 45411382 | 5423 | 1 | ENSMUST00 | 140 | Kcna7 | ENSMUSG00000038201.9 |
| chr12 | 40016757 | 40017575 | 14.73538 | 34.38367 | 31.86286 | Intron (ENSM | 12 | 40009609 | 40014484 | 4876 | 2 | ENSMUST00 | -2274 | Arl4a | ENSMUSG00000047446.13 |
| chr15 | 76205354 | 76207876 | 14.73538 | 34.38367 | 31.86286 | Promoter | 15 | 76170974 | 76206321 | 35348 | 2 | ENSMUST00 | 0 | Plec | ENSMUSG00000022565.10 |
| chr2 | 136531921 | 136532531 | 14.73538 | 34.38367 | 31.86286 | Promoter | 2 | 136532212 | 136562091 | 29880 | 1 | ENSMUST00 | 0 | Ankef1 | ENSMUSG00000074771.5 |
| chr10 | 60686537 | 60687680 | 14.333 | 34.37709 | 31.85639 | Promoter | 10 | 60657266 | 60687257 | 29992 | 2 | ENSMUST00 | 0 | Cdh23 | ENSMUSG00000012819.10 |
| chr12 | 23968295 | 23968986 | 14.333 | 34.37709 | 31.85639 | Promoter | 12 | 23968510 | 23968845 | 336 | 2 | ENSMUST00 | 0 | Gm17330 | ENSMUSG00000095329.1 |
| chr2 | 71528850 | 71530279 | 14.333 | 34.37709 | 31.85639 | Promoter | 2 | 71529101 | 71530231 | 1131 | 1 | ENSMUST00 | 0 | Dlx1 | ENSMUSG00000041911.3 |
| chr3 | 88493159 | 88493450 | 14.333 | 34.37709 | 31.85639 | Promoter | 3 | 88483206 | 88493312 | 10107 | 2 | ENSMUST00 | 0 | Lmna | ENSMUSG00000028063.10 |
| chr4 | 152119592 | 152121333 | 14.333 | 34.37709 | 31.85639 | 3' UTR | 4 | 152117971 | 152119323 | 1353 | 1 | ENSMUST00 | 1622 | Tnfrsf25 | ENSMUSG00000024793.9 |
| chr7 | 62348166 | 62348941 | 14.333 | 34.37709 | 31.85639 | Promoter | 7 | 62348277 | 62349928 | 1652 | 1 | ENSMUST00 | 0 | Ndn | ENSMUSG00000033585.4 |

| | | | | | | | | | | | | | | | |
|---|---|---|---|---|---|---|---|---|---|---|---|---|---|---|---|
| chr8 | 67974068 | 67974798 | 14.333 | 34.37709 | 31.85639 | Promoter | 8 | 67689082 | 67974574 | 285493 | 2 | ENSMUST00 | 0 | Psd3 | ENSMUSG00000030465.13 |
| chr8 | 48221368 | 48222551 | 14.333 | 34.37709 | 31.85639 | Distal Interge | 8 | 48275915 | 48300099 | 24185 | 2 | ENSMUST00 | 77548 | Tenm3 | ENSMUSG00000031561.10 |
| chr1 | 78216325 | 78217774 | 14.70535 | 34.3328 | 31.81257 | Distal Interge | 1 | 78101270 | 78197134 | 95865 | 2 | ENSMUST00 | -19192 | Pax3 | ENSMUSG00000004872.10 |
| chr12 | 85218847 | 85219349 | 14.70535 | 34.3328 | 31.81257 | Promoter | 12 | 85219481 | 85226628 | 7148 | 1 | ENSMUST00 | -132 | Eif2b2 | ENSMUSG00000004788.6 |
| chr7 | 4532056 | 4532778 | 14.70535 | 34.3328 | 31.81257 | Promoter | 7 | 4522933 | 4532453 | 9521 | 2 | ENSMUST00 | 0 | Dnaaf3 | ENSMUSG00000055809.7 |
| chr1 | 62883755 | 62885574 | 13.51752 | 34.31881 | 31.79871 | Distal Interge | 1 | 62821355 | 62832370 | 11016 | 2 | ENSMUST00 | -51386 | RP23-149A5. | ENSMUSG00000100426.1 |
| chr11 | 22617363 | 22617807 | 13.51752 | 34.31881 | 31.79871 | Promoter | 11 | 22617408 | 22618487 | 1080 | 1 | ENSMUST00 | 0 | Gm12052 | ENSMUSG00000086256.1 |
| chr13 | 114460262 | 114460704 | 13.51752 | 34.31881 | 31.79871 | Distal Interge | 13 | 114452262 | 114458730 | 6469 | 2 | ENSMUST00 | -1533 | Fst | ENSMUSG00000021765.7 |
| chr14 | 31661250 | 31662390 | 13.51752 | 34.31881 | 31.79871 | Exon (ENSMU | 14 | 31635648 | 31641286 | 5639 | 2 | ENSMUST00 | -19965 | Hacl1 | ENSMUSG00000021884.12 |
| chr14 | 122104926 | 122105742 | 13.51752 | 34.31881 | 31.79871 | Exon (ENSMU | 14 | 122103430 | 122106953 | 3524 | 2 | ENSMUST00 | 1211 | A330035P11 | ENSMUSG00000085615.2 |
| chr17 | 71526343 | 71526848 | 13.51752 | 34.31881 | 31.79871 | Promoter | 17 | 71496100 | 71526857 | 30758 | 2 | ENSMUST00 | 9 | Ndc80 | ENSMUSG00000024056.9 |
| chr2 | 104589594 | 104590306 | 13.51752 | 34.31881 | 31.79871 | Promoter | 2 | 104590523 | 104665429 | 74907 | 1 | ENSMUST00 | -217 | Cstf3 | ENSMUSG00000027176.8 |
| chr4 | 133467806 | 133468493 | 13.51752 | 34.31881 | 31.79871 | Distal Interge | 4 | 133480132 | 133487938 | 7807 | 1 | ENSMUST00 | -11639 | Fam46b | ENSMUSG00000046694.6 |
| chr4 | 133293289 | 133294010 | 13.51752 | 34.31881 | 31.79871 | 3' UTR | 4 | 133280857 | 133281312 | 456 | 2 | ENSMUST00 | -11978 | Gm13259 | ENSMUSG00000083241.1 |
| chr6 | 42245704 | 42246455 | 13.51752 | 34.31881 | 31.79871 | Promoter | 6 | 42245935 | 42250441 | 4507 | 1 | ENSMUST00 | 0 | Gstk1 | ENSMUSG00000029864.5 |
| chr6 | 83161730 | 83162987 | 13.51752 | 34.31881 | 31.79871 | 3' UTR | 6 | 83165920 | 83200116 | 34197 | 1 | ENSMUST00 | -2933 | Dctn1 | ENSMUSG00000031865.10 |
| chr6 | 140856720 | 140857690 | 13.51752 | 34.31881 | 31.79871 | Distal Interge | 6 | 140746960 | 140747172 | 213 | 2 | ENSMUST00 | -109549 | Rpl38-ps2 | ENSMUSG00000080921.4 |
| chr7 | 27408304 | 27409201 | 13.51752 | 34.31881 | 31.79871 | Exon (ENSMU | 7 | 27356383 | 27396553 | 40171 | 2 | ENSMUST00 | -11752 | Sptbn4 | ENSMUSG00000011751.11 |
| chr7 | 16129517 | 16130695 | 13.51752 | 34.31881 | 31.79871 | Promoter | 7 | 16130300 | 16160509 | 30210 | 1 | ENSMUST00 | 0 | Slc8a2 | ENSMUSG00000030376.7 |
| chr7 | 116237243 | 116237984 | 13.51752 | 34.31881 | 31.79871 | Promoter | 7 | 116123493 | 116237776 | 114284 | 2 | ENSMUST00 | 0 | Plekha7 | ENSMUSG00000045659.12 |
| chr9 | 36832334 | 36833168 | 13.51752 | 34.31881 | 31.79871 | Promoter | 9 | 36832684 | 36878924 | 46241 | 1 | ENSMUST00 | 0 | Fez1 | ENSMUSG00000032118.10 |
| chr16 | 94724026 | 94724972 | 11.73166 | 34.30278 | 31.78458 | Distal Interge | 16 | 94668896 | 94679521 | 10626 | 1 | ENSMUST00 | 55131 | Dyrk1a | ENSMUSG00000022897.9 |
| chr18 | 74215484 | 74216055 | 11.73166 | 34.30278 | 31.78458 | Promoter | 18 | 74216131 | 74221491 | 5361 | 1 | ENSMUST00 | -76 | Cxxc1 | ENSMUSG00000024560.6 |
| chr3 | 87768580 | 87769243 | 11.73166 | 34.30278 | 31.78458 | Promoter | 3 | 87747890 | 87768948 | 21059 | 2 | ENSMUST00 | 0 | Pear1 | ENSMUSG00000028073.10 |
| chr4 | 117303912 | 117304867 | 11.73166 | 34.30278 | 31.78458 | Intron (ENSM | 4 | 117285889 | 117300028 | 14140 | 2 | ENSMUST00 | -3885 | Rnf220 | ENSMUSG00000028677.13 |
| chr4 | 115657049 | 115657840 | 14.67544 | 34.28214 | 31.7644 | Distal Interge | 4 | 115652389 | 115653480 | 1092 | 1 | ENSMUST00 | 4661 | Gm12848 | ENSMUSG00000082043.1 |
| chr1 | 16092798 | 16093563 | 14.24974 | 34.23454 | 31.71693 | Promoter | 1 | 16065979 | 16093325 | 27347 | 2 | ENSMUST00 | 0 | 4930444P10 | ENSMUSG00000067795.8 |
| chr10 | 120552254 | 120553433 | 14.24974 | 34.23454 | 31.71693 | Distal Interge | 10 | 120605537 | 120605596 | 60 | 1 | ENSMUST00 | -52104 | Gm24298 | ENSMUSG00000088924.1 |
| chr11 | 47988319 | 47989395 | 14.24974 | 34.23454 | 31.71693 | Promoter | 11 | 47578805 | 47988969 | 410165 | 2 | ENSMUST00 | 0 | Sgcd | ENSMUSG00000020354.10 |
| chr11 | 51805326 | 51805880 | 14.24974 | 34.23454 | 31.71693 | Distal Interge | 11 | 51763694 | 51782801 | 19108 | 1 | ENSMUST00 | 41633 | Sar1b | ENSMUSG00000020386.5 |
| chr11 | 58008353 | 58008729 | 14.24974 | 34.23454 | 31.71693 | Promoter | 11 | 58009064 | 58062031 | 52968 | 1 | ENSMUST00 | -335 | Larp1 | ENSMUSG00000037331.10 |
| chr11 | 109331477 | 109332076 | 14.24974 | 34.23454 | 31.71693 | Distal Interge | 11 | 109346839 | 109353651 | 6813 | 1 | ENSMUST00 | -14763 | RP23-12N20. | ENSMUSG00000099630.1 |
| chr12 | 25098482 | 25099118 | 14.24974 | 34.23454 | 31.71693 | Distal Interge | 12 | 25093801 | 25096092 | 2292 | 2 | ENSMUST00 | -2391 | Id2 | ENSMUSG00000020644.8 |
| chr13 | 21981143 | 21982021 | 14.24974 | 34.23454 | 31.71693 | Promoter | 13 | 21981194 | 21988734 | 7541 | 1 | ENSMUST00 | 0 | Pom121l2 | ENSMUSG00000016982.6 |
| chr14 | 103469524 | 103469952 | 14.24974 | 34.23454 | 31.71693 | Distal Interge | 14 | 103513341 | 103613346 | 100006 | 1 | ENSMUST00 | -43389 | Scel | ENSMUSG00000022123.8 |
| chr16 | 18620708 | 18621817 | 14.24974 | 34.23454 | 31.71693 | Promoter | 16 | 18620722 | 18621366 | 645 | 2 | ENSMUST00 | 0 | Gp1bb | ENSMUSG00000050761.3 |
| chr17 | 3102250 | 3102879 | 14.24974 | 34.23454 | 31.71693 | Distal Interge | 17 | 3114972 | 3198855 | 83884 | 1 | ENSMUST00 | -12093 | Scaf8 | ENSMUSG00000046201.8 |
| chr18 | 5819564 | 5820389 | 14.24974 | 34.23454 | 31.71693 | Distal Interge | 18 | 5731081 | 5772804 | 41724 | 1 | ENSMUST00 | 88484 | Zeb1 | ENSMUSG00000024238.8 |
| chr19 | 25610467 | 25611285 | 14.24974 | 34.23454 | 31.71693 | Promoter | 19 | 25610537 | 25623920 | 13384 | 1 | ENSMUST00 | 0 | Dmrt3 | ENSMUSG00000042372.4 |
| chr4 | 126734964 | 126736321 | 14.24974 | 34.23454 | 31.71693 | Promoter | 4 | 126716006 | 126736240 | 20235 | 2 | ENSMUST00 | 0 | Tfap2e | ENSMUSG00000042477.7 |
| chr4 | 128609460 | 128610121 | 14.24974 | 34.23454 | 31.71693 | Promoter | 4 | 128583547 | 128610098 | 26552 | 2 | ENSMUST00 | 0 | Zscan20 | ENSMUSG00000061894.10 |
| chr6 | 92534427 | 92535202 | 14.24974 | 34.23454 | 31.71693 | Promoter | 6 | 92370910 | 92534861 | 163952 | 2 | ENSMUST00 | 0 | Prickle2 | ENSMUSG00000030020.8 |
| chr6 | 119451936 | 119452726 | 14.24974 | 34.23454 | 31.71693 | Intron (ENSM | 6 | 119432531 | 119449386 | 16856 | 2 | ENSMUST00 | -2551 | Wnt5b | ENSMUSG00000030170.9 |
| chr6 | 142570923 | 142572394 | 14.24974 | 34.23454 | 31.71693 | Promoter | 6 | 142564939 | 142571356 | 6418 | 2 | ENSMUST00 | 0 | Kcnj8 | ENSMUSG00000030247.7 |
| chr8 | 91134235 | 91134618 | 14.24974 | 34.23454 | 31.71693 | Promoter | 8 | 91123499 | 91134465 | 10967 | 2 | ENSMUST00 | 0 | Aktip | ENSMUSG00000031667.10 |
| chr8 | 71610623 | 71610917 | 14.24974 | 34.23454 | 31.71693 | Promoter | 8 | 71611024 | 71624909 | 13886 | 1 | ENSMUST00 | -107 | Colgalt1 | ENSMUSG00000034807.8 |
| chr8 | 124792395 | 124793359 | 14.24974 | 34.23454 | 31.71693 | Promoter | 8 | 124793019 | 124834702 | 41684 | 1 | ENSMUST00 | 0 | Trim67 | ENSMUSG00000036913.9 |
| chr8 | 31031931 | 31032847 | 14.24974 | 34.23454 | 31.71693 | Distal Interge | 8 | 31089471 | 31097047 | 7577 | 1 | ENSMUST00 | -56624 | Dusp26 | ENSMUSG00000039661.9 |
| chr8 | 110754982 | 110755873 | 14.24974 | 34.23454 | 31.71693 | Promoter | 8 | 110754992 | 110758970 | 3979 | 1 | ENSMUST00 | 0 | Gm15894 | ENSMUSG00000090039.1 |
| chr18 | 62734497 | 62735029 | 14.19614 | 34.14266 | 31.62711 | Intron (ENSM | 18 | 62607539 | 62741387 | 133849 | 2 | ENSMUST00 | 6358 | Spink13 | ENSMUSG00000073551.4 |
| chr17 | 47174701 | 47175281 | 14.58643 | 34.13128 | 31.61576 | Intron (ENSM | 17 | 47141216 | 47318790 | 177575 | 1 | ENSMUST00 | 33486 | Trerf1 | ENSMUSG00000064043.8 |
| chr8 | 84979560 | 84980539 | 14.16908 | 34.09626 | 31.58084 | Promoter | 8 | 84976909 | 84978748 | 1840 | 2 | ENSMUST00 | -813 | Junb | ENSMUSG00000052837.5 |
| chr2 | 168080352 | 168080877 | 13.37139 | 34.05986 | 31.54458 | Promoter | 2 | 168081004 | 168101203 | 20200 | 1 | ENSMUST00 | -127 | Pard6b | ENSMUSG00000044641.7 |
| chr5 | 104859665 | 104860613 | 13.37139 | 34.05986 | 31.54458 | Promoter | 5 | 104813169 | 104860070 | 46902 | 2 | ENSMUST00 | 0 | Zfp951 | ENSMUSG00000072774.5 |
| chr17 | 50698336 | 50698753 | 12.61417 | 34.02772 | 31.51277 | Promoter | 17 | 50698525 | 50699880 | 1356 | 1 | ENSMUST00 | 0 | Gm7334 | ENSMUSG00000044645.7 |
| chr14 | 64586775 | 64587966 | 10.40121 | 34.00095 | 31.48606 | Promoter | 14 | 64587331 | 64588112 | 782 | 1 | ENSMUST00 | 0 | A930011O12 | ENSMUSG00000097545.2 |
| chr5 | 92674986 | 92676099 | 14.4985 | 33.98212 | 31.46727 | Promoter | 5 | 92608253 | 92675125 | 66873 | 2 | ENSMUST00 | 0 | Ccdc158 | ENSMUSG00000050050.11 |
| chr8 | 80493131 | 80493759 | 14.4985 | 33.98212 | 31.46727 | Promoter | 8 | 80494045 | 80510785 | 16741 | 1 | ENSMUST00 | -286 | Gypa | ENSMUSG00000051839.6 |
| chr10 | 61418195 | 61418962 | 13.69809 | 33.9589 | 31.44413 | Promoter | 10 | 61417972 | 61425338 | 7367 | 1 | ENSMUST00 | 224 | Nodal | ENSMUSG00000037171.6 |
| chr2 | 121509345 | 121510085 | 13.69809 | 33.9589 | 31.44413 | Promoter | 2 | 121509848 | 121543009 | 33162 | 1 | ENSMUST00 | 0 | Wdr76 | ENSMUSG00000027242.9 |
| chr6 | 28396990 | 28398186 | 13.69809 | 33.9589 | 31.44413 | Promoter | 6 | 28240240 | 28398005 | 157766 | 2 | ENSMUST00 | 0 | Zfp800 | ENSMUSG00000039841.9 |
| chr15 | 56692783 | 56693586 | 14.46942 | 33.93277 | 31.41837 | Promoter | 15 | 56693552 | 56778399 | 84848 | 1 | ENSMUST00 | 0 | Has2os | ENSMUSG00000086541.3 |
| chr19 | 38124449 | 38125143 | 14.46942 | 33.93277 | 31.41837 | Promoter | 19 | 38116620 | 38124811 | 8192 | 2 | ENSMUST00 | 0 | Rbp4 | ENSMUSG00000024990.7 |
| chr3 | 67373820 | 67374516 | 14.46942 | 33.93277 | 31.41837 | Promoter | 3 | 67374097 | 67399998 | 25902 | 1 | ENSMUST00 | 0 | Mlf1 | ENSMUSG00000048416.10 |
| chr3 | 13471455 | 13472006 | 14.46942 | 33.93277 | 31.41837 | Exon (ENSMU | 3 | 13471906 | 13474596 | 2691 | 2 | ENSMUST00 | 2590 | Gm2464 | ENSMUSG00000097338.1 |
| chr7 | 123500216 | 123500938 | 14.46942 | 33.93277 | 31.41837 | Promoter | 7 | 123479516 | 123500449 | 20934 | 2 | ENSMUST00 | 0 | Zkscan2 | ENSMUSG00000030757.8 |
| chr8 | 79388978 | 79389788 | 14.46942 | 33.93277 | 31.41837 | Intron (ENSM | 8 | 79338395 | 79399518 | 61124 | 2 | ENSMUST00 | 9730 | Smad1 | ENSMUSG00000031681.9 |
| chr3 | 127161260 | 127162756 | 14.06187 | 33.91219 | 31.39823 | Promoter | 3 | 127031173 | 127162342 | 131170 | 2 | ENSMUST00 | 0 | Ank2 | ENSMUSG00000032826.11 |
| chr7 | 141221240 | 141221767 | 14.06187 | 33.91219 | 31.39823 | Promoter | 7 | 141221384 | 141221493 | 110 | 2 | ENSMUST00 | 0 | Mir210 | ENSMUSG00000065551.1 |
| chr7 | 18910682 | 18911164 | 14.06187 | 33.91219 | 31.39823 | Promoter | 7 | 18890883 | 18910415 | 19533 | 2 | ENSMUST00 | -268 | Ccdc61 | ENSMUSG00000074358.4 |
| chr1 | 152800193 | 152801982 | 14.44046 | 33.88361 | 31.37022 | Promoter | 1 | 152800194 | 152836991 | 36798 | 1 | ENSMUST00 | 0 | Ncf2 | ENSMUSG00000026480.7 |
| chr7 | 60243496 | 60244770 | 14.44046 | 33.88361 | 31.37022 | Distal Interge | 7 | 60155538 | 60156170 | 633 | 1 | ENSMUST00 | 87959 | Gm7367 | ENSMUSG00000047370.4 |
| chr4 | 140960684 | 140961116 | 14.03532 | 33.86656 | 31.35328 | Promoter | 4 | 140961203 | 140979193 | 17991 | 1 | ENSMUST00 | -87 | Sdhb | ENSMUSG00000009863.9 |
| chr2 | 167071380 | 167072178 | 14.41162 | 33.83462 | 31.32136 | Distal Interge | 2 | 167065276 | 167065365 | 90 | 1 | ENSMUST00 | 6105 | Snord12 | ENSMUSG00000077698.1 |
| chr1 | 90506776 | 90507928 | 12.12078 | 33.7584 | 31.24543 | Distal Interge | 1 | 90453394 | 90454501 | 1108 | 2 | ENSMUST00 | -52276 | RP24-154L11 | ENSMUSG00000101997.1 |
| chr15 | 102954459 | 102955586 | 12.12078 | 33.7584 | 31.24543 | Promoter | 15 | 102954427 | 102957708 | 3282 | 1 | ENSMUST00 | 33 | Hoxc11 | ENSMUSG00000001656.3 |
| chr15 | 78913740 | 78914615 | 12.12078 | 33.7584 | 31.24543 | Promoter | 15 | 78913919 | 78919517 | 5599 | 1 | ENSMUST00 | 0 | Sh3bp1 | ENSMUSG00000022436.10 |
| chr15 | 98895678 | 98896482 | 12.12078 | 33.7584 | 31.24543 | Intron (ENSM | 15 | 98893031 | 98898540 | 5510 | 2 | ENSMUST00 | 2058 | Dhh | ENSMUSG00000023000.4 |
| chr18 | 3280698 | 3281715 | 12.12078 | 33.7584 | 31.24543 | Promoter | 18 | 3267534 | 3281132 | 13599 | 2 | ENSMUST00 | 0 | Crem | ENSMUSG00000063889.11 |
| chr4 | 126102974 | 126103823 | 12.12078 | 33.7584 | 31.24543 | Promoter | 4 | 126103957 | 126141028 | 37072 | 1 | ENSMUST00 | -134 | Stk40 | ENSMUSG00000042608.10 |
| chr5 | 17574196 | 17575121 | 12.12078 | 33.7584 | 31.24543 | Promoter | 5 | 17574281 | 17730268 | 155988 | 1 | ENSMUST00 | 0 | Sema3c | ENSMUSG00000028780.8 |
| chr5 | 43672230 | 43673093 | 14.35427 | 33.73719 | 31.22503 | Promoter | 5 | 43672289 | 43689101 | 16813 | 1 | ENSMUST00 | 0 | Cc2d2a | ENSMUSG00000039765.10 |
| chr6 | 112490534 | 112491098 | 14.35427 | 33.73719 | 31.22503 | Promoter | 6 | 112473684 | 112489808 | 16125 | 2 | ENSMUST00 | -727 | Oxtr | ENSMUSG00000049112.8 |
| chr9 | 103108878 | 103109540 | 11.13463 | 33.72523 | 31.21317 | Distal Interge | 9 | 103111787 | 103162657 | 50871 | 1 | ENSMUST00 | -2247 | Rab6b | ENSMUSG00000032549.7 |
| chr1 | 172481777 | 172482838 | 14.70503 | 33.66069 | 31.14909 | Promoter | 1 | 172481788 | 172489593 | 7806 | 1 | ENSMUST00 | 0 | Igsf9 | ENSMUSG00000037995.10 |
| chr1 | 39720593 | 39721466 | 14.70503 | 33.66069 | 31.14909 | Promoter | 1 | 39665306 | 39720997 | 55692 | 2 | ENSMUST00 | 0 | Rfx8 | ENSMUSG00000057173.7 |
| chr10 | 67002504 | 67005089 | 14.70503 | 33.66069 | 31.14909 | Distal Interge | 10 | 66920902 | 66923599 | 2698 | 1 | ENSMUST00 | 81603 | Gm26576 | ENSMUSG00000097921.1 |
| chr12 | 34906501 | 34907530 | 14.70503 | 33.66069 | 31.14909 | Distal Interge | 12 | 34895024 | 34895155 | 132 | 1 | ENSMUST00 | 11478 | Gm24429 | ENSMUSG00000077264.1 |
| chr13 | 67844432 | 67845131 | 14.70503 | 33.66069 | 31.14909 | Distal Interge | 13 | 67833235 | 67843113 | 9879 | 1 | ENSMUST00 | 11198 | Gm10037 | ENSMUSG00000058246.6 |
| chr14 | 31220307 | 31223809 | 14.70503 | 33.66069 | 31.14909 | Exon (ENSMU | 14 | 31217860 | 31230350 | 12491 | 1 | ENSMUST00 | 2448 | Sema3g | ENSMUSG00000021904.5 |
| chr15 | 36715780 | 36716745 | 14.70503 | 33.66069 | 31.14909 | Distal Interge | 15 | 36790712 | 36792414 | 1703 | 2 | ENSMUST00 | 75669 | Ywhaz | ENSMUSG00000022285.11 |
| chr18 | 84719874 | 84720857 | 14.70503 | 33.66069 | 31.14909 | Promoter | 18 | 84720019 | 84730447 | 10429 | 1 | ENSMUST00 | 0 | Fam69c | ENSMUSG00000047992.5 |
| chr18 | 80169370 | 80170152 | 14.70503 | 33.66069 | 31.14909 | Distal Interge | 18 | 80126311 | 80151482 | 25172 | 2 | ENSMUST00 | -17889 | Adnp2 | ENSMUSG00000053950.10 |
| chr2 | 36011094 | 36011921 | 14.70503 | 33.66069 | 31.14909 | Distal Interge | 2 | 35902351 | 35979913 | 77563 | 2 | ENSMUST00 | -31182 | Ttll11 | ENSMUSG00000026885.8 |
| chr2 | 175738026 | 175738432 | 14.70503 | 33.66069 | 31.14909 | Distal Interge | 2 | 175741537 | 175748960 | 7424 | 1 | ENSMUST00 | -3105 | Gm2007 | ENSMUSG00000094932.2 |
| chr4 | 152325289 | 152325655 | 14.70503 | 33.66069 | 31.14909 | Promoter | 4 | 152325742 | 152332476 | 6735 | 1 | ENSMUST00 | -87 | Rpl22 | ENSMUSG00000028936.10 |
| chr4 | 129356376 | 129357908 | 14.70503 | 33.66069 | 31.14909 | Exon (ENSMU | 4 | 129343940 | 129346930 | 2991 | 1 | ENSMUST00 | 12437 | Zbtb8os | ENSMUSG00000057572.10 |
| chr4 | 49058681 | 49060050 | 14.70503 | 33.66069 | 31.14909 | Promoter | 4 | 49059273 | 49208600 | 149328 | 1 | ENSMUST00 | 0 | E130309F12 | ENSMUSG00000063446.4 |
| chr4 | 42172628 | 42173470 | 14.70503 | 33.66069 | 31.14909 | Distal Interge | 4 | 42158842 | 42168603 | 9762 | 2 | ENSMUST00 | -4026 | Gm13305 | ENSMUSG00000073876.3 |
| chr5 | 135903387 | 135904098 | 14.70503 | 33.66069 | 31.14909 | Distal Interge | 5 | 135888018 | 135889561 | 1544 | 1 | ENSMUST00 | 15370 | Hspb1 | ENSMUSG00000004951.10 |
| chr6 | 124769180 | 124769864 | 14.70503 | 33.66069 | 31.14909 | Promoter | 6 | 124760055 | 124769621 | 9567 | 2 | ENSMUST00 | 0 | Eno2 | ENSMUSG00000004267.11 |
| chr6 | 126643407 | 126644879 | 14.70503 | 33.66069 | 31.14909 | Promoter | 6 | 126636463 | 126645801 | 9339 | 2 | ENSMUST00 | 922 | Kcna1 | ENSMUSG00000047976.3 |
| chrX | 20961230 | 20962646 | 14.70503 | 33.66069 | 31.14909 | Promoter | 20 | 20941722 | 20961991 | 20270 | 2 | ENSMUST00 | 0 | Uxt | ENSMUSG00000001134.4 |
| chrX | 20658069 | 20659434 | 14.70503 | 33.66069 | 31.14909 | Promoter | 20 | 20658326 | 20683179 | 24854 | 1 | ENSMUST00 | 0 | Uba1 | ENSMUSG00000001924.10 |
| chrX | 85775963 | 85776674 | 14.70503 | 33.66069 | 31.14909 | Promoter | 20 | 85701937 | 85776624 | 74688 | 2 | ENSMUST00 | 0 | Gyk | ENSMUSG00000025059.11 |
| chrX | 42067931 | 42068500 | 14.70503 | 33.66069 | 31.14909 | Promoter | 20 | 42068398 | 42109656 | 41259 | 1 | ENSMUST00 | 0 | Xiap | ENSMUSG00000025860.9 |
| chrX | 7947402 | 7947948 | 14.70503 | 33.66069 | 31.14909 | Promoter | 20 | 7930120 | 7947889 | 17770 | 2 | ENSMUST00 | 0 | Hdac6 | ENSMUSG00000031161.10 |
| chrX | 160766116 | 160766999 | 14.70503 | 33.66069 | 31.14909 | Distal Interge | 20 | 160768013 | 160799663 | 31651 | 1 | ENSMUST00 | -1014 | Rs1 | ENSMUSG00000031293.8 |
| chrX | 168795149 | 168796799 | 14.70503 | 33.66069 | 31.14909 | Promoter | 20 | 168795099 | 169304435 | 509337 | 1 | ENSMUST00 | 51 | Arhgap6 | ENSMUSG00000031355.11 |
| chrX | 136740373 | 136741241 | 14.70503 | 33.66069 | 31.14909 | Promoter | 20 | 136732953 | 136741155 | 8203 | 2 | ENSMUST00 | 0 | Morf4l2 | ENSMUSG00000031422.11 |
| chrX | 169879551 | 169881706 | 14.70503 | 33.66069 | 31.14909 | Promoter | 20 | 169879622 | 169990798 | 111177 | 1 | ENSMUST00 | 0 | Mid1 | ENSMUSG00000035299.11 |
| chrX | 150458651 | 150459519 | 14.70503 | 33.66069 | 31.14909 | Promoter | 20 | 150397773 | 150459150 | 61378 | 2 | ENSMUST00 | 0 | Tmem29 | ENSMUSG00000041353.7 |
| chrX | 142227929 | 142228954 | 14.70503 | 33.66069 | 31.14909 | Promoter | 20 | 142227937 | 142239692 | 11756 | 1 | ENSMUST00 | 0 | Nxt2 | ENSMUSG00000042271.8 |
| chrX | 56374410 | 56375095 | 14.70503 | 33.66069 | 31.14909 | Promoter | 20 | 56374610 | 56377799 | 3190 | 1 | ENSMUST00 | 0 | 6330419J24 | ENSMUSG00000054850.3 |
| chr11 | 109721013 | 109722475 | 14.64785 | 33.56424 | 31.05456 | Promoter | 11 | 109669749 | 109722279 | 52531 | 2 | ENSMUST00 | 0 | Fam20a | ENSMUSG00000020614.8 |
| chr11 | 94209793 | 94210208 | 14.64785 | 33.56424 | 31.05456 | Distal Interge | 11 | 94211454 | 94215495 | 4042 | 1 | ENSMUST00 | -1246 | Tob1 | ENSMUSG00000037573.5 |
| chr2 | 163918432 | 163919307 | 14.64785 | 33.56424 | 31.05456 | Promoter | 2 | 163859751 | 163918683 | 58933 | 2 | ENSMUST00 | 0 | Rims4 | ENSMUSG00000035226.5 |
| chr11 | 62788958 | 62789669 | 13.45719 | 33.53756 | 31.02805 | Promoter | 11 | 62753377 | 62788980 | 35604 | 2 | ENSMUST00 | 0 | Zfp286 | ENSMUSG00000047342.11 |
| chr17 | 50292344 | 50293225 | 13.45719 | 33.53756 | 31.02805 | Promoter | 17 | 50279394 | 50293599 | 14206 | 2 | ENSMUST00 | 374 | Dazl | ENSMUSG00000010592.8 |
| chr8 | 70373306 | 70375073 | 13.45719 | 33.53756 | 31.02805 | Promoter | 8 | 70373548 | 70382065 | 8518 | 1 | ENSMUST00 | 0 | Comp | ENSMUSG00000031849.8 |
| chr9 | 102267473 | 102268314 | 13.45719 | 33.53756 | 31.02805 | Intron (ENSM | 9 | 102304924 | 102304975 | 52 | 2 | ENSMUST00 | 36661 | Mir7243 | ENSMUSG00000098521.1 |
| chr11 | 69757625 | 69758649 | 14.61724 | 33.51261 | 31.00336 | Promoter | 11 | 69733997 | 69758637 | 24641 | 2 | ENSMUST00 | 0 | Polr2a | ENSMUSG00000005198.10 |
| chr13 | 72628036 | 72629650 | 14.61724 | 33.51261 | 31.00336 | Promoter | 13 | 72628820 | 72634198 | 5379 | 1 | ENSMUST00 | 0 | Irx2 | ENSMUSG00000001504.10 |
| chr17 | 24151808 | 24152323 | 13.8008 | 33.46267 | 30.95374 | Promoter | 17 | 24078928 | 24150924 | 71997 | 2 | ENSMUST00 | -885 | Pdpk1 | ENSMUSG00000024122.11 |
| chr6 | 98105372 | 98105836 | 13.8008 | 33.46267 | 30.95374 | Distal Interge | 6 | 97991819 | 98018497 | 26679 | 1 | ENSMUST00 | 113554 | Mitf | ENSMUSG00000035158.10 |
| chr16 | 38483085 | 38484037 | 14.58677 | 33.46119 | 30.95275 | Intron (ENSM | 16 | 38458933 | 38486933 | 28001 | 1 | ENSMUST00 | 24153 | Cd80 | ENSMUSG00000075122.4 |
| chr1 | 50926512 | 50928267 | 12.66686 | 33.45994 | 30.9516 | Promoter | 1 | 50927519 | 51187270 | 259752 | 1 | ENSMUST00 | 0 | Tmeff2 | ENSMUSG00000026109.9 |
| chr12 | 69907977 | 69908847 | 14.18495 | 33.44911 | 30.9408 | Exon (ENSMU | 12 | 69920472 | 69921491 | 1020 | 2 | ENSMUST00 | 12644 | Gm6316 | ENSMUSG00000078370.1 |

| | | | | | | | | | | | | | | | |
|---|---|---|---|---|---|---|---|---|---|---|---|---|---|---|---|
| chr14 | 26637196 | 26637927 | 14.18495 | 33.44911 | 30.9408 | Promoter | 14 | 26638074 | 26657258 | 19185 | 1 | ENSMUST00 | -147 | Arf4 | ENSMUSG00000021877.6 |
| chr15 | 30172300 | 30173509 | 14.18495 | 33.44911 | 30.9408 | Promoter | 15 | 30172593 | 31029311 | 856719 | 1 | ENSMUST00 | 0 | Ctnnd2 | ENSMUSG00000022240.9 |
| chr4 | 117095805 | 117096454 | 14.18495 | 33.44911 | 30.9408 | Promoter | 4 | 117096075 | 117115383 | 19309 | 1 | ENSMUST00 | 0 | Ptch2 | ENSMUSG00000028681.6 |
| chr7 | 24174970 | 24175468 | 14.18495 | 33.44911 | 30.9408 | Promoter | 7 | 24176152 | 24182879 | 6728 | 1 | ENSMUST00 | -684 | Zfp114 | ENSMUSG00000068962.6 |
| chr9 | 120127025 | 120127564 | 14.15711 | 33.4017 | 30.89395 | Promoter | 9 | 120127766 | 120132369 | 4604 | 1 | ENSMUST00 | -202 | Rpsa | ENSMUSG00000032518.5 |
| chr17 | 28312972 | 28313369 | 14.5262 | 33.35895 | 30.8513 | Promoter | 17 | 28313530 | 28326567 | 13038 | 1 | ENSMUST00 | -161 | Fance | ENSMUSG00000007570.10 |
| chr3 | 13472696 | 13473450 | 14.5262 | 33.35895 | 30.8513 | Exon (ENSMU | 3 | 13471906 | 13474596 | 2691 | 2 | ENSMUST00 | 1146 | Gm2464 | ENSMUSG00000097338.1 |
| chr5 | 73338026 | 73338932 | 14.5262 | 33.35895 | 30.8513 | Promoter | 5 | 73322199 | 73338593 | 16395 | 2 | ENSMUST00 | 0 | Ociad2 | ENSMUSG00000029153.8 |
| chr7 | 25781465 | 25782062 | 14.5262 | 33.35895 | 30.8513 | Intron (ENSM | 7 | 25778680 | 25787066 | 8387 | 2 | ENSMUST00 | 5004 | Axl | ENSMUSG00000002602.11 |
| chr10 | 77339774 | 77340238 | 14.12939 | 33.35446 | 30.84692 | Promoter | 10 | 77325715 | 77340176 | 14462 | 2 | ENSMUST00 | 0 | Adarb1 | ENSMUSG00000020262.10 |
| chr9 | 108991098 | 108991644 | 14.4961 | 33.30813 | 30.80067 | Promoter | 9 | 108991902 | 109010572 | 18671 | 1 | ENSMUST00 | -258 | Pfkfb4 | ENSMUSG00000025648.12 |
| chr15 | 102671274 | 102672865 | 14.10177 | 33.30739 | 30.80003 | Promoter | 15 | 102665801 | 102672352 | 6552 | 2 | ENSMUST00 | 0 | Atp5g2 | ENSMUSG00000062683.6 |
| chr7 | 19131692 | 19132157 | 14.10177 | 33.30739 | 30.80003 | Promoter | 7 | 19131693 | 19137812 | 6120 | 1 | ENSMUST00 | 0 | Fbxo46 | ENSMUSG00000050428.6 |
| chr1 | 52844480 | 52844797 | 12.58327 | 33.30664 | 30.79935 | Promoter | 1 | 52844929 | 52848693 | 3765 | 1 | ENSMUST00 | -132 | Hibch | ENSMUSG00000041426.7 |
| chr12 | 110991791 | 110992562 | 12.58327 | 33.30664 | 30.79935 | Distal Interge | 12 | 110975353 | 110979040 | 3688 | 2 | ENSMUST00 | -12752 | Ankrd9 | ENSMUSG00000037904.9 |
| chr12 | 84175689 | 84176234 | 12.58327 | 33.30664 | 30.79935 | Intron (ENSM | 12 | 84156528 | 84160138 | 3611 | 2 | ENSMUST00 | -15552 | Elmsan1 | ENSMUSG00000042507.9 |
| chr12 | 105453638 | 105454485 | 12.58327 | 33.30664 | 30.79935 | Promoter | 12 | 105453856 | 105493095 | 39240 | 1 | ENSMUST00 | 0 | D430019H16 | ENSMUSG00000094910.1 |
| chr13 | 21233545 | 21234742 | 12.58327 | 33.30664 | 30.79935 | Distal Interge | 13 | 21190092 | 21192475 | 2384 | 1 | ENSMUST00 | 43454 | Trim27 | ENSMUSG00000021326.6 |
| chr13 | 112993511 | 112994366 | 12.58327 | 33.30664 | 30.79935 | Promoter | 13 | 112993868 | 113000394 | 6527 | 1 | ENSMUST00 | 0 | Mcidas | ENSMUSG00000074651.2 |
| chr15 | 74516050 | 74517226 | 12.58327 | 33.30664 | 30.79935 | Promoter | 15 | 74516195 | 74589464 | 73270 | 1 | ENSMUST00 | 0 | Bai1 | ENSMUSG00000034730.11 |
| chr16 | 77593935 | 77596149 | 12.58327 | 33.30664 | 30.79935 | Promoter | 16 | 77594366 | 77594967 | 602 | 1 | ENSMUST00 | 0 | 2810055G20 | ENSMUSG00000090386.4 |
| chr17 | 15536080 | 15536778 | 12.58327 | 33.30664 | 30.79935 | Distal Interge | 17 | 15521575 | 15527301 | 5727 | 2 | ENSMUST00 | -8780 | Pdcd2 | ENSMUSG00000014771.7 |
| chr17 | 37049809 | 37051239 | 12.58327 | 33.30664 | 30.79935 | Promoter | 17 | 37050631 | 37074304 | 23674 | 1 | ENSMUST00 | 0 | Gabbr1 | ENSMUSG00000024462.12 |
| chr18 | 34578611 | 34579243 | 12.58327 | 33.30664 | 30.79935 | Promoter | 18 | 34562634 | 34579100 | 16467 | 2 | ENSMUST00 | 0 | Nme5 | ENSMUSG00000035984.9 |
| chr18 | 36195978 | 36197656 | 12.58327 | 33.30664 | 30.79935 | Promoter | 18 | 36017707 | 36197380 | 179674 | 2 | ENSMUST00 | 0 | Nrg2 | ENSMUSG00000060275.6 |
| chr19 | 44314370 | 44315405 | 12.58327 | 33.30664 | 30.79935 | Distal Interge | 19 | 44303171 | 44303231 | 61 | 1 | ENSMUST00 | 11200 | Mir5114 | ENSMUSG00000093315.1 |
| chr2 | 34874154 | 34874986 | 12.58327 | 33.30664 | 30.79935 | Promoter | 2 | 34866939 | 34874968 | 8030 | 2 | ENSMUST00 | 0 | Psmd5 | ENSMUSG00000026869.7 |
| chr2 | 24367188 | 24367899 | 12.58327 | 33.30664 | 30.79935 | Promoter | 2 | 24367580 | 24394258 | 26679 | 1 | ENSMUST00 | 0 | Psd4 | ENSMUSG00000026979.11 |
| chr4 | 148742784 | 148743183 | 12.58327 | 33.30664 | 30.79935 | Distal Interge | 4 | 148713703 | 148716581 | 2879 | 1 | ENSMUST00 | 29082 | Gm13203 | ENSMUSG00000087273.1 |
| chr7 | 29309140 | 29309827 | 12.58327 | 33.30664 | 30.79935 | Promoter | 7 | 29309513 | 29313135 | 3623 | 1 | ENSMUST00 | 0 | Dpf1 | ENSMUSG00000030584.8 |
| chr7 | 104218771 | 104219653 | 12.58327 | 33.30664 | 30.79935 | Promoter | 7 | 104218793 | 104227733 | 8941 | 1 | ENSMUST00 | 0 | Trim6 | ENSMUSG00000072244.4 |
| chr8 | 110969912 | 110971162 | 12.58327 | 33.30664 | 30.79935 | 3' UTR | 8 | 110978542 | 110980634 | 2093 | 2 | ENSMUST00 | 9472 | Ddx19a | ENSMUSG00000015023.7 |
| chr9 | 45138152 | 45139055 | 12.58327 | 33.30664 | 30.79935 | Promoter | 9 | 45138437 | 45154152 | 15716 | 1 | ENSMUST00 | 0 | Scn4b | ENSMUSG00000046480.5 |
| chr9 | 60648251 | 60649883 | 12.58327 | 33.30664 | 30.79935 | Promoter | 9 | 60569172 | 60649801 | 80630 | 2 | ENSMUST00 | 0 | Lrrc49 | ENSMUSG00000047766.10 |
| chrX | 74270824 | 74272232 | 12.58327 | 33.30664 | 30.79935 | Promoter | 20 | 74270857 | 74273025 | 2169 | 1 | ENSMUST00 | 0 | Rpl10 | ENSMUSG00000008682.8 |
| chrX | 20703768 | 20705244 | 12.58327 | 33.30664 | 30.79935 | Promoter | 20 | 20703906 | 20720539 | 16634 | 1 | ENSMUST00 | 0 | Usp11 | ENSMUSG00000031066.5 |
| chr2 | 125723298 | 125724511 | 12.932 | 33.27539 | 30.76948 | Promoter | 2 | 125627447 | 125723387 | 95941 | 2 | ENSMUST00 | 0 | Shc4 | ENSMUSG00000035109.9 |
| chr4 | 136147444 | 136148224 | 14.46613 | 33.25751 | 30.75175 | Distal Interge | 4 | 136144974 | 136145755 | 782 | 1 | ENSMUST00 | 2471 | Id3 | ENSMUSG00000007872.3 |
| chr6 | 147263220 | 147264521 | 10.61933 | 33.25416 | 30.74849 | Promoter | 6 | 147252101 | 147264167 | 12067 | 2 | ENSMUST00 | 0 | Pthlh | ENSMUSG00000048776.6 |
| chr16 | 20695643 | 20696368 | 14.43627 | 33.20709 | 30.70158 | Promoter | 16 | 20696175 | 20701659 | 5485 | 1 | ENSMUST00 | 0 | Fam131a | ENSMUSG00000050821.8 |
| chr1 | 137927850 | 137928914 | 14.01957 | 33.16717 | 30.66176 | Exon (ENSMU | 1 | 137917690 | 137921312 | 3623 | 2 | ENSMUST00 | -6539 | Gm26979 | ENSMUSG00000098045.1 |
| chr10 | 86302922 | 86303457 | 14.01957 | 33.16717 | 30.66176 | Promoter | 10 | 86302854 | 86343981 | 41128 | 1 | ENSMUST00 | 69 | Timp3 | ENSMUSG00000020044.8 |
| chr11 | 117930684 | 117931423 | 14.40655 | 33.15686 | 30.65151 | Distal Interge | 11 | 117921905 | 117922328 | 424 | 1 | ENSMUST00 | 8780 | Gm11726 | ENSMUSG00000082562.1 |
| chr17 | 45544787 | 45545945 | 13.22462 | 33.12865 | 30.6234 | 3' UTR | 17 | 45541940 | 45549677 | 7738 | 2 | ENSMUST00 | 3732 | Tmem151b | ENSMUSG00000096847.1 |
| chr4 | 56989770 | 56990548 | 13.99238 | 33.12076 | 30.61571 | Promoter | 4 | 56957173 | 56990391 | 33219 | 2 | ENSMUST00 | 0 | Frrs1l | ENSMUSG00000045589.7 |
| chr1 | 56971395 | 56971814 | 14.37694 | 33.10684 | 30.60184 | Promoter | 1 | 56971412 | 56975197 | 3786 | 1 | ENSMUST00 | 0 | 9130024F11I | ENSMUSG00000087022.3 |
| chr2 | 79634694 | 79635165 | 14.37694 | 33.10684 | 30.60184 | Promoter | 2 | 79635352 | 79672966 | 37615 | 1 | ENSMUST00 | -187 | Ssfa2 | ENSMUSG00000027007.11 |
| chr4 | 151128380 | 151129581 | 14.37694 | 33.10684 | 30.60184 | Promoter | 4 | 151059814 | 151129495 | 69682 | 2 | ENSMUST00 | 0 | Camta1 | ENSMUSG00000014592.14 |
| chr13 | 113034844 | 113035598 | 11.44553 | 33.08637 | 30.58152 | Promoter | 13 | 113035111 | 113038324 | 3214 | 1 | ENSMUST00 | 0 | Cdc20b | ENSMUSG00000078926.3 |
| chr15 | 77955723 | 77957008 | 11.44553 | 33.08637 | 30.58152 | Promoter | 15 | 77940522 | 77956722 | 16201 | 2 | ENSMUST00 | 0 | Foxred2 | ENSMUSG00000016552.8 |
| chr17 | 25836594 | 25837370 | 11.44553 | 33.08637 | 30.58152 | Promoter | 17 | 25834465 | 25837295 | 2831 | 2 | ENSMUST00 | 0 | Rhbdl1 | ENSMUSG00000025735.8 |
| chr17 | 55970210 | 55970902 | 11.44553 | 33.08637 | 30.58152 | Promoter | 17 | 55970467 | 55976617 | 6151 | 1 | ENSMUST00 | 0 | Shd | ENSMUSG00000039154.6 |
| chr18 | 12333647 | 12334572 | 11.44553 | 33.08637 | 30.58152 | Promoter | 18 | 12333819 | 12583013 | 249195 | 1 | ENSMUST00 | 0 | Lama3 | ENSMUSG00000024421.11 |
| chr19 | 36736398 | 36737119 | 11.44553 | 33.08637 | 30.58152 | Promoter | 19 | 36731737 | 36736653 | 4917 | 2 | ENSMUST00 | 0 | Ppp1r3c | ENSMUSG00000067279.2 |
| chr3 | 142559873 | 142560833 | 11.44553 | 33.08637 | 30.58152 | Promoter | 3 | 142560052 | 142573202 | 13151 | 1 | ENSMUST00 | 0 | Gbp3 | ENSMUSG00000028268.9 |
| chr9 | 78520556 | 78521574 | 11.44553 | 33.08637 | 30.58152 | Distal Interge | 9 | 78479649 | 78489151 | 9503 | 2 | ENSMUST00 | -31406 | Eef1a1 | ENSMUSG00000037742.9 |
| chr1 | 153345734 | 153346574 | 14.34746 | 33.057 | 30.55262 | Distal Interge | 1 | 153218922 | 153332786 | 113865 | 2 | ENSMUST00 | -12949 | Lamc1 | ENSMUSG00000026478.9 |
| chr11 | 110398875 | 110399963 | 13.54925 | 33.02757 | 30.52334 | Promoter | 11 | 110399122 | 110513641 | 114520 | 1 | ENSMUST00 | 0 | Map2k6 | ENSMUSG00000020623.6 |
| chr6 | 37870207 | 37870570 | 13.54925 | 33.02757 | 30.52334 | Promoter | 6 | 37870811 | 37921505 | 50695 | 1 | ENSMUST00 | -241 | Trim24 | ENSMUSG00000029833.12 |
| chr7 | 6728931 | 6729561 | 13.54925 | 33.02757 | 30.52334 | Promoter | 7 | 6709395 | 6730419 | 21025 | 2 | ENSMUST00 | 858 | Peg3 | ENSMUSG00000002265.10 |
| chr8 | 48136998 | 48138471 | 13.54925 | 33.02757 | 30.52334 | 3' UTR | 8 | 48110211 | 48153233 | 43023 | 1 | ENSMUST00 | 26788 | Dctd | ENSMUSG00000031562.10 |
| chr19 | 57359910 | 57360885 | 14.3181 | 33.00736 | 30.50354 | Promoter | 19 | 57323385 | 57360775 | 37391 | 2 | ENSMUST00 | 0 | B230217O12 | ENSMUSG00000097785.1 |
| chr2 | 157130898 | 157131574 | 14.3181 | 33.00736 | 30.50354 | Promoter | 2 | 157121847 | 157130057 | 8211 | 2 | ENSMUST00 | -842 | Samhd1 | ENSMUSG00000027639.11 |
| chr10 | 120428392 | 120429562 | 13.91144 | 32.98249 | 30.47944 | Intron (ENSM | 10 | 120447991 | 120448110 | 120 | 2 | ENSMUST00 | 18548 | Mir763 | ENSMUSG00000076448.1 |
| chr11 | 102893083 | 102893719 | 13.91144 | 32.98249 | 30.47944 | Exon (ENSMU | 11 | 102887336 | 102897131 | 9796 | 2 | ENSMUST00 | 3412 | Gfap | ENSMUSG00000020932.9 |
| chr11 | 117332013 | 117332741 | 13.14203 | 32.98289 | 30.47944 | Promoter | 11 | 117332345 | 117362324 | 29980 | 1 | ENSMUST00 | 0 | Sep-09 | ENSMUSG00000059248.8 |
| chr11 | 94277276 | 94277735 | 13.14203 | 32.98289 | 30.47944 | Distal Interge | 11 | 94284652 | 94285985 | 1334 | 1 | ENSMUST00 | -6917 | Gm21885 | ENSMUSG00000094091.1 |
| chr12 | 57211543 | 57212345 | 13.14203 | 32.98289 | 30.47944 | Distal Interge | 12 | 56712634 | 57197472 | 484839 | 2 | ENSMUST00 | -14072 | Slc25a21 | ENSMUSG00000035472.8 |
| chr13 | 64343879 | 64344471 | 13.14203 | 32.98289 | 30.47944 | Distal Interge | 13 | 64363214 | 64370306 | 7093 | 2 | ENSMUST00 | 25835 | Ctsl | ENSMUSG00000021477.8 |
| chr13 | 31625127 | 31627193 | 13.14203 | 32.98289 | 30.47944 | Promoter | 13 | 31625816 | 31631403 | 5588 | 1 | ENSMUST00 | 0 | Foxf2 | ENSMUSG00000038402.2 |
| chr13 | 23432740 | 23433428 | 13.91144 | 32.98249 | 30.47944 | Distal Interge | 13 | 23427975 | 23431022 | 3048 | 2 | ENSMUST00 | -1719 | C230035I16R | ENSMUSG00000085024.1 |
| chr14 | 69823383 | 69824161 | 13.14203 | 32.98289 | 30.47944 | Distal Interge | 14 | 69840420 | 70059886 | 219467 | 1 | ENSMUST00 | -16259 | Pebp4 | ENSMUSG00000022085.3 |
| chr15 | 81936283 | 81937895 | 13.91144 | 32.98249 | 30.47944 | Promoter | 15 | 81936759 | 81950941 | 14183 | 1 | ENSMUST00 | 0 | Csdc2 | ENSMUSG00000042109.7 |
| chr15 | 103272255 | 103272634 | 13.14203 | 32.98289 | 30.47944 | Promoter | 15 | 103272911 | 103299862 | 26952 | 1 | ENSMUST00 | -277 | Copz1 | ENSMUSG00000060992.8 |
| chr15 | 38384184 | 38385474 | 13.14203 | 32.98289 | 30.47944 | Promoter | 15 | 38384981 | 38386300 | 1320 | 1 | ENSMUST00 | 0 | Gm26863 | ENSMUSG00000097645.1 |
| chr17 | 10334584 | 10336344 | 13.14203 | 32.98289 | 30.47944 | Distal Interge | 17 | 10324602 | 10329307 | 4706 | 1 | ENSMUST00 | 9983 | RP24-162O2 | ENSMUSG00000099384.1 |
| chr18 | 31938961 | 31939495 | 13.14203 | 32.98289 | 30.47944 | Intron (ENSM | 18 | 31931507 | 31958619 | 27113 | 1 | ENSMUST00 | 7455 | Lims2 | ENSMUSG00000024395.7 |
| chr18 | 23041049 | 23041930 | 13.14203 | 32.98289 | 30.47944 | Promoter | 18 | 22693634 | 23041653 | 348020 | 2 | ENSMUST00 | 0 | Nol4 | ENSMUSG00000041923.10 |
| chr18 | 76034773 | 76035255 | 13.14203 | 32.98289 | 30.47944 | Intron (ENSM | 18 | 76059458 | 76146443 | 86986 | 1 | ENSMUST00 | -24203 | Zbtb7c | ENSMUSG00000044646.9 |
| chr2 | 72242138 | 72242504 | 13.14203 | 32.98289 | 30.47944 | Promoter | 2 | 72242294 | 72256655 | 14362 | 1 | ENSMUST00 | 0 | Rapgef4 | ENSMUSG00000049044.11 |
| chr2 | 25185643 | 25186369 | 13.14203 | 32.98289 | 30.47944 | Distal Interge | 2 | 25180758 | 25183339 | 2582 | 1 | ENSMUST00 | 4886 | Nrarp | ENSMUSG00000078202.3 |
| chr2 | 152048491 | 152049467 | 13.14203 | 32.98289 | 30.47944 | Distal Interge | 2 | 152074855 | 152076808 | 1954 | 2 | ENSMUST00 | 27341 | AA387200 | ENSMUSG00000085946.1 |
| chr3 | 148959034 | 148960036 | 13.14203 | 32.98289 | 30.47944 | Intron (ENSM | 3 | 148815586 | 148954635 | 139050 | 2 | ENSMUST00 | -4400 | Lphn2 | ENSMUSG00000028184.8 |
| chr6 | 113305833 | 113306507 | 13.14203 | 32.98289 | 30.47944 | Promoter | 6 | 113307181 | 113324857 | 17677 | 1 | ENSMUST00 | -674 | Brpf1 | ENSMUSG00000001632.11 |
| chr7 | 145302113 | 145302634 | 13.14203 | 32.98289 | 30.47944 | Intron (ENSM | 7 | 145300899 | 145309349 | 8451 | 1 | ENSMUST00 | 1215 | Mrgprf | ENSMUSG00000031070.10 |
| chr7 | 99468120 | 99469149 | 13.14203 | 32.98289 | 30.47944 | Promoter | 7 | 99468963 | 99471786 | 2824 | 1 | ENSMUST00 | 0 | Klhl35 | ENSMUSG00000035298.3 |
| chr9 | 94669728 | 94670600 | 13.14203 | 32.98289 | 30.47944 | Promoter | 9 | 94669909 | 95230445 | 560537 | 1 | ENSMUST00 | 0 | Slc9a9 | ENSMUSG00000031129.9 |
| chr9 | 40241510 | 40242244 | 13.14203 | 32.98289 | 30.47944 | Distal Interge | 9 | 40269216 | 40291617 | 22402 | 1 | ENSMUST00 | -26972 | Scn3b | ENSMUSG00000049281.10 |
| chr15 | 102942884 | 102943530 | 14.28886 | 32.9579 | 30.45679 | Downstream | 15 | 102945359 | 102945427 | 69 | 1 | ENSMUST00 | -1829 | Gm27337 | ENSMUSG00000099099.1 |
| chr15 | 3356432 | 3356825 | 14.25973 | 32.90864 | 30.40768 | Intron (ENSM | 15 | 3280627 | 3303526 | 22900 | 2 | ENSMUST00 | -52907 | Ccdc152 | ENSMUSG00000091119.1 |
| chr10 | 80329422 | 80330508 | 13.83063 | 32.8443 | 30.34371 | Promoter | 10 | 80321283 | 80329453 | 8171 | 2 | ENSMUST00 | 0 | Pcsk4 | ENSMUSG00000020131.9 |
| chr11 | 96345436 | 96346158 | 13.83063 | 32.8443 | 30.34371 | Exon (ENSMU | 11 | 96343769 | 96346507 | 2739 | 1 | ENSMUST00 | 1668 | Hoxb3 | ENSMUSG00000048763.6 |
| chr11 | 101640920 | 101641727 | 13.83063 | 32.8443 | 30.34371 | Promoter | 11 | 101641748 | 101641938 | 191 | 1 | ENSMUST00 | -21 | Gm24950 | ENSMUSG00000064702.1 |
| chr13 | 95960851 | 95961885 | 13.83063 | 32.8443 | 30.34371 | 3' UTR | 13 | 95954594 | 95957171 | 2578 | 2 | ENSMUST00 | -3681 | Gm27031 | ENSMUSG00000098220.1 |
| chr14 | 50956195 | 50956465 | 13.83063 | 32.8443 | 30.34371 | Promoter | 14 | 50955992 | 50964749 | 8758 | 1 | ENSMUST00 | 204 | Pnp2 | ENSMUSG00000068417.7 |
| chr15 | 77358496 | 77358851 | 13.83063 | 32.8443 | 30.34371 | Distal Interge | 15 | 77391702 | 77392381 | 680 | 2 | ENSMUST00 | 33530 | Gm20688 | ENSMUSG00000093458.1 |
| chr16 | 22548186 | 22548727 | 13.83063 | 32.8443 | 30.34371 | Promoter | 16 | 22548553 | 22551149 | 2597 | 1 | ENSMUST00 | 0 | Gm15651 | ENSMUSG00000089663.1 |
| chr17 | 32528467 | 32528914 | 13.83063 | 32.8443 | 30.34371 | Exon (ENSMU | 17 | 32536558 | 32551798 | 15241 | 1 | ENSMUST00 | -7644 | Cyp4f16 | ENSMUSG00000048440.10 |
| chr18 | 12971989 | 12973254 | 13.83063 | 32.8443 | 30.34371 | Promoter | 18 | 12972252 | 12992948 | 20697 | 1 | ENSMUST00 | 0 | Impact | ENSMUSG00000024423.5 |
| chr19 | 18749576 | 18750221 | 13.83063 | 32.8443 | 30.34371 | Promoter | 19 | 18749983 | 18892510 | 142528 | 1 | ENSMUST00 | 0 | Trpm6 | ENSMUSG00000024727.8 |
| chr19 | 10240363 | 10240950 | 13.83063 | 32.8443 | 30.34371 | Promoter | 19 | 10208272 | 10240748 | 32477 | 2 | ENSMUST00 | 0 | Myrf | ENSMUSG00000036098.10 |
| chr2 | 147193879 | 147195631 | 13.83063 | 32.8443 | 30.34371 | Promoter | 2 | 147186305 | 147194243 | 7939 | 2 | ENSMUST00 | 0 | Nkx2-2 | ENSMUSG00000027434.11 |
| chr2 | 162943188 | 162943869 | 13.83063 | 32.8443 | 30.34371 | Promoter | 2 | 162943472 | 162972021 | 28550 | 1 | ENSMUST00 | 0 | L3mbtl1 | ENSMUSG00000035576.8 |
| chr2 | 148045749 | 148047027 | 13.83063 | 32.8443 | 30.34371 | Promoter | 2 | 148042877 | 148045948 | 3072 | 2 | ENSMUST00 | 0 | Foxa2 | ENSMUSG00000037025.6 |
| chr3 | 107517308 | 107518234 | 13.83063 | 32.8443 | 30.34371 | Promoter | 3 | 107467543 | 107518018 | 50476 | 2 | ENSMUST00 | 0 | Slc6a17 | ENSMUSG00000027894.9 |
| chr3 | 89320616 | 89323083 | 13.83063 | 32.8443 | 30.34371 | Promoter | 3 | 89313899 | 89322965 | 9067 | 2 | ENSMUST00 | 0 | Efna3 | ENSMUSG00000028039.7 |
| chr3 | 52196002 | 52196469 | 13.83063 | 32.8443 | 30.34371 | Distal Interge | 3 | 52268337 | 52350109 | 81773 | 1 | ENSMUST00 | -71868 | Foxo1 | ENSMUSG00000044167.5 |
| chr4 | 97774995 | 97776568 | 13.83063 | 32.8443 | 30.34371 | Intron (ENSM | 4 | 97777608 | 98118871 | 341264 | 1 | ENSMUST00 | -1040 | Nfia | ENSMUSG00000028565.13 |
| chr4 | 141238949 | 141240145 | 13.83063 | 32.8443 | 30.34371 | Promoter | 4 | 141239499 | 141246412 | 6914 | 1 | ENSMUST00 | 0 | Arhgef19 | ENSMUSG00000028919.6 |
| chr4 | 141247745 | 141248300 | 13.83063 | 32.8443 | 30.34371 | Promoter | 4 | 141246926 | 141247976 | 1051 | 1 | ENSMUST00 | 820 | Arhgef19 | ENSMUSG00000028919.6 |
| chr5 | 139796675 | 139797064 | 13.83063 | 32.8443 | 30.34371 | Promoter | 5 | 139797514 | 139802331 | 4818 | 1 | ENSMUST00 | -450 | Mafk | ENSMUSG00000018143.5 |
| chr5 | 148341821 | 148342904 | 13.83063 | 32.8443 | 30.34371 | Exon (ENSMU | 5 | 148352153 | 148371546 | 19394 | 2 | ENSMUST00 | 28642 | Slc7a1 | ENSMUSG00000041313.9 |
| chr5 | 110362823 | 110363863 | 13.83063 | 32.8443 | 30.34371 | 3' UTR | 5 | 110354158 | 110356031 | 1874 | 1 | ENSMUST00 | 8666 | Lrcol1 | ENSMUSG00000072754.4 |
| chr7 | 84086151 | 84086853 | 13.83063 | 32.8443 | 30.34371 | Promoter | 7 | 83932857 | 84086502 | 153646 | 2 | ENSMUST00 | 0 | Cemip | ENSMUSG00000052353.8 |
| chrX | 102070296 | 102071600 | 13.83063 | 32.8443 | 30.34371 | Promoter | 20 | 102066544 | 102071196 | 4653 | 2 | ENSMUST00 | 0 | Rgag4 | ENSMUSG00000049191.7 |
| chrY | 1244853 | 1246203 | 13.83063 | 32.8443 | 30.34371 | Promoter | 21 | 1096861 | 1245691 | 148831 | 2 | ENSMUST00 | 0 | Uty | ENSMUSG00000068457.9 |
| chr5 | 102724794 | 102726565 | 14.20184 | 32.81068 | 30.31183 | Promoter | 5 | 102724973 | 102897829 | 172857 | 1 | ENSMUST00 | 0 | Arhgap24 | ENSMUSG00000057315.9 |
| chr6 | 31542598 | 31548466 | 14.20184 | 32.81068 | 30.31183 | Intron (ENSM | 6 | 31519488 | 31563981 | 44494 | 2 | ENSMUST00 | 15515 | Podxl | ENSMUSG00000025608.9 |
| chr15 | 102224350 | 102225040 | 14.17307 | 32.76197 | 30.26328 | Exon (ENSMU | 15 | 102215995 | 102231935 | 15941 | 2 | ENSMUST00 | 6895 | Itgb7 | ENSMUSG00000001281.4 |
| chr9 | 50462691 | 50463069 | 14.17307 | 32.76197 | 30.26328 | Distal Interge | 9 | 50494525 | 50505482 | 10958 | 1 | ENSMUST00 | -31456 | Plet1 | ENSMUSG00000032068.9 |
| chr17 | 35359058 | 35359791 | 12.99996 | 32.73146 | 30.23292 | Promoter | 17 | 35358989 | 35359972 | 984 | 1 | ENSMUST00 | 70 | H2-Q3 | ENSMUSG00000092253.1 |
| chr17 | 30657942 | 30658531 | 12.99996 | 32.73146 | 30.23292 | Exon (ENSMU | 17 | 30622441 | 30626905 | 4465 | 2 | ENSMUST00 | -31038 | RP23-362B7. | ENSMUSG00000099474.1 |
| chr7 | 45277187 | 45277586 | 12.99996 | 32.73146 | 30.23292 | Promoter | 7 | 45277513 | 45288994 | 11482 | 1 | ENSMUST00 | 0 | 1700039E15I | ENSMUSG00000070568.5 |
| chr10 | 80295669 | 80296375 | 14.11588 | 32.66511 | 30.16695 | Promoter | 10 | 80295977 | 80318263 | 22287 | 1 | ENSMUST00 | 0 | Apc2 | ENSMUSG00000020135.8 |
| chr11 | 97025529 | 97026251 | 14.11588 | 32.66511 | 30.16695 | Promoter | 11 | 97020508 | 97026248 | 5741 | 2 | ENSMUST00 | 0 | Gm11532 | ENSMUSG00000087067.1 |
| chr13 | 21408148 | 21409023 | 13.70011 | 32.62073 | 30.12267 | Distal Interge | 13 | 21387013 | 21402755 | 15743 | 2 | ENSMUST00 | -5394 | Zkscan3 | ENSMUSG00000021327.13 |
| chr10 | 19185709 | 19186178 | 13.30671 | 32.60603 | 30.10809 | Distal Interge | 10 | 19008266 | 19015657 | 7392 | 2 | ENSMUST00 | -170053 | Tnfaip3 | ENSMUSG00000019850.6 |
| chr5 | 135834781 | 135835840 | 13.30671 | 32.60603 | 30.10809 | 5' UTR | 5 | 135837295 | 135837372 | 78 | 2 | ENSMUST00 | 1532 | Gm23268 | ENSMUSG00000094486.1 |

| | | | | | | | | | | | | | | | |
|---|---|---|---|---|---|---|---|---|---|---|---|---|---|---|---|
| chr7 | 143461259 | 143461604 | 13.30671 | 32.60603 | 30.10809 | Promoter | 7 | 143458350 | 143461050 | 2701 | 2 | ENSMUST00 | -210 | Cdkn1c | ENSMUSG00000037664.7 |
| chr4 | 136454867 | 136455300 | 10.86952 | 32.54407 | 30.04659 | Distal Interge | 4 | 136462250 | 136463977 | 1728 | 1 | ENSMUST00 | -6950 | 6030445D17I | ENSMUSG00000066178.2 |
| chr8 | 105268532 | 105270228 | 10.86952 | 32.54407 | 30.04659 | Promoter | 8 | 105269801 | 105275686 | 5886 | 1 | ENSMUST00 | 0 | Hsf4 | ENSMUSG00000033249.5 |
| chr1 | 5916590 | 5918001 | 13.64828 | 32.53182 | 30.03459 | Promoter | 1 | 5913707 | 5917398 | 3692 | 2 | ENSMUST00 | 0 | Npbwr1 | ENSMUSG00000033774.4 |
| chr15 | 79883113 | 79883856 | 13.64828 | 32.53182 | 30.03459 | Distal Interge | 15 | 79891659 | 79905547 | 13889 | 1 | ENSMUST00 | -7803 | Apobec3 | ENSMUSG00000009585.12 |
| chr2 | 127362704 | 127363849 | 13.64828 | 32.53182 | 30.03459 | Promoter | 2 | 127363208 | 127367221 | 4014 | 1 | ENSMUST00 | 0 | Adra2b | ENSMUSG00000058620.5 |
| chr4 | 151058172 | 151058575 | 13.64828 | 32.53182 | 30.03459 | Promoter | 4 | 151055911 | 151057963 | 2053 | 2 | ENSMUST00 | -210 | Vamp3 | ENSMUSG00000028955.3 |
| chr5 | 148923479 | 148924297 | 13.64828 | 32.53182 | 30.03459 | Intron (ENSM | 5 | 148904593 | 148928628 | 24036 | 2 | ENSMUST00 | 4331 | Katnal1 | ENSMUSG00000041298.10 |
| chr8 | 121585611 | 121586005 | 13.64828 | 32.53182 | 30.03459 | Intron (ENSM | 8 | 121581765 | 121582191 | 427 | 1 | ENSMUST00 | 3847 | Gm17786 | ENSMUSG00000097987.1 |
| chr13 | 44839940 | 44841021 | 14.03095 | 32.52118 | 30.02454 | Promoter | 13 | 44840691 | 44914319 | 73629 | 1 | ENSMUST00 | 0 | Jarid2 | ENSMUSG00000038518.10 |
| chr14 | 70553906 | 70555094 | 14.03095 | 32.52118 | 30.02454 | Promoter | 14 | 70554056 | 70573548 | 19493 | 1 | ENSMUST00 | 0 | Hr | ENSMUSG00000022096.9 |
| chr3 | 146220786 | 146221522 | 14.03095 | 32.52118 | 30.02454 | Promoter | 3 | 146220961 | 146286214 | 65254 | 1 | ENSMUST00 | 0 | Lpar3 | ENSMUSG00000036832.4 |
| chr11 | 6444643 | 6445190 | 11.80999 | 32.51257 | 30.01635 | Promoter | 11 | 6427229 | 6444443 | 17215 | 2 | ENSMUST00 | -201 | H2afv | ENSMUSG00000041126.11 |
| chr13 | 72639040 | 72640294 | 11.80999 | 32.51257 | 30.01635 | Distal Interge | 13 | 72632597 | 72633770 | 1174 | 1 | ENSMUST00 | 6444 | Irx2 | ENSMUSG00000001504.10 |
| chr13 | 89741778 | 89742945 | 11.80999 | 32.51257 | 30.01635 | Promoter | 13 | 89655312 | 89742509 | 87198 | 2 | ENSMUST00 | 0 | Vcan | ENSMUSG00000021614.11 |
| chr15 | 102404801 | 102405657 | 11.80999 | 32.51257 | 30.01635 | Promoter | 15 | 102406143 | 102408235 | 2093 | 1 | ENSMUST00 | -486 | Sp1 | ENSMUSG00000001280.7 |
| chr19 | 45076599 | 45077292 | 11.80999 | 32.51257 | 30.01635 | Promoter | 19 | 45076139 | 45079289 | 3151 | 1 | ENSMUST00 | 461 | Kazald1 | ENSMUSG00000025213.6 |
| chr19 | 42431427 | 42432531 | 11.80999 | 32.51257 | 30.01635 | Promoter | 19 | 42283037 | 42431783 | 148747 | 2 | ENSMUST00 | 0 | Crtac1 | ENSMUSG00000042401.6 |
| chr3 | 89280179 | 89281971 | 11.80999 | 32.51257 | 30.01635 | Promoter | 3 | 89271736 | 89281142 | 9407 | 2 | ENSMUST00 | 0 | Efna1 | ENSMUSG00000027954.4 |
| chr7 | 19300799 | 19301216 | 11.80999 | 32.51257 | 30.01635 | Promoter | 7 | 19298260 | 19300770 | 2511 | 2 | ENSMUST00 | -30 | Gm26802 | ENSMUSG00000097266.1 |
| chr8 | 48508648 | 48509535 | 11.80999 | 32.51257 | 30.01635 | Intron (ENSM | 8 | 48334753 | 48555693 | 220941 | 2 | ENSMUST00 | 46158 | Tenm3 | ENSMUSG00000031561.10 |
| chr15 | 77929161 | 77929701 | 14.00287 | 32.47356 | 29.97816 | Promoter | 15 | 77915047 | 77929006 | 13960 | 2 | ENSMUST00 | -156 | Txn2 | ENSMUSG00000005354.10 |
| chr2 | 131186305 | 131187478 | 14.00287 | 32.47356 | 29.97816 | Promoter | 2 | 131170261 | 131187282 | 17022 | 2 | ENSMUST00 | 0 | Spef1 | ENSMUSG00000027329.4 |
| chr2 | 118779129 | 118779887 | 14.00287 | 32.47356 | 29.97816 | Promoter | 2 | 118779719 | 118811174 | 31456 | 1 | ENSMUST00 | 0 | Disp2 | ENSMUSG00000040035.9 |
| chr6 | 108570723 | 108571632 | 14.00287 | 32.47356 | 29.97816 | Distal Interge | 6 | 108563445 | 108565693 | 2249 | 2 | ENSMUST00 | -5031 | Gm16433 | ENSMUSG00000043024.5 |
| chr9 | 102723447 | 102723832 | 14.00287 | 32.47356 | 29.97816 | Promoter | 9 | 102722534 | 102733418 | 10885 | 1 | ENSMUST00 | 914 | Amotl2 | ENSMUSG00000032531.10 |
| chr15 | 73809660 | 73809926 | 13.59684 | 32.4435 | 29.94819 | Promoter | 15 | 73809507 | 73809896 | 390 | 2 | ENSMUST00 | 0 | Gm3244 | ENSMUSG00000083380.1 |
| chr2 | 71980919 | 71981441 | 13.9749 | 32.42611 | 29.93083 | Promoter | 2 | 71981240 | 72257474 | 276235 | 1 | ENSMUST00 | 0 | Rapgef4 | ENSMUSG00000049044.11 |
| chr5 | 149006295 | 149007397 | 13.9749 | 32.42611 | 29.93083 | Promoter | 5 | 149006962 | 149012776 | 5815 | 1 | ENSMUST00 | 0 | Gm15408 | ENSMUSG00000086610.1 |
| chr11 | 77766313 | 77767112 | 13.57127 | 32.39957 | 29.9044 | Promoter | 11 | 77766736 | 77865980 | 99245 | 1 | ENSMUST00 | 0 | Myo18a | ENSMUSG00000000631.14 |
| chr11 | 100931177 | 100931983 | 13.94704 | 32.37885 | 29.8837 | Intron (ENSM | 11 | 100888236 | 100939380 | 51145 | 2 | ENSMUST00 | 7397 | Stat3 | ENSMUSG00000004040.11 |
| chr6 | 48537141 | 48538171 | 12.7828 | 32.34538 | 29.85033 | Promoter | 6 | 48537569 | 48541798 | 4230 | 1 | ENSMUST00 | 0 | Atp6v0e2 | ENSMUSG00000039347.5 |
| chr3 | 96449949 | 96450744 | 11.38976 | 32.33759 | 29.8427 | Promoter | 3 | 96450446 | 96450611 | 166 | 1 | ENSMUST00 | 0 | Gm26232 | ENSMUSG00000096838.1 |
| chr5 | 102768501 | 102770258 | 13.5204 | 32.31214 | 29.81732 | Promoter | 5 | 102768771 | 102897829 | 129059 | 1 | ENSMUST00 | 0 | Arhgap24 | ENSMUSG00000057315.9 |
| chr8 | 71405447 | 71406497 | 13.89166 | 32.28482 | 29.79003 | Promoter | 8 | 71406010 | 71409904 | 3895 | 1 | ENSMUST00 | 0 | Ankle1 | ENSMUSG00000046295.8 |
| chr10 | 103235961 | 103236447 | 14.2455 | 32.22549 | 29.73087 | Promoter | 10 | 103063198 | 103236322 | 173125 | 2 | ENSMUST00 | 0 | Lrriq1 | ENSMUSG00000019892.7 |
| chr11 | 115153806 | 115154288 | 14.2455 | 32.22549 | 29.73087 | Promoter | 11 | 115154157 | 115162236 | 8080 | 1 | ENSMUST00 | 0 | Rab37 | ENSMUSG00000020732.8 |
| chr12 | 17324274 | 17325141 | 14.2455 | 32.22549 | 29.73087 | Promoter | 12 | 17284721 | 17324712 | 39992 | 2 | ENSMUST00 | 0 | Atp6v1c2 | ENSMUSG00000020566.13 |
| chr12 | 118846716 | 118847398 | 14.2455 | 32.22549 | 29.73087 | Promoter | 12 | 118846329 | 118852578 | 6250 | 1 | ENSMUST00 | 388 | Sp8 | ENSMUSG00000048562.6 |
| chr12 | 30883955 | 30884965 | 14.2455 | 32.22549 | 29.73087 | Promoter | 12 | 30884322 | 30893854 | 9533 | 1 | ENSMUST00 | 0 | Fam150b | ENSMUSG00000054204.6 |
| chr13 | 120096030 | 120096565 | 14.2455 | 32.22549 | 29.73087 | Downstream | 13 | 120099349 | 120100332 | 984 | 2 | ENSMUST00 | 3767 | Gm21378 | ENSMUSG00000095709.1 |
| chr14 | 51358120 | 51358826 | 14.2455 | 32.22549 | 29.73087 | Downstream | 14 | 51361756 | 51569982 | 208227 | 1 | ENSMUST00 | -2930 | Gm7247 | ENSMUSG00000068399.5 |
| chr14 | 120831972 | 120832579 | 14.2455 | 32.22549 | 29.73087 | Distal Interge | 14 | 120798853 | 120798936 | 84 | 1 | ENSMUST00 | 33120 | Gm25781 | ENSMUSG00000084476.1 |
| chr17 | 46483616 | 46484335 | 14.2455 | 32.22549 | 29.73087 | 5' UTR | 17 | 46442448 | 46487675 | 45228 | 2 | ENSMUST00 | 3340 | Ttbk1 | ENSMUSG00000015599.8 |
| chr18 | 78047205 | 78047810 | 14.2455 | 32.22549 | 29.73087 | Exon (ENSMU | 18 | 78043614 | 78057395 | 13782 | 2 | ENSMUST00 | 9585 | Siglec15 | ENSMUSG00000091055.2 |
| chr19 | 32210123 | 32210979 | 14.2455 | 32.22549 | 29.73087 | Promoter | 19 | 32123479 | 32210990 | 87512 | 2 | ENSMUST00 | 11 | Sgms1 | ENSMUSG00000040451.12 |
| chr2 | 85060830 | 85061421 | 13.46992 | 32.22528 | 29.73087 | Intron (ENSM | 2 | 85058159 | 85073048 | 14890 | 1 | ENSMUST00 | 2672 | Tnks1bp1 | ENSMUSG00000033955.7 |
| chr2 | 176670467 | 176671081 | 14.2455 | 32.22549 | 29.73087 | Promoter | 2 | 176670522 | 176679874 | 9353 | 1 | ENSMUST00 | 0 | Gm14306 | ENSMUSG00000078879.5 |
| chr2 | 175965319 | 175965876 | 14.2455 | 32.22549 | 29.73087 | Promoter | 2 | 175965331 | 175977635 | 12305 | 1 | ENSMUST00 | 0 | Gm2026 | ENSMUSG00000078886.7 |
| chr3 | 137980890 | 137981715 | 14.2455 | 32.22549 | 29.73087 | Promoter | 3 | 137931007 | 137981545 | 50539 | 2 | ENSMUST00 | 0 | Dapp1 | ENSMUSG00000028159.9 |
| chr4 | 22489145 | 22490047 | 14.2455 | 32.22549 | 29.73087 | Promoter | 4 | 22482780 | 22488366 | 5587 | 2 | ENSMUST00 | -780 | Pou3f2 | ENSMUSG00000095139.2 |
| chr5 | 123038887 | 123039622 | 14.2455 | 32.22549 | 29.73087 | Exon (ENSMU | 5 | 123035862 | 123037712 | 1851 | 2 | ENSMUST00 | -1176 | Morn3 | ENSMUSG00000029477.9 |
| chr5 | 102481163 | 102482254 | 14.2455 | 32.22549 | 29.73087 | Promoter | 5 | 102481391 | 102897829 | 416439 | 1 | ENSMUST00 | 0 | Arhgap24 | ENSMUSG00000057315.9 |
| chr6 | 118011185 | 118012018 | 14.2455 | 32.22549 | 29.73087 | Distal Interge | 6 | 117997207 | 117997992 | 786 | 1 | ENSMUST00 | 13979 | Gm4875 | ENSMUSG00000082791.3 |
| chr7 | 138940359 | 138941459 | 14.2455 | 32.22549 | 29.73087 | Promoter | 7 | 138940730 | 139026888 | 86159 | 1 | ENSMUST00 | 0 | Jakmip3 | ENSMUSG00000056856.6 |
| chr8 | 88647105 | 88647818 | 14.2455 | 32.22549 | 29.73087 | Promoter | 8 | 88647315 | 88688474 | 41160 | 1 | ENSMUST00 | 0 | Nod2 | ENSMUSG00000055994.10 |
| chrX | 73659758 | 73660302 | 14.2455 | 32.22549 | 29.73087 | Promoter | 20 | 73655994 | 73659828 | 3835 | 2 | ENSMUST00 | 0 | Pnck | ENSMUSG00000002012.8 |
| chrX | 74392559 | 74393958 | 14.2455 | 32.22549 | 29.73087 | Promoter | 20 | 74393290 | 74433993 | 40704 | 1 | ENSMUST00 | 0 | Ikbkg | ENSMUSG00000004221.11 |
| chrX | 7789265 | 7789704 | 14.2455 | 32.22549 | 29.73087 | Promoter | 20 | 7789765 | 7820567 | 30803 | 1 | ENSMUST00 | -61 | Gripap1 | ENSMUSG00000031153.11 |
| chrX | 75381951 | 75382893 | 14.2455 | 32.22549 | 29.73087 | Promoter | 20 | 75172715 | 75382316 | 209602 | 2 | ENSMUST00 | 0 | F8 | ENSMUSG00000031196.8 |
| chrX | 105927957 | 105928607 | 14.2455 | 32.22549 | 29.73087 | Promoter | 20 | 105881958 | 105928577 | 46620 | 2 | ENSMUST00 | 0 | Atrx | ENSMUSG00000031229.11 |
| chrX | 73880391 | 73880858 | 14.2455 | 32.22549 | 29.73087 | Promoter | 20 | 73853955 | 73880834 | 26880 | 2 | ENSMUST00 | 0 | L1cam | ENSMUSG00000031391.13 |
| chrX | 103820678 | 103822759 | 14.2455 | 32.22549 | 29.73087 | Promoter | 20 | 103697414 | 103821965 | 124552 | 2 | ENSMUST00 | 0 | Slc16a2 | ENSMUSG00000033965.7 |
| chrX | 48034198 | 48035168 | 14.2455 | 32.22549 | 29.73087 | Promoter | 20 | 48025146 | 48034853 | 9708 | 2 | ENSMUST00 | 0 | Apln | ENSMUSG00000037010.7 |
| chrX | 36743605 | 36744128 | 14.2455 | 32.22549 | 29.73087 | Promoter | 20 | 36743659 | 36777202 | 33544 | 1 | ENSMUST00 | 0 | Slc25a43 | ENSMUSG00000037636.6 |
| chrX | 150440722 | 150441422 | 14.2455 | 32.22549 | 29.73087 | Promoter | 20 | 150397773 | 150441137 | 43365 | 2 | ENSMUST00 | 0 | Tmem29 | ENSMUSG00000041353.7 |
| chrX | 68820962 | 68821460 | 14.2455 | 32.22549 | 29.73087 | Promoter | 20 | 68821093 | 68825318 | 4226 | 1 | ENSMUST00 | 0 | Gm14698 | ENSMUSG00000071748.2 |
| chrX | 11664285 | 11665249 | 14.2455 | 32.22549 | 29.73087 | Distal Interge | 20 | 11684002 | 11684124 | 123 | 2 | ENSMUST00 | 18875 | Gm14513 | ENSMUSG00000080811.1 |
| chr14 | 118922555 | 118923223 | 13.0727 | 32.19727 | 29.70434 | Promoter | 14 | 118875520 | 118923167 | 47648 | 2 | ENSMUST00 | 0 | Dzip1 | ENSMUSG00000042156.10 |
| chr15 | 85205804 | 85206354 | 13.0727 | 32.19727 | 29.70434 | Promoter | 15 | 85205949 | 85286535 | 80587 | 1 | ENSMUST00 | 0 | Fbln1 | ENSMUSG00000006369.9 |
| chr2 | 75639299 | 75639865 | 13.0727 | 32.19727 | 29.70434 | Distal Interge | 2 | 75632638 | 75633382 | 745 | 2 | ENSMUST00 | -5918 | Gm13655 | ENSMUSG00000082152.1 |
| chr4 | 115095699 | 115096337 | 13.0727 | 32.19727 | 29.70434 | Distal Interge | 4 | 115090932 | 115093864 | 2933 | 1 | ENSMUST00 | 4768 | Pdzk1ip1 | ENSMUSG00000028716.10 |
| chr3 | 96268178 | 96268868 | 14.21987 | 32.18232 | 29.68973 | Promoter | 3 | 96268654 | 96269141 | 488 | 1 | ENSMUST00 | 0 | Hist2h3b | ENSMUSG00000074403.2 |
| chr8 | 126298356 | 126299059 | 14.21987 | 32.18232 | 29.68973 | Promoter | 8 | 126298558 | 126395482 | 96925 | 1 | ENSMUST00 | 0 | Slc35f3 | ENSMUSG00000057060.7 |
| chr1 | 170644101 | 170645106 | 12.32451 | 32.17465 | 29.68214 | Promoter | 1 | 170644532 | 170682789 | 38258 | 1 | ENSMUST00 | 0 | Olfml2b | ENSMUSG00000038463.8 |
| chr11 | 63951987 | 63952570 | 14.1901 | 32.13216 | 29.63978 | Distal Interge | 11 | 63964226 | 63968040 | 3815 | 1 | ENSMUST00 | -11656 | 9630013K17I | ENSMUSG00000086359.1 |
| chr6 | 52165559 | 52166190 | 14.1901 | 32.13216 | 29.63978 | Promoter | 6 | 52165674 | 52169451 | 3778 | 1 | ENSMUST00 | 0 | Hoxaas2 | ENSMUSG00000056445.8 |
| chr9 | 55048337 | 55049058 | 14.1901 | 32.13216 | 29.63978 | Promoter | 9 | 55028156 | 55048544 | 20389 | 2 | ENSMUST00 | 0 | Chrnb4 | ENSMUSG00000035200.6 |
| chr15 | 82354442 | 82355001 | 10.37237 | 32.10413 | 29.61181 | Promoter | 15 | 82350139 | 82354291 | 4153 | 2 | ENSMUST00 | -152 | Ndufa6 | ENSMUSG00000022450.5 |
| chr5 | 65198928 | 65199486 | 10.37237 | 32.10413 | 29.61181 | Promoter | 5 | 65199696 | 65260415 | 60720 | 1 | ENSMUST00 | -210 | Wdr19 | ENSMUSG00000037890.9 |
| chr11 | 97040971 | 97041402 | 13.3949 | 32.09606 | 29.60391 | Promoter | 11 | 97034602 | 97041407 | 6806 | 2 | ENSMUST00 | 5 | Lrrc46 | ENSMUSG00000020878.6 |
| chr11 | 44611066 | 44611887 | 13.3949 | 32.09606 | 29.60391 | Distal Interge | 11 | 44617317 | 45004995 | 387679 | 1 | ENSMUST00 | -5430 | Ebf1 | ENSMUSG00000057098.9 |
| chr17 | 34153128 | 34153765 | 13.3949 | 32.09606 | 29.60391 | Promoter | 17 | 34153072 | 34160230 | 7159 | 1 | ENSMUST00 | 57 | H2-DMb1 | ENSMUSG00000079547.3 |
| chr4 | 118446046 | 118446707 | 13.3949 | 32.09606 | 29.60391 | Exon (ENSMU | 4 | 118441013 | 118441220 | 208 | 2 | ENSMUST00 | -4827 | Gm12858 | ENSMUSG00000081921.1 |
| chr16 | 92341138 | 92341603 | 14.16046 | 32.08221 | 29.59053 | Distal Interge | 16 | 92348432 | 92358874 | 10443 | 2 | ENSMUST00 | 17271 | Kcne1 | ENSMUSG00000039639.6 |
| chr6 | 115838083 | 115838829 | 14.16046 | 32.08221 | 29.59053 | Promoter | 6 | 115810729 | 115838412 | 27684 | 2 | ENSMUST00 | 0 | Efcab12 | ENSMUSG00000030321.10 |
| chr15 | 11064632 | 11065872 | 13.7551 | 32.05274 | 29.56132 | Promoter | 15 | 11064790 | 11346867 | 282078 | 1 | ENSMUST00 | 0 | Adamts12 | ENSMUSG00000047497.9 |
| chr17 | 34933721 | 34934805 | 13.7551 | 32.05274 | 29.56132 | Exon (ENSMU | 17 | 34931863 | 34932765 | 903 | 1 | ENSMUST00 | 1859 | Neu1 | ENSMUSG00000007038.5 |
| chr1 | 135368265 | 135368833 | 14.13093 | 32.03245 | 29.5415 | Downstream | 1 | 135371060 | 135373450 | 2391 | 2 | ENSMUST00 | 4617 | Shisa4 | ENSMUSG00000041889.7 |
| chr11 | 96291649 | 96294061 | 14.13093 | 32.03245 | 29.5415 | Promoter | 11 | 96292476 | 96301569 | 9094 | 1 | ENSMUST00 | 0 | Hoxb6 | ENSMUSG00000000690.5 |
| chr15 | 99260052 | 99260513 | 14.13093 | 32.03245 | 29.5415 | Intron (ENSM | 15 | 99254882 | 99255014 | 133 | 2 | ENSMUST00 | -5039 | Gm25999 | ENSMUSG00000093139.1 |
| chr7 | 127993202 | 127994018 | 14.13093 | 32.03245 | 29.5415 | Promoter | 7 | 127992103 | 127993867 | 1765 | 2 | ENSMUST00 | 0 | Pycard | ENSMUSG00000030793.3 |
| chr1 | 135510116 | 135511017 | 12.24318 | 32.0257 | 29.53485 | Promoter | 1 | 135457782 | 135510384 | 52603 | 2 | ENSMUST00 | 0 | Nav1 | ENSMUSG00000009418.10 |
| chr1 | 167802629 | 167803394 | 12.24318 | 32.0257 | 29.53485 | Intron (ENSM | 1 | 167689557 | 167848741 | 159185 | 1 | ENSMUST00 | 113073 | Lmx1a | ENSMUSG00000026686.9 |
| chr1 | 15286407 | 15287186 | 12.24318 | 32.0257 | 29.53485 | Promoter | 1 | 15287254 | 15712548 | 425295 | 1 | ENSMUST00 | 0 | Kcnb2 | ENSMUSG00000092083.4 |
| chr10 | 128526186 | 128526660 | 12.24318 | 32.0257 | 29.53485 | Promoter | 10 | 128510248 | 128525859 | 15612 | 2 | ENSMUST00 | -328 | Esyt1 | ENSMUSG00000025366.7 |
| chr11 | 101155759 | 101156260 | 12.24318 | 32.0257 | 29.53485 | Promoter | 11 | 101155907 | 101161787 | 5881 | 1 | ENSMUST00 | 0 | Tubg2 | ENSMUSG00000045007.9 |
| chr11 | 58239296 | 58239787 | 12.24318 | 32.0257 | 29.53485 | Distal Interge | 11 | 58215028 | 58222779 | 7752 | 1 | ENSMUST00 | 24269 | Irgm2 | ENSMUSG00000069874.6 |
| chr14 | 13901512 | 13902013 | 12.24318 | 32.0257 | 29.53485 | Intron (ENSM | 14 | 13814949 | 13914478 | 99530 | 2 | ENSMUST00 | 12465 | Gm281 | ENSMUSG00000084902.1 |
| chr17 | 62784077 | 62784691 | 12.24318 | 32.0257 | 29.53485 | Intron (ENSM | 17 | 62607319 | 62881144 | 273826 | 2 | ENSMUST00 | 96453 | Efna5 | ENSMUSG00000048915.7 |
| chr19 | 3719173 | 3719935 | 12.24318 | 32.0257 | 29.53485 | Distal Interge | 19 | 3708333 | 3717881 | 9549 | 1 | ENSMUST00 | 10841 | 1810055G02 | ENSMUSG00000035372.1 |
| chr4 | 115052419 | 115053279 | 12.24318 | 32.0257 | 29.53485 | Distal Interge | 4 | 115056426 | 115071755 | 15330 | 1 | ENSMUST00 | -3147 | Tal1 | ENSMUSG00000028717.7 |
| chr5 | 116453326 | 116454089 | 12.24318 | 32.0257 | 29.53485 | Promoter | 5 | 116439275 | 116453913 | 14639 | 2 | ENSMUST00 | 0 | Srrm4 | ENSMUSG00000063919.7 |
| chr5 | 123058082 | 123058657 | 12.24318 | 32.0257 | 29.53485 | Intron (ENSM | 5 | 123066242 | 123066699 | 458 | 1 | ENSMUST00 | -7585 | Gm6444 | ENSMUSG00000080836.1 |
| chr6 | 100563040 | 100564015 | 12.24318 | 32.0257 | 29.53485 | Distal Interge | 6 | 100527400 | 100533426 | 6027 | 1 | ENSMUST00 | 35641 | RP24-114M8 | ENSMUSG00000101553.1 |
| chr7 | 79578722 | 79579171 | 12.24318 | 32.0257 | 29.53485 | Distal Interge | 7 | 79593369 | 79594924 | 1556 | 2 | ENSMUST00 | 15753 | Rhcg | ENSMUSG00000030549.5 |
| chr8 | 111061761 | 111062744 | 12.24318 | 32.0257 | 29.53485 | 3' UTR | 8 | 111047912 | 111050010 | 2099 | 1 | ENSMUST00 | 13850 | Aars | ENSMUSG00000031960.9 |
| chr9 | 75554709 | 75555949 | 12.24318 | 32.0257 | 29.53485 | Intron (ENSM | 9 | 75497796 | 75559657 | 61862 | 2 | ENSMUST00 | 3708 | Tmod3 | ENSMUSG00000058587.7 |
| chr6 | 6881365 | 6882228 | 14.10153 | 31.98288 | 29.49352 | Promoter | 6 | 6877805 | 6882068 | 4264 | 2 | ENSMUST00 | 0 | Dlx5 | ENSMUSG00000029755.9 |
| chr11 | 78261261 | 78261550 | 13.70122 | 31.96107 | 29.4719 | Promoter | 11 | 78261752 | 78264032 | 2281 | 1 | ENSMUST00 | -202 | 2610507B11I | ENSMUSG00000010277.7 |
| chr15 | 68363411 | 68363916 | 13.67444 | 31.91549 | 29.42649 | Promoter | 15 | 68340983 | 68363176 | 22194 | 2 | ENSMUST00 | -236 | Gm20732 | ENSMUSG00000093387.1 |
| chr8 | 89272345 | 89272821 | 14.0431 | 31.88435 | 29.39542 | Distal Interge | 8 | 89186946 | 89187560 | 615 | 2 | ENSMUST00 | -84786 | Gm5356 | ENSMUSG00000069962.3 |
| chr7 | 100285390 | 100285819 | 13.2717 | 31.88349 | 29.39467 | Promoter | 7 | 100285520 | 100319699 | 34180 | 1 | ENSMUST00 | 0 | P4ha3 | ENSMUSG00000051048.11 |
| chr1 | 37028830 | 37029768 | 11.15939 | 31.88065 | 29.39184 | Promoter | 1 | 37029304 | 37063649 | 34346 | 1 | ENSMUST00 | 0 | Vwa3b | ENSMUSG00000050122.13 |
| chr10 | 42761070 | 42761389 | 11.15939 | 31.88065 | 29.39184 | Promoter | 10 | 42761496 | 42832514 | 71019 | 1 | ENSMUST00 | -107 | Sec63 | ENSMUSG00000019802.8 |
| chr16 | 32419255 | 32420321 | 11.15939 | 31.88065 | 29.39184 | Promoter | 16 | 32419702 | 32429099 | 9398 | 1 | ENSMUST00 | 0 | Tctex1d2 | ENSMUSG00000014075.10 |
| chr16 | 20589454 | 20590860 | 11.15939 | 31.88065 | 29.39184 | Promoter | 16 | 20589471 | 20598467 | 8997 | 1 | ENSMUST00 | 0 | Vwa5b2 | ENSMUSG00000046613.13 |
| chr19 | 7422035 | 7423817 | 11.15939 | 31.88065 | 29.39184 | Promoter | 19 | 7422573 | 7422642 | 70 | 1 | ENSMUST00 | 0 | Mir6991 | ENSMUSG00000098293.1 |
| chr3 | 87960593 | 87961130 | 11.15939 | 31.88065 | 29.39184 | Distal Interge | 3 | 87952144 | 87952985 | 842 | 1 | ENSMUST00 | 8450 | Crabp2 | ENSMUSG00000004885.5 |
| chr7 | 38085045 | 38086196 | 11.15939 | 31.88065 | 29.39184 | Distal Interge | 7 | 38098811 | 38099369 | 559 | 2 | ENSMUST00 | 13173 | Ccne1 | ENSMUSG00000002068.11 |
| chr7 | 99267638 | 99269330 | 11.15939 | 31.88065 | 29.39184 | Promoter | 7 | 99268346 | 99337137 | 68792 | 1 | ENSMUST00 | 0 | Map6 | ENSMUSG00000055407.8 |
| chr4 | 3870692 | 3871780 | 14.01406 | 31.83537 | 29.34707 | Promoter | 4 | 3870657 | 3872105 | 1449 | 2 | ENSMUST00 | 325 | Mos | ENSMUSG00000078365.5 |
| chr9 | 95637383 | 95638160 | 14.01406 | 31.83537 | 29.34707 | Promoter | 9 | 95637601 | 95698096 | 60496 | 1 | ENSMUST00 | 0 | Pcolce2 | ENSMUSG00000015354.8 |
| chr16 | 63863789 | 63864508 | 12.84678 | 31.80056 | 29.31244 | Promoter | 16 | 63543538 | 63864158 | 320621 | 2 | ENSMUST00 | 0 | Epha3 | ENSMUSG00000052504.6 |
| chr7 | 45832670 | 45834038 | 12.84678 | 31.80056 | 29.31244 | 3' UTR | 7 | 45825232 | 45830944 | 5713 | 2 | ENSMUST00 | -1727 | Grwd1 | ENSMUSG00000053801.11 |
| chr14 | 122104452 | 122104845 | 13.98514 | 31.78658 | 29.29866 | Exon (ENSMU | 14 | 122103430 | 122106953 | 3524 | 2 | ENSMUST00 | 2108 | A330035P11I | ENSMUSG00000085615.2 |
| chr14 | 57797680 | 57798077 | 13.59473 | 31.77968 | 29.29185 | Promoter | 14 | 57798180 | 57804980 | 6801 | 1 | ENSMUST00 | -103 | Sap18 | ENSMUSG00000021963.11 |
| chr13 | 99353350 | 99353852 | 13.95634 | 31.73799 | 29.25027 | Intron (ENSM | 13 | 99344786 | 99415561 | 70776 | 1 | ENSMUST00 | 8565 | Mrps27 | ENSMUSG00000041632.5 |

| | | | | | | | | | | | | | | | |
|---|---|---|---|---|---|---|---|---|---|---|---|---|---|---|---|
| chr9 | 108048504 | 108049004 | 13.95634 | 31.73799 | 29.25027 | Promoter | 9 | 108049269 | 108049581 | 313 | 1 | ENSMUST00 | -265 | Gmppb | ENSMUSG00000070284.5 |
| chr4 | 146580154 | 146580848 | 8.34776 | 31.702 | 29.21443 | Promoter | 4 | 146580629 | 146582312 | 1684 | 1 | ENSMUST00 | 0 | Gm13233 | ENSMUSG00000083038.1 |
| chr16 | 4885493 | 4886075 | 13.92766 | 31.68958 | 29.20206 | Promoter | 16 | 4886100 | 4938296 | 52197 | 1 | ENSMUST00 | -25 | Mgrn1 | ENSMUSG00000022517.11 |
| chr11 | 30471563 | 30472399 | 13.15075 | 31.67431 | 29.18687 | Promoter | 11 | 30426006 | 30471808 | 45803 | 2 | ENSMUST00 | 0 | 4930505A04I | ENSMUSG00000040919.8 |
| chr2 | 32423247 | 32424137 | 13.15075 | 31.67431 | 29.18687 | Promoter | 2 | 32415060 | 32424005 | 8946 | 2 | ENSMUST00 | 0 | Slc25a25 | ENSMUSG00000026819.10 |
| chr1 | 180225733 | 180226564 | 12.76655 | 31.65917 | 29.17213 | Downstream | 1 | 180227468 | 180230025 | 2558 | 2 | ENSMUST00 | 3461 | Psen2 | ENSMUSG00000010609.10 |
| chr1 | 153424745 | 153425504 | 12.76655 | 31.65917 | 29.17213 | Promoter | 1 | 153425162 | 153452574 | 27413 | 1 | ENSMUST00 | 0 | Shcbp1l | ENSMUSG00000042708.7 |
| chr1 | 88639579 | 88640560 | 12.76655 | 31.65917 | 29.17213 | Distal Interge | 1 | 88656308 | 88658100 | 1793 | 1 | ENSMUST00 | -15748 | RP24-108I6.2 | ENSMUSG00000100367.1 |
| chr10 | 80264304 | 80264815 | 12.76655 | 31.65917 | 29.17213 | Promoter | 10 | 80264984 | 80288408 | 23425 | 1 | ENSMUST00 | -169 | Dazap1 | ENSMUSG00000069565.7 |
| chr11 | 96823537 | 96825054 | 12.76655 | 31.65917 | 29.17213 | Promoter | 11 | 96817414 | 96824013 | 6600 | 2 | ENSMUST00 | 0 | Nfe2l1 | ENSMUSG00000038615.12 |
| chr11 | 101173012 | 101174627 | 12.76655 | 31.65917 | 29.17213 | Promoter | 11 | 101172997 | 101175443 | 2447 | 2 | ENSMUST00 | 816 | Ccr10 | ENSMUSG00000044052.3 |
| chr11 | 97138628 | 97139531 | 12.76655 | 31.65917 | 29.17213 | 3' UTR | 11 | 97146364 | 97146419 | 56 | 2 | ENSMUST00 | 6888 | Mir7235 | ENSMUSG00000099286.1 |
| chr12 | 111274003 | 111275894 | 12.76655 | 31.65917 | 29.17213 | Exon (ENSMU | 12 | 111271111 | 111276426 | 5316 | 1 | ENSMUST00 | 2893 | Amn | ENSMUSG00000021278.6 |
| chr12 | 84053416 | 84054068 | 12.76655 | 31.65917 | 29.17213 | Promoter | 12 | 84039469 | 84053807 | 14339 | 2 | ENSMUST00 | 0 | 4732463B04I | ENSMUSG00000097762.1 |
| chr13 | 34963410 | 34964669 | 12.76655 | 31.65917 | 29.17213 | Promoter | 13 | 34946614 | 34963810 | 17197 | 2 | ENSMUST00 | 0 | Eci3 | ENSMUSG00000021416.6 |
| chr15 | 85657468 | 85658117 | 12.76655 | 31.65917 | 29.17213 | Distal Interge | 15 | 85639747 | 85645934 | 6188 | 1 | ENSMUST00 | 17722 | RP23-64J7.2 | ENSMUSG00000101828.1 |
| chr17 | 45474619 | 45475201 | 12.76655 | 31.65917 | 29.17213 | Promoter | 17 | 45448937 | 45474938 | 26002 | 2 | ENSMUST00 | 0 | Spats1 | ENSMUSG00000023935.9 |
| chr3 | 142620047 | 142621130 | 12.76655 | 31.65917 | 29.17213 | Promoter | 3 | 142620602 | 142638008 | 17407 | 1 | ENSMUST00 | 0 | Gbp2 | ENSMUSG00000028270.7 |
| chr3 | 130714841 | 130715487 | 12.76655 | 31.65917 | 29.17213 | Distal Interge | 3 | 130695921 | 130709444 | 13524 | 2 | ENSMUST00 | -5398 | Ostc | ENSMUSG00000041084.7 |
| chr4 | 41506658 | 41507586 | 12.76655 | 31.65917 | 29.17213 | Promoter | 4 | 41505009 | 41507754 | 2746 | 2 | ENSMUST00 | 168 | 1110017D15I | ENSMUSG00000028441.7 |
| chr4 | 4138361 | 4139059 | 12.76655 | 31.65917 | 29.17213 | Promoter | 4 | 4133531 | 4138477 | 4947 | 2 | ENSMUST00 | 0 | Penk | ENSMUSG00000045573.9 |
| chr4 | 127313651 | 127314447 | 12.76655 | 31.65917 | 29.17213 | Promoter | 4 | 127311421 | 127314039 | 2619 | 2 | ENSMUST00 | 0 | Gja4 | ENSMUSG00000050234.7 |
| chr6 | 122764328 | 122765197 | 12.76655 | 31.65917 | 29.17213 | Intron (ENSM | 6 | 122737278 | 122779957 | 42680 | 2 | ENSMUST00 | 14760 | Slc2a3 | ENSMUSG00000003153.5 |
| chr7 | 140057896 | 140058484 | 12.76655 | 31.65917 | 29.17213 | Distal Interge | 7 | 140046157 | 140049089 | 2933 | 2 | ENSMUST00 | -8808 | Msx3 | ENSMUSG00000025469.10 |
| chr7 | 105400400 | 105400956 | 12.76655 | 31.65917 | 29.17213 | Promoter | 7 | 105378610 | 105400054 | 21445 | 2 | ENSMUST00 | -347 | Fam160a2 | ENSMUSG00000044465.13 |
| chr7 | 4739282 | 4739778 | 12.76655 | 31.65917 | 29.17213 | Promoter | 7 | 4740115 | 4747514 | 7400 | 1 | ENSMUST00 | -337 | Suv420h2 | ENSMUSG00000059851.10 |
| chr7 | 35269265 | 35269927 | 12.76655 | 31.65917 | 29.17213 | Distal Interge | 7 | 35276544 | 35318421 | 41878 | 2 | ENSMUST00 | 48494 | Gpatch1 | ENSMUSG00000063808.9 |
| chr8 | 24823511 | 24824295 | 12.76655 | 31.65917 | 29.17213 | Promoter | 8 | 24727093 | 24824059 | 96967 | 2 | ENSMUST00 | 0 | Adam5 | ENSMUSG00000031554.12 |
| chr8 | 35375405 | 35376980 | 12.76655 | 31.65917 | 29.17213 | Promoter | 8 | 35375741 | 35388124 | 12384 | 1 | ENSMUST00 | 0 | Ppp1r3b | ENSMUSG00000046794.8 |
| chr8 | 71788919 | 71789579 | 12.76655 | 31.65917 | 29.17213 | Distal Interge | 8 | 71708394 | 71725716 | 17323 | 2 | ENSMUST00 | -63204 | Fcho1 | ENSMUSG00000070000.8 |
| chr1 | 174501636 | 174502285 | 13.48988 | 31.60083 | 29.11589 | Promoter | 1 | 174501825 | 174822717 | 320893 | 1 | ENSMUST00 | 0 | Fmn2 | ENSMUSG00000028354.8 |
| chr15 | 35326442 | 35327177 | 13.48988 | 31.60083 | 29.11589 | Distal Interge | 15 | 35296098 | 35303304 | 7207 | 1 | ENSMUST00 | 30345 | Osr2 | ENSMUSG00000022330.4 |
| chr2 | 33218133 | 33219620 | 13.48988 | 31.60083 | 29.11589 | Intron (ENSM | 2 | 33216098 | 33230124 | 14027 | 1 | ENSMUST00 | 2036 | Angptl2 | ENSMUSG00000004105.6 |
| chr4 | 149955261 | 149955826 | 13.48988 | 31.60083 | 29.11589 | Promoter | 4 | 149907142 | 149955043 | 47902 | 2 | ENSMUST00 | -219 | Spsb1 | ENSMUSG00000039911.8 |
| chr7 | 98193646 | 98194697 | 13.48988 | 31.60083 | 29.11589 | Exon (ENSMU | 7 | 98192420 | 98199481 | 7062 | 2 | ENSMUST00 | 4784 | B3gnt6 | ENSMUSG00000074004.2 |
| chr8 | 120276314 | 120277228 | 13.48988 | 31.60083 | 29.11589 | Intron (ENSM | 8 | 120230536 | 120553634 | 323099 | 1 | ENSMUST00 | 45779 | Gse1 | ENSMUSG00000031822.13 |
| chr9 | 105520778 | 105521652 | 13.48988 | 31.60083 | 29.11589 | Promoter | 9 | 105403539 | 105521147 | 117609 | 2 | ENSMUST00 | 0 | Atp2c1 | ENSMUSG00000032570.11 |
| chr12 | 4518963 | 4519538 | 13.87066 | 31.59332 | 29.10884 | Distal Interge | 12 | 4247363 | 4477182 | 229820 | 2 | ENSMUST00 | -41782 | Ncoa1 | ENSMUSG00000020647.9 |
| chr13 | 60177660 | 60178651 | 13.87066 | 31.59332 | 29.10884 | Promoter | 13 | 60174405 | 60177365 | 2961 | 2 | ENSMUST00 | -296 | Gas1 | ENSMUSG00000052957.7 |
| chr4 | 109981574 | 109983169 | 13.87066 | 31.59332 | 29.10884 | 3' UTR | 4 | 109978053 | 109983687 | 5635 | 1 | ENSMUST00 | 3522 | Dmrta2 | ENSMUSG00000047143.3 |
| chr6 | 85502675 | 85503169 | 13.87066 | 31.59332 | 29.10884 | Promoter | 6 | 85469574 | 85502994 | 33421 | 2 | ENSMUST00 | 0 | Fbxo41 | ENSMUSG00000047013.10 |
| chr8 | 94182886 | 94183576 | 13.87066 | 31.59332 | 29.10884 | Distal Interge | 8 | 94179089 | 94180325 | 1237 | 1 | ENSMUST00 | 3798 | Mt1 | ENSMUSG00000031765.7 |
| chr9 | 58805926 | 58806708 | 13.87066 | 31.59332 | 29.10884 | Distal Interge | 9 | 58823512 | 58860955 | 37444 | 1 | ENSMUST00 | -16804 | Hcn4 | ENSMUSG00000032338.8 |
| chr5 | 125386022 | 125386450 | 7.94617 | 31.58756 | 29.10344 | 3' UTR | 5 | 125387659 | 125389186 | 1528 | 2 | ENSMUST00 | 2736 | Ubc | ENSMUSG00000008348.9 |
| chr6 | 126702084 | 126702381 | 13.84233 | 31.54548 | 29.06139 | Distal Interge | 6 | 126532553 | 126698192 | 165640 | 2 | ENSMUST00 | -3893 | Kcna5 | ENSMUSG00000045534.3 |
| chr1 | 46425272 | 46425863 | 13.41152 | 31.46699 | 28.98322 | Promoter | 1 | 46425592 | 46807476 | 381885 | 1 | ENSMUST00 | 0 | Dnah7c | ENSMUSG00000101337.1 |
| chr10 | 20547795 | 20548894 | 13.41152 | 31.46699 | 28.98322 | Promoter | 10 | 20398928 | 20548328 | 149401 | 2 | ENSMUST00 | 0 | Pde7b | ENSMUSG00000019990.9 |
| chr11 | 98545525 | 98546199 | 13.41152 | 31.46699 | 28.98322 | Promoter | 11 | 98464902 | 98545800 | 80899 | 2 | ENSMUST00 | 0 | Ikzf3 | ENSMUSG00000018168.8 |
| chr11 | 58640302 | 58640910 | 13.41152 | 31.46699 | 28.98322 | Promoter | 11 | 58640465 | 58652404 | 11940 | 1 | ENSMUST00 | 0 | Trim58 | ENSMUSG00000037124.6 |
| chr11 | 53463398 | 53464100 | 13.41152 | 31.46699 | 28.98322 | Promoter | 11 | 53462379 | 53463613 | 1235 | 2 | ENSMUST00 | 0 | Gm12212 | ENSMUSG00000085707.1 |
| chr12 | 79130512 | 79131227 | 13.41152 | 31.46699 | 28.98322 | Promoter | 12 | 79130777 | 79156301 | 25525 | 1 | ENSMUST00 | 0 | Arg2 | ENSMUSG00000021125.6 |
| chr12 | 108002520 | 108003431 | 13.41152 | 31.46699 | 28.98322 | Promoter | 12 | 107915222 | 108003414 | 88193 | 2 | ENSMUST00 | 0 | Bcl11b | ENSMUSG00000048251.10 |
| chr13 | 9494583 | 9495186 | 13.41152 | 31.46699 | 28.98322 | Exon (ENSMU | 13 | 9387139 | 9391741 | 4603 | 1 | ENSMUST00 | 107445 | Gm26601 | ENSMUSG00000097859.1 |
| chr15 | 25621457 | 25622305 | 13.41152 | 31.46699 | 28.98322 | Promoter | 15 | 25622525 | 25813673 | 191149 | 1 | ENSMUST00 | -220 | Myo10 | ENSMUSG00000022272.12 |
| chr17 | 32350767 | 32351156 | 13.41152 | 31.46699 | 28.98322 | Promoter | 17 | 32321425 | 32350581 | 29157 | 2 | ENSMUST00 | -187 | Akap8l | ENSMUSG00000002625.9 |
| chr17 | 57032151 | 57032660 | 13.41152 | 31.46699 | 28.98322 | Promoter | 17 | 57021058 | 57031507 | 10450 | 2 | ENSMUST00 | -645 | Khsrp | ENSMUSG00000007670.9 |
| chr19 | 42089632 | 42090232 | 13.41152 | 31.46699 | 28.98322 | Promoter | 19 | 42090435 | 42122218 | 31784 | 1 | ENSMUST00 | -203 | Pi4k2a | ENSMUSG00000025178.8 |
| chr2 | 29819943 | 29820904 | 13.41152 | 31.46699 | 28.98322 | Promoter | 2 | 29820075 | 29843145 | 23071 | 1 | ENSMUST00 | 0 | Urm1 | ENSMUSG00000069020.4 |
| chr3 | 129590583 | 129591444 | 13.41152 | 31.46699 | 28.98322 | Intron (ENSM | 3 | 129532386 | 129638495 | 106110 | 1 | ENSMUST00 | 58198 | Elovl6 | ENSMUSG00000041220.6 |
| chr4 | 96553017 | 96553867 | 13.41152 | 31.46699 | 28.98322 | Promoter | 4 | 96516138 | 96553661 | 37524 | 2 | ENSMUST00 | 0 | Cyp2j6 | ENSMUSG00000052914.4 |
| chr5 | 90640210 | 90640875 | 13.41152 | 31.46699 | 28.98322 | Promoter | 5 | 90603076 | 90640487 | 37412 | 2 | ENSMUST00 | 0 | Rassf6 | ENSMUSG00000029370.7 |
| chr6 | 86584683 | 86585388 | 13.41152 | 31.46699 | 28.98322 | Distal Interge | 6 | 86574100 | 86574161 | 62 | 2 | ENSMUST00 | -10523 | Gm22184 | ENSMUSG00000093162.1 |
| chr7 | 143106936 | 143107840 | 13.41152 | 31.46699 | 28.98322 | Promoter | 7 | 143107254 | 143427042 | 319789 | 1 | ENSMUST00 | 0 | Kcnq1 | ENSMUSG00000009545.9 |
| chr7 | 29019305 | 29020194 | 13.41152 | 31.46699 | 28.98322 | Exon (ENSMU | 7 | 28982854 | 29003279 | 20426 | 1 | ENSMUST00 | 36452 | Map4k1 | ENSMUSG00000037337.10 |
| chr7 | 134937616 | 134938921 | 13.41152 | 31.46699 | 28.98322 | Promoter | 7 | 134881926 | 134938430 | 56505 | 2 | ENSMUST00 | 0 | Fam196a | ENSMUSG00000073805.4 |
| chr8 | 70315552 | 70316104 | 13.41152 | 31.46699 | 28.98322 | Promoter | 8 | 70315775 | 70331592 | 15818 | 1 | ENSMUST00 | 0 | Cers1 | ENSMUSG00000087408.4 |
| chr9 | 106211523 | 106212100 | 13.41152 | 31.46699 | 28.98322 | 3' UTR | 9 | 106214228 | 106215021 | 794 | 1 | ENSMUST00 | -2128 | Twf2 | ENSMUSG00000023277.6 |
| chr9 | 120539623 | 120540286 | 13.41152 | 31.46699 | 28.98322 | Promoter | 9 | 120539818 | 120568318 | 28501 | 1 | ENSMUST00 | 0 | Entpd3 | ENSMUSG00000041608.7 |
| chr9 | 22259333 | 22260055 | 13.41152 | 31.46699 | 28.98322 | Promoter | 9 | 22247430 | 22259895 | 12466 | 2 | ENSMUST00 | 0 | Zfp599 | ENSMUSG00000062794.8 |
| chr13 | 5714041 | 5714695 | 13.78602 | 31.45033 | 28.9683 | Distal Interge | 13 | 5855508 | 5857712 | 2205 | 2 | ENSMUST00 | 143017 | RP23-28J6.1 | ENSMUSG00000100625.1 |
| chr9 | 123478138 | 123478535 | 13.78602 | 31.45033 | 28.9683 | Promoter | 9 | 123478701 | 123521548 | 42848 | 1 | ENSMUST00 | -166 | Limd1 | ENSMUSG00000025239.2 |
| chr4 | 110050875 | 110051395 | 12.62853 | 31.41526 | 28.9334 | Distal Interge | 4 | 109978053 | 109983687 | 5635 | 1 | ENSMUST00 | 72823 | Dmrta2 | ENSMUSG00000047143.3 |
| chr15 | 103339596 | 103340537 | 13.75803 | 31.40302 | 28.92136 | Promoter | 15 | 103313895 | 103340086 | 26192 | 2 | ENSMUST00 | 0 | Zfp385a | ENSMUSG00000000552.9 |
| chr9 | 121313886 | 121314840 | 13.75803 | 31.40302 | 28.92136 | Distal Interge | 9 | 121102544 | 121277197 | 174654 | 2 | ENSMUST00 | -36690 | Ulk4 | ENSMUSG00000040936.9 |
| chr10 | 79716501 | 79717565 | 10.60441 | 31.37333 | 28.89181 | Promoter | 10 | 79716634 | 79736108 | 19475 | 1 | ENSMUST00 | 0 | Hcn2 | ENSMUSG00000020331.4 |
| chr9 | 37254555 | 37256102 | 10.60441 | 31.37333 | 28.89181 | Promoter | 9 | 37229149 | 37255738 | 26590 | 2 | ENSMUST00 | 0 | Slc37a2 | ENSMUSG00000032122.9 |
| chr11 | 95666123 | 95666720 | 13.73016 | 31.3559 | 28.87458 | Promoter | 11 | 95666957 | 95680773 | 13817 | 1 | ENSMUST00 | -237 | Phb | ENSMUSG00000038845.6 |
| chr7 | 63757525 | 63758834 | 13.73016 | 31.3559 | 28.87458 | 3' UTR | 7 | 63754712 | 63757925 | 3214 | 1 | ENSMUST00 | 2814 | Otud7a | ENSMUSG00000033510.9 |
| chr8 | 69271059 | 69271540 | 13.3356 | 31.33717 | 28.85595 | Distal Interge | 8 | 69300776 | 69303162 | 2387 | 1 | ENSMUST00 | -29236 | D130040H23 | ENSMUSG00000079038.2 |
| chr1 | 180696668 | 180697387 | 11.49921 | 31.27799 | 28.79692 | Promoter | 1 | 180693043 | 180697034 | 3992 | 2 | ENSMUST00 | 0 | Mixl1 | ENSMUSG00000026497.7 |
| chr1 | 89930027 | 89931593 | 11.49921 | 31.27799 | 28.79692 | Promoter | 1 | 89927956 | 89931179 | 3224 | 2 | ENSMUST00 | 0 | Gbx2 | ENSMUSG00000034486.8 |
| chr12 | 41483556 | 41484782 | 11.49921 | 31.27799 | 28.79692 | Promoter | 12 | 41451673 | 41485751 | 34079 | 2 | ENSMUST00 | 969 | Lrrn3 | ENSMUSG00000036295.4 |
| chr15 | 82987124 | 82987963 | 11.49921 | 31.27799 | 28.79692 | Promoter | 15 | 82982725 | 82987830 | 5106 | 2 | ENSMUST00 | 0 | RP23-223M2 | ENSMUSG00000100552.1 |
| chr17 | 24657817 | 24658559 | 11.49921 | 31.27799 | 28.79692 | Promoter | 17 | 24657330 | 24658457 | 1128 | 2 | ENSMUST00 | 0 | Npw | ENSMUSG00000071230.5 |
| chr18 | 55548461 | 55549338 | 11.49921 | 31.27799 | 28.79692 | Distal Interge | 18 | 55622636 | 55622920 | 285 | 2 | ENSMUST00 | 73582 | Gm24514 | ENSMUSG00000092702.2 |
| chr2 | 31298009 | 31298596 | 11.49921 | 31.27799 | 28.79692 | Distal Interge | 2 | 31314415 | 31460738 | 146324 | 1 | ENSMUST00 | -15819 | Hmcn2 | ENSMUSG00000055632.12 |
| chr7 | 39655440 | 39656122 | 11.49921 | 31.27799 | 28.79692 | Distal Interge | 7 | 39662546 | 39663009 | 464 | 2 | ENSMUST00 | 6887 | RP23-44J21.4 | ENSMUSG00000100912.1 |
| chr11 | 50851005 | 50851703 | 12.91534 | 31.26572 | 28.78545 | Promoter | 11 | 50850986 | 50866177 | 15192 | 1 | ENSMUST00 | 20 | Grm6 | ENSMUSG00000000617.9 |
| chr14 | 45690743 | 45691121 | 12.91534 | 31.26572 | 28.78545 | Distal Interge | 14 | 45713132 | 45713324 | 193 | 2 | ENSMUST00 | 22203 | Gm15516 | ENSMUSG00000082662.1 |
| chr16 | 85549733 | 85550551 | 12.91534 | 31.26572 | 28.78545 | Promoter | 16 | 85421533 | 85550417 | 128885 | 2 | ENSMUST00 | 0 | Cyyr1 | ENSMUSG00000041134.10 |
| chr19 | 42752018 | 42753504 | 12.91534 | 31.26572 | 28.78545 | Promoter | 19 | 42725858 | 42752775 | 26918 | 2 | ENSMUST00 | 0 | Pyroxd2 | ENSMUSG00000060224.3 |
| chr2 | 34671561 | 34671959 | 12.91534 | 31.26572 | 28.78545 | Distal Interge | 2 | 34677894 | 34684293 | 6400 | 2 | ENSMUST00 | 12334 | Gapvd1 | ENSMUSG00000026867.13 |
| chr6 | 50261482 | 50262180 | 12.91534 | 31.26572 | 28.78545 | Promoter | 6 | 50189369 | 50261755 | 72387 | 2 | ENSMUST00 | 0 | Dfna5 | ENSMUSG00000029821.10 |
| chr16 | 32814479 | 32815329 | 13.67476 | 31.26218 | 28.78225 | Distal Interge | 16 | 32804760 | 32810477 | 5718 | 2 | ENSMUST00 | -4003 | Gm933 | ENSMUSG00000094430.2 |
| chr15 | 103060301 | 103061298 | 13.64722 | 31.21559 | 28.73584 | Distal Interge | 15 | 103034395 | 103036852 | 2458 | 1 | ENSMUST00 | 25907 | Hoxc4 | ENSMUSG00000075394.4 |
| chr7 | 19816114 | 19817765 | 13.64722 | 31.21559 | 28.73584 | Promoter | 7 | 19816553 | 19818625 | 2073 | 1 | ENSMUST00 | 0 | Gm16174 | ENSMUSG00000087593.1 |
| chr1 | 180763520 | 180764118 | 13.2347 | 31.16438 | 28.68482 | Distal Interge | 1 | 180726043 | 180754203 | 28161 | 1 | ENSMUST00 | 37478 | Acbd3 | ENSMUSG00000026499.4 |
| chr1 | 172325902 | 172326760 | 13.2347 | 31.16438 | 28.68482 | 5' UTR | 1 | 172321033 | 172329263 | 8231 | 2 | ENSMUST00 | 2503 | Kcnj9 | ENSMUSG00000038026.7 |
| chr10 | 79322007 | 79322536 | 13.2347 | 31.16438 | 28.68482 | Distal Interge | 10 | 79356576 | 79397198 | 40623 | 1 | ENSMUST00 | -34040 | Vmn2r82 | ENSMUSG00000091468.2 |
| chr2 | 6721118 | 6722248 | 13.2347 | 31.16438 | 28.68482 | Promoter | 2 | 6539705 | 6721615 | 181911 | 2 | ENSMUST00 | 0 | Celf2 | ENSMUSG00000002107.13 |
| chr3 | 32708011 | 32708374 | 13.2347 | 31.16438 | 28.68482 | Promoter | 3 | 32708546 | 32726968 | 18423 | 1 | ENSMUST00 | -172 | Actl6a | ENSMUSG00000027671.9 |
| chr10 | 79681239 | 79682014 | 13.59249 | 31.12292 | 28.64397 | Promoter | 10 | 79682195 | 79688394 | 6200 | 1 | ENSMUST00 | -181 | Cdc34 | ENSMUSG00000020307.9 |
| chr11 | 97126901 | 97127544 | 13.59249 | 31.12292 | 28.64397 | Distal Interge | 11 | 97116796 | 97116915 | 120 | 2 | ENSMUST00 | -9987 | Gm11572 | ENSMUSG00000084368.1 |
| chr15 | 80671356 | 80672092 | 13.59249 | 31.12292 | 28.64397 | Promoter | 15 | 80671847 | 80700420 | 28574 | 1 | ENSMUST00 | 0 | Fam83f | ENSMUSG00000022408.6 |
| chr16 | 56064288 | 56064601 | 13.59249 | 31.12292 | 28.64397 | Distal Interge | 16 | 56070563 | 56070666 | 104 | 1 | ENSMUST00 | -5962 | Gm24047 | ENSMUSG00000076360.1 |
| chr12 | 86974173 | 86974966 | 13.56528 | 31.07685 | 28.5984 | Distal Interge | 12 | 86983381 | 86988676 | 5296 | 2 | ENSMUST00 | 13710 | Zdhhc22 | ENSMUSG00000048483.5 |
| chr17 | 31636392 | 31637607 | 12.41757 | 31.04075 | 28.56239 | Promoter | 17 | 31612623 | 31637199 | 24577 | 2 | ENSMUST00 | 0 | Cbs | ENSMUSG00000024039.9 |
| chr11 | 55097252 | 55097669 | 13.53819 | 31.03094 | 28.55276 | Promoter | 11 | 55098115 | 55113029 | 14915 | 1 | ENSMUST00 | -446 | Gm2a | ENSMUSG00000000594.7 |
| chr9 | 103009017 | 103009841 | 13.53819 | 31.03094 | 28.55276 | Promoter | 9 | 103008499 | 103096002 | 87504 | 1 | ENSMUST00 | 519 | Slco2a1 | ENSMUSG00000032548.9 |
| chr1 | 39761731 | 39762300 | 12.68821 | 30.86953 | 28.39186 | Distal Interge | 1 | 39665306 | 39720997 | 55692 | 2 | ENSMUST00 | -40735 | Rfx8 | ENSMUSG00000057173.7 |
| chr4 | 132767685 | 132768065 | 12.68821 | 30.86953 | 28.39186 | Promoter | 4 | 132768332 | 132776710 | 8379 | 1 | ENSMUST00 | -267 | Rpa2 | ENSMUSG00000028884.9 |
| chr4 | 45798936 | 45799475 | 12.68821 | 30.86953 | 28.39186 | Promoter | 4 | 45799022 | 45804604 | 5583 | 1 | ENSMUST00 | 0 | Aldh1b1 | ENSMUSG00000035561.5 |
| chr4 | 89524437 | 89525183 | 13.43087 | 30.849 | 28.3716 | Distal Interge | 4 | 89445219 | 89445367 | 149 | 1 | ENSMUST00 | 79219 | Gm27580 | ENSMUSG00000098669.1 |
| chr8 | 18950152 | 18951166 | 13.43087 | 30.849 | 28.3716 | Promoter | 8 | 18932729 | 18950871 | 18143 | 2 | ENSMUST00 | 0 | Xkr5 | ENSMUSG00000039814.10 |
| chr1 | 175491694 | 175492867 | 13.78596 | 30.80405 | 28.32676 | Promoter | 1 | 175059736 | 175491130 | 431395 | 2 | ENSMUST00 | -565 | Rgs7 | ENSMUSG00000026527.8 |
| chr1 | 119996980 | 119997646 | 13.78596 | 30.80405 | 28.32676 | Promoter | 1 | 119923341 | 119997190 | 73850 | 2 | ENSMUST00 | 0 | Gm101 | ENSMUSG00000036962.7 |
| chr11 | 4746001 | 4747299 | 13.78596 | 30.80405 | 28.32676 | Promoter | 11 | 4736892 | 4746778 | 9887 | 2 | ENSMUST00 | 0 | Cabp7 | ENSMUSG00000009075.2 |
| chr11 | 99243680 | 99244349 | 13.78596 | 30.80405 | 28.32676 | Promoter | 11 | 99232761 | 99244067 | 11307 | 2 | ENSMUST00 | 0 | Krt222 | ENSMUSG00000035849.9 |
| chr11 | 76666252 | 76666875 | 13.78596 | 30.80405 | 28.32676 | Distal Interge | 11 | 76672470 | 76673676 | 1207 | 1 | ENSMUST00 | -5595 | Bhlha9 | ENSMUSG00000044243.3 |
| chr11 | 89298332 | 89299444 | 13.78596 | 30.80405 | 28.32676 | Downstream | 11 | 89300638 | 89302332 | 1695 | 2 | ENSMUST00 | 2888 | Nog | ENSMUSG00000048616.4 |
| chr11 | 96875642 | 96876115 | 13.78596 | 30.80405 | 28.32676 | Distal Interge | 11 | 96863290 | 96873954 | 10665 | 2 | ENSMUST00 | -1689 | Gm11523 | ENSMUSG00000087117.3 |
| chr13 | 104863515 | 104864187 | 13.78596 | 30.80405 | 28.32676 | Promoter | 13 | 104845283 | 104863893 | 18611 | 2 | ENSMUST00 | 0 | Fam159b | ENSMUSG00000042655.4 |
| chr15 | 25414215 | 25414891 | 13.78596 | 30.80405 | 28.32676 | Promoter | 15 | 25414192 | 25414844 | 653 | 1 | ENSMUST00 | 24 | Gm5468 | ENSMUSG00000056089.5 |
| chr16 | 28850748 | 28852000 | 13.78596 | 30.80405 | 28.32676 | Intron (ENSM | 16 | 28826176 | 28929698 | 103523 | 2 | ENSMUST00 | 77698 | Mb21d2 | ENSMUSG00000051065.8 |
| chr16 | 21845852 | 21846613 | 13.78596 | 30.80405 | 28.32676 | Promoter | 16 | 21843451 | 21845776 | 2326 | 2 | ENSMUST00 | -77 | Gm26744 | ENSMUSG00000097401.1 |
| chr17 | 47394538 | 47395821 | 13.78596 | 30.80405 | 28.32676 | 3' UTR | 17 | 47394560 | 47400584 | 6025 | 2 | ENSMUST00 | 4763 | Guca1a | ENSMUSG00000023982.7 |
| chr18 | 89768845 | 89769825 | 13.78596 | 30.80405 | 28.32676 | Promoter | 18 | 89301082 | 89769528 | 468447 | 2 | ENSMUST00 | 0 | Dok6 | ENSMUSG00000073514.3 |
| chr19 | 6497332 | 6498438 | 13.78596 | 30.80405 | 28.32676 | Promoter | 19 | 6497671 | 6509865 | 12195 | 1 | ENSMUST00 | 0 | Nrxn2 | ENSMUSG00000033768.11 |

| | | | | | | | | | | | | | | | |
|---|---|---|---|---|---|---|---|---|---|---|---|---|---|---|---|
| chr2 | 109693275 | 109694248 | 13.78596 | 30.80405 | 28.32676 | Promoter | 2 | 109693563 | 109727007 | 33445 | 1 | ENSMUST00 | 0 | Bdnf | ENSMUSG00000048482.9 |
| chr3 | 9334185 | 9334555 | 13.78596 | 30.80405 | 28.32676 | Distal Interge | 3 | 9402175 | 9402485 | 311 | 1 | ENSMUST00 | -67620 | Gm22604 | ENSMUSG00000088229.1 |
| chr4 | 140905971 | 140906572 | 13.78596 | 30.80405 | 28.32676 | Promoter | 4 | 140906344 | 140952586 | 46243 | 1 | ENSMUST00 | 0 | Padi2 | ENSMUSG00000028927.6 |
| chr4 | 42669848 | 42670635 | 13.78596 | 30.80405 | 28.32676 | Distal Interge | 4 | 42656355 | 42661893 | 5539 | 2 | ENSMUST00 | -7956 | Il11ra2 | ENSMUSG00000078735.3 |
| chr5 | 30913426 | 30914740 | 13.78596 | 30.80405 | 28.32676 | Promoter | 5 | 30913739 | 30919703 | 5965 | 1 | ENSMUST00 | 0 | Emilin1 | ENSMUSG00000029163.7 |
| chr7 | 19319673 | 19320088 | 13.78596 | 30.80405 | 28.32676 | Distal Interge | 7 | 19302721 | 19310045 | 7325 | 2 | ENSMUST00 | -9629 | Fosb | ENSMUSG00000003545.2 |
| chr7 | 35185007 | 35185626 | 13.78596 | 30.80405 | 28.32676 | Promoter | 7 | 35186385 | 35195449 | 9065 | 1 | ENSMUST00 | -759 | Slc7a10 | ENSMUSG00000030495.7 |
| chr7 | 45783447 | 45784215 | 13.78596 | 30.80405 | 28.32676 | Promoter | 7 | 45783738 | 45804144 | 20407 | 1 | ENSMUST00 | 0 | Lmtk3 | ENSMUSG00000062044.8 |
| chr8 | 92827306 | 92827816 | 13.78596 | 30.80405 | 28.32676 | Promoter | 8 | 92827328 | 92853417 | 26090 | 1 | ENSMUST00 | 0 | Mmp2 | ENSMUSG00000031740.7 |
| chrX | 73715565 | 73717426 | 13.78596 | 30.80405 | 28.32676 | Promoter | 20 | 73686178 | 73716175 | 29998 | 2 | ENSMUST00 | 0 | Bcap31 | ENSMUSG00000002015.5 |
| chrX | 74297115 | 74297736 | 13.78596 | 30.80405 | 28.32676 | Promoter | 20 | 74297143 | 74302374 | 5232 | 1 | ENSMUST00 | 0 | Atp6ap1 | ENSMUSG00000019087.8 |
| chrX | 74280797 | 74281665 | 13.78596 | 30.80405 | 28.32676 | Promoter | 20 | 74273319 | 74281598 | 8280 | 2 | ENSMUST00 | 0 | Dnase1l1 | ENSMUSG00000019088.8 |
| chrX | 7728281 | 7728871 | 13.78596 | 30.80405 | 28.32676 | Promoter | 20 | 7728439 | 7731064 | 2626 | 1 | ENSMUST00 | 0 | Praf2 | ENSMUSG00000031149.7 |
| chrX | 95658077 | 95658967 | 13.78596 | 30.80405 | 28.32676 | Promoter | 20 | 95639193 | 95658408 | 19216 | 2 | ENSMUST00 | 0 | Zc4h2 | ENSMUSG00000035062.8 |
| chrX | 48054258 | 48055023 | 13.78596 | 30.80405 | 28.32676 | Distal Interge | 20 | 48025146 | 48034853 | 9708 | 2 | ENSMUST00 | -19406 | Apln | ENSMUSG00000037010.7 |
| chrX | 12159705 | 12160585 | 13.78596 | 30.80405 | 28.32676 | Promoter | 20 | 12036743 | 12160355 | 123613 | 2 | ENSMUST00 | 0 | Bcor | ENSMUSG00000040363.9 |
| chrX | 151519948 | 151520522 | 13.78596 | 30.80405 | 28.32676 | Promoter | 20 | 151520672 | 151570807 | 50136 | 1 | ENSMUST00 | -150 | Phf8 | ENSMUSG00000041229.10 |
| chrX | 153832026 | 153832855 | 13.78596 | 30.80405 | 28.32676 | Promoter | 20 | 153832225 | 153833510 | 1286 | 1 | ENSMUST00 | 0 | Spin2c | ENSMUSG00000046550.8 |
| chrX | 98936736 | 98937084 | 13.78596 | 30.80405 | 28.32676 | Promoter | 20 | 98937273 | 98943635 | 6363 | 1 | ENSMUST00 | -189 | Yipf6 | ENSMUSG00000047694.7 |
| chrX | 75475309 | 75475757 | 13.78596 | 30.80405 | 28.32676 | Promoter | 20 | 75475232 | 75475523 | 292 | 2 | ENSMUST00 | 0 | Gm8666 | ENSMUSG00000048709.6 |
| chrX | 38635344 | 38635619 | 13.78596 | 30.80405 | 28.32676 | Promoter | 20 | 38630785 | 38635087 | 4303 | 2 | ENSMUST00 | -258 | C1galt1c1 | ENSMUSG00000048970.9 |
| chrX | 136707780 | 136709072 | 13.78596 | 30.80405 | 28.32676 | Promoter | 20 | 136707982 | 136711478 | 3497 | 1 | ENSMUST00 | 0 | Tceal1 | ENSMUSG00000049536.5 |
| chrX | 136171308 | 136173104 | 13.78596 | 30.80405 | 28.32676 | Promoter | 20 | 136168984 | 136172251 | 3268 | 2 | ENSMUST00 | 0 | Tceal8 | ENSMUSG00000051579.5 |
| chrX | 10484779 | 10486109 | 13.78596 | 30.80405 | 28.32676 | Promoter | 20 | 10485158 | 10596605 | 111448 | 1 | ENSMUST00 | 0 | Tspan7 | ENSMUSG00000058254.7 |
| chrX | 162158562 | 162160279 | 13.78596 | 30.80405 | 28.32676 | Promoter | 20 | 161833296 | 162159730 | 326435 | 2 | ENSMUST00 | 0 | Nhs | ENSMUSG00000059493.8 |
| chrX | 94725350 | 94726078 | 13.78596 | 30.80405 | 28.32676 | Promoter | 20 | 94724569 | 94730187 | 5619 | 1 | ENSMUST00 | 782 | Zxdb | ENSMUSG00000073062.3 |
| chrX | 94367045 | 94367623 | 13.78596 | 30.80405 | 28.32676 | Promoter | 20 | 94367117 | 94416961 | 49845 | 1 | ENSMUST00 | 0 | Apoo | ENSMUSG00000079508.3 |
| chrX | 105069441 | 105070655 | 13.78596 | 30.80405 | 28.32676 | Promoter | 20 | 105037197 | 105070096 | 32900 | 2 | ENSMUST00 | 0 | 5530601H04 | ENSMUSG00000087174.2 |
| chr11 | 31949256 | 31949628 | 13.76117 | 30.76233 | 28.28681 | Distal Interge | 11 | 31965592 | 31979651 | 14060 | 1 | ENSMUST00 | -15964 | 4930524B15 | ENSMUSG00000020299.6 |
| chr5 | 3893085 | 3893588 | 13.76117 | 30.76233 | 28.28681 | Promoter | 5 | 3890584 | 3893907 | 3324 | 2 | ENSMUST00 | 319 | Mterf1a | ENSMUSG00000040429.5 |
| chr1 | 89927016 | 89927617 | 11.90309 | 30.75663 | 28.28125 | Downstream | 1 | 89927966 | 89929185 | 1220 | 2 | ENSMUST00 | 1568 | Gbx2 | ENSMUSG00000034486.8 |
| chr10 | 26077991 | 26079129 | 11.90309 | 30.75663 | 28.28125 | Promoter | 10 | 26078255 | 26079447 | 1193 | 1 | ENSMUST00 | 0 | Gm9767 | ENSMUSG00000039098.3 |
| chr10 | 18469401 | 18470148 | 11.90309 | 30.75663 | 28.28125 | Promoter | 10 | 18469888 | 18473427 | 3540 | 1 | ENSMUST00 | 0 | Nhsl1 | ENSMUSG00000039835.10 |
| chr11 | 120013275 | 120013740 | 11.90309 | 30.75663 | 28.28125 | Promoter | 11 | 120007313 | 120012408 | 5096 | 2 | ENSMUST00 | -868 | Aatk | ENSMUSG00000025375.10 |
| chr11 | 50887077 | 50887893 | 11.90309 | 30.75663 | 28.28125 | Promoter | 11 | 50872720 | 50887443 | 14724 | 2 | ENSMUST00 | 0 | Zfp454 | ENSMUSG00000048728.10 |
| chr12 | 54067680 | 54068821 | 11.90309 | 30.75663 | 28.28125 | Exon (ENSMU | 12 | 54062048 | 54079392 | 17345 | 2 | ENSMUST00 | 10571 | RP23-39G14. | ENSMUSG00000099407.1 |
| chr13 | 23562830 | 23563359 | 11.90309 | 30.75663 | 28.28125 | Promoter | 13 | 23561534 | 23562369 | 836 | 2 | ENSMUST00 | -462 | Hist1h3e | ENSMUSG00000069273.2 |
| chr13 | 67375112 | 67376109 | 11.90309 | 30.75663 | 28.28125 | Promoter | 13 | 67362113 | 67375810 | 13698 | 2 | ENSMUST00 | 0 | Zfp456 | ENSMUSG00000078995.4 |
| chr15 | 74954916 | 74956354 | 11.90309 | 30.75663 | 28.28125 | Promoter | 15 | 74955051 | 74959903 | 4853 | 1 | ENSMUST00 | 0 | Ly6e | ENSMUSG00000022587.9 |
| chr17 | 52600807 | 52601848 | 11.90309 | 30.75663 | 28.28125 | Promoter | 17 | 52592981 | 52602660 | 9680 | 2 | ENSMUST00 | 812 | Gm27217 | ENSMUSG00000098552.1 |
| chr19 | 47937386 | 47938109 | 11.90309 | 30.75663 | 28.28125 | Promoter | 19 | 47937712 | 48035379 | 97668 | 1 | ENSMUST00 | 0 | Ccdc147 | ENSMUSG00000046585.8 |
| chr2 | 12923666 | 12924791 | 11.90309 | 30.75663 | 28.28125 | Promoter | 2 | 12924041 | 12980404 | 56364 | 1 | ENSMUST00 | 0 | Pter | ENSMUSG00000026730.7 |
| chr2 | 32452220 | 32452803 | 11.90309 | 30.75663 | 28.28125 | Promoter | 2 | 32414490 | 32451445 | 36956 | 2 | ENSMUST00 | -776 | Slc25a25 | ENSMUSG00000026819.10 |
| chr2 | 91537762 | 91538271 | 11.90309 | 30.75663 | 28.28125 | Intron (ENSM | 2 | 91546320 | 91565168 | 18849 | 1 | ENSMUST00 | -8049 | Ckap5 | ENSMUSG00000040549.11 |
| chr4 | 124708582 | 124709363 | 11.90309 | 30.75663 | 28.28125 | 3' UTR | 4 | 124714776 | 124732460 | 17685 | 1 | ENSMUST00 | -5413 | Sf3a3 | ENSMUSG00000028902.4 |
| chr6 | 4505135 | 4506838 | 11.90309 | 30.75663 | 28.28125 | Promoter | 6 | 4505493 | 4541543 | 36051 | 1 | ENSMUST00 | 0 | Col1a2 | ENSMUSG00000029661.11 |
| chr7 | 135716654 | 135717186 | 11.90309 | 30.75663 | 28.28125 | Promoter | 7 | 135689788 | 135716379 | 26592 | 2 | ENSMUST00 | -276 | Mki67 | ENSMUSG00000031004.7 |
| chr7 | 134224073 | 134225505 | 11.90309 | 30.75663 | 28.28125 | Promoter | 7 | 133883199 | 134225097 | 341899 | 2 | ENSMUST00 | 0 | Adam12 | ENSMUSG00000054555.6 |
| chr7 | 73594570 | 73595554 | 11.90309 | 30.75663 | 28.28125 | Distal Interge | 7 | 73539801 | 73558395 | 18595 | 2 | ENSMUST00 | -36176 | 1810026B05 | ENSMUSG00000098496.2 |
| chr9 | 40344526 | 40346292 | 11.90309 | 30.75663 | 28.28125 | Promoter | 9 | 40293233 | 40346290 | 53058 | 2 | ENSMUST00 | 0 | Gramd1b | ENSMUSG00000040111.9 |
| chr14 | 84445019 | 84447421 | 12.98899 | 30.74239 | 28.2684 | 5' UTR | 14 | 84443563 | 84537060 | 93498 | 1 | ENSMUST00 | 1457 | Pcdh17 | ENSMUSG00000035566.6 |
| chr5 | 92856185 | 92858092 | 12.98899 | 30.74239 | 28.2684 | Intron (ENSM | 5 | 92897993 | 92965308 | 67316 | 1 | ENSMUST00 | -39901 | Shroom3 | ENSMUSG00000029381.10 |
| chr5 | 93005298 | 93006364 | 12.98899 | 30.74239 | 28.2684 | Distal Interge | 5 | 93013032 | 93013139 | 108 | 2 | ENSMUST00 | 6775 | Gm25521 | ENSMUSG00000089018.1 |
| chr19 | 36817962 | 36818941 | 13.73236 | 30.71385 | 28.24041 | Distal Interge | 19 | 36834232 | 36893477 | 59246 | 1 | ENSMUST00 | -15291 | Tnks2 | ENSMUSG00000024811.6 |
| chr3 | 88509595 | 88510627 | 13.73236 | 30.71385 | 28.24041 | Promoter | 3 | 88502753 | 88509956 | 7204 | 2 | ENSMUST00 | 0 | Lmna | ENSMUSG00000028063.10 |
| chr8 | 79680214 | 79681101 | 13.73236 | 30.71385 | 28.24041 | Promoter | 8 | 79680266 | 79680824 | 559 | 2 | ENSMUST00 | 0 | Tpd52-ps | ENSMUSG00000094594.1 |
| chr12 | 70346472 | 70348022 | 10.87325 | 30.68588 | 28.21252 | Promoter | 12 | 70244539 | 70347614 | 103076 | 2 | ENSMUST00 | 0 | Trim9 | ENSMUSG00000021071.10 |
| chr2 | 30441963 | 30443716 | 10.87325 | 30.68588 | 28.21252 | Promoter | 2 | 30442735 | 30446806 | 4072 | 1 | ENSMUST00 | 0 | Ppp2r4 | ENSMUSG00000039515.6 |
| chr4 | 141734708 | 141735304 | 10.87325 | 30.68588 | 28.21252 | Distal Interge | 4 | 141683563 | 141723419 | 39857 | 2 | ENSMUST00 | -11290 | Ddi2 | ENSMUSG00000078515.4 |
| chr12 | 29290671 | 29291622 | 13.32525 | 30.6697 | 28.19687 | Distal Interge | 12 | 29528384 | 29923209 | 394826 | 1 | ENSMUST00 | -236762 | Myt1l | ENSMUSG00000061911.9 |
| chr2 | 155750909 | 155751579 | 13.32525 | 30.6697 | 28.19687 | Promoter | 2 | 155751117 | 155755471 | 4355 | 1 | ENSMUST00 | 0 | Procr | ENSMUSG00000027611.7 |
| chr5 | 75574669 | 75576003 | 13.32525 | 30.6697 | 28.19687 | Promoter | 5 | 75574916 | 75656722 | 81807 | 1 | ENSMUST00 | 0 | Kit | ENSMUSG00000005672.7 |
| chr5 | 115844697 | 115845651 | 13.32525 | 30.6697 | 28.19687 | Promoter | 5 | 115845278 | 115925702 | 80425 | 1 | ENSMUST00 | 0 | Cit | ENSMUSG00000029516.14 |
| chr15 | 98001444 | 98002311 | 13.70367 | 30.66556 | 28.1932 | Promoter | 15 | 97998371 | 98001887 | 3517 | 2 | ENSMUST00 | 0 | Col2a1 | ENSMUSG00000022483.11 |
| chr4 | 44663929 | 44664792 | 13.70367 | 30.66556 | 28.1932 | Promoter | 4 | 44664415 | 44685347 | 20933 | 1 | ENSMUST00 | 0 | Gm12462 | ENSMUSG00000086211.1 |
| chr5 | 34326808 | 34327367 | 13.70367 | 30.66556 | 28.1932 | Distal Interge | 5 | 34336289 | 34353436 | 17148 | 1 | ENSMUST00 | -8922 | Rnf4 | ENSMUSG00000029110.8 |
| chr7 | 3289024 | 3290357 | 13.70367 | 30.66556 | 28.1932 | Promoter | 7 | 3289038 | 3299345 | 10308 | 1 | ENSMUST00 | 0 | Myadm | ENSMUSG00000068566.7 |
| chr7 | 66078937 | 66079487 | 13.70367 | 30.66556 | 28.1932 | Promoter | 7 | 66079649 | 66089412 | 9764 | 1 | ENSMUST00 | -162 | Vimp | ENSMUSG00000075701.8 |
| chr10 | 57485971 | 57486211 | 13.6751 | 30.61746 | 28.14547 | Promoter | 10 | 57478381 | 57486353 | 7973 | 2 | ENSMUST00 | 142 | 4930467K11 | ENSMUSG00000097370.1 |
| chr10 | 20724161 | 20725527 | 13.27306 | 30.58102 | 28.10912 | Promoter | 10 | 20398004 | 20725078 | 327075 | 2 | ENSMUST00 | 0 | Pde7b | ENSMUSG00000019990.9 |
| chr4 | 154864225 | 154864768 | 13.64664 | 30.56956 | 28.09773 | Exon (ENSMU | 4 | 154860552 | 154861637 | 1086 | 1 | ENSMUST00 | 3674 | Ttc34 | ENSMUSG00000046637.2 |
| chr6 | 52172354 | 52173164 | 13.64664 | 30.56956 | 28.09773 | Promoter | 6 | 52172881 | 52174059 | 1179 | 1 | ENSMUST00 | 0 | Gm15050 | ENSMUSG00000087626.1 |
| chr1 | 179847768 | 179848336 | 13.24712 | 30.53691 | 28.06522 | Distal Interge | 1 | 179744894 | 179803680 | 58787 | 2 | ENSMUST00 | -44089 | Ahctf1 | ENSMUSG00000026491.8 |
| chr14 | 69174392 | 69175322 | 12.46893 | 30.48504 | 28.01359 | 3' UTR | 14 | 69164797 | 69171857 | 7061 | 2 | ENSMUST00 | -2536 | 1700092C10 | ENSMUSG00000092499.1 |
| chr2 | 74749104 | 74750334 | 12.46893 | 30.48504 | 28.01359 | Promoter | 2 | 74748422 | 74750506 | 2085 | 2 | ENSMUST00 | 172 | Haglr | ENSMUSG00000075277.4 |
| chr9 | 100446056 | 100446547 | 12.46893 | 30.48504 | 28.01359 | Distal Interge | 9 | 100450322 | 100451912 | 1591 | 1 | ENSMUST00 | -3775 | RP23-368K2 | ENSMUSG00000101287.1 |
| chr2 | 105674527 | 105675806 | 13.59009 | 30.47432 | 28.00306 | Promoter | 2 | 105675109 | 105675972 | 864 | 1 | ENSMUST00 | 0 | Pax6 | ENSMUSG00000027168.16 |
| chr15 | 66285687 | 66286927 | 13.1699 | 30.40553 | 27.93451 | Promoter | 15 | 65986387 | 66286224 | 299838 | 2 | ENSMUST00 | 0 | Kcnq3 | ENSMUSG00000056258.8 |
| chr17 | 35919174 | 35919887 | 13.1699 | 30.40553 | 27.93451 | Intron (ENSM | 17 | 35917889 | 35917969 | 81 | 1 | ENSMUST00 | 1286 | Mir1894 | ENSMUSG00000084751.1 |
| chr6 | 131433403 | 131434070 | 13.1699 | 30.40553 | 27.93451 | Distal Interge | 6 | 131364858 | 131388450 | 23593 | 2 | ENSMUST00 | -44954 | Ybx3 | ENSMUSG00000030189.10 |
| chr2 | 174284180 | 174285286 | 13.53401 | 30.37983 | 27.90887 | Promoter | 2 | 174284320 | 174296895 | 12576 | 1 | ENSMUST00 | 0 | Gnas | ENSMUSG00000027523.14 |
| chr3 | 131105781 | 131106879 | 13.53401 | 30.37983 | 27.90887 | Distal Interge | 3 | 131110471 | 131224356 | 113886 | 1 | ENSMUST00 | -3592 | Lef1 | ENSMUSG00000027985.9 |
| chr3 | 51552646 | 51553566 | 13.53401 | 30.37983 | 27.90887 | Intron (ENSM | 3 | 51559757 | 51567117 | 7361 | 1 | ENSMUST00 | -6191 | 5031434O11 | ENSMUSG00000097885.2 |
| chr11 | 95120008 | 95121925 | 12.39106 | 30.348 | 27.8772 | Promoter | 11 | 95120119 | 95125296 | 5178 | 1 | ENSMUST00 | 0 | Dlx3 | ENSMUSG00000001510.8 |
| chr11 | 95981866 | 95983014 | 12.39106 | 30.348 | 27.8772 | Promoter | 11 | 95973144 | 95982594 | 9451 | 2 | ENSMUST00 | 0 | Igf2bp1 | ENSMUSG00000013415.4 |
| chr11 | 115974561 | 115975252 | 12.39106 | 30.348 | 27.8772 | Promoter | 11 | 115974709 | 116008412 | 33704 | 1 | ENSMUST00 | 0 | Itgb4 | ENSMUSG00000020758.10 |
| chr11 | 28583158 | 28584402 | 12.39106 | 30.348 | 27.8772 | Promoter | 11 | 28385685 | 28584296 | 198612 | 2 | ENSMUST00 | 0 | Ccdc85a | ENSMUSG00000032878.11 |
| chr12 | 112145337 | 112146975 | 12.39106 | 30.348 | 27.8772 | Promoter | 12 | 112146208 | 112181747 | 35540 | 1 | ENSMUST00 | 0 | Kif26a | ENSMUSG00000021294.7 |
| chr12 | 40223472 | 40224030 | 12.39106 | 30.348 | 27.8772 | Promoter | 12 | 40223429 | 40229169 | 5741 | 1 | ENSMUST00 | 44 | Gm7008 | ENSMUSG00000035983.4 |
| chr13 | 30985235 | 30986598 | 12.39106 | 30.348 | 27.8772 | Distal Interge | 13 | 30974204 | 30975976 | 1773 | 1 | ENSMUST00 | 11032 | Gm5447 | ENSMUSG00000056257.2 |
| chr16 | 45407983 | 45409040 | 12.39106 | 30.348 | 27.8772 | Promoter | 16 | 45382183 | 45408875 | 26693 | 2 | ENSMUST00 | 0 | Cd200 | ENSMUSG00000022661.9 |
| chr2 | 24048431 | 24049894 | 12.39106 | 30.348 | 27.8772 | Promoter | 2 | 24002910 | 24049394 | 46485 | 2 | ENSMUST00 | 0 | Hnmt | ENSMUSG00000026986.9 |
| chr2 | 84811070 | 84811502 | 12.39106 | 30.348 | 27.8772 | 3' UTR | 2 | 84806229 | 84809234 | 3006 | 1 | ENSMUST00 | 4842 | Ube2l6 | ENSMUSG00000027078.9 |
| chr2 | 155517341 | 155517818 | 12.39106 | 30.348 | 27.8772 | Promoter | 2 | 155517948 | 155562746 | 44799 | 1 | ENSMUST00 | -130 | Acss2 | ENSMUSG00000027605.13 |
| chr2 | 32741295 | 32741839 | 12.39106 | 30.348 | 27.8772 | Promoter | 2 | 32741490 | 32747064 | 5575 | 1 | ENSMUST00 | 0 | Sh2d3c | ENSMUSG00000059013.7 |
| chr3 | 123235775 | 123236582 | 12.39106 | 30.348 | 27.8772 | Promoter | 3 | 123076519 | 123236147 | 159629 | 2 | ENSMUST00 | 0 | Synpo2 | ENSMUSG00000050315.8 |
| chr5 | 138279680 | 138280548 | 12.39106 | 30.348 | 27.8772 | Promoter | 5 | 138264952 | 138279958 | 15007 | 2 | ENSMUST00 | 0 | Gpc2 | ENSMUSG00000029510.10 |
| chr5 | 72559849 | 72560202 | 12.39106 | 30.348 | 27.8772 | Promoter | 5 | 72513301 | 72559684 | 46384 | 2 | ENSMUST00 | -166 | Nfxl1 | ENSMUSG00000072889.4 |
| chr6 | 56900611 | 56902391 | 12.39106 | 30.348 | 27.8772 | Promoter | 6 | 56882400 | 56901886 | 19487 | 2 | ENSMUST00 | 0 | Nt5c3 | ENSMUSG00000029780.9 |
| chr8 | 70765692 | 70766954 | 12.39106 | 30.348 | 27.8772 | Promoter | 8 | 70762774 | 70766663 | 3890 | 2 | ENSMUST00 | 0 | Ifi30 | ENSMUSG00000031838.7 |
| chr8 | 124617354 | 124617742 | 12.39106 | 30.348 | 27.8772 | Exon (ENSMU | 8 | 124613440 | 124614626 | 1187 | 1 | ENSMUST00 | 3915 | Capn9 | ENSMUSG00000031981.7 |
| chr11 | 95041589 | 95042302 | 12.75224 | 30.33398 | 27.86505 | Promoter | 11 | 95026258 | 95041354 | 15097 | 2 | ENSMUST00 | -236 | Pdk2 | ENSMUSG00000038967.8 |
| chr5 | 32458307 | 32458851 | 12.75224 | 30.33398 | 27.86505 | Promoter | 5 | 32458974 | 32495523 | 36550 | 1 | ENSMUST00 | -123 | Ppp1cb | ENSMUSG00000014956.10 |
| chr7 | 126535798 | 126536407 | 12.75224 | 30.33398 | 27.86505 | Distal Interge | 7 | 126563874 | 126563980 | 107 | 2 | ENSMUST00 | 27573 | Gm24596 | ENSMUSG00000084453.1 |
| chr12 | 24571767 | 24572102 | 13.50614 | 30.33286 | 27.86432 | Promoter | 12 | 24572287 | 24617387 | 45101 | 1 | ENSMUST00 | -185 | Grhl1 | ENSMUSG00000020656.10 |
| chr19 | 48205887 | 48206719 | 13.50614 | 30.33286 | 27.86432 | Promoter | 19 | 48206025 | 48805505 | 599481 | 1 | ENSMUST00 | 0 | Sorcs3 | ENSMUSG00000063434.5 |
| chr3 | 52069672 | 52070311 | 13.50614 | 30.33286 | 27.86432 | Intron (ENSM | 3 | 51687611 | 52104891 | 417281 | 2 | ENSMUST00 | 34580 | Maml3 | ENSMUSG00000061143.10 |
| chr12 | 79965323 | 79966133 | 13.09357 | 30.27553 | 27.8073 | Distal Interge | 12 | 79963568 | 79963872 | 305 | 1 | ENSMUST00 | 1756 | RP23-179D3. | ENSMUSG00000101696.1 |
| chr2 | 35432032 | 35432943 | 13.45074 | 30.23947 | 27.77128 | Promoter | 2 | 35400179 | 35432633 | 32455 | 2 | ENSMUST00 | 0 | Ggta1 | ENSMUSG00000035778.12 |
| chr3 | 148976972 | 148978621 | 13.45074 | 30.23947 | 27.77128 | Intron (ENSM | 3 | 148888941 | 148989316 | 100376 | 2 | ENSMUST00 | 10695 | Lphn2 | ENSMUSG00000028184.8 |
| chr16 | 22008791 | 22009291 | 13.06832 | 30.2325 | 27.76443 | Promoter | 16 | 22009484 | 22049269 | 39786 | 1 | ENSMUST00 | -193 | Senp2 | ENSMUSG00000022855.8 |
| chr18 | 65083040 | 65084507 | 10.3393 | 30.21329 | 27.74566 | Intron (ENSM | 18 | 65023527 | 65217826 | 194300 | 1 | ENSMUST00 | 59514 | Nedd4l | ENSMUSG00000024589.10 |
| chr6 | 31552447 | 31557569 | 10.3393 | 30.21329 | 27.74566 | Intron (ENSM | 6 | 31519488 | 31563981 | 44494 | 2 | ENSMUST00 | 6412 | Podxl | ENSMUSG00000025608.9 |
| chr10 | 79841181 | 79841634 | 13.42322 | 30.19304 | 27.72564 | Distal Interge | 10 | 79854427 | 79864432 | 10006 | 1 | ENSMUST00 | -12793 | Ptbp1 | ENSMUSG00000006498.11 |
| chr18 | 36067903 | 36068819 | 13.42322 | 30.19304 | 27.72564 | Intron (ENSM | 18 | 35987019 | 36006084 | 19066 | 1 | ENSMUST00 | 80885 | Psd2 | ENSMUSG00000024347.11 |
| chr3 | 152667374 | 152668328 | 13.42322 | 30.19304 | 27.72564 | Promoter | 3 | 152462815 | 152668140 | 205326 | 2 | ENSMUST00 | 0 | Ak5 | ENSMUSG00000039058.7 |
| chr7 | 44527750 | 44528403 | 13.42322 | 30.19304 | 27.72564 | 3' UTR | 7 | 44501699 | 44524669 | 22971 | 2 | ENSMUST00 | -3082 | Mybpc2 | ENSMUSG00000038670.9 |
| chr14 | 68050852 | 68051801 | 13.3958 | 30.1468 | 27.67984 | Distal Interge | 14 | 68083863 | 68089095 | 5233 | 1 | ENSMUST00 | -32062 | Nefl | ENSMUSG00000022055.7 |
| chr1 | 86303076 | 86303803 | 12.99241 | 30.10301 | 27.6364 | Promoter | 1 | 86303234 | 86307305 | 4072 | 1 | ENSMUST00 | 0 | B3gnt7 | ENSMUSG00000079445.3 |
| chr11 | 100983133 | 100983588 | 12.99241 | 30.10301 | 27.6364 | Distal Interge | 11 | 100956733 | 100970887 | 14155 | 2 | ENSMUST00 | -12247 | Ptrf | ENSMUSG00000004044.9 |
| chr11 | 96005559 | 96006419 | 12.99241 | 30.10301 | 27.6364 | Promoter | 11 | 95957163 | 96005940 | 48778 | 2 | ENSMUST00 | 0 | Igf2bp1 | ENSMUSG00000013415.4 |
| chr11 | 101772347 | 101773047 | 12.99241 | 30.10301 | 27.6364 | Promoter | 11 | 101770354 | 101772181 | 1828 | 2 | ENSMUST00 | -167 | Etv4 | ENSMUSG00000017724.9 |
| chr11 | 3894800 | 3895615 | 12.99241 | 30.10301 | 27.6364 | Promoter | 11 | 3886088 | 3895098 | 9011 | 2 | ENSMUST00 | 0 | 4921536K21 | ENSMUSG00000020434.4 |
| chr11 | 117949418 | 117950173 | 12.99241 | 30.10301 | 27.6364 | Distal Interge | 11 | 117966079 | 117969186 | 3108 | 2 | ENSMUST00 | 19013 | Socs3 | ENSMUSG00000053113.3 |
| chr13 | 63627383 | 63628042 | 12.99241 | 30.10301 | 27.6364 | Distal Interge | 13 | 63511533 | 63565520 | 53988 | 2 | ENSMUST00 | -61864 | Ptch1 | ENSMUSG00000021466.6 |
| chr14 | 105328287 | 105329035 | 12.99241 | 30.10301 | 27.6364 | Distal Interge | 14 | 105294546 | 105308869 | 14324 | 1 | ENSMUST00 | 33742 | Ndfip2 | ENSMUSG00000053253.11 |
| chr15 | 51991949 | 51992294 | 12.99241 | 30.10301 | 27.6364 | Promoter | 15 | 51962605 | 51991760 | 29156 | 2 | ENSMUST00 | -190 | Rad21 | ENSMUSG00000022314.9 |
| chr15 | 3995986 | 3997027 | 12.99241 | 30.10301 | 27.6364 | Promoter | 15 | 3996039 | 4015858 | 19820 | 1 | ENSMUST00 | 0 | Gm2093 | ENSMUSG00000097662.1 |

| | | | | | | | | | | | | | | | |
|---|---|---|---|---|---|---|---|---|---|---|---|---|---|---|---|
| chr16 | 21210692 | 21211697 | 12.99241 | 30.10301 | 27.6364 | Promoter | 16 | 21211095 | 21222967 | 11873 | 1 | ENSMUST00 | 0 | Ephb3 | ENSMUSG00000005958.9 |
| chr16 | 29945587 | 29946602 | 12.99241 | 30.10301 | 27.6364 | Promoter | 16 | 29907519 | 29946517 | 38999 | 2 | ENSMUST00 | 0 | Gm26569 | ENSMUSG00000097313.1 |
| chr17 | 35840826 | 35841315 | 12.99241 | 30.10301 | 27.6364 | Promoter | 17 | 35841515 | 35859670 | 18156 | 1 | ENSMUST00 | -200 | Mdc1 | ENSMUSG00000061607.9 |
| chr18 | 21651443 | 21652355 | 12.99241 | 30.10301 | 27.6364 | Promoter | 18 | 21651448 | 21655148 | 3701 | 1 | ENSMUST00 | 0 | 4930426D05I | ENSMUSG00000043168.7 |
| chr19 | 44407790 | 44408746 | 12.99241 | 30.10301 | 27.6364 | Promoter | 19 | 44394451 | 44407709 | 13259 | 2 | ENSMUST00 | -82 | Scd1 | ENSMUSG00000037071.2 |
| chr2 | 122637598 | 122638055 | 12.99241 | 30.10301 | 27.6364 | Promoter | 2 | 122637850 | 122641191 | 3342 | 1 | ENSMUST00 | 0 | AA467197 | ENSMUSG00000033213.11 |
| chr2 | 105224122 | 105224799 | 12.99241 | 30.10301 | 27.6364 | Promoter | 2 | 105224320 | 105379796 | 155477 | 1 | ENSMUST00 | 0 | Them7 | ENSMUSG00000055312.2 |
| chr2 | 113216660 | 113217460 | 12.99241 | 30.10301 | 27.6364 | Promoter | 2 | 112631380 | 113217051 | 585672 | 2 | ENSMUST00 | 0 | Ryr3 | ENSMUSG00000057378.9 |
| chr3 | 89830800 | 89832029 | 12.99241 | 30.10301 | 27.6364 | Promoter | 3 | 89831370 | 89858834 | 27465 | 1 | ENSMUST00 | 0 | She | ENSMUSG00000046280.8 |
| chr4 | 108217439 | 108218202 | 12.99241 | 30.10301 | 27.6364 | Promoter | 4 | 108181934 | 108217897 | 35964 | 2 | ENSMUST00 | 0 | Zyg11a | ENSMUSG00000034645.8 |
| chr5 | 143387143 | 143387582 | 12.99241 | 30.10301 | 27.6364 | Exon (ENSMU | 5 | 143375461 | 143380442 | 4982 | 1 | ENSMUST00 | 11683 | Grid2ip | ENSMUSG00000010825.12 |
| chr6 | 136828404 | 136828863 | 12.99241 | 30.10301 | 27.6364 | Promoter | 6 | 136828843 | 136840536 | 11694 | 1 | ENSMUST00 | 0 | BC049715 | ENSMUSG00000047515.7 |
| chr7 | 70280086 | 70281607 | 12.99241 | 30.10301 | 27.6364 | Distal Interge | 7 | 70248973 | 70249776 | 804 | 2 | ENSMUST00 | -30311 | RP23-474J16 | ENSMUSG00000099387.1 |
| chr8 | 23157418 | 23158198 | 12.99241 | 30.10301 | 27.6364 | 3' UTR | 8 | 23153269 | 23158948 | 5680 | 1 | ENSMUST00 | 4150 | Nkx6-3 | ENSMUSG00000063672.7 |
| chr9 | 76014629 | 76015305 | 12.99241 | 30.10301 | 27.6364 | Promoter | 9 | 76014855 | 76136350 | 121496 | 1 | ENSMUST00 | 0 | Hmgcll1 | ENSMUSG00000007908.9 |
| chr9 | 50751428 | 50752783 | 12.99241 | 30.10301 | 27.6364 | Promoter | 9 | 50752758 | 50756633 | 3876 | 1 | ENSMUST00 | 0 | Cryab | ENSMUSG00000032060.9 |
| chr9 | 66158107 | 66158547 | 12.99241 | 30.10301 | 27.6364 | Promoter | 9 | 66158223 | 66272242 | 114020 | 1 | ENSMUST00 | 0 | Dapk2 | ENSMUSG00000032380.3 |
| chr10 | 81383724 | 81384197 | 12.9683 | 30.06186 | 27.59709 | Promoter | 10 | 81384428 | 81388352 | 3925 | 1 | ENSMUST00 | -231 | Dohh | ENSMUSG00000078440.4 |
| chr1 | 34677859 | 34679011 | 13.34131 | 30.05484 | 27.59022 | Promoter | 1 | 34678188 | 34812739 | 134552 | 1 | ENSMUST00 | 0 | Arhgef4 | ENSMUSG00000037509.13 |
| chr11 | 44612778 | 44613440 | 11.18842 | 30.05495 | 27.59022 | Distal Interge | 11 | 44617317 | 45004995 | 387679 | 1 | ENSMUST00 | -3877 | Ebf1 | ENSMUSG00000057098.9 |
| chr11 | 117482135 | 117483059 | 13.34131 | 30.05484 | 27.59022 | Distal Interge | 11 | 117484368 | 117489014 | 4647 | 1 | ENSMUST00 | -1309 | Gm11733 | ENSMUSG00000069588.3 |
| chr15 | 40654886 | 40656259 | 13.34131 | 30.05484 | 27.59022 | Promoter | 15 | 40655042 | 41104592 | 449551 | 1 | ENSMUST00 | 0 | Zfpm2 | ENSMUSG00000022306.8 |
| chr15 | 75886994 | 75888166 | 13.34131 | 30.05484 | 27.59022 | Promoter | 15 | 75884137 | 75888770 | 4634 | 2 | ENSMUST00 | 604 | Mroh6 | ENSMUSG00000098678.1 |
| chr19 | 21472144 | 21472759 | 11.18842 | 30.05495 | 27.59022 | Promoter | 19 | 21391307 | 21472672 | 81366 | 2 | ENSMUST00 | 0 | Gda | ENSMUSG00000058624.7 |
| chr2 | 131127148 | 131127789 | 11.18842 | 30.05495 | 27.59022 | Promoter | 2 | 131127280 | 131146321 | 19042 | 1 | ENSMUST00 | 0 | Hspa12b | ENSMUSG00000074793.5 |
| chr3 | 104961238 | 104962572 | 11.18842 | 30.05495 | 27.59022 | Promoter | 3 | 104944805 | 104961709 | 16905 | 2 | ENSMUST00 | 0 | Wnt2b | ENSMUSG00000027840.4 |
| chr3 | 130945662 | 130946563 | 11.18842 | 30.05495 | 27.59022 | Distal Interge | 3 | 130870949 | 130871076 | 128 | 2 | ENSMUST00 | -74587 | Gm26090 | ENSMUSG00000088522.1 |
| chr6 | 138140077 | 138141064 | 11.18842 | 30.05495 | 27.59022 | Promoter | 6 | 138140316 | 138156755 | 16440 | 1 | ENSMUST00 | 0 | Mgst1 | ENSMUSG00000008540.6 |
| chr7 | 127477016 | 127478026 | 11.18842 | 30.05495 | 27.59022 | Distal Interge | 7 | 127475501 | 127476757 | 1257 | 1 | ENSMUST00 | 1516 | Prr14 | ENSMUSG00000030822.10 |
| chr8 | 8334323 | 8336252 | 11.18842 | 30.05495 | 27.59022 | Distal Interge | 8 | 8620597 | 8639363 | 18767 | 2 | ENSMUST00 | 303111 | Efnb2 | ENSMUSG00000001300.10 |
| chr8 | 57530400 | 57531191 | 13.34131 | 30.05484 | 27.59022 | Intron (ENSM | 8 | 57526024 | 57538453 | 12430 | 2 | ENSMUST00 | 7262 | Galnt7 | ENSMUSG00000031608.8 |
| chr7 | 82869396 | 82870335 | 11.82498 | 29.97654 | 27.5129 | 3' UTR | 7 | 82867333 | 82871563 | 4231 | 1 | ENSMUST00 | 2064 | Mex3b | ENSMUSG00000057706.6 |
| chr19 | 55218311 | 55218652 | 12.52396 | 29.93832 | 27.47483 | Intron (ENSM | 19 | 55198297 | 55241236 | 42940 | 2 | ENSMUST00 | 22584 | Gucy2g | ENSMUSG00000055523.6 |
| chr2 | 163203837 | 163204874 | 12.52396 | 29.93832 | 27.47483 | Promoter | 2 | 163204553 | 163276098 | 71546 | 1 | ENSMUST00 | 0 | Tox2 | ENSMUSG00000074607.6 |
| chr10 | 81378743 | 81379211 | 12.89429 | 29.93542 | 27.47224 | Promoter | 10 | 81366222 | 81378516 | 12295 | 2 | ENSMUST00 | -228 | Fzr1 | ENSMUSG00000020235.11 |
| chrY | 90744229 | 90744616 | 4.53553 | 29.89161 | 27.42859 | Distal Interge | 21 | 90753057 | 90763485 | 10429 | 1 | ENSMUST00 | -8441 | Gm21857 | ENSMUSG00000095134.2 |
| chr3 | 129213717 | 129214683 | 13.23364 | 29.873 | 27.40998 | Promoter | 3 | 129213932 | 129216774 | 2843 | 1 | ENSMUST00 | 0 | Pitx2 | ENSMUSG00000028023.11 |
| chr16 | 11983270 | 11985236 | 9.87845 | 29.83486 | 27.37197 | Promoter | 16 | 11984113 | 11995390 | 11278 | 1 | ENSMUST00 | 0 | Shisa9 | ENSMUSG00000022494.8 |
| chr4 | 41773381 | 41773915 | 13.20699 | 29.82797 | 27.36516 | Promoter | 4 | 41769467 | 41773537 | 4071 | 2 | ENSMUST00 | 0 | Ccl27a | ENSMUSG00000073888.7 |
| chr10 | 18430832 | 18431419 | 12.82111 | 29.81026 | 27.34754 | Intron (ENSM | 10 | 18407675 | 18533892 | 126218 | 1 | ENSMUST00 | 23158 | Nhsl1 | ENSMUSG00000039835.10 |
| chr11 | 96332585 | 96333276 | 12.82111 | 29.81026 | 27.34754 | Intron (ENSM | 11 | 96328062 | 96344303 | 16242 | 1 | ENSMUST00 | 4524 | Hoxb3 | ENSMUSG00000048763.6 |
| chr12 | 8947654 | 8948319 | 12.82111 | 29.81026 | 27.34754 | Promoter | 12 | 8947929 | 8972028 | 24100 | 1 | ENSMUST00 | 0 | Matn3 | ENSMUSG00000020583.4 |
| chr2 | 92508250 | 92509732 | 12.82111 | 29.81026 | 27.34754 | Intron (ENSM | 2 | 92483926 | 92529809 | 45884 | 1 | ENSMUST00 | 24325 | Gm13814 | ENSMUSG00000085261.1 |
| chr12 | 69361102 | 69361383 | 13.18045 | 29.78312 | 27.32097 | Promoter | 12 | 69361182 | 69361481 | 300 | 2 | ENSMUST00 | 98 | Gm26191 | ENSMUSG00000099250.1 |
| chr16 | 92158227 | 92159042 | 13.18045 | 29.78312 | 27.32097 | Distal Interge | 16 | 92106261 | 92112227 | 5967 | 1 | ENSMUST00 | 51967 | Mrps6 | ENSMUSG00000039680.9 |
| chr12 | 36316978 | 36317582 | 13.15402 | 29.73841 | 27.27653 | 3' UTR | 12 | 36314169 | 36318452 | 4284 | 1 | ENSMUST00 | 2810 | Sostdc1 | ENSMUSG00000036169.5 |
| chr12 | 106174133 | 106175454 | 13.15402 | 29.73841 | 27.27653 | Distal Interge | 12 | 106221075 | 106221206 | 132 | 1 | ENSMUST00 | -45621 | Gm24986 | ENSMUSG00000077640.1 |
| chr19 | 45675717 | 45676617 | 13.15402 | 29.73841 | 27.27653 | Distal Interge | 19 | 45578254 | 45660312 | 82059 | 2 | ENSMUST00 | -15406 | Fbxw4 | ENSMUSG00000040913.5 |
| chr8 | 91147928 | 91148937 | 13.15402 | 29.73841 | 27.27653 | Distal Interge | 8 | 91123499 | 91134465 | 10967 | 2 | ENSMUST00 | -13464 | Aktip | ENSMUSG00000031667.10 |
| chr6 | 48629189 | 48629903 | 13.12769 | 29.69389 | 27.23246 | Exon (ENSMU | 6 | 48627935 | 48633689 | 5755 | 1 | ENSMUST00 | 1255 | AI854703 | ENSMUSG00000053297.9 |
| chr7 | 4943426 | 4943740 | 13.10147 | 29.64952 | 27.18821 | Exon (ENSMU | 7 | 4925844 | 4944821 | 18978 | 1 | ENSMUST00 | 17583 | Ssc5d | ENSMUSG00000035279.7 |
| chr17 | 70847695 | 70848485 | 12.72483 | 29.64534 | 27.18412 | Promoter | 17 | 70844981 | 70847170 | 2190 | 2 | ENSMUST00 | -526 | Tgif1 | ENSMUSG00000047407.12 |
| chr9 | 72169773 | 72170627 | 13.07535 | 29.60532 | 27.14417 | Distal Interge | 9 | 72174580 | 72174958 | 379 | 2 | ENSMUST00 | 4331 | Gm7866 | ENSMUSG00000061390.6 |
| chr2 | 34645826 | 34646740 | 12.30372 | 29.55471 | 27.09375 | Downstream | 2 | 34649514 | 34650240 | 727 | 2 | ENSMUST00 | 3500 | Gm13414 | ENSMUSG00000087635.3 |
| chr3 | 55782454 | 55783922 | 12.30372 | 29.55471 | 27.09375 | Promoter | 3 | 55782510 | 55785285 | 2776 | 1 | ENSMUST00 | 0 | Mab21l1 | ENSMUSG00000056947.4 |
| chr10 | 95394672 | 95395470 | 10.58711 | 29.50236 | 27.04178 | Intron (ENSM | 10 | 95387667 | 95392967 | 5301 | 2 | ENSMUST00 | -1706 | Socs2 | ENSMUSG00000020027.13 |
| chr16 | 23609387 | 23610459 | 10.58711 | 29.50236 | 27.04178 | Promoter | 16 | 23609919 | 23614222 | 4304 | 1 | ENSMUST00 | 0 | Rtp4 | ENSMUSG00000033355.5 |
| chr17 | 84626361 | 84626771 | 10.58711 | 29.50236 | 27.04178 | Promoter | 17 | 84626496 | 84655558 | 29063 | 1 | ENSMUST00 | 0 | Dync2li1 | ENSMUSG00000024253.8 |
| chr18 | 32816434 | 32817954 | 10.58711 | 29.50236 | 27.04178 | Exon (ENSMU | 18 | 32815383 | 32819799 | 4417 | 1 | ENSMUST00 | 1052 | Tslp | ENSMUSG00000024379.3 |
| chr2 | 91975502 | 91976270 | 10.58711 | 29.50236 | 27.04178 | Promoter | 2 | 91932824 | 91975864 | 43041 | 2 | ENSMUST00 | 0 | Dgkz | ENSMUSG00000040479.6 |
| chr6 | 52208245 | 52210010 | 10.58711 | 29.50236 | 27.04178 | Promoter | 6 | 52206288 | 52208722 | 2435 | 2 | ENSMUST00 | 0 | Hoxa6 | ENSMUSG00000043219.9 |
| chr6 | 145210438 | 145210938 | 10.58711 | 29.50236 | 27.04178 | Promoter | 6 | 145174834 | 145210791 | 35958 | 2 | ENSMUST00 | 0 | Casc1 | ENSMUSG00000043541.11 |
| chr1 | 193035390 | 193036163 | 11.56301 | 29.49975 | 27.03959 | Promoter | 1 | 192897307 | 193035698 | 138392 | 2 | ENSMUST00 | 0 | Syt14 | ENSMUSG00000016200.8 |
| chr1 | 177338537 | 177339308 | 11.56301 | 29.49975 | 27.03959 | Distal Interge | 1 | 177022116 | 177258203 | 236088 | 2 | ENSMUST00 | -80335 | Akt3 | ENSMUSG00000019699.10 |
| chr1 | 14506408 | 14507170 | 11.56301 | 29.49975 | 27.03959 | Distal Interge | 1 | 14168958 | 14310235 | 141278 | 2 | ENSMUST00 | -196174 | Eya1 | ENSMUSG00000025932.9 |
| chr10 | 84754721 | 84756301 | 11.56301 | 29.49975 | 27.03959 | Promoter | 10 | 84756062 | 84906144 | 150083 | 1 | ENSMUST00 | 0 | Rfx4 | ENSMUSG00000020037.10 |
| chr10 | 126905507 | 126906215 | 11.56301 | 29.49975 | 27.03959 | Distal Interge | 10 | 126887249 | 126901371 | 14123 | 2 | ENSMUST00 | -4137 | Xrcc6bp1 | ENSMUSG00000025436.6 |
| chr12 | 31073648 | 31075363 | 11.56301 | 29.49975 | 27.03959 | Promoter | 12 | 31073968 | 31079937 | 5970 | 1 | ENSMUST00 | 0 | Fam110c | ENSMUSG00000036136.8 |
| chr13 | 96387821 | 96388137 | 11.56301 | 29.49975 | 27.03959 | Promoter | 13 | 96388294 | 96415587 | 27294 | 1 | ENSMUST00 | -157 | Poc5 | ENSMUSG00000021671.8 |
| chr13 | 51637180 | 51637749 | 11.56301 | 29.49975 | 27.03959 | Distal Interge | 13 | 51645232 | 51650660 | 5429 | 1 | ENSMUST00 | -7483 | Cks2 | ENSMUSG00000062248.5 |
| chr15 | 89378549 | 89379545 | 11.56301 | 29.49975 | 27.03959 | Promoter | 15 | 89377450 | 89379254 | 1805 | 2 | ENSMUST00 | 0 | Odf3b | ENSMUSG00000047394.7 |
| chr16 | 4267691 | 4268465 | 11.56301 | 29.49975 | 27.03959 | Distal Interge | 16 | 4230301 | 4231099 | 799 | 1 | ENSMUST00 | 37391 | Gm5766 | ENSMUSG00000089645.3 |
| chr16 | 92605450 | 92606321 | 11.56301 | 29.49975 | 27.03959 | 3' UTR | 16 | 92612824 | 92620032 | 7209 | 1 | ENSMUST00 | -6503 | Gm26626 | ENSMUSG00000097139.2 |
| chr17 | 46705166 | 46705698 | 11.56301 | 29.49975 | 27.03959 | Promoter | 17 | 46682991 | 46705002 | 22012 | 2 | ENSMUST00 | -165 | Ppp2r5d | ENSMUSG00000059409.8 |
| chr2 | 9878010 | 9879779 | 11.56301 | 29.49975 | 27.03959 | Promoter | 2 | 9857078 | 9878600 | 21523 | 2 | ENSMUST00 | 0 | Gata3 | ENSMUSG00000015619.10 |
| chr2 | 180724765 | 180725766 | 11.56301 | 29.49975 | 27.03959 | Promoter | 2 | 180725263 | 180742280 | 17018 | 1 | ENSMUST00 | 0 | Slc17a9 | ENSMUSG00000023393.10 |
| chr4 | 134238239 | 134238620 | 11.56301 | 29.49975 | 27.03959 | Promoter | 4 | 134228041 | 134238399 | 10359 | 2 | ENSMUST00 | 0 | Cnksr1 | ENSMUSG00000028841.9 |
| chr5 | 137287934 | 137288703 | 11.56301 | 29.49975 | 27.03959 | Promoter | 5 | 137288277 | 137294466 | 6190 | 1 | ENSMUST00 | 0 | Ache | ENSMUSG00000023328.9 |
| chr5 | 105400187 | 105400832 | 11.56301 | 29.49975 | 27.03959 | Promoter | 5 | 105401245 | 105406970 | 5726 | 1 | ENSMUST00 | -413 | Gm26519 | ENSMUSG00000097237.1 |
| chr6 | 85187324 | 85188530 | 11.56301 | 29.49975 | 27.03959 | Promoter | 6 | 85187438 | 85204358 | 16921 | 1 | ENSMUST00 | 0 | Emx1 | ENSMUSG00000033726.8 |
| chr7 | 68263831 | 68264961 | 11.56301 | 29.49975 | 27.03959 | Promoter | 7 | 68236608 | 68264233 | 27626 | 2 | ENSMUST00 | 0 | Pgpep1l | ENSMUSG00000030553.2 |
| chr7 | 99248962 | 99249712 | 11.56301 | 29.49975 | 27.03959 | Distal Interge | 7 | 99219084 | 99238619 | 19536 | 2 | ENSMUST00 | -10344 | Mogat2 | ENSMUSG00000052396.7 |
| chr7 | 44589041 | 44589486 | 11.56301 | 29.49975 | 27.03959 | Intron (ENSM | 7 | 44579968 | 44590624 | 10657 | 2 | ENSMUST00 | 1138 | Gm15396 | ENSMUSG00000087560.1 |
| chr9 | 120001774 | 120002624 | 11.56301 | 29.49975 | 27.03959 | Intron (ENSM | 9 | 120013755 | 120023598 | 9844 | 2 | ENSMUST00 | 20974 | Xirp1 | ENSMUSG00000079243.2 |
| chr4 | 143325219 | 143325776 | 12.99762 | 29.47369 | 27.01492 | Distal Interge | 4 | 143340399 | 143340706 | 308 | 1 | ENSMUST00 | -14623 | Gm26313 | ENSMUSG00000088395.1 |
| chr16 | 56885993 | 56886628 | 12.97191 | 29.43014 | 26.97147 | Promoter | 16 | 56805161 | 56886166 | 81006 | 2 | ENSMUST00 | 0 | Tmem45a | ENSMUSG00000022754.5 |
| chr2 | 90744825 | 90745262 | 12.58308 | 29.40204 | 26.94344 | Promoter | 2 | 90745370 | 90781021 | 35652 | 1 | ENSMUST00 | -108 | Fnbp4 | ENSMUSG00000008200.9 |
| chr3 | 33843914 | 33844666 | 12.58308 | 29.40204 | 26.94344 | Promoter | 3 | 33812362 | 33844301 | 31940 | 2 | ENSMUST00 | 0 | Ccdc39 | ENSMUSG00000027676.6 |
| chr5 | 137610176 | 137611493 | 12.58308 | 29.40204 | 26.94344 | Promoter | 5 | 137605103 | 137611408 | 6306 | 2 | ENSMUST00 | 0 | Pcolce | ENSMUSG00000029718.9 |
| chr5 | 140505579 | 140506151 | 12.58308 | 29.40204 | 26.94344 | Promoter | 5 | 140505575 | 140524153 | 18579 | 1 | ENSMUST00 | 5 | Chst12 | ENSMUSG00000036599.10 |
| chr7 | 127362490 | 127362884 | 12.58308 | 29.40204 | 26.94344 | Promoter | 7 | 127363428 | 127363809 | 382 | 1 | ENSMUST00 | -544 | Gm10586 | ENSMUSG00000073823.2 |
| chr7 | 84749699 | 84750238 | 12.58308 | 29.40204 | 26.94344 | Intron (ENSM | 7 | 84775703 | 84776020 | 318 | 1 | ENSMUST00 | -25465 | 2610206C17I | ENSMUSG00000085236.2 |
| chr9 | 106225922 | 106226711 | 12.58308 | 29.40204 | 26.94344 | 3' UTR | 9 | 106222598 | 106226883 | 4286 | 1 | ENSMUST00 | 3325 | Tlr9 | ENSMUSG00000045322.10 |
| chr12 | 72236440 | 72237082 | 13.32643 | 29.39681 | 26.9387 | Promoter | 12 | 72211753 | 72236724 | 24972 | 2 | ENSMUST00 | 0 | Rtn1 | ENSMUSG00000021087.12 |
| chr12 | 83763561 | 83764217 | 13.32643 | 29.39681 | 26.9387 | Promoter | 12 | 83763634 | 83792382 | 28749 | 1 | ENSMUST00 | 0 | Papln | ENSMUSG00000021223.8 |
| chr12 | 61522830 | 61523339 | 13.32643 | 29.39681 | 26.9387 | Promoter | 12 | 61523150 | 61605008 | 81859 | 1 | ENSMUST00 | 0 | Lrfn5 | ENSMUSG00000035653.11 |
| chr14 | 6756382 | 6757499 | 13.32643 | 29.39681 | 26.9387 | Intron (ENSM | 14 | 6735685 | 6742332 | 6648 | 2 | ENSMUST00 | -14051 | Gm3636 | ENSMUSG00000091754.3 |
| chr15 | 61983467 | 61984614 | 13.32643 | 29.39681 | 26.9387 | Promoter | 15 | 61985341 | 61990352 | 5012 | 1 | ENSMUST00 | -727 | Myc | ENSMUSG00000022346.9 |
| chr16 | 78236753 | 78238152 | 13.32643 | 29.39681 | 26.9387 | Distal Interge | 16 | 78240258 | 78240388 | 131 | 1 | ENSMUST00 | -2106 | Gm25916 | ENSMUSG00000077451.1 |
| chr16 | 30008531 | 30009799 | 13.32643 | 29.39681 | 26.9387 | Promoter | 16 | 30008667 | 30021430 | 12764 | 1 | ENSMUST00 | 0 | 4632428C04I | ENSMUSG00000097184.1 |
| chr18 | 89837011 | 89837279 | 13.32643 | 29.39681 | 26.9387 | Distal Interge | 18 | 89301082 | 89769528 | 468447 | 2 | ENSMUST00 | -67484 | Dok6 | ENSMUSG00000073514.3 |
| chr19 | 53420993 | 53421471 | 13.32643 | 29.39681 | 26.9387 | Distal Interge | 19 | 53444014 | 53447314 | 3301 | 2 | ENSMUST00 | 25843 | 5830416P10I | ENSMUSG00000097636.2 |
| chr2 | 34904419 | 34904885 | 13.32643 | 29.39681 | 26.9387 | Promoter | 2 | 34898082 | 34904907 | 6826 | 2 | ENSMUST00 | 22 | Phf19 | ENSMUSG00000026873.4 |
| chr2 | 27441884 | 27442297 | 13.32643 | 29.39681 | 26.9387 | Distal Interge | 2 | 27450549 | 27450630 | 82 | 2 | ENSMUST00 | 8333 | Mir7578 | ENSMUSG00000098812.1 |
| chr4 | 43507955 | 43509048 | 13.32643 | 29.39681 | 26.9387 | Promoter | 4 | 43508543 | 43509035 | 493 | 1 | ENSMUST00 | 0 | Car9 | ENSMUSG00000028463.9 |
| chr4 | 88779245 | 88779530 | 13.32643 | 29.39681 | 26.9387 | Promoter | 4 | 88779850 | 88780660 | 811 | 1 | ENSMUST00 | -320 | Gm13272 | ENSMUSG00000096591.1 |
| chr5 | 31094725 | 31095687 | 13.32643 | 29.39681 | 26.9387 | Promoter | 5 | 31089341 | 31094801 | 5461 | 2 | ENSMUST00 | 0 | Slc30a3 | ENSMUSG00000029151.9 |
| chr5 | 8367573 | 8368387 | 13.32643 | 29.39681 | 26.9387 | Promoter | 5 | 8072352 | 8367982 | 295631 | 2 | ENSMUST00 | 0 | Adam22 | ENSMUSG00000040537.12 |
| chr5 | 118169271 | 118170291 | 13.32643 | 29.39681 | 26.9387 | Promoter | 5 | 118169748 | 118189478 | 19731 | 1 | ENSMUST00 | 0 | Hrk | ENSMUSG00000046607.6 |
| chr5 | 139912234 | 139913081 | 13.32643 | 29.39681 | 26.9387 | Intron (ENSM | 5 | 139907943 | 139974711 | 66769 | 1 | ENSMUST00 | 4292 | Elfn1 | ENSMUSG00000048988.7 |
| chr7 | 3301691 | 3303940 | 13.32643 | 29.39681 | 26.9387 | Promoter | 7 | 3303532 | 3331094 | 27563 | 1 | ENSMUST00 | 0 | Prkcg | ENSMUSG00000078816.4 |
| chr7 | 71411271 | 71412350 | 13.32643 | 29.39681 | 26.9387 | Distal Interge | 7 | 71370333 | 71379813 | 9481 | 1 | ENSMUST00 | 40939 | RP24-173C11 | ENSMUSG00000099853.1 |
| chr9 | 53974530 | 53975513 | 13.32643 | 29.39681 | 26.9387 | Promoter | 9 | 53911460 | 53975301 | 63842 | 2 | ENSMUST00 | 0 | Elmod1 | ENSMUSG00000041986.10 |
| chr9 | 81863347 | 81863964 | 13.32643 | 29.39681 | 26.9387 | Promoter | 9 | 81863670 | 82027026 | 163357 | 1 | ENSMUST00 | 0 | Mei4 | ENSMUSG00000043289.7 |
| chrX | 38600223 | 38601563 | 13.32643 | 29.39681 | 26.9387 | Promoter | 20 | 38600578 | 38611833 | 11256 | 1 | ENSMUST00 | 0 | Mcts1 | ENSMUSG00000000355.8 |
| chrX | 100774451 | 100774986 | 13.32643 | 29.39681 | 26.9387 | Promoter | 20 | 100774741 | 100818408 | 43668 | 1 | ENSMUST00 | 0 | Dlg3 | ENSMUSG00000000881.7 |
| chrX | 36863633 | 36864767 | 13.32643 | 29.39681 | 26.9387 | Promoter | 20 | 36823737 | 36864238 | 40502 | 2 | ENSMUST00 | 0 | C330007P06I | ENSMUSG00000006423.10 |
| chrX | 74353119 | 74353617 | 13.32643 | 29.39681 | 26.9387 | Promoter | 20 | 74351276 | 74353594 | 2319 | 2 | ENSMUST00 | 0 | Lage3 | ENSMUSG00000015289.4 |
| chrX | 150656794 | 150657499 | 13.32643 | 29.39681 | 26.9387 | Promoter | 20 | 150645304 | 150657436 | 12133 | 2 | ENSMUST00 | 0 | Tro | ENSMUSG00000025272.11 |
| chrX | 9662004 | 9663278 | 13.32643 | 29.39681 | 26.9387 | Promoter | 20 | 9654267 | 9663003 | 8737 | 2 | ENSMUST00 | 0 | Dynlt3 | ENSMUSG00000031176.3 |
| chrX | 73458435 | 73459008 | 13.32643 | 29.39681 | 26.9387 | Promoter | 20 | 73437315 | 73459044 | 21730 | 2 | ENSMUST00 | 36 | Haus7 | ENSMUSG00000031371.9 |
| chrX | 93288398 | 93289435 | 13.32643 | 29.39681 | 26.9387 | Exon (ENSMU | 20 | 93286529 | 93298357 | 11829 | 1 | ENSMUST00 | 1870 | Arx | ENSMUSG00000035277.10 |
| chrX | 77796300 | 77796830 | 13.32643 | 29.39681 | 26.9387 | Promoter | 20 | 77761411 | 77796278 | 34868 | 2 | ENSMUST00 | -23 | Prkx | ENSMUSG00000035725.8 |
| chrX | 23364568 | 23365444 | 13.32643 | 29.39681 | 26.9387 | Promoter | 20 | 23219271 | 23365082 | 145812 | 2 | ENSMUST00 | 0 | Klhl13 | ENSMUSG00000036782.8 |
| chrX | 139216832 | 139217673 | 13.32643 | 29.39681 | 26.9387 | Promoter | 20 | 139217166 | 139234802 | 17637 | 1 | ENSMUST00 | 0 | Mum1l1 | ENSMUSG00000042515.8 |
| chrX | 9199650 | 9200858 | 13.32643 | 29.39681 | 26.9387 | Promoter | 20 | 9199902 | 9268085 | 68184 | 1 | ENSMUST00 | 0 | Lancl3 | ENSMUSG00000047344.9 |
| chrX | 13461343 | 13462282 | 13.32643 | 29.39681 | 26.9387 | Distal Interge | 20 | 13466110 | 13489313 | 23204 | 1 | ENSMUST00 | -3828 | Nyx | ENSMUSG00000051228.5 |
| chrX | 51679867 | 51681677 | 13.32643 | 29.39681 | 26.9387 | Promoter | 20 | 51676783 | 51681348 | 4566 | 2 | ENSMUST00 | 0 | Hs6st2 | ENSMUSG00000062184.6 |

| | | | | | | | | | | | | | | | |
|---|---|---|---|---|---|---|---|---|---|---|---|---|---|---|---|
| chrX | 50634116 | 50635936 | 13.32643 | 29.39681 | 26.9387 | Promoter | 20 | 50555744 | 50635261 | 79518 | 2 | ENSMUST00 | 0 | Firre | ENSMUSG00000085396.2 |
| chr1 | 21271743 | 21272385 | 12.94631 | 29.38674 | 26.93043 | Promoter | 1 | 21272403 | 21272906 | 504 | 1 | ENSMUST00 | -18 | Gm7094 | ENSMUSG00000096375.2 |
| chr3 | 52257846 | 52258340 | 13.30246 | 29.35653 | 26.90032 | Distal Interge | 3 | 52268337 | 52350109 | 81773 | 1 | ENSMUST00 | -9997 | Foxo1 | ENSMUSG00000044167.5 |
| chr8 | 123628464 | 123629231 | 13.30246 | 29.35653 | 26.90032 | Intron (ENSM | 8 | 123623637 | 123623745 | 109 | 2 | ENSMUST00 | -4720 | Gm23647 | ENSMUSG00000095723.1 |
| chr13 | 58452745 | 58453502 | 12.53654 | 29.322 | 26.86593 | Distal Interge | 13 | 58391142 | 58403343 | 12202 | 2 | ENSMUST00 | -49403 | Hnrnpk | ENSMUSG00000021546.12 |
| chr15 | 83366854 | 83367307 | 12.8954 | 29.30041 | 26.84444 | Promoter | 15 | 83357526 | 83367282 | 9757 | 2 | ENSMUST00 | 0 | 1700001L05R | ENSMUSG00000075511.3 |
| chr3 | 93014414 | 93014905 | 12.8954 | 29.30041 | 26.84444 | Promoter | 3 | 93014519 | 93015669 | 1151 | 2 | ENSMUST00 | 764 | Crct1 | ENSMUSG00000027913.5 |
| chr8 | 95435015 | 95435417 | 12.8954 | 29.30041 | 26.84444 | Promoter | 8 | 95420252 | 95434869 | 14618 | 2 | ENSMUST00 | -147 | Gtl3 | ENSMUSG00000031796.6 |
| chr9 | 37529947 | 37531649 | 12.8954 | 29.30041 | 26.84444 | Promoter | 9 | 37530250 | 37531641 | 1392 | 1 | ENSMUST00 | 0 | Esam | ENSMUSG00000001946.8 |
| chr11 | 51915932 | 51916952 | 13.24688 | 29.26311 | 26.80765 | Distal Interge | 11 | 51964382 | 51964489 | 108 | 1 | ENSMUST00 | -47430 | Gm26061 | ENSMUSG00000093325.1 |
| chr12 | 111164322 | 111165013 | 13.24688 | 29.26311 | 26.80765 | Intron (ENSM | 12 | 111166370 | 111267153 | 100784 | 1 | ENSMUST00 | -1357 | Traf3 | ENSMUSG00000021277.11 |
| chr12 | 9574281 | 9574917 | 13.24688 | 29.26311 | 26.80765 | Promoter | 12 | 9574442 | 9581499 | 7058 | 1 | ENSMUST00 | 0 | Osr1 | ENSMUSG00000048387.7 |
| chr12 | 70134569 | 70135116 | 13.24688 | 29.26311 | 26.80765 | Distal Interge | 12 | 70154142 | 70183347 | 29206 | 1 | ENSMUST00 | -19026 | Abhd12b | ENSMUSG00000090121.5 |
| chr18 | 61647373 | 61648232 | 13.24688 | 29.26311 | 26.80765 | Promoter | 18 | 61647816 | 61647903 | 88 | 2 | ENSMUST00 | 0 | Mir145a | ENSMUSG00000065592.1 |
| chr9 | 119578481 | 119579407 | 13.24688 | 29.26311 | 26.80765 | Promoter | 9 | 119483410 | 119579016 | 95607 | 2 | ENSMUST00 | 0 | Scn5a | ENSMUSG00000032511.12 |
| chr12 | 111504389 | 111505149 | 13.21926 | 29.21668 | 26.76154 | Distal Interge | 12 | 111484611 | 111485823 | 1213 | 2 | ENSMUST00 | -18567 | Gm266 | ENSMUSG00000010529.5 |
| chr13 | 21222129 | 21223481 | 13.21926 | 29.21668 | 26.76154 | Distal Interge | 13 | 21190092 | 21192475 | 2384 | 1 | ENSMUST00 | 32038 | Trim27 | ENSMUSG00000021326.6 |
| chr16 | 20619732 | 20620546 | 13.21926 | 29.21668 | 26.76154 | Promoter | 16 | 20619215 | 20621287 | 2073 | 2 | ENSMUST00 | 741 | Camk2n2 | ENSMUSG00000051146.2 |
| chr13 | 64153467 | 64154294 | 12.8449 | 29.2147 | 26.75964 | Promoter | 13 | 64140468 | 64153202 | 12735 | 2 | ENSMUST00 | -266 | Zfp367 | ENSMUSG00000044934.7 |
| chr12 | 78911602 | 78912463 | 12.09108 | 29.18244 | 26.72745 | Distal Interge | 12 | 78888691 | 78906964 | 18274 | 2 | ENSMUST00 | -4639 | Plek2 | ENSMUSG00000021118.7 |
| chr16 | 50689182 | 50689895 | 12.09108 | 29.18244 | 26.72745 | Distal Interge | 16 | 50665008 | 50666346 | 1339 | 2 | ENSMUST00 | -22837 | Gm9575 | ENSMUSG00000083692.1 |
| chr14 | 105731973 | 105733249 | 12.81979 | 29.17209 | 26.71728 | Distal Interge | 14 | 105681828 | 105682211 | 384 | 1 | ENSMUST00 | 50146 | Gm10076 | ENSMUSG00000060143.6 |
| chr9 | 118487714 | 118489349 | 13.19176 | 29.17044 | 26.7157 | Distal Interge | 9 | 118478861 | 118482757 | 3897 | 1 | ENSMUST00 | 8854 | Eomes | ENSMUSG00000032446.9 |
| chr8 | 89045940 | 89046564 | 12.79478 | 29.12961 | 26.67496 | Distal Interge | 8 | 89027248 | 89044162 | 16915 | 2 | ENSMUST00 | -1779 | Sall1 | ENSMUSG00000031665.6 |
| chr17 | 46455204 | 46455661 | 13.16437 | 29.12438 | 26.66979 | Intron (ENSM | 17 | 46439578 | 46440099 | 522 | 2 | ENSMUST00 | -15106 | Gm5093 | ENSMUSG00000091742.1 |
| chr6 | 86471752 | 86472261 | 13.16437 | 29.12438 | 26.66979 | 3' UTR | 6 | 86464040 | 86469947 | 5908 | 1 | ENSMUST00 | 7713 | C87436 | ENSMUSG00000046679.11 |
| chr7 | 140922194 | 140923227 | 13.16437 | 29.12438 | 26.66979 | Promoter | 7 | 140922106 | 140922855 | 750 | 1 | ENSMUST00 | 89 | Nlrp6 | ENSMUSG00000038745.10 |
| chr2 | 74681949 | 74683155 | 13.13709 | 29.0785 | 26.62408 | Promoter | 2 | 74682323 | 74684168 | 1846 | 1 | ENSMUST00 | 0 | Hoxd11 | ENSMUSG00000042499.13 |
| chr17 | 31276940 | 31277678 | 10.07419 | 29.06419 | 26.60987 | Promoter | 17 | 31255019 | 31277356 | 22338 | 2 | ENSMUST00 | 0 | Rsph1 | ENSMUSG00000024033.9 |
| chr9 | 44066980 | 44067432 | 10.07419 | 29.06419 | 26.60987 | Promoter | 9 | 44067021 | 44095627 | 28607 | 1 | ENSMUST00 | 0 | Usp2 | ENSMUSG00000032010.9 |
| chr19 | 6362968 | 6363345 | 11.66271 | 29.05205 | 26.59792 | Promoter | 19 | 6355522 | 6363535 | 8014 | 2 | ENSMUST00 | 190 | Gm14966 | ENSMUSG00000079467.3 |
| chr8 | 70488255 | 70488918 | 11.66271 | 29.05205 | 26.59792 | Promoter | 8 | 70483876 | 70487358 | 3483 | 2 | ENSMUST00 | -898 | Tmem59l | ENSMUSG00000035964.7 |
| chr1 | 182954446 | 182955167 | 12.01557 | 29.04976 | 26.5957 | Promoter | 1 | 182954788 | 182976044 | 21257 | 1 | ENSMUST00 | 0 | Tlr5 | ENSMUSG00000079164.3 |
| chr10 | 38965270 | 38966282 | 12.01557 | 29.04976 | 26.5957 | Promoter | 10 | 38965515 | 39007396 | 41882 | 1 | ENSMUST00 | 0 | Lama4 | ENSMUSG00000019846.6 |
| chr10 | 127509147 | 127509521 | 12.01557 | 29.04976 | 26.5957 | Distal Interge | 10 | 127507225 | 127507992 | 768 | 1 | ENSMUST00 | 1923 | Stac3 | ENSMUSG00000040287.4 |
| chr12 | 113146275 | 113147394 | 12.01557 | 29.04976 | 26.5957 | Distal Interge | 12 | 113152012 | 113153877 | 1866 | 1 | ENSMUST00 | -4618 | Crip1 | ENSMUSG00000006360.7 |
| chr12 | 35038739 | 35039391 | 12.01557 | 29.04976 | 26.5957 | Distal Interge | 12 | 35047189 | 35147469 | 100281 | 1 | ENSMUST00 | -7798 | Snx13 | ENSMUSG00000020590.10 |
| chr15 | 13172727 | 13174414 | 12.01557 | 29.04976 | 26.5957 | Promoter | 15 | 13034200 | 13173639 | 139440 | 2 | ENSMUST00 | 0 | Cdh6 | ENSMUSG00000039385.4 |
| chr17 | 71869545 | 71870090 | 12.01557 | 29.04976 | 26.5957 | Exon (ENSMU | 17 | 71781947 | 71858351 | 76405 | 1 | ENSMUST00 | 87599 | Clip4 | ENSMUSG00000024059.9 |
| chr18 | 61785842 | 61787043 | 12.01557 | 29.04976 | 26.5957 | Promoter | 18 | 61730261 | 61786611 | 56351 | 2 | ENSMUST00 | 0 | Afap1l1 | ENSMUSG00000033032.10 |
| chr2 | 74668111 | 74668783 | 12.01557 | 29.04976 | 26.5957 | Promoter | 2 | 74668310 | 74671599 | 3290 | 1 | ENSMUST00 | 0 | Hoxd13 | ENSMUSG00000001819.4 |
| chr2 | 121294885 | 121296275 | 12.01557 | 29.04976 | 26.5957 | Promoter | 2 | 121295454 | 121309438 | 13985 | 1 | ENSMUST00 | 0 | Map1a | ENSMUSG00000027254.8 |
| chr2 | 90885618 | 90886097 | 12.01557 | 29.04976 | 26.5957 | Promoter | 2 | 90885860 | 90890525 | 4666 | 1 | ENSMUST00 | 0 | C1qtnf4 | ENSMUSG00000040794.5 |
| chr4 | 57781963 | 57782922 | 12.01557 | 29.04976 | 26.5957 | Promoter | 4 | 57782180 | 57855202 | 73023 | 1 | ENSMUST00 | 0 | Akap2 | ENSMUSG00000038729.15 |
| chr4 | 46344017 | 46345320 | 12.01557 | 29.04976 | 26.5957 | Promoter | 4 | 46328464 | 46345515 | 17052 | 2 | ENSMUST00 | 195 | Gm12446 | ENSMUSG00000087383.2 |
| chr4 | 136134607 | 136135552 | 12.01557 | 29.04976 | 26.5957 | Downstream | 4 | 136137496 | 136137598 | 103 | 1 | ENSMUST00 | -1944 | Gm26001 | ENSMUSG00000088614.1 |
| chr4 | 133084656 | 133086042 | 12.01557 | 29.04976 | 26.5957 | Promoter | 4 | 133085474 | 133089146 | 3673 | 1 | ENSMUST00 | 0 | Gm26615 | ENSMUSG00000097088.1 |
| chr5 | 118171301 | 118172726 | 12.01557 | 29.04976 | 26.5957 | Intron (ENSM | 5 | 118169748 | 118189478 | 19731 | 1 | ENSMUST00 | 1554 | Hrk | ENSMUSG00000046607.6 |
| chr5 | 113137726 | 113139131 | 12.01557 | 29.04976 | 26.5957 | Intron (ENSM | 5 | 113086323 | 113163351 | 77029 | 2 | ENSMUST00 | 24220 | 2900026A02R | ENSMUSG00000051339.8 |
| chr7 | 102099743 | 102100661 | 12.01557 | 29.04976 | 26.5957 | Promoter | 7 | 102096880 | 102100229 | 3350 | 2 | ENSMUST00 | 0 | Art5 | ENSMUSG00000070424.6 |
| chr6 | 108668274 | 108668838 | 13.10993 | 29.0328 | 26.58067 | Distal Interge | 6 | 108577039 | 108660934 | 83896 | 2 | ENSMUST00 | -7341 | 0610040F04R | ENSMUSG00000087341.2 |
| chr13 | 37434267 | 37435046 | 12.35373 | 29.00697 | 26.55493 | Promoter | 13 | 37421945 | 37435149 | 13205 | 2 | ENSMUST00 | 103 | RP23-21N15. | ENSMUSG00000100760.1 |
| chr19 | 45229351 | 45230057 | 12.35373 | 29.00697 | 26.55493 | Downstream | 19 | 45232684 | 45235812 | 3129 | 2 | ENSMUST00 | 5755 | Lbx1 | ENSMUSG00000025216.9 |
| chr8 | 105693784 | 105694409 | 12.72034 | 29.0031 | 26.55144 | Exon (ENSMU | 8 | 105690906 | 105698178 | 7273 | 1 | ENSMUST00 | 2879 | Rltpr | ENSMUSG00000050357.8 |
| chr8 | 108886818 | 108887329 | 12.72034 | 29.0031 | 26.55144 | Intron (ENSM | 8 | 108918264 | 108921874 | 3611 | 2 | ENSMUST00 | 34545 | Gm20686 | ENSMUSG00000093697.1 |
| chr11 | 109995458 | 109996216 | 11.28637 | 28.99159 | 26.53998 | Promoter | 11 | 109932190 | 109995845 | 63656 | 2 | ENSMUST00 | 0 | Abca8b | ENSMUSG00000020620.9 |
| chr2 | 74752685 | 74753541 | 12.69572 | 28.96122 | 26.50973 | Promoter | 2 | 74752684 | 74753744 | 1061 | 1 | ENSMUST00 | 2 | Gm14424 | ENSMUSG00000085015.1 |
| chr7 | 143367648 | 143368462 | 13.02911 | 28.89677 | 26.44547 | Intron (ENSM | 7 | 143322775 | 143330650 | 7876 | 2 | ENSMUST00 | -36999 | RP23-124B2. | ENSMUSG00000101360.1 |
| chr9 | 83447260 | 83448234 | 13.02911 | 28.89677 | 26.44547 | Intron (ENSM | 9 | 83391364 | 83441127 | 49764 | 2 | ENSMUST00 | -6134 | Lca5 | ENSMUSG00000032258.10 |
| chr19 | 45783923 | 45784485 | 12.64676 | 28.87791 | 26.4267 | Promoter | 19 | 45750261 | 45783520 | 33260 | 2 | ENSMUST00 | -404 | Mgea5 | ENSMUSG00000025220.7 |
| chr19 | 28975137 | 28975526 | 12.64676 | 28.87791 | 26.4267 | Distal Interge | 19 | 28923530 | 28967800 | 44271 | 2 | ENSMUST00 | -7338 | 4430402I18R | ENSMUSG00000064202.10 |
| chr16 | 62847165 | 62847554 | 13.00239 | 28.85178 | 26.40101 | Promoter | 16 | 62825605 | 62847040 | 21436 | 2 | ENSMUST00 | -126 | Arl13b | ENSMUSG00000022911.10 |
| chr5 | 122131704 | 122132441 | 13.00239 | 28.85178 | 26.40101 | Promoter | 5 | 122125032 | 122132003 | 6972 | 2 | ENSMUST00 | 0 | Ccdc63 | ENSMUSG00000043036.8 |
| chr5 | 150259569 | 150260125 | 13.00239 | 28.85178 | 26.40101 | Promoter | 5 | 150259930 | 150497752 | 237823 | 1 | ENSMUST00 | 0 | Fry | ENSMUSG00000056602.8 |
| chr10 | 128337105 | 128337564 | 10.87763 | 28.84376 | 26.39307 | Promoter | 10 | 128337832 | 128362479 | 24648 | 1 | ENSMUST00 | -268 | Cs | ENSMUSG00000005683.7 |
| chr10 | 72309156 | 72309427 | 10.87763 | 28.84376 | 26.39307 | Promoter | 10 | 72309225 | 72311848 | 2624 | 1 | ENSMUST00 | 0 | Gm9923 | ENSMUSG00000053830.3 |
| chr15 | 82274890 | 82275746 | 10.87763 | 28.84376 | 26.39307 | Promoter | 15 | 82274935 | 82294442 | 19508 | 1 | ENSMUST00 | 0 | Sep-03 | ENSMUSG00000022456.10 |
| chr15 | 88890596 | 88891670 | 10.87763 | 28.84376 | 26.39307 | Promoter | 15 | 88889780 | 88891516 | 1737 | 2 | ENSMUST00 | 0 | Gm20621 | ENSMUSG00000093394.1 |
| chr17 | 35064675 | 35065964 | 10.87763 | 28.84376 | 26.39307 | Promoter | 17 | 35065388 | 35069625 | 4238 | 1 | ENSMUST00 | 0 | Ly6g6c | ENSMUSG00000092586.3 |
| chr2 | 129038543 | 129039584 | 10.87763 | 28.84376 | 26.39307 | Exon (ENSMU | 2 | 129038145 | 129048172 | 10028 | 2 | ENSMUST00 | 8588 | Gm14025 | ENSMUSG00000079051.4 |
| chr8 | 95320354 | 95321027 | 10.87763 | 28.84376 | 26.39307 | Promoter | 8 | 95320438 | 95326323 | 5886 | 1 | ENSMUST00 | 0 | Tepp | ENSMUSG00000090206.2 |
| chr4 | 146503758 | 146504044 | 10.2576 | 28.84245 | 26.39254 | Intron (ENSM | 4 | 146502027 | 146539392 | 37366 | 1 | ENSMUST00 | 1732 | Gm13247 | ENSMUSG00000056300.9 |
| chr1 | 56970764 | 56971260 | 12.97578 | 28.80697 | 26.35731 | Promoter | 1 | 56971412 | 56975197 | 3786 | 1 | ENSMUST00 | -152 | 9130024F11R | ENSMUSG00000087022.3 |
| chr10 | 4338789 | 4339318 | 12.97578 | 28.80697 | 26.35731 | Intron (ENSM | 10 | 4367475 | 4387462 | 19988 | 2 | ENSMUST00 | 48144 | Zbtb2 | ENSMUSG00000075327.4 |
| chr12 | 108832249 | 108833572 | 12.97578 | 28.80697 | 26.35731 | Promoter | 12 | 108827224 | 108831617 | 4394 | 2 | ENSMUST00 | -633 | Slc25a29 | ENSMUSG00000021265.3 |
| chr16 | 35659993 | 35660721 | 12.97578 | 28.80697 | 26.35731 | Promoter | 16 | 35659435 | 35660280 | 846 | 1 | ENSMUST00 | 559 | Sema5b | ENSMUSG00000052133.11 |
| chr6 | 128285871 | 128286241 | 12.97578 | 28.80697 | 26.35731 | Intron (ENSM | 6 | 128294096 | 128294506 | 411 | 1 | ENSMUST00 | -7855 | Gm21893 | ENSMUSG00000095262.1 |
| chr7 | 64501706 | 64502548 | 12.97578 | 28.80697 | 26.35731 | Promoter | 7 | 64501949 | 64695307 | 193359 | 1 | ENSMUST00 | 0 | Apba2 | ENSMUSG00000030519.9 |
| chr7 | 143464792 | 143465507 | 12.97578 | 28.80697 | 26.35731 | Distal Interge | 7 | 143458350 | 143461050 | 2701 | 2 | ENSMUST00 | -3743 | Cdkn1c | ENSMUSG00000037664.7 |
| chr10 | 22140688 | 22141095 | 8.67572 | 28.77482 | 26.32555 | Intron (ENSM | 10 | 22117362 | 22148610 | 31249 | 2 | ENSMUST00 | 7515 | E030030I06R | ENSMUSG00000097327.2 |
| chr16 | 5306755 | 5307372 | 12.94928 | 28.76233 | 26.31305 | Distal Interge | 16 | 5324036 | 5324157 | 122 | 2 | ENSMUST00 | 16785 | n-R5s30 | ENSMUSG00000084573.1 |
| chr2 | 82053088 | 82053888 | 12.94928 | 28.76233 | 26.31305 | Promoter | 2 | 82053222 | 82259879 | 206658 | 1 | ENSMUST00 | 0 | Zfp804a | ENSMUSG00000070866.4 |
| chr1 | 74881991 | 74882898 | 12.5733 | 28.75278 | 26.30366 | 5' UTR | 1 | 74882407 | 74884608 | 2202 | 2 | ENSMUST00 | 1710 | Fev | ENSMUSG00000055197.4 |
| chr11 | 103749297 | 103750128 | 12.5733 | 28.75278 | 26.30366 | Promoter | 11 | 103727364 | 103749821 | 22458 | 2 | ENSMUST00 | 0 | Wnt9b | ENSMUSG00000018486.2 |
| chr12 | 22905921 | 22906606 | 12.5733 | 28.75278 | 26.30366 | Distal Interge | 12 | 22982386 | 23009998 | 27613 | 2 | ENSMUST00 | 103392 | 2410018L13R | ENSMUSG00000073164.6 |
| chr12 | 19288312 | 19289425 | 12.5733 | 28.75278 | 26.30366 | Distal Interge | 12 | 19387809 | 19390286 | 2478 | 1 | ENSMUST00 | -98384 | Gm5784 | ENSMUSG00000093908.1 |
| chr13 | 48389981 | 48390628 | 12.5733 | 28.75278 | 26.30366 | Distal Interge | 13 | 48272416 | 48273884 | 1469 | 2 | ENSMUST00 | -116098 | A330048O09 | ENSMUSG00000097326.1 |
| chr15 | 85673448 | 85674824 | 12.5733 | 28.75278 | 26.30366 | Distal Interge | 15 | 85706603 | 85706697 | 95 | 1 | ENSMUST00 | -31779 | Mirlet7c-2 | ENSMUSG00000065608.1 |
| chr16 | 20496816 | 20497660 | 12.5733 | 28.75278 | 26.30366 | Distal Interge | 16 | 20498817 | 20509955 | 11139 | 1 | ENSMUST00 | -1157 | Eif2b5 | ENSMUSG00000003235.7 |
| chr16 | 77399883 | 77400966 | 12.5733 | 28.75278 | 26.30366 | Promoter | 16 | 77400847 | 77401208 | 362 | 1 | ENSMUST00 | 0 | Gm21816 | ENSMUSG00000096870.1 |
| chr18 | 11351815 | 11352589 | 12.5733 | 28.75278 | 26.30366 | Distal Interge | 18 | 11633276 | 11743207 | 109932 | 1 | ENSMUST00 | -280687 | Rbbp8 | ENSMUSG00000041238.10 |
| chr18 | 64449642 | 64449991 | 12.5733 | 28.75278 | 26.30366 | Promoter | 18 | 64448519 | 64450014 | 1496 | 2 | ENSMUST00 | 23 | Amd-ps3 | ENSMUSG00000060096.4 |
| chr2 | 25242074 | 25242601 | 12.5733 | 28.75278 | 26.30366 | Promoter | 2 | 25239157 | 25242452 | 3296 | 2 | ENSMUST00 | 0 | 2310002J15R | ENSMUSG00000036731.7 |
| chr2 | 92370585 | 92371544 | 12.5733 | 28.75278 | 26.30366 | Promoter | 2 | 92365046 | 92370986 | 5941 | 2 | ENSMUST00 | 0 | Gyltl1b | ENSMUSG00000040434.11 |
| chr2 | 144172304 | 144172816 | 12.5733 | 28.75278 | 26.30366 | Downstream | 2 | 144174315 | 144179086 | 4772 | 2 | ENSMUST00 | 6270 | Gm5535 | ENSMUSG00000074758.4 |
| chr3 | 95230005 | 95230688 | 12.5733 | 28.75278 | 26.30366 | Intron (ENSM | 3 | 95228952 | 95234582 | 5631 | 1 | ENSMUST00 | 1054 | Cdc42se1 | ENSMUSG00000046722.9 |
| chr3 | 64039957 | 64040423 | 12.5733 | 28.75278 | 26.30366 | Distal Interge | 3 | 64053031 | 64055610 | 2580 | 2 | ENSMUST00 | 15187 | Vmn2r-ps3 | ENSMUSG00000090303.3 |
| chr4 | 139580247 | 139582371 | 12.5733 | 28.75278 | 26.30366 | Intron (ENSM | 4 | 139574720 | 139620382 | 45663 | 1 | ENSMUST00 | 5528 | Iffo2 | ENSMUSG00000041025.10 |
| chr4 | 154246634 | 154246904 | 12.5733 | 28.75278 | 26.30366 | Intron (ENSM | 4 | 154249296 | 154252217 | 2922 | 1 | ENSMUST00 | -2392 | Megf6 | ENSMUSG00000057751.9 |
| chr5 | 137787829 | 137788336 | 12.5733 | 28.75278 | 26.30366 | Promoter | 5 | 137787798 | 137822621 | 34824 | 1 | ENSMUST00 | 32 | Zcwpw1 | ENSMUSG00000037108.8 |
| chr5 | 117240148 | 117241406 | 12.5733 | 28.75278 | 26.30366 | Promoter | 5 | 117240667 | 117273869 | 33203 | 1 | ENSMUST00 | 0 | Taok3 | ENSMUSG00000061288.8 |
| chr6 | 56652041 | 56652641 | 12.5733 | 28.75278 | 26.30366 | Distal Interge | 6 | 56600818 | 56600949 | 132 | 2 | ENSMUST00 | -51093 | Gm22493 | ENSMUSG00000077682.1 |
| chr7 | 29247545 | 29248732 | 12.5733 | 28.75278 | 26.30366 | Promoter | 7 | 29246561 | 29248432 | 1872 | 2 | ENSMUST00 | 0 | 2200002D01R | ENSMUSG00000030587.4 |
| chr7 | 5125706 | 5126321 | 12.5733 | 28.75278 | 26.30366 | Promoter | 7 | 5124938 | 5125950 | 1013 | 2 | ENSMUST00 | 0 | Rasl2-9 | ENSMUSG00000083649.5 |
| chr9 | 98973035 | 98973450 | 12.5733 | 28.75278 | 26.30366 | Distal Interge | 9 | 98986373 | 99002021 | 15649 | 1 | ENSMUST00 | -12923 | Faim | ENSMUSG00000032463.5 |
| chr9 | 65398264 | 65398786 | 12.5733 | 28.75278 | 26.30366 | Promoter | 9 | 65398488 | 65412708 | 14221 | 1 | ENSMUST00 | 0 | Rasl12 | ENSMUSG00000041696.8 |
| chrX | 139997773 | 139998751 | 12.5733 | 28.75278 | 26.30366 | Promoter | 20 | 139889511 | 139998595 | 109085 | 2 | ENSMUST00 | 0 | Rbm41 | ENSMUSG00000031433.10 |
| chr3 | 122273780 | 122274208 | 12.92288 | 28.71786 | 26.27042 | Promoter | 3 | 122274414 | 122285269 | 10856 | 1 | ENSMUST00 | -206 | Dnttip2 | ENSMUSG00000039756.8 |
| chr7 | 45161878 | 45162477 | 12.92288 | 28.71786 | 26.27042 | Distal Interge | 7 | 45163921 | 45176138 | 12218 | 1 | ENSMUST00 | -1444 | Slc17a7 | ENSMUSG00000070570.4 |
| chr9 | 103092300 | 103092943 | 12.92288 | 28.71786 | 26.27042 | Intron (ENSM | 9 | 103111787 | 103162657 | 50871 | 1 | ENSMUST00 | -18844 | Rab6b | ENSMUSG00000032549.7 |
| chr12 | 16591340 | 16591966 | 11.47719 | 28.71741 | 26.27007 | Distal Interge | 12 | 16535669 | 16589770 | 54102 | 2 | ENSMUST00 | -1571 | Lpin1 | ENSMUSG00000020593.9 |
| chr10 | 40258450 | 40259635 | 12.8966 | 28.67356 | 26.22642 | Promoter | 10 | 40258467 | 40258680 | 214 | 1 | ENSMUST00 | 0 | Rnu3a | ENSMUSG00000065870.2 |
| chr1 | 86524733 | 86525362 | 12.526 | 28.67215 | 26.22511 | Distal Interge | 1 | 86526726 | 86530712 | 3987 | 1 | ENSMUST00 | -1364 | Ptma | ENSMUSG00000026238.9 |
| chr15 | 98408270 | 98408989 | 12.87042 | 28.62942 | 26.18246 | Intron (ENSM | 15 | 98398445 | 98416268 | 17824 | 1 | ENSMUST00 | 9826 | C12orf54 | ENSMUSG00000099353.1 |
| chr7 | 128687931 | 128688650 | 12.13259 | 28.62427 | 26.1774 | Promoter | 7 | 128688489 | 128696274 | 7786 | 1 | ENSMUST00 | 0 | Inpp5f | ENSMUSG00000042105.13 |
| chr11 | 3201976 | 3202582 | 5.64303 | 28.58646 | 26.13994 | Promoter | 11 | 3202392 | 3244581 | 42190 | 1 | ENSMUST00 | 0 | Eif4enif1 | ENSMUSG00000020454.12 |
| chr18 | 37655501 | 37656192 | 12.84435 | 28.58546 | 26.13894 | Promoter | 18 | 37655907 | 37658737 | 2831 | 1 | ENSMUST00 | 0 | 3222401L13R | ENSMUSG00000073590.2 |
| chr2 | 164404436 | 164405002 | 12.84435 | 28.58546 | 26.13894 | Promoter | 2 | 164389393 | 164404612 | 15220 | 2 | ENSMUST00 | 0 | Matn4 | ENSMUSG00000016995.12 |
| chr10 | 121930261 | 121932096 | 12.79252 | 28.49802 | 26.05175 | Exon (ENSMU | 10 | 121806640 | 121823133 | 16494 | 2 | ENSMUST00 | -107129 | Srgap1 | ENSMUSG00000020121.10 |
| chr13 | 83523402 | 83525575 | 12.79252 | 28.49802 | 26.05175 | Intron (ENSM | 13 | 83504034 | 83667079 | 163046 | 1 | ENSMUST00 | 19369 | Mef2c | ENSMUSG00000005583.11 |
| chr12 | 88354333 | 88354737 | 12.40753 | 28.46989 | 26.02371 | Distal Interge | 12 | 88360554 | 88461719 | 101166 | 1 | ENSMUST00 | -5817 | Adck1 | ENSMUSG00000021044.9 |
| chr18 | 57085470 | 57085831 | 12.40753 | 28.46989 | 26.02371 | Distal Interge | 18 | 57133090 | 57297467 | 164378 | 1 | ENSMUST00 | -47259 | Megf10 | ENSMUSG00000024593.10 |
| chr19 | 40561490 | 40561700 | 12.40753 | 28.46989 | 26.02371 | Promoter | 19 | 40560537 | 40561058 | 522 | 2 | ENSMUST00 | -433 | Gm15801 | ENSMUSG00000096592.1 |
| chr4 | 134342988 | 134343821 | 11.68713 | 28.46951 | 26.02371 | Promoter | 4 | 134343181 | 134352677 | 9497 | 1 | ENSMUST00 | 0 | Slc30a2 | ENSMUSG00000028836.9 |
| chr4 | 138943429 | 138943876 | 12.40753 | 28.46989 | 26.02371 | Distal Interge | 4 | 138967112 | 138968360 | 1249 | 1 | ENSMUST00 | -23236 | Rnf186 | ENSMUSG00000070661.6 |
| chr8 | 3400828 | 3402931 | 12.40753 | 28.46989 | 26.02371 | Intron (ENSM | 8 | 3393073 | 3429639 | 36567 | 1 | ENSMUST00 | 7756 | Arhgef18 | ENSMUSG00000004568.7 |
| chr9 | 121641939 | 121643224 | 12.40753 | 28.46989 | 26.02371 | Promoter | 9 | 121642716 | 121672954 | 30239 | 1 | ENSMUST00 | 0 | Vipr1 | ENSMUSG00000032528.4 |
| chr13 | 21172407 | 21173648 | 12.76676 | 28.45456 | 26.00909 | Distal Interge | 13 | 21179445 | 21180416 | 972 | 1 | ENSMUST00 | -5797 | Trim27 | ENSMUSG00000021326.6 |

| | | | | | | | | | | | | | | | |
|---|---|---|---|---|---|---|---|---|---|---|---|---|---|---|---|
| chr3 | 84189922 | 84191271 | 12.76676 | 28.45456 | 26.00909 | Promoter | 3 | 84177251 | 84190523 | 13273 | 2 | ENSMUST00 | 0 | Trim2 | ENSMUSG00000027993.11 |
| chr5 | 123135355 | 123136555 | 12.76676 | 28.45456 | 26.00909 | Exon (ENSMU | 5 | 123133723 | 123138109 | 4387 | 2 | ENSMUST00 | 1554 | AI480526 | ENSMUSG00000090086.2 |
| chr1 | 180936532 | 180937314 | 12.71555 | 28.36811 | 25.92295 | Promoter | 1 | 180936871 | 180937810 | 940 | 1 | ENSMUST00 | 0 | Lefty1 | ENSMUSG00000038793.8 |
| chr11 | 48788450 | 48788715 | 12.71555 | 28.36811 | 25.92295 | Distal Interge | 11 | 48800332 | 48804974 | 4643 | 1 | ENSMUST00 | -11617 | Gnb2l1 | ENSMUSG00000020372.10 |
| chr4 | 47207918 | 47208969 | 12.71555 | 28.36811 | 25.92295 | Promoter | 4 | 47208161 | 47313163 | 105003 | 1 | ENSMUST00 | 0 | Col15a1 | ENSMUSG00000028339.12 |
| chr4 | 132270224 | 132270904 | 12.71555 | 28.36811 | 25.92295 | Promoter | 4 | 132270080 | 132270213 | 134 | 2 | ENSMUST00 | -12 | Rnu11 | ENSMUSG00000077323.1 |
| chr5 | 9261125 | 9261360 | 12.71555 | 28.36811 | 25.92295 | Promoter | 5 | 9260945 | 9261388 | 444 | 2 | ENSMUST00 | 28 | Gm7332 | ENSMUSG00000080875.3 |
| chr8 | 69881191 | 69882622 | 12.71555 | 28.36811 | 25.92295 | Exon (ENSMU | 8 | 69880369 | 69887687 | 7319 | 2 | ENSMUST00 | 5065 | Cilp2 | ENSMUSG00000044006.8 |
| chr2 | 146834352 | 146835437 | 12.33751 | 28.35017 | 25.90545 | Intron (ENSM | 2 | 146828327 | 146839732 | 11406 | 2 | ENSMUST00 | 4295 | Gm14114 | ENSMUSG00000087618.1 |
| chr15 | 80083524 | 80084248 | 10.30097 | 28.33038 | 25.88568 | Promoter | 15 | 80077796 | 80083353 | 5558 | 2 | ENSMUST00 | -172 | Rpl3 | ENSMUSG00000060036.8 |
| chr17 | 6492603 | 6492890 | 10.30097 | 28.33038 | 25.88568 | Exon (ENSMU | 17 | 6441402 | 6467022 | 25621 | 2 | ENSMUST00 | -25582 | Tmem181b-p | ENSMUSG00000096780.2 |
| chr18 | 68057848 | 68058647 | 10.30097 | 28.33038 | 25.88568 | Intron (ENSM | 18 | 67933257 | 68255549 | 322293 | 1 | ENSMUST00 | 124592 | Ldlrad4 | ENSMUSG00000024544.8 |
| chr7 | 16924295 | 16924643 | 10.30097 | 28.33038 | 25.88568 | Promoter | 7 | 16915379 | 16924114 | 8736 | 2 | ENSMUST00 | -182 | Calm3 | ENSMUSG00000019370.10 |
| chr8 | 67956367 | 67957365 | 10.30097 | 28.33038 | 25.88568 | Promoter | 8 | 67956770 | 67978051 | 21282 | 1 | ENSMUST00 | 0 | Gm15991 | ENSMUSG00000087266.1 |
| chrX | 104670060 | 104671115 | 10.30097 | 28.33038 | 25.88568 | Promoter | 20 | 104536969 | 104671064 | 134096 | 2 | ENSMUST00 | 0 | Zdhhc15 | ENSMUSG00000033906.5 |
| chr13 | 23574767 | 23576227 | 12.6901 | 28.32512 | 25.88085 | Promoter | 13 | 23575763 | 23576322 | 560 | 1 | ENSMUST00 | 0 | Hist1h3d | ENSMUSG00000099583.1 |
| chr17 | 45548984 | 45549823 | 12.6901 | 28.32512 | 25.88085 | Promoter | 17 | 45541940 | 45549677 | 7738 | 2 | ENSMUST00 | 0 | Tmem151b | ENSMUSG00000096847.1 |
| chr14 | 51481628 | 51482787 | 12.66475 | 28.2823 | 25.83814 | Intron (ENSM | 14 | 51452003 | 51460723 | 8721 | 1 | ENSMUST00 | 29626 | Vmn2r89 | ENSMUSG00000070448.6 |
| chr15 | 36695381 | 36695976 | 12.66475 | 28.2823 | 25.83814 | Distal Interge | 15 | 36603496 | 36608973 | 5478 | 2 | ENSMUST00 | -86409 | Pabpc1 | ENSMUSG00000022283.9 |
| chr7 | 97748327 | 97749536 | 12.66475 | 28.2823 | 25.83814 | Distal Interge | 7 | 97726379 | 97738247 | 11869 | 2 | ENSMUST00 | -10081 | Aqp11 | ENSMUSG00000042797.8 |
| chr1 | 120602767 | 120603795 | 11.22292 | 28.25542 | 25.81138 | Promoter | 1 | 120602418 | 120607992 | 5575 | 1 | ENSMUST00 | 350 | En1 | ENSMUSG00000058665.8 |
| chr10 | 23076808 | 23077840 | 11.22292 | 28.25542 | 25.81138 | Distal Interge | 10 | 23166865 | 23166971 | 107 | 2 | ENSMUST00 | 89131 | Gm24655 | ENSMUSG00000094033.1 |
| chr11 | 84522497 | 84523298 | 11.22292 | 28.25542 | 25.81138 | Promoter | 11 | 84521896 | 84522711 | 816 | 2 | ENSMUST00 | 0 | Lhx1 | ENSMUSG00000018698.10 |
| chr14 | 68123433 | 68124023 | 11.22292 | 28.25542 | 25.81138 | Promoter | 14 | 68082590 | 68124846 | 42257 | 2 | ENSMUST00 | 823 | Nefm | ENSMUSG00000022054.6 |
| chr14 | 7657950 | 7658758 | 11.22292 | 28.25542 | 25.81138 | Distal Interge | 14 | 7640895 | 7643972 | 3078 | 2 | ENSMUST00 | -13979 | Gm10128 | ENSMUSG00000063277.7 |
| chr15 | 12718736 | 12720598 | 11.22292 | 28.25542 | 25.81138 | Intron (ENSM | 15 | 12443331 | 12739924 | 296594 | 2 | ENSMUST00 | 19326 | Pdzd2 | ENSMUSG00000022197.10 |
| chr16 | 97534754 | 97536023 | 11.22292 | 28.25542 | 25.81138 | Promoter | 16 | 97535308 | 97560339 | 25032 | 1 | ENSMUST00 | 0 | Mx2 | ENSMUSG00000023341.9 |
| chr19 | 5529510 | 5530100 | 11.22292 | 28.25542 | 25.81138 | Distal Interge | 19 | 5495278 | 5510489 | 15212 | 2 | ENSMUST00 | -19022 | Snx32 | ENSMUSG00000056185.4 |
| chr2 | 36103216 | 36104359 | 11.22292 | 28.25542 | 25.81138 | Promoter | 2 | 36081953 | 36104060 | 22108 | 2 | ENSMUST00 | 0 | Lhx6 | ENSMUSG00000026890.14 |
| chr6 | 128763655 | 128764152 | 11.22292 | 28.25542 | 25.81138 | Distal Interge | 6 | 128780043 | 128788486 | 8444 | 2 | ENSMUST00 | 24334 | Klrb1c | ENSMUSG00000030325.11 |
| chr6 | 83135625 | 83136526 | 11.22292 | 28.25542 | 25.81138 | Promoter | 6 | 83135812 | 83152579 | 16768 | 1 | ENSMUST00 | 0 | Rtkn | ENSMUSG00000034930.10 |
| chr7 | 27361634 | 27362010 | 11.22292 | 28.25542 | 25.81138 | Exon (ENSMU | 7 | 27363314 | 27363401 | 88 | 1 | ENSMUST00 | -1304 | Gm26043 | ENSMUSG00000080562.1 |
| chr8 | 114205483 | 114206138 | 11.22292 | 28.25542 | 25.81138 | Promoter | 8 | 114205612 | 114230325 | 24714 | 1 | ENSMUST00 | 0 | Vat1l | ENSMUSG00000046844.6 |
| chr8 | 122281032 | 122281852 | 11.22292 | 28.25542 | 25.81138 | Promoter | 8 | 122282141 | 122337251 | 55111 | 1 | ENSMUST00 | -289 | Zfpm1 | ENSMUSG00000049577.9 |
| chr15 | 76512105 | 76512529 | 10.79468 | 28.07545 | 25.63352 | Promoter | 15 | 76502015 | 76511951 | 9937 | 2 | ENSMUST00 | -155 | Dgat1 | ENSMUSG00000022555.6 |
| chr3 | 50381679 | 50382555 | 12.17718 | 28.07544 | 25.63351 | Exon (ENSMU | 3 | 50364936 | 50443613 | 78678 | 2 | ENSMUST00 | 61058 | Slc7a11 | ENSMUSG00000027737.5 |
| chr15 | 84663136 | 84663637 | 11.1233 | 28.07331 | 25.63184 | Distal Interge | 15 | 84669620 | 84703673 | 34054 | 1 | ENSMUST00 | -5983 | Prr5 | ENSMUSG00000036106.10 |
| chr16 | 33400159 | 33400729 | 11.1233 | 28.07331 | 25.63184 | Promoter | 16 | 33400445 | 33498643 | 98199 | 1 | ENSMUST00 | 0 | Zfp148 | ENSMUSG00000022811.11 |
| chr17 | 6601781 | 6601986 | 11.81534 | 28.07186 | 25.63041 | Promoter | 17 | 6601792 | 6603934 | 2143 | 1 | ENSMUST00 | 0 | Dynlt1c | ENSMUSG00000000579.9 |
| chr7 | 19369892 | 19370254 | 12.53952 | 28.07054 | 25.6291 | Exon (ENSMU | 7 | 19368498 | 19369911 | 1414 | 1 | ENSMUST00 | 1395 | Ppp1r13l | ENSMUSG00000040734.9 |
| chr1 | 74947926 | 74948425 | 12.8669 | 28.00426 | 25.56298 | Exon (ENSMU | 1 | 74947724 | 74951442 | 3719 | 2 | ENSMUST00 | 3017 | Ihh | ENSMUSG00000006538.10 |
| chr1 | 36303239 | 36303677 | 12.8669 | 28.00426 | 25.56298 | Distal Interge | 1 | 36307733 | 36319203 | 11471 | 1 | ENSMUST00 | -4056 | Arid5a | ENSMUSG00000037447.11 |
| chr1 | 180483082 | 180483783 | 12.8669 | 28.00426 | 25.56298 | Promoter | 1 | 180432387 | 180483467 | 51081 | 2 | ENSMUST00 | 0 | 6330403A02 | ENSMUSG00000053963.6 |
| chr10 | 128645449 | 128645963 | 12.8669 | 28.00426 | 25.56298 | Promoter | 10 | 128632415 | 128645826 | 13412 | 2 | ENSMUST00 | 0 | Ikzf4 | ENSMUSG00000002578.10 |
| chr10 | 103028287 | 103028941 | 12.8669 | 28.00426 | 25.56298 | Promoter | 10 | 103007847 | 103028777 | 20931 | 2 | ENSMUST00 | 0 | Alx1 | ENSMUSG00000036602.8 |
| chr11 | 58914353 | 58914857 | 12.8669 | 28.00426 | 25.56298 | Distal Interge | 11 | 58917908 | 58926608 | 8701 | 1 | ENSMUST00 | -3051 | Btnl10 | ENSMUSG00000020490.11 |
| chr12 | 84698611 | 84699076 | 12.8669 | 28.00426 | 25.56298 | Promoter | 12 | 84677278 | 84698807 | 21530 | 2 | ENSMUST00 | 0 | Syndig1l | ENSMUSG00000071234.2 |
| chr12 | 23800158 | 23800552 | 12.8669 | 28.00426 | 25.56298 | Distal Interge | 12 | 23778912 | 23780265 | 1354 | 2 | ENSMUST00 | -19894 | Gm10330 | ENSMUSG00000071362.2 |
| chr13 | 113031939 | 113032555 | 12.8669 | 28.00426 | 25.56298 | Distal Interge | 13 | 113035111 | 113038324 | 3214 | 1 | ENSMUST00 | -2556 | Cdc20b | ENSMUSG00000078926.3 |
| chr15 | 89210046 | 89210714 | 12.8669 | 28.00426 | 25.56298 | Downstream | 15 | 89211559 | 89215012 | 3454 | 2 | ENSMUST00 | 4298 | Gm26798 | ENSMUSG00000097578.1 |
| chr16 | 17561072 | 17561760 | 12.8669 | 28.00426 | 25.56298 | Promoter | 16 | 17544465 | 17561247 | 16783 | 2 | ENSMUST00 | 0 | 4930451C15 | ENSMUSG00000022759.8 |
| chr16 | 16828920 | 16829510 | 12.8669 | 28.00426 | 25.56298 | Promoter | 16 | 16753016 | 16829313 | 76298 | 2 | ENSMUST00 | 0 | Spag6 | ENSMUSG00000022783.6 |
| chr19 | 7182366 | 7182922 | 12.8669 | 28.00426 | 25.56298 | Intron (ENSM | 19 | 7199540 | 7200464 | 925 | 2 | ENSMUST00 | 17542 | Otub1 | ENSMUSG00000024767.6 |
| chr2 | 164880071 | 164881078 | 12.8669 | 28.00426 | 25.56298 | Promoter | 2 | 164880052 | 164883110 | 3059 | 1 | ENSMUST00 | 20 | Pcif1 | ENSMUSG00000039849.5 |
| chr2 | 169621331 | 169621927 | 12.8669 | 28.00426 | 25.56298 | Distal Interge | 2 | 169633013 | 169913093 | 280081 | 1 | ENSMUST00 | -11086 | Tshz2 | ENSMUSG00000047907.11 |
| chr2 | 175640382 | 175640925 | 12.8669 | 28.00426 | 25.56298 | Distal Interge | 2 | 175643978 | 175651401 | 7424 | 1 | ENSMUST00 | -3053 | Gm11007 | ENSMUSG00000094475.2 |
| chr3 | 41408461 | 41409150 | 12.8669 | 28.00426 | 25.56298 | Distal Interge | 3 | 41441343 | 41441467 | 125 | 2 | ENSMUST00 | 32317 | Gm25487 | ENSMUSG00000089350.1 |
| chr4 | 41097986 | 41098431 | 12.8669 | 28.00426 | 25.56298 | Promoter | 4 | 41092722 | 41098183 | 5462 | 2 | ENSMUST00 | 0 | Aqp3 | ENSMUSG00000028435.8 |
| chr4 | 120716190 | 120717054 | 12.8669 | 28.00426 | 25.56298 | Exon (ENSMU | 4 | 120696138 | 120747248 | 51111 | 2 | ENSMUST00 | 30194 | Kcnq4 | ENSMUSG00000028631.7 |
| chr5 | 36869640 | 36870277 | 12.8669 | 28.00426 | 25.56298 | Intron (ENSM | 5 | 36868513 | 36955078 | 86566 | 1 | ENSMUST00 | 1128 | Ppp2r2c | ENSMUSG00000029120.7 |
| chr5 | 64811664 | 64813679 | 12.8669 | 28.00426 | 25.56298 | Promoter | 5 | 64812339 | 64829482 | 17144 | 1 | ENSMUST00 | 0 | Klf3 | ENSMUSG00000029178.9 |
| chr6 | 17305865 | 17306709 | 12.8669 | 28.00426 | 25.56298 | Promoter | 6 | 17306335 | 17341324 | 34990 | 1 | ENSMUST00 | 0 | Cav1 | ENSMUSG00000007655.11 |
| chr7 | 30219547 | 30220166 | 12.8669 | 28.00426 | 25.56298 | Distal Interge | 7 | 30224133 | 30227000 | 2868 | 2 | ENSMUST00 | 6834 | Tbcb | ENSMUSG00000006095.7 |
| chr8 | 4165505 | 4166037 | 12.8669 | 28.00426 | 25.56298 | Promoter | 8 | 4166567 | 4186880 | 20314 | 1 | ENSMUST00 | -530 | Evi5l | ENSMUSG00000011832.10 |
| chr8 | 119895774 | 119896291 | 12.8669 | 28.00426 | 25.56298 | Distal Interge | 8 | 119910360 | 119957555 | 47196 | 1 | ENSMUST00 | -14069 | Usp10 | ENSMUSG00000031826.14 |
| chr9 | 113441834 | 113442248 | 12.8669 | 28.00426 | 25.56298 | Distal Interge | 9 | 113403475 | 113403771 | 297 | 1 | ENSMUST00 | 38360 | Gm22492 | ENSMUSG00000087817.1 |
| chrX | 20797876 | 20799031 | 12.8669 | 28.00426 | 25.56298 | Promoter | 20 | 20797814 | 20860497 | 62684 | 1 | ENSMUST00 | 63 | Araf | ENSMUSG00000001127.7 |
| chrX | 74372741 | 74373505 | 12.8669 | 28.00426 | 25.56298 | Promoter | 20 | 74367465 | 74373218 | 5754 | 2 | ENSMUST00 | 0 | Ubl4 | ENSMUSG00000015290.10 |
| chrX | 100493607 | 100494177 | 12.8669 | 28.00426 | 25.56298 | Promoter | 20 | 100494291 | 100516125 | 21835 | 1 | ENSMUST00 | -114 | Igbp1 | ENSMUSG00000031221.7 |
| chrX | 74249437 | 74250107 | 12.8669 | 28.00426 | 25.56298 | Promoter | 20 | 74223461 | 74249820 | 26360 | 2 | ENSMUST00 | 0 | Flna | ENSMUSG00000031328.10 |
| chrX | 134807945 | 134809034 | 12.8669 | 28.00426 | 25.56298 | Promoter | 20 | 134804147 | 134808996 | 4850 | 2 | ENSMUST00 | 0 | Armcx2 | ENSMUSG00000033436.8 |
| chrX | 7721487 | 7722729 | 12.8669 | 28.00426 | 25.56298 | Promoter | 20 | 7721973 | 7726292 | 4320 | 1 | ENSMUST00 | 0 | Wdr45 | ENSMUSG00000039382.6 |
| chrX | 166170399 | 166170959 | 12.8669 | 28.00426 | 25.56298 | Promoter | 20 | 166170454 | 166189804 | 19351 | 1 | ENSMUST00 | 0 | Gemin8 | ENSMUSG00000040621.9 |
| chrX | 44789565 | 44790255 | 12.8669 | 28.00426 | 25.56298 | Promoter | 20 | 44786570 | 44790161 | 3592 | 2 | ENSMUST00 | 0 | Dcaf12l1 | ENSMUSG00000045284.4 |
| chrX | 102156718 | 102157107 | 12.8669 | 28.00426 | 25.56298 | Promoter | 20 | 102141716 | 102157091 | 15376 | 2 | ENSMUST00 | 0 | Ercc6l | ENSMUSG00000051220.2 |
| chrX | 53607595 | 53608423 | 12.8669 | 28.00426 | 25.56298 | Promoter | 20 | 53608032 | 53608432 | 401 | 1 | ENSMUST00 | 0 | Cxx1c | ENSMUSG00000051851.5 |
| chrX | 95968142 | 95968722 | 12.8669 | 28.00426 | 25.56298 | Distal Interge | 20 | 95949435 | 95956962 | 7528 | 2 | ENSMUST00 | -11181 | Las1l | ENSMUSG00000057421.7 |
| chrX | 102933670 | 102934266 | 12.8669 | 28.00426 | 25.56298 | Distal Interge | 20 | 102908905 | 102909651 | 747 | 1 | ENSMUST00 | 24766 | 1700011M02 | ENSMUSG00000079476.1 |
| chrX | 12006144 | 12007515 | 12.8669 | 28.00426 | 25.56298 | Distal Interge | 20 | 11984475 | 11984650 | 176 | 1 | ENSMUST00 | 21670 | Gm14512 | ENSMUSG00000081947.1 |
| chrX | 75723585 | 75723797 | 12.8669 | 28.00426 | 25.56298 | Promoter | 20 | 75723672 | 75724126 | 455 | 1 | ENSMUST00 | 0 | Gm7153 | ENSMUSG00000081996.1 |
| chrX | 153723077 | 153724722 | 12.8669 | 28.00426 | 25.56298 | Promoter | 20 | 153723590 | 153741264 | 17675 | 1 | ENSMUST00 | 0 | 2210013O21 | ENSMUSG00000086316.2 |
| chr13 | 78194847 | 78195526 | 12.49011 | 27.98691 | 25.54734 | Promoter | 13 | 78189733 | 78196373 | 6641 | 2 | ENSMUST00 | 847 | Nr2f1 | ENSMUSG00000069171.8 |
| chr2 | 170731761 | 170732150 | 12.84376 | 27.96542 | 25.52592 | Promoter | 2 | 170731807 | 170879769 | 147963 | 1 | ENSMUST00 | 0 | Dok5 | ENSMUSG00000027560.4 |
| chr18 | 45055938 | 45056205 | 12.46556 | 27.94533 | 25.5059 | Distal Interge | 18 | 45089326 | 45090314 | 989 | 2 | ENSMUST00 | 34109 | Gm26928 | ENSMUSG00000098244.1 |
| chr2 | 79428460 | 79429179 | 12.46556 | 27.94533 | 25.5059 | Promoter | 2 | 79331752 | 79428954 | 97203 | 2 | ENSMUST00 | 0 | Cerkl | ENSMUSG00000075256.4 |
| chr7 | 118539979 | 118542150 | 12.46556 | 27.94533 | 25.5059 | Exon (ENSMU | 7 | 118525062 | 118533356 | 8295 | 2 | ENSMUST00 | -6624 | Coq7 | ENSMUSG00000030652.5 |
| chr9 | 104407784 | 104408446 | 12.46556 | 27.94533 | 25.5059 | Distal Interge | 9 | 104426113 | 104426187 | 75 | 1 | ENSMUST00 | -17667 | Mir2136 | ENSMUSG00000089406.1 |
| chr14 | 3208488 | 3209075 | 8.33652 | 27.9318 | 25.49284 | Intron (ENSM | 14 | 3210975 | 3211821 | 847 | 2 | ENSMUST00 | 2746 | Gm17222 | ENSMUSG00000095512.1 |
| chr3 | 79179754 | 79182117 | 9.80908 | 27.92633 | 25.48738 | Promoter | 3 | 79169449 | 79181340 | 11892 | 2 | ENSMUST00 | 0 | Rapgef2 | ENSMUSG00000062232.9 |
| chr7 | 126847377 | 126848103 | 9.80908 | 27.92633 | 25.48738 | Promoter | 7 | 126847416 | 126849525 | 2110 | 1 | ENSMUST00 | 0 | Doc2a | ENSMUSG00000052301.9 |
| chr10 | 127667930 | 127668854 | 12.81687 | 27.9203 | 25.48155 | Promoter | 10 | 127664984 | 127668568 | 3585 | 2 | ENSMUST00 | 0 | Nab2 | ENSMUSG00000025402.7 |
| chr11 | 102989818 | 102990461 | 11.71324 | 27.89317 | 25.45456 | Intron (ENSM | 11 | 102994049 | 102997548 | 3500 | 2 | ENSMUST00 | 7087 | Dcakd | ENSMUSG00000020935.3 |
| chr2 | 173019633 | 173020152 | 11.71324 | 27.89317 | 25.45456 | Promoter | 2 | 173020498 | 173034734 | 14237 | 1 | ENSMUST00 | -346 | Rbm38 | ENSMUSG00000027510.12 |
| chr3 | 146107243 | 146108279 | 11.71324 | 27.89317 | 25.45456 | Promoter | 3 | 146040526 | 146108130 | 67605 | 2 | ENSMUST00 | 0 | Wdr63 | ENSMUSG00000043020.8 |
| chr5 | 125140272 | 125141740 | 11.71324 | 27.89317 | 25.45456 | Intron (ENSM | 5 | 125017835 | 125124900 | 107066 | 2 | ENSMUST00 | -15373 | Ncor2 | ENSMUSG00000029478.11 |
| chr15 | 99583665 | 99584406 | 12.79009 | 27.87535 | 25.43694 | 3' UTR | 15 | 99579056 | 99584545 | 5490 | 1 | ENSMUST00 | 4610 | Aqp2 | ENSMUSG00000023013.4 |
| chr2 | 38658123 | 38658664 | 12.79009 | 27.87535 | 25.43694 | Distal Interge | 2 | 38671158 | 38687651 | 16494 | 1 | ENSMUST00 | -12494 | Gpr144-ps | ENSMUSG00000083722.1 |
| chr9 | 41890312 | 41891051 | 12.79009 | 27.87535 | 25.43694 | Distal Interge | 9 | 41969739 | 41974587 | 4849 | 2 | ENSMUST00 | 83536 | Sorl1 | ENSMUSG00000049313.8 |
| chr12 | 33175699 | 33176699 | 12.41673 | 27.8626 | 25.42443 | Intron (ENSM | 12 | 33147805 | 33328849 | 181045 | 1 | ENSMUST00 | 27895 | Atxn7l1 | ENSMUSG00000020564.11 |
| chr11 | 61618794 | 61620874 | 12.76342 | 27.83059 | 25.39249 | Distal Interge | 11 | 61653265 | 61672784 | 19520 | 1 | ENSMUST00 | -32391 | Grap | ENSMUSG00000004837.2 |
| chr3 | 66292304 | 66292928 | 12.39246 | 27.82146 | 25.38344 | Intron (ENSM | 3 | 66243998 | 66296748 | 52751 | 2 | ENSMUST00 | 3820 | Veph1 | ENSMUSG00000027831.9 |
| chr6 | 22875169 | 22876200 | 12.73687 | 27.786 | 25.34813 | Promoter | 6 | 22875502 | 23052916 | 177415 | 1 | ENSMUST00 | 0 | Ptprz1 | ENSMUSG00000068748.4 |
| chr19 | 9836505 | 9836905 | 7.87756 | 27.78156 | 25.34377 | Distal Interge | 19 | 9847562 | 9852808 | 5247 | 1 | ENSMUST00 | -10657 | Scgb2a2 | ENSMUSG00000096872.2 |
| chr7 | 125581064 | 125581546 | 12.36829 | 27.78046 | 25.34268 | Distal Interge | 7 | 125603429 | 125633570 | 30142 | 1 | ENSMUST00 | -21883 | Il21r | ENSMUSG00000030745.8 |
| chr1 | 34120825 | 34122623 | 11.64009 | 27.76484 | 25.32713 | Promoter | 1 | 34121476 | 34164237 | 42762 | 1 | ENSMUST00 | 0 | Dst | ENSMUSG00000026131.12 |
| chr10 | 52022397 | 52022699 | 11.64009 | 27.76484 | 25.32713 | Promoter | 10 | 52022502 | 52028471 | 5970 | 1 | ENSMUST00 | 0 | Vgll2 | ENSMUSG00000049641.7 |
| chr10 | 69924804 | 69925194 | 11.64009 | 27.76484 | 25.32713 | Promoter | 10 | 69925495 | 69926338 | 844 | 1 | ENSMUST00 | -301 | Ank3 | ENSMUSG00000069601.8 |
| chr11 | 70615188 | 70615679 | 11.64009 | 27.76484 | 25.32713 | Promoter | 11 | 70614885 | 70615673 | 789 | 2 | ENSMUST00 | 0 | Chrne | ENSMUSG00000014609.11 |
| chr11 | 52346514 | 52347001 | 11.64009 | 27.76484 | 25.32713 | Distal Interge | 11 | 52360860 | 52389397 | 28538 | 1 | ENSMUST00 | -13859 | Vdac1 | ENSMUSG00000020402.6 |
| chr11 | 75794874 | 75796383 | 11.64009 | 27.76484 | 25.32713 | Promoter | 11 | 75768966 | 75796049 | 27084 | 2 | ENSMUST00 | 0 | Doc2b | ENSMUSG00000020848.7 |
| chr11 | 35120870 | 35121895 | 11.64009 | 27.76484 | 25.32713 | Promoter | 11 | 35121224 | 35708507 | 587284 | 1 | ENSMUST00 | 0 | Slit3 | ENSMUSG00000056427.5 |
| chr12 | 100778728 | 100779497 | 11.64009 | 27.76484 | 25.32713 | Promoter | 12 | 100779057 | 100872610 | 93554 | 1 | ENSMUST00 | 0 | 9030617O03 | ENSMUSG00000021185.10 |
| chr12 | 55560166 | 55560772 | 11.64009 | 27.76484 | 25.32713 | Distal Interge | 12 | 55565239 | 55566896 | 1658 | 1 | ENSMUST00 | -4467 | Aldoart2 | ENSMUSG00000063129.1 |
| chr13 | 42300927 | 42302372 | 11.64009 | 27.76484 | 25.32713 | Promoter | 13 | 42301270 | 42307989 | 6720 | 1 | ENSMUST00 | 0 | Edn1 | ENSMUSG00000021367.7 |
| chr13 | 98364674 | 98365518 | 11.64009 | 27.76484 | 25.32713 | Distal Interge | 13 | 98354242 | 98356703 | 2462 | 1 | ENSMUST00 | 10433 | Foxd1 | ENSMUSG00000078302.4 |
| chr13 | 31801466 | 31802104 | 11.64009 | 27.76484 | 25.32713 | Distal Interge | 13 | 31797710 | 31800749 | 3040 | 1 | ENSMUST00 | 3757 | Gm11379 | ENSMUSG00000086144.1 |
| chr15 | 100604594 | 100605406 | 11.64009 | 27.76484 | 25.32713 | Distal Interge | 15 | 100575318 | 100599984 | 24667 | 2 | ENSMUST00 | -4611 | Pou6f1 | ENSMUSG00000009739.11 |
| chr15 | 78718334 | 78718859 | 11.64009 | 27.76484 | 25.32713 | Promoter | 15 | 78670022 | 78718113 | 48092 | 2 | ENSMUST00 | -222 | Elfn2 | ENSMUSG00000043460.6 |
| chr15 | 85821319 | 85821961 | 11.64009 | 27.76484 | 25.32713 | Promoter | 15 | 85814676 | 85821733 | 7058 | 2 | ENSMUST00 | 0 | Pkdrej | ENSMUSG00000052496.4 |
| chr16 | 20629671 | 20630130 | 11.64009 | 27.76484 | 25.32713 | Promoter | 16 | 20629828 | 20645915 | 16088 | 1 | ENSMUST00 | 0 | Ece2 | ENSMUSG00000022842.11 |
| chr18 | 43477514 | 43477852 | 11.64009 | 27.76484 | 25.32713 | Promoter | 18 | 43475418 | 43477796 | 2379 | 2 | ENSMUST00 | 0 | Eif3j2 | ENSMUSG00000043424.10 |
| chr18 | 61952994 | 61954105 | 11.64009 | 27.76484 | 25.32713 | Promoter | 18 | 61953075 | 62015715 | 62641 | 1 | ENSMUST00 | 0 | Sh3tc2 | ENSMUSG00000045629.7 |
| chr18 | 34102956 | 34103362 | 11.64009 | 27.76484 | 25.32713 | Distal Interge | 18 | 34093445 | 34093570 | 126 | 1 | ENSMUST00 | 9512 | Gm24432 | ENSMUSG00000077263.1 |
| chr2 | 181212224 | 181213431 | 11.64009 | 27.76484 | 25.32713 | Promoter | 2 | 181205562 | 181213185 | 7624 | 2 | ENSMUST00 | 0 | Srms | ENSMUSG00000027579.6 |
| chr2 | 154790229 | 154790835 | 11.64009 | 27.76484 | 25.32713 | Promoter | 2 | 154791096 | 154867245 | 76150 | 1 | ENSMUST00 | -261 | Raly | ENSMUSG00000027593.10 |
| chr2 | 157943499 | 157944396 | 11.64009 | 27.76484 | 25.32713 | 3' UTR | 2 | 157914653 | 157944719 | 30067 | 1 | ENSMUST00 | 28847 | Vstm2l | ENSMUSG00000037843.6 |
| chr3 | 97888350 | 97888893 | 11.64009 | 27.76484 | 25.32713 | Promoter | 3 | 97689824 | 97888707 | 198884 | 2 | ENSMUST00 | 0 | Pde4dip | ENSMUSG00000038170.10 |
| chr3 | 96327787 | 96328479 | 11.64009 | 27.76484 | 25.32713 | Promoter | 3 | 96328425 | 96328590 | 166 | 1 | ENSMUST00 | 0 | Gm24830 | ENSMUSG00000095701.1 |

| | | | | | | | | | | | | | | | |
|---|---|---|---|---|---|---|---|---|---|---|---|---|---|---|---|
| chr5 | 21645219 | 21645662 | 11.64009 | 27.76484 | 25.32713 | Promoter | 5 | 21483847 | 21645605 | 161759 | 2 | ENSMUST00 | 0 | Fbxl13 | ENSMUSG00000048520.11 |
| chr6 | 113392275 | 113393030 | 11.64009 | 27.76484 | 25.32713 | Promoter | 6 | 113392458 | 113414562 | 22105 | 1 | ENSMUST00 | 0 | Ttll3 | ENSMUSG00000030276.14 |
| chr6 | 37299442 | 37300233 | 11.64009 | 27.76484 | 25.32713 | Promoter | 6 | 36846022 | 37299974 | 453953 | 2 | ENSMUST00 | 0 | Dgki | ENSMUSG00000038665.10 |
| chr7 | 25379383 | 25380204 | 11.64009 | 27.76484 | 25.32713 | 3' UTR | 7 | 25384805 | 25385573 | 769 | 1 | ENSMUST00 | -4601 | 4732471J01F | ENSMUSG00000053714.7 |
| chr8 | 25465846 | 25466220 | 11.64009 | 27.76484 | 25.32713 | Distal Interge | 8 | 25454273 | 25454704 | 432 | 1 | ENSMUST00 | 11574 | Gm10689 | ENSMUSG00000074412.4 |
| chr9 | 114753006 | 114753423 | 11.64009 | 27.76484 | 25.32713 | Distal Interge | 9 | 114731203 | 114749343 | 18141 | 1 | ENSMUST00 | 21804 | Cmtm6 | ENSMUSG00000032434.8 |
| chr9 | 102721728 | 102723352 | 11.64009 | 27.76484 | 25.32713 | Promoter | 9 | 102722316 | 102728326 | 6011 | 1 | ENSMUST00 | 0 | Amotl2 | ENSMUSG00000032531.10 |
| chr19 | 47343331 | 47343841 | 12.71042 | 27.74159 | 25.30578 | Exon (ENSMU | 19 | 47273519 | 47273589 | 71 | 2 | ENSMUST00 | -69743 | Mir6995 | ENSMUSG00000098786.1 |
| chr7 | 45182073 | 45182883 | 12.71042 | 27.74159 | 25.30578 | Downstream | 7 | 45180995 | 45181838 | 844 | 1 | ENSMUST00 | 1079 | Pth2 | ENSMUSG00000038300.5 |
| chr9 | 103108276 | 103108618 | 12.71042 | 27.74159 | 25.30578 | Distal Interge | 9 | 103111787 | 103162657 | 50871 | 1 | ENSMUST00 | -3169 | Rab6b | ENSMUSG00000032549.7 |
| chr1 | 180365023 | 180365940 | 12.34421 | 27.73961 | 25.30389 | Intron (ENSM | 1 | 180326970 | 180330550 | 3581 | 2 | ENSMUST00 | -34474 | Gm5069 | ENSMUSG00000055676.3 |
| chr15 | 41163726 | 41164221 | 12.34421 | 27.73961 | 25.30389 | Distal Interge | 15 | 41173701 | 41173871 | 171 | 1 | ENSMUST00 | -9480 | 4930555K19I | ENSMUSG00000048559.4 |
| chr17 | 35635705 | 35636168 | 12.68409 | 27.69736 | 25.26176 | 3' UTR | 17 | 35635752 | 35643695 | 7944 | 2 | ENSMUST00 | 7527 | Dpcr1 | ENSMUSG00000073408.2 |
| chr11 | 43836105 | 43836689 | 12.65786 | 27.6533 | 25.21823 | Promoter | 11 | 43774606 | 43836332 | 61727 | 2 | ENSMUST00 | 0 | Adra1b | ENSMUSG00000050541.9 |
| chr7 | 101377745 | 101378703 | 12.65786 | 27.6533 | 25.21823 | Promoter | 7 | 101378190 | 101411744 | 33555 | 1 | ENSMUST00 | 0 | Arap1 | ENSMUSG00000032812.11 |
| chr10 | 127666932 | 127667699 | 10.56684 | 27.64477 | 25.20979 | Promoter | 10 | 127667123 | 127668851 | 1729 | 1 | ENSMUST00 | 0 | Tmem194 | ENSMUSG00000040195.6 |
| chr11 | 99040914 | 99041822 | 10.56684 | 27.64477 | 25.20979 | Promoter | 11 | 99041244 | 99054392 | 13149 | 1 | ENSMUST00 | 0 | Igfbp4 | ENSMUSG00000017493.7 |
| chr11 | 116004671 | 116005573 | 10.56684 | 27.64477 | 25.20979 | Exon (ENSMU | 11 | 116008457 | 116012719 | 4263 | 2 | ENSMUST00 | 7146 | Galk1 | ENSMUSG00000020766.4 |
| chr15 | 73748015 | 73749080 | 10.56684 | 27.64477 | 25.20979 | Promoter | 15 | 73748026 | 73758766 | 10741 | 1 | ENSMUST00 | 0 | Ptp4a3 | ENSMUSG00000059895.7 |
| chr19 | 56873879 | 56875052 | 10.56684 | 27.64477 | 25.20979 | Promoter | 19 | 56874249 | 56912078 | 37830 | 1 | ENSMUST00 | 0 | Vwa2 | ENSMUSG00000025082.8 |
| chr2 | 153778824 | 153779681 | 10.56684 | 27.64477 | 25.20979 | Distal Interge | 2 | 153780879 | 153795182 | 14304 | 1 | ENSMUST00 | -1198 | Efcab8 | ENSMUSG00000044083.11 |
| chr3 | 19694037 | 19694623 | 10.56684 | 27.64477 | 25.20979 | Promoter | 3 | 19693401 | 19695396 | 1996 | 2 | ENSMUST00 | 773 | Crh | ENSMUSG00000049796.4 |
| chr5 | 120711327 | 120711897 | 10.56684 | 27.64477 | 25.20979 | Promoter | 5 | 120680202 | 120711927 | 31726 | 2 | ENSMUST00 | 30 | Dtx1 | ENSMUSG00000029603.10 |
| chr7 | 43796888 | 43797863 | 10.56684 | 27.64477 | 25.20979 | Promoter | 7 | 43797577 | 43803822 | 6246 | 1 | ENSMUST00 | 0 | Klk8 | ENSMUSG00000064023.3 |
| chr8 | 22124667 | 22125183 | 10.56684 | 27.64477 | 25.20979 | Promoter | 8 | 22073616 | 22125053 | 51438 | 2 | ENSMUST00 | 0 | Nek5 | ENSMUSG00000037738.9 |
| chr8 | 123581095 | 123581369 | 10.56684 | 27.64477 | 25.20979 | Promoter | 8 | 123581155 | 123581273 | 119 | 2 | ENSMUST00 | 0 | n-R5s125 | ENSMUSG00000096092.1 |
| chr9 | 87729621 | 87731071 | 10.56684 | 27.64477 | 25.20979 | Promoter | 9 | 87724397 | 87730497 | 6101 | 2 | ENSMUST00 | 0 | Tbx18 | ENSMUSG00000032419.8 |
| chr12 | 113185641 | 113186291 | 12.63174 | 27.60941 | 25.17524 | Promoter | 12 | 113185903 | 113189520 | 3618 | 1 | ENSMUST00 | 0 | Tmem121 | ENSMUSG00000049036.6 |
| chr8 | 92460279 | 92461028 | 12.63174 | 27.60941 | 25.17524 | Distal Interge | 8 | 92365189 | 92365621 | 433 | 2 | ENSMUST00 | -94659 | RP24-69M8.4 | ENSMUSG00000100612.1 |
| chr12 | 67220564 | 67221245 | 12.24882 | 27.57765 | 25.14366 | Promoter | 12 | 66471182 | 67221221 | 750040 | 2 | ENSMUST00 | 0 | Mdga2 | ENSMUSG00000034912.11 |
| chr19 | 4711760 | 4712586 | 11.51425 | 27.5435 | 25.10965 | Promoter | 19 | 4711208 | 4752353 | 41146 | 1 | ENSMUST00 | 553 | Sptbn2 | ENSMUSG00000067889.4 |
| chr5 | 3432813 | 3433731 | 11.51425 | 27.5435 | 25.10965 | Intron (ENSM | 5 | 3473071 | 3473178 | 108 | 2 | ENSMUST00 | 39447 | Gm17590 | ENSMUSG00000091644.1 |
| chr2 | 32750916 | 32751866 | 12.2252 | 27.53752 | 25.10386 | Exon (ENSMU | 2 | 32757234 | 32762244 | 5011 | 1 | ENSMUST00 | -5368 | Tor2a | ENSMUSG00000009563.11 |
| chr2 | 148677699 | 148678246 | 12.2252 | 27.53752 | 25.10386 | Distal Interge | 2 | 148681023 | 148692949 | 11927 | 1 | ENSMUST00 | -2777 | Gzf1 | ENSMUSG00000027439.9 |
| chr11 | 16950883 | 16951157 | 12.57983 | 27.52215 | 25.08893 | Promoter | 11 | 16935154 | 16951099 | 15946 | 2 | ENSMUST00 | 0 | Eldr | ENSMUSG00000087060.2 |
| chr8 | 87961273 | 87962402 | 12.57983 | 27.52215 | 25.08893 | Distal Interge | 8 | 87661810 | 87959595 | 297786 | 2 | ENSMUST00 | -1679 | Zfp423 | ENSMUSG00000045333.10 |
| chr11 | 114791499 | 114792456 | 12.55403 | 27.47878 | 25.0457 | Exon (ENSMU | 11 | 114791217 | 114795873 | 4657 | 2 | ENSMUST00 | 3417 | Btbd17 | ENSMUSG00000000202.4 |
| chr10 | 22154145 | 22154638 | 12.52834 | 27.43557 | 25.00269 | Promoter | 10 | 22153280 | 22154416 | 1137 | 2 | ENSMUST00 | 0 | Gm4895 | ENSMUSG00000044268.4 |
| chr11 | 49794015 | 49795113 | 12.52834 | 27.43557 | 25.00269 | Promoter | 11 | 49794178 | 49807750 | 13573 | 1 | ENSMUST00 | 0 | Gfpt2 | ENSMUSG00000020363.6 |
| chr11 | 69833960 | 69834556 | 12.52834 | 27.43557 | 25.00269 | Promoter | 11 | 69827739 | 69834259 | 6521 | 2 | ENSMUST00 | 0 | Nlgn2 | ENSMUSG00000051790.10 |
| chr5 | 125532303 | 125533448 | 12.52834 | 27.43557 | 25.00269 | Promoter | 5 | 125532387 | 125792583 | 260197 | 1 | ENSMUST00 | 0 | Tmem132b | ENSMUSG00000070498.3 |
| chr5 | 149015402 | 149016094 | 12.52834 | 27.43557 | 25.00269 | Promoter | 5 | 148995651 | 149015813 | 20163 | 2 | ENSMUST00 | 0 | Gm15411 | ENSMUSG00000085971.1 |
| chr18 | 54452503 | 54453674 | 12.1549 | 27.41795 | 24.98545 | Exon (ENSMU | 18 | 54422295 | 54452840 | 30546 | 1 | ENSMUST00 | 30209 | Redrum | ENSMUSG00000096982.1 |
| chr1 | 119047870 | 119048324 | 12.15419 | 27.41675 | 24.98428 | Intron (ENSM | 1 | 118848460 | 119053339 | 204880 | 2 | ENSMUST00 | 5015 | Gli2 | ENSMUSG00000048402.9 |
| chr10 | 117063964 | 117064460 | 12.15419 | 27.41675 | 24.98428 | Promoter | 10 | 117050998 | 117063814 | 12817 | 2 | ENSMUST00 | -151 | Cct2 | ENSMUSG00000034024.7 |
| chr11 | 53636100 | 53636628 | 12.15419 | 27.41675 | 24.98428 | Distal Interge | 11 | 53631324 | 53634702 | 3379 | 2 | ENSMUST00 | -1399 | Il13 | ENSMUSG00000020383.1 |
| chr13 | 23756942 | 23757568 | 12.15419 | 27.41675 | 24.98428 | Promoter | 13 | 23757013 | 23757409 | 397 | 1 | ENSMUST00 | 0 | Hist1h4b | ENSMUSG00000069266.5 |
| chr13 | 23752169 | 23752794 | 12.15419 | 27.41675 | 24.98428 | Promoter | 13 | 23752267 | 23752886 | 620 | 1 | ENSMUST00 | 0 | Hist1h3b | ENSMUSG00000069267.2 |
| chr15 | 101976968 | 101977912 | 12.15419 | 27.41675 | 24.98428 | Distal Interge | 15 | 101946004 | 101954287 | 8284 | 2 | ENSMUST00 | -22682 | Krt78 | ENSMUSG00000050463.8 |
| chr17 | 85799516 | 85800540 | 12.15419 | 27.41675 | 24.98428 | Exon (ENSMU | 17 | 85796218 | 85824044 | 27827 | 2 | ENSMUST00 | 23504 | RP23-449G1 | ENSMUSG00000100345.1 |
| chr19 | 45293332 | 45294271 | 12.15419 | 27.41675 | 24.98428 | Distal Interge | 19 | 45300463 | 45303034 | 2572 | 2 | ENSMUST00 | 8763 | RP23-188J8.3 | ENSMUSG00000100618.1 |
| chr2 | 65668279 | 65668892 | 12.15419 | 27.41675 | 24.98428 | Intron (ENSM | 2 | 65670445 | 65767447 | 97003 | 1 | ENSMUST00 | -1553 | Scn2a1 | ENSMUSG00000075318.7 |
| chr2 | 25167864 | 25168368 | 12.15419 | 27.41675 | 24.98428 | Distal Interge | 2 | 25180758 | 25183339 | 2582 | 1 | ENSMUST00 | -12390 | Nrarp | ENSMUSG00000078202.3 |
| chr3 | 129204282 | 129205055 | 12.15419 | 27.41675 | 24.98428 | Promoter | 3 | 129203947 | 129218897 | 14951 | 1 | ENSMUST00 | 336 | Pitx2 | ENSMUSG00000028023.11 |
| chr5 | 8659973 | 8660849 | 12.15419 | 27.41675 | 24.98428 | Promoter | 5 | 8660077 | 8748575 | 88499 | 1 | ENSMUST00 | 0 | Abcb1a | ENSMUSG00000040584.8 |
| chr6 | 116673347 | 116674497 | 12.15419 | 27.41675 | 24.98428 | Promoter | 6 | 116633008 | 116673836 | 40829 | 2 | ENSMUST00 | 0 | Rassf4 | ENSMUSG00000042129.6 |
| chr8 | 71700851 | 71701735 | 12.15419 | 27.41675 | 24.98428 | Promoter | 8 | 71691719 | 71701800 | 10082 | 2 | ENSMUST00 | 65 | B3gnt3 | ENSMUSG00000031803.7 |
| chr9 | 87015352 | 87015849 | 12.15419 | 27.41675 | 24.98428 | Promoter | 9 | 87015537 | 87019917 | 4381 | 1 | ENSMUST00 | 0 | Ripply2 | ENSMUSG00000047897.5 |
| chr15 | 78982378 | 78982895 | 12.50275 | 27.39253 | 24.96176 | Promoter | 15 | 78983041 | 79005864 | 22824 | 1 | ENSMUST00 | -146 | Triobp | ENSMUSG00000033088.13 |
| chr5 | 107725401 | 107726026 | 12.50275 | 27.39253 | 24.96176 | Promoter | 5 | 107716657 | 107725805 | 9149 | 2 | ENSMUST00 | 0 | Gfi1 | ENSMUSG00000029275.12 |
| chr7 | 63858551 | 63859047 | 12.10847 | 27.33893 | 24.90845 | Distal Interge | 7 | 63896533 | 63899698 | 3166 | 1 | ENSMUST00 | -37486 | Gm27252 | ENSMUSG00000098708.2 |
| chr4 | 97868386 | 97869737 | 11.74122 | 27.32396 | 24.89351 | Intron (ENSM | 4 | 97880119 | 98065355 | 185237 | 1 | ENSMUST00 | -10382 | Nfia | ENSMUSG00000028565.13 |
| chr13 | 74121061 | 74121747 | 12.45189 | 27.30695 | 24.87681 | Promoter | 13 | 74121515 | 74166064 | 44550 | 1 | ENSMUST00 | 0 | Slc9a3 | ENSMUSG00000036123.6 |
| chr10 | 50894204 | 50895345 | 12.40144 | 27.22202 | 24.79214 | Promoter | 10 | 50895651 | 50989152 | 93502 | 1 | ENSMUST00 | -306 | Sim1 | ENSMUSG00000019913.3 |
| chr11 | 62560403 | 62561173 | 12.40144 | 27.22202 | 24.79214 | Distal Interge | 11 | 62564477 | 62566383 | 1907 | 2 | ENSMUST00 | 5210 | Gm12280 | ENSMUSG00000085748.1 |
| chr16 | 17798066 | 17798750 | 11.32191 | 27.20368 | 24.77393 | Promoter | 16 | 17797282 | 17808287 | 11006 | 1 | ENSMUST00 | 785 | Scarf2 | ENSMUSG00000012017.3 |
| chr17 | 84502135 | 84504110 | 11.32191 | 27.20368 | 24.77393 | Distal Interge | 17 | 84511895 | 84622142 | 110248 | 1 | ENSMUST00 | -7785 | Plekhh2 | ENSMUSG00000040852.5 |
| chr5 | 140610419 | 140611796 | 11.32191 | 27.20368 | 24.77393 | Intron (ENSM | 5 | 140607341 | 140615543 | 8203 | 1 | ENSMUST00 | 3079 | Lfng | ENSMUSG00000029570.1 |
| chr8 | 47974145 | 47975260 | 11.32191 | 27.20368 | 24.77393 | Intron (ENSM | 8 | 47826083 | 47990543 | 164461 | 2 | ENSMUST00 | 15283 | Wwc2 | ENSMUSG00000031563.7 |
| chr5 | 43469087 | 43471221 | 12.37637 | 27.17979 | 24.75021 | Distal Interge | 5 | 43515569 | 43616585 | 101017 | 1 | ENSMUST00 | -44348 | C1qtnf7 | ENSMUSG00000061535.9 |
| chr1 | 72870000 | 72872489 | 10.01484 | 27.17025 | 24.74075 | Intron (ENSM | 1 | 72858065 | 72874275 | 16211 | 2 | ENSMUST00 | 1786 | Igfbp5 | ENSMUSG00000026185.8 |
| chr10 | 90575990 | 90578186 | 10.01484 | 27.17025 | 24.74075 | Promoter | 10 | 90576252 | 90921313 | 345062 | 1 | ENSMUST00 | 0 | Anks1b | ENSMUSG00000058589.9 |
| chr11 | 116389519 | 116389912 | 10.01484 | 27.17025 | 24.74075 | Intron (ENSM | 11 | 116359889 | 116387405 | 27517 | 2 | ENSMUST00 | -2115 | Rnf157 | ENSMUSG00000052949.9 |
| chr16 | 21204069 | 21205484 | 10.01484 | 27.17025 | 24.74075 | Promoter | 16 | 21204755 | 21223304 | 18550 | 1 | ENSMUST00 | 0 | Ephb3 | ENSMUSG00000005958.9 |
| chr2 | 118558050 | 118558348 | 10.01484 | 27.17025 | 24.74075 | Distal Interge | 2 | 118528757 | 118549687 | 20931 | 2 | ENSMUST00 | -8364 | Bmf | ENSMUSG00000040093.10 |
| chr9 | 58581598 | 58582102 | 10.01484 | 27.17025 | 24.74075 | Promoter | 9 | 58582240 | 58651318 | 69079 | 1 | ENSMUST00 | -138 | Nptn | ENSMUSG00000032336.12 |
| chr1 | 166047761 | 166048915 | 11.99395 | 27.14372 | 24.71469 | Distal Interge | 1 | 166002960 | 166038256 | 35297 | 1 | ENSMUST00 | 44802 | Gm26665 | ENSMUSG00000097196.2 |
| chr11 | 50371346 | 50371885 | 11.99395 | 27.14372 | 24.71469 | Distal Interge | 11 | 50376990 | 50386528 | 9539 | 1 | ENSMUST00 | -5105 | Hnrnph1 | ENSMUSG00000007850.11 |
| chr2 | 105676026 | 105676722 | 11.99395 | 27.14372 | 24.71469 | Promoter | 2 | 105676128 | 105697071 | 20944 | 1 | ENSMUST00 | 0 | Pax6 | ENSMUSG00000027168.16 |
| chr12 | 33301950 | 33302416 | 12.3514 | 27.13773 | 24.70927 | Promoter | 12 | 33302515 | 33368273 | 65759 | 1 | ENSMUST00 | -99 | Atxn7l1 | ENSMUSG00000020564.11 |
| chr17 | 47975786 | 47976500 | 12.3514 | 27.13773 | 24.70927 | Distal Interge | 17 | 48003379 | 48003572 | 194 | 2 | ENSMUST00 | 27072 | Gm14871 | ENSMUSG00000082492.1 |
| chr19 | 16872263 | 16873181 | 12.3514 | 27.13773 | 24.70927 | Promoter | 19 | 16872316 | 16873830 | 1515 | 2 | ENSMUST00 | 649 | Foxb2 | ENSMUSG00000056829.3 |
| chr9 | 21936519 | 21937342 | 12.3514 | 27.13773 | 24.70927 | Promoter | 9 | 21937033 | 21948904 | 11872 | 1 | ENSMUST00 | 0 | BC018242 | ENSMUSG00000040563.8 |
| chr1 | 119052628 | 119054085 | 10.88283 | 27.02401 | 24.59589 | Promoter | 1 | 118834132 | 119053619 | 219488 | 2 | ENSMUST00 | 0 | Gli2 | ENSMUSG00000048402.9 |
| chr1 | 170907301 | 170908256 | 10.88283 | 27.02401 | 24.59589 | Exon (ENSMU | 1 | 170907273 | 170912941 | 5669 | 2 | ENSMUST00 | 4685 | Fcrlb | ENSMUSG00000070524.1 |
| chr1 | 44198730 | 44199191 | 10.88283 | 27.02401 | 24.59589 | Distal Interge | 1 | 44212162 | 44213118 | 957 | 1 | ENSMUST00 | -12971 | Gm15833 | ENSMUSG00000089703.1 |
| chr1 | 13426336 | 13427066 | 10.88283 | 27.02401 | 24.59589 | Distal Interge | 1 | 13407012 | 13407655 | 644 | 2 | ENSMUST00 | -18682 | RP23-125I14 | ENSMUSG00000101171.1 |
| chr10 | 81346400 | 81346802 | 10.88283 | 27.02401 | 24.59589 | Promoter | 10 | 81346052 | 81346730 | 679 | 2 | ENSMUST00 | 0 | Hmg20b | ENSMUSG00000020232.12 |
| chr10 | 14017825 | 14018758 | 10.88283 | 27.02401 | 24.59589 | Intron (ENSM | 10 | 13967024 | 14007405 | 40382 | 2 | ENSMUST00 | -10421 | Gm26835 | ENSMUSG00000097300.1 |
| chr13 | 33003996 | 33005121 | 10.88283 | 27.02401 | 24.59589 | Promoter | 13 | 33004541 | 33017955 | 13415 | 1 | ENSMUST00 | 0 | Serpinb9 | ENSMUSG00000045827.5 |
| chr13 | 98588608 | 98589492 | 10.88283 | 27.02401 | 24.59589 | Distal Interge | 13 | 98560354 | 98560461 | 108 | 1 | ENSMUST00 | 28255 | Gm22698 | ENSMUSG00000093116.1 |
| chr15 | 101859419 | 101860323 | 10.88283 | 27.02401 | 24.59589 | 3' UTR | 15 | 101845428 | 101850786 | 5359 | 2 | ENSMUST00 | -8634 | Krt1 | ENSMUSG00000046834.6 |
| chr19 | 5648804 | 5649331 | 10.88283 | 27.02401 | 24.59589 | Downstream | 19 | 5637483 | 5648130 | 10648 | 1 | ENSMUST00 | 11322 | Rela | ENSMUSG00000024927.7 |
| chr2 | 36091047 | 36091918 | 10.88283 | 27.02401 | 24.59589 | Exon (ENSMU | 2 | 36083684 | 36094273 | 10590 | 2 | ENSMUST00 | 2355 | Lhx6 | ENSMUSG00000026890.14 |
| chr2 | 174673312 | 174674120 | 10.88283 | 27.02401 | 24.59589 | Intron (ENSM | 2 | 174643534 | 174710832 | 67299 | 1 | ENSMUST00 | 29779 | Zfp831 | ENSMUSG00000050600.5 |
| chr3 | 60408491 | 60409375 | 10.88283 | 27.02401 | 24.59589 | Distal Interge | 3 | 60501252 | 60629748 | 128497 | 1 | ENSMUST00 | -91877 | Mbnl1 | ENSMUSG00000027763.8 |
| chr4 | 152152068 | 152152614 | 10.88283 | 27.02401 | 24.59589 | Promoter | 4 | 152120331 | 152152207 | 31877 | 2 | ENSMUST00 | 0 | Espn | ENSMUSG00000028943.12 |
| chr7 | 130692522 | 130693037 | 10.88283 | 27.02401 | 24.59589 | Promoter | 7 | 130692556 | 130764784 | 72229 | 1 | ENSMUST00 | 0 | Tacc2 | ENSMUSG00000030852.10 |
| chr10 | 4346764 | 4347001 | 12.27708 | 27.01248 | 24.58573 | Intron (ENSM | 10 | 4367475 | 4387462 | 19988 | 2 | ENSMUST00 | 40461 | Zbtb2 | ENSMUSG00000075327.4 |
| chr15 | 76987348 | 76987911 | 12.27708 | 27.01248 | 24.58573 | Distal Interge | 15 | 77015489 | 77022727 | 7239 | 2 | ENSMUST00 | 34816 | Mb | ENSMUSG00000018893.9 |
| chr17 | 34630731 | 34631194 | 12.27708 | 27.01248 | 24.58573 | Promoter | 17 | 34631404 | 34632397 | 994 | 1 | ENSMUST00 | -210 | Prrt1 | ENSMUSG00000015476.6 |
| chr17 | 86145345 | 86145870 | 12.27708 | 27.01248 | 24.58573 | Promoter | 17 | 85984665 | 86145175 | 160511 | 2 | ENSMUST00 | -171 | Srbd1 | ENSMUSG00000024135.9 |
| chr2 | 115859876 | 115861200 | 12.27708 | 27.01248 | 24.58573 | Downstream | 2 | 115864227 | 115868867 | 4641 | 2 | ENSMUST00 | 7667 | Meis2 | ENSMUSG00000027210.15 |
| chr4 | 87229895 | 87230738 | 12.27708 | 27.01248 | 24.58573 | Promoter | 4 | 86983124 | 87230477 | 247354 | 2 | ENSMUST00 | 0 | Slc24a2 | ENSMUSG00000037996.12 |
| chr8 | 70863906 | 70864357 | 12.27708 | 27.01248 | 24.58573 | Promoter | 8 | 70863127 | 70865428 | 2302 | 1 | ENSMUST00 | 780 | A230052G05 | ENSMUSG00000020887.3 |
| chr9 | 55546092 | 55546854 | 12.27708 | 27.01248 | 24.58573 | 3' UTR | 9 | 55542336 | 55544280 | 1945 | 1 | ENSMUST00 | 3757 | Isl2 | ENSMUSG00000032318.7 |
| chr2 | 104069617 | 104070899 | 11.90387 | 26.98992 | 24.5636 | Promoter | 2 | 104069849 | 104091160 | 21312 | 1 | ENSMUST00 | 0 | Cd59b | ENSMUSG00000068686.7 |
| chr16 | 38376292 | 38376903 | 11.53474 | 26.96547 | 24.53927 | Intron (ENSM | 16 | 38371998 | 38374137 | 2140 | 1 | ENSMUST00 | 4295 | Popdc2 | ENSMUSG00000022803.7 |
| chr14 | 50903886 | 50904583 | 12.22804 | 26.92976 | 24.50382 | Downstream | 14 | 50895942 | 50897456 | 1515 | 2 | ENSMUST00 | -6431 | A930018M24 | ENSMUSG00000091089.1 |
| chr16 | 89773961 | 89774496 | 12.22804 | 26.92976 | 24.50382 | Distal Interge | 16 | 89788422 | 89795282 | 6861 | 2 | ENSMUST00 | 20786 | Tiam1 | ENSMUSG00000002489.10 |
| chr7 | 127214675 | 127215431 | 12.22804 | 26.92976 | 24.50382 | Promoter | 7 | 127214447 | 127215125 | 679 | 2 | ENSMUST00 | 0 | Sep-01 | ENSMUSG00000000486.8 |
| chr16 | 30520268 | 30520789 | 12.17938 | 26.84765 | 24.42205 | Intron (ENSM | 16 | 30513970 | 30540909 | 26940 | 2 | ENSMUST00 | 20120 | Tmem44 | ENSMUSG00000022537.12 |
| chr8 | 71602475 | 71603603 | 12.17938 | 26.84765 | 24.42205 | Promoter | 8 | 71602351 | 71607924 | 5574 | 1 | ENSMUST00 | 125 | Fam129c | ENSMUSG00000043243.10 |
| chr10 | 127511667 | 127512422 | 9.54397 | 26.80001 | 24.3746 | Distal Interge | 10 | 127514939 | 127517152 | 2214 | 1 | ENSMUST00 | -2517 | Ndufa4l2 | ENSMUSG00000040280.9 |
| chr1 | 74748645 | 74749438 | 12.13111 | 26.76615 | 24.34094 | Promoter | 1 | 74738922 | 74748955 | 10034 | 2 | ENSMUST00 | 0 | Prkag3 | ENSMUSG00000006542.8 |
| chr8 | 70083311 | 70084772 | 12.13111 | 26.76615 | 24.34094 | Promoter | 8 | 70083529 | 70090862 | 7334 | 1 | ENSMUST00 | 0 | Hapln4 | ENSMUSG00000007594.9 |
| chr13 | 36578571 | 36578949 | 11.77127 | 26.76303 | 24.3379 | Distal Interge | 13 | 36654405 | 36654536 | 132 | 1 | ENSMUST00 | -75456 | Gm26395 | ENSMUSG00000077282.1 |
| chr17 | 72835821 | 72836279 | 11.77127 | 26.76303 | 24.3379 | Promoter | 17 | 72836704 | 72851195 | 14492 | 1 | ENSMUST00 | -425 | Ypel5 | ENSMUSG00000039770.5 |
| chr5 | 67230301 | 67231535 | 11.77127 | 26.76303 | 24.3379 | Distal Interge | 5 | 67260565 | 67291461 | 30897 | 1 | ENSMUST00 | -29030 | Tmem33 | ENSMUSG00000037720.11 |
| chr7 | 24637880 | 24638749 | 11.77127 | 26.76303 | 24.3379 | Promoter | 7 | 24631145 | 24637916 | 6772 | 2 | ENSMUST00 | 0 | Gm26550 | ENSMUSG00000097215.1 |
| chr9 | 37420799 | 37421297 | 11.77127 | 26.76303 | 24.3379 | Exon (ENSMU | 9 | 37416045 | 37422603 | 6559 | 2 | ENSMUST00 | 1306 | Robo3 | ENSMUSG00000032128.9 |
| chr2 | 71274009 | 71275047 | 11.72773 | 26.68839 | 24.26382 | 3' UTR | 2 | 71279561 | 71282789 | 3229 | 2 | ENSMUST00 | 7742 | Slc25a12 | ENSMUSG00000027010.11 |
| chr1 | 75458138 | 75458851 | 12.40737 | 26.6269 | 24.20249 | Intron (ENSM | 1 | 75450699 | 75474343 | 23645 | 1 | ENSMUST00 | 7440 | Asic4 | ENSMUSG00000033007.10 |
| chr1 | 74068297 | 74069096 | 12.40737 | 26.6269 | 24.20249 | Exon (ENSMU | 1 | 73910232 | 74091351 | 181120 | 2 | ENSMUST00 | 22255 | Tns1 | ENSMUSG00000055322.9 |
| chr11 | 71750829 | 71751893 | 12.40737 | 26.6269 | 24.20249 | Promoter | 11 | 71750980 | 71789647 | 38668 | 1 | ENSMUST00 | 0 | Wscd1 | ENSMUSG00000020811.11 |
| chr11 | 60509043 | 60509951 | 12.40737 | 26.6269 | 24.20249 | Promoter | 11 | 60509534 | 60521986 | 12453 | 1 | ENSMUST00 | 0 | Myo15 | ENSMUSG00000042678.11 |
| chr11 | 106500326 | 106500676 | 12.40737 | 26.6269 | 24.20249 | Promoter | 11 | 106500993 | 106501062 | 70 | 1 | ENSMUST00 | -317 | Snord104 | ENSMUSG00000065126.1 |
| chr12 | 86891801 | 86892577 | 12.40737 | 26.6269 | 24.20249 | Promoter | 12 | 86891404 | 86892567 | 1164 | 2 | ENSMUST00 | 0 | Gm26698 | ENSMUSG00000097208.1 |

| | | | | | | | | | | | | | | | |
|---|---|---|---|---|---|---|---|---|---|---|---|---|---|---|---|
| chr13 | 88821435 | 88822414 | 12.40737 | 26.6269 | 24.20249 | Promoter | 13 | 88821472 | 89200538 | 379067 | 1 | ENSMUST00 | 0 | Edil3 | ENSMUSG00000034488.9 |
| chr13 | 49015548 | 49016695 | 12.40737 | 26.6269 | 24.20249 | Distal Interge | 13 | 49037069 | 49060883 | 23275 | 2 | ENSMUST00 | 44188 | Wnk2 | ENSMUSG00000037989.10 |
| chr13 | 43864704 | 43865196 | 12.40737 | 26.6269 | 24.20249 | Distal Interge | 13 | 43872802 | 43872909 | 108 | 1 | ENSMUST00 | -7606 | Gm23117 | ENSMUSG00000089107.1 |
| chr14 | 54296944 | 54297540 | 12.40737 | 26.6269 | 24.20249 | Distal Interge | 14 | 54281896 | 54282925 | 1030 | 2 | ENSMUST00 | -14020 | Olfr49 | ENSMUSG00000048153.4 |
| chr14 | 46388600 | 46389182 | 12.40737 | 26.6269 | 24.20249 | Exon (ENSMU | 14 | 46387519 | 46389282 | 1764 | 1 | ENSMUST00 | 1082 | Gm15222 | ENSMUSG00000086746.1 |
| chr16 | 89960264 | 89960876 | 12.40737 | 26.6269 | 24.20249 | Promoter | 16 | 89787111 | 89960827 | 173717 | 2 | ENSMUST00 | 0 | Tiam1 | ENSMUSG00000002489.10 |
| chr19 | 42612614 | 42613247 | 12.40737 | 26.6269 | 24.20249 | Promoter | 19 | 42593982 | 42612813 | 18832 | 2 | ENSMUST00 | 0 | Loxl4 | ENSMUSG00000025185.9 |
| chr19 | 45544630 | 45545172 | 12.40737 | 26.6269 | 24.20249 | Distal Interge | 19 | 45552275 | 45560531 | 8257 | 2 | ENSMUST00 | 15359 | Poll | ENSMUSG00000025218.6 |
| chr2 | 38711544 | 38712323 | 12.40737 | 26.6269 | 24.20249 | Promoter | 2 | 38692656 | 38712234 | 19579 | 2 | ENSMUST00 | 0 | Nr5a1 | ENSMUSG00000026751.9 |
| chr2 | 33635272 | 33637838 | 12.40737 | 26.6269 | 24.20249 | Intron (ENSM | 2 | 33564325 | 33640480 | 76156 | 2 | ENSMUST00 | 2642 | Lmx1b | ENSMUSG00000038765.8 |
| chr2 | 70475836 | 70477119 | 12.40737 | 26.6269 | 24.20249 | Promoter | 2 | 70474923 | 70477729 | 2807 | 1 | ENSMUST00 | 914 | Sp5 | ENSMUSG00000075304.2 |
| chr3 | 33142611 | 33143454 | 12.40737 | 26.6269 | 24.20249 | Promoter | 3 | 32949954 | 33143082 | 193129 | 2 | ENSMUST00 | 0 | Pex5l | ENSMUSG00000027674.11 |
| chr3 | 89228233 | 89228799 | 12.40737 | 26.6269 | 24.20249 | Promoter | 3 | 89229057 | 89233381 | 4325 | 1 | ENSMUST00 | -258 | Muc1 | ENSMUSG00000042784.4 |
| chr3 | 45380636 | 45381645 | 12.40737 | 26.6269 | 24.20249 | Exon (ENSMU | 3 | 45378398 | 45434566 | 56169 | 1 | ENSMUST00 | 2239 | Pcdh10 | ENSMUSG00000049100.10 |
| chr4 | 89687915 | 89688981 | 12.40737 | 26.6269 | 24.20249 | Promoter | 4 | 89688198 | 89694772 | 6575 | 1 | ENSMUST00 | 0 | Dmrta1 | ENSMUSG00000043753.2 |
| chr4 | 12140702 | 12141191 | 12.40737 | 26.6269 | 24.20249 | Promoter | 4 | 12140317 | 12146731 | 6415 | 1 | ENSMUST00 | 386 | Rbm12b1 | ENSMUSG00000046667.9 |
| chr4 | 42766696 | 42766997 | 12.40737 | 26.6269 | 24.20249 | Intron (ENSM | 4 | 42772860 | 42773993 | 1134 | 2 | ENSMUST00 | 6996 | Ccl21a | ENSMUSG00000094686.1 |
| chr4 | 42651071 | 42651336 | 12.40737 | 26.6269 | 24.20249 | Distal Interge | 4 | 42655251 | 42656005 | 755 | 1 | ENSMUST00 | -3915 | Ccl27b | ENSMUSG00000096826.1 |
| chr5 | 31092608 | 31093631 | 12.40737 | 26.6269 | 24.20249 | Promoter | 5 | 31086106 | 31093615 | 7510 | 2 | ENSMUST00 | 0 | Slc30a3 | ENSMUSG00000029151.9 |
| chr5 | 139543254 | 139544446 | 12.40737 | 26.6269 | 24.20249 | Promoter | 5 | 139543494 | 139548179 | 4686 | 1 | ENSMUST00 | 0 | Uncx | ENSMUSG00000029546.11 |
| chr5 | 43947943 | 43949058 | 12.40737 | 26.6269 | 24.20249 | Distal Interge | 5 | 43943902 | 43944483 | 582 | 2 | ENSMUST00 | -3461 | Gm5292 | ENSMUSG00000059565.3 |
| chr6 | 28980783 | 28981955 | 12.40737 | 26.6269 | 24.20249 | Promoter | 6 | 28981634 | 29022165 | 40532 | 1 | ENSMUST00 | 0 | Gm3294 | ENSMUSG00000092090.1 |
| chr7 | 19092457 | 19093957 | 12.40737 | 26.6269 | 24.20249 | Promoter | 7 | 19092296 | 19093204 | 909 | 1 | ENSMUST00 | 162 | Dmpk | ENSMUSG00000030409.10 |
| chr7 | 28716492 | 28717170 | 12.40737 | 26.6269 | 24.20249 | Promoter | 7 | 28716804 | 28738144 | 21341 | 1 | ENSMUST00 | 0 | Fbxo17 | ENSMUSG00000030598.10 |
| chr7 | 44590675 | 44591741 | 12.40737 | 26.6269 | 24.20249 | Promoter | 7 | 44590886 | 44604754 | 13869 | 1 | ENSMUST00 | 0 | Kcnc3 | ENSMUSG00000062785.8 |
| chr9 | 44129765 | 44130483 | 12.40737 | 26.6269 | 24.20249 | Intron (ENSM | 9 | 44134469 | 44138934 | 4466 | 1 | ENSMUST00 | -3986 | Mcam | ENSMUSG00000032135.9 |
| chrX | 7762057 | 7762488 | 12.40737 | 26.6269 | 24.20249 | Promoter | 20 | 7762560 | 7766071 | 3512 | 1 | ENSMUST00 | -72 | Tfe3 | ENSMUSG00000000134.12 |
| chrX | 155337858 | 155338769 | 12.40737 | 26.6269 | 24.20249 | Promoter | 20 | 155323918 | 155338467 | 14550 | 2 | ENSMUST00 | 0 | Prdx4 | ENSMUSG00000025289.10 |
| chrX | 42148652 | 42149104 | 12.40737 | 26.6269 | 24.20249 | Promoter | 20 | 42149317 | 42277185 | 127869 | 1 | ENSMUST00 | -213 | Stag2 | ENSMUSG00000025862.9 |
| chrX | 48462760 | 48463231 | 12.40737 | 26.6269 | 24.20249 | Promoter | 20 | 48411049 | 48463132 | 52084 | 2 | ENSMUST00 | 0 | Elf4 | ENSMUSG00000031103.7 |
| chrX | 7948149 | 7948542 | 12.40737 | 26.6269 | 24.20249 | Promoter | 20 | 7931359 | 7947889 | 16531 | 2 | ENSMUST00 | -261 | Hdac6 | ENSMUSG00000031161.10 |
| chrX | 99135152 | 99136007 | 12.40737 | 26.6269 | 24.20249 | Promoter | 20 | 99136130 | 99148991 | 12862 | 1 | ENSMUST00 | -123 | Efnb1 | ENSMUSG00000031217.8 |
| chrX | 105763531 | 105765408 | 12.40737 | 26.6269 | 24.20249 | Promoter | 20 | 105764279 | 105774939 | 10661 | 1 | ENSMUST00 | 0 | Fgf16 | ENSMUSG00000031230.3 |
| chrX | 134600409 | 134600973 | 12.40737 | 26.6269 | 24.20249 | Promoter | 20 | 134595289 | 134600300 | 5012 | 2 | ENSMUST00 | -110 | Gla | ENSMUSG00000031266.6 |
| chrX | 48345385 | 48345904 | 12.40737 | 26.6269 | 24.20249 | Intron (ENSM | 20 | 48343770 | 48408049 | 64280 | 1 | ENSMUST00 | 1616 | Bcorl1 | ENSMUSG00000036959.10 |
| chrX | 153694747 | 153696248 | 12.40737 | 26.6269 | 24.20249 | Promoter | 20 | 153685154 | 153696280 | 11127 | 2 | ENSMUST00 | 32 | Kctd12b | ENSMUSG00000041633.9 |
| chrX | 60891891 | 60893466 | 12.40737 | 26.6269 | 24.20249 | Promoter | 20 | 60891366 | 60893430 | 2065 | 2 | ENSMUST00 | 0 | Sox3 | ENSMUSG00000045179.9 |
| chrX | 36988753 | 36989786 | 12.40737 | 26.6269 | 24.20249 | Promoter | 20 | 36911326 | 36989656 | 78331 | 2 | ENSMUST00 | 0 | Sep-06 | ENSMUSG00000050379.10 |
| chrX | 11807126 | 11808092 | 12.40737 | 26.6269 | 24.20249 | Distal Interge | 20 | 11850484 | 11850590 | 107 | 1 | ENSMUST00 | -42392 | Gm26314 | ENSMUSG00000065825.1 |
| chrX | 134276432 | 134277274 | 12.40737 | 26.6269 | 24.20249 | Promoter | 20 | 134222339 | 134276982 | 54644 | 2 | ENSMUST00 | 0 | Trmt2b | ENSMUSG00000067369.7 |
| chrX | 134585689 | 134586328 | 12.40737 | 26.6269 | 24.20249 | Promoter | 20 | 134585690 | 134588062 | 2373 | 1 | ENSMUST00 | 0 | Rpl36a | ENSMUSG00000079435.4 |
| chrX | 37083988 | 37085998 | 12.40737 | 26.6269 | 24.20249 | Promoter | 20 | 37082520 | 37085402 | 2883 | 2 | ENSMUST00 | 0 | Rpl39 | ENSMUSG00000079641.3 |
| chrX | 13346676 | 13347645 | 12.40737 | 26.6269 | 24.20249 | Distal Interge | 20 | 13326023 | 13327019 | 997 | 2 | ENSMUST00 | -19658 | Gm7129 | ENSMUSG00000080914.1 |
| chrX | 103622800 | 103624003 | 12.40737 | 26.6269 | 24.20249 | Promoter | 20 | 103560907 | 103623754 | 62848 | 2 | ENSMUST00 | 0 | Ftx | ENSMUSG00000086370.3 |
| chrX | 13042034 | 13042814 | 12.40737 | 26.6269 | 24.20249 | Promoter | 20 | 13042051 | 13045297 | 3247 | 1 | ENSMUST00 | 0 | 5730405O15 | ENSMUSG00000087424.1 |
| chrX | 13230142 | 13230793 | 12.40737 | 26.6269 | 24.20249 | Distal Interge | 20 | 13234641 | 13236283 | 1643 | 2 | ENSMUST00 | 5490 | RP23-158L19 | ENSMUSG00000100714.1 |
| chr10 | 86730848 | 86731292 | 11.33539 | 26.61763 | 24.19493 | Exon (ENSMU | 10 | 86731754 | 86732409 | 656 | 2 | ENSMUST00 | 1117 | Fabp3-ps1 | ENSMUSG00000056366.4 |
| chr12 | 3310097 | 3310877 | 11.33539 | 26.61763 | 24.19493 | Promoter | 12 | 3247430 | 3309969 | 62540 | 2 | ENSMUST00 | -129 | Rab10 | ENSMUSG00000020671.8 |
| chr10 | 81502832 | 81503214 | 12.03571 | 26.60491 | 24.1824 | 3' UTR | 10 | 81504473 | 81504766 | 294 | 1 | ENSMUST00 | -1259 | Gm16106 | ENSMUSG00000082828.1 |
| chr19 | 46151759 | 46152258 | 12.03571 | 26.60491 | 24.1824 | Promoter | 19 | 46152509 | 46286510 | 134002 | 1 | ENSMUST00 | -251 | Gbf1 | ENSMUSG00000025224.9 |
| chr8 | 11727624 | 11729032 | 12.03571 | 26.60491 | 24.1824 | Promoter | 8 | 11728105 | 11823061 | 94957 | 1 | ENSMUST00 | 0 | Arhgef7 | ENSMUSG00000031511.9 |
| chr9 | 21168109 | 21168647 | 12.03571 | 26.60491 | 24.1824 | Intron (ENSM | 9 | 21165714 | 21213248 | 47535 | 1 | ENSMUST00 | 2396 | Pde4a | ENSMUSG00000032177.11 |
| chr1 | 153748686 | 153749992 | 10.95588 | 26.55152 | 24.12963 | Promoter | 1 | 153749426 | 153762401 | 12976 | 1 | ENSMUST00 | 0 | Rnasel | ENSMUSG00000066800.6 |
| chr7 | 19011956 | 19012643 | 12.35912 | 26.54606 | 24.12426 | Downstream | 7 | 19004065 | 19006761 | 2697 | 1 | ENSMUST00 | 7892 | Irf2bp1 | ENSMUSG00000044030.3 |
| chr17 | 4907241 | 4907646 | 11.98857 | 26.52517 | 24.10345 | Distal Interge | 17 | 4994332 | 5097733 | 103402 | 1 | ENSMUST00 | -86686 | Arid1b | ENSMUSG00000069729.8 |
| chr2 | 32625128 | 32627071 | 9.13756 | 26.51177 | 24.09012 | Promoter | 2 | 32625437 | 32635054 | 9618 | 1 | ENSMUST00 | 0 | Ak1 | ENSMUSG00000026817.9 |
| chr9 | 74861447 | 74862581 | 9.13756 | 26.51177 | 24.09012 | Promoter | 9 | 74861921 | 74891781 | 29861 | 1 | ENSMUST00 | 0 | Onecut1 | ENSMUSG00000043013.10 |
| chr1 | 93176368 | 93176997 | 12.3333 | 26.50278 | 24.08123 | Distal Interge | 1 | 93151349 | 93160948 | 9600 | 2 | ENSMUST00 | -15421 | 2310007B03 | ENSMUSG00000034159.7 |
| chr14 | 61683840 | 61684700 | 12.3333 | 26.50278 | 24.08123 | Distal Interge | 14 | 61602839 | 61682373 | 79535 | 2 | ENSMUST00 | -1468 | Dleu2 | ENSMUSG00000097589.4 |
| chr2 | 166030178 | 166031780 | 12.3333 | 26.50278 | 24.08123 | Intron (ENSM | 2 | 166047918 | 166052398 | 4481 | 1 | ENSMUST00 | -16138 | Ncoa3 | ENSMUSG00000027678.12 |
| chr6 | 42373097 | 42373684 | 12.3333 | 26.50278 | 24.08123 | Promoter | 6 | 42358487 | 42373268 | 14782 | 2 | ENSMUST00 | 0 | Epha1 | ENSMUSG00000029859.4 |
| chr8 | 119585908 | 119586366 | 12.3333 | 26.50278 | 24.08123 | Intron (ENSM | 8 | 119575235 | 119598454 | 23220 | 1 | ENSMUST00 | 10674 | Dnaaf1 | ENSMUSG00000031831.6 |
| chr1 | 165840869 | 165841430 | 11.2646 | 26.49366 | 24.07236 | Intron (ENSM | 1 | 165858216 | 165861555 | 3340 | 1 | ENSMUST00 | -16786 | Cd247 | ENSMUSG00000005763.10 |
| chr1 | 100231413 | 100231844 | 11.2646 | 26.49366 | 24.07236 | Intron (ENSM | 1 | 100252629 | 100369839 | 117211 | 1 | ENSMUST00 | -20785 | Cntnap5b | ENSMUSG00000067028.6 |
| chr1 | 53706468 | 53707126 | 11.2646 | 26.49366 | 24.07236 | Promoter | 1 | 53397006 | 53706761 | 309756 | 2 | ENSMUST00 | 0 | Dnah7a | ENSMUSG00000096141.2 |
| chr1 | 65112241 | 65113318 | 11.2646 | 26.49366 | 24.07236 | Intron (ENSM | 1 | 65100619 | 65119558 | 18940 | 2 | ENSMUST00 | 6240 | RP23-348D4. | ENSMUSG00000100846.1 |
| chr10 | 99913091 | 99913737 | 11.2646 | 26.49366 | 24.07236 | Distal Interge | 10 | 100015630 | 100100413 | 84784 | 1 | ENSMUST00 | -101893 | Kitl | ENSMUSG00000019966.12 |
| chr11 | 77802210 | 77803345 | 11.2646 | 26.49366 | 24.07236 | Promoter | 11 | 77801375 | 77865056 | 63682 | 1 | ENSMUST00 | 836 | Myo18a | ENSMUSG00000000631.14 |
| chr11 | 36045828 | 36046126 | 11.2646 | 26.49366 | 24.07236 | Intron (ENSM | 11 | 36080664 | 36083854 | 3191 | 1 | ENSMUST00 | -34538 | Gm12126 | ENSMUSG00000087245.1 |
| chr12 | 84222295 | 84222901 | 11.2646 | 26.49366 | 24.07236 | Distal Interge | 12 | 84149176 | 84218881 | 69706 | 2 | ENSMUST00 | -3415 | Elmsan1 | ENSMUSG00000042507.9 |
| chr13 | 69481744 | 69482790 | 11.2646 | 26.49366 | 24.07236 | Distal Interge | 13 | 69533746 | 69615468 | 81723 | 1 | ENSMUST00 | -50956 | Nsun2 | ENSMUSG00000021595.13 |
| chr13 | 97205030 | 97205589 | 11.2646 | 26.49366 | 24.07236 | Distal Interge | 13 | 97176332 | 97198357 | 22026 | 2 | ENSMUST00 | -6674 | Hexb | ENSMUSG00000021665.7 |
| chr13 | 90886482 | 90886870 | 11.2646 | 26.49366 | 24.07236 | 3' UTR | 13 | 90883453 | 90898824 | 15372 | 2 | ENSMUST00 | 11954 | Atp6ap1l | ENSMUSG00000078958.4 |
| chr14 | 66910683 | 66911607 | 11.2646 | 26.49366 | 24.07236 | Promoter | 14 | 66911170 | 66920054 | 8885 | 1 | ENSMUST00 | 0 | Pnma2 | ENSMUSG00000046204.8 |
| chr14 | 5741286 | 5741753 | 11.2646 | 26.49366 | 24.07236 | Promoter | 14 | 5718336 | 5741577 | 23242 | 2 | ENSMUST00 | 0 | Gm3373 | ENSMUSG00000094370.2 |
| chr15 | 39659313 | 39660288 | 11.2646 | 26.49366 | 24.07236 | Intron (ENSM | 15 | 39745932 | 39760934 | 15003 | 1 | ENSMUST00 | -85644 | Dcstamp | ENSMUSG00000022303.8 |
| chr16 | 93938460 | 93938762 | 11.2646 | 26.49366 | 24.07236 | Intron (ENSM | 16 | 93919807 | 93929817 | 10011 | 2 | ENSMUST00 | -8644 | Cldn14 | ENSMUSG00000047109.7 |
| chr16 | 30187625 | 30188119 | 11.2646 | 26.49366 | 24.07236 | Distal Interge | 16 | 30195071 | 30200299 | 5229 | 1 | ENSMUST00 | -6952 | RP23-11P15. | ENSMUSG00000101888.1 |
| chr17 | 75552005 | 75552449 | 11.2646 | 26.49366 | 24.07236 | Promoter | 17 | 75537086 | 75551946 | 14861 | 2 | ENSMUST00 | -60 | Fam98a | ENSMUSG00000002017.9 |
| chr2 | 113828369 | 113829337 | 11.2646 | 26.49366 | 24.07236 | Promoter | 2 | 113776362 | 113829121 | 52760 | 2 | ENSMUST00 | 0 | Scg5 | ENSMUSG00000023236.7 |
| chr2 | 93976270 | 93976934 | 11.2646 | 26.49366 | 24.07236 | Intron (ENSM | 2 | 93980034 | 93983575 | 3542 | 1 | ENSMUST00 | -3100 | 4921507L20R | ENSMUSG00000027196.6 |
| chr2 | 128892242 | 128892677 | 11.2646 | 26.49366 | 24.07236 | Exon (ENSMU | 2 | 128863932 | 128897030 | 33099 | 1 | ENSMUST00 | 28311 | Fbln7 | ENSMUSG00000027386.4 |
| chr4 | 88794035 | 88794397 | 11.2646 | 26.49366 | 24.07236 | Promoter | 4 | 88784455 | 88794391 | 9937 | 2 | ENSMUST00 | 0 | Gm26525 | ENSMUSG00000096987.1 |
| chr5 | 43841343 | 43842217 | 11.2646 | 26.49366 | 24.07236 | 3' UTR | 5 | 43837272 | 43842242 | 4971 | 1 | ENSMUST00 | 4072 | Bst1 | ENSMUSG00000029082.12 |
| chr5 | 139546703 | 139547902 | 11.2646 | 26.49366 | 24.07236 | 3' UTR | 5 | 139543898 | 139548176 | 4279 | 1 | ENSMUST00 | 2806 | Uncx | ENSMUSG00000029546.11 |
| chr6 | 90428307 | 90429030 | 11.2646 | 26.49366 | 24.07236 | Promoter | 6 | 90403736 | 90428432 | 24697 | 2 | ENSMUST00 | 0 | Ccdc37 | ENSMUSG00000048794.9 |
| chr7 | 141099843 | 141100773 | 11.2646 | 26.49366 | 24.07236 | Promoter | 7 | 141091175 | 141100546 | 9372 | 2 | ENSMUST00 | 0 | Sigirr | ENSMUSG00000025494.7 |
| chr7 | 127712634 | 127713217 | 11.2646 | 26.49366 | 24.07236 | Promoter | 7 | 127712736 | 127718185 | 5450 | 1 | ENSMUST00 | 0 | Ctf1 | ENSMUSG00000042340.5 |
| chr8 | 4404483 | 4405330 | 11.2646 | 26.49366 | 24.07236 | Distal Interge | 8 | 4349588 | 4360020 | 10433 | 1 | ENSMUST00 | 54896 | Ccl25 | ENSMUSG00000023235.8 |
| chr8 | 120334886 | 120335828 | 11.2646 | 26.49366 | 24.07236 | Intron (ENSM | 8 | 120230536 | 120553634 | 323099 | 1 | ENSMUST00 | 104351 | Gse1 | ENSMUSG00000031822.13 |
| chr8 | 70597762 | 70598805 | 11.96514 | 26.48551 | 24.06606 | Exon (ENSMU | 8 | 70594481 | 70597288 | 2808 | 1 | ENSMUST00 | 3282 | Isyna1 | ENSMUSG00000019139.9 |
| chr15 | 76671182 | 76671490 | 12.30759 | 26.45968 | 24.0403 | Promoter | 15 | 76671615 | 76694919 | 23305 | 1 | ENSMUST00 | -125 | Ppp1r16a | ENSMUSG00000033819.10 |
| chr1 | 34432821 | 34433620 | 10.25605 | 26.45831 | 24.03902 | Promoter | 1 | 34433121 | 34433199 | 79 | 2 | ENSMUST00 | 0 | Mir5103 | ENSMUSG00000092852.1 |
| chr10 | 117393131 | 117393978 | 10.25605 | 26.45831 | 24.03902 | Distal Interge | 10 | 117366178 | 117380015 | 13838 | 2 | ENSMUST00 | -13117 | Cpsf6 | ENSMUSG00000055531.7 |
| chr11 | 70213768 | 70214799 | 10.25605 | 26.45831 | 24.03902 | Promoter | 11 | 70213910 | 70216412 | 2503 | 1 | ENSMUST00 | 0 | Slc16a11 | ENSMUSG00000040938.11 |
| chr15 | 88864126 | 88865224 | 10.25605 | 26.45831 | 24.03902 | 3' UTR | 15 | 88862186 | 88865718 | 3533 | 1 | ENSMUST00 | 1941 | Pim3 | ENSMUSG00000035828.10 |
| chr17 | 24646754 | 24648095 | 10.25605 | 26.45831 | 24.03902 | Exon (ENSMU | 17 | 24639282 | 24644957 | 5676 | 2 | ENSMUST00 | -1798 | Slc9a3r2 | ENSMUSG00000002504.9 |
| chr17 | 84096335 | 84097074 | 10.25605 | 26.45831 | 24.03902 | Distal Interge | 17 | 84183931 | 84187947 | 4017 | 2 | ENSMUST00 | 90873 | Zfp36l2 | ENSMUSG00000045817.8 |
| chr2 | 106649932 | 106650623 | 10.25605 | 26.45831 | 24.03902 | Promoter | 2 | 106642917 | 106650364 | 7448 | 2 | ENSMUST00 | 0 | BB218582 | ENSMUSG00000085218.2 |
| chr3 | 126363656 | 126364401 | 10.25605 | 26.45831 | 24.03902 | Promoter | 3 | 126363827 | 126440374 | 76548 | 1 | ENSMUST00 | 0 | Arsj | ENSMUSG00000046561.8 |
| chr4 | 154294295 | 154295715 | 10.25605 | 26.45831 | 24.03902 | Intron (ENSM | 4 | 154278486 | 154299892 | 21407 | 2 | ENSMUST00 | 4177 | Arhgef16 | ENSMUSG00000029032.7 |
| chr7 | 45361317 | 45361905 | 10.25605 | 26.45831 | 24.03902 | 5' UTR | 7 | 45339275 | 45360113 | 20839 | 2 | ENSMUST00 | -1205 | Ppfia3 | ENSMUSG00000003863.11 |
| chr7 | 131542002 | 131544690 | 10.25605 | 26.45831 | 24.03902 | Promoter | 7 | 131542867 | 131544925 | 2059 | 1 | ENSMUST00 | 0 | Hmx3 | ENSMUSG00000040148.5 |
| chr8 | 106338126 | 106338611 | 10.25605 | 26.45831 | 24.03902 | Promoter | 8 | 106252548 | 106337988 | 85441 | 2 | ENSMUST00 | -139 | Smpd3 | ENSMUSG00000031906.8 |
| chrX | 12936726 | 12937396 | 10.25605 | 26.45831 | 24.03902 | Promoter | 20 | 12936872 | 12938128 | 1257 | 1 | ENSMUST00 | 0 | AA414768 | ENSMUSG00000083307.1 |
| chr8 | 123332553 | 123333171 | 11.9418 | 26.44599 | 24.02746 | Promoter | 8 | 123332713 | 123369515 | 36803 | 1 | ENSMUST00 | 0 | Spire2 | ENSMUSG00000010154.6 |
| chr11 | 102309181 | 102309720 | 11.55672 | 26.39465 | 23.97632 | Promoter | 11 | 102308530 | 102309440 | 911 | 2 | ENSMUST00 | 0 | Ubtf | ENSMUSG00000020923.12 |
| chr16 | 58518435 | 58519923 | 11.55672 | 26.39465 | 23.97632 | Intron (ENSM | 16 | 58493745 | 58523344 | 29600 | 2 | ENSMUST00 | 3421 | St3gal6 | ENSMUSG00000022747.11 |
| chr3 | 90612830 | 90613379 | 11.55672 | 26.39465 | 23.97632 | Promoter | 3 | 90612894 | 90614414 | 1521 | 1 | ENSMUST00 | 0 | S100a6 | ENSMUSG00000001025.4 |
| chr9 | 118476911 | 118478298 | 11.55672 | 26.39465 | 23.97632 | Promoter | 9 | 118478212 | 118485366 | 7155 | 1 | ENSMUST00 | 0 | Eomes | ENSMUSG00000032446.9 |
| chr13 | 107645220 | 107646423 | 12.25648 | 26.374 | 23.95606 | Distal Interge | 13 | 107687397 | 107692162 | 4766 | 1 | ENSMUST00 | -40974 | RP23-125L8. | ENSMUSG00000102000.1 |
| chr7 | 79592005 | 79592743 | 12.25648 | 26.374 | 23.95606 | Downstream | 7 | 79593369 | 79594924 | 1556 | 2 | ENSMUST00 | 2181 | Rhcg | ENSMUSG00000030549.5 |
| chr13 | 74406413 | 74407019 | 8.78324 | 26.28941 | 23.87178 | Promoter | 13 | 74406396 | 74407317 | 922 | 1 | ENSMUST00 | 18 | Gm10116 | ENSMUSG00000062382.7 |
| chr7 | 60004176 | 60005278 | 11.84934 | 26.28932 | 23.87172 | Promoter | 7 | 59974149 | 60005064 | 30916 | 2 | ENSMUST00 | 0 | Snrpn | ENSMUSG00000000948.11 |
| chr8 | 84893970 | 84894278 | 11.84934 | 26.28932 | 23.87172 | Promoter | 8 | 84893094 | 84893921 | 828 | 2 | ENSMUST00 | -50 | Gcdh | ENSMUSG00000003809.9 |
| chr1 | 136050862 | 136051570 | 12.20579 | 26.28899 | 23.87144 | Promoter | 1 | 136050804 | 136051370 | 567 | 1 | ENSMUST00 | 59 | Ascl5 | ENSMUSG00000097918.1 |
| chr10 | 9117582 | 9118388 | 11.80365 | 26.21181 | 23.79474 | Distal Interge | 10 | 8722219 | 8886070 | 163852 | 2 | ENSMUST00 | -231513 | Sash1 | ENSMUSG00000015305.5 |
| chr17 | 21474663 | 21475289 | 11.80365 | 26.21181 | 23.79474 | Distal Interge | 17 | 21485177 | 21488875 | 3699 | 2 | ENSMUST00 | 13586 | Gm26753 | ENSMUSG00000097759.1 |
| chr19 | 16955698 | 16956843 | 11.80365 | 26.21181 | 23.79474 | Promoter | 19 | 16956118 | 17223932 | 267815 | 1 | ENSMUST00 | 0 | Prune2 | ENSMUSG00000039126.9 |
| chr2 | 24762214 | 24763518 | 11.80365 | 26.21181 | 23.79474 | Promoter | 2 | 24603887 | 24763152 | 159266 | 2 | ENSMUST00 | 0 | Cacna1b | ENSMUSG00000004113.13 |
| chr5 | 130256429 | 130256824 | 11.80365 | 26.21181 | 23.79474 | Promoter | 5 | 130257035 | 130277749 | 20715 | 1 | ENSMUST00 | -211 | Tyw1 | ENSMUSG00000056310.9 |
| chr7 | 132551335 | 132551921 | 11.80365 | 26.21181 | 23.79474 | Intron (ENSM | 7 | 132557475 | 132576398 | 18924 | 2 | ENSMUST00 | 24477 | Oat | ENSMUSG00000030934.8 |
| chr8 | 88272292 | 88272862 | 11.80365 | 26.21181 | 23.79474 | Promoter | 8 | 88272403 | 88329962 | 57560 | 1 | ENSMUST00 | 0 | Adcy7 | ENSMUSG00000031659.7 |
| chr15 | 101315990 | 101316526 | 12.15553 | 26.20465 | 23.78802 | Distal Interge | 15 | 101293232 | 101297426 | 4195 | 1 | ENSMUST00 | 22759 | 6030408B16 | ENSMUSG00000075408.5 |
| chr1 | 51333776 | 51334110 | 12.13055 | 26.16273 | 23.74622 | Promoter | 1 | 51218386 | 51333779 | 115394 | 2 | ENSMUST00 | 0 | RP24-555D5. | ENSMUSG00000101799.1 |
| chr4 | 116464329 | 116464714 | 12.13055 | 26.16273 | 23.74622 | Promoter | 4 | 116306763 | 116464183 | 157421 | 2 | ENSMUST00 | -147 | Mast2 | ENSMUSG00000003810.7 |
| chr10 | 81687366 | 81687848 | 8.47159 | 26.12089 | 23.70459 | Distal Interge | 10 | 81693718 | 81694185 | 468 | 2 | ENSMUST00 | 6337 | Gm8112 | ENSMUSG00000096623.1 |
| chr1 | 149960863 | 149961413 | 11.73508 | 26.0954 | 23.67916 | Promoter | 1 | 149829618 | 149961290 | 131673 | 2 | ENSMUST00 | 0 | Pla2g4a | ENSMUSG00000056220.9 |

| | | | | | | | | | | | | | | | |
|---|---|---|---|---|---|---|---|---|---|---|---|---|---|---|---|
| chr10 | 76906018 | 76906866 | 11.73508 | 26.0954 | 23.67916 | Promoter | 10 | 76789803 | 76905862 | 116060 | 2 | ENSMUST00 | -157 | Pcbp3 | ENSMUSG00000001120.10 |
| chr10 | 127079255 | 127080430 | 11.73508 | 26.0954 | 23.67916 | Promoter | 10 | 127078907 | 127093169 | 14263 | 1 | ENSMUST00 | 349 | Agap2 | ENSMUSG00000025422.8 |
| chr10 | 121505042 | 121505786 | 11.73508 | 26.0954 | 23.67916 | Distal Interge | 10 | 121410350 | 121476250 | 65901 | 2 | ENSMUST00 | -28793 | Rassf3 | ENSMUSG00000025795.7 |
| chr11 | 118433766 | 118434600 | 11.73508 | 26.0954 | 23.67916 | Promoter | 11 | 118433853 | 118449963 | 16111 | 1 | ENSMUST00 | 0 | C1qtnf1 | ENSMUSG00000017446.9 |
| chr11 | 98773371 | 98773844 | 11.73508 | 26.0954 | 23.67916 | Intron (ENSM | 11 | 98767932 | 98775333 | 7402 | 2 | ENSMUST00 | 1489 | Nr1d1 | ENSMUSG00000020889.11 |
| chr11 | 118908539 | 118909523 | 11.73508 | 26.0954 | 23.67916 | Promoter | 11 | 118489762 | 118909550 | 419789 | 2 | ENSMUST00 | 27 | Rbfox3 | ENSMUSG00000025576.12 |
| chr11 | 118290477 | 118290909 | 11.73508 | 26.0954 | 23.67916 | Promoter | 11 | 118259653 | 118290244 | 30592 | 2 | ENSMUST00 | -234 | Usp36 | ENSMUSG00000033909.12 |
| chr12 | 24251879 | 24252615 | 11.73508 | 26.0954 | 23.67916 | Promoter | 12 | 24251846 | 24252181 | 336 | 2 | ENSMUST00 | 0 | Gm9312 | ENSMUSG00000096279.1 |
| chr14 | 65266032 | 65267025 | 11.73508 | 26.0954 | 23.67916 | Promoter | 14 | 65266701 | 65321501 | 54801 | 1 | ENSMUST00 | 0 | Fbxo16 | ENSMUSG00000034532.8 |
| chr15 | 88954438 | 88955079 | 11.73508 | 26.0954 | 23.67916 | Promoter | 15 | 88890633 | 88954418 | 63786 | 2 | ENSMUST00 | -21 | Ttll8 | ENSMUSG00000022388.9 |
| chr15 | 93591531 | 93592123 | 11.73508 | 26.0954 | 23.67916 | Intron (ENSM | 15 | 93499114 | 93595891 | 96778 | 2 | ENSMUST00 | 3768 | Prickle1 | ENSMUSG00000036158.7 |
| chr15 | 89169012 | 89169636 | 11.73508 | 26.0954 | 23.67916 | Intron (ENSM | 15 | 89155549 | 89170688 | 15140 | 2 | ENSMUST00 | 1052 | Plxnb2 | ENSMUSG00000036606.11 |
| chr18 | 35930389 | 35930955 | 11.73508 | 26.0954 | 23.67916 | Distal Interge | 18 | 35952824 | 35952907 | 84 | 1 | ENSMUST00 | -21869 | Gm23882 | ENSMUSG00000080529.1 |
| chr2 | 34905348 | 34905705 | 11.73508 | 26.0954 | 23.67916 | Promoter | 2 | 34898082 | 34904907 | 6826 | 2 | ENSMUST00 | -442 | Phf19 | ENSMUSG00000026873.4 |
| chr2 | 24521089 | 24521948 | 11.73508 | 26.0954 | 23.67916 | Distal Interge | 2 | 24494068 | 24495011 | 944 | 1 | ENSMUST00 | 27022 | Gm13418 | ENSMUSG00000082307.3 |
| chr3 | 153973299 | 153973965 | 11.73508 | 26.0954 | 23.67916 | Promoter | 3 | 153973436 | 154271722 | 298287 | 1 | ENSMUST00 | 0 | Slc44a5 | ENSMUSG00000028360.9 |
| chr4 | 139573974 | 139574297 | 11.73508 | 26.0954 | 23.67916 | Promoter | 4 | 139574720 | 139620382 | 45663 | 1 | ENSMUST00 | -423 | Iffo2 | ENSMUSG00000041025.10 |
| chr4 | 123182218 | 123183811 | 11.73508 | 26.0954 | 23.67916 | Promoter | 4 | 123183227 | 123194701 | 11475 | 1 | ENSMUST00 | 0 | Hpcal4 | ENSMUSG00000046093.4 |
| chr5 | 84416190 | 84417807 | 11.73508 | 26.0954 | 23.67916 | Promoter | 5 | 84057791 | 84416806 | 359016 | 2 | ENSMUST00 | 0 | Epha5 | ENSMUSG00000029245.11 |
| chr5 | 77803785 | 77804100 | 11.73508 | 26.0954 | 23.67916 | Distal Interge | 5 | 77669724 | 77669846 | 123 | 2 | ENSMUST00 | -133940 | Gm22011 | ENSMUSG00000089415.1 |
| chr6 | 73221853 | 73222409 | 11.73508 | 26.0954 | 23.67916 | Promoter | 6 | 73017609 | 73221651 | 204043 | 2 | ENSMUST00 | -203 | Dnah6 | ENSMUSG00000052861.8 |
| chr7 | 48749481 | 48750128 | 11.73508 | 26.0954 | 23.67916 | Distal Interge | 7 | 48789003 | 48827440 | 38438 | 1 | ENSMUST00 | -38875 | Zdhhc13 | ENSMUSG00000030471.12 |
| chr7 | 49633370 | 49634869 | 11.73508 | 26.0954 | 23.67916 | Exon (ENSMU | 7 | 49631499 | 49636849 | 5351 | 2 | ENSMUST00 | 1980 | Dbx1 | ENSMUSG00000030507.6 |
| chr1 | 121096928 | 121097568 | 12.0809 | 26.07937 | 23.66483 | Intron (ENSM | 1 | 121087529 | 121088932 | 1404 | 1 | ENSMUST00 | 9400 | Celrr | ENSMUSG00000097881.2 |
| chr1 | 40085522 | 40086140 | 12.0809 | 26.07937 | 23.66483 | Promoter | 1 | 40076228 | 40085853 | 9626 | 2 | ENSMUST00 | 0 | Gm16894 | ENSMUSG00000097899.1 |
| chr14 | 70780922 | 70781415 | 12.0809 | 26.07937 | 23.66483 | Distal Interge | 14 | 70776710 | 70777595 | 886 | 1 | ENSMUST00 | 4213 | Dok2 | ENSMUSG00000022102.8 |
| chr19 | 11044857 | 11045546 | 12.0809 | 26.07937 | 23.66483 | Downstream | 19 | 11047617 | 11050566 | 2950 | 2 | ENSMUST00 | 5020 | AW112010 | ENSMUSG00000075010.4 |
| chr2 | 5746390 | 5746840 | 12.0809 | 26.07937 | 23.66483 | Distal Interge | 2 | 5293457 | 5714515 | 421059 | 2 | ENSMUST00 | -31876 | Camk1d | ENSMUSG00000039145.11 |
| chr2 | 167190013 | 167190424 | 12.0809 | 26.07937 | 23.66483 | Promoter | 2 | 167188422 | 167190155 | 1734 | 2 | ENSMUST00 | 0 | Kcnb1 | ENSMUSG00000050556.8 |
| chr2 | 179662183 | 179663173 | 12.0809 | 26.07937 | 23.66483 | Intron (ENSM | 2 | 179612622 | 179613567 | 946 | 2 | ENSMUST00 | -48617 | Gm14300 | ENSMUSG00000083410.1 |
| chr6 | 113166549 | 113167266 | 12.0809 | 26.07937 | 23.66483 | Downstream | 6 | 113168090 | 113195384 | 27295 | 2 | ENSMUST00 | 28118 | Lhfpl4 | ENSMUSG00000042873.10 |
| chr6 | 124512560 | 124513150 | 12.0809 | 26.07937 | 23.66483 | Promoter | 6 | 124512632 | 124523440 | 10809 | 1 | ENSMUST00 | 0 | C1ra | ENSMUSG00000055172.9 |
| chr7 | 73606918 | 73608349 | 12.0809 | 26.07937 | 23.66483 | Distal Interge | 7 | 73645985 | 73646089 | 105 | 2 | ENSMUST00 | 37740 | Gm26176 | ENSMUSG00000065729.1 |
| chr13 | 119961808 | 119962030 | 11.34984 | 26.03783 | 23.62382 | Promoter | 13 | 119962244 | 119963227 | 984 | 1 | ENSMUST00 | -214 | Gm20784 | ENSMUSG00000095367.1 |
| chr14 | 24778881 | 24779672 | 11.34984 | 26.03783 | 23.62382 | Distal Interge | 14 | 24909414 | 24913440 | 4027 | 1 | ENSMUST00 | -129742 | Gm10398 | ENSMUSG00000072686.2 |
| chr16 | 74412476 | 74413200 | 12.05622 | 26.03793 | 23.62382 | Promoter | 16 | 74349795 | 74411825 | 62031 | 2 | ENSMUST00 | -652 | Robo2 | ENSMUSG00000052516.12 |
| chr5 | 29477730 | 29478806 | 11.34984 | 26.03783 | 23.62382 | Promoter | 5 | 29473034 | 29478470 | 5437 | 2 | ENSMUST00 | 0 | Mnx1 | ENSMUSG00000001566.9 |
| chr7 | 101905384 | 101905726 | 11.34984 | 26.03783 | 23.62382 | Promoter | 7 | 101905837 | 101911901 | 6065 | 1 | ENSMUST00 | -111 | Lamtor1 | ENSMUSG00000030842.6 |
| chr1 | 63445763 | 63446570 | 9.7287 | 26.0223 | 23.6086 | Promoter | 1 | 63445891 | 63573217 | 127327 | 1 | ENSMUST00 | 0 | Adam23 | ENSMUSG00000025964.10 |
| chr10 | 59920707 | 59921675 | 9.7287 | 26.0223 | 23.6086 | Distal Interge | 10 | 59949675 | 59951770 | 2096 | 2 | ENSMUST00 | 30095 | Ddit4 | ENSMUSG00000020108.3 |
| chr2 | 92055270 | 92056117 | 9.7287 | 26.0223 | 23.6086 | Exon (ENSMU | 2 | 92054989 | 92059369 | 4381 | 2 | ENSMUST00 | 3252 | 2900072N19 | ENSMUSG00000087018.1 |
| chr5 | 131306954 | 131308205 | 9.7287 | 26.0223 | 23.6086 | Promoter | 5 | 130872082 | 131307918 | 435837 | 2 | ENSMUST00 | 0 | Wbscr17 | ENSMUSG00000034040.11 |
| chr7 | 79792598 | 79793522 | 9.7287 | 26.0223 | 23.6086 | Promoter | 7 | 79792241 | 79793788 | 1548 | 2 | ENSMUST00 | 266 | Mesp1 | ENSMUSG00000030544.5 |
| chr7 | 25305897 | 25306541 | 9.7287 | 26.0223 | 23.6086 | Promoter | 7 | 25306106 | 25308826 | 2721 | 1 | ENSMUST00 | 0 | Tmem145 | ENSMUSG00000043843.11 |
| chr8 | 25275246 | 25275700 | 9.7287 | 26.0223 | 23.6086 | Distal Interge | 8 | 25154552 | 25201449 | 46898 | 2 | ENSMUST00 | -73798 | Tacc1 | ENSMUSG00000065954.5 |
| chr8 | 19161767 | 19162352 | 9.7287 | 26.0223 | 23.6086 | Distal Interge | 8 | 19157886 | 19159578 | 1693 | 1 | ENSMUST00 | 3882 | Spag11a | ENSMUSG00000079842.1 |
| chr4 | 146270578 | 146271225 | 8.19534 | 25.99704 | 23.58378 | Promoter | 4 | 146270372 | 146278434 | 8063 | 1 | ENSMUST00 | 207 | Gm13166 | ENSMUSG00000085525.2 |
| chr13 | 71963050 | 71964051 | 12.00718 | 25.95554 | 23.54242 | Promoter | 13 | 71957921 | 71963723 | 5803 | 2 | ENSMUST00 | 0 | Irx1 | ENSMUSG00000060969.8 |
| chr7 | 19523777 | 19524693 | 10.95668 | 25.95157 | 23.53856 | Exon (ENSMU | 7 | 19518731 | 19525050 | 6320 | 1 | ENSMUST00 | 5047 | Nkpd1 | ENSMUSG00000060621.4 |
| chr13 | 21716723 | 21717258 | 11.64645 | 25.94473 | 23.53185 | Promoter | 13 | 21716422 | 21716814 | 393 | 1 | ENSMUST00 | 302 | Hist1h2ai | ENSMUSG00000071516.2 |
| chr9 | 20694637 | 20695228 | 11.9828 | 25.91458 | 23.50176 | Intron (ENSM | 9 | 20667986 | 20728219 | 60234 | 2 | ENSMUST00 | 32991 | Olfm2 | ENSMUSG00000032172.7 |
| chr7 | 87246468 | 87246885 | 11.95853 | 25.87377 | 23.46109 | Promoter | 7 | 87246649 | 87287981 | 41333 | 1 | ENSMUST00 | 0 | Nox4 | ENSMUSG00000030562.11 |
| chr14 | 103843054 | 103844298 | 11.93435 | 25.83312 | 23.42056 | Promoter | 14 | 103814625 | 103844173 | 29549 | 2 | ENSMUST00 | 0 | Ednrb | ENSMUSG00000022122.9 |
| chr3 | 97680736 | 97682016 | 11.93435 | 25.83312 | 23.42056 | Distal Interge | 3 | 97673031 | 97674694 | 1664 | 2 | ENSMUST00 | -6043 | Gm17651 | ENSMUSG00000090441.2 |
| chr11 | 95144709 | 95146433 | 11.58036 | 25.83223 | 23.41975 | Promoter | 11 | 95140447 | 95146263 | 5817 | 2 | ENSMUST00 | 0 | Dlx4 | ENSMUSG00000020871.8 |
| chr18 | 51117461 | 51118154 | 11.58036 | 25.83223 | 23.41975 | Promoter | 18 | 51117898 | 51304641 | 186744 | 1 | ENSMUST00 | 0 | Prr16 | ENSMUSG00000073565.4 |
| chr2 | 25577485 | 25578876 | 11.58036 | 25.83223 | 23.41975 | Exon (ENSMU | 2 | 25576973 | 25579897 | 2925 | 2 | ENSMUST00 | 1021 | Gm996 | ENSMUSG00000029419.8 |
| chr3 | 143100738 | 143101477 | 11.58036 | 25.83223 | 23.41975 | Distal Interge | 3 | 142962561 | 142963792 | 1232 | 1 | ENSMUST00 | 138178 | Gm2574 | ENSMUSG00000078162.5 |
| chr4 | 141010427 | 141011063 | 11.58036 | 25.83223 | 23.41975 | Promoter | 4 | 141010637 | 141015012 | 4376 | 1 | ENSMUST00 | 0 | Mfap2 | ENSMUSG00000060572.12 |
| chr1 | 93668863 | 93669336 | 10.54274 | 25.8059 | 23.39399 | Distal Interge | 1 | 93627132 | 93658659 | 31528 | 2 | ENSMUST00 | -10205 | Stk25 | ENSMUSG00000026277.8 |
| chr11 | 102218423 | 102218882 | 10.54274 | 25.8059 | 23.39399 | Promoter | 11 | 102206628 | 102218928 | 12301 | 2 | ENSMUST00 | 46 | Hdac5 | ENSMUSG00000008855.12 |
| chr11 | 59290416 | 59291053 | 10.54274 | 25.8059 | 23.39399 | Promoter | 11 | 59248033 | 59290752 | 42720 | 2 | ENSMUST00 | 0 | Wnt3a | ENSMUSG00000009900.7 |
| chr12 | 20196345 | 20197143 | 10.54274 | 25.8059 | 23.39399 | Distal Interge | 12 | 20071882 | 20072217 | 336 | 1 | ENSMUST00 | 124464 | Gm3993 | ENSMUSG00000096529.1 |
| chr14 | 65343982 | 65344530 | 10.54274 | 25.8059 | 23.39399 | Distal Interge | 14 | 65358676 | 65398930 | 40255 | 1 | ENSMUST00 | -14146 | Zfp395 | ENSMUSG00000034522.9 |
| chr15 | 44457389 | 44457746 | 10.54274 | 25.8059 | 23.39399 | Promoter | 15 | 44457553 | 44597137 | 139585 | 1 | ENSMUST00 | 0 | Pkhd1l1 | ENSMUSG00000038725.6 |
| chr15 | 13067006 | 13068064 | 10.54274 | 25.8059 | 23.39399 | Intron (ENSM | 15 | 12980829 | 12980928 | 100 | 1 | ENSMUST00 | 86178 | Gm24664 | ENSMUSG00000089564.1 |
| chr17 | 6132231 | 6132453 | 10.54274 | 25.8059 | 23.39399 | Promoter | 17 | 6131854 | 6132405 | 552 | 2 | ENSMUST00 | 0 | Gm15590 | ENSMUSG00000058542.3 |
| chr19 | 59166215 | 59166730 | 10.54274 | 25.8059 | 23.39399 | Promoter | 19 | 59166598 | 59166980 | 383 | 2 | ENSMUST00 | 250 | Vax1 | ENSMUSG00000006270.7 |
| chr3 | 37477172 | 37478054 | 10.54274 | 25.8059 | 23.39399 | Intron (ENSM | 3 | 37451721 | 37528356 | 76636 | 1 | ENSMUST00 | 25452 | Spata5 | ENSMUSG00000027722.9 |
| chr3 | 18242831 | 18243913 | 10.54274 | 25.8059 | 23.39399 | Promoter | 3 | 18071950 | 18243338 | 171389 | 2 | ENSMUST00 | 0 | Cyp7b1 | ENSMUSG00000039519.6 |
| chr4 | 21678131 | 21679690 | 10.54274 | 25.8059 | 23.39399 | 3' UTR | 4 | 21679747 | 21685781 | 6035 | 2 | ENSMUST00 | 6091 | Prdm13 | ENSMUSG00000040478.9 |
| chr4 | 107313597 | 107314233 | 10.54274 | 25.8059 | 23.39399 | Promoter | 4 | 107314363 | 107359822 | 45460 | 1 | ENSMUST00 | -130 | Yipf1 | ENSMUSG00000057375.8 |
| chr5 | 147186616 | 147187974 | 10.54274 | 25.8059 | 23.39399 | Promoter | 5 | 147188696 | 147190947 | 2252 | 1 | ENSMUST00 | -722 | Gsx1 | ENSMUSG00000053129.5 |
| chr7 | 16810940 | 16811671 | 10.54274 | 25.8059 | 23.39399 | Exon (ENSMU | 7 | 16815889 | 16840931 | 25043 | 1 | ENSMUST00 | -4218 | Strn4 | ENSMUSG00000030374.10 |
| chr7 | 118277147 | 118277690 | 10.54274 | 25.8059 | 23.39399 | Downstream | 7 | 118279861 | 118279965 | 105 | 1 | ENSMUST00 | -2171 | Gm23229 | ENSMUSG00000087922.1 |
| chr7 | 108950412 | 108951468 | 10.54274 | 25.8059 | 23.39399 | Exon (ENSMU | 7 | 108950754 | 108961535 | 10782 | 2 | ENSMUST00 | 10067 | Gm26599 | ENSMUSG00000097220.1 |
| chr7 | 138919098 | 138919991 | 10.54274 | 25.8059 | 23.39399 | Distal Interge | 7 | 138914388 | 138915314 | 927 | 1 | ENSMUST00 | 4711 | Gm9358 | ENSMUSG00000098199.1 |
| chr8 | 70353495 | 70353942 | 10.54274 | 25.8059 | 23.39399 | Promoter | 8 | 70331522 | 70353273 | 21752 | 2 | ENSMUST00 | -223 | Upf1 | ENSMUSG00000058301.6 |
| chr10 | 80165265 | 80166711 | 11.91027 | 25.79263 | 23.38204 | Promoter | 10 | 80165985 | 80170316 | 4332 | 1 | ENSMUST00 | 0 | Cirbp | ENSMUSG00000045193.8 |
| chr7 | 70325301 | 70326081 | 11.91027 | 25.79263 | 23.38204 | Distal Interge | 7 | 70351946 | 70360593 | 8648 | 2 | ENSMUST00 | 34512 | Nr2f2 | ENSMUSG00000030551.8 |
| chr2 | 105083047 | 105084091 | 11.1896 | 25.76025 | 23.34976 | Intron (ENSM | 2 | 105077895 | 105078107 | 213 | 2 | ENSMUST00 | -4941 | Wt1os | ENSMUSG00000074987.4 |
| chr14 | 60485110 | 60485927 | 11.88629 | 25.75228 | 23.34181 | Promoter | 14 | 60485208 | 60599031 | 113824 | 1 | ENSMUST00 | 0 | RP24-127N1 | ENSMUSG00000101797.1 |
| chr7 | 16844951 | 16846298 | 11.86241 | 25.71209 | 23.30173 | Exon (ENSMU | 7 | 16843065 | 16870461 | 27397 | 1 | ENSMUST00 | 1887 | Prkd2 | ENSMUSG00000041187.11 |
| chr16 | 78264802 | 78265562 | 11.15024 | 25.69191 | 23.28165 | Distal Interge | 16 | 78240258 | 78240388 | 131 | 1 | ENSMUST00 | 24545 | Gm25916 | ENSMUSG00000077451.1 |
| chr4 | 129620725 | 129621473 | 11.49339 | 25.68398 | 23.27414 | Promoter | 4 | 129619275 | 129620915 | 1641 | 2 | ENSMUST00 | 0 | Ccdc28b | ENSMUSG00000028795.10 |
| chr11 | 96314188 | 96315638 | 11.83862 | 25.67205 | 23.26222 | Distal Interge | 11 | 96317165 | 96317274 | 110 | 1 | ENSMUST00 | -1527 | Mir10a | ENSMUSG00000065519.1 |
| chr17 | 87612927 | 87613827 | 11.83862 | 25.67205 | 23.26222 | Distal Interge | 17 | 87635979 | 87651106 | 15128 | 1 | ENSMUST00 | -22152 | Epcam | ENSMUSG00000045394.8 |
| chr10 | 22842940 | 22844360 | 10.44916 | 25.63548 | 23.2261 | Distal Interge | 10 | 22817279 | 22820128 | 2850 | 2 | ENSMUST00 | -22813 | Tcf21 | ENSMUSG00000045680.7 |
| chr10 | 115817023 | 115818153 | 11.79132 | 25.59241 | 23.18327 | Promoter | 10 | 115817284 | 115849893 | 32610 | 1 | ENSMUST00 | 0 | Tspan8 | ENSMUSG00000034127.9 |
| chr17 | 22674118 | 22674681 | 11.79132 | 25.59241 | 23.18327 | Distal Interge | 17 | 22689771 | 22689941 | 171 | 1 | ENSMUST00 | -15090 | Gm9805 | ENSMUSG00000046088.8 |
| chr10 | 94378040 | 94379018 | 11.7444 | 25.51337 | 23.10443 | Intron (ENSM | 10 | 94408891 | 94410732 | 1842 | 1 | ENSMUST00 | -29873 | Gm16155 | ENSMUSG00000085409.1 |
| chr2 | 160451624 | 160452360 | 11.72109 | 25.47406 | 23.06522 | Distal Interge | 2 | 160363703 | 160367065 | 3363 | 2 | ENSMUST00 | -84560 | Mafb | ENSMUSG00000074622.4 |
| chr15 | 74752561 | 74753284 | 11.36537 | 25.4653 | 23.05655 | Promoter | 15 | 74747852 | 74753046 | 5195 | 2 | ENSMUST00 | 0 | Lynx1 | ENSMUSG00000022594.9 |
| chr17 | 34631792 | 34632576 | 11.36537 | 25.4653 | 23.05655 | Promoter | 17 | 34631404 | 34632397 | 994 | 1 | ENSMUST00 | 389 | Prrt1 | ENSMUSG00000015476.6 |
| chr2 | 116071147 | 116072197 | 11.36537 | 25.4653 | 23.05655 | Intron (ENSM | 2 | 116067968 | 116085910 | 17943 | 1 | ENSMUST00 | 3180 | G630016G05 | ENSMUSG00000086308.1 |
| chr5 | 43980573 | 43981952 | 11.36537 | 25.4653 | 23.05655 | Promoter | 5 | 43978865 | 43981762 | 2898 | 2 | ENSMUST00 | 0 | Fgfbp1 | ENSMUSG00000048373.7 |
| chr3 | 106684904 | 106685239 | 8.8906 | 25.42674 | 23.01856 | Promoter | 3 | 106684987 | 106735904 | 50918 | 1 | ENSMUST00 | 0 | Lrif1 | ENSMUSG00000056260.10 |
| chr16 | 15084586 | 15085367 | 10.95755 | 25.35628 | 22.94834 | Distal Interge | 16 | 15216859 | 15217328 | 470 | 2 | ENSMUST00 | 131961 | Gm7731 | ENSMUSG00000069315.2 |
| chr13 | 14021641 | 14022360 | 9.94526 | 25.28477 | 22.8772 | Intron (ENSM | 13 | 14005189 | 14009844 | 4656 | 2 | ENSMUST00 | -11798 | Tbce | ENSMUSG00000039233.7 |
| chr14 | 31129092 | 31129476 | 9.94526 | 25.28477 | 22.8772 | Promoter | 14 | 31124802 | 31128930 | 4129 | 2 | ENSMUST00 | -163 | Smim4 | ENSMUSG00000058351.6 |
| chr17 | 29451035 | 29451518 | 9.94526 | 25.28477 | 22.8772 | Distal Interge | 17 | 29490812 | 29496111 | 5300 | 1 | ENSMUST00 | -39294 | Pim1 | ENSMUSG00000024014.7 |
| chr17 | 56289551 | 56291172 | 9.94526 | 25.28477 | 22.8772 | Promoter | 17 | 56278962 | 56290511 | 11550 | 2 | ENSMUST00 | 0 | Plin3 | ENSMUSG00000024197.9 |
| chr2 | 154549413 | 154550003 | 9.94526 | 25.28477 | 22.8772 | Promoter | 2 | 154548927 | 154550003 | 1077 | 1 | ENSMUST00 | 487 | 1700003F12I | ENSMUSG00000038523.5 |
| chr9 | 19575345 | 19575603 | 9.94526 | 25.28477 | 22.8772 | Distal Interge | 9 | 19566435 | 19567382 | 948 | 2 | ENSMUST00 | -7964 | Olfr854 | ENSMUSG00000095667.1 |
| chrX | 6172085 | 6172489 | 9.94526 | 25.28477 | 22.8772 | Promoter | 20 | 6168696 | 6173015 | 4320 | 2 | ENSMUST00 | 526 | Nudt10 | ENSMUSG00000073293.4 |
| chr11 | 90417641 | 90418483 | 11.60586 | 25.27969 | 22.87289 | Distal Interge | 11 | 90336536 | 90390895 | 54360 | 2 | ENSMUST00 | -26747 | Hlf | ENSMUSG00000003949.11 |
| chr12 | 22622903 | 22623442 | 11.60586 | 25.27969 | 22.87289 | Promoter | 12 | 22622962 | 22623087 | 126 | 2 | ENSMUST00 | 0 | Gm17391 | ENSMUSG00000095261.2 |
| chr19 | 5050538 | 5051443 | 11.60586 | 25.27969 | 22.87289 | Promoter | 19 | 5050808 | 5057071 | 6264 | 1 | ENSMUST00 | 0 | Rin1 | ENSMUSG00000024883.6 |
| chr10 | 96751517 | 96752077 | 11.94783 | 25.26529 | 22.85895 | Distal Interge | 10 | 96855582 | 96855670 | 89 | 2 | ENSMUST00 | 103593 | Gm24587 | ENSMUSG00000080347.1 |
| chr11 | 97114422 | 97115468 | 11.94783 | 25.26529 | 22.85895 | Promoter | 11 | 97098071 | 97115331 | 17261 | 2 | ENSMUST00 | 0 | Tbx21 | ENSMUSG00000001444.2 |
| chr11 | 83852471 | 83853754 | 11.94783 | 25.26529 | 22.85895 | Promoter | 11 | 83852960 | 83905819 | 52860 | 1 | ENSMUST00 | 0 | Hnf1b | ENSMUSG00000020679.6 |
| chr11 | 74610647 | 74611319 | 11.94783 | 25.26529 | 22.85895 | Distal Interge | 11 | 74618260 | 74618513 | 254 | 2 | ENSMUST00 | 7194 | Gm22993 | ENSMUSG00000093164.2 |
| chr12 | 72441596 | 72442497 | 11.94783 | 25.26529 | 22.85895 | Promoter | 12 | 72441866 | 72510565 | 68700 | 1 | ENSMUST00 | 0 | Lrrc9 | ENSMUSG00000021090.10 |
| chr13 | 22035429 | 22036502 | 11.94783 | 25.26529 | 22.85895 | Promoter | 13 | 22035870 | 22036345 | 476 | 1 | ENSMUST00 | 0 | Hist1h2bk | ENSMUSG00000062727.4 |
| chr14 | 63244684 | 63245758 | 11.94783 | 25.26529 | 22.85895 | Promoter | 14 | 63198922 | 63245248 | 46327 | 2 | ENSMUST00 | 0 | Gata4 | ENSMUSG00000021944.10 |
| chr14 | 67344367 | 67345027 | 11.94783 | 25.26529 | 22.85895 | Intron (ENSM | 14 | 67324894 | 67335928 | 11035 | 2 | ENSMUST00 | -8440 | 1700120O09 | ENSMUSG00000093386.1 |
| chr17 | 47436434 | 47436870 | 11.94783 | 25.26529 | 22.85895 | Promoter | 17 | 47436615 | 47470144 | 33530 | 1 | ENSMUST00 | 0 | AI661453 | ENSMUSG00000034382.9 |
| chr19 | 45742386 | 45743031 | 11.94783 | 25.26529 | 22.85895 | Promoter | 19 | 45736798 | 45742874 | 6077 | 2 | ENSMUST00 | 0 | Fgf8 | ENSMUSG00000025219.8 |
| chr19 | 43614384 | 43615322 | 11.94783 | 25.26529 | 22.85895 | 3' UTR | 19 | 43612325 | 43615892 | 3568 | 1 | ENSMUST00 | 2060 | Nkx2-3 | ENSMUSG00000044220.13 |
| chr2 | 163481508 | 163482317 | 11.94783 | 25.26529 | 22.85895 | Distal Interge | 2 | 163466379 | 163472629 | 6251 | 2 | ENSMUST00 | -8880 | Fitm2 | ENSMUSG00000048486.6 |
| chr2 | 152141775 | 152142540 | 11.94783 | 25.26529 | 22.85895 | Distal Interge | 2 | 152143561 | 152149097 | 5537 | 1 | ENSMUST00 | -1021 | Tcf15 | ENSMUSG00000068079.5 |
| chr2 | 177000918 | 177001682 | 11.94783 | 25.26529 | 22.85895 | Promoter | 2 | 177000956 | 177010984 | 10029 | 1 | ENSMUST00 | 0 | Gm14419 | ENSMUSG00000078875.5 |
| chr2 | 176635839 | 176636226 | 11.94783 | 25.26529 | 22.85895 | Promoter | 2 | 176622842 | 176636318 | 13477 | 2 | ENSMUST00 | 92 | Gm14308 | ENSMUSG00000078880.5 |
| chr2 | 147184036 | 147184752 | 11.94783 | 25.26529 | 22.85895 | Promoter | 2 | 147184083 | 147331681 | 147599 | 1 | ENSMUST00 | 0 | Nkx2-2os | ENSMUSG00000086509.2 |
| chr2 | 172862620 | 172863294 | 11.94783 | 25.26529 | 22.85895 | Downstream | 2 | 172864264 | 172864341 | 78 | 2 | ENSMUST00 | 1047 | Gm22773 | ENSMUSG00000093160.1 |
| chr2 | 176520997 | 176521784 | 11.94783 | 25.26529 | 22.85895 | Promoter | 2 | 176521056 | 176531745 | 10690 | 1 | ENSMUST00 | 0 | Gm14431 | ENSMUSG00000095199.2 |
| chr3 | 88000028 | 88000453 | 11.94783 | 25.26529 | 22.85895 | Promoter | 3 | 87987531 | 88000356 | 12826 | 2 | ENSMUST00 | 0 | Bcan | ENSMUSG00000004892.8 |
| chr3 | 31149697 | 31150187 | 11.94783 | 25.26529 | 22.85895 | Promoter | 3 | 31149946 | 31164324 | 14379 | 1 | ENSMUST00 | 0 | Cldn11 | ENSMUSG00000037625.6 |

| | | | | | | | | | | | | | | | |
|---|---|---|---|---|---|---|---|---|---|---|---|---|---|---|---|
| chr5 | 115475497 | 115476366 | 11.94783 | 25.26529 | 22.85895 | Downstream | 5 | 115478013 | 115479938 | 1926 | 2 | ENSMUST00 | 3572 | Sirt4 | ENSMUSG00000029524.11 |
| chr5 | 114573244 | 114575010 | 11.94783 | 25.26529 | 22.85895 | Intron (ENSM | 5 | 114568017 | 114594678 | 26662 | 1 | ENSMUST00 | 5228 | Fam222a | ENSMUSG00000041930.7 |
| chr5 | 140735407 | 140735835 | 11.94783 | 25.26529 | 22.85895 | Promoter | 5 | 140735526 | 140754691 | 19166 | 1 | ENSMUST00 | 0 | Amz1 | ENSMUSG00000050022.11 |
| chr7 | 79710008 | 79710537 | 11.94783 | 25.26529 | 22.85895 | Promoter | 7 | 79710283 | 79712357 | 2075 | 1 | ENSMUST00 | 0 | 9330171B17 | ENSMUSG00000039133.6 |
| chr7 | 92180406 | 92181408 | 11.94783 | 25.26529 | 22.85895 | Promoter | 7 | 92180684 | 92288339 | 107656 | 1 | ENSMUST00 | 0 | Dlg2 | ENSMUSG00000052572.10 |
| chr8 | 83772603 | 83773508 | 11.94783 | 25.26529 | 22.85895 | Distal Interge | 8 | 83752918 | 83753204 | 287 | 1 | ENSMUST00 | 19686 | Gm22024 | ENSMUSG00000084682.1 |
| chr9 | 29411639 | 29412822 | 11.94783 | 25.26529 | 22.85895 | Promoter | 9 | 28996012 | 29412204 | 416193 | 2 | ENSMUST00 | 0 | Ntm | ENSMUSG00000059974.5 |
| chrX | 151802550 | 151803098 | 11.94783 | 25.26529 | 22.85895 | Promoter | 20 | 151803243 | 151873033 | 69791 | 1 | ENSMUST00 | -145 | Huwe1 | ENSMUSG00000025261.12 |
| chrX | 48452562 | 48453072 | 11.94783 | 25.26529 | 22.85895 | Intron (ENSM | 20 | 48411046 | 48454169 | 43124 | 2 | ENSMUST00 | 1097 | Elf4 | ENSMUSG00000031103.7 |
| chrX | 59566837 | 59568052 | 11.94783 | 25.26529 | 22.85895 | Promoter | 20 | 59131877 | 59567472 | 435596 | 2 | ENSMUST00 | 0 | Fgf13 | ENSMUSG00000031137.12 |
| chrX | 7679619 | 7680338 | 11.94783 | 25.26529 | 22.85895 | Promoter | 20 | 7674626 | 7681089 | 6464 | 2 | ENSMUST00 | 751 | Magix | ENSMUSG00000031147.8 |
| chrX | 99216303 | 99217267 | 11.94783 | 25.26529 | 22.85895 | Distal Interge | 20 | 99138131 | 99147230 | 9100 | 1 | ENSMUST00 | 78173 | Efnb1 | ENSMUSG00000031217.8 |
| chrX | 101332157 | 101332530 | 11.94783 | 25.26529 | 22.85895 | Distal Interge | 20 | 101314391 | 101318151 | 3761 | 1 | ENSMUST00 | 17767 | Nlgn3 | ENSMUSG00000031302.11 |
| chrX | 72274303 | 72275115 | 11.94783 | 25.26529 | 22.85895 | Promoter | 20 | 72255999 | 72274803 | 18805 | 2 | ENSMUST00 | 0 | Gabre | ENSMUSG00000031340.8 |
| chrX | 136741826 | 136742872 | 11.94783 | 25.26529 | 22.85895 | Promoter | 20 | 136733864 | 136742861 | 8998 | 2 | ENSMUST00 | 0 | Morf4l2 | ENSMUSG00000031422.11 |
| chrX | 153139583 | 153140467 | 11.94783 | 25.26529 | 22.85895 | Promoter | 20 | 153139981 | 153171943 | 31963 | 1 | ENSMUST00 | 0 | Rragb | ENSMUSG00000041658.7 |
| chrX | 98148653 | 98149162 | 11.94783 | 25.26529 | 22.85895 | Promoter | 20 | 98149721 | 98323215 | 173495 | 1 | ENSMUST00 | -559 | Ar | ENSMUSG00000046532.7 |
| chrX | 111894562 | 111895481 | 11.94783 | 25.26529 | 22.85895 | Distal Interge | 20 | 111860628 | 111861175 | 548 | 1 | ENSMUST00 | 33935 | Gm14899 | ENSMUSG00000082752.1 |
| chrX | 77810725 | 77811269 | 11.94783 | 25.26529 | 22.85895 | Distal Interge | 20 | 77802351 | 77803280 | 930 | 2 | ENSMUST00 | -7446 | Gm14742 | ENSMUSG00000085973.1 |
| chrX | 52744185 | 52745130 | 11.94783 | 25.26529 | 22.85895 | Promoter | 20 | 52742563 | 52744593 | 2031 | 2 | ENSMUST00 | 0 | Kis2 | ENSMUSG00000086173.2 |
| chrX | 103616021 | 103616916 | 11.94783 | 25.26529 | 22.85895 | Promoter | 20 | 103569568 | 103616863 | 47296 | 2 | ENSMUST00 | 0 | Ftx | ENSMUSG00000086370.3 |
| chrX | 108834591 | 108835262 | 11.94783 | 25.26529 | 22.85895 | Promoter | 20 | 108834657 | 108886683 | 52027 | 1 | ENSMUST00 | 0 | 2810403D21 | ENSMUSG00000086629.2 |
| chrX | 48314253 | 48315184 | 11.94783 | 25.26529 | 22.85895 | Distal Interge | 20 | 48307043 | 48307143 | 101 | 1 | ENSMUST00 | 7211 | Gm22612 | ENSMUSG00000088918.1 |
| chrX | 123612506 | 123612806 | 11.94783 | 25.26529 | 22.85895 | Distal Interge | 20 | 123581700 | 123602134 | 20435 | 1 | ENSMUST00 | 30807 | Gm6604 | ENSMUSG00000095474.1 |
| chr7 | 24883728 | 24884011 | 11.58309 | 25.24124 | 22.83669 | Promoter | 7 | 24884371 | 24889806 | 5436 | 1 | ENSMUST00 | -360 | Rps19 | ENSMUSG00000040952.11 |
| chr10 | 108333705 | 108333985 | 10.88911 | 25.23667 | 22.83218 | Intron (ENSM | 10 | 108332189 | 108414391 | 82203 | 1 | ENSMUST00 | 1517 | Pawr | ENSMUSG00000035873.7 |
| chr11 | 120530430 | 120531315 | 10.88911 | 25.23667 | 22.83218 | Promoter | 11 | 120530699 | 120538984 | 8286 | 1 | ENSMUST00 | 0 | Gcgr | ENSMUSG00000025127.10 |
| chr11 | 102248862 | 102249318 | 10.88911 | 25.23667 | 22.83218 | Promoter | 11 | 102248882 | 102265187 | 16306 | 1 | ENSMUST00 | 0 | BC030867 | ENSMUSG00000034773.11 |
| chr11 | 87426097 | 87426443 | 10.88911 | 25.23667 | 22.83218 | Promoter | 11 | 87426716 | 87426879 | 164 | 1 | ENSMUST00 | -273 | Gm22068 | ENSMUSG00000096205.1 |
| chr12 | 86893911 | 86894393 | 10.88911 | 25.23667 | 22.83218 | Distal Interge | 12 | 86891404 | 86892567 | 1164 | 2 | ENSMUST00 | -1345 | Gm26698 | ENSMUSG00000097208.1 |
| chr15 | 8627253 | 8627901 | 10.88911 | 25.23667 | 22.83218 | Distal Interge | 15 | 8642256 | 8643247 | 992 | 2 | ENSMUST00 | 15346 | Slc1a3 | ENSMUSG00000005360.9 |
| chr17 | 27655178 | 27655848 | 10.88911 | 25.23667 | 22.83218 | Promoter | 17 | 27655509 | 27708659 | 53151 | 1 | ENSMUST00 | 0 | Pacsin1 | ENSMUSG00000040276.9 |
| chr17 | 21370066 | 21370692 | 10.88911 | 25.23667 | 22.83218 | Distal Interge | 17 | 21383748 | 21399262 | 15515 | 1 | ENSMUST00 | -13056 | Zfp677 | ENSMUSG00000062743.4 |
| chr17 | 14075881 | 14076224 | 10.88911 | 25.23667 | 22.83218 | Distal Interge | 17 | 14058005 | 14059531 | 1527 | 2 | ENSMUST00 | -16351 | RP23-138P14 | ENSMUSG00000101307.1 |
| chr18 | 47330025 | 47330837 | 10.88911 | 25.23667 | 22.83218 | Intron (ENSM | 18 | 47291894 | 47333326 | 41433 | 2 | ENSMUST00 | 2489 | Sema6a | ENSMUSG00000019647.11 |
| chr18 | 5603509 | 5604123 | 10.88911 | 25.23667 | 22.83218 | Promoter | 18 | 5603744 | 5642873 | 39130 | 1 | ENSMUST00 | 0 | Zeb1 | ENSMUSG00000024238.8 |
| chr18 | 60911532 | 60912394 | 10.88911 | 25.23667 | 22.83218 | Promoter | 18 | 60911780 | 60918561 | 6782 | 1 | ENSMUST00 | 0 | Arsi | ENSMUSG00000036412.5 |
| chr18 | 69500166 | 69501048 | 10.88911 | 25.23667 | 22.83218 | Intron (ENSM | 18 | 69521538 | 69564687 | 43150 | 1 | ENSMUST00 | -20490 | Tcf4 | ENSMUSG00000053477.10 |
| chr19 | 8661814 | 8662583 | 10.88911 | 25.23667 | 22.83218 | Promoter | 19 | 8663789 | 8678154 | 14366 | 1 | ENSMUST00 | -1206 | Chrm1 | ENSMUSG00000032773.4 |
| chr2 | 153108245 | 153108975 | 10.88911 | 25.23667 | 22.83218 | Promoter | 2 | 153108468 | 153151439 | 42972 | 1 | ENSMUST00 | 0 | Hck | ENSMUSG00000003283.8 |
| chr2 | 152687030 | 152687782 | 10.88911 | 25.23667 | 22.83218 | Promoter | 2 | 152687035 | 152687951 | 917 | 1 | ENSMUST00 | 0 | Mcts2 | ENSMUSG00000042814.7 |
| chr2 | 157560051 | 157561310 | 10.88911 | 25.23667 | 22.83218 | Promoter | 2 | 157560078 | 157562504 | 2427 | 1 | ENSMUST00 | 0 | Nnat | ENSMUSG00000067786.11 |
| chr5 | 99339579 | 99340444 | 10.88911 | 25.23667 | 22.83218 | Intron (ENSM | 5 | 99217426 | 99252927 | 35502 | 2 | ENSMUST00 | -86653 | Rasgef1b | ENSMUSG00000029333.9 |
| chr5 | 106458042 | 106458827 | 10.88911 | 25.23667 | 22.83218 | Promoter | 5 | 106452523 | 106458166 | 5644 | 2 | ENSMUST00 | 0 | Barhl2 | ENSMUSG00000034384.11 |
| chr6 | 55117237 | 55118141 | 10.88911 | 25.23667 | 22.83218 | Promoter | 6 | 55091186 | 55117985 | 26800 | 2 | ENSMUST00 | 0 | Crhr2 | ENSMUSG00000003476.10 |
| chr6 | 115773805 | 115774364 | 10.88911 | 25.23667 | 22.83218 | Promoter | 6 | 115774538 | 115804893 | 30356 | 1 | ENSMUST00 | -174 | Cand2 | ENSMUSG00000030319.7 |
| chr7 | 141434331 | 141435371 | 10.88911 | 25.23667 | 22.83218 | Promoter | 7 | 141429739 | 141434594 | 4856 | 2 | ENSMUST00 | 0 | Slc25a22 | ENSMUSG00000019082.12 |
| chr7 | 43782678 | 43783002 | 10.88911 | 25.23667 | 22.83218 | Exon (ENSMU | 7 | 43781054 | 43785410 | 4357 | 1 | ENSMUST00 | 1625 | Klk10 | ENSMUSG00000030693.9 |
| chr7 | 120201403 | 120202069 | 10.88911 | 25.23667 | 22.83218 | Promoter | 7 | 120186380 | 120202111 | 15732 | 2 | ENSMUST00 | 42 | Crym | ENSMUSG00000030905.5 |
| chr7 | 132597221 | 132597952 | 10.88911 | 25.23667 | 22.83218 | Promoter | 7 | 132597387 | 132599637 | 2251 | 1 | ENSMUST00 | 0 | Gm16764 | ENSMUSG00000087657.3 |
| chr8 | 87656559 | 87657118 | 10.88911 | 25.23667 | 22.83218 | Distal Interge | 8 | 87636657 | 87637013 | 357 | 1 | ENSMUST00 | 19903 | Gm24841 | ENSMUSG00000089301.1 |
| chr17 | 81480190 | 81480815 | 11.56041 | 25.20292 | 22.8004 | Intron (ENSM | 17 | 81441763 | 81441923 | 161 | 1 | ENSMUST00 | 38428 | Gm11096 | ENSMUSG00000079333.3 |
| chr9 | 79977468 | 79978193 | 11.56041 | 25.20292 | 22.8004 | Promoter | 9 | 79815051 | 79977804 | 162754 | 2 | ENSMUST00 | 0 | Filip1 | ENSMUSG00000034898.10 |
| chr17 | 56268887 | 56269742 | 11.90138 | 25.18757 | 22.78511 | 3' UTR | 17 | 56269319 | 56276786 | 7468 | 2 | ENSMUST00 | 7044 | Ticam1 | ENSMUSG00000047123.8 |
| chr7 | 141047937 | 141048901 | 11.87651 | 25.14597 | 22.74365 | Distal Interge | 7 | 141061274 | 141072119 | 10846 | 1 | ENSMUST00 | -12373 | B4galnt4 | ENSMUSG00000055629.4 |
| chr19 | 61266379 | 61267256 | 4.43183 | 25.11952 | 22.71751 | Distal Interge | 19 | 61296902 | 61297069 | 168 | 2 | ENSMUST00 | 29813 | Gm21060 | ENSMUSG00000095993.1 |
| chr6 | 112273304 | 112274523 | 11.15821 | 25.11025 | 22.70824 | Promoter | 6 | 112273758 | 112330425 | 56668 | 1 | ENSMUST00 | 0 | Lmcd1 | ENSMUSG00000057604.9 |
| chr8 | 47994660 | 47995333 | 11.15821 | 25.11025 | 22.70824 | Distal Interge | 8 | 47826083 | 47990543 | 164461 | 2 | ENSMUST00 | -4118 | Wwc2 | ENSMUSG00000031563.7 |
| chr16 | 18590993 | 18592014 | 11.85175 | 25.10453 | 22.7029 | Promoter | 16 | 18591992 | 18609751 | 17760 | 1 | ENSMUST00 | 0 | 4930588K23 | ENSMUSG00000075335.3 |
| chr4 | 34049720 | 34050102 | 11.82709 | 25.06326 | 22.66179 | Promoter | 4 | 34024874 | 34050039 | 25166 | 2 | ENSMUST00 | 0 | Spaca1 | ENSMUSG00000028264.11 |
| chr11 | 52098170 | 52098567 | 10.77139 | 25.03037 | 22.62905 | Promoter | 11 | 52098681 | 52127778 | 29098 | 1 | ENSMUST00 | -114 | Ppp2ca | ENSMUSG00000020349.4 |
| chr19 | 24899494 | 24900814 | 10.77139 | 25.03037 | 22.62905 | Promoter | 19 | 24898965 | 24901309 | 2345 | 2 | ENSMUST00 | 495 | Foxd4 | ENSMUSG00000051490.2 |
| chr2 | 36203056 | 36203890 | 10.77139 | 25.03037 | 22.62905 | Promoter | 2 | 36202822 | 36212903 | 10082 | 1 | ENSMUST00 | 235 | Gm13429 | ENSMUSG00000085141.1 |
| chr4 | 57956554 | 57957472 | 10.77139 | 25.03037 | 22.62905 | Promoter | 4 | 57943373 | 57956411 | 13039 | 2 | ENSMUST00 | -144 | Txn1 | ENSMUSG00000028367.5 |
| chr3 | 55466037 | 55466697 | 11.80254 | 25.02215 | 22.62103 | 3' UTR | 3 | 55461758 | 55539064 | 77307 | 1 | ENSMUST00 | 4280 | Dclk1 | ENSMUSG00000027797.10 |
| chr7 | 104501941 | 104502390 | 11.80254 | 25.02215 | 22.62103 | Promoter | 7 | 104501055 | 104502149 | 1095 | 2 | ENSMUST00 | 0 | Gm16464 | ENSMUSG00000082154.1 |
| chr1 | 133434248 | 133434895 | 11.44831 | 25.0134 | 22.61242 | Distal Interge | 1 | 133382303 | 133424377 | 42075 | 2 | ENSMUST00 | -9872 | Sox13 | ENSMUSG00000070643.6 |
| chr3 | 132085139 | 132086948 | 11.77808 | 24.98121 | 22.58028 | Promoter | 3 | 132085292 | 132180293 | 95002 | 1 | ENSMUST00 | 0 | Dkk2 | ENSMUSG00000028031.5 |
| chr13 | 117602417 | 117602928 | 11.75373 | 24.94043 | 22.53964 | Promoter | 13 | 117602480 | 117976802 | 374323 | 1 | ENSMUST00 | 0 | Hcn1 | ENSMUSG00000021730.7 |
| chr6 | 128802547 | 128803035 | 11.40408 | 24.93852 | 22.53783 | Distal Interge | 6 | 128784764 | 128789215 | 4452 | 2 | ENSMUST00 | -13333 | Klrb1c | ENSMUSG00000030325.11 |
| chr4 | 147738528 | 147739185 | 7.02109 | 24.90988 | 22.50924 | Promoter | 4 | 147735477 | 147738426 | 2950 | 2 | ENSMUST00 | -103 | Gm13136 | ENSMUSG00000081179.1 |
| chr11 | 5061516 | 5062377 | 11.38209 | 24.90129 | 22.50064 | Exon (ENSMU | 11 | 5060470 | 5064851 | 4382 | 2 | ENSMUST00 | 2474 | Gas2l1 | ENSMUSG00000034201.8 |
| chr11 | 43775831 | 43776339 | 11.38209 | 24.90129 | 22.50064 | 3' UTR | 11 | 43729512 | 43748008 | 18497 | 2 | ENSMUST00 | -27824 | Ttc1 | ENSMUSG00000041278.5 |
| chr12 | 85177021 | 85177593 | 11.38209 | 24.90129 | 22.50064 | Promoter | 12 | 85166639 | 85177296 | 10658 | 2 | ENSMUST00 | 0 | Pgf | ENSMUSG00000004791.6 |
| chr14 | 52022604 | 52023338 | 11.38209 | 24.90129 | 22.50064 | Distal Interge | 14 | 52006077 | 52020733 | 14657 | 2 | ENSMUST00 | -1872 | Zfp219 | ENSMUSG00000049295.11 |
| chr15 | 79270308 | 79271025 | 11.38209 | 24.90129 | 22.50064 | 3' UTR | 15 | 79259716 | 79261297 | 1582 | 2 | ENSMUST00 | -9012 | Baiap2l2 | ENSMUSG00000018126.10 |
| chr18 | 46211707 | 46212795 | 11.38209 | 24.90129 | 22.50064 | Promoter | 18 | 46165302 | 46212607 | 47306 | 2 | ENSMUST00 | 0 | Trim36 | ENSMUSG00000033949.6 |
| chr7 | 139581879 | 139583098 | 11.38209 | 24.90129 | 22.50064 | Promoter | 7 | 139579376 | 139582797 | 3422 | 2 | ENSMUST00 | 0 | Nkx6-2 | ENSMUSG00000041309.12 |
| chr9 | 117872017 | 117872971 | 11.38209 | 24.90129 | 22.50064 | Promoter | 9 | 117873144 | 117881202 | 8059 | 1 | ENSMUST00 | -173 | Gm26614 | ENSMUSG00000097240.1 |
| chr16 | 13446736 | 13447202 | 11.72947 | 24.89981 | 22.4996 | Downstream | 16 | 13449523 | 13449601 | 79 | 1 | ENSMUST00 | -2321 | Mir193b | ENSMUSG00000076972.2 |
| chr1 | 89069458 | 89069983 | 9.44256 | 24.88688 | 22.48677 | Promoter | 1 | 89070415 | 89155068 | 84654 | 1 | ENSMUST00 | -432 | Sh3bp4 | ENSMUSG00000036206.7 |
| chr10 | 73826272 | 73826851 | 9.44256 | 24.88688 | 22.48677 | Intron (ENSM | 10 | 73821943 | 74507601 | 685659 | 1 | ENSMUST00 | 4330 | Pcdh15 | ENSMUSG00000052613.11 |
| chr15 | 82364587 | 82364884 | 9.44256 | 24.88688 | 22.48677 | Distal Interge | 15 | 82350139 | 82354291 | 4153 | 2 | ENSMUST00 | -10297 | Ndufa6 | ENSMUSG00000022450.5 |
| chr4 | 148447968 | 148448404 | 9.44256 | 24.88688 | 22.48677 | Promoter | 4 | 148448611 | 148454969 | 6359 | 1 | ENSMUST00 | -207 | Mtor | ENSMUSG00000028991.10 |
| chr5 | 125386989 | 125387437 | 11.36018 | 24.86417 | 22.46447 | Exon (ENSMU | 5 | 125387659 | 125389186 | 1528 | 2 | ENSMUST00 | 1749 | Ubc | ENSMUSG00000008348.9 |
| chr15 | 101052977 | 101054214 | 11.70532 | 24.85935 | 22.45968 | Promoter | 15 | 101050194 | 101054399 | 4206 | 2 | ENSMUST00 | 185 | Fignl2 | ENSMUSG00000095440.1 |
| chr2 | 69586884 | 69587570 | 11.70532 | 24.85935 | 22.45968 | Promoter | 2 | 69424340 | 69586065 | 161726 | 2 | ENSMUST00 | -820 | Lrp2 | ENSMUSG00000027070.9 |
| chr3 | 107594973 | 107595472 | 11.70532 | 24.85935 | 22.45968 | Promoter | 3 | 107595031 | 107605776 | 10746 | 1 | ENSMUST00 | 0 | Alx3 | ENSMUSG00000014603.2 |
| chr8 | 94191125 | 94191629 | 11.70532 | 24.85935 | 22.45968 | Distal Interge | 8 | 94179089 | 94180325 | 1237 | 1 | ENSMUST00 | 12037 | Mt1 | ENSMUSG00000031765.7 |
| chr2 | 32627341 | 32629460 | 11.68127 | 24.81904 | 22.41951 | Promoter | 2 | 32628390 | 32631135 | 2746 | 1 | ENSMUST00 | 0 | Ak1 | ENSMUSG00000026817.9 |
| chr1 | 34236192 | 34236931 | 11.31597 | 24.78923 | 22.38983 | Intron (ENSM | 1 | 34257836 | 34308404 | 50569 | 1 | ENSMUST00 | -20905 | Dst | ENSMUSG00000026131.12 |
| chr1 | 172520644 | 172521417 | 11.31597 | 24.78923 | 22.38983 | Promoter | 1 | 172521044 | 172545870 | 24827 | 1 | ENSMUST00 | 0 | Ccdc19 | ENSMUSG00000026546.11 |
| chr10 | 128696558 | 128696914 | 11.31597 | 24.78923 | 22.38983 | Promoter | 10 | 128677193 | 128696268 | 19076 | 2 | ENSMUST00 | -291 | Rab5b | ENSMUSG00000000711.2 |
| chr10 | 127339045 | 127340137 | 11.31597 | 24.78923 | 22.38983 | Intron (ENSM | 10 | 127329889 | 127341589 | 11701 | 2 | ENSMUST00 | 1452 | Gli1 | ENSMUSG00000025407.6 |
| chr11 | 78984490 | 78985305 | 11.31597 | 24.78923 | 22.38983 | Promoter | 11 | 78962974 | 78984946 | 21973 | 2 | ENSMUST00 | 0 | Lgals9 | ENSMUSG00000001123.10 |
| chr11 | 115628328 | 115628667 | 11.31597 | 24.78923 | 22.38983 | Promoter | 11 | 115614184 | 115628295 | 14112 | 2 | ENSMUST00 | -34 | Slc25a19 | ENSMUSG00000020744.8 |
| chr11 | 96124333 | 96124707 | 11.31597 | 24.78923 | 22.38983 | Distal Interge | 11 | 96133786 | 96165451 | 31666 | 1 | ENSMUST00 | -9079 | Ttll6 | ENSMUSG00000038756.8 |
| chr11 | 102946065 | 102947328 | 11.31597 | 24.78923 | 22.38983 | Promoter | 11 | 102939264 | 102946688 | 7425 | 2 | ENSMUST00 | 0 | C1ql1 | ENSMUSG00000045532.5 |
| chr11 | 60353345 | 60353669 | 11.31597 | 24.78923 | 22.38983 | Promoter | 11 | 60353357 | 60375139 | 21783 | 1 | ENSMUST00 | 0 | Lrrc48 | ENSMUSG00000056598.11 |
| chr11 | 6528865 | 6529218 | 11.31597 | 24.78923 | 22.38983 | Promoter | 11 | 6525591 | 6528760 | 3170 | 2 | ENSMUST00 | -106 | Gm11974 | ENSMUSG00000085156.2 |
| chr14 | 30353604 | 30354061 | 11.31597 | 24.78923 | 22.38983 | Promoter | 14 | 30039949 | 30353486 | 313538 | 2 | ENSMUST00 | -119 | Cacna1d | ENSMUSG00000015968.11 |
| chr14 | 69861560 | 69862037 | 11.31597 | 24.78923 | 22.38983 | Intron (ENSM | 14 | 69840420 | 70059886 | 219467 | 1 | ENSMUST00 | 21141 | Pebp4 | ENSMUSG00000022085.3 |
| chr14 | 21982401 | 21984208 | 11.31597 | 24.78923 | 22.38983 | 3' UTR | 14 | 21983962 | 21989601 | 5640 | 2 | ENSMUST00 | 5393 | Zfp503 | ENSMUSG00000039081.9 |
| chr15 | 101128682 | 101129529 | 11.31597 | 24.78923 | 22.38983 | Promoter | 15 | 101128922 | 101143640 | 14719 | 1 | ENSMUST00 | 0 | Acvrl1 | ENSMUSG00000000530.11 |
| chr17 | 47511325 | 47511699 | 11.31597 | 24.78923 | 22.38983 | Intron (ENSM | 17 | 47505241 | 47598649 | 93409 | 1 | ENSMUST00 | 6085 | Ccnd3 | ENSMUSG00000034165.11 |
| chr18 | 21652535 | 21654092 | 11.31597 | 24.78923 | 22.38983 | Promoter | 18 | 21652560 | 21655157 | 2598 | 1 | ENSMUST00 | 0 | 4930426D05 | ENSMUSG00000043168.7 |
| chr18 | 38209625 | 38211033 | 11.31597 | 24.78923 | 22.38983 | Promoter | 18 | 38196694 | 38209762 | 13069 | 2 | ENSMUST00 | 0 | Pcdh1 | ENSMUSG00000051375.10 |
| chr19 | 59459020 | 59459793 | 11.31597 | 24.78923 | 22.38983 | Promoter | 19 | 59459519 | 59465353 | 5835 | 1 | ENSMUST00 | 0 | Emx2 | ENSMUSG00000043969.4 |
| chr3 | 55242522 | 55243170 | 11.31597 | 24.78923 | 22.38983 | Promoter | 3 | 55242526 | 55471317 | 228792 | 1 | ENSMUST00 | 0 | Dclk1 | ENSMUSG00000027797.10 |
| chr3 | 125937937 | 125938692 | 11.31597 | 24.78923 | 22.38983 | Promoter | 3 | 125865359 | 125938550 | 73192 | 2 | ENSMUST00 | 0 | Ugt8a | ENSMUSG00000032854.8 |
| chr3 | 68902697 | 68902923 | 11.31597 | 24.78923 | 22.38983 | Promoter | 3 | 68902267 | 68902758 | 492 | 2 | ENSMUST00 | 0 | Gm18588 | ENSMUSG00000093402.1 |
| chr6 | 4003932 | 4004833 | 11.31597 | 24.78923 | 22.38983 | Promoter | 6 | 4003943 | 4008445 | 4503 | 1 | ENSMUST00 | 0 | Gng11 | ENSMUSG00000032766.8 |
| chr7 | 3644341 | 3644976 | 11.31597 | 24.78923 | 22.38983 | Promoter | 7 | 3632114 | 3644636 | 12523 | 2 | ENSMUST00 | 0 | Gm15927 | ENSMUSG00000085039.1 |
| chr7 | 39487471 | 39487744 | 11.31597 | 24.78923 | 22.38983 | Distal Interge | 7 | 39489438 | 39498801 | 9364 | 1 | ENSMUST00 | -1694 | RP23-9C7.3 | ENSMUSG00000100490.1 |
| chr8 | 70856682 | 70857339 | 11.31597 | 24.78923 | 22.38983 | Promoter | 8 | 70842049 | 70857008 | 14960 | 2 | ENSMUST00 | 0 | Kcnn1 | ENSMUSG00000002908.10 |
| chr8 | 113848116 | 113848868 | 11.31597 | 24.78923 | 22.38983 | Promoter | 8 | 113697126 | 113848615 | 151490 | 2 | ENSMUST00 | 0 | Adamts18 | ENSMUSG00000053399.8 |
| chr8 | 94704234 | 94704655 | 11.31597 | 24.78923 | 22.38983 | Distal Interge | 8 | 94698016 | 94698285 | 270 | 2 | ENSMUST00 | -5950 | Gm22846 | ENSMUSG00000088331.1 |
| chr4 | 141210577 | 141210868 | 10.95847 | 24.76638 | 22.3688 | Distal Interge | 4 | 141213956 | 141220148 | 6193 | 1 | ENSMUST00 | -3088 | Rsg1 | ENSMUSG00000073733.4 |
| chr11 | 95006674 | 95007123 | 11.63346 | 24.73891 | 22.34169 | 3' UTR | 11 | 95009879 | 95026087 | 16209 | 1 | ENSMUST00 | -2756 | Samd14 | ENSMUSG00000047181.7 |
| chr14 | 101517509 | 101518524 | 11.63346 | 24.73891 | 22.34169 | Intron (ENSM | 14 | 101442368 | 101507326 | 64959 | 2 | ENSMUST00 | -10184 | Tbc1d4 | ENSMUSG00000033083.11 |
| chr5 | 65130683 | 65131810 | 11.63346 | 24.73891 | 22.34169 | Promoter | 5 | 65131189 | 65168045 | 36857 | 1 | ENSMUST00 | 0 | Klhl5 | ENSMUSG00000054920.7 |
| chr5 | 117242619 | 117243166 | 11.63346 | 24.73891 | 22.34169 | Intron (ENSM | 5 | 117240667 | 117273869 | 33203 | 1 | ENSMUST00 | 1953 | Taok3 | ENSMUSG00000061288.8 |
| chr1 | 91249583 | 91250120 | 10.27043 | 24.72435 | 22.32749 | Promoter | 1 | 91250304 | 91275479 | 25176 | 1 | ENSMUST00 | -184 | Ube2f | ENSMUSG00000034343.9 |
| chr8 | 12384223 | 12384865 | 11.25191 | 24.68055 | 22.28399 | Promoter | 8 | 12385771 | 12436729 | 50959 | 1 | ENSMUST00 | -906 | Gm5607 | ENSMUSG00000047935.11 |
| chr11 | 118248811 | 118249207 | 11.58603 | 24.6594 | 22.26287 | Promoter | 11 | 118164166 | 118248592 | 84427 | 2 | ENSMUST00 | -220 | Cyth1 | ENSMUSG00000017132.12 |
| chr8 | 108820156 | 108820822 | 11.58603 | 24.6594 | 22.26287 | Intron (ENSM | 8 | 108918264 | 108921874 | 3611 | 2 | ENSMUST00 | 101052 | Gm20686 | ENSMUSG00000093697.1 |
| chr9 | 31912966 | 31913930 | 11.58603 | 24.6594 | 22.26287 | Promoter | 9 | 31846044 | 31913462 | 67419 | 2 | ENSMUST00 | 0 | Barx2 | ENSMUSG00000032033.11 |
| chr8 | 106337553 | 106338028 | 11.23051 | 24.64421 | 22.24776 | Promoter | 8 | 106252548 | 106337988 | 85441 | 2 | ENSMUST00 | 0 | Smpd3 | ENSMUSG00000031906.8 |

| | | | | | | | | | | | | | | | |
|---|---|---|---|---|---|---|---|---|---|---|---|---|---|---|---|
| chr9 | 123923065 | 123923929 | 11.56247 | 24.61987 | 22.22346 | Distal Interge | 9 | 123962126 | 123968692 | 6567 | 2 | ENSMUST00 | 44763 | Ccr1 | ENSMUSG00000025804.4 |
| chr1 | 89968092 | 89968890 | 10.20265 | 24.60154 | 22.20526 | 3' UTR | 1 | 89927956 | 89931179 | 3224 | 2 | ENSMUST00 | -36914 | Gbx2 | ENSMUSG00000034486.8 |
| chr10 | 87860051 | 87861040 | 10.20265 | 24.60154 | 22.20526 | Promoter | 10 | 87860606 | 87930899 | 70294 | 1 | ENSMUST00 | 0 | Igf1 | ENSMUSG00000020053.13 |
| chr12 | 110706461 | 110707262 | 10.20265 | 24.60154 | 22.20526 | Distal Interge | 12 | 110694629 | 110702728 | 8100 | 2 | ENSMUST00 | -3734 | Hsp90aa1 | ENSMUSG00000021270.8 |
| chr15 | 76069925 | 76070204 | 10.20265 | 24.60154 | 22.20526 | Promoter | 15 | 76047158 | 76069784 | 22627 | 2 | ENSMUST00 | -142 | Scrib | ENSMUSG00000022568.11 |
| chr15 | 54952181 | 54952784 | 10.20265 | 24.60154 | 22.20526 | Promoter | 15 | 54952948 | 54964642 | 11695 | 1 | ENSMUST00 | -164 | Gm26684 | ENSMUSG00000097286.1 |
| chr16 | 56478003 | 56479143 | 10.20265 | 24.60154 | 22.20526 | Promoter | 16 | 56477846 | 56690128 | 212283 | 1 | ENSMUST00 | 158 | Abi3bp | ENSMUSG00000035258.9 |
| chr16 | 21818308 | 21819072 | 10.20265 | 24.60154 | 22.20526 | Distal Interge | 16 | 21794392 | 21806123 | 11732 | 1 | ENSMUST00 | 23917 | 1300002E11I | ENSMUSG00000097290.2 |
| chr17 | 34139205 | 34140118 | 10.20265 | 24.60154 | 22.20526 | Distal Interge | 17 | 34137028 | 34138180 | 1153 | 1 | ENSMUST00 | 2178 | H2-DMa | ENSMUSG00000037649.9 |
| chr17 | 17448655 | 17449642 | 10.20265 | 24.60154 | 22.20526 | Promoter | 17 | 17448820 | 17459387 | 10568 | 1 | ENSMUST00 | 0 | Lix1 | ENSMUSG00000047786.12 |
| chr2 | 118554053 | 118554757 | 10.20265 | 24.60154 | 22.20526 | Distal Interge | 2 | 118528757 | 118549687 | 20931 | 2 | ENSMUST00 | -4367 | Bmf | ENSMUSG00000040093.10 |
| chr3 | 82073700 | 82074710 | 10.20265 | 24.60154 | 22.20526 | Promoter | 3 | 82032004 | 82074711 | 42708 | 2 | ENSMUST00 | 1 | Gucy1b3 | ENSMUSG00000028005.8 |
| chr3 | 144380865 | 144381619 | 10.20265 | 24.60154 | 22.20526 | Distal Interge | 3 | 144188530 | 144205220 | 16691 | 2 | ENSMUST00 | -175646 | Lmo4 | ENSMUSG00000028266.12 |
| chr3 | 9346706 | 9348003 | 10.20265 | 24.60154 | 22.20526 | Distal Interge | 3 | 9402175 | 9402485 | 311 | 1 | ENSMUST00 | -54172 | Gm22604 | ENSMUSG00000088229.1 |
| chr4 | 126325221 | 126325855 | 10.20265 | 24.60154 | 22.20526 | Promoter | 4 | 126322121 | 126325688 | 3568 | 2 | ENSMUST00 | 0 | Tekt2 | ENSMUSG00000028845.10 |
| chr4 | 132922832 | 132923347 | 10.20265 | 24.60154 | 22.20526 | Promoter | 4 | 132899213 | 132922558 | 23346 | 2 | ENSMUST00 | -275 | Fam76a | ENSMUSG00000028878.6 |
| chr4 | 32250084 | 32251467 | 10.20265 | 24.60154 | 22.20526 | Promoter | 4 | 32251761 | 32265835 | 14075 | 1 | ENSMUST00 | -294 | Bach2it1 | ENSMUSG00000087049.1 |
| chr6 | 70956206 | 70957076 | 10.20265 | 24.60154 | 22.20526 | Promoter | 6 | 70956531 | 70961063 | 4533 | 1 | ENSMUST00 | 0 | Foxi3 | ENSMUSG00000055874.6 |
| chr7 | 45916211 | 45917315 | 10.20265 | 24.60154 | 22.20526 | Promoter | 7 | 45915998 | 45916975 | 978 | 1 | ENSMUST00 | 214 | Tmem143 | ENSMUSG00000002781.12 |
| chr7 | 29237759 | 29238156 | 10.20265 | 24.60154 | 22.20526 | Promoter | 7 | 29238329 | 29247529 | 9201 | 1 | ENSMUST00 | -173 | Yif1b | ENSMUSG00000030588.6 |
| chr9 | 3532777 | 3533267 | 10.20265 | 24.60154 | 22.20526 | Promoter | 9 | 3532354 | 3897342 | 364989 | 1 | ENSMUST00 | 424 | Gucy1a2 | ENSMUSG00000041624.9 |
| chrX | 12087071 | 12088799 | 10.20265 | 24.60154 | 22.20526 | Intron (ENSM | 20 | 12036740 | 12080553 | 43814 | 2 | ENSMUST00 | -6519 | Bcor | ENSMUSG00000040363.9 |
| chr2 | 152739337 | 152739655 | 9.01375 | 24.58323 | 22.18825 | Distal Interge | 2 | 152736251 | 152737410 | 1160 | 1 | ENSMUST00 | 3087 | Id1 | ENSMUSG00000042745.9 |
| chr3 | 88200320 | 88201444 | 9.01375 | 24.58323 | 22.18825 | Promoter | 3 | 88200623 | 88203481 | 2859 | 1 | ENSMUST00 | 0 | AW047730 | ENSMUSG00000097428.1 |
| chr9 | 109054747 | 109055364 | 9.01375 | 24.58323 | 22.18825 | Promoter | 9 | 109054863 | 109057207 | 2345 | 1 | ENSMUST00 | 0 | Shisa5 | ENSMUSG00000025647.11 |
| chr10 | 121435512 | 121436344 | 11.51562 | 24.54127 | 22.1466 | Intron (ENSM | 10 | 121410350 | 121476250 | 65901 | 2 | ENSMUST00 | 39906 | Rassf3 | ENSMUSG00000025795.7 |
| chr5 | 114588116 | 114588585 | 11.51562 | 24.54127 | 22.1466 | Intron (ENSM | 5 | 114568017 | 114594678 | 26662 | 1 | ENSMUST00 | 20100 | Fam222a | ENSMUSG00000041930.7 |
| chr1 | 74295970 | 74296658 | 11.16678 | 24.53592 | 22.14134 | Promoter | 1 | 74288251 | 74295695 | 7445 | 2 | ENSMUST00 | -276 | Tmbim1 | ENSMUSG00000006301.12 |
| chr10 | 93594003 | 93594618 | 11.16678 | 24.53592 | 22.14134 | Intron (ENSM | 10 | 93583029 | 93589706 | 6678 | 2 | ENSMUST00 | -4298 | Snrpf | ENSMUSG00000020018.6 |
| chr10 | 43723289 | 43723948 | 11.16678 | 24.53592 | 22.14134 | Distal Interge | 10 | 43821231 | 43822235 | 1005 | 1 | ENSMUST00 | -97283 | Gm9034 | ENSMUSG00000094852.3 |
| chr11 | 69134025 | 69134665 | 11.16678 | 24.53592 | 22.14134 | Promoter | 11 | 69134142 | 69135106 | 965 | 1 | ENSMUST00 | 0 | Aloxe3 | ENSMUSG00000020892.5 |
| chr15 | 101246473 | 101246889 | 11.16678 | 24.53592 | 22.14134 | Distal Interge | 15 | 101266846 | 101274792 | 7947 | 1 | ENSMUST00 | -19957 | Nr4a1 | ENSMUSG00000023034.6 |
| chr8 | 26168841 | 26169289 | 11.16678 | 24.53592 | 22.14134 | Distal Interge | 8 | 26158255 | 26159412 | 1158 | 1 | ENSMUST00 | 10587 | Thap1 | ENSMUSG00000037214.6 |
| chr9 | 107906608 | 107907175 | 11.16678 | 24.53592 | 22.14134 | Promoter | 9 | 107906889 | 107920383 | 13495 | 1 | ENSMUST00 | 0 | Mst1r | ENSMUSG00000032584.7 |
| chr9 | 63927416 | 63928107 | 11.16678 | 24.53592 | 22.14134 | Distal Interge | 9 | 63954009 | 64016987 | 62979 | 2 | ENSMUST00 | 88880 | Smad6 | ENSMUSG00000036867.7 |
| chr15 | 25441192 | 25441567 | 11.46915 | 24.46326 | 22.06937 | Distal Interge | 15 | 25414192 | 25414844 | 653 | 1 | ENSMUST00 | 27001 | Gm5468 | ENSMUSG00000056089.5 |
| chr14 | 118713156 | 118713890 | 11.10376 | 24.42872 | 22.03518 | Distal Interge | 14 | 118482692 | 118706219 | 223528 | 2 | ENSMUST00 | -6938 | Abcc4 | ENSMUSG00000032849.8 |
| chr13 | 107469727 | 107470454 | 11.16606 | 24.42448 | 22.03097 | Distal Interge | 13 | 107413865 | 107414767 | 903 | 2 | ENSMUST00 | -54961 | Apoo-ps | ENSMUSG00000049233.6 |
| chr19 | 5349053 | 5349411 | 10.41744 | 24.40567 | 22.01225 | Promoter | 19 | 5344705 | 5349574 | 4870 | 2 | ENSMUST00 | 163 | Cst6 | ENSMUSG00000024846.5 |
| chr5 | 43354755 | 43355656 | 11.06215 | 24.35785 | 21.96462 | Distal Interge | 5 | 43237316 | 43246459 | 9144 | 1 | ENSMUST00 | 117440 | Cpeb2 | ENSMUSG00000039782.9 |
| chr2 | 18799708 | 18801068 | 8.64364 | 24.35355 | 21.96034 | Promoter | 2 | 18799245 | 18801818 | 2574 | 2 | ENSMUST00 | 750 | Carlr | ENSMUSG00000097638.1 |
| chr1 | 130731948 | 130732360 | 11.40015 | 24.34736 | 21.95423 | Promoter | 1 | 130731976 | 130744622 | 12647 | 1 | ENSMUST00 | 0 | AA986860 | ENSMUSG00000042510.7 |
| chr10 | 53630249 | 53630664 | 11.40015 | 24.34736 | 21.95423 | Promoter | 10 | 53536329 | 53630439 | 94111 | 2 | ENSMUST00 | 0 | Mcm9 | ENSMUSG00000058298.7 |
| chr10 | 117436997 | 117437413 | 11.40015 | 24.34736 | 21.95423 | Distal Interge | 10 | 117429618 | 117430116 | 499 | 1 | ENSMUST00 | 7380 | Gm9002 | ENSMUSG00000095826.1 |
| chr12 | 8301382 | 8302512 | 11.40015 | 24.34736 | 21.95423 | Promoter | 12 | 8297934 | 8301954 | 4021 | 2 | ENSMUST00 | 0 | Gdf7 | ENSMUSG00000037660.4 |
| chr17 | 45733451 | 45734265 | 11.40015 | 24.34736 | 21.95423 | Promoter | 17 | 45732864 | 45733844 | 981 | 2 | ENSMUST00 | 0 | 1600014C23I | ENSMUSG00000094690.1 |
| chr18 | 53647628 | 53648384 | 11.40015 | 24.34736 | 21.95423 | Distal Interge | 18 | 53551594 | 53575857 | 24264 | 1 | ENSMUST00 | 96035 | Prdm6 | ENSMUSG00000069378.8 |
| chr6 | 21822864 | 21823994 | 11.40015 | 24.34736 | 21.95423 | Intron (ENSM | 6 | 21799950 | 21851914 | 51965 | 2 | ENSMUST00 | 27920 | Tspan12 | ENSMUSG00000029669.9 |
| chr8 | 125254015 | 125254758 | 11.40015 | 24.34736 | 21.95423 | Intron (ENSM | 8 | 125224400 | 125227323 | 2924 | 2 | ENSMUST00 | -26693 | Gm16237 | ENSMUSG00000086694.1 |
| chr15 | 42675867 | 42676654 | 11.04146 | 24.3226 | 21.92989 | Promoter | 15 | 42676260 | 42704616 | 28357 | 1 | ENSMUST00 | 0 | Gm17473 | ENSMUSG00000097805.1 |
| chr15 | 99001428 | 99001851 | 11.37733 | 24.30901 | 21.91632 | Distal Interge | 15 | 99006056 | 99006370 | 315 | 1 | ENSMUST00 | -4205 | Gm8973 | ENSMUSG00000064081.3 |
| chr7 | 7136314 | 7136874 | 11.37733 | 24.30901 | 21.91632 | Promoter | 7 | 7130678 | 7136755 | 6078 | 2 | ENSMUST00 | 0 | Zfp773 | ENSMUSG00000063535.6 |
| chr7 | 145299716 | 145300691 | 11.33197 | 24.23275 | 21.84028 | Promoter | 7 | 145300828 | 145309557 | 8730 | 1 | ENSMUST00 | -137 | Mrgprf | ENSMUSG00000031070.10 |
| chr7 | 54835289 | 54836136 | 11.33197 | 24.23275 | 21.84028 | Promoter | 7 | 54835959 | 55265366 | 429408 | 1 | ENSMUST00 | 0 | Luzp2 | ENSMUSG00000063297.6 |
| chr9 | 78395546 | 78396375 | 11.30943 | 24.19482 | 21.80247 | Promoter | 9 | 78395777 | 78423587 | 27811 | 1 | ENSMUST00 | 0 | Ddx43 | ENSMUSG00000070291.4 |
| chr1 | 194938763 | 194939554 | 10.95946 | 24.18275 | 21.79051 | Promoter | 1 | 194938821 | 194961292 | 22472 | 1 | ENSMUST00 | 0 | Cd34 | ENSMUSG00000016494.4 |
| chr12 | 113141417 | 113143358 | 10.95946 | 24.18275 | 21.79051 | Intron (ENSM | 12 | 113140236 | 113145504 | 5269 | 1 | ENSMUST00 | 1182 | Crip2 | ENSMUSG00000006356.6 |
| chr19 | 47137816 | 47138551 | 10.95946 | 24.18275 | 21.79051 | Promoter | 19 | 47105353 | 47138294 | 32942 | 2 | ENSMUST00 | 0 | Calhm2 | ENSMUSG00000033033.5 |
| chr3 | 102011302 | 102012264 | 10.95946 | 24.18275 | 21.79051 | Intron (ENSM | 3 | 102010145 | 102015492 | 5348 | 1 | ENSMUST00 | 1158 | Nhlh2 | ENSMUSG00000048540.7 |
| chr4 | 154170264 | 154171266 | 10.95946 | 24.18275 | 21.79051 | Promoter | 4 | 154170730 | 154275713 | 104984 | 1 | ENSMUST00 | 0 | Megf6 | ENSMUSG00000057751.9 |
| chr11 | 69097223 | 69098719 | 11.28697 | 24.15705 | 21.76529 | Promoter | 11 | 69098948 | 69109960 | 11013 | 1 | ENSMUST00 | -229 | Per1 | ENSMUSG00000020893.12 |
| chr12 | 114378671 | 114379493 | 11.28697 | 24.15705 | 21.76529 | Distal Interge | 12 | 114366526 | 114366954 | 429 | 2 | ENSMUST00 | -11718 | Ighv12-3 | ENSMUSG00000076676.1 |
| chr10 | 80985613 | 80986233 | 9.63447 | 24.12454 | 21.73289 | Intron (ENSM | 10 | 80951187 | 80998190 | 47004 | 2 | ENSMUST00 | 11957 | Gng7 | ENSMUSG00000048240.9 |
| chr10 | 127676431 | 127676826 | 9.63447 | 24.12454 | 21.73289 | Promoter | 10 | 127671604 | 127676724 | 5121 | 2 | ENSMUST00 | 0 | Gm16230 | ENSMUSG00000087163.1 |
| chr11 | 117963726 | 117964052 | 9.63447 | 24.12454 | 21.73289 | Downstream | 11 | 117966079 | 117969186 | 3108 | 2 | ENSMUST00 | 5134 | Socs3 | ENSMUSG00000053113.3 |
| chr14 | 6286927 | 6287580 | 9.63447 | 24.12454 | 21.73289 | Promoter | 14 | 6259747 | 6287250 | 27504 | 2 | ENSMUST00 | 0 | Gm3411 | ENSMUSG00000079396.4 |
| chr15 | 85667175 | 85668042 | 9.63447 | 24.12454 | 21.73289 | Distal Interge | 15 | 85639747 | 85645934 | 6188 | 1 | ENSMUST00 | 27429 | RP23-64J7.2 | ENSMUSG00000101828.1 |
| chr18 | 80236468 | 80237094 | 9.63447 | 24.12454 | 21.73289 | 5' UTR | 18 | 80229758 | 80247102 | 17345 | 2 | ENSMUST00 | 10008 | Hsbp1l1 | ENSMUSG00000078963.2 |
| chr19 | 8711515 | 8712058 | 9.63447 | 24.12454 | 21.73289 | Exon (ENSMU | 19 | 8706882 | 8713897 | 7016 | 2 | ENSMUST00 | 1839 | Slc3a2 | ENSMUSG00000010095.8 |
| chr19 | 5098161 | 5098636 | 9.63447 | 24.12454 | 21.73289 | Promoter | 19 | 5092871 | 5098418 | 5548 | 2 | ENSMUST00 | 0 | Cnih2 | ENSMUSG00000024873.6 |
| chr2 | 142902407 | 142903691 | 9.63447 | 24.12454 | 21.73289 | Promoter | 2 | 142902965 | 142903572 | 608 | 1 | ENSMUST00 | 0 | Kif16bos | ENSMUSG00000086504.1 |
| chr5 | 30945147 | 30945693 | 9.63447 | 24.12454 | 21.73289 | Promoter | 5 | 30933146 | 30945480 | 12335 | 2 | ENSMUST00 | 0 | Cgref1 | ENSMUSG00000029161.6 |
| chr7 | 80904522 | 80905324 | 9.63447 | 24.12454 | 21.73289 | Promoter | 7 | 80902228 | 80905076 | 2849 | 2 | ENSMUST00 | 0 | Nmb | ENSMUSG00000025723.7 |
| chr7 | 139659948 | 139660198 | 9.63447 | 24.12454 | 21.73289 | Promoter | 7 | 139659688 | 139660155 | 468 | 2 | ENSMUST00 | 0 | Gm4459 | ENSMUSG00000083626.1 |
| chr7 | 19304324 | 19305368 | 9.63447 | 24.12454 | 21.73289 | 3' UTR | 7 | 19298260 | 19300770 | 2511 | 2 | ENSMUST00 | -3555 | Gm26802 | ENSMUSG00000097266.1 |
| chr9 | 26806024 | 26806771 | 9.63447 | 24.12454 | 21.73289 | Promoter | 9 | 26763044 | 26806432 | 43389 | 2 | ENSMUST00 | 0 | Glb1l2 | ENSMUSG00000036395.9 |
| chr2 | 30942371 | 30942883 | 10.5797 | 24.1096 | 21.71881 | Distal Interge | 2 | 30952959 | 30959015 | 6057 | 1 | ENSMUST00 | -10076 | Tor1b | ENSMUSG00000026848.9 |
| chr5 | 114647134 | 114648009 | 10.5797 | 24.1096 | 21.71881 | Intron (ENSM | 5 | 114622653 | 114644899 | 22247 | 2 | ENSMUST00 | -2236 | Trpv4 | ENSMUSG00000014158.6 |
| chr17 | 80391473 | 80392121 | 11.22013 | 24.04454 | 21.6541 | Downstream | 17 | 80373218 | 80373541 | 324 | 2 | ENSMUST00 | -17933 | Gm10190 | ENSMUSG00000066938.1 |
| chr17 | 24533282 | 24533753 | 11.22013 | 24.04454 | 21.6541 | Promoter | 17 | 24527241 | 24533726 | 6486 | 2 | ENSMUST00 | 0 | Rab26 | ENSMUSG00000079657.5 |
| chr9 | 53829457 | 53829922 | 11.22013 | 24.04454 | 21.6541 | Distal Interge | 9 | 53850251 | 53853844 | 3594 | 1 | ENSMUST00 | -20329 | Sln | ENSMUSG00000042045.5 |
| chr12 | 16800234 | 16801166 | 11.19803 | 24.00731 | 21.61695 | Promoter | 12 | 16670615 | 16800684 | 130070 | 2 | ENSMUST00 | 0 | Greb1 | ENSMUSG00000036523.11 |
| chr9 | 56796235 | 56797163 | 11.19803 | 24.00731 | 21.61695 | Distal Interge | 9 | 56848659 | 56851963 | 3305 | 2 | ENSMUST00 | 54800 | Odf3l1 | ENSMUSG00000045620.6 |
| chr11 | 75166821 | 75167496 | 10.51363 | 23.99435 | 21.60407 | Promoter | 11 | 75167153 | 75168146 | 994 | 2 | ENSMUST00 | 650 | Hic1 | ENSMUSG00000043099.4 |
| chr11 | 98035905 | 98036377 | 10.51363 | 23.99435 | 21.60407 | Downstream | 11 | 98029413 | 98029492 | 80 | 1 | ENSMUST00 | 6493 | Gm27597 | ENSMUSG00000098992.1 |
| chr12 | 9581068 | 9581483 | 10.51363 | 23.99435 | 21.60407 | 3' UTR | 12 | 9574442 | 9581499 | 7058 | 1 | ENSMUST00 | 6627 | Osr1 | ENSMUSG00000048387.7 |
| chr13 | 94173667 | 94174718 | 10.51363 | 23.99435 | 21.60407 | Promoter | 13 | 94173774 | 94174213 | 440 | 1 | ENSMUST00 | 0 | Lhfpl2 | ENSMUSG00000045312.6 |
| chr13 | 43304394 | 43305036 | 10.51363 | 23.99435 | 21.60407 | Promoter | 13 | 43195519 | 43304172 | 108654 | 2 | ENSMUST00 | -223 | Gfod1 | ENSMUSG00000051335.5 |
| chr16 | 36693807 | 36694434 | 10.51363 | 23.99435 | 21.60407 | Promoter | 16 | 36693978 | 36726804 | 32827 | 1 | ENSMUST00 | 0 | Ildr1 | ENSMUSG00000022900.9 |
| chr16 | 91830209 | 91830838 | 10.51363 | 23.99435 | 21.60407 | 3' UTR | 16 | 91827537 | 91828273 | 737 | 1 | ENSMUST00 | 2673 | Itsn1 | ENSMUSG00000022957.14 |
| chr17 | 25474478 | 25474935 | 10.51363 | 23.99435 | 21.60407 | Exon (ENSMU | 17 | 25471590 | 25476594 | 5005 | 1 | ENSMUST00 | 2889 | Tekt4 | ENSMUSG00000024175.1 |
| chr18 | 34247125 | 34248105 | 10.51363 | 23.99435 | 21.60407 | Promoter | 18 | 34247694 | 34263695 | 16002 | 1 | ENSMUST00 | 0 | Apc | ENSMUSG00000005871.9 |
| chr18 | 37742055 | 37742550 | 10.51363 | 23.99435 | 21.60407 | Promoter | 18 | 37742094 | 37841873 | 99780 | 1 | ENSMUST00 | 0 | Pcdhga12 | ENSMUSG00000023036.9 |
| chr18 | 38349111 | 38350084 | 10.51363 | 23.99435 | 21.60407 | Distal Interge | 18 | 38356578 | 38356674 | 97 | 1 | ENSMUST00 | -6494 | Gm24690 | ENSMUSG00000088384.1 |
| chr18 | 35847706 | 35848814 | 10.51363 | 23.99435 | 21.60407 | Intron (ENSM | 18 | 35830389 | 35831093 | 705 | 2 | ENSMUST00 | -16614 | RP24-326K11 | ENSMUSG00000100320.1 |
| chr2 | 145785884 | 145786836 | 10.51363 | 23.99435 | 21.60407 | Promoter | 2 | 145785980 | 145887616 | 101637 | 1 | ENSMUST00 | 0 | Rin2 | ENSMUSG00000001768.10 |
| chr2 | 57102265 | 57103597 | 10.51363 | 23.99435 | 21.60407 | Distal Interge | 2 | 57107499 | 57113072 | 5574 | 2 | ENSMUST00 | 9475 | Nr4a2 | ENSMUSG00000026826.8 |
| chr2 | 35282148 | 35282846 | 10.51363 | 23.99435 | 21.60407 | Promoter | 2 | 35282380 | 35307892 | 25513 | 1 | ENSMUST00 | 0 | Gsn | ENSMUSG00000026879.9 |
| chr2 | 124089927 | 124091059 | 10.51363 | 23.99435 | 21.60407 | Promoter | 2 | 124089969 | 124143353 | 53385 | 1 | ENSMUST00 | 0 | Sema6d | ENSMUSG00000027200.12 |
| chr2 | 131561899 | 131562811 | 10.51363 | 23.99435 | 21.60407 | Promoter | 2 | 131545850 | 131562283 | 16434 | 2 | ENSMUST00 | 0 | Adra1d | ENSMUSG00000027335.9 |
| chr2 | 156391473 | 156392336 | 10.51363 | 23.99435 | 21.60407 | Distal Interge | 2 | 156386741 | 156387222 | 482 | 1 | ENSMUST00 | 4733 | Rpl21-ps9 | ENSMUSG00000084017.1 |
| chr3 | 17786063 | 17787108 | 10.51363 | 23.99435 | 21.60407 | Promoter | 3 | 17783514 | 17786898 | 3385 | 2 | ENSMUST00 | 0 | Gm10742 | ENSMUSG00000074674.2 |
| chr4 | 116405516 | 116406243 | 10.51363 | 23.99435 | 21.60407 | Promoter | 4 | 116322434 | 116405986 | 83553 | 2 | ENSMUST00 | 0 | Mast2 | ENSMUSG00000003810.7 |
| chr5 | 73013497 | 73014356 | 10.51363 | 23.99435 | 21.60407 | Promoter | 5 | 73013499 | 73017376 | 3878 | 1 | ENSMUST00 | 0 | Gm15847 | ENSMUSG00000081849.1 |
| chr5 | 148958966 | 148959906 | 10.51363 | 23.99435 | 21.60407 | Promoter | 5 | 148958598 | 148959704 | 1107 | 2 | ENSMUST00 | 0 | 1810059H22I | ENSMUSG00000085455.2 |
| chr6 | 55617917 | 55618294 | 10.51363 | 23.99435 | 21.60407 | Distal Interge | 6 | 55677818 | 55681263 | 3446 | 2 | ENSMUST00 | 62969 | Neurod6 | ENSMUSG00000037984.8 |
| chr6 | 77242522 | 77244335 | 10.51363 | 23.99435 | 21.60407 | Promoter | 6 | 77242689 | 77257791 | 15103 | 1 | ENSMUST00 | 0 | Lrrtm1 | ENSMUSG00000060780.2 |
| chr7 | 139085887 | 139086736 | 10.51363 | 23.99435 | 21.60407 | Promoter | 7 | 139086001 | 139102704 | 16704 | 1 | ENSMUST00 | 0 | Dpysl4 | ENSMUSG00000025478.10 |
| chr7 | 49522624 | 49522988 | 10.51363 | 23.99435 | 21.60407 | Intron (ENSM | 7 | 49548192 | 49556996 | 8805 | 1 | ENSMUST00 | -25204 | Nav2 | ENSMUSG00000052512.10 |
| chr9 | 37544354 | 37545220 | 10.51363 | 23.99435 | 21.60407 | Promoter | 9 | 37544954 | 37546138 | 1185 | 2 | ENSMUST00 | 918 | Nrgn | ENSMUSG00000053310.6 |
| chr7 | 12909778 | 12910862 | 9.56443 | 23.99214 | 21.60373 | Distal Interge | 7 | 12922290 | 12926685 | 4396 | 1 | ENSMUST00 | -11428 | Rps5 | ENSMUSG00000012848.9 |
| chr11 | 89186667 | 89187483 | 11.17602 | 23.97021 | 21.58181 | Distal Interge | 11 | 89136505 | 89136766 | 262 | 2 | ENSMUST00 | -49902 | Gm24734 | ENSMUSG00000089485.1 |
| chr12 | 82470018 | 82470878 | 11.17602 | 23.97021 | 21.58181 | Distal Interge | 12 | 82495419 | 82496537 | 1119 | 2 | ENSMUST00 | 25659 | Gm5435 | ENSMUSG00000094374.1 |
| chr12 | 108327717 | 108328300 | 11.17602 | 23.97021 | 21.58181 | Promoter | 12 | 108313779 | 108328030 | 14252 | 2 | ENSMUST00 | 0 | 4930478K11I | ENSMUSG00000097847.1 |
| chr2 | 174473162 | 174473540 | 11.17602 | 23.97021 | 21.58181 | Promoter | 2 | 174465067 | 174473081 | 8015 | 2 | ENSMUST00 | -82 | Slmo2 | ENSMUSG00000016257.10 |
| chr7 | 4752032 | 4753355 | 11.17602 | 23.97021 | 21.58181 | Promoter | 7 | 4751792 | 4752972 | 1181 | 2 | ENSMUST00 | 0 | Cox6b2 | ENSMUSG00000051811.9 |
| chr9 | 107935766 | 107936195 | 11.17602 | 23.97021 | 21.58181 | Promoter | 9 | 107935920 | 107949691 | 13772 | 1 | ENSMUST00 | 0 | Camkv | ENSMUSG00000032936.8 |
| chr6 | 58596029 | 58596768 | 7.01212 | 23.94238 | 21.55444 | Promoter | 6 | 58596671 | 58692451 | 95781 | 1 | ENSMUST00 | 0 | Abcg2 | ENSMUSG00000029802.8 |
| chr1 | 189727043 | 189728064 | 11.4883 | 23.92001 | 21.53215 | Promoter | 1 | 189728268 | 189876689 | 148422 | 1 | ENSMUST00 | -204 | Ptpn14 | ENSMUSG00000026604.12 |
| chr10 | 63089664 | 63090489 | 11.4883 | 23.92001 | 21.53215 | Distal Interge | 10 | 63100156 | 63100605 | 450 | 1 | ENSMUST00 | -9667 | Atoh7 | ENSMUSG00000036816.3 |
| chr11 | 98327252 | 98328871 | 11.4883 | 23.92001 | 21.53215 | Promoter | 11 | 98325415 | 98329648 | 4234 | 2 | ENSMUST00 | 777 | Neurod2 | ENSMUSG00000038255.6 |
| chr11 | 98789230 | 98789849 | 11.4883 | 23.92001 | 21.53215 | Distal Interge | 11 | 98795516 | 98807859 | 12344 | 1 | ENSMUST00 | -5667 | Msl1 | ENSMUSG00000052915.9 |
| chr11 | 97534435 | 97535266 | 11.4883 | 23.92001 | 21.53215 | Exon (ENSMU | 11 | 97519612 | 97521895 | 2284 | 2 | ENSMUST00 | -12541 | Gm11611 | ENSMUSG00000087042.1 |
| chr14 | 79771016 | 79771889 | 11.4883 | 23.92001 | 21.53215 | Promoter | 14 | 79766772 | 79771312 | 4541 | 2 | ENSMUST00 | 0 | Pcdh8 | ENSMUSG00000036422.5 |
| chr15 | 36531830 | 36532522 | 11.4883 | 23.92001 | 21.53215 | Intron (ENSM | 15 | 36537508 | 36545687 | 8180 | 2 | ENSMUST00 | 13165 | Snx31 | ENSMUSG00000013611.9 |
| chr15 | 87544395 | 87545079 | 11.4883 | 23.92001 | 21.53215 | Distal Interge | 15 | 87625230 | 87759336 | 134107 | 1 | ENSMUST00 | -80151 | Fam19a5 | ENSMUSG00000054863.8 |
| chr17 | 47504679 | 47505280 | 11.4883 | 23.92001 | 21.53215 | Promoter | 17 | 47505051 | 47598070 | 93020 | 1 | ENSMUST00 | 0 | Ccnd3 | ENSMUSG00000034165.11 |
| chr19 | 6924622 | 6925369 | 11.4883 | 23.92001 | 21.53215 | Promoter | 19 | 6922426 | 6925380 | 2955 | 2 | ENSMUST00 | 11 | Tex40 | ENSMUSG00000050623.4 |

| | | | | | | | | | | | | | | | |
|---|---|---|---|---|---|---|---|---|---|---|---|---|---|---|---|
| chr2 | 131041102 | 131041701 | 11.4883 | 23.92001 | 21.53215 | Promoter | 2 | 131039639 | 131041457 | 1819 | 2 | ENSMUST00 | 0 | Gfra4 | ENSMUSG00000027316.10 |
| chr2 | 175703060 | 175703711 | 11.4883 | 23.92001 | 21.53215 | Promoter | 2 | 175692223 | 175703646 | 11424 | 2 | ENSMUST00 | 0 | Gm4245 | ENSMUSG00000095545.2 |
| chr3 | 73057314 | 73057778 | 11.4883 | 23.92001 | 21.53215 | Promoter | 3 | 73048126 | 73056943 | 8818 | 2 | ENSMUST00 | -372 | Slitrk3 | ENSMUSG00000048304.7 |
| chr3 | 104675153 | 104675685 | 11.4883 | 23.92001 | 21.53215 | Distal Interge | 3 | 104667032 | 104670010 | 2979 | 2 | ENSMUST00 | -5144 | RP23-179B1( | ENSMUSG00000101641.1 |
| chr6 | 128035593 | 128035915 | 11.4883 | 23.92001 | 21.53215 | Intron (ENSM | 6 | 127965747 | 128034592 | 68846 | 2 | ENSMUST00 | -1002 | Tspan9 | ENSMUSG00000030352.9 |
| chr6 | 126739429 | 126740688 | 11.4883 | 23.92001 | 21.53215 | Promoter | 6 | 126708329 | 126740662 | 32334 | 2 | ENSMUST00 | 0 | Kcna6 | ENSMUSG00000038077.7 |
| chr7 | 18872771 | 18873495 | 11.4883 | 23.92001 | 21.53215 | Distal Interge | 7 | 18884690 | 18890438 | 5749 | 1 | ENSMUST00 | -11195 | Pglyrp1 | ENSMUSG00000030413.6 |
| chr7 | 30456714 | 30457803 | 11.4883 | 23.92001 | 21.53215 | Promoter | 7 | 30447534 | 30457652 | 10119 | 2 | ENSMUST00 | 0 | Kirrel2 | ENSMUSG00000036915.11 |
| chr7 | 24944354 | 24945167 | 11.4883 | 23.92001 | 21.53215 | Distal Interge | 7 | 24923363 | 24925936 | 2574 | 1 | ENSMUST00 | 20992 | Arhgef1 | ENSMUSG00000040940.13 |
| chr8 | 121729863 | 121730496 | 11.4883 | 23.92001 | 21.53215 | Promoter | 8 | 121730563 | 121790821 | 60259 | 1 | ENSMUST00 | -67 | Jph3 | ENSMUSG00000025318.8 |
| chr8 | 119446557 | 119446960 | 11.4883 | 23.92001 | 21.53215 | Promoter | 8 | 119446719 | 119472640 | 25922 | 1 | ENSMUST00 | 0 | Necab2 | ENSMUSG00000031837.8 |
| chr9 | 89909504 | 89910526 | 11.4883 | 23.92001 | 21.53215 | Promoter | 9 | 89909908 | 89915847 | 5940 | 1 | ENSMUST00 | 0 | Rasgrf1 | ENSMUSG00000032356.7 |
| chr9 | 53301408 | 53302246 | 11.4883 | 23.92001 | 21.53215 | Promoter | 9 | 53301670 | 53377514 | 75845 | 1 | ENSMUST00 | 0 | Exph5 | ENSMUSG00000034584.3 |
| chrX | 7648486 | 7648895 | 11.4883 | 23.92001 | 21.53215 | 3' UTR | 20 | 7643909 | 7645033 | 1125 | 1 | ENSMUST00 | 4578 | Syp | ENSMUSG00000031144.10 |
| chrX | 160765752 | 160766023 | 11.4883 | 23.92001 | 21.53215 | Distal Interge | 20 | 160768013 | 160799663 | 31651 | 1 | ENSMUST00 | -1990 | Rs1 | ENSMUSG00000031293.8 |
| chrX | 160390406 | 160390984 | 11.4883 | 23.92001 | 21.53215 | Promoter | 20 | 160390690 | 160498070 | 107381 | 1 | ENSMUST00 | 0 | Gpr64 | ENSMUSG00000031298.10 |
| chrX | 72918562 | 72919107 | 11.4883 | 23.92001 | 21.53215 | Promoter | 20 | 72918557 | 72958514 | 39958 | 1 | ENSMUST00 | 6 | Nsdhl | ENSMUSG00000031349.4 |
| chrX | 74328470 | 74329456 | 11.4883 | 23.92001 | 21.53215 | Promoter | 20 | 74329066 | 74332243 | 3178 | 1 | ENSMUST00 | 0 | Plxna3 | ENSMUSG00000031398.8 |
| chrX | 99574427 | 99574928 | 11.4883 | 23.92001 | 21.53215 | Distal Interge | 20 | 99465734 | 99471273 | 5540 | 2 | ENSMUST00 | -103155 | Pja1 | ENSMUSG00000034403.11 |
| chrX | 111696510 | 111697222 | 11.4883 | 23.92001 | 21.53215 | Promoter | 20 | 111569931 | 111697079 | 127149 | 2 | ENSMUST00 | 0 | Hdx | ENSMUSG00000034551.7 |
| chrX | 23692413 | 23692873 | 11.4883 | 23.92001 | 21.53215 | Promoter | 20 | 23693051 | 23805990 | 112940 | 1 | ENSMUST00 | -178 | Wdr44 | ENSMUSG00000036769.9 |
| chrX | 134612119 | 134613107 | 11.4883 | 23.92001 | 21.53215 | Distal Interge | 20 | 134601286 | 134607054 | 5769 | 1 | ENSMUST00 | 10834 | Hnrnph2 | ENSMUSG00000045427.8 |
| chrX | 166510602 | 166510936 | 11.4883 | 23.92001 | 21.53215 | Promoter | 20 | 166499812 | 166510478 | 10667 | 2 | ENSMUST00 | -125 | Tceanc | ENSMUSG00000051224.8 |
| chrX | 36807277 | 36808144 | 11.4883 | 23.92001 | 21.53215 | Distal Interge | 20 | 36814771 | 36817200 | 2430 | 1 | ENSMUST00 | -6627 | Gm14549 | ENSMUSG00000084990.1 |
| chrX | 81071762 | 81072180 | 11.4883 | 23.92001 | 21.53215 | Promoter | 20 | 81070694 | 81073102 | 2409 | 2 | ENSMUST00 | 922 | Gm26775 | ENSMUSG00000096919.1 |
| chr6 | 99726103 | 99726707 | 11.13224 | 23.89642 | 21.51035 | Promoter | 6 | 99711299 | 99726392 | 15094 | 2 | ENSMUST00 | 0 | Prok2 | ENSMUSG00000030069.10 |
| chr2 | 84386598 | 84387927 | 11.11049 | 23.85973 | 21.47382 | Intron (ENSM | 2 | 84330659 | 84375358 | 44700 | 2 | ENSMUST00 | -11241 | Calcrl | ENSMUSG00000059588.8 |
| chr15 | 12010354 | 12011000 | 11.44363 | 23.84541 | 21.45958 | Distal Interge | 15 | 11984041 | 11996983 | 12943 | 2 | ENSMUST00 | -13372 | Sub1 | ENSMUSG00000022205.10 |
| chr11 | 97840756 | 97841660 | 10.7597 | 23.84104 | 21.45527 | Promoter | 11 | 97840780 | 97843043 | 2264 | 1 | ENSMUST00 | 0 | B230217C12 | ENSMUSG00000050538.7 |
| chr14 | 61012395 | 61012987 | 10.7597 | 23.84104 | 21.45527 | Intron (ENSM | 14 | 60963834 | 61037987 | 74154 | 2 | ENSMUST00 | 25000 | Tnfrsf19 | ENSMUSG00000060548.7 |
| chr9 | 37552212 | 37552734 | 10.7597 | 23.84104 | 21.45527 | Promoter | 9 | 37544492 | 37552904 | 8413 | 2 | ENSMUST00 | 170 | Nrgn | ENSMUSG00000053310.6 |
| chr9 | 43210225 | 43210775 | 11.08881 | 23.82316 | 21.43778 | Promoter | 9 | 43123939 | 43210369 | 86431 | 2 | ENSMUST00 | 0 | Pou2f3 | ENSMUSG00000032015.10 |
| chr18 | 82917323 | 82918110 | 11.41972 | 23.80548 | 21.42021 | Intron (ENSM | 18 | 82914632 | 83005314 | 90683 | 1 | ENSMUST00 | 2692 | Zfp516 | ENSMUSG00000058881.7 |
| chr2 | 49733538 | 49734524 | 11.41972 | 23.80548 | 21.42021 | Promoter | 2 | 49732611 | 49735575 | 2965 | 1 | ENSMUST00 | 928 | Kif5c | ENSMUSG00000026764.10 |
| chr3 | 51364956 | 51365455 | 11.41972 | 23.80548 | 21.42021 | Distal Interge | 3 | 51264869 | 51340658 | 75790 | 2 | ENSMUST00 | -24299 | Elf2 | ENSMUSG00000037174.13 |
| chr5 | 106825155 | 106825849 | 11.41972 | 23.80548 | 21.42021 | Distal Interge | 5 | 106879600 | 106879698 | 99 | 2 | ENSMUST00 | 53849 | Gm27527 | ENSMUSG00000098703.1 |
| chr8 | 126594752 | 126595283 | 11.41972 | 23.80548 | 21.42021 | Promoter | 8 | 126588296 | 126593986 | 5691 | 2 | ENSMUST00 | -767 | Irf2bp2 | ENSMUSG00000051495.7 |
| chr12 | 117053696 | 117054331 | 11.39591 | 23.76571 | 21.38092 | Intron (ENSM | 12 | 117144530 | 117151215 | 6686 | 2 | ENSMUST00 | 96884 | RP23-393F1( | ENSMUSG00000099907.1 |
| chr13 | 62812752 | 62813145 | 11.39591 | 23.76571 | 21.38092 | Distal Interge | 13 | 62791507 | 62791613 | 107 | 1 | ENSMUST00 | 21246 | Gm25448 | ENSMUSG00000095825.1 |
| chr12 | 116077410 | 116078234 | 9.15642 | 23.76437 | 21.37967 | Promoter | 12 | 116077726 | 116139972 | 62247 | 1 | ENSMUST00 | 0 | Vipr2 | ENSMUSG00000011171.6 |
| chr13 | 55189545 | 55190337 | 9.15642 | 23.76437 | 21.37967 | Distal Interge | 13 | 55209782 | 55318325 | 108544 | 1 | ENSMUST00 | -19445 | Nsd1 | ENSMUSG00000021488.7 |
| chr14 | 55788288 | 55789065 | 9.15642 | 23.76437 | 21.37967 | Promoter | 14 | 55784996 | 55788857 | 3862 | 2 | ENSMUST00 | 0 | Ripk3 | ENSMUSG00000022221.8 |
| chr15 | 98772264 | 98773130 | 9.15642 | 23.76437 | 21.37967 | 3' UTR | 15 | 98772200 | 98777261 | 5062 | 2 | ENSMUST00 | 4131 | Wnt10b | ENSMUSG00000022996.9 |
| chr2 | 118544272 | 118544761 | 9.15642 | 23.76437 | 21.37967 | Promoter | 2 | 118531974 | 118544587 | 12614 | 2 | ENSMUST00 | 0 | Bmf | ENSMUSG00000040093.10 |
| chr4 | 41730941 | 41731674 | 9.15642 | 23.76437 | 21.37967 | Promoter | 4 | 41723836 | 41731142 | 7307 | 2 | ENSMUST00 | 0 | Arid3c | ENSMUSG00000066224.8 |
| chr5 | 129941122 | 129941575 | 9.15642 | 23.76437 | 21.37967 | Promoter | 5 | 129942109 | 129984745 | 42637 | 1 | ENSMUST00 | -534 | Vkorc1l1 | ENSMUSG00000066735.5 |
| chr6 | 136834609 | 136835473 | 9.15642 | 23.76437 | 21.37967 | Promoter | 6 | 136829927 | 136835450 | 5524 | 2 | ENSMUST00 | 0 | Smco3 | ENSMUSG00000043298.7 |
| chr8 | 122229906 | 122231035 | 9.15642 | 23.76437 | 21.37967 | Intron (ENSM | 8 | 122269569 | 122272646 | 3078 | 1 | ENSMUST00 | -38534 | Gm22 | ENSMUSG00000043903.3 |
| chr8 | 105152871 | 105153368 | 9.15642 | 23.76437 | 21.37967 | Distal Interge | 8 | 105164587 | 105170367 | 5781 | 1 | ENSMUST00 | -11219 | RP23-144H1( | ENSMUSG00000099465.1 |
| chr1 | 150992911 | 150993638 | 9.43134 | 23.73935 | 21.3551 | Promoter | 1 | 150562524 | 150993051 | 430528 | 2 | ENSMUST00 | 0 | Hmcn1 | ENSMUSG00000066842.13 |
| chr4 | 155397822 | 155398317 | 11.3722 | 23.7261 | 21.34185 | Promoter | 4 | 155384980 | 155398047 | 13068 | 2 | ENSMUST00 | 0 | Gabrd | ENSMUSG00000029054.3 |
| chr8 | 109436770 | 109437328 | 11.3722 | 23.7261 | 21.34185 | Distal Interge | 8 | 109425644 | 109425744 | 101 | 2 | ENSMUST00 | -11027 | Gm23163 | ENSMUSG00000092821.1 |
| chr4 | 141599217 | 141600084 | 11.34859 | 23.68664 | 21.30253 | Promoter | 4 | 141576062 | 141599924 | 23863 | 2 | ENSMUST00 | 0 | Fblim1 | ENSMUSG00000006219.7 |
| chr11 | 77508217 | 77508676 | 11.32508 | 23.64734 | 21.26337 | 3' UTR | 11 | 77505136 | 77506039 | 904 | 1 | ENSMUST00 | 3082 | Git1 | ENSMUSG00000011877.8 |
| chr13 | 66868193 | 66868820 | 11.30166 | 23.6082 | 21.22438 | Distal Interge | 13 | 66864524 | 66865075 | 552 | 1 | ENSMUST00 | 3670 | Gm5196 | ENSMUSG00000071294.1 |
| chr7 | 64770886 | 64771580 | 11.30166 | 23.6082 | 21.22438 | Intron (ENSM | 7 | 64756113 | 64773652 | 17540 | 2 | ENSMUST00 | 2072 | Fam189a1 | ENSMUSG00000030518.12 |
| chr11 | 100761695 | 100762162 | 10.96053 | 23.6065 | 21.22278 | Promoter | 11 | 100761069 | 100762928 | 1860 | 2 | ENSMUST00 | 766 | Hcrt | ENSMUSG00000045471.4 |
| chr14 | 74640862 | 74642391 | 10.96053 | 23.6065 | 21.22278 | Promoter | 14 | 74640840 | 74706859 | 66020 | 1 | ENSMUST00 | 23 | Htr2a | ENSMUSG00000034997.4 |
| chr19 | 5406087 | 5406604 | 10.96053 | 23.6065 | 21.22278 | Promoter | 19 | 5406740 | 5415021 | 8282 | 1 | ENSMUST00 | -136 | 4930481A15 | ENSMUSG00000086938.3 |
| chr5 | 73006770 | 73007465 | 10.96053 | 23.6065 | 21.22278 | Promoter | 5 | 73006904 | 73014683 | 7780 | 1 | ENSMUST00 | 0 | Slc10a4 | ENSMUSG00000029219.7 |
| chr5 | 149439571 | 149440485 | 10.96053 | 23.6065 | 21.22278 | Promoter | 5 | 149439706 | 149470620 | 30915 | 1 | ENSMUST00 | 0 | Tex26 | ENSMUSG00000029660.7 |
| chr6 | 129049156 | 129049451 | 10.96053 | 23.6065 | 21.22278 | Intron (ENSM | 6 | 129045901 | 129057471 | 11571 | 1 | ENSMUST00 | 3256 | Klrb1f | ENSMUSG00000030154.8 |
| chr8 | 67515393 | 67515627 | 11.27834 | 23.56921 | 21.18598 | Promoter | 8 | 67514000 | 67515606 | 1607 | 2 | ENSMUST00 | 0 | Gm9755 | ENSMUSG00000030735.5 |
| chr11 | 35838072 | 35838821 | 10.5671 | 23.5101 | 21.12713 | 3' UTR | 11 | 35808381 | 35834506 | 26126 | 2 | ENSMUST00 | -3567 | Rars | ENSMUSG00000018848.4 |
| chr13 | 31800858 | 31801381 | 10.5671 | 23.5101 | 21.12713 | Distal Interge | 13 | 31797710 | 31800749 | 3040 | 1 | ENSMUST00 | 3149 | Gm11379 | ENSMUSG00000086144.1 |
| chr17 | 29841818 | 29842512 | 10.5671 | 23.5101 | 21.12713 | Promoter | 17 | 29832056 | 29842421 | 10366 | 2 | ENSMUST00 | 0 | Mdga1 | ENSMUSG00000043557.10 |
| chr9 | 107526515 | 107527076 | 10.5671 | 23.5101 | 21.12713 | Promoter | 9 | 107526657 | 107527567 | 911 | 1 | ENSMUST00 | 0 | Cacna2d2 | ENSMUSG00000010066.10 |
| chr11 | 96916724 | 96917069 | 10.89686 | 23.4988 | 21.11616 | Promoter | 11 | 96907426 | 96916496 | 9071 | 2 | ENSMUST00 | -229 | Cdk5rap3 | ENSMUSG00000018669.9 |
| chr11 | 110379745 | 110380415 | 10.89686 | 23.4988 | 21.11616 | Distal Interge | 11 | 110399122 | 110513641 | 114520 | 1 | ENSMUST00 | -18707 | Map2k6 | ENSMUSG00000020623.6 |
| chr11 | 73030715 | 73031315 | 10.89686 | 23.4988 | 21.11616 | Exon (ENSMU | 11 | 73026585 | 73040293 | 13709 | 1 | ENSMUST00 | 4131 | Camkk1 | ENSMUSG00000020785.12 |
| chr12 | 105336602 | 105337291 | 10.89686 | 23.4988 | 21.11616 | Promoter | 12 | 105336993 | 105383930 | 46938 | 1 | ENSMUST00 | 0 | Tunar | ENSMUSG00000097929.2 |
| chr13 | 37452678 | 37453217 | 10.89686 | 23.4988 | 21.11616 | Promoter | 13 | 37452212 | 37453208 | 997 | 2 | ENSMUST00 | 0 | RP23-21N15. | ENSMUSG00000099847.1 |
| chr15 | 83056402 | 83056991 | 10.89686 | 23.4988 | 21.11616 | Distal Interge | 15 | 82997726 | 83033306 | 35581 | 2 | ENSMUST00 | -23097 | Nfam1 | ENSMUSG00000058099.10 |
| chr16 | 76432922 | 76433603 | 10.89686 | 23.4988 | 21.11616 | Distal Interge | 16 | 76403209 | 76403712 | 504 | 2 | ENSMUST00 | -29211 | Gm9843 | ENSMUSG00000050299.7 |
| chr17 | 23644619 | 23645510 | 10.89686 | 23.4988 | 21.11616 | Promoter | 17 | 23629458 | 23645269 | 15812 | 2 | ENSMUST00 | 0 | Mmp25 | ENSMUSG00000023903.7 |
| chr17 | 23802526 | 23802961 | 10.89686 | 23.4988 | 21.11616 | Promoter | 17 | 23803187 | 23824736 | 21550 | 1 | ENSMUST00 | -226 | Srrm2 | ENSMUSG00000039218.11 |
| chr2 | 170205262 | 170206109 | 10.89686 | 23.4988 | 21.11616 | Distal Interge | 2 | 170192449 | 170194053 | 1605 | 2 | ENSMUST00 | -11210 | Gm17619 | ENSMUSG00000097514.1 |
| chr3 | 116419661 | 116421046 | 10.89686 | 23.4988 | 21.11616 | Intron (ENSM | 3 | 116400186 | 116423024 | 22839 | 2 | ENSMUST00 | 1978 | Cdc14a | ENSMUSG00000033502.9 |
| chr3 | 26153577 | 26154058 | 10.89686 | 23.4988 | 21.11616 | Distal Interge | 3 | 25431811 | 26133734 | 701924 | 2 | ENSMUST00 | -19844 | Nlgn1 | ENSMUSG00000063887.8 |
| chr3 | 54811143 | 54811398 | 10.89686 | 23.4988 | 21.11616 | Promoter | 3 | 54810564 | 54811250 | 687 | 2 | ENSMUST00 | 0 | Gm10254 | ENSMUSG00000069011.4 |
| chr5 | 75156238 | 75157193 | 10.89686 | 23.4988 | 21.11616 | Promoter | 5 | 75156166 | 75198204 | 42039 | 1 | ENSMUST00 | 73 | Pdgfra | ENSMUSG00000029231.10 |
| chr5 | 143253240 | 143254315 | 10.89686 | 23.4988 | 21.11616 | Exon (ENSMU | 5 | 143253209 | 143264656 | 11448 | 2 | ENSMUST00 | 10341 | Zfp316 | ENSMUSG00000046658.11 |
| chr6 | 18031627 | 18032337 | 10.89686 | 23.4988 | 21.11616 | Distal Interge | 6 | 17988940 | 18030585 | 41646 | 2 | ENSMUST00 | -1043 | Wnt2 | ENSMUSG00000010797.3 |
| chr7 | 49462163 | 49463003 | 10.89686 | 23.4988 | 21.11616 | Intron (ENSM | 7 | 49450760 | 49464580 | 13821 | 1 | ENSMUST00 | 11404 | Nav2 | ENSMUSG00000052512.10 |
| chr7 | 65704962 | 65705362 | 10.89686 | 23.4988 | 21.11616 | Distal Interge | 7 | 65693467 | 65697879 | 4413 | 1 | ENSMUST00 | 11496 | Tm2d3 | ENSMUSG00000078681.5 |
| chr8 | 85339317 | 85340239 | 10.89686 | 23.4988 | 21.11616 | Promoter | 8 | 85339343 | 85340272 | 930 | 1 | ENSMUST00 | 0 | AA672651 | ENSMUSG00000084895.1 |
| chrX | 133981299 | 133981842 | 10.89686 | 23.4988 | 21.11616 | Promoter | 20 | 133936385 | 133981783 | 45399 | 2 | ENSMUST00 | 0 | Sytl4 | ENSMUSG00000031255.9 |
| chrX | 139209935 | 139210583 | 10.89686 | 23.4988 | 21.11616 | Promoter | 20 | 139210042 | 139234975 | 24934 | 1 | ENSMUST00 | 0 | Mum1l1 | ENSMUSG00000042515.8 |
| chrX | 6172775 | 6173605 | 10.89686 | 23.4988 | 21.11616 | Promoter | 20 | 6168696 | 6173015 | 4320 | 2 | ENSMUST00 | 0 | Nudt10 | ENSMUSG00000073293.4 |
| chr10 | 11609230 | 11609459 | 8.74864 | 23.49345 | 21.11253 | Promoter | 10 | 11609257 | 11610438 | 1182 | 1 | ENSMUST00 | 0 | Gm9797 | ENSMUSG00000045455.7 |
| chr19 | 6400784 | 6402082 | 8.74864 | 23.49345 | 21.11253 | Promoter | 19 | 6401695 | 6402446 | 752 | 1 | ENSMUST00 | 0 | Rasgrp2 | ENSMUSG00000032946.11 |
| chr2 | 38350470 | 38352510 | 8.74864 | 23.49345 | 21.11253 | Promoter | 2 | 38350759 | 38369733 | 18975 | 1 | ENSMUST00 | 0 | Lhx2 | ENSMUSG00000000247.6 |
| chr5 | 7344264 | 7345100 | 8.74864 | 23.49345 | 21.11253 | Distal Interge | 5 | 7304146 | 7305809 | 1664 | 1 | ENSMUST00 | 40119 | 4921511H03 | ENSMUSG00000040514.5 |
| chr17 | 25221998 | 25222243 | 11.23199 | 23.4917 | 21.11096 | Promoter | 17 | 25222539 | 25229457 | 6919 | 1 | ENSMUST00 | -296 | Unkl | ENSMUSG00000015127.9 |
| chr8 | 10899661 | 10900229 | 11.20895 | 23.45317 | 21.07269 | Distal Interge | 8 | 10924427 | 10928457 | 4031 | 2 | ENSMUST00 | 28228 | 3930402G23 | ENSMUSG00000038917.3 |
| chr19 | 40659504 | 40660819 | 10.85587 | 23.4294 | 21.04904 | Promoter | 19 | 40659770 | 40741602 | 81833 | 1 | ENSMUST00 | 0 | Entpd1 | ENSMUSG00000048120.11 |
| chr1 | 36580125 | 36581021 | 11.18601 | 23.41478 | 21.03445 | Intron (ENSM | 1 | 36601802 | 36602146 | 345 | 1 | ENSMUST00 | -20781 | Gm6428 | ENSMUSG00000089906.1 |
| chr10 | 117149815 | 117150521 | 11.18601 | 23.41478 | 21.03445 | Distal Interge | 10 | 117070129 | 117148474 | 78346 | 2 | ENSMUST00 | -1342 | Frs2 | ENSMUSG00000020170.3 |
| chr11 | 100939794 | 100940244 | 11.18601 | 23.41478 | 21.03445 | Promoter | 11 | 100886810 | 100939540 | 52731 | 2 | ENSMUST00 | -255 | Stat3 | ENSMUSG00000004040.11 |
| chr11 | 94392506 | 94393096 | 11.18601 | 23.41478 | 21.03445 | Promoter | 11 | 94343295 | 94392976 | 49682 | 2 | ENSMUST00 | 0 | Abcc3 | ENSMUSG00000020865.11 |
| chr11 | 103158897 | 103159545 | 11.18601 | 23.41478 | 21.03445 | Distal Interge | 11 | 103171107 | 103198901 | 27795 | 1 | ENSMUST00 | -11562 | Fmnl1 | ENSMUSG00000055805.9 |
| chr11 | 63914092 | 63914926 | 11.18601 | 23.41478 | 21.03445 | Intron (ENSM | 11 | 63885792 | 63922290 | 36499 | 2 | ENSMUST00 | 7364 | Hs3st3b1 | ENSMUSG00000070407.5 |
| chr3 | 154332128 | 154332688 | 11.18601 | 23.41478 | 21.03445 | Promoter | 3 | 154331997 | 154332798 | 802 | 1 | ENSMUST00 | 132 | AI606473 | ENSMUSG00000093738.2 |
| chr9 | 107612415 | 107612844 | 11.18601 | 23.41478 | 21.03445 | Downstream | 9 | 107610287 | 107611005 | 719 | 2 | ENSMUST00 | -1411 | RP23-456B9. | ENSMUSG00000100829.1 |
| chr10 | 80939833 | 80940366 | 9.86256 | 23.41135 | 21.03137 | Distal Interge | 10 | 80930091 | 80932204 | 2114 | 1 | ENSMUST00 | 9743 | Gadd45b | ENSMUSG00000015312.8 |
| chr10 | 91124194 | 91124535 | 9.86256 | 23.41135 | 21.03137 | Promoter | 10 | 91116574 | 91124059 | 7486 | 2 | ENSMUST00 | -136 | Slc25a3 | ENSMUSG00000061904.7 |
| chr11 | 49244598 | 49245128 | 9.86256 | 23.41135 | 21.03137 | Promoter | 11 | 49244567 | 49245417 | 851 | 1 | ENSMUST00 | 32 | Mgat1 | ENSMUSG00000020346.11 |
| chr11 | 98348228 | 98349138 | 9.86256 | 23.41135 | 21.03137 | Promoter | 11 | 98348404 | 98357796 | 9393 | 1 | ENSMUST00 | 0 | Ppp1r1b | ENSMUSG00000061718.7 |
| chr11 | 63924926 | 63925627 | 9.86256 | 23.41135 | 21.03137 | Distal Interge | 11 | 63885792 | 63922290 | 36499 | 2 | ENSMUST00 | -2637 | Hs3st3b1 | ENSMUSG00000070407.5 |
| chr11 | 104571285 | 104571714 | 9.86256 | 23.41135 | 21.03137 | Promoter | 11 | 104571268 | 104571557 | 290 | 2 | ENSMUST00 | 0 | Gm11663 | ENSMUSG00000082706.1 |
| chr11 | 116221931 | 116222799 | 9.86256 | 23.41135 | 21.03137 | Exon (ENSMU | 11 | 116216540 | 116220287 | 3748 | 1 | ENSMUST00 | 5392 | Cdk3-ps | ENSMUSG00000092300.2 |
| chr15 | 99717383 | 99718192 | 9.86256 | 23.41135 | 21.03137 | Promoter | 15 | 99717515 | 99725005 | 7491 | 1 | ENSMUST00 | 0 | Gpd1 | ENSMUSG00000023019.7 |
| chr17 | 36942781 | 36943618 | 9.86256 | 23.41135 | 21.03137 | Promoter | 17 | 36942918 | 36947986 | 5069 | 1 | ENSMUST00 | 0 | Rnf39 | ENSMUSG00000036492.7 |
| chr19 | 6418302 | 6418810 | 9.86256 | 23.41135 | 21.03137 | Promoter | 19 | 6418731 | 6533217 | 114487 | 1 | ENSMUST00 | 0 | Nrxn2 | ENSMUSG00000033768.11 |
| chr2 | 128817330 | 128817853 | 9.86256 | 23.41135 | 21.03137 | Promoter | 2 | 128818118 | 128854261 | 36144 | 1 | ENSMUST00 | -265 | Tmem87b | ENSMUSG00000014353.9 |
| chr6 | 83438843 | 83440773 | 9.86256 | 23.41135 | 21.03137 | Promoter | 6 | 83362373 | 83441675 | 79303 | 2 | ENSMUST00 | 902 | Tet3 | ENSMUSG00000034832.10 |
| chr6 | 48015956 | 48016381 | 9.86256 | 23.41135 | 21.03137 | Distal Interge | 6 | 48029379 | 48044243 | 14865 | 2 | ENSMUST00 | 27862 | Zfp777 | ENSMUSG00000071477.5 |
| chr7 | 139943808 | 139944808 | 9.86256 | 23.41135 | 21.03137 | Promoter | 7 | 139943789 | 139945112 | 1324 | 1 | ENSMUST00 | 20 | Utf1 | ENSMUSG00000047751.9 |
| chr8 | 105471071 | 105471481 | 9.86256 | 23.41135 | 21.03137 | Promoter | 8 | 105467493 | 105471526 | 4034 | 2 | ENSMUST00 | 45 | Tppp3 | ENSMUSG00000014846.7 |
| chr8 | 36249520 | 36249917 | 9.86256 | 23.41135 | 21.03137 | Promoter | 8 | 36216064 | 36249516 | 33453 | 2 | ENSMUST00 | -5 | Lonrf1 | ENSMUSG00000039633.6 |
| chr9 | 8543747 | 8544818 | 9.86256 | 23.41135 | 21.03137 | Promoter | 9 | 8544196 | 8680565 | 136370 | 1 | ENSMUST00 | 0 | Trpc6 | ENSMUSG00000031997.8 |
| chr17 | 53462828 | 53463512 | 11.16317 | 23.37655 | 20.99793 | Promoter | 17 | 53398889 | 53463321 | 64433 | 2 | ENSMUST00 | 0 | Efhb | ENSMUSG00000023931.4 |
| chr3 | 88592125 | 88592728 | 11.16317 | 23.37655 | 20.99793 | Distal Interge | 3 | 88579671 | 88588397 | 8727 | 1 | ENSMUST00 | 12455 | Ssr2 | ENSMUSG00000041355.8 |
| chr12 | 77060108 | 77060915 | 11.14042 | 23.33847 | 20.95999 | Distal Interge | 12 | 77116365 | 77116432 | 68 | 2 | ENSMUST00 | 55517 | Gm22696 | ENSMUSG00000093114.1 |
| chr7 | 98341488 | 98341909 | 11.14042 | 23.33847 | 20.95999 | Distal Interge | 7 | 98350786 | 98360079 | 9294 | 2 | ENSMUST00 | 18170 | Tsku | ENSMUSG00000049580.7 |
| chr16 | 18395605 | 18396029 | 10.79403 | 23.3246 | 20.94622 | Intron (ENSM | 16 | 18397724 | 18399004 | 1281 | 1 | ENSMUST00 | -1695 | Arvcf | ENSMUSG00000000325.10 |
| chr11 | 101581839 | 101582796 | 8.39668 | 23.29253 | 20.91428 | Promoter | 11 | 101582242 | 101591788 | 9547 | 1 | ENSMUST00 | 0 | Tmem106a | ENSMUSG00000034947.8 |
| chr3 | 96161198 | 96162300 | 8.39668 | 23.29253 | 20.91428 | Promoter | 3 | 96162004 | 96171717 | 9714 | 1 | ENSMUST00 | 0 | Mtmr11 | ENSMUSG00000045934.10 |

| | | | | | | | | | | | | | | | |
|---|---|---|---|---|---|---|---|---|---|---|---|---|---|---|---|
| chr1 | 82284525 | 82285493 | 10.75319 | 23.25534 | 20.87728 | Intron (ENSM | 1 | 82233101 | 82291416 | 58316 | 2 | ENSMUST00 | 5923 | Irs1 | ENSMUSG00000055980.2 |
| chr11 | 63412682 | 63413418 | 10.75319 | 23.25534 | 20.87728 | Promoter | 11 | 63413169 | 63417888 | 4720 | 1 | ENSMUST00 | 0 | B130011K05I | ENSMUSG00000085113.1 |
| chr12 | 73428305 | 73428583 | 10.75319 | 23.25534 | 20.87728 | Distal Interge | 12 | 73364075 | 73371445 | 7371 | 2 | ENSMUST00 | -56861 | D830013O20 | ENSMUSG00000056359.5 |
| chr13 | 63254102 | 63254523 | 10.75319 | 23.25534 | 20.87728 | Intron (ENSM | 13 | 63282142 | 63326096 | 43955 | 1 | ENSMUST00 | -27619 | 2010111I01R | ENSMUSG00000021458.11 |
| chr5 | 34169672 | 34169994 | 10.75319 | 23.25534 | 20.87728 | Promoter | 5 | 34007198 | 34169448 | 162251 | 2 | ENSMUST00 | -225 | Poln | ENSMUSG00000045102.8 |
| chr8 | 24351674 | 24352423 | 10.75319 | 23.25534 | 20.87728 | Distal Interge | 8 | 24422404 | 24440008 | 17605 | 1 | ENSMUST00 | -69981 | Gm26714 | ENSMUSG00000097654.1 |
| chr2 | 79087186 | 79088133 | 10.05822 | 23.19368 | 20.81635 | Distal Interge | 2 | 79038533 | 79038734 | 202 | 2 | ENSMUST00 | -48453 | Gm14469 | ENSMUSG00000080984.1 |
| chr3 | 52400979 | 52401886 | 10.05822 | 23.19368 | 20.81635 | Distal Interge | 3 | 52268435 | 52269435 | 1001 | 2 | ENSMUST00 | -131545 | Gm20402 | ENSMUSG00000092405.1 |
| chr14 | 61440253 | 61440561 | 10.38126 | 23.18931 | 20.81208 | Promoter | 14 | 61365186 | 61439947 | 74762 | 2 | ENSMUST00 | -307 | Kpna3 | ENSMUSG00000021929.8 |
| chr11 | 85182987 | 85183499 | 11.05033 | 23.18759 | 20.81059 | Distal Interge | 11 | 85171096 | 85181145 | 10050 | 1 | ENSMUST00 | 11892 | 1700125H20I | ENSMUSG00000018479.7 |
| chr14 | 54572758 | 54573818 | 11.05033 | 23.18759 | 20.81059 | Exon (ENSMU | 14 | 54567469 | 54577661 | 10193 | 2 | ENSMUST00 | 3843 | Ajuba | ENSMUSG00000022178.10 |
| chr17 | 24642925 | 24645980 | 11.05033 | 23.18759 | 20.81059 | Promoter | 17 | 24639282 | 24644957 | 5676 | 2 | ENSMUST00 | 0 | Slc9a3r2 | ENSMUSG00000002504.9 |
| chr3 | 129199415 | 129200303 | 11.05033 | 23.18759 | 20.81059 | Promoter | 3 | 129199878 | 129219590 | 19713 | 1 | ENSMUST00 | 0 | Pitx2 | ENSMUSG00000028023.11 |
| chr16 | 92498579 | 92499155 | 11.02803 | 23.15023 | 20.77337 | Promoter | 16 | 92498134 | 92541243 | 43110 | 1 | ENSMUST00 | 446 | Clic6 | ENSMUSG00000022949.4 |
| chr1 | 55737106 | 55739417 | 8.08983 | 23.14769 | 20.77093 | Intron (ENSM | 1 | 55702025 | 55735227 | 33203 | 1 | ENSMUST00 | 35082 | Plcl1 | ENSMUSG00000038349.10 |
| chr11 | 88161085 | 88161510 | 8.08983 | 23.14769 | 20.77093 | Promoter | 11 | 88162261 | 88194140 | 31880 | 1 | ENSMUST00 | -751 | Cuedc1 | ENSMUSG00000018378.8 |
| chr13 | 67892184 | 67892453 | 8.08983 | 23.14769 | 20.77093 | Distal Interge | 13 | 67863326 | 67863923 | 598 | 1 | ENSMUST00 | 28859 | BC048507 | ENSMUSG00000064063.5 |
| chr10 | 106470046 | 106470671 | 11.00583 | 23.11301 | 20.73631 | Promoter | 10 | 106470310 | 106933467 | 463158 | 1 | ENSMUST00 | 0 | Ppfia2 | ENSMUSG00000053825.9 |
| chr5 | 24995381 | 24996261 | 11.00583 | 23.11301 | 20.73631 | Promoter | 5 | 24862748 | 24995753 | 133006 | 2 | ENSMUST00 | 0 | Prkag2 | ENSMUSG00000028944.9 |
| chr7 | 138397265 | 138397957 | 11.00583 | 23.11301 | 20.73631 | Promoter | 7 | 138208974 | 138397728 | 188755 | 2 | ENSMUST00 | 0 | Tcerg1l | ENSMUSG00000091002.2 |
| chr4 | 119359326 | 119359883 | 10.98371 | 23.07594 | 20.69934 | Promoter | 4 | 119322893 | 119359794 | 36902 | 2 | ENSMUST00 | 0 | Ccdc30 | ENSMUSG00000028637.10 |
| chr1 | 180793805 | 180794503 | 10.96168 | 23.03899 | 20.66252 | Distal Interge | 1 | 180802568 | 180809483 | 6916 | 1 | ENSMUST00 | -8065 | Gm17275 | ENSMUSG00000097347.1 |
| chr19 | 29325013 | 29325338 | 10.96168 | 23.03899 | 20.66252 | Promoter | 19 | 29321344 | 29325356 | 4013 | 2 | ENSMUST00 | 18 | Insl6 | ENSMUSG00000050957.4 |
| chr2 | 38378964 | 38379669 | 10.96168 | 23.03899 | 20.66252 | Distal Interge | 2 | 38359580 | 38363980 | 4401 | 2 | ENSMUST00 | -14985 | Gm13584 | ENSMUSG00000087539.1 |
| chr4 | 135876153 | 135877006 | 10.96168 | 23.03899 | 20.66252 | Distal Interge | 4 | 135870918 | 135873850 | 2933 | 2 | ENSMUST00 | -2304 | Pnrc2 | ENSMUSG00000028675.7 |
| chr6 | 53880742 | 53881243 | 10.96168 | 23.03899 | 20.66252 | Intron (ENSM | 6 | 53872017 | 53872148 | 132 | 2 | ENSMUST00 | -8595 | Gm22910 | ENSMUSG00000094527.1 |
| chr7 | 45857244 | 45858083 | 10.96168 | 23.03899 | 20.66252 | Exon (ENSMU | 7 | 45831883 | 45867129 | 35247 | 2 | ENSMUST00 | 9046 | Grin2d | ENSMUSG00000002771.10 |
| chr9 | 63144536 | 63145531 | 10.96168 | 23.03899 | 20.66252 | Exon (ENSMU | 9 | 63138176 | 63146980 | 8805 | 2 | ENSMUST00 | 1449 | Skor1 | ENSMUSG00000022245.10 |
| chr9 | 63146230 | 63147062 | 10.96168 | 23.03899 | 20.66252 | Promoter | 9 | 63138170 | 63147011 | 8842 | 2 | ENSMUST00 | 0 | Skor1 | ENSMUSG00000022245.10 |
| chr9 | 56118711 | 56119051 | 10.96168 | 23.03899 | 20.66252 | Intron (ENSM | 9 | 56120287 | 56125933 | 5647 | 1 | ENSMUST00 | -1236 | Pstpip1 | ENSMUSG00000032322.9 |
| chr9 | 65889984 | 65890817 | 10.96168 | 23.03899 | 20.66252 | Promoter | 9 | 65890237 | 65903266 | 13030 | 1 | ENSMUST00 | 0 | 2810417H13I | ENSMUSG00000040204.6 |
| chr7 | 79535280 | 79536335 | 10.93974 | 23.00219 | 20.62617 | Distal Interge | 7 | 79527850 | 79532044 | 4195 | 1 | ENSMUST00 | 7431 | AI854517 | ENSMUSG00000097023.3 |
| chr1 | 136139975 | 136140710 | 9.32368 | 22.97803 | 20.60213 | Intron (ENSM | 1 | 136131454 | 136149993 | 18540 | 1 | ENSMUST00 | 8522 | Kif21b | ENSMUSG00000041642.13 |
| chr15 | 38298493 | 38298933 | 9.32368 | 22.97803 | 20.60213 | Intron (ENSM | 15 | 38294413 | 38300707 | 6295 | 2 | ENSMUST00 | 1774 | Klf10 | ENSMUSG00000037465.9 |
| chr16 | 23984392 | 23985371 | 9.32368 | 22.97803 | 20.60213 | Intron (ENSM | 16 | 23965052 | 23988852 | 23801 | 2 | ENSMUST00 | 3481 | Bcl6 | ENSMUSG00000022508.5 |
| chr19 | 29948219 | 29948824 | 9.32368 | 22.97803 | 20.60213 | Intron (ENSM | 19 | 29945790 | 29960704 | 14915 | 1 | ENSMUST00 | 2430 | Il33 | ENSMUSG00000024810.11 |
| chr2 | 30544951 | 30545539 | 9.32368 | 22.97803 | 20.60213 | Distal Interge | 2 | 30534729 | 30539022 | 4294 | 1 | ENSMUST00 | 10223 | Gm14487 | ENSMUSG00000085275.1 |
| chr7 | 109438730 | 109439619 | 9.32368 | 22.97803 | 20.60213 | Promoter | 7 | 109279223 | 109438817 | 159595 | 2 | ENSMUST00 | 0 | Stk33 | ENSMUSG00000031027.10 |
| chr7 | 19789939 | 19790321 | 9.32368 | 22.97803 | 20.60213 | Promoter | 7 | 19778881 | 19789059 | 10179 | 2 | ENSMUST00 | -881 | Cblc | ENSMUSG00000040525.8 |
| chr8 | 109961662 | 109962388 | 9.32368 | 22.97803 | 20.60213 | Promoter | 8 | 109947914 | 109962168 | 14255 | 2 | ENSMUST00 | 0 | Marveld3 | ENSMUSG00000001672.9 |
| chr9 | 21901308 | 21901878 | 9.32368 | 22.97803 | 20.60213 | Distal Interge | 9 | 21914892 | 21918089 | 3198 | 2 | ENSMUST00 | 16211 | Rab3d | ENSMUSG00000019066.8 |
| chr14 | 5148916 | 5149596 | 7.36716 | 22.95781 | 20.58277 | Distal Interge | 14 | 5164535 | 5169233 | 4699 | 1 | ENSMUST00 | -14939 | Gm3317 | ENSMUSG00000095912.1 |
| chr11 | 3628859 | 3629564 | 10.89613 | 22.929 | 20.55399 | Promoter | 11 | 3628824 | 3629661 | 838 | 1 | ENSMUST00 | 36 | Gm11950 | ENSMUSG00000084310.1 |
| chr3 | 135307288 | 135307791 | 10.89613 | 22.929 | 20.55399 | Promoter | 3 | 135307700 | 135342535 | 34836 | 1 | ENSMUST00 | 0 | Slc9b2 | ENSMUSG00000037994.9 |
| chr4 | 63214071 | 63214443 | 10.89613 | 22.929 | 20.55399 | Promoter | 4 | 63214004 | 63292831 | 78828 | 1 | ENSMUST00 | 68 | Col27a1 | ENSMUSG00000045672.10 |
| chr8 | 90891574 | 90891998 | 10.89613 | 22.929 | 20.55399 | Intron (ENSM | 8 | 90907939 | 90908415 | 477 | 2 | ENSMUST00 | 16417 | Gm6658 | ENSMUSG00000074171.1 |
| chr15 | 99392412 | 99392762 | 10.55355 | 22.91593 | 20.54101 | Promoter | 15 | 99392882 | 99410049 | 17168 | 1 | ENSMUST00 | -120 | Tmbim6 | ENSMUSG00000023010.9 |
| chr19 | 5846110 | 5846457 | 10.55355 | 22.91593 | 20.54101 | Promoter | 19 | 5845623 | 5851388 | 5766 | 1 | ENSMUST00 | 488 | Frmd8os | ENSMUSG00000043488.5 |
| chr4 | 91375483 | 91376503 | 10.55355 | 22.91593 | 20.54101 | Promoter | 4 | 91250763 | 91376463 | 125701 | 2 | ENSMUST00 | 0 | Elavl2 | ENSMUSG00000008489.13 |
| chr4 | 134930127 | 134930749 | 10.55355 | 22.91593 | 20.54101 | Promoter | 4 | 134930898 | 134937548 | 6651 | 1 | ENSMUST00 | -149 | Syf2 | ENSMUSG00000028821.3 |
| chr4 | 9859139 | 9860021 | 10.55355 | 22.91593 | 20.54101 | Exon (ENSMU | 4 | 9844372 | 9862345 | 17974 | 1 | ENSMUST00 | 14768 | Gdf6 | ENSMUSG00000051279.2 |
| chr6 | 89235662 | 89236051 | 9.89563 | 22.90495 | 20.5305 | Distal Interge | 6 | 89249468 | 89249602 | 135 | 2 | ENSMUST00 | 13551 | Gm25961 | ENSMUSG00000087799.1 |
| chr13 | 98944683 | 98945247 | 10.87445 | 22.89261 | 20.51824 | Distal Interge | 13 | 98839019 | 98926384 | 87366 | 2 | ENSMUST00 | -18300 | Tnpo1 | ENSMUSG00000009470.11 |
| chr2 | 22744127 | 22745057 | 3.17916 | 22.88212 | 20.50784 | Distal Interge | 2 | 22755984 | 22762273 | 6290 | 1 | ENSMUST00 | -10927 | 1700092C17I | ENSMUSG00000085353.2 |
| chr16 | 22150021 | 22150799 | 10.20185 | 22.8781 | 20.50382 | Intron (ENSM | 16 | 22059460 | 22161450 | 101991 | 2 | ENSMUST00 | 10651 | Igf2bp2 | ENSMUSG00000033581.11 |
| chr18 | 55015467 | 55016555 | 10.20185 | 22.8781 | 20.50382 | Intron (ENSM | 18 | 54990310 | 55077337 | 87028 | 1 | ENSMUST00 | 25158 | Gm4221 | ENSMUSG00000096948.1 |
| chr5 | 24624190 | 24624742 | 10.20185 | 22.8781 | 20.50382 | Distal Interge | 5 | 24645453 | 24648860 | 3408 | 1 | ENSMUST00 | -20711 | 1700022A21I | ENSMUSG00000038165.8 |
| chr9 | 107666608 | 107667560 | 10.20185 | 22.8781 | 20.50382 | Promoter | 9 | 107651161 | 107667399 | 16239 | 2 | ENSMUST00 | 0 | Slc38a3 | ENSMUSG00000010064.10 |
| chr9 | 15238991 | 15239611 | 10.20185 | 22.8781 | 20.50382 | Promoter | 9 | 15239045 | 15259393 | 20349 | 1 | ENSMUST00 | 0 | Vstm5 | ENSMUSG00000031937.6 |
| chr6 | 117130601 | 117131095 | 10.83135 | 22.82023 | 20.44623 | Distal Interge | 6 | 117168535 | 117175069 | 6535 | 1 | ENSMUST00 | -37440 | Cxcl12 | ENSMUSG00000061353.6 |
| chr1 | 82285600 | 82287741 | 10.47576 | 22.7833 | 20.40945 | 3' UTR | 1 | 82233101 | 82291416 | 58316 | 2 | ENSMUST00 | 3675 | Irs1 | ENSMUSG00000055980.2 |
| chr1 | 44042185 | 44042566 | 10.13814 | 22.76719 | 20.39335 | Distal Interge | 1 | 44055952 | 44061936 | 5985 | 2 | ENSMUST00 | 19370 | Gm8251 | ENSMUSG00000091844.1 |
| chr10 | 118103047 | 118103667 | 10.13814 | 22.76719 | 20.39335 | Distal Interge | 10 | 118141787 | 118168997 | 27211 | 1 | ENSMUST00 | -38120 | Mdm1 | ENSMUSG00000020212.8 |
| chr10 | 53398249 | 53398793 | 10.13814 | 22.76719 | 20.39335 | Distal Interge | 10 | 53278081 | 53379851 | 101771 | 2 | ENSMUST00 | -18399 | Cep85l | ENSMUSG00000038594.8 |
| chr10 | 32889700 | 32890410 | 10.13814 | 22.76719 | 20.39335 | Promoter | 10 | 31689310 | 32890461 | 1201152 | 2 | ENSMUST00 | 51 | Nkain2 | ENSMUSG00000069670.2 |
| chr11 | 60810933 | 60811772 | 10.13814 | 22.76719 | 20.39335 | Promoter | 11 | 60788104 | 60811265 | 23162 | 2 | ENSMUST00 | 0 | Shmt1 | ENSMUSG00000020534.9 |
| chr11 | 22973245 | 22974098 | 10.13814 | 22.76719 | 20.39335 | Promoter | 11 | 22973215 | 22976496 | 3282 | 1 | ENSMUST00 | 31 | Zrsr1 | ENSMUSG00000044068.6 |
| chr11 | 22629538 | 22630091 | 10.13814 | 22.76719 | 20.39335 | Distal Interge | 11 | 22626375 | 22626664 | 290 | 2 | ENSMUST00 | -2875 | Gm12053 | ENSMUSG00000083008.1 |
| chr11 | 113172529 | 113173523 | 10.13814 | 22.76719 | 20.39335 | Promoter | 11 | 113043895 | 113173077 | 129183 | 2 | ENSMUST00 | 0 | 2610035D17I | ENSMUSG00000087259.2 |
| chr12 | 70476979 | 70477862 | 10.13814 | 22.76719 | 20.39335 | Distal Interge | 12 | 70459841 | 70460576 | 736 | 1 | ENSMUST00 | 17139 | Tmx1 | ENSMUSG00000021072.7 |
| chr13 | 21279213 | 21279697 | 10.13814 | 22.76719 | 20.39335 | Distal Interge | 13 | 21286429 | 21292686 | 6258 | 2 | ENSMUST00 | 12989 | Gpx5 | ENSMUSG00000004344.6 |
| chr13 | 51567052 | 51568071 | 10.13814 | 22.76719 | 20.39335 | Promoter | 13 | 51431041 | 51567084 | 136044 | 2 | ENSMUST00 | 0 | Shc3 | ENSMUSG00000021448.6 |
| chr13 | 28419808 | 28420458 | 10.13814 | 22.76719 | 20.39335 | Distal Interge | 13 | 28460778 | 28480841 | 20064 | 2 | ENSMUST00 | 60383 | 2610307P16I | ENSMUSG00000085936.2 |
| chr15 | 89453952 | 89454423 | 10.13814 | 22.76719 | 20.39335 | Promoter | 15 | 89453913 | 89464468 | 10556 | 1 | ENSMUST00 | 40 | Mapk8ip2 | ENSMUSG00000022619.5 |
| chr16 | 17124955 | 17125649 | 10.13814 | 22.76719 | 20.39335 | Promoter | 16 | 17113399 | 17125106 | 11708 | 2 | ENSMUST00 | 0 | 2610318N02 | ENSMUSG00000049916.9 |
| chr17 | 85621241 | 85621929 | 10.13814 | 22.76719 | 20.39335 | Promoter | 17 | 85621031 | 85629302 | 8272 | 1 | ENSMUST00 | 211 | Six3 | ENSMUSG00000038805.10 |
| chr17 | 4403276 | 4404013 | 10.13814 | 22.76719 | 20.39335 | Distal Interge | 17 | 4678020 | 4678126 | 107 | 1 | ENSMUST00 | -274007 | Gm23186 | ENSMUSG00000096200.1 |
| chr18 | 23498051 | 23498578 | 10.13814 | 22.76719 | 20.39335 | Intron (ENSM | 18 | 23415415 | 23659714 | 244300 | 1 | ENSMUST00 | 82637 | Dtna | ENSMUSG00000024302.10 |
| chr18 | 7001385 | 7002725 | 10.13814 | 22.76719 | 20.39335 | 5' UTR | 18 | 7000636 | 7003782 | 3147 | 2 | ENSMUST00 | 1057 | Mkx | ENSMUSG00000061013.5 |
| chr19 | 47837192 | 47837806 | 10.13814 | 22.76719 | 20.39335 | Promoter | 19 | 47737561 | 47837382 | 99822 | 2 | ENSMUST00 | 0 | Wdr96 | ENSMUSG00000044948.11 |
| chr19 | 6869237 | 6869914 | 10.13814 | 22.76719 | 20.39335 | Distal Interge | 19 | 6844623 | 6858211 | 13589 | 2 | ENSMUST00 | -11027 | Ccdc88b | ENSMUSG00000047810.8 |
| chr2 | 164389545 | 164389858 | 10.13814 | 22.76719 | 20.39335 | Promoter | 2 | 164355595 | 164389095 | 33501 | 2 | ENSMUST00 | -451 | Slpi | ENSMUSG00000017002.9 |
| chr2 | 93180255 | 93180601 | 10.13814 | 22.76719 | 20.39335 | Distal Interge | 2 | 93187548 | 93198919 | 11372 | 1 | ENSMUST00 | -6947 | Trp53i11 | ENSMUSG00000068735.9 |
| chr2 | 29747165 | 29747701 | 10.13814 | 22.76719 | 20.39335 | Promoter | 2 | 29744919 | 29747250 | 2332 | 2 | ENSMUST00 | 0 | Gm13420 | ENSMUSG00000079497.3 |
| chr3 | 14869373 | 14870147 | 10.13814 | 22.76719 | 20.39335 | Exon (ENSMU | 3 | 14863538 | 14872351 | 8814 | 1 | ENSMUST00 | 5836 | Car3 | ENSMUSG00000027559.4 |
| chr3 | 146945963 | 146946235 | 10.13814 | 22.76719 | 20.39335 | Intron (ENSM | 3 | 146944047 | 146984009 | 39963 | 1 | ENSMUST00 | 1917 | Ttll7 | ENSMUSG00000036745.10 |
| chr3 | 95624473 | 95624810 | 10.13814 | 22.76719 | 20.39335 | Promoter | 3 | 95624980 | 95630493 | 5514 | 1 | ENSMUST00 | -170 | Ensa | ENSMUSG00000038619.7 |
| chr3 | 84595024 | 84595686 | 10.13814 | 22.76719 | 20.39335 | 3' UTR | 3 | 84593574 | 84597032 | 3459 | 1 | ENSMUST00 | 1451 | Tigd4 | ENSMUSG00000047819.3 |
| chr3 | 108017241 | 108018783 | 10.13814 | 22.76719 | 20.39335 | Promoter | 3 | 108012255 | 108017973 | 5719 | 2 | ENSMUST00 | 0 | Gstm1 | ENSMUSG00000058135.7 |
| chr3 | 89227004 | 89227965 | 10.13814 | 22.76719 | 20.39335 | Promoter | 3 | 89209356 | 89227052 | 17697 | 2 | ENSMUST00 | 0 | Mtx1 | ENSMUSG00000064068.8 |
| chr3 | 27859203 | 27859661 | 10.13814 | 22.76719 | 20.39335 | Distal Interge | 3 | 27868113 | 27868273 | 161 | 1 | ENSMUST00 | -8462 | Gm26040 | ENSMUSG00000088772.1 |
| chr4 | 45819270 | 45820069 | 10.13814 | 22.76719 | 20.39335 | Intron (ENSM | 4 | 45813007 | 45816363 | 3357 | 2 | ENSMUST00 | -2908 | Igfbpl1 | ENSMUSG00000035551.6 |
| chr4 | 57308320 | 57309106 | 10.13814 | 22.76719 | 20.39335 | Distal Interge | 4 | 57190841 | 57301837 | 110997 | 2 | ENSMUST00 | -6484 | Ptpn3 | ENSMUSG00000038764.9 |
| chr4 | 137706974 | 137707752 | 10.13814 | 22.76719 | 20.39335 | Promoter | 4 | 137707522 | 137729861 | 22340 | 1 | ENSMUST00 | 0 | Rap1gap | ENSMUSG00000041351.11 |
| chr5 | 13399090 | 13400009 | 10.13814 | 22.76719 | 20.39335 | Promoter | 5 | 13399309 | 13602565 | 203257 | 1 | ENSMUST00 | 0 | Sema3a | ENSMUSG00000028883.12 |
| chr6 | 124779377 | 124780184 | 10.13814 | 22.76719 | 20.39335 | Promoter | 6 | 124769863 | 124779718 | 9856 | 2 | ENSMUST00 | 0 | Lrrc23 | ENSMUSG00000030125.6 |
| chr6 | 126162421 | 126164384 | 10.13814 | 22.76719 | 20.39335 | Intron (ENSM | 6 | 126101413 | 126166744 | 65332 | 2 | ENSMUST00 | 2360 | Ntf3 | ENSMUSG00000049107.8 |
| chr7 | 140047663 | 140048529 | 10.13814 | 22.76719 | 20.39335 | Promoter | 7 | 140046374 | 140049088 | 2715 | 2 | ENSMUST00 | 559 | Msx3 | ENSMUSG00000025469.10 |
| chr7 | 16891721 | 16892218 | 10.13814 | 22.76719 | 20.39335 | Promoter | 7 | 16891786 | 16895435 | 3650 | 1 | ENSMUST00 | 0 | Gng8 | ENSMUSG00000063594.4 |
| chr7 | 67849024 | 67849558 | 10.13814 | 22.76719 | 20.39335 | Distal Interge | 7 | 67784725 | 67803496 | 18772 | 2 | ENSMUST00 | -45529 | 4833412C05I | ENSMUSG00000097697.1 |
| chr8 | 120589489 | 120589886 | 10.13814 | 22.76719 | 20.39335 | Promoter | 8 | 120589531 | 120593801 | 4271 | 1 | ENSMUST00 | 0 | Gm10614 | ENSMUSG00000097773.1 |
| chr9 | 114583953 | 114584317 | 10.13814 | 22.76719 | 20.39335 | Downstream | 9 | 114602088 | 114602192 | 105 | 2 | ENSMUST00 | 17875 | Gm23232 | ENSMUSG00000087925.1 |
| chrX | 75084405 | 75085034 | 10.13814 | 22.76719 | 20.39335 | Promoter | 20 | 74966843 | 75084905 | 118063 | 2 | ENSMUST00 | 0 | Gab3 | ENSMUSG00000032750.8 |
| chr7 | 140070204 | 140070768 | 10.78859 | 22.74838 | 20.37643 | 3' UTR | 7 | 140069880 | 140082301 | 12422 | 2 | ENSMUST00 | 11533 | Caly | ENSMUSG00000025468.9 |
| chr7 | 140061165 | 140061746 | 10.78859 | 22.74838 | 20.37643 | Distal Interge | 7 | 140046157 | 140049089 | 2933 | 2 | ENSMUST00 | -12077 | Msx3 | ENSMUSG00000025469.10 |
| chr11 | 116319190 | 116319561 | 10.74617 | 22.67706 | 20.30526 | Distal Interge | 11 | 116326602 | 116326717 | 116 | 2 | ENSMUST00 | 7156 | n-R5s74 | ENSMUSG00000064588.1 |
| chr11 | 101604364 | 101605060 | 10.74617 | 22.67706 | 20.30526 | Promoter | 11 | 101604850 | 101605040 | 191 | 1 | ENSMUST00 | 0 | Gm26316 | ENSMUSG00000065820.1 |
| chr11 | 63922529 | 63922926 | 10.74617 | 22.67706 | 20.30526 | Promoter | 11 | 63885792 | 63922290 | 36499 | 2 | ENSMUST00 | -240 | Hs3st3b1 | ENSMUSG00000070407.5 |
| chr12 | 9580500 | 9580855 | 10.74617 | 22.67706 | 20.30526 | 3' UTR | 12 | 9574442 | 9581499 | 7058 | 1 | ENSMUST00 | 6059 | Osr1 | ENSMUSG00000048387.7 |
| chr15 | 73839425 | 73840056 | 10.74617 | 22.67706 | 20.30526 | Promoter | 15 | 73786412 | 73839671 | 53260 | 2 | ENSMUST00 | 0 | Mroh5 | ENSMUSG00000072487.5 |
| chr11 | 98907785 | 98908317 | 8.87028 | 22.65516 | 20.28383 | Promoter | 11 | 98907801 | 98923936 | 16136 | 1 | ENSMUST00 | 0 | Cdc6 | ENSMUSG00000017499.10 |
| chr11 | 43384963 | 43385869 | 8.87028 | 22.65516 | 20.28383 | Promoter | 11 | 43385482 | 43387183 | 1702 | 1 | ENSMUST00 | 0 | Gm12148 | ENSMUSG00000084889.1 |
| chr12 | 49401028 | 49401966 | 8.87028 | 22.65516 | 20.28383 | Promoter | 12 | 49401277 | 49401837 | 561 | 1 | ENSMUST00 | 0 | Gm9804 | ENSMUSG00000045946.3 |
| chr17 | 56119838 | 56120788 | 8.87028 | 22.65516 | 20.28383 | 3' UTR | 17 | 56119680 | 56121946 | 2267 | 2 | ENSMUST00 | 1158 | Lrg1 | ENSMUSG00000037095.7 |
| chr19 | 45079650 | 45080851 | 8.87028 | 22.65516 | 20.28383 | Distal Interge | 19 | 45076139 | 45079289 | 3151 | 1 | ENSMUST00 | 3512 | Kazald1 | ENSMUSG00000025213.6 |
| chr7 | 45920389 | 45921478 | 8.87028 | 22.65516 | 20.28383 | Promoter | 7 | 45918023 | 45920849 | 2827 | 2 | ENSMUST00 | 0 | Emp3 | ENSMUSG00000040212.6 |
| chr7 | 27688782 | 27689273 | 10.72508 | 22.64159 | 20.27066 | Promoter | 7 | 27689340 | 27706484 | 17145 | 1 | ENSMUST00 | -67 | C030039L03I | ENSMUSG00000057093.9 |
| chr13 | 112998438 | 112999258 | 9.73822 | 22.62381 | 20.25297 | 3' UTR | 13 | 112993868 | 113000394 | 6527 | 1 | ENSMUST00 | 4571 | Mcidas | ENSMUSG00000074651.2 |
| chr4 | 156191798 | 156192581 | 9.73822 | 22.62381 | 20.25297 | Intron (ENSM | 4 | 156165778 | 156197488 | 31711 | 2 | ENSMUST00 | 4907 | Agrn | ENSMUSG00000041936.13 |
| chr5 | 31596525 | 31597223 | 9.73822 | 22.62381 | 20.25297 | Promoter | 5 | 31596935 | 31654861 | 57927 | 1 | ENSMUST00 | 0 | Mrpl33 | ENSMUSG00000029142.9 |
| chr13 | 66007591 | 66007845 | 10.70408 | 22.60625 | 20.23545 | Promoter | 13 | 66007672 | 66009240 | 1569 | 1 | ENSMUST00 | 0 | Gm6910 | ENSMUSG00000055763.8 |
| chr1 | 42696429 | 42697915 | 11.02877 | 22.59171 | 20.22099 | Promoter | 1 | 42695768 | 42703176 | 7409 | 1 | ENSMUST00 | 662 | Pou3f3 | ENSMUSG00000045515.3 |
| chr10 | 80344776 | 80345635 | 11.02877 | 22.59171 | 20.22099 | Promoter | 10 | 80343006 | 80345238 | 2233 | 2 | ENSMUST00 | 0 | Adamtsl5 | ENSMUSG00000043822.13 |
| chr11 | 94937285 | 94937862 | 11.02877 | 22.59171 | 20.22099 | Promoter | 11 | 94937686 | 94938103 | 418 | 1 | ENSMUST00 | 0 | Col1a1 | ENSMUSG00000001506.10 |
| chr11 | 69948000 | 69949009 | 11.02877 | 22.59171 | 20.22099 | Promoter | 11 | 69942539 | 69948175 | 5637 | 2 | ENSMUST00 | 0 | Slc2a4 | ENSMUSG00000018566.9 |
| chr11 | 117414833 | 117415381 | 11.02877 | 22.59171 | 20.22099 | Promoter | 11 | 117414389 | 117415368 | 980 | 1 | ENSMUST00 | 445 | Gm11729 | ENSMUSG00000086588.1 |
| chr11 | 117792412 | 117792832 | 11.02877 | 22.59171 | 20.22099 | 3' UTR | 11 | 117789897 | 117791313 | 1417 | 2 | ENSMUST00 | -1100 | Gm11724 | ENSMUSG00000087669.1 |
| chr12 | 8585028 | 8585217 | 11.02877 | 22.59171 | 20.22099 | Intron (ENSM | 12 | 8528483 | 8599066 | 70584 | 2 | ENSMUST00 | 13849 | Slc7a15 | ENSMUSG00000020600.8 |

| | | | | | | | | | | | | | | | |
|---|---|---|---|---|---|---|---|---|---|---|---|---|---|---|---|
| chr12 | 76417420 | 76418032 | 11.02877 | 22.59171 | 20.22099 | Promoter | 12 | 76417599 | 76439473 | 21875 | 1 | ENSMUST00 | 0 | Ppp1r36 | ENSMUSG00000052221.7 |
| chr12 | 11436406 | 11437235 | 11.02877 | 22.59171 | 20.22099 | Promoter | 12 | 11325247 | 11436649 | 111403 | 2 | ENSMUST00 | 0 | Vsnl1 | ENSMUSG00000054459.6 |
| chr12 | 86894492 | 86895224 | 11.02877 | 22.59171 | 20.22099 | Distal Interge | 12 | 86891404 | 86892567 | 1164 | 2 | ENSMUST00 | -1926 | Gm26698 | ENSMUSG00000097208.1 |
| chr13 | 37477730 | 37478348 | 11.02877 | 22.59171 | 20.22099 | Distal Interge | 13 | 37452212 | 37453208 | 997 | 2 | ENSMUST00 | -24523 | RP23-21N15. | ENSMUSG00000099847.1 |
| chr16 | 5146399 | 5147192 | 11.02877 | 22.59171 | 20.22099 | Promoter | 16 | 5147109 | 5183934 | 36826 | 1 | ENSMUST00 | 0 | Sec14l5 | ENSMUSG00000091712.1 |
| chr16 | 50045771 | 50046261 | 11.02877 | 22.59171 | 20.22099 | Distal Interge | 16 | 50111073 | 50112062 | 990 | 1 | ENSMUST00 | -64812 | Gm8824 | ENSMUSG00000098028.1 |
| chr17 | 36178949 | 36179428 | 11.02877 | 22.59171 | 20.22099 | Promoter | 17 | 36179114 | 36179657 | 544 | 1 | ENSMUST00 | 0 | Gm6623 | ENSMUSG00000092374.1 |
| chr18 | 68058798 | 68059342 | 11.02877 | 22.59171 | 20.22099 | Intron (ENSM | 18 | 67933257 | 68255549 | 322293 | 1 | ENSMUST00 | 125542 | Ldlrad4 | ENSMUSG00000024544.8 |
| chr19 | 37177484 | 37178568 | 11.02877 | 22.59171 | 20.22099 | Promoter | 19 | 37174437 | 37178017 | 3581 | 2 | ENSMUST00 | 0 | Cpeb3 | ENSMUSG00000039652.11 |
| chr2 | 25863735 | 25864080 | 11.02877 | 22.59171 | 20.22099 | Promoter | 2 | 25863795 | 25888942 | 25148 | 1 | ENSMUST00 | 0 | Kcnt1 | ENSMUSG00000058740.9 |
| chr2 | 30718167 | 30719397 | 11.02877 | 22.59171 | 20.22099 | Promoter | 2 | 30713147 | 30720057 | 6911 | 2 | ENSMUST00 | 660 | Gm14488 | ENSMUSG00000075416.5 |
| chr3 | 121886304 | 121886967 | 11.02877 | 22.59171 | 20.22099 | Distal Interge | 3 | 121953229 | 122016677 | 63449 | 1 | ENSMUST00 | -66262 | Arhgap29 | ENSMUSG00000039831.11 |
| chr3 | 96596162 | 96596900 | 11.02877 | 22.59171 | 20.22099 | Promoter | 3 | 96596636 | 96599775 | 3140 | 1 | ENSMUST00 | 0 | Ankrd34a | ENSMUSG00000049097.9 |
| chr4 | 143238388 | 143238948 | 11.02877 | 22.59171 | 20.22099 | Distal Interge | 4 | 143242024 | 143243368 | 1345 | 2 | ENSMUST00 | 4420 | Gm13050 | ENSMUSG00000082575.1 |
| chr6 | 97278108 | 97278732 | 11.02877 | 22.59171 | 20.22099 | Distal Interge | 6 | 97238528 | 97252780 | 14253 | 2 | ENSMUST00 | -25329 | Lmod3 | ENSMUSG00000044086.7 |
| chr6 | 87730601 | 87731671 | 11.02877 | 22.59171 | 20.22099 | Promoter | 6 | 87730869 | 87734017 | 3149 | 1 | ENSMUST00 | 0 | Efcc1 | ENSMUSG00000068263.6 |
| chr6 | 83068532 | 83069355 | 11.02877 | 22.59171 | 20.22099 | Promoter | 6 | 83068591 | 83070143 | 1553 | 2 | ENSMUST00 | 788 | Tlx2 | ENSMUSG00000068327.4 |
| chr7 | 127691563 | 127692068 | 11.02877 | 22.59171 | 20.22099 | Intron (ENSM | 7 | 127704988 | 127707304 | 2317 | 2 | ENSMUST00 | 15236 | Bcl7c | ENSMUSG00000030814.12 |
| chr9 | 62026543 | 62027246 | 11.02877 | 22.59171 | 20.22099 | Promoter | 9 | 61953481 | 62026801 | 73321 | 2 | ENSMUST00 | 0 | Paqr5 | ENSMUSG00000032278.6 |
| chrX | 95165785 | 95166825 | 11.02877 | 22.59171 | 20.22099 | Promoter | 20 | 95048935 | 95166361 | 117427 | 2 | ENSMUST00 | 0 | Arhgef9 | ENSMUSG00000025656.12 |
| chrX | 159254523 | 159255547 | 11.02877 | 22.59171 | 20.22099 | Promoter | 20 | 159255782 | 159311525 | 55744 | 1 | ENSMUST00 | -235 | Rps6ka3 | ENSMUSG00000031309.10 |
| chrX | 101429020 | 101429471 | 11.02877 | 22.59171 | 20.22099 | Promoter | 20 | 101429318 | 101439621 | 10304 | 1 | ENSMUST00 | 0 | Nono | ENSMUSG00000031311.12 |
| chrX | 164980413 | 164981111 | 11.02877 | 22.59171 | 20.22099 | Promoter | 20 | 164980592 | 164997269 | 16678 | 1 | ENSMUST00 | 0 | Fancb | ENSMUSG00000047757.11 |
| chrX | 135993395 | 135993990 | 11.02877 | 22.59171 | 20.22099 | Promoter | 20 | 135993824 | 135995355 | 1532 | 1 | ENSMUST00 | 0 | Arxes2 | ENSMUSG00000048040.8 |
| chrX | 133689711 | 133690250 | 11.02877 | 22.59171 | 20.22099 | Promoter | 20 | 133582863 | 133688993 | 106131 | 2 | ENSMUST00 | -719 | Pcdh19 | ENSMUSG00000051323.11 |
| chrX | 53231690 | 53232476 | 11.02877 | 22.59171 | 20.22099 | Intron (ENSM | 20 | 53069995 | 53240111 | 170117 | 2 | ENSMUST00 | 7635 | Plac1 | ENSMUSG00000061082.6 |
| chrX | 106186491 | 106186788 | 11.02877 | 22.59171 | 20.22099 | Promoter | 20 | 106187100 | 106203699 | 16600 | 1 | ENSMUST00 | -312 | Pgk1 | ENSMUSG00000062070.7 |
| chrX | 56822099 | 56822470 | 11.02877 | 22.59171 | 20.22099 | Promoter | 20 | 56797858 | 56822326 | 24469 | 2 | ENSMUST00 | 0 | Mtap7d3 | ENSMUSG00000067878.8 |
| chrX | 38180677 | 38181621 | 11.02877 | 22.59171 | 20.22099 | Promoter | 20 | 38180728 | 38181626 | 899 | 1 | ENSMUST00 | 0 | Gm14556 | ENSMUSG00000080942.1 |
| chrX | 103694926 | 103695792 | 11.02877 | 22.59171 | 20.22099 | Downstream | 20 | 103684615 | 103685332 | 718 | 1 | ENSMUST00 | 10312 | Gm3943 | ENSMUSG00000081414.2 |
| chrX | 13241516 | 13243626 | 11.02877 | 22.59171 | 20.22099 | Intron (ENSM | 20 | 13238327 | 13239553 | 1227 | 1 | ENSMUST00 | 3190 | Gm16481 | ENSMUSG00000082090.1 |
| chrX | 12916693 | 12917211 | 11.02877 | 22.59171 | 20.22099 | Distal Interge | 20 | 12913910 | 12914592 | 683 | 2 | ENSMUST00 | -2102 | Gm14524 | ENSMUSG00000083760.1 |
| chrX | 21146819 | 21147013 | 11.02877 | 22.59171 | 20.22099 | Promoter | 20 | 21146844 | 21147640 | 797 | 1 | ENSMUST00 | 0 | Gm1848 | ENSMUSG00000083892.3 |
| chrY | 1009824 | 1011360 | 11.02877 | 22.59171 | 20.22099 | Promoter | 21 | 1010543 | 1028594 | 18052 | 1 | ENSMUST00 | 0 | Eif2s3y | ENSMUSG00000069049.6 |
| chr1 | 192190687 | 192191352 | 10.36119 | 22.58756 | 20.21866 | Promoter | 1 | 192190784 | 192510158 | 319375 | 1 | ENSMUST00 | 0 | Kcnh1 | ENSMUSG00000058248.7 |
| chr1 | 134099435 | 134099888 | 10.36119 | 22.58756 | 20.21866 | Distal Interge | 1 | 134109368 | 134109799 | 432 | 1 | ENSMUST00 | -9480 | Gm8618 | ENSMUSG00000073559.2 |
| chr1 | 184440945 | 184441651 | 10.36119 | 22.58756 | 20.21866 | Distal Interge | 1 | 184527841 | 184557691 | 29851 | 2 | ENSMUST00 | 116040 | RP23-172L16 | ENSMUSG00000101574.1 |
| chr19 | 16435137 | 16436781 | 10.36119 | 22.58756 | 20.21866 | Promoter | 19 | 16435667 | 16610818 | 175152 | 1 | ENSMUST00 | 0 | Gna14 | ENSMUSG00000024697.3 |
| chr8 | 122460368 | 122461062 | 10.36119 | 22.58756 | 20.21866 | Promoter | 8 | 122454206 | 122460692 | 6487 | 2 | ENSMUST00 | 0 | Snai3 | ENSMUSG00000006587.5 |
| chr13 | 110643419 | 110643976 | 10.02854 | 22.57593 | 20.20741 | Distal Interge | 13 | 110395044 | 110400844 | 5801 | 1 | ENSMUST00 | 248376 | Plk2 | ENSMUSG00000021701.7 |
| chr9 | 66713529 | 66714411 | 10.02854 | 22.57593 | 20.20741 | Promoter | 9 | 66713686 | 66766845 | 53160 | 1 | ENSMUST00 | 0 | Car12 | ENSMUSG00000032373.9 |
| chr16 | 4586876 | 4587231 | 11.00893 | 22.55865 | 20.19038 | Distal Interge | 16 | 4594713 | 4615957 | 21245 | 1 | ENSMUST00 | -7482 | Glis2 | ENSMUSG00000014303.8 |
| chr2 | 79453846 | 79454353 | 11.00893 | 22.55865 | 20.19038 | Promoter | 2 | 79453985 | 79461107 | 7123 | 1 | ENSMUST00 | 0 | Gm26755 | ENSMUSG00000097843.1 |
| chr4 | 145640693 | 145641355 | 8.54633 | 22.54461 | 20.17642 | Promoter | 4 | 145641357 | 145644399 | 3043 | 1 | ENSMUST00 | -2 | Gm13231 | ENSMUSG00000081885.1 |
| chr8 | 80057500 | 80058218 | 10.66232 | 22.53595 | 20.16776 | Promoter | 8 | 79970345 | 80058006 | 87662 | 2 | ENSMUST00 | 0 | Hhip | ENSMUSG00000064325.3 |
| chr19 | 44407226 | 44407665 | 10.98589 | 22.52023 | 20.15211 | Promoter | 19 | 44394451 | 44407709 | 13259 | 2 | ENSMUST00 | 44 | Scd1 | ENSMUSG00000037071.2 |
| chr11 | 87662512 | 87663092 | 10.96293 | 22.48196 | 20.11397 | Promoter | 11 | 87662722 | 87729934 | 67213 | 1 | ENSMUST00 | 0 | Rnf43 | ENSMUSG00000034177.10 |
| chr6 | 124919941 | 124920525 | 10.96293 | 22.48196 | 20.11397 | Intron (ENSM | 6 | 124913681 | 124917946 | 4266 | 2 | ENSMUST00 | -1996 | Ptms | ENSMUSG00000030122.10 |
| chr16 | 13160077 | 13160709 | 10.94008 | 22.44386 | 20.07616 | Distal Interge | 16 | 13132391 | 13144491 | 12101 | 1 | ENSMUST00 | 27687 | Ercc4 | ENSMUSG00000022545.8 |
| chr16 | 33517148 | 33517561 | 10.94008 | 22.44386 | 20.07616 | Promoter | 16 | 33517328 | 33663408 | 146081 | 1 | ENSMUST00 | 0 | Slc12a8 | ENSMUSG00000035506.11 |
| chr16 | 21995262 | 21995772 | 10.94008 | 22.44386 | 20.07616 | Promoter | 16 | 21953826 | 21995542 | 41717 | 2 | ENSMUST00 | 0 | Liph | ENSMUSG00000044626.7 |
| chr11 | 73160109 | 73161015 | 8.48353 | 22.41658 | 20.04902 | Promoter | 11 | 73160421 | 73172685 | 12265 | 1 | ENSMUST00 | 0 | P2rx5 | ENSMUSG00000005950.9 |
| chr14 | 55767672 | 55768280 | 8.48353 | 22.41658 | 20.04902 | Exon (ENSMU | 14 | 55765962 | 55768494 | 2533 | 1 | ENSMUST00 | 1711 | Ltb4r1 | ENSMUSG00000046908.5 |
| chr6 | 71012603 | 71013523 | 10.91731 | 22.4059 | 20.03854 | Distal Interge | 6 | 71039579 | 71039733 | 155 | 2 | ENSMUST00 | 26210 | Gm23485 | ENSMUSG00000077369.1 |
| chr7 | 73372401 | 73372945 | 10.91731 | 22.4059 | 20.03854 | Intron (ENSM | 7 | 73375520 | 73419899 | 44380 | 1 | ENSMUST00 | -2575 | Rgma | ENSMUSG00000070509.10 |
| chr11 | 116994126 | 116994476 | 10.89465 | 22.3681 | 20.00086 | Distal Interge | 11 | 116973073 | 116986940 | 13868 | 1 | ENSMUST00 | 21054 | Mgat5b | ENSMUSG00000043857.11 |
| chr15 | 35293045 | 35293448 | 10.89465 | 22.3681 | 20.00086 | Distal Interge | 15 | 35296098 | 35303304 | 7207 | 1 | ENSMUST00 | -2650 | Osr2 | ENSMUSG00000022330.4 |
| chr1 | 130422073 | 130422459 | 10.53897 | 22.32804 | 19.96106 | Promoter | 1 | 130419601 | 130422740 | 3140 | 2 | ENSMUST00 | 281 | Daf2 | ENSMUSG00000026401.10 |
| chr6 | 128445822 | 128446261 | 10.53897 | 22.32804 | 19.96106 | Intron (ENSM | 6 | 128438757 | 128475617 | 36861 | 1 | ENSMUST00 | 7066 | Gm10069 | ENSMUSG00000059659.5 |
| chr7 | 4470223 | 4471456 | 10.53897 | 22.32804 | 19.96106 | Promoter | 7 | 4445697 | 4471249 | 25553 | 2 | ENSMUST00 | 0 | Gm15494 | ENSMUSG00000087297.1 |
| chr9 | 44498133 | 44499362 | 10.53897 | 22.32804 | 19.96106 | Promoter | 9 | 44499136 | 44510388 | 11253 | 1 | ENSMUST00 | 0 | Bcl9l | ENSMUSG00000063382.5 |
| chr1 | 178186882 | 178187321 | 9.86102 | 22.28233 | 19.91596 | Promoter | 1 | 178187417 | 178250901 | 63485 | 1 | ENSMUST00 | -96 | Desi2 | ENSMUSG00000026502.8 |
| chr13 | 35026850 | 35027151 | 9.86102 | 22.28233 | 19.91596 | Promoter | 13 | 34986067 | 35027094 | 41028 | 2 | ENSMUST00 | 0 | Eci2 | ENSMUSG00000021417.9 |
| chr11 | 100738405 | 100738878 | 10.17572 | 22.26956 | 19.90332 | Promoter | 11 | 100715011 | 100738215 | 23205 | 2 | ENSMUST00 | -191 | Rab5c | ENSMUSG00000019173.6 |
| chr15 | 102002771 | 102003516 | 10.17572 | 22.26956 | 19.90332 | Promoter | 15 | 101996711 | 102004342 | 7632 | 2 | ENSMUST00 | 826 | Krt8 | ENSMUSG00000049382.9 |
| chr2 | 30463800 | 30464744 | 10.17572 | 22.26956 | 19.90332 | Distal Interge | 2 | 30471537 | 30474219 | 2683 | 2 | ENSMUST00 | 9475 | Ier5l | ENSMUSG00000089762.3 |
| chr5 | 53258035 | 53258786 | 10.17572 | 22.26956 | 19.90332 | Distal Interge | 5 | 53267106 | 53278487 | 11382 | 1 | ENSMUST00 | -8320 | Smim20 | ENSMUSG00000061461.7 |
| chr11 | 45926630 | 45927012 | 10.82721 | 22.25558 | 19.88968 | Downstream | 11 | 45928269 | 45944935 | 16667 | 2 | ENSMUST00 | 17923 | Lsm11 | ENSMUSG00000044847.8 |
| chr1 | 86018158 | 86018857 | 9.52248 | 22.23582 | 19.8701 | Promoter | 1 | 86018693 | 86029248 | 10556 | 1 | ENSMUST00 | 0 | Spata3 | ENSMUSG00000026226.10 |
| chr1 | 126830055 | 126831163 | 9.52248 | 22.23582 | 19.8701 | Promoter | 1 | 125913637 | 126830632 | 916996 | 2 | ENSMUST00 | 0 | Nckap5 | ENSMUSG00000049690.10 |
| chr10 | 130470229 | 130470733 | 9.52248 | 22.23582 | 19.8701 | Downstream | 10 | 130445707 | 130455894 | 10188 | 2 | ENSMUST00 | -14336 | Vmn2r86 | ENSMUSG00000092162.2 |
| chr11 | 88955066 | 88955679 | 9.52248 | 22.23582 | 19.8701 | Promoter | 11 | 88924020 | 88955465 | 31446 | 2 | ENSMUST00 | 0 | Scpep1 | ENSMUSG00000000278.10 |
| chr11 | 62746107 | 62746689 | 9.52248 | 22.23582 | 19.8701 | Distal Interge | 11 | 62749852 | 62752787 | 2936 | 1 | ENSMUST00 | -3163 | Gm27194 | ENSMUSG00000098292.1 |
| chr13 | 30748979 | 30749813 | 9.52248 | 22.23582 | 19.8701 | Promoter | 13 | 30749226 | 30766927 | 17702 | 1 | ENSMUST00 | 0 | Irf4 | ENSMUSG00000021356.4 |
| chr13 | 30336380 | 30336915 | 9.52248 | 22.23582 | 19.8701 | Promoter | 13 | 30336441 | 30345910 | 9470 | 1 | ENSMUST00 | 0 | Agtr1a | ENSMUSG00000049115.9 |
| chr14 | 44988571 | 44989670 | 9.52248 | 22.23582 | 19.8701 | Promoter | 14 | 44988195 | 45003820 | 15626 | 1 | ENSMUST00 | 377 | Ptger2 | ENSMUSG00000037759.5 |
| chr16 | 45093920 | 45094564 | 9.52248 | 22.23582 | 19.8701 | Promoter | 16 | 45094053 | 45127924 | 33872 | 1 | ENSMUST00 | 0 | Ccdc80 | ENSMUSG00000022665.9 |
| chr19 | 10634106 | 10635203 | 9.52248 | 22.23582 | 19.8701 | Promoter | 19 | 10634233 | 10665210 | 30978 | 1 | ENSMUST00 | 0 | Vwce | ENSMUSG00000043789.7 |
| chr19 | 24134630 | 24135455 | 9.52248 | 22.23582 | 19.8701 | Exon (ENSMU | 19 | 24064793 | 24066452 | 1660 | 1 | ENSMUST00 | 69838 | 1700021P04F | ENSMUSG00000096796.1 |
| chr2 | 118896252 | 118897402 | 9.52248 | 22.23582 | 19.8701 | Distal Interge | 2 | 118900377 | 118916479 | 16103 | 1 | ENSMUST00 | -2975 | Bahd1 | ENSMUSG00000040007.8 |
| chr2 | 65471159 | 65471763 | 9.52248 | 22.23582 | 19.8701 | Intron (ENSM | 2 | 65493125 | 65498599 | 5475 | 2 | ENSMUST00 | 26836 | Scn3a | ENSMUSG00000057182.9 |
| chr4 | 62582965 | 62583772 | 9.52248 | 22.23582 | 19.8701 | Promoter | 4 | 62583601 | 62657439 | 73839 | 1 | ENSMUST00 | 0 | Rgs3 | ENSMUSG00000059810.13 |
| chr5 | 66996704 | 66997500 | 9.52248 | 22.23582 | 19.8701 | Promoter | 5 | 66997092 | 66999346 | 2255 | 1 | ENSMUST00 | 0 | Limch1 | ENSMUSG00000037736.13 |
| chr5 | 134675807 | 134676758 | 9.52248 | 22.23582 | 19.8701 | Promoter | 5 | 134676490 | 134680145 | 3656 | 1 | ENSMUST00 | 0 | Gm10369 | ENSMUSG00000072573.2 |
| chr5 | 118331827 | 118332332 | 9.52248 | 22.23582 | 19.8701 | Promoter | 5 | 118331866 | 118332306 | 441 | 2 | ENSMUST00 | 0 | RP23-418E13 | ENSMUSG00000100766.1 |
| chr6 | 66534812 | 66535205 | 9.52248 | 22.23582 | 19.8701 | Promoter | 6 | 66535390 | 66541033 | 5644 | 1 | ENSMUST00 | -185 | Mad2l1 | ENSMUSG00000029910.9 |
| chr6 | 67168288 | 67168896 | 9.52248 | 22.23582 | 19.8701 | Exon (ENSMU | 6 | 67165203 | 67165616 | 414 | 1 | ENSMUST00 | 3086 | Gm27015 | ENSMUSG00000048665.4 |
| chr6 | 29473787 | 29474330 | 9.52248 | 22.23582 | 19.8701 | 3' UTR | 6 | 29471437 | 29473465 | 2029 | 1 | ENSMUST00 | 2351 | Gm9047 | ENSMUSG00000090685.1 |
| chr7 | 28693311 | 28693991 | 9.52248 | 22.23582 | 19.8701 | Promoter | 7 | 28694019 | 28699335 | 5317 | 1 | ENSMUST00 | -28 | Fbxo27 | ENSMUSG00000037463.9 |
| chr8 | 72443844 | 72444290 | 9.52248 | 22.23582 | 19.8701 | Promoter | 8 | 72424176 | 72443870 | 19695 | 2 | ENSMUST00 | 0 | Calr3 | ENSMUSG00000019732.9 |
| chr1 | 62795681 | 62796164 | 10.47775 | 22.22469 | 19.86031 | 3' UTR | 1 | 62821355 | 62832370 | 11016 | 2 | ENSMUST00 | 36206 | RP23-149A5. | ENSMUSG00000100426.1 |
| chr10 | 81997254 | 81997768 | 10.47775 | 22.22469 | 19.86031 | Distal Interge | 10 | 82058417 | 82061329 | 2913 | 1 | ENSMUST00 | -60649 | Zfp873 | ENSMUSG00000061371.6 |
| chr11 | 69765544 | 69765828 | 10.47775 | 22.22469 | 19.86031 | Promoter | 11 | 69765912 | 69780209 | 14298 | 1 | ENSMUST00 | -84 | Zbtb4 | ENSMUSG00000018750.9 |
| chr11 | 103111756 | 103112546 | 10.47775 | 22.22469 | 19.86031 | 3' UTR | 11 | 103116231 | 103119725 | 3495 | 1 | ENSMUST00 | -3685 | Hexim1 | ENSMUSG00000048878.6 |
| chr13 | 81632656 | 81633423 | 10.47775 | 22.22469 | 19.86031 | Promoter | 13 | 81095068 | 81633154 | 538087 | 2 | ENSMUST00 | 0 | Gpr98 | ENSMUSG00000069170.8 |
| chr13 | 58148364 | 58149139 | 10.47775 | 22.22469 | 19.86031 | Promoter | 13 | 58148777 | 58149692 | 916 | 1 | ENSMUST00 | 0 | 1700066J03F | ENSMUSG00000097717.1 |
| chr14 | 103513062 | 103513796 | 10.47775 | 22.22469 | 19.86031 | Promoter | 14 | 103513341 | 103613346 | 100006 | 1 | ENSMUST00 | 0 | Scel | ENSMUSG00000022123.8 |
| chr14 | 104472861 | 104473565 | 10.47775 | 22.22469 | 19.86031 | Distal Interge | 14 | 104461676 | 104467999 | 6324 | 2 | ENSMUST00 | -4863 | Pou4f1 | ENSMUSG00000048349.9 |
| chr16 | 90810646 | 90811112 | 10.47775 | 22.22469 | 19.86031 | Promoter | 16 | 90804294 | 90810413 | 6120 | 2 | ENSMUST00 | -234 | Urb1 | ENSMUSG00000039929.9 |
| chr16 | 18732913 | 18733740 | 10.47775 | 22.22469 | 19.86031 | Distal Interge | 16 | 18776847 | 18778260 | 1414 | 1 | ENSMUST00 | -43107 | Cldn5 | ENSMUSG00000041378.1 |
| chr17 | 83122743 | 83123358 | 10.47775 | 22.22469 | 19.86031 | Distal Interge | 17 | 83068905 | 83070632 | 1728 | 1 | ENSMUST00 | 53839 | RP24-320K4. | ENSMUSG00000100154.1 |
| chr18 | 43437661 | 43438573 | 10.47775 | 22.22469 | 19.86031 | Promoter | 18 | 43320979 | 43438286 | 117308 | 2 | ENSMUST00 | 0 | Dpysl3 | ENSMUSG00000024501.14 |
| chr18 | 80322367 | 80323195 | 10.47775 | 22.22469 | 19.86031 | Promoter | 18 | 80323335 | 80326605 | 3271 | 1 | ENSMUST00 | -140 | Gm26676 | ENSMUSG00000097593.2 |
| chr2 | 91194699 | 91195264 | 10.47775 | 22.22469 | 19.86031 | Promoter | 2 | 91184061 | 91195025 | 10965 | 2 | ENSMUST00 | 0 | Nr1h3 | ENSMUSG00000002108.5 |
| chr2 | 18680675 | 18681355 | 10.47775 | 22.22469 | 19.86031 | Promoter | 2 | 18680994 | 18683030 | 2037 | 1 | ENSMUST00 | 0 | Bmi1 | ENSMUSG00000026739.8 |
| chr2 | 154562854 | 154564520 | 10.47775 | 22.22469 | 19.86031 | Exon (ENSMU | 2 | 154547046 | 154558890 | 11845 | 2 | ENSMUST00 | -3965 | Necab3 | ENSMUSG00000027489.10 |
| chr2 | 165503762 | 165504477 | 10.47775 | 22.22469 | 19.86031 | Promoter | 2 | 165503787 | 165519917 | 16131 | 1 | ENSMUST00 | 0 | Slc2a10 | ENSMUSG00000027661.4 |
| chr2 | 72038378 | 72039195 | 10.47775 | 22.22469 | 19.86031 | Intron (ENSM | 2 | 72054604 | 72257472 | 202869 | 1 | ENSMUST00 | -15409 | Rapgef4 | ENSMUSG00000049044.11 |
| chr5 | 113310408 | 113310809 | 10.47775 | 22.22469 | 19.86031 | Promoter | 5 | 113243220 | 113310786 | 67567 | 2 | ENSMUST00 | 0 | Sgsm1 | ENSMUSG00000042216.8 |
| chr6 | 140337804 | 140338672 | 10.47775 | 22.22469 | 19.86031 | Distal Interge | 6 | 140319630 | 140319733 | 104 | 1 | ENSMUST00 | 18175 | n-R5s168 | ENSMUSG00000065882.1 |
| chr7 | 130577118 | 130577736 | 10.47775 | 22.22469 | 19.86031 | Promoter | 7 | 130577484 | 130764782 | 187299 | 1 | ENSMUST00 | 0 | Tacc2 | ENSMUSG00000030852.10 |
| chr7 | 60390975 | 60391849 | 10.47775 | 22.22469 | 19.86031 | Distal Interge | 7 | 60155538 | 60156170 | 633 | 1 | ENSMUST00 | 235438 | Gm7367 | ENSMUSG00000047370.4 |
| chr7 | 25476489 | 25477788 | 10.47775 | 22.22469 | 19.86031 | Promoter | 7 | 25461702 | 25477625 | 15924 | 2 | ENSMUST00 | 0 | Ceacam1 | ENSMUSG00000074272.5 |
| chr8 | 105305367 | 105306268 | 10.47775 | 22.22469 | 19.86031 | Promoter | 8 | 105305601 | 105310623 | 5023 | 1 | ENSMUST00 | 0 | Elmo3 | ENSMUSG00000014791.9 |
| chr8 | 93812990 | 93813675 | 10.47775 | 22.22469 | 19.86031 | Intron (ENSM | 8 | 93811313 | 93957497 | 146185 | 1 | ENSMUST00 | 1678 | Gnao1 | ENSMUSG00000031748.10 |
| chrX | 75143099 | 75143611 | 10.47775 | 22.22469 | 19.86031 | Promoter | 20 | 75143424 | 75143630 | 207 | 1 | ENSMUST00 | 0 | Gm15378 | ENSMUSG00000083416.1 |
| chr13 | 31803607 | 31804244 | 10.80491 | 22.21838 | 19.8557 | Distal Interge | 13 | 31806691 | 31810643 | 3953 | 1 | ENSMUST00 | -2447 | Foxc1 | ENSMUSG00000050295.3 |
| chr7 | 36167054 | 36167522 | 10.78271 | 22.18132 | 19.81877 | Distal Interge | 7 | 36118274 | 36118828 | 555 | 1 | ENSMUST00 | 48781 | Rpl17-ps9 | ENSMUSG00000066543.8 |
| chr17 | 35701709 | 35702597 | 10.7606 | 22.1444 | 19.78197 | Promoter | 17 | 35681570 | 35702044 | 20475 | 2 | ENSMUST00 | 0 | Ddr1 | ENSMUSG00000003534.12 |
| chr2 | 169524539 | 169525111 | 10.7606 | 22.1444 | 19.78197 | Distal Interge | 2 | 169480617 | 169504843 | 24227 | 1 | ENSMUST00 | 43923 | Gm14249 | ENSMUSG00000087074.1 |
| chr5 | 47985930 | 47986501 | 10.7606 | 22.1444 | 19.78197 | Promoter | 5 | 47984994 | 48173193 | 188200 | 1 | ENSMUST00 | 937 | Slit2 | ENSMUSG00000031558.10 |
| chr7 | 37563012 | 37563514 | 10.7606 | 22.1444 | 19.78197 | Intron (ENSM | 7 | 37472135 | 37479986 | 7852 | 2 | ENSMUST00 | -83027 | Zfp536 | ENSMUSG00000043456.11 |
| chr1 | 171113506 | 171114543 | 7.85869 | 22.12435 | 19.76205 | Promoter | 1 | 171113918 | 171126829 | 12912 | 1 | ENSMUST00 | 0 | 1700009P17F | ENSMUSG00000026649.9 |
| chr14 | 53328887 | 53329506 | 7.85869 | 22.12435 | 19.76205 | Distal Interge | 14 | 53324632 | 53326237 | 1606 | 1 | ENSMUST00 | 4256 | Trav7n-6 | ENSMUSG00000095415.1 |
| chr2 | 10825737 | 10825990 | 10.73857 | 22.10763 | 19.74536 | Distal Interge | 2 | 10898666 | 10899048 | 383 | 2 | ENSMUST00 | 73058 | Gm13265 | ENSMUSG00000081367.1 |
| chr6 | 18170282 | 18171015 | 10.73857 | 22.10763 | 19.74536 | Promoter | 6 | 18170687 | 18322767 | 152081 | 1 | ENSMUST00 | 0 | Cftr | ENSMUSG00000041301.10 |
| chr8 | 45884563 | 45885728 | 10.73857 | 22.10763 | 19.74536 | Promoter | 8 | 45885485 | 45919546 | 34062 | 1 | ENSMUST00 | 0 | Pdlim3 | ENSMUSG00000031636.6 |
| chr9 | 60562414 | 60563219 | 10.73857 | 22.10763 | 19.74536 | Distal Interge | 9 | 60587822 | 60598589 | 10768 | 2 | ENSMUST00 | 35370 | Lrrc49 | ENSMUSG00000047766.10 |

| | | | | | | | | | | | | | | | |
|---|---|---|---|---|---|---|---|---|---|---|---|---|---|---|---|
| chr3 | 68543028 | 68543363 | 10.39862 | 22.09095 | 19.72903 | Intron (ENSM | 3 | 68547520 | 68547708 | 189 | 2 | ENSMUST00 | 4345 | Gm22073 | ENSMUSG00000084617.1 |
| chr7 | 29303796 | 29304588 | 10.39862 | 22.09095 | 19.72903 | Promoter | 7 | 29303951 | 29317264 | 13314 | 1 | ENSMUST00 | 0 | Dpf1 | ENSMUSG00000030584.8 |
| chr12 | 17172844 | 17174021 | 10.71664 | 22.071 | 19.70914 | 3' UTR | 12 | 17172100 | 17176888 | 4789 | 2 | ENSMUST00 | 2867 | Kcnf1 | ENSMUSG00000051726.6 |
| chr4 | 95053469 | 95053993 | 10.71664 | 22.071 | 19.70914 | Promoter | 4 | 95053240 | 95159089 | 105850 | 1 | ENSMUST00 | 230 | Junos | ENSMUSG00000087366.2 |
| chr3 | 69517032 | 69517804 | 10.6948 | 22.03451 | 19.67281 | Intron (ENSM | 3 | 69542432 | 69552918 | 10487 | 1 | ENSMUST00 | -24628 | Ppm1l | ENSMUSG00000027784.9 |
| chr10 | 26107787 | 26108009 | 10.67305 | 21.99816 | 19.63659 | Distal Interge | 10 | 25991186 | 26079052 | 87867 | 2 | ENSMUST00 | -28736 | Tmem200a | ENSMUSG00000049420.8 |
| chr5 | 76869021 | 76869789 | 9.699 | 21.99685 | 19.63538 | Promoter | 5 | 76869188 | 76870674 | 1487 | 1 | ENSMUST00 | 0 | C530008M17 | ENSMUSG00000036377.13 |
| chr15 | 91191397 | 91192234 | 10.33961 | 21.99109 | 19.62971 | Promoter | 15 | 91145884 | 91191807 | 45924 | 2 | ENSMUST00 | 0 | Abcd2 | ENSMUSG00000055782.8 |
| chr2 | 93462618 | 93463003 | 10.33961 | 21.99109 | 19.62971 | Promoter | 2 | 93419116 | 93462946 | 43831 | 2 | ENSMUST00 | 0 | Cd82 | ENSMUSG00000027215.8 |
| chr3 | 121447856 | 121448677 | 10.33961 | 21.99109 | 19.62971 | Intron (ENSM | 3 | 121426541 | 121458205 | 31665 | 1 | ENSMUST00 | 21316 | Cnn3 | ENSMUSG00000053931.7 |
| chr5 | 5573722 | 5574407 | 10.33961 | 21.99109 | 19.62971 | Promoter | 5 | 5573952 | 5576203 | 2252 | 1 | ENSMUST00 | 0 | Gm8773 | ENSMUSG00000073234.2 |
| chr3 | 154328245 | 154328845 | 10.65138 | 21.96196 | 19.60116 | Promoter | 3 | 154322257 | 154328634 | 6378 | 2 | ENSMUST00 | 0 | Lhx8 | ENSMUSG00000096225.3 |
| chr4 | 129382531 | 129383931 | 10.65138 | 21.96196 | 19.60116 | Distal Interge | 4 | 129379844 | 129380179 | 336 | 1 | ENSMUST00 | 2688 | Gm12980 | ENSMUSG00000082468.1 |
| chr1 | 166256839 | 166257982 | 9.99677 | 21.96133 | 19.60061 | Intron (ENSM | 1 | 166254139 | 166316823 | 62685 | 1 | ENSMUST00 | 2701 | Ildr2 | ENSMUSG00000040612.8 |
| chr2 | 150385744 | 150385989 | 9.99677 | 21.96133 | 19.60061 | Distal Interge | 2 | 150378878 | 150379272 | 395 | 2 | ENSMUST00 | -6473 | Gm14126 | ENSMUSG00000083077.1 |
| chr6 | 93543430 | 93543904 | 10.60831 | 21.88996 | 19.52965 | Distal Interge | 6 | 93509573 | 93509702 | 130 | 1 | ENSMUST00 | 33858 | Gm22312 | ENSMUSG00000088630.1 |
| chr14 | 70889621 | 70890952 | 9.01289 | 21.84568 | 19.48558 | Promoter | 14 | 70890130 | 70979838 | 89709 | 1 | ENSMUST00 | 0 | Gfra2 | ENSMUSG00000022103.9 |
| chr14 | 36863943 | 36864341 | 9.01289 | 21.84568 | 19.48558 | Distal Interge | 14 | 36855929 | 36856044 | 116 | 2 | ENSMUST00 | -7900 | n-R5s46 | ENSMUSG00000065894.1 |
| chr15 | 99096554 | 99098767 | 9.01289 | 21.84568 | 19.48558 | Promoter | 15 | 99099412 | 99104737 | 5326 | 1 | ENSMUST00 | -645 | Dnajc22 | ENSMUSG00000038009.7 |
| chr18 | 11052342 | 11055088 | 9.01289 | 21.84568 | 19.48558 | Promoter | 18 | 11052510 | 11085635 | 33126 | 1 | ENSMUST00 | 0 | Gata6 | ENSMUSG00000005836.8 |
| chr18 | 10532418 | 10533244 | 9.01289 | 21.84568 | 19.48558 | 3' UTR | 18 | 10551289 | 10551406 | 118 | 2 | ENSMUST00 | 18162 | Gm24894 | ENSMUSG00000084693.1 |
| chr2 | 118972641 | 118973178 | 9.01289 | 21.84568 | 19.48558 | Distal Interge | 2 | 118997068 | 119000684 | 3617 | 1 | ENSMUST00 | -23890 | Gm14089 | ENSMUSG00000084802.1 |
| chr3 | 83054956 | 83055345 | 9.01289 | 21.84568 | 19.48558 | Promoter | 3 | 83055522 | 83072355 | 16834 | 1 | ENSMUST00 | -177 | Plrg1 | ENSMUSG00000027998.6 |
| chr5 | 110858984 | 110859576 | 9.01289 | 21.84568 | 19.48558 | Intron (ENSM | 5 | 110840017 | 110874133 | 34117 | 1 | ENSMUST00 | 18968 | Chek2 | ENSMUSG00000029521.3 |
| chr9 | 61369048 | 61369973 | 10.24273 | 21.8269 | 19.46754 | Downstream | 9 | 61370878 | 61371627 | 750 | 2 | ENSMUST00 | 1654 | Gm10655 | ENSMUSG00000074256.3 |
| chr1 | 79581027 | 79581671 | 10.56559 | 21.81851 | 19.45919 | Distal Interge | 1 | 79513739 | 79513864 | 126 | 2 | ENSMUST00 | -67164 | Gm24160 | ENSMUSG00000077782.1 |
| chr4 | 129491128 | 129491971 | 10.56559 | 21.81851 | 19.45919 | Promoter | 4 | 129491190 | 129491956 | 767 | 1 | ENSMUST00 | 0 | Fam229a | ENSMUSG00000078554.2 |
| chr5 | 139149210 | 139149956 | 10.20448 | 21.76199 | 19.40286 | Promoter | 5 | 139148054 | 139149957 | 1904 | 2 | ENSMUST00 | 1 | Prkar1b | ENSMUSG00000025855.8 |
| chr10 | 95264600 | 95265266 | 10.52322 | 21.7476 | 19.38849 | Intron (ENSM | 10 | 95174746 | 95324097 | 149352 | 2 | ENSMUST00 | 58831 | Cradd | ENSMUSG00000045867.9 |
| chr13 | 89539245 | 89539829 | 10.52322 | 21.7476 | 19.38849 | Promoter | 13 | 89540636 | 89611832 | 71197 | 1 | ENSMUST00 | -807 | Hapln1 | ENSMUSG00000021613.8 |
| chr15 | 99590479 | 99590942 | 10.52322 | 21.7476 | 19.38849 | Promoter | 15 | 99590849 | 99594608 | 3760 | 1 | ENSMUST00 | 0 | Aqp5 | ENSMUSG00000044217.11 |
| chr17 | 28007125 | 28009684 | 10.52322 | 21.7476 | 19.38849 | Promoter | 17 | 28007585 | 28062599 | 55015 | 1 | ENSMUST00 | 0 | Anks1 | ENSMUSG00000024219.10 |
| chr17 | 47468363 | 47468777 | 10.52322 | 21.7476 | 19.38849 | Exon (ENSMU | 17 | 47413628 | 47437376 | 23749 | 2 | ENSMUST00 | -30988 | 1700001C19 | ENSMUSG00000047150.8 |
| chr18 | 60596246 | 60596759 | 10.52322 | 21.7476 | 19.38849 | Exon (ENSMU | 18 | 60593991 | 60600497 | 6507 | 2 | ENSMUST00 | 3738 | Synpo | ENSMUSG00000043079.12 |
| chr2 | 38710319 | 38711046 | 10.52322 | 21.7476 | 19.38849 | Promoter | 2 | 38701322 | 38711188 | 9867 | 2 | ENSMUST00 | 142 | Nr5a1 | ENSMUSG00000026751.9 |
| chr7 | 79504664 | 79505220 | 10.52322 | 21.7476 | 19.38849 | Promoter | 7 | 79505264 | 79505353 | 90 | 1 | ENSMUST00 | -44 | Mir9-3 | ENSMUSG00000093238.1 |
| chr17 | 35241108 | 35241702 | 9.54222 | 21.71909 | 19.36044 | Promoter | 17 | 35241746 | 35253707 | 11962 | 1 | ENSMUST00 | -44 | Ddx39b | ENSMUSG00000019432.10 |
| chr11 | 119548812 | 119549783 | 10.50215 | 21.71234 | 19.35375 | Promoter | 11 | 119548630 | 119569046 | 20417 | 1 | ENSMUST00 | 183 | Gm11762 | ENSMUSG00000087675.1 |
| chr7 | 123982590 | 123983097 | 10.50215 | 21.71234 | 19.35375 | Promoter | 7 | 123982869 | 124398989 | 416121 | 1 | ENSMUST00 | 0 | Hs3st4 | ENSMUSG00000078591.1 |
| chr7 | 55968134 | 55968823 | 10.50215 | 21.71234 | 19.35375 | Intron (ENSM | 7 | 55962531 | 55980868 | 18338 | 1 | ENSMUST00 | 5604 | A230056P14 | ENSMUSG00000087178.2 |
| chr10 | 80161186 | 80161536 | 10.48118 | 21.67722 | 19.31875 | Distal Interge | 10 | 80165985 | 80170316 | 4332 | 1 | ENSMUST00 | -4449 | Cirbp | ENSMUSG00000045193.8 |
| chr12 | 111416976 | 111417638 | 10.48118 | 21.67722 | 19.31875 | Promoter | 12 | 111417430 | 111431182 | 13753 | 1 | ENSMUST00 | 0 | Exoc3l4 | ENSMUSG00000021280.8 |
| chr1 | 167929852 | 167930288 | 10.14765 | 21.66544 | 19.30706 | Distal Interge | 1 | 168115244 | 168122465 | 7222 | 1 | ENSMUST00 | -184956 | Gm20711 | ENSMUSG00000093538.1 |
| chr11 | 77624462 | 77625188 | 10.14765 | 21.66544 | 19.30706 | Distal Interge | 11 | 77639854 | 77640033 | 180 | 1 | ENSMUST00 | -14666 | Gm12732 | ENSMUSG00000082646.1 |
| chr18 | 84292960 | 84293447 | 10.14765 | 21.66544 | 19.30706 | Intron (ENSM | 18 | 84279820 | 84412420 | 132601 | 2 | ENSMUST00 | 118973 | Zfp407 | ENSMUSG00000048410.11 |
| chr18 | 77732034 | 77733389 | 10.14765 | 21.66544 | 19.30706 | Promoter | 18 | 77729681 | 77733470 | 3790 | 2 | ENSMUST00 | 81 | A330094K24 | ENSMUSG00000097666.1 |
| chr10 | 127177889 | 127178988 | 9.82401 | 21.66231 | 19.3044 | Exon (ENSMU | 10 | 127172426 | 127180645 | 8220 | 2 | ENSMUST00 | 1657 | Slc26a10 | ENSMUSG00000040441.8 |
| chr6 | 29507555 | 29508064 | 10.46028 | 21.64222 | 19.28449 | Promoter | 6 | 29482036 | 29507937 | 25902 | 2 | ENSMUST00 | 0 | Kcp | ENSMUSG00000059022.9 |
| chr7 | 27470330 | 27470532 | 10.46028 | 21.64222 | 19.28449 | Promoter | 7 | 27469771 | 27470535 | 765 | 2 | ENSMUST00 | 3 | Pgam1-ps2 | ENSMUSG00000082016.3 |
| chr11 | 84290445 | 84290896 | 10.43947 | 21.60736 | 19.24973 | Intron (ENSM | 11 | 84280455 | 84311401 | 30947 | 1 | ENSMUST00 | 9991 | Acaca | ENSMUSG00000020532.13 |
| chr11 | 97309796 | 97310392 | 8.58414 | 21.55966 | 19.20221 | Distal Interge | 11 | 97315716 | 97329920 | 14205 | 1 | ENSMUST00 | -5324 | Mrpl45 | ENSMUSG00000018882.7 |
| chr11 | 96341452 | 96342352 | 8.58414 | 21.55966 | 19.20221 | Promoter | 11 | 96336299 | 96342108 | 5810 | 2 | ENSMUST00 | 0 | Gm11536 | ENSMUSG00000075589.3 |
| chr16 | 85803920 | 85804421 | 8.58414 | 21.55966 | 19.20221 | Promoter | 16 | 85793827 | 85803113 | 9287 | 2 | ENSMUST00 | -808 | Adamts1 | ENSMUSG00000022893.9 |
| chr4 | 31937361 | 31937955 | 8.58414 | 21.55966 | 19.20221 | Distal Interge | 4 | 31964097 | 32023467 | 59371 | 1 | ENSMUST00 | -26142 | Map3k7 | ENSMUSG00000028284.8 |
| chr4 | 152158027 | 152159634 | 8.58414 | 21.55966 | 19.20221 | Promoter | 4 | 152158867 | 152162469 | 3603 | 1 | ENSMUST00 | 0 | Hes2 | ENSMUSG00000028940.2 |
| chr7 | 143756736 | 143757492 | 8.58414 | 21.55966 | 19.20221 | Promoter | 7 | 143709051 | 143756985 | 47935 | 2 | ENSMUST00 | 0 | Osbpl5 | ENSMUSG00000037606.12 |
| chr1 | 78196310 | 78197416 | 9.76265 | 21.55577 | 19.19873 | Promoter | 1 | 78101267 | 78196838 | 95572 | 2 | ENSMUST00 | 0 | Pax3 | ENSMUSG00000004872.10 |
| chr10 | 125968934 | 125969637 | 9.76265 | 21.55577 | 19.19873 | Intron (ENSM | 10 | 125966219 | 126015359 | 49141 | 1 | ENSMUST00 | 2716 | Lrig3 | ENSMUSG00000020105.8 |
| chr10 | 75036771 | 75037449 | 9.76265 | 21.55577 | 19.19873 | Promoter | 10 | 75037058 | 75048788 | 11731 | 1 | ENSMUST00 | 0 | Rab36 | ENSMUSG00000020175.8 |
| chr10 | 92162390 | 92163046 | 9.76265 | 21.55577 | 19.19873 | Promoter | 10 | 92162105 | 92162761 | 657 | 2 | ENSMUST00 | 0 | Rmst | ENSMUSG00000097348.3 |
| chr11 | 73323938 | 73324904 | 9.76265 | 21.55577 | 19.19873 | Promoter | 11 | 73304992 | 73324634 | 19643 | 2 | ENSMUST00 | 0 | Aspa | ENSMUSG00000020774.4 |
| chr13 | 58806475 | 58808170 | 9.76265 | 21.55577 | 19.19873 | Promoter | 13 | 58806596 | 58937030 | 130435 | 1 | ENSMUST00 | 0 | Ntrk2 | ENSMUSG00000055254.9 |
| chr13 | 110018494 | 110019269 | 9.76265 | 21.55577 | 19.19873 | Distal Interge | 13 | 109954109 | 109955963 | 1855 | 1 | ENSMUST00 | 64386 | Pde4d | ENSMUSG00000074661.2 |
| chr17 | 87473025 | 87474630 | 9.76265 | 21.55577 | 19.19873 | Distal Interge | 17 | 87490748 | 87502616 | 11869 | 1 | ENSMUST00 | -16118 | RP23-243O1 | ENSMUSG00000100847.1 |
| chr2 | 180227200 | 180228692 | 9.76265 | 21.55577 | 19.19873 | Distal Interge | 2 | 180176373 | 180225859 | 49487 | 2 | ENSMUST00 | -1342 | Lama5 | ENSMUSG00000015647.9 |
| chr2 | 172345068 | 172345396 | 9.76265 | 21.55577 | 19.19873 | Promoter | 2 | 172345565 | 172355749 | 10185 | 1 | ENSMUST00 | -169 | Fam210b | ENSMUSG00000027495.4 |
| chr2 | 61402476 | 61403089 | 9.76265 | 21.55577 | 19.19873 | Distal Interge | 2 | 61492766 | 61492894 | 129 | 1 | ENSMUST00 | -89677 | Gm22338 | ENSMUSG00000088340.1 |
| chr4 | 135152213 | 135153090 | 9.76265 | 21.55577 | 19.19873 | Promoter | 4 | 135152496 | 135176181 | 23686 | 1 | ENSMUST00 | 0 | Runx3 | ENSMUSG00000070691.5 |
| chr5 | 24598428 | 24599036 | 9.76265 | 21.55577 | 19.19873 | Promoter | 5 | 24595735 | 24598938 | 3204 | 2 | ENSMUST00 | 0 | Smarcd3 | ENSMUSG00000028949.8 |
| chr6 | 127151534 | 127155270 | 9.76265 | 21.55577 | 19.19873 | Promoter | 6 | 127125779 | 127151048 | 25270 | 2 | ENSMUST00 | -487 | Ccnd2 | ENSMUSG00000000184.9 |
| chr7 | 60491930 | 60492914 | 9.76265 | 21.55577 | 19.19873 | Distal Interge | 7 | 60155538 | 60156170 | 633 | 1 | ENSMUST00 | 336393 | Gm7367 | ENSMUSG00000047370.4 |
| chr7 | 28824653 | 28825109 | 9.76265 | 21.55577 | 19.19873 | Promoter | 7 | 28825217 | 28830142 | 4926 | 1 | ENSMUST00 | -108 | Ech1 | ENSMUSG00000053898.7 |
| chr7 | 112620319 | 112621241 | 9.76265 | 21.55577 | 19.19873 | Intron (ENSM | 7 | 112618096 | 112618225 | 130 | 2 | ENSMUST00 | -2095 | Gm24154 | ENSMUSG00000093002.1 |
| chr8 | 31033002 | 31033577 | 9.76265 | 21.55577 | 19.19873 | Distal Interge | 8 | 31089471 | 31097047 | 7577 | 1 | ENSMUST00 | -55894 | Dusp26 | ENSMUSG00000039661.9 |
| chr8 | 32883514 | 32884979 | 9.76265 | 21.55577 | 19.19873 | Distal Interge | 8 | 32890475 | 32942482 | 52008 | 2 | ENSMUST00 | 57503 | Gm3985 | ENSMUSG00000079070.5 |
| chr9 | 54691353 | 54691857 | 9.76265 | 21.55577 | 19.19873 | Distal Interge | 9 | 54698873 | 54714761 | 15889 | 1 | ENSMUST00 | -7016 | Dnaja4 | ENSMUSG00000032285.10 |
| chr9 | 37144060 | 37144640 | 9.76265 | 21.55577 | 19.19873 | Intron (ENSM | 9 | 36954772 | 37145660 | 190889 | 2 | ENSMUST00 | 1020 | Pknox2 | ENSMUSG00000035934.10 |
| chr10 | 121928991 | 121930130 | 10.3981 | 21.53802 | 19.18287 | Intron (ENSM | 10 | 121806640 | 121823133 | 16494 | 2 | ENSMUST00 | -105859 | Srgap1 | ENSMUSG00000020121.10 |
| chr14 | 52328012 | 52328474 | 10.3981 | 21.53802 | 19.18287 | Promoter | 14 | 52311172 | 52328762 | 17591 | 2 | ENSMUST00 | 288 | Sall2 | ENSMUSG00000049532.8 |
| chr10 | 76962154 | 76962538 | 10.31632 | 21.40087 | 19.04606 | Promoter | 10 | 76761860 | 76961887 | 200028 | 2 | ENSMUST00 | -268 | Pcbp3 | ENSMUSG00000001120.10 |
| chr10 | 30810781 | 30811138 | 10.31632 | 21.40087 | 19.04606 | Distal Interge | 10 | 30645584 | 30803107 | 157524 | 2 | ENSMUST00 | -7675 | Ncoa7 | ENSMUSG00000039697.10 |
| chr12 | 15808022 | 15810153 | 10.31632 | 21.40087 | 19.04606 | Exon (ENSMU | 12 | 15791727 | 15816785 | 25059 | 2 | ENSMUST00 | 6632 | Trib2 | ENSMUSG00000020601.7 |
| chr13 | 28416839 | 28417572 | 10.31632 | 21.40087 | 19.04606 | Distal Interge | 13 | 28460778 | 28480841 | 20064 | 2 | ENSMUST00 | 63269 | 2610307P16 | ENSMUSG00000085936.2 |
| chr9 | 50452004 | 50452563 | 10.31632 | 21.40087 | 19.04606 | Distal Interge | 9 | 50494525 | 50505482 | 10958 | 1 | ENSMUST00 | -41962 | Plet1 | ENSMUSG00000032068.9 |
| chr6 | 64727637 | 64728469 | 9.65711 | 21.37202 | 19.01769 | Promoter | 6 | 64729146 | 64731233 | 2088 | 1 | ENSMUST00 | -677 | Atoh1 | ENSMUSG00000073043.4 |
| chr8 | 95137212 | 95138167 | 8.21841 | 21.35299 | 18.99892 | 5' UTR | 8 | 95099832 | 95142540 | 42709 | 2 | ENSMUST00 | 4373 | Kifc3 | ENSMUSG00000031788.8 |
| chr2 | 169929617 | 169930134 | 9.96269 | 21.3504 | 18.99651 | Intron (ENSM | 2 | 169633676 | 169888503 | 254828 | 2 | ENSMUST00 | -41115 | Gm26883 | ENSMUSG00000096972.2 |
| chr8 | 85026384 | 85026721 | 9.96269 | 21.3504 | 18.99651 | Promoter | 8 | 85026833 | 85030285 | 3453 | 1 | ENSMUST00 | -112 | 2310036O22 | ENSMUSG00000041203.8 |
| chr10 | 53580520 | 53581052 | 10.27592 | 21.33303 | 18.97952 | Intron (ENSM | 10 | 53596961 | 53609213 | 12253 | 1 | ENSMUST00 | -15909 | Asf1a | ENSMUSG00000019857.6 |
| chr14 | 100749988 | 100750735 | 10.25583 | 21.2993 | 18.94588 | Distal Interge | 14 | 100760168 | 100760268 | 101 | 2 | ENSMUST00 | 9533 | Gm23538 | ENSMUSG00000088160.1 |
| chr1 | 158362405 | 158362912 | 10.56924 | 21.28106 | 18.92771 | Promoter | 1 | 158362340 | 158611018 | 248679 | 1 | ENSMUST00 | 66 | Astn1 | ENSMUSG00000026587.10 |
| chr1 | 165780251 | 165780982 | 10.56924 | 21.28106 | 18.92771 | Promoter | 1 | 165781075 | 165789367 | 8293 | 1 | ENSMUST00 | -93 | Gm16565 | ENSMUSG00000089829.1 |
| chr10 | 21188693 | 21189171 | 10.56924 | 21.28106 | 18.92771 | Distal Interge | 10 | 21125276 | 21160905 | 35630 | 2 | ENSMUST00 | -27789 | Myb | ENSMUSG00000019982.9 |
| chr12 | 72831732 | 72832306 | 10.56924 | 21.28106 | 18.92771 | Distal Interge | 12 | 72761211 | 72794940 | 33730 | 1 | ENSMUST00 | 70522 | Ppm1a | ENSMUSG00000021096.10 |
| chr13 | 73262572 | 73263926 | 10.56924 | 21.28106 | 18.92771 | Intron (ENSM | 13 | 73260497 | 73269608 | 9112 | 1 | ENSMUST00 | 2076 | Irx4 | ENSMUSG00000021604.10 |
| chr14 | 69609615 | 69610585 | 10.56924 | 21.28106 | 18.92771 | Promoter | 14 | 69609478 | 69693698 | 84221 | 1 | ENSMUST00 | 138 | Loxl2 | ENSMUSG00000034205.11 |
| chr14 | 100283191 | 100284041 | 10.56924 | 21.28106 | 18.92771 | Distal Interge | 14 | 100219486 | 100219789 | 304 | 1 | ENSMUST00 | 63706 | Gm16260 | ENSMUSG00000084991.1 |
| chr14 | 7026656 | 7027756 | 10.56924 | 21.28106 | 18.92771 | Promoter | 14 | 7006115 | 7027449 | 21335 | 2 | ENSMUST00 | 0 | Gm10406 | ENSMUSG00000093985.2 |
| chr18 | 84540707 | 84541347 | 10.56924 | 21.28106 | 18.92771 | Intron (ENSM | 18 | 84495776 | 84558375 | 62600 | 2 | ENSMUST00 | 17028 | Zfp407 | ENSMUSG00000048410.11 |
| chr19 | 11964767 | 11965263 | 10.56924 | 21.28106 | 18.92771 | Promoter | 19 | 11965844 | 11994105 | 28262 | 1 | ENSMUST00 | -581 | Osbp | ENSMUSG00000024687.10 |
| chr2 | 39325732 | 39326111 | 10.56924 | 21.28106 | 18.92771 | Distal Interge | 2 | 39274106 | 39274870 | 765 | 1 | ENSMUST00 | 51627 | Vmn2r-ps2 | ENSMUSG00000081144.1 |
| chr4 | 48491311 | 48491724 | 10.56924 | 21.28106 | 18.92771 | Distal Interge | 4 | 48430858 | 48473459 | 42602 | 2 | ENSMUST00 | -17853 | Tex10 | ENSMUSG00000028345.10 |
| chr4 | 143296548 | 143297836 | 10.56924 | 21.28106 | 18.92771 | Intron (ENSM | 4 | 143268346 | 143299463 | 31118 | 2 | ENSMUST00 | 1627 | Pdpn | ENSMUSG00000028583.9 |
| chr4 | 143349585 | 143350444 | 10.56924 | 21.28106 | 18.92771 | Promoter | 4 | 143349757 | 143371032 | 21276 | 1 | ENSMUST00 | 0 | Lrrc38 | ENSMUSG00000028584.3 |
| chr4 | 136755382 | 136756065 | 10.56924 | 21.28106 | 18.92771 | Intron (ENSM | 4 | 136661463 | 136694040 | 32578 | 2 | ENSMUST00 | -61343 | Ephb2 | ENSMUSG00000028664.9 |
| chr4 | 129107107 | 129108045 | 10.56924 | 21.28106 | 18.92771 | Promoter | 4 | 129106869 | 129110544 | 3676 | 1 | ENSMUST00 | 239 | Tmem54 | ENSMUSG00000028786.10 |
| chr4 | 114842462 | 114842938 | 10.56924 | 21.28106 | 18.92771 | Intron (ENSM | 4 | 114844985 | 114845117 | 133 | 2 | ENSMUST00 | 2179 | Gm23230 | ENSMUSG00000087923.1 |
| chr5 | 115491490 | 115492301 | 10.56924 | 21.28106 | 18.92771 | Promoter | 5 | 115491891 | 115492455 | 565 | 2 | ENSMUST00 | 154 | Gm13836 | ENSMUSG00000081282.1 |
| chr6 | 29620553 | 29621454 | 10.56924 | 21.28106 | 18.92771 | Distal Interge | 6 | 29540827 | 29609887 | 69061 | 2 | ENSMUST00 | -10667 | Tnpo3 | ENSMUSG00000012535.9 |
| chr6 | 18845430 | 18846090 | 10.56924 | 21.28106 | 18.92771 | Exon (ENSMU | 6 | 18844220 | 18847360 | 3141 | 1 | ENSMUST00 | 1211 | Gm26809 | ENSMUSG00000097815.1 |
| chr7 | 141293717 | 141294393 | 10.56924 | 21.28106 | 18.92771 | Exon (ENSMU | 7 | 141292006 | 141294999 | 2994 | 1 | ENSMUST00 | 1712 | Drd4 | ENSMUSG00000025496.4 |
| chr7 | 30362682 | 30363481 | 10.56924 | 21.28106 | 18.92771 | Promoter | 7 | 30355514 | 30362772 | 7259 | 2 | ENSMUST00 | 0 | Lrfn3 | ENSMUSG00000036957.7 |
| chr8 | 84284813 | 84285444 | 10.56924 | 21.28106 | 18.92771 | Distal Interge | 8 | 84257795 | 84270380 | 12586 | 2 | ENSMUST00 | -14434 | Ccdc130 | ENSMUSG00000004994.7 |
| chr9 | 22051740 | 22052197 | 10.56924 | 21.28106 | 18.92771 | Promoter | 9 | 22015005 | 22052023 | 37019 | 2 | ENSMUST00 | 0 | Elavl3 | ENSMUSG00000003410.7 |
| chr9 | 63148850 | 63149466 | 10.56924 | 21.28106 | 18.92771 | Promoter | 9 | 63138170 | 63148961 | 10792 | 2 | ENSMUST00 | 0 | Skor1 | ENSMUSG00000022245.10 |
| chrX | 113297496 | 113298635 | 10.56924 | 21.28106 | 18.92771 | Promoter | 20 | 113297510 | 113836386 | 538877 | 1 | ENSMUST00 | 0 | Dach2 | ENSMUSG00000025592.12 |
| chrX | 42068680 | 42069642 | 10.56924 | 21.28106 | 18.92771 | Promoter | 20 | 42068755 | 42109656 | 40902 | 1 | ENSMUST00 | 0 | Xiap | ENSMUSG00000025860.9 |
| chrX | 57393131 | 57393548 | 10.56924 | 21.28106 | 18.92771 | Promoter | 20 | 57386362 | 57393036 | 6675 | 2 | ENSMUST00 | -96 | Rbmx | ENSMUSG00000031134.11 |
| chrX | 107402586 | 107403363 | 10.56924 | 21.28106 | 18.92771 | Promoter | 20 | 107397099 | 107403376 | 6278 | 2 | ENSMUST00 | 13 | Itm2a | ENSMUSG00000031239.5 |
| chrX | 101332937 | 101333259 | 10.56924 | 21.28106 | 18.92771 | Distal Interge | 20 | 101314391 | 101318151 | 3761 | 1 | ENSMUST00 | 18547 | Nlgn3 | ENSMUSG00000031302.11 |
| chrX | 169036407 | 169037107 | 10.56924 | 21.28106 | 18.92771 | Promoter | 20 | 169036611 | 169251527 | 214917 | 1 | ENSMUST00 | 0 | Arhgap6 | ENSMUSG00000031355.11 |
| chrX | 164027459 | 164028016 | 10.56924 | 21.28106 | 18.92771 | Promoter | 20 | 163976822 | 164027997 | 51176 | 2 | ENSMUST00 | 0 | Car5b | ENSMUSG00000031373.4 |
| chrX | 20059453 | 20061117 | 10.56924 | 21.28106 | 18.92771 | Promoter | 20 | 20059560 | 20097521 | 37962 | 1 | ENSMUST00 | 0 | Chst7 | ENSMUSG00000037347.7 |
| chrX | 7922063 | 7922757 | 10.56924 | 21.28106 | 18.92771 | Exon (ENSMU | 20 | 7919822 | 7924410 | 4589 | 1 | ENSMUST00 | 2242 | Pcsk1n | ENSMUSG00000039278.10 |
| chrX | 164076632 | 164077043 | 10.56924 | 21.28106 | 18.92771 | Promoter | 20 | 164071885 | 164076493 | 4609 | 2 | ENSMUST00 | -140 | Siah1b | ENSMUSG00000040749.6 |
| chrX | 141721564 | 141722172 | 10.56924 | 21.28106 | 18.92771 | Exon (ENSMU | 20 | 141724838 | 141726163 | 1326 | 1 | ENSMUST00 | -2666 | Gm15295 | ENSMUSG00000086426.1 |
| chr11 | 70254466 | 70255539 | 10.52814 | 21.2127 | 18.86143 | Promoter | 11 | 70241457 | 70255353 | 13897 | 2 | ENSMUST00 | 0 | Alox12 | ENSMUSG00000000320.5 |
| chr17 | 5934599 | 5935485 | 10.52814 | 21.2127 | 18.86143 | Distal Interge | 17 | 5941280 | 6032793 | 91514 | 1 | ENSMUST00 | -5795 | Synj2 | ENSMUSG00000023805.11 |

| | | | | | | | | | | | | | | | |
|---|---|---|---|---|---|---|---|---|---|---|---|---|---|---|---|
| chr19 | 17136552 | 17137180 | 10.52814 | 21.2127 | 18.86143 | Intron (ENSM | 19 | 17060137 | 17060397 | 261 | 2 | ENSMUST00 | -76156 | Gm27796 | ENSMUSG00000099316.1 |
| chr15 | 76306973 | 76307820 | 7.90276 | 21.20842 | 18.85721 | Promoter | 15 | 76296597 | 76307101 | 10505 | 2 | ENSMUST00 | 0 | Oplah | ENSMUSG00000022562.8 |
| chr7 | 19212349 | 19212805 | 7.90276 | 21.20842 | 18.85721 | Promoter | 7 | 19212538 | 19224176 | 11639 | 1 | ENSMUST00 | 0 | Gpr4 | ENSMUSG00000044317.6 |
| chr10 | 99203639 | 99204461 | 10.50614 | 21.17611 | 18.82507 | Distal Interge | 10 | 99192892 | 99196850 | 3959 | 1 | ENSMUST00 | 10748 | Poc1b | ENSMUSG00000019952.9 |
| chr11 | 42182011 | 42182613 | 10.50614 | 21.17611 | 18.82507 | Promoter | 11 | 42162553 | 42182163 | 19611 | 2 | ENSMUST00 | 0 | Gabra1 | ENSMUSG00000010803.7 |
| chr11 | 118430506 | 118432556 | 10.50614 | 21.17611 | 18.82507 | Intron (ENSM | 11 | 118433853 | 118449963 | 16111 | 1 | ENSMUST00 | -1297 | C1qtnf1 | ENSMUSG00000017446.9 |
| chr4 | 141291573 | 141291937 | 10.50614 | 21.17611 | 18.82507 | Distal Interge | 4 | 141301240 | 141329384 | 28145 | 1 | ENSMUST00 | -9303 | Epha2 | ENSMUSG00000006445.3 |
| chr4 | 132756796 | 132757477 | 10.50614 | 21.17611 | 18.82507 | Promoter | 4 | 132732966 | 132757252 | 24287 | 2 | ENSMUST00 | 0 | Smpdl3b | ENSMUSG00000028885.8 |
| chr6 | 84583328 | 84584186 | 10.50614 | 21.17611 | 18.82507 | Intron (ENSM | 6 | 84571414 | 84593810 | 22397 | 2 | ENSMUST00 | 9624 | Cyp26b1 | ENSMUSG00000063415.7 |
| chr9 | 112235132 | 112235567 | 10.50614 | 21.17611 | 18.82507 | Promoter | 9 | 112155476 | 112235018 | 79543 | 2 | ENSMUST00 | -115 | Arpp21 | ENSMUSG00000032503.12 |
| chr18 | 35814542 | 35815564 | 10.17627 | 21.16558 | 18.81479 | Distal Interge | 18 | 35829818 | 35861687 | 31870 | 1 | ENSMUST00 | -14254 | Cxxc5 | ENSMUSG00000046668.8 |
| chr7 | 75642885 | 75643200 | 10.17627 | 21.16558 | 18.81479 | Promoter | 7 | 75643319 | 75704422 | 61104 | 1 | ENSMUST00 | -119 | Akap13 | ENSMUSG00000066406.9 |
| chr11 | 18871338 | 18872083 | 10.48424 | 21.13966 | 18.78892 | Promoter | 11 | 18845851 | 18872155 | 26305 | 2 | ENSMUST00 | 72 | 8430419K02I | ENSMUSG00000085743.1 |
| chr4 | 128733817 | 128734160 | 10.15658 | 21.13245 | 18.78179 | Intron (ENSM | 4 | 128727982 | 128747132 | 19151 | 1 | ENSMUST00 | 5836 | Phc2 | ENSMUSG00000028796.12 |
| chr2 | 74679439 | 74680319 | 10.46243 | 21.10336 | 18.75279 | Promoter | 2 | 74679557 | 74687016 | 7460 | 1 | ENSMUST00 | 0 | Hoxd11 | ENSMUSG00000042499.13 |
| chr4 | 133518988 | 133519505 | 9.49579 | 21.08997 | 18.73952 | Promoter | 4 | 133518992 | 133530790 | 11799 | 1 | ENSMUST00 | 0 | Kdf1 | ENSMUSG00000037600.11 |
| chr1 | 153740107 | 153740693 | 9.18239 | 21.07545 | 18.72512 | Promoter | 1 | 153740349 | 153745468 | 5120 | 1 | ENSMUST00 | 0 | Rgs16 | ENSMUSG00000026475.7 |
| chr1 | 72640932 | 72641421 | 9.18239 | 21.07545 | 18.72512 | Downstream | 1 | 72616522 | 72628299 | 11778 | 1 | ENSMUST00 | 24411 | Smarcal1 | ENSMUSG00000039354.11 |
| chr1 | 152436900 | 152437784 | 9.18239 | 21.07545 | 18.72512 | Intron (ENSM | 1 | 152407074 | 152407525 | 452 | 2 | ENSMUST00 | -29376 | Gm15618 | ENSMUSG00000080878.1 |
| chr10 | 117571175 | 117571917 | 9.18239 | 21.07545 | 18.72512 | Distal Interge | 10 | 117594185 | 117594932 | 748 | 1 | ENSMUST00 | -22268 | AC132139.1 | ENSMUSG00000099345.1 |
| chr10 | 61625384 | 61626109 | 9.18239 | 21.07545 | 18.72512 | Exon (ENSMU | 10 | 61585393 | 61611074 | 25682 | 2 | ENSMUST00 | -14311 | RP24-323H7. | ENSMUSG00000101792.1 |
| chr11 | 87763565 | 87764793 | 9.18239 | 21.07545 | 18.72512 | Promoter | 11 | 87764731 | 87766395 | 1665 | 1 | ENSMUST00 | 0 | Bzrap1 | ENSMUSG00000034156.11 |
| chr11 | 96311143 | 96312259 | 9.18239 | 21.07545 | 18.72512 | Distal Interge | 11 | 96291024 | 96306910 | 15887 | 2 | ENSMUST00 | -4234 | Hoxb5os | ENSMUSG00000085645.1 |
| chr11 | 23124303 | 23124950 | 9.18239 | 21.07545 | 18.72512 | Distal Interge | 11 | 23152140 | 23166375 | 14236 | 1 | ENSMUST00 | -27190 | 1700061J23R | ENSMUSG00000086366.1 |
| chr12 | 3981322 | 3982262 | 9.18239 | 21.07545 | 18.72512 | Exon (ENSMU | 12 | 3954951 | 3960618 | 5668 | 1 | ENSMUST00 | 26372 | Pomc | ENSMUSG00000020660.5 |
| chr12 | 107979941 | 107980966 | 9.18239 | 21.07545 | 18.72512 | Intron (ENSM | 12 | 107915222 | 108003414 | 88193 | 2 | ENSMUST00 | 22448 | Bcl11b | ENSMUSG00000048251.10 |
| chr13 | 98694767 | 98695385 | 9.18239 | 21.07545 | 18.72512 | Promoter | 13 | 98686235 | 98694834 | 8600 | 2 | ENSMUST00 | 0 | Tmem171 | ENSMUSG00000052485.7 |
| chr19 | 29066901 | 29067577 | 9.18239 | 21.07545 | 18.72512 | Distal Interge | 19 | 29020833 | 29047961 | 27129 | 2 | ENSMUST00 | -18941 | Ak3 | ENSMUSG00000024782.7 |
| chr3 | 84665977 | 84666595 | 9.18239 | 21.07545 | 18.72512 | Promoter | 3 | 84666192 | 84704575 | 38384 | 1 | ENSMUST00 | 0 | Tmem154 | ENSMUSG00000056498.8 |
| chr4 | 117189257 | 117190047 | 9.18239 | 21.07545 | 18.72512 | Promoter | 4 | 117189942 | 117190057 | 116 | 1 | ENSMUST00 | 0 | Gm25099 | ENSMUSG00000095676.1 |
| chr5 | 110135267 | 110135994 | 9.18239 | 21.07545 | 18.72512 | Promoter | 5 | 110135840 | 110171972 | 36133 | 1 | ENSMUST00 | -246 | Chfr | ENSMUSG00000014668.10 |
| chr5 | 31251470 | 31252038 | 9.18239 | 21.07545 | 18.72512 | Promoter | 5 | 31251691 | 31253197 | 1507 | 1 | ENSMUST00 | 0 | Krtcap3 | ENSMUSG00000029149.9 |
| chr5 | 64156097 | 64156907 | 9.18239 | 21.07545 | 18.72512 | Distal Interge | 5 | 64159451 | 64351480 | 192030 | 1 | ENSMUST00 | -2544 | Tbc1d1 | ENSMUSG00000029174.13 |
| chr6 | 114873653 | 114875291 | 9.18239 | 21.07545 | 18.72512 | Promoter | 6 | 114860628 | 114875107 | 14480 | 2 | ENSMUST00 | 0 | Vgll4 | ENSMUSG00000030315.10 |
| chr6 | 144208689 | 144209502 | 9.18239 | 21.07545 | 18.72512 | Promoter | 6 | 143828425 | 144209452 | 381028 | 2 | ENSMUST00 | 0 | Sox5 | ENSMUSG00000041540.11 |
| chr6 | 36387915 | 36388949 | 9.18239 | 21.07545 | 18.72512 | Promoter | 6 | 36388084 | 36528414 | 140331 | 1 | ENSMUST00 | 0 | Chrm2 | ENSMUSG00000045613.4 |
| chr7 | 73032267 | 73033031 | 9.18239 | 21.07545 | 18.72512 | Distal Interge | 7 | 72989154 | 72990153 | 1000 | 2 | ENSMUST00 | -42115 | Gm5335 | ENSMUSG00000098035.1 |
| chr8 | 94912977 | 94913712 | 9.18239 | 21.07545 | 18.72512 | 5' UTR | 8 | 94913565 | 94917521 | 3957 | 2 | ENSMUST00 | 3809 | Ccdc102a | ENSMUSG00000063605.5 |
| chr9 | 21961366 | 21963078 | 9.18239 | 21.07545 | 18.72512 | Promoter | 9 | 21958899 | 21963576 | 4678 | 2 | ENSMUST00 | 498 | Epor | ENSMUSG00000006235.5 |
| chr9 | 62677589 | 62678845 | 9.18239 | 21.07545 | 18.72512 | Promoter | 9 | 62677826 | 62783982 | 106157 | 1 | ENSMUST00 | 0 | Itga11 | ENSMUSG00000032243.8 |
| chr12 | 103060928 | 103061513 | 10.4407 | 21.06721 | 18.71816 | Exon (ENSMU | 12 | 103059079 | 103070742 | 11664 | 1 | ENSMUST00 | 1850 | Unc79 | ENSMUSG00000021198.11 |
| chr17 | 56472711 | 56474074 | 10.4407 | 21.06721 | 18.71816 | Intron (ENSM | 17 | 56422237 | 56476369 | 54133 | 2 | ENSMUST00 | 2295 | Ptprs | ENSMUSG00000013236.11 |
| chr5 | 103760006 | 103761022 | 10.4407 | 21.06721 | 18.71816 | Intron (ENSM | 5 | 103754573 | 103855322 | 100750 | 1 | ENSMUST00 | 5434 | Aff1 | ENSMUSG00000029313.13 |
| chr7 | 80192373 | 80192939 | 10.4407 | 21.06721 | 18.71816 | Intron (ENSM | 7 | 80186841 | 80226394 | 39554 | 1 | ENSMUST00 | 5533 | Sema4b | ENSMUSG00000030539.8 |
| chr7 | 79780734 | 79781201 | 10.4407 | 21.06721 | 18.71816 | Intron (ENSM | 7 | 79792241 | 79793788 | 1548 | 2 | ENSMUST00 | 12587 | Mesp1 | ENSMUSG00000030544.5 |
| chr1 | 40224965 | 40225561 | 10.11741 | 21.06654 | 18.71757 | Promoter | 1 | 40225080 | 40316177 | 91098 | 1 | ENSMUST00 | 0 | Il1r1 | ENSMUSG00000026072.7 |
| chr14 | 50896216 | 50897105 | 10.11741 | 21.06654 | 18.71757 | Promoter | 14 | 50895942 | 50897456 | 1515 | 2 | ENSMUST00 | 351 | A930018M24 | ENSMUSG00000091089.1 |
| chr4 | 101549951 | 101550895 | 10.11741 | 21.06654 | 18.71757 | Promoter | 4 | 101550480 | 101642799 | 92320 | 1 | ENSMUST00 | 0 | Dnajc6 | ENSMUSG00000028528.11 |
| chr4 | 63402627 | 63403839 | 10.11741 | 21.06654 | 18.71757 | Promoter | 4 | 63367125 | 63403354 | 36230 | 2 | ENSMUST00 | 0 | Akna | ENSMUSG00000039158.11 |
| chr7 | 112662557 | 112663376 | 10.11741 | 21.06654 | 18.71757 | Intron (ENSM | 7 | 112679318 | 112899459 | 220142 | 1 | ENSMUST00 | -15942 | Tead1 | ENSMUSG00000055320.11 |
| chr10 | 42862262 | 42862806 | 9.78435 | 21.04533 | 18.69683 | Intron (ENSM | 10 | 42860793 | 42960778 | 99986 | 1 | ENSMUST00 | 1470 | Scml4 | ENSMUSG00000044770.9 |
| chr16 | 21814859 | 21815168 | 9.78435 | 21.04533 | 18.69683 | Distal Interge | 16 | 21794392 | 21806123 | 11732 | 1 | ENSMUST00 | 20468 | 1300002E11I | ENSMUSG00000097290.2 |
| chr18 | 40219548 | 40219799 | 9.78435 | 21.04533 | 18.69683 | Promoter | 18 | 40204865 | 40219399 | 14535 | 2 | ENSMUST00 | -150 | Yipf5 | ENSMUSG00000024487.4 |
| chr19 | 3414347 | 3414746 | 9.78435 | 21.04533 | 18.69683 | Promoter | 19 | 3409919 | 3414472 | 4554 | 2 | ENSMUST00 | 0 | Gal | ENSMUSG00000024907.6 |
| chr10 | 61089183 | 61089762 | 10.09794 | 21.03375 | 18.68556 | Promoter | 10 | 61089343 | 61094328 | 4986 | 1 | ENSMUST00 | 0 | Pcbd1 | ENSMUSG00000020098.6 |
| chr7 | 142026194 | 142026806 | 10.09794 | 21.03375 | 18.68556 | Intron (ENSM | 7 | 141998661 | 142001441 | 2781 | 1 | ENSMUST00 | 27534 | Brsk2 | ENSMUSG00000053046.9 |
| chr18 | 55695094 | 55695556 | 10.41907 | 21.0312 | 18.68305 | Distal Interge | 18 | 55745251 | 55746228 | 978 | 2 | ENSMUST00 | 50672 | Gm26959 | ENSMUSG00000098119.1 |
| chr6 | 51592407 | 51592967 | 10.41907 | 21.0312 | 18.68305 | Distal Interge | 6 | 51621632 | 51621708 | 77 | 1 | ENSMUST00 | -28665 | Gm22914 | ENSMUSG00000094528.1 |
| chr8 | 48097583 | 48098044 | 10.39753 | 20.99534 | 18.64733 | Distal Interge | 8 | 48099092 | 48140609 | 41518 | 1 | ENSMUST00 | -1048 | Dctd | ENSMUSG00000031562.10 |
| chr1 | 177702638 | 177702973 | 10.05864 | 20.96754 | 18.61967 | Distal Interge | 1 | 177729814 | 177753324 | 23511 | 1 | ENSMUST00 | -26841 | 1700016C15I | ENSMUSG00000015962.5 |
| chr1 | 156305631 | 156306273 | 10.05864 | 20.96754 | 18.61967 | Distal Interge | 1 | 156255296 | 156303664 | 48369 | 2 | ENSMUST00 | -1968 | Tdrd5 | ENSMUSG00000060985.10 |
| chr1 | 63129700 | 63130479 | 10.05864 | 20.96754 | 18.61967 | Intron (ENSM | 1 | 63117118 | 63126837 | 9720 | 1 | ENSMUST00 | 12583 | RP23-13C17. | ENSMUSG00000101599.1 |
| chr11 | 102015369 | 102016894 | 10.05864 | 20.96754 | 18.61967 | Exon (ENSMU | 11 | 102008640 | 102010410 | 1771 | 2 | ENSMUST00 | -4960 | Mpp3 | ENSMUSG00000052373.9 |
| chr13 | 25115467 | 25116122 | 10.05864 | 20.96754 | 18.61967 | Intron (ENSM | 13 | 25130747 | 25131064 | 318 | 2 | ENSMUST00 | 14942 | Gm11348 | ENSMUSG00000082667.1 |
| chr14 | 8665985 | 8666397 | 10.05864 | 20.96754 | 18.61967 | Promoter | 14 | 8431192 | 8666240 | 235049 | 2 | ENSMUST00 | 0 | 4930452B06 | ENSMUSG00000021747.11 |
| chr15 | 77336286 | 77336936 | 10.05864 | 20.96754 | 18.61967 | Distal Interge | 15 | 77078991 | 77307053 | 228063 | 2 | ENSMUST00 | -29234 | Rbfox2 | ENSMUSG00000033565.11 |
| chr16 | 18590118 | 18590810 | 10.05864 | 20.96754 | 18.61967 | Distal Interge | 16 | 18591992 | 18609751 | 17760 | 1 | ENSMUST00 | -1182 | 4930588K23I | ENSMUSG00000075335.3 |
| chr16 | 84621108 | 84621678 | 10.05864 | 20.96754 | 18.61967 | Promoter | 16 | 84619807 | 84620128 | 322 | 2 | ENSMUST00 | -981 | Gm25292 | ENSMUSG00000084746.1 |
| chr17 | 33781004 | 33781640 | 10.05864 | 20.96754 | 18.61967 | Promoter | 17 | 33773750 | 33781575 | 7826 | 2 | ENSMUST00 | 0 | Angptl4 | ENSMUSG00000002289.10 |
| chr17 | 32091115 | 32091697 | 10.05864 | 20.96754 | 18.61967 | Promoter | 17 | 32076083 | 32091423 | 15341 | 2 | ENSMUST00 | 0 | Pdxk-ps | ENSMUSG00000002791.11 |
| chr18 | 11645090 | 11645938 | 10.05864 | 20.96754 | 18.61967 | Intron (ENSM | 18 | 11657349 | 11743207 | 85859 | 1 | ENSMUST00 | -11411 | Rbbp8 | ENSMUSG00000041238.10 |
| chr18 | 58836186 | 58837976 | 10.05864 | 20.96754 | 18.61967 | Promoter | 18 | 58836764 | 59053678 | 216915 | 1 | ENSMUST00 | 0 | Adamts19 | ENSMUSG00000053441.4 |
| chr2 | 155943576 | 155945070 | 10.05864 | 20.96754 | 18.61967 | Promoter | 2 | 155941023 | 155945367 | 4345 | 2 | ENSMUST00 | 297 | Gdf5 | ENSMUSG00000038259.4 |
| chr3 | 157566717 | 157567699 | 10.05864 | 20.96754 | 18.61967 | Promoter | 3 | 157566892 | 157644758 | 77867 | 1 | ENSMUST00 | 0 | Ptger3 | ENSMUSG00000040016.11 |
| chr3 | 95588336 | 95588712 | 10.05864 | 20.96754 | 18.61967 | Promoter | 3 | 95588934 | 95619242 | 30309 | 1 | ENSMUST00 | -222 | Golph3l | ENSMUSG00000046519.10 |
| chr4 | 134160878 | 134161070 | 10.05864 | 20.96754 | 18.61967 | Intron (ENSM | 4 | 134155901 | 134156921 | 1021 | 2 | ENSMUST00 | -3958 | Cep85 | ENSMUSG00000037443.8 |
| chr4 | 123322914 | 123323674 | 10.05864 | 20.96754 | 18.61967 | Promoter | 4 | 123321875 | 123323634 | 1760 | 2 | ENSMUST00 | 0 | Oxct2a | ENSMUSG00000076436.1 |
| chr4 | 133099162 | 133100276 | 10.05864 | 20.96754 | 18.61967 | Distal Interge | 4 | 133085474 | 133089146 | 3673 | 1 | ENSMUST00 | 13689 | Gm26615 | ENSMUSG00000097088.1 |
| chr5 | 28167884 | 28168883 | 10.05864 | 20.96754 | 18.61967 | Intron (ENSM | 5 | 28165694 | 28172166 | 6473 | 1 | ENSMUST00 | 2191 | En2 | ENSMUSG00000039095.8 |
| chr5 | 100155544 | 100156505 | 10.05864 | 20.96754 | 18.61967 | Intron (ENSM | 5 | 100086127 | 100159281 | 73155 | 2 | ENSMUST00 | 2776 | Tmem150c | ENSMUSG00000050640.11 |
| chr5 | 149312982 | 149313485 | 10.05864 | 20.96754 | 18.61967 | Distal Interge | 5 | 149281118 | 149281223 | 106 | 2 | ENSMUST00 | -31760 | n-R5s180 | ENSMUSG00000064704.1 |
| chr5 | 118513821 | 118514584 | 10.05864 | 20.96754 | 18.61967 | Distal Interge | 5 | 118502840 | 118503317 | 478 | 1 | ENSMUST00 | 10982 | Gm15754 | ENSMUSG00000082365.1 |
| chr5 | 107303238 | 107303864 | 10.05864 | 20.96754 | 18.61967 | Promoter | 5 | 107302946 | 107303047 | 102 | 1 | ENSMUST00 | 293 | Gm22557 | ENSMUSG00000088429.1 |
| chr6 | 121129674 | 121131182 | 10.05864 | 20.96754 | 18.61967 | Distal Interge | 6 | 121007241 | 121081609 | 74369 | 2 | ENSMUST00 | -48066 | Mical3 | ENSMUSG00000003178.7 |
| chr6 | 48626985 | 48627483 | 10.05864 | 20.96754 | 18.61967 | Promoter | 6 | 48626764 | 48633685 | 6922 | 1 | ENSMUST00 | 222 | AI854703 | ENSMUSG00000053297.9 |
| chr6 | 51843781 | 51844559 | 10.05864 | 20.96754 | 18.61967 | Distal Interge | 6 | 51859166 | 52012549 | 153384 | 2 | ENSMUST00 | 167990 | Skap2 | ENSMUSG00000059182.5 |
| chr7 | 16299552 | 16300220 | 10.05864 | 20.96754 | 18.61967 | Distal Interge | 7 | 16309616 | 16318205 | 8590 | 1 | ENSMUST00 | -9396 | Bbc3 | ENSMUSG00000002083.7 |
| chr7 | 143823026 | 143823389 | 10.05864 | 20.96754 | 18.61967 | Promoter | 7 | 143823145 | 143824215 | 1071 | 1 | ENSMUST00 | 0 | Dhcr7 | ENSMUSG00000058454.9 |
| chr7 | 46396203 | 46396773 | 10.05864 | 20.96754 | 18.61967 | Promoter | 7 | 46396497 | 46433353 | 36857 | 1 | ENSMUST00 | 0 | Kcnc1 | ENSMUSG00000058975.6 |
| chr8 | 48425003 | 48425834 | 10.05864 | 20.96754 | 18.61967 | Intron (ENSM | 8 | 48227682 | 48367363 | 139682 | 2 | ENSMUST00 | -57641 | Tenm3 | ENSMUSG00000031561.10 |
| chr8 | 30915638 | 30916582 | 10.05864 | 20.96754 | 18.61967 | Distal Interge | 8 | 31089471 | 31097047 | 7577 | 1 | ENSMUST00 | -172889 | Dusp26 | ENSMUSG00000039661.9 |
| chr9 | 40877564 | 40878072 | 10.05864 | 20.96754 | 18.61967 | 3' UTR | 9 | 40874127 | 40877972 | 3846 | 1 | ENSMUST00 | 3438 | Bsx | ENSMUSG00000054360.3 |
| chr9 | 107607689 | 107609447 | 10.05864 | 20.96754 | 18.61967 | Promoter | 9 | 107598217 | 107609229 | 11013 | 2 | ENSMUST00 | 0 | Sema3b | ENSMUSG00000057969.10 |
| chrX | 50537043 | 50537926 | 10.05864 | 20.96754 | 18.61967 | Distal Interge | 20 | 50542402 | 50542784 | 383 | 2 | ENSMUST00 | 4858 | Gm14620 | ENSMUSG00000083112.1 |
| chr13 | 23395850 | 23396815 | 10.37607 | 20.95961 | 18.61343 | Distal Interge | 13 | 23382505 | 23382660 | 156 | 1 | ENSMUST00 | 13346 | Gm11334 | ENSMUSG00000083667.1 |
| chr2 | 164431717 | 164432497 | 10.37607 | 20.95961 | 18.61343 | Intron (ENSM | 2 | 164424247 | 164443186 | 18940 | 2 | ENSMUST00 | 10689 | Sdc4 | ENSMUSG00000017009.3 |
| chr3 | 58372207 | 58373107 | 10.37607 | 20.95961 | 18.61343 | Distal Interge | 3 | 58415689 | 58466787 | 51099 | 1 | ENSMUST00 | -42582 | Tsc22d2 | ENSMUSG00000027806.5 |
| chr10 | 114070817 | 114071744 | 10.35471 | 20.92403 | 18.57802 | Distal Interge | 10 | 114757747 | 114785104 | 27358 | 1 | ENSMUST00 | -686003 | 4930473D10I | ENSMUSG00000089934.1 |
| chr13 | 95889518 | 95890487 | 10.35471 | 20.92403 | 18.57802 | Intron (ENSM | 13 | 95627177 | 95891922 | 264746 | 2 | ENSMUST00 | 1435 | Iqgap2 | ENSMUSG00000021676.9 |
| chr7 | 99483825 | 99484127 | 10.35471 | 20.92403 | 18.57802 | Promoter | 7 | 99477896 | 99483738 | 5843 | 2 | ENSMUST00 | -88 | Rps3 | ENSMUSG00000030744.7 |
| chr2 | 153740784 | 153741106 | 10.0208 | 20.90376 | 18.55785 | Promoter | 2 | 153741274 | 153773310 | 32037 | 1 | ENSMUST00 | -168 | Mapre1 | ENSMUSG00000027479.9 |
| chr17 | 67951860 | 67952768 | 10.31224 | 20.85329 | 18.50753 | Promoter | 17 | 67845110 | 67950940 | 105831 | 2 | ENSMUST00 | -921 | Arhgap28 | ENSMUSG00000024043.8 |
| chr11 | 63712180 | 63712635 | 10.29113 | 20.81812 | 18.4725 | Distal Interge | 11 | 63817019 | 63818300 | 1282 | 2 | ENSMUST00 | 105665 | Gm12288 | ENSMUSG00000083142.1 |
| chr3 | 9974195 | 9974924 | 10.29113 | 20.81812 | 18.4725 | Distal Interge | 3 | 10012548 | 10016607 | 4060 | 1 | ENSMUST00 | -37624 | Fabp5 | ENSMUSG00000027533.10 |
| chr4 | 130174513 | 130175098 | 10.29113 | 20.81812 | 18.4725 | Promoter | 4 | 130164454 | 130174802 | 10349 | 2 | ENSMUST00 | 0 | Tinagl1 | ENSMUSG00000028776.9 |
| chr8 | 46502766 | 46503229 | 10.29113 | 20.81812 | 18.4725 | Intron (ENSM | 8 | 46492832 | 46534480 | 41649 | 1 | ENSMUST00 | 9935 | Acsl1 | ENSMUSG00000018796.7 |
| chr8 | 95951984 | 95952218 | 10.29113 | 20.81812 | 18.4725 | Distal Interge | 8 | 95864134 | 95888547 | 24414 | 2 | ENSMUST00 | -63438 | Got2 | ENSMUSG00000031672.7 |
| chr12 | 86888060 | 86889010 | 10.27012 | 20.7831 | 18.43794 | Downstream | 12 | 86880703 | 86884814 | 4112 | 2 | ENSMUST00 | -3247 | Irf2bpl | ENSMUSG00000034168.6 |
| chr17 | 26885994 | 26886270 | 9.61228 | 20.74965 | 18.40462 | Promoter | 17 | 26860287 | 26886175 | 25889 | 2 | ENSMUST00 | 0 | Gm17382 | ENSMUSG00000078838.2 |
| chr5 | 5707819 | 5708647 | 9.61228 | 20.74965 | 18.40462 | Distal Interge | 5 | 5668530 | 5694578 | 26049 | 2 | ENSMUST00 | -13242 | Steap2 | ENSMUSG00000015653.8 |
| chr7 | 131553840 | 131554900 | 9.61228 | 20.74965 | 18.40462 | Promoter | 7 | 131554061 | 131556589 | 2529 | 1 | ENSMUST00 | 0 | Hmx2 | ENSMUSG00000050100.8 |
| chr7 | 140821710 | 140822263 | 9.61228 | 20.74965 | 18.40462 | Promoter | 7 | 140807449 | 140822178 | 14730 | 2 | ENSMUST00 | 0 | Zfp941 | ENSMUSG00000060314.7 |
| chr18 | 38025098 | 38026043 | 10.24918 | 20.7482 | 18.4034 | Distal Interge | 18 | 37972624 | 37997574 | 24951 | 2 | ENSMUST00 | -27525 | Arap3 | ENSMUSG00000024451.8 |
| chr7 | 27591054 | 27591412 | 10.24918 | 20.7482 | 18.4034 | Promoter | 7 | 27591552 | 27640826 | 49275 | 1 | ENSMUST00 | -140 | Akt2 | ENSMUSG00000004056.10 |
| chr10 | 121461520 | 121462180 | 9.92602 | 20.74379 | 18.3991 | Intron (ENSM | 10 | 121410350 | 121476250 | 65901 | 2 | ENSMUST00 | 14070 | Rassf3 | ENSMUSG00000025795.7 |
| chr11 | 90248686 | 90249203 | 9.92602 | 20.74379 | 18.3991 | Promoter | 11 | 90249456 | 90268465 | 19010 | 1 | ENSMUST00 | -253 | Mmd | ENSMUSG00000003948.11 |
| chr11 | 58092788 | 58093356 | 9.92602 | 20.74379 | 18.3991 | Distal Interge | 11 | 58097104 | 58097773 | 670 | 2 | ENSMUST00 | 4417 | Gm12247 | ENSMUSG00000081223.1 |
| chr13 | 5864109 | 5864976 | 9.92602 | 20.74379 | 18.3991 | Exon (ENSMU | 13 | 5861489 | 5870393 | 8905 | 1 | ENSMUST00 | 2621 | Klf6 | ENSMUSG00000000078.6 |
| chr13 | 107168546 | 107169093 | 9.92602 | 20.74379 | 18.3991 | Distal Interge | 13 | 107031963 | 107033673 | 1711 | 1 | ENSMUST00 | 136584 | 3830408C21I | ENSMUSG00000071181.1 |
| chr13 | 97435351 | 97436281 | 9.92602 | 20.74379 | 18.3991 | Promoter | 13 | 97434970 | 97497007 | 62038 | 1 | ENSMUST00 | 382 | Lincenc1 | ENSMUSG00000078952.4 |
| chr17 | 26503381 | 26504202 | 9.92602 | 20.74379 | 18.3991 | Downstream | 17 | 26506516 | 26507723 | 1208 | 2 | ENSMUST00 | 3521 | Dusp1 | ENSMUSG00000024190.6 |
| chr19 | 24301121 | 24301451 | 9.92602 | 20.74379 | 18.3991 | Intron (ENSM | 19 | 24289716 | 24290826 | 1111 | 1 | ENSMUST00 | 11406 | Gm16216 | ENSMUSG00000082107.1 |
| chr2 | 178118941 | 178119350 | 9.92602 | 20.74379 | 18.3991 | Promoter | 2 | 178118975 | 178338492 | 219518 | 1 | ENSMUST00 | 0 | Phactr3 | ENSMUSG00000027525.12 |
| chr3 | 34649716 | 34651096 | 9.92602 | 20.74379 | 18.3991 | Promoter | 3 | 34650405 | 34652461 | 2057 | 1 | ENSMUST00 | 0 | Sox2 | ENSMUSG00000074637.5 |
| chr7 | 44745024 | 44745333 | 9.92602 | 20.74379 | 18.3991 | Promoter | 7 | 44731481 | 44744230 | 12750 | 2 | ENSMUST00 | -795 | Zfp473 | ENSMUSG00000048012.13 |
| chr9 | 110953717 | 110954200 | 9.92602 | 20.74379 | 18.3991 | Intron (ENSM | 9 | 110951545 | 110984066 | 32522 | 1 | ENSMUST00 | 2173 | Lrrc2 | ENSMUSG00000032495.4 |
| chr10 | 18407193 | 18408159 | 8.7021 | 20.72798 | 18.38387 | Promoter | 10 | 18407638 | 18421140 | 13503 | 1 | ENSMUST00 | 0 | Nhsl1 | ENSMUSG00000039835.10 |
| chr12 | 12421014 | 12421383 | 8.7021 | 20.72798 | 18.38387 | Distal Interge | 12 | 12262139 | 12376359 | 114221 | 1 | ENSMUST00 | 158876 | Fam49a | ENSMUSG00000020589.11 |

| | | | | | | | | | | | | | | | |
|---|---|---|---|---|---|---|---|---|---|---|---|---|---|---|---|
| chr13 | 49247588 | 49248330 | 8.7021 | 20.72798 | 18.38387 | Promoter | 13 | 49230690 | 49248294 | 17605 | 2 | ENSMUST00 | 0 | Susd3 | ENSMUSG00000021384.9 |
| chr13 | 48546830 | 48547244 | 8.7021 | 20.72798 | 18.38387 | Distal Interge | 13 | 48538171 | 48538270 | 100 | 2 | ENSMUST00 | -8561 | Mirlet7a-1 | ENSMUSG00000065421.2 |
| chr14 | 65953477 | 65953983 | 8.7021 | 20.72798 | 18.38387 | Promoter | 14 | 65919395 | 65953744 | 34350 | 2 | ENSMUST00 | 0 | Scara3 | ENSMUSG00000034463.3 |
| chr15 | 12667662 | 12668239 | 8.7021 | 20.72798 | 18.38387 | Intron (ENSM | 15 | 12695124 | 12695224 | 101 | 1 | ENSMUST00 | -26885 | Gm24302 | ENSMUSG00000095842.1 |
| chr3 | 107582412 | 107583401 | 8.7021 | 20.72798 | 18.38387 | Distal Interge | 3 | 107595031 | 107605776 | 10746 | 1 | ENSMUST00 | -11630 | Alx3 | ENSMUSG00000014603.2 |
| chr3 | 84951948 | 84952607 | 8.7021 | 20.72798 | 18.38387 | Promoter | 3 | 84952146 | 84977645 | 25500 | 1 | ENSMUST00 | 0 | Fbxw7 | ENSMUSG00000028086.9 |
| chr4 | 129959649 | 129960632 | 8.7021 | 20.72798 | 18.38387 | Promoter | 4 | 129960374 | 129963520 | 3147 | 1 | ENSMUST00 | 0 | 1700003M07 | ENSMUSG00000085389.2 |
| chr5 | 119669890 | 119672818 | 8.7021 | 20.72798 | 18.38387 | Promoter | 5 | 119670669 | 119684561 | 13893 | 1 | ENSMUST00 | 0 | Tbx3 | ENSMUSG00000018604.13 |
| chr5 | 5261326 | 5262428 | 8.7021 | 20.72798 | 18.38387 | Intron (ENSM | 5 | 4805838 | 5265309 | 459472 | 2 | ENSMUST00 | 2881 | Cdk14 | ENSMUSG00000028926.10 |
| chr6 | 21857619 | 21858195 | 8.7021 | 20.72798 | 18.38387 | Distal Interge | 6 | 21771395 | 21852515 | 81121 | 2 | ENSMUST00 | -5105 | Tspan12 | ENSMUSG00000029669.9 |
| chr4 | 3230877 | 3231210 | 7.24074 | 20.72537 | 18.38198 | Distal Interge | 4 | 3242055 | 3247506 | 5452 | 1 | ENSMUST00 | -10845 | AI838599 | ENSMUSG00000085180.1 |
| chr5 | 110101629 | 110102192 | 9.90728 | 20.71212 | 18.36883 | Promoter | 5 | 110101204 | 110105953 | 4750 | 1 | ENSMUST00 | 426 | Plcxd1 | ENSMUSG00000064247.9 |
| chr10 | 41690070 | 41690580 | 10.20757 | 20.67883 | 18.33558 | Exon (ENSMU | 10 | 41648876 | 41694487 | 45612 | 2 | ENSMUST00 | 3907 | C6orf183 | ENSMUSG00000100855.1 |
| chr5 | 150457605 | 150458856 | 10.20757 | 20.67883 | 18.33558 | Exon (ENSMU | 5 | 150509694 | 150511338 | 1645 | 2 | ENSMUST00 | 52482 | Gm15933 | ENSMUSG00000082104.1 |
| chr13 | 42883873 | 42884612 | 8.9627 | 20.67791 | 18.33477 | Intron (ENSM | 13 | 42868338 | 42869226 | 889 | 2 | ENSMUST00 | -14648 | Gm15812 | ENSMUSG00000085276.2 |
| chr3 | 55542393 | 55543316 | 9.87001 | 20.64912 | 18.306 | Distal Interge | 3 | 55548446 | 55548552 | 107 | 1 | ENSMUST00 | -5130 | Gm25404 | ENSMUSG00000064904.1 |
| chr3 | 117575319 | 117575738 | 10.1869 | 20.64433 | 18.30124 | Promoter | 3 | 117575360 | 117689507 | 114148 | 1 | ENSMUST00 | 0 | 4833424O15 | ENSMUSG00000033342.8 |
| chr12 | 112644127 | 112644490 | 10.14579 | 20.57575 | 18.2329 | Promoter | 12 | 112644828 | 112649152 | 4325 | 1 | ENSMUST00 | -338 | Siva1 | ENSMUSG00000064326.7 |
| chr17 | 86937572 | 86938405 | 10.14579 | 20.57575 | 18.2329 | Distal Interge | 17 | 86944109 | 86947887 | 3779 | 2 | ENSMUST00 | 9482 | Atp6v1e2 | ENSMUSG00000053375.7 |
| chr19 | 57647558 | 57648330 | 10.14579 | 20.57575 | 18.2329 | Intron (ENSM | 19 | 57611034 | 58133338 | 522305 | 1 | ENSMUST00 | 36525 | Atrnl1 | ENSMUSG00000054843.8 |
| chr15 | 28026111 | 28026553 | 9.1888 | 20.54895 | 18.20621 | Promoter | 15 | 27730652 | 28025848 | 295197 | 2 | ENSMUST00 | -264 | Trio | ENSMUSG00000022263.9 |
| chr1 | 75235964 | 75236346 | 8.29801 | 20.47831 | 18.13586 | Promoter | 1 | 75236406 | 75245688 | 9283 | 1 | ENSMUST00 | -60 | Dnajb2 | ENSMUSG00000026203.11 |
| chr12 | 54162447 | 54163232 | 8.29801 | 20.47831 | 18.13586 | Distal Interge | 12 | 54178981 | 54203860 | 24880 | 2 | ENSMUST00 | 40628 | Egln3 | ENSMUSG00000035105.5 |
| chr16 | 38679316 | 38680012 | 8.29801 | 20.47831 | 18.13586 | Intron (ENSM | 16 | 38685315 | 38685869 | 555 | 2 | ENSMUST00 | 5857 | Gm15530 | ENSMUSG00000081651.1 |
| chr17 | 29436910 | 29437547 | 8.29801 | 20.47831 | 18.13586 | Distal Interge | 17 | 29490812 | 29496111 | 5300 | 1 | ENSMUST00 | -53265 | Pim1 | ENSMUSG00000024014.7 |
| chr17 | 36120710 | 36122163 | 8.29801 | 20.47831 | 18.13586 | Promoter | 17 | 36115876 | 36121435 | 5560 | 2 | ENSMUST00 | 0 | H2-T10 | ENSMUSG00000079491.4 |
| chr4 | 145246236 | 145246877 | 8.29801 | 20.47831 | 18.13586 | Promoter | 4 | 145213463 | 145246870 | 33408 | 2 | ENSMUST00 | 0 | Tnfrsf1b | ENSMUSG00000028599.5 |
| chr5 | 148265082 | 148265500 | 8.29801 | 20.47831 | 18.13586 | Promoter | 5 | 148265217 | 148314071 | 48855 | 1 | ENSMUST00 | 0 | Mtus2 | ENSMUSG00000029651.11 |
| chr6 | 29718684 | 29719630 | 8.29801 | 20.47831 | 18.13586 | Distal Interge | 6 | 29708028 | 29714822 | 6795 | 1 | ENSMUST00 | 10657 | Tspan33 | ENSMUSG00000001763.9 |
| chr8 | 105593587 | 105594870 | 8.29801 | 20.47831 | 18.13586 | Distal Interge | 8 | 105605229 | 105622194 | 16966 | 1 | ENSMUST00 | -10359 | Fam65a | ENSMUSG00000038604.8 |
| chr10 | 57631919 | 57632547 | 10.08475 | 20.47385 | 18.13181 | Promoter | 10 | 57631981 | 57738928 | 106948 | 1 | ENSMUST00 | 0 | Pkib | ENSMUSG00000019876.10 |
| chr10 | 79777080 | 79777577 | 10.08475 | 20.47385 | 18.13181 | Promoter | 10 | 79777272 | 79782630 | 5359 | 1 | ENSMUST00 | 0 | Fstl3 | ENSMUSG00000020325.10 |
| chr13 | 94709715 | 94710624 | 10.08475 | 20.47385 | 18.13181 | Distal Interge | 13 | 94788943 | 94842899 | 53957 | 1 | ENSMUST00 | -78319 | Tbca | ENSMUSG00000042043.5 |
| chr14 | 103778182 | 103778698 | 10.08475 | 20.47385 | 18.13181 | Distal Interge | 14 | 103814626 | 103843696 | 29071 | 2 | ENSMUST00 | 64998 | Ednrb | ENSMUSG00000022122.9 |
| chr14 | 118551095 | 118551960 | 10.08475 | 20.47385 | 18.13181 | Intron (ENSM | 14 | 118585039 | 118585138 | 100 | 1 | ENSMUST00 | -33079 | Gm23302 | ENSMUSG00000064644.1 |
| chr14 | 77706718 | 77707478 | 10.08475 | 20.47385 | 18.13181 | Distal Interge | 14 | 77794823 | 77798864 | 4042 | 2 | ENSMUST00 | 91386 | Gm1587 | ENSMUSG00000075512.3 |
| chr17 | 15577111 | 15577695 | 10.08475 | 20.47385 | 18.13181 | Distal Interge | 17 | 15566164 | 15566421 | 258 | 2 | ENSMUST00 | -10691 | Gm6686 | ENSMUSG00000095920.2 |
| chr7 | 144175440 | 144176046 | 10.08475 | 20.47385 | 18.13181 | Promoter | 7 | 144175529 | 144424494 | 248966 | 1 | ENSMUST00 | 0 | Shank2 | ENSMUSG00000037541.14 |
| chr16 | 4642600 | 4643276 | 9.44616 | 20.46285 | 18.12126 | Intron (ENSM | 16 | 4639989 | 4687292 | 47304 | 1 | ENSMUST00 | 2612 | Dnaja3 | ENSMUSG00000004069.11 |
| chr2 | 73273665 | 73274746 | 10.06456 | 20.44014 | 18.09874 | 3' UTR | 2 | 73271966 | 73284706 | 12741 | 1 | ENSMUST00 | 1700 | Sp9 | ENSMUSG00000068859.5 |
| chr17 | 56672778 | 56673047 | 9.74174 | 20.4319 | 18.09061 | Promoter | 17 | 56673225 | 56711769 | 38545 | 1 | ENSMUST00 | -178 | Ranbp3 | ENSMUSG00000002372.8 |
| chr3 | 63905190 | 63905659 | 9.74174 | 20.4319 | 18.09061 | Distal Interge | 3 | 63696731 | 63899472 | 202742 | 2 | ENSMUST00 | -5719 | Plch1 | ENSMUSG00000036834.11 |
| chr5 | 29568783 | 29569055 | 9.74174 | 20.4319 | 18.09061 | Promoter | 5 | 29569242 | 29676077 | 106836 | 1 | ENSMUST00 | -187 | Ube3c | ENSMUSG00000039000.4 |
| chr7 | 142660537 | 142661055 | 9.74174 | 20.4319 | 18.09061 | Promoter | 7 | 142655442 | 142661205 | 5764 | 2 | ENSMUST00 | 150 | Igf2 | ENSMUSG00000048583.11 |
| chr9 | 21204685 | 21205207 | 9.74174 | 20.4319 | 18.09061 | Exon (ENSMU | 9 | 21196705 | 21211627 | 14923 | 1 | ENSMUST00 | 7981 | Pde4a | ENSMUSG00000032177.11 |
| chr2 | 25355797 | 25356737 | 10.04446 | 20.40655 | 18.06574 | Promoter | 2 | 25352276 | 25356345 | 4070 | 2 | ENSMUST00 | 0 | Dpp7 | ENSMUSG00000026958.8 |
| chr7 | 35235387 | 35235694 | 10.04446 | 20.40655 | 18.06574 | Distal Interge | 7 | 35217235 | 35217360 | 126 | 1 | ENSMUST00 | 18153 | Gm24505 | ENSMUSG00000088307.1 |
| chr8 | 70610311 | 70610820 | 10.04446 | 20.40655 | 18.06574 | Distal Interge | 8 | 70597491 | 70608314 | 10824 | 2 | ENSMUST00 | -1998 | Ssbp4 | ENSMUSG00000070003.6 |
| chr4 | 135206065 | 135206508 | 10.02444 | 20.37309 | 18.0324 | Distal Interge | 4 | 135154436 | 135155342 | 907 | 1 | ENSMUST00 | 51630 | Runx3 | ENSMUSG00000070691.5 |
| chr10 | 59503618 | 59504435 | 9.38717 | 20.36066 | 18.02005 | Intron (ENSM | 10 | 59448597 | 59458160 | 9564 | 2 | ENSMUST00 | -45459 | Mcu | ENSMUSG00000009647.8 |
| chr10 | 94408886 | 94409414 | 9.38717 | 20.36066 | 18.02005 | Promoter | 10 | 94408891 | 94410732 | 1842 | 1 | ENSMUST00 | 0 | Gm16155 | ENSMUSG00000085409.1 |
| chr11 | 78360208 | 78360630 | 9.38717 | 20.36066 | 18.02005 | Intron (ENSM | 11 | 78347935 | 78348998 | 1064 | 1 | ENSMUST00 | 12274 | Unc119 | ENSMUSG00000002058.8 |
| chr11 | 18876720 | 18877551 | 9.38717 | 20.36066 | 18.02005 | Downstream | 11 | 18854043 | 18874309 | 20267 | 2 | ENSMUST00 | -2412 | 8430419K02I | ENSMUSG00000085743.1 |
| chr12 | 81844852 | 81845426 | 9.38717 | 20.36066 | 18.02005 | Distal Interge | 12 | 81860030 | 82000924 | 140895 | 1 | ENSMUST00 | -14604 | Pcnx | ENSMUSG00000021140.8 |
| chr12 | 17873393 | 17874325 | 9.38717 | 20.36066 | 18.02005 | Distal Interge | 12 | 17924294 | 17925668 | 1375 | 1 | ENSMUST00 | -49969 | B430203G13 | ENSMUSG00000067356.5 |
| chr14 | 111681074 | 111682118 | 9.38717 | 20.36066 | 18.02005 | 3' UTR | 14 | 111675115 | 111683134 | 8020 | 1 | ENSMUST00 | 5960 | Slitrk5 | ENSMUSG00000033214.9 |
| chr14 | 3810205 | 3810405 | 9.38717 | 20.36066 | 18.02005 | Promoter | 14 | 3810074 | 3830685 | 20612 | 1 | ENSMUST00 | 132 | Gm3002 | ENSMUSG00000096793.2 |
| chr17 | 5849210 | 5849793 | 9.38717 | 20.36066 | 18.02005 | Intron (ENSM | 17 | 5841328 | 5931033 | 89706 | 1 | ENSMUST00 | 7883 | Snx9 | ENSMUSG00000002365.9 |
| chr18 | 69428569 | 69429597 | 9.38717 | 20.36066 | 18.02005 | Intron (ENSM | 18 | 69346203 | 69683248 | 337046 | 1 | ENSMUST00 | 82367 | Tcf4 | ENSMUSG00000053477.10 |
| chr2 | 173173553 | 173174191 | 9.38717 | 20.36066 | 18.02005 | Distal Interge | 2 | 173153082 | 173159271 | 6190 | 1 | ENSMUST00 | 20472 | Pck1 | ENSMUSG00000027513.6 |
| chr2 | 118711048 | 118711563 | 9.38717 | 20.36066 | 18.02005 | Promoter | 2 | 118708180 | 118711370 | 3191 | 2 | ENSMUST00 | 0 | Plcb2 | ENSMUSG00000040061.12 |
| chr2 | 74714263 | 74715392 | 9.38717 | 20.36066 | 18.02005 | Promoter | 2 | 74710193 | 74715996 | 5804 | 2 | ENSMUST00 | 604 | Hoxd3os1 | ENSMUSG00000052371.5 |
| chr2 | 31469806 | 31470958 | 9.38717 | 20.36066 | 18.02005 | Promoter | 2 | 31470207 | 31520672 | 50466 | 1 | ENSMUST00 | 0 | Ass1 | ENSMUSG00000076441.4 |
| chr2 | 75154523 | 75155517 | 9.38717 | 20.36066 | 18.02005 | Intron (ENSM | 2 | 75138842 | 75322897 | 184056 | 1 | ENSMUST00 | 15682 | RP23-180A8. | ENSMUSG00000100303.1 |
| chr4 | 58671427 | 58671933 | 9.38717 | 20.36066 | 18.02005 | Distal Interge | 4 | 58648392 | 58658592 | 10201 | 2 | ENSMUST00 | -12836 | Gm12580 | ENSMUSG00000086470.1 |
| chr6 | 31548544 | 31549619 | 9.38717 | 20.36066 | 18.02005 | Intron (ENSM | 6 | 31519488 | 31563981 | 44494 | 2 | ENSMUST00 | 14362 | Podxl | ENSMUSG00000025608.9 |
| chr6 | 128597176 | 128597837 | 9.38717 | 20.36066 | 18.02005 | Distal Interge | 6 | 128539822 | 128581606 | 41785 | 2 | ENSMUST00 | -15571 | BC048546 | ENSMUSG00000047228.8 |
| chr6 | 125876838 | 125877186 | 9.38717 | 20.36066 | 18.02005 | Intron (ENSM | 6 | 125884415 | 125884524 | 110 | 1 | ENSMUST00 | -7229 | Gm24947 | ENSMUSG00000088959.1 |
| chr7 | 17056138 | 17056694 | 9.38717 | 20.36066 | 18.02005 | Promoter | 7 | 17035734 | 17056679 | 20946 | 2 | ENSMUST00 | 0 | Hif3a | ENSMUSG00000004328.10 |
| chr7 | 133088509 | 133089390 | 9.38717 | 20.36066 | 18.02005 | Intron (ENSM | 7 | 133025720 | 133111791 | 86072 | 2 | ENSMUST00 | 22401 | Ctbp2 | ENSMUSG00000030970.11 |
| chr8 | 71375743 | 71376320 | 9.38717 | 20.36066 | 18.02005 | Promoter | 8 | 71374628 | 71375790 | 1163 | 2 | ENSMUST00 | 0 | Nr2f6 | ENSMUSG00000002393.9 |
| chr8 | 34123882 | 34124462 | 9.38717 | 20.36066 | 18.02005 | Exon (ENSMU | 8 | 34115030 | 34125185 | 10156 | 1 | ENSMUST00 | 8853 | Mboat4 | ENSMUSG00000071113.3 |
| chr9 | 32343640 | 32344885 | 9.38717 | 20.36066 | 18.02005 | Promoter | 9 | 32314783 | 32344237 | 29455 | 2 | ENSMUST00 | 0 | Kcnj5 | ENSMUSG00000032034.5 |
| chrX | 99853235 | 99853795 | 9.38717 | 20.36066 | 18.02005 | Distal Interge | 20 | 99821021 | 99848790 | 27770 | 1 | ENSMUST00 | 32215 | Tmem28 | ENSMUSG00000071719.2 |
| chr4 | 117926605 | 117927279 | 9.98463 | 20.30655 | 17.96792 | Exon (ENSMU | 4 | 117928144 | 117929413 | 1270 | 2 | ENSMUST00 | 2134 | Artn | ENSMUSG00000028539.9 |
| chr19 | 44266714 | 44267515 | 7.9533 | 20.3031 | 17.96454 | Distal Interge | 19 | 44293676 | 44306864 | 13189 | 1 | ENSMUST00 | -26161 | Scd2 | ENSMUSG00000025203.5 |
| chr19 | 8801946 | 8802355 | 7.9533 | 20.3031 | 17.96454 | Promoter | 19 | 8802530 | 8804854 | 2325 | 1 | ENSMUST00 | -175 | Zbtb3 | ENSMUSG00000071661.6 |
| chr2 | 73272819 | 73273562 | 7.9533 | 20.3031 | 17.96454 | Promoter | 2 | 73271966 | 73284706 | 12741 | 1 | ENSMUST00 | 854 | Sp9 | ENSMUSG00000068859.5 |
| chr4 | 53779231 | 53780077 | 7.9533 | 20.3031 | 17.96454 | Promoter | 4 | 53779705 | 53788712 | 9008 | 1 | ENSMUST00 | 0 | Tal2 | ENSMUSG00000028417.3 |
| chr5 | 117364677 | 117366041 | 7.9533 | 20.3031 | 17.96454 | Exon (ENSMU | 5 | 117363522 | 117377807 | 14286 | 1 | ENSMUST00 | 1156 | Wsb2 | ENSMUSG00000029364.10 |
| chr2 | 125731853 | 125732287 | 9.96484 | 20.27346 | 17.93514 | Distal Interge | 2 | 125691910 | 125724148 | 32239 | 2 | ENSMUST00 | -7706 | Shc4 | ENSMUSG00000035109.9 |
| chr5 | 124729422 | 124730015 | 9.9255 | 20.20765 | 17.86949 | Intron (ENSM | 5 | 124731391 | 124834303 | 102913 | 1 | ENSMUST00 | -1376 | Dnah10 | ENSMUSG00000038011.10 |
| chr7 | 145302843 | 145303742 | 7.6558 | 20.18609 | 17.848 | Intron (ENSM | 7 | 145300899 | 145309349 | 8451 | 1 | ENSMUST00 | 1945 | Mrgprf | ENSMUSG00000031070.10 |
| chr3 | 89319640 | 89319954 | 9.28568 | 20.18442 | 17.84641 | Intron (ENSM | 3 | 89315859 | 89321631 | 5773 | 2 | ENSMUST00 | 1677 | Efna3 | ENSMUSG00000028039.7 |
| chr7 | 19326495 | 19327541 | 9.28568 | 20.18442 | 17.84641 | Distal Interge | 7 | 19302721 | 19310045 | 7325 | 2 | ENSMUST00 | -16451 | Fosb | ENSMUSG00000003545.2 |
| chr14 | 52327482 | 52327842 | 9.90595 | 20.17493 | 17.83711 | Promoter | 14 | 52311172 | 52328762 | 17591 | 2 | ENSMUST00 | 920 | Sall2 | ENSMUSG00000049532.8 |
| chr11 | 60209799 | 60210903 | 9.88648 | 20.14232 | 17.80457 | Promoter | 11 | 60200463 | 60210465 | 10003 | 2 | ENSMUST00 | 0 | Srebf1 | ENSMUSG00000020538.10 |
| chr13 | 45538779 | 45539436 | 9.88648 | 20.14232 | 17.80457 | Promoter | 13 | 45538809 | 45542550 | 3742 | 1 | ENSMUST00 | 0 | Gmpr | ENSMUSG00000000253.7 |
| chr15 | 89146382 | 89147003 | 9.88648 | 20.14232 | 17.80457 | Exon (ENSMU | 15 | 89142486 | 89149606 | 7121 | 2 | ENSMUST00 | 2603 | Mapk11 | ENSMUSG00000053137.6 |
| chr18 | 82869862 | 82870434 | 9.88648 | 20.14232 | 17.80457 | Exon (ENSMU | 18 | 82855897 | 82859162 | 3266 | 1 | ENSMUST00 | 13966 | 4930445N18 | ENSMUSG00000097738.1 |
| chr2 | 167924840 | 167925731 | 9.88648 | 20.14232 | 17.80457 | Distal Interge | 2 | 167932057 | 167974737 | 42681 | 1 | ENSMUST00 | -6326 | Ptpn1 | ENSMUSG00000027540.8 |
| chr2 | 119322108 | 119323062 | 9.88648 | 20.14232 | 17.80457 | Intron (ENSM | 2 | 119321187 | 119325543 | 4357 | 2 | ENSMUST00 | 2481 | Gm14207 | ENSMUSG00000074912.5 |
| chr3 | 92429132 | 92429360 | 9.88648 | 20.14232 | 17.80457 | Promoter | 3 | 92428066 | 92429423 | 1358 | 2 | ENSMUST00 | 63 | Gm9774 | ENSMUSG00000042165.7 |
| chr3 | 146682254 | 146682687 | 9.88648 | 20.14232 | 17.80457 | Promoter | 3 | 146646092 | 146682410 | 36319 | 2 | ENSMUST00 | 0 | Samd13 | ENSMUSG00000048652.7 |
| chr4 | 29133371 | 29134016 | 9.88648 | 20.14232 | 17.80457 | Distal Interge | 4 | 29125946 | 29126124 | 179 | 1 | ENSMUST00 | 7426 | Gm11918 | ENSMUSG00000080911.1 |
| chr5 | 75154016 | 75154558 | 9.88648 | 20.14232 | 17.80457 | Intron (ENSM | 5 | 75156166 | 75198204 | 42039 | 1 | ENSMUST00 | -1608 | Pdgfra | ENSMUSG00000029231.10 |
| chr6 | 37442448 | 37443022 | 9.88648 | 20.14232 | 17.80457 | Promoter | 6 | 37327255 | 37442146 | 114892 | 2 | ENSMUST00 | -303 | Creb3l2 | ENSMUSG00000038648.5 |
| chr9 | 109099985 | 109100989 | 9.88648 | 20.14232 | 17.80457 | 5' UTR | 9 | 109093844 | 109097796 | 3953 | 2 | ENSMUST00 | -2190 | Gm7628 | ENSMUSG00000025644.10 |
| chr10 | 107123341 | 107124017 | 9.56418 | 20.13019 | 17.79291 | Promoter | 10 | 106936164 | 107123668 | 187505 | 2 | ENSMUST00 | 0 | Acss3 | ENSMUSG00000035948.7 |
| chr13 | 44026481 | 44027141 | 9.56418 | 20.13019 | 17.79291 | Distal Interge | 13 | 44121167 | 44125179 | 4013 | 1 | ENSMUST00 | -94026 | Gm5083 | ENSMUSG00000089815.2 |
| chr19 | 59260814 | 59261469 | 9.56418 | 20.13019 | 17.79291 | Promoter | 19 | 59260878 | 59296012 | 35135 | 1 | ENSMUST00 | 0 | Slc18a2 | ENSMUSG00000025094.7 |
| chr4 | 136637292 | 136638034 | 9.56418 | 20.13019 | 17.79291 | Exon (ENSMU | 4 | 136622621 | 136639110 | 16490 | 1 | ENSMUST00 | 14672 | Lactbl1 | ENSMUSG00000070683.7 |
| chr8 | 35765244 | 35765973 | 9.56418 | 20.13019 | 17.79291 | Distal Interge | 8 | 35747449 | 35747562 | 114 | 1 | ENSMUST00 | 17796 | Gm22216 | ENSMUSG00000088182.1 |
| chr14 | 21659316 | 21659800 | 9.82851 | 20.04522 | 17.70854 | Intron (ENSM | 14 | 21622542 | 21634589 | 12048 | 1 | ENSMUST00 | 36775 | Kat6b | ENSMUSG00000021767.11 |
| chr5 | 24497977 | 24499046 | 8.90104 | 20.03618 | 17.69957 | Exon (ENSMU | 5 | 24502757 | 24527202 | 24446 | 2 | ENSMUST00 | 28156 | Gbx1 | ENSMUSG00000067724.5 |
| chr7 | 19342016 | 19342527 | 9.80933 | 20.01308 | 17.67651 | Distal Interge | 7 | 19344778 | 19356524 | 11747 | 1 | ENSMUST00 | -2251 | Ercc1 | ENSMUSG00000003549.4 |
| chr1 | 9237523 | 9237771 | 10.10971 | 19.98882 | 17.65231 | Intron (ENSM | 1 | 8950572 | 9298236 | 347665 | 2 | ENSMUST00 | 60465 | Sntg1 | ENSMUSG00000025909.11 |
| chr1 | 69790463 | 69790945 | 10.10971 | 19.98882 | 17.65231 | Distal Interge | 1 | 69826970 | 70267277 | 440308 | 1 | ENSMUST00 | -36025 | Spag16 | ENSMUSG00000053153.9 |
| chr10 | 80843872 | 80844268 | 10.10971 | 19.98882 | 17.65231 | Promoter | 10 | 80832807 | 80844039 | 11233 | 2 | ENSMUST00 | 0 | Lingo3 | ENSMUSG00000051067.7 |
| chr11 | 101066890 | 101067243 | 10.10971 | 19.98882 | 17.65231 | Distal Interge | 11 | 101070012 | 101077672 | 7661 | 1 | ENSMUST00 | -2769 | Naglu | ENSMUSG00000001751.9 |
| chr11 | 69559840 | 69560629 | 10.10971 | 19.98882 | 17.65231 | Promoter | 11 | 69554092 | 69560205 | 6114 | 2 | ENSMUST00 | 0 | Efnb3 | ENSMUSG00000003934.5 |
| chr12 | 84875705 | 84876980 | 10.10971 | 19.98882 | 17.65231 | Promoter | 12 | 84783212 | 84876532 | 93321 | 2 | ENSMUST00 | 0 | Ltbp2 | ENSMUSG00000002020.10 |
| chr12 | 35896375 | 35896960 | 10.10971 | 19.98882 | 17.65231 | Distal Interge | 12 | 35925620 | 35949736 | 24117 | 1 | ENSMUST00 | -28660 | Agr3 | ENSMUSG00000036231.5 |
| chr12 | 107952146 | 107952939 | 10.10971 | 19.98882 | 17.65231 | Intron (ENSM | 12 | 107915222 | 108003414 | 88193 | 2 | ENSMUST00 | 50475 | Bcl11b | ENSMUSG00000048251.10 |
| chr13 | 48228002 | 48228409 | 10.10971 | 19.98882 | 17.65231 | Intron (ENSM | 13 | 48261427 | 48264036 | 2610 | 1 | ENSMUST00 | -33018 | Id4 | ENSMUSG00000021379.1 |
| chr14 | 31323743 | 31324479 | 10.10971 | 19.98882 | 17.65231 | Promoter | 14 | 31260375 | 31323896 | 63522 | 2 | ENSMUST00 | 0 | Dnah1 | ENSMUSG00000019027.7 |
| chr14 | 32321770 | 32322381 | 10.10971 | 19.98882 | 17.65231 | Promoter | 14 | 32322019 | 32347820 | 25802 | 1 | ENSMUST00 | 0 | Ogdhl | ENSMUSG00000021913.7 |
| chr14 | 7314815 | 7315160 | 10.10971 | 19.98882 | 17.65231 | Promoter | 14 | 7293648 | 7315049 | 21402 | 2 | ENSMUST00 | 0 | Gm3739 | ENSMUSG00000091472.2 |
| chr15 | 97223525 | 97224039 | 10.10971 | 19.98882 | 17.65231 | Distal Interge | 15 | 97247107 | 97385689 | 138583 | 1 | ENSMUST00 | -23068 | Pced1b | ENSMUSG00000044250.7 |
| chr15 | 5009606 | 5010109 | 10.10971 | 19.98882 | 17.65231 | Intron (ENSM | 15 | 5035360 | 5035462 | 103 | 1 | ENSMUST00 | -25251 | Gm25652 | ENSMUSG00000065714.1 |
| chr17 | 71485589 | 71486701 | 10.10971 | 19.98882 | 17.65231 | Distal Interge | 17 | 71344489 | 71475343 | 130855 | 2 | ENSMUST00 | -10247 | Smchd1 | ENSMUSG00000024054.8 |
| chr18 | 20058796 | 20059427 | 10.10971 | 19.98882 | 17.65231 | Promoter | 18 | 20030633 | 20059478 | 28846 | 2 | ENSMUST00 | 51 | Dsc2 | ENSMUSG00000024331.6 |
| chr19 | 55895069 | 55895759 | 10.10971 | 19.98882 | 17.65231 | Promoter | 19 | 55895510 | 55933229 | 37720 | 1 | ENSMUST00 | 0 | Tcf7l2 | ENSMUSG00000024985.13 |
| chr19 | 11092230 | 11092517 | 10.10971 | 19.98882 | 17.65231 | Promoter | 19 | 11090598 | 11092077 | 1480 | 2 | ENSMUST00 | -154 | RP24-512H1 | ENSMUSG00000101439.1 |
| chr2 | 172549344 | 172550252 | 10.10971 | 19.98882 | 17.65231 | Promoter | 2 | 172549593 | 172558622 | 9030 | 1 | ENSMUST00 | 0 | Tfap2c | ENSMUSG00000028640.6 |
| chr2 | 25240589 | 25241373 | 10.10971 | 19.98882 | 17.65231 | Promoter | 2 | 25238818 | 25239988 | 1171 | 2 | ENSMUST00 | -602 | 2310002J15F | ENSMUSG00000036731.7 |
| chr2 | 169790304 | 169790965 | 10.10971 | 19.98882 | 17.65231 | Exon (ENSMU | 2 | 169884893 | 170071816 | 186924 | 1 | ENSMUST00 | -93928 | Tshz2 | ENSMUSG00000047907.11 |
| chr2 | 152406280 | 152406745 | 10.10971 | 19.98882 | 17.65231 | Distal Interge | 2 | 152393611 | 152398063 | 4453 | 2 | ENSMUST00 | -8218 | Sox12 | ENSMUSG00000051817.8 |

| | | | | | | | | | | | | | | | |
|---|---|---|---|---|---|---|---|---|---|---|---|---|---|---|---|
| chr2 | 156717214 | 156717930 | 10.10971 | 19.98882 | 17.65231 | Intron (ENSM | 2 | 156720572 | 156749091 | 28520 | 1 | ENSMUST00 | -2642 | Dlgap4 | ENSMUSG00000061689.10 |
| chr5 | 143363714 | 143364765 | 10.10971 | 19.98882 | 17.65231 | Promoter | 5 | 143363908 | 143391798 | 27891 | 1 | ENSMUST00 | 0 | Grid2ip | ENSMUSG00000010825.12 |
| chr5 | 118028927 | 118029785 | 10.10971 | 19.98882 | 17.65231 | Intron (ENSM | 5 | 118027870 | 118061870 | 34001 | 1 | ENSMUST00 | 1058 | Tesc | ENSMUSG00000029359.8 |
| chr5 | 114467046 | 114467808 | 10.10971 | 19.98882 | 17.65231 | Distal Interge | 5 | 114452420 | 114460572 | 8153 | 1 | ENSMUST00 | 14627 | Mvk | ENSMUSG00000041939.9 |
| chr5 | 124900102 | 124900516 | 10.10971 | 19.98882 | 17.65231 | Intron (ENSM | 5 | 124884569 | 124888630 | 4062 | 1 | ENSMUST00 | 15534 | Zfp664 | ENSMUSG00000079215.3 |
| chr6 | 125426535 | 125426845 | 10.10971 | 19.98882 | 17.65231 | Distal Interge | 6 | 125462116 | 125463024 | 909 | 2 | ENSMUST00 | 36179 | Cd9 | ENSMUSG00000030342.8 |
| chr6 | 52160172 | 52161353 | 10.10971 | 19.98882 | 17.65231 | Promoter | 6 | 52161042 | 52161255 | 214 | 1 | ENSMUST00 | 0 | Gm27861 | ENSMUSG00000099068.1 |
| chr7 | 18925418 | 18925797 | 10.10971 | 19.98882 | 17.65231 | Promoter | 7 | 18925888 | 18962057 | 36170 | 1 | ENSMUST00 | -91 | Nova2 | ENSMUSG00000030411.10 |
| chr7 | 142371802 | 142372653 | 10.10971 | 19.98882 | 17.65231 | Promoter | 7 | 142326110 | 142372441 | 46332 | 2 | ENSMUST00 | 0 | Ifitm10 | ENSMUSG00000045777.9 |
| chr7 | 24628757 | 24629379 | 10.10971 | 19.98882 | 17.65231 | 3' UTR | 7 | 24636570 | 24641116 | 4547 | 1 | ENSMUST00 | -7191 | Lypd3 | ENSMUSG00000057454.4 |
| chr8 | 57321366 | 57322304 | 10.10971 | 19.98882 | 17.65231 | Promoter | 8 | 57320983 | 57324517 | 3535 | 1 | ENSMUST00 | 384 | Hand2 | ENSMUSG00000038193.3 |
| chr8 | 87662542 | 87663339 | 10.10971 | 19.98882 | 17.65231 | Intron (ENSM | 8 | 87636657 | 87637013 | 357 | 1 | ENSMUST00 | 25886 | Gm24841 | ENSMUSG00000089301.1 |
| chrX | 74270333 | 74270635 | 10.10971 | 19.98882 | 17.65231 | Promoter | 20 | 74270812 | 74273135 | 2324 | 1 | ENSMUST00 | -177 | Rpl10 | ENSMUSG00000008682.8 |
| chrX | 59133972 | 59134861 | 10.10971 | 19.98882 | 17.65231 | Promoter | 20 | 59062145 | 59134436 | 72292 | 2 | ENSMUST00 | 0 | Fgf13 | ENSMUSG00000031137.12 |
| chrX | 7649150 | 7649536 | 10.10971 | 19.98882 | 17.65231 | Intron (ENSM | 20 | 7643909 | 7645033 | 1125 | 1 | ENSMUST00 | 5242 | Syp | ENSMUSG00000031144.10 |
| chrX | 99141344 | 99144749 | 10.10971 | 19.98882 | 17.65231 | Intron (ENSM | 20 | 99138131 | 99147230 | 9100 | 1 | ENSMUST00 | 3214 | Efnb1 | ENSMUSG00000031217.8 |
| chrX | 147553452 | 147554134 | 10.10971 | 19.98882 | 17.65231 | Promoter | 20 | 147470380 | 147554081 | 83702 | 2 | ENSMUST00 | 0 | Lrch2 | ENSMUSG00000031290.9 |
| chrX | 48409852 | 48410313 | 10.10971 | 19.98882 | 17.65231 | Downstream | 20 | 48369268 | 48384145 | 14878 | 1 | ENSMUST00 | 40585 | Bcorl1 | ENSMUSG00000036959.10 |
| chrX | 106920359 | 106921033 | 10.10971 | 19.98882 | 17.65231 | Promoter | 20 | 106920625 | 106933900 | 13276 | 1 | ENSMUST00 | 0 | Lpar4 | ENSMUSG00000049929.7 |
| chrX | 44367611 | 44368541 | 10.10971 | 19.98882 | 17.65231 | Promoter | 20 | 44365458 | 44368342 | 2885 | 2 | ENSMUST00 | 0 | Dcaf12l2 | ENSMUSG00000050926.5 |
| chrX | 150486140 | 150487210 | 10.10971 | 19.98882 | 17.65231 | Distal Interge | 20 | 150463726 | 150466156 | 2431 | 1 | ENSMUST00 | 22415 | 3110067C02I | ENSMUSG00000081187.1 |
| chrX | 75674480 | 75674813 | 10.10971 | 19.98882 | 17.65231 | Distal Interge | 20 | 75705409 | 75705681 | 273 | 1 | ENSMUST00 | -30596 | Gm15065 | ENSMUSG00000082551.1 |
| chrX | 53716366 | 53716859 | 10.10971 | 19.98882 | 17.65231 | Promoter | 20 | 53696922 | 53716578 | 19657 | 2 | ENSMUST00 | 0 | AW822252 | ENSMUSG00000085998.2 |
| chrX | 20819126 | 20819977 | 10.10971 | 19.98882 | 17.65231 | Intron (ENSM | 20 | 20823737 | 20823814 | 78 | 2 | ENSMUST00 | 3837 | Gm24824 | ENSMUSG00000092838.1 |
| chrX | 103447659 | 103448049 | 10.10971 | 19.98882 | 17.65231 | Exon (ENSMU | 20 | 103461218 | 103461285 | 68 | 2 | ENSMUST00 | 13236 | Gm27733 | ENSMUSG00000099312.1 |
| chr10 | 127661425 | 127662025 | 9.79024 | 19.98107 | 17.64646 | 3' UTR | 10 | 127663839 | 127665484 | 1646 | 1 | ENSMUST00 | -1814 | Gm16229 | ENSMUSG00000085731.1 |
| chr15 | 81514051 | 81514658 | 10.09152 | 19.95864 | 17.62411 | Distal Interge | 15 | 81499084 | 81499548 | 465 | 2 | ENSMUST00 | -14504 | Gm5218 | ENSMUSG00000091198.1 |
| chr2 | 73659848 | 73660560 | 9.77121 | 19.94916 | 17.6147 | Promoter | 2 | 73610745 | 73660406 | 49662 | 2 | ENSMUST00 | 0 | Chn1 | ENSMUSG00000056486.12 |
| chr1 | 125435824 | 125436528 | 8.8423 | 19.93079 | 17.59637 | Promoter | 1 | 125405880 | 125435727 | 29848 | 2 | ENSMUST00 | -98 | Actr3 | ENSMUSG00000026341.11 |
| chr1 | 176835541 | 176836044 | 8.8423 | 19.93079 | 17.59637 | Intron (ENSM | 1 | 176814660 | 177020437 | 205778 | 1 | ENSMUST00 | 20882 | Sdccag8 | ENSMUSG00000026504.12 |
| chr1 | 126737815 | 126738534 | 8.8423 | 19.93079 | 17.59637 | Promoter | 1 | 126258401 | 126738202 | 479802 | 2 | ENSMUST00 | 0 | Nckap5 | ENSMUSG00000049690.10 |
| chr11 | 74832439 | 74833147 | 8.8423 | 19.93079 | 17.59637 | Intron (ENSM | 11 | 74830920 | 74845725 | 14806 | 1 | ENSMUST00 | 1520 | Mnt | ENSMUSG00000000282.7 |
| chr11 | 48857416 | 48858774 | 8.8423 | 19.93079 | 17.59637 | Distal Interge | 11 | 48843648 | 48853493 | 9846 | 2 | ENSMUST00 | -3924 | Gm16170 | ENSMUSG00000086245.1 |
| chr14 | 20836197 | 20837085 | 8.8423 | 19.93079 | 17.59637 | Promoter | 14 | 20836662 | 20843388 | 6727 | 1 | ENSMUST00 | 0 | Plau | ENSMUSG00000021822.2 |
| chr15 | 6564944 | 6565470 | 8.8423 | 19.93079 | 17.59637 | Distal Interge | 15 | 6579871 | 6663304 | 83434 | 1 | ENSMUST00 | -14401 | Fyb | ENSMUSG00000022148.10 |
| chr15 | 11036386 | 11037536 | 8.8423 | 19.93079 | 17.59637 | Promoter | 15 | 11033717 | 11037991 | 4275 | 2 | ENSMUST00 | 455 | Rxfp3 | ENSMUSG00000060735.4 |
| chr15 | 81836244 | 81836833 | 8.8423 | 19.93079 | 17.59637 | Distal Interge | 15 | 81843343 | 81843699 | 357 | 2 | ENSMUST00 | 6866 | Gm8444 | ENSMUSG00000069439.2 |
| chr17 | 56078600 | 56078978 | 8.8423 | 19.93079 | 17.59637 | Promoter | 17 | 56079657 | 56100596 | 20940 | 1 | ENSMUST00 | -679 | Hdgfrp2 | ENSMUSG00000002833.10 |
| chr2 | 116009046 | 116009659 | 8.8423 | 19.93079 | 17.59637 | Intron (ENSM | 2 | 115864273 | 116042046 | 177774 | 2 | ENSMUST00 | 32387 | Meis2 | ENSMUSG00000027210.15 |
| chr2 | 132384739 | 132385579 | 8.8423 | 19.93079 | 17.59637 | Promoter | 2 | 132337733 | 132385427 | 47695 | 2 | ENSMUST00 | 0 | Prokr2 | ENSMUSG00000050558.8 |
| chr2 | 54084437 | 54085685 | 8.8423 | 19.93079 | 17.59637 | Promoter | 2 | 54084093 | 54085552 | 1460 | 2 | ENSMUST00 | 0 | Rprm | ENSMUSG00000075334.2 |
| chr2 | 49312796 | 49313290 | 8.8423 | 19.93079 | 17.59637 | Intron (ENSM | 2 | 49308155 | 49309462 | 1308 | 1 | ENSMUST00 | 4642 | Gm13509 | ENSMUSG00000081272.1 |
| chr2 | 168169812 | 168170726 | 8.8423 | 19.93079 | 17.59637 | Distal Interge | 2 | 168178158 | 168178252 | 95 | 2 | ENSMUST00 | 7526 | Gm24561 | ENSMUSG00000092923.1 |
| chr2 | 170193272 | 170194045 | 8.8423 | 19.93079 | 17.59637 | Promoter | 2 | 170192449 | 170194053 | 1605 | 2 | ENSMUST00 | 8 | Gm17619 | ENSMUSG00000097514.1 |
| chr3 | 89378414 | 89378892 | 8.8423 | 19.93079 | 17.59637 | 3' UTR | 3 | 89377644 | 89387189 | 9546 | 2 | ENSMUST00 | 8297 | Zbtb7b | ENSMUSG00000028042.10 |
| chr3 | 95905129 | 95905921 | 8.8423 | 19.93079 | 17.59637 | Promoter | 3 | 95897768 | 95904691 | 6924 | 2 | ENSMUST00 | -439 | Car14 | ENSMUSG00000038526.9 |
| chr3 | 96240485 | 96240743 | 8.8423 | 19.93079 | 17.59637 | Promoter | 3 | 96239749 | 96240276 | 528 | 2 | ENSMUST00 | -210 | Hist2h2aa1 | ENSMUSG00000063954.7 |
| chr3 | 9340717 | 9341155 | 8.8423 | 19.93079 | 17.59637 | Distal Interge | 3 | 9402175 | 9402485 | 311 | 1 | ENSMUST00 | -61020 | Gm22604 | ENSMUSG00000088229.1 |
| chr4 | 108383207 | 108383691 | 8.8423 | 19.93079 | 17.59637 | Promoter | 4 | 108367777 | 108383349 | 15573 | 2 | ENSMUST00 | 0 | Fam159a | ENSMUSG00000059816.5 |
| chr5 | 24266364 | 24266807 | 8.8423 | 19.93079 | 17.59637 | Distal Interge | 5 | 24256822 | 24257037 | 216 | 1 | ENSMUST00 | 9543 | Gm9836 | ENSMUSG00000081071.1 |
| chr6 | 28927814 | 28928043 | 8.8423 | 19.93079 | 17.59637 | Intron (ENSM | 6 | 28880043 | 28886527 | 6485 | 1 | ENSMUST00 | 47772 | Snd1 | ENSMUSG00000001424.9 |
| chr6 | 84018947 | 84019806 | 8.8423 | 19.93079 | 17.59637 | Promoter | 6 | 84019385 | 84211059 | 191675 | 1 | ENSMUST00 | 0 | Dysf | ENSMUSG00000033788.10 |
| chr7 | 98835654 | 98836986 | 8.8423 | 19.93079 | 17.59637 | Promoter | 7 | 98835131 | 98855195 | 20065 | 1 | ENSMUST00 | 524 | Wnt11 | ENSMUSG00000015957.6 |
| chr7 | 19522921 | 19523648 | 8.8423 | 19.93079 | 17.59637 | Exon (ENSMU | 7 | 19518731 | 19525050 | 6320 | 1 | ENSMUST00 | 4191 | Nkpd1 | ENSMUSG00000060621.4 |
| chr6 | 31938337 | 31938711 | 10.0704 | 19.92358 | 17.59046 | Distal Interge | 6 | 32050246 | 32058921 | 8676 | 1 | ENSMUST00 | -111535 | 1700012A03I | ENSMUSG00000029766.4 |
| chr7 | 141477003 | 141478311 | 10.0704 | 19.92358 | 17.59046 | Promoter | 7 | 141476400 | 141493421 | 17022 | 1 | ENSMUST00 | 604 | Tspan4 | ENSMUSG00000025511.9 |
| chr1 | 4491619 | 4493509 | 9.13057 | 19.91392 | 17.58092 | 5' UTR | 1 | 4491713 | 4496363 | 4651 | 2 | ENSMUST00 | 2854 | Sox17 | ENSMUSG00000025902.8 |
| chr12 | 58267268 | 58268926 | 9.13057 | 19.91392 | 17.58092 | Promoter | 12 | 58264720 | 58269258 | 4539 | 2 | ENSMUST00 | 332 | Clec14a | ENSMUSG00000045930.2 |
| chr19 | 25671988 | 25672887 | 9.13057 | 19.91392 | 17.58092 | Promoter | 19 | 25672420 | 25679010 | 6591 | 1 | ENSMUST00 | 0 | Dmrt2 | ENSMUSG00000048138.9 |
| chr9 | 44655380 | 44655732 | 9.13057 | 19.91392 | 17.58092 | Distal Interge | 9 | 44673233 | 44686305 | 13073 | 1 | ENSMUST00 | -17501 | Treh | ENSMUSG00000032098.8 |
| chr11 | 59538197 | 59538773 | 10.04936 | 19.88865 | 17.55579 | Distal Interge | 11 | 59541568 | 59566955 | 25388 | 1 | ENSMUST00 | -2795 | Nlrp3 | ENSMUSG00000032691.9 |
| chr13 | 97008415 | 97008776 | 10.04936 | 19.88865 | 17.55579 | Distal Interge | 13 | 97021864 | 97034362 | 12499 | 2 | ENSMUST00 | 25586 | 1700029F12I | ENSMUSG00000052075.6 |
| chr4 | 135762391 | 135762779 | 10.04936 | 19.88865 | 17.55579 | 5' UTR | 4 | 135759715 | 135815564 | 55850 | 1 | ENSMUST00 | 2677 | Myom3 | ENSMUSG00000037139.10 |
| chr6 | 85513257 | 85513937 | 10.04936 | 19.88865 | 17.55579 | Promoter | 6 | 85511131 | 85513589 | 2459 | 2 | ENSMUST00 | 0 | Egr4 | ENSMUSG00000071341.3 |
| chr6 | 48851958 | 48852493 | 10.04936 | 19.88865 | 17.55579 | Distal Interge | 6 | 48860356 | 48864297 | 3942 | 1 | ENSMUST00 | -7863 | Gm7932 | ENSMUSG00000073090.3 |
| chr8 | 57799954 | 57800512 | 10.04936 | 19.88865 | 17.55579 | Intron (ENSM | 8 | 57541995 | 57653032 | 111038 | 2 | ENSMUST00 | -146923 | Galnt7 | ENSMUSG00000031608.8 |
| chr5 | 77403825 | 77404462 | 9.40982 | 19.86685 | 17.53426 | Intron (ENSM | 5 | 77349240 | 77408045 | 58806 | 2 | ENSMUST00 | 3583 | Igfbp7 | ENSMUSG00000036256.8 |
| chr2 | 170319346 | 170320164 | 9.71458 | 19.85413 | 17.52155 | Distal Interge | 2 | 170347931 | 170349740 | 1810 | 2 | ENSMUST00 | 29576 | Bcas1 | ENSMUSG00000013523.8 |
| chr2 | 30441461 | 30441853 | 9.39297 | 19.83805 | 17.50559 | Promoter | 2 | 30441853 | 30447131 | 5279 | 1 | ENSMUST00 | -23 | Ppp2r4 | ENSMUSG00000039515.6 |
| chr10 | 73099173 | 73100007 | 9.69585 | 19.82268 | 17.49052 | Promoter | 10 | 73099342 | 74317264 | 1217923 | 1 | ENSMUST00 | 0 | Pcdh15 | ENSMUSG00000052613.11 |
| chr10 | 23236793 | 23237447 | 9.69585 | 19.82268 | 17.49052 | Intron (ENSM | 10 | 23166865 | 23166971 | 107 | 2 | ENSMUST00 | -69823 | Gm24655 | ENSMUSG00000094033.1 |
| chr15 | 27937526 | 27938266 | 9.69585 | 19.82268 | 17.49052 | Intron (ENSM | 15 | 27730652 | 28025848 | 295197 | 2 | ENSMUST00 | 87582 | Trio | ENSMUSG00000022263.9 |
| chr15 | 8967484 | 8968482 | 9.69585 | 19.82268 | 17.49052 | Promoter | 15 | 8968426 | 9002646 | 34221 | 1 | ENSMUST00 | 0 | Ranbp3l | ENSMUSG00000048424.10 |
| chr2 | 3749239 | 3749869 | 9.69585 | 19.82268 | 17.49052 | Intron (ENSM | 2 | 3752986 | 3755177 | 2192 | 2 | ENSMUST00 | 5308 | Gm13185 | ENSMUSG00000085205.1 |
| chr4 | 123104528 | 123105454 | 9.69585 | 19.82268 | 17.49052 | Promoter | 4 | 123105165 | 123126091 | 20927 | 1 | ENSMUST00 | 0 | Bmp8b | ENSMUSG00000002384.2 |
| chr7 | 127800542 | 127801396 | 9.69585 | 19.82268 | 17.49052 | Promoter | 7 | 127800604 | 127803802 | 3199 | 1 | ENSMUST00 | 0 | Hsd3b7 | ENSMUSG00000042289.6 |
| chr11 | 97994457 | 97996157 | 10.00754 | 19.81923 | 17.48753 | Promoter | 11 | 97993836 | 97995929 | 2094 | 2 | ENSMUST00 | 0 | Arl5c | ENSMUSG00000038352.10 |
| chr19 | 16772806 | 16773917 | 10.00754 | 19.81923 | 17.48753 | Intron (ENSM | 19 | 16780547 | 16781103 | 557 | 1 | ENSMUST00 | -6630 | Gm9806 | ENSMUSG00000046388.4 |
| chr6 | 110645133 | 110646357 | 10.00754 | 19.81923 | 17.48753 | Promoter | 6 | 110645581 | 111497090 | 851510 | 1 | ENSMUST00 | 0 | Grm7 | ENSMUSG00000056755.8 |
| chr4 | 105245470 | 105246115 | 9.67719 | 19.79134 | 17.45974 | Distal Interge | 4 | 105157347 | 105232764 | 75418 | 1 | ENSMUST00 | 88124 | Ppap2b | ENSMUSG00000028517.8 |
| chr5 | 148489585 | 148490197 | 9.96607 | 19.75036 | 17.41895 | Distal Interge | 5 | 148504631 | 148552788 | 48158 | 2 | ENSMUST00 | 62591 | Ubl3 | ENSMUSG00000001687.10 |
| chr1 | 39936191 | 39936699 | 9.63953 | 19.72805 | 17.39678 | Intron (ENSM | 1 | 39900913 | 40026310 | 125398 | 1 | ENSMUST00 | 35279 | Map4k4 | ENSMUSG00000026074.9 |
| chr10 | 125785480 | 125786804 | 9.63953 | 19.72805 | 17.39678 | Distal Interge | 10 | 125966219 | 126015359 | 49141 | 1 | ENSMUST00 | -179415 | Lrig3 | ENSMUSG00000020105.8 |
| chr10 | 123201979 | 123202456 | 9.63953 | 19.72805 | 17.39678 | Distal Interge | 10 | 123196876 | 123196971 | 96 | 2 | ENSMUST00 | -5009 | Gm27920 | ENSMUSG00000099088.1 |
| chr11 | 77491296 | 77492238 | 9.63953 | 19.72805 | 17.39678 | Exon (ENSMU | 11 | 77493562 | 77507786 | 14225 | 1 | ENSMUST00 | -1324 | Git1 | ENSMUSG00000011877.8 |
| chr11 | 69800732 | 69801817 | 9.63953 | 19.72805 | 17.39678 | Promoter | 11 | 69796068 | 69801716 | 5649 | 2 | ENSMUST00 | 0 | Fgf11 | ENSMUSG00000042826.8 |
| chr11 | 63264582 | 63265130 | 9.63953 | 19.72805 | 17.39678 | Distal Interge | 11 | 63241243 | 63241475 | 233 | 2 | ENSMUST00 | -23108 | Gm12285 | ENSMUSG00000083052.1 |
| chr12 | 40133800 | 40134449 | 9.63953 | 19.72805 | 17.39678 | Promoter | 12 | 40059769 | 40134183 | 74415 | 2 | ENSMUST00 | 0 | Scin | ENSMUSG00000002565.11 |
| chr12 | 21426282 | 21426888 | 9.63953 | 19.72805 | 17.39678 | Distal Interge | 12 | 21417911 | 21419803 | 1893 | 1 | ENSMUST00 | 8372 | Gm4419 | ENSMUSG00000097055.1 |
| chr13 | 21275305 | 21276605 | 9.63953 | 19.72805 | 17.39678 | Distal Interge | 13 | 21286429 | 21292686 | 6258 | 2 | ENSMUST00 | 16081 | Gpx5 | ENSMUSG00000004344.6 |
| chr13 | 21831437 | 21831753 | 9.63953 | 19.72805 | 17.39678 | Promoter | 13 | 21831767 | 21832196 | 430 | 2 | ENSMUST00 | 443 | Hist1h4n | ENSMUSG00000069305.3 |
| chr14 | 56864942 | 56865365 | 9.63953 | 19.72805 | 17.39678 | Distal Interge | 14 | 56852830 | 56852888 | 59 | 2 | ENSMUST00 | -12055 | Gm22218 | ENSMUSG00000088180.1 |
| chr15 | 81480956 | 81481535 | 9.63953 | 19.72805 | 17.39678 | Distal Interge | 15 | 81466316 | 81476369 | 10054 | 1 | ENSMUST00 | 14641 | Rbx1 | ENSMUSG00000022400.8 |
| chr17 | 43077267 | 43077841 | 9.63953 | 19.72805 | 17.39678 | Intron (ENSM | 17 | 43016555 | 43089188 | 72634 | 1 | ENSMUST00 | 60713 | Tnfrsf21 | ENSMUSG00000023915.4 |
| chr17 | 48001564 | 48002306 | 9.63953 | 19.72805 | 17.39678 | Downstream | 17 | 48003379 | 48003572 | 194 | 2 | ENSMUST00 | 1266 | Gm14871 | ENSMUSG00000082492.1 |
| chr18 | 54902209 | 54902640 | 9.63953 | 19.72805 | 17.39678 | Intron (ENSM | 18 | 54981435 | 54987689 | 6255 | 1 | ENSMUST00 | -78795 | Gm26742 | ENSMUSG00000097707.1 |
| chr2 | 61401859 | 61402201 | 9.63953 | 19.72805 | 17.39678 | Distal Interge | 2 | 61492766 | 61492894 | 129 | 1 | ENSMUST00 | -90565 | Gm22338 | ENSMUSG00000088340.1 |
| chr2 | 170219215 | 170220058 | 9.63953 | 19.72805 | 17.39678 | Distal Interge | 2 | 170192449 | 170194053 | 1605 | 2 | ENSMUST00 | -25163 | Gm17619 | ENSMUSG00000097514.1 |
| chr5 | 136102296 | 136102774 | 9.63953 | 19.72805 | 17.39678 | Exon (ENSMU | 5 | 136103894 | 136106033 | 2140 | 1 | ENSMUST00 | -1120 | Rasa4 | ENSMUSG00000004952.8 |
| chr5 | 30113981 | 30115022 | 9.63953 | 19.72805 | 17.39678 | Exon (ENSMU | 5 | 30119899 | 30120305 | 407 | 2 | ENSMUST00 | 5283 | Hadha | ENSMUSG00000025745.6 |
| chr5 | 43868753 | 43869911 | 9.63953 | 19.72805 | 17.39678 | Promoter | 5 | 43868809 | 43912375 | 43567 | 1 | ENSMUST00 | 0 | Cd38 | ENSMUSG00000029084.4 |
| chr5 | 109556662 | 109557291 | 9.63953 | 19.72805 | 17.39678 | Exon (ENSMU | 5 | 109554711 | 109558993 | 4283 | 2 | ENSMUST00 | 1702 | Crlf2 | ENSMUSG00000033467.8 |
| chr5 | 114796430 | 114796919 | 9.63953 | 19.72805 | 17.39678 | Promoter | 5 | 114795743 | 114806200 | 10458 | 1 | ENSMUST00 | 688 | Ankrd13a | ENSMUSG00000041870.11 |
| chr5 | 150952485 | 150953348 | 9.63953 | 19.72805 | 17.39678 | Promoter | 5 | 150952607 | 150993809 | 41203 | 1 | ENSMUST00 | 0 | Kl | ENSMUSG00000058488.6 |
| chr6 | 60828507 | 60829766 | 9.63953 | 19.72805 | 17.39678 | Promoter | 6 | 60731573 | 60828996 | 97424 | 2 | ENSMUST00 | 0 | Snca | ENSMUSG00000025889.8 |
| chr7 | 45163740 | 45164183 | 9.63953 | 19.72805 | 17.39678 | Promoter | 7 | 45163921 | 45176138 | 12218 | 1 | ENSMUST00 | 0 | Slc17a7 | ENSMUSG00000070570.4 |
| chr4 | 112979402 | 112979797 | 9.3262 | 19.72375 | 17.39417 | Promoter | 4 | 112978465 | 112979813 | 1349 | 2 | ENSMUST00 | 16 | Gm12818 | ENSMUSG00000082678.1 |
| chr17 | 47151601 | 47152468 | 9.94546 | 19.71614 | 17.38656 | Intron (ENSM | 17 | 47141216 | 47318790 | 177575 | 1 | ENSMUST00 | 10386 | Trerf1 | ENSMUSG00000064043.8 |
| chr19 | 25673336 | 25674140 | 9.60327 | 19.66707 | 17.33772 | Promoter | 19 | 25672420 | 25679010 | 6591 | 1 | ENSMUST00 | 917 | Dmrt2 | ENSMUSG00000048138.9 |
| chr10 | 78787730 | 78788089 | 8.98055 | 19.65095 | 17.32162 | 5' UTR | 10 | 78780496 | 78814825 | 34330 | 1 | ENSMUST00 | 7235 | Slc1a6 | ENSMUSG00000005357.8 |
| chr1 | 136948494 | 136949511 | 8.39131 | 19.62543 | 17.29633 | 5' UTR | 1 | 136843584 | 136953630 | 110047 | 2 | ENSMUST00 | 4119 | Nr5a2 | ENSMUSG00000026398.9 |
| chr11 | 37235491 | 37237012 | 8.39131 | 19.62543 | 17.29633 | Promoter | 11 | 36006656 | 37235882 | 1229227 | 2 | ENSMUST00 | 0 | Tenm2 | ENSMUSG00000049336.11 |
| chr12 | 54191776 | 54192456 | 8.39131 | 19.62543 | 17.29633 | Intron (ENSM | 12 | 54178981 | 54203860 | 24880 | 2 | ENSMUST00 | 11404 | Egln3 | ENSMUSG00000035105.5 |
| chr13 | 21832008 | 21832257 | 8.39131 | 19.62543 | 17.29633 | Promoter | 13 | 21831767 | 21832196 | 430 | 2 | ENSMUST00 | 0 | Hist1h4n | ENSMUSG00000069305.3 |
| chr15 | 37232608 | 37233471 | 8.39131 | 19.62543 | 17.29633 | Promoter | 15 | 37233036 | 37333647 | 100612 | 1 | ENSMUST00 | 0 | Grhl2 | ENSMUSG00000022286.10 |
| chr16 | 87796694 | 87797515 | 8.39131 | 19.62543 | 17.29633 | Intron (ENSM | 16 | 87787572 | 87839293 | 51722 | 1 | ENSMUST00 | 9123 | 2810407A14I | ENSMUSG00000055972.5 |
| chr17 | 34671416 | 34672526 | 8.39131 | 19.62543 | 17.29633 | Promoter | 17 | 34670535 | 34719815 | 49281 | 1 | ENSMUST00 | 882 | Tnxb | ENSMUSG00000033327.11 |
| chr5 | 35313557 | 35314341 | 8.39131 | 19.62543 | 17.29633 | Distal Interge | 5 | 35278566 | 35281761 | 3196 | 1 | ENSMUST00 | 34992 | Adra2c | ENSMUSG00000045318.5 |
| chr6 | 22205741 | 22206561 | 8.39131 | 19.62543 | 17.29633 | Intron (ENSM | 6 | 22135546 | 22145551 | 10006 | 1 | ENSMUST00 | 70196 | Cped1 | ENSMUSG00000062980.10 |
| chr6 | 115717883 | 115718429 | 8.39131 | 19.62543 | 17.29633 | Distal Interge | 6 | 115723695 | 115723832 | 138 | 2 | ENSMUST00 | 5403 | Gm24008 | ENSMUSG00000088358.1 |
| chr7 | 27329488 | 27329793 | 8.39131 | 19.62543 | 17.29633 | Exon (ENSMU | 7 | 27305998 | 27333613 | 27616 | 2 | ENSMUST00 | 3820 | Ltbp4 | ENSMUSG00000040488.11 |
| chr5 | 44799041 | 44799658 | 9.88415 | 19.61427 | 17.28588 | Promoter | 5 | 44472133 | 44799707 | 327575 | 2 | ENSMUST00 | 49 | Ldb2 | ENSMUSG00000039706.7 |
| chr5 | 100517714 | 100518159 | 9.56673 | 19.60558 | 17.27728 | Promoter | 5 | 100518309 | 100547803 | 29495 | 1 | ENSMUST00 | -150 | Cops4 | ENSMUSG00000035297.9 |
| chr13 | 40854407 | 40854922 | 9.86388 | 19.58058 | 17.25231 | Distal Interge | 13 | 40859768 | 40960891 | 101124 | 1 | ENSMUST00 | -4846 | Gcnt2 | ENSMUSG00000021360.10 |
| chr11 | 120086970 | 120087282 | 9.22779 | 19.55493 | 17.2268 | Promoter | 11 | 120064430 | 120086827 | 22398 | 2 | ENSMUST00 | -144 | Cep131 | ENSMUSG00000039781.9 |
| chr18 | 34751089 | 34751574 | 9.22779 | 19.55493 | 17.2268 | Promoter | 18 | 34732997 | 34751533 | 18537 | 2 | ENSMUST00 | 0 | Cdc25c | ENSMUSG00000044201.9 |
| chr13 | 23745133 | 23745575 | 9.84369 | 19.54702 | 17.21913 | Promoter | 13 | 23745013 | 23745501 | 489 | 2 | ENSMUST00 | 0 | Hist1h3c | ENSMUSG00000069310.3 |
| chr19 | 26748705 | 26748968 | 9.84369 | 19.54702 | 17.21913 | Promoter | 19 | 26748459 | 26778273 | 29815 | 1 | ENSMUST00 | 247 | Smarca2 | ENSMUSG00000024921.11 |

| | | | | | | | | | | | | | | | |
|---|---|---|---|---|---|---|---|---|---|---|---|---|---|---|---|
| chr3 | 95701798 | 95702505 | 9.84369 | 19.54702 | 17.21913 | Distal Interge | 3 | 95676201 | 95687917 | 11717 | 2 | ENSMUST00 | -13882 | Adamtsl4 | ENSMUSG00000015850.6 |
| chr4 | 44425090 | 44425666 | 9.84369 | 19.54702 | 17.21913 | Distal Interge | 4 | 44412469 | 44412595 | 127 | 2 | ENSMUST00 | -12496 | Gm24376 | ENSMUSG00000077587.1 |
| chr4 | 47603494 | 47604396 | 9.84369 | 19.54702 | 17.21913 | Promoter | 4 | 47603889 | 47604008 | 120 | 1 | ENSMUST00 | 0 | Gm27668 | ENSMUSG00000098518.1 |
| chr5 | 143287806 | 143288294 | 9.84369 | 19.54702 | 17.21913 | Exon (ENSMU | 5 | 143287755 | 143292356 | 4602 | 2 | ENSMUST00 | 4062 | Zfp853 | ENSMUSG00000093910.1 |
| chr6 | 31637885 | 31639087 | 9.84369 | 19.54702 | 17.21913 | Intron (ENSM | 6 | 31651041 | 31662021 | 10981 | 1 | ENSMUST00 | -11954 | Gm13846 | ENSMUSG00000086885.1 |
| chr7 | 79508499 | 79509128 | 9.84369 | 19.54702 | 17.21913 | Intron (ENSM | 7 | 79505264 | 79505353 | 90 | 1 | ENSMUST00 | 3236 | Mir9-3 | ENSMUSG00000093238.1 |
| chr7 | 91262035 | 91262875 | 9.53047 | 19.54451 | 17.21699 | Intron (ENSM | 7 | 91242995 | 91243097 | 103 | 1 | ENSMUST00 | 19041 | Gm24552 | ENSMUSG00000089271.1 |
| chr1 | 10008884 | 10009509 | 9.51244 | 19.51414 | 17.18665 | Promoter | 1 | 9968624 | 10009136 | 40513 | 2 | ENSMUST00 | 0 | Ppp1r42 | ENSMUSG00000025916.5 |
| chr1 | 180510705 | 180510981 | 9.51244 | 19.51414 | 17.18665 | Distal Interge | 1 | 180438329 | 180483504 | 45176 | 2 | ENSMUST00 | -27202 | 6330403A02 | ENSMUSG00000053963.6 |
| chr13 | 24965240 | 24965914 | 9.51244 | 19.51414 | 17.18665 | Intron (ENSM | 13 | 24955854 | 24962818 | 6965 | 1 | ENSMUST00 | 9387 | Gpld1 | ENSMUSG00000021340.7 |
| chr14 | 57910344 | 57910715 | 9.51244 | 19.51414 | 17.18665 | Distal Interge | 14 | 57832702 | 57890262 | 57561 | 2 | ENSMUST00 | -20083 | Zdhhc20 | ENSMUSG00000021969.7 |
| chr15 | 25773836 | 25774327 | 9.82359 | 19.5136 | 17.18665 | Promoter | 15 | 25773987 | 25781057 | 7071 | 1 | ENSMUST00 | 0 | Myo10 | ENSMUSG00000022272.12 |
| chr17 | 28689554 | 28689970 | 9.51244 | 19.51414 | 17.18665 | Promoter | 17 | 28637783 | 28689987 | 52205 | 2 | ENSMUST00 | 17 | Slc26a8 | ENSMUSG00000036196.10 |
| chr2 | 26132583 | 26133600 | 9.51244 | 19.51414 | 17.18665 | Distal Interge | 2 | 26138546 | 26139543 | 998 | 1 | ENSMUST00 | -4946 | 1810012K08 | ENSMUSG00000087595.1 |
| chr7 | 109010788 | 109011247 | 9.51244 | 19.51414 | 17.18665 | Promoter | 7 | 109010825 | 109034460 | 23636 | 1 | ENSMUST00 | 0 | Tub | ENSMUSG00000031028.8 |
| chr8 | 79209678 | 79210338 | 9.51244 | 19.51414 | 17.18665 | Downstream | 8 | 79178975 | 79189761 | 10787 | 1 | ENSMUST00 | 30704 | Zfp827 | ENSMUSG00000071064.8 |
| chr8 | 126516797 | 126517216 | 9.51244 | 19.51414 | 17.18665 | Distal Interge | 8 | 126476248 | 126476412 | 165 | 2 | ENSMUST00 | -40386 | Gm6091 | ENSMUSG00000074026.3 |
| chr8 | 27227299 | 27228152 | 9.80357 | 19.4803 | 17.1535 | Promoter | 8 | 27226799 | 27228494 | 1696 | 2 | ENSMUST00 | 342 | Adrb3 | ENSMUSG00000031489.9 |
| chr1 | 90438749 | 90439520 | 9.47659 | 19.4537 | 17.127 | Distal Interge | 1 | 90453394 | 90454501 | 1108 | 2 | ENSMUST00 | 14981 | RP24-154L11 | ENSMUSG00000101997.1 |
| chr11 | 58103618 | 58103927 | 9.78363 | 19.44713 | 17.12045 | Promoter | 11 | 58104153 | 58118594 | 14442 | 1 | ENSMUST00 | -226 | Cnot8 | ENSMUSG00000020515.8 |
| chr4 | 128610227 | 128610819 | 9.78363 | 19.44713 | 17.12045 | Promoter | 4 | 128583547 | 128610098 | 26552 | 2 | ENSMUST00 | -130 | Zscan20 | ENSMUSG00000061894.10 |
| chr8 | 8994672 | 8995271 | 9.78363 | 19.44713 | 17.12045 | Distal Interge | 8 | 8666577 | 8690537 | 23961 | 2 | ENSMUST00 | -304136 | Arglu1 | ENSMUSG00000040459.10 |
| chr7 | 109191384 | 109191964 | 9.76377 | 19.41409 | 17.08753 | Distal Interge | 7 | 109138572 | 109170308 | 31737 | 2 | ENSMUST00 | -21077 | Lmo1 | ENSMUSG00000036111.7 |
| chr15 | 36979672 | 36980234 | 8.01187 | 19.41161 | 17.08515 | Distal Interge | 15 | 36997031 | 37007402 | 10372 | 2 | ENSMUST00 | 27168 | Zfp706 | ENSMUSG00000062397.7 |
| chr16 | 72807879 | 72808982 | 8.01187 | 19.41161 | 17.08515 | Intron (ENSM | 16 | 72663149 | 73046095 | 382947 | 1 | ENSMUST00 | 144731 | Robo1 | ENSMUSG00000022883.10 |
| chr2 | 61815099 | 61816115 | 8.01187 | 19.41161 | 17.08515 | Distal Interge | 2 | 61806273 | 61811801 | 5529 | 1 | ENSMUST00 | 8827 | Tbr1 | ENSMUSG00000035033.10 |
| chr5 | 117360938 | 117362519 | 8.01187 | 19.41161 | 17.08515 | Promoter | 5 | 117361022 | 117370922 | 9901 | 1 | ENSMUST00 | 0 | Wsb2 | ENSMUSG00000029364.10 |
| chr6 | 52164166 | 52165356 | 8.01187 | 19.41161 | 17.08515 | Promoter | 6 | 52162511 | 52164831 | 2321 | 2 | ENSMUST00 | 0 | Hoxa2 | ENSMUSG00000014704.9 |
| chr11 | 88235516 | 88236134 | 8.83539 | 19.39511 | 17.06904 | Distal Interge | 11 | 88204444 | 88210971 | 6528 | 1 | ENSMUST00 | 31073 | Mrps23 | ENSMUSG00000023723.12 |
| chr6 | 37430134 | 37430874 | 8.83539 | 19.39511 | 17.06904 | Intron (ENSM | 6 | 37327255 | 37442146 | 114892 | 2 | ENSMUST00 | 11272 | Creb3l2 | ENSMUSG00000038648.5 |
| chr8 | 116504395 | 116505208 | 8.83539 | 19.39511 | 17.06904 | Promoter | 8 | 116505015 | 116515896 | 10882 | 1 | ENSMUST00 | 0 | Dynlrb2 | ENSMUSG00000034467.6 |
| chr10 | 112271575 | 112272169 | 9.74399 | 19.38118 | 17.0552 | Promoter | 10 | 112271123 | 112466304 | 195182 | 1 | ENSMUST00 | 453 | Kcnc2 | ENSMUSG00000035681.6 |
| chr10 | 80159097 | 80159881 | 9.74399 | 19.38118 | 17.0552 | Distal Interge | 10 | 80165985 | 80170316 | 4332 | 1 | ENSMUST00 | -6104 | Cirbp | ENSMUSG00000045193.8 |
| chr16 | 45724205 | 45724850 | 9.74399 | 19.38118 | 17.0552 | Promoter | 16 | 45711230 | 45724608 | 13379 | 2 | ENSMUST00 | 0 | Tagln3 | ENSMUSG00000022658.10 |
| chr8 | 31113109 | 31113711 | 9.74399 | 19.38118 | 17.0552 | Intron (ENSM | 8 | 31111830 | 31122085 | 10256 | 1 | ENSMUST00 | 1280 | Rnf122 | ENSMUSG00000039328.9 |
| chr4 | 140699593 | 140700679 | 9.72429 | 19.3484 | 17.02254 | Promoter | 4 | 140700541 | 140723220 | 22680 | 1 | ENSMUST00 | 0 | Rcc2 | ENSMUSG00000040945.8 |
| chr17 | 35396016 | 35396639 | 5.13602 | 19.32084 | 16.99509 | Promoter | 17 | 35397330 | 35397800 | 471 | 1 | ENSMUST00 | -691 | H2-Q5 | ENSMUSG00000055413.12 |
| chr11 | 69110491 | 69111428 | 9.70467 | 19.31574 | 16.98999 | Distal Interge | 11 | 69120404 | 69124055 | 3652 | 1 | ENSMUST00 | -8976 | Hes7 | ENSMUSG00000023781.2 |
| chr15 | 5107779 | 5108636 | 9.70467 | 19.31574 | 16.98999 | Promoter | 15 | 5095981 | 5108539 | 12559 | 2 | ENSMUST00 | 0 | Card6 | ENSMUSG00000041849.7 |
| chr7 | 3293062 | 3293649 | 9.68513 | 19.2832 | 16.95758 | 5' UTR | 7 | 3290556 | 3299345 | 8790 | 1 | ENSMUST00 | 2507 | Myadm | ENSMUSG00000068566.7 |
| chr19 | 44882414 | 44882931 | 9.06831 | 19.28032 | 16.9548 | Distal Interge | 19 | 44828493 | 44835938 | 7446 | 2 | ENSMUST00 | -46477 | 1700039E22 | ENSMUSG00000092369.1 |
| chr4 | 43266935 | 43267554 | 9.06831 | 19.28032 | 16.9548 | Promoter | 4 | 43267159 | 43344582 | 77424 | 1 | ENSMUST00 | 0 | Atp8b5 | ENSMUSG00000028457.13 |
| chr8 | 62284049 | 62284795 | 9.06831 | 19.28032 | 16.9548 | Distal Interge | 8 | 62057521 | 62123118 | 65598 | 2 | ENSMUST00 | -160932 | Anxa10 | ENSMUSG00000031635.8 |
| chr9 | 40397036 | 40397372 | 9.06831 | 19.28032 | 16.9548 | Intron (ENSM | 9 | 40317511 | 40392450 | 74940 | 2 | ENSMUST00 | -4587 | Gramd1b | ENSMUSG00000040111.9 |
| chr7 | 126704417 | 126705100 | 7.68819 | 19.26737 | 16.94203 | Promoter | 7 | 126699773 | 126704816 | 5044 | 2 | ENSMUST00 | 0 | Coro1a | ENSMUSG00000030707.10 |
| chr2 | 14873688 | 14874030 | 9.3532 | 19.24536 | 16.92031 | Promoter | 2 | 14873820 | 14986149 | 112330 | 1 | ENSMUST00 | 0 | Cacnb2 | ENSMUSG00000057914.10 |
| chr10 | 126809710 | 126810259 | 9.64628 | 19.2185 | 16.89346 | Distal Interge | 10 | 126840375 | 126840501 | 127 | 2 | ENSMUST00 | 30242 | Gm24179 | ENSMUSG00000089550.1 |
| chr13 | 53285506 | 53286294 | 9.64628 | 19.2185 | 16.89346 | Promoter | 13 | 53109312 | 53285549 | 176238 | 2 | ENSMUST00 | 0 | Ror2 | ENSMUSG00000021464.9 |
| chr13 | 83738662 | 83739103 | 9.64628 | 19.2185 | 16.89346 | Promoter | 13 | 83738863 | 83884194 | 145332 | 1 | ENSMUST00 | 0 | C130071C03 | ENSMUSG00000050334.7 |
| chr14 | 63255684 | 63256194 | 9.64628 | 19.2185 | 16.89346 | Intron (ENSM | 14 | 63198922 | 63245271 | 46350 | 2 | ENSMUST00 | -10414 | Gata4 | ENSMUSG00000021944.10 |
| chr14 | 118436770 | 118437600 | 9.64628 | 19.2185 | 16.89346 | Distal Interge | 14 | 118365878 | 118398170 | 32293 | 1 | ENSMUST00 | 70893 | RP24-66N5.1 | ENSMUSG00000099911.1 |
| chr15 | 73067144 | 73067419 | 9.64628 | 19.2185 | 16.89346 | Distal Interge | 15 | 72852169 | 73061204 | 209036 | 2 | ENSMUST00 | -5941 | Trappc9 | ENSMUSG00000047921.10 |
| chr2 | 75208826 | 75209471 | 9.64628 | 19.2185 | 16.89346 | Intron (ENSM | 2 | 75192277 | 75192759 | 483 | 1 | ENSMUST00 | 16550 | Gm13653 | ENSMUSG00000075276.5 |
| chr3 | 82144916 | 82145857 | 9.64628 | 19.2185 | 16.89346 | Promoter | 3 | 82092428 | 82145877 | 53450 | 2 | ENSMUST00 | 20 | Gucy1a3 | ENSMUSG00000033910.8 |
| chr3 | 18624678 | 18626077 | 9.64628 | 19.2185 | 16.89346 | Distal Interge | 3 | 18533861 | 18534289 | 429 | 1 | ENSMUST00 | 90818 | Gm6369 | ENSMUSG00000040217.6 |
| chr4 | 33586992 | 33587444 | 9.64628 | 19.2185 | 16.89346 | Distal Interge | 4 | 33614832 | 33615138 | 307 | 2 | ENSMUST00 | 27694 | Gm12398 | ENSMUSG00000082909.1 |
| chr7 | 35186257 | 35186627 | 9.64628 | 19.2185 | 16.89346 | Promoter | 7 | 35186385 | 35195449 | 9065 | 1 | ENSMUST00 | 0 | Slc7a10 | ENSMUSG00000030495.7 |
| chr1 | 79823721 | 79824504 | 9.33584 | 19.216 | 16.89141 | Intron (ENSM | 1 | 79796845 | 79821322 | 24478 | 2 | ENSMUST00 | -2400 | Serpine2 | ENSMUSG00000026249.5 |
| chr10 | 71024507 | 71025449 | 9.33584 | 19.216 | 16.89141 | Intron (ENSM | 10 | 71082876 | 71083651 | 776 | 1 | ENSMUST00 | -57427 | Gm15647 | ENSMUSG00000090212.1 |
| chr2 | 154603926 | 154604320 | 9.33584 | 19.216 | 16.89141 | Promoter | 2 | 154585758 | 154603708 | 17951 | 2 | ENSMUST00 | -219 | Pxmp4 | ENSMUSG00000000876.6 |
| chr9 | 58204020 | 58204775 | 9.33584 | 19.216 | 16.89141 | Promoter | 9 | 58196298 | 58204319 | 8022 | 2 | ENSMUST00 | 0 | Islr2 | ENSMUSG00000051243.8 |
| chr11 | 44232741 | 44233137 | 9.62697 | 19.18633 | 16.86223 | Distal Interge | 11 | 44265563 | 44267209 | 1647 | 2 | ENSMUST00 | 34072 | Gm12154 | ENSMUSG00000086738.1 |
| chr1 | 96871709 | 96872488 | 9.01168 | 19.18248 | 16.85847 | Promoter | 1 | 96816270 | 96872171 | 55902 | 2 | ENSMUST00 | 0 | Slco4c1 | ENSMUSG00000040693.7 |
| chr1 | 156298843 | 156299173 | 9.01168 | 19.18248 | 16.85847 | Intron (ENSM | 1 | 156255298 | 156303348 | 48051 | 2 | ENSMUST00 | 4175 | Tdrd5 | ENSMUSG00000060985.10 |
| chr10 | 39926545 | 39927243 | 9.01168 | 19.18248 | 16.85847 | Promoter | 10 | 39920382 | 39926923 | 6542 | 2 | ENSMUST00 | 0 | 2010001E11 | ENSMUSG00000039339.3 |
| chr10 | 17331474 | 17331961 | 9.01168 | 19.18248 | 16.85847 | Distal Interge | 10 | 17116804 | 17116900 | 97 | 1 | ENSMUST00 | 214671 | Gm25382 | ENSMUSG00000088436.1 |
| chr11 | 97669531 | 97669873 | 9.01168 | 19.18248 | 16.85847 | Exon (ENSMU | 11 | 97665001 | 97673548 | 8548 | 1 | ENSMUST00 | 4531 | Mllt6 | ENSMUSG00000038437.6 |
| chr11 | 48853806 | 48854227 | 9.01168 | 19.18248 | 16.85847 | Promoter | 11 | 48843648 | 48853493 | 9846 | 2 | ENSMUST00 | -314 | Gm16170 | ENSMUSG00000086245.1 |
| chr12 | 23189096 | 23189872 | 9.01168 | 19.18248 | 16.85847 | Distal Interge | 12 | 23199780 | 23199889 | 110 | 1 | ENSMUST00 | -9908 | Gm26140 | ENSMUSG00000096441.1 |
| chr13 | 21171663 | 21172300 | 9.01168 | 19.18248 | 16.85847 | Distal Interge | 13 | 21179445 | 21180416 | 972 | 1 | ENSMUST00 | -7145 | Trim27 | ENSMUSG00000021326.6 |
| chr13 | 44673213 | 44674399 | 9.01168 | 19.18248 | 16.85847 | Distal Interge | 13 | 44729474 | 44731574 | 2101 | 1 | ENSMUST00 | -55075 | Jarid2 | ENSMUSG00000038518.10 |
| chr14 | 54475812 | 54476771 | 9.01168 | 19.18248 | 16.85847 | Promoter | 14 | 54476100 | 54480431 | 4332 | 1 | ENSMUST00 | 0 | Rem2 | ENSMUSG00000022176.6 |
| chr14 | 19976398 | 19977442 | 9.01168 | 19.18248 | 16.85847 | Promoter | 14 | 19872559 | 19977249 | 104691 | 2 | ENSMUST00 | 0 | Gng2 | ENSMUSG00000043004.8 |
| chr14 | 104465701 | 104466337 | 9.01168 | 19.18248 | 16.85847 | Promoter | 14 | 104463973 | 104465365 | 1393 | 2 | ENSMUST00 | -337 | Pou4f1 | ENSMUSG00000048349.9 |
| chr15 | 44787524 | 44788224 | 9.01168 | 19.18248 | 16.85847 | Promoter | 15 | 44671859 | 44788063 | 116205 | 2 | ENSMUST00 | 0 | Sybu | ENSMUSG00000022340.9 |
| chr15 | 58869994 | 58870229 | 9.01168 | 19.18248 | 16.85847 | Distal Interge | 15 | 58872649 | 58876781 | 4133 | 1 | ENSMUST00 | -2420 | Trmt12 | ENSMUSG00000037085.7 |
| chr15 | 78927804 | 78928429 | 9.01168 | 19.18248 | 16.85847 | Exon (ENSMU | 15 | 78926725 | 78930465 | 3741 | 1 | ENSMUST00 | 1080 | Lgals1 | ENSMUSG00000068220.5 |
| chr15 | 77366951 | 77367455 | 9.01168 | 19.18248 | 16.85847 | Distal Interge | 15 | 77391702 | 77392381 | 680 | 2 | ENSMUST00 | 24926 | Gm20688 | ENSMUSG00000093458.1 |
| chr17 | 24646297 | 24646674 | 9.01168 | 19.18248 | 16.85847 | Intron (ENSM | 17 | 24639282 | 24644957 | 5676 | 2 | ENSMUST00 | -1341 | Slc9a3r2 | ENSMUSG00000002504.9 |
| chr19 | 59458190 | 59458850 | 9.01168 | 19.18248 | 16.85847 | Promoter | 19 | 59458372 | 59465357 | 6986 | 1 | ENSMUST00 | 0 | Emx2 | ENSMUSG00000043969.4 |
| chr19 | 29325446 | 29325979 | 9.01168 | 19.18248 | 16.85847 | Promoter | 19 | 29321344 | 29325356 | 4013 | 2 | ENSMUST00 | -91 | Insl6 | ENSMUSG00000050957.4 |
| chr19 | 4169169 | 4169501 | 9.01168 | 19.18248 | 16.85847 | Promoter | 19 | 4169566 | 4174284 | 4719 | 1 | ENSMUST00 | -65 | Gm17552 | ENSMUSG00000097233.1 |
| chr2 | 92140454 | 92141347 | 9.01168 | 19.18248 | 16.85847 | Intron (ENSM | 2 | 92184106 | 92361829 | 177724 | 1 | ENSMUST00 | -42759 | Phf21a | ENSMUSG00000058318.10 |
| chr3 | 108536395 | 108537046 | 9.01168 | 19.18248 | 16.85847 | Promoter | 3 | 108455694 | 108536536 | 80843 | 2 | ENSMUST00 | 0 | 5330417C22 | ENSMUSG00000040412.11 |
| chr3 | 27862881 | 27863865 | 9.01168 | 19.18248 | 16.85847 | Downstream | 3 | 27868113 | 27868273 | 161 | 1 | ENSMUST00 | -4248 | Gm26040 | ENSMUSG00000088772.1 |
| chr4 | 41124649 | 41124943 | 9.01168 | 19.18248 | 16.85847 | Promoter | 4 | 41114427 | 41124455 | 10029 | 2 | ENSMUST00 | -195 | Nol6 | ENSMUSG00000028430.9 |
| chr4 | 154855819 | 154856665 | 9.01168 | 19.18248 | 16.85847 | Promoter | 4 | 154856200 | 154867125 | 10926 | 1 | ENSMUST00 | 0 | Ttc34 | ENSMUSG00000046637.2 |
| chr5 | 33430761 | 33431807 | 9.01168 | 19.18248 | 16.85847 | Exon (ENSMU | 5 | 33430734 | 33433976 | 3243 | 2 | ENSMUST00 | 2169 | Nkx1-1 | ENSMUSG00000029112.5 |
| chr5 | 135877533 | 135878293 | 9.01168 | 19.18248 | 16.85847 | Distal Interge | 5 | 135875657 | 135875763 | 107 | 1 | ENSMUST00 | 1877 | Gm22450 | ENSMUSG00000095502.1 |
| chr6 | 83087459 | 83088057 | 9.01168 | 19.18248 | 16.85847 | Promoter | 6 | 83087338 | 83088096 | 759 | 1 | ENSMUST00 | 122 | Lbx2 | ENSMUSG00000034968.3 |
| chr7 | 140908680 | 140909095 | 9.01168 | 19.18248 | 16.85847 | Distal Interge | 7 | 140898943 | 140900446 | 1504 | 2 | ENSMUST00 | -8235 | Cox8b | ENSMUSG00000025488.8 |
| chr7 | 44310247 | 44311494 | 9.01168 | 19.18248 | 16.85847 | Promoter | 7 | 44310253 | 44358351 | 48099 | 1 | ENSMUST00 | 0 | Shank1 | ENSMUSG00000038738.10 |
| chr7 | 127716944 | 127717462 | 9.01168 | 19.18248 | 16.85847 | Exon (ENSMU | 7 | 127712736 | 127718185 | 5450 | 1 | ENSMUST00 | 4209 | Ctf1 | ENSMUSG00000042340.5 |
| chr7 | 90426051 | 90427006 | 9.01168 | 19.18248 | 16.85847 | Promoter | 7 | 90426312 | 90428660 | 2349 | 1 | ENSMUST00 | 0 | Ccdc89 | ENSMUSG00000044362.7 |
| chr8 | 104630488 | 104631317 | 9.01168 | 19.18248 | 16.85847 | Promoter | 8 | 104628066 | 104631321 | 3256 | 2 | ENSMUST00 | 4 | Rrad | ENSMUSG00000031880.6 |
| chr9 | 110735450 | 110737617 | 9.01168 | 19.18248 | 16.85847 | Intron (ENSM | 9 | 110722088 | 110742610 | 20523 | 2 | ENSMUST00 | 4993 | Pth1r | ENSMUSG00000032492.9 |
| chr1 | 74803126 | 74803703 | 7.40884 | 19.17745 | 16.8553 | 3' UTR | 1 | 74793363 | 74803495 | 10133 | 1 | ENSMUST00 | 9764 | Wnt10a | ENSMUSG00000026167.9 |
| chr17 | 70747534 | 70748306 | 9.60774 | 19.15429 | 16.83223 | Intron (ENSM | 17 | 70745329 | 70755582 | 10254 | 2 | ENSMUST00 | 7276 | 5031415H12 | ENSMUSG00000097282.1 |
| chr9 | 37527541 | 37528667 | 9.60774 | 19.15429 | 16.83223 | Promoter | 9 | 37528078 | 37538319 | 10242 | 1 | ENSMUST00 | 0 | Esam | ENSMUSG00000001946.8 |
| chr12 | 69574333 | 69574697 | 8.69484 | 19.14606 | 16.82409 | Downstream | 12 | 69577628 | 69583028 | 5401 | 2 | ENSMUST00 | 8331 | Vcpkmt | ENSMUSG00000049882.9 |
| chr4 | 129445146 | 129445784 | 8.69484 | 19.14606 | 16.82409 | Distal Interge | 4 | 129425765 | 129440853 | 15089 | 2 | ENSMUST00 | -4294 | Zbtb8b | ENSMUSG00000048485.7 |
| chr13 | 12701795 | 12702326 | 6.76115 | 19.13718 | 16.81526 | Promoter | 13 | 12702362 | 12702589 | 228 | 1 | ENSMUST00 | -36 | Gm2399 | ENSMUSG00000078141.2 |
| chr11 | 100618127 | 100618621 | 9.58859 | 19.12236 | 16.80049 | Promoter | 11 | 100619244 | 100626799 | 7556 | 1 | ENSMUST00 | -623 | Nkiras2 | ENSMUSG00000017837.7 |
| chr3 | 130690957 | 130691915 | 6.95105 | 19.11952 | 16.79773 | Distal Interge | 3 | 130695921 | 130709444 | 13524 | 2 | ENSMUST00 | 17529 | Ostc | ENSMUSG00000041084.7 |
| chr5 | 135957292 | 135958267 | 6.95105 | 19.11952 | 16.79773 | Downstream | 5 | 135960822 | 135962271 | 1450 | 2 | ENSMUST00 | 4004 | Srcrb4d | ENSMUSG00000029699.8 |
| chr11 | 100571036 | 100571772 | 9.56952 | 19.09056 | 16.76879 | 3' UTR | 11 | 100574904 | 100581728 | 6825 | 1 | ENSMUST00 | -3132 | Cnp | ENSMUSG00000006782.11 |
| chr18 | 13853881 | 13855401 | 9.56952 | 19.09056 | 16.76879 | Intron (ENSM | 18 | 13687013 | 13972733 | 285721 | 2 | ENSMUST00 | 117332 | Zfp521 | ENSMUSG00000024420.8 |
| chr16 | 23102674 | 23103467 | 9.55051 | 19.05887 | 16.73721 | Distal Interge | 16 | 23107444 | 23113947 | 6504 | 1 | ENSMUST00 | -3977 | Eif4a2 | ENSMUSG00000022884.9 |
| chr9 | 103212791 | 103213324 | 9.49396 | 18.96451 | 16.64326 | Promoter | 9 | 103209025 | 103212006 | 2982 | 2 | ENSMUST00 | -786 | Trf | ENSMUSG00000032554.10 |
| chr1 | 162005066 | 162005708 | 9.16567 | 18.92761 | 16.6065 | Promoter | 1 | 162005172 | 162016494 | 11323 | 1 | ENSMUST00 | 0 | 2810442N19 | ENSMUSG00000090260.1 |
| chr18 | 37747078 | 37747714 | 9.16567 | 18.92761 | 16.6065 | Exon (ENSMU | 18 | 37742094 | 37841873 | 99780 | 1 | ENSMUST00 | 4985 | Pcdhga12 | ENSMUSG00000023036.9 |
| chr18 | 5335054 | 5335296 | 9.16567 | 18.92761 | 16.6065 | Promoter | 18 | 5210031 | 5334439 | 124409 | 2 | ENSMUST00 | -616 | Zfp438 | ENSMUSG00000050945.7 |
| chr3 | 96526862 | 96527448 | 9.16567 | 18.92761 | 16.6065 | Exon (ENSMU | 3 | 96525185 | 96529216 | 4032 | 1 | ENSMUST00 | 1678 | Hfe2 | ENSMUSG00000038403.9 |
| chr11 | 24049418 | 24049666 | 8.55869 | 18.90347 | 16.58273 | Exon (ENSMU | 11 | 24048955 | 24075054 | 26100 | 2 | ENSMUST00 | 25388 | A830031A19 | ENSMUSG00000055010.1 |
| chr14 | 20451636 | 20452257 | 9.45663 | 18.90218 | 16.58146 | Promoter | 14 | 20394190 | 20452225 | 58036 | 2 | ENSMUST00 | 0 | Ttc18 | ENSMUSG00000039543.10 |
| chr17 | 56134411 | 56134877 | 9.45663 | 18.90218 | 16.58146 | Promoter | 17 | 56123085 | 56133852 | 10768 | 2 | ENSMUST00 | -560 | Sema6b | ENSMUSG00000001227.6 |
| chr18 | 35938309 | 35938838 | 9.45663 | 18.90218 | 16.58146 | Distal Interge | 18 | 35952824 | 35952907 | 84 | 1 | ENSMUST00 | -13986 | Gm23882 | ENSMUSG00000080529.1 |
| chr6 | 91843415 | 91844062 | 9.45663 | 18.90218 | 16.58146 | Distal Interge | 6 | 91761509 | 91827250 | 65742 | 2 | ENSMUST00 | -16166 | Grip2 | ENSMUSG00000030098.10 |
| chr7 | 60155560 | 60155798 | 9.45663 | 18.90218 | 16.58146 | Promoter | 7 | 60155538 | 60156170 | 633 | 1 | ENSMUST00 | 23 | Gm7367 | ENSMUSG00000047370.4 |
| chr4 | 126300999 | 126301721 | 8.25476 | 18.86148 | 16.54131 | Promoter | 4 | 126300530 | 126309785 | 9256 | 1 | ENSMUST00 | 470 | Col8a2 | ENSMUSG00000056174.8 |
| chr1 | 75209034 | 75209806 | 8.50221 | 18.80243 | 16.48241 | Promoter | 1 | 75199491 | 75209423 | 9933 | 2 | ENSMUST00 | 0 | Glb1l | ENSMUSG00000026200.8 |
| chr10 | 8755557 | 8757768 | 8.50221 | 18.80243 | 16.48241 | Exon (ENSMU | 10 | 8824011 | 8824147 | 137 | 2 | ENSMUST00 | 66379 | Gm25410 | ENSMUSG00000065692.1 |
| chr10 | 18197199 | 18197524 | 8.50221 | 18.80243 | 16.48241 | Intron (ENSM | 10 | 18129022 | 18210890 | 81869 | 2 | ENSMUST00 | 13366 | Ect2l | ENSMUSG00000071392.4 |
| chr11 | 98963188 | 98964126 | 8.50221 | 18.80243 | 16.48241 | Intron (ENSM | 11 | 98960412 | 98974942 | 14531 | 1 | ENSMUST00 | 2777 | Rara | ENSMUSG00000037992.11 |
| chr11 | 31169407 | 31169966 | 8.50221 | 18.80243 | 16.48241 | Distal Interge | 11 | 31182376 | 31182462 | 87 | 2 | ENSMUST00 | 12496 | Gm17332 | ENSMUSG00000091195.1 |
| chr13 | 24845035 | 24845372 | 8.50221 | 18.80243 | 16.48241 | Promoter | 13 | 24845135 | 24901263 | 56129 | 1 | ENSMUST00 | 0 | D130043K22 | ENSMUSG00000006711.10 |
| chr13 | 21940402 | 21941366 | 8.50221 | 18.80243 | 16.48241 | Exon (ENSMU | 13 | 21945094 | 21960485 | 15392 | 1 | ENSMUST00 | -3728 | Zfp184 | ENSMUSG00000006720.11 |
| chr13 | 37452098 | 37452559 | 8.50221 | 18.80243 | 16.48241 | Promoter | 13 | 37452212 | 37453208 | 997 | 2 | ENSMUST00 | 649 | RP23-21N15. | ENSMUSG00000099847.1 |

| | | | | | | | | | | | | | | | |
|---|---|---|---|---|---|---|---|---|---|---|---|---|---|---|---|
| chr16 | 61377022 | 61377301 | 8.50221 | 18.80243 | 16.48241 | Promoter | 16 | 61377256 | 61377916 | 661 | 1 | ENSMUST00 | 0 | Gm9027 | ENSMUSG00000092029.1 |
| chr2 | 165595042 | 165595754 | 8.50221 | 18.80243 | 16.48241 | Promoter | 2 | 165595032 | 165771719 | 176688 | 1 | ENSMUST00 | 11 | Eya2 | ENSMUSG00000017897.13 |
| chr2 | 132879262 | 132880027 | 8.50221 | 18.80243 | 16.48241 | Promoter | 2 | 132868305 | 132880891 | 12587 | 2 | ENSMUST00 | 864 | Lrrn4 | ENSMUSG00000043110.2 |
| chr2 | 155629170 | 155629637 | 8.50221 | 18.80243 | 16.48241 | Promoter | 2 | 155629060 | 155630384 | 1325 | 1 | ENSMUST00 | 111 | Myh7b | ENSMUSG00000074652.3 |
| chr3 | 85186707 | 85187108 | 8.50221 | 18.80243 | 16.48241 | Intron (ENSM | 3 | 85317519 | 85332030 | 14512 | 1 | ENSMUST00 | -130411 | RP23-94M23 | ENSMUSG00000099995.1 |
| chr6 | 48840058 | 48841305 | 8.50221 | 18.80243 | 16.48241 | Promoter | 6 | 48833811 | 48840988 | 7178 | 2 | ENSMUST00 | 0 | Tmem176b | ENSMUSG00000029810.10 |
| chr7 | 79810915 | 79811522 | 8.50221 | 18.80243 | 16.48241 | Promoter | 7 | 79810727 | 79813431 | 2705 | 1 | ENSMUST00 | 189 | Mesp2 | ENSMUSG00000030543.9 |
| chr7 | 127363457 | 127363955 | 8.50221 | 18.80243 | 16.48241 | Promoter | 7 | 127363428 | 127363809 | 382 | 1 | ENSMUST00 | 30 | Gm10586 | ENSMUSG00000073823.2 |
| chr9 | 56924859 | 56925753 | 8.50221 | 18.80243 | 16.48241 | Exon (ENSMU | 9 | 56917200 | 56928371 | 11172 | 2 | ENSMUST00 | 2618 | Snx33 | ENSMUSG00000032733.5 |
| chr1 | 167692097 | 167692455 | 9.65017 | 18.71579 | 16.39739 | Exon (ENSMU | 1 | 167689557 | 167848741 | 159185 | 1 | ENSMUST00 | 2541 | Lmx1a | ENSMUSG00000026686.9 |
| chr1 | 132212230 | 132212647 | 9.65017 | 18.71579 | 16.39739 | Intron (ENSM | 1 | 132203852 | 132204509 | 658 | 1 | ENSMUST00 | 8379 | Lemd1 | ENSMUSG00000079330.3 |
| chr10 | 77581193 | 77581597 | 9.65017 | 18.71579 | 16.39739 | Promoter | 10 | 77581720 | 77598732 | 17013 | 1 | ENSMUST00 | -123 | Pttg1ip | ENSMUSG00000009291.7 |
| chr10 | 80509583 | 80510090 | 9.65017 | 18.71579 | 16.39739 | Intron (ENSM | 10 | 80494835 | 80517276 | 22442 | 1 | ENSMUST00 | 14749 | Onecut3 | ENSMUSG00000045518.8 |
| chr10 | 111419043 | 111420465 | 9.65017 | 18.71579 | 16.39739 | Distal Interge | 10 | 111473192 | 111496886 | 23695 | 1 | ENSMUST00 | -52727 | Nap1l1 | ENSMUSG00000058799.9 |
| chr11 | 97685345 | 97686413 | 9.65017 | 18.71579 | 16.39739 | Promoter | 11 | 97685826 | 97688621 | 2796 | 1 | ENSMUST00 | 0 | Cisd3 | ENSMUSG00000078695.3 |
| chr12 | 81026673 | 81027354 | 9.65017 | 18.71579 | 16.39739 | Promoter | 12 | 81026808 | 81186414 | 159607 | 1 | ENSMUST00 | 0 | Smoc1 | ENSMUSG00000021136.8 |
| chr12 | 80760255 | 80760661 | 9.65017 | 18.71579 | 16.39739 | Promoter | 12 | 80754046 | 80760541 | 6496 | 2 | ENSMUST00 | 0 | Ccdc177 | ENSMUSG00000062961.6 |
| chr13 | 83518379 | 83518786 | 9.65017 | 18.71579 | 16.39739 | Intron (ENSM | 13 | 83504034 | 83667079 | 163046 | 1 | ENSMUST00 | 14346 | Mef2c | ENSMUSG00000005583.11 |
| chr13 | 36731581 | 36732397 | 9.65017 | 18.71579 | 16.39739 | Promoter | 13 | 36726458 | 36730668 | 4211 | 2 | ENSMUST00 | -914 | Nrn1 | ENSMUSG00000039114.10 |
| chr13 | 111867813 | 111868471 | 9.65017 | 18.71579 | 16.39739 | Promoter | 13 | 111867936 | 111871937 | 4002 | 1 | ENSMUST00 | 0 | Gm15326 | ENSMUSG00000086111.1 |
| chr13 | 58149283 | 58149577 | 9.65017 | 18.71579 | 16.39739 | Promoter | 13 | 58148777 | 58149692 | 916 | 1 | ENSMUST00 | 507 | 1700066J03R | ENSMUSG00000097717.1 |
| chr14 | 62292424 | 62293162 | 9.65017 | 18.71579 | 16.39739 | Promoter | 14 | 62292589 | 62353558 | 60970 | 1 | ENSMUST00 | 0 | Rnaseh2b | ENSMUSG00000021932.8 |
| chr14 | 52862568 | 52862815 | 9.65017 | 18.71579 | 16.39739 | Promoter | 14 | 52861767 | 52862630 | 864 | 2 | ENSMUST00 | 0 | Gm6740 | ENSMUSG00000095313.1 |
| chr15 | 13172083 | 13172625 | 9.65017 | 18.71579 | 16.39739 | Intron (ENSM | 15 | 13034200 | 13173639 | 139440 | 2 | ENSMUST00 | 1014 | Cdh6 | ENSMUSG00000039385.4 |
| chr17 | 45630054 | 45630652 | 9.65017 | 18.71579 | 16.39739 | 3' UTR | 17 | 45620679 | 45620759 | 81 | 1 | ENSMUST00 | 9376 | Gm25008 | ENSMUSG00000065743.1 |
| chr18 | 78048417 | 78048880 | 9.65017 | 18.71579 | 16.39739 | Exon (ENSMU | 18 | 78043614 | 78057395 | 13782 | 2 | ENSMUST00 | 8515 | Siglec15 | ENSMUSG00000091055.2 |
| chr19 | 45541444 | 45541979 | 9.65017 | 18.71579 | 16.39739 | Distal Interge | 19 | 45552275 | 45560531 | 8257 | 2 | ENSMUST00 | 18552 | Poll | ENSMUSG00000025218.6 |
| chr2 | 175469916 | 175470923 | 9.65017 | 18.71579 | 16.39739 | Promoter | 2 | 175469985 | 175479337 | 9353 | 1 | ENSMUST00 | 0 | Gm8923 | ENSMUSG00000078896.6 |
| chr4 | 42121295 | 42122124 | 9.65017 | 18.71579 | 16.39739 | Exon (ENSMU | 4 | 42114817 | 42115917 | 1101 | 1 | ENSMUST00 | 6479 | Gm13304 | ENSMUSG00000073878.3 |
| chr4 | 114843112 | 114843778 | 9.65017 | 18.71579 | 16.39739 | Intron (ENSM | 4 | 114844985 | 114845117 | 133 | 2 | ENSMUST00 | 1339 | Gm23230 | ENSMUSG00000087923.1 |
| chr4 | 42262363 | 42262822 | 9.65017 | 18.71579 | 16.39739 | Distal Interge | 4 | 42255767 | 42256432 | 666 | 1 | ENSMUST00 | 6597 | Ccl21b | ENSMUSG00000095675.1 |
| chr5 | 112394495 | 112395176 | 9.65017 | 18.71579 | 16.39739 | Distal Interge | 5 | 112385448 | 112392213 | 6766 | 2 | ENSMUST00 | -2283 | Asphd2 | ENSMUSG00000029348.7 |
| chr5 | 41763548 | 41765031 | 9.65017 | 18.71579 | 16.39739 | Promoter | 5 | 41761483 | 41764501 | 3019 | 2 | ENSMUST00 | 0 | Nkx3-2 | ENSMUSG00000049691.8 |
| chr5 | 92112124 | 92113154 | 9.65017 | 18.71579 | 16.39739 | Distal Interge | 5 | 92118177 | 92118283 | 107 | 2 | ENSMUST00 | 5129 | Gm24931 | ENSMUSG00000088949.1 |
| chr6 | 47044536 | 47045128 | 9.65017 | 18.71579 | 16.39739 | Promoter | 6 | 47045060 | 47095560 | 50501 | 1 | ENSMUST00 | 0 | Cntnap2 | ENSMUSG00000039419.11 |
| chr7 | 142481649 | 142481992 | 9.65017 | 18.71579 | 16.39739 | Intron (ENSM | 7 | 142483259 | 142489983 | 6725 | 1 | ENSMUST00 | -1267 | Lsp1 | ENSMUSG00000018819.5 |
| chr7 | 70624639 | 70625823 | 9.65017 | 18.71579 | 16.39739 | Distal Interge | 7 | 70752416 | 70752500 | 85 | 1 | ENSMUST00 | -126593 | Gm24880 | ENSMUSG00000089401.1 |
| chrX | 74269889 | 74270229 | 9.65017 | 18.71579 | 16.39739 | Promoter | 20 | 74270812 | 74273135 | 2324 | 1 | ENSMUST00 | -583 | Rpl10 | ENSMUSG00000008682.8 |
| chrX | 38565207 | 38565816 | 9.65017 | 18.71579 | 16.39739 | Promoter | 20 | 38533570 | 38564662 | 31093 | 2 | ENSMUST00 | -546 | Cul4b | ENSMUSG00000031095.10 |
| chrX | 168639404 | 168640281 | 9.65017 | 18.71579 | 16.39739 | Distal Interge | 20 | 168654122 | 168659018 | 4897 | 2 | ENSMUST00 | 18737 | Msl3 | ENSMUSG00000031358.12 |
| chrX | 74115500 | 74115824 | 9.65017 | 18.71579 | 16.39739 | Intron (ENSM | 20 | 74127464 | 74150760 | 23297 | 1 | ENSMUST00 | -11640 | Opn1mw | ENSMUSG00000031394.6 |
| chrX | 66652709 | 66653363 | 9.65017 | 18.71579 | 16.39739 | Promoter | 20 | 66653007 | 66656488 | 3482 | 1 | ENSMUST00 | 0 | Slitrk2 | ENSMUSG00000036790.5 |
| chrX | 12135593 | 12136131 | 9.65017 | 18.71579 | 16.39739 | Intron (ENSM | 20 | 12058981 | 12128400 | 69420 | 2 | ENSMUST00 | -7194 | Bcor | ENSMUSG00000040363.9 |
| chrX | 155622711 | 155623869 | 9.65017 | 18.71579 | 16.39739 | Promoter | 20 | 155573455 | 155623325 | 49871 | 2 | ENSMUST00 | 0 | Ptchd1 | ENSMUSG00000041552.7 |
| chrX | 136868050 | 136868710 | 9.65017 | 18.71579 | 16.39739 | Promoter | 20 | 136858147 | 136868755 | 10609 | 2 | ENSMUST00 | 45 | Rab9b | ENSMUSG00000043463.6 |
| chrX | 101222695 | 101223036 | 9.65017 | 18.71579 | 16.39739 | Promoter | 20 | 101211959 | 101222563 | 10605 | 2 | ENSMUST00 | -133 | Snx12 | ENSMUSG00000046032.11 |
| chrX | 142680507 | 142681054 | 9.65017 | 18.71579 | 16.39739 | Promoter | 20 | 142681406 | 142843494 | 162089 | 1 | ENSMUST00 | -352 | Tmem164 | ENSMUSG00000047045.12 |
| chrX | 103981454 | 103981958 | 9.65017 | 18.71579 | 16.39739 | Promoter | 20 | 103977795 | 103981284 | 3490 | 2 | ENSMUST00 | -171 | Rlim | ENSMUSG00000056537.6 |
| chrX | 103026287 | 103026539 | 9.65017 | 18.71579 | 16.39739 | Distal Interge | 20 | 103037624 | 103038368 | 745 | 1 | ENSMUST00 | -11085 | Gm9143 | ENSMUSG00000080774.1 |
| chrX | 93889430 | 93889846 | 9.65017 | 18.71579 | 16.39739 | Distal Interge | 20 | 93920295 | 93921962 | 1668 | 2 | ENSMUST00 | 32116 | Supt20-ps | ENSMUSG00000082728.1 |
| chrX | 103617079 | 103617732 | 9.65017 | 18.71579 | 16.39739 | Promoter | 20 | 103562897 | 103616954 | 54058 | 2 | ENSMUST00 | -126 | Ftx | ENSMUSG00000086370.3 |
| chrX | 140277527 | 140278398 | 9.65017 | 18.71579 | 16.39739 | Promoter | 20 | 140178698 | 140277862 | 99165 | 2 | ENSMUST00 | 0 | Gm15046 | ENSMUSG00000086441.1 |
| chrX | 38473221 | 38474198 | 9.65017 | 18.71579 | 16.39739 | Distal Interge | 20 | 38476461 | 38482486 | 6026 | 1 | ENSMUST00 | -2263 | Gm7598 | ENSMUSG00000087390.1 |
| chr9 | 28403098 | 28404283 | 9.32825 | 18.68756 | 16.37109 | Promoter | 9 | 28403149 | 28405951 | 2803 | 1 | ENSMUST00 | 0 | Opcml | ENSMUSG00000062257.7 |
| chr11 | 86813863 | 86814818 | 9.63282 | 18.68706 | 16.37064 | Distal Interge | 11 | 86768846 | 86807746 | 38901 | 2 | ENSMUST00 | -6118 | Dhx40 | ENSMUSG00000018425.13 |
| chr17 | 83322614 | 83323137 | 9.63282 | 18.68706 | 16.37064 | Distal Interge | 17 | 83350931 | 83480361 | 129431 | 1 | ENSMUST00 | -27794 | Eml4 | ENSMUSG00000032624.10 |
| chr7 | 45643297 | 45644009 | 9.63282 | 18.68706 | 16.37064 | Promoter | 7 | 45644384 | 45645009 | 626 | 1 | ENSMUST00 | -375 | Mamstr | ENSMUSG00000042918.10 |
| chr6 | 86480441 | 86481097 | 8.42674 | 18.66703 | 16.35069 | Promoter | 6 | 86470805 | 86480770 | 9966 | 2 | ENSMUST00 | 0 | A430078I02F | ENSMUSG00000085589.2 |
| chr3 | 65441666 | 65442179 | 9.0016 | 18.64841 | 16.33221 | Distal Interge | 3 | 65393337 | 65395184 | 1848 | 1 | ENSMUST00 | 48330 | Gm5540 | ENSMUSG00000048916.7 |
| chr10 | 128410044 | 128410678 | 9.59257 | 18.62041 | 16.30457 | Promoter | 10 | 128401910 | 128410551 | 8642 | 2 | ENSMUST00 | 0 | Nabp2 | ENSMUSG00000025374.8 |
| chr18 | 74440484 | 74441307 | 9.59257 | 18.62041 | 16.30457 | Promoter | 18 | 74440936 | 74617023 | 176088 | 1 | ENSMUST00 | 0 | Myo5b | ENSMUSG00000025885.13 |
| chr7 | 143977800 | 143978503 | 9.59257 | 18.62041 | 16.30457 | Distal Interge | 7 | 144001928 | 144424494 | 422567 | 1 | ENSMUST00 | -23425 | Shank2 | ENSMUSG00000037541.14 |
| chr3 | 96458044 | 96458592 | 6.92566 | 18.60103 | 16.28545 | Downstream | 3 | 96460024 | 96460189 | 166 | 2 | ENSMUST00 | 1597 | Rnu1b6 | ENSMUSG00000065773.1 |
| chr10 | 115586383 | 115588056 | 9.27429 | 18.59722 | 16.28163 | Promoter | 10 | 115450311 | 115587780 | 137470 | 2 | ENSMUST00 | 0 | Lgr5 | ENSMUSG00000020140.10 |
| chr11 | 78169180 | 78169588 | 9.27429 | 18.59722 | 16.28163 | Promoter | 11 | 78168972 | 78169828 | 857 | 2 | ENSMUST00 | 240 | Nek8 | ENSMUSG00000017405.9 |
| chr11 | 20561260 | 20561761 | 9.27429 | 18.59722 | 16.28163 | Intron (ENSM | 11 | 20574567 | 20575189 | 623 | 2 | ENSMUST00 | 13428 | Gm12035 | ENSMUSG00000083555.1 |
| chr14 | 72983242 | 72983749 | 9.27429 | 18.59722 | 16.28163 | Distal Interge | 14 | 73029128 | 73049114 | 19987 | 2 | ENSMUST00 | 65365 | Cysltr2 | ENSMUSG00000033470.6 |
| chr4 | 83051315 | 83052311 | 9.27429 | 18.59722 | 16.28163 | Promoter | 4 | 82897920 | 83052167 | 154248 | 2 | ENSMUST00 | 0 | Frem1 | ENSMUSG00000059049.9 |
| chr6 | 8336756 | 8337247 | 9.27429 | 18.59722 | 16.28163 | Intron (ENSM | 6 | 8354056 | 8364136 | 10081 | 2 | ENSMUST00 | 26889 | Gm16055 | ENSMUSG00000089890.1 |
| chr13 | 48663014 | 48663484 | 9.57257 | 18.5873 | 16.27215 | Promoter | 13 | 48662998 | 48666507 | 3510 | 1 | ENSMUST00 | 17 | Barx1 | ENSMUSG00000021381.4 |
| chr8 | 114707322 | 114708054 | 9.57257 | 18.5873 | 16.27215 | Promoter | 8 | 114698439 | 114708006 | 9568 | 2 | ENSMUST00 | 0 | Gm16118 | ENSMUSG00000089797.1 |
| chr7 | 99916997 | 99917670 | 9.55265 | 18.55431 | 16.2393 | 3' UTR | 7 | 99935278 | 99965734 | 30457 | 2 | ENSMUST00 | 48064 | Rnf169 | ENSMUSG00000058761.4 |
| chr1 | 86015817 | 86016003 | 8.08052 | 18.53858 | 16.22368 | Promoter | 1 | 86014403 | 86015886 | 1484 | 2 | ENSMUST00 | 0 | RP24-139E1! | ENSMUSG00000099342.1 |
| chr14 | 75828756 | 75829555 | 8.08052 | 18.53858 | 16.22368 | Distal Interge | 14 | 75843379 | 75844964 | 1586 | 2 | ENSMUST00 | 15409 | Gm4285 | ENSMUSG00000085829.1 |
| chr14 | 5879919 | 5880443 | 8.08052 | 18.53858 | 16.22368 | Promoter | 14 | 5857918 | 5880382 | 22465 | 2 | ENSMUST00 | 0 | Gm8237 | ENSMUSG00000090707.3 |
| chr15 | 37233878 | 37234588 | 8.08052 | 18.53858 | 16.22368 | Promoter | 15 | 37231939 | 37233309 | 1371 | 2 | ENSMUST00 | -570 | Gm16136 | ENSMUSG00000089804.1 |
| chr17 | 74993909 | 74994622 | 8.08052 | 18.53858 | 16.22368 | Distal Interge | 17 | 75005568 | 75391767 | 386200 | 1 | ENSMUST00 | -10946 | Ltbp1 | ENSMUSG00000001870.10 |
| chr2 | 28640640 | 28640883 | 8.08052 | 18.53858 | 16.22368 | Promoter | 2 | 28641228 | 28672507 | 31280 | 1 | ENSMUST00 | -345 | Tsc1 | ENSMUSG00000026812.11 |
| chr2 | 93645867 | 93646496 | 8.08052 | 18.53858 | 16.22368 | Intron (ENSM | 2 | 93642388 | 93681339 | 38952 | 1 | ENSMUST00 | 3480 | Alx4 | ENSMUSG00000040310.7 |
| chr2 | 148752089 | 148752484 | 8.08052 | 18.53858 | 16.22368 | Intron (ENSM | 2 | 148750396 | 148755378 | 4983 | 1 | ENSMUST00 | 1694 | Cstl1 | ENSMUSG00000055177.10 |
| chr2 | 160647655 | 160648146 | 8.08052 | 18.53858 | 16.22368 | Exon (ENSMU | 2 | 160645980 | 160650107 | 4128 | 1 | ENSMUST00 | 1676 | Top1 | ENSMUSG00000070544.6 |
| chr5 | 144545450 | 144546379 | 8.08052 | 18.53858 | 16.22368 | Promoter | 5 | 144545887 | 144557478 | 11592 | 1 | ENSMUST00 | 0 | Nptx2 | ENSMUSG00000059991.6 |
| chr6 | 128399799 | 128400399 | 8.08052 | 18.53858 | 16.22368 | Promoter | 6 | 128399970 | 128408932 | 8963 | 1 | ENSMUST00 | 0 | Nrip2 | ENSMUSG00000001520.7 |
| chr6 | 128764324 | 128764808 | 8.08052 | 18.53858 | 16.22368 | Distal Interge | 6 | 128780043 | 128788486 | 8444 | 2 | ENSMUST00 | 23678 | Klrb1c | ENSMUSG00000030325.11 |
| chr6 | 93065854 | 93066981 | 8.08052 | 18.53858 | 16.22368 | Intron (ENSM | 6 | 93137004 | 93137990 | 987 | 2 | ENSMUST00 | 71009 | Gm5313 | ENSMUSG00000097996.1 |
| chr7 | 99818089 | 99818903 | 8.08052 | 18.53858 | 16.22368 | Intron (ENSM | 7 | 99811439 | 99828417 | 16979 | 2 | ENSMUST00 | 9514 | Neu3 | ENSMUSG00000035239.6 |
| chr7 | 29000866 | 29001492 | 8.08052 | 18.53858 | 16.22368 | Exon (ENSMU | 7 | 28982854 | 29003279 | 20426 | 1 | ENSMUST00 | 18013 | Map4k1 | ENSMUSG00000037337.10 |
| chr9 | 44509623 | 44510396 | 8.08052 | 18.53858 | 16.22368 | 3' UTR | 9 | 44499136 | 44510388 | 11253 | 1 | ENSMUST00 | 10488 | Bcl9l | ENSMUSG00000063382.5 |
| chr9 | 72766897 | 72767190 | 8.08052 | 18.53858 | 16.22368 | Distal Interge | 9 | 72774063 | 72776225 | 2163 | 2 | ENSMUST00 | 9035 | Gm27204 | ENSMUSG00000098294.1 |
| chr15 | 94590042 | 94590396 | 9.53282 | 18.52146 | 16.20732 | Promoter | 15 | 94584443 | 94589889 | 5447 | 2 | ENSMUST00 | -154 | Twf1 | ENSMUSG00000022451.7 |
| chr2 | 91923700 | 91924207 | 9.53282 | 18.52146 | 16.20732 | Distal Interge | 2 | 91920528 | 91920928 | 401 | 2 | ENSMUST00 | -2773 | Gm10805 | ENSMUSG00000075036.1 |
| chr1 | 184727398 | 184728103 | 9.22042 | 18.50693 | 16.19293 | 3' UTR | 1 | 184727147 | 184732451 | 5305 | 2 | ENSMUST00 | 4348 | Hlx | ENSMUSG00000039377.6 |
| chr1 | 69241905 | 69242286 | 9.22042 | 18.50693 | 16.19293 | Distal Interge | 1 | 69130914 | 69131299 | 386 | 2 | ENSMUST00 | -110607 | Gm8840 | ENSMUSG00000084332.1 |
| chr10 | 87058104 | 87058755 | 9.22042 | 18.50693 | 16.19293 | Promoter | 10 | 87058046 | 87230588 | 172543 | 1 | ENSMUST00 | 59 | 1700113H08I | ENSMUSG00000047129.6 |
| chr10 | 49782569 | 49783705 | 9.22042 | 18.50693 | 16.19293 | Promoter | 10 | 49099843 | 49783259 | 683417 | 2 | ENSMUST00 | 0 | Grik2 | ENSMUSG00000056073.9 |
| chr10 | 90728525 | 90728900 | 9.22042 | 18.50693 | 16.19293 | Intron (ENSM | 10 | 90829111 | 90941465 | 112355 | 1 | ENSMUST00 | -100211 | Anks1b | ENSMUSG00000058589.9 |
| chr10 | 67285509 | 67285778 | 9.22042 | 18.50693 | 16.19293 | Promoter | 10 | 67266689 | 67285305 | 18617 | 2 | ENSMUST00 | -205 | Nrbf2 | ENSMUSG00000075000.6 |
| chr11 | 100692044 | 100692617 | 9.22042 | 18.50693 | 16.19293 | Promoter | 11 | 100637797 | 100692455 | 54659 | 2 | ENSMUST00 | 0 | Zfp385c | ENSMUSG00000014198.10 |
| chr11 | 6605345 | 6605981 | 9.22042 | 18.50693 | 16.19293 | Promoter | 11 | 6597823 | 6606053 | 8231 | 2 | ENSMUST00 | 72 | Nacad | ENSMUSG00000041073.5 |
| chr11 | 96282719 | 96283599 | 9.22042 | 18.50693 | 16.19293 | Promoter | 11 | 96282725 | 96284693 | 1969 | 1 | ENSMUST00 | 0 | Hoxb8 | ENSMUSG00000056648.5 |
| chr14 | 14294473 | 14294873 | 9.22042 | 18.50693 | 16.19293 | Distal Interge | 14 | 14328113 | 14329066 | 954 | 1 | ENSMUST00 | -33240 | Olfr31 | ENSMUSG00000072707.1 |
| chr15 | 43869619 | 43870189 | 9.22042 | 18.50693 | 16.19293 | Promoter | 15 | 43866695 | 43870029 | 3335 | 2 | ENSMUST00 | 0 | Tmem74 | ENSMUSG00000054409.4 |
| chr15 | 102073251 | 102073636 | 9.22042 | 18.50693 | 16.19293 | Promoter | 15 | 102073773 | 102097172 | 23400 | 1 | ENSMUST00 | -137 | Eif4b | ENSMUSG00000058655.8 |
| chr16 | 32154366 | 32154842 | 9.22042 | 18.50693 | 16.19293 | Intron (ENSM | 16 | 32144827 | 32148118 | 3292 | 2 | ENSMUST00 | -6249 | Nrros | ENSMUSG00000052384.8 |
| chr17 | 26849572 | 26850720 | 9.22042 | 18.50693 | 16.19293 | Distal Interge | 17 | 26838664 | 26841565 | 2902 | 2 | ENSMUST00 | -8008 | Nkx2-5 | ENSMUSG00000015579.4 |
| chr17 | 43534905 | 43535224 | 9.22042 | 18.50693 | 16.19293 | 3' UTR | 17 | 43534174 | 43543639 | 9466 | 2 | ENSMUST00 | 8415 | Ankrd66 | ENSMUSG00000096140.1 |
| chr19 | 43616262 | 43616660 | 9.22042 | 18.50693 | 16.19293 | Distal Interge | 19 | 43612325 | 43615892 | 3568 | 1 | ENSMUST00 | 3938 | Nkx2-3 | ENSMUSG00000044220.13 |
| chr19 | 25200266 | 25201136 | 9.22042 | 18.50693 | 16.19293 | 3' UTR | 19 | 25166085 | 25166216 | 132 | 2 | ENSMUST00 | -34051 | Gm24252 | ENSMUSG00000065634.1 |
| chr2 | 92915105 | 92915360 | 9.22042 | 18.50693 | 16.19293 | Promoter | 2 | 92915098 | 92956058 | 40961 | 1 | ENSMUST00 | 8 | Syt13 | ENSMUSG00000027220.2 |
| chr2 | 174077848 | 174078679 | 9.22042 | 18.50693 | 16.19293 | Promoter | 2 | 174077066 | 174096967 | 19902 | 1 | ENSMUST00 | 783 | Stx16 | ENSMUSG00000027522.10 |
| chr2 | 76070342 | 76070742 | 9.22042 | 18.50693 | 16.19293 | Intron (ENSM | 2 | 75989141 | 76006043 | 16903 | 2 | ENSMUST00 | -64300 | Pde11a | ENSMUSG00000075270.6 |
| chr3 | 65927064 | 65927955 | 9.22042 | 18.50693 | 16.19293 | Distal Interge | 3 | 65946782 | 65947998 | 1217 | 2 | ENSMUST00 | 20043 | Ccnl1 | ENSMUSG00000027829.10 |
| chr4 | 106827796 | 106828358 | 9.22042 | 18.50693 | 16.19293 | Distal Interge | 4 | 106841761 | 106847920 | 6160 | 2 | ENSMUST00 | 19562 | Gm12746 | ENSMUSG00000086940.1 |
| chr5 | 90246747 | 90247713 | 9.22042 | 18.50693 | 16.19293 | Intron (ENSM | 5 | 90214724 | 90224001 | 9278 | 2 | ENSMUST00 | -22747 | Cox18 | ENSMUSG00000035505.9 |
| chr6 | 119448532 | 119449103 | 9.22042 | 18.50693 | 16.19293 | Promoter | 6 | 119432531 | 119449386 | 16856 | 2 | ENSMUST00 | 283 | Wnt5b | ENSMUSG00000030170.9 |
| chr6 | 126645524 | 126645935 | 9.22042 | 18.50693 | 16.19293 | Promoter | 6 | 126636463 | 126645801 | 9339 | 2 | ENSMUST00 | 0 | Kcna1 | ENSMUSG00000047976.3 |
| chr8 | 70489360 | 70489855 | 9.22042 | 18.50693 | 16.19293 | Distal Interge | 8 | 70483876 | 70487358 | 3483 | 2 | ENSMUST00 | -2003 | Tmem59l | ENSMUSG00000035964.7 |
| chr9 | 32542145 | 32542941 | 9.22042 | 18.50693 | 16.19293 | Promoter | 9 | 32428059 | 32542861 | 114803 | 2 | ENSMUST00 | 0 | Fli1 | ENSMUSG00000016087.8 |
| chr9 | 102934291 | 102934768 | 9.22042 | 18.50693 | 16.19293 | Distal Interge | 9 | 102898555 | 102899546 | 992 | 1 | ENSMUST00 | 35737 | Ryk | ENSMUSG00000032547.7 |
| chr9 | 42520502 | 42521106 | 9.22042 | 18.50693 | 16.19293 | Promoter | 9 | 42520773 | 42520831 | 59 | 2 | ENSMUST00 | 0 | Gm24420 | ENSMUSG00000093745.1 |
| chrX | 52790905 | 52791508 | 9.22042 | 18.50693 | 16.19293 | Promoter | 20 | 52791200 | 52799468 | 8269 | 1 | ENSMUST00 | 0 | Ccdc160 | ENSMUSG00000073207.1 |
| chr4 | 88438697 | 88439007 | 8.62133 | 18.50308 | 16.19081 | Promoter | 4 | 88396144 | 88438915 | 42772 | 2 | ENSMUST00 | 0 | Ptplad2 | ENSMUSG00000028497.7 |
| chr7 | 19295677 | 19295972 | 8.62133 | 18.50308 | 16.19081 | 3' UTR | 7 | 19291099 | 19296159 | 5061 | 1 | ENSMUST00 | 4579 | Rtn2 | ENSMUSG00000030401.10 |
| chr1 | 25829852 | 25830715 | 9.47381 | 18.42369 | 16.11179 | Promoter | 1 | 25067550 | 25829707 | 762158 | 2 | ENSMUST00 | -146 | Bai3 | ENSMUSG00000033569.12 |
| chr4 | 139578786 | 139580030 | 9.47381 | 18.42369 | 16.11179 | Intron (ENSM | 4 | 139574720 | 139620382 | 45663 | 1 | ENSMUST00 | 4067 | Iffo2 | ENSMUSG00000041025.10 |
| chr4 | 133494078 | 133494613 | 9.16823 | 18.41938 | 16.10758 | Intron (ENSM | 4 | 133491118 | 133497719 | 6602 | 2 | ENSMUST00 | 3106 | Trnp1 | ENSMUSG00000056596.8 |
| chr1 | 78537071 | 78538085 | 9.4543 | 18.39136 | 16.07959 | Promoter | 1 | 78537628 | 78538173 | 546 | 1 | ENSMUST00 | 0 | Mogat1 | ENSMUSG00000012187.8 |
| chr13 | 43864248 | 43864590 | 8.8433 | 18.37784 | 16.06619 | Distal Interge | 13 | 43872802 | 43872909 | 108 | 1 | ENSMUST00 | -8212 | Gm23117 | ENSMUSG00000089107.1 |

| | | | | | | | | | | | | | | | |
|---|---|---|---|---|---|---|---|---|---|---|---|---|---|---|---|
| chr19 | 5727445 | 5727958 | 8.8433 | 18.37784 | 16.06619 | Promoter | 19 | 5727206 | 5731627 | 4422 | 1 | ENSMUST00 | 240 | Gm16538 | ENSMUSG00000089766.1 |
| chr11 | 53762730 | 53763974 | 7.72573 | 18.36004 | 16.04866 | Promoter | 11 | 53763597 | 53763654 | 58 | 2 | ENSMUST00 | 0 | Mir7671 | ENSMUSG00000099153.1 |
| chr12 | 58213585 | 58215060 | 7.72573 | 18.36004 | 16.04866 | 3' UTR | 12 | 58211804 | 58214444 | 2641 | 1 | ENSMUST00 | 1782 | Sstr1 | ENSMUSG00000035431.4 |
| chr14 | 5961139 | 5961930 | 7.72573 | 18.36004 | 16.04866 | Promoter | 14 | 5939225 | 5961745 | 22521 | 2 | ENSMUST00 | 0 | Gm3248 | ENSMUSG00000091275.1 |
| chr18 | 56754441 | 56755308 | 7.72573 | 18.36004 | 16.04866 | Distal Interge | 18 | 56769717 | 56801688 | 31972 | 1 | ENSMUST00 | -14409 | Gm15345 | ENSMUSG00000087597.1 |
| chr4 | 16168181 | 16169057 | 7.72573 | 18.36004 | 16.04866 | Intron (ENSM | 4 | 16164110 | 16266225 | 102116 | 1 | ENSMUST00 | 4072 | A530072M11 | ENSMUSG00000085112.1 |
| chr5 | 125003372 | 125003903 | 7.72573 | 18.36004 | 16.04866 | Promoter | 5 | 125003447 | 125012547 | 9101 | 1 | ENSMUST00 | 0 | Fam101a | ENSMUSG00000037962.3 |
| chr17 | 37197061 | 37197654 | 9.43487 | 18.35915 | 16.04816 | Promoter | 17 | 37196707 | 37198011 | 1305 | 2 | ENSMUST00 | 357 | Olfr94 | ENSMUSG00000043827.6 |
| chr12 | 108370838 | 108371468 | 9.41552 | 18.32708 | 16.0162 | Promoter | 12 | 108370957 | 108508538 | 137582 | 1 | ENSMUST00 | 0 | Eml1 | ENSMUSG00000058070.9 |
| chr18 | 25753044 | 25753982 | 9.41552 | 18.32708 | 16.0162 | Promoter | 18 | 25477632 | 25753983 | 276352 | 2 | ENSMUST00 | 1 | Celf4 | ENSMUSG00000024268.10 |
| chr1 | 6358937 | 6359468 | 9.09886 | 18.30288 | 15.9921 | Promoter | 1 | 6359331 | 6394731 | 35401 | 1 | ENSMUST00 | 0 | Fam150a | ENSMUSG00000087247.2 |
| chr1 | 186702429 | 186703040 | 9.09886 | 18.30288 | 15.9921 | Intron (ENSM | 1 | 186705695 | 186707033 | 1339 | 1 | ENSMUST00 | -2655 | RP24-276N2 | ENSMUSG00000100393.1 |
| chr11 | 70228749 | 70229761 | 9.09886 | 18.30288 | 15.9921 | Promoter | 11 | 70224128 | 70229739 | 5612 | 2 | ENSMUST00 | 0 | Bcl6b | ENSMUSG00000000317.6 |
| chr15 | 73067860 | 73068203 | 9.09886 | 18.30288 | 15.9921 | Distal Interge | 15 | 72852169 | 73061204 | 209036 | 2 | ENSMUST00 | -6657 | Trappc9 | ENSMUSG00000047921.10 |
| chr19 | 10818330 | 10818838 | 9.09886 | 18.30288 | 15.9921 | Intron (ENSM | 19 | 10794564 | 10829856 | 35293 | 2 | ENSMUST00 | 11018 | Cd6 | ENSMUSG00000024670.11 |
| chr19 | 44719942 | 44720590 | 9.09886 | 18.30288 | 15.9921 | Distal Interge | 19 | 44692367 | 44695275 | 2909 | 2 | ENSMUST00 | -24668 | Gm26644 | ENSMUSG00000097683.1 |
| chr2 | 27496392 | 27497064 | 9.09886 | 18.30288 | 15.9921 | Intron (ENSM | 2 | 27449269 | 27507662 | 58394 | 2 | ENSMUST00 | 10598 | Brd3 | ENSMUSG00000026918.11 |
| chr4 | 128924847 | 128925603 | 9.09886 | 18.30288 | 15.9921 | Distal Interge | 4 | 128936005 | 128940195 | 4191 | 1 | ENSMUST00 | -10402 | Gm15904 | ENSMUSG00000085961.1 |
| chr5 | 137533704 | 137533910 | 9.09886 | 18.30288 | 15.9921 | Promoter | 5 | 137528127 | 137533510 | 5384 | 2 | ENSMUST00 | -195 | Gnb2 | ENSMUSG00000029713.10 |
| chr15 | 102934973 | 102935273 | 9.39625 | 18.29513 | 15.98494 | Distal Interge | 15 | 102936757 | 102938609 | 1853 | 1 | ENSMUST00 | -1484 | Hoxc12 | ENSMUSG00000050328.2 |
| chr19 | 58970689 | 58971062 | 9.39625 | 18.29513 | 15.98494 | 3' UTR | 19 | 58943425 | 58971419 | 27995 | 1 | ENSMUST00 | 27265 | Eno4 | ENSMUSG00000048029.7 |
| chr4 | 135573178 | 135573715 | 9.39625 | 18.29513 | 15.98494 | Promoter | 4 | 135541888 | 135573630 | 31743 | 2 | ENSMUST00 | 0 | Grhl3 | ENSMUSG00000037188.7 |
| chr2 | 147084593 | 147085562 | 9.08168 | 18.274 | 15.96417 | Promoter | 2 | 147083316 | 147085445 | 2130 | 2 | ENSMUST00 | 0 | Nkx2-4 | ENSMUSG00000054160.2 |
| chr6 | 3182767 | 3183486 | 5.49417 | 18.26307 | 15.95337 | Distal Interge | 6 | 3107510 | 3108401 | 892 | 2 | ENSMUST00 | -74367 | Vmn1r-ps6 | ENSMUSG00000098936.1 |
| chr15 | 103005091 | 103006170 | 8.48197 | 18.2582 | 15.9485 | Intron (ENSM | 15 | 103009573 | 103011881 | 2309 | 1 | ENSMUST00 | -3403 | Hoxc6 | ENSMUSG00000001661.5 |
| chr11 | 80289736 | 80290384 | 7.42308 | 18.24625 | 15.93662 | Distal Interge | 11 | 80300912 | 80355955 | 55044 | 1 | ENSMUST00 | -10528 | Rhbdl3 | ENSMUSG00000017692.8 |
| chr14 | 34234723 | 34235153 | 7.42308 | 18.24625 | 15.93662 | Downstream | 14 | 34220047 | 34230421 | 10375 | 1 | ENSMUST00 | 14677 | Syt15 | ENSMUSG00000041479.8 |
| chr9 | 96258773 | 96260315 | 7.42308 | 18.24625 | 15.93662 | Promoter | 9 | 96259253 | 96318981 | 59729 | 1 | ENSMUST00 | 0 | Tfdp2 | ENSMUSG00000032411.10 |
| chr6 | 108456651 | 108456897 | 9.04751 | 18.21654 | 15.90722 | Promoter | 6 | 108456183 | 108456536 | 354 | 1 | ENSMUST00 | 469 | Gm27813 | ENSMUSG00000098674.1 |
| chr19 | 34607434 | 34608292 | 6.55627 | 18.20592 | 15.89663 | Promoter | 19 | 34607970 | 34613401 | 5432 | 1 | ENSMUST00 | 0 | I830012O16 | ENSMUSG00000062488.9 |
| chr14 | 27237740 | 27238500 | 7.16187 | 18.18295 | 15.87379 | Promoter | 14 | 27238039 | 27403904 | 165866 | 1 | ENSMUST00 | 0 | Arhgef3 | ENSMUSG00000021895.9 |
| chr6 | 48841432 | 48841971 | 7.16187 | 18.18295 | 15.87379 | Promoter | 6 | 48841483 | 48845364 | 3882 | 1 | ENSMUST00 | 0 | Tmem176a | ENSMUSG00000023367.9 |
| chr1 | 130878161 | 130878614 | 9.31996 | 18.16857 | 15.8595 | Promoter | 1 | 130878331 | 130880790 | 2460 | 1 | ENSMUST00 | 0 | Faim3 | ENSMUSG00000042474.6 |
| chr10 | 99204551 | 99205011 | 9.31996 | 18.16857 | 15.8595 | Distal Interge | 10 | 99192892 | 99196850 | 3959 | 1 | ENSMUST00 | 11660 | Poc1b | ENSMUSG00000019952.9 |
| chr19 | 46131632 | 46132138 | 9.31996 | 18.16857 | 15.8595 | Promoter | 19 | 46131897 | 46135307 | 3411 | 1 | ENSMUST00 | 0 | Elovl3 | ENSMUSG00000038754.4 |
| chr12 | 55464981 | 55465923 | 6.93414 | 18.15989 | 15.85092 | Distal Interge | 12 | 55489411 | 55492647 | 3237 | 2 | ENSMUST00 | 26724 | Nfkbia | ENSMUSG00000021025.7 |
| chr5 | 38479930 | 38480871 | 6.93414 | 18.15989 | 15.85092 | Promoter | 5 | 38349273 | 38480306 | 131034 | 2 | ENSMUST00 | 0 | Slc2a9 | ENSMUSG00000005107.8 |
| chr15 | 103002717 | 103003431 | 9.30108 | 18.13724 | 15.82832 | Intron (ENSM | 15 | 103000529 | 103010006 | 9478 | 1 | ENSMUST00 | 2189 | Hoxc6 | ENSMUSG00000001661.5 |
| chr2 | 164431125 | 164431530 | 9.30108 | 18.13724 | 15.82832 | Exon (ENSMU | 2 | 164424247 | 164443186 | 18940 | 2 | ENSMUST00 | 11656 | Sdc4 | ENSMUSG00000017009.3 |
| chr4 | 59784178 | 59784481 | 9.30108 | 18.13724 | 15.82832 | Promoter | 4 | 59769637 | 59783866 | 14230 | 2 | ENSMUST00 | -313 | Inip | ENSMUSG00000038544.9 |
| chr14 | 67233896 | 67234994 | 8.69047 | 18.11543 | 15.80663 | Promoter | 14 | 67234644 | 67428687 | 194044 | 1 | ENSMUST00 | 0 | Ebf2 | ENSMUSG00000022053.8 |
| chr10 | 78159384 | 78159690 | 9.28227 | 18.10602 | 15.7974 | Downstream | 10 | 78162067 | 78169768 | 7702 | 2 | ENSMUST00 | 10078 | D10Jhu81e | ENSMUSG00000053329.6 |
| chr14 | 19763469 | 19764453 | 9.28227 | 18.10602 | 15.7974 | Exon (ENSMU | 14 | 19751257 | 19811787 | 60531 | 1 | ENSMUST00 | 12213 | Nid2 | ENSMUSG00000021806.3 |
| chr15 | 102955674 | 102956517 | 9.24489 | 18.04396 | 15.73558 | Exon (ENSMU | 15 | 102954427 | 102957708 | 3282 | 1 | ENSMUST00 | 1248 | Hoxc11 | ENSMUSG00000001656.3 |
| chr7 | 30465371 | 30465955 | 9.24489 | 18.04396 | 15.73558 | Promoter | 7 | 30462164 | 30465591 | 3428 | 2 | ENSMUST00 | 0 | Nphs1os | ENSMUSG00000087009.1 |
| chr10 | 96620434 | 96621319 | 8.63619 | 18.02194 | 15.71366 | 3' UTR | 10 | 96617001 | 96622813 | 5813 | 1 | ENSMUST00 | 3434 | Btg1 | ENSMUSG00000036478.7 |
| chr10 | 127749979 | 127750560 | 8.63619 | 18.02194 | 15.71366 | Exon (ENSMU | 10 | 127747276 | 127751732 | 4457 | 2 | ENSMUST00 | 1172 | Gpr182 | ENSMUSG00000058396.7 |
| chr10 | 93432575 | 93432950 | 8.63619 | 18.02194 | 15.71366 | Promoter | 10 | 93433107 | 93433162 | 56 | 1 | ENSMUST00 | -157 | Mir7688 | ENSMUSG00000098679.1 |
| chr11 | 77930946 | 77931532 | 8.63619 | 18.02194 | 15.71366 | Promoter | 11 | 77931012 | 77979040 | 48029 | 1 | ENSMUST00 | 0 | Sez6 | ENSMUSG00000000632.8 |
| chr11 | 86472949 | 86473597 | 8.63619 | 18.02194 | 15.71366 | Distal Interge | 11 | 86484657 | 86499025 | 14369 | 1 | ENSMUST00 | -11060 | Rnft1 | ENSMUSG00000020521.7 |
| chr11 | 118202274 | 118203011 | 8.63619 | 18.02194 | 15.71366 | Intron (ENSM | 11 | 118204712 | 118205179 | 468 | 2 | ENSMUST00 | 2168 | Gm11737 | ENSMUSG00000080810.1 |
| chr12 | 83597366 | 83597737 | 8.63619 | 18.02194 | 15.71366 | Promoter | 12 | 83546941 | 83597147 | 50207 | 2 | ENSMUST00 | -220 | Zfyve1 | ENSMUSG00000042628.7 |
| chr12 | 54313015 | 54313524 | 8.63619 | 18.02194 | 15.71366 | Distal Interge | 12 | 54240755 | 54240900 | 146 | 2 | ENSMUST00 | -72116 | Gm24692 | ENSMUSG00000088386.1 |
| chr13 | 56765222 | 56765980 | 8.63619 | 18.02194 | 15.71366 | Distal Interge | 13 | 56733017 | 56738993 | 5977 | 1 | ENSMUST00 | 32206 | Smad5 | ENSMUSG00000021540.11 |
| chr13 | 78201704 | 78202398 | 8.63619 | 18.02194 | 15.71366 | Promoter | 13 | 78201984 | 78234713 | 32730 | 1 | ENSMUST00 | 0 | A830082K12 | ENSMUSG00000087143.4 |
| chr13 | 78209762 | 78210902 | 8.63619 | 18.02194 | 15.71366 | Intron (ENSM | 13 | 78202787 | 78279391 | 76605 | 1 | ENSMUST00 | 6976 | A830082K12 | ENSMUSG00000087143.4 |
| chr13 | 21396082 | 21396378 | 8.63619 | 18.02194 | 15.71366 | Promoter | 13 | 21396578 | 21396638 | 61 | 2 | ENSMUST00 | 260 | Mir6942 | ENSMUSG00000098790.1 |
| chr14 | 25902480 | 25903114 | 8.63619 | 18.02194 | 15.71366 | Promoter | 14 | 25887932 | 25903100 | 15169 | 2 | ENSMUST00 | 0 | Plac9a | ENSMUSG00000095304.3 |
| chr14 | 13749885 | 13750336 | 8.63619 | 18.02194 | 15.71366 | Distal Interge | 14 | 13778072 | 13778151 | 80 | 2 | ENSMUST00 | 27815 | Gm27506 | ENSMUSG00000098694.1 |
| chr15 | 36555316 | 36555968 | 8.63619 | 18.02194 | 15.71366 | Promoter | 15 | 36504062 | 36555573 | 51512 | 2 | ENSMUST00 | 0 | Snx31 | ENSMUSG00000013611.9 |
| chr15 | 101245840 | 101246344 | 8.63619 | 18.02194 | 15.71366 | Distal Interge | 15 | 101266846 | 101274792 | 7947 | 1 | ENSMUST00 | -20502 | Nr4a1 | ENSMUSG00000023034.6 |
| chr15 | 66560697 | 66561154 | 8.63619 | 18.02194 | 15.71366 | Promoter | 15 | 66526212 | 66561046 | 34835 | 2 | ENSMUST00 | 0 | Tmem71 | ENSMUSG00000036944.5 |
| chr15 | 97376804 | 97377279 | 8.63619 | 18.02194 | 15.71366 | Intron (ENSM | 15 | 97244074 | 97247287 | 3214 | 2 | ENSMUST00 | -129518 | Amigo2 | ENSMUSG00000048218.5 |
| chr17 | 7826875 | 7827726 | 8.63619 | 18.02194 | 15.71366 | Promoter | 17 | 7782444 | 7827302 | 44859 | 2 | ENSMUST00 | 0 | Fndc1 | ENSMUSG00000071984.5 |
| chr18 | 21300539 | 21300884 | 8.63619 | 18.02194 | 15.71366 | Promoter | 18 | 21127201 | 21300138 | 172938 | 2 | ENSMUST00 | -402 | Garem | ENSMUSG00000042680.7 |
| chr2 | 165857559 | 165858461 | 8.63619 | 18.02194 | 15.71366 | Promoter | 2 | 165784152 | 165858173 | 74022 | 2 | ENSMUST00 | 0 | Zmynd8 | ENSMUSG00000039671.13 |
| chr2 | 150500438 | 150500884 | 8.63619 | 18.02194 | 15.71366 | Promoter | 2 | 150499466 | 150500845 | 1380 | 2 | ENSMUST00 | 0 | Gm14121 | ENSMUSG00000083325.3 |
| chr3 | 68056547 | 68057138 | 8.63619 | 18.02194 | 15.71366 | 3' UTR | 3 | 68064802 | 68626464 | 561663 | 1 | ENSMUST00 | -7664 | Schip1 | ENSMUSG00000027777.10 |
| chr3 | 52007630 | 52007854 | 8.63619 | 18.02194 | 15.71366 | Intron (ENSM | 3 | 51687611 | 52104891 | 417281 | 2 | ENSMUST00 | 97037 | Maml3 | ENSMUSG00000061143.10 |
| chr3 | 100680065 | 100680648 | 8.63619 | 18.02194 | 15.71366 | Promoter | 3 | 100680265 | 100680555 | 291 | 1 | ENSMUST00 | 0 | Gm12485 | ENSMUSG00000083842.1 |
| chr3 | 11015023 | 11015207 | 8.63619 | 18.02194 | 15.71366 | Promoter | 3 | 11014399 | 11015163 | 765 | 2 | ENSMUST00 | 0 | Gm8910 | ENSMUSG00000094234.1 |
| chr4 | 76450773 | 76451286 | 8.63619 | 18.02194 | 15.71366 | Promoter | 4 | 75941238 | 76449980 | 508743 | 2 | ENSMUST00 | -794 | Ptprd | ENSMUSG00000028399.12 |
| chr4 | 142041621 | 142042305 | 8.63619 | 18.02194 | 15.71366 | Intron (ENSM | 4 | 142042912 | 142048578 | 5667 | 2 | ENSMUST00 | 6273 | Gm13053 | ENSMUSG00000085574.1 |
| chr4 | 114688177 | 114688804 | 8.63619 | 18.02194 | 15.71366 | Distal Interge | 4 | 114680804 | 114687595 | 6792 | 1 | ENSMUST00 | 7374 | Gm12829 | ENSMUSG00000085870.1 |
| chr5 | 77357532 | 77358996 | 8.63619 | 18.02194 | 15.71366 | Intron (ENSM | 5 | 77340368 | 77342936 | 2569 | 1 | ENSMUST00 | 17165 | Polr2b | ENSMUSG00000029250.5 |
| chr6 | 119443488 | 119443943 | 8.63619 | 18.02194 | 15.71366 | Intron (ENSM | 6 | 119432531 | 119449386 | 16856 | 2 | ENSMUST00 | 5443 | Wnt5b | ENSMUSG00000030170.9 |
| chr6 | 52260669 | 52261146 | 8.63619 | 18.02194 | 15.71366 | Promoter | 6 | 52257694 | 52260803 | 3110 | 2 | ENSMUST00 | 0 | Hoxa13 | ENSMUSG00000038203.13 |
| chr6 | 22025784 | 22026909 | 8.63619 | 18.02194 | 15.71366 | Promoter | 6 | 22025943 | 22085138 | 59196 | 1 | ENSMUST00 | 0 | Cped1 | ENSMUSG00000062980.10 |
| chr6 | 95041414 | 95041682 | 8.63619 | 18.02194 | 15.71366 | Distal Interge | 6 | 95100201 | 95100307 | 107 | 2 | ENSMUST00 | 58625 | Gm26501 | ENSMUSG00000093999.1 |
| chr7 | 143154022 | 143154318 | 8.63619 | 18.02194 | 15.71366 | Intron (ENSM | 7 | 143135031 | 143140608 | 5578 | 2 | ENSMUST00 | -13415 | RP23-101N2 | ENSMUSG00000101062.1 |
| chr8 | 84656344 | 84657189 | 8.63619 | 18.02194 | 15.71366 | Distal Interge | 8 | 84661331 | 84662852 | 1522 | 2 | ENSMUST00 | 5663 | Ier2 | ENSMUSG00000053560.4 |
| chr8 | 84192248 | 84192996 | 8.63619 | 18.02194 | 15.71366 | Exon (ENSMU | 8 | 84192774 | 84197667 | 4894 | 2 | ENSMUST00 | 4671 | Gm26887 | ENSMUSG00000097120.1 |
| chr11 | 87778206 | 87778967 | 8.92993 | 18.01848 | 15.7121 | Promoter | 11 | 87778793 | 87785928 | 7136 | 1 | ENSMUST00 | 0 | Bzrap1 | ENSMUSG00000034156.11 |
| chr12 | 4873981 | 4874676 | 8.92993 | 18.01848 | 15.7121 | Promoter | 12 | 4862440 | 4874355 | 11916 | 2 | ENSMUST00 | 0 | Mfsd2b | ENSMUSG00000037336.9 |
| chr17 | 72604317 | 72604823 | 8.92993 | 18.01848 | 15.7121 | Promoter | 17 | 71869442 | 72603709 | 734268 | 2 | ENSMUST00 | -609 | Alk | ENSMUSG00000055471.6 |
| chr9 | 21927566 | 21927885 | 8.92993 | 18.01848 | 15.7121 | Promoter | 9 | 21921010 | 21927535 | 6526 | 2 | ENSMUST00 | -32 | Tmem205 | ENSMUSG00000040883.10 |
| chr10 | 91166993 | 91167773 | 9.22632 | 18.0131 | 15.70718 | Intron (ENSM | 10 | 91147571 | 91171582 | 24012 | 2 | ENSMUST00 | 3809 | Tmpo | ENSMUSG00000019961.11 |
| chr10 | 126977989 | 126978495 | 9.20781 | 17.98236 | 15.67656 | Promoter | 10 | 126978717 | 126999975 | 21259 | 1 | ENSMUST00 | -222 | Ctdsp2 | ENSMUSG00000078429.3 |
| chr11 | 46055889 | 46057432 | 9.20781 | 17.98236 | 15.67656 | Promoter | 11 | 46055992 | 46147343 | 91352 | 1 | ENSMUST00 | 0 | Adam19 | ENSMUSG00000011256.11 |
| chr11 | 72361759 | 72362328 | 9.20781 | 17.98236 | 15.67656 | Promoter | 11 | 72344722 | 72361873 | 17152 | 2 | ENSMUST00 | 0 | Tekt1 | ENSMUSG00000020799.11 |
| chr13 | 63245914 | 63247212 | 9.20781 | 17.98236 | 15.67656 | Intron (ENSM | 13 | 63282142 | 63326096 | 43955 | 1 | ENSMUST00 | -34930 | 2010111I01R | ENSMUSG00000021458.11 |
| chr14 | 14067519 | 14068897 | 9.20781 | 17.98236 | 15.67656 | Intron (ENSM | 14 | 14113910 | 14116550 | 2641 | 2 | ENSMUST00 | 47653 | Psmd6 | ENSMUSG00000021737.3 |
| chr18 | 36328280 | 36329202 | 9.20781 | 17.98236 | 15.67656 | Distal Interge | 18 | 36345619 | 36366720 | 21102 | 1 | ENSMUST00 | -16417 | Cystm1 | ENSMUSG00000046727.7 |
| chr19 | 29355124 | 29355735 | 9.20781 | 17.98236 | 15.67656 | Intron (ENSM | 19 | 29350397 | 29351865 | 1469 | 2 | ENSMUST00 | -3260 | Plgrkt | ENSMUSG00000016495.7 |
| chr5 | 121700387 | 121700861 | 9.20781 | 17.98236 | 15.67656 | Distal Interge | 5 | 121711337 | 121749183 | 37847 | 1 | ENSMUST00 | -10476 | Atxn2 | ENSMUSG00000042605.10 |
| chr5 | 64647442 | 64648103 | 9.20781 | 17.98236 | 15.67656 | Promoter | 5 | 64647633 | 64647749 | 117 | 2 | ENSMUST00 | 0 | Gm25306 | ENSMUSG00000093029.1 |
| chr12 | 55464240 | 55464839 | 8.88047 | 17.93501 | 15.62978 | Distal Interge | 12 | 55489411 | 55492647 | 3237 | 2 | ENSMUST00 | 27808 | Nfkbia | ENSMUSG00000021025.7 |
| chr2 | 92528830 | 92529469 | 9.17103 | 17.92123 | 15.616 | Exon (ENSMU | 2 | 92529694 | 92540980 | 11287 | 2 | ENSMUST00 | 11511 | Gm13817 | ENSMUSG00000085293.2 |
| chr2 | 147876810 | 147877116 | 9.15275 | 17.89084 | 15.58571 | Distal Interge | 2 | 147874772 | 147874878 | 107 | 2 | ENSMUST00 | -1933 | Gm25516 | ENSMUSG00000065791.1 |
| chr4 | 140648533 | 140649019 | 9.15275 | 17.89084 | 15.58571 | Promoter | 4 | 140514485 | 140648750 | 134266 | 2 | ENSMUST00 | 0 | Arhgef10l | ENSMUSG00000040964.11 |
| chr1 | 106687550 | 106688390 | 8.8478 | 17.87982 | 15.5748 | Intron (ENSM | 1 | 106710339 | 106713179 | 2841 | 2 | ENSMUST00 | 24789 | Bcl2 | ENSMUSG00000057329.7 |
| chr11 | 85837738 | 85838128 | 8.54283 | 17.86073 | 15.5557 | Exon (ENSMU | 11 | 85832551 | 85841948 | 9398 | 1 | ENSMUST00 | 5188 | Tbx2 | ENSMUSG00000000093.6 |
| chr12 | 26456157 | 26456797 | 8.54283 | 17.86073 | 15.5557 | Promoter | 12 | 26442753 | 26456431 | 13679 | 2 | ENSMUST00 | 0 | Rsad2 | ENSMUSG00000020641.10 |
| chr4 | 15266992 | 15267293 | 9.1164 | 17.83039 | 15.52565 | Intron (ENSM | 4 | 15265831 | 15286753 | 20923 | 1 | ENSMUST00 | 1162 | Tmem64 | ENSMUSG00000043252.8 |
| chr13 | 48260400 | 48260882 | 9.09833 | 17.80034 | 15.4957 | Promoter | 13 | 48261427 | 48264036 | 2610 | 1 | ENSMUST00 | -545 | Id4 | ENSMUSG00000021379.1 |
| chr2 | 17852776 | 17853959 | 8.21634 | 17.78768 | 15.48312 | Distal Interge | 2 | 17796834 | 17797984 | 1151 | 2 | ENSMUST00 | -54793 | Gm13323 | ENSMUSG00000081851.1 |
| chr8 | 48482048 | 48482765 | 9.08034 | 17.7704 | 15.46587 | Intron (ENSM | 8 | 48334753 | 48555693 | 220941 | 2 | ENSMUST00 | 72928 | Tenm3 | ENSMUSG00000031561.10 |
| chr19 | 43571115 | 43571564 | 8.76716 | 17.74342 | 15.43897 | Distal Interge | 19 | 43609077 | 43610625 | 1549 | 2 | ENSMUST00 | 39061 | Gm20467 | ENSMUSG00000092250.1 |
| chr4 | 152318626 | 152319109 | 8.76716 | 17.74342 | 15.43897 | Promoter | 4 | 152314553 | 152318993 | 4441 | 2 | ENSMUST00 | 0 | Rnf207 | ENSMUSG00000058498.7 |
| chr7 | 27605855 | 27606384 | 8.76716 | 17.74342 | 15.43897 | Promoter | 7 | 27605858 | 27639453 | 33596 | 1 | ENSMUST00 | 0 | Akt2 | ENSMUSG00000004056.10 |
| chr11 | 4594963 | 4595309 | 9.06242 | 17.74057 | 15.43651 | Promoter | 11 | 4482559 | 4594863 | 112305 | 2 | ENSMUST00 | -101 | Mtmr3 | ENSMUSG00000034354.12 |
| chr13 | 28417689 | 28417950 | 9.06242 | 17.74057 | 15.43651 | Distal Interge | 13 | 28460778 | 28480841 | 20064 | 2 | ENSMUST00 | 62891 | 2610307P16 | ENSMUSG00000085936.2 |
| chr10 | 86303619 | 86305439 | 8.16212 | 17.69099 | 15.38708 | Promoter | 10 | 86302854 | 86343981 | 41128 | 1 | ENSMUST00 | 766 | Timp3 | ENSMUSG00000020044.8 |
| chr10 | 80074994 | 80075388 | 8.16212 | 17.69099 | 15.38708 | Intron (ENSM | 10 | 80055998 | 80056363 | 366 | 1 | ENSMUST00 | 18997 | Gpx4 | ENSMUSG00000075706.5 |
| chr11 | 94242877 | 94243767 | 8.16212 | 17.69099 | 15.38708 | Promoter | 11 | 94235956 | 94242707 | 6752 | 2 | ENSMUST00 | -171 | Wfikkn2 | ENSMUSG00000044177.4 |
| chr15 | 102963417 | 102964132 | 8.16212 | 17.69099 | 15.38708 | Distal Interge | 15 | 102966796 | 102971893 | 5098 | 1 | ENSMUST00 | -2664 | Hoxc10 | ENSMUSG00000022484.7 |
| chr16 | 26043961 | 26044765 | 8.16212 | 17.69099 | 15.38708 | Intron (ENSM | 16 | 26031810 | 26040870 | 9061 | 2 | ENSMUST00 | -3092 | AU015336 | ENSMUSG00000090203.1 |
| chr3 | 84220325 | 84221164 | 8.16212 | 17.69099 | 15.38708 | Promoter | 3 | 84160441 | 84220889 | 60449 | 2 | ENSMUST00 | 0 | Trim2 | ENSMUSG00000027993.11 |
| chr3 | 84304222 | 84305403 | 8.16212 | 17.69099 | 15.38708 | Promoter | 3 | 84160439 | 84304809 | 144371 | 2 | ENSMUST00 | 0 | Trim2 | ENSMUSG00000027993.11 |
| chr4 | 126296686 | 126297259 | 8.16212 | 17.69099 | 15.38708 | Intron (ENSM | 4 | 126300530 | 126309785 | 9256 | 1 | ENSMUST00 | -3271 | Col8a2 | ENSMUSG00000056174.8 |
| chr6 | 67532797 | 67533769 | 8.16212 | 17.69099 | 15.38708 | Downstream | 6 | 67534059 | 67535822 | 1764 | 2 | ENSMUST00 | 2053 | Tacstd2 | ENSMUSG00000051397.4 |
| chr6 | 145276141 | 145276778 | 8.16212 | 17.69099 | 15.38708 | Distal Interge | 6 | 145266400 | 145266777 | 378 | 1 | ENSMUST00 | 9742 | Rps25-ps1 | ENSMUSG00000067344.7 |
| chrX | 136138924 | 136139389 | 8.16212 | 17.69099 | 15.38708 | Promoter | 20 | 136138996 | 136140443 | 1448 | 1 | ENSMUST00 | 0 | Bex4 | ENSMUSG00000047844.3 |
| chr10 | 56468980 | 56469552 | 9.02678 | 17.68123 | 15.37864 | Distal Interge | 10 | 56377300 | 56390419 | 13120 | 1 | ENSMUST00 | 91681 | Gja1 | ENSMUSG00000050953.9 |
| chr11 | 50226569 | 50227061 | 9.02678 | 17.68123 | 15.37864 | Intron (ENSM | 11 | 50225335 | 50235103 | 9769 | 1 | ENSMUST00 | 1235 | Mgat4b | ENSMUSG00000036620.9 |
| chr3 | 73708021 | 73708687 | 9.02678 | 17.68123 | 15.37864 | Promoter | 3 | 73635808 | 73708415 | 72608 | 2 | ENSMUST00 | 0 | Bche | ENSMUSG00000027792.6 |
| chr7 | 16885123 | 16885857 | 9.02678 | 17.68123 | 15.37864 | Exon (ENSMU | 7 | 16891786 | 16895435 | 3650 | 1 | ENSMUST00 | -5929 | Gng8 | ENSMUSG00000063594.4 |
| chr8 | 14096666 | 14097376 | 9.02678 | 17.68123 | 15.37864 | Promoter | 8 | 14095875 | 14847680 | 751806 | 1 | ENSMUST00 | 792 | Dlgap2 | ENSMUSG00000047495.10 |

| | | | | | | | | | | | | | | | |
|---|---|---|---|---|---|---|---|---|---|---|---|---|---|---|---|
| chr8 | 89317589 | 89318015 | 9.02678 | 17.68123 | 15.37864 | Distal Interge | 8 | 89186946 | 89187560 | 615 | 2 | ENSMUST00 | -130030 | Gm5356 | ENSMUSG00000069962.3 |
| chr15 | 83268208 | 83268769 | 8.99143 | 17.62233 | 15.32028 | Distal Interge | 15 | 83226726 | 83251774 | 25049 | 2 | ENSMUST00 | -16435 | A4galt | ENSMUSG00000047878.7 |
| chr9 | 67699781 | 67700247 | 8.99143 | 17.62233 | 15.32028 | Distal Interge | 9 | 67727717 | 67727780 | 64 | 2 | ENSMUST00 | 27533 | Mir7241 | ENSMUSG00000098937.1 |
| chr10 | 96161383 | 96161548 | 8.40012 | 17.61332 | 15.31135 | Distal Interge | 10 | 96164331 | 96164431 | 101 | 1 | ENSMUST00 | -2783 | Gm22913 | ENSMUSG00000094523.1 |
| chr18 | 44543051 | 44544910 | 8.40012 | 17.61332 | 15.31135 | Intron (ENSM | 18 | 44661665 | 44676271 | 14607 | 1 | ENSMUST00 | -116755 | A930012L18I | ENSMUSG00000097610.1 |
| chr2 | 104098304 | 104098985 | 8.08967 | 17.56143 | 15.25973 | Intron (ENSM | 2 | 104095840 | 104114263 | 18424 | 1 | ENSMUST00 | 2465 | Cd59a | ENSMUSG00000032679.7 |
| chr10 | 127091314 | 127091967 | 8.61023 | 17.47713 | 15.17558 | 3' UTR | 10 | 127078907 | 127093169 | 14263 | 1 | ENSMUST00 | 12408 | Agap2 | ENSMUSG00000025422.8 |
| chr3 | 52046256 | 52046934 | 8.61023 | 17.47713 | 15.17558 | Intron (ENSM | 3 | 51687611 | 52104891 | 417281 | 2 | ENSMUST00 | 57957 | Maml3 | ENSMUSG00000061143.10 |
| chr1 | 172449957 | 172451318 | 7.76973 | 17.46802 | 15.16681 | Distal Interge | 1 | 172475358 | 172478575 | 3218 | 1 | ENSMUST00 | -24040 | Slamf9 | ENSMUSG00000026548.4 |
| chr1 | 128832356 | 128832811 | 7.76973 | 17.46802 | 15.16681 | Distal Interge | 1 | 128885909 | 128886019 | 111 | 2 | ENSMUST00 | 53208 | Gm23056 | ENSMUSG00000064737.1 |
| chr10 | 68723695 | 68724876 | 7.76973 | 17.46802 | 15.16681 | Promoter | 10 | 68723746 | 68782654 | 58909 | 1 | ENSMUST00 | 0 | Tmem26 | ENSMUSG00000060044.7 |
| chr11 | 86586399 | 86587647 | 7.76973 | 17.46802 | 15.16681 | Promoter | 11 | 86585020 | 86586994 | 1975 | 2 | ENSMUST00 | 0 | Vmp1 | ENSMUSG00000018171.4 |
| chr11 | 87442643 | 87443082 | 7.76973 | 17.46802 | 15.16681 | Promoter | 11 | 87443237 | 87443452 | 216 | 1 | ENSMUST00 | -155 | Rnu3b1 | ENSMUSG00000098943.1 |
| chr14 | 8800323 | 8800546 | 7.76973 | 17.46802 | 15.16681 | Promoter | 14 | 8799589 | 8800504 | 916 | 2 | ENSMUST00 | 0 | Gm9800 | ENSMUSG00000045799.5 |
| chr15 | 101006637 | 101007120 | 7.76973 | 17.46802 | 15.16681 | Intron (ENSM | 15 | 101050194 | 101054399 | 4206 | 2 | ENSMUST00 | 47279 | Fignl2 | ENSMUSG00000095440.1 |
| chr19 | 37233720 | 37234431 | 7.76973 | 17.46802 | 15.16681 | Promoter | 19 | 37233382 | 37233484 | 103 | 2 | ENSMUST00 | -237 | Gm25268 | ENSMUSG00000089143.2 |
| chr2 | 131171072 | 131171909 | 7.76973 | 17.46802 | 15.16681 | 3' UTR | 2 | 131171611 | 131174653 | 3043 | 2 | ENSMUST00 | 2744 | Spef1 | ENSMUSG00000027329.4 |
| chr5 | 113324062 | 113324515 | 7.76973 | 17.46802 | 15.16681 | Distal Interge | 5 | 113243220 | 113310786 | 67567 | 2 | ENSMUST00 | -13277 | Sgsm1 | ENSMUSG00000042216.8 |
| chr6 | 85189937 | 85190815 | 7.76973 | 17.46802 | 15.16681 | Intron (ENSM | 6 | 85187438 | 85204358 | 16921 | 1 | ENSMUST00 | 2500 | Emx1 | ENSMUSG00000033726.8 |
| chr7 | 141221948 | 141222415 | 7.76973 | 17.46802 | 15.16681 | Promoter | 7 | 141221384 | 141221493 | 110 | 2 | ENSMUST00 | -456 | Mir210 | ENSMUSG00000065551.1 |
| chr7 | 125393358 | 125394086 | 7.76973 | 17.46802 | 15.16681 | Distal Interge | 7 | 125368861 | 125371013 | 2153 | 1 | ENSMUST00 | 24498 | 4930571K23I | ENSMUSG00000090457.1 |
| chr7 | 37361765 | 37362385 | 7.76973 | 17.46802 | 15.16681 | Distal Interge | 7 | 37299265 | 37304034 | 4770 | 1 | ENSMUST00 | 62501 | RP23-291B1. | ENSMUSG00000099715.1 |
| chr9 | 69988988 | 69989351 | 7.76973 | 17.46802 | 15.16681 | Promoter | 9 | 69989466 | 70008317 | 18852 | 1 | ENSMUST00 | -115 | Bnip2 | ENSMUSG00000011958.12 |
| chr10 | 10546397 | 10547016 | 9.19064 | 17.46287 | 15.16242 | Exon (ENSMU | 10 | 10545011 | 10558233 | 13223 | 2 | ENSMUST00 | 11217 | Rab32 | ENSMUSG00000019832.3 |
| chr10 | 127081232 | 127081777 | 9.19064 | 17.46287 | 15.16242 | Intron (ENSM | 10 | 127078907 | 127093169 | 14263 | 1 | ENSMUST00 | 2326 | Agap2 | ENSMUSG00000025422.8 |
| chr10 | 69924266 | 69924724 | 9.19064 | 17.46287 | 15.16242 | Promoter | 10 | 69925495 | 69926338 | 844 | 1 | ENSMUST00 | -771 | Ank3 | ENSMUSG00000069601.8 |
| chr13 | 94868519 | 94869173 | 9.19064 | 17.46287 | 15.16242 | Distal Interge | 13 | 94875602 | 94885130 | 9529 | 1 | ENSMUST00 | -6429 | Otp | ENSMUSG00000021685.11 |
| chr16 | 32149405 | 32149621 | 9.19064 | 17.46287 | 15.16242 | Intron (ENSM | 16 | 32144827 | 32148118 | 3292 | 2 | ENSMUST00 | -1288 | Nrros | ENSMUSG00000052384.8 |
| chr16 | 78490297 | 78490608 | 9.19064 | 17.46287 | 15.16242 | Distal Interge | 16 | 78423730 | 78423859 | 130 | 2 | ENSMUST00 | -66439 | Gm25038 | ENSMUSG00000064862.1 |
| chr19 | 44734616 | 44734916 | 9.19064 | 17.46287 | 15.16242 | Distal Interge | 19 | 44756045 | 44837851 | 81807 | 1 | ENSMUST00 | -21129 | Pax2 | ENSMUSG00000004231.10 |
| chr3 | 97932042 | 97932510 | 9.19064 | 17.46287 | 15.16242 | Intron (ENSM | 3 | 97930173 | 97967018 | 36846 | 1 | ENSMUST00 | 1870 | Gm5544 | ENSMUSG00000074388.3 |
| chr5 | 119831097 | 119831843 | 9.19064 | 17.46287 | 15.16242 | Distal Interge | 5 | 119834663 | 119885218 | 50556 | 1 | ENSMUST00 | -2820 | Tbx5 | ENSMUSG00000018263.8 |
| chr5 | 120435845 | 120436506 | 9.19064 | 17.46287 | 15.16242 | Exon (ENSMU | 5 | 120431770 | 120441223 | 9454 | 1 | ENSMUST00 | 4076 | Lhx5 | ENSMUSG00000029595.8 |
| chr5 | 134747060 | 134747289 | 9.19064 | 17.46287 | 15.16242 | Promoter | 5 | 134703781 | 134747241 | 43461 | 2 | ENSMUST00 | 0 | Eln | ENSMUSG00000029675.9 |
| chr5 | 24329338 | 24329765 | 9.19064 | 17.46287 | 15.16242 | Promoter | 5 | 24319589 | 24329591 | 10003 | 2 | ENSMUST00 | 0 | Kcnh2 | ENSMUSG00000038319.9 |
| chr5 | 90588807 | 90589176 | 9.19064 | 17.46287 | 15.16242 | Intron (ENSM | 5 | 90561209 | 90597906 | 36698 | 1 | ENSMUST00 | 27599 | 5830473C10I | ENSMUSG00000070690.4 |
| chr6 | 144472506 | 144473052 | 9.19064 | 17.46287 | 15.16242 | Intron (ENSM | 6 | 144481225 | 144501416 | 20192 | 1 | ENSMUST00 | -8173 | Sox5os2 | ENSMUSG00000087157.1 |
| chr6 | 4747388 | 4748811 | 9.19064 | 17.46287 | 15.16242 | Promoter | 6 | 4747306 | 4760517 | 13212 | 1 | ENSMUST00 | 83 | Peg10 | ENSMUSG00000092035.3 |
| chr7 | 24954064 | 24954898 | 9.19064 | 17.46287 | 15.16242 | Distal Interge | 7 | 24970010 | 24970075 | 66 | 2 | ENSMUST00 | 15177 | Mir7046 | ENSMUSG00000098802.1 |
| chr8 | 15063503 | 15064193 | 9.19064 | 17.46287 | 15.16242 | 5' UTR | 8 | 15060190 | 15073840 | 13651 | 1 | ENSMUST00 | 3314 | Myom2 | ENSMUSG00000031461.4 |
| chr8 | 87974404 | 87975187 | 9.19064 | 17.46287 | 15.16242 | Distal Interge | 8 | 87661810 | 87959595 | 297786 | 2 | ENSMUST00 | -14810 | Zfp423 | ENSMUSG00000045333.10 |
| chr8 | 9771379 | 9772012 | 9.19064 | 17.46287 | 15.16242 | Promoter | 8 | 9206001 | 9771018 | 565018 | 2 | ENSMUST00 | -362 | Fam155a | ENSMUSG00000079157.3 |
| chr8 | 26780573 | 26780855 | 9.19064 | 17.46287 | 15.16242 | Distal Interge | 8 | 26814593 | 26824423 | 9831 | 2 | ENSMUST00 | 43568 | RP24-502D8. | ENSMUSG00000100335.1 |
| chr9 | 44486689 | 44486994 | 9.19064 | 17.46287 | 15.16242 | Distal Interge | 9 | 44478778 | 44482180 | 3403 | 1 | ENSMUST00 | 7912 | C030014I23R | ENSMUSG00000060380.2 |
| chr9 | 89066437 | 89066657 | 9.19064 | 17.46287 | 15.16242 | Distal Interge | 9 | 89075336 | 89075407 | 72 | 1 | ENSMUST00 | -8679 | Gm24463 | ENSMUSG00000096473.1 |
| chrX | 7319471 | 7319717 | 9.19064 | 17.46287 | 15.16242 | Promoter | 20 | 7171199 | 7319358 | 148160 | 2 | ENSMUST00 | -114 | Clcn5 | ENSMUSG00000004317.9 |
| chrX | 134308107 | 134308419 | 9.19064 | 17.46287 | 15.16242 | Promoter | 20 | 134308155 | 134362636 | 54482 | 1 | ENSMUST00 | 0 | Cenpi | ENSMUSG00000031262.7 |
| chrX | 53269954 | 53270284 | 9.19064 | 17.46287 | 15.16242 | Promoter | 20 | 53244864 | 53269805 | 24942 | 2 | ENSMUST00 | -150 | Fam122b | ENSMUSG00000036022.10 |
| chrX | 162829640 | 162830043 | 9.19064 | 17.46287 | 15.16242 | Promoter | 20 | 162828060 | 162829454 | 1395 | 2 | ENSMUST00 | -187 | Txlng | ENSMUSG00000038344.9 |
| chrX | 12090802 | 12091405 | 9.19064 | 17.46287 | 15.16242 | Intron (ENSM | 20 | 12036740 | 12080553 | 43814 | 2 | ENSMUST00 | -10250 | Bcor | ENSMUSG00000040363.9 |
| chrX | 12133287 | 12133788 | 9.19064 | 17.46287 | 15.16242 | Intron (ENSM | 20 | 12058981 | 12128400 | 69420 | 2 | ENSMUST00 | -4888 | Bcor | ENSMUSG00000040363.9 |
| chrX | 152178917 | 152179419 | 9.19064 | 17.46287 | 15.16242 | Promoter | 20 | 152178964 | 152194601 | 15638 | 1 | ENSMUST00 | 0 | Iqsec2 | ENSMUSG00000041115.11 |
| chrX | 56598264 | 56598585 | 9.19064 | 17.46287 | 15.16242 | Promoter | 20 | 56585512 | 56598069 | 12558 | 2 | ENSMUST00 | -196 | Mmgt1 | ENSMUSG00000061273.3 |
| chrX | 12839896 | 12840655 | 9.19064 | 17.46287 | 15.16242 | Distal Interge | 20 | 12870269 | 12870362 | 94 | 2 | ENSMUST00 | 29707 | Gm25063 | ENSMUSG00000077858.1 |
| chrX | 134585171 | 134585566 | 9.19064 | 17.46287 | 15.16242 | Promoter | 20 | 134585654 | 134588056 | 2403 | 1 | ENSMUST00 | -88 | Rpl36a | ENSMUSG00000079435.4 |
| chrX | 48285570 | 48286035 | 9.19064 | 17.46287 | 15.16242 | Promoter | 20 | 48276792 | 48285426 | 8635 | 2 | ENSMUST00 | -145 | 9530027J09R | ENSMUSG00000097197.1 |
| chr14 | 68083829 | 68084764 | 9.17411 | 17.43557 | 15.13716 | Promoter | 14 | 68083863 | 68089095 | 5233 | 1 | ENSMUST00 | 0 | Nefl | ENSMUSG00000022055.7 |
| chr8 | 70166577 | 70167056 | 9.17411 | 17.43557 | 15.13716 | 3' UTR | 8 | 70172356 | 70180930 | 8575 | 1 | ENSMUST00 | -5300 | Tmem161a | ENSMUSG00000002342.12 |
| chr9 | 57997852 | 57998445 | 9.17411 | 17.43557 | 15.13716 | Promoter | 9 | 57998041 | 58007451 | 9411 | 1 | ENSMUST00 | 0 | Gm17322 | ENSMUSG00000097604.1 |
| chr15 | 82166187 | 82166808 | 8.57951 | 17.42489 | 15.12655 | Intron (ENSM | 15 | 82147266 | 82205376 | 58111 | 1 | ENSMUST00 | 18922 | Srebf2 | ENSMUSG00000022463.7 |
| chr5 | 41556655 | 41557091 | 8.86984 | 17.41951 | 15.12118 | Distal Interge | 5 | 41624976 | 41708155 | 83180 | 2 | ENSMUST00 | 151064 | Rab28 | ENSMUSG00000029128.9 |
| chr9 | 71590190 | 71590678 | 8.86984 | 17.41951 | 15.12118 | Intron (ENSM | 9 | 71505058 | 71592265 | 87208 | 2 | ENSMUST00 | 1587 | Myzap | ENSMUSG00000041361.8 |
| chr15 | 103537548 | 103538251 | 9.1549 | 17.40386 | 15.10557 | Promoter | 15 | 103530279 | 103537992 | 7714 | 2 | ENSMUST00 | 0 | Ppp1r1a | ENSMUSG00000022490.6 |
| chr11 | 49249000 | 49249511 | 8.85273 | 17.39095 | 15.09274 | Promoter | 11 | 49248957 | 49251637 | 2681 | 1 | ENSMUST00 | 44 | Mgat1 | ENSMUSG00000020346.11 |
| chr11 | 59921070 | 59921481 | 8.85273 | 17.39095 | 15.09274 | Promoter | 11 | 59921499 | 59921889 | 391 | 1 | ENSMUST00 | -18 | 1810063I02R | ENSMUSG00000085551.1 |
| chr19 | 42045470 | 42045872 | 8.85273 | 17.39095 | 15.09274 | Promoter | 19 | 42045610 | 42052290 | 6681 | 1 | ENSMUST00 | 0 | Hoga1 | ENSMUSG00000025176.9 |
| chr5 | 123670503 | 123671429 | 8.85273 | 17.39095 | 15.09274 | Intron (ENSM | 5 | 123617706 | 123666690 | 48985 | 2 | ENSMUST00 | -3814 | Clip1 | ENSMUSG00000049550.12 |
| chr6 | 30175615 | 30176263 | 8.85273 | 17.39095 | 15.09274 | Distal Interge | 6 | 30169668 | 30169737 | 70 | 2 | ENSMUST00 | -5879 | Mir183 | ENSMUSG00000065619.2 |
| chr7 | 110003084 | 110003447 | 8.85273 | 17.39095 | 15.09274 | Promoter | 7 | 110003578 | 110003718 | 141 | 1 | ENSMUST00 | -131 | Gm23100 | ENSMUSG00000065389.1 |
| chr9 | 106418104 | 106418716 | 8.85273 | 17.39095 | 15.09274 | Distal Interge | 9 | 106429454 | 106431568 | 2115 | 1 | ENSMUST00 | -10738 | Rpl29 | ENSMUSG00000048758.8 |
| chr9 | 110994846 | 110995214 | 8.85273 | 17.39095 | 15.09274 | Distal Interge | 9 | 110985871 | 110989611 | 3741 | 2 | ENSMUST00 | -5236 | Rtp3 | ENSMUSG00000066319.5 |
| chr4 | 102978564 | 102979697 | 8.26211 | 17.37283 | 15.07506 | Intron (ENSM | 4 | 102986379 | 103002519 | 16141 | 1 | ENSMUST00 | -6682 | Tctex1d1 | ENSMUSG00000028523.8 |
| chr17 | 50469869 | 50471115 | 9.13578 | 17.37227 | 15.0746 | Distal Interge | 17 | 50509547 | 50688493 | 178947 | 1 | ENSMUST00 | -38432 | Plcl2 | ENSMUSG00000038910.6 |
| chr2 | 70704545 | 70704952 | 9.13578 | 17.37227 | 15.0746 | Intron (ENSM | 2 | 70716312 | 70717904 | 1593 | 2 | ENSMUST00 | 12952 | Tlk1 | ENSMUSG00000041997.11 |
| chr4 | 124277168 | 124278058 | 9.13578 | 17.37227 | 15.0746 | Distal Interge | 4 | 124293693 | 124299848 | 6156 | 2 | ENSMUST00 | 21790 | Gm12916 | ENSMUSG00000086087.1 |
| chr5 | 52565808 | 52566492 | 9.13578 | 17.37227 | 15.0746 | Promoter | 5 | 52537864 | 52566303 | 28440 | 2 | ENSMUST00 | 0 | Lgi2 | ENSMUSG00000039252.7 |
| chr5 | 21673031 | 21673622 | 9.13578 | 17.37227 | 15.0746 | Intron (ENSM | 5 | 21689541 | 21690175 | 635 | 1 | ENSMUST00 | -15919 | Gm15719 | ENSMUSG00000081242.1 |
| chr11 | 34845828 | 34846480 | 9.11673 | 17.34082 | 15.04346 | Distal Interge | 11 | 34842228 | 34842354 | 127 | 2 | ENSMUST00 | -3475 | Gm22022 | ENSMUSG00000077543.1 |
| chr6 | 71048320 | 71048969 | 9.11673 | 17.34082 | 15.04346 | Distal Interge | 6 | 71039579 | 71039733 | 155 | 2 | ENSMUST00 | -8588 | Gm23485 | ENSMUSG00000077369.1 |
| chr10 | 13007875 | 13008312 | 7.43959 | 17.32416 | 15.02694 | 5' UTR | 10 | 13005343 | 13009183 | 3841 | 1 | ENSMUST00 | 2533 | Sf3b5 | ENSMUSG00000078348.4 |
| chr11 | 96294720 | 96295416 | 7.43959 | 17.32416 | 15.02694 | Intron (ENSM | 11 | 96292476 | 96301569 | 9094 | 1 | ENSMUST00 | 2245 | Hoxb6 | ENSMUSG00000000690.5 |
| chr11 | 88161702 | 88162230 | 7.43959 | 17.32416 | 15.02694 | Promoter | 11 | 88162261 | 88194140 | 31880 | 1 | ENSMUST00 | -31 | Cuedc1 | ENSMUSG00000018378.8 |
| chr3 | 53630811 | 53631480 | 7.43959 | 17.32416 | 15.02694 | Intron (ENSM | 3 | 53513939 | 53657355 | 143417 | 2 | ENSMUST00 | 25875 | Frem2 | ENSMUSG00000037016.7 |
| chr9 | 111438929 | 111439460 | 7.43959 | 17.32416 | 15.02694 | Promoter | 9 | 111439081 | 111489608 | 50528 | 1 | ENSMUST00 | 0 | Dclk3 | ENSMUSG00000032500.9 |
| chr9 | 56923404 | 56924718 | 7.43959 | 17.32416 | 15.02694 | Intron (ENSM | 9 | 56917200 | 56928371 | 11172 | 2 | ENSMUST00 | 3653 | Snx33 | ENSMUSG00000032733.5 |
| chr9 | 59691603 | 59692159 | 7.43959 | 17.32416 | 15.02694 | Intron (ENSM | 9 | 59680144 | 59713923 | 33780 | 1 | ENSMUST00 | 11460 | Gramd2 | ENSMUSG00000074259.5 |
| chr11 | 96303367 | 96304335 | 9.09776 | 17.30949 | 15.01266 | Promoter | 11 | 96303512 | 96306120 | 2609 | 1 | ENSMUST00 | 0 | Hoxb5 | ENSMUSG00000038700.4 |
| chr6 | 141305022 | 141305605 | 9.09776 | 17.30949 | 15.01266 | Intron (ENSM | 6 | 141238278 | 141250561 | 12284 | 2 | ENSMUST00 | -54462 | RP23-97L11. | ENSMUSG00000102066.1 |
| chr7 | 123030639 | 123031072 | 9.09776 | 17.30949 | 15.01266 | Distal Interge | 7 | 122990130 | 122992512 | 2383 | 1 | ENSMUST00 | 40510 | Rbbp6 | ENSMUSG00000030779.10 |
| chr11 | 95071021 | 95073215 | 8.80182 | 17.30588 | 15.00915 | Intron (ENSM | 11 | 95044474 | 95069198 | 24725 | 2 | ENSMUST00 | -1824 | Itga3 | ENSMUSG00000001507.11 |
| chr1 | 84934196 | 84935125 | 8.80131 | 17.30502 | 15.00832 | Promoter | 1 | 84929334 | 84934857 | 5524 | 2 | ENSMUST00 | 0 | Slc16a14 | ENSMUSG00000026220.6 |
| chr1 | 20820517 | 20820757 | 8.80131 | 17.30502 | 15.00832 | Promoter | 1 | 20802968 | 20820312 | 17345 | 2 | ENSMUST00 | -206 | Mcm3 | ENSMUSG00000041859.10 |
| chr1 | 155530701 | 155531192 | 8.80131 | 17.30502 | 15.00832 | Distal Interge | 1 | 155520173 | 155527111 | 6939 | 2 | ENSMUST00 | -3591 | Gm5532 | ENSMUSG00000073535.4 |
| chr1 | 19241398 | 19241724 | 8.80131 | 17.30502 | 15.00832 | Downstream | 1 | 19242725 | 19243769 | 1045 | 1 | ENSMUST00 | -1001 | RP24-119I6.2 | ENSMUSG00000100538.1 |
| chr10 | 81257230 | 81257877 | 8.80131 | 17.30502 | 15.00832 | Promoter | 10 | 81257299 | 81262981 | 5683 | 1 | ENSMUST00 | 0 | Matk | ENSMUSG00000004933.12 |
| chr10 | 42863436 | 42864041 | 8.80131 | 17.30502 | 15.00832 | Intron (ENSM | 10 | 42865514 | 42930419 | 64906 | 1 | ENSMUST00 | -1473 | Scml4 | ENSMUSG00000044770.9 |
| chr11 | 114829564 | 114830177 | 8.80131 | 17.30502 | 15.00832 | Distal Interge | 11 | 114851152 | 114863733 | 12582 | 1 | ENSMUST00 | -20975 | Gprc5c | ENSMUSG00000051043.11 |
| chr11 | 20445756 | 20446213 | 8.80131 | 17.30502 | 15.00832 | Promoter | 11 | 20446412 | 20446810 | 399 | 1 | ENSMUST00 | -199 | Gm12034 | ENSMUSG00000083878.3 |
| chr13 | 37854017 | 37854797 | 8.80131 | 17.30502 | 15.00832 | Intron (ENSM | 13 | 37857934 | 37893914 | 35981 | 1 | ENSMUST00 | -3137 | Rreb1 | ENSMUSG00000039087.11 |
| chr13 | 23533489 | 23533998 | 8.80131 | 17.30502 | 15.00832 | Promoter | 13 | 23533906 | 23534304 | 399 | 1 | ENSMUST00 | 0 | Hist1h2af | ENSMUSG00000061991.2 |
| chr13 | 34148807 | 34149370 | 8.80131 | 17.30502 | 15.00832 | Distal Interge | 13 | 34162964 | 34178168 | 15205 | 1 | ENSMUST00 | -13594 | Psmg4 | ENSMUSG00000071451.4 |
| chr13 | 104518089 | 104519013 | 8.80131 | 17.30502 | 15.00832 | Distal Interge | 13 | 104684931 | 104694994 | 10064 | 1 | ENSMUST00 | -165918 | 2610204G07 | ENSMUSG00000084860.1 |
| chr14 | 105483079 | 105483514 | 8.80131 | 17.30502 | 15.00832 | Distal Interge | 14 | 105498788 | 105505622 | 6835 | 1 | ENSMUST00 | -15274 | 4930449E01I | ENSMUSG00000022116.2 |
| chr15 | 59306802 | 59307117 | 8.80131 | 17.30502 | 15.00832 | Promoter | 15 | 59306948 | 59312013 | 5066 | 1 | ENSMUST00 | 0 | Zfp572 | ENSMUSG00000098528.1 |
| chr16 | 4542080 | 4542481 | 8.80131 | 17.30502 | 15.00832 | Promoter | 16 | 4487579 | 4541816 | 54238 | 2 | ENSMUST00 | -265 | Srl | ENSMUSG00000022519.8 |
| chr17 | 5940860 | 5941188 | 8.80131 | 17.30502 | 15.00832 | Promoter | 17 | 5941280 | 6032793 | 91514 | 1 | ENSMUST00 | -92 | Synj2 | ENSMUSG00000023805.11 |
| chr17 | 56751337 | 56751843 | 8.80131 | 17.30502 | 15.00832 | 3' UTR | 17 | 56751325 | 56757530 | 6206 | 2 | ENSMUST00 | 5687 | Nrtn | ENSMUSG00000039481.5 |
| chr17 | 83267335 | 83267833 | 8.80131 | 17.30502 | 15.00832 | Downstream | 17 | 83269971 | 83270105 | 135 | 1 | ENSMUST00 | -2138 | Gm24240 | ENSMUSG00000088410.1 |
| chr2 | 153071700 | 153072185 | 8.80131 | 17.30502 | 15.00832 | Exon (ENSMU | 2 | 153066002 | 153081735 | 15734 | 1 | ENSMUST00 | 5699 | Ccm2l | ENSMUSG00000027474.7 |
| chr2 | 66409770 | 66410218 | 8.80131 | 17.30502 | 15.00832 | Promoter | 2 | 66270781 | 66410064 | 139284 | 2 | ENSMUST00 | 0 | Scn1a | ENSMUSG00000064329.8 |
| chr2 | 146218238 | 146218770 | 8.80131 | 17.30502 | 15.00832 | Distal Interge | 2 | 146221921 | 146225020 | 3100 | 1 | ENSMUST00 | -3151 | Insm1 | ENSMUSG00000068154.5 |
| chr3 | 30514283 | 30514846 | 8.80131 | 17.30502 | 15.00832 | Intron (ENSM | 3 | 29965456 | 30509822 | 544367 | 2 | ENSMUST00 | -4462 | Mecom | ENSMUSG00000027684.11 |
| chr3 | 84653748 | 84654757 | 8.80131 | 17.30502 | 15.00832 | Intron (ENSM | 3 | 84666192 | 84704575 | 38384 | 1 | ENSMUST00 | -11435 | Tmem154 | ENSMUSG00000056498.8 |
| chr4 | 102576788 | 102578153 | 8.80131 | 17.30502 | 15.00832 | Intron (ENSM | 4 | 102570095 | 102605847 | 35753 | 1 | ENSMUST00 | 6694 | Pde4b | ENSMUSG00000028525.11 |
| chr4 | 111886109 | 111886456 | 8.80131 | 17.30502 | 15.00832 | Intron (ENSM | 4 | 111875429 | 111884254 | 8826 | 2 | ENSMUST00 | -1856 | Slc5a9 | ENSMUSG00000028544.9 |
| chr5 | 151108315 | 151108823 | 8.80131 | 17.30502 | 15.00832 | Intron (ENSM | 5 | 151063593 | 151095747 | 32155 | 2 | ENSMUST00 | -12569 | Stard13 | ENSMUSG00000016128.9 |
| chr5 | 72796036 | 72796645 | 8.80131 | 17.30502 | 15.00832 | Intron (ENSM | 5 | 72755738 | 72786020 | 30283 | 2 | ENSMUST00 | -10017 | Tec | ENSMUSG00000029217.11 |
| chr5 | 111178152 | 111179082 | 8.80131 | 17.30502 | 15.00832 | Intron (ENSM | 5 | 111225458 | 111229415 | 3958 | 1 | ENSMUST00 | -46376 | Ttc28 | ENSMUSG00000033209.12 |
| chr5 | 103336102 | 103337433 | 8.80131 | 17.30502 | 15.00832 | Distal Interge | 5 | 103425192 | 103598359 | 173168 | 1 | ENSMUST00 | -87759 | Ptpn13 | ENSMUSG00000034573.10 |
| chr5 | 146584535 | 146585237 | 8.80131 | 17.30502 | 15.00832 | Promoter | 5 | 146582151 | 146585239 | 3089 | 2 | ENSMUST00 | 2 | Gpr12 | ENSMUSG00000041468.6 |
| chr5 | 36726045 | 36726829 | 8.80131 | 17.30502 | 15.00832 | Distal Interge | 5 | 36747374 | 36748679 | 1306 | 2 | ENSMUST00 | 21850 | Bloc1s4 | ENSMUSG00000060708.3 |
| chr6 | 91410984 | 91411900 | 8.80131 | 17.30502 | 15.00832 | Promoter | 6 | 91363981 | 91411363 | 47383 | 2 | ENSMUST00 | 0 | Wnt7a | ENSMUSG00000030093.6 |
| chr7 | 51879085 | 51879692 | 8.80131 | 17.30502 | 15.00832 | Promoter | 7 | 51879050 | 51994458 | 115409 | 1 | ENSMUST00 | 36 | Gas2 | ENSMUSG00000030498.9 |
| chr7 | 136894087 | 136894876 | 8.80131 | 17.30502 | 15.00832 | Promoter | 7 | 136894611 | 137128187 | 233577 | 1 | ENSMUST00 | 0 | Mgmt | ENSMUSG00000054612.3 |
| chr7 | 79496157 | 79496467 | 8.80131 | 17.30502 | 15.00832 | Distal Interge | 7 | 79500026 | 79531303 | 31278 | 1 | ENSMUST00 | -3559 | AI854517 | ENSMUSG00000097023.3 |
| chr8 | 120191035 | 120191417 | 8.80131 | 17.30502 | 15.00832 | Intron (ENSM | 8 | 120166397 | 120177466 | 11070 | 2 | ENSMUST00 | -13570 | Fam92b | ENSMUSG00000042269.10 |
| chr8 | 85013427 | 85013680 | 8.80131 | 17.30502 | 15.00832 | Promoter | 8 | 85007202 | 85014408 | 7207 | 2 | ENSMUST00 | 728 | Best2 | ENSMUSG00000052819.5 |
| chr9 | 108694114 | 108694705 | 8.80131 | 17.30502 | 15.00832 | Intron (ENSM | 9 | 108692143 | 108749512 | 57370 | 1 | ENSMUST00 | 1972 | Prkar2a | ENSMUSG00000032601.8 |

| | | | | | | | | | | | | | | | |
|---|---|---|---|---|---|---|---|---|---|---|---|---|---|---|---|
| chrX | 48284294 | 48284662 | 8.80131 | 17.30502 | 15.00832 | Promoter | 20 | 48276792 | 48285426 | 8635 | 2 | ENSMUST00 | 764 | 9530027J09F | ENSMUSG00000097197.1 |
| chr11 | 48823499 | 48824435 | 8.50367 | 17.29568 | 15.00078 | Downstream | 11 | 48826140 | 48851187 | 25048 | 1 | ENSMUST00 | -1705 | Trim7 | ENSMUSG00000040350.10 |
| chr12 | 112106900 | 112107289 | 9.07887 | 17.27829 | 14.98339 | Promoter | 12 | 112106683 | 112127573 | 20891 | 1 | ENSMUST00 | 218 | Aspg | ENSMUSG00000037686.5 |
| chr15 | 53342044 | 53343223 | 8.78498 | 17.27772 | 14.98292 | Intron (ENSM | 15 | 53068664 | 53345659 | 276996 | 2 | ENSMUST00 | 2436 | Ext1 | ENSMUSG00000061731.4 |
| chr18 | 73626639 | 73627248 | 8.7682 | 17.24965 | 14.9549 | Distal Interge | 18 | 73624928 | 73625037 | 110 | 1 | ENSMUST00 | 1712 | Gm22065 | ENSMUSG00000076941.1 |
| chr11 | 77793517 | 77793851 | 9.06006 | 17.24721 | 14.95248 | Intron (ENSM | 11 | 77785453 | 77787747 | 2295 | 2 | ENSMUST00 | -5771 | Gm10277 | ENSMUSG00000069804.2 |
| chr12 | 76195923 | 76196766 | 7.15797 | 17.24026 | 14.94564 | Downstream | 12 | 76120439 | 76177259 | 56821 | 2 | ENSMUST00 | -18665 | Esr2 | ENSMUSG00000021055.8 |
| chr14 | 70635714 | 70636321 | 8.45881 | 17.21912 | 14.92473 | Promoter | 14 | 70602185 | 70636048 | 33864 | 2 | ENSMUST00 | 0 | Dmtn | ENSMUSG00000022099.11 |
| chr16 | 78321352 | 78322028 | 8.45881 | 17.21912 | 14.92473 | Intron (ENSM | 16 | 78301756 | 78359785 | 58030 | 1 | ENSMUST00 | 19597 | Cxadr | ENSMUSG00000022865.8 |
| chr18 | 38804320 | 38804879 | 8.45881 | 17.21912 | 14.92473 | Intron (ENSM | 18 | 38771496 | 38772347 | 852 | 2 | ENSMUST00 | -31974 | Gm8302 | ENSMUSG00000081773.1 |
| chr2 | 122301296 | 122301988 | 8.45881 | 17.21912 | 14.92473 | Exon (ENSMU | 2 | 122298900 | 122302885 | 3986 | 1 | ENSMUST00 | 2397 | Duoxa2 | ENSMUSG00000027225.7 |
| chr4 | 76327299 | 76327958 | 8.45881 | 17.21912 | 14.92473 | Intron (ENSM | 4 | 76140358 | 76325288 | 184931 | 2 | ENSMUST00 | -2012 | Ptprd | ENSMUSG00000028399.12 |
| chr4 | 97707352 | 97708051 | 8.45881 | 17.21912 | 14.92473 | Intron (ENSM | 4 | 97569190 | 97690736 | 121547 | 2 | ENSMUST00 | -16617 | E130114P18I | ENSMUSG00000048747.9 |
| chr5 | 92348343 | 92349166 | 8.45881 | 17.21912 | 14.92473 | Promoter | 5 | 92346638 | 92348882 | 2245 | 2 | ENSMUST00 | 0 | Cxcl10 | ENSMUSG00000034855.8 |
| chr4 | 86053829 | 86054095 | 9.04133 | 17.21626 | 14.9221 | Promoter | 4 | 86053915 | 86087961 | 34047 | 1 | ENSMUST00 | 0 | Adamtsl1 | ENSMUSG00000066113.10 |
| chr8 | 72305094 | 72305280 | 9.04133 | 17.21626 | 14.9221 | Promoter | 8 | 72304118 | 72305276 | 1159 | 2 | ENSMUST00 | 0 | Gm10282 | ENSMUSG00000070713.4 |
| chr11 | 8467453 | 8469249 | 6.91491 | 17.20307 | 14.90901 | Promoter | 11 | 8451070 | 8468675 | 17606 | 2 | ENSMUST00 | 0 | Tns3 | ENSMUSG00000020422.8 |
| chr6 | 122742194 | 122742682 | 6.91491 | 17.20307 | 14.90901 | Promoter | 6 | 122727826 | 122742522 | 14697 | 2 | ENSMUST00 | 0 | Slc2a3 | ENSMUSG00000003153.5 |
| chr6 | 82402214 | 82403351 | 6.91491 | 17.20307 | 14.90901 | Promoter | 6 | 82402475 | 82560103 | 157629 | 1 | ENSMUST00 | 0 | Tacr1 | ENSMUSG00000030043.9 |
| chr17 | 71002765 | 71002990 | 8.73484 | 17.19383 | 14.89988 | Promoter | 17 | 70994549 | 71002878 | 8330 | 2 | ENSMUST00 | 0 | Myl12a | ENSMUSG00000024048.9 |
| chr16 | 11008077 | 11008765 | 9.02267 | 17.18543 | 14.89151 | Promoter | 16 | 11008898 | 11015186 | 6289 | 1 | ENSMUST00 | -133 | Gm4262 | ENSMUSG00000097056.1 |
| chr12 | 56695415 | 56696027 | 9.00409 | 17.15472 | 14.86093 | Promoter | 12 | 56695639 | 56712822 | 17184 | 1 | ENSMUST00 | 0 | Pax9 | ENSMUSG00000001497.13 |
| chr13 | 118714157 | 118715646 | 9.00409 | 17.15472 | 14.86093 | Promoter | 13 | 118714699 | 118792115 | 77417 | 1 | ENSMUST00 | 0 | Fgf10 | ENSMUSG00000021732.9 |
| chr1 | 157179116 | 157179842 | 8.12856 | 17.13892 | 14.84521 | Exon (ENSMU | 1 | 157179099 | 157181441 | 2343 | 2 | ENSMUST00 | 1599 | Rasal2 | ENSMUSG00000070565.6 |
| chr11 | 98750726 | 98751443 | 8.98559 | 17.12414 | 14.83055 | Intron (ENSM | 11 | 98753586 | 98768403 | 14818 | 1 | ENSMUST00 | -2143 | Thra | ENSMUSG00000058756.8 |
| chr15 | 81416774 | 81417390 | 8.98559 | 17.12414 | 14.83055 | Intron (ENSM | 15 | 81415560 | 81419188 | 3629 | 2 | ENSMUST00 | 1798 | Gm17025 | ENSMUSG00000091492.1 |
| chr8 | 54724163 | 54724903 | 8.98559 | 17.12414 | 14.83055 | Promoter | 8 | 54629233 | 54724388 | 95156 | 2 | ENSMUST00 | 0 | Wdr17 | ENSMUSG00000039375.11 |
| chr10 | 77333121 | 77334185 | 8.68527 | 17.11081 | 14.8173 | Intron (ENSM | 10 | 77325715 | 77340176 | 14462 | 2 | ENSMUST00 | 5991 | Adarb1 | ENSMUSG00000020262.10 |
| chr11 | 97315303 | 97315622 | 8.68527 | 17.11081 | 14.8173 | Promoter | 11 | 97315716 | 97329920 | 14205 | 1 | ENSMUST00 | -94 | Mrpl45 | ENSMUSG00000018882.7 |
| chr15 | 92396999 | 92397521 | 8.68527 | 17.11081 | 14.8173 | Promoter | 15 | 92396915 | 92771819 | 374905 | 1 | ENSMUST00 | 85 | Pdzrn4 | ENSMUSG00000036218.10 |
| chr4 | 41696836 | 41697076 | 8.68527 | 17.11081 | 14.8173 | Promoter | 4 | 41657498 | 41697040 | 39543 | 2 | ENSMUST00 | 0 | Cntfr | ENSMUSG00000028444.12 |
| chr4 | 142171372 | 142171665 | 8.68527 | 17.11081 | 14.8173 | Intron (ENSM | 4 | 142180602 | 142184139 | 3538 | 1 | ENSMUST00 | -8937 | Gm13062 | ENSMUSG00000087525.1 |
| chr9 | 54286374 | 54286816 | 8.68527 | 17.11081 | 14.8173 | Promoter | 9 | 54286486 | 54341786 | 55301 | 1 | ENSMUST00 | 0 | Gldn | ENSMUSG00000046167.5 |
| chr1 | 139929411 | 139929617 | 8.96716 | 17.09368 | 14.80075 | Intron (ENSM | 1 | 139862202 | 139936469 | 74268 | 2 | ENSMUST00 | 6852 | Gm16332 | ENSMUSG00000089991.1 |
| chr12 | 12323749 | 12324075 | 8.96716 | 17.09368 | 14.80075 | Intron (ENSM | 12 | 12262139 | 12376359 | 114221 | 1 | ENSMUST00 | 61611 | Fam49a | ENSMUSG00000020589.11 |
| chr11 | 68623493 | 68623933 | 8.94881 | 17.06333 | 14.77052 | Distal Interge | 11 | 68594055 | 68597954 | 3900 | 1 | ENSMUST00 | 29439 | Ccdc42 | ENSMUSG00000045915.10 |
| chr15 | 97782222 | 97782979 | 8.94881 | 17.06333 | 14.77052 | Distal Interge | 15 | 97784355 | 97792692 | 8338 | 1 | ENSMUST00 | -1376 | Slc48a1 | ENSMUSG00000081534.2 |
| chr16 | 70563636 | 70564043 | 8.94881 | 17.06333 | 14.77052 | Intron (ENSM | 16 | 70560994 | 70569498 | 8505 | 1 | ENSMUST00 | 2643 | Gbe1 | ENSMUSG00000022707.11 |
| chr17 | 8100872 | 8101493 | 8.94881 | 17.06333 | 14.77052 | Promoter | 17 | 8038510 | 8101228 | 62719 | 2 | ENSMUST00 | 0 | Gm1604A | ENSMUSG00000094083.1 |
| chr2 | 168161455 | 168162327 | 8.94881 | 17.06333 | 14.77052 | Distal Interge | 2 | 168152603 | 168154513 | 1911 | 2 | ENSMUST00 | -6943 | E130018N17 | ENSMUSG00000087648.1 |
| chr3 | 89519826 | 89521316 | 8.94881 | 17.06333 | 14.77052 | Promoter | 3 | 89520164 | 89667761 | 147598 | 1 | ENSMUST00 | 0 | Kcnn3 | ENSMUSG00000000794.8 |
| chr16 | 30070559 | 30071103 | 8.65254 | 17.05594 | 14.76351 | Distal Interge | 16 | 30066338 | 30067319 | 982 | 1 | ENSMUST00 | 4222 | Hes1 | ENSMUSG00000022528.7 |
| chr2 | 118388062 | 118388439 | 8.93054 | 17.03311 | 14.7407 | Promoter | 2 | 118388618 | 118428213 | 39596 | 1 | ENSMUST00 | -179 | Eif2ak4 | ENSMUSG00000005102.8 |
| chr4 | 9027573 | 9028722 | 8.93054 | 17.03311 | 14.7407 | Distal Interge | 4 | 8941259 | 8941656 | 398 | 1 | ENSMUST00 | 86315 | Rps18-ps2 | ENSMUSG00000083786.1 |
| chr5 | 143372867 | 143373246 | 8.93054 | 17.03311 | 14.7407 | Exon (ENSMU | 5 | 143375461 | 143380442 | 4982 | 1 | ENSMUST00 | -2215 | Grid2ip | ENSMUSG00000010825.12 |
| chr6 | 83142267 | 83142725 | 8.93054 | 17.03311 | 14.7407 | Promoter | 6 | 83142392 | 83148154 | 5763 | 1 | ENSMUST00 | 0 | Rtkn | ENSMUSG00000034930.10 |
| chr11 | 88086881 | 88087497 | 8.62005 | 17.00144 | 14.70927 | Distal Interge | 11 | 88098058 | 88100108 | 2051 | 1 | ENSMUST00 | -10561 | Cuedc1 | ENSMUSG00000018378.8 |
| chr10 | 36506707 | 36507777 | 8.89421 | 16.97301 | 14.68088 | Promoter | 10 | 36506814 | 36833773 | 326960 | 1 | ENSMUST00 | 0 | Hs3st5 | ENSMUSG00000044499.6 |
| chr2 | 130581016 | 130581325 | 8.89421 | 16.97301 | 14.68088 | Exon (ENSMU | 2 | 130580620 | 130582554 | 1935 | 2 | ENSMUST00 | 1229 | Avp | ENSMUSG00000037727.6 |
| chr4 | 121039570 | 121040059 | 8.89421 | 16.97301 | 14.68088 | Promoter | 4 | 121039748 | 121044327 | 4580 | 1 | ENSMUST00 | 0 | Col9a2 | ENSMUSG00000028626.5 |
| chr8 | 95533582 | 95534336 | 8.89421 | 16.97301 | 14.68088 | Promoter | 8 | 95534100 | 95558888 | 24789 | 1 | ENSMUST00 | 0 | Ccdc113 | ENSMUSG00000036598.3 |
| chr17 | 36157813 | 36158333 | 8.31262 | 16.9689 | 14.67687 | Promoter | 17 | 36157988 | 36158534 | 547 | 1 | ENSMUST00 | 0 | Gm6659 | ENSMUSG00000092289.1 |
| chr13 | 56221639 | 56222482 | 8.87615 | 16.94314 | 14.6513 | Exon (ENSMU | 13 | 56222224 | 56243940 | 21717 | 2 | ENSMUST00 | 21458 | 4930550C17I | ENSMUSG00000097361.1 |
| chr1 | 86059113 | 86059628 | 8.85817 | 16.91339 | 14.62167 | Distal Interge | 1 | 86064387 | 86070776 | 6390 | 1 | ENSMUST00 | -4759 | Psmd1 | ENSMUSG00000026229.12 |
| chr12 | 71577381 | 71577692 | 8.85817 | 16.91339 | 14.62167 | Distal Interge | 12 | 71724652 | 71724954 | 303 | 1 | ENSMUST00 | -146960 | Gm22959 | ENSMUSG00000084463.1 |
| chr19 | 10880177 | 10880611 | 8.85817 | 16.91339 | 14.62167 | Promoter | 19 | 10871850 | 10880370 | 8521 | 2 | ENSMUST00 | 0 | Tmem109 | ENSMUSG00000034659.9 |
| chr3 | 123690704 | 123691085 | 8.84026 | 16.88375 | 14.59221 | Promoter | 3 | 123526166 | 123690852 | 164687 | 2 | ENSMUST00 | 0 | Ndst3 | ENSMUSG00000027977.10 |
| chr1 | 170588168 | 170588755 | 8.26071 | 16.87976 | 14.58833 | Promoter | 1 | 170327613 | 170589297 | 261685 | 2 | ENSMUST00 | 542 | Nos1ap | ENSMUSG00000038473.9 |
| chr1 | 168163426 | 168164191 | 8.26071 | 16.87976 | 14.58833 | Promoter | 1 | 168158370 | 168164662 | 6293 | 2 | ENSMUST00 | 471 | Pbx1 | ENSMUSG00000052534.10 |
| chr10 | 75773574 | 75773807 | 8.26071 | 16.87976 | 14.58833 | Promoter | 10 | 75771233 | 75773414 | 2182 | 2 | ENSMUST00 | -161 | Ddt | ENSMUSG00000001666.8 |
| chr10 | 111422544 | 111422905 | 8.26071 | 16.87976 | 14.58833 | Distal Interge | 10 | 111473192 | 111496886 | 23695 | 1 | ENSMUST00 | -50287 | Nap1l1 | ENSMUSG00000058799.9 |
| chr10 | 22151351 | 22151747 | 8.26071 | 16.87976 | 14.58833 | Downstream | 10 | 22144421 | 22149270 | 4850 | 2 | ENSMUST00 | -2082 | E030030I06R | ENSMUSG00000097327.2 |
| chr11 | 96967277 | 96967816 | 8.26071 | 16.87976 | 14.58833 | Promoter | 11 | 96953473 | 96968665 | 15193 | 2 | ENSMUST00 | 849 | Sp2 | ENSMUSG00000018678.7 |
| chr11 | 96777785 | 96778193 | 8.26071 | 16.87976 | 14.58833 | Promoter | 11 | 96767877 | 96777559 | 9683 | 2 | ENSMUST00 | -227 | Snx11 | ENSMUSG00000020876.9 |
| chr11 | 75463274 | 75463896 | 8.26071 | 16.87976 | 14.58833 | Promoter | 11 | 75463716 | 75463810 | 95 | 1 | ENSMUST00 | 0 | Mir22 | ENSMUSG00000065529.1 |
| chr13 | 53475559 | 53476433 | 8.26071 | 16.87976 | 14.58833 | Distal Interge | 13 | 53471506 | 53473074 | 1569 | 2 | ENSMUST00 | -2486 | Msx2 | ENSMUSG00000021469.9 |
| chr15 | 98004179 | 98004982 | 8.26071 | 16.87976 | 14.58833 | Promoter | 15 | 97975602 | 98004657 | 29056 | 2 | ENSMUST00 | 0 | Col2a1 | ENSMUSG00000022483.11 |
| chr17 | 26438783 | 26439393 | 8.26071 | 16.87976 | 14.58833 | Exon (ENSMU | 17 | 26414965 | 26446349 | 31385 | 1 | ENSMUST00 | 23819 | Neurl1b | ENSMUSG00000034413.9 |
| chr17 | 70738102 | 70738362 | 8.26071 | 16.87976 | 14.58833 | Intron (ENSM | 17 | 70745329 | 70755582 | 10254 | 2 | ENSMUST00 | 17220 | 5031415H12I | ENSMUSG00000097282.1 |
| chr18 | 25754660 | 25755589 | 8.26071 | 16.87976 | 14.58833 | Promoter | 18 | 25477632 | 25753983 | 276352 | 2 | ENSMUST00 | -678 | Celf4 | ENSMUSG00000024268.10 |
| chr18 | 65490243 | 65490667 | 8.26071 | 16.87976 | 14.58833 | Distal Interge | 18 | 65430963 | 65478823 | 47861 | 1 | ENSMUST00 | 59281 | Malt1 | ENSMUSG00000032688.7 |
| chr2 | 76549391 | 76549798 | 8.26071 | 16.87976 | 14.58833 | Exon (ENSMU | 2 | 76548340 | 76596045 | 47706 | 1 | ENSMUST00 | 1052 | Osbpl6 | ENSMUSG00000042359.13 |
| chr2 | 38931753 | 38932320 | 8.26071 | 16.87976 | 14.58833 | Promoter | 2 | 38931978 | 38963753 | 31776 | 1 | ENSMUST00 | 0 | Olfml2a | ENSMUSG00000046618.7 |
| chr2 | 155613361 | 155613810 | 8.26071 | 16.87976 | 14.58833 | Exon (ENSMU | 2 | 155611212 | 155634307 | 23096 | 1 | ENSMUST00 | 2150 | Myh7b | ENSMUSG00000074652.3 |
| chr2 | 30693330 | 30693904 | 8.26071 | 16.87976 | 14.58833 | Distal Interge | 2 | 30658613 | 30678011 | 19399 | 2 | ENSMUST00 | -15320 | Gm14486 | ENSMUSG00000087268.1 |
| chr2 | 129947770 | 129948374 | 8.26071 | 16.87976 | 14.58833 | Promoter | 2 | 129948568 | 129949207 | 640 | 1 | ENSMUST00 | -194 | RP23-263L18 | ENSMUSG00000100255.1 |
| chr3 | 27282940 | 27283760 | 8.26071 | 16.87976 | 14.58833 | Intron (ENSM | 3 | 27317028 | 27339665 | 22638 | 1 | ENSMUST00 | -33268 | Tnfsf10 | ENSMUSG00000039304.6 |
| chr3 | 55787449 | 55788251 | 8.26071 | 16.87976 | 14.58833 | Intron (ENSM | 3 | 55782510 | 55785285 | 2776 | 1 | ENSMUST00 | 4940 | Mab21l1 | ENSMUSG00000056947.4 |
| chr4 | 8770488 | 8771068 | 8.26071 | 16.87976 | 14.58833 | Intron (ENSM | 4 | 8751438 | 8867224 | 115787 | 1 | ENSMUST00 | 19051 | Chd7 | ENSMUSG00000041235.6 |
| chr4 | 62288080 | 62288869 | 8.26071 | 16.87976 | 14.58833 | Intron (ENSM | 4 | 62286449 | 62298411 | 11963 | 1 | ENSMUST00 | 1632 | Slc31a2 | ENSMUSG00000066152.6 |
| chr5 | 65724441 | 65725032 | 8.26071 | 16.87976 | 14.58833 | Distal Interge | 5 | 65615260 | 65697856 | 82597 | 2 | ENSMUST00 | -26586 | Pds5a | ENSMUSG00000029202.9 |
| chr6 | 33591847 | 33592208 | 8.26071 | 16.87976 | 14.58833 | Intron (ENSM | 6 | 33701434 | 33862160 | 160727 | 1 | ENSMUST00 | -109226 | Exoc4 | ENSMUSG00000029763.12 |
| chr6 | 131353445 | 131353983 | 8.26071 | 16.87976 | 14.58833 | Promoter | 6 | 131311192 | 131353597 | 42406 | 2 | ENSMUST00 | 0 | Styk1 | ENSMUSG00000032899.9 |
| chr6 | 126711566 | 126713041 | 8.26071 | 16.87976 | 14.58833 | Intron (ENSM | 6 | 126718684 | 126741601 | 22918 | 1 | ENSMUST00 | -5643 | Gm26673 | ENSMUSG00000097596.1 |
| chr6 | 83762703 | 83763452 | 8.26071 | 16.87976 | 14.58833 | Promoter | 6 | 83761756 | 83762842 | 1087 | 2 | ENSMUST00 | 0 | RP23-252I12 | ENSMUSG00000099354.1 |
| chr7 | 25281477 | 25282003 | 8.26071 | 16.87976 | 14.58833 | Promoter | 7 | 25282179 | 25294145 | 11967 | 1 | ENSMUST00 | -176 | Cic | ENSMUSG00000005442.8 |
| chr7 | 19290788 | 19291589 | 8.26071 | 16.87976 | 14.58833 | Promoter | 7 | 19291099 | 19296159 | 5061 | 1 | ENSMUST00 | 0 | Rtn2 | ENSMUSG00000030401.10 |
| chr7 | 40898749 | 40899757 | 8.26071 | 16.87976 | 14.58833 | Promoter | 7 | 40899215 | 40930594 | 31380 | 1 | ENSMUST00 | 0 | Vstm2b | ENSMUSG00000039257.10 |
| chr8 | 126497748 | 126498330 | 8.26071 | 16.87976 | 14.58833 | Distal Interge | 8 | 126476248 | 126476412 | 165 | 2 | ENSMUST00 | -21337 | Gm6091 | ENSMUSG00000074026.3 |
| chr9 | 65162638 | 65163554 | 8.26071 | 16.87976 | 14.58833 | Intron (ENSM | 9 | 65141154 | 65185857 | 44704 | 1 | ENSMUST00 | 21485 | Igdcc3 | ENSMUSG00000032394.5 |
| chr9 | 51009289 | 51009537 | 8.26071 | 16.87976 | 14.58833 | Promoter | 9 | 50892801 | 51009073 | 116273 | 2 | ENSMUST00 | -217 | Sik2 | ENSMUSG00000037112.11 |
| chr9 | 66805791 | 66806244 | 8.26071 | 16.87976 | 14.58833 | Promoter | 9 | 66805306 | 66805938 | 633 | 2 | ENSMUST00 | 0 | Ppp1r2-ps4 | ENSMUSG00000059237.6 |
| chr18 | 34373623 | 34373917 | 8.82242 | 16.85422 | 14.56465 | Promoter | 18 | 34344889 | 34373415 | 28527 | 2 | ENSMUST00 | -209 | Reep5 | ENSMUSG00000005873.5 |
| chr1 | 58029703 | 58030517 | 8.52402 | 16.84016 | 14.5507 | Promoter | 1 | 58029931 | 58106413 | 76483 | 1 | ENSMUST00 | 0 | Aox1 | ENSMUSG00000063558.4 |
| chr10 | 95880231 | 95880640 | 8.52402 | 16.84016 | 14.5507 | Distal Interge | 10 | 95898603 | 95898735 | 133 | 1 | ENSMUST00 | -17963 | Gm24433 | ENSMUSG00000077268.1 |
| chr11 | 72940693 | 72940985 | 8.52402 | 16.84016 | 14.5507 | Distal Interge | 11 | 72920843 | 72927120 | 6278 | 1 | ENSMUST00 | 19851 | Zzef1 | ENSMUSG00000055670.8 |
| chr13 | 70841258 | 70841825 | 8.52402 | 16.84016 | 14.5507 | Promoter | 13 | 70727802 | 70841811 | 114010 | 2 | ENSMUST00 | 0 | Adamts16 | ENSMUSG00000049538.9 |
| chr18 | 64342377 | 64343150 | 8.52402 | 16.84016 | 14.5507 | Intron (ENSM | 18 | 64340364 | 64398488 | 58125 | 1 | ENSMUST00 | 2014 | Onecut2 | ENSMUSG00000045991.13 |
| chr19 | 44758909 | 44759336 | 8.52402 | 16.84016 | 14.5507 | Promoter | 19 | 44759915 | 44818554 | 58640 | 1 | ENSMUST00 | -579 | Pax2 | ENSMUSG00000004231.10 |
| chr7 | 145030802 | 145031199 | 8.52402 | 16.84016 | 14.5507 | Distal Interge | 7 | 144929931 | 144939869 | 9939 | 2 | ENSMUST00 | -90934 | Ccnd1 | ENSMUSG00000070348.4 |
| chr7 | 114103949 | 114104284 | 8.80466 | 16.8248 | 14.53583 | Intron (ENSM | 7 | 114046785 | 114117781 | 70997 | 2 | ENSMUST00 | 13497 | Rras2 | ENSMUSG00000055723.5 |
| chr9 | 40858586 | 40859135 | 8.80466 | 16.8248 | 14.53583 | Distal Interge | 9 | 40874127 | 40877972 | 3846 | 1 | ENSMUST00 | -14992 | Bsx | ENSMUSG00000054360.3 |
| chr1 | 166308886 | 166309682 | 8.78697 | 16.7955 | 14.50663 | Promoter | 1 | 166306741 | 166309585 | 2845 | 2 | ENSMUST00 | 0 | 5330438I03R | ENSMUSG00000091393.1 |
| chr15 | 98776332 | 98776744 | 8.78697 | 16.7955 | 14.50663 | Promoter | 15 | 98772200 | 98777261 | 5062 | 2 | ENSMUST00 | 517 | Wnt10b | ENSMUSG00000022996.9 |
| chr11 | 95587134 | 95587864 | 8.76935 | 16.76631 | 14.47755 | Promoter | 11 | 95568818 | 95587735 | 18918 | 2 | ENSMUST00 | 0 | Ngfr | ENSMUSG00000000120.6 |
| chr11 | 40733094 | 40733418 | 8.76935 | 16.76631 | 14.47755 | Promoter | 11 | 40720903 | 40733416 | 12514 | 2 | ENSMUST00 | 0 | Hmmr | ENSMUSG00000020330.11 |
| chr5 | 125393312 | 125394148 | 8.76935 | 16.76631 | 14.47755 | Distal Interge | 5 | 125385977 | 125390202 | 4226 | 2 | ENSMUST00 | -3111 | Ubc | ENSMUSG00000008348.9 |
| chr7 | 37773326 | 37773746 | 8.76935 | 16.76631 | 14.47755 | Promoter | 7 | 37569629 | 37773641 | 204013 | 2 | ENSMUST00 | 0 | Zfp536 | ENSMUSG00000043456.11 |
| chr9 | 62828063 | 62828925 | 8.76935 | 16.76631 | 14.47755 | Distal Interge | 9 | 62838785 | 62852006 | 13222 | 1 | ENSMUST00 | -9860 | Cln6 | ENSMUSG00000032245.7 |
| chr3 | 40091798 | 40092177 | 8.75179 | 16.73723 | 14.44895 | Distal Interge | 3 | 39799632 | 39799716 | 85 | 2 | ENSMUST00 | -292083 | Gm23831 | ENSMUSG00000089520.1 |
| chr11 | 97434949 | 97435291 | 8.1714 | 16.72606 | 14.43788 | Promoter | 11 | 97436285 | 97444228 | 7944 | 1 | ENSMUST00 | -994 | Arhgap23 | ENSMUSG00000049807.11 |
| chr17 | 67290633 | 67291103 | 8.73431 | 16.70826 | 14.42027 | Intron (ENSM | 17 | 66666848 | 67354459 | 687612 | 2 | ENSMUST00 | 63356 | Ptprm | ENSMUSG00000033278.9 |
| chr18 | 69346873 | 69347404 | 8.73431 | 16.70826 | 14.42027 | Promoter | 18 | 69346203 | 69683248 | 337046 | 1 | ENSMUST00 | 671 | Tcf4 | ENSMUSG00000053477.10 |
| chr9 | 107675771 | 107676736 | 8.7169 | 16.6794 | 14.39155 | 3' UTR | 9 | 107674477 | 107679592 | 5116 | 2 | ENSMUST00 | 2856 | Gnat1 | ENSMUSG00000034837.7 |
| chr1 | 183194279 | 183195192 | 7.82203 | 16.59716 | 14.30959 | Intron (ENSM | 1 | 183086264 | 183221529 | 135266 | 2 | ENSMUST00 | 26337 | Disp1 | ENSMUSG00000030768.7 |
| chr10 | 122448534 | 122449556 | 7.82203 | 16.59716 | 14.30959 | Promoter | 10 | 122448499 | 122453453 | 4955 | 1 | ENSMUST00 | 36 | Avpr1a | ENSMUSG00000020123.5 |
| chr10 | 122886388 | 122886736 | 7.82203 | 16.59716 | 14.30959 | Intron (ENSM | 10 | 122895368 | 122941449 | 46082 | 1 | ENSMUST00 | -8632 | Ppm1h | ENSMUSG00000034613.7 |
| chr10 | 81424144 | 81425098 | 7.82203 | 16.59716 | 14.30959 | Intron (ENSM | 10 | 81399138 | 81427114 | 27977 | 2 | ENSMUST00 | 2016 | Nfic | ENSMUSG00000055053.12 |
| chr11 | 62966746 | 62967589 | 7.82203 | 16.59716 | 14.30959 | Intron (ENSM | 11 | 62951193 | 62993095 | 41903 | 1 | ENSMUST00 | 15554 | Cdrt4 | ENSMUSG00000042200.3 |
| chr11 | 103142280 | 103143074 | 7.82203 | 16.59716 | 14.30959 | Distal Interge | 11 | 103133865 | 103138279 | 4415 | 1 | ENSMUST00 | 8416 | Hexim2 | ENSMUSG00000043372.7 |
| chr11 | 95796627 | 95796934 | 7.82203 | 16.59716 | 14.30959 | Intron (ENSM | 11 | 95824499 | 95830701 | 6203 | 1 | ENSMUST00 | -27565 | Phospho1 | ENSMUSG00000050860.6 |
| chr14 | 14632466 | 14633715 | 7.82203 | 16.59716 | 14.30959 | Distal Interge | 14 | 14703025 | 14799943 | 96919 | 1 | ENSMUST00 | -69310 | Slc4a7 | ENSMUSG00000021733.9 |
| chr16 | 60605788 | 60606816 | 7.82203 | 16.59716 | 14.30959 | Promoter | 16 | 60605755 | 60611236 | 5482 | 1 | ENSMUST00 | 34 | Gm9017 | ENSMUSG00000089685.2 |
| chr17 | 74159464 | 74159980 | 7.82203 | 16.59716 | 14.30959 | Distal Interge | 17 | 74017706 | 74047916 | 30211 | 2 | ENSMUST00 | -111549 | Srd5a2 | ENSMUSG00000038541.7 |
| chr17 | 27204625 | 27204859 | 7.82203 | 16.59716 | 14.30959 | Promoter | 17 | 27189601 | 27204438 | 14838 | 2 | ENSMUST00 | -188 | Lemd2 | ENSMUSG00000044857.7 |
| chr18 | 69347968 | 69348470 | 7.82203 | 16.59716 | 14.30959 | Intron (ENSM | 18 | 69346203 | 69683248 | 337046 | 1 | ENSMUST00 | 1766 | Tcf4 | ENSMUSG00000053477.10 |
| chr19 | 5846849 | 5849375 | 7.82203 | 16.59716 | 14.30959 | Promoter | 19 | 5848508 | 5851372 | 2865 | 1 | ENSMUST00 | 0 | Frmd8os | ENSMUSG00000043488.5 |
| chr2 | 180273745 | 180274071 | 7.82203 | 16.59716 | 14.30959 | Promoter | 2 | 180262132 | 180273496 | 11365 | 2 | ENSMUST00 | -250 | Cables2 | ENSMUSG00000038990.9 |

| | | | | | | | | | | | | | | | |
|---|---|---|---|---|---|---|---|---|---|---|---|---|---|---|---|
| chr2 | 144224128 | 144224938 | 7.82203 | 16.59716 | 14.30959 | Distal Interge | 2 | 144243241 | 144244849 | 1609 | 1 | ENSMUST00 | -18303 | 4930444E06l | ENSMUSG00000085127.1 |
| chr3 | 128990602 | 128991146 | 7.82203 | 16.59716 | 14.30959 | Distal Interge | 3 | 128953782 | 128955098 | 1317 | 2 | ENSMUST00 | -35505 | Gm9387 | ENSMUSG00000074236.2 |
| chr4 | 46534328 | 46534898 | 7.82203 | 16.59716 | 14.30959 | Promoter | 4 | 46494819 | 46534414 | 39596 | 2 | ENSMUST00 | 0 | Trim14 | ENSMUSG00000039853.13 |
| chr4 | 114911465 | 114913643 | 7.82203 | 16.59716 | 14.30959 | Exon (ENSMU | 4 | 114909257 | 114921967 | 12711 | 1 | ENSMUST00 | 2209 | Foxd2os | ENSMUSG00000085399.2 |
| chr4 | 41054791 | 41055340 | 7.82203 | 16.59716 | 14.30959 | Distal Interge | 4 | 41050220 | 41050326 | 107 | 1 | ENSMUST00 | 4572 | Gm25637 | ENSMUSG00000088154.1 |
| chr5 | 97935297 | 97935660 | 7.82203 | 16.59716 | 14.30959 | Intron (ENSM | 5 | 97882783 | 98030733 | 147951 | 2 | ENSMUST00 | 95073 | Antxr2 | ENSMUSG00000029338.9 |
| chr5 | 139540275 | 139541854 | 7.82203 | 16.59716 | 14.30959 | Distal Interge | 5 | 139543494 | 139548179 | 4686 | 1 | ENSMUST00 | -1640 | Uncx | ENSMUSG00000029546.11 |
| chr7 | 51510957 | 51511349 | 7.82203 | 16.59716 | 14.30959 | Promoter | 7 | 51511029 | 51598709 | 87681 | 1 | ENSMUST00 | 0 | Ano5 | ENSMUSG00000055489.7 |
| chr8 | 121665290 | 121665960 | 7.82203 | 16.59716 | 14.30959 | Promoter | 8 | 121666628 | 121666706 | 79 | 2 | ENSMUST00 | 746 | Gm23299 | ENSMUSG00000064642.1 |
| chr9 | 107377927 | 107378950 | 7.82203 | 16.59716 | 14.30959 | Distal Interge | 9 | 107399612 | 107529346 | 129735 | 1 | ENSMUST00 | -20662 | Cacna2d2 | ENSMUSG00000010066.10 |
| chr9 | 48161935 | 48162296 | 7.82203 | 16.59716 | 14.30959 | Distal Interge | 9 | 48319144 | 48340834 | 21691 | 2 | ENSMUST00 | 178538 | Nxpe2 | ENSMUSG00000032028.10 |
| chr9 | 114672779 | 114673381 | 7.82203 | 16.59716 | 14.30959 | Distal Interge | 9 | 114688790 | 114724302 | 35513 | 1 | ENSMUST00 | -15409 | Dync1li1 | ENSMUSG00000032435.8 |
| chr9 | 31688049 | 31688624 | 7.82203 | 16.59716 | 14.30959 | Distal Interge | 9 | 31426199 | 31464238 | 38040 | 2 | ENSMUST00 | -223812 | Tmem45b | ENSMUSG00000041737.7 |
| chr9 | 48521120 | 48521503 | 7.82203 | 16.59716 | 14.30959 | Distal Interge | 9 | 48506964 | 48507060 | 97 | 2 | ENSMUST00 | -14061 | Gm23653 | ENSMUSG00000080401.1 |
| chr2 | 90782528 | 90782840 | 8.66508 | 16.59347 | 14.30722 | Promoter | 2 | 90782727 | 90817244 | 34518 | 1 | ENSMUST00 | 0 | Agbl2 | ENSMUSG00000040812.9 |
| chr1 | 180847843 | 180848233 | 8.36866 | 16.57842 | 14.29226 | Distal Interge | 1 | 180851151 | 180868097 | 16947 | 1 | ENSMUST00 | -2918 | Sde2 | ENSMUSG00000038806.6 |
| chr1 | 36064634 | 36065631 | 8.36866 | 16.57842 | 14.29226 | Distal Interge | 1 | 36068400 | 36106446 | 38047 | 1 | ENSMUST00 | -2769 | Hs6st1 | ENSMUSG00000045216.6 |
| chr13 | 81691806 | 81692046 | 8.36866 | 16.57842 | 14.29226 | Intron (ENSM | 13 | 81673880 | 81687285 | 13406 | 2 | ENSMUST00 | -4522 | Polr3g | ENSMUSG00000035834.6 |
| chr6 | 99484863 | 99485599 | 8.36866 | 16.57842 | 14.29226 | Promoter | 6 | 99484591 | 99484702 | 112 | 2 | ENSMUST00 | -162 | Gm22328 | ENSMUSG00000088699.1 |
| chr15 | 78794341 | 78796577 | 8.64794 | 16.56503 | 14.27926 | Exon (ENSMU | 15 | 78787411 | 78790515 | 3105 | 2 | ENSMUST00 | -3827 | Card10 | ENSMUSG00000033170.9 |
| chr18 | 56620751 | 56621453 | 8.64794 | 16.56503 | 14.27926 | Distal Interge | 18 | 56588426 | 56594782 | 6357 | 1 | ENSMUST00 | 32326 | Tex43 | ENSMUSG00000032900.9 |
| chr9 | 46230033 | 46230457 | 8.64794 | 16.56503 | 14.27926 | 3' UTR | 9 | 46228721 | 46229973 | 1253 | 1 | ENSMUST00 | 1313 | Apoa1 | ENSMUSG00000032083.8 |
| chr15 | 75997408 | 75998865 | 8.63087 | 16.53671 | 14.25102 | Promoter | 15 | 75996713 | 75999152 | 2440 | 1 | ENSMUST00 | 696 | Mapk15 | ENSMUSG00000063704.7 |
| chr17 | 44192488 | 44195194 | 8.0349 | 16.49019 | 14.20466 | Intron (ENSM | 17 | 44188572 | 44280168 | 91597 | 1 | ENSMUST00 | 3917 | Clic5 | ENSMUSG00000023959.8 |
| chr6 | 30173073 | 30174607 | 8.0349 | 16.49019 | 14.20466 | Distal Interge | 6 | 30169668 | 30169737 | 70 | 2 | ENSMUST00 | -3337 | Mir183 | ENSMUSG00000065619.2 |
| chr1 | 7206398 | 7206966 | 8.59694 | 16.48037 | 14.19497 | Distal Interge | 1 | 7177739 | 7179037 | 1299 | 1 | ENSMUST00 | 28660 | Gm9826 | ENSMUSG00000048538.7 |
| chr12 | 103325312 | 103325641 | 8.59694 | 16.48037 | 14.19497 | Exon (ENSMU | 12 | 103321146 | 103335584 | 14439 | 2 | ENSMUST00 | 9943 | Asb2 | ENSMUSG00000021200.9 |
| chr12 | 29367971 | 29368628 | 8.59694 | 16.48037 | 14.19497 | Distal Interge | 12 | 29528384 | 29923209 | 394826 | 1 | ENSMUST00 | -159756 | Myt1l | ENSMUSG00000061911.9 |
| chr19 | 57274463 | 57275629 | 8.59694 | 16.48037 | 14.19497 | Intron (ENSM | 19 | 57036966 | 57239861 | 202896 | 2 | ENSMUST00 | -34603 | Ablim1 | ENSMUSG00000025085.11 |
| chr13 | 63901141 | 63901714 | 7.7526 | 16.47343 | 14.18852 | Distal Interge | 13 | 63872455 | 63900302 | 27848 | 1 | ENSMUST00 | 28687 | Ercc6l2 | ENSMUSG00000021470.12 |
| chr1 | 86440745 | 86441363 | 8.58007 | 16.45236 | 14.16747 | Distal Interge | 1 | 86426329 | 86428052 | 1724 | 1 | ENSMUST00 | 14417 | 1700019O17 | ENSMUSG00000036574.5 |
| chr7 | 27481298 | 27482075 | 8.58007 | 16.45236 | 14.16747 | Distal Interge | 7 | 27486910 | 27490316 | 3407 | 1 | ENSMUST00 | -4835 | Sertad1 | ENSMUSG00000008384.8 |
| chr6 | 86497025 | 86498372 | 8.56326 | 16.42445 | 14.13964 | Promoter | 6 | 86498781 | 86500537 | 1757 | 1 | ENSMUST00 | -409 | RP23-281H4. | ENSMUSG00000100174.1 |
| chr1 | 59574493 | 59575396 | 7.45894 | 16.41436 | 14.12962 | Intron (ENSM | 1 | 59558008 | 59566636 | 8629 | 1 | ENSMUST00 | 16486 | Gm973 | ENSMUSG00000047361.10 |
| chr10 | 128179852 | 128180477 | 7.45894 | 16.41436 | 14.12962 | Promoter | 10 | 128129470 | 128180297 | 50828 | 2 | ENSMUST00 | 0 | Rbms2 | ENSMUSG00000040043.10 |
| chr10 | 94919344 | 94919923 | 7.45894 | 16.41436 | 14.12962 | Intron (ENSM | 10 | 94870830 | 94922657 | 51828 | 2 | ENSMUST00 | 2734 | Plxnc1 | ENSMUSG00000074785.4 |
| chr11 | 70520099 | 70520598 | 7.45894 | 16.41436 | 14.12962 | Promoter | 11 | 70519639 | 70520519 | 881 | 1 | ENSMUST00 | 461 | Gltpd2 | ENSMUSG00000046811.3 |
| chr11 | 98746952 | 98747514 | 7.45894 | 16.41436 | 14.12962 | Intron (ENSM | 11 | 98744233 | 98753777 | 9545 | 1 | ENSMUST00 | 2720 | Thra | ENSMUSG00000058756.8 |
| chr15 | 103183422 | 103184164 | 7.45894 | 16.41436 | 14.12962 | Distal Interge | 15 | 103153307 | 103158002 | 4696 | 2 | ENSMUST00 | -25421 | Smug1 | ENSMUSG00000036061.9 |
| chr15 | 81511692 | 81512259 | 7.45894 | 16.41436 | 14.12962 | Distal Interge | 15 | 81499084 | 81499548 | 465 | 2 | ENSMUST00 | -12145 | Gm5218 | ENSMUSG00000091198.1 |
| chr16 | 60604856 | 60605684 | 7.45894 | 16.41436 | 14.12962 | Promoter | 16 | 59653483 | 60605248 | 951766 | 2 | ENSMUST00 | 0 | Epha6 | ENSMUSG00000055540.10 |
| chr17 | 8505790 | 8506275 | 7.45894 | 16.41436 | 14.12962 | Distal Interge | 17 | 8525372 | 8981671 | 456300 | 1 | ENSMUST00 | -19097 | Pde10a | ENSMUSG00000023868.10 |
| chr17 | 47039105 | 47039486 | 7.45894 | 16.41436 | 14.12962 | Distal Interge | 17 | 47042587 | 47043039 | 453 | 2 | ENSMUST00 | 3553 | Gm4945 | ENSMUSG00000062472.6 |
| chr17 | 33760632 | 33760919 | 7.45894 | 16.41436 | 14.12962 | Promoter | 17 | 33742484 | 33760530 | 18047 | 2 | ENSMUST00 | -103 | Rab11b | ENSMUSG00000077450.7 |
| chr19 | 8407368 | 8408094 | 7.45894 | 16.41436 | 14.12962 | Distal Interge | 19 | 8335371 | 8405111 | 69741 | 2 | ENSMUST00 | -2258 | Slc22a30 | ENSMUSG00000052562.10 |
| chr2 | 168765785 | 168766473 | 7.45894 | 16.41436 | 14.12962 | Promoter | 2 | 168766143 | 168768108 | 1966 | 1 | ENSMUST00 | 0 | Gm14261 | ENSMUSG00000085322.1 |
| chr4 | 130199066 | 130199520 | 7.45894 | 16.41436 | 14.12962 | Distal Interge | 4 | 130209109 | 130209856 | 748 | 2 | ENSMUST00 | 10336 | Gm853 | ENSMUSG00000023120.9 |
| chr4 | 153417347 | 153418135 | 7.45894 | 16.41436 | 14.12962 | Intron (ENSM | 4 | 153374231 | 153482179 | 107949 | 2 | ENSMUST00 | 64044 | Ajap1 | ENSMUSG00000039546.9 |
| chr4 | 133098204 | 133098494 | 7.45894 | 16.41436 | 14.12962 | Distal Interge | 4 | 133085474 | 133089146 | 3673 | 1 | ENSMUST00 | 12731 | Gm26615 | ENSMUSG00000097088.1 |
| chr7 | 109616081 | 109616620 | 7.45894 | 16.41436 | 14.12962 | Promoter | 7 | 109523920 | 109616568 | 92649 | 2 | ENSMUST00 | 0 | St5 | ENSMUSG00000031024.7 |
| chr8 | 102780817 | 102781107 | 7.45894 | 16.41436 | 14.12962 | Intron (ENSM | 8 | 102632995 | 102785111 | 152117 | 2 | ENSMUST00 | 4004 | Cdh11 | ENSMUSG00000031673.5 |
| chr1 | 37539432 | 37539696 | 8.21885 | 16.32505 | 14.04127 | Intron (ENSM | 1 | 37439340 | 37541016 | 101677 | 2 | ENSMUST00 | 1320 | Mgat4a | ENSMUSG00000026110.10 |
| chr15 | 97849374 | 97849843 | 8.21885 | 16.32505 | 14.04127 | Distal Interge | 15 | 97792666 | 97844502 | 51837 | 2 | ENSMUST00 | -4873 | Hdac7 | ENSMUSG00000022475.13 |
| chr3 | 82547923 | 82548904 | 8.21885 | 16.32505 | 14.04127 | Promoter | 3 | 82538383 | 82548084 | 9702 | 2 | ENSMUST00 | 0 | Npy2r | ENSMUSG00000028004.7 |
| chr6 | 100163172 | 100163502 | 8.21885 | 16.32505 | 14.04127 | Distal Interge | 6 | 100228566 | 100287441 | 58876 | 2 | ENSMUST00 | 123939 | Rybp | ENSMUSG00000072872.2 |
| chr8 | 110217355 | 110217761 | 8.21885 | 16.32505 | 14.04127 | Promoter | 8 | 110217989 | 110224486 | 6498 | 1 | ENSMUST00 | -228 | Cmtr2 | ENSMUSG00000046441.5 |
| chr1 | 171085567 | 171086034 | 7.15345 | 16.30454 | 14.02114 | Distal Interge | 1 | 171079840 | 171079901 | 62 | 2 | ENSMUST00 | -5667 | Gm27844 | ENSMUSG00000098965.1 |
| chr10 | 58322741 | 58323283 | 7.15345 | 16.30454 | 14.02114 | Promoter | 10 | 58323466 | 58424691 | 101226 | 1 | ENSMUST00 | -183 | Lims1 | ENSMUSG00000019920.12 |
| chr10 | 127997714 | 127997967 | 7.15345 | 16.30454 | 14.02114 | Exon (ENSMU | 10 | 127990941 | 128000678 | 9738 | 2 | ENSMUST00 | 2711 | Hsd17b6 | ENSMUSG00000025396.6 |
| chr10 | 69925327 | 69926348 | 7.15345 | 16.30454 | 14.02114 | Promoter | 10 | 69925495 | 69926338 | 844 | 1 | ENSMUST00 | 0 | Ank3 | ENSMUSG00000069601.8 |
| chr14 | 70457223 | 70458066 | 7.15345 | 16.30454 | 14.02114 | Promoter | 14 | 70457476 | 70468832 | 11357 | 1 | ENSMUST00 | 0 | Phyhip | ENSMUSG00000003469.4 |
| chr18 | 37822271 | 37823021 | 7.15345 | 16.30454 | 14.02114 | Intron (ENSM | 18 | 37819546 | 37841873 | 22328 | 1 | ENSMUST00 | 2726 | Pcdhga12 | ENSMUSG00000023036.9 |
| chr6 | 52219876 | 52221129 | 7.15345 | 16.30454 | 14.02114 | Promoter | 6 | 52214491 | 52220393 | 5903 | 2 | ENSMUST00 | 0 | Hoxa7 | ENSMUSG00000038236.7 |
| chr6 | 117717244 | 117717486 | 7.15345 | 16.30454 | 14.02114 | Promoter | 6 | 117646694 | 117717263 | 70570 | 2 | ENSMUST00 | 0 | RP24-479J14 | ENSMUSG00000099869.1 |
| chr8 | 4213700 | 4214425 | 7.15345 | 16.30454 | 14.02114 | Exon (ENSMU | 8 | 4215049 | 4216912 | 1864 | 2 | ENSMUST00 | 2487 | BC068157 | ENSMUSG00000064125.9 |
| chr7 | 145206041 | 145206738 | 7.90289 | 16.26095 | 13.97799 | Distal Interge | 7 | 145253923 | 145283927 | 30005 | 2 | ENSMUST00 | 77189 | Tpcn2 | ENSMUSG00000048677.8 |
| chr1 | 40324485 | 40325998 | 6.89286 | 16.24995 | 13.96717 | Promoter | 1 | 40324627 | 40365471 | 40845 | 1 | ENSMUST00 | 0 | Il1rl2 | ENSMUSG00000070942.3 |
| chr10 | 81328327 | 81330303 | 6.89286 | 16.24995 | 13.96717 | Promoter | 10 | 81328732 | 81335171 | 6440 | 1 | ENSMUST00 | 0 | Tbxa2r | ENSMUSG00000034881.7 |
| chr13 | 48552169 | 48552951 | 6.89286 | 16.24995 | 13.96717 | Distal Interge | 13 | 48538171 | 48538270 | 100 | 2 | ENSMUST00 | -13900 | Mirlet7a-1 | ENSMUSG00000065421.2 |
| chr17 | 25222541 | 25222838 | 6.89286 | 16.24995 | 13.96717 | Promoter | 17 | 25222582 | 25234442 | 11861 | 1 | ENSMUST00 | 0 | Unkl | ENSMUSG00000015127.9 |
| chr7 | 130646911 | 130647418 | 6.89286 | 16.24995 | 13.96717 | Intron (ENSM | 7 | 130692556 | 130764784 | 72229 | 1 | ENSMUST00 | -45138 | Tacc2 | ENSMUSG00000030852.10 |
| chr12 | 71376548 | 71377306 | 6.66795 | 16.23829 | 13.9557 | Distal Interge | 12 | 71383154 | 71384208 | 1055 | 2 | ENSMUST00 | 6902 | RP23-50H15. | ENSMUSG00000100668.1 |
| chr17 | 13007370 | 13007695 | 6.66795 | 16.23829 | 13.9557 | Promoter | 17 | 13007839 | 13018119 | 10281 | 1 | ENSMUST00 | -144 | Sod2 | ENSMUSG00000006818.4 |
| chr1 | 134092728 | 134092987 | 8.73111 | 16.23101 | 13.94854 | Distal Interge | 1 | 134075170 | 134079120 | 3951 | 2 | ENSMUST00 | -13609 | Btg2 | ENSMUSG00000020423.6 |
| chr1 | 134708855 | 134709621 | 8.73111 | 16.23101 | 13.94854 | Promoter | 1 | 134709300 | 134753149 | 43850 | 1 | ENSMUST00 | 0 | Syt2 | ENSMUSG00000026452.10 |
| chr1 | 80758116 | 80758814 | 8.73111 | 16.23101 | 13.94854 | Promoter | 1 | 80501073 | 80758448 | 257376 | 2 | ENSMUST00 | 0 | Dock10 | ENSMUSG00000038608.10 |
| chr11 | 90029518 | 90029907 | 8.73111 | 16.23101 | 13.94854 | Promoter | 11 | 90030348 | 90036508 | 6161 | 1 | ENSMUST00 | -441 | Tmem100 | ENSMUSG00000069763.3 |
| chr11 | 97536890 | 97537516 | 8.73111 | 16.23101 | 13.94854 | Exon (ENSMU | 11 | 97519612 | 97521895 | 2284 | 2 | ENSMUST00 | -14996 | Gm11611 | ENSMUSG00000087042.1 |
| chr11 | 94265397 | 94265834 | 8.73111 | 16.23101 | 13.94854 | Distal Interge | 11 | 94284652 | 94285985 | 1334 | 1 | ENSMUST00 | -18818 | Gm21885 | ENSMUSG00000094091.1 |
| chr12 | 86801398 | 86802297 | 8.73111 | 16.23101 | 13.94854 | Distal Interge | 12 | 86809957 | 86810258 | 302 | 1 | ENSMUST00 | -7660 | Gm25888 | ENSMUSG00000088403.1 |
| chr12 | 78806649 | 78807678 | 8.73111 | 16.23101 | 13.94854 | Intron (ENSM | 12 | 78752687 | 78752788 | 102 | 2 | ENSMUST00 | -53862 | Gm25225 | ENSMUSG00000088822.1 |
| chr12 | 21778557 | 21779123 | 8.73111 | 16.23101 | 13.94854 | Distal Interge | 12 | 21776159 | 21776707 | 549 | 2 | ENSMUST00 | -1851 | Gm5633 | ENSMUSG00000091105.1 |
| chr13 | 40859706 | 40860458 | 8.73111 | 16.23101 | 13.94854 | Promoter | 13 | 40859768 | 40960891 | 101124 | 1 | ENSMUST00 | 0 | Gcnt2 | ENSMUSG00000021360.10 |
| chr13 | 97007239 | 97008180 | 8.73111 | 16.23101 | 13.94854 | Distal Interge | 13 | 97021864 | 97034362 | 12499 | 2 | ENSMUST00 | 26182 | 1700029F12l | ENSMUSG00000052075.6 |
| chr14 | 67271952 | 67272421 | 8.73111 | 16.23101 | 13.94854 | Intron (ENSM | 14 | 67274869 | 67274963 | 95 | 2 | ENSMUST00 | 2542 | Gm23178 | ENSMUSG00000080432.1 |
| chr14 | 4856194 | 4856381 | 8.73111 | 16.23101 | 13.94854 | Promoter | 14 | 4855576 | 4875851 | 20276 | 1 | ENSMUST00 | 619 | Gm3264 | ENSMUSG00000094132.2 |
| chr16 | 21248347 | 21249092 | 8.73111 | 16.23101 | 13.94854 | Exon (ENSMU | 16 | 21256013 | 21256066 | 54 | 2 | ENSMUST00 | 6974 | Mir7680 | ENSMUSG00000098400.1 |
| chr19 | 25239393 | 25240384 | 8.73111 | 16.23101 | 13.94854 | Intron (ENSM | 19 | 25236975 | 25434496 | 197522 | 1 | ENSMUST00 | 2419 | Kank1 | ENSMUSG00000032702.11 |
| chr19 | 5072964 | 5073208 | 8.73111 | 16.23101 | 13.94854 | Promoter | 19 | 5070859 | 5072132 | 1274 | 2 | ENSMUST00 | -833 | Tmem151a | ENSMUSG00000061451.8 |
| chr2 | 113758247 | 113758839 | 8.73111 | 16.23101 | 13.94854 | Promoter | 2 | 113746164 | 113758646 | 12483 | 2 | ENSMUST00 | 0 | Grem1 | ENSMUSG00000074934.3 |
| chr2 | 173365434 | 173366026 | 8.73111 | 16.23101 | 13.94854 | Distal Interge | 2 | 173394566 | 173395720 | 1155 | 1 | ENSMUST00 | -28540 | Gm14642 | ENSMUSG00000086114.1 |
| chr2 | 116067660 | 116068240 | 8.73111 | 16.23101 | 13.94854 | Promoter | 2 | 116067968 | 116085910 | 17943 | 1 | ENSMUST00 | 0 | G630016G05 | ENSMUSG00000086308.1 |
| chr2 | 93954917 | 93955634 | 8.73111 | 16.23101 | 13.94854 | Downstream | 2 | 93955810 | 93957049 | 1240 | 2 | ENSMUST00 | 1415 | Gm13889 | ENSMUSG00000087006.3 |
| chr2 | 176423653 | 176424153 | 8.73111 | 16.23101 | 13.94854 | Promoter | 2 | 176423661 | 176433086 | 9426 | 1 | ENSMUST00 | 0 | Gm14435 | ENSMUSG00000089951.3 |
| chr2 | 32318161 | 32318504 | 8.73111 | 16.23101 | 13.94854 | Promoter | 2 | 32318249 | 32318356 | 108 | 1 | ENSMUST00 | 0 | Gm23363 | ENSMUSG00000092765.2 |
| chr3 | 28264063 | 28264292 | 8.73111 | 16.23101 | 13.94854 | Promoter | 3 | 28263643 | 28667982 | 404340 | 1 | ENSMUST00 | 421 | Tnik | ENSMUSG00000027692.11 |
| chr3 | 87763938 | 87764321 | 8.73111 | 16.23101 | 13.94854 | Intron (ENSM | 3 | 87749098 | 87762214 | 13117 | 2 | ENSMUST00 | -1725 | Pear1 | ENSMUSG00000028073.10 |
| chr3 | 95904558 | 95904945 | 8.73111 | 16.23101 | 13.94854 | Promoter | 3 | 95897768 | 95904691 | 6924 | 2 | ENSMUST00 | 0 | Car14 | ENSMUSG00000038526.9 |
| chr4 | 42734546 | 42735213 | 8.73111 | 16.23101 | 13.94854 | Promoter | 4 | 42735545 | 42767287 | 31743 | 1 | ENSMUST00 | -332 | 4930578G10 | ENSMUSG00000054885.6 |
| chr5 | 67401447 | 67401981 | 8.73111 | 16.23101 | 13.94854 | Intron (ENSM | 5 | 67427680 | 67437990 | 10311 | 1 | ENSMUST00 | -25699 | Gm26756 | ENSMUSG00000097840.1 |
| chr6 | 129180333 | 129181157 | 8.73111 | 16.23101 | 13.94854 | Promoter | 6 | 129180615 | 129186534 | 5920 | 1 | ENSMUST00 | 0 | Clec2d | ENSMUSG00000030157.5 |
| chr7 | 45092143 | 45092427 | 8.73111 | 16.23101 | 13.94854 | Promoter | 7 | 45082913 | 45092221 | 9309 | 2 | ENSMUST00 | 0 | Rcn3 | ENSMUSG00000019539.5 |
| chr7 | 70337629 | 70338074 | 8.73111 | 16.23101 | 13.94854 | Distal Interge | 7 | 70351946 | 70360593 | 8648 | 2 | ENSMUST00 | 22519 | Nr2f2 | ENSMUSG00000030551.8 |
| chr7 | 4690508 | 4691120 | 8.73111 | 16.23101 | 13.94854 | Promoter | 7 | 4690604 | 4715981 | 25378 | 1 | ENSMUST00 | 0 | Brsk1 | ENSMUSG00000035390.11 |
| chr8 | 56294467 | 56294818 | 8.73111 | 16.23101 | 13.94854 | Promoter | 8 | 56294552 | 56321046 | 26495 | 1 | ENSMUST00 | 0 | Hpgd | ENSMUSG00000031613.8 |
| chr8 | 84937019 | 84937389 | 8.73111 | 16.23101 | 13.94854 | Promoter | 8 | 84911903 | 84937353 | 25451 | 2 | ENSMUST00 | 0 | Mast1 | ENSMUSG00000053693.11 |
| chr8 | 121394497 | 121395306 | 8.73111 | 16.23101 | 13.94854 | Intron (ENSM | 8 | 121421345 | 121427188 | 5844 | 1 | ENSMUST00 | -26039 | Gm26747 | ENSMUSG00000097783.1 |
| chrX | 48519094 | 48519702 | 8.73111 | 16.23101 | 13.94854 | Promoter | 20 | 48519285 | 48530232 | 10948 | 1 | ENSMUST00 | 0 | Rab33a | ENSMUSG00000031104.2 |
| chrX | 73915341 | 73915717 | 8.73111 | 16.23101 | 13.94854 | Intron (ENSM | 20 | 73891442 | 73911336 | 19895 | 2 | ENSMUST00 | -4006 | Arhgap4 | ENSMUSG00000031389.12 |
| chrX | 97431492 | 97431758 | 8.73111 | 16.23101 | 13.94854 | Distal Interge | 20 | 97337791 | 97377216 | 39426 | 2 | ENSMUST00 | -54277 | Eda2r | ENSMUSG00000034457.5 |
| chrX | 8074784 | 8075293 | 8.73111 | 16.23101 | 13.94854 | Promoter | 20 | 8061171 | 8074760 | 13590 | 2 | ENSMUST00 | -25 | Suv39h1 | ENSMUSG00000039231.13 |
| chrX | 12140170 | 12140598 | 8.73111 | 16.23101 | 13.94854 | Intron (ENSM | 20 | 12058981 | 12128400 | 69420 | 2 | ENSMUST00 | -11771 | Bcor | ENSMUSG00000040363.9 |
| chrX | 136666378 | 136666737 | 8.73111 | 16.23101 | 13.94854 | Promoter | 20 | 136666403 | 136668374 | 1972 | 1 | ENSMUST00 | 0 | Tceal3 | ENSMUSG00000044550.6 |
| chrX | 133689147 | 133689454 | 8.73111 | 16.23101 | 13.94854 | Promoter | 20 | 133582863 | 133688993 | 106131 | 2 | ENSMUST00 | -155 | Pcdh19 | ENSMUSG00000051323.11 |
| chrX | 21360852 | 21361757 | 8.73111 | 16.23101 | 13.94854 | Promoter | 20 | 21360865 | 21364622 | 3758 | 1 | ENSMUST00 | 0 | Gm5124 | ENSMUSG00000051537.9 |
| chrX | 152973883 | 152974272 | 8.73111 | 16.23101 | 13.94854 | Distal Interge | 20 | 153006469 | 153009416 | 2948 | 1 | ENSMUST00 | -32197 | Usp51 | ENSMUSG00000067215.5 |
| chrX | 58030042 | 58030713 | 8.73111 | 16.23101 | 13.94854 | Promoter | 20 | 58030643 | 58036559 | 5917 | 1 | ENSMUST00 | 0 | Zic3 | ENSMUSG00000067860.6 |
| chrX | 161908297 | 161909038 | 8.73111 | 16.23101 | 13.94854 | Promoter | 20 | 161908223 | 161908777 | 555 | 1 | ENSMUST00 | 75 | Gm15202 | ENSMUSG00000087571.1 |
| chrX | 19237313 | 19238201 | 8.73111 | 16.23101 | 13.94854 | Promoter | 20 | 19232963 | 19237856 | 4894 | 2 | ENSMUST00 | 0 | Gm26652 | ENSMUSG00000097497.1 |
| chr10 | 122097451 | 122097845 | 8.71541 | 16.20516 | 13.92474 | Promoter | 10 | 122078114 | 122097371 | 19258 | 2 | ENSMUST00 | -81 | Tmem5 | ENSMUSG00000034620.12 |
| chr15 | 98792392 | 98793047 | 8.71541 | 16.20516 | 13.92474 | 3' UTR | 15 | 98789857 | 98793837 | 3981 | 1 | ENSMUST00 | 2536 | Wnt1 | ENSMUSG00000022997.9 |
| chr1 | 9544886 | 9545233 | 8.43118 | 16.20477 | 13.92443 | Promoter | 1 | 9545408 | 9547455 | 2048 | 1 | ENSMUST00 | -175 | Rrs1 | ENSMUSG00000061024.8 |
| chr13 | 48018995 | 48019644 | 8.43118 | 16.20477 | 13.92443 | Promoter | 13 | 47960330 | 48018608 | 58279 | 2 | ENSMUST00 | -388 | 4931429P17l | ENSMUSG00000047324.5 |
| chr16 | 73647566 | 73648413 | 8.43118 | 16.20477 | 13.92443 | Distal Interge | 16 | 73422793 | 73422959 | 167 | 2 | ENSMUST00 | -224608 | Gm25699 | ENSMUSG00000064830.1 |
| chr17 | 87459577 | 87460294 | 8.43118 | 16.20477 | 13.92443 | Distal Interge | 17 | 87453223 | 87453356 | 134 | 2 | ENSMUST00 | -6222 | Gm26005 | ENSMUSG00000089427.1 |
| chr5 | 67532700 | 67533248 | 8.43118 | 16.20477 | 13.92443 | Distal Interge | 5 | 67607883 | 67613992 | 6110 | 1 | ENSMUST00 | -74635 | Shisa3 | ENSMUSG00000050010.7 |
| chr5 | 126787627 | 126788384 | 8.43118 | 16.20477 | 13.92443 | Distal Interge | 5 | 126513875 | 126513981 | 107 | 2 | ENSMUST00 | -273647 | Gm24839 | ENSMUSG00000084551.1 |
| chr7 | 141278854 | 141279330 | 8.43118 | 16.20477 | 13.92443 | Promoter | 7 | 141278331 | 141279133 | 803 | 2 | ENSMUST00 | 0 | Sct | ENSMUSG00000038580.7 |

| | | | | | | | | | | | | | | | |
|---|---|---|---|---|---|---|---|---|---|---|---|---|---|---|---|
| chr7 | 143783582 | 143784666 | 8.43118 | 16.20477 | 13.92443 | Promoter | 7 | 143778365 | 143784500 | 6136 | 2 | ENSMUST00 | 0 | Mrgpre | ENSMUSG00000048965.8 |
| chr8 | 75033288 | 75033532 | 8.43118 | 16.20477 | 13.92443 | Promoter | 8 | 75033686 | 75070121 | 36436 | 1 | ENSMUST00 | -154 | Tom1 | ENSMUSG00000042870.9 |
| chr19 | 5817735 | 5818165 | 8.69716 | 16.17512 | 13.89528 | Distal Interge | 19 | 5824652 | 5824707 | 56 | 2 | ENSMUST00 | 6542 | Gm27702 | ENSMUSG00000098865.1 |
| chr1 | 12718393 | 12719676 | 8.39879 | 16.15083 | 13.87107 | Promoter | 1 | 12718459 | 12822916 | 104458 | 1 | ENSMUST00 | 0 | Sulf1 | ENSMUSG00000016918.10 |
| chr8 | 26351959 | 26352693 | 8.39879 | 16.15083 | 13.87107 | Distal Interge | 8 | 26240150 | 26240408 | 259 | 2 | ENSMUST00 | -111552 | Gm22227 | ENSMUSG00000087981.1 |
| chr10 | 117417913 | 117419059 | 8.67899 | 16.1452 | 13.86549 | Distal Interge | 10 | 117429618 | 117430116 | 499 | 1 | ENSMUST00 | -10559 | Gm9002 | ENSMUSG00000095826.1 |
| chr11 | 69613697 | 69614090 | 8.67899 | 16.1452 | 13.86549 | Downstream | 11 | 69615648 | 69616940 | 1293 | 2 | ENSMUST00 | 2850 | Shbg | ENSMUSG00000005202.2 |
| chr11 | 75599833 | 75600011 | 8.67899 | 16.1452 | 13.86549 | Promoter | 11 | 75599763 | 75599954 | 192 | 2 | ENSMUST00 | 0 | Gm12338 | ENSMUSG00000081485.5 |
| chr16 | 43709594 | 43709937 | 8.67899 | 16.1452 | 13.86549 | Distal Interge | 16 | 43648861 | 43664184 | 15324 | 2 | ENSMUST00 | -45411 | Tigit | ENSMUSG00000071552.4 |
| chr18 | 35730206 | 35730809 | 8.67899 | 16.1452 | 13.86549 | 3' UTR | 18 | 35726989 | 35730869 | 3881 | 1 | ENSMUST00 | 3218 | 1700066B19 | ENSMUSG00000073598.3 |
| chr5 | 77245469 | 77246382 | 8.67899 | 16.1452 | 13.86549 | Distal Interge | 5 | 77265491 | 77286432 | 20942 | 1 | ENSMUST00 | -19109 | Rest | ENSMUSG00000029249.10 |
| chr1 | 16294269 | 16294948 | 8.3822 | 16.12319 | 13.84377 | Intron (ENSM | 1 | 16253629 | 16258857 | 5229 | 1 | ENSMUST00 | 40641 | Gm7568 | ENSMUSG00000089982.2 |
| chr1 | 135687057 | 135687502 | 8.3822 | 16.12319 | 13.84377 | Promoter | 1 | 135606447 | 135688105 | 81659 | 2 | ENSMUST00 | 603 | Nav1 | ENSMUSG00000090399.1 |
| chr10 | 4429163 | 4430557 | 8.3822 | 16.12319 | 13.84377 | 5' UTR | 10 | 4427243 | 4432295 | 5053 | 2 | ENSMUST00 | 1738 | Rmnd1 | ENSMUSG00000019763.6 |
| chr10 | 12547901 | 12549168 | 8.3822 | 16.12319 | 13.84377 | Intron (ENSM | 10 | 12368623 | 12368832 | 210 | 1 | ENSMUST00 | 179279 | Gm25139 | ENSMUSG00000065181.1 |
| chr11 | 42420804 | 42421351 | 8.3822 | 16.12319 | 13.84377 | Promoter | 11 | 42421279 | 42487545 | 66267 | 1 | ENSMUST00 | 0 | Gabrb2 | ENSMUSG00000007653.7 |
| chr11 | 121156383 | 121156956 | 8.3822 | 16.12319 | 13.84377 | Distal Interge | 11 | 121160271 | 121161973 | 1703 | 1 | ENSMUST00 | -3315 | Uts2r | ENSMUSG00000039321.1 |
| chr11 | 104704093 | 104704387 | 8.3822 | 16.12319 | 13.84377 | Promoter | 11 | 104704358 | 104724500 | 20143 | 1 | ENSMUST00 | 0 | Gm11639 | ENSMUSG00000040838.9 |
| chr11 | 72576397 | 72577365 | 8.3822 | 16.12319 | 13.84377 | Distal Interge | 11 | 72572530 | 72573330 | 801 | 2 | ENSMUST00 | -3068 | Gm12712 | ENSMUSG00000081675.1 |
| chr11 | 87470727 | 87471143 | 8.3822 | 16.12319 | 13.84377 | Promoter | 11 | 87471368 | 87471582 | 215 | 1 | ENSMUST00 | -225 | Rnu3b2 | ENSMUSG00000098925.1 |
| chr12 | 118848411 | 118848919 | 8.3822 | 16.12319 | 13.84377 | Exon (ENSMU | 12 | 118846329 | 118852578 | 6250 | 1 | ENSMUST00 | 2083 | Sp8 | ENSMUSG00000048562.6 |
| chr13 | 73271681 | 73272357 | 8.3822 | 16.12319 | 13.84377 | Distal Interge | 13 | 73260497 | 73269608 | 9112 | 1 | ENSMUST00 | 11185 | Irx4 | ENSMUSG00000021604.10 |
| chr14 | 22147513 | 22148223 | 8.3822 | 16.12319 | 13.84377 | Intron (ENSM | 14 | 22021958 | 22022070 | 113 | 2 | ENSMUST00 | -125444 | Gm27532 | ENSMUSG00000098542.1 |
| chr15 | 80710256 | 80710596 | 8.3822 | 16.12319 | 13.84377 | Promoter | 15 | 80711319 | 80941086 | 229768 | 1 | ENSMUST00 | -723 | Tnrc6b | ENSMUSG00000047888.9 |
| chr15 | 76523174 | 76523702 | 8.3822 | 16.12319 | 13.84377 | Distal Interge | 15 | 76516203 | 76522129 | 5927 | 2 | ENSMUST00 | -1046 | Scrt1 | ENSMUSG00000048385.8 |
| chr16 | 63857125 | 63857753 | 8.3822 | 16.12319 | 13.84377 | Intron (ENSM | 16 | 63543538 | 63864158 | 320621 | 2 | ENSMUST00 | 6405 | Epha3 | ENSMUSG00000052504.6 |
| chr16 | 55951145 | 55951465 | 8.3822 | 16.12319 | 13.84377 | Distal Interge | 16 | 55966275 | 55971435 | 5161 | 1 | ENSMUST00 | -14810 | Rpl24 | ENSMUSG00000098274.2 |
| chr18 | 38483279 | 38484157 | 8.3822 | 16.12319 | 13.84377 | Distal Interge | 18 | 38418975 | 38464399 | 45425 | 1 | ENSMUST00 | 64305 | Ndfip1 | ENSMUSG00000024425.3 |
| chr18 | 77821872 | 77822250 | 8.3822 | 16.12319 | 13.84377 | Intron (ENSM | 18 | 77806314 | 77806666 | 353 | 2 | ENSMUST00 | -15207 | Gm23954 | ENSMUSG00000087838.1 |
| chr19 | 5433448 | 5433907 | 8.3822 | 16.12319 | 13.84377 | Downstream | 19 | 5435900 | 5435965 | 66 | 1 | ENSMUST00 | -1993 | Gm25432 | ENSMUSG00000088909.1 |
| chr2 | 130576468 | 130577191 | 8.3822 | 16.12319 | 13.84377 | Promoter | 2 | 130576173 | 130577054 | 882 | 1 | ENSMUST00 | 296 | Oxt | ENSMUSG00000027301.7 |
| chr2 | 152405834 | 152406136 | 8.3822 | 16.12319 | 13.84377 | Distal Interge | 2 | 152393611 | 152398063 | 4453 | 2 | ENSMUST00 | -7772 | Sox12 | ENSMUSG00000051817.8 |
| chr3 | 152763273 | 152763799 | 8.3822 | 16.12319 | 13.84377 | Promoter | 3 | 152763381 | 152766524 | 3144 | 1 | ENSMUST00 | 0 | Pigk | ENSMUSG00000039047.12 |
| chr3 | 10351564 | 10351863 | 8.3822 | 16.12319 | 13.84377 | Promoter | 3 | 10340682 | 10351317 | 10636 | 2 | ENSMUST00 | -248 | Zfand1 | ENSMUSG00000039795.9 |
| chr4 | 126110399 | 126111581 | 8.3822 | 16.12319 | 13.84377 | Intron (ENSM | 4 | 126104035 | 126117698 | 13664 | 1 | ENSMUST00 | 6365 | Stk40 | ENSMUSG00000042608.10 |
| chr5 | 139345326 | 139345581 | 8.3822 | 16.12319 | 13.84377 | Promoter | 5 | 139336189 | 139345233 | 9045 | 2 | ENSMUST00 | -94 | Cox19 | ENSMUSG00000045438.7 |
| chr6 | 127079902 | 127080527 | 8.3822 | 16.12319 | 13.84377 | 3' UTR | 6 | 127072902 | 127081408 | 8507 | 1 | ENSMUST00 | 7001 | Fgf23 | ENSMUSG00000000182.7 |
| chr7 | 141338629 | 141339191 | 8.3822 | 16.12319 | 13.84377 | Promoter | 7 | 141338880 | 141363020 | 24141 | 1 | ENSMUST00 | 0 | Eps8l2 | ENSMUSG00000025504.7 |
| chr8 | 85034880 | 85035446 | 8.3822 | 16.12319 | 13.84377 | Distal Interge | 8 | 85036915 | 85057583 | 20669 | 1 | ENSMUST00 | -1469 | Tnpo2 | ENSMUSG00000031691.8 |
| chr9 | 63059114 | 63059654 | 8.3822 | 16.12319 | 13.84377 | Distal Interge | 9 | 63027560 | 63027717 | 158 | 1 | ENSMUST00 | 31555 | Gm24526 | ENSMUSG00000064855.1 |
| chr9 | 88241399 | 88241965 | 8.3822 | 16.12319 | 13.84377 | Distal Interge | 9 | 88259895 | 88263035 | 3141 | 1 | ENSMUST00 | -17930 | RP23-118M2 | ENSMUSG00000100782.1 |
| chrX | 12135095 | 12135431 | 8.3822 | 16.12319 | 13.84377 | Intron (ENSM | 20 | 12058981 | 12128400 | 69420 | 2 | ENSMUST00 | -6696 | Bcor | ENSMUSG00000040363.9 |
| chrX | 12857929 | 12858463 | 8.3822 | 16.12319 | 13.84377 | Distal Interge | 20 | 12870269 | 12870362 | 94 | 2 | ENSMUST00 | 11899 | Gm25063 | ENSMUSG00000077858.1 |
| chr2 | 167658261 | 167658813 | 8.66089 | 16.11541 | 13.83781 | Intron (ENSM | 2 | 167607850 | 167661496 | 53647 | 2 | ENSMUST00 | 2683 | Gm20431 | ENSMUSG00000089739.2 |
| chr7 | 98729845 | 98730476 | 8.66089 | 16.11541 | 13.83781 | Distal Interge | 7 | 98709877 | 98715257 | 5381 | 1 | ENSMUST00 | 19969 | Prkrir | ENSMUSG00000030753.10 |
| chr9 | 58008807 | 58009081 | 8.66089 | 16.11541 | 13.83781 | Intron (ENSM | 9 | 58006411 | 58020742 | 14332 | 1 | ENSMUST00 | 2397 | Cyp11a1 | ENSMUSG00000032323.8 |
| chr17 | 46487176 | 46487718 | 8.64287 | 16.08574 | 13.80828 | Promoter | 17 | 46442448 | 46487675 | 45228 | 2 | ENSMUST00 | 0 | Ttbk1 | ENSMUSG00000015599.8 |
| chr4 | 78211580 | 78212236 | 8.64287 | 16.08574 | 13.80828 | Promoter | 4 | 76139382 | 78211895 | 2072514 | 2 | ENSMUST00 | 0 | Ptprd | ENSMUSG00000028399.12 |
| chr8 | 10153715 | 10154235 | 8.64287 | 16.08574 | 13.80828 | Distal Interge | 8 | 10272572 | 10634741 | 362170 | 1 | ENSMUST00 | -118337 | Myo16 | ENSMUSG00000039057.9 |
| chr10 | 12911788 | 12912272 | 8.07432 | 16.07958 | 13.80223 | Distal Interge | 10 | 12916679 | 12923090 | 6412 | 2 | ENSMUST00 | 10818 | B230208H11 | ENSMUSG00000097378.2 |
| chr15 | 102144314 | 102144805 | 8.07432 | 16.07958 | 13.80223 | Promoter | 15 | 102144362 | 102149511 | 5150 | 1 | ENSMUST00 | 0 | Igfbp6 | ENSMUSG00000023046.6 |
| chr2 | 30794723 | 30795130 | 8.07432 | 16.07958 | 13.80223 | Exon (ENSMU | 2 | 30795468 | 30796664 | 1197 | 2 | ENSMUST00 | 1534 | 1700001O22 | ENSMUSG00000044320.9 |
| chr4 | 120649674 | 120650249 | 8.07432 | 16.07958 | 13.80223 | Distal Interge | 4 | 120666572 | 120667820 | 1249 | 1 | ENSMUST00 | -16323 | Cited4 | ENSMUSG00000070803.6 |
| chr12 | 49383137 | 49384180 | 8.62493 | 16.05618 | 13.77911 | Promoter | 12 | 49383007 | 49386856 | 3850 | 1 | ENSMUST00 | 131 | Foxg1 | ENSMUSG00000020950.9 |
| chr13 | 15759032 | 15759613 | 8.62493 | 16.05618 | 13.77911 | Distal Interge | 13 | 16011851 | 16027188 | 15338 | 1 | ENSMUST00 | -252238 | Inhba | ENSMUSG00000041324.8 |
| chr1 | 185481802 | 185482467 | 8.60706 | 16.02675 | 13.74988 | Distal Interge | 1 | 185447116 | 185455568 | 8453 | 2 | ENSMUST00 | -26235 | Gm2061 | ENSMUSG00000097615.1 |
| chr2 | 84633888 | 84634308 | 8.60706 | 16.02675 | 13.74988 | Intron (ENSM | 2 | 84624378 | 84638105 | 13728 | 2 | ENSMUST00 | 3797 | Ctnnd1 | ENSMUSG00000034101.9 |
| chr6 | 141248711 | 141248952 | 8.60706 | 16.02675 | 13.74988 | Promoter | 6 | 141249269 | 141505940 | 256672 | 1 | ENSMUST00 | -317 | Pde3a | ENSMUSG00000041741.9 |
| chr4 | 54758124 | 54758708 | 8.58926 | 15.99743 | 13.72071 | Distal Interge | 4 | 54710624 | 54710912 | 289 | 1 | ENSMUST00 | 47501 | Gm24628 | ENSMUSG00000084710.1 |
| chr10 | 81545544 | 81545822 | 8.57154 | 15.96824 | 13.69164 | Promoter | 10 | 81528724 | 81545190 | 16467 | 2 | ENSMUST00 | -355 | Gna11 | ENSMUSG00000034781.5 |
| chr13 | 95824879 | 95825135 | 8.57154 | 15.96824 | 13.69164 | Intron (ENSM | 13 | 95627177 | 95891922 | 264746 | 2 | ENSMUST00 | 66787 | Iqgap2 | ENSMUSG00000021676.9 |
| chr8 | 111745562 | 111746185 | 8.57154 | 15.96824 | 13.69164 | Distal Interge | 8 | 111710485 | 111743849 | 33365 | 2 | ENSMUST00 | -1714 | Bcar1 | ENSMUSG00000031955.9 |
| chr11 | 119083935 | 119084584 | 8.55389 | 15.93915 | 13.66267 | Exon (ENSMU | 11 | 119077573 | 119086221 | 8649 | 2 | ENSMUST00 | 1637 | Cbx4 | ENSMUSG00000039989.7 |
| chr10 | 24131610 | 24131799 | 8.27169 | 15.93882 | 13.66243 | Intron (ENSM | 10 | 24126069 | 24132480 | 6412 | 1 | ENSMUST00 | 5542 | Gm15137 | ENSMUSG00000085451.1 |
| chr17 | 46460788 | 46461407 | 8.27169 | 15.93882 | 13.66243 | Intron (ENSM | 17 | 46471501 | 46825221 | 353721 | 1 | ENSMUST00 | -10094 | Gm26904 | ENSMUSG00000097560.1 |
| chr4 | 138367707 | 138368172 | 8.27169 | 15.93882 | 13.66243 | Promoter | 4 | 138338528 | 138367992 | 29465 | 2 | ENSMUST00 | 0 | Cda | ENSMUSG00000028755.3 |
| chr4 | 135570013 | 135570523 | 8.27169 | 15.93882 | 13.66243 | Intron (ENSM | 4 | 135541888 | 135573630 | 31743 | 2 | ENSMUST00 | 3107 | Grhl3 | ENSMUSG00000037188.7 |
| chr4 | 150488243 | 150488929 | 8.27169 | 15.93882 | 13.66243 | Promoter | 4 | 150488393 | 150621962 | 133570 | 1 | ENSMUST00 | 0 | Rere | ENSMUSG00000039852.11 |
| chr6 | 136793663 | 136794335 | 8.27169 | 15.93882 | 13.66243 | Distal Interge | 6 | 136801553 | 136804431 | 2879 | 2 | ENSMUST00 | 10096 | Hist4h4 | ENSMUSG00000096010.1 |
| chr7 | 64048343 | 64049162 | 8.27169 | 15.93882 | 13.66243 | Distal Interge | 7 | 64057602 | 64062958 | 5357 | 1 | ENSMUST00 | -8440 | Gm20670 | ENSMUSG00000093542.1 |
| chr9 | 63135468 | 63136009 | 8.27169 | 15.93882 | 13.66243 | Downstream | 9 | 63133283 | 63133389 | 107 | 1 | ENSMUST00 | 2186 | Gm25064 | ENSMUSG00000094201.1 |
| chr9 | 32671954 | 32672786 | 8.53631 | 15.91019 | 13.6344 | Intron (ENSM | 9 | 32696022 | 32757820 | 61799 | 1 | ENSMUST00 | -23236 | Ets1 | ENSMUSG00000032035.10 |
| chr8 | 95825286 | 95825528 | 8.5188 | 15.88133 | 13.60567 | Promoter | 8 | 95825354 | 95826133 | 780 | 1 | ENSMUST00 | 0 | Gm10094 | ENSMUSG00000061104.4 |
| chr18 | 38144815 | 38145363 | 8.50137 | 15.8526 | 13.57707 | Distal Interge | 18 | 38188888 | 38203163 | 14276 | 2 | ENSMUST00 | 57800 | Pcdh1 | ENSMUSG00000051375.10 |
| chr2 | 147365053 | 147365448 | 8.50137 | 15.8526 | 13.57707 | Promoter | 2 | 147364994 | 147375053 | 10060 | 1 | ENSMUST00 | 60 | Pax1 | ENSMUSG00000037034.10 |
| chr2 | 63701903 | 63703074 | 8.50137 | 15.8526 | 13.57707 | Distal Interge | 2 | 63432088 | 63432194 | 107 | 2 | ENSMUST00 | -269710 | Gm23503 | ENSMUSG00000065837.1 |
| chr19 | 53359715 | 53360387 | 7.93477 | 15.84155 | 13.56641 | Intron (ENSM | 19 | 53371566 | 53371766 | 201 | 2 | ENSMUST00 | 11379 | Gm10197 | ENSMUSG00000067085.1 |
| chr5 | 16047497 | 16048102 | 7.93477 | 15.84155 | 13.56641 | Intron (ENSM | 5 | 15934829 | 16370727 | 435899 | 1 | ENSMUST00 | 112669 | Cacna2d1 | ENSMUSG00000040118.10 |
| chr1 | 171262624 | 171263096 | 8.48401 | 15.82398 | 13.54903 | Distal Interge | 1 | 171270328 | 171276895 | 6568 | 1 | ENSMUST00 | -7232 | B4galt3 | ENSMUSG00000052423.9 |
| chr10 | 99599780 | 99600513 | 8.46672 | 15.79547 | 13.5207 | Distal Interge | 10 | 99443730 | 99447007 | 3278 | 1 | ENSMUST00 | 156051 | Gad1-ps | ENSMUSG00000090665.1 |
| chr11 | 44616535 | 44617386 | 8.46672 | 15.79547 | 13.5207 | Promoter | 11 | 44617317 | 45004995 | 387679 | 1 | ENSMUST00 | 0 | Ebf1 | ENSMUSG00000057098.9 |
| chr2 | 14874116 | 14874523 | 8.17886 | 15.78362 | 13.50897 | Promoter | 2 | 14873993 | 14963667 | 89675 | 1 | ENSMUST00 | 124 | Cacnb2 | ENSMUSG00000057914.10 |
| chr10 | 18742433 | 18742726 | 8.4495 | 15.76707 | 13.49243 | Intron (ENSM | 10 | 18588011 | 18743758 | 155748 | 2 | ENSMUST00 | 1032 | D10Bwg137 | ENSMUSG00000019852.6 |
| chr11 | 119215137 | 119215666 | 8.16359 | 15.75807 | 13.48353 | Intron (ENSM | 11 | 119220160 | 119222473 | 2314 | 1 | ENSMUST00 | -4494 | Gm11752 | ENSMUSG00000087404.1 |
| chr1 | 138843579 | 138844083 | 7.88522 | 15.75676 | 13.48224 | Intron (ENSM | 1 | 138825186 | 138842457 | 17272 | 2 | ENSMUST00 | -1123 | Lhx9 | ENSMUSG00000019230.9 |
| chr1 | 165718458 | 165718951 | 7.88522 | 15.75676 | 13.48224 | Distal Interge | 1 | 165648945 | 165708094 | 59150 | 2 | ENSMUST00 | -10365 | Rcsd1 | ENSMUSG00000040723.9 |
| chr1 | 182983152 | 182983426 | 7.88522 | 15.75676 | 13.48224 | Distal Interge | 1 | 182954788 | 182976044 | 21257 | 1 | ENSMUST00 | 28365 | Tlr5 | ENSMUSG00000079164.3 |
| chr10 | 45289174 | 45289829 | 7.88522 | 15.75676 | 13.48224 | Promoter | 10 | 45289305 | 45318452 | 29148 | 1 | ENSMUST00 | 0 | Popdc3 | ENSMUSG00000019848.9 |
| chr13 | 45709333 | 45709733 | 7.88522 | 15.75676 | 13.48224 | Intron (ENSM | 13 | 45538809 | 45542550 | 3742 | 1 | ENSMUST00 | 170525 | Gmpr | ENSMUSG00000000253.7 |
| chr14 | 69190476 | 69191036 | 7.88522 | 15.75676 | 13.48224 | Promoter | 14 | 69190638 | 69194662 | 4025 | 1 | ENSMUST00 | 0 | Nkx3-1 | ENSMUSG00000022061.8 |
| chr15 | 85131729 | 85132383 | 7.88522 | 15.75676 | 13.48224 | Promoter | 15 | 85132099 | 85144569 | 12471 | 1 | ENSMUST00 | 0 | Ribc2 | ENSMUSG00000022431.2 |
| chr15 | 76195069 | 76196043 | 7.88522 | 15.75676 | 13.48224 | Promoter | 15 | 76170976 | 76195710 | 24735 | 2 | ENSMUST00 | 0 | Plec | ENSMUSG00000022565.10 |
| chr15 | 38198637 | 38199902 | 7.88522 | 15.75676 | 13.48224 | Distal Interge | 15 | 38219203 | 38226735 | 7533 | 1 | ENSMUST00 | -19301 | Odf1 | ENSMUSG00000061923.4 |
| chr15 | 100443065 | 100443482 | 7.88522 | 15.75676 | 13.48224 | Distal Interge | 15 | 100449684 | 100449793 | 110 | 1 | ENSMUST00 | -6202 | n-R5s43 | ENSMUSG00000084688.1 |
| chr16 | 16559287 | 16560231 | 7.88522 | 15.75676 | 13.48224 | Promoter | 16 | 16416917 | 16560219 | 143303 | 2 | ENSMUST00 | 0 | Fgd4 | ENSMUSG00000022788.11 |
| chr16 | 31458374 | 31458750 | 7.88522 | 15.75676 | 13.48224 | Promoter | 16 | 31458116 | 31458901 | 786 | 1 | ENSMUST00 | 259 | Bdh1 | ENSMUSG00000046598.9 |
| chr18 | 76063435 | 76064324 | 7.88522 | 15.75676 | 13.48224 | Intron (ENSM | 18 | 76059458 | 76146443 | 86986 | 1 | ENSMUST00 | 3978 | Zbtb7c | ENSMUSG00000044646.9 |
| chr19 | 23143471 | 23143990 | 7.88522 | 15.75676 | 13.48224 | Intron (ENSM | 19 | 23141226 | 23166911 | 25686 | 1 | ENSMUST00 | 2246 | Klf9 | ENSMUSG00000033863.1 |
| chr2 | 11663935 | 11664152 | 7.88522 | 15.75676 | 13.48224 | Intron (ENSM | 2 | 11642850 | 11648637 | 5788 | 1 | ENSMUST00 | 21086 | Il2ra | ENSMUSG00000026770.4 |
| chr2 | 52860110 | 52860957 | 7.88522 | 15.75676 | 13.48224 | Exon (ENSMU | 2 | 52857868 | 53134202 | 276335 | 1 | ENSMUST00 | 2243 | Fmnl2 | ENSMUSG00000036053.11 |
| chr3 | 98042299 | 98042889 | 7.88522 | 15.75676 | 13.48224 | Intron (ENSM | 3 | 98013538 | 98150367 | 136830 | 1 | ENSMUST00 | 28762 | Notch2 | ENSMUSG00000027878.10 |
| chr3 | 152191014 | 152191569 | 7.88522 | 15.75676 | 13.48224 | Intron (ENSM | 3 | 152183955 | 152193803 | 9849 | 2 | ENSMUST00 | 2234 | Dnajb4 | ENSMUSG00000028035.8 |
| chr3 | 34663985 | 34665128 | 7.88522 | 15.75676 | 13.48224 | Promoter | 3 | 34665243 | 34677995 | 12753 | 1 | ENSMUST00 | -115 | Sox2ot | ENSMUSG00000090828.3 |
| chr5 | 115319420 | 115320102 | 7.88522 | 15.75676 | 13.48224 | Distal Interge | 5 | 115327177 | 115333080 | 5904 | 1 | ENSMUST00 | -7075 | Srsf9 | ENSMUSG00000029538.8 |
| chr6 | 99026693 | 99027548 | 7.88522 | 15.75676 | 13.48224 | Promoter | 6 | 99016550 | 99027991 | 11442 | 2 | ENSMUST00 | 443 | Foxp1 | ENSMUSG00000030067.12 |
| chr6 | 115991058 | 115992036 | 7.88522 | 15.75676 | 13.48224 | Intron (ENSM | 6 | 115954811 | 115995005 | 40195 | 2 | ENSMUST00 | 2969 | Plxnd1 | ENSMUSG00000030123.10 |
| chr6 | 85154976 | 85155430 | 7.88522 | 15.75676 | 13.48224 | Promoter | 6 | 85155005 | 85155968 | 964 | 1 | ENSMUST00 | 0 | Spr-ps1 | ENSMUSG00000068303.6 |
| chr7 | 141443688 | 141444335 | 7.88522 | 15.75676 | 13.48224 | Promoter | 7 | 141438113 | 141443992 | 5880 | 2 | ENSMUST00 | 0 | Pidd1 | ENSMUSG00000025507.8 |
| chr7 | 109082953 | 109083519 | 7.88522 | 15.75676 | 13.48224 | Promoter | 7 | 109034312 | 109083331 | 49020 | 2 | ENSMUST00 | 0 | Ric3 | ENSMUSG00000048330.9 |
| chr8 | 87938174 | 87938913 | 7.88522 | 15.75676 | 13.48224 | Intron (ENSM | 8 | 87783316 | 87944052 | 160737 | 2 | ENSMUST00 | 5139 | Zfp423 | ENSMUSG00000045333.10 |
| chr9 | 103366102 | 103366469 | 7.88522 | 15.75676 | 13.48224 | Promoter | 9 | 103353094 | 103365840 | 12747 | 2 | ENSMUST00 | -263 | Cdv3 | ENSMUSG00000032803.9 |
| chr9 | 60742982 | 60743346 | 7.88522 | 15.75676 | 13.48224 | Promoter | 9 | 60743120 | 60770276 | 27157 | 1 | ENSMUST00 | 0 | 1700036A12 | ENSMUSG00000095956.2 |
| chr9 | 21220721 | 21221199 | 7.88522 | 15.75676 | 13.48224 | Distal Interge | 9 | 21225371 | 21232639 | 7269 | 1 | ENSMUST00 | -4172 | Gm16754 | ENSMUSG00000097439.1 |
| chr17 | 71662526 | 71663085 | 8.43234 | 15.73878 | 13.46618 | Distal Interge | 17 | 71673261 | 71685172 | 11912 | 1 | ENSMUST00 | -10176 | Fam179a | ENSMUSG00000045761.10 |
| chr11 | 5556607 | 5556818 | 5.70353 | 15.72236 | 13.44988 | Distal Interge | 11 | 5560606 | 5560735 | 130 | 2 | ENSMUST00 | 3917 | Gm22471 | ENSMUSG00000077304.1 |
| chr4 | 147219441 | 147219917 | 5.70353 | 15.72236 | 13.44988 | Intron (ENSM | 4 | 147230284 | 147258628 | 28345 | 1 | ENSMUST00 | -10367 | Gm13147 | ENSMUSG00000087207.3 |
| chr19 | 59587925 | 59588595 | 8.39825 | 15.68253 | 13.41018 | Distal Interge | 19 | 59493136 | 59559391 | 66256 | 1 | ENSMUST00 | 94790 | RP23-103C1 | ENSMUSG00000100119.1 |
| chr10 | 111516038 | 111516501 | 8.11812 | 15.68191 | 13.40967 | Distal Interge | 10 | 111507696 | 111508165 | 470 | 1 | ENSMUST00 | 8343 | Phlda1 | ENSMUSG00000020205.8 |
| chr10 | 81563927 | 81564408 | 8.11812 | 15.68191 | 13.40967 | Exon (ENSMU | 10 | 81559494 | 81566362 | 6869 | 1 | ENSMUST00 | 4434 | Aes | ENSMUSG00000054452.9 |
| chr12 | 4509681 | 4510061 | 8.11812 | 15.68191 | 13.40967 | Distal Interge | 12 | 4247363 | 4477182 | 229820 | 2 | ENSMUST00 | -32500 | Ncoa1 | ENSMUSG00000020647.9 |
| chr13 | 74259812 | 74260370 | 8.11812 | 15.68191 | 13.40967 | Promoter | 13 | 74223029 | 74260672 | 37644 | 2 | ENSMUST00 | 302 | Ahrr | ENSMUSG00000021575.10 |
| chr15 | 78531967 | 78532263 | 8.11812 | 15.68191 | 13.40967 | Distal Interge | 15 | 78523346 | 78529625 | 6280 | 2 | ENSMUST00 | -2343 | C1qtnf6 | ENSMUSG00000022440.8 |
| chr16 | 89817773 | 89818553 | 8.11812 | 15.68191 | 13.40967 | Promoter | 16 | 89787730 | 89818352 | 30623 | 2 | ENSMUST00 | 0 | Tiam1 | ENSMUSG00000002489.10 |
| chr3 | 51708796 | 51709325 | 8.11812 | 15.68191 | 13.40967 | Intron (ENSM | 3 | 51693771 | 51695090 | 1320 | 1 | ENSMUST00 | 15026 | Gm10729 | ENSMUSG00000074603.2 |
| chr5 | 114667261 | 114667584 | 8.11812 | 15.68191 | 13.40967 | Downstream | 5 | 114622152 | 114658421 | 36270 | 2 | ENSMUST00 | -8841 | Trpv4 | ENSMUSG00000014158.6 |
| chr7 | 43606940 | 43607185 | 8.11812 | 15.68191 | 13.40967 | Promoter | 7 | 43607169 | 43618216 | 11048 | 1 | ENSMUST00 | 0 | Zfp819 | ENSMUSG00000055102.10 |
| chr9 | 110657592 | 110658259 | 8.11812 | 15.68191 | 13.40967 | Intron (ENSM | 9 | 110656503 | 110711590 | 55088 | 1 | ENSMUST00 | 1090 | Ccdc12 | ENSMUSG00000019659.4 |
| chr13 | 112119742 | 112120208 | 8.3813 | 15.65457 | 13.38283 | Distal Interge | 13 | 112005502 | 112017332 | 11831 | 1 | ENSMUST00 | 114241 | Gm15323 | ENSMUSG00000086092.1 |

| | | | | | | | | | | | | | | | |
|---|---|---|---|---|---|---|---|---|---|---|---|---|---|---|---|
| chr8 | 109105984 | 109106206 | 7.79997 | 15.61056 | 13.33905 | Intron (ENSM | 8 | 109075223 | 109231047 | 155825 | 1 | ENSMUST00 | 30762 | D030068K23I | ENSMUSG00000097393.2 |
| chr1 | 85703489 | 85703902 | 8.34762 | 15.59896 | 13.32767 | Promoter | 1 | 85703442 | 85707995 | 4554 | 1 | ENSMUST00 | 48 | Sp100 | ENSMUSG00000026222.11 |
| chr11 | 96826093 | 96826719 | 8.34762 | 15.59896 | 13.32767 | Promoter | 11 | 96817599 | 96827719 | 10121 | 2 | ENSMUST00 | 1000 | Nfe2l1 | ENSMUSG00000038615.12 |
| chr2 | 25305036 | 25305384 | 8.34762 | 15.59896 | 13.32767 | Promoter | 2 | 25303481 | 25305100 | 1620 | 2 | ENSMUST00 | 0 | Grin1 | ENSMUSG00000026959.8 |
| chr8 | 25098560 | 25099109 | 8.34762 | 15.59896 | 13.32767 | Intron (ENSM | 8 | 25042944 | 25101771 | 58828 | 2 | ENSMUST00 | 2662 | Plekha2 | ENSMUSG00000031557.10 |
| chr12 | 8892998 | 8893438 | 8.33088 | 15.57132 | 13.30013 | Distal Interge | 12 | 8873187 | 8877459 | 4273 | 2 | ENSMUST00 | -15540 | 9930038B18I | ENSMUSG00000042976.5 |
| chr13 | 95249805 | 95250468 | 8.33088 | 15.57132 | 13.30013 | Promoter | 13 | 95024454 | 95250336 | 225883 | 2 | ENSMUST00 | 0 | Pde8b | ENSMUSG00000021684.11 |
| chr13 | 48130377 | 48131082 | 8.33088 | 15.57132 | 13.30013 | Intron (ENSM | 13 | 47960330 | 48018608 | 58279 | 2 | ENSMUST00 | -111770 | 4931429P17I | ENSMUSG00000047324.5 |
| chr15 | 99890225 | 99890827 | 8.33088 | 15.57132 | 13.30013 | Distal Interge | 15 | 99898287 | 99898556 | 270 | 2 | ENSMUST00 | 7729 | Gm9763 | ENSMUSG00000037438.5 |
| chr5 | 107726329 | 107726988 | 8.33088 | 15.57132 | 13.30013 | Promoter | 5 | 107726398 | 107728801 | 2404 | 1 | ENSMUST00 | 0 | A430072P03I | ENSMUSG00000090030.1 |
| chr9 | 31645116 | 31645707 | 8.33088 | 15.57132 | 13.30013 | Distal Interge | 9 | 31426199 | 31464238 | 38040 | 2 | ENSMUST00 | -180879 | Tmem45b | ENSMUSG00000041737.7 |
| chr16 | 90197559 | 90197859 | 8.3142 | 15.54378 | 13.27308 | Distal Interge | 16 | 90220742 | 90226322 | 5581 | 1 | ENSMUST00 | -22883 | Sod1 | ENSMUSG00000022982.9 |
| chr2 | 31640482 | 31641359 | 8.3142 | 15.54378 | 13.27308 | Promoter | 2 | 31638170 | 31640589 | 2420 | 2 | ENSMUST00 | 0 | Gm13425 | ENSMUSG00000085224.1 |
| chr1 | 193160334 | 193160638 | 7.48194 | 15.52168 | 13.25109 | Intron (ENSM | 1 | 193156804 | 193164181 | 7378 | 1 | ENSMUST00 | 3531 | Irf6 | ENSMUSG00000026638.10 |
| chr10 | 10472143 | 10472522 | 7.48194 | 15.52168 | 13.25109 | Promoter | 10 | 10335703 | 10472314 | 136612 | 2 | ENSMUST00 | 0 | Adgb | ENSMUSG00000050994.14 |
| chr10 | 128212964 | 128213441 | 7.48194 | 15.52168 | 13.25109 | Distal Interge | 10 | 128209908 | 128211794 | 1887 | 2 | ENSMUST00 | -1171 | Spryd4 | ENSMUSG00000051346.8 |
| chr11 | 75467808 | 75468512 | 7.48194 | 15.52168 | 13.25109 | Promoter | 11 | 75468079 | 75470657 | 2579 | 1 | ENSMUST00 | 0 | Tlcd2 | ENSMUSG00000038217.8 |
| chr13 | 21809397 | 21809833 | 7.48194 | 15.52168 | 13.25109 | Promoter | 13 | 21806612 | 21810197 | 3586 | 2 | ENSMUST00 | 364 | Hist1h2bq | ENSMUSG00000069307.5 |
| chr13 | 66807620 | 66808221 | 7.48194 | 15.52168 | 13.25109 | Distal Interge | 13 | 66851343 | 66851513 | 171 | 2 | ENSMUST00 | 43292 | Gm17404 | ENSMUSG00000095206.1 |
| chr15 | 5262135 | 5262599 | 7.48194 | 15.52168 | 13.25109 | Distal Interge | 15 | 5233399 | 5244187 | 10789 | 2 | ENSMUST00 | -17949 | Ptger4 | ENSMUSG00000039942.10 |
| chr16 | 52454459 | 52454765 | 7.48194 | 15.52168 | 13.25109 | Promoter | 16 | 52248996 | 52454074 | 205079 | 2 | ENSMUST00 | -386 | Alcam | ENSMUSG00000022636.8 |
| chr17 | 48096864 | 48098447 | 7.48194 | 15.52168 | 13.25109 | Distal Interge | 17 | 48089632 | 48090920 | 1289 | 1 | ENSMUST00 | 7233 | 1700067P10I | ENSMUSG00000021545.4 |
| chr17 | 56162034 | 56162807 | 7.48194 | 15.52168 | 13.25109 | Promoter | 17 | 56162477 | 56173955 | 11479 | 1 | ENSMUST00 | 0 | Tnfaip8l1 | ENSMUSG00000044469.7 |
| chr17 | 70751129 | 70751782 | 7.48194 | 15.52168 | 13.25109 | Exon (ENSMU | 17 | 70745329 | 70755582 | 10254 | 2 | ENSMUST00 | 3800 | 5031415H12I | ENSMUSG00000097282.1 |
| chr19 | 45236536 | 45236958 | 7.48194 | 15.52168 | 13.25109 | Promoter | 19 | 45232684 | 45235812 | 3129 | 2 | ENSMUST00 | -725 | Lbx1 | ENSMUSG00000025216.9 |
| chr19 | 45812067 | 45812654 | 7.48194 | 15.52168 | 13.25109 | Promoter | 19 | 45793349 | 45812291 | 18943 | 2 | ENSMUST00 | 0 | Kcnip2 | ENSMUSG00000025221.10 |
| chr2 | 137082934 | 137083398 | 7.48194 | 15.52168 | 13.25109 | 3' UTR | 2 | 137082893 | 137085949 | 3057 | 2 | ENSMUST00 | 2551 | Jag1 | ENSMUSG00000027276.7 |
| chr2 | 146305500 | 146305738 | 7.48194 | 15.52168 | 13.25109 | Promoter | 2 | 146305547 | 146306389 | 843 | 1 | ENSMUST00 | 0 | Gm14117 | ENSMUSG00000042938.8 |
| chr2 | 75603345 | 75603783 | 7.48194 | 15.52168 | 13.25109 | Distal Interge | 2 | 75632638 | 75633382 | 745 | 2 | ENSMUST00 | 29599 | Gm13655 | ENSMUSG00000082152.1 |
| chr4 | 149445062 | 149446178 | 7.48194 | 15.52168 | 13.25109 | Distal Interge | 4 | 149449687 | 149453424 | 3738 | 2 | ENSMUST00 | 7246 | Rbp7 | ENSMUSG00000028996.4 |
| chr4 | 135397705 | 135398592 | 7.48194 | 15.52168 | 13.25109 | Promoter | 4 | 135369576 | 135398190 | 28615 | 2 | ENSMUST00 | 0 | Ncmap | ENSMUSG00000043924.11 |
| chr4 | 62674086 | 62674626 | 7.48194 | 15.52168 | 13.25109 | Promoter | 4 | 62674352 | 62690001 | 15650 | 1 | ENSMUST00 | 0 | Rgs3 | ENSMUSG00000059810.13 |
| chr5 | 72503612 | 72504422 | 7.48194 | 15.52168 | 13.25109 | Promoter | 5 | 72300026 | 72504208 | 204183 | 2 | ENSMUST00 | 0 | Corin | ENSMUSG00000005220.5 |
| chr5 | 96911519 | 96912088 | 7.48194 | 15.52168 | 13.25109 | Distal Interge | 5 | 96907632 | 96907825 | 194 | 1 | ENSMUST00 | 3888 | Gm24623 | ENSMUSG00000064550.1 |
| chr6 | 99094864 | 99096288 | 7.48194 | 15.52168 | 13.25109 | Promoter | 6 | 99009933 | 99096205 | 86273 | 2 | ENSMUST00 | 0 | Foxp1 | ENSMUSG00000030067.12 |
| chr6 | 51509048 | 51509496 | 7.48194 | 15.52168 | 13.25109 | Distal Interge | 6 | 51523901 | 51590679 | 66779 | 1 | ENSMUST00 | -14405 | Snx10 | ENSMUSG00000038301.10 |
| chr6 | 52176947 | 52177703 | 7.48194 | 15.52168 | 13.25109 | Promoter | 6 | 52177498 | 52180851 | 3354 | 1 | ENSMUST00 | 0 | 5730596B20I | ENSMUSG00000056468.3 |
| chr7 | 27325598 | 27325983 | 7.48194 | 15.52168 | 13.25109 | Exon (ENSMU | 7 | 27305998 | 27333613 | 27616 | 2 | ENSMUST00 | 7630 | Ltbp4 | ENSMUSG00000040488.11 |
| chr7 | 24546442 | 24546908 | 7.48194 | 15.52168 | 13.25109 | Promoter | 7 | 24547150 | 24573438 | 26289 | 1 | ENSMUST00 | -242 | Xrcc1 | ENSMUSG00000051768.8 |
| chr8 | 41016234 | 41017093 | 7.48194 | 15.52168 | 13.25109 | Promoter | 8 | 41000675 | 41016755 | 16081 | 2 | ENSMUST00 | 0 | Mtus1 | ENSMUSG00000045636.11 |
| chr2 | 155885623 | 155887460 | 8.2976 | 15.51635 | 13.24704 | 3' UTR | 2 | 155887119 | 155910604 | 23486 | 2 | ENSMUST00 | 23144 | Uqcc1 | ENSMUSG00000005882.13 |
| chr2 | 151970593 | 151971140 | 8.2976 | 15.51635 | 13.24704 | 5' UTR | 2 | 151969737 | 151973394 | 3658 | 2 | ENSMUST00 | 2254 | Fam110a | ENSMUSG00000027459.11 |
| chr4 | 62515264 | 62516219 | 8.2976 | 15.51635 | 13.24704 | Intron (ENSM | 4 | 62509169 | 62519881 | 10713 | 2 | ENSMUST00 | 3662 | Alad | ENSMUSG00000028393.5 |
| chr2 | 177253021 | 177253395 | 5.19557 | 15.51335 | 13.24413 | Distal Interge | 2 | 177264654 | 177267036 | 2383 | 2 | ENSMUST00 | 13641 | Gm14409 | ENSMUSG00000078869.2 |
| chr9 | 100505582 | 100506808 | 8.28106 | 15.48903 | 13.21982 | Promoter | 9 | 100495003 | 100506856 | 11854 | 2 | ENSMUST00 | 48 | Nck1 | ENSMUSG00000032475.10 |
| chr12 | 54835068 | 54835337 | 7.99931 | 15.48251 | 13.21341 | Distal Interge | 12 | 54858817 | 54862877 | 4061 | 2 | ENSMUST00 | 27540 | Cfl2 | ENSMUSG00000062929.7 |
| chr4 | 120813899 | 120814893 | 8.24817 | 15.43468 | 13.16567 | Promoter | 4 | 120757438 | 120815712 | 58275 | 2 | ENSMUST00 | 819 | Nfyc | ENSMUSG00000032897.12 |
| chr9 | 41611913 | 41612849 | 8.24817 | 15.43468 | 13.16567 | Exon (ENSMU | 9 | 41614237 | 41617772 | 3536 | 1 | ENSMUST00 | -1388 | 2610203C20I | ENSMUSG00000074415.5 |
| chr1 | 134381732 | 134382583 | 7.97015 | 15.43348 | 13.16457 | Distal Interge | 1 | 134385940 | 134390981 | 5042 | 1 | ENSMUST00 | -3357 | 4933406M09 | ENSMUSG00000050526.5 |
| chr10 | 25199877 | 25200479 | 7.97015 | 15.43348 | 13.16457 | Promoter | 10 | 25169090 | 25200158 | 31069 | 2 | ENSMUST00 | 0 | Akap7 | ENSMUSG00000039166.10 |
| chr14 | 79769859 | 79770446 | 7.97015 | 15.43348 | 13.16457 | Promoter | 14 | 79766772 | 79771312 | 4541 | 2 | ENSMUST00 | 866 | Pcdh8 | ENSMUSG00000036422.5 |
| chr16 | 33019115 | 33019652 | 7.97015 | 15.43348 | 13.16457 | Exon (ENSMU | 16 | 33016950 | 33023937 | 6988 | 2 | ENSMUST00 | 4285 | Iqcg | ENSMUSG00000035578.10 |
| chr18 | 80554833 | 80555396 | 7.97015 | 15.43348 | 13.16457 | Distal Interge | 18 | 80407959 | 80469695 | 61737 | 2 | ENSMUST00 | -85139 | Ctdp1 | ENSMUSG00000033323.8 |
| chr2 | 32618371 | 32618968 | 7.97015 | 15.43348 | 13.16457 | 3' UTR | 2 | 32621758 | 32635058 | 13301 | 1 | ENSMUST00 | -2790 | Ak1 | ENSMUSG00000026817.9 |
| chr11 | 115266669 | 115267381 | 8.23183 | 15.40765 | 13.13914 | Promoter | 11 | 115249169 | 115267243 | 18075 | 2 | ENSMUST00 | 0 | Grin2c | ENSMUSG00000020734.8 |
| chr4 | 129223258 | 129223802 | 8.23183 | 15.40765 | 13.13914 | Exon (ENSMU | 4 | 129225383 | 129227480 | 2098 | 2 | ENSMUST00 | 3678 | C77080 | ENSMUSG00000050390.7 |
| chr12 | 47812913 | 47813135 | 6.08217 | 15.40313 | 13.13473 | Distal Interge | 12 | 48554182 | 48559971 | 5790 | 2 | ENSMUST00 | 746836 | Gm1818 | ENSMUSG00000091277.1 |
| chr17 | 67945893 | 67947048 | 7.66968 | 15.38624 | 13.11786 | Intron (ENSM | 17 | 67845110 | 67950940 | 105831 | 2 | ENSMUST00 | 3892 | Arhgap28 | ENSMUSG00000024043.8 |
| chr17 | 56291300 | 56291623 | 7.66968 | 15.38624 | 13.11786 | Promoter | 17 | 56278962 | 56290511 | 11550 | 2 | ENSMUST00 | -790 | Plin3 | ENSMUSG00000024197.9 |
| chr5 | 130223135 | 130223615 | 7.66968 | 15.38624 | 13.11786 | Promoter | 5 | 130222335 | 130243765 | 21431 | 1 | ENSMUST00 | 801 | Tmem248 | ENSMUSG00000053094.10 |
| chr6 | 47811481 | 47812097 | 7.66968 | 15.38624 | 13.11786 | Intron (ENSM | 6 | 47796142 | 47813512 | 17371 | 2 | ENSMUST00 | 1415 | Pdia4 | ENSMUSG00000025823.8 |
| chr9 | 22095840 | 22096370 | 7.66968 | 15.38624 | 13.11786 | Distal Interge | 9 | 22099281 | 22109630 | 10350 | 1 | ENSMUST00 | -2911 | Cnn1 | ENSMUSG00000001349.4 |
| chr16 | 59604327 | 59604796 | 8.21555 | 15.38073 | 13.11249 | Distal Interge | 16 | 59558013 | 59600886 | 42874 | 2 | ENSMUST00 | -3442 | Gm9581 | ENSMUSG00000091562.3 |
| chr7 | 64839718 | 64840408 | 8.21555 | 15.38073 | 13.11249 | Intron (ENSM | 7 | 64819765 | 64856397 | 36633 | 2 | ENSMUST00 | 15989 | Fam189a1 | ENSMUSG00000030518.12 |
| chr7 | 100005758 | 100006273 | 8.21555 | 15.38073 | 13.11249 | Promoter | 7 | 100006404 | 100034726 | 28323 | 1 | ENSMUST00 | -131 | Chrdl2 | ENSMUSG00000030732.7 |
| chr10 | 111536824 | 111537354 | 7.14815 | 15.37828 | 13.11012 | Distal Interge | 10 | 111507696 | 111508165 | 470 | 1 | ENSMUST00 | 29129 | Phlda1 | ENSMUSG00000020205.8 |
| chr10 | 93430798 | 93431296 | 7.14815 | 15.37828 | 13.11012 | Downstream | 10 | 93433107 | 93433162 | 56 | 1 | ENSMUST00 | -1811 | Mir7688 | ENSMUSG00000098679.1 |
| chr11 | 33428206 | 33428445 | 7.14815 | 15.37828 | 13.11012 | Intron (ENSM | 11 | 33438449 | 33438980 | 532 | 1 | ENSMUST00 | -10004 | Gm12118 | ENSMUSG00000083471.1 |
| chr12 | 79208852 | 79209146 | 7.14815 | 15.37828 | 13.11012 | Promoter | 12 | 79208914 | 79222665 | 13752 | 1 | ENSMUST00 | 0 | Rdh12 | ENSMUSG00000021123.7 |
| chr13 | 63255485 | 63256554 | 7.14815 | 15.37828 | 13.11012 | Intron (ENSM | 13 | 63282142 | 63326096 | 43955 | 1 | ENSMUST00 | -25588 | 2010111I01R | ENSMUSG00000021458.11 |
| chr13 | 23556265 | 23556673 | 7.14815 | 15.37828 | 13.11012 | Distal Interge | 13 | 23555087 | 23555830 | 744 | 1 | ENSMUST00 | 1179 | Hist1h1d | ENSMUSG00000052565.6 |
| chr13 | 50958174 | 50958773 | 7.14815 | 15.37828 | 13.11012 | Distal Interge | 13 | 51014086 | 51014211 | 126 | 1 | ENSMUST00 | -55313 | Gm22199 | ENSMUSG00000084609.2 |
| chr16 | 38399626 | 38400302 | 7.14815 | 15.37828 | 13.11012 | Intron (ENSM | 16 | 38371998 | 38374137 | 2140 | 1 | ENSMUST00 | 27629 | Popdc2 | ENSMUSG00000022803.7 |
| chr17 | 21753795 | 21754439 | 7.14815 | 15.37828 | 13.11012 | Intron (ENSM | 17 | 21735912 | 21748969 | 13058 | 1 | ENSMUST00 | 17884 | Zfp229 | ENSMUSG00000061544.8 |
| chr2 | 74715539 | 74716285 | 7.14815 | 15.37828 | 13.11012 | Promoter | 2 | 74710193 | 74715996 | 5804 | 2 | ENSMUST00 | 0 | Hoxd3os1 | ENSMUSG00000052371.5 |
| chr2 | 30495996 | 30496430 | 7.14815 | 15.37828 | 13.11012 | Distal Interge | 2 | 30471537 | 30474219 | 2683 | 2 | ENSMUST00 | -21778 | Ier5l | ENSMUSG00000089762.3 |
| chr4 | 32531029 | 32531953 | 7.14815 | 15.37828 | 13.11012 | Intron (ENSM | 4 | 32514787 | 32515053 | 267 | 1 | ENSMUST00 | 16243 | Gm24371 | ENSMUSG00000084611.1 |
| chr6 | 34709285 | 34709814 | 7.14815 | 15.37828 | 13.11012 | Promoter | 6 | 34709442 | 34775473 | 66032 | 1 | ENSMUST00 | 0 | Cald1 | ENSMUSG00000029761.11 |
| chr6 | 25689658 | 25690479 | 7.14815 | 15.37828 | 13.11012 | Promoter | 6 | 25668607 | 25690729 | 22123 | 2 | ENSMUST00 | 250 | Gpr37 | ENSMUSG00000039904.8 |
| chr7 | 141070235 | 141070709 | 7.14815 | 15.37828 | 13.11012 | Exon (ENSMU | 7 | 141078218 | 141090510 | 12293 | 1 | ENSMUST00 | -7509 | Pkp3 | ENSMUSG00000054065.6 |
| chr7 | 126481885 | 126482537 | 7.14815 | 15.37828 | 13.11012 | Distal Interge | 7 | 126487355 | 126490730 | 3376 | 1 | ENSMUST00 | -4818 | Tufm | ENSMUSG00000073838.5 |
| chr8 | 105278971 | 105279248 | 7.14815 | 15.37828 | 13.11012 | 5' UTR | 8 | 105276447 | 105281939 | 5493 | 1 | ENSMUST00 | 2525 | Nol3 | ENSMUSG00000014776.6 |
| chr8 | 106511451 | 106512121 | 7.14815 | 15.37828 | 13.11012 | Promoter | 8 | 106510913 | 106556908 | 45996 | 1 | ENSMUST00 | 539 | Cdh3 | ENSMUSG00000061048.7 |
| chr5 | 134985958 | 134986636 | 8.19933 | 15.35391 | 13.0865 | Promoter | 5 | 134986214 | 134987472 | 1259 | 1 | ENSMUST00 | 0 | Cldn3 | ENSMUSG00000070473.4 |
| chr7 | 16417221 | 16417647 | 8.19933 | 15.35391 | 13.0865 | Exon (ENSMU | 7 | 16401196 | 16437694 | 36499 | 1 | ENSMUST00 | 16026 | Zc3h4 | ENSMUSG00000059273.7 |
| chr1 | 69826810 | 69827235 | 6.86732 | 15.30181 | 13.03456 | Promoter | 1 | 69826970 | 70267277 | 440308 | 1 | ENSMUST00 | 0 | Spag16 | ENSMUSG00000053153.9 |
| chr18 | 13786645 | 13787751 | 6.86732 | 15.30181 | 13.03456 | Intron (ENSM | 18 | 13687013 | 13972733 | 285721 | 2 | ENSMUST00 | 184982 | Zfp521 | ENSMUSG00000024420.8 |
| chr19 | 41206233 | 41207278 | 6.86732 | 15.30181 | 13.03456 | Promoter | 19 | 41083981 | 41206774 | 122794 | 2 | ENSMUST00 | 0 | Tll2 | ENSMUSG00000025013.9 |
| chr2 | 122298491 | 122299140 | 6.86732 | 15.30181 | 13.03456 | Promoter | 2 | 122298900 | 122302885 | 3986 | 1 | ENSMUST00 | 0 | Duoxa2 | ENSMUSG00000027225.7 |
| chr2 | 152081623 | 152081884 | 6.86732 | 15.30181 | 13.03456 | Promoter | 2 | 152081529 | 152095802 | 14274 | 1 | ENSMUST00 | 95 | Scrt2 | ENSMUSG00000060257.2 |
| chr5 | 129887594 | 129887923 | 6.86732 | 15.30181 | 13.03456 | Promoter | 5 | 129881156 | 129887470 | 6315 | 2 | ENSMUST00 | -125 | Chchd2 | ENSMUSG00000070493.3 |
| chr8 | 53638533 | 53639283 | 6.86732 | 15.30181 | 13.03456 | Promoter | 8 | 53586867 | 53639065 | 52199 | 2 | ENSMUST00 | 0 | Neil3 | ENSMUSG00000039396.5 |
| chr9 | 50738987 | 50739377 | 6.86732 | 15.30181 | 13.03456 | Promoter | 9 | 50709634 | 50739447 | 29814 | 2 | ENSMUST00 | 70 | Dixdc1 | ENSMUSG00000032064.12 |
| chr1 | 174503294 | 174503629 | 8.16709 | 15.30057 | 13.0337 | Exon (ENSMU | 1 | 174501825 | 174822717 | 320893 | 1 | ENSMUST00 | 1470 | Fmn2 | ENSMUSG00000028354.8 |
| chr12 | 9584583 | 9585248 | 8.16709 | 15.30057 | 13.0337 | Distal Interge | 12 | 9574442 | 9581499 | 7058 | 1 | ENSMUST00 | 10142 | Osr1 | ENSMUSG00000048387.7 |
| chr15 | 79251915 | 79252696 | 8.16709 | 15.30057 | 13.0337 | Exon (ENSMU | 15 | 79251014 | 79254761 | 3748 | 2 | ENSMUST00 | 2065 | Slc16a8 | ENSMUSG00000032988.3 |
| chr2 | 148873489 | 148873980 | 8.16709 | 15.30057 | 13.0337 | Intron (ENSM | 2 | 148872472 | 148875451 | 2980 | 2 | ENSMUST00 | 1471 | Cst3 | ENSMUSG00000027447.6 |
| chr2 | 118660044 | 118660306 | 8.16709 | 15.30057 | 13.0337 | Distal Interge | 2 | 118663303 | 118698020 | 34718 | 1 | ENSMUST00 | -2997 | Pak6 | ENSMUSG00000074923.5 |
| chr3 | 103575982 | 103576623 | 8.16709 | 15.30057 | 13.0337 | Promoter | 3 | 103576081 | 103587240 | 11160 | 1 | ENSMUST00 | 0 | Syt6 | ENSMUSG00000027849.13 |
| chr1 | 133182273 | 133182918 | 6.42099 | 15.28918 | 13.02279 | Promoter | 1 | 133181321 | 133301219 | 119899 | 1 | ENSMUST00 | 953 | Plekha6 | ENSMUSG00000041757.10 |
| chr10 | 89344535 | 89345176 | 6.62775 | 15.27588 | 13.00958 | Promoter | 10 | 89035091 | 89344645 | 309555 | 2 | ENSMUST00 | 0 | Ano4 | ENSMUSG00000035189.11 |
| chr11 | 4096193 | 4096707 | 6.62775 | 15.27588 | 13.00958 | Promoter | 11 | 4091480 | 4095445 | 3966 | 2 | ENSMUST00 | -749 | Mtfp1 | ENSMUSG00000004748.5 |
| chr11 | 57827684 | 57828247 | 6.62775 | 15.27588 | 13.00958 | Downstream | 11 | 57828707 | 57832147 | 3441 | 2 | ENSMUST00 | 3900 | Hand1 | ENSMUSG00000037335.8 |
| chr18 | 66510165 | 66510763 | 6.62775 | 15.27588 | 13.00958 | Distal Interge | 18 | 66549935 | 66550045 | 111 | 2 | ENSMUST00 | 39282 | Gm24679 | ENSMUSG00000089387.1 |
| chr19 | 34583134 | 34583387 | 6.62775 | 15.27588 | 13.00958 | Promoter | 19 | 34583531 | 34588731 | 5201 | 1 | ENSMUST00 | -144 | Ifit3 | ENSMUSG00000074896.3 |
| chr8 | 12384989 | 12386069 | 6.62775 | 15.27588 | 13.00958 | Promoter | 8 | 12385771 | 12436729 | 50959 | 1 | ENSMUST00 | 0 | Gm5607 | ENSMUSG00000047935.11 |
| chr7 | 130129103 | 130129623 | 8.1351 | 15.24762 | 12.98159 | Distal Interge | 7 | 130163608 | 130167884 | 4277 | 2 | ENSMUST00 | 38261 | Fgfr2 | ENSMUSG00000030849.12 |
| chr7 | 137309228 | 137310696 | 8.1192 | 15.22129 | 12.95534 | Exon (ENSMU | 7 | 137194698 | 137314414 | 119717 | 2 | ENSMUST00 | 3718 | Ebf3 | ENSMUSG00000010476.8 |
| chr11 | 119304895 | 119305234 | 7.82748 | 15.19303 | 12.92724 | Distal Interge | 11 | 119307768 | 119345375 | 37608 | 1 | ENSMUST00 | -2534 | Card14 | ENSMUSG00000013483.9 |
| chr5 | 135465527 | 135465869 | 7.82748 | 15.19303 | 12.92724 | Intron (ENSM | 5 | 135376141 | 135394546 | 18406 | 2 | ENSMUST00 | -70982 | Pom121 | ENSMUSG00000053293.6 |
| chr8 | 33931315 | 33931677 | 7.82748 | 15.19303 | 12.92724 | Distal Interge | 8 | 33782643 | 33929863 | 147221 | 2 | ENSMUST00 | -1453 | Rbpms | ENSMUSG00000031586.11 |
| chr16 | 22428892 | 22429554 | 7.54366 | 15.16822 | 12.9028 | Intron (ENSM | 16 | 22381316 | 22436514 | 55199 | 2 | ENSMUST00 | 6960 | Etv5 | ENSMUSG00000013089.10 |
| chr3 | 88436893 | 88437113 | 7.54366 | 15.16822 | 12.9028 | Exon (ENSMU | 3 | 88440879 | 88440951 | 73 | 2 | ENSMUST00 | 3838 | Mir7011 | ENSMUSG00000098655.1 |
| chr8 | 90889116 | 90889394 | 7.54366 | 15.16822 | 12.9028 | Intron (ENSM | 8 | 90907939 | 90908415 | 477 | 2 | ENSMUST00 | 19021 | Gm6658 | ENSMUSG00000074171.1 |
| chr10 | 20905969 | 20906735 | 8.07187 | 15.14287 | 12.87755 | Distal Interge | 10 | 20952547 | 21080428 | 127882 | 1 | ENSMUST00 | -45812 | Ahi1 | ENSMUSG00000019986.11 |
| chr11 | 120151517 | 120151860 | 8.04062 | 15.09107 | 12.82585 | Promoter | 11 | 120123844 | 120151346 | 27503 | 2 | ENSMUST00 | -172 | Slc38a10 | ENSMUSG00000061306.10 |
| chr10 | 108445568 | 108446258 | 8.02509 | 15.0653 | 12.80013 | Distal Interge | 10 | 108428067 | 108428160 | 94 | 1 | ENSMUST00 | 17502 | Gm25591 | ENSMUSG00000093762.1 |
| chr1 | 90334944 | 90335775 | 8.00962 | 15.03963 | 12.77451 | Distal Interge | 1 | 90327312 | 90327418 | 107 | 1 | ENSMUST00 | 7633 | Gm27648 | ENSMUSG00000098368.1 |
| chr10 | 95388667 | 95389229 | 8.00962 | 15.03963 | 12.77451 | 3' UTR | 10 | 95387667 | 95392967 | 5301 | 2 | ENSMUST00 | 3738 | Socs2 | ENSMUSG00000020027.13 |
| chr10 | 39614683 | 39615321 | 8.00962 | 15.03963 | 12.77451 | Intron (ENSM | 10 | 39595859 | 39613682 | 17824 | 2 | ENSMUST00 | -1002 | Gm16365 | ENSMUSG00000086915.2 |
| chr11 | 101140142 | 101140835 | 8.00962 | 15.03963 | 12.77451 | Promoter | 11 | 101140454 | 101140509 | 56 | 2 | ENSMUST00 | 0 | Gm27626 | ENSMUSG00000092762.2 |
| chr14 | 19839748 | 19840286 | 8.00962 | 15.03963 | 12.77451 | Distal Interge | 14 | 19811402 | 19823823 | 12422 | 2 | ENSMUST00 | -15926 | 2700060E02I | ENSMUSG00000021807.5 |
| chr17 | 64559153 | 64559489 | 8.00962 | 15.03963 | 12.77451 | Distal Interge | 17 | 64514137 | 64555660 | 41524 | 2 | ENSMUST00 | -3494 | AU016765 | ENSMUSG00000073377.4 |
| chr2 | 170220581 | 170221366 | 8.00962 | 15.03963 | 12.77451 | Distal Interge | 2 | 170192449 | 170194053 | 1605 | 2 | ENSMUST00 | -26529 | Gm17619 | ENSMUSG00000097514.1 |
| chr3 | 96576347 | 96576879 | 8.00962 | 15.03963 | 12.77451 | Promoter | 3 | 96576984 | 96580663 | 3680 | 1 | ENSMUST00 | -105 | Gm16253 | ENSMUSG00000087610.1 |
| chr6 | 31540972 | 31541994 | 8.00962 | 15.03963 | 12.77451 | Intron (ENSM | 6 | 31522992 | 31524311 | 1320 | 2 | ENSMUST00 | -16662 | Podxl | ENSMUSG00000025608.9 |
| chr7 | 16993771 | 16994287 | 8.00962 | 15.03963 | 12.77451 | 5' UTR | 7 | 16992708 | 16997516 | 4809 | 1 | ENSMUST00 | 1064 | Ccdc8 | ENSMUSG00000041117.8 |
| chr9 | 102505913 | 102506315 | 8.00962 | 15.03963 | 12.77451 | Promoter | 9 | 102505750 | 102546239 | 40490 | 1 | ENSMUST00 | 164 | Ky | ENSMUSG00000035606.8 |

| | | | | | | | | | | | | | | | |
|---|---|---|---|---|---|---|---|---|---|---|---|---|---|---|---|
| chr9 | 15525128 | 15526195 | 8.00962 | 15.03963 | 12.77451 | 5' UTR | 9 | 15520846 | 15545257 | 24412 | 1 | ENSMUST00 | 4283 | Smco4 | ENSMUSG00000058173.6 |
| chr1 | 124045753 | 124046211 | 8.27158 | 15.0213 | 12.75664 | Promoter | 1 | 123321471 | 124045559 | 724089 | 2 | ENSMUST00 | -195 | Dpp10 | ENSMUSG00000036815.11 |
| chr11 | 103649464 | 103649977 | 8.27158 | 15.0213 | 12.75664 | Promoter | 11 | 103649570 | 103650579 | 1010 | 1 | ENSMUST00 | 0 | Rprml | ENSMUSG00000046215.3 |
| chr11 | 28365337 | 28365878 | 8.27158 | 15.0213 | 12.75664 | Distal Interge | 11 | 28360362 | 28364339 | 3978 | 1 | ENSMUST00 | 4976 | Gm12081 | ENSMUSG00000084905.1 |
| chr12 | 109067546 | 109068486 | 8.27158 | 15.0213 | 12.75664 | Promoter | 12 | 109032187 | 109068217 | 36031 | 2 | ENSMUST00 | 0 | Begain | ENSMUSG00000040867.10 |
| chr13 | 54749513 | 54750126 | 8.27158 | 15.0213 | 12.75664 | Promoter | 13 | 54736671 | 54749874 | 13204 | 2 | ENSMUST00 | 0 | Gprin1 | ENSMUSG00000069227.4 |
| chr13 | 119661950 | 119662155 | 8.27158 | 15.0213 | 12.75664 | Distal Interge | 13 | 119680040 | 119681588 | 1549 | 1 | ENSMUST00 | -17885 | CH25-211A2 | ENSMUSG00000099872.1 |
| chr14 | 30352885 | 30353427 | 8.27158 | 15.0213 | 12.75664 | Promoter | 14 | 30039949 | 30353486 | 313538 | 2 | ENSMUST00 | 59 | Cacna1d | ENSMUSG00000015968.11 |
| chr14 | 12346088 | 12347051 | 8.27158 | 15.0213 | 12.75664 | Promoter | 14 | 12341892 | 12345865 | 3974 | 2 | ENSMUST00 | -224 | Fezf2 | ENSMUSG00000021743.5 |
| chr14 | 39472457 | 39473081 | 8.27158 | 15.0213 | 12.75664 | Promoter | 14 | 38370477 | 39472825 | 1102349 | 2 | ENSMUST00 | 0 | Nrg3 | ENSMUSG00000041014.12 |
| chr15 | 74563511 | 74563778 | 8.27158 | 15.0213 | 12.75664 | Promoter | 15 | 74563792 | 74589464 | 25673 | 1 | ENSMUST00 | -14 | Bai1 | ENSMUSG00000034730.11 |
| chr17 | 25255939 | 25256422 | 8.27158 | 15.0213 | 12.75664 | Promoter | 17 | 25242659 | 25256364 | 13706 | 2 | ENSMUST00 | 0 | Baiap3 | ENSMUSG00000047507.7 |
| chr18 | 34547196 | 34547496 | 8.27158 | 15.0213 | 12.75664 | Exon (ENSMU | 18 | 34542313 | 34548273 | 5961 | 1 | ENSMUST00 | 4884 | Wnt8a | ENSMUSG00000012282.2 |
| chr18 | 58009153 | 58009678 | 8.27158 | 15.0213 | 12.75664 | 3' UTR | 18 | 57878678 | 57946821 | 68144 | 1 | ENSMUST00 | 130476 | Slc12a2 | ENSMUSG00000024597.10 |
| chr18 | 61096192 | 61096968 | 8.27158 | 15.0213 | 12.75664 | Distal Interge | 18 | 61105572 | 61131134 | 25563 | 1 | ENSMUST00 | -8604 | Csf1r | ENSMUSG00000024621.10 |
| chr19 | 3083411 | 3083902 | 8.27158 | 15.0213 | 12.75664 | Intron (ENSM | 19 | 3136984 | 3137090 | 107 | 2 | ENSMUST00 | 53188 | Gm25431 | ENSMUSG00000093983.1 |
| chr2 | 152847572 | 152847820 | 8.27158 | 15.0213 | 12.75664 | Promoter | 2 | 152847964 | 152895321 | 47358 | 1 | ENSMUST00 | -144 | Tpx2 | ENSMUSG00000027469.11 |
| chr2 | 175930601 | 175931047 | 8.27158 | 15.0213 | 12.75664 | Promoter | 2 | 175918881 | 175931051 | 12171 | 2 | ENSMUST00 | 4 | Gm8898 | ENSMUSG00000089756.2 |
| chr2 | 172901585 | 172901973 | 8.27158 | 15.0213 | 12.75664 | Intron (ENSM | 2 | 172864264 | 172864341 | 78 | 2 | ENSMUST00 | -37245 | Gm22773 | ENSMUSG00000093160.1 |
| chr3 | 65586348 | 65586915 | 8.27158 | 15.0213 | 12.75664 | Distal Interge | 3 | 65546040 | 65552632 | 6593 | 1 | ENSMUST00 | 40309 | Tiparp | ENSMUSG00000034640.4 |
| chr4 | 111687118 | 111687803 | 8.27158 | 15.0213 | 12.75664 | Distal Interge | 4 | 111719984 | 111829142 | 109159 | 1 | ENSMUST00 | -32181 | Spata6 | ENSMUSG00000034401.11 |
| chr4 | 88805845 | 88806076 | 8.27158 | 15.0213 | 12.75664 | Promoter | 4 | 88805531 | 88808380 | 2850 | 1 | ENSMUST00 | 315 | Gm13288 | ENSMUSG00000070908.1 |
| chr5 | 34185653 | 34186033 | 8.27158 | 15.0213 | 12.75664 | Intron (ENSM | 5 | 34173887 | 34187710 | 13824 | 2 | ENSMUST00 | 1677 | Mxd4 | ENSMUSG00000037235.8 |
| chr5 | 115436452 | 115436953 | 8.27158 | 15.0213 | 12.75664 | Promoter | 5 | 115435498 | 115445694 | 10197 | 1 | ENSMUST00 | 955 | Msi1 | ENSMUSG00000054256.6 |
| chr7 | 101818037 | 101818659 | 8.27158 | 15.0213 | 12.75664 | Promoter | 7 | 101818313 | 101822726 | 4414 | 1 | ENSMUST00 | 0 | Phox2a | ENSMUSG00000007946.10 |
| chr7 | 89939996 | 89940278 | 8.27158 | 15.0213 | 12.75664 | Promoter | 7 | 89918703 | 89941066 | 22364 | 2 | ENSMUST00 | 788 | l7Rn6 | ENSMUSG00000062797.6 |
| chr7 | 19118154 | 19118592 | 8.27158 | 15.0213 | 12.75664 | Promoter | 7 | 19101551 | 19118341 | 16791 | 2 | ENSMUST00 | 0 | Gm4969 | ENSMUSG00000085601.2 |
| chr8 | 119837423 | 119838609 | 8.27158 | 15.0213 | 12.75664 | Intron (ENSM | 8 | 119809225 | 119840522 | 31298 | 2 | ENSMUST00 | 1913 | Cotl1 | ENSMUSG00000031827.7 |
| chr9 | 46328832 | 46329461 | 8.27158 | 15.0213 | 12.75664 | Promoter | 9 | 46326391 | 46329413 | 3023 | 2 | ENSMUST00 | 0 | 4930448E22I | ENSMUSG00000097463.1 |
| chrX | 164381579 | 164382466 | 8.27158 | 15.0213 | 12.75664 | Intron (ENSM | 20 | 164373378 | 164402650 | 29273 | 1 | ENSMUST00 | 8202 | Figf | ENSMUSG00000031380.10 |
| chrX | 73639196 | 73639784 | 8.27158 | 15.0213 | 12.75664 | Promoter | 20 | 73639419 | 73643514 | 4096 | 1 | ENSMUST00 | 0 | Dusp9 | ENSMUSG00000031383.8 |
| chrX | 73911012 | 73911466 | 8.27158 | 15.0213 | 12.75664 | Promoter | 20 | 73891442 | 73911336 | 19895 | 2 | ENSMUST00 | 0 | Arhgap4 | ENSMUSG00000031389.12 |
| chrX | 98149761 | 98150251 | 8.27158 | 15.0213 | 12.75664 | Promoter | 20 | 98149721 | 98323215 | 173495 | 1 | ENSMUST00 | 41 | Ar | ENSMUSG00000046532.7 |
| chrX | 71664155 | 71664674 | 8.27158 | 15.0213 | 12.75664 | Promoter | 20 | 71663966 | 71667392 | 3427 | 1 | ENSMUST00 | 190 | Gpr50 | ENSMUSG00000056380.8 |
| chrX | 159371820 | 159372033 | 8.27158 | 15.0213 | 12.75664 | Promoter | 20 | 159372178 | 159376558 | 4381 | 1 | ENSMUST00 | -145 | Eif1ax | ENSMUSG00000067194.6 |
| chrX | 56821632 | 56822014 | 8.27158 | 15.0213 | 12.75664 | Promoter | 20 | 56809655 | 56822314 | 12660 | 2 | ENSMUST00 | 300 | Mtap7d3 | ENSMUSG00000067878.8 |
| chrX | 136976504 | 136976859 | 8.27158 | 15.0213 | 12.75664 | Promoter | 20 | 136954988 | 136976869 | 21882 | 2 | ENSMUST00 | 10 | Tmsb15l | ENSMUSG00000072955.8 |
| chrX | 103186244 | 103186515 | 8.27158 | 15.0213 | 12.75664 | Promoter | 20 | 103184176 | 103186640 | 2465 | 2 | ENSMUST00 | 125 | Nap1l2 | ENSMUSG00000082229.1 |
| chrX | 152368836 | 152369430 | 8.27158 | 15.0213 | 12.75664 | Promoter | 20 | 152369028 | 152403045 | 34018 | 1 | ENSMUST00 | 0 | 3010001F23I | ENSMUSG00000084885.2 |
| chr16 | 20536420 | 20537624 | 8.2567 | 14.99688 | 12.7344 | Promoter | 16 | 20535549 | 20537839 | 2291 | 1 | ENSMUST00 | 872 | Ap2m1 | ENSMUSG00000022841.9 |
| chr10 | 116301308 | 116302192 | 7.96309 | 14.9624 | 12.70017 | Promoter | 10 | 116301374 | 116389538 | 88165 | 1 | ENSMUST00 | 0 | Ptprb | ENSMUSG00000020154.9 |
| chr10 | 116986585 | 116987184 | 7.96309 | 14.9624 | 12.70017 | Promoter | 10 | 116986314 | 117025040 | 38727 | 1 | ENSMUST00 | 272 | Best3 | ENSMUSG00000020169.4 |
| chr10 | 81346895 | 81347134 | 7.96309 | 14.9624 | 12.70017 | Promoter | 10 | 81346052 | 81346730 | 679 | 2 | ENSMUST00 | -166 | Hmg20b | ENSMUSG00000020232.12 |
| chr14 | 48133026 | 48134696 | 7.96309 | 14.9624 | 12.70017 | Intron (ENSM | 14 | 48120869 | 48260883 | 140015 | 1 | ENSMUST00 | 12158 | Peli2 | ENSMUSG00000021846.8 |
| chr14 | 72506443 | 72506830 | 7.96309 | 14.9624 | 12.70017 | Distal Interge | 14 | 72561447 | 72564718 | 3272 | 2 | ENSMUST00 | 57888 | Fndc3a | ENSMUSG00000033487.9 |
| chr14 | 76598621 | 76599033 | 7.96309 | 14.9624 | 12.70017 | Distal Interge | 14 | 76532814 | 76556687 | 23874 | 2 | ENSMUST00 | -41935 | Serp2 | ENSMUSG00000052584.9 |
| chr14 | 4855701 | 4855957 | 7.96309 | 14.9624 | 12.70017 | Promoter | 14 | 4855576 | 4875851 | 20276 | 1 | ENSMUST00 | 126 | Gm3264 | ENSMUSG00000094132.2 |
| chr15 | 95185326 | 95186361 | 7.96309 | 14.9624 | 12.70017 | Distal Interge | 15 | 95097542 | 95162394 | 64853 | 2 | ENSMUST00 | -22933 | RP24-303N1 | ENSMUSG00000100603.1 |
| chr17 | 25717084 | 25717171 | 7.96309 | 14.9624 | 12.70017 | Promoter | 17 | 25717171 | 25719406 | 2236 | 1 | ENSMUST00 | 0 | Gng13 | ENSMUSG00000025739.8 |
| chr18 | 68943381 | 68944034 | 7.96309 | 14.9624 | 12.70017 | Downstream | 18 | 68944633 | 68950141 | 5509 | 2 | ENSMUST00 | 6107 | 4930546C10I | ENSMUSG00000024546.5 |
| chr2 | 32430606 | 32431239 | 7.96309 | 14.9624 | 12.70017 | Promoter | 2 | 32414487 | 32431119 | 16633 | 2 | ENSMUST00 | 0 | Slc25a25 | ENSMUSG00000026819.10 |
| chr2 | 155774985 | 155775312 | 7.96309 | 14.9624 | 12.70017 | Promoter | 2 | 155775342 | 155818366 | 43025 | 1 | ENSMUST00 | -30 | Mmp24 | ENSMUSG00000027612.7 |
| chr4 | 43478304 | 43478742 | 7.96309 | 14.9624 | 12.70017 | Distal Interge | 4 | 43482187 | 43483222 | 1036 | 2 | ENSMUST00 | 4480 | Sit1 | ENSMUSG00000028460.6 |
| chr4 | 152125134 | 152126287 | 7.96309 | 14.9624 | 12.70017 | Intron (ENSM | 4 | 152120857 | 152128925 | 8069 | 2 | ENSMUST00 | 2638 | Espn | ENSMUSG00000028943.12 |
| chr4 | 152153765 | 152154225 | 7.96309 | 14.9624 | 12.70017 | Downstream | 4 | 152120336 | 152152371 | 32036 | 2 | ENSMUST00 | -1395 | Espn | ENSMUSG00000028943.12 |
| chr4 | 140609300 | 140609777 | 7.96309 | 14.9624 | 12.70017 | Intron (ENSM | 4 | 140593169 | 140617065 | 23897 | 2 | ENSMUST00 | 7288 | Arhgef10l | ENSMUSG00000040964.11 |
| chr5 | 139797164 | 139797689 | 7.96309 | 14.9624 | 12.70017 | Promoter | 5 | 139797514 | 139802331 | 4818 | 1 | ENSMUST00 | 0 | Mafk | ENSMUSG00000018143.5 |
| chr5 | 34287036 | 34287574 | 7.96309 | 14.9624 | 12.70017 | Promoter | 5 | 34194874 | 34288326 | 93453 | 2 | ENSMUST00 | 752 | Zfyve28 | ENSMUSG00000037224.10 |
| chr5 | 115398415 | 115398815 | 7.96309 | 14.9624 | 12.70017 | Intron (ENSM | 5 | 115429599 | 115455698 | 26100 | 1 | ENSMUST00 | -30784 | Msi1 | ENSMUSG00000054256.6 |
| chr5 | 97278248 | 97278583 | 7.96309 | 14.9624 | 12.70017 | Distal Interge | 5 | 97264828 | 97265719 | 892 | 1 | ENSMUST00 | 13421 | Gm5560 | ENSMUSG00000067161.6 |
| chr7 | 84528727 | 84529363 | 7.96309 | 14.9624 | 12.70017 | Promoter | 7 | 84528975 | 84578337 | 49363 | 1 | ENSMUST00 | 0 | Gm2115 | ENSMUSG00000097789.1 |
| chr8 | 31918531 | 31918874 | 7.96309 | 14.9624 | 12.70017 | Promoter | 8 | 31818028 | 31918203 | 100176 | 2 | ENSMUST00 | -329 | Nrg1 | ENSMUSG00000062991.6 |
| chr8 | 45917237 | 45917547 | 7.96309 | 14.9624 | 12.70017 | Exon (ENSMU | 8 | 45934619 | 45944145 | 9527 | 1 | ENSMUST00 | -17072 | Ccdc110 | ENSMUSG00000071104.4 |
| chr9 | 34485905 | 34486487 | 7.96309 | 14.9624 | 12.70017 | Promoter | 9 | 34486125 | 34510616 | 24492 | 1 | ENSMUST00 | 0 | Kirrel3 | ENSMUSG00000032036.10 |
| chr9 | 106371249 | 106372180 | 7.96309 | 14.9624 | 12.70017 | Promoter | 9 | 106370744 | 106373856 | 3113 | 1 | ENSMUST00 | 506 | Dusp7 | ENSMUSG00000053716.8 |
| chrX | 99820073 | 99820637 | 7.96309 | 14.9624 | 12.70017 | Promoter | 20 | 99821021 | 99848790 | 27770 | 1 | ENSMUST00 | -384 | Tmem28 | ENSMUSG00000071719.2 |
| chr13 | 60738123 | 60738647 | 7.68982 | 14.9601 | 12.69977 | Intron (ENSM | 13 | 60771780 | 60772145 | 366 | 1 | ENSMUST00 | -33133 | Gm10312 | ENSMUSG00000096874.1 |
| chr14 | 62457442 | 62457646 | 7.42172 | 14.95621 | 12.69611 | Distal Interge | 14 | 62392676 | 62456289 | 63614 | 2 | ENSMUST00 | -1154 | Gucy1b2 | ENSMUSG00000021933.6 |
| chr7 | 95688771 | 95689280 | 7.42172 | 14.95621 | 12.69611 | Distal Interge | 7 | 95442351 | 95442457 | 107 | 2 | ENSMUST00 | -246315 | Gm25684 | ENSMUSG00000065875.1 |
| chr12 | 69461850 | 69462301 | 8.2222 | 14.94027 | 12.68024 | Distal Interge | 12 | 69371835 | 69372458 | 624 | 2 | ENSMUST00 | -89393 | Gm9887 | ENSMUSG00000052673.1 |
| chr18 | 37998306 | 37999139 | 8.2222 | 14.94027 | 12.68024 | Promoter | 18 | 37972624 | 37997574 | 24951 | 2 | ENSMUST00 | -733 | Arap3 | ENSMUSG00000024451.8 |
| chr2 | 11100170 | 11100408 | 8.2222 | 14.94027 | 12.68024 | Promoter | 2 | 11099788 | 11100264 | 477 | 2 | ENSMUST00 | 0 | Gm13294 | ENSMUSG00000083900.1 |
| chr2 | 172728195 | 172728826 | 8.2222 | 14.94027 | 12.68024 | Distal Interge | 2 | 172864264 | 172864341 | 78 | 2 | ENSMUST00 | 135515 | Gm22773 | ENSMUSG00000093160.1 |
| chr8 | 94214073 | 94214406 | 8.2222 | 14.94027 | 12.68024 | Promoter | 8 | 94214597 | 94314857 | 100261 | 1 | ENSMUST00 | -191 | Nup93 | ENSMUSG00000032939.8 |
| chr9 | 22188041 | 22188507 | 8.2222 | 14.94027 | 12.68024 | Promoter | 9 | 22188168 | 22202123 | 13956 | 1 | ENSMUST00 | 0 | Zfp872 | ENSMUSG00000074472.4 |
| chr10 | 116330769 | 116331444 | 8.20506 | 14.91214 | 12.65245 | Exon (ENSMU | 10 | 116340583 | 116340719 | 137 | 1 | ENSMUST00 | -9139 | Gm26495 | ENSMUSG00000065086.1 |
| chr18 | 53552155 | 53552923 | 8.20506 | 14.91214 | 12.65245 | Promoter | 18 | 53551594 | 53575857 | 24264 | 1 | ENSMUST00 | 562 | Prdm6 | ENSMUSG00000069378.8 |
| chr3 | 87796610 | 87797256 | 8.20506 | 14.91214 | 12.65245 | Promoter | 3 | 87796951 | 87816101 | 19151 | 1 | ENSMUST00 | 0 | Insrr | ENSMUSG00000005640.6 |
| chr4 | 142014874 | 142015199 | 8.20506 | 14.91214 | 12.65245 | Promoter | 4 | 141890438 | 142015067 | 124630 | 2 | ENSMUST00 | 0 | Fhad1 | ENSMUSG00000051435.6 |
| chr6 | 16359991 | 16360235 | 7.91801 | 14.88752 | 12.62793 | Distal Interge | 6 | 16553905 | 16554939 | 1035 | 2 | ENSMUST00 | 194704 | RP24-434O2 | ENSMUSG00000101034.1 |
| chr10 | 86637074 | 86637384 | 8.17099 | 14.85621 | 12.59679 | Distal Interge | 10 | 86655939 | 86657381 | 1443 | 1 | ENSMUST00 | -18555 | Gm5174 | ENSMUSG00000090308.1 |
| chr17 | 27702583 | 27703041 | 8.17099 | 14.85621 | 12.59679 | Exon (ENSMU | 17 | 27714505 | 27717582 | 3078 | 2 | ENSMUST00 | 14541 | Spdef | ENSMUSG00000024215.8 |
| chr14 | 79677182 | 79677726 | 8.15406 | 14.82842 | 12.56914 | Distal Interge | 14 | 79637693 | 79662170 | 24478 | 2 | ENSMUST00 | -15013 | Lect1 | ENSMUSG00000022025.8 |
| chr1 | 165594841 | 165595745 | 7.8581 | 14.78787 | 12.52884 | Intron (ENSM | 1 | 165571850 | 165571976 | 127 | 2 | ENSMUST00 | -22866 | Gm23208 | ENSMUSG00000089291.1 |
| chr11 | 19423956 | 19424575 | 7.8581 | 14.78787 | 12.52884 | Distal Interge | 11 | 19383685 | 19384708 | 1024 | 1 | ENSMUST00 | 40272 | Gm12026 | ENSMUSG00000082737.3 |
| chr12 | 108958969 | 108959657 | 7.8581 | 14.78787 | 12.52884 | Intron (ENSM | 12 | 108894317 | 109028325 | 134009 | 1 | ENSMUST00 | 64653 | Wdr25 | ENSMUSG00000040877.10 |
| chr14 | 7791407 | 7792014 | 7.8581 | 14.78787 | 12.52884 | Promoter | 14 | 7744063 | 7791651 | 47589 | 2 | ENSMUST00 | 0 | Gm10044 | ENSMUSG00000072723.5 |
| chr3 | 51358741 | 51359388 | 7.8581 | 14.78787 | 12.52884 | Distal Interge | 3 | 51264869 | 51340658 | 75790 | 2 | ENSMUST00 | -18084 | Elf2 | ENSMUSG00000037174.13 |
| chr5 | 115515366 | 115515606 | 7.8581 | 14.78787 | 12.52884 | Promoter | 5 | 115515263 | 115515947 | 685 | 1 | ENSMUST00 | 104 | Gm13840 | ENSMUSG00000086849.1 |
| chr6 | 37464244 | 37464769 | 7.8581 | 14.78787 | 12.52884 | Distal Interge | 6 | 37327255 | 37442146 | 114892 | 2 | ENSMUST00 | -22099 | Creb3l2 | ENSMUSG00000038648.5 |
| chr7 | 45715484 | 45715770 | 7.8581 | 14.78787 | 12.52884 | Promoter | 7 | 45715457 | 45720834 | 5378 | 1 | ENSMUST00 | 28 | Rpl18 | ENSMUSG00000059070.9 |
| chr10 | 80053026 | 80053355 | 8.1204 | 14.77318 | 12.51473 | Promoter | 10 | 80053488 | 80056439 | 2952 | 1 | ENSMUST00 | -133 | Gpx4 | ENSMUSG00000075706.5 |
| chr14 | 120278129 | 120279252 | 8.10368 | 14.74572 | 12.48744 | Intron (ENSM | 14 | 120275669 | 120431697 | 156029 | 1 | ENSMUST00 | 2461 | Mbnl2 | ENSMUSG00000022139.11 |
| chr15 | 79038029 | 79038452 | 8.10368 | 14.74572 | 12.48744 | 3' UTR | 15 | 79041885 | 79043558 | 1674 | 1 | ENSMUST00 | -3433 | Gcat | ENSMUSG00000006378.8 |
| chr9 | 41273438 | 41273958 | 8.10368 | 14.74572 | 12.48744 | Distal Interge | 9 | 41216948 | 41217566 | 619 | 1 | ENSMUST00 | 56491 | RP24-531B2 | ENSMUSG00000100837.1 |
| chr2 | 92434246 | 92434768 | 7.55693 | 14.73426 | 12.4761 | Promoter | 2 | 92403646 | 92434043 | 30398 | 2 | ENSMUST00 | -204 | Cry2 | ENSMUSG00000068742.6 |
| chr7 | 27447415 | 27447729 | 7.55693 | 14.73426 | 12.4761 | Promoter | 7 | 27356387 | 27447686 | 91300 | 2 | ENSMUST00 | 0 | Sptbn4 | ENSMUSG00000011751.11 |
| chr7 | 6398523 | 6398980 | 7.55693 | 14.73426 | 12.4761 | Promoter | 7 | 6398715 | 6429450 | 30736 | 1 | ENSMUST00 | 0 | Smim17 | ENSMUSG00000093536.2 |
| chr19 | 46041290 | 46043334 | 8.08703 | 14.71837 | 12.46039 | Intron (ENSM | 19 | 46044886 | 46062197 | 17312 | 1 | ENSMUST00 | -1552 | Pprc1 | ENSMUSG00000055491.8 |
| chr1 | 61402309 | 61403031 | 8.07045 | 14.69113 | 12.43326 | Distal Interge | 1 | 61378432 | 61395904 | 17473 | 1 | ENSMUST00 | 23878 | 9530026F06I | ENSMUSG00000085447.1 |
| chr12 | 100520879 | 100521207 | 8.05393 | 14.664 | 12.40628 | Promoter | 12 | 100300771 | 100520822 | 220052 | 2 | ENSMUST00 | -58 | Ttc7b | ENSMUSG00000033530.7 |
| chr18 | 65767295 | 65767576 | 8.05393 | 14.664 | 12.40628 | Distal Interge | 18 | 65749014 | 65750390 | 1377 | 1 | ENSMUST00 | 18282 | Oacyl | ENSMUSG00000046610.9 |
| chr18 | 77806462 | 77806925 | 8.05393 | 14.664 | 12.40628 | Promoter | 18 | 77806314 | 77806666 | 353 | 2 | ENSMUST00 | 0 | Gm23954 | ENSMUSG00000087838.1 |
| chr2 | 5080374 | 5080713 | 8.05393 | 14.664 | 12.40628 | Distal Interge | 2 | 5046157 | 5064051 | 17895 | 2 | ENSMUST00 | -16324 | Optn | ENSMUSG00000026672.6 |
| chr2 | 84851073 | 84851490 | 8.05393 | 14.664 | 12.40628 | Intron (ENSM | 2 | 84840625 | 84863594 | 22970 | 1 | ENSMUST00 | 10449 | Slc43a1 | ENSMUSG00000027075.11 |
| chr6 | 145075772 | 145076204 | 8.05393 | 14.664 | 12.40628 | Promoter | 6 | 144999158 | 145076184 | 77027 | 2 | ENSMUST00 | 0 | Bcat1 | ENSMUSG00000030268.12 |
| chr8 | 57328599 | 57328910 | 8.05393 | 14.664 | 12.40628 | Distal Interge | 8 | 57332118 | 57340682 | 8565 | 1 | ENSMUST00 | -3208 | 5033428I22R | ENSMUSG00000097910.1 |
| chr10 | 93261687 | 93262161 | 7.50973 | 14.65382 | 12.39649 | Intron (ENSM | 10 | 93249323 | 93310811 | 61489 | 2 | ENSMUST00 | 48650 | Elk3 | ENSMUSG00000008398.9 |
| chr10 | 21881960 | 21882630 | 7.50973 | 14.65382 | 12.39649 | Promoter | 10 | 21882184 | 21999893 | 117710 | 1 | ENSMUST00 | 0 | Sgk1 | ENSMUSG00000019970.10 |
| chr10 | 22820971 | 22821303 | 7.50973 | 14.65382 | 12.39649 | Promoter | 10 | 22817279 | 22820128 | 2850 | 2 | ENSMUST00 | -844 | Tcf21 | ENSMUSG00000045680.7 |
| chr11 | 116237752 | 116238512 | 7.50973 | 14.65382 | 12.39649 | Promoter | 11 | 116220559 | 116238077 | 17519 | 2 | ENSMUST00 | 0 | Evpl | ENSMUSG00000034282.3 |
| chr12 | 85454794 | 85455155 | 7.50973 | 14.65382 | 12.39649 | Distal Interge | 12 | 85473890 | 85477273 | 3384 | 1 | ENSMUST00 | -18735 | Fos | ENSMUSG00000021250.8 |
| chr12 | 98270340 | 98271178 | 7.50973 | 14.65382 | 12.39649 | Intron (ENSM | 12 | 98268635 | 98276624 | 7990 | 1 | ENSMUST00 | 1706 | Gpr65 | ENSMUSG00000021886.6 |
| chr12 | 52700737 | 52701463 | 7.50973 | 14.65382 | 12.39649 | Intron (ENSM | 12 | 52699383 | 53151015 | 451633 | 1 | ENSMUST00 | 1355 | Akap6 | ENSMUSG00000061603.7 |
| chr13 | 98489659 | 98489906 | 7.50973 | 14.65382 | 12.39649 | Exon (ENSMU | 13 | 98489348 | 98492001 | 2654 | 2 | ENSMUST00 | 2095 | Gm10320 | ENSMUSG00000092116.1 |
| chr15 | 99664527 | 99665164 | 7.50973 | 14.65382 | 12.39649 | Distal Interge | 15 | 99670718 | 99701127 | 30410 | 1 | ENSMUST00 | -5554 | Asic1 | ENSMUSG00000023017.9 |
| chr15 | 102354923 | 102355382 | 7.50973 | 14.65382 | 12.39649 | Downstream | 15 | 102338247 | 102350759 | 12513 | 2 | ENSMUST00 | -4165 | Aaas | ENSMUSG00000036678.7 |
| chr16 | 33764569 | 33764799 | 7.50973 | 14.65382 | 12.39649 | Intron (ENSM | 16 | 33756523 | 33760858 | 4336 | 1 | ENSMUST00 | 8047 | Heg1 | ENSMUSG00000075254.6 |
| chr16 | 92019172 | 92019877 | 7.50973 | 14.65382 | 12.39649 | Distal Interge | 16 | 92042677 | 92042774 | 98 | 1 | ENSMUST00 | -22800 | Gm23210 | ENSMUSG00000089293.1 |
| chr17 | 56325286 | 56325641 | 7.50973 | 14.65382 | 12.39649 | Promoter | 17 | 56326062 | 56356056 | 29995 | 1 | ENSMUST00 | -421 | Kdm4b | ENSMUSG00000024201.7 |
| chr17 | 26537875 | 26538217 | 7.50973 | 14.65382 | 12.39649 | Distal Interge | 17 | 26542760 | 26543715 | 956 | 1 | ENSMUST00 | -4543 | Gm8225 | ENSMUSG00000068604.4 |
| chr18 | 69576339 | 69576674 | 7.50973 | 14.65382 | 12.39649 | Intron (ENSM | 18 | 69563335 | 69636964 | 73630 | 1 | ENSMUST00 | 13005 | Tcf4 | ENSMUSG00000053477.10 |
| chr19 | 44759527 | 44760444 | 7.50973 | 14.65382 | 12.39649 | Promoter | 19 | 44759915 | 44818554 | 58640 | 1 | ENSMUST00 | 0 | Pax2 | ENSMUSG00000004231.10 |
| chr2 | 181147843 | 181148326 | 7.50973 | 14.65382 | 12.39649 | 3' UTR | 2 | 181147653 | 181157014 | 9362 | 2 | ENSMUST00 | 8688 | Eef1a2 | ENSMUSG00000016349.10 |
| chr2 | 102549817 | 102550373 | 7.50973 | 14.65382 | 12.39649 | Promoter | 2 | 102550012 | 102643041 | 93030 | 1 | ENSMUST00 | 0 | Pamr1 | ENSMUSG00000027188.8 |
| chr2 | 107290367 | 107291116 | 7.50973 | 14.65382 | 12.39649 | Promoter | 2 | 107290639 | 107298502 | 7864 | 1 | ENSMUST00 | 0 | Kcna4 | ENSMUSG00000042604.5 |
| chr2 | 177897929 | 177898303 | 7.50973 | 14.65382 | 12.39649 | Promoter | 2 | 177897096 | 177905784 | 8689 | 1 | ENSMUST00 | 834 | Gm14327 | ENSMUSG00000074521.4 |
| chr2 | 170184421 | 170185254 | 7.50973 | 14.65382 | 12.39649 | Distal Interge | 2 | 170192449 | 170194053 | 1605 | 2 | ENSMUST00 | 8799 | Gm17619 | ENSMUSG00000097514.1 |

| | | | | | | | | | | | | | | | |
|---|---|---|---|---|---|---|---|---|---|---|---|---|---|---|---|
| chr3 | 103738023 | 103738378 | 7.50973 | 14.65382 | 12.39649 | Promoter | 3 | 103733712 | 103738001 | 4290 | 2 | ENSMUST00 | -23 | Olfml3 | ENSMUSG00000027848.11 |
| chr4 | 137370072 | 137370726 | 7.50973 | 14.65382 | 12.39649 | Distal Interge | 4 | 137319696 | 137357720 | 38025 | 2 | ENSMUST00 | -12353 | Cdc42 | ENSMUSG00000006699.12 |
| chr4 | 41774041 | 41774299 | 7.50973 | 14.65382 | 12.39649 | Promoter | 4 | 41769467 | 41774097 | 4631 | 2 | ENSMUST00 | 0 | Ccl27a | ENSMUSG00000073888.7 |
| chr4 | 117496046 | 117496934 | 7.50973 | 14.65382 | 12.39649 | Promoter | 4 | 117496452 | 117500602 | 4151 | 1 | ENSMUST00 | 0 | Gm12827 | ENSMUSG00000087228.1 |
| chr5 | 123879475 | 123880321 | 7.50973 | 14.65382 | 12.39649 | Promoter | 5 | 123876736 | 123880020 | 3285 | 2 | ENSMUST00 | 0 | Hcar1 | ENSMUSG00000049241.5 |
| chr6 | 83831934 | 83832365 | 7.50973 | 14.65382 | 12.39649 | Promoter | 6 | 83805401 | 83831741 | 26341 | 2 | ENSMUST00 | -194 | Paip2b | ENSMUSG00000045896.9 |
| chr7 | 79515287 | 79515835 | 7.50973 | 14.65382 | 12.39649 | Promoter | 7 | 79515435 | 79531259 | 15825 | 1 | ENSMUST00 | 0 | AI854517 | ENSMUSG00000097023.3 |
| chr8 | 77506813 | 77507122 | 7.50973 | 14.65382 | 12.39649 | Intron (ENSM | 8 | 77517056 | 77518579 | 1524 | 1 | ENSMUST00 | -9934 | 0610038B21 | ENSMUSG00000097882.2 |
| chr8 | 117224761 | 117225043 | 7.50973 | 14.65382 | 12.39649 | Distal Interge | 8 | 117210778 | 117215997 | 5220 | 1 | ENSMUST00 | 13984 | Gm20204 | ENSMUSG00000098148.1 |
| chr9 | 66305247 | 66305847 | 7.50973 | 14.65382 | 12.39649 | Distal Interge | 9 | 66268374 | 66271715 | 3342 | 1 | ENSMUST00 | 36874 | Dapk2 | ENSMUSG00000032380.3 |
| chr9 | 40376159 | 40376508 | 7.50973 | 14.65382 | 12.39649 | Promoter | 9 | 40333546 | 40376367 | 42822 | 2 | ENSMUST00 | 0 | Gramd1b | ENSMUSG00000040111.9 |
| chr4 | 141425649 | 141426218 | 8.03748 | 14.63698 | 12.38157 | Distal Interge | 4 | 141420779 | 141425311 | 4533 | 1 | ENSMUST00 | 4871 | Hspb7 | ENSMUSG00000006221.7 |
| chr4 | 136462027 | 136462236 | 8.03748 | 14.63698 | 12.38157 | Promoter | 4 | 136462250 | 136463977 | 1728 | 1 | ENSMUST00 | -14 | 6030445D17 | ENSMUSG00000066178.2 |
| chr7 | 70118197 | 70118909 | 8.0211 | 14.61006 | 12.35478 | Distal Interge | 7 | 70248973 | 70249776 | 804 | 2 | ENSMUST00 | 130867 | RP23-474J16 | ENSMUSG00000099387.1 |
| chr16 | 30502426 | 30502987 | 8.00478 | 14.58325 | 12.3281 | Distal Interge | 16 | 30513970 | 30540909 | 26940 | 2 | ENSMUST00 | 37922 | Tmem44 | ENSMUSG00000022537.12 |
| chr3 | 108880667 | 108881503 | 7.98854 | 14.55654 | 12.30152 | Promoter | 3 | 108876512 | 108881779 | 5268 | 2 | ENSMUST00 | 276 | Fndc7 | ENSMUSG00000045326.8 |
| chr10 | 70060088 | 70060477 | 7.71221 | 14.5447 | 12.28981 | Distal Interge | 10 | 70097121 | 70193200 | 96080 | 1 | ENSMUST00 | -36644 | Ccdc6 | ENSMUSG00000048701.8 |
| chr13 | 48977795 | 48978359 | 7.71221 | 14.5447 | 12.28981 | Distal Interge | 13 | 48968287 | 48972031 | 3745 | 1 | ENSMUST00 | 9509 | Fam120aos | ENSMUSG00000097059.1 |
| chr14 | 32492494 | 32493048 | 7.71221 | 14.5447 | 12.28981 | Distal Interge | 14 | 32513521 | 32580989 | 67469 | 1 | ENSMUST00 | -20473 | Ercc6 | ENSMUSG00000054051.6 |
| chr14 | 5388415 | 5389385 | 7.71221 | 14.5447 | 12.28981 | Promoter | 14 | 5365794 | 5389049 | 23256 | 2 | ENSMUST00 | 0 | Gm3500 | ENSMUSG00000096003.2 |
| chr16 | 78930804 | 78931226 | 7.71221 | 14.5447 | 12.28981 | Promoter | 16 | 78930948 | 78947155 | 16208 | 1 | ENSMUST00 | 0 | Chodl | ENSMUSG00000022860.9 |
| chr5 | 120648563 | 120649584 | 7.71221 | 14.5447 | 12.28981 | Promoter | 5 | 120648812 | 120654902 | 6091 | 1 | ENSMUST00 | 0 | Rasal1 | ENSMUSG00000029602.6 |
| chr5 | 147188737 | 147189288 | 7.71221 | 14.5447 | 12.28981 | Promoter | 5 | 147188696 | 147190947 | 2252 | 1 | ENSMUST00 | 42 | Gsx1 | ENSMUSG00000053129.5 |
| chr6 | 92869412 | 92870074 | 7.71221 | 14.5447 | 12.28981 | Promoter | 6 | 92869357 | 92884411 | 15055 | 1 | ENSMUST00 | 56 | Gm15737 | ENSMUSG00000079462.1 |
| chr8 | 33903501 | 33904183 | 7.71221 | 14.5447 | 12.28981 | Intron (ENSM | 8 | 33863007 | 33891764 | 28758 | 2 | ENSMUST00 | -11738 | Rbpms | ENSMUSG00000031586.11 |
| chr8 | 70672727 | 70673001 | 7.71221 | 14.5447 | 12.28981 | Promoter | 8 | 70673248 | 70678752 | 5505 | 1 | ENSMUST00 | -247 | Lsm4 | ENSMUSG00000031848.9 |
| chr14 | 21686580 | 21686975 | 7.97235 | 14.52993 | 12.27555 | Exon (ENSMU | 14 | 21676582 | 21714576 | 37995 | 2 | ENSMUST00 | 27601 | Dupd1 | ENSMUSG00000063821.5 |
| chr4 | 123406179 | 123407054 | 7.97235 | 14.52993 | 12.27555 | Promoter | 4 | 123397275 | 123407515 | 10241 | 2 | ENSMUST00 | 461 | Macf1 | ENSMUSG00000028649.12 |
| chr6 | 143143809 | 143144251 | 7.97235 | 14.52993 | 12.27555 | Distal Interge | 6 | 143167230 | 143208542 | 41313 | 1 | ENSMUST00 | -22979 | Etnk1 | ENSMUSG00000030275.4 |
| chr6 | 148354732 | 148355409 | 7.42855 | 14.51512 | 12.26086 | Promoter | 6 | 148354656 | 148355598 | 943 | 1 | ENSMUST00 | 77 | Rps4l | ENSMUSG00000063171.3 |
| chr4 | 132796069 | 132796534 | 7.94018 | 14.47704 | 12.22308 | Promoter | 4 | 132781843 | 132796387 | 14545 | 2 | ENSMUST00 | 0 | Themis2 | ENSMUSG00000037731.5 |
| chr10 | 99460783 | 99461706 | 7.14186 | 14.46535 | 12.21151 | Distal Interge | 10 | 99443730 | 99447007 | 3278 | 1 | ENSMUST00 | 17054 | Gad1-ps | ENSMUSG00000090665.1 |
| chr11 | 101176214 | 101176604 | 7.14186 | 14.46535 | 12.21151 | Promoter | 11 | 101176041 | 101190724 | 14684 | 1 | ENSMUST00 | 174 | Cntnap1 | ENSMUSG00000017167.6 |
| chr13 | 91421549 | 91422152 | 7.14186 | 14.46535 | 12.21151 | Distal Interge | 13 | 91368990 | 91388085 | 19096 | 2 | ENSMUST00 | -33465 | A830009L08 | ENSMUSG00000097182.2 |
| chr15 | 84105376 | 84105951 | 7.14186 | 14.46535 | 12.21151 | Promoter | 15 | 84076097 | 84105754 | 29658 | 2 | ENSMUST00 | 0 | Sult4a1 | ENSMUSG00000018865.8 |
| chr15 | 98812788 | 98813788 | 7.14186 | 14.46535 | 12.21151 | 3' UTR | 15 | 98803782 | 98807925 | 4144 | 2 | ENSMUST00 | -4864 | Ddn | ENSMUSG00000059213.6 |
| chr17 | 25080332 | 25081172 | 7.14186 | 14.46535 | 12.21151 | Promoter | 17 | 25057702 | 25081181 | 23480 | 2 | ENSMUST00 | 9 | Tmem204 | ENSMUSG00000024168.7 |
| chr2 | 10153562 | 10154109 | 7.14186 | 14.46535 | 12.21151 | Promoter | 2 | 10153571 | 10256529 | 102959 | 1 | ENSMUST00 | 0 | Itih5 | ENSMUSG00000025780.7 |
| chr2 | 25196158 | 25196963 | 7.14186 | 14.46535 | 12.21151 | Promoter | 2 | 25192966 | 25196886 | 3921 | 2 | ENSMUST00 | 0 | Tor4a | ENSMUSG00000059555.6 |
| chr2 | 73037974 | 73038856 | 7.14186 | 14.46535 | 12.21151 | Distal Interge | 2 | 72987241 | 72988607 | 1367 | 1 | ENSMUST00 | 50734 | Sp3os | ENSMUSG00000063714.10 |
| chr3 | 131564253 | 131564610 | 7.14186 | 14.46535 | 12.21151 | Promoter | 3 | 131564768 | 131643670 | 78903 | 1 | ENSMUST00 | -158 | Papss1 | ENSMUSG00000028032.9 |
| chr3 | 83287057 | 83287338 | 7.14186 | 14.46535 | 12.21151 | Distal Interge | 3 | 83125276 | 83129246 | 3971 | 2 | ENSMUST00 | -157812 | Gm10710 | ENSMUSG00000074517.2 |
| chr4 | 102511137 | 102512052 | 7.14186 | 14.46535 | 12.21151 | Intron (ENSM | 4 | 102470397 | 102488046 | 17650 | 1 | ENSMUST00 | 40741 | Pde4b | ENSMUSG00000028525.11 |
| chr4 | 154023440 | 154024806 | 7.14186 | 14.46535 | 12.21151 | Promoter | 4 | 154023311 | 154024355 | 1045 | 2 | ENSMUST00 | 0 | Smim1 | ENSMUSG00000078350.6 |
| chr4 | 87368252 | 87368868 | 7.14186 | 14.46535 | 12.21151 | Distal Interge | 4 | 87432272 | 87432336 | 65 | 1 | ENSMUST00 | -63404 | Gm23154 | ENSMUSG00000093065.1 |
| chr5 | 125284218 | 125284479 | 7.14186 | 14.46535 | 12.21151 | Promoter | 5 | 125277787 | 125284159 | 6373 | 2 | ENSMUST00 | -60 | Scarb1 | ENSMUSG00000037936.10 |
| chr5 | 104770417 | 104770677 | 7.14186 | 14.46535 | 12.21151 | Distal Interge | 5 | 104775779 | 104777227 | 1449 | 1 | ENSMUST00 | -5102 | Gm8258 | ENSMUSG00000048334.8 |
| chr5 | 149831564 | 149831870 | 7.14186 | 14.46535 | 12.21151 | Distal Interge | 5 | 149753942 | 149760607 | 6666 | 1 | ENSMUST00 | 77623 | B3glct | ENSMUSG00000051950.8 |
| chr7 | 30717324 | 30717912 | 7.14186 | 14.46535 | 12.21151 | Promoter | 7 | 30717832 | 30720620 | 2789 | 1 | ENSMUST00 | 0 | Atp4a | ENSMUSG00000005553.8 |
| chr7 | 68749500 | 68750022 | 7.14186 | 14.46535 | 12.21151 | Promoter | 7 | 68737045 | 68749241 | 12197 | 2 | ENSMUST00 | -260 | Arrdc4 | ENSMUSG00000042659.9 |
| chr8 | 69717215 | 69717455 | 7.14186 | 14.46535 | 12.21151 | Distal Interge | 8 | 69706332 | 69710724 | 4393 | 2 | ENSMUST00 | -6492 | Zfp869 | ENSMUSG00000054648.7 |
| chr9 | 48706987 | 48707402 | 7.14186 | 14.46535 | 12.21151 | Intron (ENSM | 9 | 48591877 | 48605153 | 13277 | 2 | ENSMUST00 | -101835 | Nnmt | ENSMUSG00000032271.8 |
| chr9 | 75552770 | 75554589 | 7.14186 | 14.46535 | 12.21151 | Intron (ENSM | 9 | 75497796 | 75559657 | 61862 | 2 | ENSMUST00 | 5068 | Tmod3 | ENSMUSG00000058587.7 |
| chrX | 38179776 | 38180114 | 7.14186 | 14.46535 | 12.21151 | Promoter | 20 | 38180728 | 38181626 | 899 | 1 | ENSMUST00 | -614 | Gm14556 | ENSMUSG00000080942.1 |
| chr10 | 120377293 | 120377839 | 7.92419 | 14.45074 | 12.19819 | Intron (ENSM | 10 | 120389523 | 120392793 | 3271 | 1 | ENSMUST00 | -11684 | 9230105E05 | ENSMUSG00000090191.1 |
| chr15 | 83121776 | 83122185 | 7.92419 | 14.45074 | 12.19819 | Promoter | 15 | 83119255 | 83122149 | 2895 | 2 | ENSMUST00 | 0 | Rrp7a | ENSMUSG00000018040.4 |
| chr17 | 34957398 | 34957803 | 7.92419 | 14.45074 | 12.19819 | Exon (ENSMU | 17 | 34956436 | 34959238 | 2803 | 2 | ENSMUST00 | 1435 | Hspa1b | ENSMUSG00000090877.3 |
| chr13 | 47148279 | 47148948 | 7.90827 | 14.42455 | 12.17211 | Intron (ENSM | 13 | 47122720 | 47247991 | 125272 | 1 | ENSMUST00 | 25560 | Rnf144b | ENSMUSG00000038068.9 |
| chr16 | 17277251 | 17278142 | 7.90827 | 14.42455 | 12.17211 | Promoter | 16 | 17277601 | 17283229 | 5629 | 1 | ENSMUST00 | 0 | Tmem191c | ENSMUSG00000055692.13 |
| chr4 | 95037791 | 95038098 | 7.90827 | 14.42455 | 12.17211 | Intron (ENSM | 4 | 95039162 | 95039625 | 464 | 2 | ENSMUST00 | 1527 | Gm12704 | ENSMUSG00000081030.1 |
| chr11 | 106031937 | 106032354 | 6.00959 | 14.42206 | 12.16971 | Intron (ENSM | 11 | 106036872 | 106059324 | 22453 | 1 | ENSMUST00 | -4518 | Dcaf7 | ENSMUSG00000049354.8 |
| chr17 | 31853458 | 31854514 | 7.89241 | 14.39845 | 12.14615 | Exon (ENSMU | 17 | 31844248 | 31855792 | 11545 | 2 | ENSMUST00 | 1278 | Sik1 | ENSMUSG00000024042.6 |
| chr3 | 27317038 | 27317331 | 7.89241 | 14.39845 | 12.14615 | Promoter | 3 | 27317061 | 27342427 | 25367 | 1 | ENSMUST00 | 0 | Tnfsf10 | ENSMUSG00000039304.6 |
| chr5 | 72706361 | 72706870 | 7.89241 | 14.39845 | 12.14615 | Intron (ENSM | 5 | 72695982 | 72736589 | 40608 | 2 | ENSMUST00 | 29719 | Txk | ENSMUSG00000054892.9 |
| chr6 | 6863228 | 6864145 | 7.89241 | 14.39845 | 12.14615 | Promoter | 6 | 6863334 | 6867970 | 4637 | 1 | ENSMUST00 | 0 | Dlx6 | ENSMUSG00000029754.8 |
| chr15 | 23036029 | 23036823 | 7.87662 | 14.37246 | 12.12064 | Promoter | 15 | 23036455 | 23175141 | 138687 | 1 | ENSMUST00 | 0 | Cdh18 | ENSMUSG00000040420.9 |
| chr3 | 87907717 | 87908191 | 7.87662 | 14.37246 | 12.12064 | Promoter | 3 | 87907534 | 87914166 | 6633 | 1 | ENSMUST00 | 184 | Hdgf | ENSMUSG00000004897.11 |
| chr1 | 36784841 | 36785614 | 6.83737 | 14.36054 | 12.10883 | Distal Interge | 1 | 36778984 | 36782815 | 3832 | 1 | ENSMUST00 | 5858 | Zap70 | ENSMUSG00000026117.9 |
| chr11 | 121706913 | 121708593 | 6.83737 | 14.36054 | 12.10883 | 5' UTR | 11 | 121702763 | 121715829 | 13067 | 1 | ENSMUST00 | 4151 | Metrnl | ENSMUSG00000039208.10 |
| chr11 | 48753499 | 48753725 | 6.83737 | 14.36054 | 12.10883 | Promoter | 11 | 48751867 | 48753714 | 1848 | 2 | ENSMUST00 | 0 | Gm12183 | ENSMUSG00000082454.3 |
| chr11 | 119159650 | 119160040 | 6.83737 | 14.36054 | 12.10883 | Intron (ENSM | 11 | 119154076 | 119154137 | 62 | 2 | ENSMUST00 | -5514 | Mir6934 | ENSMUSG00000098704.1 |
| chr14 | 66279041 | 66279703 | 6.83737 | 14.36054 | 12.10883 | Intron (ENSM | 14 | 66153844 | 66280982 | 127139 | 2 | ENSMUST00 | 1279 | Ptk2b | ENSMUSG00000059456.8 |
| chr14 | 62144438 | 62145073 | 6.83737 | 14.36054 | 12.10883 | Distal Interge | 14 | 62201915 | 62204352 | 2438 | 2 | ENSMUST00 | 59279 | Gm26969 | ENSMUSG00000098096.1 |
| chr15 | 36578323 | 36578924 | 6.83737 | 14.36054 | 12.10883 | Distal Interge | 15 | 36596024 | 36597426 | 1403 | 2 | ENSMUST00 | 18502 | Pabpc1 | ENSMUSG00000022283.9 |
| chr15 | 54278281 | 54278755 | 6.83737 | 14.36054 | 12.10883 | Promoter | 15 | 54250619 | 54278484 | 27866 | 2 | ENSMUST00 | 0 | Tnfrsf11b | ENSMUSG00000063727.3 |
| chr2 | 119804550 | 119805857 | 6.83737 | 14.36054 | 12.10883 | Exon (ENSMU | 2 | 119803087 | 119807990 | 4904 | 1 | ENSMUST00 | 1464 | Tyro3 | ENSMUSG00000027298.12 |
| chr4 | 119105671 | 119106174 | 6.83737 | 14.36054 | 12.10883 | Distal Interge | 4 | 119108711 | 119137330 | 28620 | 1 | ENSMUST00 | -2537 | Slc2a1 | ENSMUSG00000028645.5 |
| chr4 | 45763923 | 45764529 | 6.83737 | 14.36054 | 12.10883 | Promoter | 4 | 45762377 | 45764223 | 1847 | 2 | ENSMUST00 | 0 | Gm12409 | ENSMUSG00000085876.1 |
| chr5 | 125465513 | 125465775 | 6.83737 | 14.36054 | 12.10883 | Distal Interge | 5 | 125475814 | 125517412 | 41599 | 1 | ENSMUST00 | -10039 | Aacs | ENSMUSG00000029482.3 |
| chr5 | 104074594 | 104075195 | 6.83737 | 14.36054 | 12.10883 | Exon (ENSMU | 5 | 104070064 | 104077608 | 7545 | 2 | ENSMUST00 | 2413 | Gm17660 | ENSMUSG00000091034.3 |
| chr6 | 17307036 | 17307950 | 6.83737 | 14.36054 | 12.10883 | Promoter | 6 | 17307048 | 17339153 | 32106 | 1 | ENSMUST00 | 0 | Cav1 | ENSMUSG00000007655.11 |
| chr7 | 19131078 | 19131373 | 6.83737 | 14.36054 | 12.10883 | Promoter | 7 | 19131693 | 19137812 | 6120 | 1 | ENSMUST00 | -320 | Fbxo46 | ENSMUSG00000050428.6 |
| chr7 | 30183777 | 30184493 | 6.83737 | 14.36054 | 12.10883 | Promoter | 7 | 30184171 | 30186028 | 1858 | 1 | ENSMUST00 | 0 | Cox7a1 | ENSMUSG00000074218.2 |
| chr8 | 10924413 | 10925063 | 6.83737 | 14.36054 | 12.10883 | Exon (ENSMU | 8 | 10924427 | 10928457 | 4031 | 2 | ENSMUST00 | 3394 | 3930402G23 | ENSMUSG00000038917.3 |
| chr8 | 32293094 | 32293410 | 6.83737 | 14.36054 | 12.10883 | Distal Interge | 8 | 32186129 | 32186248 | 120 | 1 | ENSMUST00 | 106966 | Gm22140 | ENSMUSG00000089370.1 |
| chr9 | 107587217 | 107587526 | 6.83737 | 14.36054 | 12.10883 | Promoter | 9 | 107575820 | 107587425 | 11606 | 2 | ENSMUST00 | 0 | Gm16343 | ENSMUSG00000089935.1 |
| chr19 | 46499527 | 46499866 | 6.17403 | 14.35629 | 12.10532 | Distal Interge | 19 | 46501648 | 46516444 | 14797 | 1 | ENSMUST00 | -1782 | Trim8 | ENSMUSG00000025034.8 |
| chr8 | 121124179 | 121125173 | 6.17403 | 14.35629 | 12.10532 | Intron (ENSM | 8 | 121127940 | 121130644 | 2705 | 1 | ENSMUST00 | -2767 | Foxl1 | ENSMUSG00000097084.1 |
| chr18 | 23039180 | 23040349 | 7.86088 | 14.34657 | 12.09571 | Promoter | 18 | 22694524 | 23040078 | 345555 | 2 | ENSMUST00 | 0 | Nol4 | ENSMUSG00000041923.10 |
| chr1 | 180848318 | 180848823 | 6.36264 | 14.31869 | 12.06804 | Distal Interge | 1 | 180851151 | 180868097 | 16947 | 1 | ENSMUST00 | -2328 | Sde2 | ENSMUSG00000038806.6 |
| chr1 | 134234692 | 134235726 | 6.36264 | 14.31869 | 12.06804 | Promoter | 1 | 134199225 | 134235431 | 36207 | 2 | ENSMUST00 | 0 | Adora1 | ENSMUSG00000042429.8 |
| chr12 | 23969124 | 23969484 | 6.36264 | 14.31869 | 12.06804 | Promoter | 12 | 23968510 | 23968845 | 336 | 2 | ENSMUST00 | -280 | Gm17330 | ENSMUSG00000095329.1 |
| chr15 | 31568362 | 31568899 | 6.36264 | 14.31869 | 12.06804 | Promoter | 15 | 31568791 | 31588259 | 19469 | 1 | ENSMUST00 | 0 | Cmbl | ENSMUSG00000022235.9 |
| chr3 | 122245057 | 122245361 | 6.36264 | 14.31869 | 12.06804 | Promoter | 3 | 122245557 | 122269984 | 24428 | 1 | ENSMUST00 | -196 | Gclm | ENSMUSG00000028124.10 |
| chr6 | 117562364 | 117563386 | 6.36264 | 14.31869 | 12.06804 | Exon (ENSMU | 6 | 117559697 | 117572329 | 12633 | 1 | ENSMUST00 | 2668 | Gm9946 | ENSMUSG00000054488.9 |
| chr1 | 17727387 | 17727816 | 6.58118 | 14.31666 | 12.06619 | Promoter | 1 | 17692904 | 17727646 | 34743 | 2 | ENSMUST00 | 0 | Gm16070 | ENSMUSG00000085125.2 |
| chr10 | 17796336 | 17796712 | 6.58118 | 14.31666 | 12.06619 | Promoter | 10 | 17796256 | 17845665 | 49410 | 1 | ENSMUST00 | 81 | Txlnb | ENSMUSG00000039891.6 |
| chr12 | 8510984 | 8511787 | 6.58118 | 14.31666 | 12.06619 | Distal Interge | 12 | 8497763 | 8499985 | 2223 | 2 | ENSMUST00 | -11000 | Rhob | ENSMUSG00000054364.4 |
| chr13 | 33762395 | 33762943 | 6.58118 | 14.31666 | 12.06619 | Distal Interge | 13 | 33766920 | 33769463 | 2544 | 1 | ENSMUST00 | -3977 | Vmn2r-ps94 | ENSMUSG00000080317.2 |
| chr13 | 119836518 | 119837749 | 6.58118 | 14.31666 | 12.06619 | Promoter | 13 | 119836006 | 119846506 | 10501 | 1 | ENSMUST00 | 513 | D13Ertd608e | ENSMUSG00000095514.1 |
| chr15 | 75993569 | 75993970 | 6.58118 | 14.31666 | 12.06619 | Promoter | 15 | 75993769 | 75995824 | 2056 | 1 | ENSMUST00 | 0 | Mapk15 | ENSMUSG00000063704.7 |
| chr17 | 5770212 | 5770837 | 6.58118 | 14.31666 | 12.06619 | Distal Interge | 17 | 5798657 | 5802893 | 4237 | 1 | ENSMUST00 | -27820 | 3300005D01 | ENSMUSG00000096965.2 |
| chr18 | 37751632 | 37752081 | 6.58118 | 14.31666 | 12.06619 | Intron (ENSM | 18 | 37755773 | 37841870 | 86098 | 1 | ENSMUST00 | -3692 | Pcdhga12 | ENSMUSG00000023036.9 |
| chr18 | 73863629 | 73863995 | 6.58118 | 14.31666 | 12.06619 | Promoter | 18 | 73863631 | 73870209 | 6579 | 1 | ENSMUST00 | 0 | Mro | ENSMUSG00000064036.10 |
| chr7 | 104287340 | 104287870 | 6.58118 | 14.31666 | 12.06619 | Promoter | 7 | 104263386 | 104288094 | 24709 | 2 | ENSMUST00 | 224 | Trim5 | ENSMUSG00000060441.9 |
| chr10 | 96160979 | 96161203 | 7.57164 | 14.30958 | 12.05949 | Distal Interge | 10 | 96164331 | 96164431 | 101 | 1 | ENSMUST00 | -3128 | Gm22913 | ENSMUSG00000094523.1 |
| chr11 | 114727091 | 114727813 | 7.57164 | 14.30958 | 12.05949 | Promoter | 11 | 114727408 | 114745158 | 17751 | 1 | ENSMUST00 | 0 | Dnaic2 | ENSMUSG00000034706.11 |
| chr12 | 76694927 | 76696352 | 7.57164 | 14.30958 | 12.05949 | Intron (ENSM | 12 | 76691980 | 76692261 | 282 | 1 | ENSMUST00 | 2948 | Gm24010 | ENSMUSG00000088356.1 |
| chr13 | 31618257 | 31618684 | 7.57164 | 14.30958 | 12.05949 | Promoter | 13 | 31619168 | 31620369 | 1202 | 1 | ENSMUST00 | -484 | Gm11378 | ENSMUSG00000087276.1 |
| chr15 | 98708159 | 98708541 | 7.57164 | 14.30958 | 12.05949 | Promoter | 15 | 98708207 | 98723326 | 15120 | 1 | ENSMUST00 | 0 | Ccdc65 | ENSMUSG00000003354.5 |
| chr15 | 10952105 | 10952688 | 7.57164 | 14.30958 | 12.05949 | Promoter | 15 | 10952332 | 10980150 | 27819 | 1 | ENSMUST00 | 0 | C1qtnf3 | ENSMUSG00000058914.7 |
| chr4 | 152282969 | 152283354 | 7.30446 | 14.30232 | 12.05262 | Promoter | 4 | 152282370 | 152285336 | 2967 | 1 | ENSMUST00 | 600 | Gpr153 | ENSMUSG00000042804.8 |
| chr9 | 21029381 | 21029869 | 7.30446 | 14.30232 | 12.05262 | Promoter | 9 | 21029373 | 21030531 | 1159 | 1 | ENSMUST00 | 9 | Icam4 | ENSMUSG00000001014.5 |
| chr4 | 86632680 | 86632902 | 7.8296 | 14.29507 | 12.04551 | Distal Interge | 4 | 86627075 | 86631251 | 4177 | 2 | ENSMUST00 | -1430 | Gm12551 | ENSMUSG00000081169.1 |
| chr15 | 9094336 | 9094880 | 7.81406 | 14.26947 | 12.02001 | Intron (ENSM | 15 | 9100156 | 9107178 | 7023 | 1 | ENSMUST00 | -5276 | Nadk2 | ENSMUSG00000022253.9 |
| chr18 | 50064550 | 50064868 | 7.81406 | 14.26947 | 12.02001 | Intron (ENSM | 18 | 50052299 | 50053844 | 1546 | 2 | ENSMUST00 | -10707 | C030005K06 | ENSMUSG00000086600.3 |
| chr18 | 69666034 | 69666845 | 6.78766 | 14.26931 | 12.01994 | Intron (ENSM | 18 | 69679699 | 69684528 | 4830 | 1 | ENSMUST00 | -12854 | Tcf4 | ENSMUSG00000053477.10 |
| chr6 | 85843808 | 85844030 | 7.79857 | 14.24396 | 11.99461 | Promoter | 6 | 85843980 | 85844459 | 480 | 1 | ENSMUST00 | 0 | Gm9769 | ENSMUSG00000040078.9 |
| chr1 | 14307048 | 14308052 | 7.76779 | 14.19323 | 11.94406 | Intron (ENSM | 1 | 14270775 | 14309836 | 39062 | 2 | ENSMUST00 | 1784 | Eya1 | ENSMUSG00000025932.9 |
| chr10 | 37397243 | 37397892 | 7.76779 | 14.19323 | 11.94406 | Distal Interge | 10 | 37337486 | 37379652 | 42167 | 1 | ENSMUST00 | 59758 | Gm26535 | ENSMUSG00000097742.1 |
| chr7 | 49437771 | 49438084 | 7.76779 | 14.19323 | 11.94406 | Exon (ENSMU | 7 | 49450760 | 49464580 | 13821 | 1 | ENSMUST00 | -12676 | Nav2 | ENSMUSG00000052512.10 |
| chr7 | 69992435 | 69993144 | 7.76779 | 14.19323 | 11.94406 | Distal Interge | 7 | 69957776 | 69957904 | 129 | 2 | ENSMUST00 | -34532 | Gm24120 | ENSMUSG00000077166.1 |
| chr6 | 47778086 | 47778940 | 4.1696 | 14.1759 | 11.92681 | Distal Interge | 6 | 47781624 | 47781725 | 102 | 1 | ENSMUST00 | -2684 | Rny3 | ENSMUSG00000064945.1 |
| chr14 | 73375938 | 73376255 | 7.75248 | 14.16801 | 11.91892 | Intron (ENSM | 14 | 73362231 | 73385271 | 23041 | 2 | ENSMUST00 | 9016 | Itm2b | ENSMUSG00000022108.7 |
| chr4 | 86885852 | 86886568 | 7.75248 | 14.16801 | 11.91892 | Exon (ENSMU | 4 | 86874414 | 86932491 | 58078 | 1 | ENSMUST00 | 11439 | Acer2 | ENSMUSG00000038007.9 |
| chr16 | 92060166 | 92061297 | 6.97099 | 14.1625 | 11.91348 | Intron (ENSM | 16 | 92058322 | 92087473 | 29152 | 1 | ENSMUST00 | 1845 | Slc5a3 | ENSMUSG00000089774.2 |
| chr12 | 111071573 | 111071944 | 7.73724 | 14.14289 | 11.89389 | Intron (ENSM | 12 | 111039798 | 111113384 | 73587 | 1 | ENSMUST00 | 31776 | Rcor1 | ENSMUSG00000037896.12 |
| chr13 | 70836174 | 70836533 | 7.73724 | 14.14289 | 11.89389 | Exon (ENSMU | 13 | 70727810 | 70841775 | 113966 | 2 | ENSMUST00 | 5242 | Adamts16 | ENSMUSG00000049538.9 |

| | | | | | | | | | | | | | | | |
|---|---|---|---|---|---|---|---|---|---|---|---|---|---|---|---|
| chr13 | 21832405 | 21832594 | 7.73724 | 14.14289 | 11.89389 | Promoter | 13 | 21831767 | 21832196 | 430 | 2 | ENSMUST00 | -210 | Hist1h4n | ENSMUSG00000069305.3 |
| chr13 | 44055755 | 44056326 | 7.73724 | 14.14288 | 11.89389 | Distal Interge | 13 | 44121167 | 44125179 | 4013 | 1 | ENSMUST00 | -64841 | Gm5083 | ENSMUSG00000089815.2 |
| chr19 | 23057275 | 23057780 | 7.73724 | 14.14289 | 11.89389 | Intron (ENSM | 19 | 23037390 | 23075853 | 38464 | 2 | ENSMUST00 | 18073 | C330002G04 | ENSMUSG00000097930.1 |
| chr5 | 63649672 | 63650267 | 7.73724 | 14.14289 | 11.89389 | Promoter | 5 | 63649950 | 63650264 | 315 | 2 | ENSMUST00 | 0 | Gm9954 | ENSMUSG00000054714.3 |
| chr8 | 84811123 | 84811427 | 7.72206 | 14.11785 | 11.86935 | Distal Interge | 8 | 84708616 | 84800344 | 91729 | 2 | ENSMUST00 | -10780 | Nfix | ENSMUSG00000001911.11 |
| chr15 | 81772309 | 81773031 | 7.70694 | 14.09291 | 11.84459 | Exon (ENSMU | 15 | 81744848 | 81796269 | 51422 | 1 | ENSMUST00 | 27462 | Zc3h7b | ENSMUSG00000022390.10 |
| chr9 | 113811821 | 113812438 | 7.70694 | 14.09291 | 11.84459 | Promoter | 9 | 113812586 | 113919697 | 107112 | 1 | ENSMUST00 | -148 | Clasp2 | ENSMUSG00000033392.10 |
| chr14 | 73116970 | 73117356 | 7.4361 | 14.08204 | 11.83379 | Distal Interge | 14 | 73123037 | 73161986 | 38950 | 1 | ENSMUST00 | -5681 | Rcbtb2 | ENSMUSG00000022106.9 |
| chr14 | 55783498 | 55784324 | 7.4361 | 14.08204 | 11.83379 | Promoter | 14 | 55769058 | 55784042 | 14985 | 2 | ENSMUST00 | 0 | Adcy4 | ENSMUSG00000022220.8 |
| chr7 | 110626280 | 110626700 | 7.4361 | 14.08204 | 11.83379 | Promoter | 7 | 110627661 | 110629820 | 2160 | 1 | ENSMUST00 | -961 | Adm | ENSMUSG00000030790.9 |
| chr7 | 112958054 | 112958696 | 7.4361 | 14.08204 | 11.83379 | Distal Interge | 7 | 112953962 | 112957457 | 3496 | 1 | ENSMUST00 | 4093 | Rassf10 | ENSMUSG00000098132.1 |
| chr8 | 114351940 | 114352148 | 7.4361 | 14.08204 | 11.83379 | Promoter | 8 | 114341160 | 114351479 | 10320 | 2 | ENSMUST00 | -462 | 4933408N05 | ENSMUSG00000084848.1 |
| chr11 | 85841915 | 85842203 | 7.66192 | 14.01863 | 11.77084 | 3' UTR | 11 | 85846790 | 85850333 | 3544 | 2 | ENSMUST00 | 8130 | Gm11444 | ENSMUSG00000069785.4 |
| chr9 | 108827096 | 108827716 | 7.61743 | 13.94515 | 11.69753 | Promoter | 9 | 108826320 | 108852963 | 26644 | 1 | ENSMUST00 | 777 | Celsr3 | ENSMUSG00000023473.6 |
| chr9 | 84107313 | 84107610 | 7.61743 | 13.94515 | 11.69753 | Promoter | 9 | 84107507 | 84124240 | 16734 | 1 | ENSMUST00 | 0 | Bckdhb | ENSMUSG00000032263.9 |
| chr19 | 58918707 | 58919329 | 7.60272 | 13.92084 | 11.67327 | Distal Interge | 19 | 58911673 | 58911759 | 87 | 2 | ENSMUST00 | -6949 | Mir3086 | ENSMUSG00000092813.1 |
| chr2 | 115855759 | 115856316 | 7.60272 | 13.92084 | 11.67327 | Distal Interge | 2 | 115864227 | 115868867 | 4641 | 2 | ENSMUST00 | 12551 | Meis2 | ENSMUSG00000027210.15 |
| chr1 | 120340580 | 120341712 | 7.58806 | 13.89661 | 11.64909 | Promoter | 1 | 120340582 | 120343174 | 2593 | 1 | ENSMUST00 | 0 | C1ql2 | ENSMUSG00000036907.9 |
| chr11 | 78694398 | 78694833 | 7.58806 | 13.89661 | 11.64909 | Intron (ENSM | 11 | 78617272 | 78696494 | 79223 | 2 | ENSMUST00 | 1661 | Nlk | ENSMUSG00000017376.10 |
| chr11 | 64022653 | 64023151 | 7.58806 | 13.89661 | 11.64909 | Intron (ENSM | 11 | 64012615 | 64015224 | 2610 | 1 | ENSMUST00 | 10039 | Gm12289 | ENSMUSG00000084884.1 |
| chr13 | 43183139 | 43183465 | 7.58806 | 13.89661 | 11.64909 | Distal Interge | 13 | 43151744 | 43171501 | 19758 | 2 | ENSMUST00 | -11639 | Tbc1d7 | ENSMUSG00000021368.9 |
| chr18 | 37694435 | 37694759 | 7.58806 | 13.89661 | 11.64909 | Intron (ENSM | 18 | 37705979 | 37706069 | 91 | 1 | ENSMUST00 | -11220 | Gm24401 | ENSMUSG00000088493.1 |
| chr19 | 16267504 | 16267690 | 7.58806 | 13.89661 | 11.64909 | Promoter | 19 | 16267061 | 16267621 | 561 | 2 | ENSMUST00 | 0 | Gm10819 | ENSMUSG00000075268.2 |
| chr5 | 121948653 | 121948860 | 7.06831 | 13.89447 | 11.64743 | Intron (ENSM | 5 | 121860308 | 121924921 | 64614 | 2 | ENSMUST00 | -23733 | Cux2 | ENSMUSG00000042589.13 |
| chr4 | 127008669 | 127008920 | 5.42843 | 13.89429 | 11.6473 | Distal Interge | 4 | 127021324 | 127031236 | 9913 | 1 | ENSMUST00 | -12404 | Sfpq | ENSMUSG00000028820.8 |
| chr1 | 188198402 | 188199175 | 7.30533 | 13.86164 | 11.61471 | Exon (ENSMU | 1 | 188262023 | 188357520 | 95498 | 1 | ENSMUST00 | -62848 | Ush2a | ENSMUSG00000026609.10 |
| chr2 | 32646718 | 32647133 | 7.30533 | 13.86164 | 11.61471 | Promoter | 2 | 32646958 | 32681815 | 34858 | 1 | ENSMUST00 | 0 | Eng | ENSMUSG00000026814.11 |
| chr7 | 141435487 | 141435848 | 7.30533 | 13.86164 | 11.61471 | Promoter | 7 | 141431599 | 141435397 | 3799 | 2 | ENSMUST00 | -91 | Slc25a22 | ENSMUSG00000019082.12 |
| chr7 | 16297079 | 16297339 | 7.55891 | 13.84842 | 11.60173 | Distal Interge | 7 | 16282755 | 16286795 | 4041 | 2 | ENSMUST00 | -10285 | Ccdc9 | ENSMUSG00000041375.13 |
| chr1 | 87620399 | 87620985 | 7.81205 | 13.8349 | 11.58824 | Promoter | 1 | 87620627 | 87667992 | 47366 | 1 | ENSMUST00 | 0 | Inpp5d | ENSMUSG00000026288.9 |
| chr1 | 135104968 | 135105547 | 7.81205 | 13.8349 | 11.58824 | Promoter | 1 | 134983301 | 135105276 | 121976 | 2 | ENSMUST00 | 0 | Lgr6 | ENSMUSG00000042793.8 |
| chr10 | 62507899 | 62508643 | 7.81205 | 13.8349 | 11.58824 | Promoter | 10 | 62494684 | 62507938 | 13255 | 2 | ENSMUST00 | 0 | Srgn | ENSMUSG00000020077.9 |
| chr10 | 81596760 | 81596971 | 7.81205 | 13.8349 | 11.58824 | Promoter | 10 | 81595229 | 81597331 | 2103 | 2 | ENSMUST00 | 360 | Tle6 | ENSMUSG00000034758.7 |
| chr10 | 102512348 | 102512579 | 7.81205 | 13.8349 | 11.58824 | Promoter | 10 | 102512222 | 102546560 | 34339 | 1 | ENSMUST00 | 127 | Rassf9 | ENSMUSG00000044921.4 |
| chr11 | 5555388 | 5555569 | 7.81205 | 13.8349 | 11.58824 | Distal Interge | 11 | 5560606 | 5560735 | 130 | 2 | ENSMUST00 | 5166 | Gm22471 | ENSMUSG00000077304.1 |
| chr13 | 38145079 | 38145731 | 7.81205 | 13.8349 | 11.58824 | Distal Interge | 13 | 38151328 | 38198577 | 47250 | 1 | ENSMUST00 | -5597 | Dsp | ENSMUSG00000054889.4 |
| chr14 | 105802943 | 105803295 | 7.81205 | 13.8349 | 11.58824 | Distal Interge | 14 | 105816908 | 105817000 | 93 | 1 | ENSMUST00 | -13613 | Gm22406 | ENSMUSG00000092664.1 |
| chr14 | 69503539 | 69503826 | 7.81205 | 13.8349 | 11.58824 | Promoter | 14 | 69462309 | 69503362 | 41054 | 2 | ENSMUST00 | -178 | Gm28049 | ENSMUSG00000093954.3 |
| chr14 | 6426759 | 6427323 | 7.81205 | 13.8349 | 11.58824 | Intron (ENSM | 14 | 6405827 | 6412482 | 6656 | 2 | ENSMUST00 | -14278 | Gm3591 | ENSMUSG00000096904.2 |
| chr15 | 100870911 | 100871313 | 7.81205 | 13.8349 | 11.58824 | Promoter | 15 | 100870683 | 101045929 | 175247 | 1 | ENSMUST00 | 229 | Scn8a | ENSMUSG00000023033.9 |
| chr15 | 41154888 | 41155675 | 7.81205 | 13.8349 | 11.58824 | Distal Interge | 15 | 41173701 | 41173871 | 171 | 1 | ENSMUST00 | -18026 | 4930555K19 | ENSMUSG00000048559.4 |
| chr17 | 85618046 | 85618607 | 7.81205 | 13.8349 | 11.58824 | Promoter | 17 | 85601933 | 85618372 | 16440 | 2 | ENSMUST00 | 0 | Six3os1 | ENSMUSG00000093460.4 |
| chr2 | 164938367 | 164939099 | 7.81205 | 13.8349 | 11.58824 | Distal Interge | 2 | 164940780 | 164948894 | 8115 | 1 | ENSMUST00 | -1681 | Mmp9 | ENSMUSG00000017737.2 |
| chr4 | 125490232 | 125490973 | 7.81205 | 13.8349 | 11.58824 | Promoter | 4 | 125490700 | 125714173 | 223474 | 1 | ENSMUST00 | 0 | Grik3 | ENSMUSG00000001985.9 |
| chr4 | 130629533 | 130629813 | 7.81205 | 13.8349 | 11.58824 | Distal Interge | 4 | 130663321 | 130781564 | 118244 | 1 | ENSMUST00 | -33508 | Pum1 | ENSMUSG00000028580.10 |
| chr4 | 42154082 | 42154494 | 7.81205 | 13.8349 | 11.58824 | Promoter | 4 | 42154040 | 42158835 | 4796 | 1 | ENSMUST00 | 43 | Gm13306 | ENSMUSG00000073877.5 |
| chr7 | 49450235 | 49451151 | 7.81205 | 13.8349 | 11.58824 | Promoter | 7 | 49450760 | 49464580 | 13821 | 1 | ENSMUST00 | 0 | Nav2 | ENSMUSG00000052512.10 |
| chr9 | 59056374 | 59056971 | 7.81205 | 13.8349 | 11.58824 | Distal Interge | 9 | 58874679 | 59036441 | 161763 | 2 | ENSMUST00 | -19934 | Neo1 | ENSMUSG00000032340.7 |
| chr9 | 118453535 | 118453888 | 7.81205 | 13.8349 | 11.58824 | Distal Interge | 9 | 118466984 | 118467064 | 81 | 2 | ENSMUST00 | 13176 | Gm22479 | ENSMUSG00000084515.1 |
| chrX | 99198925 | 99199175 | 7.81205 | 13.8349 | 11.58824 | Distal Interge | 20 | 99138131 | 99147230 | 9100 | 1 | ENSMUST00 | 60795 | Efnb1 | ENSMUSG00000031217.8 |
| chrX | 48343229 | 48343507 | 7.81205 | 13.8349 | 11.58824 | Promoter | 20 | 48343770 | 48408049 | 64280 | 1 | ENSMUST00 | -263 | Bcorl1 | ENSMUSG00000036959.10 |
| chrX | 138914156 | 138914501 | 7.81205 | 13.8349 | 11.58824 | Promoter | 20 | 138914430 | 139009092 | 94663 | 1 | ENSMUST00 | 0 | Nrk | ENSMUSG00000052854.10 |
| chrX | 95957179 | 95957688 | 7.81205 | 13.8349 | 11.58824 | Promoter | 20 | 95949435 | 95956962 | 7528 | 2 | ENSMUST00 | -218 | Las1l | ENSMUSG00000057421.7 |
| chrX | 58031126 | 58032053 | 7.81205 | 13.8349 | 11.58824 | Promoter | 20 | 58031010 | 58041736 | 10727 | 1 | ENSMUST00 | 117 | Zic3 | ENSMUSG00000067860.6 |
| chrX | 12005738 | 12006002 | 7.81205 | 13.8349 | 11.58824 | Distal Interge | 20 | 11984475 | 11984650 | 176 | 1 | ENSMUST00 | 21264 | Gm14512 | ENSMUSG00000081947.1 |
| chrX | 104348236 | 104348515 | 7.81205 | 13.8349 | 11.58824 | Promoter | 20 | 104345829 | 104348234 | 2406 | 2 | ENSMUST00 | -3 | Vcp-rs | ENSMUSG00000083327.1 |
| chr1 | 151947611 | 151948316 | 7.54398 | 13.82373 | 11.57928 | Intron (ENSM | 1 | 151892903 | 151921227 | 28325 | 2 | ENSMUST00 | -26385 | 1700025G04 | ENSMUSG00000032666.11 |
| chr1 | 72536610 | 72537410 | 7.54398 | 13.82373 | 11.57928 | Promoter | 1 | 72427112 | 72536930 | 109819 | 2 | ENSMUST00 | 0 | Mar-04 | ENSMUSG00000039372.5 |
| chr1 | 84284243 | 84284833 | 7.54398 | 13.82373 | 11.57928 | Promoter | 1 | 84036296 | 84284645 | 248350 | 2 | ENSMUST00 | 0 | Pid1 | ENSMUSG00000045658.11 |
| chr10 | 127620923 | 127621130 | 7.54398 | 13.82373 | 11.57928 | Promoter | 10 | 127589156 | 127620970 | 31815 | 2 | ENSMUST00 | 0 | Lrp1 | ENSMUSG00000040249.10 |
| chr10 | 49788290 | 49788952 | 7.54398 | 13.82373 | 11.57928 | Promoter | 10 | 49099460 | 49788766 | 689307 | 2 | ENSMUST00 | 0 | Grik2 | ENSMUSG00000056073.9 |
| chr11 | 110479057 | 110479309 | 7.54398 | 13.82373 | 11.57928 | Promoter | 11 | 110479039 | 110497926 | 18888 | 1 | ENSMUST00 | 19 | Map2k6 | ENSMUSG00000020623.6 |
| chr11 | 75213701 | 75214156 | 7.54398 | 13.82373 | 11.57928 | Intron (ENSM | 11 | 75193783 | 75267769 | 73987 | 1 | ENSMUST00 | 19919 | Rtn4rl1 | ENSMUSG00000045287.6 |
| chr11 | 115250034 | 115250675 | 7.54398 | 13.82373 | 11.57928 | Exon (ENSMU | 11 | 115253278 | 115253385 | 108 | 1 | ENSMUST00 | -2603 | Gm25837 | ENSMUSG00000089544.1 |
| chr13 | 51849460 | 51849811 | 7.54398 | 13.82373 | 11.57928 | Distal Interge | 13 | 51846744 | 51848224 | 1481 | 1 | ENSMUST00 | 2717 | Gadd45g | ENSMUSG00000021453.2 |
| chr13 | 60125951 | 60126573 | 7.54398 | 13.82373 | 11.57928 | Distal Interge | 13 | 60174405 | 60177365 | 2961 | 2 | ENSMUST00 | 50792 | Gas1 | ENSMUSG00000052957.7 |
| chr14 | 33447382 | 33447720 | 7.54398 | 13.82373 | 11.57928 | Promoter | 14 | 33390938 | 33447158 | 56221 | 2 | ENSMUST00 | -225 | Mapk8 | ENSMUSG00000021936.8 |
| chr15 | 37803434 | 37804046 | 7.54398 | 13.82373 | 11.57928 | Distal Interge | 15 | 37397171 | 37792570 | 395400 | 2 | ENSMUST00 | -10865 | Ncald | ENSMUSG00000051359.9 |
| chr15 | 85666531 | 85666885 | 7.54398 | 13.82373 | 11.57928 | Distal Interge | 15 | 85639747 | 85645934 | 6188 | 1 | ENSMUST00 | 26785 | RP23-64J7.2 | ENSMUSG00000101828.1 |
| chr16 | 84875024 | 84875534 | 7.54398 | 13.82373 | 11.57928 | Distal Interge | 16 | 84860045 | 84862465 | 2421 | 1 | ENSMUST00 | 14980 | Gabpa | ENSMUSG00000008976.11 |
| chr17 | 46864321 | 46864861 | 7.54398 | 13.82373 | 11.57928 | Distal Interge | 17 | 46889216 | 46890405 | 1190 | 2 | ENSMUST00 | 25544 | A330017A19 | ENSMUSG00000073388.2 |
| chr18 | 54125549 | 54126387 | 7.54398 | 13.82373 | 11.57928 | Distal Interge | 18 | 53984321 | 53985328 | 1008 | 1 | ENSMUST00 | 141229 | Gm5507 | ENSMUSG00000069376.2 |
| chr19 | 22915965 | 22916162 | 7.54398 | 13.82373 | 11.57928 | Intron (ENSM | 19 | 23037390 | 23075853 | 38464 | 2 | ENSMUST00 | 159691 | C330002G04 | ENSMUSG00000097930.1 |
| chr2 | 102901108 | 102901573 | 7.54398 | 13.82373 | 11.57928 | Promoter | 2 | 102812530 | 102901495 | 88966 | 2 | ENSMUST00 | 0 | Cd44 | ENSMUSG00000005087.12 |
| chr3 | 82875939 | 82876578 | 7.54398 | 13.82373 | 11.57928 | Promoter | 3 | 82837228 | 82876483 | 39256 | 2 | ENSMUST00 | 0 | Rbm46 | ENSMUSG00000033882.10 |
| chr4 | 115979421 | 115979950 | 7.54398 | 13.82373 | 11.57928 | Intron (ENSM | 4 | 115998714 | 115999937 | 1224 | 2 | ENSMUST00 | 19987 | Faah | ENSMUSG00000034171.8 |
| chr4 | 11385286 | 11385726 | 7.54398 | 13.82373 | 11.57928 | Promoter | 4 | 11357586 | 11386394 | 28809 | 2 | ENSMUST00 | 668 | Esrp1 | ENSMUSG00000040728.10 |
| chr4 | 133816318 | 133816612 | 7.54398 | 13.82373 | 11.57928 | Promoter | 4 | 133816420 | 133817614 | 1195 | 1 | ENSMUST00 | 0 | Gm12974 | ENSMUSG00000084789.1 |
| chr5 | 134221760 | 134222293 | 7.54398 | 13.82373 | 11.57928 | Promoter | 5 | 134221372 | 134222222 | 851 | 2 | ENSMUST00 | 0 | Ncf1 | ENSMUSG00000015950.8 |
| chr5 | 76075137 | 76075798 | 7.54398 | 13.82373 | 11.57928 | Distal Interge | 5 | 76140271 | 76151515 | 11245 | 1 | ENSMUST00 | -64473 | Srd5a3 | ENSMUSG00000029233.10 |
| chr6 | 122826058 | 122826979 | 7.54398 | 13.82373 | 11.57928 | Intron (ENSM | 6 | 122813626 | 122820614 | 6989 | 2 | ENSMUST00 | -5445 | Gm26826 | ENSMUSG00000097635.1 |
| chr7 | 16476514 | 16476864 | 7.54398 | 13.82373 | 11.57928 | Promoter | 7 | 16455721 | 16476780 | 21060 | 2 | ENSMUST00 | 0 | Npas1 | ENSMUSG00000001988.8 |
| chr7 | 19320169 | 19320556 | 7.54398 | 13.82373 | 11.57928 | Distal Interge | 7 | 19302721 | 19310045 | 7325 | 2 | ENSMUST00 | -10125 | Fosb | ENSMUSG00000003545.2 |
| chr7 | 43283040 | 43283268 | 7.54398 | 13.82373 | 11.57928 | Distal Interge | 7 | 43296523 | 43311486 | 14964 | 2 | ENSMUST00 | 28218 | Zfp715 | ENSMUSG00000012640.11 |
| chr7 | 71456339 | 71457212 | 7.54398 | 13.82373 | 11.57928 | Distal Interge | 7 | 71370333 | 71379813 | 9481 | 1 | ENSMUST00 | 86007 | RP24-173C11 | ENSMUSG00000099853.1 |
| chr9 | 37432700 | 37433165 | 7.54398 | 13.82373 | 11.57928 | Promoter | 9 | 37416045 | 37433163 | 17119 | 2 | ENSMUST00 | 0 | Robo3 | ENSMUSG00000032128.9 |
| chr19 | 37692795 | 37693447 | 7.78167 | 13.78524 | 11.54284 | 3' UTR | 19 | 37697808 | 37701528 | 3721 | 1 | ENSMUST00 | -4361 | Cyp26a1 | ENSMUSG00000024987.5 |
| chr15 | 93708105 | 93708608 | 7.76541 | 13.75865 | 11.51639 | Distal Interge | 15 | 93499114 | 93595891 | 96778 | 2 | ENSMUST00 | -112215 | Prickle1 | ENSMUSG00000036158.7 |
| chr17 | 35868500 | 35868796 | 7.76541 | 13.75865 | 11.51639 | Promoter | 17 | 35868403 | 35875594 | 7192 | 1 | ENSMUST00 | 98 | Ppp1r18 | ENSMUSG00000034595.12 |
| chr9 | 78821557 | 78822041 | 7.76541 | 13.75865 | 11.51639 | Promoter | 9 | 78820638 | 78821838 | 1201 | 2 | ENSMUST00 | 0 | Gm8116 | ENSMUSG00000059422.5 |
| chr4 | 147275700 | 147276025 | 5.88593 | 13.7549 | 11.51293 | Promoter | 4 | 147275976 | 147279096 | 3121 | 1 | ENSMUST00 | 0 | Gm13135 | ENSMUSG00000082925.1 |
| chr2 | 91965214 | 91965682 | 7.48701 | 13.72943 | 11.48759 | Promoter | 2 | 91932824 | 91965587 | 32764 | 2 | ENSMUST00 | 0 | Dgkz | ENSMUSG00000040479.6 |
| chr1 | 179940089 | 179940732 | 7.47279 | 13.70588 | 11.46408 | Distal Interge | 1 | 179960472 | 180163156 | 202685 | 1 | ENSMUST00 | -19740 | Cdc42bpa | ENSMUSG00000026490.12 |
| chr1 | 39192448 | 39192740 | 7.7331 | 13.70582 | 11.46404 | Promoter | 1 | 39193731 | 39287615 | 93885 | 1 | ENSMUST00 | -991 | Npas2 | ENSMUSG00000026077.10 |
| chr6 | 30921301 | 30921677 | 7.71704 | 13.67956 | 11.43796 | Intron (ENSM | 6 | 30896981 | 30915573 | 18593 | 2 | ENSMUST00 | -5729 | Tsga13 | ENSMUSG00000039032.1 |
| chr11 | 103262169 | 103262814 | 7.44452 | 13.65903 | 11.41755 | Intron (ENSM | 11 | 103219762 | 103267472 | 47711 | 2 | ENSMUST00 | 4658 | Map3k14 | ENSMUSG00000020941.7 |
| chr11 | 117231879 | 117232740 | 7.44452 | 13.65903 | 11.41755 | Promoter | 11 | 117232285 | 117362313 | 130029 | 1 | ENSMUST00 | 0 | Sep-09 | ENSMUSG00000059248.8 |
| chr17 | 8372395 | 8372737 | 7.44452 | 13.65903 | 11.41755 | Promoter | 17 | 8372396 | 8422728 | 50333 | 1 | ENSMUST00 | 0 | T2 | ENSMUSG00000058159.8 |
| chr17 | 12939901 | 12940788 | 7.44452 | 13.65903 | 11.41755 | Promoter | 17 | 12923833 | 12940402 | 16570 | 2 | ENSMUST00 | 0 | Acat3 | ENSMUSG00000062480.6 |
| chr5 | 38307686 | 38307978 | 7.44452 | 13.65903 | 11.41755 | Distal Interge | 5 | 38319509 | 38322310 | 2802 | 1 | ENSMUST00 | -11531 | Drd5 | ENSMUSG00000039358.2 |
| chr7 | 97846918 | 97847585 | 7.44452 | 13.65903 | 11.41755 | Intron (ENSM | 7 | 97842935 | 97912381 | 69447 | 1 | ENSMUST00 | 3984 | Pak1 | ENSMUSG00000030774.8 |
| chr8 | 67431386 | 67431657 | 7.70105 | 13.65341 | 11.41252 | Distal Interge | 8 | 67490758 | 67492104 | 1347 | 1 | ENSMUST00 | -59101 | Nat1 | ENSMUSG00000025588.3 |
| chr19 | 24166569 | 24166928 | 7.17908 | 13.64798 | 11.4072 | Intron (ENSM | 19 | 24094523 | 24225026 | 130504 | 2 | ENSMUST00 | 58098 | Tjp2 | ENSMUSG00000024812.9 |
| chr7 | 45124565 | 45124814 | 7.17908 | 13.64798 | 11.4072 | Promoter | 7 | 45122388 | 45124389 | 2002 | 2 | ENSMUST00 | -177 | Rps11 | ENSMUSG00000003429.10 |
| chr3 | 105058545 | 105058994 | 5.64773 | 13.63357 | 11.39298 | Distal Interge | 3 | 105001915 | 105053146 | 51232 | 2 | ENSMUST00 | -5400 | Cttnbp2nl | ENSMUSG00000062127.6 |
| chr11 | 3517048 | 3517826 | 7.68513 | 13.62737 | 11.38679 | 3' UTR | 11 | 3514867 | 3517351 | 2485 | 1 | ENSMUST00 | 2182 | Selm | ENSMUSG00000075702.9 |
| chr15 | 27787827 | 27788402 | 7.41646 | 13.61251 | 11.37205 | Intron (ENSM | 15 | 27655071 | 27681542 | 26472 | 2 | ENSMUST00 | -106286 | Fam105a | ENSMUSG00000056069.8 |
| chr5 | 150258960 | 150259472 | 7.66927 | 13.60143 | 11.36099 | Promoter | 5 | 150259930 | 150497752 | 237823 | 1 | ENSMUST00 | -458 | Fry | ENSMUSG00000056602.8 |
| chr15 | 85665111 | 85666215 | 7.65348 | 13.57559 | 11.33527 | Distal Interge | 15 | 85639747 | 85645934 | 6188 | 1 | ENSMUST00 | 25365 | RP23-64J7.2 | ENSMUSG00000101828.1 |
| chr1 | 9237106 | 9237416 | 7.13425 | 13.57188 | 11.33167 | Intron (ENSM | 1 | 8950572 | 9298236 | 347665 | 2 | ENSMUST00 | 60820 | Sntg1 | ENSMUSG00000025909.11 |
| chr1 | 24586917 | 24587494 | 7.13425 | 13.57188 | 11.33167 | Promoter | 1 | 24263994 | 24587472 | 323479 | 2 | ENSMUST00 | 0 | Col19a1 | ENSMUSG00000026141.8 |
| chr1 | 195018430 | 195018802 | 7.13425 | 13.57188 | 11.33167 | Exon (ENSMU | 1 | 195017399 | 195037908 | 20510 | 1 | ENSMUST00 | 1032 | A330023F24 | ENSMUSG00000096929.2 |
| chr11 | 102276534 | 102276824 | 7.13425 | 13.57188 | 11.33167 | Exon (ENSMU | 11 | 102268743 | 102279462 | 10720 | 1 | ENSMUST00 | 7792 | Asb16 | ENSMUSG00000034768.4 |
| chr11 | 103199395 | 103199921 | 7.13425 | 13.57188 | 11.33167 | 3' UTR | 11 | 103196801 | 103197876 | 1076 | 1 | ENSMUST00 | 2595 | Fmnl1 | ENSMUSG00000055805.9 |
| chr12 | 70192259 | 70192726 | 7.13425 | 13.57188 | 11.33167 | Intron (ENSM | 12 | 70197399 | 70197841 | 443 | 2 | ENSMUST00 | 5115 | Pygl | ENSMUSG00000021069.11 |
| chr12 | 89812094 | 89813015 | 7.13425 | 13.57188 | 11.33167 | Promoter | 12 | 89812483 | 90332899 | 520417 | 1 | ENSMUST00 | 0 | Nrxn3 | ENSMUSG00000066392.6 |
| chr13 | 3636698 | 3637203 | 7.13425 | 13.57188 | 11.33167 | Intron (ENSM | 13 | 3634099 | 3642825 | 8727 | 1 | ENSMUST00 | 2600 | Asb13 | ENSMUSG00000033781.6 |
| chr13 | 101692406 | 101692748 | 7.13425 | 13.57188 | 11.33167 | Promoter | 13 | 101683664 | 101692630 | 8967 | 2 | ENSMUST00 | 0 | Pik3r1 | ENSMUSG00000041417.10 |
| chr14 | 62373664 | 62374357 | 7.13425 | 13.57188 | 11.33167 | Distal Interge | 14 | 62347299 | 62355327 | 8029 | 1 | ENSMUST00 | 26366 | Rnaseh2b | ENSMUSG00000021932.8 |
| chr14 | 22813341 | 22813713 | 7.13425 | 13.57188 | 11.33167 | Intron (ENSM | 14 | 22577727 | 22599268 | 21542 | 1 | ENSMUST00 | 235615 | 1700112E06 | ENSMUSG00000063458.7 |
| chr15 | 103069176 | 103069595 | 7.13425 | 13.57188 | 11.33167 | Distal Interge | 15 | 103096151 | 103102111 | 5961 | 2 | ENSMUST00 | 32516 | RP24-89G21. | ENSMUSG00000101356.1 |
| chr16 | 10489591 | 10489944 | 7.13425 | 13.57188 | 11.33167 | Intron (ENSM | 16 | 10488178 | 10527564 | 39387 | 1 | ENSMUST00 | 1414 | Ciita | ENSMUSG00000022504.9 |
| chr16 | 50752455 | 50753169 | 7.13425 | 13.57188 | 11.33167 | Distal Interge | 16 | 50719294 | 50732773 | 13480 | 2 | ENSMUST00 | -19683 | 5330426P16 | ENSMUSG00000022639.9 |
| chr17 | 84676024 | 84676324 | 7.13425 | 13.57188 | 11.33167 | Promoter | 17 | 84676302 | 84699333 | 23032 | 1 | ENSMUST00 | 0 | Abcg8 | ENSMUSG00000024254.10 |
| chr18 | 52830946 | 52831614 | 7.13425 | 13.57188 | 11.33167 | Intron (ENSM | 18 | 52837964 | 52915586 | 77623 | 1 | ENSMUST00 | -6350 | Sncaip | ENSMUSG00000024534.10 |
| chr18 | 36242029 | 36242495 | 7.13425 | 13.57188 | 11.33167 | Distal Interge | 18 | 36281097 | 36301521 | 20425 | 1 | ENSMUST00 | -38602 | Pura | ENSMUSG00000043991.3 |
| chr2 | 26585004 | 26585643 | 7.13425 | 13.57188 | 11.33167 | Promoter | 2 | 26586630 | 26592681 | 6052 | 1 | ENSMUST00 | -987 | Egfl7 | ENSMUSG00000026921.13 |
| chr2 | 20339638 | 20340405 | 7.13425 | 13.57188 | 11.33167 | Promoter | 2 | 20339824 | 20661918 | 322095 | 1 | ENSMUST00 | 0 | Etl4 | ENSMUSG00000036617.11 |

| | | | | | | | | | | | | | | | |
|---|---|---|---|---|---|---|---|---|---|---|---|---|---|---|---|
| chr2 | 154552164 | 154552660 | 7.13425 | 13.57188 | 11.33167 | Promoter | 2 | 154551776 | 154553276 | 1501 | 1 | ENSMUST00 | 389 | Actl10 | ENSMUSG00000078129.1 |
| chr3 | 87803918 | 87804494 | 7.13425 | 13.57188 | 11.33167 | Exon (ENSMU | 3 | 87801943 | 87802808 | 866 | 1 | ENSMUST00 | 1976 | Insrr | ENSMUSG00000005640.6 |
| chr3 | 87165808 | 87166737 | 7.13425 | 13.57188 | 11.33167 | Intron (ENSM | 3 | 87084628 | 87174550 | 89923 | 2 | ENSMUST00 | 7813 | Kirrel | ENSMUSG00000041734.10 |
| chr5 | 142439877 | 142440543 | 7.13425 | 13.57188 | 11.33167 | Intron (ENSM | 5 | 142463931 | 142478715 | 14785 | 1 | ENSMUST00 | -23388 | Ap5z1 | ENSMUSG00000039623.9 |
| chr6 | 67037596 | 67038091 | 7.13425 | 13.57188 | 11.33167 | Promoter | 6 | 67036897 | 67037457 | 561 | 2 | ENSMUST00 | -140 | Gadd45a | ENSMUSG00000036390.6 |
| chr7 | 96152930 | 96153381 | 7.13425 | 13.57188 | 11.33167 | Distal Interge | 7 | 96171244 | 96911093 | 739850 | 1 | ENSMUST00 | -17863 | Tenm4 | ENSMUSG00000048078.11 |
| chr7 | 75642319 | 75642769 | 7.13425 | 13.57188 | 11.33167 | Promoter | 7 | 75643319 | 75704422 | 61104 | 1 | ENSMUST00 | -550 | Akap13 | ENSMUSG00000066406.9 |
| chr8 | 121947767 | 121948704 | 7.13425 | 13.57188 | 11.33167 | Intron (ENSM | 8 | 121949750 | 121997183 | 47434 | 1 | ENSMUST00 | -1046 | Banp | ENSMUSG00000025316.11 |
| chr8 | 34804949 | 34805963 | 7.13425 | 13.57188 | 11.33167 | Distal Interge | 8 | 34807297 | 34819894 | 12598 | 1 | ENSMUST00 | -1334 | Dusp4 | ENSMUSG00000031530.6 |
| chr8 | 120264577 | 120265125 | 7.13425 | 13.57188 | 11.33167 | Intron (ENSM | 8 | 120230536 | 120553634 | 323099 | 1 | ENSMUST00 | 34042 | Gse1 | ENSMUSG00000031822.13 |
| chr9 | 40802583 | 40803075 | 7.13425 | 13.57188 | 11.33167 | Promoter | 9 | 40802342 | 40802912 | 571 | 1 | ENSMUST00 | 242 | Hspa8 | ENSMUSG00000015656.11 |
| chr9 | 72871557 | 72872031 | 7.13425 | 13.57188 | 11.33167 | Intron (ENSM | 9 | 72910627 | 72912305 | 1679 | 1 | ENSMUST00 | -38596 | Prtg | ENSMUSG00000036030.9 |
| chr9 | 75549201 | 75549725 | 7.13425 | 13.57188 | 11.33167 | Intron (ENSM | 9 | 75497796 | 75559657 | 61862 | 2 | ENSMUST00 | 9932 | Tmod3 | ENSMUSG00000058587.7 |
| chr9 | 56606212 | 56606736 | 7.13425 | 13.57188 | 11.33167 | Promoter | 9 | 56606693 | 56614471 | 7779 | 1 | ENSMUST00 | 0 | Gm26868 | ENSMUSG00000097870.1 |
| chr12 | 86402677 | 86402877 | 7.63775 | 13.54986 | 11.31163 | Intron (ENSM | 12 | 86421644 | 86515522 | 93879 | 1 | ENSMUST00 | -18767 | Esrrb | ENSMUSG00000021255.12 |
| chr11 | 70220520 | 70221618 | 7.36097 | 13.52042 | 11.28246 | Promoter | 11 | 70216789 | 70220994 | 4206 | 2 | ENSMUST00 | 0 | Slc16a13 | ENSMUSG00000044367.7 |
| chr11 | 87967861 | 87968198 | 7.60649 | 13.49871 | 11.2608 | Distal Interge | 11 | 87979525 | 87984012 | 4488 | 2 | ENSMUST00 | 15814 | Dynll2 | ENSMUSG00000020483.9 |
| chr12 | 104845678 | 104846338 | 7.60649 | 13.49871 | 11.2608 | Distal Interge | 12 | 104763114 | 104865076 | 101963 | 2 | ENSMUST00 | 18738 | Clmn | ENSMUSG00000021097.9 |
| chr4 | 8394774 | 8395368 | 7.60649 | 13.49871 | 11.2608 | Distal Interge | 4 | 8456166 | 8456498 | 333 | 2 | ENSMUST00 | 61130 | Gm11810 | ENSMUSG00000082151.3 |
| chr4 | 135709277 | 135709472 | 7.60649 | 13.49871 | 11.2608 | Distal Interge | 4 | 135715868 | 135716598 | 731 | 2 | ENSMUST00 | 7126 | Gm12989 | ENSMUSG00000083678.3 |
| chr6 | 85904513 | 85905000 | 7.60649 | 13.49871 | 11.2608 | Promoter | 6 | 85899051 | 85904884 | 5834 | 2 | ENSMUST00 | 0 | 1700019G17 | ENSMUSG00000068299.6 |
| chr6 | 64734297 | 64734996 | 7.60649 | 13.49871 | 11.2608 | Distal Interge | 6 | 64729146 | 64731233 | 2088 | 1 | ENSMUST00 | 5152 | Atoh1 | ENSMUSG00000073043.4 |
| chr6 | 86822508 | 86823042 | 7.60649 | 13.49871 | 11.2608 | Distal Interge | 6 | 86848399 | 86849440 | 1042 | 2 | ENSMUST00 | 26398 | RP24-467O4. | ENSMUSG00000100164.1 |
| chr7 | 137322536 | 137322926 | 7.60649 | 13.49871 | 11.2608 | Distal Interge | 7 | 137193673 | 137314445 | 120773 | 2 | ENSMUST00 | -8092 | Ebf3 | ENSMUSG00000010476.8 |
| chr9 | 111690059 | 111690550 | 7.60649 | 13.49871 | 11.2608 | Promoter | 9 | 111561437 | 111690225 | 128789 | 2 | ENSMUST00 | 0 | Stac | ENSMUSG00000032502.8 |
| chr13 | 78427286 | 78428202 | 7.57548 | 13.44796 | 11.21062 | Distal Interge | 13 | 78372987 | 78379134 | 6148 | 2 | ENSMUST00 | -48153 | Gm6317 | ENSMUSG00000098003.1 |
| chr6 | 108306279 | 108306911 | 7.57548 | 13.44796 | 11.21062 | Intron (ENSM | 6 | 108213118 | 108551109 | 337992 | 1 | ENSMUST00 | 93162 | Itpr1 | ENSMUSG00000030102.8 |
| chr6 | 146410032 | 146410364 | 7.57548 | 13.44796 | 11.21062 | Intron (ENSM | 6 | 146357700 | 146396986 | 39287 | 2 | ENSMUST00 | -13047 | Itpr2 | ENSMUSG00000030287.10 |
| chr1 | 7139266 | 7140156 | 6.57187 | 13.44444 | 11.20722 | Intron (ENSM | 1 | 7136199 | 7136586 | 388 | 2 | ENSMUST00 | -2681 | Gm26983 | ENSMUSG00000098201.1 |
| chr4 | 83284461 | 83285232 | 7.05712 | 13.44069 | 11.20347 | Promoter | 4 | 83252986 | 83285169 | 32184 | 2 | ENSMUST00 | 0 | Ttc39b | ENSMUSG00000038172.9 |
| chr5 | 38366568 | 38367437 | 7.05712 | 13.44069 | 11.20347 | Intron (ENSM | 5 | 38317333 | 38320894 | 3562 | 2 | ENSMUST00 | -45675 | Gm26560 | ENSMUSG00000097029.1 |
| chr8 | 15152251 | 15152605 | 7.05712 | 13.44069 | 11.20347 | Distal Interge | 8 | 15119007 | 15131160 | 12154 | 1 | ENSMUST00 | 33245 | Myom2 | ENSMUSG00000031461.4 |
| chr8 | 46941482 | 46941923 | 7.05712 | 13.44069 | 11.20347 | Distal Interge | 8 | 46986925 | 47030714 | 43790 | 1 | ENSMUST00 | -45002 | Enpp6 | ENSMUSG00000038173.9 |
| chr13 | 81064755 | 81065337 | 5.92707 | 13.44026 | 11.20323 | Promoter | 13 | 81064684 | 81065803 | 1120 | 1 | ENSMUST00 | 72 | Gm8399 | ENSMUSG00000043889.5 |
| chr2 | 148048602 | 148049169 | 5.92707 | 13.44026 | 11.20323 | Distal Interge | 2 | 148042877 | 148046969 | 4093 | 2 | ENSMUST00 | -1634 | Foxa2 | ENSMUSG00000037025.6 |
| chr13 | 113013171 | 113013521 | 7.30631 | 13.42958 | 11.19263 | Distal Interge | 13 | 112993868 | 113000394 | 6527 | 1 | ENSMUST00 | 19304 | Mcidas | ENSMUSG00000074651.2 |
| chr14 | 67608233 | 67608690 | 7.30631 | 13.42958 | 11.19263 | Distal Interge | 14 | 67566850 | 67566981 | 132 | 1 | ENSMUST00 | 41384 | Gm24981 | ENSMUSG00000077647.1 |
| chr16 | 30236118 | 30236495 | 7.30631 | 13.42958 | 11.19263 | Distal Interge | 16 | 30200910 | 30205879 | 4970 | 2 | ENSMUST00 | -30240 | Gm26562 | ENSMUSG00000097621.1 |
| chr5 | 43572231 | 43573230 | 7.30631 | 13.42958 | 11.19263 | Intron (ENSM | 5 | 43549930 | 43557769 | 7840 | 2 | ENSMUST00 | -14463 | Gm15865 | ENSMUSG00000085529.1 |
| chr1 | 91801422 | 91801921 | 6.80177 | 13.42905 | 11.19262 | Promoter | 1 | 91801461 | 91848028 | 46568 | 1 | ENSMUST00 | 0 | Twist2 | ENSMUSG00000007805.4 |
| chr10 | 58395768 | 58396871 | 6.80177 | 13.42905 | 11.19262 | Intron (ENSM | 10 | 58394388 | 58421578 | 27191 | 1 | ENSMUST00 | 1381 | Lims1 | ENSMUSG00000019920.12 |
| chr10 | 79637702 | 79638091 | 6.80177 | 13.42905 | 11.19262 | Promoter | 10 | 79617940 | 79637918 | 19979 | 2 | ENSMUST00 | 0 | Shc2 | ENSMUSG00000020312.6 |
| chr10 | 42591586 | 42592065 | 6.80177 | 13.42905 | 11.19262 | Intron (ENSM | 10 | 42583838 | 42700492 | 116655 | 1 | ENSMUST00 | 7749 | Ostm1 | ENSMUSG00000038280.7 |
| chr11 | 95585229 | 95585858 | 6.80177 | 13.42905 | 11.19262 | Intron (ENSM | 11 | 95568818 | 95587735 | 18918 | 2 | ENSMUST00 | 1877 | Ngfr | ENSMUSG00000000120.6 |
| chr11 | 88873176 | 88873912 | 6.80177 | 13.42905 | 11.19262 | Intron (ENSM | 11 | 88830792 | 88864586 | 33795 | 2 | ENSMUST00 | -8591 | Akap1 | ENSMUSG00000018428.10 |
| chr13 | 46776641 | 46777470 | 6.80177 | 13.42905 | 11.19262 | Exon (ENSMU | 13 | 46767520 | 46767649 | 130 | 2 | ENSMUST00 | -8993 | Gm24620 | ENSMUSG00000064558.1 |
| chr14 | 41072796 | 41073168 | 6.80177 | 13.42905 | 11.19262 | Promoter | 14 | 41072911 | 41092190 | 19280 | 1 | ENSMUST00 | 0 | Dydc1 | ENSMUSG00000021790.7 |
| chr14 | 33351074 | 33351558 | 6.80177 | 13.42905 | 11.19262 | Promoter | 14 | 33351312 | 33362575 | 11264 | 1 | ENSMUST00 | 0 | Arhgap22 | ENSMUSG00000063506.8 |
| chr14 | 58150046 | 58150663 | 6.80177 | 13.42905 | 11.19262 | Distal Interge | 14 | 58105772 | 58106129 | 358 | 2 | ENSMUST00 | -43918 | Gm17109 | ENSMUSG00000091132.1 |
| chr14 | 75456864 | 75457350 | 6.80177 | 13.42905 | 11.19262 | Promoter | 14 | 75455982 | 75526141 | 70160 | 1 | ENSMUST00 | 883 | Siah3 | ENSMUSG00000091722.1 |
| chr16 | 93079347 | 93080307 | 6.80177 | 13.42905 | 11.19262 | Distal Interge | 16 | 93072755 | 93072822 | 68 | 1 | ENSMUST00 | 6593 | Gm28003 | ENSMUSG00000098983.1 |
| chr18 | 60849135 | 60849404 | 6.80177 | 13.42905 | 11.19262 | Promoter | 18 | 60813755 | 60848971 | 35217 | 2 | ENSMUST00 | -165 | Tcof1 | ENSMUSG00000024613.11 |
| chr2 | 160887520 | 160888265 | 6.80177 | 13.42905 | 11.19262 | Promoter | 2 | 160888108 | 160906002 | 17895 | 1 | ENSMUST00 | 0 | Lpin3 | ENSMUSG00000027412.7 |
| chr2 | 25195432 | 25196064 | 6.80177 | 13.42905 | 11.19262 | Promoter | 2 | 25195057 | 25196759 | 1703 | 2 | ENSMUST00 | 695 | Tor4a | ENSMUSG00000059555.6 |
| chr2 | 168473014 | 168473234 | 6.80177 | 13.42905 | 11.19262 | Distal Interge | 2 | 168417730 | 168418574 | 845 | 2 | ENSMUST00 | -54441 | Gm14234 | ENSMUSG00000082239.1 |
| chr2 | 21034782 | 21035077 | 6.80177 | 13.42905 | 11.19262 | Distal Interge | 2 | 21097099 | 21120659 | 23561 | 1 | ENSMUST00 | -62022 | Gm13376 | ENSMUSG00000087219.1 |
| chr3 | 100705348 | 100707194 | 6.80177 | 13.42905 | 11.19262 | Distal Interge | 3 | 100562208 | 100685503 | 123296 | 2 | ENSMUST00 | -19846 | Man1a2 | ENSMUSG00000008763.11 |
| chr4 | 155469019 | 155469439 | 6.80177 | 13.42905 | 11.19262 | Promoter | 4 | 155469114 | 155470415 | 1302 | 1 | ENSMUST00 | 0 | Tmem52 | ENSMUSG00000023153.4 |
| chr4 | 152185346 | 152186529 | 6.80177 | 13.42905 | 11.19262 | Promoter | 4 | 152185765 | 152217796 | 32032 | 1 | ENSMUST00 | 0 | Acot7 | ENSMUSG00000028937.9 |
| chr4 | 150132729 | 150133176 | 6.80177 | 13.42905 | 11.19262 | Intron (ENSM | 4 | 150128946 | 150140039 | 11094 | 1 | ENSMUST00 | 3784 | Slc2a5 | ENSMUSG00000028976.5 |
| chr4 | 150373069 | 150373840 | 6.80177 | 13.42905 | 11.19262 | Intron (ENSM | 4 | 150406394 | 150470381 | 63988 | 1 | ENSMUST00 | -32554 | Rere | ENSMUSG00000039852.11 |
| chr4 | 11455125 | 11455517 | 6.80177 | 13.42905 | 11.19262 | Distal Interge | 4 | 11485958 | 11527621 | 41664 | 1 | ENSMUST00 | -30441 | 1110037F02I | ENSMUSG00000040720.10 |
| chr6 | 108709332 | 108710096 | 6.80177 | 13.42905 | 11.19262 | Distal Interge | 6 | 108577039 | 108660934 | 83896 | 2 | ENSMUST00 | -48399 | 0610040F04I | ENSMUSG00000087341.2 |
| chr8 | 66697089 | 66697635 | 6.80177 | 13.42905 | 11.19262 | Promoter | 8 | 66697422 | 66706798 | 9377 | 1 | ENSMUST00 | 0 | Npy1r | ENSMUSG00000036437.5 |
| chr9 | 69125223 | 69126025 | 6.80177 | 13.42905 | 11.19262 | Intron (ENSM | 9 | 69195541 | 69250834 | 55294 | 1 | ENSMUST00 | -69516 | Rora | ENSMUSG00000032238.12 |
| chr10 | 42055203 | 42055626 | 7.56007 | 13.42273 | 11.18766 | Promoter | 10 | 42054113 | 42055306 | 1194 | 2 | ENSMUST00 | 0 | Gm9855 | ENSMUSG00000085666.3 |
| chr12 | 111445458 | 111446258 | 7.56007 | 13.42273 | 11.18766 | Promoter | 12 | 111446084 | 111453825 | 7742 | 1 | ENSMUST00 | 0 | Tnfaip2 | ENSMUSG00000021281.10 |
| chr8 | 80482887 | 80483281 | 7.56007 | 13.42273 | 11.18766 | Distal Interge | 8 | 80494045 | 80510785 | 16741 | 1 | ENSMUST00 | -10764 | Gypa | ENSMUSG00000051839.6 |
| chr14 | 120296619 | 120297809 | 7.54473 | 13.39761 | 11.16265 | Intron (ENSM | 14 | 120275669 | 120431697 | 156029 | 1 | ENSMUST00 | 20951 | Mbnl2 | ENSMUSG00000022139.11 |
| chr5 | 123397696 | 123398378 | 7.54473 | 13.39761 | 11.16265 | Promoter | 5 | 123393649 | 123397957 | 4309 | 2 | ENSMUST00 | 0 | Gm15747 | ENSMUSG00000086700.1 |
| chr15 | 98790812 | 98791098 | 6.09753 | 13.37905 | 11.14422 | Promoter | 15 | 98789857 | 98793837 | 3981 | 1 | ENSMUST00 | 956 | Wnt1 | ENSMUSG00000022997.9 |
| chr10 | 12547161 | 12547698 | 7.52944 | 13.37258 | 11.13794 | Intron (ENSM | 10 | 12368623 | 12368832 | 210 | 1 | ENSMUST00 | 178539 | Gm25139 | ENSMUSG00000065181.1 |
| chr1 | 167237059 | 167238114 | 6.52658 | 13.36193 | 11.1274 | Exon (ENSMU | 1 | 167223827 | 167223919 | 93 | 2 | ENSMUST00 | -13141 | Uck2 | ENSMUSG00000053664.3 |
| chr10 | 85221148 | 85221663 | 6.52658 | 13.36193 | 11.1274 | Distal Interge | 10 | 85131702 | 85185054 | 53353 | 2 | ENSMUST00 | -36095 | Cry1 | ENSMUSG00000020038.9 |
| chr14 | 89898243 | 89898845 | 6.52658 | 13.36193 | 11.1274 | Promoter | 14 | 89896223 | 89898466 | 2244 | 2 | ENSMUST00 | 0 | Gm10110 | ENSMUSG00000062093.4 |
| chr15 | 51987635 | 51988331 | 6.52658 | 13.36193 | 11.1274 | Intron (ENSM | 15 | 51962605 | 51991760 | 29156 | 2 | ENSMUST00 | 3429 | Rad21 | ENSMUSG00000022314.9 |
| chr15 | 55280161 | 55280987 | 6.52658 | 13.36193 | 11.1274 | Distal Interge | 15 | 55262542 | 55262603 | 62 | 2 | ENSMUST00 | -17559 | Gm26296 | ENSMUSG00000095792.1 |
| chr17 | 83277398 | 83277913 | 6.52658 | 13.36193 | 11.1274 | Distal Interge | 17 | 83269971 | 83270105 | 135 | 1 | ENSMUST00 | 7428 | Gm24240 | ENSMUSG00000088410.1 |
| chr19 | 59469004 | 59469397 | 6.52658 | 13.36193 | 11.1274 | Distal Interge | 19 | 59462802 | 59464215 | 1414 | 1 | ENSMUST00 | 6203 | Emx2 | ENSMUSG00000043969.4 |
| chr19 | 46623471 | 46624159 | 6.52658 | 13.36193 | 11.1274 | Promoter | 19 | 46623401 | 46657385 | 33985 | 1 | ENSMUST00 | 71 | Wbp1l | ENSMUSG00000047731.12 |
| chr2 | 120076248 | 120076500 | 6.52658 | 13.36193 | 11.1274 | Exon (ENSMU | 2 | 120061586 | 120072523 | 10938 | 2 | ENSMUST00 | -3726 | Sptbn5 | ENSMUSG00000074899.4 |
| chr3 | 138197142 | 138197735 | 6.52658 | 13.36193 | 11.1274 | Distal Interge | 3 | 138164135 | 138186713 | 22579 | 2 | ENSMUST00 | -10430 | 4930579F01I | ENSMUSG00000012042.4 |
| chr3 | 68583413 | 68584351 | 6.52658 | 13.36193 | 11.1274 | Promoter | 3 | 68584179 | 68626464 | 42286 | 1 | ENSMUST00 | 0 | Schip1 | ENSMUSG00000027777.10 |
| chr5 | 130255656 | 130255914 | 6.52658 | 13.36193 | 11.1274 | Promoter | 5 | 130255619 | 130285386 | 29768 | 1 | ENSMUST00 | 38 | Tyw1 | ENSMUSG00000056310.9 |
| chr5 | 35395647 | 35395961 | 6.52658 | 13.36193 | 11.1274 | Intron (ENSM | 5 | 35389117 | 35392861 | 3745 | 1 | ENSMUST00 | 6531 | Hmx1 | ENSMUSG00000067438.4 |
| chr6 | 129476482 | 129476917 | 6.52658 | 13.36193 | 11.1274 | Distal Interge | 6 | 129463026 | 129472777 | 9752 | 2 | ENSMUST00 | -3706 | Clec7a | ENSMUSG00000079293.6 |
| chr9 | 40821440 | 40822106 | 6.52658 | 13.36193 | 11.1274 | Distal Interge | 9 | 40804748 | 40804837 | 90 | 1 | ENSMUST00 | 16693 | Snord14e | ENSMUSG00000064791.1 |
| chr9 | 63091662 | 63092777 | 6.52658 | 13.36193 | 11.1274 | Distal Interge | 9 | 63133283 | 63133389 | 107 | 1 | ENSMUST00 | -40506 | Gm25064 | ENSMUSG00000094201.1 |
| chr10 | 81363831 | 81364907 | 6.29504 | 13.3498 | 11.11603 | Promoter | 10 | 81363067 | 81364854 | 1788 | 2 | ENSMUST00 | 0 | 4930404N11 | ENSMUSG00000020234.5 |
| chr10 | 127123240 | 127123579 | 6.29504 | 13.3498 | 11.11603 | Distal Interge | 10 | 127094259 | 127121160 | 26902 | 2 | ENSMUST00 | -2081 | Os9 | ENSMUSG00000040462.7 |
| chr11 | 83849906 | 83850369 | 6.29504 | 13.3498 | 11.11603 | Promoter | 11 | 83850063 | 83856098 | 6036 | 1 | ENSMUST00 | 0 | Hnf1b | ENSMUSG00000020679.6 |
| chr2 | 161511181 | 161511758 | 6.29504 | 13.3498 | 11.11603 | Downstream | 2 | 161513690 | 161514012 | 323 | 2 | ENSMUST00 | 2254 | RP23-466K24 | ENSMUSG00000100943.1 |
| chr3 | 116968185 | 116969027 | 6.29504 | 13.3498 | 11.11603 | Promoter | 3 | 116918256 | 116968987 | 50732 | 2 | ENSMUST00 | 0 | Palmd | ENSMUSG00000033377.9 |
| chr5 | 121329725 | 121330141 | 6.29504 | 13.3498 | 11.11603 | Intron (ENSM | 5 | 121320023 | 121322779 | 2757 | 1 | ENSMUST00 | 9703 | Gm15800 | ENSMUSG00000042744.11 |
| chr9 | 57695966 | 57696757 | 6.29504 | 13.3498 | 11.11603 | Promoter | 9 | 57697613 | 57703822 | 6210 | 1 | ENSMUST00 | -856 | Cyp1a1 | ENSMUSG00000032315.5 |
| chr7 | 80387019 | 80388177 | 7.51422 | 13.34765 | 11.11425 | Promoter | 7 | 80377758 | 80387946 | 10189 | 2 | ENSMUST00 | 0 | Fes | ENSMUSG00000053158.9 |
| chr11 | 115436007 | 115436345 | 7.49906 | 13.32282 | 11.08953 | Distal Interge | 11 | 115440545 | 115448270 | 7726 | 1 | ENSMUST00 | -4200 | 4933422H20I | ENSMUSG00000070332.4 |
| chr4 | 54966022 | 54966999 | 6.74139 | 13.32281 | 11.08953 | Intron (ENSM | 4 | 54947976 | 55083563 | 135588 | 1 | ENSMUST00 | 18047 | Zfp462 | ENSMUSG00000060206.6 |
| chr12 | 100168383 | 100168840 | 7.48396 | 13.29808 | 11.06495 | Distal Interge | 12 | 100125452 | 100159653 | 34202 | 2 | ENSMUST00 | -8731 | Nrde2 | ENSMUSG00000021179.7 |
| chr3 | 75814716 | 75815020 | 7.48396 | 13.29808 | 11.06495 | Distal Interge | 3 | 75755605 | 75775808 | 20204 | 1 | ENSMUST00 | 59112 | RP23-426C11 | ENSMUSG00000100962.1 |
| chr11 | 120901830 | 120902741 | 7.46892 | 13.27344 | 11.04042 | Exon (ENSMU | 11 | 120873792 | 120885911 | 12120 | 2 | ENSMUST00 | -15920 | Ccdc57 | ENSMUSG00000048445.6 |
| chr11 | 69804833 | 69805700 | 7.46892 | 13.27344 | 11.04042 | Promoter | 11 | 69803603 | 69805624 | 2022 | 2 | ENSMUST00 | 0 | Tmem102 | ENSMUSG00000089876.3 |
| chr12 | 111514819 | 111515302 | 7.45394 | 13.24889 | 11.01597 | Distal Interge | 12 | 111538101 | 111546753 | 8653 | 1 | ENSMUST00 | -22799 | Eif5 | ENSMUSG00000021282.11 |
| chr4 | 130275062 | 130275554 | 7.45394 | 13.24889 | 11.01597 | Promoter | 4 | 130253495 | 130275218 | 21724 | 2 | ENSMUST00 | 0 | Serinc2 | ENSMUSG00000023232.12 |
| chr4 | 58034687 | 58035100 | 7.45394 | 13.24889 | 11.01597 | Distal Interge | 4 | 57984027 | 58009124 | 25098 | 2 | ENSMUST00 | -25564 | Txndc8 | ENSMUSG00000038709.9 |
| chr5 | 114897077 | 114897538 | 7.45394 | 13.24889 | 11.01597 | Promoter | 5 | 114897576 | 114900283 | 2708 | 1 | ENSMUST00 | -38 | Oasl2 | ENSMUSG00000029561.12 |
| chr5 | 72587058 | 72587721 | 7.45394 | 13.24889 | 11.01597 | Promoter | 5 | 72581639 | 72587550 | 5912 | 2 | ENSMUST00 | 0 | Gm5868 | ENSMUSG00000060204.4 |
| chr2 | 179852637 | 179852997 | 6.93923 | 13.23942 | 11.00697 | Intron (ENSM | 2 | 179893884 | 179896149 | 2266 | 1 | ENSMUST00 | -40887 | Cdh4 | ENSMUSG00000000305.7 |
| chr2 | 148406062 | 148406717 | 6.93923 | 13.23942 | 11.00697 | 3' UTR | 2 | 148404466 | 148408188 | 3723 | 2 | ENSMUST00 | 1471 | Thbd | ENSMUSG00000074743.4 |
| chr3 | 146328099 | 146328638 | 6.93923 | 13.23942 | 11.00697 | Distal Interge | 3 | 146387742 | 146387844 | 103 | 2 | ENSMUST00 | 59206 | Gm22078 | ENSMUSG00000088218.1 |
| chr9 | 107698856 | 107699487 | 6.93923 | 13.23942 | 11.00697 | Intron (ENSM | 9 | 107681502 | 107710475 | 28974 | 2 | ENSMUST00 | 10988 | Sema3f | ENSMUSG00000034684.7 |
| chr19 | 16873271 | 16874902 | 7.43903 | 13.22444 | 10.99213 | Promoter | 19 | 16872316 | 16873830 | 1515 | 2 | ENSMUST00 | 0 | Foxb2 | ENSMUSG00000056829.3 |
| chr2 | 28466056 | 28466527 | 7.43903 | 13.22444 | 10.99213 | Promoter | 2 | 28462001 | 28466332 | 4332 | 2 | ENSMUST00 | 0 | 1700007K13I | ENSMUSG00000026831.11 |
| chr5 | 132074832 | 132075369 | 7.43903 | 13.22444 | 10.99213 | Intron (ENSM | 5 | 132030909 | 132050409 | 19501 | 2 | ENSMUST00 | -24424 | Auts2 | ENSMUSG00000029673.12 |
| chr6 | 136543036 | 136543638 | 7.43903 | 13.22444 | 10.99213 | Intron (ENSM | 6 | 136551187 | 136559910 | 8724 | 1 | ENSMUST00 | -7549 | Atf7ip | ENSMUSG00000030213.7 |
| chr1 | 12875694 | 12875992 | 7.17313 | 13.20777 | 10.97558 | Intron (ENSM | 1 | 12842791 | 12861192 | 18402 | 1 | ENSMUST00 | 32904 | Sulf1 | ENSMUSG00000016918.10 |
| chr11 | 113611824 | 113612457 | 7.17313 | 13.20777 | 10.97558 | Intron (ENSM | 11 | 113619342 | 113626028 | 6687 | 1 | ENSMUST00 | -6885 | Sstr2 | ENSMUSG00000047904.6 |
| chr16 | 10394914 | 10395394 | 7.17313 | 13.20777 | 10.97558 | Promoter | 16 | 10361254 | 10395448 | 34195 | 2 | ENSMUST00 | 54 | Tekt5 | ENSMUSG00000039179.8 |
| chr17 | 75033522 | 75033986 | 7.17313 | 13.20777 | 10.97558 | Intron (ENSM | 17 | 75006294 | 75008799 | 2506 | 1 | ENSMUST00 | 27229 | Ltbp1 | ENSMUSG00000001870.10 |
| chr2 | 135717190 | 135717848 | 7.17313 | 13.20777 | 10.97558 | Intron (ENSM | 2 | 135741830 | 135910484 | 168655 | 1 | ENSMUST00 | -23982 | Plcb4 | ENSMUSG00000039943.11 |
| chr2 | 156775091 | 156775680 | 7.17313 | 13.20777 | 10.97558 | Promoter | 2 | 156775420 | 156781658 | 6239 | 1 | ENSMUST00 | 0 | Myl9 | ENSMUSG00000067818.6 |
| chr6 | 134640636 | 134640878 | 7.17313 | 13.20777 | 10.97558 | Promoter | 6 | 134640940 | 134711184 | 70245 | 1 | ENSMUST00 | -62 | Loh12cr1 | ENSMUSG00000042992.10 |
| chr15 | 99370675 | 99370916 | 7.42417 | 13.20008 | 10.96827 | Promoter | 15 | 99317225 | 99370482 | 53258 | 2 | ENSMUST00 | -194 | Fmnl3 | ENSMUSG00000023008.13 |
| chr15 | 3272432 | 3273029 | 7.42417 | 13.20008 | 10.96827 | Promoter | 15 | 3272085 | 3277271 | 5187 | 1 | ENSMUST00 | 348 | Sepp1 | ENSMUSG00000064373.6 |
| chr3 | 60124192 | 60124907 | 7.42417 | 13.20008 | 10.96827 | Downstream | 3 | 60127641 | 60127747 | 107 | 2 | ENSMUST00 | 2840 | Gm24382 | ENSMUSG00000084647.1 |
| chr3 | 102012481 | 102013129 | 7.40937 | 13.17581 | 10.94411 | 5' UTR | 3 | 102010145 | 102015492 | 5348 | 1 | ENSMUST00 | 2337 | Nhlh2 | ENSMUSG00000048540.7 |
| chr7 | 3331904 | 3332173 | 7.40937 | 13.17581 | 10.94411 | Promoter | 7 | 3332955 | 3368216 | 35262 | 1 | ENSMUST00 | -782 | Cacng7 | ENSMUSG00000069806.4 |
| chr1 | 134062917 | 134063765 | 7.39463 | 13.15164 | 10.92004 | Distal Interge | 1 | 134075170 | 134079120 | 3951 | 2 | ENSMUST00 | 15355 | Btg2 | ENSMUSG00000020423.6 |

| | | | | | | | | | | | | | | | |
|---|---|---|---|---|---|---|---|---|---|---|---|---|---|---|---|
| chr2 | 48815968 | 48816506 | 7.37994 | 13.12756 | 10.89607 | Intron (ENSM | 2 | 48814109 | 48903269 | 89161 | 1 | ENSMUST00 | 1860 | Acvr2a | ENSMUSG00000052155.5 |
| chr6 | 86547842 | 86548274 | 7.37994 | 13.12756 | 10.89607 | Intron (ENSM | 6 | 86527401 | 86564442 | 37042 | 1 | ENSMUST00 | 20442 | 1600020E01 | ENSMUSG00000097048.2 |
| chr10 | 30655302 | 30656193 | 7.36532 | 13.10357 | 10.87218 | Promoter | 10 | 30645584 | 30655867 | 10284 | 2 | ENSMUST00 | 0 | Ncoa7 | ENSMUSG00000039697.10 |
| chr19 | 22692640 | 22693337 | 7.35075 | 13.07967 | 10.84837 | Promoter | 19 | 22692619 | 22926489 | 233871 | 1 | ENSMUST00 | 22 | Trpm3 | ENSMUSG00000052387.10 |
| chr4 | 100527557 | 100527874 | 7.35075 | 13.07967 | 10.84837 | Promoter | 4 | 100523247 | 100527801 | 4555 | 2 | ENSMUST00 | 0 | Ube2uos | ENSMUSG00000085871.1 |
| chr5 | 125004082 | 125004352 | 7.35075 | 13.07967 | 10.84837 | Promoter | 5 | 125003447 | 125012547 | 9101 | 1 | ENSMUST00 | 636 | Fam101a | ENSMUSG00000037962.3 |
| chr7 | 80293964 | 80294175 | 6.82522 | 13.04386 | 10.81275 | Promoter | 7 | 80294450 | 80316259 | 21810 | 1 | ENSMUST00 | -275 | Prc1 | ENSMUSG00000038943.11 |
| chr12 | 91400435 | 91401125 | 7.32179 | 13.03213 | 10.80111 | Promoter | 12 | 91400994 | 91540509 | 139516 | 1 | ENSMUST00 | 0 | Tshr | ENSMUSG00000020963.9 |
| chr11 | 3193042 | 3193594 | 2.12844 | 13.01344 | 10.7825 | Promoter | 11 | 3131850 | 3193407 | 61558 | 2 | ENSMUST00 | 0 | Sfi1 | ENSMUSG00000023764.13 |
| chr1 | 74035844 | 74037381 | 7.3074 | 13.0085 | 10.77756 | Promoter | 1 | 73993163 | 74037133 | 43971 | 2 | ENSMUST00 | 0 | Tns1 | ENSMUSG00000055322.9 |
| chr2 | 3335686 | 3336407 | 7.3074 | 13.0085 | 10.77756 | Promoter | 2 | 3336168 | 3340993 | 4826 | 1 | ENSMUST00 | 0 | Acbd7 | ENSMUSG00000026644.6 |
| chr2 | 60901565 | 60901927 | 7.3074 | 13.0085 | 10.77756 | Intron (ENSM | 2 | 60842297 | 60889030 | 46734 | 2 | ENSMUST00 | -12536 | Rbms1 | ENSMUSG00000026970.11 |
| chr5 | 110770964 | 110771297 | 7.3074 | 13.0085 | 10.77756 | Promoter | 5 | 110770949 | 110779260 | 8312 | 1 | ENSMUST00 | 16 | Gm15559 | ENSMUSG00000086401.1 |
| chr9 | 63958739 | 63959192 | 7.3074 | 13.0085 | 10.77756 | Intron (ENSM | 9 | 63954009 | 64016987 | 62979 | 2 | ENSMUST00 | 57795 | Smad6 | ENSMUSG00000036867.7 |
| chr11 | 20400663 | 20401049 | 7.04473 | 12.99314 | 10.76275 | Distal Interge | 11 | 20446412 | 20446810 | 399 | 1 | ENSMUST00 | -45363 | Gm12034 | ENSMUSG00000083878.3 |
| chr12 | 100907908 | 100908261 | 7.04473 | 12.99314 | 10.76275 | Promoter | 12 | 100876682 | 100908198 | 31517 | 2 | ENSMUST00 | 0 | Gpr68 | ENSMUSG00000047415.6 |
| chr18 | 11239657 | 11240387 | 7.04473 | 12.99314 | 10.76275 | Distal Interge | 18 | 11049931 | 11052567 | 2637 | 2 | ENSMUST00 | -187091 | 1010001N08 | ENSMUSG00000097222.1 |
| chr10 | 39165719 | 39166083 | 7.26455 | 12.93812 | 10.70821 | Distal Interge | 10 | 39150971 | 39163794 | 12824 | 2 | ENSMUST00 | -1926 | Wisp3 | ENSMUSG00000062074.6 |
| chr13 | 94382244 | 94382851 | 7.26455 | 12.93812 | 10.70821 | Intron (ENSM | 13 | 94358960 | 94566316 | 207357 | 1 | ENSMUST00 | 23285 | Ap3b1 | ENSMUSG00000021686.4 |
| chr6 | 8363849 | 8364593 | 7.26455 | 12.93812 | 10.70821 | Promoter | 6 | 8354056 | 8364136 | 10081 | 2 | ENSMUST00 | 0 | Gm16055 | ENSMUSG00000089890.1 |
| chr2 | 68863800 | 68864109 | 7.25038 | 12.91483 | 10.685 | Intron (ENSM | 2 | 68861586 | 69108644 | 247059 | 1 | ENSMUST00 | 2215 | Cers6 | ENSMUSG00000027035.5 |
| chr4 | 66827641 | 66828067 | 7.23626 | 12.89163 | 10.66188 | Promoter | 4 | 66827619 | 66839429 | 11811 | 1 | ENSMUST00 | 23 | Tlr4 | ENSMUSG00000039005.8 |
| chr15 | 38709563 | 38710313 | 6.71489 | 12.85374 | 10.62409 | Distal Interge | 15 | 38661904 | 38692443 | 30540 | 1 | ENSMUST00 | 47660 | Atp6v1c1 | ENSMUSG00000022295.7 |
| chr13 | 23534124 | 23534308 | 7.20819 | 12.84548 | 10.6159 | Promoter | 13 | 23533906 | 23534304 | 399 | 1 | ENSMUST00 | 219 | Hist1h2af | ENSMUSG00000061991.2 |
| chr7 | 19342618 | 19342847 | 7.20819 | 12.84548 | 10.6159 | Distal Interge | 7 | 19344778 | 19356524 | 11747 | 1 | ENSMUST00 | -1931 | Ercc1 | ENSMUSG00000003549.4 |
| chr3 | 141466975 | 141467822 | 7.18034 | 12.79967 | 10.57019 | Intron (ENSM | 3 | 141465671 | 141830538 | 364868 | 1 | ENSMUST00 | 1305 | Unc5c | ENSMUSG00000059921.10 |
| chr6 | 55676842 | 55677274 | 7.18034 | 12.79967 | 10.57019 | Downstream | 6 | 55677818 | 55681263 | 3446 | 2 | ENSMUST00 | 3989 | Neurod6 | ENSMUSG00000037984.8 |
| chr1 | 59584845 | 59585072 | 6.92084 | 12.78527 | 10.55585 | Intron (ENSM | 1 | 59606782 | 59628261 | 21480 | 1 | ENSMUST00 | -21710 | Gm973 | ENSMUSG00000047361.10 |
| chr1 | 78182033 | 78182247 | 7.1665 | 12.77688 | 10.5477 | Intron (ENSM | 1 | 78101267 | 78196838 | 95572 | 2 | ENSMUST00 | 14591 | Pax3 | ENSMUSG00000004872.10 |
| chr17 | 24639673 | 24640299 | 7.1665 | 12.77688 | 10.5477 | 3' UTR | 17 | 24639282 | 24644957 | 5676 | 2 | ENSMUST00 | 4658 | Slc9a3r2 | ENSMUSG00000002504.9 |
| chr17 | 46042237 | 46042591 | 7.1665 | 12.77688 | 10.5477 | Distal Interge | 17 | 46016996 | 46032377 | 15382 | 2 | ENSMUST00 | -9861 | Vegfa | ENSMUSG00000023951.11 |
| chr17 | 24455564 | 24455892 | 7.1665 | 12.77688 | 10.5477 | Promoter | 17 | 24443792 | 24455311 | 11520 | 2 | ENSMUST00 | -254 | E4f1 | ENSMUSG00000024137.8 |
| chr3 | 55517155 | 55517941 | 7.1665 | 12.77688 | 10.5477 | Intron (ENSM | 3 | 55548446 | 55548552 | 107 | 1 | ENSMUST00 | -30505 | Gm25404 | ENSMUSG00000064904.1 |
| chr4 | 115280996 | 115281390 | 7.1665 | 12.77688 | 10.5477 | Distal Interge | 4 | 115299046 | 115332815 | 33770 | 1 | ENSMUST00 | -17656 | Cyp4a12a | ENSMUSG00000066071.6 |
| chr5 | 120439944 | 120440839 | 7.1665 | 12.77688 | 10.5477 | 3' UTR | 5 | 120431770 | 120441223 | 9454 | 1 | ENSMUST00 | 8175 | Lhx5 | ENSMUSG00000029595.8 |
| chr6 | 34903610 | 34904146 | 7.1665 | 12.77688 | 10.5477 | Intron (ENSM | 6 | 34908295 | 34910563 | 2269 | 2 | ENSMUST00 | 6417 | Wdr91 | ENSMUSG00000058486.6 |
| chr7 | 74063385 | 74064248 | 7.1665 | 12.77688 | 10.5477 | Distal Interge | 7 | 73939119 | 74013682 | 74564 | 2 | ENSMUST00 | -49704 | St8sia2 | ENSMUSG00000025789.4 |
| chr3 | 105005629 | 105006090 | 7.12528 | 12.70902 | 10.48042 | Exon (ENSMU | 3 | 104944805 | 104961709 | 16905 | 2 | ENSMUST00 | -43921 | Wnt2b | ENSMUSG00000027840.4 |
| chr1 | 184826893 | 184827200 | 7.12487 | 12.70834 | 10.47977 | Promoter | 1 | 184813760 | 184827257 | 13498 | 2 | ENSMUST00 | 57 | Mar-02 | ENSMUSG00000073481.4 |
| chr10 | 75944067 | 75944394 | 7.12487 | 12.70834 | 10.47977 | Intron (ENSM | 10 | 75943057 | 75949637 | 6581 | 1 | ENSMUST00 | 1011 | Vpreb3 | ENSMUSG00000000903.3 |
| chr10 | 75835398 | 75835702 | 7.12487 | 12.70834 | 10.47977 | Promoter | 10 | 75831845 | 75834881 | 3037 | 2 | ENSMUST00 | -518 | Gstt2 | ENSMUSG00000033318.6 |
| chr11 | 98941436 | 98942169 | 7.12487 | 12.70834 | 10.47977 | Intron (ENSM | 11 | 98939712 | 98950490 | 10779 | 1 | ENSMUST00 | 1725 | Rara | ENSMUSG00000037992.11 |
| chr11 | 116394644 | 116394835 | 7.12487 | 12.70834 | 10.47977 | Intron (ENSM | 11 | 116359889 | 116387405 | 27517 | 2 | ENSMUST00 | -7240 | Rnf157 | ENSMUSG00000052949.9 |
| chr14 | 122223648 | 122223864 | 7.12487 | 12.70834 | 10.47977 | Intron (ENSM | 14 | 122229905 | 122233654 | 3750 | 2 | ENSMUST00 | 9790 | RP23-18J12. | ENSMUSG00000100585.1 |
| chr15 | 97606661 | 97607313 | 7.12487 | 12.70834 | 10.47977 | Distal Interge | 15 | 97675105 | 97705822 | 30718 | 2 | ENSMUST00 | 98509 | Rpap3 | ENSMUSG00000022466.5 |
| chr15 | 74956532 | 74956967 | 7.12487 | 12.70834 | 10.47977 | Promoter | 15 | 74956555 | 74958826 | 2272 | 1 | ENSMUST00 | 0 | Ly6e | ENSMUSG00000022587.9 |
| chr17 | 29356723 | 29357076 | 7.12487 | 12.70834 | 10.47977 | Distal Interge | 17 | 29360914 | 29379660 | 18747 | 1 | ENSMUST00 | -3838 | Fgd2 | ENSMUSG00000024013.9 |
| chr17 | 5491201 | 5491399 | 7.12487 | 12.70834 | 10.47977 | Distal Interge | 17 | 5492600 | 5753891 | 261292 | 1 | ENSMUST00 | -1201 | Zdhhc14 | ENSMUSG00000034265.7 |
| chr18 | 86395323 | 86395752 | 7.12487 | 12.70834 | 10.47977 | Promoter | 18 | 86394952 | 86501897 | 106946 | 1 | ENSMUST00 | 372 | Neto1 | ENSMUSG00000050321.2 |
| chr19 | 32292705 | 32293187 | 7.12487 | 12.70834 | 10.47977 | Intron (ENSM | 19 | 32220587 | 32289579 | 68993 | 2 | ENSMUST00 | -3127 | Sgms1 | ENSMUSG00000040451.12 |
| chr2 | 105673607 | 105674337 | 7.12487 | 12.70834 | 10.47977 | Promoter | 2 | 105673963 | 105675876 | 1914 | 1 | ENSMUST00 | 0 | Pax6 | ENSMUSG00000027168.16 |
| chr3 | 129535964 | 129536284 | 7.12487 | 12.70834 | 10.47977 | Intron (ENSM | 3 | 129532386 | 129638495 | 106110 | 1 | ENSMUST00 | 3579 | Elovl6 | ENSMUSG00000041220.6 |
| chr4 | 83539239 | 83539763 | 7.12487 | 12.70834 | 10.47977 | Intron (ENSM | 4 | 83549660 | 83558841 | 9182 | 1 | ENSMUST00 | -9897 | Ccdc171 | ENSMUSG00000052407.11 |
| chr4 | 155408994 | 155409495 | 7.12487 | 12.70834 | 10.47977 | Promoter | 4 | 155409190 | 155415987 | 6798 | 1 | ENSMUST00 | 0 | C030017K20 | ENSMUSG00000078490.4 |
| chr4 | 99657196 | 99658141 | 7.12487 | 12.70834 | 10.47977 | Promoter | 4 | 99657742 | 99657830 | 89 | 1 | ENSMUST00 | 0 | Gm23366 | ENSMUSG00000092763.1 |
| chr5 | 138997277 | 138997786 | 7.12487 | 12.70834 | 10.47977 | Promoter | 5 | 138976971 | 138997370 | 20400 | 2 | ENSMUST00 | 0 | Pdgfa | ENSMUSG00000025856.10 |
| chr5 | 32346060 | 32346376 | 7.12487 | 12.70834 | 10.47977 | Intron (ENSM | 5 | 32353569 | 32366520 | 12952 | 1 | ENSMUST00 | -7193 | Plb1 | ENSMUSG00000029134.9 |
| chr8 | 70840003 | 70840221 | 7.12487 | 12.70834 | 10.47977 | Promoter | 8 | 70835155 | 70839720 | 4566 | 2 | ENSMUST00 | -284 | Arrdc2 | ENSMUSG00000002910.10 |
| chr8 | 116863312 | 116863842 | 7.12487 | 12.70834 | 10.47977 | Distal Interge | 8 | 116888685 | 116894524 | 5840 | 2 | ENSMUST00 | 30682 | Cmc2 | ENSMUSG00000014633.10 |
| chr8 | 109185561 | 109185939 | 7.12487 | 12.70834 | 10.47977 | Intron (ENSM | 8 | 109249866 | 109273811 | 23946 | 1 | ENSMUST00 | -63927 | D030068K23 | ENSMUSG00000097393.2 |
| chr8 | 46691543 | 46691873 | 7.12487 | 12.70834 | 10.47977 | Distal Interge | 8 | 46730972 | 46739515 | 8544 | 2 | ENSMUST00 | 47642 | Gm16675 | ENSMUSG00000097534.1 |
| chr9 | 36833324 | 36833816 | 7.12487 | 12.70834 | 10.47977 | Promoter | 9 | 36833635 | 36863499 | 29865 | 1 | ENSMUST00 | 0 | Fez1 | ENSMUSG00000032118.10 |
| chr9 | 59134085 | 59134850 | 7.12487 | 12.70834 | 10.47977 | Distal Interge | 9 | 59145677 | 59146210 | 534 | 2 | ENSMUST00 | 11360 | Gm7589 | ENSMUSG00000066592.3 |
| chr9 | 96759108 | 96759520 | 7.12487 | 12.70834 | 10.47977 | Distal Interge | 9 | 96765562 | 96773599 | 8038 | 1 | ENSMUST00 | -6042 | RP24-217M7 | ENSMUSG00000101581.1 |
| chrX | 152001301 | 152001689 | 7.12487 | 12.70834 | 10.47977 | Promoter | 20 | 152001845 | 152004442 | 2598 | 1 | ENSMUST00 | -156 | Hsd17b10 | ENSMUSG00000025260.5 |
| chr3 | 127225800 | 127226444 | 7.11165 | 12.68657 | 10.46002 | Promoter | 3 | 126922308 | 127225847 | 303540 | 2 | ENSMUST00 | 0 | Ank2 | ENSMUSG00000032826.11 |
| chr1 | 74794379 | 74795025 | 7.35251 | 12.67314 | 10.44663 | Intron (ENSM | 1 | 74793363 | 74803495 | 10133 | 1 | ENSMUST00 | 1017 | Wnt10a | ENSMUSG00000026167.9 |
| chr1 | 6441196 | 6441423 | 7.35251 | 12.67314 | 10.44663 | Distal Interge | 1 | 6487231 | 6860934 | 373704 | 1 | ENSMUST00 | -45808 | St18 | ENSMUSG00000033740.12 |
| chr1 | 172053737 | 172054259 | 7.35251 | 12.67314 | 10.44663 | 3' UTR | 1 | 172052292 | 172057573 | 5282 | 2 | ENSMUST00 | 3314 | Nhlh1 | ENSMUSG00000051251.3 |
| chr11 | 76509110 | 76509471 | 7.35251 | 12.67314 | 10.44663 | Promoter | 11 | 76416734 | 76509419 | 92686 | 2 | ENSMUST00 | 0 | Abr | ENSMUSG00000017631.13 |
| chr11 | 96285243 | 96285650 | 7.35251 | 12.67314 | 10.44663 | Promoter | 11 | 96286623 | 96290162 | 3540 | 1 | ENSMUST00 | -973 | Hoxb7 | ENSMUSG00000038721.8 |
| chr11 | 96251509 | 96251965 | 7.35251 | 12.67314 | 10.44663 | Promoter | 11 | 96251644 | 96264698 | 13055 | 1 | ENSMUST00 | 0 | Gm53 | ENSMUSG00000078706.4 |
| chr12 | 32910265 | 32910969 | 7.35251 | 12.67314 | 10.44663 | Distal Interge | 12 | 32924957 | 32953783 | 28827 | 2 | ENSMUST00 | 42814 | 4933406C10 | ENSMUSG00000097494.2 |
| chr13 | 56806744 | 56807269 | 7.35251 | 12.67314 | 10.44663 | Intron (ENSM | 13 | 56733017 | 56738993 | 5977 | 1 | ENSMUST00 | 73728 | Smad5 | ENSMUSG00000021540.11 |
| chr13 | 96132273 | 96132742 | 7.35251 | 12.67314 | 10.44663 | Promoter | 13 | 95959443 | 96132577 | 173135 | 2 | ENSMUST00 | 0 | Sv2c | ENSMUSG00000051111.10 |
| chr14 | 41053591 | 41053764 | 7.35251 | 12.67314 | 10.44663 | Intron (ENSM | 14 | 41049209 | 41069074 | 19866 | 2 | ENSMUST00 | 15310 | Dydc2 | ENSMUSG00000021791.6 |
| chr14 | 70530643 | 70531078 | 7.35251 | 12.67314 | 10.44663 | Promoter | 14 | 70530821 | 70538323 | 7503 | 1 | ENSMUST00 | 0 | Lgi3 | ENSMUSG00000033595.6 |
| chr15 | 103258367 | 103258880 | 7.35251 | 12.67314 | 10.44663 | Promoter | 15 | 103249349 | 103258403 | 9055 | 2 | ENSMUST00 | 0 | Nfe2 | ENSMUSG00000058794.7 |
| chr17 | 78202562 | 78203052 | 7.35251 | 12.67314 | 10.44663 | Intron (ENSM | 17 | 78200248 | 78376592 | 176345 | 1 | ENSMUST00 | 2315 | Crim1 | ENSMUSG00000024074.7 |
| chr17 | 32189173 | 32189746 | 7.35251 | 12.67314 | 10.44663 | Promoter | 17 | 32183770 | 32189463 | 5694 | 2 | ENSMUST00 | 0 | Ephx3 | ENSMUSG00000037577.7 |
| chr17 | 24755470 | 24756597 | 7.35251 | 12.67314 | 10.44663 | Intron (ENSM | 17 | 24753003 | 24758683 | 5681 | 1 | ENSMUST00 | 2468 | Hs3st6 | ENSMUSG00000039628.8 |
| chr17 | 27685002 | 27685451 | 7.35251 | 12.67314 | 10.44663 | Promoter | 17 | 27685197 | 27708659 | 23463 | 1 | ENSMUST00 | 0 | Pacsin1 | ENSMUSG00000040276.9 |
| chr18 | 68474898 | 68475161 | 7.35251 | 12.67314 | 10.44663 | Distal Interge | 18 | 68406907 | 68429251 | 22345 | 2 | ENSMUST00 | -45648 | Mc2r | ENSMUSG00000045569.6 |
| chr19 | 25505670 | 25506296 | 7.35251 | 12.67314 | 10.44663 | Promoter | 19 | 25505706 | 25603491 | 97786 | 1 | ENSMUST00 | 0 | Dmrt1 | ENSMUSG00000024837.10 |
| chr2 | 7405377 | 7405541 | 7.35251 | 12.67314 | 10.44663 | Intron (ENSM | 2 | 6615729 | 7396198 | 780470 | 2 | ENSMUST00 | -9180 | Celf2 | ENSMUSG00000002107.13 |
| chr2 | 91930476 | 91932210 | 7.35251 | 12.67314 | 10.44663 | Promoter | 2 | 91929805 | 91931688 | 1884 | 2 | ENSMUST00 | 0 | Mdk | ENSMUSG00000027239.9 |
| chr2 | 176927162 | 176927337 | 7.35251 | 12.67314 | 10.44663 | Promoter | 2 | 176892509 | 176927794 | 35286 | 2 | ENSMUST00 | 457 | Gm14296 | ENSMUSG00000074527.6 |
| chr3 | 88677578 | 88677789 | 7.35251 | 12.67314 | 10.44663 | Distal Interge | 3 | 88685794 | 88712933 | 27140 | 1 | ENSMUST00 | -8005 | 2810403A07 | ENSMUSG00000028060.10 |
| chr4 | 149437014 | 149437210 | 7.35251 | 12.67314 | 10.44663 | Distal Interge | 4 | 149328416 | 149426749 | 98334 | 2 | ENSMUST00 | -10266 | Ube4b | ENSMUSG00000028960.13 |
| chr5 | 115204622 | 115205180 | 7.35251 | 12.67314 | 10.44663 | Distal Interge | 5 | 115198343 | 115198705 | 363 | 1 | ENSMUST00 | 6280 | Gm13828 | ENSMUSG00000083182.1 |
| chr5 | 143293231 | 143293735 | 7.35251 | 12.67314 | 10.44663 | Promoter | 5 | 143287755 | 143292356 | 4602 | 2 | ENSMUST00 | -876 | Zfp853 | ENSMUSG00000093910.1 |
| chr7 | 122219378 | 122219615 | 7.35251 | 12.67314 | 10.44663 | Promoter | 7 | 122219496 | 122222824 | 3329 | 1 | ENSMUST00 | 0 | Chp2 | ENSMUSG00000030865.4 |
| chr7 | 127261007 | 127261368 | 7.35251 | 12.67314 | 10.44663 | Promoter | 7 | 127256959 | 127260709 | 3751 | 2 | ENSMUST00 | -299 | Dctpp1 | ENSMUSG00000042462.4 |
| chr7 | 126823269 | 126823658 | 7.35251 | 12.67314 | 10.44663 | Promoter | 7 | 126823303 | 126830217 | 6915 | 1 | ENSMUST00 | 0 | Fam57b | ENSMUSG00000058966.8 |
| chr8 | 84784076 | 84785113 | 7.35251 | 12.67314 | 10.44663 | Intron (ENSM | 8 | 84708626 | 84774369 | 65744 | 2 | ENSMUST00 | -9708 | Nfix | ENSMUSG00000001911.11 |
| chr8 | 4212898 | 4213553 | 7.35251 | 12.67314 | 10.44663 | Exon (ENSMU | 8 | 4215049 | 4216912 | 1864 | 2 | ENSMUST00 | 3359 | BC068157 | ENSMUSG00000064125.9 |
| chr8 | 25439977 | 25440949 | 7.35251 | 12.67314 | 10.44663 | Distal Interge | 8 | 25454273 | 25454704 | 432 | 1 | ENSMUST00 | -13324 | Gm10689 | ENSMUSG00000074412.4 |
| chr8 | 79501308 | 79501644 | 7.35251 | 12.67314 | 10.44663 | Distal Interge | 8 | 79549774 | 79550748 | 975 | 2 | ENSMUST00 | 49104 | Gm8079 | ENSMUSG00000097975.1 |
| chr9 | 89066752 | 89067094 | 7.35251 | 12.67314 | 10.44663 | Distal Interge | 9 | 89075336 | 89075407 | 72 | 1 | ENSMUST00 | -8242 | Gm24463 | ENSMUSG00000096473.1 |
| chrX | 100767728 | 100768093 | 7.35251 | 12.67314 | 10.44663 | Promoter | 20 | 100767749 | 100818408 | 50660 | 1 | ENSMUST00 | 0 | Dlg3 | ENSMUSG00000000881.7 |
| chrX | 59568175 | 59568432 | 7.35251 | 12.67314 | 10.44663 | Promoter | 20 | 59250808 | 59568071 | 317264 | 2 | ENSMUST00 | -105 | Fgf13 | ENSMUSG00000031137.12 |
| chrX | 75577864 | 75578251 | 7.35251 | 12.67314 | 10.44663 | Promoter | 20 | 75572046 | 75578231 | 6186 | 2 | ENSMUST00 | 0 | Rab39b | ENSMUSG00000031202.5 |
| chrX | 104856731 | 104857309 | 7.35251 | 12.67314 | 10.44663 | Promoter | 20 | 104854952 | 104857267 | 2316 | 2 | ENSMUST00 | 0 | Magee2 | ENSMUSG00000031224.6 |
| chrX | 12158400 | 12159041 | 7.35251 | 12.67314 | 10.44663 | Intron (ENSM | 20 | 12036743 | 12160355 | 123613 | 2 | ENSMUST00 | 1314 | Bcor | ENSMUSG00000040363.9 |
| chrX | 103621613 | 103622375 | 7.35251 | 12.67314 | 10.44663 | Promoter | 20 | 103607606 | 103622500 | 14895 | 2 | ENSMUST00 | 125 | Ftx | ENSMUSG00000086370.3 |
| chr1 | 163313277 | 163314513 | 7.32392 | 12.62659 | 10.40257 | Promoter | 1 | 163245119 | 163313672 | 68554 | 2 | ENSMUST00 | 0 | Prrx1 | ENSMUSG00000026586.11 |
| chr17 | 25079569 | 25079919 | 7.07106 | 12.61967 | 10.39574 | Intron (ENSM | 17 | 25057702 | 25081181 | 23480 | 2 | ENSMUST00 | 1262 | Tmem204 | ENSMUSG00000024168.7 |
| chr11 | 82845932 | 82846307 | 7.30862 | 12.60167 | 10.37778 | Promoter | 11 | 82812669 | 82846092 | 33424 | 2 | ENSMUST00 | 0 | Rffl | ENSMUSG00000020696.13 |
| chr11 | 70465727 | 70466420 | 7.30862 | 12.60167 | 10.37778 | 3' UTR | 11 | 70460565 | 70466090 | 5526 | 1 | ENSMUST00 | 5163 | Zmynd15 | ENSMUSG00000040829.9 |
| chr2 | 77768454 | 77768871 | 7.30862 | 12.60167 | 10.37778 | Intron (ENSM | 2 | 77411100 | 77816816 | 405717 | 2 | ENSMUST00 | 47945 | Zfp385b | ENSMUSG00000027016.12 |
| chr4 | 149436724 | 149436914 | 7.30862 | 12.60167 | 10.37778 | Distal Interge | 4 | 149328416 | 149426749 | 98334 | 2 | ENSMUST00 | -9976 | Ube4b | ENSMUSG00000028960.13 |
| chr11 | 120950486 | 120950962 | 6.80124 | 12.58378 | 10.36022 | Promoter | 11 | 120950688 | 120955273 | 4586 | 1 | ENSMUST00 | 0 | Slc16a3 | ENSMUSG00000025161.11 |
| chr12 | 75449665 | 75450394 | 6.80124 | 12.58378 | 10.36022 | Downstream | 12 | 75411720 | 75416781 | 5062 | 2 | ENSMUST00 | -32885 | Gphb5 | ENSMUSG00000048982.2 |
| chr15 | 50870573 | 50871205 | 7.29338 | 12.57686 | 10.35346 | Intron (ENSM | 15 | 50660435 | 50882806 | 222372 | 2 | ENSMUST00 | 11601 | Trps1 | ENSMUSG00000038679.11 |
| chr5 | 119680420 | 119681089 | 7.29338 | 12.57686 | 10.35346 | Promoter | 5 | 119680249 | 119680832 | 584 | 1 | ENSMUST00 | 172 | Tbx3 | ENSMUSG00000018604.13 |
| chr15 | 3994771 | 3995032 | 7.03093 | 12.55347 | 10.33021 | Promoter | 15 | 3982036 | 3995752 | 13717 | 2 | ENSMUST00 | 720 | AW549877 | ENSMUSG00000041935.9 |
| chr6 | 125572802 | 125573456 | 7.03093 | 12.55347 | 10.33021 | Intron (ENSM | 6 | 125566251 | 125578578 | 12328 | 1 | ENSMUST00 | 6552 | Vwf | ENSMUSG00000001930.12 |
| chr1 | 133908664 | 133908930 | 7.27821 | 12.55215 | 10.32949 | Promoter | 1 | 133897200 | 133907999 | 10800 | 2 | ENSMUST00 | -666 | Optc | ENSMUSG00000010311.10 |
| chr10 | 20528587 | 20529334 | 5.68011 | 12.54176 | 10.31921 | Intron (ENSM | 10 | 20518011 | 20519768 | 1758 | 2 | ENSMUST00 | -8820 | Gm17229 | ENSMUSG00000092051.1 |
| chr17 | 24689352 | 24690094 | 5.68011 | 12.54176 | 10.31921 | Promoter | 17 | 24685092 | 24689955 | 4864 | 2 | ENSMUST00 | 0 | Syngr3 | ENSMUSG00000007021.4 |
| chr4 | 142544906 | 142545224 | 5.68011 | 12.54176 | 10.31921 | Distal Interge | 4 | 142557390 | 142557836 | 447 | 2 | ENSMUST00 | 12612 | Gm13018 | ENSMUSG00000094224.1 |
| chr3 | 85172181 | 85172898 | 7.2631 | 12.52754 | 10.30509 | Intron (ENSM | 3 | 85317519 | 85332030 | 14512 | 1 | ENSMUST00 | -144621 | RP23-94M23 | ENSMUSG00000099995.1 |
| chr5 | 114897681 | 114897993 | 7.2631 | 12.52754 | 10.30509 | Promoter | 5 | 114897742 | 114912226 | 14485 | 1 | ENSMUST00 | 0 | Oasl2 | ENSMUSG00000029561.12 |
| chr7 | 68917383 | 68917674 | 7.2631 | 12.52754 | 10.30509 | Distal Interge | 7 | 68952100 | 68952236 | 137 | 2 | ENSMUST00 | 34562 | Gm23795 | ENSMUSG00000065940.1 |
| chr10 | 60350720 | 60351071 | 6.75876 | 12.51203 | 10.2897 | Intron (ENSM | 10 | 60349323 | 60369293 | 19971 | 1 | ENSMUST00 | 1398 | 4632428N05 | ENSMUSG00000020101.9 |
| chr10 | 122029367 | 122029824 | 6.75876 | 12.51203 | 10.2897 | Intron (ENSM | 10 | 121780991 | 122047308 | 266318 | 2 | ENSMUST00 | 17484 | Srgap1 | ENSMUSG00000020121.10 |
| chr10 | 5264188 | 5264846 | 6.75876 | 12.51203 | 10.2897 | Distal Interge | 10 | 5357173 | 5357279 | 107 | 1 | ENSMUST00 | -92327 | Gm23573 | ENSMUSG00000094906.1 |
| chr11 | 100759790 | 100760096 | 6.75876 | 12.51203 | 10.2897 | Promoter | 11 | 100740382 | 100759942 | 19561 | 2 | ENSMUST00 | 0 | Kcnh4 | ENSMUSG00000035355.10 |

| | | | | | | | | | | | | | | | |
|---|---|---|---|---|---|---|---|---|---|---|---|---|---|---|---|
| chr11 | 84642446 | 84642908 | 6.75876 | 12.51203 | 10.2897 | Distal Interge | 11 | 84716976 | 84720432 | 3457 | 2 | ENSMUST00 | 77524 | 1700109G15 | ENSMUSG00000085044.1 |
| chr13 | 63258717 | 63261057 | 6.75876 | 12.51203 | 10.2897 | Intron (ENSM | 13 | 63282142 | 63326096 | 43955 | 1 | ENSMUST00 | -21085 | 2010111I01R | ENSMUSG00000021458.11 |
| chr13 | 59971201 | 59972007 | 6.75876 | 12.51203 | 10.2897 | Distal Interge | 13 | 59771879 | 59823147 | 51269 | 2 | ENSMUST00 | -148055 | Zcchc6 | ENSMUSG00000035248.8 |
| chr13 | 115054456 | 115055124 | 6.75876 | 12.51203 | 10.2897 | Intron (ENSM | 13 | 115088357 | 115090188 | 1832 | 2 | ENSMUST00 | 35064 | Pelo | ENSMUSG00000042275.8 |
| chr15 | 101929511 | 101930156 | 6.75876 | 12.51203 | 10.2897 | 3' UTR | 15 | 101918536 | 101924735 | 6200 | 2 | ENSMUST00 | -4777 | Krt4 | ENSMUSG00000059668.5 |
| chr16 | 72999401 | 72999667 | 6.75876 | 12.51203 | 10.2897 | Intron (ENSM | 16 | 72663149 | 73046095 | 382947 | 1 | ENSMUST00 | 336253 | Robo1 | ENSMUSG00000022883.10 |
| chr17 | 75067107 | 75067527 | 6.75876 | 12.51203 | 10.2897 | Intron (ENSM | 17 | 75006294 | 75008799 | 2506 | 1 | ENSMUST00 | 60814 | Ltbp1 | ENSMUSG00000001870.10 |
| chr17 | 69120750 | 69121390 | 6.75876 | 12.51203 | 10.2897 | Distal Interge | 17 | 69156810 | 69289985 | 133176 | 1 | ENSMUST00 | -35420 | Epb4.1l3 | ENSMUSG00000024044.10 |
| chr17 | 87459188 | 87459424 | 6.75876 | 12.51203 | 10.2897 | Distal Interge | 17 | 87453223 | 87453356 | 134 | 2 | ENSMUST00 | -5833 | Gm26005 | ENSMUSG00000089427.1 |
| chr17 | 86329379 | 86329870 | 6.75876 | 12.51203 | 10.2897 | Intron (ENSM | 17 | 86285164 | 86287178 | 2015 | 2 | ENSMUST00 | -42202 | RP23-458O2 | ENSMUSG00000100738.1 |
| chr18 | 61565526 | 61565789 | 6.75876 | 12.51203 | 10.2897 | Intron (ENSM | 18 | 61575517 | 61582118 | 6602 | 1 | ENSMUST00 | -9728 | Csnk1a1 | ENSMUSG00000024576.9 |
| chr18 | 34778648 | 34779335 | 6.75876 | 12.51203 | 10.2897 | Intron (ENSM | 18 | 34777008 | 34839370 | 62363 | 1 | ENSMUST00 | 1641 | Kdm3b | ENSMUSG00000038773.10 |
| chr2 | 122377026 | 122377423 | 6.75876 | 12.51203 | 10.2897 | Distal Interge | 2 | 122349128 | 122369162 | 20035 | 2 | ENSMUST00 | -7865 | Shf | ENSMUSG00000033256.9 |
| chr2 | 170266635 | 170267352 | 6.75876 | 12.51203 | 10.2897 | Distal Interge | 2 | 170285035 | 170285880 | 846 | 1 | ENSMUST00 | -17683 | Gm14270 | ENSMUSG00000084013.1 |
| chr4 | 148163958 | 148164618 | 6.75876 | 12.51203 | 10.2897 | Exon (ENSMU | 4 | 148160621 | 148166424 | 5804 | 1 | ENSMUST00 | 3338 | Fbxo2 | ENSMUSG00000041556.8 |
| chr5 | 88796657 | 88797192 | 6.75876 | 12.51203 | 10.2897 | Distal Interge | 5 | 88764996 | 88783281 | 18286 | 1 | ENSMUST00 | 31662 | Dck | ENSMUSG00000029366.9 |
| chr5 | 116418999 | 116419736 | 6.75876 | 12.51203 | 10.2897 | Intron (ENSM | 5 | 116408940 | 116415892 | 6953 | 2 | ENSMUST00 | -3108 | Hspb8 | ENSMUSG00000041548.4 |
| chr7 | 31041678 | 31042331 | 6.75876 | 12.51203 | 10.2897 | Promoter | 7 | 31032722 | 31041839 | 9118 | 2 | ENSMUST00 | 0 | Fxyd5 | ENSMUSG00000009687.9 |
| chr7 | 17540090 | 17540275 | 6.75876 | 12.51203 | 10.2897 | Distal Interge | 7 | 17610766 | 17610875 | 110 | 1 | ENSMUST00 | -70491 | Gm24520 | ENSMUSG00000096673.1 |
| chr8 | 36548819 | 36549595 | 6.75876 | 12.51203 | 10.2897 | Distal Interge | 8 | 36572145 | 36577269 | 5125 | 2 | ENSMUST00 | 27674 | Dlc1 | ENSMUSG00000031523.11 |
| chr8 | 77506424 | 77506694 | 6.75876 | 12.51203 | 10.2897 | Intron (ENSM | 8 | 77517056 | 77518579 | 1524 | 1 | ENSMUST00 | -10362 | 0610038B21 | ENSMUSG00000097882.2 |
| chr9 | 77427113 | 77427819 | 6.75876 | 12.51203 | 10.2897 | Distal Interge | 9 | 77431089 | 77442195 | 11107 | 2 | ENSMUST00 | 14376 | Lrrc1 | ENSMUSG00000032352.10 |
| chr9 | 29962335 | 29963153 | 6.75876 | 12.51203 | 10.2897 | Promoter | 9 | 28994750 | 29963129 | 968380 | 2 | ENSMUST00 | 0 | Ntm | ENSMUSG00000059974.5 |
| chr9 | 63138558 | 63139098 | 6.75876 | 12.51203 | 10.2897 | 3' UTR | 9 | 63133283 | 63133389 | 107 | 1 | ENSMUST00 | 5276 | Gm25064 | ENSMUSG00000094201.1 |
| chr9 | 41735524 | 41736542 | 6.75876 | 12.51203 | 10.2897 | Distal Interge | 9 | 41654303 | 41655282 | 980 | 1 | ENSMUST00 | 81222 | Gm8707 | ENSMUSG00000098093.1 |
| chr9 | 50501023 | 50501474 | 6.75876 | 12.51203 | 10.2897 | Exon (ENSMU | 9 | 50488799 | 50504805 | 16007 | 2 | ENSMUST00 | 3331 | Plet1os | ENSMUSG00000101304.1 |
| chrX | 140061291 | 140061977 | 6.75876 | 12.51203 | 10.2897 | Promoter | 20 | 140022062 | 140062293 | 40232 | 2 | ENSMUST00 | 316 | Nup62cl | ENSMUSG00000072944.6 |
| chr10 | 19494625 | 19494961 | 7.23306 | 12.47862 | 10.25851 | Distal Interge | 10 | 19591949 | 19610229 | 18281 | 1 | ENSMUST00 | -96988 | Ifngr1 | ENSMUSG00000020009.7 |
| chr15 | 63895798 | 63896191 | 5.83242 | 12.45768 | 10.23771 | Intron (ENSM | 15 | 63891265 | 63902936 | 11672 | 2 | ENSMUST00 | 6745 | Gsdmc4 | ENSMUSG00000055748.7 |
| chr19 | 21714927 | 21715495 | 5.83242 | 12.45768 | 10.23771 | Distal Interge | 19 | 21653309 | 21685637 | 32329 | 1 | ENSMUST00 | 61619 | Abhd17b | ENSMUSG00000047368.3 |
| chr8 | 18740743 | 18741274 | 5.83242 | 12.45768 | 10.23771 | Promoter | 8 | 18690263 | 18741562 | 51300 | 2 | ENSMUST00 | 288 | Angpt2 | ENSMUSG00000031465.6 |
| chr9 | 72408036 | 72408475 | 5.83242 | 12.45768 | 10.23771 | Promoter | 9 | 72409268 | 72409878 | 611 | 1 | ENSMUST00 | -793 | Gm27255 | ENSMUSG00000098801.1 |
| chr3 | 152981131 | 152981491 | 7.21814 | 12.45431 | 10.23451 | Promoter | 3 | 152975374 | 152982052 | 6679 | 2 | ENSMUST00 | 561 | St6galnac5 | ENSMUSG00000039037.5 |
| chr1 | 55192080 | 55192331 | 6.46168 | 12.41373 | 10.19419 | Intron (ENSM | 1 | 55203945 | 55206489 | 2545 | 2 | ENSMUST00 | 14158 | Rftn2 | ENSMUSG00000025978.9 |
| chr1 | 90206355 | 90206995 | 6.46168 | 12.41373 | 10.19419 | Intron (ENSM | 1 | 90204006 | 90213833 | 9828 | 1 | ENSMUST00 | 2350 | Ackr3 | ENSMUSG00000044337.5 |
| chr11 | 98709428 | 98710246 | 6.46168 | 12.41373 | 10.19419 | Promoter | 11 | 98709646 | 98711066 | 1421 | 2 | ENSMUST00 | 820 | Med24 | ENSMUSG00000017210.9 |
| chr12 | 58212321 | 58213074 | 6.46168 | 12.41373 | 10.19419 | Promoter | 12 | 58211804 | 58214444 | 2641 | 1 | ENSMUST00 | 518 | Sstr1 | ENSMUSG00000035431.4 |
| chr13 | 40886799 | 40887548 | 6.46168 | 12.41373 | 10.19419 | Promoter | 13 | 40886758 | 40960891 | 74134 | 1 | ENSMUST00 | 42 | Gcnt2 | ENSMUSG00000021360.10 |
| chr14 | 15438403 | 15439088 | 6.46168 | 12.41373 | 10.19419 | Promoter | 14 | 15357519 | 15438987 | 81469 | 2 | ENSMUST00 | 0 | Lrrc3b | ENSMUSG00000045201.5 |
| chr16 | 21728087 | 21728295 | 6.46168 | 12.41373 | 10.19419 | Distal Interge | 16 | 21649045 | 21694665 | 45621 | 2 | ENSMUST00 | -33423 | 2510009E07I | ENSMUSG00000043391.9 |
| chr16 | 91917645 | 91917942 | 6.46168 | 12.41373 | 10.19419 | 3' UTR | 16 | 91921309 | 91921415 | 107 | 2 | ENSMUST00 | 3473 | Gm25245 | ENSMUSG00000096895.1 |
| chr19 | 53040211 | 53040927 | 6.46168 | 12.41373 | 10.19419 | Promoter | 19 | 52991183 | 53040214 | 49032 | 2 | ENSMUST00 | 0 | Xpnpep1 | ENSMUSG00000025027.12 |
| chr3 | 41182223 | 41182456 | 6.46168 | 12.41373 | 10.19419 | Distal Interge | 3 | 41066326 | 41083046 | 16721 | 2 | ENSMUST00 | -99178 | Pgrmc2 | ENSMUSG00000049940.7 |
| chr6 | 126164537 | 126165173 | 6.46168 | 12.41373 | 10.19419 | 5' UTR | 6 | 126101413 | 126166744 | 65332 | 2 | ENSMUST00 | 1571 | Ntf3 | ENSMUSG00000049107.8 |
| chr6 | 136807854 | 136808060 | 6.46168 | 12.41373 | 10.19419 | Promoter | 6 | 136808244 | 136810070 | 1827 | 1 | ENSMUST00 | -184 | H2afj | ENSMUSG00000060032.5 |
| chr6 | 31114573 | 31115101 | 6.46168 | 12.41373 | 10.19419 | Intron (ENSM | 6 | 31102846 | 31104646 | 1801 | 1 | ENSMUST00 | 11728 | Gm13833 | ENSMUSG00000087312.1 |
| chr8 | 67947100 | 67947355 | 6.46168 | 12.41373 | 10.19419 | Intron (ENSM | 8 | 67903648 | 67951879 | 48232 | 2 | ENSMUST00 | 4524 | Psd3 | ENSMUSG00000030465.13 |
| chr9 | 119981700 | 119982233 | 6.46168 | 12.41373 | 10.19419 | Intron (ENSM | 9 | 119971166 | 119977250 | 6085 | 2 | ENSMUST00 | -4451 | Csrnp1 | ENSMUSG00000032515.7 |
| chrX | 20687378 | 20687701 | 6.46168 | 12.41373 | 10.19419 | Promoter | 20 | 20687954 | 20699880 | 11927 | 1 | ENSMUST00 | -253 | Cdk16 | ENSMUSG00000031065.9 |
| chr10 | 99986640 | 99987193 | 6.93905 | 12.40169 | 10.18401 | Distal Interge | 10 | 100015630 | 100100413 | 84784 | 1 | ENSMUST00 | -28437 | Kitl | ENSMUSG00000019966.12 |
| chr10 | 94416164 | 94416736 | 6.0089 | 12.40205 | 10.18401 | Intron (ENSM | 10 | 94408891 | 94410732 | 1842 | 1 | ENSMUST00 | 7274 | Gm16155 | ENSMUSG00000085409.1 |
| chr13 | 37963542 | 37963910 | 6.0089 | 12.40205 | 10.18401 | Distal Interge | 13 | 37947016 | 37949206 | 2191 | 1 | ENSMUST00 | 16527 | Rreb1 | ENSMUSG00000039087.11 |
| chr14 | 54227701 | 54228492 | 6.0089 | 12.40205 | 10.18401 | Distal Interge | 14 | 54220521 | 54223097 | 2577 | 1 | ENSMUST00 | 7181 | Trac | ENSMUSG00000076928.1 |
| chr17 | 5752982 | 5753200 | 6.0089 | 12.40205 | 10.18401 | 3' UTR | 17 | 5798657 | 5802893 | 4237 | 1 | ENSMUST00 | -45457 | 3300005D01I | ENSMUSG00000096965.2 |
| chr6 | 82939378 | 82939804 | 6.0089 | 12.40205 | 10.18401 | Promoter | 6 | 82911885 | 82939769 | 27885 | 2 | ENSMUST00 | 0 | Sema4f | ENSMUSG00000000627.10 |
| chr7 | 128610821 | 128611065 | 6.0089 | 12.40205 | 10.18401 | Promoter | 7 | 128611328 | 128671622 | 60295 | 1 | ENSMUST00 | -263 | Inpp5f | ENSMUSG00000042105.13 |
| chr9 | 27191776 | 27192346 | 6.0089 | 12.40205 | 10.18401 | Downstream | 9 | 27193602 | 27194018 | 417 | 2 | ENSMUST00 | 1672 | RP24-79I19.2 | ENSMUSG00000100409.1 |
| chr8 | 122906611 | 122906820 | 6.68569 | 12.38833 | 10.17069 | Exon (ENSMU | 8 | 122884151 | 122915987 | 31837 | 2 | ENSMUST00 | 9167 | Ankrd11 | ENSMUSG00000035569.11 |
| chr16 | 22160453 | 22160894 | 6.21579 | 12.3833 | 10.16586 | Promoter | 16 | 22059460 | 22161450 | 101991 | 2 | ENSMUST00 | 556 | Igf2bp2 | ENSMUSG00000033581.11 |
| chr18 | 73762510 | 73762688 | 6.21579 | 12.3833 | 10.16586 | Distal Interge | 18 | 73735038 | 73754479 | 19442 | 2 | ENSMUST00 | -8032 | Elac1 | ENSMUSG00000036941.2 |
| chr18 | 84532099 | 84532651 | 6.21579 | 12.3833 | 10.16586 | Intron (ENSM | 18 | 84495776 | 84558375 | 62600 | 2 | ENSMUST00 | 25724 | Zfp407 | ENSMUSG00000048410.11 |
| chr19 | 24861414 | 24862114 | 6.21579 | 12.3833 | 10.16586 | Promoter | 19 | 24683016 | 24861855 | 178840 | 2 | ENSMUST00 | 0 | Pgm5 | ENSMUSG00000041731.8 |
| chr4 | 109977585 | 109978665 | 6.21579 | 12.3833 | 10.16586 | Promoter | 4 | 109978053 | 109983687 | 5635 | 1 | ENSMUST00 | 0 | Dmrta2 | ENSMUSG00000047143.3 |
| chr10 | 31312922 | 31313195 | 7.15905 | 12.35804 | 10.14148 | Promoter | 10 | 31313382 | 31328204 | 14823 | 1 | ENSMUST00 | -187 | Hddc2 | ENSMUSG00000000295.7 |
| chr10 | 18549170 | 18550005 | 7.15905 | 12.35804 | 10.14148 | Distal Interge | 10 | 18540123 | 18546076 | 5954 | 2 | ENSMUST00 | -3095 | Hebp2 | ENSMUSG00000019853.5 |
| chr15 | 82224060 | 82224522 | 7.15905 | 12.35804 | 10.14148 | Promoter | 15 | 82221744 | 82224336 | 2593 | 2 | ENSMUST00 | 0 | Tnfrsf13c | ENSMUSG00000068105.5 |
| chr18 | 23754287 | 23754562 | 7.15905 | 12.35804 | 10.14148 | Promoter | 18 | 23753748 | 23797997 | 44250 | 1 | ENSMUST00 | 540 | Mapre2 | ENSMUSG00000024277.9 |
| chr3 | 8717267 | 8717926 | 7.15905 | 12.35804 | 10.14148 | Distal Interge | 3 | 8679359 | 8679477 | 119 | 2 | ENSMUST00 | -37791 | Gm23670 | ENSMUSG00000088094.1 |
| chr4 | 115898019 | 115898568 | 7.15905 | 12.35804 | 10.14148 | Exon (ENSMU | 4 | 115890666 | 115899227 | 8562 | 1 | ENSMUST00 | 7354 | 6430628N08 | ENSMUSG00000034185.9 |
| chr5 | 107721384 | 107721992 | 7.15905 | 12.35804 | 10.14148 | Exon (ENSMU | 5 | 107717224 | 107723961 | 6738 | 2 | ENSMUST00 | 1969 | Gfi1 | ENSMUSG00000029275.12 |
| chr5 | 96429588 | 96430025 | 7.15905 | 12.35804 | 10.14148 | Intron (ENSM | 5 | 96373955 | 96784728 | 410774 | 1 | ENSMUST00 | 55634 | Fras1 | ENSMUSG00000034687.4 |
| chr6 | 7983114 | 7983614 | 7.15905 | 12.35804 | 10.14148 | Promoter | 6 | 7981829 | 7983430 | 1602 | 2 | ENSMUST00 | 0 | Gm9825 | ENSMUSG00000096403.1 |
| chr11 | 58939077 | 58939298 | 6.9004 | 12.33775 | 10.12162 | Promoter | 11 | 58932288 | 58938916 | 6629 | 2 | ENSMUST00 | -162 | Rnf187 | ENSMUSG00000020496.8 |
| chr12 | 75853474 | 75853996 | 6.9004 | 12.33775 | 10.12162 | Promoter | 12 | 75852730 | 75880328 | 27599 | 1 | ENSMUST00 | 745 | Syne2 | ENSMUSG00000063450.9 |
| chr17 | 8988327 | 8988701 | 6.9004 | 12.33775 | 10.12162 | Promoter | 17 | 8988333 | 9008319 | 19987 | 1 | ENSMUST00 | 0 | 1700010I14R | ENSMUSG00000023873.7 |
| chr17 | 35879361 | 35879596 | 6.9004 | 12.33775 | 10.12162 | Promoter | 17 | 35879819 | 35892670 | 12852 | 1 | ENSMUST00 | -223 | Dhx16 | ENSMUSG00000024422.9 |
| chr5 | 37245639 | 37246163 | 6.9004 | 12.33775 | 10.12162 | Promoter | 5 | 37245792 | 37292132 | 46341 | 1 | ENSMUST00 | 0 | Crmp1 | ENSMUSG00000029121.10 |
| chr5 | 124536865 | 124537314 | 6.9004 | 12.33775 | 10.12162 | Distal Interge | 5 | 124540695 | 124550506 | 9812 | 1 | ENSMUST00 | -3381 | Tmed2 | ENSMUSG00000029390.8 |
| chr11 | 69527111 | 69527509 | 7.11536 | 12.28684 | 10.0715 | Intron (ENSM | 11 | 69516007 | 69516755 | 749 | 2 | ENSMUST00 | -10357 | Dnah2 | ENSMUSG00000005237.8 |
| chr7 | 29956341 | 29956699 | 7.07221 | 12.21648 | 10.00152 | Distal Interge | 7 | 29937945 | 29954089 | 16145 | 2 | ENSMUST00 | -2253 | Zfp74 | ENSMUSG00000059975.9 |
| chr11 | 29718562 | 29718991 | 6.57401 | 12.19858 | 9.98374 | Promoter | 11 | 29718563 | 29742916 | 24354 | 1 | ENSMUST00 | 0 | Rtn4 | ENSMUSG00000020458.11 |
| chr10 | 71361850 | 71362500 | 7.05794 | 12.19321 | 9.97851 | Intron (ENSM | 10 | 71347843 | 71363880 | 16038 | 1 | ENSMUST00 | 14008 | Ipmk | ENSMUSG00000060733.8 |
| chr2 | 45024309 | 45024915 | 7.05794 | 12.19321 | 9.97851 | Intron (ENSM | 2 | 45023100 | 45023177 | 78 | 2 | ENSMUST00 | -1133 | Mir5129 | ENSMUSG00000092679.1 |
| chr1 | 23276484 | 23277111 | 6.77463 | 12.12924 | 9.91488 | Distal Interge | 1 | 23272264 | 23272345 | 82 | 1 | ENSMUST00 | 4221 | Mir30a | ENSMUSG00000065405.2 |
| chr18 | 15832752 | 15832916 | 6.77463 | 12.12924 | 9.91488 | Promoter | 18 | 15832794 | 15833330 | 537 | 1 | ENSMUST00 | 0 | Gm10036 | ENSMUSG00000058064.5 |
| chr5 | 141243225 | 141243997 | 6.77463 | 12.12924 | 9.91488 | Intron (ENSM | 5 | 141241918 | 141582504 | 340587 | 1 | ENSMUST00 | 1308 | Sdk1 | ENSMUSG00000039683.11 |
| chr7 | 44305663 | 44305870 | 6.77463 | 12.12924 | 9.91488 | Exon (ENSMU | 7 | 44303770 | 44306959 | 3190 | 2 | ENSMUST00 | 1089 | Clec11a | ENSMUSG00000004473.9 |
| chr7 | 100924133 | 100925359 | 6.77463 | 12.12924 | 9.91488 | Intron (ENSM | 7 | 100869746 | 100932161 | 62416 | 2 | ENSMUST00 | 6802 | Arhgef17 | ENSMUSG00000032875.7 |
| chr12 | 112588066 | 112588495 | 7.01548 | 12.12394 | 9.90998 | Promoter | 12 | 112588784 | 112615556 | 26773 | 1 | ENSMUST00 | -289 | Inf2 | ENSMUSG00000037679.8 |
| chr14 | 75834124 | 75834324 | 7.01548 | 12.12394 | 9.90998 | Distal Interge | 14 | 75843379 | 75844964 | 1586 | 2 | ENSMUST00 | 10640 | Gm4285 | ENSMUSG00000085829.1 |
| chr18 | 20566697 | 20566888 | 7.01548 | 12.12394 | 9.90998 | Intron (ENSM | 18 | 20558246 | 20582916 | 24671 | 1 | ENSMUST00 | 8452 | Dsg2 | ENSMUSG00000044393.10 |
| chr2 | 132944917 | 132945663 | 7.01548 | 12.12394 | 9.90998 | Promoter | 2 | 132904389 | 132945629 | 41241 | 2 | ENSMUST00 | 0 | Fermt1 | ENSMUSG00000027356.8 |
| chr7 | 28381008 | 28381445 | 7.01548 | 12.12394 | 9.90998 | Distal Interge | 7 | 28376784 | 28379255 | 2472 | 2 | ENSMUST00 | -1754 | Zfp36 | ENSMUSG00000044786.5 |
| chr8 | 14095628 | 14096217 | 7.01548 | 12.12394 | 9.90998 | Promoter | 8 | 14095865 | 14847108 | 751244 | 1 | ENSMUST00 | 0 | Dlgap2 | ENSMUSG00000047495.10 |
| chr2 | 68404845 | 68405276 | 6.95965 | 12.03282 | 9.81974 | Intron (ENSM | 2 | 68358986 | 68386681 | 27696 | 2 | ENSMUST00 | -18165 | Stk39 | ENSMUSG00000027030.10 |
| chr3 | 84345207 | 84345504 | 6.95965 | 12.03282 | 9.81974 | Distal Interge | 3 | 84192211 | 84305446 | 113236 | 2 | ENSMUST00 | -39762 | Trim2 | ENSMUSG00000027993.11 |
| chr19 | 46523878 | 46524562 | 6.466 | 12.01425 | 9.80126 | Distal Interge | 19 | 46501648 | 46516444 | 14797 | 1 | ENSMUST00 | 22231 | Trim8 | ENSMUSG00000025034.8 |
| chr15 | 103061395 | 103061996 | 6.94583 | 12.01026 | 9.79738 | Distal Interge | 15 | 103034395 | 103036852 | 2458 | 1 | ENSMUST00 | 27001 | Hoxc4 | ENSMUSG00000075394.4 |
| chr10 | 79746797 | 79747056 | 6.9047 | 11.94308 | 9.73049 | Promoter | 10 | 79736130 | 79746581 | 10452 | 2 | ENSMUST00 | -217 | Polrmt | ENSMUSG00000020329.7 |
| chr6 | 149565879 | 149566346 | 6.9047 | 11.94308 | 9.73049 | Distal Interge | 6 | 149576461 | 149576561 | 101 | 1 | ENSMUST00 | -10115 | Gm24307 | ENSMUSG00000094467.1 |
| chr14 | 22577859 | 22578238 | 6.65336 | 11.9275 | 9.71501 | Promoter | 14 | 22577727 | 22599268 | 21542 | 1 | ENSMUST00 | 133 | 1700112E06I | ENSMUSG00000063458.7 |
| chr15 | 74517582 | 74518319 | 6.65336 | 11.9275 | 9.71501 | Promoter | 15 | 74516831 | 74589464 | 72634 | 1 | ENSMUST00 | 752 | Bai1 | ENSMUSG00000034730.11 |
| chr19 | 34640504 | 34640895 | 6.65336 | 11.9275 | 9.71501 | Promoter | 19 | 34640871 | 34650009 | 9139 | 1 | ENSMUST00 | 0 | Ifit1 | ENSMUSG00000034459.8 |
| chr2 | 72203859 | 72204099 | 6.65336 | 11.9275 | 9.71501 | Intron (ENSM | 2 | 72198799 | 72208437 | 9639 | 1 | ENSMUST00 | 5061 | Rapgef4 | ENSMUSG00000049044.11 |
| chr2 | 152412352 | 152412869 | 6.65336 | 11.9275 | 9.71501 | 3' UTR | 2 | 152411955 | 152415027 | 3073 | 2 | ENSMUST00 | 2158 | Zcchc3 | ENSMUSG00000074682.4 |
| chr2 | 165986850 | 165987496 | 6.65336 | 11.9275 | 9.71501 | Promoter | 2 | 165976757 | 165985924 | 9168 | 2 | ENSMUST00 | -927 | Gm11463 | ENSMUSG00000084816.1 |
| chr6 | 93802385 | 93802765 | 6.65336 | 11.9275 | 9.71501 | Intron (ENSM | 6 | 93793772 | 93793877 | 106 | 1 | ENSMUST00 | 8614 | Gm23035 | ENSMUSG00000088627.1 |
| chr14 | 73374403 | 73375836 | 6.8911 | 11.92086 | 9.7087 | Intron (ENSM | 14 | 73362231 | 73385271 | 23041 | 2 | ENSMUST00 | 9435 | Itm2b | ENSMUSG00000022108.7 |
| chr2 | 4917429 | 4917962 | 6.8911 | 11.92086 | 9.7087 | Distal Interge | 2 | 4919019 | 4938730 | 19712 | 1 | ENSMUST00 | -1057 | Phyh | ENSMUSG00000026664.7 |
| chr1 | 90208030 | 90209212 | 6.87755 | 11.89872 | 9.68664 | Intron (ENSM | 1 | 90204006 | 90213833 | 9828 | 1 | ENSMUST00 | 4025 | Ackr3 | ENSMUSG00000044337.5 |
| chr11 | 86581276 | 86581767 | 6.87755 | 11.89872 | 9.68664 | Downstream | 11 | 86584067 | 86584158 | 92 | 2 | ENSMUST00 | 2391 | Mir21a | ENSMUSG00000065455.1 |
| chr12 | 41565882 | 41566506 | 6.87755 | 11.89872 | 9.68664 | Intron (ENSM | 12 | 41451673 | 41485751 | 34079 | 2 | ENSMUST00 | -80132 | Lrrn3 | ENSMUSG00000036295.4 |
| chr14 | 120137705 | 120137923 | 6.87755 | 11.89872 | 9.68664 | Distal Interge | 14 | 120019585 | 120042435 | 22851 | 2 | ENSMUST00 | -95271 | Oxgr1 | ENSMUSG00000044819.5 |
| chr6 | 8414024 | 8414334 | 6.87755 | 11.89872 | 9.68664 | Intron (ENSM | 6 | 8419678 | 8428238 | 8561 | 1 | ENSMUST00 | -5344 | Gm16039 | ENSMUSG00000089862.3 |
| chr9 | 63145628 | 63146059 | 6.87755 | 11.89872 | 9.68664 | Promoter | 9 | 63138176 | 63146980 | 8805 | 2 | ENSMUST00 | 921 | Skor1 | ENSMUSG00000022245.10 |
| chr2 | 32647512 | 32648076 | 6.86405 | 11.87666 | 9.66515 | Promoter | 2 | 32646993 | 32650367 | 3375 | 1 | ENSMUST00 | 520 | Eng | ENSMUSG00000026814.11 |
| chr6 | 34864144 | 34864641 | 6.86405 | 11.87666 | 9.66515 | Promoter | 6 | 34863146 | 34874946 | 11801 | 1 | ENSMUST00 | 999 | Tmem140 | ENSMUSG00000057137.4 |
| chr8 | 110722003 | 110722228 | 6.86405 | 11.87666 | 9.66515 | Promoter | 8 | 110721476 | 110741400 | 19925 | 1 | ENSMUST00 | 528 | Mtss1l | ENSMUSG00000033763.9 |
| chr5 | 119673187 | 119673585 | 6.36148 | 11.83505 | 9.62371 | Intron (ENSM | 5 | 119617901 | 119671365 | 53465 | 2 | ENSMUST00 | -1823 | Tbx3os1 | ENSMUSG00000087516.2 |
| chr2 | 59438126 | 59438474 | 6.83722 | 11.83278 | 9.62151 | Distal Interge | 2 | 59398986 | 59399781 | 796 | 2 | ENSMUST00 | -38346 | Gm13549 | ENSMUSG00000080982.1 |
| chr5 | 127235534 | 127235722 | 6.83722 | 11.83278 | 9.62151 | Distal Interge | 5 | 127241808 | 127565793 | 323986 | 1 | ENSMUST00 | -6086 | Tmem132c | ENSMUSG00000034324.11 |
| chr11 | 8662432 | 8662983 | 6.82388 | 11.81097 | 9.59978 | Intron (ENSM | 11 | 8431652 | 8664535 | 232884 | 2 | ENSMUST00 | 1552 | Tns3 | ENSMUSG00000020422.8 |
| chr6 | 137275578 | 137276236 | 6.8106 | 11.78924 | 9.57812 | Intron (ENSM | 6 | 137252299 | 137464629 | 212331 | 1 | ENSMUST00 | 23280 | Ptpro | ENSMUSG00000030223.9 |
| chr17 | 70997954 | 70998739 | 6.79737 | 11.76758 | 9.55656 | Promoter | 17 | 70994294 | 70998012 | 3719 | 2 | ENSMUST00 | 0 | Myl12a | ENSMUSG00000024048.9 |
| chr3 | 96245606 | 96245793 | 6.78418 | 11.74601 | 9.53503 | Promoter | 3 | 96245554 | 96248509 | 2956 | 1 | ENSMUST00 | 53 | Gm20634 | ENSMUSG00000070392.4 |
| chr1 | 74368235 | 74368582 | 6.53635 | 11.73215 | 9.52123 | Exon (ENSMU | 1 | 74364531 | 74366742 | 2212 | 1 | ENSMUST00 | 3705 | Catip | ENSMUSG00000073650.3 |
| chr10 | 44457249 | 44457941 | 6.53635 | 11.73215 | 9.52123 | Promoter | 10 | 44438293 | 44458687 | 20395 | 2 | ENSMUST00 | 746 | Prdm1 | ENSMUSG00000038151.7 |
| chr15 | 55120670 | 55121639 | 6.53635 | 11.73215 | 9.52123 | Intron (ENSM | 15 | 55110276 | 55113677 | 3402 | 2 | ENSMUST00 | -6994 | Gm9920 | ENSMUSG00000053749.7 |

| | | | | | | | | | | | | | | | |
|---|---|---|---|---|---|---|---|---|---|---|---|---|---|---|---|
| chr17 | 44614079 | 44614677 | 6.53635 | 11.73215 | 9.52123 | Intron (ENSM | 17 | 44641025 | 44641143 | 119 | 2 | ENSMUST00 | 26466 | n-R5s27 | ENSMUSG00000070209.1 |
| chr17 | 28076207 | 28076372 | 6.53635 | 11.73215 | 9.52123 | Promoter | 17 | 28075415 | 28076355 | 941 | 2 | ENSMUST00 | 0 | 4930526A20I | ENSMUSG00000085521.3 |
| chr8 | 11003695 | 11004270 | 6.53635 | 11.73215 | 9.52123 | Intron (ENSM | 8 | 11007850 | 11054536 | 46687 | 1 | ENSMUST00 | -3580 | 9530052E02I | ENSMUSG00000096938.2 |
| chr4 | 138555526 | 138555911 | 6.77105 | 11.72451 | 9.51382 | Distal Interge | 4 | 138571046 | 138581083 | 10038 | 2 | ENSMUST00 | 25172 | Vwa5b1 | ENSMUSG00000028753.7 |
| chr2 | 57123012 | 57123182 | 6.75797 | 11.70309 | 9.49245 | Promoter | 2 | 57106830 | 57124003 | 17174 | 2 | ENSMUST00 | 821 | Nr4a2 | ENSMUSG00000026826.8 |
| chr2 | 155514613 | 155515018 | 6.75797 | 11.70309 | 9.49245 | Promoter | 2 | 155490379 | 155514848 | 24470 | 2 | ENSMUST00 | 0 | Ggt7 | ENSMUSG00000027603.10 |
| chr14 | 74948471 | 74948751 | 6.74494 | 11.68175 | 9.47116 | Promoter | 14 | 74754674 | 74947877 | 193204 | 2 | ENSMUST00 | -595 | Lrch1 | ENSMUSG00000068015.6 |
| chr17 | 24642004 | 24642843 | 6.74494 | 11.68175 | 9.47116 | Exon (ENSMU | 17 | 24639282 | 24644957 | 5676 | 2 | ENSMUST00 | 2114 | Slc9a3r2 | ENSMUSG00000002504.9 |
| chr19 | 46598388 | 46598798 | 6.74494 | 11.68175 | 9.47116 | Promoter | 19 | 46599084 | 46657389 | 58306 | 1 | ENSMUST00 | -286 | Wbp1l | ENSMUSG00000047731.12 |
| chr5 | 107723080 | 107723974 | 6.74494 | 11.68175 | 9.47116 | Promoter | 5 | 107717224 | 107723961 | 6738 | 2 | ENSMUST00 | 0 | Gfi1 | ENSMUSG00000029275.12 |
| chr8 | 119344055 | 119344301 | 6.74494 | 11.68175 | 9.47116 | Promoter | 8 | 119344538 | 119348927 | 4390 | 1 | ENSMUST00 | -237 | Hsbp1 | ENSMUSG00000031839.6 |
| chr9 | 77806651 | 77806866 | 6.74494 | 11.68175 | 9.47116 | Distal Interge | 9 | 77754535 | 77794485 | 39951 | 1 | ENSMUST00 | 52117 | Gclc | ENSMUSG00000032350.8 |
| chr1 | 74950884 | 74952140 | 5.43315 | 11.66152 | 9.45144 | Promoter | 1 | 74945315 | 74951672 | 6358 | 2 | ENSMUST00 | 0 | Ihh | ENSMUSG00000006538.10 |
| chr15 | 79950550 | 79951176 | 5.43315 | 11.66152 | 9.45144 | Intron (ENSM | 15 | 79918148 | 79933899 | 15752 | 2 | ENSMUST00 | -16652 | Cbx7 | ENSMUSG00000053411.11 |
| chr9 | 30315008 | 30315316 | 6.26028 | 11.66074 | 9.45074 | Distal Interge | 9 | 30291868 | 30291974 | 107 | 2 | ENSMUST00 | -23035 | Gm26435 | ENSMUSG00000096018.1 |
| chr13 | 4942936 | 4943228 | 6.73196 | 11.66049 | 9.45054 | Distal Interge | 13 | 4771649 | 4836485 | 64837 | 1 | ENSMUST00 | 171288 | Gm5444 | ENSMUSG00000053499.3 |
| chr19 | 56544551 | 56545052 | 6.70615 | 11.61819 | 9.40834 | Exon (ENSMU | 19 | 56529169 | 56547155 | 17987 | 2 | ENSMUST00 | 2103 | Dclre1a | ENSMUSG00000025077.8 |
| chr1 | 181853153 | 181853626 | 6.70576 | 11.61755 | 9.40775 | Promoter | 1 | 181851574 | 181853189 | 1616 | 2 | ENSMUST00 | 0 | Gm5533 | ENSMUSG00000056699.7 |
| chr10 | 5594290 | 5594713 | 6.70576 | 11.61755 | 9.40775 | Promoter | 10 | 5593728 | 5606787 | 13060 | 1 | ENSMUST00 | 563 | Myct1 | ENSMUSG00000046916.8 |
| chr11 | 116657472 | 116658064 | 6.70576 | 11.61755 | 9.40775 | Promoter | 11 | 116657156 | 116660116 | 2961 | 1 | ENSMUST00 | 317 | Gm11744 | ENSMUSG00000075410.7 |
| chr12 | 71768675 | 71768873 | 6.70576 | 11.61755 | 9.40775 | Distal Interge | 12 | 71724652 | 71724954 | 303 | 1 | ENSMUST00 | 44024 | Gm22959 | ENSMUSG00000084463.1 |
| chr12 | 70148370 | 70149010 | 6.70576 | 11.61755 | 9.40775 | Distal Interge | 12 | 70154142 | 70183347 | 29206 | 1 | ENSMUST00 | -5132 | Abhd12b | ENSMUSG00000090121.5 |
| chr12 | 22745259 | 22745542 | 6.70576 | 11.61755 | 9.40775 | Distal Interge | 12 | 22622962 | 22623087 | 126 | 2 | ENSMUST00 | -122173 | Gm17391 | ENSMUSG00000095261.2 |
| chr13 | 58471905 | 58472172 | 6.70576 | 11.61755 | 9.40775 | Distal Interge | 13 | 58391142 | 58403343 | 12202 | 2 | ENSMUST00 | -68563 | Hnrnpk | ENSMUSG00000021546.12 |
| chr14 | 3333025 | 3333267 | 6.70576 | 11.61755 | 9.40775 | Promoter | 14 | 3332627 | 3354799 | 22173 | 1 | ENSMUST00 | 399 | Gm2956 | ENSMUSG00000096574.2 |
| chr15 | 75712207 | 75712608 | 6.70576 | 11.61755 | 9.40775 | 3' UTR | 15 | 75704394 | 75714225 | 9832 | 1 | ENSMUST00 | 7814 | Rhpn1 | ENSMUSG00000022580.7 |
| chr15 | 76003393 | 76003915 | 6.70576 | 11.61755 | 9.40775 | Exon (ENSMU | 15 | 76001093 | 76009498 | 8406 | 2 | ENSMUST00 | 5583 | Fam83h | ENSMUSG00000046761.7 |
| chr16 | 24919376 | 24919913 | 6.70576 | 11.61755 | 9.40775 | Intron (ENSM | 16 | 24884173 | 24884306 | 134 | 1 | ENSMUST00 | 35204 | Gm22672 | ENSMUSG00000077602.1 |
| chr17 | 8526891 | 8527339 | 6.70576 | 11.61755 | 9.40775 | Promoter | 17 | 8526801 | 8982496 | 455696 | 1 | ENSMUST00 | 91 | Pde10a | ENSMUSG00000023868.10 |
| chr17 | 87609545 | 87609920 | 6.70576 | 11.61755 | 9.40775 | Distal Interge | 17 | 87635979 | 87651106 | 15128 | 1 | ENSMUST00 | -26059 | Epcam | ENSMUSG00000045394.8 |
| chr18 | 61562155 | 61562649 | 6.70576 | 11.61755 | 9.40775 | Intron (ENSM | 18 | 61555689 | 61587531 | 31843 | 1 | ENSMUST00 | 6467 | Csnk1a1 | ENSMUSG00000024576.9 |
| chr2 | 4153085 | 4153731 | 6.70576 | 11.61755 | 9.40775 | Promoter | 2 | 4152872 | 4613942 | 461071 | 1 | ENSMUST00 | 214 | Frmd4a | ENSMUSG00000026657.11 |
| chr2 | 160360932 | 160361273 | 6.70576 | 11.61755 | 9.40775 | Downstream | 2 | 160363703 | 160367065 | 3363 | 2 | ENSMUST00 | 5792 | Mafb | ENSMUSG00000074622.4 |
| chr3 | 102723781 | 102724310 | 6.70576 | 11.61755 | 9.40775 | Intron (ENSM | 3 | 102734529 | 102772292 | 37764 | 1 | ENSMUST00 | -10219 | Tspan2 | ENSMUSG00000027858.8 |
| chr3 | 89238468 | 89239012 | 6.70576 | 11.61755 | 9.40775 | Promoter | 3 | 89234835 | 89239345 | 4511 | 2 | ENSMUST00 | 333 | Trim46 | ENSMUSG00000042766.12 |
| chr4 | 139598866 | 139599073 | 6.70576 | 11.61755 | 9.40775 | Intron (ENSM | 4 | 139603186 | 139662691 | 59506 | 1 | ENSMUST00 | -4113 | Gm21969 | ENSMUSG00000094439.1 |
| chr5 | 123674653 | 123675181 | 6.70576 | 11.61755 | 9.40775 | Intron (ENSM | 5 | 123617706 | 123666690 | 48985 | 2 | ENSMUST00 | -7964 | Clip1 | ENSMUSG00000049550.12 |
| chr5 | 120907010 | 120907651 | 6.70576 | 11.61755 | 9.40775 | Promoter | 5 | 120896256 | 120907521 | 11266 | 2 | ENSMUST00 | 0 | Oas1a | ENSMUSG00000052776.10 |
| chr6 | 37431077 | 37431538 | 6.70576 | 11.61755 | 9.40775 | Intron (ENSM | 6 | 37327255 | 37442146 | 114892 | 2 | ENSMUST00 | 10608 | Creb3l2 | ENSMUSG00000038648.5 |
| chr7 | 115217000 | 115217337 | 6.70576 | 11.61755 | 9.40775 | Distal Interge | 7 | 115172273 | 115172352 | 80 | 1 | ENSMUST00 | 44728 | Gm27684 | ENSMUSG00000099211.1 |
| chr9 | 62363939 | 62364164 | 6.70576 | 11.61755 | 9.40775 | Intron (ENSM | 9 | 62366938 | 62373676 | 6739 | 1 | ENSMUST00 | -2774 | Anp32a | ENSMUSG00000032249.9 |
| chr9 | 77426253 | 77426936 | 6.70576 | 11.61755 | 9.40775 | Distal Interge | 9 | 77431089 | 77442195 | 11107 | 2 | ENSMUST00 | 15259 | Lrrc1 | ENSMUSG00000032352.10 |
| chr9 | 121403240 | 121403706 | 6.70576 | 11.61755 | 9.40775 | Intron (ENSM | 9 | 121366958 | 121474916 | 107959 | 1 | ENSMUST00 | 36283 | Trak1 | ENSMUSG00000032536.9 |
| chrX | 136992689 | 136993612 | 6.70576 | 11.61755 | 9.40775 | Promoter | 20 | 136993155 | 136996378 | 3224 | 1 | ENSMUST00 | 0 | Zcchc18 | ENSMUSG00000031428.6 |
| chr11 | 119053225 | 119053510 | 5.56731 | 11.5554 | 9.34776 | Distal Interge | 11 | 119036305 | 119040969 | 4665 | 2 | ENSMUST00 | -12257 | Cbx8 | ENSMUSG00000025578.9 |
| chr11 | 103332926 | 103333287 | 5.56731 | 11.5554 | 9.34776 | Promoter | 11 | 103333229 | 103333395 | 167 | 2 | ENSMUST00 | 108 | Arhgap27 | ENSMUSG00000034255.12 |
| chr13 | 63247396 | 63248939 | 5.56731 | 11.5554 | 9.34776 | Intron (ENSM | 13 | 63282142 | 63326096 | 43955 | 1 | ENSMUST00 | -33203 | 2010111I01R | ENSMUSG00000021458.11 |
| chr17 | 29711613 | 29712086 | 5.56731 | 11.5554 | 9.34776 | Intron (ENSM | 17 | 29699999 | 29716963 | 16965 | 2 | ENSMUST00 | 4877 | Ccdc167 | ENSMUSG00000024018.12 |
| chr17 | 9763931 | 9764518 | 5.56731 | 11.5554 | 9.34776 | Distal Interge | 17 | 9666497 | 9669704 | 3208 | 2 | ENSMUST00 | -94228 | Pabpc6 | ENSMUSG00000046173.2 |
| chr2 | 33616045 | 33617221 | 5.56731 | 11.5554 | 9.34776 | Intron (ENSM | 2 | 33567156 | 33632283 | 65128 | 2 | ENSMUST00 | 15062 | Lmx1b | ENSMUSG00000038765.8 |
| chr3 | 5750911 | 5751085 | 5.56731 | 11.5554 | 9.34776 | Distal Interge | 3 | 5881041 | 5881150 | 110 | 1 | ENSMUST00 | -129956 | Gm25352 | ENSMUSG00000096384.1 |
| chr4 | 46563887 | 46564219 | 5.56731 | 11.5554 | 9.34776 | Intron (ENSM | 4 | 46536937 | 46566440 | 29504 | 2 | ENSMUST00 | 2221 | Coro2a | ENSMUSG00000028337.9 |
| chr7 | 131555417 | 131556092 | 5.56731 | 11.5554 | 9.34776 | 3' UTR | 7 | 131554061 | 131556589 | 2529 | 1 | ENSMUST00 | 1357 | Hmx2 | ENSMUSG00000050100.8 |
| chr8 | 92359483 | 92360639 | 5.56731 | 11.5554 | 9.34776 | Exon (ENSMU | 8 | 92357796 | 92361456 | 3661 | 1 | ENSMUST00 | 1688 | Irx5 | ENSMUSG00000031737.10 |
| chr8 | 22506165 | 22506808 | 5.56731 | 11.5554 | 9.34776 | Intron (ENSM | 8 | 22480291 | 22483330 | 3040 | 1 | ENSMUST00 | 25875 | Gm17491 | ENSMUSG00000097042.1 |
| chr13 | 113613347 | 113613736 | 6.42339 | 11.54283 | 9.33543 | Intron (ENSM | 13 | 113592180 | 113618564 | 26385 | 2 | ENSMUST00 | 4828 | Snx18 | ENSMUSG00000042364.10 |
| chr16 | 78427557 | 78428068 | 6.42339 | 11.54283 | 9.33543 | Distal Interge | 16 | 78423730 | 78423859 | 130 | 2 | ENSMUST00 | -3699 | Gm25038 | ENSMUSG00000064862.1 |
| chr2 | 152828618 | 152829229 | 6.42339 | 11.54283 | 9.33543 | Promoter | 2 | 152781411 | 152828535 | 47125 | 2 | ENSMUST00 | -84 | Bcl2l1 | ENSMUSG00000007659.13 |
| chr9 | 54676618 | 54676859 | 6.42339 | 11.54283 | 9.33543 | Distal Interge | 9 | 54604877 | 54661870 | 56994 | 2 | ENSMUST00 | -14749 | Acsbg1 | ENSMUSG00000032281.6 |
| chr1 | 144006606 | 144006786 | 6.89298 | 11.53747 | 9.33025 | Distal Interge | 1 | 143999338 | 144004161 | 4824 | 2 | ENSMUST00 | -2446 | Rgs2 | ENSMUSG00000026360.4 |
| chr1 | 170908396 | 170908729 | 6.89298 | 11.53747 | 9.33025 | Exon (ENSMU | 1 | 170907273 | 170912941 | 5669 | 2 | ENSMUST00 | 4212 | Fcrlb | ENSMUSG00000070524.1 |
| chr1 | 133492886 | 133493350 | 6.89298 | 11.53747 | 9.33025 | Distal Interge | 1 | 133529479 | 133530270 | 792 | 1 | ENSMUST00 | -36129 | RP23-78M21 | ENSMUSG00000101004.1 |
| chr10 | 109011396 | 109011737 | 6.89298 | 11.53747 | 9.33025 | Promoter | 10 | 108497652 | 109010982 | 513331 | 2 | ENSMUST00 | -415 | Syt1 | ENSMUSG00000035864.9 |
| chr11 | 5555736 | 5555907 | 6.89298 | 11.53747 | 9.33025 | Distal Interge | 11 | 5560606 | 5560735 | 130 | 2 | ENSMUST00 | 4828 | Gm22471 | ENSMUSG00000077304.1 |
| chr12 | 106479613 | 106479909 | 6.89298 | 11.53747 | 9.33025 | Distal Interge | 12 | 106459150 | 106459495 | 346 | 1 | ENSMUST00 | 20464 | Gm3191 | ENSMUSG00000091762.1 |
| chr12 | 117153914 | 117154219 | 6.89298 | 11.53747 | 9.33025 | Intron (ENSM | 12 | 117151090 | 117151651 | 562 | 2 | ENSMUST00 | -2264 | RP23-393F1( | ENSMUSG00000099907.1 |
| chr14 | 84442504 | 84442808 | 6.89298 | 11.53747 | 9.33025 | Promoter | 14 | 84443563 | 84537060 | 93498 | 1 | ENSMUST00 | -755 | Pcdh17 | ENSMUSG00000035566.6 |
| chr14 | 67613262 | 67613596 | 6.89298 | 11.53747 | 9.33025 | Distal Interge | 14 | 67566850 | 67566981 | 132 | 1 | ENSMUST00 | 46413 | Gm24981 | ENSMUSG00000077647.1 |
| chr15 | 82366154 | 82366319 | 6.89298 | 11.53747 | 9.33025 | Distal Interge | 15 | 82350139 | 82354291 | 4153 | 2 | ENSMUST00 | -11864 | Ndufa6 | ENSMUSG00000022450.5 |
| chr15 | 98002924 | 98003419 | 6.89298 | 11.53747 | 9.33025 | Intron (ENSM | 15 | 97998371 | 98001887 | 3517 | 2 | ENSMUST00 | -1038 | Col2a1 | ENSMUSG00000022483.11 |
| chr15 | 74563893 | 74564254 | 6.89298 | 11.53747 | 9.33025 | Promoter | 15 | 74563901 | 74572861 | 8961 | 1 | ENSMUST00 | 0 | Bai1 | ENSMUSG00000034730.11 |
| chr15 | 82963665 | 82963840 | 6.89298 | 11.53747 | 9.33025 | Distal Interge | 15 | 82959218 | 82959296 | 79 | 2 | ENSMUST00 | -4370 | Mir3080 | ENSMUSG00000092751.1 |
| chr19 | 29708001 | 29708455 | 6.89298 | 11.53747 | 9.33025 | 3' UTR | 19 | 29697921 | 29708448 | 10528 | 1 | ENSMUST00 | 10081 | Mlana | ENSMUSG00000024806.3 |
| chr19 | 5757068 | 5757240 | 6.89298 | 11.53747 | 9.33025 | Intron (ENSM | 19 | 5758427 | 5771401 | 12975 | 2 | ENSMUST00 | 14161 | Scyl1 | ENSMUSG00000024941.8 |
| chr19 | 55556244 | 55556509 | 6.89298 | 11.53747 | 9.33025 | Intron (ENSM | 19 | 55640589 | 55702204 | 61616 | 2 | ENSMUST00 | 145695 | 4930552P12I | ENSMUSG00000097632.1 |
| chr19 | 53421672 | 53421945 | 6.89298 | 11.53747 | 9.33025 | Distal Interge | 19 | 53444014 | 53447314 | 3301 | 2 | ENSMUST00 | 25369 | 5830416P10I | ENSMUSG00000097636.2 |
| chr2 | 4560609 | 4561536 | 6.89298 | 11.53747 | 9.33025 | Promoter | 2 | 4559775 | 4588179 | 28405 | 1 | ENSMUST00 | 835 | Frmd4a | ENSMUSG00000026657.11 |
| chr2 | 175605757 | 175606329 | 6.89298 | 11.53747 | 9.33025 | Distal Interge | 2 | 175643978 | 175651401 | 7424 | 1 | ENSMUST00 | -37649 | Gm11007 | ENSMUSG00000094475.2 |
| chr4 | 124894500 | 124895070 | 6.89298 | 11.53747 | 9.33025 | Promoter | 4 | 124890888 | 124894782 | 3895 | 2 | ENSMUST00 | 0 | 1700125G02 | ENSMUSG00000085263.1 |
| chr5 | 108677456 | 108677876 | 6.89298 | 11.53747 | 9.33025 | Intron (ENSM | 5 | 108669878 | 108675569 | 5692 | 2 | ENSMUST00 | -1888 | Slc26a1 | ENSMUSG00000046959.11 |
| chr5 | 99729354 | 99729761 | 6.89298 | 11.53747 | 9.33025 | Promoter | 5 | 99556527 | 99729065 | 172539 | 2 | ENSMUST00 | -290 | A930011G23 | ENSMUSG00000089809.3 |
| chr6 | 83038461 | 83038792 | 6.89298 | 11.53747 | 9.33025 | Exon (ENSMU | 6 | 83034173 | 83052562 | 18390 | 1 | ENSMUST00 | 4289 | Loxl3 | ENSMUSG00000000693.5 |
| chr6 | 129181737 | 129182215 | 6.89298 | 11.53747 | 9.33025 | Intron (ENSM | 6 | 129180615 | 129186534 | 5920 | 1 | ENSMUST00 | 1123 | Clec2d | ENSMUSG00000030157.5 |
| chr7 | 24462526 | 24462963 | 6.89298 | 11.53747 | 9.33025 | Promoter | 7 | 24462500 | 24475873 | 13374 | 1 | ENSMUST00 | 27 | Plaur | ENSMUSG00000046223.9 |
| chr7 | 19463932 | 19464710 | 6.89298 | 11.53747 | 9.33025 | Distal Interge | 7 | 19460797 | 19460935 | 139 | 2 | ENSMUST00 | -2998 | Gm23099 | ENSMUSG00000065388.1 |
| chr8 | 94695535 | 94696317 | 6.89298 | 11.53747 | 9.33025 | Promoter | 8 | 94674895 | 94696242 | 21348 | 2 | ENSMUST00 | 0 | Pllp | ENSMUSG00000031775.4 |
| chr8 | 71507465 | 71508004 | 6.89298 | 11.53747 | 9.33025 | Exon (ENSMU | 8 | 71497754 | 71511769 | 14016 | 2 | ENSMUST00 | 3765 | Plvap | ENSMUSG00000034845.5 |
| chr8 | 106507662 | 106508333 | 6.89298 | 11.53747 | 9.33025 | Distal Interge | 8 | 106510913 | 106556908 | 45996 | 1 | ENSMUST00 | -2580 | Cdh3 | ENSMUSG00000061048.7 |
| chr8 | 47763766 | 47764916 | 6.89298 | 11.53747 | 9.33025 | Distal Interge | 8 | 47719246 | 47719369 | 124 | 1 | ENSMUST00 | 44521 | n-R5s96 | ENSMUSG00000064508.1 |
| chr9 | 20746096 | 20746285 | 6.89298 | 11.53747 | 9.33025 | Distal Interge | 9 | 20667986 | 20728219 | 60234 | 2 | ENSMUST00 | -17878 | Olfm2 | ENSMUSG00000032172.7 |
| chr9 | 65123804 | 65124167 | 6.89298 | 11.53747 | 9.33025 | Exon (ENSMU | 9 | 65141154 | 65185857 | 44704 | 1 | ENSMUST00 | -16987 | Igdcc3 | ENSMUSG00000032394.5 |
| chrX | 74296673 | 74296866 | 6.89298 | 11.53747 | 9.33025 | Promoter | 20 | 74297097 | 74304721 | 7625 | 1 | ENSMUST00 | -231 | Atp6ap1 | ENSMUSG00000019087.8 |
| chrX | 6778348 | 6778521 | 6.89298 | 11.53747 | 9.33025 | Promoter | 20 | 6779306 | 6948362 | 169057 | 1 | ENSMUST00 | -785 | Dgkk | ENSMUSG00000062393.8 |
| chrX | 157612995 | 157613255 | 6.89298 | 11.53747 | 9.33025 | Promoter | 20 | 157612689 | 157613121 | 433 | 2 | ENSMUST00 | 0 | Gm15173 | ENSMUSG00000081222.1 |
| chrX | 94990452 | 94990862 | 6.89298 | 11.53747 | 9.33025 | Distal Interge | 20 | 94997439 | 94998683 | 1245 | 2 | ENSMUST00 | 7821 | Gm371 | ENSMUSG00000087522.3 |
| chr2 | 168766794 | 168767202 | 6.88058 | 11.51738 | 9.31251 | Promoter | 2 | 168748332 | 168767029 | 18698 | 2 | ENSMUST00 | 0 | Sall4 | ENSMUSG00000027547.12 |
| chr2 | 29967185 | 29968076 | 6.62989 | 11.4931 | 9.28844 | Intron (ENSM | 2 | 29965717 | 30031451 | 65735 | 1 | ENSMUST00 | 1469 | Sptan1 | ENSMUSG00000057738.8 |
| chr10 | 18329232 | 18330081 | 6.38327 | 11.47542 | 9.27082 | Distal Interge | 10 | 18390322 | 18524289 | 133968 | 1 | ENSMUST00 | -60241 | Nhsl1 | ENSMUSG00000039835.10 |
| chr10 | 14086008 | 14086489 | 6.38327 | 11.47542 | 9.27082 | Intron (ENSM | 10 | 14149084 | 14154446 | 5363 | 2 | ENSMUST00 | 67957 | RP23-202B1: | ENSMUSG00000101014.1 |
| chr12 | 105967577 | 105968251 | 6.38327 | 11.47542 | 9.27082 | Distal Interge | 12 | 106010263 | 106077410 | 67148 | 1 | ENSMUST00 | -42012 | Vrk1 | ENSMUSG00000021115.9 |
| chr12 | 110278510 | 110279231 | 6.38327 | 11.47542 | 9.27082 | Promoter | 12 | 110279068 | 110281097 | 2030 | 1 | ENSMUST00 | 0 | Dio3 | ENSMUSG00000075707.5 |
| chr12 | 12904538 | 12904996 | 6.38327 | 11.47542 | 9.27082 | Intron (ENSM | 12 | 12902720 | 12907251 | 4532 | 1 | ENSMUST00 | 1819 | 4930519A11I | ENSMUSG00000097627.1 |
| chr13 | 51594839 | 51595287 | 6.38327 | 11.47542 | 9.27082 | Distal Interge | 13 | 51431041 | 51567084 | 136044 | 2 | ENSMUST00 | -27756 | Shc3 | ENSMUSG00000021448.6 |
| chr14 | 16515883 | 16516149 | 6.38327 | 11.47542 | 9.27082 | Intron (ENSM | 14 | 16430842 | 16575472 | 144631 | 2 | ENSMUST00 | 59323 | Rarb | ENSMUSG00000017491.8 |
| chr14 | 4110445 | 4111239 | 5.72276 | 11.47427 | 9.27082 | Promoter | 14 | 4110526 | 4132784 | 22259 | 1 | ENSMUST00 | 0 | Gm8108 | ENSMUSG00000091563.3 |
| chr14 | 6604570 | 6605196 | 6.38327 | 11.47542 | 9.27082 | Distal Interge | 14 | 6583959 | 6590613 | 6655 | 2 | ENSMUST00 | -13958 | Gm3629 | ENSMUSG00000093898.2 |
| chr15 | 52478132 | 52478301 | 6.38327 | 11.47542 | 9.27082 | Promoter | 15 | 52477614 | 52478228 | 615 | 2 | ENSMUST00 | 0 | Gm10020 | ENSMUSG00000057262.1 |
| chr16 | 52449499 | 52449899 | 6.38327 | 11.47542 | 9.27082 | Intron (ENSM | 16 | 52251467 | 52452465 | 200999 | 2 | ENSMUST00 | 2566 | Alcam | ENSMUSG00000022636.8 |
| chr16 | 58512380 | 58514683 | 5.72276 | 11.47427 | 9.27082 | Intron (ENSM | 16 | 58502861 | 58507898 | 5038 | 2 | ENSMUST00 | -4483 | St3gal6 | ENSMUSG00000022747.11 |
| chr18 | 37761731 | 37762052 | 5.72276 | 11.47427 | 9.27082 | Intron (ENSM | 18 | 37765580 | 37841873 | 76294 | 1 | ENSMUST00 | -3528 | Pcdhga12 | ENSMUSG00000023036.9 |
| chr2 | 165055068 | 165056091 | 6.38327 | 11.47542 | 9.27082 | Promoter | 2 | 165055627 | 165062723 | 7097 | 1 | ENSMUST00 | 0 | Cd40 | ENSMUSG00000017652.11 |
| chr2 | 33629934 | 33630384 | 6.38327 | 11.47542 | 9.27082 | Intron (ENSM | 2 | 33567156 | 33632283 | 65128 | 2 | ENSMUST00 | 1899 | Lmx1b | ENSMUSG00000038765.8 |
| chr2 | 32727689 | 32728790 | 6.38327 | 11.47542 | 9.27082 | Promoter | 2 | 32727706 | 32744844 | 17139 | 1 | ENSMUST00 | 0 | Sh2d3c | ENSMUSG00000059013.7 |
| chr2 | 77863461 | 77864126 | 6.38327 | 11.47542 | 9.27082 | Distal Interge | 2 | 77879753 | 77881467 | 1715 | 2 | ENSMUST00 | 17341 | Gm13727 | ENSMUSG00000087353.1 |
| chr3 | 90509676 | 90509920 | 6.38327 | 11.47542 | 9.27082 | Promoter | 3 | 90498965 | 90509498 | 10534 | 2 | ENSMUST00 | -179 | Chtop | ENSMUSG00000001017.10 |
| chr3 | 107931624 | 107931912 | 5.72276 | 11.47427 | 9.27082 | Promoter | 3 | 107926334 | 107931817 | 5484 | 2 | ENSMUST00 | 0 | Gstm7 | ENSMUSG00000004035.7 |
| chr3 | 122635432 | 122635639 | 6.38327 | 11.47542 | 9.27082 | Distal Interge | 3 | 122538719 | 122619667 | 80949 | 2 | ENSMUST00 | -15766 | Fnbp1l | ENSMUSG00000039735.10 |
| chr4 | 107923760 | 107923964 | 6.38327 | 11.47542 | 9.27082 | Promoter | 4 | 107903981 | 107923610 | 19630 | 2 | ENSMUST00 | -151 | Cpt2 | ENSMUSG00000028607.11 |
| chr4 | 130047800 | 130048053 | 6.38327 | 11.47542 | 9.27082 | Promoter | 4 | 130047840 | 130099277 | 51438 | 1 | ENSMUST00 | 0 | Col16a1 | ENSMUSG00000040690.10 |
| chr5 | 115374070 | 115374310 | 6.38327 | 11.47542 | 9.27082 | Intron (ENSM | 5 | 115358700 | 115360123 | 1424 | 2 | ENSMUST00 | -13948 | 4930401G09 | ENSMUSG00000086352.1 |
| chr6 | 39781145 | 39781920 | 6.38327 | 11.47542 | 9.27082 | Distal Interge | 6 | 39802192 | 39810885 | 8694 | 2 | ENSMUST00 | 28965 | Mrps33 | ENSMUSG00000029918.8 |
| chr6 | 38926178 | 38926684 | 6.38327 | 11.47542 | 9.27082 | Intron (ENSM | 6 | 38919024 | 39084585 | 165562 | 1 | ENSMUST00 | 7155 | Tbxas1 | ENSMUSG00000029925.8 |
| chr6 | 64649780 | 64650109 | 6.38327 | 11.47542 | 9.27082 | Intron (ENSM | 6 | 64729146 | 64731233 | 2088 | 1 | ENSMUST00 | -79037 | Atoh1 | ENSMUSG00000073043.4 |
| chr7 | 100495153 | 100495958 | 6.38327 | 11.47542 | 9.27082 | Promoter | 7 | 100496079 | 100498256 | 2178 | 1 | ENSMUST00 | -121 | Ucp2 | ENSMUSG00000033685.7 |
| chr7 | 79275322 | 79275704 | 6.38327 | 11.47542 | 9.27082 | Intron (ENSM | 7 | 79273255 | 79345371 | 72117 | 1 | ENSMUST00 | 2068 | Abhd2 | ENSMUSG00000039202.7 |
| chr7 | 97453224 | 97454047 | 6.38327 | 11.47542 | 9.27082 | Promoter | 7 | 97453222 | 97457732 | 4511 | 1 | ENSMUST00 | 3 | Kctd14 | ENSMUSG00000051727.9 |
| chr7 | 19465259 | 19465442 | 6.38327 | 11.47542 | 9.27082 | Distal Interge | 7 | 19460797 | 19460935 | 139 | 2 | ENSMUST00 | -4325 | Gm23099 | ENSMUSG00000065388.1 |
| chr8 | 104101625 | 104102356 | 6.38327 | 11.47542 | 9.27082 | Promoter | 8 | 104101625 | 104144502 | 42878 | 1 | ENSMUST00 | 1 | Cdh5 | ENSMUSG00000031871.8 |

| | | | | | | | | | | | | | | | |
|---|---|---|---|---|---|---|---|---|---|---|---|---|---|---|---|
| chr8 | 58912468 | 58912925 | 6.38327 | 11.47542 | 9.27082 | Promoter | 8 | 58912257 | 58913196 | 940 | 1 | ENSMUST00 | 212 | BC030500 | ENSMUSG00000049946.7 |
| chr9 | 51101902 | 51102154 | 6.38327 | 11.47542 | 9.27082 | Promoter | 9 | 51081313 | 51102078 | 20766 | 2 | ENSMUST00 | 0 | 4833427G06 | ENSMUSG00000032057.8 |
| chr9 | 87144237 | 87144827 | 6.38327 | 11.47542 | 9.27082 | Promoter | 9 | 87144306 | 87184045 | 39740 | 1 | ENSMUST00 | 0 | Mrap2 | ENSMUSG00000042761.9 |
| chr9 | 104100927 | 104101613 | 6.38327 | 11.47542 | 9.27082 | Promoter | 9 | 104084157 | 104102560 | 18404 | 2 | ENSMUST00 | 947 | Ackr4 | ENSMUSG00000079355.3 |
| chr1 | 119641600 | 119643017 | 6.61735 | 11.47251 | 9.2704 | 5' UTR | 1 | 119595074 | 119648617 | 53544 | 2 | ENSMUST00 | 5600 | Epb4.1l5 | ENSMUSG00000026383.9 |
| chr13 | 51408500 | 51409140 | 6.61735 | 11.47251 | 9.2704 | Promoter | 13 | 51408618 | 51422797 | 14180 | 1 | ENSMUST00 | 0 | S1pr3 | ENSMUSG00000067586.3 |
| chr16 | 62940837 | 62941043 | 6.61735 | 11.47251 | 9.2704 | Distal Interge | 16 | 62921467 | 62928390 | 6924 | 1 | ENSMUST00 | 19371 | Pros1 | ENSMUSG00000022912.8 |
| chr2 | 17741662 | 17742280 | 6.61735 | 11.47251 | 9.2704 | Distal Interge | 2 | 17727386 | 17731464 | 4079 | 2 | ENSMUST00 | -10199 | Nebl | ENSMUSG00000053702.11 |
| chr4 | 148075215 | 148075563 | 6.61735 | 11.47251 | 9.2704 | Promoter | 4 | 148071607 | 148075420 | 3814 | 2 | ENSMUST00 | 0 | Gm13201 | ENSMUSG00000078494.5 |
| chr4 | 135312682 | 135313494 | 6.61735 | 11.47251 | 9.2704 | Distal Interge | 4 | 135307781 | 135307970 | 190 | 2 | ENSMUST00 | -4713 | Gm12982 | ENSMUSG00000082833.1 |
| chr4 | 130047399 | 130047687 | 6.61735 | 11.47251 | 9.2704 | Promoter | 4 | 130024256 | 130047774 | 23519 | 2 | ENSMUST00 | 87 | Gm12963 | ENSMUSG00000085517.2 |
| chr5 | 114379455 | 114379681 | 6.61735 | 11.47251 | 9.2704 | Promoter | 5 | 114365622 | 114380483 | 14862 | 2 | ENSMUST00 | 802 | Kctd10 | ENSMUSG00000001098.10 |
| chr5 | 125465989 | 125466160 | 6.61735 | 11.47251 | 9.2704 | Distal Interge | 5 | 125475814 | 125517412 | 41599 | 1 | ENSMUST00 | -9654 | Aacs | ENSMUSG00000029482.3 |
| chr6 | 17636619 | 17636840 | 6.61735 | 11.47251 | 9.2704 | Promoter | 6 | 17636983 | 17666972 | 29990 | 1 | ENSMUST00 | -143 | Capza2 | ENSMUSG00000015733.8 |
| chr7 | 121734302 | 121734907 | 6.61735 | 11.47251 | 9.2704 | Promoter | 7 | 121734479 | 121768475 | 33997 | 1 | ENSMUST00 | 0 | Scnn1g | ENSMUSG00000000216.8 |
| chr9 | 110880828 | 110881053 | 6.61735 | 11.47251 | 9.2704 | Promoter | 9 | 110880337 | 110900527 | 20191 | 1 | ENSMUST00 | 492 | Als2cl | ENSMUSG00000044037.10 |
| chr17 | 49485083 | 49485590 | 6.85183 | 11.47078 | 9.26929 | Exon (ENSMU | 17 | 49452284 | 49455435 | 3152 | 1 | ENSMUST00 | 32800 | Mocs1 | ENSMUSG00000064120.8 |
| chr7 | 15289003 | 15289253 | 4.84682 | 11.44916 | 9.24797 | Promoter | 7 | 15288251 | 15289190 | 940 | 2 | ENSMUST00 | 0 | Gm8086 | ENSMUSG00000098404.1 |
| chr1 | 92850025 | 92850464 | 6.83755 | 11.44763 | 9.24644 | Promoter | 1 | 92850378 | 92850443 | 66 | 1 | ENSMUST00 | 0 | Mir149 | ENSMUSG00000065470.1 |
| chr19 | 16050137 | 16050694 | 6.83755 | 11.44763 | 9.24644 | Distal Interge | 19 | 15985090 | 16010912 | 25823 | 1 | ENSMUST00 | 65048 | C130060C02I | ENSMUSG00000097557.1 |
| chr8 | 87943684 | 87943944 | 6.83755 | 11.44763 | 9.24644 | Promoter | 8 | 87783316 | 87944052 | 160737 | 2 | ENSMUST00 | 108 | Zfp423 | ENSMUSG00000045333.10 |
| chr1 | 135699188 | 135699580 | 5.905 | 11.42563 | 9.22457 | Distal Interge | 1 | 135606447 | 135688105 | 81659 | 2 | ENSMUST00 | -11084 | Nav1 | ENSMUSG00000090399.1 |
| chr10 | 45000418 | 45001238 | 5.905 | 11.42563 | 9.22457 | Distal Interge | 10 | 45006110 | 45007117 | 1008 | 1 | ENSMUST00 | -4872 | Gm4795 | ENSMUSG00000095168.1 |
| chr13 | 42103197 | 42103826 | 5.905 | 11.42563 | 9.22457 | Intron (ENSM | 13 | 42052021 | 42185026 | 133006 | 1 | ENSMUST00 | 51177 | Hivep1 | ENSMUSG00000021366.7 |
| chr18 | 80295753 | 80296313 | 5.905 | 11.42563 | 9.22457 | Exon (ENSMU | 18 | 80323335 | 80326605 | 3271 | 1 | ENSMUST00 | -27022 | Gm26676 | ENSMUSG00000097593.2 |
| chr2 | 166092157 | 166093106 | 5.905 | 11.42563 | 9.22457 | Intron (ENSM | 2 | 166074815 | 166078119 | 3305 | 2 | ENSMUST00 | -14039 | Sulf2 | ENSMUSG00000006800.8 |
| chr4 | 43453179 | 43454118 | 5.905 | 11.42563 | 9.22457 | Promoter | 4 | 43452321 | 43454323 | 2003 | 2 | ENSMUST00 | 205 | Cd72 | ENSMUSG00000028459.6 |
| chr6 | 31528147 | 31530472 | 5.905 | 11.42563 | 9.22457 | Exon (ENSMU | 6 | 31522992 | 31524311 | 1320 | 2 | ENSMUST00 | -3837 | Podxl | ENSMUSG00000025608.9 |
| chr7 | 44405386 | 44406132 | 5.905 | 11.42563 | 9.22457 | Distal Interge | 7 | 44378887 | 44385820 | 6934 | 2 | ENSMUST00 | -19567 | 1700008O03 | ENSMUSG00000008028.10 |
| chr8 | 27177237 | 27177571 | 5.905 | 11.42563 | 9.22457 | Distal Interge | 8 | 27138773 | 27174646 | 35874 | 2 | ENSMUST00 | -2592 | Rab11fip1 | ENSMUSG00000031488.8 |
| chr9 | 24774046 | 24774634 | 5.905 | 11.42563 | 9.22457 | Promoter | 9 | 24720812 | 24774303 | 53492 | 2 | ENSMUST00 | 0 | Tbx20 | ENSMUSG00000031965.9 |
| chr10 | 111247732 | 111249024 | 6.12159 | 11.42045 | 9.22029 | Intron (ENSM | 10 | 111298679 | 111301736 | 3058 | 1 | ENSMUST00 | -49655 | Bbs10 | ENSMUSG00000035759.3 |
| chr11 | 60933399 | 60934322 | 6.12159 | 11.42045 | 9.22029 | Intron (ENSM | 11 | 60932066 | 60952794 | 20729 | 1 | ENSMUST00 | 1334 | Map2k3 | ENSMUSG00000018932.4 |
| chr11 | 114065058 | 114065456 | 6.12159 | 11.42045 | 9.22029 | Promoter | 11 | 113848562 | 114065975 | 217414 | 2 | ENSMUST00 | 519 | Sdk2 | ENSMUSG00000041592.11 |
| chr13 | 53309173 | 53310170 | 6.12159 | 11.42045 | 9.22029 | Distal Interge | 13 | 53109317 | 53286124 | 176808 | 2 | ENSMUST00 | -23050 | Ror2 | ENSMUSG00000021464.9 |
| chr13 | 109775989 | 109776702 | 6.12159 | 11.42045 | 9.22029 | Intron (ENSM | 13 | 109757221 | 109761612 | 4392 | 1 | ENSMUST00 | 18769 | Pde4d | ENSMUSG00000021699.12 |
| chr15 | 27923374 | 27923933 | 6.12159 | 11.42045 | 9.22029 | Intron (ENSM | 15 | 27730652 | 28025848 | 295197 | 2 | ENSMUST00 | 101915 | Trio | ENSMUSG00000022263.9 |
| chr15 | 59201895 | 59202903 | 6.12159 | 11.42045 | 9.22029 | Distal Interge | 15 | 59210908 | 59211720 | 813 | 1 | ENSMUST00 | -8005 | Gm5045 | ENSMUSG00000025922.8 |
| chr15 | 89322680 | 89322966 | 6.12159 | 11.42045 | 9.22029 | Promoter | 15 | 89322972 | 89324728 | 1757 | 1 | ENSMUST00 | -6 | Adm2 | ENSMUSG00000054136.5 |
| chr17 | 45723872 | 45724432 | 6.12159 | 11.42045 | 9.22029 | Distal Interge | 17 | 45733864 | 45733844 | 981 | 2 | ENSMUST00 | 9412 | 1600014C23I | ENSMUSG00000094690.1 |
| chr2 | 70471950 | 70472524 | 6.12159 | 11.42045 | 9.22029 | Distal Interge | 2 | 70474923 | 70477729 | 2807 | 1 | ENSMUST00 | -2399 | Sp5 | ENSMUSG00000075304.2 |
| chr3 | 57470852 | 57471327 | 6.12159 | 11.42045 | 9.22029 | Intron (ENSM | 3 | 57425314 | 57441677 | 16364 | 1 | ENSMUST00 | 45539 | Tm4sf4 | ENSMUSG00000027801.7 |
| chr4 | 96591065 | 96591825 | 6.12159 | 11.42045 | 9.22029 | Promoter | 4 | 96568429 | 96591578 | 23150 | 2 | ENSMUST00 | 0 | Cyp2j9 | ENSMUSG00000015224.10 |
| chr4 | 134253342 | 134253866 | 6.12159 | 11.42045 | 9.22029 | Promoter | 4 | 134251110 | 134254106 | 2997 | 2 | ENSMUST00 | 240 | Grrp1 | ENSMUSG00000050105.5 |
| chr5 | 103828403 | 103829000 | 6.12159 | 11.42045 | 9.22029 | Exon (ENSMU | 5 | 103784359 | 103811595 | 27237 | 1 | ENSMUST00 | 44045 | Aff1 | ENSMUSG00000029313.13 |
| chr5 | 142961826 | 142962438 | 6.12159 | 11.42045 | 9.22029 | Promoter | 5 | 142962953 | 142970552 | 7600 | 1 | ENSMUST00 | -515 | Fscn1 | ENSMUSG00000029581.9 |
| chr6 | 30301031 | 30302224 | 6.12159 | 11.42045 | 9.22029 | Intron (ENSM | 6 | 30214861 | 30304522 | 89662 | 2 | ENSMUST00 | 2298 | Ube2h | ENSMUSG00000039159.11 |
| chr6 | 77979459 | 77979955 | 6.12159 | 11.42045 | 9.22029 | Promoter | 6 | 76881637 | 77979699 | 1098063 | 2 | ENSMUST00 | 0 | Ctnna2 | ENSMUSG00000063063.7 |
| chr7 | 144838873 | 144839414 | 6.12159 | 11.42045 | 9.22029 | Promoter | 7 | 144838595 | 144844436 | 5842 | 1 | ENSMUST00 | 279 | Fgf3 | ENSMUSG00000031074.9 |
| chr7 | 30626307 | 30626515 | 6.12159 | 11.42045 | 9.22029 | Promoter | 7 | 30616974 | 30626151 | 9178 | 2 | ENSMUST00 | -157 | Cox6b1 | ENSMUSG00000036751.6 |
| chr7 | 29965271 | 29965643 | 6.12159 | 11.42045 | 9.22029 | Promoter | 7 | 29965218 | 29965565 | 348 | 2 | ENSMUST00 | 0 | Gm10169 | ENSMUSG00000083669.5 |
| chr12 | 73537896 | 73538124 | 6.80915 | 11.40161 | 9.20291 | Distal Interge | 12 | 73543114 | 73546395 | 3282 | 2 | ENSMUST00 | 8271 | Tmem30b | ENSMUSG00000034435.5 |
| chr2 | 130580555 | 130580873 | 6.80915 | 11.40161 | 9.20291 | 3' UTR | 2 | 130580620 | 130582554 | 1935 | 2 | ENSMUST00 | 1681 | Avp | ENSMUSG00000037727.6 |
| chr10 | 77664119 | 77664484 | 6.55535 | 11.37063 | 9.17219 | Distal Interge | 10 | 77641224 | 77641307 | 84 | 1 | ENSMUST00 | 22896 | Mir1930 | ENSMUSG00000089424.1 |
| chr12 | 3307453 | 3307673 | 6.31427 | 11.35921 | 9.16078 | Intron (ENSM | 12 | 3247430 | 3309969 | 62540 | 2 | ENSMUST00 | 2296 | Rab10 | ENSMUSG00000020671.8 |
| chr13 | 63901824 | 63902011 | 6.31427 | 11.35921 | 9.16078 | Distal Interge | 13 | 63872455 | 63900302 | 27848 | 1 | ENSMUST00 | 29370 | Ercc6l2 | ENSMUSG00000021470.12 |
| chr13 | 25057376 | 25058263 | 6.31427 | 11.35921 | 9.16078 | Intron (ENSM | 13 | 25056253 | 25210706 | 154454 | 1 | ENSMUST00 | 1124 | Dcdc2a | ENSMUSG00000035910.10 |
| chr5 | 32139263 | 32139995 | 6.781 | 11.35596 | 9.15773 | Intron (ENSM | 5 | 32136472 | 32157831 | 21360 | 1 | ENSMUST00 | 2792 | Fosl2 | ENSMUSG00000029135.9 |
| chr2 | 158179002 | 158179760 | 6.767 | 11.33328 | 9.13518 | Downstream | 2 | 158192008 | 158195749 | 3742 | 1 | ENSMUST00 | -12248 | 1700060C20I | ENSMUSG00000078957.3 |
| chr16 | 34657574 | 34658006 | 6.75307 | 11.31069 | 9.11275 | Intron (ENSM | 16 | 34651188 | 34678618 | 27431 | 1 | ENSMUST00 | 6387 | Ropn1 | ENSMUSG00000022832.5 |
| chr5 | 114174612 | 114174946 | 6.75307 | 11.31069 | 9.11275 | Promoter | 5 | 114175906 | 114190338 | 14433 | 1 | ENSMUST00 | -960 | Acacb | ENSMUSG00000042010.11 |
| chr11 | 95841410 | 95842099 | 6.73919 | 11.28819 | 9.09039 | Promoter | 11 | 95837077 | 95842098 | 5022 | 2 | ENSMUST00 | 0 | Abi3 | ENSMUSG00000018381.10 |
| chr18 | 65739631 | 65740234 | 6.73919 | 11.28819 | 9.09039 | Exon (ENSMU | 18 | 65749014 | 65750390 | 1377 | 1 | ENSMUST00 | -8780 | Oacyl | ENSMUSG00000046610.9 |
| chr8 | 72443382 | 72443748 | 4.9172 | 11.27124 | 9.07355 | Promoter | 8 | 72424176 | 72443870 | 19695 | 2 | ENSMUST00 | 122 | Calr3 | ENSMUSG00000019732.9 |
| chr11 | 82845505 | 82845847 | 6.49449 | 11.2705 | 9.07282 | Promoter | 11 | 82812669 | 82846092 | 33424 | 2 | ENSMUST00 | 245 | Rffl | ENSMUSG00000020696.13 |
| chr14 | 66851840 | 66852393 | 6.49449 | 11.2705 | 9.07282 | Intron (ENSM | 14 | 66802864 | 66868688 | 65825 | 2 | ENSMUST00 | 16295 | Dpysl2 | ENSMUSG00000022048.8 |
| chr14 | 101560570 | 101561068 | 6.49449 | 11.2705 | 9.07282 | Intron (ENSM | 14 | 101443723 | 101608486 | 164764 | 2 | ENSMUST00 | 47418 | Tbc1d4 | ENSMUSG00000033083.11 |
| chr18 | 84293588 | 84293903 | 6.49449 | 11.2705 | 9.07282 | Intron (ENSM | 18 | 84279820 | 84412420 | 132601 | 2 | ENSMUST00 | 118517 | Zfp407 | ENSMUSG00000048410.11 |
| chr19 | 6321584 | 6321767 | 6.49449 | 11.2705 | 9.07282 | Promoter | 19 | 6321808 | 6322232 | 425 | 1 | ENSMUST00 | -41 | Cdc42bpg | ENSMUSG00000024769.7 |
| chr19 | 12457530 | 12457957 | 6.49449 | 11.2705 | 9.07282 | Distal Interge | 19 | 12460779 | 12465284 | 4506 | 1 | ENSMUST00 | -2822 | Mpeg1 | ENSMUSG00000046805.9 |
| chr3 | 104777291 | 104777660 | 6.49449 | 11.2705 | 9.07282 | Promoter | 3 | 104767406 | 104777547 | 10142 | 2 | ENSMUST00 | 0 | Fam19a3 | ENSMUSG00000055865.8 |
| chr5 | 3152295 | 3152650 | 6.49449 | 11.2705 | 9.07282 | Distal Interge | 5 | 3123777 | 3127383 | 3607 | 1 | ENSMUST00 | 28519 | Gm8715 | ENSMUSG00000091483.1 |
| chr7 | 75655825 | 75656075 | 6.49449 | 11.2705 | 9.07282 | Intron (ENSM | 7 | 75644185 | 75677330 | 33146 | 1 | ENSMUST00 | 11641 | Akap13 | ENSMUSG00000066406.9 |
| chr8 | 48488598 | 48489023 | 6.49449 | 11.2705 | 9.07282 | Intron (ENSM | 8 | 48334753 | 48555693 | 220941 | 2 | ENSMUST00 | 66670 | Tenm3 | ENSMUSG00000031561.10 |
| chr2 | 28917808 | 28918092 | 6.72537 | 11.26578 | 9.06861 | Distal Interge | 2 | 28908969 | 28916668 | 7700 | 2 | ENSMUST00 | -1141 | Barhl1 | ENSMUSG00000026805.9 |
| chr1 | 184851967 | 184852253 | 6.71161 | 11.24346 | 9.04642 | Distal Interge | 1 | 184822375 | 184846451 | 24077 | 2 | ENSMUST00 | -5517 | Mar-02 | ENSMUSG00000073481.4 |
| chr15 | 89765862 | 89766145 | 6.71161 | 11.24346 | 9.04642 | Distal Interge | 15 | 89782393 | 89841860 | 59468 | 2 | ENSMUST00 | 75715 | Syt10 | ENSMUSG00000063260.2 |
| chr17 | 45931643 | 45931834 | 6.71161 | 11.24346 | 9.04642 | Distal Interge | 17 | 46018712 | 46021372 | 2661 | 2 | ENSMUST00 | 89538 | Vegfa | ENSMUSG00000023951.11 |
| chr3 | 159841477 | 159841745 | 6.47047 | 11.23092 | 9.03433 | Intron (ENSM | 3 | 159839737 | 159935833 | 96097 | 1 | ENSMUST00 | 1741 | Wls | ENSMUSG00000028173.6 |
| chr2 | 169723292 | 169724242 | 6.6979 | 11.22123 | 9.02464 | Intron (ENSM | 2 | 169633676 | 169888504 | 254829 | 1 | ENSMUST00 | 89617 | Tshz2 | ENSMUSG00000047907.11 |
| chrX | 74272311 | 74272684 | 5.7805 | 11.20014 | 9.00368 | Promoter | 20 | 74272484 | 74272620 | 137 | 1 | ENSMUST00 | 0 | Snora70 | ENSMUSG00000064981.1 |
| chr7 | 27474795 | 27475287 | 6.68425 | 11.19909 | 9.00263 | Intron (ENSM | 7 | 27473768 | 27477359 | 3592 | 1 | ENSMUST00 | 1028 | Sertad3 | ENSMUSG00000055200.6 |
| chr13 | 119741790 | 119742214 | 6.20879 | 11.18097 | 8.98466 | Intron (ENSM | 13 | 119723483 | 119738394 | 14912 | 2 | ENSMUST00 | -3397 | Nim1k | ENSMUSG00000095930.1 |
| chr3 | 126692965 | 126693834 | 6.20879 | 11.18097 | 8.98466 | Intron (ENSM | 3 | 126667329 | 126669032 | 1704 | 2 | ENSMUST00 | -23934 | Gm15551 | ENSMUSG00000086679.1 |
| chr9 | 83550295 | 83550843 | 6.20879 | 11.18097 | 8.98466 | Intron (ENSM | 9 | 83548350 | 83638299 | 89950 | 1 | ENSMUST00 | 1946 | Sh3bgrl2 | ENSMUSG00000032261.11 |
| chr11 | 63918235 | 63918504 | 6.67065 | 11.17704 | 8.98086 | Intron (ENSM | 11 | 63885792 | 63922290 | 36499 | 2 | ENSMUST00 | 3786 | Hs3st3b1 | ENSMUSG00000070407.5 |
| chr18 | 25505088 | 25505483 | 6.67065 | 11.17704 | 8.98086 | Intron (ENSM | 18 | 25598145 | 25598454 | 310 | 2 | ENSMUST00 | 92971 | Gm25845 | ENSMUSG00000084537.1 |
| chr4 | 152096804 | 152096988 | 6.67065 | 11.17704 | 8.98086 | Promoter | 4 | 152096719 | 152115400 | 18682 | 1 | ENSMUST00 | 86 | Plekhg5 | ENSMUSG00000039713.11 |
| chr6 | 115730374 | 115730769 | 6.65711 | 11.15507 | 8.95905 | Exon (ENSMU | 6 | 115729141 | 115734951 | 5811 | 2 | ENSMUST00 | 4182 | Tmem40 | ENSMUSG00000059900.9 |
| chr1 | 167598247 | 167598670 | 6.6302 | 11.1114 | 8.91563 | Promoter | 1 | 167598384 | 167639623 | 41240 | 1 | ENSMUST00 | 0 | Rxrg | ENSMUSG00000015843.5 |
| chr17 | 6315885 | 6316257 | 4.99607 | 11.10239 | 8.90674 | Exon (ENSMU | 17 | 6310547 | 6317474 | 6928 | 2 | ENSMUST00 | 1217 | Dynlt1a | ENSMUSG00000092074.1 |
| chr1 | 51289439 | 51290264 | 6.37612 | 11.07527 | 8.87976 | Promoter | 1 | 51289126 | 51302960 | 13835 | 1 | ENSMUST00 | 314 | Sdpr | ENSMUSG00000045954.7 |
| chr13 | 31815598 | 31815989 | 6.37612 | 11.07527 | 8.87976 | Distal Interge | 13 | 31806691 | 31810643 | 3953 | 1 | ENSMUST00 | 8908 | Foxc1 | ENSMUSG00000050295.3 |
| chr13 | 44228751 | 44229291 | 6.37612 | 11.07527 | 8.87976 | Distal Interge | 13 | 44121294 | 44125179 | 3886 | 1 | ENSMUST00 | 107458 | Gm5083 | ENSMUSG00000089815.2 |
| chr5 | 148310922 | 148311159 | 6.37612 | 11.07527 | 8.87976 | Intron (ENSM | 5 | 148303174 | 148314000 | 10827 | 1 | ENSMUST00 | 7749 | Mtus2 | ENSMUSG00000029651.11 |
| chr1 | 92774534 | 92775195 | 6.6035 | 11.06807 | 8.87298 | Distal Interge | 1 | 92759213 | 92760480 | 1268 | 2 | ENSMUST00 | -14055 | RP23-145J9.1 | ENSMUSG00000099665.1 |
| chr11 | 69795035 | 69795318 | 6.6035 | 11.06807 | 8.87298 | Promoter | 11 | 69793520 | 69795905 | 2386 | 2 | ENSMUST00 | 587 | Chrnb1 | ENSMUSG00000041189.9 |
| chr17 | 24211932 | 24212366 | 6.59023 | 11.04653 | 8.85156 | Promoter | 17 | 24212515 | 24221730 | 9216 | 1 | ENSMUST00 | -149 | BC028777 | ENSMUSG00000085185.4 |
| chr12 | 15606607 | 15606802 | 6.57701 | 11.02508 | 8.83021 | Distal Interge | 12 | 15728097 | 15729095 | 999 | 1 | ENSMUST00 | -121295 | Gm4804 | ENSMUSG00000098158.1 |
| chr14 | 64653779 | 64653989 | 6.57701 | 11.02508 | 8.83021 | Intron (ENSM | 14 | 64652531 | 64806296 | 153766 | 1 | ENSMUST00 | 1249 | Kif13b | ENSMUSG00000060012.7 |
| chr16 | 30596816 | 30597074 | 6.57701 | 11.02508 | 8.83021 | Distal Interge | 16 | 30599723 | 30602797 | 3075 | 1 | ENSMUST00 | -2649 | Fam43a | ENSMUSG00000046546.3 |
| chr4 | 102588925 | 102589399 | 6.10678 | 11.00785 | 8.81353 | Promoter | 4 | 102589723 | 102605861 | 16139 | 1 | ENSMUST00 | -324 | Pde4b | ENSMUSG00000028525.11 |
| chr6 | 83443810 | 83444299 | 6.10678 | 11.00785 | 8.81353 | Intron (ENSM | 6 | 83441812 | 83442640 | 829 | 1 | ENSMUST00 | 1999 | B230319C09 | ENSMUSG00000051844.8 |
| chr5 | 123985626 | 123985906 | 6.56385 | 11.00371 | 8.80949 | Intron (ENSM | 5 | 123980254 | 123989984 | 9731 | 1 | ENSMUST00 | 5373 | Hip1r | ENSMUSG00000000915.10 |
| chr13 | 31812520 | 31813510 | 6.55073 | 10.98242 | 8.78832 | Distal Interge | 13 | 31806691 | 31810643 | 3953 | 1 | ENSMUST00 | 5830 | Foxc1 | ENSMUSG00000050295.3 |
| chr7 | 135465322 | 135465804 | 6.53768 | 10.96121 | 8.76724 | Distal Interge | 7 | 135410367 | 135413615 | 3249 | 1 | ENSMUST00 | 54956 | Foxi2 | ENSMUSG00000048377.5 |
| chr10 | 81978369 | 81978876 | 5.08503 | 10.94457 | 8.75072 | Distal Interge | 10 | 81905073 | 81910899 | 5827 | 2 | ENSMUST00 | -67471 | Zfp781 | ENSMUSG00000096718.1 |
| chr3 | 95997367 | 95997721 | 6.52467 | 10.94008 | 8.74627 | Downstream | 3 | 95988429 | 95996001 | 7573 | 2 | ENSMUST00 | -1367 | Plekho1 | ENSMUSG00000015745.4 |
| chr19 | 5324016 | 5324263 | 6.51171 | 10.91904 | 8.72533 | Distal Interge | 19 | 5335741 | 5344153 | 8413 | 1 | ENSMUST00 | -11478 | Catsper1 | ENSMUSG00000038498.3 |
| chr9 | 32699266 | 32700176 | 6.26198 | 10.88643 | 8.69293 | Intron (ENSM | 9 | 32696384 | 32754428 | 58045 | 1 | ENSMUST00 | 2883 | Ets1 | ENSMUSG00000032035.10 |
| chr9 | 92279127 | 92279432 | 6.26198 | 10.88643 | 8.69293 | Intron (ENSM | 9 | 92275746 | 92282192 | 6447 | 1 | ENSMUST00 | 3382 | Plscr2 | ENSMUSG00000032372.9 |
| chr10 | 59942894 | 59943438 | 6.48596 | 10.87719 | 8.68403 | Distal Interge | 10 | 59949675 | 59951770 | 2096 | 2 | ENSMUST00 | 8332 | Ddit4 | ENSMUSG00000020108.3 |
| chr14 | 31194886 | 31195731 | 6.48596 | 10.87719 | 8.68403 | Intron (ENSM | 14 | 31177501 | 31192619 | 15119 | 2 | ENSMUST00 | -2268 | Nisch | ENSMUSG00000021910.10 |
| chr6 | 89345210 | 89345510 | 6.47316 | 10.85638 | 8.66333 | Intron (ENSM | 6 | 89319015 | 89357717 | 38703 | 2 | ENSMUST00 | 12207 | Plxna1 | ENSMUSG00000030084.6 |
| chr7 | 27606528 | 27607407 | 6.4604 | 10.83565 | 8.64276 | Promoter | 7 | 27607748 | 27633246 | 25499 | 1 | ENSMUST00 | -341 | Akt2 | ENSMUSG00000004056.10 |
| chr1 | 156319641 | 156320431 | 6.4477 | 10.81501 | 8.6222 | Intron (ENSM | 1 | 156310727 | 156328035 | 17309 | 1 | ENSMUST00 | 8915 | Nphs2 | ENSMUSG00000026602.4 |
| chr4 | 139578228 | 139578599 | 6.4477 | 10.81501 | 8.6222 | Intron (ENSM | 4 | 139574720 | 139620382 | 45663 | 1 | ENSMUST00 | 3509 | Iffo2 | ENSMUSG00000041025.10 |
| chr5 | 122280859 | 122281193 | 6.4477 | 10.81501 | 8.6222 | Distal Interge | 5 | 122284365 | 122324281 | 39917 | 1 | ENSMUST00 | -3172 | Pptc7 | ENSMUSG00000038582.10 |
| chr1 | 152084550 | 152085405 | 5.18618 | 10.80032 | 8.60808 | Intron (ENSM | 1 | 151852403 | 152089970 | 237568 | 2 | ENSMUST00 | 4565 | 1700025G04 | ENSMUSG00000032666.11 |
| chr9 | 118862356 | 118862928 | 5.18618 | 10.80032 | 8.60808 | Intron (ENSM | 9 | 118872069 | 118897312 | 25244 | 1 | ENSMUST00 | -9141 | Itga9 | ENSMUSG00000039115.8 |
| chr8 | 127438311 | 127439114 | 6.43505 | 10.79443 | 8.60228 | Intron (ENSM | 8 | 127447746 | 127474513 | 26768 | 1 | ENSMUST00 | -8632 | Pard3 | ENSMUSG00000025812.12 |
| chr17 | 88629699 | 88630638 | 6.39739 | 10.73318 | 8.54127 | Intron (ENSM | 17 | 88627364 | 88637634 | 10271 | 1 | ENSMUST00 | 2336 | Ston1 | ENSMUSG00000033855.10 |
| chr11 | 99572885 | 99573116 | 6.38494 | 10.71291 | 8.52107 | Distal Interge | 11 | 99566027 | 99566630 | 604 | 2 | ENSMUST00 | -6256 | Krtap3-1 | ENSMUSG00000047564.4 |
| chr14 | 20546709 | 20546936 | 6.15186 | 10.70359 | 8.51182 | Promoter | 14 | 20499364 | 20546573 | 47210 | 2 | ENSMUST00 | -137 | Ppp3cb | ENSMUSG00000021816.5 |
| chr16 | 32165149 | 32165856 | 6.15186 | 10.70359 | 8.51182 | Promoter | 16 | 32142785 | 32165585 | 22801 | 2 | ENSMUST00 | 0 | Nrros | ENSMUSG00000052384.8 |
| chr17 | 43389657 | 43390350 | 6.15186 | 10.70359 | 8.51182 | Promoter | 17 | 43389466 | 43459557 | 70092 | 1 | ENSMUST00 | 192 | Gpr116 | ENSMUSG00000056492.5 |
| chr18 | 46849234 | 46849886 | 6.15186 | 10.70359 | 8.51182 | Promoter | 18 | 46850039 | 46905446 | 55408 | 1 | ENSMUST00 | -153 | 4833403I15R | ENSMUSG00000024481.4 |
| chr19 | 38527399 | 38528412 | 6.15186 | 10.70359 | 8.51182 | Intron (ENSM | 19 | 38524209 | 38782795 | 258587 | 1 | ENSMUST00 | 3191 | Plce1 | ENSMUSG00000024998.12 |

| | | | | | | | | | | | | | | | |
|---|---|---|---|---|---|---|---|---|---|---|---|---|---|---|---|
| chr8 | 8658358 | 8658749 | 6.15186 | 10.70359 | 8.51182 | Intron (ENSM | 8 | 8617434 | 8660773 | 43340 | 2 | ENSMUST00 | 2024 | Efnb2 | ENSMUSG00000001300.10 |
| chr18 | 43207368 | 43207822 | 5.3022 | 10.67304 | 8.48163 | Promoter | 18 | 43207697 | 43317481 | 109785 | 1 | ENSMUST00 | 0 | Stk32a | ENSMUSG00000039954.8 |
| chr8 | 105413246 | 105413983 | 4.65294 | 10.66687 | 8.4757 | Promoter | 8 | 105382807 | 105413502 | 30696 | 2 | ENSMUST00 | 0 | Kctd19 | ENSMUSG00000051648.8 |
| chr13 | 107838288 | 107838613 | 6.34786 | 10.65255 | 8.46139 | Intron (ENSM | 13 | 107829110 | 107829280 | 171 | 2 | ENSMUST00 | -9009 | Gm22118 | ENSMUSG00000093334.2 |
| chr11 | 95105181 | 95105787 | 6.3356 | 10.63258 | 8.44148 | Distal Interge | 11 | 95120119 | 95125296 | 5178 | 1 | ENSMUST00 | -14332 | Dlx3 | ENSMUSG00000001510.8 |
| chr10 | 42941757 | 42942105 | 6.32338 | 10.61268 | 8.42163 | Intron (ENSM | 10 | 42939881 | 42958721 | 18841 | 1 | ENSMUST00 | 1877 | Scml4 | ENSMUSG00000044770.9 |
| chr12 | 24975734 | 24975939 | 6.32338 | 10.61268 | 8.42163 | Promoter | 12 | 24974932 | 25059697 | 84766 | 1 | ENSMUST00 | 803 | Kidins220 | ENSMUSG00000036333.10 |
| chr14 | 54249167 | 54250077 | 6.32338 | 10.61268 | 8.42163 | Intron (ENSM | 14 | 54239661 | 54253914 | 14254 | 2 | ENSMUST00 | 3837 | Dad1 | ENSMUSG00000022174.8 |
| chr10 | 41934492 | 41935362 | 5.43663 | 10.5674 | 8.37697 | Intron (ENSM | 10 | 41916051 | 41976122 | 60072 | 2 | ENSMUST00 | 40760 | Armc2 | ENSMUSG00000071324.6 |
| chr12 | 111429436 | 111430266 | 5.43663 | 10.5674 | 8.37697 | Promoter | 12 | 111426874 | 111430946 | 4073 | 2 | ENSMUST00 | 680 | Gm10425 | ENSMUSG00000097081.1 |
| chr19 | 47319924 | 47320673 | 5.43663 | 10.5674 | 8.37697 | Exon (ENSMU | 19 | 47273519 | 47273589 | 71 | 2 | ENSMUST00 | -46336 | Mir6995 | ENSMUSG00000098786.1 |
| chr2 | 60963877 | 60964298 | 5.43663 | 10.5674 | 8.37697 | Promoter | 2 | 60750193 | 60963192 | 213000 | 2 | ENSMUST00 | -686 | Rbms1 | ENSMUSG00000026970.11 |
| chr9 | 7914276 | 7914850 | 5.43663 | 10.5674 | 8.37697 | Distal Interge | 9 | 7860810 | 7873186 | 12377 | 2 | ENSMUST00 | -41091 | Birc3 | ENSMUSG00000032000.12 |
| chr9 | 99799471 | 99800544 | 5.43663 | 10.5674 | 8.37697 | Distal Interge | 9 | 99750391 | 99750517 | 127 | 2 | ENSMUST00 | -48955 | Gm23949 | ENSMUSG00000077215.1 |
| chr11 | 96997117 | 96997561 | 6.28665 | 10.55282 | 8.36284 | Distal Interge | 11 | 96954219 | 96982959 | 28741 | 2 | ENSMUST00 | -14159 | Sp2 | ENSMUSG00000018678.7 |
| chr11 | 77685406 | 77685663 | 6.28665 | 10.55282 | 8.36284 | Promoter | 11 | 77686155 | 77712741 | 26587 | 1 | ENSMUST00 | -492 | Nufip2 | ENSMUSG00000037857.11 |
| chr11 | 90030482 | 90031070 | 6.28665 | 10.55282 | 8.36284 | Promoter | 11 | 90030348 | 90036508 | 6161 | 1 | ENSMUST00 | 135 | Tmem100 | ENSMUSG00000069763.3 |
| chr13 | 48706480 | 48706695 | 6.28665 | 10.55282 | 8.36284 | Distal Interge | 13 | 48662998 | 48666507 | 3510 | 1 | ENSMUST00 | 43483 | Barx1 | ENSMUSG00000021381.4 |
| chr13 | 102898513 | 102899308 | 6.28665 | 10.55282 | 8.36284 | Intron (ENSM | 13 | 102741969 | 102905786 | 163818 | 2 | ENSMUST00 | 6478 | Mast4 | ENSMUSG00000034751.10 |
| chr13 | 93190415 | 93191005 | 6.28665 | 10.55282 | 8.36284 | Intron (ENSM | 13 | 93147400 | 93192283 | 44884 | 2 | ENSMUST00 | 1278 | Papd4 | ENSMUSG00000042167.5 |
| chr15 | 97992801 | 97993009 | 6.28665 | 10.55282 | 8.36284 | Intron (ENSM | 15 | 97988111 | 97990476 | 2366 | 2 | ENSMUST00 | -2326 | Col2a1 | ENSMUSG00000022483.11 |
| chr15 | 30646612 | 30647240 | 6.28665 | 10.55282 | 8.36284 | Exon (ENSMU | 15 | 30601514 | 30601576 | 63 | 1 | ENSMUST00 | 45099 | Gm24487 | ENSMUSG00000094704.1 |
| chr17 | 26504706 | 26505437 | 6.28665 | 10.55282 | 8.36284 | Downstream | 17 | 26506516 | 26507723 | 1208 | 2 | ENSMUST00 | 2286 | Dusp1 | ENSMUSG00000024190.6 |
| chr17 | 69396246 | 69396431 | 6.28665 | 10.55282 | 8.36284 | Distal Interge | 17 | 69383840 | 69390750 | 6911 | 1 | ENSMUST00 | 12407 | Zbtb14 | ENSMUSG00000049672.9 |
| chr2 | 152093341 | 152093859 | 6.28665 | 10.55282 | 8.36284 | Exon (ENSMU | 2 | 152105516 | 152109211 | 3696 | 1 | ENSMUST00 | -11657 | Srxn1 | ENSMUSG00000032802.8 |
| chr4 | 127313044 | 127313512 | 6.28665 | 10.55282 | 8.36284 | Promoter | 4 | 127311421 | 127314039 | 2619 | 2 | ENSMUST00 | 527 | Gja4 | ENSMUSG00000050234.7 |
| chr4 | 57638093 | 57638356 | 6.28665 | 10.55282 | 8.36284 | Promoter | 4 | 57637975 | 57709489 | 71515 | 1 | ENSMUST00 | 119 | Palm2 | ENSMUSG00000090053.4 |
| chr6 | 125215073 | 125215389 | 6.28665 | 10.55282 | 8.36284 | Promoter | 6 | 125215551 | 125222306 | 6756 | 1 | ENSMUST00 | -162 | Vamp1 | ENSMUSG00000030337.11 |
| chr6 | 39055770 | 39056164 | 6.28665 | 10.55282 | 8.36284 | Intron (ENSM | 6 | 39063878 | 39067598 | 3721 | 2 | ENSMUST00 | 11434 | RP23-346B1 | ENSMUSG00000100121.1 |
| chr7 | 16101963 | 16102284 | 6.28665 | 10.55282 | 8.36284 | Intron (ENSM | 7 | 16098638 | 16109309 | 10672 | 1 | ENSMUST00 | 3326 | Napa | ENSMUSG00000006024.7 |
| chr7 | 36708031 | 36708378 | 6.28665 | 10.55282 | 8.36284 | Intron (ENSM | 7 | 36698217 | 36702113 | 3897 | 1 | ENSMUST00 | 9815 | Tshz3 | ENSMUSG00000021217.7 |
| chr8 | 72321766 | 72322156 | 6.28665 | 10.55282 | 8.36284 | Distal Interge | 8 | 72319033 | 72321656 | 2624 | 1 | ENSMUST00 | 2734 | Klf2 | ENSMUSG00000055148.7 |
| chr9 | 112118321 | 112118796 | 6.28665 | 10.55282 | 8.36284 | Promoter | 9 | 112118632 | 112118715 | 84 | 2 | ENSMUST00 | 0 | Mir128-2 | ENSMUSG00000065441.1 |
| chrX | 143664294 | 143664801 | 6.28665 | 10.55282 | 8.36284 | Promoter | 20 | 143664373 | 143693384 | 29012 | 1 | ENSMUST00 | 0 | Pak3 | ENSMUSG00000031284.11 |
| chr6 | 5008700 | 5009045 | 6.04554 | 10.52643 | 8.3386 | 3' UTR | 6 | 4958983 | 4959302 | 320 | 1 | ENSMUST00 | 49718 | Gm26556 | ENSMUSG00000097396.1 |
| chr9 | 107984387 | 107984724 | 6.04554 | 10.52643 | 8.3386 | Promoter | 9 | 107984633 | 107985632 | 1000 | 2 | ENSMUST00 | 908 | Fam212a | ENSMUSG00000042106.4 |
| chr8 | 27362177 | 27362458 | 5.81991 | 10.51666 | 8.32901 | Distal Interge | 8 | 27368711 | 27399729 | 31019 | 1 | ENSMUST00 | -6253 | Chrnb3 | ENSMUSG00000031492.7 |
| chr12 | 108443541 | 108443858 | 6.263 | 10.51426 | 8.32665 | Intron (ENSM | 12 | 108440530 | 108441270 | 741 | 2 | ENSMUST00 | -2272 | Gm15636 | ENSMUSG00000082203.1 |
| chr1 | 91017223 | 91017581 | 5.59421 | 10.48996 | 8.30244 | Intron (ENSM | 1 | 90998779 | 91128940 | 130162 | 1 | ENSMUST00 | 18445 | Lrrfip1 | ENSMUSG00000026305.10 |
| chr1 | 176213605 | 176213800 | 5.59421 | 10.48996 | 8.30244 | Promoter | 1 | 176042304 | 176213841 | 171538 | 2 | ENSMUST00 | 41 | Pld5 | ENSMUSG00000055214.10 |
| chr11 | 11855925 | 11856333 | 5.59421 | 10.48996 | 8.30244 | Intron (ENSM | 11 | 11844977 | 11864907 | 19931 | 2 | ENSMUST00 | 8574 | Ddc | ENSMUSG00000020182.11 |
| chr14 | 122406580 | 122407215 | 5.59421 | 10.48996 | 8.30244 | Distal Interge | 14 | 122388054 | 122388160 | 107 | 2 | ENSMUST00 | -18421 | Gm25464 | ENSMUSG00000095311.1 |
| chr15 | 59680472 | 59680879 | 5.59421 | 10.48996 | 8.30244 | Distal Interge | 15 | 59648653 | 59653689 | 5037 | 1 | ENSMUST00 | 31820 | Trib1 | ENSMUSG00000032501.8 |
| chr15 | 25469327 | 25469991 | 5.59421 | 10.48996 | 8.30244 | Distal Interge | 15 | 25414192 | 25414844 | 653 | 1 | ENSMUST00 | 55136 | Gm5468 | ENSMUSG00000056089.5 |
| chr19 | 6428734 | 6428945 | 5.59421 | 10.48996 | 8.30244 | Promoter | 19 | 6428767 | 6473339 | 44573 | 1 | ENSMUST00 | 0 | Nrxn2 | ENSMUSG00000033768.11 |
| chr3 | 28705714 | 28705963 | 5.59421 | 10.48996 | 8.30244 | Exon (ENSMU | 3 | 28698017 | 28727476 | 29460 | 1 | ENSMUST00 | 7698 | Slc2a2 | ENSMUSG00000027690.8 |
| chr3 | 89299374 | 89300027 | 5.59421 | 10.48996 | 8.30244 | Distal Interge | 3 | 89302478 | 89302729 | 252 | 1 | ENSMUST00 | -2451 | Gm22642 | ENSMUSG00000093093.2 |
| chr4 | 134187378 | 134187614 | 5.59421 | 10.48996 | 8.30244 | Promoter | 4 | 134129858 | 134187112 | 57255 | 2 | ENSMUST00 | -267 | Cep85 | ENSMUSG00000037443.8 |
| chr5 | 103376727 | 103377469 | 5.59421 | 10.48996 | 8.30244 | Distal Interge | 5 | 103425192 | 103598359 | 173168 | 1 | ENSMUST00 | -47723 | Ptpn13 | ENSMUSG00000034573.10 |
| chr9 | 45268933 | 45269302 | 5.59421 | 10.48996 | 8.30244 | Promoter | 9 | 45253837 | 45269149 | 15313 | 2 | ENSMUST00 | 0 | Il10ra | ENSMUSG00000032089.11 |
| chr1 | 77496676 | 77497279 | 6.00779 | 10.46335 | 8.27668 | Promoter | 1 | 77496653 | 77496768 | 116 | 2 | ENSMUST00 | 0 | Mir6352 | ENSMUSG00000099141.1 |
| chr10 | 111031498 | 111031923 | 6.00779 | 10.46335 | 8.27668 | Distal Interge | 10 | 110941782 | 111010066 | 68285 | 2 | ENSMUST00 | -21433 | Zdhhc17 | ENSMUSG00000035798.8 |
| chr11 | 68431830 | 68432358 | 6.00779 | 10.46335 | 8.27668 | Promoter | 11 | 68432121 | 68497849 | 65729 | 1 | ENSMUST00 | 0 | Pik3r5 | ENSMUSG00000020901.8 |
| chr11 | 68956358 | 68956756 | 6.00779 | 10.46335 | 8.27668 | Promoter | 11 | 68943157 | 68957480 | 14324 | 2 | ENSMUST00 | 724 | Arhgef15 | ENSMUSG00000052921.8 |
| chr13 | 91430808 | 91431276 | 6.00779 | 10.46335 | 8.27668 | Distal Interge | 13 | 91460283 | 91688866 | 228584 | 1 | ENSMUST00 | -29007 | Ssbp2 | ENSMUSG00000003992.9 |
| chr14 | 70167652 | 70168043 | 6.00779 | 10.46335 | 8.27668 | Promoter | 14 | 70164362 | 70167839 | 3478 | 2 | ENSMUST00 | 0 | Pdlim2 | ENSMUSG00000022090.5 |
| chr14 | 72532167 | 72532749 | 6.00779 | 10.46335 | 8.27668 | Distal Interge | 14 | 72561447 | 72564718 | 3272 | 2 | ENSMUST00 | 31969 | Fndc3a | ENSMUSG00000033487.9 |
| chr15 | 78581749 | 78582047 | 6.00779 | 10.46335 | 8.27668 | Distal Interge | 15 | 78559169 | 78572783 | 13615 | 2 | ENSMUST00 | -8967 | Rac2 | ENSMUSG00000033220.6 |
| chr18 | 70652361 | 70652858 | 6.00779 | 10.46335 | 8.27668 | Distal Interge | 18 | 70617773 | 70625777 | 8005 | 1 | ENSMUST00 | 34589 | Mbd2 | ENSMUSG00000024513.11 |
| chr19 | 47581052 | 47581630 | 6.00779 | 10.46335 | 8.27668 | Intron (ENSM | 19 | 47580019 | 47642455 | 62437 | 1 | ENSMUST00 | 1034 | Slk | ENSMUSG00000025060.9 |
| chr2 | 33215515 | 33215837 | 6.00779 | 10.46335 | 8.27668 | Promoter | 2 | 33216069 | 33247717 | 31649 | 1 | ENSMUST00 | -232 | Angptl2 | ENSMUSG00000004105.6 |
| chr3 | 125228295 | 125228542 | 6.00779 | 10.46335 | 8.27668 | Distal Interge | 3 | 125404094 | 125686644 | 282551 | 1 | ENSMUST00 | -175552 | Ndst4 | ENSMUSG00000027971.11 |
| chr3 | 141931239 | 141931804 | 6.00779 | 10.46335 | 8.27668 | Promoter | 3 | 141838102 | 141931523 | 93422 | 2 | ENSMUST00 | 0 | Bmpr1b | ENSMUSG00000052430.10 |
| chr3 | 54915521 | 54916037 | 6.00779 | 10.46335 | 8.27668 | Promoter | 3 | 54897071 | 54915887 | 18817 | 2 | ENSMUST00 | 0 | Sertm1 | ENSMUSG00000056306.5 |
| chr4 | 43558329 | 43559317 | 6.00779 | 10.46335 | 8.27668 | Promoter | 4 | 43555106 | 43558460 | 3355 | 2 | ENSMUST00 | 0 | Tln1 | ENSMUSG00000028465.11 |
| chr4 | 143294659 | 143295589 | 6.00779 | 10.46335 | 8.27668 | Intron (ENSM | 4 | 143268346 | 143299463 | 31118 | 2 | ENSMUST00 | 3874 | Pdpn | ENSMUSG00000028583.9 |
| chr4 | 123572504 | 123573089 | 6.00779 | 10.46335 | 8.27668 | Intron (ENSM | 4 | 123538880 | 123564694 | 25815 | 2 | ENSMUST00 | -7811 | Macf1 | ENSMUSG00000028649.12 |
| chr4 | 127015643 | 127016056 | 6.00779 | 10.46335 | 8.27668 | Distal Interge | 4 | 127021324 | 127031236 | 9913 | 1 | ENSMUST00 | -5268 | Sfpq | ENSMUSG00000028820.8 |
| chr4 | 133220719 | 133221105 | 6.00779 | 10.46335 | 8.27668 | Promoter | 4 | 133220810 | 133224555 | 3746 | 1 | ENSMUST00 | 0 | Cd164l2 | ENSMUSG00000028865.8 |
| chr5 | 138997875 | 138998373 | 6.00779 | 10.46335 | 8.27668 | Promoter | 5 | 138976971 | 138997370 | 20400 | 2 | ENSMUST00 | -506 | Pdgfa | ENSMUSG00000025856.10 |
| chr5 | 75835560 | 75835986 | 6.00779 | 10.46335 | 8.27668 | Distal Interge | 5 | 75849238 | 75852958 | 3721 | 2 | ENSMUST00 | 16972 | RP23-329A2 | ENSMUSG00000100936.1 |
| chr6 | 18511807 | 18512329 | 6.00779 | 10.46335 | 8.27668 | Promoter | 6 | 18435251 | 18513091 | 77841 | 2 | ENSMUST00 | 762 | Cttnbp2 | ENSMUSG00000000416.11 |
| chr7 | 28208857 | 28209308 | 6.00779 | 10.46335 | 8.27668 | Distal Interge | 7 | 28180316 | 28186943 | 6628 | 1 | ENSMUST00 | 28542 | Dyrk1b | ENSMUSG00000002409.12 |
| chr7 | 114719604 | 114719931 | 6.00779 | 10.46335 | 8.27668 | Promoter | 7 | 114719511 | 114720139 | 629 | 1 | ENSMUST00 | 94 | Calcb | ENSMUSG00000030666.6 |
| chr8 | 122564570 | 122564907 | 6.00779 | 10.46335 | 8.27668 | Intron (ENSM | 8 | 122568015 | 122573128 | 5114 | 1 | ENSMUST00 | -3108 | Cdt1 | ENSMUSG00000006585.2 |
| chr8 | 11003072 | 11003508 | 6.00779 | 10.46335 | 8.27668 | Intron (ENSM | 8 | 11007850 | 11054536 | 46687 | 1 | ENSMUST00 | -4342 | 9530052E02I | ENSMUSG00000096938.2 |
| chr10 | 69533830 | 69534337 | 5.7815 | 10.45037 | 8.26584 | Promoter | 10 | 69533843 | 70024626 | 490784 | 1 | ENSMUST00 | 0 | Ank3 | ENSMUSG00000069601.8 |
| chr12 | 33625974 | 33626252 | 5.7815 | 10.45037 | 8.26584 | Distal Interge | 12 | 33452712 | 33452843 | 132 | 2 | ENSMUST00 | -173132 | Gm23529 | ENSMUSG00000077750.1 |
| chr12 | 3359398 | 3359814 | 5.7815 | 10.45037 | 8.26584 | Distal Interge | 12 | 3343816 | 3357153 | 13338 | 2 | ENSMUST00 | -2246 | Gm26520 | ENSMUSG00000097429.1 |
| chr16 | 14292327 | 14292695 | 5.7815 | 10.45037 | 8.26584 | Promoter | 16 | 14194527 | 14291408 | 96882 | 2 | ENSMUST00 | -920 | Myh11 | ENSMUSG00000018830.8 |
| chr16 | 73529906 | 73530515 | 5.7815 | 10.45037 | 8.26584 | Distal Interge | 16 | 73422793 | 73422959 | 167 | 2 | ENSMUST00 | -106948 | Gm25699 | ENSMUSG00000064830.1 |
| chr19 | 34661965 | 34662471 | 5.7815 | 10.45037 | 8.26584 | Distal Interge | 19 | 34640871 | 34650009 | 9139 | 1 | ENSMUST00 | 21095 | Ifit1 | ENSMUSG00000034459.8 |
| chr19 | 11293156 | 11293730 | 5.7815 | 10.45037 | 8.26584 | Intron (ENSM | 19 | 11273866 | 11283873 | 10008 | 2 | ENSMUST00 | -9284 | Ms4a5 | ENSMUSG00000054523.8 |
| chr19 | 47253104 | 47253621 | 5.7815 | 10.45037 | 8.26584 | Exon (ENSMU | 19 | 47273519 | 47273589 | 71 | 2 | ENSMUST00 | 19968 | Mir6995 | ENSMUSG00000098786.1 |
| chr2 | 76571380 | 76571792 | 5.7815 | 10.45037 | 8.26584 | Intron (ENSM | 2 | 76568120 | 76580028 | 11909 | 1 | ENSMUST00 | 3261 | Osbpl6 | ENSMUSG00000042359.13 |
| chr2 | 31460114 | 31460356 | 5.7815 | 10.45037 | 8.26584 | Exon (ENSMU | 2 | 31454708 | 31457989 | 3282 | 1 | ENSMUST00 | 5407 | Hmcn2 | ENSMUSG00000055632.12 |
| chr3 | 30507042 | 30507407 | 5.7815 | 10.45037 | 8.26584 | Intron (ENSM | 3 | 30236569 | 30509487 | 272919 | 2 | ENSMUST00 | 2080 | Mecom | ENSMUSG00000027684.11 |
| chr3 | 100823672 | 100824511 | 5.7815 | 10.45037 | 8.26584 | Promoter | 3 | 100825459 | 100895679 | 70221 | 1 | ENSMUST00 | -948 | Vtcn1 | ENSMUSG00000051076.7 |
| chr4 | 123787743 | 123788348 | 5.7815 | 10.45037 | 8.26584 | Promoter | 4 | 123787874 | 123811498 | 23625 | 1 | ENSMUST00 | 0 | Rhbdl2 | ENSMUSG00000043333.7 |
| chr4 | 43209242 | 43209593 | 5.7815 | 10.45037 | 8.26584 | Intron (ENSM | 4 | 43201149 | 43201213 | 65 | 2 | ENSMUST00 | -8030 | Gm25010 | ENSMUSG00000089028.1 |
| chr5 | 67400044 | 67400554 | 5.7815 | 10.45037 | 8.26584 | Exon (ENSMU | 5 | 67427680 | 67437990 | 10311 | 1 | ENSMUST00 | -27126 | Gm26756 | ENSMUSG00000097840.1 |
| chr6 | 113194985 | 113195509 | 5.7815 | 10.45037 | 8.26584 | Promoter | 6 | 113168090 | 113195384 | 27295 | 2 | ENSMUST00 | 0 | Lhfpl4 | ENSMUSG00000042873.10 |
| chr7 | 116258615 | 116258902 | 5.7815 | 10.45037 | 8.26584 | Intron (ENSM | 7 | 116123493 | 116237776 | 114284 | 2 | ENSMUST00 | -20840 | Plekha7 | ENSMUSG00000045659.12 |
| chr8 | 23256083 | 23256337 | 5.7815 | 10.45037 | 8.26584 | Promoter | 8 | 23241353 | 23257021 | 15669 | 2 | ENSMUST00 | 684 | Golga7 | ENSMUSG00000015341.11 |
| chr8 | 60505735 | 60506168 | 5.7815 | 10.45037 | 8.26584 | Promoter | 8 | 60506124 | 60545677 | 39554 | 1 | ENSMUST00 | 0 | Aadat | ENSMUSG00000057228.5 |
| chr9 | 108111095 | 108111606 | 5.7815 | 10.45037 | 8.26584 | Promoter | 9 | 108096023 | 108110413 | 14391 | 2 | ENSMUST00 | -683 | Bsn | ENSMUSG00000032589.9 |
| chr11 | 117385100 | 117385455 | 6.43345 | 10.42957 | 8.24651 | Distal Interge | 11 | 117359360 | 117360866 | 1507 | 1 | ENSMUST00 | 25741 | Sep-09 | ENSMUSG00000059248.8 |
| chr12 | 100527655 | 100527955 | 6.43345 | 10.42957 | 8.24651 | Distal Interge | 12 | 100300771 | 100520822 | 220052 | 2 | ENSMUST00 | -6834 | Ttc7b | ENSMUSG00000033530.7 |
| chr13 | 12680329 | 12680620 | 6.43345 | 10.42957 | 8.24651 | Distal Interge | 13 | 12666051 | 12666314 | 264 | 1 | ENSMUST00 | 14279 | Gm24065 | ENSMUSG00000088112.1 |
| chr14 | 69285235 | 69285475 | 6.43345 | 10.42957 | 8.24651 | Promoter | 14 | 69241851 | 69285112 | 43262 | 2 | ENSMUST00 | -124 | Slc25a37 | ENSMUSG00000034248.7 |
| chr15 | 100729120 | 100729745 | 6.43345 | 10.42957 | 8.24651 | Promoter | 15 | 100691813 | 100729376 | 37564 | 2 | ENSMUST00 | 0 | Galnt6 | ENSMUSG00000037280.7 |
| chr17 | 56511288 | 56511664 | 6.43345 | 10.42957 | 8.24651 | Promoter | 17 | 56511248 | 56512483 | 1236 | 2 | ENSMUST00 | 819 | Znrf4 | ENSMUSG00000044526.2 |
| chr19 | 5307514 | 5307773 | 6.43345 | 10.42957 | 8.24651 | Exon (ENSMU | 19 | 5298331 | 5308738 | 10408 | 1 | ENSMUST00 | 9184 | Gal3st3 | ENSMUSG00000047658.6 |
| chr3 | 132087204 | 132087372 | 6.43345 | 10.42957 | 8.24651 | Intron (ENSM | 3 | 132085292 | 132180293 | 95002 | 1 | ENSMUST00 | 1913 | Dkk2 | ENSMUSG00000028031.5 |
| chr4 | 123116529 | 123116747 | 6.43345 | 10.42957 | 8.24651 | Promoter | 4 | 123116266 | 123118000 | 1735 | 1 | ENSMUST00 | 264 | Oxct2b | ENSMUSG00000076438.5 |
| chr5 | 149513943 | 149514458 | 6.43345 | 10.42957 | 8.24651 | Intron (ENSM | 5 | 149502507 | 149503047 | 541 | 2 | ENSMUST00 | -10897 | Gm2566 | ENSMUSG00000082959.1 |
| chr5 | 5782675 | 5782933 | 6.43345 | 10.42957 | 8.24651 | Promoter | 5 | 5781615 | 5783555 | 1941 | 2 | ENSMUST00 | 622 | Gm15459 | ENSMUSG00000100801.1 |
| chr6 | 18030409 | 18030775 | 6.43345 | 10.42957 | 8.24651 | Promoter | 6 | 17988940 | 18030585 | 41646 | 2 | ENSMUST00 | 0 | Wnt2 | ENSMUSG00000010797.3 |
| chr6 | 113891238 | 113891576 | 6.43345 | 10.42957 | 8.24651 | Promoter | 6 | 113745669 | 113891539 | 145871 | 2 | ENSMUST00 | 0 | Atp2b2 | ENSMUSG00000030302.11 |
| chr8 | 91629610 | 91630212 | 6.43345 | 10.42957 | 8.24651 | Promoter | 8 | 91621448 | 91629729 | 8282 | 2 | ENSMUST00 | 0 | Gm3235 | ENSMUSG00000090778.1 |
| chr9 | 122733211 | 122733668 | 6.43345 | 10.42957 | 8.24651 | Distal Interge | 9 | 122747346 | 122802074 | 54729 | 1 | ENSMUST00 | -13678 | Topaz1 | ENSMUSG00000094985.1 |
| chrX | 7765312 | 7765978 | 6.43345 | 10.42957 | 8.24651 | Promoter | 20 | 7764652 | 7775197 | 10546 | 1 | ENSMUST00 | 661 | Tfe3 | ENSMUSG00000000134.12 |
| chrX | 13282934 | 13283652 | 6.43345 | 10.42957 | 8.24651 | Intron (ENSM | 20 | 13281032 | 13287872 | 6841 | 1 | ENSMUST00 | 1903 | Ddx3x | ENSMUSG00000000787.7 |
| chrX | 152781864 | 152782288 | 6.43345 | 10.42957 | 8.24651 | Promoter | 20 | 152781921 | 152795615 | 13695 | 1 | ENSMUST00 | 0 | Gpr143 | ENSMUSG00000025333.5 |
| chrX | 95196365 | 95196627 | 6.43345 | 10.42957 | 8.24651 | Promoter | 20 | 95051919 | 95196505 | 144587 | 2 | ENSMUST00 | 0 | Arhgef9 | ENSMUSG00000025656.12 |
| chrX | 60147446 | 60147697 | 6.43345 | 10.42957 | 8.24651 | Promoter | 20 | 60055959 | 60147643 | 91685 | 2 | ENSMUST00 | 0 | Mcf2 | ENSMUSG00000031139.9 |
| chrX | 106012100 | 106012381 | 6.43345 | 10.42957 | 8.24651 | Promoter | 20 | 105968084 | 106011906 | 43823 | 2 | ENSMUST00 | -195 | Magt1 | ENSMUSG00000031232.11 |
| chrX | 134307737 | 134307939 | 6.43345 | 10.42957 | 8.24651 | Promoter | 20 | 134308084 | 134362639 | 54556 | 1 | ENSMUST00 | -145 | Cenpi | ENSMUSG00000031262.7 |
| chrX | 159254062 | 159254320 | 6.43345 | 10.42957 | 8.24651 | Intron (ENSM | 20 | 159255782 | 159311525 | 55744 | 1 | ENSMUST00 | -1462 | Rps6ka3 | ENSMUSG00000031309.10 |
| chrX | 72824932 | 72825516 | 6.43345 | 10.42957 | 8.24651 | Promoter | 20 | 72825178 | 72842602 | 17425 | 1 | ENSMUST00 | 0 | Gabrq | ENSMUSG00000031344.6 |
| chrX | 102157275 | 102157534 | 6.43345 | 10.42957 | 8.24651 | Promoter | 20 | 102141716 | 102157091 | 15376 | 2 | ENSMUST00 | -185 | Ercc6l | ENSMUSG00000051220.2 |
| chrX | 151169171 | 151169398 | 6.43345 | 10.42957 | 8.24651 | Promoter | 20 | 151169700 | 151171764 | 2065 | 1 | ENSMUST00 | -302 | Gm15138 | ENSMUSG00000086286.1 |
| chr10 | 80936399 | 80936845 | 6.20376 | 10.41759 | 8.23702 | Distal Interge | 10 | 80930091 | 80932204 | 2114 | 1 | ENSMUST00 | 6309 | Gadd45b | ENSMUSG00000015312.8 |
| chr12 | 24679372 | 24679611 | 6.20376 | 10.41759 | 8.23702 | Promoter | 12 | 24666554 | 24679947 | 13394 | 2 | ENSMUST00 | 336 | Cys1 | ENSMUSG00000062563.9 |
| chr15 | 31563739 | 31564155 | 6.20376 | 10.41759 | 8.23702 | Distal Interge | 15 | 31568791 | 31588259 | 19469 | 1 | ENSMUST00 | -4636 | Cmbl | ENSMUSG00000022235.9 |
| chr15 | 51591975 | 51592414 | 6.20376 | 10.41759 | 8.23702 | Distal Interge | 15 | 51786558 | 51865481 | 78924 | 2 | ENSMUST00 | 273067 | Eif3h | ENSMUSG00000022312.10 |
| chr15 | 82958320 | 82958750 | 6.20376 | 10.41759 | 8.23702 | Promoter | 15 | 82959218 | 82959296 | 79 | 2 | ENSMUST00 | 546 | Mir3080 | ENSMUSG00000092751.1 |
| chr16 | 92466432 | 92466706 | 6.20376 | 10.41759 | 8.23702 | Promoter | 16 | 92391953 | 92466146 | 74194 | 2 | ENSMUST00 | -287 | Rcan1 | ENSMUSG00000022951.10 |

| | | | | | | | | | | | | | | | |
|---|---|---|---|---|---|---|---|---|---|---|---|---|---|---|---|
| chr9 | 75544829 | 75545058 | 6.20376 | 10.41759 | 8.23702 | Intron (ENSM | 9 | 75497796 | 75559657 | 61862 | 2 | ENSMUST00 | 14599 | Tmod3 | ENSMUSG00000058587.7 |
| chr10 | 63496503 | 63497215 | 6.42188 | 10.41091 | 8.23098 | Intron (ENSM | 10 | 63504037 | 65002833 | 1498797 | 1 | ENSMUST00 | -6822 | Ctnna3 | ENSMUSG00000060843.6 |
| chr19 | 25610043 | 25610370 | 6.42188 | 10.41091 | 8.23098 | Promoter | 19 | 25610537 | 25623920 | 13384 | 1 | ENSMUST00 | -167 | Dmrt3 | ENSMUSG00000042372.4 |
| chr4 | 117929262 | 117929589 | 6.19205 | 10.39847 | 8.21863 | Promoter | 4 | 117926163 | 117929497 | 3335 | 2 | ENSMUST00 | 0 | Artn | ENSMUSG00000028539.9 |
| chr5 | 76363552 | 76364019 | 6.19205 | 10.39847 | 8.21863 | Promoter | 5 | 76333495 | 76363788 | 30294 | 2 | ENSMUST00 | 0 | Nmu | ENSMUSG00000029236.4 |
| chr16 | 31233478 | 31233700 | 6.39504 | 10.36765 | 8.18797 | Distal Interge | 16 | 31261505 | 31261599 | 95 | 1 | ENSMUST00 | -27805 | Gm27415 | ENSMUSG00000098403.1 |
| chr19 | 6321859 | 6322068 | 6.39504 | 10.36765 | 8.18797 | Promoter | 19 | 6321808 | 6322232 | 425 | 1 | ENSMUST00 | 52 | Cdc42bpg | ENSMUSG00000024769.7 |
| chr6 | 129182497 | 129183161 | 6.39504 | 10.36765 | 8.18797 | Exon (ENSMU | 6 | 129180615 | 129186534 | 5920 | 1 | ENSMUST00 | 1883 | Clec2d | ENSMUSG00000030157.5 |
| chr11 | 117316756 | 117317778 | 5.94284 | 10.35463 | 8.1753 | Intron (ENSM | 11 | 117308154 | 117352212 | 44059 | 1 | ENSMUST00 | 8603 | Sep-09 | ENSMUSG00000059248.8 |
| chr5 | 14028732 | 14028908 | 5.94284 | 10.35463 | 8.1753 | Intron (ENSM | 5 | 14025317 | 14256689 | 231373 | 1 | ENSMUST00 | 3416 | Sema3e | ENSMUSG00000063531.6 |
| chr16 | 10357887 | 10358129 | 6.38171 | 10.34615 | 8.16704 | 3' UTR | 16 | 10361254 | 10395448 | 34195 | 2 | ENSMUST00 | 37319 | Tekt5 | ENSMUSG00000039179.8 |
| chr8 | 87939025 | 87939348 | 6.38171 | 10.34615 | 8.16704 | Intron (ENSM | 8 | 87783316 | 87944052 | 160737 | 2 | ENSMUST00 | 4704 | Zfp423 | ENSMUSG00000045333.10 |
| chr17 | 56469874 | 56470166 | 6.36843 | 10.32475 | 8.14575 | Intron (ENSM | 17 | 56422237 | 56476369 | 54133 | 2 | ENSMUST00 | 6203 | Ptprs | ENSMUSG00000013236.11 |
| chr10 | 126845446 | 126845798 | 6.32893 | 10.26105 | 8.08251 | Distal Interge | 10 | 126840375 | 126840501 | 127 | 2 | ENSMUST00 | -4946 | Gm24179 | ENSMUSG00000089550.1 |
| chr18 | 61564499 | 61565431 | 6.32893 | 10.26105 | 8.08251 | Intron (ENSM | 18 | 61555689 | 61587531 | 31843 | 1 | ENSMUST00 | 8811 | Csnk1a1 | ENSMUSG00000024576.9 |
| chr3 | 84731830 | 84732550 | 6.32893 | 10.26105 | 8.08251 | Intron (ENSM | 3 | 84690223 | 84692544 | 2322 | 1 | ENSMUST00 | 41608 | Tmem154 | ENSMUSG00000056498.8 |
| chr5 | 115404396 | 115404732 | 6.32893 | 10.26105 | 8.08251 | Intron (ENSM | 5 | 115429599 | 115455698 | 26100 | 1 | ENSMUST00 | -24867 | Msi1 | ENSMUSG00000054256.6 |
| chr3 | 121390252 | 121390456 | 6.31587 | 10.24 | 8.06158 | Distal Interge | 3 | 121426541 | 121458205 | 31665 | 1 | ENSMUST00 | -36085 | Cnn3 | ENSMUSG00000053931.7 |
| chr1 | 150992058 | 150992288 | 6.08859 | 10.22929 | 8.05099 | Promoter | 1 | 150562524 | 150993051 | 430528 | 2 | ENSMUST00 | 763 | Hmcn1 | ENSMUSG00000066842.13 |
| chr10 | 85458903 | 85459076 | 6.08859 | 10.22929 | 8.05099 | Intron (ENSM | 10 | 85386827 | 85640594 | 253768 | 1 | ENSMUST00 | 72077 | Btbd11 | ENSMUSG00000020042.10 |
| chr14 | 21853640 | 21854013 | 6.08859 | 10.22929 | 8.05099 | Exon (ENSMU | 14 | 21845861 | 21848977 | 3117 | 2 | ENSMUST00 | -4664 | Comtd1 | ENSMUSG00000021773.5 |
| chr19 | 7606482 | 7606988 | 6.08859 | 10.22929 | 8.05099 | Promoter | 19 | 7596714 | 7607103 | 10390 | 2 | ENSMUST00 | 115 | Lgals12 | ENSMUSG00000024972.11 |
| chr19 | 4013028 | 4013210 | 6.08859 | 10.22929 | 8.05099 | Promoter | 19 | 4007497 | 4012806 | 5310 | 2 | ENSMUST00 | -223 | Ndufv1 | ENSMUSG00000037916.8 |
| chr7 | 82017113 | 82017513 | 6.08859 | 10.22929 | 8.05099 | Distal Interge | 7 | 81966672 | 81992299 | 25628 | 2 | ENSMUST00 | -24815 | Bnc1 | ENSMUSG00000025105.8 |
| chr6 | 141248130 | 141248620 | 6.28991 | 10.19814 | 8.02055 | Promoter | 6 | 141249269 | 141505940 | 256672 | 1 | ENSMUST00 | -649 | Pde3a | ENSMUSG00000041741.9 |
| chr9 | 48649206 | 48649674 | 6.28991 | 10.19814 | 8.02055 | Distal Interge | 9 | 48591877 | 48605153 | 13277 | 2 | ENSMUST00 | -44054 | Nnmt | ENSMUSG00000032271.8 |
| chr12 | 73846854 | 73847188 | 5.84356 | 10.1879 | 8.01044 | Distal Interge | 12 | 73901375 | 73947530 | 46156 | 1 | ENSMUST00 | -54187 | Hif1a | ENSMUSG00000021109.8 |
| chr8 | 35708111 | 35708289 | 5.84356 | 10.1879 | 8.01044 | Distal Interge | 8 | 35747449 | 35747562 | 114 | 1 | ENSMUST00 | -39160 | Gm22216 | ENSMUSG00000088182.1 |
| chr9 | 24770189 | 24770601 | 5.84356 | 10.1879 | 8.01044 | Intron (ENSM | 9 | 24720812 | 24774303 | 53492 | 2 | ENSMUST00 | 3702 | Tbx20 | ENSMUSG00000031965.9 |
| chr7 | 79052855 | 79053272 | 6.27701 | 10.17734 | 8.00004 | Promoter | 7 | 79018053 | 79053323 | 35271 | 2 | ENSMUST00 | 51 | Gm26633 | ENSMUSG00000097441.1 |
| chr13 | 34154730 | 34154995 | 6.26417 | 10.15662 | 7.97945 | Distal Interge | 13 | 34162964 | 34178168 | 15205 | 1 | ENSMUST00 | -7969 | Psmg4 | ENSMUSG00000071451.4 |
| chr16 | 95458645 | 95458925 | 6.26417 | 10.15662 | 7.97945 | Promoter | 16 | 95360806 | 95459245 | 98440 | 2 | ENSMUST00 | 320 | Erg | ENSMUSG00000040732.13 |
| chr19 | 45815556 | 45816359 | 6.26417 | 10.15662 | 7.97945 | Promoter | 19 | 45792346 | 45816022 | 23677 | 2 | ENSMUST00 | 0 | Kcnip2 | ENSMUSG00000025221.10 |
| chr2 | 26585722 | 26585924 | 6.26417 | 10.15662 | 7.97945 | Promoter | 2 | 26586630 | 26592681 | 6052 | 1 | ENSMUST00 | -706 | Egfl7 | ENSMUSG00000026921.13 |
| chr3 | 87170555 | 87170869 | 6.26417 | 10.15662 | 7.97945 | Intron (ENSM | 3 | 87084628 | 87174550 | 89923 | 2 | ENSMUST00 | 3681 | Kirrel | ENSMUSG00000041734.10 |
| chr17 | 36161228 | 36162045 | 4.8539 | 10.12085 | 7.94428 | Promoter | 17 | 36156531 | 36162079 | 5549 | 2 | ENSMUST00 | 34 | Gm17782 | ENSMUSG00000092366.2 |
| chr2 | 34359443 | 34360673 | 4.8539 | 10.12085 | 7.94428 | Intron (ENSM | 2 | 34176859 | 34370846 | 193988 | 2 | ENSMUST00 | 10173 | Pbx3 | ENSMUSG00000038718.10 |
| chr13 | 23488630 | 23488909 | 6.23863 | 10.11544 | 7.93891 | Promoter | 13 | 23477676 | 23488840 | 11165 | 2 | ENSMUST00 | 0 | Btn2a2 | ENSMUSG00000053216.9 |
| chr14 | 72708846 | 72709049 | 6.23863 | 10.11544 | 7.93891 | Promoter | 14 | 72564742 | 72709542 | 144801 | 2 | ENSMUST00 | 493 | Fndc3a | ENSMUSG00000033487.9 |
| chr8 | 46651750 | 46651962 | 6.23863 | 10.11544 | 7.93891 | Distal Interge | 8 | 46617449 | 46638706 | 21258 | 1 | ENSMUST00 | 34302 | Casp3 | ENSMUSG00000031628.8 |
| chr7 | 3292137 | 3292664 | 6.21331 | 10.07458 | 7.89833 | Intron (ENSM | 7 | 3290556 | 3299345 | 8790 | 1 | ENSMUST00 | 1582 | Myadm | ENSMUSG00000068566.7 |
| chr1 | 91078153 | 91078453 | 6.20072 | 10.05428 | 7.87816 | Exon (ENSMU | 1 | 91053586 | 91128287 | 74702 | 1 | ENSMUST00 | 24568 | Lrrfip1 | ENSMUSG00000026305.10 |
| chr1 | 97767595 | 97767854 | 5.97761 | 10.04734 | 7.87135 | Intron (ENSM | 1 | 97770172 | 97792969 | 22798 | 1 | ENSMUST00 | -2318 | Gin1 | ENSMUSG00000026333.9 |
| chr17 | 86536115 | 86536486 | 5.97761 | 10.04734 | 7.87135 | Intron (ENSM | 17 | 86497732 | 86505232 | 7501 | 2 | ENSMUST00 | -30884 | Gm10309 | ENSMUSG00000071036.2 |
| chr2 | 37742002 | 37742212 | 5.97761 | 10.04734 | 7.87135 | Distal Interge | 2 | 37776249 | 37799103 | 22855 | 1 | ENSMUST00 | -34037 | Crb2 | ENSMUSG00000035403.8 |
| chr5 | 147051077 | 147052037 | 5.97761 | 10.04734 | 7.87135 | Intron (ENSM | 5 | 147016655 | 147076586 | 59932 | 2 | ENSMUST00 | 24549 | Lnx2 | ENSMUSG00000016520.7 |
| chr11 | 116108033 | 116108364 | 6.18818 | 10.03405 | 7.85851 | Promoter | 11 | 116105752 | 116108119 | 2368 | 2 | ENSMUST00 | 0 | Trim47 | ENSMUSG00000020773.6 |
| chr17 | 57096274 | 57096690 | 6.18818 | 10.03405 | 7.85851 | Intron (ENSM | 17 | 57080066 | 57087782 | 7717 | 2 | ENSMUST00 | -8493 | Tubb4a | ENSMUSG00000062591.5 |
| chr7 | 105621527 | 105621752 | 5.74755 | 10.02598 | 7.85057 | Intron (ENSM | 7 | 105617031 | 105618026 | 996 | 2 | ENSMUST00 | -3502 | Trim3 | ENSMUSG00000036989.10 |
| chr8 | 22707129 | 22707293 | 5.74755 | 10.02598 | 7.85057 | Promoter | 8 | 22659768 | 22706589 | 46822 | 2 | ENSMUST00 | -541 | Ikbkb | ENSMUSG00000031537.10 |
| chr1 | 135686536 | 135686865 | 6.1757 | 10.01391 | 7.83861 | Intron (ENSM | 1 | 135606447 | 135688105 | 81659 | 2 | ENSMUST00 | 1240 | Nav1 | ENSMUSG00000090399.1 |
| chr10 | 121297192 | 121297630 | 6.1757 | 10.01391 | 7.83861 | Intron (ENSM | 10 | 121263824 | 121311189 | 47366 | 2 | ENSMUST00 | 13559 | Tbc1d30 | ENSMUSG00000052302.5 |
| chr9 | 98329747 | 98330138 | 6.1757 | 10.01391 | 7.83861 | Intron (ENSM | 9 | 98301650 | 98302226 | 577 | 1 | ENSMUST00 | 28098 | RP23-300D5. | ENSMUSG00000101452.1 |
| chr12 | 69537501 | 69538082 | 4.93922 | 9.95903 | 7.78408 | Distal Interge | 12 | 69561229 | 69561337 | 109 | 2 | ENSMUST00 | 23255 | Gm23018 | ENSMUSG00000095776.1 |
| chr16 | 8424475 | 8425196 | 4.93922 | 9.95903 | 7.78408 | Promoter | 16 | 8409276 | 8425136 | 15861 | 2 | ENSMUST00 | 0 | Cldn26 | ENSMUSG00000022715.2 |
| chr11 | 114795179 | 114795642 | 6.13854 | 9.95394 | 7.77908 | Promoter | 11 | 114791217 | 114795873 | 4657 | 2 | ENSMUST00 | 231 | Btbd17 | ENSMUSG00000000202.4 |
| chr14 | 121172163 | 121172507 | 6.13854 | 9.95394 | 7.77908 | Intron (ENSM | 14 | 121102079 | 121238659 | 136581 | 1 | ENSMUST00 | 70085 | Farp1 | ENSMUSG00000025555.9 |
| chr4 | 125125230 | 125125755 | 6.13854 | 9.95394 | 7.77908 | Intron (ENSM | 4 | 125118423 | 125127840 | 9418 | 2 | ENSMUST00 | 2085 | Zc3h12a | ENSMUSG00000042677.7 |
| chr5 | 143548287 | 143548588 | 6.13854 | 9.95394 | 7.77908 | Promoter | 5 | 143548706 | 143564525 | 15820 | 1 | ENSMUST00 | -118 | Fam220a | ENSMUSG00000083012.3 |
| chr5 | 106802140 | 106802497 | 6.13854 | 9.95394 | 7.77908 | Distal Interge | 5 | 106879600 | 106879698 | 99 | 2 | ENSMUST00 | 77201 | Gm27527 | ENSMUSG00000098703.1 |
| chr11 | 85832147 | 85832569 | 6.12626 | 9.93411 | 7.75979 | Promoter | 11 | 85832551 | 85841948 | 9398 | 1 | ENSMUST00 | 0 | Tbx2 | ENSMUSG00000000093.6 |
| chr4 | 62289092 | 62289666 | 6.11402 | 9.91435 | 7.74017 | Intron (ENSM | 4 | 62291547 | 62297477 | 5931 | 1 | ENSMUST00 | -1881 | Slc31a2 | ENSMUSG00000066152.6 |
| chr17 | 46084246 | 46084634 | 6.10183 | 9.89467 | 7.72061 | Distal Interge | 17 | 46110986 | 46128910 | 17925 | 1 | ENSMUST00 | -26352 | Mrps18a | ENSMUSG00000023967.4 |
| chr1 | 89750882 | 89751404 | 6.08969 | 9.87507 | 7.70114 | Intron (ENSM | 1 | 89684802 | 89685112 | 311 | 1 | ENSMUST00 | 66081 | Gm26037 | ENSMUSG00000088775.1 |
| chr11 | 5505827 | 5506262 | 6.08969 | 9.87507 | 7.70114 | Distal Interge | 11 | 5498840 | 5505699 | 6860 | 1 | ENSMUST00 | 6988 | Gm11963 | ENSMUSG00000085189.1 |
| chr6 | 21999093 | 21999507 | 6.08969 | 9.87507 | 7.70114 | Intron (ENSM | 6 | 21986892 | 22069450 | 82559 | 1 | ENSMUST00 | 12202 | Cped1 | ENSMUSG00000062980.10 |
| chr17 | 28769110 | 28769663 | 5.87061 | 9.87137 | 7.69756 | Promoter | 17 | 28769307 | 28778698 | 9392 | 1 | ENSMUST00 | 0 | Mapk13 | ENSMUSG00000004864.7 |
| chr1 | 166940081 | 166940317 | 5.65465 | 9.86863 | 7.69518 | Distal Interge | 1 | 166945597 | 166946358 | 762 | 2 | ENSMUST00 | 6041 | Gm6286 | ENSMUSG00000086925.1 |
| chr8 | 40634560 | 40634839 | 6.06556 | 9.83609 | 7.66283 | Promoter | 8 | 40551095 | 40634789 | 83695 | 2 | ENSMUST00 | 0 | Mtmr7 | ENSMUSG00000039431.11 |
| chr11 | 97804698 | 97805247 | 6.05356 | 9.81671 | 7.64356 | Promoter | 11 | 97805192 | 97836119 | 30928 | 1 | ENSMUST00 | 0 | Lasp1 | ENSMUSG00000038366.10 |
| chr11 | 31965730 | 31966157 | 5.03709 | 9.81157 | 7.63852 | Promoter | 11 | 31965592 | 31979651 | 14060 | 1 | ENSMUST00 | 139 | 4930524B15I | ENSMUSG00000020299.6 |
| chr3 | 122122216 | 122123004 | 5.03709 | 9.81157 | 7.63852 | Promoter | 3 | 122121754 | 122126467 | 4714 | 1 | ENSMUST00 | 463 | Abca4 | ENSMUSG00000028125.9 |
| chr3 | 158560987 | 158561847 | 5.03709 | 9.81157 | 7.63852 | Promoter | 3 | 158084439 | 158562221 | 477783 | 2 | ENSMUST00 | 374 | Lrrc7 | ENSMUSG00000028176.7 |
| chr7 | 133115034 | 133116382 | 5.03709 | 9.81157 | 7.63852 | Intron (ENSM | 7 | 133014731 | 133112699 | 97969 | 2 | ENSMUST00 | -2336 | Ctbp2 | ENSMUSG00000030970.11 |
| chr17 | 25078732 | 25079471 | 6.04161 | 9.7974 | 7.62455 | Promoter | 17 | 25057904 | 25078224 | 20321 | 2 | ENSMUST00 | -509 | Tmem204 | ENSMUSG00000024168.7 |
| chr4 | 15267445 | 15267622 | 6.04161 | 9.7974 | 7.62455 | Intron (ENSM | 4 | 15265831 | 15286753 | 20923 | 1 | ENSMUST00 | 1615 | Tmem64 | ENSMUSG00000043252.8 |
| chr1 | 133420822 | 133421787 | 6.02971 | 9.77817 | 7.60541 | Intron (ENSM | 1 | 133382303 | 133424377 | 42075 | 2 | ENSMUST00 | 2590 | Sox13 | ENSMUSG00000070643.6 |
| chr5 | 117313576 | 117314038 | 6.02971 | 9.77817 | 7.60541 | Exon (ENSMU | 5 | 117312520 | 117315539 | 3020 | 2 | ENSMUST00 | 1501 | BC051077 | ENSMUSG00000097163.1 |
| chr9 | 111273133 | 111273430 | 6.02971 | 9.77817 | 7.60541 | Exon (ENSMU | 9 | 111271945 | 111279097 | 7153 | 1 | ENSMUST00 | 1189 | Epm2aip1 | ENSMUSG00000046785.7 |
| chr12 | 112877424 | 112878083 | 6.01786 | 9.75901 | 7.58634 | Distal Interge | 12 | 112850876 | 112860916 | 10041 | 2 | ENSMUST00 | -16509 | Gpr132 | ENSMUSG00000021298.7 |
| chr14 | 121450830 | 121451075 | 6.01786 | 9.75901 | 7.58634 | Distal Interge | 14 | 121459621 | 121505254 | 45634 | 2 | ENSMUST00 | 54179 | Slc15a1 | ENSMUSG00000025557.9 |
| chr16 | 73679689 | 73680308 | 6.01786 | 9.75901 | 7.58634 | Distal Interge | 16 | 73894664 | 73899885 | 5222 | 2 | ENSMUST00 | 219577 | Robo2 | ENSMUSG00000052516.12 |
| chr19 | 43690902 | 43691246 | 6.01786 | 9.75901 | 7.58634 | Exon (ENSMU | 19 | 43689840 | 43722574 | 32735 | 1 | ENSMUST00 | 1063 | Entpd7 | ENSMUSG00000025192.8 |
| chr2 | 119300414 | 119300818 | 6.01786 | 9.75901 | 7.58634 | Distal Interge | 2 | 119288787 | 119298326 | 9540 | 1 | ENSMUST00 | 11628 | Vps18 | ENSMUSG00000034216.7 |
| chr2 | 23234523 | 23234782 | 6.01786 | 9.75901 | 7.58634 | Promoter | 2 | 23235236 | 23235739 | 504 | 1 | ENSMUST00 | -454 | Gm13422 | ENSMUSG00000084098.1 |
| chr6 | 85511525 | 85512330 | 6.01786 | 9.75901 | 7.58634 | 3' UTR | 6 | 85511131 | 85513589 | 2459 | 2 | ENSMUST00 | 1259 | Egr4 | ENSMUSG00000071341.3 |
| chr7 | 141900866 | 141901169 | 6.01786 | 9.75901 | 7.58634 | Intron (ENSM | 7 | 141874813 | 141902415 | 27603 | 2 | ENSMUST00 | 1246 | Tollip | ENSMUSG00000025139.8 |
| chr7 | 141010330 | 141010697 | 6.01786 | 9.75901 | 7.58634 | Promoter | 7 | 141009586 | 141010770 | 1185 | 2 | ENSMUST00 | 73 | Ifitm3 | ENSMUSG00000025492.6 |
| chr6 | 31317722 | 31318815 | 5.5647 | 9.71562 | 7.54373 | Intron (ENSM | 6 | 31321971 | 31337385 | 15415 | 1 | ENSMUST00 | -3156 | 2210408F21I | ENSMUSG00000087380.2 |
| chr9 | 47545390 | 47545941 | 5.98257 | 9.70197 | 7.53013 | Intron (ENSM | 9 | 47530373 | 47850390 | 320018 | 1 | ENSMUST00 | 15018 | Cadm1 | ENSMUSG00000032076.13 |
| chr1 | 183120557 | 183120860 | 5.76737 | 9.70104 | 7.52928 | Intron (ENSM | 1 | 183086270 | 183147461 | 61192 | 2 | ENSMUST00 | 26601 | Disp1 | ENSMUSG00000030768.7 |
| chr1 | 134098766 | 134099185 | 5.76737 | 9.70104 | 7.52928 | Distal Interge | 1 | 134109368 | 134109799 | 432 | 1 | ENSMUST00 | -10183 | Gm8618 | ENSMUSG00000073559.2 |
| chr4 | 115058229 | 115058772 | 5.76737 | 9.70104 | 7.52928 | Promoter | 4 | 115057811 | 115071228 | 13418 | 1 | ENSMUST00 | 419 | Tal1 | ENSMUSG00000028717.7 |
| chr7 | 101296863 | 101297027 | 5.76737 | 9.70104 | 7.52928 | Promoter | 7 | 101292403 | 101296782 | 4380 | 2 | ENSMUST00 | -82 | Atg16l2 | ENSMUSG00000047767.11 |
| chr8 | 70753118 | 70753812 | 5.76737 | 9.70104 | 7.52928 | Promoter | 8 | 70754679 | 70758677 | 3999 | 1 | ENSMUST00 | -867 | Rab3a | ENSMUSG00000031840.7 |
| chr17 | 34957002 | 34957243 | 5.9709 | 9.68309 | 7.51161 | 3' UTR | 17 | 34956436 | 34959238 | 2803 | 2 | ENSMUST00 | 1995 | Hspa1b | ENSMUSG00000090877.3 |
| chr5 | 65157462 | 65158077 | 5.9709 | 9.68309 | 7.51161 | Intron (ENSM | 5 | 65156070 | 65168142 | 12073 | 1 | ENSMUST00 | 1393 | Klhl5 | ENSMUSG00000054920.7 |
| chr7 | 84679004 | 84679878 | 5.9709 | 9.68309 | 7.51161 | Promoter | 7 | 84615054 | 84679351 | 64298 | 2 | ENSMUST00 | 0 | Zfand6 | ENSMUSG00000030629.9 |
| chr10 | 117209434 | 117209736 | 5.15049 | 9.68248 | 7.51107 | Distal Interge | 10 | 117215142 | 117224507 | 9366 | 2 | ENSMUST00 | 14771 | Yeats4 | ENSMUSG00000020171.7 |
| chr11 | 102025463 | 102025821 | 5.15049 | 9.68248 | 7.51107 | Promoter | 11 | 102023434 | 102025857 | 2424 | 2 | ENSMUST00 | 36 | Mpp3 | ENSMUSG00000052373.9 |
| chr5 | 81519672 | 81520026 | 5.15049 | 9.68248 | 7.51107 | Intron (ENSM | 5 | 81645989 | 81688522 | 42534 | 1 | ENSMUST00 | -125963 | Lphn3 | ENSMUSG00000037605.11 |
| chr6 | 3620329 | 3620970 | 5.15049 | 9.68248 | 7.51107 | Distal Interge | 6 | 3594018 | 3600262 | 6245 | 1 | ENSMUST00 | 26312 | Ccdc132 | ENSMUSG00000001376.12 |
| chr7 | 28290457 | 28291208 | 5.15049 | 9.68248 | 7.51107 | Promoter | 7 | 28284652 | 28291134 | 6483 | 2 | ENSMUST00 | 0 | BC089491 | ENSMUSG00000046750.11 |
| chr9 | 29771088 | 29771556 | 5.15049 | 9.68248 | 7.51107 | Intron (ENSM | 9 | 29590484 | 29592510 | 2027 | 2 | ENSMUST00 | -178579 | Gm15521 | ENSMUSG00000085638.1 |
| chr10 | 84439150 | 84439387 | 5.92467 | 9.6083 | 7.43749 | Intron (ENSM | 10 | 84370905 | 84440597 | 69693 | 2 | ENSMUST00 | 1210 | Nuak1 | ENSMUSG00000020032.9 |
| chr6 | 114738329 | 114738560 | 5.92467 | 9.6083 | 7.43749 | Intron (ENSM | 6 | 114723100 | 114859224 | 136125 | 1 | ENSMUST00 | 15230 | Atg7 | ENSMUSG00000030314.11 |
| chr1 | 162476735 | 162477096 | 5.28342 | 9.57744 | 7.40675 | Promoter | 1 | 162477680 | 162479943 | 2264 | 1 | ENSMUST00 | -584 | Gm10176 | ENSMUSG00000066752.4 |
| chr10 | 8213827 | 8214554 | 5.28342 | 9.57744 | 7.40675 | Intron (ENSM | 10 | 7905653 | 7956230 | 50578 | 2 | ENSMUST00 | -257598 | Tab2 | ENSMUSG00000015755.11 |
| chr10 | 13714661 | 13715258 | 5.28342 | 9.57744 | 7.40675 | Intron (ENSM | 10 | 13653588 | 13844286 | 190699 | 2 | ENSMUST00 | 129028 | Aig1 | ENSMUSG00000019806.8 |
| chr11 | 80680592 | 80681078 | 5.28342 | 9.57744 | 7.40675 | Intron (ENSM | 11 | 80673667 | 80780003 | 106337 | 2 | ENSMUST00 | 98925 | Myo1d | ENSMUSG00000035441.9 |
| chr14 | 77539744 | 77539909 | 5.28342 | 9.57744 | 7.40675 | Promoter | 14 | 77540642 | 77540765 | 124 | 2 | ENSMUST00 | 856 | Gm23823 | ENSMUSG00000089365.1 |
| chr15 | 102028050 | 102028578 | 5.28342 | 9.57744 | 7.40675 | Promoter | 15 | 102028216 | 102032026 | 3811 | 1 | ENSMUST00 | 0 | Krt18 | ENSMUSG00000023043.6 |
| chr17 | 75178575 | 75178791 | 5.28342 | 9.57744 | 7.40675 | Promoter | 17 | 75178810 | 75348933 | 170124 | 1 | ENSMUST00 | -19 | Ltbp1 | ENSMUSG00000001870.10 |
| chr19 | 44741085 | 44742310 | 5.28342 | 9.57744 | 7.40675 | Distal Interge | 19 | 44756045 | 44837851 | 81807 | 1 | ENSMUST00 | -13735 | Pax2 | ENSMUSG00000004231.10 |
| chr19 | 25033103 | 25033291 | 5.28342 | 9.57744 | 7.40675 | Intron (ENSM | 19 | 24999529 | 25202432 | 202904 | 1 | ENSMUST00 | 33575 | Dock8 | ENSMUSG00000052085.6 |
| chr2 | 31953965 | 31954653 | 5.28342 | 9.57744 | 7.40675 | Intron (ENSM | 2 | 31950517 | 31971338 | 20822 | 1 | ENSMUST00 | 3449 | Aif1l | ENSMUSG00000001864.8 |
| chr2 | 181219714 | 181220216 | 5.28342 | 9.57744 | 7.40675 | Promoter | 2 | 181220013 | 181222854 | 2842 | 1 | ENSMUST00 | 0 | BC051628 | ENSMUSG00000047841.8 |
| chr2 | 167129313 | 167129807 | 5.28342 | 9.57744 | 7.40675 | Intron (ENSM | 2 | 167098794 | 167101826 | 3033 | 2 | ENSMUST00 | -27488 | Gm14290 | ENSMUSG00000086605.1 |
| chr3 | 59130506 | 59130674 | 5.28342 | 9.57744 | 7.40675 | Promoter | 3 | 59114626 | 59130624 | 15999 | 2 | ENSMUST00 | 0 | P2ry14 | ENSMUSG00000036381.8 |
| chr5 | 33652798 | 33653025 | 5.28342 | 9.57744 | 7.40675 | Promoter | 5 | 33640055 | 33652574 | 12520 | 2 | ENSMUST00 | -225 | Slbp | ENSMUSG00000004642.8 |
| chr5 | 74092694 | 74092876 | 5.28342 | 9.57744 | 7.40675 | Promoter | 5 | 74093060 | 74094336 | 1277 | 1 | ENSMUST00 | -184 | Dancr | ENSMUSG00000053656.5 |
| chr6 | 142963918 | 142964846 | 5.28342 | 9.57744 | 7.40675 | Promoter | 6 | 142821545 | 142964452 | 142908 | 2 | ENSMUST00 | 0 | St8sia1 | ENSMUSG00000030283.5 |
| chr7 | 46179545 | 46180193 | 5.28342 | 9.57744 | 7.40675 | Promoter | 7 | 46104523 | 46180033 | 75511 | 2 | ENSMUST00 | 0 | Abcc8 | ENSMUSG00000040136.9 |
| chr7 | 127652150 | 127652419 | 5.28342 | 9.57744 | 7.40675 | Distal Interge | 7 | 127608756 | 127615797 | 7042 | 2 | ENSMUST00 | -36354 | Zfp629 | ENSMUSG00000045639.9 |
| chr8 | 61225383 | 61226302 | 5.28342 | 9.57744 | 7.40675 | 5' UTR | 8 | 61224171 | 61396072 | 171902 | 1 | ENSMUST00 | 1213 | Sh3rf1 | ENSMUSG00000031642.8 |
| chr9 | 14859895 | 14860359 | 5.28342 | 9.57744 | 7.40675 | Promoter | 9 | 14860210 | 14870789 | 10580 | 1 | ENSMUST00 | 0 | Gpr83 | ENSMUSG00000031932.9 |
| chr10 | 99265264 | 99265700 | 5.90182 | 9.57131 | 7.40145 | Intron (ENSM | 10 | 99263231 | 99267488 | 4258 | 1 | ENSMUST00 | 2034 | Dusp6 | ENSMUSG00000019960.7 |
| chr10 | 21998653 | 21999211 | 5.90182 | 9.57131 | 7.40145 | 3' UTR | 10 | 21996591 | 21998240 | 1650 | 1 | ENSMUST00 | 2063 | Sgk1 | ENSMUSG00000019970.10 |

| | | | | | | | | | | | | | | | |
|---|---|---|---|---|---|---|---|---|---|---|---|---|---|---|---|
| chr15 | 100489585 | 100490023 | 5.90182 | 9.57131 | 7.40145 | 5' UTR | 15 | 100479570 | 100495239 | 15670 | 2 | ENSMUST00 | 5216 | Csrnp2 | ENSMUSG00000044636.5 |
| chr16 | 33751270 | 33751495 | 5.90182 | 9.57131 | 7.40145 | Intron (ENSM | 16 | 33753156 | 33761012 | 7857 | 1 | ENSMUST00 | -1661 | Heg1 | ENSMUSG00000075254.6 |
| chr7 | 45627581 | 45627935 | 5.6677 | 9.53603 | 7.36682 | Promoter | 7 | 45627488 | 45639093 | 11606 | 1 | ENSMUST00 | 94 | Rasip1 | ENSMUSG00000044562.10 |
| chr1 | 91064586 | 91064951 | 5.86754 | 9.51579 | 7.34688 | Intron (ENSM | 1 | 91053586 | 91128287 | 74702 | 1 | ENSMUST00 | 11001 | Lrrfip1 | ENSMUSG00000026305.10 |
| chr11 | 117174661 | 117175249 | 5.86754 | 9.51579 | 7.34688 | Distal Interg | 11 | 117155069 | 117156475 | 1407 | 1 | ENSMUST00 | 19593 | Sec14l1 | ENSMUSG00000020823.11 |
| chr11 | 62921017 | 62921848 | 5.86754 | 9.51579 | 7.34688 | Promoter | 11 | 62919079 | 62921592 | 2514 | 2 | ENSMUST00 | 0 | Tvp23bos | ENSMUSG00000086677.1 |
| chr12 | 70975880 | 70976055 | 5.86754 | 9.51579 | 7.34688 | Intron (ENSM | 12 | 70974719 | 70984837 | 10119 | 1 | ENSMUST00 | 1162 | Psma3 | ENSMUSG00000060073.4 |
| chr13 | 102902043 | 102902635 | 5.86754 | 9.51579 | 7.34688 | Intron (ENSM | 13 | 102741969 | 102905786 | 163818 | 2 | ENSMUST00 | 3151 | Mast4 | ENSMUSG00000034751.10 |
| chr16 | 33178310 | 33178964 | 5.86754 | 9.51579 | 7.34688 | Distal Interg | 16 | 33185071 | 33243311 | 58241 | 1 | ENSMUST00 | -6107 | Osbpl11 | ENSMUSG00000022807.8 |
| chr17 | 8147042 | 8147215 | 5.86754 | 9.51579 | 7.34688 | Promoter | 17 | 8128610 | 8148097 | 19488 | 2 | ENSMUST00 | 882 | Rnaset2a | ENSMUSG00000095687.1 |
| chr19 | 58309736 | 58310180 | 5.86754 | 9.51579 | 7.34688 | Intron (ENSM | 19 | 58204139 | 58236047 | 31909 | 1 | ENSMUST00 | 105598 | Gm16277 | ENSMUSG00000087002.1 |
| chr2 | 58516147 | 58516393 | 5.86754 | 9.51579 | 7.34688 | Promoter | 2 | 58388644 | 58516272 | 127629 | 2 | ENSMUST00 | 0 | Acvr1 | ENSMUSG00000026836.10 |
| chr2 | 118555231 | 118555395 | 5.86754 | 9.51579 | 7.34688 | Distal Interg | 2 | 118528757 | 118549687 | 20931 | 2 | ENSMUST00 | -5545 | Bmf | ENSMUSG00000040093.10 |
| chr3 | 34663349 | 34663513 | 5.86754 | 9.51579 | 7.34688 | Intron (ENSM | 3 | 34665243 | 34677995 | 12753 | 1 | ENSMUST00 | -1730 | Sox2ot | ENSMUSG00000090828.3 |
| chr4 | 132113423 | 132113958 | 5.86754 | 9.51579 | 7.34688 | 3' UTR | 4 | 132093891 | 132094298 | 408 | 1 | ENSMUST00 | 19533 | Gm13214 | ENSMUSG00000083368.1 |
| chr4 | 83303690 | 83303937 | 5.86754 | 9.51579 | 7.34688 | Intron (ENSM | 4 | 83285804 | 83285910 | 107 | 2 | ENSMUST00 | -17781 | Gm23412 | ENSMUSG00000096801.1 |
| chr5 | 147400136 | 147400660 | 5.86754 | 9.51579 | 7.34688 | Promoter | 5 | 147330741 | 147400489 | 69749 | 2 | ENSMUST00 | 0 | Flt3 | ENSMUSG00000042817.12 |
| chr7 | 60622480 | 60622975 | 5.86754 | 9.51579 | 7.34688 | Distal Interg | 7 | 60155538 | 60156170 | 633 | 1 | ENSMUST00 | 466943 | Gm7367 | ENSMUSG00000047370.4 |
| chr8 | 110950094 | 110950489 | 5.86754 | 9.51579 | 7.34688 | Intron (ENSM | 8 | 110940904 | 110957012 | 16109 | 1 | ENSMUST00 | 9191 | St3gal2 | ENSMUSG00000031749.7 |
| chr8 | 47135549 | 47135994 | 5.86754 | 9.51579 | 7.34688 | Distal Interg | 8 | 47068931 | 47083476 | 14546 | 1 | ENSMUST00 | 66619 | Enpp6 | ENSMUSG00000038173.9 |
| chr8 | 8969125 | 8969289 | 5.86754 | 9.51579 | 7.34688 | Distal Interg | 8 | 8666577 | 8690537 | 23961 | 2 | ENSMUST00 | -278589 | Arglu1 | ENSMUSG00000040459.10 |
| chrX | 153499307 | 153499748 | 5.86754 | 9.51579 | 7.34688 | Exon (ENSMU | 20 | 153498227 | 153501570 | 3344 | 1 | ENSMUST00 | 1081 | Ubqln2 | ENSMUSG00000050148.9 |
| chr1 | 53977243 | 53977840 | 5.44141 | 9.50478 | 7.33812 | Intron (ENSM | 1 | 53886828 | 53886941 | 114 | 2 | ENSMUST00 | -90303 | Gm24251 | ENSMUSG00000087949.1 |
| chr10 | 69334521 | 69335256 | 5.44141 | 9.50478 | 7.33812 | Downstream | 10 | 69340599 | 69342729 | 2131 | 2 | ENSMUST00 | 7473 | Cdk1 | ENSMUSG00000019942.7 |
| chr13 | 28757850 | 28758518 | 5.44141 | 9.50478 | 7.33812 | Intron (ENSM | 13 | 28710781 | 28710873 | 93 | 2 | ENSMUST00 | -46978 | Mir6368 | ENSMUSG00000099158.1 |
| chr15 | 100613439 | 100613744 | 5.44141 | 9.50478 | 7.33812 | Downstream | 15 | 100614140 | 100615110 | 971 | 2 | ENSMUST00 | 1366 | C330013E15I | ENSMUSG00000097093.1 |
| chr19 | 5016611 | 5016824 | 5.44141 | 9.50478 | 7.33812 | Distal Interg | 19 | 5024006 | 5031971 | 7966 | 1 | ENSMUST00 | -7182 | Slc29a2 | ENSMUSG00000024891.5 |
| chr2 | 152437446 | 152437632 | 5.44141 | 9.50478 | 7.33812 | 3' UTR | 2 | 152427662 | 152439643 | 11982 | 1 | ENSMUST00 | 9785 | 6820408C15I | ENSMUSG00000032680.6 |
| chr3 | 87794791 | 87795223 | 5.44141 | 9.50478 | 7.33812 | Promoter | 3 | 87778244 | 87795162 | 16919 | 2 | ENSMUST00 | 0 | Ntrk1 | ENSMUSG00000028072.6 |
| chr3 | 145991241 | 145991576 | 5.44141 | 9.50478 | 7.33812 | Intron (ENSM | 3 | 145987870 | 146021720 | 33851 | 1 | ENSMUST00 | 3372 | Syde2 | ENSMUSG00000036863.7 |
| chr3 | 149444994 | 149445458 | 5.44141 | 9.50478 | 7.33812 | Distal Interg | 3 | 149340903 | 149341008 | 106 | 2 | ENSMUST00 | -103987 | Gm26468 | ENSMUSG00000089269.1 |
| chr4 | 88955655 | 88956039 | 5.44141 | 9.50478 | 7.33812 | Distal Interg | 4 | 88910557 | 88910662 | 106 | 2 | ENSMUST00 | -44994 | Mir31 | ENSMUSG00000065408.1 |
| chr4 | 138395984 | 138396571 | 5.44141 | 9.50478 | 7.33812 | Promoter | 4 | 138394092 | 138396528 | 2437 | 2 | ENSMUST00 | 0 | Fam43b | ENSMUSG00000078235.3 |
| chr5 | 144259207 | 144259440 | 5.44141 | 9.50478 | 7.33812 | Intron (ENSM | 5 | 144241777 | 144256186 | 14410 | 2 | ENSMUST00 | -3022 | 2900089D17I | ENSMUSG00000087484.1 |
| chr5 | 52254279 | 52254798 | 5.44141 | 9.50478 | 7.33812 | Distal Interg | 5 | 52190681 | 52197984 | 7304 | 1 | ENSMUST00 | 63599 | 9230114K14I | ENSMUSG00000097145.1 |
| chr7 | 27884388 | 27884607 | 5.44141 | 9.50478 | 7.33812 | Distal Interg | 7 | 27862557 | 27879567 | 17011 | 1 | ENSMUST00 | 21832 | Zfp607 | ENSMUSG00000020420.7 |
| chr7 | 120809763 | 120810066 | 5.44141 | 9.50478 | 7.33812 | Distal Interg | 7 | 120786481 | 120792762 | 6282 | 1 | ENSMUST00 | 23283 | Vwa3a | ENSMUSG00000030889.8 |
| chr7 | 70287773 | 70288401 | 5.44141 | 9.50478 | 7.33812 | Distal Interg | 7 | 70248973 | 70249776 | 804 | 2 | ENSMUST00 | -37998 | RP23-474J16 | ENSMUSG00000099387.1 |
| chr8 | 109105405 | 109105849 | 5.44141 | 9.50478 | 7.33812 | Intron (ENSM | 8 | 109075223 | 109231047 | 155825 | 1 | ENSMUST00 | 30183 | D030068K23I | ENSMUSG00000097393.2 |
| chr1 | 160351914 | 160352201 | 5.6323 | 9.47728 | 7.31216 | Promoter | 1 | 160219174 | 160351571 | 132398 | 2 | ENSMUST00 | -344 | Rabgap1l | ENSMUSG00000026721.10 |
| chr10 | 79920957 | 79921266 | 5.6323 | 9.47728 | 7.31216 | Exon (ENSMU | 10 | 79910238 | 79917006 | 6769 | 2 | ENSMUST00 | -3952 | R3hdm4 | ENSMUSG00000035781.8 |
| chr10 | 13300162 | 13300733 | 5.6323 | 9.47728 | 7.31216 | Intron (ENSM | 10 | 13207717 | 13324160 | 116444 | 2 | ENSMUST00 | 23427 | Phactr2 | ENSMUSG00000062866.10 |
| chr10 | 8760549 | 8761163 | 5.6323 | 9.47728 | 7.31216 | Intron (ENSM | 10 | 8824011 | 8824147 | 137 | 2 | ENSMUST00 | 62984 | Gm25410 | ENSMUSG00000065692.1 |
| chr11 | 9116248 | 9116505 | 5.6323 | 9.47728 | 7.31216 | Distal Interg | 11 | 9118103 | 9136164 | 18062 | 1 | ENSMUST00 | -1598 | Upp1 | ENSMUSG00000020407.8 |
| chr11 | 8494386 | 8495188 | 5.6323 | 9.47728 | 7.31216 | Intron (ENSM | 11 | 8451070 | 8468675 | 17606 | 2 | ENSMUST00 | -25712 | Tns3 | ENSMUSG00000020422.8 |
| chr11 | 69397510 | 69397960 | 5.6323 | 9.47728 | 7.31216 | Promoter | 11 | 69396519 | 69398234 | 1716 | 2 | ENSMUST00 | 274 | Tmem88 | ENSMUSG00000045377.5 |
| chr11 | 98951833 | 98952000 | 5.6323 | 9.47728 | 7.31216 | Intron (ENSM | 11 | 98946444 | 98946550 | 107 | 1 | ENSMUST00 | 5390 | Gm22061 | ENSMUSG00000064491.1 |
| chr13 | 45809709 | 45810251 | 5.6323 | 9.47728 | 7.31216 | Intron (ENSM | 13 | 45549758 | 45964991 | 415234 | 2 | ENSMUST00 | 154740 | Atxn1 | ENSMUSG00000046876.10 |
| chr13 | 22020411 | 22020846 | 5.6323 | 9.47728 | 7.31216 | Distal Interg | 13 | 22016690 | 22018249 | 1560 | 1 | ENSMUST00 | 3722 | Gm11291 | ENSMUSG00000086508.1 |
| chr15 | 85658206 | 85658609 | 5.6323 | 9.47728 | 7.31216 | Distal Interg | 15 | 85639747 | 85645934 | 6188 | 1 | ENSMUST00 | 18460 | RP23-64J7.2 | ENSMUSG00000101828.1 |
| chr17 | 32634817 | 32635209 | 5.6323 | 9.47728 | 7.31216 | Exon (ENSMU | 17 | 32621319 | 32636184 | 14866 | 1 | ENSMUST00 | 13499 | Cyp4f37 | ENSMUSG00000062464.5 |
| chr19 | 12573424 | 12574181 | 5.6323 | 9.47728 | 7.31216 | Promoter | 19 | 12573523 | 12589762 | 16240 | 1 | ENSMUST00 | 0 | Fam111a | ENSMUSG00000024691.8 |
| chr19 | 6427877 | 6428374 | 5.6323 | 9.47728 | 7.31216 | Promoter | 19 | 6427937 | 6533217 | 105281 | 1 | ENSMUST00 | 0 | Nrxn2 | ENSMUSG00000033768.11 |
| chr19 | 4191540 | 4191879 | 5.6323 | 9.47728 | 7.31216 | Promoter | 19 | 4192158 | 4195419 | 3262 | 1 | ENSMUST00 | -279 | Ppp1ca | ENSMUSG00000040385.6 |
| chr19 | 34551671 | 34551887 | 5.6323 | 9.47728 | 7.31216 | Promoter | 19 | 34550710 | 34576419 | 25710 | 1 | ENSMUST00 | 962 | Ifit2 | ENSMUSG00000045932.7 |
| chr2 | 73596734 | 73596915 | 5.6323 | 9.47728 | 7.31216 | Promoter | 2 | 73596824 | 73599200 | 2377 | 1 | ENSMUST00 | 0 | Chn1os3 | ENSMUSG00000086544.1 |
| chr3 | 84155962 | 84156227 | 5.6323 | 9.47728 | 7.31216 | Promoter | 3 | 84087934 | 84155786 | 67853 | 2 | ENSMUST00 | -177 | Mnd1 | ENSMUSG00000033752.6 |
| chr4 | 57141759 | 57142124 | 5.6323 | 9.47728 | 7.31216 | Intron (ENSM | 4 | 56991972 | 57143437 | 151466 | 2 | ENSMUST00 | 1313 | Epb4.1l4b | ENSMUSG00000028434.7 |
| chr4 | 148131571 | 148131756 | 5.6323 | 9.47728 | 7.31216 | Promoter | 4 | 148130915 | 148145699 | 14785 | 1 | ENSMUST00 | 657 | Mad2l2 | ENSMUSG00000029003.6 |
| chr4 | 114635284 | 114635626 | 5.6323 | 9.47728 | 7.31216 | Distal Interg | 4 | 114672697 | 114673340 | 644 | 2 | ENSMUST00 | 37714 | RP23-454F13 | ENSMUSG00000100998.1 |
| chr5 | 115977383 | 115977814 | 5.6323 | 9.47728 | 7.31216 | Intron (ENSM | 5 | 115968025 | 116006365 | 38341 | 1 | ENSMUST00 | 9359 | Cit | ENSMUSG00000029516.14 |
| chr5 | 146792136 | 146792704 | 5.6323 | 9.47728 | 7.31216 | Intron (ENSM | 5 | 146734809 | 146795006 | 60198 | 2 | ENSMUST00 | 2302 | Usp12 | ENSMUSG00000029640.12 |
| chr6 | 131292266 | 131292477 | 5.6323 | 9.47728 | 7.31216 | Promoter | 6 | 131284605 | 131292260 | 7656 | 2 | ENSMUST00 | -7 | Magohb | ENSMUSG00000030188.7 |
| chr6 | 54970306 | 54970894 | 5.6323 | 9.47728 | 7.31216 | Promoter | 6 | 54923942 | 54971661 | 47720 | 2 | ENSMUST00 | 767 | Nod1 | ENSMUSG00000038058.9 |
| chr8 | 12926422 | 12926874 | 5.6323 | 9.47728 | 7.31216 | Promoter | 8 | 12926632 | 13018177 | 91546 | 1 | ENSMUST00 | 0 | Mcf2l | ENSMUSG00000031442.15 |
| chr8 | 60995279 | 60995564 | 5.6323 | 9.47728 | 7.31216 | Intron (ENSM | 8 | 60993241 | 61030600 | 37360 | 1 | ENSMUST00 | 2039 | Nek1 | ENSMUSG00000031644.13 |
| chr8 | 128362422 | 128363340 | 5.6323 | 9.47728 | 7.31216 | Exon (ENSMU | 8 | 128359241 | 128359320 | 80 | 1 | ENSMUST00 | 3182 | Mir1903 | ENSMUSG00000084523.1 |
| chr8 | 26966391 | 26966705 | 5.6323 | 9.47728 | 7.31216 | Distal Interg | 8 | 26977336 | 26981461 | 4126 | 1 | ENSMUST00 | -10631 | Zfp703 | ENSMUSG00000085795.2 |
| chr9 | 15627640 | 15628058 | 5.6323 | 9.47728 | 7.31216 | Promoter | 9 | 15559870 | 15627933 | 68064 | 2 | ENSMUST00 | 0 | Ccdc67 | ENSMUSG00000039977.11 |
| chrX | 145506164 | 145506458 | 5.6323 | 9.47728 | 7.31216 | Promoter | 20 | 145475720 | 145505181 | 29462 | 2 | ENSMUST00 | -984 | Amot | ENSMUSG00000041688.11 |
| chr10 | 108446465 | 108446956 | 5.82323 | 9.44397 | 7.28107 | Distal Interg | 10 | 108428067 | 108428160 | 94 | 1 | ENSMUST00 | 18399 | Gm25591 | ENSMUSG00000093762.1 |
| chr11 | 75650646 | 75651124 | 5.79018 | 9.39037 | 7.2276 | Promoter | 11 | 75650504 | 75673910 | 23407 | 1 | ENSMUST00 | 143 | Myo1c | ENSMUSG00000017774.14 |
| chr14 | 103899206 | 103899751 | 5.79018 | 9.39037 | 7.2276 | Distal Interg | 14 | 103814625 | 103844173 | 29549 | 2 | ENSMUST00 | -55034 | Ednrb | ENSMUSG00000022122.9 |
| chr14 | 121697980 | 121698439 | 5.79018 | 9.39037 | 7.2276 | Promoter | 14 | 121542039 | 121698417 | 156379 | 2 | ENSMUST00 | 0 | Dock9 | ENSMUSG00000025558.10 |
| chr3 | 86076581 | 86076866 | 5.79018 | 9.39037 | 7.2276 | Intron (ENSM | 3 | 86070915 | 86130526 | 59612 | 1 | ENSMUST00 | 5667 | Sh3d19 | ENSMUSG00000028082.8 |
| chr5 | 125175332 | 125175506 | 5.79018 | 9.39037 | 7.2276 | Intron (ENSM | 5 | 125017162 | 125179053 | 161892 | 2 | ENSMUST00 | 3547 | Ncor2 | ENSMUSG00000029478.11 |
| chr12 | 85289617 | 85289833 | 5.57141 | 9.37604 | 7.21394 | Exon (ENSMU | 12 | 85288591 | 85299358 | 10768 | 1 | ENSMUST00 | 1027 | Zc2hc1c | ENSMUSG00000045064.4 |
| chr8 | 70001559 | 70001799 | 5.57141 | 9.37604 | 7.21394 | Distal Interg | 8 | 69907069 | 69996379 | 89311 | 2 | ENSMUST00 | -5181 | Gatad2a | ENSMUSG00000036180.9 |
| chr1 | 24586513 | 24586795 | 5.97392 | 9.35129 | 7.18948 | Promoter | 1 | 24279770 | 24587411 | 307642 | 2 | ENSMUST00 | 616 | Col19a1 | ENSMUSG00000026141.8 |
| chr1 | 173802206 | 173802475 | 5.97392 | 9.35129 | 7.18948 | Promoter | 1 | 173775837 | 173802720 | 26884 | 2 | ENSMUST00 | 245 | Gm16340 | ENSMUSG00000090222.1 |
| chr10 | 62507596 | 62507814 | 5.97392 | 9.35129 | 7.18948 | Promoter | 10 | 62494428 | 62507755 | 13328 | 2 | ENSMUST00 | 0 | Srgn | ENSMUSG00000020077.9 |
| chr10 | 33904835 | 33905470 | 5.97392 | 9.35129 | 7.18948 | Promoter | 10 | 33905111 | 33916021 | 10911 | 1 | ENSMUST00 | 0 | Rsph4a | ENSMUSG00000039552.8 |
| chr10 | 102512716 | 102513022 | 5.97392 | 9.35129 | 7.18948 | Promoter | 10 | 102512222 | 102546560 | 34339 | 1 | ENSMUST00 | 495 | Rassf9 | ENSMUSG00000044921.4 |
| chr10 | 81401906 | 81402504 | 5.97392 | 9.35129 | 7.18948 | Intron (ENSM | 10 | 81399196 | 81400554 | 1359 | 2 | ENSMUST00 | -1353 | Nfic | ENSMUSG00000055053.12 |
| chr10 | 128345191 | 128345427 | 5.97392 | 9.35129 | 7.18948 | Intron (ENSM | 10 | 128349241 | 128349375 | 135 | 1 | ENSMUST00 | -3814 | Gm23182 | ENSMUSG00000089313.2 |
| chr14 | 69768056 | 69768275 | 5.97392 | 9.35129 | 7.18948 | Promoter | 14 | 69767472 | 69784403 | 16932 | 1 | ENSMUST00 | 585 | Tnfrsf10b | ENSMUSG00000022074.5 |
| chr14 | 14295000 | 14295271 | 5.97392 | 9.35129 | 7.18948 | Distal Interg | 14 | 14328113 | 14329066 | 954 | 1 | ENSMUST00 | -32842 | Olfr31 | ENSMUSG00000072707.1 |
| chr14 | 7488354 | 7488753 | 5.97392 | 9.35129 | 7.18948 | Intron (ENSM | 14 | 7466362 | 7483762 | 17401 | 2 | ENSMUST00 | -4593 | Gm3752 | ENSMUSG00000091617.3 |
| chr17 | 12118134 | 12118437 | 5.97392 | 9.35129 | 7.18948 | Promoter | 17 | 12118704 | 12210710 | 92007 | 1 | ENSMUST00 | -267 | Agpat4 | ENSMUSG00000023827.3 |
| chr17 | 48585867 | 48586135 | 5.97392 | 9.35129 | 7.18948 | Distal Interg | 17 | 48617823 | 48617954 | 132 | 1 | ENSMUST00 | -31688 | Gm24071 | ENSMUSG00000077708.1 |
| chr18 | 54611450 | 54611844 | 5.97392 | 9.35129 | 7.18948 | Distal Interg | 18 | 54422295 | 54452840 | 30546 | 1 | ENSMUST00 | 189156 | Redrum | ENSMUSG00000096982.1 |
| chr4 | 55420280 | 55420444 | 5.97392 | 9.35129 | 7.18948 | Distal Interg | 4 | 55410443 | 55418842 | 8400 | 2 | ENSMUST00 | -1439 | Gm12505 | ENSMUSG00000087070.1 |
| chr4 | 82512626 | 82512980 | 5.97392 | 9.35129 | 7.18948 | Intron (ENSM | 4 | 82508016 | 82547292 | 39277 | 1 | ENSMUST00 | 4611 | Gm11266 | ENSMUSG00000087413.1 |
| chr6 | 142814434 | 142814814 | 5.97392 | 9.35129 | 7.18948 | Promoter | 6 | 142814385 | 142816955 | 2571 | 1 | ENSMUST00 | 50 | Gm7457 | ENSMUSG00000072683.3 |
| chr6 | 145187478 | 145187687 | 5.97392 | 9.35129 | 7.18948 | Exon (ENSMU | 6 | 145182275 | 145187630 | 5356 | 1 | ENSMUST00 | 5204 | Gm15543 | ENSMUSG00000086863.1 |
| chr7 | 116608212 | 116608439 | 5.97392 | 9.35129 | 7.18948 | Distal Interg | 7 | 116625122 | 116625242 | 121 | 2 | ENSMUST00 | 16803 | n-R5s157 | ENSMUSG00000084492.1 |
| chr7 | 38174151 | 38174607 | 5.97392 | 9.35129 | 7.18948 | Downstream | 7 | 38176614 | 38176671 | 58 | 2 | ENSMUST00 | 2064 | Gm22203 | ENSMUSG00000089230.1 |
| chr8 | 86787772 | 86788159 | 5.97392 | 9.35129 | 7.18948 | Distal Interg | 8 | 86723940 | 86746006 | 22067 | 2 | ENSMUST00 | -41767 | Siah1a | ENSMUSG00000036840.4 |
| chr8 | 13964973 | 13965430 | 5.97392 | 9.35129 | 7.18948 | Intron (ENSM | 8 | 13972616 | 13974743 | 2128 | 2 | ENSMUST00 | 9313 | Tdrp | ENSMUSG00000050052.6 |
| chr9 | 107735725 | 107736006 | 5.97392 | 9.35129 | 7.18948 | Distal Interg | 9 | 107743333 | 107744234 | 902 | 2 | ENSMUST00 | 8228 | Rbm5 | ENSMUSG00000032580.7 |
| chr9 | 67207848 | 67208595 | 5.97392 | 9.35129 | 7.18948 | Distal Interg | 9 | 67236651 | 67236735 | 85 | 2 | ENSMUST00 | 28140 | Mir190a | ENSMUSG00000076379.1 |
| chrX | 56731752 | 56732238 | 5.97392 | 9.35129 | 7.18948 | Promoter | 20 | 56731787 | 56793346 | 61560 | 1 | ENSMUST00 | 0 | Fhl1 | ENSMUSG00000023092.11 |
| chrX | 93296906 | 93297402 | 5.97392 | 9.35129 | 7.18948 | Exon (ENSMU | 20 | 93286529 | 93298357 | 11829 | 1 | ENSMUST00 | 10378 | Arx | ENSMUSG00000035277.10 |
| chrX | 136666822 | 136667111 | 5.97392 | 9.35129 | 7.18948 | Promoter | 20 | 136666611 | 136668065 | 1455 | 1 | ENSMUST00 | 212 | Tceal3 | ENSMUSG00000044550.6 |
| chrX | 141724470 | 141724905 | 5.97392 | 9.35129 | 7.18948 | Promoter | 20 | 141724838 | 141726163 | 1326 | 1 | ENSMUST00 | 0 | Gm15295 | ENSMUSG00000086426.1 |
| chrY | 10643624 | 10643797 | 5.97392 | 9.35129 | 7.18948 | Promoter | 21 | 10641257 | 10643107 | 1851 | 2 | ENSMUST00 | -518 | Gm20775 | ENSMUSG00000075359.2 |
| chr1 | 18058351 | 18058672 | 5.96317 | 9.33407 | 7.1749 | Promoter | 1 | 18048100 | 18058269 | 10170 | 2 | ENSMUST00 | -83 | RP24-571A14 | ENSMUSG00000101717.1 |
| chr15 | 78581448 | 78581629 | 5.96317 | 9.33407 | 7.1749 | Distal Interg | 15 | 78559169 | 78572783 | 13615 | 2 | ENSMUST00 | -8666 | Rac2 | ENSMUSG00000033220.6 |
| chr18 | 55040404 | 55040911 | 5.93826 | 9.29414 | 7.13524 | Intron (ENSM | 18 | 55057585 | 55079414 | 21830 | 1 | ENSMUST00 | -16674 | Gm4221 | ENSMUSG00000096948.1 |
| chr5 | 67534263 | 67534510 | 5.93826 | 9.29414 | 7.13524 | Distal Interg | 5 | 67607883 | 67613992 | 6110 | 1 | ENSMUST00 | -73373 | Shisa3 | ENSMUSG00000050010.7 |
| chr15 | 99259542 | 99259783 | 5.92587 | 9.2743 | 7.11577 | Intron (ENSM | 15 | 99254882 | 99255014 | 133 | 2 | ENSMUST00 | -4529 | Gm25999 | ENSMUSG00000093139.1 |
| chr16 | 21644103 | 21644448 | 5.90127 | 9.23487 | 7.07661 | 3' UTR | 16 | 21627631 | 21644416 | 16786 | 1 | ENSMUST00 | 16473 | Vps8 | ENSMUSG00000033653.12 |
| chr11 | 8467118 | 8467289 | 5.47834 | 9.2208 | 7.06269 | Intron (ENSM | 11 | 8451070 | 8468675 | 17606 | 2 | ENSMUST00 | 1386 | Tns3 | ENSMUSG00000020422.8 |
| chr3 | 102715193 | 102715649 | 5.47834 | 9.2208 | 7.06269 | Distal Interg | 3 | 102734529 | 102772292 | 37764 | 1 | ENSMUST00 | -18880 | Tspan2 | ENSMUSG00000027858.8 |
| chr5 | 136967757 | 136968173 | 5.47834 | 9.2208 | 7.06269 | Promoter | 5 | 136967869 | 136975844 | 7976 | 1 | ENSMUST00 | 0 | Cldn15 | ENSMUSG00000001739.9 |
| chr7 | 24377179 | 24377620 | 5.47834 | 9.2208 | 7.06269 | Exon (ENSMU | 7 | 24370263 | 24385209 | 14947 | 1 | ENSMUST00 | 6917 | Kcnn4 | ENSMUSG00000054342.7 |
| chr1 | 14308282 | 14308695 | 5.68268 | 9.21576 | 7.0578 | Intron (ENSM | 1 | 14270775 | 14309836 | 39062 | 2 | ENSMUST00 | 1141 | Eya1 | ENSMUSG00000025932.9 |
| chr1 | 53978264 | 53978534 | 5.68268 | 9.21576 | 7.0578 | Intron (ENSM | 1 | 53886828 | 53886941 | 114 | 2 | ENSMUST00 | -91324 | Gm24251 | ENSMUSG00000087949.1 |
| chr12 | 70837745 | 70838030 | 5.68268 | 9.21576 | 7.0578 | Intron (ENSM | 12 | 70825514 | 70902232 | 76719 | 1 | ENSMUST00 | 12232 | Frmd6 | ENSMUSG00000048285.8 |
| chr14 | 67070588 | 67070874 | 5.68268 | 9.21576 | 7.0578 | Intron (ENSM | 14 | 67014057 | 67072471 | 58415 | 2 | ENSMUST00 | 1597 | Ppp2r2a | ENSMUSG00000022052.8 |
| chr15 | 93628520 | 93628979 | 5.88904 | 9.21528 | 7.0578 | Distal Interg | 15 | 93499114 | 93595891 | 96778 | 2 | ENSMUST00 | -32630 | Prickle1 | ENSMUSG00000036158.7 |
| chr18 | 70564785 | 70565429 | 5.68268 | 9.21576 | 7.0578 | Distal Interg | 18 | 70568189 | 70626131 | 57943 | 1 | ENSMUST00 | -2760 | Mbd2 | ENSMUSG00000024513.11 |
| chr2 | 38196321 | 38196964 | 5.68268 | 9.21576 | 7.0578 | Intron (ENSM | 2 | 38021266 | 38166081 | 144816 | 2 | ENSMUST00 | -30241 | Dennd1a | ENSMUSG00000035392.12 |
| chr2 | 38348016 | 38348361 | 5.68268 | 9.21576 | 7.0578 | Promoter | 2 | 38347357 | 38348316 | 960 | 2 | ENSMUST00 | 0 | Gm27197 | ENSMUSG00000098383.1 |
| chr7 | 137001158 | 137001494 | 5.68268 | 9.21576 | 7.0578 | Intron (ENSM | 7 | 136894611 | 137128187 | 233577 | 1 | ENSMUST00 | 106548 | Mgmt | ENSMUSG00000054612.3 |
| chr12 | 78219108 | 78219789 | 5.86474 | 9.17633 | 7.01925 | Distal Interg | 12 | 78226655 | 78684767 | 458113 | 1 | ENSMUST00 | -6866 | Gphn | ENSMUSG00000047454.6 |
| chr3 | 58154174 | 58154409 | 5.86474 | 9.17633 | 7.01925 | Intron (ENSM | 3 | 58141707 | 58163807 | 22101 | 2 | ENSMUST00 | 9398 | RP23-375H15 | ENSMUSG00000100666.1 |
| chr14 | 57344793 | 57345318 | 5.85266 | 9.15698 | 7.00004 | Intron (ENSM | 14 | 57333366 | 57333485 | 120 | 2 | ENSMUST00 | -11309 | Gm23593 | ENSMUSG00000077497.1 |

| | | | | | | | | | | | | | | | |
|---|---|---|---|---|---|---|---|---|---|---|---|---|---|---|---|
| chr14 | 70195712 | 70196081 | 5.85266 | 9.15698 | 7.00004 | Exon (ENSMU | 14 | 70205324 | 70217659 | 12336 | 1 | ENSMUST00 | -9243 | Gm26908 | ENSMUSG00000097243.1 |
| chr6 | 86472525 | 86473392 | 5.85266 | 9.15698 | 7.00004 | 3' UTR | 6 | 86470805 | 86480770 | 9966 | 2 | ENSMUST00 | 7378 | A430078I02R | ENSMUSG00000085589.2 |
| chr4 | 140770480 | 140770838 | 4.69226 | 9.13858 | 6.9818 | Intron (ENSM | 4 | 140745865 | 140774236 | 28372 | 2 | ENSMUST00 | 3398 | Padi4 | ENSMUSG00000025330.6 |
| chr18 | 64644031 | 64644507 | 5.82866 | 9.11851 | 6.96196 | Intron (ENSM | 18 | 64528979 | 64661000 | 132022 | 2 | ENSMUST00 | 16493 | Atp8b1 | ENSMUSG00000039529.8 |
| chr4 | 132049672 | 132049836 | 5.62008 | 9.11386 | 6.95743 | Promoter | 4 | 131957620 | 132049091 | 91472 | 2 | ENSMUST00 | -582 | Epb4.1 | ENSMUSG00000028906.10 |
| chr1 | 173878794 | 173879028 | 5.81673 | 9.0994 | 6.94297 | Intron (ENSM | 1 | 173872852 | 173880111 | 7260 | 2 | ENSMUST00 | 1083 | Mndal | ENSMUSG00000090272.2 |
| chr15 | 83169965 | 83170528 | 5.81673 | 9.0994 | 6.94297 | Promoter | 15 | 83157064 | 83170506 | 13443 | 2 | ENSMUST00 | 0 | Cyb5r3 | ENSMUSG00000018042.12 |
| chr15 | 89765445 | 89765620 | 5.81673 | 9.0994 | 6.94297 | Distal Interg | 15 | 89782393 | 89841860 | 59468 | 2 | ENSMUST00 | 76240 | Syt10 | ENSMUSG00000063260.2 |
| chr2 | 26590442 | 26590696 | 5.81673 | 9.0994 | 6.94297 | Promoter | 2 | 26590520 | 26590972 | 453 | 1 | ENSMUST00 | 0 | Egfl7 | ENSMUSG00000026921.13 |
| chr5 | 136237705 | 136238071 | 5.81673 | 9.0994 | 6.94297 | Intron (ENSM | 5 | 136218149 | 136244903 | 26755 | 2 | ENSMUST00 | 6832 | Sh2b2 | ENSMUSG00000005057.9 |
| chr5 | 115347805 | 115347986 | 5.81673 | 9.0994 | 6.94297 | Promoter | 5 | 115345657 | 115348914 | 3258 | 2 | ENSMUST00 | 928 | Cox6a1 | ENSMUSG00000041697.8 |
| chr7 | 82335259 | 82335894 | 5.81673 | 9.0994 | 6.94297 | Promoter | 7 | 82335694 | 82340024 | 4331 | 1 | ENSMUST00 | 0 | Adamtsl3 | ENSMUSG00000070469.6 |
| chr8 | 123422168 | 123422676 | 5.80485 | 9.08036 | 6.92443 | Intron (ENSM | 8 | 123411424 | 123422015 | 10592 | 1 | ENSMUST00 | 10745 | Tubb3 | ENSMUSG00000062380.3 |
| chr10 | 22926260 | 22926660 | 5.38833 | 9.07008 | 6.91429 | Distal Interg | 10 | 22817279 | 22820128 | 2850 | 2 | ENSMUST00 | -106133 | Tcf21 | ENSMUSG00000045680.7 |
| chr18 | 56539781 | 56540159 | 5.5791 | 9.04708 | 6.89156 | Intron (ENSM | 18 | 56538139 | 56538198 | 60 | 1 | ENSMUST00 | 1643 | Mir1258 | ENSMUSG00000099196.1 |
| chr5 | 65955972 | 65956136 | 5.5791 | 9.04708 | 6.89156 | Intron (ENSM | 5 | 65967124 | 65977326 | 10203 | 1 | ENSMUST00 | -10988 | Chrna9 | ENSMUSG00000029205.7 |
| chr8 | 81012672 | 81013192 | 5.5791 | 9.04708 | 6.89156 | Promoter | 8 | 81013576 | 81038584 | 25009 | 1 | ENSMUST00 | -384 | Gm9725 | ENSMUSG00000037982.10 |
| chr1 | 93101661 | 93102121 | 4.77198 | 8.97201 | 6.81752 | Promoter | 1 | 93015464 | 93101822 | 86359 | 2 | ENSMUST00 | 0 | Kif1a | ENSMUSG00000014602.10 |
| chr16 | 72799976 | 72800617 | 4.77198 | 8.97201 | 6.81752 | Intron (ENSM | 16 | 72663149 | 73046095 | 382947 | 1 | ENSMUST00 | 136828 | Robo1 | ENSMUSG00000022883.10 |
| chr3 | 60489851 | 60490037 | 4.77198 | 8.97201 | 6.81752 | Distal Interg | 3 | 60501252 | 60629748 | 128497 | 1 | ENSMUST00 | -11215 | Mbnl1 | ENSMUSG00000027763.8 |
| chr7 | 16874417 | 16874875 | 4.77198 | 8.97201 | 6.81752 | Promoter | 7 | 16871354 | 16874886 | 3533 | 2 | ENSMUST00 | 11 | 9330104G04 | ENSMUSG00000097842.1 |
| chr8 | 107198122 | 107198746 | 4.77198 | 8.97201 | 6.81752 | Distal Interg | 8 | 107150661 | 107187470 | 36810 | 1 | ENSMUST00 | 47462 | Cyb5b | ENSMUSG00000031924.3 |
| chr9 | 63611389 | 63611663 | 4.77198 | 8.97201 | 6.81752 | Intron (ENSM | 9 | 63602655 | 63641889 | 39235 | 1 | ENSMUST00 | 8735 | Aagab | ENSMUSG00000037257.7 |
| chr12 | 75616233 | 75616407 | 5.73458 | 8.96772 | 6.81342 | Distal Interg | 12 | 75630290 | 75637746 | 7457 | 1 | ENSMUST00 | -13883 | Rplp2-ps1 | ENSMUSG00000100259.1 |
| chr6 | 95117488 | 95117685 | 5.73458 | 8.96772 | 6.81342 | Promoter | 6 | 95114017 | 95117631 | 3615 | 2 | ENSMUST00 | 0 | Kbtbd8os | ENSMUSG00000085413.1 |
| chr14 | 52104443 | 52104825 | 4.30255 | 8.94291 | 6.78887 | Promoter | 14 | 52073380 | 52104028 | 30649 | 2 | ENSMUST00 | -416 | Hnrnpc | ENSMUSG00000060373.9 |
| chr2 | 164430677 | 164431038 | 5.71153 | 8.93077 | 6.77673 | Intron (ENSM | 2 | 164424247 | 164443186 | 18940 | 2 | ENSMUST00 | 12148 | Sdc4 | ENSMUSG00000017009.3 |
| chr2 | 170509000 | 170509283 | 5.30123 | 8.92364 | 6.76972 | Intron (ENSM | 2 | 170511235 | 170519123 | 7889 | 1 | ENSMUST00 | -1952 | Pfdn4 | ENSMUSG00000052033.8 |
| chr10 | 21999432 | 21999658 | 5.70007 | 8.9124 | 6.75856 | 3' UTR | 10 | 21996591 | 21998240 | 1650 | 1 | ENSMUST00 | 2842 | Sgk1 | ENSMUSG00000019970.10 |
| chr11 | 47379053 | 47379403 | 5.70007 | 8.9124 | 6.75856 | Promoter | 11 | 46978785 | 47379522 | 400738 | 2 | ENSMUST00 | 119 | Sgcd | ENSMUSG00000020354.10 |
| chr16 | 33685891 | 33686306 | 5.70007 | 8.9124 | 6.75856 | Intron (ENSM | 16 | 33684545 | 33769443 | 84899 | 1 | ENSMUST00 | 1347 | Heg1 | ENSMUSG00000075254.6 |
| chr3 | 148336233 | 148336470 | 5.70007 | 8.9124 | 6.75856 | Distal Interg | 3 | 148354087 | 148354218 | 132 | 2 | ENSMUST00 | 17748 | Gm23131 | ENSMUSG00000077441.1 |
| chr5 | 144189902 | 144190189 | 5.70007 | 8.9124 | 6.75856 | Promoter | 5 | 144190286 | 144194441 | 4156 | 1 | ENSMUST00 | -97 | Bhlha15 | ENSMUSG00000052271.7 |
| chr8 | 24437689 | 24437893 | 5.70007 | 8.9124 | 6.75856 | 3' UTR | 8 | 24437180 | 24438984 | 1805 | 2 | ENSMUST00 | 1091 | 1810011O10 | ENSMUSG00000056313.5 |
| chr9 | 102352189 | 102352654 | 5.68867 | 8.8941 | 6.74086 | Intron (ENSM | 9 | 101922128 | 102354693 | 432566 | 2 | ENSMUST00 | 2039 | Ephb1 | ENSMUSG00000032537.9 |
| chr11 | 52283843 | 52284596 | 5.47923 | 8.88398 | 6.73086 | Promoter | 11 | 52253236 | 52283014 | 29779 | 2 | ENSMUST00 | -830 | Tcf7 | ENSMUSG00000000782.10 |
| chr14 | 59478486 | 59479202 | 5.47923 | 8.88398 | 6.73086 | Intron (ENSM | 14 | 59440981 | 59548903 | 107923 | 1 | ENSMUST00 | 37506 | Cab39l | ENSMUSG00000021981.9 |
| chr19 | 32211101 | 32211387 | 5.47923 | 8.88398 | 6.73086 | Promoter | 19 | 32123479 | 32210990 | 87512 | 2 | ENSMUST00 | -112 | Sgms1 | ENSMUSG00000040451.12 |
| chr2 | 33628771 | 33629521 | 5.47923 | 8.88398 | 6.73086 | Intron (ENSM | 2 | 33567156 | 33632283 | 65128 | 2 | ENSMUST00 | 2762 | Lmx1b | ENSMUSG00000038765.8 |
| chr7 | 90099165 | 90099468 | 5.47923 | 8.88398 | 6.73086 | Distal Interg | 7 | 90107477 | 90107968 | 492 | 2 | ENSMUST00 | 8500 | Gm5341 | ENSMUSG00000062235.5 |
| chr10 | 81183547 | 81183848 | 5.6773 | 8.87588 | 6.72314 | Promoter | 10 | 81182605 | 81183849 | 1245 | 2 | ENSMUST00 | 1 | 4930442H23 | ENSMUSG00000053603.4 |
| chr4 | 138471046 | 138471499 | 5.6773 | 8.87588 | 6.72314 | Distal Interg | 4 | 138454314 | 138460123 | 5810 | 1 | ENSMUST00 | 16733 | Camk2n1 | ENSMUSG00000046447.3 |
| chr13 | 21215395 | 21215908 | 5.65471 | 8.83964 | 6.68717 | Distal Interg | 13 | 21190092 | 21192475 | 2384 | 1 | ENSMUST00 | 25304 | Trim27 | ENSMUSG00000021326.6 |
| chr4 | 122884430 | 122884741 | 5.65471 | 8.83964 | 6.68717 | Intron (ENSM | 4 | 122871033 | 122885860 | 14828 | 2 | ENSMUST00 | 1119 | Cap1 | ENSMUSG00000028656.9 |
| chr10 | 17655889 | 17656181 | 4.86435 | 8.82068 | 6.66842 | Distal Interg | 10 | 17723228 | 17725673 | 2446 | 1 | ENSMUST00 | -67047 | Cited2 | ENSMUSG00000039910.9 |
| chr14 | 34219891 | 34220190 | 4.86435 | 8.82068 | 6.66842 | Promoter | 14 | 34220046 | 34227740 | 7695 | 1 | ENSMUST00 | 0 | Syt15 | ENSMUSG00000041479.8 |
| chr17 | 3611033 | 3611440 | 4.86435 | 8.82068 | 6.66842 | Distal Interg | 17 | 3519263 | 3557713 | 38451 | 2 | ENSMUST00 | -53321 | Tfb1m | ENSMUSG00000036983.6 |
| chr5 | 23831273 | 23831508 | 4.86435 | 8.82068 | 6.66842 | Promoter | 5 | 23831080 | 23831623 | 544 | 2 | ENSMUST00 | 115 | 4933427G23 | ENSMUSG00000086697.1 |
| chr9 | 118082689 | 118083635 | 4.86435 | 8.82068 | 6.66842 | Intron (ENSM | 9 | 118059056 | 118062260 | 3205 | 1 | ENSMUST00 | 23634 | Azi2 | ENSMUSG00000039285.7 |
| chr15 | 85655864 | 85656383 | 5.61007 | 8.76801 | 6.61646 | Distal Interg | 15 | 85639747 | 85645934 | 6188 | 1 | ENSMUST00 | 16118 | RP23-64J7.2 | ENSMUSG00000101828.1 |
| chr7 | 5052800 | 5053390 | 4.3444 | 8.73543 | 6.58407 | 5' UTR | 7 | 5051532 | 5053722 | 2191 | 1 | ENSMUST00 | 1269 | Zfp580 | ENSMUSG00000055633.5 |
| chr1 | 168342020 | 168342280 | 5.58801 | 8.7326 | 6.58127 | Intron (ENSM | 1 | 168120969 | 168405601 | 284633 | 2 | ENSMUST00 | 63321 | Pbx1 | ENSMUSG00000052534.10 |
| chr10 | 109136985 | 109137396 | 5.58801 | 8.7326 | 6.58127 | Distal Interg | 10 | 109089998 | 109090275 | 278 | 2 | ENSMUST00 | -46711 | Gm22097 | ENSMUSG00000088554.1 |
| chr17 | 31433568 | 31433776 | 5.58801 | 8.7326 | 6.58127 | Promoter | 17 | 31433071 | 31459129 | 26059 | 1 | ENSMUST00 | 498 | Pde9a | ENSMUSG00000041119.6 |
| chr3 | 9826209 | 9826897 | 5.58801 | 8.7326 | 6.58127 | Intron (ENSM | 3 | 9829620 | 9829728 | 109 | 2 | ENSMUST00 | 2831 | Gm24786 | ENSMUSG00000088004.1 |
| chr4 | 9136047 | 9136370 | 5.58801 | 8.7326 | 6.58127 | Distal Interg | 4 | 9210003 | 9210108 | 106 | 1 | ENSMUST00 | -73633 | Gm23423 | ENSMUSG00000075958.1 |
| chr9 | 40963964 | 40964135 | 5.58801 | 8.7326 | 6.58127 | Promoter | 9 | 40894849 | 40964118 | 69270 | 2 | ENSMUST00 | 0 | 4931429I11R | ENSMUSG00000032023.7 |
| chr3 | 62341897 | 62342088 | 5.38288 | 8.72614 | 6.57544 | Promoter | 3 | 62341101 | 62380991 | 39891 | 1 | ENSMUST00 | 797 | Arhgef26 | ENSMUSG00000036885.9 |
| chr5 | 36868327 | 36868536 | 5.38288 | 8.72614 | 6.57544 | Promoter | 5 | 36868513 | 36955078 | 86566 | 1 | ENSMUST00 | 0 | Ppp2r2c | ENSMUSG00000029120.7 |
| chr8 | 80669096 | 80669490 | 5.38288 | 8.72614 | 6.57544 | Exon (ENSMU | 8 | 80709738 | 80710861 | 1124 | 2 | ENSMUST00 | 41371 | Smarca5 | ENSMUSG00000031715.9 |
| chr1 | 89302847 | 89303210 | 4.97263 | 8.68936 | 6.53912 | Exon (ENSMU | 1 | 89256182 | 89256981 | 800 | 2 | ENSMUST00 | -45867 | RP24-217I2.2 | ENSMUSG00000102083.1 |
| chr10 | 24958973 | 24959390 | 4.97263 | 8.68936 | 6.53912 | Distal Interg | 10 | 24915208 | 24927470 | 12263 | 2 | ENSMUST00 | -31504 | Arg1 | ENSMUSG00000019987.8 |
| chr10 | 122544276 | 122544489 | 4.97263 | 8.68936 | 6.53912 | Distal Interg | 10 | 122448499 | 122453453 | 4955 | 1 | ENSMUST00 | 95778 | Avpr1a | ENSMUSG00000020123.5 |
| chr10 | 127180132 | 127180585 | 4.97263 | 8.68936 | 6.53912 | Promoter | 10 | 127172426 | 127180645 | 8220 | 2 | ENSMUST00 | 60 | Slc26a10 | ENSMUSG00000040441.8 |
| chr11 | 96988127 | 96988449 | 4.97263 | 8.68936 | 6.53912 | Distal Interg | 11 | 96954219 | 96982959 | 28741 | 2 | ENSMUST00 | -5169 | Sp2 | ENSMUSG00000018678.7 |
| chr11 | 102777657 | 102778063 | 4.97263 | 8.68936 | 6.53912 | 3' UTR | 11 | 102776554 | 102778467 | 1914 | 1 | ENSMUST00 | 1104 | Adam11 | ENSMUSG00000020926.11 |
| chr12 | 57538955 | 57539529 | 4.97263 | 8.68936 | 6.53912 | Downstream | 12 | 57540631 | 57546121 | 5491 | 2 | ENSMUST00 | 6592 | Foxa1 | ENSMUSG00000035451.6 |
| chr12 | 40381556 | 40381977 | 4.97263 | 8.68936 | 6.53912 | Intron (ENSM | 12 | 40315558 | 40445769 | 130212 | 2 | ENSMUST00 | 63792 | Zfp277 | ENSMUSG00000055917.9 |
| chr13 | 73264063 | 73264291 | 4.97263 | 8.68936 | 6.53912 | Intron (ENSM | 13 | 73260497 | 73269608 | 9112 | 1 | ENSMUST00 | 3567 | Irx4 | ENSMUSG00000021604.10 |
| chr13 | 99412495 | 99412862 | 4.97263 | 8.68936 | 6.53912 | Promoter | 13 | 99396924 | 99412621 | 15698 | 2 | ENSMUST00 | 0 | 6430562O15 | ENSMUSG00000097828.1 |
| chr14 | 22037586 | 22037901 | 4.97263 | 8.68936 | 6.53912 | Intron (ENSM | 14 | 22021958 | 22022070 | 113 | 2 | ENSMUST00 | -15517 | Gm27532 | ENSMUSG00000098542.1 |
| chr15 | 55265665 | 55265924 | 4.97263 | 8.68936 | 6.53912 | Distal Interg | 15 | 55262542 | 55262603 | 62 | 2 | ENSMUST00 | -3063 | Gm26296 | ENSMUSG00000095792.1 |
| chr16 | 32499613 | 32499780 | 4.97263 | 8.68936 | 6.53912 | Promoter | 16 | 32499611 | 32561966 | 62356 | 1 | ENSMUST00 | 3 | Zdhhc19 | ENSMUSG00000052363.7 |
| chr16 | 94086986 | 94087493 | 4.97263 | 8.68936 | 6.53912 | Intron (ENSM | 16 | 94085260 | 94127032 | 41773 | 1 | ENSMUST00 | 1727 | Sim2 | ENSMUSG00000062713.7 |
| chr2 | 12327505 | 12327772 | 4.97263 | 8.68936 | 6.53912 | Distal Interg | 2 | 12249800 | 12301920 | 52121 | 2 | ENSMUST00 | -25586 | Itga8 | ENSMUSG00000026768.5 |
| chr2 | 75964933 | 75965120 | 4.97263 | 8.68936 | 6.53912 | Distal Interg | 2 | 75975740 | 75978170 | 2431 | 2 | ENSMUST00 | 13050 | Ttc30a2 | ENSMUSG00000075272.4 |
| chr3 | 98042980 | 98043190 | 4.97263 | 8.68936 | 6.53912 | Intron (ENSM | 3 | 98013538 | 98150367 | 136830 | 1 | ENSMUST00 | 29443 | Notch2 | ENSMUSG00000027878.10 |
| chr3 | 142496678 | 142497220 | 4.97263 | 8.68936 | 6.53912 | Promoter | 3 | 142496934 | 142522344 | 25411 | 1 | ENSMUST00 | 0 | Gbp5 | ENSMUSG00000040264.9 |
| chr6 | 72813627 | 72814011 | 4.97263 | 8.68936 | 6.53912 | Distal Interg | 6 | 72626378 | 72789254 | 162877 | 2 | ENSMUST00 | -24374 | Tcf7l1 | ENSMUSG00000055799.8 |
| chr7 | 126482779 | 126483032 | 4.97263 | 8.68936 | 6.53912 | Distal Interg | 7 | 126487355 | 126490730 | 3376 | 1 | ENSMUST00 | -4323 | Tufm | ENSMUSG00000073838.5 |
| chr9 | 61405345 | 61405699 | 4.97263 | 8.68936 | 6.53912 | Exon (ENSMU | 9 | 61402461 | 61417835 | 15375 | 1 | ENSMUST00 | 2885 | Tle3 | ENSMUSG00000032280.11 |
| chr9 | 88553144 | 88553395 | 4.97263 | 8.68936 | 6.53912 | Promoter | 9 | 88553236 | 88553525 | 290 | 1 | ENSMUST00 | 0 | Gm26461 | ENSMUSG00000092805.2 |
| chr16 | 34926034 | 34926560 | 5.53361 | 8.64525 | 6.496 | Intron (ENSM | 16 | 34935854 | 34939329 | 3476 | 2 | ENSMUST00 | 12769 | RP23-80O10. | ENSMUSG00000100593.1 |
| chr1 | 55089011 | 55089192 | 5.51215 | 8.61076 | 6.4617 | Promoter | 1 | 55088314 | 55089915 | 1602 | 1 | ENSMUST00 | 698 | Hspe1 | ENSMUSG00000073676.4 |
| chr13 | 113394607 | 113394907 | 5.50148 | 8.59362 | 6.44461 | Distal Interg | 13 | 113403088 | 113403188 | 101 | 1 | ENSMUST00 | -8181 | Gm22385 | ENSMUSG00000088814.1 |
| chr12 | 56611910 | 56612223 | 5.10133 | 8.58512 | 6.43618 | Exon (ENSMU | 12 | 56611389 | 56613284 | 1896 | 2 | ENSMUST00 | 1061 | Nkx2-9 | ENSMUSG00000058669.7 |
| chr13 | 112439738 | 112440188 | 5.10133 | 8.58512 | 6.43618 | Distal Interg | 13 | 112464070 | 112510086 | 46017 | 1 | ENSMUST00 | -23882 | Il6st | ENSMUSG00000021756.7 |
| chr15 | 84924208 | 84924431 | 5.10133 | 8.58512 | 6.43618 | Promoter | 15 | 84923428 | 84942955 | 19528 | 1 | ENSMUST00 | 781 | Nup50 | ENSMUSG00000016619.9 |
| chr15 | 81235564 | 81236002 | 5.10133 | 8.58512 | 6.43618 | Promoter | 15 | 81235499 | 81238963 | 3465 | 1 | ENSMUST00 | 66 | Mchr1 | ENSMUSG00000050164.10 |
| chr18 | 61045113 | 61046136 | 5.10133 | 8.58512 | 6.43618 | Promoter | 18 | 61045150 | 61085061 | 39912 | 1 | ENSMUST00 | 0 | Pdgfrb | ENSMUSG00000024620.6 |
| chr2 | 165323665 | 165324081 | 5.10133 | 8.58512 | 6.43618 | Intron (ENSM | 2 | 165289155 | 165326479 | 37325 | 2 | ENSMUST00 | 2398 | Elmo2 | ENSMUSG00000017670.11 |
| chr3 | 146059513 | 146060072 | 5.10133 | 8.58512 | 6.43618 | Intron (ENSM | 3 | 146042781 | 146048437 | 5657 | 2 | ENSMUST00 | -11077 | Wdr63 | ENSMUSG00000043020.8 |
| chr3 | 57625556 | 57625795 | 5.10133 | 8.58512 | 6.43618 | Promoter | 3 | 57620905 | 57625818 | 4914 | 2 | ENSMUST00 | 23 | Gm26671 | ENSMUSG00000097339.1 |
| chr3 | 8594597 | 8594786 | 5.10133 | 8.58512 | 6.43618 | Distal Interg | 3 | 8615275 | 8616504 | 1230 | 2 | ENSMUST00 | 21718 | Gm18822 | ENSMUSG00000098047.1 |
| chr6 | 134036940 | 134037205 | 5.10133 | 8.58512 | 6.43618 | Promoter | 6 | 134035969 | 134248284 | 212316 | 1 | ENSMUST00 | 972 | Etv6 | ENSMUSG00000030199.11 |
| chr6 | 3398947 | 3399563 | 5.10133 | 8.58512 | 6.43618 | Promoter | 6 | 3372257 | 3399571 | 27315 | 2 | ENSMUST00 | 8 | Samd9l | ENSMUSG00000047735.12 |
| chr6 | 52316465 | 52316917 | 5.10133 | 8.58512 | 6.43618 | 3' UTR | 6 | 52308389 | 52314832 | 6444 | 2 | ENSMUST00 | -1634 | 5730457N03 | ENSMUSG00000086126.1 |
| chr7 | 45415655 | 45416165 | 5.10133 | 8.58512 | 6.43618 | 3' UTR | 7 | 45413695 | 45417178 | 3484 | 1 | ENSMUST00 | 1961 | Ntf5 | ENSMUSG00000074121.2 |
| chr8 | 119447118 | 119447316 | 5.10133 | 8.58512 | 6.43618 | Promoter | 8 | 119446761 | 119472411 | 25651 | 1 | ENSMUST00 | 358 | Necab2 | ENSMUSG00000031837.8 |
| chr5 | 124439387 | 124439751 | 5.28985 | 8.57326 | 6.42589 | Promoter | 5 | 124439930 | 124462308 | 22379 | 1 | ENSMUST00 | -179 | Setd8 | ENSMUSG00000049327.12 |
| chr11 | 44619663 | 44620194 | 5.48026 | 8.55952 | 6.41235 | Intron (ENSM | 11 | 44618179 | 44643469 | 25291 | 1 | ENSMUST00 | 1485 | Ebf1 | ENSMUSG00000057098.9 |
| chr16 | 90072371 | 90072993 | 5.48026 | 8.55952 | 6.41235 | Distal Interg | 16 | 89955965 | 89980080 | 24116 | 2 | ENSMUST00 | -92292 | Tiam1 | ENSMUSG00000002489.10 |
| chr5 | 143635357 | 143635806 | 5.48026 | 8.55952 | 6.41235 | Promoter | 5 | 143636351 | 143692554 | 56204 | 1 | ENSMUST00 | -545 | Cyth3 | ENSMUSG00000018001.13 |
| chr7 | 15963999 | 15964411 | 5.48026 | 8.55952 | 6.41235 | 5' UTR | 7 | 15957683 | 15967475 | 9793 | 2 | ENSMUST00 | 3064 | Ehd2 | ENSMUSG00000074364.5 |
| chr12 | 8571893 | 8572758 | 4.39162 | 8.53397 | 6.38747 | Intron (ENSM | 12 | 8528483 | 8599066 | 70584 | 2 | ENSMUST00 | 26308 | Slc7a15 | ENSMUSG00000020600.8 |
| chr1 | 37864981 | 37865153 | 5.25681 | 8.51884 | 6.37244 | Promoter | 1 | 37754572 | 37865079 | 110508 | 2 | ENSMUST00 | 0 | Tsga10 | ENSMUSG00000060771.9 |
| chr11 | 84520089 | 84520408 | 5.25681 | 8.51884 | 6.37244 | Exon (ENSMU | 11 | 84519706 | 84522067 | 2362 | 2 | ENSMUST00 | 1659 | Lhx1 | ENSMUSG00000018698.10 |
| chr11 | 115810395 | 115810961 | 5.25681 | 8.51884 | 6.37244 | Intron (ENSM | 11 | 115804736 | 115808102 | 3367 | 2 | ENSMUST00 | -2294 | Caskin2 | ENSMUSG00000034471.7 |
| chr11 | 95540791 | 95541295 | 5.25681 | 8.51884 | 6.37244 | Distal Interg | 11 | 95509845 | 95514570 | 4726 | 2 | ENSMUST00 | -26222 | Nxph3 | ENSMUSG00000046719.7 |
| chr18 | 65124922 | 65125119 | 5.25681 | 8.51884 | 6.37244 | Intron (ENSM | 18 | 65023527 | 65217826 | 194300 | 1 | ENSMUST00 | 101396 | Nedd4l | ENSMUSG00000024589.10 |
| chr2 | 26588475 | 26589674 | 5.25681 | 8.51884 | 6.37244 | Promoter | 2 | 26588543 | 26592443 | 3901 | 1 | ENSMUST00 | 0 | Egfl7 | ENSMUSG00000026921.13 |
| chr2 | 127729599 | 127729931 | 5.25681 | 8.51884 | 6.37244 | Promoter | 2 | 127704386 | 127729932 | 25547 | 2 | ENSMUST00 | 1 | Mall | ENSMUSG00000027377.2 |
| chr2 | 84633019 | 84633326 | 5.25681 | 8.51884 | 6.37244 | Intron (ENSM | 2 | 84624378 | 84638105 | 13728 | 2 | ENSMUST00 | 4779 | Ctnnd1 | ENSMUSG00000034101.9 |
| chr2 | 38164382 | 38165006 | 5.25681 | 8.51884 | 6.37244 | Intron (ENSM | 2 | 38021266 | 38166081 | 144816 | 2 | ENSMUST00 | 1075 | Dennd1a | ENSMUSG00000035392.12 |
| chr2 | 64074899 | 64075152 | 5.25681 | 8.51884 | 6.37244 | Intron (ENSM | 2 | 63980447 | 64097988 | 117542 | 2 | ENSMUST00 | 22836 | Fign | ENSMUSG00000075324.8 |
| chr3 | 60493615 | 60493875 | 5.25681 | 8.51884 | 6.37244 | Distal Interg | 3 | 60501252 | 60629748 | 128497 | 1 | ENSMUST00 | -7377 | Mbnl1 | ENSMUSG00000027763.8 |
| chr3 | 121887147 | 121887471 | 5.25681 | 8.51884 | 6.37244 | Distal Interg | 3 | 121953229 | 122016677 | 63449 | 1 | ENSMUST00 | -65758 | Arhgap29 | ENSMUSG00000039831.11 |
| chr3 | 149074929 | 149075687 | 5.25681 | 8.51884 | 6.37244 | Distal Interg | 3 | 149028932 | 149029038 | 107 | 1 | ENSMUST00 | 45998 | Gm25127 | ENSMUSG00000065270.1 |
| chr4 | 139833260 | 139833441 | 5.25681 | 8.51884 | 6.37244 | Promoter | 4 | 139737062 | 139833528 | 96467 | 2 | ENSMUST00 | 87 | Pax7 | ENSMUSG00000028736.8 |
| chr5 | 139793698 | 139793889 | 5.25681 | 8.51884 | 6.37244 | Intron (ENSM | 5 | 139791534 | 139799500 | 7967 | 1 | ENSMUST00 | 2165 | Mafk | ENSMUSG00000018143.5 |
| chr5 | 150673254 | 150673494 | 5.25681 | 8.51884 | 6.37244 | Promoter | 5 | 150673739 | 150810690 | 136952 | 1 | ENSMUST00 | -245 | Pds5b | ENSMUSG00000034021.10 |
| chr6 | 114131199 | 114131425 | 5.25681 | 8.51884 | 6.37244 | Promoter | 6 | 114131241 | 114249886 | 118646 | 1 | ENSMUST00 | 0 | Slc6a11 | ENSMUSG00000030307.7 |
| chr6 | 24894738 | 24894974 | 5.25681 | 8.51884 | 6.37244 | Promoter | 6 | 24894670 | 24894800 | 131 | 2 | ENSMUST00 | 0 | Gm22267 | ENSMUSG00000077196.1 |
| chr6 | 71270509 | 71270673 | 5.25681 | 8.51884 | 6.37244 | Promoter | 6 | 71271671 | 71271801 | 131 | 1 | ENSMUST00 | -998 | Mir8112 | ENSMUSG00000098623.1 |
| chr9 | 56864988 | 56865426 | 5.25681 | 8.51884 | 6.37244 | Promoter | 9 | 56865104 | 56899870 | 34767 | 1 | ENSMUST00 | 0 | Cspg4 | ENSMUSG00000032911.5 |
| chr1 | 53976769 | 53977124 | 5.44843 | 8.50832 | 6.36429 | Intron (ENSM | 1 | 53886828 | 53886941 | 114 | 2 | ENSMUST00 | -89829 | Gm24251 | ENSMUSG00000087949.1 |
| chr11 | 4188470 | 4188775 | 5.44843 | 8.50832 | 6.36429 | Intron (ENSM | 11 | 4186822 | 4215502 | 28681 | 1 | ENSMUST00 | 1649 | Tbc1d10a | ENSMUSG00000034412.8 |
| chr13 | 51401163 | 51401546 | 5.44843 | 8.50832 | 6.36429 | Distal Interg | 13 | 51408618 | 51422797 | 14180 | 1 | ENSMUST00 | -7072 | S1pr3 | ENSMUSG00000067586.3 |

| | | | | | | | | | | | | | | | |
|---|---|---|---|---|---|---|---|---|---|---|---|---|---|---|---|
| chr14 | 105259717 | 105260011 | 5.44843 | 8.50832 | 6.36429 | Promoter | 14 | 105258818 | 105294944 | 36127 | 1 | ENSMUST00 | 900 | Ndfip2 | ENSMUSG00000053253.11 |
| chr17 | 75307197 | 75307554 | 5.44843 | 8.50832 | 6.36429 | Intron (ENSM | 17 | 75289976 | 75294369 | 4394 | 1 | ENSMUST00 | 17222 | Ltbp1 | ENSMUSG00000001870.10 |
| chr19 | 5660935 | 5661316 | 5.44843 | 8.50832 | 6.36429 | 5' UTR | 19 | 5651185 | 5663707 | 12523 | 2 | ENSMUST00 | 2391 | Sipa1 | ENSMUSG00000056917.7 |
| chr2 | 23234240 | 23234404 | 5.44843 | 8.50832 | 6.36429 | Promoter | 2 | 23235236 | 23235739 | 504 | 1 | ENSMUST00 | -832 | Gm13422 | ENSMUSG00000084098.1 |
| chr3 | 148945697 | 148946095 | 5.44843 | 8.50832 | 6.36429 | Intron (ENSM | 3 | 148815586 | 148954635 | 139050 | 2 | ENSMUST00 | 8540 | Lphn2 | ENSMUSG00000028184.8 |
| chr5 | 96905975 | 96906284 | 5.44843 | 8.50832 | 6.36429 | Downstream | 5 | 96907632 | 96907825 | 194 | 1 | ENSMUST00 | -1348 | Gm24623 | ENSMUSG00000064550.1 |
| chr7 | 19463343 | 19463673 | 5.44843 | 8.50832 | 6.36429 | Distal Interge | 7 | 19460797 | 19460935 | 139 | 2 | ENSMUST00 | -2409 | Gm23099 | ENSMUSG00000065388.1 |
| chr7 | 73031981 | 73032187 | 5.44843 | 8.50832 | 6.36429 | Distal Interge | 7 | 72989154 | 72990153 | 1000 | 2 | ENSMUST00 | -41829 | Gm5335 | ENSMUSG00000098035.1 |
| chr8 | 94171450 | 94171894 | 5.44843 | 8.50832 | 6.36429 | Promoter | 8 | 94172618 | 94173567 | 950 | 1 | ENSMUST00 | -724 | Mt2 | ENSMUSG00000031762.6 |
| chr8 | 87690213 | 87690714 | 5.44843 | 8.50832 | 6.36429 | Intron (ENSM | 8 | 87636657 | 87637013 | 357 | 1 | ENSMUST00 | 53557 | Gm24841 | ENSMUSG00000089301.1 |
| chr9 | 40757741 | 40758096 | 5.44843 | 8.50832 | 6.36429 | Intron (ENSM | 9 | 40774094 | 40782471 | 8378 | 1 | ENSMUST00 | -15998 | Clmp | ENSMUSG00000032024.10 |
| chr9 | 79807055 | 79807370 | 5.44843 | 8.50832 | 6.36429 | Distal Interge | 9 | 79793641 | 79803235 | 9595 | 1 | ENSMUST00 | 13415 | 4930429F24 | ENSMUSG00000086918.1 |
| chrX | 140456114 | 140456368 | 5.44843 | 8.50832 | 6.36429 | Promoter | 20 | 140456613 | 140465442 | 8830 | 1 | ENSMUST00 | -245 | Prps1 | ENSMUSG00000031432.3 |
| chrX | 12136360 | 12136985 | 5.44843 | 8.50832 | 6.36429 | Intron (ENSM | 20 | 12058981 | 12128400 | 69420 | 2 | ENSMUST00 | -7961 | Bcor | ENSMUSG00000040363.9 |
| chrX | 11998979 | 11999498 | 5.44843 | 8.50832 | 6.36429 | Distal Interge | 20 | 11984475 | 11984650 | 176 | 1 | ENSMUST00 | 14505 | Gm14512 | ENSMUSG00000081947.1 |
| chr10 | 120915211 | 120915390 | 5.3766 | 8.3927 | 6.25156 | Distal Interge | 10 | 120928186 | 120928972 | 787 | 1 | ENSMUST00 | -12796 | Gm16166 | ENSMUSG00000082936.1 |
| chr11 | 60351067 | 60351398 | 5.3766 | 8.3927 | 6.25156 | Intron (ENSM | 11 | 60226714 | 60352905 | 126192 | 2 | ENSMUST00 | 1507 | Tom1l2 | ENSMUSG00000000538.13 |
| chr12 | 55599162 | 55599523 | 4.97928 | 8.37665 | 6.23625 | Promoter | 12 | 55598917 | 55602018 | 3102 | 1 | ENSMUST00 | 246 | Insm2 | ENSMUSG00000045440.3 |
| chr17 | 27200284 | 27200479 | 5.34626 | 8.34382 | 6.2035 | Promoter | 17 | 27201032 | 27203071 | 2040 | 1 | ENSMUST00 | -553 | Gm26724 | ENSMUSG00000097090.1 |
| chr13 | 15426617 | 15427491 | 4.4453 | 8.34 | 6.1997 | Distal Interge | 13 | 15459980 | 15461295 | 1316 | 2 | ENSMUST00 | 33804 | Gm26645 | ENSMUSG00000097684.1 |
| chr8 | 25027842 | 25028321 | 4.4453 | 8.34 | 6.1997 | Intron (ENSM | 8 | 25037052 | 25041757 | 4706 | 1 | ENSMUST00 | -8731 | Gm16933 | ENSMUSG00000086289.2 |
| chr1 | 75450453 | 75450646 | 5.51439 | 8.30479 | 6.16463 | Promoter | 1 | 75450436 | 75474340 | 23905 | 1 | ENSMUST00 | 18 | Asic4 | ENSMUSG00000033007.10 |
| chr10 | 79779949 | 79780145 | 5.51439 | 8.30479 | 6.16463 | Promoter | 10 | 79780581 | 79782607 | 2027 | 1 | ENSMUST00 | -436 | Fstl3 | ENSMUSG00000020325.10 |
| chr10 | 80495959 | 80496346 | 5.51439 | 8.30479 | 6.16463 | Exon (ENSMU | 10 | 80494835 | 80517276 | 22442 | 1 | ENSMUST00 | 1125 | Onecut3 | ENSMUSG00000045518.8 |
| chr10 | 39590550 | 39590744 | 5.51439 | 8.30479 | 6.16463 | Intron (ENSM | 10 | 39558496 | 39588452 | 29957 | 2 | ENSMUST00 | -2099 | Gm16365 | ENSMUSG00000086915.2 |
| chr13 | 13889901 | 13890165 | 5.51439 | 8.30479 | 6.16463 | Distal Interge | 13 | 13954674 | 13999068 | 44395 | 1 | ENSMUST00 | -64509 | B3galnt2 | ENSMUSG00000039242.9 |
| chr15 | 100870398 | 100870734 | 5.51439 | 8.30479 | 6.16463 | Promoter | 15 | 100870683 | 101045929 | 175247 | 1 | ENSMUST00 | 0 | Scn8a | ENSMUSG00000023033.9 |
| chr15 | 102722359 | 102722619 | 5.51439 | 8.30479 | 6.16463 | Promoter | 15 | 102706779 | 102722178 | 15400 | 2 | ENSMUST00 | -182 | Calcoco1 | ENSMUSG00000023055.4 |
| chr16 | 7149308 | 7149688 | 5.51439 | 8.30479 | 6.16463 | Intron (ENSM | 16 | 7069835 | 7412479 | 342645 | 1 | ENSMUST00 | 79474 | Rbfox1 | ENSMUSG00000008658.10 |
| chr18 | 86394597 | 86394761 | 5.51439 | 8.30479 | 6.16463 | Promoter | 18 | 86394952 | 86501897 | 106946 | 1 | ENSMUST00 | -191 | Neto1 | ENSMUSG00000050321.2 |
| chr18 | 37178296 | 37178587 | 5.51439 | 8.30479 | 6.16463 | Promoter | 18 | 37159855 | 37178522 | 18668 | 2 | ENSMUST00 | 0 | Gm10544 | ENSMUSG00000097430.1 |
| chr2 | 175435296 | 175435561 | 5.51439 | 8.30479 | 6.16463 | Promoter | 2 | 175423893 | 175435782 | 11890 | 2 | ENSMUST00 | 221 | Gm4724 | ENSMUSG00000078897.5 |
| chr4 | 136956606 | 136956785 | 5.51439 | 8.30479 | 6.16463 | Promoter | 4 | 136929419 | 136956816 | 27398 | 2 | ENSMUST00 | 31 | Epha8 | ENSMUSG00000028661.8 |
| chr6 | 17583938 | 17584308 | 5.51439 | 8.30479 | 6.16463 | Distal Interge | 6 | 17546973 | 17553430 | 6458 | 1 | ENSMUST00 | 36966 | Met | ENSMUSG00000009376.10 |
| chr6 | 120112550 | 120112767 | 5.51439 | 8.30479 | 6.16463 | Intron (ENSM | 6 | 120093350 | 120200339 | 106990 | 1 | ENSMUST00 | 19201 | Ninj2 | ENSMUSG00000041377.7 |
| chr6 | 84574188 | 84574561 | 5.51439 | 8.30479 | 6.16463 | 3' UTR | 6 | 84571414 | 84593810 | 22397 | 2 | ENSMUST00 | 19249 | Cyp26b1 | ENSMUSG00000063415.7 |
| chr7 | 44428797 | 44429185 | 5.51439 | 8.30479 | 6.16463 | Promoter | 7 | 44429018 | 44461361 | 32344 | 1 | ENSMUST00 | 0 | Lrrc4b | ENSMUSG00000047085.9 |
| chr7 | 3366680 | 3367120 | 5.51439 | 8.30479 | 6.16463 | 3' UTR | 7 | 3390683 | 3415648 | 24966 | 1 | ENSMUST00 | -23563 | Cacng8 | ENSMUSG00000053395.10 |
| chr7 | 136496346 | 136496518 | 5.51439 | 8.30479 | 6.16463 | Distal Interge | 7 | 136509026 | 136509333 | 308 | 2 | ENSMUST00 | 12815 | Gm24444 | ENSMUSG00000087869.1 |
| chr8 | 12396559 | 12397540 | 5.51439 | 8.30479 | 6.16463 | Promoter | 8 | 12396403 | 12396488 | 86 | 1 | ENSMUST00 | 157 | Gm25239 | ENSMUSG00000092656.1 |
| chrX | 20848038 | 20848277 | 5.51439 | 8.30479 | 6.16463 | Promoter | 20 | 20848543 | 20860519 | 11977 | 1 | ENSMUST00 | -266 | Araf | ENSMUSG00000001127.7 |
| chrX | 60178579 | 60178854 | 5.51439 | 8.30479 | 6.16463 | Promoter | 20 | 60055956 | 60179089 | 123134 | 2 | ENSMUST00 | 235 | Mcf2 | ENSMUSG00000031139.9 |
| chrX | 101639410 | 101639845 | 5.51439 | 8.30479 | 6.16463 | Promoter | 20 | 101640060 | 101682940 | 42881 | 1 | ENSMUST00 | -215 | Ogt | ENSMUSG00000034160.8 |
| chrX | 140062138 | 140062899 | 5.51439 | 8.30479 | 6.16463 | Promoter | 20 | 140062879 | 140106797 | 43919 | 1 | ENSMUST00 | 0 | E230019M04 | ENSMUSG00000042433.2 |
| chrX | 136203497 | 136203790 | 5.51439 | 8.30479 | 6.16463 | Promoter | 20 | 136201618 | 136203637 | 2020 | 2 | ENSMUST00 | 0 | Tceal5 | ENSMUSG00000054034.5 |
| chrX | 95967672 | 95967916 | 5.51439 | 8.30479 | 6.16463 | Distal Interge | 20 | 95949435 | 95956962 | 7528 | 2 | ENSMUST00 | -10711 | Las1l | ENSMUSG00000057421.7 |
| chrX | 103529063 | 103529362 | 5.51439 | 8.30479 | 6.16463 | Promoter | 20 | 103529483 | 103529707 | 225 | 1 | ENSMUST00 | -121 | Gm9157 | ENSMUSG00000081004.1 |
| chrX | 12252400 | 12252591 | 5.51439 | 8.30479 | 6.16463 | Distal Interge | 20 | 12179718 | 12180067 | 350 | 2 | ENSMUST00 | -72334 | Gm14521 | ENSMUSG00000083486.1 |
| chrX | 12252669 | 12252950 | 5.51439 | 8.30479 | 6.16463 | Distal Interge | 20 | 12179718 | 12180067 | 350 | 2 | ENSMUST00 | -72603 | Gm14521 | ENSMUSG00000083486.1 |
| chrX | 152327716 | 152327941 | 5.51439 | 8.30479 | 6.16463 | Promoter | 20 | 152298652 | 152327495 | 28844 | 2 | ENSMUST00 | -222 | Kantr | ENSMUSG00000087403.3 |
| chr5 | 129941943 | 129942136 | 5.50447 | 8.289 | 6.15165 | Promoter | 5 | 129942109 | 129984745 | 42637 | 1 | ENSMUST00 | 0 | Vkorc1l1 | ENSMUSG00000066735.5 |
| chr1 | 57429127 | 57429329 | 5.11312 | 8.28132 | 6.14407 | Distal Interge | 1 | 57422244 | 57422373 | 130 | 2 | ENSMUST00 | -6755 | Gm22371 | ENSMUSG00000077776.1 |
| chr5 | 114575159 | 114575480 | 5.11312 | 8.28132 | 6.14407 | Intron (ENSM | 5 | 114568017 | 114594678 | 26662 | 1 | ENSMUST00 | 7143 | Fam222a | ENSMUSG00000041930.7 |
| chr5 | 123135032 | 123135277 | 5.48147 | 8.2524 | 6.11546 | Intron (ENSM | 5 | 123118178 | 123132692 | 14515 | 2 | ENSMUST00 | -2341 | Rhof | ENSMUSG00000029449.6 |
| chr15 | 75711584 | 75711991 | 5.47004 | 8.23422 | 6.09765 | Exon (ENSMU | 15 | 75704394 | 75714225 | 9832 | 1 | ENSMUST00 | 7191 | Rhpn1 | ENSMUSG00000022580.7 |
| chr4 | 115645155 | 115645410 | 5.47004 | 8.23422 | 6.09765 | Intron (ENSM | 4 | 115624725 | 115647709 | 22985 | 2 | ENSMUST00 | 2299 | Cyp4b1 | ENSMUSG00000028713.12 |
| chr6 | 48742910 | 48743074 | 5.47004 | 8.23422 | 6.09765 | Exon (ENSMU | 6 | 48738346 | 48740621 | 2276 | 2 | ENSMUST00 | -2290 | Gimap1os | ENSMUSG00000044867.4 |
| chr12 | 117659030 | 117659562 | 5.27678 | 8.23176 | 6.09532 | Intron (ENSM | 12 | 117574342 | 117574447 | 106 | 1 | ENSMUST00 | 84689 | Gm24741 | ENSMUSG00000089482.1 |
| chr6 | 48675898 | 48676366 | 5.27678 | 8.23176 | 6.09532 | Promoter | 6 | 48676129 | 48678699 | 2571 | 1 | ENSMUST00 | 0 | Gimap9 | ENSMUSG00000051124.6 |
| chr18 | 61565898 | 61566062 | 5.45866 | 8.21611 | 6.08027 | Intron (ENSM | 18 | 61575517 | 61582118 | 6602 | 1 | ENSMUST00 | -9455 | Csnk1a1 | ENSMUSG00000024576.9 |
| chr13 | 60147217 | 60147532 | 5.44732 | 8.19807 | 6.06239 | Distal Interge | 13 | 60174405 | 60177365 | 2961 | 2 | ENSMUST00 | 29833 | Gas1 | ENSMUSG00000052957.7 |
| chr13 | 55760823 | 55761194 | 5.43604 | 8.18012 | 6.04458 | Intron (ENSM | 13 | 55775940 | 55784479 | 8540 | 2 | ENSMUST00 | 23285 | 4930451E10 | ENSMUSG00000069223.5 |
| chr16 | 76403379 | 76403656 | 5.43604 | 8.18012 | 6.04458 | Promoter | 16 | 76403209 | 76403712 | 504 | 2 | ENSMUST00 | 56 | Gm9843 | ENSMUSG00000050299.7 |
| chr1 | 94035118 | 94035626 | 4.50687 | 8.15551 | 6.02026 | Downstream | 1 | 94020493 | 94028334 | 7842 | 1 | ENSMUST00 | 14626 | Neu4 | ENSMUSG00000034000.10 |
| chr12 | 87407749 | 87408484 | 4.50687 | 8.15551 | 6.02026 | Distal Interge | 12 | 87305058 | 87388355 | 83298 | 2 | ENSMUST00 | -19395 | Sptlc2 | ENSMUSG00000021036.5 |
| chr15 | 76208303 | 76208821 | 4.50687 | 8.15551 | 6.02026 | Promoter | 15 | 76170976 | 76208653 | 37678 | 2 | ENSMUST00 | 0 | Plec | ENSMUSG00000022565.10 |
| chr10 | 86727563 | 86727736 | 5.4136 | 8.14442 | 6.00938 | Intron (ENSM | 10 | 86731754 | 86732409 | 656 | 2 | ENSMUST00 | 4673 | Fabp3-ps1 | ENSMUSG00000056366.4 |
| chr1 | 165091899 | 165092272 | 5.39135 | 8.10903 | 5.97439 | Distal Interge | 1 | 165148824 | 165148949 | 126 | 2 | ENSMUST00 | 56677 | Gm16550 | ENSMUSG00000089967.1 |
| chr11 | 58927844 | 58928012 | 5.39135 | 8.10903 | 5.97439 | Distal Interge | 11 | 58918057 | 58926965 | 8909 | 1 | ENSMUST00 | 9788 | Btnl10 | ENSMUSG00000020490.11 |
| chr3 | 78919470 | 78919634 | 5.3803 | 8.09144 | 5.95695 | Distal Interge | 3 | 78917258 | 78918259 | 1002 | 1 | ENSMUST00 | 2213 | Gm10291 | ENSMUSG00000070443.5 |
| chr11 | 118401612 | 118401825 | 5.1806 | 8.07634 | 5.94198 | Promoter | 11 | 118396607 | 118401852 | 5246 | 2 | ENSMUST00 | 27 | Lgals3bp | ENSMUSG00000033880.6 |
| chr18 | 56774403 | 56775071 | 5.36929 | 8.07392 | 5.94005 | Intron (ENSM | 18 | 56769717 | 56801688 | 31972 | 1 | ENSMUST00 | 4687 | Gm15345 | ENSMUSG00000087597.1 |
| chr8 | 116946304 | 116946544 | 5.36929 | 8.07392 | 5.94005 | Intron (ENSM | 8 | 116943393 | 116960445 | 17053 | 1 | ENSMUST00 | 2912 | Atmin | ENSMUSG00000047388.10 |
| chr13 | 112439437 | 112439617 | 5.35832 | 8.05648 | 5.92315 | Distal Interge | 13 | 112464070 | 112510086 | 46017 | 1 | ENSMUST00 | -24453 | Il6st | ENSMUSG00000021756.7 |
| chr7 | 30291531 | 30291939 | 5.35832 | 8.05648 | 5.92315 | Promoter | 7 | 30291672 | 30308367 | 16696 | 1 | ENSMUST00 | 0 | Clip3 | ENSMUSG00000013921.9 |
| chr1 | 120602405 | 120602595 | 5.3474 | 8.0391 | 5.90593 | Promoter | 1 | 120602418 | 120607992 | 5575 | 1 | ENSMUST00 | 0 | En1 | ENSMUSG00000058665.8 |
| chr19 | 45048228 | 45048617 | 5.3474 | 8.0391 | 5.90593 | Promoter | 19 | 45047576 | 45056383 | 8808 | 1 | ENSMUST00 | 653 | Sfxn3 | ENSMUSG00000025212.11 |
| chr10 | 21997324 | 21998158 | 4.57821 | 7.98325 | 5.85079 | Promoter | 10 | 21996591 | 21998240 | 1650 | 1 | ENSMUST00 | 734 | Sgk1 | ENSMUSG00000019970.10 |
| chr10 | 81414096 | 81414786 | 4.57821 | 7.98325 | 5.85079 | Intron (ENSM | 10 | 81409575 | 81411579 | 2005 | 1 | ENSMUST00 | 4522 | Gm16104 | ENSMUSG00000085327.1 |
| chr19 | 10043064 | 10043349 | 4.57821 | 7.98325 | 5.85079 | Intron (ENSM | 19 | 10041548 | 10059671 | 18124 | 1 | ENSMUST00 | 1517 | Fads3 | ENSMUSG00000024664.4 |
| chr2 | 173020239 | 173020594 | 4.57821 | 7.98325 | 5.85079 | Promoter | 2 | 173020498 | 173034734 | 14237 | 1 | ENSMUST00 | 0 | Rbm38 | ENSMUSG00000027510.12 |
| chr4 | 126227181 | 126227508 | 4.57821 | 7.98325 | 5.85079 | Distal Interge | 4 | 126232963 | 126233548 | 586 | 2 | ENSMUST00 | 6040 | Map7d1 | ENSMUSG00000028849.12 |
| chr4 | 150212761 | 150213040 | 4.57821 | 7.98325 | 5.85079 | Distal Interge | 4 | 150187015 | 150201332 | 14318 | 2 | ENSMUST00 | -11430 | Car6 | ENSMUSG00000028972.6 |
| chr5 | 36868642 | 36869253 | 4.57821 | 7.98325 | 5.85079 | Promoter | 5 | 36868513 | 36955078 | 86566 | 1 | ENSMUST00 | 130 | Ppp2r2c | ENSMUSG00000029120.7 |
| chr5 | 33534959 | 33535290 | 4.57821 | 7.98325 | 5.85079 | Distal Interge | 5 | 33600356 | 33628912 | 28557 | 2 | ENSMUST00 | 93622 | Fam53a | ENSMUSG00000037339.12 |
| chr5 | 72201807 | 72202378 | 4.57821 | 7.98325 | 5.85079 | Promoter | 5 | 72203329 | 72230475 | 27147 | 1 | ENSMUST00 | -951 | Atp10d | ENSMUSG00000046808.12 |
| chr6 | 119417167 | 119417719 | 4.57821 | 7.98325 | 5.85079 | Promoter | 6 | 119353150 | 119417704 | 64555 | 2 | ENSMUST00 | 0 | Adipor2 | ENSMUSG00000030168.8 |
| chr10 | 20902703 | 20903543 | 5.08786 | 7.92609 | 5.7945 | Distal Interge | 10 | 20952547 | 21080428 | 127882 | 1 | ENSMUST00 | -49004 | Ahi1 | ENSMUSG00000019986.11 |
| chr4 | 155235543 | 155235725 | 5.08786 | 7.92609 | 5.7945 | Distal Interge | 4 | 155154075 | 155222592 | 68518 | 2 | ENSMUST00 | -12952 | Ski | ENSMUSG00000029050.10 |
| chr8 | 70419225 | 70419465 | 5.08786 | 7.92609 | 5.7945 | Intron (ENSM | 8 | 70406127 | 70431285 | 25159 | 2 | ENSMUST00 | 11820 | Crtc1 | ENSMUSG00000003575.8 |
| chr3 | 97159776 | 97159950 | 5.26161 | 7.90259 | 5.77154 | Promoter | 3 | 97159103 | 97168310 | 9208 | 1 | ENSMUST00 | 674 | Acp6 | ENSMUSG00000028093.10 |
| chr9 | 120191320 | 120191484 | 5.26161 | 7.90259 | 5.77154 | Distal Interge | 9 | 120149742 | 120181484 | 31743 | 1 | ENSMUST00 | 41579 | Mobp | ENSMUSG00000032517.10 |
| chr2 | 62407797 | 62408068 | 5.24059 | 7.86914 | 5.73891 | Exon (ENSMU | 2 | 62407702 | 62411859 | 4158 | 2 | ENSMUST00 | 3791 | Dpp4 | ENSMUSG00000035000.8 |
| chr10 | 78124273 | 78124732 | 4.66184 | 7.82715 | 5.69732 | Distal Interge | 10 | 78162067 | 78169768 | 7702 | 2 | ENSMUST00 | 45036 | D10Jhu81e | ENSMUSG00000053329.6 |
| chr4 | 102087446 | 102088362 | 4.66184 | 7.82715 | 5.69732 | Promoter | 4 | 102087543 | 102605833 | 518291 | 1 | ENSMUST00 | 0 | Pde4b | ENSMUSG00000028525.11 |
| chr4 | 81744682 | 81745092 | 4.66184 | 7.82715 | 5.69732 | Exon (ENSMU | 4 | 81743146 | 81746701 | 3556 | 1 | ENSMUST00 | 1537 | Gm11412 | ENSMUSG00000087540.1 |
| chr5 | 52618398 | 52619007 | 4.66184 | 7.82715 | 5.69732 | Promoter | 5 | 52607550 | 52619011 | 11462 | 2 | ENSMUST00 | 4 | 8030423F21 | ENSMUSG00000052295.3 |
| chr6 | 30675367 | 30675698 | 4.66184 | 7.82715 | 5.69732 | Intron (ENSM | 6 | 30673944 | 30682266 | 8323 | 1 | ENSMUST00 | 1424 | Gm13782 | ENSMUSG00000086542.1 |
| chr8 | 70744115 | 70744662 | 4.66184 | 7.82715 | 5.69732 | Intron (ENSM | 8 | 70743079 | 70750473 | 7395 | 1 | ENSMUST00 | 1037 | Pde4c | ENSMUSG00000031842.8 |
| chr8 | 126138801 | 126139262 | 4.66184 | 7.82715 | 5.69732 | Distal Interge | 8 | 125995102 | 126030683 | 35582 | 1 | ENSMUST00 | 143700 | Kcnk1 | ENSMUSG00000033998.8 |
| chr9 | 55241948 | 55242383 | 4.66184 | 7.82715 | 5.69732 | Promoter | 9 | 55221222 | 55242461 | 21240 | 2 | ENSMUST00 | 78 | Nrg4 | ENSMUSG00000032311.12 |
| chr10 | 94020113 | 94020785 | 5.20937 | 7.81945 | 5.69051 | Promoter | 10 | 93996914 | 94020723 | 23810 | 2 | ENSMUST00 | 0 | Vezt | ENSMUSG00000036099.11 |
| chr2 | 91949972 | 91950136 | 5.20937 | 7.81945 | 5.69051 | Promoter | 2 | 91940758 | 91950311 | 9554 | 2 | ENSMUST00 | 175 | Dgkz | ENSMUSG00000040479.6 |
| chr5 | 24365320 | 24365611 | 5.20937 | 7.81945 | 5.69051 | Promoter | 5 | 24364810 | 24384474 | 19665 | 1 | ENSMUST00 | 511 | Nos3 | ENSMUSG00000028978.7 |
| chr3 | 90432026 | 90432433 | 5.19905 | 7.80301 | 5.67418 | Intron (ENSM | 3 | 90391380 | 90433636 | 42257 | 2 | ENSMUST00 | 1203 | Ints3 | ENSMUSG00000027933.6 |
| chr14 | 73372177 | 73372414 | 4.99839 | 7.78073 | 5.65212 | Intron (ENSM | 14 | 73362231 | 73385271 | 23041 | 2 | ENSMUST00 | 12857 | Itm2b | ENSMUSG00000022108.7 |
| chr3 | 79178094 | 79178421 | 4.99839 | 7.78073 | 5.65212 | Intron (ENSM | 3 | 79169449 | 79181340 | 11892 | 2 | ENSMUST00 | 2919 | Rapgef2 | ENSMUSG00000062232.9 |
| chr4 | 132462762 | 132463080 | 4.99839 | 7.78073 | 5.65212 | Promoter | 4 | 132459955 | 132463331 | 3377 | 2 | ENSMUST00 | 251 | Med18 | ENSMUSG00000066042.4 |
| chr8 | 120831968 | 120832387 | 4.16048 | 7.77275 | 5.64444 | Intron (ENSM | 8 | 120880206 | 120882481 | 2276 | 1 | ENSMUST00 | -47819 | Gm26878 | ENSMUSG00000097087.1 |
| chr10 | 76500859 | 76501338 | 4.76124 | 7.69302 | 5.56588 | Exon (ENSMU | 10 | 76531588 | 76535568 | 3981 | 1 | ENSMUST00 | -30250 | Lss | ENSMUSG00000033105.7 |
| chr11 | 117135886 | 117136235 | 4.76124 | 7.69302 | 5.56588 | Intron (ENSM | 11 | 117134598 | 117135045 | 448 | 1 | ENSMUST00 | 1289 | Gm11745 | ENSMUSG00000084052.1 |
| chr13 | 68298218 | 68298420 | 4.76124 | 7.69302 | 5.56588 | Distal Interge | 13 | 68560780 | 68582121 | 21342 | 2 | ENSMUST00 | 283701 | Mtrr | ENSMUSG00000034617.6 |
| chr13 | 25269844 | 25270178 | 4.76124 | 7.69302 | 5.56588 | Promoter | 13 | 25252040 | 25270082 | 18043 | 2 | ENSMUST00 | 0 | Nrsn1 | ENSMUSG00000048978.8 |
| chr13 | 65890516 | 65890805 | 4.76124 | 7.69302 | 5.56588 | Distal Interge | 13 | 65894209 | 65894745 | 537 | 2 | ENSMUST00 | 3940 | Gm6917 | ENSMUSG00000074841.1 |
| chr17 | 73806136 | 73807403 | 4.76124 | 7.69302 | 5.56588 | Intron (ENSM | 17 | 73804841 | 73832093 | 27253 | 1 | ENSMUST00 | 1296 | Ehd3 | ENSMUSG00000024065.7 |
| chr18 | 65175824 | 65176343 | 4.76124 | 7.69302 | 5.56588 | Intron (ENSM | 18 | 65248861 | 65248926 | 66 | 1 | ENSMUST00 | -72518 | Mir122a | ENSMUSG00000065402.1 |
| chr2 | 74655801 | 74656208 | 4.76124 | 7.69302 | 5.56588 | Exon (ENSMU | 2 | 74655616 | 74659419 | 3804 | 2 | ENSMUST00 | 3211 | Evx2 | ENSMUSG00000001815.10 |
| chr5 | 28209931 | 28210343 | 4.76124 | 7.69302 | 5.56588 | Promoter | 5 | 28202449 | 28210022 | 7574 | 2 | ENSMUST00 | 0 | Cnpy1 | ENSMUSG00000044681.13 |
| chr7 | 30553252 | 30553601 | 4.76124 | 7.69302 | 5.56588 | Promoter | 7 | 30553268 | 30555443 | 2176 | 1 | ENSMUST00 | 0 | Hspb6 | ENSMUSG00000036854.9 |
| chr9 | 101252059 | 101252248 | 4.76124 | 7.69302 | 5.56588 | Promoter | 9 | 101105084 | 101251795 | 146712 | 2 | ENSMUST00 | -265 | Ppp2r3a | ENSMUSG00000043154.10 |
| chr9 | 67192032 | 67192445 | 4.76124 | 7.69302 | 5.56588 | Distal Interge | 9 | 67236651 | 67236735 | 85 | 2 | ENSMUST00 | 44290 | Mir190a | ENSMUSG00000076379.1 |
| chr1 | 168357729 | 168357893 | 5.12792 | 7.68973 | 5.56422 | Intron (ENSM | 1 | 168120969 | 168405601 | 284633 | 2 | ENSMUST00 | 47708 | Pbx1 | ENSMUSG00000052534.10 |
| chr2 | 152827952 | 152828384 | 4.912 | 7.63997 | 5.51476 | Promoter | 2 | 152781411 | 152828535 | 47125 | 2 | ENSMUST00 | 151 | Bcl2l1 | ENSMUSG00000007659.13 |
| chr2 | 180332238 | 180332454 | 4.912 | 7.63997 | 5.51476 | Promoter | 2 | 180332857 | 180340732 | 7876 | 1 | ENSMUST00 | -403 | Gata5os | ENSMUSG00000056061.3 |
| chr10 | 97312739 | 97312904 | 4.88133 | 7.58987 | 5.46511 | Distal Interge | 10 | 97479500 | 97518162 | 38663 | 1 | ENSMUST00 | -166596 | Dcn | ENSMUSG00000019929.10 |
| chr10 | 81433158 | 81433369 | 4.88133 | 7.58987 | 5.46511 | Distal Interge | 10 | 81400953 | 81431005 | 30053 | 2 | ENSMUST00 | -2154 | Nfic | ENSMUSG00000055053.12 |
| chr13 | 12106441 | 12106991 | 4.88133 | 7.58987 | 5.46511 | Promoter | 13 | 11553103 | 12106945 | 553843 | 2 | ENSMUST00 | 0 | Ryr2 | ENSMUSG00000021313.10 |
| chr13 | 69737430 | 69737937 | 4.88133 | 7.58987 | 5.46511 | Intron (ENSM | 13 | 69702835 | 69739897 | 37063 | 2 | ENSMUST00 | 1960 | Ube2ql1 | ENSMUSG00000052981.5 |

| | | | | | | | | | | | | | | | |
|---|---|---|---|---|---|---|---|---|---|---|---|---|---|---|---|
| chr14 | 40965059 | 40965458 | 4.88133 | 7.58987 | 5.46511 | Intron (ENSM | 14 | 40906445 | 40966807 | 60363 | 2 | ENSMUST00 | 1349 | Tspan14 | ENSMUSG00000037824.5 |
| chr15 | 99890938 | 99891108 | 4.88133 | 7.58987 | 5.46511 | Distal Interge | 15 | 99898287 | 99898556 | 270 | 2 | ENSMUST00 | 7448 | Gm9763 | ENSMUSG00000037438.5 |
| chr16 | 85549211 | 85549626 | 4.88133 | 7.58987 | 5.46511 | Promoter | 16 | 85421533 | 85550417 | 128885 | 2 | ENSMUST00 | 791 | Cyyr1 | ENSMUSG00000041134.10 |
| chr17 | 10301002 | 10301198 | 4.88133 | 7.58987 | 5.46511 | Intron (ENSM | 17 | 10206471 | 10319361 | 112891 | 2 | ENSMUST00 | 18163 | Qk | ENSMUSG00000062078.9 |
| chr18 | 61639449 | 61639795 | 4.88133 | 7.58987 | 5.46511 | Promoter | 18 | 61639542 | 61647292 | 7751 | 1 | ENSMUST00 | 0 | Gm20748 | ENSMUSG00000098098.2 |
| chr19 | 37886486 | 37886921 | 4.88133 | 7.58987 | 5.46511 | Distal Interge | 19 | 37899036 | 38043577 | 144542 | 2 | ENSMUST00 | 156656 | Myof | ENSMUSG00000048612.10 |
| chr3 | 98063709 | 98064058 | 4.88133 | 7.58987 | 5.46511 | Intron (ENSM | 3 | 98013538 | 98150367 | 136830 | 1 | ENSMUST00 | 50172 | Notch2 | ENSMUSG00000027878.10 |
| chr4 | 106616991 | 106617414 | 4.88133 | 7.58987 | 5.46511 | Promoter | 4 | 106590909 | 106617238 | 26330 | 2 | ENSMUST00 | 0 | BC055111 | ENSMUSG00000054362.4 |
| chr4 | 106858151 | 106858528 | 4.88133 | 7.58987 | 5.46511 | Distal Interge | 4 | 106841761 | 106847920 | 6160 | 2 | ENSMUST00 | -10232 | Gm12746 | ENSMUSG00000086940.1 |
| chr6 | 52185214 | 52185621 | 4.88133 | 7.58987 | 5.46511 | Promoter | 6 | 52180560 | 52184874 | 4315 | 2 | ENSMUST00 | -341 | Hoxa3 | ENSMUSG00000079560.8 |
| chr7 | 100910881 | 100911312 | 4.88133 | 7.58987 | 5.46511 | Intron (ENSM | 7 | 100869746 | 100932161 | 62416 | 2 | ENSMUST00 | 20849 | Arhgef17 | ENSMUSG00000032875.7 |
| chr7 | 118712193 | 118712357 | 4.88133 | 7.58987 | 5.46511 | Promoter | 7 | 118712552 | 118737024 | 24473 | 1 | ENSMUST00 | -195 | Ccp110 | ENSMUSG00000033904.10 |
| chr8 | 83738893 | 83739343 | 4.88133 | 7.58987 | 5.46511 | Intron (ENSM | 8 | 83723362 | 83741169 | 17808 | 2 | ENSMUST00 | 1826 | Cd97 | ENSMUSG00000002885.9 |
| chr8 | 84105060 | 84105370 | 4.88133 | 7.58987 | 5.46511 | Promoter | 8 | 84097075 | 84104773 | 7699 | 2 | ENSMUST00 | -288 | Dcaf15 | ENSMUSG00000037103.7 |
| chr7 | 16475724 | 16476051 | 5.05871 | 7.57942 | 5.45715 | Promoter | 7 | 16455721 | 16476780 | 21060 | 2 | ENSMUST00 | 729 | Npas1 | ENSMUSG00000001988.8 |
| chr18 | 82954056 | 82954253 | 4.19834 | 7.56442 | 5.44278 | Intron (ENSM | 18 | 82976323 | 82976414 | 92 | 1 | ENSMUST00 | -22070 | Gm26258 | ENSMUSG00000077976.1 |
| chr1 | 91065079 | 91065244 | 5.02932 | 7.53255 | 5.41116 | Intron (ENSM | 1 | 91053586 | 91128287 | 74702 | 1 | ENSMUST00 | 11494 | Lrrfip1 | ENSMUSG00000026305.10 |
| chr1 | 144003141 | 144003334 | 5.02932 | 7.53255 | 5.41116 | Promoter | 1 | 144001883 | 144004032 | 2150 | 2 | ENSMUST00 | 698 | Rgs2 | ENSMUSG00000026360.4 |
| chr1 | 90441549 | 90441950 | 5.02932 | 7.53255 | 5.41116 | Distal Interge | 1 | 90453394 | 90454501 | 1108 | 2 | ENSMUST00 | 12551 | RP24-154L11 | ENSMUSG00000101997.1 |
| chr10 | 70521227 | 70521470 | 5.02932 | 7.53255 | 5.41116 | Intron (ENSM | 10 | 70501192 | 70501503 | 312 | 2 | ENSMUST00 | -19725 | Gm22320 | ENSMUSG00000087774.1 |
| chr11 | 95413649 | 95413960 | 5.02932 | 7.53255 | 5.41116 | Promoter | 11 | 95414080 | 95493406 | 79327 | 1 | ENSMUST00 | -120 | Spop | ENSMUSG00000057522.10 |
| chr12 | 46399482 | 46399876 | 5.02932 | 7.53255 | 5.41116 | Distal Interge | 12 | 45984343 | 45984549 | 207 | 1 | ENSMUST00 | 415140 | Gm24948 | ENSMUSG00000064800.1 |
| chr15 | 75048297 | 75048501 | 5.02932 | 7.53255 | 5.41116 | Promoter | 15 | 75045793 | 75048669 | 2877 | 2 | ENSMUST00 | 168 | Ly6c1 | ENSMUSG00000079018.5 |
| chr3 | 51226341 | 51226670 | 5.02932 | 7.53255 | 5.41116 | Intron (ENSM | 3 | 51224470 | 51251631 | 27162 | 1 | ENSMUST00 | 1872 | Ccrn4l | ENSMUSG00000023087.10 |
| chr4 | 19725042 | 19725385 | 5.02932 | 7.53255 | 5.41116 | Distal Interge | 4 | 19716732 | 19720199 | 3468 | 2 | ENSMUST00 | -4844 | Gm12354 | ENSMUSG00000086071.2 |
| chr7 | 141009879 | 141010155 | 5.02932 | 7.53255 | 5.41116 | Promoter | 7 | 141009586 | 141010770 | 1185 | 2 | ENSMUST00 | 615 | Ifitm3 | ENSMUSG00000025492.6 |
| chr7 | 78913752 | 78914062 | 5.02932 | 7.53255 | 5.41116 | Promoter | 7 | 78913765 | 78920362 | 6598 | 1 | ENSMUST00 | 0 | Isg20 | ENSMUSG00000039236.13 |
| chr8 | 27206409 | 27206710 | 5.02932 | 7.53255 | 5.41116 | Distal Interge | 8 | 27197460 | 27202547 | 5088 | 2 | ENSMUST00 | -3863 | Got1l1 | ENSMUSG00000039720.6 |
| chr9 | 90226128 | 90226432 | 5.02932 | 7.53255 | 5.41116 | Exon (ENSMU | 9 | 90221295 | 90221625 | 331 | 2 | ENSMUST00 | -4504 | Gm16200 | ENSMUSG00000081520.1 |
| chr10 | 59395646 | 59395812 | 3.67619 | 7.51343 | 5.39463 | Promoter | 10 | 59395632 | 59396757 | 1126 | 1 | ENSMUST00 | 15 | Gm10273 | ENSMUSG00000069622.4 |
| chr1 | 173942013 | 173942177 | 4.82856 | 7.50356 | 5.38477 | Promoter | 1 | 173936618 | 173942454 | 5837 | 2 | ENSMUST00 | 277 | Ifi203 | ENSMUSG00000039997.11 |
| chr8 | 36608832 | 36609258 | 4.82856 | 7.50356 | 5.38477 | Intron (ENSM | 8 | 36593231 | 36613874 | 20644 | 2 | ENSMUST00 | 4616 | Dlc1 | ENSMUSG00000031523.11 |
| chr4 | 43209681 | 43209863 | 5.0104 | 7.50237 | 5.3838 | Intron (ENSM | 4 | 43201149 | 43201213 | 65 | 2 | ENSMUST00 | -8469 | Gm25010 | ENSMUSG00000089028.1 |
| chr16 | 26102514 | 26102815 | 4.97242 | 7.44173 | 5.32336 | Intron (ENSM | 16 | 25959288 | 26105784 | 146497 | 2 | ENSMUST00 | 2969 | Leprel1 | ENSMUSG00000038168.5 |
| chr10 | 127756084 | 127756289 | 4.96301 | 7.42671 | 5.30838 | Distal Interge | 10 | 127759721 | 127768297 | 8577 | 1 | ENSMUST00 | -3432 | Rdh1 | ENSMUSG00000089789.1 |
| chr6 | 37438901 | 37439065 | 4.96301 | 7.42671 | 5.30838 | Intron (ENSM | 6 | 37327255 | 37442146 | 114892 | 2 | ENSMUST00 | 3081 | Creb3l2 | ENSMUSG00000038648.5 |
| chr7 | 130574620 | 130574963 | 4.96301 | 7.42671 | 5.30838 | Intron (ENSM | 7 | 130533770 | 130573118 | 39349 | 2 | ENSMUST00 | -1503 | Nsmce4a | ENSMUSG00000040331.10 |
| chr11 | 97149875 | 97150039 | 4.57782 | 7.38195 | 5.26448 | Promoter | 11 | 97136177 | 97150046 | 13870 | 2 | ENSMUST00 | 7 | Tbkbp1 | ENSMUSG00000038517.10 |
| chr18 | 66345036 | 66345367 | 4.57782 | 7.38195 | 5.26448 | Distal Interge | 18 | 66269028 | 66302739 | 33712 | 2 | ENSMUST00 | -42298 | Ccbe1 | ENSMUSG00000046318.10 |
| chr7 | 142019094 | 142019518 | 4.24176 | 7.36333 | 5.24607 | Intron (ENSM | 7 | 141998661 | 142001441 | 2781 | 1 | ENSMUST00 | 20434 | Brsk2 | ENSMUSG00000053046.9 |
| chr10 | 94505525 | 94505774 | 5.05485 | 7.29253 | 5.17558 | Intron (ENSM | 10 | 94514857 | 94590956 | 76100 | 1 | ENSMUST00 | -9083 | Tmcc3 | ENSMUSG00000020023.12 |
| chr10 | 40191653 | 40192185 | 5.05485 | 7.29253 | 5.17558 | Distal Interge | 10 | 40223683 | 40227798 | 4116 | 2 | ENSMUST00 | 35613 | Rpf2 | ENSMUSG00000038510.7 |
| chr11 | 98961431 | 98961754 | 5.05485 | 7.29253 | 5.17558 | Intron (ENSM | 11 | 98960412 | 98974942 | 14531 | 1 | ENSMUST00 | 1020 | Rara | ENSMUSG00000037992.11 |
| chr15 | 102921145 | 102921469 | 5.05485 | 7.29253 | 5.17558 | Promoter | 15 | 102921103 | 102928814 | 7712 | 1 | ENSMUST00 | 43 | Hoxc13 | ENSMUSG00000001655.6 |
| chr16 | 38714262 | 38714426 | 5.05485 | 7.29253 | 5.17558 | Promoter | 16 | 38598340 | 38713274 | 114935 | 2 | ENSMUST00 | -989 | Arhgap31 | ENSMUSG00000022799.4 |
| chr18 | 7608656 | 7608897 | 5.05485 | 7.29253 | 5.17558 | Intron (ENSM | 18 | 7347962 | 7626863 | 278902 | 2 | ENSMUST00 | 17966 | Mpp7 | ENSMUSG00000057440.7 |
| chr19 | 58228413 | 58228680 | 5.05485 | 7.29253 | 5.17558 | Intron (ENSM | 19 | 58204139 | 58236047 | 31909 | 1 | ENSMUST00 | 24275 | Gm16277 | ENSMUSG00000087002.1 |
| chr2 | 9882483 | 9883081 | 5.05485 | 7.29253 | 5.17558 | Promoter | 2 | 9882636 | 9883335 | 700 | 1 | ENSMUST00 | 0 | 4930412O13 | ENSMUSG00000025783.2 |
| chr2 | 176709103 | 176709267 | 5.05485 | 7.29253 | 5.17558 | Promoter | 2 | 176708355 | 176720203 | 11849 | 1 | ENSMUST00 | 749 | Gm14305 | ENSMUSG00000078878.4 |
| chr2 | 119189258 | 119189464 | 5.05485 | 7.29253 | 5.17558 | Intron (ENSM | 2 | 119187517 | 119188261 | 745 | 1 | ENSMUST00 | 1742 | Gm14138 | ENSMUSG00000081572.3 |
| chr4 | 131173889 | 131174066 | 5.05485 | 7.29253 | 5.17558 | Distal Interge | 4 | 131239277 | 131239410 | 134 | 1 | ENSMUST00 | -65211 | Gm25261 | ENSMUSG00000088041.1 |
| chr5 | 135460134 | 135460848 | 5.05485 | 7.29253 | 5.17558 | Exon (ENSMU | 5 | 135376141 | 135394546 | 18406 | 2 | ENSMUST00 | -65589 | Pom121 | ENSMUSG00000053293.6 |
| chr5 | 118458950 | 118459163 | 5.05485 | 7.29253 | 5.17558 | Distal Interge | 5 | 118418727 | 118418818 | 92 | 2 | ENSMUST00 | -40133 | Gm26455 | ENSMUSG00000093041.1 |
| chr7 | 28259443 | 28259621 | 5.05485 | 7.29253 | 5.17558 | Distal Interge | 7 | 28267881 | 28269272 | 1392 | 1 | ENSMUST00 | -8260 | Eid2 | ENSMUSG00000046058.7 |
| chr8 | 78420200 | 78420461 | 5.05485 | 7.29253 | 5.17558 | Distal Interge | 8 | 78433010 | 78436645 | 3636 | 2 | ENSMUST00 | 16184 | Pou4f2 | ENSMUSG00000031688.4 |
| chrX | 7188591 | 7188759 | 5.05485 | 7.29253 | 5.17558 | Promoter | 20 | 7153810 | 7188714 | 34905 | 2 | ENSMUST00 | 0 | Clcn5 | ENSMUSG00000004317.9 |
| chrX | 42540421 | 42540799 | 5.05485 | 7.29253 | 5.17558 | Intron (ENSM | 20 | 42526585 | 42529330 | 2746 | 1 | ENSMUST00 | 13837 | Sh2d1a | ENSMUSG00000005696.6 |
| chrX | 164269520 | 164269690 | 5.05485 | 7.29253 | 5.17558 | Promoter | 20 | 164269547 | 164344659 | 75113 | 1 | ENSMUST00 | 0 | Pir | ENSMUSG00000031379.8 |
| chrX | 73914612 | 73914942 | 5.05485 | 7.29253 | 5.17558 | Intron (ENSM | 20 | 73891442 | 73911336 | 19895 | 2 | ENSMUST00 | -3277 | Arhgap4 | ENSMUSG00000031389.12 |
| chrX | 106026889 | 106027053 | 5.05485 | 7.29253 | 5.17558 | Promoter | 20 | 106027276 | 106124917 | 97642 | 1 | ENSMUST00 | -223 | Atp7a | ENSMUSG00000033792.7 |
| chrX | 20921172 | 20921500 | 5.05485 | 7.29253 | 5.17558 | Promoter | 20 | 20860511 | 20921004 | 60494 | 2 | ENSMUST00 | -169 | Syn1 | ENSMUSG00000037217.10 |
| chrX | 138914655 | 138914853 | 5.05485 | 7.29253 | 5.17558 | Promoter | 20 | 138914431 | 139010532 | 96102 | 1 | ENSMUST00 | 225 | Nrk | ENSMUSG00000052854.10 |
| chrX | 94865846 | 94866012 | 5.05485 | 7.29253 | 5.17558 | Distal Interge | 20 | 94795563 | 94798183 | 2621 | 2 | ENSMUST00 | -67664 | Zxda | ENSMUSG00000073060.4 |
| chrX | 12886590 | 12886966 | 5.05485 | 7.29253 | 5.17558 | Distal Interge | 20 | 12870269 | 12870362 | 94 | 2 | ENSMUST00 | -16229 | Gm25063 | ENSMUSG00000077858.1 |
| chrX | 77433790 | 77433969 | 5.05485 | 7.29253 | 5.17558 | Distal Interge | 20 | 77456586 | 77457228 | 643 | 1 | ENSMUST00 | -22617 | Gm5937 | ENSMUSG00000081622.1 |
| chrX | 36817244 | 36817408 | 5.05485 | 7.29253 | 5.17558 | Promoter | 20 | 36814771 | 36817185 | 2415 | 2 | ENSMUST00 | -60 | 2310010G23 | ENSMUSG00000097468.1 |
| chr13 | 83525984 | 83526322 | 4.87087 | 7.27943 | 5.16553 | Intron (ENSM | 13 | 83504034 | 83667079 | 163046 | 1 | ENSMUST00 | 21951 | Mef2c | ENSMUSG00000005583.11 |
| chr17 | 63662388 | 63662793 | 4.87087 | 7.27943 | 5.16553 | Distal Interge | 17 | 63057452 | 63500017 | 442566 | 2 | ENSMUST00 | -162372 | Fbxl17 | ENSMUSG00000023965.8 |
| chr8 | 22303234 | 22303404 | 4.87087 | 7.27943 | 5.16553 | Intron (ENSM | 8 | 22283441 | 22371418 | 87978 | 1 | ENSMUST00 | 19794 | Tpte | ENSMUSG00000031481.7 |
| chr4 | 48051640 | 48051887 | 5.04576 | 7.27818 | 5.16492 | Promoter | 4 | 48051248 | 48056739 | 5492 | 1 | ENSMUST00 | 393 | Nr4a3 | ENSMUSG00000028341.4 |
| chr1 | 173879577 | 173879894 | 5.0352 | 7.2615 | 5.14839 | Promoter | 1 | 173872852 | 173880111 | 7260 | 2 | ENSMUST00 | 217 | Mndal | ENSMUSG00000090272.2 |
| chr14 | 31227679 | 31227897 | 4.66989 | 7.2429 | 5.13034 | Exon (ENSMU | 14 | 31217860 | 31230350 | 12491 | 1 | ENSMUST00 | 9820 | Sema3g | ENSMUSG00000021904.5 |
| chr4 | 136145337 | 136145883 | 4.66989 | 7.2429 | 5.13034 | Promoter | 4 | 136144974 | 136145755 | 782 | 1 | ENSMUST00 | 364 | Id3 | ENSMUSG00000007872.3 |
| chr15 | 89524035 | 89524289 | 5.0142 | 7.22837 | 5.11594 | Promoter | 15 | 89524322 | 89534394 | 10073 | 1 | ENSMUST00 | -33 | Shank3 | ENSMUSG00000022623.10 |
| chr9 | 83610442 | 83610749 | 4.99338 | 7.19551 | 5.0834 | Intron (ENSM | 9 | 83576831 | 83597524 | 20694 | 1 | ENSMUST00 | 33612 | Sh3bgrl2 | ENSMUSG00000032261.11 |
| chr1 | 143503168 | 143503845 | 4.29207 | 7.17162 | 5.05983 | Distal Interge | 1 | 143633851 | 143636577 | 2727 | 2 | ENSMUST00 | 132732 | Cdc73 | ENSMUSG00000026361.8 |
| chr10 | 77109671 | 77110299 | 4.29207 | 7.17162 | 5.05983 | Intron (ENSM | 10 | 77052181 | 77113676 | 61496 | 2 | ENSMUST00 | 3377 | Col18a1 | ENSMUSG00000001435.9 |
| chr11 | 120629421 | 120629732 | 4.29207 | 7.17162 | 5.05983 | Promoter | 11 | 120629232 | 120630148 | 917 | 2 | ENSMUST00 | 416 | Mafg | ENSMUSG00000051510.8 |
| chr3 | 37687154 | 37687631 | 4.29207 | 7.17162 | 5.05983 | Distal Interge | 3 | 37714717 | 37715212 | 496 | 1 | ENSMUST00 | -27086 | Rps23-ps1 | ENSMUSG00000063001.9 |
| chr5 | 74190110 | 74190403 | 4.29207 | 7.17162 | 5.05983 | Distal Interge | 5 | 74195288 | 74199481 | 4194 | 1 | ENSMUST00 | -4885 | Rasl11b | ENSMUSG00000049907.7 |
| chr7 | 45632572 | 45632800 | 4.29207 | 7.17162 | 5.05983 | Promoter | 7 | 45630664 | 45632809 | 2146 | 2 | ENSMUST00 | 9 | Gm16047 | ENSMUSG00000084882.1 |
| chr8 | 95255030 | 95255324 | 4.29207 | 7.17162 | 5.05983 | Intron (ENSM | 8 | 95240981 | 95281590 | 40610 | 2 | ENSMUST00 | 26266 | Cngb1 | ENSMUSG00000031789.11 |
| chr9 | 92207637 | 92207866 | 4.29207 | 7.17162 | 5.05983 | Exon (ENSMU | 9 | 92192936 | 92209772 | 16837 | 1 | ENSMUST00 | 14702 | Plscr5 | ENSMUSG00000095654.2 |
| chr13 | 43123703 | 43124095 | 4.97273 | 7.16293 | 5.05161 | Promoter | 13 | 43122872 | 43135359 | 12488 | 1 | ENSMUST00 | 832 | Phactr1 | ENSMUSG00000054728.11 |
| chr3 | 107758919 | 107759231 | 4.44487 | 7.15427 | 5.04311 | Promoter | 3 | 107750380 | 107760221 | 9842 | 2 | ENSMUST00 | 990 | Csf1 | ENSMUSG00000014599.5 |
| chr10 | 62578722 | 62578886 | 4.78209 | 7.13724 | 5.02624 | Promoter | 10 | 62558470 | 62578457 | 19988 | 2 | ENSMUST00 | -266 | 2510003E04 | ENSMUSG00000036955.10 |
| chr14 | 57800604 | 57800768 | 4.78209 | 7.13724 | 5.02624 | Intron (ENSM | 14 | 57798638 | 57802335 | 3698 | 1 | ENSMUST00 | 1967 | Sap18 | ENSMUSG00000021963.11 |
| chr4 | 148102198 | 148102395 | 4.78209 | 7.13724 | 5.02624 | 3' UTR | 4 | 148083915 | 148088031 | 4117 | 2 | ENSMUST00 | -14168 | Agtrap | ENSMUSG00000029007.8 |
| chr2 | 112267370 | 112267551 | 4.95225 | 7.13062 | 5.02016 | Promoter | 2 | 112266819 | 112363156 | 96338 | 1 | ENSMUST00 | 552 | Slc12a6 | ENSMUSG00000027130.10 |
| chr14 | 21981990 | 21982279 | 4.93194 | 7.09858 | 4.98851 | Downstream | 14 | 21983962 | 21989601 | 5640 | 2 | ENSMUST00 | 7322 | Zfp503 | ENSMUSG00000039081.9 |
| chr10 | 128806360 | 128806600 | 4.92185 | 7.08266 | 4.97274 | Promoter | 10 | 128805676 | 128819446 | 13771 | 1 | ENSMUST00 | 685 | Dnajc14 | ENSMUSG00000025354.4 |
| chr11 | 101247045 | 101247240 | 4.92185 | 7.08266 | 4.97274 | Promoter | 11 | 101247093 | 101248064 | 972 | 1 | ENSMUST00 | 0 | Ramp2 | ENSMUSG00000001240.8 |
| chr6 | 87495826 | 87496124 | 4.92185 | 7.08266 | 4.97274 | Promoter | 6 | 87458545 | 87496314 | 37770 | 2 | ENSMUST00 | 190 | Arhgap25 | ENSMUSG00000030047.9 |
| chr7 | 79578403 | 79578624 | 4.89181 | 7.03529 | 4.9263 | Distal Interge | 7 | 79593369 | 79594924 | 1556 | 2 | ENSMUST00 | 16300 | Rhcg | ENSMUSG00000030549.5 |
| chr3 | 9110969 | 9111240 | 4.69649 | 6.99982 | 4.89137 | Distal Interge | 3 | 8928626 | 9004723 | 76098 | 2 | ENSMUST00 | -106247 | Tpd52 | ENSMUSG00000027506.10 |
| chr5 | 24591381 | 24591702 | 4.69649 | 6.99982 | 4.89137 | Promoter | 5 | 24592114 | 24592211 | 98 | 1 | ENSMUST00 | -412 | Mir671 | ENSMUSG00000076387.1 |
| chr1 | 172217038 | 172217218 | 4.35105 | 6.99241 | 4.88445 | Intron (ENSM | 1 | 172215140 | 172219715 | 4576 | 2 | ENSMUST00 | 2497 | Casq1 | ENSMUSG00000007122.6 |
| chr1 | 52799382 | 52799815 | 4.35105 | 6.99241 | 4.88445 | Promoter | 1 | 52789668 | 52799488 | 9821 | 2 | ENSMUST00 | 0 | Inpp1 | ENSMUSG00000026102.4 |
| chr1 | 64735046 | 64735235 | 4.35105 | 6.99241 | 4.88445 | Exon (ENSMU | 1 | 64734469 | 64737749 | 3281 | 2 | ENSMUST00 | 2514 | Fzd5 | ENSMUSG00000045005.9 |
| chr1 | 72226270 | 72226566 | 4.35105 | 6.99241 | 4.88445 | Promoter | 1 | 72226240 | 72226430 | 191 | 1 | ENSMUST00 | 31 | Gm25360 | ENSMUSG00000094655.1 |
| chr1 | 81593584 | 81594044 | 4.35105 | 6.99241 | 4.88445 | Distal Interge | 1 | 81554157 | 81557483 | 3327 | 2 | ENSMUST00 | -36102 | RP23-424A2. | ENSMUSG00000101986.1 |
| chr11 | 95520674 | 95520931 | 4.35105 | 6.99241 | 4.88445 | Distal Interge | 11 | 95509845 | 95514570 | 4726 | 2 | ENSMUST00 | -6105 | Nxph3 | ENSMUSG00000046719.7 |
| chr12 | 69476645 | 69477020 | 4.35105 | 6.99241 | 4.88445 | Distal Interge | 12 | 69561229 | 69561337 | 109 | 2 | ENSMUST00 | 84317 | Gm23018 | ENSMUSG00000095776.1 |
| chr16 | 77499936 | 77500537 | 4.35105 | 6.99241 | 4.88445 | Promoter | 16 | 77500216 | 77598531 | 98316 | 1 | ENSMUST00 | 0 | 2810055G20 | ENSMUSG00000090386.4 |
| chr17 | 25556206 | 25556515 | 4.35105 | 6.99241 | 4.88445 | Distal Interge | 17 | 25567988 | 25570214 | 2227 | 2 | ENSMUST00 | 13699 | Sox8 | ENSMUSG00000024176.5 |
| chr19 | 56652107 | 56652557 | 4.35105 | 6.99241 | 4.88445 | Distal Interge | 19 | 56722372 | 56724862 | 2491 | 1 | ENSMUST00 | -69815 | Adrb1 | ENSMUSG00000035283.4 |
| chr2 | 26583972 | 26584172 | 4.35105 | 6.99241 | 4.88445 | Promoter | 2 | 26583905 | 26593120 | 9216 | 1 | ENSMUST00 | 68 | Egfl7 | ENSMUSG00000026921.13 |
| chr7 | 70822562 | 70822998 | 4.35105 | 6.99241 | 4.88445 | Distal Interge | 7 | 70752416 | 70752500 | 85 | 1 | ENSMUST00 | 70147 | Gm24880 | ENSMUSG00000089401.1 |
| chr1 | 30872076 | 30872284 | 4.86214 | 6.98849 | 4.88147 | Promoter | 1 | 30863070 | 30873252 | 10183 | 2 | ENSMUST00 | 968 | Phf3 | ENSMUSG00000048874.10 |
| chr11 | 61618055 | 61618288 | 4.82314 | 6.92697 | 4.82056 | Distal Interge | 11 | 61653265 | 61672784 | 19520 | 1 | ENSMUST00 | -34977 | Grap | ENSMUSG00000004837.2 |
| chr11 | 6695235 | 6695583 | 4.82314 | 6.92697 | 4.82056 | Distal Interge | 11 | 6685824 | 6686809 | 986 | 1 | ENSMUST00 | 9412 | Gm11980 | ENSMUSG00000083075.1 |
| chr17 | 56035377 | 56035589 | 4.82314 | 6.92697 | 4.82056 | Intron (ENSM | 17 | 56016753 | 56036637 | 19885 | 2 | ENSMUST00 | 1048 | Sh3gl1 | ENSMUSG00000003200.9 |
| chr2 | 58567976 | 58568226 | 4.82314 | 6.92697 | 4.82056 | Promoter | 2 | 58567387 | 58792971 | 225585 | 1 | ENSMUST00 | 590 | Upp2 | ENSMUSG00000026839.11 |
| chr4 | 133371995 | 133372174 | 4.82314 | 6.92697 | 4.82056 | Intron (ENSM | 4 | 133370644 | 133412136 | 41493 | 1 | ENSMUST00 | 1352 | Slc9a1 | ENSMUSG00000028854.9 |
| chr5 | 24551250 | 24551521 | 4.82314 | 6.92697 | 4.82056 | Intron (ENSM | 5 | 24543434 | 24555469 | 12036 | 2 | ENSMUST00 | 3948 | 4931409K22 | ENSMUSG00000038199.8 |
| chr16 | 92302168 | 92302332 | 4.80387 | 6.89658 | 4.79102 | Promoter | 16 | 92301296 | 92311424 | 10129 | 1 | ENSMUST00 | 873 | Smim11 | ENSMUSG00000051989.6 |
| chr3 | 104434079 | 104434526 | 4.80387 | 6.89658 | 4.79102 | Distal Interge | 3 | 104350635 | 104366613 | 15979 | 2 | ENSMUST00 | -67467 | Gm5546 | ENSMUSG00000022187.2 |
| chr18 | 60600089 | 60600295 | 4.7943 | 6.88148 | 4.77608 | Promoter | 18 | 60593991 | 60600497 | 6507 | 2 | ENSMUST00 | 202 | Synpo | ENSMUSG00000043079.12 |
| chr2 | 91068824 | 91069215 | 4.7943 | 6.88148 | 4.77608 | Promoter | 2 | 91068431 | 91068871 | 441 | 2 | ENSMUST00 | 0 | Slc39a13 | ENSMUSG00000002105.10 |
| chr11 | 36847543 | 36847866 | 4.61389 | 6.86692 | 4.76171 | Intron (ENSM | 11 | 36007662 | 36944166 | 936505 | 2 | ENSMUST00 | 96300 | Tenm2 | ENSMUSG00000049336.11 |
| chr2 | 67567140 | 67567438 | 4.61389 | 6.86692 | 4.76171 | Intron (ENSM | 2 | 67565871 | 67810894 | 245024 | 1 | ENSMUST00 | 1270 | B3galt1 | ENSMUSG00000034780.6 |
| chr10 | 70966931 | 70967191 | 4.42115 | 6.83029 | 4.72578 | Intron (ENSM | 10 | 70934502 | 70948489 | 13988 | 2 | ENSMUST00 | -18443 | Bicc1 | ENSMUSG00000014329.9 |
| chr11 | 8466768 | 8466942 | 4.42115 | 6.83029 | 4.72578 | Intron (ENSM | 11 | 8451070 | 8468675 | 17606 | 2 | ENSMUST00 | 1733 | Tns3 | ENSMUSG00000020422.8 |
| chr12 | 98577895 | 98578186 | 4.42115 | 6.83029 | 4.72578 | Promoter | 12 | 98433994 | 98577940 | 143947 | 2 | ENSMUST00 | 0 | Kcnk10 | ENSMUSG00000033854.9 |
| chr12 | 21415671 | 21415928 | 4.42115 | 6.83029 | 4.72578 | Intron (ENSM | 12 | 21395570 | 21417130 | 21561 | 2 | ENSMUST00 | 1202 | Ywhaq | ENSMUSG00000076432.7 |
| chr12 | 78089660 | 78089925 | 4.42115 | 6.83029 | 4.72578 | Distal Interge | 12 | 78165546 | 78165666 | 121 | 1 | ENSMUST00 | -75621 | Gm24994 | ENSMUSG00000088039.1 |
| chr14 | 21164096 | 21164544 | 4.42115 | 6.83029 | 4.72578 | Intron (ENSM | 14 | 21263624 | 21263859 | 236 | 1 | ENSMUST00 | -99080 | Gm26372 | ENSMUSG00000088456.1 |
| chr14 | 97588264 | 97588675 | 4.42115 | 6.83029 | 4.72578 | Distal Interge | 14 | 97336495 | 97336632 | 138 | 2 | ENSMUST00 | -251633 | Gm23276 | ENSMUSG00000088538.1 |

| | | | | | | | | | | | | | | | |
|---|---|---|---|---|---|---|---|---|---|---|---|---|---|---|---|
| chr17 | 29078123 | 29078693 | 4.42115 | 6.83029 | 4.72578 | Promoter | 17 | 29057474 | 29078961 | 21488 | 2 | ENSMUST00 | 268 | Trp53cor1 | ENSMUSG00000085912.1 |
| chr5 | 125179842 | 125180150 | 4.42115 | 6.83029 | 4.72578 | Promoter | 5 | 125017153 | 125179219 | 162067 | 2 | ENSMUST00 | -624 | Ncor2 | ENSMUSG00000029478.11 |
| chr6 | 116651521 | 116651815 | 4.42115 | 6.83029 | 4.72578 | Promoter | 6 | 116650684 | 116652847 | 2164 | 1 | ENSMUST00 | 838 | 8430408G22 | ENSMUSG00000048489.8 |
| chr9 | 37054394 | 37054779 | 4.42115 | 6.83029 | 4.72578 | Intron (ENSM | 9 | 37026399 | 37026507 | 109 | 1 | ENSMUST00 | 27996 | n-R5s82 | ENSMUSG00000077401.1 |
| chr3 | 28171269 | 28171444 | 4.75637 | 6.82164 | 4.71886 | Distal Interge | 3 | 28109759 | 28130795 | 21037 | 1 | ENSMUST00 | 61511 | Pld1 | ENSMUSG00000027695.11 |
| chr14 | 70032031 | 70032390 | 3.95138 | 6.81311 | 4.71044 | Intron (ENSM | 14 | 70077445 | 70080157 | 2713 | 1 | ENSMUST00 | -45055 | Egr3 | ENSMUSG00000033730.3 |
| chr16 | 36769341 | 36769526 | 3.95138 | 6.81311 | 4.71044 | Intron (ENSM | 16 | 36772071 | 36773278 | 1208 | 2 | ENSMUST00 | 3752 | Slc15a2 | ENSMUSG00000022899.3 |
| chr19 | 42164086 | 42164285 | 3.95138 | 6.81311 | 4.71044 | Promoter | 19 | 42163951 | 42177726 | 13776 | 1 | ENSMUST00 | 136 | Zfyve27 | ENSMUSG00000018820.8 |
| chr4 | 66414654 | 66415087 | 3.95138 | 6.81311 | 4.71044 | Distal Interge | 4 | 65380803 | 66404611 | 1023809 | 2 | ENSMUST00 | -10044 | Astn2 | ENSMUSG00000028373.11 |
| chr10 | 121926289 | 121926453 | 4.72832 | 6.77738 | 4.67505 | Intron (ENSM | 10 | 121806640 | 121823133 | 16494 | 2 | ENSMUST00 | -103157 | Srgap1 | ENSMUSG00000020121.10 |
| chr11 | 113750000 | 113750308 | 4.72832 | 6.77738 | 4.67505 | Promoter | 11 | 113729305 | 113751154 | 21850 | 2 | ENSMUST00 | 846 | Cdc42ep4 | ENSMUSG00000041598.7 |
| chr11 | 118327240 | 118327487 | 4.72831 | 6.77738 | 4.67505 | Intron (ENSM | 11 | 118332415 | 118334016 | 1602 | 2 | ENSMUST00 | 6529 | BC100451 | ENSMUSG00000076433.4 |
| chr11 | 52166568 | 52166762 | 4.72832 | 6.77738 | 4.67505 | Intron (ENSM | 11 | 52155378 | 52156283 | 906 | 2 | ENSMUST00 | -10286 | Olfr1372-ps1 | ENSMUSG00000084141.1 |
| chr11 | 71015094 | 71015305 | 4.71904 | 6.76274 | 4.6612 | Exon (ENSMU | 11 | 71014566 | 71019212 | 4647 | 2 | ENSMUST00 | 3907 | Derl2 | ENSMUSG00000018442.8 |
| chr11 | 100388449 | 100388773 | 4.7098 | 6.74815 | 4.64672 | Intron (ENSM | 11 | 100383404 | 100386770 | 3367 | 2 | ENSMUST00 | -1680 | Jup | ENSMUSG00000001552.9 |
| chr14 | 84447543 | 84447738 | 4.53416 | 6.73826 | 4.63693 | Exon (ENSMU | 14 | 84443563 | 84537060 | 93498 | 1 | ENSMUST00 | 3981 | Pcdh17 | ENSMUSG00000035566.6 |
| chr12 | 24493260 | 24493685 | 4.50584 | 6.69248 | 4.59165 | Promoter | 12 | 24493207 | 24493656 | 450 | 2 | ENSMUST00 | 0 | Gm16372 | ENSMUSG00000057160.2 |
| chr12 | 53033403 | 53033837 | 4.50584 | 6.69248 | 4.59165 | Intron (ENSM | 12 | 53046708 | 53046826 | 119 | 2 | ENSMUST00 | 12989 | n-R5s58 | ENSMUSG00000070192.1 |
| chr13 | 31820603 | 31821325 | 4.50584 | 6.69248 | 4.59165 | Intron (ENSM | 13 | 31806691 | 31810643 | 3953 | 1 | ENSMUST00 | 13913 | Foxc1 | ENSMUSG00000050295.3 |
| chr13 | 107846629 | 107846810 | 4.50584 | 6.69248 | 4.59165 | Intron (ENSM | 13 | 107829110 | 107829280 | 171 | 2 | ENSMUST00 | -17350 | Gm22118 | ENSMUSG00000093334.2 |
| chr14 | 21962606 | 21963094 | 4.50584 | 6.69248 | 4.59165 | Distal Interge | 14 | 21983962 | 21989601 | 5640 | 2 | ENSMUST00 | 26507 | Zfp503 | ENSMUSG00000039081.9 |
| chr17 | 36237553 | 36237856 | 4.50584 | 6.69248 | 4.59165 | Promoter | 17 | 36237254 | 36237359 | 106 | 2 | ENSMUST00 | -195 | Gm20391 | ENSMUSG00000092363.1 |
| chr2 | 26589865 | 26590061 | 4.50584 | 6.69248 | 4.59165 | Promoter | 2 | 26590520 | 26590972 | 453 | 1 | ENSMUST00 | -459 | Egfl7 | ENSMUSG00000026921.13 |
| chr3 | 95985612 | 95986225 | 4.50584 | 6.69248 | 4.59165 | Downstream | 3 | 95988819 | 95995668 | 6850 | 2 | ENSMUST00 | 9443 | Plekho1 | ENSMUSG00000015745.4 |
| chr4 | 123018337 | 123018625 | 4.50584 | 6.69248 | 4.59165 | Intron (ENSM | 4 | 123016597 | 123054949 | 38353 | 1 | ENSMUST00 | 1741 | Trit1 | ENSMUSG00000028653.11 |
| chr7 | 30423262 | 30423522 | 4.50584 | 6.69248 | 4.59165 | Promoter | 7 | 30423275 | 30428746 | 5472 | 1 | ENSMUST00 | 0 | Nfkbid | ENSMUSG00000036931.9 |
| chr7 | 25674977 | 25675193 | 4.50584 | 6.69248 | 4.59165 | Promoter | 7 | 25669139 | 25675166 | 6028 | 2 | ENSMUST00 | 0 | Tmem91 | ENSMUSG00000061702.5 |
| chr12 | 111711978 | 111712311 | 4.6551 | 6.66181 | 4.56385 | Promoter | 12 | 111709904 | 111712927 | 3024 | 2 | ENSMUST00 | 616 | Bag5 | ENSMUSG00000049792.6 |
| chr10 | 44380638 | 44380926 | 4.63715 | 6.63346 | 4.53566 | Distal Interge | 10 | 44359676 | 44359768 | 93 | 1 | ENSMUST00 | 20963 | Mir1929 | ENSMUSG00000089260.1 |
| chr19 | 34641023 | 34641316 | 4.63715 | 6.63346 | 4.53566 | Promoter | 19 | 34640871 | 34650009 | 9139 | 1 | ENSMUST00 | 153 | Ifit1 | ENSMUSG00000034459.8 |
| chr6 | 125099615 | 125100404 | 4.63715 | 6.63346 | 4.53566 | Intron (ENSM | 6 | 125097024 | 125097086 | 63 | 1 | ENSMUST00 | 2592 | Mir7045 | ENSMUSG00000098762.1 |
| chr9 | 96713280 | 96713460 | 4.45713 | 6.61362 | 4.51657 | Intron (ENSM | 9 | 96688477 | 96719565 | 31089 | 2 | ENSMUST00 | 6105 | Zbtb38 | ENSMUSG00000040433.11 |
| chr10 | 19936950 | 19937343 | 3.97665 | 6.59687 | 4.50014 | Intron (ENSM | 10 | 19935223 | 20100158 | 164936 | 1 | ENSMUST00 | 1728 | Map3k5 | ENSMUSG00000071369.5 |
| chr2 | 25461214 | 25461656 | 3.97665 | 6.59687 | 4.50014 | Promoter | 2 | 25459482 | 25461094 | 1613 | 2 | ENSMUST00 | -121 | BC029214 | ENSMUSG00000047617.6 |
| chr1 | 11329542 | 11329706 | 4.61021 | 6.59091 | 4.49445 | Distal Interge | 1 | 11283993 | 11298204 | 14212 | 1 | ENSMUST00 | 45550 | Prex2 | ENSMUSG00000048960.8 |
| chr18 | 37707097 | 37707487 | 4.61021 | 6.59091 | 4.49445 | Intron (ENSM | 18 | 37705979 | 37706069 | 91 | 1 | ENSMUST00 | 1119 | Gm24401 | ENSMUSG00000088493.1 |
| chr7 | 118704206 | 118704626 | 4.61021 | 6.59091 | 4.49445 | Promoter | 7 | 118691583 | 118705380 | 13798 | 2 | ENSMUST00 | 754 | Gde1 | ENSMUSG00000033917.9 |
| chr8 | 45658667 | 45659023 | 4.61021 | 6.59091 | 4.49445 | Promoter | 8 | 45658670 | 45776882 | 118213 | 1 | ENSMUST00 | 0 | Sorbs2 | ENSMUSG00000031626.11 |
| chrX | 140457653 | 140457817 | 4.61021 | 6.59091 | 4.49445 | Intron (ENSM | 20 | 140456613 | 140476140 | 19528 | 1 | ENSMUST00 | 1041 | Prps1 | ENSMUSG00000031432.3 |
| chr10 | 39629946 | 39630283 | 4.54943 | 6.49485 | 4.40153 | Intron (ENSM | 10 | 39641134 | 39645758 | 4625 | 1 | ENSMUST00 | -10851 | Traf3ip2 | ENSMUSG00000019842.6 |
| chr3 | 67553535 | 67553784 | 4.54943 | 6.49485 | 4.40153 | Distal Interge | 3 | 67573160 | 67573291 | 132 | 2 | ENSMUST00 | 19507 | Gm25919 | ENSMUSG00000077459.1 |
| chr4 | 149958450 | 149958614 | 4.54943 | 6.49485 | 4.40153 | Distal Interge | 4 | 149907142 | 149955043 | 47902 | 2 | ENSMUST00 | -3408 | Spsb1 | ENSMUSG00000039911.8 |
| chr17 | 73809790 | 73810003 | 4.53228 | 6.46773 | 4.37548 | Intron (ENSM | 17 | 73804841 | 73832093 | 27253 | 1 | ENSMUST00 | 4950 | Ehd3 | ENSMUSG00000024065.7 |
| chr15 | 95931631 | 95931827 | 4.00593 | 6.38738 | 4.29533 | Exon (ENSMU | 15 | 95879716 | 95879817 | 102 | 2 | ENSMUST00 | -51815 | Gm25070 | ENSMUSG00000092978.1 |
| chr18 | 60602248 | 60602700 | 4.00593 | 6.38738 | 4.29533 | Exon (ENSMU | 18 | 60593991 | 60600497 | 6507 | 2 | ENSMUST00 | -1752 | Synpo | ENSMUSG00000043079.12 |
| chr2 | 128428920 | 128429182 | 4.00593 | 6.38738 | 4.29533 | Promoter | 2 | 128195993 | 128429140 | 233148 | 2 | ENSMUST00 | 0 | Gm14005 | ENSMUSG00000074813.7 |
| chr3 | 107758202 | 107758390 | 4.00593 | 6.38738 | 4.29533 | Intron (ENSM | 3 | 107750380 | 107760221 | 9842 | 2 | ENSMUST00 | 1831 | Csf1 | ENSMUSG00000014599.5 |
| chr19 | 42163549 | 42163994 | 4.31067 | 6.37555 | 4.28397 | Promoter | 19 | 42163951 | 42177726 | 13776 | 1 | ENSMUST00 | 0 | Zfyve27 | ENSMUSG00000018820.8 |
| chr9 | 36768791 | 36768969 | 4.31067 | 6.37555 | 4.28397 | Promoter | 9 | 36767910 | 36782675 | 14766 | 1 | ENSMUST00 | 882 | Gm26787 | ENSMUSG00000096993.2 |
| chr1 | 165709651 | 165709945 | 4.46497 | 6.3612 | 4.26978 | Distal Interge | 1 | 165648945 | 165708094 | 59150 | 2 | ENSMUST00 | -1558 | Rcsd1 | ENSMUSG00000040723.9 |
| chr11 | 75165602 | 75165952 | 4.46497 | 6.3612 | 4.26978 | Exon (ENSMU | 11 | 75167153 | 75168146 | 994 | 2 | ENSMUST00 | 2194 | Hic1 | ENSMUSG00000043099.4 |
| chr16 | 8883870 | 8884133 | 4.46497 | 6.3612 | 4.26978 | Distal Interge | 16 | 8873079 | 8873171 | 93 | 2 | ENSMUST00 | -10700 | Gm22224 | ENSMUSG00000087982.1 |
| chr8 | 109251544 | 109251759 | 4.44845 | 6.33504 | 4.24436 | Exon (ENSMU | 8 | 109249866 | 109273811 | 23946 | 1 | ENSMUST00 | 1679 | D030068K23 | ENSMUSG00000097393.2 |
| chr15 | 31569021 | 31569229 | 3.69821 | 6.32042 | 4.22975 | Promoter | 15 | 31568905 | 31590119 | 21215 | 1 | ENSMUST00 | 117 | Cmbl | ENSMUSG00000022235.9 |
| chr12 | 84194496 | 84194660 | 4.59532 | 6.3174 | 4.22678 | Promoter | 12 | 84149278 | 84194015 | 44738 | 2 | ENSMUST00 | -482 | Elmsan1 | ENSMUSG00000042507.9 |
| chr15 | 25625526 | 25625844 | 4.59532 | 6.3174 | 4.22678 | Intron (ENSM | 15 | 25622549 | 25701950 | 79402 | 1 | ENSMUST00 | 2978 | Myo10 | ENSMUSG00000022272.12 |
| chr16 | 33687117 | 33687281 | 4.59532 | 6.3174 | 4.22678 | Intron (ENSM | 16 | 33684545 | 33769443 | 84899 | 1 | ENSMUST00 | 2573 | Heg1 | ENSMUSG00000075254.6 |
| chr6 | 65764997 | 65765388 | 4.59532 | 6.3174 | 4.22678 | Distal Interge | 6 | 65757936 | 65758321 | 386 | 1 | ENSMUST00 | 7062 | Gm15490 | ENSMUSG00000082115.1 |
| chr11 | 62005443 | 62005880 | 4.56789 | 6.27452 | 4.18762 | Intron (ENSM | 11 | 62029276 | 62223013 | 193738 | 1 | ENSMUST00 | -23396 | Specc1 | ENSMUSG00000042331.8 |
| chr11 | 96407545 | 96407720 | 4.56789 | 6.27452 | 4.18762 | Distal Interge | 11 | 96392112 | 96392666 | 555 | 1 | ENSMUST00 | 15434 | Gm11531 | ENSMUSG00000075587.5 |
| chr4 | 136956957 | 136957121 | 4.56789 | 6.27452 | 4.18762 | Promoter | 4 | 136929419 | 136956816 | 27398 | 2 | ENSMUST00 | -142 | Epha8 | ENSMUSG00000028661.8 |
| chr1 | 152589493 | 152589700 | 4.24099 | 6.26174 | 4.17529 | Intron (ENSM | 1 | 152557543 | 152592759 | 35217 | 2 | ENSMUST00 | 3059 | Rgl1 | ENSMUSG00000026482.8 |
| chr10 | 127063046 | 127063233 | 4.55837 | 6.25963 | 4.17328 | Promoter | 10 | 127063534 | 127067920 | 4387 | 1 | ENSMUST00 | -301 | Cdk4 | ENSMUSG00000006728.7 |
| chr17 | 26505609 | 26505773 | 4.55837 | 6.25963 | 4.17328 | 3' UTR | 17 | 26506516 | 26507723 | 1208 | 2 | ENSMUST00 | 1950 | Dusp1 | ENSMUSG00000024190.6 |
| chr1 | 135793742 | 135794601 | 4.04026 | 6.18694 | 4.10202 | Intron (ENSM | 1 | 135799402 | 135810989 | 11588 | 1 | ENSMUST00 | -4801 | Tnni1 | ENSMUSG00000026418.11 |
| chr10 | 22000239 | 22000416 | 4.04026 | 6.18694 | 4.10202 | Distal Interge | 10 | 21996591 | 21998240 | 1650 | 1 | ENSMUST00 | 3649 | Sgk1 | ENSMUSG00000019970.10 |
| chr12 | 33717637 | 33718380 | 4.04026 | 6.18694 | 4.10202 | Distal Interge | 12 | 33928425 | 33929310 | 886 | 1 | ENSMUST00 | -210045 | Ferd3l | ENSMUSG00000046518.3 |
| chr17 | 73808367 | 73808715 | 4.04026 | 6.18694 | 4.10202 | Intron (ENSM | 17 | 73804841 | 73832093 | 27253 | 1 | ENSMUST00 | 3527 | Ehd3 | ENSMUSG00000024065.7 |
| chr2 | 24405669 | 24405833 | 4.51134 | 6.18616 | 4.10202 | Promoter | 2 | 24405629 | 24407007 | 1379 | 1 | ENSMUST00 | 41 | Psd4 | ENSMUSG00000026979.11 |
| chr4 | 134253951 | 134254363 | 4.04026 | 6.18694 | 4.10202 | Promoter | 4 | 134251110 | 134254106 | 2997 | 2 | ENSMUST00 | 0 | Grrp1 | ENSMUSG00000050105.5 |
| chr4 | 141210041 | 141210380 | 4.04026 | 6.18694 | 4.10202 | Distal Interge | 4 | 141213956 | 141220148 | 6193 | 1 | ENSMUST00 | -3576 | Rsg1 | ENSMUSG00000073733.4 |
| chr5 | 38277178 | 38277709 | 4.04026 | 6.18694 | 4.10202 | Promoter | 5 | 38277404 | 38304208 | 26805 | 1 | ENSMUST00 | 0 | Otop1 | ENSMUSG00000051596.8 |
| chr6 | 99025931 | 99026530 | 4.04026 | 6.18694 | 4.10202 | Intron (ENSM | 6 | 99016550 | 99027991 | 11442 | 2 | ENSMUST00 | 1461 | Foxp1 | ENSMUSG00000030067.12 |
| chr9 | 57837070 | 57837375 | 4.04026 | 6.18694 | 4.10202 | Promoter | 9 | 57794687 | 57836793 | 42107 | 2 | ENSMUST00 | -278 | Arid3b | ENSMUSG00000004661.10 |
| chr10 | 20854973 | 20855279 | 4.17352 | 6.15118 | 4.06778 | Distal Interge | 10 | 20952547 | 21080428 | 127882 | 1 | ENSMUST00 | -97268 | Ahi1 | ENSMUSG00000019986.11 |
| chr10 | 51677636 | 51677913 | 4.47441 | 6.12849 | 4.04533 | Promoter | 10 | 51677756 | 51721908 | 44153 | 1 | ENSMUST00 | 0 | Rfx6 | ENSMUSG00000019900.12 |
| chr4 | 98436921 | 98437214 | 3.48302 | 6.12095 | 4.03844 | Promoter | 4 | 98436943 | 98441137 | 4195 | 1 | ENSMUST00 | 0 | Inadl | ENSMUSG00000061859.11 |
| chr1 | 91076517 | 91076791 | 4.45617 | 6.10001 | 4.01819 | Intron (ENSM | 1 | 91053586 | 91128287 | 74702 | 1 | ENSMUST00 | 22932 | Lrrfip1 | ENSMUSG00000026305.10 |
| chr4 | 136140348 | 136140740 | 3.70442 | 6.08744 | 4.00582 | Distal Interge | 4 | 136143497 | 136145392 | 1896 | 1 | ENSMUST00 | -2757 | Id3 | ENSMUSG00000007872.3 |
| chr7 | 100908338 | 100908675 | 4.42908 | 6.05774 | 3.97659 | Intron (ENSM | 7 | 100869746 | 100932161 | 62416 | 2 | ENSMUST00 | 23486 | Arhgef17 | ENSMUSG00000032875.7 |
| chr1 | 169949835 | 169950159 | 4.39348 | 6.00217 | 3.92175 | 3' UTR | 1 | 169952876 | 169965723 | 12848 | 2 | ENSMUST00 | 15564 | Hsd17b7 | ENSMUSG00000026675.7 |
| chr1 | 64289253 | 64289432 | 4.08106 | 5.99902 | 3.91878 | Intron (ENSM | 1 | 64185991 | 64303043 | 117053 | 2 | ENSMUST00 | 13611 | RP23-45M23 | ENSMUSG00000101895.1 |
| chr12 | 117658286 | 117658930 | 4.08106 | 5.99902 | 3.91878 | Exon (ENSMU | 12 | 117574342 | 117574447 | 106 | 1 | ENSMUST00 | 83945 | Gm24741 | ENSMUSG00000089482.1 |
| chr12 | 83620802 | 83621123 | 4.08106 | 5.99902 | 3.91878 | Promoter | 12 | 83620210 | 83621174 | 965 | 1 | ENSMUST00 | 593 | Gm26623 | ENSMUSG00000097703.1 |
| chr2 | 34302789 | 34302953 | 4.08106 | 5.99902 | 3.91878 | Exon (ENSMU | 2 | 34172873 | 34288852 | 115980 | 2 | ENSMUST00 | -13938 | Pbx3 | ENSMUSG00000038718.10 |
| chr3 | 27154032 | 27154373 | 4.08106 | 5.99902 | 3.91878 | Promoter | 3 | 27154026 | 27157019 | 2994 | 1 | ENSMUST00 | 7 | 1700125G22 | ENSMUSG00000097584.1 |
| chr4 | 133670218 | 133670409 | 4.08106 | 5.99902 | 3.91878 | Intron (ENSM | 4 | 133660387 | 133672647 | 12261 | 2 | ENSMUST00 | 2238 | Pigv | ENSMUSG00000043257.10 |
| chr4 | 62619715 | 62620044 | 4.08106 | 5.99902 | 3.91878 | Promoter | 4 | 62619582 | 62659840 | 40259 | 1 | ENSMUST00 | 134 | Rgs3 | ENSMUSG00000059810.13 |
| chr5 | 24364934 | 24365098 | 4.08106 | 5.99902 | 3.91878 | Promoter | 5 | 24364810 | 24384474 | 19665 | 1 | ENSMUST00 | 125 | Nos3 | ENSMUSG00000028978.7 |
| chr6 | 30300607 | 30300932 | 4.08106 | 5.99902 | 3.91878 | Intron (ENSM | 6 | 30214861 | 30304522 | 89662 | 2 | ENSMUST00 | 3590 | Ube2h | ENSMUSG00000039159.11 |
| chr6 | 48746545 | 48746835 | 4.08106 | 5.99902 | 3.91878 | Promoter | 6 | 48746199 | 48747512 | 1314 | 1 | ENSMUST00 | 347 | Gimap5 | ENSMUSG00000043505.7 |
| chr7 | 79272775 | 79272939 | 4.08106 | 5.99902 | 3.91878 | Promoter | 7 | 79273201 | 79365508 | 92308 | 1 | ENSMUST00 | -262 | Abhd2 | ENSMUSG00000039202.7 |
| chr9 | 62858110 | 62858392 | 4.08106 | 5.99902 | 3.91878 | Promoter | 9 | 62858104 | 62875918 | 17815 | 1 | ENSMUST00 | 7 | Calml4 | ENSMUSG00000032246.8 |
| chr11 | 19017878 | 19018042 | 4.38467 | 5.98842 | 3.91004 | Promoter | 11 | 18880428 | 19018714 | 138287 | 2 | ENSMUST00 | 672 | Meis1 | ENSMUSG00000020160.13 |
| chr11 | 96776812 | 96776976 | 4.2294 | 5.98714 | 3.90955 | Promoter | 11 | 96769891 | 96777492 | 7602 | 2 | ENSMUST00 | 516 | Snx11 | ENSMUSG00000020876.9 |
| chr18 | 84090281 | 84090445 | 4.3759 | 5.97472 | 3.89748 | Intron (ENSM | 18 | 84088158 | 84097514 | 9357 | 1 | ENSMUST00 | 2124 | Zadh2 | ENSMUSG00000049090.2 |
| chr2 | 91950326 | 91950586 | 4.34978 | 5.93396 | 3.85727 | Promoter | 2 | 91932827 | 91950427 | 17601 | 2 | ENSMUST00 | 0 | Dgkz | ENSMUSG00000040479.6 |
| chr3 | 9825905 | 9826069 | 4.15631 | 5.87056 | 3.79461 | Intron (ENSM | 3 | 9829620 | 9829728 | 109 | 2 | ENSMUST00 | 3659 | Gm24786 | ENSMUSG00000088004.1 |
| chr12 | 5119953 | 5120117 | 4.29847 | 5.85388 | 3.77856 | Intron (ENSM | 12 | 4999266 | 4999368 | 103 | 2 | ENSMUST00 | -120586 | Gm23167 | ENSMUSG00000087780.1 |
| chr14 | 34500259 | 34500469 | 4.29847 | 5.85388 | 3.77856 | Intron (ENSM | 14 | 34501715 | 34502684 | 970 | 1 | ENSMUST00 | -1246 | Gm17110 | ENSMUSG00000090779.1 |
| chr14 | 19041971 | 19042364 | 4.29847 | 5.85388 | 3.77856 | Distal Interge | 14 | 19127800 | 19132109 | 4310 | 1 | ENSMUST00 | -85436 | Gm8582 | ENSMUSG00000091929.2 |
| chr5 | 67845437 | 67845641 | 4.29847 | 5.85388 | 3.77856 | Intron (ENSM | 5 | 67847066 | 67909013 | 61948 | 1 | ENSMUST00 | -1425 | Gm15477 | ENSMUSG00000087261.1 |
| chr1 | 165706594 | 165706811 | 3.98342 | 5.8375 | 3.76311 | Intron (ENSM | 1 | 165648945 | 165708094 | 59150 | 2 | ENSMUST00 | 1283 | Rcsd1 | ENSMUSG00000040723.9 |
| chr5 | 109996657 | 109996916 | 3.98342 | 5.8375 | 3.76311 | Promoter | 5 | 109996563 | 110003844 | 7282 | 1 | ENSMUST00 | 95 | Zfp932 | ENSMUSG00000066613.9 |
| chr11 | 106381884 | 106382547 | 4.13035 | 5.82909 | 3.75473 | Promoter | 11 | 106377656 | 106382781 | 5126 | 2 | ENSMUST00 | 234 | Icam2 | ENSMUSG00000001029.10 |
| chr12 | 35046236 | 35046551 | 4.13035 | 5.82909 | 3.75473 | Promoter | 12 | 35047189 | 35147469 | 100281 | 1 | ENSMUST00 | -638 | Snx13 | ENSMUSG00000020590.10 |
| chr14 | 34376054 | 34376333 | 4.13035 | 5.82909 | 3.75473 | Promoter | 14 | 34375504 | 34404287 | 28784 | 1 | ENSMUST00 | 551 | Mmrn2 | ENSMUSG00000041445.8 |
| chr16 | 10358215 | 10358415 | 4.13035 | 5.82909 | 3.75473 | Exon (ENSMU | 16 | 10361254 | 10395448 | 34195 | 2 | ENSMUST00 | 37033 | Tekt5 | ENSMUSG00000039179.8 |
| chr18 | 37807818 | 37808050 | 4.13035 | 5.82909 | 3.75473 | Exon (ENSMU | 18 | 37806410 | 37841873 | 35464 | 1 | ENSMUST00 | 1409 | Pcdhga12 | ENSMUSG00000023036.9 |
| chr5 | 122145569 | 122145757 | 4.13035 | 5.82909 | 3.75473 | Distal Interge | 5 | 122134437 | 122138957 | 4521 | 2 | ENSMUST00 | -6613 | Ccdc63 | ENSMUSG00000043036.8 |
| chr7 | 28888676 | 28888895 | 4.13035 | 5.82909 | 3.75473 | Exon (ENSMU | 7 | 28894232 | 28894852 | 621 | 2 | ENSMUST00 | 5957 | Actn4 | ENSMUSG00000054808.8 |
| chr7 | 73552733 | 73553068 | 4.13035 | 5.82909 | 3.75473 | Intron (ENSM | 7 | 73544243 | 73555920 | 11678 | 2 | ENSMUST00 | 2852 | RP23-32A8.1 | ENSMUSG00000101970.1 |
| chr9 | 77529540 | 77529847 | 4.13035 | 5.82909 | 3.75473 | Intron (ENSM | 9 | 77458090 | 77532444 | 74355 | 2 | ENSMUST00 | 2597 | Lrrc1 | ENSMUSG00000032352.10 |
| chrX | 162759606 | 162759971 | 4.13035 | 5.82909 | 3.75473 | Promoter | 20 | 162760402 | 162779092 | 18691 | 1 | ENSMUST00 | -431 | Rbbp7 | ENSMUSG00000031353.8 |
| chr5 | 143525047 | 143525657 | 4.27326 | 5.81454 | 3.74324 | Intron (ENSM | 5 | 143505482 | 143527993 | 22512 | 2 | ENSMUST00 | 2336 | Rac1 | ENSMUSG00000001847.9 |
| chr17 | 71190663 | 71190997 | 4.26493 | 5.80153 | 3.73034 | Intron (ENSM | 17 | 71183978 | 71249817 | 65840 | 1 | ENSMUST00 | 6686 | Lpin2 | ENSMUSG00000024052.12 |
| chr2 | 30929804 | 30929969 | 4.21559 | 5.72451 | 3.65413 | Promoter | 2 | 30892296 | 30929863 | 37568 | 2 | ENSMUST00 | 0 | Ptges | ENSMUSG00000050737.12 |
| chr1 | 168426124 | 168426608 | 4.1911 | 5.68627 | 3.61683 | Exon (ENSMU | 1 | 168158732 | 168431504 | 272773 | 2 | ENSMUST00 | 4896 | Pbx1 | ENSMUSG00000052534.10 |
| chr17 | 31433169 | 31433409 | 4.1911 | 5.68627 | 3.61683 | Promoter | 17 | 31433071 | 31459129 | 26059 | 1 | ENSMUST00 | 99 | Pde9a | ENSMUSG00000041119.6 |
| chr19 | 37377009 | 37377227 | 4.1911 | 5.68627 | 3.61683 | Promoter | 19 | 37376403 | 37421859 | 45457 | 1 | ENSMUST00 | 607 | Kif11 | ENSMUSG00000012443.3 |
| chr2 | 11664236 | 11664607 | 4.1911 | 5.68627 | 3.61683 | Intron (ENSM | 2 | 11642850 | 11648637 | 5788 | 1 | ENSMUST00 | 21387 | Il2ra | ENSMUSG00000026770.4 |
| chrX | 74085975 | 74086195 | 4.1911 | 5.68627 | 3.61683 | Promoter | 20 | 74035288 | 74085690 | 50403 | 2 | ENSMUST00 | -286 | Mecp2 | ENSMUSG00000031393.11 |
| chr10 | 128179463 | 128179753 | 4.01745 | 5.64824 | 3.5822 | Promoter | 10 | 128129470 | 128180297 | 50828 | 2 | ENSMUST00 | 544 | Rbms2 | ENSMUSG00000040043.10 |
| chr17 | 34970586 | 34971027 | 4.01745 | 5.64824 | 3.5822 | Promoter | 17 | 34969912 | 34972136 | 2225 | 1 | ENSMUST00 | 675 | Gm20481 | ENSMUSG00000092609.1 |
| chr14 | 46060936 | 46061100 | 3.7198 | 5.63232 | 3.56653 | Distal Interge | 14 | 46084126 | 46084815 | 690 | 1 | ENSMUST00 | -23026 | Gm1821 | ENSMUSG00000044285.7 |
| chr15 | 66500816 | 66501130 | 3.7198 | 5.63232 | 3.56653 | Promoter | 15 | 66379858 | 66500910 | 121053 | 2 | ENSMUST00 | 0 | Lrrc6 | ENSMUSG00000022375.6 |
| chr1 | 165706276 | 165706464 | 3.72947 | 5.41281 | 3.35001 | Intron (ENSM | 1 | 165648945 | 165708094 | 59150 | 2 | ENSMUST00 | 1630 | Rcsd1 | ENSMUSG00000040723.9 |
| chr12 | 31350226 | 31350445 | 3.72947 | 5.41281 | 3.35001 | Promoter | 12 | 31331564 | 31351437 | 19874 | 2 | ENSMUST00 | 992 | Dld | ENSMUSG00000020664.9 |

| | | | | | | | | | | | | | | | |
|---|---|---|---|---|---|---|---|---|---|---|---|---|---|---|---|
| chr7 | 137056139 | 137056717 | 3.72947 | 5.41281 | 3.35001 | Intron (ENSM | 7 | 136894611 | 137128187 | 233577 | 1 | ENSMUST00 | 161529 | Mgmt | ENSMUSG00000054612.3 |
| chr9 | 37529328 | 37529617 | 3.72947 | 5.41281 | 3.35001 | Promoter | 9 | 37528781 | 37537112 | 8332 | 1 | ENSMUST00 | 548 | Esam | ENSMUSG00000001946.8 |
| chr1 | 55201899 | 55202197 | 3.45746 | 5.38899 | 3.32729 | 3' UTR | 1 | 55203945 | 55206489 | 2545 | 2 | ENSMUST00 | 4292 | Rftn2 | ENSMUSG00000025978.9 |
| chrX | 120292923 | 120293087 | 4.13579 | 5.38279 | 3.32121 | Intron (ENSM | 20 | 120290403 | 120906344 | 615942 | 1 | ENSMUST00 | 2521 | Pcdh11x | ENSMUSG00000034755.13 |
| chrX | 167328932 | 167329261 | 4.13579 | 5.38279 | 3.32121 | Intron (ENSM | 20 | 167317680 | 167330443 | 12764 | 2 | ENSMUST00 | 1182 | Tlr7 | ENSMUSG00000044583.8 |
| chr6 | 90325004 | 90325219 | 4.01874 | 5.20211 | 3.14917 | Promoter | 6 | 90308351 | 90325185 | 16835 | 2 | ENSMUST00 | 0 | Chst13 | ENSMUSG00000056643.4 |
| chr11 | 100640182 | 100640520 | 3.74097 | 5.20161 | 3.14887 | Intron (ENSM | 11 | 100627543 | 100650693 | 23151 | 2 | ENSMUST00 | 10173 | Zfp385c | ENSMUSG00000014198.10 |
| chr19 | 32292316 | 32292501 | 3.74097 | 5.20161 | 3.14887 | Intron (ENSM | 19 | 32220587 | 32289579 | 68993 | 2 | ENSMUST00 | -2738 | Sgms1 | ENSMUSG00000040451.12 |
| chr4 | 9874647 | 9874834 | 3.74097 | 5.20161 | 3.14887 | Distal Interge | 4 | 9844372 | 9862345 | 17974 | 1 | ENSMUST00 | 30276 | Gdf6 | ENSMUSG00000051279.2 |
| chr5 | 35728614 | 35728953 | 3.74097 | 5.20161 | 3.14887 | Promoter | 5 | 35723068 | 35729012 | 5945 | 2 | ENSMUST00 | 59 | Sh3tc1 | ENSMUSG00000036553.11 |
| chr10 | 22361626 | 22362045 | 3.44643 | 5.14763 | 3.09782 | Promoter | 10 | 22361894 | 22374139 | 12246 | 1 | ENSMUST00 | 0 | Raet1d | ENSMUSG00000078452.5 |
| chr17 | 55712888 | 55713125 | 3.84491 | 5.14421 | 3.0947 | Promoter | 17 | 55652025 | 55712628 | 60604 | 2 | ENSMUST00 | -261 | Pot1b | ENSMUSG00000024174.8 |
| chr3 | 97942653 | 97942817 | 3.97812 | 5.13948 | 3.09035 | Intron (ENSM | 3 | 97930173 | 97967018 | 36846 | 1 | ENSMUST00 | 12481 | Gm5544 | ENSMUSG00000074388.3 |
| chr2 | 30784468 | 30784901 | 3.94621 | 5.0903 | 3.04196 | Distal Interge | 2 | 30795468 | 30796664 | 1197 | 2 | ENSMUST00 | 11763 | 1700001O22 | ENSMUSG00000044320.9 |
| chr10 | 77591927 | 77592091 | 3.75487 | 5.0025 | 2.95672 | Promoter | 10 | 77592337 | 77597130 | 4794 | 1 | ENSMUST00 | -246 | Pttg1ip | ENSMUSG00000009291.7 |
| chr10 | 42864142 | 42864459 | 3.75487 | 5.0025 | 2.95672 | Intron (ENSM | 10 | 42865514 | 42930419 | 64906 | 1 | ENSMUST00 | -1055 | Scml4 | ENSMUSG00000044770.9 |
| chr11 | 68955087 | 68955309 | 3.75487 | 5.0025 | 2.95672 | Promoter | 11 | 68943155 | 68955074 | 11920 | 2 | ENSMUST00 | -14 | Arhgef15 | ENSMUSG00000052921.8 |
| chr16 | 26968032 | 26968326 | 3.75487 | 5.0025 | 2.95672 | Intron (ENSM | 16 | 26957236 | 26989974 | 32739 | 2 | ENSMUST00 | 21648 | Gmnc | ENSMUSG00000068428.6 |
| chr18 | 53649633 | 53649880 | 3.23593 | 4.97054 | 2.92865 | Distal Interge | 18 | 53681724 | 53744547 | 62824 | 2 | ENSMUST00 | 94667 | Cep120 | ENSMUSG00000048799.7 |
| chr6 | 115792386 | 115792826 | 3.23593 | 4.97054 | 2.92865 | Exon (ENSMU | 6 | 115807975 | 115808114 | 140 | 2 | ENSMUST00 | 15288 | Snora7a | ENSMUSG00000064563.1 |
| chr14 | 53032915 | 53033079 | 3.45835 | 4.95099 | 2.91005 | Promoter | 14 | 53032902 | 53033418 | 517 | 1 | ENSMUST00 | 14 | Trav13d-3 | ENSMUSG00000095643.1 |
| chr3 | 121547648 | 121547957 | 3.43366 | 4.90846 | 2.86831 | Distal Interge | 3 | 121459529 | 121532344 | 72816 | 2 | ENSMUST00 | -15305 | Slc44a3 | ENSMUSG00000039865.7 |
| chr3 | 132087896 | 132088216 | 3.77199 | 4.82198 | 2.78414 | Intron (ENSM | 3 | 132085292 | 132180293 | 95002 | 1 | ENSMUST00 | 2605 | Dkk2 | ENSMUSG00000028031.5 |
| chr6 | 29145780 | 29146159 | 3.77199 | 4.82198 | 2.78414 | Intron (ENSM | 6 | 29150554 | 29150679 | 126 | 2 | ENSMUST00 | 4520 | Gm24303 | ENSMUSG00000095596.1 |
| chr6 | 52260080 | 52260323 | 3.18132 | 4.46861 | 2.44521 | Promoter | 6 | 52258871 | 52260802 | 1932 | 2 | ENSMUST00 | 479 | Hoxa13 | ENSMUSG00000038203.13 |
| chr1 | 13293474 | 13293732 | 3.40088 | 4.4408 | 2.41859 | Intron (ENSM | 1 | 13190106 | 13299722 | 109617 | 2 | ENSMUST00 | 5990 | Ncoa2 | ENSMUSG00000005886.9 |
| chr12 | 111559899 | 111560159 | 3.40088 | 4.4408 | 2.41859 | Distal Interge | 12 | 111572321 | 111574402 | 2082 | 2 | ENSMUST00 | 14243 | 2810029C07I | ENSMUSG00000096995.1 |
| chr8 | 111741966 | 111742313 | 3.40088 | 4.4408 | 2.41859 | Intron (ENSM | 8 | 111710485 | 111743849 | 33365 | 2 | ENSMUST00 | 1536 | Bcar1 | ENSMUSG00000031955.9 |
| chr2 | 135711966 | 135712232 | 3.37938 | 4.21603 | 2.20503 | Intron (ENSM | 2 | 135686174 | 135693154 | 6981 | 2 | ENSMUST00 | -18813 | Gm14211 | ENSMUSG00000085709.1 |
| chr5 | 111200913 | 111201103 | 3.37938 | 4.21603 | 2.20503 | Intron (ENSM | 5 | 111225458 | 111229415 | 3958 | 1 | ENSMUST00 | -24355 | Ttc28 | ENSMUSG00000033209.12 |

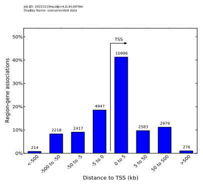

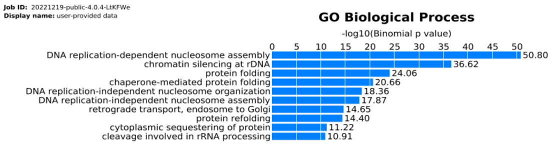

**# GREAT version 4.0.4**
**Species assembly: mm10**
**Association rule: Basal+extension: 5000 bp upstream, 1000 bp downstream, 1000000 bp max extension, curated regulatory domains included**

| # Term Name | Binom Rank | Binom Raw P-Value | Binom FDR Q-Val | Binom Fold Enri | Binom Obs | Binom Region Set C | Hyper Rank | Hyper FDR | Hyper Fold E | Hyper Observ | Hyper Tot | Hyper Gene Set Coverage |
|---|---|---|---|---|---|---|---|---|---|---|---|---|
| **abnormal DNA repair** | 2 | 8.75E-16 | 4.01E-12 | 2.010338 | 162 | 0.008596445 | 510 | 4.72E-07 | 1.336519 | 86 | 92 | 0.0057471 |
| **adipose tissue necrosis** | 4 | 3.98E-13 | 9.11E-10 | 29.10288 | 11 | 0.000583709 | 5481 | 0.8184114 | 1.429765 | 2 | 2 | 0.0001337 |
| **decreased susceptibility to type II hypersensitivity reaction** | 6 | 1.06E-10 | 1.62E-07 | 12.45616 | 13 | 0.000689838 | 5481 | 0.8184114 | 1.429765 | 2 | 2 | 0.0001337 |
| **abnormal internal anal sphincter morphology** | 7 | 1.29E-10 | 1.70E-07 | 6.925332 | 19 | 0.001008225 | 5481 | 0.8184114 | 1.429765 | 2 | 2 | 0.0001337 |
| **decreased metacarpal bone number** | 7 | 1.29E-10 | 1.70E-07 | 6.925332 | 19 | 0.001008225 | 6715 | 0.9551218 | 1.429765 | 1 | 1 | 6.683E-05 |
| **absent female preputial gland** | 7 | 1.29E-10 | 1.70E-07 | 6.925332 | 19 | 0.001008225 | 6715 | 0.9551218 | 1.429765 | 1 | 1 | 6.683E-05 |
| **abnormal female preputial gland morphology** | 12 | 1.18E-09 | 9.02E-07 | 5.39812 | 21 | 0.001114354 | 5481 | 0.8184114 | 1.429765 | 2 | 2 | 0.0001337 |
| **decreased susceptibility to autoimmune hemolytic anemia** | 13 | 6.95E-09 | 4.90174E-06 | 6.602235 | 16 | 0.000849032 | 8589 | 0.9778313 | 0.7148824 | 2 | 4 | 0.0001337 |
| **thin lung-associated mesenchyme** | 15 | 1.06E-08 | 6.49306E-06 | 7.603723 | 14 | 0.000742903 | 5481 | 0.8184114 | 1.429765 | 2 | 2 | 0.0001337 |
| **absent arcus anterior** | 18 | 1.99E-08 | 1.01537E-05 | 6.111857 | 16 | 0.000849032 | 3422 | 0.4484141 | 1.429765 | 5 | 5 | 0.0003341 |
| **increased heart right ventricle weight** | 19 | 2.53E-08 | 1.21894E-05 | 7.826249 | 13 | 0.000689838 | 3034 | 0.3537003 | 1.429765 | 6 | 6 | 0.000401 |
| **decreased mean platelet volume** | 23 | 4.09E-08 | 1.63009E-05 | 2.713317 | 40 | 0.002122579 | 1414 | 0.0148332 | 1.429765 | 17 | 17 | 0.0011361 |
| **absent palatine bone** | 25 | 7.81E-08 | 2.86589E-05 | 7.8967 | 12 | 0.000636774 | 3908 | 0.561441 | 1.429765 | 4 | 4 | 0.0002673 |
| **absent odontoid process** | 29 | 9.57E-08 | 3.02558E-05 | 12.14053 | 9 | 0.00047758 | 5481 | 0.8184114 | 1.429765 | 2 | 2 | 0.0001337 |
| **absent midbrain-hindbrain boundary** | 31 | 1.22E-07 | 3.60237E-05 | 3.789471 | 23 | 0.001220483 | 3908 | 0.561441 | 1.429765 | 4 | 4 | 0.0002673 |
| **empty decidua capsularis** | 32 | 1.29E-07 | 3.70922E-05 | 3.648492 | 24 | 0.001273547 | 3791 | 0.4864989 | 1.209801 | 11 | 13 | 0.0007351 |
| **absent proamniotic cavity** | 37 | 4.13E-07 | 0.000102286 | 3.314048 | 25 | 0.001326612 | 1986 | 0.0903649 | 1.429765 | 11 | 11 | 0.0007351 |
| **absent podocyte slit diaphragm** | 38 | 4.94E-07 | 0.000119119 | 2.964073 | 29 | 0.00153887 | 2722 | 0.2757058 | 1.429765 | 7 | 7 | 0.0004678 |
| **abnormal base-excision repair** | 39 | 5.13E-07 | 0.000120586 | 3.184072 | 26 | 0.001379676 | 2496 | 0.2102634 | 1.429765 | 8 | 8 | 0.0005346 |
| **increased cellular sensitivity to alkylating agents** | 46 | 1.81561E-06 | 0.000361938 | 2.529985 | 34 | 0.001804192 | 1986 | 0.0903649 | 1.429765 | 11 | 11 | 0.0007351 |

The test set of 18,845 genomic regions picked 14,964 (70%) of all 21,395 genes.
*Mouse Phenotype* has 9,575 terms covering 9,654 (45%) of all 21,395 genes, and 705,265 term - gene associations.
9,575 ontology terms (100%) were tested using an annotation count range of [1, Inf].

| Chrom | Start | End | -LOG10(pvalue) | fold | -LOG10(q-value) | annotation | geneChr | geneStart | geneEnd | geneLength | geneStrand | transcriptId | distanceToTSS | geneName | geneId |
|---|---|---|---|---|---|---|---|---|---|---|---|---|---|---|---|
| chr7 | 142576845 | 142577039 | 3.49516 | 4.20084 | 1.26599 | Promoter | 7 | 142576296 | 142576910 | 615 | 2 | ENSMUST00000132294.2 | 0 | H19 | ENSMUSG00000000031.10 |
| chr7 | 142581144 | 142581307 | 3.59375 | 4.57927 | 1.4998 | Distal Intergenic | 7 | 142575529 | 142578143 | 2615 | 2 | ENSMUST00000136359.2 | -3002 | H19 | ENSMUSG00000000031.10 |
| chr7 | 142613134 | 142613363 | 3.51901 | 4.19886 | 1.28809 | Distal Intergenic | 7 | 142575529 | 142578143 | 2615 | 2 | ENSMUST00000136359.2 | -34992 | H19 | ENSMUSG00000000031.10 |
| chr11 | 85831780 | 85833587 | 3.997 | 5.18468 | 1.94369 | Promoter | 11 | 85832551 | 85841948 | 9398 | 1 | ENSMUST00000000095.6 | 0 | Tbx2 | ENSMUSG00000000093.6 |
| chr11 | 85834194 | 85835342 | 4.04501 | 5.4174 | 2.05969 | Exon (ENSMUST00000000095.6/ENSMUSG00000000093.6, exon 2 of 7) | 11 | 85832551 | 85841948 | 9398 | 1 | ENSMUST00000000095.6 | 1644 | Tbx2 | ENSMUSG00000000093.6 |
| chr11 | 85836392 | 85837453 | 3.58701 | 4.79805 | 1.65738 | Exon (ENSMUST00000000095.6/ENSMUSG00000000093.6, exon 4 of 7) | 11 | 85832551 | 85841948 | 9398 | 1 | ENSMUST00000000095.6 | 3842 | Tbx2 | ENSMUSG00000000093.6 |
| chr11 | 85837816 | 85838528 | 3.95353 | 5.32827 | 1.99927 | Exon (ENSMUST00000000095.6/ENSMUSG00000000093.6, exon 6 of 7) | 11 | 85832551 | 85841948 | 9398 | 1 | ENSMUST00000000095.6 | 5266 | Tbx2 | ENSMUSG00000000093.6 |
| chr11 | 85840028 | 85840105 | 4.0707 | 5.32783 | 1.9915 | Intron (ENSMUST00000000095.6/ENSMUSG00000000093.6, intron 6 of 6) | 11 | 85832551 | 85841948 | 9398 | 1 | ENSMUST00000000095.6 | 7478 | Tbx2 | ENSMUSG00000000093.6 |
| chr11 | 85840573 | 85840897 | 4.5359 | 6.55159 | 2.95552 | Exon (ENSMUST00000000095.6/ENSMUSG00000000093.6, exon 7 of 7) | 11 | 85832551 | 85841948 | 9398 | 1 | ENSMUST00000000095.6 | 8023 | Tbx2 | ENSMUSG00000000093.6 |
| chr11 | 85879772 | 85880244 | 3.42868 | 4.03874 | 1.22628 | Distal Intergenic | 11 | 85886422 | 85916095 | 29674 | 1 | ENSMUST00000108047.2 | -6178 | Tbx4 | ENSMUSG00000000094.7 |
| chr11 | 85882025 | 85882204 | 3.86259 | 4.86659 | 1.75331 | Distal Intergenic | 11 | 85886422 | 85916095 | 29674 | 1 | ENSMUST00000108047.2 | -4218 | Tbx4 | ENSMUSG00000000094.7 |
| chr11 | 85882714 | 85882829 | 3.43049 | 4.2931 | 1.31685 | Distal Intergenic | 11 | 85886422 | 85916095 | 29674 | 1 | ENSMUST00000108047.2 | -3593 | Tbx4 | ENSMUSG00000000094.7 |
| chr11 | 85883471 | 85883549 | 3.62904 | 4.42091 | 1.42522 | Distal Intergenic | 11 | 85886422 | 85916095 | 29674 | 1 | ENSMUST00000108047.2 | -2873 | Tbx4 | ENSMUSG00000000094.7 |
| chr11 | 85884524 | 85884615 | 3.69083 | 4.58851 | 1.50563 | Distal Intergenic | 11 | 85886422 | 85916095 | 29674 | 1 | ENSMUST00000108047.2 | -1807 | Tbx4 | ENSMUSG00000000094.7 |
| chr11 | 85885826 | 85886874 | 3.83517 | 4.86277 | 1.74397 | Promoter | 11 | 85886422 | 85916095 | 29674 | 1 | ENSMUST00000108047.2 | 0 | Tbx4 | ENSMUSG00000000094.7 |
| chr11 | 85887262 | 85887641 | 3.37158 | 4.0714 | 1.15186 | Promoter | 11 | 85886422 | 85916095 | 29674 | 1 | ENSMUST00000108047.2 | 841 | Tbx4 | ENSMUSG00000000094.7 |
| chr11 | 85887954 | 85888896 | 3.62855 | 4.55239 | 1.4923 | Intron (ENSMUST00000108047.2/ENSMUSG00000000094.7, intron 1 of 8) | 11 | 85890063 | 85916097 | 26035 | 1 | ENSMUST00000000096.6 | -1167 | Tbx4 | ENSMUSG00000000094.7 |
| chr11 | 85889468 | 85890570 | 4.09046 | 5.59327 | 2.24895 | Promoter | 11 | 85890063 | 85916097 | 26035 | 1 | ENSMUST00000000096.6 | 0 | Tbx4 | ENSMUSG00000000094.7 |
| chr11 | 85890934 | 85891481 | 3.72572 | 4.5781 | 1.49203 | Promoter | 11 | 85890063 | 85916097 | 26035 | 1 | ENSMUST00000000096.6 | 872 | Tbx4 | ENSMUSG00000000094.7 |
| chr11 | 85892043 | 85892471 | 3.82716 | 4.74852 | 1.58034 | Intron (ENSMUST00000108047.2/ENSMUSG00000000094.7, intron 3 of 8) | 11 | 85890063 | 85916097 | 26035 | 1 | ENSMUST00000000096.6 | 1981 | Tbx4 | ENSMUSG00000000094.7 |
| chr11 | 85892814 | 85894260 | 3.70851 | 4.67447 | 1.57194 | Intron (ENSMUST00000108047.2/ENSMUSG00000000094.7, intron 3 of 8) | 11 | 85896765 | 85916095 | 19331 | 1 | ENSMUST00000108045.2 | -2505 | Tbx4 | ENSMUSG00000000094.7 |
| chr11 | 85894650 | 85895532 | 3.81756 | 4.74149 | 1.58335 | Intron (ENSMUST00000108047.2/ENSMUSG00000000094.7, intron 3 of 8) | 11 | 85896765 | 85916095 | 19331 | 1 | ENSMUST00000108045.2 | -1233 | Tbx4 | ENSMUSG00000000094.7 |
| chr11 | 85895890 | 85896482 | 3.65426 | 4.41257 | 1.35979 | Promoter | 11 | 85896765 | 85916095 | 19331 | 1 | ENSMUST00000108045.2 | -283 | Tbx4 | ENSMUSG00000000094.7 |
| chr11 | 85898700 | 85899104 | 3.76698 | 4.78974 | 1.63224 | Intron (ENSMUST00000108047.2/ENSMUSG00000000094.7, intron 4 of 8) | 11 | 85896765 | 85916095 | 19331 | 1 | ENSMUST00000108045.2 | 1936 | Tbx4 | ENSMUSG00000000094.7 |
| chr11 | 85899732 | 85900106 | 3.99164 | 5.05822 | 1.78959 | Exon (ENSMUST00000108047.2/ENSMUSG00000000094.7, exon 5 of 9) | 11 | 85896765 | 85916095 | 19331 | 1 | ENSMUST00000108045.2 | 2968 | Tbx4 | ENSMUSG00000000094.7 |
| chr11 | 85919073 | 85919398 | 3.48882 | 4.14636 | 1.28101 | Distal Intergenic | 11 | 85896765 | 85916095 | 19331 | 1 | ENSMUST00000108045.2 | 22309 | Tbx4 | ENSMUSG00000000094.7 |
| chr11 | 95553910 | 95554439 | 3.3416 | 3.86355 | 1.09988 | Distal Intergenic | 11 | 95568818 | 95587735 | 18918 | 2 | ENSMUST00000000122.6 | 33296 | Ngfr | ENSMUSG00000000120.6 |
| chr11 | 95571374 | 95571755 | 3.55534 | 4.21849 | 1.21339 | Intron (ENSMUST00000000122.6/ENSMUSG00000000120.6, intron 5 of 5) | 11 | 95568818 | 95587735 | 18918 | 2 | ENSMUST00000000122.6 | 15980 | Ngfr | ENSMUSG00000000120.6 |
| chr11 | 95576782 | 95576949 | 3.78442 | 4.70029 | 1.63547 | Intron (ENSMUST00000000122.6/ENSMUSG00000000120.6, intron 3 of 5) | 11 | 95568818 | 95587735 | 18918 | 2 | ENSMUST00000000122.6 | 10786 | Ngfr | ENSMUSG00000000120.6 |
| chr11 | 95577581 | 95578395 | 3.51406 | 4.27618 | 1.32822 | Exon (ENSMUST00000000122.6/ENSMUSG00000000120.6, exon 3 of 6) | 11 | 95568818 | 95587735 | 18918 | 2 | ENSMUST00000000122.6 | 9340 | Ngfr | ENSMUSG00000000120.6 |
| chr11 | 95579423 | 95580221 | 3.42072 | 4.03108 | 1.19115 | Intron (ENSMUST00000000122.6/ENSMUSG00000000120.6, intron 2 of 5) | 11 | 95568818 | 95587735 | 18918 | 2 | ENSMUST00000000122.6 | 7514 | Ngfr | ENSMUSG00000000120.6 |
| chr11 | 95580710 | 95580839 | 3.84873 | 4.89872 | 1.72644 | Intron (ENSMUST00000000122.6/ENSMUSG00000000120.6, intron 2 of 5) | 11 | 95568818 | 95587735 | 18918 | 2 | ENSMUST00000000122.6 | 6896 | Ngfr | ENSMUSG00000000120.6 |
| chr11 | 95582558 | 95583123 | 3.3948 | 4.00394 | 1.1491 | Intron (ENSMUST00000000122.6/ENSMUSG00000000120.6, intron 1 of 5) | 11 | 95568818 | 95587735 | 18918 | 2 | ENSMUST00000000122.6 | 4612 | Ngfr | ENSMUSG00000000120.6 |
| chr11 | 95583565 | 95583655 | 3.33671 | 3.8324 | 1.01733 | Intron (ENSMUST00000000122.6/ENSMUSG00000000120.6, intron 1 of 5) | 11 | 95568818 | 95587735 | 18918 | 2 | ENSMUST00000000122.6 | 4080 | Ngfr | ENSMUSG00000000120.6 |
| chr11 | 95584778 | 95585248 | 3.98655 | 5.1156 | 1.87638 | Intron (ENSMUST00000000122.6/ENSMUSG00000000120.6, intron 1 of 5) | 11 | 95568818 | 95587735 | 18918 | 2 | ENSMUST00000000122.6 | 2487 | Ngfr | ENSMUSG00000000120.6 |
| chr11 | 95586206 | 95587386 | 3.7929 | 4.78527 | 1.65211 | Promoter | 11 | 95568818 | 95587735 | 18918 | 2 | ENSMUST00000000122.6 | 349 | Ngfr | ENSMUSG00000000120.6 |
| chr11 | 95595080 | 95595238 | 3.51428 | 4.25065 | 1.26684 | Distal Intergenic | 11 | 95568818 | 95587735 | 18918 | 2 | ENSMUST00000000122.6 | -7346 | Ngfr | ENSMUSG00000000120.6 |
| chr11 | 95597389 | 95597629 | 3.44863 | 4.06319 | 1.2172 | Distal Intergenic | 11 | 95568818 | 95587735 | 18918 | 2 | ENSMUST00000000122.6 | -9655 | Ngfr | ENSMUSG00000000120.6 |
| chr11 | 103763460 | 103763642 | 3.90848 | 4.89325 | 1.67782 | Distal Intergenic | 11 | 103774150 | 103817957 | 43808 | 1 | ENSMUST00000000127.5 | -10508 | Wnt3 | ENSMUSG00000000125.5 |
| chr11 | 103765008 | 103765217 | 3.37725 | 3.89903 | 1.03675 | Distal Intergenic | 11 | 103774150 | 103817957 | 43808 | 1 | ENSMUST00000000127.5 | -8933 | Wnt3 | ENSMUSG00000000125.5 |
| chr11 | 103765981 | 103766226 | 3.35707 | 3.8631 | 1.01733 | Distal Intergenic | 11 | 103774150 | 103817957 | 43808 | 1 | ENSMUST00000000127.5 | -7924 | Wnt3 | ENSMUSG00000000125.5 |
| chr11 | 103770240 | 103771847 | 3.91871 | 4.94042 | 1.70627 | Distal Intergenic | 11 | 103774150 | 103817957 | 43808 | 1 | ENSMUST00000000127.5 | -2303 | Wnt3 | ENSMUSG00000000125.5 |
| chr11 | 103772665 | 103774989 | 3.86366 | 4.91199 | 1.70753 | Promoter | 11 | 103774150 | 103817957 | 43808 | 1 | ENSMUST00000000127.5 | 0 | Wnt3 | ENSMUSG00000000125.5 |
| chr11 | 103780624 | 103780865 | 3.55189 | 4.22164 | 1.24616 | Intron (ENSMUST00000000127.5/ENSMUSG00000000125.5, intron 1 of 4) | 11 | 103774150 | 103817957 | 43808 | 1 | ENSMUST00000000127.5 | 6475 | Wnt3 | ENSMUSG00000000125.5 |
| chr11 | 103795629 | 103795832 | 3.57775 | 4.28223 | 1.30793 | Intron (ENSMUST00000000127.5/ENSMUSG00000000125.5, intron 1 of 4) | 11 | 103774150 | 103817957 | 43808 | 1 | ENSMUST00000000127.5 | 21480 | Wnt3 | ENSMUSG00000000125.5 |
| chr11 | 103797636 | 103797820 | 3.42292 | 4.12508 | 1.2053 | Intron (ENSMUST00000000127.5/ENSMUSG00000000125.5, intron 1 of 4) | 11 | 103774150 | 103817957 | 43808 | 1 | ENSMUST00000000127.5 | 23487 | Wnt3 | ENSMUSG00000000125.5 |
| chr11 | 59308188 | 59308340 | 3.42762 | 4.16717 | 1.23647 | Intron (ENSMUST00000000128.4/ENSMUSG00000000126.6, intron 1 of 3) | 11 | 59306930 | 59333552 | 26623 | 1 | ENSMUST00000108783.3 | 1259 | Wnt9a | ENSMUSG00000000126.6 |
| chr11 | 59309027 | 59309212 | 3.2976 | 3.78975 | 1.06915 | Intron (ENSMUST00000000128.4/ENSMUSG00000000126.6, intron 1 of 3) | 11 | 59306930 | 59333552 | 26623 | 1 | ENSMUST00000108783.3 | 2098 | Wnt9a | ENSMUSG00000000126.6 |
| chr11 | 59313369 | 59313934 | 3.35956 | 3.87464 | 1.04471 | Intron (ENSMUST00000000128.4/ENSMUSG00000000126.6, intron 1 of 3) | 11 | 59306930 | 59333552 | 26623 | 1 | ENSMUST00000108783.3 | 6440 | Wnt9a | ENSMUSG00000000126.6 |
| chr11 | 108889045 | 108889160 | 3.50682 | 4.16519 | 1.26431 | Distal Intergenic | 11 | 108920349 | 108950783 | 30435 | 1 | ENSMUST00000052915.8 | -31189 | Axin2 | ENSMUSG00000000142.10 |
| chr11 | 108889523 | 108890102 | 3.85119 | 4.80217 | 1.62111 | Distal Intergenic | 11 | 108920349 | 108950783 | 30435 | 1 | ENSMUST00000052915.8 | -30247 | Axin2 | ENSMUSG00000000142.10 |
| chr11 | 108891989 | 108892229 | 3.31913 | 3.80596 | 1.01733 | Distal Intergenic | 11 | 108920349 | 108950783 | 30435 | 1 | ENSMUST00000052915.8 | -28120 | Axin2 | ENSMUSG00000000142.10 |
| chr11 | 108893296 | 108893536 | 3.37139 | 3.88775 | 1.02801 | Distal Intergenic | 11 | 108920349 | 108950783 | 30435 | 1 | ENSMUST00000052915.8 | -26813 | Axin2 | ENSMUSG00000000142.10 |
| chr11 | 108909587 | 108910177 | 3.75202 | 4.64133 | 1.59188 | Distal Intergenic | 11 | 108920349 | 108950783 | 30435 | 1 | ENSMUST00000052915.8 | -10172 | Axin2 | ENSMUSG00000000142.10 |
| chr11 | 108921742 | 108922058 | 3.67251 | 4.59262 | 1.49646 | Promoter | 11 | 108921648 | 108923494 | 1847 | 1 | ENSMUST00000144511.1 | 95 | Axin2 | ENSMUSG00000000142.10 |
| chr11 | 108923333 | 108923481 | 3.36516 | 3.8753 | 1.01733 | Promoter | 11 | 108923181 | 108949511 | 26331 | 1 | ENSMUST00000106711.1 | 153 | Axin2 | ENSMUSG00000000142.10 |
| chr5 | 140856949 | 140857255 | 3.40321 | 3.97859 | 1.14406 | Distal Intergenic | 5 | 140758408 | 140830431 | 72024 | 2 | ENSMUST00000000153.4 | -26519 | Gna12 | ENSMUSG00000000149.6 |
| chr5 | 140869549 | 140869742 | 3.37327 | 3.88754 | 1.01733 | Distal Intergenic | 5 | 140758408 | 140830431 | 72024 | 2 | ENSMUST00000000153.4 | -39119 | Gna12 | ENSMUSG00000000149.6 |
| chr5 | 140870315 | 140870553 | 3.39069 | 3.93753 | 1.08185 | Downstream (2-3kb) | 5 | 140758408 | 140830431 | 72024 | 2 | ENSMUST00000000153.4 | -39885 | Gna12 | ENSMUSG00000000149.6 |
| chr5 | 140872539 | 140872615 | 4.92545 | 6.90304 | 3.07008 | Downstream (<1kb) | 5 | 140758408 | 140830431 | 72024 | 2 | ENSMUST00000000153.4 | -42109 | Gna12 | ENSMUSG00000000149.6 |
| chr5 | 140873189 | 140873414 | 3.63015 | 4.3793 | 1.36913 | 3' UTR | 5 | 140758408 | 140830431 | 72024 | 2 | ENSMUST00000000153.4 | -42759 | Gna12 | ENSMUSG00000000149.6 |
| chr5 | 140889536 | 140889642 | 3.56062 | 4.29663 | 1.35457 | Intron (ENSMUST00000085786.5/ENSMUSG00000036526.7, intron 13 of 24) | 5 | 140758408 | 140830431 | 72024 | 2 | ENSMUST00000000153.4 | -59106 | Gna12 | ENSMUSG00000000149.6 |
| chr7 | 143480536 | 143480671 | 3.45862 | 4.04042 | 1.10208 | Intron (ENSMUST00000052348.6/ENSMUSG00000000154.10, intron 5 of 10) | 7 | 143475101 | 143499325 | 24225 | 1 | ENSMUST00000105917.2 | 5436 | Slc22a18 | ENSMUSG00000000154.10 |
| chr6 | 127079832 | 127080521 | 3.77501 | 4.74223 | 1.63047 | 3' UTR | 6 | 127072902 | 127081408 | 8507 | 1 | ENSMUST00000000186.6 | 6931 | Fgf23 | ENSMUSG00000000182.7 |
| chr6 | 127081029 | 127081428 | 3.3412 | 4.04045 | 1.14456 | 3' UTR | 6 | 127072902 | 127081408 | 8507 | 1 | ENSMUST00000000186.6 | 8128 | Fgf23 | ENSMUSG00000000182.7 |
| chr6 | 127019661 | 127020430 | 3.94277 | 4.97329 | 1.73548 | Intron (ENSMUST00000000187.5/ENSMUSG00000000183.5, intron 2 of 2) | 6 | 127015542 | 127024718 | 9177 | 1 | ENSMUST00000000187.5 | 4120 | Fgf6 | ENSMUSG00000000183.5 |
| chr6 | 127020937 | 127021240 | 3.80261 | 4.72429 | 1.54655 | Intron (ENSMUST00000000187.5/ENSMUSG00000000183.5, intron 2 of 2) | 6 | 127015542 | 127024718 | 9177 | 1 | ENSMUST00000000187.5 | 5396 | Fgf6 | ENSMUSG00000000183.5 |
| chr6 | 127150704 | 127150782 | 4.35126 | 5.86921 | 2.42216 | Promoter | 6 | 127125779 | 127151048 | 25270 | 2 | ENSMUST00000000188.8 | 266 | Ccnd2 | ENSMUSG00000000184.9 |
| chr6 | 127151600 | 127152367 | 3.63019 | 4.38601 | 1.36167 | Promoter | 6 | 127125779 | 127151048 | 25270 | 2 | ENSMUST00000000188.8 | -553 | Ccnd2 | ENSMUSG00000000184.9 |
| chr6 | 127153010 | 127153136 | 3.43641 | 4.00575 | 1.09823 | Intron (ENSMUST00000180811.1/ENSMUSG00000097166.1, intron 1 of 2) | 6 | 127125779 | 127151048 | 25270 | 2 | ENSMUST00000000188.8 | -1963 | Ccnd2 | ENSMUSG00000000184.9 |
| chr6 | 127153973 | 127154373 | 3.69759 | 4.61319 | 1.53476 | Intron (ENSMUST00000180811.1/ENSMUSG00000097166.1, intron 1 of 2) | 6 | 127125779 | 127151048 | 25270 | 2 | ENSMUST00000000188.8 | -2926 | Ccnd2 | ENSMUSG00000000184.9 |
| chr6 | 127155005 | 127156322 | 3.68446 | 4.50336 | 1.45909 | Intron (ENSMUST00000180811.1/ENSMUSG00000097166.1, intron 1 of 2) | 6 | 127125779 | 127151048 | 25270 | 2 | ENSMUST00000000188.8 | -3958 | Ccnd2 | ENSMUSG00000000184.9 |
| chr6 | 127157581 | 127157669 | 4.03556 | 5.24681 | 1.96438 | Intron (ENSMUST00000180811.1/ENSMUSG00000097166.1, intron 1 of 2) | 6 | 127125779 | 127151048 | 25270 | 2 | ENSMUST00000000188.8 | -6534 | Ccnd2 | ENSMUSG00000000184.9 |
| chr7 | 121734278 | 121734528 | 3.768 | 4.77316 | 1.63249 | Promoter | 7 | 121734479 | 121768475 | 33997 | 1 | ENSMUST00000000221.5 | 0 | Scnn1g | ENSMUSG00000000216.8 |
| chr7 | 121734902 | 121735510 | 3.52194 | 4.20204 | 1.23844 | Promoter | 7 | 121734479 | 121768475 | 33997 | 1 | ENSMUST00000000221.5 | 424 | Scnn1g | ENSMUSG00000000216.8 |
| chr7 | 121735831 | 121736289 | 4.0434 | 5.16814 | 1.87208 | Intron (ENSMUST00000000221.5/ENSMUSG00000000216.8, intron 1 of 12) | 7 | 121734479 | 121768475 | 33997 | 1 | ENSMUST00000000221.5 | 1353 | Scnn1g | ENSMUSG00000000216.8 |
| chr2 | 38315529 | 38315611 | 4.03895 | 5.63821 | 2.20853 | Distal Intergenic | 2 | 38339281 | 38369199 | 29919 | 1 | ENSMUST00000143783.3 | -23670 | Lhx2 | ENSMUSG00000000247.6 |
| chr2 | 38322607 | 38322780 | 3.45206 | 4.03245 | 1.10886 | Distal Intergenic | 2 | 38339281 | 38369199 | 29919 | 1 | ENSMUST00000143783.3 | -16501 | Lhx2 | ENSMUSG00000000247.6 |
| chr2 | 38325465 | 38325542 | 3.89744 | 4.87399 | 1.66293 | Distal Intergenic | 2 | 38339281 | 38369199 | 29919 | 1 | ENSMUST00000143783.3 | -13739 | Lhx2 | ENSMUSG00000000247.6 |
| chr2 | 38330737 | 38331229 | 3.49484 | 4.18376 | 1.28847 | Distal Intergenic | 2 | 38339281 | 38369199 | 29919 | 1 | ENSMUST00000143783.3 | -8052 | Lhx2 | ENSMUSG00000000247.6 |
| chr2 | 38332672 | 38332858 | 3.92797 | 4.99402 | 1.77444 | Distal Intergenic | 2 | 38339281 | 38369199 | 29919 | 1 | ENSMUST00000143783.3 | -6423 | Lhx2 | ENSMUSG00000000247.6 |
| chr2 | 38334225 | 38334377 | 3.78803 | 4.80384 | 1.65835 | Distal Intergenic | 2 | 38339281 | 38369199 | 29919 | 1 | ENSMUST00000143783.3 | -4904 | Lhx2 | ENSMUSG00000000247.6 |
| chr2 | 38335165 | 38335262 | 3.37284 | 4.06692 | 1.17724 | Distal Intergenic | 2 | 38339281 | 38369199 | 29919 | 1 | ENSMUST00000143783.3 | -4019 | Lhx2 | ENSMUSG00000000247.6 |
| chr2 | 38335728 | 38337230 | 4.26454 | 5.82154 | 2.36202 | Distal Intergenic | 2 | 38339281 | 38369199 | 29919 | 1 | ENSMUST00000143783.3 | -2051 | Lhx2 | ENSMUSG00000000247.6 |
| chr2 | 38337556 | 38338571 | 3.62998 | 4.37099 | 1.34944 | Promoter | 2 | 38339281 | 38369199 | 29919 | 1 | ENSMUST00000143783.3 | -710 | Lhx2 | ENSMUSG00000000247.6 |
| chr2 | 38339048 | 38339892 | 3.92121 | 5.22388 | 1.92125 | Promoter | 2 | 38339281 | 38369199 | 29919 | 1 | ENSMUST00000143783.3 | 0 | Lhx2 | ENSMUSG00000000247.6 |
| chr2 | 38340235 | 38341679 | 3.77604 | 4.65915 | 1.53711 | Promoter | 2 | 38341092 | 38354825 | 13734 | 1 | ENSMUST00000133661.2 | 0 | Lhx2 | ENSMUSG00000000247.6 |
| chr2 | 38342050 | 38342946 | 3.72571 | 4.55038 | 1.45722 | Promoter | 2 | 38341092 | 38354825 | 13734 | 1 | ENSMUST00000133661.2 | 959 | Lhx2 | ENSMUSG00000000247.6 |
| chr2 | 38343648 | 38344198 | 4.12357 | 5.36566 | 2.0166 | Intron (ENSMUST00000143783.3/ENSMUSG00000000247.6, intron 1 of 4) | 2 | 38341092 | 38354825 | 13734 | 1 | ENSMUST00000133661.2 | 2557 | Lhx2 | ENSMUSG00000000247.6 |
| chr2 | 38350417 | 38352609 | 3.88109 | 4.9568 | 1.80835 | Promoter | 2 | 38350759 | 38369733 | 18975 | 1 | ENSMUST00000000253.5 | 0 | Lhx2 | ENSMUSG00000000247.6 |
| chr2 | 38353058 | 38353542 | 3.62709 | 4.5181 | 1.50522 | Promoter | 2 | 38353802 | 38360089 | 6288 | 1 | ENSMUST00000175896.1 | -260 | Lhx2 | ENSMUSG00000000247.6 |
| chr2 | 38354253 | 38355012 | 3.68905 | 4.69367 | 1.57344 | Promoter | 2 | 38353802 | 38360089 | 6288 | 1 | ENSMUST00000175896.1 | 452 | Lhx2 | ENSMUSG00000000247.6 |
| chr2 | 38349760 | 38350046 | 3.82399 | 4.88656 | 1.72576 | Promoter | 2 | 38350759 | 38369733 | 18975 | 1 | ENSMUST00000000253.5 | -713 | Lhx2 | ENSMUSG00000000247.6 |
| chr2 | 38356164 | 38357088 | 4.0092 | 5.26237 | 1.98542 | Intron (ENSMUST00000143783.3/ENSMUSG00000000247.6, intron 3 of 4) | 2 | 38353802 | 38360089 | 6288 | 1 | ENSMUST00000175896.1 | 2363 | Lhx2 | ENSMUSG00000000247.6 |
| chr2 | 38357487 | 38358166 | 3.75446 | 5.10687 | 1.84373 | Intron (ENSMUST00000143783.3/ENSMUSG00000000247.6, intron 3 of 4) | 2 | 38353802 | 38360089 | 6288 | 1 | ENSMUST00000175896.1 | 3686 | Lhx2 | ENSMUSG00000000247.6 |
| chr2 | 38355385 | 38355602 | 3.38866 | 4.4284 | 1.40672 | Intron (ENSMUST00000143783.3/ENSMUSG00000000247.6, intron 3 of 4) | 2 | 38353802 | 38360089 | 6288 | 1 | ENSMUST00000175896.1 | 1584 | Lhx2 | ENSMUSG00000000247.6 |
| chr11 | 55608466 | 55608760 | 3.46161 | 4.12087 | 1.24155 | Promoter | 11 | 55514238 | 55608198 | 93961 | 2 | ENSMUST00000102716.4 | -269 | Glra1 | ENSMUSG00000000263.10 |
| chr11 | 55641004 | 55641275 | 3.355 | 3.85997 | 1.01733 | Distal Intergenic | 11 | 55514238 | 55608198 | 93961 | 2 | ENSMUST00000102716.4 | -32807 | Glra1 | ENSMUSG00000000263.10 |
| chr8 | 106602650 | 106603968 | 3.97748 | 5.12203 | 1.90372 | Promoter | 8 | 106603351 | 106670246 | 66896 | 1 | ENSMUST00000000312.6 | 0 | Cdh1 | ENSMUSG00000000303.7 |
| chr8 | 106604338 | 106604705 | 3.62268 | 4.45043 | 1.41792 | Promoter | 8 | 106604143 | 106653919 | 49777 | 1 | ENSMUST00000136580.1 | 196 | Cdh1 | ENSMUSG00000000303.7 |
| chr8 | 106605105 | 106605373 | 3.38717 | 4.05837 | 1.14065 | Promoter | 8 | 106604143 | 106653919 | 49777 | 1 | ENSMUST00000136580.1 | 963 | Cdh1 | ENSMUSG00000000303.7 |
| chr8 | 106605867 | 106606296 | 3.54673 | 4.44082 | 1.40983 | Intron (ENSMUST00000000312.6/ENSMUSG00000000303.7, intron 2 of 15) | 8 | 106604143 | 106653919 | 49777 | 1 | ENSMUST00000136580.1 | 1725 | Cdh1 | ENSMUSG00000000303.7 |
| chr8 | 106606712 | 106606897 | 3.53732 | 4.2298 | 1.33228 | Intron (ENSMUST00000000312.6/ENSMUSG00000000303.7, intron 2 of 15) | 8 | 106604143 | 106653919 | 49777 | 1 | ENSMUST00000136580.1 | 2570 | Cdh1 | ENSMUSG00000000303.7 |
| chr2 | 179521274 | 179521862 | 3.42828 | 4.00514 | 1.13402 | Intron (ENSMUST00000000314.7/ENSMUSG00000000305.7, intron 2 of 15) | 2 | 179444233 | 179780319 | 336087 | 1 | ENSMUST00000124708.1 | 77042 | Cdh4 | ENSMUSG00000000305.7 |
| chr2 | 179787295 | 179787600 | 3.54929 | 4.20772 | 1.2093 | Intron (ENSMUST00000000314.7/ENSMUSG00000000305.7, intron 3 of 15) | 2 | 179777163 | 179797511 | 20349 | 1 | ENSMUST00000136411.1 | 10133 | Cdh4 | ENSMUSG00000000305.7 |
| chr2 | 179905614 | 179905695 | 3.58463 | 4.28702 | 1.3031 | Distal Intergenic | 2 | 179893909 | 179894693 | 785 | 1 | ENSMUST00000098996.1 | 11706 | Cdh4 | ENSMUSG00000000305.7 |
| chr2 | 121358535 | 121359521 | 3.77586 | 4.83255 | 1.66489 | Promoter | 2 | 121358610 | 121363737 | 5128 | 1 | ENSMUST00000000317.7 | 0 | Ckmt1 | ENSMUSG00000000308.9 |
| chr11 | 70226803 | 70226926 | 3.99756 | 5.20672 | 1.9173 | Exon (ENSMUST00000000326.6/ENSMUSG00000000317.6, exon 6 of 9) | 11 | 70228540 | 70229677 | 1138 | 2 | ENSMUST00000153449.1 | 2751 | Bcl6b | ENSMUSG00000000317.6 |
| chr11 | 70228370 | 70228698 | 3.80702 | 4.7707 | 1.6442 | Promoter | 11 | 70228540 | 70229677 | 1138 | 2 | ENSMUST00000153449.1 | 979 | Bcl6b | ENSMUSG00000000317.6 |
| chr11 | 70229733 | 70230560 | 3.50937 | 4.40182 | 1.39735 | Promoter | 11 | 70224128 | 70229739 | 5612 | 2 | ENSMUST00000000326.6 | 0 | Bcl6b | ENSMUSG00000000317.6 |
| chr11 | 70166790 | 70166950 | 3.60569 | 4.33443 | 1.3064 | Promoter | 11 | 70166731 | 70168696 | 1966 | 1 | ENSMUST00000178567.1 | 60 | Clec10a | ENSMUSG00000000318.11 |
| chr11 | 70170154 | 70170229 | 3.87245 | 4.81255 | 1.601 | Promoter | 11 | 70169485 | 70170811 | 1327 | 1 | ENSMUST00000153959.1 | 670 | Clec10a | ENSMUSG00000000318.11 |
| chr11 | 70186639 | 70186748 | 3.44466 | 4.02037 | 1.10632 | Distal Intergenic | 11 | 70169485 | 70170811 | 1327 | 1 | ENSMUST00000153959.1 | 17155 | Clec10a | ENSMUSG00000000318.11 |
| chr11 | 70252999 | 70253433 | 3.4872 | 4.10878 | 1.19239 | Exon (ENSMUST00000000329.3/ENSMUSG00000000320.5, exon 3 of 14) | 11 | 70248963 | 70255341 | 6379 | 2 | ENSMUST00000108574.2 | 1908 | Alox12 | ENSMUSG00000000320.5 |
| chr11 | 70253877 | 70254791 | 3.86988 | 5.01717 | 1.79919 | Promoter | 11 | 70248963 | 70255341 | 6379 | 2 | ENSMUST00000108574.2 | 550 | Alox12 | ENSMUSG00000000320.5 |
| chr11 | 70256089 | 70256348 | 3.35406 | 3.86837 | 1.052 | Promoter | 11 | 70241457 | 70255353 | 13897 | 2 | ENSMUST00000000329.3 | -737 | Alox12 | ENSMUSG00000000320.5 |
| chr2 | 152625635 | 152625898 | 3.30476 | 3.78438 | 1.01733 | Distal Intergenic | 2 | 152626951 | 152635194 | 8244 | 1 | ENSMUST00000000369.3 | -1053 | Rem1 | ENSMUSG00000000359.3 |
| chr2 | 152626573 | 152627460 | 3.7662 | 4.75221 | 1.64346 | Promoter | 2 | 152626951 | 152635194 | 8244 | 1 | ENSMUST00000000369.3 | 0 | Rem1 | ENSMUSG00000000359.3 |
| chr16 | 97610700 | 97611797 | 3.80856 | 4.76168 | 1.59157 | Promoter | 16 | 97564684 | 97611195 | 46512 | 2 | ENSMUST00000000395.6 | 0 | Tmprss2 | ENSMUSG00000000385.7 |
| chr2 | 62573688 | 62573900 | 3.25259 | 3.70603 | 1.01733 | Promoter | 2 | 62501184 | 62573851 | 72668 | 2 | ENSMUST00000000402.10 | 0 | Fap | ENSMUSG00000000392.12 |
| chr6 | 115313619 | 115314010 | 3.53993 | 4.19587 | 1.21852 | Distal Intergenic | 6 | 115361221 | 115490401 | 129181 | 1 | ENSMUST00000171644.2 | -47211 | Pparg | ENSMUSG00000000440.7 |
| chr6 | 115344914 | 115345032 | 3.37612 | 3.94078 | 1.16614 | Distal Intergenic | 6 | 115361221 | 115490401 | 129181 | 1 | ENSMUST00000171644.2 | -16189 | Pparg | ENSMUSG00000000440.7 |
| chr6 | 115360376 | 115360544 | 3.47892 | 4.26657 | 1.25264 | Promoter | 6 | 115361221 | 115490401 | 129181 | 1 | ENSMUST00000171644.2 | -677 | Pparg | ENSMUSG00000000440.7 |
| chr6 | 115361125 | 115361311 | 3.71407 | 4.95958 | 1.77669 | Promoter | 6 | 115361221 | 115490401 | 129181 | 1 | ENSMUST00000171644.2 | 0 | Pparg | ENSMUSG00000000440.7 |
| chr6 | 115361740 | 115362358 | 4.15148 | 5.48658 | 2.12485 | Promoter | 6 | 115361221 | 115490401 | 129181 | 1 | ENSMUST00000171644.2 | 520 | Pparg | ENSMUSG00000000440.7 |
| chr15 | 101128092 | 101128238 | 3.57177 | 4.3807 | 1.41835 | Promoter | 15 | 101128522 | 101145336 | 16815 | 1 | ENSMUST00000000542.8 | -284 | Acvrl1 | ENSMUSG00000000530.11 |
| chr15 | 101128730 | 101129036 | 3.92281 | 5.35289 | 2.005 | Promoter | 15 | 101128922 | 101143640 | 14719 | 1 | ENSMUST00000119063.2 | 0 | Acvrl1 | ENSMUSG00000000530.11 |
| chr15 | 101129592 | 101129864 | 3.99469 | 5.15954 | 1.9054 | Promoter | 15 | 101128924 | 101143638 | 14715 | 1 | ENSMUST00000121718.2 | 669 | Acvrl1 | ENSMUSG00000000530.11 |
| chr15 | 101224675 | 101225313 | 3.44483 | 4.05597 | 1.20678 | Promoter | 15 | 101224207 | 101232755 | 8549 | 1 | ENSMUST00000000543.4 | 469 | Grasp | ENSMUSG00000000531.4 |
| chr1 | 166155271 | 166155636 | 3.24903 | 3.70069 | 1.01733 | Intron (ENSMUST00000060833.8/ENSMUSG00000000544.9, intron 3 of 6) | 1 | 166130467 | 166166509 | 36043 | 1 | ENSMUST00000166860.1 | 24805 | Gpa33 | ENSMUSG00000000544.9 |
| chr1 | 166158033 | 166158368 | 4.01688 | 5.25753 | 1.97584 | Intron (ENSMUST00000060833.8/ENSMUSG00000000544.9, intron 4 of 6) | 1 | 166130467 | 166166509 | 36043 | 1 | ENSMUST00000166860.1 | 27567 | Gpa33 | ENSMUSG00000000544.9 |
| chr15 | 103337135 | 103337429 | 3.4776 | 4.07502 | 1.12306 | Intron (ENSMUST00000168828.1/ENSMUSG00000000552.9, intron 1 of 7) | 15 | 103313895 | 103340086 | 26192 | 2 | ENSMUST00000168828.1 | 2657 | Zfp385a | ENSMUSG00000000552.9 |
| chr15 | 103339431 | 103339525 | 3.38409 | 3.91758 | 1.06587 | Promoter | 15 | 103313895 | 103340086 | 26192 | 2 | ENSMUST00000168828.1 | 561 | Zfp385a | ENSMUSG00000000552.9 |
| chr15 | 103339965 | 103340349 | 3.55961 | 4.35974 | 1.39737 | Promoter | 15 | 103313895 | 103340086 | 26192 | 2 | ENSMUST00000168828.1 | 0 | Zfp385a | ENSMUSG00000000552.9 |
| chr15 | 103340906 | 103341694 | 3.57321 | 4.28983 | 1.34085 | Promoter | 15 | 103313895 | 103340086 | 26192 | 2 | ENSMUST00000168828.1 | -821 | Zfp385a | ENSMUSG00000000552.9 |
| chr5 | 71092709 | 71092891 | 3.74549 | 4.587 | 1.47056 | Intron (ENSMUST00000000572.5/ENSMUSG00000000560.5, intron 2 of 9) | 5 | 70961057 | 71095849 | 134793 | 2 | ENSMUST00000000572.5 | 2958 | Gabra2 | ENSMUSG00000000560.5 |
| chr5 | 71093768 | 71094116 | 3.48161 | 4.11327 | 1.20197 | Intron (ENSMUST00000000572.5/ENSMUSG00000000560.5, intron 2 of 9) | 5 | 70961057 | 71095849 | 134793 | 2 | ENSMUST00000000572.5 | 1733 | Gabra2 | ENSMUSG00000000560.5 |
| chr5 | 71095125 | 71095275 | 4.11093 | 5.36555 | 2.04433 | Promoter | 5 | 70961057 | 71095849 | 134793 | 2 | ENSMUST00000000572.5 | 574 | Gabra2 | ENSMUSG00000000560.5 |
| chr5 | 71095788 | 71096494 | 3.89563 | 4.88891 | 1.69426 | Promoter | 5 | 70961057 | 71095849 | 134793 | 2 | ENSMUST00000000572.5 | 0 | Gabra2 | ENSMUSG00000000560.5 |
| chr11 | 112782277 | 112785470 | 3.98739 | 5.19136 | 1.89662 | Promoter | 11 | 112782224 | 112787760 | 5537 | 1 | ENSMUST00000000579.2 | 54 | Sox9 | ENSMUSG00000000567.5 |
| chr11 | 112785974 | 112787907 | 4.03218 | 5.22584 | 1.92238 | 3' UTR | 11 | 112782224 | 112787760 | 5537 | 1 | ENSMUST00000000579.2 | 3751 | Sox9 | ENSMUSG00000000567.5 |
| chr17 | 6553450 | 6554076 | 2.92049 | 4.1858 | 1.27545 | Distal Intergenic | 17 | 6601671 | 6609679 | 8009 | 1 | ENSMUST00000092966.4 | -47595 | Dynlt1c | ENSMUSG00000000579.9 |
| chr17 | 6590212 | 6590494 | 3.22175 | 4.00588 | 1.15537 | Distal Intergenic | 17 | 6601671 | 6609679 | 8009 | 1 | ENSMUST00000092966.4 | -11177 | Dynlt1c | ENSMUSG00000000579.9 |
| chr17 | 6591241 | 6591446 | 3.24825 | 4.26321 | 1.27059 | Distal Intergenic | 17 | 6601671 | 6609679 | 8009 | 1 | ENSMUST00000092966.4 | -10225 | Dynlt1c | ENSMUSG00000000579.9 |
| chr11 | 50847459 | 50848056 | 3.26738 | 3.7282 | 1.01733 | Distal Intergenic | 11 | 50850685 | 50852438 | 1754 | 1 | ENSMUST00000156890.2 | -2629 | Grm6 | ENSMUSG00000000617.9 |
| chr11 | 50849064 | 50849880 | 3.99486 | 5.11931 | 1.90045 | Promoter | 11 | 50850685 | 50852438 | 1754 | 1 | ENSMUST00000156890.2 | -805 | Grm6 | ENSMUSG00000000617.9 |
| chr11 | 50850264 | 50850796 | 3.60362 | 4.62464 | 1.54321 | Promoter | 11 | 50850685 | 50852438 | 1754 | 1 | ENSMUST00000156890.2 | 0 | Grm6 | ENSMUSG00000000617.9 |
| chr11 | 50851188 | 50851993 | 3.70641 | 4.75992 | 1.65055 | Promoter | 11 | 50850986 | 50866177 | 15192 | 1 | ENSMUST00000000631.7 | 203 | Grm6 | ENSMUSG00000000617.9 |
| chr6 | 82913737 | 82914032 | 4.16273 | 5.41233 | 2.0381 | Exon (ENSMUST00000000641.10/ENSMUSG00000000627.10, exon 14 of 14) | 6 | 82914084 | 82917586 | 3503 | 2 | ENSMUST00000125579.1 | 3554 | Sema4f | ENSMUSG00000000627.10 |
| chr6 | 82939315 | 82939987 | 3.75614 | 4.73884 | 1.59713 | Promoter | 6 | 82911885 | 82939769 | 27885 | 2 | ENSMUST00000000641.10 | 0 | Sema4f | ENSMUSG00000000627.10 |
| chr6 | 82744629 | 82744876 | 3.64644 | 4.44738 | 1.44838 | Exon (ENSMUST00000000642.5/ENSMUSG00000000628.5, exon 4 of 18) | 6 | 82725062 | 82726309 | 1248 | 2 | ENSMUST00000168725.1 | -18321 | Hk2 | ENSMUSG00000000628.5 |
| chr6 | 82774698 | 82774785 | 3.74868 | 4.58107 | 1.45693 | Promoter | 6 | 82725025 | 82774454 | 49430 | 2 | ENSMUST00000000642.5 | -245 | Hk2 | ENSMUSG00000000628.5 |
| chr6 | 82776828 | 82776953 | 3.29881 | 3.81325 | 1.06673 | Distal Intergenic | 6 | 82725025 | 82774454 | 49430 | 2 | ENSMUST00000000642.5 | -2375 | Hk2 | ENSMUSG00000000628.5 |
| chr6 | 82772164 | 82773612 | 3.63151 | 4.43069 | 1.44884 | Promoter | 6 | 82727310 | 82773363 | 46054 | 2 | ENSMUST00000170833.1 | 0 | Hk2 | ENSMUSG00000000628.5 |
| chr11 | 77930238 | 77931913 | 4.10015 | 5.36324 | 2.05917 | Promoter | 11 | 77930800 | 77979048 | 48249 | 1 | ENSMUST00000093995.4 | 0 | Sez6 | ENSMUSG00000000632.8 |
| chr11 | 77932269 | 77933548 | 3.83847 | 4.94798 | 1.76566 | Intron (ENSMUST00000093995.4/ENSMUSG00000000632.8, intron 1 of 16) | 11 | 77931039 | 77978061 | 47023 | 1 | ENSMUST00000155087.2 | 1231 | Sez6 | ENSMUSG00000000632.8 |
| chr11 | 77936825 | 77937311 | 3.53458 | 4.28212 | 1.34674 | Intron (ENSMUST00000093995.4/ENSMUSG00000000632.8, intron 1 of 16) | 11 | 77931039 | 77978061 | 47023 | 1 | ENSMUST00000155087.2 | 5787 | Sez6 | ENSMUSG00000000632.8 |
| chr11 | 77941286 | 77941654 | 3.65038 | 4.40185 | 1.34017 | Intron (ENSMUST00000093995.4/ENSMUSG00000000632.8, intron 1 of 16) | 11 | 77951305 | 77954762 | 3458 | 1 | ENSMUST00000138346.1 | -9651 | Sez6 | ENSMUSG00000000632.8 |
| chr11 | 77947563 | 77948292 | 4.10811 | 5.36422 | 2.03058 | Intron (ENSMUST00000093995.4/ENSMUSG00000000632.8, intron 1 of 16) | 11 | 77951305 | 77954762 | 3458 | 1 | ENSMUST00000138346.1 | -3013 | Sez6 | ENSMUSG00000000632.8 |
| chr11 | 77966057 | 77966150 | 3.58952 | 4.28845 | 1.2787 | Intron (ENSMUST00000093995.4/ENSMUSG00000000632.8, intron 4 of 16) | 11 | 77969011 | 77974488 | 5478 | 1 | ENSMUST00000140630.1 | -2861 | Sez6 | ENSMUSG00000000632.8 |
| chr11 | 96295594 | 96295863 | 3.66063 | 4.49953 | 1.45951 | Intron (ENSMUST00000173432.2/ENSMUSG00000000690.5, intron 1 of 2) | 11 | 96292476 | 96301569 | 9094 | 1 | ENSMUST00000173432.2 | 3119 | Hoxb6 | ENSMUSG00000000690.5 |
| chr11 | 96298269 | 96298434 | 3.31357 | 3.83196 | 1.0983 | Promoter | 11 | 96299171 | 96301568 | 2398 | 1 | ENSMUST00000000704.5 | -737 | Hoxb6 | ENSMUSG00000000690.5 |

| | | | | | | | | | | | | | | | |
|---|---|---|---|---|---|---|---|---|---|---|---|---|---|---|---|
| chr11 | 96299997 | 96300270 | 3.38418 | 3.93624 | 1.13042 | Promoter | 11 | 96299171 | 96301568 | 2398 | 1 | ENSMUST00000000704.5 | 827 | Hoxb6 | ENSMUSG00000000690.5 |
| chr13 | 23463521 | 23463600 | 3.91478 | 4.99259 | 1.79898 | Intron (ENSMUST00000041674.8/ENSMUSG00000000706.10, intron 3 of 7) | 13 | 23456992 | 23465875 | 8884 | 2 | ENSMUST00000041674.8 | 2275 | Btn1a1 | ENSMUSG00000000706.10 |
| chr13 | 23464198 | 23465117 | 3.4097 | 4.03165 | 1.18349 | Promoter | 13 | 23456992 | 23465875 | 8884 | 2 | ENSMUST00000041674.8 | 758 | Btn1a1 | ENSMUSG00000000706.10 |
| chr13 | 23465574 | 23465960 | 3.39392 | 4.02143 | 1.15113 | Promoter | 13 | 23456992 | 23465875 | 8884 | 2 | ENSMUST00000041674.8 | 0 | Btn1a1 | ENSMUSG00000000706.10 |
| chr11 | 75281103 | 75281272 | 3.34413 | 3.89738 | 1.11763 | Distal Intergenic | 11 | 75301305 | 75342188 | 40884 | 2 | ENSMUST00000135770.2 | 60916 | Rpa1 | ENSMUSG00000000751.8 |
| chr11 | 75281714 | 75282638 | 3.42413 | 4.03036 | 1.2111 | Distal Intergenic | 11 | 75301305 | 75342188 | 40884 | 2 | ENSMUST00000135770.2 | 59550 | Rpa1 | ENSMUSG00000000751.8 |
| chr11 | 75292021 | 75292472 | 3.48308 | 4.10087 | 1.18629 | Distal Intergenic | 11 | 75301305 | 75342188 | 40884 | 2 | ENSMUST00000135770.2 | 49716 | Rpa1 | ENSMUSG00000000751.8 |
| chr11 | 75292886 | 75293507 | 3.45286 | 4.05179 | 1.16852 | Distal Intergenic | 11 | 75301305 | 75342188 | 40884 | 2 | ENSMUST00000135770.2 | 48681 | Rpa1 | ENSMUSG00000000751.8 |
| chr10 | 6788605 | 6788744 | 3.31951 | 3.80651 | 1.01733 | Promoter | 10 | 6788694 | 7037177 | 248484 | 1 | ENSMUST00000105601.3 | 0 | Oprm1 | ENSMUSG00000000766.13 |
| chr8 | 70891789 | 70893449 | 3.65613 | 4.4225 | 1.35659 | Promoter | 8 | 70882889 | 70892757 | 9869 | 2 | ENSMUST00000000809.2 | 0 | Slc5a5 | ENSMUSG00000000792.2 |
| chr3 | 89490030 | 89490207 | 3.63822 | 4.38136 | 1.34257 | Distal Intergenic | 3 | 89520164 | 89667761 | 147598 | 1 | ENSMUST00000000811.7 | -29957 | Kcnn3 | ENSMUSG00000000794.8 |
| chr3 | 89506295 | 89506700 | 3.38946 | 3.94626 | 1.13868 | Distal Intergenic | 3 | 89520164 | 89667761 | 147598 | 1 | ENSMUST00000000811.7 | -13464 | Kcnn3 | ENSMUSG00000000794.8 |
| chr3 | 89508149 | 89508763 | 3.56916 | 4.33908 | 1.35392 | Distal Intergenic | 3 | 89520164 | 89667761 | 147598 | 1 | ENSMUST00000000811.7 | -11401 | Kcnn3 | ENSMUSG00000000794.8 |
| chr3 | 89519546 | 89519870 | 3.2856 | 3.75556 | 1.01733 | Promoter | 3 | 89520164 | 89667761 | 147598 | 1 | ENSMUST00000000811.7 | -294 | Kcnn3 | ENSMUSG00000000794.8 |
| chr3 | 89520940 | 89521404 | 3.48216 | 4.1135 | 1.22565 | Promoter | 3 | 89520164 | 89667761 | 147598 | 1 | ENSMUST00000000811.7 | 777 | Kcnn3 | ENSMUSG00000000794.8 |
| chr3 | 89522979 | 89523360 | 4.1533 | 5.48289 | 2.14686 | Intron (ENSMUST00000000811.7/ENSMUSG00000000794.8, intron 1 of 7) | 3 | 89520164 | 89667761 | 147598 | 1 | ENSMUST00000000811.7 | 2816 | Kcnn3 | ENSMUSG00000000794.8 |
| chr3 | 89524192 | 89524366 | 3.82303 | 4.81839 | 1.66253 | Intron (ENSMUST00000000811.7/ENSMUSG00000000794.8, intron 1 of 7) | 3 | 89520164 | 89667761 | 147598 | 1 | ENSMUST00000000811.7 | 4029 | Kcnn3 | ENSMUSG00000000794.8 |
| chr3 | 89561357 | 89561470 | 3.35317 | 3.8572 | 1.01733 | Intron (ENSMUST00000000811.7/ENSMUSG00000000794.8, intron 1 of 7) | 3 | 89520164 | 89667761 | 147598 | 1 | ENSMUST00000000811.7 | 41194 | Kcnn3 | ENSMUSG00000000794.8 |
| chr3 | 89602352 | 89602654 | 3.49447 | 4.17346 | 1.29072 | Intron (ENSMUST00000000811.7/ENSMUSG00000000794.8, intron 2 of 7) | 3 | 89662518 | 89672072 | 9555 | 1 | ENSMUST00000124584.2 | -59864 | Kcnn3 | ENSMUSG00000000794.8 |
| chr3 | 89641730 | 89642113 | 3.69934 | 4.59179 | 1.53516 | Intron (ENSMUST00000000811.7/ENSMUSG00000000794.8, intron 3 of 7) | 3 | 89662518 | 89672072 | 9555 | 1 | ENSMUST00000124584.2 | -20405 | Kcnn3 | ENSMUSG00000000794.8 |
| chr3 | 89663586 | 89663873 | 3.37091 | 3.88398 | 1.01733 | Intron (ENSMUST00000000811.7/ENSMUSG00000000794.8, intron 7 of 7) | 3 | 89662518 | 89672072 | 9555 | 1 | ENSMUST00000124584.2 | 1069 | Kcnn3 | ENSMUSG00000000794.8 |
| chr3 | 89667152 | 89667740 | 3.6928 | 4.5203 | 1.43979 | 3' UTR | 3 | 89662518 | 89672072 | 9555 | 1 | ENSMUST00000124584.2 | 4635 | Kcnn3 | ENSMUSG00000000794.8 |
| chr11 | 84942996 | 84943498 | 3.5138 | 4.18079 | 1.2954 | Distal Intergenic | 11 | 84957786 | 84962555 | 4770 | 1 | ENSMUST00000139416.1 | -14288 | Car4 | ENSMUSG00000000805.13 |
| chr11 | 84955333 | 84955647 | 3.38193 | 3.9006 | 1.01733 | Distal Intergenic | 11 | 84957786 | 84962555 | 4770 | 1 | ENSMUST00000139416.1 | -2139 | Car4 | ENSMUSG00000000805.13 |
| chr11 | 84956133 | 84956308 | 3.78425 | 4.75852 | 1.60997 | Distal Intergenic | 11 | 84957786 | 84962555 | 4770 | 1 | ENSMUST00000139416.1 | -1478 | Car4 | ENSMUSG00000000805.13 |
| chr11 | 84957091 | 84958263 | 3.9259 | 5.06757 | 1.80611 | Promoter | 11 | 84957786 | 84962555 | 4770 | 1 | ENSMUST00000139416.1 | 0 | Car4 | ENSMUSG00000000805.13 |
| chr11 | 84958743 | 84960188 | 3.72243 | 4.53555 | 1.43189 | Promoter | 11 | 84958104 | 84964228 | 6125 | 1 | ENSMUST00000127827.1 | 640 | Car4 | ENSMUSG00000000805.13 |
| chr11 | 24077951 | 24078668 | 3.9363 | 4.96042 | 1.72134 | Promoter | 11 | 24078056 | 24173194 | 95139 | 1 | ENSMUST00000000881.7 | 0 | Bcl11a | ENSMUSG00000000861.10 |
| chr11 | 24079017 | 24079256 | 3.87435 | 5.38166 | 2.04058 | Promoter | 11 | 24078173 | 24168429 | 90257 | 1 | ENSMUST00000109514.2 | 845 | Bcl11a | ENSMUSG00000000861.10 |
| chr11 | 24080290 | 24082277 | 3.86491 | 4.91497 | 1.70576 | Promoter | 11 | 24080670 | 24173558 | 92889 | 1 | ENSMUST00000118955.1 | 0 | Bcl11a | ENSMUSG00000000861.10 |
| chr11 | 24082781 | 24082856 | 3.91615 | 4.94318 | 1.71678 | Intron (ENSMUST00000000881.7/ENSMUSG00000000861.10, intron 1 of 4) | 11 | 24080670 | 24173558 | 92889 | 1 | ENSMUST00000118955.1 | 2112 | Bcl11a | ENSMUSG00000000861.10 |
| chr11 | 24083287 | 24083776 | 4.01827 | 5.19424 | 1.88749 | Intron (ENSMUST00000000881.7/ENSMUSG00000000861.10, intron 1 of 4) | 11 | 24080670 | 24173558 | 92889 | 1 | ENSMUST00000118955.1 | 2618 | Bcl11a | ENSMUSG00000000861.10 |
| chr11 | 24084582 | 24084751 | 3.68141 | 4.46036 | 1.39717 | Intron (ENSMUST00000000881.7/ENSMUSG00000000861.10, intron 1 of 4) | 11 | 24080670 | 24173558 | 92889 | 1 | ENSMUST00000118955.1 | 3913 | Bcl11a | ENSMUSG00000000861.10 |
| chr11 | 24085085 | 24085614 | 3.90581 | 5.15131 | 1.91305 | 5' UTR | 11 | 24080670 | 24173558 | 92889 | 1 | ENSMUST00000118955.1 | 4416 | Bcl11a | ENSMUSG00000000861.10 |
| chr11 | 24091504 | 24091963 | 4.19209 | 5.46132 | 2.08406 | Intron (ENSMUST00000000881.7/ENSMUSG00000000861.10, intron 2 of 4) | 11 | 24080670 | 24173558 | 92889 | 1 | ENSMUST00000118955.1 | 10835 | Bcl11a | ENSMUSG00000000861.10 |
| chr11 | 24092563 | 24093198 | 3.84217 | 4.81412 | 1.69887 | Intron (ENSMUST00000000881.7/ENSMUSG00000000861.10, intron 2 of 4) | 11 | 24080670 | 24173558 | 92889 | 1 | ENSMUST00000118955.1 | 11894 | Bcl11a | ENSMUSG00000000861.10 |
| chr11 | 24093890 | 24094335 | 3.35397 | 3.8727 | 1.0677 | Intron (ENSMUST00000000881.7/ENSMUSG00000000861.10, intron 2 of 4) | 11 | 24080670 | 24173558 | 92889 | 1 | ENSMUST00000118955.1 | 13221 | Bcl11a | ENSMUSG00000000861.10 |
| chr11 | 24094689 | 24097064 | 3.62175 | 4.39066 | 1.40038 | Intron (ENSMUST00000000881.7/ENSMUSG00000000861.10, intron 2 of 4) | 11 | 24080670 | 24173558 | 92889 | 1 | ENSMUST00000118955.1 | 14020 | Bcl11a | ENSMUSG00000000861.10 |
| chr16 | 18533220 | 18533314 | 2.87937 | 18.82721 | 13.9393 | Intron (ENSMUST00000167778.2/ENSMUSG00000000884.11, intron 2 of 7) | 16 | 18548743 | 18552584 | 3842 | 1 | ENSMUST00000151253.1 | -15429 | Gnb1l | ENSMUSG00000000884.11 |
| chr15 | 99086682 | 99088559 | 4.34103 | 6.09808 | 2.61836 | Promoter | 15 | 99084686 | 99087817 | 3132 | 2 | ENSMUST00000064462.3 | 0 | C1ql4 | ENSMUSG00000001076.6 |
| chr11 | 78421656 | 78421847 | 3.86347 | 4.8678 | 1.7051 | Promoter | 11 | 78397087 | 78422217 | 25131 | 2 | ENSMUST00000001122.5 | 370 | Slc13a2 | ENSMUSG00000001095.5 |
| chr10 | 76683520 | 76684217 | 3.36649 | 3.8984 | 1.09149 | Distal Intergenic | 10 | 76715016 | 76719236 | 4221 | 2 | ENSMUST00000137599.1 | 35019 | Col6a1 | ENSMUSG00000001119.7 |
| chr10 | 76733307 | 76733729 | 3.92496 | 4.91906 | 1.67364 | Distal Intergenic | 10 | 76708792 | 76726168 | 17377 | 2 | ENSMUST00000001147.4 | -7140 | Col6a1 | ENSMUSG00000001119.7 |
| chr10 | 76584122 | 76584227 | 3.36613 | 3.87675 | 1.01733 | Exon (ENSMUST00000001183.7/ENSMUSG00000001155.8, exon 9 of 14) | 10 | 76587133 | 76588568 | 1436 | 1 | ENSMUST00000146344.1 | -2906 | Ftcd | ENSMUSG00000001155.8 |
| chr12 | 31444219 | 31444578 | 3.63314 | 4.3912 | 1.40775 | Intron (ENSMUST00000167432.2/ENSMUSG00000001225.7, intron 2 of 6) | 12 | 31447435 | 31455079 | 7645 | 1 | ENSMUST00000165816.1 | -2857 | Slc26a3 | ENSMUSG00000001225.7 |
| chr12 | 31444921 | 31445039 | 3.50558 | 4.18346 | 1.29051 | Intron (ENSMUST00000167432.2/ENSMUSG00000001225.7, intron 2 of 6) | 12 | 31447435 | 31455079 | 7645 | 1 | ENSMUST00000165816.1 | -2396 | Slc26a3 | ENSMUSG00000001225.7 |
| chr17 | 56140254 | 56141366 | 3.52939 | 4.34966 | 1.35345 | Promoter | 17 | 56123085 | 56140343 | 17259 | 2 | ENSMUST00000001256.5 | 0 | Sema6b | ENSMUSG00000001227.6 |
| chr17 | 56141821 | 56142167 | 3.74845 | 5.05863 | 1.81418 | Distal Intergenic | 17 | 56123085 | 56140343 | 17259 | 2 | ENSMUST00000001256.5 | -1479 | Sema6b | ENSMUSG00000001227.6 |
| chr17 | 56148305 | 56148605 | 3.64699 | 4.47604 | 1.45674 | Distal Intergenic | 17 | 56123085 | 56140343 | 17259 | 2 | ENSMUST00000001256.5 | -7963 | Sema6b | ENSMUSG00000001227.6 |
| chr17 | 56149016 | 56149714 | 3.61123 | 4.36425 | 1.3997 | Distal Intergenic | 17 | 56123085 | 56140343 | 17259 | 2 | ENSMUST00000001256.5 | -8674 | Sema6b | ENSMUSG00000001227.6 |
| chr12 | 111671004 | 111671452 | 4.06536 | 5.40423 | 2.04627 | Promoter | 12 | 111669355 | 111672338 | 2984 | 2 | ENSMUST00000001304.7 | 886 | Ckb | ENSMUSG00000001270.8 |
| chr12 | 111672343 | 111673017 | 3.74493 | 4.80785 | 1.64491 | Promoter | 12 | 111669355 | 111672338 | 2984 | 2 | ENSMUST00000001304.7 | -6 | Ckb | ENSMUSG00000001270.8 |
| chr15 | 102223735 | 102223932 | 3.46705 | 4.06542 | 1.14621 | Intron (ENSMUST00000001327.4/ENSMUSG00000001281.4, intron 4 of 14) | 15 | 102215995 | 102231935 | 15941 | 2 | ENSMUST00000001327.4 | 8003 | Itgb7 | ENSMUSG00000001281.4 |
| chr9 | 22131764 | 22132221 | 3.59217 | 4.43392 | 1.4351 | Promoter | 9 | 22126731 | 22131865 | 5135 | 2 | ENSMUST00000069330.7 | 0 | Acp5 | ENSMUSG00000001348.9 |
| chr9 | 22099527 | 22100560 | 3.80914 | 4.83817 | 1.66041 | Promoter | 9 | 22099281 | 22109630 | 10350 | 1 | ENSMUST00000001384.4 | 247 | Cnn1 | ENSMUSG00000001349.4 |
| chr3 | 88253037 | 88253929 | 3.42543 | 4.04789 | 1.21931 | Promoter | 3 | 88242874 | 88254709 | 11836 | 2 | ENSMUST00000171887.2 | 780 | Rhbg | ENSMUSG00000001417.5 |
| chr3 | 88254603 | 88255034 | 3.57488 | 4.43058 | 1.38784 | Promoter | 3 | 88242874 | 88254709 | 11836 | 2 | ENSMUST00000171887.2 | 0 | Rhbg | ENSMUSG00000001417.5 |
| chr10 | 77166601 | 77167038 | 3.574 | 4.31708 | 1.30445 | Promoter | 10 | 77052178 | 77166548 | 114371 | 2 | ENSMUST00000081654.7 | -54 | Col18a1 | ENSMUSG00000001435.9 |
| chr10 | 77167356 | 77167685 | 3.69455 | 4.4764 | 1.37461 | Promoter | 10 | 77052178 | 77166548 | 114371 | 2 | ENSMUST00000081654.7 | -809 | Col18a1 | ENSMUSG00000001435.9 |
| chr11 | 97113547 | 97113797 | 3.65593 | 4.43067 | 1.37652 | Intron (ENSMUST00000001484.2/ENSMUSG00000001444.2, intron 1 of 5) | 11 | 97098071 | 97115331 | 17261 | 2 | ENSMUST00000001484.2 | 1534 | Tbx21 | ENSMUSG00000001444.2 |
| chr11 | 97114893 | 97116572 | 3.9734 | 5.10854 | 1.887 | Promoter | 11 | 97098071 | 97115331 | 17261 | 2 | ENSMUST00000001484.2 | 0 | Tbx21 | ENSMUSG00000001444.2 |
| chr18 | 67392206 | 67392458 | 3.23194 | 3.67871 | 1.02475 | Intron (ENSMUST00000001513.6/ENSMUSG00000001473.6, intron 2 of 3) | 18 | 67390717 | 67402748 | 12032 | 1 | ENSMUST00000001513.6 | 1490 | Tubb6 | ENSMUSG00000001473.6 |
| chr11 | 101891205 | 101891746 | 3.87608 | 4.91802 | 1.73682 | Exon (ENSMUST00000147438.1/ENSMUSG00000085594.1, exon 2 of 2) | 11 | 101877510 | 101894374 | 16865 | 2 | ENSMUST00000057054.7 | 2628 | Meox1 | ENSMUSG00000001493.9 |
| chr11 | 101892489 | 101892564 | 3.37278 | 3.88679 | 1.01733 | Intron (ENSMUST00000057054.7/ENSMUSG00000001493.9, intron 1 of 2) | 11 | 101877510 | 101894374 | 16865 | 2 | ENSMUST00000057054.7 | 1810 | Meox1 | ENSMUSG00000001493.9 |
| chr11 | 101892879 | 101893160 | 3.37981 | 3.89739 | 1.01733 | Intron (ENSMUST00000057054.7/ENSMUSG00000001493.9, intron 1 of 2) | 11 | 101877510 | 101894374 | 16865 | 2 | ENSMUST00000057054.7 | 1214 | Meox1 | ENSMUSG00000001493.9 |
| chr11 | 101894320 | 101894602 | 3.60024 | 4.31944 | 1.31972 | Promoter | 11 | 101877510 | 101894374 | 16865 | 2 | ENSMUST00000057054.7 | 0 | Meox1 | ENSMUSG00000001493.9 |
| chr11 | 101895116 | 101895385 | 4.18349 | 5.52725 | 2.23081 | Promoter | 11 | 101877510 | 101894374 | 16865 | 2 | ENSMUST00000057054.7 | -743 | Meox1 | ENSMUSG00000001493.9 |
| chr11 | 101896897 | 101897191 | 3.55288 | 4.28359 | 1.34476 | Distal Intergenic | 11 | 101877510 | 101894374 | 16865 | 2 | ENSMUST00000057054.7 | -2524 | Meox1 | ENSMUSG00000001493.9 |
| chr11 | 101904523 | 101904844 | 3.67338 | 4.66083 | 1.57162 | Distal Intergenic | 11 | 101877510 | 101894374 | 16865 | 2 | ENSMUST00000057054.7 | -10150 | Meox1 | ENSMUSG00000001493.9 |
| chr11 | 101906691 | 101906896 | 3.67037 | 4.43356 | 1.35269 | Distal Intergenic | 11 | 101877510 | 101894374 | 16865 | 2 | ENSMUST00000057054.7 | -12318 | Meox1 | ENSMUSG00000001493.9 |
| chr11 | 101908661 | 101908916 | 3.89512 | 5.00748 | 1.7696 | Distal Intergenic | 11 | 101877510 | 101894374 | 16865 | 2 | ENSMUST00000057054.7 | -14288 | Meox1 | ENSMUSG00000001493.9 |
| chr11 | 101916252 | 101916522 | 3.4438 | 4.10827 | 1.23833 | Distal Intergenic | 11 | 101877510 | 101894374 | 16865 | 2 | ENSMUST00000057054.7 | -21879 | Meox1 | ENSMUSG00000001493.9 |
| chr11 | 101961062 | 101961195 | 3.35912 | 3.86618 | 1.01733 | Downstream (1-2kb) | 11 | 101962458 | 101967015 | 4558 | 2 | ENSMUST00000001534.6 | 5820 | Sost | ENSMUSG00000001494.6 |
| chr11 | 101963749 | 101964903 | 4.01242 | 5.1455 | 1.87403 | 3' UTR | 11 | 101962458 | 101967015 | 4558 | 2 | ENSMUST00000001534.6 | 2112 | Sost | ENSMUSG00000001494.6 |
| chr11 | 101967023 | 101967467 | 3.7152 | 4.59397 | 1.48767 | Promoter | 11 | 101962458 | 101967015 | 4558 | 2 | ENSMUST00000001534.6 | -9 | Sost | ENSMUSG00000001494.6 |
| chr12 | 56456449 | 56456593 | 3.73507 | 4.56231 | 1.45467 | Distal Intergenic | 12 | 56531958 | 56535106 | 3149 | 2 | ENSMUST00000178477.3 | 78513 | Nkx2-1 | ENSMUSG00000001496.10 |
| chr12 | 56457810 | 56458338 | 3.84347 | 4.76656 | 1.57521 | Distal Intergenic | 12 | 56531958 | 56535106 | 3149 | 2 | ENSMUST00000178477.3 | 76768 | Nkx2-1 | ENSMUSG00000001496.10 |
| chr12 | 56458771 | 56459586 | 3.67687 | 4.46233 | 1.39969 | Distal Intergenic | 12 | 56531958 | 56535106 | 3149 | 2 | ENSMUST00000178477.3 | 75520 | Nkx2-1 | ENSMUSG00000001496.10 |
| chr12 | 56460079 | 56460329 | 3.62071 | 4.33928 | 1.29149 | Distal Intergenic | 12 | 56531958 | 56535106 | 3149 | 2 | ENSMUST00000178477.3 | 74777 | Nkx2-1 | ENSMUSG00000001496.10 |
| chr12 | 56460785 | 56460992 | 3.34153 | 3.83966 | 1.01733 | Distal Intergenic | 12 | 56531958 | 56535106 | 3149 | 2 | ENSMUST00000178477.3 | 74114 | Nkx2-1 | ENSMUSG00000001496.10 |
| chr12 | 56502074 | 56502385 | 3.31541 | 3.82012 | 1.08246 | Distal Intergenic | 12 | 56531958 | 56535106 | 3149 | 2 | ENSMUST00000178477.3 | 32721 | Nkx2-1 | ENSMUSG00000001496.10 |
| chr12 | 56503799 | 56504156 | 3.27174 | 3.73474 | 1.01733 | Distal Intergenic | 12 | 56531958 | 56535106 | 3149 | 2 | ENSMUST00000178477.3 | 30950 | Nkx2-1 | ENSMUSG00000001496.10 |
| chr12 | 56509860 | 56510450 | 4.10564 | 5.38882 | 2.0641 | Distal Intergenic | 12 | 56531958 | 56535106 | 3149 | 2 | ENSMUST00000178477.3 | 24656 | Nkx2-1 | ENSMUSG00000001496.10 |
| chr12 | 56510969 | 56511475 | 3.52003 | 4.26259 | 1.31797 | Distal Intergenic | 12 | 56531958 | 56535106 | 3149 | 2 | ENSMUST00000178477.3 | 23631 | Nkx2-1 | ENSMUSG00000001496.10 |
| chr12 | 56515005 | 56515183 | 3.43386 | 4.02941 | 1.1969 | Distal Intergenic | 12 | 56531958 | 56535106 | 3149 | 2 | ENSMUST00000178477.3 | 19923 | Nkx2-1 | ENSMUSG00000001496.10 |
| chr12 | 56517316 | 56517727 | 3.95639 | 5.09118 | 1.85994 | Distal Intergenic | 12 | 56531958 | 56535106 | 3149 | 2 | ENSMUST00000178477.3 | 17379 | Nkx2-1 | ENSMUSG00000001496.10 |
| chr12 | 56518296 | 56518788 | 3.57423 | 4.41096 | 1.4242 | Distal Intergenic | 12 | 56531958 | 56535106 | 3149 | 2 | ENSMUST00000178477.3 | 16318 | Nkx2-1 | ENSMUSG00000001496.10 |
| chr12 | 56519542 | 56520179 | 4.0004 | 5.35849 | 2.03411 | Distal Intergenic | 12 | 56531958 | 56535106 | 3149 | 2 | ENSMUST00000178477.3 | 14927 | Nkx2-1 | ENSMUSG00000001496.10 |
| chr12 | 56520923 | 56521740 | 4.32815 | 6.12697 | 2.61367 | Distal Intergenic | 12 | 56531958 | 56535106 | 3149 | 2 | ENSMUST00000178477.3 | 13366 | Nkx2-1 | ENSMUSG00000001496.10 |
| chr12 | 56523838 | 56524006 | 4.22122 | 5.64209 | 2.23609 | Distal Intergenic | 12 | 56531958 | 56535106 | 3149 | 2 | ENSMUST00000178477.3 | 11100 | Nkx2-1 | ENSMUSG00000001496.10 |
| chr12 | 56524873 | 56525185 | 3.38941 | 4.13175 | 1.24677 | Distal Intergenic | 12 | 56531958 | 56535106 | 3149 | 2 | ENSMUST00000178477.3 | 9921 | Nkx2-1 | ENSMUSG00000001496.10 |
| chr12 | 56525506 | 56525688 | 3.48994 | 4.11237 | 1.19007 | Distal Intergenic | 12 | 56531958 | 56535106 | 3149 | 2 | ENSMUST00000178477.3 | 9418 | Nkx2-1 | ENSMUSG00000001496.10 |
| chr12 | 56526234 | 56526762 | 3.51375 | 4.24347 | 1.28098 | Distal Intergenic | 12 | 56531958 | 56535106 | 3149 | 2 | ENSMUST00000178477.3 | 8344 | Nkx2-1 | ENSMUSG00000001496.10 |
| chr12 | 56527675 | 56528047 | 3.49778 | 4.16507 | 1.21474 | Distal Intergenic | 12 | 56531958 | 56535106 | 3149 | 2 | ENSMUST00000178477.3 | 7059 | Nkx2-1 | ENSMUSG00000001496.10 |
| chr12 | 56528421 | 56528669 | 4.5825 | 6.37494 | 2.87155 | Distal Intergenic | 12 | 56531958 | 56535106 | 3149 | 2 | ENSMUST00000178477.3 | 6437 | Nkx2-1 | ENSMUSG00000001496.10 |
| chr12 | 56529348 | 56530273 | 4.02517 | 5.22093 | 1.99866 | Downstream (1-2kb) | 12 | 56531958 | 56535106 | 3149 | 2 | ENSMUST00000178477.3 | 4833 | Nkx2-1 | ENSMUSG00000001496.10 |
| chr12 | 56530888 | 56531480 | 3.93751 | 5.2218 | 1.94974 | Downstream (<1kb) | 12 | 56531958 | 56535106 | 3149 | 2 | ENSMUST00000178477.3 | 3626 | Nkx2-1 | ENSMUSG00000001496.10 |
| chr12 | 56532022 | 56532556 | 4.62365 | 6.44139 | 2.89514 | 3' UTR | 12 | 56531958 | 56535106 | 3149 | 2 | ENSMUST00000178477.3 | 2550 | Nkx2-1 | ENSMUSG00000001496.10 |
| chr12 | 56533065 | 56534040 | 4.1211 | 5.82445 | 2.36303 | Exon (ENSMUST00000178477.3/ENSMUSG00000001496.10, exon 2 of 2) | 12 | 56531958 | 56535106 | 3149 | 2 | ENSMUST00000178477.3 | 1066 | Nkx2-1 | ENSMUSG00000001496.10 |
| chr12 | 56534349 | 56535381 | 4.20349 | 5.60759 | 2.22583 | Promoter | 12 | 56531958 | 56535106 | 3149 | 2 | ENSMUST00000178477.3 | 0 | Nkx2-1 | ENSMUSG00000001496.10 |
| chr12 | 56536463 | 56538222 | 4.11818 | 5.41675 | 2.07418 | Promoter | 12 | 56531958 | 56536908 | 4951 | 2 | ENSMUST00000001536.8 | 0 | Nkx2-1 | ENSMUSG00000001496.10 |
| chr12 | 56674789 | 56675083 | 3.48504 | 4.20229 | 1.268 | Distal Intergenic | 12 | 56691767 | 56712707 | 20941 | 1 | ENSMUST00000153250.3 | -16684 | Pax9 | ENSMUSG00000001497.13 |
| chr12 | 56676317 | 56676710 | 3.51592 | 4.27018 | 1.31935 | Distal Intergenic | 12 | 56691767 | 56712707 | 20941 | 1 | ENSMUST00000153250.3 | -15057 | Pax9 | ENSMUSG00000001497.13 |
| chr12 | 56677941 | 56678634 | 3.56892 | 4.44018 | 1.4134 | Distal Intergenic | 12 | 56691767 | 56712707 | 20941 | 1 | ENSMUST00000153250.3 | -13133 | Pax9 | ENSMUSG00000001497.13 |
| chr12 | 56679920 | 56680137 | 4.01109 | 5.11582 | 1.87936 | Distal Intergenic | 12 | 56691767 | 56712707 | 20941 | 1 | ENSMUST00000153250.3 | -11630 | Pax9 | ENSMUSG00000001497.13 |
| chr12 | 56681101 | 56681797 | 3.52034 | 4.19206 | 1.25092 | Distal Intergenic | 12 | 56691767 | 56712707 | 20941 | 1 | ENSMUST00000153250.3 | -9970 | Pax9 | ENSMUSG00000001497.13 |
| chr12 | 56682205 | 56683068 | 3.53748 | 4.21634 | 1.26084 | Distal Intergenic | 12 | 56691767 | 56712707 | 20941 | 1 | ENSMUST00000153250.3 | -8699 | Pax9 | ENSMUSG00000001497.13 |
| chr12 | 56683864 | 56684107 | 3.35291 | 3.86081 | 1.03138 | Distal Intergenic | 12 | 56691767 | 56712707 | 20941 | 1 | ENSMUST00000153250.3 | -7660 | Pax9 | ENSMUSG00000001497.13 |
| chr12 | 56685066 | 56687658 | 3.78863 | 4.79267 | 1.62693 | Downstream (2-3kb) | 12 | 56691767 | 56712707 | 20941 | 1 | ENSMUST00000153250.3 | -4109 | Pax9 | ENSMUSG00000001497.13 |
| chr12 | 56687982 | 56688174 | 3.38332 | 4.05574 | 1.14837 | Downstream (1-2kb) | 12 | 56691767 | 56712707 | 20941 | 1 | ENSMUST00000153250.3 | -3593 | Pax9 | ENSMUSG00000001497.13 |
| chr12 | 56688509 | 56689112 | 3.80301 | 4.76886 | 1.62689 | Downstream (<1kb) | 12 | 56691767 | 56712707 | 20941 | 1 | ENSMUST00000153250.3 | -2655 | Pax9 | ENSMUSG00000001497.13 |
| chr12 | 56689551 | 56690232 | 3.52539 | 4.17081 | 1.20425 | Exon (ENSMUST00000159874.2/ENSMUSG00000089894.2, exon 2 of 2) | 12 | 56691767 | 56712707 | 20941 | 1 | ENSMUST00000153250.3 | -1535 | Pax9 | ENSMUSG00000001497.13 |
| chr12 | 56690566 | 56691808 | 4.02383 | 5.1735 | 1.90276 | Promoter | 12 | 56691767 | 56712707 | 20941 | 1 | ENSMUST00000153250.3 | 0 | Pax9 | ENSMUSG00000001497.13 |
| chr12 | 56719557 | 56720223 | 3.62376 | 4.40791 | 1.44166 | Intron (ENSMUST00000044634.6/ENSMUSG00000035472.8, intron 7 of 9) | 12 | 56695729 | 56710346 | 14618 | 1 | ENSMUST00000152848.1 | 23829 | Pax9 | ENSMUSG00000001497.13 |
| chr12 | 56701620 | 56701747 | 4.17062 | 5.80438 | 2.32454 | Intron (ENSMUST00000153250.3/ENSMUSG00000001497.13, intron 4 of 4) | 12 | 56695729 | 56710346 | 14618 | 1 | ENSMUST00000152848.1 | 5892 | Pax9 | ENSMUSG00000001497.13 |
| chr12 | 56694344 | 56697299 | 3.93721 | 5.19817 | 1.94395 | Promoter | 12 | 56695639 | 56712822 | 17184 | 1 | ENSMUST00000001538.9 | 0 | Pax9 | ENSMUSG00000001497.13 |
| chr12 | 56697627 | 56697981 | 3.62848 | 4.4819 | 1.47764 | Intron (ENSMUST00000153250.3/ENSMUSG00000001497.13, intron 3 of 4) | 12 | 56695729 | 56710346 | 14618 | 1 | ENSMUST00000152848.1 | 1899 | Pax9 | ENSMUSG00000001497.13 |
| chr12 | 56698501 | 56701212 | 4.11259 | 5.41584 | 2.14117 | Exon (ENSMUST00000153250.3/ENSMUSG00000001497.13, exon 4 of 5) | 12 | 56695729 | 56710346 | 14618 | 1 | ENSMUST00000152848.1 | 2773 | Pax9 | ENSMUSG00000001497.13 |
| chr12 | 56706381 | 56706696 | 3.49505 | 4.12314 | 1.20087 | Intron (ENSMUST00000153250.3/ENSMUSG00000001497.13, intron 4 of 4) | 12 | 56695729 | 56710346 | 14618 | 1 | ENSMUST00000152848.1 | 10653 | Pax9 | ENSMUSG00000001497.13 |
| chr12 | 56703222 | 56703297 | 4.06753 | 5.18835 | 1.86578 | Intron (ENSMUST00000153250.3/ENSMUSG00000001497.13, intron 4 of 4) | 12 | 56695729 | 56710346 | 14618 | 1 | ENSMUST00000152848.1 | 7494 | Pax9 | ENSMUSG00000001497.13 |
| chr12 | 56704486 | 56704880 | 3.86255 | 4.84075 | 1.69844 | Intron (ENSMUST00000153250.3/ENSMUSG00000001497.13, intron 4 of 4) | 12 | 56695729 | 56710346 | 14618 | 1 | ENSMUST00000152848.1 | 8758 | Pax9 | ENSMUSG00000001497.13 |
| chr12 | 56705486 | 56705673 | 3.73092 | 4.56413 | 1.48657 | Intron (ENSMUST00000153250.3/ENSMUSG00000001497.13, intron 4 of 4) | 12 | 56695729 | 56710346 | 14618 | 1 | ENSMUST00000152848.1 | 9758 | Pax9 | ENSMUSG00000001497.13 |
| chr12 | 56709195 | 56709867 | 3.72192 | 4.58632 | 1.49158 | Exon (ENSMUST00000153250.3/ENSMUSG00000001497.13, exon 5 of 5) | 12 | 56695729 | 56710346 | 14618 | 1 | ENSMUST00000152848.1 | 13467 | Pax9 | ENSMUSG00000001497.13 |
| chr12 | 56715034 | 56715544 | 3.55888 | 4.27112 | 1.35466 | Intron (ENSMUST00000044634.6/ENSMUSG00000035472.8, intron 9 of 9) | 12 | 56695729 | 56710346 | 14618 | 1 | ENSMUST00000152848.1 | 19306 | Pax9 | ENSMUSG00000001497.13 |
| chr12 | 56716733 | 56716897 | 4.55913 | 6.32148 | 2.75928 | Intron (ENSMUST00000044634.6/ENSMUSG00000035472.8, intron 9 of 9) | 12 | 56695729 | 56710346 | 14618 | 1 | ENSMUST00000152848.1 | 21005 | Pax9 | ENSMUSG00000001497.13 |
| chr13 | 72627400 | 72629373 | 4.04397 | 5.31336 | 1.99667 | Promoter | 13 | 72628820 | 72634198 | 5379 | 1 | ENSMUST00000074372.5 | 0 | Irx2 | ENSMUSG00000001504.10 |
| chr13 | 72630345 | 72631836 | 4.0789 | 5.26596 | 1.95854 | Promoter | 13 | 72632085 | 72633613 | 1529 | 1 | ENSMUST00000163393.1 | -249 | Irx2 | ENSMUSG00000001504.10 |
| chr13 | 72633239 | 72633434 | 3.82276 | 4.75659 | 1.6375 | Promoter | 13 | 72632597 | 72633770 | 1174 | 1 | ENSMUST00000172353.2 | 643 | Irx2 | ENSMUSG00000001504.10 |
| chr13 | 72633851 | 72634379 | 4.06319 | 5.30817 | 2.03096 | 3' UTR | 13 | 72632597 | 72633770 | 1174 | 1 | ENSMUST00000172353.2 | 1255 | Irx2 | ENSMUSG00000001504.10 |
| chr13 | 72634756 | 72635177 | 3.64445 | 4.62438 | 1.50937 | Distal Intergenic | 13 | 72632597 | 72633770 | 1174 | 1 | ENSMUST00000172353.2 | 2160 | Irx2 | ENSMUSG00000001504.10 |
| chr13 | 72635553 | 72635852 | 3.46861 | 4.22582 | 1.27156 | Distal Intergenic | 13 | 72632597 | 72633770 | 1174 | 1 | ENSMUST00000172353.2 | 2957 | Irx2 | ENSMUSG00000001504.10 |
| chr13 | 72636235 | 72637706 | 3.71794 | 4.58129 | 1.5475 | Distal Intergenic | 13 | 72632597 | 72633770 | 1174 | 1 | ENSMUST00000172353.2 | 3639 | Irx2 | ENSMUSG00000001504.10 |
| chr13 | 72638448 | 72638962 | 4.34031 | 5.87575 | 2.42751 | Distal Intergenic | 13 | 72632597 | 72633770 | 1174 | 1 | ENSMUST00000172353.2 | 5852 | Irx2 | ENSMUSG00000001504.10 |
| chr13 | 72639303 | 72639917 | 3.52629 | 4.22543 | 1.34027 | Distal Intergenic | 13 | 72632597 | 72633770 | 1174 | 1 | ENSMUST00000172353.2 | 6707 | Irx2 | ENSMUSG00000001504.10 |
| chr13 | 72640249 | 72641100 | 3.46383 | 4.27181 | 1.29359 | Distal Intergenic | 13 | 72632597 | 72633770 | 1174 | 1 | ENSMUST00000172353.2 | 7653 | Irx2 | ENSMUSG00000001504.10 |
| chr13 | 72646869 | 72647093 | 3.46265 | 4.10343 | 1.26597 | Distal Intergenic | 13 | 72632597 | 72633770 | 1174 | 1 | ENSMUST00000172353.2 | 14273 | Irx2 | ENSMUSG00000001504.10 |
| chr13 | 72649639 | 72649950 | 3.59082 | 4.36282 | 1.38034 | Distal Intergenic | 13 | 72632597 | 72633770 | 1174 | 1 | ENSMUST00000172353.2 | 17043 | Irx2 | ENSMUSG00000001504.10 |
| chr13 | 72816849 | 72817255 | 3.49805 | 4.13757 | 1.19984 | Distal Intergenic | 13 | 72632597 | 72633770 | 1174 | 1 | ENSMUST00000172353.2 | 184253 | Irx2 | ENSMUSG00000001504.10 |
| chr13 | 72818224 | 72818400 | 3.50201 | 4.1296 | 1.18375 | Distal Intergenic | 13 | 72632597 | 72633770 | 1174 | 1 | ENSMUST00000172353.2 | 185628 | Irx2 | ENSMUSG00000001504.10 |
| chr13 | 72830455 | 72830643 | 3.39448 | 3.93841 | 1.08407 | Distal Intergenic | 13 | 72632597 | 72633770 | 1174 | 1 | ENSMUST00000172353.2 | 197859 | Irx2 | ENSMUSG00000001504.10 |
| chr11 | 95120229 | 95122499 | 4.02392 | 5.27522 | 1.96257 | Promoter | 11 | 95120119 | 95125296 | 5178 | 1 | ENSMUST00000092768.6 | 111 | Dlx3 | ENSMUSG00000001510.8 |
| chr11 | 95126374 | 95126609 | 3.51284 | 4.31149 | 1.31525 | Distal Intergenic | 11 | 95120119 | 95125296 | 5178 | 1 | ENSMUST00000092768.6 | 6256 | Dlx3 | ENSMUSG00000001510.8 |
| chr11 | 95128603 | 95128998 | 3.54725 | 4.21972 | 1.26452 | Distal Intergenic | 11 | 95120119 | 95125296 | 5178 | 1 | ENSMUST00000092768.6 | 8485 | Dlx3 | ENSMUSG00000001510.8 |
| chr11 | 95130897 | 95130976 | 3.75731 | 4.61704 | 1.52856 | Distal Intergenic | 11 | 95120119 | 95125296 | 5178 | 1 | ENSMUST00000092768.6 | 10779 | Dlx3 | ENSMUSG00000001510.8 |
| chr5 | 29465494 | 29465645 | 3.70139 | 4.49146 | 1.40176 | Distal Intergenic | 5 | 29473034 | 29478470 | 5437 | 2 | ENSMUST00000165512.3 | 12825 | Mnx1 | ENSMUSG00000001566.9 |
| chr5 | 29465996 | 29466432 | 4.42054 | 5.91683 | 2.39867 | Distal Intergenic | 5 | 29473034 | 29478470 | 5437 | 2 | ENSMUST00000165512.3 | 12038 | Mnx1 | ENSMUSG00000001566.9 |
| chr5 | 29466818 | 29467215 | 3.87726 | 4.8427 | 1.64114 | Distal Intergenic | 5 | 29473034 | 29478470 | 5437 | 2 | ENSMUST00000165512.3 | 11255 | Mnx1 | ENSMUSG00000001566.9 |
| chr5 | 29467598 | 29470027 | 3.9513 | 5.0392 | 1.80213 | Distal Intergenic | 5 | 29473034 | 29478470 | 5437 | 2 | ENSMUST00000165512.3 | 8443 | Mnx1 | ENSMUSG00000001566.9 |
| chr5 | 29470343 | 29473204 | 3.87864 | 4.8892 | 1.68391 | 3' UTR | 5 | 29473034 | 29478470 | 5437 | 2 | ENSMUST00000165512.3 | 5266 | Mnx1 | ENSMUSG00000001566.9 |
| chr5 | 29473599 | 29474920 | 3.66412 | 4.57442 | 1.51437 | 3' UTR | 5 | 29473034 | 29478470 | 5437 | 2 | ENSMUST00000165512.3 | 3550 | Mnx1 | ENSMUSG00000001566.9 |
| chr5 | 29475410 | 29475663 | 4.02554 | 5.12794 | 1.86145 | Intron (ENSMUST00000165512.3/ENSMUSG00000001566.9, intron 1 of 2) | 5 | 29473034 | 29478470 | 5437 | 2 | ENSMUST00000165512.3 | 2807 | Mnx1 | ENSMUSG00000001566.9 |
| chr5 | 29475975 | 29476236 | 3.8902 | 4.86355 | 1.6749 | Intron (ENSMUST00000165512.3/ENSMUSG00000001566.9, intron 1 of 2) | 5 | 29473034 | 29478470 | 5437 | 2 | ENSMUST00000165512.3 | 2234 | Mnx1 | ENSMUSG00000001566.9 |
| chr5 | 29477404 | 29478793 | 4.25921 | 5.94888 | 2.44477 | Promoter | 5 | 29473034 | 29478470 | 5437 | 2 | ENSMUST00000165512.3 | 0 | Mnx1 | ENSMUSG00000001566.9 |
| chr5 | 29479164 | 29479273 | 3.47099 | 4.18016 | 1.31026 | Promoter | 5 | 29473261 | 29478470 | 5210 | 2 | ENSMUST00000001608.7 | -695 | Mnx1 | ENSMUSG00000001566.9 |
| chr5 | 29479665 | 29480209 | 3.97319 | 5.16079 | 1.92858 | Downstream (1-2kb) | 5 | 29473261 | 29478470 | 5210 | 2 | ENSMUST00000001608.7 | -1196 | Mnx1 | ENSMUSG00000001566.9 |
| chr11 | 69858677 | 69858756 | 3.79163 | 4.67949 | 1.56185 | Promoter | 11 | 69851005 | 69858730 | 7726 | 2 | ENSMUST00000001626.4 | 0 | Tnk1 | ENSMUSG00000001583.8 |
| chr15 | 102829510 | 102829831 | 3.25757 | 3.71349 | 1.01733 | Distal Intergenic | 15 | 102921103 | 102928814 | 7712 | 1 | ENSMUST00000001700.6 | -91272 | Hoxc13 | ENSMUSG00000001655.6 |
| chr15 | 102891363 | 102891876 | 3.51443 | 4.14816 | 1.18105 | Distal Intergenic | 15 | 102921103 | 102928814 | 7712 | 1 | ENSMUST00000001700.6 | -29227 | Hoxc13 | ENSMUSG00000001655.6 |
| chr15 | 102894691 | 102895425 | 3.59373 | 4.35417 | 1.3535 | Distal Intergenic | 15 | 102921103 | 102928814 | 7712 | 1 | ENSMUST00000001700.6 | -25678 | Hoxc13 | ENSMUSG00000001655.6 |
| chr15 | 102896640 | 102897177 | 3.79182 | 4.77388 | 1.64473 | Distal Intergenic | 15 | 102921103 | 102928814 | 7712 | 1 | ENSMUST00000001700.6 | -23926 | Hoxc13 | ENSMUSG00000001655.6 |
| chr15 | 102897519 | 102897855 | 4.16936 | 5.65844 | 2.24849 | Distal Intergenic | 15 | 102921103 | 102928814 | 7712 | 1 | ENSMUST00000001700.6 | -23248 | Hoxc13 | ENSMUSG00000001655.6 |
| chr15 | 102898856 | 102900140 | 3.83906 | 4.76227 | 1.58023 | Distal Intergenic | 15 | 102921103 | 102928814 | 7712 | 1 | ENSMUST00000001700.6 | -20963 | Hoxc13 | ENSMUSG00000001655.6 |
| chr15 | 102901450 | 102902893 | 3.87464 | 4.8436 | 1.65775 | Distal Intergenic | 15 | 102921103 | 102928814 | 7712 | 1 | ENSMUST00000001700.6 | -18210 | Hoxc13 | ENSMUSG00000001655.6 |

| | | | | | | | | | | | | | | | |
|---|---|---|---|---|---|---|---|---|---|---|---|---|---|---|---|
| chr15 | 102903255 | 102903559 | 3.60985 | 4.32669 | 1.29733 | Distal Intergenic | 15 | 102921103 | 102928814 | 7712 | 1 | ENSMUST00000001700.6 | -17544 | Hoxc13 | ENSMUSG00000001655.6 |
| chr15 | 102904869 | 102904987 | 3.70688 | 4.56681 | 1.52977 | Distal Intergenic | 15 | 102921103 | 102928814 | 7712 | 1 | ENSMUST00000001700.6 | -16116 | Hoxc13 | ENSMUSG00000001655.6 |
| chr15 | 102906249 | 102906475 | 3.74944 | 4.82675 | 1.6617 | Distal Intergenic | 15 | 102921103 | 102928814 | 7712 | 1 | ENSMUST00000001700.6 | -14628 | Hoxc13 | ENSMUSG00000001655.6 |
| chr15 | 102907043 | 102907373 | 3.87606 | 5.01589 | 1.82408 | Distal Intergenic | 15 | 102921103 | 102928814 | 7712 | 1 | ENSMUST00000001700.6 | -13730 | Hoxc13 | ENSMUSG00000001655.6 |
| chr15 | 102907695 | 102908260 | 4.12922 | 5.81158 | 2.33013 | Distal Intergenic | 15 | 102921103 | 102928814 | 7712 | 1 | ENSMUST00000001700.6 | -12843 | Hoxc13 | ENSMUSG00000001655.6 |
| chr15 | 102909232 | 102909830 | 3.47293 | 4.18393 | 1.25779 | Distal Intergenic | 15 | 102921103 | 102928814 | 7712 | 1 | ENSMUST00000001700.6 | -11273 | Hoxc13 | ENSMUSG00000001655.6 |
| chr15 | 102910173 | 102911240 | 3.69046 | 4.77208 | 1.62414 | Distal Intergenic | 15 | 102921103 | 102928814 | 7712 | 1 | ENSMUST00000001700.6 | -9863 | Hoxc13 | ENSMUSG00000001655.6 |
| chr15 | 102912244 | 102912445 | 3.71587 | 4.74763 | 1.58326 | Distal Intergenic | 15 | 102921103 | 102928814 | 7712 | 1 | ENSMUST00000001700.6 | -8658 | Hoxc13 | ENSMUSG00000001655.6 |
| chr15 | 102913120 | 102913524 | 3.85596 | 4.91375 | 1.7456 | Distal Intergenic | 15 | 102921103 | 102928814 | 7712 | 1 | ENSMUST00000001700.6 | -7579 | Hoxc13 | ENSMUSG00000001655.6 |
| chr15 | 102915752 | 102915935 | 3.53499 | 4.50219 | 1.46003 | Distal Intergenic | 15 | 102921103 | 102928814 | 7712 | 1 | ENSMUST00000001700.6 | -5168 | Hoxc13 | ENSMUSG00000001655.6 |
| chr15 | 102916259 | 102917291 | 3.79118 | 4.7179 | 1.60112 | Distal Intergenic | 15 | 102921103 | 102928814 | 7712 | 1 | ENSMUST00000001700.6 | -3812 | Hoxc13 | ENSMUSG00000001655.6 |
| chr15 | 102917639 | 102919954 | 3.56737 | 4.29438 | 1.35122 | Distal Intergenic | 15 | 102921103 | 102928814 | 7712 | 1 | ENSMUST00000001700.6 | -1149 | Hoxc13 | ENSMUSG00000001655.6 |
| chr15 | 102920311 | 102922911 | 4.20126 | 5.6392 | 2.23682 | Promoter | 15 | 102921103 | 102928814 | 7712 | 1 | ENSMUST00000001700.6 | 0 | Hoxc13 | ENSMUSG00000001655.6 |
| chr15 | 102924669 | 102925245 | 3.94886 | 5.11099 | 1.87223 | Intron (ENSMUST00000001700.6/ENSMUSG00000001655.6, intron 1 of 1) | 15 | 102921103 | 102928814 | 7712 | 1 | ENSMUST00000001700.6 | 3567 | Hoxc13 | ENSMUSG00000001655.6 |
| chr15 | 102925731 | 102926507 | 3.57414 | 4.35926 | 1.37329 | Intron (ENSMUST00000001700.6/ENSMUSG00000001655.6, intron 1 of 1) | 15 | 102921103 | 102928814 | 7712 | 1 | ENSMUST00000001700.6 | 4629 | Hoxc13 | ENSMUSG00000001655.6 |
| chr15 | 102927536 | 102927756 | 3.95107 | 5.09399 | 1.85734 | 3' UTR | 15 | 102921103 | 102928814 | 7712 | 1 | ENSMUST00000001700.6 | 6434 | Hoxc13 | ENSMUSG00000001655.6 |
| chr15 | 102928087 | 102928349 | 3.7352 | 4.74432 | 1.574 | 3' UTR | 15 | 102921103 | 102928814 | 7712 | 1 | ENSMUST00000001700.6 | 6985 | Hoxc13 | ENSMUSG00000001655.6 |
| chr15 | 102928830 | 102928928 | 3.55022 | 4.30114 | 1.33615 | Distal Intergenic | 15 | 102921103 | 102928814 | 7712 | 1 | ENSMUST00000001700.6 | 7728 | Hoxc13 | ENSMUSG00000001655.6 |
| chr15 | 102952488 | 102952829 | 3.67088 | 4.52941 | 1.47209 | Distal Intergenic | 15 | 102954427 | 102957708 | 3282 | 1 | ENSMUST00000001701.3 | -1598 | Hoxc11 | ENSMUSG00000001656.3 |
| chr15 | 102953709 | 102953942 | 3.67799 | 4.58613 | 1.4921 | Promoter | 15 | 102954427 | 102957708 | 3282 | 1 | ENSMUST00000001701.3 | -485 | Hoxc11 | ENSMUSG00000001656.3 |
| chr15 | 102954781 | 102956298 | 3.95877 | 5.3983 | 2.09565 | Promoter | 15 | 102954427 | 102957708 | 3282 | 1 | ENSMUST00000001701.3 | 355 | Hoxc11 | ENSMUSG00000001656.3 |
| chr15 | 102956737 | 102957618 | 3.60062 | 4.40973 | 1.43942 | 3' UTR | 15 | 102954427 | 102957708 | 3282 | 1 | ENSMUST00000001701.3 | 2311 | Hoxc11 | ENSMUSG00000001656.3 |
| chr8 | 109946606 | 109947016 | 3.78698 | 4.86884 | 1.66203 | Downstream (<1kb) | 8 | 109959625 | 109962053 | 2429 | 2 | ENSMUST00000155052.1 | 15037 | Marveld3 | ENSMUSG00000001672.9 |
| chr8 | 109947944 | 109948666 | 3.47441 | 4.11045 | 1.23114 | 3' UTR | 8 | 109959625 | 109962053 | 2429 | 2 | ENSMUST00000155052.1 | 13387 | Marveld3 | ENSMUSG00000001672.9 |
| chr8 | 109949065 | 109949145 | 3.76443 | 4.74484 | 1.62153 | Intron (ENSMUST00000001722.8/ENSMUSG00000001672.9, intron 2 of 2) | 8 | 109959625 | 109962053 | 2429 | 2 | ENSMUST00000155052.1 | 12908 | Marveld3 | ENSMUSG00000001672.9 |
| chr8 | 109961659 | 109962724 | 3.93427 | 5.36864 | 2.05913 | Promoter | 8 | 109947914 | 109962168 | 14255 | 2 | ENSMUST00000001722.8 | 0 | Marveld3 | ENSMUSG00000001672.9 |
| chr8 | 109963213 | 109963529 | 3.21834 | 3.79841 | 1.02711 | Distal Intergenic | 8 | 109952911 | 109962203 | 9293 | 2 | ENSMUST00000051430.6 | -1011 | Marveld3 | ENSMUSG00000001672.9 |
| chr8 | 109964116 | 109964286 | 3.27107 | 3.75039 | 1.05764 | Distal Intergenic | 8 | 109952911 | 109962203 | 9293 | 2 | ENSMUST00000051430.6 | -1914 | Marveld3 | ENSMUSG00000001672.9 |
| chr8 | 109968718 | 109969725 | 3.86598 | 4.93056 | 1.72107 | Distal Intergenic | 8 | 109952911 | 109962203 | 9293 | 2 | ENSMUST00000051430.6 | -6516 | Marveld3 | ENSMUSG00000001672.9 |
| chr5 | 148478796 | 148478946 | 3.31044 | 3.79287 | 1.01733 | Distal Intergenic | 5 | 148504631 | 148552788 | 48158 | 2 | ENSMUST00000079324.8 | 73842 | Ubl3 | ENSMUSG00000001687.10 |
| chr2 | 145958753 | 145958935 | 3.44693 | 4.04464 | 1.17761 | Intron (ENSMUST00000116398.2/ENSMUSG00000037143.12, intron 6 of 6) | 2 | 145917479 | 145935014 | 17536 | 2 | ENSMUST00000001818.4 | -23740 | Crnkl1 | ENSMUSG00000001767.5 |
| chr7 | 35214816 | 35214892 | 3.55006 | 4.26072 | 1.35386 | Promoter | 7 | 35202016 | 35215292 | 13277 | 2 | ENSMUST00000118444.1 | 400 | Lrp3 | ENSMUSG00000001802.11 |
| chr2 | 74645725 | 74645929 | 3.60437 | 4.32518 | 1.31558 | Distal Intergenic | 2 | 74655616 | 74659419 | 3804 | 2 | ENSMUST00000173623.1 | 13490 | Evx2 | ENSMUSG00000001815.10 |
| chr2 | 74646292 | 74646931 | 3.78028 | 4.65063 | 1.51516 | Distal Intergenic | 2 | 74655616 | 74659419 | 3804 | 2 | ENSMUST00000173623.1 | 12488 | Evx2 | ENSMUSG00000001815.10 |
| chr2 | 74647675 | 74652717 | 3.97836 | 5.13043 | 1.86064 | Downstream (<1kb) | 2 | 74655616 | 74659419 | 3804 | 2 | ENSMUST00000173623.1 | 6702 | Evx2 | ENSMUSG00000001815.10 |
| chr2 | 74653123 | 74653367 | 3.37423 | 3.97923 | 1.12261 | 3' UTR | 2 | 74655616 | 74659419 | 3804 | 2 | ENSMUST00000173623.1 | 6052 | Evx2 | ENSMUSG00000001815.10 |
| chr2 | 74653806 | 74655178 | 3.74617 | 4.84581 | 1.66502 | 3' UTR | 2 | 74655616 | 74659419 | 3804 | 2 | ENSMUST00000173623.1 | 4241 | Evx2 | ENSMUSG00000001815.10 |
| chr2 | 74655493 | 74660053 | 4.01256 | 5.17818 | 1.92781 | Promoter | 2 | 74652991 | 74659557 | 6567 | 2 | ENSMUST00000001867.7 | 0 | Evx2 | ENSMUSG00000001815.10 |
| chr2 | 74660699 | 74662943 | 4.3036 | 5.76266 | 2.32219 | Distal Intergenic | 2 | 74652991 | 74659557 | 6567 | 2 | ENSMUST00000001867.7 | -1143 | Evx2 | ENSMUSG00000001815.10 |
| chr2 | 74663267 | 74663737 | 3.8209 | 5.0847 | 1.84267 | Distal Intergenic | 2 | 74652991 | 74659557 | 6567 | 2 | ENSMUST00000001867.7 | -3711 | Evx2 | ENSMUSG00000001815.10 |
| chr2 | 74664540 | 74666710 | 3.79639 | 4.8302 | 1.65778 | Distal Intergenic | 2 | 74668310 | 74671599 | 3290 | 1 | ENSMUST00000001872.4 | -1600 | Hoxd13 | ENSMUSG00000001819.4 |
| chr2 | 74667108 | 74668562 | 4.2881 | 5.79911 | 2.3843 | Promoter | 2 | 74668310 | 74671599 | 3290 | 1 | ENSMUST00000001872.4 | 0 | Hoxd13 | ENSMUSG00000001819.4 |
| chr2 | 74668978 | 74671256 | 3.80372 | 4.79974 | 1.63701 | Promoter | 2 | 74668310 | 74671599 | 3290 | 1 | ENSMUST00000001872.4 | 669 | Hoxd13 | ENSMUSG00000001819.4 |
| chr2 | 74672558 | 74673169 | 3.85204 | 4.90087 | 1.7189 | Distal Intergenic | 2 | 74675013 | 74677705 | 2693 | 1 | ENSMUST00000001878.5 | -1844 | Hoxd12 | ENSMUSG00000001823.5 |
| chr2 | 74673666 | 74674250 | 3.48219 | 4.12469 | 1.22618 | Promoter | 2 | 74675013 | 74677705 | 2693 | 1 | ENSMUST00000001878.5 | -763 | Hoxd12 | ENSMUSG00000001823.5 |
| chr2 | 74674599 | 74674829 | 4.04119 | 5.24646 | 1.96605 | Promoter | 2 | 74675013 | 74677705 | 2693 | 1 | ENSMUST00000001878.5 | -184 | Hoxd12 | ENSMUSG00000001823.5 |
| chr2 | 74675220 | 74675457 | 3.75583 | 4.81683 | 1.64938 | Promoter | 2 | 74675013 | 74677705 | 2693 | 1 | ENSMUST00000001878.5 | 208 | Hoxd12 | ENSMUSG00000001823.5 |
| chr2 | 74675812 | 74676104 | 3.72783 | 4.57787 | 1.52389 | Promoter | 2 | 74675013 | 74677705 | 2693 | 1 | ENSMUST00000001878.5 | 800 | Hoxd12 | ENSMUSG00000001823.5 |
| chr2 | 74676914 | 74677006 | 3.72051 | 4.52862 | 1.41986 | 3' UTR | 2 | 74675013 | 74677705 | 2693 | 1 | ENSMUST00000001878.5 | 1902 | Hoxd12 | ENSMUSG00000001823.5 |
| chr7 | 101622929 | 101623144 | 3.60773 | 4.52788 | 1.4581 | Distal Intergenic | 7 | 101663633 | 101722922 | 59290 | 1 | ENSMUST00000137214.2 | -40489 | Clpb | ENSMUSG00000001829.11 |
| chr7 | 101623766 | 101624022 | 3.5938 | 4.36738 | 1.39477 | Distal Intergenic | 7 | 101663633 | 101722922 | 59290 | 1 | ENSMUST00000137214.2 | -39611 | Clpb | ENSMUSG00000001829.11 |
| chr7 | 101624427 | 101624612 | 3.47417 | 4.32863 | 1.34475 | Distal Intergenic | 7 | 101663633 | 101722922 | 59290 | 1 | ENSMUST00000137214.2 | -39021 | Clpb | ENSMUSG00000001829.11 |
| chr11 | 106018760 | 106018933 | 3.52277 | 4.17963 | 1.23512 | Exon (ENSMUST00000106903.2/ENSMUSG00000001901.10, exon 6 of 14) | 11 | 106013773 | 106017473 | 3701 | 1 | ENSMUST00000140695.1 | 4988 | Kcnh6 | ENSMUSG00000001901.10 |
| chr11 | 106019658 | 106019891 | 3.38193 | 3.9006 | 1.01733 | Intron (ENSMUST00000106903.2/ENSMUSG00000001901.10, intron 7 of 13) | 11 | 106013773 | 106017473 | 3701 | 1 | ENSMUST00000140695.1 | 5886 | Kcnh6 | ENSMUSG00000001901.10 |
| chr11 | 106020225 | 106020907 | 3.54739 | 4.26273 | 1.27498 | Exon (ENSMUST00000106903.2/ENSMUSG00000001901.10, exon 8 of 14) | 11 | 106013773 | 106017473 | 3701 | 1 | ENSMUST00000140695.1 | 6453 | Kcnh6 | ENSMUSG00000001901.10 |
| chr8 | 84705154 | 84707297 | 4.22581 | 5.82413 | 2.41137 | Promoter | 8 | 84704712 | 84707716 | 3005 | 2 | ENSMUST00000148746.1 | 419 | Nfix | ENSMUSG00000001911.11 |
| chr8 | 84707654 | 84708353 | 3.95053 | 5.35296 | 2.02347 | Promoter | 8 | 84699876 | 84708218 | 8343 | 2 | ENSMUST00000148644.1 | 0 | Nfix | ENSMUSG00000001911.11 |
| chr8 | 84708828 | 84708942 | 3.85703 | 5.02685 | 1.79511 | Promoter | 8 | 84699876 | 84708218 | 8343 | 2 | ENSMUST00000148644.1 | -611 | Nfix | ENSMUSG00000001911.11 |
| chr8 | 84715748 | 84716174 | 3.34633 | 3.87904 | 1.12094 | Exon (ENSMUST00000109764.2/ENSMUSG00000001911.11, exon 9 of 11) | 8 | 84708634 | 84714068 | 5435 | 2 | ENSMUST00000132236.1 | -1681 | Nfix | ENSMUSG00000001911.11 |
| chr8 | 84773320 | 84773434 | 4.01864 | 5.16343 | 1.93822 | Promoter | 8 | 84704712 | 84773396 | 68685 | 2 | ENSMUST00000109764.2 | 0 | Nfix | ENSMUSG00000001911.11 |
| chr8 | 84774671 | 84775327 | 3.86257 | 4.88876 | 1.74499 | Promoter | 8 | 84708626 | 84774369 | 65744 | 2 | ENSMUST00000109762.2 | -303 | Nfix | ENSMUSG00000001911.11 |
| chr8 | 84775742 | 84775845 | 3.3263 | 3.84359 | 1.1033 | Intron (ENSMUST00000076715.7/ENSMUSG00000001911.11, intron 1 of 8) | 8 | 84708626 | 84774369 | 65744 | 2 | ENSMUST00000109762.2 | -1374 | Nfix | ENSMUSG00000001911.11 |
| chr8 | 84776276 | 84776519 | 3.35332 | 3.89932 | 1.14875 | Intron (ENSMUST00000076715.7/ENSMUSG00000001911.11, intron 1 of 8) | 8 | 84708626 | 84774369 | 65744 | 2 | ENSMUST00000109762.2 | -1908 | Nfix | ENSMUSG00000001911.11 |
| chr8 | 84779866 | 84780140 | 3.52612 | 4.26944 | 1.36111 | Intron (ENSMUST00000076715.7/ENSMUSG00000001911.11, intron 1 of 8) | 8 | 84708626 | 84774369 | 65744 | 2 | ENSMUST00000109762.2 | -5498 | Nfix | ENSMUSG00000001911.11 |
| chr8 | 84782876 | 84783118 | 3.71386 | 4.53743 | 1.47803 | Intron (ENSMUST00000076715.7/ENSMUSG00000001911.11, intron 1 of 8) | 8 | 84708626 | 84774369 | 65744 | 2 | ENSMUST00000109762.2 | -8508 | Nfix | ENSMUSG00000001911.11 |
| chr8 | 84784022 | 84785145 | 3.98869 | 5.10145 | 1.83783 | Intron (ENSMUST00000076715.7/ENSMUSG00000001911.11, intron 1 of 8) | 8 | 84708626 | 84774369 | 65744 | 2 | ENSMUST00000109762.2 | -9654 | Nfix | ENSMUSG00000001911.11 |
| chr8 | 84785594 | 84785979 | 3.38359 | 3.91043 | 1.04325 | Intron (ENSMUST00000076715.7/ENSMUSG00000001911.11, intron 1 of 8) | 8 | 84708626 | 84774369 | 65744 | 2 | ENSMUST00000109762.2 | -11226 | Nfix | ENSMUSG00000001911.11 |
| chr8 | 84786689 | 84787313 | 3.76883 | 4.76507 | 1.57391 | Intron (ENSMUST00000076715.7/ENSMUSG00000001911.11, intron 1 of 8) | 8 | 84708626 | 84774369 | 65744 | 2 | ENSMUST00000109762.2 | -12321 | Nfix | ENSMUSG00000001911.11 |
| chr8 | 84790646 | 84791222 | 3.43278 | 4.0118 | 1.13979 | Intron (ENSMUST00000076715.7/ENSMUSG00000001911.11, intron 1 of 8) | 8 | 84713795 | 84800024 | 86230 | 2 | ENSMUST00000126806.1 | 8802 | Nfix | ENSMUSG00000001911.11 |
| chr6 | 125590076 | 125590151 | 3.6235 | 4.35909 | 1.35104 | Intron (ENSMUST00000100941.3/ENSMUSG00000001930.12, intron 8 of 10) | 6 | 125603550 | 125642924 | 39375 | 1 | ENSMUST00000134073.1 | -13399 | Vwf | ENSMUSG00000001930.12 |
| chr9 | 37526717 | 37526893 | 3.22541 | 3.66532 | 1.01674 | Distal Intergenic | 9 | 37528078 | 37538319 | 10242 | 1 | ENSMUST00000002011.8 | -1185 | Esam | ENSMUSG00000001946.8 |
| chr9 | 37527206 | 37527341 | 3.62006 | 4.50795 | 1.50667 | Promoter | 9 | 37528078 | 37538319 | 10242 | 1 | ENSMUST00000002011.8 | -737 | Esam | ENSMUSG00000001946.8 |
| chr9 | 37530578 | 37531060 | 3.61339 | 4.44953 | 1.43837 | Promoter | 9 | 37530924 | 37534199 | 3276 | 1 | ENSMUST00000123198.1 | 0 | Esam | ENSMUSG00000001946.8 |
| chr4 | 125391206 | 125391900 | 3.99124 | 5.06903 | 1.80132 | Distal Intergenic | 4 | 125490700 | 125714173 | 223474 | 1 | ENSMUST00000030676.7 | -98800 | Grik3 | ENSMUSG00000001985.9 |
| chr4 | 125421314 | 125421866 | 3.47069 | 4.07319 | 1.15051 | Distal Intergenic | 4 | 125490700 | 125714173 | 223474 | 1 | ENSMUST00000030676.7 | -68834 | Grik3 | ENSMUSG00000001985.9 |
| chr4 | 125478190 | 125478422 | 3.62554 | 4.35293 | 1.31651 | Distal Intergenic | 4 | 125490700 | 125714173 | 223474 | 1 | ENSMUST00000030676.7 | -12278 | Grik3 | ENSMUSG00000001985.9 |
| chr4 | 125479356 | 125479815 | 3.4403 | 4.01526 | 1.11105 | Distal Intergenic | 4 | 125490700 | 125714173 | 223474 | 1 | ENSMUST00000030676.7 | -10885 | Grik3 | ENSMUSG00000001985.9 |
| chr4 | 125481433 | 125481984 | 3.63822 | 4.40354 | 1.37071 | Distal Intergenic | 4 | 125490700 | 125714173 | 223474 | 1 | ENSMUST00000030676.7 | -8716 | Grik3 | ENSMUSG00000001985.9 |
| chr4 | 125484359 | 125484994 | 3.79416 | 4.78594 | 1.61493 | Distal Intergenic | 4 | 125490700 | 125714173 | 223474 | 1 | ENSMUST00000030676.7 | -5706 | Grik3 | ENSMUSG00000001985.9 |
| chr4 | 125486062 | 125486156 | 3.90171 | 4.86872 | 1.64017 | Distal Intergenic | 4 | 125490700 | 125714173 | 223474 | 1 | ENSMUST00000030676.7 | -4544 | Grik3 | ENSMUSG00000001985.9 |
| chr4 | 125487533 | 125487628 | 4.5948 | 6.25948 | 2.63457 | Distal Intergenic | 4 | 125490700 | 125714173 | 223474 | 1 | ENSMUST00000030676.7 | -3072 | Grik3 | ENSMUSG00000001985.9 |
| chr4 | 125488956 | 125489031 | 3.83967 | 4.7974 | 1.6648 | Distal Intergenic | 4 | 125490700 | 125714173 | 223474 | 1 | ENSMUST00000030676.7 | -1669 | Grik3 | ENSMUSG00000001985.9 |
| chr4 | 125489771 | 125491794 | 3.93187 | 5.0498 | 1.79735 | Promoter | 4 | 125490700 | 125714173 | 223474 | 1 | ENSMUST00000030676.7 | 0 | Grik3 | ENSMUSG00000001985.9 |
| chr4 | 125492749 | 125493399 | 3.81578 | 4.81246 | 1.65772 | Intron (ENSMUST00000030676.7/ENSMUSG00000001985.9, intron 1 of 15) | 4 | 125490700 | 125714173 | 223474 | 1 | ENSMUST00000030676.7 | 2050 | Grik3 | ENSMUSG00000001985.9 |
| chr4 | 125494027 | 125494102 | 3.5844 | 4.54797 | 1.47975 | Intron (ENSMUST00000030676.7/ENSMUSG00000001985.9, intron 1 of 15) | 4 | 125490700 | 125714173 | 223474 | 1 | ENSMUST00000030676.7 | 3328 | Grik3 | ENSMUSG00000001985.9 |
| chr4 | 125494970 | 125495199 | 3.45951 | 4.11869 | 1.17107 | Intron (ENSMUST00000030676.7/ENSMUSG00000001985.9, intron 1 of 15) | 4 | 125490700 | 125714173 | 223474 | 1 | ENSMUST00000030676.7 | 4271 | Grik3 | ENSMUSG00000001985.9 |
| chr4 | 125495805 | 125496109 | 3.35707 | 3.8631 | 1.01733 | Intron (ENSMUST00000030676.7/ENSMUSG00000001985.9, intron 1 of 15) | 4 | 125490700 | 125714173 | 223474 | 1 | ENSMUST00000030676.7 | 5106 | Grik3 | ENSMUSG00000001985.9 |
| chr4 | 125497202 | 125497510 | 3.59159 | 4.31209 | 1.30327 | Intron (ENSMUST00000030676.7/ENSMUSG00000001985.9, intron 1 of 15) | 4 | 125490700 | 125714173 | 223474 | 1 | ENSMUST00000030676.7 | 6503 | Grik3 | ENSMUSG00000001985.9 |
| chr7 | 16474644 | 16474853 | 4.10134 | 5.42796 | 2.03225 | Exon (ENSMUST00000002053.8/ENSMUSG00000001988.8, exon 3 of 12) | 7 | 16455721 | 16476780 | 21060 | 2 | ENSMUST00000002053.8 | 1927 | Npas1 | ENSMUSG00000001988.8 |
| chr7 | 16475404 | 16475934 | 3.77863 | 4.73544 | 1.6068 | Promoter | 7 | 16455721 | 16476780 | 21060 | 2 | ENSMUST00000002053.8 | 846 | Npas1 | ENSMUSG00000001988.8 |
| chr7 | 16476312 | 16476966 | 3.88025 | 5.06053 | 1.81853 | Promoter | 7 | 16455721 | 16476780 | 21060 | 2 | ENSMUST00000002053.8 | 0 | Npas1 | ENSMUSG00000001988.8 |
| chr12 | 84791597 | 84791791 | 3.37772 | 3.90067 | 1.04005 | Promoter | 12 | 84785206 | 84791843 | 6638 | 2 | ENSMUST00000166383.1 | 52 | Ltbp2 | ENSMUSG00000002020.10 |
| chr12 | 84875861 | 84877904 | 4.2579 | 5.7126 | 2.36051 | Promoter | 12 | 84783212 | 84876532 | 93321 | 2 | ENSMUST00000002073.7 | 0 | Ltbp2 | ENSMUSG00000002020.10 |
| chr9 | 44970469 | 44970632 | 3.6333 | 4.40903 | 1.44097 | Promoter | 9 | 44969754 | 44970820 | 1067 | 2 | ENSMUST00000159019.1 | 188 | Cd3g | ENSMUSG00000002033.8 |
| chr9 | 44977225 | 44977333 | 3.40421 | 3.96675 | 1.13277 | Intron (ENSMUST00000002101.6/ENSMUSG00000002033.8, intron 1 of 6) | 9 | 44969572 | 44980431 | 10860 | 2 | ENSMUST00000002101.6 | 3098 | Cd3g | ENSMUSG00000002033.8 |
| chr9 | 44978646 | 44978789 | 3.28233 | 3.75331 | 1.02659 | Intron (ENSMUST00000002101.6/ENSMUSG00000002033.8, intron 1 of 6) | 9 | 44969572 | 44980431 | 10860 | 2 | ENSMUST00000002101.6 | 1642 | Cd3g | ENSMUSG00000002033.8 |
| chr8 | 83389295 | 83390064 | 3.84703 | 5.00216 | 1.78281 | Promoter | 8 | 83389867 | 83428552 | 38686 | 1 | ENSMUST00000002259.7 | 0 | Clgn | ENSMUSG00000002190.8 |
| chr17 | 25802168 | 25802340 | 3.7566 | 4.61232 | 1.50311 | Intron (ENSMUST00000180868.1/ENSMUSG00000097518.1, intron 1 of 1) | 17 | 25794949 | 25797136 | 2188 | 2 | ENSMUST00000002344.6 | -5033 | Metrn | ENSMUSG00000002274.7 |
| chr8 | 70111921 | 70112459 | 3.60729 | 4.34228 | 1.31011 | Exon (ENSMUST00000002412.8/ENSMUSG00000002341.8, exon 6 of 15) | 8 | 70093085 | 70120873 | 27789 | 2 | ENSMUST00000002412.8 | 8414 | Ncan | ENSMUSG00000002341.8 |
| chr8 | 70112798 | 70113604 | 3.58799 | 4.33349 | 1.33793 | Exon (ENSMUST00000002412.8/ENSMUSG00000002341.8, exon 4 of 15) | 8 | 70093085 | 70120873 | 27789 | 2 | ENSMUST00000002412.8 | 7269 | Ncan | ENSMUSG00000002341.8 |
| chr8 | 70114440 | 70114774 | 3.52629 | 4.20427 | 1.30867 | Intron (ENSMUST00000002412.8/ENSMUSG00000002341.8, intron 3 of 14) | 8 | 70093085 | 70120873 | 27789 | 2 | ENSMUST00000002412.8 | 6099 | Ncan | ENSMUSG00000002341.8 |
| chr8 | 70116726 | 70116901 | 3.33171 | 4.02772 | 1.17866 | Intron (ENSMUST00000002412.8/ENSMUSG00000002341.8, intron 1 of 14) | 8 | 70093085 | 70120873 | 27789 | 2 | ENSMUST00000002412.8 | 3972 | Ncan | ENSMUSG00000002341.8 |
| chr8 | 70118204 | 70118503 | 3.75119 | 4.66493 | 1.5819 | Intron (ENSMUST00000002412.8/ENSMUSG00000002341.8, intron 1 of 14) | 8 | 70093085 | 70120873 | 27789 | 2 | ENSMUST00000002412.8 | 2370 | Ncan | ENSMUSG00000002341.8 |
| chr8 | 70119205 | 70119331 | 3.28553 | 3.91671 | 1.08069 | Intron (ENSMUST00000002412.8/ENSMUSG00000002341.8, intron 1 of 14) | 8 | 70093085 | 70120873 | 27789 | 2 | ENSMUST00000002412.8 | 1542 | Ncan | ENSMUSG00000002341.8 |
| chr8 | 70119785 | 70120521 | 3.48161 | 4.16842 | 1.29707 | Promoter | 8 | 70093085 | 70120873 | 27789 | 2 | ENSMUST00000002412.8 | 352 | Ncan | ENSMUSG00000002341.8 |
| chr8 | 70121018 | 70121171 | 3.33197 | 3.8746 | 1.12039 | Promoter | 8 | 70093085 | 70120873 | 27789 | 2 | ENSMUST00000002412.8 | -146 | Ncan | ENSMUSG00000002341.8 |
| chr8 | 70166053 | 70166220 | 3.50056 | 4.2025 | 1.22872 | Exon (ENSMUST00000163756.1/ENSMUSG00000002345.12, exon 5 of 9) | 8 | 70172356 | 70180930 | 8575 | 1 | ENSMUST00000125755.2 | -6136 | Tmem161a | ENSMUSG00000002342.12 |
| chr4 | 123103291 | 123103378 | 4.09701 | 5.3494 | 2.01331 | Distal Intergenic | 4 | 123105165 | 123126091 | 20927 | 1 | ENSMUST00000002457.1 | -1787 | Bmp8b | ENSMUSG00000002384.2 |
| chr4 | 123104562 | 123105913 | 3.80837 | 4.83407 | 1.6718 | Promoter | 4 | 123105165 | 123126091 | 20927 | 1 | ENSMUST00000002457.1 | 0 | Bmp8b | ENSMUSG00000002384.2 |
| chr1 | 5013019 | 5013146 | 3.54143 | 4.26813 | 1.32976 | Intron (ENSMUST00000002533.9/ENSMUSG00000002459.12, intron 1 of 4) | 1 | 4910402 | 5018735 | 108334 | 2 | ENSMUST00000119256.2 | 5589 | Rgs20 | ENSMUSG00000002459.12 |
| chr1 | 5016844 | 5019043 | 3.69843 | 4.53805 | 1.4711 | Promoter | 1 | 4910402 | 5018735 | 108334 | 2 | ENSMUST00000119256.2 | 0 | Rgs20 | ENSMUSG00000002459.12 |
| chr1 | 5019513 | 5019694 | 3.99233 | 5.05106 | 1.78887 | Promoter | 1 | 4909576 | 5019539 | 109964 | 2 | ENSMUST00000002533.9 | 0 | Rgs20 | ENSMUSG00000002459.12 |
| chr1 | 5021010 | 5021941 | 4.03137 | 5.17481 | 1.89447 | Exon (ENSMUST00000118000.2/ENSMUSG00000002459.12, exon 2 of 6) | 1 | 4909576 | 5019539 | 109964 | 2 | ENSMUST00000002533.9 | -1472 | Rgs20 | ENSMUSG00000002459.12 |
| chr1 | 5023504 | 5023765 | 3.43554 | 4.01911 | 1.15081 | Intron (ENSMUST00000118000.2/ENSMUSG00000002459.12, intron 1 of 5) | 1 | 4909576 | 5019539 | 109964 | 2 | ENSMUST00000002533.9 | -3966 | Rgs20 | ENSMUSG00000002459.12 |
| chr1 | 5024600 | 5025211 | 3.60181 | 4.43915 | 1.42918 | Intron (ENSMUST00000118000.2/ENSMUSG00000002459.12, intron 1 of 5) | 1 | 4909576 | 5019539 | 109964 | 2 | ENSMUST00000002533.9 | -5062 | Rgs20 | ENSMUSG00000002459.12 |
| chr16 | 89796201 | 89796702 | 3.56439 | 4.24176 | 1.24947 | Promoter | 16 | 89788422 | 89795282 | 6861 | 2 | ENSMUST00000178095.1 | -920 | Tiam1 | ENSMUSG00000002489.10 |
| chr16 | 89805997 | 89806088 | 3.65886 | 4.44622 | 1.41957 | Intron (ENSMUST00000114122.2/ENSMUSG00000002489.10, intron 11 of 18) | 16 | 89791685 | 89799990 | 8306 | 2 | ENSMUST00000134021.1 | -6008 | Tiam1 | ENSMUSG00000002489.10 |
| chr16 | 89817845 | 89819054 | 3.6163 | 4.48097 | 1.43596 | Promoter | 16 | 89787730 | 89818352 | 30623 | 2 | ENSMUST00000164263.3 | 0 | Tiam1 | ENSMUSG00000002489.10 |
| chr16 | 89871087 | 89871547 | 3.6308 | 4.40222 | 1.41161 | Intron (ENSMUST00000002588.4/ENSMUSG00000002489.10, intron 6 of 27) | 16 | 89787111 | 89857817 | 70707 | 2 | ENSMUST00000114122.2 | -13271 | Tiam1 | ENSMUSG00000002489.10 |
| chr16 | 89898422 | 89898515 | 4.50363 | 6.10111 | 2.5625 | Exon (ENSMUST00000002588.4/ENSMUSG00000002489.10, exon 4 of 28) | 16 | 89898421 | 89918985 | 20565 | 2 | ENSMUST00000139544.1 | 20470 | Tiam1 | ENSMUSG00000002489.10 |
| chr16 | 89933785 | 89934038 | 3.4393 | 4.13293 | 1.21911 | Intron (ENSMUST00000002588.4/ENSMUSG00000002489.10, intron 2 of 27) | 16 | 89898421 | 89918985 | 20565 | 2 | ENSMUST00000139544.1 | -14801 | Tiam1 | ENSMUSG00000002489.10 |
| chr16 | 89945855 | 89946008 | 3.49448 | 4.1163 | 1.17983 | Intron (ENSMUST00000002588.4/ENSMUSG00000002489.10, intron 2 of 27) | 16 | 89908221 | 89956107 | 47887 | 2 | ENSMUST00000124433.2 | 10099 | Tiam1 | ENSMUSG00000002489.10 |
| chr16 | 90072588 | 90072843 | 3.46606 | 4.26788 | 1.31088 | Distal Intergenic | 16 | 89955965 | 89980080 | 24116 | 2 | ENSMUST00000136760.1 | -92509 | Tiam1 | ENSMUSG00000002489.10 |
| chr16 | 90093645 | 90093983 | 3.37637 | 3.89519 | 1.0278 | Distal Intergenic | 16 | 89955965 | 89980080 | 24116 | 2 | ENSMUST00000136760.1 | -113566 | Tiam1 | ENSMUSG00000002489.10 |
| chr16 | 90095313 | 90095397 | 4.0221 | 5.14342 | 1.88728 | Distal Intergenic | 16 | 89955965 | 89980080 | 24116 | 2 | ENSMUST00000136760.1 | -115234 | Tiam1 | ENSMUSG00000002489.10 |
| chr12 | 40131976 | 40132108 | 3.67178 | 4.50056 | 1.47815 | Intron (ENSMUST00000078481.8/ENSMUSG00000002565.11, intron 1 of 13) | 12 | 40059769 | 40134183 | 74415 | 2 | ENSMUST00000078481.8 | 2075 | Scin | ENSMUSG00000002565.11 |
| chr12 | 40133808 | 40134712 | 4.37126 | 6.04927 | 2.54336 | Promoter | 12 | 40059769 | 40134183 | 74415 | 2 | ENSMUST00000078481.8 | 0 | Scin | ENSMUSG00000002565.11 |
| chr6 | 5178021 | 5178585 | 3.33009 | 3.83062 | 1.04619 | Intron (ENSMUST00000002663.6/ENSMUSG00000002588.7, intron 5 of 8) | 6 | 5185549 | 5193763 | 8215 | 2 | ENSMUST00000176933.1 | 15178 | Pon1 | ENSMUSG00000002588.7 |
| chr5 | 28467745 | 28468083 | 3.45795 | 4.0516 | 1.14587 | Promoter | 5 | 28456815 | 28467101 | 10287 | 2 | ENSMUST00000002708.3 | -645 | Shh | ENSMUSG00000002633.3 |
| chr5 | 28469380 | 28469517 | 3.38193 | 3.9006 | 1.01733 | Intron (ENSMUST00000180878.2/ENSMUSG00000097726.2, intron 1 of 6) | 5 | 28456815 | 28467101 | 10287 | 2 | ENSMUST00000002708.3 | -2280 | Shh | ENSMUSG00000002633.3 |
| chr5 | 28472294 | 28472580 | 3.64757 | 4.3961 | 1.34057 | Intron (ENSMUST00000180878.2/ENSMUSG00000097726.2, intron 1 of 6) | 5 | 28456815 | 28467101 | 10287 | 2 | ENSMUST00000002708.3 | -5194 | Shh | ENSMUSG00000002633.3 |
| chr7 | 45857885 | 45858234 | 3.63281 | 4.49565 | 1.45079 | Exon (ENSMUST00000002848.7/ENSMUSG00000002771.10, exon 4 of 12) | 7 | 45831883 | 45867129 | 35247 | 2 | ENSMUST00000002848.7 | 8895 | Grin2d | ENSMUSG00000002771.10 |
| chr7 | 45867948 | 45868280 | 3.39087 | 3.92543 | 1.05588 | Promoter | 7 | 45831883 | 45867129 | 35247 | 2 | ENSMUST00000002848.7 | -820 | Grin2d | ENSMUSG00000002771.10 |
| chr7 | 45870724 | 45871206 | 4.16205 | 5.42468 | 2.03696 | Promoter | 7 | 45866377 | 45870928 | 4552 | 2 | ENSMUST00000146672.1 | 0 | Grin2d | ENSMUSG00000002771.10 |
| chr7 | 45871523 | 45871778 | 3.55106 | 4.26653 | 1.27288 | Promoter | 7 | 45866377 | 45870928 | 4552 | 2 | ENSMUST00000146672.1 | -596 | Grin2d | ENSMUSG00000002771.10 |
| chr12 | 112928361 | 112928627 | 3.46709 | 4.25822 | 1.30532 | Promoter | 12 | 112908590 | 112929495 | 20906 | 2 | ENSMUST00000075827.3 | 868 | Jag2 | ENSMUSG00000002799.5 |
| chr13 | 19619278 | 19619415 | 3.47228 | 4.10707 | 1.25404 | Promoter | 13 | 19591709 | 19619830 | 28122 | 2 | ENSMUST00000002885.6 | 415 | Epdr1 | ENSMUSG00000002808.6 |
| chr13 | 19619788 | 19619863 | 3.53709 | 4.55242 | 1.47595 | Promoter | 13 | 19591709 | 19619830 | 28122 | 2 | ENSMUST00000002885.6 | 0 | Epdr1 | ENSMUSG00000002808.6 |
| chr12 | 31264677 | 31265327 | 3.71071 | 4.54538 | 1.47349 | Promoter | 12 | 31265234 | 31329493 | 64260 | 1 | ENSMUST00000169088.2 | 0 | Lamb1 | ENSMUSG00000002900.10 |
| chr12 | 31265707 | 31265898 | 4.22001 | 5.75317 | 2.32915 | Promoter | 12 | 31265400 | 31329575 | 64176 | 1 | ENSMUST00000164919.1 | 308 | Lamb1 | ENSMUSG00000002900.10 |
| chr12 | 31268290 | 31268690 | 3.38264 | 3.90752 | 1.03787 | Exon (ENSMUST00000164228.1/ENSMUSG00000002900.10, exon 2 of 2) | 12 | 31265400 | 31329575 | 64176 | 1 | ENSMUST00000164919.1 | 2891 | Lamb1 | ENSMUSG00000002900.10 |
| chr12 | 31272433 | 31272586 | 4.3039 | 5.76113 | 2.36044 | Exon (ENSMUST00000169088.2/ENSMUSG00000002900.10, exon 5 of 34) | 12 | 31276656 | 31287986 | 11331 | 1 | ENSMUST00000169065.1 | -4070 | Lamb1 | ENSMUSG00000002900.10 |
| chr12 | 31296190 | 31296266 | 3.64544 | 4.80218 | 1.64117 | Intron (ENSMUST00000169088.2/ENSMUSG00000002900.10, intron 13 of 33) | 12 | 31278175 | 31278939 | 765 | 1 | ENSMUST00000164009.1 | 18016 | Lamb1 | ENSMUSG00000002900.10 |
| chr12 | 31296637 | 31297710 | 3.68227 | 4.82647 | 1.65936 | Exon (ENSMUST00000169088.2/ENSMUSG00000002900.10, exon 14 of 34) | 12 | 31278175 | 31278939 | 765 | 1 | ENSMUST00000164009.1 | 18463 | Lamb1 | ENSMUSG00000002900.10 |
| chr12 | 31298170 | 31298342 | 3.72381 | 4.716 | 1.55929 | Intron (ENSMUST00000169088.2/ENSMUSG00000002900.10, intron 14 of 33) | 12 | 31278175 | 31278939 | 765 | 1 | ENSMUST00000164009.1 | 19996 | Lamb1 | ENSMUSG00000002900.10 |
| chr12 | 31307035 | 31307203 | 3.54535 | 4.35071 | 1.37302 | Exon (ENSMUST00000169088.2/ENSMUSG00000002900.10, exon 23 of 34) | 12 | 31323521 | 31324454 | 934 | 1 | ENSMUST00000172134.1 | -16318 | Lamb1 | ENSMUSG00000002900.10 |
| chr7 | 19698968 | 19699332 | 3.69405 | 4.5469 | 1.50437 | Promoter | 7 | 19696109 | 19699166 | 3058 | 2 | ENSMUST00000174064.2 | 0 | Apoe | ENSMUSG00000002985.10 |
| chr12 | 32057896 | 32058616 | 3.54218 | 4.24387 | 1.3342 | Intron (ENSMUST00000003079.6/ENSMUSG00000002997.10, intron 1 of 11) | 12 | 31958480 | 32060589 | 102110 | 2 | ENSMUST00000146865.1 | 1973 | Prkar2b | ENSMUSG00000002997.10 |
| chr12 | 32059280 | 32059598 | 3.56895 | 4.51521 | 1.46856 | Promoter | 12 | 31958480 | 32060589 | 102110 | 2 | ENSMUST00000146865.1 | 991 | Prkar2b | ENSMUSG00000002997.10 |
| chr12 | 32060725 | 32061033 | 3.84611 | 5.36266 | 2.01939 | Promoter | 12 | 31958480 | 32060589 | 102110 | 2 | ENSMUST00000146865.1 | -137 | Prkar2b | ENSMUSG00000002997.10 |
| chr12 | 32061350 | 32061638 | 3.56833 | 4.35609 | 1.36539 | Promoter | 12 | 31958476 | 32061296 | 102821 | 2 | ENSMUST00000036497.10 | -55 | Prkar2b | ENSMUSG00000002997.10 |
| chr12 | 32127733 | 32127837 | 3.41521 | 4.29621 | 1.32534 | Distal Intergenic | 12 | 31958476 | 32061296 | 102821 | 2 | ENSMUST00000036497.10 | -66438 | Prkar2b | ENSMUSG00000002997.10 |
| chr10 | 80189261 | 80189424 | 3.71655 | 4.66264 | 1.53741 | 3' UTR | 10 | 80179482 | 80190010 | 10529 | 1 | ENSMUST00000003154.6 | 9780 | Efna2 | ENSMUSG00000003070.6 |
| chr10 | 80190664 | 80191219 | 3.82808 | 4.72993 | 1.54371 | Distal Intergenic | 10 | 80179482 | 80190010 | 10529 | 1 | ENSMUST00000003154.6 | 11183 | Efna2 | ENSMUSG00000003070.6 |
| chr6 | 122740563 | 122740662 | 3.56092 | 4.31737 | 1.34545 | Intron (ENSMUST00000032476.5/ENSMUSG00000003153.5, intron 1 of 9) | 6 | 122739833 | 122742262 | 2430 | 2 | ENSMUST00000170724.2 | 1600 | Slc2a3 | ENSMUSG00000003153.5 |
| chr6 | 122743935 | 122744030 | 4.04608 | 5.61825 | 2.19751 | Intron (ENSMUST00000165884.2/ENSMUSG00000003153.5, intron 2 of 4) | 6 | 122739892 | 122742920 | 3029 | 2 | ENSMUST00000168801.1 | -1016 | Slc2a3 | ENSMUSG00000003153.5 |
| chr6 | 122760991 | 122761280 | 3.2309 | 3.67355 | 1.01268 | Intron (ENSMUST00000165884.2/ENSMUSG00000003153.5, intron 2 of 4) | 6 | 122739892 | 122742920 | 3029 | 2 | ENSMUST00000168801.1 | -18072 | Slc2a3 | ENSMUSG00000003153.5 |
| chr6 | 122761884 | 122762095 | 3.2825 | 3.75089 | 1.01733 | Intron (ENSMUST00000165884.2/ENSMUSG00000003153.5, intron 2 of 4) | 6 | 122737278 | 122779957 | 42680 | 2 | ENSMUST00000171541.1 | 17862 | Slc2a3 | ENSMUSG00000003153.5 |
| chr6 | 122763100 | 122763517 | 3.39469 | 3.93422 | 1.06818 | Intron (ENSMUST00000165884.2/ENSMUSG00000003153.5, intron 2 of 4) | 6 | 122737278 | 122779957 | 42680 | 2 | ENSMUST00000171541.1 | 16440 | Slc2a3 | ENSMUSG00000003153.5 |
| chr6 | 122763851 | 122764104 | 3.44016 | 4.06239 | 1.21434 | Intron (ENSMUST00000165884.2/ENSMUSG00000003153.5, intron 2 of 4) | 6 | 122737278 | 122779957 | 42680 | 2 | ENSMUST00000171541.1 | 15853 | Slc2a3 | ENSMUSG00000003153.5 |
| chr6 | 122764517 | 122765080 | 3.82789 | 5.01551 | 1.78309 | Intron (ENSMUST00000165884.2/ENSMUSG00000003153.5, intron 2 of 4) | 6 | 122737278 | 122779957 | 42680 | 2 | ENSMUST00000171541.1 | 14877 | Slc2a3 | ENSMUSG00000003153.5 |
| chr6 | 122765702 | 122765946 | 3.71531 | 4.51809 | 1.41187 | Intron (ENSMUST00000165884.2/ENSMUSG00000003153.5, intron 2 of 4) | 6 | 122737278 | 122779957 | 42680 | 2 | ENSMUST00000171541.1 | 14011 | Slc2a3 | ENSMUSG00000003153.5 |
| chr10 | 58650258 | 58650346 | 3.94442 | 4.95292 | 1.70631 | Intron (ENSMUST00000003312.4/ENSMUSG00000003227.4, intron 1 of 11) | 10 | 58600789 | 58675654 | 74866 | 2 | ENSMUST00000003312.4 | 25308 | Edar | ENSMUSG00000003227.4 |

| | | | | | | | | | | | | | | | |
|---|---|---|---|---|---|---|---|---|---|---|---|---|---|---|---|
| chr7 | 45741491 | 45741776 | 3.23106 | 3.81773 | 1.04421 | Promoter | 7 | 45730000 | 45742483 | 12484 | 2 | ENSMUST00000107735.3 | 707 | Sult2b1 | ENSMUSG00000003271.10 |
| chr17 | 69968947 | 69969321 | 5.03053 | 7.39834 | 3.54805 | Promoter | 17 | 69969073 | 70821413 | 852341 | 1 | ENSMUST00000133983.2 | 0 | Dlgap1 | ENSMUSG00000003279.11 |
| chr17 | 69969663 | 69970211 | 3.94856 | 5.15955 | 1.88801 | Promoter | 17 | 69969421 | 70594408 | 624988 | 1 | ENSMUST00000130971.1 | 243 | Dlgap1 | ENSMUSG00000003279.11 |
| chr17 | 69970520 | 69970845 | 4.50699 | 6.40271 | 2.83545 | Intron (ENSMUST00000133983.2/ENSMUSG00000003279.11, intron 1 of 13) | 17 | 69969421 | 70594408 | 624988 | 1 | ENSMUST00000130971.1 | 1100 | Dlgap1 | ENSMUSG00000003279.11 |
| chr17 | 69971351 | 69971515 | 3.46675 | 4.27732 | 1.30456 | Intron (ENSMUST00000133983.2/ENSMUSG00000003279.11, intron 1 of 13) | 17 | 69969421 | 70594408 | 624988 | 1 | ENSMUST00000130971.1 | 1931 | Dlgap1 | ENSMUSG00000003279.11 |
| chr17 | 69971889 | 69972281 | 3.57974 | 4.58706 | 1.49127 | Intron (ENSMUST00000133983.2/ENSMUSG00000003279.11, intron 1 of 13) | 17 | 69969421 | 70594408 | 624988 | 1 | ENSMUST00000130971.1 | 2469 | Dlgap1 | ENSMUSG00000003279.11 |
| chr4 | 3930853 | 3931131 | 3.68707 | 4.52735 | 1.4938 | Intron (ENSMUST00000003369.4/ENSMUSG00000003282.4, intron 1 of 5) | 4 | 3900996 | 3938382 | 37387 | 2 | ENSMUST00000003369.4 | 7251 | Plag1 | ENSMUSG00000003282.4 |
| chr4 | 3931906 | 3932171 | 3.94106 | 5.29629 | 1.98718 | Intron (ENSMUST00000003369.4/ENSMUSG00000003282.4, intron 1 of 5) | 4 | 3900996 | 3938382 | 37387 | 2 | ENSMUST00000003369.4 | 6211 | Plag1 | ENSMUSG00000003282.4 |
| chr4 | 3932650 | 3932743 | 3.78452 | 4.76528 | 1.65112 | Intron (ENSMUST00000003369.4/ENSMUSG00000003282.4, intron 1 of 5) | 4 | 3900996 | 3938382 | 37387 | 2 | ENSMUST00000003369.4 | 5639 | Plag1 | ENSMUSG00000003282.4 |
| chr4 | 3933251 | 3934086 | 3.88739 | 4.90381 | 1.75987 | Intron (ENSMUST00000003369.4/ENSMUSG00000003282.4, intron 1 of 5) | 4 | 3900996 | 3938382 | 37387 | 2 | ENSMUST00000003369.4 | 4296 | Plag1 | ENSMUSG00000003282.4 |
| chr4 | 3935013 | 3935118 | 3.25799 | 3.71412 | 1.01733 | Intron (ENSMUST00000003369.4/ENSMUSG00000003282.4, intron 1 of 5) | 4 | 3900996 | 3938382 | 37387 | 2 | ENSMUST00000003369.4 | 3264 | Plag1 | ENSMUSG00000003282.4 |
| chr4 | 3936642 | 3936770 | 4.29627 | 5.82862 | 2.41376 | Intron (ENSMUST00000003369.4/ENSMUSG00000003282.4, intron 1 of 5) | 4 | 3900996 | 3938382 | 37387 | 2 | ENSMUST00000003369.4 | 1612 | Plag1 | ENSMUSG00000003282.4 |
| chr2 | 153107817 | 153108044 | 3.75471 | 4.62168 | 1.53553 | Promoter | 2 | 153108468 | 153151439 | 42972 | 1 | ENSMUST00000189688.1 | -424 | Hck | ENSMUSG00000003283.8 |
| chr2 | 153108623 | 153109564 | 3.56314 | 4.35618 | 1.40164 | Promoter | 2 | 153108468 | 153151441 | 42974 | 1 | ENSMUST00000109799.2 | 156 | Hck | ENSMUSG00000003283.8 |
| chr2 | 153110779 | 153110975 | 3.37404 | 3.96205 | 1.1544 | Intron (ENSMUST00000189688.1/ENSMUSG00000003283.8, intron 1 of 12) | 2 | 153108468 | 153151441 | 42974 | 1 | ENSMUST00000109799.2 | 2312 | Hck | ENSMUSG00000003283.8 |
| chr9 | 21242950 | 21243111 | 3.66605 | 4.45282 | 1.43038 | 3' UTR | 9 | 21229730 | 21239332 | 9603 | 2 | ENSMUST00000164812.2 | -3619 | Keap1 | ENSMUSG00000003308.9 |
| chr9 | 21312227 | 21312405 | 3.77085 | 4.92715 | 1.71577 | Promoter | 9 | 21295457 | 21312333 | 16877 | 2 | ENSMUST00000003397.7 | 0 | Ap1m2 | ENSMUSG00000003309.8 |
| chr9 | 21312930 | 21313008 | 3.86965 | 4.88906 | 1.71697 | Promoter | 9 | 21295457 | 21312333 | 16877 | 2 | ENSMUST00000115433.4 | -598 | Ap1m2 | ENSMUSG00000003309.8 |
| chr15 | 98622496 | 98622612 | 3.37701 | 3.92985 | 1.12453 | Distal Intergenic | 15 | 98632220 | 98644530 | 12311 | 1 | ENSMUST00000109150.2 | -9608 | Cacnb3 | ENSMUSG00000003352.8 |
| chr15 | 98629216 | 98629445 | 3.94855 | 5.13972 | 1.85959 | Distal Intergenic | 15 | 98632220 | 98644530 | 12311 | 1 | ENSMUST00000109150.2 | -2775 | Cacnb3 | ENSMUSG00000003352.8 |
| chr15 | 98630720 | 98631789 | 3.54876 | 4.22857 | 1.27933 | Promoter | 15 | 98632220 | 98644530 | 12311 | 1 | ENSMUST00000109150.2 | -431 | Cacnb3 | ENSMUSG00000003352.8 |
| chr15 | 98632633 | 98632983 | 4.22088 | 5.84316 | 2.35691 | Promoter | 15 | 98632220 | 98644530 | 12311 | 1 | ENSMUST00000109150.2 | 414 | Cacnb3 | ENSMUSG00000003352.8 |
| chr15 | 98633374 | 98633684 | 3.49201 | 4.10774 | 1.16194 | Intron (ENSMUST00000109150.2/ENSMUSG00000003352.8, intron 1 of 12) | 15 | 98634764 | 98644530 | 9767 | 1 | ENSMUST00000003442.7 | -1080 | Cacnb3 | ENSMUSG00000003352.8 |
| chr15 | 98634020 | 98634285 | 3.34254 | 3.84118 | 1.01733 | Promoter | 15 | 98634764 | 98644530 | 9767 | 1 | ENSMUST00000003442.7 | -479 | Cacnb3 | ENSMUSG00000003352.8 |
| chr3 | 87500341 | 87500536 | 3.30092 | 3.79295 | 1.06632 | Distal Intergenic | 3 | 87525407 | 87540156 | 14750 | 1 | ENSMUST00000119109.2 | -24871 | Etv3 | ENSMUSG00000003382.13 |
| chr3 | 87501842 | 87502094 | 3.7319 | 4.65555 | 1.58681 | Distal Intergenic | 3 | 87525407 | 87540156 | 14750 | 1 | ENSMUST00000119109.2 | -23313 | Etv3 | ENSMUSG00000003382.13 |
| chr3 | 87550171 | 87550308 | 3.29751 | 3.77344 | 1.01733 | Distal Intergenic | 3 | 87525618 | 87531034 | 5417 | 1 | ENSMUST00000152010.1 | 24554 | Etv3 | ENSMUSG00000003382.13 |
| chr9 | 22050413 | 22051183 | 3.92997 | 5.18137 | 1.89459 | Promoter | 9 | 22015005 | 22052023 | 37019 | 2 | ENSMUST00000003501.7 | 840 | Elavl3 | ENSMUSG00000003410.7 |
| chr9 | 22051579 | 22052755 | 4.28155 | 5.7362 | 2.299 | Promoter | 9 | 22015005 | 22052023 | 37019 | 2 | ENSMUST00000003501.7 | 0 | Elavl3 | ENSMUSG00000003410.7 |
| chr9 | 22053501 | 22054757 | 3.52443 | 4.17033 | 1.20992 | Downstream (<1kb) | 9 | 22015005 | 22052023 | 37019 | 2 | ENSMUST00000003501.7 | -1479 | Elavl3 | ENSMUSG00000003410.7 |
| chr2 | 13792742 | 13793739 | 4.19622 | 5.49612 | 2.10804 | Promoter | 2 | 13657154 | 13793293 | 136140 | 2 | ENSMUST00000150781.1 | 0 | St8sia6 | ENSMUSG00000003418.10 |
| chr7 | 28292520 | 28293469 | 3.77112 | 4.64993 | 1.50853 | Downstream (<1kb) | 7 | 28293599 | 28294481 | 883 | 2 | ENSMUST00000138721.1 | 1012 | Dll3 | ENSMUSG00000003436.11 |
| chr7 | 28294017 | 28295007 | 4.30991 | 5.78549 | 2.31848 | Promoter | 7 | 28293599 | 28294481 | 883 | 2 | ENSMUST00000138721.1 | 0 | Dll3 | ENSMUSG00000003436.11 |
| chr7 | 28298177 | 28298598 | 3.44996 | 4.03165 | 1.11773 | Exon (ENSMUST00000108315.3/ENSMUSG00000003436.11, exon 5 of 9) | 7 | 28298672 | 28301772 | 3101 | 2 | ENSMUST00000145512.1 | 3174 | Dll3 | ENSMUSG00000003436.11 |
| chr7 | 28298939 | 28299100 | 3.40741 | 4.23389 | 1.24614 | Exon (ENSMUST00000108315.3/ENSMUSG00000003436.11, exon 4 of 9) | 7 | 28298672 | 28301772 | 3101 | 2 | ENSMUST00000145512.1 | 2672 | Dll3 | ENSMUSG00000003436.11 |
| chr7 | 28301681 | 28301842 | 3.69845 | 4.5283 | 1.4534 | Promoter | 7 | 28298672 | 28301772 | 3101 | 2 | ENSMUST00000145512.1 | 0 | Dll3 | ENSMUSG00000003436.11 |
| chr14 | 70456137 | 70456608 | 3.65528 | 4.46724 | 1.47074 | Promoter | 14 | 70457476 | 70468832 | 11357 | 1 | ENSMUST00000003561.4 | -868 | Phyhip | ENSMUSG00000003469.4 |
| chr14 | 70467217 | 70467355 | 3.46737 | 4.05883 | 1.12037 | 3' UTR | 14 | 70458523 | 70463340 | 4818 | 1 | ENSMUST00000159180.1 | 8695 | Phyhip | ENSMUSG00000003469.4 |
| chr6 | 55088055 | 55088375 | 3.50518 | 4.16349 | 1.28272 | Downstream (1-2kb) | 6 | 55104743 | 55117771 | 13029 | 2 | ENSMUST00000169777.1 | 29396 | Crhr2 | ENSMUSG00000003476.10 |
| chr6 | 55091364 | 55091720 | 3.94395 | 4.97964 | 1.74884 | 3' UTR | 6 | 55104743 | 55117771 | 13029 | 2 | ENSMUST00000169777.1 | 26051 | Crhr2 | ENSMUSG00000003476.10 |
| chr6 | 55098238 | 55098553 | 3.51276 | 4.14139 | 1.16988 | Intron (ENSMUST00000003568.9/ENSMUSG00000003476.10, intron 9 of 12) | 6 | 55104743 | 55117771 | 13029 | 2 | ENSMUST00000169777.1 | 19218 | Crhr2 | ENSMUSG00000003476.10 |
| chr6 | 55107616 | 55107737 | 3.48428 | 4.09899 | 1.17306 | Intron (ENSMUST00000003568.9/ENSMUSG00000003476.10, intron 3 of 12) | 6 | 55104743 | 55117771 | 13029 | 2 | ENSMUST00000169777.1 | 10034 | Crhr2 | ENSMUSG00000003476.10 |
| chr6 | 55110475 | 55110877 | 3.4322 | 4.15869 | 1.25876 | Intron (ENSMUST00000003568.9/ENSMUSG00000003476.10, intron 3 of 12) | 6 | 55104743 | 55117771 | 13029 | 2 | ENSMUST00000169777.1 | 6894 | Crhr2 | ENSMUSG00000003476.10 |
| chr6 | 55111754 | 55111829 | 4.08721 | 5.22214 | 1.87432 | Intron (ENSMUST00000003568.9/ENSMUSG00000003476.10, intron 3 of 12) | 6 | 55104743 | 55117771 | 13029 | 2 | ENSMUST00000169777.1 | 5942 | Crhr2 | ENSMUSG00000003476.10 |
| chr6 | 55114997 | 55115189 | 3.63473 | 4.38589 | 1.37965 | Intron (ENSMUST00000003568.9/ENSMUSG00000003476.10, intron 3 of 12) | 6 | 55104743 | 55117771 | 13029 | 2 | ENSMUST00000169777.1 | 2582 | Crhr2 | ENSMUSG00000003476.10 |
| chr6 | 55115508 | 55115697 | 3.454 | 4.05844 | 1.19188 | Intron (ENSMUST00000003568.9/ENSMUSG00000003476.10, intron 3 of 12) | 6 | 55104743 | 55117771 | 13029 | 2 | ENSMUST00000169777.1 | 2074 | Crhr2 | ENSMUSG00000003476.10 |
| chr6 | 55116353 | 55116687 | 3.46463 | 4.32254 | 1.32199 | Intron (ENSMUST00000003568.9/ENSMUSG00000003476.10, intron 3 of 12) | 6 | 55104743 | 55117771 | 13029 | 2 | ENSMUST00000169777.1 | 1084 | Crhr2 | ENSMUSG00000003476.10 |
| chr6 | 55117422 | 55118411 | 3.86325 | 4.90691 | 1.75822 | Promoter | 6 | 55091186 | 55117985 | 26800 | 2 | ENSMUST00000095898.4 | 0 | Crhr2 | ENSMUSG00000003476.10 |
| chr7 | 19342058 | 19342364 | 4.00275 | 5.21181 | 1.91832 | Distal Intergenic | 7 | 19344778 | 19356524 | 11747 | 1 | ENSMUST00000003645.3 | -2414 | Ercc1 | ENSMUSG00000003549.4 |
| chr11 | 4119656 | 4119960 | 3.43877 | 4.00423 | 1.08015 | Promoter | 11 | 4097039 | 4118831 | 21793 | 2 | ENSMUST00000003681.7 | -826 | Sec14l2 | ENSMUSG00000003585.8 |
| chr8 | 110147190 | 110147301 | 4.62884 | 6.34385 | 2.71839 | Exon (ENSMUST00000003754.6/ENSMUSG00000003657.8, exon 9 of 11) | 8 | 110142539 | 110168206 | 25668 | 2 | ENSMUST00000003754.6 | 20905 | Calb2 | ENSMUSG00000003657.8 |
| chr8 | 110167223 | 110167334 | 3.63697 | 4.45615 | 1.41832 | Promoter | 8 | 110142539 | 110168206 | 25668 | 2 | ENSMUST00000003754.6 | 872 | Calb2 | ENSMUSG00000003657.8 |
| chr8 | 110184318 | 110184635 | 3.3013 | 3.77914 | 1.01733 | Distal Intergenic | 8 | 110142539 | 110168206 | 25668 | 2 | ENSMUST00000003754.6 | -16113 | Calb2 | ENSMUSG00000003657.8 |
| chr17 | 17850155 | 17850389 | 3.88726 | 5.18288 | 1.89342 | Exon (ENSMUST00000003762.6/ENSMUSG00000003665.6, exon 2 of 5) | 17 | 17843328 | 17855188 | 11861 | 2 | ENSMUST00000003762.6 | 4799 | Has1 | ENSMUSG00000003665.6 |
| chr1 | 121274528 | 121275018 | 3.50629 | 4.1924 | 1.32483 | Distal Intergenic | 1 | 121305893 | 121307318 | 1426 | 2 | ENSMUST00000159528.1 | 32300 | Insig2 | ENSMUSG00000003721.9 |
| chr1 | 121283397 | 121283567 | 3.26218 | 3.72042 | 1.01733 | Distal Intergenic | 1 | 121305893 | 121307318 | 1426 | 2 | ENSMUST00000159528.1 | 23751 | Insig2 | ENSMUSG00000003721.9 |
| chr8 | 84917699 | 84918047 | 3.51191 | 4.18238 | 1.28665 | Exon (ENSMUST00000109741.3/ENSMUSG00000053693.11, exon 20 of 26) | 8 | 84913391 | 84922915 | 9525 | 1 | ENSMUST00000148573.1 | 4309 | Dnase2a | ENSMUSG00000003812.8 |
| chr7 | 45355852 | 45355927 | 4.23544 | 5.5481 | 2.16749 | Intron (ENSMUST00000107779.2/ENSMUSG00000003863.11, intron 10 of 29) | 7 | 45339275 | 45360113 | 20839 | 2 | ENSMUST00000137578.1 | 4186 | Ppfia3 | ENSMUSG00000003863.11 |
| chr7 | 45356416 | 45356656 | 3.89007 | 4.91294 | 1.77298 | Exon (ENSMUST00000107779.2/ENSMUSG00000003863.11, exon 8 of 30) | 7 | 45339275 | 45360113 | 20839 | 2 | ENSMUST00000137578.1 | 3457 | Ppfia3 | ENSMUSG00000003863.11 |
| chr7 | 45360561 | 45360811 | 3.32485 | 4.06451 | 1.18811 | Promoter | 7 | 45339275 | 45360113 | 20839 | 2 | ENSMUST00000137578.1 | -449 | Ppfia3 | ENSMUSG00000003863.11 |
| chr7 | 45361788 | 45362122 | 3.38839 | 4.07135 | 1.19947 | 5' UTR | 7 | 45339275 | 45360113 | 20839 | 2 | ENSMUST00000137578.1 | -1676 | Ppfia3 | ENSMUSG00000003863.11 |
| chr7 | 45362475 | 45362725 | 3.82663 | 5.05865 | 1.83947 | Intron (ENSMUST00000107779.2/ENSMUSG00000003863.11, intron 1 of 29) | 7 | 45339275 | 45360113 | 20839 | 2 | ENSMUST00000137578.1 | -2363 | Ppfia3 | ENSMUSG00000003863.11 |
| chr11 | 69556838 | 69557559 | 3.56989 | 4.33588 | 1.37397 | Exon (ENSMUST00000004036.5/ENSMUSG00000003934.5, exon 2 of 5) | 11 | 69554092 | 69560205 | 6114 | 2 | ENSMUST00000004036.5 | 2646 | Efnb3 | ENSMUSG00000003934.5 |
| chr11 | 69559824 | 69560491 | 3.71682 | 4.52277 | 1.41245 | Promoter | 11 | 69554092 | 69560205 | 6114 | 2 | ENSMUST00000004036.5 | 0 | Efnb3 | ENSMUSG00000003934.5 |
| chr11 | 69561112 | 69561261 | 3.36987 | 3.88241 | 1.01733 | Promoter | 11 | 69554092 | 69560205 | 6114 | 2 | ENSMUST00000004036.5 | -908 | Efnb3 | ENSMUSG00000003934.5 |
| chr5 | 9725859 | 9726595 | 3.77846 | 4.84476 | 1.66918 | Promoter | 5 | 9485236 | 9725352 | 240117 | 2 | ENSMUST00000004076.3 | -508 | Grm3 | ENSMUSG00000003974.5 |
| chr5 | 9727213 | 9727407 | 3.96411 | 5.19282 | 1.91547 | Distal Intergenic | 5 | 9485236 | 9725352 | 240117 | 2 | ENSMUST00000004076.3 | -1862 | Grm3 | ENSMUSG00000003974.5 |
| chr1 | 158055015 | 158055090 | 3.57185 | 4.2912 | 1.36631 | Distal Intergenic | 1 | 158245271 | 158356260 | 110990 | 2 | ENSMUST00000004133.8 | 301170 | Brinp2 | ENSMUSG00000004031.8 |
| chr1 | 154705221 | 154705555 | 3.69291 | 4.69336 | 1.60656 | Intron (ENSMUST00000187541.1/ENSMUSG00000004110.9, intron 1 of 46) | 1 | 154392519 | 154725920 | 333402 | 2 | ENSMUST00000004214.9 | 20365 | Cacna1e | ENSMUSG00000004110.9 |
| chr1 | 154723226 | 154723940 | 3.86381 | 4.81825 | 1.62757 | Intron (ENSMUST00000187541.1/ENSMUSG00000004110.9, intron 1 of 46) | 1 | 154392519 | 154725920 | 333402 | 2 | ENSMUST00000004214.9 | 1980 | Cacna1e | ENSMUSG00000004110.9 |
| chr1 | 154724398 | 154725294 | 3.58199 | 4.2734 | 1.26332 | Promoter | 1 | 154392519 | 154725920 | 333402 | 2 | ENSMUST00000004214.9 | 626 | Cacna1e | ENSMUSG00000004110.9 |
| chr1 | 154725625 | 154726075 | 3.80243 | 4.71232 | 1.5926 | Promoter | 1 | 154392519 | 154725920 | 333402 | 2 | ENSMUST00000004214.9 | 0 | Cacna1e | ENSMUSG00000004110.9 |
| chr1 | 154726385 | 154727666 | 3.8327 | 4.77047 | 1.60698 | Promoter | 1 | 154390731 | 154726356 | 335626 | 2 | ENSMUST00000187541.1 | -30 | Cacna1e | ENSMUSG00000004110.9 |
| chr1 | 154784555 | 154784795 | 3.76925 | 4.71313 | 1.60022 | Intron (ENSMUST00000188965.1/ENSMUSG00000004110.9, intron 2 of 13) | 1 | 154481075 | 154816941 | 335867 | 2 | ENSMUST00000188965.1 | 32146 | Cacna1e | ENSMUSG00000004110.9 |
| chr1 | 154883275 | 154883813 | 3.36427 | 3.87397 | 1.01733 | Promoter | 1 | 154882166 | 154884024 | 1859 | 2 | ENSMUST00000186256.1 | 211 | Cacna1e | ENSMUSG00000004110.9 |
| chr1 | 154884325 | 154884898 | 3.4739 | 4.07297 | 1.13598 | Promoter | 1 | 154699089 | 154884501 | 185413 | 2 | ENSMUST00000188321.1 | 0 | Cacna1e | ENSMUSG00000004110.9 |
| chr2 | 24607822 | 24607955 | 3.68859 | 4.57869 | 1.49981 | Promoter | 2 | 24606379 | 24607534 | 1156 | 2 | ENSMUST00000131861.1 | -289 | Cacna1b | ENSMUSG00000004113.13 |
| chr2 | 24761752 | 24764017 | 4.3506 | 6.06938 | 2.62969 | Promoter | 2 | 24603887 | 24763152 | 159266 | 2 | ENSMUST00000070864.8 | 0 | Cacna1b | ENSMUSG00000004113.13 |
| chr2 | 24768329 | 24768645 | 3.35979 | 3.87097 | 1.03068 | Distal Intergenic | 2 | 24603887 | 24763152 | 159266 | 2 | ENSMUST00000070864.8 | -5178 | Cacna1b | ENSMUSG00000004113.13 |
| chr12 | 38780522 | 38780671 | 3.54917 | 4.25618 | 1.34042 | Promoter | 12 | 38780349 | 38835258 | 54910 | 1 | ENSMUST00000160996.2 | 174 | Etv1 | ENSMUSG00000004151.11 |
| chr12 | 38781867 | 38782216 | 3.36053 | 4.07391 | 1.20923 | Promoter | 12 | 38781129 | 38835170 | 54042 | 1 | ENSMUST00000161513.2 | 739 | Etv1 | ENSMUSG00000004151.11 |
| chr19 | 44726009 | 44726470 | 3.44035 | 4.20632 | 1.26988 | Distal Intergenic | 19 | 44756045 | 44837851 | 81807 | 1 | ENSMUST00000174490.3 | -29575 | Pax2 | ENSMUSG00000004231.10 |
| chr19 | 44732984 | 44735376 | 3.89694 | 4.95231 | 1.74372 | Distal Intergenic | 19 | 44756045 | 44837851 | 81807 | 1 | ENSMUST00000174490.3 | -20669 | Pax2 | ENSMUSG00000004231.10 |
| chr19 | 44740093 | 44740405 | 3.42599 | 4.02421 | 1.12205 | Distal Intergenic | 19 | 44756045 | 44837851 | 81807 | 1 | ENSMUST00000174490.3 | -15640 | Pax2 | ENSMUSG00000004231.10 |
| chr19 | 44741047 | 44741349 | 4.44277 | 6.04139 | 2.56205 | Distal Intergenic | 19 | 44756045 | 44837851 | 81807 | 1 | ENSMUST00000174490.3 | -14696 | Pax2 | ENSMUSG00000004231.10 |
| chr19 | 44742251 | 44742593 | 3.64379 | 4.76245 | 1.6073 | Distal Intergenic | 19 | 44756045 | 44837851 | 81807 | 1 | ENSMUST00000174490.3 | -13452 | Pax2 | ENSMUSG00000004231.10 |
| chr19 | 44744188 | 44744328 | 3.96845 | 5.24264 | 1.92439 | Distal Intergenic | 19 | 44756045 | 44837851 | 81807 | 1 | ENSMUST00000174490.3 | -11717 | Pax2 | ENSMUSG00000004231.10 |
| chr19 | 44744979 | 44745137 | 3.51391 | 4.16832 | 1.25082 | Distal Intergenic | 19 | 44756045 | 44837851 | 81807 | 1 | ENSMUST00000174490.3 | -10908 | Pax2 | ENSMUSG00000004231.10 |
| chr19 | 44746594 | 44747054 | 3.89373 | 5.11215 | 1.84446 | Distal Intergenic | 19 | 44756045 | 44837851 | 81807 | 1 | ENSMUST00000174490.3 | -8991 | Pax2 | ENSMUSG00000004231.10 |
| chr19 | 44748325 | 44749288 | 3.68812 | 4.51358 | 1.50664 | Distal Intergenic | 19 | 44756045 | 44837851 | 81807 | 1 | ENSMUST00000174490.3 | -6757 | Pax2 | ENSMUSG00000004231.10 |
| chr19 | 44749654 | 44749827 | 4.13884 | 5.42114 | 2.07069 | Distal Intergenic | 19 | 44756045 | 44837851 | 81807 | 1 | ENSMUST00000174490.3 | -6218 | Pax2 | ENSMUSG00000004231.10 |
| chr19 | 44750211 | 44751237 | 3.78781 | 4.92421 | 1.73312 | Distal Intergenic | 19 | 44756045 | 44837851 | 81807 | 1 | ENSMUST00000174490.3 | -4808 | Pax2 | ENSMUSG00000004231.10 |
| chr19 | 44751696 | 44752916 | 4.37895 | 6.22888 | 2.70394 | Distal Intergenic | 19 | 44756045 | 44837851 | 81807 | 1 | ENSMUST00000174490.3 | -3129 | Pax2 | ENSMUSG00000004231.10 |
| chr19 | 44753463 | 44753676 | 3.6657 | 4.58859 | 1.51251 | Distal Intergenic | 19 | 44756045 | 44837851 | 81807 | 1 | ENSMUST00000174490.3 | -2369 | Pax2 | ENSMUSG00000004231.10 |
| chr19 | 44754921 | 44757629 | 3.792 | 4.82307 | 1.66128 | Promoter | 19 | 44756045 | 44837851 | 81807 | 1 | ENSMUST00000174490.3 | 0 | Pax2 | ENSMUSG00000004231.10 |
| chr19 | 44757999 | 44758443 | 4.26138 | 5.58087 | 2.14177 | Promoter | 19 | 44757394 | 44837871 | 80478 | 1 | ENSMUST00000004340.5 | 606 | Pax2 | ENSMUSG00000004231.10 |
| chr19 | 44758882 | 44759195 | 3.372 | 3.91485 | 1.07216 | Promoter | 19 | 44759915 | 44818554 | 58640 | 1 | ENSMUST00000173346.3 | -720 | Pax2 | ENSMUSG00000004231.10 |
| chr19 | 44759581 | 44762011 | 3.83347 | 4.87469 | 1.68996 | Promoter | 19 | 44759915 | 44818554 | 58640 | 1 | ENSMUST00000173346.3 | 0 | Pax2 | ENSMUSG00000004231.10 |
| chr6 | 124763917 | 124764021 | 3.56207 | 4.23908 | 1.25262 | Promoter | 6 | 124762657 | 124763921 | 1265 | 2 | ENSMUST00000128598.1 | 0 | Eno2 | ENSMUSG00000004267.11 |
| chr7 | 17056309 | 17056472 | 3.49992 | 4.17734 | 1.3041 | Promoter | 7 | 17035734 | 17056679 | 20946 | 2 | ENSMUST00000037762.5 | 207 | Hif3a | ENSMUSG00000004328.10 |
| chr7 | 17062483 | 17062717 | 3.79735 | 4.73292 | 1.57576 | Promoter | 7 | 17031507 | 17062427 | 30921 | 2 | ENSMUST00000108492.3 | -57 | Hif3a | ENSMUSG00000004328.10 |
| chr13 | 21321843 | 21322034 | 3.69915 | 4.61323 | 1.52034 | Distal Intergenic | 13 | 21317258 | 21319624 | 2367 | 1 | ENSMUST00000136668.1 | 4586 | Gpx6 | ENSMUSG00000004341.7 |
| chr13 | 21243301 | 21244280 | 3.74143 | 4.59294 | 1.51641 | Distal Intergenic | 13 | 21286429 | 21292686 | 6258 | 2 | ENSMUST00000004456.4 | 48406 | Gpx5 | ENSMUSG00000004344.6 |
| chr13 | 21245387 | 21245508 | 3.32865 | 3.86937 | 1.10871 | Distal Intergenic | 13 | 21286429 | 21292686 | 6258 | 2 | ENSMUST00000004456.4 | 47178 | Gpx5 | ENSMUSG00000004344.6 |
| chr13 | 21253409 | 21254256 | 4.03502 | 5.28536 | 1.98298 | Distal Intergenic | 13 | 21286429 | 21292686 | 6258 | 2 | ENSMUST00000004456.4 | 38430 | Gpx5 | ENSMUSG00000004344.6 |
| chr13 | 21256332 | 21256599 | 3.60191 | 4.31464 | 1.30665 | Distal Intergenic | 13 | 21286429 | 21292686 | 6258 | 2 | ENSMUST00000004456.4 | 36087 | Gpx5 | ENSMUSG00000004344.6 |
| chr13 | 21257140 | 21257434 | 3.55385 | 4.40549 | 1.41028 | Distal Intergenic | 13 | 21286429 | 21292686 | 6258 | 2 | ENSMUST00000004456.4 | 35252 | Gpx5 | ENSMUSG00000004344.6 |
| chr13 | 21257861 | 21258528 | 3.4368 | 4.01827 | 1.14105 | Distal Intergenic | 13 | 21286429 | 21292686 | 6258 | 2 | ENSMUST00000004456.4 | 34158 | Gpx5 | ENSMUSG00000004344.6 |
| chr13 | 21259857 | 21260223 | 3.95173 | 5.11959 | 1.90082 | Distal Intergenic | 13 | 21286429 | 21292686 | 6258 | 2 | ENSMUST00000004456.4 | 32463 | Gpx5 | ENSMUSG00000004344.6 |
| chr13 | 21260784 | 21260984 | 4.68654 | 6.48949 | 2.8624 | Distal Intergenic | 13 | 21286429 | 21292686 | 6258 | 2 | ENSMUST00000004456.4 | 31702 | Gpx5 | ENSMUSG00000004344.6 |
| chr13 | 21261333 | 21261408 | 4.07037 | 5.20601 | 1.9036 | Distal Intergenic | 13 | 21286429 | 21292686 | 6258 | 2 | ENSMUST00000004456.4 | 31278 | Gpx5 | ENSMUSG00000004344.6 |
| chr13 | 21264007 | 21264575 | 3.42972 | 4.07329 | 1.20811 | Distal Intergenic | 13 | 21286429 | 21292686 | 6258 | 2 | ENSMUST00000004456.4 | 28111 | Gpx5 | ENSMUSG00000004344.6 |
| chr13 | 21266230 | 21266856 | 3.67249 | 4.53181 | 1.48376 | Distal Intergenic | 13 | 21286429 | 21292686 | 6258 | 2 | ENSMUST00000004456.4 | 25830 | Gpx5 | ENSMUSG00000004344.6 |
| chr13 | 21267652 | 21268155 | 3.69463 | 4.57454 | 1.54377 | Distal Intergenic | 13 | 21286429 | 21292686 | 6258 | 2 | ENSMUST00000004456.4 | 24531 | Gpx5 | ENSMUSG00000004344.6 |
| chr13 | 21273456 | 21273624 | 3.38519 | 4.03821 | 1.15189 | Distal Intergenic | 13 | 21286429 | 21292686 | 6258 | 2 | ENSMUST00000004456.4 | 19062 | Gpx5 | ENSMUSG00000004344.6 |
| chr13 | 21275157 | 21275256 | 3.66578 | 4.4291 | 1.3602 | Distal Intergenic | 13 | 21286429 | 21292686 | 6258 | 2 | ENSMUST00000004456.4 | 17430 | Gpx5 | ENSMUSG00000004344.6 |
| chr13 | 21275897 | 21276387 | 4.09881 | 5.39462 | 2.1278 | Distal Intergenic | 13 | 21286429 | 21292686 | 6258 | 2 | ENSMUST00000004456.4 | 16299 | Gpx5 | ENSMUSG00000004344.6 |
| chr13 | 21276884 | 21278552 | 3.87892 | 4.88645 | 1.72726 | Distal Intergenic | 13 | 21286429 | 21292686 | 6258 | 2 | ENSMUST00000004456.4 | 14134 | Gpx5 | ENSMUSG00000004344.6 |
| chr6 | 56361883 | 56362372 | 3.85182 | 4.90212 | 1.72823 | Promoter | 6 | 56069821 | 56362392 | 292572 | 2 | ENSMUST00000044505.8 | 20 | Pde1c | ENSMUSG00000004347.11 |
| chr16 | 23889732 | 23890236 | 3.71684 | 4.54018 | 1.46601 | Promoter | 16 | 23889581 | 23890844 | 1264 | 2 | ENSMUST00000004480.3 | 608 | Sst | ENSMUSG00000004366.3 |
| chr16 | 23897896 | 23898234 | 3.35738 | 4.28452 | 1.30826 | Distal Intergenic | 16 | 23889581 | 23890844 | 1264 | 2 | ENSMUST00000004480.3 | -7053 | Sst | ENSMUSG00000004366.3 |
| chr7 | 4777563 | 4777733 | 3.76259 | 4.61032 | 1.48352 | Promoter | 7 | 4773055 | 4778141 | 5087 | 2 | ENSMUST00000094892.6 | 408 | Il11 | ENSMUSG00000004371.9 |
| chr14 | 51913651 | 51913787 | 3.8547 | 4.96107 | 1.78183 | Promoter | 14 | 51905271 | 51913488 | 8218 | 2 | ENSMUST00000111632.3 | -164 | Ndrg2 | ENSMUSG00000004558.9 |
| chr6 | 4723665 | 4723891 | 3.43349 | 4.17034 | 1.22033 | Intron (ENSMUST00000115579.2/ENSMUSG00000004631.10, intron 1 of 10) | 6 | 4674360 | 4730126 | 55767 | 2 | ENSMUST00000153284.2 | 6235 | Sgce | ENSMUSG00000004631.10 |
| chr6 | 4738029 | 4738430 | 3.36736 | 4.12273 | 1.22811 | Intron (ENSMUST00000115579.2/ENSMUSG00000004631.10, intron 1 of 10) | 6 | 4674360 | 4730126 | 55767 | 2 | ENSMUST00000153284.2 | -7904 | Sgce | ENSMUSG00000004631.10 |
| chr6 | 4738753 | 4738868 | 3.6293 | 4.58643 | 1.48631 | Intron (ENSMUST00000115579.2/ENSMUSG00000004631.10, intron 1 of 10) | 6 | 4674418 | 4747019 | 72602 | 2 | ENSMUST00000126151.2 | 8151 | Sgce | ENSMUSG00000004631.10 |
| chr6 | 4739898 | 4740160 | 3.56158 | 4.40421 | 1.38835 | Intron (ENSMUST00000115579.2/ENSMUSG00000004631.10, intron 1 of 10) | 6 | 4674418 | 4747019 | 72602 | 2 | ENSMUST00000126151.2 | 6859 | Sgce | ENSMUSG00000004631.10 |
| chr6 | 54030362 | 54030542 | 3.51452 | 4.24604 | 1.27906 | Distal Intergenic | 6 | 54039554 | 54067798 | 28245 | 1 | ENSMUST00000138930.1 | -9012 | Chn2 | ENSMUSG00000004633.12 |
| chr6 | 54038799 | 54039170 | 3.86479 | 4.9347 | 1.77126 | Promoter | 6 | 54039554 | 54067798 | 28245 | 1 | ENSMUST00000138930.1 | -384 | Chn2 | ENSMUSG00000004633.12 |
| chr6 | 54039486 | 54040024 | 4.03788 | 5.23626 | 1.99982 | Promoter | 6 | 54039554 | 54067798 | 28245 | 1 | ENSMUST00000138930.1 | 0 | Chn2 | ENSMUSG00000004633.12 |
| chr8 | 114731787 | 114732180 | 3.85841 | 4.86105 | 1.66327 | Intron (ENSMUST00000004756.8/ENSMUSG00000004637.10, intron 8 of 8) | 8 | 114712167 | 115352143 | 639977 | 1 | ENSMUST00000162052.2 | 19621 | Wwox | ENSMUSG00000004637.10 |
| chr8 | 114848778 | 114848915 | 3.42385 | 4.10955 | 1.19715 | Intron (ENSMUST00000004756.8/ENSMUSG00000004637.10, intron 8 of 8) | 8 | 114712167 | 115352143 | 639977 | 1 | ENSMUST00000162052.2 | 136612 | Wwox | ENSMUSG00000004637.10 |
| chr6 | 55397189 | 55397404 | 3.38307 | 3.94794 | 1.16249 | Distal Intergenic | 6 | 55376295 | 55388530 | 12236 | 1 | ENSMUST00000063578.3 | 20895 | Ghrhr | ENSMUSG00000004654.4 |
| chr1 | 33714506 | 33715071 | 3.70449 | 4.53966 | 1.50645 | Distal Intergenic | 1 | 33719887 | 33726891 | 7005 | 1 | ENSMUST00000140354.2 | -4816 | Rab23 | ENSMUSG00000004768.9 |
| chr12 | 85177725 | 85177909 | 3.29362 | 3.7735 | 1.03787 | Promoter | 12 | 85166639 | 85177296 | 10658 | 2 | ENSMUST00000004913.6 | -430 | Pgf | ENSMUSG00000004791.6 |
| chr12 | 85182979 | 85183396 | 3.61453 | 4.33414 | 1.30084 | Distal Intergenic | 12 | 85166639 | 85177296 | 10658 | 2 | ENSMUST00000004913.6 | -5684 | Pgf | ENSMUSG00000004791.6 |
| chr17 | 28769145 | 28769629 | 4.1633 | 5.9209 | 2.42209 | Promoter | 17 | 28769307 | 28778698 | 9392 | 1 | ENSMUST00000004986.7 | 0 | Mapk13 | ENSMUSG00000004864.7 |
| chr17 | 28769961 | 28770902 | 3.74982 | 4.65628 | 1.59632 | Promoter | 17 | 28769372 | 28777562 | 8191 | 1 | ENSMUST00000133786.1 | 590 | Mapk13 | ENSMUSG00000004864.7 |
| chr1 | 78112936 | 78113234 | 3.61822 | 4.43465 | 1.43553 | Intron (ENSMUST00000004994.10/ENSMUSG00000004872.10, intron 7 of 8) | 1 | 78101267 | 78196838 | 95572 | 2 | ENSMUST00000004994.10 | 83604 | Pax3 | ENSMUSG00000004872.10 |
| chr1 | 78137576 | 78137696 | 3.5998 | 4.40254 | 1.40028 | Intron (ENSMUST00000004994.10/ENSMUSG00000004872.10, intron 4 of 8) | 1 | 78101267 | 78196838 | 95572 | 2 | ENSMUST00000004994.10 | 59142 | Pax3 | ENSMUSG00000004872.10 |
| chr1 | 78138644 | 78138727 | 3.54903 | 4.58816 | 1.49923 | Intron (ENSMUST00000004994.10/ENSMUSG00000004872.10, intron 4 of 8) | 1 | 78101267 | 78196838 | 95572 | 2 | ENSMUST00000004994.10 | 58111 | Pax3 | ENSMUSG00000004872.10 |
| chr1 | 78139318 | 78140190 | 3.874 | 5.04256 | 1.79873 | Intron (ENSMUST00000004994.10/ENSMUSG00000004872.10, intron 4 of 8) | 1 | 78101267 | 78196838 | 95572 | 2 | ENSMUST00000004994.10 | 56648 | Pax3 | ENSMUSG00000004872.10 |
| chr1 | 78140741 | 78141082 | 3.67449 | 4.95163 | 1.76794 | Intron (ENSMUST00000004994.10/ENSMUSG00000004872.10, intron 4 of 8) | 1 | 78101267 | 78196838 | 95572 | 2 | ENSMUST00000004994.10 | 55756 | Pax3 | ENSMUSG00000004872.10 |
| chr1 | 78142164 | 78142519 | 3.91748 | 5.07968 | 1.82764 | Intron (ENSMUST00000004994.10/ENSMUSG00000004872.10, intron 4 of 8) | 1 | 78101267 | 78196838 | 95572 | 2 | ENSMUST00000004994.10 | 54319 | Pax3 | ENSMUSG00000004872.10 |
| chr1 | 78145723 | 78145857 | 3.75603 | 4.86433 | 1.69703 | Intron (ENSMUST00000004994.10/ENSMUSG00000004872.10, intron 4 of 8) | 1 | 78101267 | 78196838 | 95572 | 2 | ENSMUST00000004994.10 | 50981 | Pax3 | ENSMUSG00000004872.10 |
| chr1 | 78146197 | 78146600 | 3.58225 | 4.28643 | 1.30117 | Intron (ENSMUST00000004994.10/ENSMUSG00000004872.10, intron 4 of 8) | 1 | 78101267 | 78196838 | 95572 | 2 | ENSMUST00000004994.10 | 50238 | Pax3 | ENSMUSG00000004872.10 |
| chr1 | 78147025 | 78147329 | 3.29688 | 3.77251 | 1.01733 | Intron (ENSMUST00000004994.10/ENSMUSG00000004872.10, intron 4 of 8) | 1 | 78101267 | 78196838 | 95572 | 2 | ENSMUST00000004994.10 | 49509 | Pax3 | ENSMUSG00000004872.10 |
| chr1 | 78147683 | 78147975 | 3.75879 | 4.67241 | 1.62616 | Intron (ENSMUST00000004994.10/ENSMUSG00000004872.10, intron 4 of 8) | 1 | 78101267 | 78196838 | 95572 | 2 | ENSMUST00000004994.10 | 48863 | Pax3 | ENSMUSG00000004872.10 |
| chr1 | 78148396 | 78148480 | 3.34061 | 3.8666 | 1.11218 | Intron (ENSMUST00000004994.10/ENSMUSG00000004872.10, intron 4 of 8) | 1 | 78101267 | 78196838 | 95572 | 2 | ENSMUST00000004994.10 | 48358 | Pax3 | ENSMUSG00000004872.10 |
| chr1 | 78153740 | 78154063 | 3.3951 | 3.96811 | 1.16909 | Intron (ENSMUST00000004994.10/ENSMUSG00000004872.10, intron 4 of 8) | 1 | 78101267 | 78196838 | 95572 | 2 | ENSMUST00000004994.10 | 42775 | Pax3 | ENSMUSG00000004872.10 |
| chr1 | 78156484 | 78156696 | 3.4607 | 4.0494 | 1.12261 | Intron (ENSMUST00000004994.10/ENSMUSG00000004872.10, intron 4 of 8) | 1 | 78101267 | 78196838 | 95572 | 2 | ENSMUST00000004994.10 | 40142 | Pax3 | ENSMUSG00000004872.10 |
| chr1 | 78161222 | 78161431 | 3.54389 | 4.20149 | 1.21167 | Intron (ENSMUST00000004994.10/ENSMUSG00000004872.10, intron 4 of 8) | 1 | 78101267 | 78196838 | 95572 | 2 | ENSMUST00000004994.10 | 35407 | Pax3 | ENSMUSG00000004872.10 |
| chr1 | 78162697 | 78162889 | 3.53334 | 4.23075 | 1.30366 | Intron (ENSMUST00000004994.10/ENSMUSG00000004872.10, intron 4 of 8) | 1 | 78101267 | 78196838 | 95572 | 2 | ENSMUST00000004994.10 | 33949 | Pax3 | ENSMUSG00000004872.10 |
| chr1 | 78168736 | 78169001 | 3.58983 | 4.53673 | 1.46936 | Intron (ENSMUST00000004994.10/ENSMUSG00000004872.10, intron 4 of 8) | 1 | 78101267 | 78196838 | 95572 | 2 | ENSMUST00000004994.10 | 27837 | Pax3 | ENSMUSG00000004872.10 |
| chr1 | 78169474 | 78169712 | 3.9328 | 5.06631 | 1.84921 | Intron (ENSMUST00000004994.10/ENSMUSG00000004872.10, intron 4 of 8) | 1 | 78101267 | 78196838 | 95572 | 2 | ENSMUST00000004994.10 | 27126 | Pax3 | ENSMUSG00000004872.10 |
| chr1 | 78170041 | 78170142 | 3.29803 | 3.77421 | 1.01733 | Intron (ENSMUST00000004994.10/ENSMUSG00000004872.10, intron 4 of 8) | 1 | 78101267 | 78196838 | 95572 | 2 | ENSMUST00000004994.10 | 26696 | Pax3 | ENSMUSG00000004872.10 |
| chr1 | 78170534 | 78171851 | 3.82185 | 4.9419 | 1.73199 | Intron (ENSMUST00000004994.10/ENSMUSG00000004872.10, intron 4 of 8) | 1 | 78101267 | 78196838 | 95572 | 2 | ENSMUST00000004994.10 | 24987 | Pax3 | ENSMUSG00000004872.10 |
| chr1 | 78172221 | 78172709 | 3.83394 | 4.83359 | 1.65514 | Intron (ENSMUST00000004994.10/ENSMUSG00000004872.10, intron 4 of 8) | 1 | 78101267 | 78196838 | 95572 | 2 | ENSMUST00000004994.10 | 24129 | Pax3 | ENSMUSG00000004872.10 |
| chr1 | 78173472 | 78173693 | 4.25494 | 6.06311 | 2.50165 | Intron (ENSMUST00000004994.10/ENSMUSG00000004872.10, intron 4 of 8) | 1 | 78101267 | 78196838 | 95572 | 2 | ENSMUST00000004994.10 | 23145 | Pax3 | ENSMUSG00000004872.10 |
| chr1 | 78175269 | 78175778 | 3.54222 | 4.2578 | 1.25497 | Intron (ENSMUST00000004994.10/ENSMUSG00000004872.10, intron 4 of 8) | 1 | 78101267 | 78196838 | 95572 | 2 | ENSMUST00000004994.10 | 21060 | Pax3 | ENSMUSG00000004872.10 |
| chr1 | 78176123 | 78176355 | 3.64066 | 4.42983 | 1.37679 | Intron (ENSMUST00000004994.10/ENSMUSG00000004872.10, intron 4 of 8) | 1 | 78101267 | 78196838 | 95572 | 2 | ENSMUST00000004994.10 | 20483 | Pax3 | ENSMUSG00000004872.10 |
| chr1 | 78176842 | 78177385 | 3.75387 | 4.59357 | 1.46146 | Intron (ENSMUST00000004994.10/ENSMUSG00000004872.10, intron 4 of 8) | 1 | 78101267 | 78196838 | 95572 | 2 | ENSMUST00000004994.10 | 19453 | Pax3 | ENSMUSG00000004872.10 |
| chr1 | 78178374 | 78179066 | 3.58763 | 4.29477 | 1.30263 | Intron (ENSMUST00000004994.10/ENSMUSG00000004872.10, intron 4 of 8) | 1 | 78101267 | 78196838 | 95572 | 2 | ENSMUST00000004994.10 | 17772 | Pax3 | ENSMUSG00000004872.10 |

| | | | | | | | | | | | | | | | |
|---|---|---|---|---|---|---|---|---|---|---|---|---|---|---|---|
| chr1 | 78179698 | 78180708 | 3.62089 | 4.49387 | 1.46742 | Intron (ENSMUST00000004994.10/ENSMUSG00000004872.10, intron 4 of 8) | 1 | 78101267 | 78196838 | 95572 | 2 | ENSMUST00000004994.10 | 16130 | Pax3 | ENSMUSG00000004872.10 |
| chr1 | 78181443 | 78182150 | 3.634 | 4.58995 | 1.50412 | Intron (ENSMUST00000004994.10/ENSMUSG00000004872.10, intron 4 of 8) | 1 | 78101267 | 78196838 | 95572 | 2 | ENSMUST00000004994.10 | 14688 | Pax3 | ENSMUSG00000004872.10 |
| chr1 | 78185318 | 78185655 | 3.43443 | 4.23605 | 1.28567 | Exon (ENSMUST00000172555.1/ENSMUSG00000004872.10, exon 5 of 5) | 1 | 78101267 | 78196838 | 95572 | 2 | ENSMUST00000004994.10 | 11183 | Pax3 | ENSMUSG00000004872.10 |
| chr1 | 78186050 | 78186374 | 3.88444 | 5.00523 | 1.77073 | Intron (ENSMUST00000004994.10/ENSMUSG00000004872.10, intron 4 of 8) | 1 | 78101267 | 78196838 | 95572 | 2 | ENSMUST00000004994.10 | 10464 | Pax3 | ENSMUSG00000004872.10 |
| chr1 | 78186762 | 78187375 | 3.68421 | 4.87745 | 1.6929 | Intron (ENSMUST00000004994.10/ENSMUSG00000004872.10, intron 4 of 8) | 1 | 78101267 | 78196838 | 95572 | 2 | ENSMUST00000004994.10 | 9463 | Pax3 | ENSMUSG00000004872.10 |
| chr1 | 78187782 | 78187991 | 3.60814 | 4.50608 | 1.45296 | Intron (ENSMUST00000004994.10/ENSMUSG00000004872.10, intron 4 of 8) | 1 | 78101267 | 78196838 | 95572 | 2 | ENSMUST00000004994.10 | 8847 | Pax3 | ENSMUSG00000004872.10 |
| chr1 | 78189056 | 78189131 | 4.35807 | 6.14991 | 2.60881 | Intron (ENSMUST00000004994.10/ENSMUSG00000004872.10, intron 4 of 8) | 1 | 78101267 | 78196838 | 95572 | 2 | ENSMUST00000004994.10 | 7707 | Pax3 | ENSMUSG00000004872.10 |
| chr1 | 78189732 | 78189917 | 3.47742 | 4.5733 | 1.49166 | Intron (ENSMUST00000004994.10/ENSMUSG00000004872.10, intron 4 of 8) | 1 | 78101267 | 78196838 | 95572 | 2 | ENSMUST00000004994.10 | 6921 | Pax3 | ENSMUSG00000004872.10 |
| chr1 | 78191670 | 78192006 | 3.53837 | 4.32874 | 1.34121 | Intron (ENSMUST00000004994.10/ENSMUSG00000004872.10, intron 4 of 8) | 1 | 78101267 | 78196838 | 95572 | 2 | ENSMUST00000004994.10 | 4832 | Pax3 | ENSMUSG00000004872.10 |
| chr1 | 78192419 | 78193421 | 3.44695 | 4.22057 | 1.27989 | Exon (ENSMUST00000004994.10/ENSMUSG00000004872.10, exon 4 of 9) | 1 | 78101267 | 78196838 | 95572 | 2 | ENSMUST00000004994.10 | 3417 | Pax3 | ENSMUSG00000004872.10 |
| chr1 | 78193783 | 78194032 | 4.17185 | 5.55357 | 2.17352 | Exon (ENSMUST00000004994.10/ENSMUSG00000004872.10, exon 3 of 9) | 1 | 78101267 | 78196838 | 95572 | 2 | ENSMUST00000004994.10 | 2806 | Pax3 | ENSMUSG00000004872.10 |
| chr1 | 78195898 | 78198125 | 4.02 | 5.37358 | 2.05722 | Promoter | 1 | 78101267 | 78196838 | 95572 | 2 | ENSMUST00000004994.10 | 0 | Pax3 | ENSMUSG00000004872.10 |
| chr1 | 78198659 | 78199883 | 3.95888 | 5.08976 | 1.8318 | Distal Intergenic | 1 | 78101270 | 78197134 | 95865 | 2 | ENSMUST00000087086.6 | -1526 | Pax3 | ENSMUSG00000004872.10 |
| chr1 | 78200451 | 78200609 | 3.46305 | 4.26513 | 1.29167 | Distal Intergenic | 1 | 78101270 | 78197134 | 95865 | 2 | ENSMUST00000087086.6 | -3318 | Pax3 | ENSMUSG00000004872.10 |
| chr1 | 78201445 | 78202191 | 3.54397 | 4.24121 | 1.25224 | Distal Intergenic | 1 | 78101270 | 78197134 | 95865 | 2 | ENSMUST00000087086.6 | -4312 | Pax3 | ENSMUSG00000004872.10 |
| chr1 | 78202848 | 78203149 | 3.37369 | 3.88816 | 1.01733 | Distal Intergenic | 1 | 78101270 | 78197134 | 95865 | 2 | ENSMUST00000087086.6 | -5715 | Pax3 | ENSMUSG00000004872.10 |
| chr1 | 78204206 | 78206033 | 4.03609 | 5.20448 | 1.91526 | Distal Intergenic | 1 | 78101270 | 78197134 | 95865 | 2 | ENSMUST00000087086.6 | -7073 | Pax3 | ENSMUSG00000004872.10 |
| chr1 | 78207081 | 78207531 | 3.76507 | 4.84847 | 1.69304 | Distal Intergenic | 1 | 78101270 | 78197134 | 95865 | 2 | ENSMUST00000087086.6 | -9948 | Pax3 | ENSMUSG00000004872.10 |
| chr1 | 78208087 | 78208891 | 3.67357 | 4.49208 | 1.4321 | Distal Intergenic | 1 | 78101270 | 78197134 | 95865 | 2 | ENSMUST00000087086.6 | -10954 | Pax3 | ENSMUSG00000004872.10 |
| chr1 | 78209904 | 78210806 | 4.19312 | 5.64822 | 2.33448 | Distal Intergenic | 1 | 78101270 | 78197134 | 95865 | 2 | ENSMUST00000087086.6 | -12771 | Pax3 | ENSMUSG00000004872.10 |
| chr1 | 78211175 | 78211293 | 3.82259 | 4.89546 | 1.6934 | Distal Intergenic | 1 | 78101270 | 78197134 | 95865 | 2 | ENSMUST00000087086.6 | -14042 | Pax3 | ENSMUSG00000004872.10 |
| chr1 | 78212691 | 78213322 | 3.90677 | 5.16234 | 1.93066 | Distal Intergenic | 1 | 78101270 | 78197134 | 95865 | 2 | ENSMUST00000087086.6 | -15558 | Pax3 | ENSMUSG00000004872.10 |
| chr1 | 78213682 | 78213859 | 3.37776 | 4.15063 | 1.25965 | Distal Intergenic | 1 | 78101270 | 78197134 | 95865 | 2 | ENSMUST00000087086.6 | -16549 | Pax3 | ENSMUSG00000004872.10 |
| chr1 | 78214174 | 78214843 | 3.55362 | 4.40411 | 1.38641 | Distal Intergenic | 1 | 78101270 | 78197134 | 95865 | 2 | ENSMUST00000087086.6 | -17041 | Pax3 | ENSMUSG00000004872.10 |
| chr1 | 78215389 | 78218776 | 3.82796 | 4.99993 | 1.83507 | Distal Intergenic | 1 | 78101270 | 78197134 | 95865 | 2 | ENSMUST00000087086.6 | -18256 | Pax3 | ENSMUSG00000004872.10 |
| chr1 | 78219279 | 78219354 | 3.93448 | 4.95274 | 1.7473 | Distal Intergenic | 1 | 78101270 | 78197134 | 95865 | 2 | ENSMUST00000087086.6 | -22146 | Pax3 | ENSMUSG00000004872.10 |
| chr1 | 78219808 | 78220135 | 3.48712 | 4.25148 | 1.29482 | Distal Intergenic | 1 | 78101270 | 78197134 | 95865 | 2 | ENSMUST00000087086.6 | -22675 | Pax3 | ENSMUSG00000004872.10 |
| chr1 | 78220579 | 78221099 | 3.77884 | 4.68946 | 1.6277 | Distal Intergenic | 1 | 78101270 | 78197134 | 95865 | 2 | ENSMUST00000087086.6 | -23446 | Pax3 | ENSMUSG00000004872.10 |
| chr1 | 78221715 | 78221843 | 4.03463 | 5.39095 | 2.03242 | Distal Intergenic | 1 | 78101270 | 78197134 | 95865 | 2 | ENSMUST00000087086.6 | -24582 | Pax3 | ENSMUSG00000004872.10 |
| chr3 | 87947724 | 87948384 | 3.43533 | 4.21343 | 1.30297 | Promoter | 3 | 87948666 | 87953376 | 4711 | 1 | ENSMUST00000005019.5 | -282 | Crabp2 | ENSMUSG00000004885.5 |
| chr3 | 87948702 | 87949666 | 4.04395 | 5.34556 | 2.0133 | Promoter | 3 | 87948666 | 87953376 | 4711 | 1 | ENSMUST00000005019.5 | 37 | Crabp2 | ENSMUSG00000004885.5 |
| chr3 | 87950252 | 87950440 | 3.8798 | 4.8387 | 1.64744 | Intron (ENSMUST00000005019.5/ENSMUSG00000004885.5, intron 1 of 3) | 3 | 87948666 | 87953376 | 4711 | 1 | ENSMUST00000005019.5 | 1587 | Crabp2 | ENSMUSG00000004885.5 |
| chr3 | 87951632 | 87951778 | 3.55506 | 4.22338 | 1.23463 | Promoter | 3 | 87952144 | 87952985 | 842 | 1 | ENSMUST00000163040.1 | -366 | Crabp2 | ENSMUSG00000004885.5 |
| chr3 | 87960659 | 87960758 | 3.54752 | 4.40542 | 1.36872 | Distal Intergenic | 3 | 87952144 | 87952985 | 842 | 1 | ENSMUST00000163040.1 | 8516 | Crabp2 | ENSMUSG00000004885.5 |
| chr3 | 87995411 | 87995555 | 3.49819 | 4.16969 | 1.21216 | Exon (ENSMUST00000090971.5/ENSMUSG00000004892.8, exon 6 of 14) | 3 | 87987531 | 88000356 | 12826 | 2 | ENSMUST00000090971.5 | 4801 | Bcan | ENSMUSG00000004892.8 |
| chr3 | 87996627 | 87997853 | 3.59058 | 4.47295 | 1.45769 | 5' UTR | 3 | 87987531 | 88000356 | 12826 | 2 | ENSMUST00000090971.5 | 2503 | Bcan | ENSMUSG00000004892.8 |
| chr3 | 87998592 | 87998851 | 3.50115 | 4.12085 | 1.15971 | Intron (ENSMUST00000090971.5/ENSMUSG00000004892.8, intron 1 of 13) | 3 | 87987531 | 88000356 | 12826 | 2 | ENSMUST00000090971.5 | 1505 | Bcan | ENSMUSG00000004892.8 |
| chr3 | 87999303 | 88000977 | 3.9798 | 5.06486 | 1.80454 | Promoter | 3 | 87987531 | 88000356 | 12826 | 2 | ENSMUST00000090971.5 | 0 | Bcan | ENSMUSG00000004892.8 |
| chr3 | 88002664 | 88002941 | 3.7921 | 4.71195 | 1.56057 | Distal Intergenic | 3 | 87987531 | 88000356 | 12826 | 2 | ENSMUST00000090971.5 | -2309 | Bcan | ENSMUSG00000004892.8 |
| chr3 | 88020236 | 88020614 | 3.41284 | 4.15577 | 1.22915 | Downstream (1-2kb) | 3 | 88022638 | 88023673 | 1036 | 2 | ENSMUST00000162352.1 | 3059 | Hapln2 | ENSMUSG00000004894.5 |
| chr3 | 88023426 | 88023925 | 3.67182 | 4.53327 | 1.4898 | Promoter | 3 | 88022638 | 88023673 | 1036 | 2 | ENSMUST00000162352.1 | 0 | Hapln2 | ENSMUSG00000004894.5 |
| chr3 | 88027843 | 88028168 | 3.29479 | 3.9323 | 1.1251 | Promoter | 3 | 88021750 | 88027583 | 5834 | 2 | ENSMUST00000160737.2 | -261 | Hapln2 | ENSMUSG00000004894.5 |
| chr10 | 81252562 | 81253313 | 3.74961 | 4.75396 | 1.61394 | Promoter | 10 | 81252935 | 81262985 | 10051 | 1 | ENSMUST00000119547.2 | 0 | Matk | ENSMUSG00000004933.12 |
| chr10 | 81254143 | 81254363 | 3.38696 | 3.93872 | 1.12566 | Intron (ENSMUST00000119547.2/ENSMUSG00000004933.12, intron 1 of 12) | 10 | 81252935 | 81262985 | 10051 | 1 | ENSMUST00000119547.2 | 1209 | Matk | ENSMUSG00000004933.12 |
| chr10 | 81256797 | 81257966 | 4.15896 | 5.52171 | 2.16488 | Promoter | 10 | 81257299 | 81262981 | 5683 | 1 | ENSMUST00000117488.2 | 0 | Matk | ENSMUSG00000004933.12 |
| chr10 | 81259504 | 81260070 | 3.49448 | 4.24844 | 1.26094 | Exon (ENSMUST00000119547.2/ENSMUSG00000004933.12, exon 5 of 13) | 10 | 81261170 | 81262667 | 1498 | 1 | ENSMUST00000148735.1 | -1100 | Matk | ENSMUSG00000004933.12 |
| chr10 | 81260522 | 81260678 | 4.13456 | 5.37286 | 2.01584 | Promoter | 10 | 81261170 | 81262667 | 1498 | 1 | ENSMUST00000148735.1 | -492 | Matk | ENSMUSG00000004933.12 |
| chr4 | 152338443 | 152340971 | 3.78628 | 4.79403 | 1.65583 | Promoter | 4 | 152338651 | 152390194 | 51544 | 1 | ENSMUST00000030775.6 | 0 | Chd5 | ENSMUSG00000005045.11 |
| chr5 | 136231606 | 136232076 | 3.49122 | 4.48409 | 1.43958 | Exon (ENSMUST00000005188.9/ENSMUSG00000005057.9, exon 2 of 9) | 5 | 136218149 | 136244903 | 26755 | 2 | ENSMUST00000005188.9 | 12827 | Sh2b2 | ENSMUSG00000005057.9 |
| chr5 | 136237820 | 136237919 | 3.83251 | 4.8099 | 1.67325 | Intron (ENSMUST00000005188.9/ENSMUSG00000005057.9, intron 1 of 8) | 5 | 136218149 | 136244903 | 26755 | 2 | ENSMUST00000005188.9 | 6984 | Sh2b2 | ENSMUSG00000005057.9 |
| chr5 | 136241365 | 136241793 | 3.83505 | 4.82076 | 1.65312 | Intron (ENSMUST00000005188.9/ENSMUSG00000005057.9, intron 1 of 8) | 5 | 136218149 | 136244903 | 26755 | 2 | ENSMUST00000005188.9 | 3110 | Sh2b2 | ENSMUSG00000005057.9 |
| chr2 | 102656360 | 102656435 | 3.53494 | 4.24841 | 1.31783 | Distal Intergenic | 2 | 102658659 | 102790784 | 132126 | 1 | ENSMUST00000080210.4 | -2224 | Slc1a2 | ENSMUSG00000005089.10 |
| chr2 | 102656776 | 102657919 | 3.50857 | 4.15548 | 1.23702 | Promoter | 2 | 102658659 | 102790784 | 132126 | 1 | ENSMUST00000080210.4 | -740 | Slc1a2 | ENSMUSG00000005089.10 |
| chr2 | 102658658 | 102659407 | 3.6823 | 4.62605 | 1.53801 | Promoter | 2 | 102658659 | 102790784 | 132126 | 1 | ENSMUST00000080210.4 | 0 | Slc1a2 | ENSMUSG00000005089.10 |
| chr2 | 102659864 | 102660165 | 3.60202 | 4.32078 | 1.32886 | Promoter | 2 | 102659281 | 102778097 | 118817 | 1 | ENSMUST00000111213.2 | 584 | Slc1a2 | ENSMUSG00000005089.10 |
| chr5 | 38486867 | 38486964 | 3.25395 | 3.70808 | 1.01733 | Intron (ENSMUST00000129099.2/ENSMUSG00000005107.8, intron 1 of 12) | 5 | 38349273 | 38483385 | 134113 | 2 | ENSMUST00000067872.6 | -3483 | Slc2a9 | ENSMUSG00000005107.8 |
| chr14 | 99298229 | 99298447 | 3.61105 | 4.45228 | 1.42468 | Promoter | 14 | 99298691 | 99313405 | 14715 | 1 | ENSMUST00000005279.6 | -244 | Klf5 | ENSMUSG00000005148.7 |
| chr14 | 99299271 | 99299502 | 3.61528 | 4.69454 | 1.59163 | Promoter | 14 | 99298691 | 99313405 | 14715 | 1 | ENSMUST00000005279.6 | 581 | Klf5 | ENSMUSG00000005148.7 |
| chr5 | 72503363 | 72504006 | 3.40415 | 3.99539 | 1.15612 | Promoter | 5 | 72300026 | 72504208 | 204183 | 2 | ENSMUST00000005352.4 | 202 | Corin | ENSMUSG00000005220.5 |
| chr11 | 69528918 | 69529214 | 3.56223 | 4.28948 | 1.36469 | Intron (ENSMUST00000035539.6/ENSMUSG00000005237.8, intron 3 of 84) | 11 | 69516007 | 69516755 | 749 | 2 | ENSMUST00000154985.1 | -12164 | Dnah2 | ENSMUSG00000005237.8 |
| chr11 | 69547559 | 69547763 | 3.56693 | 4.25616 | 1.28067 | Intron (ENSMUST00000108659.1/ENSMUSG00000005237.8, intron 1 of 85) | 11 | 69420809 | 69549108 | 128300 | 2 | ENSMUST00000108659.1 | 1345 | Dnah2 | ENSMUSG00000005237.8 |
| chr16 | 97762958 | 97764435 | 4.23898 | 5.78815 | 2.38364 | Promoter | 16 | 97741933 | 97763491 | 21559 | 2 | ENSMUST00000113743.2 | 0 | Ripk4 | ENSMUSG00000005251.9 |
| chr16 | 97764808 | 97765008 | 3.50102 | 4.35911 | 1.36019 | Distal Intergenic | 16 | 97741933 | 97763737 | 21805 | 2 | ENSMUST00000019386.8 | -1072 | Ripk4 | ENSMUSG00000005251.9 |
| chr16 | 97765663 | 97766329 | 3.65055 | 4.59457 | 1.5528 | Distal Intergenic | 16 | 97741933 | 97763737 | 21805 | 2 | ENSMUST00000019386.8 | -1927 | Ripk4 | ENSMUSG00000005251.9 |
| chr15 | 10177812 | 10177923 | 3.71399 | 4.90221 | 1.68966 | Promoter | 15 | 10177781 | 10319990 | 142210 | 1 | ENSMUST00000132361.2 | 32 | Prlr | ENSMUSG00000005268.15 |
| chr15 | 10179118 | 10179193 | 3.65338 | 4.42218 | 1.40322 | Intron (ENSMUST00000005400.7/ENSMUSG00000005268.15, intron 1 of 8) | 15 | 10177781 | 10319990 | 142210 | 1 | ENSMUST00000132361.2 | 1338 | Prlr | ENSMUSG00000005268.15 |
| chr1 | 173364091 | 173364426 | 3.36691 | 3.87794 | 1.01733 | Intron (ENSMUST00000147604.2/ENSMUSG00000037860.10, intron 1 of 11) | 1 | 173344901 | 173367633 | 22733 | 2 | ENSMUST00000126963.1 | 3207 | Cadm3 | ENSMUSG00000005338.8 |
| chr1 | 173365666 | 173366192 | 3.4133 | 3.97247 | 1.07532 | Intron (ENSMUST00000147604.2/ENSMUSG00000037860.10, intron 1 of 11) | 1 | 173344901 | 173367633 | 22733 | 2 | ENSMUST00000126963.1 | 1441 | Cadm3 | ENSMUSG00000005338.8 |
| chr1 | 173366596 | 173367031 | 4.01561 | 5.31839 | 1.96934 | Promoter | 1 | 173344901 | 173367633 | 22733 | 2 | ENSMUST00000126963.1 | 602 | Cadm3 | ENSMUSG00000005338.8 |
| chr1 | 173367348 | 173367806 | 3.82666 | 4.77321 | 1.60577 | Promoter | 1 | 173334254 | 173367695 | 33442 | 2 | ENSMUST00000111220.2 | 0 | Cadm3 | ENSMUSG00000005338.8 |
| chr1 | 173368398 | 173368569 | 3.2267 | 4.04474 | 1.15943 | Promoter | 1 | 173334254 | 173367695 | 33442 | 2 | ENSMUST00000111220.2 | -704 | Cadm3 | ENSMUSG00000005338.8 |
| chr1 | 173369146 | 173369579 | 3.41053 | 3.96672 | 1.09858 | Intron (ENSMUST00000147604.2/ENSMUSG00000037860.10, intron 1 of 11) | 1 | 173334254 | 173367695 | 33442 | 2 | ENSMUST00000111220.2 | -1452 | Cadm3 | ENSMUSG00000005338.8 |
| chr1 | 173389762 | 173390277 | 3.79879 | 4.7109 | 1.58696 | Intron (ENSMUST00000147604.2/ENSMUSG00000037860.10, intron 1 of 11) | 1 | 173334254 | 173367695 | 33442 | 2 | ENSMUST00000111220.2 | -22068 | Cadm3 | ENSMUSG00000005338.8 |
| chr10 | 78780299 | 78780733 | 3.3809 | 3.89904 | 1.01733 | Promoter | 10 | 78780496 | 78814825 | 34330 | 1 | ENSMUST00000005490.8 | 0 | Slc1a6 | ENSMUSG00000005357.8 |
| chr15 | 8628967 | 8629073 | 3.46465 | 4.13429 | 1.29009 | Distal Intergenic | 15 | 8642256 | 8643247 | 992 | 2 | ENSMUST00000128879.1 | 14174 | Slc1a3 | ENSMUSG00000005360.9 |
| chr15 | 8629468 | 8630504 | 3.48242 | 4.24165 | 1.29274 | Distal Intergenic | 15 | 8642256 | 8643247 | 992 | 2 | ENSMUST00000128879.1 | 12743 | Slc1a3 | ENSMUSG00000005360.9 |
| chr15 | 8708301 | 8710149 | 3.82738 | 4.78034 | 1.61378 | Promoter | 15 | 8708743 | 8709588 | 846 | 2 | ENSMUST00000126747.1 | 0 | Slc1a3 | ENSMUSG00000005360.9 |
| chr15 | 8710699 | 8710983 | 3.36075 | 3.96259 | 1.1404 | Promoter | 15 | 8634124 | 8710764 | 76641 | 2 | ENSMUST00000005493.8 | 0 | Slc1a3 | ENSMUSG00000005360.9 |
| chr5 | 135107674 | 135108372 | 3.6293 | 4.44079 | 1.45917 | Promoter | 5 | 135106913 | 135137922 | 31010 | 1 | ENSMUST00000153519.2 | 762 | Mlxipl | ENSMUSG00000005373.8 |
| chr5 | 135132545 | 135132888 | 3.88161 | 5.00202 | 1.77885 | Promoter | 5 | 135133610 | 135134610 | 1001 | 1 | ENSMUST00000154840.1 | -722 | Mlxipl | ENSMUSG00000005373.8 |
| chr5 | 135133639 | 135134533 | 3.62318 | 4.34375 | 1.2926 | Promoter | 5 | 135133631 | 135135738 | 2108 | 1 | ENSMUST00000123370.1 | 9 | Mlxipl | ENSMUSG00000005373.8 |
| chr5 | 135134978 | 135135273 | 3.43399 | 4.02526 | 1.16591 | Intron (ENSMUST00000005507.4/ENSMUSG00000005373.8, intron 15 of 16) | 5 | 135133631 | 135135738 | 2108 | 1 | ENSMUST00000123370.1 | 1348 | Mlxipl | ENSMUSG00000005373.8 |
| chr5 | 136714342 | 136715023 | 3.80804 | 4.76257 | 1.63464 | Distal Intergenic | 5 | 136693146 | 136701094 | 7949 | 1 | ENSMUST00000005611.5 | 21197 | Myl10 | ENSMUSG00000005474.5 |
| chr5 | 136718162 | 136718428 | 3.38995 | 3.93157 | 1.08407 | Distal Intergenic | 5 | 136693146 | 136701094 | 7949 | 1 | ENSMUST00000005611.5 | 25017 | Myl10 | ENSMUSG00000005474.5 |
| chr6 | 52290329 | 52290674 | 4.00708 | 5.2516 | 1.95001 | Distal Intergenic | 6 | 52313498 | 52318378 | 4881 | 1 | ENSMUST00000031787.7 | -22824 | Evx1 | ENSMUSG00000005503.8 |
| chr6 | 52291026 | 52292346 | 3.73516 | 4.6678 | 1.57917 | Distal Intergenic | 6 | 52313498 | 52318378 | 4881 | 1 | ENSMUST00000031787.7 | -21152 | Evx1 | ENSMUSG00000005503.8 |
| chr6 | 52292692 | 52292831 | 3.30903 | 3.79075 | 1.01733 | Distal Intergenic | 6 | 52313498 | 52318378 | 4881 | 1 | ENSMUST00000031787.7 | -20667 | Evx1 | ENSMUSG00000005503.8 |
| chr6 | 52293166 | 52293355 | 3.55947 | 4.52423 | 1.46997 | Distal Intergenic | 6 | 52313498 | 52318378 | 4881 | 1 | ENSMUST00000031787.7 | -20143 | Evx1 | ENSMUSG00000005503.8 |
| chr6 | 52294708 | 52295250 | 3.83057 | 5.03159 | 1.81912 | Distal Intergenic | 6 | 52313498 | 52318378 | 4881 | 1 | ENSMUST00000031787.7 | -18248 | Evx1 | ENSMUSG00000005503.8 |
| chr6 | 52297156 | 52297323 | 4.35421 | 5.79212 | 2.34349 | Distal Intergenic | 6 | 52313498 | 52318378 | 4881 | 1 | ENSMUST00000031787.7 | -16175 | Evx1 | ENSMUSG00000005503.8 |
| chr6 | 52300039 | 52300114 | 3.46641 | 4.08784 | 1.21951 | Distal Intergenic | 6 | 52313498 | 52318378 | 4881 | 1 | ENSMUST00000031787.7 | -13384 | Evx1 | ENSMUSG00000005503.8 |
| chr6 | 52301075 | 52301300 | 3.38193 | 3.9006 | 1.01733 | Distal Intergenic | 6 | 52313498 | 52318378 | 4881 | 1 | ENSMUST00000031787.7 | -12198 | Evx1 | ENSMUSG00000005503.8 |
| chr6 | 52302877 | 52302952 | 3.3612 | 3.95724 | 1.13433 | Distal Intergenic | 6 | 52313498 | 52318378 | 4881 | 1 | ENSMUST00000031787.7 | -10546 | Evx1 | ENSMUSG00000005503.8 |
| chr6 | 52303674 | 52303776 | 3.84857 | 4.79762 | 1.62285 | Distal Intergenic | 6 | 52313498 | 52318378 | 4881 | 1 | ENSMUST00000031787.7 | -9722 | Evx1 | ENSMUSG00000005503.8 |
| chr6 | 52305108 | 52305211 | 3.73499 | 4.55848 | 1.43564 | Distal Intergenic | 6 | 52313498 | 52318378 | 4881 | 1 | ENSMUST00000031787.7 | -8287 | Evx1 | ENSMUSG00000005503.8 |
| chr6 | 52305632 | 52307939 | 4.37848 | 5.97603 | 2.53684 | Downstream (<1kb) | 6 | 52313498 | 52318378 | 4881 | 1 | ENSMUST00000031787.7 | -5559 | Evx1 | ENSMUSG00000005503.8 |
| chr6 | 52308660 | 52314341 | 3.90833 | 5.12994 | 1.90559 | Promoter | 6 | 52313498 | 52318378 | 4881 | 1 | ENSMUST00000031787.7 | 0 | Evx1 | ENSMUSG00000005503.8 |
| chr7 | 67926614 | 67927014 | 3.24254 | 3.69098 | 1.01733 | Distal Intergenic | 7 | 67952859 | 68226780 | 273922 | 1 | ENSMUST00000005671.8 | -25845 | Igf1r | ENSMUSG00000005533.9 |
| chr7 | 67928519 | 67928656 | 4.24257 | 5.76552 | 2.30494 | Distal Intergenic | 7 | 67952859 | 68226780 | 273922 | 1 | ENSMUST00000005671.8 | -24203 | Igf1r | ENSMUSG00000005533.9 |
| chr7 | 67934066 | 67934181 | 3.77325 | 4.72402 | 1.61517 | Distal Intergenic | 7 | 67952859 | 68226780 | 273922 | 1 | ENSMUST00000005671.8 | -18678 | Igf1r | ENSMUSG00000005533.9 |
| chr7 | 30717387 | 30717545 | 3.23338 | 3.67724 | 1.01733 | Promoter | 7 | 30717832 | 30720620 | 2789 | 1 | ENSMUST00000171014.1 | -287 | Atp4a | ENSMUSG00000005553.8 |
| chr7 | 111282241 | 111282766 | 3.40026 | 4.05113 | 1.13162 | Distal Intergenic | 7 | 111070568 | 111083030 | 12463 | 2 | ENSMUST00000162415.2 | -199212 | Eif4g2 | ENSMUSG00000005610.11 |
| chr3 | 87804099 | 87804410 | 3.50536 | 4.2453 | 1.28956 | Exon (ENSMUST00000029711.3/ENSMUSG00000005640.6, exon 6 of 22) | 3 | 87801943 | 87802808 | 866 | 1 | ENSMUST00000166866.1 | 2157 | Insrr | ENSMUSG00000005640.6 |
| chr3 | 87809449 | 87809898 | 4.12749 | 5.38707 | 2.09798 | Exon (ENSMUST00000029711.3/ENSMUSG00000005640.6, exon 10 of 22) | 3 | 87801943 | 87802808 | 866 | 1 | ENSMUST00000166866.1 | 7507 | Insrr | ENSMUSG00000005640.6 |
| chr3 | 87810714 | 87811073 | 3.65075 | 4.50029 | 1.48446 | Exon (ENSMUST00000029711.3/ENSMUSG00000005640.6, exon 14 of 22) | 3 | 87801943 | 87802808 | 866 | 1 | ENSMUST00000166866.1 | 8772 | Insrr | ENSMUSG00000005640.6 |
| chr3 | 87797053 | 87797237 | 3.45648 | 4.0485 | 1.14029 | Promoter | 3 | 87797094 | 87816052 | 18959 | 1 | ENSMUST00000107582.2 | 0 | Insrr | ENSMUSG00000005640.6 |
| chr5 | 75574457 | 75575285 | 3.70093 | 4.79042 | 1.63046 | Promoter | 5 | 75574916 | 75656722 | 81807 | 1 | ENSMUST00000144270.2 | 0 | Kit | ENSMUSG00000005672.7 |
| chr5 | 75575808 | 75576141 | 3.38905 | 3.91749 | 1.03912 | Promoter | 5 | 75574955 | 75656722 | 81768 | 1 | ENSMUST00000005815.6 | 854 | Kit | ENSMUSG00000005672.7 |
| chr5 | 75576455 | 75576912 | 3.31761 | 3.80377 | 1.01733 | Intron (ENSMUST00000144270.2/ENSMUSG00000005672.7, intron 1 of 20) | 5 | 75574955 | 75656722 | 81768 | 1 | ENSMUST00000005815.6 | 1501 | Kit | ENSMUSG00000005672.7 |
| chr5 | 75578015 | 75578212 | 3.57737 | 4.26356 | 1.25396 | Intron (ENSMUST00000144270.2/ENSMUSG00000005672.7, intron 1 of 20) | 5 | 75574955 | 75656722 | 81768 | 1 | ENSMUST00000005815.6 | 3061 | Kit | ENSMUSG00000005672.7 |
| chr7 | 110768575 | 110769227 | 3.63149 | 4.38054 | 1.32982 | Promoter | 7 | 110768343 | 110791269 | 22927 | 1 | ENSMUST00000176487.2 | 233 | Ampd3 | ENSMUSG00000005686.11 |
| chr15 | 78204284 | 78204888 | 3.60632 | 4.49052 | 1.44762 | Promoter | 15 | 78191114 | 78204237 | 13124 | 2 | ENSMUST00000005860.10 | -48 | Pvalb | ENSMUSG00000005716.11 |
| chr15 | 78205659 | 78205868 | 3.48851 | 4.09856 | 1.14248 | Promoter | 15 | 78191117 | 78206400 | 15284 | 2 | ENSMUST00000120592.1 | 532 | Pvalb | ENSMUSG00000005716.11 |
| chr18 | 11048551 | 11053669 | 3.87841 | 4.97888 | 1.77108 | Promoter | 18 | 11052510 | 11085635 | 33126 | 1 | ENSMUST00000047762.7 | 0 | Gata6 | ENSMUSG00000005836.8 |
| chr6 | 92244093 | 92244417 | 3.84433 | 4.76406 | 1.58103 | Promoter | 6 | 92242061 | 92244650 | 2590 | 2 | ENSMUST00000006046.4 | 233 | Trh | ENSMUSG00000005892.4 |
| chr6 | 92244740 | 92244883 | 3.53795 | 4.23428 | 1.28618 | Promoter | 6 | 92242061 | 92244650 | 2590 | 2 | ENSMUST00000006046.4 | -91 | Trh | ENSMUSG00000005892.4 |
| chr6 | 92245243 | 92245505 | 3.86558 | 4.8189 | 1.63969 | Promoter | 6 | 92242061 | 92244650 | 2590 | 2 | ENSMUST00000006046.4 | -594 | Trh | ENSMUSG00000005892.4 |
| chr11 | 21985492 | 21986663 | 4.12461 | 5.36147 | 2.04398 | Distal Intergenic | 11 | 21994764 | 22001615 | 6852 | 2 | ENSMUST00000006071.8 | 14952 | Otx1 | ENSMUSG00000005917.10 |
| chr11 | 21987868 | 21989417 | 3.78727 | 4.89374 | 1.72865 | Distal Intergenic | 11 | 21994764 | 22001615 | 6852 | 2 | ENSMUST00000006071.8 | 12198 | Otx1 | ENSMUSG00000005917.10 |
| chr11 | 21989883 | 21990392 | 3.74996 | 4.65641 | 1.58658 | Distal Intergenic | 11 | 21994764 | 22001615 | 6852 | 2 | ENSMUST00000006071.8 | 11223 | Otx1 | ENSMUSG00000005917.10 |
| chr11 | 21990777 | 21991306 | 4.1102 | 5.40927 | 2.0864 | Distal Intergenic | 11 | 21994764 | 22001615 | 6852 | 2 | ENSMUST00000006071.8 | 10309 | Otx1 | ENSMUSG00000005917.10 |
| chr11 | 21992902 | 21994286 | 4.98002 | 7.37327 | 3.69214 | Downstream (<1kb) | 11 | 21994764 | 22001615 | 6852 | 2 | ENSMUST00000006071.8 | 7329 | Otx1 | ENSMUSG00000005917.10 |
| chr11 | 21994682 | 21994828 | 3.618 | 4.42577 | 1.41525 | 3' UTR | 11 | 21994764 | 22001615 | 6852 | 2 | ENSMUST00000006071.8 | 6787 | Otx1 | ENSMUSG00000005917.10 |
| chr11 | 21995653 | 21996935 | 3.76841 | 4.8253 | 1.66752 | 3' UTR | 11 | 21994764 | 22001615 | 6852 | 2 | ENSMUST00000006071.8 | 4680 | Otx1 | ENSMUSG00000005917.10 |
| chr11 | 21997309 | 21998828 | 4.10364 | 5.58837 | 2.23006 | Exon (ENSMUST00000006071.8/ENSMUSG00000005917.10, exon 4 of 5) | 11 | 21994764 | 22001615 | 6852 | 2 | ENSMUST00000006071.8 | 2787 | Otx1 | ENSMUSG00000005917.10 |
| chr11 | 21999386 | 22003275 | 3.9195 | 5.15192 | 1.94449 | Promoter | 11 | 21994764 | 22001615 | 6852 | 2 | ENSMUST00000006071.8 | 0 | Otx1 | ENSMUSG00000005917.10 |
| chr11 | 22003601 | 22003948 | 3.41957 | 3.98227 | 1.10545 | Promoter | 11 | 21996858 | 22002897 | 6040 | 2 | ENSMUST00000147486.2 | -705 | Otx1 | ENSMUSG00000005917.10 |
| chr11 | 22004269 | 22004997 | 3.76231 | 4.66234 | 1.5588 | Downstream (<1kb) | 11 | 21996858 | 22002897 | 6040 | 2 | ENSMUST00000147486.2 | -1373 | Otx1 | ENSMUSG00000005917.10 |
| chr11 | 22010503 | 22010855 | 3.41174 | 3.96492 | 1.08528 | Intron (ENSMUST00000045167.5/ENSMUSG00000042302.9, intron 22 of 23) | 11 | 21996858 | 22002897 | 6040 | 2 | ENSMUST00000147486.2 | -7607 | Otx1 | ENSMUSG00000005917.10 |
| chr16 | 21202563 | 21202835 | 3.81951 | 4.87276 | 1.69559 | Distal Intergenic | 16 | 21204755 | 21223304 | 18550 | 1 | ENSMUST00000006112.6 | -1920 | Ephb3 | ENSMUSG00000005958.9 |
| chr16 | 21203873 | 21204067 | 3.38337 | 3.91934 | 1.07593 | Promoter | 16 | 21204755 | 21223304 | 18550 | 1 | ENSMUST00000006112.6 | -688 | Ephb3 | ENSMUSG00000005958.9 |
| chr2 | 105311636 | 105311964 | 3.64516 | 4.42504 | 1.42323 | Intron (ENSMUST00000068813.2/ENSMUSG00000055312.2, intron 3 of 3) | 2 | 105386929 | 105395348 | 8420 | 2 | ENSMUST00000127019.1 | 83384 | Rcn1 | ENSMUSG00000005973.6 |
| chr2 | 105367571 | 105367997 | 3.52437 | 4.23176 | 1.32615 | Intron (ENSMUST00000068813.2/ENSMUSG00000055312.2, intron 3 of 3) | 2 | 105386929 | 105395348 | 8420 | 2 | ENSMUST00000127019.1 | 27351 | Rcn1 | ENSMUSG00000005973.6 |
| chr2 | 105368330 | 105368442 | 3.70663 | 4.54639 | 1.52406 | Intron (ENSMUST00000068813.2/ENSMUSG00000055312.2, intron 3 of 3) | 2 | 105386929 | 105395348 | 8420 | 2 | ENSMUST00000127019.1 | 26906 | Rcn1 | ENSMUSG00000005973.6 |
| chr2 | 105368798 | 105368970 | 3.77069 | 4.67303 | 1.59688 | Intron (ENSMUST00000068813.2/ENSMUSG00000055312.2, intron 3 of 3) | 2 | 105386929 | 105395348 | 8420 | 2 | ENSMUST00000127019.1 | 26378 | Rcn1 | ENSMUSG00000005973.6 |
| chr2 | 105370439 | 105371106 | 3.63331 | 4.48245 | 1.44 | Intron (ENSMUST00000068813.2/ENSMUSG00000055312.2, intron 3 of 3) | 2 | 105386929 | 105395348 | 8420 | 2 | ENSMUST00000127019.1 | 24242 | Rcn1 | ENSMUSG00000005973.6 |
| chr2 | 105402276 | 105402428 | 3.26174 | 3.73062 | 1.0514 | Distal Intergenic | 2 | 105386291 | 105399319 | 13029 | 2 | ENSMUST00000006128.6 | -2958 | Rcn1 | ENSMUSG00000005973.6 |
| chr2 | 105415797 | 105416267 | 3.85174 | 4.82809 | 1.70425 | Distal Intergenic | 2 | 105386291 | 105399319 | 13029 | 2 | ENSMUST00000006128.6 | -16479 | Rcn1 | ENSMUSG00000005973.6 |
| chr2 | 105417399 | 105417695 | 3.79896 | 4.78149 | 1.6427 | Distal Intergenic | 2 | 105386291 | 105399319 | 13029 | 2 | ENSMUST00000006128.6 | -18081 | Rcn1 | ENSMUSG00000005973.6 |
| chr2 | 105418171 | 105418580 | 3.86208 | 4.94265 | 1.75854 | Distal Intergenic | 2 | 105386291 | 105399319 | 13029 | 2 | ENSMUST00000006128.6 | -18853 | Rcn1 | ENSMUSG00000005973.6 |
| chr2 | 105428947 | 105429035 | 4.08624 | 5.42773 | 2.07813 | Distal Intergenic | 2 | 105386291 | 105399319 | 13029 | 2 | ENSMUST00000006128.6 | -29629 | Rcn1 | ENSMUSG00000005973.6 |
| chr2 | 105434290 | 105434365 | 3.6398 | 4.37694 | 1.32502 | Distal Intergenic | 2 | 105386291 | 105399319 | 13029 | 2 | ENSMUST00000006128.6 | -34972 | Rcn1 | ENSMUSG00000005973.6 |
| chr2 | 105436467 | 105436731 | 4.01755 | 5.13862 | 1.89493 | Distal Intergenic | 2 | 105386291 | 105399319 | 13029 | 2 | ENSMUST00000006128.6 | -37149 | Rcn1 | ENSMUSG00000005973.6 |
| chr4 | 80864618 | 80864843 | 3.70808 | 4.56708 | 1.52345 | Distal Intergenic | 4 | 80846588 | 80851000 | 4413 | 1 | ENSMUST00000133932.1 | 18031 | Tyrp1 | ENSMUSG00000005994.9 |
| chr13 | 21993690 | 21994183 | 4.30374 | 5.90138 | 2.51539 | Distal Intergenic | 13 | 22002560 | 22005222 | 2663 | 2 | ENSMUST00000129665.2 | 11039 | Prss16 | ENSMUSG00000006179.3 |
| chr13 | 21995383 | 21996415 | 3.75072 | 4.86723 | 1.69601 | Distal Intergenic | 13 | 22002560 | 22005222 | 2663 | 2 | ENSMUST00000129665.2 | 8807 | Prss16 | ENSMUSG00000006179.3 |
| chr13 | 21996736 | 21997592 | 3.95917 | 5.22425 | 1.97766 | Distal Intergenic | 13 | 22002560 | 22005222 | 2663 | 2 | ENSMUST00000129665.2 | 7630 | Prss16 | ENSMUSG00000006179.3 |
| chr13 | 21997947 | 22000604 | 3.94024 | 5.08175 | 1.93549 | Downstream (1-2kb) | 13 | 22002560 | 22005222 | 2663 | 2 | ENSMUST00000129665.2 | 4618 | Prss16 | ENSMUSG00000006179.3 |
| chr13 | 22000942 | 22001439 | 3.84768 | 4.80238 | 1.62978 | Downstream (<1kb) | 13 | 22002560 | 22005222 | 2663 | 2 | ENSMUST00000129665.2 | 3783 | Prss16 | ENSMUSG00000006179.3 |
| chr13 | 22001773 | 22002744 | 3.79416 | 4.75343 | 1.63044 | 3' UTR | 13 | 22002560 | 22005222 | 2663 | 2 | ENSMUST00000129665.2 | 2478 | Prss16 | ENSMUSG00000006179.3 |
| chr13 | 22003413 | 22003675 | 3.4952 | 4.25594 | 1.29033 | Intron (ENSMUST00000006341.2/ENSMUSG00000006179.3, intron 9 of 11) | 13 | 22002560 | 22005222 | 2663 | 2 | ENSMUST00000129665.2 | 1547 | Prss16 | ENSMUSG00000006179.3 |
| chr13 | 22004025 | 22007741 | 3.89643 | 5.02182 | 1.78571 | Promoter | 13 | 22002560 | 22005222 | 2663 | 2 | ENSMUST00000129665.2 | 0 | Prss16 | ENSMUSG00000006179.3 |
| chr13 | 22008084 | 22008805 | 3.90673 | 4.9098 | 1.70942 | Promoter | 13 | 22002176 | 22009741 | 7566 | 2 | ENSMUST00000006341.2 | 936 | Prss16 | ENSMUSG00000006179.3 |
| chr13 | 22009228 | 22009420 | 3.48386 | 4.13824 | 1.24468 | Promoter | 13 | 22002176 | 22009741 | 7566 | 2 | ENSMUST00000006341.2 | 321 | Prss16 | ENSMUSG00000006179.3 |
| chr13 | 22010413 | 22010872 | 3.94283 | 5.0673 | 1.86673 | Promoter | 13 | 22007232 | 22009742 | 2511 | 2 | ENSMUST00000130585.1 | -672 | Prss16 | ENSMUSG00000006179.3 |
| chr13 | 22011268 | 22011788 | 4.08652 | 5.56264 | 2.2286 | Distal Intergenic | 13 | 22007232 | 22009742 | 2511 | 2 | ENSMUST00000130585.1 | -1527 | Prss16 | ENSMUSG00000006179.3 |
| chr4 | 141366016 | 141366252 | 3.51371 | 4.21899 | 1.2921 | Distal Intergenic | 4 | 141368220 | 141384175 | 15956 | 1 | ENSMUST00000006380.4 | -1968 | Fam131c | ENSMUSG00000006218.4 |
| chr4 | 141368206 | 141368549 | 3.36828 | 3.89122 | 1.03682 | Promoter | 4 | 141368220 | 141384175 | 15956 | 1 | ENSMUST00000006380.4 | 0 | Fam131c | ENSMUSG00000006218.4 |
| chr4 | 141381249 | 141381558 | 3.95173 | 4.98634 | 1.73669 | Intron (ENSMUST00000006380.4/ENSMUSG00000006218.4, intron 4 of 6) | 4 | 141368220 | 141384175 | 15956 | 1 | ENSMUST00000006380.4 | 13030 | Fam131c | ENSMUSG00000006218.4 |
| chr9 | 21961525 | 21961859 | 3.76009 | 4.69758 | 1.60722 | Exon (ENSMUST00000006397.5/ENSMUSG00000006235.5, exon 3 of 8) | 9 | 21958899 | 21963576 | 4678 | 2 | ENSMUST00000006397.5 | 1717 | Epor | ENSMUSG00000006235.5 |
| chr9 | 21963107 | 21963460 | 3.53086 | 4.22274 | 1.33687 | Promoter | 9 | 21958899 | 21963576 | 4678 | 2 | ENSMUST00000006397.5 | 116 | Epor | ENSMUSG00000006235.5 |
| chr6 | 83743152 | 83743444 | 3.57938 | 4.27072 | 1.27243 | Promoter | 6 | 83743017 | 83758809 | 15793 | 1 | ENSMUST00000006431.6 | 136 | Atp6v1b1 | ENSMUSG00000006269.6 |
| chr19 | 59161546 | 59161962 | 3.56434 | 4.25946 | 1.26596 | Distal Intergenic | 19 | 59166598 | 59166980 | 383 | 2 | ENSMUST00000173710.1 | 5018 | Vax1 | ENSMUSG00000006270.7 |

| | | | | | | | | | | | | | | | |
|---|---|---|---|---|---|---|---|---|---|---|---|---|---|---|---|
| chr19 | 59163437 | 59163590 | 3.77838 | 4.67297 | 1.55872 | Downstream (2-3kb) | 19 | 59166598 | 59166980 | 383 | 2 | ENSMUST00000173710.1 | 3390 | Vax1 | ENSMUSG00000006270.7 |
| chr19 | 59164027 | 59164352 | 4.10391 | 5.53939 | 2.19417 | Downstream (1-2kb) | 19 | 59166598 | 59166980 | 383 | 2 | ENSMUST00000173710.1 | 2628 | Vax1 | ENSMUSG00000006270.7 |
| chr19 | 59164730 | 59165138 | 3.6251 | 4.35386 | 1.31827 | Downstream (1-2kb) | 19 | 59166598 | 59166980 | 383 | 2 | ENSMUST00000173710.1 | 1842 | Vax1 | ENSMUSG00000006270.7 |
| chr19 | 59165525 | 59165845 | 3.57464 | 4.26135 | 1.26462 | Downstream (<1kb) | 19 | 59166598 | 59166980 | 383 | 2 | ENSMUST00000173710.1 | 1135 | Vax1 | ENSMUSG00000006270.7 |
| chr19 | 59166176 | 59166955 | 3.43776 | 4.15106 | 1.22698 | Promoter | 19 | 59166598 | 59166980 | 383 | 2 | ENSMUST00000173710.1 | 25 | Vax1 | ENSMUSG00000006270.7 |
| chr19 | 59167288 | 59167529 | 3.82991 | 4.86323 | 1.68902 | Promoter | 19 | 59166598 | 59166980 | 383 | 2 | ENSMUST00000173710.1 | -309 | Vax1 | ENSMUSG00000006270.7 |
| chr19 | 59167906 | 59168087 | 4.16566 | 5.40866 | 2.07372 | Promoter | 19 | 59166598 | 59166980 | 383 | 2 | ENSMUST00000173710.1 | -927 | Vax1 | ENSMUSG00000006270.7 |
| chr19 | 59168544 | 59168620 | 3.7354 | 4.87929 | 1.71423 | Exon (ENSMUST00000172821.3/ENSMUSG00000006270.7, exon 2 of 3) | 19 | 59166187 | 59170056 | 3870 | 2 | ENSMUST00000172821.3 | 1436 | Vax1 | ENSMUSG00000006270.7 |
| chr19 | 59169128 | 59170327 | 4.07291 | 5.48324 | 2.12277 | Promoter | 19 | 59166187 | 59170056 | 3870 | 2 | ENSMUST00000172821.3 | 0 | Vax1 | ENSMUSG00000006270.7 |
| chr19 | 59170685 | 59170906 | 3.21222 | 3.64557 | 1.01019 | Promoter | 19 | 59166187 | 59170056 | 3870 | 2 | ENSMUST00000172821.3 | -630 | Vax1 | ENSMUSG00000006270.7 |
| chr19 | 59171228 | 59172243 | 4.40163 | 6.28979 | 2.76341 | Distal Intergenic | 19 | 59166187 | 59170056 | 3870 | 2 | ENSMUST00000172821.3 | -1173 | Vax1 | ENSMUSG00000006270.7 |
| chr19 | 59172866 | 59173456 | 3.48683 | 4.29895 | 1.31706 | Distal Intergenic | 19 | 59166187 | 59170056 | 3870 | 2 | ENSMUST00000172821.3 | -2811 | Vax1 | ENSMUSG00000006270.7 |
| chr19 | 59174679 | 59175117 | 3.73429 | 4.81415 | 1.65042 | Distal Intergenic | 19 | 59166187 | 59170056 | 3870 | 2 | ENSMUST00000172821.3 | -4624 | Vax1 | ENSMUSG00000006270.7 |
| chr19 | 59176238 | 59176328 | 3.34513 | 3.87921 | 1.12532 | Distal Intergenic | 19 | 59166187 | 59170056 | 3870 | 2 | ENSMUST00000172821.3 | -6183 | Vax1 | ENSMUSG00000006270.7 |
| chr19 | 59176783 | 59177975 | 3.62467 | 4.3878 | 1.36567 | Distal Intergenic | 19 | 59166187 | 59170056 | 3870 | 2 | ENSMUST00000172821.3 | -6728 | Vax1 | ENSMUSG00000006270.7 |
| chr19 | 59178491 | 59178692 | 3.41021 | 4.02186 | 1.18163 | Distal Intergenic | 19 | 59166187 | 59170056 | 3870 | 2 | ENSMUST00000172821.3 | -8436 | Vax1 | ENSMUSG00000006270.7 |
| chr19 | 59179061 | 59179269 | 3.60749 | 4.3639 | 1.34118 | Distal Intergenic | 19 | 59166187 | 59170056 | 3870 | 2 | ENSMUST00000172821.3 | -9006 | Vax1 | ENSMUSG00000006270.7 |
| chr19 | 59184342 | 59185173 | 3.75125 | 4.65658 | 1.56903 | Distal Intergenic | 19 | 59166187 | 59170056 | 3870 | 2 | ENSMUST00000172821.3 | -14287 | Vax1 | ENSMUSG00000006270.7 |
| chr19 | 59186956 | 59187460 | 3.91564 | 4.9916 | 1.78576 | Distal Intergenic | 19 | 59166187 | 59170056 | 3870 | 2 | ENSMUST00000172821.3 | -16901 | Vax1 | ENSMUSG00000006270.7 |
| chr19 | 59187956 | 59188068 | 3.42352 | 4.31882 | 1.33669 | Distal Intergenic | 19 | 59166187 | 59170056 | 3870 | 2 | ENSMUST00000172821.3 | -17901 | Vax1 | ENSMUSG00000006270.7 |
| chr19 | 59189273 | 59189872 | 3.85001 | 5.07252 | 1.85585 | Distal Intergenic | 19 | 59166187 | 59170056 | 3870 | 2 | ENSMUST00000172821.3 | -19218 | Vax1 | ENSMUSG00000006270.7 |
| chr19 | 59190455 | 59190824 | 3.22452 | 3.78793 | 1.01705 | Distal Intergenic | 19 | 59166187 | 59170056 | 3870 | 2 | ENSMUST00000172821.3 | -20400 | Vax1 | ENSMUSG00000006270.7 |
| chr19 | 59191343 | 59191771 | 3.82911 | 5.16544 | 1.91065 | Distal Intergenic | 19 | 59166187 | 59170056 | 3870 | 2 | ENSMUST00000172821.3 | -21288 | Vax1 | ENSMUSG00000006270.7 |
| chr19 | 59192215 | 59192518 | 3.94652 | 5.08807 | 1.84953 | Distal Intergenic | 19 | 59166187 | 59170056 | 3870 | 2 | ENSMUST00000172821.3 | -22160 | Vax1 | ENSMUSG00000006270.7 |
| chr19 | 59193958 | 59194863 | 3.64986 | 4.43257 | 1.44723 | Distal Intergenic | 19 | 59166187 | 59170056 | 3870 | 2 | ENSMUST00000172821.3 | -23903 | Vax1 | ENSMUSG00000006270.7 |
| chr7 | 30633352 | 30634232 | 4.19424 | 5.57686 | 2.16533 | 3' UTR | 7 | 30633616 | 30635852 | 2237 | 2 | ENSMUST00000108147.2 | 1620 | Etv2 | ENSMUSG00000006311.8 |
| chr7 | 30634541 | 30635414 | 3.9642 | 5.19216 | 1.89586 | Promoter | 7 | 30633616 | 30635852 | 2237 | 2 | ENSMUST00000108147.2 | 438 | Etv2 | ENSMUSG00000006311.8 |
| chr7 | 30635788 | 30636334 | 3.69328 | 4.51119 | 1.42341 | Promoter | 7 | 30633616 | 30635852 | 2237 | 2 | ENSMUST00000108147.2 | 0 | Etv2 | ENSMUSG00000006311.8 |
| chr10 | 75560706 | 75561306 | 3.60079 | 4.32155 | 1.33267 | Promoter | 10 | 75561604 | 75576255 | 14652 | 1 | ENSMUST00000145079.2 | -298 | Ggt1 | ENSMUSG00000006345.5 |
| chr8 | 122720569 | 122720644 | 3.39762 | 3.93837 | 1.06716 | Intron (ENSMUST00000127664.1/ENSMUSG00000092329.1, intron 1 of 15) | 8 | 122628739 | 122699109 | 70371 | 2 | ENSMUST00000127984.2 | -21461 | Cbfa2t3 | ENSMUSG00000006362.10 |
| chr15 | 85200630 | 85200915 | 3.42431 | 3.97787 | 1.06417 | Distal Intergenic | 15 | 85205949 | 85286535 | 80587 | 1 | ENSMUST00000057410.8 | -5034 | Fbln1 | ENSMUSG00000006369.9 |
| chr15 | 79041775 | 79041940 | 3.83952 | 4.91603 | 1.69777 | Promoter | 15 | 79041885 | 79043558 | 1674 | 1 | ENSMUST00000058004.3 | 0 | Gcat | ENSMUSG00000006378.8 |
| chr15 | 79042342 | 79042785 | 4.08012 | 5.7557 | 2.29129 | Promoter | 15 | 79041885 | 79043558 | 1674 | 1 | ENSMUST00000058004.3 | 458 | Gcat | ENSMUSG00000006378.8 |
| chr4 | 118358370 | 118358550 | 3.21895 | 3.65564 | 1.01595 | Downstream (2-3kb) | 4 | 118359999 | 118361294 | 1296 | 1 | ENSMUST00000179719.1 | -1449 | Hyi | ENSMUSG00000006395.11 |
| chr19 | 47178226 | 47178795 | 3.57005 | 4.27666 | 1.2809 | Promoter | 19 | 47178820 | 47259440 | 80621 | 1 | ENSMUST00000111808.4 | -25 | Neurl1a | ENSMUSG00000006435.10 |
| chr19 | 47179235 | 47179951 | 3.96889 | 5.10973 | 1.84119 | Promoter | 19 | 47178820 | 47259440 | 80621 | 1 | ENSMUST00000111808.4 | 416 | Neurl1a | ENSMUSG00000006435.10 |
| chr19 | 47180271 | 47180504 | 3.33249 | 3.9985 | 1.10102 | Intron (ENSMUST00000111808.4/ENSMUSG00000006435.10, intron 1 of 5) | 19 | 47178820 | 47259440 | 80621 | 1 | ENSMUST00000111808.4 | 1452 | Neurl1a | ENSMUSG00000006435.10 |
| chr19 | 47185661 | 47185926 | 3.35344 | 3.85761 | 1.01733 | Intron (ENSMUST00000111808.4/ENSMUSG00000006435.10, intron 1 of 5) | 19 | 47178820 | 47259440 | 80621 | 1 | ENSMUST00000111808.4 | 6842 | Neurl1a | ENSMUSG00000006435.10 |
| chr19 | 47201233 | 47202045 | 3.36407 | 3.88148 | 1.04448 | Intron (ENSMUST00000111808.4/ENSMUSG00000006435.10, intron 1 of 5) | 19 | 47178820 | 47259440 | 80621 | 1 | ENSMUST00000111808.4 | 22414 | Neurl1a | ENSMUSG00000006435.10 |
| chr19 | 47240085 | 47240511 | 3.43568 | 4.02863 | 1.17884 | Exon (ENSMUST00000111808.4/ENSMUSG00000006435.10, exon 3 of 6) | 19 | 47228843 | 47259440 | 30598 | 1 | ENSMUST00000111807.3 | 11243 | Neurl1a | ENSMUSG00000006435.10 |
| chr19 | 47241096 | 47241375 | 3.47748 | 4.08166 | 1.14756 | Intron (ENSMUST00000111808.4/ENSMUSG00000006435.10, intron 3 of 5) | 19 | 47228843 | 47259440 | 30598 | 1 | ENSMUST00000111807.3 | 12254 | Neurl1a | ENSMUSG00000006435.10 |
| chr19 | 47246187 | 47246325 | 3.48742 | 4.13348 | 1.21897 | Intron (ENSMUST00000111808.4/ENSMUSG00000006435.10, intron 3 of 5) | 19 | 47228843 | 47259440 | 30598 | 1 | ENSMUST00000111807.3 | 17345 | Neurl1a | ENSMUSG00000006435.10 |
| chr19 | 47250947 | 47251044 | 3.54431 | 4.48749 | 1.41903 | Intron (ENSMUST00000111808.4/ENSMUSG00000006435.10, intron 3 of 5) | 19 | 47228843 | 47259440 | 30598 | 1 | ENSMUST00000111807.3 | 22105 | Neurl1a | ENSMUSG00000006435.10 |
| chr4 | 141291708 | 141291816 | 3.52321 | 4.17381 | 1.22433 | Distal Intergenic | 4 | 141301240 | 141329384 | 28145 | 1 | ENSMUST00000006614.2 | -9424 | Epha2 | ENSMUSG00000006445.3 |
| chr1 | 74945964 | 74946409 | 3.66376 | 4.41653 | 1.33341 | 3' UTR | 1 | 74947724 | 74951442 | 3719 | 2 | ENSMUST00000189364.1 | 5033 | Ihh | ENSMUSG00000006538.10 |
| chr1 | 74947747 | 74948645 | 3.66212 | 4.45267 | 1.40824 | Exon (ENSMUST00000164097.2/ENSMUSG00000006538.10, exon 2 of 3) | 1 | 74947724 | 74951442 | 3719 | 2 | ENSMUST00000189364.1 | 2797 | Ihh | ENSMUSG00000006538.10 |
| chr1 | 74948953 | 74951987 | 4.0037 | 5.28952 | 1.99155 | Promoter | 1 | 74945315 | 74951672 | 6358 | 2 | ENSMUST00000164097.2 | 0 | Ihh | ENSMUSG00000006538.10 |
| chr1 | 74954751 | 74955282 | 3.41803 | 3.99025 | 1.14229 | Distal Intergenic | 1 | 74945315 | 74951672 | 6358 | 2 | ENSMUST00000164097.2 | -3080 | Ihh | ENSMUSG00000006538.10 |
| chr1 | 74955622 | 74955753 | 3.78578 | 4.69775 | 1.62275 | Distal Intergenic | 1 | 74945315 | 74951672 | 6358 | 2 | ENSMUST00000164097.2 | -3951 | Ihh | ENSMUSG00000006538.10 |
| chr1 | 74960523 | 74960728 | 3.21866 | 3.65522 | 1.01016 | Distal Intergenic | 1 | 74945315 | 74951672 | 6358 | 2 | ENSMUST00000164097.2 | -8852 | Ihh | ENSMUSG00000006538.10 |
| chr1 | 74890614 | 74891340 | 3.81008 | 4.72759 | 1.567 | 3' UTR | 1 | 74890531 | 74892892 | 2362 | 2 | ENSMUST00000133833.2 | 1552 | Cryba2 | ENSMUSG00000006546.3 |
| chr1 | 74892690 | 74893546 | 3.49908 | 4.47867 | 1.44621 | Promoter | 1 | 74889934 | 74893143 | 3210 | 2 | ENSMUST00000006721.2 | 0 | Cryba2 | ENSMUSG00000006546.3 |
| chr1 | 74893910 | 74894037 | 3.43645 | 4.00246 | 1.08622 | Promoter | 1 | 74889934 | 74893143 | 3210 | 2 | ENSMUST00000006721.2 | -768 | Cryba2 | ENSMUSG00000006546.3 |
| chr1 | 74895521 | 74895721 | 3.44674 | 4.03022 | 1.12974 | Distal Intergenic | 1 | 74889934 | 74893143 | 3210 | 2 | ENSMUST00000006721.2 | -2379 | Cryba2 | ENSMUSG00000006546.3 |
| chr1 | 74896353 | 74896951 | 3.81534 | 4.73718 | 1.60534 | Distal Intergenic | 1 | 74889934 | 74893143 | 3210 | 2 | ENSMUST00000006721.2 | -3211 | Cryba2 | ENSMUSG00000006546.3 |
| chr11 | 102352283 | 102352461 | 3.34826 | 3.84982 | 1.01733 | Exon (ENSMUST00000006749.9/ENSMUSG00000006574.10, exon 17 of 20) | 11 | 102359082 | 102360568 | 1487 | 2 | ENSMUST00000149993.1 | 8107 | Slc4a1 | ENSMUSG00000006574.10 |
| chr11 | 102361123 | 102361456 | 3.66362 | 4.42611 | 1.37328 | Promoter | 11 | 102359082 | 102360568 | 1487 | 2 | ENSMUST00000149993.1 | -556 | Slc4a1 | ENSMUSG00000006574.10 |
| chr11 | 102365019 | 102365395 | 3.34693 | 3.84779 | 1.01733 | Promoter | 11 | 102348824 | 102365280 | 16457 | 2 | ENSMUST00000006749.9 | 0 | Slc4a1 | ENSMUSG00000006574.10 |
| chr11 | 102390719 | 102390844 | 3.54132 | 4.19685 | 1.21447 | Distal Intergenic | 11 | 102393403 | 102401754 | 8352 | 1 | ENSMUST00000107102.2 | -2559 | Rundc3a | ENSMUSG00000006575.9 |
| chr11 | 102399263 | 102399985 | 3.83742 | 4.77163 | 1.58673 | Promoter | 11 | 102399564 | 102402226 | 2663 | 1 | ENSMUST00000142157.2 | 0 | Rundc3a | ENSMUSG00000006575.9 |
| chr4 | 13740114 | 13740225 | 3.59741 | 4.30027 | 1.28043 | Distal Intergenic | 4 | 13743436 | 13893649 | 150214 | 1 | ENSMUST00000006761.4 | -3211 | Runx1t1 | ENSMUSG00000006586.10 |
| chr4 | 13740682 | 13740936 | 3.70025 | 4.51568 | 1.43302 | Distal Intergenic | 4 | 13743436 | 13893649 | 150214 | 1 | ENSMUST00000006761.4 | -2500 | Runx1t1 | ENSMUSG00000006586.10 |
| chr4 | 13741917 | 13742373 | 3.72482 | 4.56819 | 1.46255 | Distal Intergenic | 4 | 13743436 | 13893649 | 150214 | 1 | ENSMUST00000006761.4 | -1063 | Runx1t1 | ENSMUSG00000006586.10 |
| chr4 | 13742711 | 13743786 | 4.05468 | 5.38168 | 2.04921 | Promoter | 4 | 13743436 | 13893649 | 150214 | 1 | ENSMUST00000006761.4 | 0 | Runx1t1 | ENSMUSG00000006586.10 |
| chr4 | 13744126 | 13744515 | 3.60117 | 4.46236 | 1.43082 | Promoter | 4 | 13743436 | 13893649 | 150214 | 1 | ENSMUST00000006761.4 | 691 | Runx1t1 | ENSMUSG00000006586.10 |
| chr4 | 13746157 | 13746457 | 3.63661 | 4.50597 | 1.46739 | Intron (ENSMUST00000006761.4/ENSMUSG00000006586.10, intron 1 of 10) | 4 | 13743436 | 13893649 | 150214 | 1 | ENSMUST00000006761.4 | 2722 | Runx1t1 | ENSMUSG00000006586.10 |
| chr4 | 13749099 | 13749174 | 3.87216 | 5.02108 | 1.83758 | Intron (ENSMUST00000006761.4/ENSMUSG00000006586.10, intron 1 of 10) | 4 | 13751297 | 13843027 | 91731 | 1 | ENSMUST00000150583.2 | -2123 | Runx1t1 | ENSMUSG00000006586.10 |
| chr4 | 13750137 | 13751701 | 3.59162 | 4.35225 | 1.40588 | Promoter | 4 | 13751297 | 13843027 | 91731 | 1 | ENSMUST00000150583.2 | 0 | Runx1t1 | ENSMUSG00000006586.10 |
| chr4 | 13752054 | 13752206 | 4.40654 | 5.994 | 2.51975 | Promoter | 4 | 13751297 | 13893628 | 142332 | 1 | ENSMUST00000105566.3 | 758 | Runx1t1 | ENSMUSG00000006586.10 |
| chr4 | 13752619 | 13752841 | 3.34625 | 3.85864 | 1.05921 | Intron (ENSMUST00000006761.4/ENSMUSG00000006586.10, intron 1 of 10) | 4 | 13751297 | 13893628 | 142332 | 1 | ENSMUST00000105566.3 | 1323 | Runx1t1 | ENSMUSG00000006586.10 |
| chr4 | 13753223 | 13753408 | 3.31209 | 3.90233 | 1.11221 | Intron (ENSMUST00000006761.4/ENSMUSG00000006586.10, intron 1 of 10) | 4 | 13751297 | 13893628 | 142332 | 1 | ENSMUST00000105566.3 | 1927 | Runx1t1 | ENSMUSG00000006586.10 |
| chr4 | 13785900 | 13786202 | 3.25551 | 3.71041 | 1.01733 | Intron (ENSMUST00000006761.4/ENSMUSG00000006586.10, intron 1 of 10) | 4 | 13784782 | 13891117 | 106336 | 1 | ENSMUST00000098256.3 | 1119 | Runx1t1 | ENSMUSG00000006586.10 |
| chr8 | 122458844 | 122459123 | 3.57085 | 4.28883 | 1.3561 | Intron (ENSMUST00000127664.1/ENSMUSG00000092329.1, intron 1 of 15) | 8 | 122454206 | 122460692 | 6487 | 2 | ENSMUST00000006762.5 | 1569 | Snai3 | ENSMUSG00000006587.5 |
| chr8 | 122460081 | 122460528 | 3.71808 | 4.70368 | 1.59717 | Promoter | 8 | 122454206 | 122460692 | 6487 | 2 | ENSMUST00000006762.5 | 164 | Snai3 | ENSMUSG00000006587.5 |
| chr13 | 24845255 | 24846196 | 4.05569 | 5.22932 | 1.95382 | Promoter | 13 | 24845135 | 24901270 | 56136 | 1 | ENSMUST00000006893.8 | 121 | D130043K22Rik | ENSMUSG00000006711.10 |
| chr13 | 24846538 | 24846614 | 3.80749 | 4.89511 | 1.67866 | Intron (ENSMUST00000141572.2/ENSMUSG00000006711.10, intron 1 of 19) | 13 | 24845135 | 24901270 | 56136 | 1 | ENSMUST00000006893.8 | 1404 | D130043K22Rik | ENSMUSG00000006711.10 |
| chr13 | 21926568 | 21928161 | 3.77506 | 4.74368 | 1.61134 | Distal Intergenic | 13 | 21945094 | 21960485 | 15392 | 1 | ENSMUST00000176511.2 | -16933 | Zfp184 | ENSMUSG00000006720.11 |
| chr13 | 21928606 | 21929328 | 3.84626 | 4.85149 | 1.72016 | Distal Intergenic | 13 | 21945094 | 21960485 | 15392 | 1 | ENSMUST00000176511.2 | -15766 | Zfp184 | ENSMUSG00000006720.11 |
| chr13 | 21931719 | 21932011 | 3.29114 | 3.76386 | 1.01733 | Distal Intergenic | 13 | 21945094 | 21960485 | 15392 | 1 | ENSMUST00000176511.2 | -13083 | Zfp184 | ENSMUSG00000006720.11 |
| chr13 | 21934498 | 21934796 | 3.53183 | 4.23406 | 1.3343 | Distal Intergenic | 13 | 21945094 | 21960485 | 15392 | 1 | ENSMUST00000176511.2 | -10298 | Zfp184 | ENSMUSG00000006720.11 |
| chr13 | 21935571 | 21935707 | 3.62408 | 4.41337 | 1.42309 | Distal Intergenic | 13 | 21945094 | 21960485 | 15392 | 1 | ENSMUST00000176511.2 | -9387 | Zfp184 | ENSMUSG00000006720.11 |
| chr13 | 21937197 | 21937356 | 4.28069 | 5.80428 | 2.34554 | Distal Intergenic | 13 | 21945094 | 21960485 | 15392 | 1 | ENSMUST00000176511.2 | -7738 | Zfp184 | ENSMUSG00000006720.11 |
| chr13 | 21945272 | 21945803 | 3.56553 | 4.43926 | 1.41638 | Promoter | 13 | 21945793 | 21960339 | 14547 | 1 | ENSMUST00000006903.6 | 0 | Zfp184 | ENSMUSG00000006720.11 |
| chr10 | 127047445 | 127047598 | 3.34503 | 3.87699 | 1.10901 | Promoter | 10 | 127048250 | 127052969 | 4720 | 1 | ENSMUST00000165764.2 | -652 | Cyp27b1 | ENSMUSG00000006724.7 |
| chr10 | 127047915 | 127048164 | 3.74174 | 4.75963 | 1.62571 | Promoter | 10 | 127048250 | 127052969 | 4720 | 1 | ENSMUST00000165764.2 | -86 | Cyp27b1 | ENSMUSG00000006724.7 |
| chr10 | 127049106 | 127050763 | 3.72542 | 4.72837 | 1.63126 | Promoter | 10 | 127048671 | 127049866 | 1196 | 1 | ENSMUST00000171868.1 | 436 | Cyp27b1 | ENSMUSG00000006724.7 |
| chr10 | 127165568 | 127165955 | 4.07555 | 5.84975 | 2.36498 | Promoter | 10 | 127165156 | 127172340 | 7185 | 1 | ENSMUST00000006914.9 | 413 | B4galnt1 | ENSMUSG00000006731.9 |
| chr10 | 127167819 | 127168043 | 3.38511 | 4.17918 | 1.25647 | Exon (ENSMUST00000006914.9/ENSMUSG00000006731.9, exon 6 of 11) | 10 | 127165156 | 127172340 | 7185 | 1 | ENSMUST00000006914.9 | 2664 | B4galnt1 | ENSMUSG00000006731.9 |
| chr10 | 127169322 | 127170718 | 3.67372 | 4.46777 | 1.42702 | Exon (ENSMUST00000006914.9/ENSMUSG00000006731.9, exon 7 of 11) | 10 | 127165156 | 127172340 | 7185 | 1 | ENSMUST00000006914.9 | 4167 | B4galnt1 | ENSMUSG00000006731.9 |
| chr10 | 127171077 | 127171272 | 3.36728 | 4.07297 | 1.21577 | Exon (ENSMUST00000006914.9/ENSMUSG00000006731.9, exon 10 of 11) | 10 | 127165156 | 127172340 | 7185 | 1 | ENSMUST00000006914.9 | 5922 | B4galnt1 | ENSMUSG00000006731.9 |
| chr10 | 127171620 | 127171902 | 4.35046 | 6.09451 | 2.55099 | 3' UTR | 10 | 127165156 | 127172340 | 7185 | 1 | ENSMUST00000006914.9 | 6465 | B4galnt1 | ENSMUSG00000006731.9 |
| chr11 | 99493074 | 99493202 | 3.44899 | 4.09325 | 1.24093 | Promoter | 11 | 99477974 | 99493137 | 15164 | 2 | ENSMUST00000006969.7 | 0 | Krt23 | ENSMUSG00000006777.7 |
| chr11 | 100356909 | 100357058 | 3.90145 | 4.99636 | 1.7915 | Promoter | 11 | 100347327 | 100356128 | 8802 | 2 | ENSMUST00000138603.3 | -782 | Hap1 | ENSMUSG00000006930.10 |
| chr11 | 100358578 | 100358678 | 4.2989 | 5.72249 | 2.31105 | Distal Intergenic | 11 | 100347327 | 100356128 | 8802 | 2 | ENSMUST00000138603.3 | -2451 | Hap1 | ENSMUSG00000006930.10 |
| chr17 | 24689406 | 24689571 | 3.50557 | 4.14789 | 1.23306 | Promoter | 17 | 24685092 | 24689955 | 4864 | 2 | ENSMUST00000007236.4 | 384 | Syngr3 | ENSMUSG00000007021.4 |
| chr17 | 24689999 | 24690094 | 4.37441 | 5.86577 | 2.42425 | Promoter | 17 | 24685092 | 24689955 | 4864 | 2 | ENSMUST00000007236.4 | -45 | Syngr3 | ENSMUSG00000007021.4 |
| chr1 | 172298039 | 172298327 | 3.30895 | 3.79064 | 1.01733 | Promoter | 1 | 172271709 | 172298064 | 26356 | 2 | ENSMUST00000085913.5 | 0 | Atp1a2 | ENSMUSG00000007097.9 |
| chr7 | 109864916 | 109864991 | 3.73765 | 4.77697 | 1.58842 | Promoter | 7 | 109798676 | 109865586 | 66911 | 2 | ENSMUST00000007423.6 | 595 | Scube2 | ENSMUSG00000007279.9 |
| chr7 | 109865343 | 109866026 | 4.10431 | 5.37725 | 2.1131 | Promoter | 7 | 109798676 | 109865586 | 66911 | 2 | ENSMUST00000007423.6 | 0 | Scube2 | ENSMUSG00000007279.9 |
| chr7 | 109866724 | 109866920 | 3.48844 | 4.13475 | 1.27153 | Distal Intergenic | 7 | 109798690 | 109865679 | 66990 | 2 | ENSMUST00000106729.2 | -1046 | Scube2 | ENSMUSG00000007279.9 |
| chr18 | 37011184 | 37011259 | 3.37251 | 3.92926 | 1.1536 | Promoter | 18 | 37010858 | 37187657 | 176800 | 1 | ENSMUST00000115657.4 | 327 | Pcdha10 | ENSMUSG00000007440.9 |
| chr18 | 37012013 | 37012088 | 3.89259 | 4.87898 | 1.70696 | 3' UTR | 18 | 37010858 | 37187657 | 176800 | 1 | ENSMUST00000115657.4 | 1156 | Pcdha10 | ENSMUSG00000007440.9 |
| chr18 | 37090281 | 37090551 | 3.44321 | 4.04302 | 1.19531 | Promoter | 18 | 37090136 | 37187657 | 97522 | 1 | ENSMUST00000007584.2 | 146 | Pcdha10 | ENSMUSG00000007440.9 |
| chr18 | 37142309 | 37145098 | 3.91643 | 5.13539 | 1.90744 | Promoter | 18 | 37143969 | 37187657 | 43689 | 1 | ENSMUST00000047479.1 | 0 | Pcdha10 | ENSMUSG00000007440.9 |
| chr18 | 37146255 | 37146330 | 3.88396 | 4.90996 | 1.77057 | Exon (ENSMUST00000047479.1/ENSMUSG00000007440.9, exon 1 of 4) | 18 | 37143969 | 37187657 | 43689 | 1 | ENSMUST00000047479.1 | 2287 | Pcdha10 | ENSMUSG00000007440.9 |
| chr18 | 37147066 | 37147370 | 3.29355 | 3.7767 | 1.04828 | Intron (ENSMUST00000070797.5/ENSMUSG00000007440.9, intron 1 of 3) | 18 | 37143969 | 37187657 | 43689 | 1 | ENSMUST00000047479.1 | 3098 | Pcdha10 | ENSMUSG00000007440.9 |
| chr8 | 70082873 | 70084912 | 3.7475 | 4.69965 | 1.62903 | Promoter | 8 | 70083529 | 70090862 | 7334 | 1 | ENSMUST00000007738.9 | 0 | Hapln4 | ENSMUSG00000007594.9 |
| chr8 | 70085279 | 70085420 | 3.60162 | 4.3503 | 1.41135 | Intron (ENSMUST00000007738.9/ENSMUSG00000007594.9, intron 3 of 4) | 8 | 70083529 | 70090862 | 7334 | 1 | ENSMUST00000007738.9 | 1751 | Hapln4 | ENSMUSG00000007594.9 |
| chr8 | 70085828 | 70087337 | 3.61639 | 4.44769 | 1.44299 | Exon (ENSMUST00000007738.9/ENSMUSG00000007594.9, exon 4 of 5) | 8 | 70083529 | 70090862 | 7334 | 1 | ENSMUST00000007738.9 | 2300 | Hapln4 | ENSMUSG00000007594.9 |
| chr8 | 70087995 | 70088604 | 3.86043 | 5.17408 | 1.90514 | 3' UTR | 8 | 70083529 | 70090862 | 7334 | 1 | ENSMUST00000007738.9 | 4467 | Hapln4 | ENSMUSG00000007594.9 |
| chr8 | 70089270 | 70090488 | 3.42003 | 4.00434 | 1.1665 | 3' UTR | 8 | 70083529 | 70090862 | 7334 | 1 | ENSMUST00000007738.9 | 5742 | Hapln4 | ENSMUSG00000007594.9 |
| chr4 | 47342505 | 47342981 | 3.29546 | 3.78934 | 1.07806 | Distal Intergenic | 4 | 47353222 | 47414931 | 61710 | 1 | ENSMUST00000007757.9 | -10241 | Tgfbr1 | ENSMUSG00000007613.10 |
| chr6 | 17305586 | 17305946 | 3.42526 | 3.9921 | 1.10988 | Promoter | 6 | 17306335 | 17341324 | 34990 | 1 | ENSMUST00000007799.7 | -389 | Cav1 | ENSMUSG00000007655.11 |
| chr6 | 17306343 | 17307157 | 3.52498 | 4.18483 | 1.21399 | Promoter | 6 | 17306345 | 17340429 | 34085 | 1 | ENSMUST00000115456.2 | 0 | Cav1 | ENSMUSG00000007655.11 |
| chr1 | 91728429 | 91728672 | 3.77665 | 4.67685 | 1.6031 | Distal Intergenic | 1 | 91801461 | 91848028 | 46568 | 1 | ENSMUST00000007949.3 | -72789 | Twist2 | ENSMUSG00000007805.4 |
| chr1 | 91729150 | 91729545 | 3.32849 | 3.83954 | 1.08598 | Distal Intergenic | 1 | 91801461 | 91848028 | 46568 | 1 | ENSMUST00000007949.3 | -71916 | Twist2 | ENSMUSG00000007805.4 |
| chr1 | 91730536 | 91730650 | 3.31051 | 3.80076 | 1.03442 | Distal Intergenic | 1 | 91801461 | 91848028 | 46568 | 1 | ENSMUST00000007949.3 | -70811 | Twist2 | ENSMUSG00000007805.4 |
| chr1 | 91731073 | 91731185 | 4.78352 | 6.74876 | 3.08424 | Distal Intergenic | 1 | 91801461 | 91848028 | 46568 | 1 | ENSMUST00000007949.3 | -70276 | Twist2 | ENSMUSG00000007805.4 |
| chr1 | 91731492 | 91731799 | 3.54677 | 4.21414 | 1.23065 | Distal Intergenic | 1 | 91801461 | 91848028 | 46568 | 1 | ENSMUST00000007949.3 | -69662 | Twist2 | ENSMUSG00000007805.4 |
| chr1 | 91748286 | 91748409 | 3.66299 | 4.42323 | 1.35829 | Distal Intergenic | 1 | 91801461 | 91848028 | 46568 | 1 | ENSMUST00000007949.3 | -53052 | Twist2 | ENSMUSG00000007805.4 |
| chr1 | 91757401 | 91757622 | 3.35204 | 3.93801 | 1.15326 | Distal Intergenic | 1 | 91801461 | 91848028 | 46568 | 1 | ENSMUST00000007949.3 | -43839 | Twist2 | ENSMUSG00000007805.4 |
| chr1 | 91771756 | 91771890 | 3.88461 | 4.90615 | 1.77609 | Distal Intergenic | 1 | 91801461 | 91848028 | 46568 | 1 | ENSMUST00000007949.3 | -29571 | Twist2 | ENSMUSG00000007805.4 |
| chr1 | 91799877 | 91800156 | 3.30489 | 3.78452 | 1.01733 | Exon (ENSMUST00000187564.1/ENSMUSG00000100980.1, exon 1 of 1) | 1 | 91801461 | 91848028 | 46568 | 1 | ENSMUST00000007949.3 | -1305 | Twist2 | ENSMUSG00000007805.4 |
| chr1 | 91800948 | 91801260 | 3.52779 | 4.2854 | 1.33125 | Promoter | 1 | 91801461 | 91848028 | 46568 | 1 | ENSMUST00000007949.3 | -201 | Twist2 | ENSMUSG00000007805.4 |
| chr1 | 91802017 | 91802271 | 3.56443 | 4.37793 | 1.37048 | Promoter | 1 | 91801470 | 91803994 | 2525 | 1 | ENSMUST00000186075.1 | 548 | Twist2 | ENSMUSG00000007805.4 |
| chr14 | 25607838 | 25607927 | 3.29139 | 3.78654 | 1.08714 | Promoter | 14 | 25607838 | 25645735 | 37898 | 1 | ENSMUST00000160229.1 | 1 | Zmiz1 | ENSMUSG00000007817.9 |
| chr8 | 70490846 | 70491058 | 3.65162 | 4.47686 | 1.45494 | Distal Intergenic | 8 | 70493158 | 70504081 | 10924 | 1 | ENSMUST00000008032.8 | -2100 | Crlf1 | ENSMUSG00000007888.10 |
| chr8 | 70491394 | 70494116 | 3.76843 | 4.90097 | 1.74021 | Promoter | 8 | 70493158 | 70504081 | 10924 | 1 | ENSMUST00000008032.8 | 0 | Crlf1 | ENSMUSG00000007888.10 |
| chr8 | 70494673 | 70494819 | 3.352 | 3.85544 | 1.01733 | Intron (ENSMUST00000008032.8/ENSMUSG00000007888.10, intron 1 of 8) | 8 | 70493361 | 70500930 | 7570 | 1 | ENSMUST00000132648.1 | 1313 | Crlf1 | ENSMUSG00000007888.10 |
| chr8 | 70495409 | 70495705 | 3.48541 | 4.11551 | 1.22626 | Intron (ENSMUST00000008032.8/ENSMUSG00000007888.10, intron 1 of 8) | 8 | 70493361 | 70500930 | 7570 | 1 | ENSMUST00000132648.1 | 2049 | Crlf1 | ENSMUSG00000007888.10 |
| chr8 | 70502015 | 70502284 | 3.53836 | 4.28315 | 1.33116 | Promoter | 8 | 70501133 | 70504030 | 2898 | 1 | ENSMUST00000127983.1 | 883 | Crlf1 | ENSMUSG00000007888.10 |
| chr9 | 61682696 | 61682771 | 3.60018 | 4.30444 | 1.28031 | Distal Intergenic | 9 | 61913284 | 61914538 | 1255 | 2 | ENSMUST00000008036.7 | 231767 | Rplp1 | ENSMUSG00000007892.7 |
| chr9 | 61691741 | 61692121 | 3.63741 | 4.37433 | 1.32026 | Distal Intergenic | 9 | 61913284 | 61914538 | 1255 | 2 | ENSMUST00000008036.7 | 222417 | Rplp1 | ENSMUSG00000007892.7 |
| chr9 | 61693041 | 61693197 | 3.88731 | 4.8558 | 1.65231 | Distal Intergenic | 9 | 61913284 | 61914538 | 1255 | 2 | ENSMUST00000008036.7 | 221341 | Rplp1 | ENSMUSG00000007892.7 |
| chr9 | 61698765 | 61698886 | 3.54427 | 4.2294 | 1.30266 | Distal Intergenic | 9 | 61913284 | 61914538 | 1255 | 2 | ENSMUST00000008036.7 | 215652 | Rplp1 | ENSMUSG00000007892.7 |
| chr9 | 61700173 | 61700264 | 3.7944 | 4.68498 | 1.57021 | Distal Intergenic | 9 | 61913284 | 61914538 | 1255 | 2 | ENSMUST00000008036.7 | 214274 | Rplp1 | ENSMUSG00000007892.7 |
| chr9 | 61713085 | 61713222 | 4.02511 | 5.1573 | 1.92075 | Distal Intergenic | 9 | 61913284 | 61914538 | 1255 | 2 | ENSMUST00000008036.7 | 201316 | Rplp1 | ENSMUSG00000007892.7 |
| chr9 | 61713752 | 61714030 | 3.72296 | 4.53875 | 1.44441 | Distal Intergenic | 9 | 61913284 | 61914538 | 1255 | 2 | ENSMUST00000008036.7 | 200508 | Rplp1 | ENSMUSG00000007892.7 |
| chr9 | 61714682 | 61714877 | 3.34274 | 3.84148 | 1.01733 | Distal Intergenic | 9 | 61913284 | 61914538 | 1255 | 2 | ENSMUST00000008036.7 | 199661 | Rplp1 | ENSMUSG00000007892.7 |
| chr9 | 61716919 | 61717548 | 3.44071 | 4.0393 | 1.19114 | Distal Intergenic | 9 | 61913284 | 61914538 | 1255 | 2 | ENSMUST00000008036.7 | 196990 | Rplp1 | ENSMUSG00000007892.7 |
| chr9 | 61723766 | 61724015 | 3.43697 | 4.00224 | 1.08271 | Distal Intergenic | 9 | 61913284 | 61914538 | 1255 | 2 | ENSMUST00000008036.7 | 190523 | Rplp1 | ENSMUSG00000007892.7 |
| chr9 | 61748224 | 61748469 | 3.45578 | 4.12163 | 1.22294 | Distal Intergenic | 9 | 61913284 | 61914538 | 1255 | 2 | ENSMUST00000008036.7 | 166069 | Rplp1 | ENSMUSG00000007892.7 |
| chr9 | 61756130 | 61756503 | 3.36369 | 3.87569 | 1.02654 | Distal Intergenic | 9 | 61913284 | 61914538 | 1255 | 2 | ENSMUST00000008036.7 | 158035 | Rplp1 | ENSMUSG00000007892.7 |
| chr9 | 61756949 | 61757460 | 3.9403 | 5.03477 | 1.82212 | Distal Intergenic | 9 | 61913284 | 61914538 | 1255 | 2 | ENSMUST00000008036.7 | 157078 | Rplp1 | ENSMUSG00000007892.7 |
| chr9 | 61759698 | 61759875 | 3.34153 | 3.83966 | 1.01733 | Distal Intergenic | 9 | 61913284 | 61914538 | 1255 | 2 | ENSMUST00000008036.7 | 154663 | Rplp1 | ENSMUSG00000007892.7 |
| chr9 | 61803860 | 61804186 | 3.40222 | 3.94514 | 1.06587 | Distal Intergenic | 9 | 61913284 | 61914538 | 1255 | 2 | ENSMUST00000008036.7 | 110352 | Rplp1 | ENSMUSG00000007892.7 |
| chr9 | 61804816 | 61805169 | 3.63761 | 4.37298 | 1.31616 | Distal Intergenic | 9 | 61913284 | 61914538 | 1255 | 2 | ENSMUST00000008036.7 | 109369 | Rplp1 | ENSMUSG00000007892.7 |
| chr9 | 61830115 | 61830210 | 3.50638 | 4.14252 | 1.19441 | Distal Intergenic | 9 | 61913284 | 61914538 | 1255 | 2 | ENSMUST00000008036.7 | 84328 | Rplp1 | ENSMUSG00000007892.7 |
| chr9 | 61855607 | 61855943 | 3.53125 | 4.28692 | 1.34729 | Distal Intergenic | 9 | 61913284 | 61914538 | 1255 | 2 | ENSMUST00000008036.7 | 58595 | Rplp1 | ENSMUSG00000007892.7 |
| chr9 | 61877142 | 61877487 | 3.40371 | 3.94837 | 1.07006 | Distal Intergenic | 9 | 61913284 | 61914538 | 1255 | 2 | ENSMUST00000008036.7 | 37051 | Rplp1 | ENSMUSG00000007892.7 |
| chr9 | 61884753 | 61884828 | 3.97313 | 5.00096 | 1.71471 | Distal Intergenic | 9 | 61913284 | 61914538 | 1255 | 2 | ENSMUST00000008036.7 | 29710 | Rplp1 | ENSMUSG00000007892.7 |
| chr9 | 61897852 | 61897971 | 3.71441 | 4.51238 | 1.40012 | Distal Intergenic | 9 | 61913284 | 61914538 | 1255 | 2 | ENSMUST00000008036.7 | 16567 | Rplp1 | ENSMUSG00000007892.7 |
| chr9 | 61898794 | 61899368 | 3.37225 | 3.92127 | 1.07503 | Distal Intergenic | 9 | 61913284 | 61914538 | 1255 | 2 | ENSMUST00000008036.7 | 15170 | Rplp1 | ENSMUSG00000007892.7 |
| chr9 | 76014938 | 76015040 | 3.72539 | 4.6693 | 1.58159 | Promoter | 9 | 76014984 | 76134187 | 119204 | 1 | ENSMUST00000183979.2 | 0 | Hmgcll1 | ENSMUSG00000007908.9 |
| chr7 | 27653698 | 27655073 | 3.71146 | 4.78033 | 1.64049 | Promoter | 7 | 27653915 | 27656208 | 2294 | 1 | ENSMUST00000008088.7 | 0 | Ttc9b | ENSMUSG00000007944.7 |
| chr7 | 27655481 | 27656378 | 3.65877 | 4.4276 | 1.37313 | 3' UTR | 7 | 27653915 | 27656208 | 2294 | 1 | ENSMUST00000008088.7 | 1567 | Ttc9b | ENSMUSG00000007944.7 |
| chr7 | 27658254 | 27658478 | 3.58408 | 4.27746 | 1.26917 | Exon (ENSMUST00000036453.8/ENSMUSG00000040390.8, exon 9 of 10) | 7 | 27653915 | 27656208 | 2294 | 1 | ENSMUST00000008088.7 | 4340 | Ttc9b | ENSMUSG00000007944.7 |
| chr7 | 101817965 | 101818649 | 3.83141 | 4.75553 | 1.58792 | Promoter | 7 | 101818313 | 101822726 | 4414 | 1 | ENSMUST00000008090.10 | 0 | Phox2a | ENSMUSG00000007946.10 |
| chr7 | 101818979 | 101819584 | 3.65499 | 4.43225 | 1.40142 | Promoter | 7 | 101818313 | 101822726 | 4414 | 1 | ENSMUST00000008090.10 | 667 | Phox2a | ENSMUSG00000007946.10 |
| chr7 | 101820486 | 101821383 | 3.75701 | 4.64672 | 1.56724 | Exon (ENSMUST00000008090.10/ENSMUSG00000007946.10, exon 2 of 3) | 7 | 101818313 | 101822726 | 4414 | 1 | ENSMUST00000008090.10 | 2174 | Phox2a | ENSMUSG00000007946.10 |
| chr7 | 44401878 | 44402601 | 3.37441 | 3.90222 | 1.0631 | Distal Intergenic | 7 | 44378887 | 44385820 | 6934 | 2 | ENSMUST00000134535.1 | -16059 | 1700008O03Rik | ENSMUSG00000008028.10 |
| chr1 | 43163727 | 43163945 | 3.23365 | 3.67765 | 1.01654 | Promoter | 1 | 43123074 | 43163961 | 40888 | 2 | ENSMUST00000008280.8 | 16 | Fhl2 | ENSMUSG00000008136.9 |
| chr6 | 124465453 | 124465702 | 3.54973 | 4.29715 | 1.3265 | Promoter | 6 | 124430759 | 124464794 | 34036 | 2 | ENSMUST00000008297.4 | -660 | Clstn3 | ENSMUSG00000008153.6 |

| | | | | | | | | | | | | | | | |
|---|---|---|---|---|---|---|---|---|---|---|---|---|---|---|---|
| chr10 | 93294670 | 93294988 | 3.24317 | 3.69191 | 1.01733 | Intron (ENSMUST00000008542.6/ENSMUSG00000008398.9, intron 1 of 4) | 10 | 93249323 | 93310811 | 61489 | 2 | ENSMUST00000129827.2 | 15823 | Elk3 | ENSMUSG00000008398.9 |
| chr7 | 45617375 | 45618197 | 3.67976 | 4.54539 | 1.48121 | Promoter | 7 | 45617606 | 45621059 | 3454 | 1 | ENSMUST00000008605.5 | 0 | Fut1 | ENSMUSG00000008461.5 |
| chr7 | 45618587 | 45618866 | 4.23609 | 5.70215 | 2.28529 | Promoter | 7 | 45617606 | 45621059 | 3454 | 1 | ENSMUST00000008605.5 | 982 | Fut1 | ENSMUSG00000008461.5 |
| chr4 | 91370955 | 91372100 | 4.3058 | 5.70929 | 2.28045 | Promoter | 4 | 91250763 | 91372034 | 121272 | 2 | ENSMUST00000107110.2 | 0 | Elavl2 | ENSMUSG00000008489.13 |
| chr4 | 91372606 | 91372703 | 3.5302 | 4.19596 | 1.26225 | Promoter | 4 | 91308677 | 91373030 | 64354 | 2 | ENSMUST00000147611.1 | 327 | Elavl2 | ENSMUSG00000008489.13 |
| chr4 | 91373015 | 91373260 | 3.31147 | 3.88057 | 1.09369 | Promoter | 4 | 91308677 | 91373030 | 64354 | 2 | ENSMUST00000147611.1 | 0 | Elavl2 | ENSMUSG00000008489.13 |
| chr4 | 91379840 | 91380144 | 3.35347 | 3.85766 | 1.01733 | Intron (ENSMUST00000102799.4/ENSMUSG00000008489.13, intron 2 of 7) | 4 | 91250769 | 91376496 | 125728 | 2 | ENSMUST00000107124.4 | -3345 | Elavl2 | ENSMUSG00000008489.13 |
| chr4 | 91375448 | 91375811 | 3.3946 | 3.98191 | 1.15258 | Promoter | 4 | 91258666 | 91376307 | 117642 | 2 | ENSMUST00000176469.1 | 496 | Elavl2 | ENSMUSG00000008489.13 |
| chr4 | 91376743 | 91376896 | 3.9572 | 5.07271 | 1.84686 | Promoter | 4 | 91250769 | 91376496 | 125728 | 2 | ENSMUST00000107124.4 | -248 | Elavl2 | ENSMUSG00000008489.13 |
| chr4 | 91377613 | 91377839 | 3.42372 | 4.37227 | 1.3801 | Intron (ENSMUST00000102799.4/ENSMUSG00000008489.13, intron 2 of 7) | 4 | 91250769 | 91376496 | 125728 | 2 | ENSMUST00000107124.4 | -1118 | Elavl2 | ENSMUSG00000008489.13 |
| chr4 | 91399519 | 91399594 | 3.69585 | 4.48164 | 1.39153 | Promoter | 4 | 91380432 | 91399976 | 19545 | 2 | ENSMUST00000144769.1 | 382 | Elavl2 | ENSMUSG00000008489.13 |
| chr4 | 91381588 | 91381713 | 3.37389 | 3.88849 | 1.01733 | Intron (ENSMUST00000102799.4/ENSMUSG00000008489.13, intron 2 of 7) | 4 | 91250769 | 91376496 | 125728 | 2 | ENSMUST00000107124.4 | -5093 | Elavl2 | ENSMUSG00000008489.13 |
| chr4 | 91384076 | 91384704 | 3.72151 | 4.60304 | 1.55854 | Intron (ENSMUST00000102799.4/ENSMUSG00000008489.13, intron 2 of 7) | 4 | 91250769 | 91376496 | 125728 | 2 | ENSMUST00000107124.4 | -7581 | Elavl2 | ENSMUSG00000008489.13 |
| chr4 | 91399043 | 91399118 | 3.4449 | 4.01726 | 1.09361 | Promoter | 4 | 91380432 | 91399976 | 19545 | 2 | ENSMUST00000144769.1 | 858 | Elavl2 | ENSMUSG00000008489.13 |
| chr4 | 91400697 | 91401277 | 3.63618 | 4.37682 | 1.3218 | Promoter | 4 | 91399895 | 91400785 | 891 | 2 | ENSMUST00000176151.1 | 0 | Elavl2 | ENSMUSG00000008489.13 |
| chr7 | 25129790 | 25130044 | 3.95208 | 5.02688 | 1.83811 | Intron (ENSMUST00000108415.4/ENSMUSG00000008496.13, intron 1 of 13) | 7 | 25092520 | 25132477 | 39958 | 2 | ENSMUST00000108418.5 | 2433 | Pou2f2 | ENSMUSG00000008496.13 |
| chr7 | 25131322 | 25131702 | 3.41585 | 4.10483 | 1.20594 | Promoter | 7 | 25092520 | 25132477 | 39958 | 2 | ENSMUST00000108418.5 | 775 | Pou2f2 | ENSMUSG00000008496.13 |
| chr7 | 25132423 | 25132768 | 4.4336 | 5.94497 | 2.41678 | Promoter | 7 | 25087344 | 25132488 | 45145 | 2 | ENSMUST00000108415.4 | 0 | Pou2f2 | ENSMUSG00000008496.13 |
| chr7 | 25134170 | 25134941 | 4.06649 | 5.27185 | 1.97769 | Intron (ENSMUST00000175754.2/ENSMUSG00000008496.13, intron 1 of 3) | 7 | 25093248 | 25132512 | 39265 | 2 | ENSMUST00000176408.2 | -1659 | Pou2f2 | ENSMUSG00000008496.13 |
| chr7 | 25179442 | 25180610 | 3.36446 | 4.07869 | 1.16065 | Distal Intergenic | 7 | 25120618 | 25159922 | 39305 | 2 | ENSMUST00000175754.2 | -19521 | Pou2f2 | ENSMUSG00000008496.13 |
| chr7 | 25181199 | 25182593 | 4.01453 | 5.25047 | 1.95083 | Distal Intergenic | 7 | 25120618 | 25159922 | 39305 | 2 | ENSMUST00000175754.2 | -21278 | Pou2f2 | ENSMUSG00000008496.13 |
| chr7 | 25177315 | 25177884 | 4.04856 | 5.17907 | 1.87586 | Distal Intergenic | 7 | 25120618 | 25159922 | 39305 | 2 | ENSMUST00000175754.2 | -17394 | Pou2f2 | ENSMUSG00000008496.13 |
| chr7 | 25178750 | 25179082 | 3.88439 | 4.99219 | 1.7445 | Distal Intergenic | 7 | 25120618 | 25159922 | 39305 | 2 | ENSMUST00000175754.2 | -18829 | Pou2f2 | ENSMUSG00000008496.13 |
| chr7 | 25183244 | 25183493 | 3.50233 | 4.26202 | 1.29058 | Distal Intergenic | 7 | 25120618 | 25159922 | 39305 | 2 | ENSMUST00000175754.2 | -23323 | Pou2f2 | ENSMUSG00000008496.13 |
| chr4 | 82470899 | 82471165 | 3.60013 | 4.30296 | 1.26993 | Intron (ENSMUST00000050872.9/ENSMUSG00000008575.12, intron 2 of 11) | 4 | 82498544 | 82499316 | 773 | 2 | ENSMUST00000152588.1 | 28151 | Nfib | ENSMUSG00000008575.12 |
| chr4 | 82472340 | 82472427 | 3.88037 | 4.83839 | 1.62171 | Intron (ENSMUST00000050872.9/ENSMUSG00000008575.12, intron 2 of 11) | 4 | 82498544 | 82499316 | 773 | 2 | ENSMUST00000152588.1 | 26889 | Nfib | ENSMUSG00000008575.12 |
| chr4 | 82486302 | 82486570 | 3.61155 | 4.60844 | 1.51779 | Intron (ENSMUST00000050872.9/ENSMUSG00000008575.12, intron 2 of 11) | 4 | 82498544 | 82499316 | 773 | 2 | ENSMUST00000152588.1 | 12746 | Nfib | ENSMUSG00000008575.12 |
| chr4 | 82493390 | 82493510 | 3.36652 | 3.92242 | 1.16042 | Intron (ENSMUST00000050872.9/ENSMUSG00000008575.12, intron 2 of 11) | 4 | 82498544 | 82499316 | 773 | 2 | ENSMUST00000152588.1 | 5806 | Nfib | ENSMUSG00000008575.12 |
| chr4 | 82495835 | 82495938 | 3.33741 | 3.85865 | 1.10245 | Intron (ENSMUST00000050872.9/ENSMUSG00000008575.12, intron 2 of 11) | 4 | 82498544 | 82499316 | 773 | 2 | ENSMUST00000152588.1 | 3378 | Nfib | ENSMUSG00000008575.12 |
| chr4 | 82497203 | 82497663 | 3.43275 | 4.14181 | 1.2111 | Exon (ENSMUST00000135024.1/ENSMUSG00000008575.12, exon 2 of 2) | 4 | 82498544 | 82499316 | 773 | 2 | ENSMUST00000152588.1 | 1653 | Nfib | ENSMUSG00000008575.12 |
| chr4 | 82498152 | 82498584 | 3.54972 | 4.32858 | 1.35942 | Promoter | 4 | 82498544 | 82499316 | 773 | 2 | ENSMUST00000152588.1 | 732 | Nfib | ENSMUSG00000008575.12 |
| chr4 | 82500390 | 82501036 | 3.49007 | 4.12984 | 1.24069 | Intron (ENSMUST00000050872.9/ENSMUSG00000008575.12, intron 1 of 11) | 4 | 82498544 | 82499316 | 773 | 2 | ENSMUST00000152588.1 | -1075 | Nfib | ENSMUSG00000008575.12 |
| chr4 | 82502799 | 82503012 | 3.52268 | 4.183 | 1.24813 | Intron (ENSMUST00000050872.9/ENSMUSG00000008575.12, intron 1 of 11) | 4 | 82290173 | 82505308 | 215136 | 2 | ENSMUST00000050872.9 | 2296 | Nfib | ENSMUSG00000008575.12 |
| chr4 | 82503982 | 82506836 | 3.90885 | 5.03968 | 1.84356 | Promoter | 4 | 82290173 | 82505308 | 215136 | 2 | ENSMUST00000050872.9 | 0 | Nfib | ENSMUSG00000008575.12 |
| chr9 | 48961939 | 48962464 | 3.74413 | 4.6502 | 1.58187 | Intron (ENSMUST00000008734.4/ENSMUSG00000008590.4, intron 1 of 8) | 9 | 48935008 | 48964990 | 29983 | 2 | ENSMUST00000008734.4 | 2526 | Htr3b | ENSMUSG00000008590.4 |
| chr16 | 7038675 | 7039151 | 3.93446 | 4.96289 | 1.75866 | Intron (ENSMUST00000115841.4/ENSMUSG00000008658.10, intron 1 of 12) | 16 | 7069835 | 7412479 | 342645 | 1 | ENSMUST00000056416.7 | -30684 | Rbfox1 | ENSMUSG00000008658.10 |
| chr16 | 7039963 | 7040630 | 3.52866 | 4.17291 | 1.19712 | Intron (ENSMUST00000115841.4/ENSMUSG00000008658.10, intron 1 of 12) | 16 | 7069835 | 7412479 | 342645 | 1 | ENSMUST00000056416.7 | -29205 | Rbfox1 | ENSMUSG00000008658.10 |
| chr7 | 118994548 | 118995129 | 3.75103 | 4.62086 | 1.53185 | Promoter | 7 | 118972040 | 118995211 | 23172 | 2 | ENSMUST00000008878.8 | 82 | Gprc5b | ENSMUSG00000008734.8 |
| chr7 | 118995548 | 118995795 | 3.47613 | 4.11722 | 1.25577 | Promoter | 7 | 118972040 | 118995211 | 23172 | 2 | ENSMUST00000008878.8 | -338 | Gprc5b | ENSMUSG00000008734.8 |
| chr10 | 75032371 | 75032516 | 3.44427 | 4.01531 | 1.09014 | Promoter | 10 | 75022304 | 75032528 | 10225 | 2 | ENSMUST00000159994.1 | 12 | Rtdr1 | ENSMUSG00000009070.9 |
| chr11 | 4746040 | 4746955 | 4.03911 | 5.15373 | 1.85154 | Promoter | 11 | 4736892 | 4746778 | 9887 | 2 | ENSMUST00000009219.2 | 0 | Cabp7 | ENSMUSG00000009075.2 |
| chr11 | 4747486 | 4747572 | 4.21779 | 5.52727 | 2.147 | Promoter | 11 | 4736892 | 4746778 | 9887 | 2 | ENSMUST00000009219.2 | -709 | Cabp7 | ENSMUSG00000009075.2 |
| chr11 | 4748009 | 4748204 | 3.59299 | 4.30852 | 1.32676 | Distal Intergenic | 11 | 4736892 | 4746778 | 9887 | 2 | ENSMUST00000009219.2 | -1232 | Cabp7 | ENSMUSG00000009075.2 |
| chr11 | 4749064 | 4749188 | 3.56619 | 4.30244 | 1.36455 | Distal Intergenic | 11 | 4736892 | 4746778 | 9887 | 2 | ENSMUST00000009219.2 | -2287 | Cabp7 | ENSMUSG00000009075.2 |
| chr11 | 4763917 | 4763992 | 3.35579 | 3.88447 | 1.09947 | Downstream (1-2kb) | 11 | 4736892 | 4746778 | 9887 | 2 | ENSMUST00000009219.2 | -17140 | Cabp7 | ENSMUSG00000009075.2 |
| chr16 | 18579913 | 18580946 | 3.61563 | 4.43762 | 1.45704 | Downstream (<1kb) | 16 | 18581713 | 18586969 | 5257 | 2 | ENSMUST00000009241.5 | 6023 | Tbx1 | ENSMUSG00000009097.9 |
| chr16 | 18581513 | 18583394 | 3.98878 | 5.42767 | 2.08863 | 3' UTR | 16 | 18581713 | 18586969 | 5257 | 2 | ENSMUST00000009241.5 | 3575 | Tbx1 | ENSMUSG00000009097.9 |
| chr16 | 18584618 | 18585103 | 3.65457 | 4.46344 | 1.46971 | Exon (ENSMUST00000009241.5/ENSMUSG00000009097.9, exon 2 of 7) | 16 | 18581713 | 18586969 | 5257 | 2 | ENSMUST00000009241.5 | 1866 | Tbx1 | ENSMUSG00000009097.9 |
| chr16 | 18586304 | 18587321 | 4.48052 | 6.31214 | 2.81117 | Promoter | 16 | 18581713 | 18586969 | 5257 | 2 | ENSMUST00000009241.5 | 0 | Tbx1 | ENSMUSG00000009097.9 |
| chr16 | 18587834 | 18589375 | 3.58883 | 4.34116 | 1.36952 | Promoter | 16 | 18581713 | 18586969 | 5257 | 2 | ENSMUST00000009241.5 | -866 | Tbx1 | ENSMUSG00000009097.9 |
| chr6 | 83061766 | 83062026 | 3.62249 | 4.37057 | 1.37231 | Intron (ENSMUST00000077502.2/ENSMUSG00000009145.4, intron 8 of 11) | 6 | 83057844 | 83067210 | 9367 | 1 | ENSMUST00000077502.2 | 3923 | Dqx1 | ENSMUSG00000009145.4 |
| chr6 | 83063811 | 83064068 | 3.38193 | 3.9006 | 1.01733 | Intron (ENSMUST00000077502.2/ENSMUSG00000009145.4, intron 8 of 11) | 6 | 83057844 | 83067210 | 9367 | 1 | ENSMUST00000077502.2 | 5968 | Dqx1 | ENSMUSG00000009145.4 |
| chr2 | 27141990 | 27142262 | 3.34914 | 4.15416 | 1.25676 | Promoter | 2 | 27110380 | 27142491 | 32112 | 2 | ENSMUST00000151224.2 | 229 | Fam163b | ENSMUSG00000009216.7 |
| chr7 | 142966326 | 142966635 | 3.41156 | 3.95584 | 1.05441 | Downstream (<1kb) | 7 | 142966822 | 142969264 | 2443 | 2 | ENSMUST00000009392.4 | 2629 | Ascl2 | ENSMUSG00000009248.5 |
| chr7 | 142967424 | 142968606 | 3.88041 | 4.84934 | 1.65473 | Promoter | 7 | 142966822 | 142969264 | 2443 | 2 | ENSMUST00000009392.4 | 658 | Ascl2 | ENSMUSG00000009248.5 |
| chr7 | 142969165 | 142969751 | 3.79052 | 4.7057 | 1.54556 | Promoter | 7 | 142966822 | 142969264 | 2443 | 2 | ENSMUST00000009392.4 | 0 | Ascl2 | ENSMUSG00000009248.5 |
| chr10 | 77966944 | 77967044 | 3.69355 | 4.52154 | 1.46474 | Promoter | 10 | 77909165 | 77967457 | 58293 | 2 | ENSMUST00000105400.3 | 413 | Trpm2 | ENSMUSG00000009292.12 |
| chr10 | 77968766 | 77969283 | 3.50947 | 4.13966 | 1.182 | Promoter | 10 | 77962790 | 77969804 | 7015 | 2 | ENSMUST00000105399.1 | 521 | Trpm2 | ENSMUSG00000009292.12 |
| chr10 | 77971801 | 77972284 | 3.62566 | 4.34633 | 1.29151 | Distal Intergenic | 10 | 77967045 | 77970563 | 3519 | 2 | ENSMUST00000154996.1 | -1239 | Trpm2 | ENSMUSG00000009292.12 |
| chr11 | 87816922 | 87817481 | 3.87385 | 4.85287 | 1.70584 | Promoter | 11 | 87816569 | 87817505 | 937 | 2 | ENSMUST00000157004.1 | 24 | Lpo | ENSMUSG00000009356.7 |
| chr11 | 87821739 | 87822024 | 3.45358 | 4.03719 | 1.11744 | Intron (ENSMUST00000103177.4/ENSMUSG00000009356.7, intron 2 of 12) | 11 | 87806428 | 87826113 | 19686 | 2 | ENSMUST00000103177.4 | 4089 | Lpo | ENSMUSG00000009356.7 |
| chr6 | 17461747 | 17462019 | 3.4351 | 4.02187 | 1.16341 | Distal Intergenic | 6 | 17463800 | 17573980 | 110181 | 1 | ENSMUST00000115443.2 | -1781 | Met | ENSMUSG00000009376.10 |
| chr6 | 17463373 | 17463497 | 3.65674 | 4.54212 | 1.46671 | Promoter | 6 | 17463800 | 17573980 | 110181 | 1 | ENSMUST00000115443.2 | -303 | Met | ENSMUSG00000009376.10 |
| chr6 | 17463917 | 17464529 | 4.38574 | 6.02699 | 2.5031 | Promoter | 6 | 17464038 | 17572229 | 108192 | 1 | ENSMUST00000115442.2 | 0 | Met | ENSMUSG00000009376.10 |
| chr6 | 17465334 | 17465567 | 4.13973 | 5.36531 | 2.01368 | Intron (ENSMUST00000115443.2/ENSMUSG00000009376.10, intron 1 of 21) | 6 | 17464107 | 17541412 | 77306 | 1 | ENSMUST00000145473.1 | 1228 | Met | ENSMUSG00000009376.10 |
| chr19 | 34743692 | 34743865 | 3.4396 | 4.08563 | 1.21605 | Intron (ENSMUST00000009522.3/ENSMUSG00000009378.3, intron 1 of 6) | 19 | 34668403 | 34747289 | 78887 | 2 | ENSMUST00000009522.3 | 3424 | Slc16a12 | ENSMUSG00000009378.3 |
| chr19 | 34745729 | 34746110 | 3.90246 | 4.87553 | 1.65111 | Intron (ENSMUST00000009522.3/ENSMUSG00000009378.3, intron 1 of 6) | 19 | 34668403 | 34747289 | 78887 | 2 | ENSMUST00000009522.3 | 1179 | Slc16a12 | ENSMUSG00000009378.3 |
| chr19 | 34746455 | 34747959 | 3.83249 | 4.86585 | 1.67472 | Promoter | 19 | 34668403 | 34747289 | 78887 | 2 | ENSMUST00000009522.3 | 0 | Slc16a12 | ENSMUSG00000009378.3 |
| chr6 | 115135061 | 115135657 | 3.99001 | 5.1907 | 1.94995 | Promoter | 6 | 115134902 | 115282626 | 147725 | 1 | ENSMUST00000009538.6 | 160 | Syn2 | ENSMUSG00000009394.8 |
| chr6 | 115140131 | 115140292 | 3.53984 | 4.29288 | 1.32876 | Intron (ENSMUST00000169345.1/ENSMUSG00000009394.8, intron 1 of 10) | 6 | 115134902 | 115282626 | 147725 | 1 | ENSMUST00000009538.6 | 5230 | Syn2 | ENSMUSG00000009394.8 |
| chr6 | 115144011 | 115144156 | 3.4946 | 4.11254 | 1.16529 | Intron (ENSMUST00000169345.1/ENSMUSG00000009394.8, intron 1 of 10) | 6 | 115134902 | 115282626 | 147725 | 1 | ENSMUST00000009538.6 | 9110 | Syn2 | ENSMUSG00000009394.8 |
| chr6 | 115160028 | 115160435 | 3.38464 | 3.95453 | 1.16887 | Intron (ENSMUST00000169345.1/ENSMUSG00000009394.8, intron 1 of 10) | 6 | 115134902 | 115282626 | 147725 | 1 | ENSMUST00000009538.6 | 25127 | Syn2 | ENSMUSG00000009394.8 |
| chr6 | 115164593 | 115164668 | 3.69277 | 4.60751 | 1.50308 | Intron (ENSMUST00000169345.1/ENSMUSG00000009394.8, intron 1 of 10) | 6 | 115134902 | 115282626 | 147725 | 1 | ENSMUST00000009538.6 | 29692 | Syn2 | ENSMUSG00000009394.8 |
| chr1 | 135472703 | 135473035 | 3.6994 | 4.49934 | 1.42454 | Exon (ENSMUST00000040599.9/ENSMUSG00000009418.10, exon 4 of 30) | 1 | 135455316 | 135467673 | 12358 | 2 | ENSMUST00000189252.1 | -5031 | Nav1 | ENSMUSG00000009418.10 |
| chr1 | 135510779 | 135511275 | 3.59394 | 4.42459 | 1.4298 | Promoter | 1 | 135458680 | 135510451 | 51772 | 2 | ENSMUST00000190130.1 | -329 | Nav1 | ENSMUSG00000009418.10 |
| chr7 | 46348757 | 46349033 | 3.49905 | 4.126 | 1.15655 | Distal Intergenic | 7 | 46376474 | 46379092 | 2619 | 1 | ENSMUST00000072514.1 | -27441 | Myod1 | ENSMUSG00000009471.3 |
| chr7 | 46350168 | 46350351 | 3.5099 | 4.32863 | 1.36371 | Distal Intergenic | 7 | 46376474 | 46379092 | 2619 | 1 | ENSMUST00000072514.1 | -26123 | Myod1 | ENSMUSG00000009471.3 |
| chr7 | 46376750 | 46377067 | 3.76802 | 4.65522 | 1.51684 | Promoter | 7 | 46376474 | 46379092 | 2619 | 1 | ENSMUST00000072514.1 | 277 | Myod1 | ENSMUSG00000009471.3 |
| chr7 | 46377580 | 46377847 | 3.77821 | 4.63302 | 1.47494 | Exon (ENSMUST00000072514.1/ENSMUSG00000009471.3, exon 2 of 3) | 7 | 46376474 | 46379092 | 2619 | 1 | ENSMUST00000072514.1 | 1107 | Myod1 | ENSMUSG00000009471.3 |
| chr7 | 143105439 | 143105566 | 3.91926 | 4.9975 | 1.80295 | Promoter | 7 | 143106362 | 143149537 | 43176 | 1 | ENSMUST00000186488.1 | -796 | Kcnq1 | ENSMUSG00000009545.9 |
| chr7 | 143106731 | 143107447 | 4.07399 | 5.29252 | 1.98774 | Promoter | 7 | 143106745 | 143149537 | 42793 | 1 | ENSMUST00000186798.1 | 0 | Kcnq1 | ENSMUSG00000009545.9 |
| chr7 | 143108415 | 143108578 | 3.25095 | 4.20919 | 1.23743 | Promoter | 7 | 143108351 | 143149537 | 41187 | 1 | ENSMUST00000186284.1 | 65 | Kcnq1 | ENSMUSG00000009545.9 |
| chr2 | 32741098 | 32741351 | 3.67594 | 4.55263 | 1.51554 | Promoter | 2 | 32732620 | 32741016 | 8397 | 2 | ENSMUST00000009695.2 | -83 | 6330409D20Rik | ENSMUSG00000009551.2 |
| chr8 | 33516661 | 33516755 | 3.69603 | 4.5108 | 1.4827 | Promoter | 8 | 33516738 | 33557342 | 40605 | 1 | ENSMUST00000149399.2 | 0 | Tex15 | ENSMUSG00000009628.9 |
| chr8 | 33517359 | 33517516 | 3.37545 | 3.91513 | 1.09497 | Promoter | 8 | 33517324 | 33585582 | 68259 | 1 | ENSMUST00000124501.1 | 36 | Tex15 | ENSMUSG00000009628.9 |
| chr10 | 59410003 | 59410078 | 3.40106 | 4.10772 | 1.15967 | Promoter | 10 | 59411027 | 59421970 | 10944 | 1 | ENSMUST00000162643.1 | -949 | Pla2g12b | ENSMUSG00000009646.8 |
| chr7 | 31041999 | 31042196 | 3.45933 | 4.04638 | 1.1194 | Promoter | 7 | 31032730 | 31042118 | 9389 | 2 | ENSMUST00000162733.2 | 0 | Fxyd5 | ENSMUSG00000009687.9 |
| chr11 | 59277839 | 59278030 | 3.62472 | 4.44694 | 1.43028 | Intron (ENSMUST00000010044.7/ENSMUSG00000009900.7, intron 1 of 3) | 11 | 59248033 | 59290752 | 42720 | 2 | ENSMUST00000010044.7 | 12722 | Wnt3a | ENSMUSG00000009900.7 |
| chr11 | 59288257 | 59288500 | 3.56948 | 4.2577 | 1.27992 | Intron (ENSMUST00000010044.7/ENSMUSG00000009900.7, intron 1 of 3) | 11 | 59248033 | 59290752 | 42720 | 2 | ENSMUST00000010044.7 | 2252 | Wnt3a | ENSMUSG00000009900.7 |
| chr11 | 59288837 | 59289222 | 3.58879 | 4.46215 | 1.41344 | Intron (ENSMUST00000010044.7/ENSMUSG00000009900.7, intron 1 of 3) | 11 | 59248033 | 59290752 | 42720 | 2 | ENSMUST00000010044.7 | 1530 | Wnt3a | ENSMUSG00000009900.7 |
| chr11 | 59290555 | 59290795 | 3.69467 | 4.59021 | 1.49099 | Promoter | 11 | 59248033 | 59290752 | 42720 | 2 | ENSMUST00000010044.7 | 0 | Wnt3a | ENSMUSG00000009900.7 |
| chr11 | 59291225 | 59291375 | 4.0033 | 5.09177 | 1.79769 | Promoter | 11 | 59248033 | 59290752 | 42720 | 2 | ENSMUST00000010044.7 | -474 | Wnt3a | ENSMUSG00000009900.7 |
| chr11 | 114781145 | 114782096 | 3.56398 | 4.27456 | 1.27757 | Exon (ENSMUST00000126555.2/ENSMUSG00000010021.8, exon 7 of 12) | 11 | 114784659 | 114785567 | 909 | 1 | ENSMUST00000131782.1 | -2563 | Kif19a | ENSMUSG00000010021.8 |
| chr11 | 114785411 | 114785548 | 3.7788 | 4.70992 | 1.59871 | Promoter | 11 | 114784659 | 114785567 | 909 | 1 | ENSMUST00000131782.1 | 753 | Kif19a | ENSMUSG00000010021.8 |
| chr11 | 114786340 | 114786715 | 3.50818 | 4.14084 | 1.18832 | Exon (ENSMUST00000138804.2/ENSMUSG00000010021.8, exon 12 of 16) | 11 | 114787903 | 114790739 | 2837 | 1 | ENSMUST00000138340.1 | -1188 | Kif19a | ENSMUSG00000010021.8 |
| chr11 | 114787477 | 114787576 | 3.75116 | 4.70067 | 1.57943 | Promoter | 11 | 114787903 | 114790739 | 2837 | 1 | ENSMUST00000138340.1 | -327 | Kif19a | ENSMUSG00000010021.8 |
| chr11 | 114789752 | 114789854 | 4.1987 | 5.54876 | 2.18673 | Exon (ENSMUST00000084368.6/ENSMUSG00000010021.8, exon 19 of 20) | 11 | 114787945 | 114789332 | 1388 | 1 | ENSMUST00000134165.1 | 1808 | Kif19a | ENSMUSG00000010021.8 |
| chr11 | 114790791 | 114791143 | 3.83368 | 4.77211 | 1.64741 | Downstream (<1kb) | 11 | 114787945 | 114789332 | 1388 | 1 | ENSMUST00000134165.1 | 2847 | Kif19a | ENSMUSG00000010021.8 |
| chr11 | 114791531 | 114792188 | 3.87706 | 4.92712 | 1.75167 | Exon (ENSMUST00000156053.2/ENSMUSG00000000202.4, exon 2 of 2) | 11 | 114787945 | 114789332 | 1388 | 1 | ENSMUST00000134165.1 | 3587 | Kif19a | ENSMUSG00000010021.8 |
| chr9 | 107665827 | 107666018 | 3.80346 | 4.73747 | 1.6241 | Intron (ENSMUST00000010208.8/ENSMUSG00000010064.10, intron 1 of 15) | 9 | 107651161 | 107667399 | 16239 | 2 | ENSMUST00000010208.8 | 1381 | Slc38a3 | ENSMUSG00000010064.10 |
| chr9 | 107666560 | 107667486 | 4.00028 | 5.38503 | 2.04462 | Promoter | 9 | 107651161 | 107667399 | 16239 | 2 | ENSMUST00000010208.8 | 0 | Slc38a3 | ENSMUSG00000010064.10 |
| chr9 | 107381535 | 107381665 | 3.25369 | 3.75529 | 1.0595 | Distal Intergenic | 9 | 107399612 | 107529346 | 129735 | 1 | ENSMUST00000010210.7 | -17947 | Cacna2d2 | ENSMUSG00000010066.10 |
| chr9 | 107382053 | 107382668 | 3.47974 | 4.17803 | 1.29514 | Distal Intergenic | 9 | 107399612 | 107529346 | 129735 | 1 | ENSMUST00000010210.7 | -16944 | Cacna2d2 | ENSMUSG00000010066.10 |
| chr9 | 107383739 | 107384136 | 3.46089 | 4.08663 | 1.23874 | Distal Intergenic | 9 | 107399612 | 107529346 | 129735 | 1 | ENSMUST00000010210.7 | -15476 | Cacna2d2 | ENSMUSG00000010066.10 |
| chr9 | 107385390 | 107385627 | 4.51299 | 6.30666 | 2.70329 | Distal Intergenic | 9 | 107399612 | 107529346 | 129735 | 1 | ENSMUST00000010210.7 | -13985 | Cacna2d2 | ENSMUSG00000010066.10 |
| chr9 | 107386179 | 107386523 | 3.82909 | 4.74988 | 1.60055 | Distal Intergenic | 9 | 107399612 | 107529346 | 129735 | 1 | ENSMUST00000010210.7 | -13089 | Cacna2d2 | ENSMUSG00000010066.10 |
| chr9 | 107391507 | 107391611 | 3.33846 | 3.83504 | 1.01733 | Distal Intergenic | 9 | 107399612 | 107529346 | 129735 | 1 | ENSMUST00000010210.7 | -8001 | Cacna2d2 | ENSMUSG00000010066.10 |
| chr9 | 107393404 | 107393919 | 3.64984 | 4.39123 | 1.32126 | Distal Intergenic | 9 | 107399612 | 107529346 | 129735 | 1 | ENSMUST00000010210.7 | -5693 | Cacna2d2 | ENSMUSG00000010066.10 |
| chr9 | 107401028 | 107401265 | 3.31623 | 3.80157 | 1.01733 | Promoter | 9 | 107400069 | 107529346 | 129278 | 1 | ENSMUST00000170737.1 | 960 | Cacna2d2 | ENSMUSG00000010066.10 |
| chr9 | 107401830 | 107402124 | 3.61631 | 4.39408 | 1.41999 | Intron (ENSMUST00000010210.7/ENSMUSG00000010066.10, intron 1 of 37) | 9 | 107400069 | 107529346 | 129278 | 1 | ENSMUST00000170737.1 | 1762 | Cacna2d2 | ENSMUSG00000010066.10 |
| chr9 | 107403554 | 107403729 | 3.67651 | 4.4637 | 1.44002 | Intron (ENSMUST00000010210.7/ENSMUSG00000010066.10, intron 1 of 37) | 9 | 107400069 | 107529346 | 129278 | 1 | ENSMUST00000170737.1 | 3486 | Cacna2d2 | ENSMUSG00000010066.10 |
| chr9 | 107507877 | 107508073 | 3.5173 | 4.16269 | 1.22212 | Intron (ENSMUST00000010210.7/ENSMUSG00000010066.10, intron 5 of 37) | 9 | 107509280 | 107513896 | 4617 | 1 | ENSMUST00000171809.1 | -1207 | Cacna2d2 | ENSMUSG00000010066.10 |
| chr9 | 107522445 | 107522728 | 3.45626 | 4.03611 | 1.09947 | Intron (ENSMUST00000010210.7/ENSMUSG00000010066.10, intron 23 of 37) | 9 | 107526657 | 107527567 | 911 | 1 | ENSMUST00000168959.1 | -3929 | Cacna2d2 | ENSMUSG00000010066.10 |
| chr9 | 107483015 | 107483317 | 3.36566 | 3.95847 | 1.1254 | Intron (ENSMUST00000010210.7/ENSMUSG00000010066.10, intron 3 of 37) | 9 | 107505604 | 107513534 | 7931 | 1 | ENSMUST00000169354.2 | -22287 | Cacna2d2 | ENSMUSG00000010066.10 |
| chr11 | 61452505 | 61452878 | 3.62808 | 4.45312 | 1.40241 | Exon (ENSMUST00000054927.8/ENSMUSG00000010086.12, exon 4 of 15) | 11 | 61452263 | 61453919 | 1657 | 2 | ENSMUST00000126859.2 | 1041 | Rnf112 | ENSMUSG00000010086.12 |
| chr11 | 61454035 | 61454390 | 3.41048 | 3.95875 | 1.07072 | Promoter | 11 | 61448442 | 61454106 | 5665 | 2 | ENSMUST00000102661.3 | 0 | Rnf112 | ENSMUSG00000010086.12 |
| chr11 | 61376016 | 61377039 | 3.61551 | 4.34912 | 1.31184 | Intron (ENSMUST00000010267.4/ENSMUSG00000010122.9, intron 1 of 16) | 11 | 61343402 | 61378064 | 34663 | 2 | ENSMUST00000131723.2 | 1025 | Slc47a1 | ENSMUSG00000010122.9 |
| chr11 | 61377695 | 61378124 | 4.03651 | 5.26366 | 1.93904 | Promoter | 11 | 61343401 | 61378081 | 34681 | 2 | ENSMUST00000010267.4 | 0 | Slc47a1 | ENSMUSG00000010122.9 |
| chr11 | 61379185 | 61379398 | 3.32029 | 3.80768 | 1.01733 | Promoter | 11 | 61362886 | 61378345 | 15460 | 2 | ENSMUST00000148671.1 | -841 | Slc47a1 | ENSMUSG00000010122.9 |
| chr11 | 61146480 | 61146963 | 3.53769 | 4.24388 | 1.35218 | Exon (ENSMUST00000010286.2/ENSMUSG00000010142.7, exon 4 of 5) | 11 | 61142665 | 61147759 | 5095 | 1 | ENSMUST00000123378.1 | 3816 | Tnfrsf13b | ENSMUSG00000010142.7 |
| chr8 | 123325557 | 123325831 | 3.29982 | 3.77691 | 1.01733 | Intron (ENSMUST00000127664.1/ENSMUSG00000092329.1, intron 1 of 15) | 8 | 123332713 | 123369515 | 36803 | 1 | ENSMUST00000010298.6 | -6882 | Spire2 | ENSMUSG00000010154.6 |
| chr8 | 123352493 | 123352618 | 3.61708 | 4.33576 | 1.2903 | Intron (ENSMUST00000127664.1/ENSMUSG00000092329.1, intron 1 of 15) | 8 | 123332713 | 123369515 | 36803 | 1 | ENSMUST00000010298.6 | 19781 | Spire2 | ENSMUSG00000010154.6 |
| chr1 | 190168958 | 190169529 | 5.70466 | 8.79596 | 4.74598 | Promoter | 1 | 190122453 | 190169399 | 46947 | 2 | ENSMUST00000010319.8 | 0 | Prox1 | ENSMUSG00000010175.8 |
| chr1 | 190169917 | 190170554 | 3.71004 | 4.87497 | 1.70813 | Promoter | 1 | 190121775 | 190170233 | 48459 | 2 | ENSMUST00000177288.2 | 0 | Prox1 | ENSMUSG00000010175.8 |
| chr1 | 190171004 | 190171134 | 3.27415 | 3.82468 | 1.03481 | Promoter | 1 | 190118035 | 190170714 | 52680 | 2 | ENSMUST00000175916.2 | -291 | Prox1 | ENSMUSG00000010175.8 |
| chr1 | 190171516 | 190173318 | 3.82746 | 4.82757 | 1.64867 | Promoter | 1 | 190118035 | 190170714 | 52680 | 2 | ENSMUST00000175916.2 | -803 | Prox1 | ENSMUSG00000010175.8 |
| chr1 | 190173685 | 190176328 | 3.95083 | 5.19701 | 1.91226 | Intron (ENSMUST00000110279.3/ENSMUSG00000079045.3, intron 1 of 1) | 1 | 190118035 | 190170714 | 52680 | 2 | ENSMUST00000175916.2 | -2972 | Prox1 | ENSMUSG00000010175.8 |
| chr1 | 190177006 | 190177108 | 4.12586 | 5.38215 | 2.01852 | Intron (ENSMUST00000110279.3/ENSMUSG00000079045.3, intron 1 of 1) | 1 | 190118035 | 190170714 | 52680 | 2 | ENSMUST00000175916.2 | -6293 | Prox1 | ENSMUSG00000010175.8 |
| chr1 | 190178609 | 190179049 | 3.56993 | 4.2812 | 1.33833 | Intron (ENSMUST00000110279.3/ENSMUSG00000079045.3, intron 1 of 1) | 1 | 190118035 | 190170714 | 52680 | 2 | ENSMUST00000175916.2 | -7896 | Prox1 | ENSMUSG00000010175.8 |
| chr1 | 190191672 | 190192252 | 3.68829 | 4.6715 | 1.54731 | Intron (ENSMUST00000110279.3/ENSMUSG00000079045.3, intron 1 of 1) | 1 | 190118035 | 190170714 | 52680 | 2 | ENSMUST00000175916.2 | -20959 | Prox1 | ENSMUSG00000010175.8 |
| chr1 | 133889850 | 133890007 | 4.08394 | 5.86431 | 2.39019 | Distal Intergenic | 1 | 133897199 | 133901327 | 4129 | 2 | ENSMUST00000124245.2 | 11320 | Optc | ENSMUSG00000010311.10 |
| chr1 | 133890504 | 133890655 | 3.44495 | 4.10589 | 1.17585 | Distal Intergenic | 1 | 133897199 | 133901327 | 4129 | 2 | ENSMUST00000124245.2 | 10672 | Optc | ENSMUSG00000010311.10 |
| chr1 | 133907570 | 133907721 | 3.35769 | 3.86404 | 1.01733 | Promoter | 1 | 133897200 | 133907999 | 10800 | 2 | ENSMUST00000153617.3 | 278 | Optc | ENSMUSG00000010311.10 |
| chr8 | 12495473 | 12495947 | 3.41702 | 4.07876 | 1.14073 | Distal Intergenic | 8 | 12573049 | 12600738 | 27690 | 1 | ENSMUST00000010579.6 | -77102 | Spaca7 | ENSMUSG00000010435.6 |
| chr8 | 12496522 | 12496706 | 3.61105 | 4.32864 | 1.30014 | Distal Intergenic | 8 | 12573049 | 12600738 | 27690 | 1 | ENSMUST00000010579.6 | -76343 | Spaca7 | ENSMUSG00000010435.6 |
| chr8 | 12497580 | 12497833 | 3.3672 | 3.88184 | 1.02946 | Distal Intergenic | 8 | 12573049 | 12600738 | 27690 | 1 | ENSMUST00000010579.6 | -75216 | Spaca7 | ENSMUSG00000010435.6 |
| chr8 | 12500347 | 12500668 | 3.5315 | 4.29397 | 1.30023 | Distal Intergenic | 8 | 12573049 | 12600738 | 27690 | 1 | ENSMUST00000010579.6 | -72381 | Spaca7 | ENSMUSG00000010435.6 |
| chr8 | 12501145 | 12501329 | 3.60742 | 4.32032 | 1.2952 | Distal Intergenic | 8 | 12573049 | 12600738 | 27690 | 1 | ENSMUST00000010579.6 | -71720 | Spaca7 | ENSMUSG00000010435.6 |
| chr8 | 12502166 | 12502442 | 3.43661 | 4.15028 | 1.20605 | Distal Intergenic | 8 | 12573049 | 12600738 | 27690 | 1 | ENSMUST00000010579.6 | -70607 | Spaca7 | ENSMUSG00000010435.6 |
| chr8 | 12503074 | 12505154 | 4.02329 | 5.1401 | 1.87044 | Distal Intergenic | 8 | 12573049 | 12600738 | 27690 | 1 | ENSMUST00000010579.6 | -67895 | Spaca7 | ENSMUSG00000010435.6 |
| chr8 | 12505567 | 12505842 | 3.67662 | 4.46455 | 1.4258 | Distal Intergenic | 8 | 12573049 | 12600738 | 27690 | 1 | ENSMUST00000010579.6 | -67207 | Spaca7 | ENSMUSG00000010435.6 |
| chr8 | 12548319 | 12548953 | 3.72312 | 4.53942 | 1.43313 | Distal Intergenic | 8 | 12573049 | 12600738 | 27690 | 1 | ENSMUST00000010579.6 | -24096 | Spaca7 | ENSMUSG00000010435.6 |
| chr8 | 12563158 | 12563353 | 3.36713 | 3.91191 | 1.13189 | Intron (ENSMUST00000180858.1/ENSMUSG00000097421.1, intron 1 of 1) | 8 | 12573049 | 12600738 | 27690 | 1 | ENSMUST00000010579.6 | -9696 | Spaca7 | ENSMUSG00000010435.6 |
| chr8 | 12563727 | 12563828 | 3.49934 | 4.14181 | 1.24514 | Intron (ENSMUST00000180858.1/ENSMUSG00000097421.1, intron 1 of 1) | 8 | 12573049 | 12600738 | 27690 | 1 | ENSMUST00000010579.6 | -9221 | Spaca7 | ENSMUSG00000010435.6 |
| chr7 | 137237965 | 137238335 | 3.3222 | 3.85067 | 1.07472 | Intron (ENSMUST00000033378.6/ENSMUSG00000010476.8, intron 6 of 15) | 7 | 137194698 | 137314414 | 119717 | 2 | ENSMUST00000168203.1 | 76079 | Ebf3 | ENSMUSG00000010476.8 |
| chr7 | 137253759 | 137254131 | 3.76789 | 4.71886 | 1.64117 | Intron (ENSMUST00000033378.6/ENSMUSG00000010476.8, intron 6 of 15) | 7 | 137194698 | 137314414 | 119717 | 2 | ENSMUST00000168203.1 | 60283 | Ebf3 | ENSMUSG00000010476.8 |
| chr7 | 137256150 | 137256362 | 3.85916 | 4.90635 | 1.72688 | Intron (ENSMUST00000033378.6/ENSMUSG00000010476.8, intron 6 of 15) | 7 | 137194698 | 137314414 | 119717 | 2 | ENSMUST00000168203.1 | 58052 | Ebf3 | ENSMUSG00000010476.8 |
| chr7 | 137266001 | 137266335 | 3.27066 | 3.75364 | 1.06941 | Intron (ENSMUST00000033378.6/ENSMUSG00000010476.8, intron 6 of 15) | 7 | 137194698 | 137314414 | 119717 | 2 | ENSMUST00000168203.1 | 48079 | Ebf3 | ENSMUSG00000010476.8 |
| chr7 | 137295384 | 137295514 | 3.35423 | 4.00414 | 1.11691 | Intron (ENSMUST00000033378.6/ENSMUSG00000010476.8, intron 6 of 15) | 7 | 137194698 | 137314414 | 119717 | 2 | ENSMUST00000168203.1 | 18900 | Ebf3 | ENSMUSG00000010476.8 |
| chr7 | 137301366 | 137301759 | 3.74476 | 4.66335 | 1.58581 | Intron (ENSMUST00000033378.6/ENSMUSG00000010476.8, intron 6 of 15) | 7 | 137194698 | 137314414 | 119717 | 2 | ENSMUST00000168203.1 | 12655 | Ebf3 | ENSMUSG00000010476.8 |
| chr7 | 137303738 | 137304153 | 3.68119 | 4.5722 | 1.53674 | Intron (ENSMUST00000033378.6/ENSMUSG00000010476.8, intron 6 of 15) | 7 | 137194698 | 137314414 | 119717 | 2 | ENSMUST00000168203.1 | 10261 | Ebf3 | ENSMUSG00000010476.8 |
| chr7 | 137304797 | 137305043 | 3.50712 | 4.17032 | 1.2892 | Intron (ENSMUST00000033378.6/ENSMUSG00000010476.8, intron 6 of 15) | 7 | 137194698 | 137314414 | 119717 | 2 | ENSMUST00000168203.1 | 9371 | Ebf3 | ENSMUSG00000010476.8 |
| chr7 | 137305464 | 137305581 | 3.38993 | 3.95849 | 1.16798 | Intron (ENSMUST00000033378.6/ENSMUSG00000010476.8, intron 6 of 15) | 7 | 137194698 | 137314414 | 119717 | 2 | ENSMUST00000168203.1 | 8833 | Ebf3 | ENSMUSG00000010476.8 |
| chr7 | 137307291 | 137310754 | 4.05257 | 5.35387 | 2.07445 | Exon (ENSMUST00000033378.6/ENSMUSG00000010476.8, exon 5 of 16) | 7 | 137194698 | 137314414 | 119717 | 2 | ENSMUST00000168203.1 | 3660 | Ebf3 | ENSMUSG00000010476.8 |
| chr7 | 137311182 | 137311757 | 3.92912 | 5.13328 | 1.89422 | Intron (ENSMUST00000033378.6/ENSMUSG00000010476.8, intron 4 of 15) | 7 | 137194698 | 137314414 | 119717 | 2 | ENSMUST00000168203.1 | 2657 | Ebf3 | ENSMUSG00000010476.8 |
| chr7 | 137312420 | 137312639 | 3.38777 | 4.86023 | 1.68256 | Intron (ENSMUST00000033378.6/ENSMUSG00000010476.8, intron 3 of 15) | 7 | 137194698 | 137314414 | 119717 | 2 | ENSMUST00000168203.1 | 1775 | Ebf3 | ENSMUSG00000010476.8 |
| chr7 | 137313161 | 137314785 | 4.02411 | 5.22925 | 1.99585 | Promoter | 7 | 137193673 | 137314445 | 120773 | 2 | ENSMUST00000033378.6 | 0 | Ebf3 | ENSMUSG00000010476.8 |
| chr7 | 137315131 | 137315483 | 3.70036 | 5.08937 | 1.84762 | Promoter | 7 | 137193673 | 137314445 | 120773 | 2 | ENSMUST00000169486.3 | -687 | Ebf3 | ENSMUSG00000010476.8 |
| chr7 | 137316240 | 137319894 | 3.9112 | 5.03336 | 1.84504 | Distal Intergenic | 7 | 137193673 | 137314445 | 120773 | 2 | ENSMUST00000169486.3 | -1796 | Ebf3 | ENSMUSG00000010476.8 |
| chr7 | 137320950 | 137323569 | 3.95336 | 5.20765 | 1.95048 | Distal Intergenic | 7 | 137193673 | 137314445 | 120773 | 2 | ENSMUST00000169486.3 | -6506 | Ebf3 | ENSMUSG00000010476.8 |
| chr7 | 137323977 | 137324592 | 3.84209 | 4.78011 | 1.61263 | Distal Intergenic | 7 | 137193673 | 137314445 | 120773 | 2 | ENSMUST00000169486.3 | -9533 | Ebf3 | ENSMUSG00000010476.8 |
| chr7 | 137325022 | 137325113 | 5.00704 | 7.4984 | 3.60842 | Distal Intergenic | 7 | 137193673 | 137314445 | 120773 | 2 | ENSMUST00000169486.3 | -10578 | Ebf3 | ENSMUSG00000010476.8 |
| chr7 | 137326265 | 137326720 | 3.63902 | 4.37801 | 1.33373 | Distal Intergenic | 7 | 137193673 | 137314445 | 120773 | 2 | ENSMUST00000169486.3 | -11821 | Ebf3 | ENSMUSG00000010476.8 |

| | | | | | | | | | | | | | | | |
|---|---|---|---|---|---|---|---|---|---|---|---|---|---|---|---|
| chr2 | 181763600 | 181764170 | 3.79519 | 4.67284 | 1.52011 | Promoter | 2 | 181763332 | 181827797 | 64466 | 1 | ENSMUST00000081125.5 | 269 | Myt1 | ENSMUSG00000010505.10 |
| chr2 | 181765342 | 181765628 | 3.50251 | 4.27651 | 1.29517 | Intron (ENSMUST00000081125.5/ENSMUSG00000010505.10, intron 1 of 22) | 2 | 181767029 | 181795892 | 28864 | 1 | ENSMUST00000156190.2 | -1401 | Myt1 | ENSMUSG00000010505.10 |
| chr2 | 181766679 | 181767202 | 3.40503 | 3.95524 | 1.08735 | Promoter | 2 | 181767029 | 181795892 | 28864 | 1 | ENSMUST00000156190.2 | 0 | Myt1 | ENSMUSG00000010505.10 |
| chr1 | 180263570 | 180263799 | 3.43662 | 4.18188 | 1.25104 | Promoter | 1 | 180245638 | 180263438 | 17801 | 2 | ENSMUST00000133340.1 | -133 | Psen2 | ENSMUSG00000010609.10 |
| chr1 | 180264577 | 180264707 | 4.02762 | 5.571 | 2.15143 | Distal Intergenic | 1 | 180245638 | 180263438 | 17801 | 2 | ENSMUST00000133340.1 | -1140 | Psen2 | ENSMUSG00000010609.10 |
| chr1 | 180265314 | 180265538 | 3.57365 | 4.34067 | 1.38724 | Distal Intergenic | 1 | 180245638 | 180263438 | 17801 | 2 | ENSMUST00000133340.1 | -1877 | Psen2 | ENSMUSG00000010609.10 |
| chr1 | 180282615 | 180283001 | 3.48074 | 4.08282 | 1.13394 | Distal Intergenic | 1 | 180245638 | 180263438 | 17801 | 2 | ENSMUST00000133340.1 | -19178 | Psen2 | ENSMUSG00000010609.10 |
| chr1 | 180283662 | 180284102 | 3.51166 | 4.18709 | 1.24929 | Distal Intergenic | 1 | 180245638 | 180263438 | 17801 | 2 | ENSMUST00000133340.1 | -20225 | Psen2 | ENSMUSG00000010609.10 |
| chr9 | 119154844 | 119155283 | 3.36993 | 3.93042 | 1.16289 | Intron (ENSMUST00000010795.4/ENSMUSG00000010651.4, intron 3 of 11) | 9 | 119148023 | 119157093 | 9071 | 2 | ENSMUST00000010795.4 | 1810 | Acaa1b | ENSMUSG00000010651.4 |
| chr9 | 119155896 | 119156198 | 3.95921 | 5.07469 | 1.84301 | Promoter | 9 | 119148023 | 119157093 | 9071 | 2 | ENSMUST00000010795.4 | 895 | Acaa1b | ENSMUSG00000010651.4 |
| chr7 | 143490923 | 143491144 | 3.37863 | 3.89562 | 1.01733 | 3' UTR | 7 | 143501545 | 143502541 | 997 | 2 | ENSMUST00000010904.4 | 11397 | Phlda2 | ENSMUSG00000010760.4 |
| chr7 | 143500429 | 143500567 | 3.51556 | 4.14424 | 1.16501 | Downstream (<1kb) | 7 | 143501545 | 143502541 | 997 | 2 | ENSMUST00000010904.4 | 1974 | Phlda2 | ENSMUSG00000010760.4 |
| chr7 | 143502100 | 143502472 | 3.51523 | 4.37031 | 1.34509 | Promoter | 7 | 143501545 | 143502541 | 997 | 2 | ENSMUST00000010904.4 | 69 | Phlda2 | ENSMUSG00000010760.4 |
| chr7 | 143502778 | 143503865 | 3.88843 | 4.88774 | 1.69829 | Promoter | 7 | 143501545 | 143502541 | 997 | 2 | ENSMUST00000010904.4 | -238 | Phlda2 | ENSMUSG00000010760.4 |
| chr7 | 143506202 | 143506571 | 3.72481 | 4.54333 | 1.44374 | Distal Intergenic | 7 | 143501545 | 143502541 | 997 | 2 | ENSMUST00000010904.4 | -3662 | Phlda2 | ENSMUSG00000010760.4 |
| chr6 | 18023931 | 18024195 | 3.27177 | 3.80619 | 1.08058 | Intron (ENSMUST00000010941.3/ENSMUSG00000010797.3, intron 2 of 4) | 6 | 17989215 | 18030410 | 41196 | 2 | ENSMUST00000156317.1 | 6215 | Wnt2 | ENSMUSG00000010797.3 |
| chr6 | 18030432 | 18030633 | 4.06224 | 5.19059 | 1.87522 | Promoter | 6 | 17988940 | 18030585 | 41646 | 2 | ENSMUST00000010941.3 | 0 | Wnt2 | ENSMUSG00000010797.3 |
| chr6 | 18031333 | 18031539 | 3.65646 | 4.42526 | 1.38254 | Promoter | 6 | 17988940 | 18030585 | 41646 | 2 | ENSMUST00000010941.3 | -749 | Wnt2 | ENSMUSG00000010797.3 |
| chr6 | 18032515 | 18032716 | 3.69124 | 4.4834 | 1.40364 | Distal Intergenic | 6 | 17988940 | 18030585 | 41646 | 2 | ENSMUST00000010941.3 | -1931 | Wnt2 | ENSMUSG00000010797.3 |
| chr11 | 42179712 | 42180046 | 3.59323 | 4.295 | 1.28429 | Intron (ENSMUST00000020707.6/ENSMUSG00000010803.7, intron 1 of 9) | 11 | 42162553 | 42182163 | 19611 | 2 | ENSMUST00000153147.1 | 2117 | Gabra1 | ENSMUSG00000010803.7 |
| chr11 | 42181163 | 42181259 | 3.28691 | 3.75758 | 1.01733 | Promoter | 11 | 42162553 | 42182163 | 19611 | 2 | ENSMUST00000153147.1 | 904 | Gabra1 | ENSMUSG00000010803.7 |
| chr11 | 42182609 | 42183134 | 3.84166 | 4.7932 | 1.63294 | Promoter | 11 | 42130939 | 42182930 | 51992 | 2 | ENSMUST00000020707.6 | 0 | Gabra1 | ENSMUSG00000010803.7 |
| chr5 | 143357441 | 143358164 | 3.76564 | 4.64755 | 1.51247 | Promoter | 5 | 143357338 | 143391468 | 34131 | 1 | ENSMUST00000110733.3 | 104 | Grid2ip | ENSMUSG00000010825.12 |
| chr5 | 143359310 | 143359560 | 3.84535 | 4.7678 | 1.57447 | Intron (ENSMUST00000110733.3/ENSMUSG00000010825.12, intron 1 of 21) | 5 | 143357338 | 143391468 | 34131 | 1 | ENSMUST00000110733.3 | 1973 | Grid2ip | ENSMUSG00000010825.12 |
| chr5 | 143361627 | 143361702 | 3.38193 | 3.9006 | 1.01733 | Intron (ENSMUST00000110733.3/ENSMUSG00000010825.12, intron 1 of 21) | 5 | 143363908 | 143391798 | 27891 | 1 | ENSMUST00000010969.9 | -2206 | Grid2ip | ENSMUSG00000010825.12 |
| chr5 | 143364268 | 143364995 | 3.8526 | 4.8679 | 1.66844 | Promoter | 5 | 143363908 | 143391798 | 27891 | 1 | ENSMUST00000010969.9 | 361 | Grid2ip | ENSMUSG00000010825.12 |
| chr5 | 143368434 | 143368568 | 3.65726 | 4.57307 | 1.49896 | Intron (ENSMUST00000110733.3/ENSMUSG00000010825.12, intron 2 of 21) | 5 | 143370092 | 143392152 | 22061 | 1 | ENSMUST00000120825.1 | -1524 | Grid2ip | ENSMUSG00000010825.12 |
| chr5 | 143369549 | 143372235 | 3.73447 | 4.63466 | 1.5296 | Promoter | 5 | 143370092 | 143392152 | 22061 | 1 | ENSMUST00000120825.1 | 0 | Grid2ip | ENSMUSG00000010825.12 |
| chr5 | 143373168 | 143373243 | 3.88158 | 4.89459 | 1.72966 | Exon (ENSMUST00000110733.3/ENSMUSG00000010825.12, exon 3 of 22) | 5 | 143375461 | 143380442 | 4982 | 1 | ENSMUST00000138948.1 | -2218 | Grid2ip | ENSMUSG00000010825.12 |
| chr5 | 143373665 | 143374161 | 3.8162 | 4.81261 | 1.6839 | Intron (ENSMUST00000110733.3/ENSMUSG00000010825.12, intron 4 of 21) | 5 | 143375461 | 143380442 | 4982 | 1 | ENSMUST00000138948.1 | -1300 | Grid2ip | ENSMUSG00000010825.12 |
| chr5 | 143386674 | 143387807 | 3.58823 | 4.3615 | 1.38723 | Exon (ENSMUST00000110733.3/ENSMUSG00000010825.12, exon 15 of 22) | 5 | 143375461 | 143380442 | 4982 | 1 | ENSMUST00000138948.1 | 11214 | Grid2ip | ENSMUSG00000010825.12 |
| chr15 | 79510058 | 79511685 | 3.92501 | 5.05368 | 1.8275 | Distal Intergenic | 15 | 79516408 | 79527739 | 11332 | 1 | ENSMUST00000010974.7 | -4723 | Kdelr3 | ENSMUSG00000010830.7 |
| chr15 | 79513620 | 79514545 | 3.64867 | 4.45336 | 1.43549 | Distal Intergenic | 15 | 79516408 | 79527739 | 11332 | 1 | ENSMUST00000010974.7 | -1863 | Kdelr3 | ENSMUSG00000010830.7 |
| chr15 | 79514994 | 79515145 | 3.92889 | 4.9475 | 1.73407 | Distal Intergenic | 15 | 79516408 | 79527739 | 11332 | 1 | ENSMUST00000010974.7 | -1263 | Kdelr3 | ENSMUSG00000010830.7 |
| chr15 | 79516158 | 79516284 | 3.36816 | 3.88453 | 1.03402 | Promoter | 15 | 79516408 | 79527739 | 11332 | 1 | ENSMUST00000010974.7 | -124 | Kdelr3 | ENSMUSG00000010830.7 |
| chr3 | 146149909 | 146149995 | 4.35074 | 6.04555 | 2.50639 | Promoter | 3 | 146149979 | 146172445 | 22467 | 1 | ENSMUST00000169533.1 | 0 | Mcoln2 | ENSMUSG00000011008.8 |
| chr5 | 33104139 | 33104215 | 3.90099 | 4.92999 | 1.74696 | Promoter | 5 | 33104219 | 33162872 | 58654 | 1 | ENSMUST00000011178.2 | -4 | Slc5a1 | ENSMUSG00000011034.4 |
| chr12 | 116077405 | 116077847 | 3.46342 | 4.19601 | 1.26885 | Promoter | 12 | 116077726 | 116139972 | 62247 | 1 | ENSMUST00000176078.2 | 0 | Vipr2 | ENSMUSG00000011171.6 |
| chr12 | 116078191 | 116078520 | 3.9792 | 5.16582 | 1.87857 | Promoter | 12 | 116077773 | 116133327 | 55555 | 1 | ENSMUST00000177199.1 | 419 | Vipr2 | ENSMUSG00000011171.6 |
| chr12 | 116078852 | 116078955 | 3.63348 | 4.42766 | 1.43121 | Intron (ENSMUST00000176078.2/ENSMUSG00000011171.6, intron 1 of 9) | 12 | 116077773 | 116133327 | 55555 | 1 | ENSMUST00000177199.1 | 1080 | Vipr2 | ENSMUSG00000011171.6 |
| chr11 | 46056085 | 46056211 | 3.91015 | 4.88275 | 1.64584 | Promoter | 11 | 46056058 | 46065082 | 9025 | 1 | ENSMUST00000137014.1 | 28 | Adam19 | ENSMUSG00000011256.11 |
| chr11 | 46057123 | 46057491 | 3.46329 | 4.21352 | 1.25689 | Intron (ENSMUST00000011400.7/ENSMUSG00000011256.11, intron 1 of 22) | 11 | 46056058 | 46065082 | 9025 | 1 | ENSMUST00000137014.1 | 1066 | Adam19 | ENSMUSG00000011256.11 |
| chr11 | 46057873 | 46058299 | 3.67498 | 4.54044 | 1.4528 | Intron (ENSMUST00000011400.7/ENSMUSG00000011256.11, intron 1 of 22) | 11 | 46056058 | 46065082 | 9025 | 1 | ENSMUST00000137014.1 | 1816 | Adam19 | ENSMUSG00000011256.11 |
| chr7 | 19479982 | 19480357 | 3.98512 | 5.03333 | 1.75738 | Distal Intergenic | 7 | 19489056 | 19496760 | 7705 | 1 | ENSMUST00000011407.7 | -8699 | Exoc3l2 | ENSMUSG00000011263.10 |
| chr7 | 19488148 | 19488465 | 3.35464 | 3.85943 | 1.01733 | Promoter | 7 | 19489056 | 19496760 | 7705 | 1 | ENSMUST00000011407.7 | -591 | Exoc3l2 | ENSMUSG00000011263.10 |
| chr7 | 19488865 | 19489336 | 3.44124 | 4.01605 | 1.10886 | Promoter | 7 | 19489056 | 19496760 | 7705 | 1 | ENSMUST00000011407.7 | 0 | Exoc3l2 | ENSMUSG00000011263.10 |
| chr7 | 19577520 | 19577781 | 3.37262 | 3.95891 | 1.1345 | Promoter | 7 | 19577287 | 19580656 | 3370 | 1 | ENSMUST00000108453.1 | 234 | Zfp296 | ENSMUSG00000011267.7 |
| chr7 | 27390928 | 27391003 | 4.09954 | 5.30204 | 2.01024 | Intron (ENSMUST00000108362.2/ENSMUSG00000011751.11, intron 2 of 17) | 7 | 27356383 | 27394506 | 38124 | 2 | ENSMUST00000108362.2 | 3503 | Sptbn4 | ENSMUSG00000011751.11 |
| chr7 | 27391338 | 27391661 | 3.51367 | 4.18142 | 1.27915 | Exon (ENSMUST00000108362.2/ENSMUSG00000011751.11, exon 2 of 18) | 7 | 27356383 | 27394506 | 38124 | 2 | ENSMUST00000108362.2 | 2845 | Sptbn4 | ENSMUSG00000011751.11 |
| chr7 | 27394503 | 27394782 | 3.18598 | 3.74943 | 1.01733 | Promoter | 7 | 27356383 | 27394506 | 38124 | 2 | ENSMUST00000108362.2 | 0 | Sptbn4 | ENSMUSG00000011751.11 |
| chr7 | 27395423 | 27395553 | 3.48085 | 4.09198 | 1.16652 | Promoter | 7 | 27356383 | 27396198 | 39816 | 2 | ENSMUST00000108364.2 | 645 | Sptbn4 | ENSMUSG00000011751.11 |
| chr7 | 27395949 | 27397401 | 3.71881 | 4.56933 | 1.45511 | Promoter | 7 | 27356383 | 27396198 | 39816 | 2 | ENSMUST00000108364.2 | 0 | Sptbn4 | ENSMUSG00000011751.11 |
| chr7 | 27399687 | 27399786 | 3.38193 | 3.9006 | 1.01733 | Intron (ENSMUST00000011895.8/ENSMUSG00000011751.11, intron 17 of 35) | 7 | 27356383 | 27396553 | 40171 | 2 | ENSMUST00000108363.2 | -3135 | Sptbn4 | ENSMUSG00000011751.11 |
| chr7 | 27400888 | 27401477 | 3.38008 | 3.90288 | 1.03513 | Intron (ENSMUST00000011895.8/ENSMUSG00000011751.11, intron 17 of 35) | 7 | 27356383 | 27396553 | 40171 | 2 | ENSMUST00000108363.2 | -4336 | Sptbn4 | ENSMUSG00000011751.11 |
| chr7 | 27402656 | 27402992 | 3.40747 | 4.09335 | 1.20489 | Exon (ENSMUST00000011895.8/ENSMUSG00000011751.11, exon 17 of 36) | 7 | 27356383 | 27396553 | 40171 | 2 | ENSMUST00000108363.2 | -6104 | Sptbn4 | ENSMUSG00000011751.11 |
| chr7 | 27403736 | 27405125 | 4.04876 | 5.26413 | 1.95347 | Exon (ENSMUST00000011895.8/ENSMUSG00000011751.11, exon 16 of 36) | 7 | 27356383 | 27396553 | 40171 | 2 | ENSMUST00000108363.2 | -7184 | Sptbn4 | ENSMUSG00000011751.11 |
| chr7 | 27405653 | 27405977 | 3.72613 | 4.57393 | 1.49243 | Intron (ENSMUST00000011895.8/ENSMUSG00000011751.11, intron 15 of 35) | 7 | 27356383 | 27396553 | 40171 | 2 | ENSMUST00000108363.2 | -9101 | Sptbn4 | ENSMUSG00000011751.11 |
| chr7 | 27406812 | 27407187 | 3.40771 | 3.97511 | 1.13939 | Exon (ENSMUST00000011895.8/ENSMUSG00000011751.11, exon 15 of 36) | 7 | 27356383 | 27396553 | 40171 | 2 | ENSMUST00000108363.2 | -10260 | Sptbn4 | ENSMUSG00000011751.11 |
| chr7 | 27407920 | 27408790 | 3.66191 | 4.54185 | 1.45907 | Exon (ENSMUST00000011895.8/ENSMUSG00000011751.11, exon 14 of 36) | 7 | 27356383 | 27396553 | 40171 | 2 | ENSMUST00000108363.2 | -11368 | Sptbn4 | ENSMUSG00000011751.11 |
| chr7 | 27409384 | 27409666 | 3.43898 | 4.3193 | 1.32731 | Intron (ENSMUST00000011895.8/ENSMUSG00000011751.11, intron 13 of 35) | 7 | 27356383 | 27396553 | 40171 | 2 | ENSMUST00000108363.2 | -12832 | Sptbn4 | ENSMUSG00000011751.11 |
| chr7 | 27432803 | 27433461 | 3.43751 | 4.17426 | 1.24702 | Promoter | 7 | 27433136 | 27434369 | 1234 | 2 | ENSMUST00000138087.1 | 908 | Sptbn4 | ENSMUSG00000011751.11 |
| chr5 | 107884182 | 107884431 | 3.95323 | 5.01876 | 1.78171 | Distal Intergenic | 5 | 107744795 | 107875107 | 130313 | 2 | ENSMUST00000112642.2 | -9076 | Evi5 | ENSMUSG00000011831.11 |
| chr9 | 69906863 | 69907911 | 3.75238 | 4.67054 | 1.54772 | Distal Intergenic | 9 | 69989466 | 70008317 | 18852 | 1 | ENSMUST00000034754.6 | -81555 | Bnip2 | ENSMUSG00000011958.12 |
| chr16 | 17797480 | 17797800 | 4.55061 | 6.35537 | 2.73608 | Promoter | 16 | 17797282 | 17808287 | 11006 | 1 | ENSMUST00000012161.3 | 199 | Scarf2 | ENSMUSG00000012017.3 |
| chr16 | 17798357 | 17798627 | 3.51852 | 4.38751 | 1.37771 | Intron (ENSMUST00000012161.3/ENSMUSG00000012017.3, intron 1 of 10) | 16 | 17797282 | 17808287 | 11006 | 1 | ENSMUST00000012161.3 | 1076 | Scarf2 | ENSMUSG00000012017.3 |
| chr16 | 17800685 | 17800814 | 3.7276 | 4.66573 | 1.55387 | Intron (ENSMUST00000012161.3/ENSMUSG00000012017.3, intron 1 of 10) | 16 | 17797282 | 17808287 | 11006 | 1 | ENSMUST00000012161.3 | 3404 | Scarf2 | ENSMUSG00000012017.3 |
| chr16 | 17802351 | 17802594 | 3.61589 | 4.52714 | 1.43928 | Exon (ENSMUST00000012161.3/ENSMUSG00000012017.3, exon 2 of 11) | 16 | 17797282 | 17808287 | 11006 | 1 | ENSMUST00000012161.3 | 5070 | Scarf2 | ENSMUSG00000012017.3 |
| chr16 | 17816103 | 17816243 | 3.29554 | 4.0146 | 1.17398 | Distal Intergenic | 16 | 17797282 | 17808287 | 11006 | 1 | ENSMUST00000012161.3 | 18822 | Scarf2 | ENSMUSG00000012017.3 |
| chr3 | 138196687 | 138196859 | 3.70787 | 4.83032 | 1.64812 | Distal Intergenic | 3 | 138164135 | 138186713 | 22579 | 2 | ENSMUST00000012186.4 | -9975 | 4930579F01Rik | ENSMUSG00000012042.4 |
| chr3 | 138197280 | 138197724 | 3.71122 | 4.56356 | 1.49084 | Distal Intergenic | 3 | 138164135 | 138186713 | 22579 | 2 | ENSMUST00000012186.4 | -10568 | 4930579F01Rik | ENSMUSG00000012042.4 |
| chr4 | 134064792 | 134065048 | 3.34248 | 3.84112 | 1.01733 | Promoter | 4 | 134065912 | 134072725 | 6814 | 1 | ENSMUST00000137053.1 | -864 | Aim1l | ENSMUSG00000012123.10 |
| chr4 | 134065589 | 134065850 | 3.41681 | 3.96538 | 1.06004 | Promoter | 4 | 134065912 | 134072725 | 6814 | 1 | ENSMUST00000137053.1 | -62 | Aim1l | ENSMUSG00000012123.10 |
| chr4 | 134083772 | 134083889 | 3.68082 | 4.53918 | 1.45557 | Intron (ENSMUST00000121391.2/ENSMUSG00000012123.10, intron 14 of 19) | 4 | 134082436 | 134082888 | 453 | 1 | ENSMUST00000137529.1 | 1337 | Aim1l | ENSMUSG00000012123.10 |
| chr1 | 78511700 | 78512027 | 3.59538 | 4.30589 | 1.31099 | Promoter | 1 | 78511812 | 78522837 | 11026 | 1 | ENSMUST00000152111.1 | 0 | Mogat1 | ENSMUSG00000012187.8 |
| chr18 | 34547425 | 34547739 | 3.57643 | 4.26575 | 1.27158 | 3' UTR | 18 | 34542313 | 34548273 | 5961 | 1 | ENSMUST00000012426.2 | 5113 | Wnt8a | ENSMUSG00000012282.2 |
| chr18 | 34548559 | 34548716 | 3.45495 | 4.27356 | 1.30488 | Distal Intergenic | 18 | 34542313 | 34548273 | 5961 | 1 | ENSMUST00000012426.2 | 6247 | Wnt8a | ENSMUSG00000012282.2 |
| chr18 | 34549079 | 34549505 | 3.73649 | 4.7281 | 1.5478 | Distal Intergenic | 18 | 34542313 | 34548273 | 5961 | 1 | ENSMUST00000012426.2 | 6767 | Wnt8a | ENSMUSG00000012282.2 |
| chr18 | 34551073 | 34551239 | 3.68767 | 4.55337 | 1.48586 | Distal Intergenic | 18 | 34542313 | 34548273 | 5961 | 1 | ENSMUST00000012426.2 | 8761 | Wnt8a | ENSMUSG00000012282.2 |
| chr2 | 103325081 | 103325415 | 3.39068 | 3.96972 | 1.1792 | Distal Intergenic | 2 | 103263433 | 103303278 | 39846 | 2 | ENSMUST00000090475.4 | -21804 | Ehf | ENSMUSG00000012350.10 |
| chr6 | 122707591 | 122707666 | 3.61675 | 4.33711 | 1.30669 | Promoter | 6 | 122707568 | 122714633 | 7066 | 1 | ENSMUST00000012540.4 | 24 | Nanog | ENSMUSG00000012396.7 |
| chr5 | 7960340 | 7960611 | 3.58219 | 4.52638 | 1.45581 | Promoter | 5 | 7960472 | 7982213 | 21742 | 1 | ENSMUST00000115421.1 | 0 | Steap4 | ENSMUSG00000012428.8 |
| chr19 | 37362666 | 37362936 | 3.38186 | 3.92065 | 1.08889 | Distal Intergenic | 19 | 37376403 | 37421859 | 45457 | 1 | ENSMUST00000012587.3 | -13467 | Kif11 | ENSMUSG00000012443.3 |
| chr5 | 67082812 | 67082887 | 3.43986 | 4.03138 | 1.1661 | Distal Intergenic | 5 | 67097044 | 67099057 | 2014 | 2 | ENSMUST00000174251.1 | 16170 | Phox2b | ENSMUSG00000012520.9 |
| chr5 | 67089026 | 67089104 | 3.89324 | 4.87031 | 1.67958 | Distal Intergenic | 5 | 67097044 | 67099057 | 2014 | 2 | ENSMUST00000174251.1 | 9953 | Phox2b | ENSMUSG00000012520.9 |
| chr5 | 67090708 | 67091110 | 3.6283 | 4.5857 | 1.52845 | Distal Intergenic | 5 | 67097044 | 67099057 | 2014 | 2 | ENSMUST00000174251.1 | 7947 | Phox2b | ENSMUSG00000012520.9 |
| chr5 | 67091630 | 67093021 | 3.84395 | 4.85388 | 1.74316 | Downstream (1-2kb) | 5 | 67097044 | 67099057 | 2014 | 2 | ENSMUST00000174251.1 | 6036 | Phox2b | ENSMUSG00000012520.9 |
| chr5 | 67093748 | 67094604 | 3.88666 | 5.00286 | 1.79779 | 3' UTR | 5 | 67097044 | 67099057 | 2014 | 2 | ENSMUST00000174251.1 | 4453 | Phox2b | ENSMUSG00000012520.9 |
| chr5 | 67094916 | 67095251 | 3.87535 | 5.03615 | 1.75668 | 3' UTR | 5 | 67097044 | 67099057 | 2014 | 2 | ENSMUST00000174251.1 | 3806 | Phox2b | ENSMUSG00000012520.9 |
| chr5 | 67095583 | 67097689 | 4.24189 | 5.88016 | 2.40795 | 3' UTR | 5 | 67097044 | 67099057 | 2014 | 2 | ENSMUST00000174251.1 | 1368 | Phox2b | ENSMUSG00000012520.9 |
| chr5 | 67098006 | 67098086 | 4.47758 | 6.16336 | 2.5713 | Promoter | 5 | 67097044 | 67099057 | 2014 | 2 | ENSMUST00000174251.1 | 971 | Phox2b | ENSMUSG00000012520.9 |
| chr5 | 67098534 | 67099073 | 4.5922 | 6.45071 | 2.83526 | Promoter | 5 | 67097044 | 67099057 | 2014 | 2 | ENSMUST00000174251.1 | 0 | Phox2b | ENSMUSG00000012520.9 |
| chr5 | 67099659 | 67100827 | 3.8095 | 4.83647 | 1.69884 | Promoter | 5 | 67094405 | 67099301 | 4897 | 2 | ENSMUST00000012664.9 | -359 | Phox2b | ENSMUSG00000012520.9 |
| chr5 | 67101602 | 67102318 | 3.74125 | 4.71099 | 1.60404 | Distal Intergenic | 5 | 67094405 | 67099301 | 4897 | 2 | ENSMUST00000012664.9 | -2302 | Phox2b | ENSMUSG00000012520.9 |
| chr5 | 67102627 | 67102854 | 3.51326 | 4.24505 | 1.30664 | Distal Intergenic | 5 | 67094405 | 67099301 | 4897 | 2 | ENSMUST00000012664.9 | -3327 | Phox2b | ENSMUSG00000012520.9 |
| chr5 | 67105635 | 67105829 | 3.39196 | 4.25461 | 1.30275 | Distal Intergenic | 5 | 67094405 | 67099301 | 4897 | 2 | ENSMUST00000012664.9 | -6335 | Phox2b | ENSMUSG00000012520.9 |
| chr5 | 67107302 | 67107403 | 4.17824 | 5.55703 | 2.20028 | Distal Intergenic | 5 | 67094405 | 67099301 | 4897 | 2 | ENSMUST00000012664.9 | -8002 | Phox2b | ENSMUSG00000012520.9 |
| chr5 | 67107783 | 67107858 | 4.28819 | 5.76852 | 2.31049 | Distal Intergenic | 5 | 67094405 | 67099301 | 4897 | 2 | ENSMUST00000012664.9 | -8483 | Phox2b | ENSMUSG00000012520.9 |
| chr5 | 67139131 | 67139452 | 3.44362 | 4.02947 | 1.14446 | Distal Intergenic | 5 | 67094405 | 67099301 | 4897 | 2 | ENSMUST00000012664.9 | -39831 | Phox2b | ENSMUSG00000012520.9 |
| chr5 | 67140815 | 67141661 | 3.37387 | 3.89331 | 1.03455 | Distal Intergenic | 5 | 67094405 | 67099301 | 4897 | 2 | ENSMUST00000012664.9 | -41515 | Phox2b | ENSMUSG00000012520.9 |
| chr10 | 60430925 | 60431002 | 3.98758 | 5.06796 | 1.82082 | Intron (ENSMUST00000073242.5/ENSMUSG00000012819.10, intron 16 of 66) | 10 | 60376772 | 60406453 | 29682 | 2 | ENSMUST00000144462.2 | -24473 | Cdh23 | ENSMUSG00000012819.10 |
| chr10 | 60602217 | 60602292 | 3.59233 | 4.28419 | 1.24985 | Intron (ENSMUST00000073242.5/ENSMUSG00000012819.10, intron 2 of 66) | 10 | 60628480 | 60653033 | 24554 | 2 | ENSMUST00000153677.1 | 50741 | Cdh23 | ENSMUSG00000012819.10 |
| chr10 | 60624573 | 60624813 | 3.28307 | 3.76046 | 1.04675 | Intron (ENSMUST00000073242.5/ENSMUSG00000012819.10, intron 2 of 66) | 10 | 60628480 | 60653033 | 24554 | 2 | ENSMUST00000153677.1 | 28220 | Cdh23 | ENSMUSG00000012819.10 |
| chr10 | 60685021 | 60685096 | 4.29852 | 5.73627 | 2.33196 | Intron (ENSMUST00000105461.4/ENSMUSG00000012819.10, intron 2 of 70) | 10 | 60657266 | 60687257 | 29992 | 2 | ENSMUST00000123309.1 | 2161 | Cdh23 | ENSMUSG00000012819.10 |
| chr10 | 60685569 | 60685724 | 3.30189 | 3.79688 | 1.07308 | Intron (ENSMUST00000105461.4/ENSMUSG00000012819.10, intron 2 of 70) | 10 | 60657266 | 60687257 | 29992 | 2 | ENSMUST00000123309.1 | 1533 | Cdh23 | ENSMUSG00000012819.10 |
| chr10 | 60686040 | 60687551 | 3.73552 | 4.62516 | 1.58591 | Promoter | 10 | 60657266 | 60687257 | 29992 | 2 | ENSMUST00000123309.1 | 0 | Cdh23 | ENSMUSG00000012819.10 |
| chr10 | 60697044 | 60697175 | 3.66651 | 4.46355 | 1.46101 | Promoter | 10 | 60583079 | 60696490 | 113412 | 2 | ENSMUST00000153149.2 | -555 | Cdh23 | ENSMUSG00000012819.10 |
| chr8 | 83901096 | 83901219 | 3.40877 | 3.94953 | 1.04699 | Promoter | 8 | 83900735 | 83939987 | 39253 | 1 | ENSMUST00000152978.2 | 362 | Lphn1 | ENSMUSG00000013033.11 |
| chr16 | 22441491 | 22441690 | 3.45467 | 4.03433 | 1.10163 | Distal Intergenic | 16 | 22381309 | 22439719 | 58411 | 2 | ENSMUST00000079601.7 | -1773 | Etv5 | ENSMUSG00000013089.10 |
| chr16 | 22442129 | 22442489 | 3.60638 | 4.3369 | 1.35152 | Distal Intergenic | 16 | 22381309 | 22439719 | 58411 | 2 | ENSMUST00000079601.7 | -2411 | Etv5 | ENSMUSG00000013089.10 |
| chr17 | 56473040 | 56473859 | 3.8072 | 4.73531 | 1.56758 | Intron (ENSMUST00000067538.5/ENSMUSG00000013236.11, intron 1 of 32) | 17 | 56422237 | 56476369 | 54133 | 2 | ENSMUST00000143018.1 | 2510 | Ptprs | ENSMUSG00000013236.11 |
| chr17 | 56474253 | 56476483 | 3.84946 | 4.85412 | 1.70454 | Promoter | 17 | 56412433 | 56476483 | 64051 | 2 | ENSMUST00000067538.5 | 0 | Ptprs | ENSMUSG00000013236.11 |
| chr2 | 156047604 | 156047679 | 4.02947 | 5.13941 | 1.85261 | Exon (ENSMUST00000109611.2/ENSMUSG00000013338.11, exon 11 of 45) | 2 | 156019139 | 156052947 | 33809 | 2 | ENSMUST00000109611.2 | 5268 | Fer1l4 | ENSMUSG00000013338.11 |
| chr7 | 43476905 | 43477167 | 3.62874 | 4.35935 | 1.31809 | Exon (ENSMUST00000107974.1/ENSMUSG00000013367.4, exon 5 of 8) | 7 | 43472907 | 43490075 | 17169 | 2 | ENSMUST00000107974.1 | 12908 | Iglon5 | ENSMUSG00000013367.4 |
| chr7 | 43489514 | 43490609 | 3.47564 | 4.09745 | 1.16775 | Promoter | 7 | 43472907 | 43490075 | 17169 | 2 | ENSMUST00000107974.1 | 0 | Iglon5 | ENSMUSG00000013367.4 |
| chr7 | 43491117 | 43491630 | 3.8531 | 4.99941 | 1.76575 | Downstream (<1kb) | 7 | 43472907 | 43490075 | 17169 | 2 | ENSMUST00000107974.1 | -1043 | Iglon5 | ENSMUSG00000013367.4 |
| chr7 | 43492038 | 43492113 | 4.33311 | 5.73438 | 2.25904 | 3' UTR | 7 | 43472907 | 43490075 | 17169 | 2 | ENSMUST00000107974.1 | -1964 | Iglon5 | ENSMUSG00000013367.4 |
| chr7 | 43493547 | 43493813 | 3.65929 | 4.40765 | 1.32663 | 3' UTR | 7 | 43472907 | 43490075 | 17169 | 2 | ENSMUST00000107974.1 | -3473 | Iglon5 | ENSMUSG00000013367.4 |
| chr7 | 43495056 | 43495143 | 3.50994 | 4.14087 | 1.18277 | Intron (ENSMUST00000013497.2/ENSMUSG00000013353.5, intron 7 of 7) | 7 | 43472907 | 43490075 | 17169 | 2 | ENSMUST00000107974.1 | -4982 | Iglon5 | ENSMUSG00000013367.4 |
| chr11 | 96003768 | 96006397 | 4.21502 | 5.61305 | 2.2462 | Promoter | 11 | 95957163 | 96005940 | 48778 | 2 | ENSMUST00000013559.2 | 0 | Igf2bp1 | ENSMUSG00000013415.4 |
| chr11 | 96006719 | 96009525 | 4.05775 | 5.36553 | 2.06154 | Promoter | 11 | 95957163 | 96005940 | 48778 | 2 | ENSMUST00000013559.2 | -780 | Igf2bp1 | ENSMUSG00000013415.4 |
| chr11 | 96009907 | 96012087 | 3.73753 | 4.58405 | 1.47099 | Distal Intergenic | 11 | 95957163 | 96005940 | 48778 | 2 | ENSMUST00000013559.2 | -3968 | Igf2bp1 | ENSMUSG00000013415.4 |
| chr11 | 96012399 | 96013356 | 4.11082 | 5.3571 | 2.00229 | Distal Intergenic | 11 | 95957163 | 96005940 | 48778 | 2 | ENSMUST00000013559.2 | -6460 | Igf2bp1 | ENSMUSG00000013415.4 |
| chr11 | 96013748 | 96013908 | 3.46558 | 4.09815 | 1.24365 | Distal Intergenic | 11 | 95957163 | 96005940 | 48778 | 2 | ENSMUST00000013559.2 | -7809 | Igf2bp1 | ENSMUSG00000013415.4 |
| chr11 | 96015090 | 96015199 | 3.73518 | 4.56606 | 1.45473 | Distal Intergenic | 11 | 95957163 | 96005940 | 48778 | 2 | ENSMUST00000013559.2 | -9151 | Igf2bp1 | ENSMUSG00000013415.4 |
| chr11 | 95902294 | 95902396 | 3.84383 | 4.75255 | 1.54762 | Intron (ENSMUST00000038343.6/ENSMUSG00000013418.8, intron 1 of 10) | 11 | 95865943 | 95914891 | 48949 | 2 | ENSMUST00000038343.6 | 12495 | B4galnt2 | ENSMUSG00000013418.8 |
| chr11 | 95912208 | 95912342 | 3.96083 | 5.14989 | 1.88931 | Intron (ENSMUST00000038343.6/ENSMUSG00000013418.8, intron 1 of 10) | 11 | 95865943 | 95914891 | 48949 | 2 | ENSMUST00000038343.6 | 2549 | B4galnt2 | ENSMUSG00000013418.8 |
| chr11 | 95913079 | 95914084 | 3.96753 | 5.12614 | 1.86451 | Promoter | 11 | 95865943 | 95914891 | 48949 | 2 | ENSMUST00000038343.6 | 807 | B4galnt2 | ENSMUSG00000013418.8 |
| chr11 | 95914624 | 95915497 | 3.95896 | 5.1516 | 1.90759 | Promoter | 11 | 95865943 | 95914891 | 48949 | 2 | ENSMUST00000038343.6 | 0 | B4galnt2 | ENSMUSG00000013418.8 |
| chr11 | 95916282 | 95917048 | 3.49652 | 4.73769 | 1.6216 | Distal Intergenic | 11 | 95865943 | 95914891 | 48949 | 2 | ENSMUST00000038343.6 | -1392 | B4galnt2 | ENSMUSG00000013418.8 |
| chr11 | 95917444 | 95917872 | 3.54888 | 4.34576 | 1.35287 | Distal Intergenic | 11 | 95865943 | 95914891 | 48949 | 2 | ENSMUST00000038343.6 | -2554 | B4galnt2 | ENSMUSG00000013418.8 |
| chr11 | 119310674 | 119310953 | 3.72075 | 4.55588 | 1.45993 | Intron (ENSMUST00000106250.2/ENSMUSG00000013483.9, intron 1 of 22) | 11 | 119307768 | 119345375 | 37608 | 1 | ENSMUST00000106250.2 | 2907 | Card14 | ENSMUSG00000013483.9 |
| chr2 | 170356840 | 170357175 | 3.67538 | 4.4843 | 1.47876 | Promoter | 2 | 170347945 | 170357113 | 9169 | 2 | ENSMUST00000133673.2 | 0 | Bcas1 | ENSMUSG00000013523.8 |
| chr2 | 170412069 | 170412642 | 3.54067 | 4.23198 | 1.28291 | Intron (ENSMUST00000109152.3/ENSMUSG00000013523.8, intron 3 of 11) | 2 | 170346991 | 170406509 | 59519 | 2 | ENSMUST00000154650.2 | -5561 | Bcas1 | ENSMUSG00000013523.8 |
| chr2 | 170438183 | 170438269 | 3.96324 | 5.04952 | 1.83647 | Distal Intergenic | 2 | 170397532 | 170427845 | 30314 | 2 | ENSMUST00000145920.1 | -10339 | Bcas1 | ENSMUSG00000013523.8 |
| chr2 | 170438833 | 170439086 | 3.66821 | 4.44984 | 1.41644 | Distal Intergenic | 2 | 170397532 | 170427845 | 30314 | 2 | ENSMUST00000145920.1 | -10989 | Bcas1 | ENSMUSG00000013523.8 |
| chr2 | 170406218 | 170406693 | 3.57828 | 4.56701 | 1.48981 | Promoter | 2 | 170346991 | 170406509 | 59519 | 2 | ENSMUST00000154650.2 | 0 | Bcas1 | ENSMUSG00000013523.8 |
| chr2 | 170441063 | 170441286 | 3.45501 | 4.31059 | 1.346 | Distal Intergenic | 2 | 170397532 | 170427845 | 30314 | 2 | ENSMUST00000145920.1 | -13219 | Bcas1 | ENSMUSG00000013523.8 |
| chr2 | 170439820 | 170440105 | 3.41087 | 4.04042 | 1.21432 | Distal Intergenic | 2 | 170397532 | 170427845 | 30314 | 2 | ENSMUST00000145920.1 | -11976 | Bcas1 | ENSMUSG00000013523.8 |
| chr9 | 71215282 | 71216549 | 3.43285 | 4.33092 | 1.35031 | Promoter | 9 | 71215789 | 71296243 | 80455 | 1 | ENSMUST00000034723.5 | 0 | Aldh1a2 | ENSMUSG00000013584.5 |
| chr19 | 32726660 | 32726992 | 3.61329 | 4.4561 | 1.42588 | Distal Intergenic | 19 | 32672563 | 32712298 | 39736 | 2 | ENSMUST00000070210.4 | -14363 | Atad1 | ENSMUSG00000013662.5 |
| chr19 | 32727373 | 32727546 | 3.49376 | 4.11044 | 1.16295 | Distal Intergenic | 19 | 32672563 | 32712298 | 39736 | 2 | ENSMUST00000070210.4 | -15076 | Atad1 | ENSMUSG00000013662.5 |
| chr19 | 32754472 | 32754617 | 3.96337 | 5.08253 | 1.82636 | Distal Intergenic | 19 | 32757497 | 32826160 | 68664 | 1 | ENSMUST00000013807.7 | -2880 | Pten | ENSMUSG00000013663.7 |
| chr19 | 32867633 | 32868168 | 4.06433 | 5.28196 | 1.99566 | Distal Intergenic | 19 | 32811225 | 32817977 | 6753 | 1 | ENSMUST00000154806.1 | 56409 | Pten | ENSMUSG00000013663.7 |
| chr19 | 32885659 | 32886236 | 3.2597 | 3.71669 | 1.01733 | Distal Intergenic | 19 | 32811225 | 32817977 | 6753 | 1 | ENSMUST00000154806.1 | 74435 | Pten | ENSMUSG00000013663.7 |
| chr5 | 122078382 | 122078751 | 3.78072 | 4.8527 | 1.70834 | Distal Intergenic | 5 | 122100951 | 122106854 | 5904 | 1 | ENSMUST00000014080.7 | -22200 | Myl2 | ENSMUSG00000013936.7 |
| chr5 | 122079841 | 122080216 | 3.65139 | 4.43318 | 1.43579 | Distal Intergenic | 5 | 122100951 | 122106854 | 5904 | 1 | ENSMUST00000014080.7 | -20735 | Myl2 | ENSMUSG00000013936.7 |
| chr5 | 122103628 | 122103703 | 3.51923 | 4.1616 | 1.20802 | Promoter | 5 | 122103913 | 122105003 | 1091 | 1 | ENSMUST00000126006.1 | -210 | Myl2 | ENSMUSG00000013936.7 |
| chr4 | 44692028 | 44692194 | 3.94281 | 4.94108 | 1.68012 | Exon (ENSMUST00000102932.4/ENSMUSG00000014030.10, exon 3 of 9) | 4 | 44569971 | 44683815 | 113845 | 2 | ENSMUST00000172949.2 | -8214 | Pax5 | ENSMUSG00000014030.10 |
| chr4 | 44692635 | 44693214 | 3.43018 | 4.01121 | 1.112 | Intron (ENSMUST00000102932.4/ENSMUSG00000014030.10, intron 2 of 8) | 4 | 44569971 | 44683815 | 113845 | 2 | ENSMUST00000172949.2 | -8821 | Pax5 | ENSMUSG00000014030.10 |
| chr4 | 44694583 | 44694711 | 3.5184 | 4.27888 | 1.32495 | Intron (ENSMUST00000102932.4/ENSMUSG00000014030.10, intron 2 of 8) | 4 | 44679454 | 44703413 | 23960 | 2 | ENSMUST00000174761.2 | 8702 | Pax5 | ENSMUSG00000014030.10 |
| chr4 | 44697247 | 44697634 | 3.93901 | 5.08877 | 1.81867 | Exon (ENSMUST00000102932.4/ENSMUSG00000014030.10, exon 2 of 9) | 4 | 44679454 | 44703413 | 23960 | 2 | ENSMUST00000174761.2 | 5779 | Pax5 | ENSMUSG00000014030.10 |
| chr4 | 44698204 | 44698405 | 3.41803 | 3.97684 | 1.09429 | Intron (ENSMUST00000102932.4/ENSMUSG00000014030.10, intron 1 of 8) | 4 | 44679454 | 44703413 | 23960 | 2 | ENSMUST00000174761.2 | 5008 | Pax5 | ENSMUSG00000014030.10 |
| chr4 | 44699724 | 44700241 | 3.60307 | 4.32386 | 1.32007 | Intron (ENSMUST00000102932.4/ENSMUSG00000014030.10, intron 1 of 8) | 4 | 44679454 | 44703413 | 23960 | 2 | ENSMUST00000174761.2 | 3172 | Pax5 | ENSMUSG00000014030.10 |
| chr4 | 44700727 | 44700809 | 3.79354 | 4.74853 | 1.61611 | Intron (ENSMUST00000102932.4/ENSMUSG00000014030.10, intron 1 of 8) | 4 | 44679454 | 44703413 | 23960 | 2 | ENSMUST00000174761.2 | 2604 | Pax5 | ENSMUSG00000014030.10 |
| chr4 | 44701159 | 44702570 | 3.68574 | 4.67868 | 1.56989 | Promoter | 4 | 44679454 | 44703413 | 23960 | 2 | ENSMUST00000174761.2 | 843 | Pax5 | ENSMUSG00000014030.10 |
| chr4 | 44703227 | 44705558 | 4.07616 | 5.41625 | 2.0972 | Promoter | 4 | 44679440 | 44704006 | 24567 | 2 | ENSMUST00000146335.2 | 0 | Pax5 | ENSMUSG00000014030.10 |
| chr4 | 44706010 | 44706183 | 4.23322 | 5.93213 | 2.434 | Intron (ENSMUST00000102932.4/ENSMUSG00000014030.10, intron 1 of 8) | 4 | 44679440 | 44704006 | 24567 | 2 | ENSMUST00000146335.2 | -2005 | Pax5 | ENSMUSG00000014030.10 |
| chr4 | 44707049 | 44707203 | 3.61078 | 4.39314 | 1.41373 | Intron (ENSMUST00000102932.4/ENSMUSG00000014030.10, intron 1 of 8) | 4 | 44679440 | 44704006 | 24567 | 2 | ENSMUST00000146335.2 | -3044 | Pax5 | ENSMUSG00000014030.10 |
| chr4 | 44708227 | 44708436 | 3.76806 | 4.6842 | 1.60217 | Intron (ENSMUST00000102932.4/ENSMUSG00000014030.10, intron 1 of 8) | 4 | 44531497 | 44710408 | 178912 | 2 | ENSMUST00000134968.3 | 1972 | Pax5 | ENSMUSG00000014030.10 |
| chr4 | 44710767 | 44712488 | 3.64521 | 4.79498 | 1.6487 | Promoter | 4 | 44531145 | 44710487 | 179343 | 2 | ENSMUST00000014174.8 | -281 | Pax5 | ENSMUSG00000014030.10 |
| chr4 | 44712893 | 44714393 | 3.4955 | 4.38134 | 1.38111 | Distal Intergenic | 4 | 44531145 | 44710487 | 179343 | 2 | ENSMUST00000014174.8 | -2407 | Pax5 | ENSMUSG00000014030.10 |
| chr4 | 44715090 | 44715835 | 3.43319 | 4.04156 | 1.22011 | Distal Intergenic | 4 | 44531145 | 44710487 | 179343 | 2 | ENSMUST00000014174.8 | -4604 | Pax5 | ENSMUSG00000014030.10 |
| chr4 | 44716248 | 44717187 | 3.81767 | 4.77171 | 1.60469 | Distal Intergenic | 4 | 44531145 | 44710487 | 179343 | 2 | ENSMUST00000014174.8 | -5762 | Pax5 | ENSMUSG00000014030.10 |

| | | | | | | | | | | | | | | | |
|---|---|---|---|---|---|---|---|---|---|---|---|---|---|---|---|
| chr4 | 44717792 | 44718702 | 3.48669 | 4.12729 | 1.24017 | Distal Intergenic | 4 | 44531145 | 44710487 | 179343 | 2 | ENSMUST00000014174.8 | -7306 | Pax5 | ENSMUSG00000014030.10 |
| chr11 | 96015716 | 96015791 | 4.00067 | 5.19586 | 1.92156 | Distal Intergenic | 11 | 96024545 | 96030831 | 6287 | 1 | ENSMUST00000103157.4 | -8754 | Gip | ENSMUSG00000014351.7 |
| chr4 | 151129023 | 151129615 | 3.49162 | 4.13067 | 1.2436 | Promoter | 4 | 151059814 | 151129495 | 69682 | 2 | ENSMUST00000105668.2 | 0 | Camta1 | ENSMUSG00000014592.14 |
| chr4 | 151131080 | 151131389 | 3.34685 | 3.87072 | 1.09565 | Intron (ENSMUST00000169423.3/ENSMUSG00000014592.14, intron 11 of 24) | 4 | 151059814 | 151129495 | 69682 | 2 | ENSMUST00000105668.2 | -1586 | Camta1 | ENSMUSG00000014592.14 |
| chr4 | 151139247 | 151139495 | 3.18598 | 3.74943 | 1.01733 | Promoter | 4 | 151086039 | 151139584 | 53546 | 2 | ENSMUST00000143434.1 | 89 | Camta1 | ENSMUSG00000014592.14 |
| chr4 | 151872991 | 151873167 | 3.84223 | 4.94225 | 1.72297 | Distal Intergenic | 4 | 151059525 | 151861876 | 802352 | 2 | ENSMUST00000097774.3 | -11116 | Camta1 | ENSMUSG00000014592.14 |
| chr4 | 151874333 | 151874511 | 3.70326 | 4.55861 | 1.52129 | Distal Intergenic | 4 | 151059525 | 151861876 | 802352 | 2 | ENSMUST00000097774.3 | -12458 | Camta1 | ENSMUSG00000014592.14 |
| chr4 | 151875111 | 151875309 | 3.56302 | 4.37915 | 1.36385 | Distal Intergenic | 4 | 151059525 | 151861876 | 802352 | 2 | ENSMUST00000097774.3 | -13236 | Camta1 | ENSMUSG00000014592.14 |
| chr4 | 151877022 | 151877340 | 3.4673 | 4.25148 | 1.26784 | Distal Intergenic | 4 | 151059525 | 151861876 | 802352 | 2 | ENSMUST00000097774.3 | -15147 | Camta1 | ENSMUSG00000014592.14 |
| chr4 | 151878720 | 151878798 | 3.60092 | 4.36431 | 1.42308 | Distal Intergenic | 4 | 151059525 | 151861876 | 802352 | 2 | ENSMUST00000097774.3 | -16845 | Camta1 | ENSMUSG00000014592.14 |
| chr4 | 151638665 | 151638763 | 3.74707 | 4.58485 | 1.47018 | Intron (ENSMUST00000169423.3/ENSMUSG00000014592.14, intron 3 of 24) | 4 | 151470176 | 151861650 | 391475 | 2 | ENSMUST00000155685.2 | 222887 | Camta1 | ENSMUSG00000014592.14 |
| chr4 | 151654099 | 151654174 | 4.05505 | 5.17223 | 1.86266 | Intron (ENSMUST00000169423.3/ENSMUSG00000014592.14, intron 3 of 24) | 4 | 151470176 | 151861650 | 391475 | 2 | ENSMUST00000155685.2 | 207476 | Camta1 | ENSMUSG00000014592.14 |
| chr4 | 151635320 | 151635479 | 3.31897 | 4.15971 | 1.20075 | Intron (ENSMUST00000169423.3/ENSMUSG00000014592.14, intron 3 of 24) | 4 | 151470176 | 151861650 | 391475 | 2 | ENSMUST00000155685.2 | 226171 | Camta1 | ENSMUSG00000014592.14 |
| chr4 | 151637121 | 151638048 | 3.79142 | 4.68639 | 1.54743 | Intron (ENSMUST00000169423.3/ENSMUSG00000014592.14, intron 3 of 24) | 4 | 151470176 | 151861650 | 391475 | 2 | ENSMUST00000155685.2 | 223602 | Camta1 | ENSMUSG00000014592.14 |
| chr4 | 151718313 | 151718548 | 3.34675 | 3.92256 | 1.08395 | Intron (ENSMUST00000169423.3/ENSMUSG00000014592.14, intron 3 of 24) | 4 | 151470176 | 151861650 | 391475 | 2 | ENSMUST00000155685.2 | 143102 | Camta1 | ENSMUSG00000014592.14 |
| chr4 | 151732406 | 151732933 | 3.42381 | 4.03875 | 1.2311 | Intron (ENSMUST00000169423.3/ENSMUSG00000014592.14, intron 3 of 24) | 4 | 151470176 | 151861650 | 391475 | 2 | ENSMUST00000155685.2 | 128717 | Camta1 | ENSMUSG00000014592.14 |
| chr3 | 107807960 | 107808572 | 3.38076 | 3.92794 | 1.09522 | Distal Intergenic | 3 | 107741048 | 107760469 | 19422 | 2 | ENSMUST00000014743.4 | -47492 | Csf1 | ENSMUSG00000014599.5 |
| chr3 | 107816348 | 107816433 | 3.82426 | 5.0825 | 1.87071 | Distal Intergenic | 3 | 107741048 | 107760469 | 19422 | 2 | ENSMUST00000014743.4 | -55880 | Csf1 | ENSMUSG00000014599.5 |
| chr3 | 107818108 | 107818215 | 3.5331 | 4.30996 | 1.3347 | Distal Intergenic | 3 | 107741048 | 107760469 | 19422 | 2 | ENSMUST00000014743.4 | -57640 | Csf1 | ENSMUSG00000014599.5 |
| chr1 | 93100114 | 93100683 | 4.36603 | 5.96438 | 2.44992 | Intron (ENSMUST00000171796.2/ENSMUSG00000014602.10, intron 1 of 47) | 1 | 93015464 | 93101822 | 86359 | 2 | ENSMUST00000171796.2 | 1139 | Kif1a | ENSMUSG00000014602.10 |
| chr1 | 93101432 | 93101859 | 3.74074 | 5.11134 | 1.86407 | Promoter | 1 | 93015464 | 93101822 | 86359 | 2 | ENSMUST00000171796.2 | 0 | Kif1a | ENSMUSG00000014602.10 |
| chr1 | 93102566 | 93103002 | 4.2162 | 5.63011 | 2.2515 | Promoter | 1 | 93017940 | 93101951 | 84012 | 2 | ENSMUST00000190723.1 | -616 | Kif1a | ENSMUSG00000014602.10 |
| chr3 | 107581248 | 107581350 | 3.67292 | 4.71542 | 1.58661 | Distal Intergenic | 3 | 107595031 | 107605776 | 10746 | 1 | ENSMUST00000014747.2 | -13681 | Alx3 | ENSMUSG00000014603.2 |
| chr3 | 107582342 | 107583607 | 3.8281 | 5.03533 | 1.83892 | Distal Intergenic | 3 | 107595031 | 107605776 | 10746 | 1 | ENSMUST00000014747.2 | -11424 | Alx3 | ENSMUSG00000014603.2 |
| chr3 | 107584082 | 107585403 | 4.00318 | 5.16597 | 1.93618 | Distal Intergenic | 3 | 107595031 | 107605776 | 10746 | 1 | ENSMUST00000014747.2 | -9628 | Alx3 | ENSMUSG00000014603.2 |
| chr3 | 107589964 | 107590210 | 3.49469 | 4.17176 | 1.28498 | Distal Intergenic | 3 | 107595031 | 107605776 | 10746 | 1 | ENSMUST00000014747.2 | -4821 | Alx3 | ENSMUSG00000014603.2 |
| chr3 | 107590623 | 107590764 | 3.47347 | 4.10959 | 1.24425 | Distal Intergenic | 3 | 107595031 | 107605776 | 10746 | 1 | ENSMUST00000014747.2 | -4267 | Alx3 | ENSMUSG00000014603.2 |
| chr3 | 107592472 | 107593003 | 3.51378 | 4.4194 | 1.41111 | Distal Intergenic | 3 | 107595031 | 107605776 | 10746 | 1 | ENSMUST00000014747.2 | -2028 | Alx3 | ENSMUSG00000014603.2 |
| chr3 | 107593805 | 107594424 | 3.89256 | 4.94158 | 1.77537 | Promoter | 3 | 107595031 | 107605776 | 10746 | 1 | ENSMUST00000014747.2 | -607 | Alx3 | ENSMUSG00000014603.2 |
| chr3 | 107594756 | 107595488 | 4.13307 | 5.41492 | 2.08766 | Promoter | 3 | 107595031 | 107605776 | 10746 | 1 | ENSMUST00000014747.2 | 0 | Alx3 | ENSMUSG00000014603.2 |
| chr3 | 107595834 | 107597107 | 4.22274 | 5.66477 | 2.28767 | Promoter | 3 | 107595031 | 107605776 | 10746 | 1 | ENSMUST00000014747.2 | 804 | Alx3 | ENSMUSG00000014603.2 |
| chr3 | 107597548 | 107597874 | 3.65222 | 4.43581 | 1.44162 | Intron (ENSMUST00000014747.2/ENSMUSG00000014603.2, intron 1 of 3) | 3 | 107595031 | 107605776 | 10746 | 1 | ENSMUST00000014747.2 | 2518 | Alx3 | ENSMUSG00000014603.2 |
| chr3 | 107598208 | 107598414 | 3.87859 | 4.90863 | 1.72593 | Intron (ENSMUST00000014747.2/ENSMUSG00000014603.2, intron 1 of 3) | 3 | 107595031 | 107605776 | 10746 | 1 | ENSMUST00000014747.2 | 3178 | Alx3 | ENSMUSG00000014603.2 |
| chr3 | 107598854 | 107599524 | 3.92138 | 5.1751 | 1.90178 | Intron (ENSMUST00000014747.2/ENSMUSG00000014603.2, intron 1 of 3) | 3 | 107595031 | 107605776 | 10746 | 1 | ENSMUST00000014747.2 | 3824 | Alx3 | ENSMUSG00000014603.2 |
| chr3 | 107599924 | 107600643 | 3.34742 | 3.93395 | 1.11871 | Exon (ENSMUST00000014747.2/ENSMUSG00000014603.2, exon 2 of 4) | 3 | 107595031 | 107605776 | 10746 | 1 | ENSMUST00000014747.2 | 4894 | Alx3 | ENSMUSG00000014603.2 |
| chr8 | 116863378 | 116863482 | 4.38092 | 5.92614 | 2.49368 | Distal Intergenic | 8 | 116888685 | 116894524 | 5840 | 2 | ENSMUST00000155260.1 | 31042 | Cmc2 | ENSMUSG00000014633.10 |
| chr8 | 116873725 | 116873814 | 4.04663 | 5.15237 | 1.83752 | Distal Intergenic | 8 | 116888685 | 116894524 | 5840 | 2 | ENSMUST00000155260.1 | 20710 | Cmc2 | ENSMUSG00000014633.10 |
| chr6 | 83762403 | 83763065 | 3.37115 | 4.1967 | 1.22495 | Promoter | 6 | 83762644 | 83768326 | 5683 | 1 | ENSMUST00000014891.3 | 0 | Ankrd53 | ENSMUSG00000014747.5 |
| chr8 | 105374607 | 105374781 | 3.6071 | 4.33656 | 1.3376 | Promoter | 8 | 105374904 | 105382859 | 7956 | 1 | ENSMUST00000159286.2 | -123 | Plekhg4 | ENSMUSG00000014782.10 |
| chr8 | 105349746 | 105349923 | 3.60542 | 4.5041 | 1.48096 | Intron (ENSMUST00000073149.5/ENSMUSG00000014786.7, intron 1 of 15) | 8 | 105348258 | 105369881 | 21624 | 1 | ENSMUST00000073149.5 | 1489 | Slc9a5 | ENSMUSG00000014786.7 |
| chr8 | 105350735 | 105350810 | 4.04019 | 5.13383 | 1.81024 | Intron (ENSMUST00000073149.5/ENSMUSG00000014786.7, intron 1 of 15) | 8 | 105348258 | 105369881 | 21624 | 1 | ENSMUST00000073149.5 | 2478 | Slc9a5 | ENSMUSG00000014786.7 |
| chr14 | 68898590 | 68898892 | 3.72156 | 4.56612 | 1.51366 | Distal Intergenic | 14 | 69029289 | 69041401 | 12113 | 1 | ENSMUST00000014957.8 | -130397 | Stc1 | ENSMUSG00000014813.8 |
| chr14 | 69028486 | 69028615 | 3.36236 | 3.8711 | 1.01733 | Promoter | 14 | 69029289 | 69041401 | 12113 | 1 | ENSMUST00000014957.8 | -674 | Stc1 | ENSMUSG00000014813.8 |
| chr14 | 69028999 | 69029177 | 4.16738 | 5.42357 | 2.07449 | Promoter | 14 | 69029289 | 69041401 | 12113 | 1 | ENSMUST00000014957.8 | -112 | Stc1 | ENSMUSG00000014813.8 |
| chr14 | 69030370 | 69030807 | 3.80892 | 4.8614 | 1.70472 | Intron (ENSMUST00000014957.8/ENSMUSG00000014813.8, intron 1 of 3) | 14 | 69029289 | 69041401 | 12113 | 1 | ENSMUST00000014957.8 | 1082 | Stc1 | ENSMUSG00000014813.8 |
| chr8 | 122744707 | 122744881 | 3.52605 | 4.18733 | 1.26376 | Intron (ENSMUST00000127664.1/ENSMUSG00000092329.1, intron 1 of 15) | 8 | 122775505 | 122817880 | 42376 | 1 | ENSMUST00000015160.5 | -30624 | Acsf3 | ENSMUSG00000015016.7 |
| chr6 | 88188147 | 88188904 | 3.79171 | 4.66385 | 1.50474 | Distal Intergenic | 6 | 88193891 | 88207030 | 13140 | 1 | ENSMUST00000170089.2 | -4987 | Gata2 | ENSMUSG00000015053.9 |
| chr6 | 88191998 | 88192105 | 3.41918 | 3.9746 | 1.08015 | Distal Intergenic | 6 | 88193891 | 88207030 | 13140 | 1 | ENSMUST00000170089.2 | -1786 | Gata2 | ENSMUSG00000015053.9 |
| chr6 | 88192633 | 88192884 | 3.85695 | 4.80209 | 1.63697 | Distal Intergenic | 6 | 88193891 | 88207030 | 13140 | 1 | ENSMUST00000170089.2 | -1007 | Gata2 | ENSMUSG00000015053.9 |
| chr6 | 88193201 | 88194415 | 4.00784 | 5.1726 | 1.94151 | Promoter | 6 | 88193891 | 88207030 | 13140 | 1 | ENSMUST00000170089.2 | 0 | Gata2 | ENSMUSG00000015053.9 |
| chr6 | 88194860 | 88195665 | 4.13603 | 5.47307 | 2.13769 | Promoter | 6 | 88193891 | 88207030 | 13140 | 1 | ENSMUST00000170089.2 | 970 | Gata2 | ENSMUSG00000015053.9 |
| chr6 | 88196476 | 88197835 | 3.88131 | 5.04299 | 1.81092 | Promoter | 6 | 88198664 | 88207030 | 8367 | 1 | ENSMUST00000015197.7 | -829 | Gata2 | ENSMUSG00000015053.9 |
| chr6 | 88198180 | 88199922 | 3.74543 | 4.68689 | 1.56368 | Promoter | 6 | 88198664 | 88207030 | 8367 | 1 | ENSMUST00000015197.7 | 0 | Gata2 | ENSMUSG00000015053.9 |
| chr6 | 88200238 | 88203906 | 3.73678 | 4.5735 | 1.46202 | Exon (ENSMUST00000170089.2/ENSMUSG00000015053.9, exon 3 of 6) | 6 | 88198664 | 88207030 | 8367 | 1 | ENSMUST00000015197.7 | 1575 | Gata2 | ENSMUSG00000015053.9 |
| chr6 | 88204246 | 88206645 | 3.94264 | 4.99091 | 1.7604 | 3' UTR | 6 | 88198664 | 88207030 | 8367 | 1 | ENSMUST00000015197.7 | 5583 | Gata2 | ENSMUSG00000015053.9 |
| chr6 | 88207172 | 88207407 | 3.46303 | 4.05703 | 1.13758 | Distal Intergenic | 6 | 88198664 | 88207030 | 8367 | 1 | ENSMUST00000015197.7 | 8509 | Gata2 | ENSMUSG00000015053.9 |
| chr2 | 25395868 | 25396181 | 4.17316 | 5.50067 | 2.1373 | Promoter | 2 | 25395874 | 25401321 | 5448 | 1 | ENSMUST00000028328.2 | 0 | Entpd2 | ENSMUSG00000015085.8 |
| chr2 | 25396715 | 25397043 | 3.70224 | 4.67691 | 1.57077 | Promoter | 2 | 25395916 | 25401292 | 5377 | 1 | ENSMUST00000148859.1 | 800 | Entpd2 | ENSMUSG00000015085.8 |
| chr2 | 25399600 | 25399703 | 3.69245 | 4.51255 | 1.49788 | Promoter | 2 | 25399351 | 25406340 | 6990 | 1 | ENSMUST00000124277.1 | 250 | Npdc1 | ENSMUSG00000015094.11 |
| chr2 | 25400359 | 25400460 | 3.32431 | 3.83683 | 1.09438 | Promoter | 2 | 25400469 | 25408178 | 7710 | 1 | ENSMUST00000132287.2 | -9 | Npdc1 | ENSMUSG00000015094.11 |
| chr7 | 66238314 | 66238397 | 6.01238 | 10.65645 | 6.30917 | Intron (ENSMUST00000167705.2/ENSMUSG00000015133.11, intron 2 of 2) | 7 | 66226912 | 66262470 | 35559 | 2 | ENSMUST00000167705.2 | 24073 | Lrrk1 | ENSMUSG00000015133.11 |
| chr19 | 46089391 | 46089492 | 3.7158 | 4.57315 | 1.49202 | Distal Intergenic | 19 | 46075863 | 46085528 | 9666 | 1 | ENSMUST00000165017.1 | 13529 | Nolc1 | ENSMUSG00000015176.9 |
| chr19 | 46090285 | 46090496 | 3.39841 | 3.9409 | 1.07181 | Distal Intergenic | 19 | 46075863 | 46085528 | 9666 | 1 | ENSMUST00000165017.1 | 14423 | Nolc1 | ENSMUSG00000015176.9 |
| chr1 | 66176162 | 66176723 | 3.81708 | 4.74048 | 1.59179 | Promoter | 1 | 66175550 | 66195378 | 19829 | 1 | ENSMUST00000125844.1 | 613 | Map2 | ENSMUSG00000015222.12 |
| chr4 | 52989400 | 52989585 | 3.23252 | 3.81991 | 1.04439 | Promoter | 4 | 52989418 | 53001162 | 11745 | 1 | ENSMUST00000107666.1 | 0 | Nipsnap3a | ENSMUSG00000015242.9 |
| chr9 | 95636140 | 95636666 | 3.60508 | 4.32271 | 1.30425 | Promoter | 9 | 95637601 | 95698096 | 60496 | 1 | ENSMUST00000015498.8 | -935 | Pcolce2 | ENSMUSG00000015354.8 |
| chr9 | 95637017 | 95638768 | 3.71507 | 4.60718 | 1.52353 | Promoter | 9 | 95637601 | 95698096 | 60496 | 1 | ENSMUST00000015498.8 | 0 | Pcolce2 | ENSMUSG00000015354.8 |
| chr9 | 95692889 | 95693357 | 3.24341 | 3.69227 | 1.01665 | Exon (ENSMUST00000015498.8/ENSMUSG00000015354.8, exon 7 of 9) | 9 | 95680306 | 95694683 | 14378 | 1 | ENSMUST00000151210.1 | 12584 | Pcolce2 | ENSMUSG00000015354.8 |
| chr9 | 95693955 | 95694947 | 4.04816 | 5.2877 | 2.02939 | Exon (ENSMUST00000015498.8/ENSMUSG00000015354.8, exon 8 of 9) | 9 | 95680306 | 95694683 | 14378 | 1 | ENSMUST00000151210.1 | 13650 | Pcolce2 | ENSMUSG00000015354.8 |
| chr13 | 43783900 | 43784164 | 3.4023 | 3.97488 | 1.14173 | Promoter | 13 | 43785107 | 43803130 | 18024 | 1 | ENSMUST00000015540.2 | -943 | Cd83 | ENSMUSG00000015396.3 |
| chr13 | 43785427 | 43785970 | 4.15557 | 5.39208 | 2.03155 | Promoter | 13 | 43785107 | 43803130 | 18024 | 1 | ENSMUST00000015540.2 | 321 | Cd83 | ENSMUSG00000015396.3 |
| chr1 | 156169457 | 156169931 | 3.32889 | 3.82067 | 1.01733 | Intron (ENSMUST00000015628.3/ENSMUSG00000015484.3, intron 1 of 4) | 1 | 156075966 | 156205026 | 129061 | 2 | ENSMUST00000015628.3 | 35095 | Fam163a | ENSMUSG00000015484.3 |
| chr1 | 156172349 | 156172620 | 4.14646 | 5.38064 | 2.03965 | Intron (ENSMUST00000015628.3/ENSMUSG00000015484.3, intron 1 of 4) | 1 | 156075966 | 156205026 | 129061 | 2 | ENSMUST00000015628.3 | 32406 | Fam163a | ENSMUSG00000015484.3 |
| chr1 | 156180156 | 156180235 | 3.74061 | 4.7422 | 1.57502 | Intron (ENSMUST00000015628.3/ENSMUSG00000015484.3, intron 1 of 4) | 1 | 156075966 | 156205026 | 129061 | 2 | ENSMUST00000015628.3 | 24791 | Fam163a | ENSMUSG00000015484.3 |
| chr1 | 156182208 | 156182487 | 3.26175 | 3.71976 | 1.01733 | Intron (ENSMUST00000015628.3/ENSMUSG00000015484.3, intron 1 of 4) | 1 | 156075966 | 156205026 | 129061 | 2 | ENSMUST00000015628.3 | 22539 | Fam163a | ENSMUSG00000015484.3 |
| chr1 | 156190963 | 156191189 | 3.56798 | 4.50629 | 1.43036 | Intron (ENSMUST00000015628.3/ENSMUSG00000015484.3, intron 1 of 4) | 1 | 156075966 | 156205026 | 129061 | 2 | ENSMUST00000015628.3 | 13837 | Fam163a | ENSMUSG00000015484.3 |
| chr1 | 156201278 | 156202354 | 3.44129 | 4.0677 | 1.20963 | Intron (ENSMUST00000015628.3/ENSMUSG00000015484.3, intron 1 of 4) | 1 | 156075966 | 156205026 | 129061 | 2 | ENSMUST00000015628.3 | 2672 | Fam163a | ENSMUSG00000015484.3 |
| chr1 | 156202812 | 156206478 | 3.97747 | 5.1228 | 1.87833 | Promoter | 1 | 156075966 | 156205026 | 129061 | 2 | ENSMUST00000015628.3 | 0 | Fam163a | ENSMUSG00000015484.3 |
| chr17 | 26827011 | 26827151 | 3.67161 | 4.60571 | 1.53431 | Distal Intergenic | 17 | 26838664 | 26841565 | 2902 | 2 | ENSMUST00000015723.4 | 14414 | Nkx2-5 | ENSMUSG00000015579.4 |
| chr17 | 26827961 | 26828309 | 3.50204 | 4.31465 | 1.30351 | Distal Intergenic | 17 | 26838664 | 26841565 | 2902 | 2 | ENSMUST00000015723.4 | 13256 | Nkx2-5 | ENSMUSG00000015579.4 |
| chr17 | 26835517 | 26836398 | 3.88217 | 4.90218 | 1.69303 | Downstream (2-3kb) | 17 | 26838664 | 26841565 | 2902 | 2 | ENSMUST00000015723.4 | 5167 | Nkx2-5 | ENSMUSG00000015579.4 |
| chr17 | 26836839 | 26837708 | 4.17669 | 5.57778 | 2.25334 | Downstream (<1kb) | 17 | 26838664 | 26841565 | 2902 | 2 | ENSMUST00000015723.4 | 3857 | Nkx2-5 | ENSMUSG00000015579.4 |
| chr17 | 26838361 | 26839920 | 4.09957 | 5.3317 | 2.02038 | 3' UTR | 17 | 26838664 | 26841565 | 2902 | 2 | ENSMUST00000015723.4 | 1645 | Nkx2-5 | ENSMUSG00000015579.4 |
| chr17 | 26840311 | 26840518 | 3.71234 | 4.52521 | 1.43722 | Intron (ENSMUST00000015723.4/ENSMUSG00000015579.4, intron 1 of 1) | 17 | 26838664 | 26841565 | 2902 | 2 | ENSMUST00000015723.4 | 1047 | Nkx2-5 | ENSMUSG00000015579.4 |
| chr17 | 26840932 | 26843232 | 4.33931 | 5.87617 | 2.39917 | Promoter | 17 | 26838664 | 26841565 | 2902 | 2 | ENSMUST00000015723.4 | 0 | Nkx2-5 | ENSMUSG00000015579.4 |
| chr17 | 26843715 | 26845329 | 4.44252 | 6.09366 | 2.57232 | Distal Intergenic | 17 | 26838664 | 26841565 | 2902 | 2 | ENSMUST00000015723.4 | -2151 | Nkx2-5 | ENSMUSG00000015579.4 |
| chr17 | 26845674 | 26846725 | 4.07827 | 5.25765 | 1.97771 | Distal Intergenic | 17 | 26838664 | 26841565 | 2902 | 2 | ENSMUST00000015723.4 | -4110 | Nkx2-5 | ENSMUSG00000015579.4 |
| chr17 | 26847400 | 26847888 | 4.65232 | 6.45915 | 2.89211 | Distal Intergenic | 17 | 26838664 | 26841565 | 2902 | 2 | ENSMUST00000015723.4 | -5836 | Nkx2-5 | ENSMUSG00000015579.4 |
| chr17 | 26848297 | 26848777 | 4.82216 | 7.1614 | 3.47538 | Distal Intergenic | 17 | 26838664 | 26841565 | 2902 | 2 | ENSMUST00000015723.4 | -6733 | Nkx2-5 | ENSMUSG00000015579.4 |
| chr17 | 26849289 | 26850464 | 3.76152 | 4.7369 | 1.58833 | Distal Intergenic | 17 | 26838664 | 26841565 | 2902 | 2 | ENSMUST00000015723.4 | -7725 | Nkx2-5 | ENSMUSG00000015579.4 |
| chr17 | 26850839 | 26852018 | 3.7466 | 4.66443 | 1.5834 | Distal Intergenic | 17 | 26838664 | 26841565 | 2902 | 2 | ENSMUST00000015723.4 | -9275 | Nkx2-5 | ENSMUSG00000015579.4 |
| chr17 | 46483648 | 46483896 | 3.83344 | 4.74228 | 1.56052 | Intron (ENSMUST00000181301.1/ENSMUSG00000097560.1, intron 1 of 2) | 17 | 46442448 | 46487675 | 45228 | 2 | ENSMUST00000047034.8 | 3779 | Ttbk1 | ENSMUSG00000015599.8 |
| chr17 | 46487329 | 46488425 | 3.80237 | 4.74907 | 1.61021 | Promoter | 17 | 46442448 | 46487675 | 45228 | 2 | ENSMUST00000047034.8 | 0 | Ttbk1 | ENSMUSG00000015599.8 |
| chr17 | 46488834 | 46489139 | 3.46853 | 4.21706 | 1.24487 | Intron (ENSMUST00000181301.1/ENSMUSG00000097560.1, intron 1 of 2) | 17 | 46442448 | 46487675 | 45228 | 2 | ENSMUST00000047034.8 | -1160 | Ttbk1 | ENSMUSG00000015599.8 |
| chr2 | 9853420 | 9853670 | 3.39548 | 3.92865 | 1.04403 | Distal Intergenic | 2 | 9863183 | 9873672 | 10490 | 2 | ENSMUST00000151456.1 | 20002 | Gata3 | ENSMUSG00000015619.10 |
| chr2 | 9865817 | 9866486 | 3.37802 | 3.98231 | 1.15101 | Intron (ENSMUST00000102976.3/ENSMUSG00000015619.10, intron 4 of 5) | 2 | 9863183 | 9873672 | 10490 | 2 | ENSMUST00000151456.1 | 7186 | Gata3 | ENSMUSG00000015619.10 |
| chr2 | 9868912 | 9869040 | 3.59098 | 4.36561 | 1.3927 | Intron (ENSMUST00000102976.3/ENSMUSG00000015619.10, intron 3 of 5) | 2 | 9863183 | 9873672 | 10490 | 2 | ENSMUST00000151456.1 | 4632 | Gata3 | ENSMUSG00000015619.10 |
| chr2 | 9869504 | 9870153 | 3.48671 | 4.23029 | 1.27043 | Intron (ENSMUST00000102976.3/ENSMUSG00000015619.10, intron 3 of 5) | 2 | 9863183 | 9873672 | 10490 | 2 | ENSMUST00000151456.1 | 3519 | Gata3 | ENSMUSG00000015619.10 |
| chr2 | 9871276 | 9871351 | 4.48553 | 6.65609 | 2.96036 | Intron (ENSMUST00000102976.3/ENSMUSG00000015619.10, intron 3 of 5) | 2 | 9863183 | 9873672 | 10490 | 2 | ENSMUST00000151456.1 | 2321 | Gata3 | ENSMUSG00000015619.10 |
| chr2 | 9872602 | 9874127 | 3.50773 | 4.14182 | 1.18839 | Promoter | 2 | 9863183 | 9873672 | 10490 | 2 | ENSMUST00000151456.1 | 0 | Gata3 | ENSMUSG00000015619.10 |
| chr2 | 9874458 | 9875284 | 3.64169 | 4.39142 | 1.34457 | Promoter | 2 | 9874487 | 9875399 | 913 | 2 | ENSMUST00000153509.1 | 115 | Gata3 | ENSMUSG00000015619.10 |
| chr2 | 9875749 | 9876192 | 3.83106 | 4.74899 | 1.58249 | Promoter | 2 | 9874487 | 9875399 | 913 | 2 | ENSMUST00000153509.1 | -351 | Gata3 | ENSMUSG00000015619.10 |
| chr2 | 9896188 | 9897548 | 3.98249 | 5.11527 | 1.85624 | Distal Intergenic | 2 | 9874702 | 9890034 | 15333 | 2 | ENSMUST00000130615.1 | -6155 | Gata3 | ENSMUSG00000015619.10 |
| chr2 | 9878066 | 9879029 | 4.85116 | 6.9695 | 3.23859 | Promoter | 2 | 9857078 | 9878600 | 21523 | 2 | ENSMUST00000102976.3 | 0 | Gata3 | ENSMUSG00000015619.10 |
| chr2 | 9879346 | 9879819 | 4.19043 | 5.65469 | 2.22787 | Promoter | 2 | 9857078 | 9878600 | 21523 | 2 | ENSMUST00000102976.3 | -747 | Gata3 | ENSMUSG00000015619.10 |
| chr2 | 9892420 | 9893364 | 3.91113 | 4.9124 | 1.70423 | Distal Intergenic | 2 | 9874702 | 9890034 | 15333 | 2 | ENSMUST00000130615.1 | -2387 | Gata3 | ENSMUSG00000015619.10 |
| chr2 | 9887253 | 9888292 | 3.84998 | 4.83006 | 1.67184 | Promoter | 2 | 9877835 | 9887784 | 9950 | 2 | ENSMUST00000147533.1 | 0 | Gata3 | ENSMUSG00000015619.10 |
| chr2 | 9888943 | 9891110 | 3.76969 | 4.80675 | 1.64303 | Promoter | 2 | 9874702 | 9890034 | 15333 | 2 | ENSMUST00000130615.1 | 0 | Gata3 | ENSMUSG00000015619.10 |
| chr2 | 9891965 | 9892074 | 3.99029 | 5.05943 | 1.79964 | Distal Intergenic | 2 | 9874702 | 9890034 | 15333 | 2 | ENSMUST00000130615.1 | -1932 | Gata3 | ENSMUSG00000015619.10 |
| chr2 | 9894046 | 9895733 | 4.16589 | 5.51018 | 2.13595 | Distal Intergenic | 2 | 9874702 | 9890034 | 15333 | 2 | ENSMUST00000130615.1 | -4013 | Gata3 | ENSMUSG00000015619.10 |
| chr2 | 180333675 | 180334254 | 4.40147 | 5.97326 | 2.51215 | Promoter | 2 | 180325133 | 180334699 | 9567 | 2 | ENSMUST00000015771.2 | 445 | Gata5 | ENSMUSG00000015627.5 |
| chr2 | 180334611 | 180335318 | 3.66465 | 4.65104 | 1.56929 | Promoter | 2 | 180325133 | 180334699 | 9567 | 2 | ENSMUST00000015771.2 | 0 | Gata5 | ENSMUSG00000015627.5 |
| chr2 | 180341095 | 180341211 | 4.20738 | 5.48628 | 2.08543 | Distal Intergenic | 2 | 180325133 | 180334699 | 9567 | 2 | ENSMUST00000015771.2 | -6397 | Gata5 | ENSMUSG00000015627.5 |
| chr6 | 137648142 | 137649491 | 3.873 | 5.05127 | 1.86047 | Promoter | 6 | 137477245 | 137649285 | 172041 | 2 | ENSMUST00000058210.7 | 0 | Eps8 | ENSMUSG00000015766.9 |
| chr1 | 167551785 | 167551866 | 4.04422 | 5.16312 | 1.89092 | Distal Intergenic | 1 | 167598384 | 167639623 | 41240 | 1 | ENSMUST00000015987.4 | -46518 | Rxrg | ENSMUSG00000015843.5 |
| chr1 | 167597736 | 167597899 | 3.31979 | 3.80693 | 1.01733 | Promoter | 1 | 167598384 | 167639623 | 41240 | 1 | ENSMUST00000015987.4 | -485 | Rxrg | ENSMUSG00000015843.5 |
| chr1 | 167598331 | 167598452 | 3.979 | 5.07012 | 1.86498 | Promoter | 1 | 167598384 | 167639623 | 41240 | 1 | ENSMUST00000015987.4 | 0 | Rxrg | ENSMUSG00000015843.5 |
| chr1 | 167599943 | 167600051 | 3.5856 | 4.46349 | 1.38846 | Intron (ENSMUST00000015987.4/ENSMUSG00000015843.5, intron 1 of 9) | 1 | 167598480 | 167639618 | 41139 | 1 | ENSMUST00000111384.2 | 1464 | Rxrg | ENSMUSG00000015843.5 |
| chr2 | 27645190 | 27645340 | 3.64997 | 4.40337 | 1.34263 | Distal Intergenic | 2 | 27676440 | 27741898 | 65459 | 1 | ENSMUST00000129514.2 | -31100 | Rxra | ENSMUSG00000015846.9 |
| chr2 | 27652641 | 27652754 | 3.30311 | 3.78196 | 1.01733 | Distal Intergenic | 2 | 27676440 | 27741898 | 65459 | 1 | ENSMUST00000129514.2 | -23686 | Rxra | ENSMUSG00000015846.9 |
| chr2 | 27666068 | 27666353 | 3.8389 | 5.00996 | 1.77932 | Distal Intergenic | 2 | 27676440 | 27741898 | 65459 | 1 | ENSMUST00000129514.2 | -10087 | Rxra | ENSMUSG00000015846.9 |
| chr2 | 27766166 | 27766404 | 3.67957 | 4.46995 | 1.40621 | Distal Intergenic | 2 | 27740201 | 27748671 | 8471 | 1 | ENSMUST00000148756.1 | 25966 | Rxra | ENSMUSG00000015846.9 |
| chr2 | 27769499 | 27769907 | 4.10236 | 5.38921 | 2.09153 | Distal Intergenic | 2 | 27740201 | 27748671 | 8471 | 1 | ENSMUST00000148756.1 | 29299 | Rxra | ENSMUSG00000015846.9 |
| chr3 | 87256346 | 87256597 | 3.44251 | 4.01524 | 1.09899 | Intron (ENSMUST00000090986.5/ENSMUSG00000015852.8, intron 6 of 7) | 3 | 87250758 | 87263599 | 12842 | 2 | ENSMUST00000090986.5 | 7002 | Fcrls | ENSMUSG00000015852.8 |
| chr5 | 45639408 | 45640963 | 3.78708 | 4.74133 | 1.6807 | Promoter | 5 | 45529706 | 45639501 | 109796 | 2 | ENSMUST00000016023.7 | 0 | Fam184b | ENSMUSG00000015879.7 |
| chr13 | 55960994 | 55961960 | 3.61604 | 4.37668 | 1.3496 | Distal Intergenic | 13 | 56074166 | 56084521 | 10356 | 2 | ENSMUST00000137835.2 | 122561 | H2afy | ENSMUSG00000015937.9 |
| chr13 | 55962276 | 55963315 | 3.62319 | 4.51529 | 1.46204 | Distal Intergenic | 13 | 56074166 | 56084521 | 10356 | 2 | ENSMUST00000137835.2 | 121206 | H2afy | ENSMUSG00000015937.9 |
| chr13 | 55964503 | 55964690 | 3.6194 | 4.33861 | 1.28996 | Distal Intergenic | 13 | 56074166 | 56084521 | 10356 | 2 | ENSMUST00000137835.2 | 119831 | H2afy | ENSMUSG00000015937.9 |
| chr13 | 55973856 | 55974062 | 4.39504 | 5.86128 | 2.35646 | Distal Intergenic | 13 | 56074166 | 56084521 | 10356 | 2 | ENSMUST00000137835.2 | 110459 | H2afy | ENSMUSG00000015937.9 |
| chr13 | 55984687 | 55984971 | 3.68551 | 4.53843 | 1.47815 | Distal Intergenic | 13 | 56074166 | 56084521 | 10356 | 2 | ENSMUST00000137835.2 | 99550 | H2afy | ENSMUSG00000015937.9 |
| chr13 | 56026083 | 56026316 | 3.43478 | 4.00812 | 1.11569 | Distal Intergenic | 13 | 56074166 | 56084521 | 10356 | 2 | ENSMUST00000137835.2 | 58205 | H2afy | ENSMUSG00000015937.9 |
| chr13 | 56050345 | 56050453 | 3.58263 | 4.33495 | 1.41489 | Distal Intergenic | 13 | 56074166 | 56084521 | 10356 | 2 | ENSMUST00000137835.2 | 34068 | H2afy | ENSMUSG00000015937.9 |
| chr13 | 56063196 | 56063375 | 3.79351 | 4.76139 | 1.60784 | Distal Intergenic | 13 | 56074166 | 56084521 | 10356 | 2 | ENSMUST00000137835.2 | 21146 | H2afy | ENSMUSG00000015937.9 |
| chr3 | 96284330 | 96284516 | 3.62068 | 4.70554 | 1.56824 | Exon (ENSMUST00000029748.3/ENSMUSG00000015947.6, exon 6 of 6) | 3 | 96282909 | 96293969 | 11061 | 2 | ENSMUST00000029748.3 | 9453 | Fcgr1 | ENSMUSG00000015947.6 |
| chr3 | 96288267 | 96288367 | 3.88433 | 5.03277 | 1.79025 | Intron (ENSMUST00000029748.3/ENSMUSG00000015947.6, intron 3 of 5) | 3 | 96282909 | 96293969 | 11061 | 2 | ENSMUST00000029748.3 | 5602 | Fcgr1 | ENSMUSG00000015947.6 |
| chr3 | 96289545 | 96290079 | 3.49164 | 4.15782 | 1.29849 | Intron (ENSMUST00000029748.3/ENSMUSG00000015947.6, intron 3 of 5) | 3 | 96282909 | 96293969 | 11061 | 2 | ENSMUST00000029748.3 | 3890 | Fcgr1 | ENSMUSG00000015947.6 |
| chr3 | 96304865 | 96305069 | 3.27952 | 3.74642 | 1.01733 | Distal Intergenic | 3 | 96282909 | 96293969 | 11061 | 2 | ENSMUST00000029748.3 | -10897 | Fcgr1 | ENSMUSG00000015947.6 |
| chr3 | 96308265 | 96308396 | 3.36788 | 3.8794 | 1.01733 | Distal Intergenic | 3 | 96282909 | 96293969 | 11061 | 2 | ENSMUST00000029748.3 | -14297 | Fcgr1 | ENSMUSG00000015947.6 |
| chr5 | 134221671 | 134222051 | 3.96833 | 5.01552 | 1.76112 | Promoter | 5 | 134221372 | 134222222 | 851 | 2 | ENSMUST00000182626.1 | 171 | Ncf1 | ENSMUSG00000015950.8 |
| chr5 | 134222526 | 134222620 | 3.55885 | 4.22456 | 1.21568 | Promoter | 5 | 134221372 | 134222222 | 851 | 2 | ENSMUST00000182626.1 | -305 | Ncf1 | ENSMUSG00000015950.8 |
| chr7 | 98834499 | 98836671 | 3.94391 | 5.08871 | 1.84313 | Promoter | 7 | 98835112 | 98846587 | 11476 | 1 | ENSMUST00000165122.2 | 0 | Wnt11 | ENSMUSG00000015957.6 |
| chr7 | 98839770 | 98841169 | 3.6358 | 4.54537 | 1.49772 | Promoter | 7 | 98838898 | 98853571 | 14674 | 1 | ENSMUST00000165240.1 | 873 | Wnt11 | ENSMUSG00000015957.6 |
| chr7 | 98841634 | 98842119 | 3.65601 | 4.48906 | 1.47146 | Intron (ENSMUST00000165122.2/ENSMUSG00000015957.6, intron 2 of 2) | 7 | 98838898 | 98853571 | 14674 | 1 | ENSMUST00000165240.1 | 2737 | Wnt11 | ENSMUSG00000015957.6 |
| chr14 | 30008344 | 30009011 | 3.56116 | 4.40575 | 1.39194 | Promoter | 14 | 29996141 | 30008813 | 12673 | 2 | ENSMUST00000135888.2 | 0 | Il17rb | ENSMUSG00000015966.11 |
| chr14 | 30352041 | 30352848 | 3.86036 | 4.80104 | 1.60504 | Promoter | 14 | 30039949 | 30353486 | 313538 | 2 | ENSMUST00000112249.2 | 638 | Cacna1d | ENSMUSG00000015968.11 |
| chr14 | 30353588 | 30354128 | 3.72988 | 4.82295 | 1.63974 | Promoter | 14 | 30039949 | 30353486 | 313538 | 2 | ENSMUST00000112249.2 | -103 | Cacna1d | ENSMUSG00000015968.11 |
| chr7 | 139186346 | 139186621 | 3.95793 | 5.08021 | 1.86071 | Intron (ENSMUST00000016125.6/ENSMUSG00000015981.7, intron 1 of 11) | 7 | 139103638 | 139188517 | 84880 | 2 | ENSMUST00000016125.6 | 1896 | Stk32c | ENSMUSG00000015981.7 |
| chr7 | 139187411 | 139187686 | 3.52795 | 4.26739 | 1.31409 | Promoter | 7 | 139103638 | 139188517 | 84880 | 2 | ENSMUST00000016125.6 | 831 | Stk32c | ENSMUSG00000015981.7 |
| chr15 | 86030092 | 86030225 | 3.49602 | 4.52121 | 1.44249 | Exon (ENSMUST00000016172.7/ENSMUSG00000016028.9, exon 1 of 35) | 15 | 85898758 | 86033777 | 135020 | 2 | ENSMUST00000016172.7 | 3552 | Celsr1 | ENSMUSG00000016028.9 |
| chr15 | 86031863 | 86032121 | 3.33199 | 3.9929 | 1.10718 | Exon (ENSMUST00000016172.7/ENSMUSG00000016028.9, exon 1 of 35) | 15 | 85898758 | 86033777 | 135020 | 2 | ENSMUST00000016172.7 | 1656 | Celsr1 | ENSMUSG00000016028.9 |
| chr15 | 86033168 | 86035014 | 4.00257 | 5.15891 | 1.92091 | Promoter | 15 | 85898758 | 86033777 | 135020 | 2 | ENSMUST00000016172.7 | 0 | Celsr1 | ENSMUSG00000016028.9 |
| chr15 | 86039674 | 86039822 | 3.39056 | 4.11401 | 1.23407 | Distal Intergenic | 15 | 85898758 | 86033777 | 135020 | 2 | ENSMUST00000016172.7 | -5898 | Celsr1 | ENSMUSG00000016028.9 |
| chr15 | 86040193 | 86040468 | 3.3315 | 3.83388 | 1.05002 | Distal Intergenic | 15 | 85898758 | 86033777 | 135020 | 2 | ENSMUST00000016172.7 | -6417 | Celsr1 | ENSMUSG00000016028.9 |
| chr15 | 86041279 | 86041370 | 3.46719 | 4.05604 | 1.11155 | Distal Intergenic | 15 | 85898758 | 86033777 | 135020 | 2 | ENSMUST00000016172.7 | -7503 | Celsr1 | ENSMUSG00000016028.9 |
| chr9 | 32540369 | 32542831 | 3.95961 | 5.12044 | 1.87994 | Promoter | 9 | 32422204 | 32541595 | 119392 | 2 | ENSMUST00000016231.8 | 0 | Fli1 | ENSMUSG00000016087.8 |
| chr9 | 32543223 | 32543337 | 5.12756 | 7.40768 | 3.56186 | Promoter | 9 | 32428059 | 32542861 | 114803 | 2 | ENSMUST00000183767.1 | -363 | Fli1 | ENSMUSG00000016087.8 |
| chr9 | 32543662 | 32544052 | 4.35417 | 5.81251 | 2.33899 | Promoter | 9 | 32428059 | 32542861 | 114803 | 2 | ENSMUST00000183767.1 | -802 | Fli1 | ENSMUSG00000016087.8 |
| chr9 | 32544395 | 32545014 | 3.99394 | 5.14262 | 1.87092 | Distal Intergenic | 9 | 32428059 | 32542861 | 114803 | 2 | ENSMUST00000183767.1 | -1535 | Fli1 | ENSMUSG00000016087.8 |
| chr9 | 32545332 | 32546121 | 3.63054 | 4.44961 | 1.42608 | Distal Intergenic | 9 | 32428059 | 32542861 | 114803 | 2 | ENSMUST00000183767.1 | -2472 | Fli1 | ENSMUSG00000016087.8 |
| chr5 | 151192018 | 151192221 | 4.02275 | 5.41015 | 2.05895 | Intron (ENSMUST00000129088.2/ENSMUSG00000016128.9, intron 1 of 3) | 5 | 151037628 | 151190269 | 152642 | 2 | ENSMUST00000110483.3 | -1750 | Stard13 | ENSMUSG00000016128.9 |
| chr5 | 151232941 | 151233620 | 3.46777 | 4.10598 | 1.25834 | Promoter | 5 | 151063546 | 151233774 | 170229 | 2 | ENSMUST00000129088.2 | 154 | Stard13 | ENSMUSG00000016128.9 |
| chr1 | 193368402 | 193368543 | 3.58456 | 4.27214 | 1.2487 | Intron (ENSMUST00000169907.2/ENSMUSG00000016179.6, intron 1 of 11) | 1 | 193346346 | 193370225 | 23880 | 2 | ENSMUST00000169907.2 | 1682 | Camk1g | ENSMUSG00000016179.6 |
| chr1 | 193388231 | 193388307 | 4.98208 | 7.27568 | 3.38572 | Distal Intergenic | 1 | 193346346 | 193370297 | 23952 | 2 | ENSMUST00000016323.5 | -17935 | Camk1g | ENSMUSG00000016179.6 |

| | | | | | | | | | | | | | | | |
|---|---|---|---|---|---|---|---|---|---|---|---|---|---|---|---|
| chr1 | 193412135 | 193412399 | 3.48851 | 4.0949 | 1.13511 | Distal Intergenic | 1 | 193346346 | 193370297 | 23952 | 2 | ENSMUST00000016323.5 | -41839 | Camk1g | ENSMUSG00000016179.6 |
| chr1 | 193034521 | 193035011 | 3.67903 | 4.51391 | 1.49668 | Promoter | 1 | 192897307 | 193035698 | 138392 | 2 | ENSMUST00000016344.7 | 687 | Syt14 | ENSMUSG00000016200.8 |
| chr2 | 181133302 | 181135506 | 3.77879 | 4.70652 | 1.56926 | Promoter | 2 | 181075579 | 181135103 | 59525 | 2 | ENSMUST00000149964.3 | 0 | Kcnq2 | ENSMUSG00000016346.12 |
| chr2 | 181136737 | 181136969 | 3.91403 | 4.90728 | 1.70205 | Distal Intergenic | 2 | 181075588 | 181135291 | 59704 | 2 | ENSMUST00000016491.8 | -1447 | Kcnq2 | ENSMUSG00000016346.12 |
| chr2 | 181147368 | 181147774 | 3.3888 | 3.91808 | 1.04236 | 3' UTR | 2 | 181147653 | 181157014 | 9362 | 2 | ENSMUST00000055990.7 | 9240 | Eef1a2 | ENSMUSG00000016349.10 |
| chr2 | 181148188 | 181148263 | 3.43569 | 4.00745 | 1.10427 | Intron (ENSMUST00000055990.7/ENSMUSG00000016349.10, intron 7 of 7) | 2 | 181147653 | 181157014 | 9362 | 2 | ENSMUST00000055990.7 | 8751 | Eef1a2 | ENSMUSG00000016349.10 |
| chr2 | 181149685 | 181149952 | 3.4597 | 4.07576 | 1.20413 | Exon (ENSMUST00000055990.7/ENSMUSG00000016349.10, exon 6 of 8) | 2 | 181147653 | 181157014 | 9362 | 2 | ENSMUST00000055990.7 | 7062 | Eef1a2 | ENSMUSG00000016349.10 |
| chr2 | 181156170 | 181157274 | 3.61398 | 4.39623 | 1.38833 | Promoter | 2 | 181147653 | 181157014 | 9362 | 2 | ENSMUST00000055990.7 | 0 | Eef1a2 | ENSMUSG00000016349.10 |
| chr2 | 106692039 | 106692907 | 3.81298 | 4.75352 | 1.62272 | Promoter | 2 | 106693269 | 106868356 | 175088 | 1 | ENSMUST00000111063.2 | -362 | Mpped2 | ENSMUSG00000016386.10 |
| chr2 | 106693250 | 106695230 | 4.02087 | 5.16414 | 1.8921 | Promoter | 2 | 106693269 | 106868356 | 175088 | 1 | ENSMUST00000111063.2 | 0 | Mpped2 | ENSMUSG00000016386.10 |
| chr2 | 106695729 | 106695899 | 3.50822 | 4.16189 | 1.19821 | Promoter | 2 | 106695854 | 106867031 | 171178 | 1 | ENSMUST00000125023.1 | 0 | Mpped2 | ENSMUSG00000016386.10 |
| chr2 | 106696943 | 106697253 | 3.89229 | 4.99987 | 1.78618 | Intron (ENSMUST00000111063.2/ENSMUSG00000016386.10, intron 1 of 6) | 2 | 106695854 | 106867031 | 171178 | 1 | ENSMUST00000125023.1 | 1090 | Mpped2 | ENSMUSG00000016386.10 |
| chr1 | 194938132 | 194938432 | 3.37398 | 3.92484 | 1.11888 | Promoter | 1 | 194938821 | 194961292 | 22472 | 1 | ENSMUST00000016638.2 | -389 | Cd34 | ENSMUSG00000016494.4 |
| chr1 | 194939079 | 194939491 | 3.91063 | 5.1093 | 1.86511 | Promoter | 1 | 194938821 | 194961292 | 22472 | 1 | ENSMUST00000110815.3 | 259 | Cd34 | ENSMUSG00000016494.4 |
| chr5 | 146985644 | 146985839 | 3.61401 | 4.38226 | 1.42911 | Distal Intergenic | 5 | 146958948 | 146963800 | 4853 | 2 | ENSMUST00000140526.1 | -21845 | Mtif3 | ENSMUSG00000016510.6 |
| chr1 | 131137852 | 131138150 | 3.51378 | 4.24584 | 1.30964 | Promoter | 1 | 131129875 | 131138245 | 8371 | 2 | ENSMUST00000189756.1 | 95 | Dyrk3 | ENSMUSG00000016526.8 |
| chr1 | 131021315 | 131021644 | 3.3348 | 3.83216 | 1.02638 | Exon (ENSMUST00000016673.5/ENSMUSG00000016529.5, exon 3 of 5) | 1 | 131019845 | 131024974 | 5130 | 1 | ENSMUST00000016673.5 | 1471 | Il10 | ENSMUSG00000016529.5 |
| chr15 | 77955712 | 77955965 | 3.49289 | 4.40373 | 1.40634 | Promoter | 15 | 77941019 | 77956675 | 15657 | 2 | ENSMUST00000117725.1 | 710 | Foxred2 | ENSMUSG00000016552.8 |
| chr15 | 77984008 | 77984083 | 3.75021 | 4.59185 | 1.4704 | Distal Intergenic | 15 | 77958998 | 77970824 | 11827 | 2 | ENSMUST00000100484.4 | -13185 | Eif3d | ENSMUSG00000016554.8 |
| chr15 | 78149745 | 78150196 | 3.59529 | 4.33377 | 1.31538 | Distal Intergenic | 15 | 78159465 | 78174108 | 14644 | 2 | ENSMUST00000016781.6 | 23912 | Ift27 | ENSMUSG00000016637.6 |
| chr15 | 83605752 | 83605920 | 3.38193 | 3.9006 | 1.01733 | 3' UTR | 15 | 83575644 | 83595157 | 19514 | 2 | ENSMUST00000016901.3 | -10596 | Ttll12 | ENSMUSG00000016757.5 |
| chr15 | 83677347 | 83678004 | 3.55104 | 4.21464 | 1.21998 | Intron (ENSMUST00000171496.2/ENSMUSG00000016763.9, intron 3 of 21) | 15 | 83602583 | 83725021 | 122439 | 2 | ENSMUST00000171496.2 | 47017 | Scube1 | ENSMUSG00000016763.9 |
| chr15 | 83697535 | 83697701 | 3.68466 | 4.47791 | 1.42818 | Intron (ENSMUST00000171496.2/ENSMUSG00000016763.9, intron 3 of 21) | 15 | 83602583 | 83725021 | 122439 | 2 | ENSMUST00000171496.2 | 27320 | Scube1 | ENSMUSG00000016763.9 |
| chr15 | 83723090 | 83723482 | 3.73343 | 4.7603 | 1.64447 | Intron (ENSMUST00000171496.2/ENSMUSG00000016763.9, intron 1 of 21) | 15 | 83602583 | 83725021 | 122439 | 2 | ENSMUST00000171496.2 | 1539 | Scube1 | ENSMUSG00000016763.9 |
| chr15 | 83724663 | 83725320 | 3.78125 | 4.85861 | 1.66188 | Promoter | 15 | 83602583 | 83725021 | 122439 | 2 | ENSMUST00000171496.2 | 0 | Scube1 | ENSMUSG00000016763.9 |
| chr15 | 83734405 | 83734898 | 3.36434 | 3.87406 | 1.01733 | Distal Intergenic | 15 | 83606506 | 83725021 | 118516 | 2 | ENSMUST00000076060.6 | -9385 | Scube1 | ENSMUSG00000016763.9 |
| chr13 | 21979447 | 21979882 | 3.23887 | 3.68547 | 1.01733 | Distal Intergenic | 13 | 21981194 | 21988734 | 7541 | 1 | ENSMUST00000117882.1 | -1312 | Pom121l2 | ENSMUSG00000016982.6 |
| chr13 | 21980191 | 21980583 | 3.70036 | 4.64264 | 1.5624 | Promoter | 13 | 21981194 | 21988734 | 7541 | 1 | ENSMUST00000117882.1 | -611 | Pom121l2 | ENSMUSG00000016982.6 |
| chr13 | 21981063 | 21982111 | 4.2349 | 5.731 | 2.33308 | Promoter | 13 | 21981194 | 21988734 | 7541 | 1 | ENSMUST00000117882.1 | 0 | Pom121l2 | ENSMUSG00000016982.6 |
| chr13 | 21982679 | 21982766 | 3.8511 | 5.01629 | 1.78904 | Exon (ENSMUST00000117882.1/ENSMUSG00000016982.6, exon 1 of 2) | 13 | 21981520 | 21984479 | 2960 | 1 | ENSMUST00000017126.5 | 1160 | Pom121l2 | ENSMUSG00000016982.6 |
| chr13 | 21983243 | 21983413 | 3.83517 | 4.8183 | 1.67639 | Exon (ENSMUST00000017126.5/ENSMUSG00000016982.6, exon 1 of 1) | 13 | 21981520 | 21984479 | 2960 | 1 | ENSMUST00000017126.5 | 1724 | Pom121l2 | ENSMUSG00000016982.6 |
| chr13 | 21984627 | 21984741 | 3.39984 | 4.07317 | 1.16613 | Intron (ENSMUST00000117882.1/ENSMUSG00000016982.6, intron 1 of 1) | 13 | 21981520 | 21984479 | 2960 | 1 | ENSMUST00000017126.5 | 3108 | Pom121l2 | ENSMUSG00000016982.6 |
| chr13 | 21986155 | 21986248 | 3.63722 | 4.57422 | 1.45071 | Intron (ENSMUST00000117882.1/ENSMUSG00000016982.6, intron 1 of 1) | 13 | 21981520 | 21984479 | 2960 | 1 | ENSMUST00000017126.5 | 4636 | Pom121l2 | ENSMUSG00000016982.6 |
| chr13 | 21990447 | 21990522 | 3.82062 | 4.84411 | 1.68565 | Distal Intergenic | 13 | 21981520 | 21984479 | 2960 | 1 | ENSMUST00000017126.5 | 8928 | Pom121l2 | ENSMUSG00000016982.6 |
| chr13 | 21991300 | 21992016 | 3.87059 | 4.89325 | 1.73528 | Distal Intergenic | 13 | 21981520 | 21984479 | 2960 | 1 | ENSMUST00000017126.5 | 9781 | Pom121l2 | ENSMUSG00000016982.6 |
| chr13 | 21992783 | 21993350 | 3.50211 | 4.33697 | 1.39444 | Distal Intergenic | 13 | 21981520 | 21984479 | 2960 | 1 | ENSMUST00000017126.5 | 11264 | Pom121l2 | ENSMUSG00000016982.6 |
| chr2 | 164399644 | 164400257 | 3.67688 | 4.47023 | 1.40775 | Exon (ENSMUST00000103103.4/ENSMUSG00000016995.12, exon 4 of 11) | 2 | 164403141 | 164414791 | 11651 | 1 | ENSMUST00000137427.2 | -2884 | Rbpjl | ENSMUSG00000017007.4 |
| chr2 | 164400608 | 164401165 | 3.61776 | 4.55156 | 1.51468 | Exon (ENSMUST00000109358.2/ENSMUSG00000016995.12, exon 3 of 9) | 2 | 164403141 | 164414791 | 11651 | 1 | ENSMUST00000137427.2 | -1976 | Rbpjl | ENSMUSG00000017007.4 |
| chr11 | 98548494 | 98548669 | 3.32901 | 3.85514 | 1.12573 | Distal Intergenic | 11 | 98551097 | 98558665 | 7569 | 1 | ENSMUST00000081033.7 | -2428 | Zpbp2 | ENSMUSG00000017195.10 |
| chr11 | 98549362 | 98549523 | 3.55815 | 4.27756 | 1.34444 | Distal Intergenic | 11 | 98551097 | 98558665 | 7569 | 1 | ENSMUST00000081033.7 | -1574 | Zpbp2 | ENSMUSG00000017195.10 |
| chr11 | 98666429 | 98666532 | 3.75597 | 4.59419 | 1.45382 | Exon (ENSMUST00000017348.2/ENSMUSG00000017204.4, exon 3 of 12) | 11 | 98664351 | 98677708 | 13358 | 1 | ENSMUST00000017348.2 | 2079 | Gsdma | ENSMUSG00000017204.4 |
| chr11 | 98669117 | 98669366 | 3.65127 | 4.4586 | 1.46979 | Intron (ENSMUST00000017348.2/ENSMUSG00000017204.4, intron 3 of 11) | 11 | 98664351 | 98677708 | 13358 | 1 | ENSMUST00000017348.2 | 4767 | Gsdma | ENSMUSG00000017204.4 |
| chr11 | 98648730 | 98648982 | 3.2794 | 3.90739 | 1.12486 | Intron (ENSMUST00000093938.4/ENSMUSG00000017211.9, intron 2 of 10) | 11 | 98650659 | 98652060 | 1402 | 1 | ENSMUST00000136111.1 | -1677 | Gsdma2 | ENSMUSG00000017211.9 |
| chr11 | 102104844 | 102105479 | 3.72213 | 4.60056 | 1.55551 | Downstream (1-2kb) | 11 | 102106680 | 102107416 | 737 | 2 | ENSMUST00000127381.2 | 1937 | Pyy | ENSMUSG00000017311.10 |
| chr11 | 102107157 | 102108065 | 4.31278 | 5.83765 | 2.4473 | Promoter | 11 | 102106676 | 102107822 | 1147 | 2 | ENSMUST00000177304.2 | 0 | Pyy | ENSMUSG00000017311.10 |
| chr11 | 102108683 | 102108870 | 3.3004 | 3.81 | 1.02538 | Promoter | 11 | 102106961 | 102107832 | 872 | 2 | ENSMUST00000177149.1 | -852 | Pyy | ENSMUSG00000017311.10 |
| chr11 | 102062115 | 102062409 | 3.27065 | 3.80082 | 1.03424 | Exon (ENSMUST00000017458.5/ENSMUSG00000017314.7, exon 7 of 13) | 11 | 102062089 | 102082419 | 20331 | 2 | ENSMUST00000127522.2 | 20010 | Mpp2 | ENSMUSG00000017314.7 |
| chr11 | 102069096 | 102069442 | 3.31262 | 3.87644 | 1.08849 | Intron (ENSMUST00000017458.5/ENSMUSG00000017314.7, intron 3 of 12) | 11 | 102062089 | 102082419 | 20331 | 2 | ENSMUST00000127522.2 | 12977 | Mpp2 | ENSMUSG00000017314.7 |
| chr11 | 102082353 | 102082769 | 3.74534 | 4.60873 | 1.52922 | Promoter | 11 | 102057524 | 102082484 | 24961 | 2 | ENSMUST00000100398.4 | 0 | Mpp2 | ENSMUSG00000017314.7 |
| chr11 | 102083106 | 102083356 | 3.35829 | 4.04419 | 1.19427 | Promoter | 11 | 102057524 | 102082484 | 24961 | 2 | ENSMUST00000100398.4 | -623 | Mpp2 | ENSMUSG00000017314.7 |
| chr11 | 102083994 | 102084125 | 3.48163 | 4.09362 | 1.16195 | Intron (ENSMUST00000017458.5/ENSMUSG00000017314.7, intron 2 of 12) | 11 | 102057524 | 102082484 | 24961 | 2 | ENSMUST00000100398.4 | -1511 | Mpp2 | ENSMUSG00000017314.7 |
| chr11 | 78498560 | 78498867 | 3.48442 | 4.09169 | 1.14547 | Promoter | 11 | 78499091 | 78500585 | 1495 | 1 | ENSMUST00000128479.1 | -224 | Vtn | ENSMUSG00000017344.4 |
| chr2 | 52675517 | 52675753 | 3.84198 | 4.80121 | 1.68142 | Promoter | 2 | 52675727 | 52676143 | 417 | 2 | ENSMUST00000148837.1 | 390 | Cacnb4 | ENSMUSG00000017412.10 |
| chr2 | 52676181 | 52676704 | 3.65562 | 4.4527 | 1.46369 | Promoter | 2 | 52428320 | 52676582 | 248263 | 2 | ENSMUST00000178799.2 | 0 | Cacnb4 | ENSMUSG00000017412.10 |
| chr11 | 97986123 | 97986415 | 3.38638 | 3.96845 | 1.13018 | Promoter | 11 | 97923656 | 97986399 | 62744 | 2 | ENSMUST00000107565.2 | 0 | Plxdc1 | ENSMUSG00000017417.9 |
| chr11 | 77875122 | 77875407 | 3.34506 | 3.90453 | 1.09227 | Distal Intergenic | 11 | 77881060 | 77882686 | 1627 | 2 | ENSMUST00000151383.1 | 7279 | Pipox | ENSMUSG00000017453.4 |
| chr11 | 77882979 | 77883184 | 3.33194 | 3.82523 | 1.01733 | Promoter | 11 | 77881060 | 77882686 | 1627 | 2 | ENSMUST00000151383.1 | -294 | Pipox | ENSMUSG00000017453.4 |
| chr11 | 77897029 | 77897351 | 3.62722 | 4.36901 | 1.36681 | Distal Intergenic | 11 | 77880615 | 77894096 | 13482 | 2 | ENSMUST00000017597.4 | -2934 | Pipox | ENSMUSG00000017453.4 |
| chr11 | 77899673 | 77899903 | 3.85575 | 4.82508 | 1.68198 | Distal Intergenic | 11 | 77880615 | 77894096 | 13482 | 2 | ENSMUST00000017597.4 | -5578 | Pipox | ENSMUSG00000017453.4 |
| chr11 | 99040150 | 99040350 | 3.60601 | 4.38589 | 1.40761 | Promoter | 11 | 99041244 | 99054392 | 13149 | 1 | ENSMUST00000017637.7 | -894 | Igfbp4 | ENSMUSG00000017493.7 |
| chr11 | 99040915 | 99041705 | 3.49095 | 4.5311 | 1.47492 | Promoter | 11 | 99041244 | 99054392 | 13149 | 1 | ENSMUST00000017637.7 | 0 | Igfbp4 | ENSMUSG00000017493.7 |
| chr11 | 99080055 | 99080195 | 3.27861 | 3.74506 | 1.01733 | Exon (ENSMUST00000017751.2/ENSMUSG00000017607.4, exon 3 of 13) | 11 | 99070684 | 99075341 | 4658 | 2 | ENSMUST00000153351.1 | -4715 | Tns4 | ENSMUSG00000017607.4 |
| chr11 | 76571464 | 76571755 | 4.43274 | 6.66007 | 3.0088 | Promoter | 11 | 76416734 | 76571535 | 154802 | 2 | ENSMUST00000072740.7 | 0 | Abr | ENSMUSG00000017631.13 |
| chr11 | 76644438 | 76644513 | 3.65284 | 4.40247 | 1.3459 | Distal Intergenic | 11 | 76479057 | 76622314 | 143258 | 2 | ENSMUST00000155035.2 | -22125 | Abr | ENSMUSG00000017631.13 |
| chr11 | 76645210 | 76645418 | 3.26831 | 3.72963 | 1.01733 | Distal Intergenic | 11 | 76479057 | 76622314 | 143258 | 2 | ENSMUST00000155035.2 | -22897 | Abr | ENSMUSG00000017631.13 |
| chr11 | 79589692 | 79590623 | 3.34593 | 4.10724 | 1.19403 | Promoter | 11 | 79591212 | 79698023 | 106812 | 1 | ENSMUST00000017783.7 | -589 | Rab11fip4 | ENSMUSG00000017639.8 |
| chr11 | 79591270 | 79592229 | 4.4681 | 6.17433 | 2.63996 | Promoter | 11 | 79591212 | 79698023 | 106812 | 1 | ENSMUST00000017783.7 | 59 | Rab11fip4 | ENSMUSG00000017639.8 |
| chr11 | 79592550 | 79592937 | 3.69411 | 5.05612 | 1.82998 | Intron (ENSMUST00000017783.7/ENSMUSG00000017639.8, intron 1 of 14) | 11 | 79591212 | 79698023 | 106812 | 1 | ENSMUST00000017783.7 | 1339 | Rab11fip4 | ENSMUSG00000017639.8 |
| chr11 | 79661029 | 79661222 | 3.74924 | 4.98366 | 1.76432 | Promoter | 11 | 79660562 | 79683498 | 22937 | 1 | ENSMUST00000155381.1 | 468 | Rab11fip4 | ENSMUSG00000017639.8 |
| chr2 | 165069511 | 165069628 | 3.73277 | 4.60428 | 1.563 | Intron (ENSMUST00000017799.6/ENSMUSG00000017652.11, intron 6 of 8) | 2 | 165062676 | 165066695 | 4020 | 1 | ENSMUST00000153105.1 | 6836 | Cd40 | ENSMUSG00000017652.11 |
| chr2 | 165070183 | 165070383 | 3.31442 | 3.79885 | 1.01733 | Intron (ENSMUST00000017799.6/ENSMUSG00000017652.11, intron 8 of 8) | 2 | 165062676 | 165066695 | 4020 | 1 | ENSMUST00000153105.1 | 7508 | Cd40 | ENSMUSG00000017652.11 |
| chr2 | 165070725 | 165070999 | 3.72681 | 4.57202 | 1.51255 | 3' UTR | 2 | 165062676 | 165066695 | 4020 | 1 | ENSMUST00000153105.1 | 8050 | Cd40 | ENSMUSG00000017652.11 |
| chr2 | 165075114 | 165075554 | 4.32056 | 5.74263 | 2.3096 | Distal Intergenic | 2 | 165062676 | 165066695 | 4020 | 1 | ENSMUST00000153105.1 | 12439 | Cd40 | ENSMUSG00000017652.11 |
| chr11 | 117888291 | 117888593 | 3.6406 | 4.63037 | 1.51736 | Distal Intergenic | 11 | 117867952 | 117873481 | 5530 | 2 | ENSMUST00000153989.1 | -14811 | Tha1 | ENSMUSG00000017713.9 |
| chr11 | 117889068 | 117889334 | 3.73959 | 4.68266 | 1.56179 | Distal Intergenic | 11 | 117867952 | 117873481 | 5530 | 2 | ENSMUST00000153989.1 | -15588 | Tha1 | ENSMUSG00000017713.9 |
| chr11 | 118033243 | 118033442 | 3.34366 | 3.84287 | 1.01733 | Intron (ENSMUST00000084803.6/ENSMUSG00000033987.10, intron 74 of 80) | 11 | 118012343 | 118024011 | 11669 | 1 | ENSMUST00000155507.1 | 20901 | Pgs1 | ENSMUSG00000017715.6 |
| chr2 | 164563434 | 164563892 | 3.84543 | 4.86661 | 1.73143 | Promoter | 2 | 164563019 | 164564179 | 1161 | 1 | ENSMUST00000125051.1 | 416 | Wfdc2 | ENSMUSG00000017723.6 |
| chr11 | 101774059 | 101774219 | 3.53816 | 4.25138 | 1.36457 | Exon (ENSMUST00000164750.2/ENSMUSG00000017724.9, exon 7 of 13) | 11 | 101770354 | 101772181 | 1828 | 2 | ENSMUST00000132040.1 | -1879 | Etv4 | ENSMUSG00000017724.9 |
| chr11 | 101775975 | 101776414 | 3.95316 | 5.06729 | 1.89721 | Intron (ENSMUST00000164750.2/ENSMUSG00000017724.9, intron 5 of 12) | 11 | 101770354 | 101772181 | 1828 | 2 | ENSMUST00000132040.1 | -3795 | Etv4 | ENSMUSG00000017724.9 |
| chr11 | 101781992 | 101782276 | 3.5172 | 4.26718 | 1.28548 | Exon (ENSMUST00000131117.2/ENSMUSG00000017724.9, exon 3 of 6) | 11 | 101774168 | 101784388 | 10221 | 2 | ENSMUST00000131117.2 | 2112 | Etv4 | ENSMUSG00000017724.9 |
| chr11 | 101783130 | 101783513 | 3.58243 | 4.30126 | 1.33748 | Promoter | 11 | 101774168 | 101784388 | 10221 | 2 | ENSMUST00000131117.2 | 875 | Etv4 | ENSMUSG00000017724.9 |
| chr11 | 101784030 | 101784601 | 4.13708 | 5.36343 | 2.00692 | Promoter | 11 | 101773862 | 101784491 | 10630 | 2 | ENSMUST00000129995.2 | 0 | Etv4 | ENSMUSG00000017724.9 |
| chr11 | 101785880 | 101786147 | 3.65488 | 4.85309 | 1.6866 | Promoter | 11 | 101769747 | 101785371 | 15625 | 2 | ENSMUST00000107176.2 | -510 | Etv4 | ENSMUSG00000017724.9 |
| chr11 | 101786834 | 101787141 | 3.35769 | 3.86404 | 1.01733 | Distal Intergenic | 11 | 101769747 | 101785371 | 15625 | 2 | ENSMUST00000107176.2 | -1464 | Etv4 | ENSMUSG00000017724.9 |
| chr11 | 101811784 | 101812087 | 3.51126 | 4.14934 | 1.20717 | Distal Intergenic | 11 | 101769747 | 101785371 | 15625 | 2 | ENSMUST00000107176.2 | -26414 | Etv4 | ENSMUSG00000017724.9 |
| chr11 | 101815147 | 101815523 | 3.53381 | 4.20297 | 1.25967 | Distal Intergenic | 11 | 101769747 | 101785371 | 15625 | 2 | ENSMUST00000107176.2 | -29777 | Etv4 | ENSMUSG00000017724.9 |
| chr2 | 164589903 | 164589990 | 3.75678 | 4.6701 | 1.62656 | Intron (ENSMUST00000103100.1/ENSMUSG00000017733.4, intron 2 of 3) | 2 | 164588343 | 164593594 | 5252 | 2 | ENSMUST00000103100.1 | 3604 | Eppin | ENSMUSG00000017733.4 |
| chr2 | 164934505 | 164935074 | 3.44907 | 4.0328 | 1.12685 | Distal Intergenic | 2 | 164940780 | 164948894 | 8115 | 1 | ENSMUST00000137626.1 | -5706 | Mmp9 | ENSMUSG00000017737.2 |
| chr2 | 164939515 | 164940435 | 3.72315 | 4.59134 | 1.50572 | Promoter | 2 | 164940780 | 164948894 | 8115 | 1 | ENSMUST00000137626.1 | -345 | Mmp9 | ENSMUSG00000017737.2 |
| chr2 | 164940785 | 164940918 | 3.44049 | 4.02321 | 1.13937 | Promoter | 2 | 164940780 | 164948894 | 8115 | 1 | ENSMUST00000137626.1 | 6 | Mmp9 | ENSMUSG00000017737.2 |
| chr2 | 164948451 | 164948894 | 3.46897 | 4.09173 | 1.22748 | Promoter | 2 | 164948219 | 164955850 | 7632 | 1 | ENSMUST00000017881.2 | 233 | Mmp9 | ENSMUSG00000017737.2 |
| chr2 | 164950006 | 164951230 | 3.74605 | 4.65412 | 1.56112 | Promoter | 2 | 164950696 | 164951262 | 567 | 1 | ENSMUST00000134382.1 | 0 | Mmp9 | ENSMUSG00000017737.2 |
| chr2 | 164952993 | 164953286 | 3.37516 | 3.94684 | 1.17203 | Exon (ENSMUST00000017881.2/ENSMUSG00000017737.2, exon 10 of 13) | 2 | 164954386 | 164955850 | 1465 | 1 | ENSMUST00000144917.1 | -1100 | Mmp9 | ENSMUSG00000017737.2 |
| chr2 | 164960741 | 164961219 | 3.44025 | 4.01643 | 1.1159 | Promoter | 2 | 164960816 | 164974540 | 13725 | 1 | ENSMUST00000124372.2 | 0 | Slc12a5 | ENSMUSG00000017740.11 |
| chr2 | 164966733 | 164968827 | 3.82627 | 4.90053 | 1.71385 | Promoter | 2 | 164967983 | 164999727 | 31745 | 1 | ENSMUST00000099092.4 | 0 | Slc12a5 | ENSMUSG00000017740.11 |
| chr2 | 164969352 | 164969860 | 3.75469 | 4.59518 | 1.4649 | Intron (ENSMUST00000124372.2/ENSMUSG00000017740.11, intron 1 of 3) | 2 | 164967983 | 164999727 | 31745 | 1 | ENSMUST00000099092.4 | 1370 | Slc12a5 | ENSMUSG00000017740.11 |
| chr2 | 164970854 | 164971040 | 4.00117 | 5.2214 | 1.91123 | Intron (ENSMUST00000124372.2/ENSMUSG00000017740.11, intron 1 of 3) | 2 | 164967983 | 164999727 | 31745 | 1 | ENSMUST00000099092.4 | 2872 | Slc12a5 | ENSMUSG00000017740.11 |
| chr2 | 164997889 | 164998133 | 3.64468 | 4.42309 | 1.43322 | 3' UTR | 2 | 164996591 | 164999727 | 3137 | 1 | ENSMUST00000137302.1 | 1299 | Slc12a5 | ENSMUSG00000017740.11 |
| chr2 | 164854944 | 164855044 | 3.54205 | 4.2973 | 1.33752 | Exon (ENSMUST00000109316.2/ENSMUSG00000017754.8, exon 3 of 15) | 2 | 164846813 | 164856796 | 9984 | 2 | ENSMUST00000142603.2 | 1752 | Pltp | ENSMUSG00000017754.8 |
| chr13 | 73745510 | 73745613 | 3.5735 | 4.29742 | 1.37208 | Distal Intergenic | 13 | 73763697 | 73816743 | 53047 | 1 | ENSMUST00000017900.7 | -18084 | Slc12a7 | ENSMUSG00000017756.8 |
| chr2 | 163375754 | 163376276 | 3.89787 | 5.0051 | 1.79834 | Exon (ENSMUST00000017961.5/ENSMUSG00000017817.6, exon 2 of 6) | 2 | 163336242 | 163397949 | 61708 | 2 | ENSMUST00000017961.5 | 21673 | Jph2 | ENSMUSG00000017817.6 |
| chr12 | 110385141 | 110385449 | 3.62265 | 4.52383 | 1.47197 | Distal Intergenic | 12 | 110485739 | 110583061 | 97323 | 1 | ENSMUST00000084985.4 | -100290 | Ppp2r5c | ENSMUSG00000017843.8 |
| chr12 | 110386099 | 110387760 | 3.7484 | 4.61334 | 1.51072 | Distal Intergenic | 12 | 110485739 | 110583061 | 97323 | 1 | ENSMUST00000084985.4 | -97979 | Ppp2r5c | ENSMUSG00000017843.8 |
| chr12 | 110388181 | 110388382 | 3.41776 | 3.9689 | 1.06739 | Distal Intergenic | 12 | 110485739 | 110583061 | 97323 | 1 | ENSMUST00000084985.4 | -97357 | Ppp2r5c | ENSMUSG00000017843.8 |
| chr2 | 163013411 | 163013768 | 3.51575 | 4.15275 | 1.19531 | 3' UTR | 2 | 163017354 | 163046141 | 28788 | 1 | ENSMUST00000018002.7 | -3586 | Ift52 | ENSMUSG00000017858.7 |
| chr2 | 163054751 | 163055278 | 3.85195 | 5.15079 | 1.90554 | Promoter | 2 | 163054687 | 163084688 | 30002 | 1 | ENSMUST00000018005.4 | 65 | Mybl2 | ENSMUSG00000017861.6 |
| chr2 | 163055645 | 163055735 | 3.82529 | 4.83045 | 1.70739 | Promoter | 2 | 163054687 | 163084688 | 30002 | 1 | ENSMUST00000018005.4 | 959 | Mybl2 | ENSMUSG00000017861.6 |
| chr2 | 165562592 | 165563978 | 4.17073 | 5.45188 | 2.07451 | Distal Intergenic | 2 | 165595032 | 165771719 | 176688 | 1 | ENSMUST00000088132.7 | -31054 | Eya2 | ENSMUSG00000017897.13 |
| chr2 | 165564796 | 165565112 | 3.58626 | 4.29005 | 1.291 | Distal Intergenic | 2 | 165595032 | 165771719 | 176688 | 1 | ENSMUST00000088132.7 | -29920 | Eya2 | ENSMUSG00000017897.13 |
| chr2 | 165567522 | 165567675 | 3.67355 | 4.4391 | 1.35835 | Distal Intergenic | 2 | 165595032 | 165771719 | 176688 | 1 | ENSMUST00000088132.7 | -27357 | Eya2 | ENSMUSG00000017897.13 |
| chr2 | 165593975 | 165594599 | 4.10519 | 5.29814 | 1.9661 | Promoter | 2 | 165595032 | 165771719 | 176688 | 1 | ENSMUST00000088132.7 | -433 | Eya2 | ENSMUSG00000017897.13 |
| chr2 | 165594936 | 165596441 | 3.86735 | 4.84929 | 1.65818 | Promoter | 2 | 165595032 | 165771719 | 176688 | 1 | ENSMUST00000088132.7 | 0 | Eya2 | ENSMUSG00000017897.13 |
| chr2 | 165759511 | 165759637 | 3.57469 | 4.33091 | 1.38415 | Intron (ENSMUST00000088132.7/ENSMUSG00000017897.13, intron 10 of 15) | 2 | 165761824 | 165767129 | 5306 | 1 | ENSMUST00000128696.1 | -2187 | Eya2 | ENSMUSG00000017897.13 |
| chr2 | 165726898 | 165727062 | 3.69763 | 4.4872 | 1.39584 | Intron (ENSMUST00000088132.7/ENSMUSG00000017897.13, intron 8 of 15) | 2 | 165713569 | 165766260 | 52692 | 1 | ENSMUST00000150638.2 | 13330 | Eya2 | ENSMUSG00000017897.13 |
| chr2 | 165732296 | 165732372 | 4.59204 | 6.31691 | 2.70171 | Intron (ENSMUST00000088132.7/ENSMUSG00000017897.13, intron 9 of 15) | 2 | 165713569 | 165766260 | 52692 | 1 | ENSMUST00000150638.2 | 18728 | Eya2 | ENSMUSG00000017897.13 |
| chr2 | 165758590 | 165758945 | 3.41907 | 3.9953 | 1.15442 | Intron (ENSMUST00000088132.7/ENSMUSG00000017897.13, intron 10 of 15) | 2 | 165761824 | 165767129 | 5306 | 1 | ENSMUST00000128696.1 | -2879 | Eya2 | ENSMUSG00000017897.13 |
| chr13 | 27206048 | 27206156 | 14.26252 | 33.20194 | 27.58233 | Distal Intergenic | 13 | 27199353 | 27203612 | 4260 | 1 | ENSMUST00000178072.1 | 6696 | Prl3c1 | ENSMUSG00000017922.8 |
| chr2 | 167382204 | 167382465 | 3.42615 | 4.11302 | 1.20064 | Distal Intergenic | 2 | 167298444 | 167349183 | 50740 | 2 | ENSMUST00000109221.3 | -33022 | B4galt5 | ENSMUSG00000017929.8 |
| chr2 | 163436855 | 163436951 | 3.36278 | 3.89887 | 1.11344 | Distal Intergenic | 2 | 163438476 | 163455324 | 16849 | 1 | ENSMUST00000109420.4 | -1525 | Gdap1l1 | ENSMUSG00000017943.10 |
| chr2 | 163438201 | 163438786 | 3.75747 | 4.89151 | 1.7411 | Promoter | 2 | 163438476 | 163455324 | 16849 | 1 | ENSMUST00000109420.4 | 0 | Gdap1l1 | ENSMUSG00000017943.10 |
| chr2 | 167240108 | 167240338 | 3.4357 | 4.07917 | 1.16383 | Promoter | 2 | 167191805 | 167240589 | 48785 | 2 | ENSMUST00000088041.5 | 251 | Ptgis | ENSMUSG00000017969.8 |
| chr6 | 23838493 | 23839348 | 3.93022 | 5.16033 | 1.94634 | Promoter | 6 | 23262773 | 23839137 | 576365 | 2 | ENSMUST00000166458.3 | 0 | Cadps2 | ENSMUSG00000017978.13 |
| chr15 | 78590978 | 78591053 | 4.21946 | 5.49653 | 2.07533 | Distal Intergenic | 15 | 78597047 | 78622019 | 24973 | 1 | ENSMUST00000043069.5 | -5994 | Cyth4 | ENSMUSG00000018008.7 |
| chr15 | 78621324 | 78621883 | 3.31765 | 3.83725 | 1.11426 | 3' UTR | 15 | 78597047 | 78622019 | 24973 | 1 | ENSMUST00000043069.5 | 24278 | Cyth4 | ENSMUSG00000018008.7 |
| chr15 | 78638735 | 78638885 | 3.23822 | 3.68668 | 1.02427 | Distal Intergenic | 15 | 78597047 | 78622019 | 24973 | 1 | ENSMUST00000043069.5 | 41689 | Cyth4 | ENSMUSG00000018008.7 |
| chr15 | 78639687 | 78640064 | 3.59005 | 4.2984 | 1.30192 | Distal Intergenic | 15 | 78597047 | 78622019 | 24973 | 1 | ENSMUST00000043069.5 | 42641 | Cyth4 | ENSMUSG00000018008.7 |
| chr11 | 120721622 | 120721946 | 3.41078 | 4.00515 | 1.16319 | Promoter | 11 | 120721573 | 120723606 | 2034 | 1 | ENSMUST00000142229.1 | 50 | Rac3 | ENSMUSG00000018012.6 |
| chr11 | 120722278 | 120722491 | 3.58666 | 4.34597 | 1.37981 | Promoter | 11 | 120722014 | 120723565 | 1552 | 1 | ENSMUST00000134322.1 | 265 | Rac3 | ENSMUSG00000018012.6 |
| chr5 | 118532120 | 118532363 | 3.65039 | 4.56382 | 1.51453 | Distal Intergenic | 5 | 118560719 | 118765435 | 204717 | 1 | ENSMUST00000100816.4 | -28356 | Med13l | ENSMUSG00000018076.9 |
| chr5 | 118548999 | 118549176 | 3.17865 | 3.9928 | 1.09796 | Distal Intergenic | 5 | 118560719 | 118765435 | 204717 | 1 | ENSMUST00000100816.4 | -11543 | Med13l | ENSMUSG00000018076.9 |
| chr5 | 118556172 | 118556355 | 3.34615 | 3.86441 | 1.07679 | Distal Intergenic | 5 | 118560719 | 118765435 | 204717 | 1 | ENSMUST00000100816.4 | -4364 | Med13l | ENSMUSG00000018076.9 |
| chr5 | 118831325 | 118831543 | 3.36551 | 3.87583 | 1.01733 | Distal Intergenic | 5 | 118560719 | 118765435 | 204717 | 1 | ENSMUST00000100816.4 | 270607 | Med13l | ENSMUSG00000018076.9 |
| chr15 | 79261090 | 79263537 | 3.97815 | 5.1545 | 1.90443 | Promoter | 15 | 79259716 | 79261297 | 1582 | 2 | ENSMUST00000167970.1 | 0 | Baiap2l2 | ENSMUSG00000018126.10 |
| chr11 | 98476932 | 98477007 | 3.97094 | 5.11825 | 1.86663 | Promoter | 11 | 98467258 | 98477403 | 10146 | 2 | ENSMUST00000153902.1 | 396 | Ikzf3 | ENSMUSG00000018168.8 |
| chr11 | 98493244 | 98493400 | 3.34225 | 3.87745 | 1.13292 | Intron (ENSMUST00000103141.3/ENSMUSG00000018168.8, intron 3 of 7) | 11 | 98484267 | 98489020 | 4754 | 2 | ENSMUST00000140876.1 | -4225 | Ikzf3 | ENSMUSG00000018168.8 |
| chr11 | 98494473 | 98495244 | 3.46207 | 4.15769 | 1.25522 | Intron (ENSMUST00000103141.3/ENSMUSG00000018168.8, intron 3 of 7) | 11 | 98484267 | 98489020 | 4754 | 2 | ENSMUST00000140876.1 | -5454 | Ikzf3 | ENSMUSG00000018168.8 |
| chr11 | 98505043 | 98505328 | 3.53435 | 4.21037 | 1.30209 | Intron (ENSMUST00000103141.3/ENSMUSG00000018168.8, intron 3 of 7) | 11 | 98484267 | 98489020 | 4754 | 2 | ENSMUST00000140876.1 | -16024 | Ikzf3 | ENSMUSG00000018168.8 |
| chr11 | 98510276 | 98510351 | 4.26294 | 5.63787 | 2.25944 | Intron (ENSMUST00000103141.3/ENSMUSG00000018168.8, intron 3 of 7) | 11 | 98484267 | 98489020 | 4754 | 2 | ENSMUST00000140876.1 | -21257 | Ikzf3 | ENSMUSG00000018168.8 |
| chr11 | 98541271 | 98541954 | 3.75833 | 4.64021 | 1.56167 | Intron (ENSMUST00000103141.3/ENSMUSG00000018168.8, intron 1 of 7) | 11 | 98464902 | 98545800 | 80899 | 2 | ENSMUST00000103141.3 | 3846 | Ikzf3 | ENSMUSG00000018168.8 |
| chr11 | 98542479 | 98542998 | 3.58637 | 4.29581 | 1.32238 | Intron (ENSMUST00000103141.3/ENSMUSG00000018168.8, intron 1 of 7) | 11 | 98464902 | 98545800 | 80899 | 2 | ENSMUST00000103141.3 | 2802 | Ikzf3 | ENSMUSG00000018168.8 |
| chr11 | 98544800 | 98548132 | 3.79675 | 4.79823 | 1.67375 | Promoter | 11 | 98464902 | 98545800 | 80899 | 2 | ENSMUST00000103141.3 | 0 | Ikzf3 | ENSMUSG00000018168.8 |
| chr5 | 119770877 | 119771805 | 3.43503 | 4.01548 | 1.1375 | Distal Intergenic | 5 | 119834663 | 119885218 | 50556 | 1 | ENSMUST00000018407.6 | -62858 | Tbx5 | ENSMUSG00000018263.8 |
| chr5 | 119772788 | 119773819 | 3.49364 | 4.29718 | 1.31184 | Distal Intergenic | 5 | 119834663 | 119885218 | 50556 | 1 | ENSMUST00000018407.6 | -60844 | Tbx5 | ENSMUSG00000018263.8 |
| chr5 | 119776533 | 119776626 | 3.62818 | 4.38315 | 1.39812 | Distal Intergenic | 5 | 119834663 | 119885218 | 50556 | 1 | ENSMUST00000018407.6 | -58037 | Tbx5 | ENSMUSG00000018263.8 |
| chr5 | 119780638 | 119781036 | 3.59436 | 4.33128 | 1.38636 | Distal Intergenic | 5 | 119834663 | 119885218 | 50556 | 1 | ENSMUST00000018407.6 | -53627 | Tbx5 | ENSMUSG00000018263.8 |
| chr5 | 119787274 | 119787567 | 3.42375 | 4.04899 | 1.14714 | Distal Intergenic | 5 | 119834663 | 119885218 | 50556 | 1 | ENSMUST00000018407.6 | -47096 | Tbx5 | ENSMUSG00000018263.8 |
| chr5 | 119789739 | 119790355 | 3.50798 | 4.14308 | 1.20073 | Distal Intergenic | 5 | 119834663 | 119885218 | 50556 | 1 | ENSMUST00000018407.6 | -44308 | Tbx5 | ENSMUSG00000018263.8 |
| chr5 | 119791699 | 119791986 | 3.64404 | 4.38614 | 1.33515 | Distal Intergenic | 5 | 119834663 | 119885218 | 50556 | 1 | ENSMUST00000018407.6 | -42677 | Tbx5 | ENSMUSG00000018263.8 |
| chr5 | 119794647 | 119794898 | 4.0514 | 5.20251 | 1.87615 | Distal Intergenic | 5 | 119834663 | 119885218 | 50556 | 1 | ENSMUST00000018407.6 | -39765 | Tbx5 | ENSMUSG00000018263.8 |
| chr5 | 119795983 | 119796092 | 3.34153 | 3.83966 | 1.01733 | Distal Intergenic | 5 | 119834663 | 119885218 | 50556 | 1 | ENSMUST00000018407.6 | -38571 | Tbx5 | ENSMUSG00000018263.8 |
| chr5 | 119796759 | 119796935 | 4.3228 | 5.78561 | 2.31791 | Distal Intergenic | 5 | 119834663 | 119885218 | 50556 | 1 | ENSMUST00000018407.6 | -37728 | Tbx5 | ENSMUSG00000018263.8 |
| chr5 | 119797713 | 119797817 | 3.5349 | 4.27791 | 1.31774 | Distal Intergenic | 5 | 119834663 | 119885218 | 50556 | 1 | ENSMUST00000018407.6 | -36846 | Tbx5 | ENSMUSG00000018263.8 |
| chr5 | 119799937 | 119800040 | 4.35981 | 5.78487 | 2.2947 | Distal Intergenic | 5 | 119834663 | 119885218 | 50556 | 1 | ENSMUST00000018407.6 | -34623 | Tbx5 | ENSMUSG00000018263.8 |
| chr5 | 119800427 | 119801646 | 3.59573 | 4.3374 | 1.31205 | Distal Intergenic | 5 | 119834663 | 119885218 | 50556 | 1 | ENSMUST00000018407.6 | -33017 | Tbx5 | ENSMUSG00000018263.8 |
| chr5 | 119801968 | 119802487 | 3.42268 | 4.0338 | 1.11885 | Distal Intergenic | 5 | 119834663 | 119885218 | 50556 | 1 | ENSMUST00000018407.6 | -32176 | Tbx5 | ENSMUSG00000018263.8 |
| chr5 | 119803924 | 119804510 | 3.70954 | 4.66883 | 1.56021 | Distal Intergenic | 5 | 119834663 | 119885218 | 50556 | 1 | ENSMUST00000018407.6 | -30153 | Tbx5 | ENSMUSG00000018263.8 |
| chr5 | 119804856 | 119804985 | 3.34153 | 3.83966 | 1.01733 | Distal Intergenic | 5 | 119834663 | 119885218 | 50556 | 1 | ENSMUST00000018407.6 | -29678 | Tbx5 | ENSMUSG00000018263.8 |
| chr5 | 119806018 | 119806478 | 3.81111 | 4.71096 | 1.56153 | Distal Intergenic | 5 | 119834663 | 119885218 | 50556 | 1 | ENSMUST00000018407.6 | -28185 | Tbx5 | ENSMUSG00000018263.8 |
| chr5 | 119806942 | 119808447 | 3.4892 | 4.10051 | 1.15034 | Distal Intergenic | 5 | 119834663 | 119885218 | 50556 | 1 | ENSMUST00000018407.6 | -26216 | Tbx5 | ENSMUSG00000018263.8 |
| chr5 | 119808811 | 119809033 | 3.54991 | 4.21241 | 1.22341 | Distal Intergenic | 5 | 119834663 | 119885218 | 50556 | 1 | ENSMUST00000018407.6 | -25630 | Tbx5 | ENSMUSG00000018263.8 |
| chr5 | 119811922 | 119812554 | 3.36888 | 3.88091 | 1.01733 | Distal Intergenic | 5 | 119834663 | 119885218 | 50556 | 1 | ENSMUST00000018407.6 | -22109 | Tbx5 | ENSMUSG00000018263.8 |
| chr5 | 119813724 | 119813986 | 3.67791 | 4.45172 | 1.36717 | Distal Intergenic | 5 | 119834663 | 119885218 | 50556 | 1 | ENSMUST00000018407.6 | -20677 | Tbx5 | ENSMUSG00000018263.8 |
| chr5 | 119814512 | 119814794 | 3.38095 | 3.89911 | 1.01733 | Distal Intergenic | 5 | 119834663 | 119885218 | 50556 | 1 | ENSMUST00000018407.6 | -19869 | Tbx5 | ENSMUSG00000018263.8 |

| | | | | | | | | | | | | | | | |
|---|---|---|---|---|---|---|---|---|---|---|---|---|---|---|---|
| chr5 | 119816164 | 119816503 | 3.35407 | 3.92839 | 1.09301 | Distal Intergenic | 5 | 119834663 | 119885218 | 50556 | 1 | ENSMUST00000018407.6 | -18160 | Tbx5 | ENSMUSG00000018263.8 |
| chr5 | 119818401 | 119819042 | 3.62168 | 4.37666 | 1.32878 | Distal Intergenic | 5 | 119834663 | 119885218 | 50556 | 1 | ENSMUST00000018407.6 | -15621 | Tbx5 | ENSMUSG00000018263.8 |
| chr5 | 119819693 | 119820089 | 3.81807 | 4.74194 | 1.5816 | Distal Intergenic | 5 | 119834663 | 119885218 | 50556 | 1 | ENSMUST00000018407.6 | -14574 | Tbx5 | ENSMUSG00000018263.8 |
| chr5 | 119831223 | 119831790 | 4.49611 | 6.26944 | 2.68361 | Distal Intergenic | 5 | 119834663 | 119885218 | 50556 | 1 | ENSMUST00000018407.6 | -2873 | Tbx5 | ENSMUSG00000018263.8 |
| chr5 | 119832176 | 119832647 | 3.83381 | 4.76948 | 1.61277 | Distal Intergenic | 5 | 119834663 | 119885218 | 50556 | 1 | ENSMUST00000018407.6 | -2016 | Tbx5 | ENSMUSG00000018263.8 |
| chr5 | 119833416 | 119834525 | 3.64687 | 4.41727 | 1.4058 | Promoter | 5 | 119834663 | 119885218 | 50556 | 1 | ENSMUST00000018407.6 | -138 | Tbx5 | ENSMUSG00000018263.8 |
| chr5 | 119835377 | 119835830 | 3.71816 | 4.65925 | 1.57078 | Promoter | 5 | 119834663 | 119885218 | 50556 | 1 | ENSMUST00000018407.6 | 715 | Tbx5 | ENSMUSG00000018263.8 |
| chr5 | 119836796 | 119837528 | 3.89985 | 4.94628 | 1.73738 | 5' UTR | 5 | 119834663 | 119885218 | 50556 | 1 | ENSMUST00000018407.6 | 2134 | Tbx5 | ENSMUSG00000018263.8 |
| chr5 | 119838666 | 119840081 | 3.68053 | 4.56066 | 1.49225 | Exon (ENSMUST00000018407.6/ENSMUSG00000018263.8, exon 3 of 9) | 5 | 119834663 | 119885218 | 50556 | 1 | ENSMUST00000018407.6 | 4004 | Tbx5 | ENSMUSG00000018263.8 |
| chr5 | 119840760 | 119840877 | 3.355 | 3.85997 | 1.01733 | Exon (ENSMUST00000018407.6/ENSMUSG00000018263.8, exon 4 of 9) | 5 | 119834663 | 119885218 | 50556 | 1 | ENSMUST00000018407.6 | 6098 | Tbx5 | ENSMUSG00000018263.8 |
| chr5 | 119841395 | 119843232 | 3.58864 | 4.3159 | 1.32141 | Exon (ENSMUST00000018407.6/ENSMUSG00000018263.8, exon 5 of 9) | 5 | 119834663 | 119885218 | 50556 | 1 | ENSMUST00000018407.6 | 6733 | Tbx5 | ENSMUSG00000018263.8 |
| chr5 | 119843587 | 119843854 | 3.64951 | 4.38845 | 1.31304 | Intron (ENSMUST00000018407.6/ENSMUSG00000018263.8, intron 5 of 8) | 5 | 119834663 | 119885218 | 50556 | 1 | ENSMUST00000018407.6 | 8925 | Tbx5 | ENSMUSG00000018263.8 |
| chr5 | 119844716 | 119845719 | 3.78702 | 4.8074 | 1.68004 | Exon (ENSMUST00000018407.6/ENSMUSG00000018263.8, exon 6 of 9) | 5 | 119834663 | 119885218 | 50556 | 1 | ENSMUST00000018407.6 | 10054 | Tbx5 | ENSMUSG00000018263.8 |
| chr5 | 119847534 | 119847952 | 3.35791 | 3.89784 | 1.10267 | Intron (ENSMUST00000018407.6/ENSMUSG00000018263.8, intron 6 of 8) | 5 | 119834663 | 119885218 | 50556 | 1 | ENSMUST00000018407.6 | 12872 | Tbx5 | ENSMUSG00000018263.8 |
| chr5 | 119848996 | 119849677 | 3.49724 | 4.14069 | 1.23011 | Intron (ENSMUST00000018407.6/ENSMUSG00000018263.8, intron 6 of 8) | 5 | 119834663 | 119885218 | 50556 | 1 | ENSMUST00000018407.6 | 14334 | Tbx5 | ENSMUSG00000018263.8 |
| chr5 | 119851548 | 119851780 | 3.58226 | 4.26946 | 1.24605 | Intron (ENSMUST00000018407.6/ENSMUSG00000018263.8, intron 6 of 8) | 5 | 119834663 | 119885218 | 50556 | 1 | ENSMUST00000018407.6 | 16886 | Tbx5 | ENSMUSG00000018263.8 |
| chr5 | 119856402 | 119856745 | 3.56766 | 4.29183 | 1.28521 | Intron (ENSMUST00000018407.6/ENSMUSG00000018263.8, intron 7 of 8) | 5 | 119834663 | 119885218 | 50556 | 1 | ENSMUST00000018407.6 | 21740 | Tbx5 | ENSMUSG00000018263.8 |
| chr5 | 119866028 | 119866474 | 3.89502 | 4.89047 | 1.70809 | Intron (ENSMUST00000018407.6/ENSMUSG00000018263.8, intron 7 of 8) | 5 | 119834663 | 119885218 | 50556 | 1 | ENSMUST00000018407.6 | 31366 | Tbx5 | ENSMUSG00000018263.8 |
| chr5 | 119873408 | 119873483 | 3.54295 | 4.20796 | 1.24378 | Intron (ENSMUST00000018407.6/ENSMUSG00000018263.8, intron 8 of 8) | 5 | 119834663 | 119885218 | 50556 | 1 | ENSMUST00000018407.6 | 38746 | Tbx5 | ENSMUSG00000018263.8 |
| chr5 | 119934160 | 119934345 | 3.29961 | 3.89334 | 1.12083 | Distal Intergenic | 5 | 119834663 | 119885218 | 50556 | 1 | ENSMUST00000018407.6 | 99498 | Tbx5 | ENSMUSG00000018263.8 |
| chr5 | 119943871 | 119943946 | 3.71715 | 4.5258 | 1.42658 | Distal Intergenic | 5 | 119834663 | 119885218 | 50556 | 1 | ENSMUST00000018407.6 | 109209 | Tbx5 | ENSMUSG00000018263.8 |
| chr6 | 67376177 | 67376320 | 3.68192 | 4.81691 | 1.6389 | Promoter | 6 | 67292018 | 67376188 | 84171 | 2 | ENSMUST00000018485.3 | 0 | Il12rb2 | ENSMUSG00000018341.7 |
| chr11 | 95833686 | 95834297 | 4.37369 | 6.02739 | 2.57067 | 3' UTR | 11 | 95832723 | 95835890 | 3168 | 2 | ENSMUST00000137645.1 | 1593 | Abi3 | ENSMUSG00000018381.10 |
| chr11 | 95842094 | 95842402 | 3.34779 | 3.86042 | 1.05718 | Promoter | 11 | 95837077 | 95842098 | 5022 | 2 | ENSMUST00000143492.1 | 0 | Abi3 | ENSMUSG00000018381.10 |
| chr11 | 53456470 | 53457726 | 3.87412 | 4.89782 | 1.69681 | Promoter | 11 | 53457205 | 53467766 | 10562 | 1 | ENSMUST00000109013.3 | 0 | Shroom1 | ENSMUSG00000018387.7 |
| chr11 | 53465610 | 53466024 | 3.28109 | 3.75077 | 1.02418 | Promoter | 11 | 53465496 | 53465853 | 358 | 1 | ENSMUST00000155297.1 | 115 | Shroom1 | ENSMUSG00000018387.7 |
| chr11 | 53466955 | 53467372 | 3.64037 | 4.52222 | 1.45873 | 3' UTR | 11 | 53465496 | 53465853 | 358 | 1 | ENSMUST00000155297.1 | 1460 | Shroom1 | ENSMUSG00000018387.7 |
| chr11 | 88872503 | 88872745 | 3.51402 | 4.27592 | 1.30758 | Intron (ENSMUST00000173677.1/ENSMUSG00000092399.1, intron 3 of 4) | 11 | 88830792 | 88864586 | 33795 | 2 | ENSMUST00000018572.5 | -7918 | Akap1 | ENSMUSG00000018428.10 |
| chr11 | 71032323 | 71032584 | 3.43911 | 4.0572 | 1.23866 | Promoter | 11 | 71031941 | 71033513 | 1573 | 2 | ENSMUST00000156068.2 | 929 | 6330403K07Rik | ENSMUSG00000018451.6 |
| chr11 | 71033407 | 71033490 | 3.50542 | 4.16713 | 1.2889 | Promoter | 11 | 71031941 | 71033513 | 1573 | 2 | ENSMUST00000156068.2 | 23 | 6330403K07Rik | ENSMUSG00000018451.6 |
| chr11 | 71034331 | 71034657 | 3.33003 | 3.8383 | 1.06792 | Promoter | 11 | 71031941 | 71033513 | 1573 | 2 | ENSMUST00000156068.2 | -819 | 6330403K07Rik | ENSMUSG00000018451.6 |
| chr2 | 165466101 | 165466212 | 3.38107 | 3.90241 | 1.02823 | Intron (ENSMUST00000029208.9/ENSMUSG00000018459.10, intron 1 of 12) | 2 | 165444103 | 165473199 | 29097 | 2 | ENSMUST00000147107.1 | 6987 | Slc13a3 | ENSMUSG00000018459.10 |
| chr2 | 165472062 | 165472536 | 3.43648 | 4.00135 | 1.0822 | Promoter | 2 | 165444103 | 165473199 | 29097 | 2 | ENSMUST00000147107.1 | 663 | Slc13a3 | ENSMUSG00000018459.10 |
| chr2 | 165473036 | 165473159 | 4.04351 | 5.33878 | 2.0195 | Promoter | 2 | 165444103 | 165473199 | 29097 | 2 | ENSMUST00000147107.1 | 40 | Slc13a3 | ENSMUSG00000018459.10 |
| chr11 | 103746382 | 103746457 | 3.38314 | 3.91658 | 1.06739 | Intron (ENSMUST00000018630.2/ENSMUSG00000018486.2, intron 1 of 3) | 11 | 103727364 | 103749821 | 22458 | 2 | ENSMUST00000018630.2 | 3364 | Wnt9b | ENSMUSG00000018486.2 |
| chr11 | 103749177 | 103750379 | 3.5919 | 4.35665 | 1.39031 | Promoter | 11 | 103727364 | 103749821 | 22458 | 2 | ENSMUST00000018630.2 | 0 | Wnt9b | ENSMUSG00000018486.2 |
| chr11 | 103751106 | 103751206 | 3.52941 | 4.17756 | 1.20954 | Distal Intergenic | 11 | 103727364 | 103749821 | 22458 | 2 | ENSMUST00000018630.2 | -1286 | Wnt9b | ENSMUSG00000018486.2 |
| chr11 | 103751852 | 103752009 | 3.66991 | 4.46362 | 1.45147 | Distal Intergenic | 11 | 103727364 | 103749821 | 22458 | 2 | ENSMUST00000018630.2 | -2032 | Wnt9b | ENSMUSG00000018486.2 |
| chr11 | 62246895 | 62247107 | 3.43344 | 4.00344 | 1.10632 | Distal Intergenic | 11 | 62248984 | 62266453 | 17470 | 1 | ENSMUST00000018644.2 | -1877 | Adora2b | ENSMUSG00000018500.2 |
| chr11 | 62248023 | 62249960 | 3.7054 | 4.6359 | 1.57655 | Promoter | 11 | 62248984 | 62266453 | 17470 | 1 | ENSMUST00000018644.2 | 0 | Adora2b | ENSMUSG00000018500.2 |
| chr11 | 62250360 | 62250731 | 3.52976 | 4.27026 | 1.31435 | Intron (ENSMUST00000018644.2/ENSMUSG00000018500.2, intron 1 of 1) | 11 | 62248984 | 62266453 | 17470 | 1 | ENSMUST00000018644.2 | 1377 | Adora2b | ENSMUSG00000018500.2 |
| chr11 | 69935807 | 69936800 | 3.67003 | 4.45966 | 1.40497 | Promoter | 11 | 69935914 | 69941576 | 5663 | 1 | ENSMUST00000149194.2 | 0 | Ybx2 | ENSMUSG00000018554.8 |
| chr11 | 69962892 | 69962967 | 3.60646 | 4.3263 | 1.31079 | Distal Intergenic | 11 | 69964779 | 69967810 | 3032 | 1 | ENSMUST00000108597.2 | -1812 | Cldn7 | ENSMUSG00000018569.7 |
| chr11 | 69964528 | 69966739 | 4.0361 | 5.3871 | 2.06199 | Promoter | 11 | 69964779 | 69967810 | 3032 | 1 | ENSMUST00000108597.2 | 0 | Cldn7 | ENSMUSG00000018569.7 |
| chr12 | 117989297 | 117990467 | 3.57442 | 4.29537 | 1.36223 | Intron (ENSMUST00000176756.3/ENSMUSG00000018581.10, intron 23 of 51) | 12 | 117877982 | 118060303 | 182322 | 2 | ENSMUST00000176756.3 | 69836 | Dnah11 | ENSMUSG00000018581.10 |
| chr12 | 117991395 | 117991638 | 3.31769 | 3.81596 | 1.03328 | Intron (ENSMUST00000176756.3/ENSMUSG00000018581.10, intron 23 of 51) | 12 | 117877982 | 118060303 | 182322 | 2 | ENSMUST00000176756.3 | 68665 | Dnah11 | ENSMUSG00000018581.10 |
| chr12 | 118195703 | 118195851 | 3.94959 | 5.4084 | 2.06019 | Intron (ENSMUST00000084806.6/ENSMUSG00000018581.10, intron 2 of 81) | 12 | 117877982 | 118199043 | 321062 | 2 | ENSMUST00000084806.6 | 3192 | Dnah11 | ENSMUSG00000018581.10 |
| chr12 | 118197791 | 118198200 | 4.05238 | 5.27224 | 1.98792 | Promoter | 12 | 117877982 | 118199043 | 321062 | 2 | ENSMUST00000084806.6 | 843 | Dnah11 | ENSMUSG00000018581.10 |
| chr12 | 118198514 | 118198984 | 4.06735 | 5.31041 | 2.06927 | Promoter | 12 | 117877982 | 118199043 | 321062 | 2 | ENSMUST00000084806.6 | 59 | Dnah11 | ENSMUSG00000018581.10 |
| chr5 | 119658700 | 119659192 | 3.75014 | 4.6079 | 1.51937 | Exon (ENSMUST00000154997.1/ENSMUSG00000085345.1, exon 2 of 2) | 5 | 119670669 | 119684561 | 13893 | 1 | ENSMUST00000079719.5 | -11477 | Tbx3 | ENSMUSG00000018604.13 |
| chr5 | 119660744 | 119661353 | 4.50086 | 6.14845 | 2.65103 | Intron (ENSMUST00000129108.1/ENSMUSG00000087516.2, intron 1 of 2) | 5 | 119670669 | 119684561 | 13893 | 1 | ENSMUST00000079719.5 | -9316 | Tbx3 | ENSMUSG00000018604.13 |
| chr5 | 119662319 | 119663613 | 3.89395 | 4.87478 | 1.67085 | Intron (ENSMUST00000129108.1/ENSMUSG00000087516.2, intron 1 of 2) | 5 | 119670669 | 119684561 | 13893 | 1 | ENSMUST00000079719.5 | -7056 | Tbx3 | ENSMUSG00000018604.13 |
| chr5 | 119664296 | 119664897 | 3.5654 | 4.27413 | 1.29322 | Intron (ENSMUST00000129108.1/ENSMUSG00000087516.2, intron 1 of 2) | 5 | 119670669 | 119684561 | 13893 | 1 | ENSMUST00000079719.5 | -5772 | Tbx3 | ENSMUSG00000018604.13 |
| chr5 | 119660133 | 119660310 | 3.69988 | 4.48642 | 1.38849 | Intron (ENSMUST00000129108.1/ENSMUSG00000087516.2, intron 1 of 2) | 5 | 119670669 | 119684561 | 13893 | 1 | ENSMUST00000079719.5 | -10359 | Tbx3 | ENSMUSG00000018604.13 |
| chr5 | 119667074 | 119667434 | 3.60295 | 4.45843 | 1.42198 | Intron (ENSMUST00000129108.1/ENSMUSG00000087516.2, intron 1 of 2) | 5 | 119670669 | 119684561 | 13893 | 1 | ENSMUST00000079719.5 | -3235 | Tbx3 | ENSMUSG00000018604.13 |
| chr5 | 119668326 | 119669460 | 3.56065 | 4.28397 | 1.34652 | Intron (ENSMUST00000129108.1/ENSMUSG00000087516.2, intron 1 of 2) | 5 | 119670669 | 119684561 | 13893 | 1 | ENSMUST00000079719.5 | -1209 | Tbx3 | ENSMUSG00000018604.13 |
| chr5 | 119669809 | 119670704 | 3.55279 | 4.3999 | 1.39456 | Promoter | 5 | 119670669 | 119684561 | 13893 | 1 | ENSMUST00000079719.5 | 0 | Tbx3 | ENSMUSG00000018604.13 |
| chr5 | 119665380 | 119665859 | 3.46528 | 4.07752 | 1.19029 | Intron (ENSMUST00000129108.1/ENSMUSG00000087516.2, intron 1 of 2) | 5 | 119670669 | 119684561 | 13893 | 1 | ENSMUST00000079719.5 | -4810 | Tbx3 | ENSMUSG00000018604.13 |
| chr5 | 119674015 | 119674778 | 3.52847 | 4.44871 | 1.42051 | Promoter | 5 | 119675445 | 119680630 | 5186 | 1 | ENSMUST00000145647.1 | -667 | Tbx3 | ENSMUSG00000018604.13 |
| chr5 | 119679860 | 119681029 | 4.50576 | 6.21042 | 2.75416 | Promoter | 5 | 119680249 | 119680832 | 584 | 1 | ENSMUST00000154680.1 | 0 | Tbx3 | ENSMUSG00000018604.13 |
| chr5 | 119682521 | 119683177 | 3.70234 | 4.51955 | 1.45789 | 3' UTR | 5 | 119680249 | 119680832 | 584 | 1 | ENSMUST00000154680.1 | 2273 | Tbx3 | ENSMUSG00000018604.13 |
| chr5 | 119684868 | 119685620 | 3.93454 | 4.95379 | 1.71701 | Exon (ENSMUST00000135385.1/ENSMUSG00000086847.1, exon 3 of 3) | 5 | 119680249 | 119680832 | 584 | 1 | ENSMUST00000154680.1 | 4620 | Tbx3 | ENSMUSG00000018604.13 |
| chr11 | 104099406 | 104099664 | 3.48973 | 4.1201 | 1.19738 | Distal Intergenic | 11 | 104132855 | 104175523 | 42669 | 1 | ENSMUST00000093925.4 | -33191 | Crhr1 | ENSMUSG00000018634.10 |
| chr11 | 104105055 | 104105132 | 3.6922 | 4.53257 | 1.51828 | Distal Intergenic | 11 | 104132855 | 104175523 | 42669 | 1 | ENSMUST00000093925.4 | -27723 | Crhr1 | ENSMUSG00000018634.10 |
| chr11 | 104130504 | 104131446 | 3.43541 | 4.05135 | 1.19511 | Distal Intergenic | 11 | 104132855 | 104175523 | 42669 | 1 | ENSMUST00000093925.4 | -1409 | Crhr1 | ENSMUSG00000018634.10 |
| chr11 | 104131910 | 104132064 | 3.74606 | 4.66584 | 1.57579 | Promoter | 11 | 104132855 | 104175523 | 42669 | 1 | ENSMUST00000093925.4 | -791 | Crhr1 | ENSMUSG00000018634.10 |
| chr11 | 104132624 | 104133939 | 3.5572 | 4.28543 | 1.36344 | Promoter | 11 | 104132855 | 104175523 | 42669 | 1 | ENSMUST00000093925.4 | 0 | Crhr1 | ENSMUSG00000018634.10 |
| chr11 | 104134723 | 104134798 | 4.20591 | 5.64095 | 2.20309 | Intron (ENSMUST00000093925.4/ENSMUSG00000018634.10, intron 1 of 12) | 11 | 104132855 | 104175523 | 42669 | 1 | ENSMUST00000093925.4 | 1869 | Crhr1 | ENSMUSG00000018634.10 |
| chr11 | 104135958 | 104136179 | 3.24922 | 3.701 | 1.01733 | Intron (ENSMUST00000093925.4/ENSMUSG00000018634.10, intron 1 of 12) | 11 | 104132855 | 104175523 | 42669 | 1 | ENSMUST00000093925.4 | 3104 | Crhr1 | ENSMUSG00000018634.10 |
| chr11 | 104136495 | 104136885 | 3.47575 | 4.10224 | 1.2309 | Intron (ENSMUST00000093925.4/ENSMUSG00000018634.10, intron 1 of 12) | 11 | 104132855 | 104175523 | 42669 | 1 | ENSMUST00000093925.4 | 3641 | Crhr1 | ENSMUSG00000018634.10 |
| chr11 | 104156418 | 104156640 | 4.24662 | 5.76075 | 2.30239 | Intron (ENSMUST00000093925.4/ENSMUSG00000018634.10, intron 2 of 12) | 11 | 104132855 | 104175523 | 42669 | 1 | ENSMUST00000093925.4 | 23564 | Crhr1 | ENSMUSG00000018634.10 |
| chr11 | 104157330 | 104158753 | 3.82263 | 4.73878 | 1.58952 | Intron (ENSMUST00000093925.4/ENSMUSG00000018634.10, intron 2 of 12) | 11 | 104132855 | 104175523 | 42669 | 1 | ENSMUST00000093925.4 | 24476 | Crhr1 | ENSMUSG00000018634.10 |
| chr11 | 104159329 | 104159848 | 3.97078 | 5.02233 | 1.76703 | Exon (ENSMUST00000093925.4/ENSMUSG00000018634.10, exon 3 of 13) | 11 | 104132855 | 104175523 | 42669 | 1 | ENSMUST00000093925.4 | 26475 | Crhr1 | ENSMUSG00000018634.10 |
| chr11 | 11681812 | 11681922 | 3.38078 | 3.89886 | 1.01733 | Intron (ENSMUST00000145789.1/ENSMUSG00000085652.1, intron 1 of 3) | 11 | 11685003 | 11707800 | 22798 | 1 | ENSMUST00000141436.2 | -3081 | Ikzf1 | ENSMUSG00000018654.12 |
| chr11 | 11683554 | 11683748 | 3.38193 | 3.9006 | 1.01733 | Intron (ENSMUST00000145789.1/ENSMUSG00000085652.1, intron 1 of 3) | 11 | 11685003 | 11707800 | 22798 | 1 | ENSMUST00000141436.2 | -1255 | Ikzf1 | ENSMUSG00000018654.12 |
| chr11 | 11684766 | 11687146 | 3.70701 | 4.53987 | 1.44768 | Promoter | 11 | 11685003 | 11707800 | 22798 | 1 | ENSMUST00000141436.2 | 0 | Ikzf1 | ENSMUSG00000018654.12 |
| chr11 | 84516048 | 84516309 | 3.5816 | 4.33305 | 1.37753 | Downstream (1-2kb) | 11 | 84466905 | 84513522 | 46618 | 2 | ENSMUST00000148434.1 | -2527 | Aatf | ENSMUSG00000018697.9 |
| chr11 | 84519244 | 84520362 | 3.53541 | 4.31096 | 1.35681 | 3' UTR | 11 | 84519706 | 84522067 | 2362 | 2 | ENSMUST00000176503.1 | 1705 | Lhx1 | ENSMUSG00000018698.10 |
| chr11 | 84520937 | 84521305 | 3.97939 | 5.11682 | 1.87857 | Promoter | 11 | 84519706 | 84522067 | 2362 | 2 | ENSMUST00000176503.1 | 762 | Lhx1 | ENSMUSG00000018698.10 |
| chr11 | 84521622 | 84522334 | 3.90513 | 5.04154 | 1.83091 | Promoter | 11 | 84519442 | 84522331 | 2890 | 2 | ENSMUST00000184646.2 | 0 | Lhx1 | ENSMUSG00000018698.10 |
| chr11 | 84522703 | 84524742 | 4.15985 | 5.50783 | 2.19625 | Promoter | 11 | 84521896 | 84522711 | 816 | 2 | ENSMUST00000092827.2 | 0 | Lhx1 | ENSMUSG00000018698.10 |
| chr11 | 84525150 | 84526478 | 3.52406 | 4.38502 | 1.38709 | Promoter | 11 | 84518284 | 84525535 | 7252 | 2 | ENSMUST00000018842.8 | 0 | Lhx1 | ENSMUSG00000018698.10 |
| chr19 | 42199351 | 42200408 | 3.6236 | 4.35097 | 1.30899 | Exon (ENSMUST00000018966.7/ENSMUSG00000018822.7, exon 2 of 3) | 19 | 42197971 | 42202252 | 4282 | 2 | ENSMUST00000018966.7 | 1844 | Sfrp5 | ENSMUSG00000018822.7 |
| chr19 | 42201154 | 42202928 | 3.75076 | 4.63508 | 1.54331 | Promoter | 19 | 42197971 | 42202252 | 4282 | 2 | ENSMUST00000018966.7 | 0 | Sfrp5 | ENSMUSG00000018822.7 |
| chr19 | 42203913 | 42204307 | 3.54286 | 4.2004 | 1.21755 | Distal Intergenic | 19 | 42197971 | 42202252 | 4282 | 2 | ENSMUST00000018966.7 | -1662 | Sfrp5 | ENSMUSG00000018822.7 |
| chr11 | 82925348 | 82925604 | 3.22487 | 3.6645 | 1.01593 | Exon (ENSMUST00000156395.1/ENSMUSG00000086058.1, exon 2 of 2) | 11 | 82928959 | 82929745 | 787 | 1 | ENSMUST00000125499.1 | -3355 | Unc45b | ENSMUSG00000018845.9 |
| chr11 | 82933518 | 82934422 | 3.91129 | 4.96076 | 1.8193 | Promoter | 11 | 82933602 | 82937757 | 4156 | 1 | ENSMUST00000142336.1 | 0 | Unc45b | ENSMUSG00000018845.9 |
| chr11 | 82935897 | 82936649 | 3.4465 | 4.26605 | 1.29596 | Exon (ENSMUST00000108160.2/ENSMUSG00000018845.9, exon 16 of 20) | 11 | 82933602 | 82937757 | 4156 | 1 | ENSMUST00000142336.1 | 2296 | Unc45b | ENSMUSG00000018845.9 |
| chr11 | 35921009 | 35921084 | 4.37415 | 5.81491 | 2.32161 | Intron (ENSMUST00000018993.6/ENSMUSG00000018849.6, intron 1 of 22) | 11 | 35868745 | 35875712 | 6968 | 2 | ENSMUST00000127086.1 | -45298 | Wwc1 | ENSMUSG00000018849.6 |
| chr11 | 35979786 | 35980749 | 4.37204 | 5.96088 | 2.51092 | Promoter | 11 | 35838400 | 35980527 | 142128 | 2 | ENSMUST00000018993.6 | 0 | Wwc1 | ENSMUSG00000018849.6 |
| chr11 | 35985955 | 35986231 | 3.59072 | 4.28192 | 1.24974 | Distal Intergenic | 11 | 35838400 | 35980527 | 142128 | 2 | ENSMUST00000018993.6 | -5429 | Wwc1 | ENSMUSG00000018849.6 |
| chr11 | 115331445 | 115331888 | 4.05151 | 5.20183 | 1.9363 | Exon (ENSMUST00000055490.3/ENSMUSG00000050201.4, exon 8 of 8) | 11 | 115334731 | 115346927 | 12197 | 1 | ENSMUST00000106543.2 | -2843 | Otop3 | ENSMUSG00000018862.6 |
| chr11 | 115332869 | 115333117 | 3.46559 | 4.0768 | 1.19531 | Distal Intergenic | 11 | 115334731 | 115346927 | 12197 | 1 | ENSMUST00000106543.2 | -1614 | Otop3 | ENSMUSG00000018862.6 |
| chr11 | 115334241 | 115335461 | 3.88234 | 4.92923 | 1.76361 | Promoter | 11 | 115334731 | 115346927 | 12197 | 1 | ENSMUST00000106543.2 | 0 | Otop3 | ENSMUSG00000018862.6 |
| chr11 | 115336115 | 115336900 | 3.67358 | 4.50711 | 1.44872 | Intron (ENSMUST00000106543.2/ENSMUSG00000018862.6, intron 1 of 6) | 11 | 115334734 | 115346926 | 12193 | 1 | ENSMUST00000019006.4 | 1382 | Otop3 | ENSMUSG00000018862.6 |
| chr15 | 84103138 | 84103650 | 3.78429 | 4.6904 | 1.61265 | Intron (ENSMUST00000082365.5/ENSMUSG00000018865.8, intron 1 of 6) | 15 | 84076097 | 84105754 | 29658 | 2 | ENSMUST00000082365.5 | 2104 | Sult4a1 | ENSMUSG00000018865.8 |
| chr15 | 84104948 | 84105852 | 3.72994 | 4.88219 | 1.71044 | Promoter | 15 | 84076097 | 84105754 | 29658 | 2 | ENSMUST00000082365.5 | 0 | Sult4a1 | ENSMUSG00000018865.8 |
| chr15 | 84106411 | 84106917 | 3.90642 | 5.03921 | 1.83725 | Promoter | 15 | 84076097 | 84105754 | 29658 | 2 | ENSMUST00000082365.5 | -658 | Sult4a1 | ENSMUSG00000018865.8 |
| chr15 | 84109293 | 84109535 | 3.489 | 4.15805 | 1.21759 | Distal Intergenic | 15 | 84076097 | 84105754 | 29658 | 2 | ENSMUST00000082365.5 | -3540 | Sult4a1 | ENSMUSG00000018865.8 |
| chr11 | 54140058 | 54140179 | 2.20692 | 14.08229 | 9.42704 | Distal Intergenic | 11 | 54127693 | 54131665 | 3973 | 1 | ENSMUST00000139315.1 | 12366 | P4ha2 | ENSMUSG00000018906.9 |
| chr11 | 54140552 | 54140678 | 2.02411 | 10.01662 | 5.76368 | Distal Intergenic | 11 | 54127693 | 54131665 | 3973 | 1 | ENSMUST00000139315.1 | 12860 | P4ha2 | ENSMUSG00000018906.9 |
| chr11 | 70321366 | 70322458 | 3.92625 | 4.98943 | 1.75598 | Promoter | 11 | 70319971 | 70321577 | 1607 | 2 | ENSMUST00000123752.2 | 0 | Alox12e | ENSMUSG00000018907.5 |
| chr11 | 70322769 | 70322875 | 4.06987 | 5.19432 | 1.8537 | Promoter | 11 | 70315610 | 70322628 | 7019 | 2 | ENSMUST00000019051.2 | -142 | Alox12e | ENSMUSG00000018907.5 |
| chr11 | 70324612 | 70324689 | 3.40218 | 3.97263 | 1.15353 | Distal Intergenic | 11 | 70315610 | 70322628 | 7019 | 2 | ENSMUST00000019051.2 | -1985 | Alox12e | ENSMUSG00000018907.5 |
| chr11 | 70330549 | 70331377 | 3.55802 | 4.22968 | 1.23108 | Distal Intergenic | 11 | 70315610 | 70322628 | 7019 | 2 | ENSMUST00000019051.2 | -7922 | Alox12e | ENSMUSG00000018907.5 |
| chr11 | 70350890 | 70351453 | 3.60268 | 4.34595 | 1.35182 | Promoter | 11 | 70344152 | 70352031 | 7880 | 2 | ENSMUST00000019068.6 | 578 | Alox15 | ENSMUSG00000018924.6 |
| chr11 | 70351894 | 70352258 | 3.63619 | 4.42313 | 1.38822 | Promoter | 11 | 70344152 | 70352031 | 7880 | 2 | ENSMUST00000019068.6 | 0 | Alox15 | ENSMUSG00000018924.6 |
| chr11 | 70352683 | 70353937 | 3.36947 | 3.89888 | 1.07717 | Promoter | 11 | 70344152 | 70352031 | 7880 | 2 | ENSMUST00000019068.6 | -653 | Alox15 | ENSMUSG00000018924.6 |
| chr11 | 96363285 | 96363441 | 3.3068 | 3.7986 | 1.05301 | Distal Intergenic | 11 | 96365752 | 96368256 | 2505 | 1 | ENSMUST00000019117.2 | -2311 | Hoxb1 | ENSMUSG00000018973.2 |
| chr11 | 96364070 | 96364189 | 3.29226 | 3.76554 | 1.01733 | Distal Intergenic | 11 | 96365752 | 96368256 | 2505 | 1 | ENSMUST00000019117.2 | -1563 | Hoxb1 | ENSMUSG00000018973.2 |
| chr11 | 96365096 | 96365202 | 4.17275 | 5.52402 | 2.14737 | Promoter | 11 | 96365752 | 96368256 | 2505 | 1 | ENSMUST00000019117.2 | -550 | Hoxb1 | ENSMUSG00000018973.2 |
| chr11 | 96365584 | 96366162 | 3.65928 | 4.80389 | 1.65705 | Promoter | 11 | 96365752 | 96368256 | 2505 | 1 | ENSMUST00000019117.2 | 0 | Hoxb1 | ENSMUSG00000018973.2 |
| chr11 | 96366878 | 96367150 | 4.46218 | 6.18768 | 2.62412 | Promoter | 11 | 96366258 | 96367221 | 964 | 1 | ENSMUST00000123805.1 | 621 | Hoxb1 | ENSMUSG00000018973.2 |
| chr11 | 96367505 | 96368108 | 4.21652 | 5.5648 | 2.1982 | 3' UTR | 11 | 96366258 | 96367221 | 964 | 1 | ENSMUST00000123805.1 | 1248 | Hoxb1 | ENSMUSG00000018973.2 |
| chr11 | 96368775 | 96369104 | 4.42145 | 6.05237 | 2.55855 | Distal Intergenic | 11 | 96366258 | 96367221 | 964 | 1 | ENSMUST00000123805.1 | 2518 | Hoxb1 | ENSMUSG00000018973.2 |
| chr11 | 96372982 | 96373129 | 3.36724 | 3.87844 | 1.01733 | Distal Intergenic | 11 | 96366258 | 96367221 | 964 | 1 | ENSMUST00000123805.1 | 6725 | Hoxb1 | ENSMUSG00000018973.2 |
| chr11 | 96373440 | 96373738 | 3.83447 | 4.75722 | 1.59028 | Distal Intergenic | 11 | 96366258 | 96367221 | 964 | 1 | ENSMUST00000123805.1 | 7183 | Hoxb1 | ENSMUSG00000018973.2 |
| chr11 | 96374113 | 96374386 | 3.58974 | 4.31097 | 1.35152 | Distal Intergenic | 11 | 96366258 | 96367221 | 964 | 1 | ENSMUST00000123805.1 | 7856 | Hoxb1 | ENSMUSG00000018973.2 |
| chr11 | 61212130 | 61212391 | 3.69718 | 4.7558 | 1.62653 | 5' UTR | 11 | 61208745 | 61218421 | 9677 | 1 | ENSMUST00000019246.3 | 3386 | Aldh3a1 | ENSMUSG00000019102.5 |
| chr7 | 31125354 | 31127208 | 3.64352 | 4.4787 | 1.44178 | Promoter | 7 | 31116524 | 31126844 | 10321 | 2 | ENSMUST00000188002.1 | 0 | Scn1b | ENSMUSG00000019194.9 |
| chr1 | 138826272 | 138826826 | 3.80861 | 4.91076 | 1.71203 | 3' UTR | 1 | 138830944 | 138842429 | 11486 | 2 | ENSMUST00000112026.3 | 15603 | Lhx9 | ENSMUSG00000019230.9 |
| chr1 | 138828095 | 138828187 | 4.26396 | 5.77607 | 2.32046 | 3' UTR | 1 | 138830944 | 138842429 | 11486 | 2 | ENSMUST00000112026.3 | 14242 | Lhx9 | ENSMUSG00000019230.9 |
| chr1 | 138829782 | 138830309 | 4.01281 | 5.26456 | 1.95344 | 3' UTR | 1 | 138830944 | 138842429 | 11486 | 2 | ENSMUST00000112026.3 | 12120 | Lhx9 | ENSMUSG00000019230.9 |
| chr1 | 138831423 | 138832618 | 3.7196 | 4.71136 | 1.598 | Exon (ENSMUST00000046870.7/ENSMUSG00000019230.9, exon 7 of 7) | 1 | 138830944 | 138842429 | 11486 | 2 | ENSMUST00000112026.3 | 9811 | Lhx9 | ENSMUSG00000019230.9 |
| chr1 | 138833015 | 138833249 | 3.40982 | 4.08827 | 1.17968 | Intron (ENSMUST00000019374.8/ENSMUSG00000019230.9, intron 3 of 4) | 1 | 138830944 | 138842429 | 11486 | 2 | ENSMUST00000112026.3 | 9180 | Lhx9 | ENSMUSG00000019230.9 |
| chr1 | 138833720 | 138834081 | 3.83895 | 5.07191 | 1.87091 | Intron (ENSMUST00000019374.8/ENSMUSG00000019230.9, intron 3 of 4) | 1 | 138830944 | 138842429 | 11486 | 2 | ENSMUST00000112026.3 | 8348 | Lhx9 | ENSMUSG00000019230.9 |
| chr1 | 138836268 | 138836357 | 4.34496 | 5.87232 | 2.40536 | Intron (ENSMUST00000019374.8/ENSMUSG00000019230.9, intron 3 of 4) | 1 | 138830944 | 138842429 | 11486 | 2 | ENSMUST00000112026.3 | 6072 | Lhx9 | ENSMUSG00000019230.9 |
| chr1 | 138836874 | 138837355 | 4.08373 | 5.53426 | 2.17033 | Intron (ENSMUST00000019374.8/ENSMUSG00000019230.9, intron 3 of 4) | 1 | 138830944 | 138842429 | 11486 | 2 | ENSMUST00000112026.3 | 5074 | Lhx9 | ENSMUSG00000019230.9 |
| chr1 | 138837705 | 138838915 | 4.05939 | 5.45399 | 2.13739 | Exon (ENSMUST00000019374.8/ENSMUSG00000019230.9, exon 3 of 5) | 1 | 138830944 | 138842429 | 11486 | 2 | ENSMUST00000112026.3 | 3514 | Lhx9 | ENSMUSG00000019230.9 |
| chr1 | 138839227 | 138840310 | 3.8391 | 4.88165 | 1.76073 | Exon (ENSMUST00000019374.8/ENSMUSG00000019230.9, exon 2 of 5) | 1 | 138830944 | 138842429 | 11486 | 2 | ENSMUST00000112026.3 | 2119 | Lhx9 | ENSMUSG00000019230.9 |
| chr1 | 138840881 | 138841874 | 3.68294 | 4.64198 | 1.53889 | Promoter | 1 | 138830944 | 138842429 | 11486 | 2 | ENSMUST00000112026.3 | 555 | Lhx9 | ENSMUSG00000019230.9 |
| chr1 | 138842546 | 138842870 | 4.2978 | 5.75556 | 2.36607 | Promoter | 1 | 138825186 | 138842457 | 17272 | 2 | ENSMUST00000019374.8 | -90 | Lhx9 | ENSMUSG00000019230.9 |
| chr1 | 138843370 | 138844730 | 3.68506 | 4.61781 | 1.57478 | Promoter | 1 | 138825186 | 138842457 | 17272 | 2 | ENSMUST00000019374.8 | -914 | Lhx9 | ENSMUSG00000019230.9 |
| chr1 | 138846073 | 138846422 | 3.76248 | 4.88365 | 1.69582 | Intron (ENSMUST00000093486.4/ENSMUSG00000019230.9, intron 2 of 6) | 1 | 138825186 | 138847579 | 22394 | 2 | ENSMUST00000093486.4 | 1157 | Lhx9 | ENSMUSG00000019230.9 |
| chr1 | 138846945 | 138847587 | 4.08558 | 5.25955 | 1.97789 | Promoter | 1 | 138825186 | 138847579 | 22394 | 2 | ENSMUST00000093486.4 | 0 | Lhx9 | ENSMUSG00000019230.9 |
| chr1 | 138848071 | 138849129 | 4.06591 | 5.49232 | 2.1483 | Promoter | 1 | 138827997 | 138848576 | 20580 | 2 | ENSMUST00000112030.3 | 0 | Lhx9 | ENSMUSG00000019230.9 |
| chr1 | 138849642 | 138849783 | 3.22944 | 3.67134 | 1.01733 | Downstream (2-3kb) | 1 | 138827997 | 138848576 | 20580 | 2 | ENSMUST00000112030.3 | -1067 | Lhx9 | ENSMUSG00000019230.9 |
| chr12 | 85150727 | 85151523 | 3.7197 | 4.66916 | 1.56703 | Promoter | 12 | 85135596 | 85151264 | 15669 | 2 | ENSMUST00000019379.7 | 0 | Rps6kl1 | ENSMUSG00000019235.8 |
| chr12 | 85151882 | 85152105 | 3.43008 | 4.04967 | 1.19073 | Promoter | 12 | 85135596 | 85151264 | 15669 | 2 | ENSMUST00000019379.7 | -619 | Rps6kl1 | ENSMUSG00000019235.8 |
| chr11 | 98445472 | 98445752 | 3.92683 | 4.9988 | 1.81882 | Promoter | 11 | 98446394 | 98451417 | 5024 | 1 | ENSMUST00000132771.2 | -642 | Grb7 | ENSMUSG00000019312.5 |
| chr11 | 98447106 | 98447397 | 3.40461 | 3.95015 | 1.0714 | Promoter | 11 | 98447068 | 98455373 | 8306 | 1 | ENSMUST00000156328.2 | 39 | Grb7 | ENSMUSG00000019312.5 |
| chr11 | 98450498 | 98450635 | 3.68577 | 4.47282 | 1.39095 | Intron (ENSMUST00000132771.2/ENSMUSG00000019312.5, intron 1 of 3) | 11 | 98451842 | 98453822 | 1981 | 1 | ENSMUST00000133419.1 | -1207 | Grb7 | ENSMUSG00000019312.5 |
| chr11 | 98454938 | 98455210 | 3.32465 | 4.04379 | 1.15687 | 3' UTR | 11 | 98453789 | 98455125 | 1337 | 1 | ENSMUST00000129034.1 | 1150 | Grb7 | ENSMUSG00000019312.5 |
| chr11 | 98458175 | 98458344 | 4.57677 | 6.24951 | 2.65935 | Distal Intergenic | 11 | 98453789 | 98455125 | 1337 | 1 | ENSMUST00000129034.1 | 4387 | Grb7 | ENSMUSG00000019312.5 |
| chr11 | 98459160 | 98459540 | 3.39079 | 4.01192 | 1.1238 | Distal Intergenic | 11 | 98453789 | 98455125 | 1337 | 1 | ENSMUST00000129034.1 | 5372 | Grb7 | ENSMUSG00000019312.5 |
| chr11 | 98461049 | 98461124 | 3.38193 | 3.9006 | 1.01733 | Distal Intergenic | 11 | 98453789 | 98455125 | 1337 | 1 | ENSMUST00000129034.1 | 7261 | Grb7 | ENSMUSG00000019312.5 |
| chr10 | 127187843 | 127188447 | 3.95357 | 5.01712 | 1.78448 | Intron (ENSMUST00000167353.1/ENSMUSG00000019467.8, intron 2 of 15) | 10 | 127182521 | 127189823 | 7303 | 2 | ENSMUST00000167353.1 | 1376 | Arhgef25 | ENSMUSG00000019467.8 |
| chr10 | 127188799 | 127189325 | 4.27853 | 5.68716 | 2.28841 | Promoter | 10 | 127182521 | 127189823 | 7303 | 2 | ENSMUST00000167353.1 | 498 | Arhgef25 | ENSMUSG00000019467.8 |
| chr8 | 123783759 | 123783917 | 3.34557 | 3.84575 | 1.01733 | Intron (ENSMUST00000127664.1/ENSMUSG00000092329.1, intron 1 of 15) | 8 | 123805985 | 123835287 | 29303 | 1 | ENSMUST00000118535.2 | -22068 | Rab4a | ENSMUSG00000019478.10 |
| chr17 | 57149642 | 57150267 | 3.51886 | 4.32551 | 1.36483 | Promoter | 17 | 57145997 | 57149777 | 3781 | 2 | ENSMUST00000019633.7 | 0 | Cd70 | ENSMUSG00000019489.7 |
| chr17 | 56199994 | 56200101 | 21.76834 | 62.37172 | 56.3539 | Intron (ENSMUST00000038794.4/ENSMUSG00000001229.7, intron 10 of 20) | 17 | 56175744 | 56183920 | 8177 | 2 | ENSMUST00000019723.7 | -16075 | D17Wsu104e | ENSMUSG00000019579.8 |
| chr18 | 47367219 | 47367439 | 3.32301 | 3.81516 | 1.02919 | Promoter | 18 | 47303674 | 47368268 | 64595 | 2 | ENSMUST00000144223.1 | 829 | Sema6a | ENSMUSG00000019647.11 |
| chr18 | 47537399 | 47537877 | 3.72448 | 4.96082 | 1.74036 | Distal Intergenic | 18 | 47275658 | 47368870 | 93213 | 2 | ENSMUST00000151382.2 | -168530 | Sema6a | ENSMUSG00000019647.11 |
| chr1 | 177322705 | 177323056 | 3.4075 | 3.95631 | 1.07769 | Distal Intergenic | 1 | 177022116 | 177258203 | 236088 | 2 | ENSMUST00000111159.1 | -64503 | Akt3 | ENSMUSG00000019699.10 |
| chr1 | 177331411 | 177331707 | 3.71926 | 4.59158 | 1.56344 | Distal Intergenic | 1 | 177022116 | 177258203 | 236088 | 2 | ENSMUST00000111159.1 | -73209 | Akt3 | ENSMUSG00000019699.10 |
| chr1 | 177340206 | 177340301 | 4.12482 | 5.41227 | 2.10111 | Distal Intergenic | 1 | 177022116 | 177258203 | 236088 | 2 | ENSMUST00000111159.1 | -82004 | Akt3 | ENSMUSG00000019699.10 |
| chr11 | 99398789 | 99398937 | 3.80726 | 4.76143 | 1.63212 | Distal Intergenic | 11 | 99385254 | 99389364 | 4111 | 2 | ENSMUST00000103131.4 | -9426 | Krt10 | ENSMUSG00000019761.9 |
| chr11 | 99400312 | 99400419 | 3.67436 | 4.46265 | 1.41268 | Distal Intergenic | 11 | 99385254 | 99389364 | 4111 | 2 | ENSMUST00000103131.4 | -10949 | Krt10 | ENSMUSG00000019761.9 |
| chr10 | 4710098 | 4710219 | 3.58726 | 4.28867 | 1.29418 | Promoter | 10 | 4710155 | 4967336 | 257182 | 1 | ENSMUST00000105588.2 | 0 | Esr1 | ENSMUSG00000019768.11 |
| chr10 | 4710711 | 4711272 | 3.84353 | 5.3089 | 2.01691 | Promoter | 10 | 4710155 | 5002024 | 291870 | 1 | ENSMUST00000105589.1 | 557 | Esr1 | ENSMUSG00000019768.11 |
| chr10 | 4711835 | 4712077 | 3.15025 | 3.86542 | 1.06629 | Promoter | 10 | 4712651 | 4784824 | 72174 | 1 | ENSMUST00000127145.1 | -574 | Esr1 | ENSMUSG00000019768.11 |
| chr10 | 4712620 | 4713572 | 4.20548 | 5.85012 | 2.53454 | Promoter | 10 | 4712651 | 4784824 | 72174 | 1 | ENSMUST00000127145.1 | 0 | Esr1 | ENSMUSG00000019768.11 |
| chr10 | 5921065 | 5921202 | 3.55238 | 4.74806 | 1.61225 | Intron (ENSMUST00000131996.2/ENSMUSG00000019775.12, intron 1 of 4) | 10 | 5832169 | 5922366 | 90198 | 2 | ENSMUST00000117676.2 | 1164 | Rgs17 | ENSMUSG00000019775.12 |

| | | | | | | | | | | | | | | | |
|---|---|---|---|---|---|---|---|---|---|---|---|---|---|---|---|
| chr10 | 5921791 | 5922424 | 3.78253 | 4.81952 | 1.67838 | Promoter | 10 | 5825663 | 5922386 | 96724 | 2 | ENSMUST00000131996.2 | 0 | Rgs17 | ENSMUSG00000019775.12 |
| chr10 | 33624292 | 33624756 | 3.69516 | 4.6075 | 1.53374 | Promoter | 10 | 33589912 | 33624619 | 34708 | 2 | ENSMUST00000161692.1 | 0 | Clvs2 | ENSMUSG00000019785.8 |
| chr10 | 30844511 | 30844779 | 3.57043 | 4.31612 | 1.37446 | Distal Intergenic | 10 | 30832359 | 30842801 | 10443 | 2 | ENSMUST00000019924.8 | -1711 | Hey2 | ENSMUSG00000019789.8 |
| chr10 | 7592102 | 7592637 | 4.3283 | 5.81325 | 2.37868 | Intron (ENSMUST00000134346.2/ENSMUSG00000019796.7, intron 1 of 5) | 10 | 7590127 | 7624022 | 33896 | 1 | ENSMUST00000135907.2 | 1976 | Lrp11 | ENSMUSG00000019796.7 |
| chr10 | 42562352 | 42562597 | 3.83 | 4.7607 | 1.60832 | 3' UTR | 10 | 42563272 | 42568828 | 5557 | 2 | ENSMUST00000105498.2 | 6231 | Nr2e1 | ENSMUSG00000019803.6 |
| chr10 | 42563025 | 42563673 | 3.7143 | 4.52881 | 1.43479 | 3' UTR | 10 | 42563272 | 42568828 | 5557 | 2 | ENSMUST00000105498.2 | 5155 | Nr2e1 | ENSMUSG00000019803.6 |
| chr10 | 42564362 | 42564580 | 3.5488 | 4.47 | 1.46472 | Intron (ENSMUST00000019938.5/ENSMUSG00000019803.6, intron 8 of 8) | 10 | 42563272 | 42568828 | 5557 | 2 | ENSMUST00000105498.2 | 4248 | Nr2e1 | ENSMUSG00000019803.6 |
| chr10 | 42565000 | 42565287 | 3.87233 | 4.97199 | 1.77372 | Intron (ENSMUST00000019938.5/ENSMUSG00000019803.6, intron 8 of 8) | 10 | 42563272 | 42568828 | 5557 | 2 | ENSMUST00000105498.2 | 3541 | Nr2e1 | ENSMUSG00000019803.6 |
| chr10 | 42565988 | 42566563 | 3.72357 | 4.53965 | 1.43254 | Intron (ENSMUST00000019938.5/ENSMUSG00000019803.6, intron 8 of 8) | 10 | 42563272 | 42568828 | 5557 | 2 | ENSMUST00000105498.2 | 2265 | Nr2e1 | ENSMUSG00000019803.6 |
| chr10 | 42566911 | 42567177 | 3.49982 | 4.11712 | 1.15333 | Intron (ENSMUST00000019938.5/ENSMUSG00000019803.6, intron 8 of 8) | 10 | 42563272 | 42568828 | 5557 | 2 | ENSMUST00000105498.2 | 1651 | Nr2e1 | ENSMUSG00000019803.6 |
| chr10 | 42567937 | 42568449 | 3.66823 | 4.49086 | 1.41605 | Promoter | 10 | 42563272 | 42568828 | 5557 | 2 | ENSMUST00000105498.2 | 379 | Nr2e1 | ENSMUSG00000019803.6 |
| chr10 | 42568756 | 42570149 | 3.72379 | 4.62386 | 1.52856 | Promoter | 10 | 42563272 | 42568828 | 5557 | 2 | ENSMUST00000105498.2 | 0 | Nr2e1 | ENSMUSG00000019803.6 |
| chr10 | 42570758 | 42571091 | 3.74674 | 4.58247 | 1.46652 | Intron (ENSMUST00000019938.5/ENSMUSG00000019803.6, intron 5 of 8) | 10 | 42563272 | 42568828 | 5557 | 2 | ENSMUST00000105498.2 | -1931 | Nr2e1 | ENSMUSG00000019803.6 |
| chr10 | 42571426 | 42571956 | 3.45532 | 4.11221 | 1.1787 | Exon (ENSMUST00000019938.5/ENSMUSG00000019803.6, exon 5 of 9) | 10 | 42563272 | 42568828 | 5557 | 2 | ENSMUST00000105498.2 | -2599 | Nr2e1 | ENSMUSG00000019803.6 |
| chr10 | 42572779 | 42573308 | 4.48683 | 6.11351 | 2.61899 | Exon (ENSMUST00000019938.5/ENSMUSG00000019803.6, exon 4 of 9) | 10 | 42563272 | 42568828 | 5557 | 2 | ENSMUST00000105498.2 | -3952 | Nr2e1 | ENSMUSG00000019803.6 |
| chr10 | 42573638 | 42574826 | 3.89593 | 4.92407 | 1.73643 | 5' UTR | 10 | 42563272 | 42568828 | 5557 | 2 | ENSMUST00000105498.2 | -4811 | Nr2e1 | ENSMUSG00000019803.6 |
| chr10 | 42575255 | 42575526 | 3.60234 | 4.50932 | 1.45445 | Intron (ENSMUST00000019938.5/ENSMUSG00000019803.6, intron 2 of 8) | 10 | 42571484 | 42580345 | 8862 | 2 | ENSMUST00000126848.1 | 4819 | Nr2e1 | ENSMUSG00000019803.6 |
| chr10 | 42576234 | 42576444 | 3.72276 | 4.54484 | 1.4516 | Intron (ENSMUST00000019938.5/ENSMUSG00000019803.6, intron 2 of 8) | 10 | 42571484 | 42580345 | 8862 | 2 | ENSMUST00000126848.1 | 3901 | Nr2e1 | ENSMUSG00000019803.6 |
| chr10 | 42576759 | 42576857 | 4.40059 | 5.86864 | 2.34742 | Intron (ENSMUST00000019938.5/ENSMUSG00000019803.6, intron 2 of 8) | 10 | 42571484 | 42580345 | 8862 | 2 | ENSMUST00000126848.1 | 3488 | Nr2e1 | ENSMUSG00000019803.6 |
| chr10 | 42577677 | 42581209 | 3.86282 | 4.92984 | 1.73795 | Promoter | 10 | 42571484 | 42580345 | 8862 | 2 | ENSMUST00000126848.1 | 0 | Nr2e1 | ENSMUSG00000019803.6 |
| chr10 | 42581688 | 42582025 | 3.4253 | 4.04454 | 1.1905 | Promoter | 10 | 42572858 | 42580774 | 7917 | 2 | ENSMUST00000143891.1 | -915 | Nr2e1 | ENSMUSG00000019803.6 |
| chr10 | 42582332 | 42586216 | 4.15951 | 5.52824 | 2.16326 | Promoter | 10 | 42561963 | 42583632 | 21670 | 2 | ENSMUST00000019938.5 | 0 | Nr2e1 | ENSMUSG00000019803.6 |
| chr10 | 13060834 | 13061522 | 3.82429 | 4.82509 | 1.68298 | Distal Intergenic | 10 | 13090691 | 13124860 | 34170 | 1 | ENSMUST00000123135.2 | -29169 | Plagl1 | ENSMUSG00000019817.13 |
| chr10 | 11079652 | 11080090 | 4.03659 | 5.13476 | 1.81888 | Promoter | 10 | 10688801 | 11080956 | 392156 | 2 | ENSMUST00000105560.1 | 866 | Grm1 | ENSMUSG00000019828.8 |
| chr10 | 11081153 | 11081228 | 3.39946 | 3.93483 | 1.04471 | Promoter | 10 | 10688801 | 11080956 | 392156 | 2 | ENSMUST00000105560.1 | -198 | Grm1 | ENSMUSG00000019828.8 |
| chr10 | 11082221 | 11082296 | 3.96678 | 4.98512 | 1.70024 | Promoter | 10 | 11080489 | 11082250 | 1762 | 2 | ENSMUST00000156826.1 | 0 | Grm1 | ENSMUSG00000019828.8 |
| chr10 | 45289144 | 45289241 | 3.82237 | 4.92407 | 1.71503 | Promoter | 10 | 45289305 | 45318452 | 29148 | 1 | ENSMUST00000019994.8 | -64 | Popdc3 | ENSMUSG00000019848.9 |
| chr10 | 18846575 | 18846800 | 3.8398 | 4.86026 | 1.72063 | Intron (ENSMUST00000019998.7/ENSMUSG00000019851.7, intron 1 of 2) | 10 | 18845071 | 18857073 | 12003 | 1 | ENSMUST00000019998.7 | 1505 | Perp | ENSMUSG00000019851.7 |
| chr10 | 18544817 | 18545235 | 3.84019 | 4.83523 | 1.70119 | Promoter | 10 | 18540123 | 18546076 | 5954 | 2 | ENSMUST00000020000.5 | 841 | Hebp2 | ENSMUSG00000019853.5 |
| chr10 | 14760130 | 14760379 | 3.57154 | 4.30986 | 1.31318 | Promoter | 10 | 14760157 | 14770850 | 10694 | 1 | ENSMUST00000020015.4 | 0 | Nmbr | ENSMUSG00000019865.4 |
| chr10 | 44004272 | 44004936 | 4.21942 | 5.87997 | 2.41584 | Promoter | 10 | 43950307 | 44004846 | 54540 | 2 | ENSMUST00000020017.8 | 0 | Aim1 | ENSMUSG00000019866.9 |
| chr10 | 44005308 | 44006213 | 4.31783 | 5.95522 | 2.55032 | Promoter | 10 | 43950307 | 44004846 | 54540 | 2 | ENSMUST00000020017.8 | -463 | Aim1 | ENSMUSG00000019866.9 |
| chr10 | 44008059 | 44008312 | 3.57942 | 4.34547 | 1.37172 | Distal Intergenic | 10 | 43950307 | 44004846 | 54540 | 2 | ENSMUST00000020017.8 | -3214 | Aim1 | ENSMUSG00000019866.9 |
| chr10 | 44015521 | 44015596 | 4.08044 | 5.31074 | 1.99885 | Distal Intergenic | 10 | 43950307 | 44004846 | 54540 | 2 | ENSMUST00000020017.8 | -10676 | Aim1 | ENSMUSG00000019866.9 |
| chr10 | 57630891 | 57631384 | 3.55168 | 4.21243 | 1.20967 | Promoter | 10 | 57631981 | 57738928 | 106948 | 1 | ENSMUST00000095668.4 | -597 | Pkib | ENSMUSG00000019876.10 |
| chr10 | 57632386 | 57632970 | 3.76028 | 4.71728 | 1.56579 | Promoter | 10 | 57632124 | 57728150 | 96027 | 1 | ENSMUST00000177438.2 | 263 | Pkib | ENSMUSG00000019876.10 |
| chr10 | 29533484 | 29534571 | 3.75056 | 4.63056 | 1.53357 | Intron (ENSMUST00000092623.3/ENSMUSG00000019880.9, intron 1 of 4) | 10 | 29453109 | 29535867 | 82759 | 2 | ENSMUST00000092623.3 | 1296 | Rspo3 | ENSMUSG00000019880.9 |
| chr10 | 29535422 | 29536108 | 4.0036 | 5.16175 | 1.90858 | Promoter | 10 | 29453109 | 29535867 | 82759 | 2 | ENSMUST00000092623.3 | 0 | Rspo3 | ENSMUSG00000019880.9 |
| chr10 | 29283209 | 29283543 | 3.46288 | 4.06887 | 1.18095 | Distal Intergenic | 10 | 29313166 | 29345560 | 32395 | 1 | ENSMUST00000160399.2 | -29623 | Echdc1 | ENSMUSG00000019883.5 |
| chr10 | 29284351 | 29284476 | 3.41537 | 4.00289 | 1.19161 | Distal Intergenic | 10 | 29313166 | 29345560 | 32395 | 1 | ENSMUST00000160399.2 | -28690 | Echdc1 | ENSMUSG00000019883.5 |
| chr10 | 29284929 | 29285163 | 3.58964 | 4.33342 | 1.3863 | Distal Intergenic | 10 | 29313166 | 29345560 | 32395 | 1 | ENSMUST00000160399.2 | -28003 | Echdc1 | ENSMUSG00000019883.5 |
| chr10 | 103367438 | 103367532 | 3.66423 | 4.43243 | 1.37541 | Promoter | 10 | 103367808 | 103419378 | 51571 | 1 | ENSMUST00000074204.5 | -276 | Slc6a15 | ENSMUSG00000019894.8 |
| chr10 | 51677602 | 51678078 | 3.75349 | 4.64013 | 1.53905 | Promoter | 10 | 51677756 | 51721908 | 44153 | 1 | ENSMUST00000125729.2 | 0 | Rfx6 | ENSMUSG00000019900.12 |
| chr10 | 107272126 | 107272292 | 3.7593 | 4.76247 | 1.68038 | Promoter | 10 | 107271843 | 107425143 | 153301 | 1 | ENSMUST00000105280.3 | 284 | Lin7a | ENSMUSG00000019906.9 |
| chr10 | 50885697 | 50886338 | 3.4479 | 4.1236 | 1.24179 | Distal Intergenic | 10 | 50895651 | 50989152 | 93502 | 1 | ENSMUST00000020071.3 | -9313 | Sim1 | ENSMUSG00000019913.3 |
| chr10 | 50889018 | 50890273 | 3.7555 | 4.71916 | 1.62211 | Distal Intergenic | 10 | 50895651 | 50989152 | 93502 | 1 | ENSMUST00000020071.3 | -5378 | Sim1 | ENSMUSG00000019913.3 |
| chr10 | 50890614 | 50890941 | 3.96615 | 5.38911 | 2.04978 | Distal Intergenic | 10 | 50895651 | 50989152 | 93502 | 1 | ENSMUST00000020071.3 | -4710 | Sim1 | ENSMUSG00000019913.3 |
| chr10 | 50891327 | 50891698 | 3.89829 | 4.90929 | 1.75733 | Distal Intergenic | 10 | 50895651 | 50989152 | 93502 | 1 | ENSMUST00000020071.3 | -3953 | Sim1 | ENSMUSG00000019913.3 |
| chr10 | 50892230 | 50901639 | 3.99263 | 5.31959 | 2.05447 | Promoter | 10 | 50895651 | 50989152 | 93502 | 1 | ENSMUST00000020071.3 | 0 | Sim1 | ENSMUSG00000019913.3 |
| chr10 | 50901981 | 50902093 | 3.77179 | 4.73535 | 1.6099 | Intron (ENSMUST00000020071.3/ENSMUSG00000019913.3, intron 1 of 10) | 10 | 50895651 | 50989152 | 93502 | 1 | ENSMUST00000020071.3 | 6331 | Sim1 | ENSMUSG00000019913.3 |
| chr10 | 50903089 | 50903484 | 3.67287 | 4.8323 | 1.65646 | Intron (ENSMUST00000020071.3/ENSMUSG00000019913.3, intron 1 of 10) | 10 | 50895651 | 50989152 | 93502 | 1 | ENSMUST00000020071.3 | 7439 | Sim1 | ENSMUSG00000019913.3 |
| chr10 | 50904220 | 50905162 | 3.8025 | 4.94047 | 1.7554 | Exon (ENSMUST00000020071.3/ENSMUSG00000019913.3, exon 2 of 11) | 10 | 50895651 | 50989152 | 93502 | 1 | ENSMUST00000020071.3 | 8570 | Sim1 | ENSMUSG00000019913.3 |
| chr10 | 50905609 | 50905904 | 3.55957 | 4.81132 | 1.64399 | Intron (ENSMUST00000020071.3/ENSMUSG00000019913.3, intron 2 of 10) | 10 | 50895651 | 50989152 | 93502 | 1 | ENSMUST00000020071.3 | 9959 | Sim1 | ENSMUSG00000019913.3 |
| chr10 | 50906399 | 50907157 | 3.55974 | 4.37118 | 1.38133 | Intron (ENSMUST00000020071.3/ENSMUSG00000019913.3, intron 2 of 10) | 10 | 50895651 | 50989152 | 93502 | 1 | ENSMUST00000020071.3 | 10749 | Sim1 | ENSMUSG00000019913.3 |
| chr10 | 50907786 | 50908010 | 3.43693 | 4.12933 | 1.21596 | Exon (ENSMUST00000020071.3/ENSMUSG00000019913.3, exon 3 of 11) | 10 | 50895651 | 50989152 | 93502 | 1 | ENSMUST00000020071.3 | 12136 | Sim1 | ENSMUSG00000019913.3 |
| chr10 | 50908600 | 50908923 | 4.0706 | 5.38194 | 2.03783 | Exon (ENSMUST00000020071.3/ENSMUSG00000019913.3, exon 5 of 11) | 10 | 50895651 | 50989152 | 93502 | 1 | ENSMUST00000020071.3 | 12950 | Sim1 | ENSMUSG00000019913.3 |
| chr10 | 50909870 | 50911191 | 3.89399 | 5.1524 | 1.88576 | Exon (ENSMUST00000020071.3/ENSMUSG00000019913.3, exon 7 of 11) | 10 | 50895651 | 50989152 | 93502 | 1 | ENSMUST00000020071.3 | 14220 | Sim1 | ENSMUSG00000019913.3 |
| chr10 | 50912146 | 50912622 | 4.26739 | 5.77333 | 2.38168 | Intron (ENSMUST00000020071.3/ENSMUSG00000019913.3, intron 8 of 10) | 10 | 50895651 | 50989152 | 93502 | 1 | ENSMUST00000020071.3 | 16496 | Sim1 | ENSMUSG00000019913.3 |
| chr10 | 50913411 | 50913666 | 3.31263 | 4.16218 | 1.20666 | Intron (ENSMUST00000020071.3/ENSMUSG00000019913.3, intron 8 of 10) | 10 | 50895651 | 50989152 | 93502 | 1 | ENSMUST00000020071.3 | 17761 | Sim1 | ENSMUSG00000019913.3 |
| chr10 | 50914079 | 50914575 | 3.5832 | 4.32102 | 1.39231 | Intron (ENSMUST00000020071.3/ENSMUSG00000019913.3, intron 8 of 10) | 10 | 50895651 | 50989152 | 93502 | 1 | ENSMUST00000020071.3 | 18429 | Sim1 | ENSMUSG00000019913.3 |
| chr10 | 50915329 | 50915462 | 3.28547 | 3.79252 | 1.07872 | Intron (ENSMUST00000020071.3/ENSMUSG00000019913.3, intron 8 of 10) | 10 | 50895651 | 50989152 | 93502 | 1 | ENSMUST00000020071.3 | 19679 | Sim1 | ENSMUSG00000019913.3 |
| chr10 | 50915785 | 50916263 | 3.69816 | 4.77608 | 1.63841 | Intron (ENSMUST00000020071.3/ENSMUSG00000019913.3, intron 8 of 10) | 10 | 50895651 | 50989152 | 93502 | 1 | ENSMUST00000020071.3 | 20135 | Sim1 | ENSMUSG00000019913.3 |
| chr10 | 50917872 | 50918342 | 3.38127 | 3.91745 | 1.08056 | Intron (ENSMUST00000020071.3/ENSMUSG00000019913.3, intron 8 of 10) | 10 | 50895651 | 50989152 | 93502 | 1 | ENSMUST00000020071.3 | 22222 | Sim1 | ENSMUSG00000019913.3 |
| chr10 | 50924374 | 50924675 | 3.56645 | 4.36331 | 1.37834 | Intron (ENSMUST00000020071.3/ENSMUSG00000019913.3, intron 8 of 10) | 10 | 50895651 | 50989152 | 93502 | 1 | ENSMUST00000020071.3 | 28724 | Sim1 | ENSMUSG00000019913.3 |
| chr10 | 69398952 | 69399742 | 4.28139 | 5.84687 | 2.35911 | Distal Intergenic | 10 | 69336635 | 69352938 | 16304 | 2 | ENSMUST00000020099.6 | -46015 | Cdk1 | ENSMUSG00000019942.7 |
| chr10 | 69400433 | 69400789 | 3.61295 | 4.47012 | 1.43346 | Distal Intergenic | 10 | 69336635 | 69352938 | 16304 | 2 | ENSMUST00000020099.6 | -47496 | Cdk1 | ENSMUSG00000019942.7 |
| chr10 | 69402050 | 69402144 | 4.10944 | 5.34698 | 2.03264 | Distal Intergenic | 10 | 69336635 | 69352938 | 16304 | 2 | ENSMUST00000020099.6 | -49113 | Cdk1 | ENSMUSG00000019942.7 |
| chr10 | 100017882 | 100018180 | 3.39502 | 4.06589 | 1.16133 | Intron (ENSMUST00000105283.2/ENSMUSG00000019966.12, intron 1 of 9) | 10 | 100015916 | 100096398 | 80483 | 1 | ENSMUST00000020129.7 | 1967 | Kitl | ENSMUSG00000019966.12 |
| chr10 | 21882346 | 21882600 | 3.64787 | 4.59086 | 1.50528 | Promoter | 10 | 21882369 | 21899145 | 16777 | 1 | ENSMUST00000143094.1 | 0 | Sgk1 | ENSMUSG00000019970.10 |
| chr10 | 21883481 | 21883907 | 3.74865 | 4.60925 | 1.52487 | Intron (ENSMUST00000120509.2/ENSMUSG00000019970.10, intron 1 of 13) | 10 | 21882369 | 21899145 | 16777 | 1 | ENSMUST00000143094.1 | 1113 | Sgk1 | ENSMUSG00000019970.10 |
| chr10 | 21884437 | 21884670 | 3.31478 | 3.7995 | 1.01733 | Intron (ENSMUST00000120509.2/ENSMUSG00000019970.10, intron 1 of 13) | 10 | 21882369 | 21899145 | 16777 | 1 | ENSMUST00000143094.1 | 2069 | Sgk1 | ENSMUSG00000019970.10 |
| chr10 | 21155874 | 21156108 | 3.68974 | 4.62425 | 1.51631 | Intron (ENSMUST00000188495.1/ENSMUSG00000019982.9, intron 2 of 15) | 10 | 21138893 | 21160850 | 21958 | 2 | ENSMUST00000186446.1 | 4742 | Myb | ENSMUSG00000019982.9 |
| chr10 | 21157787 | 21158494 | 3.7656 | 4.67063 | 1.5544 | Intron (ENSMUST00000188495.1/ENSMUSG00000019982.9, intron 1 of 15) | 10 | 21138893 | 21160850 | 21958 | 2 | ENSMUST00000186446.1 | 2356 | Myb | ENSMUSG00000019982.9 |
| chr10 | 21158981 | 21160066 | 4.13882 | 5.40741 | 2.07806 | Promoter | 10 | 21138893 | 21160850 | 21958 | 2 | ENSMUST00000186446.1 | 784 | Myb | ENSMUSG00000019982.9 |
| chr10 | 21161194 | 21162161 | 3.75241 | 4.66182 | 1.6015 | Promoter | 10 | 21125276 | 21160905 | 35630 | 2 | ENSMUST00000020158.7 | -290 | Myb | ENSMUSG00000019982.9 |
| chr10 | 21162812 | 21163384 | 3.4779 | 4.11383 | 1.23385 | Distal Intergenic | 10 | 21125276 | 21160905 | 35630 | 2 | ENSMUST00000020158.7 | -1908 | Myb | ENSMUSG00000019982.9 |
| chr10 | 20148349 | 20149175 | 3.28957 | 3.76796 | 1.03963 | Promoter | 10 | 20148471 | 20278699 | 130229 | 1 | ENSMUST00000020173.9 | 0 | Map7 | ENSMUSG00000019996.11 |
| chr10 | 24222167 | 24222243 | 4.21699 | 5.51924 | 2.1313 | Distal Intergenic | 10 | 24223517 | 24302790 | 79274 | 1 | ENSMUST00000095784.2 | -1274 | Moxd1 | ENSMUSG00000020000.7 |
| chr10 | 24223417 | 24224018 | 3.77899 | 5.07473 | 1.8418 | Promoter | 10 | 24223517 | 24302790 | 79274 | 1 | ENSMUST00000095784.2 | 0 | Moxd1 | ENSMUSG00000020000.7 |
| chr10 | 24224697 | 24225051 | 3.27739 | 3.89803 | 1.08598 | Intron (ENSMUST00000095784.2/ENSMUSG00000020000.7, intron 1 of 11) | 10 | 24223545 | 24231199 | 7655 | 1 | ENSMUST00000154916.1 | 1153 | Moxd1 | ENSMUSG00000020000.7 |
| chr10 | 24225373 | 24225590 | 3.63915 | 4.39812 | 1.38154 | Intron (ENSMUST00000095784.2/ENSMUSG00000020000.7, intron 1 of 11) | 10 | 24223545 | 24231199 | 7655 | 1 | ENSMUST00000154916.1 | 1829 | Moxd1 | ENSMUSG00000020000.7 |
| chr10 | 24275819 | 24276100 | 3.55852 | 4.31373 | 1.34203 | Intron (ENSMUST00000095784.2/ENSMUSG00000020000.7, intron 4 of 11) | 10 | 24223545 | 24231199 | 7655 | 1 | ENSMUST00000154916.1 | 52275 | Moxd1 | ENSMUSG00000020000.7 |
| chr10 | 93103965 | 93104095 | 3.59689 | 4.61 | 1.49969 | Distal Intergenic | 10 | 93061137 | 93081618 | 20482 | 2 | ENSMUST00000168080.2 | -22348 | 4930485B16Rik | ENSMUSG00000020014.10 |
| chr10 | 93622254 | 93622639 | 3.38263 | 3.90878 | 1.04252 | Distal Intergenic | 10 | 93640681 | 93644957 | 4277 | 1 | ENSMUST00000147080.1 | -18042 | Ntn4 | ENSMUSG00000020019.4 |
| chr10 | 93623227 | 93623541 | 3.69285 | 4.61956 | 1.52003 | Distal Intergenic | 10 | 93640681 | 93644957 | 4277 | 1 | ENSMUST00000147080.1 | -17140 | Ntn4 | ENSMUSG00000020019.4 |
| chr10 | 93641003 | 93641470 | 3.76179 | 4.69565 | 1.64363 | Promoter | 10 | 93641049 | 93747207 | 106159 | 1 | ENSMUST00000020204.4 | 0 | Ntn4 | ENSMUSG00000020019.4 |
| chr10 | 93828391 | 93828675 | 3.48724 | 4.12459 | 1.24947 | Distal Intergenic | 10 | 93831555 | 93848524 | 16970 | 1 | ENSMUST00000095333.4 | -2880 | Usp44 | ENSMUSG00000020020.7 |
| chr10 | 93829071 | 93829364 | 3.67664 | 4.57599 | 1.5047 | Distal Intergenic | 10 | 93831555 | 93848524 | 16970 | 1 | ENSMUST00000095333.4 | -2191 | Usp44 | ENSMUSG00000020020.7 |
| chr10 | 93829732 | 93830338 | 3.6435 | 4.44812 | 1.44484 | Distal Intergenic | 10 | 93831555 | 93848524 | 16970 | 1 | ENSMUST00000095333.4 | -1217 | Usp44 | ENSMUSG00000020020.7 |
| chr10 | 93831283 | 93831472 | 3.92499 | 5.13482 | 1.86066 | Promoter | 10 | 93831555 | 93848524 | 16970 | 1 | ENSMUST00000095333.4 | -83 | Usp44 | ENSMUSG00000020020.7 |
| chr10 | 93832115 | 93832247 | 3.37439 | 4.31461 | 1.31746 | Promoter | 10 | 93831555 | 93848524 | 16970 | 1 | ENSMUST00000095333.4 | 561 | Usp44 | ENSMUSG00000020020.7 |
| chr10 | 93833390 | 93835066 | 4.22447 | 5.86557 | 2.41597 | Intron (ENSMUST00000095333.4/ENSMUSG00000020020.7, intron 1 of 3) | 10 | 93831555 | 93848524 | 16970 | 1 | ENSMUST00000095333.4 | 1836 | Usp44 | ENSMUSG00000020020.7 |
| chr10 | 93835695 | 93835919 | 3.63906 | 4.43614 | 1.43859 | Intron (ENSMUST00000095333.4/ENSMUSG00000020020.7, intron 1 of 3) | 10 | 93831555 | 93848524 | 16970 | 1 | ENSMUST00000095333.4 | 4141 | Usp44 | ENSMUSG00000020020.7 |
| chr10 | 94310541 | 94310616 | 3.42858 | 4.0143 | 1.17135 | Distal Intergenic | 10 | 94311949 | 94430897 | 118949 | 1 | ENSMUST00000147869.1 | -1333 | Tmcc3 | ENSMUSG00000020023.12 |
| chr10 | 94312550 | 94312625 | 3.63048 | 4.37656 | 1.36563 | Promoter | 10 | 94312137 | 94375595 | 63459 | 1 | ENSMUST00000139230.1 | 414 | Tmcc3 | ENSMUSG00000020023.12 |
| chr10 | 84741048 | 84741339 | 3.2823 | 3.857 | 1.01924 | Distal Intergenic | 10 | 84756062 | 84906144 | 150083 | 1 | ENSMUST00000060397.7 | -14723 | Rfx4 | ENSMUSG00000020037.10 |
| chr10 | 84751890 | 84752673 | 3.4069 | 3.96257 | 1.07265 | Distal Intergenic | 10 | 84756062 | 84906144 | 150083 | 1 | ENSMUST00000060397.7 | -3389 | Rfx4 | ENSMUSG00000020037.10 |
| chr10 | 84754857 | 84756229 | 3.80344 | 4.87829 | 1.70155 | Promoter | 10 | 84756062 | 84906144 | 150083 | 1 | ENSMUST00000060397.7 | 0 | Rfx4 | ENSMUSG00000020037.10 |
| chr10 | 84757606 | 84757717 | 3.43045 | 4.14399 | 1.22512 | Intron (ENSMUST00000060397.7/ENSMUSG00000020037.10, intron 1 of 17) | 10 | 84756062 | 84906144 | 150083 | 1 | ENSMUST00000060397.7 | 1545 | Rfx4 | ENSMUSG00000020037.10 |
| chr10 | 84758725 | 84759450 | 3.97058 | 5.13021 | 1.92648 | Intron (ENSMUST00000060397.7/ENSMUSG00000020037.10, intron 1 of 17) | 10 | 84756062 | 84906144 | 150083 | 1 | ENSMUST00000060397.7 | 2664 | Rfx4 | ENSMUSG00000020037.10 |
| chr10 | 84759870 | 84760239 | 3.73363 | 4.55933 | 1.45621 | Intron (ENSMUST00000060397.7/ENSMUSG00000020037.10, intron 1 of 17) | 10 | 84762238 | 84814670 | 52433 | 1 | ENSMUST00000165868.3 | -1999 | Rfx4 | ENSMUSG00000020037.10 |
| chr10 | 84760678 | 84760829 | 4.17018 | 5.51403 | 2.17338 | Intron (ENSMUST00000060397.7/ENSMUSG00000020037.10, intron 1 of 17) | 10 | 84762238 | 84814670 | 52433 | 1 | ENSMUST00000165868.3 | -1409 | Rfx4 | ENSMUSG00000020037.10 |
| chr10 | 84761334 | 84762013 | 3.45592 | 4.15556 | 1.21337 | Promoter | 10 | 84762238 | 84814670 | 52433 | 1 | ENSMUST00000165868.3 | -225 | Rfx4 | ENSMUSG00000020037.10 |
| chr10 | 85378968 | 85379272 | 3.40103 | 4.07502 | 1.1697 | Distal Intergenic | 10 | 85386814 | 85660292 | 273479 | 1 | ENSMUST00000105307.2 | -7542 | Btbd11 | ENSMUSG00000020042.10 |
| chr10 | 85421348 | 85421423 | 3.50681 | 4.1748 | 1.28824 | Intron (ENSMUST00000105307.2/ENSMUSG00000020042.10, intron 1 of 16) | 10 | 85386827 | 85640594 | 253768 | 1 | ENSMUST00000020231.4 | 34522 | Btbd11 | ENSMUSG00000020042.10 |
| chr10 | 85598472 | 85598852 | 3.58121 | 4.32688 | 1.37939 | Promoter | 10 | 85598411 | 85606049 | 7639 | 1 | ENSMUST00000135235.1 | 62 | Btbd11 | ENSMUSG00000020042.10 |
| chr10 | 87484300 | 87485250 | 3.3485 | 3.85017 | 1.01733 | Distal Intergenic | 10 | 87490819 | 87493660 | 2842 | 2 | ENSMUST00000020243.7 | 8410 | Ascl1 | ENSMUSG00000020052.7 |
| chr10 | 87485700 | 87485871 | 3.53112 | 4.27017 | 1.28928 | Distal Intergenic | 10 | 87490819 | 87493660 | 2842 | 2 | ENSMUST00000020243.7 | 7789 | Ascl1 | ENSMUSG00000020052.7 |
| chr10 | 87486230 | 87486790 | 3.44157 | 4.08167 | 1.15789 | Distal Intergenic | 10 | 87490819 | 87493660 | 2842 | 2 | ENSMUST00000020243.7 | 6870 | Ascl1 | ENSMUSG00000020052.7 |
| chr10 | 87487679 | 87488945 | 3.85955 | 4.91036 | 1.7248 | Downstream (1-2kb) | 10 | 87490819 | 87493660 | 2842 | 2 | ENSMUST00000020243.7 | 4715 | Ascl1 | ENSMUSG00000020052.7 |
| chr10 | 87491454 | 87491560 | 3.87187 | 4.83051 | 1.64375 | 3' UTR | 10 | 87490819 | 87493660 | 2842 | 2 | ENSMUST00000020243.7 | 2100 | Ascl1 | ENSMUSG00000020052.7 |
| chr10 | 87492239 | 87492715 | 3.96747 | 5.23442 | 1.91753 | Promoter | 10 | 87490819 | 87493660 | 2842 | 2 | ENSMUST00000020243.7 | 945 | Ascl1 | ENSMUSG00000020052.7 |
| chr10 | 87493168 | 87493598 | 3.37562 | 3.89108 | 1.01733 | Promoter | 10 | 87490819 | 87493660 | 2842 | 2 | ENSMUST00000020243.7 | 62 | Ascl1 | ENSMUSG00000020052.7 |
| chr10 | 87494081 | 87494231 | 3.24358 | 3.69256 | 1.01733 | Promoter | 10 | 87490819 | 87493660 | 2842 | 2 | ENSMUST00000020243.7 | -422 | Ascl1 | ENSMUSG00000020052.7 |
| chr10 | 87494584 | 87495436 | 3.70929 | 4.67807 | 1.55056 | Promoter | 10 | 87490819 | 87493660 | 2842 | 2 | ENSMUST00000020243.7 | -925 | Ascl1 | ENSMUSG00000020052.7 |
| chr10 | 87496136 | 87496339 | 3.87056 | 4.83012 | 1.67314 | Distal Intergenic | 10 | 87490819 | 87493660 | 2842 | 2 | ENSMUST00000020243.7 | -2477 | Ascl1 | ENSMUSG00000020052.7 |
| chr10 | 87496761 | 87498513 | 3.57091 | 4.28085 | 1.3017 | Distal Intergenic | 10 | 87490819 | 87493660 | 2842 | 2 | ENSMUST00000020243.7 | -3102 | Ascl1 | ENSMUSG00000020052.7 |
| chr10 | 87500112 | 87500453 | 3.74293 | 4.83517 | 1.67037 | Distal Intergenic | 10 | 87490819 | 87493660 | 2842 | 2 | ENSMUST00000020243.7 | -6453 | Ascl1 | ENSMUSG00000020052.7 |
| chr10 | 87501138 | 87501438 | 3.94289 | 4.9602 | 1.70925 | Distal Intergenic | 10 | 87490819 | 87493660 | 2842 | 2 | ENSMUST00000020243.7 | -7479 | Ascl1 | ENSMUSG00000020052.7 |
| chr10 | 87503918 | 87504102 | 3.41026 | 4.28514 | 1.27435 | Distal Intergenic | 10 | 87490819 | 87493660 | 2842 | 2 | ENSMUST00000020243.7 | -10259 | Ascl1 | ENSMUSG00000020052.7 |
| chr10 | 87505060 | 87505135 | 3.64002 | 4.3939 | 1.36308 | Distal Intergenic | 10 | 87490819 | 87493660 | 2842 | 2 | ENSMUST00000020243.7 | -11401 | Ascl1 | ENSMUSG00000020052.7 |
| chr10 | 87858225 | 87858649 | 3.52332 | 4.38119 | 1.38054 | Promoter | 10 | 87858265 | 87930895 | 72631 | 1 | ENSMUST00000122386.2 | 0 | Igf1 | ENSMUSG00000020053.13 |
| chr10 | 87860710 | 87861012 | 3.53407 | 4.34838 | 1.37987 | Promoter | 10 | 87860606 | 87930899 | 70294 | 1 | ENSMUST00000121161.2 | 105 | Igf1 | ENSMUSG00000020053.13 |
| chr10 | 87861402 | 87861855 | 3.44999 | 4.06116 | 1.21575 | Promoter | 10 | 87861329 | 87930877 | 69549 | 1 | ENSMUST00000122100.1 | 74 | Igf1 | ENSMUSG00000020053.13 |
| chr10 | 88604236 | 88604524 | 3.51084 | 4.35393 | 1.38133 | Promoter | 10 | 88518339 | 88605025 | 86687 | 2 | ENSMUST00000119185.2 | 501 | Mybpc1 | ENSMUSG00000020061.12 |
| chr10 | 88604936 | 88605712 | 3.41629 | 4.01595 | 1.21396 | Promoter | 10 | 88518279 | 88605152 | 86874 | 2 | ENSMUST00000121629.2 | 0 | Mybpc1 | ENSMUSG00000020061.12 |
| chr10 | 88618986 | 88619162 | 3.88381 | 5.04758 | 1.81408 | Distal Intergenic | 10 | 88518279 | 88605152 | 86874 | 2 | ENSMUST00000121629.2 | -13835 | Mybpc1 | ENSMUSG00000020061.12 |
| chr10 | 88622474 | 88622826 | 3.56409 | 4.37769 | 1.35938 | Distal Intergenic | 10 | 88518279 | 88605152 | 86874 | 2 | ENSMUST00000121629.2 | -17323 | Mybpc1 | ENSMUSG00000020061.12 |
| chr10 | 88623685 | 88623807 | 3.36291 | 3.8719 | 1.01733 | Distal Intergenic | 10 | 88518279 | 88605152 | 86874 | 2 | ENSMUST00000121629.2 | -18534 | Mybpc1 | ENSMUSG00000020061.12 |
| chr10 | 88886311 | 88886535 | 3.32398 | 3.82237 | 1.04937 | Promoter | 10 | 88885992 | 88929505 | 43514 | 1 | ENSMUST00000020255.6 | 320 | Slc5a8 | ENSMUSG00000020062.6 |
| chr10 | 62507115 | 62507640 | 3.58423 | 4.27192 | 1.24985 | Promoter | 10 | 62494428 | 62507755 | 13328 | 2 | ENSMUST00000020271.7 | 115 | Srgn | ENSMUSG00000020077.9 |
| chr10 | 62405588 | 62405756 | 3.80535 | 4.92404 | 1.72374 | Exon (ENSMUST00000020277.8/ENSMUSG00000020080.8, exon 7 of 18) | 10 | 62395159 | 62400361 | 5203 | 2 | ENSMUST00000159493.1 | -5228 | Hkdc1 | ENSMUSG00000020080.8 |
| chr10 | 62110511 | 62110825 | 3.36272 | 4.07088 | 1.14736 | Promoter | 10 | 62108920 | 62110496 | 1577 | 2 | ENSMUST00000141144.1 | -16 | 2010107G23Rik | ENSMUSG00000020083.8 |
| chr10 | 62114645 | 62114830 | 3.40087 | 3.98477 | 1.19448 | Intron (ENSMUST00000027719.6/ENSMUSG00000020083.8, intron 2 of 2) | 10 | 62108210 | 62111050 | 2841 | 2 | ENSMUST00000142796.2 | -3596 | 2010107G23Rik | ENSMUSG00000020083.8 |
| chr10 | 62136141 | 62136349 | 3.35811 | 3.86466 | 1.01733 | Intron (ENSMUST00000027719.6/ENSMUSG00000020083.8, intron 1 of 2) | 10 | 62108917 | 62134706 | 25790 | 2 | ENSMUST00000156798.1 | -1436 | 2010107G23Rik | ENSMUSG00000020083.8 |
| chr10 | 62135597 | 62135815 | 3.64905 | 4.42498 | 1.37488 | Promoter | 10 | 62108917 | 62134706 | 25790 | 2 | ENSMUST00000156798.1 | -892 | 2010107G23Rik | ENSMUSG00000020083.8 |
| chr10 | 61595197 | 61595458 | 3.77013 | 4.66697 | 1.58194 | Promoter | 10 | 61595492 | 61626385 | 30894 | 1 | ENSMUST00000020287.6 | -34 | Npffr1 | ENSMUSG00000020090.6 |
| chr10 | 61175791 | 61177105 | 3.66811 | 4.46163 | 1.4429 | Promoter | 10 | 61175942 | 61188839 | 12898 | 1 | ENSMUST00000126831.1 | 0 | Tbata | ENSMUSG00000020096.14 |
| chr10 | 60346699 | 60347397 | 3.7642 | 4.6526 | 1.54286 | Promoter | 10 | 60346851 | 60372684 | 25834 | 1 | ENSMUST00000020301.8 | 0 | 4632428N05Rik | ENSMUSG00000020101.9 |
| chr10 | 125843949 | 125844396 | 3.51191 | 4.16362 | 1.21936 | Distal Intergenic | 10 | 125966219 | 126015359 | 49141 | 1 | ENSMUST00000074807.6 | -121823 | Lrig3 | ENSMUSG00000020105.8 |
| chr10 | 121532222 | 121532501 | 3.63985 | 4.62553 | 1.51952 | Distal Intergenic | 10 | 121546455 | 121586794 | 40340 | 2 | ENSMUST00000020316.2 | 54293 | Tbk1 | ENSMUSG00000020115.2 |
| chr10 | 121533412 | 121533714 | 3.5235 | 4.29819 | 1.3388 | Distal Intergenic | 10 | 121546455 | 121586794 | 40340 | 2 | ENSMUST00000020316.2 | 53080 | Tbk1 | ENSMUSG00000020115.2 |
| chr11 | 16751785 | 16753542 | 3.67799 | 4.48128 | 1.44224 | Promoter | 11 | 16752203 | 16887923 | 135721 | 1 | ENSMUST00000102884.4 | 0 | Egfr | ENSMUSG00000020122.11 |
| chr10 | 122448133 | 122449577 | 3.83115 | 4.89082 | 1.7826 | Promoter | 10 | 122448499 | 122453453 | 4955 | 1 | ENSMUST00000020323.5 | 0 | Avpr1a | ENSMUSG00000020123.5 |
| chr10 | 122450987 | 122451062 | 3.89972 | 5.1379 | 1.86009 | Intron (ENSMUST00000020323.5/ENSMUSG00000020123.5, intron 1 of 1) | 10 | 122448499 | 122453453 | 4955 | 1 | ENSMUST00000020323.5 | 2489 | Avpr1a | ENSMUSG00000020123.5 |
| chr10 | 80313332 | 80313645 | 3.30502 | 3.94308 | 1.0814 | Exon (ENSMUST00000105359.2/ENSMUSG00000020135.8, exon 15 of 15) | 10 | 80318254 | 80320537 | 2284 | 2 | ENSMUST00000020341.8 | 6892 | 2310011J03Rik | ENSMUSG00000020133.8 |
| chr10 | 80299029 | 80299342 | 3.95305 | 5.12311 | 1.85028 | Promoter | 10 | 80299091 | 80302361 | 3271 | 1 | ENSMUST00000154212.2 | 0 | Apc2 | ENSMUSG00000020135.8 |
| chr10 | 115470787 | 115471018 | 3.64367 | 4.40481 | 1.38338 | Intron (ENSMUST00000020350.9/ENSMUSG00000020140.10, intron 7 of 17) | 10 | 115457845 | 115463079 | 5235 | 2 | ENSMUST00000144732.1 | -7709 | Lgr5 | ENSMUSG00000020140.10 |
| chr10 | 115590075 | 115590289 | 3.54564 | 4.20202 | 1.209 | Distal Intergenic | 10 | 115450311 | 115587780 | 137470 | 2 | ENSMUST00000020350.9 | -2296 | Lgr5 | ENSMUSG00000020140.10 |
| chr10 | 115583812 | 115584191 | 3.69408 | 4.52022 | 1.44747 | Intron (ENSMUST00000020350.9/ENSMUSG00000020140.10, intron 1 of 17) | 10 | 115452014 | 115587493 | 135480 | 2 | ENSMUST00000172806.2 | 3302 | Lgr5 | ENSMUSG00000020140.10 |
| chr10 | 115584497 | 115588317 | 3.63484 | 4.44116 | 1.38568 | Promoter | 10 | 115450311 | 115587780 | 137470 | 2 | ENSMUST00000020350.9 | 0 | Lgr5 | ENSMUSG00000020140.10 |
| chr10 | 115588704 | 115588997 | 3.71309 | 4.5223 | 1.42627 | Promoter | 10 | 115450311 | 115587780 | 137470 | 2 | ENSMUST00000020350.9 | -925 | Lgr5 | ENSMUSG00000020140.10 |
| chr10 | 115592504 | 115592738 | 3.40616 | 4.2575 | 1.30196 | Distal Intergenic | 10 | 115450311 | 115587780 | 137470 | 2 | ENSMUST00000020350.9 | -4725 | Lgr5 | ENSMUSG00000020140.10 |
| chr10 | 115664203 | 115664390 | 3.67117 | 4.52397 | 1.50078 | Distal Intergenic | 10 | 115450311 | 115587780 | 137470 | 2 | ENSMUST00000020350.9 | -76424 | Lgr5 | ENSMUSG00000020140.10 |
| chr10 | 116016631 | 116016861 | 3.34851 | 3.86414 | 1.06662 | Distal Intergenic | 10 | 116018213 | 116274932 | 256720 | 1 | ENSMUST00000063470.5 | -1352 | Ptprr | ENSMUSG00000020151.11 |
| chr10 | 116018453 | 116018648 | 3.55035 | 4.22593 | 1.27024 | Promoter | 10 | 116018287 | 116114959 | 96673 | 1 | ENSMUST00000142617.1 | 167 | Ptprr | ENSMUSG00000020151.11 |
| chr10 | 116019307 | 116019644 | 3.87198 | 4.88316 | 1.72424 | Intron (ENSMUST00000063470.5/ENSMUSG00000020151.11, intron 1 of 13) | 10 | 116018287 | 116114959 | 96673 | 1 | ENSMUST00000142617.1 | 1021 | Ptprr | ENSMUSG00000020151.11 |
| chr10 | 116177373 | 116177493 | 3.33331 | 3.99059 | 1.12884 | Promoter | 10 | 116177411 | 116274345 | 96935 | 1 | ENSMUST00000128399.1 | 0 | Ptprr | ENSMUSG00000020151.11 |
| chr10 | 116205795 | 116206034 | 3.33488 | 3.82966 | 1.01733 | Intron (ENSMUST00000063470.5/ENSMUSG00000020151.11, intron 6 of 13) | 10 | 116196396 | 116237261 | 40866 | 1 | ENSMUST00000124369.1 | 9400 | Ptprr | ENSMUSG00000020151.11 |
| chr10 | 116300786 | 116301933 | 3.98892 | 5.12411 | 1.84472 | Promoter | 10 | 116301374 | 116389538 | 88165 | 1 | ENSMUST00000092167.5 | 0 | Ptprb | ENSMUSG00000020154.9 |
| chr10 | 116303370 | 116303849 | 3.90378 | 5.06502 | 1.81629 | Intron (ENSMUST00000092167.5/ENSMUSG00000020154.9, intron 2 of 31) | 10 | 116301374 | 116389538 | 88165 | 1 | ENSMUST00000092167.5 | 1997 | Ptprb | ENSMUSG00000020154.9 |
| chr10 | 116317463 | 116317637 | 3.41374 | 4.00235 | 1.20581 | Exon (ENSMUST00000092167.5/ENSMUSG00000020154.9, exon 5 of 32) | 10 | 116301374 | 116389538 | 88165 | 1 | ENSMUST00000092167.5 | 16090 | Ptprb | ENSMUSG00000020154.9 |
| chr11 | 33916871 | 33917188 | 3.43192 | 4.09681 | 1.20713 | Intron (ENSMUST00000154760.1/ENSMUSG00000053519.10, intron 1 of 2) | 11 | 33963013 | 33973641 | 10629 | 1 | ENSMUST00000020362.2 | -45825 | Kcnmb1 | ENSMUSG00000020155.2 |

| | | | | | | | | | | | | | | | |
|---|---|---|---|---|---|---|---|---|---|---|---|---|---|---|---|
| chr10 | 80209877 | 80210160 | 3.42551 | 3.98324 | 1.07666 | Distal Intergenic | 10 | 80226434 | 80243903 | 17470 | 1 | ENSMUST00000020365.9 | -16274 | Mum1 | ENSMUSG00000020156.11 |
| chr11 | 19025955 | 19026264 | 3.64843 | 4.4669 | 1.44649 | Distal Intergenic | 11 | 18879817 | 19018985 | 139169 | 2 | ENSMUST00000144988.2 | -6971 | Meis1 | ENSMUSG00000020160.13 |
| chr11 | 19026712 | 19027824 | 3.4323 | 4.00076 | 1.10276 | Distal Intergenic | 11 | 18879817 | 19018985 | 139169 | 2 | ENSMUST00000144988.2 | -7728 | Meis1 | ENSMUSG00000020160.13 |
| chr10 | 80449152 | 80449321 | 3.28937 | 3.79126 | 1.08797 | Distal Intergenic | 10 | 80409547 | 80433647 | 24101 | 2 | ENSMUST00000105343.2 | -15506 | Tcf3 | ENSMUSG00000020167.9 |
| chr10 | 117002878 | 117003038 | 3.59111 | 4.30991 | 1.35104 | Intron (ENSMUST00000020378.4/ENSMUSG00000020169.4, intron 5 of 9) | 10 | 116986314 | 117025040 | 38727 | 1 | ENSMUST00000020378.4 | 16565 | Best3 | ENSMUSG00000020169.4 |
| chr11 | 12464698 | 12464780 | 4.13779 | 5.43315 | 2.06077 | Promoter | 11 | 12304867 | 12464752 | 159886 | 2 | ENSMUST00000130572.1 | 0 | Cobl | ENSMUSG00000020173.12 |
| chr11 | 12036230 | 12036397 | 3.29089 | 3.76349 | 1.01733 | Intron (ENSMUST00000093321.6/ENSMUSG00000020176.12, intron 1 of 16) | 11 | 11930508 | 12037401 | 106894 | 2 | ENSMUST00000093321.6 | 1004 | Grb10 | ENSMUSG00000020176.12 |
| chr11 | 12036725 | 12037338 | 3.73374 | 4.58222 | 1.53136 | Promoter | 11 | 11930508 | 12037401 | 106894 | 2 | ENSMUST00000093321.6 | 63 | Grb10 | ENSMUSG00000020176.12 |
| chr10 | 75316441 | 75316936 | 3.36533 | 3.90332 | 1.11504 | Promoter | 10 | 75316943 | 75334788 | 17846 | 1 | ENSMUST00000105420.1 | -7 | Adora2a | ENSMUSG00000020178.5 |
| chr10 | 75317317 | 75317802 | 4.17917 | 5.61477 | 2.23771 | Promoter | 10 | 75316943 | 75334788 | 17846 | 1 | ENSMUST00000105420.1 | 375 | Adora2a | ENSMUSG00000020178.5 |
| chr10 | 110454897 | 110455250 | 3.7641 | 4.74698 | 1.63669 | Promoter | 10 | 109899103 | 110456204 | 557102 | 2 | ENSMUST00000160159.2 | 954 | Nav3 | ENSMUSG00000020181.10 |
| chr10 | 110456092 | 110456272 | 3.28157 | 3.78107 | 1.0696 | Promoter | 10 | 109899103 | 110456204 | 557102 | 2 | ENSMUST00000160159.2 | 0 | Nav3 | ENSMUSG00000020181.10 |
| chr10 | 110456812 | 110457072 | 3.4093 | 3.98407 | 1.13255 | Promoter | 10 | 109899103 | 110456204 | 557102 | 2 | ENSMUST00000160159.2 | -609 | Nav3 | ENSMUSG00000020181.10 |
| chr10 | 117627037 | 117628067 | 3.7002 | 4.52675 | 1.47734 | Distal Intergenic | 10 | 117629500 | 117635373 | 5874 | 1 | ENSMUST00000138020.2 | -1433 | Cpm | ENSMUSG00000020183.6 |
| chr10 | 117628907 | 117629581 | 3.91713 | 4.9619 | 1.77595 | Promoter | 10 | 117629500 | 117635373 | 5874 | 1 | ENSMUST00000138020.2 | 0 | Cpm | ENSMUSG00000020183.6 |
| chr10 | 117631139 | 117631466 | 3.41369 | 3.99613 | 1.12264 | Intron (ENSMUST00000138020.2/ENSMUSG00000020183.6, intron 2 of 2) | 10 | 117629500 | 117687352 | 57853 | 1 | ENSMUST00000020399.5 | 1640 | Cpm | ENSMUSG00000020183.6 |
| chr10 | 110746271 | 110746578 | 3.98779 | 5.16762 | 1.92628 | Promoter | 10 | 110745572 | 110754577 | 9006 | 1 | ENSMUST00000173634.1 | 700 | E2f7 | ENSMUSG00000020185.11 |
| chr10 | 110746942 | 110747108 | 4.46867 | 6.26247 | 2.70058 | Intron (ENSMUST00000173294.2/ENSMUSG00000020185.11, intron 2 of 2) | 10 | 110745572 | 110754577 | 9006 | 1 | ENSMUST00000173634.1 | 1371 | E2f7 | ENSMUSG00000020185.11 |
| chr10 | 110747468 | 110747583 | 3.39813 | 3.97141 | 1.17341 | Intron (ENSMUST00000173294.2/ENSMUSG00000020185.11, intron 2 of 2) | 10 | 110745572 | 110754577 | 9006 | 1 | ENSMUST00000173634.1 | 1897 | E2f7 | ENSMUSG00000020185.11 |
| chr10 | 110747950 | 110748148 | 3.62598 | 4.39539 | 1.42912 | Exon (ENSMUST00000173294.2/ENSMUSG00000020185.11, exon 3 of 3) | 10 | 110745572 | 110754577 | 9006 | 1 | ENSMUST00000173634.1 | 2379 | E2f7 | ENSMUSG00000020185.11 |
| chr10 | 118129215 | 118129457 | 3.54158 | 4.29354 | 1.33486 | Distal Intergenic | 10 | 118141787 | 118168997 | 27211 | 1 | ENSMUST00000020437.6 | -12330 | Mdm1 | ENSMUSG00000020212.8 |
| chr10 | 121022645 | 121022943 | 3.58788 | 4.28848 | 1.28573 | Distal Intergenic | 10 | 121033960 | 121100650 | 66691 | 1 | ENSMUST00000020439.5 | -11017 | Wif1 | ENSMUSG00000020218.6 |
| chr10 | 121023748 | 121023893 | 3.49327 | 4.1141 | 1.17843 | Distal Intergenic | 10 | 121033960 | 121100650 | 66691 | 1 | ENSMUST00000020439.5 | -10067 | Wif1 | ENSMUSG00000020218.6 |
| chr10 | 120271037 | 120271630 | 3.62076 | 4.34956 | 1.32187 | Distal Intergenic | 10 | 120227186 | 120231348 | 4163 | 1 | ENSMUST00000130198.1 | 43852 | Llph | ENSMUSG00000020224.9 |
| chr10 | 120272039 | 120272805 | 3.71135 | 4.67774 | 1.54906 | Distal Intergenic | 10 | 120227186 | 120231348 | 4163 | 1 | ENSMUST00000130198.1 | 44854 | Llph | ENSMUSG00000020224.9 |
| chr10 | 120273255 | 120273944 | 3.57303 | 4.30839 | 1.33867 | Distal Intergenic | 10 | 120227186 | 120231348 | 4163 | 1 | ENSMUST00000130198.1 | 46070 | Llph | ENSMUSG00000020224.9 |
| chr10 | 76117970 | 76118094 | 3.24878 | 3.70031 | 1.01733 | Distal Intergenic | 10 | 76057494 | 76110961 | 53468 | 2 | ENSMUST00000120757.1 | -7010 | Slc5a4b | ENSMUSG00000020226.9 |
| chr10 | 83533494 | 83533802 | 3.66109 | 4.556 | 1.52641 | Promoter | 10 | 83489429 | 83533395 | 43967 | 2 | ENSMUST00000146640.2 | -100 | Aldh1l2 | ENSMUSG00000020256.9 |
| chr11 | 55202030 | 55202253 | 3.3654 | 3.87567 | 1.01733 | Distal Intergenic | 11 | 55204350 | 55236330 | 31981 | 1 | ENSMUST00000108872.3 | -2097 | Slc36a1 | ENSMUSG00000020261.10 |
| chr10 | 77416965 | 77417163 | 3.39829 | 3.99145 | 1.19596 | Promoter | 10 | 77325338 | 77417854 | 92517 | 2 | ENSMUST00000150512.2 | 691 | Adarb1 | ENSMUSG00000020262.10 |
| chr11 | 32835572 | 32835871 | 3.73496 | 4.6751 | 1.57862 | Distal Intergenic | 11 | 32820376 | 32824605 | 4230 | 2 | ENSMUST00000020508.2 | -10968 | Smim23 | ENSMUSG00000020270.4 |
| chr11 | 32574473 | 32574630 | 3.57501 | 4.38698 | 1.38985 | Exon (ENSMUST00000102821.3/ENSMUSG00000020272.8, exon 3 of 19) | 11 | 32588595 | 32598528 | 9934 | 1 | ENSMUST00000143397.1 | -13965 | Stk10 | ENSMUSG00000020272.8 |
| chr11 | 32576945 | 32577541 | 3.72357 | 4.74255 | 1.61228 | Intron (ENSMUST00000102821.3/ENSMUSG00000020272.8, intron 3 of 18) | 11 | 32588595 | 32598528 | 9934 | 1 | ENSMUST00000143397.1 | -11054 | Stk10 | ENSMUSG00000020272.8 |
| chr11 | 32300005 | 32300308 | 3.44666 | 4.02586 | 1.115 | Promoter | 11 | 32300069 | 32300873 | 805 | 1 | ENSMUST00000020535.1 | 0 | Hbq1a | ENSMUSG00000020295.1 |
| chr11 | 31997616 | 31997852 | 3.63191 | 4.75796 | 1.61816 | Distal Intergenic | 11 | 32000463 | 32018007 | 17545 | 1 | ENSMUST00000109409.2 | -2611 | Nsg2 | ENSMUSG00000020297.5 |
| chr11 | 31998962 | 31999178 | 3.30676 | 3.78972 | 1.02567 | Distal Intergenic | 11 | 32000463 | 32018007 | 17545 | 1 | ENSMUST00000109409.2 | -1285 | Nsg2 | ENSMUSG00000020297.5 |
| chr11 | 32001116 | 32001805 | 3.63285 | 4.47506 | 1.41248 | Promoter | 11 | 32000502 | 32033414 | 32913 | 1 | ENSMUST00000093219.3 | 615 | Nsg2 | ENSMUSG00000020297.5 |
| chr11 | 32002599 | 32002889 | 3.45637 | 4.06811 | 1.15207 | Intron (ENSMUST00000109409.2/ENSMUSG00000020297.5, intron 2 of 2) | 11 | 32000502 | 32033414 | 32913 | 1 | ENSMUST00000093219.3 | 2098 | Nsg2 | ENSMUSG00000020297.5 |
| chr11 | 31950639 | 31950981 | 3.25776 | 3.71379 | 1.01733 | Distal Intergenic | 11 | 31965592 | 31979651 | 14060 | 1 | ENSMUST00000102829.3 | -14611 | 4930524B15Rik | ENSMUSG00000020299.6 |
| chr11 | 31367488 | 31367798 | 3.49423 | 4.1273 | 1.22073 | Promoter | 11 | 31365441 | 31368166 | 2726 | 2 | ENSMUST00000149467.1 | 368 | Stc2 | ENSMUSG00000020303.2 |
| chr11 | 31368614 | 31369079 | 3.44934 | 4.24495 | 1.28103 | Promoter | 11 | 31368003 | 31369056 | 1054 | 2 | ENSMUST00000134199.1 | 0 | Stc2 | ENSMUSG00000020303.2 |
| chr11 | 31369406 | 31370226 | 4.31436 | 5.88659 | 2.43083 | Promoter | 11 | 31357307 | 31370074 | 12768 | 2 | ENSMUST00000020546.2 | 0 | Stc2 | ENSMUSG00000020303.2 |
| chr11 | 31370932 | 31372029 | 3.83994 | 4.85669 | 1.65868 | Promoter | 11 | 31357307 | 31370074 | 12768 | 2 | ENSMUST00000020546.2 | -859 | Stc2 | ENSMUSG00000020303.2 |
| chr11 | 31372441 | 31372691 | 3.41795 | 3.97077 | 1.07295 | Distal Intergenic | 11 | 31357307 | 31370074 | 12768 | 2 | ENSMUST00000020546.2 | -2368 | Stc2 | ENSMUSG00000020303.2 |
| chr10 | 79637277 | 79637803 | 3.39237 | 4.0514 | 1.17742 | Promoter | 10 | 79617940 | 79637918 | 19979 | 2 | ENSMUST00000163867.2 | 115 | Shc2 | ENSMUSG00000020312.6 |
| chr10 | 79638290 | 79638549 | 3.30536 | 3.98327 | 1.14027 | Promoter | 10 | 79618652 | 79637918 | 19267 | 2 | ENSMUST00000020564.5 | -373 | Shc2 | ENSMUSG00000020312.6 |
| chr10 | 79639553 | 79639881 | 3.37291 | 3.88699 | 1.01733 | 3' UTR | 10 | 79618652 | 79637918 | 19267 | 2 | ENSMUST00000020564.5 | -1636 | Shc2 | ENSMUSG00000020312.6 |
| chr10 | 79593935 | 79594153 | 3.37958 | 3.90866 | 1.0584 | Distal Intergenic | 10 | 79576477 | 79587331 | 10855 | 2 | ENSMUST00000020566.6 | -6605 | Theg | ENSMUSG00000020317.9 |
| chr10 | 79594856 | 79595232 | 3.72526 | 4.60147 | 1.50627 | Distal Intergenic | 10 | 79576477 | 79587331 | 10855 | 2 | ENSMUST00000020566.6 | -7526 | Theg | ENSMUSG00000020317.9 |
| chr10 | 79714969 | 79715135 | 4.00188 | 5.25259 | 1.958 | Distal Intergenic | 10 | 79716634 | 79736108 | 19475 | 1 | ENSMUST00000020581.2 | -1499 | Hcn2 | ENSMUSG00000020331.4 |
| chr10 | 79716180 | 79716562 | 3.70023 | 4.64912 | 1.55234 | Promoter | 10 | 79716634 | 79736108 | 19475 | 1 | ENSMUST00000020581.2 | -72 | Hcn2 | ENSMUSG00000020331.4 |
| chr10 | 79717011 | 79717790 | 3.63292 | 4.57153 | 1.48058 | Promoter | 10 | 79716634 | 79736108 | 19475 | 1 | ENSMUST00000099513.2 | 378 | Hcn2 | ENSMUSG00000020331.4 |
| chr11 | 54303916 | 54304605 | 3.69493 | 4.53774 | 1.43385 | Promoter | 11 | 54304005 | 54361535 | 57531 | 1 | ENSMUST00000000145.6 | 0 | Acsl6 | ENSMUSG00000020333.12 |
| chr11 | 47379247 | 47379832 | 3.61522 | 4.42968 | 1.39776 | Promoter | 11 | 46978785 | 47379522 | 400738 | 2 | ENSMUST00000077221.5 | 0 | Sgcd | ENSMUSG00000020354.10 |
| chr11 | 47986464 | 47986577 | 3.53479 | 4.18186 | 1.19531 | Intron (ENSMUST00000128558.1/ENSMUSG00000020354.10, intron 1 of 2) | 11 | 47578805 | 47988969 | 410165 | 2 | ENSMUST00000128558.1 | 2392 | Sgcd | ENSMUSG00000020354.10 |
| chr11 | 47987745 | 47989331 | 4.13475 | 5.49811 | 2.1502 | Promoter | 11 | 47578805 | 47988969 | 410165 | 2 | ENSMUST00000128558.1 | 0 | Sgcd | ENSMUSG00000020354.10 |
| chr11 | 49602596 | 49602751 | 3.53669 | 4.18473 | 1.19531 | Distal Intergenic | 11 | 49609263 | 49628941 | 19679 | 1 | ENSMUST00000152253.1 | -6512 | Flt4 | ENSMUSG00000020357.3 |
| chr11 | 49608502 | 49608924 | 3.96238 | 5.04835 | 1.79843 | Promoter | 11 | 49609263 | 49628941 | 19679 | 1 | ENSMUST00000152253.1 | -339 | Flt4 | ENSMUSG00000020357.3 |
| chr11 | 49609519 | 49611086 | 3.65517 | 4.54183 | 1.47183 | Promoter | 11 | 49609263 | 49652739 | 43477 | 1 | ENSMUST00000020617.2 | 257 | Flt4 | ENSMUSG00000020357.3 |
| chr11 | 49611419 | 49611656 | 3.57828 | 4.26911 | 1.26598 | Intron (ENSMUST00000152253.1/ENSMUSG00000020357.3, intron 1 of 6) | 11 | 49609263 | 49652739 | 43477 | 1 | ENSMUST00000020617.2 | 2157 | Flt4 | ENSMUSG00000020357.3 |
| chr11 | 49634283 | 49634976 | 3.63319 | 4.39544 | 1.35948 | Exon (ENSMUST00000020617.2/ENSMUSG00000020357.3, exon 13 of 30) | 11 | 49609263 | 49652739 | 43477 | 1 | ENSMUST00000020617.2 | 25021 | Flt4 | ENSMUSG00000020357.3 |
| chr11 | 49635957 | 49636260 | 3.37497 | 3.89009 | 1.01733 | Exon (ENSMUST00000020617.2/ENSMUSG00000020357.3, exon 18 of 30) | 11 | 49609263 | 49652739 | 43477 | 1 | ENSMUST00000020617.2 | 26695 | Flt4 | ENSMUSG00000020357.3 |
| chr11 | 49794091 | 49794583 | 3.77905 | 4.66819 | 1.57357 | Promoter | 11 | 49794178 | 49807750 | 13573 | 1 | ENSMUST00000130129.1 | 0 | Gfpt2 | ENSMUSG00000020363.6 |
| chr11 | 49795668 | 49795757 | 3.35896 | 3.86595 | 1.01733 | Intron (ENSMUST00000130129.1/ENSMUSG00000020363.6, intron 1 of 1) | 11 | 49794187 | 49838613 | 44427 | 1 | ENSMUST00000020629.4 | 1482 | Gfpt2 | ENSMUSG00000020363.6 |
| chr11 | 49796969 | 49797044 | 4.59269 | 6.29884 | 2.71645 | Intron (ENSMUST00000130129.1/ENSMUSG00000020363.6, intron 1 of 1) | 11 | 49794187 | 49838613 | 44427 | 1 | ENSMUST00000020629.4 | 2783 | Gfpt2 | ENSMUSG00000020363.6 |
| chr11 | 49901137 | 49902698 | 4.61337 | 6.66808 | 3.0579 | Promoter | 11 | 49901835 | 49980222 | 78388 | 1 | ENSMUST00000093141.6 | 0 | Rasgef1c | ENSMUSG00000020374.11 |
| chr11 | 49917178 | 49917282 | 3.8391 | 4.76799 | 1.61152 | Intron (ENSMUST00000093141.6/ENSMUSG00000020374.11, intron 1 of 12) | 11 | 49902161 | 49980994 | 78834 | 1 | ENSMUST00000063444.2 | 15018 | Rasgef1c | ENSMUSG00000020374.11 |
| chr11 | 49926754 | 49927154 | 3.48221 | 4.10938 | 1.19869 | Intron (ENSMUST00000093141.6/ENSMUSG00000020374.11, intron 1 of 12) | 11 | 49902161 | 49980994 | 78834 | 1 | ENSMUST00000063444.2 | 24594 | Rasgef1c | ENSMUSG00000020374.11 |
| chr11 | 49928644 | 49928953 | 3.57665 | 4.27565 | 1.2969 | Intron (ENSMUST00000093141.6/ENSMUSG00000020374.11, intron 1 of 12) | 11 | 49902161 | 49980994 | 78834 | 1 | ENSMUST00000063444.2 | 26484 | Rasgef1c | ENSMUSG00000020374.11 |
| chr11 | 49961925 | 49962010 | 3.76386 | 4.62001 | 1.50337 | Intron (ENSMUST00000093141.6/ENSMUSG00000020374.11, intron 5 of 12) | 11 | 49963785 | 49975750 | 11966 | 1 | ENSMUST00000109172.3 | -1775 | Rasgef1c | ENSMUSG00000020374.11 |
| chr11 | 53636091 | 53636607 | 3.40285 | 3.94619 | 1.067 | Distal Intergenic | 11 | 53631324 | 53634702 | 3379 | 2 | ENSMUST00000020650.1 | -1390 | Il13 | ENSMUSG00000020383.1 |
| chr11 | 54068812 | 54069218 | 3.50129 | 4.14152 | 1.22471 | Promoter | 11 | 54054928 | 54069014 | 14087 | 2 | ENSMUST00000018755.4 | 0 | Pdlim4 | ENSMUSG00000020388.7 |
| chr11 | 4945954 | 4946056 | 3.67877 | 4.70798 | 1.5753 | Intron (ENSMUST00000093369.4/ENSMUSG00000020396.8, intron 1 of 3) | 11 | 4938754 | 4948064 | 9311 | 2 | ENSMUST00000093369.4 | 2008 | Nefh | ENSMUSG00000020396.8 |
| chr11 | 4946683 | 4948206 | 3.87335 | 5.16565 | 1.90842 | Promoter | 11 | 4938754 | 4948064 | 9311 | 2 | ENSMUST00000093369.4 | 0 | Nefh | ENSMUSG00000020396.8 |
| chr11 | 4948843 | 4949068 | 3.89471 | 5.11121 | 1.84587 | Promoter | 11 | 4938754 | 4948064 | 9311 | 2 | ENSMUST00000093369.4 | -780 | Nefh | ENSMUSG00000020396.8 |
| chr11 | 4949901 | 4949980 | 3.90757 | 4.99697 | 1.78369 | Distal Intergenic | 11 | 4938754 | 4948064 | 9311 | 2 | ENSMUST00000093369.4 | -1838 | Nefh | ENSMUSG00000020396.8 |
| chr11 | 4950706 | 4950781 | 3.69472 | 4.58641 | 1.51132 | Distal Intergenic | 11 | 4938754 | 4948064 | 9311 | 2 | ENSMUST00000093369.4 | -2643 | Nefh | ENSMUSG00000020396.8 |
| chr11 | 4951966 | 4952244 | 3.44869 | 4.28707 | 1.32515 | Distal Intergenic | 11 | 4938754 | 4948064 | 9311 | 2 | ENSMUST00000093369.4 | -3903 | Nefh | ENSMUSG00000020396.8 |
| chr11 | 4952972 | 4953368 | 3.49052 | 4.25223 | 1.29508 | Distal Intergenic | 11 | 4938754 | 4948064 | 9311 | 2 | ENSMUST00000093369.4 | -4909 | Nefh | ENSMUSG00000020396.8 |
| chr11 | 4954641 | 4954810 | 3.37736 | 3.89941 | 1.03748 | Distal Intergenic | 11 | 4938754 | 4948064 | 9311 | 2 | ENSMUST00000093369.4 | -6578 | Nefh | ENSMUSG00000020396.8 |
| chr11 | 46406441 | 46406769 | 3.57531 | 4.25963 | 1.25463 | Exon (ENSMUST00000139280.1/ENSMUSG00000020401.6, exon 3 of 3) | 11 | 46404730 | 46407982 | 3253 | 1 | ENSMUST00000063166.5 | 1712 | Fam71b | ENSMUSG00000020401.6 |
| chr11 | 9113488 | 9113823 | 3.85211 | 4.92135 | 1.73762 | Distal Intergenic | 11 | 9118103 | 9136164 | 18062 | 1 | ENSMUST00000020677.2 | -4280 | Upp1 | ENSMUSG00000020407.8 |
| chr11 | 9115745 | 9115882 | 3.41827 | 4.46443 | 1.4045 | Distal Intergenic | 11 | 9118103 | 9136164 | 18062 | 1 | ENSMUST00000020677.2 | -2221 | Upp1 | ENSMUSG00000020407.8 |
| chr11 | 9118857 | 9119894 | 3.51968 | 4.29129 | 1.34191 | Promoter | 11 | 9118787 | 9133251 | 14465 | 1 | ENSMUST00000172452.2 | 71 | Upp1 | ENSMUSG00000020407.8 |
| chr11 | 46165848 | 46166661 | 3.69834 | 4.72434 | 1.60201 | Promoter | 11 | 46148155 | 46166508 | 18354 | 2 | ENSMUST00000020679.2 | 0 | Nipal4 | ENSMUSG00000020411.2 |
| chr11 | 7213076 | 7213591 | 4.3134 | 6.03839 | 2.50183 | Promoter | 11 | 7206086 | 7213897 | 7812 | 2 | ENSMUST00000020702.5 | 306 | Igfbp3 | ENSMUSG00000020427.6 |
| chr11 | 7214048 | 7214519 | 3.52224 | 4.21378 | 1.32778 | Promoter | 11 | 7207686 | 7213923 | 6238 | 2 | ENSMUST00000135887.2 | -126 | Igfbp3 | ENSMUSG00000020427.6 |
| chr11 | 7197742 | 7197861 | 3.27782 | 3.74386 | 1.01733 | Promoter | 11 | 7197782 | 7202546 | 4765 | 1 | ENSMUST00000020704.7 | 0 | Igfbp1 | ENSMUSG00000020429.7 |
| chr11 | 3744051 | 3744302 | 3.2952 | 3.76997 | 1.01733 | Intron (ENSMUST00000070552.8/ENSMUSG00000020435.12, intron 2 of 13) | 11 | 3703918 | 3722216 | 18299 | 2 | ENSMUST00000101632.3 | -21836 | Osbp2 | ENSMUSG00000020435.12 |
| chr11 | 3753403 | 3753984 | 3.33488 | 4.16184 | 1.21015 | Intron (ENSMUST00000070552.8/ENSMUSG00000020435.12, intron 2 of 13) | 11 | 3717852 | 3773789 | 55938 | 2 | ENSMUST00000127371.1 | 19805 | Osbp2 | ENSMUSG00000020435.12 |
| chr11 | 3760829 | 3761526 | 3.37838 | 4.17237 | 1.27771 | Intron (ENSMUST00000070552.8/ENSMUSG00000020435.12, intron 2 of 13) | 11 | 3717852 | 3773789 | 55938 | 2 | ENSMUST00000127371.1 | 12263 | Osbp2 | ENSMUSG00000020435.12 |
| chr11 | 3761912 | 3763192 | 4.19181 | 5.75786 | 2.39713 | Intron (ENSMUST00000070552.8/ENSMUSG00000020435.12, intron 2 of 13) | 11 | 3717852 | 3773789 | 55938 | 2 | ENSMUST00000127371.1 | 10597 | Osbp2 | ENSMUSG00000020435.12 |
| chr11 | 3773411 | 3773722 | 4.08979 | 5.34078 | 2.00118 | Promoter | 11 | 3717852 | 3773789 | 55938 | 2 | ENSMUST00000127371.1 | 67 | Osbp2 | ENSMUSG00000020435.12 |
| chr11 | 3774156 | 3775387 | 3.83531 | 4.77311 | 1.63281 | Promoter | 11 | 3717912 | 3774732 | 56821 | 2 | ENSMUST00000155197.1 | 0 | Osbp2 | ENSMUSG00000020435.12 |
| chr11 | 3776914 | 3777400 | 3.9362 | 4.95041 | 1.71369 | Intron (ENSMUST00000070552.8/ENSMUSG00000020435.12, intron 2 of 13) | 11 | 3717912 | 3774732 | 56821 | 2 | ENSMUST00000155197.1 | -2183 | Osbp2 | ENSMUSG00000020435.12 |
| chr11 | 3778976 | 3779087 | 3.43311 | 4.0795 | 1.22129 | Intron (ENSMUST00000070552.8/ENSMUSG00000020435.12, intron 2 of 13) | 11 | 3717912 | 3774732 | 56821 | 2 | ENSMUST00000155197.1 | -4245 | Osbp2 | ENSMUSG00000020435.12 |
| chr11 | 57297253 | 57297739 | 3.85572 | 4.87839 | 1.71144 | Intron (ENSMUST00000036315.10/ENSMUSG00000020524.11, intron 13 of 15) | 11 | 57176518 | 57189793 | 13276 | 1 | ENSMUST00000173531.1 | 120736 | Gria1 | ENSMUSG00000020524.11 |
| chr11 | 57298159 | 57298311 | 3.40847 | 3.99043 | 1.17631 | Intron (ENSMUST00000036315.10/ENSMUSG00000020524.11, intron 13 of 15) | 11 | 57176518 | 57189793 | 13276 | 1 | ENSMUST00000173531.1 | 121642 | Gria1 | ENSMUSG00000020524.11 |
| chr11 | 57300536 | 57300963 | 3.56707 | 4.37467 | 1.38431 | Intron (ENSMUST00000036315.10/ENSMUSG00000020524.11, intron 13 of 15) | 11 | 57176518 | 57189793 | 13276 | 1 | ENSMUST00000173531.1 | 124019 | Gria1 | ENSMUSG00000020524.11 |
| chr11 | 57302198 | 57302449 | 3.4587 | 4.05262 | 1.14547 | Intron (ENSMUST00000036315.10/ENSMUSG00000020524.11, intron 13 of 15) | 11 | 57176518 | 57189793 | 13276 | 1 | ENSMUST00000173531.1 | 125681 | Gria1 | ENSMUSG00000020524.11 |
| chr11 | 65204272 | 65204573 | 3.4132 | 4.21961 | 1.27607 | Promoter | 11 | 65196102 | 65205467 | 9366 | 2 | ENSMUST00000144399.1 | 894 | Myocd | ENSMUSG00000020542.13 |
| chr11 | 65268867 | 65269050 | 3.49546 | 4.15489 | 1.28686 | Promoter | 11 | 65223734 | 65269854 | 46121 | 2 | ENSMUST00000150823.1 | 804 | Myocd | ENSMUSG00000020542.13 |
| chr11 | 65269700 | 65271510 | 3.63511 | 4.55147 | 1.48622 | Promoter | 11 | 65176561 | 65269941 | 93381 | 2 | ENSMUST00000102635.4 | 0 | Myocd | ENSMUSG00000020542.13 |
| chr11 | 65271831 | 65271988 | 3.77373 | 4.8081 | 1.65458 | Distal Intergenic | 11 | 65176574 | 65269989 | 93416 | 2 | ENSMUST00000108695.3 | -1843 | Myocd | ENSMUSG00000020542.13 |
| chr11 | 89987143 | 89987493 | 3.51941 | 4.18753 | 1.30059 | Exon (ENSMUST00000020864.8/ENSMUSG00000020553.8, exon 4 of 6) | 11 | 89982665 | 90002894 | 20230 | 2 | ENSMUST00000020864.8 | 15401 | Pctp | ENSMUSG00000020553.8 |
| chr12 | 17323686 | 17325444 | 3.98226 | 5.20031 | 1.90545 | Promoter | 12 | 17284721 | 17324712 | 39992 | 2 | ENSMUST00000095820.6 | 0 | Atp6v1c2 | ENSMUSG00000020566.13 |
| chr12 | 17329915 | 17330054 | 3.3287 | 3.82039 | 1.01733 | Promoter | 12 | 17308345 | 17329359 | 21015 | 2 | ENSMUST00000143131.1 | -557 | Atp6v1c2 | ENSMUSG00000020566.13 |
| chr12 | 16652833 | 16653932 | 3.76959 | 4.6346 | 1.50079 | Promoter | 12 | 16653470 | 16660236 | 6767 | 1 | ENSMUST00000111064.1 | 0 | Ntsr2 | ENSMUSG00000020591.10 |
| chr12 | 16654376 | 16654728 | 3.84158 | 4.75364 | 1.55854 | Promoter | 12 | 16653470 | 16660236 | 6767 | 1 | ENSMUST00000111064.1 | 907 | Ntsr2 | ENSMUSG00000020591.10 |
| chr12 | 8772552 | 8773094 | 3.76974 | 4.72884 | 1.65163 | Promoter | 12 | 8771803 | 8790448 | 18646 | 1 | ENSMUST00000160348.1 | 750 | Sdc1 | ENSMUSG00000020592.9 |
| chr12 | 44329898 | 44330020 | 4.05645 | 5.29786 | 2.02727 | Intron (ENSMUST00000020939.9/ENSMUSG00000020598.10, intron 1 of 30) | 12 | 44328885 | 44601846 | 272962 | 1 | ENSMUST00000110748.2 | 1014 | Nrcam | ENSMUSG00000020598.10 |
| chr12 | 44331674 | 44331880 | 3.33516 | 3.8364 | 1.03958 | Intron (ENSMUST00000020939.9/ENSMUSG00000020598.10, intron 1 of 30) | 12 | 44328885 | 44601846 | 272962 | 1 | ENSMUST00000110748.2 | 2790 | Nrcam | ENSMUSG00000020598.10 |
| chr12 | 44332186 | 44332318 | 3.34895 | 3.88546 | 1.09247 | Intron (ENSMUST00000020939.9/ENSMUSG00000020598.10, intron 1 of 30) | 12 | 44328885 | 44601846 | 272962 | 1 | ENSMUST00000110748.2 | 3302 | Nrcam | ENSMUSG00000020598.10 |
| chr12 | 44332898 | 44333121 | 3.49425 | 4.12575 | 1.21465 | Intron (ENSMUST00000020939.9/ENSMUSG00000020598.10, intron 1 of 30) | 12 | 44328885 | 44601846 | 272962 | 1 | ENSMUST00000110748.2 | 4014 | Nrcam | ENSMUSG00000020598.10 |
| chr11 | 109210850 | 109210936 | 3.63727 | 4.40279 | 1.35775 | Distal Intergenic | 11 | 109239511 | 109246965 | 7455 | 2 | ENSMUST00000156785.1 | 36029 | Rgs9 | ENSMUSG00000020599.8 |
| chr11 | 109238289 | 109238623 | 3.43592 | 3.99931 | 1.078 | Intron (ENSMUST00000020920.4/ENSMUSG00000020599.8, intron 15 of 18) | 11 | 109239511 | 109246965 | 7455 | 2 | ENSMUST00000156785.1 | 8342 | Rgs9 | ENSMUSG00000020599.8 |
| chr11 | 109472982 | 109474319 | 3.56198 | 4.32654 | 1.38522 | Promoter | 11 | 109473374 | 109489999 | 16626 | 1 | ENSMUST00000152252.2 | 0 | Arsg | ENSMUSG00000020604.8 |
| chr12 | 8310093 | 8310448 | 3.6582 | 4.4296 | 1.40237 | Distal Intergenic | 12 | 8313433 | 8343824 | 30392 | 1 | ENSMUST00000020927.8 | -2985 | Hs1bp3 | ENSMUSG00000020605.8 |
| chr12 | 14149034 | 14149147 | 3.33165 | 4.05731 | 1.15964 | 3' UTR | 12 | 14147599 | 14152038 | 4440 | 2 | ENSMUST00000020926.6 | 2891 | Fam84a | ENSMUSG00000020607.6 |
| chr12 | 14149566 | 14150412 | 3.45259 | 4.24389 | 1.29309 | 3' UTR | 12 | 14147599 | 14152038 | 4440 | 2 | ENSMUST00000020926.6 | 1626 | Fam84a | ENSMUSG00000020607.6 |
| chr12 | 14151289 | 14151458 | 3.5511 | 4.26055 | 1.29042 | Promoter | 12 | 14147599 | 14152038 | 4440 | 2 | ENSMUST00000020926.6 | 580 | Fam84a | ENSMUSG00000020607.6 |
| chr11 | 109721404 | 109721810 | 3.55307 | 4.22494 | 1.24049 | Promoter | 11 | 109672926 | 109722256 | 49331 | 2 | ENSMUST00000020938.7 | 446 | Fam20a | ENSMUSG00000020614.8 |
| chr11 | 109722163 | 109723151 | 3.75206 | 4.64396 | 1.51738 | Promoter | 11 | 109669749 | 109722279 | 52531 | 2 | ENSMUST00000155559.2 | 0 | Fam20a | ENSMUSG00000020614.8 |
| chr12 | 10185933 | 10186019 | 3.8748 | 7.15688 | 3.35388 | Distal Intergenic | 12 | 10369973 | 10390170 | 20198 | 1 | ENSMUST00000147323.2 | -183954 | Nt5c1b | ENSMUSG00000020622.10 |
| chr11 | 110449116 | 110449241 | 3.60168 | 4.30178 | 1.26205 | Intron (ENSMUST00000020949.6/ENSMUSG00000020623.6, intron 1 of 11) | 11 | 110477377 | 110499522 | 22146 | 1 | ENSMUST00000146540.2 | -28136 | Map2k6 | ENSMUSG00000020623.6 |
| chr11 | 112812101 | 112812267 | 3.45052 | 4.04907 | 1.16285 | Distal Intergenic | 11 | 112825154 | 112826210 | 1057 | 1 | ENSMUST00000149136.1 | -12887 | 4933434M16Rik | ENSMUSG00000020624.3 |
| chr11 | 112869754 | 112869952 | 3.32352 | 3.81261 | 1.01733 | Distal Intergenic | 11 | 112825179 | 112832469 | 7291 | 1 | ENSMUST00000153429.1 | 44576 | 4933434M16Rik | ENSMUSG00000020624.3 |
| chr11 | 112979132 | 112979273 | 3.35057 | 3.85328 | 1.01733 | Distal Intergenic | 11 | 112825179 | 112832469 | 7291 | 1 | ENSMUST00000153429.1 | 153954 | 4933434M16Rik | ENSMUSG00000020624.3 |
| chr11 | 112979691 | 112979867 | 3.45907 | 4.04913 | 1.13093 | Distal Intergenic | 11 | 112825179 | 112832469 | 7291 | 1 | ENSMUST00000153429.1 | 154513 | 4933434M16Rik | ENSMUSG00000020624.3 |
| chr11 | 112980918 | 112981333 | 3.40041 | 3.99773 | 1.1334 | Distal Intergenic | 11 | 112825179 | 112832469 | 7291 | 1 | ENSMUST00000153429.1 | 155740 | 4933434M16Rik | ENSMUSG00000020624.3 |
| chr11 | 112982636 | 112982769 | 3.90489 | 5.00601 | 1.79246 | Distal Intergenic | 11 | 112825179 | 112832469 | 7291 | 1 | ENSMUST00000153429.1 | 157458 | 4933434M16Rik | ENSMUSG00000020624.3 |
| chr11 | 112990155 | 112990501 | 3.57964 | 4.31379 | 1.38858 | Distal Intergenic | 11 | 112825179 | 112832469 | 7291 | 1 | ENSMUST00000153429.1 | 164977 | 4933434M16Rik | ENSMUSG00000020624.3 |
| chr11 | 112991124 | 112991202 | 3.64928 | 4.43278 | 1.45067 | Distal Intergenic | 11 | 112825179 | 112832469 | 7291 | 1 | ENSMUST00000153429.1 | 165946 | 4933434M16Rik | ENSMUSG00000020624.3 |
| chr12 | 25094556 | 25096058 | 3.67293 | 4.49288 | 1.44207 | Promoter | 12 | 25093801 | 25096092 | 2292 | 2 | ENSMUST00000020974.6 | 34 | Id2 | ENSMUSG00000020644.8 |
| chr12 | 25096867 | 25097537 | 3.82812 | 4.7691 | 1.62509 | Promoter | 12 | 25093801 | 25096092 | 2292 | 2 | ENSMUST00000020974.6 | -776 | Id2 | ENSMUSG00000020644.8 |
| chr12 | 25097879 | 25098008 | 3.72789 | 4.56399 | 1.49036 | Distal Intergenic | 12 | 25093801 | 25096092 | 2292 | 2 | ENSMUST00000020974.6 | -1788 | Id2 | ENSMUSG00000020644.8 |
| chr12 | 25098537 | 25099051 | 3.92497 | 4.97967 | 1.78146 | Distal Intergenic | 12 | 25093801 | 25096092 | 2292 | 2 | ENSMUST00000020974.6 | -2446 | Id2 | ENSMUSG00000020644.8 |
| chr12 | 25100761 | 25101105 | 3.68875 | 4.59352 | 1.55778 | Distal Intergenic | 12 | 25093801 | 25096092 | 2292 | 2 | ENSMUST00000020974.6 | -4670 | Id2 | ENSMUSG00000020644.8 |
| chr12 | 25102022 | 25103004 | 3.69226 | 4.499 | 1.43504 | Distal Intergenic | 12 | 25093801 | 25096092 | 2292 | 2 | ENSMUST00000020974.6 | -5931 | Id2 | ENSMUSG00000020644.8 |
| chr12 | 24706720 | 24707499 | 3.74625 | 4.72239 | 1.57936 | Promoter | 12 | 24708241 | 24714146 | 5906 | 1 | ENSMUST00000020980.6 | -742 | Rrm2 | ENSMUSG00000020649.6 |
| chr12 | 31555927 | 31556061 | 3.70745 | 4.84522 | 1.66958 | Intron (ENSMUST00000001253.7/ENSMUSG00000020651.7, intron 3 of 20) | 12 | 31519827 | 31559969 | 40143 | 2 | ENSMUST00000001253.7 | 3908 | Slc26a4 | ENSMUSG00000020651.7 |
| chr12 | 31557186 | 31557719 | 3.31972 | 3.82677 | 1.08465 | Intron (ENSMUST00000001253.7/ENSMUSG00000020651.7, intron 2 of 20) | 12 | 31519827 | 31559969 | 40143 | 2 | ENSMUST00000001253.7 | 2250 | Slc26a4 | ENSMUSG00000020651.7 |
| chr12 | 31558058 | 31558321 | 4.24602 | 5.73393 | 2.3383 | Intron (ENSMUST00000001253.7/ENSMUSG00000020651.7, intron 2 of 20) | 12 | 31519827 | 31559969 | 40143 | 2 | ENSMUST00000001253.7 | 1648 | Slc26a4 | ENSMUSG00000020651.7 |
| chr12 | 31558707 | 31561823 | 3.60143 | 4.59726 | 1.52174 | Promoter | 12 | 31519827 | 31559969 | 40143 | 2 | ENSMUST00000001253.7 | 0 | Slc26a4 | ENSMUSG00000020651.7 |
| chr12 | 4134269 | 4134817 | 3.38166 | 3.94244 | 1.14397 | Promoter | 12 | 4134153 | 4213524 | 79372 | 1 | ENSMUST00000020984.8 | 117 | Adcy3 | ENSMUSG00000020654.10 |
| chr12 | 4036080 | 4036422 | 3.2502 | 3.7025 | 1.01733 | Intron (ENSMUST00000111178.1/ENSMUSG00000020658.9, intron 1 of 22) | 12 | 3962554 | 4038915 | 76362 | 2 | ENSMUST00000111178.1 | 2493 | Efr3b | ENSMUSG00000020658.9 |
| chr12 | 4037438 | 4038258 | 3.8188 | 5.0775 | 1.83377 | Promoter | 12 | 3962554 | 4038915 | 76362 | 2 | ENSMUST00000111178.1 | 657 | Efr3b | ENSMUSG00000020658.9 |
| chr12 | 3954809 | 3955277 | 3.81713 | 4.93823 | 1.72426 | Promoter | 12 | 3954951 | 3960618 | 5668 | 1 | ENSMUST00000020990.5 | 0 | Pomc | ENSMUSG00000020660.5 |
| chr11 | 83849708 | 83850357 | 3.86348 | 4.96448 | 1.7517 | Promoter | 11 | 83850063 | 83856098 | 6036 | 1 | ENSMUST00000135975.2 | 0 | Hnf1b | ENSMUSG00000020679.6 |
| chr11 | 83851001 | 83853569 | 3.83683 | 4.9536 | 1.74006 | Promoter | 11 | 83852242 | 83864178 | 11937 | 1 | ENSMUST00000146786.2 | 0 | Hnf1b | ENSMUSG00000020679.6 |
| chr11 | 83854177 | 83855608 | 3.98121 | 5.09954 | 1.8328 | Intron (ENSMUST00000135975.2/ENSMUSG00000020679.6, intron 2 of 2) | 11 | 83852960 | 83905819 | 52860 | 1 | ENSMUST00000108113.1 | 1218 | Hnf1b | ENSMUSG00000020679.6 |
| chr11 | 83856092 | 83856467 | 3.82777 | 4.79009 | 1.67912 | Exon (ENSMUST00000135975.2/ENSMUSG00000020679.6, exon 3 of 3) | 11 | 83852960 | 83905819 | 52860 | 1 | ENSMUST00000108113.1 | 3133 | Hnf1b | ENSMUSG00000020679.6 |
| chr11 | 83857285 | 83857496 | 3.56956 | 4.25181 | 1.24876 | Intron (ENSMUST00000021016.4/ENSMUSG00000020679.6, intron 2 of 8) | 11 | 83852960 | 83905819 | 52860 | 1 | ENSMUST00000108113.1 | 4326 | Hnf1b | ENSMUSG00000020679.6 |
| chr11 | 83858525 | 83858954 | 3.42223 | 3.98589 | 1.10003 | Intron (ENSMUST00000021016.4/ENSMUSG00000020679.6, intron 2 of 8) | 11 | 83852960 | 83905819 | 52860 | 1 | ENSMUST00000108113.1 | 5566 | Hnf1b | ENSMUSG00000020679.6 |
| chr11 | 83859666 | 83859872 | 3.38136 | 3.89974 | 1.01733 | Intron (ENSMUST00000021016.4/ENSMUSG00000020679.6, intron 2 of 8) | 11 | 83852960 | 83905819 | 52860 | 1 | ENSMUST00000108113.1 | 6707 | Hnf1b | ENSMUSG00000020679.6 |
| chr11 | 105968524 | 105968694 | 3.57762 | 4.37966 | 1.39562 | Promoter | 11 | 105967945 | 105989964 | 22020 | 1 | ENSMUST00000001963.8 | 580 | Ace | ENSMUSG00000020681.9 |
| chr11 | 83461674 | 83461869 | 3.69114 | 4.70539 | 1.57597 | Intron (ENSMUST00000103209.4/ENSMUSG00000020682.12, intron 1 of 7) | 11 | 83441876 | 83462961 | 21086 | 2 | ENSMUST00000021020.7 | 1092 | Mmp28 | ENSMUSG00000020682.12 |
| chr11 | 83462555 | 83462846 | 4.01987 | 5.48614 | 2.10885 | Promoter | 11 | 83441876 | 83462961 | 21086 | 2 | ENSMUST00000021020.7 | 115 | Mmp28 | ENSMUSG00000020682.12 |
| chr11 | 83463761 | 83464134 | 3.33351 | 3.97857 | 1.10266 | Promoter | 11 | 83451386 | 83463071 | 11686 | 2 | ENSMUST00000138780.1 | -691 | Mmp28 | ENSMUSG00000020682.12 |
| chr11 | 83408814 | 83410181 | 4.20888 | 5.59014 | 2.24927 | Promoter | 11 | 83409137 | 83418834 | 9698 | 1 | ENSMUST00000021022.4 | 0 | Rasl10b | ENSMUSG00000020684.9 |
| chr11 | 83418372 | 83418631 | 3.46864 | 4.25721 | 1.29872 | 3' UTR | 11 | 83410072 | 83421038 | 10967 | 1 | ENSMUST00000164425.1 | 8301 | Rasl10b | ENSMUSG00000020684.9 |
| chr11 | 83419161 | 83419558 | 3.61023 | 4.40044 | 1.43197 | 3' UTR | 11 | 83410072 | 83421038 | 10967 | 1 | ENSMUST00000164425.1 | 9090 | Rasl10b | ENSMUSG00000020684.9 |
| chr11 | 83420777 | 83421101 | 3.42715 | 4.02894 | 1.21476 | 3' UTR | 11 | 83421902 | 83429455 | 7554 | 2 | ENSMUST00000052521.3 | 8354 | Gas2l2 | ENSMUSG00000020686.7 |

| | | | | | | | | | | | | | | | |
|---|---|---|---|---|---|---|---|---|---|---|---|---|---|---|---|
| chr11 | 83427349 | 83427492 | 3.49549 | 4.13901 | 1.24816 | Exon (ENSMUST00000052521.3/ENSMUSG00000020686.7, exon 2 of 6) | 11 | 83421902 | 83429455 | 7554 | 2 | ENSMUST00000052521.3 | 1963 | Gas2l2 | ENSMUSG00000020686.7 |
| chr11 | 83429732 | 83430290 | 3.55591 | 4.26857 | 1.28508 | Promoter | 11 | 83421902 | 83429455 | 7554 | 2 | ENSMUST00000147301.1 | -278 | Gas2l2 | ENSMUSG00000020686.7 |
| chr11 | 104517408 | 104517512 | 6.21633 | 11.83965 | 7.43285 | Exon (ENSMUST00000093923.3/ENSMUSG00000020687.8, exon 13 of 19) | 11 | 104508347 | 104513276 | 4930 | 2 | ENSMUST00000127506.1 | -4133 | Cdc27 | ENSMUSG00000020687.8 |
| chr11 | 104521501 | 104521587 | 6.33192 | 10.66604 | 6.38483 | Exon (ENSMUST00000093923.3/ENSMUSG00000020687.8, exon 11 of 19) | 11 | 104508347 | 104513276 | 4930 | 2 | ENSMUST00000127506.1 | -8226 | Cdc27 | ENSMUSG00000020687.8 |
| chr11 | 104534697 | 104534861 | 3.82326 | 5.21572 | 1.95411 | Exon (ENSMUST00000093923.3/ENSMUSG00000020687.8, exon 3 of 19) | 11 | 104504937 | 104550392 | 45456 | 2 | ENSMUST00000106962.3 | 15531 | Cdc27 | ENSMUSG00000020687.8 |
| chr11 | 104550330 | 104550503 | 4.40344 | 6.13397 | 2.56972 | Promoter | 11 | 104502745 | 104550483 | 47739 | 2 | ENSMUST00000093923.3 | 0 | Cdc27 | ENSMUSG00000020687.8 |
| chr11 | 104608410 | 104608485 | 4.75123 | 6.74283 | 3.06677 | Promoter | 11 | 104608000 | 104670476 | 62477 | 1 | ENSMUST00000021028.4 | 411 | Itgb3 | ENSMUSG00000020689.4 |
| chr11 | 82347796 | 82348042 | 3.37276 | 3.88676 | 1.01733 | Distal Intergenic | 11 | 82388900 | 82446327 | 57428 | 1 | ENSMUST00000092852.2 | -40858 | Tmem132e | ENSMUSG00000020701.7 |
| chr11 | 82353906 | 82354555 | 3.72523 | 4.55368 | 1.45665 | Distal Intergenic | 11 | 82388900 | 82446327 | 57428 | 1 | ENSMUST00000092852.2 | -34345 | Tmem132e | ENSMUSG00000020701.7 |
| chr11 | 82377156 | 82377303 | 3.29783 | 3.80229 | 1.09753 | Distal Intergenic | 11 | 82388900 | 82446327 | 57428 | 1 | ENSMUST00000092852.2 | -11597 | Tmem132e | ENSMUSG00000020701.7 |
| chr11 | 82383875 | 82384036 | 3.50057 | 4.11957 | 1.15784 | Distal Intergenic | 11 | 82388900 | 82446327 | 57428 | 1 | ENSMUST00000092852.2 | -4864 | Tmem132e | ENSMUSG00000020701.7 |
| chr11 | 82385026 | 82385101 | 3.55961 | 4.46943 | 1.42911 | Distal Intergenic | 11 | 82388900 | 82446327 | 57428 | 1 | ENSMUST00000092852.2 | -3799 | Tmem132e | ENSMUSG00000020701.7 |
| chr11 | 82387160 | 82387518 | 3.8354 | 4.7565 | 1.58232 | Distal Intergenic | 11 | 82388900 | 82446327 | 57428 | 1 | ENSMUST00000092852.2 | -1382 | Tmem132e | ENSMUSG00000020701.7 |
| chr11 | 82388231 | 82391560 | 4.03112 | 5.3141 | 2.01831 | Promoter | 11 | 82388900 | 82446327 | 57428 | 1 | ENSMUST00000092852.2 | 0 | Tmem132e | ENSMUSG00000020701.7 |
| chr11 | 82397093 | 82397168 | 4.00903 | 5.15007 | 1.89171 | Intron (ENSMUST00000092852.2/ENSMUSG00000020701.7, intron 1 of 9) | 11 | 82389123 | 82434723 | 45601 | 1 | ENSMUST00000132477.1 | 7971 | Tmem132e | ENSMUSG00000020701.7 |
| chr11 | 82433218 | 82433436 | 3.38193 | 3.9006 | 1.01733 | Intron (ENSMUST00000092852.2/ENSMUSG00000020701.7, intron 1 of 9) | 11 | 82389123 | 82434723 | 45601 | 1 | ENSMUST00000132477.1 | 44096 | Tmem132e | ENSMUSG00000020701.7 |
| chr11 | 82434297 | 82434542 | 3.74452 | 4.62682 | 1.57038 | Exon (ENSMUST00000092852.2/ENSMUSG00000020701.7, exon 2 of 10) | 11 | 82389123 | 82434723 | 45601 | 1 | ENSMUST00000132477.1 | 45175 | Tmem132e | ENSMUSG00000020701.7 |
| chr11 | 82435030 | 82435212 | 3.54902 | 4.25902 | 1.33756 | Exon (ENSMUST00000092852.2/ENSMUSG00000020701.7, exon 3 of 10) | 11 | 82389123 | 82434723 | 45601 | 1 | ENSMUST00000132477.1 | 45908 | Tmem132e | ENSMUSG00000020701.7 |
| chr11 | 82437013 | 82437412 | 3.3175 | 3.83077 | 1.10334 | Exon (ENSMUST00000092852.2/ENSMUSG00000020701.7, exon 4 of 10) | 11 | 82389123 | 82434723 | 45601 | 1 | ENSMUST00000132477.1 | 47891 | Tmem132e | ENSMUSG00000020701.7 |
| chr11 | 82473668 | 82473871 | 3.48659 | 4.12553 | 1.24934 | Distal Intergenic | 11 | 82389123 | 82434723 | 45601 | 1 | ENSMUST00000132477.1 | 84546 | Tmem132e | ENSMUSG00000020701.7 |
| chr11 | 82190073 | 82190252 | 3.48844 | 4.11286 | 1.19791 | Intron (ENSMUST00000138333.1/ENSMUSG00000020702.8, intron 9 of 10) | 11 | 82176659 | 82179812 | 3154 | 2 | ENSMUST00000021043.4 | -10262 | Ccl1 | ENSMUSG00000020702.8 |
| chr11 | 82260482 | 82260557 | 3.51721 | 4.14723 | 1.16683 | Exon (ENSMUST00000138333.1/ENSMUSG00000020702.8, exon 7 of 11) | 11 | 82176657 | 82305690 | 129034 | 2 | ENSMUST00000138333.1 | 45133 | Ccl1 | ENSMUSG00000020702.8 |
| chr11 | 82261110 | 82261445 | 3.59292 | 4.28802 | 1.26056 | Intron (ENSMUST00000138333.1/ENSMUSG00000020702.8, intron 6 of 10) | 11 | 82176657 | 82305690 | 129034 | 2 | ENSMUST00000138333.1 | 44245 | Ccl1 | ENSMUSG00000020702.8 |
| chr11 | 82327781 | 82327986 | 3.60055 | 4.4418 | 1.39939 | Distal Intergenic | 11 | 82176657 | 82305690 | 129034 | 2 | ENSMUST00000138333.1 | -22092 | Ccl1 | ENSMUSG00000020702.8 |
| chr11 | 82329644 | 82329736 | 3.49766 | 4.11965 | 1.17276 | Distal Intergenic | 11 | 82176657 | 82305690 | 129034 | 2 | ENSMUST00000138333.1 | -23955 | Ccl1 | ENSMUSG00000020702.8 |
| chr11 | 82344988 | 82345332 | 3.64574 | 4.43341 | 1.45794 | Distal Intergenic | 11 | 82176657 | 82305690 | 129034 | 2 | ENSMUST00000138333.1 | -39299 | Ccl1 | ENSMUSG00000020702.8 |
| chr11 | 106659465 | 106659540 | 3.35029 | 3.85286 | 1.01733 | Intron (ENSMUST00000103069.4/ENSMUSG00000020717.14, intron 13 of 13) | 11 | 106661185 | 106663326 | 2142 | 2 | ENSMUST00000184424.1 | 3786 | Pecam1 | ENSMUSG00000020717.14 |
| chr11 | 106659872 | 106660017 | 3.33135 | 3.82434 | 1.01733 | Intron (ENSMUST00000103069.4/ENSMUSG00000020717.14, intron 13 of 13) | 11 | 106661185 | 106663326 | 2142 | 2 | ENSMUST00000184424.1 | 3309 | Pecam1 | ENSMUSG00000020717.14 |
| chr11 | 106663897 | 106664047 | 3.24456 | 3.69401 | 1.01733 | Promoter | 11 | 106661185 | 106663326 | 2142 | 2 | ENSMUST00000184424.1 | -572 | Pecam1 | ENSMUSG00000020717.14 |
| chr11 | 106673557 | 106674033 | 3.49235 | 4.45059 | 1.45403 | Intron (ENSMUST00000103069.4/ENSMUSG00000020717.14, intron 13 of 13) | 11 | 106661185 | 106663326 | 2142 | 2 | ENSMUST00000184424.1 | -10232 | Pecam1 | ENSMUSG00000020717.14 |
| chr11 | 106674772 | 106674911 | 3.44856 | 4.18835 | 1.21836 | Intron (ENSMUST00000103069.4/ENSMUSG00000020717.14, intron 13 of 13) | 11 | 106661185 | 106663326 | 2142 | 2 | ENSMUST00000184424.1 | -11447 | Pecam1 | ENSMUSG00000020717.14 |
| chr11 | 106684911 | 106685218 | 3.85368 | 5.10105 | 1.84634 | Exon (ENSMUST00000103069.4/ENSMUSG00000020717.14, exon 8 of 14) | 11 | 106688783 | 106691347 | 2565 | 2 | ENSMUST00000135481.1 | 6129 | Pecam1 | ENSMUSG00000020717.14 |
| chr11 | 106690037 | 106690417 | 3.40236 | 4.03051 | 1.18966 | Promoter | 11 | 106688783 | 106691347 | 2565 | 2 | ENSMUST00000135481.1 | 930 | Pecam1 | ENSMUSG00000020717.14 |
| chr11 | 106691611 | 106691901 | 3.41175 | 4.00054 | 1.13277 | Promoter | 11 | 106688783 | 106691347 | 2565 | 2 | ENSMUST00000135481.1 | -265 | Pecam1 | ENSMUSG00000020717.14 |
| chr11 | 106693064 | 106693345 | 3.99422 | 5.1003 | 1.83231 | Intron (ENSMUST00000103069.4/ENSMUSG00000020717.14, intron 5 of 13) | 11 | 106688783 | 106691347 | 2565 | 2 | ENSMUST00000135481.1 | -1718 | Pecam1 | ENSMUSG00000020717.14 |
| chr11 | 106695017 | 106695352 | 3.20664 | 3.95884 | 1.08347 | Intron (ENSMUST00000103069.4/ENSMUSG00000020717.14, intron 5 of 13) | 11 | 106688783 | 106691347 | 2565 | 2 | ENSMUST00000135481.1 | -3671 | Pecam1 | ENSMUSG00000020717.14 |
| chr11 | 106696116 | 106696641 | 3.73414 | 4.58359 | 1.47301 | Intron (ENSMUST00000103069.4/ENSMUSG00000020717.14, intron 4 of 13) | 11 | 106688783 | 106691347 | 2565 | 2 | ENSMUST00000135481.1 | -4770 | Pecam1 | ENSMUSG00000020717.14 |
| chr11 | 106697411 | 106697587 | 3.68045 | 4.52986 | 1.49211 | Intron (ENSMUST00000103069.4/ENSMUSG00000020717.14, intron 3 of 13) | 11 | 106688783 | 106691347 | 2565 | 2 | ENSMUST00000135481.1 | -6065 | Pecam1 | ENSMUSG00000020717.14 |
| chr11 | 106702446 | 106702676 | 3.66416 | 4.42323 | 1.34914 | Intron (ENSMUST00000103069.4/ENSMUSG00000020717.14, intron 2 of 13) | 11 | 106688783 | 106691347 | 2565 | 2 | ENSMUST00000135481.1 | -11100 | Pecam1 | ENSMUSG00000020717.14 |
| chr11 | 106715300 | 106715490 | 3.70788 | 4.64035 | 1.52306 | Promoter | 11 | 106654218 | 106715394 | 61177 | 2 | ENSMUST00000106796.3 | 0 | Pecam1 | ENSMUSG00000020717.14 |
| chr11 | 106719709 | 106720072 | 3.68038 | 4.50622 | 1.51013 | Intron (ENSMUST00000068021.3/ENSMUSG00000020717.14, intron 1 of 15) | 11 | 106654218 | 106715394 | 61177 | 2 | ENSMUST00000106796.3 | -4316 | Pecam1 | ENSMUSG00000020717.14 |
| chr11 | 106723774 | 106724084 | 3.65466 | 4.45922 | 1.47678 | Intron (ENSMUST00000068021.3/ENSMUSG00000020717.14, intron 1 of 15) | 11 | 106654218 | 106715394 | 61177 | 2 | ENSMUST00000106796.3 | -8381 | Pecam1 | ENSMUSG00000020717.14 |
| chr11 | 106724694 | 106725056 | 3.7077 | 4.72193 | 1.59549 | Intron (ENSMUST00000068021.3/ENSMUSG00000020717.14, intron 1 of 15) | 11 | 106654218 | 106715394 | 61177 | 2 | ENSMUST00000106796.3 | -9301 | Pecam1 | ENSMUSG00000020717.14 |
| chr11 | 106725558 | 106725848 | 3.40165 | 4.15891 | 1.23397 | Intron (ENSMUST00000068021.3/ENSMUSG00000020717.14, intron 1 of 15) | 11 | 106654218 | 106715394 | 61177 | 2 | ENSMUST00000106796.3 | -10165 | Pecam1 | ENSMUSG00000020717.14 |
| chr11 | 106726251 | 106726755 | 3.6258 | 4.39027 | 1.41555 | Intron (ENSMUST00000068021.3/ENSMUSG00000020717.14, intron 1 of 15) | 11 | 106654218 | 106715394 | 61177 | 2 | ENSMUST00000106796.3 | -10858 | Pecam1 | ENSMUSG00000020717.14 |
| chr11 | 106727063 | 106727362 | 3.3963 | 3.96217 | 1.15012 | Intron (ENSMUST00000068021.3/ENSMUSG00000020717.14, intron 1 of 15) | 11 | 106654218 | 106715394 | 61177 | 2 | ENSMUST00000106796.3 | -11670 | Pecam1 | ENSMUSG00000020717.14 |
| chr11 | 106733128 | 106733592 | 3.4172 | 3.98144 | 1.115 | Intron (ENSMUST00000068021.3/ENSMUSG00000020717.14, intron 1 of 15) | 11 | 106661797 | 106750628 | 88832 | 2 | ENSMUST00000068021.3 | 17036 | Pecam1 | ENSMUSG00000020717.14 |
| chr11 | 106734073 | 106734636 | 3.6169 | 4.36259 | 1.37608 | Intron (ENSMUST00000068021.3/ENSMUSG00000020717.14, intron 1 of 15) | 11 | 106661797 | 106750628 | 88832 | 2 | ENSMUST00000068021.3 | 15992 | Pecam1 | ENSMUSG00000020717.14 |
| chr11 | 107794821 | 107794960 | 3.99107 | 5.04712 | 1.77259 | Promoter | 11 | 107732357 | 107794557 | 62201 | 2 | ENSMUST00000021066.3 | -265 | Cacng4 | ENSMUSG00000020723.3 |
| chr11 | 107798068 | 107798229 | 3.2775 | 3.74339 | 1.01733 | Distal Intergenic | 11 | 107732357 | 107794557 | 62201 | 2 | ENSMUST00000021066.3 | -3512 | Cacng4 | ENSMUSG00000020723.3 |
| chr11 | 107799646 | 107801072 | 3.82178 | 4.79946 | 1.69651 | Distal Intergenic | 11 | 107732357 | 107794557 | 62201 | 2 | ENSMUST00000021066.3 | -5090 | Cacng4 | ENSMUSG00000020723.3 |
| chr11 | 107810869 | 107810944 | 3.89956 | 4.91707 | 1.76326 | Distal Intergenic | 11 | 107732357 | 107794557 | 62201 | 2 | ENSMUST00000021066.3 | -16313 | Cacng4 | ENSMUSG00000020723.3 |
| chr11 | 115081725 | 115081932 | 3.35932 | 3.89803 | 1.11492 | Distal Intergenic | 11 | 115091431 | 115148647 | 57217 | 1 | ENSMUST00000131046.1 | -9499 | Rab37 | ENSMUSG00000020732.8 |
| chr11 | 115083616 | 115084495 | 3.4399 | 4.02397 | 1.14469 | Distal Intergenic | 11 | 115091431 | 115148647 | 57217 | 1 | ENSMUST00000131046.1 | -6936 | Rab37 | ENSMUSG00000020732.8 |
| chr11 | 115090740 | 115090959 | 3.33175 | 3.93044 | 1.147 | Promoter | 11 | 115091431 | 115148647 | 57217 | 1 | ENSMUST00000131046.1 | -472 | Rab37 | ENSMUSG00000020732.8 |
| chr11 | 115091347 | 115092032 | 3.78617 | 4.81653 | 1.66201 | Promoter | 11 | 115091431 | 115148647 | 57217 | 1 | ENSMUST00000131046.1 | 0 | Rab37 | ENSMUSG00000020732.8 |
| chr11 | 115092773 | 115093397 | 4.15996 | 5.47599 | 2.10203 | Intron (ENSMUST00000131046.1/ENSMUSG00000020732.8, intron 1 of 1) | 11 | 115091431 | 115162236 | 70806 | 1 | ENSMUST00000067754.5 | 1343 | Rab37 | ENSMUSG00000020732.8 |
| chr11 | 115153883 | 115154189 | 3.53046 | 4.17454 | 1.19328 | Promoter | 11 | 115154157 | 115162236 | 8080 | 1 | ENSMUST00000021076.5 | 0 | Rab37 | ENSMUSG00000020732.8 |
| chr11 | 115261972 | 115262219 | 3.49737 | 4.31944 | 1.31359 | Intron (ENSMUST00000003351.7/ENSMUSG00000020734.8, intron 1 of 12) | 11 | 115249169 | 115265429 | 16261 | 2 | ENSMUST00000003351.7 | 3210 | Grin2c | ENSMUSG00000020734.8 |
| chr11 | 115265383 | 115266258 | 3.66007 | 4.58016 | 1.50713 | Promoter | 11 | 115249169 | 115265429 | 16261 | 2 | ENSMUST00000003351.7 | 0 | Grin2c | ENSMUSG00000020734.8 |
| chr11 | 115266690 | 115267329 | 3.83332 | 5.02177 | 1.85266 | Promoter | 11 | 115249169 | 115267243 | 18075 | 2 | ENSMUST00000106554.1 | 0 | Grin2c | ENSMUSG00000020734.8 |
| chr11 | 115748116 | 115748445 | 3.44099 | 4.05124 | 1.2204 | Distal Intergenic | 11 | 115765433 | 115799366 | 33934 | 1 | ENSMUST00000093912.5 | -16988 | 2310067B10Rik | ENSMUSG00000020747.12 |
| chr11 | 115974183 | 115974780 | 3.52844 | 4.21521 | 1.23686 | Promoter | 11 | 115974709 | 116008412 | 33704 | 1 | ENSMUST00000021107.8 | 0 | Itgb4 | ENSMUSG00000020758.10 |
| chr11 | 115989759 | 115989835 | 3.65748 | 4.42308 | 1.37587 | Promoter | 11 | 115989938 | 115991050 | 1113 | 1 | ENSMUST00000150129.1 | -103 | Itgb4 | ENSMUSG00000020758.10 |
| chr11 | 72961230 | 72961762 | 4.6556 | 6.52161 | 2.91871 | Promoter | 11 | 72961290 | 72970760 | 9471 | 1 | ENSMUST00000149493.1 | 0 | Atp2a3 | ENSMUSG00000020788.10 |
| chr11 | 116280422 | 116282770 | 3.7171 | 4.57239 | 1.46634 | Promoter | 11 | 116280939 | 116283938 | 3000 | 1 | ENSMUST00000055872.2 | 0 | Galr2 | ENSMUSG00000020793.5 |
| chr11 | 116283240 | 116283883 | 3.68674 | 4.47309 | 1.38154 | 3' UTR | 11 | 116281486 | 116283661 | 2176 | 1 | ENSMUST00000154277.1 | 1755 | Galr2 | ENSMUSG00000020793.5 |
| chr11 | 116284582 | 116284673 | 3.35293 | 4.11451 | 1.20031 | Distal Intergenic | 11 | 116281486 | 116283661 | 2176 | 1 | ENSMUST00000154277.1 | 3097 | Galr2 | ENSMUSG00000020793.5 |
| chr11 | 72361689 | 72362665 | 3.51063 | 4.17445 | 1.23948 | Promoter | 11 | 72344722 | 72361873 | 17152 | 2 | ENSMUST00000108503.3 | 0 | Tekt1 | ENSMUSG00000020799.11 |
| chr11 | 72267411 | 72267695 | 3.20871 | 3.64033 | 1.00617 | Promoter | 11 | 72261637 | 72267222 | 5586 | 2 | ENSMUST00000137701.1 | -190 | Slc13a5 | ENSMUSG00000020805.8 |
| chr11 | 116654228 | 116654327 | 3.34153 | 3.83966 | 1.01733 | Promoter | 11 | 116645595 | 116654313 | 8719 | 2 | ENSMUST00000021166.5 | 0 | Cygb | ENSMUSG00000020810.5 |
| chr11 | 71751233 | 71751484 | 3.6587 | 4.4784 | 1.46628 | Promoter | 11 | 71751376 | 71760968 | 9593 | 1 | ENSMUST00000138339.2 | 0 | Wscd1 | ENSMUSG00000020811.11 |
| chr11 | 71752050 | 71752212 | 3.27157 | 3.74111 | 1.01898 | Promoter | 11 | 71751376 | 71789647 | 38272 | 1 | ENSMUST00000108510.1 | 675 | Wscd1 | ENSMUSG00000020811.11 |
| chr11 | 116885957 | 116886179 | 3.57277 | 4.26948 | 1.29123 | Distal Intergenic | 11 | 116868368 | 116873837 | 5470 | 1 | ENSMUST00000134106.1 | 17590 | Mfsd11 | ENSMUSG00000020818.11 |
| chr11 | 78924844 | 78925813 | 3.8977 | 4.94911 | 1.73923 | Intron (ENSMUST00000018610.6/ENSMUSG00000020826.6, intron 2 of 26) | 11 | 78920787 | 78960254 | 39468 | 1 | ENSMUST00000018610.6 | 4058 | Nos2 | ENSMUSG00000020826.6 |
| chr11 | 78927126 | 78927223 | 3.44281 | 4.17174 | 1.22572 | Intron (ENSMUST00000018610.6/ENSMUSG00000020826.6, intron 2 of 26) | 11 | 78920787 | 78960254 | 39468 | 1 | ENSMUST00000018610.6 | 6340 | Nos2 | ENSMUSG00000020826.6 |
| chr11 | 77460877 | 77461044 | 3.62645 | 4.53429 | 1.44022 | Distal Intergenic | 11 | 77462411 | 77470484 | 8074 | 1 | ENSMUST00000102493.2 | -1367 | Coro6 | ENSMUSG00000020836.10 |
| chr11 | 77466838 | 77467011 | 3.65455 | 4.46671 | 1.47882 | Intron (ENSMUST00000102493.2/ENSMUSG00000020836.10, intron 5 of 10) | 11 | 77463911 | 77469501 | 5591 | 1 | ENSMUST00000079770.2 | 2928 | Coro6 | ENSMUSG00000020836.10 |
| chr11 | 76996783 | 76997012 | 3.37688 | 3.89298 | 1.01733 | Distal Intergenic | 11 | 76998603 | 77032339 | 33737 | 1 | ENSMUST00000108402.3 | -1591 | Slc6a4 | ENSMUSG00000020838.7 |
| chr11 | 76997690 | 76998043 | 3.56422 | 4.2331 | 1.21936 | Promoter | 11 | 76998603 | 77032339 | 33737 | 1 | ENSMUST00000108402.3 | -560 | Slc6a4 | ENSMUSG00000020838.7 |
| chr11 | 76998719 | 76999234 | 3.9301 | 5.20436 | 1.94931 | Promoter | 11 | 76998616 | 77032340 | 33725 | 1 | ENSMUST00000021195.5 | 104 | Slc6a4 | ENSMUSG00000020838.7 |
| chr11 | 77000461 | 77001324 | 3.50173 | 4.33185 | 1.3563 | Intron (ENSMUST00000108402.3/ENSMUSG00000020838.7, intron 1 of 14) | 11 | 76998616 | 77032340 | 33725 | 1 | ENSMUST00000021195.5 | 1846 | Slc6a4 | ENSMUSG00000020838.7 |
| chr11 | 77005454 | 77005701 | 3.79744 | 4.82686 | 1.65988 | Intron (ENSMUST00000108402.3/ENSMUSG00000020838.7, intron 2 of 14) | 11 | 77009274 | 77013053 | 3780 | 1 | ENSMUST00000129572.2 | -3573 | Slc6a4 | ENSMUSG00000020838.7 |
| chr11 | 77006547 | 77006922 | 3.64805 | 4.43835 | 1.43542 | Intron (ENSMUST00000108402.3/ENSMUSG00000020838.7, intron 2 of 14) | 11 | 77009274 | 77013053 | 3780 | 1 | ENSMUST00000129572.2 | -2352 | Slc6a4 | ENSMUSG00000020838.7 |
| chr11 | 77007281 | 77007494 | 3.95938 | 5.00151 | 1.77182 | Intron (ENSMUST00000108402.3/ENSMUSG00000020838.7, intron 2 of 14) | 11 | 77009274 | 77013053 | 3780 | 1 | ENSMUST00000129572.2 | -1780 | Slc6a4 | ENSMUSG00000020838.7 |
| chr11 | 75889450 | 75889712 | 3.35596 | 3.86142 | 1.01733 | Intron (ENSMUST00000121287.2/ENSMUSG00000020847.10, intron 4 of 8) | 11 | 75832137 | 75911062 | 78926 | 2 | ENSMUST00000066504.5 | 21350 | Rph3al | ENSMUSG00000020847.10 |
| chr11 | 75925628 | 75926112 | 3.62249 | 4.53587 | 1.46757 | Promoter | 11 | 75899726 | 75925891 | 26166 | 2 | ENSMUST00000021208.5 | 0 | Rph3al | ENSMUSG00000020847.10 |
| chr11 | 75965759 | 75966145 | 3.71837 | 4.57862 | 1.52988 | Distal Intergenic | 11 | 75906572 | 75938429 | 31858 | 2 | ENSMUST00000152035.1 | -27331 | Rph3al | ENSMUSG00000020847.10 |
| chr11 | 75794918 | 75796710 | 3.96583 | 5.18132 | 1.90263 | Promoter | 11 | 75768966 | 75796049 | 27084 | 2 | ENSMUST00000021209.7 | 0 | Doc2b | ENSMUSG00000020848.7 |
| chr11 | 75797570 | 75797847 | 3.39343 | 3.95775 | 1.15236 | Intron (ENSMUST00000127424.2/ENSMUSG00000087059.2, intron 2 of 4) | 11 | 75768966 | 75796049 | 27084 | 2 | ENSMUST00000021209.7 | -1522 | Doc2b | ENSMUSG00000020848.7 |
| chr11 | 94392715 | 94392909 | 3.37032 | 3.90664 | 1.10076 | Promoter | 11 | 94364855 | 94392920 | 28066 | 2 | ENSMUST00000151124.2 | 11 | Abcc3 | ENSMUSG00000020865.11 |
| chr11 | 94473055 | 94473992 | 3.68585 | 4.54609 | 1.46816 | Promoter | 11 | 94426952 | 94473641 | 46690 | 2 | ENSMUST00000152811.1 | 0 | Cacna1g | ENSMUSG00000020866.12 |
| chr11 | 95145889 | 95147027 | 3.80614 | 4.84313 | 1.70152 | Promoter | 11 | 95140447 | 95146263 | 5817 | 2 | ENSMUST00000021241.7 | 0 | Dlx4 | ENSMUSG00000020871.8 |
| chr11 | 95147896 | 95148169 | 3.82908 | 4.84974 | 1.69168 | Intron (ENSMUST00000156477.3/ENSMUSG00000086552.3, intron 1 of 2) | 11 | 95140447 | 95146263 | 5817 | 2 | ENSMUST00000021241.7 | -1634 | Dlx4 | ENSMUSG00000020871.8 |
| chr11 | 95148645 | 95150928 | 4.01942 | 5.18189 | 1.92989 | Intron (ENSMUST00000156477.3/ENSMUSG00000086552.3, intron 1 of 2) | 11 | 95140447 | 95146263 | 5817 | 2 | ENSMUST00000021241.7 | -2383 | Dlx4 | ENSMUSG00000020871.8 |
| chr11 | 95151290 | 95151616 | 3.30071 | 3.77824 | 1.01733 | Intron (ENSMUST00000156477.3/ENSMUSG00000086552.3, intron 1 of 2) | 11 | 95140447 | 95146263 | 5817 | 2 | ENSMUST00000021241.7 | -5028 | Dlx4 | ENSMUSG00000020871.8 |
| chr11 | 95152051 | 95152424 | 3.65073 | 4.41208 | 1.38415 | Intron (ENSMUST00000156477.3/ENSMUSG00000086552.3, intron 1 of 2) | 11 | 95140447 | 95146263 | 5817 | 2 | ENSMUST00000021241.7 | -5789 | Dlx4 | ENSMUSG00000020871.8 |
| chr11 | 96268778 | 96269011 | 3.3945 | 3.98828 | 1.18909 | Distal Intergenic | 11 | 96271457 | 96276595 | 5139 | 1 | ENSMUST00000000010.8 | -2446 | Hoxb9 | ENSMUSG00000020875.9 |
| chr11 | 96269781 | 96270107 | 3.43966 | 4.23758 | 1.26488 | Distal Intergenic | 11 | 96271457 | 96276595 | 5139 | 1 | ENSMUST00000000010.8 | -1350 | Hoxb9 | ENSMUSG00000020875.9 |
| chr11 | 96271528 | 96271766 | 3.40457 | 4.06874 | 1.17042 | Promoter | 11 | 96271523 | 96276595 | 5073 | 1 | ENSMUST00000174042.2 | 6 | Hoxb9 | ENSMUSG00000020875.9 |
| chr11 | 96272438 | 96272707 | 3.82294 | 4.80949 | 1.69316 | Promoter | 11 | 96271523 | 96276595 | 5073 | 1 | ENSMUST00000174042.2 | 916 | Hoxb9 | ENSMUSG00000020875.9 |
| chr11 | 96275590 | 96276069 | 4.01948 | 5.15566 | 1.90404 | 3' UTR | 11 | 96271523 | 96276595 | 5073 | 1 | ENSMUST00000174042.2 | 4068 | Hoxb9 | ENSMUSG00000020875.9 |
| chr11 | 70045762 | 70046018 | 3.54845 | 4.2099 | 1.22181 | Distal Intergenic | 11 | 70044197 | 70044718 | 522 | 1 | ENSMUST00000131497.1 | 1566 | Dlg4 | ENSMUSG00000020886.11 |
| chr8 | 70862216 | 70863692 | 3.87757 | 4.90341 | 1.77164 | Promoter | 8 | 70863127 | 70865428 | 2302 | 1 | ENSMUST00000050921.2 | 0 | A230052G05Rik | ENSMUSG00000020887.3 |
| chr8 | 70864054 | 70864130 | 3.73673 | 4.6359 | 1.57092 | Promoter | 8 | 70863127 | 70865428 | 2302 | 1 | ENSMUST00000050921.2 | 928 | A230052G05Rik | ENSMUSG00000020887.3 |
| chr11 | 69236245 | 69236495 | 3.52493 | 4.44162 | 1.44249 | Promoter | 11 | 69218117 | 69237022 | 18906 | 2 | ENSMUST00000021259.3 | 527 | Gucy2e | ENSMUSG00000020890.6 |
| chr11 | 69237110 | 69237185 | 3.63945 | 4.53797 | 1.45534 | Promoter | 11 | 69221158 | 69237036 | 15879 | 2 | ENSMUST00000108664.1 | -75 | Gucy2e | ENSMUSG00000020890.6 |
| chr11 | 68451317 | 68451574 | 3.36457 | 3.8744 | 1.01733 | Intron (ENSMUST00000021283.7/ENSMUSG00000020901.8, intron 1 of 18) | 11 | 68432160 | 68440372 | 8213 | 1 | ENSMUST00000141716.1 | 19158 | Pik3r5 | ENSMUSG00000020901.8 |
| chr11 | 68379401 | 68379495 | 3.23338 | 3.67724 | 1.01733 | Intron (ENSMUST00000021284.3/ENSMUSG00000020902.7, intron 2 of 6) | 11 | 68209364 | 68386826 | 177463 | 2 | ENSMUST00000021284.3 | 7331 | Ntn1 | ENSMUSG00000020902.7 |
| chr11 | 68380126 | 68380620 | 3.54129 | 4.26796 | 1.31647 | Intron (ENSMUST00000021284.3/ENSMUSG00000020902.7, intron 2 of 6) | 11 | 68209364 | 68386826 | 177463 | 2 | ENSMUST00000021284.3 | 6206 | Ntn1 | ENSMUSG00000020902.7 |
| chr11 | 68383320 | 68383722 | 3.722 | 4.64658 | 1.59582 | Intron (ENSMUST00000021284.3/ENSMUSG00000020902.7, intron 2 of 6) | 11 | 68209364 | 68386826 | 177463 | 2 | ENSMUST00000021284.3 | 3104 | Ntn1 | ENSMUSG00000020902.7 |
| chr11 | 68384205 | 68385142 | 3.88968 | 4.88769 | 1.72135 | Exon (ENSMUST00000021284.3/ENSMUSG00000020902.7, exon 2 of 7) | 11 | 68209364 | 68386826 | 177463 | 2 | ENSMUST00000021284.3 | 1684 | Ntn1 | ENSMUSG00000020902.7 |
| chr11 | 68385516 | 68386740 | 4.07126 | 5.33594 | 2.01959 | Promoter | 11 | 68209364 | 68386826 | 177463 | 2 | ENSMUST00000021284.3 | 86 | Ntn1 | ENSMUSG00000020902.7 |
| chr11 | 68388975 | 68389411 | 3.36456 | 3.89335 | 1.08204 | Intron (ENSMUST00000108674.3/ENSMUSG00000020902.7, intron 2 of 7) | 11 | 68385644 | 68387012 | 1369 | 2 | ENSMUST00000135141.1 | -1964 | Ntn1 | ENSMUSG00000020902.7 |
| chr11 | 68389805 | 68389934 | 3.89641 | 4.89868 | 1.72792 | Intron (ENSMUST00000108674.3/ENSMUSG00000020902.7, intron 2 of 7) | 11 | 68385644 | 68387012 | 1369 | 2 | ENSMUST00000135141.1 | -2794 | Ntn1 | ENSMUSG00000020902.7 |
| chr11 | 68390553 | 68391431 | 3.78774 | 4.71534 | 1.60481 | Intron (ENSMUST00000108674.3/ENSMUSG00000020902.7, intron 2 of 7) | 11 | 68385644 | 68387012 | 1369 | 2 | ENSMUST00000135141.1 | -3542 | Ntn1 | ENSMUSG00000020902.7 |
| chr11 | 68394370 | 68394956 | 3.38767 | 3.91259 | 1.02906 | Intron (ENSMUST00000108674.3/ENSMUSG00000020902.7, intron 2 of 7) | 11 | 68209364 | 68400823 | 191460 | 2 | ENSMUST00000108674.3 | 5867 | Ntn1 | ENSMUSG00000020902.7 |
| chr11 | 68396104 | 68396297 | 3.38022 | 4.14952 | 1.20723 | Intron (ENSMUST00000108674.3/ENSMUSG00000020902.7, intron 2 of 7) | 11 | 68209364 | 68400823 | 191460 | 2 | ENSMUST00000108674.3 | 4526 | Ntn1 | ENSMUSG00000020902.7 |
| chr11 | 68397112 | 68397492 | 3.46885 | 4.11127 | 1.26649 | Intron (ENSMUST00000108674.3/ENSMUSG00000020902.7, intron 2 of 7) | 11 | 68209364 | 68400823 | 191460 | 2 | ENSMUST00000108674.3 | 3331 | Ntn1 | ENSMUSG00000020902.7 |
| chr11 | 68397951 | 68398141 | 3.46965 | 4.08587 | 1.14359 | Intron (ENSMUST00000108674.3/ENSMUSG00000020902.7, intron 2 of 7) | 11 | 68209364 | 68400823 | 191460 | 2 | ENSMUST00000108674.3 | 2682 | Ntn1 | ENSMUSG00000020902.7 |
| chr11 | 68398610 | 68400310 | 3.73214 | 4.74691 | 1.65322 | Promoter | 11 | 68209364 | 68400823 | 191460 | 2 | ENSMUST00000108674.3 | 513 | Ntn1 | ENSMUSG00000020902.7 |
| chr11 | 68400973 | 68401809 | 3.95666 | 5.09135 | 1.89299 | Promoter | 11 | 68209364 | 68400823 | 191460 | 2 | ENSMUST00000108674.3 | -151 | Ntn1 | ENSMUSG00000020902.7 |
| chr11 | 68402181 | 68402316 | 5.02055 | 7.26007 | 3.51554 | Distal Intergenic | 11 | 68209364 | 68400823 | 191460 | 2 | ENSMUST00000108674.3 | -1359 | Ntn1 | ENSMUSG00000020902.7 |
| chr11 | 68402665 | 68402931 | 3.31261 | 3.79613 | 1.01733 | Distal Intergenic | 11 | 68209364 | 68400823 | 191460 | 2 | ENSMUST00000108674.3 | -1843 | Ntn1 | ENSMUSG00000020902.7 |
| chr11 | 68404455 | 68404619 | 3.27114 | 3.73385 | 1.01733 | Distal Intergenic | 11 | 68209364 | 68400823 | 191460 | 2 | ENSMUST00000108674.3 | -3633 | Ntn1 | ENSMUSG00000020902.7 |
| chr11 | 67084215 | 67084357 | 3.79858 | 4.81142 | 1.63679 | Intron (ENSMUST00000108689.2/ENSMUSG00000020908.9, intron 8 of 40) | 11 | 67078300 | 67102291 | 23992 | 1 | ENSMUST00000165221.1 | 5916 | Myh3 | ENSMUSG00000020908.9 |
| chr11 | 100141087 | 100141392 | 3.511 | 4.26642 | 1.30361 | Exon (ENSMUST00000007317.7/ENSMUSG00000020911.9, exon 5 of 6) | 11 | 100140907 | 100143699 | 2793 | 2 | ENSMUST00000126460.2 | 2307 | Krt19 | ENSMUSG00000020911.9 |
| chr11 | 100145902 | 100146023 | 3.40506 | 4.18848 | 1.25417 | Promoter | 11 | 100140810 | 100146120 | 5311 | 2 | ENSMUST00000007317.7 | 97 | Krt19 | ENSMUSG00000020911.9 |
| chr11 | 100146336 | 100146432 | 3.51823 | 4.31406 | 1.30319 | Promoter | 11 | 100140810 | 100146120 | 5311 | 2 | ENSMUST00000007317.7 | -217 | Krt19 | ENSMUSG00000020911.9 |
| chr11 | 100152375 | 100152731 | 3.58301 | 4.32433 | 1.32115 | Distal Intergenic | 11 | 100142120 | 100148665 | 6546 | 2 | ENSMUST00000125888.1 | -3711 | Krt19 | ENSMUSG00000020911.9 |
| chr11 | 100154259 | 100154382 | 3.69007 | 4.5314 | 1.4307 | Distal Intergenic | 11 | 100142120 | 100148665 | 6546 | 2 | ENSMUST00000125888.1 | -5595 | Krt19 | ENSMUSG00000020911.9 |
| chr11 | 99426294 | 99426679 | 3.33013 | 3.8225 | 1.01733 | Downstream (1-2kb) | 11 | 99415666 | 99422259 | 6594 | 2 | ENSMUST00000017741.3 | -4036 | Krt12 | ENSMUSG00000020912.4 |
| chr11 | 102708963 | 102709094 | 3.31367 | 3.79784 | 1.01733 | Distal Intergenic | 11 | 102684686 | 102697782 | 13097 | 2 | ENSMUST00000092569.7 | -11182 | Ccdc43 | ENSMUSG00000020925.11 |
| chr11 | 102761429 | 102762291 | 4.13263 | 5.35453 | 2.01195 | Promoter | 11 | 102761439 | 102780262 | 18824 | 1 | ENSMUST00000068150.5 | 0 | Adam11 | ENSMUSG00000020926.11 |
| chr11 | 102771850 | 102772043 | 3.48911 | 4.10443 | 1.16116 | Promoter | 11 | 102771482 | 102773661 | 2180 | 1 | ENSMUST00000126024.1 | 369 | Adam11 | ENSMUSG00000020926.11 |
| chr11 | 102772995 | 102773231 | 4.01947 | 5.09607 | 1.78782 | Promoter | 11 | 102774081 | 102776124 | 2044 | 1 | ENSMUST00000134296.2 | -850 | Adam11 | ENSMUSG00000020926.11 |
| chr11 | 102775724 | 102775960 | 3.36652 | 3.87735 | 1.01733 | Promoter | 11 | 102776554 | 102778467 | 1914 | 1 | ENSMUST00000135513.1 | -594 | Adam11 | ENSMUSG00000020926.11 |
| chr11 | 102833326 | 102833630 | 3.24664 | 3.69711 | 1.01733 | Distal Intergenic | 11 | 102835849 | 102838040 | 2192 | 1 | ENSMUST00000107073.2 | -2219 | Higd1b | ENSMUSG00000020928.9 |
| chr11 | 102988610 | 102988685 | 3.97871 | 5.06959 | 1.80744 | Intron (ENSMUST00000181125.1/ENSMUSG00000097410.1, intron 1 of 1) | 11 | 102994049 | 102997548 | 3500 | 2 | ENSMUST00000134335.1 | 8863 | Dcakd | ENSMUSG00000020935.3 |
| chr11 | 102989019 | 102989257 | 3.363 | 3.87202 | 1.01733 | Intron (ENSMUST00000181125.1/ENSMUSG00000097410.1, intron 1 of 1) | 11 | 102994049 | 102997548 | 3500 | 2 | ENSMUST00000134335.1 | 8291 | Dcakd | ENSMUSG00000020935.3 |
| chr11 | 102989711 | 102990183 | 3.98463 | 5.26579 | 1.97781 | Intron (ENSMUST00000181125.1/ENSMUSG00000097410.1, intron 1 of 1) | 11 | 102994049 | 102997548 | 3500 | 2 | ENSMUST00000134335.1 | 7365 | Dcakd | ENSMUSG00000020935.3 |
| chr11 | 102991391 | 102991911 | 3.97383 | 5.20703 | 1.92932 | Intron (ENSMUST00000181125.1/ENSMUSG00000097410.1, intron 1 of 1) | 11 | 102994049 | 102997548 | 3500 | 2 | ENSMUST00000134335.1 | 5637 | Dcakd | ENSMUSG00000020935.3 |
| chr11 | 102992421 | 102992657 | 3.8006 | 4.8477 | 1.6881 | Intron (ENSMUST00000181125.1/ENSMUSG00000097410.1, intron 1 of 1) | 11 | 102994049 | 102997548 | 3500 | 2 | ENSMUST00000134335.1 | 4891 | Dcakd | ENSMUSG00000020935.3 |
| chr11 | 103639895 | 103640427 | 3.51548 | 4.14857 | 1.18121 | Distal Intergenic | 11 | 103631073 | 103638890 | 7818 | 2 | ENSMUST00000021328.7 | -1006 | Lyzl6 | ENSMUSG00000020945.8 |
| chr11 | 103641222 | 103641392 | 3.48143 | 4.10453 | 1.16991 | Distal Intergenic | 11 | 103631073 | 103638890 | 7818 | 2 | ENSMUST00000021328.7 | -2333 | Lyzl6 | ENSMUSG00000020945.8 |
| chr11 | 103643927 | 103644271 | 3.38193 | 3.9006 | 1.01733 | Distal Intergenic | 11 | 103631073 | 103638890 | 7818 | 2 | ENSMUST00000021328.7 | -5038 | Lyzl6 | ENSMUSG00000020945.8 |
| chr12 | 49370850 | 49371105 | 3.65752 | 4.46425 | 1.46291 | Distal Intergenic | 12 | 49382693 | 49384364 | 1672 | 1 | ENSMUST00000135006.1 | -11588 | Foxg1 | ENSMUSG00000020950.9 |
| chr12 | 49371656 | 49371783 | 3.62947 | 4.42155 | 1.42581 | Distal Intergenic | 12 | 49382693 | 49384364 | 1672 | 1 | ENSMUST00000135006.1 | -10910 | Foxg1 | ENSMUSG00000020950.9 |
| chr12 | 49373736 | 49373881 | 3.54641 | 4.35578 | 1.37969 | Distal Intergenic | 12 | 49382693 | 49384364 | 1672 | 1 | ENSMUST00000135006.1 | -8812 | Foxg1 | ENSMUSG00000020950.9 |
| chr12 | 49374394 | 49374659 | 3.36759 | 3.9252 | 1.16274 | Distal Intergenic | 12 | 49382693 | 49384364 | 1672 | 1 | ENSMUST00000135006.1 | -8034 | Foxg1 | ENSMUSG00000020950.9 |
| chr12 | 49375079 | 49375472 | 3.91546 | 5.16779 | 1.89729 | Distal Intergenic | 12 | 49382693 | 49384364 | 1672 | 1 | ENSMUST00000135006.1 | -7221 | Foxg1 | ENSMUSG00000020950.9 |
| chr12 | 49375870 | 49376513 | 3.80956 | 4.82147 | 1.69557 | Distal Intergenic | 12 | 49382693 | 49384364 | 1672 | 1 | ENSMUST00000135006.1 | -6180 | Foxg1 | ENSMUSG00000020950.9 |
| chr12 | 49377694 | 49379246 | 3.63606 | 4.46049 | 1.47843 | Distal Intergenic | 12 | 49382693 | 49384364 | 1672 | 1 | ENSMUST00000135006.1 | -3447 | Foxg1 | ENSMUSG00000020950.9 |
| chr12 | 49379613 | 49380276 | 3.38672 | 4.12167 | 1.20549 | Distal Intergenic | 12 | 49382693 | 49384364 | 1672 | 1 | ENSMUST00000135006.1 | -2417 | Foxg1 | ENSMUSG00000020950.9 |
| chr12 | 49381233 | 49381577 | 3.70591 | 4.5466 | 1.51295 | Distal Intergenic | 12 | 49382693 | 49384364 | 1672 | 1 | ENSMUST00000135006.1 | -1116 | Foxg1 | ENSMUSG00000020950.9 |
| chr12 | 49382507 | 49384024 | 4.09983 | 5.56124 | 2.21794 | Promoter | 12 | 49382693 | 49384364 | 1672 | 1 | ENSMUST00000135006.1 | 0 | Foxg1 | ENSMUSG00000020950.9 |
| chr12 | 51590539 | 51590807 | 3.71935 | 4.6613 | 1.55348 | Distal Intergenic | 12 | 51593341 | 51605773 | 12433 | 1 | ENSMUST00000085412.5 | -2534 | Coch | ENSMUSG00000020953.11 |
| chr12 | 51591161 | 51592683 | 3.66042 | 4.48543 | 1.46128 | Promoter | 12 | 51593341 | 51605773 | 12433 | 1 | ENSMUST00000085412.5 | -658 | Coch | ENSMUSG00000020953.11 |
| chr12 | 51593112 | 51594626 | 4.70992 | 6.6492 | 3.0591 | Promoter | 12 | 51593341 | 51605773 | 12433 | 1 | ENSMUST00000085412.5 | 0 | Coch | ENSMUSG00000020953.11 |
| chr12 | 51594937 | 51595843 | 4.0204 | 5.26399 | 1.93691 | Exon (ENSMUST00000085412.5/ENSMUSG00000020953.11, exon 4 of 12) | 12 | 51593341 | 51605773 | 12433 | 1 | ENSMUST00000164782.3 | 1597 | Coch | ENSMUSG00000020953.11 |
| chr12 | 51596394 | 51596612 | 3.46894 | 4.06307 | 1.12685 | Exon (ENSMUST00000085412.5/ENSMUSG00000020953.11, exon 5 of 12) | 12 | 51593341 | 51605773 | 12433 | 1 | ENSMUST00000164782.3 | 3054 | Coch | ENSMUSG00000020953.11 |
| chr12 | 51596992 | 51597087 | 3.79449 | 4.93491 | 1.729 | Exon (ENSMUST00000085412.5/ENSMUSG00000020953.11, exon 6 of 12) | 12 | 51593341 | 51605773 | 12433 | 1 | ENSMUST00000164782.3 | 3652 | Coch | ENSMUSG00000020953.11 |
| chr12 | 91400574 | 91401430 | 3.90636 | 4.99335 | 1.80148 | Promoter | 12 | 91400994 | 91540509 | 139516 | 1 | ENSMUST00000021346.8 | 0 | Tshr | ENSMUSG00000020963.9 |

| | | | | | | | | | | | | | | | |
|---|---|---|---|---|---|---|---|---|---|---|---|---|---|---|---|
| chr12 | 91402538 | 91402782 | 3.54984 | 4.3731 | 1.35309 | Intron (ENSMUST00000021346.8/ENSMUSG00000020963.9, intron 1 of 9) | 12 | 91401000 | 91522541 | 121542 | 1 | ENSMUST00000021343.7 | 1539 | Tshr | ENSMUSG00000020963.9 |
| chr12 | 91403758 | 91403863 | 3.54729 | 4.31117 | 1.34118 | Intron (ENSMUST00000021346.8/ENSMUSG00000020963.9, intron 1 of 9) | 12 | 91401000 | 91522541 | 121542 | 1 | ENSMUST00000021343.7 | 2759 | Tshr | ENSMUSG00000020963.9 |
| chr12 | 69791942 | 69792178 | 3.7469 | 4.73445 | 1.57725 | Intron (ENSMUST00000021378.3/ENSMUSG00000020992.4, intron 1 of 2) | 12 | 69746848 | 69790707 | 43860 | 2 | ENSMUST00000021377.4 | -1236 | Cdkl1 | ENSMUSG00000020990.4 |
| chr12 | 69792677 | 69792991 | 3.39536 | 4.1032 | 1.23909 | Intron (ENSMUST00000021378.3/ENSMUSG00000020992.4, intron 2 of 2) | 12 | 69746848 | 69790707 | 43860 | 2 | ENSMUST00000021377.4 | -1971 | Cdkl1 | ENSMUSG00000020990.4 |
| chr12 | 53244783 | 53244865 | 4.08696 | 5.47465 | 2.08933 | Distal Intergenic | 12 | 53248677 | 54072175 | 823499 | 1 | ENSMUST00000101432.2 | -3812 | Npas3 | ENSMUSG00000021010.7 |
| chr12 | 53246476 | 53247493 | 4.04228 | 5.28054 | 1.99642 | Distal Intergenic | 12 | 53248677 | 54072175 | 823499 | 1 | ENSMUST00000101432.2 | -1184 | Npas3 | ENSMUSG00000021010.7 |
| chr12 | 53247825 | 53248035 | 3.3246 | 3.82608 | 1.05904 | Promoter | 12 | 53248677 | 54072175 | 823499 | 1 | ENSMUST00000101432.2 | -642 | Npas3 | ENSMUSG00000021010.7 |
| chr12 | 53248520 | 53248622 | 4.62381 | 6.61451 | 2.91969 | Promoter | 12 | 53248677 | 54072175 | 823499 | 1 | ENSMUST00000101432.2 | -55 | Npas3 | ENSMUSG00000021010.7 |
| chr12 | 53248946 | 53249242 | 4.0969 | 5.39432 | 2.05422 | Promoter | 12 | 53248677 | 54072175 | 823499 | 1 | ENSMUST00000101432.2 | 270 | Npas3 | ENSMUSG00000021010.7 |
| chr12 | 53249723 | 53251001 | 3.91156 | 4.9448 | 1.73845 | Intron (ENSMUST00000101432.2/ENSMUSG00000021010.7, intron 1 of 11) | 12 | 53248677 | 54072175 | 823499 | 1 | ENSMUST00000101432.2 | 1047 | Npas3 | ENSMUSG00000021010.7 |
| chr12 | 53251609 | 53252066 | 3.456 | 4.0441 | 1.12974 | Intron (ENSMUST00000101432.2/ENSMUSG00000021010.7, intron 1 of 11) | 12 | 53248677 | 54072175 | 823499 | 1 | ENSMUST00000101432.2 | 2933 | Npas3 | ENSMUSG00000021010.7 |
| chr12 | 53252931 | 53254133 | 3.79096 | 4.68712 | 1.55128 | Intron (ENSMUST00000101432.2/ENSMUSG00000021010.7, intron 1 of 11) | 12 | 53248677 | 54072175 | 823499 | 1 | ENSMUST00000101432.2 | 4255 | Npas3 | ENSMUSG00000021010.7 |
| chr12 | 46818391 | 46818485 | 3.889 | 5.59664 | 2.21631 | Promoter | 12 | 46694517 | 46818775 | 124259 | 2 | ENSMUST00000021438.6 | 290 | Nova1 | ENSMUSG00000021047.6 |
| chr12 | 76177105 | 76177705 | 3.82978 | 5.0181 | 1.80375 | Promoter | 12 | 76120419 | 76177259 | 56841 | 2 | ENSMUST00000101291.4 | 0 | Esr2 | ENSMUSG00000021055.8 |
| chr12 | 76799252 | 76799566 | 3.613 | 4.36183 | 1.38982 | 3' UTR | 12 | 76799874 | 76800834 | 961 | 2 | ENSMUST00000143863.1 | 1268 | Rab15 | ENSMUSG00000021062.10 |
| chr12 | 76804861 | 76804989 | 3.48839 | 4.40356 | 1.38597 | Promoter | 12 | 76800122 | 76805478 | 5357 | 2 | ENSMUST00000122419.2 | 489 | Rab15 | ENSMUSG00000021062.10 |
| chr12 | 76806173 | 76806448 | 3.46788 | 4.06606 | 1.14379 | Promoter | 12 | 76800122 | 76805478 | 5357 | 2 | ENSMUST00000122419.2 | -696 | Rab15 | ENSMUSG00000021062.10 |
| chr12 | 76811178 | 76812333 | 3.56879 | 4.2464 | 1.24166 | Intron (ENSMUST00000125842.2/ENSMUSG00000033373.11, intron 2 of 12) | 12 | 76800122 | 76813856 | 13735 | 2 | ENSMUST00000121716.2 | 1523 | Rab15 | ENSMUSG00000021062.10 |
| chr12 | 76812737 | 76813134 | 3.56143 | 4.77183 | 1.62561 | Promoter | 12 | 76800122 | 76813856 | 13735 | 2 | ENSMUST00000121716.2 | 722 | Rab15 | ENSMUSG00000021062.10 |
| chr12 | 76813582 | 76813874 | 3.43018 | 3.99452 | 1.09208 | Promoter | 12 | 76800122 | 76813856 | 13735 | 2 | ENSMUST00000121716.2 | 0 | Rab15 | ENSMUSG00000021062.10 |
| chr12 | 76814800 | 76815196 | 3.70769 | 4.65248 | 1.55636 | Promoter | 12 | 76800122 | 76813856 | 13735 | 2 | ENSMUST00000121716.2 | -945 | Rab15 | ENSMUSG00000021062.10 |
| chr12 | 76815628 | 76815995 | 3.58542 | 4.29184 | 1.30779 | Intron (ENSMUST00000125842.2/ENSMUSG00000033373.11, intron 2 of 12) | 12 | 76800122 | 76813856 | 13735 | 2 | ENSMUST00000121716.2 | -1773 | Rab15 | ENSMUSG00000021062.10 |
| chr12 | 76817440 | 76818021 | 3.9351 | 5.00629 | 1.80772 | Promoter | 12 | 76800122 | 76818374 | 18253 | 2 | ENSMUST00000118604.2 | 353 | Rab15 | ENSMUSG00000021062.10 |
| chr12 | 76821336 | 76821503 | 3.38331 | 3.93824 | 1.1166 | Promoter | 12 | 76802880 | 76822462 | 19583 | 2 | ENSMUST00000134124.1 | 959 | Rab15 | ENSMUSG00000021062.10 |
| chr12 | 76821881 | 76823054 | 3.72594 | 4.66637 | 1.5486 | Promoter | 12 | 76797960 | 76822553 | 24594 | 2 | ENSMUST00000021459.8 | 0 | Rab15 | ENSMUSG00000021062.10 |
| chr12 | 76823392 | 76823582 | 3.38193 | 3.9006 | 1.01733 | Promoter | 12 | 76800330 | 76822908 | 22579 | 2 | ENSMUST00000154765.2 | -485 | Rab15 | ENSMUSG00000021062.10 |
| chr12 | 76824222 | 76824486 | 3.69988 | 4.48837 | 1.38784 | Intron (ENSMUST00000125842.2/ENSMUSG00000033373.11, intron 2 of 12) | 12 | 76800330 | 76822908 | 22579 | 2 | ENSMUST00000154765.2 | -1315 | Rab15 | ENSMUSG00000021062.10 |
| chr12 | 105513954 | 105514110 | 3.64975 | 4.39447 | 1.33407 | Distal Intergenic | 12 | 105563172 | 105593071 | 29900 | 1 | ENSMUST00000001652.5 | -49062 | Bdkrb2 | ENSMUSG00000021070.5 |
| chr12 | 105519413 | 105519497 | 3.8141 | 4.72293 | 1.59466 | Distal Intergenic | 12 | 105563172 | 105593071 | 29900 | 1 | ENSMUST00000001652.5 | -43675 | Bdkrb2 | ENSMUSG00000021070.5 |
| chr12 | 105520831 | 105521464 | 3.69768 | 4.4955 | 1.41215 | Distal Intergenic | 12 | 105563172 | 105593071 | 29900 | 1 | ENSMUST00000001652.5 | -41708 | Bdkrb2 | ENSMUSG00000021070.5 |
| chr12 | 105562065 | 105563159 | 3.68642 | 4.55122 | 1.52493 | Promoter | 12 | 105563172 | 105593071 | 29900 | 1 | ENSMUST00000001652.5 | -13 | Bdkrb2 | ENSMUSG00000021070.5 |
| chr12 | 105570425 | 105570713 | 3.49094 | 4.10747 | 1.1675 | Intron (ENSMUST00000001652.5/ENSMUSG00000021070.5, intron 1 of 2) | 12 | 105563172 | 105593071 | 29900 | 1 | ENSMUST00000001652.5 | 7254 | Bdkrb2 | ENSMUSG00000021070.5 |
| chr12 | 70345966 | 70346326 | 3.91663 | 5.02918 | 1.80383 | Intron (ENSMUST00000110520.3/ENSMUSG00000021071.10, intron 1 of 11) | 12 | 70244539 | 70347614 | 103076 | 2 | ENSMUST00000110520.3 | 1288 | Trim9 | ENSMUSG00000021071.10 |
| chr12 | 72235909 | 72236125 | 3.37797 | 3.90471 | 1.05293 | Promoter | 12 | 72211752 | 72235739 | 23988 | 2 | ENSMUST00000150156.1 | -171 | Rtn1 | ENSMUSG00000021087.12 |
| chr12 | 72236523 | 72237364 | 4.12822 | 5.31881 | 1.95418 | Promoter | 12 | 72211753 | 72236724 | 24972 | 2 | ENSMUST00000021497.10 | 0 | Rtn1 | ENSMUSG00000021087.12 |
| chr12 | 72237894 | 72238224 | 3.39495 | 3.92701 | 1.04124 | Intron (ENSMUST00000078505.8/ENSMUSG00000021087.12, intron 3 of 8) | 12 | 72212289 | 72236724 | 24436 | 2 | ENSMUST00000137990.1 | -1171 | Rtn1 | ENSMUSG00000021087.12 |
| chr12 | 72407301 | 72407409 | 3.43151 | 4.12112 | 1.20877 | Intron (ENSMUST00000078505.8/ENSMUSG00000021087.12, intron 1 of 8) | 12 | 72211752 | 72409054 | 197303 | 2 | ENSMUST00000078505.8 | 1645 | Rtn1 | ENSMUSG00000021087.12 |
| chr12 | 72407770 | 72408912 | 3.99498 | 5.2008 | 1.96606 | Promoter | 12 | 72211752 | 72409054 | 197303 | 2 | ENSMUST00000078505.8 | 142 | Rtn1 | ENSMUSG00000021087.12 |
| chr12 | 72440851 | 72441305 | 3.51746 | 4.15559 | 1.19531 | Promoter | 12 | 72441866 | 72510565 | 68700 | 1 | ENSMUST00000162159.2 | -561 | Lrrc9 | ENSMUSG00000021090.10 |
| chr12 | 72441862 | 72441973 | 3.37852 | 3.89546 | 1.01733 | Promoter | 12 | 72441866 | 72510565 | 68700 | 1 | ENSMUST00000162159.2 | 0 | Lrrc9 | ENSMUSG00000021090.10 |
| chr12 | 72442300 | 72442375 | 3.29078 | 3.78531 | 1.07635 | Promoter | 12 | 72442249 | 72474210 | 31962 | 1 | ENSMUST00000161957.1 | 52 | Lrrc9 | ENSMUSG00000021090.10 |
| chr12 | 104470431 | 104470859 | 3.4596 | 4.04605 | 1.11667 | Downstream (<1kb) | 12 | 104471209 | 104473330 | 2122 | 2 | ENSMUST00000021513.5 | 2471 | Gsc | ENSMUSG00000021095.5 |
| chr12 | 104471282 | 104472286 | 3.90043 | 4.96754 | 1.77304 | 3' UTR | 12 | 104471209 | 104473330 | 2122 | 2 | ENSMUST00000021513.5 | 1044 | Gsc | ENSMUSG00000021095.5 |
| chr12 | 104472650 | 104473309 | 4.02597 | 5.47313 | 2.12209 | Promoter | 12 | 104471209 | 104473330 | 2122 | 2 | ENSMUST00000021513.5 | 21 | Gsc | ENSMUSG00000021095.5 |
| chr12 | 72820902 | 72821262 | 3.80664 | 4.81628 | 1.66356 | Distal Intergenic | 12 | 72761211 | 72794940 | 33730 | 1 | ENSMUST00000021514.8 | 59692 | Ppm1a | ENSMUSG00000021096.10 |
| chr12 | 72831437 | 72831780 | 3.77134 | 4.65354 | 1.50712 | Distal Intergenic | 12 | 72761211 | 72794940 | 33730 | 1 | ENSMUST00000021514.8 | 70227 | Ppm1a | ENSMUSG00000021096.10 |
| chr12 | 72832255 | 72832719 | 4.18489 | 5.54855 | 2.14282 | Distal Intergenic | 12 | 72761211 | 72794940 | 33730 | 1 | ENSMUST00000021514.8 | 71045 | Ppm1a | ENSMUSG00000021096.10 |
| chr12 | 72833033 | 72833152 | 3.27296 | 4.14323 | 1.21703 | Distal Intergenic | 12 | 72761211 | 72794940 | 33730 | 1 | ENSMUST00000021514.8 | 71823 | Ppm1a | ENSMUSG00000021096.10 |
| chr12 | 72834685 | 72835055 | 4.31517 | 5.70805 | 2.25248 | Distal Intergenic | 12 | 72892924 | 72906271 | 13348 | 2 | ENSMUST00000132016.1 | 71216 | 4930447C04Rik | ENSMUSG00000021098.9 |
| chr12 | 72846070 | 72846991 | 3.83196 | 4.78862 | 1.62951 | Distal Intergenic | 12 | 72892924 | 72906271 | 13348 | 2 | ENSMUST00000132016.1 | 59280 | 4930447C04Rik | ENSMUSG00000021098.9 |
| chr12 | 72848654 | 72850050 | 3.57993 | 4.30059 | 1.30326 | Distal Intergenic | 12 | 72892924 | 72906271 | 13348 | 2 | ENSMUST00000132016.1 | 56221 | 4930447C04Rik | ENSMUSG00000021098.9 |
| chr12 | 72852291 | 72852888 | 3.69454 | 4.49342 | 1.42262 | Distal Intergenic | 12 | 72892924 | 72906271 | 13348 | 2 | ENSMUST00000132016.1 | 53383 | 4930447C04Rik | ENSMUSG00000021098.9 |
| chr12 | 72853287 | 72854039 | 3.5263 | 4.24931 | 1.34088 | Distal Intergenic | 12 | 72892924 | 72906271 | 13348 | 2 | ENSMUST00000132016.1 | 52232 | 4930447C04Rik | ENSMUSG00000021098.9 |
| chr12 | 72855929 | 72856192 | 3.46189 | 4.42746 | 1.38711 | Distal Intergenic | 12 | 72892924 | 72906271 | 13348 | 2 | ENSMUST00000132016.1 | 50079 | 4930447C04Rik | ENSMUSG00000021098.9 |
| chr12 | 72917106 | 72918258 | 4.11724 | 5.36059 | 2.02029 | Promoter | 12 | 72881109 | 72917760 | 36652 | 2 | ENSMUST00000110489.3 | 0 | 4930447C04Rik | ENSMUSG00000021098.9 |
| chr12 | 72919102 | 72919276 | 3.30661 | 3.78711 | 1.01733 | Intron (ENSMUST00000136075.2/ENSMUSG00000021098.9, intron 3 of 5) | 12 | 72881109 | 72917906 | 36798 | 2 | ENSMUST00000044000.6 | -1197 | 4930447C04Rik | ENSMUSG00000021098.9 |
| chr12 | 72930845 | 72931013 | 4.36936 | 5.86817 | 2.39488 | Intron (ENSMUST00000136075.2/ENSMUSG00000021098.9, intron 3 of 5) | 12 | 72913187 | 72937977 | 24791 | 2 | ENSMUST00000130635.2 | 6964 | 4930447C04Rik | ENSMUSG00000021098.9 |
| chr12 | 72932796 | 72932989 | 3.61633 | 4.37609 | 1.3913 | Intron (ENSMUST00000136075.2/ENSMUSG00000021098.9, intron 3 of 5) | 12 | 72913187 | 72937977 | 24791 | 2 | ENSMUST00000130635.2 | 4988 | 4930447C04Rik | ENSMUSG00000021098.9 |
| chr12 | 72933490 | 72934139 | 3.82628 | 4.7608 | 1.61519 | Intron (ENSMUST00000136075.2/ENSMUSG00000021098.9, intron 3 of 5) | 12 | 72913187 | 72937977 | 24791 | 2 | ENSMUST00000130635.2 | 3838 | 4930447C04Rik | ENSMUSG00000021098.9 |
| chr12 | 72935067 | 72935460 | 4.2159 | 5.5261 | 2.12569 | Intron (ENSMUST00000136075.2/ENSMUSG00000021098.9, intron 3 of 5) | 12 | 72913187 | 72937977 | 24791 | 2 | ENSMUST00000130635.2 | 2517 | 4930447C04Rik | ENSMUSG00000021098.9 |
| chr12 | 72935852 | 72936875 | 3.76732 | 4.63559 | 1.52459 | Intron (ENSMUST00000136075.2/ENSMUSG00000021098.9, intron 3 of 5) | 12 | 72913187 | 72937977 | 24791 | 2 | ENSMUST00000130635.2 | 1102 | 4930447C04Rik | ENSMUSG00000021098.9 |
| chr12 | 72937523 | 72937945 | 4.27725 | 5.66342 | 2.26583 | Promoter | 12 | 72913187 | 72937977 | 24791 | 2 | ENSMUST00000130635.2 | 32 | 4930447C04Rik | ENSMUSG00000021098.9 |
| chr12 | 72942414 | 72943984 | 3.69855 | 4.59463 | 1.55462 | 3' UTR | 12 | 72911659 | 72940774 | 29116 | 2 | ENSMUST00000136075.2 | -1641 | 4930447C04Rik | ENSMUSG00000021098.9 |
| chr12 | 72939623 | 72940485 | 4.27479 | 5.9169 | 2.49163 | Promoter | 12 | 72913191 | 72940083 | 26893 | 2 | ENSMUST00000132067.1 | 0 | 4930447C04Rik | ENSMUSG00000021098.9 |
| chr12 | 72940876 | 72941237 | 4.31181 | 5.94798 | 2.42406 | Promoter | 12 | 72911659 | 72940774 | 29116 | 2 | ENSMUST00000136075.2 | -103 | 4930447C04Rik | ENSMUSG00000021098.9 |
| chr12 | 72941559 | 72942050 | 3.59103 | 4.57236 | 1.47838 | Promoter | 12 | 72911659 | 72940774 | 29116 | 2 | ENSMUST00000136075.2 | -786 | 4930447C04Rik | ENSMUSG00000021098.9 |
| chr12 | 72938953 | 72939104 | 3.64178 | 4.69256 | 1.55633 | Promoter | 12 | 72939892 | 72944899 | 5008 | 1 | ENSMUST00000021519.6 | -788 | Six6 | ENSMUSG00000021099.6 |
| chr12 | 73581731 | 73582140 | 3.39636 | 3.98054 | 1.19766 | Distal Intergenic | 12 | 73584797 | 73778185 | 193389 | 1 | ENSMUST00000021527.9 | -2657 | Prkch | ENSMUSG00000021108.12 |
| chr12 | 73583867 | 73584155 | 3.55042 | 4.45293 | 1.43416 | Promoter | 12 | 73584797 | 73778185 | 193389 | 1 | ENSMUST00000021527.9 | -642 | Prkch | ENSMUSG00000021108.12 |
| chr12 | 73584593 | 73585252 | 3.86747 | 4.92499 | 1.73544 | Promoter | 12 | 73584797 | 73778185 | 193389 | 1 | ENSMUST00000021527.9 | 0 | Prkch | ENSMUSG00000021108.12 |
| chr12 | 73585570 | 73586061 | 3.76219 | 4.83212 | 1.67334 | Promoter | 12 | 73584843 | 73761542 | 176700 | 1 | ENSMUST00000119092.1 | 728 | Prkch | ENSMUSG00000021108.12 |
| chr12 | 105958933 | 105959104 | 3.36779 | 3.87927 | 1.01733 | Distal Intergenic | 12 | 106010263 | 106077410 | 67148 | 1 | ENSMUST00000021539.9 | -51159 | Vrk1 | ENSMUSG00000021115.9 |
| chr12 | 105963471 | 105963576 | 3.45624 | 4.16248 | 1.23904 | Distal Intergenic | 12 | 106010263 | 106077410 | 67148 | 1 | ENSMUST00000021539.9 | -46687 | Vrk1 | ENSMUSG00000021115.9 |
| chr12 | 105967958 | 105968212 | 3.68013 | 4.46898 | 1.42625 | Distal Intergenic | 12 | 106010263 | 106077410 | 67148 | 1 | ENSMUST00000021539.9 | -42051 | Vrk1 | ENSMUSG00000021115.9 |
| chr12 | 78906933 | 78907138 | 3.22102 | 3.65875 | 1.01458 | Promoter | 12 | 78888691 | 78906964 | 18274 | 2 | ENSMUST00000021544.7 | 0 | Plek2 | ENSMUSG00000021118.7 |
| chr12 | 78910606 | 78910789 | 3.61188 | 4.40875 | 1.4109 | Distal Intergenic | 12 | 78888691 | 78906964 | 18274 | 2 | ENSMUST00000021544.7 | -3643 | Plek2 | ENSMUSG00000021118.7 |
| chr12 | 79130345 | 79130602 | 3.6954 | 4.60529 | 1.54023 | Promoter | 12 | 79130777 | 79156301 | 25525 | 1 | ENSMUST00000021550.6 | -175 | Arg2 | ENSMUSG00000021125.6 |
| chr12 | 79131433 | 79132030 | 3.34167 | 3.8399 | 1.01733 | Promoter | 12 | 79130777 | 79156301 | 25525 | 1 | ENSMUST00000021550.6 | 657 | Arg2 | ENSMUSG00000021125.6 |
| chr12 | 79133783 | 79133871 | 4.30856 | 5.73566 | 2.30996 | Intron (ENSMUST00000021550.6/ENSMUSG00000021125.6, intron 2 of 7) | 12 | 79130777 | 79156301 | 25525 | 1 | ENSMUST00000021550.6 | 3007 | Arg2 | ENSMUSG00000021125.6 |
| chr12 | 80518236 | 80519479 | 3.6757 | 4.61398 | 1.52676 | Promoter | 12 | 80518990 | 80603896 | 84907 | 1 | ENSMUST00000021558.6 | 0 | Galnt16 | ENSMUSG00000021130.7 |
| chr12 | 80519813 | 80520054 | 3.42375 | 4.3992 | 1.3693 | Promoter | 12 | 80518990 | 80603896 | 84907 | 1 | ENSMUST00000021558.6 | 824 | Galnt16 | ENSMUSG00000021130.7 |
| chr12 | 80520894 | 80521208 | 3.49061 | 4.09883 | 1.13777 | Intron (ENSMUST00000021558.6/ENSMUSG00000021130.7, intron 1 of 14) | 12 | 80518990 | 80603896 | 84907 | 1 | ENSMUST00000021558.6 | 1905 | Galnt16 | ENSMUSG00000021130.7 |
| chr12 | 80522076 | 80522201 | 3.39585 | 3.93793 | 1.07505 | Intron (ENSMUST00000021558.6/ENSMUSG00000021130.7, intron 1 of 14) | 12 | 80518990 | 80603896 | 84907 | 1 | ENSMUST00000021558.6 | 3087 | Galnt16 | ENSMUSG00000021130.7 |
| chr12 | 80526380 | 80526680 | 3.36082 | 3.93833 | 1.11637 | Intron (ENSMUST00000021558.6/ENSMUSG00000021130.7, intron 1 of 14) | 12 | 80518990 | 80603896 | 84907 | 1 | ENSMUST00000021558.6 | 7391 | Galnt16 | ENSMUSG00000021130.7 |
| chr12 | 80998675 | 80998863 | 3.7554 | 4.60981 | 1.48129 | Distal Intergenic | 12 | 81026808 | 81186414 | 159607 | 1 | ENSMUST00000110347.3 | -27945 | Smoc1 | ENSMUSG00000021136.8 |
| chr12 | 81002025 | 81002199 | 3.45357 | 4.08823 | 1.22498 | Distal Intergenic | 12 | 81026808 | 81186414 | 159607 | 1 | ENSMUST00000110347.3 | -24609 | Smoc1 | ENSMUSG00000021136.8 |
| chr12 | 81023858 | 81024115 | 3.62951 | 4.51024 | 1.44504 | Distal Intergenic | 12 | 81026808 | 81186414 | 159607 | 1 | ENSMUST00000110347.3 | -2693 | Smoc1 | ENSMUSG00000021136.8 |
| chr12 | 81024515 | 81025191 | 3.8653 | 4.95542 | 1.74675 | Distal Intergenic | 12 | 81026808 | 81186414 | 159607 | 1 | ENSMUST00000110347.3 | -1617 | Smoc1 | ENSMUSG00000021136.8 |
| chr12 | 81025516 | 81025736 | 3.71626 | 4.55254 | 1.46925 | Distal Intergenic | 12 | 81026808 | 81186414 | 159607 | 1 | ENSMUST00000110347.3 | -1072 | Smoc1 | ENSMUSG00000021136.8 |
| chr12 | 81026861 | 81027116 | 3.52891 | 4.17413 | 1.19949 | Promoter | 12 | 81026828 | 81186410 | 159583 | 1 | ENSMUST00000129362.1 | 34 | Smoc1 | ENSMUSG00000021136.8 |
| chr12 | 81027666 | 81028806 | 3.69565 | 4.49109 | 1.41339 | Promoter | 12 | 81026828 | 81186410 | 159583 | 1 | ENSMUST00000129362.1 | 839 | Smoc1 | ENSMUSG00000021136.8 |
| chr12 | 81030136 | 81030211 | 3.65173 | 4.40049 | 1.33362 | Intron (ENSMUST00000110347.3/ENSMUSG00000021136.8, intron 1 of 12) | 12 | 81026828 | 81186410 | 159583 | 1 | ENSMUST00000129362.1 | 3309 | Smoc1 | ENSMUSG00000021136.8 |
| chr12 | 81033770 | 81033850 | 3.42888 | 4.03443 | 1.20962 | Intron (ENSMUST00000110347.3/ENSMUSG00000021136.8, intron 1 of 12) | 12 | 81026828 | 81186410 | 159583 | 1 | ENSMUST00000129362.1 | 6943 | Smoc1 | ENSMUSG00000021136.8 |
| chr12 | 81037389 | 81037464 | 3.51904 | 4.15941 | 1.19481 | Intron (ENSMUST00000110347.3/ENSMUSG00000021136.8, intron 1 of 12) | 12 | 81026828 | 81186410 | 159583 | 1 | ENSMUST00000129362.1 | 10562 | Smoc1 | ENSMUSG00000021136.8 |
| chr12 | 81094969 | 81095443 | 3.70626 | 4.5012 | 1.40064 | Intron (ENSMUST00000110347.3/ENSMUSG00000021136.8, intron 1 of 12) | 12 | 81026828 | 81186410 | 159583 | 1 | ENSMUST00000129362.1 | 68142 | Smoc1 | ENSMUSG00000021136.8 |
| chr13 | 9796384 | 9796591 | 3.56075 | 4.32288 | 1.38464 | Distal Intergenic | 13 | 9697644 | 9765321 | 67678 | 2 | ENSMUST00000152725.2 | -31064 | Zmynd11 | ENSMUSG00000021156.11 |
| chr12 | 117751410 | 117751658 | 3.44978 | 4.14464 | 1.22885 | Intron (ENSMUST00000109691.2/ENSMUSG00000041992.8, intron 26 of 27) | 12 | 117804289 | 117870047 | 65759 | 1 | ENSMUST00000176499.1 | -52631 | Cdca7l | ENSMUSG00000021175.10 |
| chr12 | 117788704 | 117788825 | 3.49295 | 4.14893 | 1.28441 | Distal Intergenic | 12 | 117804289 | 117870047 | 65759 | 1 | ENSMUST00000176499.1 | -15464 | Cdca7l | ENSMUSG00000021175.10 |
| chr12 | 117806391 | 117807144 | 4.0732 | 5.34223 | 2.0486 | Intron (ENSMUST00000176499.1/ENSMUSG00000021175.10, intron 1 of 4) | 12 | 117804289 | 117870047 | 65759 | 1 | ENSMUST00000176499.1 | 2103 | Cdca7l | ENSMUSG00000021175.10 |
| chr12 | 117807728 | 117807889 | 3.87352 | 4.84635 | 1.66662 | Intron (ENSMUST00000176499.1/ENSMUSG00000021175.10, intron 1 of 4) | 12 | 117804289 | 117870047 | 65759 | 1 | ENSMUST00000176499.1 | 3440 | Cdca7l | ENSMUSG00000021175.10 |
| chr12 | 117843253 | 117844157 | 3.64647 | 4.52322 | 1.49315 | Promoter | 12 | 117843515 | 117878706 | 35192 | 1 | ENSMUST00000021592.10 | 0 | Cdca7l | ENSMUSG00000021175.10 |
| chr12 | 101026558 | 101026745 | 3.26977 | 3.73179 | 1.01733 | Intron (ENSMUST00000068411.3/ENSMUSG00000021182.9, intron 2 of 29) | 12 | 100912700 | 101028983 | 116284 | 2 | ENSMUST00000068411.3 | 2238 | Ccdc88c | ENSMUSG00000021182.9 |
| chr12 | 101028133 | 101029205 | 3.57534 | 4.43443 | 1.42738 | Promoter | 12 | 100912700 | 101028983 | 116284 | 2 | ENSMUST00000068411.3 | 0 | Ccdc88c | ENSMUSG00000021182.9 |
| chr12 | 101716715 | 101718960 | 3.63709 | 4.56781 | 1.50469 | Promoter | 12 | 101645443 | 101718523 | 73081 | 2 | ENSMUST00000162735.2 | 0 | Tc2n | ENSMUSG00000021187.9 |
| chr12 | 102554665 | 102555441 | 3.43636 | 4.07723 | 1.21546 | Promoter | 12 | 102554969 | 102565027 | 10059 | 1 | ENSMUST00000021610.5 | 0 | Chga | ENSMUSG00000021194.5 |
| chr12 | 102943678 | 102943907 | 3.37141 | 3.94701 | 1.07267 | Distal Intergenic | 12 | 102948859 | 103184001 | 235143 | 1 | ENSMUST00000101099.6 | -4952 | Unc79 | ENSMUSG00000021198.11 |
| chr12 | 102947841 | 102947916 | 3.69343 | 4.47047 | 1.36553 | Promoter | 12 | 102948859 | 103184001 | 235143 | 1 | ENSMUST00000101099.6 | -943 | Unc79 | ENSMUSG00000021198.11 |
| chr12 | 102948904 | 102949291 | 3.88244 | 4.85464 | 1.65893 | Promoter | 12 | 102948859 | 103184001 | 235143 | 1 | ENSMUST00000101099.6 | 46 | Unc79 | ENSMUSG00000021198.11 |
| chr12 | 102954389 | 102954552 | 3.55263 | 4.30481 | 1.34094 | Intron (ENSMUST00000101099.6/ENSMUSG00000021198.11, intron 1 of 48) | 12 | 102949554 | 103183613 | 234060 | 1 | ENSMUST00000179002.2 | 4836 | Unc79 | ENSMUSG00000021198.11 |
| chr12 | 103325975 | 103326385 | 3.42023 | 4.10403 | 1.19333 | Intron (ENSMUST00000021617.8/ENSMUSG00000021200.9, intron 7 of 9) | 12 | 103321146 | 103335584 | 14439 | 2 | ENSMUST00000127447.1 | 9199 | Asb2 | ENSMUSG00000021200.9 |
| chr7 | 36675620 | 36675812 | 3.73749 | 4.6256 | 1.54854 | Distal Intergenic | 7 | 36698118 | 36773553 | 75436 | 1 | ENSMUST00000021641.7 | -22306 | Tshz3 | ENSMUSG00000021217.7 |
| chr7 | 36678258 | 36678456 | 3.37228 | 3.99562 | 1.15599 | Distal Intergenic | 7 | 36698118 | 36773553 | 75436 | 1 | ENSMUST00000021641.7 | -19662 | Tshz3 | ENSMUSG00000021217.7 |
| chr7 | 36679200 | 36679275 | 4.09584 | 5.33476 | 2.00214 | Distal Intergenic | 7 | 36698118 | 36773553 | 75436 | 1 | ENSMUST00000021641.7 | -18843 | Tshz3 | ENSMUSG00000021217.7 |
| chr7 | 36679788 | 36679875 | 4.19939 | 5.48317 | 2.07237 | Distal Intergenic | 7 | 36698118 | 36773553 | 75436 | 1 | ENSMUST00000021641.7 | -18243 | Tshz3 | ENSMUSG00000021217.7 |
| chr7 | 36680181 | 36680315 | 4.20891 | 5.46662 | 2.04212 | Distal Intergenic | 7 | 36698118 | 36773553 | 75436 | 1 | ENSMUST00000021641.7 | -17803 | Tshz3 | ENSMUSG00000021217.7 |
| chr7 | 36681105 | 36681635 | 3.43086 | 3.99445 | 1.08824 | Distal Intergenic | 7 | 36698118 | 36773553 | 75436 | 1 | ENSMUST00000021641.7 | -16483 | Tshz3 | ENSMUSG00000021217.7 |
| chr7 | 36689791 | 36690097 | 3.53234 | 4.40518 | 1.37012 | Distal Intergenic | 7 | 36698118 | 36773553 | 75436 | 1 | ENSMUST00000021641.7 | -8021 | Tshz3 | ENSMUSG00000021217.7 |
| chr7 | 36690782 | 36690918 | 3.71434 | 4.68871 | 1.58763 | Distal Intergenic | 7 | 36698118 | 36773553 | 75436 | 1 | ENSMUST00000021641.7 | -7200 | Tshz3 | ENSMUSG00000021217.7 |
| chr7 | 36691423 | 36691981 | 4.41411 | 6.24486 | 2.72222 | Distal Intergenic | 7 | 36698118 | 36773553 | 75436 | 1 | ENSMUST00000021641.7 | -6137 | Tshz3 | ENSMUSG00000021217.7 |
| chr7 | 36692575 | 36692917 | 3.51865 | 4.26356 | 1.31111 | Distal Intergenic | 7 | 36698118 | 36773553 | 75436 | 1 | ENSMUST00000021641.7 | -5201 | Tshz3 | ENSMUSG00000021217.7 |
| chr7 | 36694294 | 36694922 | 3.71002 | 4.60308 | 1.53193 | Distal Intergenic | 7 | 36698118 | 36773553 | 75436 | 1 | ENSMUST00000021641.7 | -3196 | Tshz3 | ENSMUSG00000021217.7 |
| chr7 | 36695368 | 36695492 | 3.31157 | 3.79457 | 1.01733 | Distal Intergenic | 7 | 36698118 | 36773553 | 75436 | 1 | ENSMUST00000021641.7 | -2626 | Tshz3 | ENSMUSG00000021217.7 |
| chr7 | 36695869 | 36696082 | 4.22994 | 5.55715 | 2.19029 | Distal Intergenic | 7 | 36698118 | 36773553 | 75436 | 1 | ENSMUST00000021641.7 | -2036 | Tshz3 | ENSMUSG00000021217.7 |
| chr7 | 36707522 | 36707806 | 3.47279 | 4.06642 | 1.11834 | Intron (ENSMUST00000021641.7/ENSMUSG00000021217.7, intron 1 of 1) | 7 | 36698217 | 36702113 | 3897 | 1 | ENSMUST00000176413.1 | 9306 | Tshz3 | ENSMUSG00000021217.7 |
| chr7 | 36710707 | 36711020 | 3.47502 | 4.18155 | 1.27265 | Intron (ENSMUST00000021641.7/ENSMUSG00000021217.7, intron 1 of 1) | 7 | 36698217 | 36702113 | 3897 | 1 | ENSMUST00000176413.1 | 12491 | Tshz3 | ENSMUSG00000021217.7 |
| chr12 | 82615244 | 82615748 | 3.54712 | 4.27345 | 1.35315 | Distal Intergenic | 12 | 82616925 | 83092731 | 475807 | 1 | ENSMUST00000161801.2 | -1177 | Rgs6 | ENSMUSG00000021219.11 |
| chr12 | 82616644 | 82617120 | 3.674 | 4.82747 | 1.66876 | Promoter | 12 | 82616925 | 83092731 | 475807 | 1 | ENSMUST00000161801.2 | 0 | Rgs6 | ENSMUSG00000021219.11 |
| chr12 | 82617447 | 82618027 | 3.81665 | 4.9323 | 1.77704 | Promoter | 12 | 82617499 | 83162056 | 544558 | 1 | ENSMUST00000186458.1 | 0 | Rgs6 | ENSMUSG00000021219.11 |
| chr12 | 82619674 | 82619939 | 3.76316 | 4.71702 | 1.61628 | Intron (ENSMUST00000161801.2/ENSMUSG00000021219.11, intron 1 of 13) | 12 | 82617499 | 83162056 | 544558 | 1 | ENSMUST00000186458.1 | 2176 | Rgs6 | ENSMUSG00000021219.11 |
| chr12 | 82997089 | 82997191 | 3.44632 | 4.08167 | 1.20287 | Intron (ENSMUST00000161801.2/ENSMUSG00000021219.11, intron 3 of 13) | 12 | 83046990 | 83048120 | 1131 | 1 | ENSMUST00000190964.1 | -49799 | Rgs6 | ENSMUSG00000021219.11 |
| chr12 | 84562962 | 84563124 | 3.35407 | 3.85857 | 1.01733 | Distal Intergenic | 12 | 84569762 | 84595457 | 25696 | 1 | ENSMUST00000021665.6 | -6638 | Vsx2 | ENSMUSG00000021239.7 |
| chr12 | 84566124 | 84566217 | 3.34153 | 3.83966 | 1.01733 | Distal Intergenic | 12 | 84569762 | 84595457 | 25696 | 1 | ENSMUST00000021665.6 | -3545 | Vsx2 | ENSMUSG00000021239.7 |
| chr12 | 84567911 | 84568421 | 3.50714 | 4.36111 | 1.36447 | Distal Intergenic | 12 | 84569762 | 84595457 | 25696 | 1 | ENSMUST00000021665.6 | -1341 | Vsx2 | ENSMUSG00000021239.7 |
| chr12 | 84569428 | 84570658 | 4.50118 | 6.42399 | 2.93839 | Promoter | 12 | 84569762 | 84595457 | 25696 | 1 | ENSMUST00000021665.6 | 0 | Vsx2 | ENSMUSG00000021239.7 |
| chr12 | 84571265 | 84572731 | 4.31803 | 6.0379 | 2.56293 | Exon (ENSMUST00000021665.6/ENSMUSG00000021239.7, exon 2 of 5) | 12 | 84569840 | 84595457 | 25618 | 1 | ENSMUST00000169934.3 | 1426 | Vsx2 | ENSMUSG00000021239.7 |
| chr12 | 84573127 | 84573481 | 3.74317 | 4.65019 | 1.6167 | Intron (ENSMUST00000021665.6/ENSMUSG00000021239.7, intron 2 of 4) | 12 | 84569840 | 84595457 | 25618 | 1 | ENSMUST00000169934.3 | 3288 | Vsx2 | ENSMUSG00000021239.7 |
| chr12 | 84575307 | 84575652 | 3.31538 | 3.80477 | 1.03303 | Intron (ENSMUST00000021665.6/ENSMUSG00000021239.7, intron 2 of 4) | 12 | 84569840 | 84595457 | 25618 | 1 | ENSMUST00000169934.3 | 5468 | Vsx2 | ENSMUSG00000021239.7 |
| chr12 | 84579963 | 84580094 | 3.35882 | 3.86574 | 1.01733 | Intron (ENSMUST00000021665.6/ENSMUSG00000021239.7, intron 3 of 4) | 12 | 84569840 | 84595457 | 25618 | 1 | ENSMUST00000169934.3 | 10124 | Vsx2 | ENSMUSG00000021239.7 |
| chr12 | 84625185 | 84625466 | 3.6304 | 4.39721 | 1.40544 | Distal Intergenic | 12 | 84602531 | 84617466 | 14936 | 2 | ENSMUST00000021666.4 | -7720 | Abcd4 | ENSMUSG00000021240.5 |
| chr12 | 84626911 | 84627109 | 3.27844 | 3.75344 | 1.04433 | Distal Intergenic | 12 | 84602531 | 84617466 | 14936 | 2 | ENSMUST00000021666.4 | -9446 | Abcd4 | ENSMUSG00000021240.5 |
| chr12 | 84751616 | 84751753 | 3.67659 | 4.50978 | 1.48411 | Downstream (2-3kb) | 12 | 84754560 | 84773112 | 18553 | 2 | ENSMUST00000021668.8 | 21359 | Npc2 | ENSMUSG00000021242.8 |
| chr12 | 85785825 | 85786246 | 3.56626 | 4.24504 | 1.25226 | Exon (ENSMUST00000040461.3/ENSMUSG00000034258.3, exon 3 of 10) | 12 | 85819332 | 85822250 | 2919 | 2 | ENSMUST00000148323.1 | 36004 | 0610007P14Rik | ENSMUSG00000021252.6 |
| chr12 | 86210579 | 86210884 | 3.81337 | 4.70657 | 1.53546 | Distal Intergenic | 12 | 86241890 | 86291414 | 49525 | 1 | ENSMUST00000071106.4 | -31006 | Gpatch2l | ENSMUSG00000021254.8 |
| chr12 | 86212053 | 86212128 | 4.16563 | 5.37053 | 1.97586 | Distal Intergenic | 12 | 86241890 | 86291414 | 49525 | 1 | ENSMUST00000071106.4 | -29762 | Gpatch2l | ENSMUSG00000021254.8 |
| chr12 | 86212494 | 86212714 | 3.42831 | 3.9957 | 1.10632 | Distal Intergenic | 12 | 86241890 | 86291414 | 49525 | 1 | ENSMUST00000071106.4 | -29176 | Gpatch2l | ENSMUSG00000021254.8 |
| chr12 | 86213343 | 86213574 | 3.48085 | 4.09593 | 1.17983 | Distal Intergenic | 12 | 86241890 | 86291414 | 49525 | 1 | ENSMUST00000071106.4 | -28316 | Gpatch2l | ENSMUSG00000021254.8 |
| chr12 | 86214064 | 86214480 | 4.00366 | 5.11264 | 1.87231 | Distal Intergenic | 12 | 86241890 | 86291414 | 49525 | 1 | ENSMUST00000071106.4 | -27410 | Gpatch2l | ENSMUSG00000021254.8 |
| chr12 | 86215273 | 86215699 | 3.74842 | 4.58976 | 1.46452 | Distal Intergenic | 12 | 86241890 | 86291414 | 49525 | 1 | ENSMUST00000071106.4 | -26191 | Gpatch2l | ENSMUSG00000021254.8 |
| chr12 | 86217041 | 86217193 | 3.68985 | 4.53028 | 1.49408 | Distal Intergenic | 12 | 86241890 | 86291414 | 49525 | 1 | ENSMUST00000071106.4 | -24697 | Gpatch2l | ENSMUSG00000021254.8 |
| chr12 | 86217834 | 86218000 | 3.48929 | 4.09634 | 1.13598 | Distal Intergenic | 12 | 86241890 | 86291414 | 49525 | 1 | ENSMUST00000071106.4 | -23890 | Gpatch2l | ENSMUSG00000021254.8 |
| chr12 | 86219774 | 86220115 | 3.44106 | 4.03207 | 1.16683 | Distal Intergenic | 12 | 86241890 | 86291414 | 49525 | 1 | ENSMUST00000071106.4 | -21775 | Gpatch2l | ENSMUSG00000021254.8 |
| chr12 | 86220801 | 86221078 | 3.36724 | 3.92008 | 1.07203 | Distal Intergenic | 12 | 86241890 | 86291414 | 49525 | 1 | ENSMUST00000071106.4 | -20812 | Gpatch2l | ENSMUSG00000021254.8 |
| chr12 | 86221438 | 86221969 | 3.3506 | 4.10619 | 1.17356 | Distal Intergenic | 12 | 86241890 | 86291414 | 49525 | 1 | ENSMUST00000071106.4 | -19921 | Gpatch2l | ENSMUSG00000021254.8 |
| chr12 | 86379618 | 86379850 | 3.36623 | 3.88529 | 1.04699 | Intron (ENSMUST00000116402.4/ENSMUSG00000021255.12, intron 1 of 6) | 12 | 86361117 | 86521628 | 160512 | 1 | ENSMUST00000116402.4 | 18502 | Esrrb | ENSMUSG00000021255.12 |
| chr12 | 86402577 | 86402656 | 4.06223 | 5.22753 | 1.96159 | Intron (ENSMUST00000116402.4/ENSMUSG00000021255.12, intron 1 of 6) | 12 | 86421644 | 86515522 | 93879 | 1 | ENSMUST00000110203.2 | -18988 | Esrrb | ENSMUSG00000021255.12 |
| chr12 | 86403048 | 86403235 | 3.78912 | 4.69628 | 1.61359 | Intron (ENSMUST00000116402.4/ENSMUSG00000021255.12, intron 1 of 6) | 12 | 86421644 | 86515522 | 93879 | 1 | ENSMUST00000110203.2 | -18409 | Esrrb | ENSMUSG00000021255.12 |
| chr12 | 86419985 | 86421220 | 3.91116 | 5.00475 | 1.76358 | Promoter | 12 | 86421644 | 86515522 | 93879 | 1 | ENSMUST00000110203.2 | -424 | Esrrb | ENSMUSG00000021255.12 |
| chr12 | 86422331 | 86423471 | 4.35718 | 6.03641 | 2.55896 | Promoter | 12 | 86421883 | 86519800 | 97918 | 1 | ENSMUST00000021680.6 | 449 | Esrrb | ENSMUSG00000021255.12 |
| chr12 | 86424026 | 86424231 | 3.71856 | 4.7704 | 1.62678 | Intron (ENSMUST00000116402.4/ENSMUSG00000021255.12, intron 1 of 6) | 12 | 86421883 | 86519800 | 97918 | 1 | ENSMUST00000021680.6 | 2144 | Esrrb | ENSMUSG00000021255.12 |
| chr12 | 86424754 | 86424975 | 3.55033 | 4.22214 | 1.24376 | Intron (ENSMUST00000116402.4/ENSMUSG00000021255.12, intron 1 of 6) | 12 | 86421883 | 86519800 | 97918 | 1 | ENSMUST00000021680.6 | 2872 | Esrrb | ENSMUSG00000021255.12 |
| chr12 | 86425534 | 86425738 | 3.50898 | 4.13974 | 1.18468 | Intron (ENSMUST00000116402.4/ENSMUSG00000021255.12, intron 1 of 6) | 12 | 86421883 | 86519800 | 97918 | 1 | ENSMUST00000021680.6 | 3652 | Esrrb | ENSMUSG00000021255.12 |
| chr12 | 86426192 | 86426412 | 3.347 | 3.84791 | 1.01733 | Intron (ENSMUST00000116402.4/ENSMUSG00000021255.12, intron 1 of 6) | 12 | 86421883 | 86519800 | 97918 | 1 | ENSMUST00000021680.6 | 4310 | Esrrb | ENSMUSG00000021255.12 |
| chr12 | 86434057 | 86434628 | 3.34627 | 3.84681 | 1.01733 | Intron (ENSMUST00000116402.4/ENSMUSG00000021255.12, intron 1 of 6) | 12 | 86421883 | 86519800 | 97918 | 1 | ENSMUST00000021680.6 | 12175 | Esrrb | ENSMUSG00000021255.12 |

| | | | | | | | | | | | | | | | |
|---|---|---|---|---|---|---|---|---|---|---|---|---|---|---|---|
| chr12 | 86546783 | 86546874 | 3.3274 | 3.81846 | 1.01733 | Distal Intergenic | 12 | 86504409 | 86509967 | 5559 | 1 | ENSMUST00000136464.2 | 42375 | Esrrb | ENSMUSG00000021255.12 |
| chr12 | 108702162 | 108702755 | 4.30761 | 5.78232 | 2.3483 | Promoter | 12 | 108686792 | 108702306 | 15515 | 2 | ENSMUST00000167978.2 | 0 | Degs2 | ENSMUSG00000021263.5 |
| chr12 | 108703489 | 108703729 | 3.3689 | 4.1574 | 1.2308 | Distal Intergenic | 12 | 108690222 | 108702306 | 12085 | 2 | ENSMUST00000021691.4 | -1184 | Degs2 | ENSMUSG00000021263.5 |
| chr12 | 109540574 | 109541101 | 3.61152 | 4.35093 | 1.32139 | Promoter | 12 | 109541001 | 109545319 | 4319 | 1 | ENSMUST00000129245.2 | 0 | Meg3 | ENSMUSG00000021268.12 |
| chr12 | 111270491 | 111270729 | 3.57489 | 4.283 | 1.32152 | Promoter | 12 | 111271111 | 111276426 | 5316 | 1 | ENSMUST00000021707.6 | -382 | Amn | ENSMUSG00000021278.6 |
| chr12 | 111271284 | 111272845 | 3.51979 | 4.24395 | 1.31169 | Promoter | 12 | 111271111 | 111276426 | 5316 | 1 | ENSMUST00000021707.6 | 174 | Amn | ENSMUSG00000021278.6 |
| chr12 | 111273514 | 111273598 | 3.28666 | 3.95919 | 1.12409 | Intron (ENSMUST00000021707.6/ENSMUSG00000021278.6, intron 3 of 11) | 12 | 111271111 | 111276426 | 5316 | 1 | ENSMUST00000021707.6 | 2404 | Amn | ENSMUSG00000021278.6 |
| chr12 | 111274381 | 111276545 | 3.92638 | 5.31575 | 2.04288 | 3' UTR | 12 | 111271111 | 111276426 | 5316 | 1 | ENSMUST00000021707.6 | 3271 | Amn | ENSMUSG00000021278.6 |
| chr12 | 111277252 | 111277412 | 3.65661 | 4.5213 | 1.46465 | Distal Intergenic | 12 | 111271111 | 111276426 | 5316 | 1 | ENSMUST00000021707.6 | 6142 | Amn | ENSMUSG00000021278.6 |
| chr12 | 111422569 | 111422943 | 3.44888 | 4.04466 | 1.11882 | Intron (ENSMUST00000072646.6/ENSMUSG00000021280.8, intron 3 of 12) | 12 | 111417430 | 111431182 | 13753 | 1 | ENSMUST00000072646.6 | 5140 | Exoc3l4 | ENSMUSG00000021280.8 |
| chr12 | 111423796 | 111424015 | 4.06282 | 5.35228 | 2.09497 | Exon (ENSMUST00000072646.6/ENSMUSG00000021280.8, exon 4 of 13) | 12 | 111417430 | 111431182 | 13753 | 1 | ENSMUST00000072646.6 | 6367 | Exoc3l4 | ENSMUSG00000021280.8 |
| chr12 | 112142856 | 112145177 | 3.7838 | 4.71965 | 1.58984 | Distal Intergenic | 12 | 112146208 | 112181747 | 35540 | 1 | ENSMUST00000128402.2 | -1031 | Kif26a | ENSMUSG00000021294.7 |
| chr12 | 112145952 | 112147740 | 4.07632 | 5.26489 | 1.96357 | Promoter | 12 | 112146208 | 112181747 | 35540 | 1 | ENSMUST00000128402.2 | 0 | Kif26a | ENSMUSG00000021294.7 |
| chr12 | 112265628 | 112265716 | 3.49104 | 4.10465 | 1.15615 | Distal Intergenic | 12 | 112179154 | 112180533 | 1380 | 1 | ENSMUST00000183816.1 | 86475 | Kif26a | ENSMUSG00000021294.7 |
| chr12 | 112870587 | 112870899 | 3.29245 | 3.76586 | 1.01733 | Distal Intergenic | 12 | 112850876 | 112860916 | 10041 | 2 | ENSMUST00000021729.7 | -9672 | Gpr132 | ENSMUSG00000021298.7 |
| chr12 | 112871210 | 112871525 | 3.26326 | 3.72202 | 1.01733 | Distal Intergenic | 12 | 112850876 | 112860916 | 10041 | 2 | ENSMUST00000021729.7 | -10295 | Gpr132 | ENSMUSG00000021298.7 |
| chr12 | 112877212 | 112877440 | 3.75837 | 4.80146 | 1.61534 | Distal Intergenic | 12 | 112850876 | 112860916 | 10041 | 2 | ENSMUST00000021729.7 | -16297 | Gpr132 | ENSMUSG00000021298.7 |
| chr12 | 112877859 | 112878847 | 3.86609 | 5.10333 | 1.87169 | Distal Intergenic | 12 | 112850876 | 112860916 | 10041 | 2 | ENSMUST00000021729.7 | -16944 | Gpr132 | ENSMUSG00000021298.7 |
| chr12 | 112886240 | 112886447 | 3.6293 | 4.45765 | 1.39568 | Distal Intergenic | 12 | 112850876 | 112860916 | 10041 | 2 | ENSMUST00000021729.7 | -25325 | Gpr132 | ENSMUSG00000021298.7 |
| chr13 | 14522351 | 14523244 | 3.93861 | 5.07969 | 1.87391 | Promoter | 13 | 14226438 | 14523226 | 296789 | 2 | ENSMUST00000110516.2 | 0 | Hecw1 | ENSMUSG00000021301.8 |
| chr13 | 14523823 | 14523918 | 3.5169 | 4.15667 | 1.20272 | Promoter | 13 | 14226438 | 14523226 | 296789 | 2 | ENSMUST00000110516.2 | -598 | Hecw1 | ENSMUSG00000021301.8 |
| chr13 | 14524652 | 14525024 | 3.83585 | 4.76915 | 1.61115 | Distal Intergenic | 13 | 14226438 | 14523226 | 296789 | 2 | ENSMUST00000110516.2 | -1427 | Hecw1 | ENSMUSG00000021301.8 |
| chr13 | 13784093 | 13784944 | 3.87084 | 4.93565 | 1.76587 | Promoter | 13 | 13784329 | 13827897 | 43569 | 1 | ENSMUST00000021734.7 | 0 | Gng4 | ENSMUSG00000021303.8 |
| chr13 | 13785395 | 13786264 | 3.48451 | 4.0987 | 1.17051 | Intron (ENSMUST00000138783.2/ENSMUSG00000021303.8, intron 1 of 2) | 13 | 13784329 | 13827897 | 43569 | 1 | ENSMUST00000021734.7 | 1067 | Gng4 | ENSMUSG00000021303.8 |
| chr13 | 12104870 | 12105423 | 3.73255 | 4.59832 | 1.55925 | Intron (ENSMUST00000021750.8/ENSMUSG00000021313.10, intron 1 of 104) | 13 | 11553103 | 12106945 | 553843 | 2 | ENSMUST00000021750.8 | 1522 | Ryr2 | ENSMUSG00000021313.10 |
| chr13 | 12106156 | 12106724 | 3.52402 | 4.22231 | 1.33679 | Promoter | 13 | 11553103 | 12106945 | 553843 | 2 | ENSMUST00000021750.8 | 221 | Ryr2 | ENSMUSG00000021313.10 |
| chr13 | 15463420 | 15464448 | 3.60812 | 4.35531 | 1.31125 | Promoter | 13 | 15463723 | 15663518 | 199796 | 1 | ENSMUST00000130535.1 | 0 | Gli3 | ENSMUSG00000021318.10 |
| chr13 | 15464977 | 15465119 | 4.32892 | 5.73008 | 2.25694 | Promoter | 13 | 15463980 | 15730026 | 266047 | 1 | ENSMUST00000110510.3 | 998 | Gli3 | ENSMUSG00000021318.10 |
| chr13 | 15465830 | 15466186 | 3.84004 | 4.99095 | 1.75257 | Intron (ENSMUST00000130065.2/ENSMUSG00000021318.10, intron 1 of 11) | 13 | 15463980 | 15730026 | 266047 | 1 | ENSMUST00000110510.3 | 1851 | Gli3 | ENSMUSG00000021318.10 |
| chr13 | 19623405 | 19623630 | 3.46285 | 4.21422 | 1.28342 | Promoter | 13 | 19623175 | 19632821 | 9647 | 1 | ENSMUST00000002883.5 | 231 | Sfrp4 | ENSMUSG00000021319.6 |
| chr13 | 19624624 | 19624728 | 3.43949 | 4.21786 | 1.26822 | Intron (ENSMUST00000002883.5/ENSMUSG00000021319.6, intron 1 of 5) | 13 | 19623175 | 19632821 | 9647 | 1 | ENSMUST00000002883.5 | 1450 | Sfrp4 | ENSMUSG00000021319.6 |
| chr13 | 21221108 | 21221301 | 3.22109 | 4.07773 | 1.16449 | Distal Intergenic | 13 | 21190092 | 21192475 | 2384 | 1 | ENSMUST00000139287.1 | 31017 | Trim27 | ENSMUSG00000021326.6 |
| chr13 | 21222321 | 21222669 | 3.42759 | 4.01046 | 1.14369 | Distal Intergenic | 13 | 21190092 | 21192475 | 2384 | 1 | ENSMUST00000139287.1 | 32230 | Trim27 | ENSMUSG00000021326.6 |
| chr13 | 21233726 | 21233952 | 3.96563 | 5.10033 | 1.84982 | Distal Intergenic | 13 | 21190092 | 21192475 | 2384 | 1 | ENSMUST00000139287.1 | 43635 | Trim27 | ENSMUSG00000021326.6 |
| chr13 | 23992153 | 23992712 | 3.63273 | 4.37836 | 1.33803 | Promoter | 13 | 23953456 | 23991214 | 37759 | 2 | ENSMUST00000021770.7 | -940 | Scgn | ENSMUSG00000021337.7 |
| chr13 | 23993240 | 23993510 | 3.4981 | 4.15703 | 1.26656 | Distal Intergenic | 13 | 23953456 | 23991214 | 37759 | 2 | ENSMUST00000021770.7 | -2027 | Scgn | ENSMUSG00000021337.7 |
| chr13 | 30746879 | 30747297 | 4.24739 | 5.76081 | 2.30554 | Distal Intergenic | 13 | 30749226 | 30766927 | 17702 | 1 | ENSMUST00000021784.3 | -1929 | Irf4 | ENSMUSG00000021356.4 |
| chr13 | 30747889 | 30748016 | 3.51665 | 4.31916 | 1.31792 | Distal Intergenic | 13 | 30749226 | 30766927 | 17702 | 1 | ENSMUST00000021784.3 | -1210 | Irf4 | ENSMUSG00000021356.4 |
| chr13 | 30749615 | 30750015 | 3.7676 | 5.07192 | 1.82335 | Promoter | 13 | 30749226 | 30766927 | 17702 | 1 | ENSMUST00000110307.1 | 390 | Irf4 | ENSMUSG00000021356.4 |
| chr13 | 30750518 | 30750593 | 3.64622 | 4.4624 | 1.45664 | 5' UTR | 13 | 30749226 | 30766927 | 17702 | 1 | ENSMUST00000110307.1 | 1293 | Irf4 | ENSMUSG00000021356.4 |
| chr13 | 40729599 | 40731015 | 3.99993 | 5.25751 | 2.02661 | Promoter | 13 | 40715675 | 40730418 | 14744 | 2 | ENSMUST00000021787.5 | 0 | Tfap2a | ENSMUSG00000021359.10 |
| chr13 | 40731359 | 40733140 | 3.70858 | 5.03807 | 1.81142 | Promoter | 13 | 40715675 | 40733823 | 18149 | 2 | ENSMUST00000110193.3 | 683 | Tfap2a | ENSMUSG00000021359.10 |
| chr13 | 40733466 | 40734097 | 3.74833 | 4.68664 | 1.59746 | Promoter | 13 | 40715675 | 40733823 | 18149 | 2 | ENSMUST00000110193.3 | 0 | Tfap2a | ENSMUSG00000021359.10 |
| chr13 | 40734495 | 40735813 | 3.71315 | 4.76087 | 1.63895 | Promoter | 13 | 40715675 | 40733823 | 18149 | 2 | ENSMUST00000110193.3 | -673 | Tfap2a | ENSMUSG00000021359.10 |
| chr13 | 40736133 | 40736310 | 3.97834 | 5.15713 | 1.87537 | Distal Intergenic | 13 | 40715675 | 40733823 | 18149 | 2 | ENSMUST00000110193.3 | -2311 | Tfap2a | ENSMUSG00000021359.10 |
| chr13 | 40737082 | 40738816 | 4.43401 | 6.27915 | 2.76015 | Distal Intergenic | 13 | 40715675 | 40733823 | 18149 | 2 | ENSMUST00000110193.3 | -3260 | Tfap2a | ENSMUSG00000021359.10 |
| chr13 | 40739174 | 40739600 | 3.84905 | 4.87928 | 1.71783 | Distal Intergenic | 13 | 40715675 | 40733823 | 18149 | 2 | ENSMUST00000110193.3 | -5352 | Tfap2a | ENSMUSG00000021359.10 |
| chr13 | 40741001 | 40741481 | 4.25925 | 5.78396 | 2.37138 | Distal Intergenic | 13 | 40715675 | 40733823 | 18149 | 2 | ENSMUST00000110193.3 | -7179 | Tfap2a | ENSMUSG00000021359.10 |
| chr13 | 40741851 | 40743285 | 3.88755 | 4.92589 | 1.78444 | Distal Intergenic | 13 | 40715675 | 40733823 | 18149 | 2 | ENSMUST00000110193.3 | -8029 | Tfap2a | ENSMUSG00000021359.10 |
| chr13 | 40744066 | 40744442 | 3.81624 | 5.02843 | 1.77955 | Distal Intergenic | 13 | 40715675 | 40733823 | 18149 | 2 | ENSMUST00000110193.3 | -10244 | Tfap2a | ENSMUSG00000021359.10 |
| chr13 | 40744752 | 40745728 | 3.77062 | 4.79182 | 1.66161 | Distal Intergenic | 13 | 40715675 | 40733823 | 18149 | 2 | ENSMUST00000110193.3 | -10930 | Tfap2a | ENSMUSG00000021359.10 |
| chr13 | 40746336 | 40746609 | 3.31029 | 3.94803 | 1.08066 | Distal Intergenic | 13 | 40715675 | 40733823 | 18149 | 2 | ENSMUST00000110193.3 | -12514 | Tfap2a | ENSMUSG00000021359.10 |
| chr13 | 40748047 | 40749079 | 3.56376 | 4.36774 | 1.4039 | Distal Intergenic | 13 | 40715675 | 40733823 | 18149 | 2 | ENSMUST00000110193.3 | -14225 | Tfap2a | ENSMUSG00000021359.10 |
| chr13 | 40886741 | 40888711 | 3.67481 | 4.48181 | 1.45597 | Promoter | 13 | 40886758 | 40960891 | 74134 | 1 | ENSMUST00000069958.8 | 0 | Gcnt2 | ENSMUSG00000021360.10 |
| chr13 | 40916373 | 40916451 | 4.27882 | 5.78364 | 2.31778 | Intron (ENSMUST00000110191.3/ENSMUSG00000021360.10, intron 1 of 2) | 13 | 40917634 | 40960891 | 43258 | 1 | ENSMUST00000067778.6 | -1183 | Gcnt2 | ENSMUSG00000021360.10 |
| chr13 | 40917496 | 40917694 | 3.57533 | 4.32765 | 1.33489 | Promoter | 13 | 40917634 | 40960891 | 43258 | 1 | ENSMUST00000067778.6 | 0 | Gcnt2 | ENSMUSG00000021360.10 |
| chr13 | 40919155 | 40919345 | 3.4135 | 3.95835 | 1.05293 | Intron (ENSMUST00000110191.3/ENSMUSG00000021360.10, intron 1 of 2) | 13 | 40917634 | 40960891 | 43258 | 1 | ENSMUST00000067778.6 | 1522 | Gcnt2 | ENSMUSG00000021360.10 |
| chr13 | 41078479 | 41078834 | 4.25116 | 5.74974 | 2.28 | Promoter | 13 | 41025120 | 41079706 | 54587 | 2 | ENSMUST00000070193.7 | 872 | Mak | ENSMUSG00000021363.8 |
| chr13 | 41079392 | 41079749 | 3.7089 | 4.54417 | 1.51785 | Promoter | 13 | 41025120 | 41079706 | 54587 | 2 | ENSMUST00000070193.7 | 0 | Mak | ENSMUSG00000021363.8 |
| chr13 | 41080472 | 41081001 | 3.48226 | 4.17373 | 1.27382 | Promoter | 13 | 41025120 | 41079706 | 54587 | 2 | ENSMUST00000165087.2 | -767 | Mak | ENSMUSG00000021363.8 |
| chr13 | 41091047 | 41091286 | 3.30032 | 3.77782 | 1.01733 | Distal Intergenic | 13 | 41025120 | 41079706 | 54587 | 2 | ENSMUST00000165087.2 | -11342 | Mak | ENSMUSG00000021363.8 |
| chr13 | 41092483 | 41092751 | 4.13376 | 5.50035 | 2.14011 | Distal Intergenic | 13 | 41025120 | 41079706 | 54587 | 2 | ENSMUST00000165087.2 | -12778 | Mak | ENSMUSG00000021363.8 |
| chr13 | 41093207 | 41093514 | 3.62631 | 4.5434 | 1.49406 | Distal Intergenic | 13 | 41025120 | 41079706 | 54587 | 2 | ENSMUST00000165087.2 | -13502 | Mak | ENSMUSG00000021363.8 |
| chr13 | 41094414 | 41094797 | 3.59098 | 4.48428 | 1.44083 | Distal Intergenic | 13 | 41025120 | 41079706 | 54587 | 2 | ENSMUST00000165087.2 | -14709 | Mak | ENSMUSG00000021363.8 |
| chr13 | 42302636 | 42303063 | 3.66653 | 4.91209 | 1.73323 | Intron (ENSMUST00000021796.7/ENSMUSG00000021367.7, intron 1 of 4) | 13 | 42301270 | 42307989 | 6720 | 1 | ENSMUST00000021796.7 | 1367 | Edn1 | ENSMUSG00000021367.7 |
| chr13 | 42351664 | 42351919 | 3.4051 | 3.98609 | 1.08041 | Distal Intergenic | 13 | 42301270 | 42307989 | 6720 | 1 | ENSMUST00000021796.7 | 50395 | Edn1 | ENSMUSG00000021367.7 |
| chr13 | 42354719 | 42355058 | 3.38797 | 3.92426 | 1.06879 | Distal Intergenic | 13 | 42301270 | 42307989 | 6720 | 1 | ENSMUST00000021796.7 | 53450 | Edn1 | ENSMUSG00000021367.7 |
| chr13 | 46633382 | 46633551 | 3.2746 | 3.75422 | 1.06325 | Intron (ENSMUST00000021802.9/ENSMUSG00000021373.10, intron 7 of 12) | 13 | 46640058 | 46646572 | 6515 | 1 | ENSMUST00000126687.1 | -6507 | Cap2 | ENSMUSG00000021373.10 |
| chr13 | 48155438 | 48155579 | 3.46546 | 4.06139 | 1.14022 | Intron (ENSMUST00000180921.1/ENSMUSG00000097622.1, intron 1 of 2) | 13 | 48261427 | 48264036 | 2610 | 1 | ENSMUST00000021810.1 | -105848 | Id4 | ENSMUSG00000021379.1 |
| chr13 | 48158197 | 48158559 | 4.20365 | 5.57806 | 2.23716 | Intron (ENSMUST00000180921.1/ENSMUSG00000097622.1, intron 1 of 2) | 13 | 48261427 | 48264036 | 2610 | 1 | ENSMUST00000021810.1 | -102868 | Id4 | ENSMUSG00000021379.1 |
| chr13 | 48167035 | 48167110 | 4.00956 | 5.07986 | 1.79026 | Intron (ENSMUST00000180921.1/ENSMUSG00000097622.1, intron 1 of 2) | 13 | 48261427 | 48264036 | 2610 | 1 | ENSMUST00000021810.1 | -94317 | Id4 | ENSMUSG00000021379.1 |
| chr13 | 48171529 | 48171731 | 3.75359 | 4.58879 | 1.45614 | Intron (ENSMUST00000180921.1/ENSMUSG00000097622.1, intron 1 of 2) | 13 | 48261427 | 48264036 | 2610 | 1 | ENSMUST00000021810.1 | -89696 | Id4 | ENSMUSG00000021379.1 |
| chr13 | 48173150 | 48173239 | 3.47512 | 4.07787 | 1.1472 | Intron (ENSMUST00000180921.1/ENSMUSG00000097622.1, intron 1 of 2) | 13 | 48261427 | 48264036 | 2610 | 1 | ENSMUST00000021810.1 | -88188 | Id4 | ENSMUSG00000021379.1 |
| chr13 | 48185940 | 48186906 | 3.49073 | 4.11062 | 1.17903 | Intron (ENSMUST00000180921.1/ENSMUSG00000097622.1, intron 1 of 2) | 13 | 48261427 | 48264036 | 2610 | 1 | ENSMUST00000021810.1 | -74521 | Id4 | ENSMUSG00000021379.1 |
| chr13 | 48200583 | 48200658 | 3.60148 | 4.35779 | 1.41235 | Intron (ENSMUST00000180921.1/ENSMUSG00000097622.1, intron 1 of 2) | 13 | 48261427 | 48264036 | 2610 | 1 | ENSMUST00000021810.1 | -60769 | Id4 | ENSMUSG00000021379.1 |
| chr13 | 48653231 | 48653888 | 3.63646 | 4.48757 | 1.42019 | Distal Intergenic | 13 | 48662998 | 48666507 | 3510 | 1 | ENSMUST00000021813.4 | -9110 | Barx1 | ENSMUSG00000021381.4 |
| chr13 | 48656152 | 48656369 | 3.58719 | 4.28318 | 1.26767 | Distal Intergenic | 13 | 48662998 | 48666507 | 3510 | 1 | ENSMUST00000021813.4 | -6629 | Barx1 | ENSMUSG00000021381.4 |
| chr13 | 48657655 | 48661922 | 3.8909 | 5.00442 | 1.78587 | Distal Intergenic | 13 | 48662998 | 48666507 | 3510 | 1 | ENSMUST00000021813.4 | -1076 | Barx1 | ENSMUSG00000021381.4 |
| chr13 | 48662340 | 48663658 | 4.16395 | 5.43021 | 2.08111 | Promoter | 13 | 48662998 | 48666507 | 3510 | 1 | ENSMUST00000021813.4 | 0 | Barx1 | ENSMUSG00000021381.4 |
| chr13 | 48663975 | 48664212 | 4.0086 | 5.7463 | 2.28104 | Promoter | 13 | 48662998 | 48666507 | 3510 | 1 | ENSMUST00000021813.4 | 978 | Barx1 | ENSMUSG00000021381.4 |
| chr13 | 48664530 | 48667082 | 4.20312 | 5.68374 | 2.30393 | 3' UTR | 13 | 48662998 | 48666507 | 3510 | 1 | ENSMUST00000021813.4 | 1533 | Barx1 | ENSMUSG00000021381.4 |
| chr13 | 48667467 | 48667599 | 3.63478 | 4.42171 | 1.37419 | Distal Intergenic | 13 | 48662998 | 48666507 | 3510 | 1 | ENSMUST00000021813.4 | 4470 | Barx1 | ENSMUSG00000021381.4 |
| chr13 | 48668754 | 48669922 | 3.92001 | 5.21052 | 1.94205 | Distal Intergenic | 13 | 48662998 | 48666507 | 3510 | 1 | ENSMUST00000021813.4 | 5757 | Barx1 | ENSMUSG00000021381.4 |
| chr13 | 48670788 | 48671730 | 4.04033 | 5.20553 | 1.91551 | Distal Intergenic | 13 | 48662998 | 48666507 | 3510 | 1 | ENSMUST00000021813.4 | 7791 | Barx1 | ENSMUSG00000021381.4 |
| chr13 | 48672391 | 48673288 | 3.66178 | 4.42769 | 1.36668 | Distal Intergenic | 13 | 48662998 | 48666507 | 3510 | 1 | ENSMUST00000021813.4 | 9394 | Barx1 | ENSMUSG00000021381.4 |
| chr13 | 48673644 | 48673725 | 4.31179 | 5.69406 | 2.22953 | Distal Intergenic | 13 | 48662998 | 48666507 | 3510 | 1 | ENSMUST00000021813.4 | 10647 | Barx1 | ENSMUSG00000021381.4 |
| chr13 | 48674378 | 48674460 | 3.32138 | 3.80936 | 1.01733 | Distal Intergenic | 13 | 48662998 | 48666507 | 3510 | 1 | ENSMUST00000021813.4 | 11381 | Barx1 | ENSMUSG00000021381.4 |
| chr13 | 48675555 | 48675716 | 3.59314 | 4.3135 | 1.31056 | Distal Intergenic | 13 | 48662998 | 48666507 | 3510 | 1 | ENSMUST00000021813.4 | 12558 | Barx1 | ENSMUSG00000021381.4 |
| chr13 | 48685028 | 48685382 | 3.57755 | 4.29145 | 1.33698 | Distal Intergenic | 13 | 48662998 | 48666507 | 3510 | 1 | ENSMUST00000021813.4 | 22031 | Barx1 | ENSMUSG00000021381.4 |
| chr13 | 48694628 | 48694703 | 3.47215 | 4.07188 | 1.1415 | Distal Intergenic | 13 | 48662998 | 48666507 | 3510 | 1 | ENSMUST00000021813.4 | 31631 | Barx1 | ENSMUSG00000021381.4 |
| chr13 | 48698922 | 48699327 | 3.64297 | 4.42954 | 1.41232 | Distal Intergenic | 13 | 48662998 | 48666507 | 3510 | 1 | ENSMUST00000021813.4 | 35925 | Barx1 | ENSMUSG00000021381.4 |
| chr13 | 48705636 | 48705892 | 3.59442 | 4.34978 | 1.38045 | Distal Intergenic | 13 | 48662998 | 48666507 | 3510 | 1 | ENSMUST00000021813.4 | 42639 | Barx1 | ENSMUSG00000021381.4 |
| chr13 | 48706302 | 48706839 | 3.43069 | 4.088 | 1.2072 | Distal Intergenic | 13 | 48662998 | 48666507 | 3510 | 1 | ENSMUST00000021813.4 | 43305 | Barx1 | ENSMUSG00000021381.4 |
| chr13 | 48723563 | 48723808 | 3.8987 | 4.89485 | 1.70201 | Distal Intergenic | 13 | 48662998 | 48666507 | 3510 | 1 | ENSMUST00000021813.4 | 60566 | Barx1 | ENSMUSG00000021381.4 |
| chr13 | 48730298 | 48730375 | 3.88365 | 4.82679 | 1.59493 | Distal Intergenic | 13 | 48662998 | 48666507 | 3510 | 1 | ENSMUST00000021813.4 | 67301 | Barx1 | ENSMUSG00000021381.4 |
| chr13 | 48731069 | 48733899 | 3.72483 | 4.57865 | 1.48545 | Distal Intergenic | 13 | 48662998 | 48666507 | 3510 | 1 | ENSMUST00000021813.4 | 68072 | Barx1 | ENSMUSG00000021381.4 |
| chr13 | 48734365 | 48735102 | 3.78809 | 4.69518 | 1.59339 | Distal Intergenic | 13 | 48662998 | 48666507 | 3510 | 1 | ENSMUST00000021813.4 | 71368 | Barx1 | ENSMUSG00000021381.4 |
| chr13 | 48748762 | 48748863 | 3.32624 | 3.81668 | 1.01733 | Distal Intergenic | 13 | 48662998 | 48666507 | 3510 | 1 | ENSMUST00000021813.4 | 85765 | Barx1 | ENSMUSG00000021381.4 |
| chr13 | 49247761 | 49248543 | 4.02775 | 5.33565 | 2.02941 | Promoter | 13 | 49230690 | 49248294 | 17605 | 2 | ENSMUST00000058196.7 | 0 | Susd3 | ENSMUSG00000021384.9 |
| chr13 | 51171291 | 51171539 | 3.71408 | 4.63236 | 1.55175 | Promoter | 13 | 51171025 | 51175187 | 4163 | 1 | ENSMUST00000021828.4 | 267 | Nxnl2 | ENSMUSG00000021396.4 |
| chr13 | 51172733 | 51173045 | 3.38699 | 3.95663 | 1.1576 | Intron (ENSMUST00000021828.4/ENSMUSG00000021396.4, intron 1 of 1) | 13 | 51171025 | 51175187 | 4163 | 1 | ENSMUST00000021828.4 | 1709 | Nxnl2 | ENSMUSG00000021396.4 |
| chr13 | 34916820 | 34917303 | 3.79109 | 4.74445 | 1.66903 | 5' UTR | 13 | 34909964 | 34919992 | 10029 | 2 | ENSMUST00000021851.6 | 2689 | Fam217a | ENSMUSG00000021414.7 |
| chr13 | 34962551 | 34962772 | 3.29609 | 3.7713 | 1.01733 | Intron (ENSMUST00000021853.6/ENSMUSG00000021416.6, intron 1 of 8) | 13 | 34948084 | 34963788 | 15705 | 2 | ENSMUST00000164155.1 | 1016 | Eci3 | ENSMUSG00000021416.6 |
| chr13 | 37311117 | 37311410 | 3.34153 | 3.83966 | 1.01733 | Distal Intergenic | 13 | 37345345 | 37419036 | 73692 | 1 | ENSMUST00000021860.5 | -33935 | Ly86 | ENSMUSG00000021423.5 |
| chr13 | 38977507 | 38977779 | 3.35914 | 3.86625 | 1.01733 | Distal Intergenic | 13 | 38932140 | 38960875 | 28736 | 2 | ENSMUST00000021870.5 | -16633 | Slc35b3 | ENSMUSG00000021432.9 |
| chr13 | 38978250 | 38978413 | 3.58234 | 4.51029 | 1.45352 | Distal Intergenic | 13 | 38932140 | 38960875 | 28736 | 2 | ENSMUST00000021870.5 | -17376 | Slc35b3 | ENSMUSG00000021432.9 |
| chr13 | 39020165 | 39020387 | 3.53761 | 4.19059 | 1.21095 | Distal Intergenic | 13 | 38932140 | 38960875 | 28736 | 2 | ENSMUST00000021870.5 | -59291 | Slc35b3 | ENSMUSG00000021432.9 |
| chr13 | 51567314 | 51568091 | 3.66666 | 4.68043 | 1.57168 | Promoter | 13 | 51431041 | 51567084 | 136044 | 2 | ENSMUST00000021898.5 | -231 | Shc3 | ENSMUSG00000021448.6 |
| chr13 | 51568468 | 51569108 | 3.52508 | 4.34933 | 1.35829 | Distal Intergenic | 13 | 51431041 | 51567084 | 136044 | 2 | ENSMUST00000021898.5 | -1385 | Shc3 | ENSMUSG00000021448.6 |
| chr13 | 51569886 | 51570309 | 3.56055 | 4.40791 | 1.37955 | Distal Intergenic | 13 | 51431041 | 51567084 | 136044 | 2 | ENSMUST00000021898.5 | -2803 | Shc3 | ENSMUSG00000021448.6 |
| chr13 | 51570824 | 51570992 | 3.48909 | 4.2197 | 1.2818 | Distal Intergenic | 13 | 51431041 | 51567084 | 136044 | 2 | ENSMUST00000021898.5 | -3741 | Shc3 | ENSMUSG00000021448.6 |
| chr13 | 51580534 | 51580879 | 3.50619 | 4.15528 | 1.24427 | Distal Intergenic | 13 | 51431041 | 51567084 | 136044 | 2 | ENSMUST00000021898.5 | -13451 | Shc3 | ENSMUSG00000021448.6 |
| chr13 | 51594043 | 51594844 | 3.5706 | 4.5162 | 1.46655 | Distal Intergenic | 13 | 51431041 | 51567084 | 136044 | 2 | ENSMUST00000021898.5 | -26960 | Shc3 | ENSMUSG00000021448.6 |
| chr13 | 51792233 | 51793452 | 4.14299 | 5.47793 | 2.12259 | Promoter | 13 | 51725423 | 51793664 | 68242 | 2 | ENSMUST00000155896.1 | 212 | Sema4d | ENSMUSG00000021451.10 |
| chr13 | 51833720 | 51833842 | 4.27237 | 5.63416 | 2.21373 | Distal Intergenic | 13 | 51846678 | 51848468 | 1791 | 1 | ENSMUST00000021903.2 | -12836 | Gadd45g | ENSMUSG00000021453.2 |
| chr13 | 51835112 | 51835456 | 3.59422 | 4.44319 | 1.41731 | Distal Intergenic | 13 | 51846678 | 51848468 | 1791 | 1 | ENSMUST00000021903.2 | -11222 | Gadd45g | ENSMUSG00000021453.2 |
| chr13 | 51838013 | 51838230 | 3.33438 | 3.82894 | 1.01733 | Distal Intergenic | 13 | 51846678 | 51848468 | 1791 | 1 | ENSMUST00000021903.2 | -8448 | Gadd45g | ENSMUSG00000021453.2 |
| chr13 | 51842015 | 51842090 | 3.93657 | 4.94607 | 1.71412 | Distal Intergenic | 13 | 51846678 | 51848468 | 1791 | 1 | ENSMUST00000021903.2 | -4588 | Gadd45g | ENSMUSG00000021453.2 |
| chr13 | 51843492 | 51843620 | 3.2396 | 3.68656 | 1.01733 | Distal Intergenic | 13 | 51846678 | 51848468 | 1791 | 1 | ENSMUST00000021903.2 | -3058 | Gadd45g | ENSMUSG00000021453.2 |
| chr13 | 51902271 | 51902756 | 3.42631 | 4.08361 | 1.20829 | Distal Intergenic | 13 | 51846744 | 51848224 | 1481 | 1 | ENSMUST00000132480.1 | 55528 | Gadd45g | ENSMUSG00000021453.2 |
| chr13 | 52583123 | 52583461 | 3.40984 | 4.22695 | 1.2442 | Promoter | 13 | 52583173 | 52648792 | 65620 | 1 | ENSMUST00000120135.2 | 0 | Syk | ENSMUSG00000021457.9 |
| chr13 | 52654875 | 52655013 | 3.42408 | 4.19064 | 1.2793 | Distal Intergenic | 13 | 52596959 | 52624339 | 27381 | 1 | ENSMUST00000140339.1 | 57917 | Syk | ENSMUSG00000021457.9 |
| chr13 | 52702513 | 52702653 | 3.34637 | 3.84695 | 1.01733 | Distal Intergenic | 13 | 52596959 | 52624339 | 27381 | 1 | ENSMUST00000140339.1 | 105555 | Syk | ENSMUSG00000021457.9 |
| chr13 | 52804897 | 52805137 | 3.62521 | 4.39815 | 1.43266 | Distal Intergenic | 13 | 52835119 | 52919232 | 84114 | 2 | ENSMUST00000137064.2 | 114095 | Auh | ENSMUSG00000021460.11 |
| chr13 | 53163909 | 53164236 | 3.67715 | 4.45507 | 1.39503 | Intron (ENSMUST00000130235.2/ENSMUSG00000021464.9, intron 1 of 8) | 13 | 53131960 | 53275658 | 143699 | 2 | ENSMUST00000149299.1 | 111422 | Ror2 | ENSMUSG00000021464.9 |
| chr13 | 53175697 | 53175823 | 3.61727 | 4.34472 | 1.3267 | Intron (ENSMUST00000130235.2/ENSMUSG00000021464.9, intron 1 of 8) | 13 | 53131960 | 53275658 | 143699 | 2 | ENSMUST00000149299.1 | 99835 | Ror2 | ENSMUSG00000021464.9 |
| chr13 | 53176317 | 53176394 | 3.74172 | 4.57686 | 1.4747 | Intron (ENSMUST00000130235.2/ENSMUSG00000021464.9, intron 1 of 8) | 13 | 53131960 | 53275658 | 143699 | 2 | ENSMUST00000149299.1 | 99264 | Ror2 | ENSMUSG00000021464.9 |
| chr13 | 53178665 | 53178949 | 3.35925 | 3.87219 | 1.03787 | Intron (ENSMUST00000130235.2/ENSMUSG00000021464.9, intron 1 of 8) | 13 | 53131960 | 53275658 | 143699 | 2 | ENSMUST00000149299.1 | 96709 | Ror2 | ENSMUSG00000021464.9 |
| chr13 | 53179318 | 53179673 | 3.57591 | 4.29186 | 1.28517 | Intron (ENSMUST00000130235.2/ENSMUSG00000021464.9, intron 1 of 8) | 13 | 53131960 | 53275658 | 143699 | 2 | ENSMUST00000149299.1 | 95985 | Ror2 | ENSMUSG00000021464.9 |
| chr13 | 53182900 | 53183253 | 3.29656 | 3.77617 | 1.03196 | Intron (ENSMUST00000130235.2/ENSMUSG00000021464.9, intron 1 of 8) | 13 | 53131960 | 53275658 | 143699 | 2 | ENSMUST00000149299.1 | 92405 | Ror2 | ENSMUSG00000021464.9 |
| chr13 | 53187578 | 53187907 | 3.44863 | 4.02648 | 1.10632 | Intron (ENSMUST00000130235.2/ENSMUSG00000021464.9, intron 1 of 8) | 13 | 53131960 | 53275658 | 143699 | 2 | ENSMUST00000149299.1 | 87751 | Ror2 | ENSMUSG00000021464.9 |
| chr13 | 53230497 | 53230797 | 3.69263 | 4.57627 | 1.52135 | Intron (ENSMUST00000130235.2/ENSMUSG00000021464.9, intron 1 of 8) | 13 | 53131960 | 53275658 | 143699 | 2 | ENSMUST00000149299.1 | 44861 | Ror2 | ENSMUSG00000021464.9 |
| chr13 | 53284741 | 53285935 | 4.24269 | 5.76434 | 2.35427 | Promoter | 13 | 53109312 | 53285549 | 176238 | 2 | ENSMUST00000130235.2 | 0 | Ror2 | ENSMUSG00000021464.9 |
| chr13 | 53307042 | 53307292 | 3.50105 | 4.63584 | 1.55594 | Distal Intergenic | 13 | 53109317 | 53286124 | 176808 | 2 | ENSMUST00000021918.8 | -20919 | Ror2 | ENSMUSG00000021464.9 |
| chr13 | 53388701 | 53388948 | 3.35885 | 3.86578 | 1.01733 | Distal Intergenic | 13 | 53332716 | 53377365 | 44650 | 2 | ENSMUST00000021920.6 | -11337 | Sptlc1 | ENSMUSG00000021468.7 |
| chr13 | 53400561 | 53400636 | 3.41231 | 3.99832 | 1.19152 | Distal Intergenic | 13 | 53332716 | 53377365 | 44650 | 2 | ENSMUST00000021920.6 | -23197 | Sptlc1 | ENSMUSG00000021468.7 |
| chr13 | 53442423 | 53442792 | 3.60811 | 4.31554 | 1.27543 | Distal Intergenic | 13 | 53466884 | 53473074 | 6191 | 2 | ENSMUST00000021922.9 | 30282 | Msx2 | ENSMUSG00000021469.9 |
| chr13 | 53447648 | 53448192 | 3.68549 | 4.48484 | 1.41696 | Distal Intergenic | 13 | 53466884 | 53473074 | 6191 | 2 | ENSMUST00000021922.9 | 24882 | Msx2 | ENSMUSG00000021469.9 |
| chr13 | 53448646 | 53448962 | 3.64264 | 4.38053 | 1.3168 | Distal Intergenic | 13 | 53466884 | 53473074 | 6191 | 2 | ENSMUST00000021922.9 | 24112 | Msx2 | ENSMUSG00000021469.9 |
| chr13 | 53452431 | 53452583 | 3.88914 | 4.85096 | 1.63599 | Distal Intergenic | 13 | 53466884 | 53473074 | 6191 | 2 | ENSMUST00000021922.9 | 20491 | Msx2 | ENSMUSG00000021469.9 |
| chr13 | 53453321 | 53453716 | 3.55432 | 4.22161 | 1.22878 | Distal Intergenic | 13 | 53466884 | 53473074 | 6191 | 2 | ENSMUST00000021922.9 | 19358 | Msx2 | ENSMUSG00000021469.9 |
| chr13 | 53454981 | 53455068 | 4.57878 | 6.21081 | 2.57911 | Distal Intergenic | 13 | 53466884 | 53473074 | 6191 | 2 | ENSMUST00000021922.9 | 18006 | Msx2 | ENSMUSG00000021469.9 |
| chr13 | 53455629 | 53455854 | 3.70998 | 4.62421 | 1.52328 | Distal Intergenic | 13 | 53466884 | 53473074 | 6191 | 2 | ENSMUST00000021922.9 | 17220 | Msx2 | ENSMUSG00000021469.9 |
| chr13 | 53461040 | 53464076 | 3.92624 | 5.00494 | 1.75986 | Downstream (2-3kb) | 13 | 53466884 | 53473074 | 6191 | 2 | ENSMUST00000021922.9 | 8998 | Msx2 | ENSMUSG00000021469.9 |
| chr13 | 53464772 | 53465167 | 3.78869 | 4.69977 | 1.584 | Downstream (1-2kb) | 13 | 53466884 | 53473074 | 6191 | 2 | ENSMUST00000021922.9 | 7907 | Msx2 | ENSMUSG00000021469.9 |
| chr13 | 53465738 | 53467063 | 3.653 | 4.44986 | 1.41552 | 3' UTR | 13 | 53466884 | 53473074 | 6191 | 2 | ENSMUST00000021922.9 | 6011 | Msx2 | ENSMUSG00000021469.9 |
| chr13 | 53467386 | 53468514 | 3.65968 | 4.42712 | 1.36802 | 3' UTR | 13 | 53466884 | 53473074 | 6191 | 2 | ENSMUST00000021922.9 | 4560 | Msx2 | ENSMUSG00000021469.9 |
| chr13 | 53469262 | 53469474 | 3.99186 | 5.08083 | 1.85891 | Intron (ENSMUST00000021922.9/ENSMUSG00000021469.9, intron 1 of 1) | 13 | 53466884 | 53473074 | 6191 | 2 | ENSMUST00000021922.9 | 3600 | Msx2 | ENSMUSG00000021469.9 |
| chr13 | 53471080 | 53472921 | 4.14392 | 5.39115 | 2.04524 | Promoter | 13 | 53466884 | 53473074 | 6191 | 2 | ENSMUST00000021922.9 | 153 | Msx2 | ENSMUSG00000021469.9 |
| chr13 | 53473413 | 53477902 | 3.85151 | 4.82325 | 1.63785 | Promoter | 13 | 53471506 | 53473074 | 1569 | 2 | ENSMUST00000188606.1 | -340 | Msx2 | ENSMUSG00000021469.9 |
| chr13 | 53481090 | 53481289 | 3.53389 | 4.28131 | 1.27971 | Distal Intergenic | 13 | 53471506 | 53473074 | 1569 | 2 | ENSMUST00000188606.1 | -8017 | Msx2 | ENSMUSG00000021469.9 |
| chr13 | 53806968 | 53807213 | 3.25238 | 3.82115 | 1.01733 | Distal Intergenic | 13 | 54051186 | 54055658 | 4473 | 2 | ENSMUST00000021932.5 | 248445 | Drd1a | ENSMUSG00000021478.5 |
| chr13 | 54055206 | 54055825 | 3.80366 | 4.75912 | 1.61112 | Promoter | 13 | 54051186 | 54055658 | 4473 | 2 | ENSMUST00000021932.5 | 0 | Drd1a | ENSMUSG00000021478.5 |
| chr13 | 55797783 | 55797978 | 3.34474 | 3.8445 | 1.01733 | Intron (ENSMUST00000021961.6/ENSMUSG00000021499.7, intron 2 of 7) | 13 | 55784579 | 55808998 | 24420 | 1 | ENSMUST00000109898.2 | 13205 | Catsper3 | ENSMUSG00000021499.7 |
| chr13 | 55817597 | 55817672 | 3.38193 | 3.9006 | 1.01733 | Distal Intergenic | 13 | 55825051 | 55831425 | 6375 | 2 | ENSMUST00000021968.6 | 13753 | Pitx1 | ENSMUSG00000021506.7 |
| chr13 | 55818150 | 55818572 | 3.77225 | 4.64081 | 1.51937 | Distal Intergenic | 13 | 55825051 | 55831425 | 6375 | 2 | ENSMUST00000021968.6 | 12853 | Pitx1 | ENSMUSG00000021506.7 |
| chr13 | 55823219 | 55824722 | 3.62389 | 4.37019 | 1.33361 | Downstream (<1kb) | 13 | 55825051 | 55831425 | 6375 | 2 | ENSMUST00000021968.6 | 6703 | Pitx1 | ENSMUSG00000021506.7 |
| chr13 | 55825079 | 55826710 | 4.4697 | 6.16058 | 2.6558 | 3' UTR | 13 | 55825051 | 55831425 | 6375 | 2 | ENSMUST00000021968.6 | 4715 | Pitx1 | ENSMUSG00000021506.7 |
| chr13 | 55827028 | 55828989 | 4.11342 | 5.45825 | 2.12522 | Exon (ENSMUST00000021968.6/ENSMUSG00000021506.7, exon 2 of 3) | 13 | 55825051 | 55831425 | 6375 | 2 | ENSMUST00000021968.6 | 2436 | Pitx1 | ENSMUSG00000021506.7 |
| chr13 | 55829407 | 55829565 | 4.08258 | 5.40502 | 2.06411 | Intron (ENSMUST00000021968.6/ENSMUSG00000021506.7, intron 1 of 2) | 13 | 55825051 | 55831425 | 6375 | 2 | ENSMUST00000021968.6 | 1860 | Pitx1 | ENSMUSG00000021506.7 |
| chr13 | 55829896 | 55830615 | 3.88568 | 4.9716 | 1.78689 | Promoter | 13 | 55825051 | 55831425 | 6375 | 2 | ENSMUST00000021968.6 | 810 | Pitx1 | ENSMUSG00000021506.7 |
| chr13 | 55831021 | 55832291 | 4.2904 | 5.974 | 2.51123 | Promoter | 13 | 55825051 | 55831425 | 6375 | 2 | ENSMUST00000021968.6 | 0 | Pitx1 | ENSMUSG00000021506.7 |
| chr13 | 55899813 | 55900090 | 3.33628 | 4.0292 | 1.16033 | Distal Intergenic | 13 | 55828474 | 55836192 | 7719 | 2 | ENSMUST00000173618.1 | -63622 | Pitx1 | ENSMUSG00000021506.7 |

| | | | | | | | | | | | | | | | |
|---|---|---|---|---|---|---|---|---|---|---|---|---|---|---|---|
| chr13 | 55835439 | 55838849 | 4.15774 | 5.5031 | 2.16148 | Promoter | 13 | 55828474 | 55836192 | 7719 | 2 | ENSMUST00000173618.1 | 0 | Pitx1 | ENSMUSG00000021506.7 |
| chr13 | 55839821 | 55840509 | 3.63703 | 4.45715 | 1.44356 | Distal Intergenic | 13 | 55828474 | 55836192 | 7719 | 2 | ENSMUST00000173618.1 | -3630 | Pitx1 | ENSMUSG00000021506.7 |
| chr13 | 55841629 | 55841952 | 3.33753 | 3.8721 | 1.05149 | Distal Intergenic | 13 | 55828474 | 55836192 | 7719 | 2 | ENSMUST00000173618.1 | -5438 | Pitx1 | ENSMUSG00000021506.7 |
| chr13 | 55842469 | 55842557 | 3.45841 | 4.05602 | 1.15926 | Distal Intergenic | 13 | 55828474 | 55836192 | 7719 | 2 | ENSMUST00000173618.1 | -6278 | Pitx1 | ENSMUSG00000021506.7 |
| chr13 | 55845091 | 55845213 | 3.59277 | 4.31286 | 1.35294 | Distal Intergenic | 13 | 55828474 | 55836192 | 7719 | 2 | ENSMUST00000173618.1 | -8900 | Pitx1 | ENSMUSG00000021506.7 |
| chr13 | 55845553 | 55845852 | 3.56982 | 4.24913 | 1.23704 | Distal Intergenic | 13 | 55828474 | 55836192 | 7719 | 2 | ENSMUST00000173618.1 | -9362 | Pitx1 | ENSMUSG00000021506.7 |
| chr13 | 55846676 | 55846819 | 4.49146 | 6.27405 | 2.66875 | Distal Intergenic | 13 | 55828474 | 55836192 | 7719 | 2 | ENSMUST00000173618.1 | -10485 | Pitx1 | ENSMUSG00000021506.7 |
| chr13 | 55847150 | 55847348 | 3.44844 | 4.09081 | 1.24738 | Distal Intergenic | 13 | 55828474 | 55836192 | 7719 | 2 | ENSMUST00000173618.1 | -10959 | Pitx1 | ENSMUSG00000021506.7 |
| chr13 | 55847791 | 55848796 | 3.57497 | 4.26168 | 1.25972 | Distal Intergenic | 13 | 55828474 | 55836192 | 7719 | 2 | ENSMUST00000173618.1 | -11600 | Pitx1 | ENSMUSG00000021506.7 |
| chr13 | 55851247 | 55851573 | 3.35745 | 3.86367 | 1.01733 | Distal Intergenic | 13 | 55828474 | 55836192 | 7719 | 2 | ENSMUST00000173618.1 | -15056 | Pitx1 | ENSMUSG00000021506.7 |
| chr13 | 55860867 | 55861048 | 3.31649 | 3.83107 | 1.10873 | Distal Intergenic | 13 | 55828474 | 55836192 | 7719 | 2 | ENSMUST00000173618.1 | -24676 | Pitx1 | ENSMUSG00000021506.7 |
| chr13 | 55898752 | 55899184 | 3.44474 | 4.03602 | 1.1569 | Distal Intergenic | 13 | 55828474 | 55836192 | 7719 | 2 | ENSMUST00000173618.1 | -62561 | Pitx1 | ENSMUSG00000021506.7 |
| chr13 | 55924880 | 55925265 | 3.45057 | 4.03282 | 1.11878 | Distal Intergenic | 13 | 55828474 | 55836192 | 7719 | 2 | ENSMUST00000173618.1 | -88689 | Pitx1 | ENSMUSG00000021506.7 |
| chr13 | 55902186 | 55902277 | 3.6966 | 4.59488 | 1.48206 | Distal Intergenic | 13 | 55828474 | 55836192 | 7719 | 2 | ENSMUST00000173618.1 | -65995 | Pitx1 | ENSMUSG00000021506.7 |
| chr13 | 55922211 | 55922377 | 4.20028 | 5.53883 | 2.15936 | Distal Intergenic | 13 | 55828474 | 55836192 | 7719 | 2 | ENSMUST00000173618.1 | -86020 | Pitx1 | ENSMUSG00000021506.7 |
| chr13 | 55923242 | 55923695 | 3.53733 | 4.20597 | 1.22724 | Distal Intergenic | 13 | 55828474 | 55836192 | 7719 | 2 | ENSMUST00000173618.1 | -87051 | Pitx1 | ENSMUSG00000021506.7 |
| chr13 | 55926614 | 55926855 | 3.76809 | 4.62171 | 1.48212 | Distal Intergenic | 13 | 55828474 | 55836192 | 7719 | 2 | ENSMUST00000173618.1 | -90423 | Pitx1 | ENSMUSG00000021506.7 |
| chr13 | 55955765 | 55956196 | 3.44951 | 4.02426 | 1.09361 | Distal Intergenic | 13 | 55828474 | 55836192 | 7719 | 2 | ENSMUST00000173618.1 | -119574 | Pitx1 | ENSMUSG00000021506.7 |
| chr13 | 56294819 | 56296694 | 4.2489 | 5.6549 | 2.28271 | Promoter | 13 | 56288647 | 56296551 | 7905 | 2 | ENSMUST00000021970.9 | 0 | Cxcl14 | ENSMUSG00000021508.10 |
| chr13 | 56297324 | 56297483 | 3.58598 | 4.31713 | 1.31922 | Promoter | 13 | 56288647 | 56296551 | 7905 | 2 | ENSMUST00000021970.9 | -774 | Cxcl14 | ENSMUSG00000021508.10 |
| chr13 | 56417995 | 56418126 | 3.52723 | 4.17726 | 1.22073 | Distal Intergenic | 13 | 56438355 | 56474239 | 35885 | 1 | ENSMUST00000021971.5 | -20229 | Slc25a48 | ENSMUSG00000021509.5 |
| chr13 | 56422730 | 56423120 | 3.41665 | 3.97486 | 1.09471 | Distal Intergenic | 13 | 56438355 | 56474239 | 35885 | 1 | ENSMUST00000021971.5 | -15235 | Slc25a48 | ENSMUSG00000021509.5 |
| chr13 | 56428917 | 56429266 | 3.76693 | 4.61184 | 1.4616 | Distal Intergenic | 13 | 56438355 | 56474239 | 35885 | 1 | ENSMUST00000021971.5 | -9089 | Slc25a48 | ENSMUSG00000021509.5 |
| chr13 | 56437597 | 56438648 | 3.93608 | 5.07402 | 1.83453 | Promoter | 13 | 56438355 | 56474239 | 35885 | 1 | ENSMUST00000021971.5 | 0 | Slc25a48 | ENSMUSG00000021509.5 |
| chr13 | 56439254 | 56439599 | 3.69116 | 4.55879 | 1.52845 | Promoter | 13 | 56438355 | 56474239 | 35885 | 1 | ENSMUST00000021971.5 | 900 | Slc25a48 | ENSMUSG00000021509.5 |
| chr13 | 56440230 | 56440309 | 3.38301 | 3.94623 | 1.16262 | Intron (ENSMUST00000021971.5/ENSMUSG00000021509.5, intron 1 of 7) | 13 | 56438355 | 56474239 | 35885 | 1 | ENSMUST00000021971.5 | 1876 | Slc25a48 | ENSMUSG00000021509.5 |
| chr13 | 56440967 | 56441067 | 4.39369 | 5.92708 | 2.46628 | Intron (ENSMUST00000021971.5/ENSMUSG00000021509.5, intron 1 of 7) | 13 | 56438355 | 56474239 | 35885 | 1 | ENSMUST00000021971.5 | 2613 | Slc25a48 | ENSMUSG00000021509.5 |
| chr13 | 68597486 | 68597670 | 3.60812 | 4.31669 | 1.281 | Promoter | 13 | 68597439 | 68614231 | 16793 | 1 | ENSMUST00000022007.6 | 48 | 1700001L19Rik | ENSMUSG00000021534.7 |
| chr13 | 68997887 | 68998319 | 3.802 | 4.86674 | 1.68835 | Intron (ENSMUST00000022013.7/ENSMUSG00000021536.7, intron 1 of 24) | 13 | 68620043 | 68999541 | 379499 | 2 | ENSMUST00000022013.7 | 1222 | Adcy2 | ENSMUSG00000021536.7 |
| chr13 | 56500579 | 56500955 | 3.34027 | 3.97256 | 1.18092 | Distal Intergenic | 13 | 56479277 | 56482246 | 2970 | 2 | ENSMUST00000022019.3 | -18334 | Il9 | ENSMUSG00000021538.3 |
| chr13 | 56553820 | 56553963 | 3.47089 | 4.1808 | 1.261 | Distal Intergenic | 13 | 56542460 | 56548538 | 6079 | 2 | ENSMUST00000062806.4 | -5283 | Lect2 | ENSMUSG00000021539.8 |
| chr13 | 56562684 | 56562871 | 3.54262 | 4.32181 | 1.35318 | Distal Intergenic | 13 | 56542460 | 56548538 | 6079 | 2 | ENSMUST00000062806.4 | -14147 | Lect2 | ENSMUSG00000021539.8 |
| chr13 | 56752166 | 56754053 | 4.14463 | 5.43772 | 2.08327 | Distal Intergenic | 13 | 56733017 | 56738993 | 5977 | 1 | ENSMUST00000136702.1 | 19150 | Smad5 | ENSMUSG00000021540.11 |
| chr13 | 56754652 | 56754924 | 3.68932 | 4.47144 | 1.3805 | Distal Intergenic | 13 | 56733017 | 56738993 | 5977 | 1 | ENSMUST00000136702.1 | 21636 | Smad5 | ENSMUSG00000021540.11 |
| chr13 | 56773591 | 56773846 | 3.81036 | 4.7369 | 1.57947 | 3' UTR | 13 | 56733017 | 56738993 | 5977 | 1 | ENSMUST00000136702.1 | 40575 | Smad5 | ENSMUSG00000021540.11 |
| chr13 | 56810620 | 56810726 | 4.33765 | 5.73462 | 2.25352 | Intron (ENSMUST00000022023.7/ENSMUSG00000021541.8, intron 5 of 11) | 13 | 56733017 | 56738993 | 5977 | 1 | ENSMUST00000136702.1 | 77604 | Smad5 | ENSMUSG00000021540.11 |
| chr17 | 48094116 | 48094542 | 3.52287 | 4.20899 | 1.27332 | Distal Intergenic | 17 | 48089632 | 48090920 | 1289 | 1 | ENSMUST00000022028.4 | 4485 | 1700067P10Rik | ENSMUSG00000021545.4 |
| chr17 | 48094913 | 48095068 | 3.82177 | 4.74165 | 1.56211 | Distal Intergenic | 17 | 48089632 | 48090920 | 1289 | 1 | ENSMUST00000022028.4 | 5282 | 1700067P10Rik | ENSMUSG00000021545.4 |
| chr17 | 48096387 | 48096911 | 3.80355 | 4.83316 | 1.64015 | Distal Intergenic | 17 | 48089632 | 48090920 | 1289 | 1 | ENSMUST00000022028.4 | 6756 | 1700067P10Rik | ENSMUSG00000021545.4 |
| chr17 | 48097334 | 48098573 | 3.89579 | 4.93671 | 1.72362 | Distal Intergenic | 17 | 48089632 | 48090920 | 1289 | 1 | ENSMUST00000022028.4 | 7703 | 1700067P10Rik | ENSMUSG00000021545.4 |
| chr17 | 48099241 | 48099418 | 3.57176 | 4.36887 | 1.38917 | Distal Intergenic | 17 | 48089632 | 48090920 | 1289 | 1 | ENSMUST00000022028.4 | 9610 | 1700067P10Rik | ENSMUSG00000021545.4 |
| chr17 | 48100122 | 48100544 | 3.40416 | 3.95002 | 1.07353 | Distal Intergenic | 17 | 48089632 | 48090920 | 1289 | 1 | ENSMUST00000022028.4 | 10491 | 1700067P10Rik | ENSMUSG00000021545.4 |
| chr13 | 58524484 | 58524757 | 3.36973 | 3.9262 | 1.15537 | Distal Intergenic | 13 | 58545399 | 58558267 | 12869 | 2 | ENSMUST00000140760.1 | 33510 | Slc28a3 | ENSMUSG00000021553.8 |
| chr13 | 58610854 | 58611181 | 3.40734 | 3.96044 | 1.09289 | Promoter | 13 | 58553305 | 58610877 | 57573 | 2 | ENSMUST00000022036.8 | 0 | Slc28a3 | ENSMUSG00000021553.8 |
| chr13 | 58637597 | 58637853 | 3.35778 | 3.86416 | 1.01733 | Distal Intergenic | 13 | 58581690 | 58610877 | 29188 | 2 | ENSMUST00000148396.1 | -26721 | Slc28a3 | ENSMUSG00000021553.8 |
| chr13 | 58668792 | 58668884 | 3.8564 | 4.80047 | 1.61723 | Distal Intergenic | 13 | 58581690 | 58610877 | 29188 | 2 | ENSMUST00000148396.1 | -57916 | Slc28a3 | ENSMUSG00000021553.8 |
| chr13 | 59181056 | 59181431 | 3.6019 | 4.55519 | 1.48383 | Distal Intergenic | 13 | 59449537 | 59450228 | 692 | 2 | ENSMUST00000166276.1 | 268797 | Agtpbp1 | ENSMUSG00000021557.9 |
| chr13 | 59182323 | 59182666 | 3.32735 | 3.92727 | 1.15325 | Distal Intergenic | 13 | 59449537 | 59450228 | 692 | 2 | ENSMUST00000166276.1 | 267562 | Agtpbp1 | ENSMUSG00000021557.9 |
| chr13 | 73721177 | 73721420 | 3.31554 | 3.80054 | 1.01733 | Distal Intergenic | 13 | 73689087 | 73704865 | 15779 | 2 | ENSMUST00000123997.2 | -16313 | Slc6a19 | ENSMUSG00000021565.11 |
| chr13 | 73733150 | 73733879 | 3.67111 | 4.51653 | 1.50888 | Distal Intergenic | 13 | 73689087 | 73704865 | 15779 | 2 | ENSMUST00000123997.2 | -28286 | Slc6a19 | ENSMUSG00000021565.11 |
| chr13 | 73846361 | 73847609 | 3.57936 | 4.33243 | 1.39406 | Promoter | 13 | 73819694 | 73847612 | 27919 | 2 | ENSMUST00000118096.1 | 3 | Nkd2 | ENSMUSG00000021567.9 |
| chr13 | 74260011 | 74260347 | 3.26049 | 3.77295 | 1.0282 | Promoter | 13 | 74223029 | 74260672 | 37644 | 2 | ENSMUST00000137538.2 | 325 | Ahrr | ENSMUSG00000021575.10 |
| chr13 | 74292209 | 74292647 | 3.36173 | 3.87013 | 1.01733 | Promoter | 13 | 74211118 | 74292331 | 81214 | 2 | ENSMUST00000022059.8 | 0 | Ahrr | ENSMUSG00000021575.10 |
| chr13 | 74292961 | 74293437 | 3.82183 | 4.73101 | 1.56038 | Promoter | 13 | 74211118 | 74292331 | 81214 | 2 | ENSMUST00000022059.8 | -631 | Ahrr | ENSMUSG00000021575.10 |
| chr13 | 75090011 | 75090369 | 3.60228 | 4.4279 | 1.41396 | Promoter | 13 | 75089916 | 75106603 | 16688 | 1 | ENSMUST00000135349.1 | 96 | Pcsk1 | ENSMUSG00000021587.4 |
| chr13 | 76382544 | 76382676 | 3.62614 | 4.42049 | 1.42153 | Distal Intergenic | 13 | 76384535 | 77031810 | 647276 | 1 | ENSMUST00000125209.2 | -1859 | Mctp1 | ENSMUSG00000021596.11 |
| chr13 | 76383461 | 76384131 | 3.48834 | 4.12143 | 1.21929 | Promoter | 13 | 76384535 | 77031810 | 647276 | 1 | ENSMUST00000125209.2 | -404 | Mctp1 | ENSMUSG00000021596.11 |
| chr13 | 76384567 | 76385183 | 3.90868 | 4.93265 | 1.731 | Promoter | 13 | 76384535 | 77031810 | 647276 | 1 | ENSMUST00000125209.2 | 33 | Mctp1 | ENSMUSG00000021596.11 |
| chr13 | 73016328 | 73016465 | 3.36107 | 3.87802 | 1.04874 | Distal Intergenic | 13 | 73260479 | 73269608 | 9130 | 1 | ENSMUST00000176684.2 | -244014 | Irx4 | ENSMUSG00000021604.10 |
| chr13 | 73035378 | 73035453 | 3.38193 | 3.9006 | 1.01733 | Distal Intergenic | 13 | 73260479 | 73269608 | 9130 | 1 | ENSMUST00000176684.2 | -225026 | Irx4 | ENSMUSG00000021604.10 |
| chr13 | 73037867 | 73038145 | 3.52974 | 4.19386 | 1.2565 | Distal Intergenic | 13 | 73260479 | 73269608 | 9130 | 1 | ENSMUST00000176684.2 | -222334 | Irx4 | ENSMUSG00000021604.10 |
| chr13 | 73043957 | 73044095 | 3.63067 | 4.40081 | 1.42033 | Distal Intergenic | 13 | 73260479 | 73269608 | 9130 | 1 | ENSMUST00000176684.2 | -216384 | Irx4 | ENSMUSG00000021604.10 |
| chr13 | 73108408 | 73108631 | 3.56505 | 4.28903 | 1.37491 | Distal Intergenic | 13 | 73260479 | 73269608 | 9130 | 1 | ENSMUST00000176684.2 | -151848 | Irx4 | ENSMUSG00000021604.10 |
| chr13 | 73117460 | 73118027 | 3.379 | 3.94605 | 1.16745 | Distal Intergenic | 13 | 73260479 | 73269608 | 9130 | 1 | ENSMUST00000176684.2 | -142452 | Irx4 | ENSMUSG00000021604.10 |
| chr13 | 73159203 | 73159559 | 3.64588 | 4.39181 | 1.34518 | Distal Intergenic | 13 | 73260479 | 73269608 | 9130 | 1 | ENSMUST00000176684.2 | -100920 | Irx4 | ENSMUSG00000021604.10 |
| chr13 | 73159924 | 73160070 | 4.22297 | 5.79535 | 2.332 | Distal Intergenic | 13 | 73260479 | 73269608 | 9130 | 1 | ENSMUST00000176684.2 | -100409 | Irx4 | ENSMUSG00000021604.10 |
| chr13 | 73257478 | 73258201 | 3.93217 | 4.99524 | 1.78205 | Distal Intergenic | 13 | 73260479 | 73269608 | 9130 | 1 | ENSMUST00000176684.2 | -2278 | Irx4 | ENSMUSG00000021604.10 |
| chr13 | 73259521 | 73259600 | 3.52056 | 4.16948 | 1.21821 | Promoter | 13 | 73260479 | 73269608 | 9130 | 1 | ENSMUST00000176684.2 | -879 | Irx4 | ENSMUSG00000021604.10 |
| chr13 | 73260355 | 73260686 | 3.79279 | 4.67039 | 1.51498 | Promoter | 13 | 73260479 | 73269608 | 9130 | 1 | ENSMUST00000176684.2 | 0 | Irx4 | ENSMUSG00000021604.10 |
| chr13 | 73261259 | 73262854 | 3.85196 | 4.87947 | 1.70298 | Promoter | 13 | 73260497 | 73269608 | 9112 | 1 | ENSMUST00000022095.9 | 763 | Irx4 | ENSMUSG00000021604.10 |
| chr13 | 73263353 | 73264037 | 4.01434 | 5.17186 | 1.88339 | Intron (ENSMUST00000176684.2/ENSMUSG00000021604.10, intron 1 of 5) | 13 | 73260497 | 73269608 | 9112 | 1 | ENSMUST00000022095.9 | 2857 | Irx4 | ENSMUSG00000021604.10 |
| chr13 | 73264542 | 73265909 | 3.90704 | 4.97435 | 1.75008 | 5' UTR | 13 | 73260497 | 73269608 | 9112 | 1 | ENSMUST00000022095.9 | 4046 | Irx4 | ENSMUSG00000021604.10 |
| chr13 | 73267502 | 73267577 | 3.8613 | 4.81883 | 1.66153 | Exon (ENSMUST00000176684.2/ENSMUSG00000021604.10, exon 5 of 6) | 13 | 73260497 | 73269608 | 9112 | 1 | ENSMUST00000022095.9 | 7006 | Irx4 | ENSMUSG00000021604.10 |
| chr13 | 73268571 | 73269056 | 3.97051 | 5.19594 | 1.91808 | 3' UTR | 13 | 73260497 | 73269608 | 9112 | 1 | ENSMUST00000022095.9 | 8075 | Irx4 | ENSMUSG00000021604.10 |
| chr13 | 73269435 | 73269993 | 3.61395 | 4.3352 | 1.30161 | 3' UTR | 13 | 73260497 | 73269608 | 9112 | 1 | ENSMUST00000022095.9 | 8939 | Irx4 | ENSMUSG00000021604.10 |
| chr13 | 73270299 | 73271551 | 3.84272 | 4.82044 | 1.663 | Distal Intergenic | 13 | 73260497 | 73269608 | 9112 | 1 | ENSMUST00000022095.9 | 9803 | Irx4 | ENSMUSG00000021604.10 |
| chr13 | 73271941 | 73272271 | 3.8189 | 4.73764 | 1.59506 | Distal Intergenic | 13 | 73260497 | 73269608 | 9112 | 1 | ENSMUST00000022095.9 | 11445 | Irx4 | ENSMUSG00000021604.10 |
| chr13 | 73272903 | 73273194 | 3.46743 | 4.05709 | 1.11395 | Distal Intergenic | 13 | 73260497 | 73269608 | 9112 | 1 | ENSMUST00000022095.9 | 12407 | Irx4 | ENSMUSG00000021604.10 |
| chr13 | 73274064 | 73274249 | 3.90133 | 5.15618 | 1.87331 | Distal Intergenic | 13 | 73260497 | 73269608 | 9112 | 1 | ENSMUST00000022095.9 | 13568 | Irx4 | ENSMUSG00000021604.10 |
| chr13 | 73535905 | 73536837 | 3.76847 | 4.68741 | 1.56075 | Promoter | 13 | 73536747 | 73578672 | 41926 | 1 | ENSMUST00000022100.6 | 0 | Slc6a3 | ENSMUSG00000021609.6 |
| chr13 | 73537453 | 73537778 | 3.5069 | 4.15554 | 1.19563 | Promoter | 13 | 73536747 | 73578672 | 41926 | 1 | ENSMUST00000022100.6 | 707 | Slc6a3 | ENSMUSG00000021609.6 |
| chr13 | 73538981 | 73539336 | 3.49653 | 4.11717 | 1.17117 | Intron (ENSMUST00000022100.6/ENSMUSG00000021609.6, intron 2 of 14) | 13 | 73536747 | 73578672 | 41926 | 1 | ENSMUST00000022100.6 | 2235 | Slc6a3 | ENSMUSG00000021609.6 |
| chr13 | 89539507 | 89539696 | 4.12461 | 5.40091 | 2.11829 | Promoter | 13 | 89540636 | 89611832 | 71197 | 1 | ENSMUST00000022108.7 | -940 | Hapln1 | ENSMUSG00000021613.8 |
| chr13 | 102538032 | 102538107 | 3.50659 | 4.33091 | 1.36608 | Distal Intergenic | 13 | 102693558 | 102706955 | 13398 | 1 | ENSMUST00000022124.4 | -155451 | Cd180 | ENSMUSG00000021624.4 |
| chr13 | 100551369 | 100551870 | 3.6913 | 4.51948 | 1.49337 | Promoter | 13 | 100511594 | 100552460 | 40867 | 2 | ENSMUST00000159515.1 | 590 | Ocln | ENSMUSG00000021638.7 |
| chr13 | 100552202 | 100553182 | 3.95151 | 5.28857 | 1.97719 | Promoter | 13 | 100496507 | 100552498 | 55992 | 2 | ENSMUST00000069756.5 | 0 | Ocln | ENSMUSG00000021638.7 |
| chr13 | 100554022 | 100554169 | 3.76436 | 4.61349 | 1.48268 | Distal Intergenic | 13 | 100497763 | 100552718 | 54956 | 2 | ENSMUST00000022140.6 | -1305 | Ocln | ENSMUSG00000021638.7 |
| chr13 | 100554575 | 100554791 | 3.57382 | 4.4484 | 1.44039 | Distal Intergenic | 13 | 100497763 | 100552718 | 54956 | 2 | ENSMUST00000022140.6 | -1858 | Ocln | ENSMUSG00000021638.7 |
| chr13 | 100555435 | 100555528 | 3.55224 | 4.27092 | 1.35136 | Distal Intergenic | 13 | 100497763 | 100552718 | 54956 | 2 | ENSMUST00000022140.6 | -2718 | Ocln | ENSMUSG00000021638.7 |
| chr13 | 99839505 | 99839858 | 3.37706 | 3.89964 | 1.03989 | Distal Intergenic | 13 | 99898484 | 99900683 | 2200 | 2 | ENSMUST00000022150.6 | 60825 | Cartpt | ENSMUSG00000021647.6 |
| chr13 | 99896328 | 99896403 | 3.42348 | 3.98313 | 1.08758 | Downstream (2-3kb) | 13 | 99898484 | 99900683 | 2200 | 2 | ENSMUST00000022150.6 | 4280 | Cartpt | ENSMUSG00000021647.6 |
| chr13 | 99897160 | 99897489 | 3.37718 | 3.89343 | 1.01733 | Downstream (<1kb) | 13 | 99898484 | 99900683 | 2200 | 2 | ENSMUST00000022150.6 | 3194 | Cartpt | ENSMUSG00000021647.6 |
| chr13 | 99898077 | 99898452 | 3.61508 | 4.33564 | 1.30259 | Downstream (<1kb) | 13 | 99898484 | 99900683 | 2200 | 2 | ENSMUST00000022150.6 | 2231 | Cartpt | ENSMUSG00000021647.6 |
| chr13 | 99898781 | 99898993 | 4.01927 | 5.11599 | 1.8334 | 3' UTR | 13 | 99898484 | 99900683 | 2200 | 2 | ENSMUST00000022150.6 | 1690 | Cartpt | ENSMUSG00000021647.6 |
| chr13 | 99899909 | 99901043 | 3.66393 | 4.57237 | 1.48324 | Promoter | 13 | 99898484 | 99900683 | 2200 | 2 | ENSMUST00000022150.6 | 0 | Cartpt | ENSMUSG00000021647.6 |
| chr13 | 97190545 | 97190624 | 3.72513 | 15.59099 | 10.74683 | Intron (ENSMUST00000022169.7/ENSMUSG00000021665.7, intron 3 of 13) | 13 | 97176332 | 97198357 | 22026 | 2 | ENSMUST00000022169.7 | 7733 | Hexb | ENSMUSG00000021665.7 |
| chr13 | 95524303 | 95525470 | 3.64716 | 4.57407 | 1.50821 | Promoter | 13 | 95511732 | 95525240 | 13509 | 2 | ENSMUST00000022185.8 | 0 | F2rl1 | ENSMUSG00000021678.8 |
| chr13 | 95443071 | 95443163 | 3.90587 | 4.87607 | 1.64061 | Intron (ENSMUST00000045583.7/ENSMUSG00000021680.7, intron 3 of 6) | 13 | 95431371 | 95444831 | 13461 | 2 | ENSMUST00000045583.7 | 1668 | Crhbp | ENSMUSG00000021680.7 |
| chr13 | 95443722 | 95444295 | 3.63911 | 4.38326 | 1.33736 | Promoter | 13 | 95431371 | 95444831 | 13461 | 2 | ENSMUST00000045583.7 | 536 | Crhbp | ENSMUSG00000021680.7 |
| chr13 | 95444929 | 95445019 | 3.97412 | 5.02534 | 1.77231 | Promoter | 13 | 95431371 | 95444831 | 13461 | 2 | ENSMUST00000045583.7 | -99 | Crhbp | ENSMUSG00000021680.7 |
| chr13 | 95446117 | 95446221 | 3.91859 | 4.96756 | 1.74123 | Distal Intergenic | 13 | 95431371 | 95444831 | 13461 | 2 | ENSMUST00000045583.7 | -1287 | Crhbp | ENSMUSG00000021680.7 |
| chr13 | 95219407 | 95219590 | 3.35104 | 3.854 | 1.01733 | Intron (ENSMUST00000162412.2/ENSMUSG00000021684.11, intron 1 of 19) | 13 | 95026120 | 95222813 | 196694 | 2 | ENSMUST00000022192.6 | 3223 | Pde8b | ENSMUSG00000021684.11 |
| chr13 | 95221008 | 95221131 | 3.36344 | 3.87271 | 1.01733 | Intron (ENSMUST00000162412.2/ENSMUSG00000021684.11, intron 1 of 19) | 13 | 95026120 | 95222813 | 196694 | 2 | ENSMUST00000022192.6 | 1682 | Pde8b | ENSMUSG00000021684.11 |
| chr13 | 95221949 | 95222154 | 3.48584 | 4.09204 | 1.13937 | Promoter | 13 | 95026120 | 95222813 | 196694 | 2 | ENSMUST00000022192.6 | 659 | Pde8b | ENSMUSG00000021684.11 |
| chr13 | 95222473 | 95223108 | 4.49972 | 6.24472 | 2.73192 | Promoter | 13 | 95024505 | 95223066 | 198562 | 2 | ENSMUST00000162292.2 | 0 | Pde8b | ENSMUSG00000021684.11 |
| chr13 | 95223870 | 95223988 | 3.9805 | 5.02739 | 1.74693 | Promoter | 13 | 95033946 | 95223555 | 189610 | 2 | ENSMUST00000159608.2 | -316 | Pde8b | ENSMUSG00000021684.11 |
| chr13 | 95249694 | 95250197 | 4.03873 | 5.22025 | 1.97527 | Promoter | 13 | 95026103 | 95250166 | 224064 | 2 | ENSMUST00000162153.2 | 0 | Pde8b | ENSMUSG00000021684.11 |
| chr13 | 94865672 | 94865786 | 3.62441 | 4.35579 | 1.33373 | Distal Intergenic | 13 | 94875602 | 94885130 | 9529 | 1 | ENSMUST00000022195.11 | -9816 | Otp | ENSMUSG00000021685.11 |
| chr13 | 94868248 | 94869128 | 3.6203 | 4.34932 | 1.32264 | Distal Intergenic | 13 | 94875602 | 94885130 | 9529 | 1 | ENSMUST00000022195.11 | -6474 | Otp | ENSMUSG00000021685.11 |
| chr13 | 94869589 | 94871070 | 3.81884 | 4.77792 | 1.60317 | Distal Intergenic | 13 | 94875602 | 94885130 | 9529 | 1 | ENSMUST00000022195.11 | -4532 | Otp | ENSMUSG00000021685.11 |
| chr13 | 94871460 | 94871933 | 4.13341 | 5.36486 | 2.03987 | Distal Intergenic | 13 | 94875602 | 94885130 | 9529 | 1 | ENSMUST00000022195.11 | -3669 | Otp | ENSMUSG00000021685.11 |
| chr13 | 94872287 | 94872753 | 3.56826 | 4.26801 | 1.31213 | Distal Intergenic | 13 | 94875602 | 94885130 | 9529 | 1 | ENSMUST00000022195.11 | -2849 | Otp | ENSMUSG00000021685.11 |
| chr13 | 94873511 | 94873795 | 3.80696 | 4.99216 | 1.77197 | Distal Intergenic | 13 | 94875602 | 94885130 | 9529 | 1 | ENSMUST00000022195.11 | -1807 | Otp | ENSMUSG00000021685.11 |
| chr13 | 94874861 | 94878302 | 4.25094 | 5.75395 | 2.35102 | Promoter | 13 | 94875602 | 94885130 | 9529 | 1 | ENSMUST00000022195.11 | 0 | Otp | ENSMUSG00000021685.11 |
| chr13 | 94878669 | 94880298 | 3.82062 | 4.76518 | 1.63813 | Intron (ENSMUST00000022195.11/ENSMUSG00000021685.11, intron 2 of 2) | 13 | 94882600 | 94883552 | 953 | 1 | ENSMUST00000184985.1 | -2302 | Otp | ENSMUSG00000021685.11 |
| chr13 | 94880725 | 94881087 | 3.40491 | 4.12122 | 1.20718 | Intron (ENSMUST00000022195.11/ENSMUSG00000021685.11, intron 2 of 2) | 13 | 94882600 | 94883552 | 953 | 1 | ENSMUST00000184985.1 | -1513 | Otp | ENSMUSG00000021685.11 |
| chr13 | 94881572 | 94882361 | 3.68188 | 4.49933 | 1.43053 | Promoter | 13 | 94882600 | 94883552 | 953 | 1 | ENSMUST00000184985.1 | -239 | Otp | ENSMUSG00000021685.11 |
| chr13 | 94883083 | 94883647 | 3.99558 | 5.13767 | 1.85428 | Promoter | 13 | 94882600 | 94883552 | 953 | 1 | ENSMUST00000184985.1 | 484 | Otp | ENSMUSG00000021685.11 |
| chr13 | 94884344 | 94884923 | 3.67648 | 4.688 | 1.57978 | 3' UTR | 13 | 94882600 | 94883552 | 953 | 1 | ENSMUST00000184985.1 | 1745 | Otp | ENSMUSG00000021685.11 |
| chr13 | 94885270 | 94886047 | 4.65931 | 6.51587 | 2.92733 | Distal Intergenic | 13 | 94882600 | 94883552 | 953 | 1 | ENSMUST00000184985.1 | 2671 | Otp | ENSMUSG00000021685.11 |
| chr13 | 94886490 | 94886662 | 3.55458 | 4.22201 | 1.22903 | Distal Intergenic | 13 | 94882600 | 94883552 | 953 | 1 | ENSMUST00000184985.1 | 3891 | Otp | ENSMUSG00000021685.11 |
| chr13 | 94887079 | 94887314 | 3.47856 | 4.07677 | 1.12412 | Distal Intergenic | 13 | 94882600 | 94883552 | 953 | 1 | ENSMUST00000184985.1 | 4480 | Otp | ENSMUSG00000021685.11 |
| chr13 | 94887673 | 94888654 | 3.90935 | 5.06273 | 1.86269 | Distal Intergenic | 13 | 94882600 | 94883552 | 953 | 1 | ENSMUST00000184985.1 | 5074 | Otp | ENSMUSG00000021685.11 |
| chr13 | 94895702 | 94896006 | 3.33016 | 3.83941 | 1.07666 | Distal Intergenic | 13 | 94882600 | 94883552 | 953 | 1 | ENSMUST00000184985.1 | 13103 | Otp | ENSMUSG00000021685.11 |
| chr13 | 94898617 | 94898831 | 3.27584 | 3.74089 | 1.01733 | Distal Intergenic | 13 | 94882600 | 94883552 | 953 | 1 | ENSMUST00000184985.1 | 16018 | Otp | ENSMUSG00000021685.11 |
| chr13 | 94911642 | 94911854 | 3.42929 | 4.01064 | 1.15141 | Distal Intergenic | 13 | 94882600 | 94883552 | 953 | 1 | ENSMUST00000184985.1 | 29043 | Otp | ENSMUSG00000021685.11 |
| chr13 | 94918098 | 94918464 | 3.29237 | 3.76584 | 1.01733 | Distal Intergenic | 13 | 94882600 | 94883552 | 953 | 1 | ENSMUST00000184985.1 | 35499 | Otp | ENSMUSG00000021685.11 |
| chr13 | 94921795 | 94921912 | 3.67061 | 4.43814 | 1.36218 | Distal Intergenic | 13 | 94882600 | 94883552 | 953 | 1 | ENSMUST00000184985.1 | 39196 | Otp | ENSMUSG00000021685.11 |
| chr13 | 94922550 | 94923068 | 3.36846 | 3.88029 | 1.01733 | Distal Intergenic | 13 | 94882600 | 94883552 | 953 | 1 | ENSMUST00000184985.1 | 39951 | Otp | ENSMUSG00000021685.11 |
| chr13 | 94924392 | 94924501 | 3.56614 | 4.2491 | 1.26333 | Distal Intergenic | 13 | 94882600 | 94883552 | 953 | 1 | ENSMUST00000184985.1 | 41793 | Otp | ENSMUSG00000021685.11 |
| chr13 | 94925260 | 94926383 | 3.42801 | 3.99807 | 1.10597 | Distal Intergenic | 13 | 94882600 | 94883552 | 953 | 1 | ENSMUST00000184985.1 | 42661 | Otp | ENSMUSG00000021685.11 |
| chr13 | 108214767 | 108214852 | 3.47391 | 4.12925 | 1.26851 | Promoter | 13 | 108214404 | 108287105 | 72702 | 1 | ENSMUST00000022207.8 | 364 | Elovl7 | ENSMUSG00000021696.8 |
| chr13 | 109116011 | 109117644 | 3.67204 | 4.494 | 1.45302 | Promoter | 13 | 109116236 | 109335063 | 218828 | 1 | ENSMUST00000129374.1 | 0 | Pde4d | ENSMUSG00000021699.12 |
| chr13 | 109229943 | 109230502 | 3.56431 | 4.23713 | 1.23337 | Intron (ENSMUST00000122041.2/ENSMUSG00000021699.12, intron 3 of 16) | 13 | 109260654 | 109951364 | 690711 | 1 | ENSMUST00000153234.2 | -30152 | Pde4d | ENSMUSG00000021699.12 |
| chr13 | 109232794 | 109233018 | 3.31111 | 3.8073 | 1.06427 | Intron (ENSMUST00000122041.2/ENSMUSG00000021699.12, intron 3 of 16) | 13 | 109260654 | 109951364 | 690711 | 1 | ENSMUST00000153234.2 | -27636 | Pde4d | ENSMUSG00000021699.12 |
| chr13 | 109235117 | 109235652 | 3.4883 | 4.09798 | 1.147 | Intron (ENSMUST00000122041.2/ENSMUSG00000021699.12, intron 3 of 16) | 13 | 109260654 | 109951364 | 690711 | 1 | ENSMUST00000153234.2 | -25002 | Pde4d | ENSMUSG00000021699.12 |
| chr13 | 109239992 | 109240212 | 3.7284 | 4.54638 | 1.44613 | Intron (ENSMUST00000122041.2/ENSMUSG00000021699.12, intron 3 of 16) | 13 | 109260654 | 109951364 | 690711 | 1 | ENSMUST00000153234.2 | -20442 | Pde4d | ENSMUSG00000021699.12 |
| chr13 | 110279060 | 110279397 | 3.40302 | 3.98517 | 1.14059 | Promoter | 13 | 110054187 | 110280206 | 226020 | 2 | ENSMUST00000167824.1 | 809 | Rab3c | ENSMUSG00000021700.9 |
| chr13 | 110280177 | 110280476 | 3.93639 | 5.03592 | 1.87004 | Promoter | 13 | 110054187 | 110280206 | 226020 | 2 | ENSMUST00000167824.1 | 0 | Rab3c | ENSMUSG00000021700.9 |
| chr13 | 92794484 | 92794935 | 3.59581 | 4.30897 | 1.30056 | Promoter | 13 | 92751590 | 92794818 | 43229 | 2 | ENSMUST00000022213.7 | 0 | Thbs4 | ENSMUSG00000021702.7 |
| chr13 | 92131471 | 92131703 | 3.83956 | 4.87729 | 1.72586 | Promoter | 13 | 91884432 | 92131494 | 247063 | 2 | ENSMUST00000099326.4 | 0 | Rasgrf2 | ENSMUSG00000021708.11 |
| chr13 | 105053033 | 105054722 | 3.71243 | 4.70506 | 1.5995 | Promoter | 13 | 104947153 | 105054930 | 107778 | 2 | ENSMUST00000063551.5 | 208 | Rgs7bp | ENSMUSG00000021719.8 |
| chr13 | 105056096 | 105056171 | 3.4223 | 4.02626 | 1.20919 | Distal Intergenic | 13 | 104947153 | 105054930 | 107778 | 2 | ENSMUST00000063551.5 | -1167 | Rgs7bp | ENSMUSG00000021719.8 |
| chr13 | 105059167 | 105059253 | 3.64481 | 4.38353 | 1.31522 | Distal Intergenic | 13 | 104947153 | 105054930 | 107778 | 2 | ENSMUST00000063551.5 | -4238 | Rgs7bp | ENSMUSG00000021719.8 |
| chr13 | 105294001 | 105294262 | 3.70908 | 4.58306 | 1.54084 | Distal Intergenic | 13 | 105149352 | 105271039 | 121688 | 2 | ENSMUST00000069686.6 | -22963 | Rnf180 | ENSMUSG00000021720.10 |
| chr13 | 105444022 | 105444199 | 3.26233 | 4.17654 | 1.21118 | Promoter | 13 | 105443693 | 105448133 | 4441 | 1 | ENSMUST00000022235.4 | 330 | Htr1a | ENSMUSG00000021721.4 |
| chr13 | 105444991 | 105445171 | 3.97237 | 5.05091 | 1.84927 | Exon (ENSMUST00000022235.4/ENSMUSG00000021721.4, exon 1 of 1) | 13 | 105443693 | 105448133 | 4441 | 1 | ENSMUST00000022235.4 | 1299 | Htr1a | ENSMUSG00000021721.4 |
| chr13 | 105445560 | 105445706 | 3.59244 | 4.36906 | 1.39127 | 3' UTR | 13 | 105443693 | 105448133 | 4441 | 1 | ENSMUST00000022235.4 | 1868 | Htr1a | ENSMUSG00000021721.4 |
| chr13 | 105446064 | 105446926 | 3.51952 | 4.18476 | 1.24443 | 3' UTR | 13 | 105443693 | 105448133 | 4441 | 1 | ENSMUST00000022235.4 | 2372 | Htr1a | ENSMUSG00000021721.4 |
| chr13 | 117220442 | 117222074 | 4.09911 | 5.43662 | 2.08402 | Promoter | 13 | 117220625 | 117274098 | 53474 | 1 | ENSMUST00000022242.7 | 0 | Emb | ENSMUSG00000021728.7 |
| chr13 | 117602286 | 117603315 | 3.61108 | 4.34688 | 1.35623 | Promoter | 13 | 117602480 | 117976802 | 374323 | 1 | ENSMUST00000006991.7 | 0 | Hcn1 | ENSMUSG00000021730.7 |
| chr13 | 117603655 | 117604696 | 4.24592 | 5.78353 | 2.41618 | Intron (ENSMUST00000006991.7/ENSMUSG00000021730.7, intron 1 of 7) | 13 | 117602480 | 117976802 | 374323 | 1 | ENSMUST00000006991.7 | 1176 | Hcn1 | ENSMUSG00000021730.7 |
| chr13 | 118713758 | 118714398 | 3.55727 | 4.24006 | 1.25937 | Promoter | 13 | 118714699 | 118792115 | 77417 | 1 | ENSMUST00000022246.8 | -301 | Fgf10 | ENSMUSG00000021732.9 |
| chr13 | 118714877 | 118715049 | 3.6008 | 4.34299 | 1.3038 | Promoter | 13 | 118714699 | 118792115 | 77417 | 1 | ENSMUST00000022246.8 | 179 | Fgf10 | ENSMUSG00000021732.9 |
| chr14 | 12341161 | 12343084 | 3.94153 | 5.07353 | 1.85465 | 3' UTR | 14 | 12341892 | 12345865 | 3974 | 2 | ENSMUST00000022262.4 | 2781 | Fezf2 | ENSMUSG00000021743.5 |
| chr14 | 12343410 | 12343927 | 3.4554 | 4.07279 | 1.22474 | Exon (ENSMUST00000022262.4/ENSMUSG00000021743.5, exon 3 of 5) | 14 | 12341892 | 12345865 | 3974 | 2 | ENSMUST00000022262.4 | 1938 | Fezf2 | ENSMUSG00000021743.5 |
| chr14 | 12344273 | 12346197 | 4.01832 | 5.12913 | 1.83324 | Promoter | 14 | 12341892 | 12345865 | 3974 | 2 | ENSMUST00000022262.4 | 0 | Fezf2 | ENSMUSG00000021743.5 |
| chr14 | 12346905 | 12347049 | 3.69978 | 4.50671 | 1.41982 | Distal Intergenic | 14 | 12341892 | 12345865 | 3974 | 2 | ENSMUST00000022262.4 | -1041 | Fezf2 | ENSMUSG00000021743.5 |
| chr14 | 12347824 | 12348975 | 3.61832 | 4.33853 | 1.29625 | Distal Intergenic | 14 | 12341892 | 12345865 | 3974 | 2 | ENSMUST00000022262.4 | -1960 | Fezf2 | ENSMUSG00000021743.5 |
| chr14 | 12349575 | 12350008 | 3.79929 | 4.76247 | 1.63683 | Distal Intergenic | 14 | 12341892 | 12345865 | 3974 | 2 | ENSMUST00000022262.4 | -3711 | Fezf2 | ENSMUSG00000021743.5 |
| chr14 | 12355201 | 12355470 | 3.51157 | 4.199 | 1.29483 | Distal Intergenic | 14 | 12341892 | 12345865 | 3974 | 2 | ENSMUST00000022262.4 | -9337 | Fezf2 | ENSMUSG00000021743.5 |
| chr14 | 12356994 | 12357297 | 4.15177 | 5.70377 | 2.28944 | Distal Intergenic | 14 | 12341892 | 12345865 | 3974 | 2 | ENSMUST00000022262.4 | -11130 | Fezf2 | ENSMUSG00000021743.5 |
| chr14 | 8286660 | 8286739 | 2.58026 | 14.49305 | 9.78724 | Distal Intergenic | 14 | 8296274 | 8309777 | 13504 | 2 | ENSMUST00000121887.2 | 23038 | Fam107a | ENSMUSG00000021750.10 |

| | | | | | | | | | | | | | | | |
|---|---|---|---|---|---|---|---|---|---|---|---|---|---|---|---|
| chr14 | 8317129 | 8317276 | 3.25027 | 3.70258 | 1.01733 | Promoter | 14 | 8297556 | 8318023 | 20468 | 2 | ENSMUST00000120411.2 | 747 | Fam107a | ENSMUSG00000021750.10 |
| chr14 | 8317664 | 8317933 | 3.95647 | 5.19411 | 1.91284 | Promoter | 14 | 8297556 | 8318023 | 20468 | 2 | ENSMUST00000120411.2 | 90 | Fam107a | ENSMUSG00000021750.10 |
| chr14 | 8318415 | 8319065 | 3.50422 | 4.16374 | 1.27272 | Promoter | 14 | 8297556 | 8318023 | 20468 | 2 | ENSMUST00000120411.2 | -393 | Fam107a | ENSMUSG00000021750.10 |
| chr13 | 113292883 | 113293663 | 3.66165 | 4.67038 | 1.52421 | Distal Intergenic | 13 | 113317084 | 113370515 | 53432 | 1 | ENSMUST00000136755.3 | -23421 | BC067074 | ENSMUSG00000021763.11 |
| chr13 | 114165098 | 114165706 | 3.38859 | 3.9335 | 1.09699 | Distal Intergenic | 13 | 114287795 | 114388094 | 100300 | 2 | ENSMUST00000022286.6 | 222388 | Ndufs4 | ENSMUSG00000021764.6 |
| chr13 | 114453390 | 114453633 | 3.68918 | 4.61015 | 1.52964 | 3' UTR | 13 | 114452262 | 114458730 | 6469 | 2 | ENSMUST00000022287.5 | 5097 | Fst | ENSMUSG00000021765.7 |
| chr13 | 114454189 | 114454321 | 3.41485 | 4.22638 | 1.29549 | Exon (ENSMUST00000022287.5/ENSMUSG00000021765.7, exon 5 of 6) | 13 | 114452262 | 114458730 | 6469 | 2 | ENSMUST00000022287.5 | 4409 | Fst | ENSMUSG00000021765.7 |
| chr13 | 114456263 | 114456500 | 3.92979 | 5.1534 | 1.93914 | Exon (ENSMUST00000022287.5/ENSMUSG00000021765.7, exon 2 of 6) | 13 | 114452262 | 114458730 | 6469 | 2 | ENSMUST00000022287.5 | 2230 | Fst | ENSMUSG00000021765.7 |
| chr13 | 114456855 | 114459748 | 4.42562 | 6.0267 | 2.52589 | Promoter | 13 | 114452262 | 114458730 | 6469 | 2 | ENSMUST00000022287.5 | 0 | Fst | ENSMUSG00000021765.7 |
| chr13 | 114460248 | 114460323 | 3.43387 | 3.99612 | 1.07761 | Distal Intergenic | 13 | 114452262 | 114458730 | 6469 | 2 | ENSMUST00000022287.5 | -1519 | Fst | ENSMUSG00000021765.7 |
| chr13 | 114461215 | 114461502 | 3.31323 | 3.79711 | 1.01733 | Distal Intergenic | 13 | 114452262 | 114458730 | 6469 | 2 | ENSMUST00000022287.5 | -2486 | Fst | ENSMUSG00000021765.7 |
| chr14 | 17665731 | 17665860 | 4.00988 | 5.17525 | 1.91325 | Intron (ENSMUST00000022303.8/ENSMUSG00000021779.11, intron 1 of 9) | 14 | 17660960 | 18038088 | 377129 | 1 | ENSMUST00000091471.5 | 4772 | Thrb | ENSMUSG00000021779.11 |
| chr14 | 17669206 | 17669342 | 3.59709 | 4.37233 | 1.39633 | Intron (ENSMUST00000022303.8/ENSMUSG00000021779.11, intron 1 of 9) | 14 | 17660960 | 18038088 | 377129 | 1 | ENSMUST00000091471.5 | 8247 | Thrb | ENSMUSG00000021779.11 |
| chr14 | 34607285 | 34607586 | 3.33356 | 3.82768 | 1.01733 | Distal Intergenic | 14 | 34592343 | 34600142 | 7800 | 2 | ENSMUST00000022331.2 | -7144 | Opn4 | ENSMUSG00000021799.3 |
| chr14 | 37098213 | 37098808 | 3.88763 | 4.93368 | 1.7974 | Promoter | 14 | 37077850 | 37098311 | 20462 | 2 | ENSMUST00000022337.9 | 0 | Cdhr1 | ENSMUSG00000021803.9 |
| chr14 | 19828518 | 19828690 | 3.58033 | 4.26504 | 1.24616 | Distal Intergenic | 14 | 19811402 | 19823823 | 12422 | 2 | ENSMUST00000022341.5 | -4696 | 2700060E02Rik | ENSMUSG00000021807.5 |
| chr14 | 20834321 | 20834942 | 3.35921 | 3.90455 | 1.14854 | Distal Intergenic | 14 | 20836662 | 20843388 | 6727 | 1 | ENSMUST00000022368.2 | -1720 | Plau | ENSMUSG00000021822.2 |
| chr14 | 46389310 | 46389752 | 3.81365 | 5.1367 | 1.86169 | Promoter | 14 | 46386996 | 46390535 | 3540 | 2 | ENSMUST00000135408.1 | 783 | Bmp4 | ENSMUSG00000021835.9 |
| chr14 | 46390095 | 46390924 | 3.69187 | 4.56638 | 1.4709 | Promoter | 14 | 46383520 | 46390669 | 7150 | 2 | ENSMUST00000074077.6 | 0 | Bmp4 | ENSMUSG00000021835.9 |
| chr14 | 46397239 | 46397589 | 4.07047 | 5.33642 | 2.00242 | Distal Intergenic | 14 | 46383520 | 46390669 | 7150 | 2 | ENSMUST00000074077.6 | -6571 | Bmp4 | ENSMUSG00000021835.9 |
| chr14 | 46398213 | 46399426 | 3.96915 | 5.05531 | 1.8097 | Distal Intergenic | 14 | 46383520 | 46390669 | 7150 | 2 | ENSMUST00000074077.6 | -7545 | Bmp4 | ENSMUSG00000021835.9 |
| chr14 | 48657357 | 48658285 | 3.85946 | 4.80239 | 1.61019 | 3' UTR | 14 | 48658916 | 48661932 | 3017 | 2 | ENSMUST00000135279.1 | 3647 | Otx2 | ENSMUSG00000021848.10 |
| chr14 | 48659072 | 48659320 | 3.91092 | 4.89877 | 1.674 | Exon (ENSMUST00000118578.2/ENSMUSG00000021848.10, exon 4 of 4) | 14 | 48658916 | 48661932 | 3017 | 2 | ENSMUST00000135279.1 | 2612 | Otx2 | ENSMUSG00000021848.10 |
| chr14 | 48660041 | 48661178 | 3.72872 | 4.65714 | 1.55446 | Promoter | 14 | 48658916 | 48661932 | 3017 | 2 | ENSMUST00000135279.1 | 754 | Otx2 | ENSMUSG00000021848.10 |
| chr14 | 48661759 | 48662548 | 4.14023 | 5.43851 | 2.06611 | Promoter | 14 | 48658916 | 48661932 | 3017 | 2 | ENSMUST00000135279.1 | 0 | Otx2 | ENSMUSG00000021848.10 |
| chr14 | 48663039 | 48663908 | 3.64027 | 4.40222 | 1.38655 | Promoter | 14 | 48658171 | 48662872 | 4702 | 2 | ENSMUST00000122009.2 | -168 | Otx2 | ENSMUSG00000021848.10 |
| chr14 | 48664232 | 48668441 | 4.02358 | 5.21951 | 1.9339 | Promoter | 14 | 48657679 | 48665246 | 7568 | 2 | ENSMUST00000118578.2 | 0 | Otx2 | ENSMUSG00000021848.10 |
| chr14 | 49524873 | 49525461 | 3.49345 | 4.1082 | 1.1563 | Promoter | 14 | 49317055 | 49525853 | 208799 | 2 | ENSMUST00000146164.1 | 392 | Slc35f4 | ENSMUSG00000021852.8 |
| chr14 | 49525789 | 49526194 | 3.6149 | 4.3876 | 1.36192 | Promoter | 14 | 49298519 | 49525865 | 227347 | 2 | ENSMUST00000138884.2 | 0 | Slc35f4 | ENSMUSG00000021852.8 |
| chr14 | 32028723 | 32029229 | 3.4421 | 4.0408 | 1.19328 | Promoter | 14 | 32028989 | 32062197 | 33209 | 1 | ENSMUST00000022460.5 | 0 | Galnt15 | ENSMUSG00000021903.6 |
| chr14 | 32465470 | 32465724 | 3.58323 | 4.26875 | 1.24385 | Promoter | 14 | 32408203 | 32465909 | 57707 | 2 | ENSMUST00000070125.4 | 185 | Chat | ENSMUSG00000021919.7 |
| chr14 | 32466483 | 32466690 | 3.82676 | 4.76349 | 1.63039 | Promoter | 14 | 32408203 | 32465909 | 57707 | 2 | ENSMUST00000070125.4 | -575 | Chat | ENSMUSG00000021919.7 |
| chr14 | 32467051 | 32467126 | 3.69774 | 4.62554 | 1.52673 | Intron (ENSMUST00000180662.1/ENSMUSG00000097794.1, intron 1 of 1) | 14 | 32408203 | 32465909 | 57707 | 2 | ENSMUST00000070125.4 | -1143 | Chat | ENSMUSG00000021919.7 |
| chr14 | 32488174 | 32488366 | 3.34153 | 3.83966 | 1.01733 | Distal Intergenic | 14 | 32408203 | 32465909 | 57707 | 2 | ENSMUST00000070125.4 | -22266 | Chat | ENSMUSG00000021919.7 |
| chr14 | 62292454 | 62293013 | 3.76444 | 4.66811 | 1.5371 | Promoter | 14 | 62292589 | 62353558 | 60970 | 1 | ENSMUST00000166879.2 | 0 | Rnaseh2b | ENSMUSG00000021932.8 |
| chr14 | 33488409 | 33488739 | 3.36462 | 3.87448 | 1.01733 | Distal Intergenic | 14 | 33390938 | 33447158 | 56221 | 2 | ENSMUST00000130539.2 | -41252 | Mapk8 | ENSMUSG00000021936.8 |
| chr14 | 33923453 | 33923738 | 3.51248 | 4.21572 | 1.27609 | Promoter | 14 | 33923587 | 33935280 | 11694 | 1 | ENSMUST00000168727.1 | 0 | Gdf10 | ENSMUSG00000021943.6 |
| chr14 | 33924083 | 33924748 | 3.71576 | 4.62936 | 1.57802 | Promoter | 14 | 33923587 | 33935280 | 11694 | 1 | ENSMUST00000168727.1 | 497 | Gdf10 | ENSMUSG00000021943.6 |
| chr14 | 33931667 | 33931742 | 3.46007 | 4.09557 | 1.26243 | Intron (ENSMUST00000168727.1/ENSMUSG00000021943.6, intron 1 of 2) | 14 | 33923587 | 33935280 | 11694 | 1 | ENSMUST00000168727.1 | 8081 | Gdf10 | ENSMUSG00000021943.6 |
| chr14 | 63217822 | 63218533 | 4.29005 | 5.80288 | 2.34316 | Intron (ENSMUST00000118022.2/ENSMUSG00000021944.10, intron 2 of 6) | 14 | 63198924 | 63213607 | 14684 | 2 | ENSMUST00000121312.1 | -4216 | Gata4 | ENSMUSG00000021944.10 |
| chr14 | 63219006 | 63219146 | 3.51066 | 4.24125 | 1.29379 | Intron (ENSMUST00000118022.2/ENSMUSG00000021944.10, intron 2 of 6) | 14 | 63198924 | 63213607 | 14684 | 2 | ENSMUST00000121312.1 | -5400 | Gata4 | ENSMUSG00000021944.10 |
| chr14 | 63230858 | 63231057 | 3.36466 | 4.17354 | 1.25913 | Intron (ENSMUST00000118022.2/ENSMUSG00000021944.10, intron 2 of 6) | 14 | 63198922 | 63245248 | 46327 | 2 | ENSMUST00000118022.2 | 14191 | Gata4 | ENSMUSG00000021944.10 |
| chr14 | 63232921 | 63233009 | 4.85402 | 6.85582 | 3.15134 | Intron (ENSMUST00000118022.2/ENSMUSG00000021944.10, intron 2 of 6) | 14 | 63198922 | 63245248 | 46327 | 2 | ENSMUST00000118022.2 | 12239 | Gata4 | ENSMUSG00000021944.10 |
| chr14 | 63234774 | 63234985 | 3.56207 | 4.2789 | 1.35018 | Intron (ENSMUST00000118022.2/ENSMUSG00000021944.10, intron 2 of 6) | 14 | 63198922 | 63245248 | 46327 | 2 | ENSMUST00000118022.2 | 10263 | Gata4 | ENSMUSG00000021944.10 |
| chr14 | 63237037 | 63237336 | 4.30709 | 5.8166 | 2.39743 | Intron (ENSMUST00000118022.2/ENSMUSG00000021944.10, intron 2 of 6) | 14 | 63198922 | 63245248 | 46327 | 2 | ENSMUST00000118022.2 | 7912 | Gata4 | ENSMUSG00000021944.10 |
| chr14 | 63238016 | 63241061 | 3.847 | 4.98392 | 1.79216 | Exon (ENSMUST00000118022.2/ENSMUSG00000021944.10, exon 2 of 7) | 14 | 63198922 | 63245248 | 46327 | 2 | ENSMUST00000118022.2 | 4187 | Gata4 | ENSMUSG00000021944.10 |
| chr14 | 63241428 | 63242240 | 3.94459 | 5.17185 | 1.90954 | 5' UTR | 14 | 63198922 | 63245248 | 46327 | 2 | ENSMUST00000118022.2 | 3008 | Gata4 | ENSMUSG00000021944.10 |
| chr14 | 63242752 | 63246259 | 4.02165 | 5.23087 | 1.93829 | Promoter | 14 | 63198922 | 63245248 | 46327 | 2 | ENSMUST00000118022.2 | 0 | Gata4 | ENSMUSG00000021944.10 |
| chr14 | 63246640 | 63247520 | 3.7947 | 4.67235 | 1.51691 | Intron (ENSMUST00000132122.1/ENSMUSG00000021944.10, intron 1 of 1) | 14 | 63198922 | 63245271 | 46350 | 2 | ENSMUST00000067417.4 | -1370 | Gata4 | ENSMUSG00000021944.10 |
| chr14 | 63248591 | 63250832 | 3.99405 | 5.14954 | 1.86141 | Intron (ENSMUST00000132122.1/ENSMUSG00000021944.10, intron 1 of 1) | 14 | 63198922 | 63245271 | 46350 | 2 | ENSMUST00000067417.4 | -3321 | Gata4 | ENSMUSG00000021944.10 |
| chr14 | 63251490 | 63253864 | 4.0654 | 5.29634 | 2.00908 | Intron (ENSMUST00000132122.1/ENSMUSG00000021944.10, intron 1 of 1) | 14 | 63198922 | 63245271 | 46350 | 2 | ENSMUST00000067417.4 | -6220 | Gata4 | ENSMUSG00000021944.10 |
| chr14 | 63254238 | 63257547 | 3.89944 | 4.97735 | 1.765 | Intron (ENSMUST00000132122.1/ENSMUSG00000021944.10, intron 1 of 1) | 14 | 63198922 | 63245271 | 46350 | 2 | ENSMUST00000067417.4 | -8968 | Gata4 | ENSMUSG00000021944.10 |
| chr14 | 63257882 | 63258271 | 3.83723 | 4.78807 | 1.63815 | Intron (ENSMUST00000132122.1/ENSMUSG00000021944.10, intron 1 of 1) | 14 | 63198922 | 63245271 | 46350 | 2 | ENSMUST00000067417.4 | -12612 | Gata4 | ENSMUSG00000021944.10 |
| chr14 | 63259321 | 63260032 | 4.08898 | 5.46749 | 2.14511 | Intron (ENSMUST00000132122.1/ENSMUSG00000021944.10, intron 1 of 1) | 14 | 63241115 | 63271124 | 30010 | 2 | ENSMUST00000137244.2 | 11092 | Gata4 | ENSMUSG00000021944.10 |
| chr14 | 63260973 | 63261227 | 3.35194 | 3.86377 | 1.04699 | Intron (ENSMUST00000132122.1/ENSMUSG00000021944.10, intron 1 of 1) | 14 | 63241115 | 63271124 | 30010 | 2 | ENSMUST00000137244.2 | 9897 | Gata4 | ENSMUSG00000021944.10 |
| chr14 | 63261761 | 63262328 | 4.08702 | 5.31661 | 2.05139 | Intron (ENSMUST00000132122.1/ENSMUSG00000021944.10, intron 1 of 1) | 14 | 63241115 | 63271124 | 30010 | 2 | ENSMUST00000137244.2 | 8796 | Gata4 | ENSMUSG00000021944.10 |
| chr14 | 63262664 | 63262904 | 3.50914 | 4.37496 | 1.34452 | Intron (ENSMUST00000132122.1/ENSMUSG00000021944.10, intron 1 of 1) | 14 | 63241115 | 63271124 | 30010 | 2 | ENSMUST00000137244.2 | 8220 | Gata4 | ENSMUSG00000021944.10 |
| chr14 | 63263740 | 63263915 | 3.54548 | 4.25547 | 1.33171 | Intron (ENSMUST00000132122.1/ENSMUSG00000021944.10, intron 1 of 1) | 14 | 63241115 | 63271124 | 30010 | 2 | ENSMUST00000137244.2 | 7209 | Gata4 | ENSMUSG00000021944.10 |
| chr14 | 63264446 | 63267172 | 3.8583 | 4.88531 | 1.74633 | Intron (ENSMUST00000132122.1/ENSMUSG00000021944.10, intron 1 of 1) | 14 | 63241115 | 63271124 | 30010 | 2 | ENSMUST00000137244.2 | 3952 | Gata4 | ENSMUSG00000021944.10 |
| chr14 | 63267547 | 63268023 | 3.47145 | 4.15541 | 1.27668 | Intron (ENSMUST00000132122.1/ENSMUSG00000021944.10, intron 1 of 1) | 14 | 63241115 | 63271124 | 30010 | 2 | ENSMUST00000137244.2 | 3101 | Gata4 | ENSMUSG00000021944.10 |
| chr14 | 63268637 | 63269806 | 4.04153 | 5.43436 | 2.1019 | Intron (ENSMUST00000132122.1/ENSMUSG00000021944.10, intron 1 of 1) | 14 | 63241115 | 63271124 | 30010 | 2 | ENSMUST00000137244.2 | 1318 | Gata4 | ENSMUSG00000021944.10 |
| chr14 | 63270351 | 63271705 | 3.93729 | 5.09364 | 1.82561 | Promoter | 14 | 63241102 | 63271692 | 30591 | 2 | ENSMUST00000132122.1 | 0 | Gata4 | ENSMUSG00000021944.10 |
| chr14 | 63272109 | 63272376 | 3.40544 | 3.95438 | 1.0814 | Promoter | 14 | 63241102 | 63271692 | 30591 | 2 | ENSMUST00000132122.1 | -418 | Gata4 | ENSMUSG00000021944.10 |
| chr14 | 63274023 | 63274243 | 3.49922 | 4.12067 | 1.16988 | Distal Intergenic | 14 | 63241102 | 63271692 | 30591 | 2 | ENSMUST00000132122.1 | -2332 | Gata4 | ENSMUSG00000021944.10 |
| chr14 | 63284633 | 63285149 | 3.43379 | 4.00241 | 1.10076 | Distal Intergenic | 14 | 63241102 | 63271692 | 30591 | 2 | ENSMUST00000132122.1 | -12942 | Gata4 | ENSMUSG00000021944.10 |
| chr14 | 63337920 | 63338012 | 3.34153 | 3.83966 | 1.01733 | Distal Intergenic | 14 | 63241102 | 63271692 | 30591 | 2 | ENSMUST00000132122.1 | -66229 | Gata4 | ENSMUSG00000021944.10 |
| chr14 | 63850884 | 63851084 | 3.2667 | 3.73058 | 1.02887 | Distal Intergenic | 14 | 63860312 | 63919859 | 59548 | 1 | ENSMUST00000022528.4 | -9228 | Pinx1 | ENSMUSG00000021958.4 |
| chr14 | 58072703 | 58073208 | 3.99969 | 5.23955 | 1.96077 | Promoter | 14 | 58072686 | 58110174 | 37489 | 1 | ENSMUST00000022545.8 | 18 | Fgf9 | ENSMUSG00000021974.9 |
| chr14 | 58073884 | 58074513 | 4.03516 | 5.37865 | 2.08008 | Promoter | 14 | 58075115 | 58110174 | 35060 | 1 | ENSMUST00000074654.6 | -602 | Fgf9 | ENSMUSG00000021974.9 |
| chr14 | 58075068 | 58075377 | 3.72661 | 4.98656 | 1.76599 | Promoter | 14 | 58075115 | 58110174 | 35060 | 1 | ENSMUST00000074654.6 | 0 | Fgf9 | ENSMUSG00000021974.9 |
| chr14 | 58076559 | 58076819 | 3.34668 | 4.16533 | 1.24857 | Promoter | 14 | 58076507 | 58109392 | 32886 | 1 | ENSMUST00000166770.1 | 53 | Fgf9 | ENSMUSG00000021974.9 |
| chr14 | 58077208 | 58077380 | 3.26164 | 3.79852 | 1.02721 | Promoter | 14 | 58076507 | 58109392 | 32886 | 1 | ENSMUST00000166770.1 | 702 | Fgf9 | ENSMUSG00000021974.9 |
| chr14 | 60085276 | 60085402 | 3.39916 | 3.97658 | 1.18263 | Intron (ENSMUST00000080368.6/ENSMUSG00000021983.10, intron 1 of 35) | 14 | 59647531 | 60086834 | 439304 | 2 | ENSMUST00000080368.6 | 1432 | Atp8a2 | ENSMUSG00000021983.10 |
| chr14 | 60085769 | 60086129 | 3.55852 | 4.5588 | 1.47107 | Promoter | 14 | 59647531 | 60086834 | 439304 | 2 | ENSMUST00000080368.6 | 705 | Atp8a2 | ENSMUSG00000021983.10 |
| chr14 | 60170726 | 60171089 | 3.54005 | 4.18887 | 1.19207 | Intron (ENSMUST00000131670.1/ENSMUSG00000021983.10, intron 1 of 6) | 14 | 60086582 | 60177539 | 90958 | 2 | ENSMUST00000140924.1 | 6450 | Atp8a2 | ENSMUSG00000021983.10 |
| chr14 | 60177086 | 60177406 | 3.96319 | 5.2173 | 1.96763 | Promoter | 14 | 60086582 | 60177539 | 90958 | 2 | ENSMUST00000140924.1 | 133 | Atp8a2 | ENSMUSG00000021983.10 |
| chr14 | 60178276 | 60178397 | 3.32532 | 3.86909 | 1.13314 | Promoter | 14 | 60086582 | 60177539 | 90958 | 2 | ENSMUST00000140924.1 | -738 | Atp8a2 | ENSMUSG00000021983.10 |
| chr14 | 60377446 | 60377601 | 3.21904 | 3.79951 | 1.03209 | Promoter | 14 | 60378286 | 60381003 | 2718 | 1 | ENSMUST00000022561.6 | -685 | Amer2 | ENSMUSG00000021986.7 |
| chr14 | 60378063 | 60378138 | 3.45878 | 4.23237 | 1.29567 | Promoter | 14 | 60378286 | 60381003 | 2718 | 1 | ENSMUST00000022561.6 | -148 | Amer2 | ENSMUSG00000021986.7 |
| chr14 | 60378722 | 60380045 | 3.62034 | 4.67418 | 1.57507 | Promoter | 14 | 60378286 | 60381003 | 2718 | 1 | ENSMUST00000022561.6 | 437 | Amer2 | ENSMUSG00000021986.7 |
| chr14 | 29706372 | 29706699 | 3.48354 | 4.3527 | 1.39059 | Intron (ENSMUST00000022567.8/ENSMUSG00000021991.8, intron 2 of 37) | 14 | 28904943 | 29721864 | 816922 | 2 | ENSMUST00000022567.8 | 15165 | Cacna2d3 | ENSMUSG00000021991.8 |
| chr14 | 29707713 | 29707917 | 3.76025 | 4.65342 | 1.59199 | Intron (ENSMUST00000022567.8/ENSMUSG00000021991.8, intron 2 of 37) | 14 | 28904943 | 29721864 | 816922 | 2 | ENSMUST00000022567.8 | 13947 | Cacna2d3 | ENSMUSG00000021991.8 |
| chr14 | 29717725 | 29718051 | 3.45411 | 4.04784 | 1.12778 | Intron (ENSMUST00000022567.8/ENSMUSG00000021991.8, intron 2 of 37) | 14 | 28904943 | 29721864 | 816922 | 2 | ENSMUST00000022567.8 | 3813 | Cacna2d3 | ENSMUSG00000021991.8 |
| chr14 | 29720863 | 29720973 | 3.57681 | 4.26779 | 1.27644 | Promoter | 14 | 28904943 | 29721864 | 816922 | 2 | ENSMUST00000022567.8 | 891 | Cacna2d3 | ENSMUSG00000021991.8 |
| chr14 | 29721359 | 29722539 | 3.72437 | 4.6551 | 1.52728 | Promoter | 14 | 28904943 | 29721864 | 816922 | 2 | ENSMUST00000022567.8 | 0 | Cacna2d3 | ENSMUSG00000021991.8 |
| chr14 | 29722874 | 29723407 | 3.82273 | 4.77131 | 1.67805 | Distal Intergenic | 14 | 28904943 | 29721864 | 816922 | 2 | ENSMUST00000022567.8 | -1011 | Cacna2d3 | ENSMUSG00000021991.8 |
| chr14 | 29723886 | 29723961 | 3.38079 | 3.93908 | 1.15254 | Distal Intergenic | 14 | 28904943 | 29721864 | 816922 | 2 | ENSMUST00000022567.8 | -2023 | Cacna2d3 | ENSMUSG00000021991.8 |
| chr14 | 29724315 | 29724663 | 3.56495 | 4.29297 | 1.36863 | Distal Intergenic | 14 | 28904943 | 29721864 | 816922 | 2 | ENSMUST00000022567.8 | -2452 | Cacna2d3 | ENSMUSG00000021991.8 |
| chr14 | 28501115 | 28501417 | 3.32716 | 3.8181 | 1.01733 | Distal Intergenic | 14 | 28504750 | 28527447 | 22698 | 1 | ENSMUST00000063465.5 | -3333 | Wnt5a | ENSMUSG00000021994.9 |
| chr14 | 28502623 | 28503519 | 3.71549 | 4.51534 | 1.40465 | Distal Intergenic | 14 | 28504750 | 28527447 | 22698 | 1 | ENSMUST00000063465.5 | -1231 | Wnt5a | ENSMUSG00000021994.9 |
| chr14 | 28503855 | 28505263 | 3.74247 | 4.57632 | 1.46108 | Promoter | 14 | 28504750 | 28527447 | 22698 | 1 | ENSMUST00000063465.5 | 0 | Wnt5a | ENSMUSG00000021994.9 |
| chr14 | 28505574 | 28506222 | 3.8021 | 4.70859 | 1.54671 | Promoter | 14 | 28505900 | 28509121 | 3222 | 1 | ENSMUST00000134766.1 | 0 | Wnt5a | ENSMUSG00000021994.9 |
| chr14 | 28506587 | 28506929 | 3.68332 | 4.46206 | 1.38126 | Promoter | 14 | 28505900 | 28509121 | 3222 | 1 | ENSMUST00000134766.1 | 688 | Wnt5a | ENSMUSG00000021994.9 |
| chr14 | 28507510 | 28507585 | 3.36462 | 3.87448 | 1.01733 | Intron (ENSMUST00000063465.5/ENSMUSG00000021994.9, intron 1 of 4) | 14 | 28505900 | 28509121 | 3222 | 1 | ENSMUST00000134766.1 | 1611 | Wnt5a | ENSMUSG00000021994.9 |
| chr14 | 28507958 | 28508308 | 3.37312 | 3.93408 | 1.14618 | Exon (ENSMUST00000180668.1/ENSMUSG00000097002.1, exon 2 of 2) | 14 | 28505900 | 28509121 | 3222 | 1 | ENSMUST00000134766.1 | 2059 | Wnt5a | ENSMUSG00000021994.9 |
| chr14 | 28508710 | 28509672 | 3.88557 | 4.90526 | 1.7081 | Promoter | 14 | 28510618 | 28511925 | 1308 | 1 | ENSMUST00000151929.1 | -946 | Wnt5a | ENSMUSG00000021994.9 |
| chr14 | 28510680 | 28511783 | 3.81757 | 4.75055 | 1.58872 | Promoter | 14 | 28511390 | 28523517 | 12128 | 1 | ENSMUST00000134163.1 | 0 | Wnt5a | ENSMUSG00000021994.9 |
| chr14 | 28513005 | 28513105 | 4.01678 | 5.17041 | 1.88681 | Intron (ENSMUST00000063465.5/ENSMUSG00000021994.9, intron 2 of 4) | 14 | 28511451 | 28511907 | 457 | 1 | ENSMUST00000146192.1 | 1555 | Wnt5a | ENSMUSG00000021994.9 |
| chr14 | 28514402 | 28514770 | 3.60733 | 4.41951 | 1.40666 | Intron (ENSMUST00000063465.5/ENSMUSG00000021994.9, intron 3 of 4) | 14 | 28511451 | 28511907 | 457 | 1 | ENSMUST00000146192.1 | 2952 | Wnt5a | ENSMUSG00000021994.9 |
| chr14 | 75129245 | 75129474 | 3.35668 | 3.86251 | 1.01733 | Exon (ENSMUST00000022574.4/ENSMUSG00000021997.4, exon 2 of 10) | 14 | 75084303 | 75130881 | 46579 | 2 | ENSMUST00000022574.4 | 1407 | Lrrc63 | ENSMUSG00000021997.4 |
| chr14 | 75192708 | 75193025 | 3.3413 | 3.83939 | 1.01733 | Intron (ENSMUST00000143539.2/ENSMUSG00000021998.11, intron 1 of 2) | 14 | 75190329 | 75230816 | 40488 | 1 | ENSMUST00000131802.1 | 2380 | Lcp1 | ENSMUSG00000021998.11 |
| chr14 | 77153449 | 77153727 | 4.13734 | 5.54254 | 2.14478 | Distal Intergenic | 14 | 77156763 | 77721760 | 564998 | 1 | ENSMUST00000022589.7 | -3036 | Enox1 | ENSMUSG00000022012.8 |
| chr14 | 77154386 | 77154848 | 3.5842 | 4.3603 | 1.35537 | Distal Intergenic | 14 | 77156763 | 77721760 | 564998 | 1 | ENSMUST00000022589.7 | -1915 | Enox1 | ENSMUSG00000022012.8 |
| chr14 | 77155243 | 77155865 | 3.88046 | 4.92359 | 1.76882 | Promoter | 14 | 77156763 | 77721760 | 564998 | 1 | ENSMUST00000022589.7 | -898 | Enox1 | ENSMUSG00000022012.8 |
| chr14 | 77156325 | 77157807 | 3.76545 | 4.90846 | 1.71973 | Promoter | 14 | 77156763 | 77721760 | 564998 | 1 | ENSMUST00000022589.7 | 0 | Enox1 | ENSMUSG00000022012.8 |
| chr14 | 78306512 | 78306712 | 3.36931 | 3.88156 | 1.01733 | Intron (ENSMUST00000022592.7/ENSMUSG00000022015.8, intron 1 of 4) | 14 | 78277445 | 78308043 | 30599 | 2 | ENSMUST00000022592.7 | 1331 | Tnfsf11 | ENSMUSG00000022015.8 |
| chr14 | 78307655 | 78307779 | 4.10104 | 5.41183 | 2.06878 | Promoter | 14 | 78277445 | 78308043 | 30599 | 2 | ENSMUST00000022592.7 | 264 | Tnfsf11 | ENSMUSG00000022015.8 |
| chr14 | 78308619 | 78308842 | 3.38131 | 3.90681 | 1.04246 | Promoter | 14 | 78277445 | 78308043 | 30599 | 2 | ENSMUST00000022592.7 | -577 | Tnfsf11 | ENSMUSG00000022015.8 |
| chr14 | 79299070 | 79299597 | 3.44398 | 4.02714 | 1.1285 | Intron (ENSMUST00000022595.7/ENSMUSG00000022018.7, intron 2 of 4) | 14 | 79288756 | 79301645 | 12890 | 2 | ENSMUST00000022595.7 | 2048 | Rgcc | ENSMUSG00000022018.7 |
| chr14 | 79300141 | 79300471 | 3.42302 | 4.10829 | 1.19493 | Intron (ENSMUST00000022595.7/ENSMUSG00000022018.7, intron 2 of 4) | 14 | 79288756 | 79301645 | 12890 | 2 | ENSMUST00000022595.7 | 1174 | Rgcc | ENSMUSG00000022018.7 |
| chr14 | 79300985 | 79301138 | 3.91958 | 4.99433 | 1.76439 | Promoter | 14 | 79288756 | 79301645 | 12890 | 2 | ENSMUST00000022595.7 | 507 | Rgcc | ENSMUSG00000022018.7 |
| chr14 | 65730950 | 65731031 | 4.0571 | 5.2055 | 1.93902 | Exon (ENSMUST00000022610.9/ENSMUSG00000022032.9, exon 4 of 9) | 14 | 65676477 | 65744791 | 68315 | 1 | ENSMUST00000154373.1 | 54474 | Scara5 | ENSMUSG00000022032.9 |
| chr14 | 66077452 | 66077902 | 3.21403 | 3.64828 | 1.01227 | Promoter | 14 | 66027329 | 66077733 | 50405 | 2 | ENSMUST00000022618.5 | 0 | Adam2 | ENSMUSG00000022039.5 |
| chr14 | 66381170 | 66381499 | 4.03579 | 5.12753 | 1.82518 | Distal Intergenic | 14 | 66344381 | 66358102 | 13722 | 1 | ENSMUST00000147477.1 | 36790 | Stmn4 | ENSMUSG00000022044.9 |
| chr14 | 67229674 | 67233591 | 3.97717 | 5.28971 | 2.00428 | Promoter | 14 | 67233292 | 67429777 | 196486 | 1 | ENSMUST00000022637.8 | 0 | Ebf2 | ENSMUSG00000022053.8 |
| chr14 | 67233907 | 67237845 | 3.97503 | 5.1785 | 1.94162 | Promoter | 14 | 67234644 | 67428687 | 194044 | 1 | ENSMUST00000176029.1 | 0 | Ebf2 | ENSMUSG00000022053.8 |
| chr14 | 67240667 | 67240957 | 3.93223 | 5.2424 | 1.97271 | Intron (ENSMUST00000022637.8/ENSMUSG00000022053.8, intron 6 of 16) | 14 | 67234644 | 67428687 | 194044 | 1 | ENSMUST00000176029.1 | 6024 | Ebf2 | ENSMUSG00000022053.8 |
| chr14 | 67242153 | 67242545 | 3.69286 | 4.51556 | 1.48645 | Intron (ENSMUST00000022637.8/ENSMUSG00000022053.8, intron 6 of 16) | 14 | 67234644 | 67428687 | 194044 | 1 | ENSMUST00000176029.1 | 7510 | Ebf2 | ENSMUSG00000022053.8 |
| chr14 | 67245141 | 67245304 | 3.50912 | 4.16264 | 1.25731 | Intron (ENSMUST00000022637.8/ENSMUSG00000022053.8, intron 6 of 16) | 14 | 67234644 | 67428687 | 194044 | 1 | ENSMUST00000176029.1 | 10498 | Ebf2 | ENSMUSG00000022053.8 |
| chr14 | 68121611 | 68122048 | 3.46474 | 4.08051 | 1.20457 | Intron (ENSMUST00000111089.2/ENSMUSG00000022054.6, intron 2 of 3) | 14 | 68082590 | 68124846 | 42257 | 2 | ENSMUST00000111089.2 | 2798 | Nefm | ENSMUSG00000022054.6 |
| chr14 | 68123125 | 68125204 | 3.79986 | 4.7415 | 1.57906 | Promoter | 14 | 68082590 | 68124846 | 42257 | 2 | ENSMUST00000111089.2 | 0 | Nefm | ENSMUSG00000022054.6 |
| chr14 | 68125831 | 68125906 | 3.36496 | 3.875 | 1.01733 | Promoter | 14 | 68119283 | 68124846 | 5564 | 2 | ENSMUST00000022638.5 | -986 | Nefm | ENSMUSG00000022054.6 |
| chr14 | 68127032 | 68127285 | 3.38286 | 4.10984 | 1.21044 | Distal Intergenic | 14 | 68119283 | 68124846 | 5564 | 2 | ENSMUST00000022638.5 | -2187 | Nefm | ENSMUSG00000022054.6 |
| chr14 | 68127682 | 68127952 | 3.35416 | 3.8587 | 1.01733 | Distal Intergenic | 14 | 68119283 | 68124846 | 5564 | 2 | ENSMUST00000022638.5 | -2837 | Nefm | ENSMUSG00000022054.6 |
| chr14 | 68039762 | 68040156 | 4.70496 | 6.69609 | 3.07144 | Distal Intergenic | 14 | 68083863 | 68089095 | 5233 | 1 | ENSMUST00000022639.7 | -43707 | Nefl | ENSMUSG00000022055.7 |
| chr14 | 68041030 | 68041233 | 3.55889 | 4.23427 | 1.25349 | Distal Intergenic | 14 | 68083863 | 68089095 | 5233 | 1 | ENSMUST00000022639.7 | -42630 | Nefl | ENSMUSG00000022055.7 |
| chr14 | 68068689 | 68068765 | 4.89203 | 6.90615 | 3.15545 | Distal Intergenic | 14 | 68083863 | 68089095 | 5233 | 1 | ENSMUST00000022639.7 | -15098 | Nefl | ENSMUSG00000022055.7 |
| chr14 | 68069110 | 68069439 | 3.39352 | 3.95892 | 1.16016 | Distal Intergenic | 14 | 68083863 | 68089095 | 5233 | 1 | ENSMUST00000022639.7 | -14424 | Nefl | ENSMUSG00000022055.7 |
| chr14 | 68073185 | 68073557 | 3.94639 | 4.99911 | 1.779 | Distal Intergenic | 14 | 68083863 | 68089095 | 5233 | 1 | ENSMUST00000022639.7 | -10306 | Nefl | ENSMUSG00000022055.7 |
| chr14 | 68074763 | 68074924 | 3.54426 | 4.25533 | 1.33916 | Distal Intergenic | 14 | 68083863 | 68089095 | 5233 | 1 | ENSMUST00000022639.7 | -8939 | Nefl | ENSMUSG00000022055.7 |
| chr14 | 68078310 | 68078738 | 3.67375 | 4.6183 | 1.5087 | Distal Intergenic | 14 | 68083863 | 68089095 | 5233 | 1 | ENSMUST00000022639.7 | -5125 | Nefl | ENSMUSG00000022055.7 |
| chr14 | 68080226 | 68080814 | 3.46 | 4.09003 | 1.22563 | Downstream (1-2kb) | 14 | 68083863 | 68089095 | 5233 | 1 | ENSMUST00000022639.7 | -3049 | Nefl | ENSMUSG00000022055.7 |
| chr14 | 68081129 | 68081628 | 3.31581 | 3.93411 | 1.12685 | Downstream (<1kb) | 14 | 68083863 | 68089095 | 5233 | 1 | ENSMUST00000022639.7 | -2235 | Nefl | ENSMUSG00000022055.7 |
| chr14 | 68082570 | 68082780 | 3.60248 | 4.33143 | 1.34649 | 3' UTR | 14 | 68083863 | 68089095 | 5233 | 1 | ENSMUST00000022639.7 | -1083 | Nefl | ENSMUSG00000022055.7 |
| chr14 | 68083733 | 68084474 | 3.96087 | 5.03723 | 1.80111 | Promoter | 14 | 68083863 | 68089095 | 5233 | 1 | ENSMUST00000022639.7 | 0 | Nefl | ENSMUSG00000022055.7 |
| chr14 | 69189905 | 69191559 | 4.23925 | 5.87885 | 2.46171 | Promoter | 14 | 69190638 | 69194662 | 4025 | 1 | ENSMUST00000022646.8 | 0 | Nkx3-1 | ENSMUSG00000022061.8 |
| chr14 | 69191907 | 69191982 | 4.06058 | 5.40268 | 2.0673 | Exon (ENSMUST00000022646.8/ENSMUSG00000022061.8, exon 2 of 2) | 14 | 69190638 | 69194662 | 4025 | 1 | ENSMUST00000022646.8 | 1270 | Nkx3-1 | ENSMUSG00000022061.8 |
| chr14 | 69193554 | 69193755 | 3.5009 | 4.24 | 1.31775 | 3' UTR | 14 | 69190638 | 69194662 | 4025 | 1 | ENSMUST00000022646.8 | 2917 | Nkx3-1 | ENSMUSG00000022061.8 |
| chr14 | 69763367 | 69763840 | 3.69001 | 4.5347 | 1.4501 | Intron (ENSMUST00000181430.1/ENSMUSG00000097284.2, intron 2 of 2) | 14 | 69767472 | 69784403 | 16932 | 1 | ENSMUST00000022663.5 | -3632 | Tnfrsf10b | ENSMUSG00000022074.5 |
| chr14 | 96517843 | 96517979 | 3.82183 | 4.72389 | 1.5573 | Exon (ENSMUST00000022666.7/ENSMUSG00000022076.9, exon 1 of 11) | 14 | 96105259 | 96519102 | 413844 | 2 | ENSMUST00000022666.7 | 1123 | Klhl1 | ENSMUSG00000022076.9 |
| chr14 | 96518463 | 96519308 | 3.90064 | 4.88565 | 1.67459 | Promoter | 14 | 96105259 | 96519102 | 413844 | 2 | ENSMUST00000022666.7 | 0 | Klhl1 | ENSMUSG00000022076.9 |
| chr14 | 70604557 | 70604632 | 3.9867 | 5.03773 | 1.77176 | 3' UTR | 14 | 70583296 | 70599835 | 16540 | 2 | ENSMUST00000022690.9 | -4723 | Fam160b2 | ENSMUSG00000022095.9 |
| chr14 | 70605333 | 70605533 | 3.36309 | 3.90769 | 1.12806 | Exon (ENSMUST00000022695.10/ENSMUSG00000022099.11, exon 12 of 15) | 14 | 70583296 | 70599835 | 16540 | 2 | ENSMUST00000022690.9 | -5499 | Fam160b2 | ENSMUSG00000022095.9 |
| chr14 | 70554147 | 70555122 | 3.71549 | 4.6513 | 1.59274 | Promoter | 14 | 70554056 | 70573548 | 19493 | 1 | ENSMUST00000022691.8 | 92 | Hr | ENSMUSG00000022096.9 |
| chr14 | 70555889 | 70556038 | 3.26895 | 3.74107 | 1.05084 | Promoter | 14 | 70555900 | 70573544 | 17645 | 1 | ENSMUST00000163060.1 | 0 | Hr | ENSMUSG00000022096.9 |
| chr14 | 70626065 | 70626314 | 3.37278 | 3.91687 | 1.12267 | Promoter | 14 | 70602185 | 70627065 | 24881 | 2 | ENSMUST00000110984.2 | 751 | Dmtn | ENSMUSG00000022099.11 |
| chr14 | 70628243 | 70628427 | 4.01179 | 5.26739 | 1.97975 | Intron (ENSMUST00000022694.10/ENSMUSG00000022099.11, intron 1 of 15) | 14 | 70602185 | 70627065 | 24881 | 2 | ENSMUST00000110984.2 | -1179 | Dmtn | ENSMUSG00000022099.11 |
| chr14 | 70635316 | 70636399 | 3.53104 | 4.48444 | 1.43978 | Promoter | 14 | 70602185 | 70636048 | 33864 | 2 | ENSMUST00000022695.10 | 0 | Dmtn | ENSMUSG00000022099.11 |
| chr14 | 70641629 | 70642457 | 3.5467 | 4.50601 | 1.45008 | Promoter | 14 | 70636205 | 70642268 | 6064 | 2 | ENSMUST00000022697.5 | 0 | Fgf17 | ENSMUSG00000022101.5 |
| chr14 | 70889111 | 70889308 | 3.38241 | 4.11918 | 1.21325 | Promoter | 14 | 70890130 | 70979838 | 89709 | 1 | ENSMUST00000022699.8 | -822 | Gfra2 | ENSMUSG00000022103.9 |
| chr14 | 70889790 | 70889961 | 3.37068 | 4.55982 | 1.46379 | Promoter | 14 | 70890130 | 70979838 | 89709 | 1 | ENSMUST00000022699.8 | -169 | Gfra2 | ENSMUSG00000022103.9 |
| chr14 | 70890495 | 70890900 | 3.67966 | 4.56255 | 1.51401 | Promoter | 14 | 70890130 | 70979838 | 89709 | 1 | ENSMUST00000022699.8 | 366 | Gfra2 | ENSMUSG00000022103.9 |
| chr14 | 70891451 | 70891752 | 3.79752 | 4.93029 | 1.70192 | Intron (ENSMUST00000022699.8/ENSMUSG00000022103.9, intron 1 of 8) | 14 | 70890130 | 70979838 | 89709 | 1 | ENSMUST00000022699.8 | 1322 | Gfra2 | ENSMUSG00000022103.9 |
| chr14 | 70892761 | 70892967 | 3.86725 | 4.91663 | 1.7443 | Intron (ENSMUST00000022699.8/ENSMUSG00000022103.9, intron 1 of 8) | 14 | 70890130 | 70979838 | 89709 | 1 | ENSMUST00000022699.8 | 2632 | Gfra2 | ENSMUSG00000022103.9 |
| chr14 | 115090639 | 115090841 | 3.31611 | 4.04819 | 1.16194 | Distal Intergenic | 14 | 115092215 | 115134210 | 41996 | 1 | ENSMUST00000176993.1 | -1374 | Gpc5 | ENSMUSG00000022112.9 |
| chr14 | 115091198 | 115091657 | 3.7011 | 4.56394 | 1.52334 | Promoter | 14 | 115092215 | 115134210 | 41996 | 1 | ENSMUST00000176993.1 | -558 | Gpc5 | ENSMUSG00000022112.9 |
| chr14 | 115092327 | 115093084 | 3.68811 | 4.71627 | 1.57689 | Promoter | 14 | 115092626 | 116524437 | 1431812 | 1 | ENSMUST00000022707.6 | 0 | Gpc5 | ENSMUSG00000022112.9 |
| chr14 | 105870600 | 105870734 | 3.46009 | 4.16458 | 1.23963 | Distal Intergenic | 14 | 105891949 | 105896819 | 4871 | 2 | ENSMUST00000022709.4 | 26085 | Spry2 | ENSMUSG00000022114.4 |
| chr14 | 105923715 | 105924039 | 3.46298 | 4.06804 | 1.17751 | Distal Intergenic | 14 | 105891949 | 105896819 | 4871 | 2 | ENSMUST00000022709.4 | -26897 | Spry2 | ENSMUSG00000022114.4 |
| chr14 | 105501243 | 105501860 | 3.84456 | 4.84888 | 1.65453 | Intron (ENSMUST00000022711.2/ENSMUSG00000022116.2, intron 1 of 2) | 14 | 105503576 | 105505629 | 2054 | 1 | ENSMUST00000188698.1 | -1716 | 4930449E01Rik | ENSMUSG00000022116.2 |
| chr14 | 105502922 | 105503068 | 3.47986 | 4.24765 | 1.30758 | Promoter | 14 | 105503576 | 105505629 | 2054 | 1 | ENSMUST00000188698.1 | -508 | 4930449E01Rik | ENSMUSG00000022116.2 |
| chr14 | 105542870 | 105543137 | 4.205 | 5.5788 | 2.19525 | Distal Intergenic | 14 | 105503576 | 105505629 | 2054 | 1 | ENSMUST00000188698.1 | 39295 | 4930449E01Rik | ENSMUSG00000022116.2 |
| chr14 | 103842230 | 103842331 | 3.62178 | 4.40603 | 1.45593 | Intron (ENSMUST00000022718.4/ENSMUSG00000022122.9, intron 2 of 7) | 14 | 103814626 | 103843696 | 29071 | 2 | ENSMUST00000172237.1 | 1365 | Ednrb | ENSMUSG00000022122.9 |

| | | | | | | | | | | | | | | | |
|---|---|---|---|---|---|---|---|---|---|---|---|---|---|---|---|
| chr14 | 118046057 | 118046661 | 3.81534 | 4.74718 | 1.57282 | Intron (ENSMUST00000022725.2/ENSMUSG00000022129.3, intron 1 of 7) | 14 | 118012792 | 118052246 | 39455 | 2 | ENSMUST00000022725.2 | 5585 | Dct | ENSMUSG00000022129.3 |
| chr14 | 118069283 | 118069483 | 3.23022 | 3.67252 | 1.0168 | Distal Intergenic | 14 | 118012792 | 118052246 | 39455 | 2 | ENSMUST00000022725.2 | -17038 | Dct | ENSMUSG00000022129.3 |
| chr14 | 118070107 | 118070212 | 3.98319 | 5.18802 | 1.92903 | Distal Intergenic | 14 | 118012792 | 118052246 | 39455 | 2 | ENSMUST00000022725.2 | -17862 | Dct | ENSMUSG00000022129.3 |
| chr14 | 118071701 | 118072093 | 3.56106 | 4.22856 | 1.22009 | Distal Intergenic | 14 | 118012792 | 118052246 | 39455 | 2 | ENSMUST00000022725.2 | -19456 | Dct | ENSMUSG00000022129.3 |
| chr14 | 118073274 | 118073424 | 4.14038 | 5.47262 | 2.11573 | Distal Intergenic | 14 | 118012792 | 118052246 | 39455 | 2 | ENSMUST00000022725.2 | -21029 | Dct | ENSMUSG00000022129.3 |
| chr14 | 118783566 | 118783641 | 3.74385 | 4.59159 | 1.51505 | Distal Intergenic | 14 | 118787908 | 118874524 | 86617 | 1 | ENSMUST00000047761.7 | -4267 | Cldn10 | ENSMUSG00000022132.10 |
| chr14 | 118854694 | 118855162 | 3.43736 | 4.01412 | 1.12315 | Promoter | 14 | 118854732 | 118875489 | 20758 | 1 | ENSMUST00000100314.3 | 0 | Cldn10 | ENSMUSG00000022132.10 |
| chr14 | 118855469 | 118855597 | 3.35495 | 3.86212 | 1.02518 | Promoter | 14 | 118854732 | 118875489 | 20758 | 1 | ENSMUST00000100314.3 | 738 | Cldn10 | ENSMUSG00000022132.10 |
| chr14 | 118857355 | 118857645 | 3.73336 | 4.63257 | 1.55227 | Intron (ENSMUST00000047761.7/ENSMUSG00000022132.10, intron 1 of 4) | 14 | 118854732 | 118875489 | 20758 | 1 | ENSMUST00000100314.3 | 2624 | Cldn10 | ENSMUSG00000022132.10 |
| chr15 | 7808531 | 7808630 | 3.56247 | 4.27637 | 1.35478 | Distal Intergenic | 15 | 7811011 | 7837575 | 26565 | 1 | ENSMUST00000022744.3 | -2381 | Gdnf | ENSMUSG00000022144.3 |
| chr15 | 7809008 | 7809131 | 3.25627 | 3.7188 | 1.04004 | Distal Intergenic | 15 | 7811011 | 7837575 | 26565 | 1 | ENSMUST00000022744.3 | -1880 | Gdnf | ENSMUSG00000022144.3 |
| chr15 | 7809498 | 7811043 | 3.86969 | 5.02498 | 1.82321 | Promoter | 15 | 7811011 | 7837575 | 26565 | 1 | ENSMUST00000022744.3 | 0 | Gdnf | ENSMUSG00000022144.3 |
| chr15 | 7813115 | 7813711 | 3.61364 | 4.40003 | 1.43009 | Intron (ENSMUST00000022744.3/ENSMUSG00000022144.3, intron 1 of 2) | 15 | 7811011 | 7837575 | 26565 | 1 | ENSMUST00000022744.3 | 2105 | Gdnf | ENSMUSG00000022144.3 |
| chr15 | 7814513 | 7815220 | 3.75354 | 4.68619 | 1.64299 | Intron (ENSMUST00000022744.3/ENSMUSG00000022144.3, intron 1 of 2) | 15 | 7811011 | 7837575 | 26565 | 1 | ENSMUST00000022744.3 | 3503 | Gdnf | ENSMUSG00000022144.3 |
| chr15 | 7815634 | 7815814 | 3.91503 | 5.00248 | 1.82284 | Exon (ENSMUST00000022744.3/ENSMUSG00000022144.3, exon 2 of 3) | 15 | 7811011 | 7837575 | 26565 | 1 | ENSMUST00000022744.3 | 4624 | Gdnf | ENSMUSG00000022144.3 |
| chr15 | 6874331 | 6874431 | 4.35992 | 5.90983 | 2.42489 | Promoter | 15 | 6854987 | 6874296 | 19310 | 2 | ENSMUST00000177263.1 | -36 | Osmr | ENSMUSG00000022146.7 |
| chr14 | 54487384 | 54487704 | 3.46615 | 4.30746 | 1.33756 | Distal Intergenic | 14 | 54477471 | 54479117 | 1647 | 1 | ENSMUST00000168930.1 | 9914 | Rem2 | ENSMUSG00000022176.6 |
| chr14 | 54912416 | 54912704 | 3.70506 | 4.5755 | 1.52026 | Promoter | 14 | 54906727 | 54913132 | 6406 | 2 | ENSMUST00000050772.7 | 428 | Slc22a17 | ENSMUSG00000022199.10 |
| chr14 | 54925593 | 54925909 | 3.52746 | 4.29679 | 1.30831 | Promoter | 14 | 54916543 | 54926788 | 10246 | 2 | ENSMUST00000022813.6 | 879 | Efs | ENSMUSG00000022203.6 |
| chr14 | 55113846 | 55114114 | 3.28589 | 4.06502 | 1.16387 | Exon (ENSMUST00000022819.6/ENSMUSG00000022208.6, exon 3 of 6) | 14 | 55107277 | 55115552 | 8276 | 2 | ENSMUST00000124493.1 | 1438 | Jph4 | ENSMUSG00000022208.6 |
| chr14 | 55115050 | 55115856 | 3.92617 | 5.008 | 1.83319 | Promoter | 14 | 55107277 | 55115552 | 8276 | 2 | ENSMUST00000124493.1 | 0 | Jph4 | ENSMUSG00000022208.6 |
| chr14 | 55117115 | 55117603 | 3.5011 | 4.16126 | 1.28836 | Promoter | 14 | 55106830 | 55116935 | 10106 | 2 | ENSMUST00000022819.6 | -181 | Jph4 | ENSMUSG00000022208.6 |
| chr14 | 55780747 | 55780842 | 3.34279 | 3.88085 | 1.14062 | Exon (ENSMUST00000170223.2/ENSMUSG00000022220.8, exon 5 of 26) | 14 | 55769093 | 55784019 | 14927 | 2 | ENSMUST00000002398.7 | 3177 | Adcy4 | ENSMUSG00000022220.8 |
| chr14 | 55783199 | 55783843 | 3.75876 | 4.8851 | 1.72012 | Promoter | 14 | 55769093 | 55784019 | 14927 | 2 | ENSMUST00000002398.7 | 176 | Adcy4 | ENSMUSG00000022220.8 |
| chr14 | 56363654 | 56363856 | 3.99547 | 5.30157 | 1.99544 | Distal Intergenic | 14 | 56365068 | 56388551 | 23484 | 1 | ENSMUST00000007340.2 | -1212 | Atp12a | ENSMUSG00000022229.2 |
| chr14 | 56364645 | 56365233 | 4.07303 | 5.30826 | 2.02047 | Promoter | 14 | 56365068 | 56388551 | 23484 | 1 | ENSMUST00000007340.2 | 0 | Atp12a | ENSMUSG00000022229.2 |
| chr15 | 30171761 | 30171952 | 3.52696 | 4.18645 | 1.25051 | Promoter | 15 | 30172593 | 31029311 | 856719 | 1 | ENSMUST00000081728.5 | -641 | Ctnnd2 | ENSMUSG00000022240.9 |
| chr15 | 30172372 | 30172628 | 3.4636 | 4.06631 | 1.16663 | Promoter | 15 | 30172593 | 31029311 | 856719 | 1 | ENSMUST00000081728.5 | 0 | Ctnnd2 | ENSMUSG00000022240.9 |
| chr15 | 30172973 | 30173250 | 4.62359 | 6.48125 | 2.90158 | Promoter | 15 | 30172593 | 31029311 | 856719 | 1 | ENSMUST00000081728.5 | 381 | Ctnnd2 | ENSMUSG00000022240.9 |
| chr15 | 30173588 | 30173976 | 3.66866 | 4.59894 | 1.52544 | Promoter | 15 | 30172593 | 31029311 | 856719 | 1 | ENSMUST00000081728.5 | 996 | Ctnnd2 | ENSMUSG00000022240.9 |
| chr15 | 30174326 | 30174895 | 3.48913 | 4.13875 | 1.27284 | Intron (ENSMUST00000081728.5/ENSMUSG00000022240.9, intron 1 of 22) | 15 | 30172593 | 31029311 | 856719 | 1 | ENSMUST00000081728.5 | 1734 | Ctnnd2 | ENSMUSG00000022240.9 |
| chr9 | 63140972 | 63141080 | 4.00082 | 5.14008 | 1.9261 | Intron (ENSMUST00000119146.2/ENSMUSG00000022245.10, intron 5 of 8) | 9 | 63138176 | 63146980 | 8805 | 2 | ENSMUST00000055281.7 | 5900 | Skor1 | ENSMUSG00000022245.10 |
| chr9 | 63141505 | 63143488 | 3.88353 | 4.98227 | 1.79199 | Exon (ENSMUST00000119146.2/ENSMUSG00000022245.10, exon 3 of 9) | 9 | 63138176 | 63146980 | 8805 | 2 | ENSMUST00000055281.7 | 3492 | Skor1 | ENSMUSG00000022245.10 |
| chr9 | 63143806 | 63148118 | 3.9314 | 5.0861 | 1.85413 | Promoter | 9 | 63138170 | 63147011 | 8842 | 2 | ENSMUST00000119146.2 | 0 | Skor1 | ENSMUSG00000022245.10 |
| chr9 | 63148483 | 63149744 | 4.18081 | 5.46938 | 2.08477 | Promoter | 9 | 63138170 | 63148961 | 10792 | 2 | ENSMUST00000116613.3 | 0 | Skor1 | ENSMUSG00000022245.10 |
| chr9 | 63150139 | 63151413 | 4.10734 | 5.45753 | 2.1283 | Distal Intergenic | 9 | 63138170 | 63148961 | 10792 | 2 | ENSMUST00000116613.3 | -1179 | Skor1 | ENSMUSG00000022245.10 |
| chr9 | 63151798 | 63153767 | 4.09306 | 5.27375 | 1.95781 | Distal Intergenic | 9 | 63138170 | 63148961 | 10792 | 2 | ENSMUST00000116613.3 | -2838 | Skor1 | ENSMUSG00000022245.10 |
| chr15 | 28154483 | 28154981 | 3.62306 | 4.44952 | 1.38044 | Distal Intergenic | 15 | 28203752 | 28472045 | 268294 | 1 | ENSMUST00000067048.6 | -48771 | Dnah5 | ENSMUSG00000022262.7 |
| chr15 | 28070671 | 28070770 | 3.41205 | 4.00526 | 1.20511 | Distal Intergenic | 15 | 27730652 | 28025848 | 295197 | 2 | ENSMUST00000090247.5 | -44824 | Trio | ENSMUSG00000022263.9 |
| chr15 | 26308646 | 26309864 | 3.96711 | 5.11655 | 1.91031 | Promoter | 15 | 26309048 | 26312371 | 3324 | 1 | ENSMUST00000126304.1 | 0 | Mar-11 | ENSMUSG00000022269.8 |
| chr15 | 37175444 | 37175520 | 3.52008 | 4.96716 | 1.7552 | Distal Intergenic | 15 | 37233036 | 37333647 | 100612 | 1 | ENSMUST00000161405.2 | -57516 | Grhl2 | ENSMUSG00000022286.10 |
| chr15 | 37176465 | 37176555 | 4.01774 | 5.0876 | 1.77836 | Distal Intergenic | 15 | 37233036 | 37333647 | 100612 | 1 | ENSMUST00000161405.2 | -56481 | Grhl2 | ENSMUSG00000022286.10 |
| chr15 | 37198139 | 37198831 | 3.44776 | 4.10297 | 1.23158 | Distal Intergenic | 15 | 37233036 | 37333647 | 100612 | 1 | ENSMUST00000161405.2 | -34205 | Grhl2 | ENSMUSG00000022286.10 |
| chr15 | 37199700 | 37199812 | 3.29511 | 4.02629 | 1.18926 | Distal Intergenic | 15 | 37233036 | 37333647 | 100612 | 1 | ENSMUST00000161405.2 | -33224 | Grhl2 | ENSMUSG00000022286.10 |
| chr15 | 37215716 | 37216087 | 3.65918 | 4.47 | 1.48027 | Distal Intergenic | 15 | 37233036 | 37333647 | 100612 | 1 | ENSMUST00000161405.2 | -16949 | Grhl2 | ENSMUSG00000022286.10 |
| chr15 | 37232851 | 37233584 | 3.60735 | 4.70177 | 1.56952 | Promoter | 15 | 37233036 | 37333647 | 100612 | 1 | ENSMUST00000161405.2 | 0 | Grhl2 | ENSMUSG00000022286.10 |
| chr15 | 38938996 | 38939200 | 3.33172 | 3.82488 | 1.01733 | Intron (ENSMUST00000163313.1/ENSMUSG00000022296.8, intron 2 of 2) | 15 | 38933142 | 38949405 | 16264 | 1 | ENSMUST00000163313.1 | 5855 | Baalc | ENSMUSG00000022296.8 |
| chr15 | 39660768 | 39660942 | 3.32465 | 3.81424 | 1.01733 | Intron (ENSMUST00000082054.6/ENSMUSG00000037386.10, intron 22 of 26) | 15 | 39745932 | 39760934 | 15003 | 1 | ENSMUST00000022913.4 | -84990 | Dcstamp | ENSMUSG00000022303.8 |
| chr15 | 39856969 | 39857213 | 3.93074 | 5.4083 | 2.05493 | Promoter | 15 | 39768485 | 39857467 | 88983 | 2 | ENSMUST00000022915.4 | 254 | Dpys | ENSMUSG00000022304.7 |
| chr15 | 40653509 | 40653714 | 3.42123 | 4.06549 | 1.21764 | Distal Intergenic | 15 | 40655042 | 41104592 | 449551 | 1 | ENSMUST00000053467.4 | -1328 | Zfpm2 | ENSMUSG00000022306.8 |
| chr15 | 40654432 | 40656487 | 3.62678 | 4.4653 | 1.46967 | Promoter | 15 | 40655042 | 41104592 | 449551 | 1 | ENSMUST00000053467.4 | 0 | Zfpm2 | ENSMUSG00000022306.8 |
| chr15 | 40656945 | 40657269 | 3.42174 | 4.00826 | 1.18662 | Intron (ENSMUST00000053467.4/ENSMUSG00000022306.8, intron 1 of 7) | 15 | 40655042 | 41104592 | 449551 | 1 | ENSMUST00000053467.4 | 1904 | Zfpm2 | ENSMUSG00000022306.8 |
| chr15 | 40658225 | 40658509 | 4.11584 | 5.36144 | 2.039 | Intron (ENSMUST00000053467.4/ENSMUSG00000022306.8, intron 1 of 7) | 15 | 40655042 | 41104592 | 449551 | 1 | ENSMUST00000053467.4 | 3184 | Zfpm2 | ENSMUSG00000022306.8 |
| chr15 | 48790060 | 48790168 | 3.59401 | 4.31173 | 1.3279 | Intron (ENSMUST00000160658.2/ENSMUSG00000022311.10, intron 1 of 68) | 15 | 47580637 | 48791933 | 1211297 | 2 | ENSMUST00000160658.2 | 1765 | Csmd3 | ENSMUSG00000022311.10 |
| chr15 | 48790708 | 48791518 | 3.61039 | 4.38578 | 1.40116 | Promoter | 15 | 47580637 | 48791933 | 1211297 | 2 | ENSMUST00000160658.2 | 415 | Csmd3 | ENSMUSG00000022311.10 |
| chr15 | 48791917 | 48792378 | 3.47229 | 4.09781 | 1.22712 | Promoter | 15 | 47580637 | 48791933 | 1211297 | 2 | ENSMUST00000160658.2 | 0 | Csmd3 | ENSMUSG00000022311.10 |
| chr15 | 48793251 | 48793557 | 3.34536 | 4.00503 | 1.18054 | Distal Intergenic | 15 | 47580722 | 48792063 | 1211342 | 2 | ENSMUST00000162830.2 | -1189 | Csmd3 | ENSMUSG00000022311.10 |
| chr15 | 18819571 | 18819691 | 3.3356 | 3.8308 | 1.01733 | Promoter | 15 | 18818977 | 18969550 | 150574 | 1 | ENSMUST00000176593.1 | 595 | Cdh10 | ENSMUSG00000022321.10 |
| chr15 | 18820089 | 18820531 | 3.36693 | 3.87796 | 1.01733 | Promoter | 15 | 18820329 | 19014236 | 193908 | 1 | ENSMUST00000040562.8 | 0 | Cdh10 | ENSMUSG00000022321.10 |
| chr15 | 34306868 | 34307023 | 3.61069 | 4.35982 | 1.38249 | Promoter | 15 | 34306681 | 34436242 | 129562 | 1 | ENSMUST00000022947.5 | 188 | Matn2 | ENSMUSG00000022324.9 |
| chr15 | 35292677 | 35295202 | 4.06955 | 5.29622 | 2.01486 | Promoter | 15 | 35296098 | 35303304 | 7207 | 1 | ENSMUST00000022952.4 | -896 | Osr2 | ENSMUSG00000022330.4 |
| chr15 | 35296043 | 35297328 | 3.88183 | 5.05518 | 1.80319 | Promoter | 15 | 35296098 | 35303304 | 7207 | 1 | ENSMUST00000022952.4 | 0 | Osr2 | ENSMUSG00000022330.4 |
| chr15 | 35297705 | 35299118 | 3.62379 | 4.36455 | 1.33852 | Intron (ENSMUST00000022952.4/ENSMUSG00000022330.4, intron 1 of 3) | 15 | 35296098 | 35303304 | 7207 | 1 | ENSMUST00000022952.4 | 1608 | Osr2 | ENSMUSG00000022330.4 |
| chr15 | 35299702 | 35300027 | 4.48208 | 6.07403 | 2.55853 | Intron (ENSMUST00000022952.4/ENSMUSG00000022330.4, intron 1 of 3) | 15 | 35296098 | 35303304 | 7207 | 1 | ENSMUST00000022952.4 | 3605 | Osr2 | ENSMUSG00000022330.4 |
| chr15 | 35300406 | 35300945 | 3.83075 | 4.91631 | 1.70796 | Exon (ENSMUST00000022952.4/ENSMUSG00000022330.4, exon 2 of 4) | 15 | 35296098 | 35303304 | 7207 | 1 | ENSMUST00000022952.4 | 4309 | Osr2 | ENSMUSG00000022330.4 |
| chr15 | 35301650 | 35301854 | 3.41239 | 4.0283 | 1.19024 | Intron (ENSMUST00000022952.4/ENSMUSG00000022330.4, intron 2 of 3) | 15 | 35296098 | 35303304 | 7207 | 1 | ENSMUST00000022952.4 | 5553 | Osr2 | ENSMUSG00000022330.4 |
| chr15 | 35304168 | 35304288 | 3.47995 | 4.09584 | 1.18452 | Distal Intergenic | 15 | 35296098 | 35303304 | 7207 | 1 | ENSMUST00000022952.4 | 8071 | Osr2 | ENSMUSG00000022330.4 |
| chr15 | 35305377 | 35305492 | 3.44832 | 4.03092 | 1.12412 | Distal Intergenic | 15 | 35296098 | 35303304 | 7207 | 1 | ENSMUST00000022952.4 | 9280 | Osr2 | ENSMUSG00000022330.4 |
| chr15 | 45114334 | 45114807 | 3.80166 | 4.76986 | 1.65953 | Promoter | 15 | 45106284 | 45114934 | 8651 | 2 | ENSMUST00000022967.5 | 127 | Kcnv1 | ENSMUSG00000022342.5 |
| chr15 | 61984086 | 61984299 | 3.86786 | 4.80421 | 1.59479 | Distal Intergenic | 15 | 61985341 | 61990352 | 5012 | 1 | ENSMUST00000188482.1 | -1042 | Myc | ENSMUSG00000022346.9 |
| chr15 | 61985077 | 61985446 | 3.44514 | 4.13088 | 1.21699 | Promoter | 15 | 61985341 | 61990352 | 5012 | 1 | ENSMUST00000188482.1 | 0 | Myc | ENSMUSG00000022346.9 |
| chr15 | 61986524 | 61986887 | 3.40421 | 4.07517 | 1.16247 | Promoter | 15 | 61987034 | 61987617 | 584 | 1 | ENSMUST00000159338.1 | -147 | Myc | ENSMUSG00000022346.9 |
| chr15 | 61987972 | 61988225 | 3.57003 | 4.35404 | 1.38288 | Promoter | 15 | 61987422 | 61990253 | 2832 | 1 | ENSMUST00000160009.1 | 551 | Myc | ENSMUSG00000022346.9 |
| chr15 | 61988887 | 61989439 | 3.85017 | 4.9861 | 1.79003 | Exon (ENSMUST00000188482.1/ENSMUSG00000022346.9, exon 3 of 3) | 15 | 61987422 | 61990253 | 2832 | 1 | ENSMUST00000160009.1 | 1466 | Myc | ENSMUSG00000022346.9 |
| chr15 | 61990573 | 61990924 | 3.70923 | 4.55506 | 1.53389 | Distal Intergenic | 15 | 61987422 | 61990253 | 2832 | 1 | ENSMUST00000160009.1 | 3152 | Myc | ENSMUSG00000022346.9 |
| chr15 | 56694363 | 56694518 | 3.69679 | 4.52458 | 1.50585 | Promoter | 15 | 56665627 | 56694539 | 28913 | 2 | ENSMUST00000050544.7 | 21 | Has2 | ENSMUSG00000022367.7 |
| chr15 | 55306581 | 55307738 | 3.68631 | 4.60154 | 1.52704 | Promoter | 15 | 55307750 | 55520802 | 213053 | 1 | ENSMUST00000110221.3 | -12 | Col14a1 | ENSMUSG00000022371.10 |
| chr15 | 64919045 | 64919122 | 4.16898 | 5.63917 | 2.20171 | Intron (ENSMUST00000180105.1/ENSMUSG00000094296.1, intron 2 of 2) | 15 | 64699042 | 64922296 | 223255 | 2 | ENSMUST00000023007.5 | 3174 | Adcy8 | ENSMUSG00000022376.7 |
| chr15 | 64919731 | 64919856 | 3.88257 | 4.87889 | 1.71715 | Intron (ENSMUST00000180105.1/ENSMUSG00000094296.1, intron 2 of 2) | 15 | 64699042 | 64922296 | 223255 | 2 | ENSMUST00000023007.5 | 2440 | Adcy8 | ENSMUSG00000022376.7 |
| chr15 | 64920177 | 64920387 | 3.38159 | 3.92972 | 1.12237 | Exon (ENSMUST00000023007.5/ENSMUSG00000022376.7, exon 1 of 18) | 15 | 64699042 | 64922296 | 223255 | 2 | ENSMUST00000023007.5 | 1909 | Adcy8 | ENSMUSG00000022376.7 |
| chr15 | 64920908 | 64922559 | 3.75934 | 4.66697 | 1.58745 | Promoter | 15 | 64699042 | 64922296 | 223255 | 2 | ENSMUST00000023007.5 | 0 | Adcy8 | ENSMUSG00000022376.7 |
| chr15 | 85575473 | 85575700 | 3.37249 | 3.88635 | 1.01733 | Intron (ENSMUST00000109424.2/ENSMUSG00000022382.9, intron 1 of 3) | 15 | 85535439 | 85578070 | 42632 | 2 | ENSMUST00000109424.2 | 2370 | Wnt7b | ENSMUSG00000022382.9 |
| chr15 | 85576162 | 85576607 | 3.90551 | 5.16417 | 1.89471 | Intron (ENSMUST00000109424.2/ENSMUSG00000022382.9, intron 1 of 3) | 15 | 85535439 | 85578070 | 42632 | 2 | ENSMUST00000109424.2 | 1463 | Wnt7b | ENSMUSG00000022382.9 |
| chr15 | 85577095 | 85578154 | 3.86385 | 4.84872 | 1.67924 | Promoter | 15 | 85535439 | 85578070 | 42632 | 2 | ENSMUST00000109424.2 | 0 | Wnt7b | ENSMUSG00000022382.9 |
| chr15 | 85579667 | 85583032 | 3.68171 | 4.52088 | 1.42467 | Promoter | 15 | 85535439 | 85580729 | 45291 | 2 | ENSMUST00000167968.2 | 0 | Wnt7b | ENSMUSG00000022382.9 |
| chr15 | 88663223 | 88663319 | 3.12291 | 4.08855 | 1.18752 | Distal Intergenic | 15 | 88696046 | 88700864 | 4819 | 2 | ENSMUST00000150004.1 | 37545 | Brd1 | ENSMUSG00000022387.11 |
| chr15 | 88954648 | 88955406 | 3.97513 | 5.09074 | 1.84153 | Promoter | 15 | 88890633 | 88954418 | 63786 | 2 | ENSMUST00000177180.2 | -231 | Ttll8 | ENSMUSG00000022388.9 |
| chr15 | 80669737 | 80669919 | 3.54249 | 4.30813 | 1.28565 | Distal Intergenic | 15 | 80671847 | 80700420 | 28574 | 1 | ENSMUST00000023044.5 | -1928 | Fam83f | ENSMUSG00000022408.6 |
| chr15 | 80671650 | 80672742 | 3.90055 | 4.94494 | 1.75358 | Promoter | 15 | 80671847 | 80700420 | 28574 | 1 | ENSMUST00000023044.5 | 0 | Fam83f | ENSMUSG00000022408.6 |
| chr15 | 80091299 | 80091681 | 3.5563 | 4.26791 | 1.34067 | Promoter | 15 | 80091334 | 80113441 | 22108 | 1 | ENSMUST00000009728.7 | 0 | Syngr1 | ENSMUSG00000022415.7 |
| chr15 | 80280954 | 80281139 | 3.93269 | 4.96854 | 1.77773 | Distal Intergenic | 15 | 80287238 | 80398279 | 111042 | 1 | ENSMUST00000160424.2 | -6099 | Cacna1i | ENSMUSG00000022416.10 |
| chr15 | 80281666 | 80281955 | 3.36477 | 3.8747 | 1.01733 | Distal Intergenic | 15 | 80287238 | 80398279 | 111042 | 1 | ENSMUST00000160424.2 | -5283 | Cacna1i | ENSMUSG00000022416.10 |
| chr15 | 80283802 | 80283970 | 3.355 | 3.85997 | 1.01733 | Distal Intergenic | 15 | 80287238 | 80398279 | 111042 | 1 | ENSMUST00000160424.2 | -3268 | Cacna1i | ENSMUSG00000022416.10 |
| chr15 | 80286790 | 80287359 | 3.81691 | 4.86276 | 1.67067 | Promoter | 15 | 80287238 | 80398279 | 111042 | 1 | ENSMUST00000160424.2 | 0 | Cacna1i | ENSMUSG00000022416.10 |
| chr15 | 80287977 | 80288425 | 4.06484 | 5.21059 | 1.89972 | Promoter | 15 | 80287534 | 80395747 | 108214 | 1 | ENSMUST00000162155.2 | 444 | Cacna1i | ENSMUSG00000022416.10 |
| chr15 | 80319242 | 80319354 | 3.54298 | 4.19421 | 1.19531 | Intron (ENSMUST00000160424.2/ENSMUSG00000022416.10, intron 2 of 36) | 15 | 80287534 | 80395747 | 108214 | 1 | ENSMUST00000162155.2 | 31709 | Cacna1i | ENSMUSG00000022416.10 |
| chr15 | 80320619 | 80320878 | 3.38193 | 3.9006 | 1.01733 | Intron (ENSMUST00000160424.2/ENSMUSG00000022416.10, intron 3 of 36) | 15 | 80287534 | 80395747 | 108214 | 1 | ENSMUST00000162155.2 | 33086 | Cacna1i | ENSMUSG00000022416.10 |
| chr15 | 80395207 | 80395987 | 3.90412 | 5.02976 | 1.81747 | 3' UTR | 15 | 80389328 | 80393820 | 4493 | 1 | ENSMUST00000160175.1 | 5880 | Cacna1i | ENSMUSG00000022416.10 |
| chr15 | 79801845 | 79802808 | 4.07307 | 5.62466 | 2.26254 | Intron (ENSMUST00000175858.3/ENSMUSG00000022421.13, intron 1 of 4) | 15 | 79786352 | 79804709 | 18358 | 2 | ENSMUST00000175858.3 | 1901 | Nptxr | ENSMUSG00000022421.13 |
| chr15 | 79803157 | 79803961 | 3.77997 | 4.81102 | 1.66245 | Promoter | 15 | 79786352 | 79804709 | 18358 | 2 | ENSMUST00000175858.3 | 748 | Nptxr | ENSMUSG00000022421.13 |
| chr15 | 79804285 | 79804649 | 3.53753 | 4.25599 | 1.31566 | Promoter | 15 | 79786352 | 79804709 | 18358 | 2 | ENSMUST00000175858.3 | 60 | Nptxr | ENSMUSG00000022421.13 |
| chr15 | 79805979 | 79806054 | 3.45903 | 4.17821 | 1.28947 | Intron (ENSMUST00000023060.7/ENSMUSG00000089837.4, intron 4 of 7) | 15 | 79786353 | 79804766 | 18414 | 2 | ENSMUST00000023057.8 | -1214 | Nptxr | ENSMUSG00000022421.13 |
| chr15 | 54919063 | 54919226 | 3.24381 | 3.83696 | 1.02404 | Promoter | 15 | 54839366 | 54919961 | 80596 | 2 | ENSMUST00000167541.2 | 735 | Enpp2 | ENSMUSG00000022425.10 |
| chr15 | 54919857 | 54919996 | 3.5887 | 4.83215 | 1.64768 | Promoter | 15 | 54839366 | 54919961 | 80596 | 2 | ENSMUST00000167541.2 | 0 | Enpp2 | ENSMUSG00000022425.10 |
| chr15 | 54920615 | 54921485 | 3.57194 | 4.26203 | 1.27585 | Promoter | 15 | 54838898 | 54920146 | 81249 | 2 | ENSMUST00000171545.2 | -470 | Enpp2 | ENSMUSG00000022425.10 |
| chr15 | 79604433 | 79605275 | 3.87083 | 4.95483 | 1.80217 | Promoter | 15 | 79561500 | 79605084 | 43585 | 2 | ENSMUST00000023065.6 | 0 | Dmc1 | ENSMUSG00000022429.10 |
| chr15 | 85163139 | 85163643 | 3.69205 | 4.53913 | 1.47322 | Distal Intergenic | 15 | 85132099 | 85144569 | 12471 | 1 | ENSMUST00000023067.2 | 31041 | Ribc2 | ENSMUSG00000022431.2 |
| chr15 | 79469534 | 79469717 | 4.28493 | 5.6353 | 2.18098 | Distal Intergenic | 15 | 79430061 | 79443919 | 13859 | 2 | ENSMUST00000156043.1 | -25616 | Csnk1e | ENSMUSG00000022433.13 |
| chr15 | 85016993 | 85017312 | 3.89632 | 5.21242 | 1.92206 | Promoter | 15 | 85017141 | 85022560 | 5420 | 1 | ENSMUST00000023070.5 | 0 | Upk3a | ENSMUSG00000022435.5 |
| chr15 | 84221223 | 84221316 | 3.61809 | 4.53719 | 1.4451 | Distal Intergenic | 15 | 84232043 | 84315688 | 83646 | 1 | ENSMUST00000023072.6 | -10727 | Parvb | ENSMUSG00000022438.6 |
| chr15 | 84224691 | 84225035 | 3.55276 | 4.28278 | 1.361 | Distal Intergenic | 15 | 84232043 | 84315688 | 83646 | 1 | ENSMUST00000023072.6 | -7008 | Parvb | ENSMUSG00000022438.6 |
| chr15 | 84230856 | 84231030 | 3.69148 | 4.48671 | 1.41145 | Distal Intergenic | 15 | 84232043 | 84315688 | 83646 | 1 | ENSMUST00000023072.6 | -1013 | Parvb | ENSMUSG00000022438.6 |
| chr15 | 84231619 | 84232504 | 3.89493 | 4.8649 | 1.65025 | Promoter | 15 | 84232043 | 84315688 | 83646 | 1 | ENSMUST00000023072.6 | 0 | Parvb | ENSMUSG00000022438.6 |
| chr15 | 84237922 | 84238115 | 3.99776 | 5.05762 | 1.76454 | Intron (ENSMUST00000023072.6/ENSMUSG00000022438.6, intron 1 of 12) | 15 | 84232066 | 84271318 | 39253 | 1 | ENSMUST00000122818.1 | 5857 | Parvb | ENSMUSG00000022438.6 |
| chr15 | 84238624 | 84239910 | 3.96539 | 5.01403 | 1.76493 | Intron (ENSMUST00000023072.6/ENSMUSG00000022438.6, intron 1 of 12) | 15 | 84232066 | 84271318 | 39253 | 1 | ENSMUST00000122818.1 | 6559 | Parvb | ENSMUSG00000022438.6 |
| chr15 | 83871149 | 83871614 | 3.45919 | 4.04961 | 1.13131 | Exon (ENSMUST00000156187.2/ENSMUSG00000022441.12, exon 29 of 32) | 15 | 83872372 | 83883641 | 11270 | 2 | ENSMUST00000130551.1 | 12027 | Efcab6 | ENSMUSG00000022441.12 |
| chr15 | 83966827 | 83966935 | 3.97919 | 5.03268 | 1.77811 | Promoter | 15 | 83965198 | 83967842 | 2645 | 2 | ENSMUST00000149563.1 | 907 | Efcab6 | ENSMUSG00000022441.12 |
| chr15 | 84054711 | 84054865 | 3.38193 | 3.9006 | 1.01733 | Intron (ENSMUST00000156187.2/ENSMUSG00000022441.12, intron 1 of 31) | 15 | 83866712 | 84065349 | 198638 | 2 | ENSMUST00000156187.2 | 10484 | Efcab6 | ENSMUSG00000022441.12 |
| chr15 | 84055181 | 84055451 | 3.56923 | 4.24438 | 1.23238 | Intron (ENSMUST00000156187.2/ENSMUSG00000022441.12, intron 1 of 31) | 15 | 83866712 | 84065349 | 198638 | 2 | ENSMUST00000156187.2 | 9898 | Efcab6 | ENSMUSG00000022441.12 |
| chr15 | 84065205 | 84065971 | 3.84647 | 4.89051 | 1.72796 | Promoter | 15 | 83866712 | 84065349 | 198638 | 2 | ENSMUST00000156187.2 | 0 | Efcab6 | ENSMUSG00000022441.12 |
| chr15 | 84068391 | 84069020 | 3.64688 | 4.50466 | 1.46631 | Distal Intergenic | 15 | 84018668 | 84065379 | 46712 | 2 | ENSMUST00000122836.1 | -3013 | Efcab6 | ENSMUSG00000022441.12 |
| chr15 | 84069381 | 84069846 | 3.67368 | 4.56127 | 1.48955 | Distal Intergenic | 15 | 84018668 | 84065379 | 46712 | 2 | ENSMUST00000122836.1 | -4003 | Efcab6 | ENSMUSG00000022441.12 |
| chr15 | 84070175 | 84070282 | 3.49337 | 4.11432 | 1.17875 | Distal Intergenic | 15 | 84018668 | 84065379 | 46712 | 2 | ENSMUST00000122836.1 | -4797 | Efcab6 | ENSMUSG00000022441.12 |
| chr15 | 84070599 | 84070964 | 3.58371 | 4.29434 | 1.31805 | Distal Intergenic | 15 | 84018668 | 84065379 | 46712 | 2 | ENSMUST00000122836.1 | -5221 | Efcab6 | ENSMUSG00000022441.12 |
| chr15 | 84071408 | 84071684 | 3.4416 | 4.02846 | 1.15081 | Distal Intergenic | 15 | 84018668 | 84065379 | 46712 | 2 | ENSMUST00000122836.1 | -6030 | Efcab6 | ENSMUSG00000022441.12 |
| chr15 | 84072685 | 84073014 | 3.39338 | 3.96981 | 1.17265 | Distal Intergenic | 15 | 84018668 | 84065379 | 46712 | 2 | ENSMUST00000122836.1 | -7307 | Efcab6 | ENSMUSG00000022441.12 |
| chr15 | 84074798 | 84074954 | 3.31081 | 3.80893 | 1.04619 | Downstream (1-2kb) | 15 | 84018668 | 84065379 | 46712 | 2 | ENSMUST00000122836.1 | -9420 | Efcab6 | ENSMUSG00000022441.12 |
| chr15 | 94401090 | 94402078 | 3.62717 | 4.38598 | 1.36947 | Intron (ENSMUST00000035342.5/ENSMUSG00000022449.9, intron 2 of 38) | 15 | 94270163 | 94404258 | 134096 | 2 | ENSMUST00000035342.5 | 2180 | Adamts20 | ENSMUSG00000022449.9 |
| chr15 | 94403287 | 94404355 | 3.79243 | 4.69473 | 1.56732 | Promoter | 15 | 94270163 | 94404258 | 134096 | 2 | ENSMUST00000035342.5 | 0 | Adamts20 | ENSMUSG00000022449.9 |
| chr15 | 94405124 | 94405317 | 3.68692 | 4.52203 | 1.43036 | Promoter | 15 | 94351432 | 94404350 | 52919 | 2 | ENSMUST00000129195.2 | -775 | Adamts20 | ENSMUSG00000022449.9 |
| chr15 | 95528269 | 95528514 | 3.28786 | 3.77639 | 1.06256 | Promoter | 15 | 95219933 | 95528252 | 308320 | 2 | ENSMUST00000075275.2 | -18 | Nell2 | ENSMUSG00000022454.11 |
| chr15 | 82272777 | 82272925 | 3.38193 | 3.9006 | 1.01733 | Distal Intergenic | 15 | 82274935 | 82294442 | 19508 | 1 | ENSMUST00000023095.7 | -2010 | Sep-03 | ENSMUSG00000022456.10 |
| chr15 | 82274602 | 82275715 | 3.91499 | 5.35553 | 2.03973 | Promoter | 15 | 82274935 | 82294442 | 19508 | 1 | ENSMUST00000023095.7 | 0 | Sep-03 | ENSMUSG00000022456.10 |
| chr15 | 97998264 | 97998439 | 3.84383 | 4.75719 | 1.55604 | Exon (ENSMUST00000088355.6/ENSMUSG00000022483.11, exon 6 of 53) | 15 | 97994868 | 97999465 | 4598 | 2 | ENSMUST00000139246.2 | 1026 | Col2a1 | ENSMUSG00000022483.11 |
| chr15 | 97998769 | 97999109 | 3.64457 | 4.38607 | 1.33009 | Promoter | 15 | 97994868 | 97999465 | 4598 | 2 | ENSMUST00000139246.2 | 356 | Col2a1 | ENSMUSG00000022483.11 |
| chr15 | 97999447 | 98000654 | 3.82936 | 4.80781 | 1.63671 | Promoter | 15 | 97994868 | 97999465 | 4598 | 2 | ENSMUST00000139246.2 | 0 | Col2a1 | ENSMUSG00000022483.11 |
| chr15 | 98001276 | 98004094 | 4.05779 | 5.30046 | 1.99024 | Promoter | 15 | 97998371 | 98001887 | 3517 | 2 | ENSMUST00000127879.1 | 0 | Col2a1 | ENSMUSG00000022483.11 |
| chr15 | 98004405 | 98005638 | 4.46969 | 6.11484 | 2.5983 | Promoter | 15 | 97975602 | 98004657 | 29056 | 2 | ENSMUST00000088355.6 | 0 | Col2a1 | ENSMUSG00000022483.11 |
| chr15 | 98006393 | 98006833 | 3.29937 | 3.77623 | 1.01733 | Distal Intergenic | 15 | 97975602 | 98004695 | 29094 | 2 | ENSMUST00000023123.9 | -1699 | Col2a1 | ENSMUSG00000022483.11 |
| chr15 | 103495346 | 103495680 | 3.48448 | 4.13458 | 1.27625 | Intron (ENSMUST00000047405.7/ENSMUSG00000022488.8, intron 30 of 30) | 15 | 103503298 | 103530052 | 26755 | 1 | ENSMUST00000023132.3 | -7618 | Pde1b | ENSMUSG00000022489.5 |
| chr15 | 103499604 | 103499837 | 3.34427 | 3.84379 | 1.01733 | Distal Intergenic | 15 | 103503298 | 103530052 | 26755 | 1 | ENSMUST00000023132.3 | -3461 | Pde1b | ENSMUSG00000022489.5 |
| chr15 | 103504021 | 103504706 | 3.34188 | 3.87177 | 1.10786 | Promoter | 15 | 103503298 | 103530052 | 26755 | 1 | ENSMUST00000023132.3 | 724 | Pde1b | ENSMUSG00000022489.5 |
| chr15 | 103505252 | 103505327 | 3.65134 | 4.81051 | 1.62031 | Intron (ENSMUST00000023132.3/ENSMUSG00000022489.5, intron 2 of 15) | 15 | 103503298 | 103530052 | 26755 | 1 | ENSMUST00000023132.3 | 1955 | Pde1b | ENSMUSG00000022489.5 |
| chr15 | 103536904 | 103537160 | 4.14612 | 5.64723 | 2.22711 | Promoter | 15 | 103530279 | 103537992 | 7714 | 2 | ENSMUST00000023133.6 | 832 | Ppp1r1a | ENSMUSG00000022490.6 |
| chr15 | 103537797 | 103538818 | 3.66526 | 4.4896 | 1.41256 | Promoter | 15 | 103530279 | 103537992 | 7714 | 2 | ENSMUST00000023133.6 | 0 | Ppp1r1a | ENSMUSG00000022490.6 |
| chr16 | 11983510 | 11983835 | 3.85801 | 4.89378 | 1.67816 | Promoter | 16 | 11984113 | 11995390 | 11278 | 1 | ENSMUST00000182889.1 | -278 | Shisa9 | ENSMUSG00000022494.8 |
| chr16 | 11984444 | 11985871 | 3.72007 | 4.58442 | 1.48659 | Promoter | 16 | 11984581 | 12269949 | 285369 | 1 | ENSMUST00000170672.3 | 0 | Shisa9 | ENSMUSG00000022494.8 |
| chr16 | 10490430 | 10490892 | 3.4356 | 4.02356 | 1.12292 | Intron (ENSMUST00000023147.6/ENSMUSG00000022504.9, intron 1 of 18) | 16 | 10488178 | 10527564 | 39387 | 1 | ENSMUST00000023147.6 | 2253 | Ciita | ENSMUSG00000022504.9 |
| chr16 | 28442760 | 28443285 | 3.28712 | 3.75784 | 1.01733 | Intron (ENSMUST00000100024.1/ENSMUSG00000022523.9, intron 1 of 4) | 16 | 28160120 | 28445227 | 285108 | 2 | ENSMUST00000100024.1 | 1942 | Fgf12 | ENSMUSG00000022523.9 |
| chr16 | 28445118 | 28445547 | 3.50418 | 4.18715 | 1.27034 | Promoter | 16 | 28160120 | 28445227 | 285108 | 2 | ENSMUST00000100024.1 | 0 | Fgf12 | ENSMUSG00000022523.9 |
| chr16 | 28446048 | 28447016 | 4.02308 | 5.21915 | 1.94403 | Promoter | 16 | 28160120 | 28445227 | 285108 | 2 | ENSMUST00000100024.1 | -822 | Fgf12 | ENSMUSG00000022523.9 |
| chr16 | 28447552 | 28448343 | 3.54954 | 4.23729 | 1.30262 | Distal Intergenic | 16 | 28160120 | 28445227 | 285108 | 2 | ENSMUST00000100024.1 | -2326 | Fgf12 | ENSMUSG00000022523.9 |
| chr16 | 28493391 | 28493550 | 3.35714 | 3.86319 | 1.01733 | Distal Intergenic | 16 | 28160120 | 28445227 | 285108 | 2 | ENSMUST00000100024.1 | -48165 | Fgf12 | ENSMUSG00000022523.9 |
| chr16 | 30037720 | 30038155 | 3.53019 | 4.2192 | 1.31788 | Distal Intergenic | 16 | 30064384 | 30065569 | 1186 | 1 | ENSMUST00000160592.1 | -26229 | Hes1 | ENSMUSG00000022528.7 |
| chr16 | 30051074 | 30051289 | 3.49709 | 4.36917 | 1.34341 | Distal Intergenic | 16 | 30064384 | 30065569 | 1186 | 1 | ENSMUST00000160592.1 | -13095 | Hes1 | ENSMUSG00000022528.7 |
| chr16 | 30051902 | 30052010 | 3.58308 | 4.30696 | 1.37289 | Distal Intergenic | 16 | 30064384 | 30065569 | 1186 | 1 | ENSMUST00000160592.1 | -12374 | Hes1 | ENSMUSG00000022528.7 |
| chr16 | 31299191 | 31299455 | 3.37734 | 3.95597 | 1.16232 | Intron (ENSMUST00000130560.2/ENSMUSG00000022548.9, intron 5 of 5) | 16 | 31297420 | 31303445 | 6026 | 2 | ENSMUST00000156456.2 | 3990 | Apod | ENSMUSG00000022548.9 |
| chr15 | 76515998 | 76516679 | 4.09436 | 5.4362 | 2.14693 | 3' UTR | 15 | 76502015 | 76511951 | 9937 | 2 | ENSMUST00000023214.5 | -4048 | Dgat1 | ENSMUSG00000022555.6 |
| chr15 | 76548521 | 76548705 | 3.5501 | 4.22295 | 1.26051 | Distal Intergenic | 15 | 76538943 | 76542130 | 3188 | 1 | ENSMUST00000023220.8 | 9579 | Slc52a2 | ENSMUSG00000022560.8 |
| chr15 | 75564803 | 75565710 | 3.66845 | 4.5341 | 1.45661 | Promoter | 15 | 75565213 | 75566632 | 1420 | 2 | ENSMUST00000150125.1 | 922 | Ly6h | ENSMUSG00000022577.11 |
| chr15 | 75566523 | 75567569 | 4.22758 | 5.65515 | 2.28679 | Promoter | 15 | 75551268 | 75566648 | 15381 | 2 | ENSMUST00000163116.2 | 0 | Ly6h | ENSMUSG00000022577.11 |
| chr15 | 74955095 | 74955837 | 3.5759 | 4.29201 | 1.27406 | Promoter | 15 | 74955106 | 74958776 | 3671 | 1 | ENSMUST00000187606.1 | 0 | Ly6e | ENSMUSG00000022587.9 |
| chr15 | 74956600 | 74956740 | 3.57716 | 4.32738 | 1.35715 | Promoter | 15 | 74956555 | 74958826 | 2272 | 1 | ENSMUST00000188042.1 | 46 | Ly6e | ENSMUSG00000022587.9 |

| | | | | | | | | | | | | | | | |
|---|---|---|---|---|---|---|---|---|---|---|---|---|---|---|---|
| chr15 | 74752093 | 74752418 | 3.62459 | 4.84245 | 1.66184 | Promoter | 15 | 74751595 | 74753014 | 1420 | 2 | ENSMUST00000189128.1 | 596 | Lynx1 | ENSMUSG00000022594.9 |
| chr15 | 74597318 | 74597462 | 3.55267 | 4.22494 | 1.25366 | Distal Intergenic | 15 | 74606415 | 74608228 | 1814 | 2 | ENSMUST00000177179.1 | 10766 | Mroh4 | ENSMUSG00000022603.4 |
| chr15 | 74652151 | 74652476 | 3.34054 | 3.87683 | 1.13129 | Distal Intergenic | 15 | 74606029 | 74636353 | 30325 | 2 | ENSMUST00000023271.2 | -15799 | Mroh4 | ENSMUSG00000022603.4 |
| chr15 | 89376173 | 89376838 | 3.256 | 3.71606 | 1.03272 | Promoter | 15 | 89371931 | 89377039 | 5109 | 2 | ENSMUST00000023285.4 | 201 | Tymp | ENSMUSG00000022615.7 |
| chr15 | 89452961 | 89453095 | 3.57519 | 4.2545 | 1.23407 | Promoter | 15 | 89453913 | 89464468 | 10556 | 1 | ENSMUST00000023291.5 | -818 | Mapk8ip2 | ENSMUSG00000022619.5 |
| chr15 | 89453688 | 89454801 | 3.91649 | 4.95765 | 1.73446 | Promoter | 15 | 89453913 | 89464468 | 10556 | 1 | ENSMUST00000023291.5 | 0 | Mapk8ip2 | ENSMUSG00000022619.5 |
| chr15 | 89557518 | 89558066 | 3.75315 | 4.63275 | 1.50969 | 3' UTR | 15 | 89568326 | 89574585 | 6260 | 1 | ENSMUST00000023295.2 | -10260 | Acr | ENSMUSG00000022622.4 |
| chr15 | 89558436 | 89559118 | 3.50912 | 4.14479 | 1.1944 | 3' UTR | 15 | 89568326 | 89574585 | 6260 | 1 | ENSMUST00000023295.2 | -9208 | Acr | ENSMUSG00000022622.4 |
| chr15 | 89572998 | 89573543 | 3.46924 | 4.08505 | 1.20126 | Exon (ENSMUST00000023295.2/ENSMUSG00000022622.4, exon 4 of 5) | 15 | 89568326 | 89574585 | 6260 | 1 | ENSMUST00000023295.2 | 4673 | Acr | ENSMUSG00000022622.4 |
| chr15 | 89497907 | 89498011 | 3.89045 | 5.01498 | 1.804 | Distal Intergenic | 15 | 89499623 | 89560261 | 60639 | 1 | ENSMUST00000109309.2 | -1612 | Shank3 | ENSMUSG00000022623.10 |
| chr15 | 89498519 | 89498938 | 3.32391 | 3.97438 | 1.12714 | Promoter | 15 | 89499623 | 89560261 | 60639 | 1 | ENSMUST00000109309.2 | -685 | Shank3 | ENSMUSG00000022623.10 |
| chr15 | 89499736 | 89500238 | 3.77459 | 4.89763 | 1.73394 | Promoter | 15 | 89499857 | 89560261 | 60405 | 1 | ENSMUST00000039074.7 | 0 | Shank3 | ENSMUSG00000022623.10 |
| chr15 | 89500816 | 89501052 | 3.68137 | 4.516 | 1.48774 | Promoter | 15 | 89501283 | 89560259 | 58977 | 1 | ENSMUST00000066545.8 | -231 | Shank3 | ENSMUSG00000022623.10 |
| chr15 | 89502674 | 89502934 | 3.36598 | 3.87654 | 1.01733 | Intron (ENSMUST00000109309.2/ENSMUSG00000022623.10, intron 3 of 21) | 15 | 89501283 | 89560259 | 58977 | 1 | ENSMUST00000066545.8 | 1392 | Shank3 | ENSMUSG00000022623.10 |
| chr15 | 89524018 | 89524235 | 4.14076 | 5.36077 | 2.02191 | Promoter | 15 | 89524322 | 89534394 | 10073 | 1 | ENSMUST00000154240.2 | -87 | Shank3 | ENSMUSG00000022623.10 |
| chr15 | 89525458 | 89525622 | 3.44231 | 4.01828 | 1.11077 | Intron (ENSMUST00000109309.2/ENSMUSG00000022623.10, intron 12 of 21) | 15 | 89524322 | 89534394 | 10073 | 1 | ENSMUST00000154240.2 | 1137 | Shank3 | ENSMUSG00000022623.10 |
| chr15 | 89548015 | 89548351 | 3.58538 | 4.30776 | 1.35387 | Exon (ENSMUST00000109309.2/ENSMUSG00000022623.10, exon 21 of 22) | 15 | 89532890 | 89543816 | 10927 | 1 | ENSMUST00000167173.1 | 15126 | Shank3 | ENSMUSG00000022623.10 |
| chr16 | 45406760 | 45407313 | 3.36818 | 3.93057 | 1.16888 | Exon (ENSMUST00000172091.1/ENSMUSG00000022661.9, exon 2 of 2) | 16 | 45399571 | 45408864 | 9294 | 2 | ENSMUST00000167552.1 | 1551 | Cd200 | ENSMUSG00000022661.9 |
| chr16 | 19761114 | 19761371 | 3.95973 | 5.37157 | 2.02374 | Promoter | 16 | 19761054 | 19770624 | 9571 | 1 | ENSMUST00000119468.1 | 61 | B3gnt5 | ENSMUSG00000022686.9 |
| chr16 | 44332978 | 44333432 | 3.86831 | 4.94713 | 1.76539 | Promoter | 16 | 44240180 | 44332991 | 92812 | 2 | ENSMUST00000136381.2 | 0 | Sidt1 | ENSMUSG00000022696.12 |
| chr16 | 8423298 | 8423428 | 3.34572 | 3.87419 | 1.10676 | Intron (ENSMUST00000023400.2/ENSMUSG00000022715.2, intron 2 of 3) | 16 | 8409276 | 8425136 | 15861 | 2 | ENSMUST00000023400.2 | 1708 | Cldn26 | ENSMUSG00000022715.2 |
| chr16 | 8423839 | 8424001 | 3.60159 | 4.309 | 1.28852 | Intron (ENSMUST00000023400.2/ENSMUSG00000022715.2, intron 2 of 3) | 16 | 8409276 | 8425136 | 15861 | 2 | ENSMUST00000023400.2 | 1135 | Cldn26 | ENSMUSG00000022715.2 |
| chr16 | 8424650 | 8424836 | 3.93776 | 5.37075 | 2.07784 | Promoter | 16 | 8409276 | 8425136 | 15861 | 2 | ENSMUST00000023400.2 | 300 | Cldn26 | ENSMUSG00000022715.2 |
| chr16 | 8425163 | 8425375 | 3.42549 | 3.98531 | 1.08407 | Promoter | 16 | 8409276 | 8425136 | 15861 | 2 | ENSMUST00000023400.2 | -28 | Cldn26 | ENSMUSG00000022715.2 |
| chr16 | 8425801 | 8425936 | 3.67353 | 4.4379 | 1.35893 | Promoter | 16 | 8409276 | 8425136 | 15861 | 2 | ENSMUST00000023400.2 | -666 | Cldn26 | ENSMUSG00000022715.2 |
| chr16 | 8426353 | 8426586 | 3.4146 | 3.98016 | 1.12412 | Distal Intergenic | 16 | 8409276 | 8425136 | 15861 | 2 | ENSMUST00000023400.2 | -1218 | Cldn26 | ENSMUSG00000022715.2 |
| chr16 | 17913194 | 17913273 | 3.79774 | 4.69931 | 1.60189 | 3' UTR | 16 | 17913107 | 17915059 | 1953 | 2 | ENSMUST00000012279.4 | 1786 | Gsc2 | ENSMUSG00000022738.6 |
| chr16 | 17914437 | 17915217 | 3.91752 | 5.02906 | 1.87699 | Promoter | 16 | 17913107 | 17915059 | 1953 | 2 | ENSMUST00000012279.4 | 0 | Gsc2 | ENSMUSG00000022738.6 |
| chr16 | 17915836 | 17916483 | 3.43285 | 4.06161 | 1.18125 | Promoter | 16 | 17913107 | 17915059 | 1953 | 2 | ENSMUST00000012279.4 | -778 | Gsc2 | ENSMUSG00000022738.6 |
| chr16 | 57241921 | 57242020 | 13.56131 | 31.22698 | 25.67596 | Distal Intergenic | 16 | 57168866 | 57231466 | 62601 | 2 | ENSMUST00000023431.6 | -10456 | Tbc1d23 | ENSMUSG00000022749.6 |
| chr16 | 17558294 | 17558520 | 3.75764 | 4.64852 | 1.59413 | Exon (ENSMUST00000023442.7/ENSMUSG00000022759.8, exon 4 of 8) | 16 | 17548811 | 17561244 | 12434 | 2 | ENSMUST00000065125.5 | 2724 | 4930451C15Rik | ENSMUSG00000022759.8 |
| chr16 | 81198502 | 81198986 | 3.37658 | 3.92403 | 1.1238 | Distal Intergenic | 16 | 81200697 | 81597805 | 397109 | 1 | ENSMUST00000037785.8 | -1711 | Ncam2 | ENSMUSG00000022762.12 |
| chr16 | 31897697 | 31898061 | 3.74463 | 4.58495 | 1.48878 | 3' UTR | 16 | 31878810 | 31899020 | 20211 | 1 | ENSMUST00000023464.5 | 18888 | Mfi2 | ENSMUSG00000022780.5 |
| chr16 | 38897851 | 38898025 | 3.21055 | 3.64308 | 1.00828 | Distal Intergenic | 16 | 38901471 | 39025361 | 123891 | 1 | ENSMUST00000114706.2 | -3446 | Igsf11 | ENSMUSG00000022790.8 |
| chr16 | 38899252 | 38899405 | 3.24972 | 3.7138 | 1.04588 | Distal Intergenic | 16 | 38901471 | 39025361 | 123891 | 1 | ENSMUST00000114706.2 | -2066 | Igsf11 | ENSMUSG00000022790.8 |
| chr16 | 38900871 | 38901078 | 3.70175 | 4.5942 | 1.51719 | Promoter | 16 | 38901471 | 39025361 | 123891 | 1 | ENSMUST00000114706.2 | -393 | Igsf11 | ENSMUSG00000022790.8 |
| chr16 | 38904722 | 38904873 | 3.21456 | 3.64908 | 1.01288 | Intron (ENSMUST00000114706.2/ENSMUSG00000022790.8, intron 1 of 6) | 16 | 38902316 | 39027159 | 124844 | 1 | ENSMUST00000023478.7 | 2407 | Igsf11 | ENSMUSG00000022790.8 |
| chr16 | 38724280 | 38724617 | 3.88152 | 4.85995 | 1.68175 | Distal Intergenic | 16 | 38598340 | 38713274 | 114935 | 2 | ENSMUST00000023487.4 | -11007 | Arhgap31 | ENSMUSG00000022799.4 |
| chr16 | 37383683 | 37384104 | 3.58388 | 4.43926 | 1.40703 | Promoter | 16 | 37114942 | 37384923 | 269982 | 2 | ENSMUST00000114787.2 | 819 | Stxbp5l | ENSMUSG00000022829.9 |
| chr16 | 34657609 | 34657862 | 3.63313 | 4.40075 | 1.41701 | Intron (ENSMUST00000148562.2/ENSMUSG00000022832.5, intron 1 of 4) | 16 | 34651188 | 34678618 | 27431 | 1 | ENSMUST00000023530.4 | 6422 | Ropn1 | ENSMUSG00000022832.5 |
| chr16 | 34658462 | 34658852 | 3.88829 | 4.85585 | 1.63847 | Intron (ENSMUST00000148562.2/ENSMUSG00000022832.5, intron 1 of 4) | 16 | 34651188 | 34678618 | 27431 | 1 | ENSMUST00000023530.4 | 7275 | Ropn1 | ENSMUSG00000022832.5 |
| chr16 | 34659485 | 34659706 | 3.38193 | 3.9006 | 1.01733 | Intron (ENSMUST00000148562.2/ENSMUSG00000022832.5, intron 1 of 4) | 16 | 34651188 | 34678618 | 27431 | 1 | ENSMUST00000023530.4 | 8298 | Ropn1 | ENSMUSG00000022832.5 |
| chr16 | 34668652 | 34668733 | 4.11752 | 5.76231 | 2.29653 | Intron (ENSMUST00000148562.2/ENSMUSG00000022832.5, intron 2 of 4) | 16 | 34651188 | 34678618 | 27431 | 1 | ENSMUST00000023530.4 | 17465 | Ropn1 | ENSMUSG00000022832.5 |
| chr16 | 35140245 | 35140463 | 3.52219 | 4.17123 | 1.21765 | Distal Intergenic | 16 | 35154877 | 35305737 | 150861 | 1 | ENSMUST00000114913.1 | -14414 | Adcy5 | ENSMUSG00000022840.8 |
| chr16 | 20629672 | 20630043 | 3.71162 | 4.75459 | 1.61496 | Promoter | 16 | 20629828 | 20645915 | 16088 | 1 | ENSMUST00000003898.6 | 0 | Ece2 | ENSMUSG00000022842.11 |
| chr16 | 78931636 | 78931846 | 3.34814 | 3.89829 | 1.08114 | Promoter | 16 | 78930948 | 78951733 | 20786 | 1 | ENSMUST00000023568.8 | 689 | Chodl | ENSMUSG00000022860.9 |
| chr16 | 22656880 | 22657095 | 3.69482 | 4.54483 | 1.52146 | Promoter | 16 | 22595628 | 22657028 | 61401 | 2 | ENSMUST00000150553.1 | 0 | Dgkg | ENSMUSG00000022861.11 |
| chr16 | 22660150 | 22661012 | 3.43988 | 4.08806 | 1.20418 | Distal Intergenic | 16 | 22469222 | 22657221 | 188000 | 2 | ENSMUST00000023578.8 | -2930 | Dgkg | ENSMUSG00000022861.11 |
| chr16 | 22894154 | 22894554 | 3.3495 | 3.89423 | 1.14649 | Exon (ENSMUST00000023583.5/ENSMUSG00000022868.5, exon 2 of 7) | 16 | 22892042 | 22899438 | 7397 | 1 | ENSMUST00000023583.5 | 2113 | Ahsg | ENSMUSG00000022868.5 |
| chr16 | 72804359 | 72804455 | 9.60347 | 23.58152 | 18.26032 | Intron (ENSMUST00000023600.7/ENSMUSG00000022883.10, intron 2 of 28) | 16 | 72663149 | 73046095 | 382947 | 1 | ENSMUST00000023600.7 | 141211 | Robo1 | ENSMUSG00000022883.10 |
| chr16 | 85899348 | 85900009 | 3.95611 | 5.03237 | 1.83522 | Exon (ENSMUST00000023611.5/ENSMUSG00000022894.5, exon 1 of 8) | 16 | 85858170 | 85901125 | 42956 | 2 | ENSMUST00000023611.5 | 1116 | Adamts5 | ENSMUSG00000022894.5 |
| chr16 | 85900988 | 85901247 | 3.96581 | 5.05291 | 1.84093 | Promoter | 16 | 85858170 | 85901125 | 42956 | 2 | ENSMUST00000023611.5 | 0 | Adamts5 | ENSMUSG00000022894.5 |
| chr16 | 85901749 | 85901866 | 3.47072 | 4.21309 | 1.27783 | Promoter | 16 | 85858170 | 85901125 | 42956 | 2 | ENSMUST00000023611.5 | -625 | Adamts5 | ENSMUSG00000022894.5 |
| chr16 | 94714469 | 94714648 | 3.86992 | 5.19969 | 1.92779 | Distal Intergenic | 16 | 94668896 | 94679521 | 10626 | 1 | ENSMUST00000134383.1 | 45574 | Dyrk1a | ENSMUSG00000022897.9 |
| chr16 | 36692987 | 36693103 | 3.37831 | 3.90244 | 1.04316 | Promoter | 16 | 36693978 | 36726804 | 32827 | 1 | ENSMUST00000023617.7 | -875 | Ildr1 | ENSMUSG00000022900.9 |
| chr16 | 36693778 | 36694772 | 3.78673 | 4.73537 | 1.57777 | Promoter | 16 | 36693978 | 36726804 | 32827 | 1 | ENSMUST00000023617.7 | 0 | Ildr1 | ENSMUSG00000022900.9 |
| chr16 | 94328517 | 94328768 | 3.40241 | 4.03515 | 1.19591 | Promoter | 16 | 94328420 | 94336935 | 8516 | 1 | ENSMUST00000023660.8 | 98 | Ripply3 | ENSMUSG00000022941.8 |
| chr16 | 93681401 | 93681623 | 3.46927 | 4.28562 | 1.3052 | Distal Intergenic | 16 | 93683215 | 93690990 | 7776 | 1 | ENSMUST00000039620.6 | -1592 | Cbr3 | ENSMUSG00000022947.7 |
| chr16 | 92498319 | 92499630 | 4.02655 | 5.28501 | 2.03868 | Promoter | 16 | 92498134 | 92541243 | 43110 | 1 | ENSMUST00000023670.3 | 186 | Clic6 | ENSMUSG00000022949.4 |
| chr16 | 92675598 | 92675948 | 3.43885 | 4.29227 | 1.31111 | Intron (ENSMUST00000023673.8/ENSMUSG00000022952.11, intron 4 of 7) | 16 | 92686956 | 92695495 | 8540 | 2 | ENSMUST00000187242.1 | 19547 | Runx1 | ENSMUSG00000022952.11 |
| chr16 | 92683901 | 92684270 | 3.42677 | 4.02075 | 1.14444 | Intron (ENSMUST00000023673.8/ENSMUSG00000022952.11, intron 4 of 7) | 16 | 92686956 | 92695495 | 8540 | 2 | ENSMUST00000187242.1 | 11225 | Runx1 | ENSMUSG00000022952.11 |
| chr16 | 92687762 | 92687994 | 3.62989 | 4.40704 | 1.4187 | Exon (ENSMUST00000189126.1/ENSMUSG00000022952.11, exon 3 of 3) | 16 | 92686956 | 92695495 | 8540 | 2 | ENSMUST00000187242.1 | 7501 | Runx1 | ENSMUSG00000022952.11 |
| chr16 | 92692658 | 92692768 | 3.54039 | 4.19138 | 1.19909 | Intron (ENSMUST00000023673.8/ENSMUSG00000022952.11, intron 3 of 7) | 16 | 92686956 | 92695495 | 8540 | 2 | ENSMUST00000187242.1 | 2727 | Runx1 | ENSMUSG00000022952.11 |
| chr16 | 92694308 | 92694848 | 4.4701 | 6.10856 | 2.64934 | Promoter | 16 | 92686956 | 92695495 | 8540 | 2 | ENSMUST00000187242.1 | 647 | Runx1 | ENSMUSG00000022952.11 |
| chr16 | 92695206 | 92696796 | 4.03846 | 5.3044 | 2.01134 | Promoter | 16 | 92605398 | 92696125 | 90728 | 2 | ENSMUST00000168195.2 | 0 | Runx1 | ENSMUSG00000022952.11 |
| chr16 | 92698984 | 92699422 | 3.81348 | 4.77853 | 1.66573 | Intron (ENSMUST00000023673.8/ENSMUSG00000022952.11, intron 2 of 7) | 16 | 92603656 | 92697328 | 93673 | 2 | ENSMUST00000113956.4 | -1657 | Runx1 | ENSMUSG00000022952.11 |
| chr16 | 92699884 | 92700325 | 3.76157 | 4.66626 | 1.57587 | Intron (ENSMUST00000023673.8/ENSMUSG00000022952.11, intron 2 of 7) | 16 | 92603656 | 92697328 | 93673 | 2 | ENSMUST00000113956.4 | -2557 | Runx1 | ENSMUSG00000022952.11 |
| chr16 | 92700678 | 92701739 | 3.69468 | 4.63529 | 1.5715 | Intron (ENSMUST00000023673.8/ENSMUSG00000022952.11, intron 2 of 7) | 16 | 92603656 | 92697328 | 93673 | 2 | ENSMUST00000113956.4 | -3351 | Runx1 | ENSMUSG00000022952.11 |
| chr16 | 92708493 | 92708582 | 3.77411 | 4.70592 | 1.61931 | Intron (ENSMUST00000023673.8/ENSMUSG00000022952.11, intron 2 of 7) | 16 | 92603656 | 92697328 | 93673 | 2 | ENSMUST00000113956.4 | -11166 | Runx1 | ENSMUSG00000022952.11 |
| chr16 | 92718666 | 92718984 | 3.21432 | 3.70266 | 1.01733 | Intron (ENSMUST00000023673.8/ENSMUSG00000022952.11, intron 2 of 7) | 16 | 92603656 | 92697328 | 93673 | 2 | ENSMUST00000113956.4 | -21339 | Runx1 | ENSMUSG00000022952.11 |
| chr16 | 92721225 | 92721663 | 3.51803 | 4.31237 | 1.35049 | Intron (ENSMUST00000023673.8/ENSMUSG00000022952.11, intron 2 of 7) | 16 | 92603656 | 92697328 | 93673 | 2 | ENSMUST00000113956.4 | -23898 | Runx1 | ENSMUSG00000022952.11 |
| chr16 | 92722012 | 92722219 | 3.39559 | 3.96654 | 1.1713 | Intron (ENSMUST00000023673.8/ENSMUSG00000022952.11, intron 2 of 7) | 16 | 92603656 | 92697328 | 93673 | 2 | ENSMUST00000113956.4 | -24685 | Runx1 | ENSMUSG00000022952.11 |
| chr16 | 92789011 | 92789349 | 3.55357 | 4.21948 | 1.22868 | Intron (ENSMUST00000023673.8/ENSMUSG00000022952.11, intron 2 of 7) | 16 | 92695592 | 92825778 | 130187 | 2 | ENSMUST00000186296.1 | 36429 | Runx1 | ENSMUSG00000022952.11 |
| chr16 | 90708875 | 90709189 | 3.50344 | 4.16579 | 1.28602 | Distal Intergenic | 16 | 90721447 | 90727139 | 5693 | 2 | ENSMUST00000147546.1 | 17950 | Mis18a | ENSMUSG00000022978.11 |
| chr16 | 90710659 | 90711453 | 3.89623 | 5.38845 | 2.05276 | Distal Intergenic | 16 | 90721447 | 90727139 | 5693 | 2 | ENSMUST00000147546.1 | 15686 | Mis18a | ENSMUSG00000022978.11 |
| chr16 | 90712684 | 90713072 | 3.38771 | 3.94881 | 1.10444 | Distal Intergenic | 16 | 90721447 | 90727139 | 5693 | 2 | ENSMUST00000147546.1 | 14067 | Mis18a | ENSMUSG00000022978.11 |
| chr16 | 90143109 | 90143237 | 4.28158 | 5.89002 | 2.38668 | Distal Intergenic | 16 | 90220742 | 90226322 | 5581 | 1 | ENSMUST00000023707.9 | -77505 | Sod1 | ENSMUSG00000022982.9 |
| chr16 | 90145195 | 90145325 | 3.37016 | 3.92519 | 1.15185 | Distal Intergenic | 16 | 90220742 | 90226322 | 5581 | 1 | ENSMUST00000023707.9 | -75417 | Sod1 | ENSMUSG00000022982.9 |
| chr16 | 90162240 | 90162545 | 3.53964 | 4.29187 | 1.33793 | Distal Intergenic | 16 | 90220742 | 90226322 | 5581 | 1 | ENSMUST00000023707.9 | -58197 | Sod1 | ENSMUSG00000022982.9 |
| chr16 | 90202444 | 90202597 | 3.27482 | 3.75939 | 1.07331 | Distal Intergenic | 16 | 90220742 | 90226322 | 5581 | 1 | ENSMUST00000023707.9 | -18145 | Sod1 | ENSMUSG00000022982.9 |
| chr16 | 90298076 | 90298179 | 3.69908 | 4.50918 | 1.44404 | Distal Intergenic | 16 | 90229156 | 90284425 | 55270 | 2 | ENSMUST00000039280.7 | -13652 | Scaf4 | ENSMUSG00000022983.10 |
| chr16 | 90331260 | 90331335 | 3.36386 | 3.87334 | 1.01733 | Distal Intergenic | 16 | 90229156 | 90284425 | 55270 | 2 | ENSMUST00000039280.7 | -46836 | Scaf4 | ENSMUSG00000022983.10 |
| chr15 | 101572779 | 101573175 | 3.51433 | 4.15292 | 1.19762 | Promoter | 15 | 101563345 | 101573904 | 10560 | 2 | ENSMUST00000042957.5 | 729 | Krt75 | ENSMUSG00000022986.5 |
| chr15 | 98776923 | 98777420 | 3.29402 | 3.81887 | 1.06628 | Promoter | 15 | 98772200 | 98777261 | 5062 | 2 | ENSMUST00000166022.1 | 0 | Wnt10b | ENSMUSG00000022996.9 |
| chr15 | 98778018 | 98778441 | 4.48438 | 6.19792 | 2.6259 | Promoter | 15 | 98771753 | 98778150 | 6398 | 2 | ENSMUST00000023732.6 | 0 | Wnt10b | ENSMUSG00000022996.9 |
| chr15 | 98780042 | 98780206 | 3.77991 | 4.84386 | 1.63677 | Distal Intergenic | 15 | 98771753 | 98778150 | 6398 | 2 | ENSMUST00000023732.6 | -1893 | Wnt10b | ENSMUSG00000022996.9 |
| chr15 | 98787870 | 98788125 | 3.69482 | 4.47458 | 1.37594 | Distal Intergenic | 15 | 98789857 | 98793837 | 3981 | 1 | ENSMUST00000023734.7 | -1732 | Wnt1 | ENSMUSG00000022997.9 |
| chr15 | 98788523 | 98788903 | 3.6614 | 4.47491 | 1.42362 | Promoter | 15 | 98789857 | 98793837 | 3981 | 1 | ENSMUST00000023734.7 | -954 | Wnt1 | ENSMUSG00000022997.9 |
| chr15 | 98789352 | 98789917 | 3.62689 | 4.37603 | 1.37503 | Promoter | 15 | 98789857 | 98793837 | 3981 | 1 | ENSMUST00000023734.7 | 0 | Wnt1 | ENSMUSG00000022997.9 |
| chr15 | 98790234 | 98790516 | 3.33349 | 3.83858 | 1.05616 | Promoter | 15 | 98789857 | 98793837 | 3981 | 1 | ENSMUST00000023734.7 | 378 | Wnt1 | ENSMUSG00000022997.9 |
| chr15 | 98790893 | 98791022 | 3.22391 | 4.09198 | 1.1942 | Exon (ENSMUST00000023734.7/ENSMUSG00000022997.9, exon 2 of 4) | 15 | 98789857 | 98793837 | 3981 | 1 | ENSMUST00000023734.7 | 1037 | Wnt1 | ENSMUSG00000022997.9 |
| chr15 | 98791520 | 98791931 | 4.05169 | 5.23847 | 1.93163 | Exon (ENSMUST00000023734.7/ENSMUSG00000022997.9, exon 3 of 4) | 15 | 98789857 | 98793837 | 3981 | 1 | ENSMUST00000023734.7 | 1664 | Wnt1 | ENSMUSG00000022997.9 |
| chr15 | 98792428 | 98792828 | 4.27478 | 5.6599 | 2.25145 | Exon (ENSMUST00000023734.7/ENSMUSG00000022997.9, exon 4 of 4) | 15 | 98789857 | 98793837 | 3981 | 1 | ENSMUST00000023734.7 | 2572 | Wnt1 | ENSMUSG00000022997.9 |
| chr15 | 98796653 | 98796950 | 3.44726 | 4.03164 | 1.13031 | Distal Intergenic | 15 | 98789857 | 98793837 | 3981 | 1 | ENSMUST00000023734.7 | 6797 | Wnt1 | ENSMUSG00000022997.9 |
| chr15 | 98893916 | 98894888 | 3.81809 | 4.84905 | 1.70493 | 3' UTR | 15 | 98893031 | 98898540 | 5510 | 2 | ENSMUST00000023737.4 | 3652 | Dhh | ENSMUSG00000023000.4 |
| chr15 | 98895899 | 98896078 | 3.69964 | 4.93439 | 1.73249 | Intron (ENSMUST00000023737.4/ENSMUSG00000023000.4, intron 1 of 2) | 15 | 98893031 | 98898540 | 5510 | 2 | ENSMUST00000023737.4 | 2462 | Dhh | ENSMUSG00000023000.4 |
| chr15 | 98898036 | 98898149 | 4.19249 | 5.63473 | 2.21616 | Promoter | 15 | 98893031 | 98898540 | 5510 | 2 | ENSMUST00000023737.4 | 391 | Dhh | ENSMUSG00000023000.4 |
| chr15 | 98898808 | 98899191 | 3.33425 | 4.03313 | 1.17334 | Promoter | 15 | 98893031 | 98898540 | 5510 | 2 | ENSMUST00000023737.4 | -269 | Dhh | ENSMUSG00000023000.4 |
| chr15 | 99515593 | 99515789 | 3.41014 | 3.95648 | 1.06417 | Intron (ENSMUST00000023750.7/ENSMUSG00000023011.7, intron 5 of 11) | 15 | 99497005 | 99528017 | 31013 | 2 | ENSMUST00000023750.7 | 12228 | Faim2 | ENSMUSG00000023011.7 |
| chr15 | 99516598 | 99517863 | 4.07597 | 5.25002 | 1.93985 | Intron (ENSMUST00000023750.7/ENSMUSG00000023011.7, intron 5 of 11) | 15 | 99497005 | 99528017 | 31013 | 2 | ENSMUST00000023750.7 | 10154 | Faim2 | ENSMUSG00000023011.7 |
| chr15 | 99526722 | 99527033 | 3.60066 | 4.31126 | 1.30292 | Promoter | 15 | 99497005 | 99528017 | 31013 | 2 | ENSMUST00000023750.7 | 984 | Faim2 | ENSMUSG00000023011.7 |
| chr15 | 99527352 | 99528500 | 3.80193 | 4.70631 | 1.57612 | Promoter | 15 | 99497005 | 99528017 | 31013 | 2 | ENSMUST00000023750.7 | 0 | Faim2 | ENSMUSG00000023011.7 |
| chr15 | 99529380 | 99529889 | 3.72544 | 4.55089 | 1.46812 | Distal Intergenic | 15 | 99497005 | 99528017 | 31013 | 2 | ENSMUST00000023750.7 | -1364 | Faim2 | ENSMUSG00000023011.7 |
| chr15 | 99576169 | 99576307 | 3.22092 | 3.80232 | 1.0327 | Distal Intergenic | 15 | 99579056 | 99584545 | 5490 | 1 | ENSMUST00000023752.4 | -2749 | Aqp2 | ENSMUSG00000023013.4 |
| chr15 | 99580290 | 99580589 | 3.38193 | 3.9006 | 1.01733 | Intron (ENSMUST00000023752.4/ENSMUSG00000023013.4, intron 1 of 3) | 15 | 99579056 | 99584545 | 5490 | 1 | ENSMUST00000023752.4 | 1235 | Aqp2 | ENSMUSG00000023013.4 |
| chr15 | 99581669 | 99581916 | 3.59154 | 4.34714 | 1.3071 | Intron (ENSMUST00000023752.4/ENSMUSG00000023013.4, intron 1 of 3) | 15 | 99579056 | 99584545 | 5490 | 1 | ENSMUST00000023752.4 | 2614 | Aqp2 | ENSMUSG00000023013.4 |
| chr15 | 99582785 | 99583154 | 3.64032 | 4.43953 | 1.41682 | Exon (ENSMUST00000023752.4/ENSMUSG00000023013.4, exon 3 of 4) | 15 | 99579056 | 99584545 | 5490 | 1 | ENSMUST00000023752.4 | 3730 | Aqp2 | ENSMUSG00000023013.4 |
| chr15 | 99583743 | 99585963 | 3.82219 | 4.78165 | 1.62207 | 3' UTR | 15 | 99579056 | 99584545 | 5490 | 1 | ENSMUST00000023752.4 | 4688 | Aqp2 | ENSMUSG00000023013.4 |
| chr15 | 99665007 | 99665188 | 4.13531 | 5.3257 | 1.95625 | Distal Intergenic | 15 | 99670718 | 99701127 | 30410 | 1 | ENSMUST00000023758.7 | -5530 | Asic1 | ENSMUSG00000023017.9 |
| chr15 | 99670433 | 99670891 | 3.55177 | 4.23345 | 1.27947 | Promoter | 15 | 99670718 | 99701127 | 30410 | 1 | ENSMUST00000023758.7 | 0 | Asic1 | ENSMUSG00000023017.9 |
| chr15 | 99673511 | 99674245 | 3.59139 | 4.31231 | 1.34053 | Intron (ENSMUST00000023758.7/ENSMUSG00000023017.9, intron 3 of 11) | 15 | 99670718 | 99701127 | 30410 | 1 | ENSMUST00000023758.7 | 2794 | Asic1 | ENSMUSG00000023017.9 |
| chr15 | 100679870 | 100680062 | 3.34441 | 3.84399 | 1.01733 | Intron (ENSMUST00000023775.7/ENSMUSG00000023031.7, intron 6 of 7) | 15 | 100674425 | 100687920 | 13496 | 2 | ENSMUST00000023775.7 | 7858 | Cela1 | ENSMUSG00000023031.7 |
| chr15 | 100685032 | 100685481 | 3.78458 | 4.66395 | 1.53508 | Exon (ENSMUST00000023775.7/ENSMUSG00000023031.7, exon 3 of 8) | 15 | 100674425 | 100687920 | 13496 | 2 | ENSMUST00000023775.7 | 2439 | Cela1 | ENSMUSG00000023031.7 |
| chr15 | 100753355 | 100753663 | 3.55393 | 4.22554 | 1.23464 | Distal Intergenic | 15 | 100761747 | 100823968 | 62222 | 1 | ENSMUST00000023776.7 | -8084 | Slc4a8 | ENSMUSG00000023032.7 |
| chr15 | 100760970 | 100761172 | 3.42682 | 3.99251 | 1.10053 | Promoter | 15 | 100761747 | 100823968 | 62222 | 1 | ENSMUST00000023776.7 | -575 | Slc4a8 | ENSMUSG00000023032.7 |
| chr15 | 100761564 | 100762410 | 3.69106 | 4.50135 | 1.44824 | Promoter | 15 | 100761747 | 100823968 | 62222 | 1 | ENSMUST00000023776.7 | 0 | Slc4a8 | ENSMUSG00000023032.7 |
| chr15 | 100824811 | 100826214 | 3.88639 | 4.90509 | 1.71343 | Distal Intergenic | 15 | 100814354 | 100815800 | 1447 | 1 | ENSMUST00000162744.1 | 10458 | Slc4a8 | ENSMUSG00000023032.7 |
| chr15 | 100870065 | 100870653 | 3.78999 | 4.76254 | 1.62126 | Promoter | 15 | 100870683 | 101045929 | 175247 | 1 | ENSMUST00000082209.7 | -30 | Scn8a | ENSMUSG00000023033.9 |
| chr15 | 100871193 | 100871400 | 4.5296 | 6.32415 | 2.74281 | Promoter | 15 | 100870683 | 101045929 | 175247 | 1 | ENSMUST00000082209.7 | 511 | Scn8a | ENSMUSG00000023033.9 |
| chr15 | 100871863 | 100873323 | 3.72086 | 4.60336 | 1.53743 | Intron (ENSMUST00000082209.7/ENSMUSG00000023033.9, intron 1 of 26) | 15 | 100870683 | 101045929 | 175247 | 1 | ENSMUST00000082209.7 | 1181 | Scn8a | ENSMUSG00000023033.9 |
| chr15 | 100914684 | 100914910 | 3.68112 | 4.62652 | 1.52655 | Intron (ENSMUST00000082209.7/ENSMUSG00000023033.9, intron 1 of 26) | 15 | 100933429 | 100940798 | 7370 | 1 | ENSMUST00000096198.4 | -18519 | Scn8a | ENSMUSG00000023033.9 |
| chr15 | 100921360 | 100921597 | 3.42551 | 4.01407 | 1.17285 | Intron (ENSMUST00000082209.7/ENSMUSG00000023033.9, intron 1 of 26) | 15 | 100933429 | 100940798 | 7370 | 1 | ENSMUST00000096198.4 | -11832 | Scn8a | ENSMUSG00000023033.9 |
| chr15 | 100923290 | 100923392 | 3.32884 | 4.01268 | 1.15822 | Intron (ENSMUST00000082209.7/ENSMUSG00000023033.9, intron 1 of 26) | 15 | 100933429 | 100940798 | 7370 | 1 | ENSMUST00000096198.4 | -10037 | Scn8a | ENSMUSG00000023033.9 |
| chr15 | 100969617 | 100970105 | 3.62543 | 4.35139 | 1.30216 | Intron (ENSMUST00000082209.7/ENSMUSG00000023033.9, intron 7 of 26) | 15 | 100955398 | 101045929 | 90532 | 1 | ENSMUST00000108909.3 | 14220 | Scn8a | ENSMUSG00000023033.9 |
| chr15 | 100982360 | 100982535 | 3.71238 | 4.63672 | 1.54552 | Intron (ENSMUST00000082209.7/ENSMUSG00000023033.9, intron 11 of 26) | 15 | 100955398 | 101045929 | 90532 | 1 | ENSMUST00000108909.3 | 26963 | Scn8a | ENSMUSG00000023033.9 |
| chr15 | 100983320 | 100983395 | 3.90573 | 5.00044 | 1.79516 | Intron (ENSMUST00000082209.7/ENSMUSG00000023033.9, intron 11 of 26) | 15 | 100955398 | 101045929 | 90532 | 1 | ENSMUST00000108909.3 | 27923 | Scn8a | ENSMUSG00000023033.9 |
| chr15 | 100983714 | 100984885 | 4.02435 | 5.35249 | 2.0813 | Exon (ENSMUST00000082209.7/ENSMUSG00000023033.9, exon 12 of 27) | 15 | 100955398 | 101045929 | 90532 | 1 | ENSMUST00000108909.3 | 28317 | Scn8a | ENSMUSG00000023033.9 |
| chr18 | 37743633 | 37744400 | 3.77127 | 4.75872 | 1.62589 | Exon (ENSMUST00000003599.7/ENSMUSG00000023036.9, exon 1 of 4) | 18 | 37742094 | 37841873 | 99780 | 1 | ENSMUST00000003599.7 | 1540 | Pcdhga12 | ENSMUSG00000023036.9 |
| chr15 | 102016342 | 102016556 | 3.33819 | 3.83463 | 1.01733 | Distal Intergenic | 15 | 102028216 | 102032026 | 3811 | 1 | ENSMUST00000023803.6 | -11660 | Krt18 | ENSMUSG00000023043.6 |
| chr15 | 102028078 | 102028447 | 3.65719 | 4.64294 | 1.5396 | Promoter | 15 | 102028216 | 102032026 | 3811 | 1 | ENSMUST00000023803.6 | 0 | Krt18 | ENSMUSG00000023043.6 |
| chr15 | 102691935 | 102692070 | 3.65818 | 4.42786 | 1.40842 | Distal Intergenic | 15 | 102706972 | 102709902 | 2931 | 2 | ENSMUST00000168112.2 | 17832 | Calcoco1 | ENSMUSG00000023055.4 |
| chr15 | 102692410 | 102693592 | 3.91014 | 4.98315 | 1.8026 | Distal Intergenic | 15 | 102706972 | 102709902 | 2931 | 2 | ENSMUST00000168112.2 | 16310 | Calcoco1 | ENSMUSG00000023055.4 |
| chr15 | 102693932 | 102694121 | 3.49162 | 4.13877 | 1.26425 | Distal Intergenic | 15 | 102706972 | 102709902 | 2931 | 2 | ENSMUST00000168112.2 | 15781 | Calcoco1 | ENSMUSG00000023055.4 |
| chr15 | 102695009 | 102695346 | 3.30852 | 3.79001 | 1.01733 | Distal Intergenic | 15 | 102706972 | 102709902 | 2931 | 2 | ENSMUST00000168112.2 | 14556 | Calcoco1 | ENSMUSG00000023055.4 |
| chr15 | 102699090 | 102699424 | 3.79131 | 4.76753 | 1.62999 | Distal Intergenic | 15 | 102706972 | 102709902 | 2931 | 2 | ENSMUST00000168112.2 | 10478 | Calcoco1 | ENSMUSG00000023055.4 |
| chr15 | 102733295 | 102733513 | 3.47942 | 4.24423 | 1.30129 | Distal Intergenic | 15 | 102706779 | 102722178 | 15400 | 2 | ENSMUST00000023818.4 | -11118 | Calcoco1 | ENSMUSG00000023055.4 |
| chr15 | 102736090 | 102736348 | 3.65068 | 4.45384 | 1.454 | Distal Intergenic | 15 | 102706779 | 102722178 | 15400 | 2 | ENSMUST00000023818.4 | -13913 | Calcoco1 | ENSMUSG00000023055.4 |
| chr15 | 102755644 | 102755893 | 3.75959 | 4.67954 | 1.59831 | Distal Intergenic | 15 | 102706779 | 102722178 | 15400 | 2 | ENSMUST00000023818.4 | -33467 | Calcoco1 | ENSMUSG00000023055.4 |
| chr15 | 102756817 | 102757052 | 3.38193 | 3.9006 | 1.01733 | Distal Intergenic | 15 | 102706779 | 102722178 | 15400 | 2 | ENSMUST00000023818.4 | -34640 | Calcoco1 | ENSMUSG00000023055.4 |
| chr15 | 102762577 | 102763531 | 3.83697 | 4.8613 | 1.65235 | Distal Intergenic | 15 | 102706779 | 102722178 | 15400 | 2 | ENSMUST00000023818.4 | -40400 | Calcoco1 | ENSMUSG00000023055.4 |
| chr15 | 102764533 | 102764693 | 3.34464 | 3.84435 | 1.01733 | Distal Intergenic | 15 | 102706779 | 102722178 | 15400 | 2 | ENSMUST00000023818.4 | -42356 | Calcoco1 | ENSMUSG00000023055.4 |
| chr15 | 102770721 | 102771950 | 3.69891 | 4.52085 | 1.45932 | Distal Intergenic | 15 | 102706779 | 102722178 | 15400 | 2 | ENSMUST00000023818.4 | -48544 | Calcoco1 | ENSMUSG00000023055.4 |
| chr15 | 102775356 | 102775578 | 3.30218 | 3.79265 | 1.06004 | Distal Intergenic | 15 | 102706779 | 102722178 | 15400 | 2 | ENSMUST00000023818.4 | -53179 | Calcoco1 | ENSMUSG00000023055.4 |
| chr15 | 102776506 | 102776707 | 3.34687 | 3.88272 | 1.12885 | Distal Intergenic | 15 | 102706779 | 102722178 | 15400 | 2 | ENSMUST00000023818.4 | -54329 | Calcoco1 | ENSMUSG00000023055.4 |
| chr15 | 102784771 | 102784846 | 3.44316 | 4.06447 | 1.24485 | Distal Intergenic | 15 | 102706779 | 102722178 | 15400 | 2 | ENSMUST00000023818.4 | -62594 | Calcoco1 | ENSMUSG00000023055.4 |
| chr15 | 102785408 | 102785642 | 3.97003 | 5.21827 | 2.00346 | Distal Intergenic | 15 | 102706779 | 102722178 | 15400 | 2 | ENSMUST00000023818.4 | -63231 | Calcoco1 | ENSMUSG00000023055.4 |
| chr15 | 102786904 | 102787644 | 3.52032 | 4.23928 | 1.30567 | Distal Intergenic | 15 | 102706779 | 102722178 | 15400 | 2 | ENSMUST00000023818.4 | -64727 | Calcoco1 | ENSMUSG00000023055.4 |
| chr15 | 102788814 | 102789206 | 3.42112 | 4.04299 | 1.19316 | Distal Intergenic | 15 | 102706779 | 102722178 | 15400 | 2 | ENSMUST00000023818.4 | -66637 | Calcoco1 | ENSMUSG00000023055.4 |
| chr15 | 102790052 | 102790364 | 3.53672 | 4.26436 | 1.32208 | Distal Intergenic | 15 | 102706779 | 102722178 | 15400 | 2 | ENSMUST00000023818.4 | -67875 | Calcoco1 | ENSMUSG00000023055.4 |
| chr15 | 102805499 | 102805629 | 3.84506 | 4.76192 | 1.57354 | Distal Intergenic | 15 | 102706779 | 102722178 | 15400 | 2 | ENSMUST00000023818.4 | -83322 | Calcoco1 | ENSMUSG00000023055.4 |
| chr15 | 102806191 | 102806653 | 3.34856 | 4.06408 | 1.20708 | Distal Intergenic | 15 | 102706779 | 102722178 | 15400 | 2 | ENSMUST00000023818.4 | -84014 | Calcoco1 | ENSMUSG00000023055.4 |
| chr15 | 102808381 | 102808679 | 3.40742 | 3.95368 | 1.069 | Distal Intergenic | 15 | 102706779 | 102722178 | 15400 | 2 | ENSMUST00000023818.4 | -86204 | Calcoco1 | ENSMUSG00000023055.4 |
| chr15 | 102814420 | 102814744 | 3.35654 | 3.88325 | 1.09149 | Distal Intergenic | 15 | 102706779 | 102722178 | 15400 | 2 | ENSMUST00000023818.4 | -92243 | Calcoco1 | ENSMUSG00000023055.4 |
| chr15 | 102815526 | 102815702 | 3.41373 | 4.12107 | 1.2041 | Distal Intergenic | 15 | 102706779 | 102722178 | 15400 | 2 | ENSMUST00000023818.4 | -93349 | Calcoco1 | ENSMUSG00000023055.4 |
| chr17 | 29094770 | 29094977 | 3.93533 | 4.99256 | 1.80905 | Promoter | 17 | 29093772 | 29100722 | 6951 | 1 | ENSMUST00000023829.6 | 999 | Cdkn1a | ENSMUSG00000023067.8 |
| chr5 | 95949820 | 95950225 | 3.42972 | 4.19548 | 1.23919 | Distal Intergenic | 5 | 95956928 | 95961067 | 4140 | 1 | ENSMUST00000023840.5 | -6703 | Cxcl13 | ENSMUSG00000023078.5 |
| chr3 | 87745891 | 87746824 | 3.67738 | 4.52955 | 1.44165 | Exon (ENSMUST00000023846.5/ENSMUSG00000023084.6, exon 2 of 15) | 3 | 87745252 | 87748594 | 3343 | 2 | ENSMUST00000174208.1 | 1770 | Lrrc71 | ENSMUSG00000023084.6 |
| chr4 | 130200653 | 130201025 | 3.51718 | 4.16797 | 1.21835 | Distal Intergenic | 4 | 130209109 | 130209856 | 748 | 2 | ENSMUST00000124528.1 | 8831 | Gm853 | ENSMUSG00000023120.9 |
| chr4 | 130240533 | 130240673 | 3.34888 | 3.85074 | 1.01733 | Distal Intergenic | 4 | 130209109 | 130222401 | 13293 | 2 | ENSMUST00000023884.6 | -18133 | Gm853 | ENSMUSG00000023120.9 |
| chr4 | 130245220 | 130245365 | 3.57901 | 4.32405 | 1.38459 | Distal Intergenic | 4 | 130209109 | 130222401 | 13293 | 2 | ENSMUST00000023884.6 | -22820 | Gm853 | ENSMUSG00000023120.9 |
| chr4 | 130245986 | 130246182 | 3.36751 | 3.87885 | 1.01733 | Distal Intergenic | 4 | 130209109 | 130222401 | 13293 | 2 | ENSMUST00000023884.6 | -23586 | Gm853 | ENSMUSG00000023120.9 |
| chr15 | 96641102 | 96642118 | 4.154 | 5.53202 | 2.17343 | Promoter | 15 | 96571418 | 96641962 | 70545 | 2 | ENSMUST00000100262.2 | 0 | Slc38a1 | ENSMUSG00000023169.9 |
| chr15 | 96642640 | 96643192 | 4.40354 | 6.03346 | 2.58824 | Promoter | 15 | 96571418 | 96642913 | 71496 | 2 | ENSMUST00000088452.5 | 0 | Slc38a1 | ENSMUSG00000023169.9 |

| | | | | | | | | | | | | | | | |
|---|---|---|---|---|---|---|---|---|---|---|---|---|---|---|---|
| chr16 | 30240130 | 30240230 | 3.37733 | 3.90498 | 1.05737 | Distal Intergenic | 16 | 30256378 | 30267532 | 11155 | 2 | ENSMUST00000064856.7 | 27302 | Cpn2 | ENSMUSG00000023176.8 |
| chr16 | 30244423 | 30245008 | 3.37035 | 3.8868 | 1.0302 | Distal Intergenic | 16 | 30256378 | 30267532 | 11155 | 2 | ENSMUST00000064856.7 | 22524 | Cpn2 | ENSMUSG00000023176.8 |
| chr16 | 30253290 | 30253551 | 3.44569 | 4.24985 | 1.27082 | Downstream (2-3kb) | 16 | 30256378 | 30267532 | 11155 | 2 | ENSMUST00000064856.7 | 13981 | Cpn2 | ENSMUSG00000023176.8 |
| chr16 | 30254842 | 30255505 | 3.6122 | 4.33091 | 1.31047 | Downstream (<1kb) | 16 | 30256378 | 30267532 | 11155 | 2 | ENSMUST00000064856.7 | 12027 | Cpn2 | ENSMUSG00000023176.8 |
| chr6 | 124855733 | 124855985 | 3.56423 | 4.25144 | 1.26831 | Promoter | 6 | 124854396 | 124855170 | 775 | 2 | ENSMUST00000135546.1 | -564 | Leprel2 | ENSMUSG00000023191.4 |
| chr6 | 124857220 | 124857665 | 3.83938 | 4.84588 | 1.67643 | Promoter | 6 | 124841206 | 124857729 | 16524 | 2 | ENSMUST00000135127.2 | 64 | Leprel2 | ENSMUSG00000023191.4 |
| chr6 | 124858910 | 124859136 | 3.45392 | 4.11199 | 1.23269 | Exon (ENSMUST00000046893.7/ENSMUSG00000038390.9, exon 5 of 5) | 6 | 124841089 | 124857752 | 16664 | 2 | ENSMUST00000023958.4 | -1159 | Leprel2 | ENSMUSG00000023191.4 |
| chr9 | 106653789 | 106654025 | 3.81447 | 4.77449 | 1.60447 | Exon (ENSMUST00000023959.8/ENSMUSG00000023192.8, exon 2 of 6) | 9 | 106643117 | 106656080 | 12964 | 2 | ENSMUST00000023959.8 | 2055 | Grm2 | ENSMUSG00000023192.8 |
| chr9 | 106654450 | 106654622 | 3.72451 | 4.58637 | 1.52812 | Intron (ENSMUST00000023959.8/ENSMUSG00000023192.8, intron 1 of 5) | 9 | 106643117 | 106656080 | 12964 | 2 | ENSMUST00000023959.8 | 1458 | Grm2 | ENSMUSG00000023192.8 |
| chr9 | 106654938 | 106655109 | 3.34061 | 4.01762 | 1.16759 | Promoter | 9 | 106643117 | 106656080 | 12964 | 2 | ENSMUST00000023959.8 | 971 | Grm2 | ENSMUSG00000023192.8 |
| chr4 | 130264034 | 130264384 | 3.62891 | 4.6291 | 1.56953 | Exon (ENSMUST00000122374.2/ENSMUSG00000023232.12, exon 3 of 10) | 4 | 130253495 | 130275218 | 21724 | 2 | ENSMUST00000122374.2 | 10834 | Serinc2 | ENSMUSG00000023232.12 |
| chr4 | 130265830 | 130266233 | 3.63893 | 4.43824 | 1.4144 | Intron (ENSMUST00000122374.2/ENSMUSG00000023232.12, intron 1 of 9) | 4 | 130253495 | 130275218 | 21724 | 2 | ENSMUST00000122374.2 | 8985 | Serinc2 | ENSMUSG00000023232.12 |
| chr4 | 130271030 | 130271240 | 3.40996 | 3.95693 | 1.06668 | Intron (ENSMUST00000122374.2/ENSMUSG00000023232.12, intron 1 of 9) | 4 | 130253495 | 130275218 | 21724 | 2 | ENSMUST00000122374.2 | 3978 | Serinc2 | ENSMUSG00000023232.12 |
| chr4 | 130273060 | 130273941 | 4.01682 | 5.16219 | 1.88285 | Intron (ENSMUST00000122374.2/ENSMUSG00000023232.12, intron 1 of 9) | 4 | 130253495 | 130275218 | 21724 | 2 | ENSMUST00000122374.2 | 1277 | Serinc2 | ENSMUSG00000023232.12 |
| chr4 | 130274607 | 130275425 | 4.33688 | 5.77489 | 2.33087 | Promoter | 4 | 130253495 | 130275218 | 21724 | 2 | ENSMUST00000122374.2 | 0 | Serinc2 | ENSMUSG00000023232.12 |
| chr4 | 130276707 | 130276964 | 3.92486 | 4.97015 | 1.74297 | Intron (ENSMUST00000120126.3/ENSMUSG00000023232.12, intron 2 of 10) | 4 | 130253495 | 130275586 | 22092 | 2 | ENSMUST00000105996.2 | -1122 | Serinc2 | ENSMUSG00000023232.12 |
| chr4 | 130279340 | 130279573 | 3.45953 | 4.04661 | 1.11903 | Promoter | 4 | 130253899 | 130279205 | 25307 | 2 | ENSMUST00000120126.3 | -136 | Serinc2 | ENSMUSG00000023232.12 |
| chr4 | 155935378 | 155936021 | 3.48684 | 4.18838 | 1.29116 | Distal Intergenic | 4 | 155943831 | 155957711 | 13881 | 1 | ENSMUST00000103175.2 | -7810 | Ube2j2 | ENSMUSG00000023286.11 |
| chr5 | 137267337 | 137267687 | 3.50846 | 4.1318 | 1.15898 | Distal Intergenic | 5 | 137287519 | 137290154 | 2636 | 1 | ENSMUST00000132191.2 | -19832 | Ache | ENSMUSG00000023328.9 |
| chr5 | 137277458 | 137277639 | 3.29351 | 3.76742 | 1.01733 | Distal Intergenic | 5 | 137287519 | 137290154 | 2636 | 1 | ENSMUST00000132191.2 | -9880 | Ache | ENSMUSG00000023328.9 |
| chr5 | 137284219 | 137284592 | 3.44861 | 4.06775 | 1.20887 | Distal Intergenic | 5 | 137287519 | 137290154 | 2636 | 1 | ENSMUST00000132191.2 | -2927 | Ache | ENSMUSG00000023328.9 |
| chr5 | 137288085 | 137288673 | 3.6859 | 4.53885 | 1.47785 | Promoter | 5 | 137288277 | 137294466 | 6190 | 1 | ENSMUST00000024099.5 | 0 | Ache | ENSMUSG00000023328.9 |
| chr5 | 137289155 | 137289247 | 3.67294 | 4.43585 | 1.35617 | Promoter | 5 | 137289247 | 137290154 | 908 | 1 | ENSMUST00000137126.1 | 0 | Ache | ENSMUSG00000023328.9 |
| chr5 | 137289984 | 137290387 | 3.41819 | 4.01464 | 1.1803 | Promoter | 5 | 137289247 | 137290154 | 908 | 1 | ENSMUST00000137126.1 | 738 | Ache | ENSMUSG00000023328.9 |
| chr8 | 119679369 | 119679518 | 3.36566 | 3.89755 | 1.09361 | Exon (ENSMUST00000024107.5/ENSMUSG00000023336.5, exon 2 of 7) | 8 | 119666523 | 119688222 | 21700 | 1 | ENSMUST00000024107.5 | 12847 | Wfdc1 | ENSMUSG00000023336.5 |
| chr9 | 106267150 | 106267225 | 3.64217 | 4.38616 | 1.3459 | Distal Intergenic | 9 | 106281061 | 106349879 | 68819 | 1 | ENSMUST00000072206.7 | -13836 | Poc1a | ENSMUSG00000023345.10 |
| chr14 | 20238365 | 20238849 | 3.35323 | 3.87262 | 1.0715 | Distal Intergenic | 14 | 20262756 | 20269162 | 6407 | 2 | ENSMUST00000024155.7 | 30313 | Kcnk16 | ENSMUSG00000023387.7 |
| chr14 | 20241420 | 20241580 | 3.53318 | 4.17944 | 1.19531 | Distal Intergenic | 14 | 20262756 | 20269162 | 6407 | 2 | ENSMUST00000024155.7 | 27582 | Kcnk16 | ENSMUSG00000023387.7 |
| chr14 | 20252102 | 20252392 | 3.39531 | 3.93789 | 1.07761 | Distal Intergenic | 14 | 20262756 | 20269162 | 6407 | 2 | ENSMUST00000024155.7 | 16770 | Kcnk16 | ENSMUSG00000023387.7 |
| chr2 | 71543032 | 71544298 | 4.17536 | 5.59299 | 2.17391 | 3' UTR | 2 | 71544866 | 71545950 | 1085 | 2 | ENSMUST00000134249.1 | 1652 | Dlx2 | ENSMUSG00000023391.8 |
| chr2 | 71544696 | 71544935 | 4.43307 | 6.29242 | 2.69913 | Exon (ENSMUST00000024159.7/ENSMUSG00000023391.8, exon 3 of 3) | 2 | 71544866 | 71545950 | 1085 | 2 | ENSMUST00000134249.1 | 1015 | Dlx2 | ENSMUSG00000023391.8 |
| chr2 | 71545304 | 71547664 | 4.02907 | 5.33739 | 2.05714 | Promoter | 2 | 71543408 | 71546754 | 3347 | 2 | ENSMUST00000024159.7 | 0 | Dlx2 | ENSMUSG00000023391.8 |
| chr2 | 71547988 | 71548240 | 3.83994 | 4.75589 | 1.57091 | Distal Intergenic | 2 | 71543408 | 71546754 | 3347 | 2 | ENSMUST00000024159.7 | -1235 | Dlx2 | ENSMUSG00000023391.8 |
| chr2 | 71548664 | 71548779 | 3.87255 | 4.838 | 1.65832 | Distal Intergenic | 2 | 71543408 | 71546754 | 3347 | 2 | ENSMUST00000024159.7 | -1911 | Dlx2 | ENSMUSG00000023391.8 |
| chr2 | 71549477 | 71549616 | 3.52946 | 4.35439 | 1.37089 | Distal Intergenic | 2 | 71543408 | 71546754 | 3347 | 2 | ENSMUST00000024159.7 | -2724 | Dlx2 | ENSMUSG00000023391.8 |
| chr2 | 71550490 | 71550949 | 3.5492 | 4.22544 | 1.26173 | Distal Intergenic | 2 | 71543408 | 71546754 | 3347 | 2 | ENSMUST00000024159.7 | -3737 | Dlx2 | ENSMUSG00000023391.8 |
| chr2 | 71551491 | 71551637 | 3.3708 | 3.89268 | 1.04874 | Distal Intergenic | 2 | 71543408 | 71546754 | 3347 | 2 | ENSMUST00000024159.7 | -4738 | Dlx2 | ENSMUSG00000023391.8 |
| chr2 | 71552220 | 71552435 | 4.30131 | 5.71099 | 2.25032 | Distal Intergenic | 2 | 71543408 | 71546754 | 3347 | 2 | ENSMUST00000024159.7 | -5467 | Dlx2 | ENSMUSG00000023391.8 |
| chr2 | 71552839 | 71553523 | 3.64481 | 4.4025 | 1.37514 | Distal Intergenic | 2 | 71543408 | 71546754 | 3347 | 2 | ENSMUST00000024159.7 | -6086 | Dlx2 | ENSMUSG00000023391.8 |
| chr2 | 71554038 | 71554261 | 3.46493 | 4.09504 | 1.14922 | Distal Intergenic | 2 | 71543408 | 71546754 | 3347 | 2 | ENSMUST00000024159.7 | -7285 | Dlx2 | ENSMUSG00000023391.8 |
| chr2 | 71574143 | 71574371 | 3.43673 | 4.13626 | 1.22612 | Distal Intergenic | 2 | 71543408 | 71546754 | 3347 | 2 | ENSMUST00000024159.7 | -27390 | Dlx2 | ENSMUSG00000023391.8 |
| chr2 | 71581122 | 71581922 | 4.10348 | 5.34069 | 2.02525 | Distal Intergenic | 2 | 71543408 | 71546754 | 3347 | 2 | ENSMUST00000024159.7 | -34369 | Dlx2 | ENSMUSG00000023391.8 |
| chr2 | 71628575 | 71628766 | 3.91596 | 5.18849 | 1.91662 | Distal Intergenic | 2 | 71543408 | 71546754 | 3347 | 2 | ENSMUST00000024159.7 | -81822 | Dlx2 | ENSMUSG00000023391.8 |
| chr2 | 71629281 | 71630359 | 3.9885 | 5.14747 | 1.952 | Distal Intergenic | 2 | 71543408 | 71546754 | 3347 | 2 | ENSMUST00000024159.7 | -82528 | Dlx2 | ENSMUSG00000023391.8 |
| chr2 | 180755994 | 180756069 | 3.67857 | 4.58384 | 1.49923 | Distal Intergenic | 2 | 180738274 | 180741416 | 3143 | 1 | ENSMUST00000140955.1 | 17721 | Slc17a9 | ENSMUSG00000023393.10 |
| chr6 | 49517622 | 49517804 | 3.31322 | 3.79707 | 1.01733 | Distal Intergenic | 6 | 49438564 | 49469500 | 30937 | 1 | ENSMUST00000167846.1 | 79059 | Stk31 | ENSMUSG00000023403.9 |
| chr14 | 55824126 | 55824486 | 3.46653 | 4.17692 | 1.26945 | Promoter | 14 | 55824795 | 55833943 | 9149 | 1 | ENSMUST00000024179.5 | -309 | Nfatc4 | ENSMUSG00000023411.6 |
| chr14 | 55824989 | 55825904 | 3.90926 | 4.95034 | 1.74439 | Promoter | 14 | 55824795 | 55833943 | 9149 | 1 | ENSMUST00000172271.2 | 195 | Nfatc4 | ENSMUSG00000023411.6 |
| chr9 | 108823551 | 108824341 | 4.06643 | 5.60609 | 2.20556 | Distal Intergenic | 9 | 108826320 | 108852963 | 26644 | 1 | ENSMUST00000024238.5 | -1979 | Celsr3 | ENSMUSG00000023473.6 |
| chr9 | 108825210 | 108826526 | 3.90822 | 5.11642 | 1.85562 | Promoter | 9 | 108826320 | 108852963 | 26644 | 1 | ENSMUST00000024238.5 | 0 | Celsr3 | ENSMUSG00000023473.6 |
| chr9 | 108826881 | 108826956 | 3.58302 | 4.35094 | 1.37676 | Promoter | 9 | 108826320 | 108852963 | 26644 | 1 | ENSMUST00000024238.5 | 562 | Celsr3 | ENSMUSG00000023473.6 |
| chr9 | 108827649 | 108827891 | 3.80868 | 4.82512 | 1.65492 | Exon (ENSMUST00000024238.5/ENSMUSG00000023473.6, exon 1 of 35) | 9 | 108826320 | 108852963 | 26644 | 1 | ENSMUST00000024238.5 | 1330 | Celsr3 | ENSMUSG00000023473.6 |
| chr9 | 108829680 | 108829917 | 4.30846 | 5.76018 | 2.33732 | Exon (ENSMUST00000024238.5/ENSMUSG00000023473.6, exon 1 of 35) | 9 | 108826320 | 108852963 | 26644 | 1 | ENSMUST00000024238.5 | 3361 | Celsr3 | ENSMUSG00000023473.6 |
| chr9 | 108830458 | 108830570 | 3.79135 | 4.76718 | 1.64247 | Intron (ENSMUST00000024238.5/ENSMUSG00000023473.6, intron 1 of 34) | 9 | 108826320 | 108852963 | 26644 | 1 | ENSMUST00000024238.5 | 4139 | Celsr3 | ENSMUSG00000023473.6 |
| chr9 | 108831939 | 108832246 | 3.57554 | 4.27087 | 1.28218 | Exon (ENSMUST00000024238.5/ENSMUSG00000023473.6, exon 2 of 35) | 9 | 108826320 | 108852963 | 26644 | 1 | ENSMUST00000024238.5 | 5620 | Celsr3 | ENSMUSG00000023473.6 |
| chr9 | 108832758 | 108832968 | 3.77919 | 4.63734 | 1.48792 | Exon (ENSMUST00000024238.5/ENSMUSG00000023473.6, exon 3 of 35) | 9 | 108826320 | 108852963 | 26644 | 1 | ENSMUST00000024238.5 | 6439 | Celsr3 | ENSMUSG00000023473.6 |
| chr15 | 99053129 | 99053256 | 3.91146 | 4.91072 | 1.69956 | Distal Intergenic | 15 | 99055174 | 99058978 | 3805 | 1 | ENSMUST00000024249.3 | -1918 | Prph | ENSMUSG00000023484.8 |
| chr15 | 99054639 | 99055930 | 3.83572 | 4.76273 | 1.59027 | Promoter | 15 | 99055174 | 99058978 | 3805 | 1 | ENSMUST00000024249.3 | 0 | Prph | ENSMUSG00000023484.8 |
| chr15 | 99056315 | 99058777 | 3.90922 | 5.07298 | 1.84903 | 3' UTR | 15 | 99055174 | 99058978 | 3805 | 1 | ENSMUST00000047104.8 | 1142 | Prph | ENSMUSG00000023484.8 |
| chr15 | 99060250 | 99060637 | 3.5617 | 4.27673 | 1.28172 | Distal Intergenic | 15 | 99055174 | 99058978 | 3805 | 1 | ENSMUST00000047104.8 | 5077 | Prph | ENSMUSG00000023484.8 |
| chr15 | 99062756 | 99062981 | 3.43486 | 3.99734 | 1.07666 | Distal Intergenic | 15 | 99055174 | 99058978 | 3805 | 1 | ENSMUST00000047104.8 | 7583 | Prph | ENSMUSG00000023484.8 |
| chr11 | 3193043 | 3193509 | 1.56843 | 3.97803 | 1.13214 | Promoter | 11 | 3131850 | 3193407 | 61558 | 2 | ENSMUST00000081318.7 | 0 | Sfi1 | ENSMUSG00000023764.13 |
| chr17 | 7117368 | 7117697 | 3.43754 | 4.05902 | 1.23975 | Distal Intergenic | 17 | 7170115 | 7303313 | 133199 | 1 | ENSMUST00000024575.6 | -52418 | Rps6ka2 | ENSMUSG00000023809.9 |
| chr17 | 11912745 | 11913384 | 3.70762 | 4.54688 | 1.46508 | Intron (ENSMUST00000191124.1/ENSMUSG00000023826.9, intron 9 of 11) | 17 | 11845210 | 11867561 | 22352 | 1 | ENSMUST00000190849.1 | 67536 | Park2 | ENSMUSG00000023826.9 |
| chr17 | 11913695 | 11914488 | 3.46302 | 4.09239 | 1.22152 | Intron (ENSMUST00000191124.1/ENSMUSG00000023826.9, intron 9 of 11) | 17 | 11845210 | 11867561 | 22352 | 1 | ENSMUST00000190849.1 | 68486 | Park2 | ENSMUSG00000023826.9 |
| chr17 | 12506599 | 12507186 | 4.06026 | 5.26541 | 1.92471 | Promoter | 17 | 12419972 | 12507704 | 87733 | 2 | ENSMUST00000024595.2 | 518 | Slc22a3 | ENSMUSG00000023828.2 |
| chr17 | 12507517 | 12507915 | 3.5759 | 4.26661 | 1.27143 | Promoter | 17 | 12419972 | 12507704 | 87733 | 2 | ENSMUST00000024595.2 | 0 | Slc22a3 | ENSMUSG00000023828.2 |
| chr17 | 12534570 | 12534645 | 3.54674 | 4.26111 | 1.32733 | Distal Intergenic | 17 | 12419972 | 12507704 | 87733 | 2 | ENSMUST00000024595.2 | -26867 | Slc22a3 | ENSMUSG00000023828.2 |
| chr17 | 12637650 | 12638086 | 3.86232 | 4.80585 | 1.62009 | Distal Intergenic | 17 | 12648874 | 12675838 | 26965 | 2 | ENSMUST00000024596.8 | 37752 | Slc22a1 | ENSMUSG00000023829.8 |
| chr17 | 9044544 | 9044691 | 4.16246 | 5.51863 | 2.12607 | Distal Intergenic | 17 | 8988383 | 9005263 | 16881 | 1 | ENSMUST00000136954.2 | 56162 | 1700010I14Rik | ENSMUSG00000023873.7 |
| chr17 | 14514600 | 14514960 | 3.60358 | 4.63052 | 1.5337 | Distal Intergenic | 17 | 14665500 | 14694262 | 28763 | 2 | ENSMUST00000170872.1 | 179302 | Thbs2 | ENSMUSG00000023885.8 |
| chr17 | 14516099 | 14516305 | 3.68619 | 4.45878 | 1.36616 | Distal Intergenic | 17 | 14665500 | 14694262 | 28763 | 2 | ENSMUST00000170872.1 | 177957 | Thbs2 | ENSMUSG00000023885.8 |
| chr17 | 14518610 | 14518778 | 3.56204 | 4.26559 | 1.33204 | Distal Intergenic | 17 | 14665500 | 14694262 | 28763 | 2 | ENSMUST00000170872.1 | 175484 | Thbs2 | ENSMUSG00000023885.8 |
| chr17 | 14627490 | 14627606 | 3.81885 | 4.72017 | 1.5641 | Distal Intergenic | 17 | 14665500 | 14694262 | 28763 | 2 | ENSMUST00000170872.1 | 66656 | Thbs2 | ENSMUSG00000023885.8 |
| chr17 | 14277301 | 14277376 | 3.501 | 4.12791 | 1.18542 | Distal Intergenic | 17 | 14279506 | 14404790 | 125285 | 1 | ENSMUST00000024660.8 | -2130 | Smoc2 | ENSMUSG00000023886.9 |
| chr17 | 14278476 | 14278762 | 3.47817 | 4.10032 | 1.21149 | Promoter | 17 | 14279506 | 14404790 | 125285 | 1 | ENSMUST00000024660.8 | -744 | Smoc2 | ENSMUSG00000023886.9 |
| chr17 | 14279789 | 14279900 | 3.79547 | 4.71533 | 1.55677 | Promoter | 17 | 14279506 | 14404790 | 125285 | 1 | ENSMUST00000024660.8 | 284 | Smoc2 | ENSMUSG00000023886.9 |
| chr17 | 14280216 | 14280900 | 3.51895 | 4.1603 | 1.1932 | Promoter | 17 | 14279506 | 14404790 | 125285 | 1 | ENSMUST00000024660.8 | 711 | Smoc2 | ENSMUSG00000023886.9 |
| chr17 | 14282470 | 14282690 | 3.42884 | 4.25752 | 1.29125 | Intron (ENSMUST00000024660.8/ENSMUSG00000023886.9, intron 1 of 12) | 17 | 14279506 | 14404790 | 125285 | 1 | ENSMUST00000024660.8 | 2965 | Smoc2 | ENSMUSG00000023886.9 |
| chr17 | 14287795 | 14288101 | 3.44336 | 4.06407 | 1.16798 | Intron (ENSMUST00000024660.8/ENSMUSG00000023886.9, intron 1 of 12) | 17 | 14279506 | 14404790 | 125285 | 1 | ENSMUST00000024660.8 | 8290 | Smoc2 | ENSMUSG00000023886.9 |
| chr17 | 14372893 | 14373060 | 3.83049 | 4.73666 | 1.55862 | Intron (ENSMUST00000024660.8/ENSMUSG00000023886.9, intron 8 of 12) | 17 | 14279506 | 14404790 | 125285 | 1 | ENSMUST00000024660.8 | 93388 | Smoc2 | ENSMUSG00000023886.9 |
| chr17 | 23630538 | 23630655 | 3.3388 | 3.84915 | 1.065 | 3' UTR | 17 | 23629458 | 23645269 | 15812 | 2 | ENSMUST00000024696.7 | 14614 | Mmp25 | ENSMUSG00000023903.7 |
| chr17 | 23643674 | 23644262 | 3.8926 | 4.93102 | 1.76151 | Exon (ENSMUST00000024696.7/ENSMUSG00000023903.7, exon 2 of 10) | 17 | 23629458 | 23645269 | 15812 | 2 | ENSMUST00000024696.7 | 1007 | Mmp25 | ENSMUSG00000023903.7 |
| chr17 | 23644746 | 23645497 | 4.06342 | 5.3555 | 2.00029 | Promoter | 17 | 23629458 | 23645269 | 15812 | 2 | ENSMUST00000024696.7 | 0 | Mmp25 | ENSMUSG00000023903.7 |
| chr17 | 23678845 | 23680420 | 3.93954 | 5.04838 | 1.85177 | Promoter | 17 | 23679365 | 23682446 | 3082 | 1 | ENSMUST00000024699.2 | 0 | Cldn6 | ENSMUSG00000023906.2 |
| chr17 | 23740816 | 23741136 | 3.2299 | 3.67204 | 1.01733 | Promoter | 17 | 23736186 | 23740330 | 4145 | 2 | ENSMUST00000024702.3 | -487 | Paqr4 | ENSMUSG00000023909.3 |
| chr17 | 51819626 | 51819916 | 3.39062 | 3.96037 | 1.17239 | Intron (ENSMUST00000144331.2/ENSMUSG00000023927.10, intron 1 of 10) | 17 | 51804366 | 51821091 | 16726 | 2 | ENSMUST00000129205.2 | 1175 | Satb1 | ENSMUSG00000023927.10 |
| chr17 | 51823940 | 51824075 | 3.31682 | 3.80809 | 1.03711 | Intron (ENSMUST00000144331.2/ENSMUSG00000023927.10, intron 1 of 10) | 17 | 51739455 | 51826566 | 87112 | 2 | ENSMUST00000129667.2 | 2491 | Satb1 | ENSMUSG00000023927.10 |
| chr17 | 51824388 | 51824803 | 3.35428 | 3.86893 | 1.05293 | Exon (ENSMUST00000156051.2/ENSMUSG00000023927.10, exon 1 of 1) | 17 | 51739455 | 51826566 | 87112 | 2 | ENSMUST00000129667.2 | 1763 | Satb1 | ENSMUSG00000023927.10 |
| chr17 | 51826365 | 51826484 | 3.3878 | 3.95178 | 1.16 | Promoter | 17 | 51739455 | 51826566 | 87112 | 2 | ENSMUST00000129667.2 | 82 | Satb1 | ENSMUSG00000023927.10 |
| chr17 | 51828469 | 51829783 | 3.72824 | 4.58584 | 1.54393 | Promoter | 17 | 51824641 | 51829497 | 4857 | 2 | ENSMUST00000156051.2 | 0 | Satb1 | ENSMUSG00000023927.10 |
| chr17 | 51831411 | 51832322 | 3.99687 | 5.2444 | 1.9936 | Promoter | 17 | 51805363 | 51832019 | 26657 | 2 | ENSMUST00000124222.1 | 0 | Satb1 | ENSMUSG00000023927.10 |
| chr17 | 51832644 | 51833348 | 3.57662 | 4.63118 | 1.56316 | Promoter | 17 | 51736187 | 51832672 | 96486 | 2 | ENSMUST00000144331.2 | 0 | Satb1 | ENSMUSG00000023927.10 |
| chr17 | 51833803 | 51834170 | 3.6918 | 4.53795 | 1.50081 | Promoter | 17 | 51738894 | 51833290 | 94397 | 2 | ENSMUST00000133574.2 | -514 | Satb1 | ENSMUSG00000023927.10 |
| chr17 | 51834803 | 51834955 | 3.64925 | 4.46082 | 1.44978 | Distal Intergenic | 17 | 51738894 | 51833290 | 94397 | 2 | ENSMUST00000133574.2 | -1514 | Satb1 | ENSMUSG00000023927.10 |
| chr17 | 51835920 | 51836041 | 3.47785 | 4.07674 | 1.1278 | Distal Intergenic | 17 | 51738894 | 51833290 | 94397 | 2 | ENSMUST00000133574.2 | -2631 | Satb1 | ENSMUSG00000023927.10 |
| chr17 | 45476804 | 45477086 | 3.49817 | 4.15986 | 1.18649 | Distal Intergenic | 17 | 45448937 | 45474938 | 26002 | 2 | ENSMUST00000024731.8 | -1867 | Spats1 | ENSMUSG00000023935.9 |
| chr17 | 54298303 | 54298955 | 3.85146 | 4.80786 | 1.66186 | Promoter | 17 | 54273594 | 54299034 | 25441 | 2 | ENSMUST00000095712.3 | 79 | Slc5a7 | ENSMUSG00000023945.6 |
| chr17 | 45522458 | 45522746 | 3.59507 | 4.30023 | 1.28286 | Promoter | 17 | 45523434 | 45542679 | 19246 | 1 | ENSMUST00000113547.1 | -688 | Tcte1 | ENSMUSG00000023949.6 |
| chr17 | 45523084 | 45523886 | 3.59619 | 4.29837 | 1.27948 | Promoter | 17 | 45523434 | 45542679 | 19246 | 1 | ENSMUST00000113547.1 | 0 | Tcte1 | ENSMUSG00000023949.6 |
| chr17 | 45524362 | 45524959 | 3.72289 | 4.57101 | 1.47421 | Promoter | 17 | 45523434 | 45542679 | 19246 | 1 | ENSMUST00000113547.1 | 929 | Tcte1 | ENSMUSG00000023949.6 |
| chr17 | 45531771 | 45532521 | 3.80337 | 4.70163 | 1.57032 | Intron (ENSMUST00000113547.1/ENSMUSG00000023949.6, intron 1 of 4) | 17 | 45523434 | 45542679 | 19246 | 1 | ENSMUST00000113547.1 | 8338 | Tcte1 | ENSMUSG00000023949.6 |
| chr17 | 45533147 | 45533398 | 3.56918 | 4.24317 | 1.22842 | Exon (ENSMUST00000113547.1/ENSMUSG00000023949.6, exon 2 of 5) | 17 | 45523434 | 45542679 | 19246 | 1 | ENSMUST00000113547.1 | 9714 | Tcte1 | ENSMUSG00000023949.6 |
| chr17 | 45535018 | 45535225 | 3.76877 | 4.65086 | 1.5229 | Exon (ENSMUST00000113547.1/ENSMUSG00000023949.6, exon 3 of 5) | 17 | 45523434 | 45542679 | 19246 | 1 | ENSMUST00000113547.1 | 11585 | Tcte1 | ENSMUSG00000023949.6 |
| chr17 | 45535553 | 45535629 | 4.58137 | 6.22638 | 2.61118 | Intron (ENSMUST00000113547.1/ENSMUSG00000023949.6, intron 3 of 4) | 17 | 45523434 | 45542679 | 19246 | 1 | ENSMUST00000113547.1 | 12120 | Tcte1 | ENSMUSG00000023949.6 |
| chr17 | 45536034 | 45536124 | 3.60392 | 4.31117 | 1.28429 | Intron (ENSMUST00000113547.1/ENSMUSG00000023949.6, intron 3 of 4) | 17 | 45523434 | 45542679 | 19246 | 1 | ENSMUST00000113547.1 | 12601 | Tcte1 | ENSMUSG00000023949.6 |
| chr17 | 44323272 | 44323505 | 3.47152 | 4.07598 | 1.15971 | Distal Intergenic | 17 | 44188572 | 44280168 | 91597 | 1 | ENSMUST00000024755.5 | 134701 | Clic5 | ENSMUSG00000023959.8 |
| chr17 | 44325993 | 44326155 | 3.355 | 3.85997 | 1.01733 | Distal Intergenic | 17 | 44188572 | 44280168 | 91597 | 1 | ENSMUST00000024755.5 | 137422 | Clic5 | ENSMUSG00000023959.8 |
| chr17 | 44326832 | 44327138 | 3.67521 | 4.44885 | 1.38033 | Distal Intergenic | 17 | 44188572 | 44280168 | 91597 | 1 | ENSMUST00000024755.5 | 138261 | Clic5 | ENSMUSG00000023959.8 |
| chr17 | 44330128 | 44330272 | 3.30921 | 3.91787 | 1.04903 | Distal Intergenic | 17 | 44188572 | 44280168 | 91597 | 1 | ENSMUST00000024755.5 | 141557 | Clic5 | ENSMUSG00000023959.8 |
| chr17 | 44351166 | 44351281 | 3.82593 | 4.79989 | 1.6707 | Distal Intergenic | 17 | 44188572 | 44280168 | 91597 | 1 | ENSMUST00000024755.5 | 162595 | Clic5 | ENSMUSG00000023959.8 |
| chr17 | 44352878 | 44353666 | 3.55077 | 4.27198 | 1.32352 | Distal Intergenic | 17 | 44188572 | 44280168 | 91597 | 1 | ENSMUST00000024755.5 | 164307 | Clic5 | ENSMUSG00000023959.8 |
| chr17 | 44356276 | 44356562 | 3.68389 | 4.45112 | 1.35104 | Distal Intergenic | 17 | 44188572 | 44280168 | 91597 | 1 | ENSMUST00000024755.5 | 167705 | Clic5 | ENSMUSG00000023959.8 |
| chr17 | 44357141 | 44357577 | 3.42319 | 4.01054 | 1.17686 | Distal Intergenic | 17 | 44188572 | 44280168 | 91597 | 1 | ENSMUST00000024755.5 | 168570 | Clic5 | ENSMUSG00000023959.8 |
| chr17 | 44361817 | 44361917 | 4.13969 | 5.37758 | 2.02019 | Distal Intergenic | 17 | 44188572 | 44280168 | 91597 | 1 | ENSMUST00000024755.5 | 173246 | Clic5 | ENSMUSG00000023959.8 |
| chr17 | 44362760 | 44363687 | 3.42011 | 3.99969 | 1.1597 | Distal Intergenic | 17 | 44188572 | 44280168 | 91597 | 1 | ENSMUST00000024755.5 | 174189 | Clic5 | ENSMUSG00000023959.8 |
| chr17 | 44364397 | 44364638 | 3.4942 | 4.12697 | 1.21313 | Distal Intergenic | 17 | 44188572 | 44280168 | 91597 | 1 | ENSMUST00000024755.5 | 175826 | Clic5 | ENSMUSG00000023959.8 |
| chr17 | 44365352 | 44365437 | 3.80112 | 4.89008 | 1.6708 | Distal Intergenic | 17 | 44188572 | 44280168 | 91597 | 1 | ENSMUST00000024755.5 | 176781 | Clic5 | ENSMUSG00000023959.8 |
| chr17 | 44367753 | 44367973 | 3.95245 | 5.3448 | 1.98081 | Distal Intergenic | 17 | 44188572 | 44280168 | 91597 | 1 | ENSMUST00000024755.5 | 179182 | Clic5 | ENSMUSG00000023959.8 |
| chr17 | 44369121 | 44369408 | 3.83754 | 4.86567 | 1.6955 | Distal Intergenic | 17 | 44188572 | 44280168 | 91597 | 1 | ENSMUST00000024755.5 | 180550 | Clic5 | ENSMUSG00000023959.8 |
| chr17 | 44372516 | 44372622 | 3.51334 | 4.37606 | 1.35672 | Distal Intergenic | 17 | 44188572 | 44280168 | 91597 | 1 | ENSMUST00000024755.5 | 183945 | Clic5 | ENSMUSG00000023959.8 |
| chr17 | 44373252 | 44373395 | 3.73081 | 4.57637 | 1.53536 | Distal Intergenic | 17 | 44188572 | 44280168 | 91597 | 1 | ENSMUST00000024755.5 | 184681 | Clic5 | ENSMUSG00000023959.8 |
| chr17 | 44373944 | 44374119 | 3.5973 | 4.35731 | 1.42832 | Distal Intergenic | 17 | 44188572 | 44280168 | 91597 | 1 | ENSMUST00000024755.5 | 185373 | Clic5 | ENSMUSG00000023959.8 |
| chr6 | 3763202 | 3763928 | 3.84775 | 4.8728 | 1.67051 | Promoter | 6 | 3685680 | 3763629 | 77950 | 2 | ENSMUST00000171613.2 | 0 | Calcr | ENSMUSG00000023964.10 |
| chr6 | 3767247 | 3767322 | 3.36357 | 3.8729 | 1.01733 | Distal Intergenic | 6 | 3685720 | 3764713 | 78994 | 2 | ENSMUST00000115622.2 | -2535 | Calcr | ENSMUSG00000023964.10 |
| chr17 | 47385268 | 47385452 | 3.43367 | 3.99713 | 1.08244 | Promoter | 17 | 47385393 | 47392967 | 7575 | 1 | ENSMUST00000024774.8 | 0 | Guca1b | ENSMUSG00000023979.9 |
| chr17 | 47394602 | 47395033 | 3.34826 | 3.84982 | 1.01733 | 3' UTR | 17 | 47394560 | 47400584 | 6025 | 2 | ENSMUST00000059348.7 | 5551 | Guca1a | ENSMUSG00000023982.7 |
| chr17 | 47801797 | 47801918 | 3.66691 | 4.42888 | 1.36554 | Distal Intergenic | 17 | 47789441 | 47791613 | 2173 | 1 | ENSMUST00000140715.1 | 12357 | Tfeb | ENSMUSG00000023990.13 |
| chr17 | 47802798 | 47803304 | 3.68253 | 4.46403 | 1.39907 | Distal Intergenic | 17 | 47789441 | 47791613 | 2173 | 1 | ENSMUST00000140715.1 | 13358 | Tfeb | ENSMUSG00000023990.13 |
| chr17 | 49731657 | 49731821 | 3.73875 | 4.7177 | 1.60679 | Intron (ENSMUST00000024798.6/ENSMUSG00000023999.9, intron 10 of 11) | 17 | 49738753 | 49755390 | 16638 | 1 | ENSMUST00000159396.1 | -6932 | Kif6 | ENSMUSG00000023999.9 |
| chr17 | 29169054 | 29169202 | 3.47894 | 4.13023 | 1.26307 | Intron (ENSMUST00000024805.8/ENSMUSG00000024008.10, intron 13 of 20) | 17 | 29169437 | 29176683 | 7247 | 2 | ENSMUST00000124917.1 | 7481 | Cpne5 | ENSMUSG00000024008.10 |
| chr17 | 29187750 | 29187864 | 3.68219 | 4.45463 | 1.37051 | Intron (ENSMUST00000024805.8/ENSMUSG00000024008.10, intron 9 of 20) | 17 | 29157916 | 29184103 | 26188 | 2 | ENSMUST00000126993.2 | -3648 | Cpne5 | ENSMUSG00000024008.10 |
| chr17 | 29188634 | 29189537 | 3.5618 | 4.41811 | 1.36827 | Intron (ENSMUST00000024805.8/ENSMUSG00000024008.10, intron 8 of 20) | 17 | 29157916 | 29184103 | 26188 | 2 | ENSMUST00000126993.2 | -4532 | Cpne5 | ENSMUSG00000024008.10 |
| chr17 | 29237091 | 29239069 | 3.89211 | 4.93207 | 1.73325 | Promoter | 17 | 29156550 | 29237797 | 81248 | 2 | ENSMUST00000024805.8 | 0 | Cpne5 | ENSMUSG00000024008.10 |
| chr17 | 29240639 | 29240875 | 3.47056 | 4.0874 | 1.20578 | Distal Intergenic | 17 | 29156550 | 29237797 | 81248 | 2 | ENSMUST00000024805.8 | -2843 | Cpne5 | ENSMUSG00000024008.10 |
| chr17 | 29242247 | 29242504 | 3.41487 | 3.96066 | 1.05373 | Distal Intergenic | 17 | 29156550 | 29237797 | 81248 | 2 | ENSMUST00000024805.8 | -4451 | Cpne5 | ENSMUSG00000024008.10 |
| chr17 | 29243303 | 29243618 | 3.662 | 4.53688 | 1.45614 | Distal Intergenic | 17 | 29156550 | 29237797 | 81248 | 2 | ENSMUST00000024805.8 | -5507 | Cpne5 | ENSMUSG00000024008.10 |
| chr17 | 29773631 | 29773987 | 3.34296 | 3.84184 | 1.01733 | Distal Intergenic | 17 | 29703332 | 29717012 | 13681 | 2 | ENSMUST00000128751.1 | -56620 | Ccdc167 | ENSMUSG00000024018.12 |
| chr17 | 31036562 | 31036872 | 3.32786 | 3.96276 | 1.16567 | Distal Intergenic | 17 | 31057698 | 31115777 | 58080 | 1 | ENSMUST00000024829.6 | -20826 | Abcg1 | ENSMUSG00000024030.6 |
| chr17 | 31193193 | 31193294 | 3.95623 | 5.40027 | 2.04528 | Intron (ENSMUST00000024833.6/ENSMUSG00000024034.7, intron 4 of 11) | 17 | 31179272 | 31197749 | 18478 | 2 | ENSMUST00000024833.6 | 4455 | Tmprss3 | ENSMUSG00000024034.7 |
| chr17 | 31196400 | 31196501 | 3.53376 | 4.23314 | 1.32666 | Intron (ENSMUST00000024833.6/ENSMUSG00000024034.7, intron 1 of 11) | 17 | 31179272 | 31197749 | 18478 | 2 | ENSMUST00000024833.6 | 1248 | Tmprss3 | ENSMUSG00000024034.7 |
| chr17 | 31635571 | 31635994 | 4.31683 | 5.78377 | 2.35414 | Promoter | 17 | 31625131 | 31636649 | 11519 | 2 | ENSMUST00000135425.2 | 655 | Cbs | ENSMUSG00000024039.9 |
| chr17 | 31637243 | 31637451 | 3.42435 | 4.19411 | 1.26324 | Promoter | 17 | 31612623 | 31637199 | 24577 | 2 | ENSMUST00000067801.7 | -45 | Cbs | ENSMUSG00000024039.9 |
| chr17 | 69075113 | 69075555 | 3.34066 | 3.83835 | 1.01733 | Distal Intergenic | 17 | 69156810 | 69289985 | 133176 | 1 | ENSMUST00000080208.5 | -81255 | Epb4.1l3 | ENSMUSG00000024044.10 |
| chr17 | 69076119 | 69076454 | 3.33452 | 3.8292 | 1.01733 | Distal Intergenic | 17 | 69156810 | 69289985 | 133176 | 1 | ENSMUST00000080208.5 | -80356 | Epb4.1l3 | ENSMUSG00000024044.10 |
| chr17 | 69077415 | 69077876 | 4.04401 | 5.18555 | 1.88175 | Distal Intergenic | 17 | 69156810 | 69289985 | 133176 | 1 | ENSMUST00000080208.5 | -78934 | Epb4.1l3 | ENSMUSG00000024044.10 |
| chr17 | 69121854 | 69121929 | 3.88843 | 4.92555 | 1.75229 | Distal Intergenic | 17 | 69156810 | 69289985 | 133176 | 1 | ENSMUST00000080208.5 | -34881 | Epb4.1l3 | ENSMUSG00000024044.10 |
| chr17 | 69122465 | 69123041 | 3.75677 | 4.74373 | 1.63739 | Distal Intergenic | 17 | 69156810 | 69289985 | 133176 | 1 | ENSMUST00000080208.5 | -33769 | Epb4.1l3 | ENSMUSG00000024044.10 |
| chr17 | 69156864 | 69157382 | 3.5971 | 4.34935 | 1.35439 | Promoter | 17 | 69156810 | 69289985 | 133176 | 1 | ENSMUST00000112680.2 | 55 | Epb4.1l3 | ENSMUSG00000024044.10 |
| chr17 | 71310444 | 71311551 | 4.0331 | 5.40618 | 2.06966 | Promoter | 17 | 71252176 | 71310965 | 58790 | 2 | ENSMUST00000024849.9 | 0 | Emilin2 | ENSMUSG00000024053.10 |
| chr17 | 71312102 | 71312354 | 3.69622 | 4.90878 | 1.74008 | Intron (ENSMUST00000180743.1/ENSMUSG00000097625.1, intron 3 of 3) | 17 | 71252176 | 71310965 | 58790 | 2 | ENSMUST00000024849.9 | -1138 | Emilin2 | ENSMUSG00000024053.10 |
| chr17 | 71332251 | 71332774 | 3.41278 | 3.98958 | 1.09107 | Intron (ENSMUST00000180743.1/ENSMUSG00000097625.1, intron 3 of 3) | 17 | 71346510 | 71349006 | 2497 | 2 | ENSMUST00000182049.1 | 16232 | Smchd1 | ENSMUSG00000024054.8 |
| chr17 | 71333352 | 71333671 | 3.35211 | 3.85562 | 1.01733 | Intron (ENSMUST00000180743.1/ENSMUSG00000097625.1, intron 3 of 3) | 17 | 71346510 | 71349006 | 2497 | 2 | ENSMUST00000182049.1 | 15335 | Smchd1 | ENSMUSG00000024054.8 |
| chr17 | 71782426 | 71783123 | 3.86629 | 5.13213 | 1.88548 | Promoter | 17 | 71781947 | 71858351 | 76405 | 1 | ENSMUST00000024854.7 | 480 | Clip4 | ENSMUSG00000024059.9 |
| chr17 | 71869029 | 71869392 | 3.38264 | 4.08225 | 1.16915 | Downstream (<1kb) | 17 | 71781947 | 71858351 | 76405 | 1 | ENSMUST00000024854.7 | 87083 | Clip4 | ENSMUSG00000024059.9 |
| chr17 | 71869875 | 71870118 | 4.1681 | 5.49501 | 2.11443 | Exon (ENSMUST00000086639.4/ENSMUSG00000055471.6, exon 29 of 29) | 17 | 71781947 | 71858351 | 76405 | 1 | ENSMUST00000024854.7 | 87929 | Clip4 | ENSMUSG00000024059.9 |
| chr17 | 71870482 | 71870713 | 3.84387 | 4.79952 | 1.65375 | Intron (ENSMUST00000086639.4/ENSMUSG00000055471.6, intron 28 of 28) | 17 | 71781947 | 71858351 | 76405 | 1 | ENSMUST00000024854.7 | 88536 | Clip4 | ENSMUSG00000024059.9 |
| chr17 | 73708617 | 73710577 | 3.79491 | 4.75653 | 1.6082 | Promoter | 17 | 73493228 | 73710451 | 217224 | 2 | ENSMUST00000024858.6 | 0 | Galnt14 | ENSMUSG00000024064.8 |
| chr17 | 73711053 | 73711294 | 3.39127 | 3.9223 | 1.04403 | Promoter | 17 | 73572784 | 73710453 | 137670 | 2 | ENSMUST00000146565.1 | -601 | Galnt14 | ENSMUSG00000024064.8 |
| chr17 | 66768255 | 66768580 | 3.29059 | 3.76907 | 1.0345 | Intron (ENSMUST00000037974.8/ENSMUSG00000033278.9, intron 17 of 30) | 17 | 66555252 | 66594621 | 39370 | 2 | ENSMUST00000024914.2 | -173635 | Themis3 | ENSMUSG00000024105.2 |
| chr17 | 88753363 | 88753580 | 3.42213 | 4.01872 | 1.20077 | Intron (ENSMUST00000024916.5/ENSMUSG00000024107.6, intron 9 of 10) | 17 | 88741549 | 88791976 | 50428 | 2 | ENSMUST00000024916.5 | 38396 | Lhcgr | ENSMUSG00000024107.6 |
| chr17 | 88755507 | 88755646 | 3.27195 | 3.83549 | 1.10036 | Exon (ENSMUST00000024916.5/ENSMUSG00000024107.6, exon 8 of 11) | 17 | 88741549 | 88791976 | 50428 | 2 | ENSMUST00000024916.5 | 36330 | Lhcgr | ENSMUSG00000024107.6 |
| chr17 | 88792266 | 88792407 | 3.6453 | 4.44734 | 1.45894 | Promoter | 17 | 88741549 | 88791976 | 50428 | 2 | ENSMUST00000024916.5 | -291 | Lhcgr | ENSMUSG00000024107.6 |

| | | | | | | | | | | | | | | | |
|---|---|---|---|---|---|---|---|---|---|---|---|---|---|---|---|
| chr17 | 90456126 | 90456445 | 4.18275 | 5.51548 | 2.10505 | Promoter | 17 | 90037077 | 90455886 | 418810 | 2 | ENSMUST00000172466.2 | -241 | Nrxn1 | ENSMUSG00000024109.13 |
| chr17 | 90456825 | 90457018 | 3.8853 | 4.85569 | 1.65302 | Promoter | 17 | 90037077 | 90455886 | 418810 | 2 | ENSMUST00000172466.2 | -940 | Nrxn1 | ENSMUSG00000024109.13 |
| chr17 | 25400421 | 25400582 | 3.64834 | 4.40436 | 1.35917 | Intron (ENSMUST00000159610.2/ENSMUSG00000024112.11, intron 2 of 34) | 17 | 25388240 | 25389472 | 1233 | 2 | ENSMUST00000162820.1 | -10950 | Cacna1h | ENSMUSG00000024112.11 |
| chr17 | 25430148 | 25430354 | 3.40859 | 4.03847 | 1.14656 | Intron (ENSMUST00000159610.2/ENSMUSG00000024112.11, intron 2 of 34) | 17 | 25397625 | 25433278 | 35654 | 2 | ENSMUST00000159623.1 | 2924 | Cacna1h | ENSMUSG00000024112.11 |
| chr17 | 25431060 | 25431832 | 3.51592 | 4.15656 | 1.19137 | Intron (ENSMUST00000159610.2/ENSMUSG00000024112.11, intron 2 of 34) | 17 | 25397625 | 25433278 | 35654 | 2 | ENSMUST00000159623.1 | 1446 | Cacna1h | ENSMUSG00000024112.11 |
| chr17 | 25433184 | 25434779 | 4.07777 | 5.42689 | 2.10069 | Promoter | 17 | 25374285 | 25433783 | 59499 | 2 | ENSMUST00000159610.2 | 0 | Cacna1h | ENSMUSG00000024112.11 |
| chr17 | 25451816 | 25451892 | 5.00817 | 14.40749 | 9.82373 | Distal Intergenic | 17 | 25431839 | 25433783 | 1945 | 2 | ENSMUST00000159052.1 | -18034 | Cacna1h | ENSMUSG00000024112.11 |
| chr17 | 84860618 | 84860810 | 3.74206 | 4.63761 | 1.55102 | Distal Intergenic | 17 | 84766428 | 84790789 | 24362 | 2 | ENSMUST00000161928.2 | -69830 | Lrpprc | ENSMUSG00000024120.7 |
| chr17 | 23981954 | 23982193 | 3.49004 | 4.15948 | 1.23193 | Distal Intergenic | 17 | 23972126 | 23975230 | 3105 | 2 | ENSMUST00000024936.4 | -6725 | Prss30 | ENSMUSG00000024124.4 |
| chr17 | 85031290 | 85031430 | 3.23338 | 3.67724 | 1.01733 | Intron (ENSMUST00000024944.7/ENSMUSG00000024131.7, intron 1 of 9) | 17 | 85028347 | 85064243 | 35897 | 1 | ENSMUST00000024944.7 | 2944 | Slc3a1 | ENSMUSG00000024131.7 |
| chr17 | 85659610 | 85659897 | 3.33615 | 3.83157 | 1.01733 | Distal Intergenic | 17 | 85684282 | 85688254 | 3973 | 2 | ENSMUST00000024947.7 | 28357 | Six2 | ENSMUSG00000024134.11 |
| chr17 | 85662577 | 85662832 | 3.5866 | 4.30482 | 1.28105 | Distal Intergenic | 17 | 85684282 | 85688254 | 3973 | 2 | ENSMUST00000024947.7 | 25422 | Six2 | ENSMUSG00000024134.11 |
| chr17 | 85663638 | 85664249 | 3.4341 | 4.00943 | 1.12223 | Distal Intergenic | 17 | 85684282 | 85688254 | 3973 | 2 | ENSMUST00000024947.7 | 24005 | Six2 | ENSMUSG00000024134.11 |
| chr17 | 85665352 | 85665827 | 4.61722 | 6.39741 | 2.84157 | Distal Intergenic | 17 | 85684282 | 85688254 | 3973 | 2 | ENSMUST00000024947.7 | 22427 | Six2 | ENSMUSG00000024134.11 |
| chr17 | 85677193 | 85677299 | 4.01731 | 5.26884 | 1.95251 | Distal Intergenic | 17 | 85684282 | 85688254 | 3973 | 2 | ENSMUST00000024947.7 | 10955 | Six2 | ENSMUSG00000024134.11 |
| chr17 | 85678608 | 85679216 | 3.53876 | 4.42858 | 1.41757 | Distal Intergenic | 17 | 85684282 | 85688254 | 3973 | 2 | ENSMUST00000024947.7 | 9038 | Six2 | ENSMUSG00000024134.11 |
| chr17 | 85679656 | 85679967 | 3.46074 | 4.06011 | 1.15689 | Distal Intergenic | 17 | 85684282 | 85688254 | 3973 | 2 | ENSMUST00000024947.7 | 8287 | Six2 | ENSMUSG00000024134.11 |
| chr17 | 85680642 | 85681184 | 4.18838 | 5.44322 | 2.0616 | Distal Intergenic | 17 | 85684282 | 85688254 | 3973 | 2 | ENSMUST00000024947.7 | 7070 | Six2 | ENSMUSG00000024134.11 |
| chr17 | 85682429 | 85682645 | 4.151 | 5.53622 | 2.1589 | Downstream (1-2kb) | 17 | 85684282 | 85688254 | 3973 | 2 | ENSMUST00000024947.7 | 5609 | Six2 | ENSMUSG00000024134.11 |
| chr17 | 85683032 | 85684407 | 3.75181 | 4.6424 | 1.52687 | 3' UTR | 17 | 85684282 | 85688254 | 3973 | 2 | ENSMUST00000024947.7 | 3847 | Six2 | ENSMUSG00000024134.11 |
| chr17 | 85684972 | 85689411 | 3.97431 | 5.17675 | 1.91117 | Promoter | 17 | 85684277 | 85688274 | 3998 | 2 | ENSMUST00000163568.3 | 0 | Six2 | ENSMUSG00000024134.11 |
| chr17 | 85690207 | 85691258 | 4.10086 | 5.33608 | 2.03419 | Distal Intergenic | 17 | 85684277 | 85688274 | 3998 | 2 | ENSMUST00000163568.3 | -1934 | Six2 | ENSMUSG00000024134.11 |
| chr17 | 85691885 | 85692771 | 3.83077 | 4.87967 | 1.73183 | Distal Intergenic | 17 | 85684277 | 85688274 | 3998 | 2 | ENSMUST00000163568.3 | -3612 | Six2 | ENSMUSG00000024134.11 |
| chr17 | 85693235 | 85693384 | 3.60123 | 4.39812 | 1.40742 | Distal Intergenic | 17 | 85684277 | 85688274 | 3998 | 2 | ENSMUST00000163568.3 | -4962 | Six2 | ENSMUSG00000024134.11 |
| chr17 | 85693933 | 85694205 | 3.886 | 4.91801 | 1.76959 | Distal Intergenic | 17 | 85684277 | 85688274 | 3998 | 2 | ENSMUST00000163568.3 | -5660 | Six2 | ENSMUSG00000024134.11 |
| chr17 | 85694558 | 85694900 | 3.85832 | 5.06576 | 1.81836 | Distal Intergenic | 17 | 85684277 | 85688274 | 3998 | 2 | ENSMUST00000163568.3 | -6285 | Six2 | ENSMUSG00000024134.11 |
| chr17 | 85737698 | 85738560 | 3.74071 | 4.62443 | 1.51587 | Distal Intergenic | 17 | 85684277 | 85688274 | 3998 | 2 | ENSMUST00000163568.3 | -49425 | Six2 | ENSMUSG00000024134.11 |
| chr17 | 85744955 | 85745494 | 3.36928 | 3.90889 | 1.11082 | Distal Intergenic | 17 | 85684277 | 85688274 | 3998 | 2 | ENSMUST00000163568.3 | -56682 | Six2 | ENSMUSG00000024134.11 |
| chr17 | 85747143 | 85748080 | 3.52956 | 4.2008 | 1.27401 | Distal Intergenic | 17 | 85684277 | 85688274 | 3998 | 2 | ENSMUST00000163568.3 | -58870 | Six2 | ENSMUSG00000024134.11 |
| chr17 | 85748625 | 85750195 | 3.83196 | 4.75924 | 1.57961 | Distal Intergenic | 17 | 85684277 | 85688274 | 3998 | 2 | ENSMUST00000163568.3 | -60352 | Six2 | ENSMUSG00000024134.11 |
| chr17 | 87745430 | 87746066 | 4.1203 | 5.46682 | 2.08226 | 3' UTR | 17 | 87696541 | 87717678 | 21138 | 1 | ENSMUST00000174703.1 | 48890 | Msh2 | ENSMUSG00000024151.8 |
| chr17 | 87746658 | 87747179 | 3.34436 | 4.08729 | 1.21798 | Exon (ENSMUST00000055221.7/ENSMUSG00000050138.7, exon 2 of 2) | 17 | 87696541 | 87717678 | 21138 | 1 | ENSMUST00000174703.1 | 50118 | Msh2 | ENSMUSG00000024151.8 |
| chr17 | 57224296 | 57224378 | 3.64866 | 4.38741 | 1.31396 | Exon (ENSMUST00000024988.9/ENSMUSG00000024164.10, exon 9 of 41) | 17 | 57204007 | 57222827 | 18821 | 2 | ENSMUST00000177425.1 | -1470 | C3 | ENSMUSG00000024164.10 |
| chr17 | 55445058 | 55447147 | 4.12753 | 5.50421 | 2.26329 | Promoter | 17 | 55445382 | 55483032 | 37651 | 1 | ENSMUST00000133899.1 | 0 | St6gal2 | ENSMUSG00000024172.5 |
| chr17 | 55447569 | 55447655 | 4.48604 | 6.05082 | 2.50125 | Intron (ENSMUST00000133899.1/ENSMUSG00000024172.5, intron 1 of 1) | 17 | 55446244 | 55482368 | 36125 | 1 | ENSMUST00000153220.1 | 1326 | St6gal2 | ENSMUSG00000024172.5 |
| chr17 | 25471498 | 25471992 | 3.60861 | 4.38985 | 1.40532 | Promoter | 17 | 25471590 | 25476594 | 5005 | 1 | ENSMUST00000025002.1 | 0 | Tekt4 | ENSMUSG00000024175.1 |
| chr17 | 26111123 | 26111556 | 3.72035 | 4.57976 | 1.46508 | Distal Intergenic | 17 | 26113316 | 26123254 | 9939 | 1 | ENSMUST00000025010.8 | -1760 | Tmem8 | ENSMUSG00000024180.8 |
| chr17 | 26204907 | 26205082 | 3.37708 | 4.06947 | 1.17388 | Exon (ENSMUST00000152676.2/ENSMUSG00000024186.10, exon 7 of 17) | 17 | 26203012 | 26204097 | 1086 | 1 | ENSMUST00000123670.1 | 1896 | Rgs11 | ENSMUSG00000024186.10 |
| chr6 | 28132131 | 28132210 | 3.90157 | 4.93465 | 1.72667 | Promoter | 6 | 28126064 | 28133129 | 7066 | 2 | ENSMUST00000132755.1 | 919 | Grm8 | ENSMUSG00000024211.10 |
| chr6 | 28133236 | 28133356 | 3.69137 | 4.52616 | 1.51803 | Promoter | 6 | 27761343 | 28133325 | 371983 | 2 | ENSMUST00000131897.1 | 0 | Grm8 | ENSMUSG00000024211.10 |
| chr6 | 28133816 | 28133917 | 3.14654 | 3.9756 | 1.10041 | Promoter | 6 | 27761343 | 28133325 | 371983 | 2 | ENSMUST00000131897.1 | -492 | Grm8 | ENSMUSG00000024211.10 |
| chr6 | 28134295 | 28134399 | 4.03418 | 5.17631 | 1.92811 | Promoter | 6 | 27275121 | 28134545 | 859425 | 2 | ENSMUST00000115323.2 | 146 | Grm8 | ENSMUSG00000024211.10 |
| chr6 | 28135112 | 28135312 | 3.73894 | 4.74097 | 1.5874 | Promoter | 6 | 27934377 | 28135178 | 200802 | 2 | ENSMUST00000146727.1 | 0 | Grm8 | ENSMUSG00000024211.10 |
| chr6 | 28135622 | 28136111 | 3.97575 | 5.20409 | 1.95212 | Promoter | 6 | 27934377 | 28135178 | 200802 | 2 | ENSMUST00000146727.1 | -445 | Grm8 | ENSMUSG00000024211.10 |
| chr6 | 28140785 | 28140956 | 3.40288 | 4.02986 | 1.19458 | Distal Intergenic | 6 | 27934377 | 28135178 | 200802 | 2 | ENSMUST00000146727.1 | -5608 | Grm8 | ENSMUSG00000024211.10 |
| chr17 | 27706899 | 27707286 | 3.48522 | 4.09246 | 1.14266 | Exon (ENSMUST00000114873.2/ENSMUSG00000040276.9, exon 7 of 10) | 17 | 27714505 | 27717582 | 3078 | 2 | ENSMUST00000127622.1 | 10296 | Spdef | ENSMUSG00000024215.8 |
| chr17 | 27708214 | 27708544 | 3.37091 | 3.88398 | 1.01733 | 3' UTR | 17 | 27714505 | 27717582 | 3078 | 2 | ENSMUST00000127622.1 | 9038 | Spdef | ENSMUSG00000024215.8 |
| chr17 | 27713630 | 27714158 | 3.56706 | 4.25694 | 1.28174 | Downstream (<1kb) | 17 | 27714505 | 27717582 | 3078 | 2 | ENSMUST00000127622.1 | 3424 | Spdef | ENSMUSG00000024215.8 |
| chr17 | 28510914 | 28511252 | 3.43375 | 4.00926 | 1.12532 | Intron (ENSMUST00000114792.2/ENSMUSG00000024222.11, intron 1 of 13) | 17 | 28497343 | 28507406 | 10064 | 2 | ENSMUST00000147716.1 | -3509 | Fkbp5 | ENSMUSG00000024222.11 |
| chr18 | 3499055 | 3499262 | 3.6875 | 4.69055 | 1.58669 | Distal Intergenic | 18 | 3507957 | 3516404 | 8448 | 1 | ENSMUST00000025075.1 | -8695 | Bambi | ENSMUSG00000024232.1 |
| chr17 | 80944866 | 80945143 | 3.42078 | 4.25052 | 1.30021 | Promoter | 17 | 80944632 | 81001816 | 57185 | 1 | ENSMUST00000025092.4 | 235 | Tmem178 | ENSMUSG00000024245.4 |
| chr17 | 84675791 | 84675866 | 3.85368 | 4.95625 | 1.74983 | Promoter | 17 | 84676302 | 84699333 | 23032 | 1 | ENSMUST00000170725.2 | -436 | Abcg8 | ENSMUSG00000024254.10 |
| chr17 | 84676248 | 84676599 | 3.63309 | 4.71627 | 1.57287 | Promoter | 17 | 84676302 | 84699333 | 23032 | 1 | ENSMUST00000170725.2 | 0 | Abcg8 | ENSMUSG00000024254.10 |
| chr17 | 93198996 | 93199393 | 3.60124 | 4.31821 | 1.31324 | Promoter | 17 | 93199422 | 93205485 | 6064 | 1 | ENSMUST00000064775.6 | -29 | Adcyap1 | ENSMUSG00000024256.6 |
| chr17 | 93199925 | 93200556 | 3.4734 | 4.3786 | 1.37437 | Promoter | 17 | 93199422 | 93205485 | 6064 | 1 | ENSMUST00000064775.6 | 504 | Adcyap1 | ENSMUSG00000024256.6 |
| chr17 | 93200973 | 93201121 | 3.35122 | 3.86099 | 1.04089 | Intron (ENSMUST00000064775.6/ENSMUSG00000024256.6, intron 2 of 4) | 17 | 93199422 | 93205485 | 6064 | 1 | ENSMUST00000064775.6 | 1552 | Adcyap1 | ENSMUSG00000024256.6 |
| chr17 | 93201513 | 93202576 | 3.86829 | 4.91878 | 1.74138 | Exon (ENSMUST00000064775.6/ENSMUSG00000024256.6, exon 3 of 5) | 17 | 93199422 | 93205485 | 6064 | 1 | ENSMUST00000064775.6 | 2092 | Adcyap1 | ENSMUSG00000024256.6 |
| chr17 | 93202918 | 93203198 | 3.85035 | 4.79888 | 1.65437 | Intron (ENSMUST00000064775.6/ENSMUSG00000024256.6, intron 4 of 4) | 17 | 93199422 | 93205485 | 6064 | 1 | ENSMUST00000064775.6 | 3497 | Adcyap1 | ENSMUSG00000024256.6 |
| chr17 | 93203664 | 93204154 | 4.01388 | 5.11679 | 1.85838 | 3' UTR | 17 | 93199422 | 93205485 | 6064 | 1 | ENSMUST00000064775.6 | 4243 | Adcyap1 | ENSMUSG00000024256.6 |
| chr17 | 93204591 | 93205391 | 3.61663 | 4.43938 | 1.42357 | 3' UTR | 17 | 93199422 | 93205485 | 6064 | 1 | ENSMUST00000064775.6 | 5170 | Adcyap1 | ENSMUSG00000024256.6 |
| chr17 | 93205819 | 93206379 | 3.92927 | 5.0083 | 1.80767 | Distal Intergenic | 17 | 93199422 | 93205485 | 6064 | 1 | ENSMUST00000064775.6 | 6398 | Adcyap1 | ENSMUSG00000024256.6 |
| chr18 | 31447407 | 31447502 | 3.63803 | 4.49858 | 1.4543 | Promoter | 18 | 31437808 | 31447415 | 9608 | 2 | ENSMUST00000025110.3 | 0 | Syt4 | ENSMUSG00000024261.5 |
| chr18 | 25676909 | 25677101 | 3.706 | 4.53276 | 1.44977 | Intron (ENSMUST00000025117.7/ENSMUSG00000024268.10, intron 2 of 12) | 18 | 25477632 | 25753983 | 276352 | 2 | ENSMUST00000025117.7 | 76882 | Celf4 | ENSMUSG00000024268.10 |
| chr18 | 25677725 | 25677816 | 3.92096 | 5.02452 | 1.759 | Intron (ENSMUST00000025117.7/ENSMUSG00000024268.10, intron 2 of 12) | 18 | 25477632 | 25753983 | 276352 | 2 | ENSMUST00000025117.7 | 76167 | Celf4 | ENSMUSG00000024268.10 |
| chr18 | 25678707 | 25679558 | 3.91667 | 4.97363 | 1.7234 | Exon (ENSMUST00000025117.7/ENSMUSG00000024268.10, exon 2 of 13) | 18 | 25477632 | 25753983 | 276352 | 2 | ENSMUST00000025117.7 | 74425 | Celf4 | ENSMUSG00000024268.10 |
| chr18 | 25682860 | 25682951 | 3.57827 | 4.37313 | 1.37904 | Intron (ENSMUST00000025117.7/ENSMUSG00000024268.10, intron 1 of 12) | 18 | 25477632 | 25753983 | 276352 | 2 | ENSMUST00000025117.7 | 71032 | Celf4 | ENSMUSG00000024268.10 |
| chr18 | 25715256 | 25715626 | 3.52123 | 4.19142 | 1.29521 | Intron (ENSMUST00000025117.7/ENSMUSG00000024268.10, intron 1 of 12) | 18 | 25477632 | 25753983 | 276352 | 2 | ENSMUST00000025117.7 | 38357 | Celf4 | ENSMUSG00000024268.10 |
| chr18 | 25753174 | 25753627 | 3.61905 | 4.45742 | 1.45628 | Promoter | 18 | 25477632 | 25753983 | 276352 | 2 | ENSMUST00000025117.7 | 356 | Celf4 | ENSMUSG00000024268.10 |
| chr18 | 25754155 | 25754364 | 3.30157 | 3.77954 | 1.01733 | Promoter | 18 | 25477632 | 25753983 | 276352 | 2 | ENSMUST00000115816.2 | -173 | Celf4 | ENSMUSG00000024268.10 |
| chr18 | 25754776 | 25755470 | 3.99235 | 5.12013 | 1.89827 | Promoter | 18 | 25477632 | 25753983 | 276352 | 2 | ENSMUST00000115816.2 | -794 | Celf4 | ENSMUSG00000024268.10 |
| chr18 | 24466370 | 24466533 | 3.31114 | 3.79393 | 1.01733 | Distal Intergenic | 18 | 24470861 | 24477765 | 6905 | 1 | ENSMUST00000097646.3 | -4328 | 2700062C07Rik | ENSMUSG00000024273.7 |
| chr18 | 23310224 | 23311368 | 4.39503 | 5.95649 | 2.5353 | Distal Intergenic | 18 | 23415415 | 23604230 | 188816 | 1 | ENSMUST00000047954.8 | -104047 | Dtna | ENSMUSG00000024302.10 |
| chr18 | 16806852 | 16807638 | 3.9994 | 5.25501 | 1.9744 | Promoter | 18 | 16771397 | 16808462 | 37066 | 2 | ENSMUST00000152779.1 | 824 | Cdh2 | ENSMUSG00000024304.9 |
| chr18 | 16808208 | 16808516 | 3.50776 | 4.19325 | 1.32357 | Promoter | 18 | 16771397 | 16808462 | 37066 | 2 | ENSMUST00000152779.1 | 0 | Cdh2 | ENSMUSG00000024304.9 |
| chr18 | 16809464 | 16809857 | 4.14111 | 5.51412 | 2.1621 | Promoter | 18 | 16588877 | 16809246 | 220370 | 2 | ENSMUST00000025166.8 | -219 | Cdh2 | ENSMUSG00000024304.9 |
| chr18 | 16810685 | 16810760 | 4.12951 | 5.45802 | 2.08377 | Distal Intergenic | 18 | 16588877 | 16809246 | 220370 | 2 | ENSMUST00000025166.8 | -1440 | Cdh2 | ENSMUSG00000024304.9 |
| chr17 | 34039319 | 34039531 | 3.35164 | 3.85505 | 1.01733 | Promoter | 17 | 34039437 | 34052517 | 13081 | 1 | ENSMUST00000131134.2 | 0 | Col11a2 | ENSMUSG00000024330.11 |
| chr17 | 34042637 | 34042894 | 3.78595 | 4.80946 | 1.64026 | Intron (ENSMUST00000131134.2/ENSMUSG00000024330.11, intron 3 of 20) | 17 | 34039437 | 34066685 | 27249 | 1 | ENSMUST00000087497.5 | 3201 | Col11a2 | ENSMUSG00000024330.11 |
| chr18 | 20059127 | 20059369 | 3.81972 | 4.79794 | 1.63193 | Promoter | 18 | 20030633 | 20059478 | 28846 | 2 | ENSMUST00000075214.3 | 109 | Dsc2 | ENSMUSG00000024331.6 |
| chr18 | 35964850 | 35965820 | 3.80755 | 4.70374 | 1.54564 | Promoter | 18 | 35965065 | 36014712 | 49648 | 1 | ENSMUST00000176873.2 | 0 | Psd2 | ENSMUSG00000024347.11 |
| chr18 | 35966370 | 35966445 | 4.0404 | 5.13883 | 1.81875 | Intron (ENSMUST00000115716.3/ENSMUSG00000024347.11, intron 1 of 14) | 18 | 35965105 | 36014674 | 49570 | 1 | ENSMUST00000175734.1 | 1266 | Psd2 | ENSMUSG00000024347.11 |
| chr18 | 36004101 | 36004234 | 3.44251 | 4.02061 | 1.11859 | Intron (ENSMUST00000115716.3/ENSMUSG00000024347.11, intron 8 of 14) | 18 | 35987019 | 36006084 | 19066 | 1 | ENSMUST00000175720.1 | 17083 | Psd2 | ENSMUSG00000024347.11 |
| chr18 | 36066296 | 36066611 | 3.48958 | 4.10224 | 1.15531 | Intron (ENSMUST00000115712.2/ENSMUSG00000060275.6, intron 1 of 9) | 18 | 35987019 | 36006084 | 19066 | 1 | ENSMUST00000175720.1 | 79278 | Psd2 | ENSMUSG00000024347.11 |
| chr18 | 36066993 | 36067189 | 3.65018 | 4.51716 | 1.47238 | Intron (ENSMUST00000115712.2/ENSMUSG00000060275.6, intron 1 of 9) | 18 | 35987019 | 36006084 | 19066 | 1 | ENSMUST00000175720.1 | 79975 | Psd2 | ENSMUSG00000024347.11 |
| chr18 | 36068192 | 36068921 | 3.42528 | 4.03571 | 1.17886 | Intron (ENSMUST00000115712.2/ENSMUSG00000060275.6, intron 1 of 9) | 18 | 35987019 | 36006084 | 19066 | 1 | ENSMUST00000175720.1 | 81174 | Psd2 | ENSMUSG00000024347.11 |
| chr18 | 36069306 | 36069405 | 4.26408 | 5.86229 | 2.34768 | Intron (ENSMUST00000115712.2/ENSMUSG00000060275.6, intron 1 of 9) | 18 | 35987019 | 36006084 | 19066 | 1 | ENSMUST00000175720.1 | 82288 | Psd2 | ENSMUSG00000024347.11 |
| chr18 | 34690844 | 34691094 | 3.3718 | 3.88848 | 1.02845 | Exon (ENSMUST00000025224.7/ENSMUSG00000024366.7, exon 7 of 8) | 18 | 34689903 | 34720387 | 30485 | 2 | ENSMUST00000025224.7 | 29293 | Gfra3 | ENSMUSG00000024366.7 |
| chr18 | 34718978 | 34719308 | 3.4481 | 4.02511 | 1.10425 | Intron (ENSMUST00000025224.7/ENSMUSG00000024366.7, intron 1 of 7) | 18 | 34689903 | 34720387 | 30485 | 2 | ENSMUST00000025224.7 | 1079 | Gfra3 | ENSMUSG00000024366.7 |
| chr18 | 34719714 | 34720381 | 3.6354 | 4.40859 | 1.358 | Promoter | 18 | 34689903 | 34720387 | 30485 | 2 | ENSMUST00000025224.7 | 6 | Gfra3 | ENSMUSG00000024366.7 |
| chr18 | 34678003 | 34678516 | 3.41808 | 4.42217 | 1.41263 | Distal Intergenic | 18 | 34631683 | 34651735 | 20053 | 2 | ENSMUST00000133181.1 | -26269 | Cdc23 | ENSMUSG00000024370.11 |
| chr18 | 34679411 | 34679829 | 3.76304 | 4.8043 | 1.63199 | Distal Intergenic | 18 | 34631683 | 34651735 | 20053 | 2 | ENSMUST00000133181.1 | -27677 | Cdc23 | ENSMUSG00000024370.11 |
| chr17 | 34881535 | 34881858 | 3.85651 | 4.80083 | 1.61993 | Promoter | 17 | 34877054 | 34881967 | 4914 | 2 | ENSMUST00000130728.1 | 109 | C2 | ENSMUSG00000024371.9 |
| chr17 | 35490246 | 35490405 | 3.65249 | 4.43775 | 1.44371 | Distal Intergenic | 17 | 35506018 | 35510772 | 4755 | 1 | ENSMUST00000025271.11 | -15613 | Pou5f1 | ENSMUSG00000024406.11 |
| chr17 | 35505674 | 35505971 | 3.6235 | 4.34457 | 1.29146 | Promoter | 17 | 35506018 | 35510772 | 4755 | 1 | ENSMUST00000025271.11 | -47 | Pou5f1 | ENSMUSG00000024406.11 |
| chr18 | 15312436 | 15312985 | 3.80772 | 4.80475 | 1.678 | Distal Intergenic | 18 | 15389394 | 15403684 | 14291 | 2 | ENSMUST00000079081.6 | 90699 | Aqp4 | ENSMUSG00000024411.9 |
| chr18 | 15313711 | 15314707 | 3.94427 | 5.08121 | 1.89417 | Distal Intergenic | 18 | 15389394 | 15403684 | 14291 | 2 | ENSMUST00000079081.6 | 88977 | Aqp4 | ENSMUSG00000024411.9 |
| chr18 | 12643973 | 12644488 | 4.28716 | 5.69578 | 2.3093 | Promoter | 18 | 12643344 | 12737049 | 93706 | 1 | ENSMUST00000025294.7 | 630 | Ttc39c | ENSMUSG00000024424.9 |
| chr18 | 38483300 | 38483408 | 4.33442 | 5.76957 | 2.34579 | Distal Intergenic | 18 | 38418975 | 38464399 | 45425 | 1 | ENSMUST00000025293.3 | 64326 | Ndfip1 | ENSMUSG00000024425.3 |
| chr18 | 37998338 | 37998910 | 3.76714 | 4.64973 | 1.53735 | Promoter | 18 | 37972624 | 37997574 | 24951 | 2 | ENSMUST00000042944.7 | -765 | Arap3 | ENSMUSG00000024451.8 |
| chr18 | 38024262 | 38024392 | 3.7169 | 4.54558 | 1.49171 | Distal Intergenic | 18 | 37972624 | 37997574 | 24951 | 2 | ENSMUST00000042944.7 | -26689 | Arap3 | ENSMUSG00000024451.8 |
| chr18 | 38025668 | 38026474 | 3.52234 | 4.24429 | 1.30774 | Distal Intergenic | 18 | 37972624 | 37997574 | 24951 | 2 | ENSMUST00000042944.7 | -28095 | Arap3 | ENSMUSG00000024451.8 |
| chr18 | 38056808 | 38056995 | 3.47709 | 4.09907 | 1.18039 | Distal Intergenic | 18 | 37972624 | 37997574 | 24951 | 2 | ENSMUST00000042944.7 | -59235 | Arap3 | ENSMUSG00000024451.8 |
| chr18 | 38058794 | 38058911 | 3.44229 | 4.09512 | 1.20619 | Distal Intergenic | 18 | 37972624 | 37997574 | 24951 | 2 | ENSMUST00000042944.7 | -61221 | Arap3 | ENSMUSG00000024451.8 |
| chr18 | 38074644 | 38074814 | 3.40994 | 4.16833 | 1.23632 | Distal Intergenic | 18 | 37972624 | 37997574 | 24951 | 2 | ENSMUST00000042944.7 | -77071 | Arap3 | ENSMUSG00000024451.8 |
| chr18 | 38098245 | 38098704 | 3.43545 | 4.016 | 1.13987 | Distal Intergenic | 18 | 37972624 | 37997574 | 24951 | 2 | ENSMUST00000042944.7 | -100672 | Arap3 | ENSMUSG00000024451.8 |
| chr17 | 36988194 | 36988937 | 3.544 | 4.25811 | 1.31396 | Promoter | 17 | 36987497 | 36989045 | 1549 | 2 | ENSMUST00000113667.3 | 108 | H2-M5 | ENSMUSG00000024459.13 |
| chr17 | 36989349 | 36989610 | 3.61319 | 4.4636 | 1.45077 | Promoter | 17 | 36984061 | 36989537 | 5477 | 2 | ENSMUST00000145260.1 | 0 | H2-M5 | ENSMUSG00000024459.13 |
| chr17 | 37049634 | 37049890 | 3.25408 | 3.70826 | 1.01733 | Promoter | 17 | 37050631 | 37074304 | 23674 | 1 | ENSMUST00000172792.2 | -741 | Gabbr1 | ENSMUSG00000024462.12 |
| chr15 | 54571225 | 54571626 | 4.04149 | 5.20045 | 1.89083 | Promoter | 15 | 54571366 | 54602846 | 31481 | 1 | ENSMUST00000025356.2 | 0 | Mal2 | ENSMUSG00000024479.2 |
| chr18 | 46848840 | 46849069 | 3.88122 | 5.14587 | 1.84952 | Promoter | 18 | 46850039 | 46905446 | 55408 | 1 | ENSMUST00000025358.2 | -970 | 4833403I15Rik | ENSMUSG00000024481.4 |
| chr18 | 46850176 | 46850314 | 3.93687 | 5.0224 | 1.80734 | Promoter | 18 | 46850039 | 46905446 | 55408 | 1 | ENSMUST00000025358.2 | 138 | 4833403I15Rik | ENSMUSG00000024481.4 |
| chr18 | 36530101 | 36530214 | 3.83413 | 4.82051 | 1.68157 | Intron (ENSMUST00000074298.7/ENSMUSG00000024485.10, intron 5 of 19) | 18 | 36528157 | 36541293 | 13137 | 1 | ENSMUST00000115694.2 | 1945 | Slc4a9 | ENSMUSG00000024485.10 |
| chr18 | 36530968 | 36531366 | 3.76326 | 4.63438 | 1.54147 | Exon (ENSMUST00000074298.7/ENSMUSG00000024485.10, exon 8 of 20) | 18 | 36528157 | 36541293 | 13137 | 1 | ENSMUST00000115694.2 | 2812 | Slc4a9 | ENSMUSG00000024485.10 |
| chr18 | 36514084 | 36514159 | 3.36164 | 3.91037 | 1.14767 | Intron (ENSMUST00000025363.5/ENSMUSG00000024486.5, intron 2 of 5) | 18 | 36504929 | 36515805 | 10877 | 2 | ENSMUST00000025363.5 | 1646 | Hbegf | ENSMUSG00000024486.5 |
| chr18 | 42381548 | 42381838 | 3.44461 | 4.05011 | 1.21207 | Distal Intergenic | 18 | 42394539 | 42397249 | 2711 | 1 | ENSMUST00000025374.3 | -12701 | Pou4f3 | ENSMUSG00000024497.3 |
| chr18 | 42385108 | 42385358 | 4.55186 | 6.18628 | 2.60271 | Distal Intergenic | 18 | 42394539 | 42397249 | 2711 | 1 | ENSMUST00000025374.3 | -9181 | Pou4f3 | ENSMUSG00000024497.3 |
| chr18 | 42386059 | 42386422 | 3.70893 | 4.55948 | 1.44426 | Distal Intergenic | 18 | 42394539 | 42397249 | 2711 | 1 | ENSMUST00000025374.3 | -8117 | Pou4f3 | ENSMUSG00000024497.3 |
| chr18 | 42388904 | 42389080 | 3.5462 | 4.20456 | 1.20789 | Distal Intergenic | 18 | 42394539 | 42397249 | 2711 | 1 | ENSMUST00000025374.3 | -5459 | Pou4f3 | ENSMUSG00000024497.3 |
| chr18 | 42392352 | 42392443 | 3.46387 | 4.05461 | 1.12412 | Distal Intergenic | 18 | 42394539 | 42397249 | 2711 | 1 | ENSMUST00000025374.3 | -2096 | Pou4f3 | ENSMUSG00000024497.3 |
| chr18 | 42394404 | 42394582 | 3.57685 | 4.26131 | 1.25226 | Promoter | 18 | 42394539 | 42397249 | 2711 | 1 | ENSMUST00000025374.3 | 0 | Pou4f3 | ENSMUSG00000024497.3 |
| chr18 | 42395415 | 42396217 | 3.79859 | 4.78217 | 1.64432 | Promoter | 18 | 42394539 | 42397249 | 2711 | 1 | ENSMUST00000025374.3 | 877 | Pou4f3 | ENSMUSG00000024497.3 |
| chr18 | 42396596 | 42397089 | 3.45158 | 4.05949 | 1.20325 | 3' UTR | 18 | 42394539 | 42397249 | 2711 | 1 | ENSMUST00000025374.3 | 2058 | Pou4f3 | ENSMUSG00000024497.3 |
| chr18 | 42397430 | 42397938 | 3.61931 | 4.43737 | 1.43327 | Distal Intergenic | 18 | 42394539 | 42397249 | 2711 | 1 | ENSMUST00000025374.3 | 2892 | Pou4f3 | ENSMUSG00000024497.3 |
| chr18 | 42398770 | 42399407 | 4.00039 | 5.09586 | 1.84113 | Distal Intergenic | 18 | 42394539 | 42397249 | 2711 | 1 | ENSMUST00000025374.3 | 4232 | Pou4f3 | ENSMUSG00000024497.3 |
| chr18 | 42400207 | 42401673 | 3.37974 | 4.35643 | 1.3393 | Distal Intergenic | 18 | 42394539 | 42397249 | 2711 | 1 | ENSMUST00000025374.3 | 5669 | Pou4f3 | ENSMUSG00000024497.3 |
| chr18 | 42402429 | 42402703 | 3.46514 | 4.09693 | 1.2535 | Distal Intergenic | 18 | 42394539 | 42397249 | 2711 | 1 | ENSMUST00000025374.3 | 7891 | Pou4f3 | ENSMUSG00000024497.3 |
| chr18 | 42403632 | 42403728 | 4.29012 | 5.66888 | 2.26339 | Distal Intergenic | 18 | 42394539 | 42397249 | 2711 | 1 | ENSMUST00000025374.3 | 9094 | Pou4f3 | ENSMUSG00000024497.3 |
| chr18 | 42416408 | 42416891 | 3.51588 | 4.16099 | 1.2157 | Distal Intergenic | 18 | 42394539 | 42397249 | 2711 | 1 | ENSMUST00000025374.3 | 21870 | Pou4f3 | ENSMUSG00000024497.3 |
| chr18 | 42417447 | 42417522 | 3.86507 | 4.80274 | 1.59422 | Distal Intergenic | 18 | 42394539 | 42397249 | 2711 | 1 | ENSMUST00000025374.3 | 22909 | Pou4f3 | ENSMUSG00000024497.3 |
| chr18 | 42898518 | 42898654 | 4.08779 | 5.33011 | 2.01732 | Promoter | 18 | 42758468 | 42898815 | 140348 | 2 | ENSMUST00000155262.1 | 161 | Ppp2r2b | ENSMUSG00000024500.13 |
| chr18 | 42899096 | 42899413 | 3.49676 | 4.34612 | 1.3559 | Promoter | 18 | 42645269 | 42899294 | 254026 | 2 | ENSMUST00000117687.2 | 0 | Ppp2r2b | ENSMUSG00000024500.13 |
| chr18 | 43390106 | 43390235 | 4.04167 | 5.26196 | 2.00197 | Intron (ENSMUST00000121805.2/ENSMUSG00000024501.14, intron 1 of 13) | 18 | 43324288 | 43393377 | 69090 | 2 | ENSMUST00000025379.8 | 3142 | Dpysl3 | ENSMUSG00000024501.14 |
| chr18 | 43390958 | 43391577 | 3.5385 | 4.20194 | 1.24822 | Intron (ENSMUST00000121805.2/ENSMUSG00000024501.14, intron 1 of 13) | 18 | 43324288 | 43393377 | 69090 | 2 | ENSMUST00000025379.8 | 1800 | Dpysl3 | ENSMUSG00000024501.14 |
| chr18 | 43392684 | 43393123 | 3.68994 | 4.62416 | 1.54062 | Promoter | 18 | 43324288 | 43393377 | 69090 | 2 | ENSMUST00000025379.8 | 254 | Dpysl3 | ENSMUSG00000024501.14 |
| chr18 | 43437705 | 43438491 | 3.58414 | 4.36415 | 1.39161 | Promoter | 18 | 43320979 | 43438286 | 117308 | 2 | ENSMUST00000121805.2 | 0 | Dpysl3 | ENSMUSG00000024501.14 |
| chr18 | 43439177 | 43439438 | 3.46086 | 4.04742 | 1.11493 | Promoter | 18 | 43320979 | 43438286 | 117308 | 2 | ENSMUST00000121805.2 | -892 | Dpysl3 | ENSMUSG00000024501.14 |
| chr18 | 70652753 | 70653173 | 3.63074 | 4.37244 | 1.35588 | Distal Intergenic | 18 | 70617773 | 70625777 | 8005 | 1 | ENSMUST00000128555.1 | 34981 | Mbd2 | ENSMUSG00000024513.11 |
| chr18 | 65871850 | 65872009 | 3.7454 | 4.57402 | 1.45047 | Promoter | 18 | 65872820 | 65886579 | 13760 | 1 | ENSMUST00000173985.3 | -811 | Grp | ENSMUSG00000024517.11 |
| chr18 | 65872555 | 65872914 | 4.67295 | 6.51543 | 2.9131 | Promoter | 18 | 65872820 | 65886579 | 13760 | 1 | ENSMUST00000173985.3 | 0 | Grp | ENSMUSG00000024517.11 |
| chr18 | 65873295 | 65873409 | 3.68138 | 4.46305 | 1.3887 | Promoter | 18 | 65873484 | 65880258 | 6775 | 1 | ENSMUST00000173530.2 | -75 | Grp | ENSMUSG00000024517.11 |
| chr18 | 65874285 | 65874409 | 3.36108 | 3.86915 | 1.01733 | Promoter | 18 | 65873494 | 65886578 | 13085 | 1 | ENSMUST00000025395.8 | 792 | Grp | ENSMUSG00000024517.11 |
| chr18 | 65938804 | 65939276 | 3.50731 | 4.36702 | 1.35518 | Promoter | 18 | 65934639 | 65939089 | 4451 | 2 | ENSMUST00000025396.3 | 0 | Rax | ENSMUSG00000024518.3 |
| chr18 | 65939608 | 65940022 | 3.56039 | 4.5511 | 1.46616 | Promoter | 18 | 65934639 | 65939089 | 4451 | 2 | ENSMUST00000025396.3 | -520 | Rax | ENSMUSG00000024518.3 |
| chr18 | 65942906 | 65943173 | 3.58386 | 4.4468 | 1.4174 | Distal Intergenic | 18 | 65934639 | 65939089 | 4451 | 2 | ENSMUST00000025396.3 | -3818 | Rax | ENSMUSG00000024518.3 |
| chr18 | 65964607 | 65964815 | 3.26835 | 3.81245 | 1.03286 | Intron (ENSMUST00000025397.5/ENSMUSG00000024519.5, intron 2 of 2) | 18 | 65955727 | 65970178 | 14452 | 2 | ENSMUST00000025397.5 | 5363 | Cplx4 | ENSMUSG00000024519.5 |
| chr18 | 66453361 | 66453931 | 3.30118 | 3.77896 | 1.01733 | Distal Intergenic | 18 | 66458604 | 66465550 | 6947 | 1 | ENSMUST00000025399.7 | -4673 | Pmaip1 | ENSMUSG00000024521.7 |
| chr18 | 66457320 | 66457649 | 3.411 | 3.99172 | 1.1803 | Promoter | 18 | 66458604 | 66465550 | 6947 | 1 | ENSMUST00000025399.7 | -955 | Pmaip1 | ENSMUSG00000024521.7 |
| chr18 | 66458465 | 66458854 | 3.80695 | 5.26614 | 2.00501 | Promoter | 18 | 66458604 | 66465550 | 6947 | 1 | ENSMUST00000025399.7 | 0 | Pmaip1 | ENSMUSG00000024521.7 |
| chr18 | 66459191 | 66459982 | 3.72688 | 4.69775 | 1.60695 | Promoter | 18 | 66458604 | 66465550 | 6947 | 1 | ENSMUST00000025399.7 | 588 | Pmaip1 | ENSMUSG00000024521.7 |
| chr18 | 66460736 | 66461225 | 3.825 | 4.89051 | 1.72241 | Exon (ENSMUST00000025399.7/ENSMUSG00000024521.7, exon 2 of 3) | 18 | 66458604 | 66465550 | 6947 | 1 | ENSMUST00000025399.7 | 2133 | Pmaip1 | ENSMUSG00000024521.7 |
| chr18 | 66461948 | 66462514 | 3.58102 | 4.37034 | 1.39995 | Intron (ENSMUST00000025399.7/ENSMUSG00000024521.7, intron 2 of 2) | 18 | 66458604 | 66465550 | 6947 | 1 | ENSMUST00000025399.7 | 3345 | Pmaip1 | ENSMUSG00000024521.7 |
| chr18 | 66463093 | 66463185 | 3.99677 | 5.16006 | 1.91813 | Intron (ENSMUST00000025399.7/ENSMUSG00000024521.7, intron 2 of 2) | 18 | 66458604 | 66465550 | 6947 | 1 | ENSMUST00000025399.7 | 4490 | Pmaip1 | ENSMUSG00000024521.7 |
| chr18 | 66464252 | 66464887 | 3.49531 | 4.30676 | 1.35449 | 3' UTR | 18 | 66458604 | 66465550 | 6947 | 1 | ENSMUST00000025399.7 | 5649 | Pmaip1 | ENSMUSG00000024521.7 |
| chr18 | 66470800 | 66470981 | 3.46517 | 4.15615 | 1.24544 | Distal Intergenic | 18 | 66458604 | 66465550 | 6947 | 1 | ENSMUST00000025399.7 | 12197 | Pmaip1 | ENSMUSG00000024521.7 |
| chr18 | 66471793 | 66472409 | 4.34334 | 5.89996 | 2.4753 | Distal Intergenic | 18 | 66458604 | 66465550 | 6947 | 1 | ENSMUST00000025399.7 | 13190 | Pmaip1 | ENSMUSG00000024521.7 |
| chr18 | 66475513 | 66475644 | 3.7862 | 4.65567 | 1.50776 | Distal Intergenic | 18 | 66458604 | 66465550 | 6947 | 1 | ENSMUST00000025399.7 | 16910 | Pmaip1 | ENSMUSG00000024521.7 |
| chr18 | 66476212 | 66476288 | 4.32272 | 5.69364 | 2.20847 | Distal Intergenic | 18 | 66458604 | 66465550 | 6947 | 1 | ENSMUST00000025399.7 | 17609 | Pmaip1 | ENSMUSG00000024521.7 |
| chr18 | 67007064 | 67007228 | 3.31633 | 3.80175 | 1.01733 | Distal Intergenic | 18 | 67088336 | 67226792 | 138457 | 1 | ENSMUST00000025402.8 | -81108 | Gnal | ENSMUSG00000024524.11 |
| chr18 | 67058902 | 67058985 | 3.71151 | 4.91938 | 1.71373 | Distal Intergenic | 18 | 67088336 | 67226792 | 138457 | 1 | ENSMUST00000025402.8 | -29351 | Gnal | ENSMUSG00000024524.11 |
| chr18 | 67086836 | 67087744 | 3.64972 | 4.58813 | 1.53327 | Promoter | 18 | 67088336 | 67226792 | 138457 | 1 | ENSMUST00000025402.8 | -592 | Gnal | ENSMUSG00000024524.11 |
| chr18 | 67088444 | 67088649 | 4.06193 | 5.35475 | 2.02062 | Promoter | 18 | 67088336 | 67226792 | 138457 | 1 | ENSMUST00000025402.8 | 109 | Gnal | ENSMUSG00000024524.11 |
| chr18 | 67089155 | 67089725 | 3.73586 | 4.79886 | 1.66751 | Promoter | 18 | 67088336 | 67226792 | 138457 | 1 | ENSMUST00000025402.8 | 820 | Gnal | ENSMUSG00000024524.11 |
| chr18 | 67121199 | 67121576 | 3.52054 | 4.16628 | 1.21755 | Intron (ENSMUST00000025402.8/ENSMUSG00000024524.11, intron 1 of 11) | 18 | 67133826 | 67226792 | 92967 | 1 | ENSMUST00000076605.7 | -12250 | Gnal | ENSMUSG00000024524.11 |

| | | | | | | | | | | | | | | | |
|---|---|---|---|---|---|---|---|---|---|---|---|---|---|---|---|
| chr18 | 67131193 | 67131545 | 3.42015 | 3.99327 | 1.13665 | Intron (ENSMUST00000025402.8/ENSMUSG00000024524.11, intron 1 of 11) | 18 | 67133826 | 67226792 | 92967 | 1 | ENSMUST00000076605.7 | -2281 | Gnal | ENSMUSG00000024524.11 |
| chr18 | 67132302 | 67135516 | 3.90043 | 4.99182 | 1.81155 | Promoter | 18 | 67133826 | 67226792 | 92967 | 1 | ENSMUST00000076605.7 | 0 | Gnal | ENSMUSG00000024524.11 |
| chr18 | 67136116 | 67136394 | 3.52195 | 4.60168 | 1.51866 | Intron (ENSMUST00000025402.8/ENSMUSG00000024524.11, intron 4 of 11) | 18 | 67133826 | 67226792 | 92967 | 1 | ENSMUST00000076605.7 | 2291 | Gnal | ENSMUSG00000024524.11 |
| chr18 | 67137613 | 67137838 | 3.25048 | 3.70286 | 1.01733 | Intron (ENSMUST00000025402.8/ENSMUSG00000024524.11, intron 4 of 11) | 18 | 67133826 | 67226792 | 92967 | 1 | ENSMUST00000076605.7 | 3788 | Gnal | ENSMUSG00000024524.11 |
| chr18 | 52528675 | 52528846 | 3.40856 | 3.9617 | 1.09126 | Promoter | 18 | 52517238 | 52529708 | 12471 | 2 | ENSMUST00000025409.7 | 862 | Lox | ENSMUSG00000024529.8 |
| chr18 | 52529166 | 52529259 | 4.57661 | 6.4969 | 2.8138 | Promoter | 18 | 52517238 | 52529708 | 12471 | 2 | ENSMUST00000025409.7 | 449 | Lox | ENSMUSG00000024529.8 |
| chr18 | 79109256 | 79109377 | 3.76384 | 4.71976 | 1.57706 | Promoter | 18 | 78750380 | 79109391 | 359012 | 2 | ENSMUST00000025430.9 | 14 | Setbp1 | ENSMUSG00000024548.10 |
| chr18 | 79111362 | 79111709 | 3.91294 | 5.02845 | 1.78481 | Distal Intergenic | 18 | 78750380 | 79109391 | 359012 | 2 | ENSMUST00000161465.2 | -1972 | Setbp1 | ENSMUSG00000024548.10 |
| chr18 | 82405524 | 82405941 | 3.36045 | 3.97173 | 1.13032 | Promoter | 18 | 82392496 | 82406777 | 14282 | 2 | ENSMUST00000065224.6 | 836 | Galr1 | ENSMUSG00000024553.7 |
| chr18 | 82406818 | 82407170 | 4.09825 | 5.2772 | 1.94578 | Promoter | 18 | 82392496 | 82406777 | 14282 | 2 | ENSMUST00000065224.6 | -42 | Galr1 | ENSMUSG00000024553.7 |
| chr18 | 82407700 | 82407834 | 3.37032 | 3.88308 | 1.01733 | Promoter | 18 | 82392496 | 82406777 | 14282 | 2 | ENSMUST00000065224.6 | -924 | Galr1 | ENSMUSG00000024553.7 |
| chr18 | 74063055 | 74063343 | 3.54429 | 4.25803 | 1.34588 | Intron (ENSMUST00000162863.2/ENSMUSG00000024558.7, intron 1 of 4) | 18 | 73928486 | 74064392 | 135907 | 2 | ENSMUST00000162863.2 | 1049 | Mapk4 | ENSMUSG00000024558.7 |
| chr18 | 74063677 | 74064843 | 3.82588 | 4.75182 | 1.61484 | Promoter | 18 | 73928486 | 74064392 | 135907 | 2 | ENSMUST00000162863.2 | 0 | Mapk4 | ENSMUSG00000024558.7 |
| chr18 | 74065230 | 74066168 | 3.68481 | 4.49478 | 1.46587 | Promoter | 18 | 73971170 | 74065359 | 94190 | 2 | ENSMUST00000160601.1 | 0 | Mapk4 | ENSMUSG00000024558.7 |
| chr18 | 76379766 | 76380248 | 3.51057 | 4.13544 | 1.16058 | Distal Intergenic | 18 | 76262492 | 76302642 | 40151 | 1 | ENSMUST00000113930.2 | 117275 | Smad2 | ENSMUSG00000024563.10 |
| chr18 | 76427707 | 76428059 | 3.50533 | 4.19047 | 1.31006 | Distal Intergenic | 18 | 76262492 | 76302642 | 40151 | 1 | ENSMUST00000113930.2 | 165216 | Smad2 | ENSMUSG00000024563.10 |
| chr18 | 76533348 | 76533981 | 3.783 | 4.71839 | 1.56594 | Distal Intergenic | 18 | 76262492 | 76302642 | 40151 | 1 | ENSMUST00000113930.2 | 270857 | Smad2 | ENSMUSG00000024563.10 |
| chr18 | 76543062 | 76543949 | 3.62214 | 4.54884 | 1.49096 | Distal Intergenic | 18 | 76262492 | 76302642 | 40151 | 1 | ENSMUST00000113930.2 | 280571 | Smad2 | ENSMUSG00000024563.10 |
| chr18 | 76545222 | 76546181 | 3.38871 | 3.93027 | 1.07954 | Distal Intergenic | 18 | 76262492 | 76302642 | 40151 | 1 | ENSMUST00000113930.2 | 282731 | Smad2 | ENSMUSG00000024563.10 |
| chr18 | 76549743 | 76549846 | 3.78121 | 4.66232 | 1.54176 | Distal Intergenic | 18 | 76262492 | 76302642 | 40151 | 1 | ENSMUST00000113930.2 | 287252 | Smad2 | ENSMUSG00000024563.10 |
| chr18 | 80973855 | 80975006 | 3.65669 | 4.4706 | 1.38649 | Exon (ENSMUST00000057950.7/ENSMUSG00000024565.8, exon 2 of 4) | 18 | 80966376 | 80986578 | 20203 | 2 | ENSMUST00000057950.7 | 11572 | Sall3 | ENSMUSG00000024565.8 |
| chr18 | 80976041 | 80976656 | 4.37804 | 5.84312 | 2.35272 | Intron (ENSMUST00000057950.7/ENSMUSG00000024565.8, intron 1 of 3) | 18 | 80966376 | 80986578 | 20203 | 2 | ENSMUST00000057950.7 | 9922 | Sall3 | ENSMUSG00000024565.8 |
| chr18 | 80977783 | 80981693 | 3.93298 | 5.07168 | 1.82591 | Exon (ENSMUST00000185180.1/ENSMUSG00000098880.1, exon 2 of 2) | 18 | 80966376 | 80986578 | 20203 | 2 | ENSMUST00000057950.7 | 4885 | Sall3 | ENSMUSG00000024565.8 |
| chr18 | 80982008 | 80983448 | 3.87963 | 4.93021 | 1.7326 | Intron (ENSMUST00000057950.7/ENSMUSG00000024565.8, intron 1 of 3) | 18 | 80966376 | 80986578 | 20203 | 2 | ENSMUST00000057950.7 | 3130 | Sall3 | ENSMUSG00000024565.8 |
| chr18 | 80983833 | 80987297 | 3.95622 | 5.06406 | 1.80314 | Promoter | 18 | 80966376 | 80986578 | 20203 | 2 | ENSMUST00000057950.7 | 0 | Sall3 | ENSMUSG00000024565.8 |
| chr18 | 80948378 | 80948453 | 4.26171 | 5.58887 | 2.17412 | Distal Intergenic | 18 | 80734143 | 80934058 | 199916 | 2 | ENSMUST00000091790.3 | -14321 | Atp9b | ENSMUSG00000024566.8 |
| chr18 | 80949262 | 80949358 | 3.41411 | 4.02533 | 1.11697 | Distal Intergenic | 18 | 80734143 | 80934058 | 199916 | 2 | ENSMUST00000091790.3 | -15205 | Atp9b | ENSMUSG00000024566.8 |
| chr18 | 80958010 | 80958148 | 3.41188 | 4.1147 | 1.25381 | Distal Intergenic | 18 | 80734143 | 80934058 | 199916 | 2 | ENSMUST00000091790.3 | -23953 | Atp9b | ENSMUSG00000024566.8 |
| chr18 | 61691417 | 61691674 | 3.47344 | 4.08253 | 1.17134 | Intron (ENSMUST00000025471.2/ENSMUSG00000024578.2, intron 2 of 2) | 18 | 61687935 | 61692537 | 4603 | 1 | ENSMUST00000025471.2 | 3483 | Il17b | ENSMUSG00000024578.2 |
| chr18 | 61692156 | 61692446 | 3.51906 | 4.30198 | 1.36932 | 3' UTR | 18 | 61687935 | 61692537 | 4603 | 1 | ENSMUST00000025471.2 | 4222 | Il17b | ENSMUSG00000024578.2 |
| chr18 | 56678803 | 56678887 | 3.40007 | 3.94339 | 1.07181 | Distal Intergenic | 18 | 56707813 | 56753424 | 45612 | 1 | ENSMUST00000025486.8 | -28926 | Lmnb1 | ENSMUSG00000024590.8 |
| chr18 | 57787490 | 57787655 | 3.39377 | 3.97768 | 1.19519 | Distal Intergenic | 18 | 57878678 | 57946821 | 68144 | 1 | ENSMUST00000115366.2 | -91023 | Slc12a2 | ENSMUSG00000024597.10 |
| chr18 | 57965866 | 57966170 | 3.35872 | 4.00137 | 1.18407 | Distal Intergenic | 18 | 57878678 | 57946821 | 68144 | 1 | ENSMUST00000115366.2 | 87189 | Slc12a2 | ENSMUSG00000024597.10 |
| chr18 | 58053221 | 58053304 | 3.7052 | 4.79211 | 1.65175 | Intron (ENSMUST00000025497.6/ENSMUSG00000024598.8, intron 40 of 64) | 18 | 58008623 | 58209926 | 201304 | 2 | ENSMUST00000025497.6 | 156622 | Fbn2 | ENSMUSG00000024598.8 |
| chr18 | 58209129 | 58209222 | 3.61978 | 4.34408 | 1.30797 | Promoter | 18 | 58008623 | 58209926 | 201304 | 2 | ENSMUST00000025497.6 | 704 | Fbn2 | ENSMUSG00000024598.8 |
| chr18 | 58209596 | 58209826 | 4.49168 | 6.03681 | 2.45921 | Promoter | 18 | 58008623 | 58209926 | 201304 | 2 | ENSMUST00000025497.6 | 100 | Fbn2 | ENSMUSG00000024598.8 |
| chr18 | 60934715 | 60935016 | 3.62704 | 4.37007 | 1.34885 | Intron (ENSMUST00000102888.4/ENSMUSG00000024617.11, intron 1 of 17) | 18 | 60926949 | 60955639 | 28691 | 1 | ENSMUST00000137805.1 | 7767 | Camk2a | ENSMUSG00000024617.11 |
| chr18 | 60987504 | 60987625 | 3.29213 | 3.76535 | 1.01733 | 3' UTR | 18 | 60963600 | 60985697 | 22098 | 1 | ENSMUST00000039904.6 | 23905 | Camk2a | ENSMUSG00000024617.11 |
| chr18 | 61025875 | 61025964 | 3.50096 | 4.14594 | 1.25092 | Intron (ENSMUST00000025521.8/ENSMUSG00000024619.8, intron 1 of 2) | 18 | 61018862 | 61036199 | 17338 | 2 | ENSMUST00000025521.8 | 10235 | Cdx1 | ENSMUSG00000024619.8 |
| chr18 | 61035275 | 61036874 | 4.28195 | 5.81823 | 2.4201 | Promoter | 18 | 61018862 | 61036199 | 17338 | 2 | ENSMUST00000025521.8 | 0 | Cdx1 | ENSMUSG00000024619.8 |
| chr18 | 86710314 | 86710762 | 4.02938 | 5.16193 | 1.86731 | Promoter | 18 | 86711110 | 86718283 | 7174 | 1 | ENSMUST00000068423.4 | -348 | Cbln2 | ENSMUSG00000024647.9 |
| chr18 | 86711257 | 86711432 | 3.92278 | 4.90493 | 1.65277 | Promoter | 18 | 86711520 | 86717589 | 6070 | 1 | ENSMUST00000122464.2 | -88 | Cbln2 | ENSMUSG00000024647.9 |
| chr18 | 86713315 | 86713410 | 3.57426 | 4.28761 | 1.34105 | Promoter | 18 | 86713048 | 86718283 | 5236 | 1 | ENSMUST00000169470.1 | 268 | Cbln2 | ENSMUSG00000024647.9 |
| chr18 | 86714625 | 86715110 | 3.29685 | 3.77248 | 1.01733 | Intron (ENSMUST00000068423.4/ENSMUSG00000024647.9, intron 3 of 4) | 18 | 86713048 | 86718283 | 5236 | 1 | ENSMUST00000169470.1 | 1578 | Cbln2 | ENSMUSG00000024647.9 |
| chr19 | 10100611 | 10101338 | 3.80504 | 4.73755 | 1.64404 | Promoter | 19 | 10064164 | 10101503 | 37340 | 2 | ENSMUST00000025567.7 | 165 | Fads2 | ENSMUSG00000024665.7 |
| chr19 | 11760649 | 11761059 | 3.65333 | 4.41611 | 1.38323 | Intron (ENSMUST00000025585.3/ENSMUSG00000024682.3, intron 8 of 8) | 19 | 11770415 | 11774944 | 4530 | 1 | ENSMUST00000167199.1 | -9356 | Mrpl16 | ENSMUSG00000024683.5 |
| chr19 | 16435236 | 16435715 | 3.84052 | 4.80123 | 1.65666 | Promoter | 19 | 16435667 | 16610818 | 175152 | 1 | ENSMUST00000025602.2 | 0 | Gna14 | ENSMUSG00000024697.3 |
| chr19 | 16436201 | 16437221 | 3.69777 | 4.62199 | 1.54613 | Promoter | 19 | 16435667 | 16610818 | 175152 | 1 | ENSMUST00000025602.2 | 535 | Gna14 | ENSMUSG00000024697.3 |
| chr19 | 17537928 | 17538003 | 3.57179 | 4.30593 | 1.30755 | Intron (ENSMUST00000025618.9/ENSMUSG00000024713.10, intron 20 of 37) | 19 | 17397437 | 17399961 | 2525 | 1 | ENSMUST00000146171.1 | 140492 | Rfk | ENSMUSG00000024712.4 |
| chr19 | 11113538 | 11113663 | 3.61767 | 4.49719 | 1.47649 | Intron (ENSMUST00000025635.7/ENSMUSG00000024729.7, intron 2 of 6) | 19 | 11109354 | 11111876 | 2523 | 2 | ENSMUST00000188756.1 | -1663 | 1700017D01Rik | ENSMUSG00000024729.7 |
| chr19 | 10387203 | 10387278 | 3.70024 | 4.53967 | 1.51846 | Distal Intergenic | 19 | 10389090 | 10453181 | 64092 | 1 | ENSMUST00000073899.4 | -1812 | Syt7 | ENSMUSG00000024743.9 |
| chr19 | 10389475 | 10389582 | 3.59939 | 4.5808 | 1.49081 | Promoter | 19 | 10389090 | 10453181 | 64092 | 1 | ENSMUST00000169121.2 | 386 | Syt7 | ENSMUSG00000024743.9 |
| chr19 | 36055793 | 36056078 | 3.39714 | 4.06917 | 1.15523 | Intron (ENSMUST00000164639.2/ENSMUSG00000024798.10, intron 1 of 2) | 19 | 35958734 | 36057340 | 98607 | 2 | ENSMUST00000164639.2 | 1262 | Htr7 | ENSMUSG00000024798.10 |
| chr19 | 36057014 | 36057285 | 4.38003 | 6.03923 | 2.54891 | Promoter | 19 | 35958734 | 36057340 | 98607 | 2 | ENSMUST00000164639.2 | 55 | Htr7 | ENSMUSG00000024798.10 |
| chr19 | 30173289 | 30173392 | 3.52608 | 4.33132 | 1.31454 | Intron (ENSMUST00000025778.7/ENSMUSG00000024827.9, intron 2 of 24) | 19 | 30098449 | 30175418 | 76970 | 2 | ENSMUST00000025778.7 | 2026 | Gldc | ENSMUSG00000024827.9 |
| chr19 | 30174493 | 30175556 | 3.6565 | 4.50234 | 1.47242 | Promoter | 19 | 30098449 | 30175418 | 76970 | 2 | ENSMUST00000025778.7 | 0 | Gldc | ENSMUSG00000024827.9 |
| chr19 | 25487623 | 25487786 | 3.5824 | 4.36974 | 1.39025 | Distal Intergenic | 19 | 25505618 | 25604329 | 98712 | 1 | ENSMUST00000025755.5 | -17832 | Dmrt1 | ENSMUSG00000024837.10 |
| chr19 | 25502633 | 25502928 | 3.79756 | 4.68083 | 1.52186 | Distal Intergenic | 19 | 25505618 | 25604329 | 98712 | 1 | ENSMUST00000025755.5 | -2690 | Dmrt1 | ENSMUSG00000024837.10 |
| chr19 | 25503663 | 25504516 | 3.35242 | 3.91751 | 1.08445 | Distal Intergenic | 19 | 25505618 | 25604329 | 98712 | 1 | ENSMUST00000025755.5 | -1102 | Dmrt1 | ENSMUSG00000024837.10 |
| chr19 | 25505436 | 25507000 | 3.97595 | 5.08473 | 1.82741 | Promoter | 19 | 25505618 | 25604329 | 98712 | 1 | ENSMUST00000025755.5 | 0 | Dmrt1 | ENSMUSG00000024837.10 |
| chr19 | 25507631 | 25507843 | 3.39983 | 3.93322 | 1.0371 | Intron (ENSMUST00000025755.5/ENSMUSG00000024837.10, intron 1 of 4) | 19 | 25505707 | 25560265 | 54559 | 1 | ENSMUST00000160814.1 | 1925 | Dmrt1 | ENSMUSG00000024837.10 |
| chr19 | 25508502 | 25508750 | 3.35963 | 3.86696 | 1.01733 | Intron (ENSMUST00000025755.5/ENSMUSG00000024837.10, intron 1 of 4) | 19 | 25505707 | 25560265 | 54559 | 1 | ENSMUST00000160814.1 | 2796 | Dmrt1 | ENSMUSG00000024837.10 |
| chr19 | 25509212 | 25510399 | 3.74927 | 4.77171 | 1.63423 | Exon (ENSMUST00000025755.5/ENSMUSG00000024837.10, exon 2 of 5) | 19 | 25505707 | 25560265 | 54559 | 1 | ENSMUST00000160814.1 | 3506 | Dmrt1 | ENSMUSG00000024837.10 |
| chr19 | 25552153 | 25552555 | 3.40572 | 4.00737 | 1.11779 | Intron (ENSMUST00000025755.5/ENSMUSG00000024837.10, intron 3 of 4) | 19 | 25505707 | 25560265 | 54559 | 1 | ENSMUST00000160814.1 | 46447 | Dmrt1 | ENSMUSG00000024837.10 |
| chr19 | 24556574 | 24556842 | 3.46057 | 4.07094 | 1.20124 | Promoter | 19 | 24294794 | 24555872 | 261079 | 2 | ENSMUST00000025800.9 | -703 | Pip5k1b | ENSMUSG00000024867.9 |
| chr19 | 24557877 | 24558142 | 3.38705 | 4.16955 | 1.2344 | Distal Intergenic | 19 | 24294794 | 24555872 | 261079 | 2 | ENSMUST00000025800.9 | -2006 | Pip5k1b | ENSMUSG00000024867.9 |
| chr19 | 30545395 | 30545526 | 3.7455 | 4.72938 | 1.58316 | Downstream (<1kb) | 19 | 30545863 | 30549665 | 3803 | 2 | ENSMUST00000025803.8 | 4139 | Dkk1 | ENSMUSG00000024868.8 |
| chr19 | 30546912 | 30547330 | 4.06637 | 5.18377 | 1.8434 | 3' UTR | 19 | 30545863 | 30549665 | 3803 | 2 | ENSMUST00000025803.8 | 2335 | Dkk1 | ENSMUSG00000024868.8 |
| chr19 | 30548100 | 30548397 | 3.54359 | 4.19727 | 1.20243 | Intron (ENSMUST00000025803.8/ENSMUSG00000024868.8, intron 2 of 3) | 19 | 30545863 | 30549665 | 3803 | 2 | ENSMUST00000025803.8 | 1268 | Dkk1 | ENSMUSG00000024868.8 |
| chr19 | 30548832 | 30550479 | 3.679 | 4.55676 | 1.46508 | Promoter | 19 | 30545863 | 30549665 | 3803 | 2 | ENSMUST00000025803.8 | 0 | Dkk1 | ENSMUSG00000024868.8 |
| chr19 | 30551436 | 30551663 | 3.62923 | 4.363 | 1.32178 | Distal Intergenic | 19 | 30545863 | 30549665 | 3803 | 2 | ENSMUST00000025803.8 | -1772 | Dkk1 | ENSMUSG00000024868.8 |
| chr19 | 30553668 | 30554421 | 3.65421 | 4.44887 | 1.41885 | Distal Intergenic | 19 | 30545863 | 30549665 | 3803 | 2 | ENSMUST00000025803.8 | -4004 | Dkk1 | ENSMUSG00000024868.8 |
| chr19 | 23891874 | 23892218 | 3.5177 | 4.17843 | 1.27331 | Intron (ENSMUST00000025830.7/ENSMUSG00000024897.8, intron 1 of 12) | 19 | 23758876 | 23949597 | 190722 | 1 | ENSMUST00000025830.7 | 132999 | Apba1 | ENSMUSG00000024897.8 |
| chr19 | 23892739 | 23892887 | 4.23127 | 5.59585 | 2.23822 | 5' UTR | 19 | 23758876 | 23949597 | 190722 | 1 | ENSMUST00000025830.7 | 133864 | Apba1 | ENSMUSG00000024897.8 |
| chr19 | 23893287 | 23893377 | 3.6512 | 4.54421 | 1.50116 | Exon (ENSMUST00000025830.7/ENSMUSG00000024897.8, exon 2 of 13) | 19 | 23758876 | 23949597 | 190722 | 1 | ENSMUST00000025830.7 | 134412 | Apba1 | ENSMUSG00000024897.8 |
| chr19 | 32595797 | 32596173 | 3.76587 | 4.67637 | 1.58175 | Distal Intergenic | 19 | 32620005 | 32667187 | 47183 | 1 | ENSMUST00000025833.6 | -23832 | Papss2 | ENSMUSG00000024899.6 |
| chr19 | 3414627 | 3414836 | 3.55834 | 4.47592 | 1.44738 | Promoter | 19 | 3409919 | 3414472 | 4554 | 2 | ENSMUST00000025842.6 | -156 | Gal | ENSMUSG00000024907.6 |
| chr19 | 3441148 | 3441555 | 3.54096 | 4.19637 | 1.21289 | Distal Intergenic | 19 | 3409919 | 3414472 | 4554 | 2 | ENSMUST00000025842.6 | -26677 | Gal | ENSMUSG00000024907.6 |
| chr19 | 5447604 | 5448028 | 3.65293 | 4.70046 | 1.57537 | Promoter | 19 | 5447703 | 5455945 | 8243 | 1 | ENSMUST00000025850.5 | 0 | Fosl1 | ENSMUSG00000024912.5 |
| chr19 | 5557473 | 5557567 | 3.71016 | 4.65995 | 1.55867 | Intron (ENSMUST00000025861.1/ENSMUSG00000024922.1, intron 1 of 3) | 19 | 5549137 | 5560575 | 11439 | 2 | ENSMUST00000025861.1 | 3008 | Ovol1 | ENSMUSG00000024922.1 |
| chr19 | 5558321 | 5558408 | 4.17665 | 5.44362 | 2.10712 | Intron (ENSMUST00000025861.1/ENSMUSG00000024922.1, intron 1 of 3) | 19 | 5549137 | 5560575 | 11439 | 2 | ENSMUST00000025861.1 | 2167 | Ovol1 | ENSMUSG00000024922.1 |
| chr19 | 5559503 | 5561391 | 4.06407 | 5.22103 | 1.9264 | Promoter | 19 | 5549137 | 5560575 | 11439 | 2 | ENSMUST00000025861.1 | 0 | Ovol1 | ENSMUSG00000024922.1 |
| chr19 | 5562483 | 5562710 | 3.41959 | 4.02784 | 1.17093 | Distal Intergenic | 19 | 5549137 | 5560575 | 11439 | 2 | ENSMUST00000025861.1 | -1909 | Ovol1 | ENSMUSG00000024922.1 |
| chr19 | 5563202 | 5563592 | 3.48165 | 4.12653 | 1.26728 | Distal Intergenic | 19 | 5549137 | 5560575 | 11439 | 2 | ENSMUST00000025861.1 | -2628 | Ovol1 | ENSMUSG00000024922.1 |
| chr19 | 27215757 | 27216204 | 3.58679 | 4.5555 | 1.49789 | Promoter | 19 | 27216484 | 27249755 | 33272 | 1 | ENSMUST00000164509.2 | -280 | Vldlr | ENSMUSG00000024924.9 |
| chr19 | 27216671 | 27217705 | 3.90202 | 4.99485 | 1.79736 | Promoter | 19 | 27217020 | 27234856 | 17837 | 1 | ENSMUST00000164746.2 | 0 | Vldlr | ENSMUSG00000024924.9 |
| chr19 | 28834362 | 28835481 | 3.65817 | 4.52068 | 1.47397 | Promoter | 19 | 28835049 | 28879301 | 44253 | 1 | ENSMUST00000161340.1 | 0 | Slc1a1 | ENSMUSG00000024935.5 |
| chr19 | 6847892 | 6849265 | 3.65765 | 4.48036 | 1.48662 | Exon (ENSMUST00000113440.1/ENSMUSG00000047810.8, exon 19 of 27) | 19 | 6829085 | 6840601 | 11517 | 2 | ENSMUST00000170516.3 | -7292 | Rps6ka4 | ENSMUSG00000024952.7 |
| chr19 | 6934278 | 6934445 | 4.2198 | 5.58543 | 2.21672 | Promoter | 19 | 6925710 | 6934515 | 8806 | 2 | ENSMUST00000025908.6 | 70 | Kcnk4 | ENSMUSG00000024957.8 |
| chr19 | 7612638 | 7612830 | 3.63978 | 4.44613 | 1.46716 | Promoter | 19 | 7612605 | 7639642 | 27038 | 1 | ENSMUST00000148558.1 | 34 | Hrasls5 | ENSMUSG00000024973.11 |
| chr19 | 56029939 | 56030084 | 4.05844 | 5.2256 | 1.964 | Distal Intergenic | 19 | 55898553 | 55919604 | 21052 | 1 | ENSMUST00000148666.1 | 131387 | Tcf7l2 | ENSMUSG00000024985.13 |
| chr19 | 37429683 | 37429868 | 3.43341 | 4.01153 | 1.13521 | Distal Intergenic | 19 | 37434810 | 37440731 | 5922 | 1 | ENSMUST00000025944.3 | -4942 | Hhex | ENSMUSG00000024986.6 |
| chr19 | 37431635 | 37432968 | 3.50001 | 4.14817 | 1.24936 | Distal Intergenic | 19 | 37434810 | 37440731 | 5922 | 1 | ENSMUST00000025944.3 | -1842 | Hhex | ENSMUSG00000024986.6 |
| chr19 | 37433323 | 37433621 | 3.5574 | 4.34609 | 1.35519 | Distal Intergenic | 19 | 37434810 | 37440731 | 5922 | 1 | ENSMUST00000025944.3 | -1189 | Hhex | ENSMUSG00000024986.6 |
| chr19 | 37434123 | 37434261 | 3.71819 | 4.78919 | 1.62243 | Promoter | 19 | 37434810 | 37440731 | 5922 | 1 | ENSMUST00000025944.3 | -549 | Hhex | ENSMUSG00000024986.6 |
| chr19 | 37434605 | 37435566 | 3.97045 | 5.19641 | 1.94053 | Promoter | 19 | 37434810 | 37440731 | 5922 | 1 | ENSMUST00000025944.3 | 0 | Hhex | ENSMUSG00000024986.6 |
| chr19 | 37436227 | 37436741 | 3.76217 | 4.84306 | 1.69522 | Promoter | 19 | 37436739 | 37440672 | 3934 | 1 | ENSMUST00000128184.2 | 0 | Hhex | ENSMUSG00000024986.6 |
| chr19 | 37437085 | 37437217 | 3.52767 | 4.29045 | 1.32496 | Promoter | 19 | 37436739 | 37440672 | 3934 | 1 | ENSMUST00000128184.2 | 347 | Hhex | ENSMUSG00000024986.6 |
| chr19 | 37437537 | 37437718 | 3.42376 | 4.09755 | 1.25633 | Promoter | 19 | 37436739 | 37440672 | 3934 | 1 | ENSMUST00000128184.2 | 799 | Hhex | ENSMUSG00000024986.6 |
| chr19 | 37439486 | 37439640 | 3.35337 | 3.87471 | 1.078 | Exon (ENSMUST00000025944.3/ENSMUSG00000024986.6, exon 4 of 4) | 19 | 37436739 | 37440672 | 3934 | 1 | ENSMUST00000128184.2 | 2748 | Hhex | ENSMUSG00000024986.6 |
| chr19 | 37441977 | 37442445 | 3.45354 | 4.0536 | 1.1737 | Distal Intergenic | 19 | 37436739 | 37440672 | 3934 | 1 | ENSMUST00000128184.2 | 5239 | Hhex | ENSMUSG00000024986.6 |
| chr19 | 37448729 | 37448959 | 3.30073 | 3.92312 | 1.06519 | Distal Intergenic | 19 | 37436739 | 37440672 | 3934 | 1 | ENSMUST00000128184.2 | 11991 | Hhex | ENSMUSG00000024986.6 |
| chr19 | 37693033 | 37693121 | 4.24422 | 5.58957 | 2.20414 | Exon (ENSMUST00000073391.4/ENSMUSG00000062432.4, exon 6 of 6) | 19 | 37697808 | 37701528 | 3721 | 1 | ENSMUST00000025946.5 | -4687 | Cyp26a1 | ENSMUSG00000024987.5 |
| chr19 | 37693472 | 37693640 | 3.84258 | 4.75421 | 1.55807 | Distal Intergenic | 19 | 37697808 | 37701528 | 3721 | 1 | ENSMUST00000025946.5 | -4168 | Cyp26a1 | ENSMUSG00000024987.5 |
| chr19 | 37695677 | 37695860 | 4.26786 | 5.61419 | 2.18924 | Distal Intergenic | 19 | 37697808 | 37701528 | 3721 | 1 | ENSMUST00000025946.5 | -1948 | Cyp26a1 | ENSMUSG00000024987.5 |
| chr19 | 37696291 | 37697247 | 3.81185 | 4.71273 | 1.55439 | Promoter | 19 | 37697808 | 37701528 | 3721 | 1 | ENSMUST00000025946.5 | -561 | Cyp26a1 | ENSMUSG00000024987.5 |
| chr19 | 37697774 | 37698553 | 3.84858 | 4.82065 | 1.66756 | Promoter | 19 | 37697808 | 37701528 | 3721 | 1 | ENSMUST00000025946.5 | 0 | Cyp26a1 | ENSMUSG00000024987.5 |
| chr19 | 37699036 | 37699628 | 3.71415 | 4.5757 | 1.50842 | Exon (ENSMUST00000025946.5/ENSMUSG00000024987.5, exon 3 of 7) | 19 | 37697808 | 37701528 | 3721 | 1 | ENSMUST00000025946.5 | 1229 | Cyp26a1 | ENSMUSG00000024987.5 |
| chr19 | 38123150 | 38123670 | 3.56183 | 4.26159 | 1.2664 | Intron (ENSMUST00000112335.2/ENSMUSG00000024990.7, intron 4 of 5) | 19 | 38116620 | 38124811 | 8192 | 2 | ENSMUST00000112335.2 | 1141 | Rbp4 | ENSMUSG00000024990.7 |
| chr19 | 38124592 | 38125534 | 4.02692 | 5.2195 | 1.95081 | Promoter | 19 | 38116620 | 38124811 | 8192 | 2 | ENSMUST00000112335.2 | 0 | Rbp4 | ENSMUSG00000024990.7 |
| chr19 | 38126798 | 38126994 | 3.37449 | 3.88937 | 1.01733 | Distal Intergenic | 19 | 38116620 | 38125321 | 8702 | 2 | ENSMUST00000025951.7 | -1478 | Rbp4 | ENSMUSG00000024990.7 |
| chr19 | 38131244 | 38131325 | 3.57321 | 4.25913 | 1.25854 | Distal Intergenic | 19 | 38132781 | 38183947 | 51167 | 1 | ENSMUST00000025956.6 | -1456 | Pde6c | ENSMUSG00000024992.7 |
| chr19 | 38133235 | 38133364 | 3.50827 | 4.17437 | 1.2897 | Promoter | 19 | 38132781 | 38183947 | 51167 | 1 | ENSMUST00000112329.1 | 455 | Pde6c | ENSMUSG00000024992.7 |
| chr19 | 41015093 | 41015645 | 3.35663 | 3.91129 | 1.15629 | Distal Intergenic | 19 | 41029275 | 41059523 | 30249 | 1 | ENSMUST00000051806.5 | -13630 | Dntt | ENSMUSG00000025014.7 |
| chr19 | 41310858 | 41311077 | 3.66546 | 4.46009 | 1.4422 | Intron (ENSMUST00000059672.7/ENSMUSG00000025017.9, intron 8 of 16) | 19 | 41210842 | 41264004 | 53163 | 2 | ENSMUST00000025989.8 | -46855 | Tm9sf3 | ENSMUSG00000025016.10 |
| chr19 | 41316247 | 41316484 | 3.42408 | 4.00482 | 1.11774 | Intron (ENSMUST00000059672.7/ENSMUSG00000025017.9, intron 8 of 16) | 19 | 41210842 | 41264004 | 53163 | 2 | ENSMUST00000025989.8 | -52244 | Tm9sf3 | ENSMUSG00000025016.10 |
| chr19 | 41316972 | 41317256 | 4.068 | 5.2165 | 1.92807 | Intron (ENSMUST00000059672.7/ENSMUSG00000025017.9, intron 8 of 16) | 19 | 41210842 | 41264004 | 53163 | 2 | ENSMUST00000025989.8 | -52969 | Tm9sf3 | ENSMUSG00000025016.10 |
| chr19 | 41384688 | 41384994 | 3.68373 | 4.55536 | 1.50226 | Promoter | 19 | 41274218 | 41385070 | 110853 | 2 | ENSMUST00000059672.7 | 76 | Pik3ap1 | ENSMUSG00000025017.9 |
| chr19 | 41615873 | 41616072 | 3.27084 | 3.9614 | 1.08044 | Intron (ENSMUST00000169141.2/ENSMUSG00000025020.6, intron 26 of 35) | 19 | 41604369 | 41608530 | 4162 | 2 | ENSMUST00000169416.1 | -7344 | Slit1 | ENSMUSG00000025020.6 |
| chr19 | 41616406 | 41616655 | 3.71589 | 4.54403 | 1.45618 | Intron (ENSMUST00000169141.2/ENSMUSG00000025020.6, intron 26 of 35) | 19 | 41604369 | 41608530 | 4162 | 2 | ENSMUST00000169416.1 | -7877 | Slit1 | ENSMUSG00000025020.6 |
| chr19 | 41618298 | 41619094 | 3.80644 | 4.78611 | 1.61396 | Intron (ENSMUST00000169141.2/ENSMUSG00000025020.6, intron 25 of 35) | 19 | 41604369 | 41608530 | 4162 | 2 | ENSMUST00000169416.1 | -9769 | Slit1 | ENSMUSG00000025020.6 |
| chr19 | 41637523 | 41637737 | 3.55741 | 4.22884 | 1.24214 | Exon (ENSMUST00000169141.2/ENSMUSG00000025020.6, exon 18 of 36) | 19 | 41604369 | 41608530 | 4162 | 2 | ENSMUST00000169416.1 | -28994 | Slit1 | ENSMUSG00000025020.6 |
| chr19 | 41716368 | 41716517 | 3.27988 | 3.75472 | 1.03801 | Intron (ENSMUST00000169141.2/ENSMUSG00000025020.6, intron 4 of 35) | 19 | 41646103 | 41743486 | 97384 | 2 | ENSMUST00000171586.1 | 26969 | Slit1 | ENSMUSG00000025020.6 |
| chr19 | 41742217 | 41742325 | 3.35974 | 3.87934 | 1.06062 | Intron (ENSMUST00000169141.2/ENSMUSG00000025020.6, intron 1 of 35) | 19 | 41646103 | 41743486 | 97384 | 2 | ENSMUST00000171586.1 | 1161 | Slit1 | ENSMUSG00000025020.6 |
| chr19 | 41742817 | 41743688 | 3.83298 | 4.87487 | 1.65587 | Promoter | 19 | 41600257 | 41743492 | 143236 | 2 | ENSMUST00000169141.2 | 0 | Slit1 | ENSMUSG00000025020.6 |
| chr19 | 53277047 | 53277147 | 3.39273 | 3.92952 | 1.06182 | Distal Intergenic | 19 | 53310506 | 53375810 | 65305 | 1 | ENSMUST00000003870.8 | -33359 | Mxi1 | ENSMUSG00000025025.8 |
| chr19 | 47014473 | 47016121 | 4.04696 | 5.54799 | 2.18182 | Promoter | 19 | 46887828 | 47015153 | 127326 | 2 | ENSMUST00000172239.2 | 0 | Nt5c2 | ENSMUSG00000025041.11 |
| chr19 | 47019378 | 47019630 | 3.47973 | 4.13524 | 1.28344 | Intron (ENSMUST00000037636.3/ENSMUSG00000034336.3, intron 1 of 2) | 19 | 46887828 | 47015153 | 127326 | 2 | ENSMUST00000172239.2 | -4226 | Nt5c2 | ENSMUSG00000025041.11 |
| chr19 | 47691562 | 47692034 | 3.6506 | 4.45426 | 1.47515 | Promoter | 19 | 47646344 | 47692094 | 45751 | 2 | ENSMUST00000026045.8 | 60 | Col17a1 | ENSMUSG00000025064.9 |
| chr19 | 56300884 | 56301006 | 3.61314 | 4.35244 | 1.36385 | Intron (ENSMUST00000078284.7/ENSMUSG00000025075.9, intron 3 of 13) | 19 | 56311814 | 56316417 | 4604 | 1 | ENSMUST00000163502.2 | -10808 | Habp2 | ENSMUSG00000025075.9 |
| chr19 | 56319591 | 56320397 | 3.7934 | 4.74299 | 1.66685 | 3' UTR | 19 | 56312948 | 56319629 | 6682 | 1 | ENSMUST00000165522.1 | 6644 | Habp2 | ENSMUSG00000025075.9 |
| chr19 | 56326095 | 56326296 | 3.60669 | 4.31936 | 1.29835 | Intron (ENSMUST00000167239.2/ENSMUSG00000049134.10, intron 35 of 36) | 19 | 56312948 | 56319629 | 6682 | 1 | ENSMUST00000165522.1 | 13148 | Habp2 | ENSMUSG00000025075.9 |
| chr19 | 56826145 | 56826768 | 3.36274 | 3.90585 | 1.13789 | Promoter | 19 | 56826209 | 56870012 | 43804 | 1 | ENSMUST00000111606.2 | 0 | Tdrd1 | ENSMUSG00000025081.8 |
| chr19 | 56830330 | 56830617 | 3.45758 | 4.04138 | 1.11126 | Promoter | 19 | 56830712 | 56870012 | 39301 | 1 | ENSMUST00000111604.1 | -95 | Tdrd1 | ENSMUSG00000025081.8 |
| chr19 | 56863853 | 56864023 | 3.28493 | 3.75454 | 1.01733 | Intron (ENSMUST00000111606.2/ENSMUSG00000025081.8, intron 21 of 25) | 19 | 56874249 | 56912078 | 37830 | 1 | ENSMUST00000026068.7 | -10226 | Vwa2 | ENSMUSG00000025082.8 |
| chr19 | 56869827 | 56870030 | 4.01853 | 5.18038 | 1.90771 | 3' UTR | 19 | 56874249 | 56912078 | 37830 | 1 | ENSMUST00000026068.7 | -4219 | Vwa2 | ENSMUSG00000025082.8 |
| chr19 | 56871009 | 56872169 | 3.67147 | 4.46707 | 1.42068 | Distal Intergenic | 19 | 56874249 | 56912078 | 37830 | 1 | ENSMUST00000026068.7 | -2080 | Vwa2 | ENSMUSG00000025082.8 |
| chr19 | 56873207 | 56873479 | 3.58164 | 4.38744 | 1.38915 | Promoter | 19 | 56874249 | 56912078 | 37830 | 1 | ENSMUST00000026068.7 | -770 | Vwa2 | ENSMUSG00000025082.8 |
| chr19 | 56873838 | 56874976 | 4.6057 | 6.57001 | 2.97091 | Promoter | 19 | 56874249 | 56912078 | 37830 | 1 | ENSMUST00000026068.7 | 0 | Vwa2 | ENSMUSG00000025082.8 |
| chr19 | 56875695 | 56875783 | 3.79249 | 4.68738 | 1.57651 | Intron (ENSMUST00000026068.7/ENSMUSG00000025082.8, intron 1 of 13) | 19 | 56874249 | 56912078 | 37830 | 1 | ENSMUST00000026068.7 | 1447 | Vwa2 | ENSMUSG00000025082.8 |
| chr19 | 56877805 | 56878026 | 3.3752 | 3.89044 | 1.01733 | Intron (ENSMUST00000026068.7/ENSMUSG00000025082.8, intron 1 of 13) | 19 | 56874249 | 56912078 | 37830 | 1 | ENSMUST00000026068.7 | 3557 | Vwa2 | ENSMUSG00000025082.8 |
| chr19 | 56878468 | 56878749 | 3.5853 | 4.27554 | 1.25542 | Intron (ENSMUST00000026068.7/ENSMUSG00000025082.8, intron 1 of 13) | 19 | 56874249 | 56912078 | 37830 | 1 | ENSMUST00000026068.7 | 4220 | Vwa2 | ENSMUSG00000025082.8 |
| chr19 | 57008069 | 57008386 | 3.46693 | 4.34911 | 1.34905 | Promoter | 19 | 56912361 | 57008198 | 95838 | 2 | ENSMUST00000118800.2 | 0 | Afap1l2 | ENSMUSG00000025083.12 |
| chr19 | 57067104 | 57067327 | 3.32172 | 3.86451 | 1.1189 | Intron (ENSMUST00000079360.6/ENSMUSG00000025085.11, intron 12 of 25) | 19 | 57039447 | 57047238 | 7792 | 2 | ENSMUST00000137389.2 | -19867 | Ablim1 | ENSMUSG00000025085.11 |
| chr19 | 57069046 | 57069121 | 3.77889 | 4.75172 | 1.67707 | Intron (ENSMUST00000079360.6/ENSMUSG00000025085.11, intron 10 of 25) | 19 | 57039447 | 57047238 | 7792 | 2 | ENSMUST00000137389.2 | -21809 | Ablim1 | ENSMUSG00000025085.11 |
| chr19 | 57079932 | 57080295 | 3.31142 | 3.83247 | 1.08108 | Intron (ENSMUST00000079360.6/ENSMUSG00000025085.11, intron 7 of 25) | 19 | 57039447 | 57047238 | 7792 | 2 | ENSMUST00000137389.2 | -32695 | Ablim1 | ENSMUSG00000025085.11 |
| chr19 | 57081140 | 57081246 | 3.87406 | 4.92913 | 1.77178 | Intron (ENSMUST00000079360.6/ENSMUSG00000025085.11, intron 7 of 25) | 19 | 57039447 | 57047238 | 7792 | 2 | ENSMUST00000137389.2 | -33903 | Ablim1 | ENSMUSG00000025085.11 |
| chr19 | 57081616 | 57081788 | 3.4095 | 3.99761 | 1.1907 | Intron (ENSMUST00000079360.6/ENSMUSG00000025085.11, intron 7 of 25) | 19 | 57039447 | 57047238 | 7792 | 2 | ENSMUST00000137389.2 | -34379 | Ablim1 | ENSMUSG00000025085.11 |
| chr19 | 57083227 | 57083311 | 4.96901 | 7.22093 | 3.38106 | Intron (ENSMUST00000079360.6/ENSMUSG00000025085.11, intron 7 of 25) | 19 | 57036479 | 57118913 | 82435 | 2 | ENSMUST00000111526.2 | 35602 | Ablim1 | ENSMUSG00000025085.11 |
| chr19 | 57084130 | 57084336 | 3.2374 | 3.68812 | 1.03214 | Intron (ENSMUST00000079360.6/ENSMUSG00000025085.11, intron 6 of 25) | 19 | 57036479 | 57118913 | 82435 | 2 | ENSMUST00000111526.2 | 34577 | Ablim1 | ENSMUSG00000025085.11 |
| chr19 | 57086838 | 57087060 | 3.55158 | 4.39542 | 1.34408 | Intron (ENSMUST00000079360.6/ENSMUSG00000025085.11, intron 6 of 25) | 19 | 57036479 | 57118913 | 82435 | 2 | ENSMUST00000111526.2 | 31853 | Ablim1 | ENSMUSG00000025085.11 |
| chr19 | 57095231 | 57095412 | 3.19984 | 4.03003 | 1.12884 | Intron (ENSMUST00000079360.6/ENSMUSG00000025085.11, intron 6 of 25) | 19 | 57036479 | 57118913 | 82435 | 2 | ENSMUST00000111526.2 | 23501 | Ablim1 | ENSMUSG00000025085.11 |
| chr19 | 57100330 | 57100709 | 3.40744 | 4.21718 | 1.29517 | Intron (ENSMUST00000079360.6/ENSMUSG00000025085.11, intron 6 of 25) | 19 | 57036479 | 57118913 | 82435 | 2 | ENSMUST00000111526.2 | 18204 | Ablim1 | ENSMUSG00000025085.11 |
| chr19 | 57103700 | 57103879 | 3.37544 | 3.92653 | 1.14009 | Intron (ENSMUST00000079360.6/ENSMUSG00000025085.11, intron 6 of 25) | 19 | 57036479 | 57118913 | 82435 | 2 | ENSMUST00000111526.2 | 15034 | Ablim1 | ENSMUSG00000025085.11 |
| chr19 | 57105270 | 57105675 | 3.51166 | 4.18321 | 1.29179 | Intron (ENSMUST00000079360.6/ENSMUSG00000025085.11, intron 6 of 25) | 19 | 57036479 | 57118913 | 82435 | 2 | ENSMUST00000111526.2 | 13238 | Ablim1 | ENSMUSG00000025085.11 |
| chr19 | 57106459 | 57106534 | 4.69066 | 6.59168 | 2.92622 | Intron (ENSMUST00000079360.6/ENSMUSG00000025085.11, intron 6 of 25) | 19 | 57036479 | 57118913 | 82435 | 2 | ENSMUST00000111526.2 | 12379 | Ablim1 | ENSMUSG00000025085.11 |
| chr19 | 57106891 | 57107079 | 3.2903 | 3.7626 | 1.01733 | Intron (ENSMUST00000079360.6/ENSMUSG00000025085.11, intron 6 of 25) | 19 | 57036479 | 57118913 | 82435 | 2 | ENSMUST00000111526.2 | 11834 | Ablim1 | ENSMUSG00000025085.11 |
| chr19 | 57114448 | 57114975 | 3.63429 | 4.4939 | 1.48357 | Intron (ENSMUST00000079360.6/ENSMUSG00000025085.11, intron 6 of 25) | 19 | 57036479 | 57118913 | 82435 | 2 | ENSMUST00000111526.2 | 3938 | Ablim1 | ENSMUSG00000025085.11 |
| chr19 | 57115581 | 57115868 | 3.35038 | 4.12188 | 1.22701 | Intron (ENSMUST00000079360.6/ENSMUSG00000025085.11, intron 6 of 25) | 19 | 57036479 | 57118913 | 82435 | 2 | ENSMUST00000111526.2 | 3045 | Ablim1 | ENSMUSG00000025085.11 |
| chr19 | 57117198 | 57117427 | 3.26948 | 3.73135 | 1.01733 | Intron (ENSMUST00000079360.6/ENSMUSG00000025085.11, intron 6 of 25) | 19 | 57036479 | 57118913 | 82435 | 2 | ENSMUST00000111526.2 | 1486 | Ablim1 | ENSMUSG00000025085.11 |
| chr19 | 57117795 | 57117995 | 3.55312 | 4.31278 | 1.36805 | Promoter | 19 | 57036479 | 57118913 | 82435 | 2 | ENSMUST00000111526.2 | 918 | Ablim1 | ENSMUSG00000025085.11 |
| chr19 | 57121952 | 57122140 | 3.62283 | 4.35962 | 1.34466 | Intron (ENSMUST00000079360.6/ENSMUSG00000025085.11, intron 6 of 25) | 19 | 57116022 | 57119024 | 3003 | 2 | ENSMUST00000150425.1 | -2929 | Ablim1 | ENSMUSG00000025085.11 |
| chr19 | 57122801 | 57123034 | 3.30366 | 3.78268 | 1.01733 | Intron (ENSMUST00000079360.6/ENSMUSG00000025085.11, intron 6 of 25) | 19 | 57116022 | 57119024 | 3003 | 2 | ENSMUST00000150425.1 | -3778 | Ablim1 | ENSMUSG00000025085.11 |

| | | | | | | | | | | | | | | | |
|---|---|---|---|---|---|---|---|---|---|---|---|---|---|---|---|
| chr19 | 57126994 | 57127213 | 3.86399 | 4.86438 | 1.69718 | Intron (ENSMUST00000079360.6/ENSMUSG00000025085.11, intron 6 of 25) | 19 | 57077302 | 57134677 | 57376 | 2 | ENSMUST00000134430.2 | 7464 | Ablim1 | ENSMUSG00000025085.11 |
| chr19 | 57128290 | 57128702 | 3.70611 | 4.57718 | 1.48245 | Intron (ENSMUST00000079360.6/ENSMUSG00000025085.11, intron 6 of 25) | 19 | 57077302 | 57134677 | 57376 | 2 | ENSMUST00000134430.2 | 5975 | Ablim1 | ENSMUSG00000025085.11 |
| chr19 | 57130180 | 57130352 | 3.34725 | 3.8483 | 1.01733 | Exon (ENSMUST00000128212.1/ENSMUSG00000025085.11, exon 4 of 4) | 19 | 57077302 | 57134677 | 57376 | 2 | ENSMUST00000134430.2 | 4325 | Ablim1 | ENSMUSG00000025085.11 |
| chr19 | 57132453 | 57133083 | 4.13002 | 5.3871 | 2.03889 | Exon (ENSMUST00000079360.6/ENSMUSG00000025085.11, exon 6 of 26) | 19 | 57077302 | 57134677 | 57376 | 2 | ENSMUST00000134430.2 | 1594 | Ablim1 | ENSMUSG00000025085.11 |
| chr19 | 57134697 | 57134772 | 3.56463 | 4.24377 | 1.25619 | Promoter | 19 | 57060964 | 57134708 | 73745 | 2 | ENSMUST00000133782.2 | 0 | Ablim1 | ENSMUSG00000025085.11 |
| chr19 | 57144292 | 57144547 | 4.44194 | 5.93831 | 2.38694 | Intron (ENSMUST00000079360.6/ENSMUSG00000025085.11, intron 4 of 25) | 19 | 57130264 | 57135725 | 5462 | 2 | ENSMUST00000128212.1 | -8568 | Ablim1 | ENSMUSG00000025085.11 |
| chr19 | 57173968 | 57174191 | 3.60458 | 4.31229 | 1.28429 | Intron (ENSMUST00000079360.6/ENSMUSG00000025085.11, intron 1 of 25) | 19 | 57130846 | 57182308 | 51463 | 2 | ENSMUST00000133369.1 | 8117 | Ablim1 | ENSMUSG00000025085.11 |
| chr19 | 57195755 | 57195851 | 3.58456 | 4.33571 | 1.40419 | Intron (ENSMUST00000079360.6/ENSMUSG00000025085.11, intron 1 of 25) | 19 | 57037140 | 57197391 | 160252 | 2 | ENSMUST00000111546.2 | 1540 | Ablim1 | ENSMUSG00000025085.11 |
| chr19 | 57283330 | 57283556 | 3.45563 | 4.03496 | 1.09878 | Intron (ENSMUST00000156316.2/ENSMUSG00000025085.11, intron 1 of 20) | 19 | 57130606 | 57314911 | 184306 | 2 | ENSMUST00000111524.2 | 31355 | Ablim1 | ENSMUSG00000025085.11 |
| chr19 | 57314834 | 57315159 | 3.66446 | 4.63094 | 1.54691 | Promoter | 19 | 57037251 | 57314919 | 277669 | 2 | ENSMUST00000156316.2 | 0 | Ablim1 | ENSMUSG00000025085.11 |
| chr19 | 57324617 | 57324955 | 3.4578 | 4.06088 | 1.17942 | Exon (ENSMUST00000181921.1/ENSMUSG00000097785.1, exon 2 of 3) | 19 | 57037251 | 57314919 | 277669 | 2 | ENSMUST00000156316.2 | -9699 | Ablim1 | ENSMUSG00000025085.11 |
| chr19 | 57418198 | 57419147 | 3.25499 | 3.71181 | 1.02463 | Distal Intergenic | 19 | 57452906 | 57488910 | 36005 | 1 | ENSMUST00000026072.3 | -33759 | Trub1 | ENSMUSG00000025086.7 |
| chr19 | 58453184 | 58454333 | 3.84503 | 4.93473 | 1.75994 | Promoter | 19 | 58236692 | 58454466 | 217775 | 2 | ENSMUST00000169850.2 | 133 | Gfra1 | ENSMUSG00000025089.10 |
| chr19 | 58454723 | 58455431 | 3.68953 | 4.74917 | 1.60946 | Promoter | 19 | 58235604 | 58455398 | 219795 | 2 | ENSMUST00000026076.8 | 0 | Gfra1 | ENSMUSG00000025089.10 |
| chr19 | 58455866 | 58456117 | 3.80394 | 4.9435 | 1.72964 | Promoter | 19 | 58453352 | 58455909 | 2558 | 2 | ENSMUST00000131877.1 | 0 | Gfra1 | ENSMUSG00000025089.10 |
| chr19 | 58457874 | 58457949 | 3.89953 | 4.96669 | 1.77913 | Distal Intergenic | 19 | 58453352 | 58455909 | 2558 | 2 | ENSMUST00000131877.1 | -1966 | Gfra1 | ENSMUSG00000025089.10 |
| chr19 | 58459120 | 58459383 | 3.34537 | 3.89552 | 1.11862 | Distal Intergenic | 19 | 58453352 | 58455909 | 2558 | 2 | ENSMUST00000131877.1 | -3212 | Gfra1 | ENSMUSG00000025089.10 |
| chr19 | 59260760 | 59261199 | 3.55448 | 4.46422 | 1.41626 | Promoter | 19 | 59260878 | 59296012 | 35135 | 1 | ENSMUST00000026084.3 | 0 | Slc18a2 | ENSMUSG00000025094.7 |
| chr7 | 81991454 | 81993559 | 3.79191 | 4.78698 | 1.66986 | Promoter | 7 | 81966672 | 81992299 | 25628 | 2 | ENSMUST00000026096.7 | 0 | Bnc1 | ENSMUSG00000025105.8 |
| chr7 | 81993867 | 81993959 | 3.48091 | 4.1223 | 1.26324 | Distal Intergenic | 7 | 81966672 | 81992299 | 25628 | 2 | ENSMUST00000026096.7 | -1569 | Bnc1 | ENSMUSG00000025105.8 |
| chr7 | 82012626 | 82012701 | 3.81171 | 4.70723 | 1.56108 | Distal Intergenic | 7 | 81966672 | 81992299 | 25628 | 2 | ENSMUST00000026096.7 | -20328 | Bnc1 | ENSMUSG00000025105.8 |
| chr7 | 82016536 | 82017045 | 3.36478 | 3.91986 | 1.15204 | Distal Intergenic | 7 | 81966672 | 81992299 | 25628 | 2 | ENSMUST00000026096.7 | -24238 | Bnc1 | ENSMUSG00000025105.8 |
| chr11 | 120530765 | 120531436 | 4.12701 | 5.5097 | 2.14836 | Promoter | 11 | 120530774 | 120538983 | 8210 | 1 | ENSMUST00000128827.2 | 0 | Gcgr | ENSMUSG00000025127.10 |
| chr3 | 18043662 | 18043999 | 3.5934 | 4.341 | 1.3403 | Distal Intergenic | 3 | 18054174 | 18057517 | 3344 | 1 | ENSMUST00000026120.7 | -10175 | Bhlhe22 | ENSMUSG00000025128.7 |
| chr3 | 18044997 | 18045220 | 3.27246 | 3.73588 | 1.01733 | Distal Intergenic | 3 | 18054174 | 18057517 | 3344 | 1 | ENSMUST00000026120.7 | -8954 | Bhlhe22 | ENSMUSG00000025128.7 |
| chr3 | 18045951 | 18046726 | 3.58228 | 4.30828 | 1.36072 | Distal Intergenic | 3 | 18054174 | 18057517 | 3344 | 1 | ENSMUST00000026120.7 | -7448 | Bhlhe22 | ENSMUSG00000025128.7 |
| chr3 | 18047857 | 18047977 | 3.51673 | 4.20064 | 1.30206 | Distal Intergenic | 3 | 18054174 | 18057517 | 3344 | 1 | ENSMUST00000026120.7 | -6197 | Bhlhe22 | ENSMUSG00000025128.7 |
| chr3 | 18048755 | 18049850 | 3.93852 | 5.02022 | 1.84944 | Distal Intergenic | 3 | 18054174 | 18057517 | 3344 | 1 | ENSMUST00000026120.7 | -4324 | Bhlhe22 | ENSMUSG00000025128.7 |
| chr3 | 18050486 | 18051857 | 3.74115 | 4.73712 | 1.63001 | Distal Intergenic | 3 | 18054174 | 18057517 | 3344 | 1 | ENSMUST00000026120.7 | -2317 | Bhlhe22 | ENSMUSG00000025128.7 |
| chr3 | 18052184 | 18052284 | 4.05788 | 5.26285 | 2.00921 | Distal Intergenic | 3 | 18054174 | 18057517 | 3344 | 1 | ENSMUST00000026120.7 | -1890 | Bhlhe22 | ENSMUSG00000025128.7 |
| chr3 | 18053291 | 18055761 | 3.82513 | 4.99005 | 1.81469 | Promoter | 3 | 18054174 | 18057517 | 3344 | 1 | ENSMUST00000026120.7 | 0 | Bhlhe22 | ENSMUSG00000025128.7 |
| chr3 | 18056085 | 18056160 | 3.9702 | 5.02277 | 1.80705 | 3' UTR | 3 | 18054174 | 18057517 | 3344 | 1 | ENSMUST00000026120.7 | 1912 | Bhlhe22 | ENSMUSG00000025128.7 |
| chr3 | 18057326 | 18058505 | 3.78732 | 4.73075 | 1.6667 | 3' UTR | 3 | 18054174 | 18057517 | 3344 | 1 | ENSMUST00000026120.7 | 3153 | Bhlhe22 | ENSMUSG00000025128.7 |
| chr3 | 18060194 | 18060459 | 4.00379 | 5.23553 | 1.92895 | Distal Intergenic | 3 | 18054174 | 18057517 | 3344 | 1 | ENSMUST00000026120.7 | 6021 | Bhlhe22 | ENSMUSG00000025128.7 |
| chr3 | 18060877 | 18061006 | 3.83028 | 5.06764 | 1.84567 | Distal Intergenic | 3 | 18054174 | 18057517 | 3344 | 1 | ENSMUST00000026120.7 | 6704 | Bhlhe22 | ENSMUSG00000025128.7 |
| chr7 | 141913391 | 141913587 | 3.31189 | 3.96402 | 1.16445 | Intron (ENSMUST00000151890.1/ENSMUSG00000025139.8, intron 1 of 5) | 7 | 141884273 | 141918507 | 34235 | 2 | ENSMUST00000151890.1 | 4920 | Tollip | ENSMUSG00000025139.8 |
| chr7 | 141914155 | 141914337 | 3.62767 | 4.41197 | 1.43724 | Intron (ENSMUST00000151890.1/ENSMUSG00000025139.8, intron 1 of 5) | 7 | 141884273 | 141918507 | 34235 | 2 | ENSMUST00000151890.1 | 4170 | Tollip | ENSMUSG00000025139.8 |
| chr19 | 42491453 | 42491592 | 3.34756 | 3.84874 | 1.01733 | Distal Intergenic | 19 | 42518759 | 42592343 | 73585 | 1 | ENSMUST00000026188.4 | -27167 | R3hcc1l | ENSMUSG00000025184.5 |
| chr19 | 42613092 | 42613382 | 3.59837 | 4.33545 | 1.32923 | Promoter | 19 | 42595106 | 42612813 | 17708 | 2 | ENSMUST00000164786.2 | -280 | Loxl4 | ENSMUSG00000025185.9 |
| chr19 | 42636650 | 42636764 | 3.35921 | 3.86632 | 1.01733 | Distal Intergenic | 19 | 42595106 | 42612813 | 17708 | 2 | ENSMUST00000164786.2 | -23838 | Loxl4 | ENSMUSG00000025185.9 |
| chr19 | 42653210 | 42653295 | 3.57137 | 4.25902 | 1.27441 | Distal Intergenic | 19 | 42595106 | 42612813 | 17708 | 2 | ENSMUST00000164786.2 | -40398 | Loxl4 | ENSMUSG00000025185.9 |
| chr19 | 42795439 | 42795706 | 3.29894 | 3.80045 | 1.04532 | Intron (ENSMUST00000099428.3/ENSMUSG00000074852.3, intron 11 of 11) | 19 | 42755105 | 42779978 | 24874 | 2 | ENSMUST00000162004.2 | -15462 | Hps1 | ENSMUSG00000025188.10 |
| chr19 | 43440654 | 43440902 | 3.66613 | 4.83246 | 1.68794 | Promoter | 19 | 43440436 | 43497212 | 56777 | 1 | ENSMUST00000165311.1 | 219 | Cnnm1 | ENSMUSG00000025189.8 |
| chr19 | 43441262 | 43441778 | 3.51704 | 4.4284 | 1.39814 | Promoter | 19 | 43440436 | 43497212 | 56777 | 1 | ENSMUST00000165311.1 | 827 | Cnnm1 | ENSMUSG00000025189.8 |
| chr19 | 43442119 | 43442506 | 3.84804 | 4.94195 | 1.76442 | Intron (ENSMUST00000165311.1/ENSMUSG00000025189.8, intron 1 of 10) | 19 | 43440436 | 43497212 | 56777 | 1 | ENSMUST00000165311.1 | 1684 | Cnnm1 | ENSMUSG00000025189.8 |
| chr19 | 43442861 | 43443167 | 3.52043 | 4.29967 | 1.33201 | Intron (ENSMUST00000165311.1/ENSMUSG00000025189.8, intron 1 of 10) | 19 | 43440436 | 43497212 | 56777 | 1 | ENSMUST00000165311.1 | 2426 | Cnnm1 | ENSMUSG00000025189.8 |
| chr19 | 43444066 | 43444534 | 4.26059 | 5.81913 | 2.36806 | Intron (ENSMUST00000165311.1/ENSMUSG00000025189.8, intron 1 of 10) | 19 | 43440436 | 43497212 | 56777 | 1 | ENSMUST00000165311.1 | 3631 | Cnnm1 | ENSMUSG00000025189.8 |
| chr19 | 43531569 | 43531951 | 3.97368 | 5.30675 | 1.9686 | Distal Intergenic | 19 | 43499752 | 43524605 | 24854 | 2 | ENSMUST00000026196.8 | -6965 | Got1 | ENSMUSG00000025190.8 |
| chr19 | 43533003 | 43533175 | 3.40789 | 4.11812 | 1.22437 | Distal Intergenic | 19 | 43499752 | 43524605 | 24854 | 2 | ENSMUST00000026196.8 | -8399 | Got1 | ENSMUSG00000025190.8 |
| chr19 | 43543567 | 43543685 | 3.38193 | 3.9006 | 1.01733 | Distal Intergenic | 19 | 43499752 | 43524605 | 24854 | 2 | ENSMUST00000026196.8 | -18963 | Got1 | ENSMUSG00000025190.8 |
| chr19 | 43546538 | 43546940 | 3.54483 | 4.20658 | 1.22618 | Distal Intergenic | 19 | 43499752 | 43524605 | 24854 | 2 | ENSMUST00000026196.8 | -21934 | Got1 | ENSMUSG00000025190.8 |
| chr19 | 43547838 | 43548332 | 3.77271 | 4.70158 | 1.58369 | Distal Intergenic | 19 | 43499752 | 43524605 | 24854 | 2 | ENSMUST00000026196.8 | -23234 | Got1 | ENSMUSG00000025190.8 |
| chr19 | 43551679 | 43552305 | 3.6941 | 4.48841 | 1.4062 | Distal Intergenic | 19 | 43499752 | 43524605 | 24854 | 2 | ENSMUST00000026196.8 | -27075 | Got1 | ENSMUSG00000025190.8 |
| chr19 | 43566714 | 43566889 | 3.54884 | 4.22416 | 1.27216 | Distal Intergenic | 19 | 43499752 | 43524605 | 24854 | 2 | ENSMUST00000026196.8 | -42110 | Got1 | ENSMUSG00000025190.8 |
| chr19 | 43787789 | 43788103 | 3.64652 | 4.3837 | 1.31219 | Intron (ENSMUST00000026208.5/ENSMUSG00000025194.5, intron 2 of 31) | 19 | 43782326 | 43784640 | 2315 | 1 | ENSMUST00000143553.1 | 5464 | Abcc2 | ENSMUSG00000025194.5 |
| chr19 | 43807620 | 43808257 | 3.53846 | 4.19307 | 1.21596 | Intron (ENSMUST00000026208.5/ENSMUSG00000025194.5, intron 10 of 31) | 19 | 43810522 | 43813401 | 2880 | 1 | ENSMUST00000146737.1 | -2265 | Abcc2 | ENSMUSG00000025194.5 |
| chr19 | 43966240 | 43966481 | 3.88407 | 4.9731 | 1.73471 | Exon (ENSMUST00000026210.4/ENSMUSG00000025196.4, exon 6 of 9) | 19 | 43962000 | 43968693 | 6694 | 2 | ENSMUST00000131882.1 | 2212 | Cpn1 | ENSMUSG00000025196.4 |
| chr19 | 43968186 | 43968492 | 3.31542 | 3.80036 | 1.01733 | Promoter | 19 | 43962000 | 43968693 | 6694 | 2 | ENSMUST00000131882.1 | 201 | Cpn1 | ENSMUSG00000025196.4 |
| chr19 | 44989128 | 44989560 | 3.58223 | 4.2814 | 1.28598 | Promoter | 19 | 44989291 | 45003397 | 14107 | 1 | ENSMUST00000130549.2 | 0 | Sema4g | ENSMUSG00000025207.10 |
| chr19 | 45064073 | 45064339 | 3.65553 | 4.44218 | 1.37327 | Distal Intergenic | 19 | 45075241 | 45079289 | 4049 | 1 | ENSMUST00000111948.2 | -10902 | Kazald1 | ENSMUSG00000025213.6 |
| chr19 | 45064901 | 45066097 | 3.98362 | 5.12595 | 1.87872 | Distal Intergenic | 19 | 45075241 | 45079289 | 4049 | 1 | ENSMUST00000111948.2 | -9144 | Kazald1 | ENSMUSG00000025213.6 |
| chr19 | 45066685 | 45066861 | 3.36555 | 4.02125 | 1.13007 | Distal Intergenic | 19 | 45075241 | 45079289 | 4049 | 1 | ENSMUST00000111948.2 | -8380 | Kazald1 | ENSMUSG00000025213.6 |
| chr19 | 45067285 | 45067571 | 4.75997 | 6.67761 | 3.04521 | Distal Intergenic | 19 | 45075241 | 45079289 | 4049 | 1 | ENSMUST00000111948.2 | -7670 | Kazald1 | ENSMUSG00000025213.6 |
| chr19 | 45085844 | 45086442 | 3.6511 | 4.52411 | 1.46628 | Distal Intergenic | 19 | 45076139 | 45079289 | 3151 | 1 | ENSMUST00000026234.4 | 9706 | Kazald1 | ENSMUSG00000025213.6 |
| chr19 | 45086810 | 45087389 | 3.56409 | 4.28703 | 1.31721 | Distal Intergenic | 19 | 45076139 | 45079289 | 3151 | 1 | ENSMUST00000026234.4 | 10672 | Kazald1 | ENSMUSG00000025213.6 |
| chr19 | 45088068 | 45088954 | 3.42887 | 3.9885 | 1.07755 | Distal Intergenic | 19 | 45076139 | 45079289 | 3151 | 1 | ENSMUST00000026234.4 | 11930 | Kazald1 | ENSMUSG00000025213.6 |
| chr19 | 45100215 | 45100543 | 3.38397 | 3.93734 | 1.0758 | Distal Intergenic | 19 | 45076139 | 45079289 | 3151 | 1 | ENSMUST00000026234.4 | 24077 | Kazald1 | ENSMUSG00000025213.6 |
| chr19 | 45100983 | 45101348 | 3.43686 | 4.18116 | 1.27003 | Distal Intergenic | 19 | 45076139 | 45079289 | 3151 | 1 | ENSMUST00000026234.4 | 24845 | Kazald1 | ENSMUSG00000025213.6 |
| chr19 | 45102173 | 45103434 | 3.70839 | 4.52467 | 1.44137 | Distal Intergenic | 19 | 45076139 | 45079289 | 3151 | 1 | ENSMUST00000026234.4 | 26035 | Kazald1 | ENSMUSG00000025213.6 |
| chr19 | 45104458 | 45104565 | 4.97896 | 7.07349 | 3.2735 | Distal Intergenic | 19 | 45076139 | 45079289 | 3151 | 1 | ENSMUST00000026234.4 | 28320 | Kazald1 | ENSMUSG00000025213.6 |
| chr19 | 45105034 | 45105747 | 3.64333 | 4.58292 | 1.48249 | Distal Intergenic | 19 | 45076139 | 45079289 | 3151 | 1 | ENSMUST00000026234.4 | 28896 | Kazald1 | ENSMUSG00000025213.6 |
| chr19 | 45110921 | 45111278 | 3.70593 | 4.50551 | 1.41853 | Distal Intergenic | 19 | 45076139 | 45079289 | 3151 | 1 | ENSMUST00000026234.4 | 34783 | Kazald1 | ENSMUSG00000025213.6 |
| chr19 | 45115341 | 45115452 | 3.48016 | 4.08761 | 1.15442 | Distal Intergenic | 19 | 45150680 | 45156943 | 6264 | 1 | ENSMUST00000026236.10 | -35228 | Tlx1 | ENSMUSG00000025215.10 |
| chr19 | 45119127 | 45119202 | 3.44547 | 4.03982 | 1.17135 | Distal Intergenic | 19 | 45150680 | 45156943 | 6264 | 1 | ENSMUST00000026236.10 | -31478 | Tlx1 | ENSMUSG00000025215.10 |
| chr19 | 45121178 | 45121875 | 3.50743 | 4.14537 | 1.20977 | Distal Intergenic | 19 | 45150680 | 45156943 | 6264 | 1 | ENSMUST00000026236.10 | -28805 | Tlx1 | ENSMUSG00000025215.10 |
| chr19 | 45125479 | 45125590 | 3.5074 | 4.1483 | 1.21403 | Distal Intergenic | 19 | 45150680 | 45156943 | 6264 | 1 | ENSMUST00000026236.10 | -25090 | Tlx1 | ENSMUSG00000025215.10 |
| chr19 | 45126175 | 45126690 | 3.42682 | 4.08675 | 1.17496 | Distal Intergenic | 19 | 45150680 | 45156943 | 6264 | 1 | ENSMUST00000026236.10 | -23990 | Tlx1 | ENSMUSG00000025215.10 |
| chr19 | 45127467 | 45127542 | 3.3058 | 3.79226 | 1.03958 | Distal Intergenic | 19 | 45150680 | 45156943 | 6264 | 1 | ENSMUST00000026236.10 | -23138 | Tlx1 | ENSMUSG00000025215.10 |
| chr19 | 45128037 | 45128796 | 3.64349 | 4.40557 | 1.392 | Distal Intergenic | 19 | 45150680 | 45156943 | 6264 | 1 | ENSMUST00000026236.10 | -21884 | Tlx1 | ENSMUSG00000025215.10 |
| chr19 | 45129858 | 45129953 | 3.8184 | 4.75313 | 1.64178 | Distal Intergenic | 19 | 45150680 | 45156943 | 6264 | 1 | ENSMUST00000026236.10 | -20727 | Tlx1 | ENSMUSG00000025215.10 |
| chr19 | 45132377 | 45132554 | 3.81635 | 4.81262 | 1.66216 | Distal Intergenic | 19 | 45150680 | 45156943 | 6264 | 1 | ENSMUST00000026236.10 | -18126 | Tlx1 | ENSMUSG00000025215.10 |
| chr19 | 45132928 | 45133076 | 3.36981 | 3.88232 | 1.01733 | Distal Intergenic | 19 | 45150680 | 45156943 | 6264 | 1 | ENSMUST00000026236.10 | -17604 | Tlx1 | ENSMUSG00000025215.10 |
| chr19 | 45133396 | 45133749 | 3.63975 | 4.3845 | 1.33731 | Distal Intergenic | 19 | 45150680 | 45156943 | 6264 | 1 | ENSMUST00000026236.10 | -16931 | Tlx1 | ENSMUSG00000025215.10 |
| chr19 | 45134084 | 45134217 | 3.5081 | 4.23726 | 1.30315 | Distal Intergenic | 19 | 45150680 | 45156943 | 6264 | 1 | ENSMUST00000026236.10 | -16463 | Tlx1 | ENSMUSG00000025215.10 |
| chr19 | 45135643 | 45135847 | 3.89079 | 4.92131 | 1.71788 | Distal Intergenic | 19 | 45150680 | 45156943 | 6264 | 1 | ENSMUST00000026236.10 | -14833 | Tlx1 | ENSMUSG00000025215.10 |
| chr19 | 45136249 | 45136835 | 3.64361 | 4.38658 | 1.32942 | Distal Intergenic | 19 | 45150680 | 45156943 | 6264 | 1 | ENSMUST00000026236.10 | -13845 | Tlx1 | ENSMUSG00000025215.10 |
| chr19 | 45137246 | 45137326 | 4.93236 | 7.27834 | 3.43317 | Distal Intergenic | 19 | 45150680 | 45156943 | 6264 | 1 | ENSMUST00000026236.10 | -13354 | Tlx1 | ENSMUSG00000025215.10 |
| chr19 | 45138139 | 45138245 | 3.47889 | 4.08278 | 1.14379 | Distal Intergenic | 19 | 45150680 | 45156943 | 6264 | 1 | ENSMUST00000026236.10 | -12435 | Tlx1 | ENSMUSG00000025215.10 |
| chr19 | 45138637 | 45138745 | 4.25469 | 5.61033 | 2.19883 | Distal Intergenic | 19 | 45150680 | 45156943 | 6264 | 1 | ENSMUST00000026236.10 | -11935 | Tlx1 | ENSMUSG00000025215.10 |
| chr19 | 45139100 | 45139335 | 3.38699 | 3.92031 | 1.06004 | Distal Intergenic | 19 | 45150680 | 45156943 | 6264 | 1 | ENSMUST00000026236.10 | -11345 | Tlx1 | ENSMUSG00000025215.10 |
| chr19 | 45139996 | 45140220 | 3.61231 | 4.32184 | 1.2762 | Distal Intergenic | 19 | 45150680 | 45156943 | 6264 | 1 | ENSMUST00000026236.10 | -10460 | Tlx1 | ENSMUSG00000025215.10 |
| chr19 | 45140715 | 45141982 | 4.01578 | 5.15368 | 1.86466 | Distal Intergenic | 19 | 45150680 | 45156943 | 6264 | 1 | ENSMUST00000026236.10 | -8698 | Tlx1 | ENSMUSG00000025215.10 |
| chr19 | 45142439 | 45144667 | 3.98776 | 5.13801 | 1.90193 | Distal Intergenic | 19 | 45150680 | 45156943 | 6264 | 1 | ENSMUST00000026236.10 | -6013 | Tlx1 | ENSMUSG00000025215.10 |
| chr19 | 45146472 | 45147614 | 3.62224 | 4.5508 | 1.49052 | Distal Intergenic | 19 | 45150680 | 45156943 | 6264 | 1 | ENSMUST00000026236.10 | -3066 | Tlx1 | ENSMUSG00000025215.10 |
| chr19 | 45148216 | 45149029 | 4.23859 | 5.57062 | 2.17928 | Distal Intergenic | 19 | 45150680 | 45156943 | 6264 | 1 | ENSMUST00000026236.10 | -1651 | Tlx1 | ENSMUSG00000025215.10 |
| chr19 | 45149522 | 45150331 | 3.83087 | 4.81005 | 1.70079 | Promoter | 19 | 45150680 | 45156943 | 6264 | 1 | ENSMUST00000026236.10 | -349 | Tlx1 | ENSMUSG00000025215.10 |
| chr19 | 45150840 | 45151833 | 4.27394 | 5.7397 | 2.41558 | Promoter | 19 | 45151469 | 45156399 | 4931 | 1 | ENSMUST00000174617.2 | 0 | Tlx1 | ENSMUSG00000025215.10 |
| chr19 | 45153258 | 45153474 | 4.1802 | 5.60725 | 2.19563 | Intron (ENSMUST00000026236.10/ENSMUSG00000025215.10, intron 1 of 2) | 19 | 45151469 | 45156399 | 4931 | 1 | ENSMUST00000174617.2 | 1790 | Tlx1 | ENSMUSG00000025215.10 |
| chr19 | 45153830 | 45154123 | 4.015 | 5.56438 | 2.17851 | Exon (ENSMUST00000173437.1/ENSMUSG00000092346.1, exon 2 of 2) | 19 | 45151469 | 45156399 | 4931 | 1 | ENSMUST00000174617.2 | 2362 | Tlx1 | ENSMUSG00000025215.10 |
| chr19 | 45208512 | 45209082 | 3.63326 | 4.36571 | 1.31642 | Distal Intergenic | 19 | 45232684 | 45235812 | 3129 | 2 | ENSMUST00000099401.5 | 26730 | Lbx1 | ENSMUSG00000025216.9 |
| chr19 | 45209912 | 45210143 | 3.64657 | 4.40359 | 1.37176 | Distal Intergenic | 19 | 45232684 | 45235812 | 3129 | 2 | ENSMUST00000099401.5 | 25669 | Lbx1 | ENSMUSG00000025216.9 |
| chr19 | 45221575 | 45222949 | 4.20486 | 5.57232 | 2.22534 | Distal Intergenic | 19 | 45232684 | 45235812 | 3129 | 2 | ENSMUST00000099401.5 | 12863 | Lbx1 | ENSMUSG00000025216.9 |
| chr19 | 45223498 | 45223590 | 3.48002 | 4.18836 | 1.24189 | Distal Intergenic | 19 | 45232684 | 45235812 | 3129 | 2 | ENSMUST00000099401.5 | 12222 | Lbx1 | ENSMUSG00000025216.9 |
| chr19 | 45223944 | 45224504 | 3.54392 | 4.53269 | 1.48875 | Distal Intergenic | 19 | 45232684 | 45235812 | 3129 | 2 | ENSMUST00000099401.5 | 11308 | Lbx1 | ENSMUSG00000025216.9 |
| chr19 | 45224973 | 45225048 | 4.04809 | 5.17614 | 1.88276 | Distal Intergenic | 19 | 45232684 | 45235812 | 3129 | 2 | ENSMUST00000099401.5 | 10764 | Lbx1 | ENSMUSG00000025216.9 |
| chr19 | 45225384 | 45225715 | 4.03238 | 5.16876 | 1.87882 | Distal Intergenic | 19 | 45232684 | 45235812 | 3129 | 2 | ENSMUST00000099401.5 | 10097 | Lbx1 | ENSMUSG00000025216.9 |
| chr19 | 45226253 | 45227175 | 4.23326 | 5.7068 | 2.28779 | Distal Intergenic | 19 | 45232684 | 45235812 | 3129 | 2 | ENSMUST00000099401.5 | 8637 | Lbx1 | ENSMUSG00000025216.9 |
| chr19 | 45227611 | 45228310 | 3.69783 | 4.50634 | 1.44129 | Distal Intergenic | 19 | 45232684 | 45235812 | 3129 | 2 | ENSMUST00000099401.5 | 7502 | Lbx1 | ENSMUSG00000025216.9 |
| chr19 | 45228701 | 45228962 | 3.43253 | 4.00051 | 1.10076 | Distal Intergenic | 19 | 45232684 | 45235812 | 3129 | 2 | ENSMUST00000099401.5 | 6850 | Lbx1 | ENSMUSG00000025216.9 |
| chr19 | 45229419 | 45230636 | 3.76827 | 4.87696 | 1.71573 | Downstream (2-3kb) | 19 | 45232684 | 45235812 | 3129 | 2 | ENSMUST00000099401.5 | 5176 | Lbx1 | ENSMUSG00000025216.9 |
| chr19 | 45230962 | 45232903 | 3.95772 | 5.06348 | 1.81916 | 3' UTR | 19 | 45232684 | 45235812 | 3129 | 2 | ENSMUST00000099401.5 | 2909 | Lbx1 | ENSMUSG00000025216.9 |
| chr19 | 45233235 | 45235766 | 3.89507 | 5.00489 | 1.81416 | Promoter | 19 | 45232684 | 45235812 | 3129 | 2 | ENSMUST00000099401.5 | 46 | Lbx1 | ENSMUSG00000025216.9 |
| chr19 | 45236164 | 45237192 | 3.9579 | 5.15384 | 1.8722 | Promoter | 19 | 45232684 | 45235812 | 3129 | 2 | ENSMUST00000099401.5 | -353 | Lbx1 | ENSMUSG00000025216.9 |
| chr19 | 45238039 | 45239707 | 3.476 | 4.1833 | 1.28925 | Distal Intergenic | 19 | 45232684 | 45235812 | 3129 | 2 | ENSMUST00000099401.5 | -2228 | Lbx1 | ENSMUSG00000025216.9 |
| chr19 | 45240284 | 45240698 | 3.77133 | 4.67232 | 1.60255 | Distal Intergenic | 19 | 45232684 | 45235812 | 3129 | 2 | ENSMUST00000099401.5 | -4473 | Lbx1 | ENSMUSG00000025216.9 |
| chr19 | 45241085 | 45241495 | 3.97198 | 5.29508 | 1.97104 | Distal Intergenic | 19 | 45232684 | 45235812 | 3129 | 2 | ENSMUST00000099401.5 | -5274 | Lbx1 | ENSMUSG00000025216.9 |
| chr19 | 45242172 | 45244516 | 3.50676 | 4.46178 | 1.44776 | Distal Intergenic | 19 | 45232684 | 45235812 | 3129 | 2 | ENSMUST00000099401.5 | -6361 | Lbx1 | ENSMUSG00000025216.9 |
| chr19 | 45244829 | 45245034 | 3.41947 | 3.99909 | 1.16169 | Distal Intergenic | 19 | 45232684 | 45235812 | 3129 | 2 | ENSMUST00000099401.5 | -9018 | Lbx1 | ENSMUSG00000025216.9 |
| chr19 | 45245668 | 45246174 | 3.36953 | 3.91551 | 1.12641 | Distal Intergenic | 19 | 45232684 | 45235812 | 3129 | 2 | ENSMUST00000099401.5 | -9857 | Lbx1 | ENSMUSG00000025216.9 |
| chr19 | 45255781 | 45255919 | 3.30977 | 3.79186 | 1.01733 | Distal Intergenic | 19 | 45232684 | 45235812 | 3129 | 2 | ENSMUST00000099401.5 | -19970 | Lbx1 | ENSMUSG00000025216.9 |
| chr19 | 45262700 | 45262822 | 3.29673 | 3.77226 | 1.01733 | Distal Intergenic | 19 | 45232684 | 45235812 | 3129 | 2 | ENSMUST00000099401.5 | -26889 | Lbx1 | ENSMUSG00000025216.9 |
| chr19 | 45540849 | 45541216 | 4.14462 | 5.34706 | 1.97316 | Distal Intergenic | 19 | 45552275 | 45560531 | 8257 | 2 | ENSMUST00000026239.6 | 19315 | Poll | ENSMUSG00000025218.6 |
| chr19 | 45541532 | 45542109 | 3.70828 | 4.50971 | 1.41149 | Distal Intergenic | 19 | 45552275 | 45560531 | 8257 | 2 | ENSMUST00000026239.6 | 18422 | Poll | ENSMUSG00000025218.6 |
| chr19 | 45542514 | 45542816 | 3.82667 | 4.73847 | 1.56167 | Distal Intergenic | 19 | 45552275 | 45560531 | 8257 | 2 | ENSMUST00000026239.6 | 17715 | Poll | ENSMUSG00000025218.6 |
| chr19 | 45543248 | 45543846 | 3.75042 | 4.61903 | 1.52282 | Distal Intergenic | 19 | 45552275 | 45560531 | 8257 | 2 | ENSMUST00000026239.6 | 16685 | Poll | ENSMUSG00000025218.6 |
| chr19 | 45545188 | 45545613 | 3.85039 | 4.8361 | 1.64858 | Distal Intergenic | 19 | 45552275 | 45560531 | 8257 | 2 | ENSMUST00000026239.6 | 14918 | Poll | ENSMUSG00000025218.6 |
| chr19 | 45545927 | 45546440 | 3.45321 | 4.15116 | 1.22693 | Distal Intergenic | 19 | 45552275 | 45560531 | 8257 | 2 | ENSMUST00000026239.6 | 14091 | Poll | ENSMUSG00000025218.6 |
| chr19 | 45547244 | 45547489 | 3.5021 | 4.13349 | 1.1922 | Distal Intergenic | 19 | 45552275 | 45560531 | 8257 | 2 | ENSMUST00000026239.6 | 13042 | Poll | ENSMUSG00000025218.6 |
| chr19 | 45548873 | 45550079 | 3.46629 | 4.08725 | 1.17484 | Downstream (2-3kb) | 19 | 45552275 | 45560531 | 8257 | 2 | ENSMUST00000026239.6 | 10452 | Poll | ENSMUSG00000025218.6 |
| chr19 | 45707293 | 45707572 | 3.38193 | 3.9006 | 1.01733 | Distal Intergenic | 19 | 45736971 | 45742684 | 5714 | 2 | ENSMUST00000111924.2 | 35112 | Fgf8 | ENSMUSG00000025219.8 |
| chr19 | 45709588 | 45709663 | 3.61876 | 4.34194 | 1.31396 | Distal Intergenic | 19 | 45736971 | 45742684 | 5714 | 2 | ENSMUST00000111924.2 | 33021 | Fgf8 | ENSMUSG00000025219.8 |
| chr19 | 45710107 | 45710291 | 3.47312 | 4.07398 | 1.14379 | Distal Intergenic | 19 | 45736971 | 45742684 | 5714 | 2 | ENSMUST00000111924.2 | 32393 | Fgf8 | ENSMUSG00000025219.8 |
| chr19 | 45712388 | 45712981 | 3.52566 | 4.1711 | 1.20122 | Distal Intergenic | 19 | 45736971 | 45742684 | 5714 | 2 | ENSMUST00000111924.2 | 29703 | Fgf8 | ENSMUSG00000025219.8 |
| chr19 | 45716901 | 45717313 | 3.86021 | 4.926 | 1.73663 | Distal Intergenic | 19 | 45736971 | 45742684 | 5714 | 2 | ENSMUST00000111924.2 | 25371 | Fgf8 | ENSMUSG00000025219.8 |
| chr19 | 45718075 | 45718329 | 4.19306 | 5.66142 | 2.2239 | Distal Intergenic | 19 | 45736971 | 45742684 | 5714 | 2 | ENSMUST00000111924.2 | 24355 | Fgf8 | ENSMUSG00000025219.8 |
| chr19 | 45721781 | 45722080 | 3.69096 | 4.53771 | 1.51371 | Distal Intergenic | 19 | 45736971 | 45742684 | 5714 | 2 | ENSMUST00000111924.2 | 20604 | Fgf8 | ENSMUSG00000025219.8 |
| chr19 | 45722979 | 45723107 | 3.78913 | 4.99095 | 1.76437 | Distal Intergenic | 19 | 45736971 | 45742684 | 5714 | 2 | ENSMUST00000111924.2 | 19577 | Fgf8 | ENSMUSG00000025219.8 |
| chr19 | 45724513 | 45724657 | 3.80644 | 4.79626 | 1.64646 | Distal Intergenic | 19 | 45736971 | 45742684 | 5714 | 2 | ENSMUST00000111924.2 | 18027 | Fgf8 | ENSMUSG00000025219.8 |
| chr19 | 45725968 | 45726357 | 4.01489 | 5.10881 | 1.83489 | Distal Intergenic | 19 | 45736971 | 45742684 | 5714 | 2 | ENSMUST00000111924.2 | 16327 | Fgf8 | ENSMUSG00000025219.8 |
| chr19 | 45728854 | 45729193 | 3.548 | 4.20803 | 1.21834 | Distal Intergenic | 19 | 45736971 | 45742684 | 5714 | 2 | ENSMUST00000111924.2 | 13491 | Fgf8 | ENSMUSG00000025219.8 |
| chr19 | 45732836 | 45733106 | 3.50292 | 4.15904 | 1.27544 | Distal Intergenic | 19 | 45736971 | 45742684 | 5714 | 2 | ENSMUST00000111924.2 | 9578 | Fgf8 | ENSMUSG00000025219.8 |
| chr19 | 45733795 | 45733910 | 3.54679 | 4.21523 | 1.23898 | Downstream (2-3kb) | 19 | 45736971 | 45742684 | 5714 | 2 | ENSMUST00000111924.2 | 8774 | Fgf8 | ENSMUSG00000025219.8 |
| chr19 | 45735650 | 45735800 | 3.37183 | 3.88536 | 1.01733 | Downstream (<1kb) | 19 | 45736971 | 45742684 | 5714 | 2 | ENSMUST00000111924.2 | 6884 | Fgf8 | ENSMUSG00000025219.8 |
| chr19 | 45736769 | 45738653 | 3.91571 | 4.94013 | 1.72512 | 3' UTR | 19 | 45736971 | 45742684 | 5714 | 2 | ENSMUST00000111924.2 | 4031 | Fgf8 | ENSMUSG00000025219.8 |
| chr19 | 45739812 | 45740019 | 3.58018 | 4.26726 | 1.25062 | Intron (ENSMUST00000026240.8/ENSMUSG00000025219.8, intron 4 of 5) | 19 | 45736971 | 45742684 | 5714 | 2 | ENSMUST00000111924.2 | 2665 | Fgf8 | ENSMUSG00000025219.8 |
| chr19 | 45740394 | 45740545 | 4.32353 | 5.71128 | 2.24214 | Intron (ENSMUST00000026240.8/ENSMUSG00000025219.8, intron 4 of 5) | 19 | 45736971 | 45742684 | 5714 | 2 | ENSMUST00000111924.2 | 2139 | Fgf8 | ENSMUSG00000025219.8 |
| chr19 | 45740894 | 45742927 | 4.21356 | 5.58732 | 2.19243 | Promoter | 19 | 45736798 | 45742874 | 6077 | 2 | ENSMUST00000026240.8 | 0 | Fgf8 | ENSMUSG00000025219.8 |
| chr19 | 45743372 | 45745772 | 3.70209 | 4.56863 | 1.49921 | Promoter | 19 | 45736813 | 45742915 | 6103 | 2 | ENSMUST00000111928.2 | -458 | Fgf8 | ENSMUSG00000025219.8 |
| chr19 | 46313943 | 46314955 | 3.57538 | 4.32348 | 1.34034 | Exon (ENSMUST00000041391.4/ENSMUSG00000037126.10, exon 13 of 18) | 19 | 46305695 | 46312090 | 6396 | 1 | ENSMUST00000111881.2 | 8249 | Nfkb2 | ENSMUSG00000025225.9 |
| chr19 | 46315395 | 46316722 | 3.71856 | 4.57036 | 1.48725 | Exon (ENSMUST00000041391.4/ENSMUSG00000037126.10, exon 12 of 18) | 19 | 46305695 | 46312090 | 6396 | 1 | ENSMUST00000111881.2 | 9701 | Nfkb2 | ENSMUSG00000025225.9 |
| chr19 | 46134318 | 46134903 | 3.71163 | 4.73843 | 1.61039 | 3' UTR | 19 | 46136495 | 46137289 | 795 | 2 | ENSMUST00000172980.1 | 2386 | Pitx3 | ENSMUSG00000025229.10 |
| chr19 | 46136935 | 46137401 | 3.82352 | 4.81688 | 1.7281 | Promoter | 19 | 46136495 | 46137289 | 795 | 2 | ENSMUST00000172980.1 | 0 | Pitx3 | ENSMUSG00000025229.10 |
| chr19 | 46147135 | 46147721 | 3.59023 | 4.33942 | 1.35009 | Promoter | 19 | 46135685 | 46148326 | 12642 | 2 | ENSMUST00000026259.10 | 605 | Pitx3 | ENSMUSG00000025229.10 |
| chr19 | 46148314 | 46148855 | 3.68525 | 4.54258 | 1.49087 | Promoter | 19 | 46135685 | 46148326 | 12642 | 2 | ENSMUST00000026259.10 | 0 | Pitx3 | ENSMUSG00000025229.10 |
| chr19 | 46149497 | 46150112 | 4.35812 | 6.28504 | 2.75028 | Distal Intergenic | 19 | 46135685 | 46148326 | 12642 | 2 | ENSMUST00000026259.10 | -1172 | Pitx3 | ENSMUSG00000025229.10 |
| chr9 | 59266064 | 59266327 | 3.37595 | 3.89157 | 1.01733 | Distal Intergenic | 9 | 59291572 | 59316199 | 24628 | 1 | ENSMUST00000026266.7 | -25245 | Adpgk | ENSMUSG00000025236.10 |
| chr9 | 123461934 | 123462167 | 2.49174 | 13.32602 | 8.80106 | 3' UTR | 9 | 123478701 | 123521548 | 42848 | 1 | ENSMUST00000026269.2 | -16534 | Limd1 | ENSMUSG00000025239.2 |
| chr3 | 5203036 | 5203271 | 3.67338 | 4.4531 | 1.39561 | Intron (ENSMUST00000128815.3/ENSMUSG00000086224.3, intron 1 of 2) | 3 | 5218526 | 5413165 | 194640 | 1 | ENSMUST00000175866.2 | -15255 | Zfhx4 | ENSMUSG00000025255.13 |
| chr3 | 5203579 | 5203798 | 3.54695 | 4.3756 | 1.33853 | Intron (ENSMUST00000128815.3/ENSMUSG00000086224.3, intron 1 of 2) | 3 | 5218526 | 5413165 | 194640 | 1 | ENSMUST00000175866.2 | -14728 | Zfhx4 | ENSMUSG00000025255.13 |
| chr3 | 5204648 | 5205024 | 3.61866 | 4.3359 | 1.28875 | Intron (ENSMUST00000128815.3/ENSMUSG00000086224.3, intron 1 of 2) | 3 | 5218526 | 5413165 | 194640 | 1 | ENSMUST00000175866.2 | -13502 | Zfhx4 | ENSMUSG00000025255.13 |
| chr3 | 5205692 | 5206123 | 3.57876 | 4.25946 | 1.23486 | Intron (ENSMUST00000128815.3/ENSMUSG00000086224.3, intron 1 of 2) | 3 | 5218526 | 5413165 | 194640 | 1 | ENSMUST00000175866.2 | -12403 | Zfhx4 | ENSMUSG00000025255.13 |
| chr3 | 5206481 | 5206556 | 3.80037 | 4.68275 | 1.53794 | Intron (ENSMUST00000128815.3/ENSMUSG00000086224.3, intron 1 of 2) | 3 | 5218526 | 5413165 | 194640 | 1 | ENSMUST00000175866.2 | -11970 | Zfhx4 | ENSMUSG00000025255.13 |
| chr3 | 5206908 | 5207505 | 4.08098 | 5.27188 | 1.96394 | Intron (ENSMUST00000128815.3/ENSMUSG00000086224.3, intron 1 of 2) | 3 | 5218526 | 5413165 | 194640 | 1 | ENSMUST00000175866.2 | -11021 | Zfhx4 | ENSMUSG00000025255.13 |

| | | | | | | | | | | | | | | | |
|---|---|---|---|---|---|---|---|---|---|---|---|---|---|---|---|
| chr3 | 5209002 | 5209077 | 4.23021 | 5.6582 | 2.2299 | Intron (ENSMUST00000128815.3/ENSMUSG00000086224.3, intron 1 of 2) | 3 | 5218526 | 5413165 | 194640 | 1 | ENSMUST00000175866.2 | -9449 | Zfhx4 | ENSMUSG00000025255.13 |
| chr3 | 5209602 | 5209947 | 3.91805 | 4.92634 | 1.7221 | Intron (ENSMUST00000128815.3/ENSMUSG00000086224.3, intron 1 of 2) | 3 | 5218526 | 5413165 | 194640 | 1 | ENSMUST00000175866.2 | -8579 | Zfhx4 | ENSMUSG00000025255.13 |
| chr3 | 5210256 | 5210665 | 4.10296 | 5.31849 | 1.97233 | Intron (ENSMUST00000128815.3/ENSMUSG00000086224.3, intron 1 of 2) | 3 | 5218526 | 5413165 | 194640 | 1 | ENSMUST00000175866.2 | -7861 | Zfhx4 | ENSMUSG00000025255.13 |
| chr3 | 5211120 | 5212087 | 3.89631 | 5.02096 | 1.8458 | Intron (ENSMUST00000128815.3/ENSMUSG00000086224.3, intron 1 of 2) | 3 | 5218526 | 5413165 | 194640 | 1 | ENSMUST00000175866.2 | -6439 | Zfhx4 | ENSMUSG00000025255.13 |
| chr3 | 5212539 | 5213088 | 3.74531 | 4.92638 | 1.74893 | Intron (ENSMUST00000128815.3/ENSMUSG00000086224.3, intron 1 of 2) | 3 | 5218526 | 5413165 | 194640 | 1 | ENSMUST00000175866.2 | -5438 | Zfhx4 | ENSMUSG00000025255.13 |
| chr3 | 5213407 | 5213675 | 3.64413 | 5.10784 | 1.8343 | Intron (ENSMUST00000128815.3/ENSMUSG00000086224.3, intron 1 of 2) | 3 | 5218526 | 5413165 | 194640 | 1 | ENSMUST00000175866.2 | -4851 | Zfhx4 | ENSMUSG00000025255.13 |
| chr3 | 5215280 | 5215362 | 3.74469 | 4.71905 | 1.56903 | Intron (ENSMUST00000128815.3/ENSMUSG00000086224.3, intron 1 of 2) | 3 | 5218526 | 5413165 | 194640 | 1 | ENSMUST00000175866.2 | -3164 | Zfhx4 | ENSMUSG00000025255.13 |
| chr3 | 5216258 | 5217227 | 3.62037 | 4.49449 | 1.48554 | Intron (ENSMUST00000128815.3/ENSMUSG00000086224.3, intron 1 of 2) | 3 | 5218526 | 5413165 | 194640 | 1 | ENSMUST00000175866.2 | -1299 | Zfhx4 | ENSMUSG00000025255.13 |
| chr3 | 5217758 | 5218700 | 4.14354 | 5.74494 | 2.31893 | Promoter | 3 | 5218526 | 5413165 | 194640 | 1 | ENSMUST00000175866.2 | 0 | Zfhx4 | ENSMUSG00000025255.13 |
| chr3 | 5226582 | 5226836 | 3.53785 | 4.23477 | 1.32824 | Intron (ENSMUST00000175866.2/ENSMUSG00000025255.13, intron 1 of 10) | 3 | 5221505 | 5242018 | 20514 | 1 | ENSMUST00000176175.1 | 5078 | Zfhx4 | ENSMUSG00000025255.13 |
| chr3 | 5221377 | 5221734 | 3.86455 | 4.94338 | 1.78707 | Promoter | 3 | 5221505 | 5242018 | 20514 | 1 | ENSMUST00000176175.1 | 0 | Zfhx4 | ENSMUSG00000025255.13 |
| chr3 | 5222220 | 5222564 | 3.95864 | 5.08095 | 1.90219 | Promoter | 3 | 5221505 | 5242018 | 20514 | 1 | ENSMUST00000176175.1 | 716 | Zfhx4 | ENSMUSG00000025255.13 |
| chr3 | 5223574 | 5224461 | 3.33718 | 3.83879 | 1.03675 | Intron (ENSMUST00000175866.2/ENSMUSG00000025255.13, intron 1 of 10) | 3 | 5221505 | 5242018 | 20514 | 1 | ENSMUST00000176175.1 | 2070 | Zfhx4 | ENSMUSG00000025255.13 |
| chr3 | 5227373 | 5227998 | 3.51444 | 4.18928 | 1.31259 | Intron (ENSMUST00000175866.2/ENSMUSG00000025255.13, intron 1 of 10) | 3 | 5221505 | 5242018 | 20514 | 1 | ENSMUST00000176175.1 | 5869 | Zfhx4 | ENSMUSG00000025255.13 |
| chr3 | 5236920 | 5237622 | 3.49868 | 4.13072 | 1.20885 | Intron (ENSMUST00000175866.2/ENSMUSG00000025255.13, intron 1 of 10) | 3 | 5241672 | 5415857 | 174186 | 1 | ENSMUST00000176383.2 | -4050 | Zfhx4 | ENSMUSG00000025255.13 |
| chr14 | 8000528 | 8000910 | 3.5856 | 4.30394 | 1.33818 | Distal Intergenic | 14 | 8002902 | 8056555 | 53654 | 1 | ENSMUST00000166497.2 | -1992 | Abhd6 | ENSMUSG00000025277.8 |
| chr14 | 8001277 | 8001484 | 3.51077 | 4.14155 | 1.18162 | Distal Intergenic | 14 | 8002902 | 8056555 | 53654 | 1 | ENSMUST00000166497.2 | -1418 | Abhd6 | ENSMUSG00000025277.8 |
| chr14 | 7817300 | 7817421 | 3.41773 | 4.02049 | 1.18035 | Promoter | 14 | 7817957 | 7951588 | 133632 | 1 | ENSMUST00000052678.8 | -536 | Flnb | ENSMUSG00000025278.8 |
| chr14 | 7818042 | 7818849 | 3.56946 | 4.31134 | 1.3588 | Promoter | 14 | 7817957 | 7951588 | 133632 | 1 | ENSMUST00000052678.8 | 86 | Flnb | ENSMUSG00000025278.8 |
| chr14 | 7829791 | 7829918 | 3.33064 | 3.85557 | 1.11663 | Intron (ENSMUST00000052678.8/ENSMUSG00000025278.8, intron 1 of 45) | 14 | 7817957 | 7951588 | 133632 | 1 | ENSMUST00000052678.8 | 11835 | Flnb | ENSMUSG00000025278.8 |
| chr14 | 7853752 | 7853944 | 3.41878 | 4.01494 | 1.20518 | Intron (ENSMUST00000052678.8/ENSMUSG00000025278.8, intron 1 of 45) | 14 | 7817957 | 7951588 | 133632 | 1 | ENSMUST00000052678.8 | 35796 | Flnb | ENSMUSG00000025278.8 |
| chr14 | 7899074 | 7899740 | 3.93512 | 5.0525 | 1.89538 | Intron (ENSMUST00000052678.8/ENSMUSG00000025278.8, intron 18 of 45) | 14 | 7975872 | 7977222 | 1351 | 2 | ENSMUST00000145234.1 | 77482 | Dnase1l3 | ENSMUSG00000025279.7 |
| chr14 | 7903773 | 7903929 | 3.5744 | 4.33786 | 1.36335 | Intron (ENSMUST00000052678.8/ENSMUSG00000025278.8, intron 19 of 45) | 14 | 7975872 | 7977222 | 1351 | 2 | ENSMUST00000145234.1 | 73293 | Dnase1l3 | ENSMUSG00000025279.7 |
| chr14 | 24508963 | 24509188 | 3.56416 | 4.26991 | 1.27648 | Distal Intergenic | 14 | 24490681 | 24496146 | 5466 | 1 | ENSMUST00000169826.1 | 18283 | Rps24 | ENSMUSG00000025290.11 |
| chr8 | 122043728 | 122043925 | 3.73656 | 4.56101 | 1.44421 | Intron (ENSMUST00000127664.1/ENSMUSG00000092329.1, intron 1 of 15) | 8 | 122000847 | 122007140 | 6294 | 1 | ENSMUST00000173201.1 | 42882 | Banp | ENSMUSG00000025316.11 |
| chr8 | 122077026 | 122077101 | 3.86401 | 4.82307 | 1.67678 | Intron (ENSMUST00000127664.1/ENSMUSG00000092329.1, intron 1 of 15) | 8 | 122000847 | 122007140 | 6294 | 1 | ENSMUST00000173201.1 | 76180 | Banp | ENSMUSG00000025316.11 |
| chr8 | 122122915 | 122123225 | 3.38307 | 3.98359 | 1.13048 | Intron (ENSMUST00000127664.1/ENSMUSG00000092329.1, intron 1 of 15) | 8 | 122000847 | 122007140 | 6294 | 1 | ENSMUST00000173201.1 | 122069 | Banp | ENSMUSG00000025316.11 |
| chr8 | 121925157 | 121925394 | 3.27822 | 3.76503 | 1.06212 | Intron (ENSMUST00000127664.1/ENSMUSG00000092329.1, intron 1 of 15) | 8 | 121918054 | 121927474 | 9421 | 2 | ENSMUST00000151462.1 | 2080 | Car5a | ENSMUSG00000025317.7 |
| chr8 | 121727588 | 121727950 | 3.5001 | 4.1918 | 1.21752 | Intron (ENSMUST00000127664.1/ENSMUSG00000092329.1, intron 1 of 15) | 8 | 121729623 | 121789617 | 59995 | 1 | ENSMUST00000169735.1 | -1673 | Jph3 | ENSMUSG00000025318.8 |
| chr8 | 121728871 | 121729429 | 3.63807 | 4.39738 | 1.35869 | Promoter | 8 | 121729623 | 121789617 | 59995 | 1 | ENSMUST00000169735.1 | -194 | Jph3 | ENSMUSG00000025318.8 |
| chr8 | 121729742 | 121730054 | 3.61869 | 4.3399 | 1.29564 | Promoter | 8 | 121729623 | 121789617 | 59995 | 1 | ENSMUST00000169735.1 | 120 | Jph3 | ENSMUSG00000025318.8 |
| chr8 | 121730363 | 121731620 | 3.81229 | 4.75878 | 1.60083 | Promoter | 8 | 121730563 | 121790821 | 60259 | 1 | ENSMUST00000026357.6 | 0 | Jph3 | ENSMUSG00000025318.8 |
| chr8 | 121733504 | 121733741 | 3.65228 | 4.45428 | 1.43801 | Intron (ENSMUST00000127664.1/ENSMUSG00000092329.1, intron 1 of 15) | 8 | 121730567 | 121794276 | 63710 | 1 | ENSMUST00000167439.1 | 2938 | Jph3 | ENSMUSG00000025318.8 |
| chr8 | 121772298 | 121772417 | 3.35203 | 3.8644 | 1.04874 | Intron (ENSMUST00000127664.1/ENSMUSG00000092329.1, intron 1 of 15) | 8 | 121753476 | 121782570 | 29095 | 1 | ENSMUST00000172209.1 | 18823 | Jph3 | ENSMUSG00000025318.8 |
| chr7 | 58658523 | 58658759 | 3.36124 | 3.86941 | 1.01733 | Promoter | 7 | 58658202 | 58829426 | 171225 | 1 | ENSMUST00000168747.1 | 322 | Atp10a | ENSMUSG00000025324.7 |
| chr4 | 140834026 | 140834324 | 3.72118 | 4.58871 | 1.54478 | Intron (ENSMUST00000026378.3/ENSMUSG00000025329.3, intron 1 of 15) | 4 | 140812983 | 140845778 | 32796 | 2 | ENSMUST00000026378.3 | 11454 | Padi1 | ENSMUSG00000025329.3 |
| chr4 | 140837791 | 140837970 | 3.92013 | 4.9013 | 1.65581 | Intron (ENSMUST00000026378.3/ENSMUSG00000025329.3, intron 1 of 15) | 4 | 140812983 | 140845778 | 32796 | 2 | ENSMUST00000026378.3 | 7808 | Padi1 | ENSMUSG00000025329.3 |
| chr4 | 140858208 | 140858430 | 3.54491 | 4.22174 | 1.27334 | Distal Intergenic | 4 | 140812983 | 140845778 | 32796 | 2 | ENSMUST00000026378.3 | -12431 | Padi1 | ENSMUSG00000025329.3 |
| chr10 | 128932312 | 128932465 | 3.34101 | 3.88049 | 1.09505 | Distal Intergenic | 10 | 128933813 | 128958277 | 24465 | 1 | ENSMUST00000099112.2 | -1348 | Itga7 | ENSMUSG00000025348.8 |
| chr10 | 128933445 | 128934162 | 4.06969 | 5.28233 | 1.99745 | Promoter | 10 | 128933813 | 128958277 | 24465 | 1 | ENSMUST00000099112.2 | 0 | Itga7 | ENSMUSG00000025348.8 |
| chr10 | 128889707 | 128889782 | 3.45645 | 4.04229 | 1.12067 | Intron (ENSMUST00000026408.6/ENSMUSG00000025352.6, intron 1 of 2) | 10 | 128882295 | 128891718 | 9424 | 2 | ENSMUST00000026408.6 | 1936 | Gdf11 | ENSMUSG00000025352.6 |
| chr10 | 128892274 | 128892718 | 3.70716 | 4.56253 | 1.51285 | Promoter | 10 | 128882295 | 128891718 | 9424 | 2 | ENSMUST00000026408.6 | -557 | Gdf11 | ENSMUSG00000025352.6 |
| chr10 | 128893150 | 128893723 | 3.36577 | 3.87622 | 1.01733 | Distal Intergenic | 10 | 128882295 | 128891718 | 9424 | 2 | ENSMUST00000026408.6 | -1433 | Gdf11 | ENSMUSG00000025352.6 |
| chr10 | 128894623 | 128894725 | 3.28441 | 3.75386 | 1.01733 | Distal Intergenic | 10 | 128882295 | 128891718 | 9424 | 2 | ENSMUST00000026408.6 | -2906 | Gdf11 | ENSMUSG00000025352.6 |
| chr11 | 119907680 | 119908147 | 3.99398 | 5.24047 | 1.948 | Distal Intergenic | 11 | 119913441 | 119917008 | 3568 | 1 | ENSMUST00000132197.2 | -5294 | Chmp6 | ENSMUSG00000025371.7 |
| chr11 | 120011883 | 120012364 | 3.57031 | 4.41433 | 1.36737 | Promoter | 11 | 120007313 | 120012408 | 5096 | 2 | ENSMUST00000142959.1 | 44 | Aatk | ENSMUSG00000025375.10 |
| chr11 | 120046006 | 120047161 | 3.93689 | 5.1448 | 1.91022 | Promoter | 11 | 120007313 | 120047070 | 39758 | 2 | ENSMUST00000064307.4 | 0 | Aatk | ENSMUSG00000025375.10 |
| chr11 | 120048272 | 120048530 | 3.83383 | 4.92745 | 1.71709 | Distal Intergenic | 11 | 120007313 | 120047167 | 39855 | 2 | ENSMUST00000150730.2 | -1106 | Aatk | ENSMUSG00000025375.10 |
| chr11 | 120361986 | 120362545 | 3.37171 | 4.05152 | 1.18822 | Promoter | 11 | 120361534 | 120368168 | 6635 | 1 | ENSMUST00000026445.2 | 453 | Fscn2 | ENSMUSG00000025380.2 |
| chr10 | 127718541 | 127718616 | 3.6235 | 4.34528 | 1.29289 | Exon (ENSMUST00000079590.5/ENSMUSG00000025401.8, exon 22 of 27) | 10 | 127724478 | 127731767 | 7290 | 1 | ENSMUST00000179960.2 | -5862 | Tac2 | ENSMUSG00000025400.5 |
| chr10 | 127721105 | 127721366 | 4.47085 | 6.03553 | 2.51091 | Distal Intergenic | 10 | 127724478 | 127731767 | 7290 | 1 | ENSMUST00000179960.2 | -3112 | Tac2 | ENSMUSG00000025400.5 |
| chr10 | 127722075 | 127722856 | 3.75674 | 4.64459 | 1.53906 | Distal Intergenic | 10 | 127724478 | 127731767 | 7290 | 1 | ENSMUST00000179960.2 | -1622 | Tac2 | ENSMUSG00000025400.5 |
| chr10 | 127524331 | 127524963 | 3.96126 | 5.09349 | 1.85301 | Downstream (<1kb) | 10 | 127517123 | 127522444 | 5322 | 2 | ENSMUST00000026470.4 | -1888 | Shmt2 | ENSMUSG00000025403.4 |
| chr10 | 127525446 | 127527577 | 3.73732 | 4.65151 | 1.5769 | 3' UTR | 10 | 127517123 | 127522444 | 5322 | 2 | ENSMUST00000026470.4 | -3003 | Shmt2 | ENSMUSG00000025403.4 |
| chr10 | 127333165 | 127333358 | 3.29886 | 3.79323 | 1.07571 | Intron (ENSMUST00000026474.3/ENSMUSG00000025407.6, intron 11 of 12) | 10 | 127329889 | 127341589 | 11701 | 2 | ENSMUST00000026474.3 | 8231 | Gli1 | ENSMUSG00000025407.6 |
| chr10 | 127340541 | 127341268 | 3.5628 | 4.2511 | 1.28996 | Promoter | 10 | 127329889 | 127341589 | 11701 | 2 | ENSMUST00000026474.3 | 321 | Gli1 | ENSMUSG00000025407.6 |
| chr4 | 106483556 | 106483782 | 3.33297 | 4.01539 | 1.1459 | 3' UTR | 4 | 106483456 | 106492283 | 8828 | 2 | ENSMUST00000054472.3 | 8501 | Bsnd | ENSMUSG00000025418.7 |
| chr4 | 106485274 | 106485648 | 3.49458 | 4.22581 | 1.26147 | Exon (ENSMUST00000054472.3/ENSMUSG00000025418.7, exon 4 of 4) | 4 | 106483456 | 106492283 | 8828 | 2 | ENSMUST00000054472.3 | 6635 | Bsnd | ENSMUSG00000025418.7 |
| chr18 | 76988898 | 76988973 | 3.21485 | 3.64951 | 1.01262 | Intron (ENSMUST00000126153.2/ENSMUSG00000025420.8, intron 13 of 15) | 18 | 76978461 | 77006872 | 28412 | 2 | ENSMUST00000154053.2 | 17899 | Katnal2 | ENSMUSG00000025420.8 |
| chr10 | 127077569 | 127078351 | 3.47789 | 4.08602 | 1.14492 | Promoter | 10 | 127078907 | 127093169 | 14263 | 1 | ENSMUST00000039259.6 | -556 | Agap2 | ENSMUSG00000025422.8 |
| chr10 | 127079196 | 127081709 | 4.00934 | 5.1816 | 1.91464 | Promoter | 10 | 127078907 | 127093169 | 14263 | 1 | ENSMUST00000039259.6 | 290 | Agap2 | ENSMUSG00000025422.8 |
| chr10 | 127082608 | 127083158 | 3.67415 | 4.53608 | 1.43658 | Exon (ENSMUST00000039259.6/ENSMUSG00000025422.8, exon 3 of 19) | 10 | 127078907 | 127093169 | 14263 | 1 | ENSMUST00000039259.6 | 3702 | Agap2 | ENSMUSG00000025422.8 |
| chr10 | 127083480 | 127083677 | 3.34153 | 3.83966 | 1.01733 | Exon (ENSMUST00000039259.6/ENSMUSG00000025422.8, exon 5 of 19) | 10 | 127078907 | 127093169 | 14263 | 1 | ENSMUST00000039259.6 | 4574 | Agap2 | ENSMUSG00000025422.8 |
| chr10 | 127084050 | 127084893 | 3.52788 | 4.19009 | 1.22763 | Intron (ENSMUST00000039259.6/ENSMUSG00000025422.8, intron 5 of 18) | 10 | 127078907 | 127093169 | 14263 | 1 | ENSMUST00000039259.6 | 5144 | Agap2 | ENSMUSG00000025422.8 |
| chr10 | 127087840 | 127088186 | 3.48036 | 4.12703 | 1.27102 | Exon (ENSMUST00000039259.6/ENSMUSG00000025422.8, exon 11 of 19) | 10 | 127078907 | 127093169 | 14263 | 1 | ENSMUST00000039259.6 | 8934 | Agap2 | ENSMUSG00000025422.8 |
| chr10 | 127089832 | 127090750 | 4.32951 | 5.98285 | 2.49617 | Exon (ENSMUST00000039259.6/ENSMUSG00000025422.8, exon 14 of 19) | 10 | 127078907 | 127093169 | 14263 | 1 | ENSMUST00000039259.6 | 10926 | Agap2 | ENSMUSG00000025422.8 |
| chr10 | 127091255 | 127092559 | 3.50942 | 4.44878 | 1.42537 | 3' UTR | 10 | 127078907 | 127093169 | 14263 | 1 | ENSMUST00000039259.6 | 12349 | Agap2 | ENSMUSG00000025422.8 |
| chr18 | 77184924 | 77185075 | 3.56353 | 4.62491 | 1.5279 | Promoter | 18 | 77185853 | 77255450 | 69598 | 1 | ENSMUST00000079618.5 | -778 | St8sia5 | ENSMUSG00000025425.12 |
| chr18 | 77185642 | 77186232 | 3.74911 | 4.63165 | 1.51254 | Promoter | 18 | 77185853 | 77255450 | 69598 | 1 | ENSMUST00000079618.5 | 0 | St8sia5 | ENSMUSG00000025425.12 |
| chr18 | 77488949 | 77489074 | 3.45101 | 4.04424 | 1.15784 | Intron (ENSMUST00000026494.8/ENSMUSG00000025427.9, intron 1 of 7) | 18 | 77468321 | 77505117 | 36797 | 2 | ENSMUST00000182348.1 | 16043 | Rnf165 | ENSMUSG00000025427.9 |
| chr18 | 77562497 | 77562593 | 3.23338 | 3.67724 | 1.01733 | Exon (ENSMUST00000182294.1/ENSMUSG00000025427.9, exon 1 of 1) | 18 | 77562093 | 77564609 | 2517 | 2 | ENSMUST00000182294.1 | 2016 | Rnf165 | ENSMUSG00000025427.9 |
| chr18 | 77563387 | 77563683 | 3.38193 | 3.9006 | 1.01733 | Promoter | 18 | 77562093 | 77564609 | 2517 | 2 | ENSMUST00000182294.1 | 926 | Rnf165 | ENSMUSG00000025427.9 |
| chr18 | 77565096 | 77565396 | 3.68042 | 4.45954 | 1.38772 | Promoter | 18 | 77456110 | 77565136 | 109027 | 2 | ENSMUST00000026494.8 | 0 | Rnf165 | ENSMUSG00000025427.9 |
| chr18 | 77565852 | 77566728 | 3.75624 | 4.76372 | 1.64788 | Promoter | 18 | 77458685 | 77565147 | 106463 | 2 | ENSMUST00000182024.1 | -706 | Rnf165 | ENSMUSG00000025427.9 |
| chr18 | 77609806 | 77610088 | 3.70477 | 4.51937 | 1.45711 | Distal Intergenic | 18 | 77458685 | 77565147 | 106463 | 2 | ENSMUST00000182024.1 | -44660 | Rnf165 | ENSMUSG00000025427.9 |
| chr7 | 140221265 | 140221406 | 6.17114 | 12.86801 | 8.34951 | Intron (ENSMUST00000084460.6/ENSMUSG00000025461.10, intron 2 of 13) | 7 | 140218267 | 140231145 | 12879 | 1 | ENSMUST00000084460.6 | 2999 | Cd163l1 | ENSMUSG00000025461.10 |
| chr7 | 140081498 | 140081868 | 3.4685 | 4.06143 | 1.12412 | Promoter | 7 | 140069880 | 140082301 | 12422 | 2 | ENSMUST00000166758.2 | 433 | Caly | ENSMUSG00000025468.9 |
| chr7 | 140046976 | 140047179 | 3.59122 | 4.39657 | 1.41538 | 3' UTR | 7 | 140046374 | 140049088 | 2715 | 2 | ENSMUST00000172775.3 | 1909 | Msx3 | ENSMUSG00000025469.10 |
| chr7 | 140047702 | 140049081 | 4.0282 | 5.28373 | 2.03968 | Promoter | 7 | 140046374 | 140049088 | 2715 | 2 | ENSMUST00000172775.3 | 7 | Msx3 | ENSMUSG00000025469.10 |
| chr7 | 140049474 | 140050380 | 4.20554 | 5.6074 | 2.21997 | Promoter | 7 | 140046157 | 140049089 | 2933 | 2 | ENSMUST00000055353.8 | -386 | Msx3 | ENSMUSG00000025469.10 |
| chr7 | 140050963 | 140051116 | 3.64728 | 4.60101 | 1.50217 | Distal Intergenic | 7 | 140046157 | 140049089 | 2933 | 2 | ENSMUST00000055353.8 | -1875 | Msx3 | ENSMUSG00000025469.10 |
| chr7 | 140051456 | 140051976 | 3.67168 | 4.62592 | 1.54908 | Distal Intergenic | 7 | 140046157 | 140049089 | 2933 | 2 | ENSMUST00000055353.8 | -2368 | Msx3 | ENSMUSG00000025469.10 |
| chr7 | 140052572 | 140053301 | 3.48973 | 4.1296 | 1.23548 | Distal Intergenic | 7 | 140046157 | 140049089 | 2933 | 2 | ENSMUST00000055353.8 | -3484 | Msx3 | ENSMUSG00000025469.10 |
| chr7 | 140054708 | 140054932 | 4.01924 | 5.14363 | 1.91088 | Distal Intergenic | 7 | 140046157 | 140049089 | 2933 | 2 | ENSMUST00000055353.8 | -5620 | Msx3 | ENSMUSG00000025469.10 |
| chr7 | 140056846 | 140059819 | 3.60795 | 4.44006 | 1.44116 | Distal Intergenic | 7 | 140046157 | 140049089 | 2933 | 2 | ENSMUST00000055353.8 | -7758 | Msx3 | ENSMUSG00000025469.10 |
| chr7 | 140060788 | 140061840 | 3.97319 | 5.16692 | 1.90715 | Distal Intergenic | 7 | 140046157 | 140049089 | 2933 | 2 | ENSMUST00000055353.8 | -11700 | Msx3 | ENSMUSG00000025469.10 |
| chr7 | 139834890 | 139835496 | 3.60873 | 4.37848 | 1.37666 | Promoter | 7 | 139834962 | 139841204 | 6243 | 1 | ENSMUST00000129454.1 | 0 | Gpr123 | ENSMUSG00000025475.12 |
| chr7 | 139085037 | 139087241 | 3.89546 | 4.93087 | 1.76374 | Promoter | 7 | 139086001 | 139102704 | 16704 | 1 | ENSMUST00000026551.9 | 0 | Dpysl4 | ENSMUSG00000025478.10 |
| chr7 | 141099720 | 141100902 | 3.71345 | 4.57945 | 1.50322 | Promoter | 7 | 141091175 | 141100546 | 9372 | 2 | ENSMUST00000097958.2 | 0 | Sigirr | ENSMUSG00000025494.7 |
| chr7 | 141290935 | 141291349 | 3.51706 | 4.15573 | 1.19876 | Promoter | 7 | 141292006 | 141294999 | 2994 | 1 | ENSMUST00000026569.4 | -657 | Drd4 | ENSMUSG00000025496.4 |
| chr7 | 141291816 | 141294779 | 3.86515 | 4.87698 | 1.69138 | Promoter | 7 | 141292006 | 141294999 | 2994 | 1 | ENSMUST00000026569.4 | 0 | Drd4 | ENSMUSG00000025496.4 |
| chr7 | 141361432 | 141362074 | 3.43124 | 3.99759 | 1.0967 | Promoter | 7 | 141361537 | 141362667 | 1131 | 1 | ENSMUST00000155729.1 | 0 | Eps8l2 | ENSMUSG00000025504.7 |
| chr14 | 124191836 | 124193093 | 3.48702 | 4.09541 | 1.14511 | Promoter | 14 | 123978291 | 124192902 | 214612 | 2 | ENSMUST00000026631.5 | 0 | Fgf14 | ENSMUSG00000025551.8 |
| chr14 | 124675048 | 124675973 | 3.73938 | 4.71344 | 1.59203 | Intron (ENSMUST00000095529.4/ENSMUSG00000025551.8, intron 1 of 4) | 14 | 123977907 | 124677127 | 699221 | 2 | ENSMUST00000095529.4 | 1154 | Fgf14 | ENSMUSG00000025551.8 |
| chr14 | 124677470 | 124677556 | 4.89925 | 6.99524 | 3.26717 | Promoter | 14 | 123977907 | 124677127 | 699221 | 2 | ENSMUST00000095529.4 | -344 | Fgf14 | ENSMUSG00000025551.8 |
| chr14 | 121504510 | 121504766 | 3.61131 | 4.4581 | 1.43956 | Promoter | 14 | 121459621 | 121505254 | 45634 | 2 | ENSMUST00000088386.6 | 488 | Slc15a1 | ENSMUSG00000025557.9 |
| chr14 | 121505146 | 121505493 | 4.33402 | 5.94833 | 2.43699 | Promoter | 14 | 121459621 | 121505254 | 45634 | 2 | ENSMUST00000088386.6 | 0 | Slc15a1 | ENSMUSG00000025557.9 |
| chr14 | 121507144 | 121507414 | 3.21704 | 3.65279 | 1.01468 | Distal Intergenic | 14 | 121459621 | 121505254 | 45634 | 2 | ENSMUST00000088386.6 | -1891 | Slc15a1 | ENSMUSG00000025557.9 |
| chr11 | 118496248 | 118496382 | 3.23338 | 3.67724 | 1.01733 | Exon (ENSMUST00000117731.2/ENSMUSG00000025576.12, exon 11 of 14) | 11 | 118496500 | 118507353 | 10854 | 2 | ENSMUST00000154746.2 | 10971 | Rbfox3 | ENSMUSG00000025576.12 |
| chr11 | 118497080 | 118497288 | 3.41304 | 3.95789 | 1.05373 | Exon (ENSMUST00000117731.2/ENSMUSG00000025576.12, exon 9 of 14) | 11 | 118496500 | 118507353 | 10854 | 2 | ENSMUST00000154746.2 | 10065 | Rbfox3 | ENSMUSG00000025576.12 |
| chr11 | 118512603 | 118512926 | 3.42655 | 3.99707 | 1.12067 | Intron (ENSMUST00000117731.2/ENSMUSG00000025576.12, intron 4 of 13) | 11 | 118496500 | 118507353 | 10854 | 2 | ENSMUST00000154746.2 | -5251 | Rbfox3 | ENSMUSG00000025576.12 |
| chr11 | 118525525 | 118525651 | 3.44817 | 4.16487 | 1.2166 | Intron (ENSMUST00000117731.2/ENSMUSG00000025576.12, intron 3 of 13) | 11 | 118496500 | 118507353 | 10854 | 2 | ENSMUST00000154746.2 | -18173 | Rbfox3 | ENSMUSG00000025576.12 |
| chr11 | 118527110 | 118527298 | 3.48393 | 4.08858 | 1.13719 | Intron (ENSMUST00000117731.2/ENSMUSG00000025576.12, intron 3 of 13) | 11 | 118496500 | 118507353 | 10854 | 2 | ENSMUST00000154746.2 | -19758 | Rbfox3 | ENSMUSG00000025576.12 |
| chr11 | 118551518 | 118551752 | 3.36904 | 3.88115 | 1.01733 | Intron (ENSMUST00000117731.2/ENSMUSG00000025576.12, intron 3 of 13) | 11 | 118502842 | 118569910 | 67069 | 2 | ENSMUST00000136551.1 | 18158 | Rbfox3 | ENSMUSG00000025576.12 |
| chr11 | 118553331 | 118553415 | 3.69252 | 4.47277 | 1.3725 | Intron (ENSMUST00000117731.2/ENSMUSG00000025576.12, intron 3 of 13) | 11 | 118502842 | 118569910 | 67069 | 2 | ENSMUST00000136551.1 | 16495 | Rbfox3 | ENSMUSG00000025576.12 |
| chr11 | 118568426 | 118569849 | 3.78238 | 4.73606 | 1.58058 | Promoter | 11 | 118502842 | 118569910 | 67069 | 2 | ENSMUST00000136551.1 | 61 | Rbfox3 | ENSMUSG00000025576.12 |
| chr11 | 118570418 | 118570539 | 3.89998 | 4.96318 | 1.75978 | Promoter | 11 | 118502842 | 118569910 | 67069 | 2 | ENSMUST00000136551.1 | -509 | Rbfox3 | ENSMUSG00000025576.12 |
| chr11 | 118572953 | 118573367 | 3.49662 | 4.34869 | 1.34819 | Intron (ENSMUST00000117731.2/ENSMUSG00000025576.12, intron 3 of 13) | 11 | 118502842 | 118569910 | 67069 | 2 | ENSMUST00000136551.1 | -3044 | Rbfox3 | ENSMUSG00000025576.12 |
| chr11 | 118609290 | 118609664 | 3.82669 | 4.75167 | 1.59182 | Intron (ENSMUST00000117731.2/ENSMUSG00000025576.12, intron 3 of 13) | 11 | 118502842 | 118569910 | 67069 | 2 | ENSMUST00000136551.1 | -39381 | Rbfox3 | ENSMUSG00000025576.12 |
| chr11 | 118696588 | 118696853 | 3.43128 | 3.99344 | 1.08244 | Intron (ENSMUST00000117731.2/ENSMUSG00000025576.12, intron 2 of 13) | 11 | 118489764 | 118761041 | 271278 | 2 | ENSMUST00000103023.2 | 64188 | Rbfox3 | ENSMUSG00000025576.12 |
| chr11 | 118697444 | 118697695 | 3.33862 | 3.83531 | 1.01733 | Intron (ENSMUST00000117731.2/ENSMUSG00000025576.12, intron 2 of 13) | 11 | 118489764 | 118761041 | 271278 | 2 | ENSMUST00000103023.2 | 63346 | Rbfox3 | ENSMUSG00000025576.12 |
| chr11 | 118740335 | 118740427 | 3.21647 | 3.65193 | 1.01352 | Intron (ENSMUST00000117731.2/ENSMUSG00000025576.12, intron 2 of 13) | 11 | 118489764 | 118761041 | 271278 | 2 | ENSMUST00000103023.2 | 20614 | Rbfox3 | ENSMUSG00000025576.12 |
| chr11 | 118746064 | 118746407 | 3.36755 | 4.14158 | 1.23342 | Intron (ENSMUST00000117731.2/ENSMUSG00000025576.12, intron 2 of 13) | 11 | 118489764 | 118761041 | 271278 | 2 | ENSMUST00000103023.2 | 14634 | Rbfox3 | ENSMUSG00000025576.12 |
| chr11 | 118747630 | 118747955 | 3.80201 | 4.72748 | 1.58331 | Intron (ENSMUST00000117731.2/ENSMUSG00000025576.12, intron 2 of 13) | 11 | 118489764 | 118761041 | 271278 | 2 | ENSMUST00000103023.2 | 13086 | Rbfox3 | ENSMUSG00000025576.12 |
| chr11 | 118748832 | 118749126 | 3.92051 | 5.02335 | 1.80737 | Intron (ENSMUST00000117731.2/ENSMUSG00000025576.12, intron 2 of 13) | 11 | 118489764 | 118761041 | 271278 | 2 | ENSMUST00000103023.2 | 11915 | Rbfox3 | ENSMUSG00000025576.12 |
| chr11 | 118750939 | 118751268 | 3.66305 | 4.41754 | 1.34221 | Intron (ENSMUST00000117731.2/ENSMUSG00000025576.12, intron 2 of 13) | 11 | 118489764 | 118761041 | 271278 | 2 | ENSMUST00000103023.2 | 9773 | Rbfox3 | ENSMUSG00000025576.12 |
| chr11 | 118773230 | 118773311 | 3.55334 | 4.21912 | 1.22842 | Intron (ENSMUST00000117731.2/ENSMUSG00000025576.12, intron 2 of 13) | 11 | 118489764 | 118761041 | 271278 | 2 | ENSMUST00000103023.2 | -12190 | Rbfox3 | ENSMUSG00000025576.12 |
| chr11 | 118907392 | 118908135 | 4.21919 | 5.55496 | 2.18731 | Promoter | 11 | 118811280 | 118908005 | 96726 | 2 | ENSMUST00000134774.1 | 0 | Rbfox3 | ENSMUSG00000025576.12 |
| chr11 | 118908584 | 118910912 | 4.18601 | 5.52359 | 2.17535 | Promoter | 11 | 118489762 | 118909550 | 419789 | 2 | ENSMUST00000117731.2 | 0 | Rbfox3 | ENSMUSG00000025576.12 |
| chr11 | 118897278 | 118897534 | 3.49684 | 4.14004 | 1.24018 | Intron (ENSMUST00000117731.2/ENSMUSG00000025576.12, intron 1 of 13) | 11 | 118811278 | 118888689 | 77412 | 2 | ENSMUST00000128863.1 | -8590 | Rbfox3 | ENSMUSG00000025576.12 |
| chr11 | 118906327 | 118906936 | 3.62891 | 4.36513 | 1.32515 | Intron (ENSMUST00000117731.2/ENSMUSG00000025576.12, intron 1 of 13) | 11 | 118811280 | 118908005 | 96726 | 2 | ENSMUST00000134774.1 | 1069 | Rbfox3 | ENSMUSG00000025576.12 |
| chr11 | 119020753 | 119021443 | 3.42291 | 4.10982 | 1.23082 | Distal Intergenic | 11 | 119022962 | 119031270 | 8309 | 1 | ENSMUST00000026662.7 | -1519 | Cbx2 | ENSMUSG00000025577.7 |
| chr11 | 119021973 | 119022748 | 3.58607 | 4.52957 | 1.4577 | Promoter | 11 | 119022962 | 119031270 | 8309 | 1 | ENSMUST00000026662.7 | -214 | Cbx2 | ENSMUSG00000025577.7 |
| chr11 | 119033495 | 119033762 | 3.41743 | 4.00081 | 1.16278 | Downstream (2-3kb) | 11 | 119039053 | 119040831 | 1779 | 2 | ENSMUST00000128019.1 | 7069 | Cbx8 | ENSMUSG00000025578.9 |
| chr11 | 119034569 | 119036161 | 4.24136 | 5.61663 | 2.19536 | Downstream (<1kb) | 11 | 119039053 | 119040831 | 1779 | 2 | ENSMUST00000128019.1 | 4670 | Cbx8 | ENSMUSG00000025578.9 |
| chr11 | 119036658 | 119037525 | 4.0785 | 5.32181 | 2.03937 | 3' UTR | 11 | 119039053 | 119040831 | 1779 | 2 | ENSMUST00000128019.1 | 3306 | Cbx8 | ENSMUSG00000025578.9 |
| chr11 | 119041552 | 119045337 | 4.05246 | 5.35689 | 2.02315 | Promoter | 11 | 119036305 | 119040969 | 4665 | 2 | ENSMUST00000026663.7 | -584 | Cbx8 | ENSMUSG00000025578.9 |
| chr11 | 119045670 | 119045805 | 3.74292 | 4.59523 | 1.49272 | Distal Intergenic | 11 | 119036305 | 119040969 | 4665 | 2 | ENSMUST00000026663.7 | -4702 | Cbx8 | ENSMUSG00000025578.9 |
| chr11 | 119046154 | 119050125 | 3.87722 | 4.99595 | 1.79677 | Distal Intergenic | 11 | 119036305 | 119040969 | 4665 | 2 | ENSMUST00000026663.7 | -5186 | Cbx8 | ENSMUSG00000025578.9 |
| chr11 | 119050794 | 119051151 | 3.60132 | 4.41397 | 1.40956 | Distal Intergenic | 11 | 119036305 | 119040969 | 4665 | 2 | ENSMUST00000026663.7 | -9826 | Cbx8 | ENSMUSG00000025578.9 |
| chr11 | 119051476 | 119051828 | 3.49188 | 4.16449 | 1.2489 | Distal Intergenic | 11 | 119036305 | 119040969 | 4665 | 2 | ENSMUST00000026663.7 | -10508 | Cbx8 | ENSMUSG00000025578.9 |
| chr11 | 119052269 | 119052423 | 3.86914 | 5.04065 | 1.81161 | Distal Intergenic | 11 | 119036305 | 119040969 | 4665 | 2 | ENSMUST00000026663.7 | -11301 | Cbx8 | ENSMUSG00000025578.9 |
| chr11 | 119052770 | 119053296 | 3.89669 | 5.09451 | 1.82321 | Distal Intergenic | 11 | 119036305 | 119040969 | 4665 | 2 | ENSMUST00000026663.7 | -11802 | Cbx8 | ENSMUSG00000025578.9 |
| chr11 | 119054164 | 119055994 | 3.96556 | 5.09515 | 1.89728 | Distal Intergenic | 11 | 119036305 | 119040969 | 4665 | 2 | ENSMUST00000026663.7 | -13196 | Cbx8 | ENSMUSG00000025578.9 |
| chr11 | 119542473 | 119542717 | 4.02047 | 5.21482 | 1.88565 | Exon (ENSMUST00000026670.4/ENSMUSG00000025582.4, exon 5 of 5) | 11 | 119538719 | 119547753 | 9035 | 2 | ENSMUST00000026670.4 | 5036 | Nptx1 | ENSMUSG00000025582.4 |
| chr11 | 119545479 | 119545841 | 3.7457 | 4.60631 | 1.5429 | Intron (ENSMUST00000026670.4/ENSMUSG00000025582.4, intron 2 of 4) | 11 | 119538719 | 119547753 | 9035 | 2 | ENSMUST00000026670.4 | 1912 | Nptx1 | ENSMUSG00000025582.4 |
| chr11 | 119546490 | 119547870 | 4.15478 | 5.71383 | 2.33785 | Promoter | 11 | 119538719 | 119547753 | 9035 | 2 | ENSMUST00000026670.4 | 0 | Nptx1 | ENSMUSG00000025582.4 |
| chr11 | 119591497 | 119591914 | 3.47215 | 4.10711 | 1.23374 | Distal Intergenic | 11 | 119602905 | 119823176 | 220272 | 1 | ENSMUST00000125583.2 | -10991 | Rptor | ENSMUSG00000025583.10 |
| chr7 | 81196199 | 81196597 | 3.64512 | 4.41044 | 1.38617 | Distal Intergenic | 7 | 81213596 | 81334533 | 120938 | 1 | ENSMUST00000026672.7 | -16999 | Pde8a | ENSMUSG00000025584.12 |
| chr7 | 81197601 | 81197923 | 3.81276 | 4.70699 | 1.55266 | Distal Intergenic | 7 | 81213596 | 81334533 | 120938 | 1 | ENSMUST00000026672.7 | -15673 | Pde8a | ENSMUSG00000025584.12 |
| chr16 | 87576627 | 87576885 | 3.36968 | 4.07902 | 1.21333 | Intron (ENSMUST00000026700.7/ENSMUSG00000025610.7, intron 3 of 4) | 16 | 87553330 | 87595336 | 42007 | 1 | ENSMUST00000026700.7 | 23298 | Map3k7cl | ENSMUSG00000025610.7 |
| chr6 | 116460569 | 116461003 | 3.5875 | 4.2979 | 1.30699 | Promoter | 6 | 116410719 | 116461024 | 50306 | 2 | ENSMUST00000164547.2 | 21 | Alox5 | ENSMUSG00000025701.7 |
| chr6 | 116461531 | 116461670 | 3.65177 | 4.65947 | 1.57057 | Promoter | 6 | 116410077 | 116461178 | 51102 | 2 | ENSMUST00000026795.7 | -354 | Alox5 | ENSMUSG00000025701.7 |
| chr17 | 25951276 | 25952623 | 3.70247 | 4.67366 | 1.56084 | Promoter | 17 | 25946387 | 25952565 | 6179 | 2 | ENSMUST00000162431.1 | 0 | A930017K11Rik | ENSMUSG00000025727.8 |
| chr17 | 25818409 | 25818573 | 3.23178 | 3.68178 | 1.03 | Exon (ENSMUST00000045692.7/ENSMUSG00000025738.7, exon 4 of 6) | 17 | 25823627 | 25828730 | 5104 | 1 | ENSMUST00000026833.5 | -5054 | Wdr24 | ENSMUSG00000025737.5 |
| chr17 | 25819292 | 25819555 | 3.57513 | 4.39094 | 1.4048 | 3' UTR | 17 | 25823627 | 25828730 | 5104 | 1 | ENSMUST00000026833.5 | -4072 | Wdr24 | ENSMUSG00000025737.5 |
| chr17 | 25716049 | 25717783 | 3.64791 | 4.41585 | 1.35566 | Promoter | 17 | 25717171 | 25719406 | 2236 | 1 | ENSMUST00000172002.2 | 0 | Gng13 | ENSMUSG00000025739.8 |
| chr17 | 25718488 | 25718751 | 3.38193 | 3.9006 | 1.01733 | Promoter | 17 | 25717528 | 25719087 | 1560 | 1 | ENSMUST00000115108.3 | 961 | Gng13 | ENSMUSG00000025739.8 |
| chr4 | 130792038 | 130793003 | 4.30772 | 5.99523 | 2.49274 | Promoter | 4 | 130792537 | 130826319 | 33783 | 1 | ENSMUST00000070478.3 | 0 | Sdc3 | ENSMUSG00000025743.9 |
| chr4 | 130799672 | 130799890 | 3.37318 | 4.04197 | 1.13565 | Intron (ENSMUST00000070478.3/ENSMUSG00000025743.9, intron 1 of 4) | 4 | 130792704 | 130817273 | 24570 | 1 | ENSMUST00000146093.1 | 6969 | Sdc3 | ENSMUSG00000025743.9 |
| chr5 | 30112830 | 30113174 | 3.30105 | 3.80827 | 1.09914 | Intron (ENSMUST00000058045.4/ENSMUSG00000044576.6, intron 2 of 5) | 5 | 30119899 | 30120305 | 407 | 2 | ENSMUST00000132486.1 | 7131 | Hadha | ENSMUSG00000025745.6 |
| chr5 | 30113967 | 30114838 | 3.92001 | 5.09787 | 1.86466 | Exon (ENSMUST00000058045.4/ENSMUSG00000044576.6, exon 4 of 6) | 5 | 30119899 | 30120305 | 407 | 2 | ENSMUST00000132486.1 | 5467 | Hadha | ENSMUSG00000025745.6 |
| chr2 | 9880152 | 9880541 | 3.576 | 4.28351 | 1.29771 | Promoter | 2 | 9881252 | 9883921 | 2670 | 1 | ENSMUST00000026889.1 | -711 | 4930412O13Rik | ENSMUSG00000025783.2 |
| chr2 | 9880957 | 9881853 | 3.6236 | 4.36206 | 1.32238 | Promoter | 2 | 9881252 | 9883921 | 2670 | 1 | ENSMUST00000026889.1 | 0 | 4930412O13Rik | ENSMUSG00000025783.2 |
| chr2 | 9882518 | 9883103 | 4.0908 | 5.3019 | 1.97235 | Promoter | 2 | 9882636 | 9883335 | 700 | 1 | ENSMUST00000114919.1 | 0 | 4930412O13Rik | ENSMUSG00000025783.2 |
| chr7 | 73942213 | 73942288 | 4.23407 | 5.58347 | 2.20436 | 3' UTR | 7 | 73939119 | 74013682 | 74564 | 2 | ENSMUST00000026896.4 | 71394 | St8sia2 | ENSMUSG00000025789.4 |
| chr7 | 74012120 | 74012864 | 3.60783 | 4.4942 | 1.43426 | Promoter | 7 | 73939119 | 74013682 | 74564 | 2 | ENSMUST00000026896.4 | 818 | St8sia2 | ENSMUSG00000025789.4 |
| chr7 | 74013223 | 74013518 | 3.64298 | 4.39999 | 1.34567 | Promoter | 7 | 73939119 | 74013682 | 74564 | 2 | ENSMUST00000026896.4 | 164 | St8sia2 | ENSMUSG00000025789.4 |
| chr7 | 81699571 | 81699910 | 3.34826 | 3.84982 | 1.01733 | Intron (ENSMUST00000026922.8/ENSMUSG00000025813.8, intron 1 of 8) | 7 | 81600481 | 81692965 | 92485 | 2 | ENSMUST00000098326.2 | -6607 | Homer2 | ENSMUSG00000025813.8 |
| chr7 | 81703877 | 81704016 | 3.70837 | 4.66574 | 1.55182 | Intron (ENSMUST00000026922.8/ENSMUSG00000025813.8, intron 1 of 8) | 7 | 81600481 | 81706925 | 106445 | 2 | ENSMUST00000026922.8 | 2909 | Homer2 | ENSMUSG00000025813.8 |
| chr7 | 81706435 | 81706831 | 3.72907 | 4.55456 | 1.44545 | Promoter | 7 | 81600481 | 81706925 | 106445 | 2 | ENSMUST00000026922.8 | 94 | Homer2 | ENSMUSG00000025813.8 |
| chr5 | 138754100 | 138754484 | 3.55482 | 4.39515 | 1.41091 | Promoter | 5 | 138754514 | 138810077 | 55564 | 1 | ENSMUST00000026972.7 | -30 | Fam20c | ENSMUSG00000025854.10 |
| chr5 | 138755595 | 138756154 | 3.98783 | 5.2897 | 1.98203 | Promoter | 5 | 138755083 | 138809381 | 54299 | 1 | ENSMUST00000160645.2 | 513 | Fam20c | ENSMUSG00000025854.10 |

| | | | | | | | | | | | | | | | |
|---|---|---|---|---|---|---|---|---|---|---|---|---|---|---|---|
| chr5 | 139027788 | 139027863 | 4.3614 | 5.86322 | 2.36785 | Intron (ENSMUST00000026973.8/ENSMUSG00000025855.8, intron 9 of 10) | 5 | 139018666 | 139023810 | 5145 | 2 | ENSMUST00000139415.1 | -3979 | Prkar1b | ENSMUSG00000025855.8 |
| chr5 | 139057172 | 139057375 | 3.53075 | 4.21124 | 1.31179 | Intron (ENSMUST00000026973.8/ENSMUSG00000025855.8, intron 6 of 10) | 5 | 139018666 | 139023810 | 5145 | 2 | ENSMUST00000139415.1 | -33363 | Prkar1b | ENSMUSG00000025855.8 |
| chr5 | 139129388 | 139129678 | 3.65813 | 4.48784 | 1.44328 | Promoter | 5 | 139017306 | 139129759 | 112454 | 2 | ENSMUST00000026973.8 | 81 | Prkar1b | ENSMUSG00000025855.8 |
| chr5 | 138932301 | 138932544 | 3.8783 | 4.82668 | 1.60846 | Distal Intergenic | 5 | 138977145 | 138994281 | 17137 | 2 | ENSMUST00000144934.2 | 61737 | Pdgfa | ENSMUSG00000025856.10 |
| chr5 | 138992241 | 138992931 | 3.55918 | 4.54399 | 1.49512 | Intron (ENSMUST00000076095.8/ENSMUSG00000025856.10, intron 2 of 5) | 5 | 138977145 | 138994281 | 17137 | 2 | ENSMUST00000144934.2 | 1350 | Pdgfa | ENSMUSG00000025856.10 |
| chr5 | 138998156 | 138999060 | 3.6154 | 4.52816 | 1.47258 | Promoter | 5 | 138976971 | 138997370 | 20400 | 2 | ENSMUST00000110896.1 | -787 | Pdgfa | ENSMUSG00000025856.10 |
| chr13 | 54370046 | 54372125 | 3.99802 | 5.11417 | 1.85672 | Promoter | 13 | 54371349 | 54383917 | 12569 | 1 | ENSMUST00000026985.8 | 0 | Cplx2 | ENSMUSG00000025867.8 |
| chr13 | 54373171 | 54373423 | 3.36032 | 3.868 | 1.01733 | Intron (ENSMUST00000026985.8/ENSMUSG00000025867.8, intron 1 of 3) | 13 | 54371349 | 54383917 | 12569 | 1 | ENSMUST00000026985.8 | 1823 | Cplx2 | ENSMUSG00000025867.8 |
| chr13 | 54805797 | 54806414 | 4.04749 | 5.24875 | 2.01179 | Distal Intergenic | 13 | 54795384 | 54796768 | 1385 | 1 | ENSMUST00000139574.1 | 10414 | Tspan17 | ENSMUSG00000025875.9 |
| chr13 | 54941577 | 54941766 | 3.32838 | 3.81985 | 1.01733 | Distal Intergenic | 13 | 54949411 | 55006018 | 56608 | 1 | ENSMUST00000026994.8 | -7645 | Unc5a | ENSMUSG00000025876.10 |
| chr13 | 54948646 | 54949304 | 3.51759 | 4.18712 | 1.22061 | Promoter | 13 | 54949411 | 55006018 | 56608 | 1 | ENSMUST00000026994.8 | -107 | Unc5a | ENSMUSG00000025876.10 |
| chr13 | 54949667 | 54949824 | 4.79656 | 6.9707 | 3.21459 | Promoter | 13 | 54949411 | 55006018 | 56608 | 1 | ENSMUST00000109994.3 | 257 | Unc5a | ENSMUSG00000025876.10 |
| chr13 | 55005711 | 55006277 | 3.35457 | 3.88358 | 1.10227 | 3' UTR | 13 | 55003680 | 55005486 | 1807 | 1 | ENSMUST00000137967.1 | 2032 | Unc5a | ENSMUSG00000025876.10 |
| chr13 | 55012150 | 55012462 | 3.51064 | 4.14654 | 1.20039 | 3' UTR | 13 | 55006067 | 55020904 | 14838 | 2 | ENSMUST00000126234.2 | 8442 | Hk3 | ENSMUSG00000025877.9 |
| chr6 | 60828648 | 60829219 | 3.48543 | 4.11014 | 1.1996 | Promoter | 6 | 60731573 | 60828996 | 97424 | 2 | ENSMUST00000114268.3 | 0 | Snca | ENSMUSG00000025889.8 |
| chr6 | 60829580 | 60829667 | 3.40603 | 3.95508 | 1.0814 | Promoter | 6 | 60731573 | 60829855 | 98283 | 2 | ENSMUST00000163779.2 | 188 | Snca | ENSMUSG00000025889.8 |
| chr9 | 4794774 | 4794849 | 4.51781 | 6.1229 | 2.57172 | Intron (ENSMUST00000027020.7/ENSMUSG00000025892.10, intron 2 of 16) | 9 | 4417899 | 4796234 | 378336 | 2 | ENSMUST00000027020.7 | 1385 | Gria4 | ENSMUSG00000025892.10 |
| chr1 | 4489246 | 4489359 | 3.68371 | 4.46025 | 1.3733 | Downstream (1-2kb) | 1 | 4491713 | 4496363 | 4651 | 2 | ENSMUST00000116652.2 | 7004 | Sox17 | ENSMUSG00000025902.8 |
| chr1 | 4490459 | 4490699 | 4.16325 | 5.65472 | 2.25218 | Downstream (<1kb) | 1 | 4491713 | 4496363 | 4651 | 2 | ENSMUST00000116652.2 | 5664 | Sox17 | ENSMUSG00000025902.8 |
| chr1 | 4491590 | 4492477 | 4.20107 | 5.91695 | 2.45379 | 3' UTR | 1 | 4491713 | 4496363 | 4651 | 2 | ENSMUST00000116652.2 | 3886 | Sox17 | ENSMUSG00000025902.8 |
| chr1 | 4493306 | 4494733 | 4.01235 | 5.32034 | 2.04893 | 5' UTR | 1 | 4491713 | 4496363 | 4651 | 2 | ENSMUST00000116652.2 | 1630 | Sox17 | ENSMUSG00000025902.8 |
| chr1 | 4495397 | 4495788 | 3.75434 | 4.70759 | 1.63741 | Promoter | 1 | 4491713 | 4496363 | 4651 | 2 | ENSMUST00000116652.2 | 575 | Sox17 | ENSMUSG00000025902.8 |
| chr1 | 4496310 | 4496802 | 3.71857 | 4.62416 | 1.59048 | Promoter | 1 | 4490928 | 4496413 | 5486 | 2 | ENSMUST00000027035.4 | 0 | Sox17 | ENSMUSG00000025902.8 |
| chr1 | 4497192 | 4498754 | 3.55865 | 4.38707 | 1.39755 | Promoter | 1 | 4490928 | 4496413 | 5486 | 2 | ENSMUST00000027035.4 | -780 | Sox17 | ENSMUSG00000025902.8 |
| chr1 | 4499533 | 4500084 | 3.47107 | 4.13525 | 1.26419 | Distal Intergenic | 1 | 4490928 | 4496413 | 5486 | 2 | ENSMUST00000027035.4 | -3121 | Sox17 | ENSMUSG00000025902.8 |
| chr1 | 4502156 | 4502431 | 3.95998 | 5.10512 | 1.85342 | Distal Intergenic | 1 | 4490928 | 4496413 | 5486 | 2 | ENSMUST00000027035.4 | -5744 | Sox17 | ENSMUSG00000025902.8 |
| chr1 | 4502758 | 4502833 | 3.86751 | 5.43576 | 2.04667 | Distal Intergenic | 1 | 4490928 | 4496413 | 5486 | 2 | ENSMUST00000027035.4 | -6346 | Sox17 | ENSMUSG00000025902.8 |
| chr1 | 4506871 | 4507276 | 3.58801 | 4.42668 | 1.42916 | Distal Intergenic | 1 | 4490928 | 4496413 | 5486 | 2 | ENSMUST00000027035.4 | -10459 | Sox17 | ENSMUSG00000025902.8 |
| chr1 | 4508661 | 4509131 | 3.85338 | 4.80319 | 1.64554 | Distal Intergenic | 1 | 4490928 | 4496413 | 5486 | 2 | ENSMUST00000027035.4 | -12249 | Sox17 | ENSMUSG00000025902.8 |
| chr1 | 9296860 | 9297060 | 3.32244 | 3.81092 | 1.01733 | Intron (ENSMUST00000132064.2/ENSMUSG00000025909.11, intron 1 of 19) | 1 | 8950572 | 9298236 | 347665 | 2 | ENSMUST00000126083.1 | 1176 | Sntg1 | ENSMUSG00000025909.11 |
| chr1 | 9299693 | 9300106 | 3.51628 | 4.28499 | 1.32725 | Promoter | 1 | 8362678 | 9299878 | 937201 | 2 | ENSMUST00000140302.2 | 0 | Sntg1 | ENSMUSG00000025909.11 |
| chr1 | 19199851 | 19199970 | 3.35163 | 3.85489 | 1.01733 | Distal Intergenic | 1 | 19208914 | 19238576 | 29663 | 1 | ENSMUST00000027059.5 | -8944 | Tfap2b | ENSMUSG00000025927.8 |
| chr1 | 19203472 | 19203929 | 3.63031 | 4.48106 | 1.46017 | Distal Intergenic | 1 | 19208914 | 19238576 | 29663 | 1 | ENSMUST00000027059.5 | -4985 | Tfap2b | ENSMUSG00000025927.8 |
| chr1 | 19204517 | 19204654 | 3.21466 | 3.64922 | 1.01299 | Distal Intergenic | 1 | 19208914 | 19238576 | 29663 | 1 | ENSMUST00000027059.5 | -4260 | Tfap2b | ENSMUSG00000025927.8 |
| chr1 | 19207037 | 19207171 | 3.51396 | 4.45901 | 1.45097 | Distal Intergenic | 1 | 19208914 | 19238576 | 29663 | 1 | ENSMUST00000027059.5 | -1743 | Tfap2b | ENSMUSG00000025927.8 |
| chr1 | 19207582 | 19207930 | 3.80108 | 4.83752 | 1.69313 | Promoter | 1 | 19208914 | 19238576 | 29663 | 1 | ENSMUST00000027059.5 | -984 | Tfap2b | ENSMUSG00000025927.8 |
| chr1 | 19208395 | 19208732 | 3.34728 | 3.84833 | 1.01733 | Promoter | 1 | 19208914 | 19238576 | 29663 | 1 | ENSMUST00000027059.5 | -182 | Tfap2b | ENSMUSG00000025927.8 |
| chr1 | 19209224 | 19210558 | 3.86563 | 4.86594 | 1.68847 | Promoter | 1 | 19208960 | 19238083 | 29124 | 1 | ENSMUST00000187754.1 | 265 | Tfap2b | ENSMUSG00000025927.8 |
| chr1 | 19210937 | 19211289 | 3.42456 | 4.02182 | 1.11595 | Promoter | 1 | 19212054 | 19234624 | 22571 | 1 | ENSMUST00000064976.5 | -765 | Tfap2b | ENSMUSG00000025927.8 |
| chr1 | 19211744 | 19212362 | 3.43989 | 4.01676 | 1.11702 | Promoter | 1 | 19212054 | 19234624 | 22571 | 1 | ENSMUST00000064976.5 | 0 | Tfap2b | ENSMUSG00000025927.8 |
| chr1 | 19212776 | 19212876 | 3.50234 | 4.25084 | 1.27879 | Promoter | 1 | 19212306 | 19214442 | 2137 | 1 | ENSMUST00000191068.1 | 471 | Tfap2b | ENSMUSG00000025927.8 |
| chr1 | 19213879 | 19215222 | 3.9034 | 5.31589 | 2.03104 | Promoter | 1 | 19213879 | 19219253 | 5375 | 1 | ENSMUST00000186972.1 | 1 | Tfap2b | ENSMUSG00000025927.8 |
| chr1 | 19215989 | 19216201 | 3.64286 | 4.38654 | 1.33542 | Intron (ENSMUST00000027059.5/ENSMUSG00000025927.8, intron 2 of 6) | 1 | 19213879 | 19219253 | 5375 | 1 | ENSMUST00000186972.1 | 2111 | Tfap2b | ENSMUSG00000025927.8 |
| chr1 | 19216788 | 19217006 | 3.35188 | 3.85526 | 1.01733 | Intron (ENSMUST00000027059.5/ENSMUSG00000025927.8, intron 2 of 6) | 1 | 19213879 | 19219253 | 5375 | 1 | ENSMUST00000186972.1 | 2910 | Tfap2b | ENSMUSG00000025927.8 |
| chr1 | 19220807 | 19221080 | 3.47512 | 4.134 | 1.22734 | Intron (ENSMUST00000027059.5/ENSMUSG00000025927.8, intron 3 of 6) | 1 | 19213879 | 19219253 | 5375 | 1 | ENSMUST00000186972.1 | 6929 | Tfap2b | ENSMUSG00000025927.8 |
| chr1 | 19222566 | 19223858 | 3.47916 | 4.12977 | 1.24655 | Intron (ENSMUST00000027059.5/ENSMUSG00000025927.8, intron 3 of 6) | 1 | 19213879 | 19219253 | 5375 | 1 | ENSMUST00000186972.1 | 8688 | Tfap2b | ENSMUSG00000025927.8 |
| chr1 | 19224654 | 19225105 | 3.33179 | 3.82516 | 1.01733 | Intron (ENSMUST00000027059.5/ENSMUSG00000025927.8, intron 3 of 6) | 1 | 19213879 | 19219253 | 5375 | 1 | ENSMUST00000186972.1 | 10776 | Tfap2b | ENSMUSG00000025927.8 |
| chr1 | 19225416 | 19225525 | 3.77488 | 5.18175 | 1.89378 | Intron (ENSMUST00000027059.5/ENSMUSG00000025927.8, intron 3 of 6) | 1 | 19213879 | 19219253 | 5375 | 1 | ENSMUST00000186972.1 | 11538 | Tfap2b | ENSMUSG00000025927.8 |
| chr1 | 19225924 | 19226109 | 3.59101 | 4.28954 | 1.27304 | Intron (ENSMUST00000027059.5/ENSMUSG00000025927.8, intron 3 of 6) | 1 | 19213879 | 19219253 | 5375 | 1 | ENSMUST00000186972.1 | 12046 | Tfap2b | ENSMUSG00000025927.8 |
| chr1 | 19226461 | 19227108 | 3.86339 | 4.84781 | 1.65999 | Exon (ENSMUST00000027059.5/ENSMUSG00000025927.8, exon 4 of 7) | 1 | 19213879 | 19219253 | 5375 | 1 | ENSMUST00000186972.1 | 12583 | Tfap2b | ENSMUSG00000025927.8 |
| chr1 | 20743923 | 20744217 | 4.15549 | 5.5216 | 2.20391 | Distal Intergenic | 1 | 20730905 | 20734496 | 3592 | 1 | ENSMUST00000027061.4 | 13019 | Il17a | ENSMUSG00000025929.4 |
| chr1 | 14755259 | 14755795 | 3.86398 | 4.92495 | 1.75241 | Promoter | 1 | 14753746 | 14755353 | 1608 | 2 | ENSMUST00000190719.1 | 0 | Msc | ENSMUSG00000025930.6 |
| chr1 | 14757770 | 14758078 | 3.80373 | 4.70831 | 1.55728 | Exon (ENSMUST00000067599.1/ENSMUSG00000054493.2, exon 1 of 1) | 1 | 14754767 | 14755992 | 1226 | 2 | ENSMUST00000190471.1 | -1779 | Msc | ENSMUSG00000025930.6 |
| chr1 | 14503097 | 14503373 | 3.60166 | 4.52504 | 1.45363 | Distal Intergenic | 1 | 14168958 | 14310235 | 141278 | 2 | ENSMUST00000168081.3 | -192863 | Eya1 | ENSMUSG00000025932.9 |
| chr1 | 14503742 | 14503868 | 3.48842 | 4.36018 | 1.3752 | Distal Intergenic | 1 | 14168958 | 14310235 | 141278 | 2 | ENSMUST00000168081.3 | -193508 | Eya1 | ENSMUSG00000025932.9 |
| chr1 | 14504400 | 14504528 | 3.19611 | 3.93484 | 1.06629 | Distal Intergenic | 1 | 14168958 | 14310235 | 141278 | 2 | ENSMUST00000168081.3 | -194166 | Eya1 | ENSMUSG00000025932.9 |
| chr1 | 14504969 | 14505532 | 3.92968 | 5.01496 | 1.8176 | Distal Intergenic | 1 | 14168958 | 14310235 | 141278 | 2 | ENSMUST00000168081.3 | -194735 | Eya1 | ENSMUSG00000025932.9 |
| chr1 | 14505864 | 14507063 | 4.00143 | 5.1974 | 1.92714 | Distal Intergenic | 1 | 14168958 | 14310235 | 141278 | 2 | ENSMUST00000168081.3 | -195630 | Eya1 | ENSMUSG00000025932.9 |
| chr1 | 14507440 | 14508118 | 3.86558 | 5.10422 | 1.85876 | Distal Intergenic | 1 | 14168958 | 14310235 | 141278 | 2 | ENSMUST00000168081.3 | -197206 | Eya1 | ENSMUSG00000025932.9 |
| chr1 | 14508437 | 14508521 | 3.75943 | 4.68325 | 1.63653 | Distal Intergenic | 1 | 14168958 | 14310235 | 141278 | 2 | ENSMUST00000168081.3 | -198203 | Eya1 | ENSMUSG00000025932.9 |
| chr1 | 13494550 | 13494958 | 3.7996 | 4.72783 | 1.5732 | Distal Intergenic | 1 | 13576144 | 13579798 | 3655 | 2 | ENSMUST00000185447.1 | 84840 | Tram1 | ENSMUSG00000025935.5 |
| chr1 | 13495649 | 13495880 | 3.73943 | 4.73525 | 1.57277 | Distal Intergenic | 1 | 13576144 | 13579798 | 3655 | 2 | ENSMUST00000185447.1 | 83918 | Tram1 | ENSMUSG00000025935.5 |
| chr1 | 12990565 | 12991493 | 3.78779 | 4.72681 | 1.60018 | Promoter | 1 | 12866552 | 12991135 | 124584 | 2 | ENSMUST00000115403.3 | 0 | Slco5a1 | ENSMUSG00000025938.11 |
| chr1 | 12992691 | 12993409 | 3.73371 | 4.63194 | 1.53067 | Promoter | 1 | 12866549 | 12992650 | 126102 | 2 | ENSMUST00000188454.1 | -42 | Slco5a1 | ENSMUSG00000025938.11 |
| chr1 | 12994291 | 12994880 | 3.77583 | 4.64482 | 1.52163 | Distal Intergenic | 1 | 12866549 | 12992650 | 126102 | 2 | ENSMUST00000188454.1 | -1642 | Slco5a1 | ENSMUSG00000025938.11 |
| chr1 | 65310290 | 65310619 | 3.56085 | 4.32401 | 1.34341 | Promoter | 1 | 65311257 | 65389244 | 77988 | 1 | ENSMUST00000027083.6 | -638 | Pth2r | ENSMUSG00000025946.8 |
| chr1 | 65311706 | 65311781 | 4.05686 | 5.4166 | 2.029 | Promoter | 1 | 65311257 | 65389244 | 77988 | 1 | ENSMUST00000027083.6 | 450 | Pth2r | ENSMUSG00000025946.8 |
| chr1 | 63444501 | 63444727 | 3.68146 | 4.46502 | 1.39116 | Distal Intergenic | 1 | 63445891 | 63573217 | 127327 | 1 | ENSMUST00000182642.2 | -1164 | Adam23 | ENSMUSG00000025964.10 |
| chr1 | 63445409 | 63446487 | 3.70075 | 4.74729 | 1.61846 | Promoter | 1 | 63445891 | 63573217 | 127327 | 1 | ENSMUST00000182642.2 | 0 | Adam23 | ENSMUSG00000025964.10 |
| chr1 | 62702029 | 62702570 | 3.6529 | 4.41961 | 1.38494 | Promoter | 1 | 62703285 | 62739932 | 36648 | 1 | ENSMUST00000126344.1 | -715 | Nrp2 | ENSMUSG00000025969.10 |
| chr1 | 62704005 | 62705719 | 3.70875 | 4.5765 | 1.51314 | Promoter | 1 | 62703702 | 62818695 | 114994 | 1 | ENSMUST00000114157.2 | 304 | Nrp2 | ENSMUSG00000025969.10 |
| chr1 | 62706052 | 62706370 | 3.38707 | 4.07622 | 1.17811 | Intron (ENSMUST00000126344.1/ENSMUSG00000025969.10, intron 1 of 2) | 1 | 62703702 | 62818695 | 114994 | 1 | ENSMUST00000114157.2 | 2351 | Nrp2 | ENSMUSG00000025969.10 |
| chr1 | 62706846 | 62708559 | 4.32869 | 5.82113 | 2.39309 | Intron (ENSMUST00000126344.1/ENSMUSG00000025969.10, intron 1 of 2) | 1 | 62703702 | 62818695 | 114994 | 1 | ENSMUST00000114157.2 | 3145 | Nrp2 | ENSMUSG00000025969.10 |
| chr1 | 55362454 | 55362726 | 3.53818 | 4.2316 | 1.318 | Promoter | 1 | 55300070 | 55362726 | 62657 | 2 | ENSMUST00000114423.4 | 0 | Boll | ENSMUSG00000025977.10 |
| chr1 | 60042466 | 60042570 | 3.67571 | 4.89193 | 1.69739 | Promoter | 1 | 59989068 | 60043087 | 54020 | 2 | ENSMUST00000027172.7 | 517 | Ica1l | ENSMUSG00000026018.7 |
| chr1 | 60043093 | 60043452 | 3.72934 | 4.66667 | 1.59251 | Promoter | 1 | 59989075 | 60043122 | 54048 | 2 | ENSMUST00000191251.1 | 0 | Ica1l | ENSMUSG00000026018.7 |
| chr1 | 59257449 | 59257589 | 3.49446 | 4.11049 | 1.15866 | Promoter | 1 | 59257128 | 59257878 | 751 | 1 | ENSMUST00000162487.1 | 322 | Cdk15 | ENSMUSG00000026023.11 |
| chr1 | 32170890 | 32171162 | 3.44761 | 4.03321 | 1.13598 | Distal Intergenic | 1 | 32172720 | 32658564 | 485845 | 1 | ENSMUST00000027226.6 | -1558 | Khdrbs2 | ENSMUSG00000026058.6 |
| chr1 | 32171637 | 32171849 | 3.36329 | 3.87247 | 1.01733 | Promoter | 1 | 32172720 | 32658564 | 485845 | 1 | ENSMUST00000027226.6 | -871 | Khdrbs2 | ENSMUSG00000026058.6 |
| chr1 | 32172716 | 32173016 | 4.34568 | 5.77617 | 2.30404 | Promoter | 1 | 32172720 | 32658564 | 485845 | 1 | ENSMUST00000027226.6 | 0 | Khdrbs2 | ENSMUSG00000026058.6 |
| chr1 | 40680359 | 40681210 | 4.02463 | 5.21486 | 1.99289 | Promoter | 1 | 40681712 | 40768885 | 87174 | 1 | ENSMUST00000027231.7 | -502 | Slc9a2 | ENSMUSG00000026062.7 |
| chr1 | 40681589 | 40682619 | 3.70615 | 4.69857 | 1.57282 | Promoter | 1 | 40681712 | 40768885 | 87174 | 1 | ENSMUST00000027231.7 | 0 | Slc9a2 | ENSMUSG00000026062.7 |
| chr1 | 40682993 | 40683329 | 3.3442 | 3.8748 | 1.08569 | Intron (ENSMUST00000027231.7/ENSMUSG00000026062.7, intron 1 of 11) | 1 | 40681712 | 40768885 | 87174 | 1 | ENSMUST00000027231.7 | 1282 | Slc9a2 | ENSMUSG00000026062.7 |
| chr1 | 38897162 | 38897515 | 3.4125 | 4.05771 | 1.14424 | Promoter | 1 | 38863877 | 38898160 | 34284 | 2 | ENSMUST00000027249.6 | 645 | Chst10 | ENSMUSG00000026080.8 |
| chr1 | 38897903 | 38898347 | 3.8892 | 4.96262 | 1.77076 | Promoter | 1 | 38863877 | 38898160 | 34284 | 2 | ENSMUST00000027249.6 | 0 | Chst10 | ENSMUSG00000026080.8 |
| chr1 | 50920473 | 50920732 | 3.33267 | 3.82632 | 1.01733 | Intron (ENSMUST00000114565.2/ENSMUSG00000026109.9, intron 3 of 9) | 1 | 50927519 | 51187270 | 259752 | 1 | ENSMUST00000081851.3 | -6787 | Tmeff2 | ENSMUSG00000026109.9 |
| chr1 | 50922461 | 50922664 | 3.33388 | 3.85561 | 1.10727 | Intron (ENSMUST00000114565.2/ENSMUSG00000026109.9, intron 3 of 9) | 1 | 50927519 | 51187270 | 259752 | 1 | ENSMUST00000081851.3 | -4855 | Tmeff2 | ENSMUSG00000026109.9 |
| chr1 | 50923393 | 50924104 | 3.77377 | 4.67816 | 1.59823 | Intron (ENSMUST00000114565.2/ENSMUSG00000026109.9, intron 3 of 9) | 1 | 50927519 | 51187270 | 259752 | 1 | ENSMUST00000081851.3 | -3415 | Tmeff2 | ENSMUSG00000026109.9 |
| chr1 | 50925034 | 50927208 | 3.81388 | 5.04693 | 1.82874 | Promoter | 1 | 50927519 | 51187270 | 259752 | 1 | ENSMUST00000081851.3 | -311 | Tmeff2 | ENSMUSG00000026109.9 |
| chr1 | 50927689 | 50928432 | 3.6154 | 4.47556 | 1.48757 | Promoter | 1 | 50927519 | 51187270 | 259752 | 1 | ENSMUST00000081851.3 | 171 | Tmeff2 | ENSMUSG00000026109.9 |
| chr1 | 50963659 | 50963873 | 3.26212 | 3.7248 | 1.03162 | Intron (ENSMUST00000114565.2/ENSMUSG00000026109.9, intron 9 of 9) | 1 | 50927519 | 51187270 | 259752 | 1 | ENSMUST00000081851.3 | 36141 | Tmeff2 | ENSMUSG00000026109.9 |
| chr1 | 36743950 | 36744568 | 3.47851 | 4.12788 | 1.2738 | Distal Intergenic | 1 | 36761798 | 36782816 | 21019 | 1 | ENSMUST00000027291.5 | -17230 | Zap70 | ENSMUSG00000026117.9 |
| chr1 | 34535403 | 34535980 | 3.59833 | 4.54277 | 1.47252 | Promoter | 1 | 34535648 | 34544311 | 8664 | 1 | ENSMUST00000027298.3 | 0 | Cfc1 | ENSMUSG00000026124.3 |
| chr1 | 34536418 | 34537206 | 4.0278 | 5.29661 | 1.97353 | Promoter | 1 | 34535648 | 34544311 | 8664 | 1 | ENSMUST00000027298.3 | 771 | Cfc1 | ENSMUSG00000026124.3 |
| chr1 | 33907570 | 33908841 | 4.2059 | 5.89084 | 2.43613 | Promoter | 1 | 33908225 | 34183904 | 275680 | 1 | ENSMUST00000182513.2 | 0 | Dst | ENSMUSG00000026131.12 |
| chr1 | 33909173 | 33909486 | 3.53878 | 4.24295 | 1.32592 | Promoter | 1 | 33908252 | 34308654 | 400403 | 1 | ENSMUST00000183034.2 | 922 | Dst | ENSMUSG00000026131.12 |
| chr1 | 24584688 | 24585414 | 4.26684 | 5.65215 | 2.23408 | Intron (ENSMUST00000115244.3/ENSMUSG00000026141.8, intron 1 of 50) | 1 | 24279770 | 24587411 | 307642 | 2 | ENSMUST00000051344.5 | 1997 | Col19a1 | ENSMUSG00000026141.8 |
| chr1 | 24586248 | 24586810 | 4.64351 | 6.46321 | 2.87559 | Promoter | 1 | 24279770 | 24587411 | 307642 | 2 | ENSMUST00000051344.5 | 601 | Col19a1 | ENSMUSG00000026141.8 |
| chr1 | 24587364 | 24587721 | 3.67674 | 4.5795 | 1.4988 | Promoter | 1 | 24263994 | 24587472 | 323479 | 2 | ENSMUST00000115244.3 | 0 | Col19a1 | ENSMUSG00000026141.8 |
| chr1 | 24589552 | 24589712 | 3.55246 | 4.23263 | 1.2748 | Distal Intergenic | 1 | 24263994 | 24587472 | 323479 | 2 | ENSMUST00000115244.3 | -2081 | Col19a1 | ENSMUSG00000026141.8 |
| chr1 | 23760945 | 23761574 | 3.5328 | 4.1777 | 1.19126 | Promoter | 1 | 23761753 | 23815740 | 53988 | 1 | ENSMUST00000144602.1 | -179 | B3gat2 | ENSMUSG00000026156.8 |
| chr1 | 23762099 | 23762857 | 3.47641 | 4.2923 | 1.32658 | Promoter | 1 | 23762011 | 23815739 | 53729 | 1 | ENSMUST00000140583.1 | 89 | B3gat2 | ENSMUSG00000026156.8 |
| chr1 | 23763290 | 23763962 | 4.33488 | 5.84756 | 2.3857 | Intron (ENSMUST00000144602.1/ENSMUSG00000026156.8, intron 2 of 2) | 1 | 23762011 | 23815739 | 53729 | 1 | ENSMUST00000140583.1 | 1280 | B3gat2 | ENSMUSG00000026156.8 |
| chr1 | 23764312 | 23764969 | 3.42771 | 3.9896 | 1.08744 | Intron (ENSMUST00000144602.1/ENSMUSG00000026156.8, intron 2 of 2) | 1 | 23762011 | 23815739 | 53729 | 1 | ENSMUST00000140583.1 | 2302 | B3gat2 | ENSMUSG00000026156.8 |
| chr1 | 23796982 | 23797332 | 3.74457 | 4.68364 | 1.5912 | Intron (ENSMUST00000144602.1/ENSMUSG00000026156.8, intron 2 of 2) | 1 | 23762011 | 23815739 | 53729 | 1 | ENSMUST00000140583.1 | 34972 | B3gat2 | ENSMUSG00000026156.8 |
| chr1 | 83406045 | 83406260 | 3.42223 | 4.01235 | 1.18158 | Intron (ENSMUST00000160953.2/ENSMUSG00000026163.12, intron 1 of 10) | 1 | 83339146 | 83408139 | 68994 | 2 | ENSMUST00000160237.1 | 1879 | Sphkap | ENSMUSG00000026163.12 |
| chr1 | 83406856 | 83407233 | 3.43561 | 3.99847 | 1.07666 | Promoter | 1 | 83339146 | 83408139 | 68994 | 2 | ENSMUST00000160237.1 | 906 | Sphkap | ENSMUSG00000026163.12 |
| chr1 | 74788405 | 74788606 | 3.3867 | 4.09095 | 1.19804 | Distal Intergenic | 1 | 74791516 | 74804179 | 12664 | 1 | ENSMUST00000006718.9 | -2910 | Wnt10a | ENSMUSG00000026167.9 |
| chr1 | 74789043 | 74789406 | 3.55905 | 4.25789 | 1.32072 | Distal Intergenic | 1 | 74791516 | 74804179 | 12664 | 1 | ENSMUST00000006718.9 | -2110 | Wnt10a | ENSMUSG00000026167.9 |
| chr1 | 74790509 | 74790760 | 3.33402 | 3.82847 | 1.01733 | Promoter | 1 | 74791516 | 74804179 | 12664 | 1 | ENSMUST00000006718.9 | -756 | Wnt10a | ENSMUSG00000026167.9 |
| chr1 | 74791690 | 74792617 | 4.36949 | 5.9285 | 2.42679 | Promoter | 1 | 74791516 | 74804179 | 12664 | 1 | ENSMUST00000006718.9 | 175 | Wnt10a | ENSMUSG00000026167.9 |
| chr1 | 74793418 | 74793791 | 3.74348 | 4.61985 | 1.51092 | Promoter | 1 | 74793363 | 74803495 | 10133 | 1 | ENSMUST00000187238.1 | 56 | Wnt10a | ENSMUSG00000026167.9 |
| chr1 | 74794260 | 74794419 | 3.46757 | 4.06385 | 1.13789 | Promoter | 1 | 74793363 | 74803495 | 10133 | 1 | ENSMUST00000187238.1 | 898 | Wnt10a | ENSMUSG00000026167.9 |
| chr1 | 74794955 | 74795649 | 3.97295 | 5.0781 | 1.81034 | Intron (ENSMUST00000006718.9/ENSMUSG00000026167.9, intron 2 of 3) | 1 | 74793363 | 74803495 | 10133 | 1 | ENSMUST00000187238.1 | 1593 | Wnt10a | ENSMUSG00000026167.9 |
| chr1 | 74797392 | 74797857 | 4.124 | 5.37494 | 2.01591 | Intron (ENSMUST00000006718.9/ENSMUSG00000026167.9, intron 2 of 3) | 1 | 74793363 | 74803495 | 10133 | 1 | ENSMUST00000187238.1 | 4030 | Wnt10a | ENSMUSG00000026167.9 |
| chr1 | 74798586 | 74798906 | 3.36239 | 4.00979 | 1.12579 | Intron (ENSMUST00000006718.9/ENSMUSG00000026167.9, intron 2 of 3) | 1 | 74793363 | 74803495 | 10133 | 1 | ENSMUST00000187238.1 | 5224 | Wnt10a | ENSMUSG00000026167.9 |
| chr1 | 74800476 | 74800950 | 3.50585 | 4.254 | 1.31819 | Exon (ENSMUST00000006718.9/ENSMUSG00000026167.9, exon 3 of 4) | 1 | 74793363 | 74803495 | 10133 | 1 | ENSMUST00000187238.1 | 7114 | Wnt10a | ENSMUSG00000026167.9 |
| chr1 | 74801665 | 74802035 | 3.59132 | 4.40375 | 1.45636 | Intron (ENSMUST00000006718.9/ENSMUSG00000026167.9, intron 3 of 3) | 1 | 74793363 | 74803495 | 10133 | 1 | ENSMUST00000187238.1 | 8303 | Wnt10a | ENSMUSG00000026167.9 |
| chr1 | 74802530 | 74803028 | 3.43628 | 4.23839 | 1.30093 | Intron (ENSMUST00000006718.9/ENSMUSG00000026167.9, intron 3 of 3) | 1 | 74793363 | 74803495 | 10133 | 1 | ENSMUST00000187238.1 | 9168 | Wnt10a | ENSMUSG00000026167.9 |
| chr1 | 74803678 | 74803825 | 3.25476 | 3.92612 | 1.11041 | 3' UTR | 1 | 74793363 | 74803495 | 10133 | 1 | ENSMUST00000187238.1 | 10316 | Wnt10a | ENSMUSG00000026167.9 |
| chr1 | 74804631 | 74804891 | 3.56735 | 4.26 | 1.29382 | Distal Intergenic | 1 | 74793363 | 74803495 | 10133 | 1 | ENSMUST00000187238.1 | 11269 | Wnt10a | ENSMUSG00000026167.9 |
| chr1 | 74805403 | 74807144 | 3.77954 | 4.78404 | 1.67267 | Distal Intergenic | 1 | 74793363 | 74803495 | 10133 | 1 | ENSMUST00000187238.1 | 12041 | Wnt10a | ENSMUSG00000026167.9 |
| chr1 | 74807481 | 74807661 | 3.79181 | 5.08702 | 1.83358 | Distal Intergenic | 1 | 74793363 | 74803495 | 10133 | 1 | ENSMUST00000187238.1 | 14119 | Wnt10a | ENSMUSG00000026167.9 |
| chr1 | 72872196 | 72872271 | 4.37309 | 5.79203 | 2.27392 | Intron (ENSMUST00000027377.8/ENSMUSG00000026185.8, intron 1 of 3) | 1 | 72858065 | 72874275 | 16211 | 2 | ENSMUST00000137339.1 | 2004 | Igfbp5 | ENSMUSG00000026185.8 |
| chr1 | 72873910 | 72874899 | 3.53824 | 4.23144 | 1.28375 | Promoter | 1 | 72857932 | 72874884 | 16953 | 2 | ENSMUST00000027377.8 | 0 | Igfbp5 | ENSMUSG00000026185.8 |
| chr1 | 72876246 | 72876488 | 3.56035 | 4.28819 | 1.35148 | Distal Intergenic | 1 | 72857932 | 72874884 | 16953 | 2 | ENSMUST00000027377.8 | -1363 | Igfbp5 | ENSMUSG00000026185.8 |
| chr1 | 72878603 | 72878938 | 3.31512 | 4.23834 | 1.29665 | Distal Intergenic | 1 | 72857932 | 72874884 | 16953 | 2 | ENSMUST00000027377.8 | -3720 | Igfbp5 | ENSMUSG00000026185.8 |
| chr1 | 72880229 | 72880345 | 4.16035 | 5.56945 | 2.17011 | Distal Intergenic | 1 | 72857932 | 72874884 | 16953 | 2 | ENSMUST00000027377.8 | -5346 | Igfbp5 | ENSMUSG00000026185.8 |
| chr1 | 75220914 | 75220998 | 3.48289 | 4.09587 | 1.16894 | Intron (ENSMUST00000179573.1/ENSMUSG00000096094.2, intron 2 of 2) | 1 | 75216857 | 75219865 | 3009 | 2 | ENSMUST00000186758.1 | -1050 | Tuba4a | ENSMUSG00000026202.8 |
| chr1 | 75263274 | 75264198 | 3.78717 | 4.80561 | 1.63109 | Promoter | 1 | 75258491 | 75263907 | 5417 | 2 | ENSMUST00000186178.1 | 0 | Ptprn | ENSMUSG00000026204.10 |
| chr1 | 75264737 | 75265038 | 3.29413 | 3.83885 | 1.09111 | Promoter | 1 | 75258559 | 75264502 | 5944 | 2 | ENSMUST00000185849.1 | -236 | Ptprn | ENSMUSG00000026204.10 |
| chr1 | 75265514 | 75265850 | 3.3651 | 3.87521 | 1.01733 | Distal Intergenic | 1 | 75258559 | 75264502 | 5944 | 2 | ENSMUST00000185849.1 | -1013 | Ptprn | ENSMUSG00000026204.10 |
| chr1 | 75266278 | 75266422 | 3.35128 | 3.85437 | 1.01733 | Distal Intergenic | 1 | 75258559 | 75264502 | 5944 | 2 | ENSMUST00000185849.1 | -1777 | Ptprn | ENSMUSG00000026204.10 |
| chr1 | 75268480 | 75268580 | 3.62884 | 4.36269 | 1.33347 | Distal Intergenic | 1 | 75258559 | 75264502 | 5944 | 2 | ENSMUST00000185849.1 | -3979 | Ptprn | ENSMUSG00000026204.10 |
| chr1 | 75485066 | 75485519 | 3.562 | 4.6984 | 1.58316 | 3' UTR | 1 | 75479310 | 75487233 | 7924 | 2 | ENSMUST00000156499.1 | 1714 | Obsl1 | ENSMUSG00000026211.12 |
| chr1 | 75486035 | 75486653 | 3.91454 | 5.15065 | 1.87337 | Promoter | 1 | 75479310 | 75487233 | 7924 | 2 | ENSMUST00000156499.1 | 580 | Obsl1 | ENSMUSG00000026211.12 |
| chr1 | 75486993 | 75487070 | 3.48799 | 4.45743 | 1.41102 | Promoter | 1 | 75479310 | 75487233 | 7924 | 2 | ENSMUST00000156499.1 | 163 | Obsl1 | ENSMUSG00000026211.12 |
| chr1 | 75505217 | 75505311 | 3.7708 | 4.63261 | 1.50709 | Promoter | 1 | 75490534 | 75505641 | 15108 | 2 | ENSMUST00000155084.2 | 330 | Obsl1 | ENSMUSG00000026211.12 |
| chr1 | 84933925 | 84935098 | 3.72535 | 4.61127 | 1.53128 | Promoter | 1 | 84929334 | 84934857 | 5524 | 2 | ENSMUST00000186980.1 | 0 | Slc16a14 | ENSMUSG00000026220.6 |
| chr1 | 84935679 | 84935765 | 3.97986 | 5.0418 | 1.80055 | Promoter | 1 | 84905898 | 84935134 | 29237 | 2 | ENSMUST00000027422.6 | -546 | Slc16a14 | ENSMUSG00000026220.6 |
| chr1 | 86018389 | 86018530 | 3.79757 | 5.09303 | 1.82837 | Promoter | 1 | 86018693 | 86029248 | 10556 | 1 | ENSMUST00000149469.1 | -163 | Spata3 | ENSMUSG00000026226.10 |
| chr1 | 86045827 | 86046213 | 4.59521 | 6.88181 | 3.19874 | Promoter | 1 | 86045863 | 86053353 | 7491 | 1 | ENSMUST00000027429.5 | 0 | 2810459M11Rik | ENSMUSG00000026227.6 |
| chr1 | 86046524 | 86046624 | 4.6968 | 6.81081 | 3.06738 | Promoter | 1 | 86045863 | 86055456 | 9594 | 1 | ENSMUST00000165824.2 | 662 | 2810459M11Rik | ENSMUSG00000026227.6 |
| chr1 | 86048442 | 86048787 | 3.24417 | 3.6934 | 1.01733 | Exon (ENSMUST00000027429.5/ENSMUSG00000026227.6, exon 2 of 3) | 1 | 86045863 | 86055456 | 9594 | 1 | ENSMUST00000165824.2 | 2580 | 2810459M11Rik | ENSMUSG00000026227.6 |
| chr1 | 77289747 | 77289822 | 3.59266 | 4.36026 | 1.43911 | Distal Intergenic | 1 | 77373613 | 77377607 | 3995 | 2 | ENSMUST00000187346.1 | 87785 | Epha4 | ENSMUSG00000026235.9 |
| chr1 | 77311303 | 77311521 | 3.29495 | 3.76963 | 1.01733 | Distal Intergenic | 1 | 77373613 | 77377607 | 3995 | 2 | ENSMUST00000187346.1 | 66086 | Epha4 | ENSMUSG00000026235.9 |
| chr1 | 77313159 | 77313530 | 3.63485 | 4.4193 | 1.43986 | Distal Intergenic | 1 | 77373613 | 77377607 | 3995 | 2 | ENSMUST00000187346.1 | 64077 | Epha4 | ENSMUSG00000026235.9 |
| chr1 | 77315528 | 77315683 | 3.40539 | 4.01503 | 1.17995 | Distal Intergenic | 1 | 77373613 | 77377607 | 3995 | 2 | ENSMUST00000187346.1 | 61924 | Epha4 | ENSMUSG00000026235.9 |
| chr1 | 77321384 | 77321521 | 3.54858 | 4.22148 | 1.25925 | Distal Intergenic | 1 | 77373613 | 77377607 | 3995 | 2 | ENSMUST00000187346.1 | 56086 | Epha4 | ENSMUSG00000026235.9 |
| chr1 | 86669038 | 86669186 | 3.91083 | 5.08008 | 1.81812 | Intron (ENSMUST00000027449.5/ENSMUSG00000026241.5, intron 2 of 2) | 1 | 86666291 | 86670571 | 4281 | 2 | ENSMUST00000027449.5 | 1385 | Nppc | ENSMUSG00000026241.5 |
| chr1 | 86669768 | 86670987 | 3.57552 | 4.42868 | 1.40037 | Promoter | 1 | 86666291 | 86670571 | 4281 | 2 | ENSMUST00000027449.5 | 0 | Nppc | ENSMUSG00000026241.5 |
| chr1 | 86671535 | 86671676 | 3.33595 | 3.83126 | 1.01733 | Promoter | 1 | 86666291 | 86670571 | 4281 | 2 | ENSMUST00000027449.5 | -965 | Nppc | ENSMUSG00000026241.5 |
| chr1 | 87152008 | 87152277 | 4.13219 | 5.44889 | 2.09401 | Exon (ENSMUST00000161002.2/ENSMUSG00000026247.8, exon 7 of 18) | 1 | 87147658 | 87155027 | 7370 | 2 | ENSMUST00000027463.5 | 2750 | Ecel1 | ENSMUSG00000026247.8 |
| chr1 | 87153473 | 87154329 | 3.4464 | 4.06071 | 1.14199 | Promoter | 1 | 87147658 | 87155027 | 7370 | 2 | ENSMUST00000027463.5 | 698 | Ecel1 | ENSMUSG00000026247.8 |
| chr1 | 87154760 | 87155332 | 3.8834 | 4.90851 | 1.70869 | Promoter | 1 | 87147658 | 87155027 | 7370 | 2 | ENSMUST00000027463.5 | 0 | Ecel1 | ENSMUSG00000026247.8 |
| chr1 | 87155732 | 87156160 | 4.24964 | 5.61707 | 2.19266 | Promoter | 1 | 87147662 | 87156165 | 8504 | 2 | ENSMUST00000160810.1 | 5 | Ecel1 | ENSMUSG00000026247.8 |
| chr1 | 87156643 | 87157433 | 3.63052 | 4.36767 | 1.32557 | Promoter | 1 | 87147655 | 87156521 | 8867 | 2 | ENSMUST00000161002.2 | -123 | Ecel1 | ENSMUSG00000026247.8 |
| chr1 | 79857980 | 79858937 | 4.0217 | 5.29237 | 1.99072 | Promoter | 1 | 79794197 | 79858696 | 64500 | 2 | ENSMUST00000027467.5 | 0 | Serpine2 | ENSMUSG00000026249.5 |
| chr1 | 79859355 | 79859645 | 3.58323 | 4.29212 | 1.27921 | Promoter | 1 | 79815108 | 79858696 | 43589 | 2 | ENSMUST00000153862.1 | -660 | Serpine2 | ENSMUSG00000026249.5 |
| chr1 | 87206165 | 87207036 | 3.68947 | 4.48482 | 1.39673 | Promoter | 1 | 87206074 | 87208595 | 2522 | 1 | ENSMUST00000189392.1 | 92 | Chrng | ENSMUSG00000026253.9 |
| chr1 | 87261487 | 87262972 | 3.64417 | 4.39456 | 1.3397 | Distal Intergenic | 1 | 87264363 | 87310836 | 46474 | 1 | ENSMUST00000118687.2 | -1391 | Efhd1 | ENSMUSG00000026255.10 |
| chr1 | 87263386 | 87264019 | 3.94998 | 4.99599 | 1.74158 | Promoter | 1 | 87264363 | 87310836 | 46474 | 1 | ENSMUST00000118687.2 | -344 | Efhd1 | ENSMUSG00000026255.10 |
| chr1 | 87264359 | 87265277 | 3.9532 | 5.07764 | 1.86436 | Promoter | 1 | 87264363 | 87310836 | 46474 | 1 | ENSMUST00000118687.2 | 0 | Efhd1 | ENSMUSG00000026255.10 |

| | | | | | | | | | | | | | | | |
|---|---|---|---|---|---|---|---|---|---|---|---|---|---|---|---|
| chr1 | 87475282 | 87475413 | 4.02999 | 5.1364 | 1.85835 | Exon (ENSMUST00000027476.5/ENSMUSG00000026258.5, exon 3 of 3) | 1 | 87472539 | 87475458 | 2920 | 1 | ENSMUST00000191095.1 | 2744 | 3110079O15Rik | ENSMUSG00000026258.5 |
| chr1 | 87508876 | 87510661 | 3.7711 | 4.67753 | 1.58221 | Promoter | 1 | 87476834 | 87510362 | 33529 | 2 | ENSMUST00000027477.9 | 0 | Ngef | ENSMUSG00000026259.9 |
| chr1 | 90118469 | 90118935 | 3.56256 | 4.41774 | 1.39196 | Intron (ENSMUST00000113094.3/ENSMUSG00000026301.10, intron 6 of 18) | 1 | 90042135 | 90090144 | 48010 | 2 | ENSMUST00000189690.1 | -28326 | Iqca | ENSMUSG00000026301.10 |
| chr1 | 90119459 | 90119534 | 3.5767 | 4.31083 | 1.38403 | Intron (ENSMUST00000113094.3/ENSMUSG00000026301.10, intron 6 of 18) | 1 | 90042135 | 90090144 | 48010 | 2 | ENSMUST00000189690.1 | -29316 | Iqca | ENSMUSG00000026301.10 |
| chr1 | 90153328 | 90153466 | 3.36257 | 3.87138 | 1.01733 | Promoter | 1 | 90042132 | 90153401 | 111270 | 2 | ENSMUST00000113094.3 | 0 | Iqca | ENSMUSG00000026301.10 |
| chr1 | 90902991 | 90903316 | 3.34267 | 3.84137 | 1.01733 | Distal Intergenic | 1 | 90915085 | 90951142 | 36058 | 1 | ENSMUST00000027528.6 | -11769 | Mlph | ENSMUSG00000026303.6 |
| chr1 | 90928116 | 90928481 | 4.79642 | 6.99894 | 3.31421 | Promoter | 1 | 90927679 | 90928432 | 754 | 1 | ENSMUST00000125149.1 | 438 | Mlph | ENSMUSG00000026303.6 |
| chr1 | 91338806 | 91338916 | 3.41655 | 3.99872 | 1.17842 | Intron (ENSMUST00000088904.4/ENSMUSG00000049515.7, intron 5 of 8) | 1 | 91351016 | 91362416 | 11401 | 1 | ENSMUST00000027533.8 | -12100 | Klhl30 | ENSMUSG00000026308.8 |
| chr1 | 91342645 | 91342979 | 3.60553 | 4.36639 | 1.41599 | Exon (ENSMUST00000088904.4/ENSMUSG00000049515.7, exon 8 of 9) | 1 | 91351016 | 91362416 | 11401 | 1 | ENSMUST00000027533.8 | -8037 | Klhl30 | ENSMUSG00000026308.8 |
| chr1 | 91344578 | 91345599 | 3.99004 | 5.24279 | 1.95 | Exon (ENSMUST00000088904.4/ENSMUSG00000049515.7, exon 9 of 9) | 1 | 91351016 | 91362416 | 11401 | 1 | ENSMUST00000027533.8 | -5417 | Klhl30 | ENSMUSG00000026308.8 |
| chr1 | 105781844 | 105782254 | 3.80519 | 4.75076 | 1.58756 | Intron (ENSMUST00000027559.7/ENSMUSG00000026321.7, intron 1 of 9) | 1 | 105780750 | 105814403 | 33654 | 1 | ENSMUST00000187812.1 | 1095 | Tnfrsf11a | ENSMUSG00000026321.7 |
| chr1 | 105819475 | 105819580 | 3.35972 | 4.03325 | 1.16491 | Promoter | 1 | 105819870 | 105825401 | 5532 | 1 | ENSMUST00000187658.1 | -290 | Tnfrsf11a | ENSMUSG00000026321.7 |
| chr1 | 105844360 | 105844810 | 3.32925 | 3.82125 | 1.01733 | Exon (ENSMUST00000027559.7/ENSMUSG00000026321.7, exon 10 of 10) | 1 | 105819870 | 105825401 | 5532 | 1 | ENSMUST00000187658.1 | 24491 | Tnfrsf11a | ENSMUSG00000026321.7 |
| chr18 | 62323595 | 62324450 | 3.61156 | 4.40474 | 1.36777 | Promoter | 18 | 62324204 | 62467802 | 143599 | 1 | ENSMUST00000027560.6 | 0 | Htr4 | ENSMUSG00000026322.8 |
| chr18 | 62324828 | 62325108 | 3.68609 | 4.61824 | 1.51349 | Promoter | 18 | 62324204 | 62467802 | 143599 | 1 | ENSMUST00000027560.6 | 625 | Htr4 | ENSMUSG00000026322.8 |
| chr18 | 62325928 | 62326064 | 3.37696 | 3.89964 | 1.04054 | Intron (ENSMUST00000027560.6/ENSMUSG00000026322.8, intron 1 of 6) | 18 | 62324204 | 62467802 | 143599 | 1 | ENSMUST00000027560.6 | 1725 | Htr4 | ENSMUSG00000026322.8 |
| chr1 | 121416414 | 121416618 | 3.40495 | 3.95984 | 1.10391 | Distal Intergenic | 1 | 121431049 | 121506460 | 75412 | 1 | ENSMUST00000036025.10 | -14431 | Ccdc93 | ENSMUSG00000026339.13 |
| chr1 | 125910371 | 125910823 | 3.49774 | 4.15965 | 1.29094 | Promoter | 1 | 125873101 | 125910908 | 37808 | 2 | ENSMUST00000159529.1 | 85 | Lypd1 | ENSMUSG00000026344.4 |
| chr1 | 125911300 | 125911557 | 3.24107 | 3.83771 | 1.07065 | Promoter | 1 | 125872284 | 125911387 | 39104 | 2 | ENSMUST00000162899.2 | 0 | Lypd1 | ENSMUSG00000026344.4 |
| chr1 | 125912515 | 125913274 | 3.78385 | 4.78452 | 1.6541 | Promoter | 1 | 125872318 | 125912928 | 40611 | 2 | ENSMUST00000027582.4 | 0 | Lypd1 | ENSMUSG00000026344.4 |
| chr1 | 127575701 | 127576060 | 3.36051 | 3.86828 | 1.01733 | Intron (ENSMUST00000027585.8/ENSMUSG00000026347.8, intron 2 of 8) | 1 | 127491545 | 127537430 | 45886 | 2 | ENSMUST00000162406.1 | -38272 | Tmem163 | ENSMUSG00000026347.8 |
| chr1 | 127675242 | 127675840 | 3.77362 | 4.70414 | 1.60802 | Intron (ENSMUST00000027585.8/ENSMUSG00000026347.8, intron 1 of 8) | 1 | 127486558 | 127678021 | 191464 | 2 | ENSMUST00000160616.2 | 2181 | Tmem163 | ENSMUSG00000026347.8 |
| chr1 | 127676286 | 127678628 | 3.74408 | 4.75046 | 1.59679 | Promoter | 1 | 127486546 | 127678096 | 191551 | 2 | ENSMUST00000027585.8 | 0 | Tmem163 | ENSMUSG00000026347.8 |
| chr1 | 127679190 | 127679573 | 3.5032 | 4.13188 | 1.18833 | Promoter | 1 | 127490898 | 127679548 | 188651 | 2 | ENSMUST00000185560.1 | 0 | Tmem163 | ENSMUSG00000026347.8 |
| chr1 | 127679880 | 127680052 | 4.00209 | 5.26124 | 1.96485 | Promoter | 1 | 127490898 | 127679548 | 188651 | 2 | ENSMUST00000185560.1 | -333 | Tmem163 | ENSMUSG00000026347.8 |
| chr1 | 118317662 | 118317774 | 3.26674 | 3.73692 | 1.04081 | Distal Intergenic | 1 | 118321839 | 118333822 | 11984 | 1 | ENSMUST00000027626.7 | -4065 | Nifk | ENSMUSG00000026377.7 |
| chr1 | 136910351 | 136911087 | 3.82995 | 4.75107 | 1.60091 | Intron (ENSMUST00000168126.1/ENSMUSG00000026398.9, intron 6 of 8) | 1 | 136843584 | 136953630 | 110047 | 2 | ENSMUST00000168126.1 | 42543 | Nr5a2 | ENSMUSG00000026398.9 |
| chr1 | 136911487 | 136912417 | 3.77132 | 4.80542 | 1.65174 | Intron (ENSMUST00000168126.1/ENSMUSG00000026398.9, intron 6 of 8) | 1 | 136843584 | 136953630 | 110047 | 2 | ENSMUST00000168126.1 | 41213 | Nr5a2 | ENSMUSG00000026398.9 |
| chr1 | 136912824 | 136912904 | 3.5116 | 4.15629 | 1.22942 | Intron (ENSMUST00000168126.1/ENSMUSG00000026398.9, intron 6 of 8) | 1 | 136843584 | 136953630 | 110047 | 2 | ENSMUST00000168126.1 | 40726 | Nr5a2 | ENSMUSG00000026398.9 |
| chr1 | 136916191 | 136916417 | 3.70052 | 4.58939 | 1.52245 | Intron (ENSMUST00000168126.1/ENSMUSG00000026398.9, intron 6 of 8) | 1 | 136843584 | 136953630 | 110047 | 2 | ENSMUST00000168126.1 | 37213 | Nr5a2 | ENSMUSG00000026398.9 |
| chr1 | 136917001 | 136917302 | 3.69917 | 4.62146 | 1.54763 | Intron (ENSMUST00000168126.1/ENSMUSG00000026398.9, intron 6 of 8) | 1 | 136843584 | 136953630 | 110047 | 2 | ENSMUST00000168126.1 | 36328 | Nr5a2 | ENSMUSG00000026398.9 |
| chr1 | 136943154 | 136943354 | 3.41409 | 4.09476 | 1.186 | Exon (ENSMUST00000168126.1/ENSMUSG00000026398.9, exon 5 of 9) | 1 | 136843584 | 136953630 | 110047 | 2 | ENSMUST00000168126.1 | 10276 | Nr5a2 | ENSMUSG00000026398.9 |
| chr1 | 136944504 | 136946463 | 3.50911 | 4.40855 | 1.41093 | Exon (ENSMUST00000168126.1/ENSMUSG00000026398.9, exon 4 of 9) | 1 | 136843584 | 136953630 | 110047 | 2 | ENSMUST00000168126.1 | 7167 | Nr5a2 | ENSMUSG00000026398.9 |
| chr1 | 136946939 | 136947825 | 3.88427 | 4.86956 | 1.69782 | Intron (ENSMUST00000168126.1/ENSMUSG00000026398.9, intron 3 of 8) | 1 | 136843584 | 136953630 | 110047 | 2 | ENSMUST00000168126.1 | 5805 | Nr5a2 | ENSMUSG00000026398.9 |
| chr1 | 136948412 | 136950634 | 3.91301 | 5.06974 | 1.84934 | 5' UTR | 1 | 136843584 | 136953630 | 110047 | 2 | ENSMUST00000168126.1 | 2996 | Nr5a2 | ENSMUSG00000026398.9 |
| chr1 | 136951029 | 136951643 | 3.92938 | 5.00831 | 1.85059 | Intron (ENSMUST00000168126.1/ENSMUSG00000026398.9, intron 2 of 8) | 1 | 136843584 | 136953630 | 110047 | 2 | ENSMUST00000168126.1 | 1987 | Nr5a2 | ENSMUSG00000026398.9 |
| chr1 | 136953017 | 136953749 | 3.52281 | 4.5453 | 1.46865 | Promoter | 1 | 136843584 | 136953630 | 110047 | 2 | ENSMUST00000168126.1 | 0 | Nr5a2 | ENSMUSG00000026398.9 |
| chr1 | 136954060 | 136954149 | 4.12423 | 5.31886 | 1.97994 | Promoter | 1 | 136843584 | 136953630 | 110047 | 2 | ENSMUST00000168126.1 | -431 | Nr5a2 | ENSMUSG00000026398.9 |
| chr1 | 136954480 | 136955054 | 4.00331 | 5.2614 | 1.94552 | Promoter | 1 | 136843584 | 136953630 | 110047 | 2 | ENSMUST00000168126.1 | -851 | Nr5a2 | ENSMUSG00000026398.9 |
| chr1 | 136956930 | 136957338 | 3.94755 | 5.04594 | 1.79475 | Intron (ENSMUST00000027649.8/ENSMUSG00000026398.9, intron 1 of 8) | 1 | 136843584 | 136960427 | 116844 | 2 | ENSMUST00000027649.8 | 3089 | Nr5a2 | ENSMUSG00000026398.9 |
| chr1 | 136958963 | 136959220 | 3.85047 | 4.81959 | 1.63838 | Intron (ENSMUST00000027649.8/ENSMUSG00000026398.9, intron 1 of 8) | 1 | 136843584 | 136960427 | 116844 | 2 | ENSMUST00000027649.8 | 1207 | Nr5a2 | ENSMUSG00000026398.9 |
| chr1 | 136960198 | 136960356 | 3.35769 | 3.86404 | 1.01733 | Promoter | 1 | 136843584 | 136960427 | 116844 | 2 | ENSMUST00000027649.8 | 71 | Nr5a2 | ENSMUSG00000026398.9 |
| chr1 | 136971387 | 136971564 | 3.57821 | 4.44215 | 1.40548 | Distal Intergenic | 1 | 136843584 | 136960427 | 116844 | 2 | ENSMUST00000027649.8 | -10961 | Nr5a2 | ENSMUSG00000026398.9 |
| chr1 | 130491072 | 130491536 | 4.15806 | 5.48186 | 2.12461 | Intron (ENSMUST00000125681.1/ENSMUSG00000085526.1, intron 1 of 4) | 1 | 130439027 | 130462744 | 23718 | 2 | ENSMUST00000027650.7 | -28329 | Cd55 | ENSMUSG00000026399.7 |
| chr1 | 135909820 | 135909971 | 3.52818 | 4.1864 | 1.24822 | Intron (ENSMUST00000027667.7/ENSMUSG00000026413.7, intron 1 of 13) | 1 | 135872950 | 135919120 | 46171 | 2 | ENSMUST00000163260.2 | 9149 | Pkp1 | ENSMUSG00000026413.7 |
| chr1 | 135917414 | 135918045 | 3.37408 | 3.88875 | 1.01733 | Intron (ENSMUST00000027667.7/ENSMUSG00000026413.7, intron 1 of 13) | 1 | 135872950 | 135919120 | 46171 | 2 | ENSMUST00000163260.2 | 1075 | Pkp1 | ENSMUSG00000026413.7 |
| chr1 | 135918627 | 135919297 | 3.846 | 4.96661 | 1.76962 | Promoter | 1 | 135871395 | 135919207 | 47813 | 2 | ENSMUST00000027667.7 | 0 | Pkp1 | ENSMUSG00000026413.7 |
| chr1 | 135793475 | 135793822 | 3.64248 | 4.54109 | 1.43798 | Intron (ENSMUST00000132795.2/ENSMUSG00000026418.11, intron 2 of 6) | 1 | 135799402 | 135810989 | 11588 | 1 | ENSMUST00000152208.2 | -5580 | Tnni1 | ENSMUSG00000026418.11 |
| chr1 | 135794875 | 135795329 | 3.35205 | 3.85552 | 1.01733 | Intron (ENSMUST00000132795.2/ENSMUSG00000026418.11, intron 2 of 6) | 1 | 135799402 | 135810989 | 11588 | 1 | ENSMUST00000152208.2 | -4073 | Tnni1 | ENSMUSG00000026418.11 |
| chr1 | 135804050 | 135804730 | 3.35742 | 3.89687 | 1.12757 | Promoter | 1 | 135805057 | 135809909 | 4853 | 1 | ENSMUST00000129217.1 | -327 | Tnni1 | ENSMUSG00000026418.11 |
| chr1 | 131579243 | 131579663 | 3.48511 | 4.14812 | 1.19335 | Distal Intergenic | 1 | 131599239 | 131612000 | 12762 | 1 | ENSMUST00000027690.6 | -19576 | Avpr1b | ENSMUSG00000026432.7 |
| chr1 | 131596007 | 131596102 | 3.30365 | 3.80391 | 1.08298 | Distal Intergenic | 1 | 131599239 | 131612000 | 12762 | 1 | ENSMUST00000027690.6 | -3137 | Avpr1b | ENSMUSG00000026432.7 |
| chr1 | 131598108 | 131598295 | 3.35493 | 3.85986 | 1.01733 | Promoter | 1 | 131599239 | 131612000 | 12762 | 1 | ENSMUST00000027690.6 | -944 | Avpr1b | ENSMUSG00000026432.7 |
| chr1 | 131963241 | 131963977 | 3.61995 | 4.48418 | 1.45856 | Promoter | 1 | 131962967 | 131982969 | 20003 | 1 | ENSMUST00000177943.2 | 275 | Slc45a3 | ENSMUSG00000026435.10 |
| chr1 | 131964634 | 131964830 | 3.34221 | 3.84289 | 1.02507 | Intron (ENSMUST00000191034.1/ENSMUSG00000100037.1, intron 1 of 2) | 1 | 131962967 | 131982969 | 20003 | 1 | ENSMUST00000177943.2 | 1668 | Slc45a3 | ENSMUSG00000026435.10 |
| chr1 | 131965688 | 131966121 | 3.73515 | 4.65537 | 1.57364 | Intron (ENSMUST00000191034.1/ENSMUSG00000100037.1, intron 1 of 2) | 1 | 131962967 | 131982969 | 20003 | 1 | ENSMUST00000177943.2 | 2722 | Slc45a3 | ENSMUSG00000026435.10 |
| chr1 | 131966461 | 131966743 | 3.39715 | 3.94812 | 1.09798 | Intron (ENSMUST00000191034.1/ENSMUSG00000100037.1, intron 1 of 2) | 1 | 131962967 | 131982969 | 20003 | 1 | ENSMUST00000177943.2 | 3495 | Slc45a3 | ENSMUSG00000026435.10 |
| chr1 | 131971012 | 131972164 | 3.72594 | 4.57174 | 1.48012 | Promoter | 1 | 131970609 | 131982964 | 12356 | 1 | ENSMUST00000027695.7 | 404 | Slc45a3 | ENSMUSG00000026435.10 |
| chr1 | 132139000 | 132139636 | 3.33576 | 3.87969 | 1.07748 | Promoter | 1 | 132114764 | 132139605 | 24842 | 2 | ENSMUST00000112362.2 | 0 | Cdk18 | ENSMUSG00000026437.6 |
| chr1 | 132165339 | 132165657 | 3.5806 | 4.27207 | 1.27024 | Distal Intergenic | 1 | 132112237 | 132139684 | 27448 | 2 | ENSMUST00000027697.6 | -25656 | Cdk18 | ENSMUSG00000026437.6 |
| chr1 | 132623102 | 132623618 | 3.53964 | 4.19686 | 1.2162 | Promoter | 1 | 132619380 | 132624512 | 5133 | 2 | ENSMUST00000191294.1 | 894 | Nfasc | ENSMUSG00000026442.9 |
| chr1 | 132624361 | 132624659 | 3.65094 | 4.40142 | 1.3364 | Promoter | 1 | 132619380 | 132624512 | 5133 | 2 | ENSMUST00000191294.1 | 0 | Nfasc | ENSMUSG00000026442.9 |
| chr1 | 132656274 | 132656466 | 3.32609 | 3.81641 | 1.01733 | Intron (ENSMUST00000094569.5/ENSMUSG00000026442.9, intron 2 of 29) | 1 | 132570423 | 132637091 | 66669 | 2 | ENSMUST00000186389.1 | -19184 | Nfasc | ENSMUSG00000026442.9 |
| chr1 | 132735873 | 132736129 | 3.3784 | 3.90706 | 1.05904 | Intron (ENSMUST00000094569.5/ENSMUSG00000026442.9, intron 1 of 29) | 1 | 132564690 | 132741757 | 177068 | 2 | ENSMUST00000094569.5 | 5628 | Nfasc | ENSMUSG00000026442.9 |
| chr1 | 132738666 | 132739489 | 3.99036 | 5.22146 | 1.96804 | Intron (ENSMUST00000094569.5/ENSMUSG00000026442.9, intron 1 of 29) | 1 | 132564690 | 132741757 | 177068 | 2 | ENSMUST00000094569.5 | 2268 | Nfasc | ENSMUSG00000026442.9 |
| chr1 | 132741133 | 132741699 | 3.89162 | 4.9469 | 1.78455 | Promoter | 1 | 132564690 | 132741757 | 177068 | 2 | ENSMUST00000094569.5 | 58 | Nfasc | ENSMUSG00000026442.9 |
| chr1 | 132765694 | 132766297 | 3.45029 | 4.06876 | 1.19014 | Distal Intergenic | 1 | 132599835 | 132741797 | 141963 | 2 | ENSMUST00000188307.1 | -23898 | Nfasc | ENSMUSG00000026442.9 |
| chr1 | 132878303 | 132878656 | 3.41388 | 4.0022 | 1.1354 | Distal Intergenic | 1 | 132880273 | 132940005 | 59733 | 1 | ENSMUST00000027706.3 | -1617 | Lrrn2 | ENSMUSG00000026443.3 |
| chr1 | 132879846 | 132880763 | 3.62601 | 4.43774 | 1.41369 | Promoter | 1 | 132880273 | 132940005 | 59733 | 1 | ENSMUST00000027706.3 | 0 | Lrrn2 | ENSMUSG00000026443.3 |
| chr1 | 132881134 | 132881681 | 4.08519 | 5.2624 | 1.97442 | Promoter | 1 | 132880355 | 132937150 | 56796 | 1 | ENSMUST00000159088.1 | 780 | Lrrn2 | ENSMUSG00000026443.3 |
| chr1 | 132917654 | 132917773 | 3.46357 | 4.06395 | 1.15971 | Intron (ENSMUST00000027706.3/ENSMUSG00000026443.3, intron 1 of 1) | 1 | 132880355 | 132937150 | 56796 | 1 | ENSMUST00000159088.1 | 37300 | Lrrn2 | ENSMUSG00000026443.3 |
| chr1 | 133046110 | 133046906 | 3.75575 | 4.66376 | 1.62773 | Promoter | 1 | 133046012 | 133108687 | 62676 | 1 | ENSMUST00000077730.5 | 99 | Pik3c2b | ENSMUSG00000026447.11 |
| chr1 | 134646296 | 134647247 | 3.98278 | 5.17404 | 1.9126 | Promoter | 1 | 134646677 | 134741829 | 95153 | 1 | ENSMUST00000187725.1 | 0 | Syt2 | ENSMUSG00000026452.10 |
| chr1 | 134680956 | 134681173 | 3.32141 | 3.80937 | 1.01733 | Intron (ENSMUST00000187725.1/ENSMUSG00000026452.10, intron 1 of 2) | 1 | 134709300 | 134753149 | 43850 | 1 | ENSMUST00000121990.1 | -28127 | Syt2 | ENSMUSG00000026452.10 |
| chr1 | 134708287 | 134710375 | 3.85573 | 4.81901 | 1.63705 | Promoter | 1 | 134709300 | 134753149 | 43850 | 1 | ENSMUST00000121990.1 | 0 | Syt2 | ENSMUSG00000026452.10 |
| chr1 | 134711842 | 134712055 | 3.44232 | 4.0107 | 1.08407 | Intron (ENSMUST00000187725.1/ENSMUSG00000026452.10, intron 1 of 2) | 1 | 134709300 | 134753149 | 43850 | 1 | ENSMUST00000121990.1 | 2543 | Syt2 | ENSMUSG00000026452.10 |
| chr1 | 134298302 | 134299421 | 3.75984 | 4.91225 | 1.71302 | 3' UTR | 1 | 134296789 | 134304808 | 8020 | 2 | ENSMUST00000189862.1 | 5387 | Ppfia4 | ENSMUSG00000026458.8 |
| chr1 | 134299973 | 134300534 | 3.40717 | 3.99906 | 1.18807 | Exon (ENSMUST00000168515.2/ENSMUSG00000026458.8, exon 26 of 29) | 1 | 134296789 | 134304808 | 8020 | 2 | ENSMUST00000189862.1 | 4274 | Ppfia4 | ENSMUSG00000026458.8 |
| chr1 | 134332593 | 134332766 | 4.21213 | 5.49696 | 2.06792 | Promoter | 1 | 134296783 | 134332928 | 36146 | 2 | ENSMUST00000168515.2 | 162 | Ppfia4 | ENSMUSG00000026458.8 |
| chr1 | 134295665 | 134295882 | 3.34153 | 3.83966 | 1.01733 | Downstream (<1kb) | 1 | 134289989 | 134292548 | 2560 | 1 | ENSMUST00000027730.5 | 5677 | Myog | ENSMUSG00000026459.5 |
| chr1 | 133805312 | 133805840 | 3.41127 | 3.96294 | 1.08122 | Distal Intergenic | 1 | 133699457 | 133801041 | 101585 | 2 | ENSMUST00000143567.2 | -4272 | Atp2b4 | ENSMUSG00000026463.12 |
| chr1 | 133810146 | 133810470 | 3.45351 | 4.03109 | 1.09643 | Distal Intergenic | 1 | 133699457 | 133801041 | 101585 | 2 | ENSMUST00000143567.2 | -9106 | Atp2b4 | ENSMUSG00000026463.12 |
| chr1 | 155704921 | 155705146 | 3.53904 | 4.33861 | 1.37493 | Intron (ENSMUST00000027740.8/ENSMUSG00000026468.9, intron 4 of 5) | 1 | 155698031 | 155751710 | 53680 | 2 | ENSMUST00000027740.8 | 46564 | Lhx4 | ENSMUSG00000026468.9 |
| chr1 | 155712518 | 155712858 | 3.59766 | 4.29487 | 1.25911 | Intron (ENSMUST00000027740.8/ENSMUSG00000026468.9, intron 2 of 5) | 1 | 155698031 | 155751710 | 53680 | 2 | ENSMUST00000027740.8 | 38852 | Lhx4 | ENSMUSG00000026468.9 |
| chr1 | 155721410 | 155721508 | 3.67035 | 4.49638 | 1.50495 | Intron (ENSMUST00000027740.8/ENSMUSG00000026468.9, intron 2 of 5) | 1 | 155698031 | 155751710 | 53680 | 2 | ENSMUST00000027740.8 | 30202 | Lhx4 | ENSMUSG00000026468.9 |
| chr1 | 155724797 | 155725293 | 4.06356 | 5.27104 | 1.97572 | Exon (ENSMUST00000027740.8/ENSMUSG00000026468.9, exon 2 of 6) | 1 | 155698031 | 155751710 | 53680 | 2 | ENSMUST00000027740.8 | 26417 | Lhx4 | ENSMUSG00000026468.9 |
| chr1 | 155726955 | 155727400 | 3.56099 | 4.24775 | 1.2894 | Intron (ENSMUST00000027740.8/ENSMUSG00000026468.9, intron 1 of 5) | 1 | 155698031 | 155751710 | 53680 | 2 | ENSMUST00000027740.8 | 24310 | Lhx4 | ENSMUSG00000026468.9 |
| chr1 | 155727732 | 155728991 | 3.61775 | 4.35291 | 1.34632 | Intron (ENSMUST00000027740.8/ENSMUSG00000026468.9, intron 1 of 5) | 1 | 155698031 | 155751710 | 53680 | 2 | ENSMUST00000027740.8 | 22719 | Lhx4 | ENSMUSG00000026468.9 |
| chr1 | 155729755 | 155730909 | 3.5964 | 4.3618 | 1.39236 | Intron (ENSMUST00000027740.8/ENSMUSG00000026468.9, intron 1 of 5) | 1 | 155698031 | 155751710 | 53680 | 2 | ENSMUST00000027740.8 | 20801 | Lhx4 | ENSMUSG00000026468.9 |
| chr1 | 155731838 | 155732600 | 3.71866 | 4.68857 | 1.59914 | Intron (ENSMUST00000027740.8/ENSMUSG00000026468.9, intron 1 of 5) | 1 | 155698031 | 155751710 | 53680 | 2 | ENSMUST00000027740.8 | 19110 | Lhx4 | ENSMUSG00000026468.9 |
| chr1 | 155732964 | 155733341 | 3.43777 | 4.02463 | 1.15866 | Intron (ENSMUST00000027740.8/ENSMUSG00000026468.9, intron 1 of 5) | 1 | 155698031 | 155751710 | 53680 | 2 | ENSMUST00000027740.8 | 18369 | Lhx4 | ENSMUSG00000026468.9 |
| chr1 | 155733734 | 155733900 | 3.44949 | 4.35564 | 1.36006 | Intron (ENSMUST00000027740.8/ENSMUSG00000026468.9, intron 1 of 5) | 1 | 155698031 | 155751710 | 53680 | 2 | ENSMUST00000027740.8 | 17810 | Lhx4 | ENSMUSG00000026468.9 |
| chr1 | 155735036 | 155736592 | 3.97059 | 5.34195 | 2.05943 | Intron (ENSMUST00000027740.8/ENSMUSG00000026468.9, intron 1 of 5) | 1 | 155698031 | 155751710 | 53680 | 2 | ENSMUST00000027740.8 | 15118 | Lhx4 | ENSMUSG00000026468.9 |
| chr1 | 155736961 | 155737345 | 3.43231 | 4.00555 | 1.11903 | Intron (ENSMUST00000027740.8/ENSMUSG00000026468.9, intron 1 of 5) | 1 | 155698031 | 155751710 | 53680 | 2 | ENSMUST00000027740.8 | 14365 | Lhx4 | ENSMUSG00000026468.9 |
| chr1 | 155737968 | 155738078 | 3.59683 | 4.30711 | 1.29893 | Intron (ENSMUST00000027740.8/ENSMUSG00000026468.9, intron 1 of 5) | 1 | 155698031 | 155751710 | 53680 | 2 | ENSMUST00000027740.8 | 13632 | Lhx4 | ENSMUSG00000026468.9 |
| chr1 | 155738613 | 155739124 | 3.50152 | 4.19619 | 1.23571 | Intron (ENSMUST00000027740.8/ENSMUSG00000026468.9, intron 1 of 5) | 1 | 155698031 | 155751710 | 53680 | 2 | ENSMUST00000027740.8 | 12586 | Lhx4 | ENSMUSG00000026468.9 |
| chr1 | 155740275 | 155744329 | 4.13583 | 5.4742 | 2.14903 | 5' UTR | 1 | 155698031 | 155751710 | 53680 | 2 | ENSMUST00000027740.8 | 7381 | Lhx4 | ENSMUSG00000026468.9 |
| chr1 | 155745092 | 155746079 | 3.6997 | 4.54096 | 1.52247 | 5' UTR | 1 | 155698031 | 155751710 | 53680 | 2 | ENSMUST00000027740.8 | 5631 | Lhx4 | ENSMUSG00000026468.9 |
| chr1 | 155748152 | 155749107 | 3.56453 | 4.31737 | 1.36652 | 5' UTR | 1 | 155698031 | 155751710 | 53680 | 2 | ENSMUST00000027740.8 | 2603 | Lhx4 | ENSMUSG00000026468.9 |
| chr1 | 155750547 | 155750751 | 3.39351 | 4.00235 | 1.18383 | Promoter | 1 | 155698031 | 155751710 | 53680 | 2 | ENSMUST00000027740.8 | 959 | Lhx4 | ENSMUSG00000026468.9 |
| chr1 | 155753693 | 155753991 | 3.38713 | 3.92842 | 1.08674 | Distal Intergenic | 1 | 155698031 | 155751710 | 53680 | 2 | ENSMUST00000027740.8 | -1984 | Lhx4 | ENSMUSG00000026468.9 |
| chr1 | 153739524 | 153739892 | 3.79641 | 4.6838 | 1.54155 | Promoter | 1 | 153740349 | 153745468 | 5120 | 1 | ENSMUST00000027748.7 | -457 | Rgs16 | ENSMUSG00000026475.7 |
| chr1 | 153740281 | 153740628 | 3.53723 | 4.19669 | 1.22586 | Promoter | 1 | 153740349 | 153745468 | 5120 | 1 | ENSMUST00000027748.7 | 0 | Rgs16 | ENSMUSG00000026475.7 |
| chr1 | 153185753 | 153186282 | 3.95932 | 5.02443 | 1.77106 | Promoter | 1 | 153122756 | 153186047 | 63292 | 2 | ENSMUST00000185356.1 | 0 | Lamc2 | ENSMUSG00000026479.8 |
| chr1 | 178527900 | 178528755 | 3.55124 | 4.29848 | 1.36768 | Promoter | 1 | 178529125 | 178932730 | 403606 | 1 | ENSMUST00000161017.2 | -370 | Kif26b | ENSMUSG00000026494.7 |
| chr1 | 178529131 | 178529369 | 3.76899 | 4.69244 | 1.61418 | Promoter | 1 | 178529125 | 178932730 | 403606 | 1 | ENSMUST00000161017.2 | 7 | Kif26b | ENSMUSG00000026494.7 |
| chr1 | 178530847 | 178531698 | 4.24477 | 5.72177 | 2.3006 | Intron (ENSMUST00000161017.2/ENSMUSG00000026494.7, intron 2 of 14) | 1 | 178529125 | 178932730 | 403606 | 1 | ENSMUST00000161017.2 | 1723 | Kif26b | ENSMUSG00000026494.7 |
| chr1 | 178532097 | 178532434 | 3.45229 | 4.07657 | 1.24157 | Intron (ENSMUST00000161017.2/ENSMUSG00000026494.7, intron 2 of 14) | 1 | 178529125 | 178932730 | 403606 | 1 | ENSMUST00000161017.2 | 2973 | Kif26b | ENSMUSG00000026494.7 |
| chr1 | 180547723 | 180547803 | 3.53512 | 4.21262 | 1.30594 | Distal Intergenic | 1 | 180568955 | 180601253 | 32299 | 1 | ENSMUST00000027777.6 | -21152 | Parp1 | ENSMUSG00000026496.6 |
| chr1 | 180548958 | 180549165 | 3.29029 | 3.76259 | 1.01733 | Distal Intergenic | 1 | 180568955 | 180601253 | 32299 | 1 | ENSMUST00000027777.6 | -19790 | Parp1 | ENSMUSG00000026496.6 |
| chr1 | 180694656 | 180694938 | 3.61753 | 4.40403 | 1.41198 | Exon (ENSMUST00000027778.7/ENSMUSG00000026497.7, exon 2 of 2) | 1 | 180693043 | 180697034 | 3992 | 2 | ENSMUST00000027778.7 | 2096 | Mixl1 | ENSMUSG00000026497.7 |
| chr1 | 180696369 | 180697459 | 4.19564 | 5.77869 | 2.3549 | Promoter | 1 | 180693043 | 180697034 | 3992 | 2 | ENSMUST00000027778.7 | 0 | Mixl1 | ENSMUSG00000026497.7 |
| chr1 | 176997733 | 176998423 | 3.63029 | 4.51823 | 1.48891 | Promoter | 1 | 176998009 | 177020437 | 22429 | 1 | ENSMUST00000139589.1 | 0 | Sdccag8 | ENSMUSG00000026504.12 |
| chr1 | 181351095 | 181352084 | 3.45415 | 4.09377 | 1.18488 | Promoter | 1 | 181352628 | 181459909 | 107282 | 1 | ENSMUST00000161880.2 | -544 | Cnih3 | ENSMUSG00000026514.8 |
| chr1 | 181352705 | 181353970 | 3.92892 | 4.94858 | 1.72594 | Promoter | 1 | 181353198 | 181459287 | 106090 | 1 | ENSMUST00000162685.1 | 0 | Cnih3 | ENSMUSG00000026514.8 |
| chr1 | 181354417 | 181354521 | 3.78776 | 4.65139 | 1.48761 | Intron (ENSMUST00000161880.2/ENSMUSG00000026514.8, intron 1 of 5) | 1 | 181353198 | 181459287 | 106090 | 1 | ENSMUST00000162685.1 | 1220 | Cnih3 | ENSMUSG00000026514.8 |
| chr1 | 175490335 | 175490937 | 3.59555 | 4.35471 | 1.34854 | Promoter | 1 | 175059736 | 175491130 | 431395 | 2 | ENSMUST00000027812.5 | 193 | Rgs7 | ENSMUSG00000026527.8 |
| chr1 | 175491991 | 175492938 | 3.61089 | 4.33259 | 1.30516 | Promoter | 1 | 175059736 | 175491130 | 431395 | 2 | ENSMUST00000027812.5 | -862 | Rgs7 | ENSMUSG00000026527.8 |
| chr1 | 172651613 | 172651844 | 3.43144 | 4.04355 | 1.23246 | Distal Intergenic | 1 | 172630769 | 172632974 | 2206 | 2 | ENSMUST00000027826.5 | -18640 | Dusp23 | ENSMUSG00000026544.5 |
| chr1 | 172652302 | 172652582 | 3.62264 | 4.53109 | 1.4437 | Distal Intergenic | 1 | 172630769 | 172632974 | 2206 | 2 | ENSMUST00000027826.5 | -19329 | Dusp23 | ENSMUSG00000026544.5 |
| chr1 | 164456775 | 164457057 | 3.4064 | 3.98746 | 1.18186 | Intron (ENSMUST00000027863.7/ENSMUSG00000026576.7, intron 1 of 5) | 1 | 164437267 | 164458355 | 21089 | 2 | ENSMUST00000027863.7 | 1298 | Atp1b1 | ENSMUSG00000026576.7 |
| chr1 | 164457372 | 164457729 | 3.31583 | 3.80766 | 1.04089 | Promoter | 1 | 164437267 | 164458355 | 21089 | 2 | ENSMUST00000027863.7 | 626 | Atp1b1 | ENSMUSG00000026576.7 |
| chr1 | 164151819 | 164151993 | 3.84683 | 4.85542 | 1.70775 | Promoter | 1 | 164151838 | 164220277 | 68440 | 1 | ENSMUST00000086040.5 | 0 | F5 | ENSMUSG00000026579.8 |
| chr1 | 163308676 | 163309097 | 3.88259 | 4.94621 | 1.75707 | Intron (ENSMUST00000075805.7/ENSMUSG00000026586.11, intron 1 of 4) | 1 | 163252866 | 163313668 | 60803 | 2 | ENSMUST00000174397.2 | 4571 | Prrx1 | ENSMUSG00000026586.11 |
| chr1 | 163309786 | 163310078 | 3.38718 | 3.9532 | 1.16448 | Intron (ENSMUST00000075805.7/ENSMUSG00000026586.11, intron 1 of 4) | 1 | 163252866 | 163313668 | 60803 | 2 | ENSMUST00000174397.2 | 3590 | Prrx1 | ENSMUSG00000026586.11 |
| chr1 | 163311585 | 163313631 | 3.83413 | 5.03749 | 1.81146 | Promoter | 1 | 163252866 | 163313668 | 60803 | 2 | ENSMUST00000174397.2 | 37 | Prrx1 | ENSMUSG00000026586.11 |
| chr1 | 163314027 | 163314153 | 3.56939 | 4.2457 | 1.23662 | Promoter | 1 | 163245119 | 163313710 | 68592 | 2 | ENSMUST00000027878.8 | -318 | Prrx1 | ENSMUSG00000026586.11 |
| chr1 | 163314536 | 163315715 | 3.68689 | 4.48278 | 1.42613 | Promoter | 1 | 163245119 | 163313710 | 68592 | 2 | ENSMUST00000027878.8 | -827 | Prrx1 | ENSMUSG00000026586.11 |
| chr1 | 163317093 | 163317279 | 3.60755 | 4.31843 | 1.28858 | Distal Intergenic | 1 | 163245119 | 163313710 | 68592 | 2 | ENSMUST00000027878.8 | -3384 | Prrx1 | ENSMUSG00000026586.11 |
| chr1 | 163357304 | 163357457 | 3.2545 | 3.70891 | 1.01733 | Distal Intergenic | 1 | 163245119 | 163313710 | 68592 | 2 | ENSMUST00000027878.8 | -43595 | Prrx1 | ENSMUSG00000026586.11 |
| chr1 | 158362547 | 158363042 | 4.00561 | 5.10013 | 1.82775 | Promoter | 1 | 158362340 | 158611018 | 248679 | 1 | ENSMUST00000170718.1 | 208 | Astn1 | ENSMUSG00000026587.10 |
| chr1 | 189947430 | 189947662 | 3.273 | 3.76601 | 1.05562 | Distal Intergenic | 1 | 189880492 | 189922363 | 41872 | 2 | ENSMUST00000027897.7 | -25068 | Smyd2 | ENSMUSG00000026603.8 |
| chr1 | 189989008 | 189989736 | 3.75841 | 4.81119 | 1.66181 | Distal Intergenic | 1 | 189880492 | 189922363 | 41872 | 2 | ENSMUST00000027897.7 | -66646 | Smyd2 | ENSMUSG00000026603.8 |
| chr1 | 189990183 | 189990359 | 3.5188 | 4.17796 | 1.25598 | Distal Intergenic | 1 | 189880492 | 189922363 | 41872 | 2 | ENSMUST00000027897.7 | -67821 | Smyd2 | ENSMUSG00000026603.8 |
| chr1 | 190019268 | 190019429 | 3.968 | 5.51851 | 2.11878 | Distal Intergenic | 1 | 189880492 | 189922363 | 41872 | 2 | ENSMUST00000027897.7 | -96906 | Smyd2 | ENSMUSG00000026603.8 |
| chr1 | 190027274 | 190027589 | 3.8661 | 4.80296 | 1.5956 | Distal Intergenic | 1 | 189880492 | 189922363 | 41872 | 2 | ENSMUST00000027897.7 | -104912 | Smyd2 | ENSMUSG00000026603.8 |
| chr1 | 188550701 | 188550791 | 3.45579 | 4.09161 | 1.26045 | Intron (ENSMUST00000060479.8/ENSMUSG00000026609.10, intron 28 of 70) | 1 | 188537807 | 188678687 | 140881 | 1 | ENSMUST00000142159.1 | 12895 | Ush2a | ENSMUSG00000026609.10 |
| chr1 | 188551117 | 188551340 | 3.34196 | 3.89752 | 1.12394 | Intron (ENSMUST00000060479.8/ENSMUSG00000026609.10, intron 28 of 70) | 1 | 188537807 | 188678687 | 140881 | 1 | ENSMUST00000142159.1 | 13311 | Ush2a | ENSMUSG00000026609.10 |
| chr1 | 188551917 | 188552090 | 3.55784 | 4.4213 | 1.42186 | Intron (ENSMUST00000060479.8/ENSMUSG00000026609.10, intron 28 of 70) | 1 | 188537807 | 188678687 | 140881 | 1 | ENSMUST00000142159.1 | 14111 | Ush2a | ENSMUSG00000026609.10 |
| chr1 | 187439319 | 187439467 | 3.41922 | 4.06108 | 1.23083 | Distal Intergenic | 1 | 187608791 | 188043502 | 434712 | 1 | ENSMUST00000127489.2 | -169324 | Esrrg | ENSMUSG00000026610.8 |
| chr1 | 187439903 | 187440084 | 3.40708 | 3.9995 | 1.19737 | Distal Intergenic | 1 | 187608791 | 188043502 | 434712 | 1 | ENSMUST00000127489.2 | -168707 | Esrrg | ENSMUSG00000026610.8 |
| chr1 | 187603316 | 187603403 | 3.88254 | 4.84078 | 1.6303 | Distal Intergenic | 1 | 187608791 | 188043502 | 434712 | 1 | ENSMUST00000127489.2 | -5388 | Esrrg | ENSMUSG00000026610.8 |
| chr1 | 187603984 | 187604116 | 3.9151 | 4.90294 | 1.67795 | Distal Intergenic | 1 | 187608791 | 188043502 | 434712 | 1 | ENSMUST00000127489.2 | -4675 | Esrrg | ENSMUSG00000026610.8 |
| chr1 | 187604458 | 187604979 | 3.7333 | 4.55371 | 1.43747 | Distal Intergenic | 1 | 187608791 | 188043502 | 434712 | 1 | ENSMUST00000127489.2 | -3812 | Esrrg | ENSMUSG00000026610.8 |
| chr1 | 187605357 | 187605837 | 3.53175 | 4.18697 | 1.21844 | Distal Intergenic | 1 | 187608791 | 188043502 | 434712 | 1 | ENSMUST00000127489.2 | -2954 | Esrrg | ENSMUSG00000026610.8 |
| chr1 | 187606320 | 187607155 | 3.70491 | 4.50389 | 1.40811 | Distal Intergenic | 1 | 187608791 | 188043502 | 434712 | 1 | ENSMUST00000127489.2 | -1636 | Esrrg | ENSMUSG00000026610.8 |
| chr1 | 187608276 | 187610420 | 3.91385 | 4.96337 | 1.74736 | Promoter | 1 | 187608791 | 188043502 | 434712 | 1 | ENSMUST00000127489.2 | 0 | Esrrg | ENSMUSG00000026610.8 |
| chr1 | 187610966 | 187611074 | 3.6328 | 4.39079 | 1.38893 | Intron (ENSMUST00000127489.2/ENSMUSG00000026610.8, intron 2 of 3) | 1 | 187609696 | 187793742 | 184047 | 1 | ENSMUST00000152927.1 | 1271 | Esrrg | ENSMUSG00000026610.8 |
| chr1 | 187651502 | 187652303 | 3.45356 | 4.0424 | 1.12221 | Intron (ENSMUST00000127489.2/ENSMUSG00000026610.8, intron 2 of 3) | 1 | 187609696 | 187793742 | 184047 | 1 | ENSMUST00000152927.1 | 41807 | Esrrg | ENSMUSG00000026610.8 |
| chr1 | 187652766 | 187652912 | 3.58013 | 4.26268 | 1.23834 | Intron (ENSMUST00000127489.2/ENSMUSG00000026610.8, intron 2 of 3) | 1 | 187609696 | 187793742 | 184047 | 1 | ENSMUST00000152927.1 | 43071 | Esrrg | ENSMUSG00000026610.8 |
| chr1 | 187661307 | 187661585 | 3.50325 | 4.21524 | 1.29737 | Intron (ENSMUST00000127489.2/ENSMUSG00000026610.8, intron 2 of 3) | 1 | 187609696 | 187793742 | 184047 | 1 | ENSMUST00000152927.1 | 51612 | Esrrg | ENSMUSG00000026610.8 |
| chr1 | 187663609 | 187663857 | 3.68441 | 4.61108 | 1.57509 | Intron (ENSMUST00000127489.2/ENSMUSG00000026610.8, intron 2 of 3) | 1 | 187609696 | 187793742 | 184047 | 1 | ENSMUST00000152927.1 | 53914 | Esrrg | ENSMUSG00000026610.8 |
| chr1 | 187997844 | 187998668 | 3.64965 | 4.49321 | 1.45909 | Promoter | 1 | 187997868 | 188214884 | 217017 | 1 | ENSMUST00000110938.1 | 0 | Esrrg | ENSMUSG00000026610.8 |
| chr1 | 187998991 | 187999282 | 3.44299 | 4.03255 | 1.15848 | Intron (ENSMUST00000127489.2/ENSMUSG00000026610.8, intron 3 of 3) | 1 | 187997868 | 188214884 | 217017 | 1 | ENSMUST00000110938.1 | 1124 | Esrrg | ENSMUSG00000026610.8 |
| chr1 | 187999996 | 188000133 | 3.5324 | 4.25374 | 1.3362 | Intron (ENSMUST00000127489.2/ENSMUSG00000026610.8, intron 3 of 3) | 1 | 187997868 | 188214884 | 217017 | 1 | ENSMUST00000110938.1 | 2129 | Esrrg | ENSMUSG00000026610.8 |
| chr1 | 188001344 | 188001466 | 3.36639 | 3.91874 | 1.15094 | Intron (ENSMUST00000127489.2/ENSMUSG00000026610.8, intron 3 of 3) | 1 | 187997868 | 188214884 | 217017 | 1 | ENSMUST00000110938.1 | 3477 | Esrrg | ENSMUSG00000026610.8 |
| chr1 | 188007061 | 188007151 | 3.74858 | 4.66494 | 1.57943 | Intron (ENSMUST00000127489.2/ENSMUSG00000026610.8, intron 3 of 3) | 1 | 187997868 | 188214884 | 217017 | 1 | ENSMUST00000110938.1 | 9194 | Esrrg | ENSMUSG00000026610.8 |
| chr1 | 188031570 | 188031898 | 4.12077 | 5.38153 | 2.07749 | Intron (ENSMUST00000127489.2/ENSMUSG00000026610.8, intron 3 of 3) | 1 | 187997868 | 188214884 | 217017 | 1 | ENSMUST00000110938.1 | 33703 | Esrrg | ENSMUSG00000026610.8 |
| chr1 | 195176571 | 195176748 | 4.04018 | 5.17041 | 1.91198 | Promoter | 1 | 195136811 | 195176715 | 39905 | 2 | ENSMUST00000082321.3 | 0 | Cr2 | ENSMUSG00000026616.9 |
| chr1 | 184999275 | 184999597 | 4.29177 | 5.64724 | 2.19066 | Promoter | 1 | 184896443 | 184999549 | 103107 | 2 | ENSMUST00000027929.4 | 0 | Mark1 | ENSMUSG00000026620.6 |
| chr1 | 185031752 | 185032082 | 3.60273 | 4.32566 | 1.34142 | Distal Intergenic | 1 | 184896443 | 184999549 | 103107 | 2 | ENSMUST00000027929.4 | -32204 | Mark1 | ENSMUSG00000026620.6 |
| chr1 | 185032698 | 185033203 | 3.4487 | 4.07985 | 1.15376 | Distal Intergenic | 1 | 184896443 | 184999549 | 103107 | 2 | ENSMUST00000027929.4 | -33150 | Mark1 | ENSMUSG00000026620.6 |
| chr1 | 185088555 | 185088916 | 3.68285 | 4.49463 | 1.46124 | Distal Intergenic | 1 | 184896443 | 184999549 | 103107 | 2 | ENSMUST00000027929.4 | -89007 | Mark1 | ENSMUSG00000026620.6 |

| | | | | | | | | | | | | | | | |
|---|---|---|---|---|---|---|---|---|---|---|---|---|---|---|---|
| chr1 | 185089534 | 185089781 | 3.71174 | 4.58828 | 1.54557 | Distal Intergenic | 1 | 184896443 | 184999549 | 103107 | 2 | ENSMUST00000027929.4 | -89986 | Mark1 | ENSMUSG00000026620.6 |
| chr1 | 184772188 | 184772408 | 3.75577 | 4.58398 | 1.43685 | Distal Intergenic | 1 | 184787289 | 184808853 | 21565 | 2 | ENSMUST00000189492.1 | 36445 | Mar-01 | ENSMUSG00000026621.8 |
| chr1 | 184775562 | 184775691 | 3.34351 | 3.87185 | 1.11148 | Distal Intergenic | 1 | 184787289 | 184808853 | 21565 | 2 | ENSMUST00000189492.1 | 33162 | Mar-01 | ENSMUSG00000026621.8 |
| chr1 | 191185302 | 191185440 | 3.44768 | 4.14566 | 1.22642 | Distal Intergenic | 1 | 191170296 | 191183340 | 13045 | 2 | ENSMUST00000027941.8 | -1963 | Atf3 | ENSMUSG00000026628.8 |
| chr1 | 191098472 | 191098980 | 4.54188 | 6.44308 | 2.8549 | Promoter | 1 | 191098414 | 191108943 | 10530 | 1 | ENSMUST00000027943.4 | 59 | Batf3 | ENSMUSG00000026630.4 |
| chr1 | 191125310 | 191125917 | 3.70938 | 4.52885 | 1.46778 | Distal Intergenic | 1 | 191098414 | 191108943 | 10530 | 1 | ENSMUST00000027943.4 | 26897 | Batf3 | ENSMUSG00000026630.4 |
| chr1 | 193153900 | 193154165 | 3.31629 | 3.80167 | 1.01733 | Promoter | 1 | 193153112 | 193172042 | 18931 | 1 | ENSMUST00000076521.5 | 789 | Irf6 | ENSMUSG00000026638.10 |
| chr1 | 193329758 | 193330139 | 4.37638 | 5.94713 | 2.54161 | Exon (ENSMUST00000016315.10/ENSMUSG00000026639.13, exon 11 of 23) | 1 | 193320695 | 193325990 | 5296 | 1 | ENSMUST00000161526.1 | 9064 | Lamb3 | ENSMUSG00000026639.13 |
| chr1 | 193330564 | 193330813 | 3.2247 | 3.66426 | 1.01733 | Exon (ENSMUST00000016315.10/ENSMUSG00000026639.13, exon 12 of 23) | 1 | 193320695 | 193325990 | 5296 | 1 | ENSMUST00000161526.1 | 9870 | Lamb3 | ENSMUSG00000026639.13 |
| chr1 | 194617017 | 194617151 | 3.50721 | 4.31181 | 1.28972 | Distal Intergenic | 1 | 194619829 | 194816869 | 197041 | 1 | ENSMUST00000027952.6 | -2678 | Plxna2 | ENSMUSG00000026640.7 |
| chr2 | 4156362 | 4156641 | 3.57024 | 4.33157 | 1.36096 | Intron (ENSMUST00000138735.1/ENSMUSG00000026657.11, intron 1 of 1) | 2 | 4152872 | 4613942 | 461071 | 1 | ENSMUST00000091497.5 | 3491 | Frmd4a | ENSMUSG00000026657.11 |
| chr2 | 4299622 | 4299824 | 3.86113 | 4.92439 | 1.72317 | Promoter | 2 | 4300518 | 4572439 | 271922 | 1 | ENSMUST00000175669.2 | -694 | Frmd4a | ENSMUSG00000026657.11 |
| chr2 | 4300253 | 4300609 | 3.72922 | 4.7692 | 1.63845 | Promoter | 2 | 4300518 | 4572439 | 271922 | 1 | ENSMUST00000175669.2 | 0 | Frmd4a | ENSMUSG00000026657.11 |
| chr2 | 4401032 | 4401210 | 3.62331 | 4.43054 | 1.43105 | Promoter | 2 | 4400975 | 4614043 | 213069 | 1 | ENSMUST00000075767.8 | 58 | Frmd4a | ENSMUSG00000026657.11 |
| chr1 | 170896813 | 170896944 | 3.26348 | 3.72235 | 1.01733 | Distal Intergenic | 1 | 170874188 | 170885540 | 11353 | 2 | ENSMUST00000027970.8 | -11274 | Dusp12 | ENSMUSG00000026659.8 |
| chr2 | 5137645 | 5137720 | 3.55039 | 4.21269 | 1.222 | Promoter | 2 | 5137776 | 5230878 | 93103 | 1 | ENSMUST00000027988.7 | -56 | Ccdc3 | ENSMUSG00000026676.7 |
| chr1 | 167657539 | 167658222 | 3.57536 | 4.25726 | 1.24391 | Distal Intergenic | 1 | 167689237 | 167848741 | 159505 | 1 | ENSMUST00000111377.2 | -31015 | Lmx1a | ENSMUSG00000026686.9 |
| chr1 | 167659187 | 167659444 | 3.97164 | 5.16635 | 1.89045 | Distal Intergenic | 1 | 167689237 | 167848741 | 159505 | 1 | ENSMUST00000111377.2 | -29793 | Lmx1a | ENSMUSG00000026686.9 |
| chr1 | 167660901 | 167662196 | 3.88238 | 4.85034 | 1.65008 | Distal Intergenic | 1 | 167689237 | 167848741 | 159505 | 1 | ENSMUST00000111377.2 | -27041 | Lmx1a | ENSMUSG00000026686.9 |
| chr1 | 167665364 | 167665734 | 3.83609 | 4.75132 | 1.57106 | Distal Intergenic | 1 | 167689237 | 167848741 | 159505 | 1 | ENSMUST00000111377.2 | -23503 | Lmx1a | ENSMUSG00000026686.9 |
| chr1 | 167671747 | 167671895 | 4.05571 | 5.18358 | 1.8782 | Distal Intergenic | 1 | 167689237 | 167848741 | 159505 | 1 | ENSMUST00000111377.2 | -17342 | Lmx1a | ENSMUSG00000026686.9 |
| chr1 | 167672737 | 167672996 | 3.38193 | 3.9006 | 1.01733 | Distal Intergenic | 1 | 167689237 | 167848741 | 159505 | 1 | ENSMUST00000111377.2 | -16241 | Lmx1a | ENSMUSG00000026686.9 |
| chr1 | 167684975 | 167685051 | 3.58971 | 4.32039 | 1.38162 | Distal Intergenic | 1 | 167689237 | 167848741 | 159505 | 1 | ENSMUST00000111377.2 | -4186 | Lmx1a | ENSMUSG00000026686.9 |
| chr1 | 167686326 | 167686730 | 3.49581 | 4.12852 | 1.21248 | Distal Intergenic | 1 | 167689237 | 167848741 | 159505 | 1 | ENSMUST00000111377.2 | -2507 | Lmx1a | ENSMUSG00000026686.9 |
| chr1 | 167687709 | 167687960 | 3.54627 | 4.21063 | 1.23526 | Distal Intergenic | 1 | 167689237 | 167848741 | 159505 | 1 | ENSMUST00000111377.2 | -1277 | Lmx1a | ENSMUSG00000026686.9 |
| chr1 | 167688308 | 167693390 | 4.05014 | 5.29256 | 1.97316 | Promoter | 1 | 167689237 | 167848741 | 159505 | 1 | ENSMUST00000111377.2 | 0 | Lmx1a | ENSMUSG00000026686.9 |
| chr1 | 167694301 | 167694407 | 4.49401 | 6.03767 | 2.45346 | Intron (ENSMUST00000111377.2/ENSMUSG00000026686.9, intron 3 of 8) | 1 | 167689557 | 167848741 | 159185 | 1 | ENSMUST00000028003.8 | 4745 | Lmx1a | ENSMUSG00000026686.9 |
| chr1 | 167694724 | 167695335 | 3.61123 | 4.3829 | 1.35059 | Intron (ENSMUST00000111377.2/ENSMUSG00000026686.9, intron 3 of 8) | 1 | 167689557 | 167848741 | 159185 | 1 | ENSMUST00000028003.8 | 5168 | Lmx1a | ENSMUSG00000026686.9 |
| chr1 | 167696291 | 167697065 | 3.51516 | 4.49601 | 1.44416 | Intron (ENSMUST00000111377.2/ENSMUSG00000026686.9, intron 3 of 8) | 1 | 167689557 | 167848741 | 159185 | 1 | ENSMUST00000028003.8 | 6735 | Lmx1a | ENSMUSG00000026686.9 |
| chr1 | 167697860 | 167698472 | 4.12758 | 5.38033 | 2.04369 | Intron (ENSMUST00000111377.2/ENSMUSG00000026686.9, intron 3 of 8) | 1 | 167689557 | 167848741 | 159185 | 1 | ENSMUST00000028003.8 | 8304 | Lmx1a | ENSMUSG00000026686.9 |
| chr1 | 167698847 | 167699433 | 3.3118 | 3.80772 | 1.06182 | Intron (ENSMUST00000111377.2/ENSMUSG00000026686.9, intron 3 of 8) | 1 | 167689557 | 167848741 | 159185 | 1 | ENSMUST00000028003.8 | 9291 | Lmx1a | ENSMUSG00000026686.9 |
| chr1 | 167699874 | 167700141 | 3.52424 | 4.18568 | 1.24321 | Intron (ENSMUST00000111377.2/ENSMUSG00000026686.9, intron 3 of 8) | 1 | 167689557 | 167848741 | 159185 | 1 | ENSMUST00000028003.8 | 10318 | Lmx1a | ENSMUSG00000026686.9 |
| chr1 | 167702103 | 167702334 | 3.32534 | 3.81762 | 1.02542 | Intron (ENSMUST00000111377.2/ENSMUSG00000026686.9, intron 3 of 8) | 1 | 167689557 | 167848741 | 159185 | 1 | ENSMUST00000028003.8 | 12547 | Lmx1a | ENSMUSG00000026686.9 |
| chr1 | 167708688 | 167708878 | 3.60221 | 4.32378 | 1.30654 | Intron (ENSMUST00000111377.2/ENSMUSG00000026686.9, intron 3 of 8) | 1 | 167689557 | 167848741 | 159185 | 1 | ENSMUST00000028003.8 | 19132 | Lmx1a | ENSMUSG00000026686.9 |
| chr1 | 167732427 | 167732702 | 3.53885 | 4.20603 | 1.26003 | Intron (ENSMUST00000111377.2/ENSMUSG00000026686.9, intron 3 of 8) | 1 | 167689557 | 167848741 | 159185 | 1 | ENSMUST00000028003.8 | 42871 | Lmx1a | ENSMUSG00000026686.9 |
| chr1 | 167734196 | 167734722 | 3.58912 | 4.34012 | 1.35895 | Intron (ENSMUST00000111377.2/ENSMUSG00000026686.9, intron 3 of 8) | 1 | 167689557 | 167848741 | 159185 | 1 | ENSMUST00000028003.8 | 44640 | Lmx1a | ENSMUSG00000026686.9 |
| chr1 | 167762553 | 167762807 | 4.47953 | 6.05321 | 2.51835 | Intron (ENSMUST00000111377.2/ENSMUSG00000026686.9, intron 3 of 8) | 1 | 167689557 | 167848741 | 159185 | 1 | ENSMUST00000028003.8 | 72997 | Lmx1a | ENSMUSG00000026686.9 |
| chr1 | 167405615 | 167405870 | 3.67053 | 4.60003 | 1.50887 | Distal Intergenic | 1 | 167371966 | 167393841 | 21876 | 2 | ENSMUST00000028005.2 | -11775 | Mgst3 | ENSMUSG00000026688.5 |
| chr1 | 167407094 | 167407177 | 3.79269 | 4.75355 | 1.63145 | Distal Intergenic | 1 | 167371966 | 167393841 | 21876 | 2 | ENSMUST00000028005.2 | -13254 | Mgst3 | ENSMUSG00000026688.5 |
| chr1 | 167412514 | 167412645 | 3.95725 | 5.01057 | 1.81347 | Distal Intergenic | 1 | 167371966 | 167393841 | 21876 | 2 | ENSMUST00000028005.2 | -18674 | Mgst3 | ENSMUSG00000026688.5 |
| chr2 | 19442765 | 19443086 | 3.39411 | 3.92711 | 1.04593 | Intron (ENSMUST00000122978.1/ENSMUSG00000085388.1, intron 2 of 3) | 2 | 19445663 | 19447501 | 1839 | 1 | ENSMUST00000028068.2 | -2577 | Ptf1a | ENSMUSG00000026735.2 |
| chr2 | 19444794 | 19444928 | 3.54423 | 4.20079 | 1.21233 | Promoter | 2 | 19445663 | 19447501 | 1839 | 1 | ENSMUST00000028068.2 | -735 | Ptf1a | ENSMUSG00000026735.2 |
| chr2 | 19445241 | 19446953 | 3.86122 | 4.90527 | 1.70712 | Promoter | 2 | 19445663 | 19447501 | 1839 | 1 | ENSMUST00000028068.2 | 0 | Ptf1a | ENSMUSG00000026735.2 |
| chr2 | 16683062 | 16683154 | 9.22048 | 18.02842 | 13.04788 | Intron (ENSMUST00000028081.7/ENSMUSG00000026748.8, intron 9 of 13) | 2 | 16638696 | 16751690 | 112995 | 1 | ENSMUST00000126173.1 | 44367 | Plxdc2 | ENSMUSG00000026748.8 |
| chr2 | 38707647 | 38708193 | 3.58232 | 4.28289 | 1.30083 | Exon (ENSMUST00000112883.2/ENSMUSG00000026751.9, exon 4 of 7) | 2 | 38701322 | 38711188 | 9867 | 2 | ENSMUST00000149690.1 | 2995 | Nr5a1 | ENSMUSG00000026751.9 |
| chr2 | 38708868 | 38709329 | 3.83133 | 4.7726 | 1.64679 | Intron (ENSMUST00000112883.2/ENSMUSG00000026751.9, intron 3 of 6) | 2 | 38701322 | 38711188 | 9867 | 2 | ENSMUST00000149690.1 | 1859 | Nr5a1 | ENSMUSG00000026751.9 |
| chr2 | 38710316 | 38711007 | 4.08422 | 5.55579 | 2.20087 | Promoter | 2 | 38701322 | 38711188 | 9867 | 2 | ENSMUST00000149690.1 | 181 | Nr5a1 | ENSMUSG00000026751.9 |
| chr2 | 38711550 | 38712584 | 4.22822 | 5.6113 | 2.19752 | Promoter | 2 | 38692656 | 38712234 | 19579 | 2 | ENSMUST00000112883.2 | 0 | Nr5a1 | ENSMUSG00000026751.9 |
| chr2 | 38714374 | 38714597 | 3.40718 | 3.9493 | 1.05458 | Promoter | 2 | 38692660 | 38714542 | 21883 | 2 | ENSMUST00000028084.4 | 0 | Nr5a1 | ENSMUSG00000026751.9 |
| chr2 | 49618375 | 49618652 | 3.48159 | 4.22783 | 1.25301 | Promoter | 2 | 49619298 | 49774778 | 155481 | 1 | ENSMUST00000028102.8 | -646 | Kif5c | ENSMUSG00000026764.10 |
| chr2 | 49619206 | 49620187 | 4.04592 | 5.19372 | 1.91135 | Promoter | 2 | 49619298 | 49774778 | 155481 | 1 | ENSMUST00000028102.8 | 0 | Kif5c | ENSMUSG00000026764.10 |
| chr2 | 49628343 | 49628681 | 4.14311 | 5.43728 | 2.12641 | Intron (ENSMUST00000028102.8/ENSMUSG00000026764.10, intron 1 of 25) | 2 | 49619298 | 49774778 | 155481 | 1 | ENSMUST00000028102.8 | 9046 | Kif5c | ENSMUSG00000026764.10 |
| chr2 | 49721681 | 49721891 | 3.38685 | 3.92441 | 1.07537 | Promoter | 2 | 49722567 | 49735584 | 13018 | 1 | ENSMUST00000138834.1 | -676 | Kif5c | ENSMUSG00000026764.10 |
| chr2 | 49787622 | 49788014 | 3.75596 | 4.68582 | 1.59757 | Promoter | 2 | 49787688 | 49948849 | 161162 | 1 | ENSMUST00000028103.7 | 0 | Lypd6b | ENSMUSG00000026765.7 |
| chr2 | 12300855 | 12301881 | 3.59741 | 4.39192 | 1.40873 | Promoter | 2 | 12286964 | 12301588 | 14625 | 2 | ENSMUST00000130548.2 | 0 | Itga8 | ENSMUSG00000026768.5 |
| chr2 | 11172199 | 11172358 | 3.59996 | 4.48171 | 1.42453 | Promoter | 2 | 11172310 | 11284369 | 112060 | 1 | ENSMUST00000102970.3 | 0 | Prkcq | ENSMUSG00000026778.8 |
| chr2 | 22619378 | 22620225 | 3.92361 | 5.02213 | 1.78525 | Distal Intergenic | 2 | 22622205 | 22693874 | 71670 | 1 | ENSMUST00000028123.3 | -1980 | Gad2 | ENSMUSG00000026787.3 |
| chr2 | 22620674 | 22621118 | 4.09706 | 5.29482 | 1.94558 | Distal Intergenic | 2 | 22622205 | 22693874 | 71670 | 1 | ENSMUST00000028123.3 | -1087 | Gad2 | ENSMUSG00000026787.3 |
| chr2 | 22621490 | 22622583 | 3.53578 | 4.1885 | 1.21004 | Promoter | 2 | 22622205 | 22693874 | 71670 | 1 | ENSMUST00000028123.3 | 0 | Gad2 | ENSMUSG00000026787.3 |
| chr2 | 22623092 | 22624097 | 3.88948 | 5.0955 | 1.83491 | Promoter | 2 | 22623304 | 22630001 | 6698 | 1 | ENSMUST00000156728.1 | 0 | Gad2 | ENSMUSG00000026787.3 |
| chr2 | 22624623 | 22624901 | 4.43773 | 6.1888 | 2.64813 | Exon (ENSMUST00000028123.3/ENSMUSG00000026787.3, exon 4 of 16) | 2 | 22623304 | 22630001 | 6698 | 1 | ENSMUST00000156728.1 | 1320 | Gad2 | ENSMUSG00000026787.3 |
| chr2 | 22625254 | 22625354 | 4.06709 | 5.27775 | 1.98587 | Intron (ENSMUST00000028123.3/ENSMUSG00000026787.3, intron 4 of 15) | 2 | 22623304 | 22630001 | 6698 | 1 | ENSMUST00000156728.1 | 1951 | Gad2 | ENSMUSG00000026787.3 |
| chr2 | 22626479 | 22626874 | 3.56508 | 4.58855 | 1.50862 | Intron (ENSMUST00000028123.3/ENSMUSG00000026787.3, intron 4 of 15) | 2 | 22623304 | 22630001 | 6698 | 1 | ENSMUST00000156728.1 | 3176 | Gad2 | ENSMUSG00000026787.3 |
| chr2 | 22627462 | 22627645 | 3.91442 | 4.93349 | 1.75561 | Intron (ENSMUST00000028123.3/ENSMUSG00000026787.3, intron 4 of 15) | 2 | 22623304 | 22630001 | 6698 | 1 | ENSMUST00000156728.1 | 4159 | Gad2 | ENSMUSG00000026787.3 |
| chr2 | 22628215 | 22628841 | 3.57524 | 4.32693 | 1.41027 | Intron (ENSMUST00000028123.3/ENSMUSG00000026787.3, intron 4 of 15) | 2 | 22623304 | 22630001 | 6698 | 1 | ENSMUST00000156728.1 | 4912 | Gad2 | ENSMUSG00000026787.3 |
| chr2 | 22630253 | 22630552 | 3.36945 | 3.93174 | 1.16997 | Intron (ENSMUST00000028123.3/ENSMUSG00000026787.3, intron 6 of 15) | 2 | 22623304 | 22630001 | 6698 | 1 | ENSMUST00000156728.1 | 6950 | Gad2 | ENSMUSG00000026787.3 |
| chr2 | 28907913 | 28908626 | 3.86694 | 4.90111 | 1.73653 | 3' UTR | 2 | 28907684 | 28912495 | 4812 | 2 | ENSMUST00000124059.1 | 3869 | Barhl1 | ENSMUSG00000026805.9 |
| chr2 | 28908987 | 28909616 | 4.27765 | 5.7135 | 2.30732 | 3' UTR | 2 | 28907684 | 28912495 | 4812 | 2 | ENSMUST00000124059.1 | 2879 | Barhl1 | ENSMUSG00000026805.9 |
| chr2 | 28910096 | 28911585 | 3.85623 | 5.01504 | 1.82747 | Promoter | 2 | 28907684 | 28912495 | 4812 | 2 | ENSMUST00000124059.1 | 910 | Barhl1 | ENSMUSG00000026805.9 |
| chr2 | 28912607 | 28913215 | 3.69385 | 4.61293 | 1.57852 | Promoter | 2 | 28907684 | 28912495 | 4812 | 2 | ENSMUST00000124059.1 | -113 | Barhl1 | ENSMUSG00000026805.9 |
| chr2 | 28914940 | 28916231 | 4.37227 | 6.09562 | 2.62221 | Promoter | 2 | 28909640 | 28915505 | 5866 | 2 | ENSMUST00000174027.1 | 0 | Barhl1 | ENSMUSG00000026805.9 |
| chr2 | 28916672 | 28916819 | 3.47118 | 4.11529 | 1.27166 | Promoter | 2 | 28908969 | 28916668 | 7700 | 2 | ENSMUST00000113847.2 | -5 | Barhl1 | ENSMUSG00000026805.9 |
| chr2 | 28917507 | 28918663 | 3.81892 | 4.79869 | 1.70461 | Promoter | 2 | 28908969 | 28916668 | 7700 | 2 | ENSMUST00000113847.2 | -840 | Barhl1 | ENSMUSG00000026805.9 |
| chr2 | 28919481 | 28919956 | 3.48896 | 4.44828 | 1.41405 | Distal Intergenic | 2 | 28908969 | 28916668 | 7700 | 2 | ENSMUST00000113847.2 | -2814 | Barhl1 | ENSMUSG00000026805.9 |
| chr2 | 28921821 | 28922268 | 3.66479 | 4.5304 | 1.50268 | Distal Intergenic | 2 | 28908969 | 28916668 | 7700 | 2 | ENSMUST00000113847.2 | -5154 | Barhl1 | ENSMUSG00000026805.9 |
| chr2 | 28750411 | 28750730 | 3.39712 | 4.24933 | 1.29573 | Intron (ENSMUST00000074156.6/ENSMUSG00000026807.8, intron 10 of 12) | 2 | 28742250 | 28760136 | 17887 | 1 | ENSMUST00000137588.1 | 8162 | Ak8 | ENSMUSG00000026807.8 |
| chr2 | 28752649 | 28753214 | 3.72229 | 4.67568 | 1.60238 | Intron (ENSMUST00000074156.6/ENSMUSG00000026807.8, intron 10 of 12) | 2 | 28742250 | 28760136 | 17887 | 1 | ENSMUST00000137588.1 | 10400 | Ak8 | ENSMUSG00000026807.8 |
| chr2 | 28756265 | 28756340 | 3.78808 | 4.67082 | 1.56148 | Intron (ENSMUST00000074156.6/ENSMUSG00000026807.8, intron 10 of 12) | 2 | 28742250 | 28760136 | 17887 | 1 | ENSMUST00000137588.1 | 14016 | Ak8 | ENSMUSG00000026807.8 |
| chr2 | 28757291 | 28757517 | 3.75584 | 4.67943 | 1.61102 | Intron (ENSMUST00000074156.6/ENSMUSG00000026807.8, intron 10 of 12) | 2 | 28742250 | 28760136 | 17887 | 1 | ENSMUST00000137588.1 | 15042 | Ak8 | ENSMUSG00000026807.8 |
| chr2 | 28620751 | 28620883 | 4.27545 | 6.26354 | 2.68433 | Intron (ENSMUST00000164290.2/ENSMUSG00000026815.9, intron 1 of 6) | 2 | 28609455 | 28621945 | 12491 | 2 | ENSMUST00000028156.7 | 1062 | Gfi1b | ENSMUSG00000026815.9 |
| chr2 | 28621717 | 28621914 | 3.87786 | 4.83724 | 1.6415 | Promoter | 2 | 28609455 | 28621945 | 12491 | 2 | ENSMUST00000028156.7 | 31 | Gfi1b | ENSMUSG00000026815.9 |
| chr2 | 55436404 | 55436628 | 3.42154 | 4.00171 | 1.16282 | Promoter | 2 | 55435970 | 55598145 | 162176 | 1 | ENSMUST00000067101.4 | 435 | Kcnj3 | ENSMUSG00000026824.6 |
| chr2 | 32334343 | 32334576 | 3.34175 | 3.96667 | 1.16475 | Intron (ENSMUST00000078352.6/ENSMUSG00000026825.12, intron 9 of 22) | 2 | 32308497 | 32320581 | 12085 | 2 | ENSMUST00000129156.1 | -13763 | Dnm1 | ENSMUSG00000026825.12 |
| chr2 | 57100948 | 57101543 | 3.65009 | 4.49764 | 1.42697 | Distal Intergenic | 2 | 57107499 | 57113072 | 5574 | 2 | ENSMUST00000112627.1 | 11529 | Nr4a2 | ENSMUSG00000026826.8 |
| chr2 | 57101878 | 57102874 | 3.82519 | 4.7712 | 1.61933 | Distal Intergenic | 2 | 57107499 | 57113072 | 5574 | 2 | ENSMUST00000112627.1 | 10198 | Nr4a2 | ENSMUSG00000026826.8 |
| chr2 | 57103224 | 57103436 | 3.5139 | 4.14935 | 1.19314 | Distal Intergenic | 2 | 57107499 | 57113072 | 5574 | 2 | ENSMUST00000112627.1 | 9636 | Nr4a2 | ENSMUSG00000026826.8 |
| chr2 | 57103978 | 57104084 | 3.72007 | 4.58029 | 1.53276 | Downstream (2-3kb) | 2 | 57107499 | 57113072 | 5574 | 2 | ENSMUST00000112627.1 | 8988 | Nr4a2 | ENSMUSG00000026826.8 |
| chr2 | 57104533 | 57104825 | 3.22922 | 3.81492 | 1.04138 | Downstream (2-3kb) | 2 | 57107499 | 57113072 | 5574 | 2 | ENSMUST00000112627.1 | 8247 | Nr4a2 | ENSMUSG00000026826.8 |
| chr2 | 57105839 | 57106311 | 4.00283 | 5.15379 | 1.88482 | Downstream (<1kb) | 2 | 57107499 | 57113072 | 5574 | 2 | ENSMUST00000112627.1 | 6761 | Nr4a2 | ENSMUSG00000026826.8 |
| chr2 | 57106763 | 57107151 | 3.50185 | 4.28885 | 1.34148 | 3' UTR | 2 | 57107499 | 57113072 | 5574 | 2 | ENSMUST00000112627.1 | 5921 | Nr4a2 | ENSMUSG00000026826.8 |
| chr2 | 57108103 | 57109062 | 3.63769 | 4.39259 | 1.35852 | 3' UTR | 2 | 57107499 | 57113072 | 5574 | 2 | ENSMUST00000112627.1 | 4010 | Nr4a2 | ENSMUSG00000026826.8 |
| chr2 | 57110323 | 57112565 | 3.72596 | 4.68925 | 1.57037 | Promoter | 2 | 57107499 | 57113072 | 5574 | 2 | ENSMUST00000112627.1 | 507 | Nr4a2 | ENSMUSG00000026826.8 |
| chr2 | 57113166 | 57113453 | 3.53429 | 4.22046 | 1.28719 | Promoter | 2 | 57107499 | 57113072 | 5574 | 2 | ENSMUST00000112627.1 | -95 | Nr4a2 | ENSMUSG00000026826.8 |
| chr2 | 28205976 | 28206735 | 4.162 | 5.67955 | 2.37879 | Promoter | 2 | 28205659 | 28230736 | 25078 | 1 | ENSMUST00000028177.5 | 318 | Olfm1 | ENSMUSG00000026833.13 |
| chr2 | 58355872 | 58356146 | 3.34153 | 3.83966 | 1.01733 | Intron (ENSMUST00000028178.8/ENSMUSG00000026834.8, intron 1 of 8) | 2 | 58274735 | 58357752 | 83018 | 2 | ENSMUST00000112607.2 | 1606 | Acvr1c | ENSMUSG00000026834.8 |
| chr2 | 27886278 | 27887224 | 4.02443 | 5.28808 | 1.99381 | Promoter | 2 | 27886425 | 28039514 | 153090 | 1 | ENSMUST00000028280.8 | 0 | Col5a1 | ENSMUSG00000026837.10 |
| chr2 | 31872375 | 31872458 | 4.20433 | 5.65279 | 2.23798 | Distal Intergenic | 2 | 31887291 | 31945246 | 57956 | 1 | ENSMUST00000138325.2 | -14833 | Lamc3 | ENSMUSG00000026840.9 |
| chr2 | 31874207 | 31874978 | 3.84854 | 4.82508 | 1.62541 | Distal Intergenic | 2 | 31887291 | 31945246 | 57956 | 1 | ENSMUST00000138325.2 | -12313 | Lamc3 | ENSMUSG00000026840.9 |
| chr2 | 31875468 | 31876212 | 3.69567 | 4.49824 | 1.41589 | Distal Intergenic | 2 | 31887291 | 31945246 | 57956 | 1 | ENSMUST00000138325.2 | -11079 | Lamc3 | ENSMUSG00000026840.9 |
| chr2 | 31876661 | 31877106 | 3.41569 | 4.16421 | 1.25639 | Distal Intergenic | 2 | 31887291 | 31945246 | 57956 | 1 | ENSMUST00000138325.2 | -10185 | Lamc3 | ENSMUSG00000026840.9 |
| chr2 | 31880412 | 31881190 | 3.28613 | 3.77463 | 1.04928 | Distal Intergenic | 2 | 31887291 | 31945246 | 57956 | 1 | ENSMUST00000138325.2 | -6101 | Lamc3 | ENSMUSG00000026840.9 |
| chr2 | 31883541 | 31883804 | 3.27673 | 3.74223 | 1.01733 | Distal Intergenic | 2 | 31887291 | 31945246 | 57956 | 1 | ENSMUST00000138325.2 | -3487 | Lamc3 | ENSMUSG00000026840.9 |
| chr2 | 31884656 | 31884919 | 3.42295 | 4.05477 | 1.215 | Downstream (2-3kb) | 2 | 31887291 | 31945246 | 57956 | 1 | ENSMUST00000138325.2 | -2372 | Lamc3 | ENSMUSG00000026840.9 |
| chr2 | 31887244 | 31887936 | 4.04694 | 5.27273 | 1.98039 | Promoter | 2 | 31887291 | 31945246 | 57956 | 1 | ENSMUST00000138325.2 | 0 | Lamc3 | ENSMUSG00000026840.9 |
| chr2 | 31888880 | 31889099 | 3.46906 | 4.17811 | 1.2424 | Intron (ENSMUST00000138325.2/ENSMUSG00000026840.9, intron 1 of 27) | 2 | 31887344 | 31946539 | 59196 | 1 | ENSMUST00000028187.6 | 1537 | Lamc3 | ENSMUSG00000026840.9 |
| chr2 | 31828197 | 31828476 | 3.55695 | 4.22947 | 1.24616 | Intron (ENSMUST00000028188.7/ENSMUSG00000026841.7, intron 4 of 6) | 2 | 31813290 | 31846005 | 32716 | 2 | ENSMUST00000028188.7 | 17529 | Fibcd1 | ENSMUSG00000026841.7 |
| chr2 | 31836338 | 31836413 | 4.05606 | 5.16931 | 1.84257 | Intron (ENSMUST00000028188.7/ENSMUSG00000026841.7, intron 2 of 6) | 2 | 31813290 | 31846005 | 32716 | 2 | ENSMUST00000028188.7 | 9592 | Fibcd1 | ENSMUSG00000026841.7 |
| chr2 | 31840238 | 31840553 | 3.22696 | 3.82845 | 1.07137 | Intron (ENSMUST00000028188.7/ENSMUSG00000026841.7, intron 1 of 6) | 2 | 31813290 | 31846005 | 32716 | 2 | ENSMUST00000028188.7 | 5452 | Fibcd1 | ENSMUSG00000026841.7 |
| chr2 | 31844182 | 31844516 | 3.61549 | 4.34257 | 1.3241 | Intron (ENSMUST00000028188.7/ENSMUSG00000026841.7, intron 1 of 6) | 2 | 31813290 | 31846005 | 32716 | 2 | ENSMUST00000028188.7 | 1489 | Fibcd1 | ENSMUSG00000026841.7 |
| chr2 | 31844908 | 31845142 | 3.45171 | 4.07471 | 1.24197 | Promoter | 2 | 31813290 | 31846005 | 32716 | 2 | ENSMUST00000028188.7 | 863 | Fibcd1 | ENSMUSG00000026841.7 |
| chr2 | 31845512 | 31847294 | 3.85126 | 5.01792 | 1.7869 | Promoter | 2 | 31813290 | 31846005 | 32716 | 2 | ENSMUST00000028188.7 | 0 | Fibcd1 | ENSMUSG00000026841.7 |
| chr2 | 31850169 | 31850554 | 3.34153 | 3.83966 | 1.01733 | Distal Intergenic | 2 | 31813290 | 31846005 | 32716 | 2 | ENSMUST00000028188.7 | -4165 | Fibcd1 | ENSMUSG00000026841.7 |
| chr2 | 31851609 | 31851840 | 3.23338 | 3.67724 | 1.01733 | Distal Intergenic | 2 | 31813290 | 31846005 | 32716 | 2 | ENSMUST00000028188.7 | -5605 | Fibcd1 | ENSMUSG00000026841.7 |
| chr2 | 31858708 | 31858863 | 3.77876 | 4.65698 | 1.52897 | Distal Intergenic | 2 | 31813290 | 31846005 | 32716 | 2 | ENSMUST00000028188.7 | -12704 | Fibcd1 | ENSMUSG00000026841.7 |
| chr2 | 31859384 | 31859459 | 4.23901 | 5.55766 | 2.15557 | Distal Intergenic | 2 | 31813290 | 31846005 | 32716 | 2 | ENSMUST00000028188.7 | -13380 | Fibcd1 | ENSMUSG00000026841.7 |
| chr2 | 34667716 | 34667942 | 3.37939 | 3.89676 | 1.01733 | Distal Intergenic | 2 | 34677894 | 34684293 | 6400 | 2 | ENSMUST00000138203.1 | 16351 | Gapvd1 | ENSMUSG00000026867.13 |
| chr2 | 35557820 | 35558106 | 3.40204 | 3.96942 | 1.13806 | Promoter | 2 | 35558266 | 35730994 | 172729 | 1 | ENSMUST00000112986.3 | -160 | Dab2ip | ENSMUSG00000026883.12 |
| chr2 | 35661589 | 35661791 | 3.4631 | 4.10612 | 1.27109 | Promoter | 2 | 35661619 | 35709933 | 48315 | 1 | ENSMUST00000138957.1 | 0 | Dab2ip | ENSMUSG00000026883.12 |
| chr2 | 36002399 | 36002774 | 3.47398 | 4.08289 | 1.17117 | Distal Intergenic | 2 | 35902351 | 35979913 | 77563 | 2 | ENSMUST00000127742.2 | -22487 | Ttll11 | ENSMUSG00000026885.8 |
| chr2 | 36011006 | 36011691 | 3.37913 | 3.91775 | 1.04044 | Distal Intergenic | 2 | 35902351 | 35979913 | 77563 | 2 | ENSMUST00000127742.2 | -31094 | Ttll11 | ENSMUSG00000026885.8 |
| chr2 | 65021946 | 65022761 | 4.06905 | 5.21604 | 1.91247 | Promoter | 2 | 64912476 | 65022782 | 110307 | 2 | ENSMUST00000028252.8 | 21 | Grb14 | ENSMUSG00000026888.9 |
| chr2 | 36091092 | 36091673 | 3.81737 | 4.98742 | 1.79092 | Exon (ENSMUST00000112961.4/ENSMUSG00000026890.14, exon 5 of 9) | 2 | 36083684 | 36094273 | 10590 | 2 | ENSMUST00000136821.3 | 2600 | Lhx6 | ENSMUSG00000026890.14 |
| chr2 | 36092014 | 36093045 | 3.49581 | 4.18723 | 1.30908 | Intron (ENSMUST00000112961.4/ENSMUSG00000026890.14, intron 4 of 8) | 2 | 36083684 | 36094273 | 10590 | 2 | ENSMUST00000136821.3 | 1228 | Lhx6 | ENSMUSG00000026890.14 |
| chr2 | 36095714 | 36096307 | 4.14789 | 5.48112 | 2.21087 | Intron (ENSMUST00000112961.4/ENSMUSG00000026890.14, intron 3 of 8) | 2 | 36083684 | 36094273 | 10590 | 2 | ENSMUST00000136821.3 | -1442 | Lhx6 | ENSMUSG00000026890.14 |
| chr2 | 36096661 | 36096931 | 3.3629 | 3.89708 | 1.10632 | Intron (ENSMUST00000112961.4/ENSMUSG00000026890.14, intron 3 of 8) | 2 | 36083684 | 36094273 | 10590 | 2 | ENSMUST00000136821.3 | -2389 | Lhx6 | ENSMUSG00000026890.14 |
| chr2 | 36097278 | 36097958 | 4.08391 | 5.54206 | 2.1781 | Intron (ENSMUST00000112961.4/ENSMUSG00000026890.14, intron 3 of 8) | 2 | 36083684 | 36094273 | 10590 | 2 | ENSMUST00000136821.3 | -3006 | Lhx6 | ENSMUSG00000026890.14 |
| chr2 | 36098303 | 36098402 | 3.70738 | 4.86087 | 1.68698 | Intron (ENSMUST00000112961.4/ENSMUSG00000026890.14, intron 3 of 8) | 2 | 36083684 | 36094273 | 10590 | 2 | ENSMUST00000136821.3 | -4031 | Lhx6 | ENSMUSG00000026890.14 |
| chr2 | 36098853 | 36099240 | 3.23001 | 3.6722 | 1.0164 | Intron (ENSMUST00000112961.4/ENSMUSG00000026890.14, intron 3 of 8) | 2 | 36094225 | 36103592 | 9368 | 2 | ENSMUST00000187180.1 | 4352 | Lhx6 | ENSMUSG00000026890.14 |
| chr2 | 36100318 | 36100622 | 3.62423 | 4.41006 | 1.43209 | Intron (ENSMUST00000112961.4/ENSMUSG00000026890.14, intron 3 of 8) | 2 | 36094225 | 36103592 | 9368 | 2 | ENSMUST00000187180.1 | 2970 | Lhx6 | ENSMUSG00000026890.14 |
| chr2 | 36101875 | 36102937 | 3.73273 | 4.68004 | 1.60976 | Promoter | 2 | 36094225 | 36103592 | 9368 | 2 | ENSMUST00000187180.1 | 655 | Lhx6 | ENSMUSG00000026890.14 |
| chr2 | 36103440 | 36104161 | 3.77606 | 4.91046 | 1.70773 | Promoter | 2 | 36081953 | 36104060 | 22108 | 2 | ENSMUST00000112961.4 | 0 | Lhx6 | ENSMUSG00000026890.14 |
| chr2 | 36104824 | 36105420 | 4.29489 | 5.92704 | 2.44344 | Promoter | 2 | 36081953 | 36105271 | 23319 | 2 | ENSMUST00000112960.2 | 0 | Lhx6 | ENSMUSG00000026890.14 |
| chr2 | 36105815 | 36106807 | 3.48853 | 4.17902 | 1.31385 | Promoter | 2 | 36081954 | 36105408 | 23455 | 2 | ENSMUST00000112963.2 | -408 | Lhx6 | ENSMUSG00000026890.14 |
| chr2 | 36108547 | 36108818 | 3.92063 | 5.04514 | 1.85959 | Distal Intergenic | 2 | 36081954 | 36105408 | 23455 | 2 | ENSMUST00000112963.2 | -3140 | Lhx6 | ENSMUSG00000026890.14 |
| chr2 | 36112070 | 36112285 | 3.99512 | 5.05547 | 1.78128 | Distal Intergenic | 2 | 36081954 | 36105408 | 23455 | 2 | ENSMUST00000112963.2 | -6663 | Lhx6 | ENSMUSG00000026890.14 |
| chr2 | 62047504 | 62047833 | 3.42491 | 3.99405 | 1.11871 | Promoter | 2 | 62046645 | 62326730 | 280086 | 1 | ENSMUST00000112480.2 | 860 | Slc4a10 | ENSMUSG00000026904.12 |
| chr2 | 62048310 | 62048515 | 3.63504 | 4.4606 | 1.44193 | Intron (ENSMUST00000102735.4/ENSMUSG00000026904.12, intron 1 of 25) | 2 | 62046645 | 62326730 | 280086 | 1 | ENSMUST00000112480.2 | 1666 | Slc4a10 | ENSMUSG00000026904.12 |
| chr2 | 62255123 | 62255198 | 4.27317 | 5.62534 | 2.22331 | Intron (ENSMUST00000102735.4/ENSMUSG00000026904.12, intron 9 of 25) | 2 | 62267043 | 62268778 | 1736 | 1 | ENSMUST00000147917.1 | -11845 | Slc4a10 | ENSMUSG00000026904.12 |
| chr2 | 26586859 | 26586934 | 3.75485 | 4.79987 | 1.60097 | Promoter | 2 | 26586689 | 26590489 | 3801 | 1 | ENSMUST00000173920.2 | 171 | Egfl7 | ENSMUSG00000026921.13 |
| chr2 | 26487792 | 26487984 | 3.35991 | 3.86742 | 1.01733 | Intron (ENSMUST00000028288.4/ENSMUSG00000026923.10, intron 2 of 33) | 2 | 26457903 | 26503822 | 45920 | 2 | ENSMUST00000028288.4 | 15838 | Notch1 | ENSMUSG00000026923.10 |
| chr2 | 26502252 | 26502655 | 4.15745 | 5.54624 | 2.1525 | Intron (ENSMUST00000028288.4/ENSMUSG00000026923.10, intron 1 of 33) | 2 | 26457903 | 26503822 | 45920 | 2 | ENSMUST00000028288.4 | 1167 | Notch1 | ENSMUSG00000026923.10 |
| chr2 | 26197909 | 26198302 | 3.82586 | 4.74149 | 1.56825 | Downstream (1-2kb) | 2 | 26200216 | 26202866 | 2651 | 2 | ENSMUST00000127035.1 | 4564 | Lhx3 | ENSMUSG00000026934.10 |
| chr2 | 26198730 | 26199423 | 3.65129 | 4.42795 | 1.40093 | Downstream (<1kb) | 2 | 26200216 | 26202866 | 2651 | 2 | ENSMUST00000127035.1 | 3443 | Lhx3 | ENSMUSG00000026934.10 |
| chr2 | 26199995 | 26200106 | 3.30945 | 3.79153 | 1.01733 | Downstream (<1kb) | 2 | 26200216 | 26202866 | 2651 | 2 | ENSMUST00000127035.1 | 2760 | Lhx3 | ENSMUSG00000026934.10 |
| chr2 | 26201070 | 26201675 | 3.87299 | 4.83764 | 1.64536 | 3' UTR | 2 | 26200216 | 26202866 | 2651 | 2 | ENSMUST00000127035.1 | 1191 | Lhx3 | ENSMUSG00000026934.10 |
| chr2 | 26202077 | 26203340 | 3.86667 | 4.85908 | 1.67138 | Promoter | 2 | 26200216 | 26202866 | 2651 | 2 | ENSMUST00000127035.1 | 0 | Lhx3 | ENSMUSG00000026934.10 |
| chr2 | 26203739 | 26204385 | 3.55466 | 4.25989 | 1.24988 | Promoter | 2 | 26202476 | 26203441 | 966 | 2 | ENSMUST00000149637.1 | -299 | Lhx3 | ENSMUSG00000026934.10 |
| chr2 | 26204742 | 26205022 | 3.80448 | 4.71633 | 1.59218 | Intron (ENSMUST00000054099.10/ENSMUSG00000026934.10, intron 1 of 5) | 2 | 26202476 | 26203441 | 966 | 2 | ENSMUST00000149637.1 | -1302 | Lhx3 | ENSMUSG00000026934.10 |
| chr2 | 26205573 | 26206942 | 3.70381 | 4.71379 | 1.60467 | Promoter | 2 | 26200216 | 26206575 | 6360 | 2 | ENSMUST00000028302.7 | 0 | Lhx3 | ENSMUSG00000026934.10 |
| chr2 | 25372927 | 25373252 | 3.89841 | 5.24118 | 1.96152 | Promoter | 2 | 25372360 | 25375142 | 2783 | 1 | ENSMUST00000155310.1 | 568 | Sapcd2 | ENSMUSG00000026955.8 |
| chr2 | 25305225 | 25305734 | 4.03599 | 5.25681 | 2.02896 | Promoter | 2 | 25303584 | 25305397 | 1814 | 2 | ENSMUST00000153551.1 | 0 | Grin1 | ENSMUSG00000026959.8 |
| chr2 | 25306768 | 25306907 | 3.79615 | 4.72325 | 1.6455 | Intron (ENSMUST00000114308.4/ENSMUSG00000026959.8, intron 5 of 19) | 2 | 25303584 | 25305397 | 1814 | 2 | ENSMUST00000153551.1 | -1372 | Grin1 | ENSMUSG00000026959.8 |
| chr2 | 25307458 | 25307770 | 3.36425 | 3.8934 | 1.08622 | Intron (ENSMUST00000114308.4/ENSMUSG00000026959.8, intron 5 of 19) | 2 | 25303584 | 25305397 | 1814 | 2 | ENSMUST00000153551.1 | -2062 | Grin1 | ENSMUSG00000026959.8 |
| chr2 | 25308139 | 25308235 | 3.47463 | 4.07929 | 1.15442 | Intron (ENSMUST00000114308.4/ENSMUSG00000026959.8, intron 5 of 19) | 2 | 25303584 | 25305397 | 1814 | 2 | ENSMUST00000153551.1 | -2743 | Grin1 | ENSMUSG00000026959.8 |
| chr2 | 25308699 | 25309051 | 3.75833 | 4.62015 | 1.51837 | Intron (ENSMUST00000114308.4/ENSMUSG00000026959.8, intron 5 of 19) | 2 | 25303584 | 25305397 | 1814 | 2 | ENSMUST00000153551.1 | -3303 | Grin1 | ENSMUSG00000026959.8 |
| chr2 | 25310393 | 25310776 | 3.48549 | 4.50345 | 1.45088 | Exon (ENSMUST00000114308.4/ENSMUSG00000026959.8, exon 5 of 20) | 2 | 25303584 | 25305397 | 1814 | 2 | ENSMUST00000153551.1 | -4997 | Grin1 | ENSMUSG00000026959.8 |
| chr2 | 25311120 | 25312663 | 3.7586 | 4.64423 | 1.53151 | Exon (ENSMUST00000114308.4/ENSMUSG00000026959.8, exon 4 of 20) | 2 | 25303584 | 25305397 | 1814 | 2 | ENSMUST00000153551.1 | -5724 | Grin1 | ENSMUSG00000026959.8 |
| chr2 | 25313065 | 25313402 | 3.4665 | 4.05653 | 1.11622 | Exon (ENSMUST00000155627.1/ENSMUSG00000026959.8, exon 3 of 3) | 2 | 25291181 | 25319095 | 27915 | 2 | ENSMUST00000114308.4 | 5693 | Grin1 | ENSMUSG00000026959.8 |
| chr2 | 25318095 | 25318170 | 3.7911 | 4.80281 | 1.64158 | Promoter | 2 | 25291181 | 25319095 | 27915 | 2 | ENSMUST00000114308.4 | 925 | Grin1 | ENSMUSG00000026959.8 |
| chr2 | 25318783 | 25318858 | 3.91418 | 5.33485 | 2.02295 | Promoter | 2 | 25291181 | 25319095 | 27915 | 2 | ENSMUST00000114308.4 | 237 | Grin1 | ENSMUSG00000026959.8 |
| chr2 | 25323953 | 25324112 | 3.87806 | 4.83932 | 1.64344 | Distal Intergenic | 2 | 25291181 | 25319187 | 28007 | 2 | ENSMUST00000114312.1 | -4767 | Grin1 | ENSMUSG00000026959.8 |
| chr2 | 25324626 | 25324790 | 3.23338 | 3.67724 | 1.01733 | Downstream (2-3kb) | 2 | 25291181 | 25319187 | 28007 | 2 | ENSMUST00000114312.1 | -5440 | Grin1 | ENSMUSG00000026959.8 |
| chr2 | 25290196 | 25290456 | 3.54555 | 4.25346 | 1.2969 | Promoter | 2 | 25289915 | 25291180 | 1266 | 1 | ENSMUST00000028337.6 | 282 | Lrrc26 | ENSMUSG00000026961.6 |

| | | | | | | | | | | | | | | | |
|---|---|---|---|---|---|---|---|---|---|---|---|---|---|---|---|
| chr2 | 24404716 | 24404859 | 3.29048 | 3.76289 | 1.01733 | Promoter | 2 | 24405629 | 24407007 | 1379 | 1 | ENSMUST00000132924.1 | -770 | Psd4 | ENSMUSG00000026979.11 |
| chr2 | 24405411 | 24405592 | 3.72114 | 4.5445 | 1.46412 | Promoter | 2 | 24405629 | 24407007 | 1379 | 1 | ENSMUST00000132924.1 | -37 | Psd4 | ENSMUSG00000026979.11 |
| chr2 | 60383286 | 60383735 | 3.45635 | 4.06182 | 1.17867 | Promoter | 2 | 60292103 | 60383303 | 91201 | 2 | ENSMUST00000112533.2 | 0 | Ly75 | ENSMUSG00000026980.10 |
| chr2 | 66122390 | 66123529 | 3.61827 | 4.34316 | 1.31565 | Intron (ENSMUST00000028378.3/ENSMUSG00000026994.4, intron 1 of 10) | 2 | 66082766 | 66124994 | 42229 | 2 | ENSMUST00000028378.3 | 1465 | Galnt3 | ENSMUSG00000026994.4 |
| chr2 | 66123960 | 66125471 | 4.12179 | 5.71348 | 2.28822 | Promoter | 2 | 66082766 | 66124994 | 42229 | 2 | ENSMUST00000028378.3 | 0 | Galnt3 | ENSMUSG00000026994.4 |
| chr2 | 80446363 | 80447815 | 3.68476 | 4.67419 | 1.56243 | Promoter | 2 | 80411970 | 80447625 | 35656 | 2 | ENSMUST00000028389.3 | 0 | Frzb | ENSMUSG00000027004.3 |
| chr2 | 80456769 | 80456954 | 3.34194 | 3.84038 | 1.01733 | Distal Intergenic | 2 | 80411970 | 80447625 | 35656 | 2 | ENSMUST00000028389.3 | -9145 | Frzb | ENSMUSG00000027004.3 |
| chr2 | 79255638 | 79255828 | 3.38299 | 4.13614 | 1.2041 | Promoter | 2 | 79255721 | 79293273 | 37553 | 1 | ENSMUST00000135919.2 | 0 | Itga4 | ENSMUSG00000027009.13 |
| chr2 | 71118227 | 71118471 | 3.68043 | 4.57378 | 1.47529 | Promoter | 2 | 71117923 | 71142926 | 25004 | 1 | ENSMUST00000028403.2 | 305 | Cybrd1 | ENSMUSG00000027015.4 |
| chr2 | 68405741 | 68405894 | 3.41904 | 4.06055 | 1.22568 | Intron (ENSMUST00000102715.3/ENSMUSG00000027030.10, intron 2 of 17) | 2 | 68358986 | 68386681 | 27696 | 2 | ENSMUST00000149581.1 | -19061 | Stk39 | ENSMUSG00000027030.10 |
| chr2 | 68437169 | 68437333 | 3.48398 | 4.18064 | 1.29016 | Intron (ENSMUST00000102715.3/ENSMUSG00000027030.10, intron 1 of 17) | 2 | 68262791 | 68471940 | 209150 | 2 | ENSMUST00000123781.2 | 34607 | Stk39 | ENSMUSG00000027030.10 |
| chr2 | 69560569 | 69560817 | 3.51718 | 4.20763 | 1.29916 | Intron (ENSMUST00000080953.6/ENSMUSG00000027070.9, intron 1 of 78) | 2 | 69506204 | 69586029 | 79826 | 2 | ENSMUST00000100051.3 | 25212 | Lrp2 | ENSMUSG00000027070.9 |
| chr2 | 69584309 | 69584672 | 3.43237 | 4.04265 | 1.22832 | Intron (ENSMUST00000080953.6/ENSMUSG00000027070.9, intron 1 of 78) | 2 | 69506204 | 69586029 | 79826 | 2 | ENSMUST00000100051.3 | 1357 | Lrp2 | ENSMUSG00000027070.9 |
| chr2 | 69585405 | 69586464 | 4.10874 | 5.45383 | 2.11192 | Promoter | 2 | 69424340 | 69586065 | 161726 | 2 | ENSMUST00000080953.6 | 0 | Lrp2 | ENSMUSG00000027070.9 |
| chr2 | 69586981 | 69587875 | 3.78166 | 4.68025 | 1.58372 | Promoter | 2 | 69424340 | 69586065 | 161726 | 2 | ENSMUST00000080953.6 | -917 | Lrp2 | ENSMUSG00000027070.9 |
| chr2 | 84839988 | 84840302 | 3.95689 | 5.14171 | 1.89985 | Promoter | 2 | 84840199 | 84863581 | 23383 | 1 | ENSMUST00000111624.2 | 0 | Slc43a1 | ENSMUSG00000027075.11 |
| chr2 | 84840727 | 84841518 | 3.93606 | 5.06398 | 1.83451 | Promoter | 2 | 84840625 | 84863594 | 22970 | 1 | ENSMUST00000111625.1 | 103 | Slc43a1 | ENSMUSG00000027075.11 |
| chr2 | 73867943 | 73868026 | 3.87754 | 15.64175 | 10.82439 | Intron (ENSMUST00000112010.3/ENSMUSG00000027104.13, intron 3 of 12) | 2 | 73816524 | 73863290 | 46767 | 2 | ENSMUST00000100009.5 | -4654 | Atf2 | ENSMUSG00000027104.13 |
| chr2 | 71812047 | 71812432 | 3.87901 | 4.8347 | 1.62643 | Intron (ENSMUST00000133124.2/ENSMUSG00000027111.10, intron 3 of 5) | 2 | 71826762 | 71854951 | 28190 | 1 | ENSMUST00000155249.1 | -14330 | Itga6 | ENSMUSG00000027111.10 |
| chr2 | 105612025 | 105612979 | 3.72406 | 4.54688 | 1.45076 | Intron (ENSMUST00000149133.2/ENSMUSG00000086029.2, intron 1 of 3) | 2 | 105668900 | 105697364 | 28465 | 1 | ENSMUST00000111086.5 | -55921 | Pax6 | ENSMUSG00000027168.16 |
| chr2 | 105615656 | 105615910 | 3.31968 | 3.80681 | 1.01733 | Intron (ENSMUST00000149133.2/ENSMUSG00000086029.2, intron 1 of 3) | 2 | 105668900 | 105697364 | 28465 | 1 | ENSMUST00000111086.5 | -52990 | Pax6 | ENSMUSG00000027168.16 |
| chr2 | 105651665 | 105651807 | 3.27112 | 3.73382 | 1.01733 | Intron (ENSMUST00000149133.2/ENSMUSG00000086029.2, intron 1 of 3) | 2 | 105668900 | 105697364 | 28465 | 1 | ENSMUST00000111086.5 | -17093 | Pax6 | ENSMUSG00000027168.16 |
| chr2 | 105654482 | 105654646 | 3.39606 | 3.93567 | 1.06587 | Intron (ENSMUST00000149133.2/ENSMUSG00000086029.2, intron 1 of 3) | 2 | 105668900 | 105697364 | 28465 | 1 | ENSMUST00000111086.5 | -14254 | Pax6 | ENSMUSG00000027168.16 |
| chr2 | 105655235 | 105656208 | 4.02892 | 5.21927 | 1.95249 | Intron (ENSMUST00000149133.2/ENSMUSG00000086029.2, intron 1 of 3) | 2 | 105668900 | 105697364 | 28465 | 1 | ENSMUST00000111086.5 | -12692 | Pax6 | ENSMUSG00000027168.16 |
| chr2 | 105657294 | 105661011 | 3.79354 | 4.74282 | 1.60512 | Intron (ENSMUST00000149133.2/ENSMUSG00000086029.2, intron 1 of 3) | 2 | 105668900 | 105697364 | 28465 | 1 | ENSMUST00000111086.5 | -7889 | Pax6 | ENSMUSG00000027168.16 |
| chr2 | 105661564 | 105662523 | 3.75854 | 4.79062 | 1.63926 | Intron (ENSMUST00000149133.2/ENSMUSG00000086029.2, intron 1 of 3) | 2 | 105668900 | 105697364 | 28465 | 1 | ENSMUST00000111086.5 | -6377 | Pax6 | ENSMUSG00000027168.16 |
| chr2 | 105663001 | 105665831 | 3.838 | 4.8336 | 1.64806 | Intron (ENSMUST00000149133.2/ENSMUSG00000086029.2, intron 1 of 3) | 2 | 105668900 | 105697364 | 28465 | 1 | ENSMUST00000111086.5 | -3069 | Pax6 | ENSMUSG00000027168.16 |
| chr2 | 105667493 | 105672512 | 4.04766 | 5.25164 | 1.96954 | Promoter | 2 | 105668900 | 105697364 | 28465 | 1 | ENSMUST00000111086.5 | 0 | Pax6 | ENSMUSG00000027168.16 |
| chr2 | 105673234 | 105677269 | 4.01934 | 5.20147 | 1.9208 | Promoter | 2 | 105673963 | 105675876 | 1914 | 1 | ENSMUST00000147923.2 | 0 | Pax6 | ENSMUSG00000027168.16 |
| chr2 | 105677685 | 105678199 | 3.70028 | 4.542 | 1.50011 | Promoter | 2 | 105678391 | 105685795 | 7405 | 1 | ENSMUST00000156216.2 | -192 | Pax6 | ENSMUSG00000027168.16 |
| chr2 | 105678517 | 105679231 | 3.64541 | 4.51019 | 1.51079 | Promoter | 2 | 105678391 | 105685795 | 7405 | 1 | ENSMUST00000156216.2 | 127 | Pax6 | ENSMUSG00000027168.16 |
| chr2 | 105680208 | 105680968 | 4.17336 | 5.51032 | 2.14487 | Promoter | 2 | 105679867 | 105692222 | 12356 | 1 | ENSMUST00000153011.2 | 342 | Pax6 | ENSMUSG00000027168.16 |
| chr2 | 105681295 | 105682211 | 3.79401 | 4.84339 | 1.69426 | Promoter | 2 | 105681712 | 105684982 | 3271 | 1 | ENSMUST00000154302.2 | 0 | Pax6 | ENSMUSG00000027168.16 |
| chr2 | 105683233 | 105683958 | 4.08878 | 5.32359 | 2.02055 | Promoter | 2 | 105683754 | 105684943 | 1190 | 1 | ENSMUST00000123000.2 | 0 | Pax6 | ENSMUSG00000027168.16 |
| chr2 | 105684272 | 105688211 | 3.87339 | 4.97372 | 1.77721 | Promoter | 2 | 105685322 | 105697364 | 12043 | 1 | ENSMUST00000146803.3 | 0 | Pax6 | ENSMUSG00000027168.16 |
| chr2 | 105689069 | 105689144 | 3.37725 | 4.42868 | 1.39746 | Intron (ENSMUST00000111086.5/ENSMUSG00000027168.16, intron 7 of 12) | 2 | 105692483 | 105693125 | 643 | 1 | ENSMUST00000147946.1 | -3339 | Pax6 | ENSMUSG00000027168.16 |
| chr2 | 105689546 | 105689740 | 3.46787 | 4.11357 | 1.27328 | Intron (ENSMUST00000111086.5/ENSMUSG00000027168.16, intron 7 of 12) | 2 | 105692483 | 105693125 | 643 | 1 | ENSMUST00000147946.1 | -2743 | Pax6 | ENSMUSG00000027168.16 |
| chr2 | 105690449 | 105691592 | 3.97758 | 5.26486 | 1.96472 | Promoter | 2 | 105692483 | 105693125 | 643 | 1 | ENSMUST00000147946.1 | -891 | Pax6 | ENSMUSG00000027168.16 |
| chr2 | 105692375 | 105692639 | 3.32544 | 4.01048 | 1.15794 | Promoter | 2 | 105692483 | 105693125 | 643 | 1 | ENSMUST00000147946.1 | 0 | Pax6 | ENSMUSG00000027168.16 |
| chr2 | 105693055 | 105693344 | 3.91068 | 5.0641 | 1.81736 | Promoter | 2 | 105692483 | 105693125 | 643 | 1 | ENSMUST00000147946.1 | 573 | Pax6 | ENSMUSG00000027168.16 |
| chr2 | 104742651 | 104742872 | 3.38709 | 4.2817 | 1.29915 | Promoter | 2 | 104721784 | 104742878 | 21095 | 2 | ENSMUST00000028595.7 | 6 | Depdc7 | ENSMUSG00000027173.7 |
| chr2 | 93676298 | 93676431 | 3.49107 | 4.10233 | 1.14829 | 3' UTR | 2 | 93703539 | 93704509 | 971 | 2 | ENSMUST00000144780.1 | 28078 | Ext2 | ENSMUSG00000027198.11 |
| chr2 | 124090093 | 124091727 | 3.50873 | 4.18151 | 1.25384 | Promoter | 2 | 124089969 | 124667769 | 577801 | 1 | ENSMUST00000103241.2 | 125 | Sema6d | ENSMUSG00000027200.12 |
| chr2 | 125505538 | 125506290 | 3.70844 | 4.62562 | 1.55011 | Promoter | 2 | 125389756 | 125506072 | 116317 | 2 | ENSMUST00000148272.1 | 0 | Fbn1 | ENSMUSG00000027204.8 |
| chr2 | 125506818 | 125506901 | 4.3183 | 5.69082 | 2.21028 | Promoter | 2 | 125300594 | 125506385 | 205792 | 2 | ENSMUST00000028633.7 | -434 | Fbn1 | ENSMUSG00000027204.8 |
| chr2 | 126147864 | 126147960 | 6.06484 | 9.86773 | 5.67923 | Intron (ENSMUST00000110446.3/ENSMUSG00000027209.12, intron 1 of 16) | 2 | 125983628 | 126146251 | 162624 | 2 | ENSMUST00000178118.2 | -1614 | Fam227b | ENSMUSG00000027209.12 |
| chr2 | 115851908 | 115852126 | 3.33854 | 4.16699 | 1.25171 | Distal Intergenic | 2 | 115864227 | 115868867 | 4641 | 2 | ENSMUST00000189640.1 | 16741 | Meis2 | ENSMUSG00000027210.15 |
| chr2 | 115853327 | 115853480 | 3.77476 | 4.73625 | 1.6134 | Distal Intergenic | 2 | 115864227 | 115868867 | 4641 | 2 | ENSMUST00000189640.1 | 15387 | Meis2 | ENSMUSG00000027210.15 |
| chr2 | 115853933 | 115854079 | 3.3054 | 3.9302 | 1.06821 | Distal Intergenic | 2 | 115864227 | 115868867 | 4641 | 2 | ENSMUST00000189640.1 | 14788 | Meis2 | ENSMUSG00000027210.15 |
| chr2 | 115854809 | 115855310 | 3.21278 | 4.14415 | 1.20359 | Distal Intergenic | 2 | 115864227 | 115868867 | 4641 | 2 | ENSMUST00000189640.1 | 13557 | Meis2 | ENSMUSG00000027210.15 |
| chr2 | 92915162 | 92915341 | 4.10034 | 5.5171 | 2.12054 | Promoter | 2 | 92915098 | 92956058 | 40961 | 1 | ENSMUST00000028648.2 | 65 | Syt13 | ENSMUSG00000027220.2 |
| chr2 | 92915968 | 92916539 | 3.87144 | 5.22007 | 1.93485 | Promoter | 2 | 92915098 | 92956058 | 40961 | 1 | ENSMUST00000028648.2 | 871 | Syt13 | ENSMUSG00000027220.2 |
| chr2 | 92917249 | 92917647 | 3.84953 | 4.99065 | 1.77268 | Intron (ENSMUST00000028648.2/ENSMUSG00000027220.2, intron 1 of 5) | 2 | 92915098 | 92956058 | 40961 | 1 | ENSMUST00000028648.2 | 2152 | Syt13 | ENSMUSG00000027220.2 |
| chr2 | 92918639 | 92918804 | 3.80867 | 4.76274 | 1.65563 | Intron (ENSMUST00000028648.2/ENSMUSG00000027220.2, intron 1 of 5) | 2 | 92915098 | 92956058 | 40961 | 1 | ENSMUST00000028648.2 | 3542 | Syt13 | ENSMUSG00000027220.2 |
| chr2 | 122301774 | 122302941 | 3.62702 | 4.55078 | 1.48518 | 3' UTR | 2 | 122303552 | 122305711 | 2160 | 2 | ENSMUST00000155561.1 | 2770 | Duoxa1 | ENSMUSG00000027224.9 |
| chr2 | 122303320 | 122303923 | 3.63145 | 4.62393 | 1.54401 | 3' UTR | 2 | 122303552 | 122305711 | 2160 | 2 | ENSMUST00000155561.1 | 1788 | Duoxa1 | ENSMUSG00000027224.9 |
| chr2 | 122306312 | 122306455 | 3.61162 | 4.39449 | 1.41168 | Promoter | 2 | 122303745 | 122306826 | 3082 | 2 | ENSMUST00000028653.8 | 371 | Duoxa1 | ENSMUSG00000027224.9 |
| chr2 | 122307436 | 122308006 | 3.34356 | 3.87349 | 1.11968 | Promoter | 2 | 122302191 | 122306848 | 4658 | 2 | ENSMUST00000110538.2 | -589 | Duoxa1 | ENSMUSG00000027224.9 |
| chr2 | 122308669 | 122309029 | 3.66177 | 4.51693 | 1.46386 | Intron (ENSMUST00000110537.2/ENSMUSG00000027224.9, intron 3 of 9) | 2 | 122302191 | 122306848 | 4658 | 2 | ENSMUST00000110538.2 | -1822 | Duoxa1 | ENSMUSG00000027224.9 |
| chr2 | 122309723 | 122309847 | 3.34812 | 3.86991 | 1.08905 | Intron (ENSMUST00000110537.2/ENSMUSG00000027224.9, intron 3 of 9) | 2 | 122302191 | 122306848 | 4658 | 2 | ENSMUST00000110538.2 | -2876 | Duoxa1 | ENSMUSG00000027224.9 |
| chr2 | 122311764 | 122311910 | 3.45623 | 4.04146 | 1.11903 | Intron (ENSMUST00000110537.2/ENSMUSG00000027224.9, intron 3 of 9) | 2 | 122305215 | 122313644 | 8430 | 2 | ENSMUST00000147788.2 | 1734 | Duoxa1 | ENSMUSG00000027224.9 |
| chr2 | 122312406 | 122312487 | 3.60213 | 4.31502 | 1.30362 | Intron (ENSMUST00000110537.2/ENSMUSG00000027224.9, intron 3 of 9) | 2 | 122305215 | 122313644 | 8430 | 2 | ENSMUST00000147788.2 | 1157 | Duoxa1 | ENSMUSG00000027224.9 |
| chr2 | 122313074 | 122314536 | 3.83219 | 4.77797 | 1.61183 | Promoter | 2 | 122303549 | 122313693 | 10145 | 2 | ENSMUST00000110537.2 | 0 | Duoxa1 | ENSMUSG00000027224.9 |
| chr2 | 122299336 | 122301279 | 3.57761 | 4.55981 | 1.4821 | Promoter | 2 | 122298900 | 122302885 | 3986 | 1 | ENSMUST00000028656.1 | 437 | Duoxa2 | ENSMUSG00000027225.7 |
| chr2 | 91930676 | 91930751 | 3.63462 | 4.3891 | 1.39069 | Promoter | 2 | 91929805 | 91931688 | 1884 | 2 | ENSMUST00000111309.2 | 937 | Mdk | ENSMUSG00000027239.9 |
| chr2 | 91931520 | 91931716 | 3.93973 | 4.95453 | 1.70502 | Promoter | 2 | 91929805 | 91931688 | 1884 | 2 | ENSMUST00000111309.2 | 0 | Mdk | ENSMUSG00000027239.9 |
| chr2 | 121290474 | 121291146 | 3.40606 | 4.01202 | 1.13287 | Promoter | 2 | 121289770 | 121291073 | 1304 | 1 | ENSMUST00000131901.1 | 705 | Map1a | ENSMUSG00000027254.8 |
| chr2 | 136058168 | 136059672 | 3.6129 | 4.3714 | 1.35725 | Promoter | 2 | 136058252 | 136059213 | 962 | 1 | ENSMUST00000154674.2 | 0 | Lamp5 | ENSMUSG00000027270.9 |
| chr2 | 136060078 | 136060365 | 3.49719 | 4.11957 | 1.17553 | Promoter | 2 | 136059431 | 136069905 | 10475 | 1 | ENSMUST00000123436.1 | 648 | Lamp5 | ENSMUSG00000027270.9 |
| chr2 | 136060913 | 136061140 | 3.38193 | 3.9006 | 1.01733 | Exon (ENSMUST00000057503.6/ENSMUSG00000027270.9, exon 5 of 6) | 2 | 136059431 | 136069905 | 10475 | 1 | ENSMUST00000123436.1 | 1483 | Lamp5 | ENSMUSG00000027270.9 |
| chr2 | 136712727 | 136712950 | 3.78137 | 4.64296 | 1.4883 | Promoter | 2 | 136713453 | 136782428 | 68976 | 1 | ENSMUST00000028727.5 | -503 | Snap25 | ENSMUSG00000027273.8 |
| chr2 | 136713520 | 136714198 | 3.60933 | 4.32755 | 1.29969 | Promoter | 2 | 136713478 | 136781351 | 67874 | 1 | ENSMUST00000110098.3 | 43 | Snap25 | ENSMUSG00000027273.8 |
| chr2 | 119743038 | 119744461 | 3.68443 | 4.5789 | 1.51018 | Promoter | 2 | 119742337 | 119751263 | 8927 | 1 | ENSMUST00000028758.7 | 702 | Itpka | ENSMUSG00000027296.7 |
| chr2 | 119746875 | 119747150 | 3.53007 | 4.18998 | 1.25092 | Intron (ENSMUST00000028758.7/ENSMUSG00000027296.7, intron 1 of 6) | 2 | 119742337 | 119751263 | 8927 | 1 | ENSMUST00000028758.7 | 4539 | Itpka | ENSMUSG00000027296.7 |
| chr2 | 119750821 | 119751007 | 3.45974 | 4.0702 | 1.19148 | 3' UTR | 2 | 119751330 | 119758519 | 7190 | 2 | ENSMUST00000134295.1 | 7512 | Ltk | ENSMUSG00000027297.9 |
| chr2 | 119751675 | 119751916 | 3.41035 | 3.95241 | 1.04874 | Exon (ENSMUST00000140224.2/ENSMUSG00000027297.9, exon 14 of 14) | 2 | 119751330 | 119758519 | 7190 | 2 | ENSMUST00000134295.1 | 6603 | Ltk | ENSMUSG00000027297.9 |
| chr2 | 119756302 | 119756540 | 3.5141 | 4.14461 | 1.17437 | Intron (ENSMUST00000140224.2/ENSMUSG00000027297.9, intron 1 of 13) | 2 | 119751330 | 119758519 | 7190 | 2 | ENSMUST00000134295.1 | 1979 | Ltk | ENSMUSG00000027297.9 |
| chr2 | 119757963 | 119758912 | 3.80335 | 4.81298 | 1.68247 | Promoter | 2 | 119751320 | 119758525 | 7206 | 2 | ENSMUST00000140224.2 | 0 | Ltk | ENSMUSG00000027297.9 |
| chr2 | 119760308 | 119760383 | 3.38193 | 3.9006 | 1.01733 | Promoter | 2 | 119751320 | 119760295 | 8976 | 2 | ENSMUST00000082130.7 | -14 | Ltk | ENSMUSG00000027297.9 |
| chr2 | 130575851 | 130577116 | 4.23819 | 5.72924 | 2.36468 | Promoter | 2 | 130576173 | 130577054 | 882 | 1 | ENSMUST00000028764.5 | 0 | Oxt | ENSMUSG00000027301.7 |
| chr2 | 130577493 | 130577667 | 3.71299 | 4.76169 | 1.62427 | Downstream (2-3kb) | 2 | 130576173 | 130577054 | 882 | 1 | ENSMUST00000028764.5 | 1321 | Oxt | ENSMUSG00000027301.7 |
| chr2 | 130578103 | 130578273 | 3.53484 | 4.24866 | 1.33872 | Downstream (2-3kb) | 2 | 130576173 | 130577054 | 882 | 1 | ENSMUST00000028764.5 | 1931 | Oxt | ENSMUSG00000027301.7 |
| chr2 | 119350416 | 119350754 | 3.55431 | 4.28452 | 1.34992 | Promoter | 2 | 119351229 | 119354381 | 3153 | 1 | ENSMUST00000028780.3 | -475 | Chac1 | ENSMUSG00000027313.3 |
| chr2 | 119325575 | 119326640 | 3.49684 | 4.56808 | 1.50663 | Promoter | 2 | 119325784 | 119335962 | 10179 | 1 | ENSMUST00000102517.3 | 0 | Dll4 | ENSMUSG00000027314.6 |
| chr2 | 119328546 | 119328805 | 3.91874 | 4.99366 | 1.82362 | Promoter | 2 | 119328538 | 119332153 | 3616 | 1 | ENSMUST00000127102.1 | 9 | Dll4 | ENSMUSG00000027314.6 |
| chr2 | 119329385 | 119329613 | 3.40758 | 3.9837 | 1.17297 | Promoter | 2 | 119328538 | 119332153 | 3616 | 1 | ENSMUST00000127102.1 | 848 | Dll4 | ENSMUSG00000027314.6 |
| chr2 | 119235568 | 119236180 | 3.79956 | 4.77233 | 1.63522 | Distal Intergenic | 2 | 119237362 | 119249514 | 12153 | 1 | ENSMUST00000028783.8 | -1182 | Spint1 | ENSMUSG00000027315.8 |
| chr2 | 119236665 | 119238205 | 4.04632 | 5.53729 | 2.15955 | Promoter | 2 | 119237362 | 119249514 | 12153 | 1 | ENSMUST00000028783.8 | 0 | Spint1 | ENSMUSG00000027315.8 |
| chr2 | 119255653 | 119255784 | 3.68451 | 4.56876 | 1.5109 | Distal Intergenic | 2 | 119248205 | 119249049 | 845 | 1 | ENSMUST00000134872.1 | 7449 | Spint1 | ENSMUSG00000027315.8 |
| chr2 | 131040976 | 131042129 | 3.74398 | 4.68756 | 1.54763 | Promoter | 2 | 131039639 | 131041457 | 1819 | 2 | ENSMUST00000156509.1 | 0 | Gfra4 | ENSMUSG00000027316.10 |
| chr2 | 131052913 | 131053345 | 3.53462 | 4.18395 | 1.20438 | Promoter | 2 | 131050591 | 131054036 | 3446 | 2 | ENSMUST00000128828.2 | 691 | Adam33 | ENSMUSG00000027318.12 |
| chr2 | 131054264 | 131055415 | 4.28961 | 5.71761 | 2.30536 | Promoter | 2 | 131053042 | 131054526 | 1485 | 2 | ENSMUST00000184921.2 | 0 | Adam33 | ENSMUSG00000027318.12 |
| chr2 | 131056400 | 131056904 | 3.55488 | 4.22227 | 1.22969 | Promoter | 2 | 131051110 | 131056481 | 5372 | 2 | ENSMUST00000134742.2 | 0 | Adam33 | ENSMUSG00000027318.12 |
| chr2 | 131063514 | 131063908 | 4.13332 | 6.00672 | 2.47934 | Promoter | 2 | 131050817 | 131063814 | 12998 | 2 | ENSMUST00000110232.3 | 0 | Adam33 | ENSMUSG00000027318.12 |
| chr2 | 131170091 | 131170399 | 3.29577 | 3.77653 | 1.03711 | 3' UTR | 2 | 131171611 | 131174653 | 3043 | 2 | ENSMUST00000127987.1 | 4254 | Spef1 | ENSMUSG00000027329.4 |
| chr2 | 118885706 | 118885977 | 3.7788 | 4.72567 | 1.62408 | Distal Intergenic | 2 | 118876433 | 118877900 | 1468 | 1 | ENSMUST00000154104.1 | 9274 | Ivd | ENSMUSG00000027332.6 |
| chr2 | 131561615 | 131562443 | 3.68845 | 4.67204 | 1.5507 | Promoter | 2 | 131545850 | 131562283 | 16434 | 2 | ENSMUST00000103184.3 | 0 | Adra1d | ENSMUSG00000027335.9 |
| chr2 | 131562760 | 131563046 | 3.62646 | 4.41101 | 1.44369 | Promoter | 2 | 131545850 | 131562283 | 16434 | 2 | ENSMUST00000103184.3 | -478 | Adra1d | ENSMUSG00000027335.9 |
| chr2 | 131564049 | 131564162 | 3.27995 | 3.74707 | 1.01733 | Distal Intergenic | 2 | 131545850 | 131562283 | 16434 | 2 | ENSMUST00000103184.3 | -1767 | Adra1d | ENSMUSG00000027335.9 |
| chr2 | 131951560 | 131951857 | 3.48197 | 4.14606 | 1.22571 | Promoter | 2 | 131951037 | 131956130 | 5094 | 1 | ENSMUST00000110169.1 | 524 | Prnd | ENSMUSG00000027338.11 |
| chr2 | 131952437 | 131953923 | 3.62847 | 4.42718 | 1.39799 | 5' UTR | 2 | 131951037 | 131956130 | 5094 | 1 | ENSMUST00000110169.1 | 1401 | Prnd | ENSMUSG00000027338.11 |
| chr2 | 131954798 | 131955089 | 3.54443 | 4.20786 | 1.23718 | 3' UTR | 2 | 131951037 | 131956130 | 5094 | 1 | ENSMUST00000110169.1 | 3762 | Prnd | ENSMUSG00000027338.11 |
| chr2 | 132449364 | 132449493 | 3.47411 | 4.12195 | 1.27689 | Intron (ENSMUST00000124095.1/ENSMUSG00000027345.6, intron 3 of 3) | 2 | 132424741 | 132452499 | 27759 | 1 | ENSMUST00000124095.1 | 24624 | 4921508D12Rik | ENSMUSG00000027345.6 |
| chr2 | 117342226 | 117343124 | 4.02708 | 5.32925 | 2.01671 | Promoter | 2 | 117279993 | 117343001 | 63009 | 2 | ENSMUST00000102534.5 | 0 | Rasgrp1 | ENSMUSG00000027347.13 |
| chr2 | 132939245 | 132939666 | 3.71727 | 4.63428 | 1.5374 | Exon (ENSMUST00000134937.1/ENSMUSG00000027356.8, exon 3 of 14) | 2 | 132939096 | 132941963 | 2868 | 2 | ENSMUST00000143981.1 | 2297 | Fermt1 | ENSMUSG00000027356.8 |
| chr2 | 132943689 | 132943802 | 4.70283 | 6.48547 | 2.80882 | Intron (ENSMUST00000134937.1/ENSMUSG00000027356.8, intron 1 of 13) | 2 | 132936113 | 132942097 | 5985 | 2 | ENSMUST00000144342.1 | -1593 | Fermt1 | ENSMUSG00000027356.8 |
| chr2 | 132944188 | 132945821 | 3.70702 | 4.55748 | 1.48311 | Promoter | 2 | 132904389 | 132945629 | 41241 | 2 | ENSMUST00000134937.1 | 0 | Fermt1 | ENSMUSG00000027356.8 |
| chr2 | 132947132 | 132947390 | 3.82695 | 5.09358 | 1.83603 | Distal Intergenic | 2 | 132904389 | 132945906 | 41518 | 2 | ENSMUST00000038280.4 | -1227 | Fermt1 | ENSMUSG00000027356.8 |
| chr2 | 132949278 | 132949525 | 3.35964 | 3.9084 | 1.02596 | Distal Intergenic | 2 | 132904389 | 132945906 | 41518 | 2 | ENSMUST00000038280.4 | -3373 | Fermt1 | ENSMUSG00000027356.8 |
| chr2 | 133552229 | 133552639 | 3.9395 | 5.10491 | 1.86242 | Promoter | 2 | 133552159 | 133562885 | 10727 | 1 | ENSMUST00000028836.6 | 71 | Bmp2 | ENSMUSG00000027358.6 |
| chr2 | 133553331 | 133553419 | 4.30622 | 5.80419 | 2.35901 | 5' UTR | 2 | 133552159 | 133562885 | 10727 | 1 | ENSMUST00000028836.6 | 1173 | Bmp2 | ENSMUSG00000027358.6 |
| chr2 | 133554112 | 133554335 | 3.53565 | 4.30444 | 1.32428 | Intron (ENSMUST00000028836.6/ENSMUSG00000027358.6, intron 1 of 2) | 2 | 133552159 | 133562885 | 10727 | 1 | ENSMUST00000028836.6 | 1954 | Bmp2 | ENSMUSG00000027358.6 |
| chr2 | 126551438 | 126551663 | 3.55804 | 4.27749 | 1.35202 | Promoter | 2 | 126552407 | 126588243 | 35837 | 1 | ENSMUST00000061491.8 | -744 | Slc27a2 | ENSMUSG00000027359.11 |
| chr2 | 126552680 | 126554098 | 4.0776 | 5.36976 | 2.17599 | Promoter | 2 | 126553590 | 126564685 | 11096 | 1 | ENSMUST00000126249.1 | 0 | Slc27a2 | ENSMUSG00000027359.11 |
| chr2 | 126606803 | 126606907 | 3.36906 | 3.88117 | 1.01733 | Promoter | 2 | 126604315 | 126607641 | 3327 | 2 | ENSMUST00000132382.1 | 734 | Hdc | ENSMUSG00000027360.5 |
| chr2 | 127451756 | 127451831 | 3.41125 | 4.00383 | 1.20507 | Distal Intergenic | 2 | 127436216 | 127444565 | 8350 | 2 | ENSMUST00000146757.2 | -7192 | Fahd2a | ENSMUSG00000027371.5 |
| chr2 | 127453611 | 127453715 | 4.38384 | 5.8482 | 2.36422 | Downstream (2-3kb) | 2 | 127436216 | 127444565 | 8350 | 2 | ENSMUST00000146757.2 | -9047 | Fahd2a | ENSMUSG00000027371.5 |
| chr2 | 127654726 | 127655472 | 3.4077 | 4.08083 | 1.21647 | Promoter | 2 | 127634874 | 127656219 | 21346 | 2 | ENSMUST00000028853.6 | 747 | Mal | ENSMUSG00000027375.9 |
| chr2 | 127656206 | 127656528 | 3.5757 | 4.53267 | 1.47526 | Promoter | 2 | 127634874 | 127656219 | 21346 | 2 | ENSMUST00000028853.6 | 0 | Mal | ENSMUSG00000027375.9 |
| chr2 | 127532668 | 127532932 | 3.36398 | 3.87352 | 1.01733 | Exon (ENSMUST00000028855.8/ENSMUSG00000027376.10, exon 15 of 24) | 2 | 127534766 | 127535197 | 432 | 2 | ENSMUST00000140156.1 | 2265 | Prom2 | ENSMUSG00000027376.10 |
| chr2 | 127539963 | 127540534 | 3.44128 | 4.00962 | 1.08578 | Promoter | 2 | 127526955 | 127541385 | 14431 | 2 | ENSMUST00000103214.2 | 851 | Prom2 | ENSMUSG00000027376.10 |
| chr2 | 127540947 | 127541979 | 3.71764 | 4.59525 | 1.48697 | Promoter | 2 | 127526473 | 127541467 | 14995 | 2 | ENSMUST00000028855.8 | 0 | Prom2 | ENSMUSG00000027376.10 |
| chr2 | 127729659 | 127730025 | 4.11631 | 5.54298 | 2.16401 | Promoter | 2 | 127704386 | 127729932 | 25547 | 2 | ENSMUST00000028856.2 | 0 | Mall | ENSMUSG00000027377.2 |
| chr2 | 127730384 | 127730740 | 3.46584 | 4.05838 | 1.12726 | Promoter | 2 | 127704386 | 127729932 | 25547 | 2 | ENSMUST00000028856.2 | -453 | Mall | ENSMUSG00000027377.2 |
| chr2 | 127839156 | 127839394 | 3.93655 | 5.01162 | 1.8347 | Promoter | 2 | 127838876 | 127877800 | 38925 | 1 | ENSMUST00000126410.1 | 281 | Acoxl | ENSMUSG00000027380.5 |
| chr2 | 127840965 | 127841179 | 3.65276 | 4.49272 | 1.4721 | Intron (ENSMUST00000126410.1/ENSMUSG00000027380.5, intron 1 of 5) | 2 | 127838876 | 127877800 | 38925 | 1 | ENSMUST00000126410.1 | 2090 | Acoxl | ENSMUSG00000027380.5 |
| chr2 | 128892446 | 128892683 | 3.46125 | 4.10121 | 1.19648 | Exon (ENSMUST00000110324.2/ENSMUSG00000027386.4, exon 6 of 9) | 2 | 128863932 | 128897030 | 33099 | 1 | ENSMUST00000028864.2 | 28515 | Fbln7 | ENSMUSG00000027386.4 |
| chr2 | 129697085 | 129697247 | 3.72758 | 4.54942 | 1.44637 | Intron (ENSMUST00000028883.6/ENSMUSG00000027400.6, intron 2 of 3) | 2 | 129688496 | 129698658 | 10163 | 2 | ENSMUST00000130608.1 | 1411 | Pdyn | ENSMUSG00000027400.6 |
| chr2 | 129697629 | 129697793 | 3.33703 | 3.83294 | 1.01733 | Promoter | 2 | 129688496 | 129698658 | 10163 | 2 | ENSMUST00000130608.1 | 865 | Pdyn | ENSMUSG00000027400.6 |
| chr2 | 129699185 | 129699976 | 3.41775 | 3.98844 | 1.13343 | Promoter | 2 | 129686565 | 129699833 | 13269 | 2 | ENSMUST00000028883.6 | 0 | Pdyn | ENSMUSG00000027400.6 |
| chr2 | 129700618 | 129700693 | 3.32371 | 3.81283 | 1.01733 | Promoter | 2 | 129697986 | 129699844 | 1859 | 2 | ENSMUST00000152113.1 | -775 | Pdyn | ENSMUSG00000027400.6 |
| chr2 | 130397264 | 130398104 | 3.74631 | 4.59991 | 1.47851 | Promoter | 2 | 130390775 | 130397574 | 6800 | 2 | ENSMUST00000028897.7 | 0 | Cpxm1 | ENSMUSG00000027408.7 |
| chr2 | 147185416 | 147186910 | 4.07472 | 5.32421 | 1.98322 | Promoter | 2 | 147177546 | 147186101 | 8556 | 2 | ENSMUST00000109970.4 | 0 | Nkx2-2 | ENSMUSG00000027434.11 |
| chr2 | 147192931 | 147193836 | 4.05949 | 5.23752 | 1.9806 | Promoter | 2 | 147186382 | 147194242 | 7861 | 2 | ENSMUST00000172627.1 | 406 | Nkx2-2 | ENSMUSG00000027434.11 |
| chr2 | 147194164 | 147195780 | 4.1059 | 5.66368 | 2.29675 | Promoter | 2 | 147186305 | 147194243 | 7939 | 2 | ENSMUST00000109969.2 | 0 | Nkx2-2 | ENSMUSG00000027434.11 |
| chr2 | 147196494 | 147196620 | 4.2553 | 5.88041 | 2.37162 | Intron (ENSMUST00000136998.2/ENSMUSG00000086509.2, intron 1 of 5) | 2 | 147186305 | 147194243 | 7939 | 2 | ENSMUST00000109969.2 | -2252 | Nkx2-2 | ENSMUSG00000027434.11 |
| chr2 | 147191833 | 147191941 | 3.97351 | 5.02676 | 1.79507 | Intron (ENSMUST00000136998.2/ENSMUSG00000086509.2, intron 1 of 5) | 2 | 147186382 | 147194242 | 7861 | 2 | ENSMUST00000172627.1 | 2301 | Nkx2-2 | ENSMUSG00000027434.11 |
| chr2 | 147202936 | 147203049 | 3.33576 | 3.89582 | 1.11394 | Intron (ENSMUST00000136998.2/ENSMUSG00000086509.2, intron 1 of 5) | 2 | 147186305 | 147194243 | 7939 | 2 | ENSMUST00000109969.2 | -8694 | Nkx2-2 | ENSMUSG00000027434.11 |
| chr2 | 147206187 | 147206428 | 3.78203 | 4.74961 | 1.62508 | Intron (ENSMUST00000136998.2/ENSMUSG00000086509.2, intron 1 of 5) | 2 | 147186305 | 147194243 | 7939 | 2 | ENSMUST00000109969.2 | -11945 | Nkx2-2 | ENSMUSG00000027434.11 |
| chr2 | 147197415 | 147197745 | 3.52702 | 4.24151 | 1.33694 | Intron (ENSMUST00000136998.2/ENSMUSG00000086509.2, intron 1 of 5) | 2 | 147186305 | 147194243 | 7939 | 2 | ENSMUST00000109969.2 | -3173 | Nkx2-2 | ENSMUSG00000027434.11 |
| chr2 | 150668159 | 150668327 | 3.96593 | 5.0213 | 1.7969 | Promoter | 2 | 150618105 | 150668258 | 50154 | 2 | ENSMUST00000028944.3 | 0 | Acss1 | ENSMUSG00000027452.6 |
| chr2 | 151973120 | 151973459 | 3.43173 | 4.00086 | 1.10632 | Promoter | 2 | 151969737 | 151973394 | 3658 | 2 | ENSMUST00000109863.1 | 0 | Fam110a | ENSMUSG00000027459.11 |
| chr2 | 151973997 | 151974073 | 4.23887 | 5.5565 | 2.17434 | Promoter | 2 | 151969398 | 151973966 | 4569 | 2 | ENSMUST00000109864.2 | -32 | Fam110a | ENSMUSG00000027459.11 |
| chr2 | 151916215 | 151916340 | 3.2754 | 3.89347 | 1.05428 | Intron (ENSMUST00000028955.5/ENSMUSG00000027460.7, intron 1 of 8) | 2 | 151911210 | 151945337 | 34128 | 1 | ENSMUST00000028955.5 | 5006 | Angpt4 | ENSMUSG00000027460.7 |
| chr2 | 151916882 | 151917337 | 3.60882 | 4.41749 | 1.45002 | Intron (ENSMUST00000028955.5/ENSMUSG00000027460.7, intron 1 of 8) | 2 | 151911210 | 151945337 | 34128 | 1 | ENSMUST00000028955.5 | 5673 | Angpt4 | ENSMUSG00000027460.7 |
| chr2 | 152909926 | 152910265 | 3.32353 | 3.83382 | 1.08888 | Distal Intergenic | 2 | 152911352 | 152923068 | 11717 | 1 | ENSMUST00000028970.7 | -1087 | Mylk2 | ENSMUSG00000027470.9 |
| chr2 | 152911322 | 152911519 | 3.24736 | 3.69819 | 1.01733 | Promoter | 2 | 152911352 | 152923068 | 11717 | 1 | ENSMUST00000028970.7 | 0 | Mylk2 | ENSMUSG00000027470.9 |
| chr2 | 153067270 | 153067750 | 3.58804 | 4.28822 | 1.28451 | Intron (ENSMUST00000127898.2/ENSMUSG00000027474.7, intron 1 of 5) | 2 | 153066002 | 153081735 | 15734 | 1 | ENSMUST00000109800.1 | 1269 | Ccm2l | ENSMUSG00000027474.7 |
| chr2 | 153070435 | 153070912 | 3.51305 | 4.17024 | 1.20797 | Exon (ENSMUST00000127898.2/ENSMUSG00000027474.7, exon 3 of 10) | 2 | 153066002 | 153081735 | 15734 | 1 | ENSMUST00000109800.1 | 4434 | Ccm2l | ENSMUSG00000027474.7 |
| chr2 | 153071508 | 153073029 | 4.01949 | 5.25944 | 1.96319 | Exon (ENSMUST00000127898.2/ENSMUSG00000027474.7, exon 5 of 10) | 2 | 153066002 | 153081735 | 15734 | 1 | ENSMUST00000109800.1 | 5507 | Ccm2l | ENSMUSG00000027474.7 |
| chr3 | 8510150 | 8510682 | 3.85123 | 4.78128 | 1.59126 | Promoter | 3 | 8509527 | 8561604 | 52078 | 1 | ENSMUST00000029002.7 | 624 | Stmn2 | ENSMUSG00000027500.9 |
| chr3 | 8511134 | 8511670 | 3.74967 | 4.60757 | 1.47583 | Intron (ENSMUST00000029002.7/ENSMUSG00000027500.9, intron 1 of 4) | 3 | 8509527 | 8561604 | 52078 | 1 | ENSMUST00000029002.7 | 1608 | Stmn2 | ENSMUSG00000027500.9 |
| chr3 | 8512190 | 8512515 | 3.38193 | 3.9006 | 1.01733 | Intron (ENSMUST00000029002.7/ENSMUSG00000027500.9, intron 1 of 4) | 3 | 8509527 | 8561604 | 52078 | 1 | ENSMUST00000029002.7 | 2664 | Stmn2 | ENSMUSG00000027500.9 |
| chr2 | 173143729 | 173144002 | 3.85682 | 5.03233 | 1.81138 | Distal Intergenic | 2 | 173153048 | 173159273 | 6226 | 1 | ENSMUST00000029017.5 | -9046 | Pck1 | ENSMUSG00000027513.6 |
| chr2 | 173145011 | 173145537 | 3.47011 | 4.06685 | 1.13442 | Distal Intergenic | 2 | 173153048 | 173159273 | 6226 | 1 | ENSMUST00000029017.5 | -7511 | Pck1 | ENSMUSG00000027513.6 |
| chr2 | 173184639 | 173184941 | 3.50913 | 4.17569 | 1.2756 | Distal Intergenic | 2 | 173153082 | 173159271 | 6190 | 1 | ENSMUST00000136885.1 | 31558 | Pck1 | ENSMUSG00000027513.6 |
| chr2 | 173202637 | 173202712 | 4.22883 | 5.51277 | 2.10159 | Distal Intergenic | 2 | 173206615 | 173218897 | 12283 | 2 | ENSMUST00000146802.1 | 16185 | Zbp1 | ENSMUSG00000027514.9 |
| chr2 | 173211994 | 173212427 | 3.30496 | 3.79029 | 1.03499 | Exon (ENSMUST00000029018.6/ENSMUSG00000027514.9, exon 4 of 8) | 2 | 173206615 | 173218897 | 12283 | 2 | ENSMUST00000146802.1 | 6470 | Zbp1 | ENSMUSG00000027514.9 |
| chr2 | 173570745 | 173570828 | 3.63848 | 4.39947 | 1.40049 | Intron (ENSMUST00000119453.2/ENSMUSG00000027517.8, intron 3 of 3) | 2 | 173568704 | 173578198 | 9495 | 2 | ENSMUST00000109112.1 | 7370 | Ankrd60 | ENSMUSG00000027517.8 |
| chr2 | 173577154 | 173577313 | 3.69932 | 4.71061 | 1.60705 | Promoter | 2 | 173568704 | 173578198 | 9495 | 2 | ENSMUST00000109112.1 | 885 | Ankrd60 | ENSMUSG00000027517.8 |
| chr1 | 63272724 | 63273336 | 4.4248 | 6.00658 | 2.53079 | Promoter | 1 | 63273265 | 63314576 | 41312 | 1 | ENSMUST00000114132.2 | 0 | Zdbf2 | ENSMUSG00000027520.10 |
| chr2 | 174297103 | 174298775 | 3.87227 | 4.93472 | 1.71598 | Promoter | 2 | 174297859 | 174302018 | 4160 | 1 | ENSMUST00000087877.5 | 0 | Gnas | ENSMUSG00000027523.14 |
| chr2 | 174299094 | 174299499 | 3.66157 | 4.41219 | 1.33076 | Exon (ENSMUST00000087877.5/ENSMUSG00000027523.14, exon 1 of 2) | 2 | 174297859 | 174346744 | 48886 | 1 | ENSMUST00000087876.5 | 1236 | Gnas | ENSMUSG00000027523.14 |

| | | | | | | | | | | | | | | | |
|---|---|---|---|---|---|---|---|---|---|---|---|---|---|---|---|
| chr2 | 174759890 | 174760246 | 3.30961 | 3.79182 | 1.01733 | Promoter | 2 | 174760619 | 174784042 | 23424 | 1 | ENSMUST00000029030.3 | -373 | Edn3 | ENSMUSG00000027524.4 |
| chr2 | 174760680 | 174760863 | 3.90076 | 4.99558 | 1.7862 | Promoter | 2 | 174760799 | 174781980 | 21182 | 1 | ENSMUST00000140908.1 | 0 | Edn3 | ENSMUSG00000027524.4 |
| chr2 | 178118968 | 178119163 | 4.41556 | 6.33687 | 2.76751 | Promoter | 2 | 178118975 | 178338492 | 219518 | 1 | ENSMUST00000108917.2 | 0 | Phactr3 | ENSMUSG00000027525.12 |
| chr2 | 178119504 | 178119797 | 3.87258 | 4.8334 | 1.65972 | Promoter | 2 | 178119179 | 178336491 | 217313 | 1 | ENSMUST00000108916.2 | 326 | Phactr3 | ENSMUSG00000027525.12 |
| chr2 | 178141640 | 178143102 | 3.69053 | 4.78364 | 1.65105 | Promoter | 2 | 178141933 | 178336072 | 194140 | 1 | ENSMUST00000103066.4 | 0 | Phactr3 | ENSMUSG00000027525.12 |
| chr2 | 168525716 | 168526079 | 3.96249 | 5.06241 | 1.79712 | Intron (ENSMUST00000074618.4/ENSMUSG00000027544.11, intron 5 of 9) | 2 | 168476413 | 168552276 | 75864 | 2 | ENSMUST00000151292.2 | 26197 | Nfatc2 | ENSMUSG00000027544.11 |
| chr2 | 168526398 | 168526547 | 3.63144 | 4.36683 | 1.32555 | Intron (ENSMUST00000074618.4/ENSMUSG00000027544.11, intron 5 of 9) | 2 | 168476413 | 168552276 | 75864 | 2 | ENSMUST00000151292.2 | 25729 | Nfatc2 | ENSMUSG00000027544.11 |
| chr2 | 168528488 | 168528755 | 3.31288 | 3.79662 | 1.01733 | Intron (ENSMUST00000074618.4/ENSMUSG00000027544.11, intron 5 of 9) | 2 | 168476413 | 168552276 | 75864 | 2 | ENSMUST00000151292.2 | 23521 | Nfatc2 | ENSMUSG00000027544.11 |
| chr2 | 168532325 | 168532586 | 3.42735 | 4.02741 | 1.19756 | Intron (ENSMUST00000074618.4/ENSMUSG00000027544.11, intron 5 of 9) | 2 | 168476413 | 168552276 | 75864 | 2 | ENSMUST00000151292.2 | 19690 | Nfatc2 | ENSMUSG00000027544.11 |
| chr2 | 168589258 | 168589528 | 3.77726 | 4.72312 | 1.64388 | Promoter | 2 | 168519098 | 168590191 | 71094 | 2 | ENSMUST00000029057.7 | 663 | Nfatc2 | ENSMUSG00000027544.11 |
| chr2 | 168589866 | 168590375 | 3.58416 | 4.29873 | 1.3325 | Promoter | 2 | 168476410 | 168590260 | 113851 | 2 | ENSMUST00000074618.4 | 0 | Nfatc2 | ENSMUSG00000027544.11 |
| chr2 | 168598128 | 168598650 | 3.34113 | 3.87868 | 1.11108 | Intron (ENSMUST00000099067.4/ENSMUSG00000027544.11, intron 1 of 6) | 2 | 168518195 | 168601632 | 83438 | 2 | ENSMUST00000099067.4 | 2982 | Nfatc2 | ENSMUSG00000027544.11 |
| chr2 | 168599544 | 168600303 | 3.93789 | 5.03567 | 1.87024 | Intron (ENSMUST00000099067.4/ENSMUSG00000027544.11, intron 1 of 6) | 2 | 168518195 | 168601632 | 83438 | 2 | ENSMUST00000099067.4 | 1329 | Nfatc2 | ENSMUSG00000027544.11 |
| chr2 | 168603134 | 168603557 | 3.7227 | 4.59506 | 1.54774 | Distal Intergenic | 2 | 168519098 | 168601657 | 82560 | 2 | ENSMUST00000137451.1 | -1478 | Nfatc2 | ENSMUSG00000027544.11 |
| chr2 | 168760734 | 168761143 | 3.49872 | 4.33672 | 1.35058 | Intron (ENSMUST00000075044.4/ENSMUSG00000027547.12, intron 1 of 2) | 2 | 168756181 | 168765246 | 9066 | 2 | ENSMUST00000137536.1 | 4103 | Sall4 | ENSMUSG00000027547.12 |
| chr2 | 168762211 | 168762355 | 3.80845 | 4.85262 | 1.66751 | Intron (ENSMUST00000075044.4/ENSMUSG00000027547.12, intron 1 of 2) | 2 | 168756181 | 168765246 | 9066 | 2 | ENSMUST00000137536.1 | 2891 | Sall4 | ENSMUSG00000027547.12 |
| chr2 | 168763292 | 168763566 | 3.82541 | 4.7474 | 1.62018 | Intron (ENSMUST00000075044.4/ENSMUSG00000027547.12, intron 1 of 2) | 2 | 168756181 | 168765246 | 9066 | 2 | ENSMUST00000137536.1 | 1680 | Sall4 | ENSMUSG00000027547.12 |
| chr2 | 168764434 | 168764663 | 3.32348 | 3.81506 | 1.02638 | Promoter | 2 | 168756181 | 168765246 | 9066 | 2 | ENSMUST00000137536.1 | 583 | Sall4 | ENSMUSG00000027547.12 |
| chr2 | 168765078 | 168767180 | 4.27308 | 5.65083 | 2.23151 | Promoter | 2 | 168748332 | 168767029 | 18698 | 2 | ENSMUST00000075044.4 | 0 | Sall4 | ENSMUSG00000027547.12 |
| chr2 | 168767746 | 168768039 | 3.4385 | 4.3289 | 1.34112 | Promoter | 2 | 168756614 | 168767943 | 11330 | 2 | ENSMUST00000125138.1 | 0 | Sall4 | ENSMUSG00000027547.12 |
| chr2 | 168768589 | 168768722 | 3.61107 | 4.39495 | 1.4273 | Promoter | 2 | 168756614 | 168767943 | 11330 | 2 | ENSMUST00000125138.1 | -647 | Sall4 | ENSMUSG00000027547.12 |
| chr2 | 168769937 | 168770012 | 3.38193 | 3.9006 | 1.01733 | Distal Intergenic | 2 | 168756614 | 168767943 | 11330 | 2 | ENSMUST00000125138.1 | -1995 | Sall4 | ENSMUSG00000027547.12 |
| chr2 | 168771886 | 168772065 | 3.31183 | 3.79495 | 1.01733 | Distal Intergenic | 2 | 168756614 | 168767943 | 11330 | 2 | ENSMUST00000125138.1 | -3944 | Sall4 | ENSMUSG00000027547.12 |
| chr2 | 168791267 | 168791574 | 3.4504 | 4.03193 | 1.11621 | Distal Intergenic | 2 | 168756614 | 168767943 | 11330 | 2 | ENSMUST00000125138.1 | -23325 | Sall4 | ENSMUSG00000027547.12 |
| chr2 | 168799418 | 168799505 | 3.64387 | 4.38175 | 1.31406 | Distal Intergenic | 2 | 168756614 | 168767943 | 11330 | 2 | ENSMUST00000125138.1 | -31476 | Sall4 | ENSMUSG00000027547.12 |
| chr2 | 168803990 | 168804100 | 3.86442 | 4.84637 | 1.69447 | Distal Intergenic | 2 | 168756614 | 168767943 | 11330 | 2 | ENSMUST00000125138.1 | -36048 | Sall4 | ENSMUSG00000027547.12 |
| chr2 | 168805948 | 168806236 | 3.41519 | 4.00098 | 1.18988 | Distal Intergenic | 2 | 168756614 | 168767943 | 11330 | 2 | ENSMUST00000125138.1 | -38006 | Sall4 | ENSMUSG00000027547.12 |
| chr2 | 168808702 | 168808965 | 3.35126 | 3.96136 | 1.12556 | Distal Intergenic | 2 | 168756614 | 168767943 | 11330 | 2 | ENSMUST00000125138.1 | -40760 | Sall4 | ENSMUSG00000027547.12 |
| chr2 | 168809405 | 168809777 | 3.47638 | 4.08536 | 1.16616 | Distal Intergenic | 2 | 168756614 | 168767943 | 11330 | 2 | ENSMUST00000125138.1 | -41463 | Sall4 | ENSMUSG00000027547.12 |
| chr2 | 168810139 | 168810360 | 3.58877 | 4.3002 | 1.32114 | Distal Intergenic | 2 | 168756614 | 168767943 | 11330 | 2 | ENSMUST00000125138.1 | -42197 | Sall4 | ENSMUSG00000027547.12 |
| chr2 | 168810845 | 168811320 | 3.71121 | 4.5174 | 1.42444 | Distal Intergenic | 2 | 168756614 | 168767943 | 11330 | 2 | ENSMUST00000125138.1 | -42903 | Sall4 | ENSMUSG00000027547.12 |
| chr2 | 168813594 | 168813685 | 3.44552 | 4.02848 | 1.13028 | Distal Intergenic | 2 | 168756614 | 168767943 | 11330 | 2 | ENSMUST00000125138.1 | -45652 | Sall4 | ENSMUSG00000027547.12 |
| chr2 | 168815406 | 168815642 | 3.85147 | 4.78563 | 1.61263 | Distal Intergenic | 2 | 168756614 | 168767943 | 11330 | 2 | ENSMUST00000125138.1 | -47464 | Sall4 | ENSMUSG00000027547.12 |
| chr2 | 168816737 | 168817070 | 3.56285 | 4.27355 | 1.28265 | Distal Intergenic | 2 | 168756614 | 168767943 | 11330 | 2 | ENSMUST00000125138.1 | -48795 | Sall4 | ENSMUSG00000027547.12 |
| chr2 | 168818603 | 168818878 | 3.50203 | 4.12444 | 1.16822 | Distal Intergenic | 2 | 168756614 | 168767943 | 11330 | 2 | ENSMUST00000125138.1 | -50661 | Sall4 | ENSMUSG00000027547.12 |
| chr2 | 168910480 | 168911302 | 3.99974 | 5.12121 | 1.85591 | Intron (ENSMUST00000109162.3/ENSMUSG00000027551.10, intron 5 of 8) | 2 | 168906842 | 168912633 | 5792 | 2 | ENSMUST00000124599.1 | 1331 | Zfp64 | ENSMUSG00000027551.10 |
| chr2 | 168911689 | 168912679 | 3.70302 | 4.68571 | 1.57586 | Promoter | 2 | 168906842 | 168912633 | 5792 | 2 | ENSMUST00000124599.1 | 0 | Zfp64 | ENSMUSG00000027551.10 |
| chr3 | 14862468 | 14863256 | 4.21204 | 5.61849 | 2.25341 | Promoter | 3 | 14863538 | 14872351 | 8814 | 1 | ENSMUST00000029076.4 | -282 | Car3 | ENSMUSG00000027559.4 |
| chr3 | 14864446 | 14864681 | 3.25773 | 3.71372 | 1.01733 | Promoter | 3 | 14863538 | 14872351 | 8814 | 1 | ENSMUST00000029076.4 | 909 | Car3 | ENSMUSG00000027559.4 |
| chr2 | 170730896 | 170731066 | 3.61548 | 4.48077 | 1.44025 | Promoter | 2 | 170731807 | 170879769 | 147963 | 1 | ENSMUST00000029075.4 | -741 | Dok5 | ENSMUSG00000027560.4 |
| chr2 | 170731415 | 170732009 | 3.58297 | 4.2852 | 1.28571 | Promoter | 2 | 170731807 | 170879769 | 147963 | 1 | ENSMUST00000029075.4 | 0 | Dok5 | ENSMUSG00000027560.4 |
| chr2 | 170772446 | 170772570 | 3.44593 | 4.04147 | 1.17141 | Intron (ENSMUST00000029075.4/ENSMUSG00000027560.4, intron 1 of 7) | 2 | 170731807 | 170879769 | 147963 | 1 | ENSMUST00000029075.4 | 40640 | Dok5 | ENSMUSG00000027560.4 |
| chr3 | 14885239 | 14885503 | 4.15559 | 5.44666 | 2.13668 | Promoter | 3 | 14886426 | 14900769 | 14344 | 1 | ENSMUST00000029078.7 | -923 | Car2 | ENSMUSG00000027562.7 |
| chr3 | 14886208 | 14887293 | 4.38601 | 6.09579 | 2.60395 | Promoter | 3 | 14886426 | 14900769 | 14344 | 1 | ENSMUST00000029078.7 | 0 | Car2 | ENSMUSG00000027562.7 |
| chr2 | 180499496 | 180499593 | 3.73997 | 4.7157 | 1.57616 | Promoter | 2 | 180499976 | 180544980 | 45005 | 1 | ENSMUST00000029084.3 | -383 | Ntsr1 | ENSMUSG00000027568.4 |
| chr2 | 180500519 | 180501638 | 3.45618 | 4.04856 | 1.14383 | Promoter | 2 | 180500227 | 180544219 | 43993 | 1 | ENSMUST00000170448.1 | 293 | Ntsr1 | ENSMUSG00000027568.4 |
| chr2 | 180506349 | 180506670 | 3.40476 | 4.02453 | 1.18486 | Intron (ENSMUST00000029084.3/ENSMUSG00000027568.4, intron 1 of 3) | 2 | 180500227 | 180544219 | 43993 | 1 | ENSMUST00000170448.1 | 6123 | Ntsr1 | ENSMUSG00000027568.4 |
| chr2 | 180955502 | 180956016 | 3.92947 | 5.00449 | 1.8099 | Promoter | 2 | 180934772 | 180954699 | 19928 | 2 | ENSMUST00000108873.3 | -804 | Nkain4 | ENSMUSG00000027574.10 |
| chr2 | 181038316 | 181038437 | 3.42489 | 3.98209 | 1.07593 | Promoter | 2 | 181018380 | 181039187 | 20808 | 2 | ENSMUST00000108851.2 | 750 | Chrna4 | ENSMUSG00000027577.9 |
| chr2 | 181038886 | 181039820 | 4.17806 | 5.52948 | 2.14871 | Promoter | 2 | 181018380 | 181039187 | 20808 | 2 | ENSMUST00000108851.2 | 0 | Chrna4 | ENSMUSG00000027577.9 |
| chr2 | 181040487 | 181040643 | 3.37315 | 3.88735 | 1.01733 | Intron (ENSMUST00000124400.1/ENSMUSG00000027577.9, intron 1 of 3) | 2 | 181022311 | 181039305 | 16995 | 2 | ENSMUST00000067120.8 | -1183 | Chrna4 | ENSMUSG00000027577.9 |
| chr2 | 181212010 | 181212369 | 3.8234 | 4.86858 | 1.70788 | Promoter | 2 | 181205562 | 181213185 | 7624 | 2 | ENSMUST00000016498.4 | 816 | Srms | ENSMUSG00000027579.6 |
| chr2 | 181212731 | 181213164 | 3.35866 | 4.09597 | 1.22657 | Promoter | 2 | 181205562 | 181213185 | 7624 | 2 | ENSMUST00000016498.4 | 21 | Srms | ENSMUSG00000027579.6 |
| chr2 | 181306229 | 181306338 | 3.54101 | 4.20331 | 1.23828 | Downstream (<1kb) | 2 | 181306459 | 181314500 | 8042 | 2 | ENSMUST00000103045.3 | 8162 | Stmn3 | ENSMUSG00000027581.12 |
| chr2 | 181312709 | 181313132 | 3.58569 | 4.40694 | 1.40773 | Intron (ENSMUST00000103045.3/ENSMUSG00000027581.12, intron 1 of 4) | 2 | 181306459 | 181314500 | 8042 | 2 | ENSMUST00000103045.3 | 1368 | Stmn3 | ENSMUSG00000027581.12 |
| chr2 | 154951959 | 154952034 | 3.5788 | 4.59659 | 1.51899 | Promoter | 2 | 154951219 | 155051011 | 99793 | 1 | ENSMUST00000109697.2 | 741 | a | ENSMUSG00000027596.5 |
| chr2 | 155514315 | 155514650 | 3.66785 | 4.53754 | 1.47893 | Promoter | 2 | 155490379 | 155514848 | 24470 | 2 | ENSMUST00000029131.5 | 198 | Ggt7 | ENSMUSG00000027603.10 |
| chr2 | 155750874 | 155750963 | 3.73757 | 4.57162 | 1.47458 | Promoter | 2 | 155751117 | 155755471 | 4355 | 1 | ENSMUST00000029140.6 | -154 | Procr | ENSMUSG00000027611.7 |
| chr2 | 155753458 | 155753636 | 3.68523 | 4.4731 | 1.39859 | Exon (ENSMUST00000029140.6/ENSMUSG00000027611.7, exon 2 of 4) | 2 | 155751344 | 155755468 | 4125 | 1 | ENSMUST00000143493.1 | 2115 | Procr | ENSMUSG00000027611.7 |
| chr2 | 155754630 | 155754849 | 3.47744 | 4.21278 | 1.26924 | Exon (ENSMUST00000029140.6/ENSMUSG00000027611.7, exon 4 of 4) | 2 | 155751344 | 155755468 | 4125 | 1 | ENSMUST00000143493.1 | 3287 | Procr | ENSMUSG00000027611.7 |
| chr2 | 155770016 | 155770275 | 3.64746 | 4.50708 | 1.45985 | Distal Intergenic | 2 | 155775342 | 155818366 | 43025 | 1 | ENSMUST00000029141.5 | -5067 | Mmp24 | ENSMUSG00000027612.7 |
| chr2 | 155772547 | 155772802 | 3.37718 | 3.89343 | 1.01733 | Distal Intergenic | 2 | 155775342 | 155818366 | 43025 | 1 | ENSMUST00000029141.5 | -2540 | Mmp24 | ENSMUSG00000027612.7 |
| chr2 | 155773564 | 155773777 | 3.55573 | 4.25125 | 1.2574 | Distal Intergenic | 2 | 155775342 | 155818366 | 43025 | 1 | ENSMUST00000029141.5 | -1565 | Mmp24 | ENSMUSG00000027612.7 |
| chr2 | 155774248 | 155775196 | 3.71391 | 4.53602 | 1.44213 | Promoter | 2 | 155775342 | 155818366 | 43025 | 1 | ENSMUST00000029141.5 | -146 | Mmp24 | ENSMUSG00000027612.7 |
| chr2 | 155776098 | 155776614 | 3.47485 | 4.345 | 1.32892 | Promoter | 2 | 155775342 | 155818366 | 43025 | 1 | ENSMUST00000029141.5 | 757 | Mmp24 | ENSMUSG00000027612.7 |
| chr2 | 155777069 | 155777175 | 4.18863 | 5.45932 | 2.08772 | Intron (ENSMUST00000029141.5/ENSMUSG00000027612.7, intron 1 of 8) | 2 | 155775342 | 155818366 | 43025 | 1 | ENSMUST00000029141.5 | 1728 | Mmp24 | ENSMUSG00000027612.7 |
| chr2 | 155778219 | 155778404 | 3.48068 | 4.10295 | 1.20737 | Intron (ENSMUST00000029141.5/ENSMUSG00000027612.7, intron 1 of 8) | 2 | 155775342 | 155818366 | 43025 | 1 | ENSMUST00000029141.5 | 2878 | Mmp24 | ENSMUSG00000027612.7 |
| chr2 | 155779845 | 155780031 | 3.91626 | 4.90174 | 1.67455 | Intron (ENSMUST00000029141.5/ENSMUSG00000027612.7, intron 1 of 8) | 2 | 155775342 | 155818366 | 43025 | 1 | ENSMUST00000029141.5 | 4504 | Mmp24 | ENSMUSG00000027612.7 |
| chr2 | 155781023 | 155781148 | 3.33705 | 3.83292 | 1.01733 | Intron (ENSMUST00000029141.5/ENSMUSG00000027612.7, intron 1 of 8) | 2 | 155775342 | 155818366 | 43025 | 1 | ENSMUST00000029141.5 | 5682 | Mmp24 | ENSMUSG00000027612.7 |
| chr2 | 155788609 | 155788748 | 3.63137 | 4.45936 | 1.44402 | Intron (ENSMUST00000029141.5/ENSMUSG00000027612.7, intron 1 of 8) | 2 | 155775342 | 155818366 | 43025 | 1 | ENSMUST00000029141.5 | 13268 | Mmp24 | ENSMUSG00000027612.7 |
| chr2 | 156420063 | 156421243 | 3.62637 | 4.40575 | 1.44181 | Promoter | 2 | 156420909 | 156543214 | 122306 | 1 | ENSMUST00000103137.4 | 0 | Epb4.1l1 | ENSMUSG00000027624.12 |
| chr2 | 156421758 | 156422177 | 3.68671 | 4.5021 | 1.47322 | Promoter | 2 | 156421112 | 156543214 | 122103 | 1 | ENSMUST00000109577.3 | 647 | Epb4.1l1 | ENSMUSG00000027624.12 |
| chr2 | 156423933 | 156424245 | 3.39231 | 3.96498 | 1.10871 | Intron (ENSMUST00000103137.4/ENSMUSG00000027624.12, intron 1 of 22) | 2 | 156421112 | 156543214 | 122103 | 1 | ENSMUST00000109577.3 | 2822 | Epb4.1l1 | ENSMUSG00000027624.12 |
| chr2 | 156429802 | 156429955 | 3.72751 | 4.69222 | 1.56045 | Intron (ENSMUST00000103137.4/ENSMUSG00000027624.12, intron 1 of 22) | 2 | 156421112 | 156543214 | 122103 | 1 | ENSMUST00000109577.3 | 8691 | Epb4.1l1 | ENSMUSG00000027624.12 |
| chr2 | 157418454 | 157418529 | 3.37304 | 3.90729 | 1.08852 | Promoter | 2 | 157418444 | 157471835 | 53392 | 1 | ENSMUST00000109533.2 | 11 | Src | ENSMUSG00000027646.10 |
| chr2 | 157970453 | 157970742 | 4.10978 | 5.34636 | 2.05903 | Distal Intergenic | 2 | 157995280 | 158007212 | 11933 | 2 | ENSMUST00000126541.1 | 36470 | Tti1 | ENSMUSG00000027650.7 |
| chr3 | 33063956 | 33064217 | 3.70341 | 4.62335 | 1.55828 | Intron (ENSMUST00000125494.1/ENSMUSG00000087625.1, intron 3 of 7) | 3 | 32949408 | 33082068 | 132661 | 2 | ENSMUST00000108225.4 | 17851 | Pex5l | ENSMUSG00000027674.11 |
| chr3 | 33139115 | 33139301 | 3.5936 | 4.33618 | 1.3821 | Intron (ENSMUST00000108226.2/ENSMUSG00000027674.11, intron 1 of 12) | 3 | 33131566 | 33143076 | 11511 | 2 | ENSMUST00000155264.1 | 3775 | Pex5l | ENSMUSG00000027674.11 |
| chr3 | 33141319 | 33142004 | 3.55095 | 4.21437 | 1.21657 | Intron (ENSMUST00000108226.2/ENSMUSG00000027674.11, intron 1 of 12) | 3 | 33131566 | 33143076 | 11511 | 2 | ENSMUST00000155264.1 | 1072 | Pex5l | ENSMUSG00000027674.11 |
| chr3 | 33142333 | 33143423 | 3.84905 | 4.81979 | 1.6452 | Promoter | 3 | 32949954 | 33143082 | 193129 | 2 | ENSMUST00000108226.2 | 0 | Pex5l | ENSMUSG00000027674.11 |
| chr3 | 33145759 | 33145964 | 3.46986 | 4.13987 | 1.24029 | Distal Intergenic | 3 | 32955866 | 33143247 | 187382 | 2 | ENSMUST00000108219.1 | -2513 | Pex5l | ENSMUSG00000027674.11 |
| chr3 | 33147166 | 33147444 | 3.73503 | 4.61364 | 1.57724 | Distal Intergenic | 3 | 32955866 | 33143247 | 187382 | 2 | ENSMUST00000108219.1 | -3920 | Pex5l | ENSMUSG00000027674.11 |
| chr3 | 30011449 | 30011635 | 3.63439 | 4.44046 | 1.43505 | Promoter | 3 | 29997729 | 30011439 | 13711 | 2 | ENSMUST00000174406.1 | -11 | Mecom | ENSMUSG00000027684.11 |
| chr3 | 30514004 | 30516485 | 4.17354 | 5.69268 | 2.35576 | Intron (ENSMUST00000172754.2/ENSMUSG00000027684.11, intron 2 of 8) | 3 | 29965456 | 30509822 | 544367 | 2 | ENSMUST00000172697.2 | -4183 | Mecom | ENSMUSG00000027684.11 |
| chr3 | 30014990 | 30015155 | 3.70312 | 4.83889 | 1.65348 | Exon (ENSMUST00000173411.2/ENSMUSG00000027684.11, exon 3 of 3) | 3 | 29951306 | 30013505 | 62200 | 2 | ENSMUST00000108270.4 | -1486 | Mecom | ENSMUSG00000027684.11 |
| chr3 | 30219983 | 30220746 | 3.65052 | 4.41415 | 1.38077 | Intron (ENSMUST00000173495.2/ENSMUSG00000027684.11, intron 1 of 15) | 3 | 29952567 | 30238162 | 285596 | 2 | ENSMUST00000173059.2 | 17416 | Mecom | ENSMUSG00000027684.11 |
| chr3 | 30519649 | 30519757 | 3.47596 | 4.07545 | 1.1334 | Intron (ENSMUST00000172754.2/ENSMUSG00000027684.11, intron 2 of 8) | 3 | 29965456 | 30509822 | 544367 | 2 | ENSMUST00000172697.2 | -9828 | Mecom | ENSMUSG00000027684.11 |
| chr3 | 30476355 | 30476430 | 3.67846 | 4.47259 | 1.444 | Intron (ENSMUST00000173495.2/ENSMUSG00000027684.11, intron 1 of 15) | 3 | 30236569 | 30509487 | 272919 | 2 | ENSMUST00000173022.1 | 33057 | Mecom | ENSMUSG00000027684.11 |
| chr3 | 30500147 | 30500319 | 3.2938 | 3.76787 | 1.01733 | Intron (ENSMUST00000173495.2/ENSMUSG00000027684.11, intron 1 of 15) | 3 | 30236569 | 30509487 | 272919 | 2 | ENSMUST00000173022.1 | 9168 | Mecom | ENSMUSG00000027684.11 |
| chr3 | 30502176 | 30502544 | 4.443 | 6.02761 | 2.56767 | Intron (ENSMUST00000173495.2/ENSMUSG00000027684.11, intron 1 of 15) | 3 | 30236569 | 30509487 | 272919 | 2 | ENSMUST00000173022.1 | 6943 | Mecom | ENSMUSG00000027684.11 |
| chr3 | 30503009 | 30503293 | 3.5737 | 4.30773 | 1.37028 | Intron (ENSMUST00000173495.2/ENSMUSG00000027684.11, intron 1 of 15) | 3 | 30236569 | 30509487 | 272919 | 2 | ENSMUST00000173022.1 | 6194 | Mecom | ENSMUSG00000027684.11 |
| chr3 | 30504569 | 30505087 | 3.64928 | 4.42182 | 1.38382 | Intron (ENSMUST00000173495.2/ENSMUSG00000027684.11, intron 1 of 15) | 3 | 30236569 | 30509487 | 272919 | 2 | ENSMUST00000173022.1 | 4400 | Mecom | ENSMUSG00000027684.11 |
| chr3 | 30505782 | 30507254 | 3.57848 | 4.42614 | 1.39704 | Intron (ENSMUST00000173495.2/ENSMUSG00000027684.11, intron 1 of 15) | 3 | 30236569 | 30509487 | 272919 | 2 | ENSMUST00000173022.1 | 2233 | Mecom | ENSMUSG00000027684.11 |
| chr3 | 30507580 | 30507996 | 3.49642 | 4.26565 | 1.32817 | Intron (ENSMUST00000173495.2/ENSMUSG00000027684.11, intron 1 of 15) | 3 | 30236569 | 30509487 | 272919 | 2 | ENSMUST00000173022.1 | 1491 | Mecom | ENSMUSG00000027684.11 |
| chr3 | 30508349 | 30509975 | 3.71305 | 4.67569 | 1.61773 | Promoter | 3 | 29951299 | 30509733 | 558435 | 2 | ENSMUST00000173495.2 | 0 | Mecom | ENSMUSG00000027684.11 |
| chr3 | 30513190 | 30513651 | 3.67345 | 4.70028 | 1.56099 | Intron (ENSMUST00000172754.2/ENSMUSG00000027684.11, intron 2 of 8) | 3 | 29965456 | 30509822 | 544367 | 2 | ENSMUST00000172697.2 | -3369 | Mecom | ENSMUSG00000027684.11 |
| chr3 | 30521046 | 30521133 | 3.86996 | 4.8544 | 1.72325 | Intron (ENSMUST00000172754.2/ENSMUSG00000027684.11, intron 2 of 8) | 3 | 29965456 | 30509822 | 544367 | 2 | ENSMUST00000172697.2 | -11225 | Mecom | ENSMUSG00000027684.11 |
| chr3 | 30516935 | 30517801 | 4.08965 | 5.35474 | 2.02017 | Intron (ENSMUST00000172754.2/ENSMUSG00000027684.11, intron 2 of 8) | 3 | 29965456 | 30509822 | 544367 | 2 | ENSMUST00000172697.2 | -7114 | Mecom | ENSMUSG00000027684.11 |
| chr3 | 30518294 | 30518880 | 3.63227 | 4.39308 | 1.36061 | Intron (ENSMUST00000172754.2/ENSMUSG00000027684.11, intron 2 of 8) | 3 | 29965456 | 30509822 | 544367 | 2 | ENSMUST00000172697.2 | -8473 | Mecom | ENSMUSG00000027684.11 |
| chr3 | 28176465 | 28176659 | 3.80657 | 4.77423 | 1.62101 | Distal Intergenic | 3 | 28109759 | 28130795 | 21037 | 1 | ENSMUST00000126594.1 | 66707 | Pld1 | ENSMUSG00000027695.11 |
| chr3 | 30649423 | 30649684 | 3.62581 | 4.37042 | 1.35269 | Intron (ENSMUST00000108267.2/ENSMUSG00000027703.11, intron 3 of 7) | 3 | 30647886 | 30672299 | 24414 | 1 | ENSMUST00000108265.2 | 1538 | Lrriq4 | ENSMUSG00000027703.11 |
| chr3 | 30650100 | 30650935 | 3.70082 | 4.63424 | 1.55898 | Exon (ENSMUST00000108267.2/ENSMUSG00000027703.11, exon 4 of 8) | 3 | 30647886 | 30672299 | 24414 | 1 | ENSMUST00000108265.2 | 2215 | Lrriq4 | ENSMUSG00000027703.11 |
| chr3 | 36062540 | 36062927 | 3.59608 | 4.35926 | 1.42039 | Distal Intergenic | 3 | 36065979 | 36094944 | 28966 | 1 | ENSMUST00000011492.9 | -3052 | Acad9 | ENSMUSG00000027710.9 |
| chr3 | 36689010 | 36689141 | 3.74564 | 4.93163 | 1.72224 | Intron (ENSMUST00000029271.4/ENSMUSG00000027716.8, intron 1 of 11) | 3 | 36620482 | 36690167 | 69686 | 2 | ENSMUST00000029271.4 | 1026 | Trpc3 | ENSMUSG00000027716.8 |
| chr3 | 36689627 | 36690129 | 4.01523 | 5.51585 | 2.11583 | Promoter | 3 | 36620482 | 36690167 | 69686 | 2 | ENSMUST00000029271.4 | 38 | Trpc3 | ENSMUSG00000027716.8 |
| chr3 | 36691471 | 36691799 | 3.53893 | 4.60559 | 1.49819 | Distal Intergenic | 3 | 36669203 | 36690167 | 20965 | 2 | ENSMUST00000146475.1 | -1305 | Trpc3 | ENSMUSG00000027716.8 |
| chr3 | 50444460 | 50444630 | 3.4222 | 3.97916 | 1.08015 | Promoter | 3 | 50415970 | 50443613 | 27644 | 2 | ENSMUST00000142932.1 | -848 | Slc7a11 | ENSMUSG00000027737.5 |
| chr3 | 54155338 | 54156211 | 4.24097 | 5.66009 | 2.27161 | Promoter | 3 | 54156057 | 54318470 | 162414 | 1 | ENSMUST00000029311.6 | 0 | Trpc4 | ENSMUSG00000027748.7 |
| chr3 | 54157359 | 54157763 | 3.48462 | 4.33823 | 1.36616 | Intron (ENSMUST00000029311.6/ENSMUSG00000027748.7, intron 1 of 10) | 3 | 54156057 | 54318470 | 162414 | 1 | ENSMUST00000029311.6 | 1303 | Trpc4 | ENSMUSG00000027748.7 |
| chr3 | 54158463 | 54158718 | 3.60414 | 4.35552 | 1.38905 | Intron (ENSMUST00000029311.6/ENSMUSG00000027748.7, intron 1 of 10) | 3 | 54156057 | 54318470 | 162414 | 1 | ENSMUST00000029311.6 | 2407 | Trpc4 | ENSMUSG00000027748.7 |
| chr3 | 55054109 | 55054405 | 3.62861 | 4.42025 | 1.42523 | Promoter | 3 | 55045469 | 55055055 | 9587 | 2 | ENSMUST00000029368.2 | 650 | Ccna1 | ENSMUSG00000027793.2 |
| chr3 | 55055044 | 55055790 | 3.70011 | 4.94099 | 1.77717 | Promoter | 3 | 55045469 | 55055055 | 9587 | 2 | ENSMUST00000029368.2 | 0 | Ccna1 | ENSMUSG00000027793.2 |
| chr3 | 55244369 | 55244461 | 3.50668 | 4.17811 | 1.302 | Intron (ENSMUST00000167204.2/ENSMUSG00000027797.10, intron 1 of 6) | 3 | 55242526 | 55539064 | 296539 | 1 | ENSMUST00000054237.8 | 1844 | Dclk1 | ENSMUSG00000027797.10 |
| chr3 | 66739230 | 66739547 | 3.52124 | 4.35692 | 1.38818 | Distal Intergenic | 3 | 66972780 | 66978447 | 5668 | 2 | ENSMUST00000162060.3 | 238900 | Shox2 | ENSMUSG00000027833.11 |
| chr3 | 66957455 | 66957678 | 3.67624 | 4.65138 | 1.54178 | Distal Intergenic | 3 | 66972780 | 66978447 | 5668 | 2 | ENSMUST00000162060.3 | 20769 | Shox2 | ENSMUSG00000027833.11 |
| chr3 | 66960685 | 66960779 | 3.84605 | 4.85262 | 1.72221 | Distal Intergenic | 3 | 66972780 | 66978447 | 5668 | 2 | ENSMUST00000162060.3 | 17668 | Shox2 | ENSMUSG00000027833.11 |
| chr3 | 66962702 | 66963074 | 3.55717 | 4.24906 | 1.31559 | Distal Intergenic | 3 | 66972780 | 66978447 | 5668 | 2 | ENSMUST00000162060.3 | 15373 | Shox2 | ENSMUSG00000027833.11 |
| chr3 | 66965068 | 66965143 | 4.33887 | 5.72966 | 2.25961 | Distal Intergenic | 3 | 66972780 | 66978447 | 5668 | 2 | ENSMUST00000162060.3 | 13304 | Shox2 | ENSMUSG00000027833.11 |
| chr3 | 66966060 | 66966803 | 3.97133 | 5.11561 | 1.87688 | Distal Intergenic | 3 | 66972780 | 66978447 | 5668 | 2 | ENSMUST00000162060.3 | 11644 | Shox2 | ENSMUSG00000027833.11 |
| chr3 | 66967616 | 66968930 | 3.56098 | 4.26793 | 1.29319 | Downstream (2-3kb) | 3 | 66972780 | 66978447 | 5668 | 2 | ENSMUST00000162060.3 | 9517 | Shox2 | ENSMUSG00000027833.11 |
| chr3 | 66969374 | 66969774 | 3.49661 | 4.15988 | 1.26976 | Downstream (1-2kb) | 3 | 66972780 | 66978447 | 5668 | 2 | ENSMUST00000162060.3 | 8673 | Shox2 | ENSMUSG00000027833.11 |
| chr3 | 66970162 | 66971719 | 4.04446 | 5.40291 | 2.0644 | Downstream (<1kb) | 3 | 66972780 | 66978447 | 5668 | 2 | ENSMUST00000162060.3 | 6728 | Shox2 | ENSMUSG00000027833.11 |
| chr3 | 66972179 | 66974121 | 4.30612 | 6.00266 | 2.52421 | 3' UTR | 3 | 66972780 | 66978447 | 5668 | 2 | ENSMUST00000162060.3 | 4326 | Shox2 | ENSMUSG00000027833.11 |
| chr3 | 66974593 | 66976499 | 3.98279 | 5.08396 | 1.84447 | Exon (ENSMUST00000162098.3/ENSMUSG00000027833.11, exon 3 of 5) | 3 | 66972780 | 66978447 | 5668 | 2 | ENSMUST00000162060.3 | 1948 | Shox2 | ENSMUSG00000027833.11 |
| chr3 | 66974593 | 66976499 | 3.98279 | 5.08396 | 1.84447 | Exon (ENSMUST00000162098.3/ENSMUSG00000027833.11, exon 3 of 5) | 3 | 66972780 | 66978447 | 5668 | 2 | ENSMUST00000162060.3 | 1948 | Shox2 | ENSMUSG00000027833.11 |
| chr3 | 66976831 | 66977527 | 4.27754 | 5.68838 | 2.22751 | Promoter | 3 | 66972780 | 66978447 | 5668 | 2 | ENSMUST00000162060.3 | 920 | Shox2 | ENSMUSG00000027833.11 |
| chr3 | 66977973 | 66978317 | 3.36 | 4.04617 | 1.15999 | Promoter | 3 | 66972780 | 66978447 | 5668 | 2 | ENSMUST00000162060.3 | 130 | Shox2 | ENSMUSG00000027833.11 |
| chr3 | 66978644 | 66979815 | 3.89848 | 5.04604 | 1.81083 | Promoter | 3 | 66972780 | 66978447 | 5668 | 2 | ENSMUST00000162060.3 | -198 | Shox2 | ENSMUSG00000027833.11 |
| chr3 | 66980133 | 66980372 | 3.70392 | 4.59455 | 1.47773 | Promoter | 3 | 66973259 | 66981141 | 7883 | 2 | ENSMUST00000162439.2 | 769 | Shox2 | ENSMUSG00000027833.11 |
| chr3 | 66980706 | 66981704 | 3.7447 | 4.70933 | 1.58126 | Promoter | 3 | 66971727 | 66981318 | 9592 | 2 | ENSMUST00000162098.3 | 0 | Shox2 | ENSMUSG00000027833.11 |
| chr3 | 66982207 | 66983093 | 3.94931 | 5.09478 | 1.80053 | Promoter | 3 | 66972790 | 66981771 | 8982 | 2 | ENSMUST00000029422.8 | -437 | Shox2 | ENSMUSG00000027833.11 |
| chr3 | 104961376 | 104962072 | 4.19313 | 5.70275 | 2.27942 | Promoter | 3 | 104944805 | 104961709 | 16905 | 2 | ENSMUST00000029429.4 | 0 | Wnt2b | ENSMUSG00000027840.4 |
| chr3 | 104962451 | 104962996 | 4.17595 | 5.60539 | 2.22399 | Promoter | 3 | 104944805 | 104961709 | 16905 | 2 | ENSMUST00000029429.4 | -743 | Wnt2b | ENSMUSG00000027840.4 |
| chr3 | 103574009 | 103575459 | 3.69553 | 4.66388 | 1.57554 | Promoter | 3 | 103575231 | 103630946 | 55716 | 1 | ENSMUST00000121834.2 | 0 | Syt6 | ENSMUSG00000027849.13 |
| chr3 | 103576068 | 103576649 | 4.02884 | 5.29492 | 1.99737 | Promoter | 3 | 103576081 | 103587240 | 11160 | 1 | ENSMUST00000183637.2 | 0 | Syt6 | ENSMUSG00000027849.13 |
| chr3 | 103577668 | 103577775 | 3.47785 | 4.11541 | 1.2574 | Intron (ENSMUST00000121834.2/ENSMUSG00000027849.13, intron 1 of 6) | 3 | 103576142 | 103645569 | 69428 | 1 | ENSMUST00000118563.2 | 1527 | Syt6 | ENSMUSG00000027849.13 |
| chr3 | 103578202 | 103578277 | 3.30265 | 3.80957 | 1.10244 | Intron (ENSMUST00000121834.2/ENSMUSG00000027849.13, intron 1 of 6) | 3 | 103576142 | 103645569 | 69428 | 1 | ENSMUST00000118563.2 | 2061 | Syt6 | ENSMUSG00000027849.13 |
| chr3 | 102469916 | 102470142 | 3.75626 | 4.66906 | 1.6266 | Promoter | 3 | 102469928 | 102521013 | 51086 | 1 | ENSMUST00000035952.3 | 0 | Ngf | ENSMUSG00000027859.5 |
| chr3 | 101072878 | 101073438 | 4.09494 | 5.33121 | 2.04012 | Exon (ENSMUST00000102694.3/ENSMUSG00000027864.8, exon 4 of 9) | 3 | 101040232 | 101110278 | 70047 | 2 | ENSMUST00000102694.3 | 36840 | Ptgfrn | ENSMUSG00000027864.8 |
| chr3 | 101108069 | 101108264 | 3.29538 | 3.7787 | 1.04674 | Intron (ENSMUST00000102694.3/ENSMUSG00000027864.8, intron 1 of 8) | 3 | 101040232 | 101110278 | 70047 | 2 | ENSMUST00000102694.3 | 2014 | Ptgfrn | ENSMUSG00000027864.8 |
| chr3 | 101109186 | 101109512 | 3.6383 | 4.43499 | 1.43761 | Promoter | 3 | 101040232 | 101110278 | 70047 | 2 | ENSMUST00000102694.3 | 766 | Ptgfrn | ENSMUSG00000027864.8 |
| chr3 | 101128975 | 101129263 | 3.32504 | 3.81483 | 1.01733 | Distal Intergenic | 3 | 101040232 | 101110278 | 70047 | 2 | ENSMUST00000102694.3 | -18698 | Ptgfrn | ENSMUSG00000027864.8 |
| chr3 | 99231292 | 99231619 | 3.4139 | 4.03407 | 1.19119 | Intron (ENSMUST00000145650.2/ENSMUSG00000004233.9, intron 4 of 8) | 3 | 99240381 | 99313065 | 72685 | 1 | ENSMUST00000151606.2 | -8762 | Tbx15 | ENSMUSG00000027868.6 |
| chr3 | 99232792 | 99235579 | 3.60301 | 4.45614 | 1.4488 | Exon (ENSMUST00000145650.2/ENSMUSG00000004233.9, exon 5 of 9) | 3 | 99240381 | 99313065 | 72685 | 1 | ENSMUST00000151606.2 | -4802 | Tbx15 | ENSMUSG00000027868.6 |
| chr3 | 99235910 | 99237143 | 3.46473 | 4.11609 | 1.22757 | Exon (ENSMUST00000145650.2/ENSMUSG00000004233.9, exon 5 of 9) | 3 | 99240381 | 99313065 | 72685 | 1 | ENSMUST00000151606.2 | -3238 | Tbx15 | ENSMUSG00000027868.6 |
| chr3 | 99238380 | 99239672 | 3.61839 | 4.40338 | 1.41341 | Promoter | 3 | 99240381 | 99313065 | 72685 | 1 | ENSMUST00000151606.2 | -709 | Tbx15 | ENSMUSG00000027868.6 |
| chr3 | 99239993 | 99241848 | 4.25563 | 5.6772 | 2.27928 | Promoter | 3 | 99240381 | 99313065 | 72685 | 1 | ENSMUST00000151606.2 | 0 | Tbx15 | ENSMUSG00000027868.6 |
| chr3 | 99242197 | 99242764 | 4.02548 | 5.17146 | 1.92358 | Intron (ENSMUST00000151606.2/ENSMUSG00000027868.6, intron 1 of 2) | 3 | 99240381 | 99313065 | 72685 | 1 | ENSMUST00000151606.2 | 1817 | Tbx15 | ENSMUSG00000027868.6 |
| chr3 | 99243577 | 99244327 | 3.68934 | 4.51822 | 1.47314 | Intron (ENSMUST00000151606.2/ENSMUSG00000027868.6, intron 1 of 2) | 3 | 99240381 | 99313065 | 72685 | 1 | ENSMUST00000151606.2 | 3197 | Tbx15 | ENSMUSG00000027868.6 |
| chr3 | 99244784 | 99245043 | 3.35408 | 3.87311 | 1.06818 | Intron (ENSMUST00000151606.2/ENSMUSG00000027868.6, intron 1 of 2) | 3 | 99240381 | 99313065 | 72685 | 1 | ENSMUST00000151606.2 | 4404 | Tbx15 | ENSMUSG00000027868.6 |
| chr3 | 99245438 | 99246303 | 3.5049 | 4.15654 | 1.25074 | Intron (ENSMUST00000151606.2/ENSMUSG00000027868.6, intron 1 of 2) | 3 | 99240381 | 99313065 | 72685 | 1 | ENSMUST00000151606.2 | 5058 | Tbx15 | ENSMUSG00000027868.6 |
| chr3 | 99247497 | 99247604 | 3.41056 | 4.26429 | 1.31376 | Intron (ENSMUST00000151606.2/ENSMUSG00000027868.6, intron 1 of 2) | 3 | 99253760 | 99354259 | 100500 | 1 | ENSMUST00000029462.5 | -6156 | Tbx15 | ENSMUSG00000027868.6 |
| chr3 | 99247980 | 99248072 | 3.72372 | 4.93437 | 1.71448 | Intron (ENSMUST00000151606.2/ENSMUSG00000027868.6, intron 1 of 2) | 3 | 99253760 | 99354259 | 100500 | 1 | ENSMUST00000029462.5 | -5688 | Tbx15 | ENSMUSG00000027868.6 |
| chr3 | 99248423 | 99249283 | 4.16619 | 5.53621 | 2.17962 | Intron (ENSMUST00000151606.2/ENSMUSG00000027868.6, intron 1 of 2) | 3 | 99253760 | 99354259 | 100500 | 1 | ENSMUST00000029462.5 | -4477 | Tbx15 | ENSMUSG00000027868.6 |
| chr3 | 99249616 | 99250508 | 4.7257 | 6.85564 | 3.21548 | Intron (ENSMUST00000151606.2/ENSMUSG00000027868.6, intron 1 of 2) | 3 | 99253760 | 99354259 | 100500 | 1 | ENSMUST00000029462.5 | -3252 | Tbx15 | ENSMUSG00000027868.6 |
| chr3 | 99252167 | 99252702 | 3.6048 | 4.59283 | 1.4893 | Intron (ENSMUST00000151606.2/ENSMUSG00000027868.6, intron 1 of 2) | 3 | 99253760 | 99354259 | 100500 | 1 | ENSMUST00000029462.5 | -1058 | Tbx15 | ENSMUSG00000027868.6 |
| chr3 | 99253156 | 99254658 | 3.84047 | 5.1614 | 1.88889 | Promoter | 3 | 99253760 | 99354259 | 100500 | 1 | ENSMUST00000029462.5 | 0 | Tbx15 | ENSMUSG00000027868.6 |
| chr3 | 99255007 | 99257817 | 3.84481 | 4.94699 | 1.7725 | Intron (ENSMUST00000151606.2/ENSMUSG00000027868.6, intron 1 of 2) | 3 | 99253760 | 99354259 | 100500 | 1 | ENSMUST00000029462.5 | 1248 | Tbx15 | ENSMUSG00000027868.6 |
| chr3 | 99258357 | 99259118 | 3.76569 | 4.64101 | 1.50464 | Intron (ENSMUST00000151606.2/ENSMUSG00000027868.6, intron 1 of 2) | 3 | 99253760 | 99354259 | 100500 | 1 | ENSMUST00000029462.5 | 4598 | Tbx15 | ENSMUSG00000027868.6 |
| chr3 | 99259465 | 99260545 | 3.81522 | 4.79769 | 1.61723 | Intron (ENSMUST00000151606.2/ENSMUSG00000027868.6, intron 1 of 2) | 3 | 99253760 | 99354259 | 100500 | 1 | ENSMUST00000029462.5 | 5706 | Tbx15 | ENSMUSG00000027868.6 |
| chr3 | 99260902 | 99264252 | 3.89376 | 4.91844 | 1.72494 | Intron (ENSMUST00000151606.2/ENSMUSG00000027868.6, intron 1 of 2) | 3 | 99253760 | 99354259 | 100500 | 1 | ENSMUST00000029462.5 | 7143 | Tbx15 | ENSMUSG00000027868.6 |
| chr3 | 99264874 | 99265407 | 3.82881 | 4.75771 | 1.57794 | Intron (ENSMUST00000151606.2/ENSMUSG00000027868.6, intron 1 of 2) | 3 | 99253760 | 99354259 | 100500 | 1 | ENSMUST00000029462.5 | 11115 | Tbx15 | ENSMUSG00000027868.6 |
| chr3 | 99265907 | 99266456 | 3.83351 | 4.78719 | 1.65237 | Intron (ENSMUST00000151606.2/ENSMUSG00000027868.6, intron 1 of 2) | 3 | 99253760 | 99354259 | 100500 | 1 | ENSMUST00000029462.5 | 12148 | Tbx15 | ENSMUSG00000027868.6 |

| | | | | | | | | | | | | | | | |
|---|---|---|---|---|---|---|---|---|---|---|---|---|---|---|---|
| chr3 | 99268143 | 99268399 | 3.36492 | 3.87494 | 1.01733 | Intron (ENSMUST00000151606.2/ENSMUSG00000027868.6, intron 1 of 2) | 3 | 99253760 | 99354259 | 100500 | 1 | ENSMUST00000029462.5 | 14384 | Tbx15 | ENSMUSG00000027868.6 |
| chr3 | 99269179 | 99269691 | 3.55628 | 4.22145 | 1.22099 | Intron (ENSMUST00000151606.2/ENSMUSG00000027868.6, intron 1 of 2) | 3 | 99253760 | 99354259 | 100500 | 1 | ENSMUST00000029462.5 | 15420 | Tbx15 | ENSMUSG00000027868.6 |
| chr3 | 97972352 | 97972752 | 3.45737 | 4.0739 | 1.22609 | Distal Intergenic | 3 | 98013538 | 98150367 | 136830 | 1 | ENSMUST00000079812.6 | -40786 | Notch2 | ENSMUSG00000027878.10 |
| chr3 | 109055637 | 109055794 | 3.33927 | 3.86051 | 1.0984 | Distal Intergenic | 3 | 109080469 | 109116687 | 36219 | 1 | ENSMUST00000029478.3 | -24675 | Slc25a54 | ENSMUSG00000027880.4 |
| chr3 | 108226038 | 108226744 | 3.71341 | 4.80927 | 1.67206 | Promoter | 3 | 108211472 | 108226648 | 15177 | 2 | ENSMUST00000141387.2 | 0 | Sypl2 | ENSMUSG00000027887.9 |
| chr3 | 108227559 | 108228178 | 3.41814 | 4.00177 | 1.15312 | Promoter | 3 | 108211472 | 108226648 | 15177 | 2 | ENSMUST00000141387.2 | -912 | Sypl2 | ENSMUSG00000027887.9 |
| chr3 | 107464948 | 107465023 | 3.55472 | 4.2218 | 1.2309 | Downstream (2-3kb) | 3 | 107467551 | 107468705 | 1155 | 2 | ENSMUST00000171029.1 | 3682 | Slc6a17 | ENSMUSG00000027894.9 |
| chr3 | 107484106 | 107484181 | 4.14302 | 5.45134 | 2.09035 | Intron (ENSMUST00000169449.2/ENSMUSG00000027894.9, intron 6 of 11) | 3 | 107489771 | 107493196 | 3426 | 2 | ENSMUST00000167975.1 | 9015 | Slc6a17 | ENSMUSG00000027894.9 |
| chr3 | 107496338 | 107496695 | 3.47597 | 4.1034 | 1.2289 | Intron (ENSMUST00000169449.2/ENSMUSG00000027894.9, intron 2 of 11) | 3 | 107489771 | 107493196 | 3426 | 2 | ENSMUST00000167975.1 | -3143 | Slc6a17 | ENSMUSG00000027894.9 |
| chr3 | 107517147 | 107517805 | 3.90043 | 4.96024 | 1.76314 | Promoter | 3 | 107482054 | 107517749 | 35696 | 2 | ENSMUST00000170429.2 | 0 | Slc6a17 | ENSMUSG00000027894.9 |
| chr3 | 107518171 | 107518474 | 3.79939 | 4.76082 | 1.64443 | Promoter | 3 | 107489773 | 107518018 | 28246 | 2 | ENSMUST00000167898.1 | -154 | Slc6a17 | ENSMUSG00000027894.9 |
| chr3 | 107526711 | 107526852 | 3.81915 | 4.79332 | 1.65723 | Distal Intergenic | 3 | 107489773 | 107518018 | 28246 | 2 | ENSMUST00000167898.1 | -8694 | Slc6a17 | ENSMUSG00000027894.9 |
| chr3 | 107527941 | 107528345 | 3.42381 | 3.98774 | 1.10208 | Distal Intergenic | 3 | 107489773 | 107518018 | 28246 | 2 | ENSMUST00000167898.1 | -9924 | Slc6a17 | ENSMUSG00000027894.9 |
| chr3 | 107456255 | 107456332 | 4.67648 | 6.40179 | 2.70962 | Intron (ENSMUST00000009617.8/ENSMUSG00000027895.8, intron 1 of 3) | 3 | 107438303 | 107458898 | 20596 | 2 | ENSMUST00000009617.8 | 2566 | Kcnc4 | ENSMUSG00000027895.8 |
| chr3 | 107456818 | 107456941 | 3.36751 | 3.97079 | 1.13786 | Intron (ENSMUST00000009617.8/ENSMUSG00000027895.8, intron 1 of 3) | 3 | 107438303 | 107458898 | 20596 | 2 | ENSMUST00000009617.8 | 1957 | Kcnc4 | ENSMUSG00000027895.8 |
| chr3 | 107458178 | 107460498 | 4.10995 | 5.35007 | 2.02258 | Promoter | 3 | 107438303 | 107458898 | 20596 | 2 | ENSMUST00000009617.8 | 0 | Kcnc4 | ENSMUSG00000027895.8 |
| chr3 | 107460942 | 107461243 | 3.6304 | 4.35786 | 1.30051 | Distal Intergenic | 3 | 107438303 | 107458898 | 20596 | 2 | ENSMUST00000009617.8 | -2045 | Kcnc4 | ENSMUSG00000027895.8 |
| chr3 | 107461888 | 107462439 | 3.72845 | 4.59492 | 1.49984 | Distal Intergenic | 3 | 107438303 | 107458898 | 20596 | 2 | ENSMUST00000009617.8 | -2991 | Kcnc4 | ENSMUSG00000027895.8 |
| chr3 | 93013748 | 93013963 | 3.43316 | 4.06982 | 1.18453 | Downstream (<1kb) | 3 | 93014519 | 93015669 | 1151 | 2 | ENSMUST00000107301.1 | 1706 | Crct1 | ENSMUSG00000027913.5 |
| chr3 | 93014706 | 93015019 | 3.48945 | 4.11371 | 1.19771 | Promoter | 3 | 93014519 | 93015669 | 1151 | 2 | ENSMUST00000107301.1 | 650 | Crct1 | ENSMUSG00000027913.5 |
| chr3 | 89763520 | 89764361 | 3.428 | 3.99041 | 1.08889 | Promoter | 3 | 89753448 | 89764632 | 11185 | 2 | ENSMUST00000029562.3 | 271 | Chrnb2 | ENSMUSG00000027950.3 |
| chr3 | 89486202 | 89486891 | 3.53838 | 4.21533 | 1.23578 | Distal Intergenic | 3 | 89459571 | 89469006 | 9436 | 1 | ENSMUST00000184515.1 | 26632 | Pmvk | ENSMUSG00000027952.11 |
| chr3 | 89487234 | 89487377 | 3.44125 | 4.03156 | 1.13075 | Distal Intergenic | 3 | 89459571 | 89469006 | 9436 | 1 | ENSMUST00000184515.1 | 27664 | Pmvk | ENSMUSG00000027952.11 |
| chr3 | 79886952 | 79887073 | 3.73557 | 4.56271 | 1.44123 | Exon (ENSMUST00000118853.2/ENSMUSG00000027955.11, exon 2 of 5) | 3 | 79885930 | 79946278 | 60349 | 1 | ENSMUST00000029567.8 | 1023 | Fam198b | ENSMUSG00000027955.11 |
| chr3 | 114030474 | 114030908 | 3.69603 | 4.58438 | 1.51247 | Promoter | 3 | 114030540 | 114108324 | 77785 | 1 | ENSMUST00000123619.2 | 0 | Col11a1 | ENSMUSG00000027966.14 |
| chr3 | 127621307 | 127621571 | 3.38977 | 4.12377 | 1.18777 | Distal Intergenic | 3 | 127633136 | 127635631 | 2496 | 1 | ENSMUST00000029587.7 | -11565 | Neurog2 | ENSMUSG00000027967.7 |
| chr3 | 127623113 | 127623312 | 3.99854 | 5.17335 | 1.91206 | Distal Intergenic | 3 | 127633136 | 127635631 | 2496 | 1 | ENSMUST00000029587.7 | -9824 | Neurog2 | ENSMUSG00000027967.7 |
| chr3 | 127624267 | 127625987 | 4.02843 | 5.2738 | 2.00402 | Distal Intergenic | 3 | 127633136 | 127635631 | 2496 | 1 | ENSMUST00000029587.7 | -7149 | Neurog2 | ENSMUSG00000027967.7 |
| chr3 | 127626439 | 127629103 | 4.13324 | 5.47255 | 2.1863 | Distal Intergenic | 3 | 127633136 | 127635631 | 2496 | 1 | ENSMUST00000029587.7 | -4033 | Neurog2 | ENSMUSG00000027967.7 |
| chr3 | 127630315 | 127631676 | 3.92109 | 5.03496 | 1.83779 | Distal Intergenic | 3 | 127633136 | 127635631 | 2496 | 1 | ENSMUST00000029587.7 | -1460 | Neurog2 | ENSMUSG00000027967.7 |
| chr3 | 127632108 | 127632782 | 4.16863 | 5.65388 | 2.2272 | Promoter | 3 | 127633136 | 127635631 | 2496 | 1 | ENSMUST00000029587.7 | -354 | Neurog2 | ENSMUSG00000027967.7 |
| chr3 | 127633176 | 127634471 | 4.05027 | 5.38988 | 2.11016 | Promoter | 3 | 127633136 | 127635631 | 2496 | 1 | ENSMUST00000029587.7 | 41 | Neurog2 | ENSMUSG00000027967.7 |
| chr3 | 127634872 | 127635094 | 3.68432 | 4.60428 | 1.52788 | 3' UTR | 3 | 127633136 | 127635631 | 2496 | 1 | ENSMUST00000029587.7 | 1737 | Neurog2 | ENSMUSG00000027967.7 |
| chr3 | 127636116 | 127636603 | 4.0972 | 5.49861 | 2.11325 | Distal Intergenic | 3 | 127633136 | 127635631 | 2496 | 1 | ENSMUST00000029587.7 | 2981 | Neurog2 | ENSMUSG00000027967.7 |
| chr3 | 127638213 | 127638383 | 3.55639 | 4.22816 | 1.23593 | Distal Intergenic | 3 | 127633136 | 127635631 | 2496 | 1 | ENSMUST00000029587.7 | 5078 | Neurog2 | ENSMUSG00000027967.7 |
| chr3 | 127638958 | 127639179 | 3.49106 | 4.35566 | 1.34594 | Distal Intergenic | 3 | 127633136 | 127635631 | 2496 | 1 | ENSMUST00000029587.7 | 5823 | Neurog2 | ENSMUSG00000027967.7 |
| chr3 | 127639706 | 127641037 | 4.36877 | 5.92061 | 2.41505 | Distal Intergenic | 3 | 127633136 | 127635631 | 2496 | 1 | ENSMUST00000029587.7 | 6571 | Neurog2 | ENSMUSG00000027967.7 |
| chr3 | 127641380 | 127642573 | 3.85128 | 4.81362 | 1.63377 | Distal Intergenic | 3 | 127633136 | 127635631 | 2496 | 1 | ENSMUST00000029587.7 | 8245 | Neurog2 | ENSMUSG00000027967.7 |
| chr3 | 127642947 | 127643628 | 4.15584 | 5.45942 | 2.07697 | Distal Intergenic | 3 | 127633136 | 127635631 | 2496 | 1 | ENSMUST00000029587.7 | 9812 | Neurog2 | ENSMUSG00000027967.7 |
| chr3 | 127644322 | 127644686 | 3.62121 | 4.34931 | 1.31584 | Distal Intergenic | 3 | 127633136 | 127635631 | 2496 | 1 | ENSMUST00000029587.7 | 11187 | Neurog2 | ENSMUSG00000027967.7 |
| chr3 | 127646670 | 127646885 | 3.84802 | 5.2247 | 1.90658 | Distal Intergenic | 3 | 127633136 | 127635631 | 2496 | 1 | ENSMUST00000029587.7 | 13535 | Neurog2 | ENSMUSG00000027967.7 |
| chr3 | 125228370 | 125228467 | 11.98384 | 30.11829 | 24.61983 | Distal Intergenic | 3 | 125404094 | 125686644 | 282551 | 1 | ENSMUST00000147016.2 | -175627 | Ndst4 | ENSMUSG00000027971.11 |
| chr3 | 123677396 | 123677825 | 3.95161 | 5.23782 | 1.93409 | Intron (ENSMUST00000137404.2/ENSMUSG00000027977.10, intron 1 of 13) | 3 | 123526169 | 123672392 | 146224 | 2 | ENSMUST00000029602.7 | -5005 | Ndst3 | ENSMUSG00000027977.10 |
| chr3 | 123680027 | 123680441 | 3.26593 | 3.76026 | 1.05621 | Intron (ENSMUST00000137404.2/ENSMUSG00000027977.10, intron 1 of 13) | 3 | 123526169 | 123672392 | 146224 | 2 | ENSMUST00000029602.7 | -7636 | Ndst3 | ENSMUSG00000027977.10 |
| chr3 | 123680887 | 123680962 | 3.62963 | 4.47119 | 1.43848 | Intron (ENSMUST00000137404.2/ENSMUSG00000027977.10, intron 1 of 13) | 3 | 123526169 | 123672392 | 146224 | 2 | ENSMUST00000029602.7 | -8496 | Ndst3 | ENSMUSG00000027977.10 |
| chr3 | 123688507 | 123688739 | 3.40604 | 3.95736 | 1.08948 | Intron (ENSMUST00000137404.2/ENSMUSG00000027977.10, intron 1 of 13) | 3 | 123526168 | 123690817 | 164650 | 2 | ENSMUST00000154668.2 | 2078 | Ndst3 | ENSMUSG00000027977.10 |
| chr3 | 123690408 | 123690636 | 3.51552 | 4.17568 | 1.27431 | Promoter | 3 | 123526168 | 123690817 | 164650 | 2 | ENSMUST00000154668.2 | 181 | Ndst3 | ENSMUSG00000027977.10 |
| chr3 | 123446233 | 123446754 | 3.52214 | 4.21685 | 1.3018 | Promoter | 3 | 123446913 | 123506602 | 59690 | 1 | ENSMUST00000029603.8 | -159 | Prss12 | ENSMUSG00000027978.8 |
| chr3 | 123447109 | 123447305 | 3.72641 | 4.60957 | 1.53899 | Promoter | 3 | 123446913 | 123506602 | 59690 | 1 | ENSMUST00000029603.8 | 197 | Prss12 | ENSMUSG00000027978.8 |
| chr3 | 131067571 | 131067864 | 3.38001 | 3.8977 | 1.01733 | Distal Intergenic | 3 | 131110471 | 131224356 | 113886 | 1 | ENSMUST00000029611.8 | -42607 | Lef1 | ENSMUSG00000027985.9 |
| chr3 | 131084149 | 131084276 | 3.3445 | 3.8442 | 1.01733 | Distal Intergenic | 3 | 131110471 | 131224356 | 113886 | 1 | ENSMUST00000029611.8 | -26195 | Lef1 | ENSMUSG00000027985.9 |
| chr3 | 131084840 | 131085371 | 3.70644 | 4.50642 | 1.41134 | Distal Intergenic | 3 | 131110471 | 131224356 | 113886 | 1 | ENSMUST00000029611.8 | -25100 | Lef1 | ENSMUSG00000027985.9 |
| chr3 | 131086167 | 131086542 | 3.57516 | 4.36306 | 1.39383 | Distal Intergenic | 3 | 131110471 | 131224356 | 113886 | 1 | ENSMUST00000029611.8 | -23929 | Lef1 | ENSMUSG00000027985.9 |
| chr3 | 131093442 | 131094401 | 3.6255 | 4.37095 | 1.33335 | Distal Intergenic | 3 | 131110471 | 131224356 | 113886 | 1 | ENSMUST00000029611.8 | -16070 | Lef1 | ENSMUSG00000027985.9 |
| chr3 | 131103223 | 131103461 | 3.92837 | 4.93184 | 1.71271 | Distal Intergenic | 3 | 131110471 | 131224356 | 113886 | 1 | ENSMUST00000029611.8 | -7010 | Lef1 | ENSMUSG00000027985.9 |
| chr3 | 131104866 | 131105250 | 4.09498 | 5.38219 | 2.12858 | Distal Intergenic | 3 | 131110471 | 131224356 | 113886 | 1 | ENSMUST00000029611.8 | -5221 | Lef1 | ENSMUSG00000027985.9 |
| chr3 | 131106256 | 131108234 | 3.91822 | 5.05446 | 1.83 | Distal Intergenic | 3 | 131110471 | 131224356 | 113886 | 1 | ENSMUST00000029611.8 | -2237 | Lef1 | ENSMUSG00000027985.9 |
| chr3 | 131108918 | 131109820 | 3.53473 | 4.23037 | 1.28696 | Promoter | 3 | 131110471 | 131224356 | 113886 | 1 | ENSMUST00000029611.8 | -651 | Lef1 | ENSMUSG00000027985.9 |
| chr3 | 131110302 | 131110527 | 5.26654 | 7.83836 | 4.0112 | Promoter | 3 | 131110471 | 131224356 | 113886 | 1 | ENSMUST00000029611.8 | 0 | Lef1 | ENSMUSG00000027985.9 |
| chr3 | 131110962 | 131112521 | 3.87161 | 5.09179 | 1.84836 | Promoter | 3 | 131112797 | 131117387 | 4591 | 1 | ENSMUST00000136147.2 | -276 | Lef1 | ENSMUSG00000027985.9 |
| chr3 | 131113681 | 131113963 | 3.76903 | 4.61462 | 1.47651 | Promoter | 3 | 131112806 | 131188082 | 75277 | 1 | ENSMUST00000132737.1 | 876 | Lef1 | ENSMUSG00000027985.9 |
| chr3 | 84190443 | 84190592 | 3.90134 | 4.92579 | 1.76242 | Promoter | 3 | 84177251 | 84190523 | 13273 | 2 | ENSMUST00000140568.1 | 0 | Trim2 | ENSMUSG00000027993.11 |
| chr3 | 84191206 | 84191775 | 3.5565 | 4.31246 | 1.36655 | Promoter | 3 | 84190662 | 84191548 | 887 | 2 | ENSMUST00000156347.1 | 0 | Trim2 | ENSMUSG00000027993.11 |
| chr3 | 84258789 | 84259071 | 3.79884 | 4.67293 | 1.50428 | Promoter | 3 | 84193359 | 84259369 | 66011 | 2 | ENSMUST00000147901.2 | 298 | Trim2 | ENSMUSG00000027993.11 |
| chr3 | 84259414 | 84259524 | 3.55065 | 4.28921 | 1.29613 | Promoter | 3 | 84257683 | 84259369 | 1687 | 2 | ENSMUST00000152953.1 | -46 | Trim2 | ENSMUSG00000027993.11 |
| chr3 | 84260368 | 84260669 | 3.39971 | 3.94214 | 1.06816 | Promoter | 3 | 84160440 | 84259812 | 99373 | 2 | ENSMUST00000107691.2 | -557 | Trim2 | ENSMUSG00000027993.11 |
| chr3 | 84269631 | 84269776 | 3.42335 | 4.15368 | 1.25578 | Intron (ENSMUST00000107692.2/ENSMUSG00000027993.11, intron 1 of 11) | 3 | 84163722 | 84270791 | 107070 | 2 | ENSMUST00000054990.5 | 1015 | Trim2 | ENSMUSG00000027993.11 |
| chr3 | 84270122 | 84270374 | 3.90191 | 5.11773 | 1.83043 | Promoter | 3 | 84163722 | 84270791 | 107070 | 2 | ENSMUST00000054990.5 | 417 | Trim2 | ENSMUSG00000027993.11 |
| chr3 | 84301991 | 84302256 | 3.44462 | 4.06709 | 1.24612 | Intron (ENSMUST00000107692.2/ENSMUSG00000027993.11, intron 1 of 11) | 3 | 84299962 | 84304741 | 4780 | 2 | ENSMUST00000136805.1 | 2485 | Trim2 | ENSMUSG00000027993.11 |
| chr3 | 84303515 | 84303701 | 3.74758 | 4.67647 | 1.60788 | Intron (ENSMUST00000107692.2/ENSMUSG00000027993.11, intron 1 of 11) | 3 | 84299962 | 84304741 | 4780 | 2 | ENSMUST00000136805.1 | 1040 | Trim2 | ENSMUSG00000027993.11 |
| chr3 | 84304354 | 84305280 | 3.86763 | 4.92105 | 1.75384 | Promoter | 3 | 84160439 | 84304809 | 144371 | 2 | ENSMUST00000107692.2 | 0 | Trim2 | ENSMUSG00000027993.11 |
| chr3 | 84305877 | 84306185 | 3.5635 | 4.25497 | 1.28944 | Promoter | 3 | 84192211 | 84305446 | 113236 | 2 | ENSMUST00000132283.2 | -432 | Trim2 | ENSMUSG00000027993.11 |
| chr3 | 84345792 | 84345886 | 4.21911 | 5.5951 | 2.23939 | Distal Intergenic | 3 | 84192211 | 84305446 | 113236 | 2 | ENSMUST00000132283.2 | -40347 | Trim2 | ENSMUSG00000027993.11 |
| chr3 | 83840798 | 83841685 | 3.72166 | 4.86214 | 1.68333 | Promoter | 3 | 83836273 | 83841767 | 5495 | 2 | ENSMUST00000029623.9 | 82 | Tlr2 | ENSMUSG00000027995.9 |
| chr3 | 83760334 | 83760470 | 3.34153 | 3.83966 | 1.01733 | Distal Intergenic | 3 | 83766321 | 83774316 | 7996 | 1 | ENSMUST00000029625.7 | -5851 | Sfrp2 | ENSMUSG00000027996.8 |
| chr3 | 83764177 | 83765102 | 3.74836 | 4.6794 | 1.55137 | Distal Intergenic | 3 | 83766321 | 83774316 | 7996 | 1 | ENSMUST00000029625.7 | -1219 | Sfrp2 | ENSMUSG00000027996.8 |
| chr3 | 83765437 | 83766131 | 4.04851 | 5.35244 | 2.00272 | Promoter | 3 | 83766321 | 83774316 | 7996 | 1 | ENSMUST00000029625.7 | -190 | Sfrp2 | ENSMUSG00000027996.8 |
| chr3 | 83766485 | 83767088 | 3.93962 | 4.96949 | 1.73529 | Promoter | 3 | 83766565 | 83770143 | 3579 | 1 | ENSMUST00000140261.2 | 0 | Sfrp2 | ENSMUSG00000027996.8 |
| chr3 | 83767615 | 83768000 | 3.52494 | 4.17403 | 1.22162 | Promoter | 3 | 83766853 | 83772149 | 5297 | 1 | ENSMUST00000154794.1 | 763 | Sfrp2 | ENSMUSG00000027996.8 |
| chr3 | 83768758 | 83768843 | 3.28805 | 4.06572 | 1.16509 | Intron (ENSMUST00000029625.7/ENSMUSG00000027996.8, intron 1 of 2) | 3 | 83766853 | 83772149 | 5297 | 1 | ENSMUST00000154794.1 | 1906 | Sfrp2 | ENSMUSG00000027996.8 |
| chr3 | 83032823 | 83033016 | 3.38125 | 3.90203 | 1.02594 | 3' UTR | 3 | 83026153 | 83033615 | 7463 | 1 | ENSMUST00000166581.2 | 6671 | Fga | ENSMUSG00000028001.11 |
| chr3 | 82902534 | 82903977 | 3.70318 | 4.51794 | 1.43199 | Promoter | 3 | 82892579 | 82903973 | 11395 | 2 | ENSMUST00000029632.6 | 0 | Lrat | ENSMUSG00000028003.6 |
| chr3 | 82548029 | 82549052 | 3.88448 | 4.98755 | 1.80697 | Promoter | 3 | 82538383 | 82548084 | 9702 | 2 | ENSMUST00000029633.4 | 0 | Npy2r | ENSMUSG00000028004.7 |
| chr3 | 82073882 | 82073979 | 3.59661 | 4.33122 | 1.38441 | Promoter | 3 | 82032004 | 82074711 | 42708 | 2 | ENSMUST00000029635.8 | 732 | Gucy1b3 | ENSMUSG00000028005.8 |
| chr3 | 81036173 | 81037023 | 3.50747 | 4.33979 | 1.34214 | Promoter | 3 | 81036416 | 81214040 | 177625 | 1 | ENSMUST00000029652.3 | 0 | Pdgfc | ENSMUSG00000028019.4 |
| chr3 | 81037346 | 81037629 | 3.74211 | 4.64241 | 1.60721 | Promoter | 3 | 81037424 | 81210770 | 173347 | 1 | ENSMUST00000143721.1 | 0 | Pdgfc | ENSMUSG00000028019.4 |
| chr3 | 80912983 | 80913870 | 4.38878 | 5.92294 | 2.4255 | Promoter | 3 | 80843599 | 80913614 | 70016 | 2 | ENSMUST00000135043.2 | 0 | Glrb | ENSMUSG00000028020.11 |
| chr3 | 129158071 | 129158258 | 3.92219 | 5.07467 | 1.8496 | Distal Intergenic | 3 | 129199878 | 129219590 | 19713 | 1 | ENSMUST00000174661.3 | -41620 | Pitx2 | ENSMUSG00000028023.11 |
| chr3 | 129184080 | 129184295 | 3.84734 | 4.83366 | 1.71471 | Distal Intergenic | 3 | 129199878 | 129219590 | 19713 | 1 | ENSMUST00000174661.3 | -15583 | Pitx2 | ENSMUSG00000028023.11 |
| chr3 | 129187913 | 129188026 | 3.2431 | 3.69181 | 1.01733 | Distal Intergenic | 3 | 129199878 | 129219590 | 19713 | 1 | ENSMUST00000174661.3 | -11852 | Pitx2 | ENSMUSG00000028023.11 |
| chr3 | 129189069 | 129190393 | 3.66523 | 4.43936 | 1.37739 | Distal Intergenic | 3 | 129199878 | 129219590 | 19713 | 1 | ENSMUST00000174661.3 | -9485 | Pitx2 | ENSMUSG00000028023.11 |
| chr3 | 129190878 | 129191225 | 3.53639 | 4.29809 | 1.335 | Distal Intergenic | 3 | 129199878 | 129219590 | 19713 | 1 | ENSMUST00000174661.3 | -8653 | Pitx2 | ENSMUSG00000028023.11 |
| chr3 | 129192475 | 129192623 | 3.51976 | 4.52116 | 1.4511 | Distal Intergenic | 3 | 129199878 | 129219590 | 19713 | 1 | ENSMUST00000174661.3 | -7255 | Pitx2 | ENSMUSG00000028023.11 |
| chr3 | 129193381 | 129194519 | 3.85688 | 4.81478 | 1.64609 | Distal Intergenic | 3 | 129199878 | 129219590 | 19713 | 1 | ENSMUST00000174661.3 | -5359 | Pitx2 | ENSMUSG00000028023.11 |
| chr3 | 129194908 | 129195889 | 3.8881 | 5.14715 | 1.88535 | Distal Intergenic | 3 | 129199878 | 129219590 | 19713 | 1 | ENSMUST00000174661.3 | -3989 | Pitx2 | ENSMUSG00000028023.11 |
| chr3 | 129196905 | 129197329 | 3.43532 | 4.27591 | 1.27435 | Distal Intergenic | 3 | 129199878 | 129219590 | 19713 | 1 | ENSMUST00000174661.3 | -2549 | Pitx2 | ENSMUSG00000028023.11 |
| chr3 | 129197692 | 129198751 | 3.80844 | 4.82088 | 1.67472 | Distal Intergenic | 3 | 129199878 | 129219590 | 19713 | 1 | ENSMUST00000174661.3 | -1127 | Pitx2 | ENSMUSG00000028023.11 |
| chr3 | 129199080 | 129208309 | 4.01139 | 5.19573 | 1.92629 | Promoter | 3 | 129199878 | 129219590 | 19713 | 1 | ENSMUST00000174661.3 | 0 | Pitx2 | ENSMUSG00000028023.11 |
| chr3 | 129230840 | 129230984 | 3.40182 | 4.14783 | 1.21075 | Distal Intergenic | 3 | 129214275 | 129216624 | 2350 | 1 | ENSMUST00000174623.2 | 16566 | Pitx2 | ENSMUSG00000028023.11 |
| chr3 | 129235770 | 129236006 | 3.61462 | 4.38196 | 1.42206 | Distal Intergenic | 3 | 129214275 | 129216624 | 2350 | 1 | ENSMUST00000174623.2 | 21496 | Pitx2 | ENSMUSG00000028023.11 |
| chr3 | 129212945 | 129215115 | 3.92387 | 5.03455 | 1.85257 | Promoter | 3 | 129213932 | 129216774 | 2843 | 1 | ENSMUST00000187145.1 | 0 | Pitx2 | ENSMUSG00000028023.11 |
| chr3 | 129215494 | 129216418 | 3.94971 | 5.24336 | 1.946 | 3' UTR | 3 | 129214275 | 129216624 | 2350 | 1 | ENSMUST00000174623.2 | 1220 | Pitx2 | ENSMUSG00000028023.11 |
| chr3 | 129216758 | 129217130 | 3.52696 | 4.5161 | 1.47425 | Exon (ENSMUST00000187145.1/ENSMUSG00000028023.11, exon 1 of 1) | 3 | 129214275 | 129216624 | 2350 | 1 | ENSMUST00000174623.2 | 2484 | Pitx2 | ENSMUSG00000028023.11 |
| chr3 | 129217570 | 129221152 | 3.84447 | 4.95953 | 1.75147 | 3' UTR | 3 | 129214275 | 129216624 | 2350 | 1 | ENSMUST00000174623.2 | 3296 | Pitx2 | ENSMUSG00000028023.11 |
| chr3 | 129221762 | 129222189 | 3.60173 | 4.32353 | 1.3078 | Distal Intergenic | 3 | 129214275 | 129216624 | 2350 | 1 | ENSMUST00000174623.2 | 7488 | Pitx2 | ENSMUSG00000028023.11 |
| chr3 | 129222660 | 129224459 | 3.93767 | 5.06175 | 1.81777 | Distal Intergenic | 3 | 129214275 | 129216624 | 2350 | 1 | ENSMUST00000174623.2 | 8386 | Pitx2 | ENSMUSG00000028023.11 |
| chr3 | 129224784 | 129224870 | 3.61314 | 4.62853 | 1.53417 | Distal Intergenic | 3 | 129214275 | 129216624 | 2350 | 1 | ENSMUST00000174623.2 | 10510 | Pitx2 | ENSMUSG00000028023.11 |
| chr3 | 129225190 | 129225772 | 3.89608 | 5.02298 | 1.8039 | Distal Intergenic | 3 | 129214275 | 129216624 | 2350 | 1 | ENSMUST00000174623.2 | 10916 | Pitx2 | ENSMUSG00000028023.11 |
| chr3 | 129226173 | 129226947 | 4.04343 | 5.24152 | 2.01814 | Distal Intergenic | 3 | 129214275 | 129216624 | 2350 | 1 | ENSMUST00000174623.2 | 11899 | Pitx2 | ENSMUSG00000028023.11 |
| chr3 | 129228523 | 129229096 | 3.92121 | 5.08141 | 1.83633 | Distal Intergenic | 3 | 129214275 | 129216624 | 2350 | 1 | ENSMUST00000174623.2 | 14249 | Pitx2 | ENSMUSG00000028023.11 |
| chr3 | 129229446 | 129229693 | 3.84376 | 5.00026 | 1.78025 | Distal Intergenic | 3 | 129214275 | 129216624 | 2350 | 1 | ENSMUST00000174623.2 | 15172 | Pitx2 | ENSMUSG00000028023.11 |
| chr3 | 129238935 | 129239696 | 3.64004 | 4.46024 | 1.43377 | Distal Intergenic | 3 | 129214275 | 129216624 | 2350 | 1 | ENSMUST00000174623.2 | 24661 | Pitx2 | ENSMUSG00000028023.11 |
| chr3 | 132084427 | 132084517 | 3.56333 | 4.57419 | 1.52693 | Promoter | 3 | 132085292 | 132180293 | 95002 | 1 | ENSMUST00000029665.5 | -775 | Dkk2 | ENSMUSG00000028031.5 |
| chr3 | 132085663 | 132086152 | 3.42778 | 4.00225 | 1.13249 | Promoter | 3 | 132085292 | 132180293 | 95002 | 1 | ENSMUST00000029665.5 | 372 | Dkk2 | ENSMUSG00000028031.5 |
| chr3 | 131489493 | 131490101 | 3.59482 | 4.77627 | 1.64089 | Distal Intergenic | 3 | 131564768 | 131643670 | 78903 | 1 | ENSMUST00000029666.9 | -74667 | Papss1 | ENSMUSG00000028032.9 |
| chr3 | 131490975 | 131491050 | 3.41209 | 4.32301 | 1.32266 | Distal Intergenic | 3 | 131564768 | 131643670 | 78903 | 1 | ENSMUST00000029666.9 | -73718 | Papss1 | ENSMUSG00000028032.9 |
| chr3 | 131492261 | 131492584 | 3.41533 | 4.15321 | 1.20602 | Distal Intergenic | 3 | 131564768 | 131643670 | 78903 | 1 | ENSMUST00000029666.9 | -72184 | Papss1 | ENSMUSG00000028032.9 |
| chr3 | 131496486 | 131497026 | 3.72721 | 4.68949 | 1.60958 | Distal Intergenic | 3 | 131564768 | 131643670 | 78903 | 1 | ENSMUST00000029666.9 | -67742 | Papss1 | ENSMUSG00000028032.9 |
| chr1 | 21960617 | 21962609 | 3.81433 | 5.06728 | 1.82436 | Promoter | 1 | 21398403 | 21961581 | 563179 | 2 | ENSMUST00000029667.7 | 0 | Kcnq5 | ENSMUSG00000028033.11 |
| chr3 | 89318928 | 89319130 | 3.55656 | 4.23329 | 1.22955 | Intron (ENSMUST00000029673.5/ENSMUSG00000028039.7, intron 1 of 4) | 3 | 89315859 | 89321631 | 5773 | 2 | ENSMUST00000139439.1 | 2501 | Efna3 | ENSMUSG00000028039.7 |
| chr3 | 89319521 | 89319887 | 3.91584 | 5.48847 | 2.12634 | Intron (ENSMUST00000029673.5/ENSMUSG00000028039.7, intron 1 of 4) | 3 | 89315859 | 89321631 | 5773 | 2 | ENSMUST00000139439.1 | 1744 | Efna3 | ENSMUSG00000028039.7 |
| chr3 | 89320230 | 89322965 | 4.26845 | 5.71622 | 2.28526 | Promoter | 3 | 89313899 | 89322965 | 9067 | 2 | ENSMUST00000029673.5 | 0 | Efna3 | ENSMUSG00000028039.7 |
| chr3 | 89327675 | 89327959 | 3.24506 | 3.69474 | 1.01733 | Distal Intergenic | 3 | 89313899 | 89322965 | 9067 | 2 | ENSMUST00000029673.5 | -4711 | Efna3 | ENSMUSG00000028039.7 |
| chr3 | 89328807 | 89329189 | 3.76421 | 4.64708 | 1.57204 | Distal Intergenic | 3 | 89313899 | 89322965 | 9067 | 2 | ENSMUST00000029673.5 | -5843 | Efna3 | ENSMUSG00000028039.7 |
| chr3 | 89339047 | 89339684 | 3.54315 | 4.21968 | 1.27623 | Promoter | 3 | 89339745 | 89340352 | 608 | 2 | ENSMUST00000139705.1 | 668 | Adam15 | ENSMUSG00000028041.12 |
| chr3 | 89339990 | 89340394 | 3.51931 | 4.17158 | 1.24252 | Promoter | 3 | 89339745 | 89340352 | 608 | 2 | ENSMUST00000139705.1 | 0 | Adam15 | ENSMUSG00000028041.12 |
| chr3 | 89340934 | 89341169 | 3.50503 | 4.12807 | 1.16462 | Promoter | 3 | 89339641 | 89341157 | 1517 | 2 | ENSMUST00000148068.2 | 0 | Adam15 | ENSMUSG00000028041.12 |
| chr3 | 89144525 | 89144807 | 3.40485 | 3.9499 | 1.06942 | Intron (ENSMUST00000047111.7/ENSMUSG00000041237.7, intron 9 of 10) | 3 | 89148849 | 89151214 | 2366 | 2 | ENSMUST00000127654.1 | 6407 | Hcn3 | ENSMUSG00000028051.5 |
| chr3 | 89146137 | 89146286 | 3.79729 | 4.672 | 1.5091 | 3' UTR | 3 | 89148849 | 89151214 | 2366 | 2 | ENSMUST00000127654.1 | 4928 | Hcn3 | ENSMUSG00000028051.5 |
| chr3 | 89147424 | 89147917 | 3.56323 | 4.23959 | 1.23774 | 3' UTR | 3 | 89148849 | 89151214 | 2366 | 2 | ENSMUST00000127654.1 | 3297 | Hcn3 | ENSMUSG00000028051.5 |
| chr3 | 89159050 | 89159180 | 3.91152 | 4.8776 | 1.62573 | Promoter | 3 | 89151481 | 89159980 | 8500 | 2 | ENSMUST00000133368.1 | 800 | Hcn3 | ENSMUSG00000028051.5 |
| chr3 | 87792216 | 87792499 | 3.54043 | 4.20257 | 1.2398 | Intron (ENSMUST00000029714.9/ENSMUSG00000028073.10, intron 1 of 22) | 3 | 87778244 | 87795162 | 16919 | 2 | ENSMUST00000029712.4 | 2663 | Ntrk1 | ENSMUSG00000028072.6 |
| chr3 | 87793018 | 87793897 | 3.56049 | 4.25959 | 1.27287 | Intron (ENSMUST00000029714.9/ENSMUSG00000028073.10, intron 1 of 22) | 3 | 87778244 | 87795162 | 16919 | 2 | ENSMUST00000029712.4 | 1265 | Ntrk1 | ENSMUSG00000028072.6 |
| chr3 | 87794776 | 87795437 | 4.33183 | 5.94194 | 2.43125 | Promoter | 3 | 87778244 | 87795162 | 16919 | 2 | ENSMUST00000029712.4 | 0 | Ntrk1 | ENSMUSG00000028072.6 |
| chr3 | 87791243 | 87791749 | 3.49726 | 4.12524 | 1.19583 | Exon (ENSMUST00000029712.4/ENSMUSG00000028072.6, exon 2 of 17) | 3 | 87778244 | 87795162 | 16919 | 2 | ENSMUST00000029712.4 | 3413 | Ntrk1 | ENSMUSG00000028072.6 |
| chr3 | 87756636 | 87756998 | 3.45171 | 4.04711 | 1.16295 | Promoter | 3 | 87752317 | 87756853 | 4537 | 2 | ENSMUST00000174219.1 | 0 | Pear1 | ENSMUSG00000028073.10 |
| chr3 | 87771881 | 87771956 | 4.09516 | 5.27031 | 1.97502 | Intron (ENSMUST00000029714.9/ENSMUSG00000028073.10, intron 1 of 22) | 3 | 87758358 | 87768966 | 10609 | 2 | ENSMUST00000174776.2 | -2916 | Pear1 | ENSMUSG00000028073.10 |
| chr3 | 87778831 | 87779458 | 3.36496 | 3.87501 | 1.01733 | Intron (ENSMUST00000029714.9/ENSMUSG00000028073.10, intron 1 of 22) | 3 | 87758358 | 87768966 | 10609 | 2 | ENSMUST00000174776.2 | -9866 | Pear1 | ENSMUSG00000028073.10 |
| chr3 | 87813062 | 87813290 | 3.45305 | 4.08835 | 1.2618 | Exon (ENSMUST00000029711.3/ENSMUSG00000005640.6, exon 17 of 22) | 3 | 87749097 | 87824255 | 75159 | 2 | ENSMUST00000029714.9 | 10965 | Pear1 | ENSMUSG00000028073.10 |
| chr3 | 86864466 | 86864595 | 3.41662 | 3.97401 | 1.09183 | Intron (ENSMUST00000029719.8/ENSMUSG00000028078.9, intron 2 of 15) | 3 | 86786150 | 86920884 | 134735 | 2 | ENSMUST00000029719.8 | 56289 | Dclk2 | ENSMUSG00000028078.9 |
| chr3 | 86875574 | 86875769 | 3.45644 | 4.05593 | 1.16988 | Intron (ENSMUST00000029719.8/ENSMUSG00000028078.9, intron 2 of 15) | 3 | 86786150 | 86920884 | 134735 | 2 | ENSMUST00000029719.8 | 45115 | Dclk2 | ENSMUSG00000028078.9 |
| chr3 | 86883815 | 86884026 | 3.29178 | 3.76483 | 1.01733 | Intron (ENSMUST00000029719.8/ENSMUSG00000028078.9, intron 2 of 15) | 3 | 86786150 | 86920884 | 134735 | 2 | ENSMUST00000029719.8 | 36858 | Dclk2 | ENSMUSG00000028078.9 |
| chr3 | 86922165 | 86922273 | 3.23572 | 3.684 | 1.02576 | Distal Intergenic | 3 | 86786150 | 86920884 | 134735 | 2 | ENSMUST00000029719.8 | -1282 | Dclk2 | ENSMUSG00000028078.9 |
| chr3 | 97189248 | 97189367 | 3.37837 | 3.89522 | 1.01733 | Distal Intergenic | 3 | 97172642 | 97176518 | 3877 | 1 | ENSMUST00000143234.1 | 16607 | Acp6 | ENSMUSG00000028093.10 |
| chr3 | 122408963 | 122409343 | 3.55462 | 4.22558 | 1.23481 | Distal Intergenic | 3 | 122419780 | 122530180 | 110401 | 1 | ENSMUST00000029766.4 | -10437 | Bcar3 | ENSMUSG00000028121.4 |
| chr3 | 122024923 | 122025190 | 3.35027 | 3.85286 | 1.01733 | Distal Intergenic | 3 | 122044443 | 122180061 | 135619 | 1 | ENSMUST00000013995.8 | -19253 | Abca4 | ENSMUSG00000028125.9 |
| chr3 | 122034166 | 122034275 | 3.3633 | 3.90727 | 1.13518 | Distal Intergenic | 3 | 122044443 | 122180061 | 135619 | 1 | ENSMUST00000013995.8 | -10168 | Abca4 | ENSMUSG00000028125.9 |
| chr3 | 121262310 | 121262651 | 3.71398 | 4.65136 | 1.55157 | Promoter | 3 | 121202008 | 121263159 | 61152 | 2 | ENSMUST00000128909.2 | 508 | Tmem56 | ENSMUSG00000028132.10 |
| chr3 | 121263298 | 121263619 | 3.86663 | 4.87958 | 1.72451 | Promoter | 3 | 121202008 | 121263340 | 61333 | 2 | ENSMUST00000029777.8 | 0 | Tmem56 | ENSMUSG00000028132.10 |
| chr3 | 121265520 | 121265640 | 3.42285 | 4.22998 | 1.30149 | Intron (ENSMUST00000135818.2/ENSMUSG00000028132.10, intron 2 of 10) | 3 | 121202008 | 121263340 | 61333 | 2 | ENSMUST00000029777.8 | -2181 | Tmem56 | ENSMUSG00000028132.10 |
| chr3 | 121268601 | 121268759 | 3.593 | 4.34494 | 1.41013 | Intron (ENSMUST00000135818.2/ENSMUSG00000028132.10, intron 2 of 10) | 3 | 121202008 | 121263340 | 61333 | 2 | ENSMUST00000029777.8 | -5262 | Tmem56 | ENSMUSG00000028132.10 |
| chr3 | 134829110 | 134829315 | 3.34609 | 3.9918 | 1.09682 | Promoter | 3 | 134829007 | 134934581 | 105575 | 1 | ENSMUST00000029822.4 | 104 | Tacr3 | ENSMUSG00000028172.4 |
| chr3 | 158560624 | 158561102 | 3.78421 | 4.70475 | 1.5793 | Intron (ENSMUST00000106044.1/ENSMUSG00000028176.7, intron 1 of 27) | 3 | 158084439 | 158562221 | 477783 | 2 | ENSMUST00000106044.1 | 1119 | Lrrc7 | ENSMUSG00000028176.7 |
| chr3 | 158561487 | 158563027 | 3.97445 | 5.1754 | 1.9233 | Promoter | 3 | 158084439 | 158562221 | 477783 | 2 | ENSMUST00000106044.1 | 0 | Lrrc7 | ENSMUSG00000028176.7 |
| chr3 | 157523956 | 157524031 | 3.56108 | 4.29694 | 1.36447 | Distal Intergenic | 3 | 157534160 | 157548390 | 14231 | 1 | ENSMUST00000106058.2 | -10129 | Zranb2 | ENSMUSG00000028180.13 |
| chr3 | 157525103 | 157525275 | 3.35653 | 3.88871 | 1.11019 | Distal Intergenic | 3 | 157534160 | 157548390 | 14231 | 1 | ENSMUST00000106058.2 | -8885 | Zranb2 | ENSMUSG00000028180.13 |
| chr3 | 157527854 | 157528174 | 3.78204 | 4.78046 | 1.6357 | Distal Intergenic | 3 | 157534160 | 157548390 | 14231 | 1 | ENSMUST00000106058.2 | -5986 | Zranb2 | ENSMUSG00000028180.13 |
| chr3 | 157533090 | 157533251 | 3.60448 | 4.49937 | 1.45392 | Promoter | 3 | 157534160 | 157548390 | 14231 | 1 | ENSMUST00000106058.2 | -909 | Zranb2 | ENSMUSG00000028180.13 |
| chr3 | 148728600 | 148728675 | 3.4281 | 4.13876 | 1.21409 | Distal Intergenic | 3 | 148815856 | 148822979 | 7124 | 2 | ENSMUST00000168352.1 | 94304 | Lphn2 | ENSMUSG00000028184.8 |
| chr3 | 145293013 | 145293565 | 3.6337 | 4.39539 | 1.35883 | Promoter | 3 | 145292527 | 145314047 | 21521 | 1 | ENSMUST00000139001.1 | 487 | Col24a1 | ENSMUSG00000028197.4 |
| chr3 | 145294100 | 145294290 | 3.74158 | 4.67662 | 1.58258 | Intron (ENSMUST00000029848.4/ENSMUSG00000028197.4, intron 1 of 59) | 3 | 145292527 | 145314047 | 21521 | 1 | ENSMUST00000139001.1 | 1574 | Col24a1 | ENSMUSG00000028197.4 |
| chr4 | 15880708 | 15880959 | 3.34406 | 3.85416 | 1.04426 | Promoter | 4 | 15881264 | 15908064 | 26801 | 1 | ENSMUST00000029876.1 | -305 | Calb1 | ENSMUSG00000028222.2 |
| chr4 | 15881288 | 15881954 | 3.62018 | 4.36923 | 1.38537 | Promoter | 4 | 15881266 | 15895777 | 14512 | 1 | ENSMUST00000136266.1 | 23 | Calb1 | ENSMUSG00000028222.2 |

| | | | | | | | | | | | | | | | |
|---|---|---|---|---|---|---|---|---|---|---|---|---|---|---|---|
| chr4 | 17853722 | 17854185 | 3.33129 | 3.841 | 1.07287 | Promoter | 4 | 17853593 | 18054681 | 201089 | 1 | ENSMUST00000133416.1 | 130 | Mmp16 | ENSMUSG00000028226.10 |
| chr4 | 21929525 | 21929817 | 3.52413 | 4.28821 | 1.31966 | Distal Intergenic | 4 | 21931329 | 21994146 | 62818 | 1 | ENSMUST00000124440.2 | -1512 | Faxc | ENSMUSG00000028246.8 |
| chr4 | 21931091 | 21932123 | 3.85945 | 4.8259 | 1.65135 | Promoter | 4 | 21931329 | 21994146 | 62818 | 1 | ENSMUST00000124440.2 | 0 | Faxc | ENSMUSG00000028246.8 |
| chr4 | 21715607 | 21715690 | 3.45591 | 4.07723 | 1.23739 | Distal Intergenic | 4 | 21727701 | 21750546 | 22846 | 1 | ENSMUST00000065928.5 | -12011 | Ccnc | ENSMUSG00000028252.15 |
| chr3 | 144086901 | 144087198 | 3.40574 | 4.09004 | 1.17931 | Distal Intergenic | 3 | 144189062 | 144194818 | 5757 | 2 | ENSMUST00000148128.1 | 107620 | Lmo4 | ENSMUSG00000028266.12 |
| chr3 | 144096443 | 144096947 | 3.67229 | 4.61345 | 1.55404 | Distal Intergenic | 3 | 144189062 | 144194818 | 5757 | 2 | ENSMUST00000148128.1 | 97871 | Lmo4 | ENSMUSG00000028266.12 |
| chr3 | 144097635 | 144098133 | 3.28032 | 3.74964 | 1.0243 | Distal Intergenic | 3 | 144189062 | 144194818 | 5757 | 2 | ENSMUST00000148128.1 | 96685 | Lmo4 | ENSMUSG00000028266.12 |
| chr3 | 144098497 | 144098572 | 4.48776 | 6.3145 | 2.66323 | Distal Intergenic | 3 | 144189062 | 144194818 | 5757 | 2 | ENSMUST00000148128.1 | 96246 | Lmo4 | ENSMUSG00000028266.12 |
| chr3 | 144223152 | 144223323 | 3.45005 | 4.0628 | 1.20197 | Distal Intergenic | 3 | 144188530 | 144205220 | 16691 | 2 | ENSMUST00000120539.2 | -17933 | Lmo4 | ENSMUSG00000028266.12 |
| chr3 | 144367778 | 144368007 | 3.27847 | 3.74484 | 1.01733 | Distal Intergenic | 3 | 144188530 | 144205220 | 16691 | 2 | ENSMUST00000120539.2 | -162559 | Lmo4 | ENSMUSG00000028266.12 |
| chr4 | 31822517 | 31822832 | 3.32473 | 3.81436 | 1.01733 | Distal Intergenic | 4 | 31964097 | 32023467 | 59371 | 1 | ENSMUST00000037607.5 | -141265 | Map3k7 | ENSMUSG00000028284.8 |
| chr4 | 31824695 | 31824826 | 3.41428 | 3.96132 | 1.05921 | Distal Intergenic | 4 | 31964097 | 32023467 | 59371 | 1 | ENSMUST00000037607.5 | -139271 | Map3k7 | ENSMUSG00000028284.8 |
| chr4 | 31825671 | 31825815 | 3.35901 | 3.86601 | 1.01733 | Distal Intergenic | 4 | 31964097 | 32023467 | 59371 | 1 | ENSMUST00000037607.5 | -138282 | Map3k7 | ENSMUSG00000028284.8 |
| chr4 | 31835868 | 31835991 | 3.82284 | 4.85887 | 1.71034 | Distal Intergenic | 4 | 31964097 | 32023467 | 59371 | 1 | ENSMUST00000037607.5 | -128106 | Map3k7 | ENSMUSG00000028284.8 |
| chr4 | 31937305 | 31937734 | 3.52241 | 4.37479 | 1.36059 | Distal Intergenic | 4 | 31964097 | 32023467 | 59371 | 1 | ENSMUST00000037607.5 | -26363 | Map3k7 | ENSMUSG00000028284.8 |
| chr4 | 31938158 | 31938560 | 3.65931 | 4.44254 | 1.44358 | Distal Intergenic | 4 | 31964097 | 32023467 | 59371 | 1 | ENSMUST00000037607.5 | -25537 | Map3k7 | ENSMUSG00000028284.8 |
| chr4 | 34728805 | 34728900 | 4.99416 | 7.41631 | 3.56281 | Exon (ENSMUST00000029971.6/ENSMUSG00000028294.10, exon 2 of 13) | 4 | 34711332 | 34730178 | 18847 | 2 | ENSMUST00000108136.2 | 1278 | 1700003M02Rik | ENSMUSG00000028294.10 |
| chr4 | 45849034 | 45849142 | 3.79966 | 4.74214 | 1.65441 | Promoter | 4 | 45848997 | 45860016 | 11020 | 1 | ENSMUST00000128947.1 | 38 | 1300002K09Rik | ENSMUSG00000028327.9 |
| chr4 | 45853893 | 45854212 | 3.41417 | 3.99169 | 1.16414 | Intron (ENSMUST00000107783.2/ENSMUSG00000028327.9, intron 2 of 18) | 4 | 45848997 | 45860016 | 11020 | 1 | ENSMUST00000128947.1 | 4897 | 1300002K09Rik | ENSMUSG00000028327.9 |
| chr4 | 45869181 | 45869402 | 3.27032 | 3.74692 | 1.05993 | Exon (ENSMUST00000107783.2/ENSMUSG00000028327.9, exon 8 of 19) | 4 | 45874211 | 45886618 | 12408 | 1 | ENSMUST00000165478.1 | -4809 | 1300002K09Rik | ENSMUSG00000028327.9 |
| chr4 | 46039086 | 46039821 | 3.25842 | 4.23454 | 1.28094 | Promoter | 4 | 46039209 | 46090897 | 51689 | 1 | ENSMUST00000156200.2 | 0 | Tmod1 | ENSMUSG00000028328.8 |
| chr4 | 46074645 | 46075027 | 3.57555 | 4.27737 | 1.31314 | Intron (ENSMUST00000107773.2/ENSMUSG00000028328.8, intron 2 of 9) | 4 | 46077127 | 46099708 | 22582 | 1 | ENSMUST00000128689.2 | -2100 | Tmod1 | ENSMUSG00000028328.8 |
| chr4 | 46202690 | 46203169 | 3.94226 | 5.05209 | 1.83865 | Distal Intergenic | 4 | 46175222 | 46196311 | 21090 | 2 | ENSMUST00000030013.6 | -6380 | Xpa | ENSMUSG00000028329.8 |
| chr4 | 46371334 | 46371950 | 3.88242 | 4.86298 | 1.65206 | Distal Intergenic | 4 | 46376505 | 46389418 | 12914 | 2 | ENSMUST00000151903.2 | 17468 | 5830415F09Rik | ENSMUSG00000028331.7 |
| chr4 | 46600606 | 46600912 | 3.34153 | 3.83966 | 1.01733 | Promoter | 4 | 46543001 | 46601891 | 58891 | 2 | ENSMUST00000139179.1 | 979 | Coro2a | ENSMUSG00000028337.9 |
| chr4 | 46601345 | 46601747 | 3.94688 | 5.01605 | 1.78869 | Promoter | 4 | 46543001 | 46601891 | 58891 | 2 | ENSMUST00000139179.1 | 144 | Coro2a | ENSMUSG00000028337.9 |
| chr4 | 46620504 | 46620811 | 3.82494 | 4.75823 | 1.63234 | Exon (ENSMUST00000084621.6/ENSMUSG00000039813.9, exon 6 of 13) | 4 | 46536943 | 46602202 | 65260 | 2 | ENSMUST00000107756.2 | -18303 | Coro2a | ENSMUSG00000028337.9 |
| chr4 | 47207734 | 47209143 | 3.67682 | 4.57509 | 1.52343 | Promoter | 4 | 47208161 | 47313163 | 105003 | 1 | ENSMUST00000082303.7 | 0 | Col15a1 | ENSMUSG00000028339.12 |
| chr4 | 48047535 | 48048379 | 3.71749 | 4.6718 | 1.59658 | Promoter | 4 | 48049114 | 48086447 | 37334 | 1 | ENSMUST00000153369.1 | -735 | Nr4a3 | ENSMUSG00000028341.4 |
| chr4 | 48048929 | 48049106 | 3.78087 | 4.7844 | 1.65013 | Promoter | 4 | 48049114 | 48086447 | 37334 | 1 | ENSMUST00000153369.1 | -8 | Nr4a3 | ENSMUSG00000028341.4 |
| chr4 | 48051777 | 48052215 | 4.29063 | 5.81562 | 2.36199 | Promoter | 4 | 48051248 | 48056739 | 5492 | 1 | ENSMUST00000133539.1 | 530 | Nr4a3 | ENSMUSG00000028341.4 |
| chr4 | 48054614 | 48054866 | 3.45729 | 4.26051 | 1.33372 | Intron (ENSMUST00000030025.4/ENSMUSG00000028341.4, intron 3 of 7) | 4 | 48051248 | 48056739 | 5492 | 1 | ENSMUST00000133539.1 | 3367 | Nr4a3 | ENSMUSG00000028341.4 |
| chr4 | 68953865 | 68954354 | 3.82451 | 4.81875 | 1.73311 | Promoter | 4 | 68761514 | 68954397 | 192884 | 2 | ENSMUST00000030036.5 | 43 | Brinp1 | ENSMUSG00000028351.5 |
| chr1 | 174501094 | 174501169 | 3.47207 | 4.08585 | 1.19167 | Promoter | 1 | 174501825 | 174822717 | 320893 | 1 | ENSMUST00000030039.7 | -656 | Fmn2 | ENSMUSG00000028354.8 |
| chr1 | 174501812 | 174503488 | 3.87519 | 4.99762 | 1.81991 | Promoter | 1 | 174501825 | 174822717 | 320893 | 1 | ENSMUST00000030039.7 | 0 | Fmn2 | ENSMUSG00000028354.8 |
| chr3 | 153972988 | 153973905 | 3.8525 | 4.87779 | 1.65763 | Promoter | 3 | 153973436 | 154271722 | 298287 | 1 | ENSMUST00000089948.5 | 0 | Slc44a5 | ENSMUSG00000028360.9 |
| chr3 | 153974277 | 153974806 | 3.72248 | 4.53689 | 1.42158 | Promoter | 3 | 153973436 | 154271722 | 298287 | 1 | ENSMUST00000089948.5 | 842 | Slc44a5 | ENSMUSG00000028360.9 |
| chr4 | 64046674 | 64046924 | 3.48957 | 4.10529 | 1.1646 | Promoter | 4 | 63959785 | 64047015 | 87231 | 2 | ENSMUST00000107377.4 | 91 | Tnc | ENSMUSG00000028364.10 |
| chr4 | 65124601 | 65125138 | 3.80371 | 5.03765 | 1.79591 | Promoter | 4 | 65124174 | 65357509 | 233336 | 1 | ENSMUST00000084501.3 | 428 | Pappa | ENSMUSG00000028370.7 |
| chr4 | 78210210 | 78210850 | 3.63432 | 4.37895 | 1.34863 | Promoter | 4 | 76593551 | 78211739 | 1618189 | 2 | ENSMUST00000138955.2 | 889 | Ptprd | ENSMUSG00000028399.12 |
| chr4 | 78211567 | 78211858 | 3.79845 | 4.70036 | 1.59107 | Promoter | 4 | 76593551 | 78211739 | 1618189 | 2 | ENSMUST00000138955.2 | 0 | Ptprd | ENSMUSG00000028399.12 |
| chr4 | 78212173 | 78213092 | 3.57105 | 4.25989 | 1.27177 | Promoter | 4 | 77351290 | 78211961 | 860672 | 2 | ENSMUST00000140422.2 | -213 | Ptprd | ENSMUSG00000028399.12 |
| chr4 | 78213405 | 78213622 | 3.37682 | 3.90179 | 1.04874 | Intron (ENSMUST00000173049.1/ENSMUSG00000092492.1, intron 1 of 1) | 4 | 77351290 | 78211961 | 860672 | 2 | ENSMUST00000140422.2 | -1445 | Ptprd | ENSMUSG00000028399.12 |
| chr4 | 81442536 | 81442878 | 3.61324 | 4.35573 | 1.38646 | Promoter | 4 | 81278500 | 81442805 | 164306 | 2 | ENSMUST00000107262.2 | 0 | Mpdz | ENSMUSG00000028402.12 |
| chr4 | 81443633 | 81444275 | 3.44915 | 4.11367 | 1.25697 | Promoter | 4 | 81416540 | 81442815 | 26276 | 2 | ENSMUST00000142273.1 | -819 | Mpdz | ENSMUSG00000028402.12 |
| chr4 | 57092780 | 57093039 | 3.42415 | 3.98585 | 1.09361 | Intron (ENSMUST00000095076.4/ENSMUSG00000028434.7, intron 2 of 25) | 4 | 57073300 | 57103453 | 30154 | 2 | ENSMUST00000149109.2 | 10414 | Epb4.1l4b | ENSMUSG00000028434.7 |
| chr4 | 57094224 | 57094511 | 3.35452 | 3.85925 | 1.01733 | Intron (ENSMUST00000095076.4/ENSMUSG00000028434.7, intron 2 of 25) | 4 | 57073300 | 57103453 | 30154 | 2 | ENSMUST00000149109.2 | 8942 | Epb4.1l4b | ENSMUSG00000028434.7 |
| chr4 | 57141933 | 57143102 | 3.62705 | 4.43024 | 1.40057 | Promoter | 4 | 56991972 | 57143437 | 151466 | 2 | ENSMUST00000095076.4 | 335 | Epb4.1l4b | ENSMUSG00000028434.7 |
| chr4 | 41513805 | 41513907 | 3.36254 | 3.87135 | 1.01733 | Intron (ENSMUST00000030152.7/ENSMUSG00000028441.7, intron 1 of 6) | 4 | 41505009 | 41511467 | 6459 | 2 | ENSMUST00000125303.2 | -2339 | 1110017D15Rik | ENSMUSG00000028441.7 |
| chr4 | 41681931 | 41682006 | 3.43314 | 4.00262 | 1.10486 | Intron (ENSMUST00000102961.4/ENSMUSG00000028444.12, intron 2 of 9) | 4 | 41657498 | 41695442 | 37945 | 2 | ENSMUST00000102961.4 | 13436 | Cntfr | ENSMUSG00000028444.12 |
| chr4 | 41683003 | 41684209 | 4.21867 | 5.92452 | 2.44108 | Intron (ENSMUST00000102961.4/ENSMUSG00000028444.12, intron 2 of 9) | 4 | 41657498 | 41695442 | 37945 | 2 | ENSMUST00000102961.4 | 11233 | Cntfr | ENSMUSG00000028444.12 |
| chr4 | 41684669 | 41685577 | 3.41198 | 4.14094 | 1.23071 | Intron (ENSMUST00000102961.4/ENSMUSG00000028444.12, intron 2 of 9) | 4 | 41657498 | 41695442 | 37945 | 2 | ENSMUST00000102961.4 | 9865 | Cntfr | ENSMUSG00000028444.12 |
| chr4 | 41693976 | 41694329 | 3.60537 | 4.31358 | 1.27652 | Intron (ENSMUST00000102961.4/ENSMUSG00000028444.12, intron 1 of 9) | 4 | 41657498 | 41695442 | 37945 | 2 | ENSMUST00000102961.4 | 1113 | Cntfr | ENSMUSG00000028444.12 |
| chr4 | 41694649 | 41694954 | 3.46054 | 4.05238 | 1.13453 | Promoter | 4 | 41657498 | 41695442 | 37945 | 2 | ENSMUST00000102961.4 | 488 | Cntfr | ENSMUSG00000028444.12 |
| chr4 | 41725713 | 41726307 | 4.28484 | 5.79921 | 2.39358 | Exon (ENSMUST00000171251.2/ENSMUSG00000066224.8, exon 4 of 8) | 4 | 41714798 | 41723170 | 8373 | 2 | ENSMUST00000030158.5 | -2544 | Dctn3 | ENSMUSG00000028447.6 |
| chr4 | 42964622 | 42965041 | 3.58633 | 4.32481 | 1.34408 | Distal Intergenic | 4 | 42969604 | 42971332 | 1729 | 1 | ENSMUST00000189414.1 | -4563 | 1700022I11Rik | ENSMUSG00000028451.7 |
| chr4 | 43454253 | 43454580 | 3.4651 | 4.05643 | 1.12412 | Promoter | 4 | 43447724 | 43454563 | 6840 | 2 | ENSMUST00000107926.2 | 0 | Cd72 | ENSMUSG00000028459.6 |
| chr4 | 43519085 | 43519410 | 3.90872 | 4.89806 | 1.67235 | Promoter | 4 | 43515682 | 43519669 | 3988 | 2 | ENSMUST00000133355.1 | 259 | Tpm2 | ENSMUSG00000028464.11 |
| chr4 | 44084520 | 44084875 | 3.82631 | 4.81328 | 1.64227 | Promoter | 4 | 44034075 | 44084177 | 50103 | 2 | ENSMUST00000030201.8 | -344 | Gne | ENSMUSG00000028479.13 |
| chr4 | 43956680 | 43956965 | 3.39229 | 3.94354 | 1.11441 | Promoter | 4 | 43957401 | 43979118 | 21718 | 1 | ENSMUST00000030202.8 | -436 | Glipr2 | ENSMUSG00000028480.9 |
| chr4 | 43957438 | 43958676 | 4.05404 | 5.38547 | 2.06455 | Promoter | 4 | 43957692 | 43979118 | 21427 | 1 | ENSMUST00000107855.1 | 0 | Glipr2 | ENSMUSG00000028480.9 |
| chr4 | 43959556 | 43959961 | 3.44461 | 4.04651 | 1.19834 | Intron (ENSMUST00000030202.8/ENSMUSG00000028480.9, intron 1 of 4) | 4 | 43957692 | 43979118 | 21427 | 1 | ENSMUST00000107855.1 | 1865 | Glipr2 | ENSMUSG00000028480.9 |
| chr4 | 84543796 | 84544832 | 3.99443 | 5.39748 | 2.08424 | Intron (ENSMUST00000102820.3/ENSMUSG00000028487.13, intron 3 of 7) | 4 | 84411154 | 84546290 | 135137 | 2 | ENSMUST00000177040.1 | 1458 | Bnc2 | ENSMUSG00000028487.13 |
| chr4 | 84545278 | 84545502 | 4.62071 | 6.55739 | 2.9169 | Promoter | 4 | 84411154 | 84546290 | 135137 | 2 | ENSMUST00000177040.1 | 788 | Bnc2 | ENSMUSG00000028487.13 |
| chr4 | 84546398 | 84546914 | 3.70662 | 4.82137 | 1.67705 | Promoter | 4 | 84390959 | 84546619 | 155661 | 2 | ENSMUST00000177256.2 | 0 | Bnc2 | ENSMUSG00000028487.13 |
| chr4 | 84674160 | 84676063 | 4.0024 | 5.2705 | 1.97171 | Promoter | 4 | 84275095 | 84675034 | 399940 | 2 | ENSMUST00000102820.3 | 0 | Bnc2 | ENSMUSG00000028487.13 |
| chr4 | 85203576 | 85203993 | 3.75509 | 4.65357 | 1.60922 | Distal Intergenic | 4 | 85205126 | 85399529 | 194404 | 1 | ENSMUST00000107188.4 | -1133 | Sh3gl2 | ENSMUSG00000028488.10 |
| chr4 | 85205045 | 85206371 | 4.56538 | 6.47672 | 2.89127 | Promoter | 4 | 85205126 | 85399529 | 194404 | 1 | ENSMUST00000107188.4 | 0 | Sh3gl2 | ENSMUSG00000028488.10 |
| chr4 | 105137344 | 105137431 | 4.2604 | 5.59202 | 2.19536 | Distal Intergenic | 4 | 105157347 | 105232764 | 75418 | 1 | ENSMUST00000064139.7 | -19916 | Ppap2b | ENSMUSG00000028517.8 |
| chr4 | 105120424 | 105120569 | 3.46245 | 4.04741 | 1.10632 | Distal Intergenic | 4 | 105029874 | 105109890 | 80017 | 2 | ENSMUST00000030243.7 | -10535 | Prkaa2 | ENSMUSG00000028518.8 |
| chr4 | 105125082 | 105125202 | 3.31167 | 3.79472 | 1.01733 | Distal Intergenic | 4 | 105029874 | 105109890 | 80017 | 2 | ENSMUST00000030243.7 | -15193 | Prkaa2 | ENSMUSG00000028518.8 |
| chr4 | 105127625 | 105127714 | 3.88951 | 4.89241 | 1.74411 | Distal Intergenic | 4 | 105029874 | 105109890 | 80017 | 2 | ENSMUST00000030243.7 | -17736 | Prkaa2 | ENSMUSG00000028518.8 |
| chr4 | 105130377 | 105130764 | 4.10443 | 5.35793 | 2.07597 | Distal Intergenic | 4 | 105029874 | 105109890 | 80017 | 2 | ENSMUST00000030243.7 | -20488 | Prkaa2 | ENSMUSG00000028518.8 |
| chr4 | 105131247 | 105131656 | 3.67188 | 4.53451 | 1.49837 | Distal Intergenic | 4 | 105029874 | 105109890 | 80017 | 2 | ENSMUST00000030243.7 | -21358 | Prkaa2 | ENSMUSG00000028518.8 |
| chr4 | 103620139 | 103620587 | 3.568 | 4.28279 | 1.33616 | Promoter | 4 | 103619665 | 104680382 | 1060718 | 1 | ENSMUST00000106827.2 | 475 | Dab1 | ENSMUSG00000028519.11 |
| chr4 | 103621368 | 103621612 | 3.7633 | 4.63591 | 1.50614 | Intron (ENSMUST00000143294.2/ENSMUSG00000028519.11, intron 1 of 3) | 4 | 103619665 | 104680382 | 1060718 | 1 | ENSMUST00000106827.2 | 1704 | Dab1 | ENSMUSG00000028519.11 |
| chr4 | 104366664 | 104367744 | 3.77434 | 4.80864 | 1.64846 | Promoter | 4 | 104367113 | 104679496 | 312384 | 1 | ENSMUST00000143644.2 | 0 | Dab1 | ENSMUSG00000028519.11 |
| chr4 | 104368201 | 104368987 | 3.77003 | 4.71675 | 1.61929 | Promoter | 4 | 104367590 | 104744844 | 377255 | 1 | ENSMUST00000106830.3 | 612 | Dab1 | ENSMUSG00000028519.11 |
| chr4 | 104369306 | 104369733 | 3.3475 | 3.84869 | 1.01733 | Intron (ENSMUST00000106827.2/ENSMUSG00000028519.11, intron 3 of 9) | 4 | 104367590 | 104744844 | 377255 | 1 | ENSMUST00000106830.3 | 1717 | Dab1 | ENSMUSG00000028519.11 |
| chr4 | 102086339 | 102086655 | 3.89018 | 4.90816 | 1.75615 | Promoter | 4 | 102087543 | 102605833 | 518291 | 1 | ENSMUST00000106911.2 | -888 | Pde4b | ENSMUSG00000028525.11 |
| chr4 | 102087751 | 102088140 | 3.84931 | 4.91472 | 1.753 | Promoter | 4 | 102087741 | 102557597 | 469857 | 1 | ENSMUST00000173119.2 | 11 | Pde4b | ENSMUSG00000028525.11 |
| chr4 | 102088531 | 102088660 | 3.43531 | 4.02858 | 1.18173 | Promoter | 4 | 102087741 | 102557597 | 469857 | 1 | ENSMUST00000173119.2 | 791 | Pde4b | ENSMUSG00000028525.11 |
| chr4 | 102089310 | 102089647 | 3.91373 | 5.00258 | 1.79684 | Intron (ENSMUST00000106911.2/ENSMUSG00000028525.11, intron 1 of 16) | 4 | 102087741 | 102557597 | 469857 | 1 | ENSMUST00000173119.2 | 1570 | Pde4b | ENSMUSG00000028525.11 |
| chr4 | 101495732 | 101496064 | 3.52614 | 4.1729 | 1.20758 | Promoter | 4 | 101496648 | 101642799 | 146152 | 1 | ENSMUST00000106930.2 | -584 | Dnajc6 | ENSMUSG00000028528.11 |
| chr4 | 101496432 | 101496955 | 4.13202 | 5.67735 | 2.24969 | Promoter | 4 | 101496648 | 101642799 | 146152 | 1 | ENSMUST00000106930.2 | 0 | Dnajc6 | ENSMUSG00000028528.11 |
| chr4 | 101497352 | 101497427 | 4.32277 | 5.68294 | 2.18664 | Promoter | 4 | 101496685 | 101606603 | 109919 | 1 | ENSMUST00000154120.2 | 668 | Dnajc6 | ENSMUSG00000028528.11 |
| chr4 | 101507193 | 101507728 | 3.47265 | 4.27606 | 1.34924 | Promoter | 4 | 101507871 | 101642799 | 134929 | 1 | ENSMUST00000038207.6 | -143 | Dnajc6 | ENSMUSG00000028528.11 |
| chr4 | 101508576 | 101509501 | 3.5574 | 4.30447 | 1.38108 | Promoter | 4 | 101508030 | 101599104 | 91075 | 1 | ENSMUST00000146489.1 | 547 | Dnajc6 | ENSMUSG00000028528.11 |
| chr4 | 117926638 | 117927199 | 3.6075 | 4.37828 | 1.42603 | Exon (ENSMUST00000070816.3/ENSMUSG00000028539.9, exon 4 of 5) | 4 | 117928144 | 117929413 | 1270 | 2 | ENSMUST00000150612.1 | 2214 | Artn | ENSMUSG00000028539.9 |
| chr4 | 117927686 | 117927767 | 3.45718 | 4.36323 | 1.35342 | 5' UTR | 4 | 117928144 | 117929413 | 1270 | 2 | ENSMUST00000150612.1 | 1646 | Artn | ENSMUSG00000028539.9 |
| chr4 | 110225560 | 110225750 | 3.75513 | 4.66087 | 1.58797 | Exon (ENSMUST00000144554.1/ENSMUSG00000028546.12, exon 4 of 4) | 4 | 110206359 | 110223148 | 16790 | 2 | ENSMUST00000153200.1 | -2413 | Elavl4 | ENSMUSG00000028546.12 |
| chr4 | 110285467 | 110286187 | 3.58571 | 4.27641 | 1.25552 | Promoter | 4 | 110213217 | 110286616 | 73400 | 2 | ENSMUST00000138972.2 | 429 | Elavl4 | ENSMUSG00000028546.12 |
| chr4 | 110286558 | 110286635 | 4.37917 | 5.79309 | 2.26213 | Promoter | 4 | 110213217 | 110286616 | 73400 | 2 | ENSMUST00000138972.2 | 0 | Elavl4 | ENSMUSG00000028546.12 |
| chr4 | 110351447 | 110351921 | 3.3909 | 3.94337 | 1.11215 | Promoter | 4 | 110203737 | 110351909 | 148173 | 2 | ENSMUST00000106603.3 | 0 | Elavl4 | ENSMUSG00000028546.12 |
| chr4 | 109660962 | 109663594 | 3.94698 | 5.05632 | 1.86391 | 3' UTR | 4 | 109660876 | 109665372 | 4497 | 2 | ENSMUST00000063531.4 | 1778 | Cdkn2c | ENSMUSG00000028551.9 |
| chr4 | 109664186 | 109666056 | 3.86026 | 4.88408 | 1.76247 | Promoter | 4 | 109660876 | 109665372 | 4497 | 2 | ENSMUST00000063531.4 | 0 | Cdkn2c | ENSMUSG00000028551.9 |
| chr4 | 109406891 | 109407732 | 3.42245 | 3.99374 | 1.108 | Promoter | 4 | 109407087 | 109444745 | 37659 | 1 | ENSMUST00000064129.8 | 0 | Ttc39a | ENSMUSG00000028555.10 |
| chr4 | 130910550 | 130910769 | 3.49174 | 4.10331 | 1.14756 | Distal Intergenic | 4 | 130913125 | 130936141 | 23017 | 1 | ENSMUST00000151698.2 | -2356 | Laptm5 | ENSMUSG00000028581.12 |
| chr4 | 143348103 | 143348716 | 3.52971 | 4.18386 | 1.22271 | Distal Intergenic | 4 | 143349757 | 143371032 | 21276 | 1 | ENSMUST00000052458.2 | -1041 | Lrrc38 | ENSMUSG00000028584.3 |
| chr4 | 143349186 | 143350749 | 4.13847 | 5.35365 | 1.993 | Promoter | 4 | 143349757 | 143371032 | 21276 | 1 | ENSMUST00000052458.2 | 0 | Lrrc38 | ENSMUSG00000028584.3 |
| chr4 | 145245728 | 145245890 | 3.52134 | 4.1685 | 1.22119 | Promoter | 4 | 145213463 | 145246870 | 33408 | 2 | ENSMUST00000030336.5 | 980 | Tnfrsf1b | ENSMUSG00000028599.5 |
| chr4 | 108031691 | 108032048 | 3.7663 | 4.69308 | 1.56295 | Promoter | 4 | 108014851 | 108031953 | 17103 | 2 | ENSMUST00000106708.2 | 0 | Podn | ENSMUSG00000028600.10 |
| chr4 | 108032388 | 108033251 | 3.59072 | 4.31347 | 1.33252 | Promoter | 4 | 108027193 | 108032337 | 5145 | 2 | ENSMUST00000124265.1 | -52 | Podn | ENSMUSG00000028600.10 |
| chr4 | 108165039 | 108165686 | 4.16428 | 5.46038 | 2.10344 | Promoter | 4 | 108165437 | 108179308 | 13872 | 1 | ENSMUST00000052999.7 | 0 | Echdc2 | ENSMUSG00000028601.13 |
| chr4 | 107653167 | 107653247 | 3.97451 | 5.14527 | 1.8487 | Distal Intergenic | 4 | 107676904 | 107680960 | 4057 | 2 | ENSMUST00000131508.1 | 27713 | Dmrtb1 | ENSMUSG00000028610.11 |
| chr4 | 107671779 | 107672067 | 3.38193 | 3.9006 | 1.01733 | Distal Intergenic | 4 | 107676904 | 107680960 | 4057 | 2 | ENSMUST00000131508.1 | 8893 | Dmrtb1 | ENSMUSG00000028610.11 |
| chr4 | 107672980 | 107674598 | 3.69443 | 4.56828 | 1.49261 | Downstream (1-2kb) | 4 | 107676904 | 107680960 | 4057 | 2 | ENSMUST00000131508.1 | 6362 | Dmrtb1 | ENSMUSG00000028610.11 |
| chr4 | 107802574 | 107802747 | 3.98556 | 5.62344 | 2.21233 | Promoter | 4 | 107802842 | 107851362 | 48521 | 1 | ENSMUST00000147319.2 | -95 | Lrp8 | ENSMUSG00000028613.9 |
| chr4 | 107803072 | 107803631 | 3.5156 | 4.17822 | 1.26935 | Promoter | 4 | 107803251 | 107872816 | 69566 | 1 | ENSMUST00000146552.2 | 0 | Lrp8 | ENSMUSG00000028613.9 |
| chr4 | 107804826 | 107805161 | 3.49527 | 4.25137 | 1.32326 | Intron (ENSMUST00000143601.2/ENSMUSG00000028613.9, intron 2 of 20) | 4 | 107803251 | 107872816 | 69566 | 1 | ENSMUST00000146552.2 | 1576 | Lrp8 | ENSMUSG00000028613.9 |
| chr4 | 107846257 | 107846603 | 3.47088 | 4.17179 | 1.23811 | Promoter | 4 | 107846201 | 107848295 | 2095 | 1 | ENSMUST00000135591.1 | 57 | Lrp8 | ENSMUSG00000028613.9 |
| chr4 | 107847658 | 107847933 | 3.23338 | 3.67724 | 1.01733 | Intron (ENSMUST00000143601.2/ENSMUSG00000028613.9, intron 6 of 20) | 4 | 107846201 | 107848295 | 2095 | 1 | ENSMUST00000135591.1 | 1458 | Lrp8 | ENSMUSG00000028613.9 |
| chr4 | 121039249 | 121040519 | 3.87661 | 4.89925 | 1.75819 | Promoter | 4 | 121039385 | 121055322 | 15938 | 1 | ENSMUST00000030372.5 | 0 | Col9a2 | ENSMUSG00000028626.5 |
| chr4 | 121040851 | 121041478 | 3.55812 | 4.28103 | 1.37234 | Exon (ENSMUST00000030372.5/ENSMUSG00000028626.5, exon 2 of 32) | 4 | 121039748 | 121044327 | 4580 | 1 | ENSMUST00000151987.1 | 1104 | Col9a2 | ENSMUSG00000028626.5 |
| chr4 | 121052780 | 121053162 | 3.46765 | 4.06604 | 1.14267 | Promoter | 4 | 121052999 | 121054819 | 1821 | 1 | ENSMUST00000140119.1 | 0 | Col9a2 | ENSMUSG00000028626.5 |
| chr4 | 121053743 | 121054085 | 3.26822 | 3.7295 | 1.01733 | Promoter | 4 | 121052999 | 121054819 | 1821 | 1 | ENSMUST00000140119.1 | 745 | Col9a2 | ENSMUSG00000028626.5 |
| chr10 | 118655581 | 118656301 | 3.52247 | 4.18534 | 1.2669 | Distal Intergenic | 10 | 118859349 | 118868903 | 9555 | 2 | ENSMUST00000004281.8 | 212602 | Dyrk2 | ENSMUSG00000028630.8 |
| chr10 | 118676126 | 118676261 | 3.59777 | 4.35493 | 1.40178 | Distal Intergenic | 10 | 118859349 | 118868903 | 9555 | 2 | ENSMUST00000004281.8 | 192642 | Dyrk2 | ENSMUSG00000028630.8 |
| chr10 | 118708960 | 118709313 | 3.84976 | 4.80566 | 1.63925 | Distal Intergenic | 10 | 118859349 | 118868903 | 9555 | 2 | ENSMUST00000004281.8 | 159590 | Dyrk2 | ENSMUSG00000028630.8 |
| chr10 | 118711097 | 118711203 | 3.60707 | 4.57803 | 1.51399 | Distal Intergenic | 10 | 118859349 | 118868903 | 9555 | 2 | ENSMUST00000004281.8 | 157700 | Dyrk2 | ENSMUSG00000028630.8 |
| chr10 | 118711684 | 118712340 | 3.77403 | 4.63663 | 1.49495 | Distal Intergenic | 10 | 118859349 | 118868903 | 9555 | 2 | ENSMUST00000004281.8 | 156563 | Dyrk2 | ENSMUSG00000028630.8 |
| chr10 | 118713123 | 118713637 | 3.359 | 3.86601 | 1.01733 | Distal Intergenic | 10 | 118859349 | 118868903 | 9555 | 2 | ENSMUST00000004281.8 | 155266 | Dyrk2 | ENSMUSG00000028630.8 |
| chr10 | 118717018 | 118717242 | 3.38193 | 3.9006 | 1.01733 | Distal Intergenic | 10 | 118859349 | 118868903 | 9555 | 2 | ENSMUST00000004281.8 | 151661 | Dyrk2 | ENSMUSG00000028630.8 |
| chr10 | 118722245 | 118722495 | 3.36981 | 3.88232 | 1.01733 | Distal Intergenic | 10 | 118859349 | 118868903 | 9555 | 2 | ENSMUST00000004281.8 | 146408 | Dyrk2 | ENSMUSG00000028630.8 |
| chr10 | 118769866 | 118770237 | 3.46641 | 4.10701 | 1.2376 | Distal Intergenic | 10 | 118859349 | 118868903 | 9555 | 2 | ENSMUST00000004281.8 | 98666 | Dyrk2 | ENSMUSG00000028630.8 |
| chr10 | 118794485 | 118794660 | 3.55521 | 4.28205 | 1.37592 | Distal Intergenic | 10 | 118859349 | 118868903 | 9555 | 2 | ENSMUST00000004281.8 | 74243 | Dyrk2 | ENSMUSG00000028630.8 |
| chr10 | 118803853 | 118804238 | 3.79747 | 4.86413 | 1.64989 | Distal Intergenic | 10 | 118859349 | 118868903 | 9555 | 2 | ENSMUST00000004281.8 | 64665 | Dyrk2 | ENSMUSG00000028630.8 |
| chr10 | 118804647 | 118804900 | 3.50094 | 4.3422 | 1.35982 | Distal Intergenic | 10 | 118859349 | 118868903 | 9555 | 2 | ENSMUST00000004281.8 | 64003 | Dyrk2 | ENSMUSG00000028630.8 |
| chr10 | 118805313 | 118805520 | 3.62542 | 4.41573 | 1.4135 | Distal Intergenic | 10 | 118859349 | 118868903 | 9555 | 2 | ENSMUST00000004281.8 | 63383 | Dyrk2 | ENSMUSG00000028630.8 |
| chr10 | 118805902 | 118806508 | 3.69017 | 4.52772 | 1.47256 | Distal Intergenic | 10 | 118859349 | 118868903 | 9555 | 2 | ENSMUST00000004281.8 | 62395 | Dyrk2 | ENSMUSG00000028630.8 |
| chr10 | 118807230 | 118807630 | 3.69997 | 4.49721 | 1.40512 | Distal Intergenic | 10 | 118859349 | 118868903 | 9555 | 2 | ENSMUST00000004281.8 | 61273 | Dyrk2 | ENSMUSG00000028630.8 |
| chr10 | 118809469 | 118809730 | 3.42168 | 4.00286 | 1.16437 | Distal Intergenic | 10 | 118859349 | 118868903 | 9555 | 2 | ENSMUST00000004281.8 | 59173 | Dyrk2 | ENSMUSG00000028630.8 |
| chr10 | 118820137 | 118820370 | 3.2711 | 3.73378 | 1.01733 | Distal Intergenic | 10 | 118859349 | 118868903 | 9555 | 2 | ENSMUST00000004281.8 | 48533 | Dyrk2 | ENSMUSG00000028630.8 |
| chr10 | 118821473 | 118821944 | 3.30724 | 3.79533 | 1.03803 | Distal Intergenic | 10 | 118859349 | 118868903 | 9555 | 2 | ENSMUST00000004281.8 | 46959 | Dyrk2 | ENSMUSG00000028630.8 |
| chr10 | 118822945 | 118823227 | 3.80405 | 4.78706 | 1.64246 | Distal Intergenic | 10 | 118859349 | 118868903 | 9555 | 2 | ENSMUST00000004281.8 | 45676 | Dyrk2 | ENSMUSG00000028630.8 |
| chr10 | 118824771 | 118824846 | 3.29129 | 3.76415 | 1.01733 | Distal Intergenic | 10 | 118859349 | 118868903 | 9555 | 2 | ENSMUST00000004281.8 | 44057 | Dyrk2 | ENSMUSG00000028630.8 |
| chr10 | 118827170 | 118827482 | 3.60147 | 4.31537 | 1.31281 | Distal Intergenic | 10 | 118859349 | 118868903 | 9555 | 2 | ENSMUST00000004281.8 | 41421 | Dyrk2 | ENSMUSG00000028630.8 |
| chr10 | 118828593 | 118828819 | 3.48413 | 4.08693 | 1.13028 | Distal Intergenic | 10 | 118859349 | 118868903 | 9555 | 2 | ENSMUST00000004281.8 | 40084 | Dyrk2 | ENSMUSG00000028630.8 |
| chr10 | 118830414 | 118831225 | 3.51319 | 4.14816 | 1.19234 | Distal Intergenic | 10 | 118859349 | 118868903 | 9555 | 2 | ENSMUST00000004281.8 | 37678 | Dyrk2 | ENSMUSG00000028630.8 |
| chr10 | 118838416 | 118838597 | 3.56484 | 4.23908 | 1.23413 | Distal Intergenic | 10 | 118859349 | 118868903 | 9555 | 2 | ENSMUST00000004281.8 | 30306 | Dyrk2 | ENSMUSG00000028630.8 |
| chr10 | 118839858 | 118840134 | 3.34153 | 3.83966 | 1.01733 | Distal Intergenic | 10 | 118859349 | 118868903 | 9555 | 2 | ENSMUST00000004281.8 | 28769 | Dyrk2 | ENSMUSG00000028630.8 |
| chr10 | 118840530 | 118840638 | 3.72308 | 4.53306 | 1.41649 | Distal Intergenic | 10 | 118859349 | 118868903 | 9555 | 2 | ENSMUST00000004281.8 | 28265 | Dyrk2 | ENSMUSG00000028630.8 |
| chr10 | 118846626 | 118846707 | 3.58017 | 4.35545 | 1.37319 | Distal Intergenic | 10 | 118859349 | 118868903 | 9555 | 2 | ENSMUST00000004281.8 | 22196 | Dyrk2 | ENSMUSG00000028630.8 |
| chr10 | 118850373 | 118851186 | 3.70657 | 4.59417 | 1.53348 | Distal Intergenic | 10 | 118859349 | 118868903 | 9555 | 2 | ENSMUST00000004281.8 | 17717 | Dyrk2 | ENSMUSG00000028630.8 |
| chr10 | 118851679 | 118851849 | 3.43041 | 3.99062 | 1.07666 | Distal Intergenic | 10 | 118859349 | 118868903 | 9555 | 2 | ENSMUST00000004281.8 | 17054 | Dyrk2 | ENSMUSG00000028630.8 |
| chr10 | 118852662 | 118852764 | 3.81903 | 4.8028 | 1.65796 | Distal Intergenic | 10 | 118859349 | 118868903 | 9555 | 2 | ENSMUST00000004281.8 | 16139 | Dyrk2 | ENSMUSG00000028630.8 |
| chr10 | 118853962 | 118854736 | 3.83564 | 4.77371 | 1.61358 | Distal Intergenic | 10 | 118859349 | 118868903 | 9555 | 2 | ENSMUST00000004281.8 | 14167 | Dyrk2 | ENSMUSG00000028630.8 |
| chr10 | 118855080 | 118855155 | 3.96113 | 5.02279 | 1.81124 | Distal Intergenic | 10 | 118859349 | 118868903 | 9555 | 2 | ENSMUST00000004281.8 | 13748 | Dyrk2 | ENSMUSG00000028630.8 |
| chr10 | 118858119 | 118859164 | 3.54771 | 4.38199 | 1.38715 | Downstream (<1kb) | 10 | 118859349 | 118868903 | 9555 | 2 | ENSMUST00000004281.8 | 9739 | Dyrk2 | ENSMUSG00000028630.8 |
| chr10 | 118860323 | 118860945 | 3.57178 | 4.7065 | 1.58505 | Exon (ENSMUST00000004281.8/ENSMUSG00000028630.8, exon 3 of 3) | 10 | 118859349 | 118868903 | 9555 | 2 | ENSMUST00000004281.8 | 7958 | Dyrk2 | ENSMUSG00000028630.8 |
| chr10 | 118862110 | 118862746 | 4.04999 | 5.46656 | 2.14907 | Intron (ENSMUST00000004281.8/ENSMUSG00000028630.8, intron 2 of 2) | 10 | 118859349 | 118868903 | 9555 | 2 | ENSMUST00000004281.8 | 6157 | Dyrk2 | ENSMUSG00000028630.8 |
| chr10 | 118863154 | 118863522 | 3.63496 | 4.52137 | 1.46178 | Intron (ENSMUST00000004281.8/ENSMUSG00000028630.8, intron 2 of 2) | 10 | 118859349 | 118868903 | 9555 | 2 | ENSMUST00000004281.8 | 5381 | Dyrk2 | ENSMUSG00000028630.8 |
| chr10 | 118864135 | 118864526 | 3.76145 | 4.92942 | 1.7207 | Intron (ENSMUST00000004281.8/ENSMUSG00000028630.8, intron 2 of 2) | 10 | 118859349 | 118868903 | 9555 | 2 | ENSMUST00000004281.8 | 4377 | Dyrk2 | ENSMUSG00000028630.8 |
| chr10 | 118866038 | 118866387 | 4.00976 | 5.23484 | 1.94878 | Intron (ENSMUST00000004281.8/ENSMUSG00000028630.8, intron 2 of 2) | 10 | 118859349 | 118868903 | 9555 | 2 | ENSMUST00000004281.8 | 2516 | Dyrk2 | ENSMUSG00000028630.8 |
| chr10 | 118867926 | 118869082 | 4.19612 | 5.79647 | 2.42047 | Promoter | 10 | 118859349 | 118868903 | 9555 | 2 | ENSMUST00000004281.8 | 0 | Dyrk2 | ENSMUSG00000028630.8 |
| chr10 | 118870210 | 118871071 | 3.469 | 4.13462 | 1.25892 | Distal Intergenic | 10 | 118859349 | 118868903 | 9555 | 2 | ENSMUST00000004281.8 | -1308 | Dyrk2 | ENSMUSG00000028630.8 |
| chr10 | 118872051 | 118872387 | 3.46097 | 4.21949 | 1.24972 | Distal Intergenic | 10 | 118859349 | 118868903 | 9555 | 2 | ENSMUST00000004281.8 | -3149 | Dyrk2 | ENSMUSG00000028630.8 |
| chr10 | 118874233 | 118874410 | 3.43675 | 4.0278 | 1.17553 | Distal Intergenic | 10 | 118859349 | 118868903 | 9555 | 2 | ENSMUST00000004281.8 | -5331 | Dyrk2 | ENSMUSG00000028630.8 |
| chr10 | 118876298 | 118876487 | 3.32107 | 3.83835 | 1.11039 | Distal Intergenic | 10 | 118859349 | 118868903 | 9555 | 2 | ENSMUST00000004281.8 | -7396 | Dyrk2 | ENSMUSG00000028630.8 |
| chr10 | 118878999 | 118879258 | 3.52476 | 4.21025 | 1.32042 | Distal Intergenic | 10 | 118859349 | 118868903 | 9555 | 2 | ENSMUST00000004281.8 | -10097 | Dyrk2 | ENSMUSG00000028630.8 |
| chr4 | 120717456 | 120717670 | 4.54601 | 6.31616 | 2.76725 | Exon (ENSMUST00000030376.7/ENSMUSG00000028631.7, exon 2 of 14) | 4 | 120696138 | 120747248 | 51111 | 2 | ENSMUST00000030376.7 | 29578 | Kcnq4 | ENSMUSG00000028631.7 |
| chr4 | 120732962 | 120733037 | 3.30245 | 3.78087 | 1.01733 | Intron (ENSMUST00000030376.7/ENSMUSG00000028631.7, intron 1 of 13) | 4 | 120696138 | 120747248 | 51111 | 2 | ENSMUST00000030376.7 | 14211 | Kcnq4 | ENSMUSG00000028631.7 |
| chr4 | 120738526 | 120738621 | 3.93569 | 5.02539 | 1.80649 | Intron (ENSMUST00000030376.7/ENSMUSG00000028631.7, intron 1 of 13) | 4 | 120696138 | 120747248 | 51111 | 2 | ENSMUST00000030376.7 | 8627 | Kcnq4 | ENSMUSG00000028631.7 |

| | | | | | | | | | | | | | | | |
|---|---|---|---|---|---|---|---|---|---|---|---|---|---|---|---|
| chr4 | 120746674 | 120747005 | 3.91011 | 5.14969 | 1.88947 | Promoter | 4 | 120696138 | 120747248 | 51111 | 2 | ENSMUST00000030376.7 | 243 | Kcnq4 | ENSMUSG00000028631.7 |
| chr4 | 120034356 | 120034589 | 3.99064 | 5.16298 | 1.89674 | Intron (ENSMUST00000106307.3/ENSMUSG00000028634.11, intron 2 of 8) | 4 | 120094489 | 120134557 | 40069 | 1 | ENSMUST00000166542.2 | -59900 | Hivep3 | ENSMUSG00000028634.11 |
| chr4 | 120183451 | 120183667 | 3.23998 | 3.68713 | 1.01733 | Distal Intergenic | 4 | 120161206 | 120167360 | 6155 | 1 | ENSMUST00000030384.4 | 22246 | Edn2 | ENSMUSG00000028635.7 |
| chr4 | 120188781 | 120189183 | 3.34866 | 3.8504 | 1.01733 | Distal Intergenic | 4 | 120161206 | 120167360 | 6155 | 1 | ENSMUST00000030384.4 | 27576 | Edn2 | ENSMUSG00000028635.7 |
| chr2 | 172544685 | 172545575 | 3.67649 | 4.46019 | 1.3988 | Distal Intergenic | 2 | 172549593 | 172558622 | 9030 | 1 | ENSMUST00000030391.3 | -4018 | Tfap2c | ENSMUSG00000028640.6 |
| chr2 | 172545945 | 172547279 | 3.98021 | 5.13449 | 1.87467 | Distal Intergenic | 2 | 172549593 | 172558622 | 9030 | 1 | ENSMUST00000030391.3 | -2314 | Tfap2c | ENSMUSG00000028640.6 |
| chr2 | 172547595 | 172551868 | 4.4997 | 6.1827 | 2.65549 | Promoter | 2 | 172549593 | 172558622 | 9030 | 1 | ENSMUST00000030391.3 | 0 | Tfap2c | ENSMUSG00000028640.6 |
| chr2 | 172552181 | 172552999 | 3.80256 | 4.7847 | 1.60186 | Promoter | 2 | 172552116 | 172553518 | 1403 | 1 | ENSMUST00000142633.1 | 66 | Tfap2c | ENSMUSG00000028640.6 |
| chr2 | 172553641 | 172553730 | 3.48018 | 4.09044 | 1.16462 | Intron (ENSMUST00000030391.3/ENSMUSG00000028640.6, intron 4 of 6) | 2 | 172552116 | 172553518 | 1403 | 1 | ENSMUST00000142633.1 | 1526 | Tfap2c | ENSMUSG00000028640.6 |
| chr2 | 172554245 | 172554460 | 3.97779 | 5.01805 | 1.73581 | Intron (ENSMUST00000030391.3/ENSMUSG00000028640.6, intron 5 of 6) | 2 | 172552116 | 172553518 | 1403 | 1 | ENSMUST00000142633.1 | 2130 | Tfap2c | ENSMUSG00000028640.6 |
| chr2 | 172555291 | 172555489 | 4.25118 | 5.57116 | 2.14151 | Intron (ENSMUST00000030391.3/ENSMUSG00000028640.6, intron 5 of 6) | 2 | 172552116 | 172553518 | 1403 | 1 | ENSMUST00000142633.1 | 3176 | Tfap2c | ENSMUSG00000028640.6 |
| chr2 | 172558735 | 172559015 | 4.21331 | 5.49943 | 2.09587 | Distal Intergenic | 2 | 172552116 | 172553518 | 1403 | 1 | ENSMUST00000142633.1 | 6620 | Tfap2c | ENSMUSG00000028640.6 |
| chr2 | 172613395 | 172613497 | 3.83336 | 4.89304 | 1.71397 | Distal Intergenic | 2 | 172552116 | 172553518 | 1403 | 1 | ENSMUST00000142633.1 | 61280 | Tfap2c | ENSMUSG00000028640.6 |
| chr2 | 172613922 | 172614261 | 3.8321 | 4.78347 | 1.6691 | Distal Intergenic | 2 | 172552116 | 172553518 | 1403 | 1 | ENSMUST00000142633.1 | 61807 | Tfap2c | ENSMUSG00000028640.6 |
| chr2 | 172623379 | 172623465 | 3.51873 | 4.39044 | 1.36536 | Distal Intergenic | 2 | 172552116 | 172553518 | 1403 | 1 | ENSMUST00000142633.1 | 71264 | Tfap2c | ENSMUSG00000028640.6 |
| chr2 | 172623874 | 172624396 | 3.63525 | 4.38791 | 1.33589 | Distal Intergenic | 2 | 172552116 | 172553518 | 1403 | 1 | ENSMUST00000142633.1 | 71759 | Tfap2c | ENSMUSG00000028640.6 |
| chr2 | 172661225 | 172661939 | 3.62895 | 4.44633 | 1.4238 | Distal Intergenic | 2 | 172552116 | 172553518 | 1403 | 1 | ENSMUST00000142633.1 | 109110 | Tfap2c | ENSMUSG00000028640.6 |
| chr4 | 122982606 | 122982786 | 3.277 | 3.77055 | 1.06264 | Distal Intergenic | 4 | 122995652 | 122996739 | 1088 | 1 | ENSMUST00000144998.1 | -12866 | Mycl | ENSMUSG00000028654.8 |
| chr4 | 122983286 | 122983744 | 3.33286 | 3.83656 | 1.05226 | Distal Intergenic | 4 | 122995652 | 122996739 | 1088 | 1 | ENSMUST00000144998.1 | -11908 | Mycl | ENSMUSG00000028654.8 |
| chr4 | 122960022 | 122960183 | 3.80618 | 4.69314 | 1.52724 | Promoter | 4 | 122956732 | 122959990 | 3259 | 2 | ENSMUST00000138964.1 | -33 | Mfsd2a | ENSMUSG00000028655.6 |
| chr4 | 136942130 | 136942490 | 3.43353 | 4.00078 | 1.09643 | Intron (ENSMUST00000030420.8/ENSMUSG00000028661.8, intron 3 of 16) | 4 | 136929419 | 136956816 | 27398 | 2 | ENSMUST00000030420.8 | 14326 | Epha8 | ENSMUSG00000028661.8 |
| chr4 | 136946049 | 136946957 | 3.54121 | 4.22617 | 1.26821 | Exon (ENSMUST00000030420.8/ENSMUSG00000028661.8, exon 3 of 17) | 4 | 136929419 | 136956816 | 27398 | 2 | ENSMUST00000030420.8 | 9859 | Epha8 | ENSMUSG00000028661.8 |
| chr4 | 136953962 | 136954279 | 3.46352 | 4.05003 | 1.10988 | Intron (ENSMUST00000030420.8/ENSMUSG00000028661.8, intron 1 of 16) | 4 | 136929419 | 136956816 | 27398 | 2 | ENSMUST00000030420.8 | 2537 | Epha8 | ENSMUSG00000028661.8 |
| chr4 | 136956516 | 136957354 | 3.48826 | 4.18632 | 1.20758 | Promoter | 4 | 136929419 | 136956816 | 27398 | 2 | ENSMUST00000030420.8 | 0 | Epha8 | ENSMUSG00000028661.8 |
| chr4 | 136958447 | 136958671 | 3.92309 | 4.95381 | 1.73404 | Distal Intergenic | 4 | 136929419 | 136956816 | 27398 | 2 | ENSMUST00000030420.8 | -1632 | Epha8 | ENSMUSG00000028661.8 |
| chr4 | 136959102 | 136959205 | 3.88811 | 4.87106 | 1.682 | Distal Intergenic | 4 | 136929419 | 136956816 | 27398 | 2 | ENSMUST00000030420.8 | -2287 | Epha8 | ENSMUSG00000028661.8 |
| chr4 | 136959957 | 136960113 | 3.49688 | 4.14564 | 1.2132 | Distal Intergenic | 4 | 136929419 | 136956816 | 27398 | 2 | ENSMUST00000030420.8 | -3142 | Epha8 | ENSMUSG00000028661.8 |
| chr4 | 136965485 | 136965770 | 3.43523 | 4.03919 | 1.13133 | Distal Intergenic | 4 | 136929419 | 136956816 | 27398 | 2 | ENSMUST00000030420.8 | -8670 | Epha8 | ENSMUSG00000028661.8 |
| chr4 | 136966401 | 136967248 | 3.89111 | 4.90954 | 1.7134 | Distal Intergenic | 4 | 136929419 | 136956816 | 27398 | 2 | ENSMUST00000030420.8 | -9586 | Epha8 | ENSMUSG00000028661.8 |
| chr4 | 136967583 | 136967727 | 3.50272 | 4.12081 | 1.15081 | Distal Intergenic | 4 | 136929419 | 136956816 | 27398 | 2 | ENSMUST00000030420.8 | -10768 | Epha8 | ENSMUSG00000028661.8 |
| chr4 | 136969014 | 136969523 | 3.65087 | 4.62682 | 1.53536 | Distal Intergenic | 4 | 136929419 | 136956816 | 27398 | 2 | ENSMUST00000030420.8 | -12199 | Epha8 | ENSMUSG00000028661.8 |
| chr4 | 136969958 | 136970068 | 3.91138 | 4.90133 | 1.6606 | Distal Intergenic | 4 | 136929419 | 136956816 | 27398 | 2 | ENSMUST00000030420.8 | -13143 | Epha8 | ENSMUSG00000028661.8 |
| chr4 | 136971237 | 136971362 | 3.475 | 4.18313 | 1.26605 | Distal Intergenic | 4 | 136929419 | 136956816 | 27398 | 2 | ENSMUST00000030420.8 | -14422 | Epha8 | ENSMUSG00000028661.8 |
| chr4 | 136972446 | 136972569 | 3.35817 | 3.86475 | 1.01733 | Distal Intergenic | 4 | 136929419 | 136956816 | 27398 | 2 | ENSMUST00000030420.8 | -15631 | Epha8 | ENSMUSG00000028661.8 |
| chr4 | 136974185 | 136974802 | 3.36197 | 3.87049 | 1.01733 | Distal Intergenic | 4 | 136929419 | 136956816 | 27398 | 2 | ENSMUST00000030420.8 | -17370 | Epha8 | ENSMUSG00000028661.8 |
| chr4 | 136976135 | 136976277 | 3.40181 | 4.22906 | 1.28002 | Distal Intergenic | 4 | 136929419 | 136956816 | 27398 | 2 | ENSMUST00000030420.8 | -19320 | Epha8 | ENSMUSG00000028661.8 |
| chr4 | 136739240 | 136739456 | 3.69444 | 4.54078 | 1.50116 | Intron (ENSMUST00000105845.3/ENSMUSG00000028664.9, intron 3 of 15) | 4 | 136661463 | 136694040 | 32578 | 2 | ENSMUST00000156558.1 | -45201 | Ephb2 | ENSMUSG00000028664.9 |
| chr4 | 136755310 | 136755828 | 3.76568 | 4.69305 | 1.54177 | Intron (ENSMUST00000105845.3/ENSMUSG00000028664.9, intron 3 of 15) | 4 | 136661463 | 136694040 | 32578 | 2 | ENSMUST00000156558.1 | -61271 | Ephb2 | ENSMUSG00000028664.9 |
| chr4 | 136756724 | 136757115 | 3.85924 | 4.78145 | 1.56916 | Intron (ENSMUST00000105845.3/ENSMUSG00000028664.9, intron 3 of 15) | 4 | 136661463 | 136694040 | 32578 | 2 | ENSMUST00000156558.1 | -62685 | Ephb2 | ENSMUSG00000028664.9 |
| chr4 | 136783867 | 136783971 | 3.64091 | 4.48756 | 1.45508 | Intron (ENSMUST00000105845.3/ENSMUSG00000028664.9, intron 1 of 15) | 4 | 136654646 | 136835843 | 181198 | 2 | ENSMUST00000105846.3 | 51872 | Ephb2 | ENSMUSG00000028664.9 |
| chr4 | 136785092 | 136785370 | 3.49282 | 4.10646 | 1.15324 | Intron (ENSMUST00000105845.3/ENSMUSG00000028664.9, intron 1 of 15) | 4 | 136654646 | 136835843 | 181198 | 2 | ENSMUST00000105846.3 | 50473 | Ephb2 | ENSMUSG00000028664.9 |
| chr4 | 136786585 | 136786672 | 3.6375 | 4.38159 | 1.34283 | Intron (ENSMUST00000105845.3/ENSMUSG00000028664.9, intron 1 of 15) | 4 | 136654646 | 136835843 | 181198 | 2 | ENSMUST00000105846.3 | 49171 | Ephb2 | ENSMUSG00000028664.9 |
| chr4 | 136797873 | 136798126 | 3.55813 | 4.23017 | 1.24214 | Intron (ENSMUST00000105845.3/ENSMUSG00000028664.9, intron 1 of 15) | 4 | 136654646 | 136835843 | 181198 | 2 | ENSMUST00000105846.3 | 37717 | Ephb2 | ENSMUSG00000028664.9 |
| chr4 | 136817544 | 136817795 | 3.59524 | 4.29133 | 1.26003 | Intron (ENSMUST00000105845.3/ENSMUSG00000028664.9, intron 1 of 15) | 4 | 136654646 | 136835843 | 181198 | 2 | ENSMUST00000105846.3 | 18048 | Ephb2 | ENSMUSG00000028664.9 |
| chr4 | 136828246 | 136828827 | 3.53293 | 4.1982 | 1.23633 | Intron (ENSMUST00000105845.3/ENSMUSG00000028664.9, intron 1 of 15) | 4 | 136654646 | 136835843 | 181198 | 2 | ENSMUST00000105846.3 | 7016 | Ephb2 | ENSMUSG00000028664.9 |
| chr4 | 136833239 | 136833392 | 4.35092 | 5.75154 | 2.25282 | Intron (ENSMUST00000105845.3/ENSMUSG00000028664.9, intron 1 of 15) | 4 | 136654646 | 136835843 | 181198 | 2 | ENSMUST00000105846.3 | 2451 | Ephb2 | ENSMUSG00000028664.9 |
| chr4 | 136835026 | 136836007 | 4.01934 | 5.15117 | 1.88612 | Promoter | 4 | 136647539 | 136835988 | 188450 | 2 | ENSMUST00000105845.3 | 0 | Ephb2 | ENSMUSG00000028664.9 |
| chr4 | 136836898 | 136837093 | 3.75321 | 4.73969 | 1.61606 | Promoter | 4 | 136647539 | 136835988 | 188450 | 2 | ENSMUST00000105845.3 | -911 | Ephb2 | ENSMUSG00000028664.9 |
| chr4 | 136837713 | 136838951 | 3.52155 | 4.18429 | 1.2698 | Distal Intergenic | 4 | 136647539 | 136835988 | 188450 | 2 | ENSMUST00000105845.3 | -1726 | Ephb2 | ENSMUSG00000028664.9 |
| chr4 | 136839873 | 136840455 | 3.72393 | 4.56993 | 1.48981 | Distal Intergenic | 4 | 136647539 | 136835988 | 188450 | 2 | ENSMUST00000105845.3 | -3886 | Ephb2 | ENSMUSG00000028664.9 |
| chr4 | 136850359 | 136850499 | 3.49321 | 4.11036 | 1.16529 | Distal Intergenic | 4 | 136647539 | 136835988 | 188450 | 2 | ENSMUST00000105845.3 | -14372 | Ephb2 | ENSMUSG00000028664.9 |
| chr4 | 117303030 | 117304691 | 3.90083 | 5.05969 | 1.83805 | Intron (ENSMUST00000030439.9/ENSMUSG00000028677.13, intron 3 of 14) | 4 | 117285889 | 117300028 | 14140 | 2 | ENSMUST00000150148.1 | -3003 | Rnf220 | ENSMUSG00000028677.13 |
| chr4 | 117305740 | 117305939 | 3.38607 | 3.94071 | 1.13746 | Intron (ENSMUST00000030439.9/ENSMUSG00000028677.13, intron 3 of 14) | 4 | 117285889 | 117300028 | 14140 | 2 | ENSMUST00000150148.1 | -5713 | Rnf220 | ENSMUSG00000028677.13 |
| chr4 | 117315245 | 117315633 | 3.65634 | 4.50398 | 1.47013 | Intron (ENSMUST00000030439.9/ENSMUSG00000028677.13, intron 2 of 14) | 4 | 117285889 | 117300028 | 14140 | 2 | ENSMUST00000150148.1 | -15218 | Rnf220 | ENSMUSG00000028677.13 |
| chr4 | 117317470 | 117317826 | 3.93457 | 5.01955 | 1.82418 | Intron (ENSMUST00000030439.9/ENSMUSG00000028677.13, intron 2 of 14) | 4 | 117285889 | 117300028 | 14140 | 2 | ENSMUST00000150148.1 | -17443 | Rnf220 | ENSMUSG00000028677.13 |
| chr4 | 117318853 | 117319674 | 3.91201 | 5.01663 | 1.84605 | Intron (ENSMUST00000030439.9/ENSMUSG00000028677.13, intron 2 of 14) | 4 | 117285889 | 117300028 | 14140 | 2 | ENSMUST00000150148.1 | -18826 | Rnf220 | ENSMUSG00000028677.13 |
| chr4 | 117321429 | 117322571 | 3.9239 | 5.06442 | 1.89894 | Intron (ENSMUST00000030439.9/ENSMUSG00000028677.13, intron 2 of 14) | 4 | 117285889 | 117300028 | 14140 | 2 | ENSMUST00000150148.1 | -21402 | Rnf220 | ENSMUSG00000028677.13 |
| chr4 | 117323466 | 117324245 | 3.5446 | 4.33648 | 1.38761 | Intron (ENSMUST00000030439.9/ENSMUSG00000028677.13, intron 2 of 14) | 4 | 117285889 | 117300028 | 14140 | 2 | ENSMUST00000150148.1 | -23439 | Rnf220 | ENSMUSG00000028677.13 |
| chr4 | 117357213 | 117357473 | 3.36108 | 3.86915 | 1.01733 | Intron (ENSMUST00000030439.9/ENSMUSG00000028677.13, intron 2 of 14) | 4 | 117271927 | 117375457 | 103531 | 2 | ENSMUST00000094853.3 | 17984 | Rnf220 | ENSMUSG00000028677.13 |
| chr4 | 117382191 | 117382709 | 3.35564 | 3.93005 | 1.14147 | Intron (ENSMUST00000030439.9/ENSMUSG00000028677.13, intron 2 of 14) | 4 | 117271927 | 117375457 | 103531 | 2 | ENSMUST00000094853.3 | -6735 | Rnf220 | ENSMUSG00000028677.13 |
| chr4 | 117391101 | 117391181 | 3.6056 | 4.31024 | 1.26823 | Intron (ENSMUST00000030439.9/ENSMUSG00000028677.13, intron 2 of 14) | 4 | 117271927 | 117375457 | 103531 | 2 | ENSMUST00000094853.3 | -15645 | Rnf220 | ENSMUSG00000028677.13 |
| chr4 | 117391488 | 117391612 | 3.26901 | 3.73069 | 1.01733 | Intron (ENSMUST00000030439.9/ENSMUSG00000028677.13, intron 2 of 14) | 4 | 117271927 | 117375457 | 103531 | 2 | ENSMUST00000094853.3 | -16032 | Rnf220 | ENSMUSG00000028677.13 |
| chr4 | 117392008 | 117392346 | 3.81419 | 4.70792 | 1.54569 | Intron (ENSMUST00000030439.9/ENSMUSG00000028677.13, intron 2 of 14) | 4 | 117271927 | 117375457 | 103531 | 2 | ENSMUST00000094853.3 | -16552 | Rnf220 | ENSMUSG00000028677.13 |
| chr4 | 117393694 | 117393954 | 3.51227 | 4.15584 | 1.22497 | Intron (ENSMUST00000030439.9/ENSMUSG00000028677.13, intron 2 of 14) | 4 | 117271927 | 117375457 | 103531 | 2 | ENSMUST00000094853.3 | -18238 | Rnf220 | ENSMUSG00000028677.13 |
| chr4 | 117404689 | 117404872 | 3.37866 | 3.90998 | 1.0677 | Intron (ENSMUST00000030439.9/ENSMUSG00000028677.13, intron 2 of 14) | 4 | 117271927 | 117375457 | 103531 | 2 | ENSMUST00000094853.3 | -29233 | Rnf220 | ENSMUSG00000028677.13 |
| chr4 | 117407833 | 117408042 | 3.27297 | 3.73663 | 1.01733 | Intron (ENSMUST00000030439.9/ENSMUSG00000028677.13, intron 2 of 14) | 4 | 117271927 | 117375457 | 103531 | 2 | ENSMUST00000094853.3 | -32377 | Rnf220 | ENSMUSG00000028677.13 |
| chr4 | 117408763 | 117409091 | 4.03517 | 5.36919 | 2.07139 | Intron (ENSMUST00000030439.9/ENSMUSG00000028677.13, intron 2 of 14) | 4 | 117271927 | 117375457 | 103531 | 2 | ENSMUST00000094853.3 | -33307 | Rnf220 | ENSMUSG00000028677.13 |
| chr4 | 117485672 | 117486434 | 3.54104 | 4.2294 | 1.29644 | Intron (ENSMUST00000030439.9/ENSMUSG00000028677.13, intron 2 of 14) | 4 | 117490235 | 117495011 | 4777 | 2 | ENSMUST00000123222.1 | 8577 | Rnf220 | ENSMUSG00000028677.13 |
| chr4 | 117486892 | 117487572 | 3.69466 | 4.49218 | 1.41055 | Intron (ENSMUST00000030439.9/ENSMUSG00000028677.13, intron 2 of 14) | 4 | 117490235 | 117495011 | 4777 | 2 | ENSMUST00000123222.1 | 7439 | Rnf220 | ENSMUSG00000028677.13 |
| chr4 | 117489293 | 117489684 | 3.50095 | 4.15472 | 1.26995 | Exon (ENSMUST00000030439.9/ENSMUSG00000028677.13, exon 2 of 15) | 4 | 117490235 | 117495011 | 4777 | 2 | ENSMUST00000123222.1 | 5327 | Rnf220 | ENSMUSG00000028677.13 |
| chr4 | 117490105 | 117490358 | 3.4908 | 4.1701 | 1.27389 | 5' UTR | 4 | 117490235 | 117495011 | 4777 | 2 | ENSMUST00000123222.1 | 4653 | Rnf220 | ENSMUSG00000028677.13 |
| chr4 | 117490990 | 117493499 | 3.74722 | 4.70733 | 1.59803 | Intron (ENSMUST00000030439.9/ENSMUSG00000028677.13, intron 1 of 14) | 4 | 117490235 | 117495011 | 4777 | 2 | ENSMUST00000123222.1 | 1512 | Rnf220 | ENSMUSG00000028677.13 |
| chr4 | 117493839 | 117496205 | 3.83862 | 4.86829 | 1.73383 | Promoter | 4 | 117490235 | 117495011 | 4777 | 2 | ENSMUST00000123222.1 | 0 | Rnf220 | ENSMUSG00000028677.13 |
| chr4 | 117472210 | 117472316 | 3.33321 | 3.93401 | 1.11221 | Intron (ENSMUST00000030439.9/ENSMUSG00000028677.13, intron 2 of 14) | 4 | 117490235 | 117495011 | 4777 | 2 | ENSMUST00000123222.1 | 22695 | Rnf220 | ENSMUSG00000028677.13 |
| chr4 | 117472677 | 117472911 | 3.78642 | 4.65548 | 1.52052 | Intron (ENSMUST00000030439.9/ENSMUSG00000028677.13, intron 2 of 14) | 4 | 117490235 | 117495011 | 4777 | 2 | ENSMUST00000123222.1 | 22100 | Rnf220 | ENSMUSG00000028677.13 |
| chr4 | 117473657 | 117474003 | 3.86907 | 4.83 | 1.64615 | Intron (ENSMUST00000030439.9/ENSMUSG00000028677.13, intron 2 of 14) | 4 | 117490235 | 117495011 | 4777 | 2 | ENSMUST00000123222.1 | 21008 | Rnf220 | ENSMUSG00000028677.13 |
| chr4 | 117475398 | 117475724 | 3.66345 | 4.42518 | 1.36331 | Intron (ENSMUST00000030439.9/ENSMUSG00000028677.13, intron 2 of 14) | 4 | 117490235 | 117495011 | 4777 | 2 | ENSMUST00000123222.1 | 19287 | Rnf220 | ENSMUSG00000028677.13 |
| chr4 | 117476832 | 117477119 | 3.62049 | 4.38479 | 1.34821 | Intron (ENSMUST00000030439.9/ENSMUSG00000028677.13, intron 2 of 14) | 4 | 117490235 | 117495011 | 4777 | 2 | ENSMUST00000123222.1 | 17892 | Rnf220 | ENSMUSG00000028677.13 |
| chr4 | 117478125 | 117478200 | 3.60548 | 4.55883 | 1.47984 | Intron (ENSMUST00000030439.9/ENSMUSG00000028677.13, intron 2 of 14) | 4 | 117490235 | 117495011 | 4777 | 2 | ENSMUST00000123222.1 | 16811 | Rnf220 | ENSMUSG00000028677.13 |
| chr4 | 117479855 | 117480229 | 3.58677 | 4.34558 | 1.36844 | Intron (ENSMUST00000030439.9/ENSMUSG00000028677.13, intron 2 of 14) | 4 | 117490235 | 117495011 | 4777 | 2 | ENSMUST00000123222.1 | 14782 | Rnf220 | ENSMUSG00000028677.13 |
| chr4 | 117500778 | 117501135 | 3.38643 | 3.93897 | 1.12908 | Distal Intergenic | 4 | 117271463 | 117497052 | 225590 | 2 | ENSMUST00000030439.9 | -3727 | Rnf220 | ENSMUSG00000028677.13 |
| chr4 | 117496752 | 117497208 | 3.58822 | 4.58554 | 1.50973 | Promoter | 4 | 117271463 | 117497052 | 225590 | 2 | ENSMUST00000030439.9 | 0 | Rnf220 | ENSMUSG00000028677.13 |
| chr4 | 117497642 | 117497877 | 3.94581 | 5.07518 | 1.85225 | Promoter | 4 | 117271463 | 117497052 | 225590 | 2 | ENSMUST00000030439.9 | -591 | Rnf220 | ENSMUSG00000028677.13 |
| chr4 | 117105425 | 117105500 | 3.48511 | 4.09218 | 1.14353 | Promoter | 4 | 117105238 | 117108514 | 3277 | 1 | ENSMUST00000137209.1 | 188 | Ptch2 | ENSMUSG00000028681.6 |
| chr4 | 115908470 | 115908560 | 3.46832 | 4.10863 | 1.26139 | Distal Intergenic | 4 | 115915119 | 115925638 | 10520 | 2 | ENSMUST00000064806.7 | 17078 | Dmbx1 | ENSMUSG00000028707.10 |
| chr4 | 115910965 | 115911209 | 3.4701 | 4.30325 | 1.34709 | Distal Intergenic | 4 | 115915119 | 115925638 | 10520 | 2 | ENSMUST00000064806.7 | 14429 | Dmbx1 | ENSMUSG00000028707.10 |
| chr4 | 115917794 | 115919391 | 3.76723 | 4.84744 | 1.72068 | 3' UTR | 4 | 115915119 | 115925638 | 10520 | 2 | ENSMUST00000064806.7 | 6247 | Dmbx1 | ENSMUSG00000028707.10 |
| chr4 | 115922648 | 115922842 | 3.64461 | 4.40688 | 1.40236 | Intron (ENSMUST00000064806.7/ENSMUSG00000028707.10, intron 1 of 3) | 4 | 115915119 | 115925638 | 10520 | 2 | ENSMUST00000064806.7 | 2796 | Dmbx1 | ENSMUSG00000028707.10 |
| chr4 | 115927394 | 115927770 | 3.31334 | 3.89632 | 1.09808 | Intron (ENSMUST00000124071.3/ENSMUSG00000028707.10, intron 1 of 4) | 4 | 115915119 | 115925638 | 10520 | 2 | ENSMUST00000064806.7 | -1757 | Dmbx1 | ENSMUSG00000028707.10 |
| chr4 | 115928169 | 115929064 | 3.92759 | 5.0132 | 1.79244 | Intron (ENSMUST00000124071.3/ENSMUSG00000028707.10, intron 1 of 4) | 4 | 115915119 | 115925638 | 10520 | 2 | ENSMUST00000064806.7 | -2532 | Dmbx1 | ENSMUSG00000028707.10 |
| chr4 | 115929439 | 115929884 | 3.56374 | 4.29867 | 1.29948 | Intron (ENSMUST00000124071.3/ENSMUSG00000028707.10, intron 1 of 4) | 4 | 115915119 | 115925638 | 10520 | 2 | ENSMUST00000064806.7 | -3802 | Dmbx1 | ENSMUSG00000028707.10 |
| chr4 | 115931170 | 115932209 | 3.54508 | 4.33662 | 1.37218 | Intron (ENSMUST00000124071.3/ENSMUSG00000028707.10, intron 1 of 4) | 4 | 115915119 | 115925638 | 10520 | 2 | ENSMUST00000064806.7 | -5533 | Dmbx1 | ENSMUSG00000028707.10 |
| chr4 | 115933365 | 115933545 | 3.37152 | 3.89118 | 1.03958 | Intron (ENSMUST00000124071.3/ENSMUSG00000028707.10, intron 1 of 4) | 4 | 115915119 | 115939926 | 24808 | 2 | ENSMUST00000124071.3 | 6381 | Dmbx1 | ENSMUSG00000028707.10 |
| chr4 | 115934199 | 115935671 | 3.97572 | 5.13002 | 1.89621 | Intron (ENSMUST00000124071.3/ENSMUSG00000028707.10, intron 1 of 4) | 4 | 115915119 | 115939926 | 24808 | 2 | ENSMUST00000124071.3 | 4255 | Dmbx1 | ENSMUSG00000028707.10 |
| chr4 | 115936004 | 115936271 | 4.42122 | 6.21474 | 2.65612 | Intron (ENSMUST00000124071.3/ENSMUSG00000028707.10, intron 1 of 4) | 4 | 115915119 | 115939926 | 24808 | 2 | ENSMUST00000124071.3 | 3655 | Dmbx1 | ENSMUSG00000028707.10 |
| chr4 | 115936712 | 115941110 | 3.74362 | 4.72619 | 1.58642 | Promoter | 4 | 115915119 | 115939926 | 24808 | 2 | ENSMUST00000124071.3 | 0 | Dmbx1 | ENSMUSG00000028707.10 |
| chr4 | 115942288 | 115943457 | 3.8689 | 4.80766 | 1.59515 | Distal Intergenic | 4 | 115915120 | 115939926 | 24807 | 2 | ENSMUST00000084338.6 | -2363 | Dmbx1 | ENSMUSG00000028707.10 |
| chr4 | 115943992 | 115944235 | 3.50709 | 4.20617 | 1.22386 | Distal Intergenic | 4 | 115915120 | 115939926 | 24807 | 2 | ENSMUST00000084338.6 | -4067 | Dmbx1 | ENSMUSG00000028707.10 |
| chr4 | 115946609 | 115946758 | 4.37751 | 6.29925 | 2.71898 | Distal Intergenic | 4 | 115915120 | 115939926 | 24807 | 2 | ENSMUST00000084338.6 | -6684 | Dmbx1 | ENSMUSG00000028707.10 |
| chr4 | 115947538 | 115948120 | 3.70687 | 4.54507 | 1.46601 | Distal Intergenic | 4 | 115915120 | 115939926 | 24807 | 2 | ENSMUST00000084338.6 | -7613 | Dmbx1 | ENSMUSG00000028707.10 |
| chr4 | 115948794 | 115949530 | 3.55366 | 4.26551 | 1.3589 | Distal Intergenic | 4 | 115915120 | 115939926 | 24807 | 2 | ENSMUST00000084338.6 | -8869 | Dmbx1 | ENSMUSG00000028707.10 |
| chr4 | 115950082 | 115950271 | 3.54728 | 4.23229 | 1.30201 | Distal Intergenic | 4 | 115915120 | 115939926 | 24807 | 2 | ENSMUST00000084338.6 | -10157 | Dmbx1 | ENSMUSG00000028707.10 |
| chr4 | 115954393 | 115954797 | 3.37398 | 3.90156 | 1.06336 | Distal Intergenic | 4 | 115915120 | 115939926 | 24807 | 2 | ENSMUST00000084338.6 | -14468 | Dmbx1 | ENSMUSG00000028707.10 |
| chr4 | 115955219 | 115955577 | 3.48581 | 4.09256 | 1.14141 | Distal Intergenic | 4 | 115915120 | 115939926 | 24807 | 2 | ENSMUST00000084338.6 | -15294 | Dmbx1 | ENSMUSG00000028707.10 |
| chr4 | 115956408 | 115956584 | 3.46405 | 4.06924 | 1.17607 | Distal Intergenic | 4 | 115915120 | 115939926 | 24807 | 2 | ENSMUST00000084338.6 | -16483 | Dmbx1 | ENSMUSG00000028707.10 |
| chr4 | 115959426 | 115959607 | 3.38193 | 3.9006 | 1.01733 | Distal Intergenic | 4 | 115915120 | 115939926 | 24807 | 2 | ENSMUST00000084338.6 | -19501 | Dmbx1 | ENSMUSG00000028707.10 |
| chr4 | 115959992 | 115960070 | 3.67411 | 4.51478 | 1.48002 | Distal Intergenic | 4 | 115915120 | 115939926 | 24807 | 2 | ENSMUST00000084338.6 | -20067 | Dmbx1 | ENSMUSG00000028707.10 |
| chr4 | 115961410 | 115963689 | 3.91413 | 5.10722 | 1.87028 | Distal Intergenic | 4 | 115915120 | 115939926 | 24807 | 2 | ENSMUST00000084338.6 | -21485 | Dmbx1 | ENSMUSG00000028707.10 |
| chr4 | 115964603 | 115965356 | 3.55987 | 4.27965 | 1.33681 | Downstream (1-2kb) | 4 | 115915120 | 115939926 | 24807 | 2 | ENSMUST00000084338.6 | -24678 | Dmbx1 | ENSMUSG00000028707.10 |
| chr4 | 115966254 | 115966847 | 3.36819 | 3.89313 | 1.05319 | Downstream (<1kb) | 4 | 115915120 | 115939926 | 24807 | 2 | ENSMUST00000084338.6 | -26329 | Dmbx1 | ENSMUSG00000028707.10 |
| chr4 | 115095710 | 115096199 | 3.76793 | 4.80907 | 1.6411 | Distal Intergenic | 4 | 115090932 | 115093864 | 2933 | 1 | ENSMUST00000146578.1 | 4779 | Pdzk1ip1 | ENSMUSG00000028716.10 |
| chr4 | 115052723 | 115053311 | 3.61166 | 4.59571 | 1.48608 | Distal Intergenic | 4 | 115056426 | 115071755 | 15330 | 1 | ENSMUST00000030489.3 | -3115 | Tal1 | ENSMUSG00000028717.7 |
| chr4 | 115056540 | 115057034 | 3.77053 | 4.76018 | 1.62052 | Promoter | 4 | 115056426 | 115071755 | 15330 | 1 | ENSMUST00000030489.3 | 115 | Tal1 | ENSMUSG00000028717.7 |
| chr4 | 115057411 | 115058136 | 3.66226 | 4.87014 | 1.6965 | Promoter | 4 | 115057438 | 115063570 | 6133 | 1 | ENSMUST00000136946.2 | 0 | Tal1 | ENSMUSG00000028717.7 |
| chr4 | 115058873 | 115059145 | 3.54184 | 4.42827 | 1.40721 | Promoter | 4 | 115059508 | 115070469 | 10962 | 1 | ENSMUST00000162489.1 | -363 | Tal1 | ENSMUSG00000028717.7 |
| chr4 | 115059465 | 115060855 | 3.64746 | 4.48517 | 1.49748 | Promoter | 4 | 115059508 | 115070469 | 10962 | 1 | ENSMUST00000162489.1 | 0 | Tal1 | ENSMUSG00000028717.7 |
| chr4 | 115061567 | 115064105 | 3.77664 | 4.93542 | 1.76537 | 5' UTR | 4 | 115059508 | 115070469 | 10962 | 1 | ENSMUST00000162489.1 | 2060 | Tal1 | ENSMUSG00000028717.7 |
| chr4 | 115064524 | 115064737 | 3.39087 | 4.30263 | 1.31366 | Exon (ENSMUST00000030489.3/ENSMUSG00000028717.7, exon 4 of 5) | 4 | 115059508 | 115070469 | 10962 | 1 | ENSMUST00000162489.1 | 5017 | Tal1 | ENSMUSG00000028717.7 |
| chr4 | 115065968 | 115066365 | 3.58677 | 4.55197 | 1.46358 | Intron (ENSMUST00000030489.3/ENSMUSG00000028717.7, intron 4 of 4) | 4 | 115059508 | 115070469 | 10962 | 1 | ENSMUST00000162489.1 | 6461 | Tal1 | ENSMUSG00000028717.7 |
| chr4 | 115066951 | 115067249 | 3.3555 | 3.86073 | 1.01733 | Intron (ENSMUST00000030489.3/ENSMUSG00000028717.7, intron 4 of 4) | 4 | 115059508 | 115070469 | 10962 | 1 | ENSMUST00000162489.1 | 7444 | Tal1 | ENSMUSG00000028717.7 |
| chr4 | 118609790 | 118609865 | 3.49895 | 4.12983 | 1.19976 | Intron (ENSMUST00000071972.5/ENSMUSG00000028730.9, intron 5 of 22) | 4 | 118554696 | 118620405 | 65710 | 2 | ENSMUST00000081921.6 | 10540 | Wdr65 | ENSMUSG00000028730.9 |
| chr4 | 139755380 | 139755703 | 3.35051 | 3.8532 | 1.01733 | Intron (ENSMUST00000030508.8/ENSMUSG00000028736.8, intron 7 of 8) | 4 | 139740676 | 139833007 | 92332 | 2 | ENSMUST00000174681.1 | 77304 | Pax7 | ENSMUSG00000028736.8 |
| chr4 | 139757119 | 139757674 | 3.68228 | 4.46556 | 1.38787 | Intron (ENSMUST00000030508.8/ENSMUSG00000028736.8, intron 7 of 8) | 4 | 139740676 | 139833007 | 92332 | 2 | ENSMUST00000174681.1 | 75333 | Pax7 | ENSMUSG00000028736.8 |
| chr4 | 139767084 | 139767243 | 3.58548 | 4.43343 | 1.38503 | Intron (ENSMUST00000030508.8/ENSMUSG00000028736.8, intron 7 of 8) | 4 | 139740676 | 139833007 | 92332 | 2 | ENSMUST00000174681.1 | 65764 | Pax7 | ENSMUSG00000028736.8 |
| chr4 | 139771277 | 139771368 | 3.36981 | 3.88232 | 1.01733 | Intron (ENSMUST00000030508.8/ENSMUSG00000028736.8, intron 7 of 8) | 4 | 139740676 | 139833007 | 92332 | 2 | ENSMUST00000174681.1 | 61639 | Pax7 | ENSMUSG00000028736.8 |
| chr4 | 139777377 | 139777685 | 3.42977 | 3.99563 | 1.09823 | Intron (ENSMUST00000030508.8/ENSMUSG00000028736.8, intron 7 of 8) | 4 | 139740676 | 139833007 | 92332 | 2 | ENSMUST00000174681.1 | 55322 | Pax7 | ENSMUSG00000028736.8 |
| chr4 | 139779626 | 139779851 | 3.3957 | 3.92891 | 1.04403 | Exon (ENSMUST00000030508.8/ENSMUSG00000028736.8, exon 7 of 9) | 4 | 139740676 | 139833007 | 92332 | 2 | ENSMUST00000174681.1 | 53156 | Pax7 | ENSMUSG00000028736.8 |
| chr4 | 139791478 | 139791568 | 3.48249 | 4.08784 | 1.14049 | Intron (ENSMUST00000030508.8/ENSMUSG00000028736.8, intron 4 of 8) | 4 | 139740676 | 139833007 | 92332 | 2 | ENSMUST00000174681.1 | 41439 | Pax7 | ENSMUSG00000028736.8 |
| chr4 | 139794766 | 139794999 | 3.5664 | 4.23692 | 1.22119 | Intron (ENSMUST00000030508.8/ENSMUSG00000028736.8, intron 4 of 8) | 4 | 139740676 | 139833007 | 92332 | 2 | ENSMUST00000174681.1 | 38008 | Pax7 | ENSMUSG00000028736.8 |
| chr4 | 139798126 | 139798572 | 4.17255 | 5.40865 | 2.01526 | Intron (ENSMUST00000030508.8/ENSMUSG00000028736.8, intron 4 of 8) | 4 | 139740676 | 139833007 | 92332 | 2 | ENSMUST00000174681.1 | 34435 | Pax7 | ENSMUSG00000028736.8 |
| chr4 | 139799737 | 139800267 | 3.48729 | 4.10311 | 1.15217 | Intron (ENSMUST00000030508.8/ENSMUSG00000028736.8, intron 4 of 8) | 4 | 139740676 | 139833007 | 92332 | 2 | ENSMUST00000174681.1 | 32740 | Pax7 | ENSMUSG00000028736.8 |
| chr4 | 139800589 | 139800824 | 3.42978 | 3.99507 | 1.09547 | Intron (ENSMUST00000030508.8/ENSMUSG00000028736.8, intron 4 of 8) | 4 | 139740676 | 139833007 | 92332 | 2 | ENSMUST00000174681.1 | 32183 | Pax7 | ENSMUSG00000028736.8 |
| chr4 | 139801581 | 139801999 | 3.4384 | 4.01838 | 1.12914 | Intron (ENSMUST00000030508.8/ENSMUSG00000028736.8, intron 4 of 8) | 4 | 139740676 | 139833007 | 92332 | 2 | ENSMUST00000174681.1 | 31008 | Pax7 | ENSMUSG00000028736.8 |
| chr4 | 139804186 | 139804269 | 3.3628 | 3.8744 | 1.0267 | Intron (ENSMUST00000030508.8/ENSMUSG00000028736.8, intron 4 of 8) | 4 | 139740676 | 139833007 | 92332 | 2 | ENSMUST00000174681.1 | 28738 | Pax7 | ENSMUSG00000028736.8 |
| chr4 | 139805604 | 139805872 | 3.43531 | 4.00626 | 1.10632 | Intron (ENSMUST00000030508.8/ENSMUSG00000028736.8, intron 4 of 8) | 4 | 139740676 | 139833007 | 92332 | 2 | ENSMUST00000174681.1 | 27135 | Pax7 | ENSMUSG00000028736.8 |
| chr4 | 139807338 | 139807637 | 3.49636 | 4.10922 | 1.14379 | Intron (ENSMUST00000030508.8/ENSMUSG00000028736.8, intron 4 of 8) | 4 | 139740676 | 139833007 | 92332 | 2 | ENSMUST00000174681.1 | 25370 | Pax7 | ENSMUSG00000028736.8 |
| chr4 | 139809359 | 139809872 | 3.74034 | 4.56853 | 1.45769 | Intron (ENSMUST00000030508.8/ENSMUSG00000028736.8, intron 4 of 8) | 4 | 139740676 | 139833007 | 92332 | 2 | ENSMUST00000174681.1 | 23135 | Pax7 | ENSMUSG00000028736.8 |
| chr4 | 139810761 | 139810864 | 3.46655 | 4.06597 | 1.15081 | Intron (ENSMUST00000030508.8/ENSMUSG00000028736.8, intron 4 of 8) | 4 | 139740676 | 139833007 | 92332 | 2 | ENSMUST00000174681.1 | 22143 | Pax7 | ENSMUSG00000028736.8 |
| chr4 | 139811199 | 139811274 | 3.97949 | 5.02047 | 1.73792 | Intron (ENSMUST00000030508.8/ENSMUSG00000028736.8, intron 4 of 8) | 4 | 139740676 | 139833007 | 92332 | 2 | ENSMUST00000174681.1 | 21733 | Pax7 | ENSMUSG00000028736.8 |
| chr4 | 139816907 | 139817772 | 3.53639 | 4.19342 | 1.22473 | Intron (ENSMUST00000030508.8/ENSMUSG00000028736.8, intron 4 of 8) | 4 | 139740676 | 139833007 | 92332 | 2 | ENSMUST00000174681.1 | 15235 | Pax7 | ENSMUSG00000028736.8 |
| chr4 | 139818277 | 139818525 | 4.26098 | 5.65204 | 2.21594 | Intron (ENSMUST00000030508.8/ENSMUSG00000028736.8, intron 4 of 8) | 4 | 139740676 | 139833007 | 92332 | 2 | ENSMUST00000174681.1 | 14482 | Pax7 | ENSMUSG00000028736.8 |
| chr4 | 139819756 | 139819834 | 4.06189 | 5.17248 | 1.83646 | Intron (ENSMUST00000030508.8/ENSMUSG00000028736.8, intron 4 of 8) | 4 | 139740676 | 139833007 | 92332 | 2 | ENSMUST00000174681.1 | 13173 | Pax7 | ENSMUSG00000028736.8 |
| chr4 | 139820184 | 139820756 | 3.61548 | 4.34815 | 1.31752 | Intron (ENSMUST00000030508.8/ENSMUSG00000028736.8, intron 4 of 8) | 4 | 139740676 | 139833007 | 92332 | 2 | ENSMUST00000174681.1 | 12251 | Pax7 | ENSMUSG00000028736.8 |
| chr4 | 139821125 | 139821966 | 3.79756 | 4.70388 | 1.57495 | Intron (ENSMUST00000030508.8/ENSMUSG00000028736.8, intron 4 of 8) | 4 | 139740676 | 139833007 | 92332 | 2 | ENSMUST00000174681.1 | 11041 | Pax7 | ENSMUSG00000028736.8 |
| chr4 | 139822493 | 139822956 | 3.57453 | 4.26264 | 1.26038 | Intron (ENSMUST00000030508.8/ENSMUSG00000028736.8, intron 4 of 8) | 4 | 139740676 | 139833007 | 92332 | 2 | ENSMUST00000174681.1 | 10051 | Pax7 | ENSMUSG00000028736.8 |
| chr4 | 139823620 | 139825663 | 4.05868 | 5.3521 | 2.02069 | Intron (ENSMUST00000030508.8/ENSMUSG00000028736.8, intron 4 of 8) | 4 | 139740676 | 139833007 | 92332 | 2 | ENSMUST00000174681.1 | 7344 | Pax7 | ENSMUSG00000028736.8 |
| chr4 | 139826028 | 139826753 | 3.84847 | 4.78417 | 1.59997 | Intron (ENSMUST00000030508.8/ENSMUSG00000028736.8, intron 4 of 8) | 4 | 139740676 | 139833007 | 92332 | 2 | ENSMUST00000174681.1 | 6254 | Pax7 | ENSMUSG00000028736.8 |
| chr4 | 139827666 | 139827966 | 3.57998 | 4.27221 | 1.26964 | Intron (ENSMUST00000030508.8/ENSMUSG00000028736.8, intron 4 of 8) | 4 | 139740676 | 139833007 | 92332 | 2 | ENSMUST00000174681.1 | 5041 | Pax7 | ENSMUSG00000028736.8 |
| chr4 | 139828452 | 139828687 | 3.70627 | 4.5034 | 1.4097 | Exon (ENSMUST00000030508.8/ENSMUSG00000028736.8, exon 4 of 9) | 4 | 139740676 | 139833007 | 92332 | 2 | ENSMUST00000174681.1 | 4320 | Pax7 | ENSMUSG00000028736.8 |
| chr4 | 139829593 | 139830143 | 3.57455 | 4.28434 | 1.30551 | Exon (ENSMUST00000030508.8/ENSMUSG00000028736.8, exon 2 of 9) | 4 | 139740676 | 139833007 | 92332 | 2 | ENSMUST00000174681.1 | 2864 | Pax7 | ENSMUSG00000028736.8 |
| chr4 | 139830551 | 139830747 | 3.47645 | 4.18368 | 1.25884 | Intron (ENSMUST00000030508.8/ENSMUSG00000028736.8, intron 1 of 8) | 4 | 139740676 | 139833007 | 92332 | 2 | ENSMUST00000174681.1 | 2260 | Pax7 | ENSMUSG00000028736.8 |
| chr4 | 139831703 | 139834189 | 3.92563 | 5.06497 | 1.82377 | Promoter | 4 | 139737062 | 139833528 | 96467 | 2 | ENSMUST00000030508.8 | 0 | Pax7 | ENSMUSG00000028736.8 |
| chr4 | 139835518 | 139836456 | 3.7014 | 4.49991 | 1.41502 | Distal Intergenic | 4 | 139737062 | 139833528 | 96467 | 2 | ENSMUST00000030508.8 | -1991 | Pax7 | ENSMUSG00000028736.8 |
| chr4 | 139839712 | 139840410 | 3.69076 | 4.51573 | 1.46145 | Distal Intergenic | 4 | 139737062 | 139833528 | 96467 | 2 | ENSMUST00000030508.8 | -6185 | Pax7 | ENSMUSG00000028736.8 |
| chr4 | 139841034 | 139841768 | 4.32673 | 5.72187 | 2.25468 | Distal Intergenic | 4 | 139737062 | 139833528 | 96467 | 2 | ENSMUST00000030508.8 | -7507 | Pax7 | ENSMUSG00000028736.8 |
| chr4 | 139842619 | 139843141 | 3.35847 | 3.86521 | 1.01733 | Distal Intergenic | 4 | 139737062 | 139833528 | 96467 | 2 | ENSMUST00000030508.8 | -9092 | Pax7 | ENSMUSG00000028736.8 |
| chr4 | 139843907 | 139844093 | 3.73055 | 4.57288 | 1.44786 | Distal Intergenic | 4 | 139737062 | 139833528 | 96467 | 2 | ENSMUST00000030508.8 | -10380 | Pax7 | ENSMUSG00000028736.8 |
| chr4 | 139848803 | 139849161 | 3.50836 | 4.14501 | 1.20274 | Distal Intergenic | 4 | 139737062 | 139833528 | 96467 | 2 | ENSMUST00000030508.8 | -15276 | Pax7 | ENSMUSG00000028736.8 |
| chr4 | 139850212 | 139852353 | 3.8073 | 4.74175 | 1.59927 | Distal Intergenic | 4 | 139737062 | 139833528 | 96467 | 2 | ENSMUST00000030508.8 | -16685 | Pax7 | ENSMUSG00000028736.8 |
| chr4 | 139852796 | 139853253 | 4.13644 | 5.63382 | 2.22519 | Distal Intergenic | 4 | 139737062 | 139833528 | 96467 | 2 | ENSMUST00000030508.8 | -19269 | Pax7 | ENSMUSG00000028736.8 |
| chr4 | 139854906 | 139855155 | 3.35019 | 3.85272 | 1.01733 | Distal Intergenic | 4 | 139737062 | 139833528 | 96467 | 2 | ENSMUST00000030508.8 | -21379 | Pax7 | ENSMUSG00000028736.8 |
| chr4 | 139855573 | 139855648 | 3.80785 | 4.68955 | 1.51309 | Distal Intergenic | 4 | 139737062 | 139833528 | 96467 | 2 | ENSMUST00000030508.8 | -22046 | Pax7 | ENSMUSG00000028736.8 |
| chr4 | 139856242 | 139856539 | 3.36173 | 3.87013 | 1.01733 | Distal Intergenic | 4 | 139737062 | 139833528 | 96467 | 2 | ENSMUST00000030508.8 | -22715 | Pax7 | ENSMUSG00000028736.8 |

| | | | | | | | | | | | | | | | |
|---|---|---|---|---|---|---|---|---|---|---|---|---|---|---|---|
| chr4 | 139861565 | 139862060 | 3.76271 | 4.68897 | 1.61473 | Distal Intergenic | 4 | 139737062 | 139833528 | 96467 | 2 | ENSMUST00000030508.8 | -28038 | Pax7 | ENSMUSG00000028736.8 |
| chr4 | 139869674 | 139870405 | 4.18263 | 5.51941 | 2.21549 | Distal Intergenic | 4 | 139737062 | 139833528 | 96467 | 2 | ENSMUST00000030508.8 | -36147 | Pax7 | ENSMUSG00000028736.8 |
| chr4 | 139871616 | 139871818 | 3.29673 | 3.77226 | 1.01733 | Distal Intergenic | 4 | 139737062 | 139833528 | 96467 | 2 | ENSMUST00000030508.8 | -38089 | Pax7 | ENSMUSG00000028736.8 |
| chr4 | 139697827 | 139698186 | 3.64717 | 4.39493 | 1.33299 | Distal Intergenic | 4 | 139653549 | 139669855 | 16307 | 1 | ENSMUST00000166773.1 | 44279 | Tas1r2 | ENSMUSG00000028738.9 |
| chr4 | 139068550 | 139068625 | 3.57349 | 4.38728 | 1.37479 | Intron (ENSMUST00000105802.2/ENSMUSG00000028747.5, intron 2 of 3) | 4 | 139061409 | 139074789 | 13381 | 2 | ENSMUST00000068036.1 | 6164 | Htr6 | ENSMUSG00000028747.5 |
| chr4 | 139073914 | 139075675 | 3.80683 | 4.85192 | 1.70443 | Promoter | 4 | 139061108 | 139075570 | 14463 | 2 | ENSMUST00000105802.2 | 0 | Htr6 | ENSMUSG00000028747.5 |
| chr4 | 139076595 | 139076805 | 3.24828 | 3.6996 | 1.01733 | Distal Intergenic | 4 | 139061108 | 139075570 | 14463 | 2 | ENSMUST00000105802.2 | -1026 | Htr6 | ENSMUSG00000028747.5 |
| chr4 | 138748563 | 138748793 | 3.47294 | 4.08199 | 1.17395 | Downstream (1-2kb) | 4 | 138750533 | 138757626 | 7094 | 2 | ENSMUST00000030526.6 | 8833 | Pla2g2f | ENSMUSG00000028749.7 |
| chr4 | 138749236 | 138749346 | 3.77646 | 4.64453 | 1.51581 | Downstream (1-2kb) | 4 | 138750533 | 138757626 | 7094 | 2 | ENSMUST00000030526.6 | 8280 | Pla2g2f | ENSMUSG00000028749.7 |
| chr4 | 138697006 | 138697400 | 3.63495 | 4.40951 | 1.39362 | Distal Intergenic | 4 | 138724792 | 138746132 | 21341 | 1 | ENSMUST00000105808.2 | -27392 | Pla2g2c | ENSMUSG00000028750.7 |
| chr4 | 138709785 | 138710061 | 3.37782 | 3.8944 | 1.01733 | 3' UTR | 4 | 138724792 | 138746132 | 21341 | 1 | ENSMUST00000105808.2 | -14731 | Pla2g2c | ENSMUSG00000028750.7 |
| chr4 | 138712814 | 138713078 | 3.57471 | 4.33834 | 1.361 | Intron (ENSMUST00000146415.1/ENSMUSG00000043621.8, intron 3 of 3) | 4 | 138724792 | 138746132 | 21341 | 1 | ENSMUST00000105808.2 | -11714 | Pla2g2c | ENSMUSG00000028750.7 |
| chr4 | 138720943 | 138721422 | 3.76496 | 4.6609 | 1.59512 | 5' UTR | 4 | 138724792 | 138746132 | 21341 | 1 | ENSMUST00000105808.2 | -3370 | Pla2g2c | ENSMUSG00000028750.7 |
| chr4 | 138723787 | 138724492 | 3.54829 | 4.24302 | 1.33244 | Promoter | 4 | 138724792 | 138746132 | 21341 | 1 | ENSMUST00000105808.2 | -300 | Pla2g2c | ENSMUSG00000028750.7 |
| chr4 | 138881842 | 138881976 | 3.58155 | 4.26508 | 1.2398 | Intron (ENSMUST00000030531.8/ENSMUSG00000028751.8, intron 4 of 4) | 4 | 138879360 | 138882817 | 3458 | 1 | ENSMUST00000105804.1 | 2483 | Pla2g2e | ENSMUSG00000028751.8 |
| chr4 | 138603197 | 138603365 | 4.25102 | 5.57689 | 2.16175 | Intron (ENSMUST00000030533.6/ENSMUSG00000028753.7, intron 7 of 21) | 4 | 138594340 | 138606530 | 12191 | 2 | ENSMUST00000154312.2 | 3165 | Vwa5b1 | ENSMUSG00000028753.7 |
| chr4 | 138341520 | 138341842 | 3.36914 | 3.88398 | 1.0267 | Intron (ENSMUST00000144986.1/ENSMUSG00000028755.3, intron 1 of 1) | 4 | 138338424 | 138343649 | 5226 | 2 | ENSMUST00000144986.1 | 1807 | Cda | ENSMUSG00000028755.3 |
| chr4 | 138362282 | 138362477 | 3.36038 | 3.8681 | 1.01733 | Intron (ENSMUST00000030535.3/ENSMUSG00000028755.3, intron 1 of 3) | 4 | 138338528 | 138367992 | 29465 | 2 | ENSMUST00000030535.3 | 5515 | Cda | ENSMUSG00000028755.3 |
| chr4 | 138368031 | 138368533 | 3.34304 | 3.87287 | 1.11409 | Promoter | 4 | 138338528 | 138367992 | 29465 | 2 | ENSMUST00000030535.3 | -40 | Cda | ENSMUSG00000028755.3 |
| chr4 | 138330581 | 138330877 | 3.58429 | 4.4217 | 1.40456 | Distal Intergenic | 4 | 138313412 | 138326307 | 12896 | 2 | ENSMUST00000030536.7 | -4275 | Pink1 | ENSMUSG00000028756.7 |
| chr4 | 137487282 | 137487582 | 3.63951 | 4.40696 | 1.41974 | Intron (ENSMUST00000030547.9/ENSMUSG00000028763.12, intron 1 of 96) | 4 | 137468803 | 137570629 | 101827 | 1 | ENSMUST00000171332.1 | 18480 | Hspg2 | ENSMUSG00000028763.12 |
| chr4 | 137569526 | 137569795 | 3.46036 | 4.06329 | 1.16579 | Promoter | 4 | 137569171 | 137570117 | 947 | 1 | ENSMUST00000155648.1 | 356 | Hspg2 | ENSMUSG00000028763.12 |
| chr4 | 137748739 | 137749068 | 3.74722 | 4.5933 | 1.46612 | Intron (ENSMUST00000030551.5/ENSMUSG00000028766.5, intron 7 of 11) | 4 | 137746420 | 137747147 | 728 | 2 | ENSMUST00000141451.1 | -1593 | Alpl | ENSMUSG00000028766.5 |
| chr4 | 137784741 | 137784915 | 3.43512 | 4.00229 | 1.09143 | Promoter | 4 | 137755086 | 137785371 | 30286 | 2 | ENSMUST00000133473.1 | 456 | Alpl | ENSMUSG00000028766.5 |
| chr4 | 137796060 | 137796348 | 3.86167 | 4.95387 | 1.74107 | Promoter | 4 | 137755630 | 137796230 | 40601 | 2 | ENSMUST00000138463.1 | 0 | Alpl | ENSMUSG00000028766.5 |
| chr4 | 130307237 | 130307592 | 3.85572 | 4.86383 | 1.73876 | Downstream (<1kb) | 4 | 130308595 | 130315463 | 6869 | 1 | ENSMUST00000070532.7 | -1003 | Fabp3 | ENSMUSG00000028773.8 |
| chr4 | 130162936 | 130163011 | 4.03018 | 5.14896 | 1.87586 | Downstream (1-2kb) | 4 | 130165604 | 130166611 | 1008 | 2 | ENSMUST00000145000.2 | 3600 | Tinagl1 | ENSMUSG00000028776.9 |
| chr4 | 130163560 | 130164022 | 3.40499 | 3.9433 | 1.04514 | Downstream (<1kb) | 4 | 130165604 | 130166611 | 1008 | 2 | ENSMUST00000145000.2 | 2589 | Tinagl1 | ENSMUSG00000028776.9 |
| chr4 | 130166799 | 130166898 | 3.40877 | 3.94954 | 1.04699 | Promoter | 4 | 130165604 | 130166611 | 1008 | 2 | ENSMUST00000145000.2 | -189 | Tinagl1 | ENSMUSG00000028776.9 |
| chr4 | 130168887 | 130169130 | 3.43564 | 4.03082 | 1.15175 | Promoter | 4 | 130166977 | 130169484 | 2508 | 2 | ENSMUST00000145774.2 | 354 | Tinagl1 | ENSMUSG00000028776.9 |
| chr4 | 130174123 | 130174266 | 3.45236 | 4.28651 | 1.31851 | Promoter | 4 | 130169113 | 130174691 | 5579 | 2 | ENSMUST00000132545.2 | 425 | Tinagl1 | ENSMUSG00000028776.9 |
| chr4 | 130174592 | 130174751 | 3.72489 | 4.75154 | 1.59824 | Promoter | 4 | 130169113 | 130174691 | 5579 | 2 | ENSMUST00000132545.2 | 0 | Tinagl1 | ENSMUSG00000028776.9 |
| chr4 | 130137182 | 130137413 | 3.53139 | 4.23259 | 1.29056 | Promoter | 4 | 130130841 | 130137628 | 6788 | 2 | ENSMUST00000164887.2 | 215 | Hcrtr1 | ENSMUSG00000028778.10 |
| chr4 | 130138002 | 130139937 | 3.99851 | 5.17461 | 1.91528 | Promoter | 4 | 130130217 | 130138551 | 8335 | 2 | ENSMUST00000120154.3 | 0 | Hcrtr1 | ENSMUSG00000028778.10 |
| chr5 | 17574563 | 17574742 | 3.95239 | 5.28586 | 1.94386 | Promoter | 5 | 17574649 | 17682083 | 107435 | 1 | ENSMUST00000170181.2 | 0 | Sema3c | ENSMUSG00000028780.8 |
| chr4 | 129980472 | 129980868 | 3.65717 | 4.42917 | 1.38873 | Distal Intergenic | 4 | 129984870 | 130022633 | 37764 | 1 | ENSMUST00000120204.2 | -4002 | Bai2 | ENSMUSG00000028782.9 |
| chr4 | 129981322 | 129981404 | 3.66657 | 4.58579 | 1.48681 | Distal Intergenic | 4 | 129984870 | 130022633 | 37764 | 1 | ENSMUST00000120204.2 | -3466 | Bai2 | ENSMUSG00000028782.9 |
| chr4 | 129982256 | 129982333 | 3.78447 | 4.81846 | 1.63024 | Distal Intergenic | 4 | 129984870 | 130022633 | 37764 | 1 | ENSMUST00000120204.2 | -2537 | Bai2 | ENSMUSG00000028782.9 |
| chr4 | 129982739 | 129984928 | 3.79608 | 4.71264 | 1.57328 | Promoter | 4 | 129984870 | 130022633 | 37764 | 1 | ENSMUST00000120204.2 | 0 | Bai2 | ENSMUSG00000028782.9 |
| chr4 | 129985255 | 129986406 | 4.24813 | 5.89276 | 2.48836 | Promoter | 4 | 129985268 | 130022633 | 37366 | 1 | ENSMUST00000121049.2 | 0 | Bai2 | ENSMUSG00000028782.9 |
| chr4 | 129987211 | 129988303 | 3.7517 | 4.63641 | 1.51271 | Intron (ENSMUST00000120204.2/ENSMUSG00000028782.9, intron 1 of 27) | 4 | 129985268 | 130022633 | 37366 | 1 | ENSMUST00000121049.2 | 1944 | Bai2 | ENSMUSG00000028782.9 |
| chr4 | 129998454 | 129998780 | 3.44148 | 4.01273 | 1.09564 | Intron (ENSMUST00000120204.2/ENSMUSG00000028782.9, intron 2 of 27) | 4 | 129992069 | 130022338 | 30270 | 1 | ENSMUST00000106018.3 | 6386 | Bai2 | ENSMUSG00000028782.9 |
| chr4 | 130006470 | 130006931 | 3.94692 | 4.97259 | 1.72337 | Exon (ENSMUST00000106017.2/ENSMUSG00000028782.9, exon 5 of 30) | 4 | 130009876 | 130011866 | 1991 | 1 | ENSMUST00000149282.2 | -2945 | Bai2 | ENSMUSG00000028782.9 |
| chr4 | 130007293 | 130007601 | 3.37026 | 3.88299 | 1.01733 | Exon (ENSMUST00000120204.2/ENSMUSG00000028782.9, exon 5 of 28) | 4 | 130009876 | 130011866 | 1991 | 1 | ENSMUST00000149282.2 | -2275 | Bai2 | ENSMUSG00000028782.9 |
| chr4 | 129943342 | 129943595 | 3.37286 | 3.88692 | 1.01733 | Distal Intergenic | 4 | 129947547 | 129953562 | 6016 | 1 | ENSMUST00000128007.2 | -3952 | Spocd1 | ENSMUSG00000028784.8 |
| chr4 | 129950074 | 129950649 | 3.51227 | 4.26128 | 1.32067 | Intron (ENSMUST00000128007.2/ENSMUSG00000028784.8, intron 3 of 4) | 4 | 129947721 | 129957115 | 9395 | 1 | ENSMUST00000084263.4 | 2354 | Spocd1 | ENSMUSG00000028784.8 |
| chr4 | 127985771 | 127986293 | 3.49048 | 4.11433 | 1.16586 | Distal Intergenic | 4 | 127987857 | 128339657 | 351801 | 1 | ENSMUST00000129619.2 | -1564 | Csmd2 | ENSMUSG00000028804.13 |
| chr4 | 127987852 | 127987927 | 4.10027 | 5.24519 | 1.88028 | Promoter | 4 | 127987857 | 128339657 | 351801 | 1 | ENSMUST00000129619.2 | 0 | Csmd2 | ENSMUSG00000028804.13 |
| chr4 | 127988310 | 127989156 | 4.33732 | 5.80644 | 2.33603 | Promoter | 4 | 127988044 | 128419617 | 431574 | 1 | ENSMUST00000144298.1 | 267 | Csmd2 | ENSMUSG00000028804.13 |
| chr4 | 127970231 | 127970418 | 3.37117 | 3.91877 | 1.13868 | Promoter | 4 | 127927592 | 127970951 | 43360 | 2 | ENSMUST00000030614.2 | 533 | CK137956 | ENSMUSG00000028813.2 |
| chr4 | 127970791 | 127971841 | 3.68776 | 4.64433 | 1.57013 | Promoter | 4 | 127927592 | 127970951 | 43360 | 2 | ENSMUST00000030614.2 | 0 | CK137956 | ENSMUSG00000028813.2 |
| chr4 | 134967881 | 134968265 | 4.01681 | 5.10762 | 1.81499 | Distal Intergenic | 4 | 134931010 | 134936078 | 5069 | 1 | ENSMUST00000144037.1 | 36872 | Syf2 | ENSMUSG00000028821.3 |
| chr4 | 134968708 | 134968971 | 3.42758 | 4.00437 | 1.1412 | Distal Intergenic | 4 | 134931010 | 134936078 | 5069 | 1 | ENSMUST00000144037.1 | 37699 | Syf2 | ENSMUSG00000028821.3 |
| chr4 | 134969415 | 134969936 | 3.42989 | 4.08606 | 1.2225 | Distal Intergenic | 4 | 134931010 | 134936078 | 5069 | 1 | ENSMUST00000144037.1 | 38406 | Syf2 | ENSMUSG00000028821.3 |
| chr4 | 134983207 | 134983282 | 3.40804 | 3.97705 | 1.14805 | Distal Intergenic | 4 | 134931010 | 134936078 | 5069 | 1 | ENSMUST00000144037.1 | 52198 | Syf2 | ENSMUSG00000028821.3 |
| chr4 | 134994129 | 134994305 | 3.4272 | 3.98722 | 1.08177 | Distal Intergenic | 4 | 134931010 | 134936078 | 5069 | 1 | ENSMUST00000144037.1 | 63120 | Syf2 | ENSMUSG00000028821.3 |
| chr4 | 135025575 | 135025774 | 3.42522 | 4.25998 | 1.29522 | Distal Intergenic | 4 | 134931010 | 134936078 | 5069 | 1 | ENSMUST00000144037.1 | 94566 | Syf2 | ENSMUSG00000028821.3 |
| chr4 | 134868643 | 134868733 | 3.37135 | 4.03 | 1.12641 | Intron (ENSMUST00000030627.7/ENSMUSG00000028825.7, intron 1 of 9) | 4 | 134864536 | 134896172 | 31637 | 1 | ENSMUST00000030627.7 | 4108 | Rhd | ENSMUSG00000028825.7 |
| chr4 | 134343101 | 134343246 | 3.93968 | 5.16679 | 1.87829 | Promoter | 4 | 134343181 | 134352677 | 9497 | 1 | ENSMUST00000105873.2 | 0 | Slc30a2 | ENSMUSG00000028836.9 |
| chr4 | 134343916 | 134344080 | 3.59229 | 4.43242 | 1.41698 | Promoter | 4 | 134343709 | 134353186 | 9478 | 1 | ENSMUST00000081094.5 | 208 | Slc30a2 | ENSMUSG00000028836.9 |
| chr4 | 133432463 | 133432621 | 3.31179 | 3.79491 | 1.01733 | Distal Intergenic | 4 | 133419972 | 133422320 | 2349 | 1 | ENSMUST00000141658.1 | 12492 | Slc9a1 | ENSMUSG00000028854.9 |
| chr4 | 125990550 | 125991085 | 3.43086 | 3.99213 | 1.07981 | Distal Intergenic | 4 | 126024550 | 126044440 | 19891 | 1 | ENSMUST00000106162.2 | -33465 | Csf3r | ENSMUSG00000028859.9 |
| chr4 | 125994196 | 125994271 | 3.6479 | 4.40161 | 1.35567 | Distal Intergenic | 4 | 126024550 | 126044440 | 19891 | 1 | ENSMUST00000106162.2 | -30279 | Csf3r | ENSMUSG00000028859.9 |
| chr4 | 133249999 | 133250074 | 3.34038 | 3.88453 | 1.13824 | Promoter | 4 | 133250849 | 133252928 | 2080 | 1 | ENSMUST00000134895.1 | -775 | Map3k6 | ENSMUSG00000028862.6 |
| chr4 | 124984635 | 124984946 | 3.44852 | 4.02886 | 1.1154 | Intron (ENSMUST00000146679.1/ENSMUSG00000087011.1, intron 1 of 1) | 4 | 124986430 | 125009099 | 22670 | 1 | ENSMUST00000030687.7 | -1484 | Rspo1 | ENSMUSG00000028871.7 |
| chr4 | 124985548 | 124985925 | 4.09346 | 5.38573 | 2.03576 | Promoter | 4 | 124986430 | 125009099 | 22670 | 1 | ENSMUST00000030687.7 | -505 | Rspo1 | ENSMUSG00000028871.7 |
| chr4 | 124986350 | 124986796 | 3.97186 | 5.08157 | 1.89765 | Promoter | 4 | 124986430 | 125009099 | 22670 | 1 | ENSMUST00000030687.7 | 0 | Rspo1 | ENSMUSG00000028871.7 |
| chr4 | 132973438 | 132973525 | 3.95621 | 5.00887 | 1.77984 | Promoter | 4 | 132974095 | 132984360 | 10266 | 1 | ENSMUST00000124451.1 | -570 | Fgr | ENSMUSG00000028874.9 |
| chr4 | 132973869 | 132973982 | 3.98425 | 5.12563 | 1.92911 | Promoter | 4 | 132974095 | 132984360 | 10266 | 1 | ENSMUST00000124451.1 | -113 | Fgr | ENSMUSG00000028874.9 |
| chr4 | 132974634 | 132975230 | 3.85983 | 4.82715 | 1.66565 | Promoter | 4 | 132974102 | 133001910 | 27809 | 1 | ENSMUST00000030693.7 | 533 | Fgr | ENSMUSG00000028874.9 |
| chr4 | 124879125 | 124879395 | 3.6298 | 4.39242 | 1.39645 | Distal Intergenic | 4 | 124880899 | 124886930 | 6032 | 1 | ENSMUST00000059343.6 | -1504 | Epha10 | ENSMUSG00000028876.7 |
| chr4 | 124879725 | 124879985 | 3.58456 | 4.29964 | 1.28164 | Promoter | 4 | 124880899 | 124886930 | 6032 | 1 | ENSMUST00000059343.6 | -914 | Epha10 | ENSMUSG00000028876.7 |
| chr4 | 124881350 | 124882997 | 3.84868 | 4.79908 | 1.61937 | Promoter | 4 | 124880899 | 124886930 | 6032 | 1 | ENSMUST00000059343.6 | 452 | Epha10 | ENSMUSG00000028876.7 |
| chr4 | 124883335 | 124883440 | 3.36009 | 3.86766 | 1.01733 | Intron (ENSMUST00000059343.6/ENSMUSG00000028876.7, intron 1 of 2) | 4 | 124885799 | 124917800 | 32002 | 1 | ENSMUST00000149146.1 | -2359 | Epha10 | ENSMUSG00000028876.7 |
| chr4 | 124883762 | 124884388 | 3.70152 | 4.4949 | 1.3994 | Exon (ENSMUST00000059343.6/ENSMUSG00000028876.7, exon 2 of 3) | 4 | 124885799 | 124917800 | 32002 | 1 | ENSMUST00000149146.1 | -1411 | Epha10 | ENSMUSG00000028876.7 |
| chr4 | 124886197 | 124886408 | 3.30486 | 3.93897 | 1.11098 | Promoter | 4 | 124885799 | 124917800 | 32002 | 1 | ENSMUST00000149146.1 | 399 | Epha10 | ENSMUSG00000028876.7 |
| chr4 | 124887702 | 124887977 | 3.37777 | 3.89432 | 1.01733 | Intron (ENSMUST00000149146.1/ENSMUSG00000028876.7, intron 1 of 14) | 4 | 124885799 | 124917800 | 32002 | 1 | ENSMUST00000149146.1 | 1904 | Epha10 | ENSMUSG00000028876.7 |
| chr4 | 124888328 | 124888763 | 3.54381 | 4.21116 | 1.23309 | Intron (ENSMUST00000149146.1/ENSMUSG00000028876.7, intron 1 of 14) | 4 | 124885799 | 124917800 | 32002 | 1 | ENSMUST00000149146.1 | 2530 | Epha10 | ENSMUSG00000028876.7 |
| chr5 | 13398420 | 13398527 | 3.29735 | 3.84868 | 1.1045 | Promoter | 5 | 13398688 | 13451207 | 52520 | 1 | ENSMUST00000125629.2 | -161 | Sema3a | ENSMUSG00000028883.12 |
| chr5 | 13399289 | 13399369 | 4.0146 | 5.27469 | 1.97766 | Promoter | 5 | 13399309 | 13602565 | 203257 | 1 | ENSMUST00000030714.7 | 0 | Sema3a | ENSMUSG00000028883.12 |
| chr4 | 132756783 | 132757260 | 3.37038 | 3.88318 | 1.01733 | Promoter | 4 | 132732966 | 132757252 | 24287 | 2 | ENSMUST00000030709.8 | 0 | Smpdl3b | ENSMUSG00000028885.8 |
| chr4 | 131835204 | 131835293 | 3.9139 | 4.91451 | 1.7074 | Intron (ENSMUST00000105987.3/ENSMUSG00000028909.12, intron 1 of 29) | 4 | 131769498 | 131838231 | 68734 | 2 | ENSMUST00000030741.3 | 2938 | Ptpru | ENSMUSG00000028909.12 |
| chr4 | 131836057 | 131836649 | 3.47432 | 4.07281 | 1.13296 | Intron (ENSMUST00000105987.3/ENSMUSG00000028909.12, intron 1 of 29) | 4 | 131769498 | 131838231 | 68734 | 2 | ENSMUST00000030741.3 | 1582 | Ptpru | ENSMUSG00000028909.12 |
| chr4 | 141240918 | 141241643 | 3.6149 | 4.35879 | 1.35174 | Intron (ENSMUST00000141834.3/ENSMUSG00000028919.6, intron 2 of 3) | 4 | 141242855 | 141246356 | 3502 | 1 | ENSMUST00000138096.2 | -1212 | Arhgef19 | ENSMUSG00000028919.6 |
| chr4 | 141247156 | 141249387 | 3.82243 | 4.79271 | 1.61876 | Promoter | 4 | 141246926 | 141247976 | 1051 | 1 | ENSMUST00000184162.1 | 231 | Arhgef19 | ENSMUSG00000028919.6 |
| chr4 | 141250822 | 141251170 | 3.37202 | 3.88565 | 1.01733 | Exon (ENSMUST00000006618.3/ENSMUSG00000028919.6, exon 13 of 17) | 4 | 141246926 | 141247976 | 1051 | 1 | ENSMUST00000184162.1 | 3897 | Arhgef19 | ENSMUSG00000028919.6 |
| chr5 | 5380549 | 5380948 | 3.41845 | 4.14258 | 1.21322 | Promoter | 5 | 5091614 | 5380251 | 288638 | 2 | ENSMUST00000132390.2 | -299 | Cdk14 | ENSMUSG00000028926.10 |
| chr4 | 140901244 | 140901595 | 3.42512 | 3.9873 | 1.09361 | Distal Intergenic | 4 | 140906344 | 140952586 | 46243 | 1 | ENSMUST00000030765.6 | -4749 | Padi2 | ENSMUSG00000028927.6 |
| chr4 | 140904150 | 140904273 | 3.36594 | 3.87648 | 1.01733 | Distal Intergenic | 4 | 140906344 | 140952586 | 46243 | 1 | ENSMUST00000030765.6 | -2071 | Padi2 | ENSMUSG00000028927.6 |
| chr4 | 140906258 | 140906953 | 4.14194 | 5.42064 | 2.11238 | Promoter | 4 | 140906344 | 140952586 | 46243 | 1 | ENSMUST00000030765.6 | 0 | Padi2 | ENSMUSG00000028927.6 |
| chr4 | 140922126 | 140922255 | 3.56311 | 4.23091 | 1.21755 | Intron (ENSMUST00000030765.6/ENSMUSG00000028927.6, intron 3 of 15) | 4 | 140924159 | 140932787 | 8629 | 1 | ENSMUST00000148160.1 | -1904 | Padi2 | ENSMUSG00000028927.6 |
| chr4 | 140931424 | 140931717 | 3.37295 | 3.88706 | 1.01733 | Intron (ENSMUST00000030765.6/ENSMUSG00000028927.6, intron 6 of 15) | 4 | 140924159 | 140932787 | 8629 | 1 | ENSMUST00000148160.1 | 7266 | Padi2 | ENSMUSG00000028927.6 |
| chr4 | 152418003 | 152418361 | 3.35377 | 3.85813 | 1.01733 | Promoter | 4 | 152412827 | 152418407 | 5581 | 2 | ENSMUST00000161844.1 | 46 | Kcnab2 | ENSMUSG00000028931.6 |
| chr4 | 152418926 | 152419172 | 3.54296 | 4.19564 | 1.20039 | Promoter | 4 | 152412827 | 152418407 | 5581 | 2 | ENSMUST00000161844.1 | -520 | Kcnab2 | ENSMUSG00000028931.6 |
| chr4 | 152456627 | 152456963 | 3.51456 | 4.1849 | 1.28219 | Intron (ENSMUST00000160884.2/ENSMUSG00000028931.6, intron 1 of 15) | 4 | 152401938 | 152455043 | 53106 | 2 | ENSMUST00000161236.2 | -1585 | Kcnab2 | ENSMUSG00000028931.6 |
| chr4 | 152157709 | 152158889 | 3.54506 | 4.35255 | 1.37456 | Promoter | 4 | 152158867 | 152162469 | 3603 | 1 | ENSMUST00000030782.1 | 0 | Hes2 | ENSMUSG00000028940.2 |
| chr4 | 152159457 | 152160872 | 3.50337 | 4.2505 | 1.30646 | Promoter | 4 | 152159549 | 152160600 | 1052 | 1 | ENSMUST00000138991.1 | 0 | Hes2 | ENSMUSG00000028940.2 |
| chr4 | 152131398 | 152131864 | 3.62427 | 4.43404 | 1.50083 | Promoter | 4 | 152121132 | 152131841 | 10710 | 2 | ENSMUST00000136627.2 | 0 | Espn | ENSMUSG00000028943.12 |
| chr4 | 152288989 | 152289230 | 3.417 | 4.54661 | 1.48862 | Intron (ENSMUST00000094438.1/ENSMUSG00000028946.6, intron 1 of 4) | 4 | 152285972 | 152291662 | 5691 | 2 | ENSMUST00000094438.1 | 2432 | Hes3 | ENSMUSG00000028946.6 |
| chr4 | 152289588 | 152289667 | 3.85367 | 5.41318 | 2.0586 | Intron (ENSMUST00000094438.1/ENSMUSG00000028946.6, intron 1 of 4) | 4 | 152285972 | 152291662 | 5691 | 2 | ENSMUST00000094438.1 | 1995 | Hes3 | ENSMUSG00000028946.6 |
| chr4 | 152289973 | 152290892 | 4.04692 | 5.30093 | 2.04683 | Promoter | 4 | 152285972 | 152291662 | 5691 | 2 | ENSMUST00000094438.1 | 770 | Hes3 | ENSMUSG00000028946.6 |
| chr4 | 152291652 | 152293049 | 3.65728 | 4.42615 | 1.37652 | Promoter | 4 | 152285972 | 152291662 | 5691 | 2 | ENSMUST00000094438.1 | 0 | Hes3 | ENSMUSG00000028946.6 |
| chr5 | 8796955 | 8797579 | 3.26326 | 3.73504 | 1.05544 | Promoter | 5 | 8798147 | 8866314 | 68168 | 1 | ENSMUST00000009058.5 | -568 | Abcb1b | ENSMUSG00000028970.5 |
| chr5 | 8798042 | 8798205 | 3.57843 | 4.35354 | 1.40624 | Promoter | 5 | 8798147 | 8866314 | 68168 | 1 | ENSMUST00000009058.5 | 0 | Abcb1b | ENSMUSG00000028970.5 |
| chr4 | 150119767 | 150120396 | 3.76606 | 4.70817 | 1.5988 | Promoter | 4 | 150119283 | 150144169 | 24887 | 1 | ENSMUST00000030826.3 | 485 | Slc2a5 | ENSMUSG00000028976.5 |
| chr4 | 148771316 | 148772642 | 3.61382 | 4.40006 | 1.41461 | Distal Intergenic | 4 | 148804429 | 148945129 | 140701 | 1 | ENSMUST00000094464.4 | -31787 | Casz1 | ENSMUSG00000028977.11 |
| chr4 | 148115862 | 148115941 | 3.94356 | 4.99183 | 1.8053 | Exon (ENSMUST00000030862.4/ENSMUSG00000029005.4, exon 2 of 7) | 4 | 148130384 | 148140791 | 10408 | 1 | ENSMUST00000126261.1 | -14443 | Mad2l2 | ENSMUSG00000029003.6 |
| chr4 | 148106825 | 148107147 | 3.31795 | 3.82943 | 1.09977 | Intron (ENSMUST00000030862.4/ENSMUSG00000029005.4, intron 6 of 6) | 4 | 148083915 | 148088031 | 4117 | 2 | ENSMUST00000137393.1 | -18795 | Agtrap | ENSMUSG00000029007.8 |
| chr4 | 148107831 | 148108051 | 3.58148 | 4.37358 | 1.37855 | Exon (ENSMUST00000030862.4/ENSMUSG00000029005.4, exon 6 of 7) | 4 | 148083915 | 148088031 | 4117 | 2 | ENSMUST00000137393.1 | -19801 | Agtrap | ENSMUSG00000029007.8 |
| chr4 | 148108799 | 148108923 | 3.57905 | 4.34964 | 1.3421 | Intron (ENSMUST00000030862.4/ENSMUSG00000029005.4, intron 5 of 6) | 4 | 148083915 | 148088031 | 4117 | 2 | ENSMUST00000137393.1 | -20769 | Agtrap | ENSMUSG00000029007.8 |
| chr5 | 21865378 | 21865677 | 4.0833 | 5.25336 | 1.91255 | Promoter | 5 | 21810655 | 21865604 | 54950 | 2 | ENSMUST00000030878.2 | 0 | Slc26a5 | ENSMUSG00000029015.4 |
| chr5 | 21866514 | 21866888 | 3.87497 | 4.84735 | 1.66776 | Promoter | 5 | 21810655 | 21865604 | 54950 | 2 | ENSMUST00000030878.2 | -911 | Slc26a5 | ENSMUSG00000029015.4 |
| chr4 | 154071041 | 154071524 | 3.44272 | 4.04917 | 1.15489 | Intron (ENSMUST00000133533.2/ENSMUSG00000029026.11, intron 2 of 11) | 4 | 154064445 | 154085322 | 20878 | 2 | ENSMUST00000155642.1 | 13798 | Trp73 | ENSMUSG00000029026.11 |
| chr4 | 154072472 | 154072665 | 4.104 | 5.484 | 2.10977 | Intron (ENSMUST00000133533.2/ENSMUSG00000029026.11, intron 2 of 11) | 4 | 154064445 | 154085322 | 20878 | 2 | ENSMUST00000155642.1 | 12657 | Trp73 | ENSMUSG00000029026.11 |
| chr4 | 154080451 | 154080866 | 3.40535 | 3.94805 | 1.06004 | Intron (ENSMUST00000133533.2/ENSMUSG00000029026.11, intron 2 of 11) | 4 | 154064445 | 154085322 | 20878 | 2 | ENSMUST00000155642.1 | 4456 | Trp73 | ENSMUSG00000029026.11 |
| chr4 | 154081389 | 154081476 | 4.79186 | 6.68993 | 2.97101 | Exon (ENSMUST00000133533.2/ENSMUSG00000029026.11, exon 2 of 12) | 4 | 154064445 | 154085322 | 20878 | 2 | ENSMUST00000155642.1 | 3846 | Trp73 | ENSMUSG00000029026.11 |
| chr4 | 154137687 | 154137983 | 3.41676 | 4.18644 | 1.25005 | Intron (ENSMUST00000105644.3/ENSMUSG00000029026.11, intron 1 of 13) | 4 | 154056253 | 154140208 | 83956 | 2 | ENSMUST00000105644.3 | 2225 | Trp73 | ENSMUSG00000029026.11 |
| chr4 | 154138356 | 154140172 | 3.95157 | 5.10839 | 1.90007 | Promoter | 4 | 154056253 | 154140208 | 83956 | 2 | ENSMUST00000105644.3 | 36 | Trp73 | ENSMUSG00000029026.11 |
| chr4 | 154140512 | 154140947 | 3.88576 | 4.95611 | 1.82547 | Promoter | 4 | 154056253 | 154140208 | 83956 | 2 | ENSMUST00000105644.3 | -305 | Trp73 | ENSMUSG00000029026.11 |
| chr4 | 154141335 | 154142035 | 3.52672 | 4.21282 | 1.30665 | Promoter | 4 | 154142372 | 154156816 | 14445 | 1 | ENSMUST00000030895.6 | -337 | Wrap73 | ENSMUSG00000029029.9 |
| chr5 | 26515098 | 26515173 | 3.73792 | 4.55552 | 1.42262 | Distal Intergenic | 5 | 26460074 | 26493278 | 33205 | 2 | ENSMUST00000030910.5 | -21821 | 4930584F24Rik | ENSMUSG00000029044.5 |
| chr4 | 155344805 | 155344956 | 4.02379 | 5.14697 | 1.8712 | Promoter | 4 | 155294353 | 155344740 | 50388 | 2 | ENSMUST00000145373.1 | -66 | Prkcz | ENSMUSG00000029053.11 |
| chr4 | 155364695 | 155364842 | 3.35075 | 3.85355 | 1.01733 | Distal Intergenic | 4 | 155261721 | 155361361 | 99641 | 2 | ENSMUST00000030922.9 | -3335 | Prkcz | ENSMUSG00000029053.11 |
| chr4 | 155367505 | 155368139 | 3.67644 | 4.48493 | 1.47491 | Distal Intergenic | 4 | 155261721 | 155361361 | 99641 | 2 | ENSMUST00000030922.9 | -6145 | Prkcz | ENSMUSG00000029053.11 |
| chr4 | 155381754 | 155382405 | 3.54209 | 4.19891 | 1.21755 | Downstream (2-3kb) | 4 | 155388565 | 155398036 | 9472 | 2 | ENSMUST00000129892.1 | 15631 | Gabrd | ENSMUSG00000029054.3 |
| chr4 | 155382868 | 155382990 | 3.94525 | 4.9647 | 1.72557 | Downstream (1-2kb) | 4 | 155388565 | 155398036 | 9472 | 2 | ENSMUST00000129892.1 | 15046 | Gabrd | ENSMUSG00000029054.3 |
| chr4 | 155386265 | 155386435 | 3.35528 | 3.86041 | 1.01733 | Exon (ENSMUST00000150423.2/ENSMUSG00000029054.3, exon 6 of 7) | 4 | 155388565 | 155398036 | 9472 | 2 | ENSMUST00000129892.1 | 11601 | Gabrd | ENSMUSG00000029054.3 |
| chr4 | 155396900 | 155397130 | 3.4467 | 4.32319 | 1.3302 | Promoter | 4 | 155388565 | 155398036 | 9472 | 2 | ENSMUST00000129892.1 | 906 | Gabrd | ENSMUSG00000029054.3 |
| chr4 | 155397828 | 155398582 | 3.72504 | 4.57912 | 1.52764 | Promoter | 4 | 155384980 | 155398047 | 13068 | 2 | ENSMUST00000150423.2 | 0 | Gabrd | ENSMUSG00000029054.3 |
| chr4 | 155014549 | 155014881 | 3.41938 | 4.21323 | 1.26799 | Intron (ENSMUST00000176194.2/ENSMUSG00000029055.12, intron 1 of 21) | 4 | 154998144 | 155010984 | 12841 | 2 | ENSMUST00000145662.3 | -3566 | Plch2 | ENSMUSG00000029055.12 |
| chr4 | 155015662 | 155016012 | 3.32775 | 3.85864 | 1.06923 | Intron (ENSMUST00000176194.2/ENSMUSG00000029055.12, intron 1 of 21) | 4 | 155009077 | 155019428 | 10352 | 2 | ENSMUST00000126098.1 | 3416 | Plch2 | ENSMUSG00000029055.12 |
| chr4 | 155018430 | 155018543 | 3.43311 | 3.99492 | 1.07749 | Promoter | 4 | 155009077 | 155019428 | 10352 | 2 | ENSMUST00000126098.1 | 885 | Plch2 | ENSMUSG00000029055.12 |
| chr4 | 155019196 | 155019363 | 3.58552 | 4.31184 | 1.35139 | Promoter | 4 | 155009077 | 155019428 | 10352 | 2 | ENSMUST00000126098.1 | 65 | Plch2 | ENSMUSG00000029055.12 |
| chr4 | 155042467 | 155044417 | 3.61354 | 4.37759 | 1.35119 | Promoter | 4 | 154983662 | 155043154 | 59493 | 2 | ENSMUST00000135665.3 | 0 | Plch2 | ENSMUSG00000029055.12 |
| chr5 | 43868672 | 43868827 | 3.76348 | 4.85881 | 1.66684 | Promoter | 5 | 43868809 | 43912375 | 43567 | 1 | ENSMUST00000030964.4 | 0 | Cd38 | ENSMUSG00000029084.4 |
| chr5 | 44099450 | 44099602 | 3.56813 | 4.25961 | 1.28759 | Promoter | 5 | 43994304 | 44099224 | 104921 | 2 | ENSMUST00000165909.2 | -227 | Prom1 | ENSMUSG00000029086.10 |
| chr5 | 44101918 | 44102498 | 3.77572 | 4.78386 | 1.60469 | Promoter | 5 | 43993625 | 44102032 | 108408 | 2 | ENSMUST00000171543.2 | 0 | Prom1 | ENSMUSG00000029086.10 |
| chr5 | 49524564 | 49525257 | 3.73437 | 4.6988 | 1.63179 | Promoter | 5 | 48389502 | 49524904 | 1135403 | 2 | ENSMUST00000087395.5 | 0 | Kcnip4 | ENSMUSG00000029088.11 |
| chr5 | 36278139 | 36278214 | 3.67859 | 4.45367 | 1.3698 | Intron (ENSMUST00000037370.8/ENSMUSG00000029093.9, intron 1 of 26) | 5 | 36065376 | 36268479 | 203104 | 2 | ENSMUST00000135324.2 | -9661 | Sorcs2 | ENSMUSG00000029093.9 |
| chr5 | 36397512 | 36397845 | 3.61171 | 4.52894 | 1.44477 | Promoter | 5 | 36061081 | 36398107 | 337027 | 2 | ENSMUST00000137040.2 | 262 | Sorcs2 | ENSMUSG00000029093.9 |
| chr5 | 36398750 | 36398838 | 3.54407 | 4.22317 | 1.29497 | Promoter | 5 | 36150202 | 36398139 | 247938 | 2 | ENSMUST00000070720.6 | -612 | Sorcs2 | ENSMUSG00000029093.9 |
| chr5 | 35679366 | 35679522 | 3.5256 | 4.33381 | 1.36499 | Promoter | 5 | 35661559 | 35679472 | 17914 | 2 | ENSMUST00000114233.2 | 0 | Htra3 | ENSMUSG00000029096.10 |
| chr5 | 35120757 | 35120856 | 3.28417 | 3.75344 | 1.01733 | Distal Intergenic | 5 | 35091490 | 35105766 | 14277 | 2 | ENSMUST00000030986.9 | -14992 | Lrpap1 | ENSMUSG00000029103.11 |
| chr5 | 57718427 | 57720256 | 3.86026 | 4.83709 | 1.67974 | Promoter | 5 | 57718021 | 58129351 | 411331 | 1 | ENSMUST00000094783.4 | 407 | Pcdh7 | ENSMUSG00000029108.9 |
| chr5 | 57722675 | 57722921 | 3.43884 | 4.04207 | 1.20968 | Intron (ENSMUST00000068110.7/ENSMUSG00000029108.9, intron 1 of 1) | 5 | 57718021 | 58129351 | 411331 | 1 | ENSMUST00000094783.4 | 4655 | Pcdh7 | ENSMUSG00000029108.9 |
| chr5 | 33428912 | 33429495 | 3.29922 | 3.77603 | 1.01733 | Downstream (1-2kb) | 5 | 33430734 | 33433976 | 3243 | 2 | ENSMUST00000173348.1 | 4481 | Nkx1-1 | ENSMUSG00000029112.5 |
| chr5 | 33430257 | 33432399 | 4.07735 | 5.31454 | 2.01223 | Exon (ENSMUST00000173348.1/ENSMUSG00000029112.5, exon 2 of 2) | 5 | 33430734 | 33433976 | 3243 | 2 | ENSMUST00000173348.1 | 1577 | Nkx1-1 | ENSMUSG00000029112.5 |
| chr5 | 33432790 | 33434900 | 4.32307 | 5.81561 | 2.34754 | Promoter | 5 | 33430734 | 33433976 | 3243 | 2 | ENSMUST00000173348.1 | 0 | Nkx1-1 | ENSMUSG00000029112.5 |
| chr5 | 33435341 | 33435708 | 3.41794 | 3.98961 | 1.14054 | Distal Intergenic | 5 | 33430734 | 33433976 | 3243 | 2 | ENSMUST00000173348.1 | -1366 | Nkx1-1 | ENSMUSG00000029112.5 |
| chr5 | 33436128 | 33436488 | 3.81128 | 4.71738 | 1.56627 | Distal Intergenic | 5 | 33430734 | 33433976 | 3243 | 2 | ENSMUST00000173348.1 | -2153 | Nkx1-1 | ENSMUSG00000029112.5 |
| chr5 | 33437188 | 33437458 | 4.02991 | 5.3295 | 1.99485 | Distal Intergenic | 5 | 33430734 | 33433976 | 3243 | 2 | ENSMUST00000173348.1 | -3213 | Nkx1-1 | ENSMUSG00000029112.5 |
| chr5 | 33438581 | 33439158 | 3.39587 | 3.93413 | 1.05168 | Distal Intergenic | 5 | 33430734 | 33433976 | 3243 | 2 | ENSMUST00000173348.1 | -4606 | Nkx1-1 | ENSMUSG00000029112.5 |
| chr5 | 33440640 | 33440964 | 3.63953 | 4.41794 | 1.34772 | Distal Intergenic | 5 | 33430734 | 33433976 | 3243 | 2 | ENSMUST00000173348.1 | -6665 | Nkx1-1 | ENSMUSG00000029112.5 |
| chr5 | 33441928 | 33442669 | 3.59183 | 4.28703 | 1.25927 | Distal Intergenic | 5 | 33430734 | 33433976 | 3243 | 2 | ENSMUST00000173348.1 | -7953 | Nkx1-1 | ENSMUSG00000029112.5 |
| chr5 | 33443007 | 33444098 | 4.04737 | 5.82937 | 2.41479 | Distal Intergenic | 5 | 33430734 | 33433976 | 3243 | 2 | ENSMUST00000173348.1 | -9032 | Nkx1-1 | ENSMUSG00000029112.5 |
| chr5 | 33444503 | 33444791 | 3.35358 | 3.86035 | 1.02623 | Distal Intergenic | 5 | 33430734 | 33433976 | 3243 | 2 | ENSMUST00000173348.1 | -10528 | Nkx1-1 | ENSMUSG00000029112.5 |
| chr5 | 33501592 | 33501872 | 3.48166 | 4.12425 | 1.24892 | Distal Intergenic | 5 | 33430734 | 33433976 | 3243 | 2 | ENSMUST00000173348.1 | -67617 | Nkx1-1 | ENSMUSG00000029112.5 |
| chr5 | 33502714 | 33503024 | 3.44822 | 4.04391 | 1.17135 | Distal Intergenic | 5 | 33430734 | 33433976 | 3243 | 2 | ENSMUST00000173348.1 | -68739 | Nkx1-1 | ENSMUSG00000029112.5 |
| chr5 | 33529087 | 33529674 | 3.62707 | 4.42907 | 1.38196 | Distal Intergenic | 5 | 33430734 | 33433976 | 3243 | 2 | ENSMUST00000173348.1 | -95112 | Nkx1-1 | ENSMUSG00000029112.5 |
| chr5 | 36857789 | 36858108 | 3.48795 | 4.11895 | 1.22497 | Distal Intergenic | 5 | 36868513 | 36955078 | 86566 | 1 | ENSMUST00000031003.7 | -10405 | Ppp2r2c | ENSMUSG00000029120.7 |
| chr5 | 36862028 | 36862277 | 3.29063 | 3.76323 | 1.01733 | Distal Intergenic | 5 | 36868513 | 36955078 | 86566 | 1 | ENSMUST00000031003.7 | -6236 | Ppp2r2c | ENSMUSG00000029120.7 |
| chr5 | 36866473 | 36867862 | 3.55602 | 4.27397 | 1.28286 | Promoter | 5 | 36868513 | 36955078 | 86566 | 1 | ENSMUST00000031003.7 | -651 | Ppp2r2c | ENSMUSG00000029120.7 |
| chr5 | 36868335 | 36868430 | 3.52336 | 4.32428 | 1.31622 | Promoter | 5 | 36868513 | 36955078 | 86566 | 1 | ENSMUST00000031003.7 | -83 | Ppp2r2c | ENSMUSG00000029120.7 |

| | | | | | | | | | | | | | | | |
|---|---|---|---|---|---|---|---|---|---|---|---|---|---|---|---|
| chr5 | 36868898 | 36870134 | 3.50423 | 4.15711 | 1.20242 | Promoter | 5 | 36868513 | 36955078 | 86566 | 1 | ENSMUST00000031003.7 | 386 | Ppp2r2c | ENSMUSG00000029120.7 |
| chr5 | 36871240 | 36871688 | 3.86415 | 4.92701 | 1.74244 | Intron (ENSMUST00000031003.7/ENSMUSG00000029120.7, intron 1 of 8) | 5 | 36868513 | 36955078 | 86566 | 1 | ENSMUST00000031003.7 | 2728 | Ppp2r2c | ENSMUSG00000029120.7 |
| chr5 | 36911854 | 36912046 | 3.39222 | 4.17336 | 1.22928 | Intron (ENSMUST00000031003.7/ENSMUSG00000029120.7, intron 1 of 8) | 5 | 36939430 | 36940187 | 758 | 1 | ENSMUST00000138124.1 | -27384 | Ppp2r2c | ENSMUSG00000029120.7 |
| chr5 | 36949385 | 36949460 | 3.68637 | 4.45973 | 1.36202 | Intron (ENSMUST00000031003.7/ENSMUSG00000029120.7, intron 7 of 8) | 5 | 36939430 | 36940187 | 758 | 1 | ENSMUST00000138124.1 | 9956 | Ppp2r2c | ENSMUSG00000029120.7 |
| chr5 | 37239985 | 37240339 | 3.59218 | 4.337 | 1.37501 | Distal Intergenic | 5 | 37242080 | 37292131 | 50052 | 1 | ENSMUST00000114158.3 | -1741 | Crmp1 | ENSMUSG00000029121.10 |
| chr5 | 37241416 | 37241530 | 3.71678 | 4.54887 | 1.50378 | Promoter | 5 | 37242080 | 37292131 | 50052 | 1 | ENSMUST00000114158.3 | -550 | Crmp1 | ENSMUSG00000029121.10 |
| chr5 | 37241856 | 37242913 | 3.52154 | 4.18962 | 1.25774 | Promoter | 5 | 37242080 | 37292131 | 50052 | 1 | ENSMUST00000114158.3 | 0 | Crmp1 | ENSMUSG00000029121.10 |
| chr5 | 37243892 | 37244357 | 3.65433 | 4.41467 | 1.35444 | Intron (ENSMUST00000114158.3/ENSMUSG00000029121.10, intron 1 of 13) | 5 | 37245792 | 37292132 | 46341 | 1 | ENSMUST00000031004.7 | -1435 | Crmp1 | ENSMUSG00000029121.10 |
| chr5 | 37244740 | 37244815 | 3.67612 | 4.44954 | 1.37164 | Promoter | 5 | 37245792 | 37292132 | 46341 | 1 | ENSMUST00000031004.7 | -977 | Crmp1 | ENSMUSG00000029121.10 |
| chr5 | 37245812 | 37246509 | 3.82307 | 4.90695 | 1.70416 | Promoter | 5 | 37245792 | 37292132 | 46341 | 1 | ENSMUST00000031004.7 | 21 | Crmp1 | ENSMUSG00000029121.10 |
| chr5 | 37336528 | 37337478 | 4.12551 | 5.43515 | 2.13464 | Promoter | 5 | 37299171 | 37336894 | 37724 | 2 | ENSMUST00000031005.5 | 0 | Evc | ENSMUSG00000029122.6 |
| chr5 | 37716939 | 37717247 | 3.40585 | 3.98231 | 1.17668 | Promoter | 5 | 37446828 | 37717153 | 270326 | 2 | ENSMUST00000094836.4 | 0 | Stk32b | ENSMUSG00000029123.7 |
| chr5 | 37966858 | 37967901 | 3.8579 | 4.90213 | 1.74731 | Distal Intergenic | 5 | 38038742 | 38137765 | 99024 | 1 | ENSMUST00000114126.3 | -70841 | Stx18 | ENSMUSG00000029125.9 |
| chr5 | 37968315 | 37969465 | 4.06647 | 5.27111 | 1.98867 | Distal Intergenic | 5 | 38038742 | 38137765 | 99024 | 1 | ENSMUST00000114126.3 | -69277 | Stx18 | ENSMUSG00000029125.9 |
| chr5 | 37970038 | 37970504 | 3.50911 | 4.28979 | 1.32464 | Distal Intergenic | 5 | 38038742 | 38137765 | 99024 | 1 | ENSMUST00000114126.3 | -68238 | Stx18 | ENSMUSG00000029125.9 |
| chr5 | 37971732 | 37971911 | 3.3958 | 3.95738 | 1.14296 | Distal Intergenic | 5 | 38038742 | 38137765 | 99024 | 1 | ENSMUST00000114126.3 | -66831 | Stx18 | ENSMUSG00000029125.9 |
| chr5 | 38159058 | 38159478 | 3.44401 | 4.02991 | 1.14257 | Promoter | 5 | 38137193 | 38159467 | 22275 | 2 | ENSMUST00000031009.4 | 0 | Nsg1 | ENSMUSG00000029126.4 |
| chr5 | 29197485 | 29197898 | 3.45134 | 4.07375 | 1.24078 | Promoter | 5 | 29198006 | 29225291 | 27286 | 1 | ENSMUST00000168460.2 | -108 | Rnf32 | ENSMUSG00000029130.7 |
| chr5 | 32121622 | 32121948 | 3.37105 | 3.88419 | 1.01733 | Distal Intergenic | 5 | 32136472 | 32157831 | 21360 | 1 | ENSMUST00000031017.9 | -14524 | Fosl2 | ENSMUSG00000029135.9 |
| chr5 | 31083151 | 31083703 | 3.50416 | 4.32754 | 1.38254 | Downstream (2-3kb) | 5 | 31087399 | 31088356 | 958 | 2 | ENSMUST00000182290.1 | 4653 | Slc30a3 | ENSMUSG00000029151.9 |
| chr5 | 31089503 | 31089732 | 3.91424 | 5.08077 | 1.86325 | Exon (ENSMUST00000031037.8/ENSMUSG00000029151.9, exon 3 of 8) | 5 | 31087399 | 31088356 | 958 | 2 | ENSMUST00000182290.1 | -1148 | Slc30a3 | ENSMUSG00000029151.9 |
| chr5 | 31090066 | 31090488 | 3.61735 | 4.64787 | 1.51488 | 5' UTR | 5 | 31087399 | 31088356 | 958 | 2 | ENSMUST00000182290.1 | -1711 | Slc30a3 | ENSMUSG00000029151.9 |
| chr5 | 31090795 | 31091136 | 3.67368 | 4.5798 | 1.51219 | Intron (ENSMUST00000031037.8/ENSMUSG00000029151.9, intron 1 of 7) | 5 | 31088782 | 31093469 | 4688 | 2 | ENSMUST00000182364.1 | 2333 | Slc30a3 | ENSMUSG00000029151.9 |
| chr5 | 31091993 | 31095844 | 4.16039 | 5.49121 | 2.1328 | Promoter | 5 | 31086106 | 31093615 | 7510 | 2 | ENSMUST00000031037.8 | 0 | Slc30a3 | ENSMUSG00000029151.9 |
| chr5 | 31096169 | 31096521 | 3.6105 | 4.47434 | 1.46012 | Intron (ENSMUST00000182444.2/ENSMUSG00000029151.9, intron 1 of 8) | 5 | 31089341 | 31094801 | 5461 | 2 | ENSMUST00000182672.2 | -1369 | Slc30a3 | ENSMUSG00000029151.9 |
| chr5 | 31098206 | 31098304 | 3.28555 | 3.75549 | 1.01733 | Intron (ENSMUST00000182444.2/ENSMUSG00000029151.9, intron 1 of 8) | 5 | 31089341 | 31094801 | 5461 | 2 | ENSMUST00000182672.2 | -3406 | Slc30a3 | ENSMUSG00000029151.9 |
| chr5 | 31098716 | 31099155 | 3.38063 | 3.89863 | 1.01733 | Intron (ENSMUST00000182444.2/ENSMUSG00000029151.9, intron 1 of 8) | 5 | 31089341 | 31094801 | 5461 | 2 | ENSMUST00000182672.2 | -3916 | Slc30a3 | ENSMUSG00000029151.9 |
| chr5 | 73350196 | 73350359 | 3.38531 | 3.92613 | 1.08948 | Distal Intergenic | 5 | 73332128 | 73338947 | 6820 | 2 | ENSMUST00000133164.1 | -11250 | Ociad2 | ENSMUSG00000029153.8 |
| chr5 | 73404701 | 73405444 | 3.28625 | 3.75662 | 1.01733 | Promoter | 5 | 73406076 | 73453435 | 47360 | 1 | ENSMUST00000031040.7 | -632 | Cwh43 | ENSMUSG00000029154.8 |
| chr5 | 73405833 | 73407019 | 3.96598 | 5.02807 | 1.77641 | Promoter | 5 | 73406076 | 73453435 | 47360 | 1 | ENSMUST00000031040.7 | 0 | Cwh43 | ENSMUSG00000029154.8 |
| chr5 | 73442159 | 73442536 | 3.73739 | 4.58656 | 1.47436 | Intron (ENSMUST00000031040.7/ENSMUSG00000029154.8, intron 15 of 15) | 5 | 73430331 | 73434476 | 4146 | 1 | ENSMUST00000135820.1 | 11829 | Cwh43 | ENSMUSG00000029154.8 |
| chr5 | 73651000 | 73651813 | 4.19494 | 5.71432 | 2.32572 | Promoter | 5 | 73651380 | 73679096 | 27717 | 1 | ENSMUST00000041422.9 | 0 | Spata18 | ENSMUSG00000029155.11 |
| chr5 | 73720434 | 73720578 | 3.32476 | 3.83602 | 1.08926 | Distal Intergenic | 5 | 73657535 | 73679482 | 21948 | 1 | ENSMUST00000113548.2 | 62900 | Spata18 | ENSMUSG00000029155.11 |
| chr5 | 73649125 | 73649676 | 3.47981 | 4.27668 | 1.31131 | Distal Intergenic | 5 | 73632749 | 73647713 | 14965 | 2 | ENSMUST00000081170.7 | -1413 | Sgcb | ENSMUSG00000029156.10 |
| chr5 | 30707008 | 30707295 | 3.38193 | 3.9006 | 1.01733 | Distal Intergenic | 5 | 30711564 | 30799375 | 87812 | 1 | ENSMUST00000114729.2 | -4269 | Dpysl5 | ENSMUSG00000029168.9 |
| chr5 | 30713353 | 30713801 | 3.76156 | 4.64428 | 1.56712 | Intron (ENSMUST00000114729.2/ENSMUSG00000029168.9, intron 1 of 12) | 5 | 30711901 | 30748014 | 36114 | 1 | ENSMUST00000101442.3 | 1453 | Dpysl5 | ENSMUSG00000029168.9 |
| chr5 | 64047780 | 64047882 | 3.78951 | 4.6953 | 1.60967 | Distal Intergenic | 5 | 64092936 | 64128355 | 35420 | 1 | ENSMUST00000087324.5 | -45054 | Pgm1 | ENSMUSG00000029171.8 |
| chr5 | 30618912 | 30618987 | 3.57554 | 4.26256 | 1.26265 | Intron (ENSMUST00000066295.2/ENSMUSG00000049265.6, intron 1 of 1) | 5 | 30647933 | 30659731 | 11799 | 1 | ENSMUST00000062962.6 | -28946 | Slc35f6 | ENSMUSG00000029175.12 |
| chr5 | 30620762 | 30621090 | 3.52967 | 4.52925 | 1.46695 | Intron (ENSMUST00000066295.2/ENSMUSG00000049265.6, intron 1 of 1) | 5 | 30647933 | 30659731 | 11799 | 1 | ENSMUST00000062962.6 | -26843 | Slc35f6 | ENSMUSG00000029175.12 |
| chr5 | 30621855 | 30623010 | 3.73747 | 4.6386 | 1.54365 | 3' UTR | 5 | 30647933 | 30659731 | 11799 | 1 | ENSMUST00000062962.6 | -24923 | Slc35f6 | ENSMUSG00000029175.12 |
| chr5 | 30623756 | 30623831 | 3.38193 | 3.9006 | 1.01733 | 3' UTR | 5 | 30647933 | 30659731 | 11799 | 1 | ENSMUST00000062962.6 | -24102 | Slc35f6 | ENSMUSG00000029175.12 |
| chr5 | 30465483 | 30466131 | 3.65723 | 4.43784 | 1.41548 | Promoter | 5 | 30466047 | 30480906 | 14860 | 1 | ENSMUST00000127815.1 | 0 | 1700001C02Rik | ENSMUSG00000029182.5 |
| chr5 | 53048940 | 53049400 | 4.06 | 5.2747 | 1.99977 | Promoter | 5 | 53049353 | 53066300 | 16948 | 1 | ENSMUST00000147243.2 | 0 | Slc34a2 | ENSMUSG00000029188.9 |
| chr5 | 53210276 | 53210486 | 3.50311 | 4.15929 | 1.18101 | Intron (ENSMUST00000031090.6/ENSMUSG00000029189.6, intron 1 of 23) | 5 | 53107084 | 53213452 | 106369 | 2 | ENSMUST00000031090.6 | 2966 | Sel1l3 | ENSMUSG00000029189.6 |
| chr5 | 53211307 | 53211563 | 3.76643 | 5.20199 | 1.92051 | Intron (ENSMUST00000031090.6/ENSMUSG00000029189.6, intron 1 of 23) | 5 | 53107084 | 53213452 | 106369 | 2 | ENSMUST00000031090.6 | 1889 | Sel1l3 | ENSMUSG00000029189.6 |
| chr5 | 53211959 | 53212211 | 3.74097 | 4.58214 | 1.46662 | Intron (ENSMUST00000031090.6/ENSMUSG00000029189.6, intron 1 of 23) | 5 | 53107084 | 53213452 | 106369 | 2 | ENSMUST00000031090.6 | 1241 | Sel1l3 | ENSMUSG00000029189.6 |
| chr5 | 53212867 | 53213226 | 3.48421 | 4.272 | 1.29085 | Promoter | 5 | 53107084 | 53213452 | 106369 | 2 | ENSMUST00000031090.6 | 226 | Sel1l3 | ENSMUSG00000029189.6 |
| chr5 | 66259566 | 66260273 | 3.63222 | 4.43665 | 1.37994 | Promoter | 5 | 66259912 | 66297062 | 37151 | 1 | ENSMUST00000065530.7 | 0 | Nsun7 | ENSMUSG00000029206.8 |
| chr5 | 73005485 | 73005565 | 3.70075 | 4.74902 | 1.61651 | Distal Intergenic | 5 | 73006904 | 73014683 | 7780 | 1 | ENSMUST00000031127.7 | -1339 | Slc10a4 | ENSMUSG00000029219.7 |
| chr5 | 73006070 | 73006175 | 3.29675 | 3.77238 | 1.01733 | Promoter | 5 | 73006904 | 73014683 | 7780 | 1 | ENSMUST00000031127.7 | -729 | Slc10a4 | ENSMUSG00000029219.7 |
| chr5 | 73007058 | 73009514 | 4.2659 | 5.77229 | 2.43063 | Promoter | 5 | 73006904 | 73014683 | 7780 | 1 | ENSMUST00000031127.7 | 155 | Slc10a4 | ENSMUSG00000029219.7 |
| chr5 | 66676776 | 66677021 | 3.40391 | 4.08697 | 1.18607 | Promoter | 5 | 66676893 | 66687229 | 10337 | 1 | ENSMUST00000176015.1 | 0 | Uchl1 | ENSMUSG00000029223.9 |
| chr5 | 66677965 | 66678292 | 3.78662 | 4.93932 | 1.72369 | Intron (ENSMUST00000031131.9/ENSMUSG00000029223.9, intron 3 of 8) | 5 | 66676893 | 66687229 | 10337 | 1 | ENSMUST00000176015.1 | 1073 | Uchl1 | ENSMUSG00000029223.9 |
| chr5 | 74631194 | 74631371 | 3.57252 | 4.28013 | 1.32161 | Intron (ENSMUST00000039744.7/ENSMUSG00000029228.10, intron 1 of 10) | 5 | 74606037 | 74623521 | 17485 | 2 | ENSMUST00000140084.1 | -7674 | Lnx1 | ENSMUSG00000029228.10 |
| chr5 | 74632429 | 74632504 | 3.355 | 3.85997 | 1.01733 | Intron (ENSMUST00000039744.7/ENSMUSG00000029228.10, intron 1 of 10) | 5 | 74606037 | 74623521 | 17485 | 2 | ENSMUST00000140084.1 | -8909 | Lnx1 | ENSMUSG00000029228.10 |
| chr5 | 74632911 | 74633756 | 3.59377 | 4.29665 | 1.27895 | Intron (ENSMUST00000039744.7/ENSMUSG00000029228.10, intron 1 of 10) | 5 | 74606037 | 74623521 | 17485 | 2 | ENSMUST00000140084.1 | -9391 | Lnx1 | ENSMUSG00000029228.10 |
| chr5 | 75128426 | 75128635 | 3.74892 | 4.60404 | 1.50233 | Distal Intergenic | 5 | 75152291 | 75198204 | 45914 | 1 | ENSMUST00000000476.9 | -23656 | Pdgfra | ENSMUSG00000029231.10 |
| chr5 | 75142627 | 75143218 | 3.69279 | 4.60262 | 1.52475 | Distal Intergenic | 5 | 75152291 | 75198204 | 45914 | 1 | ENSMUST00000000476.9 | -9073 | Pdgfra | ENSMUSG00000029231.10 |
| chr5 | 75143752 | 75143954 | 3.34571 | 4.08946 | 1.2137 | Distal Intergenic | 5 | 75152291 | 75198204 | 45914 | 1 | ENSMUST00000000476.9 | -8337 | Pdgfra | ENSMUSG00000029231.10 |
| chr5 | 75144335 | 75144729 | 3.66896 | 4.53331 | 1.48697 | Distal Intergenic | 5 | 75152291 | 75198204 | 45914 | 1 | ENSMUST00000000476.9 | -7562 | Pdgfra | ENSMUSG00000029231.10 |
| chr5 | 75146321 | 75146990 | 3.76125 | 4.73065 | 1.62024 | Distal Intergenic | 5 | 75152291 | 75198204 | 45914 | 1 | ENSMUST00000000476.9 | -5301 | Pdgfra | ENSMUSG00000029231.10 |
| chr5 | 75147762 | 75148289 | 3.95521 | 5.16072 | 1.89233 | Distal Intergenic | 5 | 75152291 | 75198204 | 45914 | 1 | ENSMUST00000000476.9 | -4002 | Pdgfra | ENSMUSG00000029231.10 |
| chr5 | 75148799 | 75151539 | 3.97508 | 5.05253 | 1.79179 | Promoter | 5 | 75152291 | 75198204 | 45914 | 1 | ENSMUST00000000476.9 | -752 | Pdgfra | ENSMUSG00000029231.10 |
| chr5 | 75151884 | 75151974 | 3.81728 | 4.71994 | 1.54804 | Promoter | 5 | 75152291 | 75198204 | 45914 | 1 | ENSMUST00000000476.9 | -317 | Pdgfra | ENSMUSG00000029231.10 |
| chr5 | 75152298 | 75152502 | 3.60665 | 4.34738 | 1.33032 | Promoter | 5 | 75152291 | 75198204 | 45914 | 1 | ENSMUST00000000476.9 | 8 | Pdgfra | ENSMUSG00000029231.10 |
| chr5 | 75152890 | 75154433 | 4.27365 | 5.83944 | 2.37292 | Promoter | 5 | 75152291 | 75198204 | 45914 | 1 | ENSMUST00000000476.9 | 600 | Pdgfra | ENSMUSG00000029231.10 |
| chr5 | 75154804 | 75155299 | 3.52374 | 4.188 | 1.26396 | Promoter | 5 | 75156166 | 75198204 | 42039 | 1 | ENSMUST00000168162.1 | -867 | Pdgfra | ENSMUSG00000029231.10 |
| chr5 | 75155643 | 75156958 | 3.96198 | 5.02164 | 1.77202 | Promoter | 5 | 75156166 | 75198204 | 42039 | 1 | ENSMUST00000168162.1 | 0 | Pdgfra | ENSMUSG00000029231.10 |
| chr5 | 75158185 | 75158602 | 3.66277 | 4.42501 | 1.36945 | Intron (ENSMUST00000000476.9/ENSMUSG00000029231.10, intron 1 of 22) | 5 | 75156166 | 75198204 | 42039 | 1 | ENSMUST00000168162.1 | 2020 | Pdgfra | ENSMUSG00000029231.10 |
| chr5 | 75161721 | 75161897 | 3.48644 | 4.09974 | 1.16416 | Intron (ENSMUST00000000476.9/ENSMUSG00000029231.10, intron 1 of 22) | 5 | 75156166 | 75198204 | 42039 | 1 | ENSMUST00000168162.1 | 5556 | Pdgfra | ENSMUSG00000029231.10 |
| chr5 | 75162473 | 75162809 | 3.39652 | 3.946 | 1.10039 | Intron (ENSMUST00000000476.9/ENSMUSG00000029231.10, intron 2 of 22) | 5 | 75156166 | 75198204 | 42039 | 1 | ENSMUST00000168162.1 | 6308 | Pdgfra | ENSMUSG00000029231.10 |
| chr5 | 76361566 | 76361825 | 3.68901 | 4.56247 | 1.50933 | Intron (ENSMUST00000031146.2/ENSMUSG00000029236.4, intron 1 of 9) | 5 | 76333495 | 76363788 | 30294 | 2 | ENSMUST00000031146.2 | 1963 | Nmu | ENSMUSG00000029236.4 |
| chr5 | 76363175 | 76363918 | 4.01375 | 5.38874 | 2.04735 | Promoter | 5 | 76333495 | 76363788 | 30294 | 2 | ENSMUST00000031146.2 | 0 | Nmu | ENSMUSG00000029236.4 |
| chr5 | 84411947 | 84412237 | 3.41657 | 4.06907 | 1.22389 | Intron (ENSMUST00000113403.2/ENSMUSG00000029245.11, intron 1 of 17) | 5 | 84057791 | 84416806 | 359016 | 2 | ENSMUST00000113403.2 | 4569 | Epha5 | ENSMUSG00000029245.11 |
| chr5 | 84412986 | 84413406 | 3.45239 | 4.26578 | 1.29899 | Intron (ENSMUST00000113403.2/ENSMUSG00000029245.11, intron 1 of 17) | 5 | 84057791 | 84416806 | 359016 | 2 | ENSMUST00000113403.2 | 3400 | Epha5 | ENSMUSG00000029245.11 |
| chr5 | 84415467 | 84415547 | 3.76449 | 4.66813 | 1.60437 | Intron (ENSMUST00000113403.2/ENSMUSG00000029245.11, intron 1 of 17) | 5 | 84057791 | 84416806 | 359016 | 2 | ENSMUST00000113403.2 | 1259 | Epha5 | ENSMUSG00000029245.11 |
| chr5 | 84416020 | 84416857 | 4.02916 | 5.18883 | 1.95309 | Promoter | 5 | 84057791 | 84416806 | 359016 | 2 | ENSMUST00000113403.2 | 0 | Epha5 | ENSMUSG00000029245.11 |
| chr5 | 84417295 | 84417773 | 3.55905 | 4.36123 | 1.38985 | Promoter | 5 | 84065512 | 84417359 | 351848 | 2 | ENSMUST00000113401.2 | 0 | Epha5 | ENSMUSG00000029245.11 |
| chr5 | 77016653 | 77017006 | 3.62968 | 4.49277 | 1.44344 | Promoter | 5 | 77016023 | 77061529 | 45507 | 1 | ENSMUST00000117880.2 | 631 | 1700023E05Rik | ENSMUSG00000029248.9 |
| chr5 | 108304545 | 108304822 | 3.37295 | 3.88706 | 1.01733 | Distal Intergenic | 5 | 108312609 | 108348233 | 35625 | 1 | ENSMUST00000031189.8 | -7787 | Pigg | ENSMUSG00000029263.10 |
| chr5 | 107707972 | 107708234 | 3.3501 | 3.85259 | 1.01733 | Distal Intergenic | 5 | 107717224 | 107723961 | 6738 | 2 | ENSMUST00000165344.1 | 15727 | Gfi1 | ENSMUSG00000029275.12 |
| chr5 | 107711567 | 107711813 | 3.39959 | 3.93985 | 1.06182 | Distal Intergenic | 5 | 107717224 | 107723961 | 6738 | 2 | ENSMUST00000165344.1 | 12148 | Gfi1 | ENSMUSG00000029275.12 |
| chr5 | 107712495 | 107712591 | 3.60407 | 4.50557 | 1.41581 | Distal Intergenic | 5 | 107717224 | 107723961 | 6738 | 2 | ENSMUST00000165344.1 | 11370 | Gfi1 | ENSMUSG00000029275.12 |
| chr5 | 107714959 | 107715141 | 3.38345 | 3.91762 | 1.06942 | Downstream (1-2kb) | 5 | 107717224 | 107723961 | 6738 | 2 | ENSMUST00000165344.1 | 8820 | Gfi1 | ENSMUSG00000029275.12 |
| chr5 | 107717820 | 107720338 | 3.92238 | 4.96999 | 1.77793 | Exon (ENSMUST00000031205.10/ENSMUSG00000029275.12, exon 5 of 6) | 5 | 107717224 | 107723961 | 6738 | 2 | ENSMUST00000165344.1 | 3623 | Gfi1 | ENSMUSG00000029275.12 |
| chr5 | 107720715 | 107722554 | 4.41895 | 6.08604 | 2.59889 | 5' UTR | 5 | 107717224 | 107723961 | 6738 | 2 | ENSMUST00000165344.1 | 1407 | Gfi1 | ENSMUSG00000029275.12 |
| chr5 | 107723096 | 107725989 | 3.94059 | 5.11173 | 1.87374 | Promoter | 5 | 107716657 | 107724343 | 7687 | 2 | ENSMUST00000031205.10 | 0 | Gfi1 | ENSMUSG00000029275.12 |
| chr5 | 106963556 | 106963825 | 3.3569 | 3.87893 | 1.07413 | Promoter | 5 | 106964322 | 106984431 | 20110 | 1 | ENSMUST00000117196.3 | -497 | Cdc7 | ENSMUSG00000029283.12 |
| chr5 | 104114071 | 104114249 | 3.30845 | 3.79788 | 1.04393 | Promoter | 5 | 104079111 | 104114088 | 34978 | 2 | ENSMUST00000031249.3 | 0 | Sparcl1 | ENSMUSG00000029309.3 |
| chr5 | 100582130 | 100582345 | 3.4972 | 4.11775 | 1.16861 | Distal Intergenic | 5 | 100553725 | 100572245 | 18521 | 2 | ENSMUST00000031264.6 | -9886 | Plac8 | ENSMUSG00000029322.7 |
| chr5 | 99174906 | 99175325 | 3.3695 | 3.88185 | 1.01733 | Distal Intergenic | 5 | 99227794 | 99234066 | 6273 | 2 | ENSMUST00000166632.1 | 58741 | Rasgef1b | ENSMUSG00000029333.9 |
| chr5 | 99191207 | 99191375 | 3.91643 | 5.07369 | 1.81736 | Distal Intergenic | 5 | 99227794 | 99234066 | 6273 | 2 | ENSMUST00000166632.1 | 42691 | Rasgef1b | ENSMUSG00000029333.9 |
| chr5 | 99276317 | 99276653 | 3.5309 | 4.1917 | 1.24802 | Distal Intergenic | 5 | 99217426 | 99252927 | 35502 | 2 | ENSMUST00000168092.2 | -23391 | Rasgef1b | ENSMUSG00000029333.9 |
| chr5 | 99034607 | 99034821 | 3.31144 | 3.7944 | 1.01733 | Intron (ENSMUST00000161490.2/ENSMUSG00000029334.9, intron 1 of 18) | 5 | 98931720 | 99037035 | 105316 | 2 | ENSMUST00000031277.6 | 2214 | Prkg2 | ENSMUSG00000029334.9 |
| chr5 | 99036676 | 99037244 | 4.1244 | 5.32312 | 1.97223 | Promoter | 5 | 98931720 | 99037035 | 105316 | 2 | ENSMUST00000031277.6 | 0 | Prkg2 | ENSMUSG00000029334.9 |
| chr5 | 99037602 | 99037677 | 3.5978 | 4.29417 | 1.25589 | Promoter | 5 | 98929773 | 99037351 | 107579 | 2 | ENSMUST00000161490.2 | -252 | Prkg2 | ENSMUSG00000029334.9 |
| chr5 | 99038142 | 99038502 | 4.00153 | 5.07152 | 1.79454 | Promoter | 5 | 98929773 | 99037351 | 107579 | 2 | ENSMUST00000161490.2 | -792 | Prkg2 | ENSMUSG00000029334.9 |
| chr5 | 98853135 | 98853415 | 3.63902 | 4.42163 | 1.40245 | Distal Intergenic | 5 | 98854439 | 98880960 | 26522 | 1 | ENSMUST00000031278.4 | -1024 | Bmp3 | ENSMUSG00000029335.4 |
| chr5 | 98854120 | 98854490 | 3.53554 | 4.18677 | 1.20913 | Promoter | 5 | 98854439 | 98880960 | 26522 | 1 | ENSMUST00000031278.4 | 0 | Bmp3 | ENSMUSG00000029335.4 |
| chr5 | 98854891 | 98855132 | 3.95056 | 5.05722 | 1.8313 | Promoter | 5 | 98854439 | 98880960 | 26522 | 1 | ENSMUST00000031278.4 | 453 | Bmp3 | ENSMUSG00000029335.4 |
| chr5 | 98252275 | 98252470 | 4.30925 | 5.82671 | 2.4466 | Distal Intergenic | 5 | 98254184 | 98277030 | 22847 | 1 | ENSMUST00000031280.1 | -1714 | Fgf5 | ENSMUSG00000029337.1 |
| chr5 | 98252836 | 98253057 | 3.69911 | 4.78146 | 1.60257 | Distal Intergenic | 5 | 98254184 | 98277030 | 22847 | 1 | ENSMUST00000031280.1 | -1127 | Fgf5 | ENSMUSG00000029337.1 |
| chr5 | 98253906 | 98255826 | 3.76179 | 4.75519 | 1.636 | Promoter | 5 | 98254184 | 98277030 | 22847 | 1 | ENSMUST00000031280.1 | 0 | Fgf5 | ENSMUSG00000029337.1 |
| chr5 | 98256231 | 98257132 | 3.81835 | 4.78434 | 1.67521 | Intron (ENSMUST00000031280.1/ENSMUSG00000029337.1, intron 1 of 2) | 5 | 98254184 | 98277030 | 22847 | 1 | ENSMUST00000031280.1 | 2048 | Fgf5 | ENSMUSG00000029337.1 |
| chr5 | 98257691 | 98258217 | 3.54012 | 4.22838 | 1.31185 | Intron (ENSMUST00000031280.1/ENSMUSG00000029337.1, intron 1 of 2) | 5 | 98254184 | 98277030 | 22847 | 1 | ENSMUST00000031280.1 | 3508 | Fgf5 | ENSMUSG00000029337.1 |
| chr5 | 98290529 | 98290732 | 3.35092 | 3.85382 | 1.01733 | Distal Intergenic | 5 | 98254184 | 98277030 | 22847 | 1 | ENSMUST00000031280.1 | 36346 | Fgf5 | ENSMUSG00000029337.1 |
| chr5 | 97935405 | 97935590 | 3.68066 | 5.12732 | 1.87069 | Intron (ENSMUST00000031281.9/ENSMUSG00000029338.9, intron 16 of 16) | 5 | 97882783 | 98030733 | 147951 | 2 | ENSMUST00000031281.9 | 95143 | Antxr2 | ENSMUSG00000029338.9 |
| chr5 | 112268411 | 112269431 | 3.75923 | 4.61949 | 1.49111 | Exon (ENSMUST00000031286.7/ENSMUSG00000029343.12, exon 6 of 6) | 5 | 112276691 | 112315361 | 38671 | 1 | ENSMUST00000031287.5 | -7260 | Tpst2 | ENSMUSG00000029344.9 |
| chr5 | 112391897 | 112392082 | 3.59772 | 4.29592 | 1.26265 | Promoter | 5 | 112385448 | 112392213 | 6766 | 2 | ENSMUST00000031291.7 | 131 | Asphd2 | ENSMUSG00000029348.7 |
| chr5 | 112393341 | 112394016 | 3.54916 | 4.26785 | 1.29149 | Distal Intergenic | 5 | 112385448 | 112392213 | 6766 | 2 | ENSMUST00000031291.7 | -1129 | Asphd2 | ENSMUSG00000029348.7 |
| chr5 | 112394501 | 112394925 | 3.48314 | 4.09369 | 1.16006 | Distal Intergenic | 5 | 112385448 | 112392213 | 6766 | 2 | ENSMUST00000031291.7 | -2289 | Asphd2 | ENSMUSG00000029348.7 |
| chr5 | 112395346 | 112395621 | 3.83434 | 4.74313 | 1.55957 | Distal Intergenic | 5 | 112385448 | 112392213 | 6766 | 2 | ENSMUST00000031291.7 | -3134 | Asphd2 | ENSMUSG00000029348.7 |
| chr5 | 112396189 | 112396801 | 3.72668 | 4.53335 | 1.407 | Distal Intergenic | 5 | 112385448 | 112392213 | 6766 | 2 | ENSMUST00000031291.7 | -3977 | Asphd2 | ENSMUSG00000029348.7 |
| chr5 | 113121465 | 113121770 | 3.3428 | 3.85686 | 1.07072 | Intron (ENSMUST00000050125.8/ENSMUSG00000051339.8, intron 1 of 7) | 5 | 113075839 | 113081584 | 5746 | 2 | ENSMUST00000119627.2 | -39882 | Crybb3 | ENSMUSG00000029352.9 |
| chr5 | 118027073 | 118029328 | 4.05595 | 5.54111 | 2.18891 | Promoter | 5 | 118027743 | 118061878 | 34136 | 1 | ENSMUST00000031304.8 | 0 | Tesc | ENSMUSG00000029359.8 |
| chr5 | 118029685 | 118030166 | 3.54312 | 4.22869 | 1.24418 | Intron (ENSMUST00000031304.8/ENSMUSG00000029359.8, intron 1 of 7) | 5 | 118027870 | 118061870 | 34001 | 1 | ENSMUST00000135540.2 | 1816 | Tesc | ENSMUSG00000029359.8 |
| chr5 | 118037079 | 118037701 | 3.46529 | 4.14134 | 1.19203 | Intron (ENSMUST00000031304.8/ENSMUSG00000029359.8, intron 1 of 7) | 5 | 118036031 | 118061773 | 25743 | 1 | ENSMUST00000151792.2 | 1049 | Tesc | ENSMUSG00000029359.8 |
| chr5 | 118038099 | 118038659 | 3.64107 | 4.53144 | 1.45465 | Intron (ENSMUST00000031304.8/ENSMUSG00000029359.8, intron 1 of 7) | 5 | 118036031 | 118061773 | 25743 | 1 | ENSMUST00000151792.2 | 2069 | Tesc | ENSMUSG00000029359.8 |
| chr5 | 117790766 | 117790905 | 3.63041 | 4.38367 | 1.39175 | Intron (ENSMUST00000138579.2/ENSMUSG00000029361.13, intron 1 of 1) | 5 | 117781055 | 117867086 | 86032 | 1 | ENSMUST00000124194.1 | 9712 | Nos1 | ENSMUSG00000029361.13 |
| chr5 | 117801526 | 117801672 | 3.36462 | 3.87448 | 1.01733 | Intron (ENSMUST00000138579.2/ENSMUSG00000029361.13, intron 1 of 1) | 5 | 117781055 | 117867086 | 86032 | 1 | ENSMUST00000124194.1 | 20472 | Nos1 | ENSMUSG00000029361.13 |
| chr5 | 117837429 | 117838245 | 3.52923 | 4.28928 | 1.32813 | Intron (ENSMUST00000138579.2/ENSMUSG00000029361.13, intron 1 of 1) | 5 | 117841850 | 117958840 | 116991 | 1 | ENSMUST00000142742.3 | -3605 | Nos1 | ENSMUSG00000029361.13 |
| chr5 | 117841838 | 117842042 | 3.56951 | 4.65322 | 1.56405 | Promoter | 5 | 117841850 | 117958840 | 116991 | 1 | ENSMUST00000142742.3 | 0 | Nos1 | ENSMUSG00000029361.13 |
| chr5 | 117842401 | 117842531 | 3.72804 | 4.5946 | 1.54981 | Promoter | 5 | 117842186 | 117954364 | 112179 | 1 | ENSMUST00000086451.6 | 216 | Nos1 | ENSMUSG00000029361.13 |
| chr5 | 117842882 | 117843143 | 4.04364 | 5.22872 | 1.91943 | Promoter | 5 | 117842186 | 117954364 | 112179 | 1 | ENSMUST00000086451.6 | 697 | Nos1 | ENSMUSG00000029361.13 |
| chr5 | 117844074 | 117844247 | 3.82596 | 4.73249 | 1.55984 | Intron (ENSMUST00000138579.2/ENSMUSG00000029361.13, intron 1 of 1) | 5 | 117842186 | 117954364 | 112179 | 1 | ENSMUST00000086451.6 | 1889 | Nos1 | ENSMUSG00000029361.13 |
| chr5 | 117856421 | 117856807 | 3.43929 | 4.16902 | 1.28811 | Intron (ENSMUST00000138579.2/ENSMUSG00000029361.13, intron 1 of 1) | 5 | 117867114 | 117953909 | 86796 | 1 | ENSMUST00000102557.4 | -10307 | Nos1 | ENSMUSG00000029361.13 |
| chr5 | 117900450 | 117900525 | 4.12798 | 5.31625 | 1.96941 | Exon (ENSMUST00000142742.3/ENSMUSG00000029361.13, exon 8 of 29) | 5 | 117919175 | 117925731 | 6557 | 1 | ENSMUST00000138554.1 | -18650 | Nos1 | ENSMUSG00000029361.13 |
| chr5 | 88791327 | 88791456 | 3.65399 | 4.40384 | 1.33259 | Distal Intergenic | 5 | 88764996 | 88783281 | 18286 | 1 | ENSMUST00000031311.9 | 26332 | Dck | ENSMUSG00000029366.9 |
| chr5 | 90759359 | 90760175 | 4.04282 | 5.28232 | 2.02091 | Promoter | 5 | 90759299 | 90761625 | 2327 | 1 | ENSMUST00000031318.4 | 61 | Cxcl5 | ENSMUSG00000029371.6 |
| chr5 | 91074337 | 91074504 | 3.39312 | 4.25157 | 1.2945 | Promoter | 5 | 91074617 | 91093649 | 19033 | 1 | ENSMUST00000031324.4 | -113 | Ereg | ENSMUSG00000029377.4 |
| chr5 | 91139795 | 91140022 | 3.8415 | 4.99716 | 1.79528 | Promoter | 5 | 91139615 | 91148432 | 8818 | 1 | ENSMUST00000031325.4 | 181 | Areg | ENSMUSG00000029378.4 |
| chr5 | 91140566 | 91141076 | 3.67532 | 4.63742 | 1.50784 | Promoter | 5 | 91139615 | 91148432 | 8818 | 1 | ENSMUST00000031325.4 | 952 | Areg | ENSMUSG00000029378.4 |
| chr5 | 90784991 | 90785256 | 3.95098 | 4.96459 | 1.70381 | Promoter | 5 | 90786103 | 90788086 | 1984 | 1 | ENSMUST00000031326.8 | -847 | Cxcl3 | ENSMUSG00000029379.9 |
| chr5 | 90786269 | 90786417 | 3.48099 | 4.14007 | 1.21334 | Promoter | 5 | 90786103 | 90788086 | 1984 | 1 | ENSMUST00000031326.8 | 167 | Cxcl3 | ENSMUSG00000029379.9 |
| chr5 | 127632346 | 127632818 | 3.94916 | 4.97431 | 1.72182 | Promoter | 5 | 127601930 | 127632880 | 30951 | 2 | ENSMUST00000144603.1 | 62 | Slc15a4 | ENSMUSG00000029416.12 |
| chr5 | 127633236 | 127633707 | 3.75688 | 4.6081 | 1.49675 | Promoter | 5 | 127608793 | 127632897 | 24105 | 2 | ENSMUST00000155321.1 | -340 | Slc15a4 | ENSMUSG00000029416.12 |
| chr5 | 92327579 | 92328153 | 3.33469 | 3.85397 | 1.10009 | Promoter | 5 | 92321331 | 92328079 | 6749 | 2 | ENSMUST00000113093.3 | 0 | Cxcl9 | ENSMUSG00000029417.8 |
| chr2 | 25578704 | 25580144 | 4.06286 | 5.39913 | 2.05882 | Promoter | 2 | 25576973 | 25579897 | 2925 | 2 | ENSMUST00000188161.1 | 0 | Gm996 | ENSMUSG00000029419.8 |
| chr2 | 25580478 | 25580866 | 3.75804 | 4.62425 | 1.53728 | Promoter | 2 | 25579811 | 25580263 | 453 | 2 | ENSMUST00000187017.1 | -216 | Gm996 | ENSMUSG00000029419.8 |
| chr5 | 128877568 | 128877662 | 3.45362 | 4.03668 | 1.11569 | Intron (ENSMUST00000111346.1/ENSMUSG00000029420.8, intron 1 of 21) | 5 | 128760401 | 128953362 | 192962 | 2 | ENSMUST00000111346.1 | 75700 | Rimbp2 | ENSMUSG00000029420.8 |
| chr5 | 128904315 | 128904561 | 3.64092 | 4.40417 | 1.39802 | Intron (ENSMUST00000111346.1/ENSMUSG00000029420.8, intron 1 of 21) | 5 | 128760401 | 128953362 | 192962 | 2 | ENSMUST00000111346.1 | 48801 | Rimbp2 | ENSMUSG00000029420.8 |
| chr5 | 128905000 | 128905217 | 3.89025 | 4.8463 | 1.62636 | Intron (ENSMUST00000111346.1/ENSMUSG00000029420.8, intron 1 of 21) | 5 | 128760401 | 128953362 | 192962 | 2 | ENSMUST00000111346.1 | 48145 | Rimbp2 | ENSMUSG00000029420.8 |
| chr5 | 128950730 | 128951029 | 3.59803 | 4.30644 | 1.29717 | Intron (ENSMUST00000111346.1/ENSMUSG00000029420.8, intron 1 of 21) | 5 | 128760401 | 128953362 | 192962 | 2 | ENSMUST00000111346.1 | 2333 | Rimbp2 | ENSMUSG00000029420.8 |
| chr5 | 128952103 | 128952178 | 3.77529 | 4.63138 | 1.49288 | Intron (ENSMUST00000111346.1/ENSMUSG00000029420.8, intron 1 of 21) | 5 | 128760401 | 128953362 | 192962 | 2 | ENSMUST00000111346.1 | 1184 | Rimbp2 | ENSMUSG00000029420.8 |
| chr5 | 128685113 | 128685188 | 3.40655 | 3.95871 | 1.09149 | Distal Intergenic | 5 | 128736246 | 128755474 | 19229 | 1 | ENSMUST00000086056.3 | -51058 | Piwil1 | ENSMUSG00000029423.6 |
| chr5 | 128704661 | 128704968 | 3.57694 | 4.25654 | 1.23424 | Distal Intergenic | 5 | 128736246 | 128755474 | 19229 | 1 | ENSMUST00000086056.3 | -31278 | Piwil1 | ENSMUSG00000029423.6 |
| chr5 | 128741299 | 128741390 | 3.3655 | 3.87582 | 1.01733 | Intron (ENSMUST00000086056.3/ENSMUSG00000029423.6, intron 5 of 21) | 5 | 128736246 | 128755474 | 19229 | 1 | ENSMUST00000086056.3 | 5054 | Piwil1 | ENSMUSG00000029423.6 |
| chr5 | 129380814 | 129381103 | 3.38967 | 4.00027 | 1.18181 | Distal Intergenic | 5 | 129501231 | 129571383 | 70153 | 1 | ENSMUST00000053737.7 | -120128 | Sfswap | ENSMUSG00000029439.10 |
| chr5 | 123294406 | 123294666 | 3.77057 | 4.64067 | 1.52531 | Promoter | 5 | 123293901 | 123326630 | 32730 | 1 | ENSMUST00000069311.4 | 506 | Wdr66 | ENSMUSG00000029442.13 |
| chr5 | 123300163 | 123300666 | 3.50322 | 4.13338 | 1.19376 | Intron (ENSMUST00000121964.2/ENSMUSG00000029442.13, intron 17 of 21) | 5 | 123302516 | 123326649 | 24134 | 1 | ENSMUST00000125979.1 | -1850 | Wdr66 | ENSMUSG00000029442.13 |
| chr5 | 122734840 | 122735061 | 3.35703 | 3.86724 | 1.03216 | Intron (ENSMUST00000111668.3/ENSMUSG00000029471.9, intron 15 of 15) | 5 | 122727664 | 122728585 | 922 | 1 | ENSMUST00000139712.1 | 7177 | P2rx4 | ENSMUSG00000029470.10 |
| chr5 | 110341876 | 110342054 | 3.90032 | 5.00499 | 1.7994 | Promoter | 5 | 110339812 | 110343035 | 3224 | 2 | ENSMUST00000058016.10 | 981 | P2rx2 | ENSMUSG00000029503.11 |
| chr5 | 110343649 | 110343761 | 4.18722 | 5.48335 | 2.14948 | Promoter | 5 | 110339812 | 110343035 | 3224 | 2 | ENSMUST00000112478.3 | -615 | P2rx2 | ENSMUSG00000029503.11 |
| chr5 | 115906363 | 115906734 | 3.36208 | 3.87877 | 1.04604 | Promoter | 5 | 115907470 | 115925600 | 18131 | 1 | ENSMUST00000128702.1 | -736 | Cit | ENSMUSG00000029516.14 |
| chr5 | 115907269 | 115907865 | 3.97222 | 5.19081 | 1.91666 | Promoter | 5 | 115907470 | 115925600 | 18131 | 1 | ENSMUST00000128702.1 | 0 | Cit | ENSMUSG00000029516.14 |
| chr5 | 115926636 | 115926716 | 3.7941 | 4.67526 | 1.54095 | Promoter | 5 | 115926086 | 116006368 | 80283 | 1 | ENSMUST00000147479.2 | 551 | Cit | ENSMUSG00000029516.14 |
| chr5 | 115928537 | 115928839 | 3.23896 | 3.6856 | 1.01733 | Intron (ENSMUST00000051704.9/ENSMUSG00000029516.14, intron 12 of 46) | 5 | 115926086 | 116006368 | 80283 | 1 | ENSMUST00000147479.2 | 2452 | Cit | ENSMUSG00000029516.14 |
| chr9 | 123677895 | 123679398 | 4.23426 | 5.86142 | 2.42935 | Promoter | 9 | 123678439 | 123781965 | 103527 | 1 | ENSMUST00000163559.2 | 0 | Ccr9 | ENSMUSG00000029530.10 |
| chr5 | 139352617 | 139352898 | 3.6568 | 4.42373 | 1.38706 | Promoter | 5 | 139352617 | 139357033 | 4417 | 1 | ENSMUST00000031521.8 | 1 | Cyp2w1 | ENSMUSG00000029541.8 |
| chr5 | 115190099 | 115190419 | 3.3305 | 4.15415 | 1.21842 | Intron (ENSMUST00000112113.2/ENSMUSG00000029544.11, intron 1 of 5) | 5 | 115168689 | 115186308 | 17620 | 2 | ENSMUST00000031519.8 | -3792 | Cabp1 | ENSMUSG00000029544.11 |
| chr5 | 115191884 | 115193286 | 3.55833 | 4.33419 | 1.39037 | Intron (ENSMUST00000112113.2/ENSMUSG00000029544.11, intron 1 of 5) | 5 | 115168689 | 115194381 | 25693 | 2 | ENSMUST00000112113.2 | 1095 | Cabp1 | ENSMUSG00000029544.11 |
| chr5 | 115193624 | 115194527 | 4.10674 | 5.51487 | 2.13555 | Promoter | 5 | 115168689 | 115194381 | 25693 | 2 | ENSMUST00000112113.2 | 0 | Cabp1 | ENSMUSG00000029544.11 |
| chr5 | 139531981 | 139532134 | 3.31273 | 3.79633 | 1.01733 | Distal Intergenic | 5 | 139543494 | 139548179 | 4686 | 1 | ENSMUST00000172997.1 | -11360 | Uncx | ENSMUSG00000029546.11 |

| | | | | | | | | | | | | | | | |
|---|---|---|---|---|---|---|---|---|---|---|---|---|---|---|---|
| chr5 | 139534438 | 139534753 | 3.61536 | 4.44318 | 1.38721 | Distal Intergenic | 5 | 139543494 | 139548179 | 4686 | 1 | ENSMUST00000172997.1 | -8741 | Uncx | ENSMUSG00000029546.11 |
| chr5 | 139535128 | 139536238 | 4.46156 | 6.44267 | 2.82948 | Distal Intergenic | 5 | 139543494 | 139548179 | 4686 | 1 | ENSMUST00000172997.1 | -7256 | Uncx | ENSMUSG00000029546.11 |
| chr5 | 139537103 | 139537383 | 3.61251 | 4.34152 | 1.31911 | Distal Intergenic | 5 | 139543494 | 139548179 | 4686 | 1 | ENSMUST00000172997.1 | -6111 | Uncx | ENSMUSG00000029546.11 |
| chr5 | 139538429 | 139538644 | 3.97557 | 5.03735 | 1.78156 | Distal Intergenic | 5 | 139543494 | 139548179 | 4686 | 1 | ENSMUST00000172997.1 | -4850 | Uncx | ENSMUSG00000029546.11 |
| chr5 | 139539347 | 139540510 | 3.99292 | 5.15289 | 1.88622 | Distal Intergenic | 5 | 139543494 | 139548179 | 4686 | 1 | ENSMUST00000172997.1 | -2984 | Uncx | ENSMUSG00000029546.11 |
| chr5 | 139541157 | 139541956 | 4.13944 | 5.43218 | 2.06576 | Distal Intergenic | 5 | 139543494 | 139548179 | 4686 | 1 | ENSMUST00000172997.1 | -1538 | Uncx | ENSMUSG00000029546.11 |
| chr5 | 139542625 | 139547784 | 3.89262 | 4.99411 | 1.77512 | Promoter | 5 | 139543494 | 139548179 | 4686 | 1 | ENSMUST00000172997.1 | 0 | Uncx | ENSMUSG00000029546.11 |
| chr5 | 139548610 | 139549166 | 3.72102 | 4.54995 | 1.45976 | Distal Intergenic | 5 | 139543898 | 139548176 | 4279 | 1 | ENSMUST00000031523.8 | 4713 | Uncx | ENSMUSG00000029546.11 |
| chr5 | 139549510 | 139549936 | 3.84088 | 4.7819 | 1.59311 | Distal Intergenic | 5 | 139543898 | 139548176 | 4279 | 1 | ENSMUST00000031523.8 | 5613 | Uncx | ENSMUSG00000029546.11 |
| chr5 | 139550498 | 139551194 | 4.22422 | 5.57206 | 2.20609 | Distal Intergenic | 5 | 139543898 | 139548176 | 4279 | 1 | ENSMUST00000031523.8 | 6601 | Uncx | ENSMUSG00000029546.11 |
| chr5 | 139552525 | 139553470 | 3.75977 | 4.66695 | 1.57025 | Distal Intergenic | 5 | 139543898 | 139548176 | 4279 | 1 | ENSMUST00000031523.8 | 8628 | Uncx | ENSMUSG00000029546.11 |
| chr5 | 139553863 | 139554441 | 3.90582 | 4.93627 | 1.71784 | Distal Intergenic | 5 | 139543898 | 139548176 | 4279 | 1 | ENSMUST00000031523.8 | 9966 | Uncx | ENSMUSG00000029546.11 |
| chr5 | 139555568 | 139556091 | 3.78259 | 4.72663 | 1.57331 | Distal Intergenic | 5 | 139543898 | 139548176 | 4279 | 1 | ENSMUST00000031523.8 | 11671 | Uncx | ENSMUSG00000029546.11 |
| chr5 | 139556517 | 139556831 | 3.42361 | 4.10654 | 1.19293 | Distal Intergenic | 5 | 139543898 | 139548176 | 4279 | 1 | ENSMUST00000031523.8 | 12620 | Uncx | ENSMUSG00000029546.11 |
| chr5 | 139557182 | 139557446 | 3.95325 | 5.35124 | 2.03026 | Distal Intergenic | 5 | 139543898 | 139548176 | 4279 | 1 | ENSMUST00000031523.8 | 13285 | Uncx | ENSMUSG00000029546.11 |
| chr5 | 139557832 | 139558713 | 3.98323 | 5.0778 | 1.81777 | Distal Intergenic | 5 | 139543898 | 139548176 | 4279 | 1 | ENSMUST00000031523.8 | 13935 | Uncx | ENSMUSG00000029546.11 |
| chr5 | 139559585 | 139560276 | 3.49997 | 4.19734 | 1.26917 | Distal Intergenic | 5 | 139543898 | 139548176 | 4279 | 1 | ENSMUST00000031523.8 | 15688 | Uncx | ENSMUSG00000029546.11 |
| chr5 | 139833143 | 139833388 | 3.37905 | 3.89625 | 1.01733 | Distal Intergenic | 5 | 139823592 | 139826885 | 3294 | 2 | ENSMUST00000031531.8 | -6259 | Psmg3 | ENSMUSG00000029551.8 |
| chr5 | 114952466 | 114952627 | 3.68791 | 4.48067 | 1.41615 | 3' UTR | 5 | 114945805 | 114947784 | 1980 | 1 | ENSMUST00000131771.2 | 6662 | 2210016L21Rik | ENSMUSG00000029559.10 |
| chr6 | 14895287 | 14895472 | 3.38271 | 3.95526 | 1.16251 | Distal Intergenic | 6 | 14901349 | 15441977 | 540629 | 1 | ENSMUST00000115477.2 | -5877 | Foxp2 | ENSMUSG00000029563.11 |
| chr6 | 14896235 | 14896601 | 3.43637 | 4.41845 | 1.38524 | Distal Intergenic | 6 | 14901349 | 15441977 | 540629 | 1 | ENSMUST00000115477.2 | -4748 | Foxp2 | ENSMUSG00000029563.11 |
| chr6 | 14897437 | 14898742 | 3.91967 | 5.05285 | 1.86956 | Distal Intergenic | 6 | 14901349 | 15441977 | 540629 | 1 | ENSMUST00000115477.2 | -2607 | Foxp2 | ENSMUSG00000029563.11 |
| chr6 | 14899536 | 14899795 | 3.29079 | 3.79085 | 1.08983 | Distal Intergenic | 6 | 14901349 | 15441977 | 540629 | 1 | ENSMUST00000115477.2 | -1554 | Foxp2 | ENSMUSG00000029563.11 |
| chr6 | 14900155 | 14900933 | 3.90035 | 4.95607 | 1.79019 | Promoter | 6 | 14901349 | 15441977 | 540629 | 1 | ENSMUST00000115477.2 | -416 | Foxp2 | ENSMUSG00000029563.11 |
| chr6 | 14901251 | 14902048 | 3.74797 | 4.88726 | 1.7247 | Promoter | 6 | 14901349 | 15441977 | 540629 | 1 | ENSMUST00000115477.2 | 0 | Foxp2 | ENSMUSG00000029563.11 |
| chr6 | 14902683 | 14903122 | 3.85684 | 4.98987 | 1.73024 | Promoter | 6 | 14903457 | 15196959 | 293503 | 1 | ENSMUST00000156883.1 | -335 | Foxp2 | ENSMUSG00000029563.11 |
| chr6 | 14903469 | 14903550 | 3.98831 | 5.13755 | 1.94648 | Promoter | 6 | 14903457 | 15196959 | 293503 | 1 | ENSMUST00000156883.1 | 13 | Foxp2 | ENSMUSG00000029563.11 |
| chr6 | 15183668 | 15184716 | 4.12839 | 5.44424 | 2.10385 | Promoter | 6 | 15184647 | 15199344 | 14698 | 1 | ENSMUST00000115470.2 | 0 | Foxp2 | ENSMUSG00000029563.11 |
| chr6 | 15187150 | 15187299 | 3.41517 | 4.41283 | 1.3655 | Intron (ENSMUST00000115477.2/ENSMUSG00000029563.11, intron 3 of 18) | 6 | 15185563 | 15394815 | 209253 | 1 | ENSMUST00000137628.2 | 1588 | Foxp2 | ENSMUSG00000029563.11 |
| chr6 | 15188133 | 15188208 | 4.08425 | 5.42976 | 2.07065 | Intron (ENSMUST00000115477.2/ENSMUSG00000029563.11, intron 3 of 18) | 6 | 15185563 | 15394815 | 209253 | 1 | ENSMUST00000137628.2 | 2571 | Foxp2 | ENSMUSG00000029563.11 |
| chr5 | 114857417 | 114857696 | 3.39916 | 3.94588 | 1.08513 | Intron (ENSMUST00000031547.6/ENSMUSG00000029564.7, intron 2 of 11) | 5 | 114859710 | 114876702 | 16993 | 1 | ENSMUST00000112153.2 | -2014 | 4930519G04Rik | ENSMUSG00000029564.7 |
| chr5 | 142960362 | 142960447 | 3.4393 | 4.17738 | 1.26643 | Promoter | 5 | 142960343 | 142973185 | 12843 | 1 | ENSMUST00000031565.9 | 20 | Fscn1 | ENSMUSG00000029581.9 |
| chr5 | 142960927 | 142961968 | 4.1762 | 5.68685 | 2.34794 | Promoter | 5 | 142960843 | 142972170 | 11328 | 1 | ENSMUST00000129306.2 | 85 | Fscn1 | ENSMUSG00000029581.9 |
| chr5 | 142974899 | 142975154 | 3.39394 | 3.9519 | 1.07202 | Distal Intergenic | 5 | 142970085 | 142971989 | 1905 | 1 | ENSMUST00000135293.1 | 4815 | Fscn1 | ENSMUSG00000029581.9 |
| chr5 | 120090606 | 120090681 | 3.3774 | 3.8999 | 1.03869 | Distal Intergenic | 5 | 120116510 | 120197979 | 81470 | 1 | ENSMUST00000031590.8 | -25829 | Rbm19 | ENSMUSG00000029594.9 |
| chr5 | 120092305 | 120092438 | 3.64705 | 4.42098 | 1.42233 | Distal Intergenic | 5 | 120116510 | 120197979 | 81470 | 1 | ENSMUST00000031590.8 | -24072 | Rbm19 | ENSMUSG00000029594.9 |
| chr5 | 120093306 | 120093631 | 3.34034 | 3.83787 | 1.01733 | Distal Intergenic | 5 | 120116510 | 120197979 | 81470 | 1 | ENSMUST00000031590.8 | -22879 | Rbm19 | ENSMUSG00000029594.9 |
| chr5 | 120432127 | 120432450 | 3.55422 | 4.24462 | 1.27603 | Promoter | 5 | 120431770 | 120441223 | 9454 | 1 | ENSMUST00000031591.8 | 358 | Lhx5 | ENSMUSG00000029595.8 |
| chr5 | 120432818 | 120433000 | 3.8828 | 5.03077 | 1.80533 | Intron (ENSMUST00000031591.8/ENSMUSG00000029595.8, intron 1 of 4) | 5 | 120431770 | 120441223 | 9454 | 1 | ENSMUST00000031591.8 | 1049 | Lhx5 | ENSMUSG00000029595.8 |
| chr5 | 120435093 | 120436368 | 4.05352 | 5.3159 | 1.99359 | Exon (ENSMUST00000031591.8/ENSMUSG00000029595.8, exon 3 of 5) | 5 | 120431770 | 120441223 | 9454 | 1 | ENSMUST00000031591.8 | 3324 | Lhx5 | ENSMUSG00000029595.8 |
| chr5 | 120436916 | 120437317 | 3.52141 | 4.18136 | 1.20665 | Intron (ENSMUST00000031591.8/ENSMUSG00000029595.8, intron 4 of 4) | 5 | 120431770 | 120441223 | 9454 | 1 | ENSMUST00000031591.8 | 5147 | Lhx5 | ENSMUSG00000029595.8 |
| chr5 | 120437659 | 120438836 | 4.06601 | 5.25235 | 1.951 | Intron (ENSMUST00000031591.8/ENSMUSG00000029595.8, intron 4 of 4) | 5 | 120431770 | 120441223 | 9454 | 1 | ENSMUST00000031591.8 | 5890 | Lhx5 | ENSMUSG00000029595.8 |
| chr5 | 120439532 | 120439664 | 3.68672 | 4.60722 | 1.54044 | Intron (ENSMUST00000031591.8/ENSMUSG00000029595.8, intron 4 of 4) | 5 | 120431770 | 120441223 | 9454 | 1 | ENSMUST00000031591.8 | 7763 | Lhx5 | ENSMUSG00000029595.8 |
| chr5 | 120440027 | 120440470 | 3.33252 | 3.99491 | 1.15077 | 3' UTR | 5 | 120431770 | 120441223 | 9454 | 1 | ENSMUST00000031591.8 | 8258 | Lhx5 | ENSMUSG00000029595.8 |
| chr5 | 120448007 | 120448099 | 3.41273 | 3.97977 | 1.13151 | Distal Intergenic | 5 | 120431770 | 120441223 | 9454 | 1 | ENSMUST00000031591.8 | 16238 | Lhx5 | ENSMUSG00000029595.8 |
| chr5 | 120454212 | 120454321 | 4.27934 | 5.72583 | 2.27471 | Distal Intergenic | 5 | 120460688 | 120467360 | 6673 | 2 | ENSMUST00000132916.1 | 13039 | Sdsl | ENSMUSG00000029596.8 |
| chr5 | 120454735 | 120454917 | 3.32477 | 3.82363 | 1.02546 | Distal Intergenic | 5 | 120460688 | 120467360 | 6673 | 2 | ENSMUST00000132916.1 | 12443 | Sdsl | ENSMUSG00000029596.8 |
| chr5 | 120480194 | 120480625 | 3.6533 | 4.40196 | 1.33575 | Promoter | 5 | 120480363 | 120480888 | 526 | 1 | ENSMUST00000124978.1 | 0 | Sds | ENSMUSG00000029597.9 |
| chr5 | 120648745 | 120650446 | 4.09571 | 5.40051 | 2.09962 | Promoter | 5 | 120648812 | 120654902 | 6091 | 1 | ENSMUST00000125804.2 | 0 | Rasal1 | ENSMUSG00000029602.6 |
| chr5 | 120652457 | 120652532 | 3.44849 | 4.07689 | 1.23893 | Promoter | 5 | 120653367 | 120659650 | 6284 | 1 | ENSMUST00000148695.1 | -835 | Rasal1 | ENSMUSG00000029602.6 |
| chr5 | 120674730 | 120674986 | 3.41815 | 3.98594 | 1.12518 | Exon (ENSMUST00000031606.4/ENSMUSG00000029602.6, exon 17 of 22) | 5 | 120678593 | 120679306 | 714 | 1 | ENSMUST00000154759.1 | -3607 | Rasal1 | ENSMUSG00000029602.6 |
| chr5 | 120679014 | 120679274 | 3.37011 | 3.88276 | 1.01733 | Promoter | 5 | 120678593 | 120679306 | 714 | 1 | ENSMUST00000154759.1 | 422 | Rasal1 | ENSMUSG00000029602.6 |
| chr5 | 120680275 | 120680516 | 3.65293 | 4.44006 | 1.40483 | 3' UTR | 5 | 120678593 | 120679306 | 714 | 1 | ENSMUST00000154759.1 | 1683 | Rasal1 | ENSMUSG00000029602.6 |
| chr5 | 120680928 | 120681310 | 3.43172 | 4.01137 | 1.13759 | 3' UTR | 5 | 120681452 | 120682813 | 1362 | 2 | ENSMUST00000151562.1 | 1503 | Dtx1 | ENSMUSG00000029603.10 |
| chr5 | 120682355 | 120682844 | 3.69721 | 4.52822 | 1.49057 | Promoter | 5 | 120681452 | 120682813 | 1362 | 2 | ENSMUST00000151562.1 | 0 | Dtx1 | ENSMUSG00000029603.10 |
| chr5 | 120683401 | 120683630 | 3.45867 | 4.04537 | 1.11522 | Promoter | 5 | 120681202 | 120682915 | 1714 | 2 | ENSMUST00000144889.2 | -487 | Dtx1 | ENSMUSG00000029603.10 |
| chr5 | 120694149 | 120694709 | 3.38192 | 3.90145 | 1.02036 | Promoter | 5 | 120680203 | 120695403 | 15201 | 2 | ENSMUST00000145174.2 | 694 | Dtx1 | ENSMUSG00000029603.10 |
| chr5 | 120695505 | 120695673 | 3.75478 | 4.59592 | 1.46919 | Promoter | 5 | 120680203 | 120695403 | 15201 | 2 | ENSMUST00000145174.2 | -103 | Dtx1 | ENSMUSG00000029603.10 |
| chr5 | 120695982 | 120696057 | 3.47356 | 4.06766 | 1.11859 | Promoter | 5 | 120680203 | 120695403 | 15201 | 2 | ENSMUST00000145174.2 | -580 | Dtx1 | ENSMUSG00000029603.10 |
| chr5 | 120696997 | 120697174 | 3.32622 | 3.81684 | 1.01733 | Intron (ENSMUST00000031607.6/ENSMUSG00000029603.10, intron 2 of 9) | 5 | 120680203 | 120695403 | 15201 | 2 | ENSMUST00000145174.2 | -1595 | Dtx1 | ENSMUSG00000029603.10 |
| chr5 | 120699138 | 120699311 | 3.68565 | 4.54263 | 1.50407 | Intron (ENSMUST00000031607.6/ENSMUSG00000029603.10, intron 2 of 9) | 5 | 120680203 | 120695403 | 15201 | 2 | ENSMUST00000145174.2 | -3736 | Dtx1 | ENSMUSG00000029603.10 |
| chr5 | 120708200 | 120708530 | 3.42737 | 4.03279 | 1.20009 | Intron (ENSMUST00000031607.6/ENSMUSG00000029603.10, intron 2 of 9) | 5 | 120694721 | 120705889 | 11169 | 2 | ENSMUST00000124079.1 | -2312 | Dtx1 | ENSMUSG00000029603.10 |
| chr5 | 120710153 | 120711893 | 4.09333 | 5.49618 | 2.13499 | Promoter | 5 | 120680202 | 120711927 | 31726 | 2 | ENSMUST00000031607.6 | 34 | Dtx1 | ENSMUSG00000029603.10 |
| chr5 | 120712823 | 120712903 | 3.89612 | 4.86981 | 1.6593 | Promoter | 5 | 120680202 | 120711927 | 31726 | 2 | ENSMUST00000031607.6 | -897 | Dtx1 | ENSMUSG00000029603.10 |
| chr5 | 120714795 | 120715107 | 3.57107 | 4.25238 | 1.2437 | Distal Intergenic | 5 | 120680202 | 120711927 | 31726 | 2 | ENSMUST00000031607.6 | -2869 | Dtx1 | ENSMUSG00000029603.10 |
| chr5 | 120967876 | 120968170 | 3.53515 | 4.19044 | 1.21273 | Intron (ENSMUST00000079204.5/ENSMUSG00000029608.7, intron 5 of 21) | 5 | 120940502 | 121009518 | 69017 | 2 | ENSMUST00000079204.5 | 41348 | Rph3a | ENSMUSG00000029608.7 |
| chr5 | 121007674 | 121007916 | 3.54823 | 4.20748 | 1.21404 | Intron (ENSMUST00000079204.5/ENSMUSG00000029608.7, intron 2 of 21) | 5 | 120940502 | 121009518 | 69017 | 2 | ENSMUST00000079204.5 | 1602 | Rph3a | ENSMUSG00000029608.7 |
| chr5 | 121008366 | 121008524 | 3.41342 | 4.15995 | 1.21782 | Promoter | 5 | 120940502 | 121009518 | 69017 | 2 | ENSMUST00000079204.5 | 994 | Rph3a | ENSMUSG00000029608.7 |
| chr5 | 121039687 | 121039984 | 3.72784 | 4.53596 | 1.4091 | Distal Intergenic | 5 | 120940502 | 121009518 | 69017 | 2 | ENSMUST00000079204.5 | -30170 | Rph3a | ENSMUSG00000029608.7 |
| chr5 | 121041318 | 121041587 | 3.50313 | 4.13963 | 1.21755 | Distal Intergenic | 5 | 120940502 | 121009518 | 69017 | 2 | ENSMUST00000079204.5 | -31801 | Rph3a | ENSMUSG00000029608.7 |
| chr5 | 146351052 | 146351367 | 3.47072 | 4.06861 | 1.137 | Distal Intergenic | 5 | 146384985 | 146435471 | 50487 | 1 | ENSMUST00000110600.2 | -33618 | Wasf3 | ENSMUSG00000029636.8 |
| chr5 | 146384978 | 146385068 | 3.22653 | 3.66698 | 1.01733 | Promoter | 5 | 146384985 | 146435471 | 50487 | 1 | ENSMUST00000110600.2 | 0 | Wasf3 | ENSMUSG00000029636.8 |
| chr5 | 146385433 | 146385682 | 3.37031 | 4.24134 | 1.30235 | Promoter | 5 | 146384985 | 146472411 | 87427 | 1 | ENSMUST00000016143.7 | 449 | Wasf3 | ENSMUSG00000029636.8 |
| chr5 | 146386287 | 146386475 | 3.36965 | 4.04892 | 1.21012 | Intron (ENSMUST00000110600.2/ENSMUSG00000029636.8, intron 1 of 2) | 5 | 146384985 | 146472411 | 87427 | 1 | ENSMUST00000016143.7 | 1303 | Wasf3 | ENSMUSG00000029636.8 |
| chr5 | 147237188 | 147237263 | 4.36818 | 6.00714 | 2.50018 | Intron (ENSMUST00000174285.2/ENSMUSG00000092570.2, intron 3 of 5) | 5 | 147269959 | 147278008 | 8050 | 1 | ENSMUST00000085591.5 | -32696 | Pdx1 | ENSMUSG00000029644.6 |
| chr5 | 147238976 | 147239148 | 3.60311 | 4.34796 | 1.35505 | Exon (ENSMUST00000172461.1/ENSMUSG00000092570.2, exon 4 of 4) | 5 | 147269959 | 147278008 | 8050 | 1 | ENSMUST00000085591.5 | -30811 | Pdx1 | ENSMUSG00000029644.6 |
| chr5 | 147239907 | 147240244 | 3.29681 | 3.89607 | 1.03922 | Intron (ENSMUST00000174285.2/ENSMUSG00000092570.2, intron 2 of 5) | 5 | 147269959 | 147278008 | 8050 | 1 | ENSMUST00000085591.5 | -29715 | Pdx1 | ENSMUSG00000029644.6 |
| chr5 | 147242899 | 147243150 | 3.58227 | 4.27666 | 1.27429 | Intron (ENSMUST00000174285.2/ENSMUSG00000092570.2, intron 1 of 5) | 5 | 147269959 | 147278008 | 8050 | 1 | ENSMUST00000085591.5 | -26809 | Pdx1 | ENSMUSG00000029644.6 |
| chr5 | 147258492 | 147258583 | 3.72056 | 4.70098 | 1.58675 | Intron (ENSMUST00000174285.2/ENSMUSG00000092570.2, intron 1 of 5) | 5 | 147269959 | 147278008 | 8050 | 1 | ENSMUST00000085591.5 | -11376 | Pdx1 | ENSMUSG00000029644.6 |
| chr5 | 147258993 | 147259362 | 4.07325 | 5.54875 | 2.18238 | Intron (ENSMUST00000174285.2/ENSMUSG00000092570.2, intron 1 of 5) | 5 | 147269959 | 147278008 | 8050 | 1 | ENSMUST00000085591.5 | -10597 | Pdx1 | ENSMUSG00000029644.6 |
| chr5 | 147261763 | 147262361 | 3.60228 | 4.34769 | 1.40018 | Intron (ENSMUST00000174285.2/ENSMUSG00000092570.2, intron 1 of 5) | 5 | 147269959 | 147278008 | 8050 | 1 | ENSMUST00000085591.5 | -7598 | Pdx1 | ENSMUSG00000029644.6 |
| chr5 | 147262836 | 147262960 | 3.64214 | 4.49583 | 1.50007 | Intron (ENSMUST00000174285.2/ENSMUSG00000092570.2, intron 1 of 5) | 5 | 147269959 | 147278008 | 8050 | 1 | ENSMUST00000085591.5 | -6999 | Pdx1 | ENSMUSG00000029644.6 |
| chr5 | 147263903 | 147264189 | 3.52958 | 4.43017 | 1.40486 | Intron (ENSMUST00000174285.2/ENSMUSG00000092570.2, intron 1 of 5) | 5 | 147269959 | 147278008 | 8050 | 1 | ENSMUST00000085591.5 | -5770 | Pdx1 | ENSMUSG00000029644.6 |
| chr5 | 147265271 | 147265648 | 3.35633 | 4.02014 | 1.18171 | Intron (ENSMUST00000174285.2/ENSMUSG00000092570.2, intron 1 of 5) | 5 | 147269959 | 147278008 | 8050 | 1 | ENSMUST00000085591.5 | -4311 | Pdx1 | ENSMUSG00000029644.6 |
| chr5 | 147266020 | 147266867 | 3.54188 | 4.60808 | 1.51495 | Intron (ENSMUST00000174285.2/ENSMUSG00000092570.2, intron 1 of 5) | 5 | 147269959 | 147278008 | 8050 | 1 | ENSMUST00000085591.5 | -3092 | Pdx1 | ENSMUSG00000029644.6 |
| chr5 | 147267439 | 147267605 | 4.25493 | 5.73848 | 2.33929 | Intron (ENSMUST00000174285.2/ENSMUSG00000092570.2, intron 1 of 5) | 5 | 147269959 | 147278008 | 8050 | 1 | ENSMUST00000085591.5 | -2354 | Pdx1 | ENSMUSG00000029644.6 |
| chr5 | 147267965 | 147269077 | 3.45669 | 4.24289 | 1.32567 | Promoter | 5 | 147269959 | 147278008 | 8050 | 1 | ENSMUST00000085591.5 | -882 | Pdx1 | ENSMUSG00000029644.6 |
| chr5 | 147269690 | 147269825 | 3.80249 | 4.95148 | 1.74511 | Promoter | 5 | 147269959 | 147278008 | 8050 | 1 | ENSMUST00000085591.5 | -134 | Pdx1 | ENSMUSG00000029644.6 |
| chr5 | 147295941 | 147296524 | 3.75711 | 4.66683 | 1.54706 | Distal Intergenic | 5 | 147300805 | 147307270 | 6466 | 2 | ENSMUST00000031650.3 | 10746 | Cdx2 | ENSMUSG00000029646.3 |
| chr5 | 147296906 | 147297325 | 3.57175 | 4.29936 | 1.34121 | Distal Intergenic | 5 | 147300805 | 147307270 | 6466 | 2 | ENSMUST00000031650.3 | 9945 | Cdx2 | ENSMUSG00000029646.3 |
| chr5 | 147298680 | 147298801 | 3.79219 | 4.73576 | 1.64491 | Downstream (2-3kb) | 5 | 147300805 | 147307270 | 6466 | 2 | ENSMUST00000031650.3 | 8469 | Cdx2 | ENSMUSG00000029646.3 |
| chr5 | 147299256 | 147302016 | 4.06945 | 5.23695 | 1.9354 | 3' UTR | 5 | 147300805 | 147307270 | 6466 | 2 | ENSMUST00000031650.3 | 5254 | Cdx2 | ENSMUSG00000029646.3 |
| chr5 | 147302423 | 147307398 | 4.21881 | 5.65405 | 2.26177 | Promoter | 5 | 147300805 | 147307270 | 6466 | 2 | ENSMUST00000031650.3 | 0 | Cdx2 | ENSMUSG00000029646.3 |
| chr5 | 147308263 | 147308660 | 4.02796 | 5.59651 | 2.19765 | Promoter | 5 | 147300805 | 147307270 | 6466 | 2 | ENSMUST00000031650.3 | -994 | Cdx2 | ENSMUSG00000029646.3 |
| chr5 | 147309182 | 147309316 | 3.4391 | 4.04218 | 1.15812 | Distal Intergenic | 5 | 147300805 | 147307270 | 6466 | 2 | ENSMUST00000031650.3 | -1913 | Cdx2 | ENSMUSG00000029646.3 |
| chr5 | 147309979 | 147310201 | 3.68501 | 4.59435 | 1.49532 | Distal Intergenic | 5 | 147300805 | 147307270 | 6466 | 2 | ENSMUST00000031650.3 | -2710 | Cdx2 | ENSMUSG00000029646.3 |
| chr5 | 147310690 | 147311751 | 3.8827 | 4.85757 | 1.67075 | Distal Intergenic | 5 | 147300805 | 147307270 | 6466 | 2 | ENSMUST00000031650.3 | -3421 | Cdx2 | ENSMUSG00000029646.3 |
| chr5 | 147312072 | 147312249 | 3.98444 | 5.04934 | 1.81234 | Downstream (2-3kb) | 5 | 147300805 | 147307270 | 6466 | 2 | ENSMUST00000031650.3 | -4803 | Cdx2 | ENSMUSG00000029646.3 |
| chr5 | 147312981 | 147313318 | 3.49194 | 4.10591 | 1.15595 | Downstream (1-2kb) | 5 | 147300805 | 147307270 | 6466 | 2 | ENSMUST00000031650.3 | -5712 | Cdx2 | ENSMUSG00000029646.3 |
| chr5 | 147560522 | 147560773 | 3.45594 | 4.03715 | 1.10499 | Downstream (<1kb) | 5 | 147539277 | 147548493 | 9217 | 1 | ENSMUST00000176553.1 | 21246 | Pan3 | ENSMUSG00000029647.10 |
| chr5 | 147637719 | 147637994 | 3.22918 | 3.6764 | 1.02698 | 3' UTR | 5 | 147604660 | 147634418 | 29759 | 2 | ENSMUST00000125272.1 | -3302 | Flt1 | ENSMUSG00000029648.8 |
| chr5 | 147652409 | 147652492 | 3.95877 | 5.05029 | 1.85495 | Intron (ENSMUST00000031653.6/ENSMUSG00000029648.8, intron 11 of 29) | 5 | 147604660 | 147634418 | 29759 | 2 | ENSMUST00000125272.1 | -17992 | Flt1 | ENSMUSG00000029648.8 |
| chr5 | 147653068 | 147653338 | 3.39529 | 4.31825 | 1.32714 | Intron (ENSMUST00000031653.6/ENSMUSG00000029648.8, intron 11 of 29) | 5 | 147604660 | 147634418 | 29759 | 2 | ENSMUST00000125272.1 | -18651 | Flt1 | ENSMUSG00000029648.8 |
| chr5 | 147653825 | 147653996 | 3.41381 | 4.03554 | 1.20448 | Intron (ENSMUST00000031653.6/ENSMUSG00000029648.8, intron 11 of 29) | 5 | 147604660 | 147634418 | 29759 | 2 | ENSMUST00000125272.1 | -19408 | Flt1 | ENSMUSG00000029648.8 |
| chr5 | 147682218 | 147682586 | 3.2748 | 3.76502 | 1.0401 | Intron (ENSMUST00000031653.6/ENSMUSG00000029648.8, intron 4 of 29) | 5 | 147635159 | 147725988 | 90830 | 2 | ENSMUST00000110529.2 | 43402 | Flt1 | ENSMUSG00000029648.8 |
| chr5 | 147695450 | 147696042 | 3.60367 | 4.44467 | 1.41861 | Intron (ENSMUST00000031653.6/ENSMUSG00000029648.8, intron 3 of 29) | 5 | 147635159 | 147725988 | 90830 | 2 | ENSMUST00000110529.2 | 29946 | Flt1 | ENSMUSG00000029648.8 |
| chr5 | 147698252 | 147698441 | 4.02639 | 5.14066 | 1.86003 | Intron (ENSMUST00000031653.6/ENSMUSG00000029648.8, intron 3 of 29) | 5 | 147635159 | 147725988 | 90830 | 2 | ENSMUST00000110529.2 | 27547 | Flt1 | ENSMUSG00000029648.8 |
| chr5 | 147714934 | 147715145 | 3.41106 | 3.96203 | 1.07876 | Intron (ENSMUST00000031653.6/ENSMUSG00000029648.8, intron 1 of 29) | 5 | 147635159 | 147725988 | 90830 | 2 | ENSMUST00000110529.2 | 10843 | Flt1 | ENSMUSG00000029648.8 |
| chr5 | 147721265 | 147722123 | 4.12742 | 5.35227 | 2.02885 | Intron (ENSMUST00000031653.6/ENSMUSG00000029648.8, intron 1 of 29) | 5 | 147635159 | 147725988 | 90830 | 2 | ENSMUST00000110529.2 | 3865 | Flt1 | ENSMUSG00000029648.8 |
| chr5 | 147722818 | 147723398 | 3.88104 | 4.85783 | 1.66095 | Intron (ENSMUST00000031653.6/ENSMUSG00000029648.8, intron 1 of 29) | 5 | 147635159 | 147725988 | 90830 | 2 | ENSMUST00000110529.2 | 2590 | Flt1 | ENSMUSG00000029648.8 |
| chr5 | 147723742 | 147726695 | 3.73071 | 4.64151 | 1.51876 | Promoter | 5 | 147561604 | 147726011 | 164408 | 2 | ENSMUST00000031653.6 | 0 | Flt1 | ENSMUSG00000029648.8 |
| chr5 | 147728071 | 147728943 | 3.96745 | 5.05193 | 1.82483 | Distal Intergenic | 5 | 147561604 | 147726011 | 164408 | 2 | ENSMUST00000031653.6 | -2061 | Flt1 | ENSMUSG00000029648.8 |
| chr5 | 147747608 | 147748491 | 3.46747 | 4.1085 | 1.23587 | Distal Intergenic | 5 | 147561604 | 147726011 | 164408 | 2 | ENSMUST00000031653.6 | -21598 | Flt1 | ENSMUSG00000029648.8 |
| chr5 | 147750737 | 147750936 | 3.39275 | 3.97371 | 1.18148 | Distal Intergenic | 5 | 147561604 | 147726011 | 164408 | 2 | ENSMUST00000031653.6 | -24727 | Flt1 | ENSMUSG00000029648.8 |
| chr5 | 147751489 | 147752095 | 3.51103 | 4.20408 | 1.32166 | Distal Intergenic | 5 | 147561604 | 147726011 | 164408 | 2 | ENSMUST00000031653.6 | -25479 | Flt1 | ENSMUSG00000029648.8 |
| chr5 | 147753105 | 147753455 | 3.76397 | 4.81921 | 1.6437 | Distal Intergenic | 5 | 147561604 | 147726011 | 164408 | 2 | ENSMUST00000031653.6 | -27095 | Flt1 | ENSMUSG00000029648.8 |
| chr5 | 147753994 | 147754127 | 3.31833 | 4.00457 | 1.17093 | Distal Intergenic | 5 | 147561604 | 147726011 | 164408 | 2 | ENSMUST00000031653.6 | -27984 | Flt1 | ENSMUSG00000029648.8 |
| chr5 | 147847535 | 147847610 | 4.02611 | 5.10872 | 1.79668 | Distal Intergenic | 5 | 147860628 | 147875778 | 15151 | 1 | ENSMUST00000031654.6 | -13018 | Pomp | ENSMUSG00000029649.7 |
| chr5 | 147957465 | 147957837 | 4.84771 | 6.97865 | 3.34247 | Promoter | 5 | 147957342 | 148076537 | 119196 | 1 | ENSMUST00000129092.1 | 124 | Mtus2 | ENSMUSG00000029651.11 |
| chr5 | 147958537 | 147958612 | 3.56356 | 4.24707 | 1.27002 | Intron (ENSMUST00000085558.5/ENSMUSG00000029651.11, intron 1 of 14) | 5 | 147957342 | 148076537 | 119196 | 1 | ENSMUST00000129092.1 | 1196 | Mtus2 | ENSMUSG00000029651.11 |
| chr5 | 147958925 | 147959121 | 3.62369 | 4.35629 | 1.33536 | Intron (ENSMUST00000085558.5/ENSMUSG00000029651.11, intron 1 of 14) | 5 | 147957342 | 148076537 | 119196 | 1 | ENSMUST00000129092.1 | 1584 | Mtus2 | ENSMUSG00000029651.11 |
| chr5 | 148263397 | 148263647 | 3.36251 | 3.90731 | 1.13698 | Intron (ENSMUST00000085558.5/ENSMUSG00000029651.11, intron 7 of 14) | 5 | 148265217 | 148314071 | 48855 | 1 | ENSMUST00000110515.3 | -1570 | Mtus2 | ENSMUSG00000029651.11 |
| chr5 | 148264684 | 148265073 | 4.28187 | 5.63994 | 2.21709 | Promoter | 5 | 148265217 | 148314071 | 48855 | 1 | ENSMUST00000110515.3 | -144 | Mtus2 | ENSMUSG00000029651.11 |
| chr5 | 148265468 | 148265745 | 3.36917 | 4.10808 | 1.23455 | Promoter | 5 | 148265345 | 148314038 | 48694 | 1 | ENSMUST00000110514.2 | 124 | Mtus2 | ENSMUSG00000029651.11 |
| chr5 | 149527430 | 149527729 | 3.39416 | 3.9285 | 1.0507 | Promoter | 5 | 149528679 | 149611894 | 83216 | 1 | ENSMUST00000110502.2 | -950 | Wdr95 | ENSMUSG00000029658.12 |
| chr5 | 149528141 | 149528829 | 3.47694 | 4.0798 | 1.14197 | Promoter | 5 | 149528679 | 149611894 | 83216 | 1 | ENSMUST00000110502.2 | 0 | Wdr95 | ENSMUSG00000029658.12 |
| chr5 | 149411261 | 149411416 | 3.57954 | 4.4924 | 1.4113 | Promoter | 5 | 149411749 | 149431723 | 19975 | 1 | ENSMUST00000093110.6 | -333 | Medag | ENSMUSG00000029659.11 |
| chr5 | 149411745 | 149412158 | 3.66292 | 4.58814 | 1.49721 | Promoter | 5 | 149411749 | 149431723 | 19975 | 1 | ENSMUST00000093110.6 | 0 | Medag | ENSMUSG00000029659.11 |
| chr5 | 149440204 | 149440384 | 3.839 | 4.89901 | 1.70417 | Promoter | 5 | 149439706 | 149470620 | 30915 | 1 | ENSMUST00000031667.4 | 499 | Tex26 | ENSMUSG00000029660.7 |
| chr5 | 149441366 | 149441881 | 3.40206 | 3.99704 | 1.19848 | Intron (ENSMUST00000031667.4/ENSMUSG00000029660.7, intron 1 of 6) | 5 | 149439706 | 149470620 | 30915 | 1 | ENSMUST00000031667.4 | 1661 | Tex26 | ENSMUSG00000029660.7 |
| chr5 | 149464160 | 149464235 | 3.87409 | 4.9183 | 1.69836 | Intron (ENSMUST00000031667.4/ENSMUSG00000029660.7, intron 6 of 6) | 5 | 149439706 | 149470620 | 30915 | 1 | ENSMUST00000031667.4 | 24455 | Tex26 | ENSMUSG00000029660.7 |
| chr6 | 22287060 | 22287220 | 3.43195 | 4.00032 | 1.10253 | Distal Intergenic | 6 | 22288227 | 22293579 | 5353 | 1 | ENSMUST00000128245.2 | -1007 | Wnt16 | ENSMUSG00000029671.10 |
| chr6 | 22287633 | 22287767 | 3.46118 | 4.05158 | 1.12857 | Promoter | 6 | 22288227 | 22293579 | 5353 | 1 | ENSMUST00000128245.2 | -460 | Wnt16 | ENSMUSG00000029671.10 |
| chr6 | 22288436 | 22289639 | 3.62822 | 4.3655 | 1.32993 | Promoter | 6 | 22288861 | 22297827 | 8967 | 1 | ENSMUST00000176681.1 | 0 | Wnt16 | ENSMUSG00000029671.10 |
| chr6 | 22290015 | 22290388 | 4.15048 | 5.38691 | 2.01447 | Intron (ENSMUST00000128245.2/ENSMUSG00000029671.10, intron 2 of 2) | 6 | 22288861 | 22297827 | 8967 | 1 | ENSMUST00000176681.1 | 1155 | Wnt16 | ENSMUSG00000029671.10 |
| chr6 | 22290747 | 22291310 | 3.73678 | 4.5761 | 1.47736 | 3' UTR | 6 | 22288861 | 22297827 | 8967 | 1 | ENSMUST00000176681.1 | 1887 | Wnt16 | ENSMUSG00000029671.10 |
| chr6 | 22292233 | 22293230 | 3.52633 | 4.23927 | 1.29514 | 3' UTR | 6 | 22288861 | 22297827 | 8967 | 1 | ENSMUST00000176681.1 | 3373 | Wnt16 | ENSMUSG00000029671.10 |
| chr6 | 22293555 | 22294214 | 3.75988 | 4.63815 | 1.55414 | 3' UTR | 6 | 22288861 | 22297827 | 8967 | 1 | ENSMUST00000176681.1 | 4695 | Wnt16 | ENSMUSG00000029671.10 |
| chr5 | 131519716 | 131519812 | 5.35511 | 8.77284 | 4.73513 | Intron (ENSMUST00000161226.3/ENSMUSG00000029673.12, intron 6 of 18) | 5 | 131476742 | 131534397 | 57656 | 2 | ENSMUST00000183153.1 | 14585 | Auts2 | ENSMUSG00000029673.12 |
| chr5 | 134746171 | 134746482 | 3.75099 | 4.57885 | 1.4367 | Promoter | 5 | 134703781 | 134747241 | 43461 | 2 | ENSMUST00000015138.9 | 759 | Eln | ENSMUSG00000029675.9 |
| chr5 | 134747012 | 134747237 | 3.82923 | 4.82891 | 1.63401 | Promoter | 5 | 134703781 | 134747241 | 43461 | 2 | ENSMUST00000015138.9 | 4 | Eln | ENSMUSG00000029675.9 |
| chr6 | 23132863 | 23132978 | 4.18682 | 5.75617 | 2.28282 | Promoter | 6 | 23109477 | 23132916 | 23440 | 2 | ENSMUST00000152280.1 | 0 | Aass | ENSMUSG00000029695.8 |
| chr6 | 23133810 | 23134316 | 3.69123 | 4.61536 | 1.52933 | Promoter | 6 | 23072173 | 23132986 | 60814 | 2 | ENSMUST00000031707.8 | -825 | Aass | ENSMUSG00000029695.8 |
| chr6 | 23234418 | 23234917 | 3.52886 | 4.22348 | 1.31734 | Distal Intergenic | 6 | 23245047 | 23248264 | 3218 | 2 | ENSMUST00000031709.5 | 13347 | Fezf1 | ENSMUSG00000029697.5 |
| chr6 | 23235932 | 23236018 | 3.83589 | 4.76417 | 1.59239 | Distal Intergenic | 6 | 23245047 | 23248264 | 3218 | 2 | ENSMUST00000031709.5 | 12246 | Fezf1 | ENSMUSG00000029697.5 |
| chr6 | 23238764 | 23239161 | 3.78245 | 4.66933 | 1.55841 | Distal Intergenic | 6 | 23245047 | 23248264 | 3218 | 2 | ENSMUST00000031709.5 | 9103 | Fezf1 | ENSMUSG00000029697.5 |
| chr6 | 23240161 | 23240236 | 4.1641 | 5.41472 | 2.07524 | Distal Intergenic | 6 | 23245047 | 23248264 | 3218 | 2 | ENSMUST00000031709.5 | 8028 | Fezf1 | ENSMUSG00000029697.5 |
| chr6 | 23243624 | 23243708 | 4.46401 | 5.97581 | 2.41026 | Downstream (1-2kb) | 6 | 23245047 | 23248264 | 3218 | 2 | ENSMUST00000031709.5 | 4556 | Fezf1 | ENSMUSG00000029697.5 |
| chr6 | 23244033 | 23250997 | 3.9641 | 5.0713 | 1.81387 | Promoter | 6 | 23245047 | 23248264 | 3218 | 2 | ENSMUST00000031709.5 | 0 | Fezf1 | ENSMUSG00000029697.5 |
| chr6 | 23251362 | 23254255 | 4.09581 | 5.49659 | 2.14532 | Distal Intergenic | 6 | 23245047 | 23248264 | 3218 | 2 | ENSMUST00000031709.5 | -3099 | Fezf1 | ENSMUSG00000029697.5 |
| chr6 | 23254832 | 23255374 | 3.65664 | 4.53105 | 1.50844 | Distal Intergenic | 6 | 23245047 | 23248264 | 3218 | 2 | ENSMUST00000031709.5 | -6569 | Fezf1 | ENSMUSG00000029697.5 |
| chr6 | 23256468 | 23256697 | 3.81709 | 4.84333 | 1.67501 | Distal Intergenic | 6 | 23245047 | 23248264 | 3218 | 2 | ENSMUST00000031709.5 | -8205 | Fezf1 | ENSMUSG00000029697.5 |
| chr6 | 23257485 | 23257839 | 4.0329 | 5.2576 | 1.932 | Distal Intergenic | 6 | 23245047 | 23248264 | 3218 | 2 | ENSMUST00000031709.5 | -9222 | Fezf1 | ENSMUSG00000029697.5 |
| chr6 | 23259033 | 23259345 | 3.34683 | 3.84767 | 1.01733 | Distal Intergenic | 6 | 23245047 | 23248264 | 3218 | 2 | ENSMUST00000031709.5 | -10770 | Fezf1 | ENSMUSG00000029697.5 |
| chr6 | 23259876 | 23260054 | 3.35051 | 3.8532 | 1.01733 | Downstream (2-3kb) | 6 | 23245047 | 23248264 | 3218 | 2 | ENSMUST00000031709.5 | -11613 | Fezf1 | ENSMUSG00000029697.5 |
| chr6 | 23260387 | 23260490 | 3.56895 | 4.24157 | 1.22401 | Downstream (2-3kb) | 6 | 23245047 | 23248264 | 3218 | 2 | ENSMUST00000031709.5 | -12124 | Fezf1 | ENSMUSG00000029697.5 |
| chr6 | 23260999 | 23261242 | 3.69764 | 4.62213 | 1.51998 | Downstream (1-2kb) | 6 | 23245047 | 23248264 | 3218 | 2 | ENSMUST00000031709.5 | -12736 | Fezf1 | ENSMUSG00000029697.5 |
| chr6 | 23261717 | 23261810 | 3.49292 | 4.46182 | 1.42132 | Downstream (<1kb) | 6 | 23245047 | 23248264 | 3218 | 2 | ENSMUST00000031709.5 | -13454 | Fezf1 | ENSMUSG00000029697.5 |
| chr6 | 23262874 | 23263101 | 3.4408 | 4.01566 | 1.11019 | 3' UTR | 6 | 23245047 | 23248264 | 3218 | 2 | ENSMUST00000031709.5 | -14611 | Fezf1 | ENSMUSG00000029697.5 |
| chr6 | 23263508 | 23263763 | 3.65981 | 4.4156 | 1.3488 | 3' UTR | 6 | 23245047 | 23248264 | 3218 | 2 | ENSMUST00000031709.5 | -15245 | Fezf1 | ENSMUSG00000029697.5 |
| chr6 | 23264947 | 23265320 | 3.38379 | 3.94538 | 1.14014 | Intron (ENSMUST00000166458.3/ENSMUSG00000017978.13, intron 29 of 29) | 6 | 23245047 | 23248264 | 3218 | 2 | ENSMUST00000031709.5 | -16684 | Fezf1 | ENSMUSG00000029697.5 |
| chr6 | 23269618 | 23269752 | 3.38566 | 4.22785 | 1.26043 | Intron (ENSMUST00000166458.3/ENSMUSG00000017978.13, intron 28 of 29) | 6 | 23245047 | 23248264 | 3218 | 2 | ENSMUST00000031709.5 | -21355 | Fezf1 | ENSMUSG00000029697.5 |
| chr6 | 23271579 | 23272469 | 3.59909 | 4.39365 | 1.42707 | Intron (ENSMUST00000166458.3/ENSMUSG00000017978.13, intron 28 of 29) | 6 | 23245047 | 23248264 | 3218 | 2 | ENSMUST00000031709.5 | -23316 | Fezf1 | ENSMUSG00000029697.5 |

| | | | | | | | | | | | | | | | |
|---|---|---|---|---|---|---|---|---|---|---|---|---|---|---|---|
| chr6 | 23273466 | 23274258 | 3.65754 | 4.47627 | 1.42504 | Intron (ENSMUST00000166458.3/ENSMUSG00000017978.13, intron 28 of 29) | 6 | 23245047 | 23248264 | 3218 | 2 | ENSMUST00000031709.5 | -25203 | Fezf1 | ENSMUSG00000029697.5 |
| chr6 | 23274621 | 23275043 | 3.83702 | 4.7686 | 1.61643 | Intron (ENSMUST00000166458.3/ENSMUSG00000017978.13, intron 28 of 29) | 6 | 23245047 | 23248264 | 3218 | 2 | ENSMUST00000031709.5 | -26358 | Fezf1 | ENSMUSG00000029697.5 |
| chr6 | 24309308 | 24309804 | 3.79259 | 4.7199 | 1.64538 | Distal Intergenic | 6 | 24088283 | 24168092 | 79810 | 2 | ENSMUST00000031713.8 | -141217 | Slc13a1 | ENSMUSG00000029700.11 |
| chr6 | 24314894 | 24315036 | 3.73906 | 4.71774 | 1.58968 | Distal Intergenic | 6 | 24088283 | 24168092 | 79810 | 2 | ENSMUST00000031713.8 | -146803 | Slc13a1 | ENSMUSG00000029700.11 |
| chr6 | 24318737 | 24319019 | 3.35342 | 4.00291 | 1.15559 | Distal Intergenic | 6 | 24088283 | 24168092 | 79810 | 2 | ENSMUST00000031713.8 | -150646 | Slc13a1 | ENSMUSG00000029700.11 |
| chr6 | 24320055 | 24320185 | 3.75999 | 4.74527 | 1.5903 | Distal Intergenic | 6 | 24088283 | 24168092 | 79810 | 2 | ENSMUST00000031713.8 | -151964 | Slc13a1 | ENSMUSG00000029700.11 |
| chr6 | 24322750 | 24322953 | 3.31186 | 3.80985 | 1.06942 | Distal Intergenic | 6 | 24088283 | 24168092 | 79810 | 2 | ENSMUST00000031713.8 | -154659 | Slc13a1 | ENSMUSG00000029700.11 |
| chr6 | 24327171 | 24327265 | 3.8501 | 4.82954 | 1.70063 | Distal Intergenic | 6 | 24088283 | 24168092 | 79810 | 2 | ENSMUST00000031713.8 | -159080 | Slc13a1 | ENSMUSG00000029700.11 |
| chr6 | 24328819 | 24329143 | 3.79824 | 4.75899 | 1.68883 | Distal Intergenic | 6 | 24088283 | 24168092 | 79810 | 2 | ENSMUST00000031713.8 | -160728 | Slc13a1 | ENSMUSG00000029700.11 |
| chr6 | 24330324 | 24330519 | 3.88916 | 4.94192 | 1.79708 | Distal Intergenic | 6 | 24088283 | 24168092 | 79810 | 2 | ENSMUST00000031713.8 | -162233 | Slc13a1 | ENSMUSG00000029700.11 |
| chr6 | 24334820 | 24335467 | 3.69466 | 4.6251 | 1.5106 | Distal Intergenic | 6 | 24088283 | 24168092 | 79810 | 2 | ENSMUST00000031713.8 | -166729 | Slc13a1 | ENSMUSG00000029700.11 |
| chr6 | 29170375 | 29170840 | 3.73989 | 4.73302 | 1.61974 | Exon (ENSMUST00000159200.1/ENSMUSG00000079654.3, exon 5 of 5) | 6 | 29123573 | 29165006 | 41434 | 2 | ENSMUST00000007993.10 | -5370 | Rbm28 | ENSMUSG00000029701.10 |
| chr6 | 29171290 | 29171466 | 3.44632 | 4.17321 | 1.21939 | Exon (ENSMUST00000159200.1/ENSMUSG00000079654.3, exon 5 of 5) | 6 | 29123573 | 29165006 | 41434 | 2 | ENSMUST00000007993.10 | -6285 | Rbm28 | ENSMUSG00000029701.10 |
| chr6 | 29172002 | 29172178 | 3.34454 | 3.8442 | 1.01733 | Intron (ENSMUST00000159200.1/ENSMUSG00000079654.3, intron 4 of 4) | 6 | 29123573 | 29165006 | 41434 | 2 | ENSMUST00000007993.10 | -6997 | Rbm28 | ENSMUSG00000029701.10 |
| chr5 | 136581353 | 136581845 | 3.68598 | 4.55015 | 1.47725 | Distal Intergenic | 5 | 136482875 | 136567490 | 84616 | 2 | ENSMUST00000177305.1 | -13864 | Cux1 | ENSMUSG00000029705.12 |
| chr6 | 28450658 | 28451164 | 3.57199 | 4.52332 | 1.49721 | Distal Intergenic | 6 | 28442334 | 28449353 | 7020 | 2 | ENSMUST00000031718.8 | -1306 | Pax4 | ENSMUSG00000029706.10 |
| chr5 | 137348728 | 137349287 | 3.49373 | 4.16677 | 1.21023 | Promoter | 5 | 137350109 | 137374528 | 24420 | 1 | ENSMUST00000061244.9 | -822 | Ephb4 | ENSMUSG00000029710.10 |
| chr5 | 137350052 | 137350630 | 4.00408 | 5.0852 | 1.81299 | Promoter | 5 | 137350109 | 137374528 | 24420 | 1 | ENSMUST00000061244.9 | 0 | Ephb4 | ENSMUSG00000029710.10 |
| chr5 | 137486399 | 137486727 | 3.53616 | 4.18389 | 1.19291 | Promoter | 5 | 137483026 | 137485816 | 2791 | 2 | ENSMUST00000111039.2 | -584 | Epo | ENSMUSG00000029711.8 |
| chr5 | 137553862 | 137554439 | 3.66639 | 4.44701 | 1.40929 | Promoter | 5 | 137553557 | 137569557 | 16001 | 1 | ENSMUST00000031725.8 | 306 | Actl6b | ENSMUSG00000029712.9 |
| chr5 | 137574727 | 137574898 | 3.57343 | 4.34263 | 1.38576 | Exon (ENSMUST00000031729.8/ENSMUSG00000029716.9, exon 8 of 20) | 5 | 137569851 | 137587481 | 17631 | 1 | ENSMUST00000031729.8 | 4877 | Tfr2 | ENSMUSG00000029716.9 |
| chr5 | 137583039 | 137583687 | 3.46661 | 4.06143 | 1.13413 | Exon (ENSMUST00000031729.8/ENSMUSG00000029716.9, exon 16 of 20) | 5 | 137569851 | 137587481 | 17631 | 1 | ENSMUST00000031729.8 | 13189 | Tfr2 | ENSMUSG00000029716.9 |
| chr6 | 6862671 | 6865388 | 3.84618 | 4.88749 | 1.72261 | Promoter | 6 | 6863334 | 6867970 | 4637 | 1 | ENSMUST00000171311.2 | 0 | Dlx6 | ENSMUSG00000029754.8 |
| chr6 | 6877169 | 6877443 | 3.62505 | 4.35403 | 1.31466 | Downstream (<1kb) | 6 | 6877805 | 6882068 | 4264 | 2 | ENSMUST00000142635.1 | 4625 | Dlx5 | ENSMUSG00000029755.9 |
| chr6 | 6878098 | 6878951 | 3.77482 | 5.03523 | 1.80449 | 3' UTR | 6 | 6877805 | 6882068 | 4264 | 2 | ENSMUST00000142635.1 | 3117 | Dlx5 | ENSMUSG00000029755.9 |
| chr6 | 6879380 | 6880168 | 3.97097 | 5.07406 | 1.81929 | 3' UTR | 6 | 6877805 | 6882068 | 4264 | 2 | ENSMUST00000142635.1 | 1900 | Dlx5 | ENSMUSG00000029755.9 |
| chr6 | 6880498 | 6880902 | 4.53103 | 6.18978 | 2.66337 | Intron (ENSMUST00000142635.1/ENSMUSG00000029755.9, intron 1 of 2) | 6 | 6877805 | 6882068 | 4264 | 2 | ENSMUST00000142635.1 | 1166 | Dlx5 | ENSMUSG00000029755.9 |
| chr6 | 6881234 | 6882156 | 4.14109 | 5.46588 | 2.11466 | Promoter | 6 | 6877805 | 6882068 | 4264 | 2 | ENSMUST00000142635.1 | 0 | Dlx5 | ENSMUSG00000029755.9 |
| chr6 | 6882743 | 6882923 | 3.41025 | 4.08658 | 1.22847 | Promoter | 6 | 6877805 | 6882085 | 4281 | 2 | ENSMUST00000052609.8 | -659 | Dlx5 | ENSMUSG00000029755.9 |
| chr6 | 6884315 | 6885023 | 3.90173 | 5.09333 | 1.86714 | Distal Intergenic | 6 | 6877805 | 6882085 | 4281 | 2 | ENSMUST00000052609.8 | -2231 | Dlx5 | ENSMUSG00000029755.9 |
| chr6 | 32563555 | 32563799 | 3.2323 | 3.68282 | 1.02718 | Intron (ENSMUST00000115096.3/ENSMUSG00000029765.11, intron 1 of 31) | 6 | 32144268 | 32588192 | 443925 | 2 | ENSMUST00000115096.3 | 24393 | Plxna4 | ENSMUSG00000029765.11 |
| chr6 | 32583653 | 32584478 | 3.40366 | 4.00033 | 1.19974 | Intron (ENSMUST00000115096.3/ENSMUSG00000029765.11, intron 1 of 31) | 6 | 32144268 | 32588192 | 443925 | 2 | ENSMUST00000115096.3 | 3714 | Plxna4 | ENSMUSG00000029765.11 |
| chr6 | 32585268 | 32585813 | 3.86451 | 4.86125 | 1.7096 | Intron (ENSMUST00000115096.3/ENSMUSG00000029765.11, intron 1 of 31) | 6 | 32144268 | 32588192 | 443925 | 2 | ENSMUST00000115096.3 | 2379 | Plxna4 | ENSMUSG00000029765.11 |
| chr6 | 32586326 | 32586935 | 3.35081 | 3.92176 | 1.12667 | Intron (ENSMUST00000115096.3/ENSMUSG00000029765.11, intron 1 of 31) | 6 | 32144268 | 32588192 | 443925 | 2 | ENSMUST00000115096.3 | 1257 | Plxna4 | ENSMUSG00000029765.11 |
| chr6 | 32587270 | 32587978 | 3.66097 | 4.61595 | 1.54171 | Promoter | 6 | 32144268 | 32588192 | 443925 | 2 | ENSMUST00000115096.3 | 214 | Plxna4 | ENSMUSG00000029765.11 |
| chr6 | 31936665 | 31936970 | 3.40731 | 3.95777 | 1.08407 | Distal Intergenic | 6 | 32050246 | 32058921 | 8676 | 1 | ENSMUST00000031778.4 | -113276 | 1700012A03Rik | ENSMUSG00000029766.4 |
| chr6 | 29394511 | 29395488 | 3.56056 | 4.24711 | 1.26528 | Promoter | 6 | 29396309 | 29399449 | 3141 | 1 | ENSMUST00000132232.1 | -821 | Ccdc136 | ENSMUSG00000029769.11 |
| chr6 | 29402501 | 29402994 | 3.70833 | 4.94001 | 1.74307 | Promoter | 6 | 29402905 | 29410138 | 7234 | 1 | ENSMUST00000129846.1 | 0 | Ccdc136 | ENSMUSG00000029769.11 |
| chr6 | 29525878 | 29527648 | 4.18637 | 5.4736 | 2.09807 | Promoter | 6 | 29526625 | 29537319 | 10695 | 1 | ENSMUST00000004392.6 | 0 | Irf5 | ENSMUSG00000029771.7 |
| chr6 | 29528001 | 29528136 | 3.5819 | 4.32942 | 1.39717 | Promoter | 6 | 29528270 | 29533997 | 5728 | 1 | ENSMUST00000164626.2 | -134 | Irf5 | ENSMUSG00000029771.7 |
| chr6 | 29529322 | 29529397 | 3.61809 | 4.34477 | 1.32284 | Promoter | 6 | 29529282 | 29537319 | 8038 | 1 | ENSMUST00000163511.2 | 41 | Irf5 | ENSMUSG00000029771.7 |
| chr6 | 29530448 | 29530760 | 3.3903 | 3.92266 | 1.0507 | Promoter | 6 | 29531248 | 29535346 | 4099 | 1 | ENSMUST00000164922.1 | -488 | Irf5 | ENSMUSG00000029771.7 |
| chr6 | 29531207 | 29531742 | 3.63734 | 4.38208 | 1.34314 | Promoter | 6 | 29531248 | 29535346 | 4099 | 1 | ENSMUST00000164922.1 | 0 | Irf5 | ENSMUSG00000029771.7 |
| chr6 | 55451663 | 55452949 | 4.23847 | 5.63579 | 2.25412 | Promoter | 6 | 55451978 | 55498905 | 46928 | 1 | ENSMUST00000070736.6 | 0 | Adcyap1r1 | ENSMUSG00000029778.7 |
| chr6 | 55453482 | 55453689 | 3.62939 | 4.36352 | 1.31327 | Intron (ENSMUST00000070736.6/ENSMUSG00000029778.7, intron 1 of 16) | 6 | 55452177 | 55460245 | 8069 | 1 | ENSMUST00000167484.1 | 1306 | Adcyap1r1 | ENSMUSG00000029778.7 |
| chr6 | 55454486 | 55454768 | 3.49824 | 4.1165 | 1.15938 | Intron (ENSMUST00000070736.6/ENSMUSG00000029778.7, intron 1 of 16) | 6 | 55452177 | 55460245 | 8069 | 1 | ENSMUST00000167484.1 | 2310 | Adcyap1r1 | ENSMUSG00000029778.7 |
| chr6 | 30600178 | 30600436 | 3.30269 | 3.78124 | 1.01733 | Distal Intergenic | 6 | 30611010 | 30631745 | 20736 | 1 | ENSMUST00000062758.5 | -10574 | Cpa5 | ENSMUSG00000029788.8 |
| chr6 | 30615890 | 30616043 | 3.82009 | 4.81527 | 1.66521 | Intron (ENSMUST00000062758.5/ENSMUSG00000029788.8, intron 4 of 11) | 6 | 30612555 | 30631436 | 18882 | 1 | ENSMUST00000165949.1 | 3336 | Cpa5 | ENSMUSG00000029788.8 |
| chr6 | 30617517 | 30617734 | 3.28472 | 3.75422 | 1.01733 | Intron (ENSMUST00000062758.5/ENSMUSG00000029788.8, intron 4 of 11) | 6 | 30612555 | 30631436 | 18882 | 1 | ENSMUST00000165949.1 | 4963 | Cpa5 | ENSMUSG00000029788.8 |
| chr6 | 30618257 | 30618477 | 3.49618 | 4.15606 | 1.28964 | Intron (ENSMUST00000062758.5/ENSMUSG00000029788.8, intron 4 of 11) | 6 | 30612555 | 30631436 | 18882 | 1 | ENSMUST00000165949.1 | 5703 | Cpa5 | ENSMUSG00000029788.8 |
| chr6 | 48823650 | 48823872 | 3.26621 | 3.89391 | 1.11088 | Distal Intergenic | 6 | 48833811 | 48840988 | 7178 | 2 | ENSMUST00000164733.1 | 17116 | Tmem176b | ENSMUSG00000029810.10 |
| chr6 | 48825440 | 48825578 | 4.23821 | 5.56893 | 2.17973 | Distal Intergenic | 6 | 48833811 | 48840988 | 7178 | 2 | ENSMUST00000164733.1 | 15410 | Tmem176b | ENSMUSG00000029810.10 |
| chr6 | 48826247 | 48826429 | 3.56932 | 4.24593 | 1.23758 | Distal Intergenic | 6 | 48833811 | 48840988 | 7178 | 2 | ENSMUST00000164733.1 | 14559 | Tmem176b | ENSMUSG00000029810.10 |
| chr6 | 49210358 | 49211020 | 3.61375 | 4.44306 | 1.43347 | Intron (ENSMUST00000031838.7/ENSMUSG00000029814.8, intron 2 of 14) | 6 | 49085221 | 49214954 | 129734 | 2 | ENSMUST00000031838.7 | 3934 | Igf2bp3 | ENSMUSG00000029814.8 |
| chr6 | 49212674 | 49212968 | 3.73747 | 4.70181 | 1.59784 | Intron (ENSMUST00000031838.7/ENSMUSG00000029814.8, intron 2 of 14) | 6 | 49085221 | 49214954 | 129734 | 2 | ENSMUST00000031838.7 | 1986 | Igf2bp3 | ENSMUSG00000029814.8 |
| chr6 | 49213700 | 49215847 | 4.04908 | 5.68017 | 2.28519 | Promoter | 6 | 49085221 | 49214954 | 129734 | 2 | ENSMUST00000031838.7 | 0 | Igf2bp3 | ENSMUSG00000029814.8 |
| chr6 | 49216174 | 49221038 | 4.00008 | 5.252 | 1.95826 | Distal Intergenic | 6 | 49085221 | 49214954 | 129734 | 2 | ENSMUST00000031838.7 | -1221 | Igf2bp3 | ENSMUSG00000029814.8 |
| chr6 | 49221483 | 49222268 | 4.13823 | 5.48651 | 2.10915 | Distal Intergenic | 6 | 49085221 | 49214954 | 129734 | 2 | ENSMUST00000031838.7 | -6530 | Igf2bp3 | ENSMUSG00000029814.8 |
| chr6 | 49222823 | 49223443 | 3.61052 | 4.56742 | 1.48413 | Distal Intergenic | 6 | 49085221 | 49214954 | 129734 | 2 | ENSMUST00000031838.7 | -7870 | Igf2bp3 | ENSMUSG00000029814.8 |
| chr6 | 49223991 | 49224564 | 3.92548 | 4.95341 | 1.72133 | Distal Intergenic | 6 | 49085221 | 49214954 | 129734 | 2 | ENSMUST00000031838.7 | -9038 | Igf2bp3 | ENSMUSG00000029814.8 |
| chr6 | 49224973 | 49226940 | 3.84919 | 4.91497 | 1.72117 | Distal Intergenic | 6 | 49085221 | 49214954 | 129734 | 2 | ENSMUST00000031838.7 | -10020 | Igf2bp3 | ENSMUSG00000029814.8 |
| chr6 | 49227321 | 49228629 | 3.62776 | 4.50337 | 1.45616 | Distal Intergenic | 6 | 49085221 | 49214954 | 129734 | 2 | ENSMUST00000031838.7 | -12368 | Igf2bp3 | ENSMUSG00000029814.8 |
| chr6 | 49236470 | 49236592 | 2.39614 | 13.62381 | 9.03613 | Distal Intergenic | 6 | 49085221 | 49214954 | 129734 | 2 | ENSMUST00000031838.7 | -21517 | Igf2bp3 | ENSMUSG00000029814.8 |
| chr6 | 49808469 | 49809000 | 3.70476 | 4.52075 | 1.4125 | Distal Intergenic | 6 | 49822710 | 49829507 | 6798 | 1 | ENSMUST00000031843.6 | -13710 | Npy | ENSMUSG00000029819.6 |
| chr6 | 49819529 | 49819604 | 3.78357 | 4.77427 | 1.66624 | Distal Intergenic | 6 | 49822710 | 49829507 | 6798 | 1 | ENSMUST00000031843.6 | -3106 | Npy | ENSMUSG00000029819.6 |
| chr6 | 49820491 | 49821793 | 4.29295 | 5.93591 | 2.45693 | Promoter | 6 | 49822710 | 49829507 | 6798 | 1 | ENSMUST00000031843.6 | -917 | Npy | ENSMUSG00000029819.6 |
| chr6 | 49822131 | 49824222 | 4.08947 | 5.39028 | 2.09022 | Promoter | 6 | 49822710 | 49829507 | 6798 | 1 | ENSMUST00000031843.6 | 0 | Npy | ENSMUSG00000029819.6 |
| chr6 | 50330425 | 50330560 | 3.3707 | 3.94922 | 1.15602 | Intron (ENSMUST00000114468.3/ENSMUSG00000029822.10, intron 11 of 21) | 6 | 50295183 | 50300354 | 5172 | 2 | ENSMUST00000141466.1 | -30072 | Osbpl3 | ENSMUSG00000029822.10 |
| chr6 | 50361843 | 50362099 | 3.35435 | 3.93256 | 1.15234 | Intron (ENSMUST00000114468.3/ENSMUSG00000029822.10, intron 2 of 21) | 6 | 50297065 | 50370215 | 73151 | 2 | ENSMUST00000090019.4 | 8116 | Osbpl3 | ENSMUSG00000029822.10 |
| chr6 | 50382623 | 50382776 | 3.49198 | 4.10681 | 1.15885 | Promoter | 6 | 50293330 | 50382837 | 89508 | 2 | ENSMUST00000114468.3 | 61 | Osbpl3 | ENSMUSG00000029822.10 |
| chr6 | 50453303 | 50453403 | 3.78985 | 4.73234 | 1.64284 | Intron (ENSMUST00000071728.5/ENSMUSG00000029822.10, intron 1 of 21) | 6 | 50327412 | 50456145 | 128734 | 2 | ENSMUST00000147621.1 | 2742 | Osbpl3 | ENSMUSG00000029822.10 |
| chr6 | 50455301 | 50455623 | 3.44672 | 4.26905 | 1.31088 | Promoter | 6 | 50327412 | 50456145 | 128734 | 2 | ENSMUST00000147621.1 | 522 | Osbpl3 | ENSMUSG00000029822.10 |
| chr6 | 50456767 | 50457668 | 3.72527 | 4.62822 | 1.52078 | Promoter | 6 | 50411569 | 50456201 | 44633 | 2 | ENSMUST00000154333.1 | -567 | Osbpl3 | ENSMUSG00000029822.10 |
| chr6 | 50459205 | 50459490 | 3.25527 | 3.71009 | 1.01733 | Distal Intergenic | 6 | 50411569 | 50456201 | 44633 | 2 | ENSMUST00000154333.1 | -3005 | Osbpl3 | ENSMUSG00000029822.10 |
| chr6 | 51433030 | 51433484 | 4.03974 | 5.23709 | 2.01197 | Promoter | 6 | 51432749 | 51456876 | 24128 | 1 | ENSMUST00000160133.1 | 282 | Nfe2l3 | ENSMUSG00000029832.11 |
| chr6 | 42323707 | 42325259 | 3.70697 | 4.62358 | 1.52529 | Promoter | 6 | 42315312 | 42324640 | 9329 | 2 | ENSMUST00000031891.9 | 0 | Fam131b | ENSMUSG00000029861.12 |
| chr15 | 98167085 | 98167444 | 3.65667 | 4.69861 | 1.59519 | Promoter | 15 | 98167806 | 98170134 | 2329 | 1 | ENSMUST00000031914.4 | -362 | Ccdc184 | ENSMUSG00000029875.4 |
| chr15 | 98167851 | 98168679 | 3.76618 | 4.67351 | 1.62446 | Promoter | 15 | 98167806 | 98170134 | 2329 | 1 | ENSMUST00000031914.4 | 46 | Ccdc184 | ENSMUSG00000029875.4 |
| chr15 | 98172449 | 98172549 | 3.4289 | 4.02762 | 1.20753 | Distal Intergenic | 15 | 98167806 | 98170134 | 2329 | 1 | ENSMUST00000031914.4 | 4644 | Ccdc184 | ENSMUSG00000029875.4 |
| chr15 | 98172866 | 98172970 | 3.31192 | 3.82033 | 1.09623 | Distal Intergenic | 15 | 98167806 | 98170134 | 2329 | 1 | ENSMUST00000031914.4 | 5061 | Ccdc184 | ENSMUSG00000029875.4 |
| chr6 | 65381418 | 65381574 | 3.50923 | 4.1521 | 1.18219 | Promoter | 6 | 65381294 | 65458150 | 76857 | 1 | ENSMUST00000170608.2 | 125 | C130060K24Rik | ENSMUSG00000029917.10 |
| chr6 | 85837443 | 85837555 | 3.34364 | 3.84285 | 1.01733 | Distal Intergenic | 6 | 85830388 | 85832082 | 1695 | 2 | ENSMUST00000032073.6 | -5362 | Nat8 | ENSMUSG00000030004.6 |
| chr6 | 82971869 | 82972150 | 3.32743 | 3.83086 | 1.05653 | Intron (ENSMUST00000113980.3/ENSMUSG00000030041.8, intron 3 of 9) | 6 | 82946902 | 83030183 | 83282 | 1 | ENSMUST00000113980.3 | 24968 | M1ap | ENSMUSG00000030041.8 |
| chr6 | 81991429 | 81991568 | 3.43778 | 4.03869 | 1.20802 | Distal Intergenic | 6 | 81957851 | 81965958 | 8108 | 2 | ENSMUST00000032124.8 | -25472 | Mrpl19 | ENSMUSG00000030045.10 |
| chr6 | 87384196 | 87384581 | 3.65999 | 4.44008 | 1.4189 | Exon (ENSMUST00000032127.4/ENSMUSG00000030048.4, exon 5 of 6) | 6 | 87383268 | 87388935 | 5668 | 2 | ENSMUST00000032127.4 | 4354 | Gkn3 | ENSMUSG00000030048.4 |
| chr6 | 97425396 | 97425527 | 3.90177 | 5.0683 | 1.84294 | Intron (ENSMUST00000113355.3/ENSMUSG00000030064.11, intron 2 of 23) | 6 | 97396257 | 97431429 | 35173 | 2 | ENSMUST00000124050.1 | 5902 | Frmd4b | ENSMUSG00000030064.11 |
| chr6 | 99725875 | 99726350 | 3.77853 | 4.73319 | 1.62525 | Promoter | 6 | 99711299 | 99726392 | 15094 | 2 | ENSMUST00000008273.6 | 42 | Prok2 | ENSMUSG00000030069.10 |
| chr6 | 99726787 | 99727526 | 3.74128 | 5.06185 | 1.84474 | Promoter | 6 | 99711299 | 99726392 | 15094 | 2 | ENSMUST00000101120.5 | -396 | Prok2 | ENSMUSG00000030069.10 |
| chr6 | 102563706 | 102564411 | 3.58767 | 4.66721 | 1.55431 | Distal Intergenic | 6 | 102163306 | 102464667 | 301362 | 2 | ENSMUST00000032159.6 | -99040 | Cntn3 | ENSMUSG00000030075.8 |
| chr6 | 103510269 | 103510422 | 3.56515 | 4.30052 | 1.38537 | Promoter | 6 | 103510876 | 103733035 | 222160 | 1 | ENSMUST00000066905.6 | -454 | Chl1 | ENSMUSG00000030077.9 |
| chr6 | 103510816 | 103512140 | 4.06005 | 5.34592 | 2.09924 | Promoter | 6 | 103510876 | 103733035 | 222160 | 1 | ENSMUST00000066905.6 | 0 | Chl1 | ENSMUSG00000030077.9 |
| chr6 | 91118039 | 91118356 | 3.34638 | 3.84697 | 1.01733 | Distal Intergenic | 6 | 91013068 | 91116829 | 103762 | 2 | ENSMUST00000032179.8 | -1211 | Nup210 | ENSMUSG00000030091.12 |
| chr6 | 91408788 | 91409664 | 3.5935 | 4.30115 | 1.29405 | Exon (ENSMUST00000032180.6/ENSMUSG00000030093.6, exon 2 of 4) | 6 | 91363981 | 91411363 | 47383 | 2 | ENSMUST00000032180.6 | 1699 | Wnt7a | ENSMUSG00000030093.6 |
| chr6 | 91409996 | 91410374 | 3.65615 | 4.41855 | 1.36956 | Promoter | 6 | 91363981 | 91411363 | 47383 | 2 | ENSMUST00000032180.6 | 989 | Wnt7a | ENSMUSG00000030093.6 |
| chr6 | 91410709 | 91412523 | 3.75419 | 4.66003 | 1.5578 | Promoter | 6 | 91363981 | 91411363 | 47383 | 2 | ENSMUST00000032180.6 | 0 | Wnt7a | ENSMUSG00000030093.6 |
| chr6 | 91412835 | 91413273 | 3.39075 | 3.93518 | 1.08547 | Distal Intergenic | 6 | 91363981 | 91411363 | 47383 | 2 | ENSMUST00000032180.6 | -1473 | Wnt7a | ENSMUSG00000030093.6 |
| chr6 | 91417646 | 91417740 | 3.29782 | 3.77398 | 1.01733 | Distal Intergenic | 6 | 91363981 | 91411363 | 47383 | 2 | ENSMUST00000032180.6 | -6284 | Wnt7a | ENSMUSG00000030093.6 |
| chr6 | 91843230 | 91844235 | 3.94729 | 5.28204 | 1.98399 | Distal Intergenic | 6 | 91761509 | 91827250 | 65742 | 2 | ENSMUST00000162293.2 | -15981 | Grip2 | ENSMUSG00000030098.10 |
| chr6 | 91844564 | 91844775 | 3.75127 | 4.77999 | 1.63877 | Distal Intergenic | 6 | 91761509 | 91827250 | 65742 | 2 | ENSMUST00000162293.2 | -17315 | Grip2 | ENSMUSG00000030098.10 |
| chr6 | 121412436 | 121413045 | 3.94099 | 4.97027 | 1.74207 | 3' UTR | 6 | 121358965 | 121363856 | 4892 | 1 | ENSMUST00000170339.1 | 53472 | Slc6a12 | ENSMUSG00000030109.10 |
| chr6 | 121413495 | 121413746 | 3.41108 | 3.99199 | 1.16254 | 3' UTR | 6 | 121358965 | 121363856 | 4892 | 1 | ENSMUST00000170339.1 | 54531 | Slc6a12 | ENSMUSG00000030109.10 |
| chr6 | 121415234 | 121415542 | 3.36376 | 3.87319 | 1.01733 | Intron (ENSMUST00000129815.2/ENSMUSG00000040797.11, intron 2 of 13) | 6 | 121358965 | 121363856 | 4892 | 1 | ENSMUST00000170339.1 | 56270 | Slc6a12 | ENSMUSG00000030109.10 |
| chr6 | 118195614 | 118196223 | 3.48671 | 4.09537 | 1.14636 | Intron (ENSMUST00000032201.5/ENSMUSG00000030110.8, intron 1 of 19) | 6 | 118151748 | 118197744 | 45997 | 2 | ENSMUST00000032201.5 | 1521 | Ret | ENSMUSG00000030110.8 |
| chr6 | 118197051 | 118197297 | 3.95728 | 5.08615 | 1.85485 | Promoter | 6 | 118151748 | 118197744 | 45997 | 2 | ENSMUST00000032201.5 | 447 | Ret | ENSMUSG00000030110.8 |
| chr6 | 118197608 | 118197978 | 3.57686 | 4.26769 | 1.26145 | Promoter | 6 | 118151748 | 118197744 | 45997 | 2 | ENSMUST00000032201.5 | 0 | Ret | ENSMUSG00000030110.8 |
| chr6 | 118198337 | 118198841 | 4.01069 | 5.09551 | 1.79759 | Promoter | 6 | 118151748 | 118197744 | 45997 | 2 | ENSMUST00000088790.4 | -594 | Ret | ENSMUSG00000030110.8 |
| chr6 | 121636201 | 121636410 | 3.89507 | 4.91102 | 1.75089 | Promoter | 6 | 121636173 | 121679237 | 43065 | 1 | ENSMUST00000032203.7 | 29 | A2m | ENSMUSG00000030111.8 |
| chr6 | 122270417 | 122270500 | 4.10959 | 5.30657 | 2.01018 | Downstream (<1kb) | 6 | 122270596 | 122282882 | 12287 | 2 | ENSMUST00000032207.8 | 12382 | Klrg1 | ENSMUSG00000030114.8 |
| chr6 | 121210807 | 121210929 | 3.36921 | 3.88967 | 1.04645 | Promoter | 6 | 121210734 | 121226854 | 16121 | 1 | ENSMUST00000032233.7 | 74 | Tuba8 | ENSMUSG00000030137.7 |
| chr6 | 121211326 | 121211666 | 3.44393 | 4.01523 | 1.09149 | Promoter | 6 | 121210734 | 121226854 | 16121 | 1 | ENSMUST00000032233.7 | 593 | Tuba8 | ENSMUSG00000030137.7 |
| chr6 | 132463701 | 132463820 | 24.45408 | 68.22082 | 62.22096 | Distal Intergenic | 6 | 132361041 | 132364134 | 3094 | 2 | ENSMUST00000080849.5 | -99568 | Gm8882 | ENSMUSG00000030143.7 |
| chr6 | 131433420 | 131434063 | 3.62598 | 4.55371 | 1.47041 | Distal Intergenic | 6 | 131364858 | 131388450 | 23593 | 2 | ENSMUST00000087865.2 | -44971 | Ybx3 | ENSMUSG00000030189.10 |
| chr6 | 134388432 | 134388574 | 3.36579 | 3.89736 | 1.09208 | Distal Intergenic | 6 | 134396318 | 134438736 | 42419 | 1 | ENSMUST00000032321.5 | -7744 | Bcl2l14 | ENSMUSG00000030200.8 |
| chr6 | 134389193 | 134389425 | 3.49192 | 4.14262 | 1.2706 | Distal Intergenic | 6 | 134396318 | 134438736 | 42419 | 1 | ENSMUST00000032321.5 | -6893 | Bcl2l14 | ENSMUSG00000030200.8 |
| chr6 | 135251731 | 135251996 | 4.1062 | 5.40534 | 2.06965 | Intron (ENSMUST00000111909.2/ENSMUSG00000030206.8, intron 1 of 5) | 6 | 135237329 | 135254170 | 16842 | 2 | ENSMUST00000111909.2 | 2174 | Gsg1 | ENSMUSG00000030206.8 |
| chr6 | 135183063 | 135183191 | 3.30889 | 3.79054 | 1.01733 | Distal Intergenic | 6 | 135197977 | 135236242 | 38266 | 1 | ENSMUST00000111915.2 | -14786 | 8430419L09Rik | ENSMUSG00000030207.10 |
| chr6 | 135376828 | 135377866 | 3.75929 | 4.68098 | 1.59085 | 5' UTR | 6 | 135367493 | 135383173 | 15681 | 1 | ENSMUST00000111907.1 | 9336 | Emp1 | ENSMUSG00000030208.10 |
| chr6 | 136171855 | 136173035 | 3.57841 | 4.3015 | 1.29285 | Promoter | 6 | 136044203 | 136171889 | 127687 | 2 | ENSMUST00000188999.1 | 0 | Grin2b | ENSMUSG00000030209.9 |
| chr6 | 136173626 | 136173918 | 3.45155 | 4.02779 | 1.09519 | Promoter | 6 | 136044196 | 136173511 | 129316 | 2 | ENSMUST00000143943.2 | -116 | Grin2b | ENSMUSG00000030209.9 |
| chr6 | 136661391 | 136662299 | 3.64715 | 4.87814 | 1.69022 | Promoter | 6 | 136612070 | 136661928 | 49859 | 2 | ENSMUST00000032336.4 | 0 | Plbd1 | ENSMUSG00000030214.4 |
| chr6 | 138421158 | 138421438 | 3.3327 | 3.99118 | 1.11875 | Promoter | 6 | 138364196 | 138421452 | 57257 | 2 | ENSMUST00000163065.2 | 14 | Lmo3 | ENSMUSG00000030226.7 |
| chr6 | 138422647 | 138423210 | 3.72426 | 4.56315 | 1.45866 | Promoter | 6 | 138364439 | 138422831 | 58393 | 2 | ENSMUST00000067437.5 | 0 | Lmo3 | ENSMUSG00000030226.7 |
| chr6 | 138424578 | 138424873 | 3.37599 | 3.89164 | 1.01733 | Promoter | 6 | 138364962 | 138424615 | 59654 | 2 | ENSMUST00000160050.1 | 0 | Lmo3 | ENSMUSG00000030226.7 |
| chr6 | 138426519 | 138426675 | 3.36669 | 3.88986 | 1.06004 | Promoter | 6 | 138416415 | 138426816 | 10402 | 2 | ENSMUST00000162932.1 | 141 | Lmo3 | ENSMUSG00000030226.7 |
| chr6 | 139587737 | 139587941 | 3.45024 | 4.04547 | 1.16564 | Promoter | 6 | 139587237 | 139636536 | 49300 | 1 | ENSMUST00000190962.1 | 501 | Pik3c2g | ENSMUSG00000030228.10 |
| chr6 | 142510773 | 142511067 | 3.50047 | 4.45653 | 1.41934 | Distal Intergenic | 6 | 142490249 | 142507957 | 17709 | 2 | ENSMUST00000032373.6 | -2817 | Ldhb | ENSMUSG00000030246.6 |
| chr6 | 142511681 | 142511860 | 3.21779 | 3.6539 | 1.01337 | Distal Intergenic | 6 | 142490249 | 142507957 | 17709 | 2 | ENSMUST00000032373.6 | -3725 | Ldhb | ENSMUSG00000030246.6 |
| chr6 | 142512283 | 142512451 | 3.48396 | 4.13553 | 1.28216 | Distal Intergenic | 6 | 142490249 | 142507957 | 17709 | 2 | ENSMUST00000032373.6 | -4327 | Ldhb | ENSMUSG00000030246.6 |
| chr6 | 142569182 | 142569295 | 3.68878 | 4.53144 | 1.49941 | Intron (ENSMUST00000032374.7/ENSMUSG00000030247.7, intron 2 of 2) | 6 | 142564939 | 142571356 | 6418 | 2 | ENSMUST00000032374.7 | 2061 | Kcnj8 | ENSMUSG00000030247.7 |
| chr6 | 142569628 | 142570320 | 4.04773 | 5.322 | 2.04585 | Exon (ENSMUST00000032374.7/ENSMUSG00000030247.7, exon 2 of 3) | 6 | 142564939 | 142571356 | 6418 | 2 | ENSMUST00000032374.7 | 1036 | Kcnj8 | ENSMUSG00000030247.7 |
| chr6 | 142571014 | 142571591 | 4.1052 | 5.30814 | 1.99865 | Promoter | 6 | 142564939 | 142571356 | 6418 | 2 | ENSMUST00000032374.7 | 0 | Kcnj8 | ENSMUSG00000030247.7 |
| chr6 | 142572045 | 142572134 | 5.12242 | 7.39075 | 3.52681 | Promoter | 6 | 142564939 | 142571356 | 6418 | 2 | ENSMUST00000032374.7 | -690 | Kcnj8 | ENSMUSG00000030247.7 |
| chr6 | 142572446 | 142572696 | 3.81548 | 4.92108 | 1.73863 | Distal Intergenic | 6 | 142564939 | 142571356 | 6418 | 2 | ENSMUST00000032374.7 | -1091 | Kcnj8 | ENSMUSG00000030247.7 |
| chr6 | 142702277 | 142702921 | 3.59646 | 4.50771 | 1.4641 | Promoter | 6 | 142587862 | 142702274 | 114413 | 2 | ENSMUST00000111771.2 | -4 | Abcc9 | ENSMUSG00000030249.10 |
| chr6 | 145174828 | 145175422 | 4.14649 | 5.46496 | 2.13425 | 3' UTR | 6 | 145169505 | 145171989 | 2485 | 1 | ENSMUST00000135191.1 | 5324 | Lrmp | ENSMUSG00000030263.8 |
| chr6 | 145047107 | 145048105 | 3.6777 | 4.56109 | 1.45743 | Promoter | 6 | 145019192 | 145047853 | 28662 | 2 | ENSMUST00000123930.2 | 0 | Bcat1 | ENSMUSG00000030268.12 |
| chr6 | 145048513 | 145049082 | 3.79085 | 4.8438 | 1.65706 | Promoter | 6 | 144993835 | 145048812 | 54978 | 2 | ENSMUST00000032402.6 | 0 | Bcat1 | ENSMUSG00000030268.12 |
| chr6 | 145074109 | 145075401 | 3.92795 | 4.99913 | 1.83477 | Promoter | 6 | 144999158 | 145076184 | 77027 | 2 | ENSMUST00000111742.2 | 783 | Bcat1 | ENSMUSG00000030268.12 |
| chr6 | 145075890 | 145075965 | 3.93063 | 5.00125 | 1.82985 | Promoter | 6 | 144999158 | 145076184 | 77027 | 2 | ENSMUST00000111742.2 | 219 | Bcat1 | ENSMUSG00000030268.12 |
| chr6 | 145076342 | 145076460 | 3.67693 | 4.69775 | 1.57728 | Promoter | 6 | 145004118 | 145076184 | 72067 | 2 | ENSMUST00000048252.5 | -159 | Bcat1 | ENSMUSG00000030268.12 |
| chr6 | 145077126 | 145077353 | 3.35988 | 3.91227 | 1.11615 | Promoter | 6 | 145004118 | 145076184 | 72067 | 2 | ENSMUST00000048252.5 | -943 | Bcat1 | ENSMUSG00000030268.12 |
| chr6 | 113282125 | 113283249 | 3.93656 | 5.03575 | 1.80123 | Promoter | 6 | 113282307 | 113305569 | 23263 | 1 | ENSMUST00000041203.5 | 0 | Cpne9 | ENSMUSG00000030270.6 |
| chr6 | 142735722 | 142736136 | 3.6033 | 4.31052 | 1.28114 | Distal Intergenic | 6 | 142756686 | 142775714 | 19029 | 1 | ENSMUST00000032419.8 | -20550 | Cmas | ENSMUSG00000030282.9 |
| chr6 | 142962533 | 142962769 | 3.35101 | 3.85395 | 1.01733 | Intron (ENSMUST00000032421.3/ENSMUSG00000030283.5, intron 1 of 4) | 6 | 142821545 | 142964452 | 142908 | 2 | ENSMUST00000032421.3 | 1683 | St8sia1 | ENSMUSG00000030283.5 |
| chr6 | 142963865 | 142964594 | 3.81748 | 4.77916 | 1.61311 | Promoter | 6 | 142821545 | 142964452 | 142908 | 2 | ENSMUST00000032421.3 | 0 | St8sia1 | ENSMUSG00000030283.5 |
| chr6 | 146190603 | 146190977 | 3.58173 | 4.31225 | 1.37777 | Exon (ENSMUST00000053273.9/ENSMUSG00000030287.10, exon 47 of 57) | 6 | 146180336 | 146196960 | 16625 | 2 | ENSMUST00000155339.1 | 5983 | Itpr2 | ENSMUSG00000030287.10 |
| chr6 | 146504160 | 146504355 | 3.30861 | 3.93505 | 1.06319 | Distal Intergenic | 6 | 146373123 | 146502223 | 129101 | 2 | ENSMUST00000139732.1 | -1938 | Itpr2 | ENSMUSG00000030287.10 |
| chr6 | 113748130 | 113748828 | 3.51827 | 4.28946 | 1.31915 | 3' UTR | 6 | 113728067 | 113740743 | 12677 | 2 | ENSMUST00000032440.4 | -7388 | Sec13 | ENSMUSG00000030298.5 |
| chr6 | 113890519 | 113890773 | 3.66312 | 4.42338 | 1.35966 | Promoter | 6 | 113803338 | 113891370 | 88033 | 2 | ENSMUST00000135199.1 | 597 | Atp2b2 | ENSMUSG00000030302.11 |
| chr6 | 113891571 | 113892350 | 3.60149 | 4.30379 | 1.26801 | Promoter | 6 | 113745669 | 113891539 | 145871 | 2 | ENSMUST00000089003.6 | -33 | Atp2b2 | ENSMUSG00000030302.11 |
| chr6 | 114040065 | 114040318 | 3.48467 | 4.10086 | 1.17751 | Intron (ENSMUST00000138670.1/ENSMUSG00000087083.1, intron 1 of 2) | 6 | 113745670 | 114042015 | 296346 | 2 | ENSMUST00000152831.2 | 1697 | Atp2b2 | ENSMUSG00000030302.11 |
| chr6 | 114041297 | 114041868 | 4.02812 | 5.18765 | 1.89475 | Promoter | 6 | 113745670 | 114042015 | 296346 | 2 | ENSMUST00000152831.2 | 147 | Atp2b2 | ENSMUSG00000030302.11 |
| chr6 | 114042300 | 114042852 | 3.77297 | 4.62904 | 1.48269 | Promoter | 6 | 113842549 | 114042613 | 200065 | 2 | ENSMUST00000154738.1 | 0 | Atp2b2 | ENSMUSG00000030302.11 |
| chr6 | 148047175 | 148047556 | 3.48065 | 4.08844 | 1.15478 | Promoter | 6 | 148047259 | 148176849 | 129591 | 1 | ENSMUST00000032443.8 | 0 | Far2 | ENSMUSG00000030303.10 |
| chr6 | 114129823 | 114130026 | 3.37793 | 3.93915 | 1.13831 | Distal Intergenic | 6 | 114131241 | 114249886 | 118646 | 1 | ENSMUST00000032451.7 | -1215 | Slc6a11 | ENSMUSG00000030307.7 |
| chr6 | 114131013 | 114131526 | 3.81295 | 5.1028 | 1.85429 | Promoter | 6 | 114131241 | 114249886 | 118646 | 1 | ENSMUST00000032451.7 | 0 | Slc6a11 | ENSMUSG00000030307.7 |
| chr6 | 114173573 | 114173813 | 3.51779 | 4.19504 | 1.22754 | Intron (ENSMUST00000032451.7/ENSMUSG00000030307.7, intron 5 of 13) | 6 | 114131241 | 114249886 | 118646 | 1 | ENSMUST00000032451.7 | 42333 | Slc6a11 | ENSMUSG00000030307.7 |
| chr6 | 114282924 | 114283249 | 4.03024 | 5.29296 | 1.99567 | Promoter | 6 | 114282635 | 114317522 | 34888 | 1 | ENSMUST00000032454.5 | 290 | Slc6a1 | ENSMUSG00000030310.8 |
| chr6 | 114284004 | 114284495 | 3.57802 | 4.3803 | 1.41303 | Intron (ENSMUST00000032454.5/ENSMUSG00000030310.8, intron 1 of 15) | 6 | 114282635 | 114317522 | 34888 | 1 | ENSMUST00000032454.5 | 1370 | Slc6a1 | ENSMUSG00000030310.8 |
| chr6 | 114286259 | 114286358 | 3.4174 | 4.04924 | 1.20944 | Intron (ENSMUST00000032454.5/ENSMUSG00000030310.8, intron 1 of 15) | 6 | 114282635 | 114317522 | 34888 | 1 | ENSMUST00000032454.5 | 3625 | Slc6a1 | ENSMUSG00000030310.8 |
| chr6 | 114286680 | 114287354 | 3.55657 | 4.47269 | 1.46684 | Intron (ENSMUST00000032454.5/ENSMUSG00000030310.8, intron 1 of 15) | 6 | 114282635 | 114317522 | 34888 | 1 | ENSMUST00000032454.5 | 4046 | Slc6a1 | ENSMUSG00000030310.8 |
| chr6 | 115251606 | 115251810 | 3.71157 | 4.59267 | 1.51195 | Promoter | 6 | 115245616 | 115251849 | 6234 | 2 | ENSMUST00000032462.6 | 39 | Timp4 | ENSMUSG00000030317.6 |
| chr6 | 124995050 | 124995400 | 3.60665 | 4.36714 | 1.39612 | Distal Intergenic | 6 | 124996694 | 125003096 | 6403 | 1 | ENSMUST00000032479.5 | -1294 | Pianp | ENSMUSG00000030329.6 |

| | | | | | | | | | | | | | | | |
|---|---|---|---|---|---|---|---|---|---|---|---|---|---|---|---|
| chr6 | 124997028 | 124997879 | 3.83958 | 4.90629 | 1.75375 | Promoter | 6 | 124997075 | 124999731 | 2657 | 1 | ENSMUST00000159391.2 | 0 | Pianp | ENSMUSG00000030329.6 |
| chr6 | 124998875 | 124999099 | 3.83084 | 5.00904 | 1.76094 | Promoter | 6 | 124998639 | 124999445 | 807 | 1 | ENSMUST00000161292.1 | 237 | Pianp | ENSMUSG00000030329.6 |
| chr6 | 125321145 | 125321465 | 3.61817 | 4.36178 | 1.38078 | Promoter | 6 | 125321205 | 125330973 | 9769 | 1 | ENSMUST00000176365.1 | 0 | Scnn1a | ENSMUSG00000030340.11 |
| chr6 | 125322045 | 125322204 | 3.4636 | 4.06437 | 1.16077 | Promoter | 6 | 125321521 | 125344612 | 23092 | 1 | ENSMUST00000177329.1 | 525 | Scnn1a | ENSMUSG00000030340.11 |
| chr6 | 125366154 | 125366308 | 3.5142 | 4.14356 | 1.16988 | Intron (ENSMUST00000042647.6/ENSMUSG00000038167.6, intron 14 of 16) | 6 | 125360427 | 125361874 | 1448 | 1 | ENSMUST00000134803.1 | 5728 | Tnfrsf1a | ENSMUSG00000030341.12 |
| chr6 | 125495021 | 125495170 | 3.59982 | 4.44709 | 1.4515 | Promoter | 6 | 125460266 | 125494791 | 34526 | 2 | ENSMUST00000032492.8 | -231 | Cd9 | ENSMUSG00000030342.8 |
| chr6 | 127766680 | 127767281 | 3.60242 | 4.31409 | 1.29899 | Intron (ENSMUST00000032500.8/ENSMUSG00000030350.8, intron 1 of 9) | 6 | 127689011 | 127769472 | 80462 | 2 | ENSMUST00000032500.8 | 2191 | Prmt8 | ENSMUSG00000030350.8 |
| chr6 | 127767598 | 127769358 | 4.04677 | 5.28332 | 1.99661 | Promoter | 6 | 127689011 | 127769472 | 80462 | 2 | ENSMUST00000032500.8 | 114 | Prmt8 | ENSMUSG00000030350.8 |
| chr6 | 127770138 | 127770841 | 3.66241 | 4.45508 | 1.40777 | Promoter | 6 | 127689011 | 127769472 | 80462 | 2 | ENSMUST00000032500.8 | -667 | Prmt8 | ENSMUSG00000030350.8 |
| chr6 | 127771862 | 127772160 | 3.56186 | 4.29665 | 1.33326 | Distal Intergenic | 6 | 127689011 | 127769472 | 80462 | 2 | ENSMUST00000032500.8 | -2391 | Prmt8 | ENSMUSG00000030350.8 |
| chr6 | 127887978 | 127888385 | 3.85742 | 4.81896 | 1.67106 | Promoter | 6 | 127887772 | 127923881 | 36110 | 1 | ENSMUST00000149801.1 | 207 | Tspan11 | ENSMUSG00000030351.4 |
| chr6 | 128033259 | 128033473 | 3.40179 | 4.26355 | 1.29494 | Intron (ENSMUST00000112173.2/ENSMUSG00000030352.9, intron 2 of 7) | 6 | 127965747 | 128034592 | 68846 | 2 | ENSMUST00000123786.2 | 1119 | Tspan9 | ENSMUSG00000030352.9 |
| chr6 | 128035416 | 128035806 | 3.50755 | 4.12962 | 1.15615 | Promoter | 6 | 127965747 | 128034592 | 68846 | 2 | ENSMUST00000123786.2 | -825 | Tspan9 | ENSMUSG00000030352.9 |
| chr6 | 128037772 | 128037893 | 3.36705 | 3.87815 | 1.01733 | Intron (ENSMUST00000112173.2/ENSMUSG00000030352.9, intron 1 of 7) | 6 | 127965747 | 128034592 | 68846 | 2 | ENSMUST00000123786.2 | -3181 | Tspan9 | ENSMUSG00000030352.9 |
| chr7 | 16130996 | 16131157 | 3.44981 | 4.05385 | 1.14516 | Promoter | 7 | 16130300 | 16160509 | 30210 | 1 | ENSMUST00000168693.1 | 697 | Slc8a2 | ENSMUSG00000030376.7 |
| chr7 | 12464084 | 12464201 | 3.53502 | 4.27812 | 1.32051 | Intron (ENSMUST00000173045.1/ENSMUSG00000092583.1, intron 4 of 4) | 7 | 12478293 | 12496235 | 17943 | 1 | ENSMUST00000098822.4 | -14092 | Zfp606 | ENSMUSG00000030386.12 |
| chr7 | 12473061 | 12473603 | 3.6498 | 4.49517 | 1.4917 | Distal Intergenic | 7 | 12478293 | 12496235 | 17943 | 1 | ENSMUST00000098822.4 | -4690 | Zfp606 | ENSMUSG00000030386.12 |
| chr7 | 19414325 | 19414519 | 3.50264 | 4.14633 | 1.24385 | Exon (ENSMUST00000003643.1/ENSMUSG00000030399.1, exon 2 of 8) | 7 | 19411094 | 19421583 | 10490 | 1 | ENSMUST00000003643.1 | 3232 | Ckm | ENSMUSG00000030399.1 |
| chr7 | 19415421 | 19415759 | 3.59184 | 4.29174 | 1.27977 | Exon (ENSMUST00000003643.1/ENSMUSG00000030399.1, exon 4 of 8) | 7 | 19411094 | 19421583 | 10490 | 1 | ENSMUST00000003643.1 | 4328 | Ckm | ENSMUSG00000030399.1 |
| chr7 | 19277804 | 19278593 | 3.6071 | 4.41622 | 1.40039 | Exon (ENSMUST00000032560.4/ENSMUSG00000030402.4, exon 2 of 5) | 7 | 19276807 | 19280049 | 3243 | 2 | ENSMUST00000032560.4 | 1456 | Ppm1n | ENSMUSG00000030402.4 |
| chr7 | 19279292 | 19280086 | 3.65506 | 4.5552 | 1.49246 | Promoter | 7 | 19276807 | 19280049 | 3243 | 2 | ENSMUST00000032560.4 | 0 | Ppm1n | ENSMUSG00000030402.4 |
| chr7 | 19159349 | 19159805 | 3.52839 | 4.19887 | 1.28582 | Exon (ENSMUST00000094790.3/ENSMUSG00000030406.6, exon 11 of 14) | 7 | 19157125 | 19166127 | 9003 | 2 | ENSMUST00000094790.3 | 6322 | Gipr | ENSMUSG00000030406.6 |
| chr7 | 19165054 | 19165401 | 4.64026 | 6.37833 | 2.76135 | Promoter | 7 | 19157125 | 19166127 | 9003 | 2 | ENSMUST00000094790.3 | 726 | Gipr | ENSMUSG00000030406.6 |
| chr7 | 19067310 | 19067570 | 3.53434 | 4.29776 | 1.35841 | Intron (ENSMUST00000035521.5/ENSMUSG00000040866.8, intron 4 of 5) | 7 | 19076249 | 19082776 | 6528 | 1 | ENSMUST00000032570.8 | -8679 | Dmwd | ENSMUSG00000030410.11 |
| chr7 | 18919501 | 18919872 | 3.51552 | 4.15002 | 1.18683 | Distal Intergenic | 7 | 18925888 | 18962057 | 36170 | 1 | ENSMUST00000172835.2 | -6016 | Nova2 | ENSMUSG00000030411.10 |
| chr7 | 18924587 | 18924683 | 3.7775 | 4.64031 | 1.49474 | Distal Intergenic | 7 | 18925888 | 18962057 | 36170 | 1 | ENSMUST00000172835.2 | -1205 | Nova2 | ENSMUSG00000030411.10 |
| chr7 | 18924998 | 18925693 | 3.57519 | 4.26026 | 1.24592 | Promoter | 7 | 18925888 | 18962057 | 36170 | 1 | ENSMUST00000172835.2 | -195 | Nova2 | ENSMUSG00000030411.10 |
| chr7 | 18926028 | 18927006 | 3.65102 | 4.40292 | 1.34277 | Promoter | 7 | 18925888 | 18965319 | 39432 | 1 | ENSMUST00000032571.8 | 141 | Nova2 | ENSMUSG00000030411.10 |
| chr7 | 18927313 | 18927503 | 4.09855 | 5.431 | 2.05278 | Intron (ENSMUST00000172835.2/ENSMUSG00000030411.10, intron 1 of 3) | 7 | 18925888 | 18965319 | 39432 | 1 | ENSMUST00000032571.8 | 1426 | Nova2 | ENSMUSG00000030411.10 |
| chr7 | 18942564 | 18942737 | 3.34538 | 3.84546 | 1.01733 | Intron (ENSMUST00000172835.2/ENSMUSG00000030411.10, intron 2 of 3) | 7 | 18925888 | 18965319 | 39432 | 1 | ENSMUST00000032571.8 | 16677 | Nova2 | ENSMUSG00000030411.10 |
| chr7 | 18878267 | 18878472 | 3.66507 | 4.4465 | 1.42325 | Distal Intergenic | 7 | 18884690 | 18890438 | 5749 | 1 | ENSMUST00000032573.6 | -6218 | Pglyrp1 | ENSMUSG00000030413.6 |
| chr7 | 4118444 | 4118861 | 3.43739 | 4.02415 | 1.15898 | Promoter | 7 | 4119408 | 4125588 | 6181 | 1 | ENSMUST00000130150.2 | -547 | Ttyh1 | ENSMUSG00000030428.11 |
| chr7 | 4119223 | 4119864 | 4.31379 | 5.90532 | 2.45691 | Promoter | 7 | 4119408 | 4125588 | 6181 | 1 | ENSMUST00000130150.2 | 0 | Ttyh1 | ENSMUSG00000030428.11 |
| chr7 | 4120217 | 4120366 | 3.31329 | 3.79715 | 1.01733 | Promoter | 7 | 4120220 | 4125560 | 5341 | 1 | ENSMUST00000151959.2 | 0 | Ttyh1 | ENSMUSG00000030428.11 |
| chr7 | 35184919 | 35185999 | 4.05111 | 5.28109 | 2.00505 | Promoter | 7 | 35186385 | 35195449 | 9065 | 1 | ENSMUST00000135452.2 | -386 | Slc7a10 | ENSMUSG00000030495.7 |
| chr7 | 35186322 | 35187173 | 3.91951 | 5.0131 | 1.78328 | Promoter | 7 | 35186385 | 35195449 | 9065 | 1 | ENSMUST00000135452.2 | 0 | Slc7a10 | ENSMUSG00000030495.7 |
| chr7 | 51619356 | 51619647 | 3.43661 | 4.00568 | 1.09685 | Distal Intergenic | 7 | 51621830 | 51671125 | 49296 | 1 | ENSMUST00000032710.5 | -2183 | Slc17a6 | ENSMUSG00000030500.5 |
| chr7 | 51621880 | 51622698 | 3.73144 | 4.57308 | 1.46207 | Promoter | 7 | 51621830 | 51671125 | 49296 | 1 | ENSMUST00000032710.5 | 51 | Slc17a6 | ENSMUSG00000030500.5 |
| chr7 | 51623356 | 51623937 | 3.75865 | 4.6173 | 1.50812 | Intron (ENSMUST00000032710.5/ENSMUSG00000030500.5, intron 1 of 11) | 7 | 51621830 | 51671125 | 49296 | 1 | ENSMUST00000032710.5 | 1527 | Slc17a6 | ENSMUSG00000030500.5 |
| chr7 | 51624338 | 51625365 | 3.8701 | 4.81775 | 1.61764 | Exon (ENSMUST00000032710.5/ENSMUSG00000030500.5, exon 2 of 12) | 7 | 51621830 | 51671125 | 49296 | 1 | ENSMUST00000032710.5 | 2509 | Slc17a6 | ENSMUSG00000030500.5 |
| chr7 | 51625736 | 51626384 | 3.70752 | 4.5052 | 1.40557 | Intron (ENSMUST00000032710.5/ENSMUSG00000030500.5, intron 2 of 11) | 7 | 51621830 | 51671125 | 49296 | 1 | ENSMUST00000032710.5 | 3907 | Slc17a6 | ENSMUSG00000030500.5 |
| chr7 | 51627221 | 51628539 | 3.72576 | 4.60636 | 1.48796 | Intron (ENSMUST00000032710.5/ENSMUSG00000030500.5, intron 3 of 11) | 7 | 51621830 | 51671125 | 49296 | 1 | ENSMUST00000032710.5 | 5392 | Slc17a6 | ENSMUSG00000030500.5 |
| chr7 | 51628936 | 51629078 | 3.59584 | 4.41199 | 1.3588 | Intron (ENSMUST00000032710.5/ENSMUSG00000030500.5, intron 3 of 11) | 7 | 51621830 | 51671125 | 49296 | 1 | ENSMUST00000032710.5 | 7107 | Slc17a6 | ENSMUSG00000030500.5 |
| chr7 | 51629389 | 51630630 | 3.51252 | 4.18362 | 1.23586 | Intron (ENSMUST00000032710.5/ENSMUSG00000030500.5, intron 3 of 11) | 7 | 51621830 | 51671125 | 49296 | 1 | ENSMUST00000032710.5 | 7560 | Slc17a6 | ENSMUSG00000030500.5 |
| chr7 | 51631975 | 51632163 | 3.39231 | 3.93544 | 1.05542 | Intron (ENSMUST00000032710.5/ENSMUSG00000030500.5, intron 3 of 11) | 7 | 51621830 | 51671125 | 49296 | 1 | ENSMUST00000032710.5 | 10146 | Slc17a6 | ENSMUSG00000030500.5 |
| chr7 | 51632638 | 51633182 | 4.02498 | 5.19638 | 1.894 | Intron (ENSMUST00000032710.5/ENSMUSG00000030500.5, intron 3 of 11) | 7 | 51621830 | 51671125 | 49296 | 1 | ENSMUST00000032710.5 | 10809 | Slc17a6 | ENSMUSG00000030500.5 |
| chr7 | 49627206 | 49627523 | 3.78598 | 4.68359 | 1.59676 | Distal Intergenic | 7 | 49631499 | 49636849 | 5351 | 2 | ENSMUST00000032717.6 | 9326 | Dbx1 | ENSMUSG00000030507.6 |
| chr7 | 49629878 | 49630254 | 3.64859 | 4.42596 | 1.37847 | Downstream (1-2kb) | 7 | 49631499 | 49636849 | 5351 | 2 | ENSMUST00000032717.6 | 6595 | Dbx1 | ENSMUSG00000030507.6 |
| chr7 | 49630640 | 49630984 | 3.86981 | 4.8207 | 1.61781 | Downstream (<1kb) | 7 | 49631499 | 49636849 | 5351 | 2 | ENSMUST00000032717.6 | 5865 | Dbx1 | ENSMUSG00000030507.6 |
| chr7 | 49632370 | 49632623 | 3.64239 | 4.52623 | 1.44101 | 3' UTR | 7 | 49631499 | 49636849 | 5351 | 2 | ENSMUST00000032717.6 | 4226 | Dbx1 | ENSMUSG00000030507.6 |
| chr7 | 49633445 | 49633653 | 4.55141 | 6.25048 | 2.67967 | Exon (ENSMUST00000032717.6/ENSMUSG00000030507.6, exon 3 of 4) | 7 | 49631499 | 49636849 | 5351 | 2 | ENSMUST00000032717.6 | 3196 | Dbx1 | ENSMUSG00000030507.6 |
| chr7 | 49634170 | 49634459 | 3.72552 | 4.53413 | 1.41434 | Intron (ENSMUST00000032717.6/ENSMUSG00000030507.6, intron 2 of 3) | 7 | 49631499 | 49636849 | 5351 | 2 | ENSMUST00000032717.6 | 2390 | Dbx1 | ENSMUSG00000030507.6 |
| chr7 | 49634803 | 49635735 | 3.8185 | 4.73281 | 1.56757 | Exon (ENSMUST00000032717.6/ENSMUSG00000030507.6, exon 2 of 4) | 7 | 49631499 | 49636849 | 5351 | 2 | ENSMUST00000032717.6 | 1114 | Dbx1 | ENSMUSG00000030507.6 |
| chr7 | 49636305 | 49637217 | 3.62571 | 4.64117 | 1.53078 | Promoter | 7 | 49631499 | 49636849 | 5351 | 2 | ENSMUST00000032717.6 | 0 | Dbx1 | ENSMUSG00000030507.6 |
| chr7 | 49637970 | 49638649 | 4.51987 | 6.16486 | 2.60955 | Distal Intergenic | 7 | 49631499 | 49636849 | 5351 | 2 | ENSMUST00000032717.6 | -1122 | Dbx1 | ENSMUSG00000030507.6 |
| chr7 | 49639060 | 49640052 | 3.9709 | 5.10598 | 1.86423 | Distal Intergenic | 7 | 49631499 | 49636849 | 5351 | 2 | ENSMUST00000032717.6 | -2212 | Dbx1 | ENSMUSG00000030507.6 |
| chr7 | 49642580 | 49643239 | 3.66467 | 4.49415 | 1.46297 | Distal Intergenic | 7 | 49631499 | 49636849 | 5351 | 2 | ENSMUST00000032717.6 | -5732 | Dbx1 | ENSMUSG00000030507.6 |
| chr7 | 49645463 | 49645679 | 3.68148 | 4.49799 | 1.47076 | Distal Intergenic | 7 | 49631499 | 49636849 | 5351 | 2 | ENSMUST00000032717.6 | -8615 | Dbx1 | ENSMUSG00000030507.6 |
| chr7 | 49661793 | 49662201 | 3.57565 | 4.26303 | 1.25934 | Distal Intergenic | 7 | 49631499 | 49636849 | 5351 | 2 | ENSMUST00000032717.6 | -24945 | Dbx1 | ENSMUSG00000030507.6 |
| chr7 | 49662534 | 49662730 | 3.34153 | 3.83966 | 1.01733 | Distal Intergenic | 7 | 49631499 | 49636849 | 5351 | 2 | ENSMUST00000032717.6 | -25686 | Dbx1 | ENSMUSG00000030507.6 |
| chr7 | 49664296 | 49664520 | 3.33588 | 3.83123 | 1.01733 | Distal Intergenic | 7 | 49631499 | 49636849 | 5351 | 2 | ENSMUST00000032717.6 | -27448 | Dbx1 | ENSMUSG00000030507.6 |
| chr7 | 49674763 | 49675466 | 3.33594 | 3.85598 | 1.09552 | Distal Intergenic | 7 | 49631499 | 49636849 | 5351 | 2 | ENSMUST00000032717.6 | -37915 | Dbx1 | ENSMUSG00000030507.6 |
| chr7 | 49676024 | 49679283 | 3.94903 | 5.12835 | 1.88572 | Distal Intergenic | 7 | 49631499 | 49636849 | 5351 | 2 | ENSMUST00000032717.6 | -39176 | Dbx1 | ENSMUSG00000030507.6 |
| chr7 | 49679840 | 49680395 | 3.78205 | 4.75766 | 1.61112 | Distal Intergenic | 7 | 49631499 | 49636849 | 5351 | 2 | ENSMUST00000032717.6 | -42992 | Dbx1 | ENSMUSG00000030507.6 |
| chr7 | 49681640 | 49681923 | 3.52867 | 4.17037 | 1.18707 | Distal Intergenic | 7 | 49631499 | 49636849 | 5351 | 2 | ENSMUST00000032717.6 | -44792 | Dbx1 | ENSMUSG00000030507.6 |
| chr7 | 49684052 | 49684192 | 3.43649 | 4.04692 | 1.22174 | Distal Intergenic | 7 | 49631499 | 49636849 | 5351 | 2 | ENSMUST00000032717.6 | -47204 | Dbx1 | ENSMUSG00000030507.6 |
| chr7 | 49685571 | 49685839 | 3.31977 | 3.96737 | 1.15964 | Distal Intergenic | 7 | 49631499 | 49636849 | 5351 | 2 | ENSMUST00000032717.6 | -48723 | Dbx1 | ENSMUSG00000030507.6 |
| chr7 | 49686540 | 49687316 | 3.71469 | 4.53832 | 1.46223 | Distal Intergenic | 7 | 49631499 | 49636849 | 5351 | 2 | ENSMUST00000032717.6 | -49692 | Dbx1 | ENSMUSG00000030507.6 |
| chr7 | 49687849 | 49688001 | 4.16726 | 5.42191 | 2.07848 | Distal Intergenic | 7 | 49631499 | 49636849 | 5351 | 2 | ENSMUST00000032717.6 | -51001 | Dbx1 | ENSMUSG00000030507.6 |
| chr7 | 65847916 | 65848225 | 3.39107 | 3.94298 | 1.1174 | Distal Intergenic | 7 | 65861734 | 65952984 | 91251 | 1 | ENSMUST00000176199.2 | -13509 | Pcsk6 | ENSMUSG00000030513.9 |
| chr7 | 65155479 | 65155907 | 3.78989 | 4.7656 | 1.63134 | Promoter | 7 | 64786690 | 65156416 | 369727 | 2 | ENSMUST00000148459.1 | 509 | Fam189a1 | ENSMUSG00000030518.12 |
| chr7 | 65156285 | 65156363 | 3.96402 | 5.10323 | 1.87153 | Promoter | 7 | 64786690 | 65156416 | 369727 | 2 | ENSMUST00000148459.1 | 53 | Fam189a1 | ENSMUSG00000030518.12 |
| chr7 | 64501721 | 64502350 | 3.62434 | 4.36657 | 1.35121 | Promoter | 7 | 64501949 | 64695307 | 193359 | 1 | ENSMUST00000138829.2 | 0 | Apba2 | ENSMUSG00000030519.9 |
| chr7 | 63210617 | 63210692 | 3.95633 | 5.23944 | 1.91924 | Intron (ENSMUST00000032738.5/ENSMUSG00000030525.7, intron 2 of 9) | 7 | 63098692 | 63212526 | 113835 | 2 | ENSMUST00000032738.5 | 1834 | Chrna7 | ENSMUSG00000030525.7 |
| chr7 | 63211651 | 63211959 | 3.30778 | 3.79899 | 1.05251 | Promoter | 7 | 63098692 | 63212526 | 113835 | 2 | ENSMUST00000032738.5 | 567 | Chrna7 | ENSMUSG00000030525.7 |
| chr7 | 79791516 | 79792243 | 3.92355 | 5.02182 | 1.80421 | 3' UTR | 7 | 79792241 | 79793788 | 1548 | 2 | ENSMUST00000032760.5 | 1545 | Mesp1 | ENSMUSG00000030544.5 |
| chr7 | 79792608 | 79793538 | 3.82316 | 4.99518 | 1.7858 | Promoter | 7 | 79792241 | 79793788 | 1548 | 2 | ENSMUST00000032760.5 | 250 | Mesp1 | ENSMUSG00000030544.5 |
| chr7 | 79561377 | 79561610 | 3.49396 | 4.25025 | 1.28788 | Distal Intergenic | 7 | 79593369 | 79594924 | 1556 | 2 | ENSMUST00000172788.1 | 33314 | Rhcg | ENSMUSG00000030549.5 |
| chr7 | 79563500 | 79564195 | 3.94854 | 4.99138 | 1.75192 | Distal Intergenic | 7 | 79593369 | 79594924 | 1556 | 2 | ENSMUST00000172788.1 | 30729 | Rhcg | ENSMUSG00000030549.5 |
| chr7 | 79565283 | 79565730 | 3.66909 | 4.44825 | 1.38457 | Distal Intergenic | 7 | 79593369 | 79594924 | 1556 | 2 | ENSMUST00000172788.1 | 29194 | Rhcg | ENSMUSG00000030549.5 |
| chr7 | 79588647 | 79588769 | 3.64281 | 4.40074 | 1.38709 | Distal Intergenic | 7 | 79593369 | 79594924 | 1556 | 2 | ENSMUST00000172788.1 | 6155 | Rhcg | ENSMUSG00000030549.5 |
| chr7 | 79590413 | 79592309 | 4.09438 | 5.30748 | 2.01739 | Downstream (1-2kb) | 7 | 79593369 | 79594924 | 1556 | 2 | ENSMUST00000172788.1 | 2615 | Rhcg | ENSMUSG00000030549.5 |
| chr7 | 79602224 | 79602541 | 3.79537 | 4.67706 | 1.53666 | Intron (ENSMUST00000032766.4/ENSMUSG00000030549.5, intron 3 of 10) | 7 | 79603766 | 79607874 | 4109 | 2 | ENSMUST00000161084.1 | 5333 | Rhcg | ENSMUSG00000030549.5 |
| chr7 | 79604636 | 79604852 | 3.38732 | 3.91691 | 1.04619 | Intron (ENSMUST00000032766.4/ENSMUSG00000030549.5, intron 2 of 10) | 7 | 79603766 | 79607874 | 4109 | 2 | ENSMUST00000161084.1 | 3022 | Rhcg | ENSMUSG00000030549.5 |
| chr7 | 79606878 | 79606953 | 3.39 | 4.15304 | 1.2308 | Promoter | 7 | 79603766 | 79607874 | 4109 | 2 | ENSMUST00000161084.1 | 921 | Rhcg | ENSMUSG00000030549.5 |
| chr7 | 79613377 | 79613559 | 4.17879 | 5.41402 | 2.0281 | Intron (ENSMUST00000032766.4/ENSMUSG00000030549.5, intron 1 of 10) | 7 | 79593363 | 79617657 | 24295 | 2 | ENSMUST00000032766.4 | 4098 | Rhcg | ENSMUSG00000030549.5 |
| chr7 | 79616409 | 79616919 | 3.48771 | 4.18119 | 1.24445 | Promoter | 7 | 79593363 | 79617657 | 24295 | 2 | ENSMUST00000032766.4 | 738 | Rhcg | ENSMUSG00000030549.5 |
| chr7 | 79631815 | 79632129 | 3.79073 | 4.65816 | 1.49534 | Distal Intergenic | 7 | 79593363 | 79617657 | 24295 | 2 | ENSMUST00000032766.4 | -14159 | Rhcg | ENSMUSG00000030549.5 |
| chr7 | 70325971 | 70326179 | 3.54061 | 4.26761 | 1.32807 | Distal Intergenic | 7 | 70351946 | 70360593 | 8648 | 2 | ENSMUST00000032768.8 | 34414 | Nr2f2 | ENSMUSG00000030551.8 |
| chr7 | 67761616 | 67761870 | 3.39482 | 3.96913 | 1.15561 | Distal Intergenic | 7 | 67730161 | 67759742 | 29582 | 2 | ENSMUST00000074233.5 | -1875 | Synm | ENSMUSG00000030554.10 |
| chr7 | 67736549 | 67736806 | 3.48242 | 4.22267 | 1.26723 | Exon (ENSMUST00000051389.8/ENSMUSG00000030554.10, exon 4 of 5) | 7 | 67725261 | 67726014 | 754 | 1 | ENSMUST00000154697.1 | 11289 | Ttc23 | ENSMUSG00000030555.10 |
| chr7 | 67737358 | 67737799 | 3.29625 | 3.79292 | 1.08451 | Intron (ENSMUST00000051389.8/ENSMUSG00000030554.10, intron 3 of 4) | 7 | 67725261 | 67726014 | 754 | 1 | ENSMUST00000154697.1 | 12098 | Ttc23 | ENSMUSG00000030555.10 |
| chr7 | 87246596 | 87246948 | 3.48441 | 4.13129 | 1.27002 | Promoter | 7 | 87246649 | 87287981 | 41333 | 1 | ENSMUST00000124269.2 | 0 | Nox4 | ENSMUSG00000030562.11 |
| chr7 | 30873510 | 30874243 | 3.52036 | 4.17555 | 1.24176 | Promoter | 7 | 30870492 | 30873419 | 2928 | 2 | ENSMUST00000190455.1 | -92 | Cd22 | ENSMUSG00000030577.8 |
| chr7 | 29304179 | 29305300 | 4.17099 | 5.63037 | 2.22287 | Promoter | 7 | 29303976 | 29317588 | 13613 | 1 | ENSMUST00000049977.7 | 204 | Dpf1 | ENSMUSG00000030584.8 |
| chr7 | 29317587 | 29318361 | 3.56597 | 4.34432 | 1.36252 | 3' UTR | 7 | 29314143 | 29316563 | 2421 | 1 | ENSMUST00000128145.1 | 3445 | Dpf1 | ENSMUSG00000030584.8 |
| chr7 | 29318763 | 29319201 | 3.21042 | 3.82138 | 1.07426 | Downstream (1-2kb) | 7 | 29314143 | 29316563 | 2421 | 1 | ENSMUST00000128145.1 | 4621 | Dpf1 | ENSMUSG00000030584.8 |
| chr7 | 29135490 | 29135752 | 3.58016 | 4.32353 | 1.35608 | Promoter | 7 | 29135073 | 29151499 | 16427 | 1 | ENSMUST00000159351.1 | 418 | Rasgrp4 | ENSMUSG00000030589.10 |
| chr7 | 29158630 | 29158986 | 3.47882 | 4.08054 | 1.13598 | 3' UTR | 7 | 29153393 | 29156234 | 2842 | 2 | ENSMUST00000136256.2 | -2397 | Fam98c | ENSMUSG00000030590.9 |
| chr7 | 29119767 | 29119850 | 3.44392 | 4.11992 | 1.2533 | Intron (ENSMUST00000179893.2/ENSMUSG00000030592.11, intron 1 of 105) | 7 | 29003340 | 29125151 | 121812 | 2 | ENSMUST00000179893.2 | 5301 | Ryr1 | ENSMUSG00000030592.11 |
| chr7 | 29124882 | 29125273 | 4.18458 | 5.4784 | 2.12256 | Promoter | 7 | 29003340 | 29125151 | 121812 | 2 | ENSMUST00000179893.2 | 0 | Ryr1 | ENSMUSG00000030592.11 |
| chr7 | 28451726 | 28452313 | 3.36954 | 3.88193 | 1.01733 | Promoter | 7 | 28451980 | 28459285 | 7306 | 1 | ENSMUST00000190954.1 | 0 | Lrfn1 | ENSMUSG00000030600.10 |
| chr7 | 28467737 | 28468015 | 3.81325 | 4.89677 | 1.70396 | 3' UTR | 7 | 28458658 | 28468242 | 9585 | 1 | ENSMUST00000108288.4 | 9080 | Lrfn1 | ENSMUSG00000030600.10 |
| chr7 | 28468808 | 28468923 | 3.87197 | 4.8794 | 1.71501 | Intron (ENSMUST00000040531.8/ENSMUSG00000030602.7, intron 1 of 12) | 7 | 28458658 | 28468242 | 9585 | 1 | ENSMUST00000108288.4 | 10151 | Lrfn1 | ENSMUSG00000030600.10 |
| chr7 | 79052637 | 79053693 | 4.27656 | 5.72186 | 2.35715 | Promoter | 7 | 79053483 | 79115099 | 61617 | 1 | ENSMUST00000032835.5 | 0 | Acan | ENSMUSG00000030607.6 |
| chr7 | 79054173 | 79054579 | 3.78808 | 5.03463 | 1.80224 | Promoter | 7 | 79053483 | 79115099 | 61617 | 1 | ENSMUST00000032835.5 | 691 | Acan | ENSMUSG00000030607.6 |
| chr7 | 89497099 | 89497407 | 3.23338 | 3.67724 | 1.01733 | Distal Intergenic | 7 | 89510007 | 89511368 | 1362 | 1 | ENSMUST00000179361.1 | -12600 | Prss23os | ENSMUSG00000030623.3 |
| chr7 | 89497866 | 89497983 | 3.3653 | 3.87551 | 1.01733 | Distal Intergenic | 7 | 89510007 | 89511368 | 1362 | 1 | ENSMUST00000179361.1 | -12024 | Prss23os | ENSMUSG00000030623.3 |
| chr7 | 82175478 | 82175952 | 3.80432 | 4.97429 | 1.7703 | Promoter | 7 | 82175234 | 82176292 | 1059 | 1 | ENSMUST00000179150.1 | 245 | Sh3gl3 | ENSMUSG00000030638.8 |
| chr7 | 82206358 | 82206720 | 3.38288 | 3.91915 | 1.07761 | Intron (ENSMUST00000179318.2/ENSMUSG00000030638.8, intron 1 of 8) | 7 | 82175234 | 82176292 | 1059 | 1 | ENSMUST00000179150.1 | 31125 | Sh3gl3 | ENSMUSG00000030638.8 |
| chr7 | 118597254 | 118597528 | 4.07605 | 5.50348 | 2.15135 | Promoter | 7 | 118597297 | 118675085 | 77789 | 1 | ENSMUST00000098088.5 | 0 | Tmc5 | ENSMUSG00000030650.12 |
| chr7 | 118597869 | 118598809 | 3.7588 | 4.61959 | 1.50542 | Promoter | 7 | 118597297 | 118675086 | 77790 | 1 | ENSMUST00000121715.2 | 573 | Tmc5 | ENSMUSG00000030650.12 |
| chr7 | 101420457 | 101420782 | 3.36521 | 3.87538 | 1.01733 | Promoter | 7 | 101421691 | 101512829 | 91139 | 1 | ENSMUST00000084894.8 | -909 | Pde2a | ENSMUSG00000030653.11 |
| chr7 | 101421687 | 101422339 | 3.45593 | 4.04823 | 1.14474 | Promoter | 7 | 101421691 | 101512829 | 91139 | 1 | ENSMUST00000084894.8 | 0 | Pde2a | ENSMUSG00000030653.11 |
| chr7 | 101425570 | 101426018 | 3.69677 | 4.50663 | 1.46374 | Intron (ENSMUST00000084894.8/ENSMUSG00000030653.11, intron 1 of 30) | 7 | 101421709 | 101512814 | 91106 | 1 | ENSMUST00000166652.1 | 3862 | Pde2a | ENSMUSG00000030653.11 |
| chr7 | 116035146 | 116035501 | 3.5125 | 4.24402 | 1.29325 | Intron (ENSMUST00000166877.2/ENSMUSG00000051910.8, intron 1 of 15) | 7 | 116031979 | 116032550 | 572 | 1 | ENSMUST00000032900.3 | 3168 | Sox6os | ENSMUSG00000030664.3 |
| chr7 | 114716663 | 114716896 | 3.32444 | 3.82285 | 1.04874 | Distal Intergenic | 7 | 114718643 | 114723365 | 4723 | 1 | ENSMUST00000032902.5 | -1747 | Calcb | ENSMUSG00000030666.6 |
| chr7 | 114718867 | 114719022 | 3.53198 | 4.2439 | 1.34279 | Promoter | 7 | 114718675 | 114723063 | 4389 | 1 | ENSMUST00000182816.1 | 193 | Calcb | ENSMUSG00000030666.6 |
| chr7 | 114719486 | 114719664 | 3.47538 | 4.1307 | 1.27684 | Promoter | 7 | 114719511 | 114720139 | 629 | 1 | ENSMUST00000182044.1 | 0 | Calcb | ENSMUSG00000030666.6 |
| chr7 | 127121967 | 127122467 | 3.37147 | 3.91728 | 1.12209 | Promoter | 7 | 127107114 | 127122226 | 15113 | 2 | ENSMUST00000032912.5 | 0 | Qprt | ENSMUSG00000030674.5 |
| chr7 | 126950139 | 126951147 | 3.63463 | 4.42659 | 1.39131 | Promoter | 7 | 126950563 | 126970606 | 20044 | 1 | ENSMUST00000106335.2 | 0 | Sez6l2 | ENSMUSG00000030683.6 |
| chr7 | 126951922 | 126952205 | 4.2284 | 5.53305 | 2.11183 | Promoter | 7 | 126950966 | 126970270 | 19305 | 1 | ENSMUST00000106332.1 | 957 | Sez6l2 | ENSMUSG00000030683.6 |
| chr7 | 126963307 | 126963387 | 3.92227 | 4.9288 | 1.71947 | Exon (ENSMUST00000106335.2/ENSMUSG00000030683.6, exon 11 of 17) | 7 | 126967743 | 126970270 | 2528 | 1 | ENSMUST00000134471.1 | -4356 | Sez6l2 | ENSMUSG00000030683.6 |
| chr7 | 43782814 | 43783786 | 3.92098 | 4.98538 | 1.78019 | Exon (ENSMUST00000014058.9/ENSMUSG00000030693.9, exon 3 of 6) | 7 | 43781054 | 43785410 | 4357 | 1 | ENSMUST00000014058.9 | 1761 | Klk10 | ENSMUSG00000030693.9 |
| chr7 | 126703495 | 126703581 | 3.83659 | 4.83233 | 1.68698 | Promoter | 7 | 126699881 | 126703173 | 3293 | 2 | ENSMUST00000173108.2 | -323 | Coro1a | ENSMUSG00000030707.10 |
| chr7 | 126704123 | 126704482 | 3.53134 | 4.1849 | 1.22535 | Promoter | 7 | 126699773 | 126704220 | 4448 | 2 | ENSMUST00000106364.2 | 0 | Coro1a | ENSMUSG00000030707.10 |
| chr7 | 100205247 | 100205440 | 3.34153 | 3.83966 | 1.01733 | Distal Intergenic | 7 | 100227394 | 100275358 | 47965 | 1 | ENSMUST00000084935.4 | -21954 | Pgm2l1 | ENSMUSG00000030729.11 |
| chr7 | 100216776 | 100217098 | 3.35295 | 3.8621 | 1.03574 | Distal Intergenic | 7 | 100227394 | 100275358 | 47965 | 1 | ENSMUST00000084935.4 | -10296 | Pgm2l1 | ENSMUSG00000030729.11 |
| chr7 | 100004928 | 100006212 | 3.85008 | 4.99129 | 1.83006 | Promoter | 7 | 100006404 | 100034726 | 28323 | 1 | ENSMUST00000032977.5 | -192 | Chrdl2 | ENSMUSG00000030732.7 |
| chr7 | 100007185 | 100007649 | 3.58138 | 4.36211 | 1.40959 | Promoter | 7 | 100006404 | 100034726 | 28323 | 1 | ENSMUST00000032977.5 | 782 | Chrdl2 | ENSMUSG00000030732.7 |
| chr7 | 44602756 | 44602936 | 3.35292 | 3.86342 | 1.04054 | 3' UTR | 7 | 44605805 | 44614125 | 8321 | 2 | ENSMUST00000125870.1 | 11189 | Myh14 | ENSMUSG00000030739.12 |
| chr7 | 44603244 | 44604633 | 4.12403 | 5.44986 | 2.11316 | 3' UTR | 7 | 44605805 | 44614125 | 8321 | 2 | ENSMUST00000125870.1 | 9492 | Myh14 | ENSMUSG00000030739.12 |
| chr7 | 44606300 | 44606611 | 4.10851 | 5.28463 | 1.94178 | Exon (ENSMUST00000107899.3/ENSMUSG00000030739.12, exon 40 of 40) | 7 | 44605805 | 44614125 | 8321 | 2 | ENSMUST00000125870.1 | 7514 | Myh14 | ENSMUSG00000030739.12 |
| chr7 | 44607950 | 44608137 | 3.35002 | 3.88284 | 1.12184 | Intron (ENSMUST00000107899.3/ENSMUSG00000030739.12, intron 38 of 39) | 7 | 44605805 | 44614125 | 8321 | 2 | ENSMUST00000125870.1 | 5988 | Myh14 | ENSMUSG00000030739.12 |
| chr7 | 44615192 | 44615772 | 3.39671 | 3.94135 | 1.08271 | Exon (ENSMUST00000107899.3/ENSMUSG00000030739.12, exon 33 of 40) | 7 | 44605805 | 44614125 | 8321 | 2 | ENSMUST00000125870.1 | -1068 | Myh14 | ENSMUSG00000030739.12 |
| chr7 | 44638239 | 44638570 | 3.57432 | 4.25622 | 1.24178 | Exon (ENSMUST00000107899.3/ENSMUSG00000030739.12, exon 13 of 40) | 7 | 44629966 | 44632785 | 2820 | 2 | ENSMUST00000139186.1 | -5455 | Myh14 | ENSMUSG00000030739.12 |
| chr7 | 44640244 | 44640853 | 3.66527 | 4.44971 | 1.41539 | Intron (ENSMUST00000107899.3/ENSMUSG00000030739.12, intron 10 of 39) | 7 | 44629966 | 44632785 | 2820 | 2 | ENSMUST00000139186.1 | -7460 | Myh14 | ENSMUSG00000030739.12 |
| chr7 | 44641644 | 44641745 | 3.40938 | 3.98415 | 1.16611 | Intron (ENSMUST00000107899.3/ENSMUSG00000030739.12, intron 9 of 39) | 7 | 44629966 | 44632785 | 2820 | 2 | ENSMUST00000139186.1 | -8860 | Myh14 | ENSMUSG00000030739.12 |
| chr7 | 44645070 | 44645223 | 3.64871 | 4.48825 | 1.45314 | Intron (ENSMUST00000107899.3/ENSMUSG00000030739.12, intron 8 of 39) | 7 | 44629966 | 44632785 | 2820 | 2 | ENSMUST00000139186.1 | -12286 | Myh14 | ENSMUSG00000030739.12 |
| chr7 | 44646166 | 44646515 | 3.45176 | 4.39116 | 1.39812 | Intron (ENSMUST00000107899.3/ENSMUSG00000030739.12, intron 8 of 39) | 7 | 44629966 | 44632785 | 2820 | 2 | ENSMUST00000139186.1 | -13382 | Myh14 | ENSMUSG00000030739.12 |
| chr7 | 44647721 | 44647877 | 3.60634 | 4.36957 | 1.40229 | Intron (ENSMUST00000107899.3/ENSMUSG00000030739.12, intron 8 of 39) | 7 | 44629966 | 44632785 | 2820 | 2 | ENSMUST00000139186.1 | -14937 | Myh14 | ENSMUSG00000030739.12 |
| chr7 | 125581396 | 125581485 | 4.22093 | 5.62884 | 2.23078 | Distal Intergenic | 7 | 125603429 | 125633570 | 30142 | 1 | ENSMUST00000033000.6 | -21944 | Il21r | ENSMUSG00000030745.8 |
| chr7 | 98755020 | 98755336 | 3.47446 | 4.07198 | 1.12908 | Distal Intergenic | 7 | 98709877 | 98715257 | 5381 | 1 | ENSMUST00000146473.1 | 45144 | Prkrir | ENSMUSG00000030753.10 |
| chr7 | 98756763 | 98757279 | 3.42794 | 4.01493 | 1.17637 | Distal Intergenic | 7 | 98709877 | 98715257 | 5381 | 1 | ENSMUST00000146473.1 | 46887 | Prkrir | ENSMUSG00000030753.10 |
| chr7 | 98758020 | 98758245 | 3.31973 | 3.81063 | 1.03051 | Distal Intergenic | 7 | 98709877 | 98715257 | 5381 | 1 | ENSMUST00000146473.1 | 48144 | Prkrir | ENSMUSG00000030753.10 |
| chr7 | 98763794 | 98764057 | 3.35121 | 3.86747 | 1.06017 | Distal Intergenic | 7 | 98709877 | 98715257 | 5381 | 1 | ENSMUST00000146473.1 | 53918 | Prkrir | ENSMUSG00000030753.10 |
| chr7 | 123499693 | 123499804 | 4.13708 | 5.38772 | 2.07579 | Promoter | 7 | 123479516 | 123500449 | 20934 | 2 | ENSMUST00000042470.8 | 645 | Zkscan2 | ENSMUSG00000030757.8 |
| chr7 | 123500406 | 123500514 | 3.85859 | 4.86097 | 1.68344 | Promoter | 7 | 123479516 | 123500449 | 20934 | 2 | ENSMUST00000042470.8 | 0 | Zkscan2 | ENSMUSG00000030757.8 |
| chr7 | 98060013 | 98060111 | 15.29837 | 36.60757 | 30.86012 | Intron (ENSMUST00000107128.2/ENSMUSG00000030761.10, intron 39 of 48) | 7 | 98062643 | 98065202 | 2560 | 2 | ENSMUST00000124787.1 | 5091 | Myo7a | ENSMUSG00000030761.10 |
| chr7 | 98111377 | 98111589 | 4.04771 | 5.2006 | 1.85712 | Intron (ENSMUST00000107128.2/ENSMUSG00000030761.10, intron 2 of 48) | 7 | 98097613 | 98112662 | 15050 | 2 | ENSMUST00000153657.2 | 1073 | Myo7a | ENSMUSG00000030761.10 |
| chr1 | 183249521 | 183249596 | 3.86506 | 4.79583 | 1.58028 | Distal Intergenic | 1 | 183086264 | 183221529 | 135266 | 2 | ENSMUST00000003035.6 | -27993 | Disp1 | ENSMUSG00000030768.7 |
| chr7 | 112158854 | 112159436 | 4.07615 | 5.37375 | 2.07313 | Promoter | 7 | 112116019 | 112159057 | 43039 | 2 | ENSMUST00000033036.5 | 0 | Dkk3 | ENSMUSG00000030772.5 |
| chr7 | 112168195 | 112168291 | 3.99523 | 5.06029 | 1.7835 | Distal Intergenic | 7 | 112116019 | 112159057 | 43039 | 2 | ENSMUST00000033036.5 | -9139 | Dkk3 | ENSMUSG00000030772.5 |
| chr7 | 110794337 | 110794487 | 3.34629 | 3.85877 | 1.05921 | Exon (ENSMUST00000106686.1/ENSMUSG00000005686.11, exon 5 of 5) | 7 | 110800432 | 110801599 | 1168 | 2 | ENSMUST00000148367.1 | 7112 | Rnf141 | ENSMUSG00000030788.10 |
| chr7 | 45211240 | 45211780 | 3.85961 | 4.96329 | 1.78822 | Promoter | 7 | 45207525 | 45211883 | 4359 | 2 | ENSMUST00000033057.7 | 103 | Dkkl1 | ENSMUSG00000030792.7 |
| chr7 | 45212251 | 45212566 | 3.52062 | 4.19971 | 1.30697 | Promoter | 7 | 45207525 | 45211883 | 4359 | 2 | ENSMUST00000033057.7 | -369 | Dkkl1 | ENSMUSG00000030792.7 |
| chr7 | 45216006 | 45216254 | 3.41448 | 4.00277 | 1.15235 | Promoter | 7 | 45215852 | 45233619 | 17768 | 1 | ENSMUST00000107801.2 | 155 | Tead2 | ENSMUSG00000030796.10 |
| chr7 | 45233850 | 45233978 | 3.88761 | 5.00555 | 1.76582 | 3' UTR | 7 | 45234350 | 45235458 | 1109 | 2 | ENSMUST00000154014.1 | 1480 | Cd37 | ENSMUSG00000030798.8 |
| chr7 | 45615925 | 45616235 | 3.3402 | 3.83766 | 1.01733 | Promoter | 7 | 45613907 | 45615490 | 1584 | 2 | ENSMUST00000033099.4 | -436 | Fgf21 | ENSMUSG00000030827.4 |
| chr7 | 128416238 | 128417030 | 3.62148 | 4.40089 | 1.38744 | Intron (ENSMUST00000033133.6/ENSMUSG00000030844.6, intron 1 of 4) | 7 | 128402910 | 128418163 | 15254 | 2 | ENSMUST00000145739.1 | 1133 | Rgs10 | ENSMUSG00000030844.6 |
| chr7 | 128417346 | 128417950 | 3.46876 | 4.16538 | 1.27323 | Promoter | 7 | 128402910 | 128418163 | 15254 | 2 | ENSMUST00000145739.1 | 213 | Rgs10 | ENSMUSG00000030844.6 |
| chr7 | 128422679 | 128422772 | 3.32819 | 3.81959 | 1.01733 | Distal Intergenic | 7 | 128403181 | 128418758 | 15578 | 2 | ENSMUST00000147840.1 | -3922 | Rgs10 | ENSMUSG00000030844.6 |
| chr7 | 130290815 | 130291399 | 3.38263 | 3.93047 | 1.11364 | Intron (ENSMUST00000124096.2/ENSMUSG00000030849.12, intron 2 of 6) | 7 | 130163709 | 130266263 | 102555 | 2 | ENSMUST00000120187.2 | -24553 | Fgfr2 | ENSMUSG00000030849.12 |

| | | | | | | | | | | | | | | | |
|---|---|---|---|---|---|---|---|---|---|---|---|---|---|---|---|
| chr7 | 47131694 | 47133140 | 4.01455 | 5.14666 | 1.88209 | Promoter | 7 | 47077798 | 47132326 | 54529 | 2 | ENSMUST00000126370.2 | 0 | Ptpn5 | ENSMUSG00000030854.10 |
| chr7 | 47133620 | 47134007 | 3.65191 | 4.43714 | 1.39036 | Promoter | 7 | 47077802 | 47133684 | 55883 | 2 | ENSMUST00000102626.2 | 0 | Ptpn5 | ENSMUSG00000030854.10 |
| chr7 | 132153629 | 132155216 | 3.89372 | 5.01051 | 1.82905 | Promoter | 7 | 132042816 | 132154739 | 111924 | 2 | ENSMUST00000033149.4 | 0 | Cpxm2 | ENSMUSG00000030862.7 |
| chr7 | 122219269 | 122219382 | 3.64257 | 4.38233 | 1.31856 | Promoter | 7 | 122219496 | 122222824 | 3329 | 1 | ENSMUST00000033152.4 | -114 | Chp2 | ENSMUSG00000030865.4 |
| chr7 | 122220528 | 122220749 | 3.70731 | 4.50259 | 1.40974 | Promoter | 7 | 122221573 | 122222183 | 611 | 1 | ENSMUST00000160828.1 | -824 | Chp2 | ENSMUSG00000030865.4 |
| chr7 | 121864767 | 121865183 | 3.4401 | 4.04147 | 1.14121 | Promoter | 7 | 121865111 | 121918515 | 53405 | 1 | ENSMUST00000033161.5 | 0 | Scnn1b | ENSMUSG00000030873.8 |
| chr7 | 121899065 | 121899289 | 3.73159 | 4.54838 | 1.42848 | 5' UTR | 7 | 121865111 | 121918515 | 53405 | 1 | ENSMUST00000033161.5 | 33955 | Scnn1b | ENSMUSG00000030873.8 |
| chr7 | 120809771 | 120810050 | 3.80745 | 4.7096 | 1.57726 | Distal Intergenic | 7 | 120786481 | 120792762 | 6282 | 1 | ENSMUST00000166083.1 | 23291 | Vwa3a | ENSMUSG00000030889.8 |
| chr7 | 105425134 | 105425501 | 3.56825 | 4.25534 | 1.27515 | Promoter | 7 | 105425817 | 105470898 | 45082 | 1 | ENSMUST00000181339.2 | -316 | Cckbr | ENSMUSG00000030898.9 |
| chr7 | 105425921 | 105426297 | 3.61823 | 4.34282 | 1.30413 | Promoter | 7 | 105425820 | 105436339 | 10520 | 1 | ENSMUST00000033189.4 | 102 | Cckbr | ENSMUSG00000030898.9 |
| chr7 | 133205416 | 133205625 | 3.55984 | 4.28701 | 1.33995 | Distal Intergenic | 7 | 133025702 | 133124354 | 98653 | 2 | ENSMUST00000171882.2 | -81063 | Ctbp2 | ENSMUSG00000030970.11 |
| chr7 | 133245374 | 133245690 | 3.27242 | 3.73577 | 1.01733 | Distal Intergenic | 7 | 133025702 | 133124354 | 98653 | 2 | ENSMUST00000171882.2 | -121021 | Ctbp2 | ENSMUSG00000030970.11 |
| chr7 | 133324104 | 133324286 | 3.35769 | 3.86404 | 1.01733 | Distal Intergenic | 7 | 133025702 | 133124354 | 98653 | 2 | ENSMUST00000171882.2 | -199751 | Ctbp2 | ENSMUSG00000030970.11 |
| chr7 | 133605212 | 133605405 | 3.37555 | 3.89098 | 1.01733 | Distal Intergenic | 7 | 133587024 | 133602115 | 15092 | 2 | ENSMUST00000033275.2 | -3098 | Tex36 | ENSMUSG00000030976.2 |
| chr7 | 133607165 | 133607424 | 3.59955 | 4.40149 | 1.3982 | Distal Intergenic | 7 | 133587024 | 133602115 | 15092 | 2 | ENSMUST00000033275.2 | -5051 | Tex36 | ENSMUSG00000030976.2 |
| chr7 | 134290051 | 134290339 | 3.23338 | 3.67724 | 1.01733 | Intron (ENSMUST00000172947.2/ENSMUSG00000030994.10, intron 6 of 8) | 7 | 134266275 | 134341986 | 75712 | 2 | ENSMUST00000106129.3 | 51647 | D7Ertd443e | ENSMUSG00000030994.10 |
| chr7 | 135715916 | 135716123 | 3.449 | 4.33222 | 1.35192 | Promoter | 7 | 135689788 | 135716379 | 26592 | 2 | ENSMUST00000033310.7 | 256 | Mki67 | ENSMUSG00000031004.7 |
| chr7 | 109493416 | 109493572 | 3.46161 | 4.25619 | 1.30059 | Promoter | 7 | 109449006 | 109493738 | 44733 | 2 | ENSMUST00000106739.2 | 166 | Trim66 | ENSMUSG00000031026.9 |
| chr7 | 109438256 | 109439232 | 3.55489 | 4.7535 | 1.62862 | Promoter | 7 | 109279223 | 109438817 | 159595 | 2 | ENSMUST00000121748.2 | 0 | Stk33 | ENSMUSG00000031027.10 |
| chr7 | 109439635 | 109440387 | 3.33572 | 3.85368 | 1.05987 | Promoter | 7 | 109279223 | 109439081 | 159859 | 2 | ENSMUST00000141210.1 | -555 | Stk33 | ENSMUSG00000031027.10 |
| chr7 | 109441091 | 109441207 | 3.23338 | 3.67724 | 1.01733 | Intron (ENSMUST00000143715.1/ENSMUSG00000085407.1, intron 1 of 1) | 7 | 109279223 | 109439081 | 159859 | 2 | ENSMUST00000141210.1 | -2011 | Stk33 | ENSMUSG00000031027.10 |
| chr7 | 109010485 | 109011393 | 3.80255 | 5.01929 | 1.81592 | Promoter | 7 | 109010825 | 109034460 | 23636 | 1 | ENSMUST00000033341.6 | 0 | Tub | ENSMUSG00000031028.8 |
| chr7 | 109011963 | 109012133 | 3.62815 | 4.59024 | 1.49146 | Intron (ENSMUST00000033341.6/ENSMUSG00000031028.8, intron 1 of 11) | 7 | 109010825 | 109034460 | 23636 | 1 | ENSMUST00000033341.6 | 1139 | Tub | ENSMUSG00000031028.8 |
| chr7 | 108950358 | 108950884 | 3.69102 | 4.71581 | 1.57762 | Exon (ENSMUST00000180590.1/ENSMUSG00000097220.1, exon 2 of 2) | 7 | 108940308 | 108941008 | 701 | 1 | ENSMUST00000147369.1 | 10051 | Eif3f | ENSMUSG00000031029.6 |
| chr7 | 137475114 | 137475335 | 3.27668 | 3.80392 | 1.07207 | Distal Intergenic | 7 | 137437648 | 137468594 | 30947 | 1 | ENSMUST00000064404.6 | 37467 | Glrx3 | ENSMUSG00000031068.7 |
| chr7 | 137475798 | 137475905 | 3.44304 | 4.23159 | 1.28018 | Distal Intergenic | 7 | 137437648 | 137468594 | 30947 | 1 | ENSMUST00000064404.6 | 38151 | Glrx3 | ENSMUSG00000031068.7 |
| chr7 | 144888273 | 144888514 | 3.3449 | 3.84474 | 1.01733 | Distal Intergenic | 7 | 144896532 | 144900953 | 4422 | 1 | ENSMUST00000033389.5 | -8018 | Fgf15 | ENSMUSG00000031073.5 |
| chr7 | 144894290 | 144894667 | 4.08574 | 5.39421 | 2.04075 | Exon (ENSMUST00000180930.1/ENSMUSG00000096999.1, exon 2 of 2) | 7 | 144896532 | 144900953 | 4422 | 1 | ENSMUST00000033389.5 | -1865 | Fgf15 | ENSMUSG00000031073.5 |
| chr7 | 144895077 | 144895737 | 3.8789 | 4.95746 | 1.77999 | Promoter | 7 | 144896532 | 144900953 | 4422 | 1 | ENSMUST00000033389.5 | -795 | Fgf15 | ENSMUSG00000031073.5 |
| chr7 | 144896188 | 144896505 | 3.57406 | 4.46629 | 1.40317 | Promoter | 7 | 144896532 | 144900953 | 4422 | 1 | ENSMUST00000033389.5 | -27 | Fgf15 | ENSMUSG00000031073.5 |
| chr7 | 144828494 | 144828677 | 3.44969 | 4.27356 | 1.31341 | Distal Intergenic | 7 | 144838083 | 144842719 | 4637 | 1 | ENSMUST00000155320.2 | -9406 | Fgf3 | ENSMUSG00000031074.9 |
| chr7 | 144838310 | 144838648 | 3.42619 | 4.03026 | 1.21586 | Promoter | 7 | 144838595 | 144844436 | 5842 | 1 | ENSMUST00000105898.1 | 0 | Fgf3 | ENSMUSG00000031074.9 |
| chr7 | 144839545 | 144839859 | 3.53327 | 4.53819 | 1.47476 | Promoter | 7 | 144838595 | 144844436 | 5842 | 1 | ENSMUST00000105898.1 | 951 | Fgf3 | ENSMUSG00000031074.9 |
| chr7 | 144840211 | 144840473 | 3.29594 | 4.14767 | 1.20785 | Intron (ENSMUST00000155320.2/ENSMUSG00000031074.9, intron 2 of 3) | 7 | 144838595 | 144844436 | 5842 | 1 | ENSMUST00000105898.1 | 1617 | Fgf3 | ENSMUSG00000031074.9 |
| chr7 | 144622751 | 144623077 | 3.30973 | 3.79682 | 1.03484 | Intron (ENSMUST00000121758.2/ENSMUSG00000031075.12, intron 12 of 25) | 7 | 144595442 | 144608295 | 12854 | 2 | ENSMUST00000131571.1 | -14457 | Ano1 | ENSMUSG00000031075.12 |
| chr7 | 144678356 | 144678906 | 3.99107 | 5.10222 | 1.82382 | Promoter | 7 | 144655601 | 144678885 | 23285 | 2 | ENSMUST00000131731.1 | 0 | Ano1 | ENSMUSG00000031075.12 |
| chr7 | 144738054 | 144738748 | 3.82883 | 4.74997 | 1.59546 | Promoter | 7 | 144588549 | 144738543 | 149995 | 2 | ENSMUST00000121758.2 | 0 | Ano1 | ENSMUSG00000031075.12 |
| chr8 | 12872348 | 12872525 | 3.44631 | 4.21347 | 1.27913 | Intron (ENSMUST00000145372.1/ENSMUSG00000085465.1, intron 1 of 2) | 8 | 12873806 | 12880146 | 6341 | 1 | ENSMUST00000156560.1 | -1281 | Mcf2l | ENSMUSG00000031442.15 |
| chr8 | 13248129 | 13248585 | 3.35529 | 3.88192 | 1.06137 | Exon (ENSMUST00000033825.5/ENSMUSG00000031448.7, exon 2 of 7) | 8 | 13242262 | 13243085 | 824 | 2 | ENSMUST00000166438.1 | -5045 | Adprhl1 | ENSMUSG00000031448.7 |
| chr8 | 13493708 | 13494165 | 3.50088 | 4.19374 | 1.30929 | Promoter | 8 | 13465374 | 13494535 | 29162 | 2 | ENSMUST00000033828.5 | 370 | Gas6 | ENSMUSG00000031451.5 |
| chr8 | 13678481 | 13678647 | 3.45866 | 4.04877 | 1.13174 | Promoter | 8 | 13566950 | 13677603 | 110654 | 2 | ENSMUST00000117551.2 | -879 | Rasa3 | ENSMUSG00000031453.10 |
| chr8 | 18739067 | 18739463 | 4.82936 | 7.16291 | 3.41437 | Intron (ENSMUST00000039412.9/ENSMUSG00000039842.10, intron 12 of 13) | 8 | 18690263 | 18741562 | 51300 | 2 | ENSMUST00000033846.6 | 2099 | Angpt2 | ENSMUSG00000031465.6 |
| chr8 | 18740209 | 18740369 | 3.4383 | 4.20513 | 1.25403 | Intron (ENSMUST00000039412.9/ENSMUSG00000039842.10, intron 12 of 13) | 8 | 18690263 | 18741562 | 51300 | 2 | ENSMUST00000033846.6 | 1193 | Angpt2 | ENSMUSG00000031465.6 |
| chr8 | 18740833 | 18740994 | 3.34084 | 4.27294 | 1.2938 | Promoter | 8 | 18690263 | 18741562 | 51300 | 2 | ENSMUST00000033846.6 | 568 | Angpt2 | ENSMUSG00000031465.6 |
| chr8 | 18741322 | 18741681 | 3.90219 | 5.23928 | 1.91941 | Promoter | 8 | 18690263 | 18741562 | 51300 | 2 | ENSMUST00000033846.6 | 0 | Angpt2 | ENSMUSG00000031465.6 |
| chr8 | 27226140 | 27226954 | 3.68572 | 4.49713 | 1.41041 | 3' UTR | 8 | 27226799 | 27228494 | 1696 | 2 | ENSMUST00000117565.1 | 1540 | Adrb3 | ENSMUSG00000031489.9 |
| chr8 | 27227279 | 27228505 | 3.85057 | 5.001 | 1.79471 | Promoter | 8 | 27226799 | 27228494 | 1696 | 2 | ENSMUST00000117565.1 | 0 | Adrb3 | ENSMUSG00000031489.9 |
| chr8 | 27228884 | 27229128 | 3.51896 | 4.18593 | 1.28098 | Promoter | 8 | 27226001 | 27228583 | 2583 | 2 | ENSMUST00000121838.1 | -302 | Adrb3 | ENSMUSG00000031489.9 |
| chr8 | 27281521 | 27281685 | 3.8509 | 4.84553 | 1.71074 | Distal Intergenic | 8 | 27260327 | 27275656 | 15330 | 1 | ENSMUST00000033880.5 | 21195 | Eif4ebp1 | ENSMUSG00000031490.5 |
| chr8 | 27282114 | 27282292 | 3.65814 | 4.47341 | 1.46761 | Distal Intergenic | 8 | 27260327 | 27275656 | 15330 | 1 | ENSMUST00000033880.5 | 21788 | Eif4ebp1 | ENSMUSG00000031490.5 |
| chr7 | 29171203 | 29171523 | 3.41762 | 4.12968 | 1.17416 | Promoter | 7 | 29170390 | 29173912 | 3523 | 1 | ENSMUST00000033886.7 | 814 | Ggn | ENSMUSG00000031493.9 |
| chr7 | 29171958 | 29172471 | 3.41897 | 3.98175 | 1.10202 | Exon (ENSMUST00000098609.2/ENSMUSG00000031493.9, exon 2 of 3) | 7 | 29170390 | 29173912 | 3523 | 1 | ENSMUST00000033886.7 | 1569 | Ggn | ENSMUSG00000031493.9 |
| chr8 | 3754408 | 3754483 | 3.51031 | 12.83576 | 8.24823 | Distal Intergenic | 8 | 3743395 | 3748984 | 5590 | 2 | ENSMUST00000012847.1 | -5425 | Cd209a | ENSMUSG00000031494.6 |
| chr8 | 11217426 | 11217542 | 3.56845 | 4.32881 | 1.35614 | Promoter | 8 | 11212401 | 11216801 | 4401 | 2 | ENSMUST00000130488.1 | -626 | Col4a1 | ENSMUSG00000031502.9 |
| chr8 | 11219688 | 11219763 | 3.7473 | 4.84971 | 1.68577 | Intron (ENSMUST00000033898.9/ENSMUSG00000031502.9, intron 33 of 51) | 8 | 11212401 | 11216801 | 4401 | 2 | ENSMUST00000130488.1 | -2888 | Col4a1 | ENSMUSG00000031502.9 |
| chr8 | 11226377 | 11226495 | 3.30193 | 3.78007 | 1.01733 | Exon (ENSMUST00000033898.9/ENSMUSG00000031502.9, exon 26 of 52) | 8 | 11212401 | 11216801 | 4401 | 2 | ENSMUST00000130488.1 | -9577 | Col4a1 | ENSMUSG00000031502.9 |
| chr8 | 11230062 | 11230303 | 3.48745 | 4.12554 | 1.25174 | Intron (ENSMUST00000033898.9/ENSMUSG00000031502.9, intron 24 of 51) | 8 | 11212401 | 11216801 | 4401 | 2 | ENSMUST00000130488.1 | -13262 | Col4a1 | ENSMUSG00000031502.9 |
| chr8 | 11312518 | 11313047 | 4.20031 | 5.50549 | 2.1544 | Promoter | 8 | 11198423 | 11312826 | 114404 | 2 | ENSMUST00000033898.9 | 0 | Col4a1 | ENSMUSG00000031502.9 |
| chr8 | 11699873 | 11700315 | 3.44149 | 4.20603 | 1.27736 | Promoter | 8 | 11655479 | 11701063 | 45585 | 2 | ENSMUST00000190741.1 | 748 | 1700016D06Rik | ENSMUSG00000031509.5 |
| chr8 | 11725390 | 11726024 | 3.30579 | 3.80057 | 1.06675 | Distal Intergenic | 8 | 11728105 | 11823061 | 94957 | 1 | ENSMUST00000033908.7 | -2081 | Arhgef7 | ENSMUSG00000031511.9 |
| chr8 | 11726521 | 11726925 | 3.425 | 4.02238 | 1.17495 | Distal Intergenic | 8 | 11728105 | 11823061 | 94957 | 1 | ENSMUST00000033908.7 | -1180 | Arhgef7 | ENSMUSG00000031511.9 |
| chr8 | 11727309 | 11728599 | 3.95761 | 5.07282 | 1.82229 | Promoter | 8 | 11728105 | 11823061 | 94957 | 1 | ENSMUST00000033908.7 | 0 | Arhgef7 | ENSMUSG00000031511.9 |
| chr8 | 11915111 | 11915418 | 3.71765 | 4.54302 | 1.47279 | Distal Intergenic | 8 | 11842061 | 11855757 | 13697 | 1 | ENSMUST00000132536.1 | 73051 | Tex29 | ENSMUSG00000031512.9 |
| chr8 | 11957373 | 11957448 | 4.30885 | 5.75783 | 2.33473 | Distal Intergenic | 8 | 11842061 | 11855757 | 13697 | 1 | ENSMUST00000132536.1 | 115313 | Tex29 | ENSMUSG00000031512.9 |
| chr8 | 12030272 | 12030806 | 3.32531 | 3.86479 | 1.11812 | Distal Intergenic | 8 | 11842061 | 11855757 | 13697 | 1 | ENSMUST00000132536.1 | 188212 | Tex29 | ENSMUSG00000031512.9 |
| chr8 | 12041555 | 12041729 | 3.57161 | 4.26143 | 1.27412 | Distal Intergenic | 8 | 11842061 | 11855757 | 13697 | 1 | ENSMUST00000132536.1 | 199495 | Tex29 | ENSMUSG00000031512.9 |
| chr8 | 34079305 | 34079478 | 3.66217 | 4.61527 | 1.49172 | Distal Intergenic | 8 | 34090607 | 34092925 | 2319 | 2 | ENSMUST00000123113.1 | 13447 | Dctn6 | ENSMUSG00000031516.6 |
| chr8 | 34079887 | 34080104 | 3.27828 | 3.97654 | 1.15964 | Distal Intergenic | 8 | 34090607 | 34092925 | 2319 | 2 | ENSMUST00000123113.1 | 12821 | Dctn6 | ENSMUSG00000031516.6 |
| chr8 | 54600638 | 54601039 | 3.73221 | 4.54965 | 1.42907 | Promoter | 8 | 54600781 | 54610098 | 9318 | 1 | ENSMUST00000033917.6 | 0 | Spata4 | ENSMUSG00000031518.6 |
| chr8 | 54077546 | 54077753 | 3.58254 | 4.39451 | 1.38514 | Promoter | 8 | 54077532 | 54186454 | 108923 | 1 | ENSMUST00000033919.4 | 15 | Vegfc | ENSMUSG00000031520.5 |
| chr8 | 34680062 | 34680166 | 3.62396 | 4.3601 | 1.3326 | Distal Intergenic | 8 | 34807297 | 34819894 | 12598 | 1 | ENSMUST00000033930.4 | -127131 | Dusp4 | ENSMUSG00000031530.6 |
| chr8 | 34792947 | 34793526 | 3.45355 | 4.13141 | 1.18803 | Distal Intergenic | 8 | 34807297 | 34819894 | 12598 | 1 | ENSMUST00000033930.4 | -13771 | Dusp4 | ENSMUSG00000031530.6 |
| chr8 | 34805226 | 34805383 | 3.82341 | 5.10634 | 1.85652 | Distal Intergenic | 8 | 34807297 | 34819894 | 12598 | 1 | ENSMUST00000033930.4 | -1914 | Dusp4 | ENSMUSG00000031530.6 |
| chr8 | 34805737 | 34805974 | 3.63973 | 4.50946 | 1.46411 | Distal Intergenic | 8 | 34807297 | 34819894 | 12598 | 1 | ENSMUST00000033930.4 | -1323 | Dusp4 | ENSMUSG00000031530.6 |
| chr8 | 34809072 | 34810026 | 3.82009 | 4.97118 | 1.75703 | Intron (ENSMUST00000033930.4/ENSMUSG00000031530.6, intron 1 of 3) | 8 | 34807297 | 34819894 | 12598 | 1 | ENSMUST00000033930.4 | 1776 | Dusp4 | ENSMUSG00000031530.6 |
| chr8 | 34814213 | 34815334 | 3.4829 | 4.10898 | 1.20985 | Intron (ENSMUST00000033930.4/ENSMUSG00000031530.6, intron 1 of 3) | 8 | 34807297 | 34819894 | 12598 | 1 | ENSMUST00000033930.4 | 6917 | Dusp4 | ENSMUSG00000031530.6 |
| chr8 | 22616842 | 22617204 | 4.21538 | 5.58166 | 2.25553 | Distal Intergenic | 8 | 22624043 | 22627536 | 3494 | 1 | ENSMUST00000033936.6 | -6839 | Dkk4 | ENSMUSG00000031535.6 |
| chr8 | 22957966 | 22958159 | 4.10768 | 5.34573 | 2.04115 | Distal Intergenic | 8 | 22974844 | 23149933 | 175090 | 1 | ENSMUST00000110688.3 | -16685 | Ank1 | ENSMUSG00000031543.13 |
| chr8 | 22959130 | 22959246 | 3.65074 | 4.45203 | 1.47893 | Distal Intergenic | 8 | 22974844 | 23149933 | 175090 | 1 | ENSMUST00000110688.3 | -15598 | Ank1 | ENSMUSG00000031543.13 |
| chr8 | 22974124 | 22974671 | 3.52337 | 4.19651 | 1.26216 | Promoter | 8 | 22974844 | 23149933 | 175090 | 1 | ENSMUST00000110688.3 | -173 | Ank1 | ENSMUSG00000031543.13 |
| chr8 | 22975128 | 22976019 | 4.02475 | 5.41673 | 2.08503 | Promoter | 8 | 22975136 | 23148307 | 173172 | 1 | ENSMUST00000117270.3 | 0 | Ank1 | ENSMUSG00000031543.13 |
| chr8 | 22976788 | 22976867 | 4.04624 | 5.14294 | 1.81267 | Intron (ENSMUST00000110688.3/ENSMUSG00000031543.13, intron 1 of 42) | 8 | 22975136 | 23148307 | 173172 | 1 | ENSMUST00000117270.3 | 1653 | Ank1 | ENSMUSG00000031543.13 |
| chr8 | 22978692 | 22979067 | 3.78101 | 4.6741 | 1.55505 | Intron (ENSMUST00000110688.3/ENSMUSG00000031543.13, intron 1 of 42) | 8 | 22975136 | 23148307 | 173172 | 1 | ENSMUST00000117270.3 | 3557 | Ank1 | ENSMUSG00000031543.13 |
| chr8 | 22979823 | 22979910 | 3.56471 | 4.25272 | 1.28164 | Intron (ENSMUST00000110688.3/ENSMUSG00000031543.13, intron 1 of 42) | 8 | 22975136 | 23148307 | 173172 | 1 | ENSMUST00000117270.3 | 4688 | Ank1 | ENSMUSG00000031543.13 |
| chr8 | 23351257 | 23351635 | 3.4822 | 4.27625 | 1.28777 | Distal Intergenic | 8 | 23411502 | 23449632 | 38131 | 1 | ENSMUST00000033952.7 | -59867 | Sfrp1 | ENSMUSG00000031548.7 |
| chr8 | 23386910 | 23387107 | 4.06255 | 5.20896 | 1.92342 | Distal Intergenic | 8 | 23411502 | 23449632 | 38131 | 1 | ENSMUST00000033952.7 | -24395 | Sfrp1 | ENSMUSG00000031548.7 |
| chr8 | 23394372 | 23394478 | 3.33999 | 3.83734 | 1.01733 | Distal Intergenic | 8 | 23411502 | 23449632 | 38131 | 1 | ENSMUST00000033952.7 | -17024 | Sfrp1 | ENSMUSG00000031548.7 |
| chr8 | 23409757 | 23410489 | 3.58737 | 4.35765 | 1.40266 | Distal Intergenic | 8 | 23411502 | 23449632 | 38131 | 1 | ENSMUST00000033952.7 | -1013 | Sfrp1 | ENSMUSG00000031548.7 |
| chr8 | 23411448 | 23412694 | 3.87891 | 4.89313 | 1.72036 | Promoter | 8 | 23411502 | 23449632 | 38131 | 1 | ENSMUST00000033952.7 | 0 | Sfrp1 | ENSMUSG00000031548.7 |
| chr8 | 24554489 | 24554574 | 4.4583 | 6.11279 | 2.55446 | Promoter | 8 | 24550770 | 24554554 | 3785 | 2 | ENSMUST00000138155.1 | 0 | Ido2 | ENSMUSG00000031549.10 |
| chr8 | 24555110 | 24555314 | 3.66014 | 4.49729 | 1.47125 | Promoter | 8 | 24531894 | 24554597 | 22704 | 2 | ENSMUST00000140417.2 | -514 | Ido2 | ENSMUSG00000031549.10 |
| chr8 | 24595521 | 24595748 | 3.61244 | 4.70065 | 1.57187 | Promoter | 8 | 24584136 | 24595666 | 11531 | 2 | ENSMUST00000110667.2 | 0 | Ido1 | ENSMUSG00000031551.7 |
| chr5 | 47985211 | 47985441 | 3.88667 | 4.91851 | 1.77103 | Promoter | 5 | 47984994 | 48173193 | 188200 | 1 | ENSMUST00000173926.2 | 218 | Slit2 | ENSMUSG00000031558.10 |
| chr5 | 47985918 | 47986437 | 3.80459 | 4.8041 | 1.67118 | Promoter | 5 | 47984994 | 48173193 | 188200 | 1 | ENSMUST00000173926.2 | 925 | Slit2 | ENSMUSG00000031558.10 |
| chr8 | 49222605 | 49222694 | 3.35162 | 3.85487 | 1.01733 | Distal Intergenic | 8 | 49370886 | 49377214 | 6329 | 1 | ENSMUST00000143556.2 | -148192 | 4930555F03Rik | ENSMUSG00000031559.9 |
| chr8 | 49901764 | 49902040 | 3.31848 | 3.80508 | 1.01733 | Distal Intergenic | 8 | 49485481 | 49521095 | 35615 | 1 | ENSMUST00000126285.1 | 416284 | 4930555F03Rik | ENSMUSG00000031559.9 |
| chr8 | 48041607 | 48041753 | 3.51577 | 4.15656 | 1.20802 | Distal Intergenic | 8 | 47826083 | 47990543 | 164461 | 2 | ENSMUST00000057561.7 | -51065 | Wwc2 | ENSMUSG00000031563.7 |
| chr8 | 33972842 | 33972970 | 3.36316 | 3.87228 | 1.01733 | Distal Intergenic | 8 | 33782643 | 33929863 | 147221 | 2 | ENSMUST00000053251.6 | -42980 | Rbpms | ENSMUSG00000031586.11 |
| chr8 | 33974042 | 33974278 | 3.39241 | 4.02464 | 1.1199 | Distal Intergenic | 8 | 33782643 | 33929863 | 147221 | 2 | ENSMUST00000053251.6 | -44180 | Rbpms | ENSMUSG00000031586.11 |
| chr8 | 40926288 | 40926377 | 4.3167 | 5.8468 | 2.41215 | Promoter | 8 | 40926233 | 40990785 | 64553 | 1 | ENSMUST00000034004.9 | 56 | Pdgfrl | ENSMUSG00000031595.9 |
| chr8 | 40927370 | 40927589 | 3.71316 | 4.64648 | 1.56834 | Intron (ENSMUST00000143216.1/ENSMUSG00000031595.9, intron 1 of 1) | 8 | 40926233 | 40990785 | 64553 | 1 | ENSMUST00000034004.9 | 1138 | Pdgfrl | ENSMUSG00000031595.9 |
| chr8 | 40284735 | 40285037 | 3.78416 | 4.71521 | 1.65495 | Intron (ENSMUST00000034014.8/ENSMUSG00000031603.9, intron 1 of 2) | 8 | 40279166 | 40286953 | 7788 | 2 | ENSMUST00000034014.8 | 1916 | Fgf20 | ENSMUSG00000031603.9 |
| chr8 | 40285654 | 40287142 | 4.04303 | 5.25511 | 1.9809 | Promoter | 8 | 40279166 | 40286953 | 7788 | 2 | ENSMUST00000034014.8 | 0 | Fgf20 | ENSMUSG00000031603.9 |
| chr8 | 40287512 | 40287818 | 3.42857 | 4.03186 | 1.13156 | Promoter | 8 | 40279166 | 40286953 | 7788 | 2 | ENSMUST00000034014.8 | -560 | Fgf20 | ENSMUSG00000031603.9 |
| chr8 | 40289130 | 40290007 | 3.81524 | 4.79059 | 1.63258 | Intron (ENSMUST00000118639.1/ENSMUSG00000031603.9, intron 1 of 2) | 8 | 40279166 | 40286953 | 7788 | 2 | ENSMUST00000034014.8 | -2178 | Fgf20 | ENSMUSG00000031603.9 |
| chr8 | 57399136 | 57399459 | 3.97346 | 5.04718 | 1.80136 | Distal Intergenic | 8 | 57455923 | 57477585 | 21663 | 1 | ENSMUST00000034023.3 | -56464 | Scrg1 | ENSMUSG00000031610.3 |
| chr8 | 57400160 | 57400241 | 3.51721 | 4.17679 | 1.26599 | Distal Intergenic | 8 | 57455923 | 57477585 | 21663 | 1 | ENSMUST00000034023.3 | -55682 | Scrg1 | ENSMUSG00000031610.3 |
| chr8 | 57400660 | 57401047 | 3.55752 | 4.48786 | 1.44119 | Distal Intergenic | 8 | 57455923 | 57477585 | 21663 | 1 | ENSMUST00000034023.3 | -54876 | Scrg1 | ENSMUSG00000031610.3 |
| chr8 | 57402591 | 57402962 | 3.28566 | 3.75565 | 1.01733 | Distal Intergenic | 8 | 57455923 | 57477585 | 21663 | 1 | ENSMUST00000034023.3 | -52961 | Scrg1 | ENSMUSG00000031610.3 |
| chr8 | 57403729 | 57403959 | 3.24735 | 3.78306 | 1.01733 | Distal Intergenic | 8 | 57455923 | 57477585 | 21663 | 1 | ENSMUST00000034023.3 | -51964 | Scrg1 | ENSMUSG00000031610.3 |
| chr8 | 57404733 | 57405591 | 3.55291 | 4.31395 | 1.33751 | Distal Intergenic | 8 | 57455923 | 57477585 | 21663 | 1 | ENSMUST00000034023.3 | -50332 | Scrg1 | ENSMUSG00000031610.3 |
| chr8 | 57406181 | 57406979 | 3.66679 | 4.70876 | 1.57474 | Distal Intergenic | 8 | 57455923 | 57477585 | 21663 | 1 | ENSMUST00000034023.3 | -48944 | Scrg1 | ENSMUSG00000031610.3 |
| chr8 | 57407758 | 57407947 | 3.72405 | 4.71637 | 1.59055 | Distal Intergenic | 8 | 57455923 | 57477585 | 21663 | 1 | ENSMUST00000034023.3 | -47976 | Scrg1 | ENSMUSG00000031610.3 |
| chr8 | 57410115 | 57411224 | 3.93487 | 5.12295 | 1.90568 | Distal Intergenic | 8 | 57455923 | 57477585 | 21663 | 1 | ENSMUST00000034023.3 | -44699 | Scrg1 | ENSMUSG00000031610.3 |
| chr8 | 57412979 | 57413137 | 3.59369 | 4.53851 | 1.49084 | Distal Intergenic | 8 | 57455923 | 57477585 | 21663 | 1 | ENSMUST00000034023.3 | -42786 | Scrg1 | ENSMUSG00000031610.3 |
| chr8 | 57413814 | 57413998 | 3.66425 | 4.48127 | 1.47025 | Distal Intergenic | 8 | 57455923 | 57477585 | 21663 | 1 | ENSMUST00000034023.3 | -41925 | Scrg1 | ENSMUSG00000031610.3 |
| chr8 | 57414481 | 57414776 | 3.28785 | 3.96514 | 1.16384 | Distal Intergenic | 8 | 57455923 | 57477585 | 21663 | 1 | ENSMUST00000034023.3 | -41147 | Scrg1 | ENSMUSG00000031610.3 |
| chr8 | 57423151 | 57423358 | 3.21938 | 3.65629 | 1.01523 | Distal Intergenic | 8 | 57455923 | 57477585 | 21663 | 1 | ENSMUST00000034023.3 | -32565 | Scrg1 | ENSMUSG00000031610.3 |
| chr8 | 57426658 | 57427145 | 3.83011 | 4.76843 | 1.64853 | Distal Intergenic | 8 | 57455923 | 57477585 | 21663 | 1 | ENSMUST00000034023.3 | -28778 | Scrg1 | ENSMUSG00000031610.3 |
| chr8 | 57427537 | 57428506 | 3.72749 | 4.77274 | 1.63599 | Distal Intergenic | 8 | 57455923 | 57477585 | 21663 | 1 | ENSMUST00000034023.3 | -27417 | Scrg1 | ENSMUSG00000031610.3 |
| chr8 | 56294911 | 56295137 | 3.46011 | 4.04485 | 1.10974 | Promoter | 8 | 56294552 | 56321046 | 26495 | 1 | ENSMUST00000034026.8 | 360 | Hpgd | ENSMUSG00000031613.8 |
| chr8 | 45884999 | 45885219 | 3.46118 | 4.06668 | 1.17172 | Promoter | 8 | 45885485 | 45919546 | 34062 | 1 | ENSMUST00000034053.5 | -266 | Pdlim3 | ENSMUSG00000031636.6 |
| chr8 | 45885531 | 45885778 | 3.88734 | 4.84462 | 1.62202 | Promoter | 8 | 45885485 | 45919546 | 34062 | 1 | ENSMUST00000034053.5 | 47 | Pdlim3 | ENSMUSG00000031636.6 |
| chr8 | 60633254 | 60633459 | 3.24813 | 3.69934 | 1.01733 | Promoter | 8 | 60632860 | 60676729 | 43870 | 1 | ENSMUST00000160719.2 | 395 | Mfap3l | ENSMUSG00000031647.5 |
| chr8 | 87466369 | 87466764 | 3.58447 | 4.27873 | 1.26786 | Downstream (2-3kb) | 8 | 87469238 | 87472386 | 3149 | 2 | ENSMUST00000169693.1 | 5622 | Cbln1 | ENSMUSG00000031654.10 |
| chr8 | 87467648 | 87470753 | 3.99515 | 5.1191 | 1.86723 | 3' UTR | 8 | 87469238 | 87472386 | 3149 | 2 | ENSMUST00000169693.1 | 1633 | Cbln1 | ENSMUSG00000031654.10 |
| chr8 | 87471350 | 87472485 | 3.89359 | 5.07519 | 1.87327 | Promoter | 8 | 87469238 | 87472386 | 3149 | 2 | ENSMUST00000169693.1 | 0 | Cbln1 | ENSMUSG00000031654.10 |
| chr8 | 88272087 | 88272771 | 3.7568 | 4.83778 | 1.66668 | Promoter | 8 | 88272403 | 88329962 | 57560 | 1 | ENSMUST00000169037.2 | 0 | Adcy7 | ENSMUSG00000031659.7 |
| chr8 | 88451632 | 88451760 | 3.51819 | 4.31617 | 1.34883 | Distal Intergenic | 8 | 88521344 | 88594887 | 73544 | 1 | ENSMUST00000034086.6 | -69584 | Nkd1 | ENSMUSG00000031661.6 |
| chr8 | 88452648 | 88452936 | 3.77648 | 4.77179 | 1.61657 | Distal Intergenic | 8 | 88521344 | 88594887 | 73544 | 1 | ENSMUST00000034086.6 | -68408 | Nkd1 | ENSMUSG00000031661.6 |
| chr8 | 88454174 | 88454684 | 3.8715 | 4.89627 | 1.73057 | Distal Intergenic | 8 | 88521344 | 88594887 | 73544 | 1 | ENSMUST00000034086.6 | -66660 | Nkd1 | ENSMUSG00000031661.6 |
| chr8 | 88455157 | 88456304 | 3.5598 | 4.53219 | 1.45911 | Distal Intergenic | 8 | 88521344 | 88594887 | 73544 | 1 | ENSMUST00000034086.6 | -65040 | Nkd1 | ENSMUSG00000031661.6 |
| chr8 | 88495994 | 88496215 | 3.55271 | 4.21394 | 1.21333 | Distal Intergenic | 8 | 88521344 | 88594887 | 73544 | 1 | ENSMUST00000034086.6 | -25129 | Nkd1 | ENSMUSG00000031661.6 |
| chr8 | 88512119 | 88513182 | 3.67318 | 4.44428 | 1.37444 | Distal Intergenic | 8 | 88521344 | 88594887 | 73544 | 1 | ENSMUST00000034086.6 | -8162 | Nkd1 | ENSMUSG00000031661.6 |
| chr8 | 88602990 | 88603542 | 4.09301 | 5.25167 | 1.92536 | Distal Intergenic | 8 | 88626828 | 88636128 | 9301 | 2 | ENSMUST00000034087.7 | 32586 | Snx20 | ENSMUSG00000031662.7 |
| chr8 | 88627149 | 88627535 | 3.76929 | 4.62148 | 1.48326 | 3' UTR | 8 | 88626828 | 88636128 | 9301 | 2 | ENSMUST00000034087.7 | 8593 | Snx20 | ENSMUSG00000031662.7 |
| chr8 | 88627898 | 88628395 | 3.66148 | 4.42516 | 1.36133 | Intron (ENSMUST00000034087.7/ENSMUSG00000031662.7, intron 3 of 3) | 8 | 88626828 | 88636128 | 9301 | 2 | ENSMUST00000034087.7 | 7733 | Snx20 | ENSMUSG00000031662.7 |
| chr8 | 78419577 | 78419768 | 3.68114 | 4.55758 | 1.50085 | Distal Intergenic | 8 | 78433010 | 78436645 | 3636 | 2 | ENSMUST00000034115.4 | 16877 | Pou4f2 | ENSMUSG00000031688.4 |
| chr8 | 78420209 | 78420437 | 3.37183 | 3.88536 | 1.01733 | Distal Intergenic | 8 | 78433010 | 78436645 | 3636 | 2 | ENSMUST00000034115.4 | 16208 | Pou4f2 | ENSMUSG00000031688.4 |
| chr8 | 78421445 | 78421949 | 4.31433 | 5.83475 | 2.37633 | Distal Intergenic | 8 | 78433010 | 78436645 | 3636 | 2 | ENSMUST00000034115.4 | 14696 | Pou4f2 | ENSMUSG00000031688.4 |
| chr8 | 78424926 | 78425047 | 4.26033 | 5.5819 | 2.14367 | Distal Intergenic | 8 | 78433010 | 78436645 | 3636 | 2 | ENSMUST00000034115.4 | 11598 | Pou4f2 | ENSMUSG00000031688.4 |
| chr8 | 78426095 | 78426278 | 3.41181 | 3.9767 | 1.1263 | Distal Intergenic | 8 | 78433010 | 78436645 | 3636 | 2 | ENSMUST00000034115.4 | 10367 | Pou4f2 | ENSMUSG00000031688.4 |
| chr8 | 78426608 | 78426683 | 3.40016 | 3.93384 | 1.03748 | Distal Intergenic | 8 | 78433010 | 78436645 | 3636 | 2 | ENSMUST00000034115.4 | 9962 | Pou4f2 | ENSMUSG00000031688.4 |
| chr8 | 78427042 | 78427919 | 3.94657 | 4.97932 | 1.73953 | Distal Intergenic | 8 | 78433010 | 78436645 | 3636 | 2 | ENSMUST00000034115.4 | 8726 | Pou4f2 | ENSMUSG00000031688.4 |
| chr8 | 78429098 | 78429644 | 3.40398 | 3.94449 | 1.0547 | Distal Intergenic | 8 | 78433010 | 78436645 | 3636 | 2 | ENSMUST00000034115.4 | 7001 | Pou4f2 | ENSMUSG00000031688.4 |
| chr8 | 78429963 | 78430256 | 3.69742 | 4.5865 | 1.53332 | Downstream (2-3kb) | 8 | 78433010 | 78436645 | 3636 | 2 | ENSMUST00000034115.4 | 6389 | Pou4f2 | ENSMUSG00000031688.4 |
| chr8 | 78431870 | 78433467 | 3.63429 | 4.43661 | 1.4109 | 3' UTR | 8 | 78433010 | 78436645 | 3636 | 2 | ENSMUST00000034115.4 | 3178 | Pou4f2 | ENSMUSG00000031688.4 |
| chr8 | 78434094 | 78434505 | 3.69235 | 4.59763 | 1.54129 | 3' UTR | 8 | 78433010 | 78436645 | 3636 | 2 | ENSMUST00000034115.4 | 2140 | Pou4f2 | ENSMUSG00000031688.4 |
| chr8 | 78434820 | 78435406 | 3.5314 | 4.34471 | 1.3706 | Exon (ENSMUST00000034115.4/ENSMUSG00000031688.4, exon 2 of 2) | 8 | 78433010 | 78436645 | 3636 | 2 | ENSMUST00000034115.4 | 1239 | Pou4f2 | ENSMUSG00000031688.4 |
| chr8 | 78435814 | 78436581 | 3.67211 | 4.4781 | 1.47492 | Promoter | 8 | 78433010 | 78436645 | 3636 | 2 | ENSMUST00000034115.4 | 64 | Pou4f2 | ENSMUSG00000031688.4 |
| chr8 | 78437091 | 78437468 | 3.85411 | 4.85441 | 1.74052 | Promoter | 8 | 78433010 | 78436645 | 3636 | 2 | ENSMUST00000034115.4 | -447 | Pou4f2 | ENSMUSG00000031688.4 |
| chr8 | 78438486 | 78439601 | 3.85975 | 4.91272 | 1.75126 | Distal Intergenic | 8 | 78433010 | 78436645 | 3636 | 2 | ENSMUST00000034115.4 | -1842 | Pou4f2 | ENSMUSG00000031688.4 |
| chr8 | 78440155 | 78440824 | 3.38434 | 3.94812 | 1.12474 | Distal Intergenic | 8 | 78433010 | 78436645 | 3636 | 2 | ENSMUST00000034115.4 | -3511 | Pou4f2 | ENSMUSG00000031688.4 |
| chr8 | 83290019 | 83290433 | 3.40648 | 3.951 | 1.06444 | Promoter | 8 | 83290348 | 83298456 | 8109 | 1 | ENSMUST00000034146.3 | 0 | Ucp1 | ENSMUSG00000031710.3 |
| chr8 | 109464574 | 109464658 | 3.44271 | 4.02197 | 1.12202 | Distal Intergenic | 8 | 109494027 | 109542642 | 48616 | 1 | ENSMUST00000034162.6 | -29369 | Pmfbp1 | ENSMUSG00000031727.7 |
| chr8 | 109467129 | 109467229 | 3.42653 | 3.98191 | 1.06662 | Distal Intergenic | 8 | 109494027 | 109542642 | 48616 | 1 | ENSMUST00000034162.6 | -26798 | Pmfbp1 | ENSMUSG00000031727.7 |
| chr8 | 109468246 | 109468520 | 3.31445 | 3.79906 | 1.01733 | Distal Intergenic | 8 | 109494027 | 109542642 | 48616 | 1 | ENSMUST00000034162.6 | -25507 | Pmfbp1 | ENSMUSG00000031727.7 |
| chr8 | 92353458 | 92353690 | 3.31971 | 3.98667 | 1.14693 | Intron (ENSMUST00000179421.2/ENSMUSG00000031736.5, intron 1 of 3) | 8 | 92336211 | 92355764 | 19554 | 2 | ENSMUST00000180102.2 | 2074 | Crnde | ENSMUSG00000031736.5 |
| chr8 | 92356616 | 92357374 | 3.9263 | 5.01619 | 1.84189 | Promoter | 8 | 92357796 | 92361456 | 3661 | 1 | ENSMUST00000034184.10 | -422 | Irx5 | ENSMUSG00000031737.10 |
| chr8 | 92357725 | 92361112 | 3.96518 | 5.26522 | 1.9878 | Promoter | 8 | 92357796 | 92361456 | 3661 | 1 | ENSMUST00000034184.10 | 0 | Irx5 | ENSMUSG00000031737.10 |
| chr8 | 92669338 | 92669655 | 3.48467 | 4.09183 | 1.14501 | Distal Intergenic | 8 | 92674288 | 92680956 | 6669 | 1 | ENSMUST00000034185.7 | -4633 | Irx6 | ENSMUSG00000031738.9 |
| chr8 | 92673841 | 92674994 | 3.74571 | 4.82651 | 1.6837 | Promoter | 8 | 92674288 | 92680956 | 6669 | 1 | ENSMUST00000034185.7 | 0 | Irx6 | ENSMUSG00000031738.9 |
| chr8 | 92675832 | 92676073 | 3.57306 | 4.29697 | 1.36377 | Exon (ENSMUST00000034185.7/ENSMUSG00000031738.9, exon 2 of 6) | 8 | 92674826 | 92680956 | 6131 | 1 | ENSMUST00000167261.2 | 1007 | Irx6 | ENSMUSG00000031738.9 |

| | | | | | | | | | | | | | | | |
|---|---|---|---|---|---|---|---|---|---|---|---|---|---|---|---|
| chr8 | 92827609 | 92827763 | 3.36579 | 3.87976 | 1.02975 | Promoter | 8 | 92827328 | 92853417 | 26090 | 1 | ENSMUST00000034187.7 | 282 | Mmp2 | ENSMUSG00000031740.7 |
| chr8 | 94152170 | 94152828 | 3.76233 | 4.71618 | 1.60367 | Promoter | 8 | 94152607 | 94154148 | 1542 | 1 | ENSMUST00000034211.8 | 0 | Mt3 | ENSMUSG00000031760.8 |
| chr8 | 94153757 | 94153854 | 3.72501 | 4.54551 | 1.44451 | Intron (ENSMUST00000034211.8/ENSMUSG00000031760.8, intron 2 of 2) | 8 | 94152607 | 94154148 | 1542 | 1 | ENSMUST00000034211.8 | 1151 | Mt3 | ENSMUSG00000031760.8 |
| chr8 | 94166332 | 94166518 | 3.53576 | 4.20831 | 1.28525 | Distal Intergenic | 8 | 94172618 | 94173567 | 950 | 1 | ENSMUST00000034214.6 | -6100 | Mt2 | ENSMUSG00000031762.6 |
| chr8 | 94166858 | 94167041 | 3.89315 | 4.90861 | 1.76238 | Distal Intergenic | 8 | 94172618 | 94173567 | 950 | 1 | ENSMUST00000034214.6 | -5577 | Mt2 | ENSMUSG00000031762.6 |
| chr8 | 94172464 | 94173400 | 3.54225 | 4.20606 | 1.22165 | Promoter | 8 | 94172618 | 94173567 | 950 | 1 | ENSMUST00000034214.6 | 0 | Mt2 | ENSMUSG00000031762.6 |
| chr8 | 112570506 | 112570722 | 3.81096 | 4.72644 | 1.60104 | Promoter | 8 | 112570055 | 112786578 | 216524 | 1 | ENSMUST00000125196.2 | 452 | Cntnap4 | ENSMUSG00000031772.12 |
| chr8 | 94694916 | 94695002 | 3.6275 | 4.41213 | 1.43961 | Intron (ENSMUST00000034227.4/ENSMUSG00000031775.4, intron 1 of 3) | 8 | 94674895 | 94696242 | 21348 | 2 | ENSMUST00000034227.4 | 1240 | Pllp | ENSMUSG00000031775.4 |
| chr8 | 94695473 | 94696088 | 3.94045 | 4.97029 | 1.72421 | Promoter | 8 | 94674895 | 94696242 | 21348 | 2 | ENSMUST00000034227.4 | 154 | Pllp | ENSMUSG00000031775.4 |
| chr8 | 94696814 | 94697449 | 3.53894 | 4.19898 | 1.21063 | Promoter | 8 | 94674895 | 94696242 | 21348 | 2 | ENSMUST00000034227.4 | -573 | Pllp | ENSMUSG00000031775.4 |
| chr8 | 94977148 | 94977389 | 3.2762 | 4.08435 | 1.20267 | Promoter | 8 | 94977109 | 95014208 | 37100 | 1 | ENSMUST00000179619.2 | 40 | Gpr56 | ENSMUSG00000031785.10 |
| chr8 | 94996171 | 94996493 | 3.5881 | 4.29479 | 1.30095 | Intron (ENSMUST00000179619.2/ENSMUSG00000031785.10, intron 1 of 13) | 8 | 94984399 | 95014208 | 29810 | 1 | ENSMUST00000093271.6 | 11773 | Gpr56 | ENSMUSG00000031785.10 |
| chr8 | 95184421 | 95184729 | 3.43274 | 4.00498 | 1.1154 | Distal Intergenic | 8 | 95099832 | 95142540 | 42709 | 2 | ENSMUST00000169748.2 | -41882 | Kifc3 | ENSMUSG00000031788.8 |
| chr8 | 71699513 | 71699773 | 3.34174 | 3.85519 | 1.07072 | Intron (ENSMUST00000034260.7/ENSMUSG00000031803.7, intron 1 of 2) | 8 | 71691719 | 71701800 | 10082 | 2 | ENSMUST00000034260.7 | 2027 | B3gnt3 | ENSMUSG00000031803.7 |
| chr8 | 71700904 | 71701573 | 3.54787 | 4.3067 | 1.37324 | Promoter | 8 | 71691719 | 71701800 | 10082 | 2 | ENSMUST00000034260.7 | 227 | B3gnt3 | ENSMUSG00000031803.7 |
| chr8 | 71704258 | 71704553 | 4.19489 | 5.60308 | 2.20071 | Distal Intergenic | 8 | 71691719 | 71701800 | 10082 | 2 | ENSMUST00000034260.7 | -2459 | B3gnt3 | ENSMUSG00000031803.7 |
| chr8 | 71677077 | 71677546 | 3.45607 | 4.04219 | 1.12261 | Promoter | 8 | 71677551 | 71688558 | 11008 | 1 | ENSMUST00000110012.1 | -5 | Jak3 | ENSMUSG00000031805.12 |
| chr8 | 71678701 | 71679917 | 3.72782 | 4.57199 | 1.47369 | Promoter | 8 | 71678335 | 71690224 | 11890 | 1 | ENSMUST00000130624.1 | 367 | Jak3 | ENSMUSG00000031805.12 |
| chr8 | 71680525 | 71681121 | 3.76738 | 4.6989 | 1.58463 | Exon (ENSMUST00000110013.4/ENSMUSG00000031805.12, exon 7 of 24) | 8 | 71678335 | 71690224 | 11890 | 1 | ENSMUST00000130624.1 | 2191 | Jak3 | ENSMUSG00000031805.12 |
| chr8 | 71681564 | 71682291 | 4.13606 | 5.41077 | 2.05178 | Exon (ENSMUST00000110013.4/ENSMUSG00000031805.12, exon 10 of 24) | 8 | 71678335 | 71690224 | 11890 | 1 | ENSMUST00000130624.1 | 3230 | Jak3 | ENSMUSG00000031805.12 |
| chr8 | 119992276 | 119992653 | 3.75486 | 4.80876 | 1.64327 | Promoter | 8 | 119992438 | 120052793 | 60356 | 1 | ENSMUST00000132583.2 | 0 | Crispld2 | ENSMUSG00000031825.11 |
| chr8 | 119897467 | 119897611 | 3.68494 | 4.45371 | 1.35455 | Distal Intergenic | 8 | 119910360 | 119957555 | 47196 | 1 | ENSMUST00000144458.2 | -12749 | Usp10 | ENSMUSG00000031826.14 |
| chr8 | 119446045 | 119447954 | 3.78551 | 4.71804 | 1.57735 | Promoter | 8 | 119446719 | 119472640 | 25922 | 1 | ENSMUST00000098363.4 | 0 | Necab2 | ENSMUSG00000031837.8 |
| chr8 | 119449596 | 119449864 | 3.35411 | 3.85869 | 1.01733 | Intron (ENSMUST00000098363.4/ENSMUSG00000031837.8, intron 1 of 12) | 8 | 119452432 | 119470945 | 18514 | 1 | ENSMUST00000148328.1 | -2568 | Necab2 | ENSMUSG00000031837.8 |
| chr8 | 119460026 | 119460198 | 3.37071 | 3.88367 | 1.01733 | Intron (ENSMUST00000098363.4/ENSMUSG00000031837.8, intron 5 of 12) | 8 | 119452432 | 119470945 | 18514 | 1 | ENSMUST00000148328.1 | 7595 | Necab2 | ENSMUSG00000031837.8 |
| chr8 | 119464422 | 119464567 | 3.34153 | 3.83966 | 1.01733 | Intron (ENSMUST00000098363.4/ENSMUSG00000031837.8, intron 6 of 12) | 8 | 119452432 | 119470945 | 18514 | 1 | ENSMUST00000148328.1 | 11991 | Necab2 | ENSMUSG00000031837.8 |
| chr8 | 119465079 | 119465879 | 3.55867 | 4.25105 | 1.29509 | Exon (ENSMUST00000098363.4/ENSMUSG00000031837.8, exon 8 of 13) | 8 | 119452432 | 119470945 | 18514 | 1 | ENSMUST00000148328.1 | 12648 | Necab2 | ENSMUSG00000031837.8 |
| chr8 | 119466220 | 119466435 | 3.36917 | 3.88136 | 1.01733 | Intron (ENSMUST00000098363.4/ENSMUSG00000031837.8, intron 8 of 12) | 8 | 119452432 | 119470945 | 18514 | 1 | ENSMUST00000148328.1 | 13789 | Necab2 | ENSMUSG00000031837.8 |
| chr8 | 118274435 | 118274512 | 4.61619 | 6.30982 | 2.68443 | Distal Intergenic | 8 | 118283733 | 119324921 | 1041189 | 1 | ENSMUST00000117160.1 | -9221 | Cdh13 | ENSMUSG00000031841.13 |
| chr8 | 118282055 | 118282411 | 3.58292 | 4.29566 | 1.32453 | Distal Intergenic | 8 | 118283733 | 119324921 | 1041189 | 1 | ENSMUST00000117160.1 | -1322 | Cdh13 | ENSMUSG00000031841.13 |
| chr8 | 118284253 | 118284804 | 3.39712 | 4.02669 | 1.17194 | Promoter | 8 | 118283797 | 119202402 | 918606 | 1 | ENSMUST00000148836.2 | 457 | Cdh13 | ENSMUSG00000031841.13 |
| chr8 | 118286156 | 118286401 | 3.55639 | 4.31509 | 1.3977 | Intron (ENSMUST00000117160.1/ENSMUSG00000031841.13, intron 1 of 13) | 8 | 118283797 | 119202402 | 918606 | 1 | ENSMUST00000148836.2 | 2360 | Cdh13 | ENSMUSG00000031841.13 |
| chr8 | 118308961 | 118309134 | 3.52849 | 4.17833 | 1.21755 | Intron (ENSMUST00000117160.1/ENSMUSG00000031841.13, intron 1 of 13) | 8 | 118314413 | 118856206 | 541794 | 1 | ENSMUST00000123567.1 | -5279 | Cdh13 | ENSMUSG00000031841.13 |
| chr8 | 118310345 | 118310907 | 3.35583 | 3.86124 | 1.01733 | Intron (ENSMUST00000117160.1/ENSMUSG00000031841.13, intron 1 of 13) | 8 | 118314413 | 118856206 | 541794 | 1 | ENSMUST00000123567.1 | -3506 | Cdh13 | ENSMUSG00000031841.13 |
| chr8 | 118355174 | 118355566 | 3.72855 | 4.62281 | 1.55252 | Intron (ENSMUST00000117160.1/ENSMUSG00000031841.13, intron 1 of 13) | 8 | 118314413 | 118856206 | 541794 | 1 | ENSMUST00000123567.1 | 40762 | Cdh13 | ENSMUSG00000031841.13 |
| chr8 | 70375902 | 70376215 | 3.74162 | 4.68285 | 1.56912 | Exon (ENSMUST00000003659.7/ENSMUSG00000031849.8, exon 6 of 19) | 8 | 70373548 | 70382065 | 8518 | 1 | ENSMUST00000003659.7 | 2355 | Comp | ENSMUSG00000031849.8 |
| chr8 | 125910685 | 125911656 | 4.17597 | 5.48484 | 2.1139 | Promoter | 8 | 125910450 | 125947439 | 36990 | 1 | ENSMUST00000034316.4 | 236 | BC021891 | ENSMUSG00000031853.4 |
| chr8 | 69833984 | 69834452 | 3.85073 | 4.94693 | 1.71692 | Intron (ENSMUST00000081503.7/ENSMUSG00000031860.12, intron 1 of 7) | 8 | 69832690 | 69872292 | 39603 | 1 | ENSMUST00000134777.4 | 1295 | Pbx4 | ENSMUSG00000031860.12 |
| chr6 | 83161971 | 83162257 | 3.76561 | 4.80623 | 1.63728 | Exon (ENSMUST00000032106.4/ENSMUSG00000030030.4, exon 4 of 4) | 6 | 83165920 | 83200116 | 34197 | 1 | ENSMUST00000077407.6 | -3663 | Dctn1 | ENSMUSG00000031865.10 |
| chr9 | 8900359 | 8900998 | 3.72896 | 4.78035 | 1.63707 | Promoter | 9 | 8900419 | 8948607 | 48189 | 1 | ENSMUST00000151080.1 | 0 | Pgr | ENSMUSG00000031870.11 |
| chr9 | 8901327 | 8902204 | 4.02272 | 5.21683 | 1.95455 | Promoter | 9 | 8900468 | 8965425 | 64958 | 1 | ENSMUST00000098986.3 | 860 | Pgr | ENSMUSG00000031870.11 |
| chr8 | 104062577 | 104062680 | 3.37941 | 3.89679 | 1.01733 | Distal Intergenic | 8 | 104101625 | 104144502 | 42878 | 1 | ENSMUST00000034339.8 | -38945 | Cdh5 | ENSMUSG00000031871.8 |
| chr8 | 104169446 | 104169707 | 3.96994 | 5.08618 | 1.90276 | Promoter | 8 | 104170513 | 104219092 | 48580 | 1 | ENSMUST00000093245.6 | -806 | Bean1 | ENSMUSG00000031872.8 |
| chr8 | 104170327 | 104171349 | 3.55865 | 4.27498 | 1.35258 | Promoter | 8 | 104170513 | 104219092 | 48580 | 1 | ENSMUST00000093245.6 | 0 | Bean1 | ENSMUSG00000031872.8 |
| chr8 | 104171758 | 104172079 | 3.44439 | 4.0714 | 1.20849 | Intron (ENSMUST00000093245.6/ENSMUSG00000031872.8, intron 1 of 4) | 8 | 104170513 | 104219092 | 48580 | 1 | ENSMUST00000171018.2 | 1246 | Bean1 | ENSMUSG00000031872.8 |
| chr8 | 104630780 | 104631150 | 3.52129 | 4.32877 | 1.38195 | Promoter | 8 | 104628066 | 104631321 | 3256 | 2 | ENSMUST00000034351.6 | 171 | Rrad | ENSMUSG00000031880.6 |
| chr8 | 104540372 | 104541483 | 4.09614 | 5.39556 | 2.06197 | Promoter | 8 | 104540807 | 104550343 | 9537 | 1 | ENSMUST00000056051.5 | 0 | Car7 | ENSMUSG00000031883.7 |
| chr8 | 104541809 | 104542652 | 3.75121 | 4.60686 | 1.49418 | Promoter | 8 | 104541309 | 104550337 | 9029 | 1 | ENSMUST00000162761.1 | 501 | Car7 | ENSMUSG00000031883.7 |
| chr8 | 105258927 | 105259245 | 3.39485 | 3.93848 | 1.08244 | Promoter | 8 | 105259190 | 105260120 | 931 | 2 | ENSMUST00000136239.1 | 875 | Tradd | ENSMUSG00000031887.8 |
| chr8 | 106334734 | 106335072 | 4.02808 | 5.36809 | 2.06958 | Intron (ENSMUST00000180410.1/ENSMUSG00000097026.1, intron 1 of 1) | 8 | 106252548 | 106337988 | 85441 | 2 | ENSMUST00000067512.7 | 2916 | Smpd3 | ENSMUSG00000031906.8 |
| chr8 | 106336010 | 106337065 | 3.79955 | 5.20679 | 1.93654 | Promoter | 8 | 106252548 | 106337988 | 85441 | 2 | ENSMUST00000067512.7 | 923 | Smpd3 | ENSMUSG00000031906.8 |
| chr8 | 106871956 | 106872286 | 3.3308 | 4.09247 | 1.18668 | Intron (ENSMUST00000034385.6/ENSMUSG00000031910.9, intron 1 of 3) | 8 | 106870891 | 106884566 | 13676 | 1 | ENSMUST00000175987.1 | 1066 | Has3 | ENSMUSG00000031910.9 |
| chr9 | 14770655 | 14771030 | 3.89487 | 4.94328 | 1.73068 | Promoter | 9 | 14756593 | 14771013 | 14421 | 2 | ENSMUST00000188350.1 | 0 | 1700012B09Rik | ENSMUSG00000031927.4 |
| chr9 | 14859533 | 14860984 | 3.61202 | 4.49728 | 1.44199 | Promoter | 9 | 14860210 | 14870789 | 10580 | 1 | ENSMUST00000034408.6 | 0 | Gpr83 | ENSMUSG00000031932.9 |
| chr9 | 14861895 | 14862268 | 3.37022 | 3.93624 | 1.16692 | Intron (ENSMUST00000034408.6/ENSMUSG00000031932.9, intron 1 of 3) | 9 | 14860254 | 14870310 | 10057 | 1 | ENSMUST00000115624.2 | 1642 | Gpr83 | ENSMUSG00000031932.9 |
| chr9 | 15239344 | 15239570 | 3.82484 | 5.12822 | 1.87762 | Promoter | 9 | 15239045 | 15259393 | 20349 | 1 | ENSMUST00000034413.6 | 300 | Vstm5 | ENSMUSG00000031937.6 |
| chr8 | 111909146 | 111909221 | 3.47679 | 4.09752 | 1.19804 | Promoter | 8 | 111889135 | 111910199 | 21065 | 2 | ENSMUST00000034430.4 | 978 | Chst5 | ENSMUSG00000031952.4 |
| chr8 | 111909548 | 111909885 | 3.4161 | 4.00625 | 1.18718 | Promoter | 8 | 111889135 | 111910199 | 21065 | 2 | ENSMUST00000034430.4 | 314 | Chst5 | ENSMUSG00000031952.4 |
| chr8 | 111910196 | 111910394 | 3.56397 | 4.36724 | 1.35888 | Promoter | 8 | 111889135 | 111910199 | 21065 | 2 | ENSMUST00000034430.4 | 0 | Chst5 | ENSMUSG00000031952.4 |
| chr8 | 122853701 | 122853798 | 3.37293 | 3.92546 | 1.13962 | Intron (ENSMUST00000127664.1/ENSMUSG00000092329.1, intron 1 of 15) | 8 | 122848374 | 122867397 | 19024 | 1 | ENSMUST00000034443.5 | 5328 | Cdh15 | ENSMUSG00000031962.5 |
| chr9 | 23035998 | 23036456 | 3.54811 | 4.25057 | 1.31269 | Distal Intergenic | 9 | 23223076 | 23485202 | 262127 | 1 | ENSMUST00000071982.5 | -186620 | Bmper | ENSMUSG00000031963.7 |
| chr9 | 23037271 | 23037346 | 3.64674 | 4.47839 | 1.48147 | Distal Intergenic | 9 | 23223076 | 23485202 | 262127 | 1 | ENSMUST00000071982.5 | -185730 | Bmper | ENSMUSG00000031963.7 |
| chr9 | 23037795 | 23037968 | 3.21794 | 3.65414 | 1.01175 | Distal Intergenic | 9 | 23223076 | 23485202 | 262127 | 1 | ENSMUST00000071982.5 | -185108 | Bmper | ENSMUSG00000031963.7 |
| chr9 | 23223235 | 23223514 | 3.69603 | 4.88217 | 1.70663 | Promoter | 9 | 23223076 | 23485202 | 262127 | 1 | ENSMUST00000071982.5 | 160 | Bmper | ENSMUSG00000031963.7 |
| chr9 | 23224041 | 23224317 | 3.63644 | 4.43217 | 1.4351 | Promoter | 9 | 23223076 | 23485202 | 262127 | 1 | ENSMUST00000071982.5 | 966 | Bmper | ENSMUSG00000031963.7 |
| chr9 | 24746693 | 24747107 | 3.66657 | 4.43478 | 1.37764 | Intron (ENSMUST00000052946.5/ENSMUSG00000031965.9, intron 6 of 7) | 9 | 24720812 | 24774303 | 53492 | 2 | ENSMUST00000052946.5 | 27196 | Tbx20 | ENSMUSG00000031965.9 |
| chr9 | 24761846 | 24761936 | 3.63991 | 4.50729 | 1.48898 | Intron (ENSMUST00000052946.5/ENSMUSG00000031965.9, intron 4 of 7) | 9 | 24720812 | 24774303 | 53492 | 2 | ENSMUST00000052946.5 | 12367 | Tbx20 | ENSMUSG00000031965.9 |
| chr9 | 24763896 | 24764050 | 3.61409 | 4.40719 | 1.41611 | Intron (ENSMUST00000052946.5/ENSMUSG00000031965.9, intron 4 of 7) | 9 | 24720812 | 24774303 | 53492 | 2 | ENSMUST00000052946.5 | 10253 | Tbx20 | ENSMUSG00000031965.9 |
| chr9 | 24764618 | 24765692 | 4.01504 | 5.47282 | 2.0989 | Exon (ENSMUST00000052946.5/ENSMUSG00000031965.9, exon 4 of 8) | 9 | 24720812 | 24774303 | 53492 | 2 | ENSMUST00000052946.5 | 8611 | Tbx20 | ENSMUSG00000031965.9 |
| chr9 | 24766093 | 24766541 | 4.17682 | 5.57372 | 2.18279 | Intron (ENSMUST00000052946.5/ENSMUSG00000031965.9, intron 3 of 7) | 9 | 24720812 | 24774303 | 53492 | 2 | ENSMUST00000052946.5 | 7762 | Tbx20 | ENSMUSG00000031965.9 |
| chr9 | 24767017 | 24767261 | 4.13635 | 5.8084 | 2.34013 | Intron (ENSMUST00000052946.5/ENSMUSG00000031965.9, intron 3 of 7) | 9 | 24720812 | 24774303 | 53492 | 2 | ENSMUST00000052946.5 | 7042 | Tbx20 | ENSMUSG00000031965.9 |
| chr9 | 24767643 | 24767761 | 3.84541 | 5.14246 | 1.8677 | Intron (ENSMUST00000052946.5/ENSMUSG00000031965.9, intron 3 of 7) | 9 | 24720812 | 24774303 | 53492 | 2 | ENSMUST00000052946.5 | 6542 | Tbx20 | ENSMUSG00000031965.9 |
| chr9 | 24768096 | 24769888 | 3.9025 | 5.18431 | 1.93144 | Exon (ENSMUST00000052946.5/ENSMUSG00000031965.9, exon 3 of 8) | 9 | 24720812 | 24774303 | 53492 | 2 | ENSMUST00000052946.5 | 4415 | Tbx20 | ENSMUSG00000031965.9 |
| chr9 | 24770738 | 24770941 | 3.58947 | 4.79207 | 1.6266 | Exon (ENSMUST00000052946.5/ENSMUSG00000031965.9, exon 2 of 8) | 9 | 24720812 | 24774303 | 53492 | 2 | ENSMUST00000052946.5 | 3362 | Tbx20 | ENSMUSG00000031965.9 |
| chr9 | 24771762 | 24771889 | 3.65768 | 4.53552 | 1.48022 | Intron (ENSMUST00000052946.5/ENSMUSG00000031965.9, intron 1 of 7) | 9 | 24720812 | 24774303 | 53492 | 2 | ENSMUST00000052946.5 | 2414 | Tbx20 | ENSMUSG00000031965.9 |
| chr9 | 24772576 | 24775152 | 3.8749 | 5.21227 | 1.94336 | Promoter | 9 | 24720812 | 24774303 | 53492 | 2 | ENSMUST00000052946.5 | 0 | Tbx20 | ENSMUSG00000031965.9 |
| chr9 | 24776321 | 24777276 | 3.61609 | 4.53716 | 1.47193 | Distal Intergenic | 9 | 24740256 | 24774303 | 34048 | 2 | ENSMUST00000166018.2 | -2019 | Tbx20 | ENSMUSG00000031965.9 |
| chr9 | 24777605 | 24779070 | 4.06494 | 5.42217 | 2.09786 | Distal Intergenic | 9 | 24740256 | 24774303 | 34048 | 2 | ENSMUST00000166018.2 | -3303 | Tbx20 | ENSMUSG00000031965.9 |
| chr9 | 24779460 | 24779717 | 3.51001 | 4.50055 | 1.44118 | Distal Intergenic | 9 | 24740256 | 24774303 | 34048 | 2 | ENSMUST00000166018.2 | -5158 | Tbx20 | ENSMUSG00000031965.9 |
| chr9 | 24780273 | 24781424 | 3.84673 | 4.92714 | 1.76118 | Distal Intergenic | 9 | 24740256 | 24774303 | 34048 | 2 | ENSMUST00000166018.2 | -5971 | Tbx20 | ENSMUSG00000031965.9 |
| chr9 | 24781775 | 24782044 | 3.54396 | 4.30816 | 1.34743 | Distal Intergenic | 9 | 24740256 | 24774303 | 34048 | 2 | ENSMUST00000166018.2 | -7473 | Tbx20 | ENSMUSG00000031965.9 |
| chr9 | 24782594 | 24783113 | 3.71638 | 4.67653 | 1.57495 | Distal Intergenic | 9 | 24740256 | 24774303 | 34048 | 2 | ENSMUST00000166018.2 | -8292 | Tbx20 | ENSMUSG00000031965.9 |
| chr9 | 24783535 | 24784259 | 3.78583 | 4.73601 | 1.58868 | Distal Intergenic | 9 | 24740256 | 24774303 | 34048 | 2 | ENSMUST00000166018.2 | -9233 | Tbx20 | ENSMUSG00000031965.9 |
| chr8 | 123513523 | 123515328 | 4.20079 | 5.63318 | 2.26994 | Promoter | 8 | 123505687 | 123515390 | 9704 | 2 | ENSMUST00000177240.2 | 62 | Dbndd1 | ENSMUSG00000031970.11 |
| chr8 | 123515638 | 123515713 | 3.76931 | 4.63991 | 1.53537 | Promoter | 8 | 123505687 | 123515463 | 9777 | 2 | ENSMUST00000176155.1 | -176 | Dbndd1 | ENSMUSG00000031970.11 |
| chr8 | 123516128 | 123516451 | 3.33187 | 3.82511 | 1.01733 | Promoter | 8 | 123505687 | 123515463 | 9777 | 2 | ENSMUST00000176155.1 | -666 | Dbndd1 | ENSMUSG00000031970.11 |
| chr8 | 123864165 | 123864387 | 3.37775 | 3.8943 | 1.01733 | Intron (ENSMUST00000127664.1/ENSMUSG00000092329.1, intron 1 of 15) | 8 | 123840844 | 123860209 | 19366 | 2 | ENSMUST00000122421.1 | -3957 | Ccsap | ENSMUSG00000031971.10 |
| chr8 | 123867325 | 123867420 | 3.87126 | 4.83109 | 1.63151 | Intron (ENSMUST00000127664.1/ENSMUSG00000092329.1, intron 1 of 15) | 8 | 123840844 | 123860209 | 19366 | 2 | ENSMUST00000122421.1 | -7117 | Ccsap | ENSMUSG00000031971.10 |
| chr8 | 123891644 | 123892450 | 4.09401 | 5.39716 | 2.12938 | 3' UTR | 8 | 123891767 | 123894736 | 2970 | 2 | ENSMUST00000034453.4 | 2286 | Acta1 | ENSMUSG00000031972.4 |
| chr8 | 123894189 | 123894440 | 3.61642 | 4.34373 | 1.32402 | Promoter | 8 | 123891767 | 123894736 | 2970 | 2 | ENSMUST00000034453.4 | 296 | Acta1 | ENSMUSG00000031972.4 |
| chr8 | 124515810 | 124516129 | 3.65166 | 4.40984 | 1.37257 | Distal Intergenic | 8 | 124520767 | 124552008 | 31242 | 1 | ENSMUST00000034460.5 | -4638 | Cog2 | ENSMUSG00000031979.12 |
| chr8 | 124517614 | 124517689 | 3.88298 | 4.87075 | 1.71993 | Distal Intergenic | 8 | 124520767 | 124552008 | 31242 | 1 | ENSMUST00000034460.5 | -3078 | Cog2 | ENSMUSG00000031979.12 |
| chr8 | 124616828 | 124616943 | 4.25101 | 5.55922 | 2.12466 | Intron (ENSMUST00000093033.5/ENSMUSG00000031981.7, intron 18 of 19) | 8 | 124613440 | 124614626 | 1187 | 1 | ENSMUST00000133583.1 | 3389 | Capn9 | ENSMUSG00000031981.7 |
| chr8 | 124968431 | 124968521 | 3.43907 | 4.01623 | 1.12151 | Distal Intergenic | 8 | 124908596 | 124949254 | 40659 | 2 | ENSMUST00000034469.5 | -19178 | Egln1 | ENSMUSG00000031987.5 |
| chr9 | 27155856 | 27156106 | 3.56496 | 4.36302 | 1.37855 | Promoter | 9 | 27097385 | 27155421 | 58037 | 2 | ENSMUST00000034472.9 | -436 | Jam3 | ENSMUSG00000031990.9 |
| chr9 | 30941828 | 30943028 | 3.67335 | 4.51636 | 1.50476 | Promoter | 9 | 30942562 | 30962856 | 20295 | 1 | ENSMUST00000068135.7 | 0 | Adamts8 | ENSMUSG00000031994.8 |
| chr9 | 31131093 | 31132702 | 3.73384 | 5.09901 | 1.86416 | Promoter | 9 | 31089402 | 31131853 | 42452 | 2 | ENSMUST00000034478.2 | 0 | St14 | ENSMUSG00000031995.3 |
| chr9 | 31134019 | 31134110 | 3.2173 | 3.65318 | 1.01485 | Distal Intergenic | 9 | 31089402 | 31131853 | 42452 | 2 | ENSMUST00000034478.2 | -2167 | St14 | ENSMUSG00000031995.3 |
| chr9 | 8544855 | 8544975 | 4.06104 | 5.45916 | 2.06487 | Promoter | 9 | 8544196 | 8680565 | 136370 | 1 | ENSMUST00000050433.6 | 660 | Trpc6 | ENSMUSG00000031997.8 |
| chr9 | 44043446 | 44043850 | 4.01574 | 5.09704 | 1.78905 | Promoter | 9 | 44043384 | 44048579 | 5196 | 1 | ENSMUST00000114840.1 | 63 | Thy1 | ENSMUSG00000032011.4 |
| chr9 | 44044405 | 44044821 | 4.74732 | 6.65394 | 3.02632 | Intron (ENSMUST00000114840.1/ENSMUSG00000032011.4, intron 1 of 3) | 9 | 44043384 | 44048579 | 5196 | 1 | ENSMUST00000114840.1 | 1022 | Thy1 | ENSMUSG00000032011.4 |
| chr9 | 44045191 | 44045377 | 3.37718 | 3.90568 | 1.06062 | Intron (ENSMUST00000114840.1/ENSMUSG00000032011.4, intron 1 of 3) | 9 | 44043384 | 44048579 | 5196 | 1 | ENSMUST00000114840.1 | 1808 | Thy1 | ENSMUSG00000032011.4 |
| chr9 | 43668573 | 43668839 | 3.75981 | 4.5986 | 1.45403 | Distal Intergenic | 9 | 43744576 | 43807454 | 62879 | 1 | ENSMUST00000034510.7 | -75737 | Pvrl1 | ENSMUSG00000032012.7 |
| chr9 | 43723644 | 43723865 | 3.43428 | 4.00571 | 1.10939 | Distal Intergenic | 9 | 43744576 | 43807454 | 62879 | 1 | ENSMUST00000034510.7 | -20711 | Pvrl1 | ENSMUSG00000032012.7 |
| chr9 | 43744326 | 43745732 | 4.05157 | 5.22025 | 1.93547 | Promoter | 9 | 43744576 | 43807454 | 62879 | 1 | ENSMUST00000034510.7 | 0 | Pvrl1 | ENSMUSG00000032012.7 |
| chr9 | 43746038 | 43746115 | 4.13102 | 5.4198 | 2.07691 | Intron (ENSMUST00000034510.7/ENSMUSG00000032012.7, intron 1 of 5) | 9 | 43744576 | 43807454 | 62879 | 1 | ENSMUST00000034510.7 | 1463 | Pvrl1 | ENSMUSG00000032012.7 |
| chr9 | 43358729 | 43359304 | 3.66134 | 4.46189 | 1.41708 | Distal Intergenic | 9 | 43310763 | 43336125 | 25363 | 1 | ENSMUST00000034511.5 | 47967 | Trim29 | ENSMUSG00000032013.5 |
| chr9 | 43203920 | 43204217 | 3.36223 | 3.91093 | 1.05291 | Intron (ENSMUST00000034513.7/ENSMUSG00000032015.10, intron 1 of 12) | 9 | 43124540 | 43205755 | 81216 | 2 | ENSMUST00000176636.3 | 1538 | Pou2f3 | ENSMUSG00000032015.10 |
| chr9 | 43205098 | 43206186 | 3.90865 | 5.17284 | 1.93463 | Promoter | 9 | 43124540 | 43205755 | 81216 | 2 | ENSMUST00000176636.3 | 0 | Pou2f3 | ENSMUSG00000032015.10 |
| chr9 | 42683164 | 42683467 | 3.46726 | 4.26345 | 1.28489 | Intron (ENSMUST00000114865.2/ENSMUSG00000032017.9, intron 4 of 20) | 9 | 42520412 | 42808699 | 288288 | 2 | ENSMUST00000034515.6 | 125232 | Grik4 | ENSMUSG00000032017.9 |
| chr9 | 42703173 | 42703418 | 3.55253 | 4.48537 | 1.44073 | Intron (ENSMUST00000114865.2/ENSMUSG00000032017.9, intron 3 of 20) | 9 | 42520412 | 42808699 | 288288 | 2 | ENSMUST00000034515.6 | 105281 | Grik4 | ENSMUSG00000032017.9 |
| chr9 | 42754054 | 42754334 | 3.33537 | 3.83049 | 1.01733 | Intron (ENSMUST00000114865.2/ENSMUSG00000032017.9, intron 3 of 20) | 9 | 42520412 | 42808699 | 288288 | 2 | ENSMUST00000034515.6 | 54365 | Grik4 | ENSMUSG00000032017.9 |
| chr9 | 42774240 | 42774364 | 3.31829 | 3.80467 | 1.01733 | Intron (ENSMUST00000114865.2/ENSMUSG00000032017.9, intron 3 of 20) | 9 | 42520412 | 42808699 | 288288 | 2 | ENSMUST00000034515.6 | 34335 | Grik4 | ENSMUSG00000032017.9 |
| chr9 | 42944357 | 42944484 | 3.33757 | 3.89259 | 1.12699 | Promoter | 9 | 42518135 | 42944493 | 426359 | 2 | ENSMUST00000114865.2 | 9 | Grik4 | ENSMUSG00000032017.9 |
| chr9 | 42946650 | 42946895 | 3.3431 | 3.84207 | 1.01733 | Distal Intergenic | 9 | 42518135 | 42944493 | 426359 | 2 | ENSMUST00000114865.2 | -2158 | Grik4 | ENSMUSG00000032017.9 |
| chr9 | 42938744 | 42938875 | 3.64785 | 4.40019 | 1.35356 | Intron (ENSMUST00000114865.2/ENSMUSG00000032017.9, intron 1 of 20) | 9 | 42518135 | 42944493 | 426359 | 2 | ENSMUST00000114865.2 | 5618 | Grik4 | ENSMUSG00000032017.9 |
| chr9 | 42943284 | 42944026 | 3.58919 | 4.37156 | 1.36007 | Promoter | 9 | 42518135 | 42944493 | 426359 | 2 | ENSMUST00000114865.2 | 467 | Grik4 | ENSMUSG00000032017.9 |
| chr9 | 42957507 | 42957808 | 3.45175 | 4.27002 | 1.2776 | Distal Intergenic | 9 | 42518135 | 42944493 | 426359 | 2 | ENSMUST00000114865.2 | -13015 | Grik4 | ENSMUSG00000032017.9 |
| chr9 | 42220386 | 42220484 | 3.30973 | 3.7918 | 1.01733 | Distal Intergenic | 9 | 42254733 | 42264200 | 9468 | 2 | ENSMUST00000169609.1 | 43716 | Sc5d | ENSMUSG00000032018.8 |
| chr9 | 42228933 | 42229115 | 3.37645 | 3.91942 | 1.11296 | Distal Intergenic | 9 | 42254733 | 42264200 | 9468 | 2 | ENSMUST00000169609.1 | 35085 | Sc5d | ENSMUSG00000032018.8 |
| chr9 | 42240581 | 42240683 | 3.70204 | 4.60158 | 1.52148 | Distal Intergenic | 9 | 42254733 | 42264200 | 9468 | 2 | ENSMUST00000169609.1 | 23517 | Sc5d | ENSMUSG00000032018.8 |
| chr9 | 40888598 | 40888927 | 3.68671 | 4.52478 | 1.49815 | Distal Intergenic | 9 | 40894851 | 40897618 | 2768 | 2 | ENSMUST00000162122.1 | 8691 | 4931429I11Rik | ENSMUSG00000032023.7 |
| chr9 | 40889442 | 40889699 | 4.08542 | 5.33983 | 2.02181 | Distal Intergenic | 9 | 40894851 | 40897618 | 2768 | 2 | ENSMUST00000162122.1 | 7919 | 4931429I11Rik | ENSMUSG00000032023.7 |
| chr9 | 40891041 | 40891219 | 3.36004 | 4.09398 | 1.22114 | Distal Intergenic | 9 | 40894851 | 40897618 | 2768 | 2 | ENSMUST00000162122.1 | 6399 | 4931429I11Rik | ENSMUSG00000032023.7 |
| chr9 | 40891872 | 40891970 | 3.84981 | 5.03943 | 1.81318 | Downstream (2-3kb) | 9 | 40894851 | 40897618 | 2768 | 2 | ENSMUST00000162122.1 | 5648 | 4931429I11Rik | ENSMUSG00000032023.7 |
| chr9 | 40897981 | 40898097 | 3.41565 | 3.96994 | 1.08271 | Promoter | 9 | 40894851 | 40897618 | 2768 | 2 | ENSMUST00000162122.1 | -364 | 4931429I11Rik | ENSMUSG00000032023.7 |
| chr9 | 40960694 | 40960850 | 3.42582 | 4.11265 | 1.19634 | Intron (ENSMUST00000034521.7/ENSMUSG00000032023.7, intron 2 of 8) | 9 | 40960886 | 40962333 | 1448 | 2 | ENSMUST00000160120.1 | 1483 | 4931429I11Rik | ENSMUSG00000032023.7 |
| chr9 | 40963810 | 40964390 | 4.00747 | 5.2629 | 1.9737 | Promoter | 9 | 40894849 | 40964118 | 69270 | 2 | ENSMUST00000034521.7 | 0 | 4931429I11Rik | ENSMUSG00000032023.7 |
| chr9 | 40684421 | 40684504 | 3.89002 | 4.9018 | 1.74898 | Distal Intergenic | 9 | 40685962 | 40761060 | 75099 | 1 | ENSMUST00000139577.1 | -1458 | Clmp | ENSMUSG00000032024.10 |
| chr9 | 40685823 | 40685923 | 3.77199 | 4.65008 | 1.55175 | Promoter | 9 | 40685962 | 40761060 | 75099 | 1 | ENSMUST00000139577.1 | -39 | Clmp | ENSMUSG00000032024.10 |
| chr9 | 40686248 | 40686419 | 4.41191 | 5.94491 | 2.46857 | Promoter | 9 | 40686035 | 40761183 | 75149 | 1 | ENSMUST00000141759.1 | 214 | Clmp | ENSMUSG00000032024.10 |
| chr9 | 48253002 | 48253116 | 3.68451 | 4.48482 | 1.45008 | Distal Intergenic | 9 | 48319144 | 48340834 | 21691 | 2 | ENSMUST00000135457.2 | 87718 | Nxpe2 | ENSMUSG00000032028.10 |
| chr9 | 31858979 | 31859105 | 3.51546 | 4.29559 | 1.35641 | Exon (ENSMUST00000116615.4/ENSMUSG00000032033.11, exon 2 of 4) | 9 | 31846044 | 31913462 | 67419 | 2 | ENSMUST00000116615.4 | 54357 | Barx2 | ENSMUSG00000032033.11 |
| chr9 | 31859467 | 31859828 | 4.0308 | 5.28223 | 1.94996 | Intron (ENSMUST00000116615.4/ENSMUSG00000032033.11, intron 1 of 3) | 9 | 31846044 | 31913462 | 67419 | 2 | ENSMUST00000116615.4 | 53634 | Barx2 | ENSMUSG00000032033.11 |
| chr9 | 31908457 | 31908901 | 3.48498 | 4.16771 | 1.2641 | Intron (ENSMUST00000116615.4/ENSMUSG00000032033.11, intron 1 of 3) | 9 | 31846044 | 31913462 | 67419 | 2 | ENSMUST00000116615.4 | 4561 | Barx2 | ENSMUSG00000032033.11 |
| chr9 | 31912609 | 31913926 | 3.78801 | 4.71133 | 1.6173 | Promoter | 9 | 31846044 | 31913462 | 67419 | 2 | ENSMUST00000116615.4 | 0 | Barx2 | ENSMUSG00000032033.11 |
| chr9 | 31914362 | 31915564 | 4.17051 | 5.62221 | 2.25925 | Promoter | 9 | 31846044 | 31913462 | 67419 | 2 | ENSMUST00000116615.4 | -901 | Barx2 | ENSMUSG00000032033.11 |
| chr9 | 31916081 | 31916395 | 3.3176 | 3.81448 | 1.05525 | Distal Intergenic | 9 | 31846044 | 31913462 | 67419 | 2 | ENSMUST00000116615.4 | -2620 | Barx2 | ENSMUSG00000032033.11 |
| chr9 | 31917298 | 31917895 | 4.10208 | 5.36976 | 2.06558 | Distal Intergenic | 9 | 31846044 | 31913462 | 67419 | 2 | ENSMUST00000116615.4 | -3837 | Barx2 | ENSMUSG00000032033.11 |
| chr9 | 31918446 | 31918691 | 3.87591 | 4.86296 | 1.72652 | Distal Intergenic | 9 | 31846044 | 31913462 | 67419 | 2 | ENSMUST00000116615.4 | -4985 | Barx2 | ENSMUSG00000032033.11 |
| chr9 | 31919801 | 31919876 | 4.36973 | 6.16755 | 2.63387 | Distal Intergenic | 9 | 31846044 | 31913462 | 67419 | 2 | ENSMUST00000116615.4 | -6340 | Barx2 | ENSMUSG00000032033.11 |
| chr9 | 31921403 | 31921478 | 4.27231 | 5.63269 | 2.23645 | Distal Intergenic | 9 | 31846044 | 31913462 | 67419 | 2 | ENSMUST00000116615.4 | -7942 | Barx2 | ENSMUSG00000032033.11 |
| chr9 | 32312788 | 32313147 | 3.38634 | 4.08287 | 1.22489 | Downstream (1-2kb) | 9 | 32314783 | 32344237 | 29455 | 2 | ENSMUST00000034533.5 | 31090 | Kcnj5 | ENSMUSG00000032034.5 |
| chr9 | 32343678 | 32343791 | 3.92883 | 5.2693 | 1.95725 | Promoter | 9 | 32314783 | 32344237 | 29455 | 2 | ENSMUST00000034533.5 | 446 | Kcnj5 | ENSMUSG00000032034.5 |
| chr9 | 32345893 | 32346040 | 3.68487 | 4.46689 | 1.38857 | Distal Intergenic | 9 | 32314783 | 32344237 | 29455 | 2 | ENSMUST00000034533.5 | -1657 | Kcnj5 | ENSMUSG00000032034.5 |
| chr9 | 34485002 | 34485567 | 3.4944 | 4.12439 | 1.2033 | Promoter | 9 | 34485894 | 34489158 | 3265 | 1 | ENSMUST00000184203.2 | -327 | Kirrel3 | ENSMUSG00000032036.10 |
| chr9 | 34486006 | 34486905 | 4.06171 | 5.28469 | 1.99289 | Promoter | 9 | 34486125 | 34510616 | 24492 | 1 | ENSMUST00000183580.1 | 0 | Kirrel3 | ENSMUSG00000032036.10 |
| chr9 | 34487686 | 34488093 | 4.29626 | 5.67845 | 2.24589 | Promoter | 9 | 34488733 | 35036716 | 547984 | 1 | ENSMUST00000187625.1 | -640 | Kirrel3 | ENSMUSG00000032036.10 |
| chr9 | 34488487 | 34488869 | 3.4278 | 3.9991 | 1.12101 | Promoter | 9 | 34488733 | 35036716 | 547984 | 1 | ENSMUST00000187625.1 | 0 | Kirrel3 | ENSMUSG00000032036.10 |
| chr9 | 34489198 | 34489305 | 3.51266 | 4.16365 | 1.24449 | Promoter | 9 | 34488936 | 35036310 | 547375 | 1 | ENSMUST00000187182.1 | 263 | Kirrel3 | ENSMUSG00000032036.10 |
| chr9 | 34489617 | 34490181 | 4.19229 | 5.89379 | 2.46713 | Promoter | 9 | 34488936 | 35036310 | 547375 | 1 | ENSMUST00000187182.1 | 682 | Kirrel3 | ENSMUSG00000032036.10 |
| chr9 | 50698941 | 50699155 | 3.76418 | 4.70826 | 1.5777 | Promoter | 9 | 50662756 | 50699362 | 36607 | 2 | ENSMUST00000141919.2 | 207 | Dixdc1 | ENSMUSG00000032064.12 |
| chr9 | 50727596 | 50727709 | 3.42721 | 4.11461 | 1.20288 | Promoter | 9 | 50710137 | 50727938 | 17802 | 2 | ENSMUST00000118707.1 | 229 | Dixdc1 | ENSMUSG00000032064.12 |
| chr9 | 50421976 | 50422107 | 3.18598 | 3.74943 | 1.01733 | Distal Intergenic | 9 | 50494525 | 50505482 | 10958 | 1 | ENSMUST00000114474.2 | -72418 | Plet1 | ENSMUSG00000032068.9 |
| chr9 | 50425223 | 50425355 | 3.69099 | 4.486 | 1.41258 | Distal Intergenic | 9 | 50494525 | 50505482 | 10958 | 1 | ENSMUST00000114474.2 | -69170 | Plet1 | ENSMUSG00000032068.9 |
| chr9 | 50475894 | 50476324 | 3.46527 | 4.10859 | 1.21617 | Distal Intergenic | 9 | 50494525 | 50505482 | 10958 | 1 | ENSMUST00000114474.2 | -18201 | Plet1 | ENSMUSG00000032068.9 |
| chr9 | 50485763 | 50485930 | 3.30475 | 3.80598 | 1.08673 | Downstream (2-3kb) | 9 | 50494525 | 50505482 | 10958 | 1 | ENSMUST00000114474.2 | -8595 | Plet1 | ENSMUSG00000032068.9 |
| chr9 | 47530747 | 47531255 | 3.73596 | 4.70382 | 1.61467 | Promoter | 9 | 47530373 | 47850390 | 320018 | 1 | ENSMUST00000034581.3 | 375 | Cadm1 | ENSMUSG00000032076.13 |
| chr9 | 47535640 | 47535735 | 3.52859 | 4.19806 | 1.26982 | Intron (ENSMUST00000114548.2/ENSMUSG00000032076.13, intron 1 of 9) | 9 | 47530373 | 47850390 | 320018 | 1 | ENSMUST00000034581.3 | 5268 | Cadm1 | ENSMUSG00000032076.13 |
| chr9 | 47543693 | 47544120 | 3.89731 | 4.99569 | 1.79366 | Intron (ENSMUST00000114548.2/ENSMUSG00000032076.13, intron 1 of 9) | 9 | 47530373 | 47850390 | 320018 | 1 | ENSMUST00000034581.3 | 13321 | Cadm1 | ENSMUSG00000032076.13 |
| chr9 | 47619427 | 47619666 | 3.60385 | 4.30691 | 1.26859 | Intron (ENSMUST00000114548.2/ENSMUSG00000032076.13, intron 1 of 9) | 9 | 47530373 | 47850390 | 320018 | 1 | ENSMUST00000034581.3 | 89055 | Cadm1 | ENSMUSG00000032076.13 |
| chr9 | 47770909 | 47771100 | 3.39748 | 4.07739 | 1.15696 | Intron (ENSMUST00000114548.2/ENSMUSG00000032076.13, intron 1 of 9) | 9 | 47530373 | 47850390 | 320018 | 1 | ENSMUST00000034581.3 | 240537 | Cadm1 | ENSMUSG00000032076.13 |
| chr9 | 45429631 | 45429779 | 3.38193 | 3.9006 | 1.01733 | Promoter | 9 | 45430293 | 45753164 | 322872 | 1 | ENSMUST00000034592.8 | -514 | Dscaml1 | ENSMUSG00000032087.9 |

| | | | | | | | | | | | | | | | |
|---|---|---|---|---|---|---|---|---|---|---|---|---|---|---|---|
| chr9 | 45430090 | 45430366 | 4.19854 | 5.48675 | 2.10608 | Promoter | 9 | 45430293 | 45753164 | 322872 | 1 | ENSMUST00000034592.8 | 0 | Dscaml1 | ENSMUSG00000032087.9 |
| chr9 | 45268804 | 45269001 | 4.20248 | 5.48827 | 2.1128 | Promoter | 9 | 45254527 | 45269146 | 14620 | 2 | ENSMUST00000176222.1 | 145 | Il10ra | ENSMUSG00000032089.11 |
| chr9 | 45028750 | 45028873 | 3.26803 | 3.73342 | 1.03152 | Distal Intergenic | 9 | 45042425 | 45054015 | 11591 | 1 | ENSMUST00000034600.4 | -13552 | Mpzl2 | ENSMUSG00000032092.4 |
| chr9 | 45037468 | 45037630 | 3.17761 | 3.73677 | 1.01733 | Distal Intergenic | 9 | 45042425 | 45054015 | 11591 | 1 | ENSMUST00000034600.4 | -4795 | Mpzl2 | ENSMUSG00000032092.4 |
| chr9 | 45039738 | 45040007 | 3.75067 | 4.80643 | 1.64824 | Distal Intergenic | 9 | 45042425 | 45054015 | 11591 | 1 | ENSMUST00000034600.4 | -2418 | Mpzl2 | ENSMUSG00000032092.4 |
| chr9 | 45040585 | 45040660 | 3.61628 | 4.39154 | 1.43939 | Distal Intergenic | 9 | 45042425 | 45054015 | 11591 | 1 | ENSMUST00000034600.4 | -1765 | Mpzl2 | ENSMUSG00000032092.4 |
| chr9 | 45042406 | 45043405 | 3.56908 | 4.28098 | 1.29268 | Promoter | 9 | 45042425 | 45054015 | 11591 | 1 | ENSMUST00000034600.4 | 0 | Mpzl2 | ENSMUSG00000032092.4 |
| chr9 | 45043726 | 45043801 | 4.05578 | 5.16452 | 1.83135 | Exon (ENSMUST00000034600.4/ENSMUSG00000032092.4, exon 2 of 6) | 9 | 45042425 | 45054015 | 11591 | 1 | ENSMUST00000034600.4 | 1302 | Mpzl2 | ENSMUSG00000032092.4 |
| chr9 | 45044158 | 45044237 | 3.82171 | 4.71322 | 1.52519 | Exon (ENSMUST00000034600.4/ENSMUSG00000032092.4, exon 3 of 6) | 9 | 45042425 | 45054015 | 11591 | 1 | ENSMUST00000034600.4 | 1734 | Mpzl2 | ENSMUSG00000032092.4 |
| chr9 | 35558275 | 35559024 | 3.89837 | 4.94165 | 1.77037 | Promoter | 9 | 35541847 | 35558528 | 16682 | 2 | ENSMUST00000034612.5 | 0 | Ddx25 | ENSMUSG00000032101.5 |
| chr9 | 37406993 | 37407155 | 3.68359 | 4.52632 | 1.49377 | Intron (ENSMUST00000156972.1/ENSMUSG00000032125.14, intron 9 of 16) | 9 | 37402046 | 37414022 | 11977 | 1 | ENSMUST00000102895.4 | 4948 | Robo4 | ENSMUSG00000032125.14 |
| chr9 | 37407604 | 37407894 | 3.50672 | 4.17411 | 1.29889 | Intron (ENSMUST00000156972.1/ENSMUSG00000032125.14, intron 9 of 16) | 9 | 37402046 | 37414022 | 11977 | 1 | ENSMUST00000102895.4 | 5559 | Robo4 | ENSMUSG00000032125.14 |
| chr9 | 37409211 | 37409316 | 3.37377 | 4.03371 | 1.14667 | Intron (ENSMUST00000156972.1/ENSMUSG00000032125.14, intron 11 of 16) | 9 | 37402046 | 37414022 | 11977 | 1 | ENSMUST00000102895.4 | 7166 | Robo4 | ENSMUSG00000032125.14 |
| chr9 | 37430991 | 37431229 | 3.75051 | 5.24957 | 1.94952 | Promoter | 9 | 37416162 | 37430134 | 13973 | 2 | ENSMUST00000170512.2 | -858 | Robo3 | ENSMUSG00000032128.9 |
| chr9 | 37432480 | 37433120 | 4.26787 | 5.68438 | 2.31131 | Promoter | 9 | 37416045 | 37433163 | 17119 | 2 | ENSMUST00000115038.1 | 43 | Robo3 | ENSMUSG00000032128.9 |
| chr9 | 37434284 | 37434574 | 3.81569 | 4.84137 | 1.69035 | Distal Intergenic | 9 | 37416045 | 37433163 | 17119 | 2 | ENSMUST00000115038.1 | -1122 | Robo3 | ENSMUSG00000032128.9 |
| chr9 | 37435084 | 37435299 | 3.89092 | 4.98407 | 1.79819 | Distal Intergenic | 9 | 37416045 | 37433163 | 17119 | 2 | ENSMUST00000115038.1 | -1922 | Robo3 | ENSMUSG00000032128.9 |
| chr9 | 37435718 | 37436027 | 4.28158 | 5.75876 | 2.39575 | Distal Intergenic | 9 | 37416045 | 37433163 | 17119 | 2 | ENSMUST00000115038.1 | -2556 | Robo3 | ENSMUSG00000032128.9 |
| chr9 | 37436340 | 37436613 | 3.31489 | 3.82568 | 1.10109 | Distal Intergenic | 9 | 37416045 | 37433163 | 17119 | 2 | ENSMUST00000115038.1 | -3178 | Robo3 | ENSMUSG00000032128.9 |
| chr9 | 37437002 | 37437093 | 3.57528 | 4.30755 | 1.38364 | Distal Intergenic | 9 | 37416045 | 37433163 | 17119 | 2 | ENSMUST00000115038.1 | -3840 | Robo3 | ENSMUSG00000032128.9 |
| chr9 | 37450764 | 37450913 | 3.36906 | 3.88118 | 1.01733 | Distal Intergenic | 9 | 37416045 | 37433163 | 17119 | 2 | ENSMUST00000115038.1 | -17602 | Robo3 | ENSMUSG00000032128.9 |
| chr9 | 37451298 | 37452104 | 3.59843 | 4.41371 | 1.40212 | Distal Intergenic | 9 | 37416045 | 37433163 | 17119 | 2 | ENSMUST00000115038.1 | -18136 | Robo3 | ENSMUSG00000032128.9 |
| chr9 | 37453014 | 37453484 | 3.50347 | 4.19405 | 1.26285 | Distal Intergenic | 9 | 37416045 | 37433163 | 17119 | 2 | ENSMUST00000115038.1 | -19852 | Robo3 | ENSMUSG00000032128.9 |
| chr9 | 37453996 | 37454448 | 3.75959 | 4.64999 | 1.52586 | Distal Intergenic | 9 | 37416045 | 37433163 | 17119 | 2 | ENSMUST00000115038.1 | -20834 | Robo3 | ENSMUSG00000032128.9 |
| chr9 | 37454858 | 37455129 | 3.71781 | 4.52428 | 1.41585 | Distal Intergenic | 9 | 37416045 | 37433163 | 17119 | 2 | ENSMUST00000115038.1 | -21696 | Robo3 | ENSMUSG00000032128.9 |
| chr9 | 37455977 | 37456925 | 3.62166 | 4.34771 | 1.31008 | Distal Intergenic | 9 | 37416045 | 37433163 | 17119 | 2 | ENSMUST00000115038.1 | -22815 | Robo3 | ENSMUSG00000032128.9 |
| chr9 | 37457559 | 37459711 | 3.54898 | 4.24959 | 1.2883 | Distal Intergenic | 9 | 37416045 | 37433163 | 17119 | 2 | ENSMUST00000115038.1 | -24397 | Robo3 | ENSMUSG00000032128.9 |
| chr9 | 37460128 | 37461051 | 4.27723 | 5.67869 | 2.28975 | Distal Intergenic | 9 | 37416045 | 37433163 | 17119 | 2 | ENSMUST00000115038.1 | -26966 | Robo3 | ENSMUSG00000032128.9 |
| chr9 | 44287218 | 44288132 | 4.39381 | 6.43474 | 2.90992 | Promoter | 9 | 44273188 | 44287544 | 14357 | 2 | ENSMUST00000034648.10 | 0 | Abcg4 | ENSMUSG00000032131.10 |
| chr9 | 44134446 | 44135314 | 3.41613 | 4.18549 | 1.27219 | Promoter | 9 | 44134469 | 44138934 | 4466 | 1 | ENSMUST00000149241.2 | 0 | Mcam | ENSMUSG00000032135.9 |
| chr9 | 20693920 | 20694023 | 3.47282 | 4.22611 | 1.30637 | Intron (ENSMUST00000034692.7/ENSMUSG00000032172.7, intron 1 of 5) | 9 | 20667986 | 20728219 | 60234 | 2 | ENSMUST00000034692.7 | 34196 | Olfm2 | ENSMUSG00000032172.7 |
| chr9 | 20695030 | 20695311 | 3.50539 | 4.1668 | 1.28623 | Intron (ENSMUST00000034692.7/ENSMUSG00000032172.7, intron 1 of 5) | 9 | 20667986 | 20728219 | 60234 | 2 | ENSMUST00000034692.7 | 32908 | Olfm2 | ENSMUSG00000032172.7 |
| chr9 | 20726288 | 20727050 | 3.98045 | 5.29872 | 1.98847 | Intron (ENSMUST00000034692.7/ENSMUSG00000032172.7, intron 1 of 5) | 9 | 20667986 | 20728219 | 60234 | 2 | ENSMUST00000034692.7 | 1169 | Olfm2 | ENSMUSG00000032172.7 |
| chr9 | 20727357 | 20727593 | 3.78767 | 4.94566 | 1.73526 | Promoter | 9 | 20667986 | 20728219 | 60234 | 2 | ENSMUST00000034692.7 | 626 | Olfm2 | ENSMUSG00000032172.7 |
| chr9 | 20744268 | 20745600 | 3.75402 | 4.72729 | 1.59957 | Distal Intergenic | 9 | 20667986 | 20728219 | 60234 | 2 | ENSMUST00000034692.7 | -16050 | Olfm2 | ENSMUSG00000032172.7 |
| chr9 | 20745934 | 20746296 | 4.12967 | 5.35762 | 2.01012 | Distal Intergenic | 9 | 20667986 | 20728219 | 60234 | 2 | ENSMUST00000034692.7 | -17716 | Olfm2 | ENSMUSG00000032172.7 |
| chr9 | 21165031 | 21165296 | 3.64532 | 4.4095 | 1.39732 | Promoter | 9 | 21165714 | 21213248 | 47535 | 1 | ENSMUST00000039413.9 | -418 | Pde4a | ENSMUSG00000032177.11 |
| chr9 | 21165741 | 21166258 | 4.07369 | 5.51807 | 2.12519 | Promoter | 9 | 21165714 | 21213248 | 47535 | 1 | ENSMUST00000039413.9 | 28 | Pde4a | ENSMUSG00000032177.11 |
| chr9 | 21166714 | 21167270 | 3.55041 | 4.44916 | 1.43634 | Intron (ENSMUST00000039413.9/ENSMUSG00000032177.11, intron 1 of 14) | 9 | 21165714 | 21213248 | 47535 | 1 | ENSMUST00000039413.9 | 1001 | Pde4a | ENSMUSG00000032177.11 |
| chr9 | 75683921 | 75684333 | 4.02169 | 5.26509 | 1.96975 | Promoter | 9 | 75643264 | 75683996 | 40733 | 2 | ENSMUST00000034699.6 | 0 | Scg3 | ENSMUSG00000032181.6 |
| chr9 | 75610777 | 75610935 | 3.6487 | 4.50163 | 1.46307 | Promoter | 9 | 75565621 | 75611325 | 45705 | 2 | ENSMUST00000064433.4 | 390 | Tmod2 | ENSMUSG00000032186.9 |
| chr9 | 69507178 | 69507523 | 3.36368 | 3.88332 | 1.05373 | Distal Intergenic | 9 | 69486783 | 69488108 | 1326 | 1 | ENSMUST00000154591.1 | 20396 | Anxa2 | ENSMUSG00000032231.9 |
| chr9 | 69540356 | 69540445 | 4.28194 | 5.74699 | 2.32048 | Distal Intergenic | 9 | 69486783 | 69488108 | 1326 | 1 | ENSMUST00000154591.1 | 53574 | Anxa2 | ENSMUSG00000032231.9 |
| chr9 | 62608584 | 62608673 | 12.44241 | 32.85821 | 27.17674 | Distal Intergenic | 9 | 62677826 | 62783982 | 106157 | 1 | ENSMUST00000034774.8 | -69153 | Itga11 | ENSMUSG00000032243.8 |
| chr9 | 62676979 | 62677376 | 3.48241 | 4.08957 | 1.14944 | Promoter | 9 | 62677826 | 62783982 | 106157 | 1 | ENSMUST00000034774.8 | -450 | Itga11 | ENSMUSG00000032243.8 |
| chr9 | 62678486 | 62679041 | 3.62721 | 4.59896 | 1.52037 | Promoter | 9 | 62677860 | 62777111 | 99252 | 1 | ENSMUST00000159012.1 | 627 | Itga11 | ENSMUSG00000032243.8 |
| chr9 | 62679388 | 62679856 | 3.41029 | 3.95263 | 1.04983 | Intron (ENSMUST00000034774.8/ENSMUSG00000032243.8, intron 1 of 29) | 9 | 62677860 | 62777111 | 99252 | 1 | ENSMUST00000159012.1 | 1529 | Itga11 | ENSMUSG00000032243.8 |
| chr9 | 62140514 | 62140809 | 3.3433 | 3.8845 | 1.14384 | Distal Intergenic | 9 | 62070468 | 62122655 | 52188 | 2 | ENSMUST00000185873.1 | -17860 | Glce | ENSMUSG00000032252.9 |
| chr9 | 49338708 | 49339128 | 3.5005 | 4.26038 | 1.29828 | Distal Intergenic | 9 | 49340662 | 49407214 | 66553 | 1 | ENSMUST00000075764.6 | -1534 | Drd2 | ENSMUSG00000032259.7 |
| chr9 | 49339968 | 49341330 | 3.67794 | 4.57931 | 1.53529 | Promoter | 9 | 49340662 | 49407214 | 66553 | 1 | ENSMUST00000075764.6 | 0 | Drd2 | ENSMUSG00000032259.7 |
| chr9 | 83804270 | 83804859 | 3.62818 | 4.37887 | 1.36768 | Intron (ENSMUST00000034796.8/ENSMUSG00000032262.8, intron 1 of 5) | 9 | 83778692 | 83806252 | 27561 | 2 | ENSMUST00000034796.8 | 1393 | Elovl4 | ENSMUSG00000032262.8 |
| chr9 | 83805587 | 83806475 | 4.29784 | 5.73349 | 2.29863 | Promoter | 9 | 83778692 | 83806252 | 27561 | 2 | ENSMUST00000034796.8 | 0 | Elovl4 | ENSMUSG00000032262.8 |
| chr9 | 49126814 | 49127007 | 3.2886 | 3.76008 | 1.01733 | Distal Intergenic | 9 | 49109002 | 49117585 | 8584 | 1 | ENSMUST00000170426.1 | 17813 | Tmprss5 | ENSMUSG00000032268.8 |
| chr9 | 49128238 | 49128357 | 3.57848 | 4.41295 | 1.42189 | Distal Intergenic | 9 | 49109002 | 49117585 | 8584 | 1 | ENSMUST00000170426.1 | 19237 | Tmprss5 | ENSMUSG00000032268.8 |
| chr9 | 49128977 | 49129244 | 3.70937 | 4.6362 | 1.56176 | Distal Intergenic | 9 | 49109002 | 49117585 | 8584 | 1 | ENSMUST00000170426.1 | 19976 | Tmprss5 | ENSMUSG00000032268.8 |
| chr9 | 48933212 | 48933539 | 3.46776 | 4.06251 | 1.13123 | Downstream (1-2kb) | 9 | 48899214 | 48911099 | 11886 | 2 | ENSMUST00000003826.7 | -22114 | Htr3a | ENSMUSG00000032269.7 |
| chr9 | 48662896 | 48662971 | 3.59683 | 4.31238 | 1.32775 | Intron (ENSMUST00000093852.3/ENSMUSG00000066687.4, intron 5 of 6) | 9 | 48591877 | 48605153 | 13277 | 2 | ENSMUST00000119426.1 | -57744 | Nnmt | ENSMUSG00000032271.8 |
| chr9 | 62026667 | 62027491 | 3.63522 | 4.45187 | 1.39836 | Promoter | 9 | 61953481 | 62026801 | 73321 | 2 | ENSMUST00000135050.1 | 0 | Paqr5 | ENSMUSG00000032278.6 |
| chr9 | 61381650 | 61381828 | 3.31715 | 3.84259 | 1.1004 | Intron (ENSMUST00000178113.2/ENSMUSG00000032280.11, intron 4 of 19) | 9 | 61375529 | 61416976 | 41448 | 1 | ENSMUST00000162127.1 | 6122 | Tle3 | ENSMUSG00000032280.11 |
| chr9 | 61384235 | 61384310 | 4.03452 | 5.23861 | 1.93113 | Intron (ENSMUST00000178113.2/ENSMUSG00000032280.11, intron 4 of 19) | 9 | 61375529 | 61416976 | 41448 | 1 | ENSMUST00000162127.1 | 8707 | Tle3 | ENSMUSG00000032280.11 |
| chr9 | 61390778 | 61391319 | 3.41657 | 3.98811 | 1.09658 | Intron (ENSMUST00000178113.2/ENSMUSG00000032280.11, intron 4 of 19) | 9 | 61401859 | 61410134 | 8276 | 1 | ENSMUST00000162962.2 | -10540 | Tle3 | ENSMUSG00000032280.11 |
| chr9 | 61406668 | 61407418 | 3.68918 | 4.56909 | 1.509 | Exon (ENSMUST00000178113.2/ENSMUSG00000032280.11, exon 8 of 20) | 9 | 61409937 | 61418419 | 8483 | 1 | ENSMUST00000162281.1 | -2519 | Tle3 | ENSMUSG00000032280.11 |
| chr9 | 61425327 | 61425516 | 3.52974 | 4.24264 | 1.31593 | Distal Intergenic | 9 | 61409937 | 61418419 | 8483 | 1 | ENSMUST00000162281.1 | 15391 | Tle3 | ENSMUSG00000032280.11 |
| chr9 | 61426611 | 61426721 | 3.58832 | 4.33662 | 1.334 | Distal Intergenic | 9 | 61409937 | 61418419 | 8483 | 1 | ENSMUST00000162281.1 | 16675 | Tle3 | ENSMUSG00000032280.11 |
| chr9 | 61429447 | 61429717 | 3.61808 | 4.35108 | 1.34947 | Distal Intergenic | 9 | 61409937 | 61418419 | 8483 | 1 | ENSMUST00000162281.1 | 19511 | Tle3 | ENSMUSG00000032280.11 |
| chr9 | 61432288 | 61432560 | 3.51797 | 4.17957 | 1.27944 | Distal Intergenic | 9 | 61409937 | 61418419 | 8483 | 1 | ENSMUST00000162281.1 | 22352 | Tle3 | ENSMUSG00000032280.11 |
| chr9 | 61433333 | 61433563 | 3.48762 | 4.27051 | 1.31131 | Distal Intergenic | 9 | 61409937 | 61418419 | 8483 | 1 | ENSMUST00000162281.1 | 23397 | Tle3 | ENSMUSG00000032280.11 |
| chr9 | 61435358 | 61435445 | 3.81385 | 4.84913 | 1.66613 | Distal Intergenic | 9 | 61409937 | 61418419 | 8483 | 1 | ENSMUST00000162281.1 | 25422 | Tle3 | ENSMUSG00000032280.11 |
| chr9 | 61435951 | 61436463 | 3.78724 | 4.86609 | 1.68039 | Distal Intergenic | 9 | 61409937 | 61418419 | 8483 | 1 | ENSMUST00000162281.1 | 26015 | Tle3 | ENSMUSG00000032280.11 |
| chr9 | 61438146 | 61438221 | 3.58024 | 4.31525 | 1.38627 | Distal Intergenic | 9 | 61409937 | 61418419 | 8483 | 1 | ENSMUST00000162281.1 | 28210 | Tle3 | ENSMUSG00000032280.11 |
| chr9 | 61441368 | 61442066 | 3.79874 | 4.68547 | 1.53524 | Distal Intergenic | 9 | 61409937 | 61418419 | 8483 | 1 | ENSMUST00000162281.1 | 31432 | Tle3 | ENSMUSG00000032280.11 |
| chr9 | 61444713 | 61444901 | 3.37625 | 3.90108 | 1.04937 | Distal Intergenic | 9 | 61409937 | 61418419 | 8483 | 1 | ENSMUST00000162281.1 | 34777 | Tle3 | ENSMUSG00000032280.11 |
| chr9 | 61448419 | 61448652 | 3.30145 | 3.77935 | 1.01733 | Distal Intergenic | 9 | 61409937 | 61418419 | 8483 | 1 | ENSMUST00000162281.1 | 38483 | Tle3 | ENSMUSG00000032280.11 |
| chr9 | 61449537 | 61449664 | 3.33746 | 3.83353 | 1.01733 | Distal Intergenic | 9 | 61409937 | 61418419 | 8483 | 1 | ENSMUST00000162281.1 | 39601 | Tle3 | ENSMUSG00000032280.11 |
| chr9 | 61453693 | 61454339 | 3.41777 | 3.96773 | 1.06321 | Distal Intergenic | 9 | 61409937 | 61418419 | 8483 | 1 | ENSMUST00000162281.1 | 43757 | Tle3 | ENSMUSG00000032280.11 |
| chr9 | 61455715 | 61455947 | 3.67301 | 4.44884 | 1.39071 | Distal Intergenic | 9 | 61409937 | 61418419 | 8483 | 1 | ENSMUST00000162281.1 | 45779 | Tle3 | ENSMUSG00000032280.11 |
| chr9 | 61457026 | 61457316 | 3.32867 | 3.82452 | 1.03216 | Distal Intergenic | 9 | 61409937 | 61418419 | 8483 | 1 | ENSMUST00000162281.1 | 47090 | Tle3 | ENSMUSG00000032280.11 |
| chr9 | 61457760 | 61457909 | 3.71956 | 4.53498 | 1.43742 | Distal Intergenic | 9 | 61409937 | 61418419 | 8483 | 1 | ENSMUST00000162281.1 | 47824 | Tle3 | ENSMUSG00000032280.11 |
| chr9 | 61458357 | 61458561 | 3.86955 | 4.80432 | 1.58902 | Distal Intergenic | 9 | 61409937 | 61418419 | 8483 | 1 | ENSMUST00000162281.1 | 48421 | Tle3 | ENSMUSG00000032280.11 |
| chr9 | 61459506 | 61459830 | 3.4618 | 4.08059 | 1.21794 | Distal Intergenic | 9 | 61409937 | 61418419 | 8483 | 1 | ENSMUST00000162281.1 | 49570 | Tle3 | ENSMUSG00000032280.11 |
| chr9 | 61462903 | 61463067 | 3.53589 | 4.18352 | 1.19531 | Distal Intergenic | 9 | 61409937 | 61418419 | 8483 | 1 | ENSMUST00000162281.1 | 52967 | Tle3 | ENSMUSG00000032280.11 |
| chr9 | 61465757 | 61466162 | 4.2317 | 5.69609 | 2.28186 | Distal Intergenic | 9 | 61409937 | 61418419 | 8483 | 1 | ENSMUST00000162281.1 | 55821 | Tle3 | ENSMUSG00000032280.11 |
| chr9 | 61469244 | 61469354 | 3.24162 | 3.68958 | 1.01733 | Distal Intergenic | 9 | 61409937 | 61418419 | 8483 | 1 | ENSMUST00000162281.1 | 59308 | Tle3 | ENSMUSG00000032280.11 |
| chr9 | 61474285 | 61474399 | 4.05881 | 5.2405 | 1.98549 | Distal Intergenic | 9 | 61409937 | 61418419 | 8483 | 1 | ENSMUST00000162281.1 | 64349 | Tle3 | ENSMUSG00000032280.11 |
| chr9 | 61478939 | 61479437 | 3.39292 | 3.97138 | 1.18926 | Distal Intergenic | 9 | 61409937 | 61418419 | 8483 | 1 | ENSMUST00000162281.1 | 69003 | Tle3 | ENSMUSG00000032280.11 |
| chr9 | 61482754 | 61483211 | 3.53253 | 4.27813 | 1.31784 | Distal Intergenic | 9 | 61409937 | 61418419 | 8483 | 1 | ENSMUST00000162281.1 | 72818 | Tle3 | ENSMUSG00000032280.11 |
| chr9 | 61483754 | 61483935 | 3.51833 | 4.32165 | 1.32067 | Distal Intergenic | 9 | 61409937 | 61418419 | 8483 | 1 | ENSMUST00000162281.1 | 73818 | Tle3 | ENSMUSG00000032280.11 |
| chr9 | 61489317 | 61489592 | 3.62091 | 4.53718 | 1.4541 | Distal Intergenic | 9 | 61409937 | 61418419 | 8483 | 1 | ENSMUST00000162281.1 | 79381 | Tle3 | ENSMUSG00000032280.11 |
| chr9 | 61501399 | 61501597 | 3.83087 | 4.79659 | 1.66811 | Distal Intergenic | 9 | 61409937 | 61418419 | 8483 | 1 | ENSMUST00000162281.1 | 91463 | Tle3 | ENSMUSG00000032280.11 |
| chr9 | 61507859 | 61507998 | 3.33669 | 3.83242 | 1.01733 | Distal Intergenic | 9 | 61409937 | 61418419 | 8483 | 1 | ENSMUST00000162281.1 | 97923 | Tle3 | ENSMUSG00000032280.11 |
| chr9 | 61508888 | 61509122 | 3.37789 | 3.89451 | 1.01733 | Distal Intergenic | 9 | 61409937 | 61418419 | 8483 | 1 | ENSMUST00000162281.1 | 98952 | Tle3 | ENSMUSG00000032280.11 |
| chr9 | 61509481 | 61509996 | 3.55097 | 4.44348 | 1.41169 | Distal Intergenic | 9 | 61409937 | 61418419 | 8483 | 1 | ENSMUST00000162281.1 | 99545 | Tle3 | ENSMUSG00000032280.11 |
| chr9 | 61510490 | 61510769 | 3.78131 | 4.70997 | 1.63461 | Distal Intergenic | 9 | 61409937 | 61418419 | 8483 | 1 | ENSMUST00000162281.1 | 100554 | Tle3 | ENSMUSG00000032280.11 |
| chr9 | 61511076 | 61511292 | 3.34966 | 3.89412 | 1.14738 | Distal Intergenic | 9 | 61409937 | 61418419 | 8483 | 1 | ENSMUST00000162281.1 | 101140 | Tle3 | ENSMUSG00000032280.11 |
| chr9 | 61524715 | 61525031 | 3.65489 | 4.41221 | 1.36788 | Distal Intergenic | 9 | 61409937 | 61418419 | 8483 | 1 | ENSMUST00000162281.1 | 114779 | Tle3 | ENSMUSG00000032280.11 |
| chr9 | 61528372 | 61528571 | 3.72939 | 4.57697 | 1.50564 | Distal Intergenic | 9 | 61409937 | 61418419 | 8483 | 1 | ENSMUST00000162281.1 | 118436 | Tle3 | ENSMUSG00000032280.11 |
| chr9 | 61530407 | 61530542 | 3.48026 | 4.09246 | 1.17135 | Distal Intergenic | 9 | 61409937 | 61418419 | 8483 | 1 | ENSMUST00000162281.1 | 120471 | Tle3 | ENSMUSG00000032280.11 |
| chr9 | 61541360 | 61541550 | 3.50436 | 4.14709 | 1.22599 | Distal Intergenic | 9 | 61409937 | 61418419 | 8483 | 1 | ENSMUST00000162281.1 | 131424 | Tle3 | ENSMUSG00000032280.11 |
| chr9 | 61592934 | 61593018 | 4.01223 | 5.1799 | 1.86761 | Distal Intergenic | 9 | 61409937 | 61418419 | 8483 | 1 | ENSMUST00000162281.1 | 182998 | Tle3 | ENSMUSG00000032280.11 |
| chr9 | 61596971 | 61597155 | 3.4331 | 3.99519 | 1.07851 | Distal Intergenic | 9 | 61409937 | 61418419 | 8483 | 1 | ENSMUST00000162281.1 | 187035 | Tle3 | ENSMUSG00000032280.11 |
| chr9 | 61597699 | 61597909 | 3.78815 | 4.71324 | 1.55266 | Distal Intergenic | 9 | 61409937 | 61418419 | 8483 | 1 | ENSMUST00000162281.1 | 187763 | Tle3 | ENSMUSG00000032280.11 |
| chr9 | 61620078 | 61620345 | 3.50048 | 4.19706 | 1.30538 | Distal Intergenic | 9 | 61409937 | 61418419 | 8483 | 1 | ENSMUST00000162281.1 | 210142 | Tle3 | ENSMUSG00000032280.11 |
| chr9 | 61623662 | 61623782 | 3.42501 | 3.9932 | 1.11522 | Distal Intergenic | 9 | 61409937 | 61418419 | 8483 | 1 | ENSMUST00000162281.1 | 213726 | Tle3 | ENSMUSG00000032280.11 |
| chr9 | 61644980 | 61645283 | 4.45325 | 5.98371 | 2.45436 | Distal Intergenic | 9 | 61409937 | 61418419 | 8483 | 1 | ENSMUST00000162281.1 | 235044 | Tle3 | ENSMUSG00000032280.11 |
| chr9 | 61659068 | 61659486 | 4.16057 | 5.43058 | 2.10959 | Distal Intergenic | 9 | 61409937 | 61418419 | 8483 | 1 | ENSMUST00000162281.1 | 249132 | Tle3 | ENSMUSG00000032280.11 |
| chr9 | 54691651 | 54691730 | 4.06517 | 5.18351 | 1.86403 | Distal Intergenic | 9 | 54698873 | 54714761 | 15889 | 1 | ENSMUST00000120452.2 | -7143 | Dnaja4 | ENSMUSG00000032285.10 |
| chr9 | 60510182 | 60510378 | 3.44025 | 4.13903 | 1.241 | Promoter | 9 | 60509875 | 60511007 | 1133 | 2 | ENSMUST00000146099.1 | 629 | Thsd4 | ENSMUSG00000032289.10 |
| chr9 | 60519496 | 60519890 | 3.35468 | 3.8595 | 1.01733 | Intron (ENSMUST00000140824.1/ENSMUSG00000032289.10, intron 1 of 3) | 9 | 60428226 | 60522046 | 93821 | 2 | ENSMUST00000140824.1 | 2156 | Thsd4 | ENSMUSG00000032289.10 |
| chr9 | 60520633 | 60521177 | 3.54003 | 4.2128 | 1.26166 | Promoter | 9 | 60428226 | 60522046 | 93821 | 2 | ENSMUST00000140824.1 | 869 | Thsd4 | ENSMUSG00000032289.10 |
| chr9 | 60522002 | 60522376 | 3.61932 | 4.68654 | 1.57389 | Promoter | 9 | 60428226 | 60522046 | 93821 | 2 | ENSMUST00000140824.1 | 0 | Thsd4 | ENSMUSG00000032289.10 |
| chr9 | 54762531 | 54763069 | 3.87041 | 4.8656 | 1.74189 | Distal Intergenic | 9 | 54764748 | 54773110 | 8363 | 1 | ENSMUST00000034830.8 | -1679 | Crabp1 | ENSMUSG00000032291.8 |
| chr9 | 54764431 | 54764952 | 3.82249 | 4.82527 | 1.73223 | Promoter | 9 | 54764748 | 54773110 | 8363 | 1 | ENSMUST00000034830.8 | 0 | Crabp1 | ENSMUSG00000032291.8 |
| chr9 | 54765467 | 54766012 | 3.5305 | 4.37955 | 1.37693 | Promoter | 9 | 54764748 | 54773110 | 8363 | 1 | ENSMUST00000034830.8 | 720 | Crabp1 | ENSMUSG00000032291.8 |
| chr9 | 54766561 | 54767064 | 3.52569 | 4.25555 | 1.36078 | Intron (ENSMUST00000034830.8/ENSMUSG00000032291.8, intron 2 of 3) | 9 | 54764748 | 54773110 | 8363 | 1 | ENSMUST00000034830.8 | 1814 | Crabp1 | ENSMUSG00000032291.8 |
| chr9 | 54769461 | 54769849 | 3.54201 | 4.45784 | 1.41569 | Intron (ENSMUST00000034830.8/ENSMUSG00000032291.8, intron 3 of 3) | 9 | 54764748 | 54773110 | 8363 | 1 | ENSMUST00000034830.8 | 4714 | Crabp1 | ENSMUSG00000032291.8 |
| chr9 | 54770797 | 54771053 | 4.00798 | 5.18035 | 1.964 | Intron (ENSMUST00000034830.8/ENSMUSG00000032291.8, intron 3 of 3) | 9 | 54764748 | 54773110 | 8363 | 1 | ENSMUST00000034830.8 | 6050 | Crabp1 | ENSMUSG00000032291.8 |
| chr9 | 59583728 | 59583954 | 3.29186 | 3.76495 | 1.01733 | Intron (ENSMUST00000143916.2/ENSMUSG00000032297.6, intron 2 of 10) | 9 | 59589288 | 59607291 | 18004 | 1 | ENSMUST00000121266.2 | -5334 | Celf6 | ENSMUSG00000032297.6 |
| chr9 | 59588640 | 59588983 | 3.56905 | 4.24422 | 1.23296 | Promoter | 9 | 59589288 | 59607291 | 18004 | 1 | ENSMUST00000121266.2 | -305 | Celf6 | ENSMUSG00000032297.6 |
| chr9 | 59603556 | 59604211 | 3.39511 | 3.96225 | 1.16234 | Promoter | 9 | 59604033 | 59606101 | 2069 | 1 | ENSMUST00000129728.1 | 0 | Celf6 | ENSMUSG00000032297.6 |
| chr9 | 57281726 | 57281801 | 3.59628 | 4.57656 | 1.50859 | Distal Intergenic | 9 | 57252322 | 57262599 | 10278 | 2 | ENSMUST00000034846.5 | -19128 | 1700017B05Rik | ENSMUSG00000032300.6 |
| chr9 | 55023774 | 55023950 | 3.76582 | 4.64134 | 1.56291 | Intron (ENSMUST00000034851.4/ENSMUSG00000032303.6, intron 3 of 5) | 9 | 55011343 | 55026559 | 15217 | 2 | ENSMUST00000034851.4 | 2609 | Chrna3 | ENSMUSG00000032303.6 |
| chr9 | 55024320 | 55024462 | 3.69887 | 4.6308 | 1.55471 | Exon (ENSMUST00000034851.4/ENSMUSG00000032303.6, exon 2 of 6) | 9 | 55011343 | 55026559 | 15217 | 2 | ENSMUST00000034851.4 | 2097 | Chrna3 | ENSMUSG00000032303.6 |
| chr9 | 55024903 | 55025471 | 3.472 | 4.10426 | 1.23202 | Intron (ENSMUST00000034851.4/ENSMUSG00000032303.6, intron 1 of 5) | 9 | 55011343 | 55026559 | 15217 | 2 | ENSMUST00000034851.4 | 1088 | Chrna3 | ENSMUSG00000032303.6 |
| chr9 | 55026023 | 55026687 | 3.43618 | 4.23739 | 1.30976 | Promoter | 9 | 55011343 | 55026559 | 15217 | 2 | ENSMUST00000034851.4 | 0 | Chrna3 | ENSMUSG00000032303.6 |
| chr9 | 55027102 | 55027386 | 3.58027 | 4.28654 | 1.32611 | Promoter | 9 | 55011343 | 55026559 | 15217 | 2 | ENSMUST00000034851.4 | -544 | Chrna3 | ENSMUSG00000032303.6 |
| chr9 | 55029570 | 55029715 | 3.6723 | 4.45586 | 1.43287 | 3' UTR | 9 | 55011343 | 55026559 | 15217 | 2 | ENSMUST00000034851.4 | -3012 | Chrna3 | ENSMUSG00000032303.6 |
| chr9 | 55240842 | 55241076 | 3.4805 | 4.19537 | 1.27375 | 3' UTR | 9 | 55221222 | 55242461 | 21240 | 2 | ENSMUST00000144939.2 | 1385 | Nrg4 | ENSMUSG00000032311.12 |
| chr9 | 55241742 | 55242128 | 3.79165 | 4.90235 | 1.74866 | Promoter | 9 | 55221222 | 55242461 | 21240 | 2 | ENSMUST00000144939.2 | 333 | Nrg4 | ENSMUSG00000032311.12 |
| chr9 | 55283635 | 55284158 | 3.68801 | 4.49755 | 1.40986 | Promoter | 9 | 55220223 | 55283625 | 63403 | 2 | ENSMUST00000164721.2 | -11 | Nrg4 | ENSMUSG00000032311.12 |
| chr9 | 55318647 | 55318751 | 3.47854 | 4.29236 | 1.32923 | Intron (ENSMUST00000130158.2/ENSMUSG00000032311.12, intron 2 of 8) | 9 | 55306204 | 55325187 | 18984 | 2 | ENSMUST00000133659.1 | 6436 | Nrg4 | ENSMUSG00000032311.12 |
| chr9 | 55321683 | 55322080 | 3.68065 | 4.56657 | 1.50677 | Intron (ENSMUST00000130158.2/ENSMUSG00000032311.12, intron 1 of 8) | 9 | 55306204 | 55325187 | 18984 | 2 | ENSMUST00000133659.1 | 3107 | Nrg4 | ENSMUSG00000032311.12 |
| chr9 | 55322630 | 55322880 | 3.86787 | 4.84706 | 1.71517 | Intron (ENSMUST00000130158.2/ENSMUSG00000032311.12, intron 1 of 8) | 9 | 55306204 | 55325187 | 18984 | 2 | ENSMUST00000133659.1 | 2307 | Nrg4 | ENSMUSG00000032311.12 |
| chr9 | 55325047 | 55325277 | 4.23932 | 5.62772 | 2.2335 | Promoter | 9 | 55306204 | 55325187 | 18984 | 2 | ENSMUST00000133659.1 | 0 | Nrg4 | ENSMUSG00000032311.12 |
| chr9 | 55326523 | 55327792 | 3.74105 | 4.65696 | 1.62763 | Promoter | 9 | 55221301 | 55326749 | 105449 | 2 | ENSMUST00000130158.2 | 0 | Nrg4 | ENSMUSG00000032311.12 |
| chr9 | 55331696 | 55331953 | 3.66644 | 4.49311 | 1.47074 | Intron (ENSMUST00000085754.4/ENSMUSG00000032313.6, intron 1 of 10) | 9 | 55333201 | 55437358 | 104158 | 1 | ENSMUST00000130465.1 | -1248 | AI118078 | ENSMUSG00000032313.6 |
| chr9 | 55335592 | 55335779 | 3.63316 | 4.3844 | 1.36621 | Intron (ENSMUST00000085754.4/ENSMUSG00000032313.6, intron 1 of 10) | 9 | 55333201 | 55437358 | 104158 | 1 | ENSMUST00000130465.1 | 2392 | AI118078 | ENSMUSG00000032313.6 |
| chr9 | 55337642 | 55337847 | 3.78341 | 4.80852 | 1.63537 | Intron (ENSMUST00000085754.4/ENSMUSG00000032313.6, intron 1 of 10) | 9 | 55333201 | 55437358 | 104158 | 1 | ENSMUST00000130465.1 | 4442 | AI118078 | ENSMUSG00000032313.6 |
| chr9 | 55346263 | 55347043 | 3.64502 | 4.3863 | 1.32377 | Intron (ENSMUST00000085754.4/ENSMUSG00000032313.6, intron 1 of 10) | 9 | 55333201 | 55437358 | 104158 | 1 | ENSMUST00000130465.1 | 13063 | AI118078 | ENSMUSG00000032313.6 |
| chr9 | 55360282 | 55360571 | 3.221 | 3.65871 | 1.01733 | Intron (ENSMUST00000085754.4/ENSMUSG00000032313.6, intron 1 of 10) | 9 | 55333201 | 55437358 | 104158 | 1 | ENSMUST00000130465.1 | 27082 | AI118078 | ENSMUSG00000032313.6 |
| chr9 | 55360910 | 55361185 | 3.84021 | 5.10882 | 1.8377 | Intron (ENSMUST00000085754.4/ENSMUSG00000032313.6, intron 1 of 10) | 9 | 55333201 | 55437358 | 104158 | 1 | ENSMUST00000130465.1 | 27710 | AI118078 | ENSMUSG00000032313.6 |
| chr9 | 55362669 | 55362873 | 3.44714 | 4.07352 | 1.24885 | Intron (ENSMUST00000085754.4/ENSMUSG00000032313.6, intron 1 of 10) | 9 | 55333201 | 55437358 | 104158 | 1 | ENSMUST00000130465.1 | 29469 | AI118078 | ENSMUSG00000032313.6 |
| chr9 | 55366977 | 55367203 | 3.30588 | 3.81628 | 1.10625 | Intron (ENSMUST00000085754.4/ENSMUSG00000032313.6, intron 1 of 10) | 9 | 55333201 | 55437358 | 104158 | 1 | ENSMUST00000130465.1 | 33777 | AI118078 | ENSMUSG00000032313.6 |
| chr9 | 55369492 | 55369601 | 3.67127 | 4.57658 | 1.51006 | Intron (ENSMUST00000085754.4/ENSMUSG00000032313.6, intron 1 of 10) | 9 | 55333201 | 55437358 | 104158 | 1 | ENSMUST00000130465.1 | 36292 | AI118078 | ENSMUSG00000032313.6 |
| chr9 | 55384066 | 55384575 | 3.69967 | 4.58256 | 1.51494 | Intron (ENSMUST00000085754.4/ENSMUSG00000032313.6, intron 3 of 10) | 9 | 55333201 | 55437358 | 104158 | 1 | ENSMUST00000130465.1 | 50866 | AI118078 | ENSMUSG00000032313.6 |
| chr9 | 55392135 | 55392353 | 3.56801 | 4.33111 | 1.37808 | Intron (ENSMUST00000085754.4/ENSMUSG00000032313.6, intron 3 of 10) | 9 | 55333201 | 55437358 | 104158 | 1 | ENSMUST00000130465.1 | 58935 | AI118078 | ENSMUSG00000032313.6 |
| chr9 | 55393481 | 55393664 | 3.61832 | 4.36805 | 1.39041 | Intron (ENSMUST00000085754.4/ENSMUSG00000032313.6, intron 3 of 10) | 9 | 55333201 | 55437358 | 104158 | 1 | ENSMUST00000130465.1 | 60281 | AI118078 | ENSMUSG00000032313.6 |
| chr9 | 55409879 | 55410058 | 3.34586 | 3.84619 | 1.01733 | Intron (ENSMUST00000085754.4/ENSMUSG00000032313.6, intron 6 of 10) | 9 | 55333201 | 55437358 | 104158 | 1 | ENSMUST00000130465.1 | 76679 | AI118078 | ENSMUSG00000032313.6 |
| chr9 | 55428662 | 55429253 | 3.44114 | 4.08035 | 1.17916 | Intron (ENSMUST00000085754.4/ENSMUSG00000032313.6, intron 9 of 10) | 9 | 55474537 | 55487513 | 12977 | 2 | ENSMUST00000139342.1 | 58260 | Etfa | ENSMUSG00000032314.9 |
| chr9 | 55430121 | 55430413 | 3.34901 | 3.87518 | 1.08298 | Exon (ENSMUST00000085754.4/ENSMUSG00000032313.6, exon 10 of 11) | 9 | 55474537 | 55487513 | 12977 | 2 | ENSMUST00000139342.1 | 57100 | Etfa | ENSMUSG00000032314.9 |
| chr9 | 57692911 | 57693234 | 3.62243 | 4.44875 | 1.42769 | Distal Intergenic | 9 | 57697613 | 57703822 | 6210 | 1 | ENSMUST00000034865.4 | -4379 | Cyp1a1 | ENSMUSG00000032315.5 |
| chr9 | 57696380 | 57697157 | 3.75074 | 4.62762 | 1.54939 | Promoter | 9 | 57697613 | 57703822 | 6210 | 1 | ENSMUST00000034865.4 | -456 | Cyp1a1 | ENSMUSG00000032315.5 |
| chr9 | 57697467 | 57697595 | 3.34827 | 3.84984 | 1.01733 | Promoter | 9 | 57697613 | 57703822 | 6210 | 1 | ENSMUST00000034865.4 | -18 | Cyp1a1 | ENSMUSG00000032315.5 |
| chr9 | 55525728 | 55525829 | 3.89685 | 4.93834 | 1.79604 | Distal Intergenic | 9 | 55538672 | 55542570 | 3899 | 1 | ENSMUST00000175950.2 | -12843 | Isl2 | ENSMUSG00000032318.7 |
| chr9 | 55535166 | 55535449 | 3.46914 | 4.06368 | 1.12815 | Distal Intergenic | 9 | 55538672 | 55542570 | 3899 | 1 | ENSMUST00000175950.2 | -3223 | Isl2 | ENSMUSG00000032318.7 |
| chr9 | 55536484 | 55537481 | 3.63596 | 4.48277 | 1.45884 | Distal Intergenic | 9 | 55538672 | 55542570 | 3899 | 1 | ENSMUST00000175950.2 | -1191 | Isl2 | ENSMUSG00000032318.7 |
| chr9 | 55537856 | 55546938 | 4.28852 | 5.86884 | 2.44772 | Promoter | 9 | 55538672 | 55542570 | 3899 | 1 | ENSMUST00000175950.2 | 0 | Isl2 | ENSMUSG00000032318.7 |
| chr9 | 55547374 | 55548535 | 3.83089 | 4.76189 | 1.60549 | Downstream (1-2kb) | 9 | 55542336 | 55544280 | 1945 | 1 | ENSMUST00000165302.1 | 5039 | Isl2 | ENSMUSG00000032318.7 |
| chr9 | 58055012 | 58055172 | 3.44 | 4.04168 | 1.20447 | Intron (ENSMUST00000042205.7/ENSMUSG00000037716.9, intron 11 of 18) | 9 | 58063788 | 58142031 | 78244 | 1 | ENSMUST00000145886.1 | -8616 | Stra6 | ENSMUSG00000032327.9 |
| chr9 | 58064228 | 58064314 | 3.90043 | 4.86304 | 1.62639 | Promoter | 9 | 58063788 | 58142031 | 78244 | 1 | ENSMUST00000145886.1 | 441 | Stra6 | ENSMUSG00000032327.9 |
| chr9 | 58129463 | 58129586 | 3.54097 | 4.22595 | 1.25524 | Promoter | 9 | 58129501 | 58140871 | 11371 | 1 | ENSMUST00000133287.2 | 0 | Stra6 | ENSMUSG00000032327.9 |
| chr9 | 58131302 | 58131429 | 4.05831 | 5.26883 | 1.98208 | Promoter | 9 | 58131897 | 58140444 | 8548 | 1 | ENSMUST00000136338.2 | -468 | Stra6 | ENSMUSG00000032327.9 |
| chr9 | 58135315 | 58135551 | 3.34765 | 3.84889 | 1.01733 | Promoter | 9 | 58134810 | 58139366 | 4557 | 1 | ENSMUST00000134450.1 | 506 | Stra6 | ENSMUSG00000032327.9 |
| chr9 | 58136047 | 58136312 | 3.72031 | 4.53266 | 1.42373 | Intron (ENSMUST00000145886.1/ENSMUSG00000032327.9, intron 2 of 4) | 9 | 58134810 | 58139366 | 4557 | 1 | ENSMUST00000134450.1 | 1238 | Stra6 | ENSMUSG00000032327.9 |

| | | | | | | | | | | | | | | | |
|---|---|---|---|---|---|---|---|---|---|---|---|---|---|---|---|
| chr9 | 58140405 | 58140535 | 3.64292 | 4.54841 | 1.46625 | Exon (ENSMUST00000145886.1/ENSMUSG00000032327.9, exon 5 of 5) | 9 | 58134810 | 58139366 | 4557 | 1 | ENSMUST00000134450.1 | 5596 | Stra6 | ENSMUSG00000032327.9 |
| chr9 | 79714012 | 79714479 | 4.09939 | 5.32604 | 2.04113 | Exon (ENSMUST00000122827.1/ENSMUSG00000032332.12, exon 2 of 2) | 9 | 79714905 | 79718494 | 3590 | 2 | ENSMUST00000141667.1 | 4015 | Col12a1 | ENSMUSG00000032332.12 |
| chr9 | 79717290 | 79717494 | 3.2547 | 3.70919 | 1.01733 | Promoter | 9 | 79714905 | 79718494 | 3590 | 2 | ENSMUST00000141667.1 | 1000 | Col12a1 | ENSMUSG00000032332.12 |
| chr9 | 79717830 | 79718116 | 3.55604 | 4.2542 | 1.3306 | Promoter | 9 | 79714905 | 79718494 | 3590 | 2 | ENSMUST00000141667.1 | 378 | Col12a1 | ENSMUSG00000032332.12 |
| chr9 | 79718452 | 79718952 | 4.10526 | 5.3028 | 1.98791 | Promoter | 9 | 79598991 | 79718518 | 119528 | 2 | ENSMUST00000071750.7 | 0 | Col12a1 | ENSMUSG00000032332.12 |
| chr9 | 79720005 | 79720629 | 3.65165 | 4.66267 | 1.5377 | Distal Intergenic | 9 | 79712534 | 79718831 | 6298 | 2 | ENSMUST00000122827.1 | -1175 | Col12a1 | ENSMUSG00000032332.12 |
| chr9 | 79721048 | 79721195 | 3.64049 | 4.37843 | 1.32685 | Distal Intergenic | 9 | 79712534 | 79718831 | 6298 | 2 | ENSMUST00000122827.1 | -2218 | Col12a1 | ENSMUSG00000032332.12 |
| chr9 | 58813663 | 58813835 | 3.53955 | 4.20192 | 1.23751 | Distal Intergenic | 9 | 58823512 | 58860955 | 37444 | 1 | ENSMUST00000034889.8 | -9677 | Hcn4 | ENSMUSG00000032338.8 |
| chr9 | 58822249 | 58824055 | 3.67155 | 4.49712 | 1.49371 | Promoter | 9 | 58823512 | 58860955 | 37444 | 1 | ENSMUST00000034889.8 | 0 | Hcn4 | ENSMUSG00000032338.8 |
| chr9 | 59056979 | 59057388 | 3.49151 | 4.11907 | 1.20519 | Distal Intergenic | 9 | 58874679 | 59036441 | 161763 | 2 | ENSMUST00000068664.5 | -20539 | Neo1 | ENSMUSG00000032340.7 |
| chr9 | 59071375 | 59071597 | 3.54354 | 4.27992 | 1.3231 | Distal Intergenic | 9 | 58874679 | 59036441 | 161763 | 2 | ENSMUST00000068664.5 | -34935 | Neo1 | ENSMUSG00000032340.7 |
| chr9 | 59072702 | 59072792 | 3.81423 | 4.72543 | 1.58473 | Distal Intergenic | 9 | 58874679 | 59036441 | 161763 | 2 | ENSMUST00000068664.5 | -36262 | Neo1 | ENSMUSG00000032340.7 |
| chr9 | 78441068 | 78441160 | 3.74416 | 4.62465 | 1.5769 | Intron (ENSMUST00000070742.8/ENSMUSG00000032344.12, intron 1 of 4) | 9 | 78431789 | 78442885 | 11097 | 2 | ENSMUST00000127190.1 | 1725 | Mb21d1 | ENSMUSG00000032344.12 |
| chr9 | 78442729 | 78443120 | 3.18789 | 4.16529 | 1.23435 | Promoter | 9 | 78431789 | 78442885 | 11097 | 2 | ENSMUST00000127190.1 | 0 | Mb21d1 | ENSMUSG00000032344.12 |
| chr9 | 89874412 | 89874579 | 3.23368 | 3.67769 | 1.01733 | Distal Intergenic | 9 | 89909908 | 89915847 | 5940 | 1 | ENSMUST00000189545.1 | -35329 | Rasgrf1 | ENSMUSG00000032356.7 |
| chr9 | 89881705 | 89882137 | 3.40388 | 3.95282 | 1.08407 | Distal Intergenic | 9 | 89909908 | 89915847 | 5940 | 1 | ENSMUST00000189545.1 | -27771 | Rasgrf1 | ENSMUSG00000032356.7 |
| chr9 | 89882956 | 89883561 | 4.16426 | 5.4152 | 2.0515 | Distal Intergenic | 9 | 89909908 | 89915847 | 5940 | 1 | ENSMUST00000189545.1 | -26347 | Rasgrf1 | ENSMUSG00000032356.7 |
| chr9 | 89884295 | 89884672 | 3.64407 | 4.47422 | 1.44336 | Distal Intergenic | 9 | 89909908 | 89915847 | 5940 | 1 | ENSMUST00000189545.1 | -25236 | Rasgrf1 | ENSMUSG00000032356.7 |
| chr9 | 89885580 | 89885815 | 3.64815 | 4.39154 | 1.32279 | Distal Intergenic | 9 | 89909908 | 89915847 | 5940 | 1 | ENSMUST00000189545.1 | -24093 | Rasgrf1 | ENSMUSG00000032356.7 |
| chr9 | 89903623 | 89904424 | 3.66577 | 4.44466 | 1.38434 | Distal Intergenic | 9 | 89909908 | 89915847 | 5940 | 1 | ENSMUST00000189545.1 | -5484 | Rasgrf1 | ENSMUSG00000032356.7 |
| chr9 | 89906979 | 89907147 | 3.48557 | 4.10457 | 1.18018 | Distal Intergenic | 9 | 89909908 | 89915847 | 5940 | 1 | ENSMUST00000189545.1 | -2761 | Rasgrf1 | ENSMUSG00000032356.7 |
| chr9 | 89907461 | 89908273 | 3.53571 | 4.19015 | 1.21614 | Distal Intergenic | 9 | 89909908 | 89915847 | 5940 | 1 | ENSMUST00000189545.1 | -1635 | Rasgrf1 | ENSMUSG00000032356.7 |
| chr9 | 89908671 | 89909185 | 3.65517 | 4.47844 | 1.46872 | Promoter | 9 | 89909908 | 89915847 | 5940 | 1 | ENSMUST00000189545.1 | -723 | Rasgrf1 | ENSMUSG00000032356.7 |
| chr9 | 89909590 | 89910488 | 3.94959 | 5.0097 | 1.77175 | Promoter | 9 | 89909908 | 89915847 | 5940 | 1 | ENSMUST00000189545.1 | 0 | Rasgrf1 | ENSMUSG00000032356.7 |
| chr9 | 89911033 | 89911469 | 3.50677 | 4.1776 | 1.27054 | Exon (ENSMUST00000189545.1/ENSMUSG00000032356.7, exon 2 of 2) | 9 | 89909912 | 89926740 | 16829 | 1 | ENSMUST00000190073.1 | 1122 | Rasgrf1 | ENSMUSG00000032356.7 |
| chr9 | 89912282 | 89912454 | 3.7885 | 4.66077 | 1.51415 | 3' UTR | 9 | 89909912 | 89926740 | 16829 | 1 | ENSMUST00000190073.1 | 2371 | Rasgrf1 | ENSMUSG00000032356.7 |
| chr9 | 76567053 | 76567338 | 3.45349 | 4.20248 | 1.24853 | Promoter | 9 | 76490054 | 76567116 | 77063 | 2 | ENSMUST00000183437.2 | 0 | Fam83b | ENSMUSG00000032358.7 |
| chr9 | 76321357 | 76321857 | 3.54395 | 4.22399 | 1.29579 | Exon (ENSMUST00000184200.1/ENSMUSG00000032360.11, exon 1 of 1) | 9 | 76321676 | 76323322 | 1647 | 2 | ENSMUST00000184200.1 | 1465 | Hcrtr2 | ENSMUSG00000032360.11 |
| chr9 | 76322177 | 76322452 | 3.59082 | 4.33903 | 1.40284 | Promoter | 9 | 76321676 | 76323322 | 1647 | 2 | ENSMUST00000184200.1 | 870 | Hcrtr2 | ENSMUSG00000032360.11 |
| chr9 | 90163568 | 90163791 | 3.48842 | 4.30647 | 1.314 | Promoter | 9 | 90163226 | 90200102 | 36877 | 1 | ENSMUST00000113060.2 | 343 | Adamts7 | ENSMUSG00000032363.10 |
| chr9 | 91361560 | 91366010 | 4.26797 | 5.70329 | 2.27489 | Promoter | 9 | 91358058 | 91365768 | 7711 | 2 | ENSMUST00000034927.7 | 0 | Zic1 | ENSMUSG00000032368.9 |
| chr9 | 91366748 | 91367275 | 4.24986 | 5.64776 | 2.23651 | Promoter | 9 | 91360500 | 91365810 | 5311 | 2 | ENSMUST00000065360.4 | -939 | Zic1 | ENSMUSG00000032368.9 |
| chr9 | 66152319 | 66152624 | 3.55858 | 4.33412 | 1.38092 | Distal Intergenic | 9 | 66158223 | 66272242 | 114020 | 1 | ENSMUST00000034944.3 | -5599 | Dapk2 | ENSMUSG00000032380.3 |
| chr9 | 66158270 | 66158365 | 3.97603 | 5.09943 | 1.86898 | Promoter | 9 | 66158235 | 66222080 | 63846 | 1 | ENSMUST00000129442.1 | 36 | Dapk2 | ENSMUSG00000032380.3 |
| chr9 | 65608834 | 65609160 | 4.12595 | 5.44809 | 2.11205 | Distal Intergenic | 9 | 65629648 | 65660528 | 30881 | 1 | ENSMUST00000169003.2 | -20488 | Rbpms2 | ENSMUSG00000032387.9 |
| chr9 | 65613221 | 65613626 | 3.62816 | 4.45745 | 1.44236 | Distal Intergenic | 9 | 65629648 | 65660528 | 30881 | 1 | ENSMUST00000169003.2 | -16022 | Rbpms2 | ENSMUSG00000032387.9 |
| chr9 | 65619202 | 65619315 | 3.24143 | 3.83336 | 1.01733 | Distal Intergenic | 9 | 65629648 | 65660528 | 30881 | 1 | ENSMUST00000169003.2 | -10333 | Rbpms2 | ENSMUSG00000032387.9 |
| chr9 | 65622854 | 65623379 | 3.88708 | 4.99253 | 1.81246 | Distal Intergenic | 9 | 65629648 | 65660528 | 30881 | 1 | ENSMUST00000169003.2 | -6269 | Rbpms2 | ENSMUSG00000032387.9 |
| chr9 | 65623979 | 65624540 | 3.59175 | 4.32391 | 1.37136 | Distal Intergenic | 9 | 65629648 | 65660528 | 30881 | 1 | ENSMUST00000169003.2 | -5108 | Rbpms2 | ENSMUSG00000032387.9 |
| chr9 | 65625078 | 65625162 | 3.88541 | 4.83207 | 1.60233 | Distal Intergenic | 9 | 65629648 | 65660528 | 30881 | 1 | ENSMUST00000169003.2 | -4486 | Rbpms2 | ENSMUSG00000032387.9 |
| chr9 | 65627465 | 65627763 | 4.43327 | 5.93933 | 2.41692 | Distal Intergenic | 9 | 65629648 | 65660528 | 30881 | 1 | ENSMUST00000169003.2 | -1885 | Rbpms2 | ENSMUSG00000032387.9 |
| chr9 | 65628916 | 65629523 | 4.02737 | 5.39646 | 2.08807 | Promoter | 9 | 65629648 | 65660528 | 30881 | 1 | ENSMUST00000169003.2 | -125 | Rbpms2 | ENSMUSG00000032387.9 |
| chr9 | 65630133 | 65630448 | 3.54155 | 4.30486 | 1.32285 | Promoter | 9 | 65630582 | 65660505 | 29924 | 1 | ENSMUST00000055844.8 | -134 | Rbpms2 | ENSMUSG00000032387.9 |
| chr9 | 65631042 | 65631297 | 4.21155 | 5.51069 | 2.13713 | Promoter | 9 | 65630582 | 65660505 | 29924 | 1 | ENSMUST00000055844.8 | 461 | Rbpms2 | ENSMUSG00000032387.9 |
| chr9 | 65636403 | 65636719 | 3.41179 | 3.95855 | 1.06291 | Intron (ENSMUST00000169003.2/ENSMUSG00000032387.9, intron 2 of 8) | 9 | 65630582 | 65660505 | 29924 | 1 | ENSMUST00000055844.8 | 5822 | Rbpms2 | ENSMUSG00000032387.9 |
| chr9 | 65640925 | 65641055 | 3.39993 | 3.99294 | 1.09788 | Intron (ENSMUST00000169003.2/ENSMUSG00000032387.9, intron 2 of 8) | 9 | 65630582 | 65660505 | 29924 | 1 | ENSMUST00000055844.8 | 10344 | Rbpms2 | ENSMUSG00000032387.9 |
| chr9 | 65641562 | 65641873 | 3.44448 | 4.62071 | 1.52984 | Intron (ENSMUST00000169003.2/ENSMUSG00000032387.9, intron 2 of 8) | 9 | 65630582 | 65660505 | 29924 | 1 | ENSMUST00000055844.8 | 10981 | Rbpms2 | ENSMUSG00000032387.9 |
| chr9 | 65642303 | 65642746 | 3.50295 | 4.22141 | 1.26988 | Intron (ENSMUST00000169003.2/ENSMUSG00000032387.9, intron 2 of 8) | 9 | 65630582 | 65660505 | 29924 | 1 | ENSMUST00000055844.8 | 11722 | Rbpms2 | ENSMUSG00000032387.9 |
| chr9 | 65121932 | 65122224 | 3.69429 | 4.4844 | 1.39585 | Intron (ENSMUST00000035499.4/ENSMUSG00000032816.9, intron 5 of 19) | 9 | 65141154 | 65185857 | 44704 | 1 | ENSMUST00000034961.4 | -18930 | Igdcc3 | ENSMUSG00000032394.5 |
| chr9 | 65122538 | 65122803 | 3.51041 | 4.14054 | 1.17983 | Exon (ENSMUST00000035499.4/ENSMUSG00000032816.9, exon 6 of 20) | 9 | 65141154 | 65185857 | 44704 | 1 | ENSMUST00000034961.4 | -18351 | Igdcc3 | ENSMUSG00000032394.5 |
| chr9 | 65123989 | 65124591 | 3.76006 | 4.67188 | 1.53134 | Exon (ENSMUST00000035499.4/ENSMUSG00000032816.9, exon 7 of 20) | 9 | 65141154 | 65185857 | 44704 | 1 | ENSMUST00000034961.4 | -16563 | Igdcc3 | ENSMUSG00000032394.5 |
| chr9 | 65128408 | 65129117 | 3.51295 | 4.15377 | 1.20682 | Exon (ENSMUST00000035499.4/ENSMUSG00000032816.9, exon 11 of 20) | 9 | 65141154 | 65185857 | 44704 | 1 | ENSMUST00000034961.4 | -12037 | Igdcc3 | ENSMUSG00000032394.5 |
| chr9 | 65140345 | 65141218 | 3.8888 | 4.95064 | 1.80614 | Promoter | 9 | 65141154 | 65185857 | 44704 | 1 | ENSMUST00000034961.4 | 0 | Igdcc3 | ENSMUSG00000032394.5 |
| chr9 | 65142114 | 65142566 | 3.27396 | 4.0748 | 1.16726 | Promoter | 9 | 65141154 | 65185857 | 44704 | 1 | ENSMUST00000034961.4 | 961 | Igdcc3 | ENSMUSG00000032394.5 |
| chr9 | 65143578 | 65143782 | 3.7613 | 4.63455 | 1.54949 | Intron (ENSMUST00000034961.4/ENSMUSG00000032394.5, intron 1 of 13) | 9 | 65141154 | 65185857 | 44704 | 1 | ENSMUST00000034961.4 | 2425 | Igdcc3 | ENSMUSG00000032394.5 |
| chr9 | 65145226 | 65145384 | 3.35173 | 3.85504 | 1.01733 | Intron (ENSMUST00000034961.4/ENSMUSG00000032394.5, intron 2 of 13) | 9 | 65141154 | 65185857 | 44704 | 1 | ENSMUST00000034961.4 | 4073 | Igdcc3 | ENSMUSG00000032394.5 |
| chr9 | 65153212 | 65153287 | 3.70477 | 4.53311 | 1.49075 | Intron (ENSMUST00000034961.4/ENSMUSG00000032394.5, intron 2 of 13) | 9 | 65141154 | 65185857 | 44704 | 1 | ENSMUST00000034961.4 | 12059 | Igdcc3 | ENSMUSG00000032394.5 |
| chr9 | 65156467 | 65158275 | 3.94305 | 5.1423 | 1.9438 | Intron (ENSMUST00000034961.4/ENSMUSG00000032394.5, intron 2 of 13) | 9 | 65141154 | 65185857 | 44704 | 1 | ENSMUST00000034961.4 | 15314 | Igdcc3 | ENSMUSG00000032394.5 |
| chr9 | 65161981 | 65163027 | 3.56604 | 4.32508 | 1.38512 | Intron (ENSMUST00000034961.4/ENSMUSG00000032394.5, intron 2 of 13) | 9 | 65141154 | 65185857 | 44704 | 1 | ENSMUST00000034961.4 | 20828 | Igdcc3 | ENSMUSG00000032394.5 |
| chr9 | 65163388 | 65163837 | 3.49931 | 4.25251 | 1.30457 | Intron (ENSMUST00000034961.4/ENSMUSG00000032394.5, intron 2 of 13) | 9 | 65141154 | 65185857 | 44704 | 1 | ENSMUST00000034961.4 | 22235 | Igdcc3 | ENSMUSG00000032394.5 |
| chr9 | 65165418 | 65165987 | 3.55246 | 4.28148 | 1.35479 | Intron (ENSMUST00000034961.4/ENSMUSG00000032394.5, intron 2 of 13) | 9 | 65141154 | 65185857 | 44704 | 1 | ENSMUST00000034961.4 | 24265 | Igdcc3 | ENSMUSG00000032394.5 |
| chr9 | 96386162 | 96386452 | 3.29673 | 3.77226 | 1.01733 | Distal Intergenic | 9 | 96332655 | 96364442 | 31788 | 2 | ENSMUST00000034983.6 | -21721 | Atp1b3 | ENSMUSG00000032412.8 |
| chr9 | 96663777 | 96664076 | 3.76496 | 4.65313 | 1.52544 | Distal Intergenic | 9 | 96539300 | 96631617 | 92318 | 2 | ENSMUST00000034984.7 | -32161 | Rasa2 | ENSMUSG00000032413.7 |
| chr9 | 87710125 | 87710520 | 3.67984 | 4.67588 | 1.55755 | Intron (ENSMUST00000034991.7/ENSMUSG00000032419.8, intron 6 of 7) | 9 | 87724397 | 87730497 | 6101 | 2 | ENSMUST00000186698.1 | 19977 | Tbx18 | ENSMUSG00000032419.8 |
| chr9 | 87719392 | 87719567 | 3.7015 | 4.63809 | 1.52997 | Intron (ENSMUST00000034991.7/ENSMUSG00000032419.8, intron 4 of 7) | 9 | 87724397 | 87730497 | 6101 | 2 | ENSMUST00000186698.1 | 10930 | Tbx18 | ENSMUSG00000032419.8 |
| chr9 | 87723072 | 87723547 | 3.91315 | 5.00622 | 1.82581 | Intron (ENSMUST00000034991.7/ENSMUSG00000032419.8, intron 4 of 7) | 9 | 87724397 | 87730497 | 6101 | 2 | ENSMUST00000186698.1 | 6950 | Tbx18 | ENSMUSG00000032419.8 |
| chr9 | 87724367 | 87724482 | 3.33051 | 4.10962 | 1.22915 | Exon (ENSMUST00000034991.7/ENSMUSG00000032419.8, exon 4 of 8) | 9 | 87724397 | 87730497 | 6101 | 2 | ENSMUST00000186698.1 | 6015 | Tbx18 | ENSMUSG00000032419.8 |
| chr9 | 87725328 | 87725657 | 3.8994 | 5.08014 | 1.8127 | Intron (ENSMUST00000034991.7/ENSMUSG00000032419.8, intron 3 of 7) | 9 | 87724397 | 87730497 | 6101 | 2 | ENSMUST00000186698.1 | 4840 | Tbx18 | ENSMUSG00000032419.8 |
| chr9 | 87727191 | 87728143 | 3.95237 | 5.11078 | 1.88384 | Exon (ENSMUST00000034991.7/ENSMUSG00000032419.8, exon 3 of 8) | 9 | 87724397 | 87730497 | 6101 | 2 | ENSMUST00000186698.1 | 2354 | Tbx18 | ENSMUSG00000032419.8 |
| chr9 | 87729461 | 87731064 | 4.12271 | 5.96143 | 2.50795 | Promoter | 9 | 87724397 | 87730497 | 6101 | 2 | ENSMUST00000186698.1 | 0 | Tbx18 | ENSMUSG00000032419.8 |
| chr9 | 87731375 | 87732029 | 4.14402 | 5.46858 | 2.14252 | Promoter | 9 | 87702800 | 87731260 | 28461 | 2 | ENSMUST00000034991.7 | -116 | Tbx18 | ENSMUSG00000032419.8 |
| chr9 | 87732366 | 87732876 | 4.55374 | 6.43251 | 2.85833 | Distal Intergenic | 9 | 87702800 | 87731260 | 28461 | 2 | ENSMUST00000034991.7 | -1107 | Tbx18 | ENSMUSG00000032419.8 |
| chr9 | 87733318 | 87734027 | 3.86122 | 5.00837 | 1.7717 | Distal Intergenic | 9 | 87702800 | 87731260 | 28461 | 2 | ENSMUST00000034991.7 | -2059 | Tbx18 | ENSMUSG00000032419.8 |
| chr9 | 87734750 | 87734938 | 3.73568 | 4.7545 | 1.62705 | Distal Intergenic | 9 | 87702800 | 87731260 | 28461 | 2 | ENSMUST00000034991.7 | -3491 | Tbx18 | ENSMUSG00000032419.8 |
| chr9 | 87735278 | 87737194 | 3.83061 | 4.94929 | 1.75651 | Distal Intergenic | 9 | 87702800 | 87731260 | 28461 | 2 | ENSMUST00000034991.7 | -4019 | Tbx18 | ENSMUSG00000032419.8 |
| chr9 | 87737591 | 87738916 | 3.50685 | 4.44284 | 1.43477 | Distal Intergenic | 9 | 87702800 | 87731260 | 28461 | 2 | ENSMUST00000034991.7 | -6332 | Tbx18 | ENSMUSG00000032419.8 |
| chr9 | 87739296 | 87740100 | 4.29658 | 5.78341 | 2.37046 | Distal Intergenic | 9 | 87702800 | 87731260 | 28461 | 2 | ENSMUST00000034991.7 | -8037 | Tbx18 | ENSMUSG00000032419.8 |
| chr9 | 87740572 | 87741002 | 3.54849 | 4.33029 | 1.3861 | Distal Intergenic | 9 | 87702800 | 87731260 | 28461 | 2 | ENSMUST00000034991.7 | -9313 | Tbx18 | ENSMUSG00000032419.8 |
| chr9 | 87741393 | 87742482 | 3.40063 | 3.97366 | 1.16457 | Distal Intergenic | 9 | 87702800 | 87731260 | 28461 | 2 | ENSMUST00000034991.7 | -10134 | Tbx18 | ENSMUSG00000032419.8 |
| chr9 | 87743093 | 87743390 | 3.59464 | 4.33141 | 1.36676 | Distal Intergenic | 9 | 87702800 | 87731260 | 28461 | 2 | ENSMUST00000034991.7 | -11834 | Tbx18 | ENSMUSG00000032419.8 |
| chr9 | 87744092 | 87744221 | 4.10876 | 5.29427 | 1.96967 | Distal Intergenic | 9 | 87702800 | 87731260 | 28461 | 2 | ENSMUST00000034991.7 | -12833 | Tbx18 | ENSMUSG00000032419.8 |
| chr9 | 88327369 | 88328108 | 3.73606 | 4.56938 | 1.47178 | Promoter | 9 | 88327618 | 88331219 | 3602 | 1 | ENSMUST00000186789.1 | 0 | Nt5e | ENSMUSG00000032420.7 |
| chr9 | 118472861 | 118473497 | 3.84762 | 4.80922 | 1.62702 | Distal Intergenic | 9 | 118478212 | 118485366 | 7155 | 1 | ENSMUST00000111763.2 | -4715 | Eomes | ENSMUSG00000032446.9 |
| chr9 | 118473990 | 118475620 | 3.88576 | 4.88072 | 1.68173 | Distal Intergenic | 9 | 118478212 | 118485366 | 7155 | 1 | ENSMUST00000111763.2 | -2592 | Eomes | ENSMUSG00000032446.9 |
| chr9 | 118475955 | 118478155 | 4.1043 | 5.49246 | 2.1409 | Promoter | 9 | 118478212 | 118485366 | 7155 | 1 | ENSMUST00000111763.2 | -57 | Eomes | ENSMUSG00000032446.9 |
| chr9 | 118478515 | 118480081 | 4.15743 | 5.48505 | 2.17076 | Promoter | 9 | 118478861 | 118482757 | 3897 | 1 | ENSMUST00000150633.1 | 0 | Eomes | ENSMUSG00000032446.9 |
| chr9 | 118480579 | 118480925 | 3.57993 | 4.36381 | 1.40435 | Exon (ENSMUST00000111763.2/ENSMUSG00000032446.9, exon 2 of 6) | 9 | 118478861 | 118482757 | 3897 | 1 | ENSMUST00000150633.1 | 1719 | Eomes | ENSMUSG00000032446.9 |
| chr9 | 118482537 | 118482673 | 3.43448 | 4.05933 | 1.2061 | 3' UTR | 9 | 118478861 | 118482757 | 3897 | 1 | ENSMUST00000150633.1 | 3677 | Eomes | ENSMUSG00000032446.9 |
| chr9 | 118483042 | 118483250 | 3.43586 | 4.0674 | 1.20378 | Intron (ENSMUST00000111763.2/ENSMUSG00000032446.9, intron 5 of 5) | 9 | 118478861 | 118482757 | 3897 | 1 | ENSMUST00000150633.1 | 4182 | Eomes | ENSMUSG00000032446.9 |
| chr9 | 118484376 | 118484591 | 3.36564 | 3.89233 | 1.05341 | Exon (ENSMUST00000111763.2/ENSMUSG00000032446.9, exon 6 of 6) | 9 | 118478861 | 118482757 | 3897 | 1 | ENSMUST00000150633.1 | 5516 | Eomes | ENSMUSG00000032446.9 |
| chr9 | 118486624 | 118488322 | 4.25194 | 5.7198 | 2.32898 | Distal Intergenic | 9 | 118478861 | 118482757 | 3897 | 1 | ENSMUST00000150633.1 | 7764 | Eomes | ENSMUSG00000032446.9 |
| chr9 | 118488862 | 118490064 | 3.7081 | 4.55847 | 1.50064 | Distal Intergenic | 9 | 118478861 | 118482757 | 3897 | 1 | ENSMUST00000150633.1 | 10002 | Eomes | ENSMUSG00000032446.9 |
| chr9 | 118491250 | 118491594 | 3.34208 | 3.8405 | 1.01733 | Distal Intergenic | 9 | 118478861 | 118482757 | 3897 | 1 | ENSMUST00000150633.1 | 12390 | Eomes | ENSMUSG00000032446.9 |
| chr9 | 98031482 | 98032620 | 3.8482 | 4.92039 | 1.76249 | Promoter | 9 | 97453257 | 98032983 | 579727 | 2 | ENSMUST00000162295.1 | 363 | Clstn2 | ENSMUSG00000032452.7 |
| chr9 | 98033337 | 98033492 | 3.24102 | 3.89758 | 1.06617 | Promoter | 9 | 97444395 | 98033181 | 588787 | 2 | ENSMUST00000035027.7 | -157 | Clstn2 | ENSMUSG00000032452.7 |
| chr9 | 98033900 | 98033991 | 3.43374 | 4.17218 | 1.21146 | Promoter | 9 | 97444395 | 98033181 | 588787 | 2 | ENSMUST00000035027.7 | -720 | Clstn2 | ENSMUSG00000032452.7 |
| chr9 | 98035054 | 98035204 | 3.65215 | 4.47185 | 1.46256 | Distal Intergenic | 9 | 97444395 | 98033181 | 588787 | 2 | ENSMUST00000035027.7 | -1874 | Clstn2 | ENSMUSG00000032452.7 |
| chr9 | 98604311 | 98604634 | 3.39654 | 3.93053 | 1.04507 | Distal Intergenic | 9 | 98588730 | 98601660 | 12931 | 2 | ENSMUST00000035034.9 | -2652 | Mrps22 | ENSMUSG00000032459.9 |
| chr9 | 98612862 | 98613158 | 3.89947 | 4.90674 | 1.71411 | Distal Intergenic | 9 | 98588730 | 98601660 | 12931 | 2 | ENSMUST00000035034.9 | -11203 | Mrps22 | ENSMUSG00000032459.9 |
| chr9 | 98613501 | 98613579 | 3.55546 | 4.40709 | 1.37677 | Distal Intergenic | 9 | 98588730 | 98601660 | 12931 | 2 | ENSMUST00000035034.9 | -11842 | Mrps22 | ENSMUSG00000032459.9 |
| chr9 | 98650586 | 98650826 | 3.59569 | 4.30069 | 1.27971 | Distal Intergenic | 9 | 98588730 | 98601660 | 12931 | 2 | ENSMUST00000035034.9 | -48927 | Mrps22 | ENSMUSG00000032459.9 |
| chr9 | 99171272 | 99171347 | 4.04576 | 5.6119 | 2.16898 | Distal Intergenic | 9 | 99101350 | 99140621 | 39272 | 2 | ENSMUST00000124723.1 | -30652 | Pik3cb | ENSMUSG00000032462.9 |
| chr9 | 98970845 | 98971816 | 3.9534 | 5.12627 | 1.86221 | Distal Intergenic | 9 | 98986373 | 99002021 | 15649 | 1 | ENSMUST00000035038.2 | -14557 | Faim | ENSMUSG00000032463.5 |
| chr9 | 98972313 | 98972502 | 3.76994 | 4.61938 | 1.471 | Distal Intergenic | 9 | 98986373 | 99002021 | 15649 | 1 | ENSMUST00000035038.2 | -13871 | Faim | ENSMUSG00000032463.5 |
| chr9 | 98973060 | 98973732 | 4.02807 | 5.30087 | 2.01971 | Distal Intergenic | 9 | 98986373 | 99002021 | 15649 | 1 | ENSMUST00000035038.2 | -12641 | Faim | ENSMUSG00000032463.5 |
| chr9 | 98974093 | 98974424 | 3.47621 | 4.07322 | 1.12412 | Distal Intergenic | 9 | 98986373 | 99002021 | 15649 | 1 | ENSMUST00000035038.2 | -11949 | Faim | ENSMUSG00000032463.5 |
| chr9 | 98974732 | 98975170 | 3.68666 | 4.53242 | 1.4757 | Distal Intergenic | 9 | 98986373 | 99002021 | 15649 | 1 | ENSMUST00000035038.2 | -11203 | Faim | ENSMUSG00000032463.5 |
| chr9 | 98976417 | 98977312 | 3.82303 | 4.9135 | 1.71539 | Distal Intergenic | 9 | 98986373 | 99002021 | 15649 | 1 | ENSMUST00000035038.2 | -9061 | Faim | ENSMUSG00000032463.5 |
| chr9 | 98978006 | 98978085 | 3.75533 | 4.69594 | 1.58157 | Distal Intergenic | 9 | 98986373 | 99002021 | 15649 | 1 | ENSMUST00000035038.2 | -8288 | Faim | ENSMUSG00000032463.5 |
| chr9 | 98979235 | 98979531 | 3.51291 | 4.29215 | 1.30592 | Distal Intergenic | 9 | 98986373 | 99002021 | 15649 | 1 | ENSMUST00000035038.2 | -6842 | Faim | ENSMUSG00000032463.5 |
| chr9 | 110249837 | 110250210 | 3.54075 | 4.19407 | 1.20395 | Intron (ENSMUST00000035058.5/ENSMUSG00000032482.5, intron 2 of 4) | 9 | 110243783 | 110262575 | 18793 | 1 | ENSMUST00000035058.5 | 6055 | Cspg5 | ENSMUSG00000032482.5 |
| chr9 | 110250543 | 110250686 | 3.36235 | 3.87105 | 1.01733 | Intron (ENSMUST00000035058.5/ENSMUSG00000032482.5, intron 2 of 4) | 9 | 110243783 | 110262575 | 18793 | 1 | ENSMUST00000035058.5 | 6761 | Cspg5 | ENSMUSG00000032482.5 |
| chr9 | 110944449 | 110944678 | 3.27723 | 3.74965 | 1.04054 | Intron (ENSMUST00000035075.8/ENSMUSG00000032494.8, intron 1 of 5) | 9 | 110939610 | 110946158 | 6549 | 2 | ENSMUST00000035075.8 | 1480 | Tdgf1 | ENSMUSG00000032494.8 |
| chr9 | 110953884 | 110954117 | 3.43182 | 4.20899 | 1.27259 | Intron (ENSMUST00000035076.4/ENSMUSG00000032495.4, intron 1 of 8) | 9 | 110951545 | 110984066 | 32522 | 1 | ENSMUST00000035076.4 | 2340 | Lrrc2 | ENSMUSG00000032495.4 |
| chr9 | 110954521 | 110954896 | 3.38143 | 4.03692 | 1.19435 | Intron (ENSMUST00000035076.4/ENSMUSG00000032495.4, intron 1 of 8) | 9 | 110951545 | 110984066 | 32522 | 1 | ENSMUST00000035076.4 | 2977 | Lrrc2 | ENSMUSG00000032495.4 |
| chr9 | 110955655 | 110956376 | 3.48206 | 4.13631 | 1.24399 | Intron (ENSMUST00000035076.4/ENSMUSG00000032495.4, intron 1 of 8) | 9 | 110951545 | 110984066 | 32522 | 1 | ENSMUST00000035076.4 | 4111 | Lrrc2 | ENSMUSG00000032495.4 |
| chr9 | 111439187 | 111439595 | 3.69134 | 4.68016 | 1.56833 | Promoter | 9 | 111439081 | 111489608 | 50528 | 1 | ENSMUST00000111879.3 | 107 | Dclk3 | ENSMUSG00000032500.9 |
| chr9 | 111491321 | 111491597 | 4.01222 | 5.15679 | 1.90002 | Distal Intergenic | 9 | 111439081 | 111489608 | 50528 | 1 | ENSMUST00000111879.3 | 52241 | Dclk3 | ENSMUSG00000032500.9 |
| chr15 | 59655772 | 59655909 | 3.43141 | 4.02923 | 1.20427 | 3' UTR | 15 | 59648653 | 59653689 | 5037 | 1 | ENSMUST00000118228.1 | 7120 | Trib1 | ENSMUSG00000032501.8 |
| chr15 | 59679766 | 59679862 | 4.23295 | 5.65083 | 2.23442 | Distal Intergenic | 15 | 59648653 | 59653689 | 5037 | 1 | ENSMUST00000118228.1 | 31114 | Trib1 | ENSMUSG00000032501.8 |
| chr9 | 111689180 | 111690365 | 3.85052 | 4.84961 | 1.70496 | Promoter | 9 | 111561437 | 111690225 | 128789 | 2 | ENSMUST00000161995.1 | 0 | Stac | ENSMUSG00000032502.8 |
| chr9 | 112232420 | 112232716 | 3.50057 | 4.15316 | 1.26433 | Promoter | 9 | 112065260 | 112232501 | 167242 | 2 | ENSMUST00000035085.6 | 0 | Arpp21 | ENSMUSG00000032503.12 |
| chr9 | 112233052 | 112233162 | 3.34138 | 3.83944 | 1.01733 | Promoter | 9 | 112175580 | 112233626 | 58047 | 2 | ENSMUST00000161138.2 | 464 | Arpp21 | ENSMUSG00000032503.12 |
| chr9 | 112233514 | 112235661 | 3.82786 | 4.80554 | 1.64383 | Promoter | 9 | 112065236 | 112234423 | 169188 | 2 | ENSMUST00000159432.2 | 0 | Arpp21 | ENSMUSG00000032503.12 |
| chr9 | 112236219 | 112236486 | 3.85537 | 4.91057 | 1.70832 | Promoter | 9 | 112187629 | 112235938 | 48310 | 2 | ENSMUST00000161945.1 | -282 | Arpp21 | ENSMUSG00000032503.12 |
| chr9 | 112238860 | 112239301 | 3.51001 | 4.17599 | 1.24712 | Intron (ENSMUST00000144424.3/ENSMUSG00000087038.4, intron 1 of 3) | 9 | 112187629 | 112235938 | 48310 | 2 | ENSMUST00000161945.1 | -2923 | Arpp21 | ENSMUSG00000032503.12 |
| chr9 | 113629955 | 113630070 | 3.57773 | 4.29765 | 1.35194 | Distal Intergenic | 9 | 113655570 | 113657294 | 1725 | 2 | ENSMUST00000142256.1 | 27224 | Pdcd6ip | ENSMUSG00000032504.10 |
| chr9 | 119518881 | 119519185 | 3.38193 | 3.9006 | 1.01733 | Intron (ENSMUST00000065196.7/ENSMUSG00000032511.12, intron 15 of 26) | 9 | 119483408 | 119562678 | 79271 | 2 | ENSMUST00000065196.7 | 43493 | Scn5a | ENSMUSG00000032511.12 |
| chr9 | 119554385 | 119554706 | 3.37504 | 3.91248 | 1.09313 | Intron (ENSMUST00000065196.7/ENSMUSG00000032511.12, intron 2 of 26) | 9 | 119483408 | 119562678 | 79271 | 2 | ENSMUST00000065196.7 | 7972 | Scn5a | ENSMUSG00000032511.12 |
| chr9 | 119571153 | 119571333 | 3.38114 | 3.89941 | 1.01733 | Intron (ENSMUST00000117911.2/ENSMUSG00000032511.12, intron 1 of 27) | 9 | 119562537 | 119563503 | 967 | 2 | ENSMUST00000138934.1 | -7651 | Scn5a | ENSMUSG00000032511.12 |
| chr9 | 119571693 | 119571768 | 4.1871 | 5.62222 | 2.18966 | Intron (ENSMUST00000117911.2/ENSMUSG00000032511.12, intron 1 of 27) | 9 | 119483410 | 119579016 | 95607 | 2 | ENSMUST00000117911.2 | 7248 | Scn5a | ENSMUSG00000032511.12 |
| chr9 | 119575848 | 119575985 | 3.90812 | 4.89762 | 1.67903 | Intron (ENSMUST00000117911.2/ENSMUSG00000032511.12, intron 1 of 27) | 9 | 119483410 | 119579016 | 95607 | 2 | ENSMUST00000117911.2 | 3031 | Scn5a | ENSMUSG00000032511.12 |
| chr9 | 119576395 | 119576566 | 4.3321 | 5.75621 | 2.30409 | Intron (ENSMUST00000117911.2/ENSMUSG00000032511.12, intron 1 of 27) | 9 | 119483410 | 119579016 | 95607 | 2 | ENSMUST00000117911.2 | 2450 | Scn5a | ENSMUSG00000032511.12 |
| chr9 | 119578427 | 119579595 | 4.13575 | 5.7491 | 2.32093 | Promoter | 9 | 119483410 | 119579016 | 95607 | 2 | ENSMUST00000117911.2 | 0 | Scn5a | ENSMUSG00000032511.12 |
| chr9 | 119579957 | 119580383 | 3.60677 | 4.34295 | 1.30775 | Promoter | 9 | 119484324 | 119579016 | 94693 | 2 | ENSMUST00000120420.1 | -942 | Scn5a | ENSMUSG00000032511.12 |
| chr9 | 119586377 | 119586533 | 3.79054 | 4.66382 | 1.51456 | Distal Intergenic | 9 | 119484324 | 119579016 | 94693 | 2 | ENSMUST00000120420.1 | -7362 | Scn5a | ENSMUSG00000032511.12 |
| chr9 | 119590609 | 119590940 | 3.73503 | 4.55651 | 1.43907 | Distal Intergenic | 9 | 119484324 | 119579016 | 94693 | 2 | ENSMUST00000120420.1 | -11594 | Scn5a | ENSMUSG00000032511.12 |
| chr9 | 119597632 | 119597871 | 3.71485 | 4.64806 | 1.5791 | Distal Intergenic | 9 | 119484324 | 119579016 | 94693 | 2 | ENSMUST00000120420.1 | -18617 | Scn5a | ENSMUSG00000032511.12 |
| chr9 | 119599021 | 119599134 | 3.60212 | 4.35473 | 1.35613 | Distal Intergenic | 9 | 119484324 | 119579016 | 94693 | 2 | ENSMUST00000120420.1 | -20006 | Scn5a | ENSMUSG00000032511.12 |
| chr9 | 119603778 | 119603869 | 3.6783 | 4.46315 | 1.40232 | Distal Intergenic | 9 | 119484324 | 119579016 | 94693 | 2 | ENSMUST00000120420.1 | -24763 | Scn5a | ENSMUSG00000032511.12 |
| chr9 | 119604407 | 119604555 | 3.34493 | 3.86943 | 1.09552 | Distal Intergenic | 9 | 119484324 | 119579016 | 94693 | 2 | ENSMUST00000120420.1 | -25392 | Scn5a | ENSMUSG00000032511.12 |
| chr9 | 119606333 | 119606737 | 3.47754 | 4.08756 | 1.16885 | Downstream (1-2kb) | 9 | 119484324 | 119579016 | 94693 | 2 | ENSMUST00000120420.1 | -27318 | Scn5a | ENSMUSG00000032511.12 |
| chr9 | 119614052 | 119614780 | 3.84116 | 4.79551 | 1.65159 | Intron (ENSMUST00000084787.5/ENSMUSG00000034533.9, intron 26 of 27) | 9 | 119484324 | 119579016 | 94693 | 2 | ENSMUST00000120420.1 | -35037 | Scn5a | ENSMUSG00000032511.12 |
| chr9 | 119617350 | 119618127 | 3.4407 | 4.15567 | 1.23732 | Exon (ENSMUST00000084787.5/ENSMUSG00000034533.9, exon 25 of 28) | 9 | 119484324 | 119579016 | 94693 | 2 | ENSMUST00000120420.1 | -38335 | Scn5a | ENSMUSG00000032511.12 |
| chr9 | 119623344 | 119623526 | 3.43518 | 3.99963 | 1.08304 | Intron (ENSMUST00000084787.5/ENSMUSG00000034533.9, intron 22 of 27) | 9 | 119484324 | 119579016 | 94693 | 2 | ENSMUST00000120420.1 | -44329 | Scn5a | ENSMUSG00000032511.12 |
| chr9 | 119625565 | 119625784 | 4.38372 | 5.87361 | 2.37208 | Intron (ENSMUST00000084787.5/ENSMUSG00000034533.9, intron 21 of 27) | 9 | 119484324 | 119579016 | 94693 | 2 | ENSMUST00000120420.1 | -46550 | Scn5a | ENSMUSG00000032511.12 |
| chr9 | 119627982 | 119628107 | 3.66842 | 4.55455 | 1.4796 | Intron (ENSMUST00000084787.5/ENSMUSG00000034533.9, intron 20 of 27) | 9 | 119484324 | 119579016 | 94693 | 2 | ENSMUST00000120420.1 | -48967 | Scn5a | ENSMUSG00000032511.12 |
| chr9 | 119629977 | 119630291 | 3.3964 | 4.09029 | 1.1959 | Exon (ENSMUST00000084787.5/ENSMUSG00000034533.9, exon 20 of 28) | 9 | 119484324 | 119579016 | 94693 | 2 | ENSMUST00000120420.1 | -50962 | Scn5a | ENSMUSG00000032511.12 |
| chr9 | 119645439 | 119645796 | 3.43647 | 4.0231 | 1.16013 | Intron (ENSMUST00000084787.5/ENSMUSG00000034533.9, intron 15 of 27) | 9 | 119484324 | 119579016 | 94693 | 2 | ENSMUST00000120420.1 | -66424 | Scn5a | ENSMUSG00000032511.12 |
| chr9 | 119647754 | 119648123 | 3.55982 | 4.41363 | 1.39305 | Exon (ENSMUST00000084787.5/ENSMUSG00000034533.9, exon 15 of 28) | 9 | 119484324 | 119579016 | 94693 | 2 | ENSMUST00000120420.1 | -68739 | Scn5a | ENSMUSG00000032511.12 |
| chr9 | 120184590 | 120184714 | 3.32372 | 3.81287 | 1.01733 | Distal Intergenic | 9 | 120149742 | 120181484 | 31743 | 1 | ENSMUST00000174193.2 | 34849 | Mobp | ENSMUSG00000032517.10 |
| chr9 | 121642806 | 121643263 | 4.04934 | 5.31686 | 2.05239 | Promoter | 9 | 121643051 | 121660225 | 17175 | 1 | ENSMUST00000139189.1 | 0 | Vipr1 | ENSMUSG00000032528.4 |
| chr9 | 121643652 | 121644929 | 3.70175 | 4.52777 | 1.45616 | Promoter | 9 | 121643051 | 121660225 | 17175 | 1 | ENSMUST00000139189.1 | 602 | Vipr1 | ENSMUSG00000032528.4 |
| chr9 | 121669300 | 121669536 | 3.9745 | 5.30755 | 2.00263 | Promoter | 9 | 121668713 | 121669509 | 797 | 1 | ENSMUST00000149959.1 | 588 | Vipr1 | ENSMUSG00000032528.4 |
| chr9 | 121641041 | 121641397 | 3.28127 | 3.74906 | 1.01733 | Promoter | 9 | 121584087 | 121641526 | 57440 | 2 | ENSMUST00000125075.1 | 129 | Lyzl4 | ENSMUSG00000032530.8 |
| chr9 | 121641707 | 121641923 | 3.38906 | 3.93348 | 1.09585 | Promoter | 9 | 121577843 | 121641566 | 63724 | 2 | ENSMUST00000077706.4 | -142 | Lyzl4 | ENSMUSG00000032530.8 |
| chr9 | 121491906 | 121491981 | 3.33962 | 3.8564 | 1.08654 | Intron (ENSMUST00000035120.4/ENSMUSG00000032532.6, intron 2 of 2) | 9 | 121489825 | 121495689 | 5865 | 2 | ENSMUST00000035120.4 | 3708 | Cck | ENSMUSG00000032532.6 |
| chr9 | 121493310 | 121494734 | 3.50864 | 4.23263 | 1.29962 | Promoter | 9 | 121489825 | 121495689 | 5865 | 2 | ENSMUST00000035120.4 | 955 | Cck | ENSMUSG00000032532.6 |
| chr9 | 121495732 | 121495910 | 3.91703 | 5.33872 | 2.01557 | Promoter | 9 | 121489825 | 121495689 | 5865 | 2 | ENSMUST00000035120.4 | -44 | Cck | ENSMUSG00000032532.6 |
| chr9 | 104282851 | 104283044 | 3.4889 | 4.15089 | 1.27403 | Distal Intergenic | 9 | 104291049 | 104301225 | 10177 | 2 | ENSMUST00000131084.1 | 18181 | Acpp | ENSMUSG00000032561.9 |
| chr9 | 104569600 | 104569683 | 3.25013 | 3.70234 | 1.01733 | Promoter | 9 | 104569787 | 105034544 | 464758 | 1 | ENSMUST00000077190.6 | -104 | Cpne4 | ENSMUSG00000032564.10 |
| chr9 | 104570246 | 104570689 | 4.04443 | 5.54279 | 2.15393 | Promoter | 9 | 104569787 | 105034544 | 464758 | 1 | ENSMUST00000077190.6 | 460 | Cpne4 | ENSMUSG00000032564.10 |
| chr9 | 105990023 | 105990170 | 3.67202 | 4.46188 | 1.43459 | Exon (ENSMUST00000149207.1/ENSMUSG00000032572.7, exon 4 of 4) | 9 | 105989454 | 105998322 | 8869 | 2 | ENSMUST00000149207.1 | 8152 | Col6a4 | ENSMUSG00000032572.7 |

| | | | | | | | | | | | | | | | |
|---|---|---|---|---|---|---|---|---|---|---|---|---|---|---|---|
| chr9 | 105999318 | 105999624 | 3.32066 | 3.83096 | 1.09225 | Promoter | 9 | 105989454 | 105998322 | 8869 | 2 | ENSMUST00000149207.1 | -997 | Col6a4 | ENSMUSG00000032572.7 |
| chr9 | 106100075 | 106100420 | 3.55584 | 4.28531 | 1.35556 | Distal Intergenic | 9 | 105990461 | 106096783 | 106323 | 2 | ENSMUST00000121963.1 | -3293 | Col6a4 | ENSMUSG00000032572.7 |
| chr9 | 107917022 | 107917182 | 3.38193 | 3.9006 | 1.01733 | Intron (ENSMUST00000035203.7/ENSMUSG00000032584.7, intron 16 of 18) | 9 | 107906889 | 107920383 | 13495 | 1 | ENSMUST00000035203.7 | 10134 | Mst1r | ENSMUSG00000032584.7 |
| chr9 | 108109432 | 108109507 | 3.58899 | 4.2781 | 1.24616 | Promoter | 9 | 108096023 | 108110413 | 14391 | 2 | ENSMUST00000124763.1 | 906 | Bsn | ENSMUSG00000032589.9 |
| chr9 | 108110052 | 108111686 | 3.69593 | 4.51109 | 1.4382 | Promoter | 9 | 108096023 | 108110413 | 14391 | 2 | ENSMUST00000124763.1 | 0 | Bsn | ENSMUSG00000032589.9 |
| chr9 | 108189919 | 108190093 | 3.30169 | 3.79853 | 1.07825 | Promoter | 9 | 108096022 | 108190384 | 94363 | 2 | ENSMUST00000035208.8 | 291 | Bsn | ENSMUSG00000032589.9 |
| chr13 | 103710666 | 103710749 | 3.86942 | 4.81086 | 1.60243 | Distal Intergenic | 13 | 103741615 | 103764504 | 22890 | 2 | ENSMUST00000074616.5 | 53755 | Srek1 | ENSMUSG00000032621.7 |
| chr6 | 134886994 | 134888176 | 3.4134 | 3.97851 | 1.12367 | Promoter | 6 | 134869157 | 134887832 | 18676 | 2 | ENSMUST00000066107.6 | 0 | Gpr19 | ENSMUSG00000032641.12 |
| chr18 | 65497763 | 65497905 | 3.5442 | 4.20508 | 1.22515 | Distal Intergenic | 18 | 65430963 | 65478823 | 47861 | 1 | ENSMUST00000049248.5 | 66801 | Malt1 | ENSMUSG00000032688.7 |
| chr5 | 120726408 | 120726605 | 3.63968 | 4.38699 | 1.34041 | Distal Intergenic | 5 | 120730336 | 120735575 | 5240 | 2 | ENSMUST00000147941.1 | 8970 | Oas2 | ENSMUSG00000032690.11 |
| chr2 | 103967458 | 103967689 | 3.38048 | 3.90577 | 1.04338 | Intron (ENSMUST00000156813.2/ENSMUSG00000032698.10, intron 1 of 3) | 2 | 103969528 | 103981092 | 11565 | 2 | ENSMUST00000123437.2 | -1839 | Lmo2 | ENSMUSG00000032698.10 |
| chr2 | 103969336 | 103969520 | 3.37022 | 3.88292 | 1.01733 | Promoter | 2 | 103969528 | 103981092 | 11565 | 1 | ENSMUST00000123437.2 | -8 | Lmo2 | ENSMUSG00000032698.10 |
| chr17 | 47815579 | 47815795 | 4.05221 | 5.29949 | 1.99992 | 3' UTR | 17 | 47816039 | 47833169 | 17131 | 2 | ENSMUST00000131971.1 | 17374 | Mdfi | ENSMUSG00000032717.9 |
| chr17 | 47831111 | 47831292 | 3.36398 | 3.87352 | 1.01733 | Intron (ENSMUST00000113280.2/ENSMUSG00000032717.9, intron 2 of 3) | 17 | 47816039 | 47833169 | 17131 | 2 | ENSMUST00000131971.1 | 1877 | Mdfi | ENSMUSG00000032717.9 |
| chr17 | 47832941 | 47833681 | 3.96655 | 5.02677 | 1.79179 | Promoter | 17 | 47815328 | 47833241 | 17914 | 2 | ENSMUST00000113280.2 | 0 | Mdfi | ENSMUSG00000032717.9 |
| chr17 | 47834045 | 47834501 | 3.66447 | 4.52846 | 1.46628 | Promoter | 17 | 47815328 | 47834691 | 19364 | 2 | ENSMUST00000066368.7 | 190 | Mdfi | ENSMUSG00000032717.9 |
| chr1 | 15892385 | 15893554 | 4.3904 | 6.00717 | 2.56353 | Promoter | 1 | 15853862 | 15892722 | 38861 | 2 | ENSMUST00000040695.4 | 0 | Sbspon | ENSMUSG00000032719.4 |
| chr7 | 125887035 | 125887137 | 3.9136 | 4.92351 | 1.69711 | Intron (ENSMUST00000073935.5/ENSMUSG00000046182.7, intron 6 of 6) | 7 | 125865271 | 125868064 | 2794 | 1 | ENSMUST00000155059.1 | 21765 | D430042O09Rik | ENSMUSG00000032743.10 |
| chr7 | 125887483 | 125887733 | 3.42757 | 3.99572 | 1.10653 | Intron (ENSMUST00000073935.5/ENSMUSG00000046182.7, intron 6 of 6) | 7 | 125865271 | 125868064 | 2794 | 1 | ENSMUST00000155059.1 | 22213 | D430042O09Rik | ENSMUSG00000032743.10 |
| chr7 | 125959191 | 125959388 | 3.62328 | 4.41589 | 1.43045 | Intron (ENSMUST00000073935.5/ENSMUSG00000046182.7, intron 2 of 6) | 7 | 125865271 | 125868064 | 2794 | 1 | ENSMUST00000155059.1 | 93921 | D430042O09Rik | ENSMUSG00000032743.10 |
| chr4 | 123230969 | 123231251 | 3.22428 | 3.66363 | 1.01509 | Distal Intergenic | 4 | 123233556 | 123249875 | 16320 | 1 | ENSMUST00000040821.4 | -2305 | Heyl | ENSMUSG00000032744.4 |
| chr4 | 123232963 | 123233248 | 3.33355 | 3.82764 | 1.01733 | Promoter | 4 | 123233556 | 123249875 | 16320 | 1 | ENSMUST00000040821.4 | -308 | Heyl | ENSMUSG00000032744.4 |
| chr4 | 123233606 | 123233927 | 3.7428 | 4.9613 | 1.75836 | Promoter | 4 | 123233853 | 123240659 | 6807 | 1 | ENSMUST00000151870.1 | 0 | Heyl | ENSMUSG00000032744.4 |
| chr7 | 72305457 | 72306075 | 3.68639 | 4.68083 | 1.53831 | Promoter | 7 | 72077830 | 72306595 | 228766 | 2 | ENSMUST00000079323.6 | 520 | Mctp2 | ENSMUSG00000032776.8 |
| chr15 | 99066425 | 99066706 | 3.37474 | 4.152 | 1.25021 | Distal Intergenic | 15 | 99074973 | 99083407 | 8435 | 1 | ENSMUST00000039665.6 | -8267 | Troap | ENSMUSG00000032783.8 |
| chr17 | 67697660 | 67698074 | 3.89712 | 5.23054 | 1.96373 | Promoter | 17 | 67697265 | 67822645 | 125381 | 1 | ENSMUST00000035471.7 | 396 | Lama1 | ENSMUSG00000032796.7 |
| chr17 | 67701625 | 67701700 | 3.26318 | 3.73088 | 1.04591 | Intron (ENSMUST00000035471.7/ENSMUSG00000032796.7, intron 1 of 62) | 17 | 67697265 | 67822645 | 125381 | 1 | ENSMUST00000035471.7 | 4361 | Lama1 | ENSMUSG00000032796.7 |
| chr2 | 152092896 | 152094870 | 3.85653 | 4.84081 | 1.70946 | 3' UTR | 2 | 152105516 | 152109211 | 3696 | 1 | ENSMUST00000137751.1 | -10646 | Srxn1 | ENSMUSG00000032802.8 |
| chr2 | 152095187 | 152095417 | 5.01231 | 7.36507 | 3.53974 | 3' UTR | 2 | 152105516 | 152109211 | 3696 | 1 | ENSMUST00000137751.1 | -10099 | Srxn1 | ENSMUSG00000032802.8 |
| chr2 | 152095976 | 152096146 | 3.29801 | 3.98214 | 1.12811 | Distal Intergenic | 2 | 152105516 | 152109211 | 3696 | 1 | ENSMUST00000137751.1 | -9370 | Srxn1 | ENSMUSG00000032802.8 |
| chr2 | 152114178 | 152114926 | 3.96708 | 5.18616 | 1.91898 | Distal Intergenic | 2 | 152105524 | 152111376 | 5853 | 1 | ENSMUST00000041500.7 | 8655 | Srxn1 | ENSMUSG00000032802.8 |
| chr2 | 152120401 | 152120477 | 3.87716 | 4.83022 | 1.63997 | Distal Intergenic | 2 | 152105524 | 152111376 | 5853 | 1 | ENSMUST00000041500.7 | 14878 | Srxn1 | ENSMUSG00000032802.8 |
| chr2 | 152121148 | 152121272 | 3.66337 | 4.43508 | 1.38575 | Distal Intergenic | 2 | 152105524 | 152111376 | 5853 | 1 | ENSMUST00000041500.7 | 15625 | Srxn1 | ENSMUSG00000032802.8 |
| chr9 | 65101508 | 65101647 | 3.47859 | 4.09583 | 1.19265 | Promoter | 9 | 65101535 | 65135580 | 34046 | 1 | ENSMUST00000166273.1 | 0 | Igdcc4 | ENSMUSG00000032816.9 |
| chr9 | 65102464 | 65103151 | 3.53742 | 4.22567 | 1.30543 | Promoter | 9 | 65101535 | 65135580 | 34046 | 1 | ENSMUST00000166273.1 | 930 | Igdcc4 | ENSMUSG00000032816.9 |
| chr18 | 77277381 | 77277550 | 3.55334 | 4.2183 | 1.22569 | Distal Intergenic | 18 | 77281958 | 77442257 | 160300 | 1 | ENSMUST00000096547.4 | -4408 | Loxhd1 | ENSMUSG00000032818.10 |
| chr18 | 77366302 | 77366625 | 3.48833 | 4.10671 | 1.17875 | Exon (ENSMUST00000096547.4/ENSMUSG00000032818.10, exon 17 of 39) | 18 | 77375065 | 77442341 | 67277 | 1 | ENSMUST00000123410.2 | -8440 | Loxhd1 | ENSMUSG00000032818.10 |
| chr3 | 127161282 | 127163168 | 3.56713 | 4.42629 | 1.42185 | Promoter | 3 | 127031173 | 127162342 | 131170 | 2 | ENSMUST00000182994.2 | 0 | Ank2 | ENSMUSG00000032826.11 |
| chr3 | 127224704 | 127224878 | 3.602 | 4.37979 | 1.40194 | Promoter | 3 | 126922308 | 127225847 | 303540 | 2 | ENSMUST00000182726.2 | 969 | Ank2 | ENSMUSG00000032826.11 |
| chr3 | 127407629 | 127408227 | 3.60292 | 4.59217 | 1.53981 | Promoter | 3 | 127079943 | 127408952 | 329010 | 2 | ENSMUST00000183095.1 | 725 | Ank2 | ENSMUSG00000032826.11 |
| chr3 | 127459077 | 127459271 | 3.35464 | 4.04331 | 1.15311 | Intron (ENSMUST00000182594.2/ENSMUSG00000032826.11, intron 1 of 45) | 3 | 127336394 | 127499001 | 162608 | 2 | ENSMUST00000182316.1 | 39730 | Ank2 | ENSMUSG00000032826.11 |
| chr3 | 127498194 | 127499374 | 3.69677 | 4.67385 | 1.57851 | Promoter | 3 | 126923957 | 127499115 | 575159 | 2 | ENSMUST00000182594.2 | 0 | Ank2 | ENSMUSG00000032826.11 |
| chr9 | 95751645 | 95751850 | 4.687 | 6.55206 | 2.95213 | Downstream (<1kb) | 9 | 95749038 | 95750375 | 1338 | 2 | ENSMUST00000190205.1 | -1271 | Trpc1 | ENSMUSG00000032839.8 |
| chr14 | 118704810 | 118705323 | 3.96899 | 5.08117 | 1.85312 | Promoter | 14 | 118482692 | 118706219 | 223528 | 2 | ENSMUST00000036554.7 | 896 | Abcc4 | ENSMUSG00000032849.8 |
| chr14 | 118705691 | 118706008 | 3.75231 | 4.64869 | 1.57847 | Promoter | 14 | 118482692 | 118706219 | 223528 | 2 | ENSMUST00000036554.7 | 211 | Abcc4 | ENSMUSG00000032849.8 |
| chr2 | 151842610 | 151843225 | 3.53108 | 4.37533 | 1.37421 | Promoter | 2 | 151842927 | 151874668 | 31742 | 1 | ENSMUST00000042217.3 | 0 | Rspo4 | ENSMUSG00000032852.3 |
| chr3 | 125937339 | 125939057 | 4.00655 | 5.2319 | 1.97667 | Promoter | 3 | 125865359 | 125938550 | 73192 | 2 | ENSMUST00000057944.7 | 0 | Ugt8a | ENSMUSG00000032854.8 |
| chr3 | 125940288 | 125940671 | 3.64743 | 4.52318 | 1.4676 | Distal Intergenic | 3 | 125865359 | 125938550 | 73192 | 2 | ENSMUST00000057944.7 | -1739 | Ugt8a | ENSMUSG00000032854.8 |
| chr7 | 101011795 | 101012071 | 3.69321 | 4.66445 | 1.53564 | Promoter | 7 | 100996572 | 101011991 | 15420 | 2 | ENSMUST00000037540.3 | 0 | P2ry2 | ENSMUSG00000032860.4 |
| chr7 | 101012955 | 101013161 | 3.79976 | 4.93608 | 1.72534 | Promoter | 7 | 100996572 | 101011991 | 15420 | 2 | ENSMUST00000037540.3 | -965 | P2ry2 | ENSMUSG00000032860.4 |
| chr7 | 101014233 | 101014455 | 3.76245 | 4.68105 | 1.62303 | Distal Intergenic | 7 | 100996572 | 101011991 | 15420 | 2 | ENSMUST00000037540.3 | -2243 | P2ry2 | ENSMUSG00000032860.4 |
| chr7 | 101033249 | 101033360 | 3.33504 | 3.97504 | 1.08088 | Distal Intergenic | 7 | 100996572 | 101011991 | 15420 | 2 | ENSMUST00000037540.3 | -21259 | P2ry2 | ENSMUSG00000032860.4 |
| chr2 | 151746186 | 151746261 | 3.86861 | 4.85562 | 1.71941 | Downstream (1-2kb) | 2 | 151718265 | 151744186 | 25922 | 2 | ENSMUST00000109869.1 | -2001 | Psmf1 | ENSMUSG00000032869.10 |
| chr11 | 28582047 | 28582279 | 3.32621 | 3.8166 | 1.01733 | Intron (ENSMUST00000093253.4/ENSMUSG00000032878.11, intron 1 of 4) | 11 | 28387152 | 28583995 | 196844 | 2 | ENSMUST00000146385.3 | 1716 | Ccdc85a | ENSMUSG00000032878.11 |
| chr11 | 28583400 | 28583781 | 3.55741 | 4.23907 | 1.27926 | Promoter | 11 | 28387152 | 28583995 | 196844 | 2 | ENSMUST00000146385.3 | 214 | Ccdc85a | ENSMUSG00000032878.11 |
| chr11 | 28584313 | 28584607 | 3.66283 | 4.44791 | 1.43634 | Promoter | 11 | 28385692 | 28584324 | 198633 | 2 | ENSMUST00000042534.9 | 0 | Ccdc85a | ENSMUSG00000032878.11 |
| chr4 | 120856234 | 120856543 | 3.42696 | 3.98398 | 1.07163 | Intron (ENSMUST00000132895.2/ENSMUSG00000032890.12, intron 1 of 3) | 4 | 120854819 | 120896579 | 41761 | 1 | ENSMUST00000071093.3 | 1416 | Rims3 | ENSMUSG00000032890.12 |
| chr6 | 131353497 | 131353640 | 3.44458 | 4.10478 | 1.22294 | Promoter | 6 | 131311192 | 131353597 | 42406 | 2 | ENSMUST00000135859.1 | 0 | Styk1 | ENSMUSG00000032899.9 |
| chr1 | 78309891 | 78310633 | 3.98992 | 5.17137 | 1.9255 | Promoter | 1 | 78310345 | 78420289 | 109945 | 1 | ENSMUST00000036172.9 | 0 | Sgpp2 | ENSMUSG00000032908.9 |
| chr1 | 78310946 | 78311313 | 4.1193 | 5.46078 | 2.15374 | Promoter | 1 | 78310345 | 78420289 | 109945 | 1 | ENSMUST00000036172.9 | 602 | Sgpp2 | ENSMUSG00000032908.9 |
| chr1 | 78313736 | 78314529 | 3.48569 | 4.11002 | 1.19962 | Intron (ENSMUST00000036172.9/ENSMUSG00000032908.9, intron 1 of 4) | 1 | 78310345 | 78420289 | 109945 | 1 | ENSMUST00000036172.9 | 3392 | Sgpp2 | ENSMUSG00000032908.9 |
| chr9 | 56863914 | 56864181 | 3.37572 | 3.89122 | 1.01733 | Promoter | 9 | 56865104 | 56899870 | 34767 | 1 | ENSMUST00000035661.5 | -923 | Cspg4 | ENSMUSG00000032911.5 |
| chr9 | 56885095 | 56886098 | 3.45028 | 4.03232 | 1.11834 | Exon (ENSMUST00000035661.5/ENSMUSG00000032911.5, exon 3 of 10) | 9 | 56865104 | 56899870 | 34767 | 1 | ENSMUST00000035661.5 | 19992 | Cspg4 | ENSMUSG00000032911.5 |
| chr9 | 107935783 | 107936404 | 3.89939 | 5.04751 | 1.8366 | Promoter | 9 | 107935920 | 107949691 | 13772 | 1 | ENSMUST00000035700.8 | 0 | Camkv | ENSMUSG00000032936.8 |
| chr9 | 107936719 | 107936956 | 3.49419 | 4.14824 | 1.28571 | Promoter | 9 | 107935920 | 107949691 | 13772 | 1 | ENSMUST00000035700.8 | 800 | Camkv | ENSMUSG00000032936.8 |
| chr19 | 6399426 | 6399581 | 3.57404 | 4.26456 | 1.2781 | Promoter | 19 | 6399456 | 6403633 | 4178 | 1 | ENSMUST00000113475.2 | 0 | Rasgrp2 | ENSMUSG00000032946.11 |
| chr19 | 6401186 | 6401321 | 3.58301 | 4.29983 | 1.30762 | Promoter | 19 | 6401695 | 6402446 | 752 | 1 | ENSMUST00000122930.1 | -374 | Rasgrp2 | ENSMUSG00000032946.11 |
| chr19 | 6401870 | 6402076 | 3.16261 | 3.71406 | 1.01733 | Promoter | 19 | 6401719 | 6403638 | 1920 | 1 | ENSMUST00000133968.1 | 152 | Rasgrp2 | ENSMUSG00000032946.11 |
| chr11 | 26210527 | 26210602 | 3.31358 | 3.81446 | 1.07501 | Promoter | 11 | 25617668 | 26210569 | 592902 | 2 | ENSMUST00000125773.2 | 0 | 5730522E02Rik | ENSMUSG00000032985.10 |
| chr15 | 79251696 | 79252831 | 4.02168 | 5.28002 | 2.00883 | Exon (ENSMUST00000039752.3/ENSMUSG00000032988.3, exon 5 of 6) | 15 | 79251014 | 79254761 | 3748 | 2 | ENSMUST00000039752.3 | 1930 | Slc16a8 | ENSMUSG00000032988.3 |
| chr15 | 79253576 | 79254096 | 3.56248 | 4.28301 | 1.36941 | Promoter | 15 | 79251014 | 79254761 | 3748 | 2 | ENSMUST00000039752.3 | 665 | Slc16a8 | ENSMUSG00000032988.3 |
| chr15 | 79254736 | 79255005 | 3.57942 | 4.35357 | 1.41228 | Promoter | 15 | 79251014 | 79254761 | 3748 | 2 | ENSMUST00000039752.3 | 0 | Slc16a8 | ENSMUSG00000032988.3 |
| chr15 | 79162428 | 79162503 | 3.43315 | 4.22069 | 1.25896 | Intron (ENSMUST00000040019.4/ENSMUSG00000033006.9, intron 2 of 3) | 15 | 79154913 | 79164490 | 9578 | 2 | ENSMUST00000040019.4 | 1987 | Sox10 | ENSMUSG00000033006.9 |
| chr15 | 79163302 | 79164427 | 3.64937 | 4.55461 | 1.47355 | Promoter | 15 | 79154913 | 79164490 | 9578 | 2 | ENSMUST00000040019.4 | 63 | Sox10 | ENSMUSG00000033006.9 |
| chr1 | 75446327 | 75447158 | 3.83899 | 4.81499 | 1.66464 | Distal Intergenic | 1 | 75450436 | 75474340 | 23905 | 1 | ENSMUST00000113577.2 | -3278 | Asic4 | ENSMUSG00000033007.10 |
| chr1 | 75447731 | 75447938 | 5.28501 | 7.8087 | 3.92327 | Distal Intergenic | 1 | 75450436 | 75474340 | 23905 | 1 | ENSMUST00000113577.2 | -2498 | Asic4 | ENSMUSG00000033007.10 |
| chr1 | 75448839 | 75449492 | 4.10445 | 5.39701 | 2.0161 | Promoter | 1 | 75450436 | 75474340 | 23905 | 1 | ENSMUST00000113577.2 | -944 | Asic4 | ENSMUSG00000033007.10 |
| chr1 | 75449871 | 75449963 | 3.97944 | 6.11757 | 2.56443 | Promoter | 1 | 75450436 | 75474340 | 23905 | 1 | ENSMUST00000113577.2 | -473 | Asic4 | ENSMUSG00000033007.10 |
| chr1 | 75450276 | 75451780 | 3.86883 | 4.91927 | 1.71419 | Promoter | 1 | 75450436 | 75474340 | 23905 | 1 | ENSMUST00000113577.2 | 0 | Asic4 | ENSMUSG00000033007.10 |
| chr1 | 75452749 | 75452930 | 3.29758 | 3.77365 | 1.01733 | Intron (ENSMUST00000113577.2/ENSMUSG00000033007.10, intron 1 of 8) | 1 | 75450699 | 75474343 | 23645 | 1 | ENSMUST00000037708.9 | 2051 | Asic4 | ENSMUSG00000033007.10 |
| chr11 | 25100057 | 25100219 | 18.21335 | 75.047 | 68.93687 | Distal Intergenic | 11 | 25232626 | 25367133 | 134508 | 1 | ENSMUST00000191131.1 | -132407 | 4933427E13Rik | ENSMUSG00000033015.11 |
| chr18 | 80711001 | 80711591 | 3.74077 | 4.65372 | 1.594 | Intron (ENSMUST00000078049.5/ENSMUSG00000033016.10, intron 1 of 9) | 18 | 80606205 | 80713071 | 106867 | 2 | ENSMUST00000078049.5 | 1480 | Nfatc1 | ENSMUSG00000033016.10 |
| chr18 | 80712638 | 80713952 | 3.66959 | 4.53214 | 1.49244 | Promoter | 18 | 80606205 | 80713071 | 106867 | 2 | ENSMUST00000078049.5 | 0 | Nfatc1 | ENSMUSG00000033016.10 |
| chr18 | 80725278 | 80725610 | 3.38193 | 3.9006 | 1.01733 | Distal Intergenic | 18 | 80606205 | 80713071 | 106867 | 2 | ENSMUST00000170905.2 | -12208 | Nfatc1 | ENSMUSG00000033016.10 |
| chr18 | 61762127 | 61762413 | 3.38193 | 3.9006 | 1.01733 | Intron (ENSMUST00000154876.2/ENSMUSG00000033032.10, intron 1 of 15) | 18 | 61730261 | 61786611 | 56351 | 2 | ENSMUST00000154876.2 | 24198 | Afap1l1 | ENSMUSG00000033032.10 |
| chr18 | 61785813 | 61786003 | 3.64095 | 4.4896 | 1.50103 | Promoter | 18 | 61730261 | 61786611 | 56351 | 2 | ENSMUST00000154876.2 | 608 | Afap1l1 | ENSMUSG00000033032.10 |
| chr14 | 101730144 | 101730728 | 3.35262 | 3.85648 | 1.01733 | Promoter | 14 | 101729957 | 101934710 | 204754 | 1 | ENSMUST00000100337.4 | 188 | Lmo7 | ENSMUSG00000033060.10 |
| chr14 | 101731709 | 101731993 | 3.38193 | 3.9006 | 1.01733 | Intron (ENSMUST00000100337.4/ENSMUSG00000033060.10, intron 1 of 30) | 14 | 101729957 | 101934710 | 204754 | 1 | ENSMUST00000100337.4 | 1753 | Lmo7 | ENSMUSG00000033060.10 |
| chr1 | 75277608 | 75277763 | 3.33514 | 3.84869 | 1.0829 | Promoter | 1 | 75274070 | 75278284 | 4215 | 2 | ENSMUST00000189203.1 | 521 | Resp18 | ENSMUSG00000033061.10 |
| chr1 | 75278256 | 75278361 | 3.28089 | 3.74851 | 1.01733 | Promoter | 1 | 75272311 | 75278316 | 6006 | 2 | ENSMUST00000186229.1 | 0 | Resp18 | ENSMUSG00000033061.10 |
| chr2 | 150687635 | 150689342 | 3.71328 | 4.64235 | 1.55015 | Promoter | 2 | 150680702 | 150689360 | 8659 | 2 | ENSMUST00000046095.9 | 18 | Vsx1 | ENSMUSG00000033080.9 |
| chr2 | 150690047 | 150690327 | 3.62253 | 4.40997 | 1.44108 | Promoter | 2 | 150680702 | 150689360 | 8659 | 2 | ENSMUST00000046095.9 | -688 | Vsx1 | ENSMUSG00000033080.9 |
| chr2 | 150694069 | 150694212 | 3.48031 | 4.19507 | 1.27349 | Distal Intergenic | 2 | 150680702 | 150689360 | 8659 | 2 | ENSMUST00000046095.9 | -4710 | Vsx1 | ENSMUSG00000033080.9 |
| chr6 | 88873609 | 88873929 | 3.62632 | 4.36056 | 1.3283 | Promoter | 6 | 88842565 | 88874045 | 31481 | 2 | ENSMUST00000038409.6 | 116 | Podxl2 | ENSMUSG00000033152.8 |
| chr6 | 88874300 | 88874672 | 3.40852 | 3.95102 | 1.05353 | Promoter | 6 | 88842565 | 88874597 | 32033 | 2 | ENSMUST00000061262.5 | 0 | Podxl2 | ENSMUSG00000033152.8 |
| chr6 | 88875614 | 88876112 | 3.64275 | 4.48825 | 1.45621 | Promoter | 6 | 88849467 | 88875044 | 25578 | 2 | ENSMUST00000145944.1 | -571 | Podxl2 | ENSMUSG00000033152.8 |
| chr3 | 101494289 | 101494674 | 3.3771 | 3.92451 | 1.07575 | Distal Intergenic | 3 | 101576316 | 101577560 | 1245 | 2 | ENSMUST00000130649.1 | 82886 | Atp1a1 | ENSMUSG00000033161.9 |
| chr16 | 45653835 | 45654210 | 3.66439 | 4.46225 | 1.4657 | Promoter | 16 | 45626848 | 45654118 | 27271 | 2 | ENSMUST00000036732.7 | 0 | BC016579 | ENSMUSG00000033187.7 |
| chr2 | 122637972 | 122638237 | 3.4664 | 4.17421 | 1.24358 | Promoter | 2 | 122637915 | 122641085 | 3171 | 1 | ENSMUST00000110512.3 | 58 | AA467197 | ENSMUSG00000033213.11 |
| chr14 | 111675399 | 111677232 | 4.23118 | 5.65556 | 2.25339 | Promoter | 14 | 111675115 | 111683134 | 8020 | 1 | ENSMUST00000042767.7 | 285 | Slitrk5 | ENSMUSG00000033214.9 |
| chr14 | 111680081 | 111680274 | 3.62388 | 4.36147 | 1.35338 | Exon (ENSMUST00000042767.7/ENSMUSG00000033214.9, exon 2 of 2) | 14 | 111675115 | 111683134 | 8020 | 1 | ENSMUST00000042767.7 | 4967 | Slitrk5 | ENSMUSG00000033214.9 |
| chr14 | 111680582 | 111680928 | 3.51541 | 4.21634 | 1.27518 | Exon (ENSMUST00000042767.7/ENSMUSG00000033214.9, exon 2 of 2) | 14 | 111675115 | 111683134 | 8020 | 1 | ENSMUST00000042767.7 | 5468 | Slitrk5 | ENSMUSG00000033214.9 |
| chr14 | 111681329 | 111681578 | 4.35397 | 5.81128 | 2.36439 | Exon (ENSMUST00000042767.7/ENSMUSG00000033214.9, exon 2 of 2) | 14 | 111675115 | 111683134 | 8020 | 1 | ENSMUST00000042767.7 | 6215 | Slitrk5 | ENSMUSG00000033214.9 |
| chr1 | 74770871 | 74771079 | 3.77331 | 4.70602 | 1.61273 | Promoter | 1 | 74771892 | 74785322 | 13431 | 1 | ENSMUST00000006716.7 | -813 | Wnt6 | ENSMUSG00000033227.7 |
| chr1 | 74772151 | 74772433 | 3.91419 | 5.09894 | 1.90613 | Promoter | 1 | 74771892 | 74785322 | 13431 | 1 | ENSMUST00000006716.7 | 260 | Wnt6 | ENSMUSG00000033227.7 |
| chr1 | 74782642 | 74782737 | 3.59401 | 4.61228 | 1.5067 | Promoter | 1 | 74782439 | 74784602 | 2164 | 1 | ENSMUST00000189544.1 | 204 | Wnt6 | ENSMUSG00000033227.7 |
| chr1 | 74783101 | 74784359 | 3.78722 | 4.69927 | 1.5829 | Promoter | 1 | 74782439 | 74784602 | 2164 | 1 | ENSMUST00000189544.1 | 663 | Wnt6 | ENSMUSG00000033227.7 |
| chr15 | 96488648 | 96489018 | 3.38153 | 3.92608 | 1.10441 | Distal Intergenic | 15 | 96411698 | 96460843 | 49146 | 2 | ENSMUST00000047835.6 | -27806 | Scaf11 | ENSMUSG00000033228.7 |
| chr15 | 96542184 | 96542421 | 3.74873 | 4.58552 | 1.46463 | Distal Intergenic | 15 | 96411698 | 96460843 | 49146 | 2 | ENSMUST00000047835.6 | -81342 | Scaf11 | ENSMUSG00000033228.7 |
| chr2 | 122338217 | 122338364 | 3.27809 | 3.74428 | 1.01733 | Exon (ENSMUST00000099461.3/ENSMUSG00000033268.8, exon 25 of 33) | 2 | 122349504 | 122354131 | 4628 | 2 | ENSMUST00000135848.1 | 15767 | Shf | ENSMUSG00000033256.9 |
| chr17 | 67289798 | 67289964 | 3.86446 | 4.88381 | 1.71225 | Intron (ENSMUST00000037974.8/ENSMUSG00000033278.9, intron 1 of 30) | 17 | 66666848 | 67354459 | 687612 | 2 | ENSMUST00000037974.8 | 64495 | Ptprm | ENSMUSG00000033278.9 |
| chr17 | 67353151 | 67353453 | 3.96602 | 5.2067 | 1.90894 | Intron (ENSMUST00000037974.8/ENSMUSG00000033278.9, intron 1 of 30) | 17 | 66666848 | 67354459 | 687612 | 2 | ENSMUST00000037974.8 | 1006 | Ptprm | ENSMUSG00000033278.9 |
| chr17 | 67353841 | 67354305 | 3.40307 | 4.74832 | 1.62382 | Promoter | 17 | 66666848 | 67354459 | 687612 | 2 | ENSMUST00000037974.8 | 154 | Ptprm | ENSMUSG00000033278.9 |
| chr15 | 78428606 | 78428954 | 4.04417 | 5.2533 | 1.9995 | Promoter | 15 | 78428628 | 78438909 | 10282 | 1 | ENSMUST00000089414.5 | 0 | Kctd17 | ENSMUSG00000033287.10 |
| chr15 | 78429316 | 78429529 | 3.30358 | 3.79728 | 1.06757 | Promoter | 15 | 78428796 | 78438647 | 9852 | 1 | ENSMUST00000162321.2 | 521 | Kctd17 | ENSMUSG00000033287.10 |
| chr15 | 78437918 | 78438199 | 3.36864 | 4.03279 | 1.16866 | Intron (ENSMUST00000162517.2/ENSMUSG00000033287.10, intron 7 of 7) | 15 | 78433982 | 78437579 | 3598 | 1 | ENSMUST00000162605.1 | 3937 | Kctd17 | ENSMUSG00000033287.10 |
| chr15 | 78439113 | 78439465 | 3.33498 | 3.8378 | 1.04543 | Downstream (<1kb) | 15 | 78433982 | 78437579 | 3598 | 1 | ENSMUST00000162605.1 | 5132 | Kctd17 | ENSMUSG00000033287.10 |
| chr15 | 78439779 | 78440035 | 3.39523 | 4.0662 | 1.16133 | 3' UTR | 15 | 78433982 | 78437579 | 3598 | 1 | ENSMUST00000162605.1 | 5798 | Kctd17 | ENSMUSG00000033287.10 |
| chr15 | 78440799 | 78440966 | 3.30978 | 3.79188 | 1.01733 | Intron (ENSMUST00000017086.3/ENSMUSG00000016942.5, intron 16 of 17) | 15 | 78433982 | 78437579 | 3598 | 1 | ENSMUST00000162605.1 | 6818 | Kctd17 | ENSMUSG00000033287.10 |
| chr5 | 110628433 | 110628540 | 3.49084 | 4.17804 | 1.30229 | Distal Intergenic | 5 | 110648847 | 110650017 | 1171 | 2 | ENSMUST00000123380.1 | 21477 | Noc4l | ENSMUSG00000033294.6 |
| chr4 | 118290760 | 118291243 | 3.46003 | 4.06895 | 1.19531 | Promoter | 4 | 118208213 | 118291405 | 83193 | 2 | ENSMUST00000049074.7 | 162 | Ptprf | ENSMUSG00000033295.8 |
| chr4 | 118294106 | 118294434 | 3.432 | 4.00356 | 1.11441 | Distal Intergenic | 4 | 118208213 | 118291405 | 83193 | 2 | ENSMUST00000049074.7 | -2702 | Ptprf | ENSMUSG00000033295.8 |
| chr4 | 118294999 | 118295609 | 3.6364 | 4.37557 | 1.32292 | Distal Intergenic | 4 | 118208213 | 118291405 | 83193 | 2 | ENSMUST00000049074.7 | -3595 | Ptprf | ENSMUSG00000033295.8 |
| chr4 | 118297343 | 118297440 | 3.73133 | 4.55798 | 1.45617 | Distal Intergenic | 4 | 118208213 | 118291405 | 83193 | 2 | ENSMUST00000049074.7 | -5939 | Ptprf | ENSMUSG00000033295.8 |
| chr16 | 24353249 | 24353769 | 3.79831 | 4.6813 | 1.52505 | Distal Intergenic | 16 | 24393350 | 24992576 | 599227 | 1 | ENSMUST00000038053.7 | -39581 | Lpp | ENSMUSG00000033306.9 |
| chr5 | 110541565 | 110541756 | 3.64209 | 4.38805 | 1.35351 | Distal Intergenic | 5 | 110544345 | 110621382 | 77038 | 1 | ENSMUST00000040001.8 | -2589 | Galnt9 | ENSMUSG00000033316.9 |
| chr5 | 110543624 | 110545711 | 3.78818 | 4.70426 | 1.54457 | Promoter | 5 | 110544345 | 110621382 | 77038 | 1 | ENSMUST00000040001.8 | 0 | Galnt9 | ENSMUSG00000033316.9 |
| chr17 | 34671203 | 34672987 | 3.96785 | 5.26481 | 1.96408 | Promoter | 17 | 34670535 | 34719815 | 49281 | 1 | ENSMUST00000087399.4 | 669 | Tnxb | ENSMUSG00000033327.11 |
| chr3 | 117575048 | 117575583 | 3.96858 | 5.06016 | 1.86039 | Promoter | 3 | 117575227 | 117689507 | 114281 | 1 | ENSMUST00000106473.3 | 0 | 4833424O15Rik | ENSMUSG00000033342.8 |
| chr3 | 117575928 | 117576013 | 4.11021 | 5.46235 | 2.07241 | Promoter | 3 | 117575360 | 117689507 | 114148 | 1 | ENSMUST00000039564.6 | 569 | 4833424O15Rik | ENSMUSG00000033342.8 |
| chr9 | 95322533 | 95322751 | 3.42602 | 4.01183 | 1.17637 | Distal Intergenic | 9 | 95399292 | 95406722 | 7431 | 2 | ENSMUST00000036267.7 | 83971 | Chst2 | ENSMUSG00000033350.7 |
| chr9 | 95405034 | 95405938 | 3.68978 | 4.63819 | 1.57308 | Promoter | 9 | 95399292 | 95406722 | 7431 | 2 | ENSMUST00000036267.7 | 784 | Chst2 | ENSMUSG00000033350.7 |
| chr9 | 95406561 | 95406784 | 3.87976 | 4.9822 | 1.78348 | Promoter | 9 | 95399292 | 95406722 | 7431 | 2 | ENSMUST00000036267.7 | 0 | Chst2 | ENSMUSG00000033350.7 |
| chr15 | 94519802 | 94519978 | 3.35353 | 3.85782 | 1.01733 | Downstream (2-3kb) | 15 | 94540735 | 94543486 | 2752 | 2 | ENSMUST00000134061.1 | 23508 | Pus7l | ENSMUSG00000033356.3 |
| chr8 | 113688407 | 113688697 | 3.41527 | 4.02206 | 1.21396 | Distal Intergenic | 8 | 113643213 | 113655533 | 12321 | 1 | ENSMUST00000095173.1 | 45195 | Syce1l | ENSMUSG00000033409.8 |
| chr9 | 86877553 | 86877760 | 3.26659 | 3.7415 | 1.06254 | Intron (ENSMUST00000098495.4/ENSMUSG00000033419.10, intron 2 of 25) | 9 | 86835109 | 86880397 | 45289 | 2 | ENSMUST00000191290.1 | 2637 | Snap91 | ENSMUSG00000033419.10 |
| chr9 | 86878971 | 86880807 | 4.045 | 5.29033 | 2.01959 | Promoter | 9 | 86765923 | 86880439 | 114517 | 2 | ENSMUST00000098495.4 | 0 | Snap91 | ENSMUSG00000033419.10 |
| chr6 | 87335004 | 87335448 | 3.83439 | 4.83238 | 1.69085 | Promoter | 6 | 87133854 | 87335775 | 201922 | 2 | ENSMUST00000042025.9 | 327 | Antxr1 | ENSMUSG00000033420.10 |
| chr9 | 30920615 | 30921643 | 3.7754 | 4.64006 | 1.50454 | Promoter | 9 | 30899155 | 30922452 | 23298 | 2 | ENSMUST00000065112.6 | 809 | Adamts15 | ENSMUSG00000033453.7 |
| chr14 | 72531571 | 72533004 | 3.91026 | 4.97485 | 1.77718 | Distal Intergenic | 14 | 72561447 | 72564718 | 3272 | 2 | ENSMUST00000159144.1 | 31714 | Fndc3a | ENSMUSG00000033487.9 |
| chr14 | 72533534 | 72533828 | 3.35756 | 3.86382 | 1.01733 | Distal Intergenic | 14 | 72561447 | 72564718 | 3272 | 2 | ENSMUST00000159144.1 | 30890 | Fndc3a | ENSMUSG00000033487.9 |
| chr14 | 72534392 | 72534711 | 3.76565 | 4.80765 | 1.6202 | Distal Intergenic | 14 | 72561447 | 72564718 | 3272 | 2 | ENSMUST00000159144.1 | 30007 | Fndc3a | ENSMUSG00000033487.9 |
| chr9 | 86743650 | 86743740 | 3.85491 | 4.84523 | 1.71859 | Promoter | 9 | 86743649 | 86758443 | 14795 | 1 | ENSMUST00000036426.7 | 2 | Prss35 | ENSMUSG00000033491.8 |
| chr13 | 9187233 | 9187335 | 3.93838 | 5.00582 | 1.83523 | Distal Intergenic | 13 | 9168361 | 9173121 | 4761 | 1 | ENSMUST00000189330.1 | 18873 | Larp4b | ENSMUSG00000033499.8 |
| chr13 | 9199869 | 9199955 | 3.51861 | 4.16833 | 1.23576 | Distal Intergenic | 13 | 9168361 | 9173121 | 4761 | 1 | ENSMUST00000189330.1 | 31509 | Larp4b | ENSMUSG00000033499.8 |
| chr16 | 22807781 | 22807856 | 3.55289 | 4.21228 | 1.20626 | Intron (ENSMUST00000040592.4/ENSMUSG00000033501.4, intron 1 of 2) | 16 | 22805203 | 22811410 | 6208 | 2 | ENSMUST00000040592.4 | 3554 | Crygs | ENSMUSG00000033501.4 |
| chr7 | 63445018 | 63445771 | 3.92027 | 4.97949 | 1.80854 | Promoter | 7 | 63444777 | 63617900 | 173124 | 1 | ENSMUST00000176734.1 | 242 | Otud7a | ENSMUSG00000033510.9 |
| chr7 | 63446602 | 63446992 | 3.33473 | 3.82942 | 1.01733 | Intron (ENSMUST00000058476.8/ENSMUSG00000033510.9, intron 1 of 13) | 7 | 63444777 | 63617900 | 173124 | 1 | ENSMUST00000176734.1 | 1826 | Otud7a | ENSMUSG00000033510.9 |
| chr7 | 63757491 | 63758661 | 3.67368 | 4.5012 | 1.51042 | 3' UTR | 7 | 63754712 | 63757925 | 3214 | 1 | ENSMUST00000177511.1 | 2780 | Otud7a | ENSMUSG00000033510.9 |
| chr7 | 63759606 | 63760251 | 4.13289 | 5.42834 | 2.13226 | Distal Intergenic | 7 | 63754712 | 63757925 | 3214 | 1 | ENSMUST00000177511.1 | 4895 | Otud7a | ENSMUSG00000033510.9 |
| chr1 | 25228390 | 25228730 | 3.91122 | 5.15501 | 1.86301 | Promoter | 1 | 25067476 | 25228826 | 161351 | 2 | ENSMUST00000126626.2 | 96 | Bai3 | ENSMUSG00000033569.12 |
| chr1 | 25828173 | 25828392 | 3.51106 | 4.26807 | 1.28285 | Intron (ENSMUST00000135518.2/ENSMUSG00000033569.12, intron 1 of 31) | 1 | 25067550 | 25829511 | 761962 | 2 | ENSMUST00000135518.2 | 1119 | Bai3 | ENSMUSG00000033569.12 |
| chr1 | 25830004 | 25830138 | 3.63729 | 4.38288 | 1.35624 | Promoter | 1 | 25067550 | 25829707 | 762158 | 2 | ENSMUST00000151309.2 | -298 | Bai3 | ENSMUSG00000033569.12 |
| chr1 | 25830514 | 25830765 | 4.01676 | 5.11535 | 1.85101 | Promoter | 1 | 25067550 | 25829707 | 762158 | 2 | ENSMUST00000151309.2 | -808 | Bai3 | ENSMUSG00000033569.12 |
| chr1 | 25831338 | 25831441 | 3.26294 | 3.72158 | 1.01733 | Distal Intergenic | 1 | 25067550 | 25829707 | 762158 | 2 | ENSMUST00000151309.2 | -1632 | Bai3 | ENSMUSG00000033569.12 |
| chr1 | 25832152 | 25832432 | 3.62855 | 4.6609 | 1.55032 | Distal Intergenic | 1 | 25067550 | 25829707 | 762158 | 2 | ENSMUST00000151309.2 | -2446 | Bai3 | ENSMUSG00000033569.12 |
| chr9 | 80351825 | 80351999 | 3.30172 | 3.79147 | 1.05793 | Intron (ENSMUST00000085289.6/ENSMUSG00000032343.10, intron 11 of 15) | 9 | 80306914 | 80308602 | 1689 | 1 | ENSMUST00000138650.2 | 44912 | Myo6 | ENSMUSG00000033577.13 |
| chr9 | 75231952 | 75233218 | 3.97248 | 5.14547 | 1.962 | Promoter | 9 | 75232014 | 75305451 | 73438 | 1 | ENSMUST00000036555.6 | 0 | Myo5c | ENSMUSG00000033590.7 |
| chr9 | 75253108 | 75253403 | 3.34153 | 3.83966 | 1.01733 | Intron (ENSMUST00000036555.6/ENSMUSG00000033590.7, intron 8 of 40) | 9 | 75232014 | 75305451 | 73438 | 1 | ENSMUST00000036555.6 | 21095 | Myo5c | ENSMUSG00000033590.7 |
| chr14 | 70529228 | 70529459 | 4.81175 | 6.73713 | 3.0373 | Distal Intergenic | 14 | 70530821 | 70538323 | 7503 | 1 | ENSMUST00000047331.6 | -1362 | Lgi3 | ENSMUSG00000033595.6 |
| chr14 | 70530848 | 70531121 | 3.42046 | 3.97371 | 1.06997 | Promoter | 14 | 70530821 | 70538323 | 7503 | 1 | ENSMUST00000047331.6 | 28 | Lgi3 | ENSMUSG00000033595.6 |
| chr14 | 70532041 | 70532360 | 3.89931 | 4.86239 | 1.63092 | Exon (ENSMUST00000047331.6/ENSMUSG00000033595.6, exon 2 of 8) | 14 | 70530821 | 70538323 | 7503 | 1 | ENSMUST00000047331.6 | 1221 | Lgi3 | ENSMUSG00000033595.6 |
| chr5 | 108549902 | 108551467 | 3.90971 | 4.95657 | 1.75193 | Promoter | 5 | 108518554 | 108550024 | 31471 | 2 | ENSMUST00000046892.9 | 0 | Cplx1 | ENSMUSG00000033615.9 |

| | | | | | | | | | | | | | | | |
|---|---|---|---|---|---|---|---|---|---|---|---|---|---|---|---|
| chr18 | 30268123 | 30268288 | 3.52507 | 4.29004 | 1.30699 | Distal Intergenic | 18 | 30272747 | 30348126 | 75380 | 1 | ENSMUST00000115812.4 | -4459 | Pik3c3 | ENSMUSG00000033628.10 |
| chr7 | 57590533 | 57591238 | 3.77138 | 4.71152 | 1.60359 | Promoter | 7 | 57591162 | 57765781 | 174620 | 1 | ENSMUST00000156375.1 | 0 | Gabrb3 | ENSMUSG00000033676.8 |
| chr7 | 57591671 | 57592025 | 3.9045 | 4.95272 | 1.7782 | Promoter | 7 | 57591530 | 57816370 | 224841 | 1 | ENSMUST00000138350.1 | 142 | Gabrb3 | ENSMUSG00000033676.8 |
| chr1 | 155825498 | 155826038 | 3.50311 | 4.15298 | 1.22226 | Distal Intergenic | 1 | 155778155 | 155812883 | 34729 | 2 | ENSMUST00000035325.9 | -12616 | Qsox1 | ENSMUSG00000033684.9 |
| chr1 | 155829327 | 155829402 | 3.79411 | 4.66518 | 1.50584 | Distal Intergenic | 1 | 155778155 | 155812883 | 34729 | 2 | ENSMUST00000035325.9 | -16445 | Qsox1 | ENSMUSG00000033684.9 |
| chr1 | 155830018 | 155830116 | 3.46821 | 4.05789 | 1.11267 | Distal Intergenic | 1 | 155778155 | 155812883 | 34729 | 2 | ENSMUST00000035325.9 | -17136 | Qsox1 | ENSMUSG00000033684.9 |
| chr15 | 76718236 | 76718882 | 3.48344 | 4.32521 | 1.34997 | Promoter | 15 | 76716207 | 76718416 | 2210 | 2 | ENSMUST00000146855.1 | 0 | Lrrc24 | ENSMUSG00000033707.8 |
| chr19 | 54043283 | 54043617 | 3.60302 | 4.54097 | 1.48305 | Distal Intergenic | 19 | 54045182 | 54048982 | 3801 | 1 | ENSMUST00000036700.5 | -1565 | Adra2a | ENSMUSG00000033717.5 |
| chr19 | 54044254 | 54045249 | 3.51683 | 4.29651 | 1.34105 | Promoter | 19 | 54045182 | 54048982 | 3801 | 1 | ENSMUST00000036700.5 | 0 | Adra2a | ENSMUSG00000033717.5 |
| chr19 | 54045579 | 54046296 | 3.74489 | 4.6335 | 1.57784 | Promoter | 19 | 54045182 | 54048982 | 3801 | 1 | ENSMUST00000036700.5 | 398 | Adra2a | ENSMUSG00000033717.5 |
| chr19 | 54046804 | 54047761 | 3.66569 | 4.56569 | 1.51533 | 3' UTR | 19 | 54045182 | 54048982 | 3801 | 1 | ENSMUST00000036700.5 | 1623 | Adra2a | ENSMUSG00000033717.5 |
| chr19 | 54052255 | 54052353 | 3.83468 | 4.76005 | 1.60128 | Distal Intergenic | 19 | 54045182 | 54048982 | 3801 | 1 | ENSMUST00000036700.5 | 7074 | Adra2a | ENSMUSG00000033717.5 |
| chr19 | 54073731 | 54073871 | 3.4831 | 4.36043 | 1.35716 | Distal Intergenic | 19 | 54045182 | 54048982 | 3801 | 1 | ENSMUST00000036700.5 | 28550 | Adra2a | ENSMUSG00000033717.5 |
| chr19 | 54114238 | 54114951 | 3.48577 | 4.14248 | 1.23345 | Distal Intergenic | 19 | 54045182 | 54048982 | 3801 | 1 | ENSMUST00000036700.5 | 69057 | Adra2a | ENSMUSG00000033717.5 |
| chr19 | 54117117 | 54117565 | 3.37224 | 3.97775 | 1.10958 | Distal Intergenic | 19 | 54045182 | 54048982 | 3801 | 1 | ENSMUST00000036700.5 | 71936 | Adra2a | ENSMUSG00000033717.5 |
| chr19 | 54244035 | 54244147 | 3.24243 | 3.69081 | 1.01733 | Distal Intergenic | 19 | 54045182 | 54048982 | 3801 | 1 | ENSMUST00000036700.5 | 198854 | Adra2a | ENSMUSG00000033717.5 |
| chr19 | 54244550 | 54245004 | 3.5988 | 4.30539 | 1.28773 | Distal Intergenic | 19 | 54045182 | 54048982 | 3801 | 1 | ENSMUST00000036700.5 | 199369 | Adra2a | ENSMUSG00000033717.5 |
| chr3 | 109340396 | 109341135 | 3.9001 | 5.02811 | 1.84749 | Promoter | 3 | 109340653 | 109685698 | 345046 | 1 | ENSMUST00000046864.8 | 0 | Vav3 | ENSMUSG00000033721.11 |
| chr6 | 85183649 | 85184027 | 3.39655 | 4.01481 | 1.19064 | Distal Intergenic | 6 | 85187438 | 85204358 | 16921 | 1 | ENSMUST00000045942.8 | -3411 | Emx1 | ENSMUSG00000033726.8 |
| chr6 | 85185263 | 85185530 | 4.02791 | 5.28846 | 1.9818 | Distal Intergenic | 6 | 85187438 | 85204358 | 16921 | 1 | ENSMUST00000045942.8 | -1908 | Emx1 | ENSMUSG00000033726.8 |
| chr6 | 85186421 | 85188241 | 4.07216 | 5.45902 | 2.10313 | Promoter | 6 | 85187438 | 85204358 | 16921 | 1 | ENSMUST00000045942.8 | 0 | Emx1 | ENSMUSG00000033726.8 |
| chr6 | 85188688 | 85189002 | 3.5854 | 4.44509 | 1.45047 | Intron (ENSMUST00000045942.8/ENSMUSG00000033726.8, intron 1 of 2) | 6 | 85187438 | 85204358 | 16921 | 1 | ENSMUST00000045942.8 | 1251 | Emx1 | ENSMUSG00000033726.8 |
| chr6 | 85189388 | 85189672 | 3.52251 | 4.2104 | 1.3113 | Intron (ENSMUST00000045942.8/ENSMUSG00000033726.8, intron 1 of 2) | 6 | 85187438 | 85204358 | 16921 | 1 | ENSMUST00000045942.8 | 1951 | Emx1 | ENSMUSG00000033726.8 |
| chr6 | 85190008 | 85190545 | 3.44587 | 4.15373 | 1.25727 | Intron (ENSMUST00000045942.8/ENSMUSG00000033726.8, intron 1 of 2) | 6 | 85187438 | 85204358 | 16921 | 1 | ENSMUST00000045942.8 | 2571 | Emx1 | ENSMUSG00000033726.8 |
| chr6 | 85191246 | 85192225 | 3.69768 | 4.61027 | 1.56344 | Intron (ENSMUST00000045942.8/ENSMUSG00000033726.8, intron 1 of 2) | 6 | 85193519 | 85204462 | 10944 | 1 | ENSMUST00000173919.1 | -1294 | Emx1 | ENSMUSG00000033726.8 |
| chr6 | 85194693 | 85195280 | 3.81878 | 4.78248 | 1.63564 | Intron (ENSMUST00000045942.8/ENSMUSG00000033726.8, intron 2 of 2) | 6 | 85193519 | 85204462 | 10944 | 1 | ENSMUST00000173919.1 | 1175 | Emx1 | ENSMUSG00000033726.8 |
| chr6 | 85206487 | 85206785 | 4.15827 | 5.38824 | 2.02884 | Distal Intergenic | 6 | 85193519 | 85204462 | 10944 | 1 | ENSMUST00000173919.1 | 12969 | Emx1 | ENSMUSG00000033726.8 |
| chr14 | 69976795 | 69977007 | 3.50784 | 4.21222 | 1.30016 | Intron (ENSMUST00000022678.3/ENSMUSG00000022085.3, intron 4 of 6) | 14 | 70077445 | 70080157 | 2713 | 1 | ENSMUST00000035908.1 | -100438 | Egr3 | ENSMUSG00000033730.3 |
| chr14 | 70075739 | 70075959 | 3.47081 | 4.08028 | 1.17913 | Distal Intergenic | 14 | 70077445 | 70080157 | 2713 | 1 | ENSMUST00000035908.1 | -1486 | Egr3 | ENSMUSG00000033730.3 |
| chr14 | 70077459 | 70079671 | 4.08014 | 5.41048 | 2.10468 | Promoter | 14 | 70077445 | 70080157 | 2713 | 1 | ENSMUST00000035908.1 | 15 | Egr3 | ENSMUSG00000033730.3 |
| chr14 | 70080688 | 70080826 | 3.93408 | 5.01662 | 1.84438 | Distal Intergenic | 14 | 70077445 | 70080157 | 2713 | 1 | ENSMUST00000035908.1 | 3244 | Egr3 | ENSMUSG00000033730.3 |
| chr1 | 6441110 | 6441534 | 3.49113 | 4.10717 | 1.16473 | Distal Intergenic | 1 | 6487231 | 6860934 | 373704 | 1 | ENSMUST00000140079.2 | -45697 | St18 | ENSMUSG00000033740.12 |
| chr1 | 6442698 | 6442920 | 3.33836 | 3.83489 | 1.01733 | Distal Intergenic | 1 | 6487231 | 6860934 | 373704 | 1 | ENSMUST00000140079.2 | -44311 | St18 | ENSMUSG00000033740.12 |
| chr19 | 6418716 | 6418845 | 3.64638 | 4.52227 | 1.45745 | Promoter | 19 | 6418731 | 6533217 | 114487 | 1 | ENSMUST00000113462.2 | 0 | Nrxn2 | ENSMUSG00000033768.11 |
| chr19 | 6419246 | 6419784 | 3.62328 | 4.4603 | 1.4279 | Promoter | 19 | 6418765 | 6532278 | 113514 | 1 | ENSMUST00000113461.2 | 482 | Nrxn2 | ENSMUSG00000033768.11 |
| chr19 | 6420931 | 6421185 | 3.65399 | 4.40315 | 1.33544 | Intron (ENSMUST00000113462.2/ENSMUSG00000033768.11, intron 1 of 22) | 19 | 6418765 | 6532278 | 113514 | 1 | ENSMUST00000113461.2 | 2167 | Nrxn2 | ENSMUSG00000033768.11 |
| chr19 | 6424133 | 6424235 | 3.35634 | 4.0073 | 1.10827 | Intron (ENSMUST00000113462.2/ENSMUSG00000033768.11, intron 1 of 22) | 19 | 6427937 | 6533217 | 105281 | 1 | ENSMUST00000137166.2 | -3702 | Nrxn2 | ENSMUSG00000033768.11 |
| chr19 | 6426923 | 6429578 | 3.80496 | 4.99754 | 1.7899 | Promoter | 19 | 6427937 | 6533217 | 105281 | 1 | ENSMUST00000137166.2 | 0 | Nrxn2 | ENSMUSG00000033768.11 |
| chr19 | 6430028 | 6430136 | 4.51502 | 6.19834 | 2.62159 | Intron (ENSMUST00000113462.2/ENSMUSG00000033768.11, intron 2 of 22) | 19 | 6428767 | 6473339 | 44573 | 1 | ENSMUST00000155158.1 | 1262 | Nrxn2 | ENSMUSG00000033768.11 |
| chr19 | 6464094 | 6464169 | 3.49671 | 4.21999 | 1.27972 | Exon (ENSMUST00000128272.2/ENSMUSG00000033768.11, exon 5 of 5) | 19 | 6473109 | 6482696 | 9588 | 1 | ENSMUST00000140435.1 | -8940 | Nrxn2 | ENSMUSG00000033768.11 |
| chr19 | 6487553 | 6487672 | 3.64941 | 4.42266 | 1.41144 | Intron (ENSMUST00000113462.2/ENSMUSG00000033768.11, intron 11 of 22) | 19 | 6497671 | 6509865 | 12195 | 1 | ENSMUST00000148391.1 | -9999 | Nrxn2 | ENSMUSG00000033768.11 |
| chr19 | 6495064 | 6495827 | 3.3607 | 3.86863 | 1.01733 | Intron (ENSMUST00000113462.2/ENSMUSG00000033768.11, intron 17 of 22) | 19 | 6497671 | 6509865 | 12195 | 1 | ENSMUST00000148391.1 | -1844 | Nrxn2 | ENSMUSG00000033768.11 |
| chr19 | 6496598 | 6497099 | 3.77861 | 4.63392 | 1.47845 | Promoter | 19 | 6497671 | 6509865 | 12195 | 1 | ENSMUST00000148391.1 | -572 | Nrxn2 | ENSMUSG00000033768.11 |
| chr19 | 6497835 | 6498885 | 3.67397 | 4.44541 | 1.37276 | Promoter | 19 | 6497835 | 6544169 | 46335 | 1 | ENSMUST00000113459.1 | 1 | Nrxn2 | ENSMUSG00000033768.11 |
| chr19 | 6499191 | 6499497 | 3.35668 | 3.86251 | 1.01733 | Intron (ENSMUST00000113462.2/ENSMUSG00000033768.11, intron 17 of 22) | 19 | 6497835 | 6544169 | 46335 | 1 | ENSMUST00000113459.1 | 1357 | Nrxn2 | ENSMUSG00000033768.11 |
| chr19 | 6501282 | 6501510 | 3.68964 | 4.54166 | 1.50578 | Intron (ENSMUST00000113462.2/ENSMUSG00000033768.11, intron 17 of 22) | 19 | 6497835 | 6544169 | 46335 | 1 | ENSMUST00000113459.1 | 3448 | Nrxn2 | ENSMUSG00000033768.11 |
| chr19 | 6501874 | 6503213 | 3.76105 | 4.63485 | 1.506 | Intron (ENSMUST00000113462.2/ENSMUSG00000033768.11, intron 17 of 22) | 19 | 6497835 | 6544169 | 46335 | 1 | ENSMUST00000113459.1 | 4040 | Nrxn2 | ENSMUSG00000033768.11 |
| chr19 | 6503887 | 6504012 | 3.90403 | 4.88119 | 1.64924 | Intron (ENSMUST00000113462.2/ENSMUSG00000033768.11, intron 17 of 22) | 19 | 6497835 | 6544169 | 46335 | 1 | ENSMUST00000113459.1 | 6053 | Nrxn2 | ENSMUSG00000033768.11 |
| chr19 | 6504387 | 6504545 | 3.36799 | 3.883 | 1.02946 | Intron (ENSMUST00000113462.2/ENSMUSG00000033768.11, intron 17 of 22) | 19 | 6497835 | 6544169 | 46335 | 1 | ENSMUST00000113459.1 | 6553 | Nrxn2 | ENSMUSG00000033768.11 |
| chr19 | 6506626 | 6507012 | 3.36821 | 3.8799 | 1.01733 | Intron (ENSMUST00000113462.2/ENSMUSG00000033768.11, intron 18 of 22) | 19 | 6497835 | 6544169 | 46335 | 1 | ENSMUST00000113459.1 | 8792 | Nrxn2 | ENSMUSG00000033768.11 |
| chr19 | 6507334 | 6507642 | 3.40708 | 3.96635 | 1.11621 | Intron (ENSMUST00000113462.2/ENSMUSG00000033768.11, intron 18 of 22) | 19 | 6497835 | 6544169 | 46335 | 1 | ENSMUST00000113459.1 | 9500 | Nrxn2 | ENSMUSG00000033768.11 |
| chr19 | 6510916 | 6511328 | 3.69065 | 4.48895 | 1.42694 | Intron (ENSMUST00000113462.2/ENSMUSG00000033768.11, intron 19 of 22) | 19 | 6517205 | 6532296 | 15092 | 1 | ENSMUST00000154580.1 | -5877 | Nrxn2 | ENSMUSG00000033768.11 |
| chr19 | 6514278 | 6514812 | 3.7745 | 4.74747 | 1.61598 | Intron (ENSMUST00000113462.2/ENSMUSG00000033768.11, intron 20 of 22) | 19 | 6517205 | 6532296 | 15092 | 1 | ENSMUST00000154580.1 | -2393 | Nrxn2 | ENSMUSG00000033768.11 |
| chr4 | 141382064 | 141382326 | 3.72818 | 4.65579 | 1.54015 | Intron (ENSMUST00000006380.4/ENSMUSG00000006218.4, intron 4 of 6) | 4 | 141391396 | 141394827 | 3432 | 2 | ENSMUST00000150334.1 | 12501 | Clcnka | ENSMUSG00000033770.8 |
| chr4 | 141392696 | 141392771 | 3.99953 | 5.04985 | 1.74935 | Exon (ENSMUST00000105790.2/ENSMUSG00000033770.8, exon 8 of 19) | 4 | 141391396 | 141394827 | 3432 | 2 | ENSMUST00000150334.1 | 2056 | Clcnka | ENSMUSG00000033770.8 |
| chr1 | 5918549 | 5918653 | 3.43702 | 4.23036 | 1.29169 | Distal Intergenic | 1 | 5913707 | 5917398 | 3692 | 2 | ENSMUST00000044180.4 | -1152 | Npbwr1 | ENSMUSG00000033774.4 |
| chr6 | 84008407 | 84008533 | 3.53551 | 4.43635 | 1.42219 | Promoter | 6 | 84008590 | 84211042 | 202453 | 1 | ENSMUST00000113823.2 | -57 | Dysf | ENSMUSG00000033788.10 |
| chr6 | 84017772 | 84018107 | 3.81975 | 4.95627 | 1.78952 | Intron (ENSMUST00000113823.2/ENSMUSG00000033788.10, intron 1 of 54) | 6 | 84019385 | 84211059 | 191675 | 1 | ENSMUST00000089595.6 | -1278 | Dysf | ENSMUSG00000033788.10 |
| chr6 | 84018596 | 84018809 | 3.90224 | 4.97346 | 1.78058 | Promoter | 6 | 84019385 | 84211059 | 191675 | 1 | ENSMUST00000089595.6 | -576 | Dysf | ENSMUSG00000033788.10 |
| chr6 | 84020262 | 84020394 | 4.00967 | 5.30061 | 1.988 | Promoter | 6 | 84019389 | 84211059 | 191671 | 1 | ENSMUST00000081904.6 | 874 | Dysf | ENSMUSG00000033788.10 |
| chr6 | 84102061 | 84102332 | 3.77671 | 4.68372 | 1.5567 | Exon (ENSMUST00000137673.1/ENSMUSG00000087001.1, exon 2 of 3) | 6 | 84105373 | 84112199 | 6827 | 1 | ENSMUST00000127459.2 | -3041 | Dysf | ENSMUSG00000033788.10 |
| chr6 | 84104229 | 84104545 | 3.83136 | 4.73328 | 1.54082 | Promoter | 6 | 84105373 | 84112199 | 6827 | 1 | ENSMUST00000127459.2 | -828 | Dysf | ENSMUSG00000033788.10 |
| chr6 | 84105043 | 84105992 | 3.76107 | 4.75002 | 1.58134 | Promoter | 6 | 84105373 | 84112199 | 6827 | 1 | ENSMUST00000127459.2 | 0 | Dysf | ENSMUSG00000033788.10 |
| chr6 | 84314344 | 84314555 | 3.62063 | 4.34526 | 1.31633 | Distal Intergenic | 6 | 84210484 | 84211041 | 558 | 1 | ENSMUST00000128965.1 | 103861 | Dysf | ENSMUSG00000033788.10 |
| chr6 | 84197930 | 84198117 | 3.30525 | 3.7851 | 1.01733 | Intron (ENSMUST00000113823.2/ENSMUSG00000033788.10, intron 50 of 54) | 6 | 84210484 | 84211041 | 558 | 1 | ENSMUST00000128965.1 | -12367 | Dysf | ENSMUSG00000033788.10 |
| chr6 | 84263917 | 84264035 | 3.43578 | 4.05436 | 1.15806 | Distal Intergenic | 6 | 84210484 | 84211041 | 558 | 1 | ENSMUST00000128965.1 | 53434 | Dysf | ENSMUSG00000033788.10 |
| chr6 | 84276913 | 84277429 | 3.5132 | 4.18769 | 1.24948 | Distal Intergenic | 6 | 84210484 | 84211041 | 558 | 1 | ENSMUST00000128965.1 | 66430 | Dysf | ENSMUSG00000033788.10 |
| chr6 | 84375905 | 84376016 | 3.73577 | 4.57848 | 1.50603 | Distal Intergenic | 6 | 84210484 | 84211041 | 558 | 1 | ENSMUST00000128965.1 | 165422 | Dysf | ENSMUSG00000033788.10 |
| chr6 | 84315290 | 84315495 | 3.56219 | 4.24335 | 1.26428 | Distal Intergenic | 6 | 84210484 | 84211041 | 558 | 1 | ENSMUST00000128965.1 | 104807 | Dysf | ENSMUSG00000033788.10 |
| chr6 | 84371380 | 84371455 | 3.68219 | 4.46164 | 1.38381 | Distal Intergenic | 6 | 84210484 | 84211041 | 558 | 1 | ENSMUST00000128965.1 | 160897 | Dysf | ENSMUSG00000033788.10 |
| chr6 | 84371975 | 84372246 | 3.38846 | 3.9201 | 1.05141 | Distal Intergenic | 6 | 84210484 | 84211041 | 558 | 1 | ENSMUST00000128965.1 | 161492 | Dysf | ENSMUSG00000033788.10 |
| chr6 | 84376698 | 84377261 | 3.5647 | 4.262 | 1.30996 | Distal Intergenic | 6 | 84210484 | 84211041 | 558 | 1 | ENSMUST00000128965.1 | 166215 | Dysf | ENSMUSG00000033788.10 |
| chr5 | 107403304 | 107403774 | 3.93574 | 5.339 | 2.01934 | Promoter | 5 | 107403496 | 107430035 | 26540 | 1 | ENSMUST00000049146.6 | 0 | Ephx4 | ENSMUSG00000033805.7 |
| chr5 | 107404368 | 107404622 | 3.43419 | 4.04376 | 1.22339 | Promoter | 5 | 107403775 | 107413528 | 9754 | 1 | ENSMUST00000161246.2 | 594 | Ephx4 | ENSMUSG00000033805.7 |
| chr12 | 98574579 | 98575450 | 3.74455 | 4.68634 | 1.64617 | Intron (ENSMUST00000110113.1/ENSMUSG00000033854.9, intron 1 of 6) | 12 | 98433994 | 98577940 | 143947 | 2 | ENSMUST00000110113.1 | 2490 | Kcnk10 | ENSMUSG00000033854.9 |
| chr12 | 98575998 | 98576368 | 3.56937 | 4.43395 | 1.40496 | Intron (ENSMUST00000110113.1/ENSMUSG00000033854.9, intron 1 of 6) | 12 | 98433994 | 98577940 | 143947 | 2 | ENSMUST00000110113.1 | 1572 | Kcnk10 | ENSMUSG00000033854.9 |
| chr12 | 98577033 | 98578270 | 3.74225 | 4.87643 | 1.73147 | Promoter | 12 | 98433994 | 98577940 | 143947 | 2 | ENSMUST00000110113.1 | 0 | Kcnk10 | ENSMUSG00000033854.9 |
| chr12 | 98578775 | 98579485 | 3.45398 | 4.06394 | 1.21194 | Promoter | 12 | 98433994 | 98577940 | 143947 | 2 | ENSMUST00000110113.1 | -836 | Kcnk10 | ENSMUSG00000033854.9 |
| chr3 | 82875748 | 82877241 | 4.54534 | 6.46534 | 2.91662 | Promoter | 3 | 82837228 | 82876483 | 39256 | 2 | ENSMUST00000048647.8 | 0 | Rbm46 | ENSMUSG00000033882.10 |
| chr3 | 82145276 | 82145520 | 3.56975 | 4.25409 | 1.26542 | Promoter | 3 | 82092428 | 82145877 | 53450 | 2 | ENSMUST00000048976.6 | 357 | Gucy1a3 | ENSMUSG00000033910.8 |
| chr8 | 127006478 | 127006844 | 3.87922 | 5.16366 | 1.9106 | Distal Intergenic | 8 | 126947175 | 126971079 | 23905 | 2 | ENSMUST00000045994.6 | -35400 | Rbm34 | ENSMUSG00000033931.8 |
| chr18 | 46195290 | 46196553 | 4.03261 | 5.2034 | 1.95119 | Exon (ENSMUST00000167364.2/ENSMUSG00000033949.6, exon 2 of 10) | 18 | 46165302 | 46198818 | 33517 | 2 | ENSMUST00000167364.2 | 2265 | Trim36 | ENSMUSG00000033949.6 |
| chr18 | 46197534 | 46198260 | 3.47186 | 4.26823 | 1.33372 | Promoter | 18 | 46165302 | 46198818 | 33517 | 2 | ENSMUST00000167364.2 | 558 | Trim36 | ENSMUSG00000033949.6 |
| chr18 | 46200412 | 46200928 | 3.31974 | 3.8127 | 1.03787 | Intron (ENSMUST00000037011.4/ENSMUSG00000033949.6, intron 1 of 9) | 18 | 46165302 | 46198818 | 33517 | 2 | ENSMUST00000167364.2 | -1595 | Trim36 | ENSMUSG00000033949.6 |
| chr18 | 46207793 | 46208505 | 3.65615 | 4.41873 | 1.36436 | Intron (ENSMUST00000037011.4/ENSMUSG00000033949.6, intron 1 of 9) | 18 | 46165302 | 46212607 | 47306 | 2 | ENSMUST00000037011.4 | 4102 | Trim36 | ENSMUSG00000033949.6 |
| chr18 | 46208949 | 46210563 | 3.75413 | 4.69985 | 1.58265 | Intron (ENSMUST00000037011.4/ENSMUSG00000033949.6, intron 1 of 9) | 18 | 46165302 | 46212607 | 47306 | 2 | ENSMUST00000037011.4 | 2044 | Trim36 | ENSMUSG00000033949.6 |
| chr18 | 46210919 | 46212579 | 4.04494 | 5.35109 | 2.01655 | Promoter | 18 | 46165302 | 46212607 | 47306 | 2 | ENSMUST00000037011.4 | 28 | Trim36 | ENSMUSG00000033949.6 |
| chr18 | 46213424 | 46213816 | 3.25773 | 3.72369 | 1.01733 | Promoter | 18 | 46165302 | 46212607 | 47306 | 2 | ENSMUST00000037011.4 | -818 | Trim36 | ENSMUSG00000033949.6 |
| chr18 | 46214626 | 46214821 | 3.35617 | 3.89306 | 1.12397 | Distal Intergenic | 18 | 46165302 | 46212607 | 47306 | 2 | ENSMUST00000037011.4 | -2020 | Trim36 | ENSMUSG00000033949.6 |
| chr18 | 46215953 | 46216301 | 3.50677 | 4.2065 | 1.31327 | Distal Intergenic | 18 | 46165302 | 46212607 | 47306 | 2 | ENSMUST00000037011.4 | -3347 | Trim36 | ENSMUSG00000033949.6 |
| chr1 | 139454892 | 139455214 | 3.5005 | 4.28532 | 1.35713 | Promoter | 1 | 139454819 | 139457134 | 2316 | 1 | ENSMUST00000128413.1 | 74 | Aspm | ENSMUSG00000033952.9 |
| chr1 | 139455543 | 139455687 | 3.54827 | 4.3717 | 1.39034 | Promoter | 1 | 139454819 | 139457134 | 2316 | 1 | ENSMUST00000128413.1 | 725 | Aspm | ENSMUSG00000033952.9 |
| chr17 | 80576863 | 80577280 | 3.92734 | 5.0116 | 1.80257 | Distal Intergenic | 17 | 80523550 | 80563834 | 40285 | 2 | ENSMUST00000086545.4 | -13030 | Cdkl4 | ENSMUSG00000033966.7 |
| chr7 | 12927864 | 12928015 | 3.70788 | 4.55858 | 1.4445 | Promoter | 7 | 12927416 | 12931072 | 3657 | 1 | ENSMUST00000045870.4 | 449 | 2310014L17Rik | ENSMUSG00000033967.4 |
| chr3 | 80800485 | 80801368 | 3.82966 | 4.77536 | 1.63081 | Intron (ENSMUST00000075316.4/ENSMUSG00000033981.9, intron 2 of 15) | 3 | 80684936 | 80802791 | 117856 | 2 | ENSMUST00000075316.4 | 1423 | Gria2 | ENSMUSG00000033981.9 |
| chr11 | 88970204 | 88970512 | 2.24599 | 4.11929 | 1.22451 | Promoter | 11 | 88970252 | 88982645 | 12394 | 1 | ENSMUST00000135325.2 | 0 | Coil | ENSMUSG00000033983.8 |
| chr8 | 125963907 | 125964092 | 3.45318 | 4.2645 | 1.32624 | Distal Intergenic | 8 | 125995102 | 126030683 | 35582 | 1 | ENSMUST00000046765.8 | -31010 | Kcnk1 | ENSMUSG00000033998.8 |
| chr8 | 125995261 | 125996383 | 3.74679 | 4.62185 | 1.51551 | Promoter | 8 | 125995102 | 126030683 | 35582 | 1 | ENSMUST00000046765.8 | 160 | Kcnk1 | ENSMUSG00000033998.8 |
| chr8 | 126054059 | 126054353 | 3.41081 | 3.9564 | 1.06062 | Distal Intergenic | 8 | 125995102 | 126030683 | 35582 | 1 | ENSMUST00000046765.8 | 58958 | Kcnk1 | ENSMUSG00000033998.8 |
| chr8 | 126056438 | 126056806 | 3.27564 | 3.95203 | 1.08164 | Distal Intergenic | 8 | 125995102 | 126030683 | 35582 | 1 | ENSMUST00000046765.8 | 61337 | Kcnk1 | ENSMUSG00000033998.8 |
| chr8 | 126066054 | 126066158 | 3.67587 | 4.53946 | 1.47957 | Distal Intergenic | 8 | 125995102 | 126030683 | 35582 | 1 | ENSMUST00000046765.8 | 70953 | Kcnk1 | ENSMUSG00000033998.8 |
| chr8 | 126067191 | 126067494 | 3.28123 | 3.74898 | 1.01733 | Distal Intergenic | 8 | 125995102 | 126030683 | 35582 | 1 | ENSMUST00000046765.8 | 72090 | Kcnk1 | ENSMUSG00000033998.8 |
| chr8 | 126138700 | 126139368 | 3.76604 | 4.87457 | 1.66832 | Distal Intergenic | 8 | 125995102 | 126030683 | 35582 | 1 | ENSMUST00000046765.8 | 143599 | Kcnk1 | ENSMUSG00000033998.8 |
| chr8 | 126140038 | 126140128 | 4.28674 | 5.73245 | 2.30302 | Distal Intergenic | 8 | 125995102 | 126030683 | 35582 | 1 | ENSMUST00000046765.8 | 144937 | Kcnk1 | ENSMUSG00000033998.8 |
| chr8 | 126141530 | 126141605 | 4.06729 | 5.27825 | 1.98204 | Distal Intergenic | 8 | 125995102 | 126030683 | 35582 | 1 | ENSMUST00000046765.8 | 146429 | Kcnk1 | ENSMUSG00000033998.8 |
| chr3 | 79783681 | 79783756 | 3.71866 | 4.51678 | 1.39402 | Distal Intergenic | 3 | 79644112 | 79737880 | 93769 | 2 | ENSMUST00000182491.1 | -45802 | Rxfp1 | ENSMUSG00000034009.9 |
| chr11 | 88274750 | 88275309 | 3.34435 | 3.84393 | 1.01733 | Distal Intergenic | 11 | 88294058 | 88295261 | 1204 | 1 | ENSMUST00000037268.5 | -18749 | 1700106J16Rik | ENSMUSG00000034031.5 |
| chr6 | 91978418 | 91978864 | 3.84939 | 4.99341 | 1.76384 | Promoter | 6 | 91978878 | 92076004 | 97127 | 1 | ENSMUST00000089334.3 | -14 | Fgd5 | ENSMUSG00000034037.9 |
| chr5 | 131178193 | 131178534 | 3.44691 | 4.03349 | 1.14007 | Intron (ENSMUST00000086023.6/ENSMUSG00000034040.11, intron 1 of 10) | 5 | 131085272 | 131306915 | 221644 | 2 | ENSMUST00000160807.1 | 128381 | Wbscr17 | ENSMUSG00000034040.11 |
| chr5 | 131237511 | 131237789 | 3.57391 | 4.35184 | 1.3767 | Intron (ENSMUST00000086023.6/ENSMUSG00000034040.11, intron 1 of 10) | 5 | 131085272 | 131306915 | 221644 | 2 | ENSMUST00000160807.1 | 69126 | Wbscr17 | ENSMUSG00000034040.11 |
| chr5 | 131306010 | 131307543 | 3.78493 | 4.71838 | 1.60355 | Promoter | 5 | 131085272 | 131306915 | 221644 | 2 | ENSMUST00000160807.1 | 0 | Wbscr17 | ENSMUSG00000034040.11 |
| chr5 | 131319452 | 131319624 | 3.32645 | 3.81698 | 1.01733 | Distal Intergenic | 5 | 131306850 | 131308078 | 1229 | 2 | ENSMUST00000162966.1 | -11375 | Wbscr17 | ENSMUSG00000034040.11 |
| chr8 | 84701660 | 84701887 | 3.86948 | 4.86569 | 1.70492 | Promoter | 8 | 84701457 | 84704715 | 3259 | 1 | ENSMUST00000037165.4 | 204 | Lyl1 | ENSMUSG00000034041.4 |
| chr8 | 84703718 | 84704697 | 3.71617 | 4.6186 | 1.54437 | 3' UTR | 8 | 84701457 | 84704715 | 3259 | 1 | ENSMUST00000037165.4 | 2262 | Lyl1 | ENSMUSG00000034041.4 |
| chr10 | 116815278 | 116815488 | 3.80676 | 4.73114 | 1.63401 | Intron (ENSMUST00000048229.7/ENSMUSG00000034057.7, intron 13 of 24) | 10 | 116776545 | 116896879 | 120335 | 2 | ENSMUST00000048229.7 | 81391 | Myrfl | ENSMUSG00000034057.7 |
| chr3 | 75857792 | 75858026 | 3.28238 | 3.77466 | 1.08124 | Distal Intergenic | 3 | 75876479 | 75879008 | 2530 | 2 | ENSMUST00000134219.1 | 20982 | Golim4 | ENSMUSG00000034109.10 |
| chr8 | 119699065 | 119699260 | 3.26887 | 3.73045 | 1.01733 | Promoter | 8 | 119700009 | 119757718 | 57710 | 1 | ENSMUST00000095171.3 | -749 | Atp2c2 | ENSMUSG00000034112.8 |
| chr8 | 119699571 | 119700594 | 3.87095 | 4.95944 | 1.77933 | Promoter | 8 | 119700009 | 119757718 | 57710 | 1 | ENSMUST00000095171.3 | 0 | Atp2c2 | ENSMUSG00000034112.8 |
| chr9 | 119805691 | 119805766 | 3.44824 | 4.10763 | 1.1769 | Exon (ENSMUST00000070617.7/ENSMUSG00000034115.9, exon 9 of 27) | 9 | 119753763 | 119825456 | 71694 | 2 | ENSMUST00000070617.7 | 19690 | Scn11a | ENSMUSG00000034115.9 |
| chr12 | 87010660 | 87010769 | 3.66769 | 4.6733 | 1.54929 | Distal Intergenic | 12 | 87021340 | 87090043 | 68704 | 1 | ENSMUST00000154801.2 | -10571 | Tmem63c | ENSMUSG00000034145.9 |
| chr12 | 87013577 | 87013796 | 3.34637 | 3.84695 | 1.01733 | Distal Intergenic | 12 | 87021340 | 87090043 | 68704 | 1 | ENSMUST00000154801.2 | -7544 | Tmem63c | ENSMUSG00000034145.9 |
| chr12 | 87014402 | 87014831 | 3.3752 | 3.91381 | 1.10018 | Distal Intergenic | 12 | 87021340 | 87090043 | 68704 | 1 | ENSMUST00000154801.2 | -6509 | Tmem63c | ENSMUSG00000034145.9 |
| chr12 | 87026487 | 87027062 | 3.59212 | 4.50455 | 1.43362 | Promoter | 12 | 87026564 | 87090033 | 63470 | 1 | ENSMUST00000110187.2 | 0 | Tmem63c | ENSMUSG00000034145.9 |
| chr12 | 87028194 | 87028496 | 3.81893 | 4.79208 | 1.70478 | Intron (ENSMUST00000154801.2/ENSMUSG00000034145.9, intron 1 of 25) | 12 | 87026616 | 87057670 | 31055 | 1 | ENSMUST00000156162.2 | 1579 | Tmem63c | ENSMUSG00000034145.9 |
| chr11 | 87760386 | 87760839 | 3.84087 | 4.78433 | 1.6316 | Promoter | 11 | 87760541 | 87785928 | 25388 | 1 | ENSMUST00000039627.6 | 0 | Bzrap1 | ENSMUSG00000034156.11 |
| chr11 | 87761487 | 87761665 | 3.5175 | 4.15561 | 1.19531 | Promoter | 11 | 87760587 | 87785928 | 25342 | 1 | ENSMUST00000100644.4 | 901 | Bzrap1 | ENSMUSG00000034156.11 |
| chr11 | 87762617 | 87762810 | 3.87361 | 4.86399 | 1.65771 | Exon (ENSMUST00000039627.6/ENSMUSG00000034156.11, exon 3 of 32) | 11 | 87764731 | 87766395 | 1665 | 1 | ENSMUST00000148422.1 | -1921 | Bzrap1 | ENSMUSG00000034156.11 |
| chr11 | 87763303 | 87763389 | 4.36822 | 5.92543 | 2.41927 | Intron (ENSMUST00000039627.6/ENSMUSG00000034156.11, intron 3 of 31) | 11 | 87764731 | 87766395 | 1665 | 1 | ENSMUST00000148422.1 | -1342 | Bzrap1 | ENSMUSG00000034156.11 |
| chr11 | 87763748 | 87765184 | 4.12075 | 5.375 | 2.06212 | Promoter | 11 | 87764731 | 87766395 | 1665 | 1 | ENSMUST00000148422.1 | 0 | Bzrap1 | ENSMUSG00000034156.11 |
| chr11 | 87767104 | 87767329 | 3.5547 | 4.24622 | 1.32037 | Intron (ENSMUST00000039627.6/ENSMUSG00000034156.11, intron 10 of 31) | 11 | 87764731 | 87766395 | 1665 | 1 | ENSMUST00000148422.1 | 2374 | Bzrap1 | ENSMUSG00000034156.11 |
| chr11 | 87777724 | 87778564 | 3.65168 | 4.40713 | 1.35411 | Promoter | 11 | 87778793 | 87785928 | 7136 | 1 | ENSMUST00000148814.2 | -229 | Bzrap1 | ENSMUSG00000034156.11 |
| chr11 | 5152302 | 5152536 | 3.3032 | 3.78199 | 1.01733 | Promoter | 11 | 5106265 | 5152257 | 45993 | 2 | ENSMUST00000062821.7 | -46 | Emid1 | ENSMUSG00000034164.12 |
| chr4 | 115971122 | 115971286 | 3.40626 | 3.95776 | 1.08973 | Intron (ENSMUST00000125761.2/ENSMUSG00000034171.8, intron 13 of 14) | 4 | 115998714 | 115999937 | 1224 | 2 | ENSMUST00000156126.1 | 28651 | Faah | ENSMUSG00000034171.8 |
| chr4 | 115972248 | 115972829 | 3.61852 | 4.55476 | 1.50258 | Intron (ENSMUST00000125761.2/ENSMUSG00000034171.8, intron 13 of 14) | 4 | 115998714 | 115999937 | 1224 | 2 | ENSMUST00000156126.1 | 27108 | Faah | ENSMUSG00000034171.8 |
| chr4 | 115977445 | 115977521 | 4.75185 | 6.56039 | 2.8405 | Intron (ENSMUST00000125761.2/ENSMUSG00000034171.8, intron 13 of 14) | 4 | 115998714 | 115999937 | 1224 | 2 | ENSMUST00000156126.1 | 22416 | Faah | ENSMUSG00000034171.8 |
| chr4 | 115977948 | 115978023 | 3.33257 | 3.82618 | 1.01733 | Intron (ENSMUST00000125761.2/ENSMUSG00000034171.8, intron 13 of 14) | 4 | 115998714 | 115999937 | 1224 | 2 | ENSMUST00000156126.1 | 21914 | Faah | ENSMUSG00000034171.8 |
| chr4 | 115979040 | 115980761 | 3.60911 | 4.34002 | 1.34088 | Intron (ENSMUST00000125761.2/ENSMUSG00000034171.8, intron 13 of 14) | 4 | 115998714 | 115999937 | 1224 | 2 | ENSMUST00000156126.1 | 19176 | Faah | ENSMUSG00000034171.8 |
| chr4 | 115981352 | 115981462 | 3.60285 | 4.40787 | 1.38407 | Intron (ENSMUST00000125761.2/ENSMUSG00000034171.8, intron 13 of 14) | 4 | 115998714 | 115999937 | 1224 | 2 | ENSMUST00000156126.1 | 18475 | Faah | ENSMUSG00000034171.8 |
| chr4 | 115982183 | 115982336 | 3.56568 | 4.25388 | 1.28172 | Intron (ENSMUST00000125761.2/ENSMUSG00000034171.8, intron 13 of 14) | 4 | 115998714 | 115999937 | 1224 | 2 | ENSMUST00000156126.1 | 17601 | Faah | ENSMUSG00000034171.8 |
| chr4 | 115998830 | 115999097 | 3.61061 | 4.34197 | 1.34846 | Promoter | 4 | 115998714 | 115999937 | 1224 | 2 | ENSMUST00000156126.1 | 840 | Faah | ENSMUSG00000034171.8 |
| chr4 | 116010238 | 116010438 | 3.3591 | 3.86616 | 1.01733 | Intron (ENSMUST00000125761.2/ENSMUSG00000034171.8, intron 1 of 14) | 4 | 115967145 | 116017675 | 50531 | 2 | ENSMUST00000125761.2 | 7237 | Faah | ENSMUSG00000034171.8 |
| chr4 | 116014849 | 116015034 | 3.33224 | 3.82568 | 1.01733 | Intron (ENSMUST00000125761.2/ENSMUSG00000034171.8, intron 1 of 14) | 4 | 115967145 | 116017675 | 50531 | 2 | ENSMUST00000125761.2 | 2641 | Faah | ENSMUSG00000034171.8 |
| chr4 | 116016087 | 116016529 | 3.55505 | 4.2538 | 1.2973 | Intron (ENSMUST00000125761.2/ENSMUSG00000034171.8, intron 1 of 14) | 4 | 115967145 | 116017675 | 50531 | 2 | ENSMUST00000125761.2 | 1146 | Faah | ENSMUSG00000034171.8 |
| chr11 | 87663070 | 87663246 | 3.49986 | 4.16424 | 1.27513 | Promoter | 11 | 87663087 | 87718459 | 55373 | 1 | ENSMUST00000123658.3 | 0 | Rnf43 | ENSMUSG00000034177.10 |
| chr4 | 115890524 | 115892151 | 3.94354 | 5.11884 | 1.89647 | Promoter | 4 | 115890665 | 115899227 | 8563 | 1 | ENSMUST00000132221.2 | 0 | 6430628N08Rik | ENSMUSG00000034185.9 |
| chr4 | 115896937 | 115897143 | 4.10984 | 5.2818 | 1.93012 | Intron (ENSMUST00000132221.2/ENSMUSG00000034185.9, intron 1 of 1) | 4 | 115890666 | 115899227 | 8562 | 1 | ENSMUST00000165938.1 | 6272 | 6430628N08Rik | ENSMUSG00000034185.9 |
| chr4 | 115897595 | 115898613 | 4.16609 | 5.45457 | 2.11593 | Exon (ENSMUST00000132221.2/ENSMUSG00000034185.9, exon 2 of 2) | 4 | 115890666 | 115899227 | 8562 | 1 | ENSMUST00000165938.1 | 6930 | 6430628N08Rik | ENSMUSG00000034185.9 |
| chr11 | 103808033 | 103808359 | 3.4629 | 4.08318 | 1.13675 | Exon (ENSMUST00000000127.5/ENSMUSG00000000125.5, exon 2 of 5) | 11 | 103822934 | 103824100 | 1167 | 2 | ENSMUST00000040443.10 | 15741 | Nsf | ENSMUSG00000034187.13 |
| chr11 | 103809036 | 103809223 | 3.78673 | 4.80581 | 1.64615 | Intron (ENSMUST00000000127.5/ENSMUSG00000000125.5, intron 2 of 4) | 11 | 103822934 | 103824100 | 1167 | 2 | ENSMUST00000040443.10 | 14877 | Nsf | ENSMUSG00000034187.13 |
| chr11 | 103809708 | 103811346 | 4.04376 | 5.27299 | 1.97789 | Exon (ENSMUST00000000127.5/ENSMUSG00000000125.5, exon 3 of 5) | 11 | 103822934 | 103824100 | 1167 | 2 | ENSMUST00000040443.10 | 12754 | Nsf | ENSMUSG00000034187.13 |
| chr11 | 103811895 | 103812661 | 3.83011 | 4.75927 | 1.59935 | Exon (ENSMUST00000000127.5/ENSMUSG00000000125.5, exon 4 of 5) | 11 | 103822934 | 103824100 | 1167 | 2 | ENSMUST00000040443.10 | 11439 | Nsf | ENSMUSG00000034187.13 |
| chr11 | 5057913 | 5058125 | 3.6095 | 4.32642 | 1.30856 | Promoter | 11 | 5058128 | 5060385 | 2258 | 1 | ENSMUST00000037218.1 | -3 | Rasl10a | ENSMUSG00000034209.4 |
| chr11 | 5058448 | 5058591 | 3.63693 | 4.36901 | 1.3124 | Promoter | 11 | 5058128 | 5060385 | 2258 | 1 | ENSMUST00000037218.1 | 321 | Rasl10a | ENSMUSG00000034209.4 |
| chr11 | 5059134 | 5059638 | 3.58833 | 4.31117 | 1.32183 | Promoter | 11 | 5058966 | 5060062 | 1097 | 1 | ENSMUST00000126974.1 | 169 | Rasl10a | ENSMUSG00000034209.4 |
| chr5 | 129683191 | 129683519 | 3.46949 | 4.08778 | 1.17733 | 3' UTR | 5 | 129683391 | 129693232 | 9842 | 2 | ENSMUST00000182381.1 | 9713 | Sep-14 | ENSMUSG00000034219.10 |
| chr5 | 129703328 | 129703642 | 3.60352 | 4.38213 | 1.39752 | Promoter | 5 | 129683402 | 129704259 | 20858 | 2 | ENSMUST00000042266.7 | 617 | Sep-14 | ENSMUSG00000034219.10 |
| chr1 | 92877468 | 92877630 | 3.64762 | 4.44918 | 1.44995 | Intron (ENSMUST00000112998.1/ENSMUSG00000034212.8, intron 13 of 17) | 1 | 92856768 | 92857595 | 828 | 1 | ENSMUST00000190215.1 | 20701 | Gpc1 | ENSMUSG00000034220.7 |
| chr8 | 119492715 | 119492791 | 4.7482 | 6.56359 | 2.85536 | Intron (ENSMUST00000036748.9/ENSMUSG00000034224.9, intron 5 of 10) | 8 | 119486099 | 119494895 | 8797 | 2 | ENSMUST00000125594.1 | 2104 | Slc38a8 | ENSMUSG00000034224.9 |
| chr2 | 119270735 | 119271320 | 3.64016 | 4.47129 | 1.44399 | Promoter | 2 | 119269201 | 119271272 | 2072 | 2 | ENSMUST00000037360.7 | 0 | Rhov | ENSMUSG00000034226.7 |
| chr11 | 116334045 | 116335239 | 3.91939 | 4.9646 | 1.74126 | Promoter | 11 | 116330704 | 116335399 | 4696 | 2 | ENSMUST00000036215.7 | 160 | Foxj1 | ENSMUSG00000034227.7 |
| chr11 | 103323695 | 103324036 | 3.37555 | 3.89098 | 1.01733 | Distal Intergenic | 11 | 103333229 | 103333395 | 167 | 2 | ENSMUST00000163250.1 | 9359 | Arhgap27 | ENSMUSG00000034255.12 |
| chr11 | 103324940 | 103325028 | 3.82091 | 4.71944 | 1.54214 | Distal Intergenic | 11 | 103333229 | 103333395 | 167 | 2 | ENSMUST00000163250.1 | 8367 | Arhgap27 | ENSMUSG00000034255.12 |
| chr11 | 103325754 | 103326038 | 3.74674 | 4.64386 | 1.52824 | Distal Intergenic | 11 | 103333229 | 103333395 | 167 | 2 | ENSMUST00000163250.1 | 7357 | Arhgap27 | ENSMUSG00000034255.12 |

| | | | | | | | | | | | | | | |
|---|---|---|---|---|---|---|---|---|---|---|---|---|---|---|
| chr11 | 103333175 | 103333902 | 3.57983 | 4.44535 | 1.42936 | Promoter | 11 | 103333229 | 103333395 | 167 | 2 | ENSMUST00000163250.1 | 0 | Arhgap27 | ENSMUSG00000034255.12 |
| chr11 | 103339435 | 103339550 | 3.71043 | 4.51693 | 1.43448 | Promoter | 11 | 103337777 | 103339303 | 1527 | 2 | ENSMUST00000124490.1 | -133 | Arhgap27 | ENSMUSG00000034255.12 |
| chr11 | 103340210 | 103340383 | 3.75682 | 4.7256 | 1.61391 | Promoter | 11 | 103335268 | 103339719 | 4452 | 2 | ENSMUST00000139830.2 | -492 | Arhgap27 | ENSMUSG00000034255.12 |
| chr11 | 103360389 | 103361106 | 3.4063 | 4.25196 | 1.31562 | Promoter | 11 | 103331497 | 103360900 | 29404 | 2 | ENSMUST00000107024.2 | 0 | Arhgap27 | ENSMUSG00000034255.12 |
| chr12 | 85744460 | 85744605 | 3.35163 | 3.85489 | 1.01733 | Distal Intergenic | 12 | 85746539 | 85813585 | 67047 | 1 | ENSMUST00000040461.3 | -1934 | Mfsd7c | ENSMUSG00000034258.3 |
| chr12 | 85746597 | 85747083 | 4.25713 | 5.59794 | 2.17464 | Promoter | 12 | 85746539 | 85813585 | 67047 | 1 | ENSMUST00000040461.3 | 59 | Mfsd7c | ENSMUSG00000034258.3 |
| chr12 | 85581980 | 85582204 | 3.31325 | 3.94206 | 1.05356 | Distal Intergenic | 12 | 85599105 | 85639878 | 40774 | 1 | ENSMUST00000177587.2 | -16901 | Jdp2 | ENSMUSG00000034271.10 |
| chr9 | 27299623 | 27299882 | 3.34579 | 3.88153 | 1.07325 | Promoter | 9 | 27299230 | 27334763 | 35534 | 1 | ENSMUST00000133213.1 | 394 | Igsf9b | ENSMUSG00000034275.12 |
| chr9 | 27300300 | 27300549 | 3.30726 | 3.78822 | 1.01733 | Intron (ENSMUST00000115247.2/ENSMUSG00000034275.12, intron 1 of 14) | 9 | 27299230 | 27334763 | 35534 | 1 | ENSMUST00000133213.1 | 1071 | Igsf9b | ENSMUSG00000034275.12 |
| chr18 | 24709468 | 24710776 | 3.72778 | 4.61434 | 1.57785 | Promoter | 18 | 24709445 | 25133500 | 424056 | 1 | ENSMUST00000037097.7 | 24 | Fhod3 | ENSMUSG00000034295.9 |
| chr8 | 117683497 | 117684324 | 3.42903 | 4.01266 | 1.16267 | Distal Intergenic | 8 | 117663333 | 117673689 | 10357 | 2 | ENSMUST00000174450.1 | -9809 | Sdr42e1 | ENSMUSG00000034308.8 |
| chr5 | 128387390 | 128387612 | 3.48515 | 4.24867 | 1.30598 | Intron (ENSMUST00000044441.6/ENSMUSG00000034310.7, intron 1 of 8) | 5 | 127783491 | 128433077 | 649587 | 2 | ENSMUST00000044441.6 | 45465 | Tmem132d | ENSMUSG00000034310.7 |
| chr5 | 128432370 | 128432948 | 3.61873 | 4.45674 | 1.46231 | Promoter | 5 | 127783491 | 128433077 | 649587 | 2 | ENSMUST00000044441.6 | 129 | Tmem132d | ENSMUSG00000034310.7 |
| chr5 | 127241258 | 127243298 | 4.0598 | 5.39189 | 2.08788 | Promoter | 5 | 127241808 | 127565793 | 323986 | 1 | ENSMUST00000119026.2 | 0 | Tmem132c | ENSMUSG00000034324.11 |
| chr5 | 127243738 | 127244035 | 4.95242 | 7.13425 | 3.37421 | Intron (ENSMUST00000119026.2/ENSMUSG00000034324.11, intron 1 of 8) | 5 | 127241826 | 127287336 | 45511 | 1 | ENSMUST00000134780.1 | 1913 | Tmem132c | ENSMUSG00000034324.11 |
| chr5 | 127244485 | 127244915 | 3.54745 | 4.21839 | 1.24617 | Intron (ENSMUST00000119026.2/ENSMUSG00000034324.11, intron 1 of 8) | 5 | 127241826 | 127287336 | 45511 | 1 | ENSMUST00000134780.1 | 2660 | Tmem132c | ENSMUSG00000034324.11 |
| chr5 | 127484626 | 127484988 | 3.67573 | 4.50495 | 1.49305 | Intron (ENSMUST00000119026.2/ENSMUSG00000034324.11, intron 3 of 8) | 5 | 127538260 | 127564866 | 26607 | 1 | ENSMUST00000128068.2 | -53272 | Tmem132c | ENSMUSG00000034324.11 |
| chr8 | 117498116 | 117499586 | 3.48736 | 4.10868 | 1.15518 | Promoter | 8 | 117498291 | 117635140 | 136850 | 1 | ENSMUST00000081232.7 | 0 | Plcg2 | ENSMUSG00000034330.9 |
| chr19 | 47008052 | 47008379 | 3.86763 | 5.08138 | 1.84678 | Intron (ENSMUST00000172239.2/ENSMUSG00000025041.11, intron 1 of 17) | 19 | 47014698 | 47025327 | 10630 | 1 | ENSMUST00000037636.3 | -6319 | Ina | ENSMUSG00000034336.3 |
| chr19 | 47012406 | 47012615 | 5.12427 | 8.04483 | 4.198 | Intron (ENSMUST00000172239.2/ENSMUSG00000025041.11, intron 1 of 17) | 19 | 47014698 | 47025327 | 10630 | 1 | ENSMUST00000037636.3 | -2083 | Ina | ENSMUSG00000034336.3 |
| chr1 | 91180140 | 91180531 | 3.76516 | 4.66557 | 1.60576 | Promoter | 1 | 91180046 | 91225196 | 45151 | 1 | ENSMUST00000097648.5 | 95 | Ramp1 | ENSMUSG00000034353.9 |
| chr8 | 94531961 | 94534095 | 3.73143 | 4.681 | 1.57387 | Promoter | 8 | 94532990 | 94570529 | 37540 | 1 | ENSMUST00000048653.3 | 0 | Cpne2 | ENSMUSG00000034361.4 |
| chr17 | 47436677 | 47436997 | 4.076 | 5.26366 | 1.96642 | Promoter | 17 | 47436805 | 47470638 | 33834 | 1 | ENSMUST00000150819.2 | 0 | AI661453 | ENSMUSG00000034382.9 |
| chr5 | 106292319 | 106292583 | 3.36173 | 3.87013 | 1.01733 | Distal Intergenic | 5 | 106452523 | 106458166 | 5644 | 2 | ENSMUST00000086795.7 | 165583 | Barhl2 | ENSMUSG00000034384.11 |
| chr5 | 106343942 | 106344171 | 3.32358 | 3.82673 | 1.02319 | Distal Intergenic | 5 | 106452523 | 106458166 | 5644 | 2 | ENSMUST00000086795.7 | 113995 | Barhl2 | ENSMUSG00000034384.11 |
| chr5 | 106364460 | 106364535 | 3.37083 | 3.88934 | 1.03675 | Distal Intergenic | 5 | 106452523 | 106458166 | 5644 | 2 | ENSMUST00000086795.7 | 93631 | Barhl2 | ENSMUSG00000034384.11 |
| chr5 | 106420608 | 106420747 | 3.60655 | 4.42234 | 1.41881 | Distal Intergenic | 5 | 106452523 | 106458166 | 5644 | 2 | ENSMUST00000086795.7 | 37419 | Barhl2 | ENSMUSG00000034384.11 |
| chr5 | 106426553 | 106426651 | 3.47825 | 4.07847 | 1.13174 | Distal Intergenic | 5 | 106452523 | 106458166 | 5644 | 2 | ENSMUST00000086795.7 | 31515 | Barhl2 | ENSMUSG00000034384.11 |
| chr5 | 106428717 | 106429034 | 3.55169 | 4.2124 | 1.21333 | Distal Intergenic | 5 | 106452523 | 106458166 | 5644 | 2 | ENSMUST00000086795.7 | 29132 | Barhl2 | ENSMUSG00000034384.11 |
| chr5 | 106429732 | 106429943 | 3.55174 | 4.21609 | 1.22709 | Distal Intergenic | 5 | 106452523 | 106458166 | 5644 | 2 | ENSMUST00000086795.7 | 28223 | Barhl2 | ENSMUSG00000034384.11 |
| chr5 | 106437811 | 106438011 | 3.31302 | 3.79677 | 1.01733 | Distal Intergenic | 5 | 106452523 | 106458166 | 5644 | 2 | ENSMUST00000086795.7 | 20155 | Barhl2 | ENSMUSG00000034384.11 |
| chr5 | 106438841 | 106439036 | 3.43766 | 4.12885 | 1.24343 | Distal Intergenic | 5 | 106452523 | 106458166 | 5644 | 2 | ENSMUST00000086795.7 | 19130 | Barhl2 | ENSMUSG00000034384.11 |
| chr5 | 106440747 | 106441813 | 3.39401 | 4.01911 | 1.16498 | Distal Intergenic | 5 | 106452523 | 106458166 | 5644 | 2 | ENSMUST00000086795.7 | 16353 | Barhl2 | ENSMUSG00000034384.11 |
| chr5 | 106442389 | 106443227 | 3.59388 | 4.47731 | 1.4703 | Distal Intergenic | 5 | 106452523 | 106458166 | 5644 | 2 | ENSMUST00000086795.7 | 14939 | Barhl2 | ENSMUSG00000034384.11 |
| chr5 | 106444074 | 106444251 | 3.47672 | 4.10993 | 1.25114 | Distal Intergenic | 5 | 106452523 | 106458166 | 5644 | 2 | ENSMUST00000086795.7 | 13915 | Barhl2 | ENSMUSG00000034384.11 |
| chr5 | 106444593 | 106445210 | 3.52552 | 4.26004 | 1.34279 | Distal Intergenic | 5 | 106452523 | 106458166 | 5644 | 2 | ENSMUST00000086795.7 | 12956 | Barhl2 | ENSMUSG00000034384.11 |
| chr5 | 106445970 | 106446225 | 3.25808 | 3.71426 | 1.01733 | Distal Intergenic | 5 | 106452523 | 106458166 | 5644 | 2 | ENSMUST00000086795.7 | 11941 | Barhl2 | ENSMUSG00000034384.11 |
| chr5 | 106446585 | 106446984 | 3.82926 | 4.86637 | 1.72303 | Distal Intergenic | 5 | 106452523 | 106458166 | 5644 | 2 | ENSMUST00000086795.7 | 11182 | Barhl2 | ENSMUSG00000034384.11 |
| chr5 | 106447961 | 106451094 | 3.98381 | 5.17949 | 1.9496 | Downstream (1-2kb) | 5 | 106452523 | 106458166 | 5644 | 2 | ENSMUST00000086795.7 | 7072 | Barhl2 | ENSMUSG00000034384.11 |
| chr5 | 106452014 | 106453745 | 4.01592 | 5.39809 | 2.08104 | 3' UTR | 5 | 106452523 | 106458166 | 5644 | 2 | ENSMUST00000086795.7 | 4421 | Barhl2 | ENSMUSG00000034384.11 |
| chr5 | 106454204 | 106454451 | 3.78903 | 4.68274 | 1.52643 | Intron (ENSMUST00000086795.7/ENSMUSG00000034384.11, intron 2 of 2) | 5 | 106452523 | 106458166 | 5644 | 2 | ENSMUST00000086795.7 | 3715 | Barhl2 | ENSMUSG00000034384.11 |
| chr5 | 106454785 | 106455707 | 3.97484 | 5.05883 | 1.80412 | Exon (ENSMUST00000086795.7/ENSMUSG00000034384.11, exon 2 of 3) | 5 | 106452523 | 106458166 | 5644 | 2 | ENSMUST00000086795.7 | 2459 | Barhl2 | ENSMUSG00000034384.11 |
| chr5 | 106456251 | 106459872 | 4.01669 | 5.19857 | 1.90869 | Promoter | 5 | 106452523 | 106458166 | 5644 | 2 | ENSMUST00000086795.7 | 0 | Barhl2 | ENSMUSG00000034384.11 |
| chr5 | 106460210 | 106461908 | 3.99483 | 5.314 | 1.99113 | Distal Intergenic | 5 | 106452523 | 106458166 | 5644 | 2 | ENSMUST00000086795.7 | -2045 | Barhl2 | ENSMUSG00000034384.11 |
| chr5 | 106462310 | 106463020 | 3.86873 | 4.88501 | 1.66788 | Downstream (2-3kb) | 5 | 106452523 | 106458166 | 5644 | 2 | ENSMUST00000086795.7 | -4145 | Barhl2 | ENSMUSG00000034384.11 |
| chr5 | 106463834 | 106464306 | 4.67588 | 6.62535 | 3.00899 | Downstream (1-2kb) | 5 | 106452523 | 106458166 | 5644 | 2 | ENSMUST00000086795.7 | -5669 | Barhl2 | ENSMUSG00000034384.11 |
| chr12 | 75175208 | 75175543 | 3.70338 | 4.56835 | 1.53396 | Intron (ENSMUST00000042299.2/ENSMUSG00000034402.2, intron 1 of 10) | 12 | 74897217 | 75177332 | 280116 | 2 | ENSMUST00000042299.2 | 1789 | Kcnh5 | ENSMUSG00000034402.2 |
| chr12 | 75176001 | 75177959 | 4.01145 | 5.23207 | 1.94304 | Promoter | 12 | 74897217 | 75177332 | 280116 | 2 | ENSMUST00000042299.2 | 0 | Kcnh5 | ENSMUSG00000034402.2 |
| chr12 | 75178746 | 75179125 | 3.67198 | 4.49393 | 1.45457 | Distal Intergenic | 12 | 74897217 | 75177332 | 280116 | 2 | ENSMUST00000042299.2 | -1415 | Kcnh5 | ENSMUSG00000034402.2 |
| chr17 | 26412819 | 26413691 | 3.42633 | 4.00815 | 1.11901 | Distal Intergenic | 17 | 26414829 | 26441795 | 26967 | 1 | ENSMUST00000182897.2 | -1138 | Neurl1b | ENSMUSG00000034413.9 |
| chr17 | 26414693 | 26415409 | 3.53404 | 4.24704 | 1.24506 | Promoter | 17 | 26414829 | 26441795 | 26967 | 1 | ENSMUST00000182897.2 | 0 | Neurl1b | ENSMUSG00000034413.9 |
| chr17 | 26438903 | 26439261 | 4.33875 | 5.83308 | 2.39132 | Exon (ENSMUST00000182897.2/ENSMUSG00000034413.9, exon 2 of 4) | 17 | 26414965 | 26446349 | 31385 | 1 | ENSMUST00000053020.7 | 23939 | Neurl1b | ENSMUSG00000034413.9 |
| chr17 | 26439571 | 26440309 | 3.81695 | 4.78509 | 1.65555 | Intron (ENSMUST00000182897.2/ENSMUSG00000034413.9, intron 2 of 3) | 17 | 26414965 | 26446349 | 31385 | 1 | ENSMUST00000053020.7 | 24607 | Neurl1b | ENSMUSG00000034413.9 |
| chr11 | 115858162 | 115858540 | 3.61612 | 4.4269 | 1.44044 | Distal Intergenic | 11 | 115861612 | 115892592 | 30981 | 1 | ENSMUST00000093911.5 | -3072 | Myo15b | ENSMUSG00000034427.11 |
| chr11 | 115858938 | 115860398 | 3.4468 | 4.40546 | 1.3961 | Distal Intergenic | 11 | 115861612 | 115892592 | 30981 | 1 | ENSMUST00000093911.5 | -1214 | Myo15b | ENSMUSG00000034427.11 |
| chr11 | 115886153 | 115886456 | 3.38092 | 3.89908 | 1.01733 | Promoter | 11 | 115886436 | 115887446 | 1011 | 1 | ENSMUST00000125835.1 | 0 | Myo15b | ENSMUSG00000034427.11 |
| chr12 | 73467271 | 73467407 | 3.45827 | 4.23554 | 1.27848 | Distal Intergenic | 12 | 73543114 | 73546395 | 3282 | 2 | ENSMUST00000042975.5 | 78988 | Tmem30b | ENSMUSG00000034435.5 |
| chr12 | 73468165 | 73468593 | 3.4466 | 4.02424 | 1.10866 | Distal Intergenic | 12 | 73543114 | 73546395 | 3282 | 2 | ENSMUST00000042975.5 | 77802 | Tmem30b | ENSMUSG00000034435.5 |
| chr12 | 73506061 | 73506139 | 4.04214 | 5.30986 | 1.98561 | Distal Intergenic | 12 | 73543114 | 73546395 | 3282 | 2 | ENSMUST00000042975.5 | 40256 | Tmem30b | ENSMUSG00000034435.5 |
| chr12 | 73517473 | 73517780 | 3.4269 | 4.03969 | 1.22471 | Distal Intergenic | 12 | 73543114 | 73546395 | 3282 | 2 | ENSMUST00000042975.5 | 28615 | Tmem30b | ENSMUSG00000034435.5 |
| chr12 | 73521089 | 73521374 | 3.45961 | 4.15386 | 1.24207 | Distal Intergenic | 12 | 73543114 | 73546395 | 3282 | 2 | ENSMUST00000042975.5 | 25021 | Tmem30b | ENSMUSG00000034435.5 |
| chr12 | 73529204 | 73529522 | 3.44897 | 4.02901 | 1.11316 | Distal Intergenic | 12 | 73543114 | 73546395 | 3282 | 2 | ENSMUST00000042975.5 | 16873 | Tmem30b | ENSMUSG00000034435.5 |
| chr12 | 73529895 | 73530107 | 3.53864 | 4.23766 | 1.26633 | Distal Intergenic | 12 | 73543114 | 73546395 | 3282 | 2 | ENSMUST00000042975.5 | 16288 | Tmem30b | ENSMUSG00000034435.5 |
| chr12 | 73532475 | 73532584 | 3.94006 | 5.06849 | 1.8137 | Distal Intergenic | 12 | 73543114 | 73546395 | 3282 | 2 | ENSMUST00000042975.5 | 13811 | Tmem30b | ENSMUSG00000034435.5 |
| chr12 | 73537947 | 73538189 | 3.57611 | 4.32246 | 1.3135 | Distal Intergenic | 12 | 73543114 | 73546395 | 3282 | 2 | ENSMUST00000042975.5 | 8206 | Tmem30b | ENSMUSG00000034435.5 |
| chr12 | 73541052 | 73541297 | 3.34579 | 3.99131 | 1.09436 | Downstream (1-2kb) | 12 | 73543114 | 73546395 | 3282 | 2 | ENSMUST00000042975.5 | 5098 | Tmem30b | ENSMUSG00000034435.5 |
| chr12 | 73541997 | 73542082 | 3.76494 | 5.02998 | 1.80374 | Downstream (1-2kb) | 12 | 73543114 | 73546395 | 3282 | 2 | ENSMUST00000042975.5 | 4313 | Tmem30b | ENSMUSG00000034435.5 |
| chr12 | 73542532 | 73543079 | 3.5881 | 4.44614 | 1.40076 | Downstream (<1kb) | 12 | 73543114 | 73546395 | 3282 | 2 | ENSMUST00000042975.5 | 3316 | Tmem30b | ENSMUSG00000034435.5 |
| chr12 | 73544214 | 73544916 | 3.903 | 5.27739 | 1.99021 | 3' UTR | 12 | 73543114 | 73546395 | 3282 | 2 | ENSMUST00000042975.5 | 1479 | Tmem30b | ENSMUSG00000034435.5 |
| chr12 | 73545316 | 73546812 | 3.61178 | 4.68387 | 1.55659 | Promoter | 12 | 73543114 | 73546395 | 3282 | 2 | ENSMUST00000042975.5 | 0 | Tmem30b | ENSMUSG00000034435.5 |
| chr12 | 73547836 | 73547969 | 3.52122 | 4.20533 | 1.31894 | Distal Intergenic | 12 | 73543114 | 73546395 | 3282 | 2 | ENSMUST00000042975.5 | -1442 | Tmem30b | ENSMUSG00000034435.5 |
| chr12 | 73548302 | 73548848 | 3.52192 | 4.2123 | 1.33068 | Distal Intergenic | 12 | 73543114 | 73546395 | 3282 | 2 | ENSMUST00000042975.5 | -1908 | Tmem30b | ENSMUSG00000034435.5 |
| chr12 | 73549797 | 73550761 | 4.117 | 5.51424 | 2.22568 | Distal Intergenic | 12 | 73543114 | 73546395 | 3282 | 2 | ENSMUST00000042975.5 | -3403 | Tmem30b | ENSMUSG00000034435.5 |
| chr12 | 73551951 | 73552239 | 3.98627 | 5.04845 | 1.79141 | Distal Intergenic | 12 | 73543114 | 73546395 | 3282 | 2 | ENSMUST00000042975.5 | -5557 | Tmem30b | ENSMUSG00000034435.5 |
| chr12 | 73563017 | 73563102 | 3.70922 | 4.54118 | 1.50664 | Distal Intergenic | 12 | 73543114 | 73546395 | 3282 | 2 | ENSMUST00000042975.5 | -16623 | Tmem30b | ENSMUSG00000034435.5 |
| chr12 | 73563966 | 73564348 | 3.64159 | 4.46554 | 1.47051 | Distal Intergenic | 12 | 73543114 | 73546395 | 3282 | 2 | ENSMUST00000042975.5 | -17572 | Tmem30b | ENSMUSG00000034435.5 |
| chr12 | 73110844 | 73111491 | 3.9843 | 5.08445 | 1.86113 | Intron (ENSMUST00000043208.7/ENSMUSG00000034460.9, intron 1 of 2) | 12 | 73099609 | 73113425 | 13817 | 2 | ENSMUST00000043208.7 | 1934 | Six4 | ENSMUSG00000034460.9 |
| chr12 | 73111847 | 73112475 | 3.71004 | 4.59433 | 1.47701 | Promoter | 12 | 73099609 | 73113425 | 13817 | 2 | ENSMUST00000043208.7 | 950 | Six4 | ENSMUSG00000034460.9 |
| chr12 | 73112806 | 73112887 | 4.08687 | 6.01886 | 2.51287 | Promoter | 12 | 73099609 | 73113425 | 13817 | 2 | ENSMUST00000043208.7 | 538 | Six4 | ENSMUSG00000034460.9 |
| chr12 | 73113442 | 73113726 | 3.47925 | 4.084 | 1.14621 | Promoter | 12 | 73103441 | 73113456 | 10016 | 2 | ENSMUST00000175693.2 | 0 | Six4 | ENSMUSG00000034460.9 |
| chr14 | 65952776 | 65953437 | 3.64795 | 4.41618 | 1.35378 | Promoter | 14 | 65919395 | 65953744 | 34350 | 2 | ENSMUST00000042046.3 | 307 | Scara3 | ENSMUSG00000034463.3 |
| chr14 | 65953774 | 65953963 | 3.53123 | 4.35303 | 1.32228 | Promoter | 14 | 65919395 | 65953744 | 34350 | 2 | ENSMUST00000042046.3 | -31 | Scara3 | ENSMUSG00000034463.3 |
| chr8 | 75213457 | 75214122 | 3.66786 | 4.83733 | 1.66734 | Promoter | 8 | 75213944 | 75224113 | 10170 | 1 | ENSMUST00000139848.2 | 0 | Rasd2 | ENSMUSG00000034472.8 |
| chr1 | 89917804 | 89917980 | 3.47709 | 4.07455 | 1.12412 | Distal Intergenic | 1 | 89927966 | 89929185 | 1220 | 2 | ENSMUST00000187633.1 | 11205 | Gbx2 | ENSMUSG00000034486.8 |
| chr1 | 89923291 | 89923654 | 4.0167 | 5.16523 | 1.91332 | Distal Intergenic | 1 | 89927966 | 89929185 | 1220 | 2 | ENSMUST00000187633.1 | 5531 | Gbx2 | ENSMUSG00000034486.8 |
| chr1 | 89924202 | 89925107 | 3.5624 | 4.3132 | 1.31725 | Downstream (2-3kb) | 1 | 89927966 | 89929185 | 1220 | 2 | ENSMUST00000187633.1 | 4078 | Gbx2 | ENSMUSG00000034486.8 |
| chr1 | 89926935 | 89929299 | 3.95869 | 5.16612 | 1.93492 | Promoter | 1 | 89927966 | 89929185 | 1220 | 2 | ENSMUST00000187633.1 | 0 | Gbx2 | ENSMUSG00000034486.8 |
| chr1 | 89930065 | 89931771 | 3.86833 | 5.10251 | 1.86583 | Promoter | 1 | 89927956 | 89931179 | 3224 | 2 | ENSMUST00000036954.8 | 0 | Gbx2 | ENSMUSG00000034486.8 |
| chr1 | 89932440 | 89933345 | 3.58818 | 4.5271 | 1.45058 | Exon (ENSMUST00000189763.1/ENSMUSG00000092627.2, exon 1 of 1) | 1 | 89927956 | 89931179 | 3224 | 2 | ENSMUST00000036954.8 | -1262 | Gbx2 | ENSMUSG00000034486.8 |
| chr1 | 89933851 | 89935602 | 3.85419 | 5.23413 | 1.95808 | Distal Intergenic | 1 | 89927956 | 89931179 | 3224 | 2 | ENSMUST00000036954.8 | -2673 | Gbx2 | ENSMUSG00000034486.8 |
| chr1 | 89936750 | 89937403 | 3.84803 | 5.04806 | 1.79638 | Distal Intergenic | 1 | 89927956 | 89931179 | 3224 | 2 | ENSMUST00000036954.8 | -5572 | Gbx2 | ENSMUSG00000034486.8 |
| chr1 | 89939703 | 89941150 | 4.19198 | 5.79132 | 2.35214 | Distal Intergenic | 1 | 89927956 | 89931179 | 3224 | 2 | ENSMUST00000036954.8 | -8525 | Gbx2 | ENSMUSG00000034486.8 |
| chr1 | 89941578 | 89942396 | 3.68099 | 4.67966 | 1.55971 | Distal Intergenic | 1 | 89927956 | 89931179 | 3224 | 2 | ENSMUST00000036954.8 | -10400 | Gbx2 | ENSMUSG00000034486.8 |
| chr1 | 89944010 | 89944420 | 3.6972 | 4.64838 | 1.55006 | Distal Intergenic | 1 | 89927956 | 89931179 | 3224 | 2 | ENSMUST00000036954.8 | -12832 | Gbx2 | ENSMUSG00000034486.8 |
| chr1 | 89948115 | 89948316 | 3.51 | 4.17769 | 1.29919 | Downstream (1-2kb) | 1 | 89927956 | 89931179 | 3224 | 2 | ENSMUST00000036954.8 | -16937 | Gbx2 | ENSMUSG00000034486.8 |
| chr1 | 89953095 | 89953547 | 3.40782 | 4.00359 | 1.20777 | Intron (ENSMUST00000086882.2/ENSMUSG00000067081.7, intron 5 of 5) | 1 | 89927956 | 89931179 | 3224 | 2 | ENSMUST00000036954.8 | -21917 | Gbx2 | ENSMUSG00000034486.8 |
| chr1 | 89959218 | 89959590 | 3.31389 | 3.79823 | 1.01733 | Intron (ENSMUST00000086882.2/ENSMUSG00000067081.7, intron 4 of 5) | 1 | 89927956 | 89931179 | 3224 | 2 | ENSMUST00000036954.8 | -28040 | Gbx2 | ENSMUSG00000034486.8 |
| chr1 | 89961238 | 89961422 | 3.45174 | 4.03121 | 1.10632 | Intron (ENSMUST00000086882.2/ENSMUSG00000067081.7, intron 4 of 5) | 1 | 89927956 | 89931179 | 3224 | 2 | ENSMUST00000036954.8 | -30060 | Gbx2 | ENSMUSG00000034486.8 |
| chr1 | 89962362 | 89962646 | 3.86505 | 4.80813 | 1.605 | Intron (ENSMUST00000086882.2/ENSMUSG00000067081.7, intron 4 of 5) | 1 | 89927956 | 89931179 | 3224 | 2 | ENSMUST00000036954.8 | -31184 | Gbx2 | ENSMUSG00000034486.8 |
| chr1 | 89964228 | 89964334 | 3.37295 | 3.88706 | 1.01733 | Intron (ENSMUST00000086882.2/ENSMUSG00000067081.7, intron 4 of 5) | 1 | 89927956 | 89931179 | 3224 | 2 | ENSMUST00000036954.8 | -33050 | Gbx2 | ENSMUSG00000034486.8 |
| chr1 | 89964736 | 89965039 | 3.51123 | 4.14259 | 1.18303 | Intron (ENSMUST00000086882.2/ENSMUSG00000067081.7, intron 4 of 5) | 1 | 89927956 | 89931179 | 3224 | 2 | ENSMUST00000036954.8 | -33558 | Gbx2 | ENSMUSG00000034486.8 |
| chr1 | 89965358 | 89965440 | 4.7201 | 6.63205 | 2.93863 | Intron (ENSMUST00000086882.2/ENSMUSG00000067081.7, intron 4 of 5) | 1 | 89927956 | 89931179 | 3224 | 2 | ENSMUST00000036954.8 | -34180 | Gbx2 | ENSMUSG00000034486.8 |
| chr1 | 89967157 | 89967408 | 3.47164 | 4.07001 | 1.137 | Intron (ENSMUST00000086882.2/ENSMUSG00000067081.7, intron 4 of 5) | 1 | 89927956 | 89931179 | 3224 | 2 | ENSMUST00000036954.8 | -35979 | Gbx2 | ENSMUSG00000034486.8 |
| chr1 | 89967771 | 89968758 | 3.94389 | 5.13833 | 1.89517 | 3' UTR | 1 | 89927956 | 89931179 | 3224 | 2 | ENSMUST00000036954.8 | -36593 | Gbx2 | ENSMUSG00000034486.8 |
| chr1 | 89969472 | 89969667 | 3.8829 | 4.84292 | 1.63716 | Intron (ENSMUST00000086882.2/ENSMUSG00000067081.7, intron 3 of 5) | 1 | 89927956 | 89931179 | 3224 | 2 | ENSMUST00000036954.8 | -38294 | Gbx2 | ENSMUSG00000034486.8 |
| chr1 | 89970047 | 89970122 | 4.10512 | 5.27094 | 1.9334 | Intron (ENSMUST00000086882.2/ENSMUSG00000067081.7, intron 3 of 5) | 1 | 89927956 | 89931179 | 3224 | 2 | ENSMUST00000036954.8 | -38869 | Gbx2 | ENSMUSG00000034486.8 |
| chr13 | 88821892 | 88822173 | 3.53645 | 4.1979 | 1.24425 | Promoter | 13 | 88821642 | 89323223 | 501582 | 1 | ENSMUST00000043111.6 | 251 | Edil3 | ENSMUSG00000034488.9 |
| chr13 | 88822844 | 88823084 | 3.74108 | 4.55816 | 1.42043 | Intron (ENSMUST00000118731.2/ENSMUSG00000034488.9, intron 1 of 9) | 13 | 88821642 | 89323223 | 501582 | 1 | ENSMUST00000043111.6 | 1203 | Edil3 | ENSMUSG00000034488.9 |
| chr11 | 102815805 | 102817335 | 3.56255 | 4.28625 | 1.29162 | Intron (ENSMUST00000068933.6/ENSMUSG00000034520.9, intron 1 of 2) | 11 | 102799579 | 102819179 | 19601 | 2 | ENSMUST00000068933.6 | 1844 | Gjc1 | ENSMUSG00000034520.9 |
| chr11 | 102817716 | 102818663 | 3.56267 | 4.4327 | 1.43702 | Promoter | 11 | 102799579 | 102819179 | 19601 | 2 | ENSMUST00000068933.6 | 516 | Gjc1 | ENSMUSG00000034520.9 |
| chr11 | 102819008 | 102820131 | 4.0148 | 5.15246 | 1.90321 | Promoter | 11 | 102799579 | 102819179 | 19601 | 2 | ENSMUST00000068933.6 | 0 | Gjc1 | ENSMUSG00000034520.9 |
| chr11 | 102820439 | 102821473 | 4.23294 | 5.62064 | 2.20898 | Promoter | 11 | 102799579 | 102819700 | 20122 | 2 | ENSMUST00000092567.5 | -740 | Gjc1 | ENSMUSG00000034520.9 |
| chr11 | 102824070 | 102824400 | 3.3435 | 3.84262 | 1.01733 | Distal Intergenic | 11 | 102799579 | 102819700 | 20122 | 2 | ENSMUST00000092567.5 | -4371 | Gjc1 | ENSMUSG00000034520.9 |
| chr9 | 119650335 | 119650459 | 3.36705 | 3.87815 | 1.01733 | Intron (ENSMUST00000084787.5/ENSMUSG00000034533.9, intron 14 of 27) | 9 | 119608456 | 119719032 | 110577 | 2 | ENSMUST00000084787.5 | 68573 | Scn10a | ENSMUSG00000034533.9 |
| chr9 | 119651066 | 119651196 | 4.20563 | 5.48588 | 2.09623 | Exon (ENSMUST00000084787.5/ENSMUSG00000034533.9, exon 14 of 28) | 9 | 119608456 | 119719032 | 110577 | 2 | ENSMUST00000084787.5 | 67836 | Scn10a | ENSMUSG00000034533.9 |
| chr9 | 119656333 | 119656461 | 3.37252 | 4.09005 | 1.17185 | Intron (ENSMUST00000084787.5/ENSMUSG00000034533.9, intron 13 of 27) | 9 | 119608456 | 119719032 | 110577 | 2 | ENSMUST00000084787.5 | 62571 | Scn10a | ENSMUSG00000034533.9 |
| chr9 | 119665614 | 119665966 | 3.28351 | 3.75243 | 1.01733 | Exon (ENSMUST00000084787.5/ENSMUSG00000034533.9, exon 10 of 28) | 9 | 119608456 | 119719032 | 110577 | 2 | ENSMUST00000084787.5 | 53066 | Scn10a | ENSMUSG00000034533.9 |
| chr9 | 119675903 | 119676429 | 3.75964 | 4.78241 | 1.64336 | Intron (ENSMUST00000084787.5/ENSMUSG00000034533.9, intron 4 of 27) | 9 | 119608456 | 119719032 | 110577 | 2 | ENSMUST00000084787.5 | 42603 | Scn10a | ENSMUSG00000034533.9 |
| chr9 | 119677552 | 119677729 | 3.79929 | 4.81234 | 1.65031 | Intron (ENSMUST00000084787.5/ENSMUSG00000034533.9, intron 4 of 27) | 9 | 119608456 | 119719032 | 110577 | 2 | ENSMUST00000084787.5 | 41303 | Scn10a | ENSMUSG00000034533.9 |
| chr9 | 119684315 | 119684573 | 3.65044 | 4.44977 | 1.45669 | Intron (ENSMUST00000084787.5/ENSMUSG00000034533.9, intron 4 of 27) | 9 | 119608456 | 119719032 | 110577 | 2 | ENSMUST00000084787.5 | 34459 | Scn10a | ENSMUSG00000034533.9 |
| chr9 | 119684949 | 119685469 | 3.50578 | 4.24084 | 1.33253 | Intron (ENSMUST00000084787.5/ENSMUSG00000034533.9, intron 4 of 27) | 9 | 119608456 | 119719032 | 110577 | 2 | ENSMUST00000084787.5 | 33563 | Scn10a | ENSMUSG00000034533.9 |
| chr9 | 119686106 | 119686251 | 4.19082 | 5.5064 | 2.15387 | Intron (ENSMUST00000084787.5/ENSMUSG00000034533.9, intron 4 of 27) | 9 | 119608456 | 119719032 | 110577 | 2 | ENSMUST00000084787.5 | 32781 | Scn10a | ENSMUSG00000034533.9 |
| chr9 | 119687358 | 119687711 | 3.98671 | 5.19946 | 1.93758 | Intron (ENSMUST00000084787.5/ENSMUSG00000034533.9, intron 4 of 27) | 9 | 119608456 | 119719032 | 110577 | 2 | ENSMUST00000084787.5 | 31321 | Scn10a | ENSMUSG00000034533.9 |
| chr9 | 119688065 | 119688724 | 3.4867 | 4.16404 | 1.31066 | Intron (ENSMUST00000084787.5/ENSMUSG00000034533.9, intron 3 of 27) | 9 | 119608456 | 119719032 | 110577 | 2 | ENSMUST00000084787.5 | 30308 | Scn10a | ENSMUSG00000034533.9 |
| chr9 | 119690658 | 119690858 | 3.30611 | 3.83161 | 1.10854 | Intron (ENSMUST00000084787.5/ENSMUSG00000034533.9, intron 3 of 27) | 9 | 119608456 | 119719032 | 110577 | 2 | ENSMUST00000084787.5 | 28174 | Scn10a | ENSMUSG00000034533.9 |
| chr9 | 119693818 | 119693919 | 3.38376 | 3.96457 | 1.19615 | Intron (ENSMUST00000084787.5/ENSMUSG00000034533.9, intron 2 of 27) | 9 | 119608456 | 119719032 | 110577 | 2 | ENSMUST00000084787.5 | 25113 | Scn10a | ENSMUSG00000034533.9 |
| chr9 | 119694334 | 119694608 | 3.77126 | 4.70755 | 1.60946 | 5' UTR | 9 | 119608456 | 119719032 | 110577 | 2 | ENSMUST00000084787.5 | 24424 | Scn10a | ENSMUSG00000034533.9 |
| chr9 | 119701537 | 119701957 | 3.31362 | 3.79767 | 1.01733 | Intron (ENSMUST00000084787.5/ENSMUSG00000034533.9, intron 1 of 27) | 9 | 119608456 | 119719032 | 110577 | 2 | ENSMUST00000084787.5 | 17075 | Scn10a | ENSMUSG00000034533.9 |
| chr9 | 119704993 | 119705100 | 3.29338 | 3.76723 | 1.01733 | Intron (ENSMUST00000084787.5/ENSMUSG00000034533.9, intron 1 of 27) | 9 | 119608456 | 119719032 | 110577 | 2 | ENSMUST00000084787.5 | 13932 | Scn10a | ENSMUSG00000034533.9 |
| chr9 | 119707050 | 119707467 | 3.88938 | 4.87498 | 1.67213 | Intron (ENSMUST00000084787.5/ENSMUSG00000034533.9, intron 1 of 27) | 9 | 119608456 | 119719032 | 110577 | 2 | ENSMUST00000084787.5 | 11565 | Scn10a | ENSMUSG00000034533.9 |
| chr9 | 119711102 | 119711306 | 3.55539 | 4.22524 | 1.23944 | Intron (ENSMUST00000084787.5/ENSMUSG00000034533.9, intron 1 of 27) | 9 | 119608456 | 119719032 | 110577 | 2 | ENSMUST00000084787.5 | 7726 | Scn10a | ENSMUSG00000034533.9 |
| chr9 | 119714846 | 119715178 | 3.44744 | 4.0705 | 1.23876 | Intron (ENSMUST00000084787.5/ENSMUSG00000034533.9, intron 1 of 27) | 9 | 119608456 | 119719032 | 110577 | 2 | ENSMUST00000084787.5 | 3854 | Scn10a | ENSMUSG00000034533.9 |
| chr9 | 119731063 | 119731369 | 3.36072 | 3.86861 | 1.01733 | Distal Intergenic | 9 | 119608456 | 119719032 | 110577 | 2 | ENSMUST00000084787.5 | -12032 | Scn10a | ENSMUSG00000034533.9 |
| chr9 | 119732232 | 119732383 | 3.65178 | 4.39384 | 1.32037 | Distal Intergenic | 9 | 119608456 | 119719032 | 110577 | 2 | ENSMUST00000084787.5 | -13201 | Scn10a | ENSMUSG00000034533.9 |
| chr9 | 119739873 | 119739948 | 3.5899 | 4.29314 | 1.2959 | Distal Intergenic | 9 | 119608456 | 119719032 | 110577 | 2 | ENSMUST00000084787.5 | -20842 | Scn10a | ENSMUSG00000034533.9 |
| chr9 | 119748975 | 119749491 | 3.56854 | 4.30373 | 1.39194 | Distal Intergenic | 9 | 119608456 | 119719032 | 110577 | 2 | ENSMUST00000084787.5 | -29944 | Scn10a | ENSMUSG00000034533.9 |
| chr3 | 66983649 | 66984253 | 3.5396 | 4.67052 | 1.54522 | Intron (ENSMUST00000162036.2/ENSMUSG00000034544.12, intron 1 of 9) | 3 | 66985670 | 66996473 | 10804 | 1 | ENSMUST00000162362.2 | -1417 | Rsrc1 | ENSMUSG00000034544.12 |
| chr11 | 3488154 | 3488318 | 3.27438 | 3.7387 | 1.01733 | Promoter | 11 | 3488284 | 3494166 | 5883 | 1 | ENSMUST00000064265.7 | 0 | Pla2g3 | ENSMUSG00000034579.12 |
| chr9 | 53340438 | 53340754 | 3.36766 | 3.909 | 1.10822 | Promoter | 9 | 53341084 | 53368955 | 27872 | 1 | ENSMUST00000132410.1 | -330 | Exph5 | ENSMUSG00000034584.3 |
| chr9 | 53341182 | 53341611 | 3.48799 | 4.16006 | 1.27067 | Promoter | 9 | 53341189 | 53355634 | 14446 | 1 | ENSMUST00000139096.1 | 0 | Exph5 | ENSMUSG00000034584.3 |
| chr9 | 75070800 | 75071050 | 3.34741 | 3.85218 | 1.02975 | Promoter | 9 | 75071015 | 75117308 | 46294 | 1 | ENSMUST00000149823.1 | 0 | Myo5a | ENSMUSG00000034593.11 |
| chr10 | 82983476 | 82983719 | 3.72919 | 4.90978 | 1.69597 | Distal Intergenic | 10 | 82985498 | 83195900 | 210403 | 1 | ENSMUST00000040110.7 | -1779 | Chst11 | ENSMUSG00000034612.7 |
| chr10 | 82985395 | 82986283 | 3.44324 | 4.10287 | 1.24672 | Promoter | 10 | 82985498 | 83195900 | 210403 | 1 | ENSMUST00000040110.7 | 0 | Chst11 | ENSMUSG00000034612.7 |
| chr4 | 108217350 | 108218216 | 4.09732 | 5.57207 | 2.19281 | Promoter | 4 | 108181934 | 108217897 | 35964 | 2 | ENSMUST00000106690.2 | 0 | Zyg11a | ENSMUSG00000034645.8 |
| chr6 | 107529544 | 107530239 | 3.66786 | 4.44102 | 1.37373 | Promoter | 6 | 107529726 | 107570228 | 40503 | 1 | ENSMUST00000049285.8 | 0 | Lrrn1 | ENSMUSG00000034648.8 |
| chr8 | 84414342 | 84414847 | 3.70418 | 4.88212 | 1.69505 | Promoter | 8 | 84415364 | 84640246 | 224883 | 1 | ENSMUST00000121390.2 | -517 | Cacna1a | ENSMUSG00000034656.11 |
| chr8 | 84415199 | 84415499 | 3.85903 | 4.9942 | 1.79931 | Promoter | 8 | 84415364 | 84640246 | 224883 | 1 | ENSMUST00000121390.2 | 0 | Cacna1a | ENSMUSG00000034656.11 |
| chr8 | 84416372 | 84416471 | 4.25142 | 5.65569 | 2.29656 | Promoter | 8 | 84415377 | 84640063 | 224687 | 1 | ENSMUST00000122053.1 | 996 | Cacna1a | ENSMUSG00000034656.11 |
| chr8 | 84554281 | 84554446 | 3.3767 | 3.89271 | 1.01733 | Intron (ENSMUST00000121390.2/ENSMUSG00000034656.11, intron 17 of 46) | 8 | 84547094 | 84554084 | 6991 | 1 | ENSMUST00000129620.1 | 7188 | Cacna1a | ENSMUSG00000034656.11 |
| chr8 | 84559004 | 84559645 | 3.46488 | 4.10553 | 1.19172 | Exon (ENSMUST00000121390.2/ENSMUSG00000034656.11, exon 19 of 47) | 8 | 84547094 | 84554084 | 6991 | 1 | ENSMUST00000129620.1 | 11911 | Cacna1a | ENSMUSG00000034656.11 |
| chr8 | 84567005 | 84567234 | 3.38193 | 3.9006 | 1.01733 | Intron (ENSMUST00000121390.2/ENSMUSG00000034656.11, intron 20 of 46) | 8 | 84571797 | 84580462 | 8666 | 1 | ENSMUST00000126302.1 | -4563 | Cacna1a | ENSMUSG00000034656.11 |
| chr8 | 84569567 | 84570499 | 3.55191 | 4.22672 | 1.23899 | Exon (ENSMUST00000121390.2/ENSMUSG00000034656.11, exon 22 of 47) | 8 | 84571797 | 84580462 | 8666 | 1 | ENSMUST00000126302.1 | -1298 | Cacna1a | ENSMUSG00000034656.11 |
| chr8 | 84600283 | 84601286 | 3.63727 | 4.37246 | 1.31767 | Promoter | 8 | 84601431 | 84612080 | 10650 | 1 | ENSMUST00000143215.2 | -145 | Cacna1a | ENSMUSG00000034656.11 |
| chr7 | 6197070 | 6197271 | 3.45187 | 4.03145 | 1.10632 | Promoter | 7 | 6197090 | 6216137 | 19048 | 1 | ENSMUST00000037553.5 | 0 | Galp | ENSMUSG00000034660.8 |
| chr13 | 55464310 | 55465009 | 4.14 | 5.63718 | 2.2674 | Promoter | 13 | 55464267 | 55473155 | 8889 | 1 | ENSMUST00000046533.7 | 44 | Prr7 | ENSMUSG00000034686.7 |
| chr5 | 96374303 | 96374689 | 3.41405 | 3.96408 | 1.07019 | Promoter | 5 | 96373955 | 96784728 | 410774 | 1 | ENSMUST00000036019.4 | 349 | Fras1 | ENSMUSG00000034687.4 |
| chr5 | 96375835 | 96375931 | 4.5085 | 6.18112 | 2.59592 | Intron (ENSMUST00000036019.4/ENSMUSG00000034687.4, intron 1 of 73) | 5 | 96373955 | 96784728 | 410774 | 1 | ENSMUST00000036019.4 | 1881 | Fras1 | ENSMUSG00000034687.4 |
| chr2 | 79455927 | 79457625 | 3.65858 | 4.49613 | 1.47216 | Promoter | 2 | 79452521 | 79456751 | 4231 | 2 | ENSMUST00000041099.4 | 0 | Neurod1 | ENSMUSG00000034701.9 |
| chr11 | 114727489 | 114727792 | 3.91879 | 5.23286 | 1.99201 | Promoter | 11 | 114727416 | 114754352 | 26937 | 1 | ENSMUST00000136403.2 | 74 | Dnaic2 | ENSMUSG00000034706.11 |
| chr10 | 121347025 | 121347100 | 3.58028 | 4.29683 | 1.34684 | Distal Intergenic | 10 | 121365090 | 121397245 | 32156 | 1 | ENSMUST00000040344.6 | -17990 | Gns | ENSMUSG00000034707.6 |
| chr10 | 121350404 | 121351928 | 3.75445 | 4.72974 | 1.62212 | Distal Intergenic | 10 | 121365090 | 121397245 | 32156 | 1 | ENSMUST00000040344.6 | -13162 | Gns | ENSMUSG00000034707.6 |
| chr5 | 96035200 | 96035279 | 3.66488 | 4.46922 | 1.47881 | Distal Intergenic | 5 | 96075576 | 96086933 | 11358 | 2 | ENSMUST00000152490.2 | 51654 | Cnot6l | ENSMUSG00000034724.12 |
| chr15 | 74487221 | 74487397 | 3.34412 | 3.90183 | 1.06908 | Distal Intergenic | 15 | 74516195 | 74589464 | 73270 | 1 | ENSMUST00000042035.10 | -28798 | Bai1 | ENSMUSG00000034730.11 |

| | | | | | | | | | | | | | | | |
|---|---|---|---|---|---|---|---|---|---|---|---|---|---|---|---|
| chr15 | 74506577 | 74507045 | 3.87062 | 4.80984 | 1.59427 | Distal Intergenic | 15 | 74516195 | 74589464 | 73270 | 1 | ENSMUST00000042035.10 | -9150 | Bai1 | ENSMUSG00000034730.11 |
| chr15 | 74515966 | 74516708 | 3.6386 | 4.3991 | 1.34287 | Promoter | 15 | 74516195 | 74589464 | 73270 | 1 | ENSMUST00000042035.10 | 0 | Bai1 | ENSMUSG00000034730.11 |
| chr15 | 74517579 | 74520225 | 3.81681 | 4.8387 | 1.70476 | Promoter | 15 | 74516831 | 74589464 | 72634 | 1 | ENSMUST00000187485.1 | 749 | Bai1 | ENSMUSG00000034730.11 |
| chr15 | 74529291 | 74529477 | 4.05771 | 5.34106 | 2.02537 | Exon (ENSMUST00000042035.10/ENSMUSG00000034730.11, exon 2 of 31) | 15 | 74540397 | 74543522 | 3126 | 1 | ENSMUST00000189200.1 | -10920 | Bai1 | ENSMUSG00000034730.11 |
| chr15 | 74538697 | 74539164 | 4.17665 | 5.41509 | 2.01574 | Intron (ENSMUST00000042035.10/ENSMUSG00000034730.11, intron 3 of 30) | 15 | 74540397 | 74543522 | 3126 | 1 | ENSMUST00000189200.1 | -1233 | Bai1 | ENSMUSG00000034730.11 |
| chr15 | 74539658 | 74539826 | 3.88407 | 4.87601 | 1.68832 | Promoter | 15 | 74540397 | 74543522 | 3126 | 1 | ENSMUST00000189200.1 | -571 | Bai1 | ENSMUSG00000034730.11 |
| chr15 | 74559528 | 74560106 | 3.4544 | 4.03273 | 1.09742 | Intron (ENSMUST00000042035.10/ENSMUSG00000034730.11, intron 17 of 30) | 15 | 74561284 | 74574119 | 12836 | 1 | ENSMUST00000189353.1 | -1178 | Bai1 | ENSMUSG00000034730.11 |
| chr15 | 74561629 | 74561890 | 3.37466 | 3.88963 | 1.01733 | Promoter | 15 | 74561284 | 74574119 | 12836 | 1 | ENSMUST00000189353.1 | 346 | Bai1 | ENSMUSG00000034730.11 |
| chr15 | 74563138 | 74564729 | 3.94465 | 5.02901 | 1.78598 | Promoter | 15 | 74563378 | 74572207 | 8830 | 1 | ENSMUST00000190524.1 | 0 | Bai1 | ENSMUSG00000034730.11 |
| chr15 | 74565474 | 74565872 | 3.69922 | 4.486 | 1.38942 | Intron (ENSMUST00000042035.10/ENSMUSG00000034730.11, intron 18 of 30) | 15 | 74563901 | 74572861 | 8961 | 1 | ENSMUST00000187599.1 | 1574 | Bai1 | ENSMUSG00000034730.11 |
| chr13 | 102958080 | 102958631 | 3.33668 | 3.87314 | 1.13564 | Promoter | 13 | 102735002 | 102958467 | 223466 | 2 | ENSMUST00000099202.4 | 0 | Mast4 | ENSMUSG00000034751.10 |
| chr13 | 103027686 | 103027936 | 3.60752 | 4.43613 | 1.43178 | Intron (ENSMUST00000166726.2/ENSMUSG00000034751.10, intron 3 of 26) | 13 | 102794128 | 103096828 | 302701 | 2 | ENSMUST00000166336.2 | 68892 | Mast4 | ENSMUSG00000034751.10 |
| chr13 | 103096265 | 103097128 | 3.74927 | 4.77668 | 1.64861 | Promoter | 13 | 102794128 | 103096828 | 302701 | 2 | ENSMUST00000166336.2 | 0 | Mast4 | ENSMUSG00000034751.10 |
| chr13 | 103097529 | 103097920 | 3.56131 | 4.33333 | 1.34696 | Promoter | 13 | 103094177 | 103096844 | 2668 | 2 | ENSMUST00000185314.1 | -686 | Mast4 | ENSMUSG00000034751.10 |
| chr13 | 103098312 | 103098387 | 3.70579 | 4.60963 | 1.49091 | Intron (ENSMUST00000166726.2/ENSMUSG00000034751.10, intron 3 of 26) | 13 | 103094177 | 103096844 | 2668 | 2 | ENSMUST00000185314.1 | -1469 | Mast4 | ENSMUSG00000034751.10 |
| chr13 | 103334546 | 103334904 | 3.95846 | 5.26692 | 1.96004 | Promoter | 13 | 103058696 | 103334497 | 275802 | 2 | ENSMUST00000171358.2 | -50 | Mast4 | ENSMUSG00000034751.10 |
| chr13 | 103331377 | 103331662 | 3.59051 | 4.39227 | 1.40691 | Intron (ENSMUST00000166726.2/ENSMUSG00000034751.10, intron 1 of 26) | 13 | 102732486 | 103334192 | 601707 | 2 | ENSMUST00000166726.2 | 2530 | Mast4 | ENSMUSG00000034751.10 |
| chr13 | 103331974 | 103332158 | 4.19009 | 5.74496 | 2.29779 | Intron (ENSMUST00000166726.2/ENSMUSG00000034751.10, intron 1 of 26) | 13 | 102732486 | 103334192 | 601707 | 2 | ENSMUST00000166726.2 | 2034 | Mast4 | ENSMUSG00000034751.10 |
| chr13 | 103333329 | 103334035 | 4.34313 | 5.87116 | 2.44847 | Promoter | 13 | 102732486 | 103334192 | 601707 | 2 | ENSMUST00000166726.2 | 157 | Mast4 | ENSMUSG00000034751.10 |
| chr10 | 81591952 | 81592350 | 3.48702 | 4.10597 | 1.18158 | Promoter | 10 | 81590905 | 81592715 | 1811 | 2 | ENSMUST00000129282.2 | 365 | Tle6 | ENSMUSG00000034758.7 |
| chr10 | 81592657 | 81592981 | 3.42901 | 3.98708 | 1.07163 | Promoter | 10 | 81590905 | 81592715 | 1811 | 2 | ENSMUST00000129282.2 | 0 | Tle6 | ENSMUSG00000034758.7 |
| chr10 | 81594445 | 81594635 | 3.80847 | 4.77105 | 1.65698 | Promoter | 10 | 81593919 | 81595244 | 1326 | 2 | ENSMUST00000134457.1 | 609 | Tle6 | ENSMUSG00000034758.7 |
| chr4 | 107434793 | 107435100 | 4.46334 | 6.03458 | 2.51525 | Promoter | 4 | 107434719 | 107635061 | 200343 | 1 | ENSMUST00000046005.3 | 75 | Glis1 | ENSMUSG00000034762.4 |
| chr4 | 107444375 | 107444728 | 3.86932 | 4.84251 | 1.69096 | Intron (ENSMUST00000135835.2/ENSMUSG00000034762.4, intron 2 of 2) | 4 | 107438998 | 107635061 | 196064 | 1 | ENSMUST00000106738.1 | 5378 | Glis1 | ENSMUSG00000034762.4 |
| chr10 | 120361918 | 120362174 | 3.32507 | 3.97376 | 1.1039 | 3' UTR | 10 | 120364157 | 120382836 | 18680 | 1 | ENSMUST00000041178.6 | -1983 | 1700006J14Rik | ENSMUSG00000034764.10 |
| chr10 | 120362586 | 120363338 | 3.68314 | 4.55974 | 1.50499 | Promoter | 10 | 120364157 | 120382836 | 18680 | 1 | ENSMUST00000041178.6 | -819 | 1700006J14Rik | ENSMUSG00000034764.10 |
| chr10 | 120364250 | 120365333 | 3.90927 | 5.02519 | 1.83498 | Promoter | 10 | 120364213 | 120383829 | 19617 | 1 | ENSMUST00000161511.1 | 38 | 1700006J14Rik | ENSMUSG00000034764.10 |
| chr10 | 120366534 | 120366821 | 3.41131 | 4.17947 | 1.27699 | Intron (ENSMUST00000041178.6/ENSMUSG00000034764.10, intron 1 of 2) | 10 | 120364213 | 120383829 | 19617 | 1 | ENSMUST00000161511.1 | 2322 | 1700006J14Rik | ENSMUSG00000034764.10 |
| chr10 | 120369351 | 120369824 | 3.68586 | 4.53486 | 1.5016 | Intron (ENSMUST00000041178.6/ENSMUSG00000034764.10, intron 1 of 2) | 10 | 120364213 | 120383829 | 19617 | 1 | ENSMUST00000161511.1 | 5139 | 1700006J14Rik | ENSMUSG00000034764.10 |
| chr10 | 120370647 | 120370939 | 3.78317 | 4.75528 | 1.65157 | Exon (ENSMUST00000159699.1/ENSMUSG00000056758.9, exon 5 of 6) | 10 | 120364213 | 120383829 | 19617 | 1 | ENSMUST00000161511.1 | 6435 | 1700006J14Rik | ENSMUSG00000034764.10 |
| chr10 | 120371292 | 120371432 | 3.67781 | 4.69629 | 1.62154 | Exon (ENSMUST00000160091.2/ENSMUSG00000034764.10, exon 3 of 4) | 10 | 120364213 | 120383829 | 19617 | 1 | ENSMUST00000161511.1 | 7080 | 1700006J14Rik | ENSMUSG00000034764.10 |
| chr10 | 120375992 | 120376330 | 3.69463 | 4.59779 | 1.55266 | Intron (ENSMUST00000041178.6/ENSMUSG00000034764.10, intron 2 of 2) | 10 | 120364213 | 120383829 | 19617 | 1 | ENSMUST00000161511.1 | 11780 | 1700006J14Rik | ENSMUSG00000034764.10 |
| chr19 | 53544495 | 53544710 | 3.29104 | 3.76383 | 1.01733 | Distal Intergenic | 19 | 53529109 | 53542431 | 13323 | 1 | ENSMUST00000038287.6 | 15387 | Dusp5 | ENSMUSG00000034765.6 |
| chr11 | 102275413 | 102275869 | 3.80132 | 4.77466 | 1.67704 | Intron (ENSMUST00000036467.4/ENSMUSG00000034768.4, intron 2 of 4) | 11 | 102268743 | 102279462 | 10720 | 1 | ENSMUST00000036467.4 | 6671 | Asb16 | ENSMUSG00000034768.4 |
| chr11 | 102276234 | 102276775 | 4.08021 | 5.4835 | 2.16965 | Exon (ENSMUST00000036467.4/ENSMUSG00000034768.4, exon 3 of 5) | 11 | 102268743 | 102279462 | 10720 | 1 | ENSMUST00000036467.4 | 7492 | Asb16 | ENSMUSG00000034768.4 |
| chr11 | 102244699 | 102245558 | 3.51746 | 4.16662 | 1.22984 | Distal Intergenic | 11 | 102248882 | 102265187 | 16306 | 1 | ENSMUST00000100392.4 | -3324 | BC030867 | ENSMUSG00000034773.11 |
| chr11 | 102263444 | 102263774 | 3.80427 | 4.82386 | 1.67613 | Intron (ENSMUST00000100392.4/ENSMUSG00000034773.11, intron 9 of 9) | 11 | 102261964 | 102265187 | 3224 | 1 | ENSMUST00000140040.1 | 1481 | BC030867 | ENSMUSG00000034773.11 |
| chr6 | 83711017 | 83711415 | 3.76598 | 4.7427 | 1.62683 | Promoter | 6 | 83711264 | 83738313 | 27050 | 1 | ENSMUST00000037807.2 | 0 | Vax2 | ENSMUSG00000034777.2 |
| chr2 | 67565835 | 67566162 | 3.60468 | 4.5468 | 1.46528 | Promoter | 2 | 67565871 | 67810894 | 245024 | 1 | ENSMUST00000188614.1 | 0 | B3galt1 | ENSMUSG00000034780.6 |
| chr2 | 67566677 | 67568709 | 3.51449 | 4.29808 | 1.34099 | Promoter | 2 | 67565871 | 67810894 | 245024 | 1 | ENSMUST00000188614.1 | 807 | B3galt1 | ENSMUSG00000034780.6 |
| chr6 | 83680090 | 83680430 | 3.41832 | 3.96799 | 1.06121 | Distal Intergenic | 6 | 83671207 | 83677857 | 6651 | 2 | ENSMUST00000037882.6 | -2234 | Cd207 | ENSMUSG00000034783.6 |
| chr10 | 81511938 | 81512299 | 3.71925 | 4.53063 | 1.43174 | Promoter | 10 | 81503087 | 81512639 | 9553 | 2 | ENSMUST00000150586.1 | 340 | Gna15 | ENSMUSG00000034792.8 |
| chr8 | 123115766 | 123116274 | 3.75759 | 4.69056 | 1.63406 | Intron (ENSMUST00000127664.1/ENSMUSG00000092329.1, intron 1 of 15) | 8 | 123117374 | 123135180 | 17807 | 1 | ENSMUST00000134235.2 | -1100 | Cpne7 | ENSMUSG00000034796.9 |
| chr8 | 123116611 | 123118104 | 3.83701 | 4.89709 | 1.73368 | Promoter | 8 | 123117374 | 123135180 | 17807 | 1 | ENSMUST00000134235.2 | 0 | Cpne7 | ENSMUSG00000034796.9 |
| chr8 | 123126534 | 123128306 | 3.43111 | 3.99873 | 1.10144 | Promoter | 8 | 123126517 | 123127373 | 857 | 1 | ENSMUST00000123191.1 | 18 | Cpne7 | ENSMUSG00000034796.9 |
| chr8 | 71654574 | 71654968 | 3.53164 | 4.23668 | 1.27836 | Exon (ENSMUST00000030170.9/ENSMUSG00000034799.11, exon 16 of 44) | 8 | 71653607 | 71658495 | 4889 | 2 | ENSMUST00000176426.1 | 3527 | Unc13a | ENSMUSG00000034799.11 |
| chr8 | 71657352 | 71658275 | 3.76937 | 4.66957 | 1.59581 | Promoter | 8 | 71653607 | 71658495 | 4889 | 2 | ENSMUST00000176426.1 | 220 | Unc13a | ENSMUSG00000034799.11 |
| chr8 | 71658975 | 71659164 | 3.38329 | 3.9562 | 1.1354 | Promoter | 8 | 71653607 | 71658495 | 4889 | 2 | ENSMUST00000176426.1 | -481 | Unc13a | ENSMUSG00000034799.11 |
| chr8 | 71660519 | 71660811 | 3.82459 | 4.94212 | 1.73137 | Exon (ENSMUST00000030170.9/ENSMUSG00000034799.11, exon 9 of 44) | 8 | 71653607 | 71658495 | 4889 | 2 | ENSMUST00000176426.1 | -2025 | Unc13a | ENSMUSG00000034799.11 |
| chr8 | 71666787 | 71667095 | 3.38193 | 3.9006 | 1.01733 | Promoter | 8 | 71660346 | 71668082 | 7737 | 2 | ENSMUST00000175780.1 | 987 | Unc13a | ENSMUSG00000034799.11 |
| chr8 | 71670615 | 71670803 | 3.71792 | 4.52363 | 1.41162 | Promoter | 8 | 71629054 | 71671736 | 42683 | 2 | ENSMUST00000177517.2 | 933 | Unc13a | ENSMUSG00000034799.11 |
| chr10 | 119452751 | 119452894 | 3.34153 | 3.83966 | 1.01733 | Intron (ENSMUST00000154566.1/ENSMUSG00000085454.1, intron 1 of 2) | 10 | 119454034 | 120087260 | 633227 | 1 | ENSMUST00000147356.2 | -1140 | Grip1 | ENSMUSG00000034813.11 |
| chr10 | 119453356 | 119454551 | 3.83362 | 4.76437 | 1.59197 | Promoter | 10 | 119454034 | 120087260 | 633227 | 1 | ENSMUST00000147356.2 | 0 | Grip1 | ENSMUSG00000034813.11 |
| chr10 | 81472002 | 81472233 | 3.72414 | 4.53784 | 1.42377 | Promoter | 10 | 81469445 | 81473002 | 3558 | 2 | ENSMUST00000147524.1 | 769 | Celf5 | ENSMUSG00000034818.11 |
| chr10 | 81473700 | 81474010 | 3.78021 | 4.74492 | 1.63017 | Promoter | 10 | 81467507 | 81473378 | 5872 | 2 | ENSMUST00000141207.2 | -323 | Celf5 | ENSMUSG00000034818.11 |
| chr10 | 81474507 | 81474879 | 3.66427 | 4.44589 | 1.40413 | Promoter | 10 | 81460930 | 81474844 | 13915 | 2 | ENSMUST00000120856.2 | 0 | Celf5 | ENSMUSG00000034818.11 |
| chr10 | 81480870 | 81481088 | 3.36161 | 3.86994 | 1.01733 | Intron (ENSMUST00000118763.2/ENSMUSG00000034818.11, intron 1 of 11) | 10 | 81467759 | 81482672 | 14914 | 2 | ENSMUST00000128494.1 | 1584 | Celf5 | ENSMUSG00000034818.11 |
| chr10 | 81481860 | 81482229 | 3.77091 | 4.63896 | 1.51992 | Promoter | 10 | 81467759 | 81482672 | 14914 | 2 | ENSMUST00000128494.1 | 443 | Celf5 | ENSMUSG00000034818.11 |
| chr7 | 109778764 | 109779400 | 3.69621 | 4.67572 | 1.54429 | Intron (ENSMUST00000041460.7/ENSMUSG00000034825.8, intron 2 of 7) | 7 | 109758059 | 109781545 | 23487 | 2 | ENSMUST00000033331.6 | 2145 | Nrip3 | ENSMUSG00000034825.8 |
| chr7 | 109780371 | 109781350 | 3.67855 | 4.64612 | 1.57759 | Promoter | 7 | 109758059 | 109781545 | 23487 | 2 | ENSMUST00000033331.6 | 195 | Nrip3 | ENSMUSG00000034825.8 |
| chr7 | 109781728 | 109781943 | 4.16522 | 5.5774 | 2.16152 | Promoter | 7 | 109758055 | 109782025 | 23971 | 2 | ENSMUST00000041460.7 | 82 | Nrip3 | ENSMUSG00000034825.8 |
| chr7 | 109783695 | 109783816 | 3.56669 | 4.31825 | 1.35427 | Distal Intergenic | 7 | 109758055 | 109782025 | 23971 | 2 | ENSMUST00000041460.7 | -1671 | Nrip3 | ENSMUSG00000034825.8 |
| chr8 | 71562747 | 71562822 | 3.69112 | 4.67613 | 1.5371 | Exon (ENSMUST00000048243.6/ENSMUSG00000034829.7, exon 2 of 3) | 8 | 71560556 | 71566601 | 6046 | 2 | ENSMUST00000048243.6 | 3779 | Nxnl1 | ENSMUSG00000034829.7 |
| chr9 | 107677045 | 107677369 | 3.81569 | 4.82357 | 1.67459 | Exon (ENSMUST00000010205.7/ENSMUSG00000034837.7, exon 4 of 9) | 9 | 107674477 | 107679592 | 5116 | 2 | ENSMUST00000010205.7 | 2223 | Gnat1 | ENSMUSG00000034837.7 |
| chr5 | 92390977 | 92391132 | 3.45518 | 4.24023 | 1.29887 | Intron (ENSMUST00000154245.2/ENSMUSG00000034842.11, intron 3 of 3) | 5 | 92392585 | 92414622 | 22038 | 1 | ENSMUST00000126281.2 | -1453 | Art3 | ENSMUSG00000034842.11 |
| chr5 | 92391704 | 92392114 | 3.72807 | 4.57775 | 1.52328 | Promoter | 5 | 92392585 | 92414622 | 22038 | 1 | ENSMUST00000126281.2 | -471 | Art3 | ENSMUSG00000034842.11 |
| chr8 | 71507568 | 71508188 | 3.6668 | 4.44906 | 1.37755 | Exon (ENSMUST00000048452.4/ENSMUSG00000034845.5, exon 3 of 6) | 8 | 71497754 | 71511769 | 14016 | 2 | ENSMUST00000048452.4 | 3581 | Plvap | ENSMUSG00000034845.5 |
| chr8 | 71509006 | 71509168 | 3.48001 | 4.08634 | 1.15081 | Intron (ENSMUST00000048452.4/ENSMUSG00000034845.5, intron 1 of 5) | 8 | 71497754 | 71511769 | 14016 | 2 | ENSMUST00000048452.4 | 2601 | Plvap | ENSMUSG00000034845.5 |
| chr8 | 71509623 | 71509990 | 3.38864 | 3.91592 | 1.03574 | Intron (ENSMUST00000048452.4/ENSMUSG00000034845.5, intron 1 of 5) | 8 | 71497754 | 71511769 | 14016 | 2 | ENSMUST00000048452.4 | 1779 | Plvap | ENSMUSG00000034845.5 |
| chr8 | 71511494 | 71511838 | 3.40713 | 3.95643 | 1.08015 | Promoter | 8 | 71497754 | 71511769 | 14016 | 2 | ENSMUST00000048452.4 | 0 | Plvap | ENSMUSG00000034845.5 |
| chr17 | 70963702 | 70963781 | 3.04546 | 15.53772 | 10.73667 | Intron (ENSMUST00000180743.1/ENSMUSG00000097625.1, intron 1 of 3) | 17 | 70974271 | 70975966 | 1696 | 2 | ENSMUST00000137537.1 | 12185 | Myl12b | ENSMUSG00000034868.8 |
| chr13 | 54757781 | 54757856 | 3.63304 | 4.36265 | 1.30972 | Downstream (1-2kb) | 13 | 54759428 | 54763067 | 3640 | 2 | ENSMUST00000153829.1 | 5211 | Sncb | ENSMUSG00000034891.8 |
| chr13 | 54758819 | 54758961 | 4.06859 | 5.54954 | 2.17419 | 3' UTR | 13 | 54759428 | 54763067 | 3640 | 2 | ENSMUST00000153829.1 | 4106 | Sncb | ENSMUSG00000034891.8 |
| chr13 | 54764515 | 54765006 | 3.55882 | 4.23641 | 1.24022 | Intron (ENSMUST00000036825.8/ENSMUSG00000034891.8, intron 2 of 5) | 13 | 54759381 | 54766097 | 6717 | 2 | ENSMUST00000134110.1 | 1091 | Sncb | ENSMUSG00000034891.8 |
| chr13 | 54765388 | 54765889 | 3.66174 | 4.46212 | 1.42322 | Promoter | 13 | 54759381 | 54766097 | 6717 | 2 | ENSMUST00000134110.1 | 208 | Sncb | ENSMUSG00000034891.8 |
| chr13 | 54766265 | 54766907 | 3.77929 | 5.04991 | 1.82191 | Promoter | 13 | 54758860 | 54766595 | 7736 | 2 | ENSMUST00000036825.8 | 0 | Sncb | ENSMUSG00000034891.8 |
| chr2 | 65238850 | 65239019 | 3.43912 | 4.05345 | 1.14455 | Promoter | 2 | 65125972 | 65239044 | 113073 | 2 | ENSMUST00000155916.2 | 25 | Cobll1 | ENSMUSG00000034903.13 |
| chr2 | 127132346 | 127132512 | 3.4234 | 4.00511 | 1.16602 | Intron (ENSMUST00000110387.3/ENSMUSG00000034906.10, intron 1 of 17) | 2 | 127127661 | 127133897 | 6237 | 2 | ENSMUST00000156625.2 | 1385 | Ncaph | ENSMUSG00000034906.10 |
| chr12 | 67059184 | 67059652 | 1.93646 | 8.96652 | 4.93204 | Intron (ENSMUST00000037181.9/ENSMUSG00000034912.11, intron 1 of 16) | 12 | 66471469 | 67221202 | 749734 | 2 | ENSMUST00000101379.3 | 161550 | Mdga2 | ENSMUSG00000034912.11 |
| chr12 | 67220340 | 67220989 | 3.86624 | 4.92864 | 1.76383 | Promoter | 12 | 66471469 | 67221202 | 749734 | 2 | ENSMUST00000101379.3 | 213 | Mdga2 | ENSMUSG00000034912.11 |
| chr12 | 67222091 | 67222333 | 3.67289 | 4.67411 | 1.55222 | Promoter | 12 | 67212530 | 67222381 | 9852 | 2 | ENSMUST00000179025.1 | 48 | Mdga2 | ENSMUSG00000034912.11 |
| chr12 | 67222766 | 67222967 | 4.59312 | 6.55732 | 2.89359 | Promoter | 12 | 66469568 | 67222549 | 752982 | 2 | ENSMUST00000037181.9 | -218 | Mdga2 | ENSMUSG00000034912.11 |
| chr13 | 54714344 | 54714419 | 3.38193 | 3.9006 | 1.01733 | Intron (ENSMUST00000037145.7/ENSMUSG00000034918.8, intron 5 of 31) | 13 | 54701461 | 54736662 | 35202 | 1 | ENSMUST00000037145.7 | 12884 | Cdhr2 | ENSMUSG00000034918.8 |
| chr4 | 106622490 | 106622719 | 3.42849 | 4.06265 | 1.20715 | Promoter | 4 | 106622432 | 106640189 | 17758 | 1 | ENSMUST00000047922.2 | 59 | Ttc22 | ENSMUSG00000034919.2 |
| chr10 | 81240199 | 81240591 | 3.9955 | 5.12605 | 1.84638 | Exon (ENSMUST00000117798.2/ENSMUSG00000034949.13, exon 4 of 18) | 10 | 81233187 | 81252121 | 18935 | 1 | ENSMUST00000132174.2 | 7013 | Zfr2 | ENSMUSG00000034949.13 |
| chr7 | 35119629 | 35119741 | 3.51624 | 4.24958 | 1.30897 | Promoter | 7 | 35119293 | 35121926 | 2634 | 1 | ENSMUST00000042985.9 | 337 | Cebpa | ENSMUSG00000034957.9 |
| chr7 | 35120182 | 35120259 | 5.49758 | 8.73804 | 4.61431 | Promoter | 7 | 35119293 | 35121926 | 2634 | 1 | ENSMUST00000042985.9 | 890 | Cebpa | ENSMUSG00000034957.9 |
| chr7 | 35121194 | 35121493 | 3.23183 | 3.67492 | 1.01733 | 3' UTR | 7 | 35119293 | 35121926 | 2634 | 1 | ENSMUST00000042985.9 | 1902 | Cebpa | ENSMUSG00000034957.9 |
| chr7 | 35149300 | 35149482 | 3.70346 | 4.52875 | 1.49454 | Distal Intergenic | 7 | 35119293 | 35121926 | 2634 | 1 | ENSMUST00000042985.9 | 30008 | Cebpa | ENSMUSG00000034957.9 |
| chr10 | 81228670 | 81228879 | 3.35389 | 3.88108 | 1.0896 | Intron (ENSMUST00000146030.2/ENSMUSG00000034958.6, intron 1 of 12) | 10 | 81204508 | 81230785 | 26278 | 2 | ENSMUST00000146030.2 | 1906 | Atcay | ENSMUSG00000034958.6 |
| chr10 | 81230559 | 81231508 | 3.86059 | 4.82872 | 1.66157 | Promoter | 10 | 81204508 | 81230785 | 26278 | 2 | ENSMUST00000146030.2 | 0 | Atcay | ENSMUSG00000034958.6 |
| chr6 | 83084449 | 83084650 | 3.36363 | 3.88243 | 1.0507 | Distal Intergenic | 6 | 83086367 | 83088243 | 1877 | 1 | ENSMUST00000041265.3 | -1717 | Lbx2 | ENSMUSG00000034968.3 |
| chr6 | 83086821 | 83087053 | 4.09493 | 5.24743 | 1.90543 | Promoter | 6 | 83087338 | 83088096 | 759 | 1 | ENSMUST00000151508.1 | -285 | Lbx2 | ENSMUSG00000034968.3 |
| chr6 | 83087993 | 83089505 | 3.66396 | 4.56462 | 1.46528 | Promoter | 6 | 83087338 | 83088096 | 759 | 1 | ENSMUST00000151508.1 | 656 | Lbx2 | ENSMUSG00000034968.3 |
| chr5 | 91518047 | 91518635 | 3.70628 | 4.50492 | 1.40144 | Promoter | 5 | 91517615 | 91626894 | 109280 | 1 | ENSMUST00000040576.9 | 433 | Parm1 | ENSMUSG00000034981.9 |
| chr5 | 91519018 | 91519330 | 3.62973 | 4.363 | 1.32473 | Intron (ENSMUST00000040576.9/ENSMUSG00000034981.9, intron 1 of 3) | 5 | 91517615 | 91626894 | 109280 | 1 | ENSMUST00000040576.9 | 1404 | Parm1 | ENSMUSG00000034981.9 |
| chr14 | 74640837 | 74642109 | 3.96214 | 5.143 | 1.9044 | Promoter | 14 | 74640840 | 74706859 | 66020 | 1 | ENSMUST00000036653.3 | 0 | Htr2a | ENSMUSG00000034997.4 |
| chr14 | 74642538 | 74642714 | 3.75472 | 4.655 | 1.5772 | Intron (ENSMUST00000036653.3/ENSMUSG00000034997.4, intron 1 of 2) | 14 | 74640840 | 74706859 | 66020 | 1 | ENSMUST00000036653.3 | 1699 | Htr2a | ENSMUSG00000034997.4 |
| chr2 | 61802584 | 61803076 | 4.04588 | 5.19227 | 1.91584 | Promoter | 2 | 61802930 | 61804558 | 1629 | 1 | ENSMUST00000136867.1 | 0 | Tbr1 | ENSMUSG00000035033.10 |
| chr2 | 61803579 | 61803815 | 3.4156 | 4.44778 | 1.42251 | Promoter | 2 | 61804453 | 61814114 | 9662 | 1 | ENSMUST00000048934.9 | -638 | Tbr1 | ENSMUSG00000035033.10 |
| chr2 | 61805192 | 61806258 | 4.00228 | 5.14967 | 1.85763 | Promoter | 2 | 61806262 | 61812531 | 6270 | 1 | ENSMUST00000102737.4 | -4 | Tbr1 | ENSMUSG00000035033.10 |
| chr2 | 61806892 | 61809820 | 3.81158 | 4.95854 | 1.78423 | Promoter | 2 | 61806273 | 61811801 | 5529 | 1 | ENSMUST00000131538.1 | 620 | Tbr1 | ENSMUSG00000035033.10 |
| chr2 | 61810173 | 61812958 | 3.87801 | 5.00564 | 1.82373 | 3' UTR | 2 | 61806273 | 61811801 | 5529 | 1 | ENSMUST00000131538.1 | 3901 | Tbr1 | ENSMUSG00000035033.10 |
| chr2 | 61813407 | 61813798 | 3.88831 | 4.9486 | 1.7522 | 3' UTR | 2 | 61806273 | 61811801 | 5529 | 1 | ENSMUST00000131538.1 | 7135 | Tbr1 | ENSMUSG00000035033.10 |
| chr2 | 61814153 | 61816906 | 4.13099 | 5.58862 | 2.21396 | Distal Intergenic | 2 | 61806273 | 61811801 | 5529 | 1 | ENSMUST00000131538.1 | 7881 | Tbr1 | ENSMUSG00000035033.10 |
| chr2 | 61817373 | 61817466 | 3.46109 | 4.20969 | 1.27161 | Distal Intergenic | 2 | 61806273 | 61811801 | 5529 | 1 | ENSMUST00000131538.1 | 11101 | Tbr1 | ENSMUSG00000035033.10 |
| chr2 | 61818301 | 61818510 | 3.69354 | 4.85822 | 1.66996 | Distal Intergenic | 2 | 61806273 | 61811801 | 5529 | 1 | ENSMUST00000131538.1 | 12029 | Tbr1 | ENSMUSG00000035033.10 |
| chr2 | 61819953 | 61820532 | 3.74867 | 4.71653 | 1.6128 | Distal Intergenic | 2 | 61806273 | 61811801 | 5529 | 1 | ENSMUST00000131538.1 | 13681 | Tbr1 | ENSMUSG00000035033.10 |
| chr2 | 61821268 | 61821755 | 3.52405 | 4.21508 | 1.33049 | Distal Intergenic | 2 | 61806273 | 61811801 | 5529 | 1 | ENSMUST00000131538.1 | 14996 | Tbr1 | ENSMUSG00000035033.10 |
| chr2 | 61824347 | 61824793 | 4.05656 | 5.21905 | 1.89202 | Distal Intergenic | 2 | 61806273 | 61811801 | 5529 | 1 | ENSMUST00000131538.1 | 18075 | Tbr1 | ENSMUSG00000035033.10 |
| chr2 | 61825246 | 61825451 | 4.12541 | 5.4048 | 2.08789 | Distal Intergenic | 2 | 61806273 | 61811801 | 5529 | 1 | ENSMUST00000131538.1 | 18974 | Tbr1 | ENSMUSG00000035033.10 |
| chr2 | 61826387 | 61826882 | 3.58874 | 4.2937 | 1.29125 | Distal Intergenic | 2 | 61806273 | 61811801 | 5529 | 1 | ENSMUST00000131538.1 | 20115 | Tbr1 | ENSMUSG00000035033.10 |
| chr2 | 61827270 | 61827544 | 3.72853 | 4.54774 | 1.43808 | Distal Intergenic | 2 | 61806273 | 61811801 | 5529 | 1 | ENSMUST00000131538.1 | 20998 | Tbr1 | ENSMUSG00000035033.10 |
| chr2 | 61828655 | 61829050 | 3.41743 | 3.96757 | 1.06444 | Distal Intergenic | 2 | 61806273 | 61811801 | 5529 | 1 | ENSMUST00000131538.1 | 22383 | Tbr1 | ENSMUSG00000035033.10 |
| chr2 | 61830850 | 61831636 | 3.38536 | 3.91173 | 1.03841 | Distal Intergenic | 2 | 61806273 | 61811801 | 5529 | 1 | ENSMUST00000131538.1 | 24578 | Tbr1 | ENSMUSG00000035033.10 |
| chr2 | 61833545 | 61833724 | 4.07752 | 5.26792 | 1.94883 | Distal Intergenic | 2 | 61806273 | 61811801 | 5529 | 1 | ENSMUST00000131538.1 | 27273 | Tbr1 | ENSMUSG00000035033.10 |
| chr2 | 61834249 | 61834643 | 3.59078 | 4.3344 | 1.33198 | Distal Intergenic | 2 | 61806273 | 61811801 | 5529 | 1 | ENSMUST00000131538.1 | 27977 | Tbr1 | ENSMUSG00000035033.10 |
| chr2 | 61837263 | 61837598 | 3.54627 | 4.22583 | 1.29339 | Distal Intergenic | 2 | 61806273 | 61811801 | 5529 | 1 | ENSMUST00000131538.1 | 30991 | Tbr1 | ENSMUSG00000035033.10 |
| chr14 | 63578304 | 63578503 | 3.5258 | 4.17311 | 1.2131 | Distal Intergenic | 14 | 63606503 | 63820809 | 214307 | 1 | ENSMUST00000119973.2 | -28000 | Xkr6 | ENSMUSG00000035067.9 |
| chr14 | 63583248 | 63583610 | 3.69475 | 4.50975 | 1.46788 | Distal Intergenic | 14 | 63606503 | 63820809 | 214307 | 1 | ENSMUST00000119973.2 | -22893 | Xkr6 | ENSMUSG00000035067.9 |
| chr14 | 63584103 | 63584279 | 3.858 | 4.82523 | 1.69279 | Distal Intergenic | 14 | 63606503 | 63820809 | 214307 | 1 | ENSMUST00000119973.2 | -22224 | Xkr6 | ENSMUSG00000035067.9 |
| chr14 | 63584691 | 63584817 | 4.19164 | 5.78009 | 2.29669 | Distal Intergenic | 14 | 63606503 | 63820809 | 214307 | 1 | ENSMUST00000119973.2 | -21686 | Xkr6 | ENSMUSG00000035067.9 |
| chr14 | 63585970 | 63586414 | 3.65565 | 4.45047 | 1.4642 | Distal Intergenic | 14 | 63606503 | 63820809 | 214307 | 1 | ENSMUST00000119973.2 | -20089 | Xkr6 | ENSMUSG00000035067.9 |
| chr14 | 63586765 | 63586905 | 3.32013 | 4.02478 | 1.16257 | Distal Intergenic | 14 | 63606503 | 63820809 | 214307 | 1 | ENSMUST00000119973.2 | -19598 | Xkr6 | ENSMUSG00000035067.9 |
| chr14 | 63587367 | 63587843 | 3.49653 | 4.14113 | 1.24319 | Distal Intergenic | 14 | 63606503 | 63820809 | 214307 | 1 | ENSMUST00000119973.2 | -18660 | Xkr6 | ENSMUSG00000035067.9 |
| chr14 | 63589611 | 63589823 | 3.30953 | 3.80036 | 1.04105 | Distal Intergenic | 14 | 63606503 | 63820809 | 214307 | 1 | ENSMUST00000119973.2 | -16680 | Xkr6 | ENSMUSG00000035067.9 |
| chr14 | 63593721 | 63593847 | 3.3652 | 3.91289 | 1.12939 | Distal Intergenic | 14 | 63606503 | 63820809 | 214307 | 1 | ENSMUST00000119973.2 | -12656 | Xkr6 | ENSMUSG00000035067.9 |
| chr14 | 63599824 | 63599899 | 3.4875 | 4.12498 | 1.24947 | Downstream (2-3kb) | 14 | 63606503 | 63820809 | 214307 | 1 | ENSMUST00000119973.2 | -6604 | Xkr6 | ENSMUSG00000035067.9 |
| chr14 | 63601899 | 63602231 | 3.30837 | 3.79849 | 1.04784 | Downstream (<1kb) | 14 | 63606503 | 63820809 | 214307 | 1 | ENSMUST00000119973.2 | -4272 | Xkr6 | ENSMUSG00000035067.9 |
| chr14 | 63603150 | 63603240 | 3.49412 | 4.11637 | 1.18162 | Intron (ENSMUST00000137259.1/ENSMUSG00000087652.1, intron 1 of 1) | 14 | 63606503 | 63820809 | 214307 | 1 | ENSMUST00000119973.2 | -3263 | Xkr6 | ENSMUSG00000035067.9 |
| chr14 | 63547593 | 63547677 | 4.07695 | 5.27144 | 1.9971 | Distal Intergenic | 14 | 63523616 | 63543953 | 20338 | 2 | ENSMUST00000058679.5 | -3641 | Mtmr9 | ENSMUSG00000035078.5 |
| chr14 | 63549710 | 63549895 | 3.82659 | 4.85532 | 1.69688 | Distal Intergenic | 14 | 63523616 | 63543953 | 20338 | 2 | ENSMUST00000058679.5 | -5758 | Mtmr9 | ENSMUSG00000035078.5 |
| chr14 | 63551578 | 63553175 | 3.50948 | 4.20486 | 1.31011 | Distal Intergenic | 14 | 63523616 | 63543953 | 20338 | 2 | ENSMUST00000058679.5 | -7626 | Mtmr9 | ENSMUSG00000035078.5 |
| chr14 | 63554136 | 63554354 | 3.31462 | 3.79916 | 1.01733 | Distal Intergenic | 14 | 63523616 | 63543953 | 20338 | 2 | ENSMUST00000058679.5 | -10184 | Mtmr9 | ENSMUSG00000035078.5 |
| chr14 | 63563701 | 63563992 | 3.49515 | 4.17431 | 1.29843 | Distal Intergenic | 14 | 63523616 | 63543953 | 20338 | 2 | ENSMUST00000058679.5 | -19749 | Mtmr9 | ENSMUSG00000035078.5 |
| chr14 | 63571877 | 63571975 | 3.7358 | 4.61161 | 1.57564 | Distal Intergenic | 14 | 63523616 | 63543953 | 20338 | 2 | ENSMUST00000058679.5 | -27925 | Mtmr9 | ENSMUSG00000035078.5 |
| chr14 | 63572962 | 63573106 | 3.41074 | 3.96373 | 1.08697 | Distal Intergenic | 14 | 63523616 | 63543953 | 20338 | 2 | ENSMUST00000058679.5 | -29010 | Mtmr9 | ENSMUSG00000035078.5 |
| chr11 | 83442084 | 83442303 | 3.36002 | 3.87294 | 1.0364 | 3' UTR | 11 | 83440131 | 83440828 | 698 | 1 | ENSMUST00000188702.1 | 1954 | 1700020L24Rik | ENSMUSG00000035085.5 |
| chr11 | 83442763 | 83442904 | 4.01762 | 5.18341 | 1.91554 | Exon (ENSMUST00000103209.4/ENSMUSG00000020682.12, exon 8 of 8) | 11 | 83440131 | 83440828 | 698 | 1 | ENSMUST00000188702.1 | 2633 | 1700020L24Rik | ENSMUSG00000035085.5 |
| chr11 | 83444519 | 83444986 | 3.56591 | 4.30856 | 1.32626 | Exon (ENSMUST00000103209.4/ENSMUSG00000020682.12, exon 5 of 8) | 11 | 83440131 | 83440828 | 698 | 1 | ENSMUST00000188702.1 | 4389 | 1700020L24Rik | ENSMUSG00000035085.5 |
| chr14 | 63434084 | 63434364 | 3.71083 | 4.77223 | 1.62942 | Distal Intergenic | 14 | 63436394 | 63465498 | 29105 | 1 | ENSMUST00000121288.1 | -2030 | Fam167a | ENSMUSG00000035095.11 |
| chr14 | 63435731 | 63436632 | 3.75992 | 4.66586 | 1.56569 | Promoter | 14 | 63436394 | 63465498 | 29105 | 1 | ENSMUST00000121288.1 | 0 | Fam167a | ENSMUSG00000035095.11 |
| chr14 | 63449819 | 63450113 | 3.46963 | 4.06831 | 1.14229 | Intron (ENSMUST00000121288.1/ENSMUSG00000035095.11, intron 2 of 3) | 14 | 63436409 | 63442056 | 5648 | 1 | ENSMUST00000128313.1 | 13411 | Fam167a | ENSMUSG00000035095.11 |
| chr2 | 125722852 | 125723285 | 3.46258 | 4.09766 | 1.24978 | Promoter | 2 | 125627447 | 125723387 | 95941 | 2 | ENSMUST00000042246.8 | 102 | Shc4 | ENSMUSG00000035109.9 |
| chr2 | 125723623 | 125723737 | 3.50361 | 4.63701 | 1.51275 | Promoter | 2 | 125627447 | 125723387 | 95941 | 2 | ENSMUST00000042246.8 | -237 | Shc4 | ENSMUSG00000035109.9 |
| chr11 | 101265041 | 101265316 | 3.93087 | 4.96136 | 1.75615 | Promoter | 11 | 101265308 | 101274129 | 8822 | 1 | ENSMUST00000170056.1 | 0 | Wnk4 | ENSMUSG00000035112.12 |
| chr1 | 146494805 | 146494880 | 3.86655 | 4.99695 | 1.73314 | Promoter | 1 | 146494760 | 146902471 | 407712 | 1 | ENSMUST00000074622.5 | 46 | Brinp3 | ENSMUSG00000035131.9 |
| chr1 | 146497512 | 146497740 | 3.8238 | 4.97442 | 1.77229 | Promoter | 1 | 146497614 | 146751780 | 254167 | 1 | ENSMUST00000132847.2 | 0 | Brinp3 | ENSMUSG00000035131.9 |
| chr1 | 146498410 | 146498986 | 3.69026 | 4.50453 | 1.44232 | Promoter | 1 | 146497738 | 146517554 | 19817 | 1 | ENSMUST00000125331.1 | 673 | Brinp3 | ENSMUSG00000035131.9 |
| chr12 | 52222811 | 52222901 | 4.99539 | 7.331 | 3.55339 | Intron (ENSMUST00000040090.10/ENSMUSG00000035142.13, intron 6 of 10) | 12 | 52176087 | 52271079 | 94993 | 1 | ENSMUST00000162111.1 | 46725 | Nubpl | ENSMUSG00000035142.13 |
| chr7 | 100178698 | 100178970 | 3.53804 | 4.33397 | 1.34855 | Promoter | 7 | 100178683 | 100184869 | 6187 | 1 | ENSMUST00000179842.1 | 16 | Kcne3 | ENSMUSG00000035165.8 |
| chr7 | 100179735 | 100180196 | 3.60948 | 4.38181 | 1.3982 | Intron (ENSMUST00000178946.2/ENSMUSG00000035165.8, intron 2 of 4) | 7 | 100178683 | 100184869 | 6187 | 1 | ENSMUST00000179842.1 | 1053 | Kcne3 | ENSMUSG00000035165.8 |
| chr19 | 56775589 | 56776003 | 3.74637 | 4.7931 | 1.62304 | Distal Intergenic | 19 | 56790531 | 56795155 | 4625 | 2 | ENSMUST00000135666.2 | 19152 | A630007B06Rik | ENSMUSG00000035173.9 |
| chr19 | 56776321 | 56776782 | 3.42052 | 4.06162 | 1.21082 | Distal Intergenic | 19 | 56790531 | 56795155 | 4625 | 2 | ENSMUST00000135666.2 | 18373 | A630007B06Rik | ENSMUSG00000035173.9 |
| chr19 | 10486822 | 10487388 | 3.76622 | 4.75198 | 1.6018 | Distal Intergenic | 19 | 10474257 | 10482897 | 8641 | 2 | ENSMUST00000038842.3 | -3926 | Ppp1r32 | ENSMUSG00000035179.3 |
| chr5 | 101641072 | 101641244 | 3.22079 | 3.65841 | 1.01502 | Distal Intergenic | 5 | 101659177 | 101664996 | 5820 | 2 | ENSMUST00000044125.9 | 23752 | Nkx6-1 | ENSMUSG00000035187.9 |
| chr5 | 101644953 | 101645635 | 3.60172 | 4.36891 | 1.41999 | Distal Intergenic | 5 | 101659177 | 101664996 | 5820 | 2 | ENSMUST00000044125.9 | 19361 | Nkx6-1 | ENSMUSG00000035187.9 |
| chr5 | 101646391 | 101646684 | 3.60641 | 4.47517 | 1.44133 | Distal Intergenic | 5 | 101659177 | 101664996 | 5820 | 2 | ENSMUST00000044125.9 | 18312 | Nkx6-1 | ENSMUSG00000035187.9 |
| chr5 | 101647041 | 101647472 | 4.26998 | 5.74078 | 2.33994 | Distal Intergenic | 5 | 101659177 | 101664996 | 5820 | 2 | ENSMUST00000044125.9 | 17524 | Nkx6-1 | ENSMUSG00000035187.9 |
| chr5 | 101649191 | 101649438 | 3.54714 | 4.38257 | 1.375 | Distal Intergenic | 5 | 101659177 | 101664996 | 5820 | 2 | ENSMUST00000044125.9 | 15558 | Nkx6-1 | ENSMUSG00000035187.9 |
| chr5 | 101650651 | 101650805 | 3.73555 | 4.63946 | 1.56518 | Distal Intergenic | 5 | 101659177 | 101664996 | 5820 | 2 | ENSMUST00000044125.9 | 14191 | Nkx6-1 | ENSMUSG00000035187.9 |
| chr5 | 101652401 | 101652814 | 3.33908 | 3.86481 | 1.11488 | Distal Intergenic | 5 | 101659177 | 101664996 | 5820 | 2 | ENSMUST00000044125.9 | 12182 | Nkx6-1 | ENSMUSG00000035187.9 |
| chr5 | 101656100 | 101656194 | 3.51263 | 4.32862 | 1.36005 | Downstream (2-3kb) | 5 | 101659177 | 101664996 | 5820 | 2 | ENSMUST00000044125.9 | 8802 | Nkx6-1 | ENSMUSG00000035187.9 |

| | | | | | | | | | | | | | | |
|---|---|---|---|---|---|---|---|---|---|---|---|---|---|---|
| chr5 | 101657561 | 101660177 | 3.87748 | 5.19325 | 1.95257 | 3' UTR | 5 | 101659177 | 101664996 | 5820 | 2 | ENSMUST00000044125.9 | 4819 | Nkx6-1 | ENSMUSG00000035187.9 |
| chr5 | 101661038 | 101661218 | 3.86306 | 4.83948 | 1.70978 | Intron (ENSMUST00000044125.9/ENSMUSG00000035187.9, intron 2 of 2) | 5 | 101659177 | 101664996 | 5820 | 2 | ENSMUST00000044125.9 | 3778 | Nkx6-1 | ENSMUSG00000035187.9 |
| chr5 | 101661579 | 101666361 | 4.01381 | 5.25184 | 1.98167 | Promoter | 5 | 101659177 | 101664996 | 5820 | 2 | ENSMUST00000044125.9 | 0 | Nkx6-1 | ENSMUSG00000035187.9 |
| chr5 | 101666698 | 101668929 | 3.96625 | 5.15389 | 1.90172 | Distal Intergenic | 5 | 101659177 | 101664996 | 5820 | 2 | ENSMUST00000044125.9 | -1703 | Nkx6-1 | ENSMUSG00000035187.9 |
| chr5 | 101670955 | 101671135 | 3.38493 | 3.91884 | 1.06587 | Distal Intergenic | 5 | 101659177 | 101664996 | 5820 | 2 | ENSMUST00000044125.9 | -5960 | Nkx6-1 | ENSMUSG00000035187.9 |
| chr5 | 101673175 | 101673504 | 3.69383 | 4.48138 | 1.38765 | Distal Intergenic | 5 | 101659177 | 101664996 | 5820 | 2 | ENSMUST00000044125.9 | -8180 | Nkx6-1 | ENSMUSG00000035187.9 |
| chr10 | 89256009 | 89256340 | 3.73931 | 4.59737 | 1.47189 | Promoter | 10 | 89067732 | 89257136 | 189405 | 2 | ENSMUST00000181976.2 | 796 | Ano4 | ENSMUSG00000035189.11 |
| chr10 | 89256939 | 89257383 | 3.44255 | 4.01701 | 1.10426 | Promoter | 10 | 88949002 | 89257300 | 308299 | 2 | ENSMUST00000182790.2 | 0 | Ano4 | ENSMUSG00000035189.11 |
| chr10 | 89343566 | 89343702 | 3.67919 | 4.50173 | 1.49113 | Promoter | 10 | 89035091 | 89344645 | 309555 | 2 | ENSMUST00000182419.2 | 943 | Ano4 | ENSMUSG00000035189.11 |
| chr10 | 89344193 | 89344335 | 3.31338 | 3.79747 | 1.01733 | Promoter | 10 | 89035091 | 89344645 | 309555 | 2 | ENSMUST00000182419.2 | 310 | Ano4 | ENSMUSG00000035189.11 |
| chr10 | 89344854 | 89345094 | 3.35932 | 4.1494 | 1.23359 | Promoter | 10 | 89335560 | 89344762 | 9203 | 2 | ENSMUST00000180843.1 | -93 | Ano4 | ENSMUSG00000035189.11 |
| chr9 | 55044615 | 55045125 | 3.49754 | 4.11917 | 1.16916 | Intron (ENSMUST00000034854.6/ENSMUSG00000035200.6, intron 1 of 5) | 9 | 55028156 | 55048544 | 20389 | 2 | ENSMUST00000034854.6 | 3419 | Chrnb4 | ENSMUSG00000035200.6 |
| chr9 | 55048151 | 55049504 | 4.02842 | 5.20046 | 1.91968 | Promoter | 9 | 55028156 | 55048544 | 20389 | 2 | ENSMUST00000034854.6 | 0 | Chrnb4 | ENSMUSG00000035200.6 |
| chr2 | 163914110 | 163914481 | 3.50271 | 4.16487 | 1.24235 | Intron (ENSMUST00000044734.2/ENSMUSG00000035226.5, intron 1 of 5) | 2 | 163859751 | 163918683 | 58933 | 2 | ENSMUST00000044734.2 | 4202 | Rims4 | ENSMUSG00000035226.5 |
| chr2 | 163917879 | 163918003 | 3.78584 | 4.66425 | 1.52395 | Promoter | 2 | 163859751 | 163918683 | 58933 | 2 | ENSMUST00000044734.2 | 680 | Rims4 | ENSMUSG00000035226.5 |
| chr2 | 163918724 | 163919846 | 3.80163 | 4.70116 | 1.56269 | Promoter | 2 | 163859751 | 163918683 | 58933 | 2 | ENSMUST00000044734.2 | -42 | Rims4 | ENSMUSG00000035226.5 |
| chr7 | 5054471 | 5054814 | 3.77743 | 4.94828 | 1.784 | Intron (ENSMUST00000123644.1/ENSMUSG00000086604.1, intron 1 of 1) | 7 | 5056152 | 5060782 | 4631 | 1 | ENSMUST00000108571.2 | -1338 | Ccdc106 | ENSMUSG00000035228.9 |
| chr7 | 5055771 | 5056471 | 3.75378 | 4.70734 | 1.63316 | Promoter | 7 | 5056152 | 5060782 | 4631 | 1 | ENSMUST00000108571.2 | 0 | Ccdc106 | ENSMUSG00000035228.9 |
| chr7 | 5056811 | 5057228 | 3.70299 | 4.79367 | 1.66499 | Promoter | 7 | 5056856 | 5058085 | 1230 | 1 | ENSMUST00000131368.2 | 0 | Ccdc106 | ENSMUSG00000035228.9 |
| chr2 | 163851751 | 163852079 | 3.41235 | 4.02437 | 1.20193 | Distal Intergenic | 2 | 163853651 | 163858942 | 5292 | 1 | ENSMUST00000044798.3 | -1572 | Kcnk15 | ENSMUSG00000035238.4 |
| chr2 | 163855459 | 163855739 | 3.27414 | 3.93612 | 1.13388 | Intron (ENSMUST00000044798.3/ENSMUSG00000035238.4, intron 1 of 1) | 2 | 163853733 | 163859746 | 6014 | 1 | ENSMUST00000109396.1 | 1727 | Kcnk15 | ENSMUSG00000035238.4 |
| chr2 | 163857404 | 163857694 | 3.4881 | 4.15338 | 1.30126 | Intron (ENSMUST00000044798.3/ENSMUSG00000035238.4, intron 1 of 1) | 2 | 163853733 | 163859746 | 6014 | 1 | ENSMUST00000109396.1 | 3672 | Kcnk15 | ENSMUSG00000035238.4 |
| chr2 | 163858115 | 163858479 | 3.88958 | 5.08511 | 1.85094 | Exon (ENSMUST00000044798.3/ENSMUSG00000035238.4, exon 2 of 2) | 2 | 163853733 | 163859746 | 6014 | 1 | ENSMUST00000109396.1 | 4383 | Kcnk15 | ENSMUSG00000035238.4 |
| chr6 | 97102527 | 97102688 | 3.93267 | 5.0905 | 1.835 | Distal Intergenic | 6 | 97112186 | 97118754 | 6569 | 2 | ENSMUST00000130562.1 | 16066 | Eogt | ENSMUSG00000035245.7 |
| chr5 | 100719311 | 100719647 | 4.07668 | 5.33218 | 2.02763 | Promoter | 5 | 100684581 | 100719705 | 35125 | 2 | ENSMUST00000112908.1 | 58 | Hpse | ENSMUSG00000035273.9 |
| chr19 | 56719241 | 56719499 | 3.40027 | 3.95433 | 1.10939 | Distal Intergenic | 19 | 56722372 | 56724862 | 2491 | 1 | ENSMUST00000038949.4 | -2873 | Adrb1 | ENSMUSG00000035283.4 |
| chr19 | 56721316 | 56721726 | 3.59588 | 4.34485 | 1.38082 | Promoter | 19 | 56722372 | 56724862 | 2491 | 1 | ENSMUST00000038949.4 | -646 | Adrb1 | ENSMUSG00000035283.4 |
| chr19 | 56722140 | 56723983 | 3.69974 | 4.70131 | 1.60988 | Promoter | 19 | 56722372 | 56724862 | 2491 | 1 | ENSMUST00000038949.4 | 0 | Adrb1 | ENSMUSG00000035283.4 |
| chr19 | 56724324 | 56724514 | 3.83529 | 5.18584 | 1.91547 | 3' UTR | 19 | 56722372 | 56724862 | 2491 | 1 | ENSMUST00000038949.4 | 1953 | Adrb1 | ENSMUSG00000035283.4 |
| chr19 | 56725186 | 56725906 | 3.5349 | 4.2316 | 1.30485 | Distal Intergenic | 19 | 56722372 | 56724862 | 2491 | 1 | ENSMUST00000038949.4 | 2815 | Adrb1 | ENSMUSG00000035283.4 |
| chr19 | 56729765 | 56730075 | 3.34153 | 3.83966 | 1.01733 | Distal Intergenic | 19 | 56722372 | 56724862 | 2491 | 1 | ENSMUST00000038949.4 | 7394 | Adrb1 | ENSMUSG00000035283.4 |
| chr14 | 61274578 | 61274888 | 3.71726 | 4.53231 | 1.43338 | Distal Intergenic | 14 | 61221059 | 61258490 | 37432 | 2 | ENSMUST00000077954.6 | -16089 | Sgcg | ENSMUSG00000035296.8 |
| chr14 | 61275461 | 61275679 | 4.26262 | 5.61115 | 2.19148 | Distal Intergenic | 14 | 61221059 | 61258490 | 37432 | 2 | ENSMUST00000077954.6 | -16972 | Sgcg | ENSMUSG00000035296.8 |
| chr11 | 100759460 | 100760072 | 3.96511 | 5.01461 | 1.77508 | Promoter | 11 | 100740376 | 100759740 | 19365 | 2 | ENSMUST00000107361.3 | 0 | Kcnh4 | ENSMUSG00000035355.10 |
| chr6 | 101199405 | 101199953 | 3.66704 | 4.44341 | 1.39726 | Promoter | 6 | 101169856 | 101199200 | 29345 | 2 | ENSMUST00000144505.1 | -206 | Pdzrn3 | ENSMUSG00000035357.11 |
| chr6 | 101214937 | 101215114 | 3.31707 | 3.84627 | 1.07504 | Intron (ENSMUST00000075994.7/ENSMUSG00000035357.11, intron 3 of 9) | 6 | 101169856 | 101199200 | 29345 | 2 | ENSMUST00000144505.1 | -15738 | Pdzrn3 | ENSMUSG00000035357.11 |
| chr6 | 101235582 | 101235676 | 3.24678 | 3.69732 | 1.01733 | Intron (ENSMUST00000075994.7/ENSMUSG00000035357.11, intron 3 of 9) | 6 | 101169856 | 101199200 | 29345 | 2 | ENSMUST00000144505.1 | -36383 | Pdzrn3 | ENSMUSG00000035357.11 |
| chr6 | 101376795 | 101377256 | 3.77058 | 4.76658 | 1.63393 | Promoter | 6 | 101179878 | 101377342 | 197465 | 2 | ENSMUST00000151175.1 | 86 | Pdzrn3 | ENSMUSG00000035357.11 |
| chr6 | 101377822 | 101377951 | 4.19712 | 5.6508 | 2.22201 | Promoter | 6 | 101149609 | 101377897 | 228289 | 2 | ENSMUST00000075994.7 | 0 | Pdzrn3 | ENSMUSG00000035357.11 |
| chr2 | 38312888 | 38313514 | 3.95569 | 5.29779 | 2.01191 | Distal Intergenic | 2 | 37816916 | 38287390 | 470475 | 2 | ENSMUST00000140552.2 | -25499 | Dennd1a | ENSMUSG00000035392.12 |
| chr4 | 98778681 | 98778778 | 3.9137 | 5.3461 | 2.01156 | Exon (ENSMUST00000102790.3/ENSMUSG00000035407.8, exon 3 of 10) | 4 | 98756309 | 98771475 | 15167 | 2 | ENSMUST00000137270.1 | -7207 | Kank4 | ENSMUSG00000035407.8 |
| chr4 | 98780146 | 98780312 | 3.22704 | 3.67463 | 1.02476 | Exon (ENSMUST00000102790.3/ENSMUSG00000035407.8, exon 3 of 10) | 4 | 98756309 | 98771475 | 15167 | 2 | ENSMUST00000137270.1 | -8672 | Kank4 | ENSMUSG00000035407.8 |
| chr4 | 98808987 | 98809141 | 3.81647 | 4.72444 | 1.57386 | Intron (ENSMUST00000102790.3/ENSMUSG00000035407.8, intron 1 of 9) | 4 | 98754898 | 98817537 | 62640 | 2 | ENSMUST00000102790.3 | 8396 | Kank4 | ENSMUSG00000035407.8 |
| chr4 | 98817224 | 98818043 | 3.64352 | 4.64703 | 1.54353 | Promoter | 4 | 98754898 | 98817537 | 62640 | 2 | ENSMUST00000102790.3 | 0 | Kank4 | ENSMUSG00000035407.8 |
| chr12 | 58212487 | 58213914 | 3.56167 | 4.29991 | 1.34037 | Promoter | 12 | 58211804 | 58214444 | 2641 | 1 | ENSMUST00000044299.2 | 684 | Sstr1 | ENSMUSG00000035431.4 |
| chr8 | 71188444 | 71188673 | 3.08413 | 5.64045 | 2.22949 | Distal Intergenic | 8 | 71256038 | 71256653 | 616 | 2 | ENSMUST00000129455.1 | 67980 | Haus8 | ENSMUSG00000035439.8 |
| chr12 | 57533668 | 57534211 | 3.3389 | 3.90897 | 1.12135 | Distal Intergenic | 12 | 57540631 | 57546121 | 5491 | 2 | ENSMUST00000044380.6 | 11910 | Foxa1 | ENSMUSG00000035451.6 |
| chr12 | 57534525 | 57536508 | 4.10776 | 5.3985 | 2.09918 | Distal Intergenic | 12 | 57540631 | 57546121 | 5491 | 2 | ENSMUST00000044380.6 | 9613 | Foxa1 | ENSMUSG00000035451.6 |
| chr12 | 57537215 | 57538169 | 4.25643 | 5.86824 | 2.43415 | Downstream (2-3kb) | 12 | 57540631 | 57546121 | 5491 | 2 | ENSMUST00000044380.6 | 7952 | Foxa1 | ENSMUSG00000035451.6 |
| chr12 | 57538490 | 57539407 | 4.7046 | 7.40382 | 3.73688 | Downstream (1-2kb) | 12 | 57540631 | 57546121 | 5491 | 2 | ENSMUST00000044380.6 | 6714 | Foxa1 | ENSMUSG00000035451.6 |
| chr12 | 57539981 | 57540429 | 3.50314 | 4.27291 | 1.32521 | Downstream (<1kb) | 12 | 57540631 | 57546121 | 5491 | 2 | ENSMUST00000044380.6 | 5692 | Foxa1 | ENSMUSG00000035451.6 |
| chr12 | 57540907 | 57544031 | 4.06789 | 5.41894 | 2.10852 | 3' UTR | 12 | 57540631 | 57546121 | 5491 | 2 | ENSMUST00000044380.6 | 2090 | Foxa1 | ENSMUSG00000035451.6 |
| chr12 | 57544421 | 57546745 | 4.38437 | 5.94767 | 2.46703 | Promoter | 12 | 57540631 | 57546121 | 5491 | 2 | ENSMUST00000044380.6 | 0 | Foxa1 | ENSMUSG00000035451.6 |
| chr12 | 57547083 | 57547209 | 3.75984 | 4.63709 | 1.54986 | Promoter | 12 | 57540631 | 57546121 | 5491 | 2 | ENSMUST00000044380.6 | -963 | Foxa1 | ENSMUSG00000035451.6 |
| chr12 | 57547957 | 57548873 | 3.81552 | 4.78494 | 1.61067 | Distal Intergenic | 12 | 57540631 | 57546121 | 5491 | 2 | ENSMUST00000044380.6 | -1837 | Foxa1 | ENSMUSG00000035451.6 |
| chr12 | 57549453 | 57550283 | 3.8075 | 4.70441 | 1.54332 | Distal Intergenic | 12 | 57540631 | 57546121 | 5491 | 2 | ENSMUST00000044380.6 | -3333 | Foxa1 | ENSMUSG00000035451.6 |
| chr12 | 57550605 | 57550708 | 3.61824 | 4.34756 | 1.33206 | Distal Intergenic | 12 | 57540631 | 57546121 | 5491 | 2 | ENSMUST00000044380.6 | -4485 | Foxa1 | ENSMUSG00000035451.6 |
| chr12 | 57551017 | 57551567 | 4.44588 | 6.10634 | 2.55278 | Distal Intergenic | 12 | 57540631 | 57546121 | 5491 | 2 | ENSMUST00000044380.6 | -4897 | Foxa1 | ENSMUSG00000035451.6 |
| chr12 | 57552498 | 57553772 | 3.78592 | 4.7536 | 1.62561 | Distal Intergenic | 12 | 57540631 | 57546121 | 5491 | 2 | ENSMUST00000044380.6 | -6378 | Foxa1 | ENSMUSG00000035451.6 |
| chr12 | 57554198 | 57554353 | 3.77801 | 4.6375 | 1.48486 | Distal Intergenic | 12 | 57540631 | 57546121 | 5491 | 2 | ENSMUST00000044380.6 | -8078 | Foxa1 | ENSMUSG00000035451.6 |
| chr5 | 98166441 | 98170689 | 3.8455 | 4.91143 | 1.73343 | Distal Intergenic | 5 | 98180869 | 98187448 | 6580 | 1 | ENSMUST00000112959.1 | -10180 | Prdm8 | ENSMUSG00000035456.6 |
| chr5 | 98171117 | 98171931 | 3.64055 | 4.77158 | 1.60685 | Distal Intergenic | 5 | 98180869 | 98187448 | 6580 | 1 | ENSMUST00000112959.1 | -8938 | Prdm8 | ENSMUSG00000035456.6 |
| chr5 | 98172309 | 98172593 | 3.86604 | 5.33018 | 1.97529 | Distal Intergenic | 5 | 98180869 | 98187448 | 6580 | 1 | ENSMUST00000112959.1 | -8276 | Prdm8 | ENSMUSG00000035456.6 |
| chr5 | 98172943 | 98174103 | 3.93883 | 5.27625 | 1.97871 | Distal Intergenic | 5 | 98180869 | 98187448 | 6580 | 1 | ENSMUST00000112959.1 | -6766 | Prdm8 | ENSMUSG00000035456.6 |
| chr5 | 98177797 | 98177872 | 3.24221 | 3.83455 | 1.01733 | Distal Intergenic | 5 | 98180869 | 98187448 | 6580 | 1 | ENSMUST00000112959.1 | -2997 | Prdm8 | ENSMUSG00000035456.6 |
| chr5 | 98178285 | 98178486 | 3.71996 | 4.6 | 1.57045 | Distal Intergenic | 5 | 98180869 | 98187448 | 6580 | 1 | ENSMUST00000112959.1 | -2383 | Prdm8 | ENSMUSG00000035456.6 |
| chr5 | 98178892 | 98179208 | 4.24677 | 5.68736 | 2.24881 | Distal Intergenic | 5 | 98180869 | 98187448 | 6580 | 1 | ENSMUST00000112959.1 | -1661 | Prdm8 | ENSMUSG00000035456.6 |
| chr5 | 98179520 | 98179894 | 3.83881 | 4.8924 | 1.71025 | Promoter | 5 | 98180869 | 98187448 | 6580 | 1 | ENSMUST00000112959.1 | -975 | Prdm8 | ENSMUSG00000035456.6 |
| chr5 | 98180671 | 98181660 | 3.68352 | 4.57431 | 1.53037 | Promoter | 5 | 98180869 | 98187448 | 6580 | 1 | ENSMUST00000112959.1 | 0 | Prdm8 | ENSMUSG00000035456.6 |
| chr5 | 98182440 | 98183370 | 4.0736 | 5.37577 | 2.08657 | 5' UTR | 5 | 98180869 | 98187448 | 6580 | 1 | ENSMUST00000112959.1 | 1572 | Prdm8 | ENSMUSG00000035456.6 |
| chr7 | 4520584 | 4520692 | 4.0755 | 5.55476 | 2.1753 | Promoter | 7 | 4518308 | 4520567 | 2260 | 2 | ENSMUST00000126873.1 | -18 | Tnni3 | ENSMUSG00000035458.9 |
| chr7 | 4521055 | 4521764 | 3.52996 | 4.20691 | 1.29827 | Promoter | 7 | 4521042 | 4521984 | 943 | 2 | ENSMUST00000127736.1 | 220 | Tnni3 | ENSMUSG00000035458.9 |
| chr10 | 86934733 | 86934942 | 3.45401 | 4.04947 | 1.15915 | Exon (ENSMUST00000035288.9/ENSMUSG00000035459.10, exon 24 of 69) | 10 | 86918662 | 86940640 | 21979 | 2 | ENSMUST00000160580.1 | 5698 | Stab2 | ENSMUSG00000035459.10 |
| chr12 | 56824643 | 56825156 | 3.65228 | 4.57876 | 1.5001 | Intron (ENSMUST00000044634.6/ENSMUSG00000035472.8, intron 3 of 9) | 12 | 56727127 | 56775360 | 48234 | 2 | ENSMUST00000160709.1 | -49284 | Slc25a21 | ENSMUSG00000035472.8 |
| chr12 | 56956993 | 56957251 | 3.88 | 4.87193 | 1.7278 | Intron (ENSMUST00000044634.6/ENSMUSG00000035472.8, intron 1 of 9) | 12 | 56773512 | 56980393 | 206882 | 2 | ENSMUST00000160892.1 | 23142 | Slc25a21 | ENSMUSG00000035472.8 |
| chr12 | 56957662 | 56957857 | 3.73871 | 4.6543 | 1.56199 | Intron (ENSMUST00000044634.6/ENSMUSG00000035472.8, intron 1 of 9) | 12 | 56773512 | 56980393 | 206882 | 2 | ENSMUST00000160892.1 | 22536 | Slc25a21 | ENSMUSG00000035472.8 |
| chr12 | 56958253 | 56958675 | 3.32076 | 4.0302 | 1.19884 | Intron (ENSMUST00000044634.6/ENSMUSG00000035472.8, intron 1 of 9) | 12 | 56773512 | 56980393 | 206882 | 2 | ENSMUST00000160892.1 | 21718 | Slc25a21 | ENSMUSG00000035472.8 |
| chr12 | 56959415 | 56959675 | 3.82618 | 4.79506 | 1.69096 | Intron (ENSMUST00000044634.6/ENSMUSG00000035472.8, intron 1 of 9) | 12 | 56773512 | 56980393 | 206882 | 2 | ENSMUST00000160892.1 | 20718 | Slc25a21 | ENSMUSG00000035472.8 |
| chr12 | 56984534 | 56984830 | 3.53489 | 4.30831 | 1.33917 | Intron (ENSMUST00000044634.6/ENSMUSG00000035472.8, intron 1 of 9) | 12 | 56773512 | 56980393 | 206882 | 2 | ENSMUST00000160892.1 | -4142 | Slc25a21 | ENSMUSG00000035472.8 |
| chr12 | 56985636 | 56985951 | 3.41519 | 4.12626 | 1.18614 | Intron (ENSMUST00000044634.6/ENSMUSG00000035472.8, intron 1 of 9) | 12 | 56773512 | 56980393 | 206882 | 2 | ENSMUST00000160892.1 | -5244 | Slc25a21 | ENSMUSG00000035472.8 |
| chr12 | 56987906 | 56988377 | 3.66517 | 4.53449 | 1.47838 | Intron (ENSMUST00000044634.6/ENSMUSG00000035472.8, intron 1 of 9) | 12 | 56773512 | 56980393 | 206882 | 2 | ENSMUST00000160892.1 | -7514 | Slc25a21 | ENSMUSG00000035472.8 |
| chr12 | 57196826 | 57197341 | 3.91765 | 5.12267 | 1.88061 | Promoter | 12 | 56712634 | 57197472 | 484839 | 2 | ENSMUST00000044634.6 | 131 | Slc25a21 | ENSMUSG00000035472.8 |
| chr10 | 80356874 | 80357099 | 3.97838 | 5.256 | 1.95408 | Promoter | 10 | 80356459 | 80365489 | 9031 | 1 | ENSMUST00000105351.1 | 416 | Plk5 | ENSMUSG00000035486.9 |
| chr13 | 56605550 | 56605737 | 3.34961 | 3.85185 | 1.01733 | Distal Intergenic | 13 | 56609603 | 56639339 | 29737 | 1 | ENSMUST00000045173.8 | -3866 | Tgfbi | ENSMUSG00000035493.9 |
| chr13 | 56606064 | 56606816 | 3.65764 | 4.43051 | 1.39528 | Distal Intergenic | 13 | 56609603 | 56639339 | 29737 | 1 | ENSMUST00000045173.8 | -2787 | Tgfbi | ENSMUSG00000035493.9 |
| chr13 | 56607419 | 56608120 | 3.4501 | 4.0355 | 1.13059 | Distal Intergenic | 13 | 56609603 | 56639339 | 29737 | 1 | ENSMUST00000045173.8 | -1483 | Tgfbi | ENSMUSG00000035493.9 |
| chr13 | 56608936 | 56609124 | 3.54598 | 4.25734 | 1.29067 | Promoter | 13 | 56609603 | 56639339 | 29737 | 1 | ENSMUST00000045173.8 | -479 | Tgfbi | ENSMUSG00000035493.9 |
| chr13 | 56609604 | 56610313 | 3.96384 | 5.00972 | 1.74434 | Promoter | 13 | 56609603 | 56639339 | 29737 | 1 | ENSMUST00000045173.8 | 2 | Tgfbi | ENSMUSG00000035493.9 |
| chr13 | 56633369 | 56633512 | 3.58448 | 4.28968 | 1.31396 | Intron (ENSMUST00000045173.8/ENSMUSG00000035493.9, intron 12 of 16) | 13 | 56609603 | 56639339 | 29737 | 1 | ENSMUST00000045173.8 | 23767 | Tgfbi | ENSMUSG00000035493.9 |
| chr9 | 123215156 | 123215460 | 3.20377 | 3.77636 | 1.01733 | Promoter | 9 | 123180763 | 123215999 | 35237 | 2 | ENSMUST00000140915.1 | 539 | Cdcp1 | ENSMUSG00000035498.9 |
| chr9 | 123215819 | 123216164 | 3.78333 | 4.74476 | 1.62667 | Promoter | 9 | 123170824 | 123216038 | 45215 | 2 | ENSMUST00000039229.7 | 0 | Cdcp1 | ENSMUSG00000035498.9 |
| chr16 | 33554847 | 33555032 | 3.37261 | 3.88654 | 1.01733 | Intron (ENSMUST00000121925.2/ENSMUSG00000035506.11, intron 5 of 13) | 16 | 33518536 | 33663408 | 144873 | 1 | ENSMUST00000119173.2 | 36312 | Slc12a8 | ENSMUSG00000035506.11 |
| chr16 | 33576616 | 33576891 | 3.32802 | 3.84142 | 1.08354 | Intron (ENSMUST00000121925.2/ENSMUSG00000035506.11, intron 5 of 13) | 16 | 33596562 | 33664135 | 67574 | 1 | ENSMUST00000122314.2 | -19671 | Slc12a8 | ENSMUSG00000035506.11 |
| chr16 | 33607003 | 33607100 | 4.18401 | 5.54245 | 2.15499 | Intron (ENSMUST00000121925.2/ENSMUSG00000035506.11, intron 7 of 13) | 16 | 33616390 | 33624602 | 8213 | 1 | ENSMUST00000149291.1 | -9290 | Slc12a8 | ENSMUSG00000035506.11 |
| chr13 | 56502469 | 56502840 | 3.54947 | 4.26034 | 1.34838 | Distal Intergenic | 13 | 56522472 | 56527467 | 4996 | 1 | ENSMUST00000121095.2 | -19632 | Fbxl21 | ENSMUSG00000035509.12 |
| chr5 | 89524906 | 89525080 | 3.38193 | 3.9006 | 1.01733 | Distal Intergenic | 5 | 89527429 | 89583740 | 56312 | 1 | ENSMUST00000048557.2 | -2349 | Npffr2 | ENSMUSG00000035528.4 |
| chr5 | 89528724 | 89528836 | 3.71683 | 4.51457 | 1.39428 | Intron (ENSMUST00000048557.2/ENSMUSG00000035528.4, intron 1 of 3) | 5 | 89527429 | 89583740 | 56312 | 1 | ENSMUST00000048557.2 | 1296 | Npffr2 | ENSMUSG00000035528.4 |
| chr5 | 89536180 | 89536264 | 6.37818 | 10.64604 | 6.32218 | Intron (ENSMUST00000048557.2/ENSMUSG00000035528.4, intron 1 of 3) | 5 | 89527429 | 89583740 | 56312 | 1 | ENSMUST00000048557.2 | 8752 | Npffr2 | ENSMUSG00000035528.4 |
| chr4 | 45824378 | 45824720 | 3.54516 | 4.20811 | 1.2279 | Intron (ENSMUST00000044297.6/ENSMUSG00000035551.6, intron 1 of 4) | 4 | 45809468 | 45826923 | 17456 | 2 | ENSMUST00000044297.6 | 2203 | Igfbpl1 | ENSMUSG00000035551.6 |
| chr4 | 45825094 | 45825238 | 3.4217 | 3.9765 | 1.07353 | Intron (ENSMUST00000044297.6/ENSMUSG00000035551.6, intron 1 of 4) | 4 | 45809468 | 45826923 | 17456 | 2 | ENSMUST00000044297.6 | 1685 | Igfbpl1 | ENSMUSG00000035551.6 |
| chr4 | 45826236 | 45826414 | 3.75625 | 4.59017 | 1.44521 | Promoter | 4 | 45809468 | 45826923 | 17456 | 2 | ENSMUST00000044297.6 | 509 | Igfbpl1 | ENSMUSG00000035551.6 |
| chr4 | 45826771 | 45826988 | 4.34473 | 5.85134 | 2.3598 | Promoter | 4 | 45809468 | 45826923 | 17456 | 2 | ENSMUST00000044297.6 | 0 | Igfbpl1 | ENSMUSG00000035551.6 |
| chr4 | 45827391 | 45828029 | 3.64252 | 4.47081 | 1.4342 | Promoter | 4 | 45809468 | 45826923 | 17456 | 2 | ENSMUST00000044297.6 | -469 | Igfbpl1 | ENSMUSG00000035551.6 |
| chr4 | 45830475 | 45830699 | 3.42091 | 3.98353 | 1.10276 | Distal Intergenic | 4 | 45809468 | 45826923 | 17456 | 2 | ENSMUST00000044297.6 | -3553 | Igfbpl1 | ENSMUSG00000035551.6 |
| chr14 | 84438141 | 84438228 | 3.72112 | 4.56875 | 1.48529 | Distal Intergenic | 14 | 84443563 | 84537060 | 93498 | 1 | ENSMUST00000071370.5 | -5335 | Pcdh17 | ENSMUSG00000035566.6 |
| chr14 | 84440054 | 84440367 | 3.36613 | 3.87675 | 1.01733 | Distal Intergenic | 14 | 84443563 | 84537060 | 93498 | 1 | ENSMUST00000071370.5 | -3196 | Pcdh17 | ENSMUSG00000035566.6 |
| chr14 | 84441254 | 84441507 | 3.53952 | 4.19155 | 1.20045 | Distal Intergenic | 14 | 84443563 | 84537060 | 93498 | 1 | ENSMUST00000071370.5 | -2056 | Pcdh17 | ENSMUSG00000035566.6 |
| chr14 | 84441965 | 84442487 | 3.50698 | 4.13564 | 1.17585 | Distal Intergenic | 14 | 84443563 | 84537060 | 93498 | 1 | ENSMUST00000071370.5 | -1076 | Pcdh17 | ENSMUSG00000035566.6 |
| chr14 | 84443705 | 84443877 | 3.36846 | 3.88029 | 1.01733 | Promoter | 14 | 84443563 | 84537060 | 93498 | 1 | ENSMUST00000071370.5 | 143 | Pcdh17 | ENSMUSG00000035566.6 |
| chr14 | 84444401 | 84444600 | 4.02972 | 5.13266 | 1.84231 | Promoter | 14 | 84443563 | 84537060 | 93498 | 1 | ENSMUST00000071370.5 | 839 | Pcdh17 | ENSMUSG00000035566.6 |
| chr14 | 84445160 | 84445317 | 3.41535 | 3.9719 | 1.09075 | 5' UTR | 14 | 84443563 | 84537060 | 93498 | 1 | ENSMUST00000071370.5 | 1598 | Pcdh17 | ENSMUSG00000035566.6 |
| chr14 | 84446005 | 84447334 | 3.96593 | 5.18888 | 1.93648 | 5' UTR | 14 | 84443563 | 84537060 | 93498 | 1 | ENSMUST00000071370.5 | 2443 | Pcdh17 | ENSMUSG00000035566.6 |
| chr14 | 84447644 | 84448848 | 3.80788 | 4.91174 | 1.72007 | Exon (ENSMUST00000071370.5/ENSMUSG00000035566.6, exon 1 of 4) | 14 | 84443563 | 84537060 | 93498 | 1 | ENSMUST00000071370.5 | 4082 | Pcdh17 | ENSMUSG00000035566.6 |
| chr11 | 79973279 | 79973452 | 3.43069 | 4.01435 | 1.15971 | Distal Intergenic | 11 | 79933953 | 79962390 | 28438 | 2 | ENSMUST00000043152.5 | -10890 | Utp6 | ENSMUSG00000035575.6 |
| chr11 | 79974017 | 79974423 | 3.59383 | 4.30475 | 1.31026 | Distal Intergenic | 11 | 79933953 | 79962390 | 28438 | 2 | ENSMUST00000043152.5 | -11628 | Utp6 | ENSMUSG00000035575.6 |
| chr2 | 162947838 | 162948047 | 3.38193 | 3.9006 | 1.01733 | Promoter | 2 | 162948005 | 162949539 | 1535 | 1 | ENSMUST00000124264.1 | 0 | L3mbtl1 | ENSMUSG00000035576.8 |
| chr17 | 52602243 | 52602843 | 4.34884 | 5.99555 | 2.58611 | Promoter | 17 | 52602709 | 52979194 | 376486 | 1 | ENSMUST00000039366.10 | 0 | Kcnh8 | ENSMUSG00000035580.10 |
| chr17 | 52603280 | 52603622 | 3.43041 | 4.01258 | 1.15576 | Promoter | 17 | 52602709 | 52979194 | 376486 | 1 | ENSMUST00000039366.10 | 572 | Kcnh8 | ENSMUSG00000035580.10 |
| chr17 | 52603934 | 52604306 | 3.74348 | 4.60451 | 1.5432 | Intron (ENSMUST00000039366.10/ENSMUSG00000035580.10, intron 1 of 15) | 17 | 52602709 | 52979194 | 376486 | 1 | ENSMUST00000039366.10 | 1226 | Kcnh8 | ENSMUSG00000035580.10 |
| chr9 | 54978410 | 54978617 | 3.54176 | 4.22595 | 1.30867 | Distal Intergenic | 9 | 54980880 | 55007779 | 26900 | 1 | ENSMUST00000093844.3 | -2263 | Chrna5 | ENSMUSG00000035594.9 |
| chr9 | 54980662 | 54981038 | 3.58896 | 4.36831 | 1.41079 | Promoter | 9 | 54980880 | 55007779 | 26900 | 1 | ENSMUST00000093844.3 | 0 | Chrna5 | ENSMUSG00000035594.9 |
| chr9 | 54981402 | 54982387 | 3.79168 | 4.77456 | 1.63628 | Promoter | 9 | 54980880 | 55007779 | 26900 | 1 | ENSMUST00000093844.3 | 523 | Chrna5 | ENSMUSG00000035594.9 |
| chr9 | 54983684 | 54983879 | 3.30121 | 3.779 | 1.01733 | Intron (ENSMUST00000093844.3/ENSMUSG00000035594.9, intron 1 of 5) | 9 | 54980880 | 55007779 | 26900 | 1 | ENSMUST00000093844.3 | 2805 | Chrna5 | ENSMUSG00000035594.9 |
| chr9 | 102505871 | 102506666 | 3.58882 | 4.3595 | 1.38374 | Promoter | 9 | 102505750 | 102546239 | 40490 | 1 | ENSMUST00000039390.5 | 122 | Ky | ENSMUSG00000035606.8 |
| chr9 | 102515868 | 102516108 | 3.47737 | 4.09164 | 1.18418 | Intron (ENSMUST00000039390.5/ENSMUSG00000035606.8, intron 3 of 10) | 9 | 102521336 | 102528059 | 6724 | 1 | ENSMUST00000190609.1 | -5228 | Ky | ENSMUSG00000035606.8 |
| chr9 | 102531488 | 102531860 | 3.39148 | 3.96991 | 1.18831 | Intron (ENSMUST00000039390.5/ENSMUSG00000035606.8, intron 8 of 10) | 9 | 102521336 | 102528059 | 6724 | 1 | ENSMUST00000190609.1 | 10153 | Ky | ENSMUSG00000035606.8 |
| chr4 | 45186098 | 45186246 | 3.33281 | 3.83143 | 1.03455 | Intron (ENSMUST00000134280.2/ENSMUSG00000035615.7, intron 1 of 6) | 4 | 45184906 | 45285936 | 101031 | 1 | ENSMUST00000044773.6 | 1193 | Frmpd1 | ENSMUSG00000035615.7 |
| chr4 | 45186607 | 45187124 | 3.58346 | 4.28432 | 1.29905 | Intron (ENSMUST00000134280.2/ENSMUSG00000035615.7, intron 1 of 6) | 4 | 45184906 | 45285936 | 101031 | 1 | ENSMUST00000044773.6 | 1702 | Frmpd1 | ENSMUSG00000035615.7 |
| chr12 | 61519856 | 61520066 | 3.62988 | 4.37576 | 1.36375 | Distal Intergenic | 12 | 61523150 | 61605008 | 81859 | 1 | ENSMUST00000130887.1 | -3084 | Lrfn5 | ENSMUSG00000035653.11 |
| chr12 | 61521661 | 61521736 | 3.77449 | 4.63565 | 1.50627 | Distal Intergenic | 12 | 61523150 | 61605008 | 81859 | 1 | ENSMUST00000130887.1 | -1414 | Lrfn5 | ENSMUSG00000035653.11 |
| chr12 | 61523738 | 61523852 | 3.81242 | 4.7155 | 1.55829 | Promoter | 12 | 61523948 | 61845258 | 321311 | 1 | ENSMUST00000055815.8 | -96 | Lrfn5 | ENSMUSG00000035653.11 |
| chr12 | 61524163 | 61524805 | 3.84693 | 5.01442 | 1.77322 | Promoter | 12 | 61523948 | 61857928 | 333981 | 1 | ENSMUST00000119481.1 | 216 | Lrfn5 | ENSMUSG00000035653.11 |
| chr12 | 61526088 | 61526190 | 3.55806 | 4.35517 | 1.37519 | Intron (ENSMUST00000130887.1/ENSMUSG00000035653.11, intron 1 of 1) | 12 | 61523948 | 61857928 | 333981 | 1 | ENSMUST00000119481.1 | 2141 | Lrfn5 | ENSMUSG00000035653.11 |
| chr17 | 57104607 | 57104782 | 4.30338 | 5.66632 | 2.20802 | Promoter | 17 | 57105385 | 57107757 | 2373 | 1 | ENSMUST00000039490.7 | -603 | Tnfsf9 | ENSMUSG00000035678.7 |
| chr17 | 57105496 | 57105769 | 3.58017 | 4.27177 | 1.26441 | Promoter | 17 | 57105385 | 57107757 | 2373 | 1 | ENSMUST00000039490.7 | 112 | Tnfsf9 | ENSMUSG00000035678.7 |
| chr10 | 112270466 | 112270813 | 3.22474 | 4.09823 | 1.16756 | Promoter | 10 | 112271123 | 112466304 | 195182 | 1 | ENSMUST00000092175.2 | -310 | Kcnc2 | ENSMUSG00000035681.6 |
| chr10 | 112271871 | 112272151 | 3.7432 | 4.61662 | 1.57229 | Promoter | 10 | 112271123 | 112466304 | 195182 | 1 | ENSMUST00000092175.2 | 749 | Kcnc2 | ENSMUSG00000035681.6 |
| chr10 | 112272830 | 112273016 | 3.59498 | 4.35425 | 1.39455 | Intron (ENSMUST00000092175.2/ENSMUSG00000035681.6, intron 2 of 4) | 10 | 112271123 | 112466304 | 195182 | 1 | ENSMUST00000092175.2 | 1708 | Kcnc2 | ENSMUSG00000035681.6 |
| chr4 | 44247383 | 44247747 | 3.53417 | 4.20895 | 1.27728 | Distal Intergenic | 4 | 44152434 | 44233789 | 81356 | 2 | ENSMUST00000153116.1 | -13595 | Rnf38 | ENSMUSG00000035696.10 |
| chr7 | 97325120 | 97325398 | 3.71798 | 4.5193 | 1.40577 | Promoter | 7 | 97309380 | 97325964 | 16585 | 2 | ENSMUST00000168435.2 | 566 | Usp35 | ENSMUSG00000035713.10 |
| chr7 | 97325988 | 97326270 | 3.49288 | 4.1039 | 1.14267 | Promoter | 7 | 97309380 | 97325964 | 16585 | 2 | ENSMUST00000168435.2 | -25 | Usp35 | ENSMUSG00000035713.10 |
| chr19 | 10340906 | 10341093 | 3.59459 | 4.29977 | 1.29456 | Distal Intergenic | 19 | 10245265 | 10304877 | 59613 | 2 | ENSMUST00000039327.5 | -36030 | Dagla | ENSMUSG00000035735.5 |
| chr10 | 79970560 | 79971365 | 3.78872 | 4.66278 | 1.51499 | Promoter | 10 | 79970715 | 79977190 | 6476 | 1 | ENSMUST00000045085.2 | 0 | Grin3b | ENSMUSG00000035745.3 |
| chr10 | 79956865 | 79956940 | 3.80449 | 4.69951 | 1.5611 | Distal Intergenic | 10 | 79960152 | 79969246 | 9095 | 1 | ENSMUST00000045247.7 | -3212 | Wdr18 | ENSMUSG00000035754.7 |
| chr11 | 99461820 | 99462095 | 3.41453 | 4.0962 | 1.24176 | Distal Intergenic | 11 | 99428403 | 99438150 | 9748 | 2 | ENSMUST00000017743.2 | -23671 | Krt20 | ENSMUSG00000035775.2 |
| chr10 | 79918503 | 79918835 | 3.38921 | 3.92985 | 1.08205 | 5' UTR | 10 | 79910238 | 79917006 | 6769 | 2 | ENSMUST00000171416.1 | -1498 | R3hdm4 | ENSMUSG00000035781.8 |
| chr10 | 79920150 | 79920264 | 3.76862 | 4.7028 | 1.59787 | Intron (ENSMUST00000045529.2/ENSMUSG00000035773.5, intron 2 of 4) | 10 | 79910238 | 79917006 | 6769 | 2 | ENSMUST00000171416.1 | -3145 | R3hdm4 | ENSMUSG00000035781.8 |
| chr10 | 79921504 | 79922147 | 4.17373 | 5.45757 | 2.11141 | 3' UTR | 10 | 79910238 | 79917006 | 6769 | 2 | ENSMUST00000171416.1 | -4499 | R3hdm4 | ENSMUSG00000035781.8 |
| chr19 | 34252377 | 34253145 | 3.43149 | 4.01545 | 1.15971 | 5' UTR | 19 | 34241090 | 34255336 | 14247 | 2 | ENSMUST00000039631.8 | 2191 | Acta2 | ENSMUSG00000035783.8 |
| chr19 | 34253906 | 34254127 | 3.3486 | 3.86717 | 1.07666 | Intron (ENSMUST00000039631.8/ENSMUSG00000035783.8, intron 1 of 8) | 19 | 34241090 | 34255336 | 14247 | 2 | ENSMUST00000039631.8 | 1209 | Acta2 | ENSMUSG00000035783.8 |
| chr12 | 33943713 | 33943788 | 4.31997 | 6.00146 | 2.47043 | Distal Intergenic | 12 | 33957671 | 33959817 | 2147 | 1 | ENSMUST00000049089.5 | -13883 | Twist1 | ENSMUSG00000035799.5 |
| chr12 | 33950075 | 33950176 | 3.58573 | 4.35505 | 1.38059 | Distal Intergenic | 12 | 33957671 | 33959817 | 2147 | 1 | ENSMUST00000049089.5 | -7495 | Twist1 | ENSMUSG00000035799.5 |
| chr12 | 33953941 | 33954315 | 4.08907 | 5.33921 | 2.02766 | Distal Intergenic | 12 | 33957671 | 33959817 | 2147 | 1 | ENSMUST00000049089.5 | -3356 | Twist1 | ENSMUSG00000035799.5 |
| chr12 | 33954803 | 33955121 | 3.78528 | 4.85361 | 1.69583 | Distal Intergenic | 12 | 33957671 | 33959817 | 2147 | 1 | ENSMUST00000049089.5 | -2550 | Twist1 | ENSMUSG00000035799.5 |
| chr12 | 33955497 | 33956266 | 3.75607 | 4.74964 | 1.63473 | Distal Intergenic | 12 | 33957671 | 33959817 | 2147 | 1 | ENSMUST00000049089.5 | -1405 | Twist1 | ENSMUSG00000035799.5 |
| chr12 | 33956851 | 33958793 | 4.36595 | 6.101 | 2.64006 | Promoter | 12 | 33957671 | 33959817 | 2147 | 1 | ENSMUST00000049089.5 | 0 | Twist1 | ENSMUSG00000035799.5 |
| chr12 | 33959714 | 33960172 | 4.43822 | 5.98829 | 2.482 | 3' UTR | 12 | 33957671 | 33959817 | 2147 | 1 | ENSMUST00000049089.5 | 2044 | Twist1 | ENSMUSG00000035799.5 |
| chr12 | 33961200 | 33962529 | 4.43459 | 6.00975 | 2.51217 | Distal Intergenic | 12 | 33957671 | 33959817 | 2147 | 1 | ENSMUST00000049089.5 | 3530 | Twist1 | ENSMUSG00000035799.5 |
| chr12 | 33962928 | 33965225 | 3.90172 | 5.019 | 1.79013 | Distal Intergenic | 12 | 33957671 | 33959817 | 2147 | 1 | ENSMUST00000049089.5 | 5258 | Twist1 | ENSMUSG00000035799.5 |
| chr12 | 33965725 | 33965986 | 3.66127 | 4.61749 | 1.51749 | Distal Intergenic | 12 | 33957671 | 33959817 | 2147 | 1 | ENSMUST00000049089.5 | 8055 | Twist1 | ENSMUSG00000035799.5 |
| chr12 | 33966663 | 33967587 | 3.91094 | 4.96635 | 1.72696 | Distal Intergenic | 12 | 33957671 | 33959817 | 2147 | 1 | ENSMUST00000049089.5 | 8993 | Twist1 | ENSMUSG00000035799.5 |
| chr12 | 33968253 | 33969271 | 3.87063 | 4.84062 | 1.63961 | Distal Intergenic | 12 | 33957671 | 33959817 | 2147 | 1 | ENSMUST00000049089.5 | 10583 | Twist1 | ENSMUSG00000035799.5 |
| chr12 | 33970193 | 33970691 | 3.53938 | 4.20943 | 1.26799 | Distal Intergenic | 12 | 33957671 | 33959817 | 2147 | 1 | ENSMUST00000049089.5 | 12523 | Twist1 | ENSMUSG00000035799.5 |

| | | | | | | | | | | | | | | | |
|---|---|---|---|---|---|---|---|---|---|---|---|---|---|---|---|
| chr12 | 33971074 | 33971382 | 3.4726 | 4.10967 | 1.17457 | Distal Intergenic | 12 | 33957671 | 33959817 | 2147 | 1 | ENSMUST00000049089.5 | 13404 | Twist1 | ENSMUSG00000035799.5 |
| chr12 | 33973093 | 33973252 | 3.5476 | 4.31147 | 1.3443 | Distal Intergenic | 12 | 33957671 | 33959817 | 2147 | 1 | ENSMUST00000049089.5 | 15423 | Twist1 | ENSMUSG00000035799.5 |
| chr12 | 33973588 | 33974218 | 3.96665 | 5.12688 | 1.89389 | Distal Intergenic | 12 | 33957671 | 33959817 | 2147 | 1 | ENSMUST00000049089.5 | 15918 | Twist1 | ENSMUSG00000035799.5 |
| chr12 | 33975405 | 33975679 | 3.71382 | 4.57959 | 1.55751 | Distal Intergenic | 12 | 33957671 | 33959817 | 2147 | 1 | ENSMUST00000049089.5 | 17735 | Twist1 | ENSMUSG00000035799.5 |
| chr12 | 33977948 | 33978187 | 4.1263 | 5.55515 | 2.23778 | Distal Intergenic | 12 | 33957671 | 33959817 | 2147 | 1 | ENSMUST00000049089.5 | 20278 | Twist1 | ENSMUSG00000035799.5 |
| chr12 | 33978771 | 33978988 | 3.40752 | 4.03837 | 1.223 | Distal Intergenic | 12 | 33957671 | 33959817 | 2147 | 1 | ENSMUST00000049089.5 | 21101 | Twist1 | ENSMUSG00000035799.5 |
| chr12 | 33979357 | 33979551 | 3.53643 | 4.23338 | 1.33576 | Distal Intergenic | 12 | 33957671 | 33959817 | 2147 | 1 | ENSMUST00000049089.5 | 21687 | Twist1 | ENSMUSG00000035799.5 |
| chr12 | 33981211 | 33981764 | 3.31439 | 3.82229 | 1.09347 | Distal Intergenic | 12 | 33957671 | 33959817 | 2147 | 1 | ENSMUST00000049089.5 | 23541 | Twist1 | ENSMUSG00000035799.5 |
| chr12 | 33987768 | 33987843 | 3.36523 | 4.02074 | 1.11874 | Distal Intergenic | 12 | 33957671 | 33959817 | 2147 | 1 | ENSMUST00000049089.5 | 30098 | Twist1 | ENSMUSG00000035799.5 |
| chr10 | 79874855 | 79875089 | 3.57789 | 4.30668 | 1.30608 | Promoter | 10 | 79864781 | 79874634 | 9854 | 2 | ENSMUST00000092325.5 | -222 | BC005764 | ENSMUSG00000035835.9 |
| chr10 | 109008316 | 109008425 | 3.5022 | 4.13271 | 1.19531 | Promoter | 10 | 108899432 | 109009100 | 109669 | 2 | ENSMUST00000126278.1 | 675 | Syt1 | ENSMUSG00000035864.9 |
| chr10 | 109010450 | 109010812 | 3.72249 | 4.62458 | 1.57973 | Promoter | 10 | 108899215 | 109010962 | 111748 | 2 | ENSMUST00000146618.1 | 150 | Syt1 | ENSMUSG00000035864.9 |
| chr15 | 86194888 | 86195250 | 3.80198 | 12.62475 | 8.25723 | Distal Intergenic | 15 | 86139129 | 86186141 | 47013 | 2 | ENSMUST00000044332.10 | -8748 | Cerk | ENSMUSG00000035891.11 |
| chr5 | 86181564 | 86181697 | 3.33727 | 3.8333 | 1.01733 | 3' UTR | 5 | 86110720 | 86172803 | 62084 | 2 | ENSMUST00000039373.8 | -8762 | Uba6 | ENSMUSG00000035898.8 |
| chr9 | 58554061 | 58554398 | 3.31342 | 3.97915 | 1.10611 | Promoter | 9 | 58524302 | 58555132 | 30831 | 2 | ENSMUST00000165365.1 | 734 | Cd276 | ENSMUSG00000035914.5 |
| chr9 | 58555515 | 58555625 | 3.43261 | 4.01576 | 1.15137 | Promoter | 9 | 58524302 | 58555132 | 30831 | 2 | ENSMUST00000165365.1 | -384 | Cd276 | ENSMUSG00000035914.5 |
| chr10 | 107493892 | 107494126 | 3.57585 | 4.28239 | 1.32895 | Promoter | 10 | 107492860 | 107494729 | 1870 | 2 | ENSMUST00000044210.3 | 603 | Myf6 | ENSMUSG00000035923.3 |
| chr10 | 107494561 | 107495094 | 3.46629 | 4.10088 | 1.22362 | Promoter | 10 | 107492860 | 107494729 | 1870 | 2 | ENSMUST00000044210.3 | 0 | Myf6 | ENSMUSG00000035923.3 |
| chr5 | 75065001 | 75065300 | 3.48476 | 4.15369 | 1.28617 | Distal Intergenic | 5 | 75075464 | 75077625 | 2162 | 1 | ENSMUST00000160104.1 | -10164 | Gsx2 | ENSMUSG00000035946.6 |
| chr5 | 75065854 | 75066243 | 3.30199 | 3.80943 | 1.10133 | Distal Intergenic | 5 | 75075464 | 75077625 | 2162 | 1 | ENSMUST00000160104.1 | -9221 | Gsx2 | ENSMUSG00000035946.6 |
| chr5 | 75066582 | 75066682 | 4.22699 | 5.6131 | 2.24122 | Distal Intergenic | 5 | 75075464 | 75077625 | 2162 | 1 | ENSMUST00000160104.1 | -8782 | Gsx2 | ENSMUSG00000035946.6 |
| chr5 | 75066995 | 75067661 | 3.94807 | 5.22157 | 1.9344 | Distal Intergenic | 5 | 75075464 | 75077625 | 2162 | 1 | ENSMUST00000160104.1 | -7803 | Gsx2 | ENSMUSG00000035946.6 |
| chr5 | 75068484 | 75069087 | 3.87105 | 4.99572 | 1.77239 | Distal Intergenic | 5 | 75075464 | 75077625 | 2162 | 1 | ENSMUST00000160104.1 | -6377 | Gsx2 | ENSMUSG00000035946.6 |
| chr5 | 75069448 | 75069523 | 3.95832 | 5.62508 | 2.22038 | Distal Intergenic | 5 | 75075464 | 75077625 | 2162 | 1 | ENSMUST00000160104.1 | -5941 | Gsx2 | ENSMUSG00000035946.6 |
| chr5 | 75070227 | 75071304 | 3.72823 | 4.66197 | 1.59104 | Distal Intergenic | 5 | 75075464 | 75077625 | 2162 | 1 | ENSMUST00000160104.1 | -4160 | Gsx2 | ENSMUSG00000035946.6 |
| chr5 | 75071663 | 75073530 | 3.89651 | 5.06723 | 1.83251 | Distal Intergenic | 5 | 75075464 | 75077625 | 2162 | 1 | ENSMUST00000160104.1 | -1934 | Gsx2 | ENSMUSG00000035946.6 |
| chr5 | 75074791 | 75074930 | 3.75471 | 4.70415 | 1.60348 | Promoter | 5 | 75075464 | 75077625 | 2162 | 1 | ENSMUST00000160104.1 | -534 | Gsx2 | ENSMUSG00000035946.6 |
| chr5 | 75075839 | 75076310 | 3.50706 | 4.55353 | 1.47184 | Promoter | 5 | 75075601 | 75077893 | 2293 | 1 | ENSMUST00000040477.3 | 239 | Gsx2 | ENSMUSG00000035946.6 |
| chr5 | 75077035 | 75077402 | 3.99476 | 5.22808 | 1.96653 | 3' UTR | 5 | 75075601 | 75077893 | 2293 | 1 | ENSMUST00000040477.3 | 1435 | Gsx2 | ENSMUSG00000035946.6 |
| chr5 | 75077980 | 75078076 | 4.15014 | 5.49739 | 2.13731 | Distal Intergenic | 5 | 75075601 | 75077893 | 2293 | 1 | ENSMUST00000040477.3 | 2380 | Gsx2 | ENSMUSG00000035946.6 |
| chr5 | 75078772 | 75078885 | 3.5138 | 4.40611 | 1.40343 | Distal Intergenic | 5 | 75075601 | 75077893 | 2293 | 1 | ENSMUST00000040477.3 | 3172 | Gsx2 | ENSMUSG00000035946.6 |
| chr5 | 75079259 | 75079881 | 3.72332 | 4.6943 | 1.60267 | Distal Intergenic | 5 | 75075601 | 75077893 | 2293 | 1 | ENSMUST00000040477.3 | 3659 | Gsx2 | ENSMUSG00000035946.6 |
| chr5 | 75080239 | 75080939 | 3.32797 | 3.85747 | 1.12729 | Distal Intergenic | 5 | 75075601 | 75077893 | 2293 | 1 | ENSMUST00000040477.3 | 4639 | Gsx2 | ENSMUSG00000035946.6 |
| chr5 | 75081886 | 75082739 | 3.5588 | 4.39075 | 1.41276 | Distal Intergenic | 5 | 75075601 | 75077893 | 2293 | 1 | ENSMUST00000040477.3 | 6286 | Gsx2 | ENSMUSG00000035946.6 |
| chr5 | 75083314 | 75084375 | 4.05144 | 5.42875 | 2.11973 | Distal Intergenic | 5 | 75075601 | 75077893 | 2293 | 1 | ENSMUST00000040477.3 | 7714 | Gsx2 | ENSMUSG00000035946.6 |
| chr5 | 75084841 | 75086213 | 4.28852 | 5.78712 | 2.34281 | Distal Intergenic | 5 | 75075601 | 75077893 | 2293 | 1 | ENSMUST00000040477.3 | 9241 | Gsx2 | ENSMUSG00000035946.6 |
| chr5 | 75086918 | 75086993 | 4.03545 | 5.57297 | 2.18684 | Distal Intergenic | 5 | 75075601 | 75077893 | 2293 | 1 | ENSMUST00000040477.3 | 11318 | Gsx2 | ENSMUSG00000035946.6 |
| chr5 | 75087587 | 75087827 | 3.2467 | 3.75567 | 1.06235 | Distal Intergenic | 5 | 75075601 | 75077893 | 2293 | 1 | ENSMUST00000040477.3 | 11987 | Gsx2 | ENSMUSG00000035946.6 |
| chr5 | 75088418 | 75088636 | 3.57731 | 4.31351 | 1.38967 | Distal Intergenic | 5 | 75075601 | 75077893 | 2293 | 1 | ENSMUST00000040477.3 | 12818 | Gsx2 | ENSMUSG00000035946.6 |
| chr10 | 79654269 | 79654824 | 3.40724 | 3.95865 | 1.08758 | Distal Intergenic | 10 | 79639526 | 79645738 | 6213 | 2 | ENSMUST00000095464.2 | -8532 | Odf3l2 | ENSMUSG00000035963.8 |
| chr8 | 70487086 | 70487342 | 3.79563 | 5.11058 | 1.84136 | Promoter | 8 | 70483876 | 70487358 | 3483 | 2 | ENSMUST00000045286.7 | 16 | Tmem59l | ENSMUSG00000035964.7 |
| chr8 | 70487972 | 70488223 | 3.6316 | 4.7121 | 1.58565 | Promoter | 8 | 70483876 | 70487358 | 3483 | 2 | ENSMUST00000045286.7 | -615 | Tmem59l | ENSMUSG00000035964.7 |
| chr8 | 70488562 | 70488934 | 3.69814 | 4.82406 | 1.65131 | Distal Intergenic | 8 | 70483876 | 70487358 | 3483 | 2 | ENSMUST00000045286.7 | -1205 | Tmem59l | ENSMUSG00000035964.7 |
| chr8 | 70489495 | 70490209 | 4.18225 | 5.65902 | 2.25941 | Distal Intergenic | 8 | 70483876 | 70487358 | 3483 | 2 | ENSMUST00000045286.7 | -2138 | Tmem59l | ENSMUSG00000035964.7 |
| chr18 | 34560706 | 34560854 | 3.37493 | 3.89004 | 1.01733 | Downstream (1-2kb) | 18 | 34567038 | 34579083 | 12046 | 2 | ENSMUST00000154342.1 | 18229 | Nme5 | ENSMUSG00000035984.9 |
| chr13 | 24551795 | 24552146 | 3.36462 | 3.87448 | 1.01733 | Distal Intergenic | 13 | 24582189 | 24733816 | 151628 | 1 | ENSMUST00000110383.2 | -30043 | Fam65b | ENSMUSG00000036006.14 |
| chr13 | 24555312 | 24555390 | 3.52831 | 4.17184 | 1.19531 | Distal Intergenic | 13 | 24582189 | 24733816 | 151628 | 1 | ENSMUST00000110383.2 | -26799 | Fam65b | ENSMUSG00000036006.14 |
| chr13 | 24638332 | 24639041 | 3.66423 | 4.55941 | 1.51199 | Promoter | 13 | 24638648 | 24733816 | 95169 | 1 | ENSMUST00000110384.3 | 0 | Fam65b | ENSMUSG00000036006.14 |
| chr13 | 24713029 | 24713315 | 3.26808 | 3.72925 | 1.01733 | Intron (ENSMUST00000110383.2/ENSMUSG00000036006.14, intron 16 of 22) | 13 | 24710063 | 24715684 | 5622 | 1 | ENSMUST00000177489.1 | 2967 | Fam65b | ENSMUSG00000036006.14 |
| chr9 | 72798929 | 72799004 | 3.47722 | 4.17829 | 1.27371 | Distal Intergenic | 9 | 72806874 | 72917291 | 110418 | 1 | ENSMUST00000055535.8 | -7870 | Prtg | ENSMUSG00000036030.9 |
| chr9 | 72806819 | 72807184 | 3.574 | 4.48679 | 1.43807 | Promoter | 9 | 72806874 | 72917291 | 110418 | 1 | ENSMUST00000055535.8 | 0 | Prtg | ENSMUSG00000036030.9 |
| chr9 | 72913472 | 72913719 | 3.46524 | 4.28094 | 1.31916 | 3' UTR | 9 | 72910627 | 72912305 | 1679 | 1 | ENSMUST00000184274.1 | 2846 | Prtg | ENSMUSG00000036030.9 |
| chr17 | 37002628 | 37002866 | 3.35112 | 3.85411 | 1.01733 | Promoter | 17 | 37002670 | 37009746 | 7077 | 1 | ENSMUST00000172580.2 | 0 | Zfp57 | ENSMUSG00000036036.10 |
| chr4 | 42914368 | 42914595 | 3.4561 | 4.0781 | 1.23652 | Distal Intergenic | 4 | 42916660 | 42937900 | 21241 | 1 | ENSMUST00000151824.1 | -2065 | N28178 | ENSMUSG00000036062.8 |
| chr4 | 42916639 | 42917704 | 4.28847 | 5.67306 | 2.23806 | Promoter | 4 | 42916660 | 42937900 | 21241 | 1 | ENSMUST00000151824.1 | 0 | N28178 | ENSMUSG00000036062.8 |
| chr4 | 42918714 | 42919029 | 3.42324 | 4.14202 | 1.24603 | Intron (ENSMUST00000151824.1/ENSMUSG00000036062.8, intron 1 of 1) | 4 | 42917334 | 42940357 | 23024 | 1 | ENSMUST00000124380.1 | 1381 | N28178 | ENSMUSG00000036062.8 |
| chr4 | 42933362 | 42933750 | 3.60942 | 4.3255 | 1.30221 | Promoter | 4 | 42933919 | 42939513 | 5595 | 1 | ENSMUST00000138425.2 | -169 | N28178 | ENSMUSG00000036062.8 |
| chr4 | 42938096 | 42938349 | 3.70476 | 4.53243 | 1.49206 | Promoter | 4 | 42938254 | 42939833 | 1580 | 1 | ENSMUST00000131234.1 | 0 | N28178 | ENSMUSG00000036062.8 |
| chr12 | 37880569 | 37880862 | 3.44835 | 4.14678 | 1.21713 | Promoter | 12 | 37880705 | 38633410 | 752706 | 1 | ENSMUST00000040500.7 | 0 | Dgkb | ENSMUSG00000036095.10 |
| chr19 | 44883747 | 44884420 | 3.53993 | 4.20291 | 1.21997 | Distal Intergenic | 19 | 44931119 | 44983787 | 52669 | 1 | ENSMUST00000096053.3 | -46699 | Fam178a | ENSMUSG00000036097.7 |
| chr19 | 44886939 | 44887723 | 3.55724 | 4.23793 | 1.26345 | Distal Intergenic | 19 | 44931119 | 44983787 | 52669 | 1 | ENSMUST00000096053.3 | -43396 | Fam178a | ENSMUSG00000036097.7 |
| chr19 | 44889538 | 44889613 | 3.68753 | 4.47191 | 1.39234 | Distal Intergenic | 19 | 44931119 | 44983787 | 52669 | 1 | ENSMUST00000096053.3 | -41506 | Fam178a | ENSMUSG00000036097.7 |
| chr19 | 44890512 | 44890805 | 3.3266 | 3.81719 | 1.01733 | Distal Intergenic | 19 | 44931119 | 44983787 | 52669 | 1 | ENSMUST00000096053.3 | -40314 | Fam178a | ENSMUSG00000036097.7 |
| chr19 | 10224716 | 10224996 | 3.39789 | 4.02076 | 1.1555 | Intron (ENSMUST00000088013.6/ENSMUSG00000036098.10, intron 4 of 26) | 19 | 10210300 | 10217528 | 7229 | 2 | ENSMUST00000186854.1 | -7189 | Myrf | ENSMUSG00000036098.10 |
| chr19 | 10231127 | 10231339 | 3.87732 | 4.86478 | 1.71209 | Intron (ENSMUST00000088013.6/ENSMUSG00000036098.10, intron 1 of 26) | 19 | 10210385 | 10240604 | 30220 | 2 | ENSMUST00000189897.1 | 9265 | Myrf | ENSMUSG00000036098.10 |
| chr15 | 84640933 | 84641291 | 3.52809 | 4.16707 | 1.17885 | Distal Intergenic | 15 | 84669620 | 84703673 | 34054 | 1 | ENSMUST00000171460.2 | -28329 | Prr5 | ENSMUSG00000036106.10 |
| chr7 | 109168774 | 109170435 | 3.70191 | 4.60979 | 1.54997 | Promoter | 7 | 109138572 | 109170308 | 31737 | 2 | ENSMUST00000036992.7 | 0 | Lmo1 | ENSMUSG00000036111.7 |
| chr7 | 109176281 | 109176478 | 3.76721 | 4.61882 | 1.48597 | Distal Intergenic | 7 | 109138572 | 109170308 | 31737 | 2 | ENSMUST00000036992.7 | -5974 | Lmo1 | ENSMUSG00000036111.7 |
| chr7 | 109187308 | 109187653 | 3.53866 | 4.21329 | 1.27127 | Distal Intergenic | 7 | 109138572 | 109170308 | 31737 | 2 | ENSMUST00000036992.7 | -17001 | Lmo1 | ENSMUSG00000036111.7 |
| chr7 | 109196635 | 109196874 | 3.27927 | 3.74605 | 1.01733 | Distal Intergenic | 7 | 109138572 | 109170308 | 31737 | 2 | ENSMUST00000036992.7 | -26328 | Lmo1 | ENSMUSG00000036111.7 |
| chr13 | 74121148 | 74121291 | 4.2931 | 5.7208 | 2.33065 | Promoter | 13 | 74121515 | 74166064 | 44550 | 1 | ENSMUST00000036208.6 | -224 | Slc9a3 | ENSMUSG00000036123.6 |
| chr13 | 74122140 | 74122426 | 3.33473 | 3.85991 | 1.08649 | Promoter | 13 | 74121515 | 74166064 | 44550 | 1 | ENSMUST00000036208.6 | 626 | Slc9a3 | ENSMUSG00000036123.6 |
| chr12 | 31073632 | 31074151 | 4.07271 | 5.68803 | 2.26856 | Promoter | 12 | 31073968 | 31079937 | 5970 | 1 | ENSMUST00000041133.8 | 0 | Fam110c | ENSMUSG00000036136.8 |
| chr12 | 31074519 | 31075798 | 3.94219 | 5.21845 | 1.94116 | Promoter | 12 | 31073968 | 31079937 | 5970 | 1 | ENSMUST00000041133.8 | 552 | Fam110c | ENSMUSG00000036136.8 |
| chr12 | 31076199 | 31076279 | 4.0668 | 5.29685 | 2.02287 | Intron (ENSMUST00000041133.8/ENSMUSG00000036136.8, intron 1 of 1) | 12 | 31073968 | 31079937 | 5970 | 1 | ENSMUST00000041133.8 | 2232 | Fam110c | ENSMUSG00000036136.8 |
| chr12 | 31076739 | 31077022 | 3.6578 | 4.46468 | 1.46325 | Intron (ENSMUST00000041133.8/ENSMUSG00000036136.8, intron 1 of 1) | 12 | 31073968 | 31079937 | 5970 | 1 | ENSMUST00000041133.8 | 2772 | Fam110c | ENSMUSG00000036136.8 |
| chr12 | 37108614 | 37109378 | 3.90116 | 4.92912 | 1.76453 | Promoter | 12 | 37108540 | 37179534 | 70995 | 1 | ENSMUST00000041183.6 | 75 | Meox2 | ENSMUSG00000036144.6 |
| chr12 | 37109724 | 37109834 | 3.63952 | 4.59023 | 1.51588 | Intron (ENSMUST00000041183.6/ENSMUSG00000036144.6, intron 1 of 2) | 12 | 37108540 | 37179534 | 70995 | 1 | ENSMUST00000041183.6 | 1185 | Meox2 | ENSMUSG00000036144.6 |
| chr12 | 37111045 | 37111702 | 3.59633 | 4.41138 | 1.42223 | Intron (ENSMUST00000041183.6/ENSMUSG00000036144.6, intron 1 of 2) | 12 | 37108540 | 37179534 | 70995 | 1 | ENSMUST00000041183.6 | 2506 | Meox2 | ENSMUSG00000036144.6 |
| chr15 | 93763699 | 93763998 | 3.36294 | 3.87194 | 1.01733 | Distal Intergenic | 15 | 93499114 | 93595891 | 96778 | 2 | ENSMUST00000048982.5 | -167809 | Prickle1 | ENSMUSG00000036158.7 |
| chr15 | 93764357 | 93764433 | 4.35635 | 6.00034 | 2.46326 | Distal Intergenic | 15 | 93499114 | 93595891 | 96778 | 2 | ENSMUST00000048982.5 | -168467 | Prickle1 | ENSMUSG00000036158.7 |
| chr12 | 36314133 | 36314697 | 3.55456 | 4.32302 | 1.35086 | Promoter | 12 | 36314169 | 36318452 | 4284 | 1 | ENSMUST00000041407.5 | 0 | Sostdc1 | ENSMUSG00000036169.5 |
| chr19 | 19108100 | 19108738 | 3.3499 | 3.86303 | 1.0552 | Intron (ENSMUST00000112832.2/ENSMUSG00000036192.10, intron 1 of 9) | 19 | 18930605 | 19111196 | 180592 | 2 | ENSMUST00000112832.2 | 2458 | Rorb | ENSMUSG00000036192.10 |
| chr16 | 44678830 | 44678908 | 4.07375 | 5.24043 | 1.95895 | Distal Intergenic | 16 | 44724301 | 44737284 | 12984 | 1 | ENSMUST00000048788.8 | -45393 | BC027231 | ENSMUSG00000036208.8 |
| chr16 | 44685538 | 44685667 | 3.96224 | 5.0681 | 1.8628 | Distal Intergenic | 16 | 44724301 | 44737284 | 12984 | 1 | ENSMUST00000048788.8 | -38634 | BC027231 | ENSMUSG00000036208.8 |
| chr16 | 44702807 | 44702995 | 3.37651 | 3.89241 | 1.01733 | Distal Intergenic | 16 | 44724301 | 44737284 | 12984 | 1 | ENSMUST00000048788.8 | -21306 | BC027231 | ENSMUSG00000036208.8 |
| chr16 | 44703583 | 44703893 | 3.69925 | 4.48257 | 1.37519 | Distal Intergenic | 16 | 44724301 | 44737284 | 12984 | 1 | ENSMUST00000048788.8 | -20408 | BC027231 | ENSMUSG00000036208.8 |
| chr15 | 92395741 | 92396608 | 3.33483 | 3.84313 | 1.065 | Promoter | 15 | 92396915 | 92771819 | 374905 | 1 | ENSMUST00000169942.3 | -307 | Pdzrn4 | ENSMUSG00000036218.10 |
| chr15 | 92396937 | 92397509 | 4.10971 | 5.53117 | 2.156 | Promoter | 15 | 92396915 | 92771819 | 374905 | 1 | ENSMUST00000169942.3 | 23 | Pdzrn4 | ENSMUSG00000036218.10 |
| chr15 | 92520056 | 92520131 | 3.78773 | 4.81051 | 1.65118 | Intron (ENSMUST00000169942.3/ENSMUSG00000036218.10, intron 3 of 9) | 15 | 92597109 | 92771819 | 174711 | 1 | ENSMUST00000035399.8 | -76978 | Pdzrn4 | ENSMUSG00000036218.10 |
| chr15 | 92597172 | 92597279 | 3.54205 | 4.21469 | 1.27241 | Promoter | 15 | 92597109 | 92771819 | 174711 | 1 | ENSMUST00000035399.8 | 64 | Pdzrn4 | ENSMUSG00000036218.10 |
| chr15 | 92597946 | 92598521 | 4.21658 | 5.56029 | 2.16272 | Promoter | 15 | 92597109 | 92771819 | 174711 | 1 | ENSMUST00000035399.8 | 838 | Pdzrn4 | ENSMUSG00000036218.10 |
| chr15 | 92599144 | 92599513 | 3.58451 | 4.40278 | 1.36862 | Intron (ENSMUST00000169942.3/ENSMUSG00000036218.10, intron 3 of 9) | 15 | 92597109 | 92771819 | 174711 | 1 | ENSMUST00000035399.8 | 2036 | Pdzrn4 | ENSMUSG00000036218.10 |
| chr18 | 15064092 | 15064203 | 3.62561 | 4.41569 | 1.4206 | Promoter | 18 | 14968685 | 15063582 | 94898 | 2 | ENSMUST00000168989.2 | -511 | Kctd1 | ENSMUSG00000036225.9 |
| chr18 | 15064555 | 15065130 | 3.73794 | 4.73456 | 1.6074 | Promoter | 18 | 14968685 | 15063582 | 94898 | 2 | ENSMUST00000168989.2 | -974 | Kctd1 | ENSMUSG00000036225.9 |
| chr18 | 15066698 | 15067058 | 3.66927 | 4.53833 | 1.47693 | Intron (ENSMUST00000025992.6/ENSMUSG00000036225.9, intron 1 of 4) | 18 | 14968685 | 15063582 | 94898 | 2 | ENSMUST00000168989.2 | -3117 | Kctd1 | ENSMUSG00000036225.9 |
| chr18 | 15148857 | 15148992 | 3.4135 | 3.97086 | 1.09742 | Intron (ENSMUST00000025992.6/ENSMUSG00000036225.9, intron 1 of 4) | 18 | 14968685 | 15151446 | 182762 | 2 | ENSMUST00000025992.6 | 2454 | Kctd1 | ENSMUSG00000036225.9 |
| chr18 | 15150402 | 15150509 | 3.54451 | 4.42842 | 1.40391 | Promoter | 18 | 14968685 | 15151446 | 182762 | 2 | ENSMUST00000025992.6 | 937 | Kctd1 | ENSMUSG00000036225.9 |
| chr18 | 15151037 | 15151195 | 3.29255 | 3.94951 | 1.1265 | Promoter | 18 | 14968685 | 15151446 | 182762 | 2 | ENSMUST00000025992.6 | 251 | Kctd1 | ENSMUSG00000036225.9 |
| chr18 | 15151961 | 15152422 | 3.84373 | 4.97046 | 1.76254 | Promoter | 18 | 14968685 | 15151446 | 182762 | 2 | ENSMUST00000025992.6 | -516 | Kctd1 | ENSMUSG00000036225.9 |
| chr18 | 15152791 | 15153150 | 3.57223 | 4.26567 | 1.28969 | Distal Intergenic | 18 | 14968685 | 15151446 | 182762 | 2 | ENSMUST00000025992.6 | -1346 | Kctd1 | ENSMUSG00000036225.9 |
| chr11 | 52763032 | 52763470 | 3.59913 | 4.3106 | 1.29502 | Distal Intergenic | 11 | 52764634 | 53188538 | 423905 | 1 | ENSMUST00000036796.7 | -1164 | Fstl4 | ENSMUSG00000036264.9 |
| chr11 | 52764380 | 52765047 | 4.29093 | 5.69158 | 2.27629 | Promoter | 11 | 52764634 | 53188538 | 423905 | 1 | ENSMUST00000036796.7 | 0 | Fstl4 | ENSMUSG00000036264.9 |
| chr11 | 52765688 | 52765988 | 3.85501 | 4.86833 | 1.71928 | Intron (ENSMUST00000036796.7/ENSMUSG00000036264.9, intron 1 of 15) | 11 | 52764634 | 53188538 | 423905 | 1 | ENSMUST00000036796.7 | 1055 | Fstl4 | ENSMUSG00000036264.9 |
| chr11 | 52766791 | 52767135 | 3.31072 | 3.80779 | 1.06818 | Intron (ENSMUST00000036796.7/ENSMUSG00000036264.9, intron 1 of 15) | 11 | 52764634 | 53188538 | 423905 | 1 | ENSMUST00000036796.7 | 2158 | Fstl4 | ENSMUSG00000036264.9 |
| chr11 | 52767923 | 52768021 | 3.56318 | 4.36429 | 1.38702 | Intron (ENSMUST00000036796.7/ENSMUSG00000036264.9, intron 1 of 15) | 11 | 52764634 | 53188538 | 423905 | 1 | ENSMUST00000036796.7 | 3290 | Fstl4 | ENSMUSG00000036264.9 |
| chr12 | 41483062 | 41483274 | 3.60937 | 4.42205 | 1.40898 | Intron (ENSMUST00000132121.2/ENSMUSG00000056899.5, intron 4 of 6) | 12 | 41451673 | 41485751 | 34079 | 2 | ENSMUST00000043884.4 | 2477 | Lrrn3 | ENSMUSG00000036295.4 |
| chr12 | 41485082 | 41485293 | 3.69129 | 4.58024 | 1.50444 | Promoter | 12 | 41451673 | 41485751 | 34079 | 2 | ENSMUST00000043884.4 | 458 | Lrrn3 | ENSMUSG00000036295.4 |
| chr15 | 91570571 | 91570682 | 3.43505 | 4.04121 | 1.21576 | Intron (ENSMUST00000109283.1/ENSMUSG00000036298.10, intron 1 of 9) | 15 | 91542225 | 91573194 | 30970 | 2 | ENSMUST00000109282.1 | 2512 | Slc2a13 | ENSMUSG00000036298.10 |
| chr15 | 91572603 | 91572799 | 3.23314 | 3.6769 | 1.01646 | Promoter | 15 | 91542225 | 91573194 | 30970 | 2 | ENSMUST00000109282.1 | 395 | Slc2a13 | ENSMUSG00000036298.10 |
| chr16 | 43979268 | 43979669 | 4.37509 | 6.15495 | 2.63828 | Promoter | 16 | 43969078 | 43979093 | 10016 | 2 | ENSMUST00000036321.8 | -176 | Zdhhc23 | ENSMUSG00000036304.8 |
| chr8 | 69185070 | 69185479 | 3.43744 | 4.03688 | 1.20127 | Promoter | 8 | 69132669 | 69184225 | 51557 | 2 | ENSMUST00000185176.2 | -846 | Lzts1 | ENSMUSG00000036306.6 |
| chr8 | 69186467 | 69186930 | 3.4827 | 4.10922 | 1.21834 | Distal Intergenic | 8 | 69132669 | 69184225 | 51557 | 2 | ENSMUST00000185176.2 | -2243 | Lzts1 | ENSMUSG00000036306.6 |
| chr14 | 48537970 | 48538578 | 3.55503 | 4.31485 | 1.3092 | Distal Intergenic | 14 | 48446398 | 48508372 | 61975 | 1 | ENSMUST00000156480.1 | 91573 | Tmem260 | ENSMUSG00000036339.12 |
| chr14 | 48543754 | 48543963 | 3.57716 | 4.31969 | 1.37834 | Distal Intergenic | 14 | 48446398 | 48508372 | 61975 | 1 | ENSMUST00000156480.1 | 97357 | Tmem260 | ENSMUSG00000036339.12 |
| chr13 | 23404296 | 23404584 | 3.51941 | 4.3097 | 1.3105 | Distal Intergenic | 13 | 23421539 | 23423866 | 2328 | 2 | ENSMUST00000041782.2 | 19282 | Abt1 | ENSMUSG00000036376.2 |
| chr13 | 23408201 | 23408332 | 3.39671 | 3.948 | 1.10632 | Distal Intergenic | 13 | 23421539 | 23423866 | 2328 | 2 | ENSMUST00000041782.2 | 15534 | Abt1 | ENSMUSG00000036376.2 |
| chr13 | 23411389 | 23413152 | 3.80531 | 4.85447 | 1.67094 | Distal Intergenic | 13 | 23421539 | 23423866 | 2328 | 2 | ENSMUST00000041782.2 | 10714 | Abt1 | ENSMUSG00000036376.2 |
| chr5 | 76657079 | 76658497 | 4.35961 | 6.13096 | 2.67521 | Promoter | 5 | 76657683 | 76840881 | 183199 | 1 | ENSMUST00000128112.2 | 0 | C530008M17Rik | ENSMUSG00000036377.13 |
| chr5 | 76763363 | 76763493 | 3.47241 | 4.25305 | 1.30196 | Intron (ENSMUST00000128112.2/ENSMUSG00000036377.13, intron 2 of 3) | 5 | 76752368 | 76873330 | 120963 | 1 | ENSMUST00000121160.2 | 10996 | C530008M17Rik | ENSMUSG00000036377.13 |
| chr5 | 76856207 | 76856317 | 3.36462 | 3.87448 | 1.01733 | Intron (ENSMUST00000121160.2/ENSMUSG00000036377.13, intron 7 of 10) | 5 | 76858842 | 76866491 | 7650 | 1 | ENSMUST00000152373.1 | -2525 | C530008M17Rik | ENSMUSG00000036377.13 |
| chr5 | 76856710 | 76857051 | 4.20986 | 5.58439 | 2.19059 | Exon (ENSMUST00000121160.2/ENSMUSG00000036377.13, exon 8 of 11) | 5 | 76858842 | 76866491 | 7650 | 1 | ENSMUST00000152373.1 | -1791 | C530008M17Rik | ENSMUSG00000036377.13 |
| chr5 | 76857548 | 76857831 | 3.8645 | 5.10735 | 1.8549 | Exon (ENSMUST00000121160.2/ENSMUSG00000036377.13, exon 8 of 11) | 5 | 76858842 | 76866491 | 7650 | 1 | ENSMUST00000152373.1 | -1011 | C530008M17Rik | ENSMUSG00000036377.13 |
| chr5 | 76858771 | 76859046 | 3.5175 | 4.38223 | 1.37932 | Promoter | 5 | 76858842 | 76866491 | 7650 | 1 | ENSMUST00000152373.1 | 0 | C530008M17Rik | ENSMUSG00000036377.13 |
| chr18 | 60911938 | 60912138 | 3.40302 | 3.95043 | 1.07981 | Promoter | 18 | 60911780 | 60918561 | 6782 | 1 | ENSMUST00000040359.5 | 159 | Arsi | ENSMUSG00000036412.5 |
| chr14 | 79767349 | 79767860 | 3.62569 | 4.35952 | 1.3276 | 3' UTR | 14 | 79766772 | 79771312 | 4541 | 2 | ENSMUST00000039568.5 | 3452 | Pcdh8 | ENSMUSG00000036422.5 |
| chr14 | 79768603 | 79772965 | 4.08285 | 5.35734 | 2.03418 | Promoter | 14 | 79766772 | 79771312 | 4541 | 2 | ENSMUST00000039568.5 | 0 | Pcdh8 | ENSMUSG00000036422.5 |
| chr14 | 79773940 | 79774310 | 3.75884 | 4.63336 | 1.52878 | Distal Intergenic | 14 | 79766772 | 79771312 | 4541 | 2 | ENSMUST00000039568.5 | -2629 | Pcdh8 | ENSMUSG00000036422.5 |
| chr14 | 79774770 | 79775175 | 4.03603 | 5.25064 | 1.96638 | Distal Intergenic | 14 | 79766772 | 79771312 | 4541 | 2 | ENSMUST00000039568.5 | -3459 | Pcdh8 | ENSMUSG00000036422.5 |
| chr8 | 66697672 | 66697836 | 3.58857 | 4.39892 | 1.39865 | Promoter | 8 | 66697422 | 66706798 | 9377 | 1 | ENSMUST00000039303.5 | 251 | Npy1r | ENSMUSG00000036437.5 |
| chr19 | 44608361 | 44608498 | 3.3585 | 3.86957 | 1.03258 | Distal Intergenic | 19 | 44562850 | 44576274 | 13425 | 1 | ENSMUST00000040455.4 | 45512 | Hif1an | ENSMUSG00000036450.4 |
| chr9 | 64383305 | 64383527 | 3.34671 | 3.84751 | 1.01733 | Distal Intergenic | 9 | 64385626 | 64700975 | 315350 | 1 | ENSMUST00000093829.3 | -2099 | Megf11 | ENSMUSG00000036466.12 |
| chr9 | 64384344 | 64384861 | 3.71419 | 4.52107 | 1.41846 | Promoter | 9 | 64385626 | 64700975 | 315350 | 1 | ENSMUST00000093829.3 | -765 | Megf11 | ENSMUSG00000036466.12 |
| chr9 | 64385660 | 64387428 | 4.17775 | 5.50777 | 2.13668 | Promoter | 9 | 64385683 | 64649845 | 264163 | 1 | ENSMUST00000148449.2 | 0 | Megf11 | ENSMUSG00000036466.12 |
| chr9 | 64387838 | 64387989 | 3.55409 | 4.29273 | 1.29494 | Intron (ENSMUST00000093829.3/ENSMUSG00000036466.12, intron 1 of 22) | 9 | 64385686 | 64709205 | 323520 | 1 | ENSMUST00000068967.5 | 2153 | Megf11 | ENSMUSG00000036466.12 |
| chr9 | 64475974 | 64476086 | 3.29587 | 3.78078 | 1.05082 | Intron (ENSMUST00000093829.3/ENSMUSG00000036466.12, intron 1 of 22) | 9 | 64501336 | 64509072 | 7737 | 1 | ENSMUST00000148963.2 | -25250 | Megf11 | ENSMUSG00000036466.12 |
| chr9 | 64508633 | 64508932 | 3.36931 | 3.88156 | 1.01733 | Exon (ENSMUST00000093829.3/ENSMUSG00000036466.12, exon 4 of 23) | 9 | 64501601 | 64687753 | 186153 | 1 | ENSMUST00000144383.1 | 7033 | Megf11 | ENSMUSG00000036466.12 |
| chr9 | 64509772 | 64509891 | 3.51612 | 4.1535 | 1.19531 | Intron (ENSMUST00000093829.3/ENSMUSG00000036466.12, intron 4 of 22) | 9 | 64501601 | 64687753 | 186153 | 1 | ENSMUST00000144383.1 | 8172 | Megf11 | ENSMUSG00000036466.12 |
| chr9 | 64511656 | 64511827 | 3.43744 | 4.00308 | 1.08325 | Intron (ENSMUST00000093829.3/ENSMUSG00000036466.12, intron 4 of 22) | 9 | 64501601 | 64687753 | 186153 | 1 | ENSMUST00000144383.1 | 10056 | Megf11 | ENSMUSG00000036466.12 |
| chr9 | 64560182 | 64560519 | 3.38193 | 3.9006 | 1.01733 | Intron (ENSMUST00000093829.3/ENSMUSG00000036466.12, intron 4 of 22) | 9 | 64546899 | 64638466 | 91568 | 1 | ENSMUST00000147388.1 | 13284 | Megf11 | ENSMUSG00000036466.12 |
| chr8 | 66312617 | 66312722 | 13.80489 | 32.10509 | 26.5475 | Intron (ENSMUST00000152320.2/ENSMUSG00000036469.11, intron 5 of 8) | 8 | 66386294 | 66470421 | 84128 | 1 | ENSMUST00000039540.6 | -73572 | Mar-01 | ENSMUSG00000036469.11 |
| chr17 | 24206152 | 24206334 | 4.04864 | 5.18677 | 1.89784 | Promoter | 17 | 24175431 | 24205562 | 30132 | 2 | ENSMUST00000168410.3 | -591 | Tbc1d24 | ENSMUSG00000036473.10 |
| chr17 | 24206787 | 24206976 | 4.22095 | 5.59463 | 2.19841 | Exon (ENSMUST00000024931.4/ENSMUSG00000079662.1, exon 6 of 6) | 17 | 24175431 | 24205562 | 30132 | 2 | ENSMUST00000168410.3 | -1226 | Tbc1d24 | ENSMUSG00000036473.10 |
| chr1 | 87169825 | 87170874 | 3.87223 | 4.88377 | 1.71008 | Distal Intergenic | 1 | 87183313 | 87188405 | 5093 | 1 | ENSMUST00000044533.8 | -12439 | Prss56 | ENSMUSG00000036480.9 |
| chr1 | 87172290 | 87172402 | 3.56459 | 4.29294 | 1.33513 | Distal Intergenic | 1 | 87183313 | 87188405 | 5093 | 1 | ENSMUST00000044533.8 | -10911 | Prss56 | ENSMUSG00000036480.9 |
| chr1 | 87174576 | 87174686 | 3.96138 | 5.37828 | 2.04412 | Distal Intergenic | 1 | 87183313 | 87188405 | 5093 | 1 | ENSMUST00000044533.8 | -8627 | Prss56 | ENSMUSG00000036480.9 |
| chr1 | 87176003 | 87176128 | 3.52891 | 4.21245 | 1.26377 | Distal Intergenic | 1 | 87183313 | 87188405 | 5093 | 1 | ENSMUST00000044533.8 | -7185 | Prss56 | ENSMUSG00000036480.9 |
| chr1 | 87182938 | 87183192 | 3.38193 | 3.9006 | 1.01733 | Promoter | 1 | 87183313 | 87188405 | 5093 | 1 | ENSMUST00000044533.8 | -121 | Prss56 | ENSMUSG00000036480.9 |
| chr1 | 87183704 | 87184237 | 3.76602 | 4.6866 | 1.53558 | Promoter | 1 | 87183313 | 87188405 | 5093 | 1 | ENSMUST00000044533.8 | 392 | Prss56 | ENSMUSG00000036480.9 |
| chr1 | 87184762 | 87185427 | 3.8477 | 4.96131 | 1.75701 | Exon (ENSMUST00000044533.8/ENSMUSG00000036480.9, exon 4 of 13) | 1 | 87183313 | 87188405 | 5093 | 1 | ENSMUST00000044533.8 | 1450 | Prss56 | ENSMUSG00000036480.9 |
| chr1 | 87185805 | 87187081 | 3.73301 | 4.65852 | 1.56989 | Exon (ENSMUST00000044533.8/ENSMUSG00000036480.9, exon 7 of 13) | 1 | 87183313 | 87188405 | 5093 | 1 | ENSMUST00000044533.8 | 2493 | Prss56 | ENSMUSG00000036480.9 |
| chr17 | 36943141 | 36943492 | 4.1202 | 5.53576 | 2.14329 | Promoter | 17 | 36943055 | 36945615 | 2561 | 1 | ENSMUST00000173072.1 | 87 | Rnf39 | ENSMUSG00000036492.7 |
| chr1 | 87128604 | 87128697 | 3.39536 | 3.93598 | 1.07072 | Distal Intergenic | 1 | 87125008 | 87127912 | 2905 | 1 | ENSMUST00000044878.3 | 3597 | Akp3 | ENSMUSG00000036500.4 |
| chr8 | 99415311 | 99415548 | 3.41353 | 4.04399 | 1.17569 | Promoter | 8 | 99028961 | 99416319 | 387359 | 2 | ENSMUST00000145601.2 | 771 | Cdh8 | ENSMUSG00000036510.12 |
| chr12 | 16753093 | 16753351 | 3.61894 | 4.36729 | 1.40087 | Intron (ENSMUST00000048064.10/ENSMUSG00000036523.11, intron 1 of 32) | 12 | 16672079 | 16757195 | 85117 | 2 | ENSMUST00000159120.2 | 3844 | Greb1 | ENSMUSG00000036523.11 |
| chr12 | 16800339 | 16800521 | 4.68408 | 6.59916 | 2.95291 | Promoter | 12 | 16670615 | 16800684 | 130070 | 2 | ENSMUST00000162112.2 | 163 | Greb1 | ENSMUSG00000036523.11 |
| chr11 | 50586108 | 50586294 | 3.71431 | 4.57738 | 1.55653 | Distal Intergenic | 11 | 50602084 | 50807573 | 205490 | 1 | ENSMUST00000040523.8 | -15790 | Adamts2 | ENSMUSG00000036545.8 |
| chr11 | 50593052 | 50593326 | 3.66735 | 4.43413 | 1.38176 | Distal Intergenic | 11 | 50602084 | 50807573 | 205490 | 1 | ENSMUST00000040523.8 | -8758 | Adamts2 | ENSMUSG00000036545.8 |
| chr11 | 50594056 | 50594391 | 3.8686 | 4.87522 | 1.70634 | Distal Intergenic | 11 | 50602084 | 50807573 | 205490 | 1 | ENSMUST00000040523.8 | -7693 | Adamts2 | ENSMUSG00000036545.8 |
| chr11 | 50595769 | 50596034 | 3.22893 | 3.79809 | 1.04319 | Distal Intergenic | 11 | 50602084 | 50807573 | 205490 | 1 | ENSMUST00000040523.8 | -6050 | Adamts2 | ENSMUSG00000036545.8 |
| chr11 | 50596423 | 50596702 | 3.25709 | 3.71276 | 1.01733 | Downstream (2-3kb) | 11 | 50602084 | 50807573 | 205490 | 1 | ENSMUST00000040523.8 | -5382 | Adamts2 | ENSMUSG00000036545.8 |
| chr11 | 50600835 | 50601154 | 3.64874 | 4.58561 | 1.49872 | Promoter | 11 | 50602084 | 50807573 | 205490 | 1 | ENSMUST00000040523.8 | -930 | Adamts2 | ENSMUSG00000036545.8 |
| chr11 | 50763189 | 50763357 | 3.62864 | 4.35039 | 1.28997 | Intron (ENSMUST00000040523.8/ENSMUSG00000036545.8, intron 5 of 21) | 11 | 50776179 | 50777015 | 837 | 1 | ENSMUST00000125988.1 | -12822 | Adamts2 | ENSMUSG00000036545.8 |

| | | | | | | | | | | | | | | | |
|---|---|---|---|---|---|---|---|---|---|---|---|---|---|---|---|
| chr8 | 95677037 | 95679148 | 4.14628 | 5.53757 | 2.19233 | Promoter | 8 | 95678085 | 95714956 | 36872 | 1 | ENSMUST00000041318.8 | 0 | Ndrg4 | ENSMUSG00000036564.11 |
| chr8 | 95695041 | 95695574 | 3.612 | 4.32477 | 1.2884 | Intron (ENSMUST00000160964.2/ENSMUSG00000036564.11, intron 2 of 10) | 8 | 95703037 | 95715119 | 12083 | 1 | ENSMUST00000073139.8 | -7463 | Ndrg4 | ENSMUSG00000036564.11 |
| chr8 | 95697198 | 95697484 | 3.61179 | 4.33714 | 1.33515 | 5' UTR | 8 | 95703037 | 95715119 | 12083 | 1 | ENSMUST00000073139.8 | -5553 | Ndrg4 | ENSMUSG00000036564.11 |
| chr7 | 31053843 | 31054100 | 3.61356 | 4.34235 | 1.33302 | Promoter | 7 | 31051681 | 31054853 | 3173 | 2 | ENSMUST00000071697.5 | 753 | Fxyd1 | ENSMUSG00000036570.9 |
| chr7 | 31049100 | 31050991 | 3.7573 | 4.6755 | 1.54457 | Promoter | 7 | 31042517 | 31051462 | 8946 | 2 | ENSMUST00000073892.4 | 471 | Fxyd7 | ENSMUSG00000036578.6 |
| chr7 | 31051843 | 31053010 | 3.51601 | 4.21573 | 1.24112 | Promoter | 7 | 31042517 | 31051462 | 8946 | 2 | ENSMUST00000073892.4 | -382 | Fxyd7 | ENSMUSG00000036578.6 |
| chr18 | 38930017 | 38930098 | 4.23208 | 5.5857 | 2.23054 | Promoter | 18 | 38838675 | 38929404 | 90730 | 2 | ENSMUST00000131348.2 | -614 | Fgf1 | ENSMUSG00000036585.10 |
| chr2 | 20899893 | 20900204 | 4.38176 | 11.36285 | 7.1396 | Intron (ENSMUST00000154230.3/ENSMUSG00000036591.10, intron 3 of 24) | 2 | 20886358 | 20914694 | 28337 | 2 | ENSMUST00000156142.2 | 14490 | Arhgap21 | ENSMUSG00000036591.10 |
| chr5 | 35465487 | 35465849 | 3.69217 | 4.48669 | 1.41784 | Distal Intergenic | 5 | 35517565 | 35525571 | 8007 | 2 | ENSMUST00000132959.1 | 59722 | Cpz | ENSMUSG00000036596.5 |
| chr5 | 35467699 | 35467774 | 3.60996 | 4.33713 | 1.3351 | Distal Intergenic | 5 | 35517565 | 35525571 | 8007 | 2 | ENSMUST00000132959.1 | 57797 | Cpz | ENSMUSG00000036596.5 |
| chr5 | 35489028 | 35489333 | 3.68 | 4.49692 | 1.43589 | Distal Intergenic | 5 | 35517565 | 35525571 | 8007 | 2 | ENSMUST00000132959.1 | 36238 | Cpz | ENSMUSG00000036596.5 |
| chr8 | 95528570 | 95528930 | 3.41599 | 3.96426 | 1.06048 | Distal Intergenic | 8 | 95534100 | 95558888 | 24789 | 1 | ENSMUST00000041569.3 | -5170 | Ccdc113 | ENSMUSG00000036598.3 |
| chr5 | 140508377 | 140508708 | 3.32679 | 3.85489 | 1.11815 | Intron (ENSMUST00000043050.8/ENSMUSG00000036599.10, intron 1 of 1) | 5 | 140505575 | 140524153 | 18579 | 1 | ENSMUST00000124142.1 | 2803 | Chst12 | ENSMUSG00000036599.10 |
| chr10 | 103022136 | 103022316 | 3.73279 | 4.61785 | 1.5775 | Promoter | 10 | 103008151 | 103022574 | 14424 | 2 | ENSMUST00000164226.2 | 258 | Alx1 | ENSMUSG00000036602.8 |
| chr10 | 103023760 | 103025076 | 4.29906 | 5.85283 | 2.40655 | Intron (ENSMUST00000167156.3/ENSMUSG00000036602.8, intron 3 of 4) | 10 | 103008151 | 103022574 | 14424 | 2 | ENSMUST00000164226.2 | -1187 | Alx1 | ENSMUSG00000036602.8 |
| chr10 | 103025897 | 103026131 | 3.82578 | 4.77352 | 1.61255 | Intron (ENSMUST00000167156.3/ENSMUSG00000036602.8, intron 2 of 4) | 10 | 103007847 | 103028777 | 20931 | 2 | ENSMUST00000040859.5 | 2646 | Alx1 | ENSMUSG00000036602.8 |
| chr10 | 103026971 | 103028763 | 4.26762 | 5.71866 | 2.30566 | Promoter | 10 | 103007847 | 103028777 | 20931 | 2 | ENSMUST00000040859.5 | 14 | Alx1 | ENSMUSG00000036602.8 |
| chr2 | 19909667 | 19910540 | 4.02117 | 5.25921 | 1.96135 | Promoter | 2 | 19909780 | 20744816 | 835037 | 1 | ENSMUST00000114610.2 | 0 | Etl4 | ENSMUSG00000036617.11 |
| chr15 | 73489628 | 73490059 | 3.34216 | 3.88043 | 1.12487 | Distal Intergenic | 15 | 73512560 | 73572242 | 59683 | 1 | ENSMUST00000043414.6 | -22501 | Dennd3 | ENSMUSG00000036661.8 |
| chr15 | 102354763 | 102355269 | 3.37226 | 3.88601 | 1.01733 | Downstream (1-2kb) | 15 | 102338247 | 102350759 | 12513 | 2 | ENSMUST00000041208.7 | -4005 | Aaas | ENSMUSG00000036678.7 |
| chr15 | 102355747 | 102356185 | 4.09037 | 5.29322 | 1.94266 | Downstream (<1kb) | 15 | 102338247 | 102350759 | 12513 | 2 | ENSMUST00000041208.7 | -4989 | Aaas | ENSMUSG00000036678.7 |
| chr15 | 102356743 | 102357216 | 3.7071 | 4.54601 | 1.47034 | 3' UTR | 15 | 102338247 | 102350759 | 12513 | 2 | ENSMUST00000041208.7 | -5985 | Aaas | ENSMUSG00000036678.7 |
| chr5 | 139658799 | 139659033 | 3.71507 | 4.56002 | 1.46139 | Distal Intergenic | 5 | 139706886 | 139710227 | 3342 | 2 | ENSMUST00000164584.2 | 51194 | Micall2 | ENSMUSG00000036718.12 |
| chr5 | 139659370 | 139659756 | 3.51794 | 4.36432 | 1.33679 | Distal Intergenic | 5 | 139706886 | 139710227 | 3342 | 2 | ENSMUST00000164584.2 | 50471 | Micall2 | ENSMUSG00000036718.12 |
| chr5 | 139660497 | 139660688 | 3.52965 | 4.20751 | 1.29515 | Distal Intergenic | 5 | 139706886 | 139710227 | 3342 | 2 | ENSMUST00000164584.2 | 49539 | Micall2 | ENSMUSG00000036718.12 |
| chr5 | 139668105 | 139669501 | 3.80112 | 4.70701 | 1.5593 | Distal Intergenic | 5 | 139706886 | 139710227 | 3342 | 2 | ENSMUST00000164584.2 | 40726 | Micall2 | ENSMUSG00000036718.12 |
| chr5 | 139669892 | 139670232 | 3.54686 | 4.24101 | 1.2926 | Distal Intergenic | 5 | 139706886 | 139710227 | 3342 | 2 | ENSMUST00000164584.2 | 39995 | Micall2 | ENSMUSG00000036718.12 |
| chr5 | 139675389 | 139675464 | 3.72826 | 4.5413 | 1.42649 | Distal Intergenic | 5 | 139706886 | 139710227 | 3342 | 2 | ENSMUST00000164584.2 | 34763 | Micall2 | ENSMUSG00000036718.12 |
| chr5 | 139680691 | 139681090 | 3.51915 | 4.21331 | 1.24137 | Distal Intergenic | 5 | 139706886 | 139710227 | 3342 | 2 | ENSMUST00000164584.2 | 29137 | Micall2 | ENSMUSG00000036718.12 |
| chr5 | 139682595 | 139682790 | 3.39368 | 3.92587 | 1.04403 | Distal Intergenic | 5 | 139706886 | 139710227 | 3342 | 2 | ENSMUST00000164584.2 | 27437 | Micall2 | ENSMUSG00000036718.12 |
| chr5 | 139686905 | 139687045 | 3.41086 | 3.98443 | 1.16035 | Distal Intergenic | 5 | 139706886 | 139710227 | 3342 | 2 | ENSMUST00000164584.2 | 23182 | Micall2 | ENSMUSG00000036718.12 |
| chr5 | 139687705 | 139687780 | 3.90174 | 4.94396 | 1.75885 | Distal Intergenic | 5 | 139706886 | 139710227 | 3342 | 2 | ENSMUST00000164584.2 | 22447 | Micall2 | ENSMUSG00000036718.12 |
| chr2 | 25241560 | 25242891 | 3.52544 | 4.20539 | 1.22839 | Promoter | 2 | 25239157 | 25242452 | 3296 | 2 | ENSMUST00000114356.2 | 0 | 2310002J15Rik | ENSMUSG00000036731.7 |
| chr3 | 147030168 | 147030350 | 3.29561 | 3.77058 | 1.01733 | Distal Intergenic | 3 | 146944047 | 146984009 | 39963 | 1 | ENSMUST00000037942.6 | 86122 | Ttll7 | ENSMUSG00000036745.10 |
| chr3 | 147098127 | 147098300 | 3.42025 | 3.98126 | 1.09823 | Distal Intergenic | 3 | 146944047 | 146984009 | 39963 | 1 | ENSMUST00000037942.6 | 154081 | Ttll7 | ENSMUSG00000036745.10 |
| chr3 | 147182516 | 147182740 | 3.56062 | 4.29749 | 1.31999 | Distal Intergenic | 3 | 146944047 | 146984009 | 39963 | 1 | ENSMUST00000037942.6 | 238470 | Ttll7 | ENSMUSG00000036745.10 |
| chr15 | 72546107 | 72547224 | 4.09706 | 5.30037 | 1.97532 | Promoter | 15 | 72512119 | 72546279 | 34161 | 2 | ENSMUST00000044624.6 | 0 | Kcnk9 | ENSMUSG00000036760.6 |
| chr15 | 72547591 | 72547863 | 4.57446 | 6.24559 | 2.62647 | Distal Intergenic | 15 | 72512119 | 72546279 | 34161 | 2 | ENSMUST00000044624.6 | -1313 | Kcnk9 | ENSMUSG00000036760.6 |
| chr1 | 84710385 | 84710634 | 3.35854 | 3.86532 | 1.01733 | Distal Intergenic | 1 | 84369839 | 84696221 | 326383 | 2 | ENSMUST00000049126.7 | -14165 | Dner | ENSMUSG00000036766.7 |
| chr15 | 71729071 | 71729272 | 3.27808 | 3.74426 | 1.01733 | Distal Intergenic | 15 | 71445678 | 71727838 | 282161 | 2 | ENSMUST00000022953.8 | -1234 | Fam135b | ENSMUSG00000036800.7 |
| chr1 | 124045928 | 124046440 | 3.87425 | 4.82556 | 1.61732 | Promoter | 1 | 123321471 | 124045559 | 724089 | 2 | ENSMUST00000112606.2 | -370 | Dpp10 | ENSMUSG00000036815.11 |
| chr10 | 63099852 | 63100054 | 3.3616 | 3.87354 | 1.03004 | Promoter | 10 | 63100156 | 63100605 | 450 | 1 | ENSMUST00000044059.3 | -102 | Atoh7 | ENSMUSG00000036816.3 |
| chr10 | 63100457 | 63100792 | 4.70919 | 6.71591 | 3.03647 | Promoter | 10 | 63100156 | 63100605 | 450 | 1 | ENSMUST00000044059.3 | 302 | Atoh7 | ENSMUSG00000036816.3 |
| chr3 | 146218637 | 146218741 | 3.73212 | 4.54797 | 1.4339 | Distal Intergenic | 3 | 146220961 | 146286214 | 65254 | 1 | ENSMUST00000039164.2 | -2220 | Lpar3 | ENSMUSG00000036832.4 |
| chr3 | 146220988 | 146221194 | 3.45654 | 4.08649 | 1.17828 | Promoter | 3 | 146220961 | 146286214 | 65254 | 1 | ENSMUST00000039164.2 | 28 | Lpar3 | ENSMUSG00000036832.4 |
| chr3 | 146221961 | 146222216 | 3.71031 | 4.52284 | 1.4524 | Intron (ENSMUST00000039164.2/ENSMUSG00000036832.4, intron 1 of 2) | 3 | 146220961 | 146286214 | 65254 | 1 | ENSMUST00000039164.2 | 1001 | Lpar3 | ENSMUSG00000036832.4 |
| chr3 | 146222852 | 146223355 | 3.71226 | 4.65635 | 1.5449 | Intron (ENSMUST00000039164.2/ENSMUSG00000036832.4, intron 1 of 2) | 3 | 146220961 | 146286214 | 65254 | 1 | ENSMUST00000039164.2 | 1892 | Lpar3 | ENSMUSG00000036832.4 |
| chr8 | 86789802 | 86790486 | 3.74425 | 4.74538 | 1.6197 | Distal Intergenic | 8 | 86723940 | 86746006 | 22067 | 2 | ENSMUST00000045296.4 | -43797 | Siah1a | ENSMUSG00000036840.4 |
| chr3 | 146117388 | 146118223 | 3.49779 | 4.14472 | 1.23934 | Promoter | 3 | 146117459 | 146124806 | 7348 | 1 | ENSMUST00000140214.1 | 0 | Mcoln3 | ENSMUSG00000036853.4 |
| chr4 | 137277233 | 137277328 | 3.86571 | 4.80474 | 1.5998 | Promoter | 4 | 137277489 | 137299726 | 22238 | 1 | ENSMUST00000045747.4 | -161 | Wnt4 | ENSMUSG00000036856.4 |
| chr4 | 137277760 | 137278593 | 3.54201 | 4.2313 | 1.24129 | Promoter | 4 | 137277489 | 137299726 | 22238 | 1 | ENSMUST00000045747.4 | 272 | Wnt4 | ENSMUSG00000036856.4 |
| chr7 | 105786786 | 105786961 | 3.70929 | 4.57191 | 1.55573 | Promoter | 7 | 105752989 | 105787550 | 34562 | 2 | ENSMUST00000078482.6 | 589 | Dchs1 | ENSMUSG00000036862.9 |
| chr7 | 105787338 | 105787846 | 4.02363 | 5.27853 | 2.00345 | Promoter | 7 | 105752989 | 105787550 | 34562 | 2 | ENSMUST00000078482.6 | 0 | Dchs1 | ENSMUSG00000036862.9 |
| chr18 | 74822959 | 74823514 | 3.71075 | 4.50871 | 1.39932 | Distal Intergenic | 18 | 74779197 | 74806207 | 27011 | 1 | ENSMUST00000041053.9 | 43763 | Acaa2 | ENSMUSG00000036880.9 |
| chr4 | 136896027 | 136896626 | 3.85358 | 4.88666 | 1.69388 | 3' UTR | 4 | 136895917 | 136898803 | 2887 | 2 | ENSMUST00000046285.5 | 2177 | C1qa | ENSMUSG00000036887.5 |
| chr4 | 136897488 | 136897802 | 3.44052 | 4.01883 | 1.12287 | Exon (ENSMUST00000046285.5/ENSMUSG00000036887.5, exon 2 of 3) | 4 | 136895917 | 136898803 | 2887 | 2 | ENSMUST00000046285.5 | 1001 | C1qa | ENSMUSG00000036887.5 |
| chr4 | 136899106 | 136899589 | 3.69171 | 4.49785 | 1.45275 | Promoter | 4 | 136895917 | 136898803 | 2887 | 2 | ENSMUST00000046285.5 | -304 | C1qa | ENSMUSG00000036887.5 |
| chr4 | 136900172 | 136900278 | 3.15995 | 3.71003 | 1.01733 | Distal Intergenic | 4 | 136895917 | 136898803 | 2887 | 2 | ENSMUST00000046285.5 | -1370 | C1qa | ENSMUSG00000036887.5 |
| chr4 | 136903731 | 136903837 | 3.30251 | 3.78096 | 1.01733 | Distal Intergenic | 4 | 136895917 | 136898803 | 2887 | 2 | ENSMUST00000046285.5 | -4929 | C1qa | ENSMUSG00000036887.5 |
| chr4 | 136906566 | 136907189 | 3.64203 | 4.39391 | 1.35878 | Distal Intergenic | 4 | 136895917 | 136898803 | 2887 | 2 | ENSMUST00000046285.5 | -7764 | C1qa | ENSMUSG00000036887.5 |
| chr4 | 136911328 | 136911567 | 3.5507 | 4.21299 | 1.21308 | Distal Intergenic | 4 | 136895917 | 136898803 | 2887 | 2 | ENSMUST00000046285.5 | -12526 | C1qa | ENSMUSG00000036887.5 |
| chr4 | 136911977 | 136912250 | 3.68011 | 4.45271 | 1.36996 | Distal Intergenic | 4 | 136895917 | 136898803 | 2887 | 2 | ENSMUST00000046285.5 | -13175 | C1qa | ENSMUSG00000036887.5 |
| chr4 | 136912608 | 136913489 | 3.45034 | 4.08597 | 1.20642 | Distal Intergenic | 4 | 136895917 | 136898803 | 2887 | 2 | ENSMUST00000046285.5 | -13806 | C1qa | ENSMUSG00000036887.5 |
| chr4 | 136914110 | 136914598 | 3.61651 | 4.35311 | 1.35026 | Distal Intergenic | 4 | 136895917 | 136898803 | 2887 | 2 | ENSMUST00000046285.5 | -15308 | C1qa | ENSMUSG00000036887.5 |
| chr4 | 136917198 | 136917537 | 3.68357 | 4.46112 | 1.37738 | Distal Intergenic | 4 | 136895917 | 136898803 | 2887 | 2 | ENSMUST00000046285.5 | -18396 | C1qa | ENSMUSG00000036887.5 |
| chr4 | 136926900 | 136926975 | 3.67289 | 4.45706 | 1.41095 | Downstream (2-3kb) | 4 | 136895917 | 136898803 | 2887 | 2 | ENSMUST00000046285.5 | -28098 | C1qa | ENSMUSG00000036887.5 |
| chr2 | 44557085 | 44557375 | 3.66653 | 4.50713 | 1.48062 | Distal Intergenic | 2 | 44565133 | 44571347 | 6215 | 2 | ENSMUST00000134813.1 | 13972 | Gtdc1 | ENSMUSG00000036890.8 |
| chr4 | 136892226 | 136892622 | 3.48196 | 4.09612 | 1.16765 | Promoter | 4 | 136891651 | 136892876 | 1226 | 2 | ENSMUST00000153104.1 | 254 | C1qc | ENSMUSG00000036896.5 |
| chr4 | 136895278 | 136895649 | 4.02619 | 5.13042 | 1.82852 | Downstream (<1kb) | 4 | 136889804 | 136893065 | 3262 | 2 | ENSMUST00000046332.5 | -2214 | C1qc | ENSMUSG00000036896.5 |
| chr4 | 136861949 | 136862107 | 3.44957 | 4.06213 | 1.15673 | Distal Intergenic | 4 | 136880129 | 136886187 | 6059 | 2 | ENSMUST00000046384.8 | 24080 | C1qb | ENSMUSG00000036905.8 |
| chr4 | 136865170 | 136865646 | 3.96394 | 5.0002 | 1.73914 | Distal Intergenic | 4 | 136880129 | 136886187 | 6059 | 2 | ENSMUST00000046384.8 | 20541 | C1qb | ENSMUSG00000036905.8 |
| chr4 | 136867411 | 136868061 | 3.55527 | 4.21814 | 1.21195 | Distal Intergenic | 4 | 136880129 | 136886187 | 6059 | 2 | ENSMUST00000046384.8 | 18126 | C1qb | ENSMUSG00000036905.8 |
| chr4 | 136878799 | 136878917 | 4.28091 | 5.69031 | 2.27425 | Downstream (1-2kb) | 4 | 136880129 | 136886187 | 6059 | 2 | ENSMUST00000046384.8 | 7270 | C1qb | ENSMUSG00000036905.8 |
| chr1 | 120333234 | 120333625 | 3.82133 | 4.76656 | 1.63933 | Distal Intergenic | 1 | 120340582 | 120343174 | 2593 | 1 | ENSMUST00000037286.9 | -6957 | C1ql2 | ENSMUSG00000036907.9 |
| chr1 | 120336164 | 120336430 | 3.43837 | 4.06366 | 1.18718 | Distal Intergenic | 1 | 120340582 | 120343174 | 2593 | 1 | ENSMUST00000037286.9 | -4152 | C1ql2 | ENSMUSG00000036907.9 |
| chr1 | 120336890 | 120337402 | 3.34089 | 3.8387 | 1.01733 | Distal Intergenic | 1 | 120340582 | 120343174 | 2593 | 1 | ENSMUST00000037286.9 | -3180 | C1ql2 | ENSMUSG00000036907.9 |
| chr1 | 120337971 | 120338095 | 3.34632 | 3.84688 | 1.01733 | Distal Intergenic | 1 | 120340582 | 120343174 | 2593 | 1 | ENSMUST00000037286.9 | -2487 | C1ql2 | ENSMUSG00000036907.9 |
| chr1 | 120338929 | 120339363 | 3.33141 | 3.83905 | 1.0673 | Distal Intergenic | 1 | 120340582 | 120343174 | 2593 | 1 | ENSMUST00000037286.9 | -1219 | C1ql2 | ENSMUSG00000036907.9 |
| chr1 | 120340448 | 120340832 | 3.29296 | 4.33032 | 1.31473 | Promoter | 1 | 120340582 | 120343174 | 2593 | 1 | ENSMUST00000037286.9 | 0 | C1ql2 | ENSMUSG00000036907.9 |
| chr1 | 120341173 | 120343240 | 3.52431 | 4.24625 | 1.31011 | Promoter | 1 | 120340582 | 120343174 | 2593 | 1 | ENSMUST00000037286.9 | 592 | C1ql2 | ENSMUSG00000036907.9 |
| chr1 | 120343637 | 120343726 | 3.41794 | 4.20065 | 1.27105 | Distal Intergenic | 1 | 120340582 | 120343174 | 2593 | 1 | ENSMUST00000037286.9 | 3056 | C1ql2 | ENSMUSG00000036907.9 |
| chr1 | 120344749 | 120345119 | 3.34506 | 3.95754 | 1.12351 | Distal Intergenic | 1 | 120340582 | 120343174 | 2593 | 1 | ENSMUST00000037286.9 | 4168 | C1ql2 | ENSMUSG00000036907.9 |
| chr8 | 124775196 | 124775544 | 3.501 | 4.27563 | 1.31162 | Distal Intergenic | 8 | 124793019 | 124834702 | 41684 | 1 | ENSMUST00000167588.2 | -17475 | Trim67 | ENSMUSG00000036913.9 |
| chr8 | 124790725 | 124791249 | 3.37399 | 4.02766 | 1.16123 | Distal Intergenic | 8 | 124793019 | 124834702 | 41684 | 1 | ENSMUST00000167588.2 | -1770 | Trim67 | ENSMUSG00000036913.9 |
| chr8 | 124791624 | 124791723 | 3.76196 | 4.68786 | 1.63926 | Distal Intergenic | 8 | 124793019 | 124834702 | 41684 | 1 | ENSMUST00000167588.2 | -1296 | Trim67 | ENSMUSG00000036913.9 |
| chr8 | 124793005 | 124793233 | 3.76937 | 4.69355 | 1.62336 | Promoter | 8 | 124793019 | 124834702 | 41684 | 1 | ENSMUST00000167588.2 | 0 | Trim67 | ENSMUSG00000036913.9 |
| chr8 | 124793546 | 124795179 | 3.83756 | 5.09588 | 1.88553 | Promoter | 8 | 124793092 | 124834713 | 41622 | 1 | ENSMUST00000041106.7 | 455 | Trim67 | ENSMUSG00000036913.9 |
| chr5 | 138280559 | 138280644 | 3.50502 | 4.15253 | 1.25046 | Promoter | 5 | 138280556 | 138290406 | 9851 | 1 | ENSMUST00000161691.1 | 4 | Stag3 | ENSMUSG00000036928.9 |
| chr9 | 91350253 | 91353198 | 4.07595 | 5.42366 | 2.07635 | Distal Intergenic | 9 | 91362413 | 91378915 | 16503 | 1 | ENSMUST00000173342.2 | -9215 | Zic4 | ENSMUSG00000036972.9 |
| chr9 | 91353722 | 91354889 | 3.71326 | 4.58896 | 1.50158 | Distal Intergenic | 9 | 91362413 | 91378915 | 16503 | 1 | ENSMUST00000173342.2 | -7524 | Zic4 | ENSMUSG00000036972.9 |
| chr9 | 91355246 | 91355335 | 4.316 | 5.692 | 2.23672 | Downstream (2-3kb) | 9 | 91362413 | 91378915 | 16503 | 1 | ENSMUST00000173342.2 | -7078 | Zic4 | ENSMUSG00000036972.9 |
| chr9 | 91355674 | 91356524 | 3.58217 | 4.30311 | 1.28911 | Downstream (1-2kb) | 9 | 91362413 | 91378915 | 16503 | 1 | ENSMUST00000173342.2 | -5889 | Zic4 | ENSMUSG00000036972.9 |
| chr9 | 91356954 | 91358378 | 4.27698 | 5.7223 | 2.33611 | 3' UTR | 9 | 91362413 | 91378915 | 16503 | 1 | ENSMUST00000173342.2 | -4035 | Zic4 | ENSMUSG00000036972.9 |
| chr9 | 91358848 | 91360923 | 3.87131 | 4.89484 | 1.70677 | 3' UTR | 9 | 91362413 | 91378915 | 16503 | 1 | ENSMUST00000173342.2 | -1490 | Zic4 | ENSMUSG00000036972.9 |
| chr9 | 91390964 | 91391055 | 4.07514 | 5.21503 | 1.8952 | Distal Intergenic | 9 | 91382410 | 91388044 | 5635 | 1 | ENSMUST00000085242.4 | 8555 | Zic4 | ENSMUSG00000036972.9 |
| chr9 | 91391919 | 91392076 | 4.00814 | 5.07716 | 1.79137 | Distal Intergenic | 9 | 91382410 | 91388044 | 5635 | 1 | ENSMUST00000085242.4 | 9510 | Zic4 | ENSMUSG00000036972.9 |
| chr9 | 91367700 | 91367893 | 3.36379 | 3.87324 | 1.01733 | Promoter | 9 | 91368367 | 91369133 | 767 | 1 | ENSMUST00000172978.1 | -474 | Zic4 | ENSMUSG00000036972.9 |
| chr9 | 91368494 | 91371992 | 3.81937 | 4.82744 | 1.67639 | Promoter | 9 | 91368812 | 91379257 | 10446 | 1 | ENSMUST00000173054.2 | 0 | Zic4 | ENSMUSG00000036972.9 |
| chr9 | 91373708 | 91374243 | 3.38561 | 3.96251 | 1.17029 | Exon (ENSMUST00000174802.1/ENSMUSG00000036972.9, exon 3 of 3) | 9 | 91371350 | 91373923 | 2574 | 1 | ENSMUST00000174802.1 | 2359 | Zic4 | ENSMUSG00000036972.9 |
| chr9 | 91374562 | 91375161 | 3.45042 | 4.17569 | 1.24352 | Intron (ENSMUST00000173342.2/ENSMUSG00000036972.9, intron 2 of 2) | 9 | 91378197 | 91384398 | 6202 | 1 | ENSMUST00000172858.1 | -3036 | Zic4 | ENSMUSG00000036972.9 |
| chr9 | 91375601 | 91375697 | 4.40165 | 5.93785 | 2.46618 | Intron (ENSMUST00000173342.2/ENSMUSG00000036972.9, intron 2 of 2) | 9 | 91378197 | 91384398 | 6202 | 1 | ENSMUST00000172858.1 | -2500 | Zic4 | ENSMUSG00000036972.9 |
| chr9 | 91377009 | 91377530 | 3.86303 | 4.83237 | 1.68495 | Promoter | 9 | 91378197 | 91384398 | 6202 | 1 | ENSMUST00000172858.1 | -667 | Zic4 | ENSMUSG00000036972.9 |
| chr9 | 91378414 | 91382508 | 4.15811 | 5.55871 | 2.19138 | Promoter | 9 | 91378642 | 91389348 | 10707 | 1 | ENSMUST00000066384.6 | 0 | Zic4 | ENSMUSG00000036972.9 |
| chr9 | 91383608 | 91384922 | 4.08939 | 5.4437 | 2.10128 | Exon (ENSMUST00000172646.2/ENSMUSG00000036972.9, exon 4 of 5) | 9 | 91382410 | 91388044 | 5635 | 1 | ENSMUST00000085242.4 | 1199 | Zic4 | ENSMUSG00000036972.9 |
| chr9 | 91385407 | 91385490 | 4.04334 | 5.16239 | 1.87643 | Intron (ENSMUST00000172646.2/ENSMUSG00000036972.9, intron 4 of 4) | 9 | 91382410 | 91388044 | 5635 | 1 | ENSMUST00000085242.4 | 2998 | Zic4 | ENSMUSG00000036972.9 |
| chr9 | 91386025 | 91387737 | 4.1773 | 5.4937 | 2.14676 | 3' UTR | 9 | 91382410 | 91388044 | 5635 | 1 | ENSMUST00000085242.4 | 3616 | Zic4 | ENSMUSG00000036972.9 |
| chr9 | 91388710 | 91390320 | 3.73973 | 4.5929 | 1.49566 | 3' UTR | 9 | 91382410 | 91388044 | 5635 | 1 | ENSMUST00000085242.4 | 6301 | Zic4 | ENSMUSG00000036972.9 |
| chr9 | 91392461 | 91392771 | 3.48359 | 4.08625 | 1.13079 | Distal Intergenic | 9 | 91382410 | 91388044 | 5635 | 1 | ENSMUST00000085242.4 | 10052 | Zic4 | ENSMUSG00000036972.9 |
| chr2 | 148394998 | 148396207 | 4.11387 | 5.51791 | 2.23281 | Promoter | 2 | 148395344 | 148396767 | 1424 | 1 | ENSMUST00000109962.3 | 0 | Sstr4 | ENSMUSG00000037014.4 |
| chr2 | 148396661 | 148397040 | 3.38818 | 4.32937 | 1.33094 | 3' UTR | 2 | 148395344 | 148396767 | 1424 | 1 | ENSMUST00000109962.3 | 1318 | Sstr4 | ENSMUSG00000037014.4 |
| chr3 | 53655225 | 53657107 | 3.8435 | 4.85184 | 1.70968 | Promoter | 3 | 53513939 | 53657355 | 143417 | 2 | ENSMUST00000091137.4 | 248 | Frem2 | ENSMUSG00000037016.7 |
| chr3 | 53658167 | 53658463 | 4.21437 | 5.49977 | 2.11126 | Promoter | 3 | 53513939 | 53657355 | 143417 | 2 | ENSMUST00000091137.4 | -813 | Frem2 | ENSMUSG00000037016.7 |
| chr2 | 148043116 | 148043422 | 3.71917 | 4.64893 | 1.56701 | 3' UTR | 2 | 148044240 | 148045464 | 1225 | 2 | ENSMUST00000146242.1 | 2042 | Foxa2 | ENSMUSG00000037025.6 |
| chr2 | 148043846 | 148045334 | 3.73463 | 4.83772 | 1.68919 | Promoter | 2 | 148044240 | 148045464 | 1225 | 2 | ENSMUST00000146242.1 | 130 | Foxa2 | ENSMUSG00000037025.6 |
| chr2 | 148046028 | 148047800 | 3.87485 | 5.06745 | 1.83094 | Promoter | 2 | 148042877 | 148046969 | 4093 | 2 | ENSMUST00000047315.4 | 0 | Foxa2 | ENSMUSG00000037025.6 |
| chr2 | 148049159 | 148049426 | 3.36463 | 4.31695 | 1.32208 | Distal Intergenic | 2 | 148042877 | 148046969 | 4093 | 2 | ENSMUST00000047315.4 | -2191 | Foxa2 | ENSMUSG00000037025.6 |
| chr2 | 147350420 | 147350495 | 3.5538 | 4.26626 | 1.33836 | Distal Intergenic | 2 | 147361925 | 147367878 | 5954 | 1 | ENSMUST00000126068.2 | -11430 | Pax1 | ENSMUSG00000037034.10 |
| chr2 | 147354007 | 147354381 | 3.69334 | 4.48693 | 1.41495 | Distal Intergenic | 2 | 147361925 | 147367878 | 5954 | 1 | ENSMUST00000126068.2 | -7544 | Pax1 | ENSMUSG00000037034.10 |
| chr2 | 147356334 | 147356927 | 3.62712 | 4.38816 | 1.38395 | Distal Intergenic | 2 | 147361925 | 147367878 | 5954 | 1 | ENSMUST00000126068.2 | -4998 | Pax1 | ENSMUSG00000037034.10 |
| chr2 | 147357240 | 147357562 | 3.47011 | 4.06894 | 1.1415 | Distal Intergenic | 2 | 147361925 | 147367878 | 5954 | 1 | ENSMUST00000126068.2 | -4363 | Pax1 | ENSMUSG00000037034.10 |
| chr2 | 147358609 | 147358781 | 3.54021 | 4.25038 | 1.26651 | Downstream (2-3kb) | 2 | 147361925 | 147367878 | 5954 | 1 | ENSMUST00000126068.2 | -3144 | Pax1 | ENSMUSG00000037034.10 |
| chr2 | 147359340 | 147359598 | 3.55242 | 4.34471 | 1.35074 | Downstream (1-2kb) | 2 | 147361925 | 147367878 | 5954 | 1 | ENSMUST00000126068.2 | -2327 | Pax1 | ENSMUSG00000037034.10 |
| chr2 | 147360343 | 147361971 | 3.96699 | 5.16406 | 1.91825 | Promoter | 2 | 147361925 | 147367878 | 5954 | 1 | ENSMUST00000126068.2 | 0 | Pax1 | ENSMUSG00000037034.10 |
| chr2 | 147382096 | 147382217 | 3.45987 | 4.13693 | 1.25119 | Intron (ENSMUST00000156584.1/ENSMUSG00000037034.10, intron 1 of 1) | 2 | 147368282 | 147393295 | 25014 | 1 | ENSMUST00000156584.1 | 13815 | Pax1 | ENSMUSG00000037034.10 |
| chr2 | 147369689 | 147371063 | 3.58934 | 4.33289 | 1.35729 | Intron (ENSMUST00000109968.2/ENSMUSG00000037034.10, intron 4 of 4) | 2 | 147368282 | 147393295 | 25014 | 1 | ENSMUST00000156584.1 | 1408 | Pax1 | ENSMUSG00000037034.10 |
| chr2 | 147364273 | 147365359 | 3.75493 | 4.65587 | 1.56078 | Promoter | 2 | 147364994 | 147375053 | 10060 | 1 | ENSMUST00000109968.2 | 0 | Pax1 | ENSMUSG00000037034.10 |
| chr2 | 147365769 | 147366791 | 3.55298 | 4.23264 | 1.25015 | Promoter | 2 | 147364994 | 147375053 | 10060 | 1 | ENSMUST00000109968.2 | 776 | Pax1 | ENSMUSG00000037034.10 |
| chr2 | 147367136 | 147368552 | 4.34638 | 5.89532 | 2.43999 | Promoter | 2 | 147368282 | 147393295 | 25014 | 1 | ENSMUST00000156584.1 | 0 | Pax1 | ENSMUSG00000037034.10 |
| chr2 | 147375006 | 147375315 | 3.5613 | 4.29148 | 1.38492 | 3' UTR | 2 | 147368282 | 147393295 | 25014 | 1 | ENSMUST00000156584.1 | 6725 | Pax1 | ENSMUSG00000037034.10 |
| chr2 | 147371685 | 147371947 | 3.54848 | 4.38279 | 1.38527 | Intron (ENSMUST00000109968.2/ENSMUSG00000037034.10, intron 4 of 4) | 2 | 147368282 | 147393295 | 25014 | 1 | ENSMUST00000156584.1 | 3404 | Pax1 | ENSMUSG00000037034.10 |
| chr2 | 147372456 | 147372531 | 3.32928 | 4.01338 | 1.15122 | Intron (ENSMUST00000109968.2/ENSMUSG00000037034.10, intron 4 of 4) | 2 | 147368282 | 147393295 | 25014 | 1 | ENSMUST00000156584.1 | 4175 | Pax1 | ENSMUSG00000037034.10 |
| chr2 | 147373172 | 147373967 | 3.74853 | 4.68492 | 1.58464 | 3' UTR | 2 | 147368282 | 147393295 | 25014 | 1 | ENSMUST00000156584.1 | 4891 | Pax1 | ENSMUSG00000037034.10 |
| chr2 | 147375829 | 147376229 | 3.44486 | 4.06103 | 1.23284 | Intron (ENSMUST00000156584.1/ENSMUSG00000037034.10, intron 1 of 1) | 2 | 147368282 | 147393295 | 25014 | 1 | ENSMUST00000156584.1 | 7548 | Pax1 | ENSMUSG00000037034.10 |
| chr2 | 147376859 | 147377126 | 3.66072 | 4.77727 | 1.64268 | Intron (ENSMUST00000156584.1/ENSMUSG00000037034.10, intron 1 of 1) | 2 | 147368282 | 147393295 | 25014 | 1 | ENSMUST00000156584.1 | 8578 | Pax1 | ENSMUSG00000037034.10 |
| chr2 | 147377477 | 147378021 | 3.73142 | 4.65322 | 1.58943 | Intron (ENSMUST00000156584.1/ENSMUSG00000037034.10, intron 1 of 1) | 2 | 147368282 | 147393295 | 25014 | 1 | ENSMUST00000156584.1 | 9196 | Pax1 | ENSMUSG00000037034.10 |
| chr1 | 119417317 | 119417779 | 3.66857 | 4.48929 | 1.45618 | 3' UTR | 1 | 119415465 | 119422248 | 6784 | 2 | ENSMUST00000038765.5 | 4469 | Inhbb | ENSMUSG00000037035.5 |
| chr1 | 119418646 | 119418916 | 3.54178 | 4.2261 | 1.2952 | Intron (ENSMUST00000038765.5/ENSMUSG00000037035.5, intron 1 of 1) | 1 | 119415465 | 119422248 | 6784 | 2 | ENSMUST00000038765.5 | 3332 | Inhbb | ENSMUSG00000037035.5 |
| chr1 | 119420277 | 119421186 | 3.85373 | 4.82936 | 1.63922 | 5' UTR | 1 | 119415465 | 119422248 | 6784 | 2 | ENSMUST00000038765.5 | 1062 | Inhbb | ENSMUSG00000037035.5 |
| chr1 | 119421495 | 119421727 | 3.89656 | 4.87848 | 1.68887 | Promoter | 1 | 119415465 | 119422248 | 6784 | 2 | ENSMUST00000038765.5 | 521 | Inhbb | ENSMUSG00000037035.5 |
| chr1 | 119447893 | 119447997 | 3.43916 | 4.02587 | 1.13924 | Distal Intergenic | 1 | 119415465 | 119422248 | 6784 | 2 | ENSMUST00000038765.5 | -25646 | Inhbb | ENSMUSG00000037035.5 |
| chr19 | 44406690 | 44407102 | 3.36963 | 3.88442 | 1.02567 | Promoter | 19 | 44394451 | 44407709 | 13259 | 2 | ENSMUST00000041331.2 | 607 | Scd1 | ENSMUSG00000037071.2 |
| chr19 | 44439298 | 44439373 | 3.65766 | 4.43049 | 1.41358 | Distal Intergenic | 19 | 44394451 | 44407709 | 13259 | 2 | ENSMUST00000041331.2 | -31590 | Scd1 | ENSMUSG00000037071.2 |
| chr17 | 56120570 | 56121625 | 4.1267 | 5.40377 | 2.06864 | Promoter | 17 | 56119680 | 56121946 | 2267 | 2 | ENSMUST00000041357.7 | 321 | Lrg1 | ENSMUSG00000037095.7 |
| chr17 | 56122476 | 56122861 | 3.60658 | 4.32701 | 1.31829 | Promoter | 17 | 56119680 | 56121946 | 2267 | 2 | ENSMUST00000041357.7 | -531 | Lrg1 | ENSMUSG00000037095.7 |
| chr17 | 56123921 | 56124915 | 3.8399 | 4.93267 | 1.7612 | 3' UTR | 17 | 56119680 | 56121946 | 2267 | 2 | ENSMUST00000041357.7 | -1976 | Lrg1 | ENSMUSG00000037095.7 |
| chr3 | 78966791 | 78967086 | 3.29167 | 3.77602 | 1.05544 | Promoter | 3 | 78966356 | 78966799 | 444 | 2 | ENSMUST00000036976.4 | 0 | Gm9762 | ENSMUSG00000037096.4 |
| chr8 | 78067822 | 78068504 | 3.53275 | 4.21439 | 1.31358 | Distal Intergenic | 8 | 78213297 | 78220975 | 7679 | 1 | ENSMUST00000117845.2 | -144793 | Ttc29 | ENSMUSG00000037101.11 |
| chr11 | 58640604 | 58640738 | 3.99421 | 5.14631 | 1.94291 | Promoter | 11 | 58640465 | 58652404 | 11940 | 1 | ENSMUST00000075084.4 | 140 | Trim58 | ENSMUSG00000037124.6 |
| chr19 | 46317471 | 46318432 | 4.29147 | 5.68295 | 2.2487 | Intron (ENSMUST00000041391.4/ENSMUSG00000037126.10, intron 11 of 17) | 19 | 46312087 | 46327156 | 15070 | 2 | ENSMUST00000041391.4 | 8724 | Psd | ENSMUSG00000037126.10 |
| chr19 | 46318982 | 46319641 | 3.72527 | 4.57821 | 1.47444 | Exon (ENSMUST00000041391.4/ENSMUSG00000037126.10, exon 11 of 18) | 19 | 46312087 | 46327156 | 15070 | 2 | ENSMUST00000041391.4 | 7515 | Psd | ENSMUSG00000037126.10 |
| chr19 | 46319973 | 46320048 | 3.43945 | 4.03456 | 1.18269 | Intron (ENSMUST00000041391.4/ENSMUSG00000037126.10, intron 9 of 17) | 19 | 46312087 | 46327156 | 15070 | 2 | ENSMUST00000041391.4 | 7108 | Psd | ENSMUSG00000037126.10 |
| chr19 | 46324873 | 46325486 | 3.37891 | 3.90522 | 1.04921 | 5' UTR | 19 | 46312087 | 46327156 | 15070 | 2 | ENSMUST00000041391.4 | 1670 | Psd | ENSMUSG00000037126.10 |
| chr19 | 46326019 | 46326162 | 3.37849 | 4.02355 | 1.13118 | Promoter | 19 | 46312087 | 46327156 | 15070 | 2 | ENSMUST00000041391.4 | 994 | Psd | ENSMUSG00000037126.10 |
| chr2 | 146209839 | 146210215 | 3.51274 | 4.20137 | 1.32932 | Intron (ENSMUST00000133433.2/ENSMUSG00000037143.12, intron 26 of 26) | 2 | 146214265 | 146215035 | 771 | 1 | ENSMUST00000146891.1 | -4050 | 4930529M08Rik | ENSMUSG00000037143.12 |
| chr2 | 146214549 | 146214656 | 3.7547 | 4.68663 | 1.61275 | Promoter | 2 | 146214613 | 146214795 | 183 | 1 | ENSMUST00000167179.1 | 0 | 4930529M08Rik | ENSMUSG00000037143.12 |
| chr2 | 146216045 | 146216390 | 4.08976 | 5.30096 | 2.02785 | Distal Intergenic | 2 | 146214613 | 146214795 | 183 | 1 | ENSMUST00000167179.1 | 1433 | 4930529M08Rik | ENSMUSG00000037143.12 |
| chr3 | 51371598 | 51371991 | 3.71427 | 4.55726 | 1.44679 | Distal Intergenic | 3 | 51391261 | 51396488 | 5228 | 2 | ENSMUST00000145938.1 | 24497 | Mgarp | ENSMUSG00000037161.9 |
| chr3 | 51376697 | 51376978 | 3.55991 | 4.35939 | 1.37215 | Distal Intergenic | 3 | 51391261 | 51396488 | 5228 | 2 | ENSMUST00000145938.1 | 19510 | Mgarp | ENSMUSG00000037161.9 |
| chr3 | 51378721 | 51378951 | 3.37385 | 3.935 | 1.16535 | Distal Intergenic | 3 | 51391261 | 51396488 | 5228 | 2 | ENSMUST00000145938.1 | 17537 | Mgarp | ENSMUSG00000037161.9 |
| chr3 | 51383678 | 51384062 | 3.45253 | 4.04664 | 1.15738 | Distal Intergenic | 3 | 51391261 | 51396488 | 5228 | 2 | ENSMUST00000145938.1 | 12426 | Mgarp | ENSMUSG00000037161.9 |
| chr3 | 51387292 | 51387415 | 3.5221 | 4.23003 | 1.31414 | Downstream (<1kb) | 3 | 51391261 | 51396488 | 5228 | 2 | ENSMUST00000145938.1 | 9073 | Mgarp | ENSMUSG00000037161.9 |
| chr3 | 51388432 | 51389457 | 3.54006 | 4.31716 | 1.33156 | 3' UTR | 3 | 51391261 | 51396488 | 5228 | 2 | ENSMUST00000145938.1 | 7031 | Mgarp | ENSMUSG00000037161.9 |
| chr3 | 51391695 | 51392107 | 3.86117 | 4.87521 | 1.74823 | Exon (ENSMUST00000108046.2/ENSMUSG00000037161.9, exon 2 of 4) | 3 | 51391261 | 51396488 | 5228 | 2 | ENSMUST00000145938.1 | 4381 | Mgarp | ENSMUSG00000037161.9 |

| | | | | | | | | | | | | | | | |
|---|---|---|---|---|---|---|---|---|---|---|---|---|---|---|---|
| chr3 | 51394659 | 51395023 | 3.37101 | 3.924 | 1.14737 | Intron (ENSMUST00000108046.2/ENSMUSG00000037161.9, intron 1 of 3) | 3 | 51391261 | 51396488 | 5228 | 2 | ENSMUST00000145938.1 | 1465 | Mgarp | ENSMUSG00000037161.9 |
| chr3 | 51395988 | 51396648 | 3.89132 | 5.298 | 1.99622 | Promoter | 3 | 51388415 | 51396502 | 8088 | 2 | ENSMUST00000108046.2 | 0 | Mgarp | ENSMUSG00000037161.9 |
| chr3 | 51398957 | 51399032 | 3.66328 | 4.46821 | 1.47202 | Distal Intergenic | 3 | 51388772 | 51396738 | 7967 | 2 | ENSMUST00000141156.1 | -2220 | Mgarp | ENSMUSG00000037161.9 |
| chr10 | 61405963 | 61406419 | 3.50905 | 4.19044 | 1.25797 | Distal Intergenic | 10 | 61417972 | 61425338 | 7367 | 1 | ENSMUST00000049339.6 | -11553 | Nodal | ENSMUSG00000037171.6 |
| chr10 | 61418413 | 61418617 | 3.78142 | 4.94338 | 1.70721 | Promoter | 10 | 61417972 | 61425338 | 7367 | 1 | ENSMUST00000049339.6 | 442 | Nodal | ENSMUSG00000037171.6 |
| chr10 | 61419190 | 61420267 | 3.78976 | 4.66557 | 1.51999 | Intron (ENSMUST00000049339.6/ENSMUSG00000037171.6, intron 1 of 2) | 10 | 61417972 | 61425338 | 7367 | 1 | ENSMUST00000049339.6 | 1219 | Nodal | ENSMUSG00000037171.6 |
| chr3 | 51356714 | 51356789 | 4.00406 | 5.18239 | 1.89843 | Distal Intergenic | 3 | 51264869 | 51340658 | 75790 | 2 | ENSMUST00000183338.1 | -16057 | Elf2 | ENSMUSG00000037174.13 |
| chr3 | 51357681 | 51358516 | 3.43085 | 4.04877 | 1.22446 | Distal Intergenic | 3 | 51264869 | 51340658 | 75790 | 2 | ENSMUST00000183338.1 | -17024 | Elf2 | ENSMUSG00000037174.13 |
| chr3 | 51358970 | 51360321 | 3.67967 | 4.61645 | 1.53061 | Distal Intergenic | 3 | 51264869 | 51340658 | 75790 | 2 | ENSMUST00000183338.1 | -18313 | Elf2 | ENSMUSG00000037174.13 |
| chr3 | 51361485 | 51361717 | 3.48851 | 4.10364 | 1.16633 | Distal Intergenic | 3 | 51264869 | 51340658 | 75790 | 2 | ENSMUST00000183338.1 | -20828 | Elf2 | ENSMUSG00000037174.13 |
| chr3 | 51362097 | 51362876 | 3.7365 | 4.69076 | 1.59145 | Distal Intergenic | 3 | 51264869 | 51340658 | 75790 | 2 | ENSMUST00000183338.1 | -21440 | Elf2 | ENSMUSG00000037174.13 |
| chr3 | 51363738 | 51364037 | 3.32237 | 3.81082 | 1.01733 | Distal Intergenic | 3 | 51264869 | 51340658 | 75790 | 2 | ENSMUST00000183338.1 | -23081 | Elf2 | ENSMUSG00000037174.13 |
| chr3 | 51364640 | 51364733 | 3.75387 | 4.78014 | 1.62051 | Distal Intergenic | 3 | 51264869 | 51340658 | 75790 | 2 | ENSMUST00000183338.1 | -23983 | Elf2 | ENSMUSG00000037174.13 |
| chr3 | 51366728 | 51368610 | 3.8032 | 4.73723 | 1.61637 | Distal Intergenic | 3 | 51264869 | 51340658 | 75790 | 2 | ENSMUST00000183338.1 | -26071 | Elf2 | ENSMUSG00000037174.13 |
| chr4 | 135548859 | 135549030 | 3.25216 | 3.70539 | 1.01733 | Intron (ENSMUST00000105855.1/ENSMUSG00000037188.7, intron 13 of 15) | 4 | 135541888 | 135573630 | 31743 | 2 | ENSMUST00000105855.1 | 24600 | Grhl3 | ENSMUSG00000037188.7 |
| chr4 | 135568988 | 135569588 | 3.37781 | 3.94068 | 1.16363 | Intron (ENSMUST00000105855.1/ENSMUSG00000037188.7, intron 1 of 15) | 4 | 135541888 | 135573630 | 31743 | 2 | ENSMUST00000105855.1 | 4042 | Grhl3 | ENSMUSG00000037188.7 |
| chr4 | 135570333 | 135570964 | 3.67192 | 4.59088 | 1.52536 | Intron (ENSMUST00000105855.1/ENSMUSG00000037188.7, intron 1 of 15) | 4 | 135541888 | 135573630 | 31743 | 2 | ENSMUST00000105855.1 | 2666 | Grhl3 | ENSMUSG00000037188.7 |
| chr4 | 135572068 | 135572143 | 3.63255 | 4.36136 | 1.30856 | Intron (ENSMUST00000105855.1/ENSMUSG00000037188.7, intron 1 of 15) | 4 | 135541888 | 135573630 | 31743 | 2 | ENSMUST00000105855.1 | 1487 | Grhl3 | ENSMUSG00000037188.7 |
| chr4 | 135572738 | 135574176 | 3.58043 | 4.50171 | 1.44955 | Promoter | 4 | 135541888 | 135573630 | 31743 | 2 | ENSMUST00000105855.1 | 0 | Grhl3 | ENSMUSG00000037188.7 |
| chr5 | 34286518 | 34286740 | 3.27301 | 3.78474 | 1.06849 | Intron (ENSMUST00000094868.4/ENSMUSG00000037224.10, intron 1 of 12) | 5 | 34194874 | 34288326 | 93453 | 2 | ENSMUST00000094868.4 | 1586 | Zfyve28 | ENSMUSG00000037224.10 |
| chr5 | 34287286 | 34288160 | 3.67212 | 4.68531 | 1.56811 | Promoter | 5 | 34194874 | 34288326 | 93453 | 2 | ENSMUST00000094868.4 | 166 | Zfyve28 | ENSMUSG00000037224.10 |
| chr5 | 34288512 | 34288829 | 3.61977 | 4.81691 | 1.63562 | Promoter | 5 | 34226588 | 34288449 | 61862 | 2 | ENSMUST00000114369.1 | -64 | Zfyve28 | ENSMUSG00000037224.10 |
| chr5 | 34289339 | 34289636 | 4.02308 | 5.24397 | 1.95396 | Promoter | 5 | 34226588 | 34288449 | 61862 | 2 | ENSMUST00000114369.1 | -891 | Zfyve28 | ENSMUSG00000037224.10 |
| chr3 | 37348093 | 37349467 | 4.11666 | 5.74471 | 2.34207 | Promoter | 3 | 37348346 | 37405377 | 57032 | 1 | ENSMUST00000138563.3 | 0 | Fgf2 | ENSMUSG00000037225.8 |
| chr10 | 60945934 | 60946082 | 3.50228 | 4.17441 | 1.26937 | Distal Intergenic | 10 | 60928226 | 60938132 | 9907 | 2 | ENSMUST00000036304.3 | -7803 | Pldi | ENSMUSG00000037247.3 |
| chr2 | 144330551 | 144333258 | 3.85346 | 5.01461 | 1.81097 | Promoter | 2 | 144305175 | 144331695 | 26521 | 2 | ENSMUST00000103171.4 | 0 | Ovol2 | ENSMUSG00000037279.8 |
| chr2 | 144334289 | 144334679 | 4.46272 | 6.03411 | 2.49678 | Distal Intergenic | 2 | 144305175 | 144332146 | 26972 | 2 | ENSMUST00000037423.3 | -2144 | Ovol2 | ENSMUSG00000037279.8 |
| chr15 | 100728341 | 100730305 | 4.34438 | 5.83297 | 2.41203 | Promoter | 15 | 100691813 | 100729376 | 37564 | 2 | ENSMUST00000159715.2 | 0 | Galnt6 | ENSMUSG00000037280.7 |
| chr15 | 100731762 | 100732062 | 3.8948 | 4.86109 | 1.63789 | Distal Intergenic | 15 | 100691815 | 100729376 | 37562 | 2 | ENSMUST00000052069.6 | -2387 | Galnt6 | ENSMUSG00000037280.7 |
| chr4 | 134675982 | 134676057 | 3.55748 | 4.23367 | 1.25887 | Intron (ENSMUST00000038628.3/ENSMUSG00000037306.8, intron 2 of 12) | 4 | 134567889 | 134703767 | 135879 | 2 | ENSMUST00000176606.1 | 27710 | Man1c1 | ENSMUSG00000037306.8 |
| chr1 | 193186826 | 193187102 | 3.36311 | 3.90937 | 1.11815 | Promoter | 1 | 193184663 | 193187140 | 2478 | 2 | ENSMUST00000159666.1 | 38 | Traf3ip3 | ENSMUSG00000037318.5 |
| chr11 | 57825807 | 57826191 | 3.56651 | 4.29995 | 1.36968 | Downstream (2-3kb) | 11 | 57828707 | 57832147 | 3441 | 2 | ENSMUST00000036917.2 | 5956 | Hand1 | ENSMUSG00000037335.8 |
| chr11 | 57826806 | 57828034 | 3.73259 | 4.70774 | 1.62024 | Downstream (<1kb) | 11 | 57828707 | 57832147 | 3441 | 2 | ENSMUST00000036917.2 | 4113 | Hand1 | ENSMUSG00000037335.8 |
| chr11 | 57828389 | 57828520 | 3.5741 | 4.39128 | 1.37107 | Downstream (<1kb) | 11 | 57828707 | 57832147 | 3441 | 2 | ENSMUST00000036917.2 | 3627 | Hand1 | ENSMUSG00000037335.8 |
| chr11 | 57828850 | 57830955 | 4.14187 | 5.50739 | 2.17408 | 3' UTR | 11 | 57828707 | 57832147 | 3441 | 2 | ENSMUST00000036917.2 | 1192 | Hand1 | ENSMUSG00000037335.8 |
| chr11 | 57831275 | 57832053 | 3.79906 | 4.72948 | 1.61388 | Promoter | 11 | 57828707 | 57832147 | 3441 | 2 | ENSMUST00000036917.2 | 94 | Hand1 | ENSMUSG00000037335.8 |
| chr11 | 57832537 | 57833935 | 3.71032 | 4.63567 | 1.55526 | Promoter | 11 | 57828705 | 57832818 | 4114 | 2 | ENSMUST00000160392.3 | 0 | Hand1 | ENSMUSG00000037335.8 |
| chr11 | 57834268 | 57834738 | 3.54349 | 4.23316 | 1.2439 | Distal Intergenic | 11 | 57831694 | 57832818 | 1125 | 2 | ENSMUST00000108845.2 | -1451 | Hand1 | ENSMUSG00000037335.8 |
| chr11 | 57836013 | 57836502 | 3.79722 | 4.74102 | 1.58323 | Distal Intergenic | 11 | 57831694 | 57832818 | 1125 | 2 | ENSMUST00000108845.2 | -3196 | Hand1 | ENSMUSG00000037335.8 |
| chr11 | 57837012 | 57837697 | 4.23857 | 5.58351 | 2.17318 | Distal Intergenic | 11 | 57831694 | 57832818 | 1125 | 2 | ENSMUST00000108845.2 | -4195 | Hand1 | ENSMUSG00000037335.8 |
| chr11 | 57838093 | 57838693 | 4.12317 | 5.36095 | 2.05014 | Distal Intergenic | 11 | 57831694 | 57832818 | 1125 | 2 | ENSMUST00000108845.2 | -5276 | Hand1 | ENSMUSG00000037335.8 |
| chr11 | 57839627 | 57839780 | 3.7059 | 4.49859 | 1.39794 | Distal Intergenic | 11 | 57831694 | 57832818 | 1125 | 2 | ENSMUST00000108845.2 | -6810 | Hand1 | ENSMUSG00000037335.8 |
| chr11 | 57840934 | 57841175 | 4.40591 | 5.87861 | 2.3676 | Distal Intergenic | 11 | 57831694 | 57832818 | 1125 | 2 | ENSMUST00000108845.2 | -8117 | Hand1 | ENSMUSG00000037335.8 |
| chr11 | 57844116 | 57844191 | 3.95198 | 5.01804 | 1.75485 | Distal Intergenic | 11 | 57831694 | 57832818 | 1125 | 2 | ENSMUST00000108845.2 | -11299 | Hand1 | ENSMUSG00000037335.8 |
| chr7 | 29001575 | 29001650 | 3.85908 | 4.89934 | 1.70668 | Exon (ENSMUST00000085835.6/ENSMUSG00000037337.10, exon 28 of 32) | 7 | 28982854 | 29003279 | 20426 | 1 | ENSMUST00000085835.6 | 18722 | Map4k1 | ENSMUSG00000037337.10 |
| chr7 | 29002060 | 29002577 | 3.68719 | 4.54028 | 1.44236 | Exon (ENSMUST00000085835.6/ENSMUSG00000037337.10, exon 31 of 32) | 7 | 28982854 | 29003279 | 20426 | 1 | ENSMUST00000085835.6 | 19207 | Map4k1 | ENSMUSG00000037337.10 |
| chr7 | 29019371 | 29019915 | 4.658 | 6.42732 | 2.80787 | Exon (ENSMUST00000179893.2/ENSMUSG00000030592.11, exon 91 of 106) | 7 | 28982854 | 29003279 | 20426 | 1 | ENSMUST00000085835.6 | 36518 | Map4k1 | ENSMUSG00000037337.10 |
| chr5 | 33540370 | 33540726 | 3.61055 | 4.31825 | 1.27308 | Distal Intergenic | 5 | 33600356 | 33628912 | 28557 | 2 | ENSMUST00000065162.5 | 88186 | Fam53a | ENSMUSG00000037339.12 |
| chr5 | 33542393 | 33542578 | 3.52897 | 4.16868 | 1.17983 | Distal Intergenic | 5 | 33600356 | 33628912 | 28557 | 2 | ENSMUST00000065162.5 | 86334 | Fam53a | ENSMUSG00000037339.12 |
| chr5 | 33563185 | 33563543 | 3.54338 | 4.20138 | 1.21834 | Distal Intergenic | 5 | 33600356 | 33628912 | 28557 | 2 | ENSMUST00000065162.5 | 65369 | Fam53a | ENSMUSG00000037339.12 |
| chr15 | 54745898 | 54746006 | 3.60223 | 4.36998 | 1.43377 | Promoter | 15 | 54745702 | 54754039 | 8338 | 1 | ENSMUST00000050027.8 | 197 | Nov | ENSMUSG00000037362.8 |
| chr15 | 54746319 | 54746635 | 3.28985 | 3.84219 | 1.08849 | Promoter | 15 | 54745702 | 54754039 | 8338 | 1 | ENSMUST00000050027.8 | 618 | Nov | ENSMUSG00000037362.8 |
| chr10 | 24639454 | 24639727 | 4.09421 | 5.33748 | 2.06504 | 3' UTR | 10 | 24668486 | 24674258 | 5773 | 2 | ENSMUST00000142974.1 | 34531 | Enpp1 | ENSMUSG00000037370.8 |
| chr10 | 24679608 | 24679756 | 3.36136 | 3.88517 | 1.07256 | Intron (ENSMUST00000105520.2/ENSMUSG00000037370.8, intron 1 of 24) | 10 | 24668486 | 24674258 | 5773 | 2 | ENSMUST00000142974.1 | -5351 | Enpp1 | ENSMUSG00000037370.8 |
| chr10 | 24681671 | 24681917 | 3.35128 | 3.86645 | 1.06004 | Intron (ENSMUST00000105520.2/ENSMUSG00000037370.8, intron 1 of 24) | 10 | 24668486 | 24674258 | 5773 | 2 | ENSMUST00000142974.1 | -7414 | Enpp1 | ENSMUSG00000037370.8 |
| chr10 | 24708380 | 24708510 | 3.35459 | 3.85935 | 1.01733 | Intron (ENSMUST00000105520.2/ENSMUSG00000037370.8, intron 1 of 24) | 10 | 24641806 | 24711991 | 70186 | 2 | ENSMUST00000135846.1 | 3481 | Enpp1 | ENSMUSG00000037370.8 |
| chr10 | 24710259 | 24710334 | 3.70639 | 4.51435 | 1.44205 | Intron (ENSMUST00000105520.2/ENSMUSG00000037370.8, intron 1 of 24) | 10 | 24641806 | 24711991 | 70186 | 2 | ENSMUST00000135846.1 | 1657 | Enpp1 | ENSMUSG00000037370.8 |
| chr10 | 24711340 | 24711872 | 3.49239 | 4.20741 | 1.26405 | Promoter | 10 | 24641806 | 24711991 | 70186 | 2 | ENSMUST00000135846.1 | 119 | Enpp1 | ENSMUSG00000037370.8 |
| chr15 | 39193546 | 39193626 | 3.81973 | 4.71507 | 1.53861 | Distal Intergenic | 15 | 39198332 | 39681940 | 483609 | 1 | ENSMUST00000082054.6 | -4706 | Rims2 | ENSMUSG00000037386.10 |
| chr15 | 39196891 | 39197930 | 3.86963 | 4.97825 | 1.864 | Promoter | 15 | 39198332 | 39681940 | 483609 | 1 | ENSMUST00000082054.6 | -402 | Rims2 | ENSMUSG00000037386.10 |
| chr15 | 39198308 | 39199125 | 4.06179 | 5.30736 | 2.03954 | Promoter | 15 | 39198332 | 39681940 | 483609 | 1 | ENSMUST00000082054.6 | 0 | Rims2 | ENSMUSG00000037386.10 |
| chr5 | 137130542 | 137130617 | 4.11405 | 5.2753 | 1.9185 | Distal Intergenic | 5 | 137134961 | 137139127 | 4167 | 2 | ENSMUST00000160630.1 | 8510 | Muc3 | ENSMUSG00000037390.10 |
| chr5 | 137136831 | 137136939 | 3.54896 | 4.31029 | 1.31621 | Intron (ENSMUST00000160533.2/ENSMUSG00000037390.10, intron 6 of 6) | 5 | 137134961 | 137139127 | 4167 | 2 | ENSMUST00000160630.1 | 2188 | Muc3 | ENSMUSG00000037390.10 |
| chr9 | 21015556 | 21016211 | 3.76366 | 4.60716 | 1.46199 | Promoter | 9 | 21015960 | 21028797 | 12838 | 1 | ENSMUST00000086399.4 | 0 | Icam1 | ENSMUSG00000037405.7 |
| chr9 | 21016819 | 21016998 | 4.22835 | 5.77123 | 2.28911 | Promoter | 9 | 21015960 | 21028797 | 12838 | 1 | ENSMUST00000086399.4 | 860 | Icam1 | ENSMUSG00000037405.7 |
| chr9 | 21017474 | 21017812 | 3.62045 | 4.43482 | 1.41813 | Intron (ENSMUST00000086399.4/ENSMUSG00000037405.7, intron 1 of 6) | 9 | 21015960 | 21028797 | 12838 | 1 | ENSMUST00000086399.4 | 1515 | Icam1 | ENSMUSG00000037405.7 |
| chr9 | 21018393 | 21018838 | 3.33617 | 3.8409 | 1.04937 | Intron (ENSMUST00000086399.4/ENSMUSG00000037405.7, intron 1 of 6) | 9 | 21015960 | 21028797 | 12838 | 1 | ENSMUST00000086399.4 | 2434 | Icam1 | ENSMUSG00000037405.7 |
| chr8 | 25037324 | 25037780 | 3.69711 | 4.64593 | 1.52163 | Promoter | 8 | 25029654 | 25037330 | 7677 | 2 | ENSMUST00000176814.1 | 0 | Htra4 | ENSMUSG00000037406.6 |
| chr8 | 25038493 | 25038648 | 3.4937 | 4.40863 | 1.42236 | Promoter | 8 | 25025451 | 25038962 | 13512 | 2 | ENSMUST00000084031.4 | 314 | Htra4 | ENSMUSG00000037406.6 |
| chr5 | 137024655 | 137024791 | 3.59713 | 4.30197 | 1.27527 | Distal Intergenic | 5 | 137026392 | 137033351 | 6960 | 1 | ENSMUST00000190827.1 | -1601 | Vgf | ENSMUSG00000037428.9 |
| chr5 | 137025692 | 137025807 | 3.37102 | 3.88858 | 1.03303 | Promoter | 5 | 137026392 | 137033351 | 6960 | 1 | ENSMUST00000190827.1 | -585 | Vgf | ENSMUSG00000037428.9 |
| chr5 | 137029398 | 137029689 | 3.81725 | 4.72585 | 1.57068 | Promoter | 5 | 137030295 | 137033351 | 3057 | 1 | ENSMUST00000041543.8 | -606 | Vgf | ENSMUSG00000037428.9 |
| chr5 | 137030583 | 137030741 | 3.50333 | 4.30592 | 1.33002 | Promoter | 5 | 137030701 | 137031407 | 707 | 1 | ENSMUST00000187382.1 | 0 | Vgf | ENSMUSG00000037428.9 |
| chr5 | 137031104 | 137031769 | 4.42062 | 6.18401 | 2.64651 | Promoter | 5 | 137030701 | 137031407 | 707 | 1 | ENSMUST00000187382.1 | 404 | Vgf | ENSMUSG00000037428.9 |
| chr17 | 28362303 | 28362677 | 3.67086 | 4.4721 | 1.42674 | Promoter | 17 | 28358450 | 28362432 | 3983 | 2 | ENSMUST00000131604.1 | 0 | Tulp1 | ENSMUSG00000037446.8 |
| chr17 | 28363313 | 28364484 | 3.72345 | 4.71538 | 1.59724 | Promoter | 17 | 28361650 | 28362957 | 1308 | 2 | ENSMUST00000125876.1 | -357 | Tulp1 | ENSMUSG00000037446.8 |
| chr17 | 28364842 | 28365256 | 3.90134 | 4.90929 | 1.73454 | Promoter | 17 | 28351515 | 28365148 | 13634 | 2 | ENSMUST00000041819.7 | 0 | Tulp1 | ENSMUSG00000037446.8 |
| chr17 | 28374799 | 28375308 | 3.92465 | 4.93943 | 1.71609 | Distal Intergenic | 17 | 28355817 | 28365182 | 9366 | 2 | ENSMUST00000114794.1 | -9618 | Tulp1 | ENSMUSG00000037446.8 |
| chr17 | 28375865 | 28375992 | 3.98903 | 5.30444 | 1.97771 | Distal Intergenic | 17 | 28355817 | 28365182 | 9366 | 2 | ENSMUST00000114794.1 | -10684 | Tulp1 | ENSMUSG00000037446.8 |
| chr17 | 28379858 | 28380012 | 3.45085 | 4.03634 | 1.12974 | Distal Intergenic | 17 | 28355817 | 28365182 | 9366 | 2 | ENSMUST00000114794.1 | -14677 | Tulp1 | ENSMUSG00000037446.8 |
| chr17 | 28388678 | 28388753 | 3.83001 | 4.80657 | 1.64294 | Distal Intergenic | 17 | 28355817 | 28365182 | 9366 | 2 | ENSMUST00000114794.1 | -23497 | Tulp1 | ENSMUSG00000037446.8 |
| chr17 | 28391445 | 28392250 | 3.7225 | 4.56598 | 1.50432 | Distal Intergenic | 17 | 28355817 | 28365182 | 9366 | 2 | ENSMUST00000114794.1 | -26264 | Tulp1 | ENSMUSG00000037446.8 |
| chr7 | 28664802 | 28665147 | 3.61316 | 4.43957 | 1.42991 | Distal Intergenic | 7 | 28692849 | 28699338 | 6490 | 1 | ENSMUST00000039998.5 | -27702 | Fbxo27 | ENSMUSG00000037463.9 |
| chr7 | 28665959 | 28666054 | 3.55528 | 4.24475 | 1.31168 | Distal Intergenic | 7 | 28692849 | 28699338 | 6490 | 1 | ENSMUST00000039998.5 | -26795 | Fbxo27 | ENSMUSG00000037463.9 |
| chr7 | 28695102 | 28695177 | 3.51931 | 4.1607 | 1.20378 | Intron (ENSMUST00000039998.5/ENSMUSG00000037463.9, intron 4 of 5) | 7 | 28694019 | 28699335 | 5317 | 1 | ENSMUST00000108280.1 | 1084 | Fbxo27 | ENSMUSG00000037463.9 |
| chr7 | 28628966 | 28629052 | 3.29896 | 3.78806 | 1.06062 | Intron (ENSMUST00000040112.3/ENSMUSG00000037469.4, intron 2 of 12) | 7 | 28629491 | 28630500 | 1010 | 2 | ENSMUST00000159095.1 | 1448 | Papl | ENSMUSG00000037469.4 |
| chr7 | 28630250 | 28631387 | 3.92633 | 5.08711 | 1.84887 | Promoter | 7 | 28607634 | 28631067 | 23434 | 2 | ENSMUST00000040112.3 | 0 | Papl | ENSMUSG00000037469.4 |
| chr8 | 23669602 | 23670420 | 3.94728 | 5.03228 | 1.82859 | Promoter | 8 | 23669658 | 24063105 | 393448 | 1 | ENSMUST00000042352.5 | 0 | Zmat4 | ENSMUSG00000037492.10 |
| chr1 | 34678143 | 34678230 | 3.4443 | 4.04796 | 1.20702 | Promoter | 1 | 34678188 | 34812739 | 134552 | 1 | ENSMUST00000159747.3 | 0 | Arhgef4 | ENSMUSG00000037509.13 |
| chr1 | 34678561 | 34678925 | 3.37048 | 3.90393 | 1.09014 | Promoter | 1 | 34678188 | 34812739 | 134552 | 1 | ENSMUST00000159747.3 | 374 | Arhgef4 | ENSMUSG00000037509.13 |
| chr1 | 34679264 | 34679760 | 3.89867 | 4.93953 | 1.80084 | Intron (ENSMUST00000159747.3/ENSMUSG00000037509.13, intron 1 of 13) | 1 | 34678188 | 34812739 | 134552 | 1 | ENSMUST00000159747.3 | 1077 | Arhgef4 | ENSMUSG00000037509.13 |
| chr1 | 34680143 | 34680475 | 3.3969 | 4.1624 | 1.25271 | Intron (ENSMUST00000159747.3/ENSMUSG00000037509.13, intron 1 of 13) | 1 | 34678188 | 34812739 | 134552 | 1 | ENSMUST00000159747.3 | 1956 | Arhgef4 | ENSMUSG00000037509.13 |
| chr1 | 34748894 | 34749085 | 3.37948 | 3.89691 | 1.01733 | Intron (ENSMUST00000159747.3/ENSMUSG00000037509.13, intron 4 of 13) | 1 | 34750881 | 34807144 | 56264 | 1 | ENSMUST00000159021.2 | -1796 | Arhgef4 | ENSMUSG00000037509.13 |
| chr1 | 34750290 | 34751032 | 4.55268 | 6.32131 | 2.72441 | Promoter | 1 | 34750881 | 34807144 | 56264 | 1 | ENSMUST00000159021.2 | 0 | Arhgef4 | ENSMUSG00000037509.13 |
| chr1 | 34751675 | 34751796 | 3.54934 | 4.2974 | 1.29397 | Promoter | 1 | 34750881 | 34807144 | 56264 | 1 | ENSMUST00000159021.2 | 795 | Arhgef4 | ENSMUSG00000037509.13 |
| chr1 | 34752501 | 34752601 | 3.77415 | 4.80098 | 1.65772 | Intron (ENSMUST00000159747.3/ENSMUSG00000037509.13, intron 4 of 13) | 1 | 34750881 | 34807144 | 56264 | 1 | ENSMUST00000159021.2 | 1621 | Arhgef4 | ENSMUSG00000037509.13 |
| chr15 | 99498119 | 99498457 | 3.39422 | 3.92552 | 1.03989 | 3' UTR | 15 | 99470084 | 99474730 | 4647 | 2 | ENSMUST00000040313.4 | -23390 | Bcdin3d | ENSMUSG00000037525.4 |
| chr10 | 21465984 | 21466234 | 3.28343 | 3.89696 | 1.04779 | Distal Intergenic | 10 | 21425342 | 21425650 | 309 | 1 | ENSMUST00000042671.4 | 40643 | 1700021A07Rik | ENSMUSG00000037535.5 |
| chr7 | 143977080 | 143979506 | 3.76623 | 4.65276 | 1.53362 | Distal Intergenic | 7 | 144001928 | 144424494 | 422567 | 1 | ENSMUST00000105902.2 | -22422 | Shank2 | ENSMUSG00000037541.14 |
| chr7 | 143980222 | 143980633 | 3.90887 | 5.10311 | 1.83672 | Distal Intergenic | 7 | 144001928 | 144424494 | 422567 | 1 | ENSMUST00000105902.2 | -21295 | Shank2 | ENSMUSG00000037541.14 |
| chr7 | 144153861 | 144153936 | 3.85022 | 4.77336 | 1.57414 | Intron (ENSMUST00000105902.2/ENSMUSG00000037541.14, intron 10 of 22) | 7 | 144175529 | 144424494 | 248966 | 1 | ENSMUST00000105900.2 | -21593 | Shank2 | ENSMUSG00000037541.14 |
| chr7 | 144175494 | 144175784 | 3.41958 | 4.13402 | 1.21879 | Promoter | 7 | 144175529 | 144424494 | 248966 | 1 | ENSMUST00000105900.2 | 0 | Shank2 | ENSMUSG00000037541.14 |
| chr7 | 144178043 | 144178311 | 3.44814 | 4.03211 | 1.12928 | Intron (ENSMUST00000105902.2/ENSMUSG00000037541.14, intron 10 of 22) | 7 | 144175529 | 144424494 | 248966 | 1 | ENSMUST00000105900.2 | 2515 | Shank2 | ENSMUSG00000037541.14 |
| chr7 | 144284077 | 144284526 | 3.70261 | 4.50183 | 1.41224 | Promoter | 7 | 144284385 | 144422675 | 138291 | 1 | ENSMUST00000097929.2 | 0 | Shank2 | ENSMUSG00000037541.14 |
| chr7 | 144284953 | 144285028 | 3.65505 | 4.39889 | 1.32037 | Promoter | 7 | 144284439 | 144424482 | 140044 | 1 | ENSMUST00000146006.1 | 515 | Shank2 | ENSMUSG00000037541.14 |
| chr7 | 144417409 | 144417778 | 3.3522 | 3.85578 | 1.01733 | Promoter | 7 | 144417825 | 144420664 | 2840 | 1 | ENSMUST00000136979.1 | -47 | Shank2 | ENSMUSG00000037541.14 |
| chr1 | 190976716 | 190976791 | 3.75609 | 4.71095 | 1.58733 | Intron (ENSMUST00000047409.7/ENSMUSG00000037568.7, intron 2 of 7) | 1 | 190947648 | 190978998 | 31351 | 2 | ENSMUST00000047409.7 | 2207 | Vash2 | ENSMUSG00000037568.7 |
| chr1 | 190978112 | 190978559 | 3.74218 | 4.70629 | 1.59356 | Promoter | 1 | 190947648 | 190978998 | 31351 | 2 | ENSMUST00000047409.7 | 439 | Vash2 | ENSMUSG00000037568.7 |
| chr1 | 190978942 | 190979120 | 3.9468 | 5.04062 | 1.84875 | Promoter | 1 | 190947648 | 190978998 | 31351 | 2 | ENSMUST00000047409.7 | 0 | Vash2 | ENSMUSG00000037568.7 |
| chr1 | 190981760 | 190982013 | 3.47004 | 4.22445 | 1.24222 | Distal Intergenic | 1 | 190949573 | 190979296 | 29724 | 2 | ENSMUST00000166139.1 | -2465 | Vash2 | ENSMUSG00000037568.7 |
| chr17 | 32188838 | 32189548 | 3.66629 | 4.44253 | 1.39075 | Promoter | 17 | 32183770 | 32189463 | 5694 | 2 | ENSMUST00000087721.4 | 0 | Ephx3 | ENSMUSG00000037577.7 |
| chr15 | 99223154 | 99223396 | 3.81377 | 4.76972 | 1.676 | Distal Intergenic | 15 | 99224976 | 99242817 | 17842 | 1 | ENSMUST00000041415.3 | -1580 | Kcnh3 | ENSMUSG00000037579.6 |
| chr14 | 47188508 | 47188984 | 3.32002 | 3.83804 | 1.11087 | Promoter | 14 | 47153895 | 47189413 | 35519 | 2 | ENSMUST00000089959.6 | 429 | Gch1 | ENSMUSG00000037580.9 |
| chr4 | 133518115 | 133518389 | 3.32674 | 3.85446 | 1.12429 | Promoter | 4 | 133518963 | 133528768 | 9806 | 1 | ENSMUST00000149807.1 | -574 | Kdf1 | ENSMUSG00000037600.11 |
| chr4 | 133518921 | 133519734 | 3.96678 | 5.31391 | 2.02748 | Promoter | 4 | 133518963 | 133528768 | 9806 | 1 | ENSMUST00000149807.1 | 0 | Kdf1 | ENSMUSG00000037600.11 |
| chr4 | 133520579 | 133520880 | 3.67875 | 4.58786 | 1.521 | Intron (ENSMUST00000149807.1/ENSMUSG00000037600.11, intron 1 of 1) | 4 | 133519083 | 133530787 | 11705 | 1 | ENSMUST00000121797.1 | 1497 | Kdf1 | ENSMUSG00000037600.11 |
| chr5 | 81019678 | 81020655 | 3.66237 | 4.51827 | 1.44042 | Promoter | 5 | 81020138 | 81388784 | 368647 | 1 | ENSMUST00000147226.2 | 0 | Lphn3 | ENSMUSG00000037605.11 |
| chr5 | 81021000 | 81022475 | 3.80227 | 4.87924 | 1.71013 | Promoter | 5 | 81021199 | 81467610 | 446412 | 1 | ENSMUST00000140356.2 | 0 | Lphn3 | ENSMUSG00000037605.11 |
| chr5 | 81022851 | 81023347 | 4.39652 | 5.92218 | 2.45199 | Intron (ENSMUST00000147226.2/ENSMUSG00000037605.11, intron 1 of 5) | 5 | 81021810 | 81561051 | 539242 | 1 | ENSMUST00000127046.1 | 1042 | Lphn3 | ENSMUSG00000037605.11 |
| chr5 | 81024574 | 81024896 | 3.51555 | 4.15264 | 1.19531 | Intron (ENSMUST00000147226.2/ENSMUSG00000037605.11, intron 1 of 5) | 5 | 81021810 | 81561051 | 539242 | 1 | ENSMUST00000127046.1 | 2765 | Lphn3 | ENSMUSG00000037605.11 |
| chr3 | 31901555 | 31901698 | 3.57544 | 4.27064 | 1.29456 | Promoter | 3 | 31902507 | 32199765 | 297259 | 1 | ENSMUST00000119310.2 | -809 | Kcnmb2 | ENSMUSG00000037610.10 |
| chr3 | 31903035 | 31903217 | 3.74375 | 4.64008 | 1.60373 | Promoter | 3 | 31902703 | 32200180 | 297478 | 1 | ENSMUST00000119970.2 | 333 | Kcnmb2 | ENSMUSG00000037610.10 |
| chr3 | 31903647 | 31904461 | 3.64743 | 4.54725 | 1.49572 | Promoter | 3 | 31902703 | 32200180 | 297478 | 1 | ENSMUST00000119970.2 | 945 | Kcnmb2 | ENSMUSG00000037610.10 |
| chr1 | 189341286 | 189341821 | 3.75167 | 4.65605 | 1.49562 | Intron (ENSMUST00000079451.7/ENSMUSG00000037624.10, intron 1 of 6) | 1 | 189207931 | 189343353 | 135423 | 2 | ENSMUST00000079451.7 | 1532 | Kcnk2 | ENSMUSG00000037624.10 |
| chr1 | 189343538 | 189343731 | 3.50153 | 4.40971 | 1.41806 | Promoter | 1 | 189207931 | 189343704 | 135774 | 2 | ENSMUST00000180044.2 | 0 | Kcnk2 | ENSMUSG00000037624.10 |
| chr1 | 189344226 | 189344487 | 3.3752 | 3.89044 | 1.01733 | Promoter | 1 | 189207937 | 189343832 | 135896 | 2 | ENSMUST00000110920.1 | -395 | Kcnk2 | ENSMUSG00000037624.10 |
| chr3 | 31149814 | 31149981 | 3.82004 | 4.72979 | 1.57555 | Promoter | 3 | 31149946 | 31164324 | 14379 | 1 | ENSMUST00000046174.6 | 0 | Cldn11 | ENSMUSG00000037625.6 |
| chr15 | 36139651 | 36140643 | 4.75111 | 6.9004 | 3.31991 | Promoter | 15 | 36009479 | 36140400 | 130922 | 2 | ENSMUST00000172831.2 | 0 | Rgs22 | ENSMUSG00000037627.10 |
| chr12 | 8294489 | 8294655 | 3.67224 | 4.48061 | 1.45154 | Distal Intergenic | 12 | 8297934 | 8301954 | 4021 | 2 | ENSMUST00000037313.4 | 7299 | Gdf7 | ENSMUSG00000037660.4 |
| chr12 | 8295636 | 8296224 | 3.43114 | 4.00785 | 1.13461 | Downstream (1-2kb) | 12 | 8297934 | 8301954 | 4021 | 2 | ENSMUST00000037313.4 | 5730 | Gdf7 | ENSMUSG00000037660.4 |
| chr12 | 8297215 | 8302524 | 4.07978 | 5.32473 | 2.04701 | Promoter | 12 | 8297934 | 8301954 | 4021 | 2 | ENSMUST00000037313.4 | 0 | Gdf7 | ENSMUSG00000037660.4 |
| chr12 | 8305047 | 8305278 | 3.36653 | 3.87736 | 1.01733 | Distal Intergenic | 12 | 8297934 | 8301954 | 4021 | 2 | ENSMUST00000037313.4 | -3094 | Gdf7 | ENSMUSG00000037660.4 |
| chr9 | 99358049 | 99358865 | 3.90149 | 5.08874 | 1.83189 | Promoter | 9 | 99309963 | 99358518 | 48556 | 2 | ENSMUST00000042158.7 | 0 | Esyt3 | ENSMUSG00000037681.8 |
| chr2 | 18696223 | 18696983 | 3.58255 | 4.38573 | 1.3888 | Intron (ENSMUST00000137796.2/ENSMUSG00000087182.2, intron 1 of 2) | 2 | 18699023 | 18750413 | 51391 | 1 | ENSMUST00000095132.4 | -2040 | BC061194 | ENSMUSG00000037708.11 |
| chr2 | 18697521 | 18697969 | 3.60221 | 4.35367 | 1.39976 | Intron (ENSMUST00000137796.2/ENSMUSG00000087182.2, intron 1 of 2) | 2 | 18699023 | 18750413 | 51391 | 1 | ENSMUST00000095132.4 | -1054 | BC061194 | ENSMUSG00000037708.11 |
| chr2 | 18698325 | 18699710 | 3.99991 | 5.20076 | 1.93009 | Promoter | 2 | 18699023 | 18750413 | 51391 | 1 | ENSMUST00000095132.4 | 0 | BC061194 | ENSMUSG00000037708.11 |
| chr2 | 18700718 | 18701104 | 3.56184 | 4.47494 | 1.44452 | Intron (ENSMUST00000095132.4/ENSMUSG00000037708.11, intron 2 of 10) | 2 | 18699084 | 18732034 | 32951 | 1 | ENSMUST00000173763.1 | 1635 | BC061194 | ENSMUSG00000037708.11 |
| chr2 | 18702134 | 18702634 | 3.8374 | 5.04756 | 1.81466 | Intron (ENSMUST00000095132.4/ENSMUSG00000037708.11, intron 2 of 10) | 2 | 18699084 | 18732034 | 32951 | 1 | ENSMUST00000173763.1 | 3051 | BC061194 | ENSMUSG00000037708.11 |
| chr2 | 18705718 | 18705821 | 3.49223 | 4.15223 | 1.28868 | Intron (ENSMUST00000095132.4/ENSMUSG00000037708.11, intron 2 of 10) | 2 | 18699084 | 18732034 | 32951 | 1 | ENSMUST00000173763.1 | 6635 | BC061194 | ENSMUSG00000037708.11 |
| chr2 | 18711616 | 18711847 | 4.4189 | 5.92331 | 2.42298 | Intron (ENSMUST00000095132.4/ENSMUSG00000037708.11, intron 3 of 10) | 2 | 18699084 | 18732034 | 32951 | 1 | ENSMUST00000173763.1 | 12533 | BC061194 | ENSMUSG00000037708.11 |
| chr9 | 58037869 | 58038327 | 3.47788 | 4.08305 | 1.15081 | Intron (ENSMUST00000153106.1/ENSMUSG00000086624.1, intron 4 of 4) | 9 | 58033232 | 58042036 | 8805 | 2 | ENSMUST00000143797.1 | 3709 | Ccdc33 | ENSMUSG00000037716.9 |
| chr9 | 58113655 | 58113770 | 3.83986 | 4.75908 | 1.56891 | Promoter | 9 | 58067503 | 58114360 | 46858 | 2 | ENSMUST00000146741.1 | 590 | Ccdc33 | ENSMUSG00000037716.9 |
| chr9 | 58114172 | 58114260 | 3.30793 | 3.78911 | 1.01733 | Promoter | 9 | 58067503 | 58114360 | 46858 | 2 | ENSMUST00000146741.1 | 100 | Ccdc33 | ENSMUSG00000037716.9 |
| chr9 | 58085056 | 58085216 | 3.36678 | 3.87775 | 1.01733 | Intron (ENSMUST00000145886.1/ENSMUSG00000032327.9, intron 1 of 4) | 9 | 58077013 | 58082069 | 5057 | 2 | ENSMUST00000151404.1 | -2988 | Ccdc33 | ENSMUSG00000037716.9 |
| chr9 | 58112255 | 58112487 | 3.45522 | 4.0383 | 1.11267 | Intron (ENSMUST00000145886.1/ENSMUSG00000032327.9, intron 1 of 4) | 9 | 58067503 | 58114360 | 46858 | 2 | ENSMUST00000146741.1 | 1873 | Ccdc33 | ENSMUSG00000037716.9 |
| chr9 | 58114597 | 58114735 | 3.49293 | 4.12659 | 1.18693 | Promoter | 9 | 58067503 | 58114360 | 46858 | 2 | ENSMUST00000146741.1 | -238 | Ccdc33 | ENSMUSG00000037716.9 |
| chr5 | 67184659 | 67185131 | 3.34253 | 3.88571 | 1.12074 | Distal Intergenic | 5 | 67260565 | 67291461 | 30897 | 1 | ENSMUST00000037918.6 | -75434 | Tmem33 | ENSMUSG00000037720.11 |
| chr5 | 67208082 | 67208376 | 3.6832 | 4.52117 | 1.5134 | Distal Intergenic | 5 | 67260565 | 67291461 | 30897 | 1 | ENSMUST00000037918.6 | -52189 | Tmem33 | ENSMUSG00000037720.11 |
| chr5 | 67211889 | 67212537 | 3.84582 | 4.83225 | 1.66985 | Distal Intergenic | 5 | 67260565 | 67291461 | 30897 | 1 | ENSMUST00000037918.6 | -48028 | Tmem33 | ENSMUSG00000037720.11 |
| chr5 | 67213435 | 67213756 | 3.34602 | 3.84643 | 1.01733 | Distal Intergenic | 5 | 67260565 | 67291461 | 30897 | 1 | ENSMUST00000037918.6 | -46809 | Tmem33 | ENSMUSG00000037720.11 |
| chr5 | 67214086 | 67214161 | 3.74284 | 4.55901 | 1.41778 | Distal Intergenic | 5 | 67260565 | 67291461 | 30897 | 1 | ENSMUST00000037918.6 | -46404 | Tmem33 | ENSMUSG00000037720.11 |
| chr5 | 67214605 | 67214912 | 3.584 | 4.33111 | 1.34437 | Distal Intergenic | 5 | 67260565 | 67291461 | 30897 | 1 | ENSMUST00000037918.6 | -45653 | Tmem33 | ENSMUSG00000037720.11 |
| chr5 | 67215606 | 67215734 | 4.51959 | 6.12852 | 2.56372 | Distal Intergenic | 5 | 67260565 | 67291461 | 30897 | 1 | ENSMUST00000037918.6 | -44831 | Tmem33 | ENSMUSG00000037720.11 |
| chr5 | 67216340 | 67217161 | 3.9403 | 5.02305 | 1.77403 | Distal Intergenic | 5 | 67260565 | 67291461 | 30897 | 1 | ENSMUST00000037918.6 | -43404 | Tmem33 | ENSMUSG00000037720.11 |
| chr5 | 67219586 | 67219702 | 3.52604 | 4.36911 | 1.37299 | Distal Intergenic | 5 | 67260565 | 67291461 | 30897 | 1 | ENSMUST00000037918.6 | -40863 | Tmem33 | ENSMUSG00000037720.11 |
| chr5 | 67220071 | 67221088 | 3.70135 | 4.67601 | 1.56687 | Distal Intergenic | 5 | 67260565 | 67291461 | 30897 | 1 | ENSMUST00000037918.6 | -39477 | Tmem33 | ENSMUSG00000037720.11 |
| chr5 | 67222216 | 67223084 | 3.9534 | 5.34006 | 2.03764 | Distal Intergenic | 5 | 67260565 | 67291461 | 30897 | 1 | ENSMUST00000037918.6 | -37481 | Tmem33 | ENSMUSG00000037720.11 |
| chr5 | 67223625 | 67223785 | 3.43915 | 4.39871 | 1.39069 | Distal Intergenic | 5 | 67260565 | 67291461 | 30897 | 1 | ENSMUST00000037918.6 | -36780 | Tmem33 | ENSMUSG00000037720.11 |
| chr5 | 67224733 | 67225906 | 3.92769 | 5.05276 | 1.85421 | Distal Intergenic | 5 | 67260565 | 67291461 | 30897 | 1 | ENSMUST00000037918.6 | -34659 | Tmem33 | ENSMUSG00000037720.11 |
| chr5 | 67226235 | 67226368 | 3.25246 | 3.71172 | 1.03635 | Distal Intergenic | 5 | 67260565 | 67291461 | 30897 | 1 | ENSMUST00000037918.6 | -34197 | Tmem33 | ENSMUSG00000037720.11 |
| chr5 | 67226829 | 67226932 | 5.12861 | 7.5677 | 3.75774 | Distal Intergenic | 5 | 67260565 | 67291461 | 30897 | 1 | ENSMUST00000037918.6 | -33633 | Tmem33 | ENSMUSG00000037720.11 |
| chr5 | 67227360 | 67228668 | 4.03001 | 5.25621 | 1.94596 | Distal Intergenic | 5 | 67260565 | 67291461 | 30897 | 1 | ENSMUST00000037918.6 | -31897 | Tmem33 | ENSMUSG00000037720.11 |
| chr5 | 67229069 | 67230402 | 4.01386 | 5.24095 | 1.93128 | Distal Intergenic | 5 | 67260565 | 67291461 | 30897 | 1 | ENSMUST00000037918.6 | -30163 | Tmem33 | ENSMUSG00000037720.11 |

| | | | | | | | | | | | | | | | |
|---|---|---|---|---|---|---|---|---|---|---|---|---|---|---|---|
| chr5 | 67230726 | 67232005 | 3.87534 | 4.91626 | 1.71075 | Distal Intergenic | 5 | 67260565 | 67291461 | 30897 | 1 | ENSMUST00000037918.6 | -28560 | Tmem33 | ENSMUSG00000037720.11 |
| chr5 | 67232649 | 67233006 | 3.41689 | 4.20956 | 1.24634 | Distal Intergenic | 5 | 67260565 | 67291461 | 30897 | 1 | ENSMUST00000037918.6 | -27559 | Tmem33 | ENSMUSG00000037720.11 |
| chr5 | 67233324 | 67233922 | 3.81602 | 4.74471 | 1.63142 | Distal Intergenic | 5 | 67260565 | 67291461 | 30897 | 1 | ENSMUST00000037918.6 | -26643 | Tmem33 | ENSMUSG00000037720.11 |
| chr5 | 67234601 | 67235246 | 3.37397 | 3.9023 | 1.06587 | Distal Intergenic | 5 | 67260565 | 67291461 | 30897 | 1 | ENSMUST00000037918.6 | -25319 | Tmem33 | ENSMUSG00000037720.11 |
| chr2 | 130578895 | 130581376 | 4.02439 | 5.29085 | 2.05376 | 3' UTR | 2 | 130580620 | 130582554 | 1935 | 2 | ENSMUST00000046001.6 | 1178 | Avp | ENSMUSG00000037727.6 |
| chr4 | 132793899 | 132794260 | 3.65707 | 4.5272 | 1.50509 | Intron (ENSMUST00000045154.5/ENSMUSG00000037731.5, intron 1 of 5) | 4 | 132781843 | 132796387 | 14545 | 2 | ENSMUST00000045154.5 | 2127 | Themis2 | ENSMUSG00000037731.5 |
| chr4 | 132794914 | 132795389 | 3.34877 | 4.23182 | 1.27728 | Promoter | 4 | 132781843 | 132796387 | 14545 | 2 | ENSMUST00000045154.5 | 998 | Themis2 | ENSMUSG00000037731.5 |
| chr4 | 132795735 | 132796037 | 3.51769 | 4.24823 | 1.32418 | Promoter | 4 | 132781843 | 132796387 | 14545 | 2 | ENSMUST00000045154.5 | 350 | Themis2 | ENSMUSG00000037731.5 |
| chr5 | 66747000 | 66747147 | 3.43153 | 4.13696 | 1.20143 | Intron (ENSMUST00000101164.5/ENSMUSG00000037736.13, intron 1 of 25) | 5 | 66745889 | 66953678 | 207790 | 1 | ENSMUST00000130228.1 | 1112 | Limch1 | ENSMUSG00000037736.13 |
| chr5 | 66747759 | 66748166 | 3.43447 | 4.06093 | 1.23928 | Intron (ENSMUST00000101164.5/ENSMUSG00000037736.13, intron 1 of 25) | 5 | 66745889 | 66953678 | 207790 | 1 | ENSMUST00000130228.1 | 1871 | Limch1 | ENSMUSG00000037736.13 |
| chr5 | 66969915 | 66970218 | 3.7262 | 4.73085 | 1.61279 | Promoter | 5 | 66968961 | 66998686 | 29726 | 1 | ENSMUST00000132991.1 | 955 | Limch1 | ENSMUSG00000037736.13 |
| chr10 | 70597399 | 70597718 | 3.64485 | 4.44173 | 1.47085 | Intron (ENSMUST00000046513.4/ENSMUSG00000037747.4, intron 1 of 4) | 10 | 70563865 | 70599082 | 35218 | 2 | ENSMUST00000162470.1 | 1364 | Phyhipl | ENSMUSG00000037747.4 |
| chr10 | 70598156 | 70598251 | 4.08373 | 5.53929 | 2.14189 | Promoter | 10 | 70563865 | 70599082 | 35218 | 2 | ENSMUST00000162470.1 | 831 | Phyhipl | ENSMUSG00000037747.4 |
| chr10 | 70598582 | 70599256 | 3.58278 | 4.46434 | 1.4225 | Promoter | 10 | 70563865 | 70599082 | 35218 | 2 | ENSMUST00000162470.1 | 0 | Phyhipl | ENSMUSG00000037747.4 |
| chr10 | 70599825 | 70600384 | 4.25957 | 5.93385 | 2.47085 | Promoter | 10 | 70557682 | 70599291 | 41610 | 2 | ENSMUST00000046513.4 | -535 | Phyhipl | ENSMUSG00000037747.4 |
| chr2 | 158665362 | 158665832 | 3.41019 | 3.95293 | 1.05141 | Promoter | 2 | 158665398 | 158759334 | 93937 | 1 | ENSMUST00000129902.1 | 0 | Ppp1r16b | ENSMUSG00000037754.8 |
| chr2 | 158666371 | 158666446 | 3.54548 | 4.19958 | 1.20087 | Promoter | 2 | 158666733 | 158766334 | 99602 | 1 | ENSMUST00000103116.4 | -287 | Ppp1r16b | ENSMUSG00000037754.8 |
| chr2 | 158666854 | 158668449 | 4.07006 | 5.26602 | 1.97054 | Promoter | 2 | 158667119 | 158766334 | 99216 | 1 | ENSMUST00000045503.5 | 0 | Ppp1r16b | ENSMUSG00000037754.8 |
| chr2 | 158669621 | 158670011 | 3.83661 | 4.75884 | 1.59343 | Intron (ENSMUST00000129902.1/ENSMUSG00000037754.8, intron 2 of 10) | 2 | 158667119 | 158766334 | 99216 | 1 | ENSMUST00000045503.5 | 2503 | Ppp1r16b | ENSMUSG00000037754.8 |
| chr2 | 158670616 | 158670916 | 3.33491 | 3.84282 | 1.06277 | Intron (ENSMUST00000129902.1/ENSMUSG00000037754.8, intron 2 of 10) | 2 | 158667119 | 158766334 | 99216 | 1 | ENSMUST00000045503.5 | 3498 | Ppp1r16b | ENSMUSG00000037754.8 |
| chr2 | 158681853 | 158682117 | 3.57156 | 4.25165 | 1.24616 | Intron (ENSMUST00000129902.1/ENSMUSG00000037754.8, intron 2 of 10) | 2 | 158696135 | 158766334 | 70200 | 1 | ENSMUST00000145073.1 | -14018 | Ppp1r16b | ENSMUSG00000037754.8 |
| chr14 | 44989663 | 44990201 | 3.72957 | 4.83249 | 1.64995 | Exon (ENSMUST00000046891.5/ENSMUSG00000037759.5, exon 1 of 2) | 14 | 44988195 | 45003820 | 15626 | 1 | ENSMUST00000046891.5 | 1469 | Ptger2 | ENSMUSG00000037759.5 |
| chr2 | 158618777 | 158620201 | 3.54032 | 4.19656 | 1.21115 | Distal Intergenic | 2 | 158624888 | 158629157 | 4270 | 1 | ENSMUST00000125390.2 | -4687 | Actr5 | ENSMUSG00000037761.11 |
| chr10 | 70244536 | 70244793 | 3.75795 | 4.6013 | 1.46675 | Promoter | 10 | 70245100 | 70285968 | 40869 | 1 | ENSMUST00000046807.6 | -307 | Slc16a9 | ENSMUSG00000037762.6 |
| chr10 | 70245437 | 70245756 | 3.34488 | 3.87832 | 1.09113 | Promoter | 10 | 70245100 | 70285968 | 40869 | 1 | ENSMUST00000046807.6 | 338 | Slc16a9 | ENSMUSG00000037762.6 |
| chr10 | 70246168 | 70246558 | 3.48469 | 4.13878 | 1.17926 | Intron (ENSMUST00000046807.6/ENSMUSG00000037762.6, intron 1 of 5) | 10 | 70245100 | 70285968 | 40869 | 1 | ENSMUST00000046807.6 | 1069 | Slc16a9 | ENSMUSG00000037762.6 |
| chr2 | 158609815 | 158611557 | 3.9796 | 5.10916 | 1.84371 | Promoter | 2 | 158610767 | 158615748 | 4982 | 1 | ENSMUST00000045738.4 | 0 | Slc32a1 | ENSMUSG00000037771.11 |
| chr2 | 158228319 | 158228417 | 4.13207 | 5.3823 | 2.02625 | Promoter | 2 | 158182533 | 158229222 | 46690 | 2 | ENSMUST00000103121.4 | 805 | D630003M21Rik | ENSMUSG00000037813.8 |
| chr2 | 158229198 | 158229875 | 3.56402 | 4.29002 | 1.31289 | Promoter | 2 | 158182533 | 158229222 | 46690 | 2 | ENSMUST00000103121.4 | 0 | D630003M21Rik | ENSMUSG00000037813.8 |
| chr14 | 40895525 | 40895803 | 3.48187 | 4.26248 | 1.33581 | Distal Intergenic | 14 | 40815868 | 40893266 | 77399 | 2 | ENSMUST00000096000.3 | -2260 | Sh2d4b | ENSMUSG00000037833.8 |
| chr2 | 157914322 | 157914922 | 4.36908 | 5.872 | 2.4246 | Promoter | 2 | 157914653 | 157944719 | 30067 | 1 | ENSMUST00000109523.1 | 0 | Vstm2l | ENSMUSG00000037843.6 |
| chr2 | 157920888 | 157921252 | 3.34975 | 3.85206 | 1.01733 | Intron (ENSMUST00000109523.1/ENSMUSG00000037843.6, intron 1 of 3) | 2 | 157914653 | 157944719 | 30067 | 1 | ENSMUST00000109523.1 | 6236 | Vstm2l | ENSMUSG00000037843.6 |
| chr2 | 157921802 | 157922210 | 3.28805 | 3.90397 | 1.09297 | Intron (ENSMUST00000109523.1/ENSMUSG00000037843.6, intron 1 of 3) | 2 | 157914653 | 157944719 | 30067 | 1 | ENSMUST00000109523.1 | 7150 | Vstm2l | ENSMUSG00000037843.6 |
| chr2 | 157923114 | 157923344 | 3.31913 | 3.80596 | 1.01733 | Intron (ENSMUST00000109523.1/ENSMUSG00000037843.6, intron 1 of 3) | 2 | 157914653 | 157944719 | 30067 | 1 | ENSMUST00000109523.1 | 8462 | Vstm2l | ENSMUSG00000037843.6 |
| chr2 | 157923937 | 157924189 | 3.94717 | 5.17659 | 1.88332 | Intron (ENSMUST00000109523.1/ENSMUSG00000037843.6, intron 1 of 3) | 2 | 157914653 | 157944719 | 30067 | 1 | ENSMUST00000109523.1 | 9285 | Vstm2l | ENSMUSG00000037843.6 |
| chr2 | 157926273 | 157926645 | 3.95735 | 4.97372 | 1.7045 | Intron (ENSMUST00000109523.1/ENSMUSG00000037843.6, intron 1 of 3) | 2 | 157914653 | 157944719 | 30067 | 1 | ENSMUST00000109523.1 | 11621 | Vstm2l | ENSMUSG00000037843.6 |
| chr2 | 157929638 | 157929854 | 3.63687 | 4.38752 | 1.3513 | Intron (ENSMUST00000109523.1/ENSMUSG00000037843.6, intron 1 of 3) | 2 | 157914653 | 157944719 | 30067 | 1 | ENSMUST00000109523.1 | 14986 | Vstm2l | ENSMUSG00000037843.6 |
| chr2 | 157941191 | 157942130 | 3.54661 | 4.26221 | 1.36217 | Intron (ENSMUST00000109523.1/ENSMUSG00000037843.6, intron 3 of 3) | 2 | 157914653 | 157944719 | 30067 | 1 | ENSMUST00000109523.1 | 26539 | Vstm2l | ENSMUSG00000037843.6 |
| chr2 | 157943269 | 157944474 | 4.05564 | 5.35101 | 2.05895 | 3' UTR | 2 | 157914653 | 157944719 | 30067 | 1 | ENSMUST00000109523.1 | 28617 | Vstm2l | ENSMUSG00000037843.6 |
| chr10 | 67911054 | 67911172 | 3.32504 | 3.81483 | 1.01733 | Intron (ENSMUST00000064656.7/ENSMUSG00000037855.10, intron 1 of 4) | 10 | 67896694 | 67912435 | 15742 | 2 | ENSMUST00000138543.1 | 1263 | Zfp365 | ENSMUSG00000037855.10 |
| chr10 | 67534476 | 67534650 | 3.44613 | 4.18433 | 1.28893 | Promoter | 10 | 67535475 | 67540246 | 4772 | 1 | ENSMUST00000145754.2 | -825 | Egr2 | ENSMUSG00000037868.10 |
| chr10 | 67535352 | 67536137 | 3.66403 | 4.78276 | 1.63997 | Promoter | 10 | 67535475 | 67540246 | 4772 | 1 | ENSMUST00000145754.2 | 0 | Egr2 | ENSMUSG00000037868.10 |
| chr3 | 40397875 | 40398123 | 3.52444 | 4.28171 | 1.32142 | Distal Intergenic | 3 | 40508430 | 40509770 | 1341 | 2 | ENSMUST00000179966.1 | 111647 | 1700017G19Rik | ENSMUSG00000037884.4 |
| chr11 | 99188552 | 99188686 | 3.34346 | 3.84256 | 1.01733 | Distal Intergenic | 11 | 99209992 | 99213443 | 3452 | 2 | ENSMUST00000156160.1 | 24757 | Smarce1 | ENSMUSG00000037935.11 |
| chr11 | 99189075 | 99189186 | 3.32777 | 3.81895 | 1.01733 | Distal Intergenic | 11 | 99209992 | 99213443 | 3452 | 2 | ENSMUST00000156160.1 | 24257 | Smarce1 | ENSMUSG00000037935.11 |
| chr11 | 99189662 | 99189852 | 3.60069 | 4.41055 | 1.40758 | Distal Intergenic | 11 | 99209992 | 99213443 | 3452 | 2 | ENSMUST00000156160.1 | 23591 | Smarce1 | ENSMUSG00000037935.11 |
| chr5 | 125245801 | 125246016 | 3.39885 | 3.93413 | 1.04543 | Distal Intergenic | 5 | 125277787 | 125284159 | 6373 | 2 | ENSMUST00000124582.1 | 38143 | Scarb1 | ENSMUSG00000037936.10 |
| chr8 | 81342541 | 81342733 | 3.33188 | 3.84439 | 1.08372 | Promoter | 8 | 81342556 | 81745890 | 403335 | 1 | ENSMUST00000172167.2 | 0 | Inpp4b | ENSMUSG00000037940.10 |
| chr11 | 99134083 | 99134444 | 3.45495 | 4.05819 | 1.14695 | Distal Intergenic | 11 | 99144196 | 99155077 | 10882 | 2 | ENSMUST00000103134.3 | 20633 | Ccr7 | ENSMUSG00000037944.8 |
| chr5 | 124964011 | 124964571 | 3.40862 | 4.12865 | 1.21922 | Distal Intergenic | 5 | 125003447 | 125012547 | 9101 | 1 | ENSMUST00000036109.3 | -38876 | Fam101a | ENSMUSG00000037962.3 |
| chr5 | 125003572 | 125003909 | 3.23092 | 3.67901 | 1.02909 | Promoter | 5 | 125003447 | 125012547 | 9101 | 1 | ENSMUST00000036109.3 | 126 | Fam101a | ENSMUSG00000037962.3 |
| chr13 | 49025500 | 49025736 | 3.38049 | 3.89842 | 1.01733 | Distal Intergenic | 13 | 49037609 | 49060883 | 23275 | 2 | ENSMUST00000159196.2 | 35147 | Wnk2 | ENSMUSG00000037989.10 |
| chr13 | 49146470 | 49146901 | 3.47254 | 4.30146 | 1.31707 | Promoter | 13 | 49036309 | 49147232 | 110924 | 2 | ENSMUST00000162403.2 | 331 | Wnk2 | ENSMUSG00000037989.10 |
| chr13 | 49147292 | 49147432 | 3.96463 | 5.2767 | 1.98505 | Promoter | 13 | 49036310 | 49147232 | 110923 | 2 | ENSMUST00000159559.2 | -61 | Wnk2 | ENSMUSG00000037989.10 |
| chr13 | 49148105 | 49148374 | 3.25387 | 4.09172 | 1.17488 | Promoter | 13 | 49038874 | 49148014 | 109141 | 2 | ENSMUST00000091623.4 | -92 | Wnk2 | ENSMUSG00000037989.10 |
| chr13 | 49148982 | 49149338 | 3.84024 | 4.8555 | 1.71036 | Promoter | 13 | 49038874 | 49148014 | 109141 | 2 | ENSMUST00000091623.4 | -969 | Wnk2 | ENSMUSG00000037989.10 |
| chr10 | 58814060 | 58814431 | 3.56131 | 4.47357 | 1.4355 | Promoter | 10 | 58814059 | 59007193 | 193135 | 1 | ENSMUST00000133151.1 | 2 | Sh3rf3 | ENSMUSG00000037990.13 |
| chr4 | 87227682 | 87227904 | 3.33701 | 3.83285 | 1.01733 | Promoter | 4 | 86991173 | 87227873 | 236701 | 2 | ENSMUST00000044990.5 | 0 | Slc24a2 | ENSMUSG00000037996.12 |
| chr4 | 87229345 | 87229420 | 4.34427 | 5.84418 | 2.36063 | Intron (ENSMUST00000107157.3/ENSMUSG00000037996.12, intron 1 of 10) | 4 | 87072830 | 87230422 | 157593 | 2 | ENSMUST00000134643.2 | 1002 | Slc24a2 | ENSMUSG00000037996.12 |
| chr4 | 87230064 | 87230665 | 3.61826 | 4.43382 | 1.41874 | Promoter | 4 | 86983124 | 87230477 | 247354 | 2 | ENSMUST00000107157.3 | 0 | Slc24a2 | ENSMUSG00000037996.12 |
| chr6 | 127377779 | 127377891 | 3.80029 | 4.70596 | 1.58536 | Distal Intergenic | 6 | 127446840 | 127494240 | 47401 | 1 | ENSMUST00000112191.2 | -68949 | Parp11 | ENSMUSG00000037997.7 |
| chr6 | 127439808 | 127439930 | 3.4234 | 3.97698 | 1.06587 | Distal Intergenic | 6 | 127446840 | 127494240 | 47401 | 1 | ENSMUST00000112191.2 | -6910 | Parp11 | ENSMUSG00000037997.7 |
| chr6 | 127441546 | 127441771 | 3.54878 | 4.20582 | 1.20542 | Distal Intergenic | 6 | 127446840 | 127494240 | 47401 | 1 | ENSMUST00000112191.2 | -5069 | Parp11 | ENSMUSG00000037997.7 |
| chr15 | 99093736 | 99093920 | 3.88853 | 5.02307 | 1.77163 | Promoter | 15 | 99093170 | 99100917 | 7748 | 1 | ENSMUST00000146173.1 | 567 | Dnajc22 | ENSMUSG00000038009.7 |
| chr15 | 99094666 | 99095275 | 3.54946 | 4.38762 | 1.45097 | Intron (ENSMUST00000146173.1/ENSMUSG00000038009.7, intron 1 of 2) | 15 | 99093170 | 99100917 | 7748 | 1 | ENSMUST00000146173.1 | 1497 | Dnajc22 | ENSMUSG00000038009.7 |
| chr15 | 99096387 | 99096866 | 4.17114 | 5.66985 | 2.24688 | Intron (ENSMUST00000146173.1/ENSMUSG00000038009.7, intron 1 of 2) | 15 | 99099412 | 99104737 | 5326 | 1 | ENSMUST00000061295.6 | -2546 | Dnajc22 | ENSMUSG00000038009.7 |
| chr15 | 99097998 | 99098495 | 4.44146 | 6.35429 | 2.82889 | Promoter | 15 | 99099412 | 99104737 | 5326 | 1 | ENSMUST00000061295.6 | -917 | Dnajc22 | ENSMUSG00000038009.7 |
| chr1 | 172325676 | 172326690 | 3.93266 | 5.18194 | 1.90293 | 5' UTR | 1 | 172321033 | 172329263 | 8231 | 2 | ENSMUST00000062387.2 | 2573 | Kcnj9 | ENSMUSG00000038026.7 |
| chr1 | 172327397 | 172327641 | 3.72153 | 4.77881 | 1.62832 | Intron (ENSMUST00000062387.2/ENSMUSG00000038026.7, intron 2 of 3) | 1 | 172321033 | 172329263 | 8231 | 2 | ENSMUST00000062387.2 | 1622 | Kcnj9 | ENSMUSG00000038026.7 |
| chr1 | 172333531 | 172333864 | 3.79159 | 4.72098 | 1.58762 | Distal Intergenic | 1 | 172321033 | 172329263 | 8231 | 2 | ENSMUST00000062387.2 | -4269 | Kcnj9 | ENSMUSG00000038026.7 |
| chr13 | 47452640 | 47452753 | 3.37925 | 3.94511 | 1.16604 | Distal Intergenic | 13 | 47194003 | 47247991 | 53989 | 1 | ENSMUST00000110111.2 | 258638 | Rnf144b | ENSMUSG00000038068.9 |
| chr6 | 126737377 | 126737713 | 3.3699 | 3.88245 | 1.01733 | Promoter | 6 | 126708331 | 126736811 | 28481 | 2 | ENSMUST00000128137.1 | -567 | Kcna6 | ENSMUSG00000038077.7 |
| chr6 | 126738476 | 126738706 | 4.15099 | 5.40207 | 2.05568 | Exon (ENSMUST00000181126.1/ENSMUSG00000097596.1, exon 2 of 2) | 6 | 126736688 | 126740151 | 3464 | 2 | ENSMUST00000112242.1 | 1445 | Kcna6 | ENSMUSG00000038077.7 |
| chr6 | 126739907 | 126740308 | 4.67899 | 6.46487 | 2.82589 | Promoter | 6 | 126736688 | 126740151 | 3464 | 2 | ENSMUST00000112242.1 | 0 | Kcna6 | ENSMUSG00000038077.7 |
| chr6 | 126740648 | 126741019 | 3.51848 | 4.16028 | 1.20173 | Promoter | 6 | 126708329 | 126740662 | 32334 | 2 | ENSMUST00000040751.5 | 0 | Kcna6 | ENSMUSG00000038077.7 |
| chr16 | 29657707 | 29657887 | 3.35416 | 3.8587 | 1.01733 | Distal Intergenic | 16 | 29616625 | 29618707 | 2083 | 1 | ENSMUST00000160101.1 | 41083 | Opa1 | ENSMUSG00000038084.11 |
| chr16 | 29663031 | 29663243 | 3.69288 | 4.48433 | 1.40831 | Distal Intergenic | 16 | 29616625 | 29618707 | 2083 | 1 | ENSMUST00000160101.1 | 46407 | Opa1 | ENSMUSG00000038084.11 |
| chr9 | 39603401 | 39603651 | 3.28696 | 3.75759 | 1.01733 | Promoter | 9 | 39600295 | 39603627 | 3333 | 2 | ENSMUST00000136814.1 | 0 | AW551984 | ENSMUSG00000038112.10 |
| chr9 | 39604429 | 39604890 | 3.59335 | 4.41161 | 1.40047 | Promoter | 9 | 39599658 | 39604403 | 4746 | 2 | ENSMUST00000141370.2 | -27 | AW551984 | ENSMUSG00000038112.10 |
| chr6 | 125690226 | 125690458 | 3.46585 | 4.12431 | 1.25634 | Promoter | 6 | 125690419 | 125957021 | 266603 | 1 | ENSMUST00000159111.2 | 0 | Ano2 | ENSMUSG00000038115.11 |
| chr6 | 125803985 | 125804201 | 3.51123 | 4.14741 | 1.20039 | Intron (ENSMUST00000159111.2/ENSMUSG00000038115.11, intron 7 of 15) | 6 | 125815534 | 125903085 | 87552 | 1 | ENSMUST00000160317.1 | -11333 | Ano2 | ENSMUSG00000038115.11 |
| chr9 | 35361673 | 35361789 | 3.38304 | 3.9154 | 1.06376 | Distal Intergenic | 9 | 35421128 | 35455365 | 34238 | 1 | ENSMUST00000151682.2 | -59339 | Cdon | ENSMUSG00000038119.9 |
| chr9 | 35399759 | 35400071 | 3.34153 | 3.83966 | 1.01733 | Distal Intergenic | 9 | 35421128 | 35455365 | 34238 | 1 | ENSMUST00000151682.2 | -21057 | Cdon | ENSMUSG00000038119.9 |
| chr9 | 35400608 | 35400735 | 3.4237 | 4.10932 | 1.19481 | Distal Intergenic | 9 | 35421128 | 35455365 | 34238 | 1 | ENSMUST00000151682.2 | -20393 | Cdon | ENSMUSG00000038119.9 |
| chr9 | 35401583 | 35401801 | 3.41915 | 3.99592 | 1.14284 | Distal Intergenic | 9 | 35421128 | 35455365 | 34238 | 1 | ENSMUST00000151682.2 | -19327 | Cdon | ENSMUSG00000038119.9 |
| chr18 | 32938479 | 32939906 | 3.90431 | 5.05053 | 1.81671 | Promoter | 18 | 32939041 | 33195767 | 256727 | 1 | ENSMUST00000042868.4 | 0 | Camk4 | ENSMUSG00000038128.6 |
| chr13 | 46418181 | 46419413 | 3.78621 | 4.76998 | 1.65674 | Promoter | 13 | 46418300 | 46431090 | 12791 | 1 | ENSMUST00000037923.3 | 0 | Rbm24 | ENSMUSG00000038132.5 |
| chr5 | 24778939 | 24779084 | 3.50733 | 4.16336 | 1.26564 | Distal Intergenic | 5 | 24751003 | 24758008 | 7006 | 2 | ENSMUST00000047119.4 | -20932 | Crygn | ENSMUSG00000038135.4 |
| chr17 | 6268389 | 6268627 | 3.29249 | 4.58534 | 1.52608 | Distal Intergenic | 17 | 6270475 | 6305783 | 35309 | 1 | ENSMUST00000088940.4 | -1848 | Tmem181a | ENSMUSG00000038141.8 |
| chr17 | 32162183 | 32162911 | 3.92234 | 4.93243 | 1.71132 | Intron (ENSMUST00000087723.3/ENSMUSG00000038146.7, intron 2 of 32) | 17 | 32120820 | 32166880 | 46061 | 2 | ENSMUST00000087723.3 | 3969 | Notch3 | ENSMUSG00000038146.7 |
| chr17 | 32166680 | 32167021 | 3.99474 | 5.17779 | 1.93133 | Promoter | 17 | 32120820 | 32166880 | 46061 | 2 | ENSMUST00000087723.3 | 0 | Notch3 | ENSMUSG00000038146.7 |
| chr10 | 44455930 | 44456930 | 3.73415 | 4.86753 | 1.68458 | Exon (ENSMUST00000039174.5/ENSMUSG00000038151.7, exon 3 of 8) | 10 | 44438293 | 44458687 | 20395 | 2 | ENSMUST00000105490.2 | 1757 | Prdm1 | ENSMUSG00000038151.7 |
| chr10 | 44457733 | 44458243 | 3.39019 | 4.14532 | 1.23372 | Promoter | 10 | 44438293 | 44458687 | 20395 | 2 | ENSMUST00000105490.2 | 444 | Prdm1 | ENSMUSG00000038151.7 |
| chr10 | 44458586 | 44458842 | 3.81259 | 5.12957 | 1.8731 | Promoter | 10 | 44437177 | 44458748 | 21572 | 2 | ENSMUST00000039174.5 | 0 | Prdm1 | ENSMUSG00000038151.7 |
| chr10 | 44459284 | 44459568 | 3.62202 | 4.40541 | 1.4349 | Promoter | 10 | 44437177 | 44458748 | 21572 | 2 | ENSMUST00000039174.5 | -537 | Prdm1 | ENSMUSG00000038151.7 |
| chr10 | 44468632 | 44468707 | 3.65137 | 4.59724 | 1.51994 | Intron (ENSMUST00000167340.1/ENSMUSG00000038151.7, intron 1 of 3) | 10 | 44437177 | 44458748 | 21572 | 2 | ENSMUST00000039174.5 | -9885 | Prdm1 | ENSMUSG00000038151.7 |
| chr10 | 44477285 | 44477470 | 3.70252 | 4.52881 | 1.50292 | Intron (ENSMUST00000167340.1/ENSMUSG00000038151.7, intron 1 of 3) | 10 | 44437177 | 44458748 | 21572 | 2 | ENSMUST00000039174.5 | -18538 | Prdm1 | ENSMUSG00000038151.7 |
| chr10 | 44478121 | 44478550 | 3.74923 | 4.63735 | 1.58852 | Intron (ENSMUST00000167340.1/ENSMUSG00000038151.7, intron 1 of 3) | 10 | 44437177 | 44458748 | 21572 | 2 | ENSMUST00000039174.5 | -19374 | Prdm1 | ENSMUSG00000038151.7 |
| chr10 | 44479116 | 44479473 | 3.32371 | 3.95909 | 1.14652 | Intron (ENSMUST00000167340.1/ENSMUSG00000038151.7, intron 1 of 3) | 10 | 44437177 | 44458748 | 21572 | 2 | ENSMUST00000039174.5 | -20369 | Prdm1 | ENSMUSG00000038151.7 |
| chr10 | 44491114 | 44491262 | 3.72343 | 4.53782 | 1.44902 | Intron (ENSMUST00000167340.1/ENSMUSG00000038151.7, intron 1 of 3) | 10 | 44437177 | 44458748 | 21572 | 2 | ENSMUST00000039174.5 | -32367 | Prdm1 | ENSMUSG00000038151.7 |
| chr10 | 44499000 | 44500760 | 3.72116 | 4.66872 | 1.57625 | Intron (ENSMUST00000167340.1/ENSMUSG00000038151.7, intron 1 of 3) | 10 | 44446717 | 44528501 | 81785 | 2 | ENSMUST00000167340.1 | 27741 | Prdm1 | ENSMUSG00000038151.7 |
| chr10 | 44525447 | 44525537 | 4.54089 | 6.65374 | 2.94047 | Intron (ENSMUST00000167340.1/ENSMUSG00000038151.7, intron 1 of 3) | 10 | 44446717 | 44528501 | 81785 | 2 | ENSMUST00000167340.1 | 2964 | Prdm1 | ENSMUSG00000038151.7 |
| chr10 | 44525971 | 44526238 | 4.14227 | 5.5028 | 2.13579 | Intron (ENSMUST00000167340.1/ENSMUSG00000038151.7, intron 1 of 3) | 10 | 44446717 | 44528501 | 81785 | 2 | ENSMUST00000167340.1 | 2263 | Prdm1 | ENSMUSG00000038151.7 |
| chr10 | 44526686 | 44527430 | 4.13719 | 5.50641 | 2.17069 | Intron (ENSMUST00000167340.1/ENSMUSG00000038151.7, intron 1 of 3) | 10 | 44446717 | 44528501 | 81785 | 2 | ENSMUST00000167340.1 | 1071 | Prdm1 | ENSMUSG00000038151.7 |
| chr10 | 44527757 | 44527942 | 3.49473 | 4.37058 | 1.36335 | Promoter | 10 | 44446717 | 44528501 | 81785 | 2 | ENSMUST00000167340.1 | 559 | Prdm1 | ENSMUSG00000038151.7 |
| chr10 | 44528263 | 44528795 | 3.83957 | 4.95569 | 1.74555 | Promoter | 10 | 44446717 | 44528501 | 81785 | 2 | ENSMUST00000167340.1 | 0 | Prdm1 | ENSMUSG00000038151.7 |
| chr10 | 44531687 | 44531835 | 3.63432 | 4.37355 | 1.33316 | Distal Intergenic | 10 | 44446717 | 44528501 | 81785 | 2 | ENSMUST00000167340.1 | -3187 | Prdm1 | ENSMUSG00000038151.7 |
| chr10 | 44533288 | 44533374 | 3.79352 | 4.67305 | 1.51712 | Distal Intergenic | 10 | 44446717 | 44528501 | 81785 | 2 | ENSMUST00000167340.1 | -4788 | Prdm1 | ENSMUSG00000038151.7 |
| chr10 | 44533759 | 44533959 | 3.5222 | 4.15633 | 1.17234 | Distal Intergenic | 10 | 44446717 | 44528501 | 81785 | 2 | ENSMUST00000167340.1 | -5259 | Prdm1 | ENSMUSG00000038151.7 |
| chr10 | 44535799 | 44536638 | 3.80483 | 4.72797 | 1.59686 | Distal Intergenic | 10 | 44446717 | 44528501 | 81785 | 2 | ENSMUST00000167340.1 | -7299 | Prdm1 | ENSMUSG00000038151.7 |
| chr10 | 44538853 | 44538949 | 3.46287 | 4.3452 | 1.38042 | Distal Intergenic | 10 | 44446717 | 44528501 | 81785 | 2 | ENSMUST00000167340.1 | -10353 | Prdm1 | ENSMUSG00000038151.7 |
| chr6 | 125381913 | 125382296 | 3.60799 | 4.36784 | 1.42072 | Distal Intergenic | 6 | 125362660 | 125380793 | 18134 | 2 | ENSMUST00000042647.6 | -1121 | Plekhg6 | ENSMUSG00000038167.6 |
| chr3 | 97888108 | 97889239 | 3.80142 | 4.91284 | 1.72369 | Promoter | 3 | 97689824 | 97888707 | 198884 | 2 | ENSMUST00000168438.3 | 0 | Pde4dip | ENSMUSG00000038170.10 |
| chr3 | 97890064 | 97890194 | 3.49412 | 4.30351 | 1.33872 | Downstream (2-3kb) | 3 | 97689865 | 97888707 | 198843 | 2 | ENSMUST00000090750.6 | -1358 | Pde4dip | ENSMUSG00000038170.10 |
| chr8 | 46989631 | 46989914 | 3.93087 | 5.04871 | 1.82844 | Intron (ENSMUST00000140149.1/ENSMUSG00000038173.9, intron 1 of 1) | 8 | 46986954 | 47082794 | 95841 | 1 | ENSMUST00000119686.1 | 2678 | Enpp6 | ENSMUSG00000038173.9 |
| chr8 | 57320444 | 57322495 | 4.015 | 5.21097 | 1.90999 | Promoter | 8 | 57320983 | 57324517 | 3535 | 1 | ENSMUST00000040104.3 | 0 | Hand2 | ENSMUSG00000038193.3 |
| chr7 | 45418017 | 45418501 | 3.53729 | 4.1971 | 1.23175 | Promoter | 7 | 45418353 | 45421845 | 3493 | 1 | ENSMUST00000127256.2 | 0 | Lhb | ENSMUSG00000038194.8 |
| chr7 | 45405756 | 45405913 | 3.77846 | 4.84709 | 1.65688 | Promoter | 7 | 45405960 | 45411382 | 5423 | 1 | ENSMUST00000107774.1 | -47 | Kcna7 | ENSMUSG00000038201.9 |
| chr7 | 45406311 | 45406707 | 4.32051 | 6.36068 | 2.81634 | Promoter | 7 | 45405960 | 45411382 | 5423 | 1 | ENSMUST00000107774.1 | 352 | Kcna7 | ENSMUSG00000038201.9 |
| chr6 | 52255602 | 52256977 | 3.8734 | 4.90395 | 1.78478 | Downstream (<1kb) | 6 | 52258871 | 52260802 | 1932 | 2 | ENSMUST00000114416.1 | 3825 | Hoxa13 | ENSMUSG00000038203.13 |
| chr6 | 52257341 | 52257815 | 3.6645 | 4.56411 | 1.49058 | 3' UTR | 6 | 52258871 | 52260802 | 1932 | 2 | ENSMUST00000114416.1 | 2987 | Hoxa13 | ENSMUSG00000038203.13 |
| chr6 | 52258268 | 52258618 | 3.54232 | 4.38854 | 1.40787 | 3' UTR | 6 | 52258871 | 52260802 | 1932 | 2 | ENSMUST00000114416.1 | 2184 | Hoxa13 | ENSMUSG00000038203.13 |
| chr6 | 52259360 | 52260869 | 3.72718 | 4.78976 | 1.66891 | Promoter | 6 | 52257694 | 52260803 | 3110 | 2 | ENSMUST00000147595.3 | 0 | Hoxa13 | ENSMUSG00000038203.13 |
| chr6 | 52244505 | 52246708 | 4.14934 | 5.50013 | 2.11961 | Promoter | 6 | 52242106 | 52245810 | 3705 | 2 | ENSMUST00000048026.9 | 0 | Hoxa11 | ENSMUSG00000038210.10 |
| chr11 | 98386483 | 98386790 | 3.6339 | 4.37117 | 1.3288 | Promoter | 11 | 98386712 | 98387482 | 771 | 1 | ENSMUST00000124527.1 | 0 | Pnmt | ENSMUSG00000038216.7 |
| chr11 | 98388367 | 98388605 | 3.8611 | 4.80345 | 1.59092 | Downstream (<1kb) | 11 | 98386712 | 98387482 | 771 | 1 | ENSMUST00000124527.1 | 1656 | Pnmt | ENSMUSG00000038216.7 |
| chr7 | 45337669 | 45338247 | 3.37831 | 3.89513 | 1.01733 | Exon (ENSMUST00000085351.4/ENSMUSG00000038239.9, exon 3 of 6) | 7 | 45335290 | 45338973 | 3684 | 1 | ENSMUST00000085351.4 | 2380 | Hrc | ENSMUSG00000038239.9 |
| chr11 | 98311116 | 98311644 | 3.37475 | 3.88977 | 1.01733 | Distal Intergenic | 11 | 98325415 | 98329648 | 4234 | 2 | ENSMUST00000041685.6 | 18004 | Neurod2 | ENSMUSG00000038255.6 |
| chr11 | 98313601 | 98313676 | 4.19815 | 5.46455 | 2.06897 | Distal Intergenic | 11 | 98325415 | 98329648 | 4234 | 2 | ENSMUST00000041685.6 | 15972 | Neurod2 | ENSMUSG00000038255.6 |
| chr11 | 98315112 | 98316041 | 3.84419 | 4.82007 | 1.67431 | Distal Intergenic | 11 | 98325415 | 98329648 | 4234 | 2 | ENSMUST00000041685.6 | 13607 | Neurod2 | ENSMUSG00000038255.6 |
| chr11 | 98317712 | 98318028 | 4.09818 | 5.29049 | 1.94214 | Distal Intergenic | 11 | 98325415 | 98329648 | 4234 | 2 | ENSMUST00000041685.6 | 11620 | Neurod2 | ENSMUSG00000038255.6 |
| chr11 | 98320008 | 98320147 | 3.57971 | 4.52791 | 1.45416 | Distal Intergenic | 11 | 98325415 | 98329648 | 4234 | 2 | ENSMUST00000041685.6 | 9501 | Neurod2 | ENSMUSG00000038255.6 |
| chr11 | 98320569 | 98320701 | 3.61892 | 4.58847 | 1.50588 | Distal Intergenic | 11 | 98325415 | 98329648 | 4234 | 2 | ENSMUST00000041685.6 | 8947 | Neurod2 | ENSMUSG00000038255.6 |
| chr11 | 98321203 | 98321800 | 3.78788 | 4.74956 | 1.58325 | Distal Intergenic | 11 | 98325415 | 98329648 | 4234 | 2 | ENSMUST00000041685.6 | 7848 | Neurod2 | ENSMUSG00000038255.6 |
| chr11 | 98322274 | 98323063 | 3.56225 | 4.24105 | 1.24544 | Downstream (2-3kb) | 11 | 98325415 | 98329648 | 4234 | 2 | ENSMUST00000041685.6 | 6585 | Neurod2 | ENSMUSG00000038255.6 |
| chr11 | 98323478 | 98323885 | 3.66723 | 4.44084 | 1.39311 | Downstream (1-2kb) | 11 | 98325415 | 98329648 | 4234 | 2 | ENSMUST00000041685.6 | 5763 | Neurod2 | ENSMUSG00000038255.6 |
| chr11 | 98324255 | 98325122 | 3.59355 | 4.34824 | 1.32966 | Downstream (<1kb) | 11 | 98325415 | 98329648 | 4234 | 2 | ENSMUST00000041685.6 | 4526 | Neurod2 | ENSMUSG00000038255.6 |
| chr11 | 98325463 | 98326484 | 3.728 | 4.54621 | 1.43329 | 3' UTR | 11 | 98325415 | 98329648 | 4234 | 2 | ENSMUST00000041685.6 | 3164 | Neurod2 | ENSMUSG00000038255.6 |
| chr11 | 98326803 | 98330179 | 4.24141 | 5.62348 | 2.20562 | Promoter | 11 | 98325415 | 98329648 | 4234 | 2 | ENSMUST00000041685.6 | 0 | Neurod2 | ENSMUSG00000038255.6 |
| chr11 | 98330786 | 98331489 | 3.76117 | 4.61114 | 1.47888 | Distal Intergenic | 11 | 98325415 | 98329648 | 4234 | 2 | ENSMUST00000041685.6 | -1139 | Neurod2 | ENSMUSG00000038255.6 |
| chr11 | 98333895 | 98334700 | 3.93139 | 5.10247 | 1.84757 | Distal Intergenic | 11 | 98325415 | 98329648 | 4234 | 2 | ENSMUST00000041685.6 | -4248 | Neurod2 | ENSMUSG00000038255.6 |
| chr11 | 98335352 | 98337640 | 3.85494 | 4.90732 | 1.71987 | Distal Intergenic | 11 | 98325415 | 98329648 | 4234 | 2 | ENSMUST00000041685.6 | -5705 | Neurod2 | ENSMUSG00000038255.6 |
| chr11 | 98338066 | 98338921 | 3.81185 | 4.75096 | 1.57604 | Distal Intergenic | 11 | 98325415 | 98329648 | 4234 | 2 | ENSMUST00000041685.6 | -8419 | Neurod2 | ENSMUSG00000038255.6 |
| chr9 | 57940608 | 57941170 | 3.8288 | 4.76227 | 1.64361 | Promoter | 9 | 57940113 | 57962864 | 22752 | 1 | ENSMUST00000043059.7 | 496 | Sema7a | ENSMUSG00000038264.7 |
| chr13 | 34343933 | 34344285 | 3.81007 | 4.79328 | 1.67855 | Promoter | 13 | 34221370 | 34344203 | 122834 | 2 | ENSMUST00000143353.1 | 0 | Slc22a23 | ENSMUSG00000038267.9 |
| chr13 | 34344598 | 34345045 | 3.49411 | 4.28025 | 1.3184 | Promoter | 13 | 34204020 | 34344595 | 140576 | 2 | ENSMUST00000145038.2 | -4 | Slc22a23 | ENSMUSG00000038267.9 |
| chr10 | 42586559 | 42586785 | 3.57805 | 4.39638 | 1.38785 | Intron (ENSMUST00000105497.2/ENSMUSG00000038280.7, intron 2 of 5) | 10 | 42583838 | 42700492 | 116655 | 1 | ENSMUST00000144806.2 | 2722 | Ostm1 | ENSMUSG00000038280.7 |
| chr10 | 42587826 | 42588283 | 3.9741 | 5.1733 | 1.92688 | Intron (ENSMUST00000105497.2/ENSMUSG00000038280.7, intron 2 of 5) | 10 | 42583838 | 42700492 | 116655 | 1 | ENSMUST00000144806.2 | 3989 | Ostm1 | ENSMUSG00000038280.7 |
| chr10 | 42588662 | 42589286 | 3.83547 | 4.84887 | 1.72404 | Intron (ENSMUST00000105497.2/ENSMUSG00000038280.7, intron 2 of 5) | 10 | 42583838 | 42700492 | 116655 | 1 | ENSMUST00000144806.2 | 4825 | Ostm1 | ENSMUSG00000038280.7 |
| chr10 | 42589628 | 42591012 | 4.17181 | 5.65479 | 2.2981 | Intron (ENSMUST00000105497.2/ENSMUSG00000038280.7, intron 2 of 5) | 10 | 42583838 | 42700492 | 116655 | 1 | ENSMUST00000144806.2 | 5791 | Ostm1 | ENSMUSG00000038280.7 |
| chr10 | 42591365 | 42592627 | 3.72383 | 4.87413 | 1.68984 | Intron (ENSMUST00000105497.2/ENSMUSG00000038280.7, intron 2 of 5) | 10 | 42583838 | 42700492 | 116655 | 1 | ENSMUST00000144806.2 | 7528 | Ostm1 | ENSMUSG00000038280.7 |
| chr10 | 42593191 | 42593698 | 4.02078 | 5.18059 | 1.88973 | Intron (ENSMUST00000105497.2/ENSMUSG00000038280.7, intron 2 of 5) | 10 | 42583838 | 42700492 | 116655 | 1 | ENSMUST00000144806.2 | 9354 | Ostm1 | ENSMUSG00000038280.7 |
| chr10 | 42594347 | 42594892 | 3.41447 | 4.01758 | 1.15466 | Intron (ENSMUST00000105497.2/ENSMUSG00000038280.7, intron 2 of 5) | 10 | 42583838 | 42700492 | 116655 | 1 | ENSMUST00000144806.2 | 10510 | Ostm1 | ENSMUSG00000038280.7 |
| chr10 | 42596649 | 42596757 | 3.77469 | 4.68495 | 1.6183 | Intron (ENSMUST00000105497.2/ENSMUSG00000038280.7, intron 2 of 5) | 10 | 42583838 | 42700492 | 116655 | 1 | ENSMUST00000144806.2 | 12812 | Ostm1 | ENSMUSG00000038280.7 |
| chr10 | 42601518 | 42601843 | 3.49706 | 4.11808 | 1.17234 | Intron (ENSMUST00000105497.2/ENSMUSG00000038280.7, intron 2 of 5) | 10 | 42583838 | 42700492 | 116655 | 1 | ENSMUST00000144806.2 | 17681 | Ostm1 | ENSMUSG00000038280.7 |
| chr10 | 42602325 | 42602472 | 3.73219 | 4.67863 | 1.56371 | Intron (ENSMUST00000105497.2/ENSMUSG00000038280.7, intron 2 of 5) | 10 | 42583838 | 42700492 | 116655 | 1 | ENSMUST00000144806.2 | 18488 | Ostm1 | ENSMUSG00000038280.7 |
| chr10 | 42608884 | 42609210 | 3.71546 | 4.65915 | 1.54316 | Intron (ENSMUST00000105497.2/ENSMUSG00000038280.7, intron 2 of 5) | 10 | 42583838 | 42700492 | 116655 | 1 | ENSMUST00000144806.2 | 25047 | Ostm1 | ENSMUSG00000038280.7 |
| chr10 | 42611274 | 42611614 | 3.61344 | 4.58738 | 1.47637 | Intron (ENSMUST00000105497.2/ENSMUSG00000038280.7, intron 2 of 5) | 10 | 42583838 | 42700492 | 116655 | 1 | ENSMUST00000144806.2 | 27437 | Ostm1 | ENSMUSG00000038280.7 |
| chr10 | 42612760 | 42613553 | 3.95157 | 5.28993 | 1.98618 | Intron (ENSMUST00000105497.2/ENSMUSG00000038280.7, intron 2 of 5) | 10 | 42583838 | 42700492 | 116655 | 1 | ENSMUST00000144806.2 | 28923 | Ostm1 | ENSMUSG00000038280.7 |
| chr10 | 42614025 | 42616369 | 4.06243 | 5.47787 | 2.14773 | Intron (ENSMUST00000105497.2/ENSMUSG00000038280.7, intron 2 of 5) | 10 | 42583838 | 42700492 | 116655 | 1 | ENSMUST00000144806.2 | 30188 | Ostm1 | ENSMUSG00000038280.7 |
| chr10 | 42616798 | 42618328 | 4.00154 | 5.18728 | 1.99455 | Intron (ENSMUST00000105497.2/ENSMUSG00000038280.7, intron 2 of 5) | 10 | 42583838 | 42700492 | 116655 | 1 | ENSMUST00000144806.2 | 32961 | Ostm1 | ENSMUSG00000038280.7 |
| chr10 | 42618741 | 42619168 | 3.43379 | 4.05085 | 1.20194 | Intron (ENSMUST00000105497.2/ENSMUSG00000038280.7, intron 2 of 5) | 10 | 42583838 | 42700492 | 116655 | 1 | ENSMUST00000144806.2 | 34904 | Ostm1 | ENSMUSG00000038280.7 |
| chr10 | 42620019 | 42620260 | 3.48298 | 4.10988 | 1.20621 | Intron (ENSMUST00000105497.2/ENSMUSG00000038280.7, intron 2 of 5) | 10 | 42583838 | 42700492 | 116655 | 1 | ENSMUST00000144806.2 | 36182 | Ostm1 | ENSMUSG00000038280.7 |

| | | | | | | | | | | | | | | | |
|---|---|---|---|---|---|---|---|---|---|---|---|---|---|---|---|
| chr10 | 42623624 | 42623720 | 4.13885 | 5.44962 | 2.07234 | Intron (ENSMUST00000105497.2/ENSMUSG00000038280.7, intron 2 of 5) | 10 | 42583838 | 42700492 | 116655 | 1 | ENSMUST00000144806.2 | 39787 | Ostm1 | ENSMUSG00000038280.7 |
| chr10 | 42626937 | 42627407 | 3.46757 | 4.05811 | 1.11679 | Intron (ENSMUST00000105497.2/ENSMUSG00000038280.7, intron 2 of 5) | 10 | 42583838 | 42700492 | 116655 | 1 | ENSMUST00000144806.2 | 43100 | Ostm1 | ENSMUSG00000038280.7 |
| chr10 | 42628831 | 42629189 | 3.60094 | 4.31427 | 1.30361 | Intron (ENSMUST00000105497.2/ENSMUSG00000038280.7, intron 2 of 5) | 10 | 42583838 | 42700492 | 116655 | 1 | ENSMUST00000144806.2 | 44994 | Ostm1 | ENSMUSG00000038280.7 |
| chr10 | 42629509 | 42629726 | 3.66663 | 4.45501 | 1.43359 | Intron (ENSMUST00000105497.2/ENSMUSG00000038280.7, intron 2 of 5) | 10 | 42583838 | 42700492 | 116655 | 1 | ENSMUST00000144806.2 | 45672 | Ostm1 | ENSMUSG00000038280.7 |
| chr10 | 42630047 | 42630122 | 3.46574 | 4.06229 | 1.14167 | Intron (ENSMUST00000105497.2/ENSMUSG00000038280.7, intron 2 of 5) | 10 | 42583838 | 42700492 | 116655 | 1 | ENSMUST00000144806.2 | 46210 | Ostm1 | ENSMUSG00000038280.7 |
| chr10 | 42630470 | 42632444 | 3.79441 | 4.86567 | 1.71659 | Intron (ENSMUST00000105497.2/ENSMUSG00000038280.7, intron 2 of 5) | 10 | 42678916 | 42702459 | 23544 | 1 | ENSMUST00000040718.5 | -46472 | Ostm1 | ENSMUSG00000038280.7 |
| chr10 | 42632754 | 42633781 | 4.06482 | 5.52382 | 2.16583 | Intron (ENSMUST00000105497.2/ENSMUSG00000038280.7, intron 2 of 5) | 10 | 42678916 | 42702459 | 23544 | 1 | ENSMUST00000040718.5 | -45135 | Ostm1 | ENSMUSG00000038280.7 |
| chr10 | 42634110 | 42634660 | 3.75855 | 4.80145 | 1.67259 | Intron (ENSMUST00000105497.2/ENSMUSG00000038280.7, intron 2 of 5) | 10 | 42678916 | 42702459 | 23544 | 1 | ENSMUST00000040718.5 | -44256 | Ostm1 | ENSMUSG00000038280.7 |
| chr10 | 42636080 | 42636235 | 3.56033 | 4.31704 | 1.36949 | Intron (ENSMUST00000105497.2/ENSMUSG00000038280.7, intron 2 of 5) | 10 | 42678916 | 42702459 | 23544 | 1 | ENSMUST00000040718.5 | -42681 | Ostm1 | ENSMUSG00000038280.7 |
| chr10 | 42636741 | 42637055 | 3.96794 | 5.06161 | 1.85987 | Intron (ENSMUST00000105497.2/ENSMUSG00000038280.7, intron 2 of 5) | 10 | 42678916 | 42702459 | 23544 | 1 | ENSMUST00000040718.5 | -41861 | Ostm1 | ENSMUSG00000038280.7 |
| chr7 | 45189433 | 45189747 | 4.18261 | 5.50225 | 2.16324 | Promoter | 7 | 45183928 | 45190066 | 6139 | 2 | ENSMUST00000147253.1 | 319 | Ccdc155 | ENSMUSG00000038292.12 |
| chr7 | 45181114 | 45181934 | 4.0855 | 5.50095 | 2.13689 | Promoter | 7 | 45180995 | 45181838 | 844 | 1 | ENSMUST00000042754.5 | 120 | Pth2 | ENSMUSG00000038300.5 |
| chr17 | 36144610 | 36145125 | 3.64932 | 4.72126 | 1.59823 | Promoter | 17 | 36145018 | 36162374 | 17357 | 1 | ENSMUST00000056774.10 | 0 | 2410017I17Rik | ENSMUSG00000038311.8 |
| chr17 | 36152014 | 36152210 | 3.60425 | 4.43009 | 1.42826 | Intron (ENSMUST00000056774.10/ENSMUSG00000038311.8, intron 1 of 4) | 17 | 36154847 | 36163176 | 8330 | 1 | ENSMUST00000090537.6 | -2637 | 2410017I17Rik | ENSMUSG00000038311.8 |
| chr17 | 36152622 | 36152845 | 3.59243 | 4.31631 | 1.36772 | Intron (ENSMUST00000056774.10/ENSMUSG00000038311.8, intron 1 of 4) | 17 | 36154847 | 36163176 | 8330 | 1 | ENSMUST00000090537.6 | -2002 | 2410017I17Rik | ENSMUSG00000038311.8 |
| chr17 | 36153353 | 36153428 | 3.74826 | 4.60148 | 1.5229 | Intron (ENSMUST00000056774.10/ENSMUSG00000038311.8, intron 1 of 4) | 17 | 36154847 | 36163176 | 8330 | 1 | ENSMUST00000090537.6 | -1419 | 2410017I17Rik | ENSMUSG00000038311.8 |
| chr17 | 36151419 | 36151538 | 4.13501 | 5.35484 | 2.03732 | Intron (ENSMUST00000056774.10/ENSMUSG00000038311.8, intron 1 of 4) | 17 | 36154847 | 36163176 | 8330 | 1 | ENSMUST00000090537.6 | -3309 | 2410017I17Rik | ENSMUSG00000038311.8 |
| chr17 | 36154935 | 36155362 | 3.65966 | 4.57522 | 1.53514 | Promoter | 17 | 36154847 | 36163176 | 8330 | 1 | ENSMUST00000090537.6 | 89 | 2410017I17Rik | ENSMUSG00000038311.8 |
| chr5 | 24328838 | 24329577 | 3.74441 | 4.57531 | 1.44947 | Promoter | 5 | 24319589 | 24329591 | 10003 | 2 | ENSMUST00000115098.2 | 14 | Kcnh2 | ENSMUSG00000038319.9 |
| chr5 | 24349154 | 24349539 | 3.48416 | 4.09198 | 1.14801 | Intron (ENSMUST00000036092.8/ENSMUSG00000038319.9, intron 1 of 14) | 5 | 24319589 | 24351604 | 32016 | 2 | ENSMUST00000036092.8 | 2065 | Kcnh2 | ENSMUSG00000038319.9 |
| chr5 | 24350501 | 24350873 | 3.7134 | 4.53716 | 1.45072 | Promoter | 5 | 24319589 | 24351604 | 32016 | 2 | ENSMUST00000036092.8 | 731 | Kcnh2 | ENSMUSG00000038319.9 |
| chr5 | 24351353 | 24351602 | 3.6257 | 4.77454 | 1.63403 | Promoter | 5 | 24319589 | 24351604 | 32016 | 2 | ENSMUST00000036092.8 | 2 | Kcnh2 | ENSMUSG00000038319.9 |
| chr5 | 24352018 | 24352213 | 3.47439 | 4.48419 | 1.4284 | Promoter | 5 | 24319589 | 24351604 | 32016 | 2 | ENSMUST00000036092.8 | -415 | Kcnh2 | ENSMUSG00000038319.9 |
| chr5 | 24352703 | 24353035 | 3.43186 | 4.1406 | 1.21651 | Distal Intergenic | 5 | 24319589 | 24351604 | 32016 | 2 | ENSMUST00000036092.8 | -1100 | Kcnh2 | ENSMUSG00000038319.9 |
| chr1 | 56967767 | 56968577 | 4.26233 | 5.73534 | 2.39824 | Intron (ENSMUST00000042857.8/ENSMUSG00000038331.10, intron 2 of 9) | 1 | 56948134 | 56969840 | 21707 | 2 | ENSMUST00000176759.1 | 1263 | Satb2 | ENSMUSG00000038331.10 |
| chr1 | 56969151 | 56971238 | 3.80995 | 4.89072 | 1.73831 | Promoter | 1 | 56793988 | 56969872 | 175885 | 2 | ENSMUST00000177424.1 | 0 | Satb2 | ENSMUSG00000038331.10 |
| chr1 | 56971604 | 56972723 | 3.43594 | 4.03024 | 1.17727 | Promoter | 1 | 56793981 | 56972502 | 178522 | 2 | ENSMUST00000042857.8 | 0 | Satb2 | ENSMUSG00000038331.10 |
| chr1 | 56973759 | 56973961 | 4.0148 | 5.18127 | 1.96878 | Intron (ENSMUST00000135238.3/ENSMUSG00000087022.3, intron 1 of 1) | 1 | 56793981 | 56972502 | 178522 | 2 | ENSMUST00000042857.8 | -1258 | Satb2 | ENSMUSG00000038331.10 |
| chr1 | 56974336 | 56974565 | 3.55261 | 4.24904 | 1.33657 | Exon (ENSMUST00000135238.3/ENSMUSG00000087022.3, exon 2 of 2) | 1 | 56793981 | 56972502 | 178522 | 2 | ENSMUST00000042857.8 | -1835 | Satb2 | ENSMUSG00000038331.10 |
| chr1 | 56974910 | 56975017 | 4.00617 | 5.28429 | 1.99 | Exon (ENSMUST00000135238.3/ENSMUSG00000087022.3, exon 2 of 2) | 1 | 56793981 | 56972502 | 178522 | 2 | ENSMUST00000042857.8 | -2409 | Satb2 | ENSMUSG00000038331.10 |
| chr1 | 56975683 | 56976807 | 3.91583 | 5.09014 | 1.86138 | Intron (ENSMUST00000185204.1/ENSMUSG00000087022.3, intron 1 of 4) | 1 | 56969740 | 56978650 | 8911 | 2 | ENSMUST00000177282.1 | 1843 | Satb2 | ENSMUSG00000038331.10 |
| chr1 | 56977933 | 56980394 | 3.72986 | 4.71241 | 1.58603 | Promoter | 1 | 56969740 | 56978650 | 8911 | 2 | ENSMUST00000177282.1 | 0 | Satb2 | ENSMUSG00000038331.10 |
| chr1 | 56980742 | 56981413 | 3.83988 | 4.98015 | 1.77094 | Intron (ENSMUST00000185204.1/ENSMUSG00000087022.3, intron 1 of 4) | 1 | 56969740 | 56978650 | 8911 | 2 | ENSMUST00000177282.1 | -2093 | Satb2 | ENSMUSG00000038331.10 |
| chr1 | 56981747 | 56982356 | 4.28052 | 6.0535 | 2.65807 | Intron (ENSMUST00000185204.1/ENSMUSG00000087022.3, intron 1 of 4) | 1 | 56969740 | 56978650 | 8911 | 2 | ENSMUST00000177282.1 | -3098 | Satb2 | ENSMUSG00000038331.10 |
| chr1 | 56982679 | 56982931 | 3.98349 | 5.11822 | 1.8594 | Intron (ENSMUST00000185204.1/ENSMUSG00000087022.3, intron 1 of 4) | 1 | 56969740 | 56978650 | 8911 | 2 | ENSMUST00000177282.1 | -4030 | Satb2 | ENSMUSG00000038331.10 |
| chr1 | 56983449 | 56984404 | 3.85487 | 4.85855 | 1.6903 | Intron (ENSMUST00000185204.1/ENSMUSG00000087022.3, intron 1 of 4) | 1 | 56969740 | 56978650 | 8911 | 2 | ENSMUST00000177282.1 | -4800 | Satb2 | ENSMUSG00000038331.10 |
| chr1 | 56984791 | 56985587 | 3.63289 | 4.37748 | 1.32744 | Intron (ENSMUST00000185204.1/ENSMUSG00000087022.3, intron 1 of 4) | 1 | 56969740 | 56978650 | 8911 | 2 | ENSMUST00000177282.1 | -6142 | Satb2 | ENSMUSG00000038331.10 |
| chr1 | 56986081 | 56986331 | 3.41942 | 4.18931 | 1.26049 | Intron (ENSMUST00000185204.1/ENSMUSG00000087022.3, intron 1 of 4) | 1 | 56969740 | 56978650 | 8911 | 2 | ENSMUST00000177282.1 | -7432 | Satb2 | ENSMUSG00000038331.10 |
| chr17 | 13668296 | 13668930 | 3.48023 | 4.19007 | 1.268 | Promoter | 17 | 13489273 | 13668891 | 179619 | 2 | ENSMUST00000185626.1 | 0 | Tcte2 | ENSMUSG00000038347.9 |
| chr17 | 13764357 | 13764521 | 3.28421 | 4.06835 | 1.16408 | Intron (ENSMUST00000139347.2/ENSMUSG00000068036.9, intron 1 of 4) | 17 | 13716436 | 13761825 | 45390 | 2 | ENSMUST00000148430.1 | -2533 | Tcte2 | ENSMUSG00000038347.9 |
| chr17 | 13716872 | 13717131 | 3.54044 | 4.23774 | 1.31475 | 3' UTR | 17 | 13714049 | 13732898 | 18850 | 2 | ENSMUST00000142863.1 | 15767 | Tcte2 | ENSMUSG00000038347.9 |
| chr17 | 13737481 | 13737633 | 3.65801 | 4.40393 | 1.32243 | Intron (ENSMUST00000135850.2/ENSMUSG00000038347.9, intron 1 of 4) | 17 | 13714049 | 13732898 | 18850 | 2 | ENSMUST00000142863.1 | -4584 | Tcte2 | ENSMUSG00000038347.9 |
| chr17 | 13740099 | 13740174 | 3.30772 | 3.78877 | 1.01733 | Intron (ENSMUST00000135850.2/ENSMUSG00000038347.9, intron 1 of 4) | 17 | 13714049 | 13732898 | 18850 | 2 | ENSMUST00000142863.1 | -7202 | Tcte2 | ENSMUSG00000038347.9 |
| chr17 | 13742796 | 13743042 | 3.86453 | 4.88824 | 1.72382 | Intron (ENSMUST00000135850.2/ENSMUSG00000038347.9, intron 1 of 4) | 17 | 13714049 | 13732898 | 18850 | 2 | ENSMUST00000142863.1 | -9899 | Tcte2 | ENSMUSG00000038347.9 |
| chr17 | 13744382 | 13744521 | 3.55903 | 4.36998 | 1.35827 | Intron (ENSMUST00000135850.2/ENSMUSG00000038347.9, intron 1 of 4) | 17 | 13714049 | 13732898 | 18850 | 2 | ENSMUST00000142863.1 | -11485 | Tcte2 | ENSMUSG00000038347.9 |
| chr17 | 13745870 | 13746142 | 3.59118 | 4.50966 | 1.45992 | Intron (ENSMUST00000135850.2/ENSMUSG00000038347.9, intron 1 of 4) | 17 | 13714049 | 13732898 | 18850 | 2 | ENSMUST00000142863.1 | -12973 | Tcte2 | ENSMUSG00000038347.9 |
| chr17 | 13761119 | 13761980 | 3.97468 | 5.32172 | 2.00472 | Promoter | 17 | 13489273 | 13761378 | 272106 | 2 | ENSMUST00000135850.2 | 0 | Tcte2 | ENSMUSG00000038347.9 |
| chr17 | 13764914 | 13765106 | 3.83043 | 4.84389 | 1.67903 | Intron (ENSMUST00000139347.2/ENSMUSG00000068036.9, intron 1 of 4) | 17 | 13716436 | 13761825 | 45390 | 2 | ENSMUST00000148430.1 | -3090 | Tcte2 | ENSMUSG00000038347.9 |
| chr3 | 96683378 | 96683623 | 3.47644 | 4.18149 | 1.2378 | 3' UTR | 3 | 96680093 | 96683202 | 3110 | 1 | ENSMUST00000130429.1 | 3286 | Ankrd35 | ENSMUSG00000038354.8 |
| chr1 | 171191922 | 171192011 | 3.75506 | 4.62745 | 1.55663 | Intron (ENSMUST00000111332.1/ENSMUSG00000038370.6, intron 1 of 2) | 1 | 171173262 | 171196268 | 23007 | 2 | ENSMUST00000111332.1 | 4257 | Pcp4l1 | ENSMUSG00000038370.6 |
| chr1 | 171193388 | 171193573 | 3.41547 | 4.012 | 1.20379 | Intron (ENSMUST00000111332.1/ENSMUSG00000038370.6, intron 1 of 2) | 1 | 171173262 | 171196268 | 23007 | 2 | ENSMUST00000111332.1 | 2695 | Pcp4l1 | ENSMUSG00000038370.6 |
| chr1 | 171195100 | 171196271 | 3.9674 | 5.20439 | 1.9994 | Promoter | 1 | 171173262 | 171196268 | 23007 | 2 | ENSMUST00000111332.1 | 0 | Pcp4l1 | ENSMUSG00000038370.6 |
| chr1 | 171196857 | 171197062 | 3.41601 | 4.16163 | 1.2151 | Promoter | 1 | 171173262 | 171196268 | 23007 | 2 | ENSMUST00000111332.1 | -590 | Pcp4l1 | ENSMUSG00000038370.6 |
| chr6 | 50219887 | 50220176 | 3.35014 | 3.88581 | 1.12891 | Intron (ENSMUST00000165099.2/ENSMUSG00000029821.10, intron 7 of 9) | 6 | 50194416 | 50198327 | 3912 | 1 | ENSMUST00000167319.1 | 25472 | Mpp6 | ENSMUSG00000038388.9 |
| chr6 | 124861540 | 124862035 | 3.3965 | 3.96 | 1.11198 | 5' UTR | 6 | 124858444 | 124863917 | 5474 | 2 | ENSMUST00000046893.7 | 1882 | Gpr162 | ENSMUSG00000038390.9 |
| chr6 | 124863332 | 124863545 | 3.30268 | 3.7812 | 1.01733 | Promoter | 6 | 124858444 | 124863917 | 5474 | 2 | ENSMUST00000046893.7 | 372 | Gpr162 | ENSMUSG00000038390.9 |
| chr13 | 31622225 | 31622791 | 4.21429 | 5.57529 | 2.1665 | Distal Intergenic | 13 | 31625816 | 31631403 | 5588 | 1 | ENSMUST00000042054.2 | -3025 | Foxf2 | ENSMUSG00000038402.2 |
| chr13 | 31623170 | 31623917 | 3.64016 | 4.52148 | 1.44391 | Distal Intergenic | 13 | 31625816 | 31631403 | 5588 | 1 | ENSMUST00000042054.2 | -1899 | Foxf2 | ENSMUSG00000038402.2 |
| chr13 | 31624245 | 31624627 | 4.11161 | 5.38102 | 2.03567 | Distal Intergenic | 13 | 31625816 | 31631403 | 5588 | 1 | ENSMUST00000042054.2 | -1189 | Foxf2 | ENSMUSG00000038402.2 |
| chr13 | 31625354 | 31626446 | 4.42728 | 5.95164 | 2.44789 | Promoter | 13 | 31625816 | 31631403 | 5588 | 1 | ENSMUST00000042054.2 | 0 | Foxf2 | ENSMUSG00000038402.2 |
| chr3 | 96526796 | 96527485 | 3.66601 | 4.58199 | 1.48832 | Exon (ENSMUST00000049208.9/ENSMUSG00000038403.9, exon 3 of 4) | 3 | 96525185 | 96529216 | 4032 | 1 | ENSMUST00000049208.9 | 1612 | Hfe2 | ENSMUSG00000038403.9 |
| chr3 | 96527845 | 96528461 | 3.51464 | 4.15744 | 1.21755 | Exon (ENSMUST00000049208.9/ENSMUSG00000038403.9, exon 4 of 4) | 3 | 96525185 | 96529216 | 4032 | 1 | ENSMUST00000049208.9 | 2661 | Hfe2 | ENSMUSG00000038403.9 |
| chr13 | 31576813 | 31577029 | 3.51371 | 4.1444 | 1.17607 | Intron (ENSMUST00000150418.2/ENSMUSG00000038408.8, intron 3 of 5) | 13 | 31565492 | 31582505 | 17014 | 2 | ENSMUST00000150418.2 | 5476 | 1700018A04Rik | ENSMUSG00000038408.8 |
| chr13 | 31505476 | 31505561 | 4.08267 | 5.45746 | 2.05424 | Distal Intergenic | 13 | 31558324 | 31560976 | 2653 | 1 | ENSMUST00000042118.9 | -52763 | Foxq1 | ENSMUSG00000038415.9 |
| chr13 | 31556033 | 31556116 | 3.47599 | 4.09357 | 1.19887 | Downstream (<1kb) | 13 | 31558324 | 31560976 | 2653 | 1 | ENSMUST00000042118.9 | -2208 | Foxq1 | ENSMUSG00000038415.9 |
| chr13 | 31557206 | 31557564 | 3.18189 | 3.74325 | 1.01733 | Promoter | 13 | 31558324 | 31560976 | 2653 | 1 | ENSMUST00000042118.9 | -760 | Foxq1 | ENSMUSG00000038415.9 |
| chr13 | 31558095 | 31560861 | 4.20023 | 5.73216 | 2.34312 | Promoter | 13 | 31558324 | 31560976 | 2653 | 1 | ENSMUST00000042118.9 | 0 | Foxq1 | ENSMUSG00000038415.9 |
| chr11 | 97570875 | 97570990 | 3.83066 | 4.85303 | 1.65479 | Intron (ENSMUST00000145845.1/ENSMUSG00000086394.1, intron 1 of 1) | 11 | 97534887 | 97573929 | 39043 | 2 | ENSMUST00000126287.1 | 2939 | Srcin1 | ENSMUSG00000038453.10 |
| chr11 | 97572086 | 97572226 | 3.50501 | 4.12536 | 1.15463 | Intron (ENSMUST00000145845.1/ENSMUSG00000086394.1, intron 1 of 1) | 11 | 97534887 | 97573929 | 39043 | 2 | ENSMUST00000126287.1 | 1703 | Srcin1 | ENSMUSG00000038453.10 |
| chr11 | 97573045 | 97573348 | 3.43776 | 4.034 | 1.18272 | Promoter | 11 | 97534887 | 97573929 | 39043 | 2 | ENSMUST00000126287.1 | 581 | Srcin1 | ENSMUSG00000038453.10 |
| chr11 | 97574137 | 97574484 | 3.54404 | 4.21278 | 1.24402 | Promoter | 11 | 97512859 | 97574053 | 61195 | 2 | ENSMUST00000107593.2 | -85 | Srcin1 | ENSMUSG00000038453.10 |
| chr11 | 97574874 | 97576682 | 3.75151 | 4.69501 | 1.5597 | Promoter | 11 | 97509340 | 97575215 | 65876 | 2 | ENSMUST00000107596.3 | 0 | Srcin1 | ENSMUSG00000038453.10 |
| chr8 | 13435614 | 13435699 | 3.77501 | 4.63552 | 1.49133 | Promoter | 8 | 13435459 | 13461452 | 25994 | 1 | ENSMUST00000167505.1 | 156 | Tmem255b | ENSMUSG00000038457.11 |
| chr8 | 13436185 | 13436358 | 4.7912 | 6.84236 | 3.15117 | Promoter | 8 | 13435459 | 13461452 | 25994 | 1 | ENSMUST00000167505.1 | 727 | Tmem255b | ENSMUSG00000038457.11 |
| chr1 | 170643273 | 170643348 | 3.69835 | 4.61979 | 1.5208 | Distal Intergenic | 1 | 170644532 | 170682789 | 38258 | 1 | ENSMUST00000046792.8 | -1184 | Olfml2b | ENSMUSG00000038463.8 |
| chr1 | 170644327 | 170644612 | 3.32579 | 4.16952 | 1.23684 | Promoter | 1 | 170644532 | 170682789 | 38258 | 1 | ENSMUST00000046792.8 | 0 | Olfml2b | ENSMUSG00000038463.8 |
| chr10 | 40191934 | 40192696 | 3.60127 | 4.30742 | 1.28243 | Distal Intergenic | 10 | 40223683 | 40227798 | 4116 | 2 | ENSMUST00000182287.1 | 35102 | Rpf2 | ENSMUSG00000038510.7 |
| chr8 | 13200390 | 13200480 | 3.36604 | 4.0265 | 1.11882 | Promoter | 8 | 13176869 | 13200624 | 23756 | 2 | ENSMUST00000165605.2 | 144 | Grtp1 | ENSMUSG00000038515.9 |
| chr11 | 97150148 | 97151229 | 4.08484 | 5.4125 | 2.13534 | Promoter | 11 | 97137099 | 97151036 | 13938 | 2 | ENSMUST00000107614.2 | 0 | Tbkbp1 | ENSMUSG00000038517.10 |
| chr11 | 97151746 | 97151836 | 4.01272 | 5.17677 | 1.97349 | Promoter | 11 | 97148629 | 97151495 | 2867 | 2 | ENSMUST00000146374.1 | -252 | Tbkbp1 | ENSMUSG00000038517.10 |
| chr11 | 97152261 | 97152545 | 3.28546 | 3.75534 | 1.01733 | Promoter | 11 | 97148629 | 97151495 | 2867 | 2 | ENSMUST00000146374.1 | -767 | Tbkbp1 | ENSMUSG00000038517.10 |
| chr13 | 44869570 | 44870002 | 3.75644 | 16.72723 | 11.88095 | Intron (ENSMUST00000173704.2/ENSMUSG00000038518.10, intron 4 of 18) | 13 | 44840783 | 44896481 | 55699 | 1 | ENSMUST00000173003.1 | 28788 | Jarid2 | ENSMUSG00000038518.10 |
| chr1 | 169747576 | 169748290 | 3.41927 | 3.97154 | 1.06867 | Promoter | 1 | 169741477 | 169747642 | 6166 | 2 | ENSMUST00000027991.6 | 0 | Rgs4 | ENSMUSG00000038530.6 |
| chr2 | 172246709 | 172246876 | 3.25883 | 3.71546 | 1.01733 | Distal Intergenic | 2 | 172248492 | 172251114 | 2623 | 1 | ENSMUST00000038532.1 | -1616 | Mc3r | ENSMUSG00000038537.1 |
| chr17 | 74045662 | 74046899 | 3.93056 | 5.08049 | 1.86469 | Intron (ENSMUST00000043458.7/ENSMUSG00000038541.7, intron 1 of 4) | 17 | 74017706 | 74047916 | 30211 | 2 | ENSMUST00000043458.7 | 1017 | Srd5a2 | ENSMUSG00000038541.7 |
| chr17 | 74047495 | 74048600 | 3.69331 | 4.55087 | 1.52021 | Promoter | 17 | 74017706 | 74047916 | 30211 | 2 | ENSMUST00000043458.7 | 0 | Srd5a2 | ENSMUSG00000038541.7 |
| chr17 | 74049788 | 74050542 | 3.8243 | 4.85281 | 1.69776 | Distal Intergenic | 17 | 74017706 | 74047916 | 30211 | 2 | ENSMUST00000043458.7 | -1873 | Srd5a2 | ENSMUSG00000038541.7 |
| chr11 | 97016172 | 97016404 | 3.74161 | 4.59728 | 1.48124 | Intron (ENSMUST00000047997.7/ENSMUSG00000038560.7, intron 1 of 1) | 11 | 97018733 | 97024731 | 5999 | 1 | ENSMUST00000107622.1 | -2329 | Sp6 | ENSMUSG00000038560.7 |
| chr11 | 97018696 | 97019287 | 3.81868 | 4.86792 | 1.6928 | Promoter | 11 | 97018733 | 97024731 | 5999 | 1 | ENSMUST00000107622.1 | 0 | Sp6 | ENSMUSG00000038560.7 |
| chr2 | 170499020 | 170499261 | 3.81463 | 4.7391 | 1.6133 | Intron (ENSMUST00000075087.2/ENSMUSG00000052033.8, intron 1 of 3) | 2 | 170482708 | 170497145 | 14438 | 2 | ENSMUST00000038824.5 | -1876 | Cyp24a1 | ENSMUSG00000038567.5 |
| chr2 | 170495175 | 170497529 | 3.72224 | 4.64453 | 1.54921 | Promoter | 2 | 170482708 | 170497145 | 14438 | 2 | ENSMUST00000038824.5 | 0 | Cyp24a1 | ENSMUSG00000038567.5 |
| chr2 | 170497854 | 170498585 | 3.69227 | 4.84366 | 1.66255 | Promoter | 2 | 170482708 | 170497145 | 14438 | 2 | ENSMUST00000038824.5 | -710 | Cyp24a1 | ENSMUSG00000038567.5 |
| chr2 | 170503414 | 170503653 | 3.5045 | 4.16893 | 1.22818 | Exon (ENSMUST00000126086.1/ENSMUSG00000085495.1, exon 2 of 3) | 2 | 170482708 | 170497145 | 14438 | 2 | ENSMUST00000038824.5 | -6270 | Cyp24a1 | ENSMUSG00000038567.5 |
| chr2 | 170500773 | 170500848 | 3.72393 | 4.56037 | 1.51302 | Intron (ENSMUST00000075087.2/ENSMUSG00000052033.8, intron 1 of 3) | 2 | 170482708 | 170497145 | 14438 | 2 | ENSMUST00000038824.5 | -3629 | Cyp24a1 | ENSMUSG00000038567.5 |
| chr2 | 170501513 | 170501588 | 4.23765 | 5.5586 | 2.16555 | Intron (ENSMUST00000075087.2/ENSMUSG00000052033.8, intron 1 of 3) | 2 | 170482708 | 170497145 | 14438 | 2 | ENSMUST00000038824.5 | -4369 | Cyp24a1 | ENSMUSG00000038567.5 |
| chr2 | 170501995 | 170502972 | 3.58 | 4.41272 | 1.39817 | Exon (ENSMUST00000126086.1/ENSMUSG00000085495.1, exon 2 of 3) | 2 | 170482708 | 170497145 | 14438 | 2 | ENSMUST00000038824.5 | -4851 | Cyp24a1 | ENSMUSG00000038567.5 |
| chr1 | 182763314 | 182764306 | 3.57102 | 4.26536 | 1.25554 | Promoter | 1 | 182763860 | 182854040 | 90181 | 1 | ENSMUST00000155229.1 | 0 | Susd4 | ENSMUSG00000038576.10 |
| chr1 | 182764986 | 182765708 | 3.28728 | 3.95964 | 1.07733 | Promoter | 1 | 182764895 | 182896598 | 131704 | 1 | ENSMUST00000085724.3 | 92 | Susd4 | ENSMUSG00000038576.10 |
| chr1 | 182778282 | 182778439 | 3.21528 | 3.65015 | 1.01079 | Intron (ENSMUST00000155229.1/ENSMUSG00000038576.10, intron 2 of 3) | 1 | 182764895 | 182896598 | 131704 | 1 | ENSMUST00000085724.3 | 13388 | Susd4 | ENSMUSG00000038576.10 |
| chr1 | 182814131 | 182814336 | 3.59245 | 4.41441 | 1.42787 | Intron (ENSMUST00000155229.1/ENSMUSG00000038576.10, intron 2 of 3) | 1 | 182764895 | 182896598 | 131704 | 1 | ENSMUST00000085724.3 | 49237 | Susd4 | ENSMUSG00000038576.10 |
| chr1 | 182832189 | 182832307 | 3.31054 | 3.79302 | 1.01733 | Intron (ENSMUST00000155229.1/ENSMUSG00000038576.10, intron 2 of 3) | 1 | 182764895 | 182896598 | 131704 | 1 | ENSMUST00000085724.3 | 67295 | Susd4 | ENSMUSG00000038576.10 |
| chr1 | 182837656 | 182837938 | 3.52126 | 4.17043 | 1.22855 | Intron (ENSMUST00000155229.1/ENSMUSG00000038576.10, intron 3 of 3) | 1 | 182764895 | 182896598 | 131704 | 1 | ENSMUST00000085724.3 | 72762 | Susd4 | ENSMUSG00000038576.10 |
| chr1 | 182838755 | 182839303 | 3.50987 | 4.18465 | 1.26731 | Intron (ENSMUST00000155229.1/ENSMUSG00000038576.10, intron 3 of 3) | 1 | 182764895 | 182896598 | 131704 | 1 | ENSMUST00000085724.3 | 73861 | Susd4 | ENSMUSG00000038576.10 |
| chr7 | 141279237 | 141279402 | 3.43922 | 4.52386 | 1.43524 | Promoter | 7 | 141278331 | 141279133 | 803 | 2 | ENSMUST00000046156.7 | -105 | Sct | ENSMUSG00000038580.7 |
| chr10 | 4265867 | 4266167 | 3.60393 | 4.45041 | 1.44521 | Promoter | 10 | 4266329 | 4359468 | 93140 | 1 | ENSMUST00000045730.5 | -162 | Akap12 | ENSMUSG00000038587.8 |
| chr10 | 4266969 | 4267184 | 3.53092 | 4.64714 | 1.50953 | Promoter | 10 | 4266329 | 4359468 | 93140 | 1 | ENSMUST00000045730.5 | 641 | Akap12 | ENSMUSG00000038587.8 |
| chr1 | 182642123 | 182642356 | 3.58445 | 4.27059 | 1.24385 | Distal Intergenic | 1 | 182565007 | 182632352 | 67346 | 1 | ENSMUST00000048941.8 | 77117 | Capn8 | ENSMUSG00000038599.9 |
| chr10 | 52689743 | 52690408 | 4.1041 | 5.40196 | 2.0977 | Promoter | 10 | 52690533 | 53111622 | 421090 | 1 | ENSMUST00000105473.2 | -125 | Slc35f1 | ENSMUSG00000038602.7 |
| chr10 | 52691077 | 52691823 | 3.50251 | 4.47126 | 1.43779 | Promoter | 10 | 52690533 | 53111622 | 421090 | 1 | ENSMUST00000105473.2 | 545 | Slc35f1 | ENSMUSG00000038602.7 |
| chr10 | 52692614 | 52693514 | 3.49248 | 4.16613 | 1.27301 | Intron (ENSMUST00000105473.2/ENSMUSG00000038602.7, intron 1 of 7) | 10 | 52690533 | 53111622 | 421090 | 1 | ENSMUST00000105473.2 | 2082 | Slc35f1 | ENSMUSG00000038602.7 |
| chr10 | 52704740 | 52704971 | 3.34433 | 3.89207 | 1.13087 | Intron (ENSMUST00000105473.2/ENSMUSG00000038602.7, intron 1 of 7) | 10 | 52690533 | 53111622 | 421090 | 1 | ENSMUST00000105473.2 | 14208 | Slc35f1 | ENSMUSG00000038602.7 |
| chr1 | 80757718 | 80758383 | 3.83228 | 4.74167 | 1.56353 | Promoter | 1 | 80501073 | 80758448 | 257376 | 2 | ENSMUST00000077946.6 | 65 | Dock10 | ENSMUSG00000038608.10 |
| chr7 | 81859185 | 81859284 | 4.16579 | 5.40588 | 2.04985 | Promoter | 7 | 81859151 | 81884434 | 25284 | 1 | ENSMUST00000126334.2 | 35 | Tm6sf1 | ENSMUSG00000038623.4 |
| chr4 | 58942765 | 58942899 | 3.28108 | 3.90412 | 1.04874 | Promoter | 4 | 58943628 | 58958355 | 14728 | 1 | ENSMUST00000107554.1 | -729 | Zkscan16 | ENSMUSG00000038630.8 |
| chr4 | 58944291 | 58944571 | 3.36703 | 3.89665 | 1.0829 | Promoter | 4 | 58943628 | 58958355 | 14728 | 1 | ENSMUST00000107554.1 | 664 | Zkscan16 | ENSMUSG00000038630.8 |
| chr4 | 58945109 | 58945184 | 3.34524 | 3.84525 | 1.01733 | Intron (ENSMUST00000107554.1/ENSMUSG00000038630.8, intron 1 of 5) | 4 | 58943628 | 58958355 | 14728 | 1 | ENSMUST00000107554.1 | 1482 | Zkscan16 | ENSMUSG00000038630.8 |
| chr4 | 58945653 | 58945768 | 3.55199 | 4.21686 | 1.22206 | Intron (ENSMUST00000107554.1/ENSMUSG00000038630.8, intron 1 of 5) | 4 | 58943628 | 58958355 | 14728 | 1 | ENSMUST00000107554.1 | 2026 | Zkscan16 | ENSMUSG00000038630.8 |
| chr13 | 41114648 | 41114849 | 3.74829 | 4.62102 | 1.54375 | Promoter | 13 | 41114307 | 41146658 | 32352 | 1 | ENSMUST00000124093.1 | 342 | Sycp2l | ENSMUSG00000038651.9 |
| chr13 | 41115188 | 41115386 | 3.66672 | 4.64631 | 1.54727 | Promoter | 13 | 41114307 | 41146658 | 32352 | 1 | ENSMUST00000124093.1 | 882 | Sycp2l | ENSMUSG00000038651.9 |
| chr6 | 37096370 | 37097200 | 3.51404 | 4.21894 | 1.32651 | Intron (ENSMUST00000101532.4/ENSMUSG00000038665.10, intron 4 of 33) | 6 | 37034608 | 37064510 | 29903 | 2 | ENSMUST00000146656.1 | -31861 | Dgki | ENSMUSG00000038665.10 |
| chr6 | 37299109 | 37301044 | 4.30429 | 5.80142 | 2.41307 | Promoter | 6 | 36846022 | 37299974 | 453953 | 2 | ENSMUST00000101532.4 | 0 | Dgki | ENSMUSG00000038665.10 |
| chr6 | 37301552 | 37301695 | 3.52804 | 4.18876 | 1.22562 | Distal Intergenic | 6 | 36846981 | 37300184 | 453204 | 2 | ENSMUST00000150300.3 | -1369 | Dgki | ENSMUSG00000038665.10 |
| chr6 | 37302076 | 37302183 | 3.27116 | 3.87836 | 1.02639 | Distal Intergenic | 6 | 36846981 | 37300184 | 453204 | 2 | ENSMUST00000150300.3 | -1893 | Dgki | ENSMUSG00000038665.10 |
| chr6 | 37303254 | 37303386 | 3.38193 | 3.9006 | 1.01733 | Distal Intergenic | 6 | 36846981 | 37300184 | 453204 | 2 | ENSMUST00000150300.3 | -3071 | Dgki | ENSMUSG00000038665.10 |
| chr5 | 31137772 | 31139335 | 3.76215 | 4.64272 | 1.50541 | Promoter | 5 | 31138063 | 31138829 | 767 | 2 | ENSMUST00000043475.2 | 0 | Ucn | ENSMUSG00000038676.2 |
| chr17 | 28141758 | 28143074 | 3.63778 | 4.76783 | 1.6425 | Promoter | 17 | 28142316 | 28174852 | 32537 | 1 | ENSMUST00000043503.4 | 0 | Scube3 | ENSMUSG00000038677.8 |
| chr17 | 28143936 | 28144753 | 3.57631 | 4.29805 | 1.33686 | Intron (ENSMUST00000043503.4/ENSMUSG00000038677.8, intron 1 of 21) | 17 | 28142316 | 28174852 | 32537 | 1 | ENSMUST00000043503.4 | 1621 | Scube3 | ENSMUSG00000038677.8 |
| chr17 | 28155045 | 28155129 | 3.72594 | 4.55705 | 1.47852 | Intron (ENSMUST00000043503.4/ENSMUSG00000038677.8, intron 3 of 21) | 17 | 28153602 | 28170145 | 16544 | 1 | ENSMUST00000132670.1 | 1444 | Scube3 | ENSMUSG00000038677.8 |
| chr17 | 28164287 | 28164478 | 3.43018 | 3.9945 | 1.09208 | Exon (ENSMUST00000043503.4/ENSMUSG00000038677.8, exon 12 of 22) | 17 | 28165809 | 28167648 | 1840 | 1 | ENSMUST00000142170.1 | -1331 | Scube3 | ENSMUSG00000038677.8 |
| chr7 | 44477903 | 44478996 | 3.52757 | 4.38853 | 1.39466 | Distal Intergenic | 7 | 44470925 | 44471651 | 727 | 1 | ENSMUST00000134398.1 | 6979 | Josd2 | ENSMUSG00000038695.7 |
| chr11 | 96303788 | 96303865 | 3.44068 | 4.01818 | 1.11957 | Promoter | 11 | 96303512 | 96306120 | 2609 | 1 | ENSMUST00000049272.4 | 277 | Hoxb5 | ENSMUSG00000038700.4 |
| chr7 | 44460449 | 44460642 | 3.6262 | 4.34719 | 1.29171 | Intron (ENSMUST00000127790.2/ENSMUSG00000047085.9, intron 2 of 2) | 7 | 44465391 | 44467757 | 2367 | 1 | ENSMUST00000035929.5 | -4749 | Aspdh | ENSMUSG00000038704.5 |
| chr7 | 44461403 | 44461576 | 3.60143 | 4.48035 | 1.41793 | Exon (ENSMUST00000058667.9/ENSMUSG00000047085.9, exon 3 of 3) | 7 | 44465391 | 44467757 | 2367 | 1 | ENSMUST00000035929.5 | -3815 | Aspdh | ENSMUSG00000038704.5 |
| chr7 | 44461984 | 44463347 | 4.10978 | 5.30362 | 1.95797 | 3' UTR | 7 | 44465391 | 44467757 | 2367 | 1 | ENSMUST00000035929.5 | -2044 | Aspdh | ENSMUSG00000038704.5 |
| chr11 | 96287156 | 96287253 | 3.43231 | 4.06981 | 1.22706 | Promoter | 11 | 96286623 | 96290162 | 3540 | 1 | ENSMUST00000049352.7 | 534 | Hoxb7 | ENSMUSG00000038721.8 |
| chr15 | 44457499 | 44457719 | 3.61963 | 4.40496 | 1.44293 | Promoter | 15 | 44457553 | 44597137 | 139585 | 1 | ENSMUST00000038336.6 | 0 | Pkhd1l1 | ENSMUSG00000038725.6 |
| chr7 | 44310707 | 44312461 | 3.91309 | 4.96126 | 1.75086 | Promoter | 7 | 44312688 | 44357487 | 44800 | 1 | ENSMUST00000107935.2 | -227 | Shank1 | ENSMUSG00000038738.10 |
| chr7 | 44312774 | 44312893 | 3.32103 | 3.82048 | 1.05788 | Promoter | 7 | 44312707 | 44357337 | 44631 | 1 | ENSMUST00000107934.1 | 68 | Shank1 | ENSMUSG00000038738.10 |
| chr7 | 44335491 | 44335566 | 3.59451 | 4.29082 | 1.25761 | Promoter | 7 | 44336016 | 44343738 | 7723 | 1 | ENSMUST00000134470.2 | -450 | Shank1 | ENSMUSG00000038738.10 |
| chr7 | 44336030 | 44336420 | 3.91603 | 4.91516 | 1.69278 | Promoter | 7 | 44336017 | 44342509 | 6493 | 1 | ENSMUST00000134205.1 | 14 | Shank1 | ENSMUSG00000038738.10 |
| chr7 | 44336861 | 44337329 | 3.9538 | 4.99108 | 1.74825 | Promoter | 7 | 44336017 | 44342509 | 6493 | 1 | ENSMUST00000134205.1 | 845 | Shank1 | ENSMUSG00000038738.10 |
| chr7 | 44347952 | 44348327 | 3.35622 | 3.86182 | 1.01733 | Intron (ENSMUST00000107938.2/ENSMUSG00000038738.10, intron 22 of 24) | 7 | 44350762 | 44352057 | 1296 | 1 | ENSMUST00000127164.1 | -2435 | Shank1 | ENSMUSG00000038738.10 |
| chr7 | 44349847 | 44350178 | 3.7077 | 4.60037 | 1.54771 | Promoter | 7 | 44350762 | 44352057 | 1296 | 1 | ENSMUST00000127164.1 | -584 | Shank1 | ENSMUSG00000038738.10 |
| chr7 | 44351004 | 44351138 | 4.03653 | 5.41184 | 2.06024 | Promoter | 7 | 44350762 | 44352057 | 1296 | 1 | ENSMUST00000127164.1 | 243 | Shank1 | ENSMUSG00000038738.10 |
| chr7 | 44351779 | 44354872 | 3.8144 | 4.86162 | 1.69689 | Exon (ENSMUST00000107938.2/ENSMUSG00000038738.10, exon 24 of 25) | 7 | 44350762 | 44352057 | 1296 | 1 | ENSMUST00000127164.1 | 1018 | Shank1 | ENSMUSG00000038738.10 |
| chr7 | 44356739 | 44356898 | 3.94479 | 4.95503 | 1.6994 | Exon (ENSMUST00000107938.2/ENSMUSG00000038738.10, exon 25 of 25) | 7 | 44350762 | 44352057 | 1296 | 1 | ENSMUST00000127164.1 | 5978 | Shank1 | ENSMUSG00000038738.10 |
| chr7 | 44357709 | 44358044 | 3.63015 | 4.36444 | 1.32298 | 3' UTR | 7 | 44350762 | 44352057 | 1296 | 1 | ENSMUST00000127164.1 | 6948 | Shank1 | ENSMUSG00000038738.10 |
| chr7 | 44359047 | 44359588 | 3.81118 | 4.73413 | 1.57028 | Downstream (<1kb) | 7 | 44350762 | 44352057 | 1296 | 1 | ENSMUST00000127164.1 | 8286 | Shank1 | ENSMUSG00000038738.10 |
| chr7 | 140919294 | 140919700 | 3.48802 | 4.23299 | 1.33655 | Distal Intergenic | 7 | 140920902 | 140928540 | 7639 | 1 | ENSMUST00000106045.2 | -1202 | Nlrp6 | ENSMUSG00000038745.10 |
| chr7 | 140920552 | 140920939 | 3.69532 | 4.58671 | 1.51344 | Promoter | 7 | 140920902 | 140928540 | 7639 | 1 | ENSMUST00000106045.2 | 0 | Nlrp6 | ENSMUSG00000038745.10 |
| chr7 | 140921313 | 140924097 | 4.08419 | 5.34501 | 2.0784 | Promoter | 7 | 140922106 | 140922855 | 750 | 1 | ENSMUST00000183761.1 | 0 | Nlrp6 | ENSMUSG00000038745.10 |
| chr19 | 46131760 | 46132069 | 3.55627 | 4.25578 | 1.33628 | Promoter | 19 | 46131897 | 46135307 | 3411 | 1 | ENSMUST00000043739.4 | 0 | Elovl3 | ENSMUSG00000038754.4 |
| chr19 | 46133044 | 46133934 | 3.91871 | 5.01138 | 1.82206 | Exon (ENSMUST00000043739.4/ENSMUSG00000038754.4, exon 2 of 4) | 19 | 46131897 | 46135307 | 3411 | 1 | ENSMUST00000043739.4 | 1148 | Elovl3 | ENSMUSG00000038754.4 |
| chr11 | 96123373 | 96125110 | 3.76036 | 4.68612 | 1.55801 | Distal Intergenic | 11 | 96133786 | 96165451 | 31666 | 1 | ENSMUST00000167258.2 | -8676 | Ttll6 | ENSMUSG00000038756.8 |
| chr11 | 96126367 | 96126697 | 3.36678 | 3.87775 | 1.01733 | Distal Intergenic | 11 | 96133786 | 96165451 | 31666 | 1 | ENSMUST00000167258.2 | -7089 | Ttll6 | ENSMUSG00000038756.8 |
| chr11 | 96134289 | 96134364 | 3.60481 | 4.3773 | 1.3571 | Promoter | 11 | 96134219 | 96165451 | 31233 | 1 | ENSMUST00000107680.1 | 71 | Ttll6 | ENSMUSG00000038756.8 |
| chr11 | 96135220 | 96135309 | 3.42905 | 3.99934 | 1.11522 | Intron (ENSMUST00000167258.2/ENSMUSG00000038756.8, intron 2 of 14) | 11 | 96134219 | 96165451 | 31233 | 1 | ENSMUST00000107680.1 | 1002 | Ttll6 | ENSMUSG00000038756.8 |
| chr11 | 96137833 | 96138094 | 4.47597 | 6.02721 | 2.47153 | Intron (ENSMUST00000167258.2/ENSMUSG00000038756.8, intron 4 of 14) | 11 | 96134219 | 96165451 | 31233 | 1 | ENSMUST00000107680.1 | 3615 | Ttll6 | ENSMUSG00000038756.8 |
| chr11 | 96140249 | 96140324 | 3.44427 | 4.01426 | 1.08622 | Intron (ENSMUST00000167258.2/ENSMUSG00000038756.8, intron 6 of 14) | 11 | 96134219 | 96165451 | 31233 | 1 | ENSMUST00000107680.1 | 6031 | Ttll6 | ENSMUSG00000038756.8 |
| chr11 | 96143534 | 96143889 | 3.53822 | 4.18704 | 1.19531 | Intron (ENSMUST00000167258.2/ENSMUSG00000038756.8, intron 7 of 14) | 11 | 96134219 | 96165451 | 31233 | 1 | ENSMUST00000107680.1 | 9316 | Ttll6 | ENSMUSG00000038756.8 |
| chr11 | 96145101 | 96147087 | 4.10935 | 5.38802 | 2.09072 | Exon (ENSMUST00000167258.2/ENSMUSG00000038756.8, exon 8 of 15) | 11 | 96134219 | 96165451 | 31233 | 1 | ENSMUST00000107680.1 | 10883 | Ttll6 | ENSMUSG00000038756.8 |
| chr11 | 96148832 | 96148963 | 3.28596 | 3.75615 | 1.01733 | Intron (ENSMUST00000167258.2/ENSMUSG00000038756.8, intron 9 of 14) | 11 | 96134219 | 96165451 | 31233 | 1 | ENSMUST00000107680.1 | 14614 | Ttll6 | ENSMUSG00000038756.8 |
| chr11 | 96151884 | 96152172 | 3.54541 | 4.20526 | 1.21729 | Exon (ENSMUST00000167258.2/ENSMUSG00000038756.8, exon 10 of 15) | 11 | 96134219 | 96165451 | 31233 | 1 | ENSMUST00000107680.1 | 17666 | Ttll6 | ENSMUSG00000038756.8 |

| | | | | | | | | | | | | | | | |
|---|---|---|---|---|---|---|---|---|---|---|---|---|---|---|---|
| chr11 | 96153272 | 96153992 | 3.71984 | 4.54422 | 1.46129 | Intron (ENSMUST00000167258.2/ENSMUSG00000038756.8, intron 11 of 14) | 11 | 96134219 | 96165451 | 31233 | 1 | ENSMUST00000107680.1 | 19054 | Ttll6 | ENSMUSG00000038756.8 |
| chr4 | 57322010 | 57322166 | 3.31066 | 3.7932 | 1.01733 | Distal Intergenic | 4 | 57190841 | 57301837 | 110997 | 2 | ENSMUST00000075637.5 | -20174 | Ptpn3 | ENSMUSG00000038764.9 |
| chr4 | 57324088 | 57324626 | 3.41318 | 3.96474 | 1.07749 | Distal Intergenic | 4 | 57190841 | 57301837 | 110997 | 2 | ENSMUST00000075637.5 | -22252 | Ptpn3 | ENSMUSG00000038764.9 |
| chr14 | 24618509 | 24618750 | 3.3254 | 3.82108 | 1.03735 | Promoter | 14 | 24617568 | 24621693 | 4126 | 1 | ENSMUST00000038250.3 | 942 | 4931403M11Rik | ENSMUSG00000038792.3 |
| chr7 | 140841772 | 140841953 | 3.25308 | 3.70675 | 1.01733 | Exon (ENSMUST00000190581.1/ENSMUSG00000099590.1, exon 2 of 4) | 7 | 140845547 | 140846769 | 1223 | 1 | ENSMUST00000035300.6 | -3594 | Scgb1c1 | ENSMUSG00000038801.6 |
| chr17 | 85621187 | 85622439 | 3.97856 | 5.24335 | 1.96822 | Promoter | 17 | 85621031 | 85629302 | 8272 | 1 | ENSMUST00000162695.2 | 157 | Six3 | ENSMUSG00000038805.10 |
| chr17 | 85623299 | 85624459 | 4.2271 | 5.62044 | 2.23324 | 3' UTR | 17 | 85621031 | 85629302 | 8272 | 1 | ENSMUST00000162695.2 | 2269 | Six3 | ENSMUSG00000038805.10 |
| chr17 | 85624807 | 85625642 | 3.95784 | 5.05641 | 1.79804 | 3' UTR | 17 | 85621031 | 85629302 | 8272 | 1 | ENSMUST00000162695.2 | 3777 | Six3 | ENSMUSG00000038805.10 |
| chr17 | 85626027 | 85626490 | 4.16332 | 5.42316 | 2.06587 | 3' UTR | 17 | 85621031 | 85629302 | 8272 | 1 | ENSMUST00000162695.2 | 4997 | Six3 | ENSMUSG00000038805.10 |
| chr17 | 85626969 | 85627325 | 3.63276 | 4.362 | 1.30299 | Intron (ENSMUST00000162695.2/ENSMUSG00000038805.10, intron 2 of 2) | 17 | 85621031 | 85629302 | 8272 | 1 | ENSMUST00000162695.2 | 5939 | Six3 | ENSMUSG00000038805.10 |
| chr17 | 85628476 | 85628886 | 3.37674 | 3.98297 | 1.11721 | 3' UTR | 17 | 85621031 | 85629302 | 8272 | 1 | ENSMUST00000162695.2 | 7446 | Six3 | ENSMUSG00000038805.10 |
| chr17 | 85631289 | 85631485 | 3.35149 | 3.88823 | 1.09637 | Distal Intergenic | 17 | 85621031 | 85629302 | 8272 | 1 | ENSMUST00000162695.2 | 10259 | Six3 | ENSMUSG00000038805.10 |
| chr17 | 85632671 | 85632746 | 3.37433 | 3.944 | 1.17482 | Distal Intergenic | 17 | 85621031 | 85629302 | 8272 | 1 | ENSMUST00000162695.2 | 11641 | Six3 | ENSMUSG00000038805.10 |
| chr17 | 85634108 | 85634290 | 3.91495 | 5.0092 | 1.83754 | Distal Intergenic | 17 | 85621031 | 85629302 | 8272 | 1 | ENSMUST00000162695.2 | 13078 | Six3 | ENSMUSG00000038805.10 |
| chr17 | 85637168 | 85637482 | 3.75455 | 4.69403 | 1.59642 | Distal Intergenic | 17 | 85621031 | 85629302 | 8272 | 1 | ENSMUST00000162695.2 | 16138 | Six3 | ENSMUSG00000038805.10 |
| chr17 | 85637892 | 85638074 | 3.34679 | 3.85218 | 1.03351 | Distal Intergenic | 17 | 85621031 | 85629302 | 8272 | 1 | ENSMUST00000162695.2 | 16862 | Six3 | ENSMUSG00000038805.10 |
| chr17 | 85639521 | 85639648 | 3.44908 | 4.04843 | 1.18222 | Distal Intergenic | 17 | 85621031 | 85629302 | 8272 | 1 | ENSMUST00000162695.2 | 18491 | Six3 | ENSMUSG00000038805.10 |
| chr17 | 85639999 | 85640194 | 3.53663 | 4.18795 | 1.2075 | Distal Intergenic | 17 | 85621031 | 85629302 | 8272 | 1 | ENSMUST00000162695.2 | 18969 | Six3 | ENSMUSG00000038805.10 |
| chr17 | 85641895 | 85641997 | 3.45845 | 4.0445 | 1.11744 | Distal Intergenic | 17 | 85621031 | 85629302 | 8272 | 1 | ENSMUST00000162695.2 | 20865 | Six3 | ENSMUSG00000038805.10 |
| chr17 | 85643842 | 85643979 | 3.44365 | 4.0483 | 1.21075 | Distal Intergenic | 17 | 85621031 | 85629302 | 8272 | 1 | ENSMUST00000162695.2 | 22812 | Six3 | ENSMUSG00000038805.10 |
| chr17 | 85644461 | 85644688 | 3.38892 | 4.0998 | 1.15425 | Distal Intergenic | 17 | 85621031 | 85629302 | 8272 | 1 | ENSMUST00000162695.2 | 23431 | Six3 | ENSMUSG00000038805.10 |
| chr17 | 85645206 | 85645379 | 3.73123 | 4.60029 | 1.56797 | Distal Intergenic | 17 | 85621031 | 85629302 | 8272 | 1 | ENSMUST00000162695.2 | 24176 | Six3 | ENSMUSG00000038805.10 |
| chr17 | 85645980 | 85646691 | 3.78561 | 4.69486 | 1.61748 | Distal Intergenic | 17 | 85621031 | 85629302 | 8272 | 1 | ENSMUST00000162695.2 | 24950 | Six3 | ENSMUSG00000038805.10 |
| chr11 | 74496052 | 74496257 | 3.60269 | 4.33226 | 1.31773 | Intron (ENSMUST00000102521.2/ENSMUSG00000038807.12, intron 2 of 24) | 11 | 74383484 | 74408025 | 24542 | 2 | ENSMUST00000123440.1 | -88028 | Rap1gap2 | ENSMUSG00000038807.12 |
| chr11 | 74497105 | 74497442 | 3.60446 | 4.31098 | 1.28064 | Intron (ENSMUST00000102521.2/ENSMUSG00000038807.12, intron 2 of 24) | 11 | 74383484 | 74408025 | 24542 | 2 | ENSMUST00000123440.1 | -89081 | Rap1gap2 | ENSMUSG00000038807.12 |
| chr11 | 74521996 | 74522488 | 3.56285 | 4.35994 | 1.39831 | Intron (ENSMUST00000102521.2/ENSMUSG00000038807.12, intron 2 of 24) | 11 | 74437043 | 74590124 | 153082 | 2 | ENSMUST00000145524.1 | 67636 | Rap1gap2 | ENSMUSG00000038807.12 |
| chr11 | 74527620 | 74527785 | 3.96256 | 5.0319 | 1.80175 | Intron (ENSMUST00000102521.2/ENSMUSG00000038807.12, intron 2 of 24) | 11 | 74437043 | 74590124 | 153082 | 2 | ENSMUST00000145524.1 | 62339 | Rap1gap2 | ENSMUSG00000038807.12 |
| chr11 | 74530918 | 74530993 | 3.51001 | 4.20584 | 1.29867 | Intron (ENSMUST00000102521.2/ENSMUSG00000038807.12, intron 2 of 24) | 11 | 74437043 | 74590124 | 153082 | 2 | ENSMUST00000145524.1 | 59131 | Rap1gap2 | ENSMUSG00000038807.12 |
| chr11 | 74589364 | 74589439 | 4.11663 | 5.28895 | 1.93698 | Promoter | 11 | 74437043 | 74590124 | 153082 | 2 | ENSMUST00000145524.1 | 685 | Rap1gap2 | ENSMUSG00000038807.12 |
| chr11 | 74590508 | 74590704 | 3.3845 | 3.9139 | 1.0507 | Promoter | 11 | 74383356 | 74590186 | 206831 | 2 | ENSMUST00000102521.2 | -323 | Rap1gap2 | ENSMUSG00000038807.12 |
| chr11 | 74592498 | 74592901 | 3.43888 | 4.00845 | 1.09499 | Distal Intergenic | 11 | 74383356 | 74590186 | 206831 | 2 | ENSMUST00000102521.2 | -2313 | Rap1gap2 | ENSMUSG00000038807.12 |
| chr19 | 17356446 | 17356649 | 3.90526 | 4.99021 | 1.80182 | Promoter | 19 | 17326141 | 17356667 | 30527 | 2 | ENSMUST00000174236.2 | 18 | Gcnt1 | ENSMUSG00000038843.12 |
| chr11 | 95645205 | 95645779 | 3.91401 | 4.94795 | 1.70608 | Distal Intergenic | 11 | 95666957 | 95680773 | 13817 | 1 | ENSMUST00000125172.2 | -21178 | Phb | ENSMUSG00000038845.6 |
| chr11 | 95646946 | 95647885 | 3.70701 | 4.52675 | 1.43649 | Distal Intergenic | 11 | 95666957 | 95680773 | 13817 | 1 | ENSMUST00000125172.2 | -19072 | Phb | ENSMUSG00000038845.6 |
| chr1 | 180312958 | 180313254 | 3.35399 | 3.90162 | 1.11345 | Distal Intergenic | 1 | 180330485 | 180424802 | 94318 | 1 | ENSMUST00000070181.6 | -17231 | Itpkb | ENSMUSG00000038855.9 |
| chr1 | 180313568 | 180313815 | 3.93414 | 5.07797 | 1.84967 | Distal Intergenic | 1 | 180330485 | 180424802 | 94318 | 1 | ENSMUST00000070181.6 | -16670 | Itpkb | ENSMUSG00000038855.9 |
| chr1 | 180314893 | 180315037 | 3.29828 | 3.7746 | 1.01733 | Distal Intergenic | 1 | 180330485 | 180424802 | 94318 | 1 | ENSMUST00000070181.6 | -15448 | Itpkb | ENSMUSG00000038855.9 |
| chr1 | 180315780 | 180316202 | 3.97077 | 5.0588 | 1.86372 | Distal Intergenic | 1 | 180330485 | 180424802 | 94318 | 1 | ENSMUST00000070181.6 | -14283 | Itpkb | ENSMUSG00000038855.9 |
| chr1 | 180329350 | 180329618 | 3.26983 | 3.75756 | 1.0705 | Promoter | 1 | 180330485 | 180424802 | 94318 | 1 | ENSMUST00000070181.6 | -867 | Itpkb | ENSMUSG00000038855.9 |
| chr8 | 108301017 | 108301321 | 3.69482 | 4.67224 | 1.5446 | Distal Intergenic | 8 | 108703100 | 108800270 | 97171 | 1 | ENSMUST00000188994.1 | -401779 | Zfhx3 | ENSMUSG00000038872.9 |
| chr8 | 108549808 | 108550031 | 3.34497 | 3.84485 | 1.01733 | Distal Intergenic | 8 | 108703100 | 108800270 | 97171 | 1 | ENSMUST00000188994.1 | -153069 | Zfhx3 | ENSMUSG00000038872.9 |
| chr8 | 108566327 | 108566739 | 3.55599 | 4.3475 | 1.34777 | Distal Intergenic | 8 | 108703100 | 108800270 | 97171 | 1 | ENSMUST00000188994.1 | -136361 | Zfhx3 | ENSMUSG00000038872.9 |
| chr8 | 108567222 | 108567479 | 3.49322 | 4.12331 | 1.2035 | Distal Intergenic | 8 | 108703100 | 108800270 | 97171 | 1 | ENSMUST00000188994.1 | -135621 | Zfhx3 | ENSMUSG00000038872.9 |
| chr8 | 108581967 | 108582042 | 3.90778 | 4.91487 | 1.73507 | Distal Intergenic | 8 | 108703100 | 108800270 | 97171 | 1 | ENSMUST00000188994.1 | -121058 | Zfhx3 | ENSMUSG00000038872.9 |
| chr8 | 108596490 | 108596565 | 4.0124 | 5.11801 | 1.87566 | Distal Intergenic | 8 | 108703100 | 108800270 | 97171 | 1 | ENSMUST00000188994.1 | -106535 | Zfhx3 | ENSMUSG00000038872.9 |
| chr8 | 108622409 | 108622484 | 4.08916 | 5.60111 | 2.18531 | Distal Intergenic | 8 | 108703100 | 108800270 | 97171 | 1 | ENSMUST00000188994.1 | -80616 | Zfhx3 | ENSMUSG00000038872.9 |
| chr8 | 108674849 | 108675087 | 3.30825 | 3.81902 | 1.10722 | Distal Intergenic | 8 | 108703100 | 108800270 | 97171 | 1 | ENSMUST00000188994.1 | -28013 | Zfhx3 | ENSMUSG00000038872.9 |
| chr8 | 108697247 | 108697572 | 3.76037 | 4.75576 | 1.60872 | Distal Intergenic | 8 | 108703100 | 108800270 | 97171 | 1 | ENSMUST00000188994.1 | -5528 | Zfhx3 | ENSMUSG00000038872.9 |
| chr8 | 108698452 | 108698573 | 3.56053 | 4.35681 | 1.3682 | Distal Intergenic | 8 | 108703100 | 108800270 | 97171 | 1 | ENSMUST00000188994.1 | -4527 | Zfhx3 | ENSMUSG00000038872.9 |
| chr8 | 108698911 | 108699027 | 4.74052 | 6.98735 | 3.22134 | Distal Intergenic | 8 | 108703100 | 108800270 | 97171 | 1 | ENSMUST00000188994.1 | -4073 | Zfhx3 | ENSMUSG00000038872.9 |
| chr8 | 108699666 | 108700712 | 4.11677 | 5.59345 | 2.24021 | Distal Intergenic | 8 | 108703100 | 108800270 | 97171 | 1 | ENSMUST00000188994.1 | -2388 | Zfhx3 | ENSMUSG00000038872.9 |
| chr8 | 108701022 | 108701231 | 3.58206 | 4.45236 | 1.43498 | Distal Intergenic | 8 | 108703100 | 108800270 | 97171 | 1 | ENSMUST00000188994.1 | -1869 | Zfhx3 | ENSMUSG00000038872.9 |
| chr8 | 108701605 | 108701780 | 4.13033 | 5.64219 | 2.22791 | Distal Intergenic | 8 | 108703100 | 108800270 | 97171 | 1 | ENSMUST00000188994.1 | -1320 | Zfhx3 | ENSMUSG00000038872.9 |
| chr8 | 108702363 | 108703126 | 3.61011 | 4.44575 | 1.40913 | Promoter | 8 | 108703100 | 108800270 | 97171 | 1 | ENSMUST00000188994.1 | 0 | Zfhx3 | ENSMUSG00000038872.9 |
| chr15 | 34678303 | 34679468 | 4.13546 | 5.61518 | 2.24734 | Promoter | 15 | 34572799 | 34678706 | 105908 | 2 | ENSMUST00000040791.7 | 0 | Nipal2 | ENSMUSG00000038879.7 |
| chr15 | 34679805 | 34680159 | 3.61699 | 4.38115 | 1.37111 | Distal Intergenic | 15 | 34572799 | 34678706 | 105908 | 2 | ENSMUST00000040791.7 | -1100 | Nipal2 | ENSMUSG00000038879.7 |
| chr15 | 34681452 | 34681607 | 3.38828 | 3.9541 | 1.16628 | Distal Intergenic | 15 | 34572799 | 34678706 | 105908 | 2 | ENSMUST00000040791.7 | -2747 | Nipal2 | ENSMUSG00000038879.7 |
| chr10 | 29142469 | 29143064 | 3.57896 | 4.34883 | 1.38349 | Promoter | 10 | 29143996 | 29199630 | 55635 | 1 | ENSMUST00000092629.2 | -932 | Soga3 | ENSMUSG00000038916.6 |
| chr10 | 29143549 | 29144876 | 3.80048 | 4.69735 | 1.53982 | Promoter | 10 | 29143996 | 29199630 | 55635 | 1 | ENSMUST00000092629.2 | 0 | Soga3 | ENSMUSG00000038916.6 |
| chr8 | 10879123 | 10879461 | 3.28819 | 3.75951 | 1.01733 | Distal Intergenic | 8 | 10924427 | 10928457 | 4031 | 2 | ENSMUST00000040608.3 | 48996 | 3930402G23Rik | ENSMUSG00000038917.3 |
| chr2 | 180630090 | 180630262 | 3.40828 | 3.98042 | 1.15866 | Intron (ENSMUST00000037877.10/ENSMUSG00000038932.13, intron 5 of 5) | 2 | 180637466 | 180642347 | 4882 | 2 | ENSMUST00000131881.1 | 12085 | Tcfl5 | ENSMUSG00000038932.13 |
| chr2 | 180640917 | 180641388 | 3.86286 | 4.96403 | 1.76789 | Promoter | 2 | 180637466 | 180642347 | 4882 | 2 | ENSMUST00000131881.1 | 959 | Tcfl5 | ENSMUSG00000038932.13 |
| chr2 | 180641736 | 180642504 | 3.96064 | 5.08677 | 1.85224 | Promoter | 2 | 180637466 | 180642347 | 4882 | 2 | ENSMUST00000131881.1 | 0 | Tcfl5 | ENSMUSG00000038932.13 |
| chr2 | 180643015 | 180643643 | 3.33874 | 4.14449 | 1.24107 | Promoter | 2 | 180621956 | 180642708 | 20753 | 2 | ENSMUST00000037877.10 | -308 | Tcfl5 | ENSMUSG00000038932.13 |
| chr2 | 180644129 | 180644328 | 3.35713 | 4.05352 | 1.15288 | Distal Intergenic | 2 | 180621956 | 180642708 | 20753 | 2 | ENSMUST00000037877.10 | -1422 | Tcfl5 | ENSMUSG00000038932.13 |
| chr2 | 180456130 | 180457055 | 4.12916 | 5.46076 | 2.14452 | Promoter | 2 | 180456245 | 180474867 | 18623 | 1 | ENSMUST00000038259.7 | 0 | Slco4a1 | ENSMUSG00000038963.10 |
| chr2 | 180472250 | 180472653 | 3.45083 | 4.11291 | 1.22686 | Promoter | 2 | 180472429 | 180474852 | 2424 | 1 | ENSMUST00000128367.1 | 0 | Slco4a1 | ENSMUSG00000038963.10 |
| chr5 | 144098048 | 144098195 | 3.42833 | 4.05787 | 1.17359 | Distal Intergenic | 5 | 144100436 | 144188204 | 87769 | 1 | ENSMUST00000041804.7 | -2241 | Lmtk2 | ENSMUSG00000038970.7 |
| chr13 | 38344059 | 38344387 | 3.354 | 4.0464 | 1.21021 | Distal Intergenic | 13 | 38345716 | 38499728 | 154013 | 1 | ENSMUST00000171970.1 | -1329 | Bmp6 | ENSMUSG00000039004.5 |
| chr13 | 38345610 | 38347303 | 4.08016 | 5.45673 | 2.14658 | Promoter | 13 | 38345716 | 38499728 | 154013 | 1 | ENSMUST00000171970.1 | 0 | Bmp6 | ENSMUSG00000039004.5 |
| chr13 | 38347630 | 38347888 | 3.48743 | 4.24561 | 1.26677 | Intron (ENSMUST00000171970.1/ENSMUSG00000039004.5, intron 1 of 6) | 13 | 38345716 | 38499728 | 154013 | 1 | ENSMUST00000171970.1 | 1915 | Bmp6 | ENSMUSG00000039004.5 |
| chr13 | 38354828 | 38355135 | 3.46253 | 4.13565 | 1.27212 | Intron (ENSMUST00000171970.1/ENSMUSG00000039004.5, intron 1 of 6) | 13 | 38345716 | 38499728 | 154013 | 1 | ENSMUST00000171970.1 | 9113 | Bmp6 | ENSMUSG00000039004.5 |
| chr13 | 38409460 | 38409787 | 3.22223 | 3.66056 | 1.01463 | Intron (ENSMUST00000171970.1/ENSMUSG00000039004.5, intron 1 of 6) | 13 | 38345716 | 38499728 | 154013 | 1 | ENSMUST00000171970.1 | 63745 | Bmp6 | ENSMUSG00000039004.5 |
| chr6 | 30920626 | 30920876 | 3.41402 | 3.96445 | 1.07181 | Intron (ENSMUST00000139006.1/ENSMUSG00000085321.1, intron 2 of 2) | 6 | 30896981 | 30915573 | 18593 | 2 | ENSMUST00000048580.1 | -5054 | Tsga13 | ENSMUSG00000039032.1 |
| chr6 | 30921962 | 30922816 | 3.48563 | 4.1057 | 1.18802 | Intron (ENSMUST00000139006.1/ENSMUSG00000085321.1, intron 2 of 2) | 6 | 30896981 | 30915573 | 18593 | 2 | ENSMUST00000048580.1 | -6390 | Tsga13 | ENSMUSG00000039032.1 |
| chr3 | 152981210 | 152982195 | 4.37307 | 5.91628 | 2.48436 | Promoter | 3 | 152975374 | 152982052 | 6679 | 2 | ENSMUST00000161757.1 | 0 | St6galnac5 | ENSMUSG00000039037.5 |
| chr3 | 153123746 | 153123850 | 15.55999 | 39.30928 | 33.60904 | Distal Intergenic | 3 | 152817081 | 152982246 | 165166 | 2 | ENSMUST00000044278.5 | -141501 | St6galnac5 | ENSMUSG00000039037.5 |
| chr8 | 10153552 | 10154770 | 4.01049 | 5.11108 | 1.84817 | Distal Intergenic | 8 | 10272572 | 10634741 | 362170 | 1 | ENSMUST00000042103.8 | -117802 | Myo16 | ENSMUSG00000039057.9 |
| chr8 | 10418652 | 10418796 | 3.35139 | 3.85452 | 1.01733 | Intron (ENSMUST00000042103.8/ENSMUSG00000039057.9, intron 15 of 34) | 8 | 10272572 | 10634741 | 362170 | 1 | ENSMUST00000042103.8 | 146081 | Myo16 | ENSMUSG00000039057.9 |
| chr8 | 10450565 | 10450732 | 3.30933 | 3.84607 | 1.10846 | Intron (ENSMUST00000042103.8/ENSMUSG00000039057.9, intron 19 of 34) | 8 | 10272572 | 10634741 | 362170 | 1 | ENSMUST00000042103.8 | 177994 | Myo16 | ENSMUSG00000039057.9 |
| chr8 | 10453325 | 10453588 | 3.52121 | 4.21972 | 1.32343 | Intron (ENSMUST00000042103.8/ENSMUSG00000039057.9, intron 19 of 34) | 8 | 10272572 | 10634741 | 362170 | 1 | ENSMUST00000042103.8 | 180754 | Myo16 | ENSMUSG00000039057.9 |
| chr8 | 10454671 | 10454856 | 3.42442 | 4.02967 | 1.22358 | Intron (ENSMUST00000042103.8/ENSMUSG00000039057.9, intron 19 of 34) | 8 | 10272572 | 10634741 | 362170 | 1 | ENSMUST00000042103.8 | 182100 | Myo16 | ENSMUSG00000039057.9 |
| chr3 | 152667461 | 152668062 | 3.89751 | 4.91598 | 1.75924 | Promoter | 3 | 152462815 | 152668140 | 205326 | 2 | ENSMUST00000045262.6 | 78 | Ak5 | ENSMUSG00000039058.7 |
| chr3 | 152668979 | 152669091 | 3.42126 | 4.00632 | 1.18236 | Promoter | 3 | 152462815 | 152668140 | 205326 | 2 | ENSMUST00000045262.6 | -840 | Ak5 | ENSMUSG00000039058.7 |
| chr2 | 180102115 | 180102468 | 3.55816 | 4.2667 | 1.2832 | Promoter | 2 | 180100984 | 180103335 | 2352 | 2 | ENSMUST00000166392.1 | 867 | Hrh3 | ENSMUSG00000039059.6 |
| chr2 | 180102834 | 180104540 | 4.11474 | 5.34181 | 2.00542 | Promoter | 2 | 180099465 | 180104488 | 5024 | 2 | ENSMUST00000056480.4 | 0 | Hrh3 | ENSMUSG00000039059.6 |
| chr2 | 180104906 | 180105105 | 3.57681 | 4.33191 | 1.36034 | Promoter | 2 | 180099465 | 180104488 | 5024 | 2 | ENSMUST00000056480.4 | -419 | Hrh3 | ENSMUSG00000039059.6 |
| chr2 | 180105775 | 180106133 | 3.36077 | 3.86868 | 1.01733 | Distal Intergenic | 2 | 180099465 | 180104488 | 5024 | 2 | ENSMUST00000056480.4 | -1288 | Hrh3 | ENSMUSG00000039059.6 |
| chr7 | 79847797 | 79848083 | 3.70198 | 4.68996 | 1.56943 | Promoter | 7 | 79842614 | 79848193 | 5580 | 2 | ENSMUST00000149164.1 | 110 | Anpep | ENSMUSG00000039062.10 |
| chr2 | 6195721 | 6196143 | 3.64009 | 4.37876 | 1.32563 | Promoter | 2 | 6189509 | 6196208 | 6700 | 2 | ENSMUST00000132810.1 | 65 | Echdc3 | ENSMUSG00000039063.4 |
| chr2 | 6219107 | 6219310 | 3.23118 | 3.67395 | 1.01549 | Intron (ENSMUST00000151782.1/ENSMUSG00000087125.1, intron 1 of 6) | 2 | 6188465 | 6213033 | 24569 | 2 | ENSMUST00000042658.3 | -6075 | Echdc3 | ENSMUSG00000039063.4 |
| chr5 | 28119931 | 28120187 | 3.36449 | 3.87428 | 1.01733 | Distal Intergenic | 5 | 28165694 | 28172166 | 6473 | 1 | ENSMUST00000036177.8 | -45507 | En2 | ENSMUSG00000039095.8 |
| chr5 | 28120634 | 28121437 | 3.61847 | 4.41169 | 1.39055 | Distal Intergenic | 5 | 28165694 | 28172166 | 6473 | 1 | ENSMUST00000036177.8 | -44257 | En2 | ENSMUSG00000039095.8 |
| chr5 | 28122352 | 28122429 | 3.83225 | 4.73965 | 1.55768 | Distal Intergenic | 5 | 28165694 | 28172166 | 6473 | 1 | ENSMUST00000036177.8 | -43265 | En2 | ENSMUSG00000039095.8 |
| chr5 | 28122991 | 28123523 | 3.61946 | 4.63704 | 1.53743 | Distal Intergenic | 5 | 28165694 | 28172166 | 6473 | 1 | ENSMUST00000036177.8 | -42171 | En2 | ENSMUSG00000039095.8 |
| chr5 | 28124481 | 28124572 | 3.31079 | 3.82004 | 1.09716 | Distal Intergenic | 5 | 28165694 | 28172166 | 6473 | 1 | ENSMUST00000036177.8 | -41122 | En2 | ENSMUSG00000039095.8 |
| chr5 | 28124878 | 28125152 | 4.03494 | 5.26869 | 1.95101 | Distal Intergenic | 5 | 28165694 | 28172166 | 6473 | 1 | ENSMUST00000036177.8 | -40542 | En2 | ENSMUSG00000039095.8 |
| chr5 | 28127566 | 28127874 | 3.61586 | 4.67095 | 1.54423 | Distal Intergenic | 5 | 28165694 | 28172166 | 6473 | 1 | ENSMUST00000036177.8 | -37820 | En2 | ENSMUSG00000039095.8 |
| chr5 | 28131201 | 28131463 | 3.97854 | 5.07657 | 1.82485 | Distal Intergenic | 5 | 28165694 | 28172166 | 6473 | 1 | ENSMUST00000036177.8 | -34231 | En2 | ENSMUSG00000039095.8 |
| chr5 | 28134565 | 28134926 | 3.36342 | 3.91902 | 1.1624 | Distal Intergenic | 5 | 28165694 | 28172166 | 6473 | 1 | ENSMUST00000036177.8 | -30768 | En2 | ENSMUSG00000039095.8 |
| chr5 | 28152181 | 28152974 | 3.36103 | 3.86907 | 1.01733 | Distal Intergenic | 5 | 28165694 | 28172166 | 6473 | 1 | ENSMUST00000036177.8 | -12720 | En2 | ENSMUSG00000039095.8 |
| chr5 | 28154107 | 28154262 | 3.60006 | 4.30657 | 1.28388 | Distal Intergenic | 5 | 28165694 | 28172166 | 6473 | 1 | ENSMUST00000036177.8 | -11432 | En2 | ENSMUSG00000039095.8 |
| chr5 | 28155682 | 28156238 | 3.44601 | 4.02576 | 1.11744 | Distal Intergenic | 5 | 28165694 | 28172166 | 6473 | 1 | ENSMUST00000036177.8 | -9456 | En2 | ENSMUSG00000039095.8 |
| chr5 | 28156663 | 28157468 | 3.86043 | 4.84475 | 1.66737 | Distal Intergenic | 5 | 28165694 | 28172166 | 6473 | 1 | ENSMUST00000036177.8 | -8226 | En2 | ENSMUSG00000039095.8 |
| chr5 | 28157798 | 28157919 | 3.67863 | 4.63671 | 1.50339 | Distal Intergenic | 5 | 28165694 | 28172166 | 6473 | 1 | ENSMUST00000036177.8 | -7775 | En2 | ENSMUSG00000039095.8 |
| chr5 | 28158472 | 28160117 | 4.06006 | 5.29717 | 1.97861 | Distal Intergenic | 5 | 28165694 | 28172166 | 6473 | 1 | ENSMUST00000036177.8 | -5577 | En2 | ENSMUSG00000039095.8 |
| chr5 | 28160554 | 28160629 | 4.52978 | 6.32615 | 2.69747 | Distal Intergenic | 5 | 28165694 | 28172166 | 6473 | 1 | ENSMUST00000036177.8 | -5065 | En2 | ENSMUSG00000039095.8 |
| chr5 | 28161027 | 28162716 | 3.99359 | 5.12656 | 1.8687 | Distal Intergenic | 5 | 28165694 | 28172166 | 6473 | 1 | ENSMUST00000036177.8 | -2978 | En2 | ENSMUSG00000039095.8 |
| chr5 | 28163172 | 28163353 | 3.3535 | 3.85772 | 1.01733 | Distal Intergenic | 5 | 28165694 | 28172166 | 6473 | 1 | ENSMUST00000036177.8 | -2341 | En2 | ENSMUSG00000039095.8 |
| chr5 | 28163691 | 28164389 | 4.15338 | 5.69526 | 2.2992 | Distal Intergenic | 5 | 28165694 | 28172166 | 6473 | 1 | ENSMUST00000036177.8 | -1305 | En2 | ENSMUSG00000039095.8 |
| chr5 | 28165222 | 28167827 | 3.83932 | 4.95417 | 1.74342 | Promoter | 5 | 28165694 | 28172166 | 6473 | 1 | ENSMUST00000036177.8 | 0 | En2 | ENSMUSG00000039095.8 |
| chr5 | 28168213 | 28169726 | 3.74655 | 4.65308 | 1.54651 | Intron (ENSMUST00000036177.8/ENSMUSG00000039095.8, intron 1 of 1) | 5 | 28165694 | 28172166 | 6473 | 1 | ENSMUST00000036177.8 | 2520 | En2 | ENSMUSG00000039095.8 |
| chr5 | 28170098 | 28170639 | 3.68376 | 4.4947 | 1.43749 | 3' UTR | 5 | 28165694 | 28172166 | 6473 | 1 | ENSMUST00000036177.8 | 4405 | En2 | ENSMUSG00000039095.8 |
| chr5 | 28170982 | 28171494 | 4.04025 | 5.1481 | 1.84304 | 3' UTR | 5 | 28165694 | 28172166 | 6473 | 1 | ENSMUST00000036177.8 | 5289 | En2 | ENSMUSG00000039095.8 |
| chr5 | 28171932 | 28173155 | 3.79737 | 4.88489 | 1.70735 | 3' UTR | 5 | 28165694 | 28172166 | 6473 | 1 | ENSMUST00000036177.8 | 6239 | En2 | ENSMUSG00000039095.8 |
| chr5 | 28173487 | 28175336 | 3.77758 | 4.69415 | 1.54695 | Distal Intergenic | 5 | 28165694 | 28172166 | 6473 | 1 | ENSMUST00000036177.8 | 7794 | En2 | ENSMUSG00000039095.8 |
| chr5 | 28176098 | 28176177 | 3.99113 | 5.05722 | 1.80079 | Distal Intergenic | 5 | 28165694 | 28172166 | 6473 | 1 | ENSMUST00000036177.8 | 10405 | En2 | ENSMUSG00000039095.8 |
| chr5 | 28176606 | 28176951 | 3.46755 | 4.11896 | 1.23867 | Distal Intergenic | 5 | 28165694 | 28172166 | 6473 | 1 | ENSMUST00000036177.8 | 10913 | En2 | ENSMUSG00000039095.8 |
| chr5 | 28177312 | 28177481 | 3.33622 | 3.83187 | 1.01733 | Distal Intergenic | 5 | 28165694 | 28172166 | 6473 | 1 | ENSMUST00000036177.8 | 11619 | En2 | ENSMUSG00000039095.8 |
| chr5 | 28178739 | 28179621 | 3.45112 | 4.04539 | 1.13596 | Distal Intergenic | 5 | 28165694 | 28172166 | 6473 | 1 | ENSMUST00000036177.8 | 13046 | En2 | ENSMUSG00000039095.8 |
| chr5 | 28182940 | 28183114 | 3.36372 | 3.88712 | 1.06587 | Distal Intergenic | 5 | 28165694 | 28172166 | 6473 | 1 | ENSMUST00000036177.8 | 17247 | En2 | ENSMUSG00000039095.8 |
| chr10 | 26078044 | 26078607 | 3.68842 | 4.54422 | 1.50725 | Promoter | 10 | 26078255 | 26079447 | 1193 | 1 | ENSMUST00000041011.3 | 0 | Gm9767 | ENSMUSG00000039098.3 |
| chr5 | 27841988 | 27842330 | 3.33519 | 3.83011 | 1.01733 | Promoter | 5 | 27841947 | 27855086 | 13140 | 1 | ENSMUST00000036227.5 | 42 | Htr5a | ENSMUSG00000039106.5 |
| chr13 | 36722832 | 36723375 | 3.64409 | 4.42571 | 1.43252 | Downstream (2-3kb) | 13 | 36726458 | 36730668 | 4211 | 2 | ENSMUST00000122286.1 | 7293 | Nrn1 | ENSMUSG00000039114.10 |
| chr13 | 36723728 | 36725482 | 3.69734 | 4.7063 | 1.59352 | Downstream (<1kb) | 13 | 36726458 | 36730668 | 4211 | 2 | ENSMUST00000122286.1 | 5186 | Nrn1 | ENSMUSG00000039114.10 |
| chr13 | 36726030 | 36727235 | 4.27034 | 5.95151 | 2.47575 | 3' UTR | 13 | 36726458 | 36730668 | 4211 | 2 | ENSMUST00000122286.1 | 3433 | Nrn1 | ENSMUSG00000039114.10 |
| chr13 | 36727556 | 36729121 | 3.89619 | 4.89725 | 1.71527 | Intron (ENSMUST00000037623.9/ENSMUSG00000039114.10, intron 2 of 2) | 13 | 36726458 | 36730668 | 4211 | 2 | ENSMUST00000122286.1 | 1547 | Nrn1 | ENSMUSG00000039114.10 |
| chr13 | 36729566 | 36729940 | 3.37669 | 3.90276 | 1.05293 | Promoter | 13 | 36726458 | 36730668 | 4211 | 2 | ENSMUST00000122286.1 | 728 | Nrn1 | ENSMUSG00000039114.10 |
| chr13 | 36730563 | 36730724 | 3.65992 | 4.50858 | 1.43594 | Promoter | 13 | 36726458 | 36730668 | 4211 | 2 | ENSMUST00000122286.1 | 0 | Nrn1 | ENSMUSG00000039114.10 |
| chr13 | 36731194 | 36732692 | 4.05858 | 5.33614 | 2.01425 | Promoter | 13 | 36726458 | 36730668 | 4211 | 2 | ENSMUST00000122286.1 | -527 | Nrn1 | ENSMUSG00000039114.10 |
| chr13 | 36733086 | 36734204 | 4.36516 | 5.91868 | 2.43977 | Promoter | 13 | 36725625 | 36734477 | 8853 | 2 | ENSMUST00000037623.9 | 273 | Nrn1 | ENSMUSG00000039114.10 |
| chr13 | 36734624 | 36735019 | 3.80938 | 4.71057 | 1.55988 | Promoter | 13 | 36725625 | 36734477 | 8853 | 2 | ENSMUST00000037623.9 | -148 | Nrn1 | ENSMUSG00000039114.10 |
| chr13 | 36735457 | 36736754 | 4.23485 | 5.62196 | 2.25777 | Promoter | 13 | 36725625 | 36734477 | 8853 | 2 | ENSMUST00000037623.9 | -981 | Nrn1 | ENSMUSG00000039114.10 |
| chr13 | 36737791 | 36738210 | 3.80871 | 4.73319 | 1.62141 | Distal Intergenic | 13 | 36725625 | 36734477 | 8853 | 2 | ENSMUST00000037623.9 | -3315 | Nrn1 | ENSMUSG00000039114.10 |
| chr13 | 36738984 | 36739108 | 4.11463 | 5.44363 | 2.1191 | Distal Intergenic | 13 | 36725625 | 36734477 | 8853 | 2 | ENSMUST00000037623.9 | -4508 | Nrn1 | ENSMUSG00000039114.10 |
| chr9 | 118606309 | 118606413 | 3.34153 | 3.83966 | 1.01733 | Promoter | 9 | 118606690 | 118901003 | 294314 | 1 | ENSMUST00000044165.8 | -277 | Itga9 | ENSMUSG00000039115.8 |
| chr9 | 118606782 | 118607686 | 4.35938 | 5.83887 | 2.37477 | Promoter | 9 | 118606690 | 118901003 | 294314 | 1 | ENSMUST00000044165.8 | 93 | Itga9 | ENSMUSG00000039115.8 |
| chr9 | 118608605 | 118608927 | 3.89345 | 4.89309 | 1.66641 | Intron (ENSMUST00000044165.8/ENSMUSG00000039115.8, intron 1 of 27) | 9 | 118606690 | 118901003 | 294314 | 1 | ENSMUST00000044165.8 | 1916 | Itga9 | ENSMUSG00000039115.8 |
| chr9 | 118611413 | 118611604 | 3.59915 | 4.32279 | 1.34042 | Intron (ENSMUST00000044165.8/ENSMUSG00000039115.8, intron 1 of 27) | 9 | 118606690 | 118901003 | 294314 | 1 | ENSMUST00000044165.8 | 4724 | Itga9 | ENSMUSG00000039115.8 |
| chr9 | 118814251 | 118815050 | 3.38159 | 3.97324 | 1.16219 | Intron (ENSMUST00000044165.8/ENSMUSG00000039115.8, intron 18 of 27) | 9 | 118807284 | 118843848 | 36565 | 1 | ENSMUST00000149150.1 | 6968 | Itga9 | ENSMUSG00000039115.8 |
| chr9 | 118817325 | 118817484 | 3.31092 | 3.79377 | 1.01733 | Intron (ENSMUST00000044165.8/ENSMUSG00000039115.8, intron 18 of 27) | 9 | 118807284 | 118843848 | 36565 | 1 | ENSMUST00000149150.1 | 10042 | Itga9 | ENSMUSG00000039115.8 |
| chr10 | 14545137 | 14545402 | 3.72796 | 4.64269 | 1.5968 | Promoter | 10 | 14402585 | 14545036 | 142452 | 2 | ENSMUST00000041168.4 | -102 | Gpr126 | ENSMUSG00000039116.4 |
| chr19 | 16958892 | 16959181 | 3.51597 | 4.24915 | 1.30196 | Intron (ENSMUST00000087689.4/ENSMUSG00000039126.9, intron 1 of 19) | 19 | 16956118 | 17223932 | 267815 | 1 | ENSMUST00000087689.4 | 2775 | Prune2 | ENSMUSG00000039126.9 |
| chr3 | 152066168 | 152066451 | 3.7678 | 4.6899 | 1.59877 | Distal Intergenic | 3 | 152094019 | 152102854 | 8836 | 2 | ENSMUST00000152680.1 | 36403 | Gipc2 | ENSMUSG00000039131.10 |
| chr3 | 152165707 | 152166043 | 4.09534 | 5.41118 | 2.05584 | Promoter | 3 | 152130434 | 152165750 | 35317 | 2 | ENSMUST00000133102.1 | 0 | Gipc2 | ENSMUSG00000039131.10 |
| chr3 | 152167295 | 152167433 | 3.78844 | 4.76064 | 1.65145 | Distal Intergenic | 3 | 152093839 | 152166230 | 72392 | 2 | ENSMUST00000046614.9 | -1066 | Gipc2 | ENSMUSG00000039131.10 |
| chr7 | 79710142 | 79710447 | 3.35868 | 3.86553 | 1.01733 | Promoter | 7 | 79710283 | 79712357 | 2075 | 1 | ENSMUST00000035939.6 | 0 | 9330171B17Rik | ENSMUSG00000039133.6 |
| chr4 | 63418015 | 63418307 | 3.61834 | 4.43235 | 1.41693 | Exon (ENSMUST00000145133.2/ENSMUSG00000039137.13, exon 5 of 8) | 4 | 63419469 | 63431929 | 12461 | 2 | ENSMUST00000145630.2 | 13622 | Whrn | ENSMUSG00000039137.13 |
| chr4 | 63494628 | 63494982 | 4.01834 | 5.11693 | 1.82582 | Promoter | 4 | 63460163 | 63494759 | 34597 | 2 | ENSMUST00000133425.1 | 0 | Whrn | ENSMUSG00000039137.13 |
| chr4 | 63495435 | 63496013 | 3.5828 | 4.2773 | 1.27127 | Promoter | 4 | 63414910 | 63495951 | 81042 | 2 | ENSMUST00000063650.4 | 0 | Whrn | ENSMUSG00000039137.13 |
| chr4 | 63500732 | 63500937 | 3.49259 | 4.19829 | 1.2706 | Distal Intergenic | 4 | 63431433 | 63495991 | 64559 | 2 | ENSMUST00000063672.4 | -4742 | Whrn | ENSMUSG00000039137.13 |
| chr4 | 63501262 | 63501337 | 3.78928 | 4.64327 | 1.46751 | Distal Intergenic | 4 | 63431433 | 63495991 | 64559 | 2 | ENSMUST00000063672.4 | -5272 | Whrn | ENSMUSG00000039137.13 |
| chr2 | 5712272 | 5712347 | 4.07485 | 5.20337 | 1.85938 | Intron (ENSMUST00000044009.8/ENSMUSG00000039145.11, intron 1 of 10) | 2 | 5565486 | 5714503 | 149018 | 2 | ENSMUST00000155927.1 | 2156 | Camk1d | ENSMUSG00000039145.11 |
| chr2 | 5712795 | 5713012 | 3.41909 | 3.97495 | 1.08177 | Intron (ENSMUST00000044009.8/ENSMUSG00000039145.11, intron 1 of 10) | 2 | 5565486 | 5714503 | 149018 | 2 | ENSMUST00000155927.1 | 1491 | Camk1d | ENSMUSG00000039145.11 |
| chr2 | 5725773 | 5725906 | 3.29903 | 3.77573 | 1.01733 | Distal Intergenic | 2 | 5293457 | 5714515 | 421059 | 2 | ENSMUST00000044009.8 | -11259 | Camk1d | ENSMUSG00000039145.11 |
| chr17 | 44729703 | 44729810 | 3.7292 | 4.54999 | 1.44032 | Intron (ENSMUST00000162816.2/ENSMUSG00000039153.11, intron 1 of 5) | 17 | 44639681 | 44734693 | 95013 | 2 | ENSMUST00000160199.1 | 4883 | Runx2 | ENSMUSG00000039153.11 |
| chr17 | 44730399 | 44731192 | 3.87484 | 4.82501 | 1.61513 | Intron (ENSMUST00000162816.2/ENSMUSG00000039153.11, intron 1 of 5) | 17 | 44639681 | 44734693 | 95013 | 2 | ENSMUST00000160199.1 | 3501 | Runx2 | ENSMUSG00000039153.11 |
| chr17 | 44732011 | 44732203 | 3.5107 | 4.16125 | 1.21542 | Intron (ENSMUST00000162816.2/ENSMUSG00000039153.11, intron 1 of 5) | 17 | 44639681 | 44734693 | 95013 | 2 | ENSMUST00000160199.1 | 2490 | Runx2 | ENSMUSG00000039153.11 |
| chr17 | 44732518 | 44732861 | 3.497 | 4.14732 | 1.26597 | Intron (ENSMUST00000162816.2/ENSMUSG00000039153.11, intron 1 of 5) | 17 | 44639681 | 44734693 | 95013 | 2 | ENSMUST00000160199.1 | 1832 | Runx2 | ENSMUSG00000039153.11 |
| chr17 | 44733210 | 44734606 | 4.38707 | 6.10744 | 2.62367 | Promoter | 17 | 44639681 | 44734693 | 95013 | 2 | ENSMUST00000160199.1 | 87 | Runx2 | ENSMUSG00000039153.11 |
| chr17 | 44735018 | 44736325 | 4.13788 | 5.47697 | 2.13794 | Promoter | 17 | 44495987 | 44736176 | 240190 | 2 | ENSMUST00000162816.2 | 0 | Runx2 | ENSMUSG00000039153.11 |
| chr17 | 44737313 | 44738043 | 3.88991 | 5.05524 | 1.81347 | Promoter | 17 | 44604001 | 44736648 | 132648 | 2 | ENSMUST00000113572.3 | -666 | Runx2 | ENSMUSG00000039153.11 |
| chr17 | 44738673 | 44738835 | 3.89307 | 5.0147 | 1.76428 | Intron (ENSMUST00000113571.4/ENSMUSG00000039153.11, intron 2 of 8) | 17 | 44604001 | 44736648 | 132648 | 2 | ENSMUST00000113572.3 | -2026 | Runx2 | ENSMUSG00000039153.11 |

| | | | | | | | | | | | | | | | |
|---|---|---|---|---|---|---|---|---|---|---|---|---|---|---|---|
| chr17 | 44739191 | 44739734 | 3.35168 | 3.87511 | 1.08816 | Intron (ENSMUST00000113571.4/ENSMUSG00000039153.11, intron 2 of 8) | 17 | 44604001 | 44736648 | 132648 | 2 | ENSMUST00000113572.3 | -2544 | Runx2 | ENSMUSG00000039153.11 |
| chr17 | 44740406 | 44740701 | 3.35134 | 3.85445 | 1.01733 | Intron (ENSMUST00000113571.4/ENSMUSG00000039153.11, intron 2 of 8) | 17 | 44604001 | 44736648 | 132648 | 2 | ENSMUST00000113572.3 | -3759 | Runx2 | ENSMUSG00000039153.11 |
| chr17 | 44741184 | 44741514 | 3.3991 | 3.95368 | 1.11267 | Intron (ENSMUST00000113571.4/ENSMUSG00000039153.11, intron 2 of 8) | 17 | 44604001 | 44736648 | 132648 | 2 | ENSMUST00000113572.3 | -4537 | Runx2 | ENSMUSG00000039153.11 |
| chr17 | 55970648 | 55970737 | 4.2257 | 5.98346 | 2.46172 | Promoter | 17 | 55970467 | 55976617 | 6151 | 1 | ENSMUST00000044216.6 | 182 | Shd | ENSMUSG00000039154.6 |
| chr17 | 55971172 | 55971285 | 4.33514 | 5.90486 | 2.43014 | Promoter | 17 | 55970467 | 55976617 | 6151 | 1 | ENSMUST00000044216.6 | 706 | Shd | ENSMUSG00000039154.6 |
| chr2 | 178610528 | 178610833 | 3.65331 | 4.50753 | 1.50046 | Distal Intergenic | 2 | 178460630 | 178466572 | 5943 | 1 | ENSMUST00000176869.1 | 149899 | Cdh26 | ENSMUSG00000039155.10 |
| chr2 | 178612880 | 178613032 | 3.48257 | 4.22077 | 1.26863 | Distal Intergenic | 2 | 178460630 | 178466572 | 5943 | 1 | ENSMUST00000176869.1 | 152251 | Cdh26 | ENSMUSG00000039155.10 |
| chr2 | 178623038 | 178623255 | 3.28708 | 3.75777 | 1.01733 | Distal Intergenic | 2 | 178460630 | 178466572 | 5943 | 1 | ENSMUST00000176869.1 | 162409 | Cdh26 | ENSMUSG00000039155.10 |
| chr2 | 178626599 | 178626895 | 3.55234 | 4.21489 | 1.21834 | Distal Intergenic | 2 | 178460630 | 178466572 | 5943 | 1 | ENSMUST00000176869.1 | 165970 | Cdh26 | ENSMUSG00000039155.10 |
| chr2 | 178627928 | 178628319 | 3.77018 | 4.6231 | 1.47654 | Distal Intergenic | 2 | 178460630 | 178466572 | 5943 | 1 | ENSMUST00000176869.1 | 167299 | Cdh26 | ENSMUSG00000039155.10 |
| chr2 | 178670984 | 178671095 | 3.44323 | 4.02103 | 1.11621 | Distal Intergenic | 2 | 178460630 | 178466572 | 5943 | 1 | ENSMUST00000176869.1 | 210355 | Cdh26 | ENSMUSG00000039155.10 |
| chr2 | 178674090 | 178674319 | 3.4178 | 3.96792 | 1.06376 | Distal Intergenic | 2 | 178460630 | 178466572 | 5943 | 1 | ENSMUST00000176869.1 | 213461 | Cdh26 | ENSMUSG00000039155.10 |
| chr2 | 178687471 | 178687650 | 3.42436 | 4.05495 | 1.21794 | Distal Intergenic | 2 | 178460630 | 178466572 | 5943 | 1 | ENSMUST00000176869.1 | 226842 | Cdh26 | ENSMUSG00000039155.10 |
| chr2 | 178713929 | 178714296 | 3.55391 | 4.254 | 1.24363 | Distal Intergenic | 2 | 178460630 | 178466572 | 5943 | 1 | ENSMUST00000176869.1 | 253300 | Cdh26 | ENSMUSG00000039155.10 |
| chr16 | 10192993 | 10193107 | 3.48142 | 4.11663 | 1.21846 | Promoter | 16 | 10192947 | 10214231 | 21285 | 1 | ENSMUST00000133674.1 | 47 | Atf7ip2 | ENSMUSG00000039200.10 |
| chr16 | 10193794 | 10194257 | 3.70551 | 4.51822 | 1.45475 | Promoter | 16 | 10192947 | 10214231 | 21285 | 1 | ENSMUST00000133674.1 | 848 | Atf7ip2 | ENSMUSG00000039200.10 |
| chr16 | 10194662 | 10194833 | 3.41364 | 4.02509 | 1.11989 | Intron (ENSMUST00000044005.8/ENSMUSG00000039200.10, intron 1 of 10) | 16 | 10192947 | 10214231 | 21285 | 1 | ENSMUST00000133674.1 | 1716 | Atf7ip2 | ENSMUSG00000039200.10 |
| chr5 | 52534195 | 52534290 | 3.75604 | 4.63652 | 1.55905 | Distal Intergenic | 5 | 52537864 | 52566303 | 28440 | 2 | ENSMUST00000039750.5 | 32013 | Lgi2 | ENSMUSG00000039252.7 |
| chr5 | 52564729 | 52565828 | 3.54935 | 4.26922 | 1.29297 | Promoter | 5 | 52537864 | 52566303 | 28440 | 2 | ENSMUST00000039750.5 | 475 | Lgi2 | ENSMUSG00000039252.7 |
| chr7 | 40899204 | 40901789 | 3.8734 | 4.89831 | 1.70717 | Promoter | 7 | 40899215 | 40930594 | 31380 | 1 | ENSMUST00000044705.10 | 0 | Vstm2b | ENSMUSG00000039257.10 |
| chr17 | 35568970 | 35569277 | 3.42742 | 4.02281 | 1.18854 | Distal Intergenic | 17 | 35567485 | 35568930 | 1446 | 1 | ENSMUST00000174246.1 | 1486 | 2300002M23Rik | ENSMUSG00000039269.5 |
| chr4 | 70378133 | 70378233 | 2.07019 | 12.9751 | 8.51941 | Intron (ENSMUST00000144099.2/ENSMUSG00000039298.11, intron 4 of 36) | 4 | 70398886 | 70403541 | 4656 | 2 | ENSMUST00000126416.1 | 25308 | Cdk5rap2 | ENSMUSG00000039298.11 |
| chr9 | 89619606 | 89620633 | 4.04753 | 5.22308 | 1.92046 | Intron (ENSMUST00000044491.7/ENSMUSG00000039313.9, intron 1 of 3) | 9 | 89587216 | 89623125 | 35910 | 2 | ENSMUST00000044491.7 | 2492 | AF529169 | ENSMUSG00000039313.9 |
| chr9 | 89621113 | 89621908 | 3.86151 | 4.83573 | 1.6459 | Intron (ENSMUST00000044491.7/ENSMUSG00000039313.9, intron 1 of 3) | 9 | 89587216 | 89623125 | 35910 | 2 | ENSMUST00000044491.7 | 1217 | AF529169 | ENSMUSG00000039313.9 |
| chr9 | 89622405 | 89623674 | 3.80641 | 4.72676 | 1.55992 | Promoter | 9 | 89587216 | 89623125 | 35910 | 2 | ENSMUST00000044491.7 | 0 | AF529169 | ENSMUSG00000039313.9 |
| chr5 | 38812432 | 38812621 | 3.44401 | 4.2538 | 1.304 | Intron (ENSMUST00000169819.1/ENSMUSG00000039315.10, intron 2 of 18) | 5 | 38706462 | 38876693 | 170232 | 2 | ENSMUST00000169819.1 | 64072 | Clnk | ENSMUSG00000039315.10 |
| chr17 | 50190087 | 50190896 | 3.88905 | 5.23531 | 1.99354 | Promoter | 17 | 49992257 | 50190674 | 198418 | 2 | ENSMUST00000044503.8 | 0 | Rftn1 | ENSMUSG00000039316.9 |
| chr11 | 121156272 | 121156676 | 3.92123 | 4.95618 | 1.76212 | Distal Intergenic | 11 | 121160271 | 121161973 | 1703 | 1 | ENSMUST00000039044.1 | -3595 | Uts2r | ENSMUSG00000039321.1 |
| chr11 | 121157499 | 121157667 | 3.4806 | 4.11008 | 1.2309 | Distal Intergenic | 11 | 121160271 | 121161973 | 1703 | 1 | ENSMUST00000039044.1 | -2604 | Uts2r | ENSMUSG00000039321.1 |
| chr1 | 72824583 | 72825888 | 3.65693 | 4.46217 | 1.42223 | Promoter | 1 | 72825322 | 72852467 | 27146 | 1 | ENSMUST00000120564.1 | 0 | Igfbp2 | ENSMUSG00000039323.13 |
| chr1 | 72826265 | 72826452 | 3.96852 | 5.004 | 1.74688 | Promoter | 1 | 72825322 | 72852467 | 27146 | 1 | ENSMUST00000120564.1 | 944 | Igfbp2 | ENSMUSG00000039323.13 |
| chr1 | 72827079 | 72827268 | 3.39231 | 3.92471 | 1.04699 | Intron (ENSMUST00000047328.5/ENSMUSG00000039323.13, intron 1 of 3) | 1 | 72825322 | 72852467 | 27146 | 1 | ENSMUST00000120564.1 | 1758 | Igfbp2 | ENSMUSG00000039323.13 |
| chr1 | 72827714 | 72828033 | 3.5564 | 4.22803 | 1.23703 | Intron (ENSMUST00000047328.5/ENSMUSG00000039323.13, intron 1 of 3) | 1 | 72825322 | 72852467 | 27146 | 1 | ENSMUST00000120564.1 | 2393 | Igfbp2 | ENSMUSG00000039323.13 |
| chr10 | 39926710 | 39927072 | 5.24312 | 8.30907 | 4.3642 | Promoter | 10 | 39920382 | 39926923 | 6542 | 2 | ENSMUST00000045526.3 | 0 | 2010001E11Rik | ENSMUSG00000039339.3 |
| chr10 | 39927439 | 39927668 | 3.59107 | 4.36001 | 1.43358 | Promoter | 10 | 39920382 | 39926923 | 6542 | 2 | ENSMUST00000045526.3 | -517 | 2010001E11Rik | ENSMUSG00000039339.3 |
| chr1 | 72576043 | 72576453 | 3.95429 | 5.04848 | 1.82624 | Distal Intergenic | 1 | 72583251 | 72585991 | 2741 | 1 | ENSMUST00000145868.2 | -6798 | Smarcal1 | ENSMUSG00000039354.11 |
| chr5 | 38317922 | 38318409 | 3.43697 | 4.00892 | 1.10632 | Exon (ENSMUST00000181240.1/ENSMUSG00000097029.1, exon 2 of 3) | 5 | 38319509 | 38322310 | 2802 | 1 | ENSMUST00000041646.2 | -1100 | Drd5 | ENSMUSG00000039358.2 |
| chr5 | 38318970 | 38320183 | 4.24518 | 5.63732 | 2.23669 | Promoter | 5 | 38319509 | 38322310 | 2802 | 1 | ENSMUST00000041646.2 | 0 | Drd5 | ENSMUSG00000039358.2 |
| chr11 | 121064091 | 121064273 | 3.41988 | 4.01735 | 1.21052 | Promoter | 11 | 121053458 | 121063569 | 10112 | 2 | ENSMUST00000039309.6 | -523 | Sectm1b | ENSMUSG00000039364.8 |
| chr1 | 72534726 | 72535033 | 3.55887 | 4.28211 | 1.33922 | Exon (ENSMUST00000047786.5/ENSMUSG00000039372.5, exon 1 of 4) | 1 | 72427112 | 72536930 | 109819 | 2 | ENSMUST00000047786.5 | 1897 | Mar-04 | ENSMUSG00000039372.5 |
| chr1 | 72536119 | 72536682 | 3.66493 | 4.64773 | 1.55816 | Promoter | 1 | 72427112 | 72536930 | 109819 | 2 | ENSMUST00000047786.5 | 248 | Mar-04 | ENSMUSG00000039372.5 |
| chr1 | 72536997 | 72537424 | 3.62115 | 4.44217 | 1.3865 | Promoter | 1 | 72427112 | 72536930 | 109819 | 2 | ENSMUST00000047786.5 | -68 | Mar-04 | ENSMUSG00000039372.5 |
| chr1 | 72537972 | 72538165 | 3.67557 | 4.44513 | 1.37583 | Distal Intergenic | 1 | 72427112 | 72536930 | 109819 | 2 | ENSMUST00000047786.5 | -1043 | Mar-04 | ENSMUSG00000039372.5 |
| chr8 | 54779669 | 54779747 | 3.39832 | 3.9571 | 1.13028 | Intron (ENSMUST00000176866.2/ENSMUSG00000039375.11, intron 2 of 4) | 8 | 54629247 | 54724504 | 95258 | 2 | ENSMUST00000175915.2 | -55166 | Wdr17 | ENSMUSG00000039375.11 |
| chr1 | 184723391 | 184723604 | 3.3413 | 4.07234 | 1.19958 | Distal Intergenic | 1 | 184727147 | 184732451 | 5305 | 2 | ENSMUST00000174257.1 | 8847 | Hlx | ENSMUSG00000039377.6 |
| chr1 | 184723915 | 184724404 | 3.91867 | 4.9425 | 1.7506 | Downstream (2-3kb) | 1 | 184727147 | 184732451 | 5305 | 2 | ENSMUST00000174257.1 | 8047 | Hlx | ENSMUSG00000039377.6 |
| chr1 | 184725252 | 184726140 | 3.85365 | 4.87635 | 1.71778 | Downstream (<1kb) | 1 | 184727147 | 184732451 | 5305 | 2 | ENSMUST00000174257.1 | 6311 | Hlx | ENSMUSG00000039377.6 |
| chr1 | 184726599 | 184727096 | 3.4475 | 4.07075 | 1.19967 | Downstream (<1kb) | 1 | 184727147 | 184732451 | 5305 | 2 | ENSMUST00000174257.1 | 5355 | Hlx | ENSMUSG00000039377.6 |
| chr1 | 184727427 | 184728113 | 3.65436 | 4.45875 | 1.42644 | 3' UTR | 1 | 184727147 | 184732451 | 5305 | 2 | ENSMUST00000174257.1 | 4338 | Hlx | ENSMUSG00000039377.6 |
| chr1 | 184728516 | 184729068 | 3.60447 | 4.38709 | 1.33691 | Intron (ENSMUST00000048572.6/ENSMUSG00000039377.6, intron 3 of 3) | 1 | 184727147 | 184732451 | 5305 | 2 | ENSMUST00000174257.1 | 3383 | Hlx | ENSMUSG00000039377.6 |
| chr1 | 184729513 | 184730043 | 3.36627 | 3.89111 | 1.067 | Exon (ENSMUST00000048572.6/ENSMUSG00000039377.6, exon 3 of 4) | 1 | 184727147 | 184732451 | 5305 | 2 | ENSMUST00000174257.1 | 2408 | Hlx | ENSMUSG00000039377.6 |
| chr1 | 184730621 | 184731080 | 3.77787 | 4.63773 | 1.49528 | Exon (ENSMUST00000048572.6/ENSMUSG00000039377.6, exon 2 of 4) | 1 | 184727147 | 184732451 | 5305 | 2 | ENSMUST00000174257.1 | 1371 | Hlx | ENSMUSG00000039377.6 |
| chr1 | 184731682 | 184735586 | 3.84 | 4.92119 | 1.74274 | Promoter | 1 | 184727140 | 184732619 | 5480 | 2 | ENSMUST00000048572.6 | 0 | Hlx | ENSMUSG00000039377.6 |
| chr1 | 184735956 | 184736052 | 3.58748 | 4.40856 | 1.39611 | Distal Intergenic | 1 | 184727140 | 184732619 | 5480 | 2 | ENSMUST00000048572.6 | -3338 | Hlx | ENSMUSG00000039377.6 |
| chr1 | 184737187 | 184737675 | 3.71993 | 4.8372 | 1.67067 | Distal Intergenic | 1 | 184727140 | 184732619 | 5480 | 2 | ENSMUST00000048572.6 | -4569 | Hlx | ENSMUSG00000039377.6 |
| chr1 | 184739311 | 184739743 | 3.46081 | 4.06489 | 1.17444 | Distal Intergenic | 1 | 184727140 | 184732619 | 5480 | 2 | ENSMUST00000048572.6 | -6693 | Hlx | ENSMUSG00000039377.6 |
| chr1 | 184740918 | 184741364 | 3.76075 | 4.62024 | 1.50213 | Distal Intergenic | 1 | 184727140 | 184732619 | 5480 | 2 | ENSMUST00000048572.6 | -8300 | Hlx | ENSMUSG00000039377.6 |
| chr1 | 184742076 | 184742915 | 3.35633 | 3.95085 | 1.11022 | Distal Intergenic | 1 | 184727140 | 184732619 | 5480 | 2 | ENSMUST00000048572.6 | -9458 | Hlx | ENSMUSG00000039377.6 |
| chr1 | 184745550 | 184745890 | 3.36883 | 3.88084 | 1.01733 | Distal Intergenic | 1 | 184727140 | 184732619 | 5480 | 2 | ENSMUST00000048572.6 | -12932 | Hlx | ENSMUSG00000039377.6 |
| chr1 | 184746789 | 184746877 | 4.06344 | 5.32029 | 2.01844 | Distal Intergenic | 1 | 184727140 | 184732619 | 5480 | 2 | ENSMUST00000048572.6 | -14171 | Hlx | ENSMUSG00000039377.6 |
| chr1 | 184748128 | 184748274 | 3.55859 | 4.31512 | 1.36728 | Distal Intergenic | 1 | 184727140 | 184732619 | 5480 | 2 | ENSMUST00000048572.6 | -15510 | Hlx | ENSMUSG00000039377.6 |
| chr1 | 184748926 | 184749516 | 3.68268 | 4.47489 | 1.42389 | Distal Intergenic | 1 | 184727140 | 184732619 | 5480 | 2 | ENSMUST00000048572.6 | -16308 | Hlx | ENSMUSG00000039377.6 |
| chr1 | 184274782 | 184274912 | 3.28577 | 3.7558 | 1.01733 | Distal Intergenic | 1 | 184034381 | 184075636 | 41256 | 1 | ENSMUST00000048655.7 | 240402 | Dusp10 | ENSMUSG00000039384.8 |
| chr15 | 13171649 | 13171790 | 3.37654 | 3.89247 | 1.01733 | Intron (ENSMUST00000036439.4/ENSMUSG00000039385.4, intron 1 of 11) | 15 | 13034200 | 13173639 | 139440 | 2 | ENSMUST00000036439.4 | 1849 | Cdh6 | ENSMUSG00000039385.4 |
| chr15 | 13173226 | 13173623 | 3.59922 | 4.44516 | 1.38685 | Promoter | 15 | 13034200 | 13173639 | 139440 | 2 | ENSMUST00000036439.4 | 16 | Cdh6 | ENSMUSG00000039385.4 |
| chr15 | 13174051 | 13174870 | 3.58403 | 4.36334 | 1.37172 | Promoter | 15 | 13034200 | 13173639 | 139440 | 2 | ENSMUST00000036439.4 | -413 | Cdh6 | ENSMUSG00000039385.4 |
| chr15 | 13175959 | 13176377 | 3.39076 | 3.954 | 1.0865 | Distal Intergenic | 15 | 13034200 | 13173639 | 139440 | 2 | ENSMUST00000036439.4 | -2321 | Cdh6 | ENSMUSG00000039385.4 |
| chr1 | 72188588 | 72188739 | 3.84081 | 4.90524 | 1.71851 | Intron (ENSMUST00000048860.8/ENSMUSG00000039395.8, intron 2 of 4) | 1 | 72159442 | 72212307 | 52866 | 2 | ENSMUST00000048860.8 | 23568 | Mreg | ENSMUSG00000039395.8 |
| chr4 | 154322790 | 154323156 | 3.5303 | 4.3347 | 1.35027 | Intron (ENSMUST00000030902.7/ENSMUSG00000039410.11, intron 15 of 16) | 4 | 154323331 | 154334852 | 11522 | 2 | ENSMUST00000145217.2 | 11696 | Prdm16 | ENSMUSG00000039410.11 |
| chr4 | 154330214 | 154330401 | 3.37163 | 3.88505 | 1.01733 | Intron (ENSMUST00000030902.7/ENSMUSG00000039410.11, intron 12 of 16) | 4 | 154323331 | 154334852 | 11522 | 2 | ENSMUST00000145217.2 | 4451 | Prdm16 | ENSMUSG00000039410.11 |
| chr4 | 154331165 | 154331955 | 3.356 | 3.88469 | 1.09223 | Intron (ENSMUST00000030902.7/ENSMUSG00000039410.11, intron 12 of 16) | 4 | 154323331 | 154334852 | 11522 | 2 | ENSMUST00000145217.2 | 2897 | Prdm16 | ENSMUSG00000039410.11 |
| chr4 | 154332262 | 154332514 | 3.72486 | 4.74076 | 1.62222 | Intron (ENSMUST00000030902.7/ENSMUSG00000039410.11, intron 12 of 16) | 4 | 154323331 | 154334852 | 11522 | 2 | ENSMUST00000145217.2 | 2338 | Prdm16 | ENSMUSG00000039410.11 |
| chr4 | 154332876 | 154333363 | 3.64535 | 4.45463 | 1.46718 | Intron (ENSMUST00000030902.7/ENSMUSG00000039410.11, intron 12 of 16) | 4 | 154323331 | 154334852 | 11522 | 2 | ENSMUST00000145217.2 | 1489 | Prdm16 | ENSMUSG00000039410.11 |
| chr4 | 154334759 | 154335118 | 3.5523 | 4.27204 | 1.36561 | Promoter | 4 | 154323331 | 154334852 | 11522 | 2 | ENSMUST00000145217.2 | 0 | Prdm16 | ENSMUSG00000039410.11 |
| chr4 | 154499902 | 154500349 | 3.36173 | 3.87013 | 1.01733 | Intron (ENSMUST00000030902.7/ENSMUSG00000039410.11, intron 2 of 16) | 4 | 154476469 | 154528765 | 52297 | 2 | ENSMUST00000135064.1 | 28416 | Prdm16 | ENSMUSG00000039410.11 |
| chr4 | 154501461 | 154501846 | 3.39049 | 3.95894 | 1.11981 | Intron (ENSMUST00000030902.7/ENSMUSG00000039410.11, intron 2 of 16) | 4 | 154476469 | 154528765 | 52297 | 2 | ENSMUST00000135064.1 | 26919 | Prdm16 | ENSMUSG00000039410.11 |
| chr4 | 154343735 | 154344179 | 3.51821 | 4.32385 | 1.34821 | Intron (ENSMUST00000181143.1/ENSMUSG00000097871.1, intron 2 of 3) | 4 | 154346139 | 154348457 | 2319 | 2 | ENSMUST00000131762.1 | 4278 | Prdm16 | ENSMUSG00000039410.11 |
| chr4 | 154348108 | 154348307 | 3.47887 | 4.10111 | 1.20306 | Promoter | 4 | 154346139 | 154348457 | 2319 | 2 | ENSMUST00000131762.1 | 150 | Prdm16 | ENSMUSG00000039410.11 |
| chr4 | 154348646 | 154348770 | 3.91395 | 5.254 | 1.94582 | Promoter | 4 | 154346139 | 154348457 | 2319 | 2 | ENSMUST00000131762.1 | -190 | Prdm16 | ENSMUSG00000039410.11 |
| chr4 | 154349408 | 154349501 | 3.53862 | 4.30832 | 1.31499 | Promoter | 4 | 154346139 | 154348457 | 2319 | 2 | ENSMUST00000131762.1 | -952 | Prdm16 | ENSMUSG00000039410.11 |
| chr4 | 154351507 | 154352279 | 3.99115 | 5.14091 | 1.85129 | Intron (ENSMUST00000030902.7/ENSMUSG00000039410.11, intron 5 of 16) | 4 | 154346139 | 154348457 | 2319 | 2 | ENSMUST00000131762.1 | -3051 | Prdm16 | ENSMUSG00000039410.11 |
| chr4 | 154353217 | 154353335 | 3.53095 | 4.1798 | 1.20874 | Intron (ENSMUST00000030902.7/ENSMUSG00000039410.11, intron 5 of 16) | 4 | 154346139 | 154348457 | 2319 | 2 | ENSMUST00000131762.1 | -4761 | Prdm16 | ENSMUSG00000039410.11 |
| chr4 | 154374033 | 154374361 | 3.38151 | 3.93339 | 1.12839 | Intron (ENSMUST00000030902.7/ENSMUSG00000039410.11, intron 3 of 16) | 4 | 154346139 | 154348457 | 2319 | 2 | ENSMUST00000131762.1 | -25577 | Prdm16 | ENSMUSG00000039410.11 |
| chr4 | 154377193 | 154377268 | 3.26154 | 3.71947 | 1.01733 | Intron (ENSMUST00000030902.7/ENSMUSG00000039410.11, intron 3 of 16) | 4 | 154346139 | 154348457 | 2319 | 2 | ENSMUST00000131762.1 | -28737 | Prdm16 | ENSMUSG00000039410.11 |
| chr4 | 154381470 | 154381570 | 3.43294 | 4.00644 | 1.11957 | Intron (ENSMUST00000030902.7/ENSMUSG00000039410.11, intron 3 of 16) | 4 | 154346139 | 154348457 | 2319 | 2 | ENSMUST00000131762.1 | -33014 | Prdm16 | ENSMUSG00000039410.11 |
| chr4 | 154382814 | 154383003 | 3.37688 | 3.89298 | 1.01733 | Intron (ENSMUST00000030902.7/ENSMUSG00000039410.11, intron 3 of 16) | 4 | 154346139 | 154348457 | 2319 | 2 | ENSMUST00000131762.1 | -34358 | Prdm16 | ENSMUSG00000039410.11 |
| chr4 | 154393063 | 154393177 | 3.41525 | 3.96135 | 1.05415 | Intron (ENSMUST00000030902.7/ENSMUSG00000039410.11, intron 3 of 16) | 4 | 154346139 | 154348457 | 2319 | 2 | ENSMUST00000131762.1 | -44607 | Prdm16 | ENSMUSG00000039410.11 |
| chr4 | 154394026 | 154394194 | 3.50773 | 4.13154 | 1.16194 | Intron (ENSMUST00000030902.7/ENSMUSG00000039410.11, intron 3 of 16) | 4 | 154346139 | 154348457 | 2319 | 2 | ENSMUST00000131762.1 | -45570 | Prdm16 | ENSMUSG00000039410.11 |
| chr4 | 154399834 | 154400013 | 3.3384 | 3.83499 | 1.01733 | Intron (ENSMUST00000030902.7/ENSMUSG00000039410.11, intron 3 of 16) | 4 | 154346139 | 154348457 | 2319 | 2 | ENSMUST00000131762.1 | -51378 | Prdm16 | ENSMUSG00000039410.11 |
| chr4 | 154400566 | 154401796 | 3.57625 | 4.2852 | 1.2789 | Intron (ENSMUST00000030902.7/ENSMUSG00000039410.11, intron 3 of 16) | 4 | 154346139 | 154348457 | 2319 | 2 | ENSMUST00000131762.1 | -52110 | Prdm16 | ENSMUSG00000039410.11 |
| chr4 | 154406933 | 154407077 | 3.51944 | 4.17132 | 1.22303 | Intron (ENSMUST00000030902.7/ENSMUSG00000039410.11, intron 3 of 16) | 4 | 154346139 | 154348457 | 2319 | 2 | ENSMUST00000131762.1 | -58477 | Prdm16 | ENSMUSG00000039410.11 |
| chr4 | 154407443 | 154408377 | 3.72772 | 4.55835 | 1.4612 | Intron (ENSMUST00000030902.7/ENSMUSG00000039410.11, intron 3 of 16) | 4 | 154346139 | 154348457 | 2319 | 2 | ENSMUST00000131762.1 | -58987 | Prdm16 | ENSMUSG00000039410.11 |
| chr4 | 154409393 | 154410169 | 3.72945 | 4.54916 | 1.43946 | Intron (ENSMUST00000030902.7/ENSMUSG00000039410.11, intron 3 of 16) | 4 | 154346139 | 154348457 | 2319 | 2 | ENSMUST00000131762.1 | -60937 | Prdm16 | ENSMUSG00000039410.11 |
| chr4 | 154415183 | 154415258 | 3.48508 | 4.10997 | 1.20802 | Intron (ENSMUST00000030902.7/ENSMUSG00000039410.11, intron 3 of 16) | 4 | 154346139 | 154348457 | 2319 | 2 | ENSMUST00000131762.1 | -66727 | Prdm16 | ENSMUSG00000039410.11 |
| chr4 | 154420997 | 154421343 | 3.7419 | 4.58577 | 1.46042 | Intron (ENSMUST00000030902.7/ENSMUSG00000039410.11, intron 3 of 16) | 4 | 154346139 | 154348457 | 2319 | 2 | ENSMUST00000131762.1 | -72541 | Prdm16 | ENSMUSG00000039410.11 |
| chr4 | 154424807 | 154424926 | 3.63709 | 4.41793 | 1.38163 | Intron (ENSMUST00000030902.7/ENSMUSG00000039410.11, intron 3 of 16) | 4 | 154346139 | 154348457 | 2319 | 2 | ENSMUST00000131762.1 | -76351 | Prdm16 | ENSMUSG00000039410.11 |
| chr4 | 154426455 | 154426608 | 3.51787 | 4.23417 | 1.30144 | Intron (ENSMUST00000030902.7/ENSMUSG00000039410.11, intron 3 of 16) | 4 | 154346139 | 154348457 | 2319 | 2 | ENSMUST00000131762.1 | -77999 | Prdm16 | ENSMUSG00000039410.11 |
| chr4 | 154427945 | 154428170 | 3.3349 | 4.19166 | 1.24977 | Intron (ENSMUST00000030902.7/ENSMUSG00000039410.11, intron 3 of 16) | 4 | 154346139 | 154348457 | 2319 | 2 | ENSMUST00000131762.1 | -79489 | Prdm16 | ENSMUSG00000039410.11 |
| chr4 | 154437613 | 154438851 | 3.62874 | 4.38922 | 1.38969 | Intron (ENSMUST00000030902.7/ENSMUSG00000039410.11, intron 3 of 16) | 4 | 154346139 | 154348457 | 2319 | 2 | ENSMUST00000131762.1 | -89157 | Prdm16 | ENSMUSG00000039410.11 |
| chr4 | 154446670 | 154447207 | 4.09969 | 5.30598 | 1.98029 | Intron (ENSMUST00000030902.7/ENSMUSG00000039410.11, intron 3 of 16) | 4 | 154476469 | 154528765 | 52297 | 2 | ENSMUST00000135064.1 | 81558 | Prdm16 | ENSMUSG00000039410.11 |
| chr4 | 154447702 | 154447821 | 3.48127 | 4.21181 | 1.26063 | Intron (ENSMUST00000030902.7/ENSMUSG00000039410.11, intron 3 of 16) | 4 | 154476469 | 154528765 | 52297 | 2 | ENSMUST00000135064.1 | 80944 | Prdm16 | ENSMUSG00000039410.11 |
| chr4 | 154453639 | 154453871 | 3.86372 | 4.90858 | 1.7294 | Intron (ENSMUST00000030902.7/ENSMUSG00000039410.11, intron 3 of 16) | 4 | 154476469 | 154528765 | 52297 | 2 | ENSMUST00000135064.1 | 74894 | Prdm16 | ENSMUSG00000039410.11 |
| chr4 | 154465303 | 154465390 | 4.0579 | 5.20938 | 1.90552 | Intron (ENSMUST00000030902.7/ENSMUSG00000039410.11, intron 3 of 16) | 4 | 154476469 | 154528765 | 52297 | 2 | ENSMUST00000135064.1 | 63375 | Prdm16 | ENSMUSG00000039410.11 |
| chr4 | 154466752 | 154466859 | 3.29718 | 3.77305 | 1.01733 | Intron (ENSMUST00000030902.7/ENSMUSG00000039410.11, intron 3 of 16) | 4 | 154476469 | 154528765 | 52297 | 2 | ENSMUST00000135064.1 | 61906 | Prdm16 | ENSMUSG00000039410.11 |
| chr4 | 154471743 | 154472121 | 3.71149 | 4.78722 | 1.64674 | Intron (ENSMUST00000030902.7/ENSMUSG00000039410.11, intron 3 of 16) | 4 | 154476469 | 154528765 | 52297 | 2 | ENSMUST00000135064.1 | 56644 | Prdm16 | ENSMUSG00000039410.11 |
| chr4 | 154473704 | 154474454 | 3.46665 | 4.07014 | 1.15786 | Intron (ENSMUST00000030902.7/ENSMUSG00000039410.11, intron 3 of 16) | 4 | 154476469 | 154528765 | 52297 | 2 | ENSMUST00000135064.1 | 54311 | Prdm16 | ENSMUSG00000039410.11 |
| chr4 | 154477131 | 154477224 | 4.02239 | 5.10241 | 1.8098 | Intron (ENSMUST00000030902.7/ENSMUSG00000039410.11, intron 2 of 16) | 4 | 154476469 | 154528765 | 52297 | 2 | ENSMUST00000135064.1 | 51541 | Prdm16 | ENSMUSG00000039410.11 |
| chr4 | 154480920 | 154481025 | 3.37134 | 3.8846 | 1.01733 | Intron (ENSMUST00000030902.7/ENSMUSG00000039410.11, intron 2 of 16) | 4 | 154476469 | 154528765 | 52297 | 2 | ENSMUST00000135064.1 | 47740 | Prdm16 | ENSMUSG00000039410.11 |
| chr4 | 154482736 | 154482968 | 3.68431 | 4.72895 | 1.6051 | Intron (ENSMUST00000030902.7/ENSMUSG00000039410.11, intron 2 of 16) | 4 | 154476469 | 154528765 | 52297 | 2 | ENSMUST00000135064.1 | 45797 | Prdm16 | ENSMUSG00000039410.11 |
| chr4 | 154487833 | 154488034 | 3.711 | 4.52372 | 1.43526 | Intron (ENSMUST00000030902.7/ENSMUSG00000039410.11, intron 2 of 16) | 4 | 154476469 | 154528765 | 52297 | 2 | ENSMUST00000135064.1 | 40731 | Prdm16 | ENSMUSG00000039410.11 |
| chr4 | 154489430 | 154489692 | 3.94961 | 4.95348 | 1.68036 | Intron (ENSMUST00000030902.7/ENSMUSG00000039410.11, intron 2 of 16) | 4 | 154476469 | 154528765 | 52297 | 2 | ENSMUST00000135064.1 | 39073 | Prdm16 | ENSMUSG00000039410.11 |
| chr4 | 154490381 | 154490600 | 3.49007 | 4.11837 | 1.18891 | Intron (ENSMUST00000030902.7/ENSMUSG00000039410.11, intron 2 of 16) | 4 | 154476469 | 154528765 | 52297 | 2 | ENSMUST00000135064.1 | 38165 | Prdm16 | ENSMUSG00000039410.11 |
| chr4 | 154497171 | 154497262 | 3.60723 | 4.49243 | 1.46278 | Intron (ENSMUST00000030902.7/ENSMUSG00000039410.11, intron 2 of 16) | 4 | 154476469 | 154528765 | 52297 | 2 | ENSMUST00000135064.1 | 31503 | Prdm16 | ENSMUSG00000039410.11 |
| chr4 | 154499036 | 154499111 | 3.89262 | 4.85747 | 1.64878 | Intron (ENSMUST00000030902.7/ENSMUSG00000039410.11, intron 2 of 16) | 4 | 154476469 | 154528765 | 52297 | 2 | ENSMUST00000135064.1 | 29654 | Prdm16 | ENSMUSG00000039410.11 |
| chr4 | 154631847 | 154632009 | 3.84371 | 4.76238 | 1.5761 | Intron (ENSMUST00000030902.7/ENSMUSG00000039410.11, intron 1 of 16) | 4 | 154476500 | 154636797 | 160298 | 2 | ENSMUST00000149605.1 | 4788 | Prdm16 | ENSMUSG00000039410.11 |
| chr4 | 154632937 | 154634463 | 4.11651 | 5.37272 | 2.07811 | Intron (ENSMUST00000030902.7/ENSMUSG00000039410.11, intron 1 of 16) | 4 | 154476500 | 154636797 | 160298 | 2 | ENSMUST00000149605.1 | 2334 | Prdm16 | ENSMUSG00000039410.11 |
| chr4 | 154506323 | 154506864 | 3.48285 | 4.17255 | 1.29182 | Intron (ENSMUST00000030902.7/ENSMUSG00000039410.11, intron 2 of 16) | 4 | 154476469 | 154528765 | 52297 | 2 | ENSMUST00000135064.1 | 21901 | Prdm16 | ENSMUSG00000039410.11 |
| chr4 | 154512581 | 154512741 | 3.31375 | 3.82686 | 1.10832 | Intron (ENSMUST00000030902.7/ENSMUSG00000039410.11, intron 2 of 16) | 4 | 154476469 | 154528765 | 52297 | 2 | ENSMUST00000135064.1 | 16024 | Prdm16 | ENSMUSG00000039410.11 |
| chr4 | 154514229 | 154514675 | 3.31644 | 3.83022 | 1.09977 | Intron (ENSMUST00000030902.7/ENSMUSG00000039410.11, intron 2 of 16) | 4 | 154476469 | 154528765 | 52297 | 2 | ENSMUST00000135064.1 | 14090 | Prdm16 | ENSMUSG00000039410.11 |
| chr4 | 154517365 | 154517539 | 3.31116 | 3.79395 | 1.01733 | Intron (ENSMUST00000030902.7/ENSMUSG00000039410.11, intron 2 of 16) | 4 | 154476469 | 154528765 | 52297 | 2 | ENSMUST00000135064.1 | 11226 | Prdm16 | ENSMUSG00000039410.11 |
| chr4 | 154523748 | 154524122 | 3.54533 | 4.20483 | 1.22107 | Intron (ENSMUST00000030902.7/ENSMUSG00000039410.11, intron 2 of 16) | 4 | 154476469 | 154528765 | 52297 | 2 | ENSMUST00000135064.1 | 4643 | Prdm16 | ENSMUSG00000039410.11 |
| chr4 | 154528629 | 154528914 | 3.56182 | 4.28701 | 1.37155 | Promoter | 4 | 154476469 | 154528765 | 52297 | 2 | ENSMUST00000135064.1 | 0 | Prdm16 | ENSMUSG00000039410.11 |
| chr4 | 154533974 | 154534657 | 3.49675 | 4.11764 | 1.17234 | Intron (ENSMUST00000030902.7/ENSMUSG00000039410.11, intron 1 of 16) | 4 | 154476469 | 154528765 | 52297 | 2 | ENSMUST00000135064.1 | -5210 | Prdm16 | ENSMUSG00000039410.11 |
| chr4 | 154536493 | 154536889 | 3.38614 | 4.11332 | 1.22249 | Intron (ENSMUST00000030902.7/ENSMUSG00000039410.11, intron 1 of 16) | 4 | 154476469 | 154528765 | 52297 | 2 | ENSMUST00000135064.1 | -7729 | Prdm16 | ENSMUSG00000039410.11 |
| chr4 | 154627699 | 154627890 | 3.61767 | 4.3487 | 1.33106 | Intron (ENSMUST00000030902.7/ENSMUSG00000039410.11, intron 1 of 16) | 4 | 154476500 | 154636797 | 160298 | 2 | ENSMUST00000149605.1 | 8907 | Prdm16 | ENSMUSG00000039410.11 |
| chr4 | 154631036 | 154631264 | 3.57162 | 4.33545 | 1.32869 | Intron (ENSMUST00000030902.7/ENSMUSG00000039410.11, intron 1 of 16) | 4 | 154476500 | 154636797 | 160298 | 2 | ENSMUST00000149605.1 | 5533 | Prdm16 | ENSMUSG00000039410.11 |
| chr4 | 154634847 | 154636504 | 4.06563 | 5.43877 | 2.11794 | Promoter | 4 | 154476500 | 154636797 | 160298 | 2 | ENSMUST00000149605.1 | 293 | Prdm16 | ENSMUSG00000039410.11 |
| chr4 | 154636827 | 154636962 | 3.81343 | 4.71007 | 1.55317 | Promoter | 4 | 154316125 | 154636866 | 320742 | 2 | ENSMUST00000030902.7 | 0 | Prdm16 | ENSMUSG00000039410.11 |
| chr6 | 45060047 | 45060565 | 3.84476 | 4.83083 | 1.68458 | Promoter | 6 | 45060061 | 47301369 | 2241309 | 1 | ENSMUST00000114641.2 | 0 | Cntnap2 | ENSMUSG00000039419.11 |
| chr6 | 47044702 | 47045191 | 3.81828 | 4.75117 | 1.60107 | Promoter | 6 | 47045060 | 47095560 | 50501 | 1 | ENSMUST00000150737.1 | 0 | Cntnap2 | ENSMUSG00000039419.11 |
| chr7 | 89062525 | 89062916 | 3.56909 | 4.30417 | 1.37392 | Distal Intergenic | 7 | 89141856 | 89154300 | 12445 | 2 | ENSMUST00000156153.1 | 91384 | Tmem135 | ENSMUSG00000039428.5 |
| chr8 | 40590603 | 40590680 | 3.60266 | 4.31875 | 1.31756 | Promoter | 8 | 40553858 | 40590514 | 36657 | 2 | ENSMUST00000173487.2 | -90 | Mtmr7 | ENSMUSG00000039431.11 |
| chr16 | 5120333 | 5120429 | 3.64362 | 4.39714 | 1.3589 | Intron (ENSMUST00000035672.3/ENSMUSG00000039457.3, intron 1 of 21) | 16 | 5086291 | 5132481 | 46191 | 2 | ENSMUST00000035672.3 | 12052 | Ppl | ENSMUSG00000039457.3 |
| chr16 | 5130969 | 5131199 | 3.79571 | 4.76042 | 1.66906 | Intron (ENSMUST00000035672.3/ENSMUSG00000039457.3, intron 1 of 21) | 16 | 5086291 | 5132481 | 46191 | 2 | ENSMUST00000035672.3 | 1282 | Ppl | ENSMUSG00000039457.3 |
| chr16 | 5131707 | 5132396 | 3.38352 | 3.95955 | 1.1269 | Promoter | 16 | 5086291 | 5132481 | 46191 | 2 | ENSMUST00000035672.3 | 85 | Ppl | ENSMUSG00000039457.3 |
| chr16 | 5132734 | 5132924 | 3.96616 | 5.25183 | 1.92926 | Promoter | 16 | 5086291 | 5132481 | 46191 | 2 | ENSMUST00000035672.3 | -254 | Ppl | ENSMUSG00000039457.3 |
| chr2 | 30844313 | 30844531 | 3.355 | 3.85997 | 1.01733 | Promoter | 2 | 30845059 | 30881251 | 36193 | 1 | ENSMUST00000041659.6 | -528 | Prrx2 | ENSMUSG00000039476.8 |
| chr2 | 30845819 | 30846334 | 3.99292 | 5.14657 | 1.88454 | Promoter | 2 | 30845444 | 30878657 | 33214 | 1 | ENSMUST00000150299.1 | 376 | Prrx2 | ENSMUSG00000039476.8 |
| chr2 | 30846748 | 30846915 | 3.4201 | 3.99777 | 1.15784 | Intron (ENSMUST00000113592.3/ENSMUSG00000039476.8, intron 1 of 3) | 2 | 30845444 | 30878657 | 33214 | 1 | ENSMUST00000150299.1 | 1305 | Prrx2 | ENSMUSG00000039476.8 |
| chr5 | 142607967 | 142609100 | 4.01653 | 5.11544 | 1.83788 | Promoter | 5 | 142562365 | 142608800 | 46436 | 2 | ENSMUST00000037048.7 | 0 | Mmd2 | ENSMUSG00000039533.7 |
| chr8 | 38658038 | 38658240 | 3.2443 | 3.6936 | 1.01733 | Intron (ENSMUST00000135764.2/ENSMUSG00000039539.8, intron 1 of 9) | 8 | 37522298 | 38661508 | 1139211 | 2 | ENSMUST00000135764.2 | 3268 | Sgcz | ENSMUSG00000039539.8 |
| chr8 | 38660024 | 38660885 | 3.89813 | 5.06657 | 1.89095 | Promoter | 8 | 37522298 | 38661508 | 1139211 | 2 | ENSMUST00000135764.2 | 623 | Sgcz | ENSMUSG00000039539.8 |
| chr8 | 38661457 | 38661698 | 3.35048 | 3.89243 | 1.14057 | Promoter | 8 | 37522298 | 38661508 | 1139211 | 2 | ENSMUST00000135764.2 | 0 | Sgcz | ENSMUSG00000039539.8 |
| chr4 | 153479956 | 153480266 | 3.34337 | 3.84243 | 1.01733 | Intron (ENSMUST00000105646.2/ENSMUSG00000039546.9, intron 1 of 5) | 4 | 153374231 | 153482179 | 107949 | 2 | ENSMUST00000149177.1 | 1913 | Ajap1 | ENSMUSG00000039546.9 |
| chr4 | 153481384 | 153481828 | 3.77721 | 4.65915 | 1.53764 | Promoter | 4 | 153374231 | 153482179 | 107949 | 2 | ENSMUST00000149177.1 | 351 | Ajap1 | ENSMUSG00000039546.9 |
| chr4 | 153482308 | 153482683 | 3.44975 | 4.0539 | 1.14937 | Promoter | 4 | 153373221 | 153482811 | 109591 | 2 | ENSMUST00000105646.2 | 128 | Ajap1 | ENSMUSG00000039546.9 |
| chr4 | 153530462 | 153530769 | 3.4191 | 3.96836 | 1.0584 | Distal Intergenic | 4 | 153373221 | 153482811 | 109591 | 2 | ENSMUST00000105646.2 | -47652 | Ajap1 | ENSMUSG00000039546.9 |
| chr4 | 153543067 | 153543189 | 3.40246 | 3.99565 | 1.20954 | Distal Intergenic | 4 | 153373221 | 153482811 | 109591 | 2 | ENSMUST00000105646.2 | -60257 | Ajap1 | ENSMUSG00000039546.9 |
| chr4 | 153544685 | 153544854 | 3.49943 | 4.23954 | 1.2998 | Distal Intergenic | 4 | 153373221 | 153482811 | 109591 | 2 | ENSMUST00000105646.2 | -61875 | Ajap1 | ENSMUSG00000039546.9 |
| chr4 | 153545348 | 153545986 | 3.39157 | 3.96803 | 1.18418 | Distal Intergenic | 4 | 153373221 | 153482811 | 109591 | 2 | ENSMUST00000105646.2 | -62538 | Ajap1 | ENSMUSG00000039546.9 |
| chr4 | 153557834 | 153558147 | 3.37018 | 3.88287 | 1.01733 | Distal Intergenic | 4 | 153373221 | 153482811 | 109591 | 2 | ENSMUST00000105646.2 | -75024 | Ajap1 | ENSMUSG00000039546.9 |
| chr4 | 153633385 | 153633593 | 3.76477 | 4.67371 | 1.601 | Distal Intergenic | 4 | 153373221 | 153482811 | 109591 | 2 | ENSMUST00000105646.2 | -150575 | Ajap1 | ENSMUSG00000039546.9 |
| chr4 | 153634147 | 153634328 | 3.38058 | 3.99008 | 1.09865 | Distal Intergenic | 4 | 153373221 | 153482811 | 109591 | 2 | ENSMUST00000105646.2 | -151337 | Ajap1 | ENSMUSG00000039546.9 |
| chr4 | 153636982 | 153637256 | 3.34268 | 3.8414 | 1.01733 | Distal Intergenic | 4 | 153373221 | 153482811 | 109591 | 2 | ENSMUST00000105646.2 | -154172 | Ajap1 | ENSMUSG00000039546.9 |
| chr4 | 152496368 | 152496489 | 3.23338 | 3.67724 | 1.01733 | Intron (ENSMUST00000142027.2/ENSMUSG00000039577.12, intron 2 of 28) | 4 | 152502914 | 152518257 | 15344 | 1 | ENSMUST00000123635.1 | -6425 | Nphp4 | ENSMUSG00000039577.12 |
| chr4 | 49843726 | 49843801 | 3.33571 | 3.83652 | 1.0371 | Intron (ENSMUST00000076674.3/ENSMUSG00000039579.10, intron 1 of 8) | 4 | 49661611 | 49845549 | 183939 | 2 | ENSMUST00000076674.3 | 1748 | Grin3a | ENSMUSG00000039579.10 |
| chr4 | 49844393 | 49845614 | 3.86018 | 5.06291 | 1.8194 | Promoter | 4 | 49661611 | 49845549 | 183939 | 2 | ENSMUST00000076674.3 | 0 | Grin3a | ENSMUSG00000039579.10 |
| chr8 | 36459218 | 36459384 | 3.23151 | 3.6801 | 1.03006 | Intron (ENSMUST00000135373.2/ENSMUSG00000039620.11, intron 2 of 6) | 8 | 36458182 | 36511591 | 53410 | 1 | ENSMUST00000152039.3 | 1037 | 6430573F11Rik | ENSMUSG00000039620.11 |
| chr8 | 36522565 | 36522772 | 3.5157 | 4.14542 | 1.16768 | Distal Intergenic | 8 | 36505489 | 36512593 | 7105 | 1 | ENSMUST00000125492.1 | 17077 | 6430573F11Rik | ENSMUSG00000039620.11 |
| chr2 | 166544704 | 166544807 | 3.26993 | 3.74467 | 1.05503 | Distal Intergenic | 2 | 166568317 | 166572977 | 4661 | 2 | ENSMUST00000136974.1 | 28170 | Prex1 | ENSMUSG00000039621.8 |
| chr2 | 166580317 | 166580481 | 3.34153 | 3.83966 | 1.01733 | Exon (ENSMUST00000036719.6/ENSMUSG00000039621.8, exon 28 of 40) | 2 | 166566966 | 166582071 | 15106 | 2 | ENSMUST00000109246.1 | 1590 | Prex1 | ENSMUSG00000039621.8 |
| chr2 | 166605850 | 166606051 | 3.36038 | 3.8681 | 1.01733 | Intron (ENSMUST00000036719.6/ENSMUSG00000039621.8, intron 10 of 39) | 2 | 166599678 | 166603087 | 3410 | 2 | ENSMUST00000136564.1 | -2764 | Prex1 | ENSMUSG00000039621.8 |
| chr6 | 29918437 | 29918683 | 3.54854 | 4.32155 | 1.34411 | Intron (ENSMUST00000137068.2/ENSMUSG00000039629.9, intron 1 of 17) | 6 | 29917208 | 29959675 | 42468 | 1 | ENSMUST00000151738.1 | 1230 | Strip2 | ENSMUSG00000039629.9 |

| | | | | | | | | | | | | | | | |
|---|---|---|---|---|---|---|---|---|---|---|---|---|---|---|---|
| chr6 | 29919141 | 29919638 | 3.65276 | 4.48255 | 1.46401 | Intron (ENSMUST00000137068.2/ENSMUSG00000039629.9, intron 1 of 17) | 6 | 29917208 | 29959675 | 42468 | 1 | ENSMUST00000151738.1 | 1934 | Strip2 | ENSMUSG00000039629.9 |
| chr8 | 36201541 | 36201675 | 3.67329 | 4.48157 | 1.47157 | Distal Intergenic | 8 | 36216064 | 36249516 | 33453 | 2 | ENSMUST00000065297.5 | 47841 | Lonrf1 | ENSMUSG00000039633.6 |
| chr16 | 92358662 | 92358875 | 3.4291 | 4.0272 | 1.20711 | Promoter | 16 | 92348432 | 92358874 | 10443 | 2 | ENSMUST00000166707.1 | 0 | Kcne1 | ENSMUSG00000039639.6 |
| chr8 | 30959457 | 30959663 | 3.3842 | 3.91123 | 1.04276 | Distal Intergenic | 8 | 31089471 | 31097047 | 7577 | 1 | ENSMUST00000036631.8 | -129808 | Dusp26 | ENSMUSG00000039661.9 |
| chr8 | 31088842 | 31088958 | 3.72059 | 4.62341 | 1.51752 | Promoter | 8 | 31089471 | 31097047 | 7577 | 1 | ENSMUST00000036631.8 | -513 | Dusp26 | ENSMUSG00000039661.9 |
| chr8 | 31089795 | 31089963 | 3.58677 | 4.27772 | 1.25574 | Promoter | 8 | 31089804 | 31092232 | 2429 | 1 | ENSMUST00000162551.1 | 0 | Dusp26 | ENSMUSG00000039661.9 |
| chr5 | 141197003 | 141197645 | 3.58546 | 4.27986 | 1.26706 | Distal Intergenic | 5 | 141241490 | 142215586 | 974097 | 1 | ENSMUST00000085774.5 | -43845 | Sdk1 | ENSMUSG00000039683.11 |
| chr5 | 141241094 | 141241233 | 3.58496 | 4.35302 | 1.37921 | Promoter | 5 | 141241490 | 142215586 | 974097 | 1 | ENSMUST00000085774.5 | -257 | Sdk1 | ENSMUSG00000039683.11 |
| chr5 | 141241735 | 141242241 | 3.8916 | 4.96271 | 1.75966 | Promoter | 5 | 141241918 | 141582504 | 340587 | 1 | ENSMUST00000129831.1 | 0 | Sdk1 | ENSMUSG00000039683.11 |
| chr5 | 141242558 | 141243774 | 3.78708 | 4.78812 | 1.6534 | Promoter | 5 | 141241918 | 141582504 | 340587 | 1 | ENSMUST00000129831.1 | 641 | Sdk1 | ENSMUSG00000039683.11 |
| chr5 | 141751680 | 141751919 | 3.28177 | 3.76404 | 1.06204 | Intron (ENSMUST00000085774.5/ENSMUSG00000039683.11, intron 4 of 44) | 5 | 141856787 | 142213194 | 356408 | 1 | ENSMUST00000074546.6 | -104868 | Sdk1 | ENSMUSG00000039683.11 |
| chr5 | 141753450 | 141753633 | 3.42489 | 4.03003 | 1.15029 | Intron (ENSMUST00000085774.5/ENSMUSG00000039683.11, intron 4 of 44) | 5 | 141856787 | 142213194 | 356408 | 1 | ENSMUST00000074546.6 | -103154 | Sdk1 | ENSMUSG00000039683.11 |
| chr5 | 141755150 | 141755251 | 3.6542 | 4.53775 | 1.46937 | Intron (ENSMUST00000085774.5/ENSMUSG00000039683.11, intron 4 of 44) | 5 | 141856787 | 142213194 | 356408 | 1 | ENSMUST00000074546.6 | -101536 | Sdk1 | ENSMUSG00000039683.11 |
| chr5 | 141755859 | 141756133 | 3.4679 | 4.11295 | 1.27489 | Intron (ENSMUST00000085774.5/ENSMUSG00000039683.11, intron 4 of 44) | 5 | 141856787 | 142213194 | 356408 | 1 | ENSMUST00000074546.6 | -100654 | Sdk1 | ENSMUSG00000039683.11 |
| chr5 | 141756779 | 141757061 | 3.81415 | 4.77182 | 1.67766 | Intron (ENSMUST00000085774.5/ENSMUSG00000039683.11, intron 4 of 44) | 5 | 141856787 | 142213194 | 356408 | 1 | ENSMUST00000074546.6 | -99726 | Sdk1 | ENSMUSG00000039683.11 |
| chr5 | 141758346 | 141758514 | 3.24505 | 3.82066 | 1.04287 | Intron (ENSMUST00000085774.5/ENSMUSG00000039683.11, intron 4 of 44) | 5 | 141856787 | 142213194 | 356408 | 1 | ENSMUST00000074546.6 | -98273 | Sdk1 | ENSMUSG00000039683.11 |
| chr5 | 141866133 | 141866444 | 3.33235 | 3.82584 | 1.01733 | Intron (ENSMUST00000085774.5/ENSMUSG00000039683.11, intron 5 of 44) | 5 | 141856787 | 142213194 | 356408 | 1 | ENSMUST00000074546.6 | 9347 | Sdk1 | ENSMUSG00000039683.11 |
| chr9 | 57606747 | 57606892 | 3.60999 | 4.32203 | 1.29023 | Promoter | 9 | 57600019 | 57606281 | 6263 | 2 | ENSMUST00000045068.8 | -467 | Cplx3 | ENSMUSG00000039714.8 |
| chr9 | 107223853 | 107224123 | 3.531 | 4.18437 | 1.22082 | Intron (ENSMUST00000044532.5/ENSMUSG00000039716.7, intron 1 of 52) | 9 | 107108220 | 107231582 | 123363 | 2 | ENSMUST00000165182.1 | 7459 | Dock3 | ENSMUSG00000039716.7 |
| chr9 | 107224507 | 107224795 | 3.31919 | 3.95112 | 1.12907 | Intron (ENSMUST00000044532.5/ENSMUSG00000039716.7, intron 1 of 52) | 9 | 107108220 | 107231582 | 123363 | 2 | ENSMUST00000165182.1 | 6787 | Dock3 | ENSMUSG00000039716.7 |
| chr9 | 107226435 | 107226789 | 3.45466 | 4.09906 | 1.25989 | Intron (ENSMUST00000044532.5/ENSMUSG00000039716.7, intron 1 of 52) | 9 | 107108220 | 107231582 | 123363 | 2 | ENSMUST00000165182.1 | 4793 | Dock3 | ENSMUSG00000039716.7 |
| chr9 | 107227153 | 107227972 | 3.53177 | 4.34669 | 1.34156 | Intron (ENSMUST00000044532.5/ENSMUSG00000039716.7, intron 1 of 52) | 9 | 107108220 | 107231582 | 123363 | 2 | ENSMUST00000165182.1 | 3610 | Dock3 | ENSMUSG00000039716.7 |
| chr9 | 107228334 | 107229586 | 3.99674 | 5.33842 | 2.01708 | Intron (ENSMUST00000044532.5/ENSMUSG00000039716.7, intron 1 of 52) | 9 | 107108220 | 107231582 | 123363 | 2 | ENSMUST00000165182.1 | 1996 | Dock3 | ENSMUSG00000039716.7 |
| chr9 | 107229912 | 107231043 | 3.62167 | 4.533 | 1.48366 | Promoter | 9 | 107108220 | 107231582 | 123363 | 2 | ENSMUST00000165182.1 | 539 | Dock3 | ENSMUSG00000039716.7 |
| chr9 | 107231959 | 107232514 | 3.30323 | 3.82036 | 1.0815 | Promoter | 9 | 106892834 | 107231909 | 339076 | 2 | ENSMUST00000044532.5 | -51 | Dock3 | ENSMUSG00000039716.7 |
| chr9 | 107233266 | 107233686 | 3.63211 | 4.45537 | 1.44146 | Distal Intergenic | 9 | 106892834 | 107231909 | 339076 | 2 | ENSMUST00000044532.5 | -1358 | Dock3 | ENSMUSG00000039716.7 |
| chr9 | 107235047 | 107235209 | 3.42216 | 3.99781 | 1.147 | Distal Intergenic | 9 | 106892834 | 107231909 | 339076 | 2 | ENSMUST00000044532.5 | -3139 | Dock3 | ENSMUSG00000039716.7 |
| chr9 | 107235527 | 107235746 | 3.34251 | 3.84114 | 1.01733 | Distal Intergenic | 9 | 106892834 | 107231909 | 339076 | 2 | ENSMUST00000044532.5 | -3619 | Dock3 | ENSMUSG00000039716.7 |
| chr9 | 107236288 | 107236561 | 3.54985 | 4.29553 | 1.32512 | Distal Intergenic | 9 | 106892834 | 107231909 | 339076 | 2 | ENSMUST00000044532.5 | -4380 | Dock3 | ENSMUSG00000039716.7 |
| chr9 | 107236885 | 107237177 | 3.31985 | 3.80714 | 1.01733 | Distal Intergenic | 9 | 106892834 | 107231909 | 339076 | 2 | ENSMUST00000044532.5 | -4977 | Dock3 | ENSMUSG00000039716.7 |
| chr9 | 107240468 | 107240829 | 3.91651 | 4.92722 | 1.7415 | Distal Intergenic | 9 | 106892834 | 107231909 | 339076 | 2 | ENSMUST00000044532.5 | -8560 | Dock3 | ENSMUSG00000039716.7 |
| chr9 | 107242350 | 107242666 | 3.45051 | 4.07191 | 1.17539 | Distal Intergenic | 9 | 106892834 | 107231909 | 339076 | 2 | ENSMUST00000044532.5 | -10442 | Dock3 | ENSMUSG00000039716.7 |
| chr8 | 27206046 | 27206757 | 3.73594 | 4.65464 | 1.61796 | Distal Intergenic | 8 | 27197460 | 27202547 | 5088 | 2 | ENSMUST00000038174.6 | -3500 | Got1l1 | ENSMUSG00000039720.6 |
| chr7 | 49906404 | 49906529 | 3.46475 | 4.05496 | 1.12067 | Distal Intergenic | 7 | 49910146 | 49963856 | 53711 | 1 | ENSMUST00000056442.6 | -3617 | Slc6a5 | ENSMUSG00000039728.9 |
| chr7 | 49908080 | 49908213 | 4.21921 | 5.48834 | 2.06002 | Distal Intergenic | 7 | 49910146 | 49963856 | 53711 | 1 | ENSMUST00000056442.6 | -1933 | Slc6a5 | ENSMUSG00000039728.9 |
| chr7 | 49908681 | 49909259 | 3.7266 | 4.55392 | 1.45831 | Promoter | 7 | 49910146 | 49963856 | 53711 | 1 | ENSMUST00000056442.6 | -887 | Slc6a5 | ENSMUSG00000039728.9 |
| chr7 | 49909736 | 49910282 | 3.59665 | 4.53058 | 1.45365 | Promoter | 7 | 49910146 | 49963856 | 53711 | 1 | ENSMUST00000056442.6 | 0 | Slc6a5 | ENSMUSG00000039728.9 |
| chr7 | 49911151 | 49912345 | 4.04979 | 5.28714 | 1.99845 | Promoter | 7 | 49911700 | 49959493 | 47794 | 1 | ENSMUST00000107605.2 | 0 | Slc6a5 | ENSMUSG00000039728.9 |
| chr7 | 49912717 | 49912912 | 3.43262 | 4.00095 | 1.10163 | Intron (ENSMUST00000056442.6/ENSMUSG00000039728.9, intron 2 of 15) | 7 | 49911700 | 49959493 | 47794 | 1 | ENSMUST00000107605.2 | 1018 | Slc6a5 | ENSMUSG00000039728.9 |
| chr7 | 49914540 | 49914850 | 3.56326 | 4.3347 | 1.32564 | Exon (ENSMUST00000056442.6/ENSMUSG00000039728.9, exon 3 of 16) | 7 | 49911700 | 49959493 | 47794 | 1 | ENSMUST00000107605.2 | 2841 | Slc6a5 | ENSMUSG00000039728.9 |
| chr7 | 49701290 | 49701614 | 3.64371 | 4.42831 | 1.38125 | Distal Intergenic | 7 | 49759106 | 49773999 | 14894 | 1 | ENSMUST00000085272.5 | -57492 | Htatip2 | ENSMUSG00000039745.7 |
| chr7 | 49751492 | 49751780 | 4.24495 | 5.59593 | 2.20217 | Distal Intergenic | 7 | 49759106 | 49773999 | 14894 | 1 | ENSMUST00000085272.5 | -7326 | Htatip2 | ENSMUSG00000039745.7 |
| chr7 | 49752488 | 49752986 | 3.64251 | 4.42092 | 1.43021 | Distal Intergenic | 7 | 49759106 | 49773999 | 14894 | 1 | ENSMUST00000085272.5 | -6120 | Htatip2 | ENSMUSG00000039745.7 |
| chr4 | 151905777 | 151906172 | 3.80275 | 4.72963 | 1.54901 | Distal Intergenic | 4 | 151933691 | 151982137 | 48447 | 1 | ENSMUST00000062904.5 | -27519 | Dnajc11 | ENSMUSG00000039768.11 |
| chr4 | 47091804 | 47092206 | 3.6776 | 4.55297 | 1.49819 | Promoter | 4 | 47091909 | 47123070 | 31162 | 1 | ENSMUST00000045041.6 | 0 | Galnt12 | ENSMUSG00000039774.7 |
| chr4 | 47092875 | 47093432 | 3.64621 | 4.78663 | 1.62684 | Promoter | 4 | 47091923 | 47123043 | 31121 | 1 | ENSMUST00000107744.1 | 953 | Galnt12 | ENSMUSG00000039774.7 |
| chr4 | 47094335 | 47095273 | 3.48463 | 4.23139 | 1.32076 | Intron (ENSMUST00000045041.6/ENSMUSG00000039774.7, intron 1 of 9) | 4 | 47091923 | 47123043 | 31121 | 1 | ENSMUST00000107744.1 | 2413 | Galnt12 | ENSMUSG00000039774.7 |
| chr4 | 46688977 | 46689081 | 4.02865 | 5.2367 | 1.94543 | Intron (ENSMUST00000107749.2/ENSMUSG00000039809.9, intron 13 of 18) | 4 | 46681131 | 46684494 | 3364 | 2 | ENSMUST00000129328.1 | -4484 | Gabbr2 | ENSMUSG00000039809.9 |
| chr4 | 46719629 | 46719873 | 3.51191 | 4.14471 | 1.18722 | Intron (ENSMUST00000107749.2/ENSMUSG00000039809.9, intron 12 of 18) | 4 | 46681131 | 46684494 | 3364 | 2 | ENSMUST00000129328.1 | -35136 | Gabbr2 | ENSMUSG00000039809.9 |
| chr4 | 46747387 | 46747643 | 3.31156 | 3.79471 | 1.01733 | Intron (ENSMUST00000107749.2/ENSMUSG00000039809.9, intron 8 of 18) | 4 | 46681131 | 46684494 | 3364 | 2 | ENSMUST00000129328.1 | -62894 | Gabbr2 | ENSMUSG00000039809.9 |
| chr4 | 46988162 | 46988396 | 3.34604 | 3.84649 | 1.01733 | Intron (ENSMUST00000107749.2/ENSMUSG00000039809.9, intron 1 of 18) | 4 | 46662318 | 46991873 | 329556 | 2 | ENSMUST00000107749.2 | 3477 | Gabbr2 | ENSMUSG00000039809.9 |
| chr4 | 46989750 | 46990227 | 3.34055 | 3.88577 | 1.08621 | Intron (ENSMUST00000107749.2/ENSMUSG00000039809.9, intron 1 of 18) | 4 | 46662318 | 46991873 | 329556 | 2 | ENSMUST00000107749.2 | 1646 | Gabbr2 | ENSMUSG00000039809.9 |
| chr4 | 46990633 | 46992248 | 4.11824 | 5.43671 | 2.09406 | Promoter | 4 | 46662318 | 46991873 | 329556 | 2 | ENSMUST00000107749.2 | 0 | Gabbr2 | ENSMUSG00000039809.9 |
| chr4 | 46992701 | 46992815 | 3.69939 | 4.61883 | 1.56309 | Promoter | 4 | 46662318 | 46991873 | 329556 | 2 | ENSMUST00000107749.2 | -829 | Gabbr2 | ENSMUSG00000039809.9 |
| chr4 | 46994031 | 46994376 | 3.6148 | 4.49357 | 1.45113 | Distal Intergenic | 4 | 46662318 | 46991873 | 329556 | 2 | ENSMUST00000107749.2 | -2159 | Gabbr2 | ENSMUSG00000039809.9 |
| chr16 | 91205548 | 91205753 | 3.72637 | 4.58574 | 1.5485 | Distal Intergenic | 16 | 91225550 | 91228677 | 3128 | 1 | ENSMUST00000035608.8 | -19797 | Olig2 | ENSMUSG00000039830.8 |
| chr16 | 91206165 | 91206291 | 3.30159 | 3.77956 | 1.01733 | Distal Intergenic | 16 | 91225550 | 91228677 | 3128 | 1 | ENSMUST00000035608.8 | -19259 | Olig2 | ENSMUSG00000039830.8 |
| chr16 | 91211050 | 91211178 | 3.78469 | 4.7878 | 1.66966 | Distal Intergenic | 16 | 91225550 | 91228677 | 3128 | 1 | ENSMUST00000035608.8 | -14372 | Olig2 | ENSMUSG00000039830.8 |
| chr16 | 91212334 | 91212617 | 3.44991 | 4.10936 | 1.25613 | Distal Intergenic | 16 | 91225550 | 91228677 | 3128 | 1 | ENSMUST00000035608.8 | -12933 | Olig2 | ENSMUSG00000039830.8 |
| chr16 | 91213800 | 91214111 | 3.43901 | 4.0167 | 1.12412 | Distal Intergenic | 16 | 91225550 | 91228677 | 3128 | 1 | ENSMUST00000035608.8 | -11439 | Olig2 | ENSMUSG00000039830.8 |
| chr16 | 91216855 | 91218341 | 3.6022 | 4.37137 | 1.33969 | Distal Intergenic | 16 | 91225550 | 91228677 | 3128 | 1 | ENSMUST00000035608.8 | -7209 | Olig2 | ENSMUSG00000039830.8 |
| chr16 | 91218697 | 91220341 | 3.79245 | 4.67691 | 1.53215 | Distal Intergenic | 16 | 91225550 | 91228677 | 3128 | 1 | ENSMUST00000035608.8 | -5209 | Olig2 | ENSMUSG00000039830.8 |
| chr16 | 91220846 | 91221565 | 3.94429 | 4.95606 | 1.7068 | Distal Intergenic | 16 | 91225550 | 91228677 | 3128 | 1 | ENSMUST00000035608.8 | -3985 | Olig2 | ENSMUSG00000039830.8 |
| chr16 | 91222245 | 91223157 | 4.15939 | 5.45615 | 2.09471 | Distal Intergenic | 16 | 91225550 | 91228677 | 3128 | 1 | ENSMUST00000035608.8 | -2393 | Olig2 | ENSMUSG00000039830.8 |
| chr16 | 91223784 | 91223864 | 3.80795 | 4.68773 | 1.51124 | Distal Intergenic | 16 | 91225550 | 91228677 | 3128 | 1 | ENSMUST00000035608.8 | -1686 | Olig2 | ENSMUSG00000039830.8 |
| chr16 | 91224785 | 91225725 | 3.85402 | 4.8164 | 1.63254 | Promoter | 16 | 91225550 | 91228677 | 3128 | 1 | ENSMUST00000035608.8 | 0 | Olig2 | ENSMUSG00000039830.8 |
| chr16 | 91226137 | 91227466 | 3.76092 | 4.65572 | 1.53085 | Promoter | 16 | 91225550 | 91228677 | 3128 | 1 | ENSMUST00000035608.8 | 588 | Olig2 | ENSMUSG00000039830.8 |
| chr16 | 91227788 | 91228048 | 3.51142 | 4.13702 | 1.16164 | 3' UTR | 16 | 91225550 | 91228677 | 3128 | 1 | ENSMUST00000035608.8 | 2239 | Olig2 | ENSMUSG00000039830.8 |
| chr16 | 91228458 | 91229278 | 3.95969 | 5.06865 | 1.83365 | 3' UTR | 16 | 91225550 | 91228677 | 3128 | 1 | ENSMUST00000035608.8 | 2909 | Olig2 | ENSMUSG00000039830.8 |
| chr16 | 91230806 | 91231027 | 3.38193 | 3.9006 | 1.01733 | Distal Intergenic | 16 | 91225550 | 91228677 | 3128 | 1 | ENSMUST00000035608.8 | 5257 | Olig2 | ENSMUSG00000039830.8 |
| chr16 | 91232371 | 91234019 | 3.73695 | 4.62709 | 1.52164 | Distal Intergenic | 16 | 91225550 | 91228677 | 3128 | 1 | ENSMUST00000035608.8 | 6822 | Olig2 | ENSMUSG00000039830.8 |
| chr16 | 91234395 | 91234644 | 3.40143 | 4.07556 | 1.16759 | Distal Intergenic | 16 | 91225550 | 91228677 | 3128 | 1 | ENSMUST00000035608.8 | 8846 | Olig2 | ENSMUSG00000039830.8 |
| chr16 | 91244649 | 91244727 | 3.76585 | 4.75602 | 1.62736 | Distal Intergenic | 16 | 91225550 | 91228677 | 3128 | 1 | ENSMUST00000035608.8 | 19100 | Olig2 | ENSMUSG00000039830.8 |
| chr16 | 91245633 | 91245792 | 3.28625 | 3.75669 | 1.01733 | Distal Intergenic | 16 | 91225550 | 91228677 | 3128 | 1 | ENSMUST00000035608.8 | 20084 | Olig2 | ENSMUSG00000039830.8 |
| chr4 | 150651578 | 150652236 | 3.63541 | 4.48928 | 1.44051 | Promoter | 4 | 150628572 | 150652174 | 23603 | 2 | ENSMUST00000037827.4 | 0 | Slc45a1 | ENSMUSG00000039838.6 |
| chr4 | 150652576 | 150652976 | 3.75257 | 4.7305 | 1.59367 | Promoter | 4 | 150628572 | 150652174 | 23603 | 2 | ENSMUST00000037827.4 | -403 | Slc45a1 | ENSMUSG00000039838.6 |
| chr4 | 150661378 | 150662027 | 3.47896 | 4.09902 | 1.18097 | Distal Intergenic | 4 | 150628572 | 150652174 | 23603 | 2 | ENSMUST00000037827.4 | -9205 | Slc45a1 | ENSMUSG00000039838.6 |
| chr6 | 28395140 | 28395397 | 3.28062 | 3.75247 | 1.03258 | Intron (ENSMUST00000035930.5/ENSMUSG00000039841.9, intron 1 of 6) | 6 | 28240240 | 28398005 | 157766 | 2 | ENSMUST00000035930.5 | 2608 | Zfp800 | ENSMUSG00000039841.9 |
| chr6 | 28395819 | 28395975 | 3.67826 | 4.50931 | 1.52202 | Intron (ENSMUST00000035930.5/ENSMUSG00000039841.9, intron 1 of 6) | 6 | 28240240 | 28398005 | 157766 | 2 | ENSMUST00000035930.5 | 2030 | Zfp800 | ENSMUSG00000039841.9 |
| chr6 | 28396657 | 28397490 | 3.73545 | 4.60843 | 1.57079 | Promoter | 6 | 28240240 | 28398005 | 157766 | 2 | ENSMUST00000035930.5 | 515 | Zfp800 | ENSMUSG00000039841.9 |
| chr6 | 28398324 | 28398500 | 3.70562 | 4.63849 | 1.53749 | Promoter | 6 | 28240240 | 28398005 | 157766 | 2 | ENSMUST00000035930.5 | -320 | Zfp800 | ENSMUSG00000039841.9 |
| chr6 | 28399824 | 28400195 | 3.56065 | 4.32526 | 1.37831 | Distal Intergenic | 6 | 28240240 | 28398005 | 157766 | 2 | ENSMUST00000035930.5 | -1820 | Zfp800 | ENSMUSG00000039841.9 |
| chr8 | 18563801 | 18563878 | 4.1007 | 5.52941 | 2.08959 | Distal Intergenic | 8 | 18595131 | 18803189 | 208059 | 1 | ENSMUST00000039412.9 | -31253 | Mcph1 | ENSMUSG00000039842.10 |
| chr2 | 29588180 | 29588324 | 3.62912 | 4.39715 | 1.39343 | Distal Intergenic | 2 | 29619720 | 29740353 | 120634 | 1 | ENSMUST00000095087.5 | -31396 | Rapgef1 | ENSMUSG00000039844.14 |
| chr2 | 29605725 | 29606056 | 3.60725 | 4.32712 | 1.32419 | Distal Intergenic | 2 | 29619720 | 29740353 | 120634 | 1 | ENSMUST00000095087.5 | -13664 | Rapgef1 | ENSMUSG00000039844.14 |
| chr11 | 119514149 | 119515339 | 3.97378 | 5.14211 | 1.83537 | Distal Intergenic | 11 | 119501978 | 119507389 | 5412 | 1 | ENSMUST00000132015.1 | 12172 | Endov | ENSMUSG00000039850.11 |
| chr16 | 91084176 | 91084509 | 3.53393 | 4.20458 | 1.27625 | Intron (ENSMUST00000146047.1/ENSMUSG00000039851.6, intron 1 of 1) | 16 | 91055838 | 91095122 | 39285 | 2 | ENSMUST00000146047.1 | 10613 | 4932438H23Rik | ENSMUSG00000039851.6 |
| chr16 | 91104216 | 91104356 | 3.77322 | 4.68731 | 1.57134 | Distal Intergenic | 16 | 91055838 | 91095122 | 39285 | 2 | ENSMUST00000146047.1 | -9095 | 4932438H23Rik | ENSMUSG00000039851.6 |
| chr4 | 46534496 | 46534583 | 3.66725 | 4.67989 | 1.56007 | Promoter | 4 | 46523576 | 46534604 | 11029 | 2 | ENSMUST00000142606.1 | 21 | Trim14 | ENSMUSG00000039853.13 |
| chr4 | 46536705 | 46537077 | 3.60247 | 4.41297 | 1.40435 | Promoter | 4 | 46506847 | 46536141 | 29295 | 2 | ENSMUST00000046897.7 | -565 | Trim14 | ENSMUSG00000039853.13 |
| chr4 | 46537729 | 46538042 | 3.27561 | 3.74059 | 1.01733 | 3' UTR | 4 | 46506847 | 46536141 | 29295 | 2 | ENSMUST00000046897.7 | -1589 | Trim14 | ENSMUSG00000039853.13 |
| chr5 | 135850882 | 135851072 | 3.59621 | 4.3072 | 1.3124 | Intron (ENSMUST00000126232.2/ENSMUSG00000039860.14, intron 3 of 3) | 5 | 135854414 | 135857509 | 3096 | 1 | ENSMUST00000135615.1 | -3342 | Srrm3 | ENSMUSG00000039860.14 |
| chr5 | 135852013 | 135852521 | 4.94458 | 7.34684 | 3.68231 | Exon (ENSMUST00000126232.2/ENSMUSG00000039860.14, exon 4 of 4) | 5 | 135854414 | 135857509 | 3096 | 1 | ENSMUST00000135615.1 | -1893 | Srrm3 | ENSMUSG00000039860.14 |
| chr5 | 135852977 | 135853226 | 3.5523 | 4.22015 | 1.22832 | Intron (ENSMUST00000013615.8/ENSMUSG00000039860.14, intron 6 of 15) | 5 | 135854414 | 135857509 | 3096 | 1 | ENSMUST00000135615.1 | -1188 | Srrm3 | ENSMUSG00000039860.14 |
| chr5 | 135855960 | 135857831 | 4.02526 | 5.15528 | 1.87237 | Exon (ENSMUST00000013615.8/ENSMUSG00000039860.14, exon 10 of 16) | 5 | 135854414 | 135857509 | 3096 | 1 | ENSMUST00000135615.1 | 1547 | Srrm3 | ENSMUSG00000039860.14 |
| chr5 | 135866802 | 135866877 | 3.77369 | 4.63657 | 1.51562 | Promoter | 5 | 135866617 | 135868209 | 1593 | 1 | ENSMUST00000150566.1 | 186 | Srrm3 | ENSMUSG00000039860.14 |
| chr5 | 135867683 | 135869540 | 3.82731 | 4.76704 | 1.59713 | 3' UTR | 5 | 135866617 | 135868209 | 1593 | 1 | ENSMUST00000150566.1 | 1067 | Srrm3 | ENSMUSG00000039860.14 |
| chr3 | 121533701 | 121533870 | 3.62738 | 4.34933 | 1.29291 | Exon (ENSMUST00000180804.1/ENSMUSG00000097124.1, exon 2 of 2) | 3 | 121459529 | 121532344 | 72816 | 2 | ENSMUST00000039197.7 | -1358 | Slc44a3 | ENSMUSG00000039865.7 |
| chr16 | 90831156 | 90831340 | 3.8599 | 4.92162 | 1.74224 | Promoter | 16 | 90831113 | 90904695 | 73583 | 1 | ENSMUST00000099543.4 | 44 | Eva1c | ENSMUSG00000039903.11 |
| chr4 | 149588237 | 149588529 | 3.44523 | 4.07828 | 1.23957 | Promoter | 4 | 149587248 | 149627325 | 40078 | 1 | ENSMUST00000137232.1 | 990 | Clstn1 | ENSMUSG00000039953.8 |
| chr8 | 123638881 | 123639168 | 3.33654 | 3.955 | 1.07785 | Intron (ENSMUST00000127664.1/ENSMUSG00000092329.1, intron 1 of 15) | 8 | 123653929 | 123655985 | 2057 | 1 | ENSMUST00000136615.1 | -14761 | Rhou | ENSMUSG00000039960.5 |
| chr8 | 123645246 | 123645422 | 3.63252 | 4.39086 | 1.40112 | Intron (ENSMUST00000127664.1/ENSMUSG00000092329.1, intron 1 of 15) | 8 | 123653929 | 123655985 | 2057 | 1 | ENSMUST00000136615.1 | -8507 | Rhou | ENSMUSG00000039960.5 |
| chr8 | 123649150 | 123649707 | 3.51558 | 4.14845 | 1.18022 | Intron (ENSMUST00000127664.1/ENSMUSG00000092329.1, intron 1 of 15) | 8 | 123653929 | 123655985 | 2057 | 1 | ENSMUST00000136615.1 | -4222 | Rhou | ENSMUSG00000039960.5 |
| chr8 | 123653341 | 123653459 | 4.03606 | 5.13903 | 1.83803 | Promoter | 8 | 123653929 | 123655985 | 2057 | 1 | ENSMUST00000136615.1 | -470 | Rhou | ENSMUSG00000039960.5 |
| chr8 | 123654118 | 123654412 | 4.01165 | 5.12425 | 1.87401 | Promoter | 8 | 123653929 | 123663884 | 9956 | 1 | ENSMUST00000045487.3 | 190 | Rhou | ENSMUSG00000039960.5 |
| chr8 | 123682853 | 123682928 | 3.48393 | 4.10663 | 1.20257 | Intron (ENSMUST00000127664.1/ENSMUSG00000092329.1, intron 1 of 15) | 8 | 123653929 | 123663884 | 9956 | 1 | ENSMUST00000045487.3 | 28925 | Rhou | ENSMUSG00000039960.5 |
| chr8 | 123701186 | 123701330 | 3.34615 | 3.84662 | 1.01733 | Intron (ENSMUST00000127664.1/ENSMUSG00000092329.1, intron 1 of 15) | 8 | 123653929 | 123663884 | 9956 | 1 | ENSMUST00000045487.3 | 47258 | Rhou | ENSMUSG00000039960.5 |
| chr11 | 119251881 | 119252256 | 3.40025 | 3.94751 | 1.08549 | Exon (ENSMUST00000035935.3/ENSMUSG00000039963.13, exon 13 of 19) | 11 | 119253553 | 119264436 | 10884 | 1 | ENSMUST00000133788.2 | -1297 | Ccdc40 | ENSMUSG00000039963.13 |
| chr3 | 157940502 | 157940620 | 4.42195 | 5.97213 | 2.51105 | Distal Intergenic | 3 | 157947414 | 158005910 | 58497 | 1 | ENSMUST00000164582.2 | -6794 | Ankrd13c | ENSMUSG00000039988.9 |
| chr11 | 119087296 | 119088009 | 4.0482 | 5.19859 | 1.88991 | Distal Intergenic | 11 | 119081373 | 119086221 | 4849 | 2 | ENSMUST00000145058.1 | -1076 | Cbx4 | ENSMUSG00000039989.7 |
| chr11 | 119088444 | 119090298 | 4.17799 | 5.57686 | 2.2466 | Distal Intergenic | 11 | 119081373 | 119086221 | 4849 | 2 | ENSMUST00000145058.1 | -2224 | Cbx4 | ENSMUSG00000039989.7 |
| chr2 | 118894004 | 118894421 | 4.0439 | 5.15603 | 1.84695 | Distal Intergenic | 2 | 118900377 | 118916479 | 16103 | 1 | ENSMUST00000151162.1 | -5956 | Bahd1 | ENSMUSG00000040007.8 |
| chr2 | 118896878 | 118897520 | 3.71065 | 4.67245 | 1.5672 | Distal Intergenic | 2 | 118900377 | 118916479 | 16103 | 1 | ENSMUST00000151162.1 | -2857 | Bahd1 | ENSMUSG00000040007.8 |
| chr2 | 118897935 | 118898065 | 3.23128 | 3.93421 | 1.13177 | Distal Intergenic | 2 | 118900377 | 118916479 | 16103 | 1 | ENSMUST00000151162.1 | -2312 | Bahd1 | ENSMUSG00000040007.8 |
| chr2 | 118898845 | 118899062 | 3.62568 | 4.37377 | 1.3831 | Distal Intergenic | 2 | 118900377 | 118916479 | 16103 | 1 | ENSMUST00000151162.1 | -1315 | Bahd1 | ENSMUSG00000040007.8 |
| chr10 | 74990448 | 74990641 | 3.38193 | 3.9006 | 1.01733 | Promoter | 10 | 74990841 | 75016898 | 26058 | 1 | ENSMUST00000159991.1 | -200 | Gnaz | ENSMUSG00000040009.5 |
| chr10 | 74991345 | 74991879 | 3.39201 | 3.92862 | 1.06258 | Promoter | 10 | 74990841 | 75016898 | 26058 | 1 | ENSMUST00000159991.1 | 505 | Gnaz | ENSMUSG00000040009.5 |
| chr10 | 74995558 | 74995704 | 3.54088 | 4.23197 | 1.31794 | Intron (ENSMUST00000037813.4/ENSMUSG00000040009.5, intron 3 of 3) | 10 | 74990841 | 75016898 | 26058 | 1 | ENSMUST00000159991.1 | 4718 | Gnaz | ENSMUSG00000040009.5 |
| chr8 | 121871163 | 121871368 | 3.41081 | 3.99504 | 1.1797 | Intron (ENSMUST00000127664.1/ENSMUSG00000092329.1, intron 1 of 15) | 8 | 121882802 | 121883914 | 1113 | 2 | ENSMUST00000182165.1 | 12546 | Slc7a5 | ENSMUSG00000040010.10 |
| chr8 | 121872694 | 121874139 | 3.59793 | 4.35722 | 1.36293 | Intron (ENSMUST00000127664.1/ENSMUSG00000092329.1, intron 1 of 15) | 8 | 121882802 | 121883914 | 1113 | 2 | ENSMUST00000182165.1 | 9775 | Slc7a5 | ENSMUSG00000040010.10 |
| chr8 | 121878548 | 121878813 | 4.24848 | 6.15166 | 2.65298 | Intron (ENSMUST00000127664.1/ENSMUSG00000092329.1, intron 1 of 15) | 8 | 121882802 | 121883914 | 1113 | 2 | ENSMUST00000182165.1 | 5101 | Slc7a5 | ENSMUSG00000040010.10 |
| chr8 | 121893069 | 121893314 | 3.52029 | 4.15849 | 1.1882 | Intron (ENSMUST00000127664.1/ENSMUSG00000092329.1, intron 1 of 15) | 8 | 121888374 | 121896502 | 8129 | 2 | ENSMUST00000183015.1 | 3188 | Slc7a5 | ENSMUSG00000040010.10 |
| chr8 | 121896108 | 121897071 | 3.67857 | 4.49551 | 1.4176 | Promoter | 8 | 121888374 | 121896502 | 8129 | 2 | ENSMUST00000183015.1 | 0 | Slc7a5 | ENSMUSG00000040010.10 |
| chr8 | 121898341 | 121898635 | 3.88916 | 4.89705 | 1.71559 | Intron (ENSMUST00000127664.1/ENSMUSG00000092329.1, intron 1 of 15) | 8 | 121888374 | 121896502 | 8129 | 2 | ENSMUST00000183015.1 | -1840 | Slc7a5 | ENSMUSG00000040010.10 |
| chr8 | 121906779 | 121907633 | 3.53472 | 4.31423 | 1.35126 | Promoter | 8 | 121881150 | 121907694 | 26545 | 2 | ENSMUST00000045557.9 | 61 | Slc7a5 | ENSMUSG00000040010.10 |
| chr3 | 157566650 | 157568274 | 3.98187 | 5.1091 | 1.84906 | Promoter | 3 | 157566892 | 157644758 | 77867 | 1 | ENSMUST00000041175.7 | 0 | Ptger3 | ENSMUSG00000040016.11 |
| chr2 | 118790732 | 118790973 | 3.5039 | 4.20885 | 1.2925 | Exon (ENSMUST00000037547.3/ENSMUSG00000040035.9, exon 8 of 8) | 2 | 118779759 | 118790048 | 10290 | 1 | ENSMUST00000142072.1 | 10974 | Disp2 | ENSMUSG00000040035.9 |
| chr3 | 156561613 | 156561887 | 3.32212 | 3.81044 | 1.01733 | Promoter | 3 | 156561794 | 157316445 | 754652 | 1 | ENSMUST00000074015.5 | 0 | Negr1 | ENSMUSG00000040037.8 |
| chr3 | 156562241 | 156562367 | 3.46786 | 4.10407 | 1.2509 | Promoter | 3 | 156562150 | 157016244 | 454095 | 1 | ENSMUST00000175773.1 | 92 | Negr1 | ENSMUSG00000040037.8 |
| chr7 | 46672181 | 46672726 | 3.63673 | 4.3744 | 1.32253 | Promoter | 7 | 46647161 | 46672537 | 25377 | 2 | ENSMUST00000049298.8 | 0 | Tph1 | ENSMUSG00000040046.8 |
| chr14 | 57130153 | 57130228 | 3.90137 | 4.89948 | 1.7092 | Intron (ENSMUST00000039380.8/ENSMUSG00000040055.8, intron 2 of 2) | 14 | 57130837 | 57132059 | 1223 | 2 | ENSMUST00000162931.1 | 1831 | Gjb6 | ENSMUSG00000040055.8 |
| chr14 | 57133161 | 57133378 | 3.86314 | 4.85161 | 1.72044 | Promoter | 14 | 57124282 | 57133281 | 9000 | 2 | ENSMUST00000160703.1 | 0 | Gjb6 | ENSMUSG00000040055.8 |
| chr14 | 57133688 | 57134116 | 4.2366 | 5.56876 | 2.1796 | Promoter | 14 | 57123303 | 57133611 | 10309 | 2 | ENSMUST00000039380.8 | -78 | Gjb6 | ENSMUSG00000040055.8 |
| chr2 | 118706986 | 118707287 | 3.55479 | 4.30808 | 1.34206 | Downstream (<1kb) | 2 | 118708699 | 118709741 | 1043 | 2 | ENSMUST00000126907.1 | 2454 | Plcb2 | ENSMUSG00000040061.12 |
| chr2 | 118710239 | 118711957 | 3.68062 | 4.5418 | 1.46069 | Promoter | 2 | 118708180 | 118711370 | 3191 | 2 | ENSMUST00000129153.1 | 0 | Plcb2 | ENSMUSG00000040061.12 |
| chr5 | 15935362 | 15935634 | 3.29908 | 3.92075 | 1.11474 | Promoter | 5 | 15934829 | 16370727 | 435899 | 1 | ENSMUST00000115281.2 | 534 | Cacna2d1 | ENSMUSG00000040118.10 |
| chr7 | 131800745 | 131800936 | 3.32014 | 3.81236 | 1.03402 | Intron (ENSMUST00000124096.2/ENSMUSG00000030849.12, intron 2 of 6) | 7 | 131966460 | 131994405 | 27946 | 1 | ENSMUST00000045840.4 | -165524 | Gpr26 | ENSMUSG00000040125.4 |
| chr7 | 131801945 | 131802204 | 3.71392 | 4.51727 | 1.42709 | Intron (ENSMUST00000124096.2/ENSMUSG00000030849.12, intron 2 of 6) | 7 | 131966460 | 131994405 | 27946 | 1 | ENSMUST00000045840.4 | -164256 | Gpr26 | ENSMUSG00000040125.4 |
| chr7 | 131883225 | 131883324 | 3.9301 | 4.96691 | 1.79218 | Intron (ENSMUST00000124096.2/ENSMUSG00000030849.12, intron 2 of 6) | 7 | 131966460 | 131994405 | 27946 | 1 | ENSMUST00000045840.4 | -83136 | Gpr26 | ENSMUSG00000040125.4 |
| chr7 | 131966798 | 131967243 | 3.41134 | 3.96622 | 1.09253 | Promoter | 7 | 131966460 | 131994405 | 27946 | 1 | ENSMUST00000045840.4 | 339 | Gpr26 | ENSMUSG00000040125.4 |
| chr2 | 118370810 | 118371103 | 3.97614 | 5.11407 | 1.82973 | Intron (ENSMUST00000039160.2/ENSMUSG00000040133.2, intron 1 of 2) | 2 | 118277110 | 118373419 | 96310 | 2 | ENSMUST00000039160.2 | 2316 | Gpr176 | ENSMUSG00000040133.2 |
| chr2 | 118371577 | 118372177 | 3.84411 | 4.79392 | 1.65034 | Intron (ENSMUST00000039160.2/ENSMUSG00000040133.2, intron 1 of 2) | 2 | 118277110 | 118373419 | 96310 | 2 | ENSMUST00000039160.2 | 1242 | Gpr176 | ENSMUSG00000040133.2 |
| chr2 | 118372489 | 118374246 | 4.04213 | 5.28538 | 1.98825 | Promoter | 2 | 118277110 | 118373419 | 96310 | 2 | ENSMUST00000039160.2 | 0 | Gpr176 | ENSMUSG00000040133.2 |
| chr7 | 46179081 | 46179197 | 3.71601 | 4.52889 | 1.44365 | Promoter | 7 | 46104523 | 46180033 | 75511 | 2 | ENSMUST00000033123.6 | 836 | Abcc8 | ENSMUSG00000040136.9 |
| chr7 | 46179524 | 46179988 | 3.45978 | 4.07613 | 1.20608 | Promoter | 7 | 46104523 | 46180033 | 75511 | 2 | ENSMUST00000033123.6 | 45 | Abcc8 | ENSMUSG00000040136.9 |
| chr7 | 46180779 | 46181021 | 3.54517 | 4.20387 | 1.21755 | Promoter | 7 | 46104523 | 46180033 | 75511 | 2 | ENSMUST00000033123.6 | -747 | Abcc8 | ENSMUSG00000040136.9 |
| chr17 | 43629329 | 43630769 | 3.8319 | 4.93811 | 1.73138 | Promoter | 17 | 43615335 | 43630299 | 14965 | 2 | ENSMUST00000168073.2 | 0 | Tdrd6 | ENSMUSG00000040140.9 |
| chr7 | 131535061 | 131535204 | 3.22312 | 3.66189 | 1.01674 | Intron (ENSMUST00000124096.2/ENSMUSG00000030849.12, intron 2 of 6) | 7 | 131542867 | 131544925 | 2059 | 1 | ENSMUST00000046093.5 | -7663 | Hmx3 | ENSMUSG00000040148.5 |
| chr7 | 131538711 | 131539217 | 3.70741 | 4.90254 | 1.71201 | Intron (ENSMUST00000124096.2/ENSMUSG00000030849.12, intron 2 of 6) | 7 | 131542867 | 131544925 | 2059 | 1 | ENSMUST00000046093.5 | -3650 | Hmx3 | ENSMUSG00000040148.5 |
| chr7 | 131539935 | 131540248 | 3.88284 | 4.92171 | 1.79605 | Intron (ENSMUST00000124096.2/ENSMUSG00000030849.12, intron 2 of 6) | 7 | 131542867 | 131544925 | 2059 | 1 | ENSMUST00000046093.5 | -2619 | Hmx3 | ENSMUSG00000040148.5 |
| chr7 | 131541140 | 131542656 | 4.0101 | 5.32288 | 2.04488 | Promoter | 7 | 131542867 | 131544925 | 2059 | 1 | ENSMUST00000046093.5 | -211 | Hmx3 | ENSMUSG00000040148.5 |
| chr7 | 131543105 | 131546229 | 3.86264 | 5.06161 | 1.88012 | Promoter | 7 | 131542867 | 131544925 | 2059 | 1 | ENSMUST00000046093.5 | 239 | Hmx3 | ENSMUSG00000040148.5 |
| chr2 | 164167440 | 164167757 | 3.98352 | 5.14034 | 1.91405 | Intron (ENSMUST00000045196.3/ENSMUSG00000040164.3, intron 3 of 3) | 2 | 164163619 | 164171113 | 7495 | 2 | ENSMUST00000045196.3 | 3356 | Kcns1 | ENSMUSG00000040164.3 |
| chr2 | 164168240 | 164168557 | 3.72006 | 4.52346 | 1.41224 | Exon (ENSMUST00000045196.3/ENSMUSG00000040164.3, exon 3 of 4) | 2 | 164163619 | 164171113 | 7495 | 2 | ENSMUST00000045196.3 | 2556 | Kcns1 | ENSMUSG00000040164.3 |
| chr2 | 164170081 | 164170507 | 3.74563 | 4.59191 | 1.49389 | Promoter | 2 | 164163619 | 164171113 | 7495 | 2 | ENSMUST00000045196.3 | 606 | Kcns1 | ENSMUSG00000040164.3 |
| chr2 | 164170968 | 164172478 | 3.89269 | 4.95769 | 1.77085 | Promoter | 2 | 164163619 | 164171113 | 7495 | 2 | ENSMUST00000045196.3 | 0 | Kcns1 | ENSMUSG00000040164.3 |
| chr2 | 164172945 | 164173277 | 3.72527 | 4.55847 | 1.48348 | Distal Intergenic | 2 | 164163619 | 164171113 | 7495 | 2 | ENSMUST00000045196.3 | -1833 | Kcns1 | ENSMUSG00000040164.3 |
| chr2 | 94017901 | 94018256 | 3.54755 | 4.26184 | 1.30123 | Distal Intergenic | 2 | 93980550 | 94010807 | 30258 | 2 | ENSMUST00000111240.2 | -7095 | Alkbh3 | ENSMUSG00000040174.9 |
| chr4 | 32952880 | 32953119 | 3.61331 | 4.5753 | 1.49012 | Distal Intergenic | 4 | 32804035 | 32950841 | 146807 | 2 | ENSMUST00000084748.3 | -2040 | Ankrd6 | ENSMUSG00000040183.8 |
| chr6 | 146795356 | 146795673 | 3.4217 | 4.00559 | 1.1712 | Promoter | 6 | 146796055 | 146805578 | 9524 | 1 | ENSMUST00000123444.1 | -382 | Arntl2 | ENSMUSG00000040187.10 |
| chr3 | 9629757 | 9630254 | 3.38965 | 3.992 | 1.13563 | Distal Intergenic | 3 | 9427011 | 9610085 | 183075 | 2 | ENSMUST00000041124.7 | -19673 | Zfp704 | ENSMUSG00000040209.7 |
| chr3 | 9631169 | 9631545 | 3.33557 | 3.83073 | 1.01733 | Distal Intergenic | 3 | 9427011 | 9610085 | 183075 | 2 | ENSMUST00000041124.7 | -21085 | Zfp704 | ENSMUSG00000040209.7 |
| chr10 | 127549367 | 127549794 | 3.63132 | 4.42078 | 1.44082 | Promoter | 10 | 127546395 | 127549917 | 3523 | 2 | ENSMUST00000129727.1 | 123 | Lrp1 | ENSMUSG00000040249.10 |

| | | | | | | | | | | | | | | | |
|---|---|---|---|---|---|---|---|---|---|---|---|---|---|---|---|
| chr10 | 127577938 | 127578274 | 3.35594 | 3.90033 | 1.13884 | Exon (ENSMUST00000049149.9/ENSMUSG00000040249.10, exon 22 of 89) | 10 | 127578329 | 127581885 | 3557 | 2 | ENSMUST00000129877.1 | 3611 | Lrp1 | ENSMUSG00000040249.10 |
| chr10 | 127613996 | 127614250 | 3.80822 | 4.73581 | 1.64259 | Intron (ENSMUST00000049149.9/ENSMUSG00000040249.10, intron 1 of 88) | 10 | 127599884 | 127620922 | 21039 | 2 | ENSMUST00000118455.1 | 6672 | Lrp1 | ENSMUSG00000040249.10 |
| chr5 | 12383468 | 12383700 | 3.87022 | 5.02482 | 1.78334 | Promoter | 5 | 12383166 | 12588943 | 205778 | 1 | ENSMUST00000030868.6 | 303 | Sema3d | ENSMUSG00000040254.7 |
| chr10 | 127528417 | 127529200 | 3.93679 | 5.05364 | 1.82936 | Intron (ENSMUST00000095266.2/ENSMUSG00000040258.6, intron 1 of 1) | 10 | 127525473 | 127534559 | 9087 | 2 | ENSMUST00000095266.2 | 5359 | Nxph4 | ENSMUSG00000040258.6 |
| chr10 | 127529533 | 127530180 | 3.43679 | 4.01794 | 1.136 | Intron (ENSMUST00000095266.2/ENSMUSG00000040258.6, intron 1 of 1) | 10 | 127525473 | 127534559 | 9087 | 2 | ENSMUST00000095266.2 | 4379 | Nxph4 | ENSMUSG00000040258.6 |
| chr10 | 127530523 | 127531004 | 3.78539 | 4.81305 | 1.66561 | Intron (ENSMUST00000095266.2/ENSMUSG00000040258.6, intron 1 of 1) | 10 | 127525473 | 127534559 | 9087 | 2 | ENSMUST00000095266.2 | 3555 | Nxph4 | ENSMUSG00000040258.6 |
| chr10 | 127531563 | 127531704 | 3.41963 | 3.96954 | 1.05971 | Intron (ENSMUST00000095266.2/ENSMUSG00000040258.6, intron 1 of 1) | 10 | 127525473 | 127534559 | 9087 | 2 | ENSMUST00000095266.2 | 2855 | Nxph4 | ENSMUSG00000040258.6 |
| chr10 | 127532071 | 127534679 | 4.16427 | 5.5677 | 2.22624 | Promoter | 10 | 127525473 | 127534559 | 9087 | 2 | ENSMUST00000095266.2 | 0 | Nxph4 | ENSMUSG00000040258.6 |
| chr10 | 127535070 | 127535795 | 3.66787 | 4.43985 | 1.37145 | Promoter | 10 | 127525473 | 127534559 | 9087 | 2 | ENSMUST00000095266.2 | -512 | Nxph4 | ENSMUSG00000040258.6 |
| chr10 | 127536149 | 127536455 | 3.36846 | 3.88029 | 1.01733 | Downstream (1-2kb) | 10 | 127525473 | 127534559 | 9087 | 2 | ENSMUST00000095266.2 | -1591 | Nxph4 | ENSMUSG00000040258.6 |
| chr8 | 121784528 | 121784855 | 3.7994 | 4.89032 | 1.73491 | Exon (ENSMUST00000169735.1/ENSMUSG00000025318.8, exon 4 of 5) | 8 | 121796344 | 121797428 | 1085 | 2 | ENSMUST00000174206.1 | 12573 | Klhdc4 | ENSMUSG00000040263.11 |
| chr8 | 121785777 | 121785917 | 3.96621 | 5.14451 | 1.89643 | Intron (ENSMUST00000127664.1/ENSMUSG00000092329.1, intron 1 of 15) | 8 | 121796344 | 121797428 | 1085 | 2 | ENSMUST00000174206.1 | 11511 | Klhdc4 | ENSMUSG00000040263.11 |
| chr7 | 130834524 | 130834606 | 6.23422 | 19.18786 | 14.19755 | Intron (ENSMUST00000124096.2/ENSMUSG00000030849.12, intron 2 of 6) | 7 | 130865756 | 130913312 | 47557 | 1 | ENSMUST00000120441.2 | -31150 | Plekha1 | ENSMUSG00000040268.11 |
| chr5 | 3341296 | 3341593 | 3.55144 | 4.44758 | 1.41294 | Distal Intergenic | 5 | 3343893 | 3523218 | 179326 | 1 | ENSMUST00000165117.2 | -2300 | Cdk6 | ENSMUSG00000040274.6 |
| chr17 | 27655345 | 27655754 | 3.641 | 4.40579 | 1.35275 | Promoter | 17 | 27655509 | 27708659 | 53151 | 1 | ENSMUST00000114873.2 | 0 | Pacsin1 | ENSMUSG00000040276.9 |
| chr17 | 27656118 | 27656235 | 3.53333 | 4.17964 | 1.19362 | Promoter | 17 | 27655643 | 27708677 | 53035 | 1 | ENSMUST00000114872.2 | 476 | Pacsin1 | ENSMUSG00000040276.9 |
| chr17 | 27683369 | 27683670 | 3.31125 | 3.80921 | 1.06502 | Intron (ENSMUST00000114873.2/ENSMUSG00000040276.9, intron 1 of 9) | 17 | 27685197 | 27708659 | 23463 | 1 | ENSMUST00000097360.2 | -1527 | Pacsin1 | ENSMUSG00000040276.9 |
| chr17 | 27684926 | 27686117 | 4.04621 | 5.23919 | 1.98024 | Promoter | 17 | 27685197 | 27708659 | 23463 | 1 | ENSMUST00000097360.2 | 0 | Pacsin1 | ENSMUSG00000040276.9 |
| chr10 | 127511218 | 127512311 | 3.67578 | 4.67674 | 1.57342 | Distal Intergenic | 10 | 127514939 | 127517152 | 2214 | 1 | ENSMUST00000035735.9 | -2628 | Ndufa4l2 | ENSMUSG00000040280.9 |
| chr10 | 127514049 | 127514496 | 4.08612 | 5.35124 | 2.03118 | Promoter | 10 | 127514939 | 127517152 | 2214 | 1 | ENSMUST00000035735.9 | -443 | Ndufa4l2 | ENSMUSG00000040280.9 |
| chr10 | 127514892 | 127514979 | 4.51559 | 6.22735 | 2.66326 | Promoter | 10 | 127514939 | 127517152 | 2214 | 1 | ENSMUST00000035735.9 | 0 | Ndufa4l2 | ENSMUSG00000040280.9 |
| chr10 | 127515606 | 127516049 | 3.65569 | 4.74447 | 1.60625 | Promoter | 10 | 127514939 | 127517152 | 2214 | 1 | ENSMUST00000035735.9 | 668 | Ndufa4l2 | ENSMUSG00000040280.9 |
| chr10 | 127516729 | 127516886 | 3.57587 | 4.32382 | 1.4037 | 3' UTR | 10 | 127514939 | 127517152 | 2214 | 1 | ENSMUST00000035735.9 | 1791 | Ndufa4l2 | ENSMUSG00000040280.9 |
| chr11 | 49169151 | 49169522 | 4.01761 | 5.37825 | 2.00701 | Exon (ENSMUST00000046522.7/ENSMUSG00000040283.9, exon 10 of 10) | 11 | 49169322 | 49170675 | 1354 | 2 | ENSMUST00000131363.1 | 1153 | Btnl9 | ENSMUSG00000040283.9 |
| chr10 | 127509320 | 127509753 | 3.76114 | 4.89035 | 1.70511 | Distal Intergenic | 10 | 127507225 | 127507992 | 768 | 1 | ENSMUST00000162302.1 | 2096 | Stac3 | ENSMUSG00000040287.4 |
| chr3 | 8665230 | 8665799 | 3.8205 | 4.79247 | 1.69461 | Intron (ENSMUST00000042412.3/ENSMUSG00000040289.3, intron 4 of 4) | 3 | 8663362 | 8667039 | 3678 | 2 | ENSMUST00000042412.3 | 1240 | Hey1 | ENSMUSG00000040289.3 |
| chr2 | 93618111 | 93618190 | 3.64772 | 4.64109 | 1.52993 | Distal Intergenic | 2 | 93642384 | 93681339 | 38956 | 1 | ENSMUST00000042078.4 | -24194 | Alx4 | ENSMUSG00000040310.7 |
| chr2 | 93618529 | 93618941 | 3.35605 | 4.15262 | 1.22537 | Distal Intergenic | 2 | 93642384 | 93681339 | 38956 | 1 | ENSMUST00000042078.4 | -23443 | Alx4 | ENSMUSG00000040310.7 |
| chr2 | 93619380 | 93619707 | 4.21539 | 5.60646 | 2.20267 | Distal Intergenic | 2 | 93642384 | 93681339 | 38956 | 1 | ENSMUST00000042078.4 | -22677 | Alx4 | ENSMUSG00000040310.7 |
| chr2 | 93628878 | 93629577 | 3.61291 | 4.36306 | 1.40726 | Distal Intergenic | 2 | 93642384 | 93681339 | 38956 | 1 | ENSMUST00000042078.4 | -12807 | Alx4 | ENSMUSG00000040310.7 |
| chr2 | 93630148 | 93630337 | 3.29673 | 3.77226 | 1.01733 | Distal Intergenic | 2 | 93642384 | 93681339 | 38956 | 1 | ENSMUST00000042078.4 | -12047 | Alx4 | ENSMUSG00000040310.7 |
| chr2 | 93632909 | 93633138 | 3.44061 | 4.0149 | 1.10821 | Distal Intergenic | 2 | 93642384 | 93681339 | 38956 | 1 | ENSMUST00000042078.4 | -9246 | Alx4 | ENSMUSG00000040310.7 |
| chr2 | 93633819 | 93635146 | 3.60887 | 4.35521 | 1.38059 | Distal Intergenic | 2 | 93642384 | 93681339 | 38956 | 1 | ENSMUST00000042078.4 | -7238 | Alx4 | ENSMUSG00000040310.7 |
| chr2 | 93636814 | 93637058 | 3.37736 | 4.16079 | 1.25452 | Distal Intergenic | 2 | 93642384 | 93681339 | 38956 | 1 | ENSMUST00000042078.4 | -5326 | Alx4 | ENSMUSG00000040310.7 |
| chr2 | 93637926 | 93638042 | 3.57885 | 4.27031 | 1.2671 | Distal Intergenic | 2 | 93642384 | 93681339 | 38956 | 1 | ENSMUST00000042078.4 | -4342 | Alx4 | ENSMUSG00000040310.7 |
| chr2 | 93638666 | 93638855 | 3.32656 | 3.81712 | 1.01733 | Distal Intergenic | 2 | 93642384 | 93681339 | 38956 | 1 | ENSMUST00000042078.4 | -3529 | Alx4 | ENSMUSG00000040310.7 |
| chr2 | 93639572 | 93642092 | 3.64807 | 4.47306 | 1.46266 | Promoter | 2 | 93642384 | 93681339 | 38956 | 1 | ENSMUST00000042078.4 | -292 | Alx4 | ENSMUSG00000040310.7 |
| chr2 | 93642844 | 93643592 | 4.05392 | 5.41902 | 2.09692 | Promoter | 2 | 93642388 | 93681339 | 38952 | 1 | ENSMUST00000111254.1 | 457 | Alx4 | ENSMUSG00000040310.7 |
| chr2 | 93644386 | 93648733 | 4.00223 | 5.28794 | 2.03872 | Intron (ENSMUST00000042078.4/ENSMUSG00000040310.7, intron 1 of 3) | 2 | 93642388 | 93681339 | 38952 | 1 | ENSMUST00000111254.1 | 1999 | Alx4 | ENSMUSG00000040310.7 |
| chr2 | 93649491 | 93649841 | 3.45794 | 4.25884 | 1.30685 | Intron (ENSMUST00000042078.4/ENSMUSG00000040310.7, intron 1 of 3) | 2 | 93642388 | 93681339 | 38952 | 1 | ENSMUST00000111254.1 | 7104 | Alx4 | ENSMUSG00000040310.7 |
| chr2 | 93651179 | 93651325 | 3.43356 | 4.22289 | 1.28133 | Intron (ENSMUST00000042078.4/ENSMUSG00000040310.7, intron 1 of 3) | 2 | 93642388 | 93681339 | 38952 | 1 | ENSMUST00000111254.1 | 8792 | Alx4 | ENSMUSG00000040310.7 |
| chr2 | 93657469 | 93657777 | 3.3922 | 3.95956 | 1.16513 | Intron (ENSMUST00000042078.4/ENSMUSG00000040310.7, intron 1 of 3) | 2 | 93642388 | 93681339 | 38952 | 1 | ENSMUST00000111254.1 | 15082 | Alx4 | ENSMUSG00000040310.7 |
| chr1 | 161693173 | 161693321 | 3.72272 | 4.53999 | 1.44582 | Distal Intergenic | 1 | 161726242 | 161727489 | 1248 | 2 | ENSMUST00000151268.1 | 34168 | Gm15429 | ENSMUSG00000040323.13 |
| chr1 | 161734475 | 161735384 | 3.40472 | 3.98408 | 1.1553 | Distal Intergenic | 1 | 161726242 | 161727489 | 1248 | 2 | ENSMUST00000151268.1 | -6987 | Gm15429 | ENSMUSG00000040323.13 |
| chr1 | 161741242 | 161741439 | 3.7731 | 4.85114 | 1.65401 | Distal Intergenic | 1 | 161726242 | 161727489 | 1248 | 2 | ENSMUST00000151268.1 | -13754 | Gm15429 | ENSMUSG00000040323.13 |
| chr1 | 161741818 | 161742450 | 3.34012 | 3.90996 | 1.15337 | Distal Intergenic | 1 | 161726242 | 161727489 | 1248 | 2 | ENSMUST00000151268.1 | -14330 | Gm15429 | ENSMUSG00000040323.13 |
| chr1 | 161745893 | 161746518 | 4.14656 | 5.63115 | 2.22083 | Distal Intergenic | 1 | 161726242 | 161727489 | 1248 | 2 | ENSMUST00000151268.1 | -18405 | Gm15429 | ENSMUSG00000040323.13 |
| chr1 | 161754479 | 161754831 | 3.41862 | 3.97459 | 1.08325 | Distal Intergenic | 1 | 161726242 | 161727489 | 1248 | 2 | ENSMUST00000151268.1 | -26991 | Gm15429 | ENSMUSG00000040323.13 |
| chr10 | 127325077 | 127325304 | 3.45483 | 4.07051 | 1.22843 | 5' UTR | 10 | 127323727 | 127329943 | 6217 | 1 | ENSMUST00000069548.5 | 1351 | Arhgap9 | ENSMUSG00000040345.9 |
| chr10 | 127326264 | 127326832 | 3.55034 | 4.21897 | 1.24436 | Exon (ENSMUST00000069548.5/ENSMUSG00000040345.9, exon 6 of 15) | 10 | 127323727 | 127329943 | 6217 | 1 | ENSMUST00000069548.5 | 2538 | Arhgap9 | ENSMUSG00000040345.9 |
| chr10 | 127332357 | 127332505 | 3.6562 | 4.45387 | 1.46406 | Exon (ENSMUST00000026474.3/ENSMUSG00000025407.6, exon 12 of 13) | 10 | 127323727 | 127329943 | 6217 | 1 | ENSMUST00000069548.5 | 8631 | Arhgap9 | ENSMUSG00000040345.9 |
| chr11 | 107914964 | 107915073 | 3.53747 | 4.19117 | 1.20594 | Promoter | 11 | 107874671 | 107915055 | 40385 | 2 | ENSMUST00000039071.2 | 0 | Cacng5 | ENSMUSG00000040373.7 |
| chr11 | 107915612 | 107915827 | 3.5115 | 4.14295 | 1.18285 | Promoter | 11 | 107874671 | 107915055 | 40385 | 2 | ENSMUST00000039071.2 | -558 | Cacng5 | ENSMUSG00000040373.7 |
| chr11 | 107917146 | 107917294 | 3.54108 | 4.19767 | 1.21842 | Distal Intergenic | 11 | 107874671 | 107915055 | 40385 | 2 | ENSMUST00000039071.2 | -2092 | Cacng5 | ENSMUSG00000040373.7 |
| chr19 | 43653674 | 43654003 | 4.12486 | 5.35545 | 2.0421 | Distal Intergenic | 19 | 43663822 | 43674881 | 11060 | 2 | ENSMUST00000046038.7 | 20878 | Slc25a28 | ENSMUSG00000040414.7 |
| chr15 | 23037097 | 23037283 | 3.72358 | 4.72678 | 1.63273 | Promoter | 15 | 23036463 | 23474418 | 437956 | 1 | ENSMUST00000164787.2 | 635 | Cdh18 | ENSMUSG00000040420.9 |
| chr14 | 55761722 | 55762883 | 4.04686 | 5.37901 | 2.07881 | Promoter | 14 | 55761428 | 55763229 | 1802 | 1 | ENSMUST00000044554.4 | 295 | Ltb4r2 | ENSMUSG00000040432.4 |
| chr14 | 55763370 | 55763998 | 3.50212 | 4.16104 | 1.28273 | Distal Intergenic | 14 | 55761428 | 55763229 | 1802 | 1 | ENSMUST00000044554.4 | 1943 | Ltb4r2 | ENSMUSG00000040432.4 |
| chr10 | 127173724 | 127174032 | 3.49811 | 4.1172 | 1.16295 | Exon (ENSMUST00000095270.2/ENSMUSG00000040441.8, exon 13 of 18) | 10 | 127172426 | 127180645 | 8220 | 2 | ENSMUST00000095270.2 | 6613 | Slc26a10 | ENSMUSG00000040441.8 |
| chr10 | 127174444 | 127175252 | 3.5006 | 4.32612 | 1.36941 | Exon (ENSMUST00000095270.2/ENSMUSG00000040441.8, exon 9 of 18) | 10 | 127172426 | 127180645 | 8220 | 2 | ENSMUST00000095270.2 | 5393 | Slc26a10 | ENSMUSG00000040441.8 |
| chr10 | 127175628 | 127176096 | 3.41847 | 4.09137 | 1.23459 | Intron (ENSMUST00000095270.2/ENSMUSG00000040441.8, intron 8 of 17) | 10 | 127172426 | 127180645 | 8220 | 2 | ENSMUST00000095270.2 | 4549 | Slc26a10 | ENSMUSG00000040441.8 |
| chr10 | 127176435 | 127176943 | 3.79014 | 4.74589 | 1.65517 | Exon (ENSMUST00000095270.2/ENSMUSG00000040441.8, exon 7 of 18) | 10 | 127172426 | 127180645 | 8220 | 2 | ENSMUST00000095270.2 | 3702 | Slc26a10 | ENSMUSG00000040441.8 |
| chr10 | 127178152 | 127178524 | 3.80047 | 4.96603 | 1.80568 | Exon (ENSMUST00000095270.2/ENSMUSG00000040441.8, exon 4 of 18) | 10 | 127172426 | 127180645 | 8220 | 2 | ENSMUST00000095270.2 | 2121 | Slc26a10 | ENSMUSG00000040441.8 |
| chr10 | 127180176 | 127180712 | 3.89199 | 4.97582 | 1.79271 | Promoter | 10 | 127172426 | 127180645 | 8220 | 2 | ENSMUST00000095270.2 | 0 | Slc26a10 | ENSMUSG00000040441.8 |
| chr10 | 127181946 | 127182191 | 3.48765 | 4.15329 | 1.25544 | Downstream (<1kb) | 10 | 127172426 | 127180645 | 8220 | 2 | ENSMUST00000095270.2 | -1302 | Slc26a10 | ENSMUSG00000040441.8 |
| chr10 | 127182745 | 127183320 | 3.65559 | 4.52061 | 1.48186 | 3' UTR | 10 | 127172426 | 127180645 | 8220 | 2 | ENSMUST00000095270.2 | -2101 | Slc26a10 | ENSMUSG00000040441.8 |
| chr10 | 127184404 | 127184479 | 3.82594 | 4.72904 | 1.55656 | Exon (ENSMUST00000167353.1/ENSMUSG00000019467.8, exon 12 of 16) | 10 | 127172426 | 127180645 | 8220 | 2 | ENSMUST00000095270.2 | -3760 | Slc26a10 | ENSMUSG00000040441.8 |
| chr15 | 21110871 | 21111629 | 3.62884 | 4.47707 | 1.45004 | Promoter | 15 | 21111452 | 21589533 | 478082 | 1 | ENSMUST00000075132.6 | 0 | Cdh12 | ENSMUSG00000040452.8 |
| chr5 | 5563524 | 5563838 | 3.24848 | 3.69987 | 1.01733 | Distal Intergenic | 5 | 5537454 | 5559538 | 22085 | 2 | ENSMUST00000115441.3 | -3987 | Gtpbp10 | ENSMUSG00000040464.10 |
| chr4 | 21678127 | 21680148 | 4.49347 | 6.29302 | 2.73894 | 3' UTR | 4 | 21679747 | 21685781 | 6035 | 2 | ENSMUST00000149516.2 | 5633 | Prdm13 | ENSMUSG00000040478.9 |
| chr4 | 21682292 | 21682826 | 3.69624 | 4.48762 | 1.39588 | Exon (ENSMUST00000139568.1/ENSMUSG00000040478.9, exon 3 of 3) | 4 | 21679747 | 21685781 | 6035 | 2 | ENSMUST00000149516.2 | 2955 | Prdm13 | ENSMUSG00000040478.9 |
| chr4 | 21685269 | 21685625 | 4.26101 | 5.73378 | 2.28845 | Promoter | 4 | 21679747 | 21685781 | 6035 | 2 | ENSMUST00000149516.2 | 156 | Prdm13 | ENSMUSG00000040478.9 |
| chr4 | 21686564 | 21688080 | 3.74612 | 4.67989 | 1.54772 | Promoter | 4 | 21677481 | 21685963 | 8483 | 2 | ENSMUST00000095141.3 | -602 | Prdm13 | ENSMUSG00000040478.9 |
| chr4 | 21688471 | 21689122 | 3.92995 | 5.46412 | 2.12301 | Distal Intergenic | 4 | 21677481 | 21685963 | 8483 | 2 | ENSMUST00000095141.3 | -2509 | Prdm13 | ENSMUSG00000040478.9 |
| chr4 | 21689794 | 21690829 | 4.2788 | 5.75252 | 2.34978 | Distal Intergenic | 4 | 21677481 | 21685963 | 8483 | 2 | ENSMUST00000095141.3 | -3832 | Prdm13 | ENSMUSG00000040478.9 |
| chr4 | 21691398 | 21691724 | 3.4068 | 3.99196 | 1.18236 | Distal Intergenic | 4 | 21677481 | 21685963 | 8483 | 2 | ENSMUST00000095141.3 | -5436 | Prdm13 | ENSMUSG00000040478.9 |
| chr4 | 21692072 | 21692551 | 4.00471 | 5.1222 | 1.87538 | Distal Intergenic | 4 | 21677481 | 21685963 | 8483 | 2 | ENSMUST00000095141.3 | -6110 | Prdm13 | ENSMUSG00000040478.9 |
| chr4 | 21693469 | 21694229 | 3.71862 | 4.54597 | 1.47602 | Distal Intergenic | 4 | 21677481 | 21685963 | 8483 | 2 | ENSMUST00000095141.3 | -7507 | Prdm13 | ENSMUSG00000040478.9 |
| chr4 | 21705295 | 21705689 | 3.56569 | 4.2914 | 1.33401 | Distal Intergenic | 4 | 21677481 | 21685963 | 8483 | 2 | ENSMUST00000095141.3 | -19333 | Prdm13 | ENSMUSG00000040478.9 |
| chr2 | 91975565 | 91976388 | 3.52808 | 4.52449 | 1.47907 | Promoter | 2 | 91932824 | 91975864 | 43041 | 2 | ENSMUST00000099709.4 | 0 | Dgkz | ENSMUSG00000040479.6 |
| chr1 | 167440428 | 167440579 | 3.3489 | 3.85077 | 1.01733 | Distal Intergenic | 1 | 167445675 | 167466780 | 21106 | 2 | ENSMUST00000036643.4 | 26201 | Lrrc52 | ENSMUSG00000040485.4 |
| chr7 | 27336473 | 27337588 | 4.00236 | 5.13634 | 1.88186 | Promoter | 7 | 27305136 | 27337692 | 32557 | 2 | ENSMUST00000038618.7 | 104 | Ltbp4 | ENSMUSG00000040488.11 |
| chr17 | 48930714 | 48931277 | 3.76999 | 4.66389 | 1.54253 | Distal Intergenic | 17 | 48932379 | 49097588 | 165210 | 1 | ENSMUST00000046254.2 | -1102 | Lrfn2 | ENSMUSG00000040490.4 |
| chr17 | 48931606 | 48931724 | 3.34952 | 3.8517 | 1.01733 | Promoter | 17 | 48932379 | 49097588 | 165210 | 1 | ENSMUST00000046254.2 | -655 | Lrfn2 | ENSMUSG00000040490.4 |
| chr17 | 48932754 | 48932923 | 3.73117 | 4.68121 | 1.59976 | Promoter | 17 | 48932379 | 49097588 | 165210 | 1 | ENSMUST00000046254.2 | 376 | Lrfn2 | ENSMUSG00000040490.4 |
| chr17 | 48933349 | 48933569 | 3.49338 | 4.10939 | 1.16077 | Promoter | 17 | 48932379 | 49097588 | 165210 | 1 | ENSMUST00000046254.2 | 971 | Lrfn2 | ENSMUSG00000040490.4 |
| chr2 | 91924478 | 91924631 | 3.69101 | 4.46624 | 1.36525 | Distal Intergenic | 2 | 91927249 | 91928688 | 1440 | 1 | ENSMUST00000045537.3 | -2618 | Chrm4 | ENSMUSG00000040495.4 |
| chr17 | 84674559 | 84674842 | 3.46354 | 4.12667 | 1.2554 | Promoter | 17 | 84669917 | 84674370 | 4454 | 2 | ENSMUST00000171544.2 | -190 | Abcg5 | ENSMUSG00000040505.8 |
| chr7 | 19796676 | 19796890 | 3.99477 | 5.09251 | 1.82459 | Promoter | 7 | 19779718 | 19796793 | 17076 | 2 | ENSMUST00000108449.3 | 0 | Cblc | ENSMUSG00000040525.8 |
| chr4 | 15147892 | 15148003 | 3.40888 | 4.06289 | 1.19065 | Promoter | 4 | 15110962 | 15148961 | 38000 | 2 | ENSMUST00000141105.1 | 958 | Necab1 | ENSMUSG00000040536.10 |
| chr5 | 8367173 | 8367354 | 3.80852 | 4.71324 | 1.56136 | Promoter | 5 | 8072352 | 8367982 | 295631 | 2 | ENSMUST00000050166.8 | 628 | Adam22 | ENSMUSG00000040537.12 |
| chr5 | 8367780 | 8368219 | 3.83557 | 4.91406 | 1.68816 | Promoter | 5 | 8072352 | 8367982 | 295631 | 2 | ENSMUST00000050166.8 | 0 | Adam22 | ENSMUSG00000040537.12 |
| chr11 | 106635790 | 106635921 | 3.32436 | 3.8138 | 1.01733 | Distal Intergenic | 11 | 106568098 | 106613423 | 45326 | 2 | ENSMUST00000128933.1 | -22368 | Tex2 | ENSMUSG00000040548.10 |
| chr11 | 106637715 | 106637790 | 3.92722 | 5.02129 | 1.82625 | Distal Intergenic | 11 | 106568098 | 106613423 | 45326 | 2 | ENSMUST00000128933.1 | -24293 | Tex2 | ENSMUSG00000040548.10 |
| chr11 | 72039464 | 72039539 | 3.5755 | 4.27404 | 1.30654 | Distal Intergenic | 11 | 72028717 | 72037509 | 8793 | 2 | ENSMUST00000059082.3 | -1956 | Aipl1 | ENSMUSG00000040554.5 |
| chr9 | 21943334 | 21943543 | 3.26077 | 3.71986 | 1.02234 | Exon (ENSMUST00000046371.7/ENSMUSG00000040563.8, exon 6 of 10) | 9 | 21938453 | 21948017 | 9565 | 1 | ENSMUST00000188468.1 | 4882 | BC018242 | ENSMUSG00000040563.8 |
| chr5 | 8621312 | 8621618 | 3.90176 | 4.97703 | 1.77121 | Intron (ENSMUST00000047485.9/ENSMUSG00000040570.9, intron 1 of 9) | 5 | 8490334 | 8622952 | 132619 | 2 | ENSMUST00000047485.9 | 1334 | Rundc3b | ENSMUSG00000040570.9 |
| chr5 | 8622314 | 8622533 | 3.23366 | 3.68259 | 1.03099 | Promoter | 5 | 8490334 | 8622952 | 132619 | 2 | ENSMUST00000047485.9 | 419 | Rundc3b | ENSMUSG00000040570.9 |
| chr5 | 8660256 | 8660532 | 3.90494 | 4.96642 | 1.77671 | Promoter | 5 | 8660077 | 8748575 | 88499 | 1 | ENSMUST00000047753.4 | 180 | Abcb1a | ENSMUSG00000040584.8 |
| chr3 | 107829640 | 107829870 | 3.39423 | 3.94125 | 1.09547 | Distal Intergenic | 3 | 107877230 | 107892899 | 15670 | 1 | ENSMUST00000037375.8 | -47360 | Eps8l3 | ENSMUSG00000040600.8 |
| chr16 | 97293075 | 97293350 | 3.30312 | 3.78987 | 1.04543 | Distal Intergenic | 16 | 97356728 | 97439012 | 82285 | 1 | ENSMUST00000047275.6 | -63378 | Bace2 | ENSMUSG00000040605.6 |
| chr4 | 142254123 | 142254360 | 3.37006 | 3.88268 | 1.01733 | Distal Intergenic | 4 | 142139240 | 142239401 | 100162 | 2 | ENSMUST00000036476.4 | -14723 | Kazn | ENSMUSG00000040606.8 |
| chr4 | 142255512 | 142255829 | 3.89863 | 4.93149 | 1.74252 | Distal Intergenic | 4 | 142139240 | 142239401 | 100162 | 2 | ENSMUST00000036476.4 | -16112 | Kazn | ENSMUSG00000040606.8 |
| chr4 | 142274335 | 142274469 | 4.11487 | 5.29701 | 1.9405 | Distal Intergenic | 4 | 142139240 | 142239401 | 100162 | 2 | ENSMUST00000036476.4 | -34935 | Kazn | ENSMUSG00000040606.8 |
| chr11 | 33192941 | 33193214 | 3.36529 | 3.88021 | 1.03402 | Distal Intergenic | 11 | 33200752 | 33203589 | 2838 | 2 | ENSMUST00000037746.7 | 10375 | Tlx3 | ENSMUSG00000040610.7 |
| chr11 | 33193670 | 33194112 | 3.66773 | 4.43537 | 1.36913 | Distal Intergenic | 11 | 33200752 | 33203589 | 2838 | 2 | ENSMUST00000037746.7 | 9477 | Tlx3 | ENSMUSG00000040610.7 |
| chr11 | 33195365 | 33195497 | 3.43692 | 4.02024 | 1.14623 | Distal Intergenic | 11 | 33200752 | 33203589 | 2838 | 2 | ENSMUST00000037746.7 | 8092 | Tlx3 | ENSMUSG00000040610.7 |
| chr11 | 33195858 | 33196956 | 3.65121 | 4.48555 | 1.45698 | Distal Intergenic | 11 | 33200752 | 33203589 | 2838 | 2 | ENSMUST00000037746.7 | 6633 | Tlx3 | ENSMUSG00000040610.7 |
| chr11 | 33197269 | 33198961 | 4.00819 | 5.32482 | 2.03031 | Downstream (1-2kb) | 11 | 33200752 | 33203589 | 2838 | 2 | ENSMUST00000037746.7 | 4628 | Tlx3 | ENSMUSG00000040610.7 |
| chr11 | 33199414 | 33201735 | 4.12721 | 5.55672 | 2.19661 | 3' UTR | 11 | 33200752 | 33203589 | 2838 | 2 | ENSMUST00000037746.7 | 1854 | Tlx3 | ENSMUSG00000040610.7 |
| chr11 | 33202052 | 33204619 | 3.96199 | 5.42101 | 2.08919 | Promoter | 11 | 33200752 | 33203589 | 2838 | 2 | ENSMUST00000037746.7 | 0 | Tlx3 | ENSMUSG00000040610.7 |
| chr4 | 142084040 | 142084398 | 3.45986 | 4.08859 | 1.24791 | Promoter | 4 | 142030992 | 142084304 | 53313 | 2 | ENSMUST00000036572.3 | 0 | Tmem51 | ENSMUSG00000040616.3 |
| chr6 | 122555361 | 122555445 | 3.52021 | 4.16139 | 1.20087 | Intron (ENSMUST00000043301.8/ENSMUSG00000040627.9, intron 1 of 4) | 6 | 122553801 | 122564180 | 10380 | 1 | ENSMUST00000043301.8 | 1561 | Aicda | ENSMUSG00000040627.9 |
| chr14 | 55519178 | 55519296 | 3.56976 | 4.25736 | 1.27054 | 3' UTR | 14 | 55518978 | 55524191 | 5214 | 2 | ENSMUST00000111404.2 | 4895 | Nrl | ENSMUSG00000040632.10 |
| chr14 | 55520695 | 55520926 | 3.92005 | 5.26416 | 1.94907 | Exon (ENSMUST00000111404.2/ENSMUSG00000040632.10, exon 3 of 3) | 14 | 55518978 | 55524191 | 5214 | 2 | ENSMUST00000111404.2 | 3265 | Nrl | ENSMUSG00000040632.10 |
| chr14 | 55524121 | 55524334 | 3.34115 | 3.86236 | 1.09808 | Promoter | 14 | 55518978 | 55524191 | 5214 | 2 | ENSMUST00000111404.2 | 0 | Nrl | ENSMUSG00000040632.10 |
| chr14 | 27622331 | 27622579 | 3.26822 | 3.94315 | 1.15141 | Promoter | 14 | 27622442 | 28478537 | 856096 | 1 | ENSMUST00000090302.5 | 0 | Erc2 | ENSMUSG00000040640.9 |
| chr14 | 27623288 | 27623578 | 3.99224 | 5.17323 | 1.93895 | Promoter | 14 | 27622442 | 28478537 | 856096 | 1 | ENSMUST00000090302.5 | 847 | Erc2 | ENSMUSG00000040640.9 |
| chr6 | 122485450 | 122486258 | 3.98825 | 5.16254 | 1.92464 | Promoter | 6 | 122453101 | 122486313 | 33213 | 2 | ENSMUST00000068242.6 | 55 | Rimklb | ENSMUSG00000040649.10 |
| chr6 | 122486715 | 122487455 | 4.12053 | 5.47631 | 2.14435 | Promoter | 6 | 122453101 | 122486313 | 33213 | 2 | ENSMUST00000068242.6 | -403 | Rimklb | ENSMUSG00000040649.10 |
| chr6 | 122488705 | 122489011 | 3.48371 | 4.18303 | 1.26095 | Distal Intergenic | 6 | 122453101 | 122486313 | 33213 | 2 | ENSMUST00000068242.6 | -2393 | Rimklb | ENSMUSG00000040649.10 |
| chr6 | 122490576 | 122490910 | 3.32617 | 3.81654 | 1.01733 | Distal Intergenic | 6 | 122453101 | 122486313 | 33213 | 2 | ENSMUST00000068242.6 | -4264 | Rimklb | ENSMUSG00000040649.10 |
| chr6 | 122491388 | 122491692 | 3.75847 | 4.69617 | 1.58764 | Distal Intergenic | 6 | 122453101 | 122486313 | 33213 | 2 | ENSMUST00000068242.6 | -5076 | Rimklb | ENSMUSG00000040649.10 |
| chr6 | 122492507 | 122493311 | 3.55653 | 4.41423 | 1.39226 | Distal Intergenic | 6 | 122453101 | 122486313 | 33213 | 2 | ENSMUST00000068242.6 | -6195 | Rimklb | ENSMUSG00000040649.10 |
| chr6 | 122495778 | 122496160 | 3.8979 | 4.99235 | 1.79265 | Distal Intergenic | 6 | 122453101 | 122486313 | 33213 | 2 | ENSMUST00000068242.6 | -9466 | Rimklb | ENSMUSG00000040649.10 |
| chr17 | 23744716 | 23746296 | 3.57178 | 4.33806 | 1.38014 | Promoter | 17 | 23741200 | 23745829 | 4630 | 2 | ENSMUST00000046525.8 | 0 | Kremen2 | ENSMUSG00000040680.8 |
| chr17 | 23747063 | 23747632 | 3.72118 | 4.69531 | 1.58192 | Intron (ENSMUST00000181291.2/ENSMUSG00000097006.2, intron 3 of 4) | 17 | 23741200 | 23745829 | 4630 | 2 | ENSMUST00000046525.8 | -1235 | Kremen2 | ENSMUSG00000040680.8 |
| chr4 | 130047787 | 130047929 | 3.59837 | 4.7454 | 1.60942 | Promoter | 4 | 130047840 | 130099277 | 51438 | 1 | ENSMUST00000044565.9 | 0 | Col16a1 | ENSMUSG00000040690.10 |
| chr1 | 96871729 | 96872515 | 3.63062 | 4.37615 | 1.35896 | Promoter | 1 | 96816270 | 96872171 | 55902 | 2 | ENSMUST00000071985.5 | 0 | Slco4c1 | ENSMUSG00000040693.7 |
| chr7 | 19420113 | 19420450 | 3.38455 | 4.32107 | 1.34287 | Promoter | 7 | 19418812 | 19421417 | 2606 | 2 | ENSMUST00000047020.1 | 967 | A930016O22Rik | ENSMUSG00000040705.1 |
| chr4 | 141746728 | 141746811 | 3.63442 | 4.58319 | 1.48769 | Promoter | 4 | 141746672 | 141759263 | 12592 | 1 | ENSMUST00000038161.4 | 57 | Agmat | ENSMUSG00000040706.4 |
| chr1 | 95666024 | 95666690 | 3.63998 | 4.5105 | 1.47775 | Promoter | 1 | 95618551 | 95667535 | 48985 | 2 | ENSMUST00000189556.1 | 845 | St8sia4 | ENSMUSG00000040710.5 |
| chr1 | 95668523 | 95668995 | 3.85072 | 4.94719 | 1.74933 | Promoter | 1 | 95587682 | 95667571 | 79890 | 2 | ENSMUST00000043336.5 | -953 | St8sia4 | ENSMUSG00000040710.5 |
| chr7 | 19398500 | 19398651 | 3.41158 | 3.97197 | 1.11155 | Promoter | 7 | 19394564 | 19399550 | 4987 | 2 | ENSMUST00000108457.1 | 899 | Klc3 | ENSMUSG00000040714.8 |
| chr7 | 19402763 | 19403220 | 3.63221 | 4.42218 | 1.46097 | Promoter | 7 | 19394437 | 19404104 | 9668 | 2 | ENSMUST00000108458.4 | 884 | Klc3 | ENSMUSG00000040714.8 |
| chr7 | 19403833 | 19404128 | 4.0359 | 5.27801 | 2.04219 | Promoter | 7 | 19394437 | 19404104 | 9668 | 2 | ENSMUST00000108458.4 | 0 | Klc3 | ENSMUSG00000040714.8 |
| chr14 | 55078048 | 55078845 | 3.35633 | 3.86198 | 1.01733 | Exon (ENSMUST00000185430.1/ENSMUSG00000040721.9, exon 1 of 1) | 14 | 55074569 | 55080452 | 5884 | 2 | ENSMUST00000185430.1 | 1607 | Zfhx2 | ENSMUSG00000040721.9 |
| chr3 | 107099426 | 107099628 | 3.76816 | 4.64038 | 1.5521 | Distal Intergenic | 3 | 107101567 | 107106879 | 5313 | 1 | ENSMUST00000038695.1 | -1939 | Kcna2 | ENSMUSG00000040724.1 |
| chr3 | 107100479 | 107101847 | 3.9426 | 5.04195 | 1.82554 | Promoter | 3 | 107101567 | 107106879 | 5313 | 1 | ENSMUST00000038695.1 | 0 | Kcna2 | ENSMUSG00000040724.1 |
| chr3 | 107102209 | 107102438 | 3.59644 | 4.50533 | 1.46511 | Promoter | 3 | 107101567 | 107106879 | 5313 | 1 | ENSMUST00000038695.1 | 643 | Kcna2 | ENSMUSG00000040724.1 |
| chr14 | 26999811 | 27000095 | 3.54226 | 4.24517 | 1.32497 | Promoter | 14 | 27000362 | 27002329 | 1968 | 1 | ENSMUST00000035433.9 | -267 | Hesx1 | ENSMUSG00000040726.9 |
| chr14 | 27000547 | 27000899 | 3.41989 | 3.9783 | 1.08844 | Promoter | 14 | 27000362 | 27002329 | 1968 | 1 | ENSMUST00000035433.9 | 186 | Hesx1 | ENSMUSG00000040726.9 |
| chr4 | 11378341 | 11378429 | 3.32303 | 3.84774 | 1.12255 | Intron (ENSMUST00000043781.8/ENSMUSG00000040728.10, intron 4 of 14) | 4 | 11357586 | 11386394 | 28809 | 2 | ENSMUST00000155519.1 | 7965 | Esrp1 | ENSMUSG00000040728.10 |
| chr4 | 11381621 | 11382069 | 3.28628 | 4.17588 | 1.2455 | Intron (ENSMUST00000043781.8/ENSMUSG00000040728.10, intron 3 of 14) | 4 | 11357586 | 11386394 | 28809 | 2 | ENSMUST00000155519.1 | 4325 | Esrp1 | ENSMUSG00000040728.10 |
| chr4 | 11383197 | 11383682 | 3.89312 | 4.99028 | 1.80187 | Intron (ENSMUST00000043781.8/ENSMUSG00000040728.10, intron 3 of 14) | 4 | 11357586 | 11386394 | 28809 | 2 | ENSMUST00000155519.1 | 2712 | Esrp1 | ENSMUSG00000040728.10 |
| chr4 | 11384094 | 11389863 | 3.9987 | 5.16032 | 1.9099 | Promoter | 4 | 11331933 | 11386679 | 54747 | 2 | ENSMUST00000043781.8 | 0 | Esrp1 | ENSMUSG00000040728.10 |
| chr16 | 95553422 | 95553634 | 3.48033 | 4.08834 | 1.14928 | Intron (ENSMUST00000077773.7/ENSMUSG00000040732.13, intron 1 of 10) | 16 | 95530265 | 95700420 | 170156 | 2 | ENSMUST00000113848.3 | -23058 | Erg | ENSMUSG00000040732.13 |
| chr16 | 95579882 | 95579957 | 3.36837 | 3.88014 | 1.01733 | Intron (ENSMUST00000077773.7/ENSMUSG00000040732.13, intron 1 of 10) | 16 | 95360210 | 95586593 | 226384 | 2 | ENSMUST00000077773.7 | 6636 | Erg | ENSMUSG00000040732.13 |
| chr16 | 95582663 | 95582971 | 3.59401 | 4.29283 | 1.26464 | Intron (ENSMUST00000077773.7/ENSMUSG00000040732.13, intron 1 of 10) | 16 | 95360210 | 95586593 | 226384 | 2 | ENSMUST00000077773.7 | 3622 | Erg | ENSMUSG00000040732.13 |
| chr16 | 95583321 | 95584308 | 3.47247 | 4.07184 | 1.13852 | Intron (ENSMUST00000077773.7/ENSMUSG00000040732.13, intron 1 of 10) | 16 | 95360210 | 95586593 | 226384 | 2 | ENSMUST00000077773.7 | 2285 | Erg | ENSMUSG00000040732.13 |
| chr16 | 95585309 | 95585974 | 3.91793 | 4.98424 | 1.77276 | Promoter | 16 | 95360210 | 95586593 | 226384 | 2 | ENSMUST00000077773.7 | 619 | Erg | ENSMUSG00000040732.13 |
| chr16 | 95586765 | 95587044 | 3.46278 | 4.17379 | 1.24466 | Promoter | 16 | 95360210 | 95586593 | 226384 | 2 | ENSMUST00000077773.7 | -173 | Erg | ENSMUSG00000040732.13 |
| chr16 | 95588093 | 95588379 | 3.54309 | 4.27472 | 1.28616 | Distal Intergenic | 16 | 95360210 | 95586593 | 226384 | 2 | ENSMUST00000077773.7 | -1501 | Erg | ENSMUSG00000040732.13 |
| chr16 | 95588807 | 95589093 | 3.75087 | 4.58952 | 1.47743 | Distal Intergenic | 16 | 95360210 | 95586593 | 226384 | 2 | ENSMUST00000077773.7 | -2215 | Erg | ENSMUSG00000040732.13 |
| chr2 | 90885500 | 90886302 | 4.57587 | 6.5806 | 2.99059 | Promoter | 2 | 90885860 | 90890525 | 4666 | 1 | ENSMUST00000111466.2 | 0 | C1qtnf4 | ENSMUSG00000040794.5 |
| chr2 | 90886625 | 90888781 | 3.5663 | 4.3419 | 1.38967 | Promoter | 2 | 90885860 | 90890525 | 4666 | 1 | ENSMUST00000111466.2 | 766 | C1qtnf4 | ENSMUSG00000040794.5 |
| chr2 | 90889256 | 90889932 | 3.76175 | 4.94126 | 1.75026 | 5' UTR | 2 | 90885860 | 90890525 | 4666 | 1 | ENSMUST00000111466.2 | 3397 | C1qtnf4 | ENSMUSG00000040794.5 |
| chr6 | 121462849 | 121463184 | 3.87855 | 4.86891 | 1.69734 | Intron (ENSMUST00000129815.2/ENSMUSG00000040797.11, intron 1 of 13) | 6 | 121372933 | 121473623 | 100691 | 2 | ENSMUST00000129815.2 | 10439 | Iqsec3 | ENSMUSG00000040797.11 |
| chr6 | 121473603 | 121473779 | 3.3081 | 4.15565 | 1.24425 | Promoter | 6 | 121372933 | 121473623 | 100691 | 2 | ENSMUST00000129815.2 | 0 | Iqsec3 | ENSMUSG00000040797.11 |
| chr6 | 121474517 | 121474869 | 3.69459 | 4.6604 | 1.55986 | Promoter | 6 | 121404801 | 121473678 | 68878 | 2 | ENSMUST00000151397.1 | -840 | Iqsec3 | ENSMUSG00000040797.11 |
| chr7 | 19173019 | 19173303 | 3.41599 | 4.08158 | 1.19056 | Distal Intergenic | 7 | 19176421 | 19206257 | 29837 | 1 | ENSMUST00000117338.2 | -3118 | Eml2 | ENSMUSG00000040811.10 |
| chr7 | 19176503 | 19176718 | 4.4127 | 6.16308 | 2.61349 | Promoter | 7 | 19176421 | 19206257 | 29837 | 1 | ENSMUST00000117338.2 | 83 | Eml2 | ENSMUSG00000040811.10 |
| chr7 | 19177062 | 19177300 | 3.94148 | 5.08932 | 1.86766 | Promoter | 7 | 19176421 | 19206257 | 29837 | 1 | ENSMUST00000117338.2 | 642 | Eml2 | ENSMUSG00000040811.10 |

| | | | | | | | | | | | | | | | |
|---|---|---|---|---|---|---|---|---|---|---|---|---|---|---|---|
| chr11 | 70465160 | 70465264 | 3.43675 | 4.14983 | 1.2118 | Exon (ENSMUST00000108563.3/ENSMUSG00000040829.9, exon 12 of 14) | 11 | 70460565 | 70466090 | 5526 | 1 | ENSMUST00000092958.3 | 4596 | Zmynd15 | ENSMUSG00000040829.9 |
| chr1 | 165281417 | 165282699 | 3.97602 | 5.05772 | 1.83758 | Distal Intergenic | 1 | 165301863 | 165322930 | 21068 | 1 | ENSMUST00000178700.2 | -19164 | Gpr161 | ENSMUSG00000040836.10 |
| chr11 | 104685901 | 104686121 | 3.45682 | 4.1595 | 1.24191 | Promoter | 11 | 104685707 | 104753962 | 68256 | 1 | ENSMUST00000148007.2 | 195 | Gm11639 | ENSMUSG00000040838.9 |
| chr12 | 109439024 | 109439144 | 3.71771 | 4.57725 | 1.52052 | Distal Intergenic | 12 | 109452823 | 109453782 | 960 | 1 | ENSMUST00000174539.1 | -13679 | Dlk1 | ENSMUSG00000040856.12 |
| chr12 | 109444783 | 109445077 | 3.50189 | 4.29872 | 1.28672 | Distal Intergenic | 12 | 109452823 | 109453782 | 960 | 1 | ENSMUST00000174539.1 | -7746 | Dlk1 | ENSMUSG00000040856.12 |
| chr12 | 109449959 | 109450165 | 3.35558 | 3.86086 | 1.01733 | Distal Intergenic | 12 | 109452823 | 109453782 | 960 | 1 | ENSMUST00000174539.1 | -2658 | Dlk1 | ENSMUSG00000040856.12 |
| chr12 | 109450527 | 109451153 | 3.44065 | 4.01058 | 1.09248 | Distal Intergenic | 12 | 109452823 | 109453782 | 960 | 1 | ENSMUST00000174539.1 | -1670 | Dlk1 | ENSMUSG00000040856.12 |
| chr12 | 109451543 | 109451618 | 4.03917 | 5.1523 | 1.85609 | Downstream (2-3kb) | 12 | 109452823 | 109453782 | 960 | 1 | ENSMUST00000174539.1 | -1205 | Dlk1 | ENSMUSG00000040856.12 |
| chr12 | 109452449 | 109453132 | 3.80504 | 4.69696 | 1.5459 | Promoter | 12 | 109452823 | 109453782 | 960 | 1 | ENSMUST00000174539.1 | 0 | Dlk1 | ENSMUSG00000040856.12 |
| chr12 | 109453484 | 109453777 | 4.26776 | 5.62892 | 2.21722 | Promoter | 12 | 109453491 | 109460381 | 6891 | 1 | ENSMUST00000109842.3 | 0 | Dlk1 | ENSMUSG00000040856.12 |
| chr12 | 109459551 | 109460131 | 3.80614 | 4.77639 | 1.67426 | Promoter | 12 | 109459843 | 109460475 | 633 | 1 | ENSMUST00000173812.1 | 0 | Dlk1 | ENSMUSG00000040856.12 |
| chr12 | 109477522 | 109478092 | 3.26321 | 3.88558 | 1.0721 | Distal Intergenic | 12 | 109459843 | 109460475 | 633 | 1 | ENSMUST00000173812.1 | 17680 | Dlk1 | ENSMUSG00000040856.12 |
| chr12 | 109050512 | 109050714 | 3.49758 | 4.11559 | 1.16021 | Intron (ENSMUST00000073156.7/ENSMUSG00000040867.10, intron 2 of 6) | 12 | 109037652 | 109053581 | 15930 | 2 | ENSMUST00000185745.1 | 2867 | Begain | ENSMUSG00000040867.10 |
| chr12 | 109052387 | 109054573 | 4.093 | 5.44408 | 2.09916 | Promoter | 12 | 109037652 | 109053581 | 15930 | 2 | ENSMUST00000185745.1 | 0 | Begain | ENSMUSG00000040867.10 |
| chr12 | 109055083 | 109055201 | 3.38193 | 3.9006 | 1.01733 | Intron (ENSMUST00000073156.7/ENSMUSG00000040867.10, intron 1 of 6) | 12 | 109037652 | 109053581 | 15930 | 2 | ENSMUST00000185745.1 | -1503 | Begain | ENSMUSG00000040867.10 |
| chr12 | 109055509 | 109055913 | 3.76955 | 4.62933 | 1.49346 | Intron (ENSMUST00000073156.7/ENSMUSG00000040867.10, intron 1 of 6) | 12 | 109037652 | 109053581 | 15930 | 2 | ENSMUST00000185745.1 | -1929 | Begain | ENSMUSG00000040867.10 |
| chr12 | 109067440 | 109068352 | 3.44634 | 4.03587 | 1.14674 | Promoter | 12 | 109032187 | 109068217 | 36031 | 2 | ENSMUST00000073156.7 | 0 | Begain | ENSMUSG00000040867.10 |
| chr12 | 109073272 | 109073347 | 4.21283 | 5.51402 | 2.12556 | Distal Intergenic | 12 | 109032187 | 109068217 | 36031 | 2 | ENSMUST00000190647.1 | -5056 | Begain | ENSMUSG00000040867.10 |
| chr9 | 114977602 | 114978141 | 3.3857 | 3.92251 | 1.07454 | Promoter | 9 | 114978569 | 115232223 | 253655 | 1 | ENSMUST00000183104.2 | -428 | Osbpl10 | ENSMUSG00000040875.7 |
| chr7 | 19021024 | 19021165 | 3.60602 | 4.33225 | 1.35104 | Intron (ENSMUST00000153321.1/ENSMUSG00000040891.5, intron 1 of 2) | 7 | 19013282 | 19023538 | 10257 | 2 | ENSMUST00000153321.1 | 2373 | Foxa3 | ENSMUSG00000040891.5 |
| chr7 | 19021550 | 19022384 | 3.35222 | 3.85579 | 1.01733 | Intron (ENSMUST00000153321.1/ENSMUSG00000040891.5, intron 1 of 2) | 7 | 19013282 | 19023538 | 10257 | 2 | ENSMUST00000153321.1 | 1154 | Foxa3 | ENSMUSG00000040891.5 |
| chr7 | 19023510 | 19023691 | 3.69711 | 4.73372 | 1.61063 | Promoter | 7 | 19013282 | 19023538 | 10257 | 2 | ENSMUST00000153321.1 | 0 | Foxa3 | ENSMUSG00000040891.5 |
| chr3 | 105452823 | 105453108 | 3.58458 | 4.69954 | 1.56194 | Promoter | 3 | 105453086 | 105669460 | 216375 | 1 | ENSMUST00000118360.2 | 0 | Kcnd3 | ENSMUSG00000040896.11 |
| chr3 | 105453851 | 105454009 | 3.66411 | 4.55291 | 1.49352 | Promoter | 3 | 105453086 | 105669460 | 216375 | 1 | ENSMUST00000118360.2 | 766 | Kcnd3 | ENSMUSG00000040896.11 |
| chr3 | 105454523 | 105454747 | 3.80463 | 4.82182 | 1.6612 | Intron (ENSMUST00000098761.4/ENSMUSG00000040896.11, intron 1 of 6) | 3 | 105453086 | 105669460 | 216375 | 1 | ENSMUST00000118360.2 | 1438 | Kcnd3 | ENSMUSG00000040896.11 |
| chr3 | 105531988 | 105532124 | 3.40544 | 3.98758 | 1.19288 | Intron (ENSMUST00000098761.4/ENSMUSG00000040896.11, intron 2 of 6) | 3 | 105458782 | 105669309 | 210528 | 1 | ENSMUST00000079169.4 | 73207 | Kcnd3 | ENSMUSG00000040896.11 |
| chr19 | 59195251 | 59195839 | 3.49476 | 4.15559 | 1.2938 | Distal Intergenic | 19 | 59219648 | 59237370 | 17723 | 1 | ENSMUST00000065204.6 | -23809 | Kcnk18 | ENSMUSG00000040901.7 |
| chr19 | 59196198 | 59196690 | 4.16224 | 5.52101 | 2.16847 | Distal Intergenic | 19 | 59219648 | 59237370 | 17723 | 1 | ENSMUST00000065204.6 | -22958 | Kcnk18 | ENSMUSG00000040901.7 |
| chr19 | 59197134 | 59197818 | 3.83406 | 5.07123 | 1.83981 | Distal Intergenic | 19 | 59219648 | 59237370 | 17723 | 1 | ENSMUST00000065204.6 | -21830 | Kcnk18 | ENSMUSG00000040901.7 |
| chr19 | 59198917 | 59199021 | 3.5455 | 4.23337 | 1.32218 | Distal Intergenic | 19 | 59219648 | 59237370 | 17723 | 1 | ENSMUST00000065204.6 | -20627 | Kcnk18 | ENSMUSG00000040901.7 |
| chr7 | 24994337 | 24994596 | 3.51455 | 4.16189 | 1.23486 | Exon (ENSMUST00000102858.4/ENSMUSG00000040907.10, exon 9 of 22) | 7 | 24998659 | 25001197 | 2539 | 2 | ENSMUST00000154077.2 | 6601 | Atp1a3 | ENSMUSG00000040907.10 |
| chr7 | 25003372 | 25003493 | 3.64205 | 4.37857 | 1.3169 | Intron (ENSMUST00000102858.4/ENSMUSG00000040907.10, intron 1 of 21) | 7 | 24998659 | 25001197 | 2539 | 2 | ENSMUST00000154077.2 | -2176 | Atp1a3 | ENSMUSG00000040907.10 |
| chr7 | 25003820 | 25004037 | 3.34153 | 3.83966 | 1.01733 | Intron (ENSMUST00000102858.4/ENSMUSG00000040907.10, intron 1 of 21) | 7 | 24978167 | 25005895 | 27729 | 2 | ENSMUST00000102858.4 | 1858 | Atp1a3 | ENSMUSG00000040907.10 |
| chr7 | 25005200 | 25006000 | 3.84171 | 4.78291 | 1.60957 | Promoter | 7 | 24978167 | 25005895 | 27729 | 2 | ENSMUST00000102858.4 | 0 | Atp1a3 | ENSMUSG00000040907.10 |
| chr7 | 25007833 | 25008173 | 3.26014 | 3.92908 | 1.13318 | Downstream (1-2kb) | 7 | 25000571 | 25005957 | 5387 | 2 | ENSMUST00000125444.1 | -1877 | Atp1a3 | ENSMUSG00000040907.10 |
| chr7 | 25010212 | 25010852 | 4.25647 | 5.70923 | 2.35466 | 3' UTR | 7 | 25000571 | 25005957 | 5387 | 2 | ENSMUST00000125444.1 | -4256 | Atp1a3 | ENSMUSG00000040907.10 |
| chr19 | 45649973 | 45650233 | 2.26526 | 10.77831 | 6.43646 | Intron (ENSMUST00000046869.5/ENSMUSG00000040913.5, intron 1 of 8) | 19 | 45630875 | 45640803 | 9929 | 2 | ENSMUST00000159590.1 | -9171 | Fbxw4 | ENSMUSG00000040913.5 |
| chr19 | 45671994 | 45672506 | 3.68867 | 4.63445 | 1.52137 | Distal Intergenic | 19 | 45578254 | 45660312 | 82059 | 2 | ENSMUST00000046869.5 | -11683 | Fbxw4 | ENSMUSG00000040913.5 |
| chr19 | 45674139 | 45674249 | 3.82279 | 4.72292 | 1.54185 | Distal Intergenic | 19 | 45578254 | 45660312 | 82059 | 2 | ENSMUST00000046869.5 | -13828 | Fbxw4 | ENSMUSG00000040913.5 |
| chr19 | 45674905 | 45676532 | 3.61789 | 4.4613 | 1.43346 | Distal Intergenic | 19 | 45578254 | 45660312 | 82059 | 2 | ENSMUST00000046869.5 | -14594 | Fbxw4 | ENSMUSG00000040913.5 |
| chr19 | 45677481 | 45677990 | 3.41792 | 4.07945 | 1.17851 | Distal Intergenic | 19 | 45578254 | 45660312 | 82059 | 2 | ENSMUST00000046869.5 | -17170 | Fbxw4 | ENSMUSG00000040913.5 |
| chr19 | 45678371 | 45678745 | 3.36713 | 4.03205 | 1.15395 | Distal Intergenic | 19 | 45578254 | 45660312 | 82059 | 2 | ENSMUST00000046869.5 | -18060 | Fbxw4 | ENSMUSG00000040913.5 |
| chr19 | 45686322 | 45686825 | 3.75061 | 4.65046 | 1.57639 | Distal Intergenic | 19 | 45578254 | 45660312 | 82059 | 2 | ENSMUST00000046869.5 | -26011 | Fbxw4 | ENSMUSG00000040913.5 |
| chr19 | 45688343 | 45688540 | 3.63444 | 4.61506 | 1.54226 | Distal Intergenic | 19 | 45578254 | 45660312 | 82059 | 2 | ENSMUST00000046869.5 | -28032 | Fbxw4 | ENSMUSG00000040913.5 |
| chr19 | 45689394 | 45689678 | 3.61171 | 4.39053 | 1.38956 | Distal Intergenic | 19 | 45578254 | 45660312 | 82059 | 2 | ENSMUST00000046869.5 | -29083 | Fbxw4 | ENSMUSG00000040913.5 |
| chr19 | 45691488 | 45691972 | 3.58699 | 4.32108 | 1.38468 | Distal Intergenic | 19 | 45578254 | 45660312 | 82059 | 2 | ENSMUST00000046869.5 | -31177 | Fbxw4 | ENSMUSG00000040913.5 |
| chr1 | 164208086 | 164208161 | 4.57817 | 6.59151 | 2.91494 | Intron (ENSMUST00000086040.5/ENSMUSG00000026579.8, intron 19 of 24) | 1 | 164249046 | 164265385 | 16340 | 1 | ENSMUST00000044021.6 | -40885 | Slc19a2 | ENSMUSG00000040918.7 |
| chr1 | 164209733 | 164210133 | 3.68662 | 4.78264 | 1.61024 | Intron (ENSMUST00000086040.5/ENSMUSG00000026579.8, intron 21 of 24) | 1 | 164249046 | 164265385 | 16340 | 1 | ENSMUST00000044021.6 | -38913 | Slc19a2 | ENSMUSG00000040918.7 |
| chr1 | 164214373 | 164214778 | 3.52298 | 4.17122 | 1.22001 | Exon (ENSMUST00000086040.5/ENSMUSG00000026579.8, exon 23 of 25) | 1 | 164249046 | 164265385 | 16340 | 1 | ENSMUST00000044021.6 | -34268 | Slc19a2 | ENSMUSG00000040918.7 |
| chr1 | 164216513 | 164216696 | 3.44041 | 4.0934 | 1.17221 | Intron (ENSMUST00000086040.5/ENSMUSG00000026579.8, intron 23 of 24) | 1 | 164249046 | 164265385 | 16340 | 1 | ENSMUST00000044021.6 | -32350 | Slc19a2 | ENSMUSG00000040918.7 |
| chr1 | 164217423 | 164217686 | 3.62969 | 4.35359 | 1.29371 | Exon (ENSMUST00000086040.5/ENSMUSG00000026579.8, exon 24 of 25) | 1 | 164249046 | 164265385 | 16340 | 1 | ENSMUST00000044021.6 | -31360 | Slc19a2 | ENSMUSG00000040918.7 |
| chr1 | 164220097 | 164220804 | 3.35431 | 4.02416 | 1.15616 | 3' UTR | 1 | 164249046 | 164265385 | 16340 | 1 | ENSMUST00000044021.6 | -28242 | Slc19a2 | ENSMUSG00000040918.7 |
| chr9 | 121125241 | 121125533 | 3.37459 | 3.88952 | 1.01733 | Intron (ENSMUST00000171923.2/ENSMUSG00000040936.9, intron 31 of 36) | 9 | 120960268 | 121108525 | 148258 | 2 | ENSMUST00000170406.2 | -16717 | Ulk4 | ENSMUSG00000040936.9 |
| chr11 | 70208583 | 70208658 | 3.63519 | 4.45104 | 1.48532 | Distal Intergenic | 11 | 70212752 | 70216413 | 3662 | 1 | ENSMUST00000126388.2 | -4094 | Slc16a11 | ENSMUSG00000040938.11 |
| chr11 | 70212784 | 70213333 | 3.47162 | 4.31064 | 1.34775 | Promoter | 11 | 70212752 | 70216413 | 3662 | 1 | ENSMUST00000126388.2 | 33 | Slc16a11 | ENSMUSG00000040938.11 |
| chr11 | 70213669 | 70214828 | 3.90467 | 5.0055 | 1.79301 | Promoter | 11 | 70213910 | 70216412 | 2503 | 1 | ENSMUST00000171032.2 | 0 | Slc16a11 | ENSMUSG00000040938.11 |
| chr11 | 70215717 | 70215893 | 4.26199 | 5.83318 | 2.37524 | Promoter | 11 | 70214818 | 70216413 | 1596 | 1 | ENSMUST00000149880.1 | 900 | Slc16a11 | ENSMUSG00000040938.11 |
| chr7 | 24931273 | 24931359 | 4.01169 | 5.08267 | 1.78076 | Distal Intergenic | 7 | 24923363 | 24925936 | 2574 | 1 | ENSMUST00000132786.1 | 7911 | Arhgef1 | ENSMUSG00000040940.13 |
| chr7 | 24932013 | 24932233 | 3.78969 | 4.66496 | 1.51632 | Distal Intergenic | 7 | 24923363 | 24925936 | 2574 | 1 | ENSMUST00000132786.1 | 8651 | Arhgef1 | ENSMUSG00000040940.13 |
| chr7 | 24932600 | 24932896 | 3.34239 | 3.84096 | 1.01733 | Distal Intergenic | 7 | 24923363 | 24925936 | 2574 | 1 | ENSMUST00000132786.1 | 9238 | Arhgef1 | ENSMUSG00000040940.13 |
| chr7 | 24933217 | 24934501 | 3.6434 | 4.40104 | 1.36165 | Distal Intergenic | 7 | 24923363 | 24925936 | 2574 | 1 | ENSMUST00000132786.1 | 9855 | Arhgef1 | ENSMUSG00000040940.13 |
| chr7 | 24935328 | 24935958 | 3.41704 | 4.04635 | 1.13668 | Distal Intergenic | 7 | 24923363 | 24925936 | 2574 | 1 | ENSMUST00000132786.1 | 11966 | Arhgef1 | ENSMUSG00000040940.13 |
| chr7 | 24941952 | 24942296 | 4.06971 | 5.28312 | 1.95856 | Distal Intergenic | 7 | 24923363 | 24925936 | 2574 | 1 | ENSMUST00000132786.1 | 18590 | Arhgef1 | ENSMUSG00000040940.13 |
| chr7 | 24942851 | 24943608 | 4.11014 | 5.30218 | 1.97151 | Distal Intergenic | 7 | 24923363 | 24925936 | 2574 | 1 | ENSMUST00000132786.1 | 19489 | Arhgef1 | ENSMUSG00000040940.13 |
| chr7 | 24944219 | 24946012 | 3.81785 | 4.78868 | 1.64756 | Distal Intergenic | 7 | 24923363 | 24925936 | 2574 | 1 | ENSMUST00000132786.1 | 20857 | Arhgef1 | ENSMUSG00000040940.13 |
| chr4 | 140504455 | 140504753 | 3.30772 | 3.80217 | 1.04225 | Distal Intergenic | 4 | 140544314 | 140568381 | 24068 | 2 | ENSMUST00000140403.1 | 63628 | Arhgef10l | ENSMUSG00000040964.11 |
| chr4 | 140506417 | 140506649 | 3.72175 | 4.53515 | 1.44064 | Distal Intergenic | 4 | 140544314 | 140568381 | 24068 | 2 | ENSMUST00000140403.1 | 61732 | Arhgef10l | ENSMUSG00000040964.11 |
| chr4 | 140648807 | 140649002 | 3.94366 | 5.17764 | 1.91738 | Promoter | 4 | 140514485 | 140648756 | 134272 | 2 | ENSMUST00000097820.3 | -52 | Arhgef10l | ENSMUSG00000040964.11 |
| chr4 | 140655621 | 140655884 | 3.87864 | 4.83522 | 1.64561 | Intron (ENSMUST00000069623.6/ENSMUSG00000040964.11, intron 1 of 27) | 4 | 140591932 | 140649360 | 57429 | 2 | ENSMUST00000143614.2 | -6262 | Arhgef10l | ENSMUSG00000040964.11 |
| chr3 | 133234790 | 133235289 | 3.33345 | 3.86921 | 1.12898 | Promoter | 3 | 133112278 | 133234889 | 122612 | 2 | ENSMUST00000147041.4 | 0 | Arhgef38 | ENSMUSG00000040969.9 |
| chr4 | 140199994 | 140200221 | 3.34557 | 3.84575 | 1.01733 | Intron (ENSMUST00000039331.8/ENSMUSG00000040972.8, intron 1 of 9) | 4 | 140026846 | 140246784 | 219939 | 2 | ENSMUST00000039331.8 | 46563 | Igsf21 | ENSMUSG00000040972.8 |
| chr4 | 140204904 | 140205604 | 3.52812 | 4.19381 | 1.2648 | Intron (ENSMUST00000039331.8/ENSMUSG00000040972.8, intron 1 of 9) | 4 | 140026846 | 140246784 | 219939 | 2 | ENSMUST00000039331.8 | 41180 | Igsf21 | ENSMUSG00000040972.8 |
| chr4 | 140206066 | 140206244 | 3.42972 | 4.01768 | 1.17751 | Intron (ENSMUST00000039331.8/ENSMUSG00000040972.8, intron 1 of 9) | 4 | 140026846 | 140246784 | 219939 | 2 | ENSMUST00000039331.8 | 40540 | Igsf21 | ENSMUSG00000040972.8 |
| chr4 | 140220713 | 140221350 | 3.50277 | 4.28895 | 1.32477 | Intron (ENSMUST00000039331.8/ENSMUSG00000040972.8, intron 1 of 9) | 4 | 140026846 | 140246784 | 219939 | 2 | ENSMUST00000039331.8 | 25434 | Igsf21 | ENSMUSG00000040972.8 |
| chr4 | 140221812 | 140222512 | 3.58639 | 4.29574 | 1.30751 | Intron (ENSMUST00000039331.8/ENSMUSG00000040972.8, intron 1 of 9) | 4 | 140026846 | 140246784 | 219939 | 2 | ENSMUST00000039331.8 | 24272 | Igsf21 | ENSMUSG00000040972.8 |
| chr4 | 140223460 | 140223627 | 3.7485 | 4.59376 | 1.4998 | Intron (ENSMUST00000039331.8/ENSMUSG00000040972.8, intron 1 of 9) | 4 | 140026846 | 140246784 | 219939 | 2 | ENSMUST00000039331.8 | 23157 | Igsf21 | ENSMUSG00000040972.8 |
| chr4 | 140224845 | 140225050 | 3.51243 | 4.14776 | 1.19531 | Intron (ENSMUST00000039331.8/ENSMUSG00000040972.8, intron 1 of 9) | 4 | 140026846 | 140246784 | 219939 | 2 | ENSMUST00000039331.8 | 21734 | Igsf21 | ENSMUSG00000040972.8 |
| chr4 | 140237576 | 140237672 | 4.13901 | 5.33357 | 1.96399 | Intron (ENSMUST00000039331.8/ENSMUSG00000040972.8, intron 1 of 9) | 4 | 140026846 | 140246784 | 219939 | 2 | ENSMUST00000039331.8 | 9112 | Igsf21 | ENSMUSG00000040972.8 |
| chr4 | 140243192 | 140243661 | 3.71241 | 4.52028 | 1.42515 | Intron (ENSMUST00000039331.8/ENSMUSG00000040972.8, intron 1 of 9) | 4 | 140026846 | 140246784 | 219939 | 2 | ENSMUST00000039331.8 | 3123 | Igsf21 | ENSMUSG00000040972.8 |
| chr4 | 140244005 | 140244728 | 3.69639 | 4.50454 | 1.41019 | Intron (ENSMUST00000039331.8/ENSMUSG00000040972.8, intron 1 of 9) | 4 | 140026846 | 140246784 | 219939 | 2 | ENSMUST00000039331.8 | 2056 | Igsf21 | ENSMUSG00000040972.8 |
| chr4 | 140245546 | 140245781 | 3.68567 | 4.70554 | 1.56989 | Intron (ENSMUST00000039331.8/ENSMUSG00000040972.8, intron 1 of 9) | 4 | 140026846 | 140246784 | 219939 | 2 | ENSMUST00000039331.8 | 1003 | Igsf21 | ENSMUSG00000040972.8 |
| chr4 | 140246187 | 140246714 | 3.49947 | 4.19695 | 1.24074 | Promoter | 4 | 140026846 | 140246784 | 219939 | 2 | ENSMUST00000039331.8 | 70 | Igsf21 | ENSMUSG00000040972.8 |
| chr4 | 140247206 | 140248960 | 3.72515 | 4.55502 | 1.46061 | Promoter | 4 | 140026846 | 140246784 | 219939 | 2 | ENSMUST00000039331.8 | -423 | Igsf21 | ENSMUSG00000040972.8 |
| chr4 | 140252135 | 140252333 | 3.42981 | 3.99231 | 1.08578 | Distal Intergenic | 4 | 140026846 | 140246784 | 219939 | 2 | ENSMUST00000039331.8 | -5352 | Igsf21 | ENSMUSG00000040972.8 |
| chr4 | 140255432 | 140255554 | 3.38193 | 3.9006 | 1.01733 | Distal Intergenic | 4 | 140026846 | 140246784 | 219939 | 2 | ENSMUST00000039331.8 | -8649 | Igsf21 | ENSMUSG00000040972.8 |
| chr4 | 140256076 | 140256242 | 3.37146 | 3.8848 | 1.01733 | Distal Intergenic | 4 | 140026846 | 140246784 | 219939 | 2 | ENSMUST00000039331.8 | -9293 | Igsf21 | ENSMUSG00000040972.8 |
| chr4 | 140256831 | 140256931 | 3.65318 | 4.40261 | 1.34095 | Distal Intergenic | 4 | 140026846 | 140246784 | 219939 | 2 | ENSMUST00000039331.8 | -10048 | Igsf21 | ENSMUSG00000040972.8 |
| chr7 | 18833725 | 18834011 | 3.2637 | 3.72963 | 1.03901 | Distal Intergenic | 7 | 18839966 | 18858653 | 18688 | 1 | ENSMUST00000072415.6 | -5955 | Mill2 | ENSMUSG00000040987.9 |
| chr4 | 128879532 | 128879635 | 3.87867 | 4.8403 | 1.66377 | Distal Intergenic | 4 | 128883580 | 128903565 | 19986 | 1 | ENSMUST00000147852.1 | -3945 | Trim62 | ENSMUSG00000041000.8 |
| chr14 | 39471790 | 39472183 | 4.03917 | 5.16383 | 1.85907 | Promoter | 14 | 39009187 | 39472666 | 463480 | 2 | ENSMUST00000176122.1 | 483 | Nrg3 | ENSMUSG00000041014.12 |
| chr14 | 39472635 | 39474426 | 3.82006 | 4.76794 | 1.5969 | Promoter | 14 | 38368952 | 39473088 | 1104137 | 2 | ENSMUST00000166968.3 | 0 | Nrg3 | ENSMUSG00000041014.12 |
| chr14 | 39475081 | 39475347 | 3.59882 | 4.43251 | 1.43388 | Distal Intergenic | 14 | 38368952 | 39473088 | 1104137 | 2 | ENSMUST00000166968.3 | -1994 | Nrg3 | ENSMUSG00000041014.12 |
| chr9 | 65588971 | 65589605 | 4.04159 | 5.2837 | 2.05989 | Exon (ENSMUST00000134538.2/ENSMUSG00000041064.8, exon 3 of 13) | 9 | 65587226 | 65588280 | 1055 | 1 | ENSMUST00000141046.1 | 1746 | Pif1 | ENSMUSG00000041064.8 |
| chr9 | 65596516 | 65596591 | 3.7713 | 4.62173 | 1.47083 | Distal Intergenic | 9 | 65587226 | 65588280 | 1055 | 1 | ENSMUST00000141046.1 | 9291 | Pif1 | ENSMUSG00000041064.8 |
| chr11 | 6606249 | 6606555 | 3.35673 | 3.87316 | 1.0547 | Promoter | 11 | 6597823 | 6606053 | 8231 | 2 | ENSMUST00000045713.3 | -197 | Nacad | ENSMUSG00000041073.5 |
| chr11 | 6606872 | 6607094 | 3.50061 | 4.15609 | 1.23584 | Promoter | 11 | 6597823 | 6606053 | 8231 | 2 | ENSMUST00000045713.3 | -820 | Nacad | ENSMUSG00000041073.5 |
| chr11 | 6607404 | 6607650 | 3.47576 | 4.11701 | 1.24485 | Distal Intergenic | 11 | 6597823 | 6606053 | 8231 | 2 | ENSMUST00000045713.3 | -1352 | Nacad | ENSMUSG00000041073.5 |
| chr14 | 34811142 | 34811235 | 3.54667 | 4.2214 | 1.26545 | Distal Intergenic | 14 | 34819825 | 35583379 | 763555 | 1 | ENSMUST00000043349.5 | -8590 | Grid1 | ENSMUSG00000041078.6 |
| chr14 | 34815248 | 34815462 | 4.05223 | 5.30107 | 2.01141 | Distal Intergenic | 14 | 34819825 | 35583379 | 763555 | 1 | ENSMUST00000043349.5 | -4363 | Grid1 | ENSMUSG00000041078.6 |
| chr14 | 34816485 | 34817052 | 3.65489 | 4.57429 | 1.53201 | Distal Intergenic | 14 | 34819825 | 35583379 | 763555 | 1 | ENSMUST00000043349.5 | -2773 | Grid1 | ENSMUSG00000041078.6 |
| chr14 | 34817621 | 34817723 | 3.52618 | 4.17535 | 1.21958 | Distal Intergenic | 14 | 34819825 | 35583379 | 763555 | 1 | ENSMUST00000043349.5 | -2102 | Grid1 | ENSMUSG00000041078.6 |
| chr14 | 34818493 | 34818616 | 3.65312 | 4.4852 | 1.45759 | Distal Intergenic | 14 | 34819825 | 35583379 | 763555 | 1 | ENSMUST00000043349.5 | -1209 | Grid1 | ENSMUSG00000041078.6 |
| chr14 | 34819148 | 34820779 | 4.13729 | 5.44692 | 2.12113 | Promoter | 14 | 34819825 | 35583379 | 763555 | 1 | ENSMUST00000043349.5 | 0 | Grid1 | ENSMUSG00000041078.6 |
| chr14 | 34821101 | 34821605 | 4.41343 | 5.96766 | 2.51817 | Intron (ENSMUST00000043349.5/ENSMUSG00000041078.6, intron 1 of 15) | 14 | 34819825 | 35583379 | 763555 | 1 | ENSMUST00000043349.5 | 1277 | Grid1 | ENSMUSG00000041078.6 |
| chr14 | 34822633 | 34823410 | 3.68056 | 4.50222 | 1.42741 | Exon (ENSMUST00000043349.5/ENSMUSG00000041078.6, exon 2 of 16) | 14 | 34819825 | 35583379 | 763555 | 1 | ENSMUST00000043349.5 | 2809 | Grid1 | ENSMUSG00000041078.6 |
| chr14 | 34824704 | 34825019 | 3.44705 | 4.15765 | 1.20629 | Intron (ENSMUST00000043349.5/ENSMUSG00000041078.6, intron 2 of 15) | 14 | 34819825 | 35583379 | 763555 | 1 | ENSMUST00000043349.5 | 4880 | Grid1 | ENSMUSG00000041078.6 |
| chr7 | 16983087 | 16983201 | 3.49824 | 4.15798 | 1.27965 | Distal Intergenic | 7 | 16992708 | 16997516 | 4809 | 1 | ENSMUST00000094805.4 | -9507 | Ccdc8 | ENSMUSG00000041117.8 |
| chr7 | 16993975 | 16994258 | 3.54682 | 4.27815 | 1.31156 | 5' UTR | 7 | 16992708 | 16997516 | 4809 | 1 | ENSMUST00000094805.4 | 1268 | Ccdc8 | ENSMUSG00000041117.8 |
| chr7 | 16995710 | 16996060 | 3.73085 | 4.63354 | 1.54462 | Exon (ENSMUST00000094805.4/ENSMUSG00000041117.8, exon 1 of 1) | 7 | 16992708 | 16997516 | 4809 | 1 | ENSMUST00000094805.4 | 3003 | Ccdc8 | ENSMUSG00000041117.8 |
| chr17 | 31386213 | 31386345 | 3.64922 | 4.4007 | 1.34798 | Promoter | 17 | 31386234 | 31476310 | 90077 | 1 | ENSMUST00000047168.6 | 0 | Pde9a | ENSMUSG00000041119.6 |
| chr17 | 31388332 | 31388433 | 3.58533 | 4.27334 | 1.24917 | Intron (ENSMUST00000047168.6/ENSMUSG00000041119.6, intron 1 of 18) | 17 | 31386294 | 31455122 | 68829 | 1 | ENSMUST00000141314.2 | 2039 | Pde9a | ENSMUSG00000041119.6 |
| chr17 | 31433634 | 31434067 | 3.55409 | 4.30921 | 1.35761 | Promoter | 17 | 31433071 | 31459129 | 26059 | 1 | ENSMUST00000136384.2 | 564 | Pde9a | ENSMUSG00000041119.6 |
| chr17 | 31457397 | 31457631 | 3.24362 | 3.69258 | 1.01733 | Intron (ENSMUST00000047168.6/ENSMUSG00000041119.6, intron 9 of 18) | 17 | 31461628 | 31476141 | 14514 | 1 | ENSMUST00000154392.1 | -3997 | Pde9a | ENSMUSG00000041119.6 |
| chr17 | 31458461 | 31458885 | 4.00279 | 5.12733 | 1.90931 | Intron (ENSMUST00000047168.6/ENSMUSG00000041119.6, intron 9 of 18) | 17 | 31461628 | 31476141 | 14514 | 1 | ENSMUST00000154392.1 | -2743 | Pde9a | ENSMUSG00000041119.6 |
| chr5 | 150595183 | 150595461 | 3.58325 | 4.3513 | 1.37676 | Promoter | 5 | 150571643 | 150594525 | 22883 | 2 | ENSMUST00000016279.7 | -659 | N4bp2l1 | ENSMUSG00000041132.8 |
| chr7 | 16959797 | 16960267 | 3.79827 | 4.69581 | 1.56697 | Promoter | 7 | 16959795 | 16962319 | 2525 | 1 | ENSMUST00000038163.6 | 3 | Pnmal1 | ENSMUSG00000041141.6 |
| chr16 | 20239106 | 20239181 | 3.49209 | 4.12043 | 1.20737 | Intron (ENSMUST00000040880.7/ENSMUSG00000041205.7, intron 1 of 2) | 16 | 20233310 | 20241358 | 8049 | 2 | ENSMUST00000040880.7 | 2177 | Map6d1 | ENSMUSG00000041205.7 |
| chr16 | 20240566 | 20241550 | 3.88357 | 4.96677 | 1.74877 | Promoter | 16 | 20233310 | 20241358 | 8049 | 2 | ENSMUST00000040880.7 | 0 | Map6d1 | ENSMUSG00000041205.7 |
| chr4 | 9265352 | 9265477 | 3.32724 | 3.82448 | 1.03958 | Downstream (1-2kb) | 4 | 9269293 | 9282272 | 12980 | 1 | ENSMUST00000147285.1 | -3816 | Clvs1 | ENSMUSG00000041216.10 |
| chr4 | 9267770 | 9267889 | 3.63624 | 4.49456 | 1.44446 | Exon (ENSMUST00000181165.1/ENSMUSG00000097094.1, exon 2 of 2) | 4 | 9269293 | 9282272 | 12980 | 1 | ENSMUST00000147285.1 | -1404 | Clvs1 | ENSMUSG00000041216.10 |
| chr4 | 9269192 | 9269969 | 4.02719 | 5.28347 | 1.97403 | Promoter | 4 | 9269293 | 9282272 | 12980 | 1 | ENSMUST00000147285.1 | 0 | Clvs1 | ENSMUSG00000041216.10 |
| chr4 | 9307876 | 9308129 | 3.35899 | 4.0113 | 1.11181 | Intron (ENSMUST00000128117.2/ENSMUSG00000041216.10, intron 2 of 2) | 4 | 9311756 | 9429803 | 118048 | 1 | ENSMUST00000141085.2 | -3627 | Clvs1 | ENSMUSG00000041216.10 |
| chr4 | 9337405 | 9337481 | 3.60995 | 4.59738 | 1.52263 | Intron (ENSMUST00000128117.2/ENSMUSG00000041216.10, intron 2 of 2) | 4 | 9335609 | 9429840 | 94232 | 1 | ENSMUST00000135864.1 | 1797 | Clvs1 | ENSMUSG00000041216.10 |
| chr18 | 11350423 | 11350725 | 3.60931 | 4.3332 | 1.32757 | Distal Intergenic | 18 | 11633276 | 11743207 | 109932 | 1 | ENSMUST00000115861.3 | -282551 | Rbbp8 | ENSMUSG00000041238.10 |
| chr18 | 11352124 | 11352568 | 3.73675 | 4.56765 | 1.45064 | Distal Intergenic | 18 | 11633276 | 11743207 | 109932 | 1 | ENSMUST00000115861.3 | -280708 | Rbbp8 | ENSMUSG00000041238.10 |
| chr18 | 11471909 | 11472295 | 3.29922 | 3.81297 | 1.09838 | Distal Intergenic | 18 | 11633276 | 11743207 | 109932 | 1 | ENSMUST00000115861.3 | -160981 | Rbbp8 | ENSMUSG00000041238.10 |
| chr9 | 32407486 | 32407760 | 3.95435 | 5.28467 | 1.98067 | Distal Intergenic | 9 | 32393985 | 32399192 | 5208 | 1 | ENSMUST00000172015.1 | 13502 | Kcnj1 | ENSMUSG00000041248.8 |
| chr9 | 32408285 | 32408693 | 3.99961 | 5.22162 | 1.97372 | Distal Intergenic | 9 | 32393985 | 32399192 | 5208 | 1 | ENSMUST00000172015.1 | 14301 | Kcnj1 | ENSMUSG00000041248.8 |
| chr9 | 32409243 | 32409535 | 3.92551 | 5.39552 | 2.08923 | Distal Intergenic | 9 | 32393985 | 32399192 | 5208 | 1 | ENSMUST00000172015.1 | 15259 | Kcnj1 | ENSMUSG00000041248.8 |
| chr4 | 8238620 | 8239118 | 4.16774 | 5.42773 | 2.06201 | Promoter | 4 | 8143362 | 8239041 | 95680 | 2 | ENSMUST00000066674.7 | 0 | Car8 | ENSMUSG00000041261.9 |
| chr3 | 89092265 | 89092859 | 3.36538 | 3.96275 | 1.14986 | Promoter | 3 | 89083981 | 89093363 | 9383 | 2 | ENSMUST00000052539.7 | 504 | Rusc1 | ENSMUSG00000041263.9 |
| chr4 | 6989304 | 6991542 | 3.82058 | 4.83073 | 1.71055 | Promoter | 4 | 6686353 | 6990799 | 304447 | 2 | ENSMUST00000039987.3 | 0 | Tox | ENSMUSG00000041272.6 |
| chr11 | 43768110 | 43768369 | 3.46228 | 4.19693 | 1.25981 | Distal Intergenic | 11 | 43729512 | 43748008 | 18497 | 2 | ENSMUST00000109278.2 | -20103 | Ttc1 | ENSMUSG00000041278.5 |
| chr11 | 43775857 | 43776715 | 3.55338 | 4.25669 | 1.34341 | 3' UTR | 11 | 43729512 | 43748008 | 18497 | 2 | ENSMUST00000109278.2 | -27850 | Ttc1 | ENSMUSG00000041278.5 |
| chr6 | 18170022 | 18170995 | 3.98686 | 5.17069 | 1.91025 | Promoter | 6 | 18170687 | 18322767 | 152081 | 1 | ENSMUST00000115405.3 | 0 | Cftr | ENSMUSG00000041301.10 |
| chr1 | 54446026 | 54446308 | 3.36538 | 3.87562 | 1.01733 | Distal Intergenic | 1 | 54396004 | 54438971 | 42968 | 2 | ENSMUST00000041638.7 | -7056 | Gtf3c3 | ENSMUSG00000041303.7 |
| chr7 | 139582013 | 139582624 | 3.79323 | 4.90759 | 1.72476 | Promoter | 7 | 139581220 | 139582790 | 1571 | 2 | ENSMUST00000106095.2 | 166 | Nkx6-2 | ENSMUSG00000041309.12 |
| chr7 | 139585127 | 139585236 | 4.21272 | 5.58564 | 2.19556 | Distal Intergenic | 7 | 139579376 | 139582797 | 3422 | 2 | ENSMUST00000097974.3 | -2331 | Nkx6-2 | ENSMUSG00000041309.12 |
| chr7 | 139585549 | 139585638 | 3.5839 | 4.39288 | 1.38245 | Distal Intergenic | 7 | 139579376 | 139582797 | 3422 | 2 | ENSMUST00000097974.3 | -2753 | Nkx6-2 | ENSMUSG00000041309.12 |
| chr7 | 139586439 | 139586838 | 3.66247 | 4.47976 | 1.48719 | Distal Intergenic | 7 | 139579376 | 139582797 | 3422 | 2 | ENSMUST00000097974.3 | -3643 | Nkx6-2 | ENSMUSG00000041309.12 |
| chr7 | 139587513 | 139587714 | 3.69237 | 4.51992 | 1.51089 | Distal Intergenic | 7 | 139579376 | 139582797 | 3422 | 2 | ENSMUST00000097974.3 | -4717 | Nkx6-2 | ENSMUSG00000041309.12 |
| chr7 | 139588524 | 139588879 | 3.56532 | 4.25587 | 1.29662 | Distal Intergenic | 7 | 139579376 | 139582797 | 3422 | 2 | ENSMUST00000097974.3 | -5728 | Nkx6-2 | ENSMUSG00000041309.12 |
| chr13 | 16006212 | 16006517 | 3.31582 | 3.82262 | 1.08926 | Distal Intergenic | 13 | 16011851 | 16027188 | 15338 | 1 | ENSMUST00000042603.7 | -5334 | Inhba | ENSMUSG00000041324.8 |
| chr13 | 16009930 | 16010182 | 3.43879 | 4.0656 | 1.23748 | Distal Intergenic | 13 | 16011851 | 16027188 | 15338 | 1 | ENSMUST00000042603.7 | -1669 | Inhba | ENSMUSG00000041324.8 |
| chr13 | 16011966 | 16012072 | 4.43791 | 6.09023 | 2.59172 | Promoter | 13 | 16011851 | 16027188 | 15338 | 1 | ENSMUST00000042603.7 | 116 | Inhba | ENSMUSG00000041324.8 |
| chr13 | 16015833 | 16016677 | 3.45121 | 4.09887 | 1.19298 | Intron (ENSMUST00000042603.7/ENSMUSG00000041324.8, intron 3 of 4) | 13 | 16014475 | 16027211 | 12737 | 1 | ENSMUST00000164993.1 | 1359 | Inhba | ENSMUSG00000041324.8 |
| chr13 | 16017977 | 16019064 | 3.64995 | 4.47318 | 1.44903 | Intron (ENSMUST00000042603.7/ENSMUSG00000041324.8, intron 4 of 4) | 13 | 16014475 | 16027211 | 12737 | 1 | ENSMUST00000164993.1 | 3503 | Inhba | ENSMUSG00000041324.8 |
| chr12 | 105584498 | 105584646 | 3.50967 | 4.23965 | 1.29078 | Intron (ENSMUST00000001652.5/ENSMUSG00000021070.5, intron 1 of 2) | 12 | 105603085 | 105605339 | 2255 | 1 | ENSMUST00000182899.1 | -18439 | Bdkrb1 | ENSMUSG00000041347.5 |
| chr12 | 105586369 | 105586687 | 3.65158 | 4.39585 | 1.32685 | Intron (ENSMUST00000001652.5/ENSMUSG00000021070.5, intron 1 of 2) | 12 | 105603085 | 105605339 | 2255 | 1 | ENSMUST00000182899.1 | -16398 | Bdkrb1 | ENSMUSG00000041347.5 |
| chr4 | 137664682 | 137665244 | 4.00413 | 5.13891 | 1.93284 | Promoter | 4 | 137664726 | 137716570 | 51845 | 1 | ENSMUST00000139831.2 | 0 | Rap1gap | ENSMUSG00000041351.11 |
| chr4 | 137676570 | 137676724 | 3.46852 | 4.08128 | 1.19531 | Intron (ENSMUST00000139831.2/ENSMUSG00000041351.11, intron 1 of 8) | 4 | 137680753 | 137715790 | 35038 | 1 | ENSMUST00000152567.2 | -4029 | Rap1gap | ENSMUSG00000041351.11 |
| chr4 | 137680492 | 137680678 | 3.34408 | 3.86806 | 1.10345 | Promoter | 4 | 137680753 | 137715790 | 35038 | 1 | ENSMUST00000152567.2 | -75 | Rap1gap | ENSMUSG00000041351.11 |
| chr4 | 137681704 | 137681927 | 3.78086 | 4.70155 | 1.61327 | Promoter | 4 | 137681667 | 137729860 | 48194 | 1 | ENSMUST00000047243.6 | 38 | Rap1gap | ENSMUSG00000041351.11 |
| chr4 | 137683188 | 137683469 | 3.61941 | 4.34817 | 1.33373 | Exon (ENSMUST00000150928.2/ENSMUSG00000041351.11, exon 2 of 11) | 4 | 137681667 | 137729860 | 48194 | 1 | ENSMUST00000047243.6 | 1522 | Rap1gap | ENSMUSG00000041351.11 |
| chr4 | 137707375 | 137707675 | 3.29609 | 3.91608 | 1.0491 | Promoter | 4 | 137707522 | 137729861 | 22340 | 1 | ENSMUST00000097837.5 | 0 | Rap1gap | ENSMUSG00000041351.11 |
| chr4 | 137710554 | 137710771 | 3.54043 | 4.20517 | 1.24947 | Exon (ENSMUST00000139831.2/ENSMUSG00000041351.11, exon 4 of 9) | 4 | 137707522 | 137729861 | 22340 | 1 | ENSMUST00000097837.5 | 3033 | Rap1gap | ENSMUSG00000041351.11 |
| chr4 | 137719451 | 137719534 | 4.53747 | 6.25137 | 2.68385 | Promoter | 4 | 137718872 | 137728358 | 9487 | 1 | ENSMUST00000141306.1 | 580 | Rap1gap | ENSMUSG00000041351.11 |
| chr4 | 137723769 | 137723909 | 3.85266 | 4.87319 | 1.71367 | Promoter | 4 | 137723989 | 137729861 | 5873 | 1 | ENSMUST00000148054.2 | -80 | Rap1gap | ENSMUSG00000041351.11 |
| chr4 | 137728178 | 137729938 | 4.13893 | 5.58233 | 2.22899 | Promoter | 4 | 137728181 | 137728819 | 639 | 1 | ENSMUST00000123134.1 | 0 | Rap1gap | ENSMUSG00000041351.11 |
| chr4 | 137730980 | 137731616 | 3.45304 | 4.10908 | 1.22681 | Distal Intergenic | 4 | 137728211 | 137729702 | 1492 | 1 | ENSMUST00000144059.1 | 2770 | Rap1gap | ENSMUSG00000041351.11 |
| chr12 | 105271266 | 105271528 | 3.51983 | 4.17583 | 1.25319 | Distal Intergenic | 12 | 105217498 | 105222793 | 5296 | 2 | ENSMUST00000175652.1 | -48474 | Tcl1 | ENSMUSG00000041359.9 |
| chr19 | 59075292 | 59075985 | 3.43069 | 4.02769 | 1.18899 | Promoter | 19 | 58973358 | 59076069 | 102712 | 2 | ENSMUST00000047511.8 | 84 | 4930506M07Rik | ENSMUSG00000041362.11 |
| chr19 | 59076584 | 59076674 | 3.60087 | 4.34455 | 1.39877 | Promoter | 19 | 58973358 | 59076069 | 102712 | 2 | ENSMUST00000163821.1 | -516 | 4930506M07Rik | ENSMUSG00000041362.11 |

| | | | | | | | | | | | | | | | |
|---|---|---|---|---|---|---|---|---|---|---|---|---|---|---|---|
| chr19 | 59079343 | 59079598 | 4.37969 | 5.9378 | 2.47895 | Distal Intergenic | 19 | 58973358 | 59076069 | 102712 | 2 | ENSMUST00000163821.1 | -3275 | 4930506M07Rik | ENSMUSG00000041362.11 |
| chr6 | 120294692 | 120295675 | 3.69541 | 4.57429 | 1.48667 | Promoter | 6 | 120203094 | 120294559 | 91466 | 2 | ENSMUST00000057283.7 | -134 | B4galnt3 | ENSMUSG00000041372.9 |
| chr6 | 120296488 | 120297024 | 3.50935 | 4.18006 | 1.23637 | Distal Intergenic | 6 | 120203094 | 120294559 | 91466 | 2 | ENSMUST00000057283.7 | -1930 | B4galnt3 | ENSMUSG00000041372.9 |
| chr6 | 120303051 | 120303378 | 4.05686 | 5.18247 | 1.84754 | Distal Intergenic | 6 | 120203094 | 120294559 | 91466 | 2 | ENSMUST00000057283.7 | -8493 | B4galnt3 | ENSMUSG00000041372.9 |
| chr14 | 34661142 | 34661335 | 3.31383 | 3.81343 | 1.07067 | Distal Intergenic | 14 | 34673928 | 34747983 | 74056 | 1 | ENSMUST00000048263.8 | -12593 | Wapal | ENSMUSG00000041408.11 |
| chr12 | 104637201 | 104637373 | 3.77401 | 4.75183 | 1.65768 | Distal Intergenic | 12 | 104687742 | 104751952 | 64211 | 2 | ENSMUST00000041987.6 | 114579 | Dicer1 | ENSMUSG00000041415.9 |
| chr12 | 104637753 | 104637863 | 3.49923 | 4.16934 | 1.30302 | Distal Intergenic | 12 | 104687742 | 104751952 | 64211 | 2 | ENSMUST00000041987.6 | 114089 | Dicer1 | ENSMUSG00000041415.9 |
| chr13 | 101800976 | 101801051 | 3.55387 | 4.23145 | 1.261 | Distal Intergenic | 13 | 101680563 | 101768217 | 87655 | 2 | ENSMUST00000055518.7 | -32760 | Pik3r1 | ENSMUSG00000041417.10 |
| chr13 | 101801637 | 101801980 | 3.58446 | 4.28219 | 1.27597 | Distal Intergenic | 13 | 101680563 | 101768217 | 87655 | 2 | ENSMUST00000055518.7 | -33421 | Pik3r1 | ENSMUSG00000041417.10 |
| chr7 | 16158962 | 16159632 | 3.71359 | 4.64843 | 1.52902 | 3' UTR | 7 | 16175090 | 16178793 | 3704 | 1 | ENSMUST00000176342.2 | -15458 | Meis3 | ENSMUSG00000041420.13 |
| chr3 | 88365590 | 88365904 | 3.5253 | 4.23393 | 1.32349 | Promoter | 3 | 88366722 | 88368541 | 1820 | 1 | ENSMUST00000147818.1 | -818 | Paqr6 | ENSMUSG00000041423.11 |
| chr3 | 88366235 | 88366363 | 3.61074 | 4.56817 | 1.51759 | Promoter | 3 | 88366722 | 88368541 | 1820 | 1 | ENSMUST00000147818.1 | -359 | Paqr6 | ENSMUSG00000041423.11 |
| chr9 | 32032843 | 32032923 | 3.89027 | 4.87331 | 1.69296 | Distal Intergenic | 9 | 32116136 | 32268446 | 152311 | 1 | ENSMUST00000174641.2 | -83213 | Arhgap32 | ENSMUSG00000041444.9 |
| chr5 | 146585074 | 146585149 | 3.55013 | 4.26618 | 1.36083 | Promoter | 5 | 146582151 | 146585239 | 3089 | 2 | ENSMUST00000036211.6 | 90 | Gpr12 | ENSMUSG00000041468.6 |
| chr5 | 146586421 | 146587100 | 3.3221 | 3.84003 | 1.11075 | Distal Intergenic | 5 | 146582151 | 146585239 | 3089 | 2 | ENSMUST00000036211.6 | -1183 | Gpr12 | ENSMUSG00000041468.6 |
| chr5 | 146644739 | 146645075 | 3.46493 | 4.09223 | 1.24241 | Distal Intergenic | 5 | 146582151 | 146585239 | 3089 | 2 | ENSMUST00000036211.6 | -59501 | Gpr12 | ENSMUSG00000041468.6 |
| chr6 | 119224959 | 119225120 | 3.86092 | 4.89907 | 1.70934 | Distal Intergenic | 6 | 119217544 | 119221614 | 4071 | 1 | ENSMUST00000127059.1 | 7416 | Dcp1b | ENSMUSG00000041477.9 |
| chr18 | 63386048 | 63386167 | 3.557 | 4.23503 | 1.2665 | Promoter | 18 | 63059718 | 63386457 | 326740 | 2 | ENSMUST00000183217.2 | 290 | Piezo2 | ENSMUSG00000041482.11 |
| chr18 | 63386478 | 63387257 | 3.92535 | 5.10157 | 1.87226 | Promoter | 18 | 63010213 | 63387183 | 376971 | 2 | ENSMUST00000047480.7 | 0 | Piezo2 | ENSMUSG00000041482.11 |
| chr5 | 146515915 | 146516079 | 3.44902 | 4.06321 | 1.21372 | Exon (ENSMUST00000036715.10/ENSMUSG00000041505.10, exon 4 of 5) | 5 | 146514151 | 146516724 | 2574 | 1 | ENSMUST00000077133.6 | 1765 | Gm3402 | ENSMUSG00000041505.10 |
| chr8 | 120735031 | 120735402 | 3.38756 | 3.91923 | 1.05293 | Promoter | 8 | 120736358 | 120756694 | 20337 | 1 | ENSMUST00000047737.4 | -956 | Irf8 | ENSMUSG00000041515.4 |
| chr8 | 120735901 | 120736431 | 4.39629 | 5.89282 | 2.41202 | Promoter | 8 | 120736358 | 120756694 | 20337 | 1 | ENSMUST00000047737.4 | 0 | Irf8 | ENSMUSG00000041515.4 |
| chr8 | 120736762 | 120737121 | 4.28584 | 5.65544 | 2.22098 | Promoter | 8 | 120736385 | 120753478 | 17094 | 1 | ENSMUST00000162658.2 | 378 | Irf8 | ENSMUSG00000041515.4 |
| chr8 | 120737502 | 120738023 | 3.66496 | 4.66853 | 1.57213 | Promoter | 8 | 120738639 | 120755581 | 16943 | 1 | ENSMUST00000162001.2 | -616 | Irf8 | ENSMUSG00000041515.4 |
| chr8 | 120768783 | 120768858 | 3.31017 | 3.80647 | 1.06149 | Intron (ENSMUST00000127664.1/ENSMUSG00000092329.1, intron 1 of 15) | 8 | 120752209 | 120753711 | 1503 | 1 | ENSMUST00000160594.1 | 16575 | Irf8 | ENSMUSG00000041515.4 |
| chr8 | 120800832 | 120801182 | 3.98416 | 5.04325 | 1.79911 | Intron (ENSMUST00000127664.1/ENSMUSG00000092329.1, intron 1 of 15) | 8 | 120752209 | 120753711 | 1503 | 1 | ENSMUST00000160594.1 | 48624 | Irf8 | ENSMUSG00000041515.4 |
| chr4 | 148220702 | 148220914 | 3.80034 | 4.68756 | 1.54038 | Distal Intergenic | 4 | 148258216 | 148260751 | 2536 | 2 | ENSMUST00000143851.1 | 39837 | Ptchd2 | ENSMUSG00000041544.11 |
| chr4 | 148221737 | 148222001 | 3.58989 | 4.33707 | 1.40233 | Distal Intergenic | 4 | 148258216 | 148260751 | 2536 | 2 | ENSMUST00000143851.1 | 38750 | Ptchd2 | ENSMUSG00000041544.11 |
| chr4 | 148222984 | 148223219 | 4.067 | 5.4031 | 2.02947 | Distal Intergenic | 4 | 148258216 | 148260751 | 2536 | 2 | ENSMUST00000143851.1 | 37532 | Ptchd2 | ENSMUSG00000041544.11 |
| chr4 | 148271694 | 148272572 | 3.60481 | 4.39531 | 1.36667 | 5' UTR | 4 | 148259879 | 148261231 | 1353 | 2 | ENSMUST00000129734.1 | -10464 | Ptchd2 | ENSMUSG00000041544.11 |
| chr4 | 148284707 | 148285103 | 3.82028 | 4.82581 | 1.67085 | Intron (ENSMUST00000047720.8/ENSMUSG00000041544.11, intron 1 of 20) | 4 | 148240264 | 148287965 | 47702 | 2 | ENSMUST00000047720.8 | 2862 | Ptchd2 | ENSMUSG00000041544.11 |
| chr4 | 148285597 | 148288628 | 3.67334 | 4.50958 | 1.47134 | Promoter | 4 | 148240264 | 148287965 | 47702 | 2 | ENSMUST00000047720.8 | 0 | Ptchd2 | ENSMUSG00000041544.11 |
| chr4 | 148293886 | 148293966 | 4.02611 | 5.11794 | 1.81129 | Distal Intergenic | 4 | 148240264 | 148287965 | 47702 | 2 | ENSMUST00000047720.8 | -5922 | Ptchd2 | ENSMUSG00000041544.11 |
| chr4 | 148164985 | 148165249 | 3.41123 | 4.04658 | 1.13544 | Exon (ENSMUST00000047951.8/ENSMUSG00000041556.8, exon 4 of 6) | 4 | 148160621 | 148166424 | 5804 | 1 | ENSMUST00000047951.8 | 4365 | Fbxo2 | ENSMUSG00000041556.8 |
| chr4 | 148165625 | 148166714 | 3.89539 | 4.87229 | 1.66164 | 3' UTR | 4 | 148160621 | 148166424 | 5804 | 1 | ENSMUST00000047951.8 | 5005 | Fbxo2 | ENSMUSG00000041556.8 |
| chr4 | 148169363 | 148169680 | 3.32753 | 3.82215 | 1.02919 | Distal Intergenic | 4 | 148160621 | 148166424 | 5804 | 1 | ENSMUST00000047951.8 | 8743 | Fbxo2 | ENSMUSG00000041556.8 |
| chr4 | 148172563 | 148173402 | 3.54827 | 4.28956 | 1.34113 | Distal Intergenic | 4 | 148160621 | 148166424 | 5804 | 1 | ENSMUST00000047951.8 | 11943 | Fbxo2 | ENSMUSG00000041556.8 |
| chr4 | 148173823 | 148175098 | 3.94849 | 5.12711 | 1.89391 | Distal Intergenic | 4 | 148160621 | 148166424 | 5804 | 1 | ENSMUST00000047951.8 | 13203 | Fbxo2 | ENSMUSG00000041556.8 |
| chr4 | 148175663 | 148176708 | 3.45612 | 4.0545 | 1.16393 | Distal Intergenic | 4 | 148160621 | 148166424 | 5804 | 1 | ENSMUST00000047951.8 | 15043 | Fbxo2 | ENSMUSG00000041556.8 |
| chr17 | 68273837 | 68274323 | 3.60878 | 4.39042 | 1.4122 | Promoter | 17 | 68273797 | 68777961 | 504165 | 1 | ENSMUST00000093007.4 | 41 | L3mbtl4 | ENSMUSG00000041565.11 |
| chr17 | 68274683 | 68275005 | 3.95653 | 5.09224 | 1.87911 | Promoter | 17 | 68273797 | 68777961 | 504165 | 1 | ENSMUST00000093007.4 | 887 | L3mbtl4 | ENSMUSG00000041565.11 |
| chr7 | 15877610 | 15877833 | 3.17115 | 3.691 | 1.01733 | Intron (ENSMUST00000044434.7/ENSMUSG00000041578.10, intron 1 of 3) | 7 | 15867226 | 15879844 | 12619 | 2 | ENSMUST00000172758.2 | 2011 | Crx | ENSMUSG00000041578.10 |
| chr11 | 113923198 | 113923694 | 3.57063 | 4.25105 | 1.2487 | Intron (ENSMUST00000041627.8/ENSMUSG00000041592.11, intron 2 of 44) | 11 | 113806341 | 113826219 | 19879 | 2 | ENSMUST00000125879.1 | -96980 | Sdk2 | ENSMUSG00000041592.11 |
| chr11 | 113945642 | 113946144 | 3.64876 | 4.44372 | 1.40025 | Intron (ENSMUST00000041627.8/ENSMUSG00000041592.11, intron 1 of 44) | 11 | 113806341 | 113826219 | 19879 | 2 | ENSMUST00000125879.1 | -119424 | Sdk2 | ENSMUSG00000041592.11 |
| chr11 | 113991308 | 113991464 | 3.38316 | 3.92647 | 1.10163 | Intron (ENSMUST00000041627.8/ENSMUSG00000041592.11, intron 1 of 44) | 11 | 113848562 | 114065975 | 217414 | 2 | ENSMUST00000141943.1 | 74511 | Sdk2 | ENSMUSG00000041592.11 |
| chr11 | 114041865 | 114042021 | 3.32968 | 3.82186 | 1.01733 | Intron (ENSMUST00000041627.8/ENSMUSG00000041592.11, intron 1 of 44) | 11 | 113848562 | 114065975 | 217414 | 2 | ENSMUST00000141943.1 | 23954 | Sdk2 | ENSMUSG00000041592.11 |
| chr11 | 114048097 | 114048221 | 3.45021 | 4.31329 | 1.33456 | Intron (ENSMUST00000041627.8/ENSMUSG00000041592.11, intron 1 of 44) | 11 | 113848562 | 114065975 | 217414 | 2 | ENSMUST00000141943.1 | 17754 | Sdk2 | ENSMUSG00000041592.11 |
| chr11 | 114048909 | 114049359 | 3.77287 | 4.65551 | 1.56049 | Intron (ENSMUST00000041627.8/ENSMUSG00000041592.11, intron 1 of 44) | 11 | 113848562 | 114065975 | 217414 | 2 | ENSMUST00000141943.1 | 16616 | Sdk2 | ENSMUSG00000041592.11 |
| chr11 | 114050572 | 114051289 | 3.41651 | 3.998 | 1.13725 | Intron (ENSMUST00000041627.8/ENSMUSG00000041592.11, intron 1 of 44) | 11 | 113848562 | 114065975 | 217414 | 2 | ENSMUST00000141943.1 | 14686 | Sdk2 | ENSMUSG00000041592.11 |
| chr11 | 114064073 | 114064415 | 3.56355 | 4.31501 | 1.34969 | Intron (ENSMUST00000041627.8/ENSMUSG00000041592.11, intron 1 of 44) | 11 | 113848562 | 114065975 | 217414 | 2 | ENSMUST00000141943.1 | 1560 | Sdk2 | ENSMUSG00000041592.11 |
| chr11 | 114064853 | 114065141 | 3.50137 | 4.28557 | 1.27741 | Promoter | 11 | 113848562 | 114065975 | 217414 | 2 | ENSMUST00000141943.1 | 834 | Sdk2 | ENSMUSG00000041592.11 |
| chr11 | 114065835 | 114066906 | 3.42507 | 4.06807 | 1.1949 | Promoter | 11 | 113776374 | 114066218 | 289845 | 2 | ENSMUST00000041627.8 | 0 | Sdk2 | ENSMUSG00000041592.11 |
| chr11 | 114068837 | 114069004 | 3.75022 | 4.58933 | 1.47883 | Distal Intergenic | 11 | 113865600 | 114067046 | 201447 | 2 | ENSMUST00000155651.1 | -1792 | Sdk2 | ENSMUSG00000041592.11 |
| chr11 | 114071053 | 114071338 | 3.27561 | 3.74059 | 1.01733 | Distal Intergenic | 11 | 113865600 | 114067046 | 201447 | 2 | ENSMUST00000155651.1 | -4008 | Sdk2 | ENSMUSG00000041592.11 |
| chr11 | 114072142 | 114072559 | 3.64646 | 4.53285 | 1.48004 | Distal Intergenic | 11 | 113865600 | 114067046 | 201447 | 2 | ENSMUST00000155651.1 | -5097 | Sdk2 | ENSMUSG00000041592.11 |
| chr7 | 14623279 | 14623507 | 3.39083 | 3.9634 | 1.17912 | Promoter | 7 | 14530652 | 14622479 | 91828 | 2 | ENSMUST00000044683.6 | -801 | Nlrp5-ps | ENSMUSG00000041596.7 |
| chr1 | 136227965 | 136228585 | 3.95862 | 5.08009 | 1.89928 | Intron (ENSMUST00000120339.2/ENSMUSG00000041605.11, intron 1 of 9) | 1 | 136220278 | 136230289 | 10012 | 2 | ENSMUST00000150163.2 | 1704 | 5730559C18Rik | ENSMUSG00000041605.11 |
| chr1 | 136230190 | 136230418 | 3.63655 | 4.42295 | 1.44329 | Promoter | 1 | 136220278 | 136230289 | 10012 | 2 | ENSMUST00000150163.2 | 0 | 5730559C18Rik | ENSMUSG00000041605.11 |
| chr1 | 136231728 | 136231870 | 4.21781 | 5.64321 | 2.22288 | Intron (ENSMUST00000120339.2/ENSMUSG00000041605.11, intron 1 of 9) | 1 | 136220316 | 136230310 | 9995 | 2 | ENSMUST00000144464.1 | -1419 | 5730559C18Rik | ENSMUSG00000041605.11 |
| chr1 | 136232901 | 136233355 | 3.83117 | 4.85503 | 1.68521 | Promoter | 1 | 136226193 | 136234119 | 7927 | 2 | ENSMUST00000153910.1 | 764 | 5730559C18Rik | ENSMUSG00000041605.11 |
| chr1 | 136233922 | 136234001 | 3.83151 | 4.87518 | 1.68543 | Promoter | 1 | 136226193 | 136234119 | 7927 | 2 | ENSMUST00000153910.1 | 118 | 5730559C18Rik | ENSMUSG00000041605.11 |
| chr1 | 136234395 | 136234621 | 3.51745 | 4.46602 | 1.44929 | Promoter | 1 | 136213531 | 136234264 | 20734 | 2 | ENSMUST00000120339.2 | -132 | 5730559C18Rik | ENSMUSG00000041605.11 |
| chr18 | 82475205 | 82475692 | 3.66477 | 4.63469 | 1.53047 | Promoter | 18 | 82475370 | 82584217 | 108848 | 1 | ENSMUST00000143506.2 | 0 | Mbp | ENSMUSG00000041607.11 |
| chr18 | 82476078 | 82476361 | 3.70209 | 4.62821 | 1.54826 | Promoter | 18 | 82475370 | 82584217 | 108848 | 1 | ENSMUST00000143506.2 | 709 | Mbp | ENSMUSG00000041607.11 |
| chr18 | 82545994 | 82546069 | 3.68175 | 4.46982 | 1.42184 | Intron (ENSMUST00000091789.5/ENSMUSG00000041607.11, intron 3 of 3) | 18 | 82554463 | 82585637 | 31175 | 1 | ENSMUST00000062446.8 | -8394 | Mbp | ENSMUSG00000041607.11 |
| chr18 | 82556693 | 82556935 | 3.38193 | 3.9006 | 1.01733 | 3' UTR | 18 | 82554583 | 82585626 | 31044 | 1 | ENSMUST00000132369.1 | 2111 | Mbp | ENSMUSG00000041607.11 |
| chr18 | 82559171 | 82559374 | 3.44209 | 4.01398 | 1.09742 | Intron (ENSMUST00000047865.8/ENSMUSG00000041607.11, intron 4 of 6) | 18 | 82563262 | 82584287 | 21026 | 1 | ENSMUST00000150952.1 | -3888 | Mbp | ENSMUSG00000041607.11 |
| chr18 | 82560327 | 82560699 | 3.70515 | 4.50224 | 1.41324 | Intron (ENSMUST00000047865.8/ENSMUSG00000041607.11, intron 4 of 6) | 18 | 82563262 | 82584287 | 21026 | 1 | ENSMUST00000150952.1 | -2563 | Mbp | ENSMUSG00000041607.11 |
| chr18 | 82561197 | 82561565 | 3.66525 | 4.42067 | 1.34362 | Intron (ENSMUST00000047865.8/ENSMUSG00000041607.11, intron 4 of 6) | 18 | 82563262 | 82584287 | 21026 | 1 | ENSMUST00000150952.1 | -1697 | Mbp | ENSMUSG00000041607.11 |
| chr9 | 120540537 | 120540899 | 3.50383 | 4.24541 | 1.31584 | Promoter | 9 | 120539818 | 120568318 | 28501 | 1 | ENSMUST00000047687.7 | 720 | Entpd3 | ENSMUSG00000041608.7 |
| chr5 | 115728595 | 115728804 | 3.50823 | 4.23742 | 1.30271 | Intron (ENSMUST00000055408.7/ENSMUSG00000041609.11, intron 1 of 8) | 5 | 115670185 | 115731367 | 61183 | 2 | ENSMUST00000141950.1 | 2563 | Ccdc64 | ENSMUSG00000041609.11 |
| chr5 | 115730637 | 115731066 | 3.84945 | 4.86203 | 1.70118 | Promoter | 5 | 115670185 | 115731367 | 61183 | 2 | ENSMUST00000141950.1 | 301 | Ccdc64 | ENSMUSG00000041609.11 |
| chr9 | 3531092 | 3531485 | 3.69538 | 4.48958 | 1.41519 | Promoter | 9 | 3532354 | 3897342 | 364989 | 1 | ENSMUST00000115733.1 | -869 | Gucy1a2 | ENSMUSG00000041624.9 |
| chr9 | 3532071 | 3532146 | 3.42475 | 4.00571 | 1.11661 | Promoter | 9 | 3532354 | 3897342 | 364989 | 1 | ENSMUST00000115733.1 | -208 | Gucy1a2 | ENSMUSG00000041624.9 |
| chr1 | 136131534 | 136131709 | 3.96203 | 5.26399 | 1.93931 | Promoter | 1 | 136131454 | 136149993 | 18540 | 1 | ENSMUST00000122892.1 | 81 | Kif21b | ENSMUSG00000041642.13 |
| chr12 | 103242093 | 103242198 | 3.60958 | 4.38392 | 1.40318 | Promoter | 12 | 103196908 | 103242146 | 45239 | 2 | ENSMUST00000127843.2 | 0 | Prima1 | ENSMUSG00000041669.10 |
| chr12 | 103242642 | 103242979 | 3.70406 | 4.68175 | 1.58156 | Promoter | 12 | 103196908 | 103242150 | 45243 | 2 | ENSMUST00000074416.4 | -493 | Prima1 | ENSMUSG00000041669.10 |
| chr12 | 103243699 | 103244137 | 3.53624 | 4.3149 | 1.29404 | Distal Intergenic | 12 | 103196908 | 103242150 | 45243 | 2 | ENSMUST00000074416.4 | -1550 | Prima1 | ENSMUSG00000041669.10 |
| chr1 | 22533019 | 22533987 | 3.89951 | 5.13257 | 1.86755 | Exon (ENSMUST00000081544.7/ENSMUSG00000041670.10, exon 8 of 32) | 1 | 22288422 | 22432843 | 144422 | 2 | ENSMUST00000185942.1 | -100177 | Rims1 | ENSMUSG00000041670.10 |
| chr1 | 22805545 | 22806285 | 3.66928 | 4.61865 | 1.55248 | Promoter | 1 | 22286251 | 22805724 | 519474 | 2 | ENSMUST00000081544.7 | 0 | Rims1 | ENSMUSG00000041670.10 |
| chr1 | 22806658 | 22806974 | 3.72776 | 4.60307 | 1.56885 | Promoter | 1 | 22468191 | 22805724 | 337534 | 2 | ENSMUST00000097808.5 | -935 | Rims1 | ENSMUSG00000041670.10 |
| chr11 | 112620992 | 112621284 | 3.54781 | 4.21852 | 1.25564 | Distal Intergenic | 11 | 112664839 | 112669398 | 4560 | 2 | ENSMUST00000141399.1 | 48114 | BC006965 | ENSMUSG00000041674.11 |
| chr11 | 112679646 | 112679855 | 3.34348 | 3.84259 | 1.01733 | Intron (ENSMUST00000124028.2/ENSMUSG00000041674.11, intron 2 of 5) | 11 | 112669201 | 112677220 | 8020 | 2 | ENSMUST00000133231.2 | -2427 | BC006965 | ENSMUSG00000041674.11 |
| chr11 | 112773666 | 112774062 | 3.66986 | 4.62922 | 1.52493 | Intron (ENSMUST00000137081.2/ENSMUSG00000041674.11, intron 3 of 5) | 11 | 112687645 | 112781976 | 94332 | 2 | ENSMUST00000137081.2 | 7914 | BC006965 | ENSMUSG00000041674.11 |
| chr11 | 112775325 | 112777195 | 3.42587 | 4.07371 | 1.16061 | Exon (ENSMUST00000137081.2/ENSMUSG00000041674.11, exon 2 of 6) | 11 | 112687645 | 112781976 | 94332 | 2 | ENSMUST00000137081.2 | 4781 | BC006965 | ENSMUSG00000041674.11 |
| chr11 | 112777883 | 112778290 | 3.46566 | 4.05856 | 1.12857 | Intron (ENSMUST00000137081.2/ENSMUSG00000041674.11, intron 1 of 5) | 11 | 112687645 | 112781976 | 94332 | 2 | ENSMUST00000137081.2 | 3686 | BC006965 | ENSMUSG00000041674.11 |
| chr11 | 112779241 | 112781594 | 3.87624 | 4.92617 | 1.74766 | Promoter | 11 | 112687645 | 112781976 | 94332 | 2 | ENSMUST00000137081.2 | 382 | BC006965 | ENSMUSG00000041674.11 |
| chr11 | 111067005 | 111067126 | 3.58076 | 4.27974 | 1.28775 | Promoter | 11 | 111066164 | 111076821 | 10658 | 1 | ENSMUST00000042970.2 | 842 | Kcnj2 | ENSMUSG00000041695.2 |
| chr11 | 111067980 | 111068645 | 4.15057 | 5.48781 | 2.17412 | Intron (ENSMUST00000042970.2/ENSMUSG00000041695.2, intron 1 of 1) | 11 | 111066164 | 111076821 | 10658 | 1 | ENSMUST00000042970.2 | 1817 | Kcnj2 | ENSMUSG00000041695.2 |
| chr11 | 111088356 | 111089261 | 3.6216 | 4.39491 | 1.39747 | Distal Intergenic | 11 | 111066164 | 111076821 | 10658 | 1 | ENSMUST00000042970.2 | 22193 | Kcnj2 | ENSMUSG00000041695.2 |
| chr14 | 122457331 | 122458283 | 3.72286 | 4.61194 | 1.57522 | 3' UTR | 14 | 122459193 | 122460021 | 829 | 2 | ENSMUST00000143084.1 | 1738 | Zic5 | ENSMUSG00000041703.7 |
| chr14 | 122458933 | 122459600 | 3.61068 | 4.53391 | 1.48931 | Promoter | 14 | 122459193 | 122460021 | 829 | 2 | ENSMUST00000143084.1 | 421 | Zic5 | ENSMUSG00000041703.7 |
| chr14 | 122460446 | 122460521 | 3.47585 | 4.11697 | 1.26566 | Promoter | 14 | 122459193 | 122460021 | 829 | 2 | ENSMUST00000143084.1 | -426 | Zic5 | ENSMUSG00000041703.7 |
| chr14 | 122460907 | 122461065 | 4.13337 | 5.38941 | 2.05817 | Promoter | 14 | 122459193 | 122460021 | 829 | 2 | ENSMUST00000143084.1 | -887 | Zic5 | ENSMUSG00000041703.7 |
| chr14 | 122461727 | 122462873 | 3.70137 | 4.66219 | 1.59593 | Intron (ENSMUST00000039118.6/ENSMUSG00000041703.7, intron 1 of 1) | 14 | 122459193 | 122460021 | 829 | 2 | ENSMUST00000143084.1 | -1707 | Zic5 | ENSMUSG00000041703.7 |
| chr14 | 122463464 | 122465701 | 4.35093 | 6.02562 | 2.54789 | Promoter | 14 | 122456795 | 122465677 | 8883 | 2 | ENSMUST00000039118.6 | 0 | Zic5 | ENSMUSG00000041703.7 |
| chr14 | 122466178 | 122467736 | 4.14017 | 5.50431 | 2.15938 | Promoter | 14 | 122456795 | 122465677 | 8883 | 2 | ENSMUST00000039118.6 | -502 | Zic5 | ENSMUSG00000041703.7 |
| chr14 | 122468137 | 122469515 | 4.43416 | 6.07962 | 2.6718 | Downstream (<1kb) | 14 | 122456795 | 122465677 | 8883 | 2 | ENSMUST00000039118.6 | -2461 | Zic5 | ENSMUSG00000041703.7 |
| chr15 | 83771656 | 83771788 | 3.73411 | 4.56838 | 1.47297 | Distal Intergenic | 15 | 83779467 | 83792131 | 12665 | 1 | ENSMUST00000109469.1 | -7679 | Mpped1 | ENSMUSG00000041708.7 |
| chr15 | 83778946 | 83779704 | 3.87532 | 4.87914 | 1.65672 | Promoter | 15 | 83779467 | 83792131 | 12665 | 1 | ENSMUST00000109469.1 | 0 | Mpped1 | ENSMUSG00000041708.7 |
| chr15 | 83781319 | 83781537 | 3.99786 | 5.12664 | 1.83827 | Intron (ENSMUST00000109469.1/ENSMUSG00000041708.7, intron 1 of 1) | 15 | 83780023 | 83858484 | 78462 | 1 | ENSMUST00000046168.6 | 1297 | Mpped1 | ENSMUSG00000041708.7 |
| chr14 | 32588735 | 32588885 | 3.64951 | 4.45206 | 1.45246 | Distal Intergenic | 14 | 32599407 | 32649204 | 49798 | 1 | ENSMUST00000187377.1 | -10522 | Prrxl1 | ENSMUSG00000041730.8 |
| chr14 | 32593482 | 32593677 | 3.63192 | 4.42529 | 1.42908 | Distal Intergenic | 14 | 32599407 | 32649204 | 49798 | 1 | ENSMUST00000187377.1 | -5730 | Prrxl1 | ENSMUSG00000041730.8 |
| chr14 | 32596708 | 32596867 | 3.5063 | 4.23475 | 1.28616 | Distal Intergenic | 14 | 32599407 | 32649204 | 49798 | 1 | ENSMUST00000187377.1 | -2540 | Prrxl1 | ENSMUSG00000041730.8 |
| chr14 | 32597619 | 32598439 | 3.86216 | 4.93185 | 1.74088 | Promoter | 14 | 32599407 | 32649204 | 49798 | 1 | ENSMUST00000187377.1 | -968 | Prrxl1 | ENSMUSG00000041730.8 |
| chr14 | 32598943 | 32600746 | 4.24172 | 5.75092 | 2.39104 | Promoter | 14 | 32599407 | 32649204 | 49798 | 1 | ENSMUST00000187377.1 | 0 | Prrxl1 | ENSMUSG00000041730.8 |
| chr19 | 24814577 | 24815055 | 3.29599 | 3.77116 | 1.01733 | Intron (ENSMUST00000047666.4/ENSMUSG00000041731.8, intron 6 of 10) | 19 | 24816231 | 24861652 | 45422 | 2 | ENSMUST00000129127.1 | 46597 | Pgm5 | ENSMUSG00000041731.8 |
| chr19 | 24816298 | 24816428 | 3.66559 | 4.68268 | 1.54083 | Exon (ENSMUST00000047666.4/ENSMUSG00000041731.8, exon 5 of 11) | 19 | 24816231 | 24861652 | 45422 | 2 | ENSMUST00000129127.1 | 45224 | Pgm5 | ENSMUSG00000041731.8 |
| chr19 | 24860011 | 24860763 | 3.82104 | 4.93375 | 1.72097 | Promoter | 19 | 24816231 | 24861652 | 45422 | 2 | ENSMUST00000129127.1 | 889 | Pgm5 | ENSMUSG00000041731.8 |
| chr19 | 24861212 | 24861461 | 3.64839 | 4.41947 | 1.40835 | Promoter | 19 | 24816231 | 24861652 | 45422 | 2 | ENSMUST00000129127.1 | 191 | Pgm5 | ENSMUSG00000041731.8 |
| chr19 | 24861825 | 24862333 | 3.6963 | 4.49522 | 1.42482 | Promoter | 19 | 24683016 | 24861855 | 178840 | 2 | ENSMUST00000047666.4 | 0 | Pgm5 | ENSMUSG00000041731.8 |
| chr9 | 31462818 | 31462893 | 3.48161 | 4.19705 | 1.27497 | Intron (ENSMUST00000048050.7/ENSMUSG00000041737.7, intron 1 of 5) | 9 | 31426199 | 31464238 | 38040 | 2 | ENSMUST00000048050.7 | 1345 | Tmem45b | ENSMUSG00000041737.7 |
| chr9 | 31463937 | 31464409 | 3.50121 | 4.2712 | 1.33066 | Promoter | 9 | 31426199 | 31464238 | 38040 | 2 | ENSMUST00000048050.7 | 0 | Tmem45b | ENSMUSG00000041737.7 |
| chr9 | 31681391 | 31681590 | 3.28502 | 3.75468 | 1.01733 | Distal Intergenic | 9 | 31426199 | 31464238 | 38040 | 2 | ENSMUST00000048050.7 | -217154 | Tmem45b | ENSMUSG00000041737.7 |
| chr1 | 133162550 | 133162772 | 3.84799 | 5.01897 | 1.78356 | Distal Intergenic | 1 | 133181321 | 133301219 | 119899 | 1 | ENSMUST00000187285.1 | -18549 | Plekha6 | ENSMUSG00000041757.10 |
| chr1 | 133164188 | 133164263 | 3.69463 | 4.7245 | 1.58998 | Distal Intergenic | 1 | 133181321 | 133301219 | 119899 | 1 | ENSMUST00000187285.1 | -17058 | Plekha6 | ENSMUSG00000041757.10 |
| chr1 | 133298377 | 133298776 | 3.42932 | 4.01197 | 1.15776 | Promoter | 1 | 133297951 | 133301617 | 3667 | 1 | ENSMUST00000190208.1 | 427 | Plekha6 | ENSMUSG00000041757.10 |
| chr12 | 102128048 | 102128495 | 3.6024 | 4.3145 | 1.29684 | Promoter | 12 | 102128733 | 102267091 | 138359 | 1 | ENSMUST00000159329.2 | -238 | Slc24a4 | ENSMUSG00000041771.8 |
| chr12 | 102128843 | 102130436 | 3.74049 | 4.58474 | 1.48179 | Promoter | 12 | 102129174 | 102266749 | 137576 | 1 | ENSMUST00000079020.5 | 0 | Slc24a4 | ENSMUSG00000041771.8 |
| chr13 | 97243152 | 97243320 | 3.59097 | 4.34949 | 1.34877 | Intron (ENSMUST00000041623.7/ENSMUSG00000041773.7, intron 1 of 2) | 13 | 97241105 | 97253039 | 11935 | 1 | ENSMUST00000041623.7 | 2048 | Enc1 | ENSMUSG00000041773.7 |
| chr13 | 97244992 | 97245223 | 3.47027 | 4.19303 | 1.26256 | Exon (ENSMUST00000041623.7/ENSMUSG00000041773.7, exon 2 of 3) | 13 | 97241105 | 97253039 | 11935 | 1 | ENSMUST00000041623.7 | 3888 | Enc1 | ENSMUSG00000041773.7 |
| chr13 | 97257351 | 97257506 | 4.03155 | 5.14715 | 1.89569 | Distal Intergenic | 13 | 97241105 | 97253039 | 11935 | 1 | ENSMUST00000041623.7 | 16247 | Enc1 | ENSMUSG00000041773.7 |
| chr1 | 135817916 | 135818258 | 3.26627 | 3.89288 | 1.09648 | Promoter | 1 | 135818598 | 135833342 | 14745 | 1 | ENSMUST00000038760.9 | -340 | Lad1 | ENSMUSG00000041782.9 |
| chr1 | 135818634 | 135819126 | 4.18726 | 5.71476 | 2.31642 | Promoter | 1 | 135818598 | 135833342 | 14745 | 1 | ENSMUST00000038760.9 | 37 | Lad1 | ENSMUSG00000041782.9 |
| chr1 | 135819721 | 135820236 | 3.65602 | 4.49336 | 1.42589 | Intron (ENSMUST00000038760.9/ENSMUSG00000041782.9, intron 1 of 9) | 1 | 135818598 | 135833342 | 14745 | 1 | ENSMUST00000038760.9 | 1124 | Lad1 | ENSMUSG00000041782.9 |
| chr1 | 135820826 | 135821067 | 3.67504 | 4.46485 | 1.4367 | Intron (ENSMUST00000038760.9/ENSMUSG00000041782.9, intron 1 of 9) | 1 | 135818598 | 135833342 | 14745 | 1 | ENSMUST00000038760.9 | 2229 | Lad1 | ENSMUSG00000041782.9 |
| chr9 | 120301209 | 120301816 | 3.378 | 3.91163 | 1.03006 | Distal Intergenic | 9 | 120303936 | 120474841 | 170906 | 1 | ENSMUST00000048121.7 | -2120 | Myrip | ENSMUSG00000041794.7 |
| chr9 | 120302307 | 120302718 | 3.97325 | 5.02341 | 1.76403 | Distal Intergenic | 9 | 120303936 | 120474841 | 170906 | 1 | ENSMUST00000048121.7 | -1218 | Myrip | ENSMUSG00000041794.7 |
| chr9 | 120303026 | 120303386 | 3.90374 | 5.20348 | 1.91415 | Promoter | 9 | 120303936 | 120474841 | 170906 | 1 | ENSMUST00000048121.7 | -550 | Myrip | ENSMUSG00000041794.7 |
| chr9 | 120303706 | 120304666 | 3.99509 | 5.07268 | 1.80526 | Promoter | 9 | 120303936 | 120474841 | 170906 | 1 | ENSMUST00000048121.7 | 0 | Myrip | ENSMUSG00000041794.7 |
| chr13 | 97065087 | 97066229 | 3.45365 | 4.03697 | 1.11688 | Distal Intergenic | 13 | 97067287 | 97129521 | 62235 | 1 | ENSMUST00000169863.2 | -1058 | Fam169a | ENSMUSG00000041817.8 |
| chr13 | 97069988 | 97070113 | 3.57184 | 4.24823 | 1.23206 | Intron (ENSMUST00000169863.2/ENSMUSG00000041817.8, intron 1 of 12) | 13 | 97071643 | 97129521 | 57879 | 1 | ENSMUST00000042517.6 | -1530 | Fam169a | ENSMUSG00000041817.8 |
| chr13 | 97070586 | 97070974 | 4.03173 | 5.13745 | 1.83977 | Promoter | 13 | 97071643 | 97129521 | 57879 | 1 | ENSMUST00000042517.6 | -669 | Fam169a | ENSMUSG00000041817.8 |
| chr13 | 97071355 | 97072000 | 4.08452 | 5.40347 | 2.05384 | Promoter | 13 | 97071643 | 97129521 | 57879 | 1 | ENSMUST00000042517.6 | 0 | Fam169a | ENSMUSG00000041817.8 |
| chr17 | 6700309 | 6700401 | 3.29795 | 4.50121 | 1.45284 | Intron (ENSMUST00000115763.4/ENSMUSG00000041831.10, intron 3 of 7) | 17 | 6705830 | 6737836 | 32007 | 1 | ENSMUST00000159880.1 | -5429 | Sytl3 | ENSMUSG00000041831.10 |
| chr17 | 6702816 | 6703302 | 3.46736 | 4.58386 | 1.48995 | 5' UTR | 17 | 6705830 | 6737836 | 32007 | 1 | ENSMUST00000159880.1 | -2528 | Sytl3 | ENSMUSG00000041831.10 |
| chr3 | 84461571 | 84461765 | 3.56485 | 4.56177 | 1.47973 | Intron (ENSMUST00000091002.2/ENSMUSG00000041842.10, intron 4 of 11) | 3 | 84448809 | 84453763 | 4955 | 2 | ENSMUST00000125643.1 | -7809 | Fhdc1 | ENSMUSG00000041842.10 |
| chr3 | 84478082 | 84478959 | 3.87513 | 5.09268 | 1.85517 | Promoter | 3 | 84474523 | 84478982 | 4460 | 2 | ENSMUST00000131065.1 | 23 | Fhdc1 | ENSMUSG00000041842.10 |
| chr3 | 84479285 | 84479874 | 3.61475 | 4.61822 | 1.52735 | Promoter | 3 | 84444035 | 84479491 | 35457 | 2 | ENSMUST00000091002.2 | 0 | Fhdc1 | ENSMUSG00000041842.10 |
| chr3 | 84480506 | 84480788 | 3.37645 | 4.12648 | 1.22533 | Promoter | 3 | 84444038 | 84480429 | 36392 | 2 | ENSMUST00000107689.1 | -78 | Fhdc1 | ENSMUSG00000041842.10 |
| chr15 | 82907403 | 82907599 | 3.84807 | 4.88013 | 1.71717 | Intron (ENSMUST00000048966.5/ENSMUSG00000041852.9, intron 1 of 4) | 15 | 82808629 | 82912134 | 103506 | 2 | ENSMUST00000048966.5 | 4535 | Tcf20 | ENSMUSG00000041852.9 |
| chr15 | 82908462 | 82908887 | 3.39546 | 4.05458 | 1.19199 | Intron (ENSMUST00000048966.5/ENSMUSG00000041852.9, intron 1 of 4) | 15 | 82808629 | 82912134 | 103506 | 2 | ENSMUST00000048966.5 | 3247 | Tcf20 | ENSMUSG00000041852.9 |
| chr15 | 82909848 | 82910060 | 3.43856 | 4.02155 | 1.14229 | Intron (ENSMUST00000048966.5/ENSMUSG00000041852.9, intron 1 of 4) | 15 | 82808629 | 82912134 | 103506 | 2 | ENSMUST00000048966.5 | 2074 | Tcf20 | ENSMUSG00000041852.9 |
| chr15 | 82911598 | 82912143 | 3.7884 | 4.7077 | 1.6387 | Promoter | 15 | 82808629 | 82912134 | 103506 | 2 | ENSMUST00000048966.5 | 0 | Tcf20 | ENSMUSG00000041852.9 |
| chr15 | 82912458 | 82912535 | 3.98944 | 5.07931 | 1.86045 | Promoter | 15 | 82808629 | 82912134 | 103506 | 2 | ENSMUST00000109510.3 | -325 | Tcf20 | ENSMUSG00000041852.9 |
| chr15 | 82912874 | 82913110 | 3.27949 | 3.78158 | 1.05213 | Promoter | 15 | 82808629 | 82912134 | 103506 | 2 | ENSMUST00000109510.3 | -741 | Tcf20 | ENSMUSG00000041852.9 |
| chr15 | 82913655 | 82913871 | 3.45601 | 4.08642 | 1.25259 | Distal Intergenic | 15 | 82808629 | 82912134 | 103506 | 2 | ENSMUST00000109510.3 | -1522 | Tcf20 | ENSMUSG00000041852.9 |
| chr15 | 82916295 | 82916473 | 3.42246 | 4.22924 | 1.32369 | Distal Intergenic | 15 | 82808629 | 82912134 | 103506 | 2 | ENSMUST00000109510.3 | -4162 | Tcf20 | ENSMUSG00000041852.9 |
| chr15 | 82917434 | 82917785 | 4.07556 | 5.22167 | 1.90259 | Distal Intergenic | 15 | 82808629 | 82912134 | 103506 | 2 | ENSMUST00000109510.3 | -5301 | Tcf20 | ENSMUSG00000041852.9 |
| chr15 | 82918818 | 82919034 | 3.41987 | 4.04205 | 1.21088 | Distal Intergenic | 15 | 82808629 | 82912134 | 103506 | 2 | ENSMUST00000109510.3 | -6685 | Tcf20 | ENSMUSG00000041852.9 |
| chr15 | 82919350 | 82919671 | 3.52823 | 4.18454 | 1.20521 | Distal Intergenic | 15 | 82808629 | 82912134 | 103506 | 2 | ENSMUST00000109510.3 | -7217 | Tcf20 | ENSMUSG00000041852.9 |
| chr15 | 82921848 | 82922899 | 4.32799 | 5.78599 | 2.36125 | Distal Intergenic | 15 | 82808629 | 82912134 | 103506 | 2 | ENSMUST00000109510.3 | -9715 | Tcf20 | ENSMUSG00000041852.9 |
| chr15 | 82923791 | 82924231 | 3.7328 | 4.59992 | 1.49929 | Distal Intergenic | 15 | 82808629 | 82912134 | 103506 | 2 | ENSMUST00000109510.3 | -11658 | Tcf20 | ENSMUSG00000041852.9 |
| chr15 | 82925266 | 82925509 | 3.71924 | 4.52357 | 1.41142 | Distal Intergenic | 15 | 82808629 | 82912134 | 103506 | 2 | ENSMUST00000109510.3 | -13133 | Tcf20 | ENSMUSG00000041852.9 |
| chr15 | 82930326 | 82930717 | 3.4779 | 4.13769 | 1.26119 | Distal Intergenic | 15 | 82808629 | 82912134 | 103506 | 2 | ENSMUST00000109510.3 | -18193 | Tcf20 | ENSMUSG00000041852.9 |
| chr15 | 82931442 | 82931765 | 4.02033 | 5.35297 | 2.05161 | Distal Intergenic | 15 | 82808629 | 82912134 | 103506 | 2 | ENSMUST00000109510.3 | -19309 | Tcf20 | ENSMUSG00000041852.9 |
| chr15 | 82934703 | 82935949 | 3.51986 | 4.39627 | 1.40919 | Distal Intergenic | 15 | 82808629 | 82912134 | 103506 | 2 | ENSMUST00000109510.3 | -22570 | Tcf20 | ENSMUSG00000041852.9 |
| chr1 | 135368944 | 135369423 | 3.53312 | 4.20382 | 1.28429 | Downstream (1-2kb) | 1 | 135371060 | 135373450 | 2391 | 2 | ENSMUST00000159918.1 | 4027 | Shisa4 | ENSMUSG00000041889.7 |
| chr1 | 42918916 | 42919233 | 3.38496 | 3.94953 | 1.16274 | Distal Intergenic | 1 | 42952872 | 42955155 | 2284 | 1 | ENSMUST00000188621.1 | -33639 | Gpr45 | ENSMUSG00000041907.9 |
| chr1 | 42951027 | 42951127 | 4.51661 | 6.09602 | 2.51259 | Distal Intergenic | 1 | 42952872 | 42955155 | 2284 | 1 | ENSMUST00000188621.1 | -1745 | Gpr45 | ENSMUSG00000041907.9 |
| chr1 | 42952659 | 42953931 | 4.18166 | 5.48436 | 2.13016 | Promoter | 1 | 42952872 | 42955155 | 2284 | 1 | ENSMUST00000188621.1 | 0 | Gpr45 | ENSMUSG00000041907.9 |
| chr1 | 42954327 | 42954965 | 3.67042 | 4.50948 | 1.43952 | Exon (ENSMUST00000188621.1/ENSMUSG00000041907.9, exon 2 of 2) | 1 | 42953182 | 42953878 | 697 | 1 | ENSMUST00000187998.1 | 1146 | Gpr45 | ENSMUSG00000041907.9 |
| chr2 | 71528248 | 71529857 | 4.05253 | 5.37108 | 2.04667 | Promoter | 2 | 71528683 | 71530065 | 1383 | 1 | ENSMUST00000126400.1 | 0 | Dlx1 | ENSMUSG00000041911.3 |

| | | | | | | | | | | | | | | | |
|---|---|---|---|---|---|---|---|---|---|---|---|---|---|---|---|
| chr2 | 71530168 | 71530308 | 3.94911 | 5.46299 | 2.11119 | Promoter | 2 | 71529947 | 71533981 | 4035 | 1 | ENSMUST00000145088.1 | 222 | Dlx1 | ENSMUSG00000041911.3 |
| chr2 | 71532134 | 71533067 | 4.05174 | 5.31716 | 2.01481 | 3' UTR | 2 | 71530780 | 71532689 | 1910 | 1 | ENSMUST00000172629.1 | 1355 | Dlx1 | ENSMUSG00000041911.3 |
| chr2 | 71533427 | 71533610 | 3.79004 | 4.87162 | 1.67876 | 3' UTR | 2 | 71530780 | 71532689 | 1910 | 1 | ENSMUST00000172629.1 | 2648 | Dlx1 | ENSMUSG00000041911.3 |
| chr2 | 71530798 | 71531069 | 3.37323 | 3.92408 | 1.13977 | Promoter | 2 | 71530780 | 71532689 | 1910 | 1 | ENSMUST00000172629.1 | 19 | Dlx1 | ENSMUSG00000041911.3 |
| chr2 | 71531407 | 71531697 | 3.29283 | 3.77348 | 1.04188 | Promoter | 2 | 71530780 | 71532689 | 1910 | 1 | ENSMUST00000172629.1 | 628 | Dlx1 | ENSMUSG00000041911.3 |
| chr11 | 109470369 | 109471102 | 3.56279 | 4.23995 | 1.24797 | Intron (ENSMUST00000070152.6/ENSMUSG00000041920.9, intron 1 of 5) | 11 | 109459799 | 109472688 | 12890 | 2 | ENSMUST00000168740.1 | 1586 | Slc16a6 | ENSMUSG00000041920.9 |
| chr11 | 109472248 | 109472503 | 4.10508 | 5.73665 | 2.30466 | Promoter | 11 | 109459799 | 109472688 | 12890 | 2 | ENSMUST00000168740.1 | 185 | Slc16a6 | ENSMUSG00000041920.9 |
| chr11 | 109477133 | 109477319 | 3.43042 | 4.01576 | 1.16652 | Intron (ENSMUST00000152252.2/ENSMUSG00000020604.8, intron 1 of 3) | 11 | 109450855 | 109473598 | 22744 | 2 | ENSMUST00000070152.6 | -3536 | Slc16a6 | ENSMUSG00000041920.9 |
| chr2 | 71521366 | 71521453 | 4.63356 | 6.55126 | 2.85266 | Intron (ENSMUST00000037210.8/ENSMUSG00000041921.11, intron 6 of 9) | 2 | 71515412 | 71516284 | 873 | 1 | ENSMUST00000148164.1 | 5955 | Metap1d | ENSMUSG00000041921.11 |
| chr18 | 23036633 | 23036904 | 3.35371 | 3.85802 | 1.01733 | Intron (ENSMUST00000081423.7/ENSMUSG00000041923.10, intron 1 of 10) | 18 | 22693381 | 23038656 | 345276 | 2 | ENSMUST00000081423.7 | 1752 | Nol4 | ENSMUSG00000041923.10 |
| chr18 | 23038318 | 23038428 | 3.92622 | 5.3963 | 2.05949 | Promoter | 18 | 22693381 | 23038656 | 345276 | 2 | ENSMUST00000081423.7 | 228 | Nol4 | ENSMUSG00000041923.10 |
| chr18 | 23038951 | 23040584 | 3.86485 | 4.89751 | 1.76594 | Promoter | 18 | 22694524 | 23040078 | 345555 | 2 | ENSMUST00000164893.2 | 0 | Nol4 | ENSMUSG00000041923.10 |
| chr18 | 23041333 | 23043368 | 3.75237 | 4.65883 | 1.56474 | Promoter | 18 | 22693634 | 23041653 | 348020 | 2 | ENSMUST00000097651.4 | 0 | Nol4 | ENSMUSG00000041923.10 |
| chr18 | 23046154 | 23046229 | 3.48241 | 4.11477 | 1.2398 | Distal Intergenic | 18 | 22693634 | 23041653 | 348020 | 2 | ENSMUST00000097651.4 | -4502 | Nol4 | ENSMUSG00000041923.10 |
| chr5 | 114474518 | 114474979 | 3.58027 | 4.2903 | 1.33029 | Distal Intergenic | 5 | 114452420 | 114460572 | 8153 | 1 | ENSMUST00000139420.1 | 22099 | Mvk | ENSMUSG00000041939.9 |
| chr3 | 93362173 | 93362879 | 3.5849 | 4.30732 | 1.35306 | Distal Intergenic | 3 | 93393699 | 93399442 | 5744 | 1 | ENSMUST00000045912.2 | -30820 | Rptn | ENSMUSG00000041984.2 |
| chr9 | 53934713 | 53934837 | 3.64578 | 4.41119 | 1.40185 | Intron (ENSMUST00000166580.1/ENSMUSG00000041986.10, intron 3 of 9) | 9 | 53911460 | 53936834 | 25375 | 2 | ENSMUST00000166580.1 | 1997 | Elmod1 | ENSMUSG00000041986.10 |
| chr9 | 53973305 | 53973624 | 3.92521 | 5.10069 | 1.85172 | Intron (ENSMUST00000048409.8/ENSMUSG00000041986.10, intron 1 of 10) | 9 | 53911460 | 53975301 | 63842 | 2 | ENSMUST00000048409.8 | 1677 | Elmod1 | ENSMUSG00000041986.10 |
| chr9 | 53973961 | 53975935 | 4.0423 | 5.22299 | 1.92515 | Promoter | 9 | 53911460 | 53975301 | 63842 | 2 | ENSMUST00000048409.8 | 0 | Elmod1 | ENSMUSG00000041986.10 |
| chr9 | 53977099 | 53977242 | 4.05374 | 5.18968 | 1.85983 | Distal Intergenic | 9 | 53911460 | 53975301 | 63842 | 2 | ENSMUST00000048409.8 | -1799 | Elmod1 | ENSMUSG00000041986.10 |
| chr9 | 53986989 | 53987250 | 4.00274 | 5.07103 | 1.7817 | Distal Intergenic | 9 | 53911460 | 53975301 | 63842 | 2 | ENSMUST00000048409.8 | -11689 | Elmod1 | ENSMUSG00000041986.10 |
| chr12 | 117534158 | 117534726 | 4.11186 | 5.72917 | 2.30136 | Intron (ENSMUST00000109691.2/ENSMUSG00000041992.8, intron 3 of 27) | 12 | 117516479 | 117756978 | 240500 | 1 | ENSMUST00000109691.2 | 17680 | Rapgef5 | ENSMUSG00000041992.8 |
| chr5 | 114269450 | 114270180 | 3.7761 | 4.64719 | 1.50354 | Intron (ENSMUST00000044790.6/ENSMUSG00000042002.6, intron 2 of 9) | 5 | 114260356 | 114273702 | 13347 | 2 | ENSMUST00000144050.1 | 3522 | Foxn4 | ENSMUSG00000042002.6 |
| chr5 | 114270874 | 114271059 | 3.42671 | 3.99839 | 1.12412 | Intron (ENSMUST00000044790.6/ENSMUSG00000042002.6, intron 2 of 9) | 5 | 114260356 | 114273702 | 13347 | 2 | ENSMUST00000144050.1 | 2643 | Foxn4 | ENSMUSG00000042002.6 |
| chr5 | 114271412 | 114271707 | 3.55289 | 4.21266 | 1.20703 | Intron (ENSMUST00000044790.6/ENSMUSG00000042002.6, intron 2 of 9) | 5 | 114260356 | 114273702 | 13347 | 2 | ENSMUST00000144050.1 | 1995 | Foxn4 | ENSMUSG00000042002.6 |
| chr5 | 114272673 | 114273940 | 3.73127 | 4.61994 | 1.50655 | Promoter | 5 | 114254164 | 114273807 | 19644 | 2 | ENSMUST00000044790.6 | 0 | Foxn4 | ENSMUSG00000042002.6 |
| chr5 | 114274306 | 114274967 | 4.26338 | 5.6303 | 2.22327 | Promoter | 5 | 114254164 | 114273807 | 19644 | 2 | ENSMUST00000044790.6 | -500 | Foxn4 | ENSMUSG00000042002.6 |
| chr5 | 114275435 | 114276102 | 3.55455 | 4.28081 | 1.3 | Distal Intergenic | 5 | 114254164 | 114273807 | 19644 | 2 | ENSMUST00000044790.6 | -1629 | Foxn4 | ENSMUSG00000042002.6 |
| chr5 | 114173077 | 114173183 | 10.58331 | 22.7139 | 17.50692 | Intron (ENSMUST00000102582.2/ENSMUSG00000042010.11, intron 2 of 52) | 5 | 114175906 | 114190338 | 14433 | 1 | ENSMUST00000146841.1 | -2723 | Acacb | ENSMUSG00000042010.11 |
| chr13 | 94941907 | 94942256 | 3.46892 | 4.06103 | 1.12001 | Distal Intergenic | 13 | 94976344 | 95014531 | 38188 | 1 | ENSMUST00000160115.2 | -34088 | Wdr41 | ENSMUSG00000042015.11 |
| chr3 | 101378000 | 101378194 | 3.74595 | 4.57908 | 1.45337 | Promoter | 3 | 101377125 | 101463060 | 85936 | 1 | ENSMUST00000043983.5 | 876 | Igsf3 | ENSMUSG00000042035.6 |
| chr13 | 94666135 | 94666763 | 3.93502 | 4.94 | 1.69591 | Distal Intergenic | 13 | 94788943 | 94842899 | 53957 | 1 | ENSMUST00000046644.5 | -122180 | Tbca | ENSMUSG00000042043.5 |
| chr13 | 94686084 | 94686402 | 3.34153 | 3.83966 | 1.01733 | Distal Intergenic | 13 | 94788943 | 94842899 | 53957 | 1 | ENSMUST00000046644.5 | -102541 | Tbca | ENSMUSG00000042043.5 |
| chr13 | 94696883 | 94696983 | 3.69517 | 4.48354 | 1.38684 | Distal Intergenic | 13 | 94788943 | 94842899 | 53957 | 1 | ENSMUST00000046644.5 | -91960 | Tbca | ENSMUSG00000042043.5 |
| chr13 | 94701202 | 94701449 | 3.41742 | 3.97569 | 1.09361 | Distal Intergenic | 13 | 94788943 | 94842899 | 53957 | 1 | ENSMUST00000046644.5 | -87494 | Tbca | ENSMUSG00000042043.5 |
| chr13 | 94702066 | 94703333 | 3.96505 | 5.11256 | 1.8665 | Distal Intergenic | 13 | 94788943 | 94842899 | 53957 | 1 | ENSMUST00000046644.5 | -85610 | Tbca | ENSMUSG00000042043.5 |
| chr13 | 94703916 | 94704036 | 3.80211 | 4.69828 | 1.56118 | Distal Intergenic | 13 | 94788943 | 94842899 | 53957 | 1 | ENSMUST00000046644.5 | -84907 | Tbca | ENSMUSG00000042043.5 |
| chr13 | 94706452 | 94707693 | 3.74657 | 4.58947 | 1.4717 | Distal Intergenic | 13 | 94788943 | 94842899 | 53957 | 1 | ENSMUST00000046644.5 | -81250 | Tbca | ENSMUSG00000042043.5 |
| chr13 | 94708002 | 94708342 | 3.63064 | 4.37341 | 1.35526 | Distal Intergenic | 13 | 94788943 | 94842899 | 53957 | 1 | ENSMUST00000046644.5 | -80601 | Tbca | ENSMUSG00000042043.5 |
| chr13 | 94708711 | 94708896 | 3.43049 | 3.99331 | 1.08578 | Distal Intergenic | 13 | 94788943 | 94842899 | 53957 | 1 | ENSMUST00000046644.5 | -80047 | Tbca | ENSMUSG00000042043.5 |
| chr13 | 94709636 | 94710634 | 3.9695 | 5.29798 | 2.022 | Distal Intergenic | 13 | 94788943 | 94842899 | 53957 | 1 | ENSMUST00000046644.5 | -78309 | Tbca | ENSMUSG00000042043.5 |
| chr13 | 94711384 | 94712018 | 3.4848 | 4.09883 | 1.16661 | Distal Intergenic | 13 | 94788943 | 94842899 | 53957 | 1 | ENSMUST00000046644.5 | -76925 | Tbca | ENSMUSG00000042043.5 |
| chr13 | 94712439 | 94712532 | 3.8418 | 4.91201 | 1.72689 | Distal Intergenic | 13 | 94788943 | 94842899 | 53957 | 1 | ENSMUST00000046644.5 | -76411 | Tbca | ENSMUSG00000042043.5 |
| chr9 | 53825904 | 53826044 | 3.55275 | 4.37382 | 1.39863 | Distal Intergenic | 9 | 53850251 | 53853844 | 3594 | 1 | ENSMUST00000048485.5 | -24207 | Sln | ENSMUSG00000042045.5 |
| chr9 | 53826579 | 53826913 | 3.4652 | 4.10959 | 1.25197 | Distal Intergenic | 9 | 53850251 | 53853844 | 3594 | 1 | ENSMUST00000048485.5 | -23338 | Sln | ENSMUSG00000042045.5 |
| chr2 | 70126789 | 70126864 | 3.65927 | 4.64533 | 1.55513 | Exon (ENSMUST00000060208.5/ENSMUSG00000042064.8, exon 8 of 35) | 2 | 70096298 | 70183267 | 86970 | 1 | ENSMUST00000112241.1 | 30492 | Myo3b | ENSMUSG00000042064.8 |
| chr2 | 70127181 | 70127920 | 3.75087 | 4.6007 | 1.48976 | Intron (ENSMUST00000060208.5/ENSMUSG00000042064.8, intron 8 of 34) | 2 | 70096298 | 70183267 | 86970 | 1 | ENSMUST00000112241.1 | 30884 | Myo3b | ENSMUSG00000042064.8 |
| chr2 | 70128474 | 70128942 | 3.50371 | 4.17373 | 1.25851 | Intron (ENSMUST00000060208.5/ENSMUSG00000042064.8, intron 8 of 34) | 2 | 70096298 | 70183267 | 86970 | 1 | ENSMUST00000112241.1 | 32177 | Myo3b | ENSMUSG00000042064.8 |
| chr2 | 70129420 | 70129803 | 4.01914 | 5.15935 | 1.92683 | Intron (ENSMUST00000060208.5/ENSMUSG00000042064.8, intron 8 of 34) | 2 | 70096298 | 70183267 | 86970 | 1 | ENSMUST00000112241.1 | 33123 | Myo3b | ENSMUSG00000042064.8 |
| chr2 | 70130844 | 70131133 | 3.45657 | 4.06664 | 1.2075 | Intron (ENSMUST00000060208.5/ENSMUSG00000042064.8, intron 8 of 34) | 2 | 70096298 | 70183267 | 86970 | 1 | ENSMUST00000112241.1 | 34547 | Myo3b | ENSMUSG00000042064.8 |
| chr2 | 70135200 | 70135570 | 3.32996 | 3.83692 | 1.069 | Intron (ENSMUST00000060208.5/ENSMUSG00000042064.8, intron 8 of 34) | 2 | 70096298 | 70183267 | 86970 | 1 | ENSMUST00000112241.1 | 38903 | Myo3b | ENSMUSG00000042064.8 |
| chr2 | 70136083 | 70136406 | 3.36631 | 3.89944 | 1.09643 | Intron (ENSMUST00000060208.5/ENSMUSG00000042064.8, intron 8 of 34) | 2 | 70096298 | 70183267 | 86970 | 1 | ENSMUST00000112241.1 | 39786 | Myo3b | ENSMUSG00000042064.8 |
| chr2 | 70137312 | 70137635 | 3.38022 | 4.06222 | 1.21397 | Intron (ENSMUST00000060208.5/ENSMUSG00000042064.8, intron 8 of 34) | 2 | 70096298 | 70183267 | 86970 | 1 | ENSMUST00000112241.1 | 41015 | Myo3b | ENSMUSG00000042064.8 |
| chr2 | 70138785 | 70139124 | 3.67359 | 4.44949 | 1.40295 | Intron (ENSMUST00000060208.5/ENSMUSG00000042064.8, intron 8 of 34) | 2 | 70096298 | 70183267 | 86970 | 1 | ENSMUST00000112241.1 | 42488 | Myo3b | ENSMUSG00000042064.8 |
| chr2 | 70140967 | 70141279 | 3.40003 | 4.0638 | 1.16581 | Intron (ENSMUST00000060208.5/ENSMUSG00000042064.8, intron 8 of 34) | 2 | 70096298 | 70183267 | 86970 | 1 | ENSMUST00000112241.1 | 44670 | Myo3b | ENSMUSG00000042064.8 |
| chr2 | 70143822 | 70144068 | 3.51066 | 4.33283 | 1.35845 | Intron (ENSMUST00000060208.5/ENSMUSG00000042064.8, intron 8 of 34) | 2 | 70096298 | 70183267 | 86970 | 1 | ENSMUST00000112241.1 | 47525 | Myo3b | ENSMUSG00000042064.8 |
| chr2 | 70146968 | 70147140 | 3.60518 | 4.52025 | 1.4529 | Intron (ENSMUST00000060208.5/ENSMUSG00000042064.8, intron 8 of 34) | 2 | 70096298 | 70183267 | 86970 | 1 | ENSMUST00000112241.1 | 50671 | Myo3b | ENSMUSG00000042064.8 |
| chr2 | 70156475 | 70156611 | 4.10703 | 5.36671 | 2.09322 | Intron (ENSMUST00000060208.5/ENSMUSG00000042064.8, intron 8 of 34) | 2 | 70096298 | 70183267 | 86970 | 1 | ENSMUST00000112241.1 | 60178 | Myo3b | ENSMUSG00000042064.8 |
| chr2 | 70158198 | 70159180 | 3.42718 | 4.04512 | 1.22976 | Intron (ENSMUST00000060208.5/ENSMUSG00000042064.8, intron 8 of 34) | 2 | 70096298 | 70183267 | 86970 | 1 | ENSMUST00000112241.1 | 61901 | Myo3b | ENSMUSG00000042064.8 |
| chr2 | 70159608 | 70159943 | 4.07273 | 5.76476 | 2.29608 | Intron (ENSMUST00000060208.5/ENSMUSG00000042064.8, intron 8 of 34) | 2 | 70096298 | 70183267 | 86970 | 1 | ENSMUST00000112241.1 | 63311 | Myo3b | ENSMUSG00000042064.8 |
| chr2 | 70162791 | 70163051 | 3.5504 | 4.23672 | 1.30176 | Intron (ENSMUST00000060208.5/ENSMUSG00000042064.8, intron 8 of 34) | 2 | 70096298 | 70183267 | 86970 | 1 | ENSMUST00000112241.1 | 66494 | Myo3b | ENSMUSG00000042064.8 |
| chr2 | 70163711 | 70163786 | 3.48548 | 4.09768 | 1.16164 | Intron (ENSMUST00000060208.5/ENSMUSG00000042064.8, intron 8 of 34) | 2 | 70096298 | 70183267 | 86970 | 1 | ENSMUST00000112241.1 | 67414 | Myo3b | ENSMUSG00000042064.8 |
| chr2 | 70167366 | 70167691 | 4.29652 | 6.07052 | 2.54026 | Intron (ENSMUST00000060208.5/ENSMUSG00000042064.8, intron 8 of 34) | 2 | 70096298 | 70183267 | 86970 | 1 | ENSMUST00000112241.1 | 71069 | Myo3b | ENSMUSG00000042064.8 |
| chr2 | 70168079 | 70168812 | 3.7298 | 4.54882 | 1.43609 | Intron (ENSMUST00000060208.5/ENSMUSG00000042064.8, intron 8 of 34) | 2 | 70096298 | 70183267 | 86970 | 1 | ENSMUST00000112241.1 | 71782 | Myo3b | ENSMUSG00000042064.8 |
| chr2 | 70179999 | 70180279 | 3.5958 | 4.46477 | 1.43406 | Intron (ENSMUST00000060208.5/ENSMUSG00000042064.8, intron 8 of 34) | 2 | 70096298 | 70183267 | 86970 | 1 | ENSMUST00000112241.1 | 83702 | Myo3b | ENSMUSG00000042064.8 |
| chr2 | 70202013 | 70202401 | 3.73269 | 4.54885 | 1.43548 | Intron (ENSMUST00000060208.5/ENSMUSG00000042064.8, intron 9 of 34) | 2 | 70096298 | 70183267 | 86970 | 1 | ENSMUST00000112241.1 | 105716 | Myo3b | ENSMUSG00000042064.8 |
| chr2 | 70202729 | 70203008 | 3.41102 | 3.97318 | 1.11903 | Intron (ENSMUST00000060208.5/ENSMUSG00000042064.8, intron 9 of 34) | 2 | 70096298 | 70183267 | 86970 | 1 | ENSMUST00000112241.1 | 106432 | Myo3b | ENSMUSG00000042064.8 |
| chr2 | 70216214 | 70216390 | 3.36953 | 4.09278 | 1.18663 | Intron (ENSMUST00000060208.5/ENSMUSG00000042064.8, intron 9 of 34) | 2 | 70096298 | 70183267 | 86970 | 1 | ENSMUST00000112241.1 | 119917 | Myo3b | ENSMUSG00000042064.8 |
| chr2 | 70217211 | 70217445 | 3.30253 | 3.81698 | 1.10145 | Intron (ENSMUST00000060208.5/ENSMUSG00000042064.8, intron 10 of 34) | 2 | 70096298 | 70183267 | 86970 | 1 | ENSMUST00000112241.1 | 120914 | Myo3b | ENSMUSG00000042064.8 |
| chr2 | 70222119 | 70222290 | 3.38193 | 3.9006 | 1.01733 | Intron (ENSMUST00000060208.5/ENSMUSG00000042064.8, intron 10 of 34) | 2 | 70096298 | 70183267 | 86970 | 1 | ENSMUST00000112241.1 | 125822 | Myo3b | ENSMUSG00000042064.8 |
| chr2 | 70223438 | 70223804 | 3.54695 | 4.21025 | 1.2309 | Intron (ENSMUST00000060208.5/ENSMUSG00000042064.8, intron 10 of 34) | 2 | 70096298 | 70183267 | 86970 | 1 | ENSMUST00000112241.1 | 127141 | Myo3b | ENSMUSG00000042064.8 |
| chr13 | 93674269 | 93674587 | 3.71439 | 4.69996 | 1.59275 | Promoter | 13 | 93674433 | 93752831 | 78399 | 1 | ENSMUST00000048001.6 | 0 | Dmgdh | ENSMUSG00000042102.6 |
| chr14 | 119099230 | 119099352 | 3.38193 | 3.9006 | 1.01733 | Promoter | 14 | 118985039 | 119099430 | 114392 | 2 | ENSMUST00000156203.2 | 78 | Uggt2 | ENSMUSG00000042104.13 |
| chr6 | 116673553 | 116673657 | 3.85733 | 4.95349 | 1.72838 | Promoter | 6 | 116633008 | 116673836 | 40829 | 2 | ENSMUST00000035842.4 | 179 | Rassf4 | ENSMUSG00000042129.6 |
| chr6 | 116674072 | 116674294 | 3.53734 | 4.18617 | 1.19731 | Promoter | 6 | 116633008 | 116673836 | 40829 | 2 | ENSMUST00000035842.4 | -237 | Rassf4 | ENSMUSG00000042129.6 |
| chr9 | 37462058 | 37462234 | 3.38758 | 4.19441 | 1.25228 | Distal Intergenic | 9 | 37489321 | 37524148 | 34828 | 1 | ENSMUST00000048604.6 | -27087 | Msantd2 | ENSMUSG00000042138.6 |
| chr14 | 118922107 | 118923231 | 3.4838 | 4.25576 | 1.28347 | Promoter | 14 | 118875520 | 118923167 | 47648 | 2 | ENSMUST00000047208.6 | 0 | Dzip1 | ENSMUSG00000042156.10 |
| chr14 | 118923650 | 118924507 | 3.59575 | 4.30243 | 1.28906 | Promoter | 14 | 118875520 | 118923167 | 47648 | 2 | ENSMUST00000047208.6 | -484 | Dzip1 | ENSMUSG00000042156.10 |
| chr11 | 63061856 | 63061966 | 3.48859 | 4.22428 | 1.29488 | Promoter | 11 | 63061654 | 63094964 | 33311 | 1 | ENSMUST00000035732.4 | 203 | Tekt3 | ENSMUSG00000042189.5 |
| chr11 | 63090765 | 63090840 | 3.67764 | 4.46652 | 1.42402 | Intron (ENSMUST00000035732.4/ENSMUSG00000042189.5, intron 7 of 7) | 11 | 63061654 | 63094964 | 33311 | 1 | ENSMUST00000035732.4 | 29112 | Tekt3 | ENSMUSG00000042189.5 |
| chr5 | 113610592 | 113610667 | 3.95151 | 5.02044 | 1.75403 | Downstream (1-2kb) | 5 | 113614036 | 113627615 | 13580 | 2 | ENSMUST00000132065.2 | 16948 | Cmklr1 | ENSMUSG00000042190.7 |
| chr5 | 113649548 | 113649802 | 3.29548 | 3.77039 | 1.01733 | Promoter | 5 | 113631306 | 113650396 | 19091 | 2 | ENSMUST00000151460.1 | 594 | Cmklr1 | ENSMUSG00000042190.7 |
| chr9 | 53754061 | 53754350 | 3.65432 | 4.45442 | 1.42012 | Distal Intergenic | 9 | 53771535 | 53818161 | 46627 | 1 | ENSMUST00000048670.8 | -17185 | Slc35f2 | ENSMUSG00000042195.8 |
| chr9 | 53755194 | 53755680 | 3.78236 | 4.80403 | 1.67688 | Distal Intergenic | 9 | 53771535 | 53818161 | 46627 | 1 | ENSMUST00000048670.8 | -15855 | Slc35f2 | ENSMUSG00000042195.8 |
| chr9 | 53756056 | 53756436 | 3.68196 | 4.66287 | 1.56378 | Distal Intergenic | 9 | 53771535 | 53818161 | 46627 | 1 | ENSMUST00000048670.8 | -15099 | Slc35f2 | ENSMUSG00000042195.8 |
| chr9 | 53771598 | 53771692 | 4.19339 | 5.91952 | 2.46043 | Promoter | 9 | 53771535 | 53818161 | 46627 | 1 | ENSMUST00000048670.8 | 64 | Slc35f2 | ENSMUSG00000042195.8 |
| chr9 | 53772379 | 53772714 | 3.56842 | 4.45337 | 1.41037 | Promoter | 9 | 53771535 | 53818161 | 46627 | 1 | ENSMUST00000048670.8 | 845 | Slc35f2 | ENSMUSG00000042195.8 |
| chr5 | 113286828 | 113286903 | 3.40413 | 3.94608 | 1.05948 | Exon (ENSMUST00000048112.7/ENSMUSG00000042216.8, exon 5 of 25) | 5 | 113279610 | 113282892 | 3283 | 2 | ENSMUST00000138266.1 | -3937 | Sgsm1 | ENSMUSG00000042216.8 |
| chr5 | 113287888 | 113288488 | 3.84548 | 4.90164 | 1.72681 | Intron (ENSMUST00000048112.7/ENSMUSG00000042216.8, intron 3 of 24) | 5 | 113279610 | 113282892 | 3283 | 2 | ENSMUST00000138266.1 | -4997 | Sgsm1 | ENSMUSG00000042216.8 |
| chr5 | 113291246 | 113291346 | 4.27552 | 5.62669 | 2.19963 | Intron (ENSMUST00000048112.7/ENSMUSG00000042216.8, intron 2 of 24) | 5 | 113279610 | 113282892 | 3283 | 2 | ENSMUST00000138266.1 | -8355 | Sgsm1 | ENSMUSG00000042216.8 |
| chr5 | 113301540 | 113302305 | 3.51778 | 4.25831 | 1.27138 | Intron (ENSMUST00000048112.7/ENSMUSG00000042216.8, intron 2 of 24) | 5 | 113266287 | 113310697 | 44411 | 2 | ENSMUST00000154248.2 | 8392 | Sgsm1 | ENSMUSG00000042216.8 |
| chr5 | 113308695 | 113308867 | 3.47933 | 4.09283 | 1.17751 | Intron (ENSMUST00000048112.7/ENSMUSG00000042216.8, intron 2 of 24) | 5 | 113266287 | 113310697 | 44411 | 2 | ENSMUST00000154248.2 | 1830 | Sgsm1 | ENSMUSG00000042216.8 |
| chr5 | 113309850 | 113310572 | 3.64337 | 4.48868 | 1.42042 | Promoter | 5 | 113266287 | 113310697 | 44411 | 2 | ENSMUST00000154248.2 | 125 | Sgsm1 | ENSMUSG00000042216.8 |
| chr5 | 113311582 | 113311793 | 3.5564 | 4.22188 | 1.22181 | Promoter | 5 | 113243220 | 113310786 | 67567 | 2 | ENSMUST00000048112.7 | -797 | Sgsm1 | ENSMUSG00000042216.8 |
| chr5 | 113313483 | 113313817 | 3.95046 | 5.05641 | 1.8439 | Distal Intergenic | 5 | 113243220 | 113310786 | 67567 | 2 | ENSMUST00000048112.7 | -2698 | Sgsm1 | ENSMUSG00000042216.8 |
| chr5 | 113314192 | 113314372 | 3.87084 | 4.83611 | 1.65151 | Distal Intergenic | 5 | 113243220 | 113310786 | 67567 | 2 | ENSMUST00000048112.7 | -3407 | Sgsm1 | ENSMUSG00000042216.8 |
| chr5 | 113319547 | 113319860 | 3.4049 | 3.95459 | 1.08598 | Distal Intergenic | 5 | 113243220 | 113310786 | 67567 | 2 | ENSMUST00000048112.7 | -8762 | Sgsm1 | ENSMUSG00000042216.8 |
| chr5 | 113323880 | 113324071 | 3.99576 | 5.18358 | 1.95229 | Distal Intergenic | 5 | 113243220 | 113310786 | 67567 | 2 | ENSMUST00000048112.7 | -13095 | Sgsm1 | ENSMUSG00000042216.8 |
| chr5 | 113332395 | 113332831 | 3.46635 | 4.30204 | 1.34142 | Distal Intergenic | 5 | 113243220 | 113310786 | 67567 | 2 | ENSMUST00000048112.7 | -21610 | Sgsm1 | ENSMUSG00000042216.8 |
| chr5 | 113014655 | 113015004 | 3.61977 | 4.34859 | 1.32385 | Promoter | 5 | 112910478 | 113015514 | 105037 | 2 | ENSMUST00000065167.4 | 510 | Adrbk2 | ENSMUSG00000042249.7 |
| chr5 | 113024928 | 113025111 | 3.54485 | 4.19763 | 1.19738 | Distal Intergenic | 5 | 112910478 | 113015514 | 105037 | 2 | ENSMUST00000065167.4 | -9415 | Adrbk2 | ENSMUSG00000042249.7 |
| chr1 | 131797405 | 131797525 | 3.62615 | 4.34586 | 1.28723 | Promoter | 1 | 131797398 | 131821473 | 24076 | 1 | ENSMUST00000048660.6 | 8 | Pm20d1 | ENSMUSG00000042251.7 |
| chr1 | 131798267 | 131798645 | 3.66158 | 4.43179 | 1.38297 | Promoter | 1 | 131797398 | 131821473 | 24076 | 1 | ENSMUST00000048660.6 | 870 | Pm20d1 | ENSMUSG00000042251.7 |
| chr13 | 116290481 | 116291262 | 3.63216 | 4.47203 | 1.45438 | Distal Intergenic | 13 | 116299395 | 116303351 | 3957 | 2 | ENSMUST00000176444.1 | 12089 | Isl1 | ENSMUSG00000042258.8 |
| chr13 | 116292154 | 116292558 | 3.31769 | 3.80379 | 1.01733 | Distal Intergenic | 13 | 116299395 | 116303351 | 3957 | 2 | ENSMUST00000176444.1 | 10793 | Isl1 | ENSMUSG00000042258.8 |
| chr13 | 116293003 | 116294403 | 3.73835 | 4.79226 | 1.64631 | Distal Intergenic | 13 | 116299395 | 116303351 | 3957 | 2 | ENSMUST00000176444.1 | 8948 | Isl1 | ENSMUSG00000042258.8 |
| chr13 | 116294869 | 116295558 | 3.66001 | 4.51048 | 1.4819 | Downstream (2-3kb) | 13 | 116299395 | 116303351 | 3957 | 2 | ENSMUST00000176444.1 | 7793 | Isl1 | ENSMUSG00000042258.8 |
| chr13 | 116295980 | 116296615 | 3.46857 | 4.09546 | 1.22873 | Downstream (1-2kb) | 13 | 116299395 | 116303351 | 3957 | 2 | ENSMUST00000176444.1 | 6736 | Isl1 | ENSMUSG00000042258.8 |
| chr13 | 116297142 | 116300146 | 3.99228 | 5.14739 | 1.8848 | 3' UTR | 13 | 116299395 | 116303351 | 3957 | 2 | ENSMUST00000176444.1 | 3205 | Isl1 | ENSMUSG00000042258.8 |
| chr13 | 116297142 | 116300146 | 3.99228 | 5.14739 | 1.8848 | 3' UTR | 13 | 116299395 | 116303351 | 3957 | 2 | ENSMUST00000176444.1 | 3205 | Isl1 | ENSMUSG00000042258.8 |
| chr13 | 116300736 | 116300977 | 3.72621 | 4.60938 | 1.49084 | Intron (ENSMUST00000036060.7/ENSMUSG00000042258.8, intron 5 of 5) | 13 | 116299395 | 116303351 | 3957 | 2 | ENSMUST00000176444.1 | 2374 | Isl1 | ENSMUSG00000042258.8 |
| chr13 | 116301373 | 116301734 | 4.62841 | 6.3743 | 2.77618 | Exon (ENSMUST00000036060.7/ENSMUSG00000042258.8, exon 5 of 6) | 13 | 116299395 | 116303351 | 3957 | 2 | ENSMUST00000176444.1 | 1617 | Isl1 | ENSMUSG00000042258.8 |
| chr13 | 116302283 | 116302493 | 3.50856 | 4.13672 | 1.17482 | Promoter | 13 | 116299395 | 116303351 | 3957 | 2 | ENSMUST00000176444.1 | 858 | Isl1 | ENSMUSG00000042258.8 |
| chr13 | 116302809 | 116303435 | 4.95565 | 7.14254 | 3.43566 | Promoter | 13 | 116299395 | 116303351 | 3957 | 2 | ENSMUST00000176444.1 | 0 | Isl1 | ENSMUSG00000042258.8 |
| chr13 | 116304110 | 116304601 | 3.79974 | 4.68225 | 1.51894 | Promoter | 13 | 116299395 | 116303351 | 3957 | 2 | ENSMUST00000176444.1 | -760 | Isl1 | ENSMUSG00000042258.8 |
| chr13 | 116305002 | 116306354 | 3.76549 | 4.64212 | 1.51106 | Promoter | 13 | 116303046 | 116306397 | 3352 | 2 | ENSMUST00000176812.2 | 43 | Isl1 | ENSMUSG00000042258.8 |
| chr13 | 116306731 | 116307495 | 3.42343 | 4.52803 | 1.46151 | Promoter | 13 | 116303046 | 116306397 | 3352 | 2 | ENSMUST00000176812.2 | -335 | Isl1 | ENSMUSG00000042258.8 |
| chr13 | 116307806 | 116307997 | 3.53349 | 4.27983 | 1.28604 | Exon (ENSMUST00000177469.2/ENSMUSG00000042258.8, exon 2 of 2) | 13 | 116303046 | 116306397 | 3352 | 2 | ENSMUST00000176812.2 | -1410 | Isl1 | ENSMUSG00000042258.8 |
| chr13 | 116308307 | 116310559 | 4.13178 | 5.47108 | 2.10077 | Promoter | 13 | 116298281 | 116309689 | 11409 | 2 | ENSMUST00000036060.7 | 0 | Isl1 | ENSMUSG00000042258.8 |
| chr13 | 116311113 | 116312396 | 3.64357 | 4.38397 | 1.32315 | Distal Intergenic | 13 | 116298281 | 116309689 | 11409 | 2 | ENSMUST00000036060.7 | -1425 | Isl1 | ENSMUSG00000042258.8 |
| chr13 | 116313731 | 116313958 | 3.6597 | 4.81436 | 1.65393 | Distal Intergenic | 13 | 116298281 | 116309689 | 11409 | 2 | ENSMUST00000036060.7 | -4043 | Isl1 | ENSMUSG00000042258.8 |
| chr13 | 116314819 | 116315407 | 3.72536 | 4.55949 | 1.44656 | Distal Intergenic | 13 | 116298281 | 116309689 | 11409 | 2 | ENSMUST00000036060.7 | -5131 | Isl1 | ENSMUSG00000042258.8 |
| chr13 | 116315715 | 116315907 | 3.36372 | 3.87313 | 1.01733 | Distal Intergenic | 13 | 116298281 | 116309689 | 11409 | 2 | ENSMUST00000036060.7 | -6027 | Isl1 | ENSMUSG00000042258.8 |
| chr13 | 116317201 | 116317502 | 3.34336 | 3.87832 | 1.13089 | Distal Intergenic | 13 | 116298281 | 116309689 | 11409 | 2 | ENSMUST00000036060.7 | -7513 | Isl1 | ENSMUSG00000042258.8 |
| chr13 | 116318705 | 116319051 | 5.13777 | 7.60114 | 3.69147 | Distal Intergenic | 13 | 116298281 | 116309689 | 11409 | 2 | ENSMUST00000036060.7 | -9017 | Isl1 | ENSMUSG00000042258.8 |
| chr8 | 120177447 | 120177793 | 3.68986 | 4.52186 | 1.43488 | Promoter | 8 | 120166397 | 120177466 | 11070 | 2 | ENSMUST00000048786.7 | 0 | Fam92b | ENSMUSG00000042269.10 |
| chr11 | 61956913 | 61957274 | 4.45291 | 6.24638 | 2.73303 | Promoter | 11 | 61956810 | 62045484 | 88675 | 1 | ENSMUST00000136971.1 | 104 | Specc1 | ENSMUSG00000042331.8 |
| chr11 | 61957917 | 61958396 | 3.85929 | 4.86115 | 1.69839 | Intron (ENSMUST00000049836.8/ENSMUSG00000042331.8, intron 1 of 7) | 11 | 61956810 | 62045484 | 88675 | 1 | ENSMUST00000136971.1 | 1108 | Specc1 | ENSMUSG00000042331.8 |
| chr11 | 61958944 | 61959188 | 3.77999 | 5.0381 | 1.82169 | Intron (ENSMUST00000049836.8/ENSMUSG00000042331.8, intron 1 of 7) | 11 | 61956810 | 62045484 | 88675 | 1 | ENSMUST00000136971.1 | 2135 | Specc1 | ENSMUSG00000042331.8 |
| chr11 | 61959644 | 61959819 | 3.43685 | 4.06104 | 1.23151 | Intron (ENSMUST00000049836.8/ENSMUSG00000042331.8, intron 1 of 7) | 11 | 61956810 | 62045484 | 88675 | 1 | ENSMUST00000136971.1 | 2835 | Specc1 | ENSMUSG00000042331.8 |
| chr11 | 62005286 | 62005622 | 3.78884 | 4.6592 | 1.50904 | Intron (ENSMUST00000049836.8/ENSMUSG00000042331.8, intron 1 of 7) | 11 | 62029276 | 62223013 | 193738 | 1 | ENSMUST00000108709.2 | -23654 | Specc1 | ENSMUSG00000042331.8 |
| chr7 | 127717253 | 127717527 | 3.44807 | 4.09847 | 1.2338 | 3' UTR | 7 | 127712736 | 127718185 | 5450 | 1 | ENSMUST00000047393.5 | 4518 | Ctf1 | ENSMUSG00000042340.5 |
| chr13 | 113888286 | 113888902 | 3.69403 | 4.48784 | 1.41015 | Intron (ENSMUST00000091201.6/ENSMUSG00000042348.9, intron 1 of 4) | 13 | 113794505 | 114157461 | 362957 | 1 | ENSMUST00000091201.6 | 93782 | Arl15 | ENSMUSG00000042348.9 |
| chr15 | 80616082 | 80616431 | 3.44772 | 4.05152 | 1.19938 | Distal Intergenic | 15 | 80623505 | 80650559 | 27055 | 1 | ENSMUST00000043149.7 | -7074 | Grap2 | ENSMUSG00000042351.8 |
| chr15 | 80616956 | 80617188 | 3.33399 | 3.82832 | 1.01733 | Distal Intergenic | 15 | 80623505 | 80650559 | 27055 | 1 | ENSMUST00000043149.7 | -6317 | Grap2 | ENSMUSG00000042351.8 |
| chr4 | 127327671 | 127328041 | 3.60589 | 4.36045 | 1.37037 | Intron (ENSMUST00000106091.3/ENSMUSG00000042367.7, intron 1 of 1) | 4 | 127325235 | 127329161 | 3927 | 2 | ENSMUST00000106091.3 | 1120 | Gjb3 | ENSMUSG00000042367.7 |
| chr4 | 127329999 | 127330125 | 3.48543 | 4.11867 | 1.23041 | Promoter | 4 | 127325235 | 127330844 | 5610 | 2 | ENSMUST00000046532.3 | 719 | Gjb3 | ENSMUSG00000042367.7 |
| chr4 | 127330517 | 127331089 | 3.89493 | 4.91574 | 1.6846 | Promoter | 4 | 127325235 | 127330844 | 5610 | 2 | ENSMUST00000046532.3 | 0 | Gjb3 | ENSMUSG00000042367.7 |
| chr4 | 127331857 | 127332445 | 3.48741 | 4.09861 | 1.15436 | Distal Intergenic | 4 | 127325235 | 127330844 | 5610 | 2 | ENSMUST00000046532.3 | -1014 | Gjb3 | ENSMUSG00000042367.7 |
| chr4 | 127332790 | 127332983 | 3.48966 | 4.10604 | 1.16218 | Distal Intergenic | 4 | 127325235 | 127330844 | 5610 | 2 | ENSMUST00000046532.3 | -1947 | Gjb3 | ENSMUSG00000042367.7 |
| chr19 | 25566523 | 25567075 | 3.81929 | 4.85441 | 1.69962 | Intron (ENSMUST00000025755.5/ENSMUSG00000024837.10, intron 4 of 4) | 19 | 25610537 | 25623920 | 13384 | 1 | ENSMUST00000048935.4 | -43462 | Dmrt3 | ENSMUSG00000042372.4 |
| chr19 | 25568276 | 25568621 | 4.14431 | 5.40549 | 2.09345 | Intron (ENSMUST00000025755.5/ENSMUSG00000024837.10, intron 4 of 4) | 19 | 25610537 | 25623920 | 13384 | 1 | ENSMUST00000048935.4 | -41916 | Dmrt3 | ENSMUSG00000042372.4 |
| chr19 | 25570322 | 25570640 | 3.4117 | 3.99132 | 1.17596 | Intron (ENSMUST00000025755.5/ENSMUSG00000024837.10, intron 4 of 4) | 19 | 25610537 | 25623920 | 13384 | 1 | ENSMUST00000048935.4 | -39897 | Dmrt3 | ENSMUSG00000042372.4 |
| chr19 | 25572152 | 25572447 | 3.60078 | 4.35417 | 1.41519 | Intron (ENSMUST00000025755.5/ENSMUSG00000024837.10, intron 4 of 4) | 19 | 25610537 | 25623920 | 13384 | 1 | ENSMUST00000048935.4 | -38090 | Dmrt3 | ENSMUSG00000042372.4 |
| chr19 | 25572964 | 25573261 | 3.52331 | 4.26281 | 1.33544 | Intron (ENSMUST00000025755.5/ENSMUSG00000024837.10, intron 4 of 4) | 19 | 25610537 | 25623920 | 13384 | 1 | ENSMUST00000048935.4 | -37276 | Dmrt3 | ENSMUSG00000042372.4 |
| chr19 | 25585308 | 25585840 | 3.34898 | 4.06758 | 1.15991 | Intron (ENSMUST00000025755.5/ENSMUSG00000024837.10, intron 4 of 4) | 19 | 25610537 | 25623920 | 13384 | 1 | ENSMUST00000048935.4 | -24697 | Dmrt3 | ENSMUSG00000042372.4 |
| chr19 | 25586203 | 25586294 | 3.41214 | 4.00167 | 1.16856 | Intron (ENSMUST00000025755.5/ENSMUSG00000024837.10, intron 4 of 4) | 19 | 25610537 | 25623920 | 13384 | 1 | ENSMUST00000048935.4 | -24243 | Dmrt3 | ENSMUSG00000042372.4 |
| chr19 | 25586914 | 25587184 | 3.87889 | 4.93462 | 1.77153 | Intron (ENSMUST00000025755.5/ENSMUSG00000024837.10, intron 4 of 4) | 19 | 25610537 | 25623920 | 13384 | 1 | ENSMUST00000048935.4 | -23353 | Dmrt3 | ENSMUSG00000042372.4 |
| chr19 | 25587645 | 25587946 | 3.63523 | 4.41565 | 1.44798 | Intron (ENSMUST00000025755.5/ENSMUSG00000024837.10, intron 4 of 4) | 19 | 25610537 | 25623920 | 13384 | 1 | ENSMUST00000048935.4 | -22591 | Dmrt3 | ENSMUSG00000042372.4 |
| chr19 | 25589025 | 25589288 | 3.91057 | 4.95627 | 1.80663 | Intron (ENSMUST00000025755.5/ENSMUSG00000024837.10, intron 4 of 4) | 19 | 25610537 | 25623920 | 13384 | 1 | ENSMUST00000048935.4 | -21249 | Dmrt3 | ENSMUSG00000042372.4 |
| chr19 | 25597667 | 25598249 | 3.48622 | 4.15744 | 1.25744 | Intron (ENSMUST00000025755.5/ENSMUSG00000024837.10, intron 4 of 4) | 19 | 25610537 | 25623920 | 13384 | 1 | ENSMUST00000048935.4 | -12288 | Dmrt3 | ENSMUSG00000042372.4 |
| chr19 | 25598769 | 25600008 | 3.79035 | 4.72216 | 1.59524 | Intron (ENSMUST00000025755.5/ENSMUSG00000024837.10, intron 4 of 4) | 19 | 25610537 | 25623920 | 13384 | 1 | ENSMUST00000048935.4 | -10529 | Dmrt3 | ENSMUSG00000042372.4 |
| chr19 | 25600441 | 25601060 | 3.9708 | 5.0455 | 1.81115 | Intron (ENSMUST00000025755.5/ENSMUSG00000024837.10, intron 4 of 4) | 19 | 25610537 | 25623920 | 13384 | 1 | ENSMUST00000048935.4 | -9477 | Dmrt3 | ENSMUSG00000042372.4 |
| chr19 | 25601615 | 25602776 | 3.78397 | 4.8015 | 1.69166 | Intron (ENSMUST00000025755.5/ENSMUSG00000024837.10, intron 4 of 4) | 19 | 25610537 | 25623920 | 13384 | 1 | ENSMUST00000048935.4 | -7761 | Dmrt3 | ENSMUSG00000042372.4 |
| chr19 | 25603402 | 25604034 | 3.83666 | 4.84907 | 1.66119 | 3' UTR | 19 | 25610537 | 25623920 | 13384 | 1 | ENSMUST00000048935.4 | -6503 | Dmrt3 | ENSMUSG00000042372.4 |
| chr19 | 25604592 | 25604819 | 3.46538 | 4.27595 | 1.29015 | Distal Intergenic | 19 | 25610537 | 25623920 | 13384 | 1 | ENSMUST00000048935.4 | -5718 | Dmrt3 | ENSMUSG00000042372.4 |
| chr19 | 25605156 | 25605878 | 3.77277 | 4.81101 | 1.65891 | Distal Intergenic | 19 | 25610537 | 25623920 | 13384 | 1 | ENSMUST00000048935.4 | -4659 | Dmrt3 | ENSMUSG00000042372.4 |
| chr19 | 25606340 | 25607267 | 3.69537 | 4.54855 | 1.4796 | Distal Intergenic | 19 | 25610537 | 25623920 | 13384 | 1 | ENSMUST00000048935.4 | -3270 | Dmrt3 | ENSMUSG00000042372.4 |
| chr19 | 25608120 | 25614242 | 4.07115 | 5.47037 | 2.14442 | Promoter | 19 | 25610537 | 25623920 | 13384 | 1 | ENSMUST00000048935.4 | 0 | Dmrt3 | ENSMUSG00000042372.4 |
| chr19 | 25615140 | 25616919 | 3.59635 | 4.43444 | 1.39441 | Intron (ENSMUST00000048935.4/ENSMUSG00000042372.4, intron 1 of 1) | 19 | 25610537 | 25623920 | 13384 | 1 | ENSMUST00000048935.4 | 4604 | Dmrt3 | ENSMUSG00000042372.4 |
| chr19 | 25618170 | 25619111 | 3.39498 | 3.99006 | 1.10331 | Intron (ENSMUST00000048935.4/ENSMUSG00000042372.4, intron 1 of 1) | 19 | 25610537 | 25623920 | 13384 | 1 | ENSMUST00000048935.4 | 7634 | Dmrt3 | ENSMUSG00000042372.4 |
| chr19 | 25619429 | 25619505 | 4.69914 | 6.50699 | 2.8331 | Intron (ENSMUST00000048935.4/ENSMUSG00000042372.4, intron 1 of 1) | 19 | 25610537 | 25623920 | 13384 | 1 | ENSMUST00000048935.4 | 8893 | Dmrt3 | ENSMUSG00000042372.4 |
| chr19 | 25619919 | 25620013 | 3.3765 | 3.91295 | 1.08948 | Intron (ENSMUST00000048935.4/ENSMUSG00000042372.4, intron 1 of 1) | 19 | 25610537 | 25623920 | 13384 | 1 | ENSMUST00000048935.4 | 9383 | Dmrt3 | ENSMUSG00000042372.4 |

| | | | | | | | | | | | | | | | |
|---|---|---|---|---|---|---|---|---|---|---|---|---|---|---|---|
| chr19 | 25620533 | 25620720 | 3.59998 | 4.58527 | 1.49039 | Intron (ENSMUST00000048935.4/ENSMUSG00000042372.4, intron 1 of 1) | 19 | 25610537 | 25623920 | 13384 | 1 | ENSMUST00000048935.4 | 9997 | Dmrt3 | ENSMUSG00000042372.4 |
| chr19 | 25621145 | 25621544 | 3.5539 | 4.27521 | 1.3485 | Intron (ENSMUST00000048935.4/ENSMUSG00000042372.4, intron 1 of 1) | 19 | 25610537 | 25623920 | 13384 | 1 | ENSMUST00000048935.4 | 10609 | Dmrt3 | ENSMUSG00000042372.4 |
| chr19 | 25622322 | 25623297 | 4.144 | 5.60153 | 2.29554 | 3' UTR | 19 | 25610537 | 25623920 | 13384 | 1 | ENSMUST00000048935.4 | 11786 | Dmrt3 | ENSMUSG00000042372.4 |
| chr19 | 25623714 | 25623993 | 3.51574 | 4.14846 | 1.17913 | 3' UTR | 19 | 25610537 | 25623920 | 13384 | 1 | ENSMUST00000048935.4 | 13178 | Dmrt3 | ENSMUSG00000042372.4 |
| chr19 | 25624324 | 25624410 | 3.95112 | 4.97348 | 1.72657 | Distal Intergenic | 19 | 25610537 | 25623920 | 13384 | 1 | ENSMUST00000048935.4 | 13788 | Dmrt3 | ENSMUSG00000042372.4 |
| chr19 | 25625134 | 25625505 | 3.72126 | 4.57329 | 1.50735 | Distal Intergenic | 19 | 25610537 | 25623920 | 13384 | 1 | ENSMUST00000048935.4 | 14598 | Dmrt3 | ENSMUSG00000042372.4 |
| chr19 | 25626068 | 25626312 | 3.48594 | 4.13094 | 1.2026 | Distal Intergenic | 19 | 25610537 | 25623920 | 13384 | 1 | ENSMUST00000048935.4 | 15532 | Dmrt3 | ENSMUSG00000042372.4 |
| chr19 | 25627824 | 25628473 | 4.11264 | 5.36288 | 2.02562 | Distal Intergenic | 19 | 25610537 | 25623920 | 13384 | 1 | ENSMUST00000048935.4 | 17288 | Dmrt3 | ENSMUSG00000042372.4 |
| chr13 | 113209940 | 113210021 | 3.71935 | 4.70417 | 1.56407 | Promoter | 13 | 113209659 | 113218098 | 8440 | 1 | ENSMUST00000038144.8 | 282 | Esm1 | ENSMUSG00000042379.8 |
| chr4 | 127223536 | 127224195 | 3.88428 | 4.87748 | 1.6636 | Intron (ENSMUST00000046659.8/ENSMUSG00000042388.9, intron 8 of 11) | 4 | 127243784 | 127247809 | 4026 | 1 | ENSMUST00000142029.1 | -19589 | Smim12 | ENSMUSG00000042380.8 |
| chr4 | 127226064 | 127226479 | 3.38188 | 3.91939 | 1.08407 | Intron (ENSMUST00000046659.8/ENSMUSG00000042388.9, intron 8 of 11) | 4 | 127243784 | 127247809 | 4026 | 1 | ENSMUST00000142029.1 | -17305 | Smim12 | ENSMUSG00000042380.8 |
| chr4 | 127227081 | 127227626 | 3.47123 | 4.08498 | 1.1902 | Intron (ENSMUST00000046659.8/ENSMUSG00000042388.9, intron 8 of 11) | 4 | 127243784 | 127247809 | 4026 | 1 | ENSMUST00000142029.1 | -16158 | Smim12 | ENSMUSG00000042380.8 |
| chr4 | 127228987 | 127229109 | 3.77143 | 4.64309 | 1.55043 | Intron (ENSMUST00000046659.8/ENSMUSG00000042388.9, intron 8 of 11) | 4 | 127243784 | 127247809 | 4026 | 1 | ENSMUST00000142029.1 | -14675 | Smim12 | ENSMUSG00000042380.8 |
| chr4 | 127236731 | 127237294 | 3.90197 | 4.95069 | 1.80831 | 3' UTR | 4 | 127243784 | 127247809 | 4026 | 1 | ENSMUST00000142029.1 | -6490 | Smim12 | ENSMUSG00000042380.8 |
| chr4 | 127166337 | 127166412 | 4.08108 | 5.25718 | 1.98277 | Distal Intergenic | 4 | 127169204 | 127237022 | 67819 | 1 | ENSMUST00000046659.8 | -2792 | Dlgap3 | ENSMUSG00000042388.9 |
| chr4 | 127166764 | 127167208 | 3.54859 | 4.22596 | 1.27342 | Distal Intergenic | 4 | 127169204 | 127237022 | 67819 | 1 | ENSMUST00000046659.8 | -1996 | Dlgap3 | ENSMUSG00000042388.9 |
| chr4 | 127167863 | 127169549 | 4.02569 | 5.35432 | 2.05837 | Promoter | 4 | 127169204 | 127237022 | 67819 | 1 | ENSMUST00000046659.8 | 0 | Dlgap3 | ENSMUSG00000042388.9 |
| chr4 | 127171194 | 127171400 | 3.4566 | 4.22122 | 1.28596 | Intron (ENSMUST00000046659.8/ENSMUSG00000042388.9, intron 1 of 11) | 4 | 127172880 | 127237022 | 64143 | 1 | ENSMUST00000106094.3 | -1480 | Dlgap3 | ENSMUSG00000042388.9 |
| chr4 | 127172603 | 127172828 | 3.59217 | 4.28766 | 1.2634 | Promoter | 4 | 127172880 | 127237022 | 64143 | 1 | ENSMUST00000106094.3 | -52 | Dlgap3 | ENSMUSG00000042388.9 |
| chr4 | 127175263 | 127175715 | 3.67984 | 4.46441 | 1.39675 | Intron (ENSMUST00000046659.8/ENSMUSG00000042388.9, intron 1 of 11) | 4 | 127172880 | 127237022 | 64143 | 1 | ENSMUST00000106094.3 | 2384 | Dlgap3 | ENSMUSG00000042388.9 |
| chr4 | 127176708 | 127176964 | 3.49432 | 4.11554 | 1.17751 | Intron (ENSMUST00000046659.8/ENSMUSG00000042388.9, intron 1 of 11) | 4 | 127172880 | 127237022 | 64143 | 1 | ENSMUST00000106094.3 | 3829 | Dlgap3 | ENSMUSG00000042388.9 |
| chr4 | 127178838 | 127179112 | 3.42277 | 4.06811 | 1.1495 | Intron (ENSMUST00000046659.8/ENSMUSG00000042388.9, intron 1 of 11) | 4 | 127172880 | 127237022 | 64143 | 1 | ENSMUST00000106094.3 | 5959 | Dlgap3 | ENSMUSG00000042388.9 |
| chr4 | 127179466 | 127179813 | 3.36833 | 3.88008 | 1.01733 | Intron (ENSMUST00000046659.8/ENSMUSG00000042388.9, intron 1 of 11) | 4 | 127172880 | 127237022 | 64143 | 1 | ENSMUST00000106094.3 | 6587 | Dlgap3 | ENSMUSG00000042388.9 |
| chr4 | 127214386 | 127215358 | 3.98253 | 5.09059 | 1.85247 | Exon (ENSMUST00000046659.8/ENSMUSG00000042388.9, exon 6 of 12) | 4 | 127203306 | 127214754 | 11449 | 1 | ENSMUST00000137062.1 | 11081 | Dlgap3 | ENSMUSG00000042388.9 |
| chr4 | 127222209 | 127222323 | 3.45318 | 4.22531 | 1.26939 | Intron (ENSMUST00000046659.8/ENSMUSG00000042388.9, intron 8 of 11) | 4 | 127203306 | 127214754 | 11449 | 1 | ENSMUST00000137062.1 | 18904 | Dlgap3 | ENSMUSG00000042388.9 |
| chr6 | 115519567 | 115519884 | 3.54544 | 4.20313 | 1.21481 | Distal Intergenic | 6 | 115544664 | 115554272 | 9609 | 1 | ENSMUST00000130425.2 | -24780 | Tsen2 | ENSMUSG00000042389.8 |
| chr19 | 42380006 | 42380504 | 3.52751 | 4.17856 | 1.21482 | Intron (ENSMUST00000048630.6/ENSMUSG00000042401.6, intron 2 of 14) | 19 | 42283037 | 42431783 | 148747 | 2 | ENSMUST00000048630.6 | 51279 | Crtac1 | ENSMUSG00000042401.6 |
| chr19 | 42381047 | 42381122 | 4.47534 | 5.99892 | 2.42175 | Intron (ENSMUST00000048630.6/ENSMUSG00000042401.6, intron 2 of 14) | 19 | 42283037 | 42431783 | 148747 | 2 | ENSMUST00000048630.6 | 50661 | Crtac1 | ENSMUSG00000042401.6 |
| chr19 | 42381500 | 42382398 | 3.72619 | 4.54379 | 1.42953 | Intron (ENSMUST00000048630.6/ENSMUSG00000042401.6, intron 2 of 14) | 19 | 42283037 | 42431783 | 148747 | 2 | ENSMUST00000048630.6 | 49385 | Crtac1 | ENSMUSG00000042401.6 |
| chr19 | 42408698 | 42409064 | 3.63936 | 4.46142 | 1.45427 | Intron (ENSMUST00000048630.6/ENSMUSG00000042401.6, intron 2 of 14) | 19 | 42283037 | 42431783 | 148747 | 2 | ENSMUST00000048630.6 | 22719 | Crtac1 | ENSMUSG00000042401.6 |
| chr19 | 42409755 | 42409908 | 3.59447 | 4.43843 | 1.44609 | Intron (ENSMUST00000048630.6/ENSMUSG00000042401.6, intron 2 of 14) | 19 | 42283037 | 42431783 | 148747 | 2 | ENSMUST00000048630.6 | 21875 | Crtac1 | ENSMUSG00000042401.6 |
| chr19 | 42429412 | 42429888 | 3.683 | 4.72013 | 1.59106 | Intron (ENSMUST00000048630.6/ENSMUSG00000042401.6, intron 1 of 14) | 19 | 42283037 | 42431783 | 148747 | 2 | ENSMUST00000048630.6 | 1895 | Crtac1 | ENSMUSG00000042401.6 |
| chr19 | 42430200 | 42430373 | 3.26005 | 3.71725 | 1.01733 | Intron (ENSMUST00000048630.6/ENSMUSG00000042401.6, intron 1 of 14) | 19 | 42283037 | 42431783 | 148747 | 2 | ENSMUST00000048630.6 | 1410 | Crtac1 | ENSMUSG00000042401.6 |
| chr19 | 42430755 | 42431152 | 4.09074 | 5.473 | 2.14664 | Promoter | 19 | 42283037 | 42431783 | 148747 | 2 | ENSMUST00000048630.6 | 631 | Crtac1 | ENSMUSG00000042401.6 |
| chr19 | 42431562 | 42433165 | 4.16052 | 5.69991 | 2.29876 | Promoter | 19 | 42283037 | 42431783 | 148747 | 2 | ENSMUST00000048630.6 | 0 | Crtac1 | ENSMUSG00000042401.6 |
| chr1 | 13103529 | 13104366 | 3.75013 | 4.74411 | 1.63633 | Distal Intergenic | 1 | 13113457 | 13127163 | 13707 | 2 | ENSMUST00000047577.6 | 22797 | Prdm14 | ENSMUSG00000042414.7 |
| chr1 | 13119535 | 13119804 | 3.45262 | 4.11359 | 1.21321 | Intron (ENSMUST00000047577.6/ENSMUSG00000042414.7, intron 5 of 7) | 1 | 13113457 | 13127163 | 13707 | 2 | ENSMUST00000047577.6 | 7359 | Prdm14 | ENSMUSG00000042414.7 |
| chr1 | 13123439 | 13124462 | 3.71721 | 4.57689 | 1.53385 | Exon (ENSMUST00000047577.6/ENSMUSG00000042414.7, exon 4 of 8) | 1 | 13113457 | 13127163 | 13707 | 2 | ENSMUST00000047577.6 | 2701 | Prdm14 | ENSMUSG00000042414.7 |
| chr1 | 13124947 | 13127698 | 3.97024 | 5.15381 | 1.91174 | Promoter | 1 | 13113457 | 13127163 | 13707 | 2 | ENSMUST00000047577.6 | 0 | Prdm14 | ENSMUSG00000042414.7 |
| chr1 | 13128064 | 13128181 | 3.75528 | 4.61947 | 1.51486 | Promoter | 1 | 13113457 | 13127163 | 13707 | 2 | ENSMUST00000047577.6 | -902 | Prdm14 | ENSMUSG00000042414.7 |
| chr1 | 13128654 | 13129270 | 3.86015 | 4.92359 | 1.76649 | Distal Intergenic | 1 | 13113457 | 13127163 | 13707 | 2 | ENSMUST00000047577.6 | -1492 | Prdm14 | ENSMUSG00000042414.7 |
| chr1 | 13129819 | 13130039 | 3.54605 | 4.24633 | 1.33837 | Distal Intergenic | 1 | 13113457 | 13127163 | 13707 | 2 | ENSMUST00000047577.6 | -2657 | Prdm14 | ENSMUSG00000042414.7 |
| chr1 | 13130606 | 13130869 | 3.36169 | 3.91774 | 1.13447 | Distal Intergenic | 1 | 13113457 | 13127163 | 13707 | 2 | ENSMUST00000047577.6 | -3444 | Prdm14 | ENSMUSG00000042414.7 |
| chr13 | 112976593 | 112976914 | 3.28423 | 3.75348 | 1.01733 | Distal Intergenic | 13 | 112987802 | 112990778 | 2977 | 1 | ENSMUST00000038404.4 | -10888 | Ccno | ENSMUSG00000042417.4 |
| chr13 | 112983502 | 112983577 | 3.38607 | 3.91318 | 1.03958 | Distal Intergenic | 13 | 112987802 | 112990778 | 2977 | 1 | ENSMUST00000038404.4 | -4225 | Ccno | ENSMUSG00000042417.4 |
| chr13 | 112984062 | 112984664 | 3.80771 | 4.73654 | 1.60027 | Distal Intergenic | 13 | 112987802 | 112990778 | 2977 | 1 | ENSMUST00000038404.4 | -3138 | Ccno | ENSMUSG00000042417.4 |
| chr13 | 112986584 | 112986887 | 3.42957 | 3.98776 | 1.07104 | Promoter | 13 | 112987802 | 112990778 | 2977 | 1 | ENSMUST00000038404.4 | -915 | Ccno | ENSMUSG00000042417.4 |
| chr13 | 112988823 | 112989144 | 3.60572 | 4.39771 | 1.36297 | Exon (ENSMUST00000038404.4/ENSMUSG00000042417.4, exon 2 of 3) | 13 | 112987802 | 112990778 | 2977 | 1 | ENSMUST00000038404.4 | 1022 | Ccno | ENSMUSG00000042417.4 |
| chr13 | 112989549 | 112989653 | 3.90705 | 4.92851 | 1.76801 | Exon (ENSMUST00000038404.4/ENSMUSG00000042417.4, exon 3 of 3) | 13 | 112987802 | 112990778 | 2977 | 1 | ENSMUST00000038404.4 | 1748 | Ccno | ENSMUSG00000042417.4 |
| chr15 | 80172934 | 80174576 | 3.59684 | 4.3818 | 1.37894 | Promoter | 15 | 80173721 | 80215519 | 41799 | 1 | ENSMUST00000044970.6 | 0 | Mgat3 | ENSMUSG00000042428.6 |
| chr15 | 80175288 | 80175408 | 3.31772 | 3.94883 | 1.05916 | Intron (ENSMUST00000044970.6/ENSMUSG00000042428.6, intron 1 of 1) | 15 | 80173721 | 80215519 | 41799 | 1 | ENSMUST00000044970.6 | 1568 | Mgat3 | ENSMUSG00000042428.6 |
| chr1 | 134234385 | 134235201 | 4.2299 | 5.76515 | 2.35565 | Promoter | 1 | 134199223 | 134234578 | 35356 | 2 | ENSMUST00000169927.1 | 0 | Adora1 | ENSMUSG00000042429.8 |
| chr18 | 65549835 | 65550108 | 3.66079 | 4.48756 | 1.46674 | Distal Intergenic | 18 | 65580230 | 65689443 | 109214 | 1 | ENSMUST00000049016.6 | -30122 | Zfp532 | ENSMUSG00000042439.7 |
| chr2 | 74763261 | 74763414 | 3.56425 | 4.53193 | 1.43477 | Promoter | 2 | 74762980 | 74765142 | 2163 | 1 | ENSMUST00000047793.5 | 282 | Hoxd1 | ENSMUSG00000042448.5 |
| chr5 | 22341201 | 22341326 | 3.30993 | 3.96404 | 1.10803 | Intron (ENSMUST00000161356.2/ENSMUSG00000042453.9, intron 1 of 64) | 5 | 21885250 | 22344690 | 459441 | 2 | ENSMUST00000062372.8 | 3364 | Reln | ENSMUSG00000042453.9 |
| chr5 | 22343667 | 22344668 | 3.95474 | 5.05238 | 1.85502 | Promoter | 5 | 21885250 | 22344690 | 459441 | 2 | ENSMUST00000062372.8 | 22 | Reln | ENSMUSG00000042453.9 |
| chr5 | 22345026 | 22346749 | 3.70019 | 4.64921 | 1.57809 | Promoter | 5 | 22342071 | 22344702 | 2632 | 2 | ENSMUST00000161782.1 | -325 | Reln | ENSMUSG00000042453.9 |
| chr1 | 130877695 | 130878413 | 4.22865 | 5.63927 | 2.27741 | Promoter | 1 | 130878158 | 130880779 | 2622 | 1 | ENSMUST00000126821.1 | 0 | Faim3 | ENSMUSG00000042474.6 |
| chr1 | 130878731 | 130879225 | 3.92541 | 5.01737 | 1.83598 | Promoter | 1 | 130878331 | 130880790 | 2460 | 1 | ENSMUST00000145446.1 | 401 | Faim3 | ENSMUSG00000042474.6 |
| chr5 | 8893316 | 8893711 | 3.61305 | 4.46109 | 1.41834 | Promoter | 5 | 8893721 | 8959225 | 65505 | 1 | ENSMUST00000003717.8 | -10 | Abcb4 | ENSMUSG00000042476.8 |
| chr4 | 126729560 | 126729949 | 3.35536 | 4.00513 | 1.11995 | Intron (ENSMUST00000048194.7/ENSMUSG00000042477.7, intron 3 of 6) | 4 | 126716006 | 126736240 | 20235 | 2 | ENSMUST00000048194.7 | 6291 | Tfap2e | ENSMUSG00000042477.7 |
| chr4 | 126731536 | 126731646 | 3.62634 | 4.36047 | 1.315 | Intron (ENSMUST00000048194.7/ENSMUSG00000042477.7, intron 3 of 6) | 4 | 126716006 | 126736240 | 20235 | 2 | ENSMUST00000048194.7 | 4594 | Tfap2e | ENSMUSG00000042477.7 |
| chr4 | 126732222 | 126732589 | 4.2099 | 5.60747 | 2.22152 | Intron (ENSMUST00000048194.7/ENSMUSG00000042477.7, intron 3 of 6) | 4 | 126716006 | 126736240 | 20235 | 2 | ENSMUST00000048194.7 | 3651 | Tfap2e | ENSMUSG00000042477.7 |
| chr4 | 126733415 | 126733753 | 3.56082 | 4.5942 | 1.49256 | Intron (ENSMUST00000048194.7/ENSMUSG00000042477.7, intron 3 of 6) | 4 | 126716006 | 126736240 | 20235 | 2 | ENSMUST00000048194.7 | 2487 | Tfap2e | ENSMUSG00000042477.7 |
| chr4 | 126734107 | 126735960 | 4.15349 | 5.50418 | 2.15529 | Promoter | 4 | 126716006 | 126736240 | 20235 | 2 | ENSMUST00000048194.7 | 280 | Tfap2e | ENSMUSG00000042477.7 |
| chr4 | 126736307 | 126736641 | 3.55501 | 4.23753 | 1.28118 | Promoter | 4 | 126716006 | 126736240 | 20235 | 2 | ENSMUST00000048194.7 | -68 | Tfap2e | ENSMUSG00000042477.7 |
| chr4 | 126738185 | 126738441 | 4.0663 | 5.29579 | 2.05425 | Distal Intergenic | 4 | 126716006 | 126736240 | 20235 | 2 | ENSMUST00000048194.7 | -1946 | Tfap2e | ENSMUSG00000042477.7 |
| chr2 | 74677438 | 74677831 | 4.18709 | 5.46739 | 2.10392 | 3' UTR | 2 | 74679557 | 74687016 | 7460 | 1 | ENSMUST00000048086.8 | -1726 | Hoxd11 | ENSMUSG00000042499.13 |
| chr2 | 74678808 | 74680295 | 3.83735 | 4.94861 | 1.75741 | Promoter | 2 | 74679557 | 74687016 | 7460 | 1 | ENSMUST00000048086.8 | 0 | Hoxd11 | ENSMUSG00000042499.13 |
| chr2 | 74680632 | 74680941 | 3.85671 | 5.09879 | 1.85548 | Intron (ENSMUST00000048086.8/ENSMUSG00000042499.13, intron 1 of 1) | 2 | 74679557 | 74687016 | 7460 | 1 | ENSMUST00000048086.8 | 1076 | Hoxd11 | ENSMUSG00000042499.13 |
| chr2 | 74681279 | 74683370 | 4.47571 | 6.20024 | 2.68232 | Promoter | 2 | 74682323 | 74684168 | 1846 | 1 | ENSMUST00000142312.2 | 0 | Hoxd11 | ENSMUSG00000042499.13 |
| chr19 | 42225972 | 42226164 | 3.85571 | 4.84479 | 1.70058 | Distal Intergenic | 19 | 42247574 | 42270348 | 22775 | 1 | ENSMUST00000122375.2 | -21410 | Golga7b | ENSMUSG00000042532.9 |
| chr19 | 42254475 | 42254683 | 3.6633 | 4.62669 | 1.50951 | Intron (ENSMUST00000122375.2/ENSMUSG00000042532.9, intron 2 of 6) | 19 | 42255739 | 42270348 | 14610 | 1 | ENSMUST00000087123.5 | -1056 | Golga7b | ENSMUSG00000042532.9 |
| chr19 | 42255062 | 42256168 | 3.92576 | 4.98283 | 1.80236 | Promoter | 19 | 42255739 | 42270348 | 14610 | 1 | ENSMUST00000087123.5 | 0 | Golga7b | ENSMUSG00000042532.9 |
| chr12 | 84068276 | 84068543 | 3.81692 | 4.93674 | 1.69978 | Promoter | 12 | 84069325 | 84076019 | 6695 | 1 | ENSMUST00000046422.5 | -782 | Acot5 | ENSMUSG00000042540.7 |
| chr12 | 84069048 | 84070023 | 4.06664 | 5.4393 | 2.073 | Promoter | 12 | 84069325 | 84076019 | 6695 | 1 | ENSMUST00000046422.5 | 0 | Acot5 | ENSMUSG00000042540.7 |
| chr1 | 129273497 | 129273753 | 3.49466 | 4.16425 | 1.24663 | Promoter | 1 | 129273485 | 129629012 | 355528 | 1 | ENSMUST00000140629.1 | 13 | Thsd7b | ENSMUSG00000042581.9 |
| chr5 | 121894780 | 121895059 | 4.45458 | 5.97878 | 2.43643 | Promoter | 5 | 121879821 | 121895381 | 15561 | 2 | ENSMUST00000144069.1 | 322 | Cux2 | ENSMUSG00000042589.13 |
| chr5 | 121895546 | 121896039 | 3.91931 | 4.90654 | 1.66675 | Promoter | 5 | 121879821 | 121895381 | 15561 | 2 | ENSMUST00000144069.1 | -166 | Cux2 | ENSMUSG00000042589.13 |
| chr5 | 121925506 | 121926060 | 3.63721 | 4.37448 | 1.32134 | Promoter | 5 | 121860308 | 121924921 | 64614 | 2 | ENSMUST00000168288.3 | -586 | Cux2 | ENSMUSG00000042589.13 |
| chr5 | 121948560 | 121948808 | 3.3511 | 4.03699 | 1.13028 | Intron (ENSMUST00000111752.4/ENSMUSG00000042589.13, intron 2 of 22) | 5 | 121860308 | 121924921 | 64614 | 2 | ENSMUST00000168288.3 | -23640 | Cux2 | ENSMUSG00000042589.13 |
| chr5 | 122049920 | 122050114 | 3.60712 | 4.32155 | 1.30024 | Promoter | 5 | 121884754 | 122050102 | 165349 | 2 | ENSMUST00000154139.2 | 0 | Cux2 | ENSMUSG00000042589.13 |
| chr5 | 122053364 | 122053627 | 3.55586 | 4.21805 | 1.21078 | Intron (ENSMUST00000135544.1/ENSMUSG00000087386.1, intron 2 of 2) | 5 | 121884754 | 122050102 | 165349 | 2 | ENSMUST00000154139.2 | -3263 | Cux2 | ENSMUSG00000042589.13 |
| chr1 | 19097236 | 19097311 | 3.53248 | 4.18916 | 1.2258 | Intron (ENSMUST00000159108.2/ENSMUSG00000089787.2, intron 1 of 3) | 1 | 19103022 | 19166346 | 63325 | 1 | ENSMUST00000037294.7 | -5711 | Tfap2d | ENSMUSG00000042596.7 |
| chr1 | 19109919 | 19110612 | 3.64055 | 4.40598 | 1.35843 | Intron (ENSMUST00000037294.7/ENSMUSG00000042596.7, intron 3 of 7) | 1 | 19107181 | 19148367 | 41187 | 1 | ENSMUST00000149505.1 | 2739 | Tfap2d | ENSMUSG00000042596.7 |
| chr1 | 19111283 | 19111451 | 3.99425 | 5.06011 | 1.78572 | Intron (ENSMUST00000037294.7/ENSMUSG00000042596.7, intron 3 of 7) | 1 | 19107181 | 19148367 | 41187 | 1 | ENSMUST00000149505.1 | 4103 | Tfap2d | ENSMUSG00000042596.7 |
| chr2 | 107290993 | 107291215 | 3.61246 | 4.42101 | 1.4221 | Promoter | 2 | 107291395 | 107294472 | 3078 | 1 | ENSMUST00000173191.1 | -180 | Kcna4 | ENSMUSG00000042604.5 |
| chr6 | 5380383 | 5381693 | 3.82144 | 4.82178 | 1.65282 | Distal Intergenic | 6 | 5383386 | 5433022 | 49637 | 1 | ENSMUST00000043294.6 | -1693 | Asb4 | ENSMUSG00000042607.10 |
| chr6 | 5383935 | 5384058 | 4.19761 | 5.63495 | 2.19303 | Promoter | 6 | 5383386 | 5433022 | 49637 | 1 | ENSMUST00000043294.6 | 550 | Asb4 | ENSMUSG00000042607.10 |
| chr6 | 5384796 | 5385416 | 3.77306 | 4.66378 | 1.57997 | Intron (ENSMUST00000043294.6/ENSMUSG00000042607.10, intron 1 of 5) | 6 | 5383386 | 5433022 | 49637 | 1 | ENSMUST00000043294.6 | 1411 | Asb4 | ENSMUSG00000042607.10 |
| chr6 | 5385728 | 5386190 | 3.33453 | 3.83722 | 1.03913 | Intron (ENSMUST00000043294.6/ENSMUSG00000042607.10, intron 1 of 5) | 6 | 5383386 | 5433022 | 49637 | 1 | ENSMUST00000043294.6 | 2343 | Asb4 | ENSMUSG00000042607.10 |
| chr6 | 5388007 | 5388465 | 3.56395 | 4.26816 | 1.3033 | Intron (ENSMUST00000043294.6/ENSMUSG00000042607.10, intron 1 of 5) | 6 | 5390427 | 5402249 | 11823 | 1 | ENSMUST00000183358.1 | -1962 | Asb4 | ENSMUSG00000042607.10 |
| chr6 | 5391015 | 5392867 | 3.99857 | 5.13218 | 1.92405 | Promoter | 6 | 5390427 | 5402249 | 11823 | 1 | ENSMUST00000183358.1 | 589 | Asb4 | ENSMUSG00000042607.10 |
| chr6 | 5395136 | 5395266 | 3.74276 | 4.70316 | 1.60684 | Intron (ENSMUST00000043294.6/ENSMUSG00000042607.10, intron 2 of 5) | 6 | 5390427 | 5402249 | 11823 | 1 | ENSMUST00000183358.1 | 4710 | Asb4 | ENSMUSG00000042607.10 |
| chr2 | 153030307 | 153030679 | 3.76098 | 4.6967 | 1.63476 | Distal Intergenic | 2 | 153031852 | 153055775 | 23924 | 1 | ENSMUST00000037235.6 | -1173 | Xkr7 | ENSMUSG00000042631.7 |
| chr2 | 153031099 | 153031400 | 4.12141 | 5.43108 | 2.11828 | Promoter | 2 | 153031852 | 153055775 | 23924 | 1 | ENSMUST00000037235.6 | -452 | Xkr7 | ENSMUSG00000042631.7 |
| chr2 | 153031720 | 153032409 | 4.78568 | 6.83157 | 3.21496 | Promoter | 2 | 153031852 | 153055775 | 23924 | 1 | ENSMUST00000037235.6 | 0 | Xkr7 | ENSMUSG00000042631.7 |
| chr2 | 153032760 | 153032884 | 3.9354 | 5.03826 | 1.813 | Promoter | 2 | 153031852 | 153055775 | 23924 | 1 | ENSMUST00000037235.6 | 909 | Xkr7 | ENSMUSG00000042631.7 |
| chr2 | 153033238 | 153033398 | 3.7788 | 4.7829 | 1.62171 | Intron (ENSMUST00000037235.6/ENSMUSG00000042631.7, intron 1 of 2) | 2 | 153031852 | 153055775 | 23924 | 1 | ENSMUST00000037235.6 | 1387 | Xkr7 | ENSMUSG00000042631.7 |
| chr13 | 104863582 | 104864011 | 4.1717 | 5.42282 | 2.05183 | Promoter | 13 | 104845283 | 104863893 | 18611 | 2 | ENSMUST00000043061.4 | 0 | Fam159b | ENSMUSG00000042655.4 |
| chr1 | 153641719 | 153641865 | 4.03264 | 5.43459 | 2.06309 | Distal Intergenic | 1 | 153653025 | 153700323 | 47299 | 1 | ENSMUST00000041776.6 | -11160 | Rgs8 | ENSMUSG00000042671.7 |
| chr1 | 153650260 | 153650365 | 3.66813 | 4.49711 | 1.48702 | Distal Intergenic | 1 | 153653025 | 153700323 | 47299 | 1 | ENSMUST00000041776.6 | -2660 | Rgs8 | ENSMUSG00000042671.7 |
| chr1 | 153651089 | 153651318 | 3.63618 | 4.37567 | 1.34092 | Distal Intergenic | 1 | 153653025 | 153700323 | 47299 | 1 | ENSMUST00000041776.6 | -1707 | Rgs8 | ENSMUSG00000042671.7 |
| chr1 | 153652449 | 153652601 | 3.74387 | 4.571 | 1.44255 | Promoter | 1 | 153653025 | 153700323 | 47299 | 1 | ENSMUST00000041776.6 | -424 | Rgs8 | ENSMUSG00000042671.7 |
| chr1 | 153653118 | 153653386 | 3.71966 | 4.53897 | 1.45169 | Promoter | 1 | 153653025 | 153700323 | 47299 | 1 | ENSMUST00000041776.6 | 94 | Rgs8 | ENSMUSG00000042671.7 |
| chr1 | 153655795 | 153656210 | 3.27287 | 3.7365 | 1.01733 | Intron (ENSMUST00000041776.6/ENSMUSG00000042671.7, intron 1 of 6) | 1 | 153653025 | 153700323 | 47299 | 1 | ENSMUST00000041776.6 | 2771 | Rgs8 | ENSMUSG00000042671.7 |
| chr1 | 153660559 | 153660640 | 4.07919 | 5.40424 | 2.03058 | Promoter | 1 | 153661081 | 153691000 | 29920 | 1 | ENSMUST00000124500.2 | -441 | Rgs8 | ENSMUSG00000042671.7 |
| chr1 | 153661565 | 153662001 | 3.72592 | 4.55548 | 1.46768 | Promoter | 1 | 153661081 | 153691000 | 29920 | 1 | ENSMUST00000124500.2 | 485 | Rgs8 | ENSMUSG00000042671.7 |
| chr1 | 153665230 | 153665372 | 3.42526 | 3.98233 | 1.07483 | Promoter | 1 | 153665274 | 153692777 | 27504 | 1 | ENSMUST00000152114.2 | 0 | Rgs8 | ENSMUSG00000042671.7 |
| chr1 | 153668069 | 153668144 | 3.53377 | 4.17885 | 1.188 | Promoter | 1 | 153667489 | 153670839 | 3351 | 1 | ENSMUST00000147794.1 | 581 | Rgs8 | ENSMUSG00000042671.7 |
| chr4 | 125146094 | 125146513 | 4.09059 | 5.24495 | 1.90525 | Distal Intergenic | 4 | 125118423 | 125127840 | 9418 | 2 | ENSMUST00000036188.7 | -18255 | Zc3h12a | ENSMUSG00000042677.7 |
| chr4 | 125151957 | 125152032 | 3.24987 | 3.7074 | 1.03436 | Distal Intergenic | 4 | 125118423 | 125127840 | 9418 | 2 | ENSMUST00000036188.7 | -24118 | Zc3h12a | ENSMUSG00000042677.7 |
| chr4 | 125160848 | 125161170 | 3.56308 | 4.32951 | 1.34039 | Distal Intergenic | 4 | 125118423 | 125127840 | 9418 | 2 | ENSMUST00000036188.7 | -33009 | Zc3h12a | ENSMUSG00000042677.7 |
| chr4 | 125169577 | 125170113 | 4.0773 | 5.29431 | 2.04723 | Distal Intergenic | 4 | 125118423 | 125127840 | 9418 | 2 | ENSMUST00000036188.7 | -41738 | Zc3h12a | ENSMUSG00000042677.7 |
| chr4 | 125176955 | 125177039 | 4.27781 | 5.64451 | 2.21596 | Distal Intergenic | 4 | 125118423 | 125127840 | 9418 | 2 | ENSMUST00000036188.7 | -49116 | Zc3h12a | ENSMUSG00000042677.7 |
| chr11 | 60509831 | 60510200 | 3.50934 | 4.13372 | 1.16108 | Promoter | 11 | 60509534 | 60521986 | 12453 | 1 | ENSMUST00000122825.1 | 298 | Myo15 | ENSMUSG00000042678.11 |
| chr1 | 153422915 | 153423029 | 4.11597 | 5.4359 | 2.08589 | Distal Intergenic | 1 | 153425162 | 153452574 | 27413 | 1 | ENSMUST00000042373.6 | -2133 | Shcbp1l | ENSMUSG00000042708.7 |
| chr1 | 153425100 | 153425532 | 4.08437 | 5.50359 | 2.16028 | Promoter | 1 | 153425162 | 153452574 | 27413 | 1 | ENSMUST00000042373.6 | 0 | Shcbp1l | ENSMUSG00000042708.7 |
| chr11 | 60385215 | 60385360 | 3.59686 | 4.31131 | 1.3138 | Intron (ENSMUST00000108723.3/ENSMUSG00000056598.11, intron 10 of 13) | 11 | 60401478 | 60417018 | 15541 | 2 | ENSMUST00000145532.2 | 31658 | Atpaf2 | ENSMUSG00000042709.9 |
| chr12 | 81779172 | 81779263 | 3.95434 | 4.96311 | 1.68806 | Intron (ENSMUST00000035987.7/ENSMUSG00000042724.7, intron 1 of 10) | 12 | 81714950 | 81781170 | 66221 | 2 | ENSMUST00000035987.7 | 1907 | Map3k9 | ENSMUSG00000042724.7 |
| chr12 | 81780531 | 81781062 | 3.60232 | 4.45085 | 1.41594 | Promoter | 12 | 81714950 | 81781170 | 66221 | 2 | ENSMUST00000035987.7 | 108 | Map3k9 | ENSMUSG00000042724.7 |
| chr12 | 81631346 | 81632418 | 3.93794 | 5.14627 | 1.90496 | Promoter | 12 | 81631369 | 81664941 | 33573 | 1 | ENSMUST00000036116.5 | 0 | Ttc9 | ENSMUSG00000042734.5 |
| chr1 | 152953933 | 152954297 | 3.52972 | 4.27393 | 1.31898 | Promoter | 1 | 152954993 | 153119164 | 164172 | 1 | ENSMUST00000186368.1 | -696 | Nmnat2 | ENSMUSG00000042751.9 |
| chr1 | 152954869 | 152956261 | 3.54388 | 4.24795 | 1.32352 | Promoter | 1 | 152954993 | 153119164 | 164172 | 1 | ENSMUST00000186368.1 | 0 | Nmnat2 | ENSMUSG00000042751.9 |
| chr9 | 87143858 | 87144943 | 3.71903 | 4.65328 | 1.58336 | Promoter | 9 | 87144306 | 87184045 | 39740 | 1 | ENSMUST00000049457.8 | 0 | Mrap2 | ENSMUSG00000042761.9 |
| chr9 | 87145709 | 87145901 | 4.35117 | 5.79032 | 2.33418 | Intron (ENSMUST00000049457.8/ENSMUSG00000042761.9, intron 1 of 3) | 9 | 87144343 | 87184045 | 39703 | 1 | ENSMUST00000179313.2 | 1367 | Mrap2 | ENSMUSG00000042761.9 |
| chr4 | 124861174 | 124862134 | 3.71821 | 4.55609 | 1.45017 | Promoter | 4 | 124855239 | 124862171 | 6933 | 2 | ENSMUST00000064444.7 | 37 | Maneal | ENSMUSG00000042763.9 |
| chr4 | 124862490 | 124862845 | 3.35671 | 3.86254 | 1.01733 | Promoter | 4 | 124855239 | 124862171 | 6933 | 2 | ENSMUST00000064444.7 | -320 | Maneal | ENSMUSG00000042763.9 |
| chr3 | 89243732 | 89243966 | 3.39994 | 3.96313 | 1.14296 | Exon (ENSMUST00000107464.2/ENSMUSG00000042766.12, exon 3 of 10) | 3 | 89234177 | 89245013 | 10837 | 2 | ENSMUST00000107464.2 | 1047 | Trim46 | ENSMUSG00000042766.12 |
| chr3 | 89244837 | 89245121 | 3.43133 | 3.99671 | 1.09174 | Promoter | 3 | 89234177 | 89245013 | 10837 | 2 | ENSMUST00000107464.2 | 0 | Trim46 | ENSMUSG00000042766.12 |
| chr6 | 135179920 | 135180020 | 3.76439 | 4.70106 | 1.61219 | Distal Intergenic | 6 | 135137519 | 135168215 | 30697 | 2 | ENSMUST00000045855.7 | -11706 | Hebp1 | ENSMUSG00000042770.7 |
| chr9 | 119479154 | 119479406 | 3.56328 | 4.29749 | 1.34817 | Distal Intergenic | 9 | 119444951 | 119464230 | 19280 | 1 | ENSMUST00000164213.2 | 34204 | Exog | ENSMUSG00000042787.8 |
| chr9 | 119483618 | 119483942 | 3.39687 | 3.95148 | 1.11783 | 3' UTR | 9 | 119444951 | 119464230 | 19280 | 1 | ENSMUST00000164213.2 | 38668 | Exog | ENSMUSG00000042787.8 |
| chr1 | 135017742 | 135017999 | 3.36814 | 3.8798 | 1.01733 | Promoter | 1 | 134986355 | 135017779 | 31425 | 2 | ENSMUST00000137968.2 | 0 | Lgr6 | ENSMUSG00000042793.8 |
| chr1 | 135018823 | 135019147 | 3.40963 | 4.00883 | 1.19925 | Intron (ENSMUST00000044828.8/ENSMUSG00000042793.8, intron 6 of 17) | 1 | 134986355 | 135017779 | 31425 | 2 | ENSMUST00000137968.2 | -1045 | Lgr6 | ENSMUSG00000042793.8 |
| chr1 | 135084910 | 135085116 | 3.45169 | 4.0369 | 1.12726 | Intron (ENSMUST00000044828.8/ENSMUSG00000042793.8, intron 1 of 17) | 1 | 134983301 | 135105276 | 121976 | 2 | ENSMUST00000044828.8 | 20160 | Lgr6 | ENSMUSG00000042793.8 |
| chr1 | 135086113 | 135086440 | 3.9482 | 4.98698 | 1.74663 | Intron (ENSMUST00000044828.8/ENSMUSG00000042793.8, intron 1 of 17) | 1 | 134983301 | 135105276 | 121976 | 2 | ENSMUST00000044828.8 | 18836 | Lgr6 | ENSMUSG00000042793.8 |
| chr1 | 135099360 | 135099790 | 3.46616 | 4.11049 | 1.18161 | Intron (ENSMUST00000044828.8/ENSMUSG00000042793.8, intron 1 of 17) | 1 | 134983301 | 135105276 | 121976 | 2 | ENSMUST00000044828.8 | 5486 | Lgr6 | ENSMUSG00000042793.8 |
| chr1 | 135100295 | 135100871 | 3.48093 | 4.1031 | 1.20311 | Intron (ENSMUST00000044828.8/ENSMUSG00000042793.8, intron 1 of 17) | 1 | 134983301 | 135105276 | 121976 | 2 | ENSMUST00000044828.8 | 4405 | Lgr6 | ENSMUSG00000042793.8 |
| chr1 | 135103620 | 135104377 | 3.68178 | 4.46031 | 1.38374 | Promoter | 1 | 134983301 | 135105276 | 121976 | 2 | ENSMUST00000044828.8 | 899 | Lgr6 | ENSMUSG00000042793.8 |
| chr1 | 135104834 | 135105260 | 3.96285 | 5.02938 | 1.78721 | Promoter | 1 | 134983301 | 135105276 | 121976 | 2 | ENSMUST00000044828.8 | 16 | Lgr6 | ENSMUSG00000042793.8 |
| chr4 | 152275383 | 152275732 | 3.51194 | 4.45991 | 1.42097 | Intron (ENSMUST00000105650.2/ENSMUSG00000042804.8, intron 1 of 5) | 4 | 152274362 | 152285337 | 10976 | 1 | ENSMUST00000105651.2 | 1022 | Gpr153 | ENSMUSG00000042804.8 |
| chr4 | 152281469 | 152281767 | 3.24043 | 3.68781 | 1.01733 | Promoter | 4 | 152282370 | 152285336 | 2967 | 1 | ENSMUST00000144035.1 | -603 | Gpr153 | ENSMUSG00000042804.8 |
| chr4 | 152282737 | 152283197 | 4.00078 | 5.32873 | 1.9921 | Promoter | 4 | 152282370 | 152285336 | 2967 | 1 | ENSMUST00000144035.1 | 368 | Gpr153 | ENSMUSG00000042804.8 |
| chr4 | 152286205 | 152287713 | 3.64057 | 4.64257 | 1.56437 | 3' UTR | 4 | 152282370 | 152285336 | 2967 | 1 | ENSMUST00000144035.1 | 3836 | Gpr153 | ENSMUSG00000042804.8 |
| chr1 | 54192915 | 54193239 | 3.664 | 4.45942 | 1.43733 | Promoter | 1 | 53806876 | 54194048 | 387173 | 2 | ENSMUST00000120904.2 | 809 | Hecw2 | ENSMUSG00000042807.10 |
| chr1 | 54194157 | 54194647 | 3.84797 | 4.80361 | 1.62559 | Promoter | 1 | 53806876 | 54194048 | 387173 | 2 | ENSMUST00000120904.2 | -110 | Hecw2 | ENSMUSG00000042807.10 |
| chr1 | 54195100 | 54195418 | 4.1733 | 5.45682 | 2.09775 | Promoter | 1 | 53941496 | 54195168 | 253673 | 2 | ENSMUST00000146850.1 | 0 | Hecw2 | ENSMUSG00000042807.10 |
| chr12 | 76796358 | 76796521 | 3.30741 | 3.78832 | 1.01733 | Promoter | 12 | 76792335 | 76795554 | 3220 | 2 | ENSMUST00000082431.3 | -805 | Gpx2 | ENSMUSG00000042808.7 |
| chr8 | 121084293 | 121085418 | 4.42528 | 6.32486 | 2.81197 | Promoter | 8 | 121084386 | 121088144 | 3759 | 1 | ENSMUST00000181504.1 | 0 | Foxf1 | ENSMUSG00000042812.5 |
| chr18 | 42691328 | 42691684 | 3.78698 | 4.6676 | 1.53554 | Intron (ENSMUST00000136118.2/ENSMUSG00000024500.13, intron 5 of 8) | 18 | 42578020 | 42579652 | 1633 | 2 | ENSMUST00000054738.3 | -111677 | Gpr151 | ENSMUSG00000042816.3 |
| chr5 | 147399939 | 147400593 | 4.33454 | 5.84043 | 2.36649 | Promoter | 5 | 147330741 | 147400489 | 69749 | 2 | ENSMUST00000049324.9 | 0 | Flt3 | ENSMUSG00000042817.12 |
| chr10 | 64082372 | 64082497 | 3.30495 | 3.78468 | 1.01733 | Intron (ENSMUST00000105441.2/ENSMUSG00000060843.6, intron 7 of 17) | 10 | 64087496 | 64090247 | 2752 | 2 | ENSMUST00000148712.1 | 7750 | Lrrtm3 | ENSMUSG00000042846.8 |
| chr10 | 64083779 | 64083988 | 3.46853 | 4.11186 | 1.25519 | Intron (ENSMUST00000105441.2/ENSMUSG00000060843.6, intron 7 of 17) | 10 | 64087496 | 64090247 | 2752 | 2 | ENSMUST00000148712.1 | 6259 | Lrrtm3 | ENSMUSG00000042846.8 |
| chr10 | 64085040 | 64085708 | 3.58285 | 4.33417 | 1.39273 | Intron (ENSMUST00000105441.2/ENSMUSG00000060843.6, intron 7 of 17) | 10 | 64087496 | 64090247 | 2752 | 2 | ENSMUST00000148712.1 | 4539 | Lrrtm3 | ENSMUSG00000042846.8 |
| chr10 | 64086713 | 64087036 | 3.73341 | 4.59992 | 1.56601 | Intron (ENSMUST00000105441.2/ENSMUSG00000060843.6, intron 7 of 17) | 10 | 64087496 | 64090247 | 2752 | 2 | ENSMUST00000148712.1 | 3211 | Lrrtm3 | ENSMUSG00000042846.8 |
| chr10 | 64088751 | 64088946 | 3.98697 | 5.36096 | 2.04829 | Exon (ENSMUST00000105439.1/ENSMUSG00000042846.8, exon 2 of 3) | 10 | 64087496 | 64090247 | 2752 | 2 | ENSMUST00000148712.1 | 1301 | Lrrtm3 | ENSMUSG00000042846.8 |
| chr10 | 64089332 | 64090350 | 3.68544 | 4.55962 | 1.52673 | Promoter | 10 | 63928472 | 64090274 | 161803 | 2 | ENSMUST00000105439.1 | 0 | Lrrtm3 | ENSMUSG00000042846.8 |
| chr10 | 64091041 | 64091252 | 3.6033 | 4.53133 | 1.45524 | Promoter | 10 | 63929890 | 64090277 | 160388 | 2 | ENSMUST00000131945.1 | -765 | Lrrtm3 | ENSMUSG00000042846.8 |
| chr6 | 113193400 | 113196172 | 4.16461 | 5.45908 | 2.10287 | Promoter | 6 | 113168090 | 113195384 | 27295 | 2 | ENSMUST00000162280.1 | 0 | Lhfpl4 | ENSMUSG00000042873.10 |
| chr18 | 10324335 | 10325005 | 3.71328 | 4.70328 | 1.56513 | Promoter | 18 | 10325177 | 10502412 | 177236 | 1 | ENSMUST00000173356.2 | -172 | Greb1l | ENSMUSG00000042942.10 |
| chr3 | 154652971 | 154653164 | 3.93495 | 5.01375 | 1.83647 | Distal Intergenic | 3 | 154664096 | 154757527 | 93432 | 1 | ENSMUST00000172865.2 | -10932 | Erich3 | ENSMUSG00000042943.9 |
| chr3 | 154662742 | 154663062 | 4.05658 | 5.60515 | 2.23183 | Distal Intergenic | 3 | 154664096 | 154757527 | 93432 | 1 | ENSMUST00000172865.2 | -1034 | Erich3 | ENSMUSG00000042943.9 |

| | | | | | | | | | | | | | | | |
|---|---|---|---|---|---|---|---|---|---|---|---|---|---|---|---|
| chr3 | 154704862 | 154704944 | 3.94207 | 5.03835 | 1.82265 | Exon (ENSMUST00000172865.2/ENSMUSG00000042943.9, exon 5 of 13) | 3 | 154711133 | 154749012 | 37880 | 1 | ENSMUST00000051862.6 | -6189 | Erich3 | ENSMUSG00000042943.9 |
| chr15 | 7397431 | 7398435 | 4.26193 | 5.72618 | 2.37299 | Promoter | 15 | 7206120 | 7398304 | 192185 | 2 | ENSMUST00000058593.4 | 0 | Egflam | ENSMUSG00000042961.8 |
| chr15 | 7398778 | 7399498 | 4.13587 | 5.36498 | 2.03315 | Promoter | 15 | 7206583 | 7398395 | 191813 | 2 | ENSMUST00000160207.2 | -384 | Egflam | ENSMUSG00000042961.8 |
| chr15 | 7460088 | 7460356 | 3.45516 | 4.07007 | 1.17864 | Distal Intergenic | 15 | 7206583 | 7398395 | 191813 | 2 | ENSMUST00000160207.2 | -61694 | Egflam | ENSMUSG00000042961.8 |
| chr15 | 7462889 | 7463224 | 3.4976 | 4.11873 | 1.17117 | Distal Intergenic | 15 | 7206583 | 7398395 | 191813 | 2 | ENSMUST00000160207.2 | -64495 | Egflam | ENSMUSG00000042961.8 |
| chr15 | 7463544 | 7463931 | 3.56582 | 4.25206 | 1.27024 | Distal Intergenic | 15 | 7206583 | 7398395 | 191813 | 2 | ENSMUST00000160207.2 | -65150 | Egflam | ENSMUSG00000042961.8 |
| chr12 | 8856295 | 8856503 | 3.37868 | 3.93326 | 1.14487 | Distal Intergenic | 12 | 8873187 | 8877459 | 4273 | 2 | ENSMUST00000151816.1 | 20956 | 9930038B18Rik | ENSMUSG00000042976.5 |
| chr7 | 126248369 | 126248526 | 3.50328 | 4.17742 | 1.24298 | Distal Intergenic | 7 | 126272619 | 126294999 | 22381 | 1 | ENSMUST00000056028.9 | -24093 | Sbk1 | ENSMUSG00000042978.9 |
| chr11 | 120658276 | 120658351 | 3.22044 | 3.65787 | 1.01677 | Promoter | 11 | 120653845 | 120658044 | 4200 | 2 | ENSMUST00000055439.3 | -233 | Notum | ENSMUSG00000042988.5 |
| chr11 | 120660871 | 120662028 | 3.72745 | 4.76978 | 1.62606 | Promoter | 11 | 120657563 | 120661175 | 3613 | 2 | ENSMUST00000150458.1 | 0 | Notum | ENSMUSG00000042988.5 |
| chr11 | 120662375 | 120662735 | 3.90677 | 5.00472 | 1.79604 | Distal Intergenic | 11 | 120657563 | 120661175 | 3613 | 2 | ENSMUST00000150458.1 | -1201 | Notum | ENSMUSG00000042988.5 |
| chr4 | 35121089 | 35121194 | 3.41202 | 3.96107 | 1.07072 | Intron (ENSMUST00000102975.4/ENSMUSG00000073910.5, intron 1 of 3) | 4 | 35152056 | 35154005 | 1950 | 1 | ENSMUST00000058595.6 | -30862 | Ifnk | ENSMUSG00000042993.6 |
| chr4 | 35121785 | 35122675 | 3.56667 | 4.24633 | 1.25056 | Intron (ENSMUST00000102975.4/ENSMUSG00000073910.5, intron 1 of 3) | 4 | 35152056 | 35154005 | 1950 | 1 | ENSMUST00000058595.6 | -29381 | Ifnk | ENSMUSG00000042993.6 |
| chr4 | 73790263 | 73791413 | 3.89195 | 4.94215 | 1.7603 | Promoter | 4 | 73714579 | 73790602 | 76024 | 2 | ENSMUST00000058292.7 | 0 | Rasef | ENSMUSG00000043003.8 |
| chr14 | 19974591 | 19975279 | 3.54054 | 4.31591 | 1.38008 | Intron (ENSMUST00000160013.2/ENSMUSG00000043004.8, intron 1 of 3) | 14 | 19875793 | 19976693 | 100901 | 2 | ENSMUST00000161247.1 | 1414 | Gng2 | ENSMUSG00000043004.8 |
| chr14 | 19975677 | 19975988 | 3.41905 | 4.06831 | 1.21107 | Promoter | 14 | 19875793 | 19976693 | 100901 | 2 | ENSMUST00000161247.1 | 705 | Gng2 | ENSMUSG00000043004.8 |
| chr14 | 19986750 | 19986963 | 3.43624 | 4.05815 | 1.23678 | Distal Intergenic | 14 | 19875655 | 19977627 | 101973 | 2 | ENSMUST00000159073.2 | -9124 | Gng2 | ENSMUSG00000043004.8 |
| chr9 | 74860652 | 74862077 | 4.15747 | 5.47266 | 2.12407 | Promoter | 9 | 74861921 | 74891781 | 29861 | 1 | ENSMUST00000056006.10 | 0 | Onecut1 | ENSMUSG00000043013.10 |
| chr9 | 74862478 | 74862556 | 4.18251 | 5.60385 | 2.19624 | Promoter | 9 | 74861921 | 74891781 | 29861 | 1 | ENSMUST00000056006.10 | 558 | Onecut1 | ENSMUSG00000043013.10 |
| chr9 | 74862931 | 74864269 | 4.01243 | 5.28625 | 1.97536 | Exon (ENSMUST00000056006.10/ENSMUSG00000043013.10, exon 1 of 2) | 9 | 74861921 | 74891781 | 29861 | 1 | ENSMUST00000056006.10 | 1011 | Onecut1 | ENSMUSG00000043013.10 |
| chr9 | 74864619 | 74865638 | 3.74138 | 4.5692 | 1.44455 | Promoter | 9 | 74866484 | 74890360 | 23877 | 1 | ENSMUST00000160499.2 | -846 | Onecut1 | ENSMUSG00000043013.10 |
| chr9 | 74866383 | 74867256 | 4.34757 | 5.98751 | 2.48332 | Promoter | 9 | 74866484 | 74890360 | 23877 | 1 | ENSMUST00000160499.2 | 0 | Onecut1 | ENSMUSG00000043013.10 |
| chr9 | 74867700 | 74868943 | 3.66594 | 4.57079 | 1.48894 | Intron (ENSMUST00000160950.2/ENSMUSG00000093662.2, intron 2 of 2) | 9 | 74866484 | 74890360 | 23877 | 1 | ENSMUST00000160499.2 | 1217 | Onecut1 | ENSMUSG00000043013.10 |
| chr9 | 74869304 | 74869598 | 3.38116 | 3.9213 | 1.09471 | Intron (ENSMUST00000160950.2/ENSMUSG00000093662.2, intron 2 of 2) | 9 | 74866484 | 74890360 | 23877 | 1 | ENSMUST00000160499.2 | 2821 | Onecut1 | ENSMUSG00000043013.10 |
| chr9 | 74870654 | 74870892 | 3.35951 | 4.16421 | 1.24813 | Intron (ENSMUST00000160950.2/ENSMUSG00000093662.2, intron 2 of 2) | 9 | 74866484 | 74890360 | 23877 | 1 | ENSMUST00000160499.2 | 4171 | Onecut1 | ENSMUSG00000043013.10 |
| chr8 | 125053126 | 125053341 | 3.28507 | 3.75492 | 1.01733 | Promoter | 8 | 125054195 | 125261151 | 206957 | 1 | ENSMUST00000074562.5 | -854 | Disc1 | ENSMUSG00000043051.12 |
| chr8 | 125053647 | 125053755 | 4.0329 | 5.14242 | 1.85952 | Promoter | 8 | 125054195 | 125261151 | 206957 | 1 | ENSMUST00000074562.5 | -440 | Disc1 | ENSMUSG00000043051.12 |
| chr8 | 125054130 | 125054529 | 4.28332 | 5.73128 | 2.31668 | Promoter | 8 | 125054195 | 125261151 | 206957 | 1 | ENSMUST00000074562.5 | 0 | Disc1 | ENSMUSG00000043051.12 |
| chr8 | 125055112 | 125055294 | 3.81336 | 4.76978 | 1.62983 | Promoter | 8 | 125054230 | 125261148 | 206919 | 1 | ENSMUST00000121953.1 | 883 | Disc1 | ENSMUSG00000043051.12 |
| chr8 | 125055745 | 125055939 | 3.38925 | 3.99421 | 1.188 | Intron (ENSMUST00000074562.5/ENSMUSG00000043051.12, intron 1 of 12) | 8 | 125054230 | 125261148 | 206919 | 1 | ENSMUST00000121953.1 | 1516 | Disc1 | ENSMUSG00000043051.12 |
| chr8 | 125134376 | 125134451 | 3.42312 | 3.99086 | 1.11688 | Intron (ENSMUST00000074562.5/ENSMUSG00000043051.12, intron 5 of 12) | 8 | 125087987 | 125102878 | 14892 | 1 | ENSMUST00000126111.1 | 46390 | Disc1 | ENSMUSG00000043051.12 |
| chr8 | 125134947 | 125135496 | 3.73893 | 4.63416 | 1.5469 | Exon (ENSMUST00000074562.5/ENSMUSG00000043051.12, exon 6 of 13) | 8 | 125087987 | 125102878 | 14892 | 1 | ENSMUST00000126111.1 | 46961 | Disc1 | ENSMUSG00000043051.12 |
| chr9 | 24542052 | 24542142 | 2.3155 | 15.11302 | 10.38442 | Distal Intergenic | 9 | 24411803 | 24503140 | 91338 | 2 | ENSMUST00000170356.1 | -38913 | Dpy19l1 | ENSMUSG00000043067.10 |
| chr8 | 124750550 | 124750810 | 3.23008 | 3.84128 | 1.06325 | Promoter | 8 | 124740257 | 124751809 | 11553 | 2 | ENSMUST00000055257.5 | 999 | Fam89a | ENSMUSG00000043068.5 |
| chr8 | 124751469 | 124751735 | 3.81516 | 4.92057 | 1.70476 | Promoter | 8 | 124740257 | 124751809 | 11553 | 2 | ENSMUST00000055257.5 | 74 | Fam89a | ENSMUSG00000043068.5 |
| chr4 | 141617906 | 141618306 | 4.32953 | 5.91693 | 2.42052 | Promoter | 4 | 141614233 | 141618238 | 4006 | 2 | ENSMUST00000053263.8 | 0 | Tmem82 | ENSMUSG00000043085.9 |
| chr15 | 99040925 | 99041235 | 3.47036 | 4.15204 | 1.28125 | Distal Intergenic | 15 | 99029891 | 99038105 | 8215 | 1 | ENSMUST00000058914.8 | 11035 | Tuba1c | ENSMUSG00000043091.8 |
| chr11 | 75165885 | 75166687 | 4.1625 | 5.57027 | 2.29262 | Exon (ENSMUST00000055619.4/ENSMUSG00000043099.4, exon 2 of 2) | 11 | 75167153 | 75168146 | 994 | 2 | ENSMUST00000131720.1 | 1459 | Hic1 | ENSMUSG00000043099.4 |
| chr11 | 75167121 | 75167412 | 3.98653 | 5.21911 | 1.95826 | Promoter | 11 | 75167153 | 75168146 | 994 | 2 | ENSMUST00000131720.1 | 734 | Hic1 | ENSMUSG00000043099.4 |
| chr11 | 75168120 | 75168839 | 3.71356 | 4.8758 | 1.7036 | Promoter | 11 | 75167153 | 75168146 | 994 | 2 | ENSMUST00000131720.1 | 0 | Hic1 | ENSMUSG00000043099.4 |
| chr2 | 31812854 | 31813093 | 3.9762 | 5.15028 | 1.90713 | Downstream (<1kb) | 2 | 31806166 | 31810580 | 4415 | 2 | ENSMUST00000057407.2 | -2275 | Qrfp | ENSMUSG00000043102.2 |
| chr2 | 31821032 | 31821473 | 3.3711 | 3.9168 | 1.12101 | Exon (ENSMUST00000028188.7/ENSMUSG00000026841.7, exon 5 of 7) | 2 | 31806166 | 31810580 | 4415 | 2 | ENSMUST00000057407.2 | -10453 | Qrfp | ENSMUSG00000043102.2 |
| chr2 | 31821804 | 31821940 | 3.48425 | 4.10921 | 1.20639 | Intron (ENSMUST00000028188.7/ENSMUSG00000026841.7, intron 4 of 6) | 2 | 31806166 | 31810580 | 4415 | 2 | ENSMUST00000057407.2 | -11225 | Qrfp | ENSMUSG00000043102.2 |
| chr2 | 132877816 | 132878419 | 3.57998 | 4.26606 | 1.25226 | Exon (ENSMUST00000049787.2/ENSMUSG00000043110.2, exon 3 of 5) | 2 | 132868305 | 132880891 | 12587 | 2 | ENSMUST00000049787.2 | 2472 | Lrrn4 | ENSMUSG00000043110.2 |
| chr2 | 132878927 | 132879875 | 3.9943 | 5.13097 | 1.86779 | Exon (ENSMUST00000049787.2/ENSMUSG00000043110.2, exon 2 of 5) | 2 | 132868305 | 132880891 | 12587 | 2 | ENSMUST00000049787.2 | 1016 | Lrrn4 | ENSMUSG00000043110.2 |
| chr2 | 132881421 | 132881947 | 3.42992 | 4.00158 | 1.11475 | Promoter | 2 | 132868305 | 132880891 | 12587 | 2 | ENSMUST00000049787.2 | -531 | Lrrn4 | ENSMUSG00000043110.2 |
| chr12 | 112467471 | 112468041 | 3.58593 | 4.28215 | 1.27794 | Distal Intergenic | 12 | 112489448 | 112499927 | 10480 | 1 | ENSMUST00000057465.6 | -21407 | A530016L24Rik | ENSMUSG00000043122.6 |
| chr15 | 99599368 | 99599515 | 4.37763 | 5.84014 | 2.34892 | Distal Intergenic | 15 | 99601400 | 99605477 | 4078 | 1 | ENSMUST00000023754.5 | -1885 | Aqp6 | ENSMUSG00000043144.5 |
| chr15 | 99599928 | 99600730 | 3.80172 | 4.69478 | 1.54899 | Promoter | 15 | 99601400 | 99605477 | 4078 | 1 | ENSMUST00000023754.5 | -670 | Aqp6 | ENSMUSG00000043144.5 |
| chr15 | 99601115 | 99601556 | 3.45944 | 4.0522 | 1.13979 | Promoter | 15 | 99601400 | 99605477 | 4078 | 1 | ENSMUST00000023754.5 | 0 | Aqp6 | ENSMUSG00000043144.5 |
| chr15 | 99606548 | 99606680 | 3.56109 | 4.24577 | 1.26866 | Distal Intergenic | 15 | 99601400 | 99605477 | 4078 | 1 | ENSMUST00000023754.5 | 5149 | Aqp6 | ENSMUSG00000043144.5 |
| chr15 | 99608142 | 99608339 | 3.30908 | 3.79093 | 1.01733 | Distal Intergenic | 15 | 99601400 | 99605477 | 4078 | 1 | ENSMUST00000023754.5 | 6743 | Aqp6 | ENSMUSG00000043144.5 |
| chr15 | 99610741 | 99611113 | 3.47207 | 4.09247 | 1.216 | Distal Intergenic | 15 | 99601400 | 99605477 | 4078 | 1 | ENSMUST00000023754.5 | 9342 | Aqp6 | ENSMUSG00000043144.5 |
| chr15 | 99613707 | 99613832 | 3.7626 | 4.68246 | 1.60939 | Distal Intergenic | 15 | 99601400 | 99605477 | 4078 | 1 | ENSMUST00000023754.5 | 12308 | Aqp6 | ENSMUSG00000043144.5 |
| chr13 | 22023471 | 22023839 | 3.6447 | 4.48788 | 1.44872 | Distal Intergenic | 13 | 22026699 | 22031610 | 4912 | 1 | ENSMUST00000146121.1 | -2860 | Gm11292 | ENSMUSG00000043145.6 |
| chr13 | 22024176 | 22024983 | 3.47491 | 4.11111 | 1.19479 | Distal Intergenic | 13 | 22026699 | 22031610 | 4912 | 1 | ENSMUST00000146121.1 | -1716 | Gm11292 | ENSMUSG00000043145.6 |
| chr13 | 22025336 | 22025541 | 4.1358 | 5.38167 | 2.07254 | Distal Intergenic | 13 | 22026699 | 22031610 | 4912 | 1 | ENSMUST00000146121.1 | -1158 | Gm11292 | ENSMUSG00000043145.6 |
| chr13 | 22026796 | 22027272 | 3.54245 | 4.30894 | 1.3317 | Promoter | 13 | 22026699 | 22031610 | 4912 | 1 | ENSMUST00000146121.1 | 98 | Gm11292 | ENSMUSG00000043145.6 |
| chr13 | 22028058 | 22028389 | 3.63413 | 4.5303 | 1.47596 | Intron (ENSMUST00000146121.1/ENSMUSG00000043145.6, intron 1 of 1) | 13 | 22026699 | 22031610 | 4912 | 1 | ENSMUST00000146121.1 | 1360 | Gm11292 | ENSMUSG00000043145.6 |
| chr18 | 21651633 | 21652165 | 3.71352 | 4.55377 | 1.44845 | Promoter | 18 | 21651581 | 21655046 | 3466 | 1 | ENSMUST00000127684.2 | 53 | 4930426D05Rik | ENSMUSG00000043168.7 |
| chr18 | 21652566 | 21654139 | 4.02557 | 5.26223 | 1.97771 | Promoter | 18 | 21652563 | 21656105 | 3543 | 1 | ENSMUST00000139804.1 | 4 | 4930426D05Rik | ENSMUSG00000043168.7 |
| chr2 | 5997067 | 5997422 | 3.54565 | 4.22936 | 1.29431 | Intron (ENSMUST00000060092.7/ENSMUSG00000043241.9, intron 8 of 21) | 2 | 6023339 | 6056698 | 33360 | 1 | ENSMUST00000128200.1 | -25917 | Upf2 | ENSMUSG00000043241.9 |
| chr2 | 6063760 | 6064746 | 3.58823 | 4.35089 | 1.42543 | Distal Intergenic | 2 | 6050427 | 6055502 | 5076 | 1 | ENSMUST00000144901.1 | 13334 | Upf2 | ENSMUSG00000043241.9 |
| chr2 | 6066901 | 6067002 | 3.38441 | 3.96122 | 1.1598 | Distal Intergenic | 2 | 6050427 | 6055502 | 5076 | 1 | ENSMUST00000144901.1 | 16475 | Upf2 | ENSMUSG00000043241.9 |
| chr2 | 6067958 | 6068593 | 3.43629 | 4.25181 | 1.31979 | Distal Intergenic | 2 | 6050427 | 6055502 | 5076 | 1 | ENSMUST00000144901.1 | 17532 | Upf2 | ENSMUSG00000043241.9 |
| chr2 | 6069974 | 6070500 | 3.86105 | 4.86638 | 1.74256 | Distal Intergenic | 2 | 6050427 | 6055502 | 5076 | 1 | ENSMUST00000144901.1 | 19548 | Upf2 | ENSMUSG00000043241.9 |
| chr2 | 6071883 | 6072332 | 3.82676 | 4.77127 | 1.6516 | Distal Intergenic | 2 | 6050427 | 6055502 | 5076 | 1 | ENSMUST00000144901.1 | 21457 | Upf2 | ENSMUSG00000043241.9 |
| chr2 | 6075029 | 6075794 | 3.83696 | 4.94632 | 1.72782 | Distal Intergenic | 2 | 6050427 | 6055502 | 5076 | 1 | ENSMUST00000144901.1 | 24603 | Upf2 | ENSMUSG00000043241.9 |
| chr2 | 6076165 | 6076348 | 3.74591 | 4.59573 | 1.5101 | Distal Intergenic | 2 | 6050427 | 6055502 | 5076 | 1 | ENSMUST00000144901.1 | 25739 | Upf2 | ENSMUSG00000043241.9 |
| chr2 | 6076703 | 6077013 | 3.46407 | 4.09774 | 1.25147 | Distal Intergenic | 2 | 6050427 | 6055502 | 5076 | 1 | ENSMUST00000144901.1 | 26277 | Upf2 | ENSMUSG00000043241.9 |
| chr2 | 6080088 | 6080388 | 3.4684 | 4.17712 | 1.23821 | Distal Intergenic | 2 | 6050427 | 6055502 | 5076 | 1 | ENSMUST00000144901.1 | 29662 | Upf2 | ENSMUSG00000043241.9 |
| chr2 | 6081718 | 6082017 | 3.67382 | 4.47179 | 1.42041 | Distal Intergenic | 2 | 6050427 | 6055502 | 5076 | 1 | ENSMUST00000144901.1 | 31292 | Upf2 | ENSMUSG00000043241.9 |
| chr2 | 6082528 | 6082708 | 4.16073 | 5.3752 | 1.99564 | Distal Intergenic | 2 | 6050427 | 6055502 | 5076 | 1 | ENSMUST00000144901.1 | 32102 | Upf2 | ENSMUSG00000043241.9 |
| chr2 | 6083248 | 6083911 | 3.93554 | 4.99568 | 1.78757 | Distal Intergenic | 2 | 6050427 | 6055502 | 5076 | 1 | ENSMUST00000144901.1 | 32822 | Upf2 | ENSMUSG00000043241.9 |
| chr2 | 6084621 | 6084789 | 3.3614 | 3.88889 | 1.07947 | Distal Intergenic | 2 | 6050427 | 6055502 | 5076 | 1 | ENSMUST00000144901.1 | 34195 | Upf2 | ENSMUSG00000043241.9 |
| chr2 | 6085234 | 6085359 | 3.34153 | 3.83966 | 1.01733 | Distal Intergenic | 2 | 6050427 | 6055502 | 5076 | 1 | ENSMUST00000144901.1 | 34808 | Upf2 | ENSMUSG00000043241.9 |
| chr2 | 6085902 | 6086594 | 4.13393 | 5.4147 | 2.05992 | Distal Intergenic | 2 | 6050427 | 6055502 | 5076 | 1 | ENSMUST00000144901.1 | 35476 | Upf2 | ENSMUSG00000043241.9 |
| chr2 | 6087003 | 6087270 | 3.98069 | 5.08815 | 1.82688 | Distal Intergenic | 2 | 6050427 | 6055502 | 5076 | 1 | ENSMUST00000144901.1 | 36577 | Upf2 | ENSMUSG00000043241.9 |
| chr2 | 6090143 | 6090307 | 3.68267 | 4.51881 | 1.44441 | Distal Intergenic | 2 | 6050427 | 6055502 | 5076 | 1 | ENSMUST00000144901.1 | 39717 | Upf2 | ENSMUSG00000043241.9 |
| chr9 | 81862807 | 81863913 | 3.63942 | 4.39355 | 1.34749 | Promoter | 9 | 81863670 | 82027026 | 163357 | 1 | ENSMUST00000057067.4 | 0 | Mei4 | ENSMUSG00000043289.7 |
| chr3 | 69599357 | 69599527 | 3.52872 | 4.26851 | 1.31562 | Promoter | 3 | 69574186 | 69598861 | 24676 | 2 | ENSMUST00000061826.1 | -497 | B3galnt1 | ENSMUSG00000043300.1 |
| chr16 | 94996041 | 94996177 | 3.67667 | 4.5545 | 1.50699 | Intron (ENSMUST00000095873.5/ENSMUSG00000043301.10, intron 1 of 4) | 16 | 94749266 | 94997696 | 248431 | 2 | ENSMUST00000095873.5 | 1519 | Kcnj6 | ENSMUSG00000043301.10 |
| chr16 | 94997816 | 94997914 | 3.75744 | 4.66645 | 1.61915 | Promoter | 16 | 94761080 | 94997696 | 236617 | 2 | ENSMUST00000099508.2 | -121 | Kcnj6 | ENSMUSG00000043301.10 |
| chr16 | 57391455 | 57391671 | 2.41039 | 17.64875 | 12.65958 | Intron (ENSMUST00000159816.2/ENSMUSG00000043336.9, intron 1 of 4) | 16 | 57353277 | 57515690 | 162414 | 1 | ENSMUST00000099667.2 | 38179 | Filip1l | ENSMUSG00000043336.9 |
| chr5 | 144760909 | 144762002 | 3.80437 | 4.74649 | 1.63834 | Promoter | 5 | 144735915 | 144761649 | 25735 | 2 | ENSMUST00000061446.7 | 0 | Tmem130 | ENSMUSG00000043388.7 |
| chr1 | 182663933 | 182664040 | 3.15839 | 3.8976 | 1.07224 | Distal Intergenic | 1 | 182747126 | 182749180 | 2055 | 2 | ENSMUST00000060041.6 | 85140 | Ccdc185 | ENSMUSG00000043429.7 |
| chr1 | 182670424 | 182670523 | 3.32315 | 3.81201 | 1.01733 | Distal Intergenic | 1 | 182747126 | 182749180 | 2055 | 2 | ENSMUST00000060041.6 | 78657 | Ccdc185 | ENSMUSG00000043429.7 |
| chr1 | 182683058 | 182683428 | 3.33661 | 3.83226 | 1.01733 | Distal Intergenic | 1 | 182747126 | 182749180 | 2055 | 2 | ENSMUST00000060041.6 | 65752 | Ccdc185 | ENSMUSG00000043429.7 |
| chr1 | 182727962 | 182728130 | 3.70859 | 4.58159 | 1.53601 | Distal Intergenic | 1 | 182747126 | 182749180 | 2055 | 2 | ENSMUST00000060041.6 | 21050 | Ccdc185 | ENSMUSG00000043429.7 |
| chr11 | 97627193 | 97627279 | 4.01401 | 5.23226 | 1.93081 | Downstream (<1kb) | 11 | 97627389 | 97629702 | 2314 | 2 | ENSMUST00000052281.5 | 2423 | E130012A19Rik | ENSMUSG00000043439.5 |
| chr11 | 97628266 | 97630370 | 3.8722 | 5.04738 | 1.8282 | Promoter | 11 | 97627389 | 97629702 | 2314 | 2 | ENSMUST00000052281.5 | 0 | E130012A19Rik | ENSMUSG00000043439.5 |
| chr11 | 97631118 | 97631562 | 3.82108 | 4.86307 | 1.7021 | Distal Intergenic | 11 | 97627389 | 97629702 | 2314 | 2 | ENSMUST00000052281.5 | -1417 | E130012A19Rik | ENSMUSG00000043439.5 |
| chr3 | 62603775 | 62604270 | 4.38502 | 5.93059 | 2.46071 | Promoter | 3 | 62529077 | 62605140 | 76064 | 2 | ENSMUST00000058535.5 | 870 | Gpr149 | ENSMUSG00000043441.5 |
| chr11 | 59176136 | 59176433 | 3.48941 | 4.21337 | 1.28341 | 3' UTR | 11 | 59175568 | 59182826 | 7259 | 2 | ENSMUST00000108793.3 | 6393 | Gjc2 | ENSMUSG00000043448.8 |
| chr11 | 59177226 | 59177327 | 4.6674 | 6.98722 | 3.2312 | Exon (ENSMUST00000108793.3/ENSMUSG00000043448.8, exon 2 of 2) | 11 | 59175568 | 59182826 | 7259 | 2 | ENSMUST00000108793.3 | 5499 | Gjc2 | ENSMUSG00000043448.8 |
| chr7 | 37563345 | 37563551 | 3.51306 | 4.16836 | 1.2665 | Intron (ENSMUST00000175941.2/ENSMUSG00000043456.11, intron 5 of 6) | 7 | 37472135 | 37479986 | 7852 | 2 | ENSMUST00000176129.1 | -83360 | Zfp536 | ENSMUSG00000043456.11 |
| chr7 | 37565985 | 37566101 | 3.23338 | 3.67724 | 1.01733 | Intron (ENSMUST00000175941.2/ENSMUSG00000043456.11, intron 5 of 6) | 7 | 37472135 | 37479986 | 7852 | 2 | ENSMUST00000176129.1 | -86000 | Zfp536 | ENSMUSG00000043456.11 |
| chr7 | 37770678 | 37771284 | 3.6136 | 4.337 | 1.31261 | Promoter | 7 | 37479122 | 37770980 | 291859 | 2 | ENSMUST00000176114.2 | 0 | Zfp536 | ENSMUSG00000043456.11 |
| chr7 | 37745805 | 37746173 | 3.63011 | 4.3816 | 1.37452 | Intron (ENSMUST00000175941.2/ENSMUSG00000043456.11, intron 1 of 6) | 7 | 37479104 | 37769624 | 290521 | 2 | ENSMUST00000175941.2 | 23451 | Zfp536 | ENSMUSG00000043456.11 |
| chr7 | 37767519 | 37767798 | 3.70684 | 4.50299 | 1.39825 | Intron (ENSMUST00000175941.2/ENSMUSG00000043456.11, intron 1 of 6) | 7 | 37479104 | 37769624 | 290521 | 2 | ENSMUST00000175941.2 | 1826 | Zfp536 | ENSMUSG00000043456.11 |
| chr7 | 37768709 | 37769851 | 4.34015 | 5.86206 | 2.39442 | Promoter | 7 | 37479104 | 37769624 | 290521 | 2 | ENSMUST00000175941.2 | 0 | Zfp536 | ENSMUSG00000043456.11 |
| chr7 | 37771675 | 37772262 | 3.77806 | 4.68974 | 1.54181 | Promoter | 7 | 37479177 | 37772875 | 293699 | 2 | ENSMUST00000176205.2 | 613 | Zfp536 | ENSMUSG00000043456.11 |
| chr7 | 37772741 | 37773155 | 3.69266 | 4.79801 | 1.65304 | Promoter | 7 | 37479177 | 37772875 | 293699 | 2 | ENSMUST00000176205.2 | 0 | Zfp536 | ENSMUSG00000043456.11 |
| chr7 | 37775438 | 37775971 | 3.64071 | 4.42744 | 1.38649 | Distal Intergenic | 7 | 37569629 | 37773641 | 204013 | 2 | ENSMUST00000176534.2 | -1798 | Zfp536 | ENSMUSG00000043456.11 |
| chr15 | 78657574 | 78657867 | 3.60041 | 4.30562 | 1.28429 | Distal Intergenic | 15 | 78670022 | 78718113 | 48092 | 2 | ENSMUST00000088592.4 | 60246 | Elfn2 | ENSMUSG00000043460.6 |
| chr15 | 78710485 | 78710986 | 3.80134 | 4.77921 | 1.65539 | Intron (ENSMUST00000088592.4/ENSMUSG00000043460.6, intron 1 of 1) | 15 | 78670022 | 78718113 | 48092 | 2 | ENSMUST00000088592.4 | 7127 | Elfn2 | ENSMUSG00000043460.6 |
| chr15 | 78711294 | 78711776 | 3.36614 | 3.91896 | 1.14866 | Intron (ENSMUST00000088592.4/ENSMUSG00000043460.6, intron 1 of 1) | 15 | 78670022 | 78718113 | 48092 | 2 | ENSMUST00000088592.4 | 6337 | Elfn2 | ENSMUSG00000043460.6 |
| chr15 | 78712357 | 78713622 | 3.50581 | 4.27651 | 1.29901 | Intron (ENSMUST00000088592.4/ENSMUSG00000043460.6, intron 1 of 1) | 15 | 78670022 | 78718113 | 48092 | 2 | ENSMUST00000088592.4 | 4491 | Elfn2 | ENSMUSG00000043460.6 |
| chr15 | 78715340 | 78715566 | 4.00686 | 5.18136 | 1.93387 | Intron (ENSMUST00000088592.4/ENSMUSG00000043460.6, intron 1 of 1) | 15 | 78670022 | 78718113 | 48092 | 2 | ENSMUST00000088592.4 | 2547 | Elfn2 | ENSMUSG00000043460.6 |
| chr15 | 78718349 | 78719155 | 3.8592 | 4.93504 | 1.75072 | Promoter | 15 | 78670022 | 78718113 | 48092 | 2 | ENSMUST00000088592.4 | -237 | Elfn2 | ENSMUSG00000043460.6 |
| chr15 | 78719519 | 78719912 | 4.21056 | 5.71755 | 2.26909 | Distal Intergenic | 15 | 78670022 | 78718113 | 48092 | 2 | ENSMUST00000088592.4 | -1407 | Elfn2 | ENSMUSG00000043460.6 |
| chr15 | 78726259 | 78726380 | 3.34615 | 3.88551 | 1.14041 | Distal Intergenic | 15 | 78670022 | 78718113 | 48092 | 2 | ENSMUST00000088592.4 | -8147 | Elfn2 | ENSMUSG00000043460.6 |
| chr15 | 78735575 | 78735921 | 3.54251 | 4.28942 | 1.33067 | Distal Intergenic | 15 | 78670022 | 78718113 | 48092 | 2 | ENSMUST00000088592.4 | -17463 | Elfn2 | ENSMUSG00000043460.6 |
| chr3 | 69859449 | 69860920 | 3.63995 | 4.54374 | 1.4749 | Promoter | 3 | 69819538 | 69859896 | 40359 | 2 | ENSMUST00000051239.7 | 0 | Sptssb | ENSMUSG00000043461.8 |
| chr3 | 69861470 | 69861633 | 3.32518 | 3.81504 | 1.01733 | Distal Intergenic | 3 | 69819538 | 69859896 | 40359 | 2 | ENSMUST00000171529.2 | -1575 | Sptssb | ENSMUSG00000043461.8 |
| chr6 | 53819965 | 53821920 | 3.91765 | 5.09915 | 1.84322 | Promoter | 6 | 53815466 | 53820830 | 5365 | 2 | ENSMUST00000127748.3 | 0 | Tril | ENSMUSG00000043496.6 |
| chr6 | 48743572 | 48743741 | 3.94838 | 5.47878 | 2.12043 | 3' UTR | 6 | 48746197 | 48754210 | 8014 | 1 | ENSMUST00000055558.5 | -2456 | Gimap5 | ENSMUSG00000043505.7 |
| chr6 | 48746226 | 48746577 | 3.62374 | 4.57342 | 1.49369 | Promoter | 6 | 48746199 | 48747512 | 1314 | 1 | ENSMUST00000135174.1 | 28 | Gimap5 | ENSMUSG00000043505.7 |
| chr11 | 94883148 | 94883245 | 3.4965 | 4.36462 | 1.35541 | Distal Intergenic | 11 | 94861018 | 94861759 | 742 | 2 | ENSMUST00000124176.1 | -21390 | A430060F13Rik | ENSMUSG00000043522.8 |
| chr19 | 50678111 | 50678962 | 3.53814 | 4.32379 | 1.3574 | Promoter | 19 | 50143301 | 50678646 | 535346 | 2 | ENSMUST00000072685.7 | 0 | Sorcs1 | ENSMUSG00000043531.10 |
| chr17 | 29794560 | 29796025 | 3.81611 | 4.94741 | 1.75793 | Distal Intergenic | 17 | 29832266 | 29835358 | 3093 | 2 | ENSMUST00000170786.1 | 39333 | Mdga1 | ENSMUSG00000043557.10 |
| chr17 | 29814071 | 29814276 | 3.64624 | 4.44707 | 1.44812 | Distal Intergenic | 17 | 29832266 | 29835358 | 3093 | 2 | ENSMUST00000170786.1 | 21082 | Mdga1 | ENSMUSG00000043557.10 |
| chr17 | 29849067 | 29849349 | 3.359 | 3.86601 | 1.01733 | Exon (ENSMUST00000171691.2/ENSMUSG00000043557.10, exon 6 of 17) | 17 | 29832056 | 29842421 | 10366 | 2 | ENSMUST00000165528.2 | -6647 | Mdga1 | ENSMUSG00000043557.10 |
| chr17 | 29852234 | 29852748 | 3.81538 | 4.80795 | 1.63006 | Exon (ENSMUST00000171691.2/ENSMUSG00000043557.10, exon 3 of 17) | 17 | 29832056 | 29842421 | 10366 | 2 | ENSMUST00000165528.2 | -9814 | Mdga1 | ENSMUSG00000043557.10 |
| chr17 | 29883523 | 29883700 | 3.76983 | 4.73249 | 1.61778 | Intron (ENSMUST00000171691.2/ENSMUSG00000043557.10, intron 1 of 16) | 17 | 29849683 | 29887482 | 37800 | 2 | ENSMUST00000165211.1 | 3782 | Mdga1 | ENSMUSG00000043557.10 |
| chr17 | 29886634 | 29888173 | 3.55088 | 4.50188 | 1.43876 | Promoter | 17 | 29827958 | 29887882 | 59925 | 2 | ENSMUST00000073556.6 | 0 | Mdga1 | ENSMUSG00000043557.10 |
| chr17 | 29888764 | 29890507 | 3.64568 | 4.71727 | 1.60519 | Promoter | 17 | 29850562 | 29889950 | 39389 | 2 | ENSMUST00000167102.1 | 0 | Mdga1 | ENSMUSG00000043557.10 |
| chr17 | 29964695 | 29965008 | 3.54529 | 4.20511 | 1.22269 | Intron (ENSMUST00000167190.2/ENSMUSG00000043557.10, intron 1 of 12) | 17 | 29842119 | 29970087 | 127969 | 2 | ENSMUST00000167190.2 | 5079 | Mdga1 | ENSMUSG00000043557.10 |
| chr17 | 29971397 | 29971511 | 3.2874 | 3.75835 | 1.01733 | Distal Intergenic | 17 | 29842119 | 29970087 | 127969 | 2 | ENSMUST00000167190.2 | -1311 | Mdga1 | ENSMUSG00000043557.10 |
| chr17 | 48467362 | 48467452 | 3.43614 | 4.05081 | 1.20681 | 3' UTR | 17 | 48462404 | 48539607 | 77204 | 1 | ENSMUST00000162467.1 | 4959 | Unc5cl | ENSMUSG00000043592.10 |
| chr17 | 48469753 | 48469828 | 3.33662 | 3.84392 | 1.0584 | Intron (ENSMUST00000162313.2/ENSMUSG00000043592.10, intron 4 of 5) | 17 | 48462404 | 48539607 | 77204 | 1 | ENSMUST00000162467.1 | 7350 | Unc5cl | ENSMUSG00000043592.10 |
| chr17 | 48538713 | 48539019 | 3.46344 | 4.13396 | 1.17748 | Intron (ENSMUST00000162313.2/ENSMUSG00000043592.10, intron 5 of 5) | 17 | 48462404 | 48539607 | 77204 | 1 | ENSMUST00000162467.1 | 76310 | Unc5cl | ENSMUSG00000043592.10 |
| chr5 | 90162625 | 90162840 | 3.33184 | 3.97021 | 1.09731 | Distal Intergenic | 5 | 90132229 | 90132675 | 447 | 1 | ENSMUST00000135410.1 | 30397 | Eif5al3-ps | ENSMUSG00000043618.9 |
| chr1 | 52951613 | 52951688 | 3.21556 | 3.65056 | 1.01346 | Intron (ENSMUST00000050567.5/ENSMUSG00000043629.7, intron 1 of 1) | 1 | 52922324 | 52952844 | 30521 | 2 | ENSMUST00000050567.5 | 1156 | 1700019D03Rik | ENSMUSG00000043629.7 |
| chr1 | 52952221 | 52952951 | 4.00114 | 5.25875 | 1.98529 | Promoter | 1 | 52922324 | 52952844 | 30521 | 2 | ENSMUST00000050567.5 | 0 | 1700019D03Rik | ENSMUSG00000043629.7 |
| chr1 | 52953277 | 52953410 | 3.41149 | 4.21754 | 1.27287 | Promoter | 1 | 52922324 | 52953350 | 31027 | 2 | ENSMUST00000114492.2 | 0 | 1700019D03Rik | ENSMUSG00000043629.7 |
| chr19 | 53676356 | 53677654 | 4.04438 | 5.44894 | 2.10836 | Promoter | 19 | 53677306 | 53843763 | 166458 | 1 | ENSMUST00000095969.4 | 0 | Rbm20 | ENSMUSG00000043639.9 |
| chr19 | 53678077 | 53678552 | 3.58863 | 4.28565 | 1.27122 | Promoter | 19 | 53677306 | 53867080 | 189775 | 1 | ENSMUST00000164202.2 | 772 | Rbm20 | ENSMUSG00000043639.9 |
| chr19 | 53703850 | 53703955 | 3.38193 | 3.9006 | 1.01733 | Intron (ENSMUST00000095969.4/ENSMUSG00000043639.9, intron 1 of 8) | 19 | 53677306 | 53867080 | 189775 | 1 | ENSMUST00000164202.2 | 26545 | Rbm20 | ENSMUSG00000043639.9 |
| chr19 | 53806870 | 53807082 | 3.37539 | 3.90664 | 1.07353 | Intron (ENSMUST00000095969.4/ENSMUSG00000043639.9, intron 1 of 8) | 19 | 53793308 | 53813573 | 20266 | 1 | ENSMUST00000162910.1 | 13563 | Rbm20 | ENSMUSG00000043639.9 |
| chr8 | 71558541 | 71558687 | 3.57807 | 4.42289 | 1.3761 | Promoter | 8 | 71554303 | 71558871 | 4569 | 2 | ENSMUST00000052072.6 | 184 | Tmem221 | ENSMUSG00000043664.6 |
| chr8 | 71560975 | 71561471 | 3.58118 | 4.32325 | 1.37213 | 3' UTR | 8 | 71554303 | 71558871 | 4569 | 2 | ENSMUST00000052072.6 | -2105 | Tmem221 | ENSMUSG00000043664.6 |
| chr8 | 90347206 | 90347594 | 4.30108 | 5.6761 | 2.22459 | Promoter | 8 | 90257946 | 90348126 | 90181 | 2 | ENSMUST00000176034.1 | 532 | Tox3 | ENSMUSG00000043668.8 |
| chr8 | 90348665 | 90348768 | 4.17816 | 5.72099 | 2.25803 | Promoter | 8 | 90247040 | 90348343 | 101304 | 2 | ENSMUST00000109621.4 | -323 | Tox3 | ENSMUSG00000043668.8 |
| chr10 | 81024969 | 81025560 | 3.73738 | 4.56452 | 1.44808 | Promoter | 10 | 81019589 | 81025409 | 5821 | 2 | ENSMUST00000055125.4 | 0 | Diras1 | ENSMUSG00000043670.4 |
| chr12 | 11148322 | 11148566 | 3.36245 | 3.89384 | 1.09742 | Intron (ENSMUST00000055673.1/ENSMUSG00000043673.5, intron 1 of 2) | 12 | 11090202 | 11150373 | 60172 | 2 | ENSMUST00000055673.1 | 1807 | Kcns3 | ENSMUSG00000043673.5 |
| chr12 | 11149953 | 11150143 | 4.12866 | 5.40271 | 2.1058 | Promoter | 12 | 11090202 | 11150373 | 60172 | 2 | ENSMUST00000055673.1 | 230 | Kcns3 | ENSMUSG00000043673.5 |
| chr17 | 73487716 | 73487927 | 3.86127 | 4.83458 | 1.674 | Distal Intergenic | 17 | 73306464 | 73399296 | 92833 | 2 | ENSMUST00000095208.3 | -88421 | Capn13 | ENSMUSG00000043705.6 |
| chr9 | 105827964 | 105828100 | 3.321 | 3.84355 | 1.0481 | Promoter | 9 | 105689417 | 105828085 | 138669 | 2 | ENSMUST00000166431.3 | 0 | Col6a6 | ENSMUSG00000043719.9 |
| chr9 | 105834631 | 105835027 | 3.61426 | 4.32694 | 1.28429 | Distal Intergenic | 9 | 105687809 | 105828160 | 140352 | 2 | ENSMUST00000098441.4 | -6472 | Col6a6 | ENSMUSG00000043719.9 |
| chr9 | 105835612 | 105835857 | 3.54274 | 4.2028 | 1.2219 | Distal Intergenic | 9 | 105687809 | 105828160 | 140352 | 2 | ENSMUST00000098441.4 | -7453 | Col6a6 | ENSMUSG00000043719.9 |
| chr5 | 121112153 | 121112384 | 3.61549 | 4.33822 | 1.30566 | Distal Intergenic | 5 | 121133840 | 121143635 | 9796 | 2 | ENSMUST00000148871.1 | 31251 | Ptpn11 | ENSMUSG00000043733.9 |
| chr4 | 89687636 | 89688988 | 3.96996 | 5.03428 | 1.77298 | Promoter | 4 | 89688198 | 89694772 | 6575 | 1 | ENSMUST00000052478.2 | 0 | Dmrta1 | ENSMUSG00000043753.2 |
| chr4 | 43663311 | 43663539 | 3.65342 | 4.54459 | 1.47337 | Promoter | 4 | 43664043 | 43664339 | 297 | 1 | ENSMUST00000102482.4 | -504 | Gm12481 | ENSMUSG00000043770.10 |
| chr17 | 23660130 | 23661187 | 3.54716 | 4.35492 | 1.34235 | Promoter | 17 | 23660523 | 23668610 | 8088 | 1 | ENSMUST00000062967.8 | 0 | Ccdc64b | ENSMUSG00000043782.8 |
| chr17 | 23661497 | 23661710 | 3.41413 | 4.22305 | 1.27197 | Promoter | 17 | 23660523 | 23668610 | 8088 | 1 | ENSMUST00000062967.8 | 975 | Ccdc64b | ENSMUSG00000043782.8 |
| chr17 | 23662916 | 23663047 | 3.57445 | 4.28413 | 1.33248 | Intron (ENSMUST00000062967.8/ENSMUSG00000043782.8, intron 2 of 9) | 17 | 23660523 | 23668610 | 8088 | 1 | ENSMUST00000062967.8 | 2394 | Ccdc64b | ENSMUSG00000043782.8 |
| chr16 | 18126403 | 18126704 | 3.37662 | 3.94717 | 1.07368 | Promoter | 16 | 18127642 | 18152408 | 24767 | 1 | ENSMUST00000059589.5 | -938 | Rtn4r | ENSMUSG00000043811.5 |
| chr16 | 18127830 | 18128594 | 3.42166 | 4.08574 | 1.21335 | Promoter | 16 | 18127642 | 18152408 | 24767 | 1 | ENSMUST00000059589.5 | 189 | Rtn4r | ENSMUSG00000043811.5 |
| chr17 | 37196609 | 37198384 | 3.70491 | 4.65168 | 1.56287 | Promoter | 17 | 37196707 | 37198011 | 1305 | 2 | ENSMUST00000055324.6 | 0 | Olfr94 | ENSMUSG00000043827.6 |
| chr11 | 116905651 | 116906268 | 3.91206 | 4.93782 | 1.72587 | Distal Intergenic | 11 | 116918863 | 116986948 | 68086 | 1 | ENSMUST00000103027.4 | -12595 | Mgat5b | ENSMUSG00000043857.11 |
| chr11 | 116996620 | 116996881 | 3.46511 | 4.06421 | 1.15293 | Distal Intergenic | 11 | 116973073 | 116986940 | 13868 | 1 | ENSMUST00000126757.1 | 23548 | Mgat5b | ENSMUSG00000043857.11 |
| chr11 | 117003576 | 117003858 | 3.7031 | 4.60749 | 1.52235 | Distal Intergenic | 11 | 116973073 | 116986940 | 13868 | 1 | ENSMUST00000126757.1 | 30504 | Mgat5b | ENSMUSG00000043857.11 |

| | | | | | | | | | | | | | | | |
|---|---|---|---|---|---|---|---|---|---|---|---|---|---|---|---|
| chr11 | 117004607 | 117005388 | 3.87447 | 4.83389 | 1.63712 | Distal Intergenic | 11 | 116973073 | 116986940 | 13868 | 1 | ENSMUST00000126757.1 | 31535 | Mgat5b | ENSMUSG00000043857.11 |
| chr11 | 117006150 | 117006432 | 3.48292 | 4.09556 | 1.1675 | Distal Intergenic | 11 | 116973073 | 116986940 | 13868 | 1 | ENSMUST00000126757.1 | 33078 | Mgat5b | ENSMUSG00000043857.11 |
| chr8 | 122178946 | 122179324 | 4.02699 | 5.14614 | 1.86652 | Intron (ENSMUST00000127664.1/ENSMUSG00000092329.1, intron 1 of 15) | 8 | 122269569 | 122272646 | 3078 | 1 | ENSMUST00000055537.2 | -90245 | Gm22 | ENSMUSG00000043903.3 |
| chr8 | 122229788 | 122229930 | 4.2192 | 5.54624 | 2.16776 | Intron (ENSMUST00000127664.1/ENSMUSG00000092329.1, intron 1 of 15) | 8 | 122269569 | 122272646 | 3078 | 1 | ENSMUST00000055537.2 | -39639 | Gm22 | ENSMUSG00000043903.3 |
| chr8 | 122230266 | 122231367 | 4.06143 | 5.30345 | 1.99286 | Intron (ENSMUST00000127664.1/ENSMUSG00000092329.1, intron 1 of 15) | 8 | 122269569 | 122272646 | 3078 | 1 | ENSMUST00000055537.2 | -38202 | Gm22 | ENSMUSG00000043903.3 |
| chr8 | 122232951 | 122233045 | 3.8738 | 4.86282 | 1.72445 | Intron (ENSMUST00000127664.1/ENSMUSG00000092329.1, intron 1 of 15) | 8 | 122269569 | 122272646 | 3078 | 1 | ENSMUST00000055537.2 | -36524 | Gm22 | ENSMUSG00000043903.3 |
| chr5 | 116288168 | 116288442 | 3.40727 | 3.98961 | 1.18964 | Promoter | 5 | 116169225 | 116288963 | 119739 | 2 | ENSMUST00000086483.3 | 521 | Ccdc60 | ENSMUSG00000043913.9 |
| chr5 | 116290088 | 116290174 | 3.5446 | 4.46632 | 1.43589 | Distal Intergenic | 5 | 116124641 | 116288985 | 164345 | 2 | ENSMUST00000050178.7 | -1104 | Ccdc60 | ENSMUSG00000043913.9 |
| chr4 | 135381477 | 135381602 | 3.29775 | 3.77385 | 1.01733 | Intron (ENSMUST00000105858.2/ENSMUSG00000043924.11, intron 1 of 4) | 4 | 135370227 | 135385645 | 15419 | 2 | ENSMUST00000105857.2 | 4043 | Ncmap | ENSMUSG00000043924.11 |
| chr4 | 135396277 | 135396369 | 3.4939 | 4.14857 | 1.27178 | Intron (ENSMUST00000105859.2/ENSMUSG00000043924.11, intron 1 of 4) | 4 | 135369576 | 135398190 | 28615 | 2 | ENSMUST00000105859.2 | 1821 | Ncmap | ENSMUSG00000043924.11 |
| chr4 | 135397993 | 135398338 | 3.61803 | 4.49917 | 1.46188 | Promoter | 4 | 135369576 | 135398190 | 28615 | 2 | ENSMUST00000105859.2 | 0 | Ncmap | ENSMUSG00000043924.11 |
| chr19 | 59413552 | 59413752 | 3.27843 | 3.74478 | 1.01733 | Distal Intergenic | 19 | 59458372 | 59465357 | 6986 | 1 | ENSMUST00000062216.3 | -44620 | Emx2 | ENSMUSG00000043969.4 |
| chr19 | 59417472 | 59417679 | 3.34689 | 3.88295 | 1.12882 | Distal Intergenic | 19 | 59458372 | 59465357 | 6986 | 1 | ENSMUST00000062216.3 | -40693 | Emx2 | ENSMUSG00000043969.4 |
| chr19 | 59446208 | 59446703 | 3.38378 | 3.97814 | 1.17443 | Intron (ENSMUST00000136990.2/ENSMUSG00000087095.2, intron 1 of 3) | 19 | 59458372 | 59465357 | 6986 | 1 | ENSMUST00000062216.3 | -11669 | Emx2 | ENSMUSG00000043969.4 |
| chr19 | 59448008 | 59448220 | 3.38254 | 3.94261 | 1.14826 | Intron (ENSMUST00000136990.2/ENSMUSG00000087095.2, intron 1 of 3) | 19 | 59458372 | 59465357 | 6986 | 1 | ENSMUST00000062216.3 | -10152 | Emx2 | ENSMUSG00000043969.4 |
| chr19 | 59449001 | 59449414 | 3.6151 | 4.36652 | 1.31491 | Intron (ENSMUST00000136990.2/ENSMUSG00000087095.2, intron 1 of 3) | 19 | 59458372 | 59465357 | 6986 | 1 | ENSMUST00000062216.3 | -8958 | Emx2 | ENSMUSG00000043969.4 |
| chr19 | 59449982 | 59450755 | 3.58503 | 4.28455 | 1.28223 | Intron (ENSMUST00000136990.2/ENSMUSG00000087095.2, intron 1 of 3) | 19 | 59458372 | 59465357 | 6986 | 1 | ENSMUST00000062216.3 | -7617 | Emx2 | ENSMUSG00000043969.4 |
| chr19 | 59451671 | 59452579 | 4.87003 | 7.00451 | 3.33874 | Intron (ENSMUST00000136990.2/ENSMUSG00000087095.2, intron 1 of 3) | 19 | 59458372 | 59465357 | 6986 | 1 | ENSMUST00000062216.3 | -5793 | Emx2 | ENSMUSG00000043969.4 |
| chr19 | 59453485 | 59454720 | 3.78085 | 4.69266 | 1.55788 | Intron (ENSMUST00000136990.2/ENSMUSG00000087095.2, intron 1 of 3) | 19 | 59458372 | 59465357 | 6986 | 1 | ENSMUST00000062216.3 | -3652 | Emx2 | ENSMUSG00000043969.4 |
| chr19 | 59455425 | 59457172 | 3.70481 | 4.59272 | 1.49305 | Intron (ENSMUST00000136990.2/ENSMUSG00000087095.2, intron 1 of 3) | 19 | 59458372 | 59465357 | 6986 | 1 | ENSMUST00000062216.3 | -1200 | Emx2 | ENSMUSG00000043969.4 |
| chr19 | 59457631 | 59458469 | 3.72661 | 4.55231 | 1.45894 | Promoter | 19 | 59458372 | 59465357 | 6986 | 1 | ENSMUST00000062216.3 | 0 | Emx2 | ENSMUSG00000043969.4 |
| chr19 | 59459231 | 59459755 | 4.46474 | 6.25462 | 2.73722 | Promoter | 19 | 59459519 | 59465353 | 5835 | 1 | ENSMUST00000174353.1 | 0 | Emx2 | ENSMUSG00000043969.4 |
| chr19 | 59460117 | 59462383 | 4.09049 | 5.42883 | 2.11196 | Promoter | 19 | 59462802 | 59464215 | 1414 | 1 | ENSMUST00000174573.1 | -419 | Emx2 | ENSMUSG00000043969.4 |
| chr19 | 59462968 | 59463266 | 3.62655 | 4.36565 | 1.34407 | Promoter | 19 | 59462802 | 59464215 | 1414 | 1 | ENSMUST00000174573.1 | 167 | Emx2 | ENSMUSG00000043969.4 |
| chr19 | 59463621 | 59464144 | 3.58718 | 4.30563 | 1.30815 | Promoter | 19 | 59462802 | 59464215 | 1414 | 1 | ENSMUST00000174573.1 | 820 | Emx2 | ENSMUSG00000043969.4 |
| chr19 | 59464790 | 59467742 | 3.84243 | 4.92262 | 1.73432 | 3' UTR | 19 | 59462802 | 59464215 | 1414 | 1 | ENSMUST00000174573.1 | 1989 | Emx2 | ENSMUSG00000043969.4 |
| chr19 | 59468341 | 59468703 | 3.54568 | 4.21292 | 1.23846 | Distal Intergenic | 19 | 59462802 | 59464215 | 1414 | 1 | ENSMUST00000174573.1 | 5540 | Emx2 | ENSMUSG00000043969.4 |
| chr19 | 59469218 | 59469771 | 3.73132 | 4.60005 | 1.51476 | Distal Intergenic | 19 | 59462802 | 59464215 | 1414 | 1 | ENSMUST00000174573.1 | 6417 | Emx2 | ENSMUSG00000043969.4 |
| chr19 | 59470213 | 59470301 | 3.41969 | 3.9716 | 1.06662 | Distal Intergenic | 19 | 59462802 | 59464215 | 1414 | 1 | ENSMUST00000174573.1 | 7412 | Emx2 | ENSMUSG00000043969.4 |
| chr19 | 59470892 | 59472118 | 3.57099 | 4.28131 | 1.30567 | Distal Intergenic | 19 | 59462802 | 59464215 | 1414 | 1 | ENSMUST00000174573.1 | 8091 | Emx2 | ENSMUSG00000043969.4 |
| chr8 | 69879470 | 69879726 | 3.35607 | 3.90261 | 1.14741 | Downstream (<1kb) | 8 | 69880369 | 69887687 | 7319 | 2 | ENSMUST00000057831.7 | 7961 | Cilp2 | ENSMUSG00000044006.8 |
| chr8 | 69880771 | 69882853 | 4.11173 | 5.5507 | 2.18569 | 3' UTR | 8 | 69880369 | 69887687 | 7319 | 2 | ENSMUST00000057831.7 | 4834 | Cilp2 | ENSMUSG00000044006.8 |
| chr8 | 69884781 | 69886998 | 3.88464 | 4.8999 | 1.74418 | Promoter | 8 | 69880369 | 69887687 | 7319 | 2 | ENSMUST00000057831.7 | 689 | Cilp2 | ENSMUSG00000044006.8 |
| chr8 | 66688443 | 66689918 | 3.53901 | 4.22353 | 1.25843 | Promoter | 8 | 66679965 | 66688094 | 8130 | 2 | ENSMUST00000070810.7 | -350 | Npy5r | ENSMUSG00000044014.8 |
| chr5 | 129063774 | 129063933 | 3.38499 | 3.90989 | 1.03376 | Distal Intergenic | 5 | 129096750 | 129204599 | 107850 | 1 | ENSMUST00000056617.8 | -32817 | Gpr133 | ENSMUSG00000044017.11 |
| chr5 | 129096952 | 129097099 | 3.37243 | 3.88626 | 1.01733 | Promoter | 5 | 129096757 | 129115095 | 18339 | 1 | ENSMUST00000156437.1 | 196 | Gpr133 | ENSMUSG00000044017.11 |
| chr5 | 129109978 | 129110140 | 3.63822 | 4.40544 | 1.42086 | Intron (ENSMUST00000056617.8/ENSMUSG00000044017.11, intron 3 of 25) | 5 | 129096757 | 129115095 | 18339 | 1 | ENSMUST00000156437.1 | 13222 | Gpr133 | ENSMUSG00000044017.11 |
| chr2 | 77177076 | 77177224 | 3.76158 | 4.6162 | 1.48593 | Distal Intergenic | 2 | 77044971 | 77170636 | 125666 | 2 | ENSMUST00000028406.5 | -6441 | Ccdc141 | ENSMUSG00000044033.11 |
| chr9 | 110873663 | 110874412 | 3.78774 | 4.72604 | 1.607 | 5' UTR | 9 | 110879870 | 110882722 | 2853 | 1 | ENSMUST00000143416.1 | -5458 | Als2cl | ENSMUSG00000044037.10 |
| chr11 | 100123940 | 100124015 | 3.35498 | 3.89628 | 1.1346 | Distal Intergenic | 11 | 100117327 | 100121566 | 4240 | 2 | ENSMUST00000007275.2 | -2375 | Krt13 | ENSMUSG00000044041.4 |
| chr11 | 59614643 | 59614805 | 3.34498 | 4.16301 | 1.22951 | Distal Intergenic | 11 | 59612966 | 59614225 | 1260 | 1 | ENSMUST00000055276.4 | 1678 | Olfr225 | ENSMUSG00000044061.5 |
| chr12 | 80692367 | 80693017 | 3.58787 | 4.72174 | 1.60412 | Promoter | 12 | 80686370 | 80692466 | 6097 | 2 | ENSMUST00000153297.1 | 0 | Plekhd1os | ENSMUSG00000044062.7 |
| chr19 | 41441639 | 41442443 | 3.91337 | 4.9261 | 1.70449 | Distal Intergenic | 19 | 41459813 | 41460718 | 906 | 1 | ENSMUST00000057038.3 | -17370 | Gm9788 | ENSMUSG00000044065.3 |
| chr10 | 123264479 | 123265850 | 3.94365 | 5.06276 | 1.86039 | Promoter | 10 | 123264076 | 123741204 | 477129 | 1 | ENSMUST00000050756.7 | 404 | Fam19a2 | ENSMUSG00000044071.7 |
| chr10 | 123266386 | 123266585 | 3.3213 | 3.87215 | 1.08526 | Intron (ENSMUST00000050756.7/ENSMUSG00000044071.7, intron 1 of 5) | 10 | 123264076 | 123741204 | 477129 | 1 | ENSMUST00000050756.7 | 2311 | Fam19a2 | ENSMUSG00000044071.7 |
| chr10 | 123267579 | 123267804 | 3.96371 | 5.13584 | 1.91496 | Intron (ENSMUST00000050756.7/ENSMUSG00000044071.7, intron 1 of 5) | 10 | 123264076 | 123741204 | 477129 | 1 | ENSMUST00000050756.7 | 3504 | Fam19a2 | ENSMUSG00000044071.7 |
| chr10 | 123268531 | 123268861 | 3.7635 | 4.64979 | 1.57664 | Intron (ENSMUST00000050756.7/ENSMUSG00000044071.7, intron 1 of 5) | 10 | 123264076 | 123741204 | 477129 | 1 | ENSMUST00000050756.7 | 4456 | Fam19a2 | ENSMUSG00000044071.7 |
| chr7 | 120597480 | 120598015 | 3.6585 | 4.41205 | 1.34336 | Promoter | 7 | 120597625 | 120598475 | 851 | 1 | ENSMUST00000123576.1 | 0 | E130201H02Rik | ENSMUSG00000044141.8 |
| chr6 | 3487138 | 3487514 | 3.7781 | 4.72132 | 1.59983 | Promoter | 6 | 3457395 | 3487380 | 29986 | 2 | ENSMUST00000184466.1 | 0 | Hepacam2 | ENSMUSG00000044156.9 |
| chr13 | 43616193 | 43616804 | 3.88827 | 4.92363 | 1.74953 | Promoter | 13 | 43615983 | 43668066 | 52084 | 1 | ENSMUST00000161817.1 | 211 | Rnf182 | ENSMUSG00000044164.2 |
| chr14 | 69168228 | 69169556 | 4.14809 | 5.4246 | 2.0909 | Intron (ENSMUST00000172547.1/ENSMUSG00000092499.1, intron 1 of 2) | 14 | 69171802 | 69175518 | 3717 | 1 | ENSMUST00000062437.9 | -2246 | Nkx2-6 | ENSMUSG00000044186.10 |
| chr14 | 69170477 | 69171680 | 4.54014 | 6.31982 | 2.80644 | Promoter | 14 | 69171802 | 69175518 | 3717 | 1 | ENSMUST00000062437.9 | -122 | Nkx2-6 | ENSMUSG00000044186.10 |
| chr15 | 79479237 | 79479405 | 3.47474 | 4.1051 | 1.23415 | Distal Intergenic | 15 | 79483718 | 79505241 | 21524 | 2 | ENSMUST00000057801.6 | 25836 | Kcnj4 | ENSMUSG00000044216.6 |
| chr15 | 79481237 | 79481414 | 3.53827 | 4.19627 | 1.22889 | Downstream (2-3kb) | 15 | 79483718 | 79505241 | 21524 | 2 | ENSMUST00000057801.6 | 23827 | Kcnj4 | ENSMUSG00000044216.6 |
| chr15 | 79491585 | 79491660 | 3.49265 | 4.10246 | 1.13969 | Intron (ENSMUST00000057801.6/ENSMUSG00000044216.6, intron 1 of 1) | 15 | 79483718 | 79505241 | 21524 | 2 | ENSMUST00000057801.6 | 13581 | Kcnj4 | ENSMUSG00000044216.6 |
| chr15 | 79492748 | 79493084 | 3.46197 | 4.05327 | 1.13013 | Intron (ENSMUST00000057801.6/ENSMUSG00000044216.6, intron 1 of 1) | 15 | 79483718 | 79505241 | 21524 | 2 | ENSMUST00000057801.6 | 12157 | Kcnj4 | ENSMUSG00000044216.6 |
| chr15 | 79493837 | 79494151 | 4.53073 | 6.23964 | 2.68301 | Intron (ENSMUST00000057801.6/ENSMUSG00000044216.6, intron 1 of 1) | 15 | 79483718 | 79505241 | 21524 | 2 | ENSMUST00000057801.6 | 11090 | Kcnj4 | ENSMUSG00000044216.6 |
| chr15 | 79497778 | 79497853 | 3.59563 | 4.2902 | 1.25349 | Intron (ENSMUST00000057801.6/ENSMUSG00000044216.6, intron 1 of 1) | 15 | 79483718 | 79505241 | 21524 | 2 | ENSMUST00000057801.6 | 7388 | Kcnj4 | ENSMUSG00000044216.6 |
| chr15 | 79504107 | 79506936 | 3.73643 | 4.60691 | 1.51002 | Promoter | 15 | 79483718 | 79505241 | 21524 | 2 | ENSMUST00000057801.6 | 0 | Kcnj4 | ENSMUSG00000044216.6 |
| chr15 | 79509665 | 79509740 | 3.79648 | 5.08409 | 1.83206 | Distal Intergenic | 15 | 79483718 | 79505241 | 21524 | 2 | ENSMUST00000057801.6 | -4425 | Kcnj4 | ENSMUSG00000044216.6 |
| chr15 | 99586937 | 99587623 | 3.53697 | 4.18972 | 1.21177 | Distal Intergenic | 15 | 99590849 | 99594608 | 3760 | 1 | ENSMUST00000088200.7 | -3226 | Aqp5 | ENSMUSG00000044217.11 |
| chr15 | 99589862 | 99591907 | 3.78997 | 4.76309 | 1.62079 | Promoter | 15 | 99590849 | 99594608 | 3760 | 1 | ENSMUST00000088200.7 | 0 | Aqp5 | ENSMUSG00000044217.11 |
| chr15 | 99592335 | 99592565 | 3.73502 | 4.55352 | 1.42676 | Intron (ENSMUST00000088200.7/ENSMUSG00000044217.11, intron 2 of 4) | 15 | 99591028 | 99594829 | 3802 | 1 | ENSMUST00000169082.1 | 1308 | Aqp5 | ENSMUSG00000044217.11 |
| chr15 | 99595551 | 99595922 | 3.49049 | 4.11451 | 1.19531 | Distal Intergenic | 15 | 99591028 | 99594829 | 3802 | 1 | ENSMUST00000169082.1 | 4524 | Aqp5 | ENSMUSG00000044217.11 |
| chr19 | 43611207 | 43617523 | 4.18612 | 5.52378 | 2.15685 | Promoter | 19 | 43612325 | 43615892 | 3568 | 1 | ENSMUST00000057178.10 | 0 | Nkx2-3 | ENSMUSG00000044220.13 |
| chr5 | 88684907 | 88685577 | 3.83863 | 4.77077 | 1.60085 | Distal Intergenic | 5 | 88659448 | 88676171 | 16724 | 2 | ENSMUST00000078945.6 | -8737 | Grsf1 | ENSMUSG00000044221.9 |
| chr11 | 76665982 | 76666467 | 3.52793 | 4.19306 | 1.215 | Distal Intergenic | 11 | 76672470 | 76673676 | 1207 | 1 | ENSMUST00000056184.1 | -6003 | Bhlha9 | ENSMUSG00000044243.3 |
| chr11 | 76671842 | 76672146 | 3.39007 | 3.92703 | 1.06739 | Promoter | 11 | 76672470 | 76673676 | 1207 | 1 | ENSMUST00000056184.1 | -324 | Bhlha9 | ENSMUSG00000044243.3 |
| chr11 | 76672494 | 76673294 | 4.14852 | 5.53259 | 2.17663 | Promoter | 11 | 76672470 | 76673676 | 1207 | 1 | ENSMUST00000056184.1 | 25 | Bhlha9 | ENSMUSG00000044243.3 |
| chr11 | 76674257 | 76674366 | 4.33556 | 5.73263 | 2.24933 | Distal Intergenic | 11 | 76672470 | 76673676 | 1207 | 1 | ENSMUST00000056184.1 | 1788 | Bhlha9 | ENSMUSG00000044243.3 |
| chr4 | 106462816 | 106463720 | 3.71292 | 4.62827 | 1.52402 | Promoter | 4 | 106442329 | 106464329 | 22001 | 2 | ENSMUST00000049507.5 | 609 | Pcsk9 | ENSMUSG00000044254.6 |
| chr4 | 106464947 | 106465237 | 3.64661 | 4.39325 | 1.34711 | Promoter | 4 | 106442329 | 106464329 | 22001 | 2 | ENSMUST00000049507.5 | -619 | Pcsk9 | ENSMUSG00000044254.6 |
| chr10 | 22152646 | 22153041 | 3.43653 | 4.04296 | 1.21689 | Downstream (<1kb) | 10 | 22153280 | 22154416 | 1137 | 2 | ENSMUST00000181646.1 | 1375 | Gm4895 | ENSMUSG00000044268.4 |
| chr10 | 22153877 | 22154948 | 4.36897 | 5.97964 | 2.57617 | Promoter | 10 | 22153280 | 22154416 | 1137 | 2 | ENSMUST00000181646.1 | 0 | Gm4895 | ENSMUSG00000044268.4 |
| chr10 | 22156297 | 22156380 | 3.29278 | 3.94303 | 1.10723 | Distal Intergenic | 10 | 22153280 | 22154416 | 1137 | 2 | ENSMUST00000181646.1 | -1882 | Gm4895 | ENSMUSG00000044268.4 |
| chr4 | 33920895 | 33921064 | 3.26976 | 3.73176 | 1.01733 | Distal Intergenic | 4 | 33924593 | 33935316 | 10724 | 1 | ENSMUST00000133462.1 | -3529 | Cnr1 | ENSMUSG00000044288.6 |
| chr4 | 33923598 | 33925349 | 3.97139 | 5.23486 | 1.96358 | Promoter | 4 | 33924593 | 33935316 | 10724 | 1 | ENSMUST00000133462.1 | 0 | Cnr1 | ENSMUSG00000044288.6 |
| chr4 | 33926719 | 33927063 | 3.63939 | 4.44983 | 1.44435 | Promoter | 4 | 33926202 | 33948819 | 22618 | 1 | ENSMUST00000084736.2 | 518 | Cnr1 | ENSMUSG00000044288.6 |
| chr15 | 101537677 | 101537892 | 3.37158 | 3.88822 | 1.02869 | Distal Intergenic | 15 | 101525026 | 101532820 | 7795 | 2 | ENSMUST00000023720.7 | -4858 | Krt84 | ENSMUSG00000044294.7 |
| chr4 | 89278117 | 89279176 | 3.82056 | 4.95156 | 1.75573 | Intron (ENSMUST00000060501.4/ENSMUSG00000044303.6, intron 1 of 2) | 4 | 89274471 | 89282192 | 7722 | 2 | ENSMUST00000060501.4 | 3016 | Cdkn2a | ENSMUSG00000044303.6 |
| chr4 | 89279609 | 89279886 | 3.5843 | 4.37073 | 1.38714 | Intron (ENSMUST00000060501.4/ENSMUSG00000044303.6, intron 1 of 2) | 4 | 89274471 | 89282192 | 7722 | 2 | ENSMUST00000060501.4 | 2306 | Cdkn2a | ENSMUSG00000044303.6 |
| chr4 | 89280365 | 89283175 | 3.78076 | 4.70472 | 1.58844 | Promoter | 4 | 89274471 | 89282192 | 7722 | 2 | ENSMUST00000060501.4 | 0 | Cdkn2a | ENSMUSG00000044303.6 |
| chr4 | 89283634 | 89283709 | 3.96528 | 4.99928 | 1.74949 | Intron (ENSMUST00000107131.1/ENSMUSG00000044303.6, intron 1 of 2) | 4 | 89274471 | 89282192 | 7722 | 2 | ENSMUST00000060501.4 | -1443 | Cdkn2a | ENSMUSG00000044303.6 |
| chr4 | 89284183 | 89285866 | 3.92991 | 4.97467 | 1.76943 | Intron (ENSMUST00000107131.1/ENSMUSG00000044303.6, intron 1 of 2) | 4 | 89274471 | 89282192 | 7722 | 2 | ENSMUST00000060501.4 | -1992 | Cdkn2a | ENSMUSG00000044303.6 |
| chr4 | 89286207 | 89286847 | 3.46705 | 4.35974 | 1.35121 | Intron (ENSMUST00000107131.1/ENSMUSG00000044303.6, intron 1 of 2) | 4 | 89274471 | 89282192 | 7722 | 2 | ENSMUST00000060501.4 | -4016 | Cdkn2a | ENSMUSG00000044303.6 |
| chr4 | 89287277 | 89289737 | 3.78753 | 4.80612 | 1.66841 | Intron (ENSMUST00000107131.1/ENSMUSG00000044303.6, intron 1 of 2) | 4 | 89274541 | 89294653 | 20113 | 2 | ENSMUST00000107131.1 | 4916 | Cdkn2a | ENSMUSG00000044303.6 |
| chr4 | 89290276 | 89291423 | 3.93548 | 5.22607 | 1.95729 | Intron (ENSMUST00000107131.1/ENSMUSG00000044303.6, intron 1 of 2) | 4 | 89274541 | 89294653 | 20113 | 2 | ENSMUST00000107131.1 | 3230 | Cdkn2a | ENSMUSG00000044303.6 |
| chr4 | 89292353 | 89292511 | 4.07229 | 5.4743 | 2.06997 | Intron (ENSMUST00000107131.1/ENSMUSG00000044303.6, intron 1 of 2) | 4 | 89274541 | 89294653 | 20113 | 2 | ENSMUST00000107131.1 | 2142 | Cdkn2a | ENSMUSG00000044303.6 |
| chr4 | 89292849 | 89293052 | 3.42869 | 4.13627 | 1.21912 | Intron (ENSMUST00000107131.1/ENSMUSG00000044303.6, intron 1 of 2) | 4 | 89274541 | 89294653 | 20113 | 2 | ENSMUST00000107131.1 | 1601 | Cdkn2a | ENSMUSG00000044303.6 |
| chr4 | 89294153 | 89295289 | 3.84655 | 4.88846 | 1.73113 | Promoter | 4 | 89274541 | 89294653 | 20113 | 2 | ENSMUST00000107131.1 | 0 | Cdkn2a | ENSMUSG00000044303.6 |
| chr4 | 89297140 | 89297215 | 3.80017 | 4.80498 | 1.65707 | Distal Intergenic | 4 | 89274541 | 89294653 | 20113 | 2 | ENSMUST00000107131.1 | -2488 | Cdkn2a | ENSMUSG00000044303.6 |
| chr4 | 89300113 | 89300251 | 3.6516 | 4.50116 | 1.46207 | Distal Intergenic | 4 | 89274541 | 89294653 | 20113 | 2 | ENSMUST00000107131.1 | -5461 | Cdkn2a | ENSMUSG00000044303.6 |
| chr4 | 89302137 | 89302520 | 3.91984 | 5.28288 | 1.94708 | Distal Intergenic | 4 | 89274541 | 89294653 | 20113 | 2 | ENSMUST00000107131.1 | -7485 | Cdkn2a | ENSMUSG00000044303.6 |
| chr10 | 62132607 | 62132702 | 3.33468 | 3.82936 | 1.01733 | Promoter | 10 | 62133090 | 62134763 | 1674 | 1 | ENSMUST00000050103.1 | -388 | Neurog3 | ENSMUSG00000044312.3 |
| chr10 | 62133043 | 62133230 | 3.767 | 4.7097 | 1.62193 | Promoter | 10 | 62133090 | 62134763 | 1674 | 1 | ENSMUST00000050103.1 | 0 | Neurog3 | ENSMUSG00000044312.3 |
| chr10 | 62130516 | 62130741 | 3.7926 | 4.67887 | 1.55395 | Intron (ENSMUST00000027719.6/ENSMUSG00000020083.8, intron 2 of 2) | 10 | 62133090 | 62134763 | 1674 | 1 | ENSMUST00000050103.1 | -2349 | Neurog3 | ENSMUSG00000044312.3 |
| chr3 | 101780943 | 101781419 | 3.36043 | 3.87073 | 1.02638 | Distal Intergenic | 3 | 101813076 | 101836223 | 23148 | 2 | ENSMUST00000061831.5 | 54804 | Mab21l3 | ENSMUSG00000044313.8 |
| chr3 | 101800337 | 101800414 | 3.81346 | 4.72077 | 1.59904 | Distal Intergenic | 3 | 101813076 | 101836223 | 23148 | 2 | ENSMUST00000061831.5 | 35809 | Mab21l3 | ENSMUSG00000044313.8 |
| chr2 | 30784516 | 30784861 | 3.61276 | 4.33761 | 1.31897 | Distal Intergenic | 2 | 30795468 | 30796664 | 1197 | 2 | ENSMUST00000130396.2 | 11803 | 1700001O22Rik | ENSMUSG00000044320.9 |
| chr2 | 158375400 | 158376365 | 3.4963 | 4.2859 | 1.32673 | Promoter | 2 | 158375638 | 158386145 | 10508 | 1 | ENSMUST00000109488.2 | 0 | Snhg11 | ENSMUSG00000044349.10 |
| chr11 | 53486397 | 53486888 | 3.41286 | 3.97043 | 1.09947 | Distal Intergenic | 11 | 53476573 | 53480274 | 3702 | 2 | ENSMUST00000104955.3 | -6124 | Sowaha | ENSMUSG00000044352.6 |
| chr11 | 53487244 | 53487646 | 3.55037 | 4.38775 | 1.38018 | Distal Intergenic | 11 | 53476573 | 53480274 | 3702 | 2 | ENSMUST00000104955.3 | -6971 | Sowaha | ENSMUSG00000044352.6 |
| chr11 | 53488377 | 53488729 | 3.3551 | 3.88258 | 1.09327 | Distal Intergenic | 11 | 53476573 | 53480274 | 3702 | 2 | ENSMUST00000104955.3 | -8104 | Sowaha | ENSMUSG00000044352.6 |
| chr2 | 151701984 | 151703240 | 3.57686 | 4.26906 | 1.27355 | Promoter | 2 | 151702008 | 151707310 | 5303 | 1 | ENSMUST00000060196.2 | 0 | Tmem74b | ENSMUSG00000044364.5 |
| chr2 | 151703816 | 151704238 | 3.56359 | 4.27112 | 1.26517 | Intron (ENSMUST00000060196.2/ENSMUSG00000044364.5, intron 1 of 1) | 2 | 151702311 | 151712157 | 9847 | 1 | ENSMUST00000109872.1 | 1506 | Tmem74b | ENSMUSG00000044364.5 |
| chr3 | 134145478 | 134145583 | 4.03515 | 5.16432 | 1.90921 | Intron (ENSMUST00000181952.1/ENSMUSG00000097911.1, intron 1 of 2) | 3 | 134236491 | 134262090 | 25600 | 1 | ENSMUST00000181904.1 | -90908 | Cxxc4 | ENSMUSG00000044365.11 |
| chr3 | 134190196 | 134190372 | 3.50454 | 4.1437 | 1.22378 | Intron (ENSMUST00000181952.1/ENSMUSG00000097911.1, intron 1 of 2) | 3 | 134236491 | 134262090 | 25600 | 1 | ENSMUST00000181904.1 | -46119 | Cxxc4 | ENSMUSG00000044365.11 |
| chr11 | 70220393 | 70220685 | 3.75846 | 4.82341 | 1.65212 | Promoter | 11 | 70219289 | 70220784 | 1496 | 2 | ENSMUST00000141290.1 | 99 | Slc16a13 | ENSMUSG00000044367.7 |
| chr11 | 70223606 | 70223718 | 3.80167 | 4.82979 | 1.66174 | Downstream (<1kb) | 11 | 70219176 | 70221064 | 1889 | 2 | ENSMUST00000190533.1 | -2543 | Slc16a13 | ENSMUSG00000044367.7 |
| chr6 | 138006977 | 138007144 | 3.74673 | 4.63582 | 1.58296 | Intron (ENSMUST00000111873.2/ENSMUSG00000044378.7, intron 4 of 5) | 6 | 138017385 | 138018104 | 720 | 2 | ENSMUST00000150278.1 | 10960 | Slc15a5 | ENSMUSG00000044378.7 |
| chr18 | 20557528 | 20558652 | 3.98245 | 5.16041 | 1.96491 | Promoter | 18 | 20558074 | 20604521 | 46448 | 1 | ENSMUST00000059787.9 | 0 | Dsg2 | ENSMUSG00000044393.10 |
| chr18 | 20559217 | 20559346 | 3.52677 | 4.25527 | 1.31201 | Promoter | 18 | 20558246 | 20582916 | 24671 | 1 | ENSMUST00000121837.1 | 972 | Dsg2 | ENSMUSG00000044393.10 |
| chr12 | 54487064 | 54487202 | 3.24255 | 3.83506 | 1.03496 | Distal Intergenic | 12 | 54645374 | 54656572 | 11199 | 2 | ENSMUST00000056228.6 | 169370 | Sptssa | ENSMUSG00000044408.6 |
| chr13 | 55414986 | 55415145 | 3.61355 | 4.3444 | 1.33104 | Promoter | 13 | 55414753 | 55415166 | 414 | 2 | ENSMUST00000054146.3 | 21 | Pfn3 | ENSMUSG00000044444.5 |
| chr13 | 55419882 | 55420586 | 3.85612 | 4.82233 | 1.67933 | Exon (ENSMUST00000021948.9/ENSMUSG00000021492.10, exon 11 of 14) | 13 | 55414753 | 55415166 | 414 | 2 | ENSMUST00000054146.3 | -4717 | Pfn3 | ENSMUSG00000044444.5 |
| chr14 | 59612836 | 59613116 | 3.28997 | 3.98632 | 1.11767 | Distal Intergenic | 14 | 59625281 | 59631658 | 6378 | 1 | ENSMUST00000053949.5 | -12165 | Shisa2 | ENSMUSG00000044461.5 |
| chr14 | 59625267 | 59626555 | 4.09224 | 5.41949 | 2.0991 | Promoter | 14 | 59625281 | 59631658 | 6378 | 1 | ENSMUST00000053949.5 | 0 | Shisa2 | ENSMUSG00000044461.5 |
| chr14 | 59627290 | 59627387 | 3.68084 | 4.57308 | 1.52083 | Intron (ENSMUST00000053949.5/ENSMUSG00000044461.5, intron 1 of 1) | 14 | 59625281 | 59631658 | 6378 | 1 | ENSMUST00000053949.5 | 2010 | Shisa2 | ENSMUSG00000044461.5 |
| chr14 | 59630890 | 59631172 | 3.45148 | 4.03407 | 1.11783 | 3' UTR | 14 | 59625281 | 59631658 | 6378 | 1 | ENSMUST00000053949.5 | 5610 | Shisa2 | ENSMUSG00000044461.5 |
| chr14 | 59633355 | 59633654 | 3.37327 | 3.88754 | 1.01733 | Distal Intergenic | 14 | 59625281 | 59631658 | 6378 | 1 | ENSMUST00000053949.5 | 8075 | Shisa2 | ENSMUSG00000044461.5 |
| chr14 | 59643547 | 59643843 | 3.34961 | 3.85185 | 1.01733 | Distal Intergenic | 14 | 59625281 | 59631658 | 6378 | 1 | ENSMUST00000053949.5 | 18267 | Shisa2 | ENSMUSG00000044461.5 |
| chr14 | 59644727 | 59644985 | 3.53491 | 4.20223 | 1.25913 | Downstream (2-3kb) | 14 | 59625281 | 59631658 | 6378 | 1 | ENSMUST00000053949.5 | 19447 | Shisa2 | ENSMUSG00000044461.5 |
| chr14 | 59648647 | 59648924 | 3.37385 | 3.88841 | 1.01733 | Intron (ENSMUST00000080368.6/ENSMUSG00000021983.10, intron 35 of 35) | 14 | 59625281 | 59631658 | 6378 | 1 | ENSMUST00000053949.5 | 23367 | Shisa2 | ENSMUSG00000044461.5 |
| chr14 | 59686558 | 59686633 | 3.23575 | 3.68398 | 1.02727 | Intron (ENSMUST00000080368.6/ENSMUSG00000021983.10, intron 33 of 35) | 14 | 59625281 | 59631658 | 6378 | 1 | ENSMUST00000053949.5 | 61278 | Shisa2 | ENSMUSG00000044461.5 |
| chr14 | 59703882 | 59703957 | 3.5263 | 4.17561 | 1.21985 | Intron (ENSMUST00000080368.6/ENSMUSG00000021983.10, intron 32 of 35) | 14 | 59625281 | 59631658 | 6378 | 1 | ENSMUST00000053949.5 | 78602 | Shisa2 | ENSMUSG00000044461.5 |
| chr14 | 59714689 | 59714851 | 3.28628 | 3.75656 | 1.01733 | Intron (ENSMUST00000080368.6/ENSMUSG00000021983.10, intron 32 of 35) | 14 | 59625281 | 59631658 | 6378 | 1 | ENSMUST00000053949.5 | 89409 | Shisa2 | ENSMUSG00000044461.5 |
| chr14 | 59721533 | 59721857 | 3.40086 | 3.96886 | 1.15779 | Intron (ENSMUST00000080368.6/ENSMUSG00000021983.10, intron 32 of 35) | 14 | 59625281 | 59631658 | 6378 | 1 | ENSMUST00000053949.5 | 96253 | Shisa2 | ENSMUSG00000044461.5 |
| chr14 | 59763103 | 59763178 | 3.43804 | 4.02731 | 1.16683 | Intron (ENSMUST00000080368.6/ENSMUSG00000021983.10, intron 32 of 35) | 14 | 59625281 | 59631658 | 6378 | 1 | ENSMUST00000053949.5 | 137823 | Shisa2 | ENSMUSG00000044461.5 |
| chr14 | 59763658 | 59763813 | 3.34594 | 3.85203 | 1.03748 | Intron (ENSMUST00000080368.6/ENSMUSG00000021983.10, intron 32 of 35) | 14 | 59625281 | 59631658 | 6378 | 1 | ENSMUST00000053949.5 | 138378 | Shisa2 | ENSMUSG00000044461.5 |
| chr14 | 59764136 | 59764876 | 3.8352 | 4.81445 | 1.67221 | Intron (ENSMUST00000080368.6/ENSMUSG00000021983.10, intron 32 of 35) | 14 | 59625281 | 59631658 | 6378 | 1 | ENSMUST00000053949.5 | 138856 | Shisa2 | ENSMUSG00000044461.5 |
| chr14 | 59767123 | 59767306 | 3.30807 | 3.92969 | 1.10479 | Intron (ENSMUST00000080368.6/ENSMUSG00000021983.10, intron 32 of 35) | 14 | 59625281 | 59631658 | 6378 | 1 | ENSMUST00000053949.5 | 141843 | Shisa2 | ENSMUSG00000044461.5 |
| chr14 | 59768453 | 59768574 | 3.76324 | 4.60984 | 1.48006 | Intron (ENSMUST00000080368.6/ENSMUSG00000021983.10, intron 32 of 35) | 14 | 59625281 | 59631658 | 6378 | 1 | ENSMUST00000053949.5 | 143173 | Shisa2 | ENSMUSG00000044461.5 |
| chr14 | 59773608 | 59774331 | 3.92652 | 4.95668 | 1.73545 | Exon (ENSMUST00000080368.6/ENSMUSG00000021983.10, exon 30 of 36) | 14 | 59625281 | 59631658 | 6378 | 1 | ENSMUST00000053949.5 | 148328 | Shisa2 | ENSMUSG00000044461.5 |
| chr14 | 59774786 | 59775353 | 3.72914 | 4.57548 | 1.48287 | Intron (ENSMUST00000080368.6/ENSMUSG00000021983.10, intron 29 of 35) | 14 | 59625281 | 59631658 | 6378 | 1 | ENSMUST00000053949.5 | 149506 | Shisa2 | ENSMUSG00000044461.5 |
| chr14 | 59777744 | 59778239 | 3.60479 | 4.4807 | 1.42191 | Intron (ENSMUST00000080368.6/ENSMUSG00000021983.10, intron 29 of 35) | 14 | 59625281 | 59631658 | 6378 | 1 | ENSMUST00000053949.5 | 152464 | Shisa2 | ENSMUSG00000044461.5 |
| chr14 | 59779265 | 59779783 | 4.50232 | 6.3657 | 2.76678 | Intron (ENSMUST00000080368.6/ENSMUSG00000021983.10, intron 29 of 35) | 14 | 59625281 | 59631658 | 6378 | 1 | ENSMUST00000053949.5 | 153985 | Shisa2 | ENSMUSG00000044461.5 |
| chr14 | 59780198 | 59780465 | 3.83467 | 4.8703 | 1.67631 | Intron (ENSMUST00000080368.6/ENSMUSG00000021983.10, intron 29 of 35) | 14 | 59625281 | 59631658 | 6378 | 1 | ENSMUST00000053949.5 | 154918 | Shisa2 | ENSMUSG00000044461.5 |
| chr14 | 59781295 | 59781903 | 4.14034 | 5.338 | 1.95802 | Intron (ENSMUST00000080368.6/ENSMUSG00000021983.10, intron 29 of 35) | 14 | 59625281 | 59631658 | 6378 | 1 | ENSMUST00000053949.5 | 156015 | Shisa2 | ENSMUSG00000044461.5 |
| chr14 | 59783912 | 59784373 | 3.50411 | 4.14128 | 1.21642 | Intron (ENSMUST00000080368.6/ENSMUSG00000021983.10, intron 29 of 35) | 14 | 59625281 | 59631658 | 6378 | 1 | ENSMUST00000053949.5 | 158632 | Shisa2 | ENSMUSG00000044461.5 |
| chr14 | 59785879 | 59785995 | 3.53239 | 4.19453 | 1.24481 | Intron (ENSMUST00000080368.6/ENSMUSG00000021983.10, intron 29 of 35) | 14 | 59625281 | 59631658 | 6378 | 1 | ENSMUST00000053949.5 | 160599 | Shisa2 | ENSMUSG00000044461.5 |
| chr14 | 59787419 | 59787898 | 3.73879 | 4.6475 | 1.57461 | Intron (ENSMUST00000080368.6/ENSMUSG00000021983.10, intron 29 of 35) | 14 | 59625281 | 59631658 | 6378 | 1 | ENSMUST00000053949.5 | 162139 | Shisa2 | ENSMUSG00000044461.5 |
| chr14 | 59789013 | 59789360 | 3.29098 | 3.80552 | 1.06393 | Intron (ENSMUST00000080368.6/ENSMUSG00000021983.10, intron 29 of 35) | 14 | 59625281 | 59631658 | 6378 | 1 | ENSMUST00000053949.5 | 163733 | Shisa2 | ENSMUSG00000044461.5 |
| chr14 | 59790934 | 59791065 | 3.82152 | 5.08444 | 1.81707 | Intron (ENSMUST00000080368.6/ENSMUSG00000021983.10, intron 29 of 35) | 14 | 59625281 | 59631658 | 6378 | 1 | ENSMUST00000053949.5 | 165654 | Shisa2 | ENSMUSG00000044461.5 |
| chr14 | 59792678 | 59792799 | 3.54431 | 4.23824 | 1.33273 | Intron (ENSMUST00000080368.6/ENSMUSG00000021983.10, intron 28 of 35) | 14 | 59625281 | 59631658 | 6378 | 1 | ENSMUST00000053949.5 | 167398 | Shisa2 | ENSMUSG00000044461.5 |
| chr14 | 59808002 | 59808077 | 3.45828 | 4.18564 | 1.25896 | Intron (ENSMUST00000080368.6/ENSMUSG00000021983.10, intron 26 of 35) | 14 | 59625281 | 59631658 | 6378 | 1 | ENSMUST00000053949.5 | 182722 | Shisa2 | ENSMUSG00000044461.5 |
| chr17 | 56159308 | 56159383 | 3.77607 | 4.62702 | 1.46873 | Distal Intergenic | 17 | 56162477 | 56173955 | 11479 | 1 | ENSMUST00000077788.6 | -3094 | Tnfaip8l1 | ENSMUSG00000044469.7 |
| chr17 | 56159785 | 56159879 | 3.55462 | 4.24433 | 1.29661 | Distal Intergenic | 17 | 56162477 | 56173955 | 11479 | 1 | ENSMUST00000077788.6 | -2598 | Tnfaip8l1 | ENSMUSG00000044469.7 |
| chr10 | 36505933 | 36506296 | 3.52369 | 4.21383 | 1.31854 | Promoter | 10 | 36506814 | 36833773 | 326960 | 1 | ENSMUST00000167191.2 | -518 | Hs3st5 | ENSMUSG00000044499.6 |
| chr10 | 36506718 | 36506811 | 4.10539 | 5.33454 | 2.05794 | Promoter | 10 | 36506814 | 36833773 | 326960 | 1 | ENSMUST00000167191.2 | -3 | Hs3st5 | ENSMUSG00000044499.6 |
| chr10 | 36507503 | 36507643 | 3.42353 | 4.00198 | 1.15261 | Promoter | 10 | 36507407 | 36810038 | 302632 | 1 | ENSMUST00000171521.1 | 97 | Hs3st5 | ENSMUSG00000044499.6 |
| chr10 | 36508029 | 36508555 | 3.71265 | 4.56942 | 1.55443 | Promoter | 10 | 36507407 | 36810038 | 302632 | 1 | ENSMUST00000171521.1 | 623 | Hs3st5 | ENSMUSG00000044499.6 |
| chr10 | 36509947 | 36510234 | 3.51207 | 4.43199 | 1.43276 | Intron (ENSMUST00000167191.2/ENSMUSG00000044499.6, intron 1 of 4) | 10 | 36507407 | 36810038 | 302632 | 1 | ENSMUST00000171521.1 | 2541 | Hs3st5 | ENSMUSG00000044499.6 |
| chr3 | 94399152 | 94399390 | 3.46998 | 4.09098 | 1.22221 | Promoter | 3 | 94398517 | 94404501 | 5985 | 1 | ENSMUST00000050975.3 | 636 | Lingo4 | ENSMUSG00000044505.3 |
| chr4 | 114925747 | 114926423 | 3.47481 | 4.12018 | 1.2709 | Promoter | 4 | 114925147 | 114926013 | 867 | 2 | ENSMUST00000050940.3 | 0 | Foxe3 | ENSMUSG00000044518.3 |
| chr4 | 114926799 | 114927087 | 3.68873 | 4.53663 | 1.539 | Promoter | 4 | 114925147 | 114926013 | 867 | 2 | ENSMUST00000050940.3 | -787 | Foxe3 | ENSMUSG00000044518.3 |
| chr17 | 56511345 | 56512053 | 3.85727 | 4.80257 | 1.61409 | Promoter | 17 | 56511248 | 56512483 | 1236 | 2 | ENSMUST00000052211.2 | 430 | Znrf4 | ENSMUSG00000044526.2 |
| chr3 | 124318207 | 124318524 | 3.33873 | 3.98067 | 1.1058 | Distal Intergenic | 3 | 124321037 | 124323260 | 2224 | 1 | ENSMUST00000058994.4 | -2513 | Tram1l1 | ENSMUSG00000044528.4 |
| chr3 | 124321019 | 124321612 | 3.49119 | 4.26183 | 1.31272 | Promoter | 3 | 124321037 | 124323260 | 2224 | 1 | ENSMUST00000058994.4 | 0 | Tram1l1 | ENSMUSG00000044528.4 |
| chr10 | 120694588 | 120694917 | 3.38141 | 4.11782 | 1.19456 | Distal Intergenic | 10 | 120765774 | 120778886 | 13113 | 1 | ENSMUST00000056994.3 | -70857 | 4921513I03Rik | ENSMUSG00000044544.3 |
| chr10 | 120720778 | 120721095 | 3.8766 | 4.90254 | 1.76131 | Distal Intergenic | 10 | 120765774 | 120778886 | 13113 | 1 | ENSMUST00000056994.3 | -44679 | 4921513I03Rik | ENSMUSG00000044544.3 |
| chr10 | 120722104 | 120722562 | 3.98779 | 5.24111 | 1.97538 | Distal Intergenic | 10 | 120765774 | 120778886 | 13113 | 1 | ENSMUST00000056994.3 | -43212 | 4921513I03Rik | ENSMUSG00000044544.3 |
| chr10 | 120734788 | 120735106 | 3.36678 | 3.87775 | 1.01733 | Distal Intergenic | 10 | 120765774 | 120778886 | 13113 | 1 | ENSMUST00000056994.3 | -30668 | 4921513I03Rik | ENSMUSG00000044544.3 |
| chr10 | 120744223 | 120744480 | 3.59895 | 4.49209 | 1.43483 | Distal Intergenic | 10 | 120765774 | 120778886 | 13113 | 1 | ENSMUST00000056994.3 | -21294 | 4921513I03Rik | ENSMUSG00000044544.3 |

| | | | | | | | | | | | | | | | |
|---|---|---|---|---|---|---|---|---|---|---|---|---|---|---|---|
| chr12 | 71309587 | 71309875 | 4.04118 | 5.20615 | 1.91565 | Promoter | 12 | 71309884 | 71320107 | 10224 | 1 | ENSMUST00000061273.6 | -9 | Dact1 | ENSMUSG00000044548.6 |
| chr12 | 71310265 | 71310488 | 3.44358 | 4.17831 | 1.21571 | Promoter | 12 | 71309984 | 71318894 | 8911 | 1 | ENSMUST00000150639.1 | 282 | Dact1 | ENSMUSG00000044548.6 |
| chr5 | 30105004 | 30106237 | 4.21457 | 5.78489 | 2.37701 | Promoter | 5 | 30105161 | 30118378 | 13218 | 1 | ENSMUST00000058045.4 | 0 | Gareml | ENSMUSG00000044576.6 |
| chr2 | 65845573 | 65846001 | 3.63681 | 4.67526 | 1.55668 | Promoter | 2 | 65845767 | 66031546 | 185780 | 1 | ENSMUST00000122912.2 | 0 | Csrnp3 | ENSMUSG00000044647.11 |
| chr2 | 65846414 | 65846509 | 4.05944 | 5.36044 | 2.00692 | Promoter | 2 | 65845855 | 66023279 | 177425 | 1 | ENSMUST00000112397.2 | 560 | Csrnp3 | ENSMUSG00000044647.11 |
| chr2 | 65847457 | 65847597 | 4.68875 | 6.76611 | 3.05297 | Intron (ENSMUST00000122912.2/ENSMUSG00000044647.11, intron 1 of 6) | 2 | 65845855 | 66023279 | 177425 | 1 | ENSMUST00000112397.2 | 1603 | Csrnp3 | ENSMUSG00000044647.11 |
| chr2 | 65930201 | 65930425 | 3.3097 | 3.79176 | 1.01733 | Promoter | 2 | 65930137 | 66023946 | 93810 | 1 | ENSMUST00000176109.2 | 65 | Csrnp3 | ENSMUSG00000044647.11 |
| chr2 | 65932180 | 65932255 | 3.65233 | 4.42383 | 1.42872 | Promoter | 2 | 65931865 | 66023484 | 91620 | 1 | ENSMUST00000145598.3 | 316 | Csrnp3 | ENSMUSG00000044647.11 |
| chr3 | 117361021 | 117361153 | 3.38442 | 3.90752 | 1.02845 | Promoter | 3 | 117321109 | 117360876 | 39768 | 2 | ENSMUST00000125664.1 | -146 | D3Bwg0562e | ENSMUSG00000044667.8 |
| chr5 | 4666482 | 4666687 | 3.56149 | 4.39495 | 1.35561 | Distal Intergenic | 5 | 4753873 | 4758035 | 4163 | 2 | ENSMUST00000054294.5 | 91348 | Fzd1 | ENSMUSG00000044674.5 |
| chr5 | 4756448 | 4756703 | 3.83505 | 5.09724 | 1.84995 | Exon (ENSMUST00000054294.5/ENSMUSG00000044674.5, exon 1 of 1) | 5 | 4753873 | 4758035 | 4163 | 2 | ENSMUST00000054294.5 | 1332 | Fzd1 | ENSMUSG00000044674.5 |
| chr5 | 28188582 | 28188912 | 3.88429 | 4.9081 | 1.75685 | Distal Intergenic | 5 | 28202449 | 28210022 | 7574 | 2 | ENSMUST00000118882.1 | 21110 | Cnpy1 | ENSMUSG00000044681.13 |
| chr5 | 28198553 | 28198907 | 3.77162 | 4.96281 | 1.73367 | Downstream (1-2kb) | 5 | 28202449 | 28210022 | 7574 | 2 | ENSMUST00000118882.1 | 11115 | Cnpy1 | ENSMUSG00000044681.13 |
| chr5 | 28202844 | 28203294 | 3.40568 | 4.07111 | 1.16548 | 3' UTR | 5 | 28202449 | 28210022 | 7574 | 2 | ENSMUST00000118882.1 | 6728 | Cnpy1 | ENSMUSG00000044681.13 |
| chr5 | 28203652 | 28203951 | 3.45496 | 4.07 | 1.18976 | Intron (ENSMUST00000141601.2/ENSMUSG00000044681.13, intron 3 of 3) | 5 | 28202449 | 28210022 | 7574 | 2 | ENSMUST00000118882.1 | 6071 | Cnpy1 | ENSMUSG00000044681.13 |
| chr5 | 28204793 | 28205319 | 4.13035 | 5.62423 | 2.23524 | Intron (ENSMUST00000141601.2/ENSMUSG00000044681.13, intron 3 of 3) | 5 | 28202449 | 28210022 | 7574 | 2 | ENSMUST00000118882.1 | 4703 | Cnpy1 | ENSMUSG00000044681.13 |
| chr5 | 28205676 | 28205776 | 3.47032 | 4.22866 | 1.28796 | 3' UTR | 5 | 28202449 | 28210022 | 7574 | 2 | ENSMUST00000118882.1 | 4246 | Cnpy1 | ENSMUSG00000044681.13 |
| chr5 | 28206084 | 28206292 | 3.46663 | 4.06799 | 1.15621 | 3' UTR | 5 | 28202449 | 28210022 | 7574 | 2 | ENSMUST00000118882.1 | 3730 | Cnpy1 | ENSMUSG00000044681.13 |
| chr5 | 28207094 | 28207419 | 4.00175 | 5.08037 | 1.80923 | Exon (ENSMUST00000141601.2/ENSMUSG00000044681.13, exon 3 of 4) | 5 | 28202449 | 28210022 | 7574 | 2 | ENSMUST00000118882.1 | 2603 | Cnpy1 | ENSMUSG00000044681.13 |
| chr5 | 28208592 | 28209276 | 3.65579 | 4.41746 | 1.3611 | Promoter | 5 | 28202449 | 28210022 | 7574 | 2 | ENSMUST00000118882.1 | 746 | Cnpy1 | ENSMUSG00000044681.13 |
| chr5 | 28209591 | 28211008 | 3.85413 | 4.92477 | 1.72618 | Promoter | 5 | 28202449 | 28210022 | 7574 | 2 | ENSMUST00000118882.1 | 0 | Cnpy1 | ENSMUSG00000044681.13 |
| chr5 | 28211779 | 28211854 | 3.99317 | 5.28287 | 1.92139 | Intron (ENSMUST00000141601.2/ENSMUSG00000044681.13, intron 1 of 3) | 5 | 28209245 | 28210363 | 1119 | 2 | ENSMUST00000141196.1 | -1417 | Cnpy1 | ENSMUSG00000044681.13 |
| chr5 | 28212639 | 28213832 | 3.64362 | 4.39544 | 1.35713 | Intron (ENSMUST00000141601.2/ENSMUSG00000044681.13, intron 1 of 3) | 5 | 28209245 | 28210363 | 1119 | 2 | ENSMUST00000141196.1 | -2277 | Cnpy1 | ENSMUSG00000044681.13 |
| chr5 | 28214151 | 28214283 | 4.29431 | 5.64963 | 2.21122 | Intron (ENSMUST00000141601.2/ENSMUSG00000044681.13, intron 1 of 3) | 5 | 28209245 | 28210363 | 1119 | 2 | ENSMUST00000141196.1 | -3789 | Cnpy1 | ENSMUSG00000044681.13 |
| chr5 | 28215016 | 28215347 | 3.64837 | 4.46476 | 1.44674 | Intron (ENSMUST00000141601.2/ENSMUSG00000044681.13, intron 1 of 3) | 5 | 28209245 | 28210363 | 1119 | 2 | ENSMUST00000141196.1 | -4654 | Cnpy1 | ENSMUSG00000044681.13 |
| chr5 | 28228231 | 28228306 | 3.5794 | 4.26753 | 1.25041 | Intron (ENSMUST00000141601.2/ENSMUSG00000044681.13, intron 1 of 3) | 5 | 28205460 | 28237873 | 32414 | 2 | ENSMUST00000120068.1 | 9567 | Cnpy1 | ENSMUSG00000044681.13 |
| chr1 | 172337950 | 172338119 | 3.52319 | 4.17611 | 1.23192 | Distal Intergenic | 1 | 172341210 | 172374085 | 32876 | 1 | ENSMUST00000056136.3 | -3091 | Kcnj10 | ENSMUSG00000044708.5 |
| chr1 | 172340409 | 172341881 | 3.66694 | 4.57513 | 1.5005 | Promoter | 1 | 172341210 | 172374085 | 32876 | 1 | ENSMUST00000056136.3 | 0 | Kcnj10 | ENSMUSG00000044708.5 |
| chr1 | 172342204 | 172342279 | 3.98967 | 5.08803 | 1.87261 | Promoter | 1 | 172341210 | 172374085 | 32876 | 1 | ENSMUST00000056136.3 | 995 | Kcnj10 | ENSMUSG00000044708.5 |
| chr1 | 172350660 | 172350949 | 3.63463 | 4.41512 | 1.42713 | Intron (ENSMUST00000056136.3/ENSMUSG00000044708.5, intron 1 of 1) | 1 | 172341210 | 172374085 | 32876 | 1 | ENSMUST00000056136.3 | 9451 | Kcnj10 | ENSMUSG00000044708.5 |
| chr5 | 35056336 | 35056763 | 3.31885 | 3.80552 | 1.01733 | Promoter | 5 | 35056813 | 35080964 | 24152 | 1 | ENSMUST00000101298.3 | -50 | Dok7 | ENSMUSG00000044716.7 |
| chr5 | 35057127 | 35057259 | 3.77821 | 4.68868 | 1.595 | Promoter | 5 | 35057077 | 35087839 | 30763 | 1 | ENSMUST00000050709.3 | 51 | Dok7 | ENSMUSG00000044716.7 |
| chr5 | 35058487 | 35059009 | 4.16778 | 5.44033 | 2.11891 | Intron (ENSMUST00000101298.3/ENSMUSG00000044716.7, intron 2 of 4) | 5 | 35057077 | 35087839 | 30763 | 1 | ENSMUST00000050709.3 | 1411 | Dok7 | ENSMUSG00000044716.7 |
| chr15 | 34453450 | 34454052 | 3.98717 | 5.1038 | 1.85365 | Promoter | 15 | 34453312 | 34473892 | 20581 | 1 | ENSMUST00000060894.7 | 139 | Erich5 | ENSMUSG00000044726.7 |
| chr10 | 42853093 | 42853471 | 3.50336 | 4.1284 | 1.16834 | Distal Intergenic | 10 | 42860370 | 42960780 | 100411 | 1 | ENSMUST00000063063.8 | -6899 | Scml4 | ENSMUSG00000044770.9 |
| chr10 | 42951719 | 42951904 | 3.66092 | 4.53232 | 1.4667 | Intron (ENSMUST00000063063.8/ENSMUSG00000044770.9, intron 7 of 8) | 10 | 42939881 | 42958721 | 18841 | 1 | ENSMUST00000127675.1 | 11839 | Scml4 | ENSMUSG00000044770.9 |
| chr10 | 42957529 | 42957923 | 4.17891 | 5.48751 | 2.15984 | 3' UTR | 10 | 42939881 | 42958721 | 18841 | 1 | ENSMUST00000127675.1 | 17649 | Scml4 | ENSMUSG00000044770.9 |
| chr10 | 42958851 | 42959270 | 3.63831 | 4.39656 | 1.39311 | 3' UTR | 10 | 42939881 | 42958721 | 18841 | 1 | ENSMUST00000127675.1 | 18971 | Scml4 | ENSMUSG00000044770.9 |
| chr10 | 42994647 | 42995120 | 3.53148 | 4.22767 | 1.34057 | Distal Intergenic | 10 | 42939881 | 42958721 | 18841 | 1 | ENSMUST00000127675.1 | 54767 | Scml4 | ENSMUSG00000044770.9 |
| chr10 | 42995655 | 42996141 | 3.27805 | 3.76445 | 1.07731 | Distal Intergenic | 10 | 42939881 | 42958721 | 18841 | 1 | ENSMUST00000127675.1 | 55775 | Scml4 | ENSMUSG00000044770.9 |
| chr10 | 43001599 | 43001674 | 3.35206 | 3.88348 | 1.11347 | Downstream (<1kb) | 10 | 42939881 | 42958721 | 18841 | 1 | ENSMUST00000127675.1 | 61719 | Scml4 | ENSMUSG00000044770.9 |
| chr10 | 43021331 | 43021943 | 4.33293 | 5.8739 | 2.42813 | Exon (ENSMUST00000040275.8/ENSMUSG00000038248.8, exon 6 of 7) | 10 | 42939881 | 42958721 | 18841 | 1 | ENSMUST00000127675.1 | 81451 | Scml4 | ENSMUSG00000044770.9 |
| chr10 | 43023444 | 43023565 | 3.49173 | 4.1412 | 1.27324 | Intron (ENSMUST00000040275.8/ENSMUSG00000038248.8, intron 5 of 6) | 10 | 42939881 | 42958721 | 18841 | 1 | ENSMUST00000127675.1 | 83564 | Scml4 | ENSMUSG00000044770.9 |
| chr10 | 43028038 | 43028164 | 3.67102 | 4.47762 | 1.46797 | Intron (ENSMUST00000040275.8/ENSMUSG00000038248.8, intron 5 of 6) | 10 | 42939881 | 42958721 | 18841 | 1 | ENSMUST00000127675.1 | 88158 | Scml4 | ENSMUSG00000044770.9 |
| chr1 | 161142085 | 161142274 | 3.46675 | 4.10802 | 1.26739 | Promoter | 1 | 161142691 | 161163295 | 20605 | 1 | ENSMUST00000125018.2 | -417 | Ankrd45 | ENSMUSG00000044835.10 |
| chr6 | 48742407 | 48743065 | 3.4452 | 4.05978 | 1.15052 | Exon (ENSMUST00000054368.4/ENSMUSG00000090019.3, exon 3 of 3) | 6 | 48738346 | 48740621 | 2276 | 2 | ENSMUST00000155906.1 | -1787 | Gimap1os | ENSMUSG00000044867.4 |
| chr12 | 73997374 | 73998284 | 3.72586 | 4.60641 | 1.53383 | Promoter | 12 | 73997761 | 74267916 | 270156 | 1 | ENSMUST00000110451.2 | 0 | Syt16 | ENSMUSG00000044912.9 |
| chr12 | 73999268 | 73999582 | 3.5641 | 4.26369 | 1.33197 | Intron (ENSMUST00000110451.2/ENSMUSG00000044912.9, intron 1 of 6) | 12 | 73997761 | 74267916 | 270156 | 1 | ENSMUST00000110451.2 | 1508 | Syt16 | ENSMUSG00000044912.9 |
| chr6 | 87981369 | 87982329 | 3.63087 | 4.403 | 1.38858 | Promoter | 6 | 87980421 | 87981637 | 1217 | 2 | ENSMUST00000056403.6 | 0 | H1fx | ENSMUSG00000044927.6 |
| chr15 | 78543933 | 78544277 | 3.41952 | 3.98586 | 1.11654 | Promoter | 15 | 78537015 | 78544345 | 7331 | 2 | ENSMUST00000053239.2 | 68 | Sstr3 | ENSMUSG00000044933.2 |
| chr15 | 78544845 | 78545252 | 4.69723 | 6.56234 | 2.94338 | Promoter | 15 | 78537015 | 78544345 | 7331 | 2 | ENSMUST00000053239.2 | -501 | Sstr3 | ENSMUSG00000044933.2 |
| chr9 | 77654837 | 77655630 | 3.82193 | 4.76772 | 1.61334 | Exon (ENSMUST00000057781.7/ENSMUSG00000044938.8, exon 3 of 3) | 9 | 77636500 | 77660127 | 23628 | 1 | ENSMUST00000057781.7 | 18338 | Klhl31 | ENSMUSG00000044938.8 |
| chr19 | 47822876 | 47823057 | 3.3939 | 3.9329 | 1.0677 | Exon (ENSMUST00000160247.1/ENSMUSG00000044948.11, exon 5 of 38) | 19 | 47794851 | 47812146 | 17296 | 2 | ENSMUST00000160970.1 | -10731 | Wdr96 | ENSMUSG00000044948.11 |
| chr19 | 47837206 | 47837526 | 3.61979 | 4.40876 | 1.36407 | Promoter | 19 | 47737561 | 47837382 | 99822 | 2 | ENSMUST00000160247.1 | 0 | Wdr96 | ENSMUSG00000044948.11 |
| chr9 | 74110395 | 74110639 | 3.47481 | 4.3429 | 1.35536 | Promoter | 9 | 74110404 | 74113679 | 3276 | 1 | ENSMUST00000184725.1 | 0 | Wdr72 | ENSMUSG00000044976.11 |
| chr9 | 74186786 | 74187004 | 3.68757 | 4.70544 | 1.56008 | Intron (ENSMUST00000055879.8/ENSMUSG00000044976.11, intron 14 of 19) | 9 | 74110404 | 74113679 | 3276 | 1 | ENSMUST00000184725.1 | 76383 | Wdr72 | ENSMUSG00000044976.11 |
| chr6 | 113501667 | 113502331 | 3.89322 | 4.89343 | 1.67686 | Promoter | 6 | 113494095 | 113501818 | 7724 | 2 | ENSMUST00000101059.1 | 0 | Prrt3 | ENSMUSG00000045009.7 |
| chr13 | 92425625 | 92427942 | 3.81755 | 4.86211 | 1.72427 | Promoter | 13 | 92425896 | 92441658 | 15763 | 1 | ENSMUST00000061594.7 | 0 | Ankrd34b | ENSMUSG00000045034.7 |
| chr7 | 25313460 | 25313619 | 3.56555 | 4.29285 | 1.35964 | Intron (ENSMUST00000119703.2/ENSMUSG00000043843.11, intron 13 of 14) | 7 | 25317164 | 25365917 | 48754 | 1 | ENSMUST00000128119.1 | -3545 | Megf8 | ENSMUSG00000045039.9 |
| chr7 | 25314269 | 25314406 | 3.58094 | 4.26967 | 1.25374 | Intron (ENSMUST00000119703.2/ENSMUSG00000043843.11, intron 13 of 14) | 7 | 25317164 | 25365917 | 48754 | 1 | ENSMUST00000128119.1 | -2758 | Megf8 | ENSMUSG00000045039.9 |
| chr7 | 25314715 | 25315884 | 4.28983 | 5.71256 | 2.26388 | 3' UTR | 7 | 25317164 | 25365917 | 48754 | 1 | ENSMUST00000128119.1 | -1280 | Megf8 | ENSMUSG00000045039.9 |
| chr19 | 60464555 | 60464907 | 3.65027 | 4.42002 | 1.41763 | Downstream (1-2kb) | 19 | 60466733 | 60468304 | 1572 | 2 | ENSMUST00000051277.2 | 3397 | Prlhr | ENSMUSG00000045052.2 |
| chr19 | 60465482 | 60466292 | 3.5273 | 4.23363 | 1.31296 | Downstream (<1kb) | 19 | 60466733 | 60468304 | 1572 | 2 | ENSMUST00000051277.2 | 2012 | Prlhr | ENSMUSG00000045052.2 |
| chr19 | 60466994 | 60467931 | 3.94823 | 5.22838 | 1.96032 | Promoter | 19 | 60466733 | 60468304 | 1572 | 2 | ENSMUST00000051277.2 | 373 | Prlhr | ENSMUSG00000045052.2 |
| chr19 | 60468279 | 60468927 | 3.86486 | 5.17675 | 1.90329 | Promoter | 19 | 60466733 | 60468304 | 1572 | 2 | ENSMUST00000051277.2 | 0 | Prlhr | ENSMUSG00000045052.2 |
| chr17 | 83629880 | 83632059 | 3.66791 | 4.69202 | 1.58051 | Promoter | 17 | 83585957 | 83631895 | 45939 | 2 | ENSMUST00000051482.1 | 0 | Kcng3 | ENSMUSG00000045053.1 |
| chr17 | 83632468 | 83632711 | 4.82136 | 6.80384 | 3.13198 | Promoter | 17 | 83585957 | 83631895 | 45939 | 2 | ENSMUST00000051482.1 | -574 | Kcng3 | ENSMUSG00000045053.1 |
| chr4 | 36951430 | 36952014 | 3.94787 | 5.06839 | 1.8576 | Promoter | 4 | 36715820 | 36951744 | 235925 | 2 | ENSMUST00000145615.1 | 0 | Lingo2 | ENSMUSG00000045083.9 |
| chr9 | 21244173 | 21245323 | 3.81264 | 4.88353 | 1.73299 | 5' UTR | 9 | 21242912 | 21248443 | 5532 | 2 | ENSMUST00000122088.1 | 3120 | S1pr5 | ENSMUSG00000045087.8 |
| chr9 | 21245649 | 21246455 | 3.60334 | 4.37124 | 1.43652 | 5' UTR | 9 | 21242912 | 21248443 | 5532 | 2 | ENSMUST00000122088.1 | 1988 | S1pr5 | ENSMUSG00000045087.8 |
| chr9 | 21247938 | 21248839 | 3.68269 | 4.65055 | 1.54789 | Promoter | 9 | 21242912 | 21248443 | 5532 | 2 | ENSMUST00000122088.1 | 0 | S1pr5 | ENSMUSG00000045087.8 |
| chr9 | 21251134 | 21251410 | 3.44929 | 4.02625 | 1.10227 | Distal Intergenic | 9 | 21242912 | 21248443 | 5532 | 2 | ENSMUST00000122088.1 | -2692 | S1pr5 | ENSMUSG00000045087.8 |
| chr3 | 115714466 | 115715355 | 4.11672 | 5.51801 | 2.15239 | Promoter | 3 | 115710433 | 115715055 | 4623 | 2 | ENSMUST00000055676.2 | 0 | S1pr1 | ENSMUSG00000045092.7 |
| chr18 | 61533484 | 61533584 | 3.77268 | 4.74684 | 1.64292 | Intron (ENSMUST00000171629.1/ENSMUSG00000045094.7, intron 1 of 12) | 18 | 61493794 | 61536536 | 42743 | 2 | ENSMUST00000171629.1 | 2952 | Arhgef37 | ENSMUSG00000045094.7 |
| chr6 | 94282707 | 94282863 | 3.7125 | 4.57671 | 1.53821 | Promoter | 6 | 93675455 | 94283322 | 607868 | 2 | ENSMUST00000055224.9 | 459 | Magi1 | ENSMUSG00000045095.11 |
| chr14 | 20034416 | 20035140 | 3.49023 | 4.10341 | 1.15507 | Distal Intergenic | 14 | 20075646 | 20083172 | 7527 | 2 | ENSMUST00000059666.4 | 48032 | Saysd1 | ENSMUSG00000045107.4 |
| chr14 | 20113465 | 20113553 | 3.51338 | 4.15321 | 1.20954 | Distal Intergenic | 14 | 20075646 | 20083172 | 7527 | 2 | ENSMUST00000059666.4 | -30294 | Saysd1 | ENSMUSG00000045107.4 |
| chr14 | 20116438 | 20116676 | 3.86435 | 4.82297 | 1.63981 | Distal Intergenic | 14 | 20075646 | 20083172 | 7527 | 2 | ENSMUST00000059666.4 | -33267 | Saysd1 | ENSMUSG00000045107.4 |
| chr1 | 34579923 | 34580140 | 3.8889 | 4.89285 | 1.73444 | Promoter | 1 | 34579727 | 34590944 | 11218 | 1 | ENSMUST00000052670.9 | 197 | Amer3 | ENSMUSG00000045174.9 |
| chr8 | 84043638 | 84044059 | 4.22622 | 5.77806 | 2.32281 | Promoter | 8 | 84042929 | 84045000 | 2072 | 2 | ENSMUST00000061923.4 | 941 | Rln3 | ENSMUSG00000045232.4 |
| chr8 | 84044526 | 84044818 | 3.57626 | 4.34086 | 1.38675 | Promoter | 8 | 84042929 | 84045000 | 2072 | 2 | ENSMUST00000061923.4 | 182 | Rln3 | ENSMUSG00000045232.4 |
| chr7 | 142301252 | 142301375 | 3.52946 | 4.18652 | 1.23607 | Distal Intergenic | 7 | 142303502 | 142304503 | 1002 | 1 | ENSMUST00000061403.4 | -2127 | Krtap5-4 | ENSMUSG00000045236.4 |
| chr7 | 142303598 | 142303837 | 3.48887 | 4.10269 | 1.16131 | Promoter | 7 | 142303502 | 142304503 | 1002 | 1 | ENSMUST00000061403.4 | 97 | Krtap5-4 | ENSMUSG00000045236.4 |
| chr5 | 98183966 | 98184719 | 4.1414 | 5.6538 | 2.30722 | Exon (ENSMUST00000112959.1/ENSMUSG00000035456.6, exon 3 of 4) | 5 | 98185296 | 98185984 | 689 | 2 | ENSMUST00000057889.6 | 1265 | A730035I17Rik | ENSMUSG00000045238.7 |
| chr5 | 98185127 | 98185351 | 4.59232 | 6.3455 | 2.76251 | Promoter | 5 | 98185296 | 98185984 | 689 | 2 | ENSMUST00000057889.6 | 633 | A730035I17Rik | ENSMUSG00000045238.7 |
| chr5 | 98185677 | 98186443 | 3.87647 | 5.15963 | 1.89079 | Promoter | 5 | 98185296 | 98185984 | 689 | 2 | ENSMUST00000057889.6 | 0 | A730035I17Rik | ENSMUSG00000045238.7 |
| chr5 | 98187101 | 98187360 | 3.60136 | 4.34033 | 1.39073 | 3' UTR | 5 | 98185296 | 98185984 | 689 | 2 | ENSMUST00000057889.6 | -1118 | A730035I17Rik | ENSMUSG00000045238.7 |
| chr5 | 98187785 | 98187886 | 3.46723 | 4.10364 | 1.25337 | Distal Intergenic | 5 | 98185296 | 98185984 | 689 | 2 | ENSMUST00000057889.6 | -1802 | A730035I17Rik | ENSMUSG00000045238.7 |
| chr5 | 98188427 | 98188917 | 3.56105 | 4.24336 | 1.26981 | Distal Intergenic | 5 | 98185296 | 98185984 | 689 | 2 | ENSMUST00000057889.6 | -2444 | A730035I17Rik | ENSMUSG00000045238.7 |
| chr5 | 98189404 | 98189834 | 3.79878 | 4.73735 | 1.63368 | Distal Intergenic | 5 | 98185296 | 98185984 | 689 | 2 | ENSMUST00000057889.6 | -3421 | A730035I17Rik | ENSMUSG00000045238.7 |
| chr5 | 98191245 | 98191804 | 3.49764 | 4.44596 | 1.41522 | Distal Intergenic | 5 | 98185296 | 98185984 | 689 | 2 | ENSMUST00000057889.6 | -5262 | A730035I17Rik | ENSMUSG00000045238.7 |
| chr5 | 98193558 | 98193981 | 3.40297 | 3.94748 | 1.07072 | Distal Intergenic | 5 | 98185296 | 98185984 | 689 | 2 | ENSMUST00000057889.6 | -7575 | A730035I17Rik | ENSMUSG00000045238.7 |
| chr5 | 98198388 | 98199078 | 3.44036 | 4.02189 | 1.1086 | Distal Intergenic | 5 | 98185296 | 98185984 | 689 | 2 | ENSMUST00000057889.6 | -12405 | A730035I17Rik | ENSMUSG00000045238.7 |
| chr5 | 98200205 | 98200445 | 3.35377 | 3.86199 | 1.03102 | Distal Intergenic | 5 | 98185296 | 98185984 | 689 | 2 | ENSMUST00000057889.6 | -14222 | A730035I17Rik | ENSMUSG00000045238.7 |
| chr8 | 119633129 | 119633290 | 3.44277 | 4.02079 | 1.11744 | Exon (ENSMUST00000164382.1/ENSMUSG00000045246.6, exon 2 of 4) | 8 | 119623854 | 119635346 | 11493 | 2 | ENSMUST00000164382.1 | 2056 | Kcng4 | ENSMUSG00000045246.6 |
| chr11 | 75194104 | 75194447 | 3.59839 | 4.56143 | 1.48013 | Promoter | 11 | 75193783 | 75267769 | 73987 | 1 | ENSMUST00000102514.3 | 322 | Rtn4rl1 | ENSMUSG00000045287.6 |
| chr11 | 75195238 | 75195313 | 3.90321 | 4.86275 | 1.61715 | Intron (ENSMUST00000102514.3/ENSMUSG00000045287.6, intron 1 of 1) | 11 | 75193783 | 75267769 | 73987 | 1 | ENSMUST00000102514.3 | 1456 | Rtn4rl1 | ENSMUSG00000045287.6 |
| chr11 | 75196737 | 75197086 | 3.46877 | 4.06513 | 1.13598 | Intron (ENSMUST00000102514.3/ENSMUSG00000045287.6, intron 1 of 1) | 11 | 75193783 | 75267769 | 73987 | 1 | ENSMUST00000102514.3 | 2955 | Rtn4rl1 | ENSMUSG00000045287.6 |
| chr11 | 75199935 | 75200309 | 3.6107 | 4.38242 | 1.3219 | Intron (ENSMUST00000102514.3/ENSMUSG00000045287.6, intron 1 of 1) | 11 | 75193783 | 75267769 | 73987 | 1 | ENSMUST00000102514.3 | 6153 | Rtn4rl1 | ENSMUSG00000045287.6 |
| chr11 | 115318130 | 115319110 | 3.55867 | 4.28592 | 1.34714 | Exon (ENSMUST00000103037.4/ENSMUSG00000045288.10, exon 2 of 3) | 11 | 115315192 | 115322041 | 6850 | 2 | ENSMUST00000103037.4 | 2931 | Ush1g | ENSMUSG00000045288.10 |
| chr11 | 115321580 | 115322128 | 3.93021 | 4.9886 | 1.77563 | Promoter | 11 | 115315192 | 115322041 | 6850 | 2 | ENSMUST00000103037.4 | 0 | Ush1g | ENSMUSG00000045288.10 |
| chr5 | 28116263 | 28116567 | 3.95601 | 4.99023 | 1.73228 | Distal Intergenic | 5 | 28071363 | 28078662 | 7300 | 1 | ENSMUST00000059155.10 | 44901 | Insig1 | ENSMUSG00000045294.10 |
| chr5 | 28116921 | 28117180 | 3.34153 | 3.83966 | 1.01733 | Distal Intergenic | 5 | 28071363 | 28078662 | 7300 | 1 | ENSMUST00000059155.10 | 45559 | Insig1 | ENSMUSG00000045294.10 |
| chr5 | 28117712 | 28118803 | 4.0521 | 5.22465 | 1.93297 | Distal Intergenic | 5 | 28071363 | 28078662 | 7300 | 1 | ENSMUST00000059155.10 | 46350 | Insig1 | ENSMUSG00000045294.10 |
| chr13 | 94057461 | 94058730 | 3.77436 | 4.87659 | 1.69487 | Promoter | 13 | 94057796 | 94195409 | 137614 | 1 | ENSMUST00000054274.2 | 0 | Lhfpl2 | ENSMUSG00000045312.6 |
| chr13 | 94059091 | 94059244 | 3.51273 | 4.2351 | 1.30348 | Intron (ENSMUST00000054274.2/ENSMUSG00000045312.6, intron 1 of 3) | 13 | 94057832 | 94123974 | 66143 | 1 | ENSMUST00000144838.1 | 1260 | Lhfpl2 | ENSMUSG00000045312.6 |
| chr13 | 94059759 | 94060300 | 3.62039 | 4.46013 | 1.43112 | Intron (ENSMUST00000054274.2/ENSMUSG00000045312.6, intron 1 of 3) | 13 | 94057832 | 94123974 | 66143 | 1 | ENSMUST00000144838.1 | 1928 | Lhfpl2 | ENSMUSG00000045312.6 |
| chr13 | 94060807 | 94060982 | 3.55552 | 4.30918 | 1.34367 | Intron (ENSMUST00000054274.2/ENSMUSG00000045312.6, intron 1 of 3) | 13 | 94057832 | 94123974 | 66143 | 1 | ENSMUST00000144838.1 | 2976 | Lhfpl2 | ENSMUSG00000045312.6 |
| chr13 | 94176120 | 94176682 | 4.11182 | 5.4411 | 2.11212 | Intron (ENSMUST00000054274.2/ENSMUSG00000045312.6, intron 3 of 3) | 13 | 94173992 | 94195409 | 21418 | 1 | ENSMUST00000121618.1 | 2129 | Lhfpl2 | ENSMUSG00000045312.6 |
| chr13 | 94173663 | 94174178 | 3.76152 | 4.85063 | 1.65916 | Promoter | 13 | 94173774 | 94174213 | 440 | 1 | ENSMUST00000131595.1 | 0 | Lhfpl2 | ENSMUSG00000045312.6 |
| chr5 | 93045026 | 93045578 | 4.03766 | 5.51513 | 2.13703 | Promoter | 5 | 93041123 | 93045022 | 3900 | 2 | ENSMUST00000061328.5 | -5 | Sowahb | ENSMUSG00000045314.5 |
| chr2 | 6097674 | 6098348 | 3.65084 | 4.44713 | 1.42471 | Downstream (1-2kb) | 2 | 6101190 | 6130139 | 28950 | 2 | ENSMUST00000126551.1 | 31791 | Proser2 | ENSMUSG00000045319.8 |
| chr2 | 6100224 | 6101515 | 3.91665 | 4.9467 | 1.74098 | 3' UTR | 2 | 6101190 | 6130139 | 28950 | 2 | ENSMUST00000126551.1 | 28624 | Proser2 | ENSMUSG00000045319.8 |
| chr2 | 6102225 | 6102316 | 3.84477 | 4.94383 | 1.77327 | Intron (ENSMUST00000054254.6/ENSMUSG00000045319.8, intron 3 of 3) | 2 | 6101190 | 6130139 | 28950 | 2 | ENSMUST00000126551.1 | 27823 | Proser2 | ENSMUSG00000045319.8 |
| chr2 | 6102753 | 6102960 | 3.24856 | 3.69999 | 1.01733 | Exon (ENSMUST00000054254.6/ENSMUSG00000045319.8, exon 3 of 4) | 2 | 6101190 | 6130139 | 28950 | 2 | ENSMUST00000126551.1 | 27179 | Proser2 | ENSMUSG00000045319.8 |
| chr2 | 6106197 | 6106657 | 3.41015 | 4.06744 | 1.20523 | Intron (ENSMUST00000054254.6/ENSMUSG00000045319.8, intron 2 of 3) | 2 | 6101190 | 6130139 | 28950 | 2 | ENSMUST00000126551.1 | 23482 | Proser2 | ENSMUSG00000045319.8 |
| chr2 | 6112872 | 6112947 | 3.35326 | 3.85735 | 1.01733 | Intron (ENSMUST00000054254.6/ENSMUSG00000045319.8, intron 2 of 3) | 2 | 6101190 | 6130139 | 28950 | 2 | ENSMUST00000126551.1 | 17192 | Proser2 | ENSMUSG00000045319.8 |
| chr2 | 6114106 | 6114490 | 3.52175 | 4.17292 | 1.22332 | 5' UTR | 2 | 6101190 | 6130139 | 28950 | 2 | ENSMUST00000126551.1 | 15649 | Proser2 | ENSMUSG00000045319.8 |
| chr2 | 6114843 | 6114958 | 3.3941 | 3.93087 | 1.05948 | Intron (ENSMUST00000054254.6/ENSMUSG00000045319.8, intron 1 of 3) | 2 | 6101190 | 6130139 | 28950 | 2 | ENSMUST00000126551.1 | 15181 | Proser2 | ENSMUSG00000045319.8 |
| chr2 | 6116251 | 6116363 | 3.469 | 4.07133 | 1.15662 | Intron (ENSMUST00000054254.6/ENSMUSG00000045319.8, intron 1 of 3) | 2 | 6101190 | 6130139 | 28950 | 2 | ENSMUST00000126551.1 | 13776 | Proser2 | ENSMUSG00000045319.8 |
| chr2 | 6118323 | 6118950 | 3.46017 | 4.05515 | 1.14656 | Intron (ENSMUST00000054254.6/ENSMUSG00000045319.8, intron 1 of 3) | 2 | 6101190 | 6130139 | 28950 | 2 | ENSMUST00000126551.1 | 11189 | Proser2 | ENSMUSG00000045319.8 |
| chr2 | 6121651 | 6121780 | 3.24197 | 3.69011 | 1.01733 | Intron (ENSMUST00000054254.6/ENSMUSG00000045319.8, intron 1 of 3) | 2 | 6101190 | 6130139 | 28950 | 2 | ENSMUST00000126551.1 | 8359 | Proser2 | ENSMUSG00000045319.8 |
| chr2 | 6127343 | 6128035 | 3.88713 | 4.99205 | 1.7932 | Intron (ENSMUST00000054254.6/ENSMUSG00000045319.8, intron 1 of 3) | 2 | 6101190 | 6130139 | 28950 | 2 | ENSMUST00000126551.1 | 2104 | Proser2 | ENSMUSG00000045319.8 |
| chr2 | 6128390 | 6128741 | 3.313 | 3.9669 | 1.14929 | Intron (ENSMUST00000054254.6/ENSMUSG00000045319.8, intron 1 of 3) | 2 | 6101190 | 6130139 | 28950 | 2 | ENSMUST00000126551.1 | 1398 | Proser2 | ENSMUSG00000045319.8 |
| chr2 | 6129549 | 6131001 | 3.95141 | 5.10721 | 1.88751 | Promoter | 2 | 6100025 | 6130146 | 30122 | 2 | ENSMUST00000054254.6 | 0 | Proser2 | ENSMUSG00000045319.8 |
| chr2 | 6131519 | 6132248 | 4.07076 | 5.25809 | 1.95235 | Exon (ENSMUST00000117166.2/ENSMUSG00000082332.2, exon 2 of 2) | 2 | 6100045 | 6130211 | 30167 | 2 | ENSMUST00000114942.3 | -1309 | Proser2 | ENSMUSG00000045319.8 |
| chr5 | 137731648 | 137731911 | 3.34962 | 3.85442 | 1.02638 | 3' UTR | 5 | 137730883 | 137739998 | 9116 | 2 | ENSMUST00000118326.2 | 8087 | Nyap1 | ENSMUSG00000045348.10 |
| chr4 | 138250641 | 138251094 | 3.87774 | 5.08786 | 1.83653 | Promoter | 4 | 138250464 | 138261332 | 10869 | 1 | ENSMUST00000105823.1 | 178 | Sh2d5 | ENSMUSG00000045349.10 |
| chr4 | 138252620 | 138253168 | 4.21261 | 5.78829 | 2.37116 | Intron (ENSMUST00000105824.2/ENSMUSG00000045349.10, intron 1 of 9) | 4 | 138250464 | 138261332 | 10869 | 1 | ENSMUST00000105823.1 | 2157 | Sh2d5 | ENSMUSG00000045349.10 |
| chr4 | 138256823 | 138257158 | 3.42687 | 4.0053 | 1.14829 | Promoter | 4 | 138256735 | 138259111 | 2377 | 1 | ENSMUST00000154421.1 | 89 | Sh2d5 | ENSMUSG00000045349.10 |
| chr1 | 128590061 | 128590372 | 3.77951 | 4.76978 | 1.6138 | Intron (ENSMUST00000052172.6/ENSMUSG00000045382.6, intron 1 of 1) | 1 | 128588199 | 128592290 | 4092 | 2 | ENSMUST00000052172.6 | 1918 | Cxcr4 | ENSMUSG00000045382.6 |
| chr1 | 128590794 | 128591022 | 3.80144 | 5.07145 | 1.82634 | Intron (ENSMUST00000052172.6/ENSMUSG00000045382.6, intron 1 of 1) | 1 | 128588199 | 128592290 | 4092 | 2 | ENSMUST00000052172.6 | 1268 | Cxcr4 | ENSMUSG00000045382.6 |
| chr1 | 128591459 | 128591828 | 3.71263 | 4.84835 | 1.67869 | Promoter | 1 | 128588199 | 128592290 | 4092 | 2 | ENSMUST00000052172.6 | 462 | Cxcr4 | ENSMUSG00000045382.6 |
| chr1 | 128592319 | 128592549 | 3.43964 | 4.04221 | 1.19182 | Promoter | 1 | 128589099 | 128592293 | 3195 | 2 | ENSMUST00000142893.1 | -27 | Cxcr4 | ENSMUSG00000045382.6 |
| chr1 | 128593215 | 128593754 | 3.6588 | 4.41703 | 1.34762 | Promoter | 1 | 128589099 | 128592293 | 3195 | 2 | ENSMUST00000142893.1 | -923 | Cxcr4 | ENSMUSG00000045382.6 |
| chr1 | 128595077 | 128595774 | 4.27931 | 5.63091 | 2.1878 | Distal Intergenic | 1 | 128589099 | 128592293 | 3195 | 2 | ENSMUST00000142893.1 | -2785 | Cxcr4 | ENSMUSG00000045382.6 |
| chr1 | 128596985 | 128597120 | 3.32851 | 3.8201 | 1.01733 | Distal Intergenic | 1 | 128589099 | 128592293 | 3195 | 2 | ENSMUST00000142893.1 | -4693 | Cxcr4 | ENSMUSG00000045382.6 |
| chr1 | 128653259 | 128653675 | 3.35264 | 3.86115 | 1.03351 | Distal Intergenic | 1 | 128589099 | 128592293 | 3195 | 2 | ENSMUST00000142893.1 | -60967 | Cxcr4 | ENSMUSG00000045382.6 |
| chr17 | 87635812 | 87636063 | 3.95274 | 4.97992 | 1.73958 | Promoter | 17 | 87635979 | 87651106 | 15128 | 1 | ENSMUST00000053577.8 | 0 | Epcam | ENSMUSG00000045394.8 |
| chr17 | 87636694 | 87637232 | 3.79667 | 5.10185 | 1.86994 | Promoter | 17 | 87635979 | 87651106 | 15128 | 1 | ENSMUST00000053577.8 | 716 | Epcam | ENSMUSG00000045394.8 |
| chr17 | 87637934 | 87638266 | 4.05295 | 5.36648 | 2.04139 | Intron (ENSMUST00000053577.8/ENSMUSG00000045394.8, intron 2 of 9) | 17 | 87635979 | 87651106 | 15128 | 1 | ENSMUST00000053577.8 | 1956 | Epcam | ENSMUSG00000045394.8 |
| chr12 | 99964632 | 99965298 | 4.10449 | 5.46352 | 2.13389 | Promoter | 12 | 99964525 | 100062674 | 98150 | 1 | ENSMUST00000049788.8 | 108 | Kcnk13 | ENSMUSG00000045404.10 |
| chr12 | 99965648 | 99966018 | 3.50277 | 4.51577 | 1.46231 | 5' UTR | 12 | 99964525 | 100062674 | 98150 | 1 | ENSMUST00000049788.8 | 1124 | Kcnk13 | ENSMUSG00000045404.10 |
| chr12 | 99967153 | 99968010 | 3.67 | 4.59024 | 1.53652 | Intron (ENSMUST00000160413.2/ENSMUSG00000045404.10, intron 2 of 2) | 12 | 99964525 | 100062674 | 98150 | 1 | ENSMUST00000049788.8 | 2629 | Kcnk13 | ENSMUSG00000045404.10 |
| chr12 | 55597617 | 55597958 | 3.23652 | 3.68195 | 1.01733 | Promoter | 12 | 55598917 | 55602018 | 3102 | 1 | ENSMUST00000051857.3 | -959 | Insm2 | ENSMUSG00000045440.3 |
| chr12 | 55598885 | 55600345 | 3.68462 | 4.9158 | 1.7352 | Promoter | 12 | 55598917 | 55602018 | 3102 | 1 | ENSMUST00000051857.3 | 0 | Insm2 | ENSMUSG00000045440.3 |
| chr12 | 55600774 | 55600926 | 3.61973 | 4.42776 | 1.41816 | Exon (ENSMUST00000051857.3/ENSMUSG00000045440.3, exon 1 of 1) | 12 | 55598917 | 55602018 | 3102 | 1 | ENSMUST00000051857.3 | 1858 | Insm2 | ENSMUSG00000045440.3 |
| chr11 | 100762788 | 100763117 | 3.82287 | 4.76132 | 1.59253 | Promoter | 11 | 100761069 | 100762928 | 1860 | 2 | ENSMUST00000107360.2 | 0 | Hcrt | ENSMUSG00000045471.4 |
| chr2 | 180776101 | 180776311 | 4.05184 | 5.21013 | 1.9511 | Promoter | 2 | 180774381 | 180776900 | 2520 | 2 | ENSMUST00000108878.1 | 589 | Bhlhe23 | ENSMUSG00000045493.4 |
| chr2 | 180776818 | 180776994 | 3.47404 | 4.07647 | 1.14756 | Promoter | 2 | 180774381 | 180776900 | 2520 | 2 | ENSMUST00000108878.1 | 0 | Bhlhe23 | ENSMUSG00000045493.4 |
| chr2 | 180777595 | 180777798 | 3.9155 | 4.96869 | 1.79182 | Promoter | 2 | 180774381 | 180776900 | 2520 | 2 | ENSMUST00000108878.1 | -696 | Bhlhe23 | ENSMUSG00000045493.4 |
| chr2 | 180778706 | 180778839 | 4.06165 | 5.32313 | 2.00338 | Distal Intergenic | 2 | 180774381 | 180776900 | 2520 | 2 | ENSMUST00000108878.1 | -1807 | Bhlhe23 | ENSMUSG00000045493.4 |
| chr2 | 180786570 | 180786950 | 3.6284 | 4.43898 | 1.43495 | Distal Intergenic | 2 | 180774381 | 180776900 | 2520 | 2 | ENSMUST00000108878.1 | -9671 | Bhlhe23 | ENSMUSG00000045493.4 |
| chr2 | 180790853 | 180791056 | 3.5366 | 4.25431 | 1.33015 | Distal Intergenic | 2 | 180774381 | 180776900 | 2520 | 2 | ENSMUST00000108878.1 | -13954 | Bhlhe23 | ENSMUSG00000045493.4 |
| chr13 | 76052608 | 76052789 | 3.25436 | 3.72349 | 1.05282 | Downstream (2-3kb) | 13 | 76054851 | 76056996 | 2146 | 2 | ENSMUST00000056130.5 | 4207 | Gpr150 | ENSMUSG00000045509.5 |
| chr13 | 76055903 | 76056634 | 3.65217 | 4.746 | 1.59687 | Promoter | 13 | 76054851 | 76056996 | 2146 | 2 | ENSMUST00000056130.5 | 362 | Gpr150 | ENSMUSG00000045509.5 |
| chr1 | 42695742 | 42702776 | 4.03897 | 5.32314 | 2.03485 | Promoter | 1 | 42695768 | 42703176 | 7409 | 1 | ENSMUST00000054883.3 | 0 | Pou3f3 | ENSMUSG00000045515.3 |
| chr1 | 42703192 | 42703340 | 3.42001 | 4.02787 | 1.20927 | Intron (ENSMUST00000176807.2/ENSMUSG00000093436.2, intron 1 of 1) | 1 | 42695768 | 42703176 | 7409 | 1 | ENSMUST00000054883.3 | 7425 | Pou3f3 | ENSMUSG00000045515.3 |
| chr1 | 42703661 | 42703896 | 3.24586 | 3.69594 | 1.01733 | Intron (ENSMUST00000176807.2/ENSMUSG00000093436.2, intron 1 of 1) | 1 | 42695768 | 42703176 | 7409 | 1 | ENSMUST00000054883.3 | 7894 | Pou3f3 | ENSMUSG00000045515.3 |
| chr10 | 80487080 | 80487588 | 3.46621 | 4.05623 | 1.11744 | Distal Intergenic | 10 | 80494835 | 80517276 | 22442 | 1 | ENSMUST00000051773.8 | -7247 | Onecut3 | ENSMUSG00000045518.8 |
| chr10 | 80488609 | 80488739 | 3.47442 | 4.07088 | 1.12539 | Distal Intergenic | 10 | 80494835 | 80517276 | 22442 | 1 | ENSMUST00000051773.8 | -6096 | Onecut3 | ENSMUSG00000045518.8 |
| chr10 | 80489184 | 80489717 | 3.40997 | 4.00154 | 1.10683 | Distal Intergenic | 10 | 80494835 | 80517276 | 22442 | 1 | ENSMUST00000051773.8 | -5118 | Onecut3 | ENSMUSG00000045518.8 |
| chr10 | 80490036 | 80490469 | 3.51605 | 4.43378 | 1.39457 | Distal Intergenic | 10 | 80494835 | 80517276 | 22442 | 1 | ENSMUST00000051773.8 | -4366 | Onecut3 | ENSMUSG00000045518.8 |

| | | | | | | | | | | | | | | | |
|---|---|---|---|---|---|---|---|---|---|---|---|---|---|---|---|
| chr10 | 80490918 | 80491399 | 3.4445 | 4.30613 | 1.3455 | Distal Intergenic | 10 | 80494835 | 80517276 | 22442 | 1 | ENSMUST00000051773.8 | -3436 | Onecut3 | ENSMUSG00000045518.8 |
| chr10 | 80492455 | 80492934 | 3.89753 | 4.85535 | 1.61814 | Distal Intergenic | 10 | 80494835 | 80517276 | 22442 | 1 | ENSMUST00000051773.8 | -1901 | Onecut3 | ENSMUSG00000045518.8 |
| chr10 | 80493437 | 80493725 | 3.66871 | 4.43912 | 1.37222 | Distal Intergenic | 10 | 80494835 | 80517276 | 22442 | 1 | ENSMUST00000051773.8 | -1110 | Onecut3 | ENSMUSG00000045518.8 |
| chr10 | 80494076 | 80494516 | 3.90039 | 4.89718 | 1.68898 | Promoter | 10 | 80494835 | 80517276 | 22442 | 1 | ENSMUST00000051773.8 | -319 | Onecut3 | ENSMUSG00000045518.8 |
| chr10 | 80494854 | 80497512 | 3.88621 | 4.89785 | 1.70576 | Promoter | 10 | 80494835 | 80517276 | 22442 | 1 | ENSMUST00000051773.8 | 20 | Onecut3 | ENSMUSG00000045518.8 |
| chr10 | 80498221 | 80498514 | 3.48817 | 4.13857 | 1.22958 | Intron (ENSMUST00000051773.8/ENSMUSG00000045518.8, intron 1 of 1) | 10 | 80494835 | 80517276 | 22442 | 1 | ENSMUST00000051773.8 | 3387 | Onecut3 | ENSMUSG00000045518.8 |
| chr10 | 80501010 | 80501203 | 3.53414 | 4.25501 | 1.29632 | Intron (ENSMUST00000051773.8/ENSMUSG00000045518.8, intron 1 of 1) | 10 | 80494835 | 80517276 | 22442 | 1 | ENSMUST00000051773.8 | 6176 | Onecut3 | ENSMUSG00000045518.8 |
| chr10 | 80501562 | 80501853 | 4.05268 | 5.2055 | 1.89117 | Intron (ENSMUST00000051773.8/ENSMUSG00000045518.8, intron 1 of 1) | 10 | 80494835 | 80517276 | 22442 | 1 | ENSMUST00000051773.8 | 6728 | Onecut3 | ENSMUSG00000045518.8 |
| chr10 | 80502332 | 80503127 | 4.35667 | 5.92609 | 2.48127 | Intron (ENSMUST00000051773.8/ENSMUSG00000045518.8, intron 1 of 1) | 10 | 80494835 | 80517276 | 22442 | 1 | ENSMUST00000051773.8 | 7498 | Onecut3 | ENSMUSG00000045518.8 |
| chr10 | 80503473 | 80503697 | 4.15972 | 5.74115 | 2.28218 | Intron (ENSMUST00000051773.8/ENSMUSG00000045518.8, intron 1 of 1) | 10 | 80494835 | 80517276 | 22442 | 1 | ENSMUST00000051773.8 | 8639 | Onecut3 | ENSMUSG00000045518.8 |
| chr10 | 80504177 | 80504375 | 3.75067 | 4.60191 | 1.47108 | Intron (ENSMUST00000051773.8/ENSMUSG00000045518.8, intron 1 of 1) | 10 | 80494835 | 80517276 | 22442 | 1 | ENSMUST00000051773.8 | 9343 | Onecut3 | ENSMUSG00000045518.8 |
| chr10 | 80504971 | 80506386 | 3.77272 | 4.72219 | 1.57735 | Intron (ENSMUST00000051773.8/ENSMUSG00000045518.8, intron 1 of 1) | 10 | 80494835 | 80517276 | 22442 | 1 | ENSMUST00000051773.8 | 10137 | Onecut3 | ENSMUSG00000045518.8 |
| chr10 | 80507807 | 80508075 | 3.64378 | 4.3899 | 1.35104 | Intron (ENSMUST00000051773.8/ENSMUSG00000045518.8, intron 1 of 1) | 10 | 80494835 | 80517276 | 22442 | 1 | ENSMUST00000051773.8 | 12973 | Onecut3 | ENSMUSG00000045518.8 |
| chr10 | 80510675 | 80511080 | 3.55816 | 4.23101 | 1.24404 | Intron (ENSMUST00000051773.8/ENSMUSG00000045518.8, intron 1 of 1) | 10 | 80494835 | 80517276 | 22442 | 1 | ENSMUST00000051773.8 | 15841 | Onecut3 | ENSMUSG00000045518.8 |
| chr10 | 80511633 | 80511921 | 3.99049 | 5.03892 | 1.75423 | Intron (ENSMUST00000051773.8/ENSMUSG00000045518.8, intron 1 of 1) | 10 | 80494835 | 80517276 | 22442 | 1 | ENSMUST00000051773.8 | 16799 | Onecut3 | ENSMUSG00000045518.8 |
| chr10 | 80513942 | 80514150 | 3.88309 | 4.83133 | 1.60539 | 3' UTR | 10 | 80494835 | 80517276 | 22442 | 1 | ENSMUST00000051773.8 | 19108 | Onecut3 | ENSMUSG00000045518.8 |
| chr10 | 80514566 | 80514935 | 3.47234 | 4.14557 | 1.20584 | 3' UTR | 10 | 80494835 | 80517276 | 22442 | 1 | ENSMUST00000051773.8 | 19732 | Onecut3 | ENSMUSG00000045518.8 |
| chr10 | 80515260 | 80515623 | 3.53762 | 4.23687 | 1.26919 | 3' UTR | 10 | 80494835 | 80517276 | 22442 | 1 | ENSMUST00000051773.8 | 20426 | Onecut3 | ENSMUSG00000045518.8 |
| chr11 | 102936603 | 102936887 | 3.62876 | 4.42049 | 1.42484 | Intron (ENSMUST00000181125.1/ENSMUSG00000097410.1, intron 1 of 1) | 11 | 102939264 | 102946688 | 7425 | 2 | ENSMUST00000057849.5 | 9801 | C1ql1 | ENSMUSG00000045532.5 |
| chr11 | 102939459 | 102939616 | 3.46908 | 4.06594 | 1.13687 | 3' UTR | 11 | 102939264 | 102946688 | 7425 | 2 | ENSMUST00000057849.5 | 7072 | C1ql1 | ENSMUSG00000045532.5 |
| chr11 | 102943305 | 102943786 | 3.64958 | 4.46004 | 1.48322 | Intron (ENSMUST00000181125.1/ENSMUSG00000097410.1, intron 1 of 1) | 11 | 102939264 | 102946688 | 7425 | 2 | ENSMUST00000057849.5 | 2902 | C1ql1 | ENSMUSG00000045532.5 |
| chr11 | 102944259 | 102944654 | 3.22539 | 3.66528 | 1.01527 | Intron (ENSMUST00000181125.1/ENSMUSG00000097410.1, intron 1 of 1) | 11 | 102939264 | 102946688 | 7425 | 2 | ENSMUST00000057849.5 | 2034 | C1ql1 | ENSMUSG00000045532.5 |
| chr11 | 102945115 | 102946477 | 3.79416 | 4.7726 | 1.64916 | Promoter | 11 | 102939264 | 102946688 | 7425 | 2 | ENSMUST00000057849.5 | 211 | C1ql1 | ENSMUSG00000045532.5 |
| chr11 | 102954052 | 102954376 | 3.58465 | 4.34592 | 1.37842 | Intron (ENSMUST00000181125.1/ENSMUSG00000097410.1, intron 1 of 1) | 11 | 102939264 | 102946688 | 7425 | 2 | ENSMUST00000057849.5 | -7365 | C1ql1 | ENSMUSG00000045532.5 |
| chr10 | 40674153 | 40674397 | 3.42601 | 4.01579 | 1.18948 | Distal Intergenic | 10 | 40683282 | 40811083 | 127802 | 1 | ENSMUST00000058747.3 | -8885 | Mettl24 | ENSMUSG00000045555.3 |
| chr4 | 4137833 | 4138911 | 4.04584 | 5.4213 | 2.06339 | Promoter | 4 | 4133531 | 4138477 | 4947 | 2 | ENSMUST00000070375.7 | 0 | Penk | ENSMUSG00000045573.9 |
| chr4 | 56989902 | 56990535 | 3.61436 | 4.48318 | 1.44975 | Promoter | 4 | 56957173 | 56990391 | 33219 | 2 | ENSMUST00000053681.5 | 0 | Frrs1l | ENSMUSG00000045589.7 |
| chr10 | 19337965 | 19338146 | 3.30588 | 3.78604 | 1.01733 | Distal Intergenic | 10 | 19356560 | 19358606 | 2047 | 1 | ENSMUST00000053225.5 | -18414 | Olig3 | ENSMUSG00000045591.5 |
| chr10 | 19338969 | 19339764 | 3.5037 | 4.26094 | 1.32078 | Distal Intergenic | 10 | 19356560 | 19358606 | 2047 | 1 | ENSMUST00000053225.5 | -16796 | Olig3 | ENSMUSG00000045591.5 |
| chr10 | 19340457 | 19340689 | 3.35468 | 3.88732 | 1.1154 | Distal Intergenic | 10 | 19356560 | 19358606 | 2047 | 1 | ENSMUST00000053225.5 | -15871 | Olig3 | ENSMUSG00000045591.5 |
| chr10 | 19341038 | 19341237 | 3.4254 | 4.00497 | 1.15512 | Distal Intergenic | 10 | 19356560 | 19358606 | 2047 | 1 | ENSMUST00000053225.5 | -15323 | Olig3 | ENSMUSG00000045591.5 |
| chr10 | 19341548 | 19341821 | 3.43355 | 4.018 | 1.15784 | Distal Intergenic | 10 | 19356560 | 19358606 | 2047 | 1 | ENSMUST00000053225.5 | -14739 | Olig3 | ENSMUSG00000045591.5 |
| chr10 | 19346551 | 19346699 | 3.29527 | 3.77007 | 1.01733 | Distal Intergenic | 10 | 19356560 | 19358606 | 2047 | 1 | ENSMUST00000053225.5 | -9861 | Olig3 | ENSMUSG00000045591.5 |
| chr10 | 19348183 | 19348486 | 3.49315 | 4.13397 | 1.24153 | Distal Intergenic | 10 | 19356560 | 19358606 | 2047 | 1 | ENSMUST00000053225.5 | -8074 | Olig3 | ENSMUSG00000045591.5 |
| chr10 | 19350598 | 19351428 | 3.80235 | 5.0346 | 1.82214 | Distal Intergenic | 10 | 19356560 | 19358606 | 2047 | 1 | ENSMUST00000053225.5 | -5132 | Olig3 | ENSMUSG00000045591.5 |
| chr10 | 19352255 | 19352529 | 3.56104 | 4.34076 | 1.36048 | Distal Intergenic | 10 | 19356560 | 19358606 | 2047 | 1 | ENSMUST00000053225.5 | -4031 | Olig3 | ENSMUSG00000045591.5 |
| chr10 | 19353708 | 19355485 | 3.74395 | 4.69596 | 1.58562 | Distal Intergenic | 10 | 19356560 | 19358606 | 2047 | 1 | ENSMUST00000053225.5 | -1075 | Olig3 | ENSMUSG00000045591.5 |
| chr10 | 19355843 | 19356328 | 3.69162 | 4.60161 | 1.51983 | Promoter | 10 | 19356560 | 19358606 | 2047 | 1 | ENSMUST00000053225.5 | -232 | Olig3 | ENSMUSG00000045591.5 |
| chr10 | 19356801 | 19357741 | 4.43383 | 6.12404 | 2.63457 | Promoter | 10 | 19356560 | 19358606 | 2047 | 1 | ENSMUST00000053225.5 | 242 | Olig3 | ENSMUSG00000045591.5 |
| chr10 | 19358168 | 19359907 | 3.58252 | 4.32312 | 1.30694 | 3' UTR | 10 | 19356560 | 19358606 | 2047 | 1 | ENSMUST00000053225.5 | 1609 | Olig3 | ENSMUSG00000045591.5 |
| chr10 | 19360993 | 19362950 | 3.93191 | 4.97242 | 1.73915 | Distal Intergenic | 10 | 19356560 | 19358606 | 2047 | 1 | ENSMUST00000053225.5 | 4434 | Olig3 | ENSMUSG00000045591.5 |
| chr10 | 19363660 | 19364077 | 4.15554 | 5.42097 | 2.06419 | Distal Intergenic | 10 | 19356560 | 19358606 | 2047 | 1 | ENSMUST00000053225.5 | 7101 | Olig3 | ENSMUSG00000045591.5 |
| chr10 | 19364886 | 19365026 | 3.26167 | 3.89005 | 1.07379 | Distal Intergenic | 10 | 19356560 | 19358606 | 2047 | 1 | ENSMUST00000053225.5 | 8327 | Olig3 | ENSMUSG00000045591.5 |
| chr15 | 95639725 | 95639800 | 3.74597 | 4.58181 | 1.4585 | Intron (ENSMUST00000054244.6/ENSMUSG00000045608.6, intron 2 of 3) | 15 | 95623563 | 95655960 | 32398 | 2 | ENSMUST00000054244.6 | 16160 | Dbx2 | ENSMUSG00000045608.6 |
| chr15 | 95640127 | 95640722 | 3.60501 | 4.32759 | 1.29205 | Exon (ENSMUST00000054244.6/ENSMUSG00000045608.6, exon 2 of 4) | 15 | 95623563 | 95655960 | 32398 | 2 | ENSMUST00000054244.6 | 15238 | Dbx2 | ENSMUSG00000045608.6 |
| chr15 | 95651109 | 95652049 | 3.61957 | 4.41312 | 1.4592 | Intron (ENSMUST00000054244.6/ENSMUSG00000045608.6, intron 1 of 3) | 15 | 95623563 | 95655960 | 32398 | 2 | ENSMUST00000054244.6 | 3911 | Dbx2 | ENSMUSG00000045608.6 |
| chr15 | 95652826 | 95654615 | 3.83721 | 4.88709 | 1.75768 | Exon (ENSMUST00000054244.6/ENSMUSG00000045608.6, exon 1 of 4) | 15 | 95623563 | 95655960 | 32398 | 2 | ENSMUST00000054244.6 | 1345 | Dbx2 | ENSMUSG00000045608.6 |
| chr15 | 95655058 | 95655428 | 3.58919 | 4.49021 | 1.46332 | Promoter | 15 | 95623563 | 95655960 | 32398 | 2 | ENSMUST00000054244.6 | 532 | Dbx2 | ENSMUSG00000045608.6 |
| chr6 | 36387991 | 36388226 | 3.79937 | 5.03864 | 1.79661 | Promoter | 6 | 36388084 | 36528414 | 140331 | 1 | ENSMUST00000172278.2 | 0 | Chrm2 | ENSMUSG00000045613.4 |
| chr9 | 56771501 | 56771799 | 3.29436 | 3.7687 | 1.01733 | Distal Intergenic | 9 | 56848659 | 56851963 | 3305 | 2 | ENSMUST00000055036.6 | 80164 | Odf3l1 | ENSMUSG00000045620.6 |
| chr9 | 56793612 | 56793924 | 3.24126 | 3.68904 | 1.01733 | Distal Intergenic | 9 | 56848659 | 56851963 | 3305 | 2 | ENSMUST00000055036.6 | 58039 | Odf3l1 | ENSMUSG00000045620.6 |
| chr9 | 56795287 | 56796327 | 4.01039 | 5.2823 | 1.98036 | Distal Intergenic | 9 | 56848659 | 56851963 | 3305 | 2 | ENSMUST00000055036.6 | 55636 | Odf3l1 | ENSMUSG00000045620.6 |
| chr9 | 56796762 | 56797537 | 4.16101 | 5.7996 | 2.34187 | Distal Intergenic | 9 | 56848659 | 56851963 | 3305 | 2 | ENSMUST00000055036.6 | 54426 | Odf3l1 | ENSMUSG00000045620.6 |
| chr9 | 56798495 | 56798849 | 3.50216 | 4.16839 | 1.29659 | Distal Intergenic | 9 | 56848659 | 56851963 | 3305 | 2 | ENSMUST00000055036.6 | 53114 | Odf3l1 | ENSMUSG00000045620.6 |
| chr9 | 56799265 | 56799476 | 3.57386 | 4.34264 | 1.37422 | Distal Intergenic | 9 | 56848659 | 56851963 | 3305 | 2 | ENSMUST00000055036.6 | 52487 | Odf3l1 | ENSMUSG00000045620.6 |
| chr9 | 56815045 | 56815288 | 3.26026 | 3.71752 | 1.01733 | Distal Intergenic | 9 | 56848659 | 56851963 | 3305 | 2 | ENSMUST00000055036.6 | 36675 | Odf3l1 | ENSMUSG00000045620.6 |
| chr9 | 56825708 | 56826560 | 3.51007 | 4.1645 | 1.25489 | Distal Intergenic | 9 | 56848659 | 56851963 | 3305 | 2 | ENSMUST00000055036.6 | 25403 | Odf3l1 | ENSMUSG00000045620.6 |
| chr16 | 31933776 | 31935950 | 3.75104 | 4.80099 | 1.6577 | Promoter | 16 | 31933851 | 31946046 | 12196 | 1 | ENSMUST00000052174.2 | 0 | Pigz | ENSMUSG00000045625.4 |
| chr16 | 31936525 | 31936634 | 3.39999 | 3.98951 | 1.20185 | Intron (ENSMUST00000052174.2/ENSMUSG00000045625.4, intron 1 of 2) | 16 | 31933982 | 31942623 | 8642 | 1 | ENSMUST00000134928.1 | 2544 | Pigz | ENSMUSG00000045625.4 |
| chr8 | 41133156 | 41133668 | 3.37838 | 4.09863 | 1.21867 | Promoter | 8 | 41084563 | 41133718 | 49156 | 2 | ENSMUST00000155055.1 | 50 | Mtus1 | ENSMUSG00000045636.11 |
| chr7 | 127644998 | 127645386 | 3.47766 | 4.09512 | 1.19531 | Distal Intergenic | 7 | 127608756 | 127615797 | 7042 | 2 | ENSMUST00000132524.2 | -29202 | Zfp629 | ENSMUSG00000045639.9 |
| chr1 | 84284550 | 84284818 | 3.90285 | 5.02568 | 1.77329 | Promoter | 1 | 84036296 | 84284645 | 248350 | 2 | ENSMUST00000168574.3 | 0 | Pid1 | ENSMUSG00000045658.11 |
| chr3 | 108940394 | 108940600 | 3.27406 | 3.73833 | 1.01733 | Promoter | 3 | 108940500 | 108960773 | 20274 | 1 | ENSMUST00000106586.2 | 0 | Henmt1 | ENSMUSG00000045662.11 |
| chr4 | 63213824 | 63214286 | 3.63988 | 4.577 | 1.54599 | Promoter | 4 | 63214004 | 63292831 | 78828 | 1 | ENSMUST00000125504.2 | 0 | Col27a1 | ENSMUSG00000045672.10 |
| chr4 | 63215696 | 63216398 | 4.02484 | 5.24068 | 1.96763 | Promoter | 4 | 63215435 | 63334991 | 119557 | 1 | ENSMUST00000036300.7 | 262 | Col27a1 | ENSMUSG00000045672.10 |
| chr4 | 63307185 | 63307424 | 3.52914 | 4.19592 | 1.26164 | Intron (ENSMUST00000036300.7/ENSMUSG00000045672.10, intron 36 of 60) | 4 | 63330993 | 63334138 | 3146 | 1 | ENSMUST00000151992.1 | -23569 | Col27a1 | ENSMUSG00000045672.10 |
| chr6 | 29974588 | 29975174 | 4.20055 | 5.79162 | 2.36803 | Distal Intergenic | 6 | 29985329 | 29993530 | 8202 | 1 | ENSMUST00000124522.1 | -10155 | 1700023L04Rik | ENSMUSG00000045709.10 |
| chr6 | 29979529 | 29979914 | 3.55912 | 4.28744 | 1.35211 | Distal Intergenic | 6 | 29985329 | 29993530 | 8202 | 1 | ENSMUST00000124522.1 | -5415 | 1700023L04Rik | ENSMUSG00000045709.10 |
| chr6 | 29984761 | 29984987 | 3.36639 | 3.87716 | 1.01733 | Promoter | 6 | 29985329 | 29993530 | 8202 | 1 | ENSMUST00000124522.1 | -342 | 1700023L04Rik | ENSMUSG00000045709.10 |
| chr6 | 29986632 | 29986726 | 3.66332 | 4.42126 | 1.34372 | Intron (ENSMUST00000124522.1/ENSMUSG00000045709.10, intron 1 of 1) | 6 | 29985331 | 29993776 | 8446 | 1 | ENSMUST00000141679.1 | 1302 | 1700023L04Rik | ENSMUSG00000045709.10 |
| chr14 | 65422328 | 65422659 | 3.49879 | 4.15521 | 1.19883 | Intron (ENSMUST00000059339.5/ENSMUSG00000045731.5, intron 1 of 3) | 14 | 65400673 | 65425472 | 24800 | 2 | ENSMUST00000059339.5 | 2813 | Pnoc | ENSMUSG00000045731.5 |
| chr14 | 65423092 | 65423349 | 3.43673 | 4.00647 | 1.09947 | Intron (ENSMUST00000059339.5/ENSMUSG00000045731.5, intron 1 of 3) | 14 | 65400673 | 65425472 | 24800 | 2 | ENSMUST00000059339.5 | 2123 | Pnoc | ENSMUSG00000045731.5 |
| chr14 | 65424415 | 65424558 | 3.52761 | 4.18043 | 1.22279 | Promoter | 14 | 65400673 | 65425472 | 24800 | 2 | ENSMUST00000059339.5 | 914 | Pnoc | ENSMUSG00000045731.5 |
| chr14 | 65424938 | 65425029 | 3.81452 | 4.72559 | 1.58325 | Promoter | 14 | 65400673 | 65425472 | 24800 | 2 | ENSMUST00000059339.5 | 443 | Pnoc | ENSMUSG00000045731.5 |
| chr14 | 65425995 | 65426619 | 3.76737 | 4.68087 | 1.56649 | Promoter | 14 | 65400673 | 65425472 | 24800 | 2 | ENSMUST00000059339.5 | -524 | Pnoc | ENSMUSG00000045731.5 |
| chr7 | 140153117 | 140153790 | 3.55715 | 4.22394 | 1.22312 | Promoter | 7 | 140150964 | 140153541 | 2578 | 2 | ENSMUST00000156791.1 | 0 | Sprn | ENSMUSG00000045733.7 |
| chr7 | 140154122 | 140154359 | 3.51755 | 4.15047 | 1.17248 | Promoter | 7 | 140150628 | 140154877 | 4250 | 2 | ENSMUST00000059241.7 | 518 | Sprn | ENSMUSG00000045733.7 |
| chr15 | 25284637 | 25284720 | 3.7114 | 4.51487 | 1.41947 | Distal Intergenic | 15 | 25363285 | 25413764 | 50480 | 2 | ENSMUST00000058845.7 | 129044 | Basp1 | ENSMUSG00000045763.7 |
| chr15 | 25360151 | 25360265 | 3.39841 | 3.93293 | 1.04359 | Distal Intergenic | 15 | 25363285 | 25413764 | 50480 | 2 | ENSMUST00000058845.7 | 53499 | Basp1 | ENSMUSG00000045763.7 |
| chr15 | 25361728 | 25362204 | 3.48446 | 4.09578 | 1.16058 | Downstream (1-2kb) | 15 | 25363285 | 25413764 | 50480 | 2 | ENSMUST00000058845.7 | 51560 | Basp1 | ENSMUSG00000045763.7 |
| chr15 | 25363419 | 25365471 | 3.96003 | 5.04812 | 1.80421 | 5' UTR | 15 | 25363285 | 25413764 | 50480 | 2 | ENSMUST00000058845.7 | 48293 | Basp1 | ENSMUSG00000045763.7 |
| chr15 | 25412593 | 25412809 | 3.3281 | 3.81946 | 1.01733 | Promoter | 15 | 25363285 | 25413764 | 50480 | 2 | ENSMUST00000058845.7 | 955 | Basp1 | ENSMUSG00000045763.7 |
| chr15 | 25413150 | 25413887 | 4.31493 | 5.71124 | 2.2648 | Promoter | 15 | 25363285 | 25413764 | 50480 | 2 | ENSMUST00000058845.7 | 0 | Basp1 | ENSMUSG00000045763.7 |
| chr7 | 142349618 | 142349722 | 3.51672 | 4.21712 | 1.32507 | Intron (ENSMUST00000059223.9/ENSMUSG00000045777.9, intron 3 of 3) | 7 | 142325837 | 142338022 | 12186 | 2 | ENSMUST00000131791.1 | -11597 | Ifitm10 | ENSMUSG00000045777.9 |
| chr7 | 142372063 | 142372496 | 3.5867 | 4.28713 | 1.28625 | Promoter | 7 | 142326110 | 142372441 | 46332 | 2 | ENSMUST00000059223.9 | 0 | Ifitm10 | ENSMUSG00000045777.9 |
| chr5 | 52471199 | 52471636 | 3.50642 | 4.13941 | 1.18964 | Promoter | 5 | 52374651 | 52471543 | 96893 | 2 | ENSMUST00000059428.5 | 0 | Ccdc149 | ENSMUSG00000045790.5 |
| chr5 | 52516262 | 52516459 | 3.33935 | 3.87657 | 1.14037 | Distal Intergenic | 5 | 52374651 | 52471543 | 96893 | 2 | ENSMUST00000059428.5 | -44720 | Ccdc149 | ENSMUSG00000045790.5 |
| chr7 | 62230907 | 62231205 | 4.53716 | 6.1397 | 2.5573 | Distal Intergenic | 7 | 62114641 | 62118987 | 4347 | 2 | ENSMUST00000054860.6 | -111921 | Gm9801 | ENSMUSG00000045813.6 |
| chr7 | 62232200 | 62232275 | 3.66804 | 4.42276 | 1.33542 | Distal Intergenic | 7 | 62114641 | 62118987 | 4347 | 2 | ENSMUST00000054860.6 | -113214 | Gm9801 | ENSMUSG00000045813.6 |
| chr1 | 34579439 | 34579601 | 3.2729 | 3.82806 | 1.06511 | Promoter | 1 | 34579224 | 34579613 | 390 | 2 | ENSMUST00000062557.3 | 12 | 1700101I19Rik | ENSMUSG00000045815.4 |
| chr17 | 84096309 | 84096658 | 3.26913 | 3.73092 | 1.01733 | Distal Intergenic | 17 | 84183931 | 84187947 | 4017 | 2 | ENSMUST00000060366.6 | 91289 | Zfp36l2 | ENSMUSG00000045817.8 |
| chr17 | 84097512 | 84097773 | 3.56258 | 4.25567 | 1.30856 | Distal Intergenic | 17 | 84183931 | 84187947 | 4017 | 2 | ENSMUST00000060366.6 | 90174 | Zfp36l2 | ENSMUSG00000045817.8 |
| chr17 | 84143188 | 84143776 | 3.89666 | 4.92982 | 1.76541 | Distal Intergenic | 17 | 84183931 | 84187947 | 4017 | 2 | ENSMUST00000060366.6 | 44171 | Zfp36l2 | ENSMUSG00000045817.8 |
| chr14 | 66635402 | 66635700 | 3.9082 | 5.02282 | 1.80713 | Promoter | 14 | 66635556 | 66732741 | 97186 | 1 | ENSMUST00000161339.1 | 0 | Adra1a | ENSMUSG00000045875.7 |
| chr14 | 66636461 | 66637855 | 3.79408 | 4.69878 | 1.58704 | Promoter | 14 | 66635556 | 66732741 | 97186 | 1 | ENSMUST00000161339.1 | 906 | Adra1a | ENSMUSG00000045875.7 |
| chr19 | 4984258 | 4984679 | 4.39212 | 5.97944 | 2.49954 | 3' UTR | 19 | 4984355 | 4989971 | 5617 | 2 | ENSMUST00000056129.7 | 5292 | Npas4 | ENSMUSG00000045903.8 |
| chr19 | 4989498 | 4990063 | 4.29661 | 5.77383 | 2.42688 | Promoter | 19 | 4984355 | 4989971 | 5617 | 2 | ENSMUST00000056129.7 | 0 | Npas4 | ENSMUSG00000045903.8 |
| chr19 | 4990405 | 4991120 | 3.9752 | 5.1363 | 1.92422 | Promoter | 19 | 4984355 | 4989971 | 5617 | 2 | ENSMUST00000056129.7 | -435 | Npas4 | ENSMUSG00000045903.8 |
| chr19 | 5001846 | 5001950 | 3.6491 | 4.87097 | 1.68094 | Distal Intergenic | 19 | 4984355 | 4989971 | 5617 | 2 | ENSMUST00000056129.7 | -11876 | Npas4 | ENSMUSG00000045903.8 |
| chr10 | 79609497 | 79609582 | 3.72953 | 5.08555 | 1.80404 | 3' UTR | 10 | 79606857 | 79614018 | 7162 | 2 | ENSMUST00000059699.7 | 4436 | C2cd4c | ENSMUSG00000045912.8 |
| chr10 | 79611993 | 79614367 | 3.8923 | 4.92537 | 1.72437 | Promoter | 10 | 79606857 | 79614018 | 7162 | 2 | ENSMUST00000059699.7 | 0 | C2cd4c | ENSMUSG00000045912.8 |
| chr12 | 49397226 | 49397380 | 3.65317 | 4.45762 | 1.45729 | Intron (ENSMUST00000127041.1/ENSMUSG00000089922.1, intron 2 of 2) | 12 | 49401277 | 49401837 | 561 | 1 | ENSMUST00000056234.3 | -3897 | Gm9804 | ENSMUSG00000045946.3 |
| chr12 | 49397892 | 49398189 | 3.46153 | 4.16347 | 1.2887 | Intron (ENSMUST00000127041.1/ENSMUSG00000089922.1, intron 2 of 2) | 12 | 49401277 | 49401837 | 561 | 1 | ENSMUST00000056234.3 | -3088 | Gm9804 | ENSMUSG00000045946.3 |
| chr12 | 49398683 | 49398792 | 4.1931 | 5.52803 | 2.18731 | Intron (ENSMUST00000127041.1/ENSMUSG00000089922.1, intron 2 of 2) | 12 | 49401277 | 49401837 | 561 | 1 | ENSMUST00000056234.3 | -2485 | Gm9804 | ENSMUSG00000045946.3 |
| chr12 | 49399574 | 49399742 | 3.48681 | 4.12467 | 1.25226 | Intron (ENSMUST00000127041.1/ENSMUSG00000089922.1, intron 2 of 2) | 12 | 49401277 | 49401837 | 561 | 1 | ENSMUST00000056234.3 | -1535 | Gm9804 | ENSMUSG00000045946.3 |
| chr12 | 49400101 | 49400393 | 3.96867 | 5.15214 | 1.88256 | Promoter | 12 | 49401277 | 49401837 | 561 | 1 | ENSMUST00000056234.3 | -884 | Gm9804 | ENSMUSG00000045946.3 |
| chr12 | 49401256 | 49401724 | 3.74125 | 5.20463 | 1.9264 | Promoter | 12 | 49401277 | 49401837 | 561 | 1 | ENSMUST00000056234.3 | 0 | Gm9804 | ENSMUSG00000045946.3 |
| chr12 | 49402785 | 49402937 | 3.53825 | 4.30434 | 1.35204 | Intron (ENSMUST00000127041.1/ENSMUSG00000089922.1, intron 2 of 2) | 12 | 49401277 | 49401837 | 561 | 1 | ENSMUST00000056234.3 | 1509 | Gm9804 | ENSMUSG00000045946.3 |
| chr12 | 49403491 | 49404039 | 3.39293 | 3.96532 | 1.17336 | Intron (ENSMUST00000127041.1/ENSMUSG00000089922.1, intron 2 of 2) | 12 | 49401277 | 49401837 | 561 | 1 | ENSMUST00000056234.3 | 2215 | Gm9804 | ENSMUSG00000045946.3 |
| chr12 | 49404689 | 49405192 | 3.46382 | 4.14501 | 1.27322 | Intron (ENSMUST00000127041.1/ENSMUSG00000089922.1, intron 2 of 2) | 12 | 49401277 | 49401837 | 561 | 1 | ENSMUST00000056234.3 | 3413 | Gm9804 | ENSMUSG00000045946.3 |
| chr12 | 49407403 | 49407495 | 3.38756 | 4.22393 | 1.27464 | Distal Intergenic | 12 | 49401277 | 49401837 | 561 | 1 | ENSMUST00000056234.3 | 6127 | Gm9804 | ENSMUSG00000045946.3 |
| chr12 | 49409329 | 49409404 | 4.38862 | 5.97592 | 2.47688 | Distal Intergenic | 12 | 49401277 | 49401837 | 561 | 1 | ENSMUST00000056234.3 | 8053 | Gm9804 | ENSMUSG00000045946.3 |
| chr12 | 49417907 | 49418149 | 3.97396 | 5.17574 | 1.89498 | Distal Intergenic | 12 | 49401277 | 49401837 | 561 | 1 | ENSMUST00000056234.3 | 16631 | Gm9804 | ENSMUSG00000045946.3 |
| chr18 | 64318554 | 64319388 | 3.30211 | 3.92525 | 1.11164 | Distal Intergenic | 18 | 64340364 | 64398486 | 58123 | 1 | ENSMUST00000115145.3 | -20976 | Onecut2 | ENSMUSG00000045991.13 |
| chr18 | 64323264 | 64323444 | 3.2508 | 3.70334 | 1.01733 | Distal Intergenic | 18 | 64340364 | 64398486 | 58123 | 1 | ENSMUST00000115145.3 | -16920 | Onecut2 | ENSMUSG00000045991.13 |
| chr18 | 64323919 | 64324156 | 3.32402 | 3.8133 | 1.01733 | Distal Intergenic | 18 | 64340364 | 64398486 | 58123 | 1 | ENSMUST00000115145.3 | -16208 | Onecut2 | ENSMUSG00000045991.13 |
| chr18 | 64326892 | 64326990 | 3.56887 | 4.28044 | 1.33985 | Distal Intergenic | 18 | 64340364 | 64398486 | 58123 | 1 | ENSMUST00000115145.3 | -13374 | Onecut2 | ENSMUSG00000045991.13 |
| chr18 | 64327562 | 64327708 | 3.5544 | 4.22058 | 1.2206 | Distal Intergenic | 18 | 64340364 | 64398486 | 58123 | 1 | ENSMUST00000115145.3 | -12656 | Onecut2 | ENSMUSG00000045991.13 |
| chr18 | 64328110 | 64328216 | 3.41787 | 3.98147 | 1.11155 | Distal Intergenic | 18 | 64340364 | 64398486 | 58123 | 1 | ENSMUST00000115145.3 | -12148 | Onecut2 | ENSMUSG00000045991.13 |
| chr18 | 64329507 | 64329729 | 3.74745 | 4.63744 | 1.58584 | Distal Intergenic | 18 | 64340364 | 64398486 | 58123 | 1 | ENSMUST00000115145.3 | -10635 | Onecut2 | ENSMUSG00000045991.13 |
| chr18 | 64330251 | 64330650 | 4.02434 | 5.11996 | 1.83707 | Distal Intergenic | 18 | 64340364 | 64398486 | 58123 | 1 | ENSMUST00000115145.3 | -9714 | Onecut2 | ENSMUSG00000045991.13 |
| chr18 | 64331263 | 64334647 | 4.09055 | 5.43718 | 2.0945 | Distal Intergenic | 18 | 64340364 | 64398486 | 58123 | 1 | ENSMUST00000115145.3 | -5717 | Onecut2 | ENSMUSG00000045991.13 |
| chr18 | 64334999 | 64336026 | 4.08234 | 5.41056 | 2.04983 | Distal Intergenic | 18 | 64340364 | 64398486 | 58123 | 1 | ENSMUST00000115145.3 | -4338 | Onecut2 | ENSMUSG00000045991.13 |
| chr18 | 64336348 | 64337369 | 4.08735 | 5.55424 | 2.17477 | Distal Intergenic | 18 | 64340364 | 64398486 | 58123 | 1 | ENSMUST00000115145.3 | -2995 | Onecut2 | ENSMUSG00000045991.13 |
| chr18 | 64337716 | 64342937 | 4.01277 | 5.2025 | 1.95489 | Promoter | 18 | 64340364 | 64398486 | 58123 | 1 | ENSMUST00000115145.3 | 0 | Onecut2 | ENSMUSG00000045991.13 |
| chr18 | 64343257 | 64343332 | 3.46257 | 4.22747 | 1.24284 | Intron (ENSMUST00000115145.3/ENSMUSG00000045991.13, intron 1 of 1) | 18 | 64340364 | 64398488 | 58125 | 1 | ENSMUST00000175965.3 | 2894 | Onecut2 | ENSMUSG00000045991.13 |
| chr18 | 64343939 | 64344221 | 3.3715 | 4.11051 | 1.23505 | Intron (ENSMUST00000115145.3/ENSMUSG00000045991.13, intron 1 of 1) | 18 | 64340364 | 64398488 | 58125 | 1 | ENSMUST00000175965.3 | 3576 | Onecut2 | ENSMUSG00000045991.13 |
| chr18 | 64345163 | 64345474 | 3.62803 | 4.69623 | 1.60564 | Intron (ENSMUST00000115145.3/ENSMUSG00000045991.13, intron 1 of 1) | 18 | 64340364 | 64398488 | 58125 | 1 | ENSMUST00000175965.3 | 4800 | Onecut2 | ENSMUSG00000045991.13 |
| chr18 | 64347265 | 64347427 | 3.60836 | 4.42647 | 1.42872 | Intron (ENSMUST00000115145.3/ENSMUSG00000045991.13, intron 1 of 1) | 18 | 64340364 | 64398488 | 58125 | 1 | ENSMUST00000175965.3 | 6902 | Onecut2 | ENSMUSG00000045991.13 |
| chr18 | 64348155 | 64350011 | 3.70907 | 4.59908 | 1.56326 | Intron (ENSMUST00000115145.3/ENSMUSG00000045991.13, intron 1 of 1) | 18 | 64340364 | 64398488 | 58125 | 1 | ENSMUST00000175965.3 | 7792 | Onecut2 | ENSMUSG00000045991.13 |
| chr18 | 64350376 | 64350723 | 4.18948 | 6.23087 | 2.69507 | Intron (ENSMUST00000115145.3/ENSMUSG00000045991.13, intron 1 of 1) | 18 | 64340364 | 64398488 | 58125 | 1 | ENSMUST00000175965.3 | 10013 | Onecut2 | ENSMUSG00000045991.13 |
| chr18 | 64351252 | 64351941 | 4.45614 | 6.41307 | 2.81927 | Intron (ENSMUST00000115145.3/ENSMUSG00000045991.13, intron 1 of 1) | 18 | 64340364 | 64398488 | 58125 | 1 | ENSMUST00000175965.3 | 10889 | Onecut2 | ENSMUSG00000045991.13 |
| chr18 | 64354142 | 64354369 | 3.24389 | 3.7025 | 1.04144 | Intron (ENSMUST00000115145.3/ENSMUSG00000045991.13, intron 1 of 1) | 18 | 64340364 | 64398488 | 58125 | 1 | ENSMUST00000175965.3 | 13779 | Onecut2 | ENSMUSG00000045991.13 |
| chr18 | 64391862 | 64392012 | 3.33301 | 3.82696 | 1.01733 | 3' UTR | 18 | 64340364 | 64398488 | 58125 | 1 | ENSMUST00000175965.3 | 51499 | Onecut2 | ENSMUSG00000045991.13 |
| chr18 | 64392638 | 64392827 | 3.57747 | 4.26815 | 1.25436 | 3' UTR | 18 | 64340364 | 64398488 | 58125 | 1 | ENSMUST00000175965.3 | 52275 | Onecut2 | ENSMUSG00000045991.13 |
| chr18 | 64393435 | 64394331 | 3.46753 | 4.06221 | 1.13142 | 3' UTR | 18 | 64340364 | 64398488 | 58125 | 1 | ENSMUST00000175965.3 | 53072 | Onecut2 | ENSMUSG00000045991.13 |
| chr9 | 26730800 | 26731036 | 3.51181 | 4.18492 | 1.24635 | Distal Intergenic | 9 | 26733728 | 26761350 | 27623 | 1 | ENSMUST00000160899.2 | -2692 | B3gat1 | ENSMUSG00000045994.11 |
| chr9 | 26733121 | 26734172 | 3.62658 | 4.35293 | 1.30482 | Promoter | 9 | 26733728 | 26761350 | 27623 | 1 | ENSMUST00000160899.2 | 0 | B3gat1 | ENSMUSG00000045994.11 |
| chr9 | 26734496 | 26734678 | 3.34153 | 3.83966 | 1.01733 | Promoter | 9 | 26733851 | 26757982 | 24132 | 1 | ENSMUST00000115269.3 | 646 | B3gat1 | ENSMUSG00000045994.11 |
| chr1 | 156780163 | 156780624 | 3.45467 | 4.08309 | 1.24944 | Distal Intergenic | 1 | 156793512 | 156793844 | 333 | 1 | ENSMUST00000121352.1 | -12888 | Gm15428 | ENSMUSG00000046057.4 |
| chr1 | 156781534 | 156781826 | 3.44726 | 4.2151 | 1.26526 | Distal Intergenic | 1 | 156793512 | 156793844 | 333 | 1 | ENSMUST00000121352.1 | -11686 | Gm15428 | ENSMUSG00000046057.4 |
| chr4 | 123175607 | 123175795 | 3.58715 | 4.34579 | 1.40863 | Distal Intergenic | 4 | 123183227 | 123194701 | 11475 | 1 | ENSMUST00000059667.3 | -7432 | Hpcal4 | ENSMUSG00000046093.4 |
| chr4 | 123179480 | 123180024 | 3.32518 | 3.82532 | 1.05353 | Distal Intergenic | 4 | 123183227 | 123194701 | 11475 | 1 | ENSMUST00000059667.3 | -3203 | Hpcal4 | ENSMUSG00000046093.4 |
| chr4 | 123180565 | 123181120 | 3.86784 | 5.06183 | 1.84865 | Distal Intergenic | 4 | 123183227 | 123194701 | 11475 | 1 | ENSMUST00000059667.3 | -2107 | Hpcal4 | ENSMUSG00000046093.4 |
| chr4 | 123181561 | 123181699 | 3.25427 | 4.07339 | 1.18946 | Distal Intergenic | 4 | 123183227 | 123194701 | 11475 | 1 | ENSMUST00000059667.3 | -1528 | Hpcal4 | ENSMUSG00000046093.4 |
| chr4 | 123182172 | 123183887 | 3.5195 | 4.222 | 1.31998 | Promoter | 4 | 123183227 | 123194701 | 11475 | 1 | ENSMUST00000059667.3 | 0 | Hpcal4 | ENSMUSG00000046093.4 |
| chr4 | 123184261 | 123184336 | 3.76826 | 4.8021 | 1.65117 | Promoter | 4 | 123183729 | 123190744 | 7016 | 1 | ENSMUST00000152194.1 | 533 | Hpcal4 | ENSMUSG00000046093.4 |
| chr4 | 138394254 | 138396322 | 3.92907 | 5.12854 | 1.92677 | Promoter | 4 | 138395198 | 138397714 | 2517 | 1 | ENSMUST00000062902.6 | 0 | AB041806 | ENSMUSG00000046109.6 |
| chr13 | 10360389 | 10360473 | 4.21166 | 6.10928 | 2.5253 | Promoter | 13 | 10215694 | 10360563 | 144870 | 2 | ENSMUST00000190395.1 | 90 | Chrm3 | ENSMUSG00000046159.10 |
| chr16 | 91257265 | 91257708 | 4.05488 | 5.21709 | 1.92635 | Distal Intergenic | 16 | 91269772 | 91271933 | 2162 | 1 | ENSMUST00000056882.6 | -12064 | Olig1 | ENSMUSG00000046160.6 |
| chr16 | 91258804 | 91259087 | 3.54996 | 4.21578 | 1.23541 | Distal Intergenic | 16 | 91269772 | 91271933 | 2162 | 1 | ENSMUST00000056882.6 | -10685 | Olig1 | ENSMUSG00000046160.6 |
| chr16 | 91266095 | 91266468 | 3.53112 | 4.2801 | 1.32504 | Distal Intergenic | 16 | 91269772 | 91271933 | 2162 | 1 | ENSMUST00000056882.6 | -3304 | Olig1 | ENSMUSG00000046160.6 |
| chr16 | 91266957 | 91267470 | 3.68863 | 4.71859 | 1.58515 | Distal Intergenic | 16 | 91269772 | 91271933 | 2162 | 1 | ENSMUST00000056882.6 | -2302 | Olig1 | ENSMUSG00000046160.6 |
| chr16 | 91269791 | 91270604 | 3.97909 | 5.1115 | 1.89889 | Promoter | 16 | 91269772 | 91271933 | 2162 | 1 | ENSMUST00000056882.6 | 20 | Olig1 | ENSMUSG00000046160.6 |
| chr16 | 91271182 | 91271901 | 3.76017 | 4.86145 | 1.67497 | 3' UTR | 16 | 91269772 | 91271933 | 2162 | 1 | ENSMUST00000056882.6 | 1411 | Olig1 | ENSMUSG00000046160.6 |
| chr16 | 91272524 | 91272976 | 4.13809 | 5.48062 | 2.09989 | Distal Intergenic | 16 | 91269772 | 91271933 | 2162 | 1 | ENSMUST00000056882.6 | 2753 | Olig1 | ENSMUSG00000046160.6 |
| chr16 | 91273924 | 91274792 | 3.92264 | 4.94405 | 1.73261 | Distal Intergenic | 16 | 91269772 | 91271933 | 2162 | 1 | ENSMUST00000056882.6 | 4153 | Olig1 | ENSMUSG00000046160.6 |
| chr16 | 91275820 | 91275944 | 3.35579 | 3.86117 | 1.01733 | Distal Intergenic | 16 | 91269772 | 91271933 | 2162 | 1 | ENSMUST00000056882.6 | 6049 | Olig1 | ENSMUSG00000046160.6 |
| chr16 | 91276512 | 91276866 | 3.98068 | 5.03783 | 1.77411 | Distal Intergenic | 16 | 91269772 | 91271933 | 2162 | 1 | ENSMUST00000056882.6 | 6741 | Olig1 | ENSMUSG00000046160.6 |
| chr16 | 91277419 | 91277683 | 3.26615 | 3.72639 | 1.01733 | Distal Intergenic | 16 | 91269772 | 91271933 | 2162 | 1 | ENSMUST00000056882.6 | 7648 | Olig1 | ENSMUSG00000046160.6 |
| chr16 | 91279626 | 91279973 | 3.4217 | 3.98144 | 1.09075 | Distal Intergenic | 16 | 91269772 | 91271933 | 2162 | 1 | ENSMUST00000056882.6 | 9855 | Olig1 | ENSMUSG00000046160.6 |
| chr16 | 91284137 | 91284212 | 3.63512 | 4.38533 | 1.35371 | Distal Intergenic | 16 | 91269772 | 91271933 | 2162 | 1 | ENSMUST00000056882.6 | 14366 | Olig1 | ENSMUSG00000046160.6 |
| chr9 | 54286339 | 54286872 | 3.34353 | 4.12737 | 1.20675 | Promoter | 9 | 54286486 | 54341786 | 55301 | 1 | ENSMUST00000056740.5 | 0 | Gldn | ENSMUSG00000046167.5 |

| | | | | | | | | | | | | | | | |
|---|---|---|---|---|---|---|---|---|---|---|---|---|---|---|---|
| chr6 | 8944710 | 8945044 | 3.70381 | 4.5658 | 1.54559 | Distal Intergenic | 6 | 8948431 | 8952430 | 4000 | 1 | ENSMUST00000162942.1 | -3387 | Nxph1 | ENSMUSG00000046178.3 |
| chr6 | 8945674 | 8945749 | 3.44737 | 4.24819 | 1.31348 | Distal Intergenic | 6 | 8948431 | 8952430 | 4000 | 1 | ENSMUST00000162942.1 | -2682 | Nxph1 | ENSMUSG00000046178.3 |
| chr6 | 8947682 | 8947833 | 3.91755 | 5.11515 | 1.84547 | Promoter | 6 | 8948431 | 8952430 | 4000 | 1 | ENSMUST00000162942.1 | -598 | Nxph1 | ENSMUSG00000046178.3 |
| chr6 | 8948140 | 8952038 | 4.06918 | 5.44348 | 2.12952 | Promoter | 6 | 8948431 | 8952430 | 4000 | 1 | ENSMUST00000162942.1 | 0 | Nxph1 | ENSMUSG00000046178.3 |
| chr6 | 8953553 | 8954036 | 3.52074 | 4.21522 | 1.27848 | Intron (ENSMUST00000160300.1/ENSMUSG00000046178.3, intron 2 of 2) | 6 | 8949676 | 9249030 | 299355 | 1 | ENSMUST00000060369.3 | 3878 | Nxph1 | ENSMUSG00000046178.3 |
| chr6 | 8954439 | 8955169 | 4.10627 | 5.31624 | 2.00296 | Intron (ENSMUST00000160300.1/ENSMUSG00000046178.3, intron 2 of 2) | 6 | 8949676 | 9249030 | 299355 | 1 | ENSMUST00000060369.3 | 4764 | Nxph1 | ENSMUSG00000046178.3 |
| chr6 | 8955486 | 8957807 | 3.70337 | 4.57688 | 1.47798 | Intron (ENSMUST00000160300.1/ENSMUSG00000046178.3, intron 2 of 2) | 6 | 8949676 | 9249030 | 299355 | 1 | ENSMUST00000060369.3 | 5811 | Nxph1 | ENSMUSG00000046178.3 |
| chr6 | 8958185 | 8958413 | 3.44121 | 4.01057 | 1.08948 | Intron (ENSMUST00000160300.1/ENSMUSG00000046178.3, intron 2 of 2) | 6 | 8949676 | 9249030 | 299355 | 1 | ENSMUST00000060369.3 | 8510 | Nxph1 | ENSMUSG00000046178.3 |
| chr6 | 8958812 | 8959046 | 3.48723 | 4.09434 | 1.13987 | Intron (ENSMUST00000160300.1/ENSMUSG00000046178.3, intron 2 of 2) | 6 | 8949676 | 9249030 | 299355 | 1 | ENSMUST00000060369.3 | 9137 | Nxph1 | ENSMUSG00000046178.3 |
| chr6 | 8960044 | 8960151 | 3.36398 | 3.87352 | 1.01733 | Intron (ENSMUST00000160300.1/ENSMUSG00000046178.3, intron 2 of 2) | 6 | 8949676 | 9249030 | 299355 | 1 | ENSMUST00000060369.3 | 10369 | Nxph1 | ENSMUSG00000046178.3 |
| chr7 | 48881152 | 48882078 | 3.64133 | 4.47557 | 1.46185 | Promoter | 7 | 48866429 | 48881596 | 15168 | 2 | ENSMUST00000119223.1 | 0 | E2f8 | ENSMUSG00000046179.12 |
| chr7 | 126001697 | 126001778 | 4.14581 | 5.34501 | 1.97245 | Intron (ENSMUST00000073935.5/ENSMUSG00000046182.7, intron 2 of 6) | 7 | 125878419 | 126082411 | 203993 | 2 | ENSMUST00000073935.5 | 80633 | Gsg1l | ENSMUSG00000046182.7 |
| chr7 | 126012693 | 126012997 | 3.38193 | 3.9006 | 1.01733 | Intron (ENSMUST00000073935.5/ENSMUSG00000046182.7, intron 2 of 6) | 7 | 125878419 | 126082411 | 203993 | 2 | ENSMUST00000073935.5 | 69414 | Gsg1l | ENSMUSG00000046182.7 |
| chr7 | 126081019 | 126081104 | 4.25224 | 5.75799 | 2.30031 | Intron (ENSMUST00000073935.5/ENSMUSG00000046182.7, intron 1 of 6) | 7 | 125878419 | 126082411 | 203993 | 2 | ENSMUST00000073935.5 | 1307 | Gsg1l | ENSMUSG00000046182.7 |
| chr7 | 126081452 | 126083172 | 3.68851 | 4.60923 | 1.50739 | Promoter | 7 | 125878419 | 126082411 | 203993 | 2 | ENSMUST00000073935.5 | 0 | Gsg1l | ENSMUSG00000046182.7 |
| chr7 | 126083532 | 126084451 | 3.49052 | 4.18008 | 1.22715 | Distal Intergenic | 7 | 125878419 | 126082411 | 203993 | 2 | ENSMUST00000073935.5 | -1122 | Gsg1l | ENSMUSG00000046182.7 |
| chr9 | 78614510 | 78614585 | 3.43077 | 4.05414 | 1.2162 | Promoter | 9 | 78615546 | 78716253 | 100708 | 1 | ENSMUST00000093812.4 | -961 | Cd109 | ENSMUSG00000046186.8 |
| chr9 | 78615586 | 78616401 | 3.73997 | 4.61637 | 1.55362 | Promoter | 9 | 78615763 | 78618089 | 2327 | 1 | ENSMUST00000163608.1 | 0 | Cd109 | ENSMUSG00000046186.8 |
| chr9 | 78616721 | 78616818 | 4.15254 | 5.4256 | 2.11894 | Promoter | 9 | 78615763 | 78618089 | 2327 | 1 | ENSMUST00000163608.1 | 959 | Cd109 | ENSMUSG00000046186.8 |
| chr9 | 78618217 | 78618357 | 3.15415 | 3.70126 | 1.01733 | Intron (ENSMUST00000093812.4/ENSMUSG00000046186.8, intron 2 of 32) | 9 | 78615763 | 78618089 | 2327 | 1 | ENSMUST00000163608.1 | 2455 | Cd109 | ENSMUSG00000046186.8 |
| chr6 | 24343109 | 24343553 | 3.42911 | 4.04994 | 1.21184 | Distal Intergenic | 6 | 24444865 | 24515067 | 70203 | 2 | ENSMUST00000052277.4 | 171514 | Iqub | ENSMUSG00000046192.4 |
| chr6 | 24349250 | 24349611 | 3.55656 | 4.2319 | 1.25597 | Distal Intergenic | 6 | 24444865 | 24515067 | 70203 | 2 | ENSMUST00000052277.4 | 165456 | Iqub | ENSMUSG00000046192.4 |
| chr6 | 24349970 | 24350566 | 3.80658 | 4.78124 | 1.64342 | Distal Intergenic | 6 | 24444865 | 24515067 | 70203 | 2 | ENSMUST00000052277.4 | 164501 | Iqub | ENSMUSG00000046192.4 |
| chr14 | 66908592 | 66908721 | 3.30709 | 3.79533 | 1.04359 | Distal Intergenic | 14 | 66911170 | 66920054 | 8885 | 1 | ENSMUST00000089236.4 | -2449 | Pnma2 | ENSMUSG00000046204.8 |
| chr14 | 66909567 | 66909923 | 4.42692 | 6.1201 | 2.58437 | Distal Intergenic | 14 | 66911170 | 66920054 | 8885 | 1 | ENSMUST00000089236.4 | -1247 | Pnma2 | ENSMUSG00000046204.8 |
| chr14 | 66910254 | 66911330 | 3.97576 | 5.2135 | 1.95025 | Promoter | 14 | 66911170 | 66920054 | 8885 | 1 | ENSMUST00000089236.4 | 0 | Pnma2 | ENSMUSG00000046204.8 |
| chr14 | 66912052 | 66912613 | 3.46847 | 4.07845 | 1.18531 | Promoter | 14 | 66911242 | 66914726 | 3485 | 1 | ENSMUST00000168010.1 | 811 | Pnma2 | ENSMUSG00000046204.8 |
| chr11 | 68531216 | 68531542 | 3.36061 | 3.86842 | 1.01733 | Exon (ENSMUST00000060441.6/ENSMUSG00000046207.9, exon 9 of 20) | 11 | 68536200 | 68544383 | 8184 | 1 | ENSMUST00000126069.1 | -4658 | Pik3r6 | ENSMUSG00000046207.9 |
| chr11 | 103644675 | 103644777 | 4.3757 | 5.80866 | 2.30586 | Distal Intergenic | 11 | 103649570 | 103650579 | 1010 | 1 | ENSMUST00000057870.3 | -4793 | Rprml | ENSMUSG00000046215.3 |
| chr11 | 103647915 | 103648503 | 3.43849 | 4.01435 | 1.11793 | Downstream (2-3kb) | 11 | 103649570 | 103650579 | 1010 | 1 | ENSMUST00000057870.3 | -1067 | Rprml | ENSMUSG00000046215.3 |
| chr11 | 103648882 | 103649216 | 3.6845 | 4.45907 | 1.36878 | Promoter | 11 | 103649570 | 103650579 | 1010 | 1 | ENSMUST00000057870.3 | -354 | Rprml | ENSMUSG00000046215.3 |
| chr11 | 103649603 | 103650312 | 3.87886 | 4.87723 | 1.67162 | Promoter | 11 | 103649570 | 103650579 | 1010 | 1 | ENSMUST00000057870.3 | 34 | Rprml | ENSMUSG00000046215.3 |
| chr11 | 103650820 | 103651432 | 3.81222 | 4.72535 | 1.57744 | Distal Intergenic | 11 | 103649570 | 103650579 | 1010 | 1 | ENSMUST00000057870.3 | 1251 | Rprml | ENSMUSG00000046215.3 |
| chr11 | 103652507 | 103652854 | 3.56744 | 4.24251 | 1.23576 | Distal Intergenic | 11 | 103649570 | 103650579 | 1010 | 1 | ENSMUST00000057870.3 | 2938 | Rprml | ENSMUSG00000046215.3 |
| chr11 | 103655054 | 103655129 | 3.40205 | 3.94224 | 1.05718 | Distal Intergenic | 11 | 103649570 | 103650579 | 1010 | 1 | ENSMUST00000057870.3 | 5485 | Rprml | ENSMUSG00000046215.3 |
| chr11 | 103658301 | 103658409 | 3.5746 | 4.2806 | 1.32693 | Distal Intergenic | 11 | 103649570 | 103650579 | 1010 | 1 | ENSMUST00000057870.3 | 8732 | Rprml | ENSMUSG00000046215.3 |
| chr11 | 103660823 | 103660898 | 3.93087 | 5.03096 | 1.80139 | Distal Intergenic | 11 | 103649570 | 103650579 | 1010 | 1 | ENSMUST00000057870.3 | 11254 | Rprml | ENSMUSG00000046215.3 |
| chr11 | 103667972 | 103668296 | 3.99619 | 5.09371 | 1.86935 | Distal Intergenic | 11 | 103649570 | 103650579 | 1010 | 1 | ENSMUST00000057870.3 | 18403 | Rprml | ENSMUSG00000046215.3 |
| chr19 | 16825165 | 16825435 | 3.28603 | 3.77169 | 1.06853 | Distal Intergenic | 19 | 16616763 | 16780822 | 164060 | 2 | ENSMUST00000068156.6 | -44344 | Vps13a | ENSMUSG00000046230.9 |
| chr9 | 99450761 | 99450885 | 3.56345 | 4.37295 | 1.3633 | Distal Intergenic | 9 | 99456207 | 99470932 | 14726 | 1 | ENSMUST00000163199.3 | -5322 | 1600029I14Rik | ENSMUSG00000046242.7 |
| chr9 | 99471609 | 99471949 | 3.36364 | 3.87303 | 1.01733 | Intron (ENSMUST00000056103.5/ENSMUSG00000046242.7, intron 1 of 1) | 9 | 99470421 | 99474751 | 4331 | 1 | ENSMUST00000056103.5 | 1189 | 1600029I14Rik | ENSMUSG00000046242.7 |
| chr3 | 89829120 | 89829351 | 3.32215 | 4.18331 | 1.25102 | Distal Intergenic | 3 | 89831370 | 89858834 | 27465 | 1 | ENSMUST00000050401.5 | -2019 | She | ENSMUSG00000046280.8 |
| chr3 | 89830455 | 89831904 | 3.81797 | 4.92688 | 1.78889 | Promoter | 3 | 89831370 | 89858834 | 27465 | 1 | ENSMUST00000050401.5 | 0 | She | ENSMUSG00000046280.8 |
| chr3 | 89832236 | 89832425 | 3.26575 | 3.72576 | 1.01733 | Promoter | 3 | 89831370 | 89858834 | 27465 | 1 | ENSMUST00000050401.5 | 867 | She | ENSMUSG00000046280.8 |
| chr3 | 89833110 | 89833185 | 3.30837 | 3.80124 | 1.05666 | Intron (ENSMUST00000050401.5/ENSMUSG00000046280.8, intron 1 of 5) | 3 | 89831370 | 89858834 | 27465 | 1 | ENSMUST00000050401.5 | 1741 | She | ENSMUSG00000046280.8 |
| chr4 | 41498041 | 41498901 | 3.56178 | 4.24098 | 1.25577 | Exon (ENSMUST00000054920.4/ENSMUSG00000046312.4, exon 2 of 2) | 4 | 41499494 | 41502668 | 3175 | 2 | ENSMUST00000149596.1 | 3767 | AI464131 | ENSMUSG00000046312.4 |
| chr12 | 45071950 | 45072080 | 3.51807 | 4.19528 | 1.3035 | Intron (ENSMUST00000120531.2/ENSMUSG00000046314.10, intron 1 of 5) | 12 | 44852488 | 45074112 | 221625 | 2 | ENSMUST00000120531.2 | 2032 | Stxbp6 | ENSMUSG00000046314.10 |
| chr12 | 45073807 | 45074250 | 3.60645 | 4.38774 | 1.37639 | Promoter | 12 | 44852488 | 45074112 | 221625 | 2 | ENSMUST00000120531.2 | 0 | Stxbp6 | ENSMUSG00000046314.10 |
| chr12 | 45074564 | 45074699 | 3.47972 | 4.10549 | 1.18492 | Promoter | 12 | 44855221 | 45074493 | 219273 | 2 | ENSMUST00000053768.7 | -72 | Stxbp6 | ENSMUSG00000046314.10 |
| chr12 | 45075821 | 45076123 | 3.25591 | 3.71099 | 1.01733 | Distal Intergenic | 12 | 44855221 | 45074493 | 219273 | 2 | ENSMUST00000053768.7 | -1329 | Stxbp6 | ENSMUSG00000046314.10 |
| chr3 | 96454648 | 96454872 | 3.97014 | 5.31462 | 1.9701 | Distal Intergenic | 3 | 96433788 | 96452306 | 18519 | 2 | ENSMUST00000098841.3 | -2343 | BC107364 | ENSMUSG00000046317.9 |
| chr3 | 96455599 | 96455815 | 4.37359 | 5.90281 | 2.43652 | Distal Intergenic | 3 | 96433788 | 96452306 | 18519 | 2 | ENSMUST00000098841.3 | -3294 | BC107364 | ENSMUSG00000046317.9 |
| chr7 | 121392257 | 121393325 | 3.6743 | 4.48708 | 1.46984 | Promoter | 7 | 121392296 | 121501768 | 109473 | 1 | ENSMUST00000084628.3 | 0 | Hs3st2 | ENSMUSG00000046321.7 |
| chr6 | 122628019 | 122628278 | 3.37806 | 3.90257 | 1.04495 | Intron (ENSMUST00000049644.3/ENSMUSG00000046323.3, intron 1 of 3) | 6 | 122626753 | 122630272 | 3520 | 1 | ENSMUST00000123429.1 | 1267 | Dppa3 | ENSMUSG00000046323.3 |
| chr6 | 122628595 | 122628748 | 3.50334 | 4.12518 | 1.16295 | Exon (ENSMUST00000049644.3/ENSMUSG00000046323.3, exon 2 of 4) | 6 | 122626753 | 122630272 | 3520 | 1 | ENSMUST00000123429.1 | 1843 | Dppa3 | ENSMUSG00000046323.3 |
| chr14 | 57105291 | 57105413 | 3.47254 | 4.11356 | 1.25847 | Promoter | 14 | 57098600 | 57104702 | 6103 | 2 | ENSMUST00000055698.7 | -590 | Gjb2 | ENSMUSG00000046352.7 |
| chr9 | 98424875 | 98425117 | 3.73722 | 4.95607 | 1.74526 | 5' UTR | 9 | 98422961 | 98446575 | 23615 | 1 | ENSMUST00000052068.10 | 1915 | Rbp1 | ENSMUSG00000046402.10 |
| chr7 | 29232412 | 29232487 | 3.75483 | 4.65883 | 1.58009 | Promoter | 7 | 29221928 | 29232522 | 10595 | 2 | ENSMUST00000085818.4 | 35 | Kcnk6 | ENSMUSG00000046410.9 |
| chr8 | 91798131 | 91798547 | 3.92025 | 4.94198 | 1.71177 | 3' UTR | 8 | 91800415 | 91807539 | 7125 | 1 | ENSMUST00000062522.6 | -1868 | Irx3os | ENSMUSG00000046413.7 |
| chr8 | 91798858 | 91801321 | 4.09689 | 5.41242 | 2.0939 | Promoter | 8 | 91800415 | 91807539 | 7125 | 1 | ENSMUST00000062522.6 | 0 | Irx3os | ENSMUSG00000046413.7 |
| chr8 | 91803958 | 91805794 | 4.01245 | 5.31811 | 2.0412 | Exon (ENSMUST00000062522.6/ENSMUSG00000046413.7, exon 2 of 5) | 8 | 91802175 | 91805215 | 3041 | 1 | ENSMUST00000133528.1 | 1784 | Irx3os | ENSMUSG00000046413.7 |
| chr8 | 91806199 | 91807165 | 3.8907 | 5.28295 | 2.0292 | Exon (ENSMUST00000062522.6/ENSMUSG00000046413.7, exon 5 of 5) | 8 | 91802175 | 91805215 | 3041 | 1 | ENSMUST00000133528.1 | 4025 | Irx3os | ENSMUSG00000046413.7 |
| chr8 | 91808472 | 91808574 | 4.67346 | 6.52 | 2.89948 | Distal Intergenic | 8 | 91802175 | 91805215 | 3041 | 1 | ENSMUST00000133528.1 | 6298 | Irx3os | ENSMUSG00000046413.7 |
| chr8 | 91816359 | 91816612 | 3.32794 | 3.81924 | 1.01733 | Distal Intergenic | 8 | 91802175 | 91805215 | 3041 | 1 | ENSMUST00000133528.1 | 14185 | Irx3os | ENSMUSG00000046413.7 |
| chr8 | 91821689 | 91821764 | 4.24649 | 5.53562 | 2.08662 | Distal Intergenic | 8 | 91802175 | 91805215 | 3041 | 1 | ENSMUST00000133528.1 | 19515 | Irx3os | ENSMUSG00000046413.7 |
| chr8 | 91822072 | 91822267 | 3.7704 | 4.63836 | 1.50769 | Distal Intergenic | 8 | 91802175 | 91805215 | 3041 | 1 | ENSMUST00000133528.1 | 19898 | Irx3os | ENSMUSG00000046413.7 |
| chr8 | 91823054 | 91823184 | 3.36392 | 3.87342 | 1.01733 | Distal Intergenic | 8 | 91802175 | 91805215 | 3041 | 1 | ENSMUST00000133528.1 | 20880 | Irx3os | ENSMUSG00000046413.7 |
| chr8 | 91826533 | 91826635 | 3.38282 | 4.06957 | 1.22221 | Distal Intergenic | 8 | 91802175 | 91805215 | 3041 | 1 | ENSMUST00000133528.1 | 24359 | Irx3os | ENSMUSG00000046413.7 |
| chr8 | 91831690 | 91831819 | 3.58156 | 4.34404 | 1.37083 | Distal Intergenic | 8 | 91802175 | 91805215 | 3041 | 1 | ENSMUST00000133528.1 | 29516 | Irx3os | ENSMUSG00000046413.7 |
| chr18 | 67345156 | 67345311 | 3.58829 | 4.38175 | 1.38577 | Intron (ENSMUST00000025404.8/ENSMUSG00000024526.8, intron 1 of 4) | 18 | 67321209 | 67344103 | 22895 | 2 | ENSMUST00000145081.2 | -1054 | B430212C06Rik | ENSMUSG00000046415.6 |
| chr18 | 67347110 | 67347423 | 3.47643 | 4.21226 | 1.29388 | Intron (ENSMUST00000025404.8/ENSMUSG00000024526.8, intron 1 of 4) | 18 | 67321209 | 67344103 | 22895 | 2 | ENSMUST00000145081.2 | -3008 | B430212C06Rik | ENSMUSG00000046415.6 |
| chr4 | 138448037 | 138448269 | 3.23373 | 3.67777 | 1.01733 | Distal Intergenic | 4 | 138454314 | 138460123 | 5810 | 1 | ENSMUST00000050918.3 | -6045 | Camk2n1 | ENSMUSG00000046447.3 |
| chr4 | 138496120 | 138496195 | 3.38012 | 3.89787 | 1.01733 | Distal Intergenic | 4 | 138454314 | 138460123 | 5810 | 1 | ENSMUST00000050918.3 | 41807 | Camk2n1 | ENSMUSG00000046447.3 |
| chr4 | 138498647 | 138498766 | 3.78906 | 4.72773 | 1.55534 | Distal Intergenic | 4 | 138454314 | 138460123 | 5810 | 1 | ENSMUST00000050918.3 | 44334 | Camk2n1 | ENSMUSG00000046447.3 |
| chr4 | 138502179 | 138502309 | 3.49947 | 4.30416 | 1.31098 | Distal Intergenic | 4 | 138454314 | 138460123 | 5810 | 1 | ENSMUST00000050918.3 | 47866 | Camk2n1 | ENSMUSG00000046447.3 |
| chr4 | 138504457 | 138505376 | 3.48423 | 4.13361 | 1.20788 | Distal Intergenic | 4 | 138454314 | 138460123 | 5810 | 1 | ENSMUST00000050918.3 | 50144 | Camk2n1 | ENSMUSG00000046447.3 |
| chr4 | 138511324 | 138512125 | 3.55751 | 4.29549 | 1.29908 | Distal Intergenic | 4 | 138454314 | 138460123 | 5810 | 1 | ENSMUST00000050918.3 | 57011 | Camk2n1 | ENSMUSG00000046447.3 |
| chr10 | 37140037 | 37140834 | 3.40343 | 4.01574 | 1.17646 | Promoter | 10 | 37139914 | 37151315 | 11402 | 1 | ENSMUST00000191067.1 | 124 | 5930403N24Rik | ENSMUSG00000046463.4 |
| chr2 | 181670160 | 181671840 | 4.04405 | 5.34143 | 2.06365 | Promoter | 2 | 181669836 | 181671640 | 1805 | 2 | ENSMUST00000054491.5 | 0 | Sox18 | ENSMUSG00000046470.5 |
| chr9 | 45137024 | 45137238 | 3.63567 | 4.43661 | 1.42043 | Distal Intergenic | 9 | 45138437 | 45154152 | 15716 | 1 | ENSMUST00000060125.5 | -1199 | Scn4b | ENSMUSG00000046480.5 |
| chr9 | 45138218 | 45139595 | 3.5672 | 4.26946 | 1.29235 | Promoter | 9 | 45138437 | 45154152 | 15716 | 1 | ENSMUST00000060125.5 | 0 | Scn4b | ENSMUSG00000046480.5 |
| chr9 | 45140102 | 45140371 | 3.47671 | 4.2449 | 1.27891 | Intron (ENSMUST00000060125.5/ENSMUSG00000046480.5, intron 1 of 4) | 9 | 45138437 | 45154152 | 15716 | 1 | ENSMUST00000060125.5 | 1666 | Scn4b | ENSMUSG00000046480.5 |
| chr9 | 45141147 | 45141405 | 3.49146 | 4.27041 | 1.29394 | Intron (ENSMUST00000060125.5/ENSMUSG00000046480.5, intron 1 of 4) | 9 | 45138437 | 45154152 | 15716 | 1 | ENSMUST00000060125.5 | 2711 | Scn4b | ENSMUSG00000046480.5 |
| chr9 | 45141756 | 45141831 | 3.8478 | 4.77534 | 1.59434 | Intron (ENSMUST00000060125.5/ENSMUSG00000046480.5, intron 1 of 4) | 9 | 45138437 | 45154152 | 15716 | 1 | ENSMUST00000060125.5 | 3320 | Scn4b | ENSMUSG00000046480.5 |
| chr9 | 45142786 | 45143071 | 3.50166 | 4.29575 | 1.28553 | Intron (ENSMUST00000060125.5/ENSMUSG00000046480.5, intron 1 of 4) | 9 | 45138437 | 45154152 | 15716 | 1 | ENSMUST00000060125.5 | 4350 | Scn4b | ENSMUSG00000046480.5 |
| chr9 | 45144088 | 45144621 | 3.90645 | 4.92859 | 1.74274 | Intron (ENSMUST00000060125.5/ENSMUSG00000046480.5, intron 1 of 4) | 9 | 45138437 | 45154152 | 15716 | 1 | ENSMUST00000060125.5 | 5652 | Scn4b | ENSMUSG00000046480.5 |
| chr9 | 45151343 | 45151654 | 3.46698 | 4.09324 | 1.23096 | 3' UTR | 9 | 45138437 | 45154152 | 15716 | 1 | ENSMUST00000060125.5 | 12907 | Scn4b | ENSMUSG00000046480.5 |
| chr18 | 46464030 | 46464207 | 3.57959 | 4.62244 | 1.52024 | Downstream (1-2kb) | 18 | 46465215 | 46465790 | 576 | 2 | ENSMUST00000180618.1 | 1583 | Mospd4 | ENSMUSG00000046487.6 |
| chr18 | 46465580 | 46465904 | 3.69058 | 4.55994 | 1.53658 | Promoter | 18 | 46465215 | 46465790 | 576 | 2 | ENSMUST00000180618.1 | 0 | Mospd4 | ENSMUSG00000046487.6 |
| chr6 | 97058953 | 97059234 | 3.45089 | 4.15048 | 1.24018 | Intron (ENSMUST00000089295.4/ENSMUSG00000046500.7, intron 1 of 5) | 6 | 96831209 | 97060411 | 229203 | 2 | ENSMUST00000089295.4 | 1177 | Fam19a4 | ENSMUSG00000046500.7 |
| chr12 | 33923058 | 33923229 | 3.39226 | 4.15955 | 1.22695 | Distal Intergenic | 12 | 33928425 | 33929310 | 886 | 1 | ENSMUST00000061035.3 | -5196 | Ferd3l | ENSMUSG00000046518.3 |
| chr12 | 33927323 | 33927723 | 4.3317 | 5.76905 | 2.34449 | Promoter | 12 | 33928425 | 33929310 | 886 | 1 | ENSMUST00000061035.3 | -702 | Ferd3l | ENSMUSG00000046518.3 |
| chr12 | 33928090 | 33929058 | 3.55123 | 4.22378 | 1.23873 | Promoter | 12 | 33928425 | 33929310 | 886 | 1 | ENSMUST00000061035.3 | 0 | Ferd3l | ENSMUSG00000046518.3 |
| chr12 | 33929597 | 33930131 | 3.52895 | 4.30734 | 1.32924 | Distal Intergenic | 12 | 33928425 | 33929310 | 886 | 1 | ENSMUST00000061035.3 | 1173 | Ferd3l | ENSMUSG00000046518.3 |
| chr16 | 30695368 | 30695938 | 3.63034 | 4.3889 | 1.38337 | Distal Intergenic | 16 | 30599723 | 30602797 | 3075 | 1 | ENSMUST00000059078.3 | 95646 | Fam43a | ENSMUSG00000046546.3 |
| chr12 | 76152898 | 76153145 | 3.62738 | 4.39003 | 1.41065 | Promoter | 12 | 76152336 | 76152627 | 292 | 2 | ENSMUST00000121461.1 | -272 | Gm7862 | ENSMUSG00000046580.10 |
| chr19 | 47938458 | 47938544 | 3.35567 | 3.94707 | 1.12705 | Promoter | 19 | 47937712 | 48035379 | 97668 | 1 | ENSMUST00000066308.7 | 747 | Ccdc147 | ENSMUSG00000046585.8 |
| chr19 | 47938899 | 47939836 | 3.77618 | 4.71716 | 1.62015 | Intron (ENSMUST00000066308.7/ENSMUSG00000046585.8, intron 1 of 17) | 19 | 47937712 | 48035379 | 97668 | 1 | ENSMUST00000066308.7 | 1188 | Ccdc147 | ENSMUSG00000046585.8 |
| chr19 | 47997879 | 47998058 | 3.94835 | 5.04757 | 1.8067 | Intron (ENSMUST00000066308.7/ENSMUSG00000046585.8, intron 15 of 17) | 19 | 47937712 | 48035379 | 97668 | 1 | ENSMUST00000066308.7 | 60168 | Ccdc147 | ENSMUSG00000046585.8 |
| chr8 | 4226357 | 4226681 | 3.88561 | 4.86448 | 1.69516 | Promoter | 8 | 4226827 | 4237470 | 10644 | 1 | ENSMUST00000053035.6 | -146 | Lrrc8e | ENSMUSG00000046589.6 |
| chr4 | 40472762 | 40472904 | 3.54236 | 4.28922 | 1.33198 | Promoter | 4 | 40473130 | 40475653 | 2524 | 1 | ENSMUST00000179526.1 | -226 | Tmem215 | ENSMUSG00000046593.3 |
| chr4 | 40473992 | 40474241 | 3.49503 | 4.2903 | 1.32533 | Promoter | 4 | 40473130 | 40475653 | 2524 | 1 | ENSMUST00000179526.1 | 863 | Tmem215 | ENSMUSG00000046593.3 |
| chr16 | 31428161 | 31429315 | 4.08022 | 5.22926 | 1.89904 | Promoter | 16 | 31428748 | 31458901 | 30154 | 1 | ENSMUST00000115227.4 | 0 | Bdh1 | ENSMUSG00000046598.9 |
| chr16 | 31436114 | 31436664 | 3.66227 | 4.42087 | 1.34689 | Promoter | 16 | 31436091 | 31456750 | 20660 | 1 | ENSMUST00000149039.1 | 24 | Bdh1 | ENSMUSG00000046598.9 |
| chr16 | 31453353 | 31453558 | 3.42691 | 4.0176 | 1.13614 | Intron (ENSMUST00000089759.3/ENSMUSG00000046598.9, intron 5 of 6) | 16 | 31458116 | 31458901 | 786 | 1 | ENSMUST00000126825.1 | -4558 | Bdh1 | ENSMUSG00000046598.9 |
| chr5 | 118160938 | 118161070 | 3.5629 | 4.23361 | 1.22842 | Distal Intergenic | 5 | 118164648 | 118184828 | 20181 | 1 | ENSMUST00000184679.1 | -3578 | Hrk | ENSMUSG00000046607.6 |
| chr5 | 118163274 | 118163648 | 3.57584 | 4.26664 | 1.27096 | Promoter | 5 | 118164648 | 118184828 | 20181 | 1 | ENSMUST00000184679.1 | -1000 | Hrk | ENSMUSG00000046607.6 |
| chr5 | 118164159 | 118164439 | 3.98545 | 5.1724 | 1.89388 | Promoter | 5 | 118164648 | 118184828 | 20181 | 1 | ENSMUST00000184679.1 | -209 | Hrk | ENSMUSG00000046607.6 |
| chr5 | 118167552 | 118168148 | 4.04751 | 5.27583 | 1.9814 | Intron (ENSMUST00000184679.1/ENSMUSG00000046607.6, intron 1 of 1) | 5 | 118169748 | 118189478 | 19731 | 1 | ENSMUST00000054836.6 | -1600 | Hrk | ENSMUSG00000046607.6 |
| chr5 | 118168490 | 118169482 | 3.90931 | 4.89414 | 1.66488 | Promoter | 5 | 118169748 | 118189478 | 19731 | 1 | ENSMUST00000054836.6 | -266 | Hrk | ENSMUSG00000046607.6 |
| chr5 | 118169865 | 118170424 | 3.48463 | 4.08895 | 1.13424 | Promoter | 5 | 118169748 | 118189478 | 19731 | 1 | ENSMUST00000054836.6 | 118 | Hrk | ENSMUSG00000046607.6 |
| chr5 | 118170765 | 118171104 | 4.03371 | 5.14009 | 1.8332 | Intron (ENSMUST00000184679.1/ENSMUSG00000046607.6, intron 1 of 1) | 5 | 118169748 | 118189478 | 19731 | 1 | ENSMUST00000054836.6 | 1018 | Hrk | ENSMUSG00000046607.6 |
| chr5 | 118171724 | 118171865 | 3.71518 | 4.64241 | 1.55622 | Intron (ENSMUST00000184679.1/ENSMUSG00000046607.6, intron 1 of 1) | 5 | 118169748 | 118189478 | 19731 | 1 | ENSMUST00000054836.6 | 1977 | Hrk | ENSMUSG00000046607.6 |
| chr5 | 118172258 | 118174031 | 3.81562 | 4.78 | 1.62498 | Intron (ENSMUST00000184679.1/ENSMUSG00000046607.6, intron 1 of 1) | 5 | 118169748 | 118189478 | 19731 | 1 | ENSMUST00000054836.6 | 2511 | Hrk | ENSMUSG00000046607.6 |
| chr5 | 118174693 | 118174768 | 3.36983 | 3.93744 | 1.08165 | Intron (ENSMUST00000184679.1/ENSMUSG00000046607.6, intron 1 of 1) | 5 | 118169748 | 118189478 | 19731 | 1 | ENSMUST00000054836.6 | 4946 | Hrk | ENSMUSG00000046607.6 |
| chr18 | 65736187 | 65736873 | 3.41882 | 4.09902 | 1.19816 | Intron (ENSMUST00000115097.2/ENSMUSG00000046610.9, intron 8 of 14) | 18 | 65749014 | 65750390 | 1377 | 1 | ENSMUST00000134343.1 | -12141 | Oacyl | ENSMUSG00000046610.9 |
| chr16 | 20588185 | 20588828 | 4.44881 | 6.0137 | 2.52308 | Promoter | 16 | 20589471 | 20598467 | 8997 | 1 | ENSMUST00000100074.4 | -643 | Vwa5b2 | ENSMUSG00000046613.13 |
| chr16 | 20589208 | 20589634 | 3.57259 | 4.25645 | 1.24884 | Promoter | 16 | 20589471 | 20598467 | 8997 | 1 | ENSMUST00000100074.4 | 0 | Vwa5b2 | ENSMUSG00000046613.13 |
| chr16 | 20590025 | 20590418 | 3.81348 | 4.71903 | 1.57546 | Promoter | 16 | 20590059 | 20590531 | 473 | 1 | ENSMUST00000161715.1 | 0 | Vwa5b2 | ENSMUSG00000046613.13 |
| chr16 | 20590762 | 20591685 | 3.62959 | 4.36448 | 1.31765 | Promoter | 16 | 20591156 | 20605374 | 14219 | 1 | ENSMUST00000159780.1 | 0 | Vwa5b2 | ENSMUSG00000046613.13 |
| chr4 | 154854880 | 154855123 | 3.78267 | 4.66703 | 1.54655 | Distal Intergenic | 4 | 154856200 | 154867125 | 10926 | 1 | ENSMUST00000050220.2 | -1077 | Ttc34 | ENSMUSG00000046637.2 |
| chr4 | 154855552 | 154856745 | 3.91991 | 5.04723 | 1.80251 | Promoter | 4 | 154856200 | 154867125 | 10926 | 1 | ENSMUST00000050220.2 | 0 | Ttc34 | ENSMUSG00000046637.2 |
| chr5 | 143253372 | 143253479 | 3.5971 | 4.38492 | 1.39284 | Exon (ENSMUST00000161448.2/ENSMUSG00000046658.11, exon 7 of 7) | 5 | 143253209 | 143264656 | 11448 | 2 | ENSMUST00000051665.8 | 11177 | Zfp316 | ENSMUSG00000046658.11 |
| chr5 | 143254269 | 143255356 | 4.04513 | 5.40259 | 2.06714 | Exon (ENSMUST00000161448.2/ENSMUSG00000046658.11, exon 7 of 7) | 5 | 143253209 | 143264656 | 11448 | 2 | ENSMUST00000051665.8 | 9300 | Zfp316 | ENSMUSG00000046658.11 |
| chr4 | 133480897 | 133481265 | 3.36157 | 3.86988 | 1.01733 | Promoter | 4 | 133480132 | 133487938 | 7807 | 1 | ENSMUST00000051676.6 | 766 | Fam46b | ENSMUSG00000046694.6 |
| chr11 | 118972619 | 118972759 | 3.55179 | 4.21245 | 1.21267 | Distal Intergenic | 11 | 118988188 | 118992841 | 4654 | 1 | ENSMUST00000092373.7 | -15429 | Enpp7 | ENSMUSG00000046697.11 |
| chr11 | 118994014 | 118994290 | 3.26958 | 3.73154 | 1.01733 | Distal Intergenic | 11 | 118988216 | 118992665 | 4450 | 1 | ENSMUST00000106273.1 | 5799 | Enpp7 | ENSMUSG00000046697.11 |
| chr11 | 118998211 | 118998341 | 3.28353 | 3.75247 | 1.01733 | Distal Intergenic | 11 | 118988216 | 118992665 | 4450 | 1 | ENSMUST00000106273.1 | 9996 | Enpp7 | ENSMUSG00000046697.11 |
| chr11 | 118999260 | 119000423 | 4.09044 | 5.29676 | 1.98982 | Distal Intergenic | 11 | 118988216 | 118992665 | 4450 | 1 | ENSMUST00000106273.1 | 11045 | Enpp7 | ENSMUSG00000046697.11 |
| chr11 | 119000740 | 119000958 | 3.65883 | 4.42245 | 1.36308 | Distal Intergenic | 11 | 118988216 | 118992665 | 4450 | 1 | ENSMUST00000106273.1 | 12525 | Enpp7 | ENSMUSG00000046697.11 |
| chr8 | 71525140 | 71525697 | 3.71311 | 4.5731 | 1.49886 | Distal Intergenic | 8 | 71534264 | 71537437 | 3174 | 2 | ENSMUST00000051672.7 | 11740 | Bst2 | ENSMUSG00000046718.7 |
| chr18 | 36324197 | 36324320 | 3.45377 | 4.42209 | 1.38845 | Distal Intergenic | 18 | 36345619 | 36366720 | 21102 | 1 | ENSMUST00000139727.1 | -21299 | Cystm1 | ENSMUSG00000046727.7 |
| chr3 | 38883913 | 38884272 | 3.58968 | 4.30432 | 1.32052 | Intron (ENSMUST00000153659.1/ENSMUSG00000086181.1, intron 2 of 3) | 3 | 38886940 | 39011985 | 125046 | 1 | ENSMUST00000061260.7 | -2668 | Fat4 | ENSMUSG00000046743.6 |
| chr3 | 38886095 | 38886551 | 3.85127 | 4.94374 | 1.74771 | Promoter | 3 | 38886940 | 39011985 | 125046 | 1 | ENSMUST00000061260.7 | -389 | Fat4 | ENSMUSG00000046743.6 |
| chr3 | 38887215 | 38887641 | 3.5446 | 4.42273 | 1.4049 | Promoter | 3 | 38886940 | 39011985 | 125046 | 1 | ENSMUST00000061260.7 | 276 | Fat4 | ENSMUSG00000046743.6 |
| chr3 | 38894747 | 38894898 | 3.29837 | 4.0372 | 1.12977 | Intron (ENSMUST00000061260.7/ENSMUSG00000046743.6, intron 1 of 16) | 3 | 38890942 | 38952429 | 61488 | 1 | ENSMUST00000129182.1 | 3806 | Fat4 | ENSMUSG00000046743.6 |
| chr7 | 28288862 | 28289714 | 3.55428 | 4.36437 | 1.39839 | Promoter | 7 | 28287060 | 28288182 | 1123 | 2 | ENSMUST00000156408.1 | -681 | BC089491 | ENSMUSG00000046750.11 |
| chr7 | 28290727 | 28291157 | 3.70853 | 4.69258 | 1.57494 | Promoter | 7 | 28284652 | 28291134 | 6483 | 2 | ENSMUST00000108316.3 | 0 | BC089491 | ENSMUSG00000046750.11 |
| chr15 | 76004527 | 76004733 | 3.89714 | 5.00127 | 1.7964 | Exon (ENSMUST00000170153.1/ENSMUSG00000046761.7, exon 5 of 5) | 15 | 76001093 | 76009498 | 8406 | 2 | ENSMUST00000170153.1 | 4765 | Fam83h | ENSMUSG00000046761.7 |
| chr15 | 76005041 | 76005345 | 3.8589 | 4.87663 | 1.75448 | Exon (ENSMUST00000170153.1/ENSMUSG00000046761.7, exon 3 of 5) | 15 | 76001093 | 76009498 | 8406 | 2 | ENSMUST00000170153.1 | 4153 | Fam83h | ENSMUSG00000046761.7 |
| chr15 | 76005678 | 76005753 | 3.52582 | 4.27982 | 1.31592 | Intron (ENSMUST00000170153.1/ENSMUSG00000046761.7, intron 2 of 4) | 15 | 76001093 | 76009498 | 8406 | 2 | ENSMUST00000170153.1 | 3745 | Fam83h | ENSMUSG00000046761.7 |
| chr12 | 75308801 | 75309399 | 3.71499 | 4.82788 | 1.63966 | Promoter | 12 | 75308881 | 75400212 | 91332 | 1 | ENSMUST00000172981.1 | 0 | Rhoj | ENSMUSG00000046768.8 |
| chr12 | 57555381 | 57555692 | 3.66666 | 4.45848 | 1.4486 | Distal Intergenic | 12 | 57564113 | 57622112 | 58000 | 1 | ENSMUST00000101398.4 | -8421 | Ttc6 | ENSMUSG00000046782.9 |
| chr12 | 57556888 | 57557182 | 3.6802 | 4.4853 | 1.47913 | Distal Intergenic | 12 | 57564113 | 57622112 | 58000 | 1 | ENSMUST00000101398.4 | -6931 | Ttc6 | ENSMUSG00000046782.9 |
| chr12 | 57563231 | 57563462 | 3.37871 | 3.92923 | 1.13454 | Promoter | 12 | 57564113 | 57622112 | 58000 | 1 | ENSMUST00000101398.4 | -651 | Ttc6 | ENSMUSG00000046782.9 |
| chr12 | 57569426 | 57569746 | 3.61623 | 4.44768 | 1.42657 | Intron (ENSMUST00000101398.4/ENSMUSG00000046782.9, intron 1 of 4) | 12 | 57564113 | 57622112 | 58000 | 1 | ENSMUST00000101398.4 | 5314 | Ttc6 | ENSMUSG00000046782.9 |
| chr12 | 57570524 | 57570893 | 3.37613 | 3.89184 | 1.01733 | Intron (ENSMUST00000101398.4/ENSMUSG00000046782.9, intron 1 of 4) | 12 | 57575817 | 57737928 | 162112 | 1 | ENSMUST00000172939.2 | -4924 | Ttc6 | ENSMUSG00000046782.9 |
| chr12 | 57571872 | 57572059 | 3.46829 | 4.17307 | 1.27185 | Intron (ENSMUST00000101398.4/ENSMUSG00000046782.9, intron 1 of 4) | 12 | 57575817 | 57737928 | 162112 | 1 | ENSMUST00000172939.2 | -3758 | Ttc6 | ENSMUSG00000046782.9 |
| chr12 | 57574006 | 57574180 | 3.36026 | 4.04434 | 1.18282 | Intron (ENSMUST00000101398.4/ENSMUSG00000046782.9, intron 1 of 4) | 12 | 57575817 | 57737928 | 162112 | 1 | ENSMUST00000172939.2 | -1637 | Ttc6 | ENSMUSG00000046782.9 |
| chr12 | 57574640 | 57575222 | 4.03321 | 5.20557 | 1.98454 | Promoter | 12 | 57575817 | 57737928 | 162112 | 1 | ENSMUST00000172939.2 | -595 | Ttc6 | ENSMUSG00000046782.9 |
| chr12 | 57577309 | 57577894 | 3.5864 | 4.81948 | 1.65298 | Intron (ENSMUST00000101398.4/ENSMUSG00000046782.9, intron 2 of 4) | 12 | 57575817 | 57737928 | 162112 | 1 | ENSMUST00000172939.2 | 1493 | Ttc6 | ENSMUSG00000046782.9 |
| chr12 | 57776348 | 57776423 | 3.47605 | 4.10562 | 1.23505 | Distal Intergenic | 12 | 57649415 | 57651719 | 2305 | 1 | ENSMUST00000172852.1 | 126934 | Ttc6 | ENSMUSG00000046782.9 |
| chr8 | 35351675 | 35351849 | 3.37805 | 4.00941 | 1.21088 | Distal Intergenic | 8 | 35375741 | 35388124 | 12384 | 1 | ENSMUST00000070481.6 | -23892 | Ppp1r3b | ENSMUSG00000046794.8 |
| chr10 | 75559938 | 75560059 | 3.48506 | 4.24525 | 1.28933 | Promoter | 10 | 75550125 | 75560330 | 10206 | 2 | ENSMUST00000051129.9 | 271 | Lrrc75b | ENSMUSG00000046807.9 |
| chr3 | 137623604 | 137624694 | 3.32299 | 3.82672 | 1.06575 | Promoter | 3 | 137623672 | 137628333 | 4662 | 1 | ENSMUST00000053855.7 | 0 | Ddit4l | ENSMUSG00000046818.7 |
| chr3 | 137625046 | 137625277 | 3.8038 | 4.76287 | 1.65697 | Intron (ENSMUST00000167564.1/ENSMUSG00000046818.7, intron 2 of 2) | 3 | 137624037 | 137626174 | 2138 | 1 | ENSMUST00000165845.1 | 1010 | Ddit4l | ENSMUSG00000046818.7 |
| chr15 | 101844254 | 101844537 | 3.28279 | 3.75133 | 1.01733 | Downstream (<1kb) | 15 | 101845428 | 101850786 | 5359 | 2 | ENSMUST00000023790.3 | 6249 | Krt1 | ENSMUSG00000046834.6 |
| chr15 | 101845554 | 101846198 | 3.71366 | 4.59229 | 1.54091 | 3' UTR | 15 | 101845428 | 101850786 | 5359 | 2 | ENSMUST00000023790.3 | 4588 | Krt1 | ENSMUSG00000046834.6 |
| chr15 | 101851431 | 101851561 | 3.89052 | 4.85462 | 1.64847 | Promoter | 15 | 101845428 | 101850786 | 5359 | 2 | ENSMUST00000023790.3 | -646 | Krt1 | ENSMUSG00000046834.6 |
| chr1 | 183299161 | 183299579 | 1.86248 | 9.28565 | 5.14653 | Intron (ENSMUST00000109166.2/ENSMUSG00000042901.5, intron 1 of 9) | 1 | 183277402 | 183297256 | 19855 | 2 | ENSMUST00000163528.2 | -1906 | Brox | ENSMUSG00000046836.8 |
| chr8 | 114206072 | 114206254 | 3.66326 | 4.43298 | 1.37875 | Promoter | 8 | 114205634 | 114230319 | 24686 | 1 | ENSMUST00000150963.1 | 439 | Vat1l | ENSMUSG00000046844.6 |
| chr8 | 114206965 | 114207244 | 3.8115 | 4.81763 | 1.65466 | Intron (ENSMUST00000142591.1/ENSMUSG00000046844.6, intron 1 of 1) | 8 | 114205634 | 114230319 | 24686 | 1 | ENSMUST00000150963.1 | 1332 | Vat1l | ENSMUSG00000046844.6 |
| chr1 | 63213362 | 63213676 | 3.32968 | 3.82186 | 1.01733 | Promoter | 1 | 63208742 | 63214223 | 5482 | 2 | ENSMUST00000187533.1 | 547 | Gpr1 | ENSMUSG00000046856.8 |

| | | | | | | | | | | | | | | | |
|---|---|---|---|---|---|---|---|---|---|---|---|---|---|---|---|
| chr1 | 63214331 | 63214602 | 3.37637 | 3.92391 | 1.12974 | Promoter | 1 | 63182691 | 63214543 | 31853 | 2 | ENSMUST00000050536.8 | 0 | Gpr1 | ENSMUSG00000046856.8 |
| chr14 | 55765709 | 55765800 | 3.6318 | 4.41316 | 1.43504 | Promoter | 14 | 55765962 | 55768494 | 2533 | 1 | ENSMUST00000057569.3 | -162 | Ltb4r1 | ENSMUSG00000046908.5 |
| chr14 | 55766713 | 55766899 | 3.35711 | 3.86315 | 1.01733 | Promoter | 14 | 55765962 | 55768494 | 2533 | 1 | ENSMUST00000057569.3 | 752 | Ltb4r1 | ENSMUSG00000046908.5 |
| chr14 | 55767233 | 55767645 | 3.22427 | 4.03733 | 1.13801 | 5' UTR | 14 | 55765962 | 55768494 | 2533 | 1 | ENSMUST00000057569.3 | 1272 | Ltb4r1 | ENSMUSG00000046908.5 |
| chr10 | 41069811 | 41069951 | 3.77138 | 4.62194 | 1.4745 | Downstream (<1kb) | 10 | 41069989 | 41072285 | 2297 | 2 | ENSMUST00000061796.6 | 2334 | Gpr6 | ENSMUSG00000046922.6 |
| chr10 | 41070634 | 41072643 | 3.62073 | 4.44206 | 1.40955 | Promoter | 10 | 41069989 | 41072285 | 2297 | 2 | ENSMUST00000061796.6 | 0 | Gpr6 | ENSMUSG00000046922.6 |
| chr16 | 37914994 | 37915318 | 3.69971 | 4.51454 | 1.44339 | Distal Intergenic | 16 | 37916496 | 38007527 | 91032 | 1 | ENSMUST00000061274.7 | -1178 | Gpr156 | ENSMUSG00000046961.7 |
| chr16 | 37916338 | 37916631 | 3.63322 | 4.3724 | 1.33627 | Promoter | 16 | 37916496 | 38007527 | 91032 | 1 | ENSMUST00000061274.7 | 0 | Gpr156 | ENSMUSG00000046961.7 |
| chr16 | 37917068 | 37917192 | 3.48753 | 4.09718 | 1.14847 | Promoter | 16 | 37916496 | 38007527 | 91032 | 1 | ENSMUST00000061274.7 | 573 | Gpr156 | ENSMUSG00000046961.7 |
| chr3 | 68867851 | 68867927 | 4.17238 | 5.39175 | 1.99471 | 3' UTR | 3 | 68869586 | 68872163 | 2578 | 1 | ENSMUST00000054551.2 | -1659 | 1110032F04Rik | ENSMUSG00000046999.2 |
| chr3 | 68869008 | 68870871 | 4.11026 | 5.42387 | 2.09592 | Promoter | 3 | 68869586 | 68872163 | 2578 | 1 | ENSMUST00000054551.2 | 0 | 1110032F04Rik | ENSMUSG00000046999.2 |
| chr6 | 85483763 | 85484136 | 3.59142 | 4.29947 | 1.30334 | Exon (ENSMUST00000159062.2/ENSMUSG00000047013.10, exon 2 of 13) | 6 | 85469580 | 85502858 | 33279 | 2 | ENSMUST00000161546.1 | 18722 | Fbxo41 | ENSMUSG00000047013.10 |
| chr6 | 85484605 | 85484792 | 3.43045 | 4.25257 | 1.27848 | 5' UTR | 6 | 85469580 | 85502858 | 33279 | 2 | ENSMUST00000161546.1 | 18066 | Fbxo41 | ENSMUSG00000047013.10 |
| chr6 | 85496166 | 85496468 | 3.48993 | 4.15789 | 1.29406 | Intron (ENSMUST00000159062.2/ENSMUSG00000047013.10, intron 1 of 12) | 6 | 85469580 | 85502858 | 33279 | 2 | ENSMUST00000161546.1 | 6390 | Fbxo41 | ENSMUSG00000047013.10 |
| chr6 | 85501546 | 85501868 | 3.50696 | 4.1888 | 1.2999 | Promoter | 6 | 85469580 | 85502858 | 33279 | 2 | ENSMUST00000161546.1 | 990 | Fbxo41 | ENSMUSG00000047013.10 |
| chr6 | 85502502 | 85503926 | 3.87848 | 4.9544 | 1.77461 | Promoter | 6 | 85469574 | 85502994 | 33421 | 2 | ENSMUST00000159062.2 | 0 | Fbxo41 | ENSMUSG00000047013.10 |
| chr6 | 85504977 | 85505052 | 3.67436 | 4.43962 | 1.35237 | Distal Intergenic | 6 | 85469574 | 85502994 | 33421 | 2 | ENSMUST00000159062.2 | -1984 | Fbxo41 | ENSMUSG00000047013.10 |
| chr6 | 85505368 | 85505596 | 3.44036 | 4.00836 | 1.08622 | Distal Intergenic | 6 | 85469574 | 85502994 | 33421 | 2 | ENSMUST00000159062.2 | -2375 | Fbxo41 | ENSMUSG00000047013.10 |
| chr6 | 85507073 | 85507325 | 3.3452 | 3.8452 | 1.01733 | Distal Intergenic | 6 | 85469574 | 85502994 | 33421 | 2 | ENSMUST00000159062.2 | -4080 | Fbxo41 | ENSMUSG00000047013.10 |
| chr1 | 74926978 | 74927739 | 3.74015 | 5.02137 | 1.77771 | Exon (ENSMUST00000094844.3/ENSMUSG00000047021.9, exon 7 of 33) | 1 | 74923392 | 74935592 | 12201 | 2 | ENSMUST00000130489.1 | 7853 | Ccdc108 | ENSMUSG00000047021.9 |
| chr1 | 74929998 | 74930120 | 3.35004 | 3.85249 | 1.01733 | Intron (ENSMUST00000094844.3/ENSMUSG00000047021.9, intron 3 of 32) | 1 | 74923392 | 74935592 | 12201 | 2 | ENSMUST00000130489.1 | 5472 | Ccdc108 | ENSMUSG00000047021.9 |
| chr1 | 74931652 | 74932449 | 3.56438 | 4.24131 | 1.24397 | Exon (ENSMUST00000094844.3/ENSMUSG00000047021.9, exon 3 of 33) | 1 | 74923392 | 74935592 | 12201 | 2 | ENSMUST00000130489.1 | 3143 | Ccdc108 | ENSMUSG00000047021.9 |
| chr1 | 74932840 | 74933596 | 3.83965 | 4.75375 | 1.57004 | 5' UTR | 1 | 74923392 | 74935592 | 12201 | 2 | ENSMUST00000130489.1 | 1996 | Ccdc108 | ENSMUSG00000047021.9 |
| chr1 | 74934380 | 74934961 | 3.64852 | 4.44923 | 1.40127 | Promoter | 1 | 74923392 | 74935592 | 12201 | 2 | ENSMUST00000130489.1 | 631 | Ccdc108 | ENSMUSG00000047021.9 |
| chr1 | 74936302 | 74936579 | 4.36817 | 6.02132 | 2.54527 | Promoter | 1 | 74902071 | 74935599 | 33529 | 2 | ENSMUST00000094844.3 | -704 | Ccdc108 | ENSMUSG00000047021.9 |
| chr1 | 74937849 | 74938088 | 3.09477 | 3.79432 | 1.03782 | Distal Intergenic | 1 | 74902071 | 74935599 | 33529 | 2 | ENSMUST00000094844.3 | -2251 | Ccdc108 | ENSMUSG00000047021.9 |
| chr1 | 74941783 | 74941890 | 3.36628 | 3.87698 | 1.01733 | Distal Intergenic | 1 | 74902071 | 74935599 | 33529 | 2 | ENSMUST00000094844.3 | -6185 | Ccdc108 | ENSMUSG00000047021.9 |
| chr6 | 88235645 | 88236027 | 4.00306 | 5.09788 | 1.81543 | Distal Intergenic | 6 | 88222602 | 88225678 | 3077 | 2 | ENSMUST00000151901.1 | -9968 | 1700031F10Rik | ENSMUSG00000047046.7 |
| chr6 | 88236495 | 88236659 | 3.54706 | 4.30563 | 1.33636 | Distal Intergenic | 6 | 88222602 | 88225678 | 3077 | 2 | ENSMUST00000151901.1 | -10818 | 1700031F10Rik | ENSMUSG00000047046.7 |
| chr7 | 44423296 | 44423444 | 3.43375 | 4.04522 | 1.1805 | Distal Intergenic | 7 | 44429018 | 44461361 | 32344 | 1 | ENSMUST00000127790.2 | -5574 | Lrrc4b | ENSMUSG00000047085.9 |
| chr7 | 44427798 | 44427982 | 4.20647 | 5.46322 | 2.04637 | Distal Intergenic | 7 | 44429018 | 44461361 | 32344 | 1 | ENSMUST00000127790.2 | -1036 | Lrrc4b | ENSMUSG00000047085.9 |
| chr7 | 44429080 | 44429605 | 3.99445 | 5.06836 | 1.79611 | Promoter | 7 | 44429018 | 44461361 | 32344 | 1 | ENSMUST00000127790.2 | 63 | Lrrc4b | ENSMUSG00000047085.9 |
| chr7 | 44433564 | 44433956 | 3.59838 | 4.3087 | 1.29581 | Intron (ENSMUST00000127790.2/ENSMUSG00000047085.9, intron 1 of 2) | 7 | 44429018 | 44461361 | 32344 | 1 | ENSMUST00000127790.2 | 4547 | Lrrc4b | ENSMUSG00000047085.9 |
| chr7 | 44440324 | 44440549 | 3.52003 | 4.15582 | 1.18303 | Intron (ENSMUST00000127790.2/ENSMUSG00000047085.9, intron 1 of 2) | 7 | 44442485 | 44463351 | 20867 | 1 | ENSMUST00000058667.9 | -1936 | Lrrc4b | ENSMUSG00000047085.9 |
| chr7 | 44443375 | 44443550 | 3.42343 | 3.98255 | 1.08528 | Promoter | 7 | 44442737 | 44461407 | 18671 | 1 | ENSMUST00000156093.1 | 639 | Lrrc4b | ENSMUSG00000047085.9 |
| chr7 | 44443963 | 44444259 | 3.7243 | 4.77322 | 1.61005 | Intron (ENSMUST00000127790.2/ENSMUSG00000047085.9, intron 1 of 2) | 7 | 44442737 | 44461407 | 18671 | 1 | ENSMUST00000156093.1 | 1227 | Lrrc4b | ENSMUSG00000047085.9 |
| chr7 | 44444909 | 44445133 | 3.49202 | 4.11285 | 1.16091 | Exon (ENSMUST00000127790.2/ENSMUSG00000047085.9, exon 2 of 3) | 7 | 44442737 | 44461407 | 18671 | 1 | ENSMUST00000156093.1 | 2173 | Lrrc4b | ENSMUSG00000047085.9 |
| chr7 | 44445647 | 44446009 | 3.70487 | 4.63798 | 1.55279 | Intron (ENSMUST00000127790.2/ENSMUSG00000047085.9, intron 2 of 2) | 7 | 44442737 | 44461407 | 18671 | 1 | ENSMUST00000156093.1 | 2911 | Lrrc4b | ENSMUSG00000047085.9 |
| chr13 | 40285121 | 40285802 | 3.31052 | 3.79299 | 1.01733 | Intron (ENSMUST00000054635.7/ENSMUSG00000047094.8, intron 1 of 18) | 13 | 40001882 | 40288011 | 286130 | 2 | ENSMUST00000054635.7 | 2209 | Ofcc1 | ENSMUSG00000047094.8 |
| chr13 | 40286828 | 40288070 | 3.74602 | 4.61371 | 1.52999 | Promoter | 13 | 40001882 | 40288011 | 286130 | 2 | ENSMUST00000054635.7 | 0 | Ofcc1 | ENSMUSG00000047094.8 |
| chr16 | 93934189 | 93934522 | 3.34787 | 3.99449 | 1.10568 | Intron (ENSMUST00000050962.4/ENSMUSG00000047109.7, intron 2 of 2) | 16 | 93919807 | 93929817 | 10011 | 2 | ENSMUST00000137163.1 | -4373 | Cldn14 | ENSMUSG00000047109.7 |
| chr16 | 93934856 | 93935152 | 3.17264 | 3.95349 | 1.06448 | Intron (ENSMUST00000050962.4/ENSMUSG00000047109.7, intron 2 of 2) | 16 | 93919807 | 93929817 | 10011 | 2 | ENSMUST00000137163.1 | -5040 | Cldn14 | ENSMUSG00000047109.7 |
| chr16 | 93935592 | 93936388 | 3.87249 | 4.91233 | 1.75182 | Intron (ENSMUST00000050962.4/ENSMUSG00000047109.7, intron 2 of 2) | 16 | 93919807 | 93929817 | 10011 | 2 | ENSMUST00000137163.1 | -5776 | Cldn14 | ENSMUSG00000047109.7 |
| chr16 | 93937177 | 93937308 | 3.22649 | 3.66693 | 1.01733 | Intron (ENSMUST00000050962.4/ENSMUSG00000047109.7, intron 2 of 2) | 16 | 93919807 | 93929817 | 10011 | 2 | ENSMUST00000137163.1 | -7361 | Cldn14 | ENSMUSG00000047109.7 |
| chr16 | 93957287 | 93957454 | 3.2501 | 3.7023 | 1.01733 | Intron (ENSMUST00000050962.4/ENSMUSG00000047109.7, intron 1 of 2) | 16 | 93919807 | 93929817 | 10011 | 2 | ENSMUST00000137163.1 | -27471 | Cldn14 | ENSMUSG00000047109.7 |
| chr16 | 93958141 | 93958291 | 3.30233 | 3.78069 | 1.01733 | Intron (ENSMUST00000050962.4/ENSMUSG00000047109.7, intron 1 of 2) | 16 | 93919807 | 93929817 | 10011 | 2 | ENSMUST00000137163.1 | -28325 | Cldn14 | ENSMUSG00000047109.7 |
| chr16 | 94038921 | 94039192 | 3.52759 | 4.37509 | 1.38219 | Distal Intergenic | 16 | 93919032 | 94008837 | 89806 | 2 | ENSMUST00000050962.4 | -30085 | Cldn14 | ENSMUSG00000047109.7 |
| chr16 | 94039665 | 94039888 | 3.43492 | 4.02065 | 1.15971 | Distal Intergenic | 16 | 93919032 | 94008837 | 89806 | 2 | ENSMUST00000050962.4 | -30829 | Cldn14 | ENSMUSG00000047109.7 |
| chr16 | 94040893 | 94040987 | 3.43265 | 4.01365 | 1.14641 | Distal Intergenic | 16 | 93919032 | 94008837 | 89806 | 2 | ENSMUST00000050962.4 | -32057 | Cldn14 | ENSMUSG00000047109.7 |
| chr16 | 94041536 | 94041775 | 3.61917 | 4.35302 | 1.35294 | Distal Intergenic | 16 | 93919032 | 94008837 | 89806 | 2 | ENSMUST00000050962.4 | -32700 | Cldn14 | ENSMUSG00000047109.7 |
| chr10 | 87057595 | 87058805 | 4.01194 | 5.30253 | 2.02003 | Promoter | 10 | 87058045 | 87230589 | 172545 | 1 | ENSMUST00000189456.1 | 0 | 1700113H08Rik | ENSMUSG00000047129.6 |
| chr10 | 43578433 | 43579446 | 3.84102 | 4.84983 | 1.77028 | Promoter | 10 | 43579169 | 43584262 | 5094 | 1 | ENSMUST00000058714.8 | 0 | Cd24a | ENSMUSG00000047139.8 |
| chr10 | 43579958 | 43580111 | 4.02759 | 5.22345 | 1.92549 | Promoter | 10 | 43579169 | 43584262 | 5094 | 1 | ENSMUST00000058714.8 | 790 | Cd24a | ENSMUSG00000047139.8 |
| chr10 | 43580927 | 43581048 | 3.55418 | 4.4235 | 1.38177 | Intron (ENSMUST00000058714.8/ENSMUSG00000047139.8, intron 1 of 1) | 10 | 43579169 | 43584262 | 5094 | 1 | ENSMUST00000058714.8 | 1759 | Cd24a | ENSMUSG00000047139.8 |
| chr4 | 109977552 | 109978703 | 4.01155 | 5.43656 | 2.10772 | Promoter | 4 | 109978053 | 109983687 | 5635 | 1 | ENSMUST00000061187.3 | 0 | Dmrta2 | ENSMUSG00000047143.3 |
| chr4 | 109979084 | 109980417 | 3.99573 | 5.23833 | 1.96528 | 5' UTR | 4 | 109978053 | 109983687 | 5635 | 1 | ENSMUST00000061187.3 | 1032 | Dmrta2 | ENSMUSG00000047143.3 |
| chr4 | 109980783 | 109981173 | 3.61944 | 4.40857 | 1.4104 | Intron (ENSMUST00000061187.3/ENSMUSG00000047143.3, intron 2 of 2) | 4 | 109978053 | 109983687 | 5635 | 1 | ENSMUST00000061187.3 | 2731 | Dmrta2 | ENSMUSG00000047143.3 |
| chr4 | 109981635 | 109984519 | 4.29677 | 5.84002 | 2.3926 | 3' UTR | 4 | 109978053 | 109983687 | 5635 | 1 | ENSMUST00000061187.3 | 3583 | Dmrta2 | ENSMUSG00000047143.3 |
| chr4 | 109984954 | 109985587 | 4.37568 | 5.99914 | 2.49874 | Distal Intergenic | 4 | 109978053 | 109983687 | 5635 | 1 | ENSMUST00000061187.3 | 6902 | Dmrta2 | ENSMUSG00000047143.3 |
| chr4 | 109985947 | 109986553 | 3.87621 | 5.02359 | 1.8088 | Distal Intergenic | 4 | 109978053 | 109983687 | 5635 | 1 | ENSMUST00000061187.3 | 7895 | Dmrta2 | ENSMUSG00000047143.3 |
| chr4 | 109987732 | 109988054 | 3.77031 | 4.69782 | 1.62533 | Distal Intergenic | 4 | 109978053 | 109983687 | 5635 | 1 | ENSMUST00000061187.3 | 9680 | Dmrta2 | ENSMUSG00000047143.3 |
| chr4 | 109989285 | 109989632 | 4.10577 | 5.47627 | 2.10556 | Distal Intergenic | 4 | 109978053 | 109983687 | 5635 | 1 | ENSMUST00000061187.3 | 11233 | Dmrta2 | ENSMUSG00000047143.3 |
| chr4 | 109989977 | 109991736 | 3.70469 | 4.66191 | 1.55381 | Distal Intergenic | 4 | 109978053 | 109983687 | 5635 | 1 | ENSMUST00000061187.3 | 11925 | Dmrta2 | ENSMUSG00000047143.3 |
| chr4 | 109992081 | 109992442 | 4.16487 | 5.63105 | 2.20282 | Distal Intergenic | 4 | 109978053 | 109983687 | 5635 | 1 | ENSMUST00000061187.3 | 14029 | Dmrta2 | ENSMUSG00000047143.3 |
| chr4 | 110051170 | 110052018 | 3.58563 | 4.57904 | 1.52042 | Distal Intergenic | 4 | 109978053 | 109983687 | 5635 | 1 | ENSMUST00000061187.3 | 73118 | Dmrta2 | ENSMUSG00000047143.3 |
| chr4 | 115133718 | 115134593 | 3.92111 | 5.04022 | 1.83113 | Promoter | 4 | 115106323 | 115134281 | 27959 | 2 | ENSMUST00000106545.2 | 0 | Cyp4x1 | ENSMUSG00000047155.8 |
| chr4 | 115138806 | 115139062 | 3.81084 | 4.81225 | 1.65414 | Intron (ENSMUST00000121884.1/ENSMUSG00000081823.1, intron 11 of 11) | 4 | 115106323 | 115134281 | 27959 | 2 | ENSMUST00000106545.2 | -4526 | Cyp4x1 | ENSMUSG00000047155.8 |
| chr18 | 15759998 | 15760621 | 3.83849 | 4.7618 | 1.59466 | Promoter | 18 | 15451924 | 15760157 | 308234 | 2 | ENSMUST00000130553.1 | 0 | Chst9 | ENSMUSG00000047161.7 |
| chr8 | 46290849 | 46291553 | 4.02234 | 5.12919 | 1.84703 | Downstream (<1kb) | 8 | 46292050 | 46294671 | 2622 | 2 | ENSMUST00000058636.7 | 3118 | Helt | ENSMUSG00000047171.7 |
| chr8 | 46292477 | 46293435 | 3.68739 | 4.48871 | 1.40342 | Exon (ENSMUST00000058636.7/ENSMUSG00000047171.7, exon 3 of 4) | 8 | 46292050 | 46294671 | 2622 | 2 | ENSMUST00000058636.7 | 1236 | Helt | ENSMUSG00000047171.7 |
| chr8 | 46293900 | 46294583 | 3.60247 | 4.32194 | 1.3089 | Promoter | 8 | 46292050 | 46294671 | 2622 | 2 | ENSMUST00000058636.7 | 88 | Helt | ENSMUSG00000047171.7 |
| chr8 | 46296186 | 46296332 | 3.52903 | 4.16928 | 1.18162 | Distal Intergenic | 8 | 46292050 | 46294671 | 2622 | 2 | ENSMUST00000058636.7 | -1516 | Helt | ENSMUSG00000047171.7 |
| chr8 | 46297992 | 46298122 | 3.55589 | 4.21985 | 1.21755 | Distal Intergenic | 8 | 46292050 | 46294671 | 2622 | 2 | ENSMUST00000058636.7 | -3322 | Helt | ENSMUSG00000047171.7 |
| chr1 | 36269465 | 36269655 | 3.64206 | 4.38089 | 1.32701 | Exon (ENSMUST00000056946.7/ENSMUSG00000047180.7, exon 2 of 4) | 1 | 36264597 | 36273435 | 8839 | 2 | ENSMUST00000056946.7 | 3780 | Neurl3 | ENSMUSG00000047180.7 |
| chr11 | 98978324 | 98978730 | 3.50629 | 4.14533 | 1.22022 | Distal Intergenic | 11 | 98982180 | 98983016 | 837 | 2 | ENSMUST00000062931.1 | 4286 | Gjd3 | ENSMUSG00000047197.1 |
| chr11 | 98981897 | 98982094 | 3.8018 | 4.89722 | 1.7262 | Promoter | 11 | 98982180 | 98983016 | 837 | 2 | ENSMUST00000062931.1 | 922 | Gjd3 | ENSMUSG00000047197.1 |
| chr11 | 98982408 | 98983016 | 3.70924 | 4.60984 | 1.54242 | Promoter | 11 | 98982180 | 98983016 | 837 | 2 | ENSMUST00000062931.1 | 0 | Gjd3 | ENSMUSG00000047197.1 |
| chr18 | 66833464 | 66833707 | 3.46781 | 4.16178 | 1.26406 | Distal Intergenic | 18 | 66857715 | 66860472 | 2758 | 2 | ENSMUST00000057942.2 | 26765 | Mc4r | ENSMUSG00000047259.2 |
| chr18 | 66859967 | 66860194 | 3.62841 | 4.51991 | 1.44964 | Promoter | 18 | 66857715 | 66860472 | 2758 | 2 | ENSMUST00000057942.2 | 278 | Mc4r | ENSMUSG00000047259.2 |
| chr18 | 66860599 | 66861348 | 3.5866 | 4.37707 | 1.3861 | Promoter | 18 | 66857715 | 66860472 | 2758 | 2 | ENSMUST00000057942.2 | -128 | Mc4r | ENSMUSG00000047259.2 |
| chr16 | 42339228 | 42340627 | 4.01091 | 5.30751 | 2.05882 | Promoter | 16 | 42248442 | 42340651 | 92210 | 2 | ENSMUST00000102817.4 | 24 | Gap43 | ENSMUSG00000047261.9 |
| chr19 | 27317193 | 27317454 | 3.35434 | 3.86664 | 1.04403 | Distal Intergenic | 19 | 27322588 | 27337179 | 14592 | 1 | ENSMUST00000056708.3 | -5134 | Kcnv2 | ENSMUSG00000047298.3 |
| chr13 | 47649708 | 47649946 | 3.43198 | 4.12184 | 1.20884 | Distal Intergenic | 13 | 47960330 | 48018608 | 58279 | 2 | ENSMUST00000061899.5 | 368662 | 4931429P17Rik | ENSMUSG00000047324.5 |
| chr13 | 47856262 | 47856468 | 4.13188 | 5.61071 | 2.2542 | Distal Intergenic | 13 | 47960330 | 48018608 | 58279 | 2 | ENSMUST00000061899.5 | 162140 | 4931429P17Rik | ENSMUSG00000047324.5 |
| chr13 | 47940784 | 47941248 | 3.5194 | 4.28689 | 1.34524 | Distal Intergenic | 13 | 47960330 | 48018608 | 58279 | 2 | ENSMUST00000061899.5 | 77360 | 4931429P17Rik | ENSMUSG00000047324.5 |
| chr13 | 47948042 | 47948358 | 3.32111 | 3.80892 | 1.01733 | Distal Intergenic | 13 | 47960330 | 48018608 | 58279 | 2 | ENSMUST00000061899.5 | 70250 | 4931429P17Rik | ENSMUSG00000047324.5 |
| chr13 | 47951434 | 47951517 | 3.6157 | 4.41472 | 1.41422 | Distal Intergenic | 13 | 47960330 | 48018608 | 58279 | 2 | ENSMUST00000061899.5 | 67091 | 4931429P17Rik | ENSMUSG00000047324.5 |
| chr13 | 47954120 | 47954371 | 3.52377 | 4.19394 | 1.2856 | Distal Intergenic | 13 | 47960330 | 48018608 | 58279 | 2 | ENSMUST00000061899.5 | 64237 | 4931429P17Rik | ENSMUSG00000047324.5 |
| chr13 | 47954763 | 47955340 | 3.45583 | 4.05466 | 1.16766 | Distal Intergenic | 13 | 47960330 | 48018608 | 58279 | 2 | ENSMUST00000061899.5 | 63268 | 4931429P17Rik | ENSMUSG00000047324.5 |
| chr13 | 47956617 | 47957732 | 3.39812 | 3.94104 | 1.07396 | Downstream (2-3kb) | 13 | 47960330 | 48018608 | 58279 | 2 | ENSMUST00000061899.5 | 60876 | 4931429P17Rik | ENSMUSG00000047324.5 |
| chr13 | 47964891 | 47965256 | 3.68535 | 4.58535 | 1.52743 | Intron (ENSMUST00000061899.5/ENSMUSG00000047324.5, intron 7 of 7) | 13 | 47960330 | 48018608 | 58279 | 2 | ENSMUST00000061899.5 | 53352 | 4931429P17Rik | ENSMUSG00000047324.5 |
| chr13 | 47967870 | 47968472 | 3.41868 | 4.01954 | 1.21591 | Intron (ENSMUST00000061899.5/ENSMUSG00000047324.5, intron 5 of 7) | 13 | 47960330 | 48018608 | 58279 | 2 | ENSMUST00000061899.5 | 50136 | 4931429P17Rik | ENSMUSG00000047324.5 |
| chr13 | 47970258 | 47970565 | 3.45309 | 4.18351 | 1.23099 | Exon (ENSMUST00000061899.5/ENSMUSG00000047324.5, exon 4 of 8) | 13 | 47960330 | 48018608 | 58279 | 2 | ENSMUST00000061899.5 | 48043 | 4931429P17Rik | ENSMUSG00000047324.5 |
| chr13 | 47977752 | 47978103 | 3.37894 | 3.9308 | 1.13912 | Intron (ENSMUST00000061899.5/ENSMUSG00000047324.5, intron 2 of 7) | 13 | 47960330 | 48018608 | 58279 | 2 | ENSMUST00000061899.5 | 40505 | 4931429P17Rik | ENSMUSG00000047324.5 |
| chr13 | 47978602 | 47978677 | 3.53332 | 4.18361 | 1.21014 | Intron (ENSMUST00000061899.5/ENSMUSG00000047324.5, intron 2 of 7) | 13 | 47960330 | 48018608 | 58279 | 2 | ENSMUST00000061899.5 | 39931 | 4931429P17Rik | ENSMUSG00000047324.5 |
| chr13 | 47979146 | 47979588 | 3.96542 | 5.12382 | 1.91137 | Intron (ENSMUST00000061899.5/ENSMUSG00000047324.5, intron 2 of 7) | 13 | 47960330 | 48018608 | 58279 | 2 | ENSMUST00000061899.5 | 39020 | 4931429P17Rik | ENSMUSG00000047324.5 |
| chr13 | 47981668 | 47982120 | 3.32662 | 3.81721 | 1.01733 | Intron (ENSMUST00000061899.5/ENSMUSG00000047324.5, intron 2 of 7) | 13 | 47960330 | 48018608 | 58279 | 2 | ENSMUST00000061899.5 | 36488 | 4931429P17Rik | ENSMUSG00000047324.5 |
| chr13 | 47985211 | 47985667 | 3.41251 | 4.11906 | 1.22597 | Intron (ENSMUST00000061899.5/ENSMUSG00000047324.5, intron 2 of 7) | 13 | 47960330 | 48018608 | 58279 | 2 | ENSMUST00000061899.5 | 32941 | 4931429P17Rik | ENSMUSG00000047324.5 |
| chr13 | 47997130 | 47997438 | 3.26379 | 3.72283 | 1.01733 | Intron (ENSMUST00000061899.5/ENSMUSG00000047324.5, intron 2 of 7) | 13 | 47960330 | 48018608 | 58279 | 2 | ENSMUST00000061899.5 | 21170 | 4931429P17Rik | ENSMUSG00000047324.5 |
| chr13 | 47998492 | 47998970 | 3.60521 | 4.31704 | 1.28546 | Intron (ENSMUST00000061899.5/ENSMUSG00000047324.5, intron 2 of 7) | 13 | 47960330 | 48018608 | 58279 | 2 | ENSMUST00000061899.5 | 19638 | 4931429P17Rik | ENSMUSG00000047324.5 |
| chr13 | 48007248 | 48007329 | 4.60155 | 6.23406 | 2.57488 | Intron (ENSMUST00000061899.5/ENSMUSG00000047324.5, intron 2 of 7) | 13 | 47960330 | 48018608 | 58279 | 2 | ENSMUST00000061899.5 | 11279 | 4931429P17Rik | ENSMUSG00000047324.5 |
| chr13 | 48008222 | 48008398 | 3.25061 | 3.71584 | 1.03398 | Intron (ENSMUST00000061899.5/ENSMUSG00000047324.5, intron 2 of 7) | 13 | 47960330 | 48018608 | 58279 | 2 | ENSMUST00000061899.5 | 10210 | 4931429P17Rik | ENSMUSG00000047324.5 |
| chr13 | 48013805 | 48013937 | 3.60859 | 4.38919 | 1.44098 | Intron (ENSMUST00000061899.5/ENSMUSG00000047324.5, intron 2 of 7) | 13 | 47960330 | 48018608 | 58279 | 2 | ENSMUST00000061899.5 | 4671 | 4931429P17Rik | ENSMUSG00000047324.5 |
| chr13 | 48044093 | 48044389 | 3.97148 | 5.01473 | 1.76125 | Exon (ENSMUST00000180921.1/ENSMUSG00000097622.1, exon 3 of 3) | 13 | 47960330 | 48018608 | 58279 | 2 | ENSMUST00000061899.5 | -25486 | 4931429P17Rik | ENSMUSG00000047324.5 |
| chr13 | 48077968 | 48078540 | 3.62129 | 4.37311 | 1.37291 | Intron (ENSMUST00000180921.1/ENSMUSG00000097622.1, intron 1 of 2) | 13 | 47960330 | 48018608 | 58279 | 2 | ENSMUST00000061899.5 | -59361 | 4931429P17Rik | ENSMUSG00000047324.5 |
| chr13 | 48101765 | 48102159 | 3.52165 | 4.19313 | 1.28338 | Intron (ENSMUST00000180921.1/ENSMUSG00000097622.1, intron 1 of 2) | 13 | 47960330 | 48018608 | 58279 | 2 | ENSMUST00000061899.5 | -83158 | 4931429P17Rik | ENSMUSG00000047324.5 |
| chr13 | 48108177 | 48108796 | 3.36908 | 3.88121 | 1.01733 | Intron (ENSMUST00000180921.1/ENSMUSG00000097622.1, intron 1 of 2) | 13 | 47960330 | 48018608 | 58279 | 2 | ENSMUST00000061899.5 | -89570 | 4931429P17Rik | ENSMUSG00000047324.5 |
| chr1 | 181743014 | 181743807 | 3.00197 | 6.95602 | 3.37644 | Promoter | 1 | 181742740 | 181747214 | 4475 | 1 | ENSMUST00000160793.1 | 275 | Dnah14 | ENSMUSG00000047369.9 |
| chr1 | 192834424 | 192834648 | 3.78167 | 4.90382 | 1.72417 | Promoter | 1 | 192833169 | 192834719 | 1551 | 2 | ENSMUST00000057543.2 | 71 | A730013G03Rik | ENSMUSG00000047384.2 |
| chr15 | 89377721 | 89378886 | 3.90532 | 4.92038 | 1.7349 | Promoter | 15 | 89377450 | 89379254 | 1805 | 2 | ENSMUST00000049968.7 | 368 | Odf3b | ENSMUSG00000047394.7 |
| chr12 | 100896813 | 100897193 | 3.69213 | 4.48192 | 1.40568 | 3' UTR | 12 | 100879314 | 100899498 | 20185 | 2 | ENSMUST00000135684.1 | 2305 | Gpr68 | ENSMUSG00000047415.6 |
| chr12 | 100897901 | 100899683 | 4.20085 | 5.56288 | 2.22273 | Promoter | 12 | 100876685 | 100899524 | 22840 | 2 | ENSMUST00000053668.4 | 0 | Gpr68 | ENSMUSG00000047415.6 |
| chr12 | 100908823 | 100909054 | 3.53145 | 4.17578 | 1.19167 | Promoter | 12 | 100887684 | 100908198 | 20515 | 2 | ENSMUST00000124459.1 | -626 | Gpr68 | ENSMUSG00000047415.6 |
| chr12 | 78350926 | 78351139 | 2.88904 | 11.37897 | 7.04337 | Intron (ENSMUST00000052472.4/ENSMUSG00000047454.6, intron 2 of 23) | 12 | 78226655 | 78684767 | 458113 | 1 | ENSMUST00000110388.3 | 124272 | Gphn | ENSMUSG00000047454.6 |
| chr11 | 89447671 | 89447797 | 3.41061 | 3.96495 | 1.09197 | Intron (ENSMUST00000128717.2/ENSMUSG00000047773.7, intron 8 of 15) | 11 | 89390223 | 89418948 | 28726 | 2 | ENSMUST00000050983.1 | -28724 | 4932411E22Rik | ENSMUSG00000047490.1 |
| chr10 | 127350971 | 127351420 | 3.50626 | 4.35098 | 1.3885 | Promoter | 10 | 127349402 | 127351772 | 2371 | 2 | ENSMUST00000059718.4 | 352 | Inhbe | ENSMUSG00000047492.4 |
| chr10 | 127351819 | 127352002 | 3.48415 | 4.13259 | 1.22603 | Promoter | 10 | 127349402 | 127351772 | 2371 | 2 | ENSMUST00000059718.4 | -48 | Inhbe | ENSMUSG00000047492.4 |
| chr8 | 14094969 | 14095330 | 3.46966 | 4.17941 | 1.29697 | Promoter | 8 | 14095865 | 14847108 | 751244 | 1 | ENSMUST00000152652.2 | -535 | Dlgap2 | ENSMUSG00000047495.10 |
| chr8 | 14095703 | 14096639 | 4.18305 | 5.61426 | 2.31925 | Promoter | 8 | 14095865 | 14847108 | 751244 | 1 | ENSMUST00000152652.2 | 0 | Dlgap2 | ENSMUSG00000047495.10 |
| chr8 | 14096953 | 14097221 | 4.12426 | 5.39609 | 2.07457 | Intron (ENSMUST00000152652.2/ENSMUSG00000047495.10, intron 1 of 14) | 8 | 14095875 | 14847680 | 751806 | 1 | ENSMUST00000133298.2 | 1079 | Dlgap2 | ENSMUSG00000047495.10 |
| chr8 | 14098019 | 14098873 | 3.42284 | 4.03473 | 1.16097 | Intron (ENSMUST00000152652.2/ENSMUSG00000047495.10, intron 1 of 14) | 8 | 14095875 | 14847680 | 751806 | 1 | ENSMUST00000133298.2 | 2145 | Dlgap2 | ENSMUSG00000047495.10 |
| chr8 | 14099269 | 14100122 | 3.63972 | 4.42932 | 1.39259 | Intron (ENSMUST00000152652.2/ENSMUSG00000047495.10, intron 1 of 14) | 8 | 14095875 | 14847680 | 751806 | 1 | ENSMUST00000133298.2 | 3395 | Dlgap2 | ENSMUSG00000047495.10 |
| chr8 | 14594709 | 14594784 | 4.24292 | 5.5584 | 2.17066 | Intron (ENSMUST00000152652.2/ENSMUSG00000047495.10, intron 3 of 14) | 8 | 14742753 | 14760902 | 18150 | 1 | ENSMUST00000137130.1 | -147969 | Dlgap2 | ENSMUSG00000047495.10 |
| chr5 | 134945558 | 134945651 | 4.9181 | 6.98017 | 3.20726 | 3' UTR | 5 | 134945126 | 134946934 | 1809 | 2 | ENSMUST00000051401.2 | 1283 | Cldn4 | ENSMUSG00000047501.2 |
| chr5 | 134946020 | 134946641 | 3.55552 | 4.2429 | 1.24066 | Promoter | 5 | 134945126 | 134946934 | 1809 | 2 | ENSMUST00000051401.2 | 293 | Cldn4 | ENSMUSG00000047501.2 |
| chr5 | 134947072 | 134947351 | 3.80402 | 4.78286 | 1.61629 | Promoter | 5 | 134945126 | 134946934 | 1809 | 2 | ENSMUST00000051401.2 | -139 | Cldn4 | ENSMUSG00000047501.2 |
| chr17 | 25253533 | 25253608 | 4.32277 | 5.67732 | 2.1762 | Promoter | 17 | 25243355 | 25253947 | 10593 | 2 | ENSMUST00000182825.2 | 339 | Baiap3 | ENSMUSG00000047507.7 |
| chr17 | 25254149 | 25254301 | 3.73275 | 4.55921 | 1.47331 | Promoter | 17 | 25243355 | 25253947 | 10593 | 2 | ENSMUST00000182825.2 | -203 | Baiap3 | ENSMUSG00000047507.7 |
| chr17 | 25255627 | 25256600 | 3.62204 | 4.41258 | 1.37967 | Promoter | 17 | 25242659 | 25256364 | 13706 | 2 | ENSMUST00000182056.2 | 0 | Baiap3 | ENSMUSG00000047507.7 |
| chr3 | 154545556 | 154546044 | 3.7141 | 4.5392 | 1.4918 | Distal Intergenic | 3 | 154579845 | 154593755 | 13911 | 2 | ENSMUST00000148680.1 | 47711 | Tyw3 | ENSMUSG00000047583.4 |
| chr3 | 154547012 | 154547183 | 3.4533 | 4.22392 | 1.27464 | Distal Intergenic | 3 | 154579845 | 154593755 | 13911 | 2 | ENSMUST00000148680.1 | 46572 | Tyw3 | ENSMUSG00000047583.4 |
| chr3 | 154550175 | 154550395 | 3.39165 | 4.12139 | 1.2125 | Distal Intergenic | 3 | 154579845 | 154593755 | 13911 | 2 | ENSMUST00000148680.1 | 43360 | Tyw3 | ENSMUSG00000047583.4 |
| chr3 | 154568039 | 154568200 | 3.31355 | 3.79756 | 1.01733 | Distal Intergenic | 3 | 154579845 | 154593755 | 13911 | 2 | ENSMUST00000148680.1 | 25555 | Tyw3 | ENSMUSG00000047583.4 |
| chr3 | 154569267 | 154569490 | 3.61521 | 4.37429 | 1.4179 | Distal Intergenic | 3 | 154579845 | 154593755 | 13911 | 2 | ENSMUST00000148680.1 | 24265 | Tyw3 | ENSMUSG00000047583.4 |
| chr3 | 154570221 | 154570461 | 3.66488 | 4.45215 | 1.44494 | Distal Intergenic | 3 | 154579845 | 154593755 | 13911 | 2 | ENSMUST00000148680.1 | 23294 | Tyw3 | ENSMUSG00000047583.4 |
| chr3 | 154571293 | 154571595 | 3.53268 | 4.39176 | 1.35926 | Distal Intergenic | 3 | 154579845 | 154593755 | 13911 | 2 | ENSMUST00000148680.1 | 22160 | Tyw3 | ENSMUSG00000047583.4 |
| chr3 | 154572855 | 154573264 | 3.37941 | 3.92899 | 1.12974 | Distal Intergenic | 3 | 154579845 | 154593755 | 13911 | 2 | ENSMUST00000148680.1 | 20491 | Tyw3 | ENSMUSG00000047583.4 |
| chr7 | 28544224 | 28544451 | 3.76562 | 4.81319 | 1.64001 | Exon (ENSMUST00000057974.3/ENSMUSG00000047586.3, exon 5 of 6) | 7 | 28543596 | 28547254 | 3659 | 2 | ENSMUST00000057974.3 | 2803 | Nccrp1 | ENSMUSG00000047586.3 |
| chr7 | 28546159 | 28547000 | 4.33264 | 5.86437 | 2.44657 | Promoter | 7 | 28543596 | 28547254 | 3659 | 2 | ENSMUST00000057974.3 | 254 | Nccrp1 | ENSMUSG00000047586.3 |
| chr19 | 41841550 | 41841672 | 3.35893 | 3.86591 | 1.01733 | Distal Intergenic | 19 | 41845978 | 41848132 | 2155 | 2 | ENSMUST00000059231.2 | 6460 | Frat2 | ENSMUSG00000047604.2 |
| chr9 | 89738441 | 89738527 | 3.93888 | 4.94845 | 1.69589 | Promoter | 9 | 89728249 | 89738475 | 10227 | 2 | ENSMUST00000060700.1 | 0 | Ankrd34c | ENSMUSG00000047606.2 |
| chr15 | 101438501 | 101439098 | 3.84867 | 5.04534 | 1.80619 | Promoter | 15 | 101431490 | 101438804 | 7315 | 2 | ENSMUST00000081945.3 | 0 | Krt83 | ENSMUSG00000047641.7 |
| chr15 | 101440891 | 101441100 | 3.77924 | 4.84305 | 1.69134 | Distal Intergenic | 15 | 101431490 | 101438804 | 7315 | 2 | ENSMUST00000081945.3 | -2088 | Krt83 | ENSMUSG00000047641.7 |
| chr10 | 121641610 | 121642634 | 3.78834 | 4.72818 | 1.64167 | Promoter | 10 | 121641701 | 121692892 | 51192 | 1 | ENSMUST00000140802.2 | 0 | D930020B18Rik | ENSMUSG00000047642.9 |
| chr10 | 121643425 | 121644077 | 3.63149 | 4.41162 | 1.4277 | Intron (ENSMUST00000142501.2/ENSMUSG00000047642.9, intron 1 of 6) | 10 | 121641872 | 121692621 | 50750 | 1 | ENSMUST00000132744.1 | 1554 | D930020B18Rik | ENSMUSG00000047642.9 |
| chr5 | 77581104 | 77581473 | 3.99683 | 5.05472 | 1.76069 | Distal Intergenic | 5 | 77511194 | 77511748 | 555 | 2 | ENSMUST00000183185.1 | -69357 | Pea15b | ENSMUSG00000047652.4 |
| chr3 | 149845099 | 149845357 | 3.55116 | 4.27748 | 1.34869 | Distal Intergenic | 3 | 150072590 | 150073620 | 1031 | 2 | ENSMUST00000057740.5 | 228263 | Rpsa-ps10 | ENSMUSG00000047676.5 |
| chr7 | 28066437 | 28066748 | 3.41466 | 3.96817 | 1.08117 | Distal Intergenic | 7 | 28071236 | 28120860 | 49625 | 1 | ENSMUST00000138392.2 | -4488 | Fcgbp | ENSMUSG00000047730.12 |
| chr7 | 139943517 | 139945026 | 4.42503 | 6.06171 | 2.58853 | Promoter | 7 | 139943789 | 139945112 | 1324 | 1 | ENSMUST00000168457.2 | 0 | Utf1 | ENSMUSG00000047751.9 |
| chr7 | 139945944 | 139946026 | 4.60035 | 6.28364 | 2.67716 | Distal Intergenic | 7 | 139943789 | 139945112 | 1324 | 1 | ENSMUST00000168457.2 | 2156 | Utf1 | ENSMUSG00000047751.9 |
| chr11 | 89545226 | 89545494 | 3.38688 | 4.03213 | 1.19876 | Intron (ENSMUST00000128717.2/ENSMUSG00000047773.7, intron 1 of 15) | 11 | 89421086 | 89538556 | 117471 | 2 | ENSMUST00000169201.1 | -6671 | Ankfn1 | ENSMUSG00000047773.7 |
| chr11 | 89624248 | 89624547 | 3.38193 | 3.9006 | 1.01733 | Intron (ENSMUST00000128717.2/ENSMUSG00000047773.7, intron 1 of 15) | 11 | 89420987 | 89639631 | 218645 | 2 | ENSMUST00000128717.2 | 15084 | Ankfn1 | ENSMUSG00000047773.7 |
| chr19 | 6849946 | 6850035 | 3.37022 | 3.9358 | 1.17108 | Exon (ENSMUST00000113440.1/ENSMUSG00000047810.8, exon 17 of 27) | 19 | 6844623 | 6858211 | 13589 | 2 | ENSMUST00000113440.1 | 8176 | Ccdc88b | ENSMUSG00000047810.8 |
| chr3 | 84596076 | 84596354 | 3.67307 | 4.67077 | 1.54996 | 3' UTR | 3 | 84593574 | 84597032 | 3459 | 1 | ENSMUST00000062623.3 | 2503 | Tigd4 | ENSMUSG00000047819.3 |
| chr11 | 62821944 | 62822157 | 3.72558 | 4.5859 | 1.53835 | Intron (ENSMUST00000055006.6/ENSMUSG00000047821.11, intron 1 of 5) | 11 | 62820469 | 62842936 | 22468 | 1 | ENSMUST00000108703.1 | 1476 | Trim16 | ENSMUSG00000047821.11 |
| chr12 | 112301551 | 112301771 | 3.73821 | 4.83793 | 1.66092 | Distal Intergenic | 12 | 112413915 | 112423212 | 9298 | 2 | ENSMUST00000050754.1 | 121441 | A730018C14Rik | ENSMUSG00000047828.3 |
| chr12 | 112309378 | 112309458 | 3.74428 | 4.56561 | 1.42662 | Distal Intergenic | 12 | 112413915 | 112423212 | 9298 | 2 | ENSMUST00000050754.1 | 113754 | A730018C14Rik | ENSMUSG00000047828.3 |
| chr12 | 112328846 | 112328921 | 3.73417 | 4.6147 | 1.5645 | Distal Intergenic | 12 | 112413915 | 112423212 | 9298 | 2 | ENSMUST00000050754.1 | 94291 | A730018C14Rik | ENSMUSG00000047828.3 |
| chr12 | 112329535 | 112329834 | 3.97031 | 5.16498 | 1.89303 | Distal Intergenic | 12 | 112413915 | 112423212 | 9298 | 2 | ENSMUST00000050754.1 | 93378 | A730018C14Rik | ENSMUSG00000047828.3 |
| chr12 | 112337547 | 112337810 | 3.84582 | 4.80903 | 1.68552 | Distal Intergenic | 12 | 112413915 | 112423212 | 9298 | 2 | ENSMUST00000050754.1 | 85402 | A730018C14Rik | ENSMUSG00000047828.3 |
| chr12 | 112405960 | 112406042 | 3.78371 | 4.66001 | 1.53214 | Distal Intergenic | 12 | 112413915 | 112423212 | 9298 | 2 | ENSMUST00000050754.1 | 17170 | A730018C14Rik | ENSMUSG00000047828.3 |
| chr13 | 52261291 | 52261643 | 3.3958 | 3.93158 | 1.05293 | Distal Intergenic | 13 | 52504375 | 52530836 | 26462 | 2 | ENSMUST00000057442.6 | 269193 | Diras2 | ENSMUSG00000047842.6 |
| chr13 | 52460654 | 52460767 | 3.30311 | 3.7914 | 1.0507 | Distal Intergenic | 13 | 52504375 | 52530836 | 26462 | 2 | ENSMUST00000057442.6 | 70069 | Diras2 | ENSMUSG00000047842.6 |

| | | | | | | | | | | | | | | | |
|---|---|---|---|---|---|---|---|---|---|---|---|---|---|---|---|
| chr13 | 52529720 | 52529795 | 3.84749 | 4.7752 | 1.60584 | Intron (ENSMUST00000057442.6/ENSMUSG00000047842.6, intron 1 of 1) | 13 | 52504375 | 52530836 | 26462 | 2 | ENSMUST00000057442.6 | 1041 | Diras2 | ENSMUSG00000047842.6 |
| chr13 | 52530195 | 52530903 | 3.66716 | 4.44983 | 1.41491 | Promoter | 13 | 52504375 | 52530836 | 26462 | 2 | ENSMUST00000057442.6 | 0 | Diras2 | ENSMUSG00000047842.6 |
| chr15 | 83251406 | 83251764 | 3.54101 | 4.45519 | 1.44628 | Promoter | 15 | 83226726 | 83251731 | 25006 | 2 | ENSMUST00000164614.1 | 0 | A4galt | ENSMUSG00000047878.7 |
| chr9 | 86984872 | 86985494 | 3.44713 | 4.05665 | 1.20339 | Distal Intergenic | 9 | 87015537 | 87019917 | 4381 | 1 | ENSMUST00000058846.5 | -30043 | Ripply2 | ENSMUSG00000047897.5 |
| chr9 | 87014876 | 87016080 | 3.80514 | 4.71503 | 1.55771 | Promoter | 9 | 87015537 | 87019917 | 4381 | 1 | ENSMUST00000058846.5 | 0 | Ripply2 | ENSMUSG00000047897.5 |
| chr9 | 87017279 | 87017661 | 3.58506 | 4.27584 | 1.25404 | Intron (ENSMUST00000058846.5/ENSMUSG00000047897.5, intron 3 of 3) | 9 | 87015636 | 87019916 | 4281 | 1 | ENSMUST00000188187.1 | 1644 | Ripply2 | ENSMUSG00000047897.5 |
| chr9 | 114498994 | 114499469 | 3.46168 | 4.06815 | 1.1749 | Distal Intergenic | 9 | 114490316 | 114496544 | 6229 | 2 | ENSMUST00000054414.3 | -2451 | Ccr4 | ENSMUSG00000047898.6 |
| chr11 | 113619412 | 113619991 | 3.93454 | 4.99008 | 1.75502 | Promoter | 11 | 113619476 | 113625895 | 6420 | 1 | ENSMUST00000106630.1 | 0 | Sstr2 | ENSMUSG00000047904.6 |
| chr11 | 113620432 | 113620838 | 3.44618 | 4.04436 | 1.17463 | Promoter | 11 | 113619476 | 113625895 | 6420 | 1 | ENSMUST00000106630.1 | 957 | Sstr2 | ENSMUSG00000047904.6 |
| chr6 | 67230209 | 67230414 | 3.87092 | 4.81675 | 1.61142 | Distal Intergenic | 6 | 67205619 | 67206083 | 465 | 1 | ENSMUST00000120174.1 | 24591 | Gm8566 | ENSMUSG00000047905.10 |
| chr8 | 12379651 | 12379987 | 3.71556 | 4.61888 | 1.54118 | Distal Intergenic | 8 | 12385771 | 12436729 | 50959 | 1 | ENSMUST00000080795.8 | -5784 | Gm5607 | ENSMUSG00000047935.11 |
| chr8 | 12380935 | 12381461 | 3.54075 | 4.54164 | 1.46742 | Distal Intergenic | 8 | 12385771 | 12436729 | 50959 | 1 | ENSMUST00000080795.8 | -4310 | Gm5607 | ENSMUSG00000047935.11 |
| chr8 | 12382803 | 12383035 | 3.45875 | 4.30023 | 1.32793 | Distal Intergenic | 8 | 12385771 | 12436729 | 50959 | 1 | ENSMUST00000080795.8 | -2736 | Gm5607 | ENSMUSG00000047935.11 |
| chr8 | 12383848 | 12387577 | 3.92702 | 5.32379 | 2.01062 | Promoter | 8 | 12385771 | 12436729 | 50959 | 1 | ENSMUST00000080795.8 | 0 | Gm5607 | ENSMUSG00000047935.11 |
| chr8 | 12388451 | 12388530 | 4.16961 | 5.58103 | 2.15176 | Intron (ENSMUST00000080795.8/ENSMUSG00000047935.11, intron 1 of 8) | 8 | 12385771 | 12436750 | 50980 | 1 | ENSMUST00000186174.1 | 2681 | Gm5607 | ENSMUSG00000047935.11 |
| chr8 | 12388963 | 12391417 | 4.08971 | 5.68415 | 2.27011 | Intron (ENSMUST00000080795.8/ENSMUSG00000047935.11, intron 1 of 8) | 8 | 12385771 | 12436750 | 50980 | 1 | ENSMUST00000186174.1 | 3193 | Gm5607 | ENSMUSG00000047935.11 |
| chr15 | 81212609 | 81212996 | 3.41735 | 3.9733 | 1.08472 | Distal Intergenic | 15 | 81190853 | 81192756 | 1904 | 1 | ENSMUST00000138715.1 | 21757 | 4930483J18Rik | ENSMUSG00000047938.7 |
| chr4 | 129513879 | 129514284 | 3.7548 | 4.74924 | 1.6261 | Promoter | 4 | 129513581 | 129515985 | 2405 | 1 | ENSMUST00000062356.6 | 299 | Marcksl1 | ENSMUSG00000047945.6 |
| chr4 | 129514591 | 129514776 | 3.3977 | 4.10708 | 1.24953 | Exon (ENSMUST00000062356.6/ENSMUSG00000047945.6, exon 2 of 2) | 4 | 129513581 | 129515985 | 2405 | 1 | ENSMUST00000062356.6 | 1011 | Marcksl1 | ENSMUSG00000047945.6 |
| chr16 | 30300478 | 30300765 | 3.57269 | 4.32021 | 1.37344 | Distal Intergenic | 16 | 30307686 | 30310779 | 3094 | 2 | ENSMUST00000061190.6 | 10014 | Gp5 | ENSMUSG00000047953.6 |
| chr16 | 30301819 | 30302233 | 3.35348 | 3.8897 | 1.12375 | Distal Intergenic | 16 | 30307686 | 30310779 | 3094 | 2 | ENSMUST00000061190.6 | 8546 | Gp5 | ENSMUSG00000047953.6 |
| chr16 | 30304406 | 30304497 | 3.53184 | 4.1957 | 1.26071 | Distal Intergenic | 16 | 30307686 | 30310779 | 3094 | 2 | ENSMUST00000061190.6 | 6282 | Gp5 | ENSMUSG00000047953.6 |
| chr16 | 30306306 | 30306621 | 3.60248 | 4.32952 | 1.34869 | Downstream (1-2kb) | 16 | 30307686 | 30310779 | 3094 | 2 | ENSMUST00000061190.6 | 4158 | Gp5 | ENSMUSG00000047953.6 |
| chr16 | 30307484 | 30307591 | 4.40128 | 6.08733 | 2.53291 | Downstream (<1kb) | 16 | 30307686 | 30310779 | 3094 | 2 | ENSMUST00000061190.6 | 3188 | Gp5 | ENSMUSG00000047953.6 |
| chr16 | 30308010 | 30308795 | 4.1482 | 5.86001 | 2.41056 | 3' UTR | 16 | 30307686 | 30310779 | 3094 | 2 | ENSMUST00000061190.6 | 1984 | Gp5 | ENSMUSG00000047953.6 |
| chr16 | 30309514 | 30310122 | 3.45449 | 4.10578 | 1.21359 | Promoter | 16 | 30307686 | 30310779 | 3094 | 2 | ENSMUST00000061190.6 | 657 | Gp5 | ENSMUSG00000047953.6 |
| chr3 | 107036154 | 107037466 | 3.68396 | 4.86051 | 1.69778 | Promoter | 3 | 107036162 | 107038129 | 1968 | 1 | ENSMUST00000052718.3 | 0 | Kcna3 | ENSMUSG00000047959.3 |
| chr2 | 70312562 | 70312940 | 3.36874 | 3.8807 | 1.01733 | Intron (ENSMUST00000060208.5/ENSMUSG00000042064.8, intron 26 of 34) | 2 | 70458982 | 70459560 | 579 | 1 | ENSMUST00000140507.1 | -146042 | Rpl9-ps7 | ENSMUSG00000047965.9 |
| chr2 | 70327386 | 70327762 | 3.39629 | 4.07905 | 1.18729 | Intron (ENSMUST00000060208.5/ENSMUSG00000042064.8, intron 29 of 34) | 2 | 70458982 | 70459560 | 579 | 1 | ENSMUST00000140507.1 | -131220 | Rpl9-ps7 | ENSMUSG00000047965.9 |
| chr2 | 70330194 | 70330604 | 3.58679 | 4.55234 | 1.46536 | Intron (ENSMUST00000060208.5/ENSMUSG00000042064.8, intron 29 of 34) | 2 | 70458982 | 70459560 | 579 | 1 | ENSMUST00000140507.1 | -128378 | Rpl9-ps7 | ENSMUSG00000047965.9 |
| chr2 | 70335521 | 70335596 | 3.49779 | 4.29278 | 1.27207 | Intron (ENSMUST00000060208.5/ENSMUSG00000042064.8, intron 30 of 34) | 2 | 70458982 | 70459560 | 579 | 1 | ENSMUST00000140507.1 | -123386 | Rpl9-ps7 | ENSMUSG00000047965.9 |
| chr2 | 70370663 | 70370843 | 3.40218 | 3.94373 | 1.06182 | Intron (ENSMUST00000060208.5/ENSMUSG00000042064.8, intron 32 of 34) | 2 | 70458982 | 70459560 | 579 | 1 | ENSMUST00000140507.1 | -88139 | Rpl9-ps7 | ENSMUSG00000047965.9 |
| chr2 | 70378881 | 70379092 | 3.43272 | 4.30005 | 1.32749 | Intron (ENSMUST00000060208.5/ENSMUSG00000042064.8, intron 32 of 34) | 2 | 70458982 | 70459560 | 579 | 1 | ENSMUST00000140507.1 | -79890 | Rpl9-ps7 | ENSMUSG00000047965.9 |
| chr2 | 70380101 | 70380472 | 3.35593 | 4.30156 | 1.29218 | Intron (ENSMUST00000060208.5/ENSMUSG00000042064.8, intron 32 of 34) | 2 | 70458982 | 70459560 | 579 | 1 | ENSMUST00000140507.1 | -78510 | Rpl9-ps7 | ENSMUSG00000047965.9 |
| chr2 | 70386181 | 70386355 | 3.60193 | 4.37971 | 1.39287 | Intron (ENSMUST00000060208.5/ENSMUSG00000042064.8, intron 32 of 34) | 2 | 70458982 | 70459560 | 579 | 1 | ENSMUST00000140507.1 | -72627 | Rpl9-ps7 | ENSMUSG00000047965.9 |
| chr2 | 70400197 | 70400446 | 3.46772 | 4.0924 | 1.23503 | Intron (ENSMUST00000060208.5/ENSMUSG00000042064.8, intron 32 of 34) | 2 | 70458982 | 70459560 | 579 | 1 | ENSMUST00000140507.1 | -58536 | Rpl9-ps7 | ENSMUSG00000047965.9 |
| chr2 | 70412304 | 70413282 | 3.87712 | 4.84865 | 1.65138 | Intron (ENSMUST00000060208.5/ENSMUSG00000042064.8, intron 32 of 34) | 2 | 70458982 | 70459560 | 579 | 1 | ENSMUST00000140507.1 | -45700 | Rpl9-ps7 | ENSMUSG00000047965.9 |
| chr2 | 70443926 | 70445733 | 3.82571 | 4.78781 | 1.64395 | Distal Intergenic | 2 | 70458982 | 70459560 | 579 | 1 | ENSMUST00000140507.1 | -13249 | Rpl9-ps7 | ENSMUSG00000047965.9 |
| chr2 | 70446397 | 70446597 | 3.36251 | 3.8713 | 1.01733 | Distal Intergenic | 2 | 70458982 | 70459560 | 579 | 1 | ENSMUST00000140507.1 | -12385 | Rpl9-ps7 | ENSMUSG00000047965.9 |
| chr2 | 70456311 | 70456457 | 4.07229 | 5.25585 | 1.98549 | Distal Intergenic | 2 | 70458982 | 70459560 | 579 | 1 | ENSMUST00000140507.1 | -2525 | Rpl9-ps7 | ENSMUSG00000047965.9 |
| chr6 | 126534881 | 126535436 | 3.57042 | 4.42259 | 1.40377 | 5' UTR | 6 | 126636463 | 126645801 | 9339 | 2 | ENSMUST00000055168.3 | 110365 | Kcna1 | ENSMUSG00000047976.3 |
| chr6 | 126640101 | 126640239 | 3.54677 | 4.29594 | 1.33158 | 3' UTR | 6 | 126636463 | 126645801 | 9339 | 2 | ENSMUST00000055168.3 | 5562 | Kcna1 | ENSMUSG00000047976.3 |
| chr6 | 126642408 | 126642596 | 3.44791 | 4.10821 | 1.22575 | Exon (ENSMUST00000055168.3/ENSMUSG00000047976.3, exon 2 of 2) | 6 | 126636463 | 126645801 | 9339 | 2 | ENSMUST00000055168.3 | 3205 | Kcna1 | ENSMUSG00000047976.3 |
| chr6 | 126642992 | 126643861 | 3.74491 | 4.59135 | 1.48499 | 5' UTR | 6 | 126636463 | 126645801 | 9339 | 2 | ENSMUST00000055168.3 | 1940 | Kcna1 | ENSMUSG00000047976.3 |
| chr6 | 126644265 | 126644749 | 3.89564 | 4.92205 | 1.71097 | 5' UTR | 6 | 126636463 | 126645801 | 9339 | 2 | ENSMUST00000055168.3 | 1052 | Kcna1 | ENSMUSG00000047976.3 |
| chr6 | 126645625 | 126646300 | 3.79572 | 4.69502 | 1.5887 | Promoter | 6 | 126636463 | 126645801 | 9339 | 2 | ENSMUST00000055168.3 | 0 | Kcna1 | ENSMUSG00000047976.3 |
| chr6 | 126647060 | 126647153 | 3.43368 | 4.56409 | 1.47758 | Intron (ENSMUST00000060972.3/ENSMUSG00000045534.3, intron 1 of 1) | 6 | 126636463 | 126645801 | 9339 | 2 | ENSMUST00000055168.3 | -1260 | Kcna1 | ENSMUSG00000047976.3 |
| chr8 | 84021080 | 84021486 | 3.58453 | 4.28008 | 1.27148 | Promoter | 8 | 84021471 | 84029188 | 7718 | 1 | ENSMUST00000142367.2 | 0 | Palm3 | ENSMUSG00000047986.7 |
| chr8 | 84026003 | 84026387 | 3.54071 | 4.19804 | 1.22127 | Promoter | 8 | 84026075 | 84027049 | 975 | 1 | ENSMUST00000154171.1 | 0 | Palm3 | ENSMUSG00000047986.7 |
| chr9 | 67829971 | 67830310 | 3.83834 | 5.12682 | 1.84661 | Downstream (<1kb) | 9 | 67830532 | 67832330 | 1799 | 2 | ENSMUST00000054500.5 | 2020 | C2cd4a | ENSMUSG00000047990.5 |
| chr9 | 67830809 | 67831935 | 4.36535 | 5.96479 | 2.45306 | Promoter | 9 | 67830532 | 67832330 | 1799 | 2 | ENSMUST00000054500.5 | 395 | C2cd4a | ENSMUSG00000047990.5 |
| chr9 | 67832370 | 67832490 | 3.84649 | 4.76927 | 1.58423 | Promoter | 9 | 67830532 | 67832330 | 1799 | 2 | ENSMUST00000054500.5 | -41 | C2cd4a | ENSMUSG00000047990.5 |
| chr9 | 67832919 | 67833074 | 3.31072 | 3.79328 | 1.01733 | Promoter | 9 | 67830532 | 67832330 | 1799 | 2 | ENSMUST00000054500.5 | -590 | C2cd4a | ENSMUSG00000047990.5 |
| chr18 | 84719318 | 84719756 | 4.69953 | 6.61061 | 3.04022 | Promoter | 18 | 84720019 | 84730447 | 10429 | 1 | ENSMUST00000161429.1 | -263 | Fam69c | ENSMUSG00000047992.5 |
| chr18 | 84720073 | 84721152 | 3.70436 | 4.54467 | 1.44807 | Promoter | 18 | 84720242 | 84740436 | 20195 | 1 | ENSMUST00000052501.2 | 0 | Fam69c | ENSMUSG00000047992.5 |
| chr18 | 84721641 | 84722049 | 3.69908 | 4.4884 | 1.39364 | Promoter | 18 | 84720883 | 84730224 | 9342 | 1 | ENSMUST00000161033.1 | 759 | Fam69c | ENSMUSG00000047992.5 |
| chr18 | 84722530 | 84723177 | 3.84043 | 4.92881 | 1.72292 | Intron (ENSMUST00000161429.1/ENSMUSG00000047992.5, intron 1 of 1) | 18 | 84720883 | 84730224 | 9342 | 1 | ENSMUST00000161033.1 | 1648 | Fam69c | ENSMUSG00000047992.5 |
| chr4 | 154948510 | 154949907 | 3.86266 | 4.84552 | 1.67828 | Distal Intergenic | 4 | 154960923 | 154962371 | 1449 | 1 | ENSMUST00000049621.6 | -11016 | Hes5 | ENSMUSG00000048001.7 |
| chr4 | 154950244 | 154950379 | 3.38193 | 3.9006 | 1.01733 | Distal Intergenic | 4 | 154960923 | 154962371 | 1449 | 1 | ENSMUST00000049621.6 | -10544 | Hes5 | ENSMUSG00000048001.7 |
| chr4 | 154950879 | 154951064 | 3.34826 | 3.84982 | 1.01733 | Distal Intergenic | 4 | 154960923 | 154962371 | 1449 | 1 | ENSMUST00000049621.6 | -9859 | Hes5 | ENSMUSG00000048001.7 |
| chr4 | 154952884 | 154953246 | 3.58716 | 4.30826 | 1.32015 | Distal Intergenic | 4 | 154960923 | 154962371 | 1449 | 1 | ENSMUST00000049621.6 | -7677 | Hes5 | ENSMUSG00000048001.7 |
| chr4 | 154953688 | 154953925 | 3.37756 | 3.89401 | 1.01733 | Distal Intergenic | 4 | 154960923 | 154962371 | 1449 | 1 | ENSMUST00000049621.6 | -6998 | Hes5 | ENSMUSG00000048001.7 |
| chr4 | 154955785 | 154955897 | 3.44085 | 4.00802 | 1.08244 | Distal Intergenic | 4 | 154960923 | 154962371 | 1449 | 1 | ENSMUST00000049621.6 | -5026 | Hes5 | ENSMUSG00000048001.7 |
| chr4 | 154957055 | 154957259 | 3.3799 | 3.99471 | 1.1358 | Distal Intergenic | 4 | 154960923 | 154962371 | 1449 | 1 | ENSMUST00000049621.6 | -3664 | Hes5 | ENSMUSG00000048001.7 |
| chr4 | 154958241 | 154958887 | 3.91118 | 4.9226 | 1.72313 | Downstream (1-2kb) | 4 | 154960923 | 154962371 | 1449 | 1 | ENSMUST00000049621.6 | -2036 | Hes5 | ENSMUSG00000048001.7 |
| chr4 | 154959387 | 154959967 | 3.65378 | 4.52269 | 1.46394 | Promoter | 4 | 154960923 | 154962371 | 1449 | 1 | ENSMUST00000049621.6 | -956 | Hes5 | ENSMUSG00000048001.7 |
| chr4 | 154960346 | 154962219 | 3.96118 | 5.07324 | 1.8677 | Promoter | 4 | 154960923 | 154962371 | 1449 | 1 | ENSMUST00000049621.6 | 0 | Hes5 | ENSMUSG00000048001.7 |
| chr12 | 119945335 | 119946153 | 3.24694 | 3.77761 | 1.04813 | Promoter | 12 | 119945957 | 120019211 | 73255 | 1 | ENSMUST00000058644.9 | 0 | Tmem196 | ENSMUSG00000048004.9 |
| chr11 | 100095839 | 100096029 | 4.0404 | 5.14506 | 1.83523 | Promoter | 11 | 100092193 | 100096225 | 4033 | 2 | ENSMUST00000103127.3 | 196 | Krt35 | ENSMUSG00000048013.9 |
| chr6 | 24953149 | 24953542 | 3.59624 | 4.29932 | 1.28429 | 3' UTR | 6 | 24951141 | 24956297 | 5157 | 2 | ENSMUST00000127247.3 | 2755 | Tmem229a | ENSMUSG00000048022.11 |
| chr6 | 24954555 | 24956669 | 4.48942 | 6.31352 | 2.80162 | Promoter | 6 | 24951141 | 24956297 | 5157 | 2 | ENSMUST00000127247.3 | 0 | Tmem229a | ENSMUSG00000048022.11 |
| chr19 | 58933834 | 58934189 | 3.71784 | 4.55243 | 1.48977 | Distal Intergenic | 19 | 58943425 | 58971419 | 27995 | 1 | ENSMUST00000054280.6 | -9236 | Eno4 | ENSMUSG00000048029.7 |
| chr19 | 58953096 | 58953345 | 3.32024 | 3.80761 | 1.01733 | Exon (ENSMUST00000054280.6/ENSMUSG00000048029.7, exon 4 of 14) | 19 | 58943425 | 58971419 | 27995 | 1 | ENSMUST00000054280.6 | 9672 | Eno4 | ENSMUSG00000048029.7 |
| chr19 | 58966761 | 58967612 | 3.68589 | 4.65898 | 1.53959 | Intron (ENSMUST00000054280.6/ENSMUSG00000048029.7, intron 11 of 13) | 19 | 58943425 | 58971419 | 27995 | 1 | ENSMUST00000054280.6 | 23337 | Eno4 | ENSMUSG00000048029.7 |
| chr19 | 58968842 | 58969207 | 3.60653 | 4.32094 | 1.30295 | Exon (ENSMUST00000054280.6/ENSMUSG00000048029.7, exon 13 of 14) | 19 | 58943425 | 58971419 | 27995 | 1 | ENSMUST00000054280.6 | 25418 | Eno4 | ENSMUSG00000048029.7 |
| chr19 | 58970277 | 58970943 | 3.65919 | 4.50851 | 1.48512 | 3' UTR | 19 | 58943425 | 58971419 | 27995 | 1 | ENSMUST00000054280.6 | 26853 | Eno4 | ENSMUSG00000048029.7 |
| chr19 | 58974379 | 58974466 | 3.23338 | 3.67724 | 1.01733 | 3' UTR | 19 | 58943425 | 58971419 | 27995 | 1 | ENSMUST00000054280.6 | 30955 | Eno4 | ENSMUSG00000048029.7 |
| chr19 | 58979878 | 58980151 | 3.37633 | 4.05971 | 1.22453 | Intron (ENSMUST00000047511.8/ENSMUSG00000041362.11, intron 16 of 16) | 19 | 58943425 | 58971419 | 27995 | 1 | ENSMUST00000054280.6 | 36454 | Eno4 | ENSMUSG00000048029.7 |
| chr2 | 102119876 | 102120236 | 3.49516 | 4.16329 | 1.30311 | Intron (ENSMUST00000111222.1/ENSMUSG00000048058.12, intron 1 of 5) | 2 | 102111569 | 102138119 | 26551 | 2 | ENSMUST00000127888.1 | 17883 | Ldlrad3 | ENSMUSG00000048058.12 |
| chr7 | 96170707 | 96171044 | 3.26126 | 3.72655 | 1.03749 | Promoter | 7 | 96171244 | 96911093 | 739850 | 1 | ENSMUST00000107162.2 | -200 | Tenm4 | ENSMUSG00000048078.11 |
| chr7 | 96207281 | 96207694 | 3.46776 | 4.06251 | 1.13123 | Intron (ENSMUST00000107162.2/ENSMUSG00000048078.11, intron 1 of 32) | 7 | 96209663 | 96437138 | 227476 | 1 | ENSMUST00000144263.2 | -1969 | Tenm4 | ENSMUSG00000048078.11 |
| chr7 | 96209750 | 96209999 | 3.50588 | 4.14421 | 1.21682 | Promoter | 7 | 96209713 | 96498917 | 289205 | 1 | ENSMUST00000133770.2 | 38 | Tenm4 | ENSMUSG00000048078.11 |
| chr7 | 96210369 | 96211984 | 3.91365 | 4.97569 | 1.76013 | Promoter | 7 | 96210637 | 96911093 | 700457 | 1 | ENSMUST00000107166.2 | 0 | Tenm4 | ENSMUSG00000048078.11 |
| chr7 | 96213049 | 96214327 | 3.73453 | 4.56113 | 1.44497 | Intron (ENSMUST00000107162.2/ENSMUSG00000048078.11, intron 1 of 32) | 7 | 96211671 | 96911093 | 699423 | 1 | ENSMUST00000107165.2 | 1379 | Tenm4 | ENSMUSG00000048078.11 |
| chr7 | 96642811 | 96643531 | 3.67325 | 4.46975 | 1.4299 | Promoter | 7 | 96644172 | 96732280 | 88109 | 1 | ENSMUST00000131541.1 | -641 | Tenm4 | ENSMUSG00000048078.11 |
| chr7 | 96644166 | 96644347 | 4.22437 | 5.7166 | 2.27179 | Promoter | 7 | 96644172 | 96732280 | 88109 | 1 | ENSMUST00000131541.1 | 0 | Tenm4 | ENSMUSG00000048078.11 |
| chr7 | 96645731 | 96646088 | 3.38974 | 4.01621 | 1.15034 | Intron (ENSMUST00000107162.2/ENSMUSG00000048078.11, intron 7 of 32) | 7 | 96644172 | 96732280 | 88109 | 1 | ENSMUST00000131541.1 | 1560 | Tenm4 | ENSMUSG00000048078.11 |
| chr7 | 96647484 | 96648009 | 3.26998 | 3.74657 | 1.06254 | Intron (ENSMUST00000107162.2/ENSMUSG00000048078.11, intron 7 of 32) | 7 | 96644172 | 96732280 | 88109 | 1 | ENSMUST00000131541.1 | 3313 | Tenm4 | ENSMUSG00000048078.11 |
| chr7 | 96794621 | 96794778 | 3.48518 | 4.2534 | 1.26665 | Intron (ENSMUST00000107162.2/ENSMUSG00000048078.11, intron 16 of 32) | 7 | 96796641 | 96798576 | 1936 | 1 | ENSMUST00000130299.2 | -1863 | Tenm4 | ENSMUSG00000048078.11 |
| chr7 | 96797413 | 96797543 | 3.70223 | 4.59932 | 1.52061 | Promoter | 7 | 96796641 | 96843440 | 46800 | 1 | ENSMUST00000147659.1 | 773 | Tenm4 | ENSMUSG00000048078.11 |
| chr1 | 135324675 | 135324884 | 3.88772 | 4.87961 | 1.71769 | Promoter | 1 | 135324807 | 135368065 | 43259 | 1 | ENSMUST00000059352.2 | 0 | Lmod1 | ENSMUSG00000048096.7 |
| chr19 | 40773999 | 40774074 | 3.96967 | 4.99698 | 1.72457 | Distal Intergenic | 19 | 40736184 | 40736865 | 682 | 1 | ENSMUST00000151726.1 | 37816 | Entpd1 | ENSMUSG00000048120.11 |
| chr1 | 90737605 | 90737917 | 3.56988 | 4.24483 | 1.2309 | Distal Intergenic | 1 | 90767389 | 90773215 | 5827 | 2 | ENSMUST00000150769.1 | 35298 | Col6a3 | ENSMUSG00000048126.11 |
| chr1 | 90813509 | 90813658 | 3.75519 | 4.59935 | 1.47727 | Exon (ENSMUST00000056925.10/ENSMUSG00000048126.11, exon 6 of 44) | 1 | 90791467 | 90807982 | 16516 | 2 | ENSMUST00000136916.2 | -5528 | Col6a3 | ENSMUSG00000048126.11 |
| chr1 | 90852397 | 90852669 | 3.68413 | 4.56744 | 1.50476 | Distal Intergenic | 1 | 90822271 | 90843971 | 21701 | 2 | ENSMUST00000187753.1 | -8427 | Col6a3 | ENSMUSG00000048126.11 |
| chr19 | 25672462 | 25672849 | 4.47852 | 6.21587 | 2.66338 | Promoter | 19 | 25672420 | 25679010 | 6591 | 1 | ENSMUST00000053068.5 | 43 | Dmrt2 | ENSMUSG00000048138.9 |
| chr19 | 25673472 | 25674087 | 4.47166 | 6.31669 | 2.75305 | Exon (ENSMUST00000053068.5/ENSMUSG00000048138.9, exon 2 of 4) | 19 | 25672420 | 25679010 | 6591 | 1 | ENSMUST00000053068.5 | 1053 | Dmrt2 | ENSMUSG00000048138.9 |
| chr19 | 25676036 | 25676330 | 3.45153 | 4.21832 | 1.27872 | Intron (ENSMUST00000053068.5/ENSMUSG00000048138.9, intron 3 of 3) | 19 | 25672420 | 25679010 | 6591 | 1 | ENSMUST00000053068.5 | 3617 | Dmrt2 | ENSMUSG00000048138.9 |
| chr19 | 25677508 | 25677984 | 3.49263 | 4.40573 | 1.38451 | Exon (ENSMUST00000053068.5/ENSMUSG00000048138.9, exon 4 of 4) | 19 | 25672420 | 25679010 | 6591 | 1 | ENSMUST00000053068.5 | 5089 | Dmrt2 | ENSMUSG00000048138.9 |
| chr19 | 25688870 | 25688968 | 3.35585 | 4.05852 | 1.16745 | Distal Intergenic | 19 | 25672420 | 25679010 | 6591 | 1 | ENSMUST00000053068.5 | 16451 | Dmrt2 | ENSMUSG00000048138.9 |
| chr5 | 33995360 | 33997676 | 3.63415 | 4.47664 | 1.42414 | Promoter | 5 | 33995984 | 34005916 | 9933 | 1 | ENSMUST00000056355.8 | 0 | Nat8l | ENSMUSG00000048142.8 |
| chr5 | 33998023 | 33998430 | 3.89153 | 4.93457 | 1.75648 | Exon (ENSMUST00000056355.8/ENSMUSG00000048142.8, exon 2 of 3) | 5 | 33995984 | 34005916 | 9933 | 1 | ENSMUST00000056355.8 | 2040 | Nat8l | ENSMUSG00000048142.8 |
| chr5 | 33999643 | 34000170 | 3.37003 | 3.88265 | 1.01733 | Intron (ENSMUST00000056355.8/ENSMUSG00000048142.8, intron 2 of 2) | 5 | 33995984 | 34005916 | 9933 | 1 | ENSMUST00000056355.8 | 3660 | Nat8l | ENSMUSG00000048142.8 |
| chr5 | 113825501 | 113825780 | 3.40571 | 3.95288 | 1.07537 | Intron (ENSMUST00000100874.4/ENSMUSG00000048163.9, intron 1 of 1) | 5 | 113817798 | 113830501 | 12704 | 2 | ENSMUST00000100874.4 | 4721 | Selplg | ENSMUSG00000048163.9 |
| chr7 | 128699985 | 128700064 | 3.7163 | 15.10774 | 10.34703 | Promoter | 7 | 128698187 | 128699219 | 1033 | 2 | ENSMUST00000140714.1 | -767 | Mcmbp | ENSMUSG00000048170.9 |
| chr15 | 98165697 | 98166090 | 3.81137 | 4.71782 | 1.59256 | Promoter | 15 | 98134637 | 98165625 | 30989 | 2 | ENSMUST00000143400.2 | -73 | Asb8 | ENSMUSG00000048175.8 |
| chr6 | 13835796 | 13836231 | 3.58094 | 4.29964 | 1.35868 | Exon (ENSMUST00000060442.8/ENSMUSG00000048216.8, exon 4 of 4) | 6 | 13835077 | 13837241 | 2165 | 2 | ENSMUST00000115491.1 | 1010 | Gpr85 | ENSMUSG00000048216.8 |
| chr6 | 13838328 | 13839265 | 3.61552 | 4.56264 | 1.52973 | Promoter | 6 | 13835068 | 13838454 | 3387 | 2 | ENSMUST00000115492.1 | 0 | Gpr85 | ENSMUSG00000048216.8 |
| chr11 | 102145152 | 102148278 | 3.82291 | 4.80804 | 1.67459 | Promoter | 11 | 102145513 | 102149477 | 3965 | 1 | ENSMUST00000129799.2 | 0 | Nags | ENSMUSG00000048217.6 |
| chr11 | 102149019 | 102149338 | 3.42515 | 4.21342 | 1.27182 | 3' UTR | 11 | 102146366 | 102149477 | 3112 | 1 | ENSMUST00000147252.1 | 2654 | Nags | ENSMUSG00000048217.6 |
| chr15 | 97469019 | 97469655 | 3.66104 | 4.41621 | 1.34408 | Distal Intergenic | 15 | 97244074 | 97247287 | 3214 | 2 | ENSMUST00000053106.5 | -221733 | Amigo2 | ENSMUSG00000048218.5 |
| chr10 | 80985517 | 80985744 | 3.50921 | 4.36628 | 1.38208 | Intron (ENSMUST00000099462.2/ENSMUSG00000048240.9, intron 2 of 3) | 10 | 80951187 | 80998190 | 47004 | 2 | ENSMUST00000118465.2 | 12446 | Gng7 | ENSMUSG00000048240.9 |
| chr10 | 80999578 | 80999679 | 3.57757 | 4.33033 | 1.37069 | Intron (ENSMUST00000099462.2/ENSMUSG00000048240.9, intron 1 of 3) | 10 | 80951187 | 80998190 | 47004 | 2 | ENSMUST00000118465.2 | -1389 | Gng7 | ENSMUSG00000048240.9 |
| chr12 | 107928404 | 107928775 | 3.78054 | 4.71724 | 1.64159 | Intron (ENSMUST00000066060.5/ENSMUSG00000048251.10, intron 3 of 3) | 12 | 107915222 | 108003414 | 88193 | 2 | ENSMUST00000109887.2 | 74639 | Bcl11b | ENSMUSG00000048251.10 |
| chr12 | 107952971 | 107953124 | 3.80392 | 4.70207 | 1.55273 | Intron (ENSMUST00000066060.5/ENSMUSG00000048251.10, intron 3 of 3) | 12 | 107915222 | 108003414 | 88193 | 2 | ENSMUST00000109887.2 | 50290 | Bcl11b | ENSMUSG00000048251.10 |
| chr12 | 107959838 | 107960126 | 3.65491 | 4.41601 | 1.37271 | Intron (ENSMUST00000066060.5/ENSMUSG00000048251.10, intron 3 of 3) | 12 | 107915222 | 108003414 | 88193 | 2 | ENSMUST00000109887.2 | 43288 | Bcl11b | ENSMUSG00000048251.10 |
| chr12 | 107961445 | 107961552 | 4.27671 | 5.67226 | 2.29215 | Intron (ENSMUST00000066060.5/ENSMUSG00000048251.10, intron 3 of 3) | 12 | 107915222 | 108003414 | 88193 | 2 | ENSMUST00000109887.2 | 41862 | Bcl11b | ENSMUSG00000048251.10 |
| chr12 | 107969811 | 107970501 | 3.37426 | 3.89935 | 1.05341 | Intron (ENSMUST00000066060.5/ENSMUSG00000048251.10, intron 2 of 3) | 12 | 107915222 | 108003414 | 88193 | 2 | ENSMUST00000109887.2 | 32913 | Bcl11b | ENSMUSG00000048251.10 |
| chr12 | 107975620 | 107976729 | 4.03361 | 5.16025 | 1.86762 | Intron (ENSMUST00000066060.5/ENSMUSG00000048251.10, intron 2 of 3) | 12 | 107915222 | 108003414 | 88193 | 2 | ENSMUST00000109887.2 | 26685 | Bcl11b | ENSMUSG00000048251.10 |
| chr12 | 107977036 | 107977264 | 3.6031 | 4.31358 | 1.29622 | Intron (ENSMUST00000066060.5/ENSMUSG00000048251.10, intron 2 of 3) | 12 | 107915222 | 108003414 | 88193 | 2 | ENSMUST00000109887.2 | 26150 | Bcl11b | ENSMUSG00000048251.10 |
| chr12 | 107977594 | 107977863 | 3.60541 | 4.32108 | 1.31099 | Intron (ENSMUST00000066060.5/ENSMUSG00000048251.10, intron 2 of 3) | 12 | 107915222 | 108003414 | 88193 | 2 | ENSMUST00000109887.2 | 25551 | Bcl11b | ENSMUSG00000048251.10 |
| chr12 | 107979763 | 107981130 | 3.79603 | 4.83823 | 1.70724 | Intron (ENSMUST00000066060.5/ENSMUSG00000048251.10, intron 2 of 3) | 12 | 107915222 | 108003414 | 88193 | 2 | ENSMUST00000109887.2 | 22284 | Bcl11b | ENSMUSG00000048251.10 |
| chr12 | 107981579 | 107981894 | 3.26261 | 3.72104 | 1.01733 | Intron (ENSMUST00000066060.5/ENSMUSG00000048251.10, intron 2 of 3) | 12 | 107915222 | 108003414 | 88193 | 2 | ENSMUST00000109887.2 | 21520 | Bcl11b | ENSMUSG00000048251.10 |
| chr12 | 107988748 | 107988905 | 3.355 | 3.85997 | 1.01733 | Intron (ENSMUST00000066060.5/ENSMUSG00000048251.10, intron 2 of 3) | 12 | 107915222 | 108003414 | 88193 | 2 | ENSMUST00000109887.2 | 14509 | Bcl11b | ENSMUSG00000048251.10 |
| chr12 | 107991393 | 107991649 | 3.45308 | 4.03249 | 1.10362 | Intron (ENSMUST00000066060.5/ENSMUSG00000048251.10, intron 1 of 3) | 12 | 107915222 | 108003414 | 88193 | 2 | ENSMUST00000109887.2 | 11765 | Bcl11b | ENSMUSG00000048251.10 |
| chr12 | 107998358 | 107998725 | 3.55455 | 4.5212 | 1.4722 | Intron (ENSMUST00000066060.5/ENSMUSG00000048251.10, intron 1 of 3) | 12 | 107915222 | 108003414 | 88193 | 2 | ENSMUST00000109887.2 | 4689 | Bcl11b | ENSMUSG00000048251.10 |
| chr12 | 108000868 | 108001970 | 3.67835 | 4.52193 | 1.50326 | Intron (ENSMUST00000066060.5/ENSMUSG00000048251.10, intron 1 of 3) | 12 | 107915222 | 108003414 | 88193 | 2 | ENSMUST00000109887.2 | 1444 | Bcl11b | ENSMUSG00000048251.10 |
| chr12 | 108002900 | 108003062 | 3.8121 | 4.87044 | 1.69519 | Promoter | 12 | 107915222 | 108003414 | 88193 | 2 | ENSMUST00000109887.2 | 352 | Bcl11b | ENSMUSG00000048251.10 |
| chr12 | 108003385 | 108004330 | 3.77322 | 4.89731 | 1.70949 | Promoter | 12 | 107910403 | 108003602 | 93200 | 2 | ENSMUST00000066060.5 | 0 | Bcl11b | ENSMUSG00000048251.10 |
| chr12 | 108005041 | 108005636 | 4.38179 | 5.91865 | 2.46197 | Distal Intergenic | 12 | 107910403 | 108003602 | 93200 | 2 | ENSMUST00000066060.5 | -1440 | Bcl11b | ENSMUSG00000048251.10 |
| chr12 | 108006039 | 108006298 | 3.38901 | 3.93708 | 1.10447 | Distal Intergenic | 12 | 107910403 | 108003602 | 93200 | 2 | ENSMUST00000066060.5 | -2438 | Bcl11b | ENSMUSG00000048251.10 |
| chr12 | 108009148 | 108009352 | 3.90199 | 4.87777 | 1.65566 | Distal Intergenic | 12 | 107910403 | 108003602 | 93200 | 2 | ENSMUST00000066060.5 | -5547 | Bcl11b | ENSMUSG00000048251.10 |
| chr12 | 108010051 | 108010876 | 3.87537 | 4.91949 | 1.71544 | Distal Intergenic | 12 | 107910403 | 108003602 | 93200 | 2 | ENSMUST00000066060.5 | -6450 | Bcl11b | ENSMUSG00000048251.10 |
| chr5 | 28293084 | 28293321 | 3.28557 | 3.75552 | 1.01733 | Distal Intergenic | 5 | 28317121 | 28352868 | 35748 | 1 | ENSMUST00000090355.5 | -23800 | Rbm33 | ENSMUSG00000048271.9 |
| chr14 | 61135265 | 61135354 | 3.32747 | 3.81849 | 1.01733 | Distal Intergenic | 14 | 61138457 | 61240695 | 102239 | 1 | ENSMUST00000089394.4 | -3103 | Sacs | ENSMUSG00000048279.12 |
| chr14 | 61137589 | 61137877 | 3.45751 | 4.09921 | 1.24777 | Promoter | 14 | 61138457 | 61240695 | 102239 | 1 | ENSMUST00000089394.4 | -580 | Sacs | ENSMUSG00000048279.12 |
| chr14 | 61138627 | 61138871 | 3.34578 | 3.88022 | 1.12921 | Promoter | 14 | 61138491 | 61199666 | 61176 | 1 | ENSMUST00000119509.2 | 137 | Sacs | ENSMUSG00000048279.12 |
| chr14 | 61139244 | 61139936 | 3.62426 | 4.39636 | 1.43411 | Promoter | 14 | 61138491 | 61199666 | 61176 | 1 | ENSMUST00000119509.2 | 754 | Sacs | ENSMUSG00000048279.12 |
| chr3 | 73053180 | 73053630 | 4.19063 | 5.44834 | 2.06256 | Intron (ENSMUST00000059407.7/ENSMUSG00000048304.7, intron 1 of 1) | 3 | 73048126 | 73056943 | 8818 | 2 | ENSMUST00000059407.7 | 3313 | Slitrk3 | ENSMUSG00000048304.7 |
| chr3 | 73055506 | 73055678 | 4.56538 | 6.28638 | 2.68132 | Intron (ENSMUST00000059407.7/ENSMUSG00000048304.7, intron 1 of 1) | 3 | 73048126 | 73056943 | 8818 | 2 | ENSMUST00000059407.7 | 1265 | Slitrk3 | ENSMUSG00000048304.7 |
| chr3 | 73056564 | 73056957 | 4.04535 | 5.1748 | 1.87578 | Promoter | 3 | 73048126 | 73056943 | 8818 | 2 | ENSMUST00000059407.7 | 0 | Slitrk3 | ENSMUSG00000048304.7 |
| chr3 | 73057340 | 73057653 | 3.96772 | 5.11455 | 1.85708 | Promoter | 3 | 73048126 | 73056943 | 8818 | 2 | ENSMUST00000059407.7 | -398 | Slitrk3 | ENSMUSG00000048304.7 |
| chr3 | 73058512 | 73058587 | 3.51283 | 4.27846 | 1.29186 | Distal Intergenic | 3 | 73048126 | 73056943 | 8818 | 2 | ENSMUST00000059407.7 | -1570 | Slitrk3 | ENSMUSG00000048304.7 |
| chr11 | 68555764 | 68556315 | 3.53145 | 4.31115 | 1.32751 | Promoter | 11 | 68556186 | 68558245 | 2060 | 1 | ENSMUST00000053211.7 | 0 | Mfsd6l | ENSMUSG00000048329.7 |
| chr7 | 109083027 | 109083630 | 3.56272 | 4.23884 | 1.24333 | Promoter | 7 | 109034312 | 109083331 | 49020 | 2 | ENSMUST00000055993.7 | 0 | Ric3 | ENSMUSG00000048330.9 |
| chr7 | 109083979 | 109084221 | 3.48799 | 4.09395 | 1.13453 | Promoter | 7 | 109034312 | 109083331 | 49020 | 2 | ENSMUST00000055993.7 | -649 | Ric3 | ENSMUSG00000048330.9 |
| chr7 | 109085119 | 109085404 | 3.9715 | 5.00588 | 1.74671 | Distal Intergenic | 7 | 109034312 | 109083331 | 49020 | 2 | ENSMUST00000055993.7 | -1789 | Ric3 | ENSMUSG00000048330.9 |
| chr3 | 53041445 | 53041784 | 3.60334 | 4.46448 | 1.42368 | Promoter | 3 | 53041528 | 53261679 | 220152 | 1 | ENSMUST00000059562.8 | 0 | Lhfp | ENSMUSG00000048332.8 |
| chr3 | 53042168 | 53043013 | 4.11996 | 5.38726 | 2.09125 | Promoter | 3 | 53041583 | 53261679 | 220097 | 1 | ENSMUST00000147139.1 | 586 | Lhfp | ENSMUSG00000048332.8 |
| chr3 | 53043461 | 53043969 | 3.59443 | 4.40521 | 1.41675 | Exon (ENSMUST00000059562.8/ENSMUSG00000048332.8, exon 2 of 4) | 3 | 53041583 | 53261679 | 220097 | 1 | ENSMUST00000147139.1 | 1879 | Lhfp | ENSMUSG00000048332.8 |
| chr14 | 34152031 | 34152215 | 4.0758 | 5.38283 | 2.06921 | Promoter | 14 | 34143992 | 34152489 | 8498 | 2 | ENSMUST00000052164.3 | 274 | Npy4r | ENSMUSG00000048337.3 |
| chr14 | 104336928 | 104337059 | 23.79681 | 82.5297 | 76.29892 | Distal Intergenic | 14 | 104463973 | 104465365 | 1393 | 2 | ENSMUST00000139505.1 | 128306 | Pou4f1 | ENSMUSG00000048349.9 |
| chr14 | 104454228 | 104454677 | 4.44912 | 6.31196 | 2.70583 | Distal Intergenic | 14 | 104463973 | 104465365 | 1393 | 2 | ENSMUST00000139505.1 | 10688 | Pou4f1 | ENSMUSG00000048349.9 |
| chr14 | 104456497 | 104456694 | 3.4575 | 4.28702 | 1.33119 | Distal Intergenic | 14 | 104463973 | 104465365 | 1393 | 2 | ENSMUST00000139505.1 | 8671 | Pou4f1 | ENSMUSG00000048349.9 |
| chr14 | 104457241 | 104457607 | 4.16673 | 5.76668 | 2.35363 | Distal Intergenic | 14 | 104463973 | 104465365 | 1393 | 2 | ENSMUST00000139505.1 | 7758 | Pou4f1 | ENSMUSG00000048349.9 |
| chr14 | 104458001 | 104458793 | 3.71676 | 4.63762 | 1.54019 | Downstream (2-3kb) | 14 | 104463973 | 104465365 | 1393 | 2 | ENSMUST00000139505.1 | 6572 | Pou4f1 | ENSMUSG00000048349.9 |
| chr14 | 104459198 | 104459749 | 4.1347 | 5.45887 | 2.13966 | Downstream (1-2kb) | 14 | 104463973 | 104465365 | 1393 | 2 | ENSMUST00000139505.1 | 5616 | Pou4f1 | ENSMUSG00000048349.9 |
| chr14 | 104460521 | 104461465 | 4.29783 | 5.90193 | 2.41379 | Downstream (<1kb) | 14 | 104463973 | 104465365 | 1393 | 2 | ENSMUST00000139505.1 | 3900 | Pou4f1 | ENSMUSG00000048349.9 |
| chr14 | 104461888 | 104465204 | 3.9233 | 5.06175 | 1.82442 | Promoter | 14 | 104463973 | 104465365 | 1393 | 2 | ENSMUST00000139505.1 | 161 | Pou4f1 | ENSMUSG00000048349.9 |
| chr14 | 104465533 | 104469810 | 3.86818 | 4.94791 | 1.7519 | Promoter | 14 | 104461676 | 104467999 | 6324 | 2 | ENSMUST00000053016.9 | 0 | Pou4f1 | ENSMUSG00000048349.9 |
| chr14 | 104470718 | 104470951 | 4.11971 | 5.38104 | 2.07135 | Distal Intergenic | 14 | 104461676 | 104467999 | 6324 | 2 | ENSMUST00000053016.9 | -2720 | Pou4f1 | ENSMUSG00000048349.9 |
| chr14 | 104471340 | 104471602 | 3.83168 | 5.02912 | 1.7846 | Distal Intergenic | 14 | 104461676 | 104467999 | 6324 | 2 | ENSMUST00000053016.9 | -3342 | Pou4f1 | ENSMUSG00000048349.9 |
| chr14 | 104471964 | 104472329 | 3.65032 | 4.7482 | 1.61375 | Distal Intergenic | 14 | 104461676 | 104467999 | 6324 | 2 | ENSMUST00000053016.9 | -3966 | Pou4f1 | ENSMUSG00000048349.9 |

| | | | | | | | | | | | | | | | |
|---|---|---|---|---|---|---|---|---|---|---|---|---|---|---|---|
| chr14 | 104472931 | 104474426 | 3.69339 | 4.58405 | 1.51693 | Distal Intergenic | 14 | 104461676 | 104467999 | 6324 | 2 | ENSMUST00000053016.9 | -4933 | Pou4f1 | ENSMUSG00000048349.9 |
| chr14 | 104475622 | 104475794 | 3.48837 | 4.37252 | 1.3536 | Downstream (1-2kb) | 14 | 104461676 | 104467999 | 6324 | 2 | ENSMUST00000053016.9 | -7624 | Pou4f1 | ENSMUSG00000048349.9 |
| chr8 | 104570903 | 104571129 | 3.40373 | 3.94431 | 1.05547 | Distal Intergenic | 8 | 104591468 | 104596836 | 5369 | 1 | ENSMUST00000059588.6 | -20339 | Pdp2 | ENSMUSG00000048371.7 |
| chr7 | 135411337 | 135411989 | 3.62347 | 4.39576 | 1.38705 | Promoter | 7 | 135410367 | 135413615 | 3249 | 1 | ENSMUST00000060356.5 | 971 | Foxi2 | ENSMUSG00000048377.5 |
| chr7 | 135412949 | 135413216 | 3.78429 | 4.70869 | 1.55478 | 3' UTR | 7 | 135410367 | 135413615 | 3249 | 1 | ENSMUST00000060356.5 | 2583 | Foxi2 | ENSMUSG00000048377.5 |
| chr15 | 76517071 | 76518748 | 3.64455 | 4.47008 | 1.42033 | 3' UTR | 15 | 76516213 | 76521902 | 5690 | 2 | ENSMUST00000164703.1 | 3154 | Scrt1 | ENSMUSG00000048385.8 |
| chr15 | 76519114 | 76520056 | 4.21575 | 5.63016 | 2.22065 | Exon (ENSMUST00000096365.3/ENSMUSG00000048385.8, exon 2 of 2) | 15 | 76516213 | 76521902 | 5690 | 2 | ENSMUST00000164703.1 | 1846 | Scrt1 | ENSMUSG00000048385.8 |
| chr15 | 76520409 | 76521575 | 3.79788 | 4.86567 | 1.70924 | Promoter | 15 | 76516213 | 76521902 | 5690 | 2 | ENSMUST00000164703.1 | 327 | Scrt1 | ENSMUSG00000048385.8 |
| chr15 | 76521908 | 76523673 | 3.77081 | 4.73266 | 1.66039 | Promoter | 15 | 76516203 | 76522129 | 5927 | 2 | ENSMUST00000096365.3 | 0 | Scrt1 | ENSMUSG00000048385.8 |
| chr12 | 9572953 | 9573894 | 3.91901 | 5.06103 | 1.83637 | Promoter | 12 | 9574442 | 9581499 | 7058 | 1 | ENSMUST00000057021.7 | -548 | Osr1 | ENSMUSG00000048387.7 |
| chr12 | 9574557 | 9575123 | 4.16372 | 5.42383 | 2.05705 | Promoter | 12 | 9574442 | 9581499 | 7058 | 1 | ENSMUST00000057021.7 | 116 | Osr1 | ENSMUSG00000048387.7 |
| chr12 | 9575755 | 9576891 | 4.10753 | 5.317 | 2.01281 | Intron (ENSMUST00000057021.7/ENSMUSG00000048387.7, intron 1 of 2) | 12 | 9574442 | 9581499 | 7058 | 1 | ENSMUST00000057021.7 | 1314 | Osr1 | ENSMUSG00000048387.7 |
| chr12 | 9577612 | 9578555 | 3.63775 | 4.45843 | 1.46137 | Intron (ENSMUST00000057021.7/ENSMUSG00000048387.7, intron 1 of 2) | 12 | 9574442 | 9581499 | 7058 | 1 | ENSMUST00000057021.7 | 3171 | Osr1 | ENSMUSG00000048387.7 |
| chr12 | 9579047 | 9580237 | 4.11505 | 5.40313 | 2.09286 | 5' UTR | 12 | 9574442 | 9581499 | 7058 | 1 | ENSMUST00000057021.7 | 4606 | Osr1 | ENSMUSG00000048387.7 |
| chr12 | 9581201 | 9581829 | 4.31788 | 5.90881 | 2.48395 | 3' UTR | 12 | 9574442 | 9581499 | 7058 | 1 | ENSMUST00000057021.7 | 6760 | Osr1 | ENSMUSG00000048387.7 |
| chr12 | 9582146 | 9583060 | 3.60584 | 4.36582 | 1.38588 | Distal Intergenic | 12 | 9574442 | 9581499 | 7058 | 1 | ENSMUST00000057021.7 | 7705 | Osr1 | ENSMUSG00000048387.7 |
| chr12 | 9584398 | 9584800 | 3.88803 | 4.88788 | 1.70193 | Distal Intergenic | 12 | 9574442 | 9581499 | 7058 | 1 | ENSMUST00000057021.7 | 9957 | Osr1 | ENSMUSG00000048387.7 |
| chr2 | 83812283 | 83812378 | 4.30933 | 5.83842 | 2.35009 | Promoter | 2 | 83812636 | 83883486 | 70851 | 1 | ENSMUST00000051454.3 | -258 | Fam171b | ENSMUSG00000048388.3 |
| chr2 | 83812740 | 83813140 | 3.79193 | 4.84722 | 1.70064 | Promoter | 2 | 83812636 | 83883486 | 70851 | 1 | ENSMUST00000051454.3 | 105 | Fam171b | ENSMUSG00000048388.3 |
| chr1 | 118982585 | 118982711 | 3.37587 | 3.89146 | 1.01733 | Promoter | 1 | 118868042 | 118982588 | 114547 | 2 | ENSMUST00000159678.2 | 0 | Gli2 | ENSMUSG00000048402.9 |
| chr1 | 119048558 | 119049412 | 3.64149 | 4.44889 | 1.39652 | Intron (ENSMUST00000062483.9/ENSMUSG00000048402.9, intron 1 of 13) | 1 | 118848460 | 119053339 | 204880 | 2 | ENSMUST00000159839.2 | 3927 | Gli2 | ENSMUSG00000048402.9 |
| chr1 | 119049773 | 119050543 | 3.38668 | 3.93215 | 1.09471 | Intron (ENSMUST00000062483.9/ENSMUSG00000048402.9, intron 1 of 13) | 1 | 118848460 | 119053339 | 204880 | 2 | ENSMUST00000159839.2 | 2796 | Gli2 | ENSMUSG00000048402.9 |
| chr1 | 119051498 | 119051787 | 3.39537 | 3.94582 | 1.10519 | Intron (ENSMUST00000062483.9/ENSMUSG00000048402.9, intron 1 of 13) | 1 | 118848460 | 119053339 | 204880 | 2 | ENSMUST00000159839.2 | 1552 | Gli2 | ENSMUSG00000048402.9 |
| chr1 | 119052641 | 119054678 | 4.20459 | 5.64651 | 2.28073 | Promoter | 1 | 118834132 | 119053619 | 219488 | 2 | ENSMUST00000062483.9 | 0 | Gli2 | ENSMUSG00000048402.9 |
| chr15 | 8968339 | 8968507 | 3.61853 | 4.41542 | 1.42147 | Promoter | 15 | 8968426 | 9002646 | 34221 | 1 | ENSMUST00000166524.2 | 0 | Ranbp3l | ENSMUSG00000048424.10 |
| chr5 | 37790313 | 37790667 | 3.89129 | 4.8761 | 1.68262 | Distal Intergenic | 5 | 37822703 | 37824582 | 1880 | 2 | ENSMUST00000183013.1 | 33915 | Msx1 | ENSMUSG00000048450.10 |
| chr5 | 37816100 | 37816254 | 3.62867 | 4.6084 | 1.51296 | Distal Intergenic | 5 | 37822703 | 37824582 | 1880 | 2 | ENSMUST00000183013.1 | 8328 | Msx1 | ENSMUSG00000048450.10 |
| chr5 | 37816944 | 37817247 | 3.76907 | 4.89912 | 1.71563 | Distal Intergenic | 5 | 37822703 | 37824582 | 1880 | 2 | ENSMUST00000183013.1 | 7335 | Msx1 | ENSMUSG00000048450.10 |
| chr5 | 37818301 | 37819427 | 3.90626 | 4.99678 | 1.83002 | Downstream (1-2kb) | 5 | 37822703 | 37824582 | 1880 | 2 | ENSMUST00000183013.1 | 5155 | Msx1 | ENSMUSG00000048450.10 |
| chr5 | 37819856 | 37820226 | 3.66648 | 4.51728 | 1.47394 | Downstream (<1kb) | 5 | 37822703 | 37824582 | 1880 | 2 | ENSMUST00000183013.1 | 4356 | Msx1 | ENSMUSG00000048450.10 |
| chr5 | 37821190 | 37821656 | 4.53248 | 6.23072 | 2.71015 | 3' UTR | 5 | 37822703 | 37824582 | 1880 | 2 | ENSMUST00000183013.1 | 2926 | Msx1 | ENSMUSG00000048450.10 |
| chr5 | 37822008 | 37822302 | 3.53016 | 4.17979 | 1.21404 | Intron (ENSMUST00000063116.9/ENSMUSG00000048450.10, intron 1 of 1) | 5 | 37822703 | 37824582 | 1880 | 2 | ENSMUST00000183013.1 | 2280 | Msx1 | ENSMUSG00000048450.10 |
| chr5 | 37823572 | 37824344 | 4.42174 | 5.95779 | 2.45722 | Promoter | 5 | 37822703 | 37824582 | 1880 | 2 | ENSMUST00000183013.1 | 238 | Msx1 | ENSMUSG00000048450.10 |
| chr5 | 37824945 | 37825603 | 3.81463 | 4.78309 | 1.64794 | Promoter | 5 | 37820485 | 37824583 | 4099 | 2 | ENSMUST00000063116.9 | -363 | Msx1 | ENSMUSG00000048450.10 |
| chr5 | 37825922 | 37827818 | 3.78035 | 4.78542 | 1.63647 | Distal Intergenic | 5 | 37820485 | 37824583 | 4099 | 2 | ENSMUST00000063116.9 | -1340 | Msx1 | ENSMUSG00000048450.10 |
| chr5 | 37828146 | 37828555 | 3.64388 | 4.67696 | 1.54654 | Distal Intergenic | 5 | 37820485 | 37824583 | 4099 | 2 | ENSMUST00000063116.9 | -3564 | Msx1 | ENSMUSG00000048450.10 |
| chr5 | 37829024 | 37830182 | 4.32957 | 5.83468 | 2.37708 | Distal Intergenic | 5 | 37820485 | 37824583 | 4099 | 2 | ENSMUST00000063116.9 | -4442 | Msx1 | ENSMUSG00000048450.10 |
| chr5 | 37830639 | 37831657 | 3.94524 | 4.98555 | 1.75213 | Distal Intergenic | 5 | 37820485 | 37824583 | 4099 | 2 | ENSMUST00000063116.9 | -6057 | Msx1 | ENSMUSG00000048450.10 |
| chr5 | 37831965 | 37832611 | 3.79837 | 4.69306 | 1.55744 | Distal Intergenic | 5 | 37820485 | 37824583 | 4099 | 2 | ENSMUST00000063116.9 | -7383 | Msx1 | ENSMUSG00000048450.10 |
| chr5 | 37832958 | 37833229 | 3.32031 | 3.80773 | 1.01733 | Distal Intergenic | 5 | 37820485 | 37824583 | 4099 | 2 | ENSMUST00000063116.9 | -8376 | Msx1 | ENSMUSG00000048450.10 |
| chr5 | 37836678 | 37836810 | 3.45791 | 4.10929 | 1.22029 | Distal Intergenic | 5 | 37820485 | 37824583 | 4099 | 2 | ENSMUST00000063116.9 | -12096 | Msx1 | ENSMUSG00000048450.10 |
| chr5 | 37837159 | 37837487 | 3.38078 | 3.89886 | 1.01733 | Distal Intergenic | 5 | 37820485 | 37824583 | 4099 | 2 | ENSMUST00000063116.9 | -12577 | Msx1 | ENSMUSG00000048450.10 |
| chr5 | 37837793 | 37837924 | 3.55841 | 4.23646 | 1.26376 | Distal Intergenic | 5 | 37820485 | 37824583 | 4099 | 2 | ENSMUST00000063116.9 | -13211 | Msx1 | ENSMUSG00000048450.10 |
| chr5 | 37838409 | 37839559 | 3.51561 | 4.15091 | 1.18662 | Distal Intergenic | 5 | 37820485 | 37824583 | 4099 | 2 | ENSMUST00000063116.9 | -13827 | Msx1 | ENSMUSG00000048450.10 |
| chr5 | 37870878 | 37871080 | 3.56337 | 4.25912 | 1.31955 | Distal Intergenic | 5 | 37820485 | 37824583 | 4099 | 2 | ENSMUST00000063116.9 | -46296 | Msx1 | ENSMUSG00000048450.10 |
| chr5 | 37891051 | 37891269 | 3.63265 | 4.66238 | 1.5458 | Distal Intergenic | 5 | 37820485 | 37824583 | 4099 | 2 | ENSMUST00000063116.9 | -66469 | Msx1 | ENSMUSG00000048450.10 |
| chr5 | 37893695 | 37895668 | 3.88039 | 4.85472 | 1.66363 | Distal Intergenic | 5 | 37820485 | 37824583 | 4099 | 2 | ENSMUST00000063116.9 | -69113 | Msx1 | ENSMUSG00000048450.10 |
| chr5 | 37898514 | 37898818 | 3.35418 | 3.85873 | 1.01733 | Distal Intergenic | 5 | 37820485 | 37824583 | 4099 | 2 | ENSMUST00000063116.9 | -73932 | Msx1 | ENSMUSG00000048450.10 |
| chr5 | 37900883 | 37901171 | 3.3967 | 3.9359 | 1.06336 | Distal Intergenic | 5 | 37820485 | 37824583 | 4099 | 2 | ENSMUST00000063116.9 | -76301 | Msx1 | ENSMUSG00000048450.10 |
| chr5 | 37916710 | 37916875 | 3.44629 | 4.02303 | 1.10632 | Distal Intergenic | 5 | 37820485 | 37824583 | 4099 | 2 | ENSMUST00000063116.9 | -92128 | Msx1 | ENSMUSG00000048450.10 |
| chr5 | 37917353 | 37917549 | 3.45457 | 4.03332 | 1.09858 | Distal Intergenic | 5 | 37820485 | 37824583 | 4099 | 2 | ENSMUST00000063116.9 | -92771 | Msx1 | ENSMUSG00000048450.10 |
| chr5 | 37918228 | 37918715 | 3.42691 | 3.98731 | 1.08407 | Distal Intergenic | 5 | 37820485 | 37824583 | 4099 | 2 | ENSMUST00000063116.9 | -93646 | Msx1 | ENSMUSG00000048450.10 |
| chr5 | 37919062 | 37919252 | 3.60732 | 4.31759 | 1.27946 | Distal Intergenic | 5 | 37820485 | 37824583 | 4099 | 2 | ENSMUST00000063116.9 | -94480 | Msx1 | ENSMUSG00000048450.10 |
| chr6 | 142800955 | 142801442 | 3.85592 | 4.89273 | 1.71298 | Intron (ENSMUST00000156662.2/ENSMUSG00000048473.10, intron 2 of 6) | 6 | 142786307 | 142804390 | 18084 | 2 | ENSMUST00000111768.1 | 2948 | Gm766 | ENSMUSG00000048473.10 |
| chr6 | 142802073 | 142802148 | 4.06986 | 5.33909 | 2.01308 | Intron (ENSMUST00000156662.2/ENSMUSG00000048473.10, intron 1 of 6) | 6 | 142786307 | 142804390 | 18084 | 2 | ENSMUST00000111768.1 | 2242 | Gm766 | ENSMUSG00000048473.10 |
| chr2 | 109673887 | 109674240 | 3.89571 | 4.92721 | 1.72089 | Promoter | 2 | 109674700 | 109727007 | 52308 | 1 | ENSMUST00000053317.6 | -460 | Bdnf | ENSMUSG00000048482.9 |
| chr2 | 109674716 | 109675750 | 3.45755 | 4.10391 | 1.22459 | Promoter | 2 | 109674700 | 109727007 | 52308 | 1 | ENSMUST00000053317.6 | 17 | Bdnf | ENSMUSG00000048482.9 |
| chr2 | 109676081 | 109677769 | 3.99205 | 5.1835 | 1.93255 | Promoter | 2 | 109677032 | 109727007 | 49976 | 1 | ENSMUST00000111046.3 | 0 | Bdnf | ENSMUSG00000048482.9 |
| chr2 | 109678653 | 109678764 | 3.45537 | 4.04371 | 1.13174 | Intron (ENSMUST00000053317.6/ENSMUSG00000048482.9, intron 1 of 1) | 2 | 109677032 | 109727007 | 49976 | 1 | ENSMUST00000111046.3 | 1622 | Bdnf | ENSMUSG00000048482.9 |
| chr2 | 109679130 | 109679217 | 3.93172 | 5.06374 | 1.83175 | Intron (ENSMUST00000053317.6/ENSMUSG00000048482.9, intron 1 of 1) | 2 | 109677032 | 109727007 | 49976 | 1 | ENSMUST00000111046.3 | 2099 | Bdnf | ENSMUSG00000048482.9 |
| chr2 | 109691931 | 109692258 | 3.7082 | 4.51105 | 1.41424 | Promoter | 2 | 109692436 | 109727007 | 34572 | 1 | ENSMUST00000111050.4 | -178 | Bdnf | ENSMUSG00000048482.9 |
| chr2 | 109692565 | 109692672 | 3.36112 | 3.86921 | 1.01733 | Promoter | 2 | 109692436 | 109727007 | 34572 | 1 | ENSMUST00000111050.4 | 130 | Bdnf | ENSMUSG00000048482.9 |
| chr2 | 109693359 | 109694073 | 4.81923 | 6.81274 | 3.14029 | Promoter | 2 | 109693563 | 109727007 | 33445 | 1 | ENSMUST00000111045.3 | 0 | Bdnf | ENSMUSG00000048482.9 |
| chr2 | 109694455 | 109694659 | 4.16156 | 5.75426 | 2.35546 | Promoter | 2 | 109694469 | 109727007 | 32539 | 1 | ENSMUST00000111044.2 | 0 | Bdnf | ENSMUSG00000048482.9 |
| chr2 | 109697568 | 109698042 | 3.48115 | 4.08743 | 1.14704 | Intron (ENSMUST00000053317.6/ENSMUSG00000048482.9, intron 1 of 1) | 2 | 109694469 | 109727007 | 32539 | 1 | ENSMUST00000111044.2 | 3100 | Bdnf | ENSMUSG00000048482.9 |
| chr12 | 86982817 | 86983416 | 3.67102 | 4.50425 | 1.508 | Exon (ENSMUST00000095521.2/ENSMUSG00000048483.5, exon 2 of 2) | 12 | 86983381 | 86988676 | 5296 | 2 | ENSMUST00000095521.2 | 5260 | Zdhhc22 | ENSMUSG00000048483.5 |
| chr12 | 86985100 | 86985242 | 3.30674 | 3.80283 | 1.06832 | Intron (ENSMUST00000095521.2/ENSMUSG00000048483.5, intron 1 of 1) | 12 | 86983381 | 86988676 | 5296 | 2 | ENSMUST00000095521.2 | 3434 | Zdhhc22 | ENSMUSG00000048483.5 |
| chr12 | 86987141 | 86987216 | 3.42963 | 4.01303 | 1.16108 | Intron (ENSMUST00000095521.2/ENSMUSG00000048483.5, intron 1 of 1) | 12 | 86983381 | 86988676 | 5296 | 2 | ENSMUST00000095521.2 | 1460 | Zdhhc22 | ENSMUSG00000048483.5 |
| chr12 | 86988348 | 86988721 | 3.69509 | 4.57345 | 1.57427 | Promoter | 12 | 86983381 | 86988676 | 5296 | 2 | ENSMUST00000095521.2 | 0 | Zdhhc22 | ENSMUSG00000048483.5 |
| chr12 | 86989495 | 86990612 | 3.83763 | 4.95795 | 1.7559 | Promoter | 12 | 86983381 | 86988676 | 5296 | 2 | ENSMUST00000095521.2 | -820 | Zdhhc22 | ENSMUSG00000048483.5 |
| chr7 | 132599370 | 132600032 | 4.37142 | 5.85858 | 2.385 | Promoter | 7 | 132594878 | 132599637 | 4760 | 2 | ENSMUST00000054562.3 | 0 | Nkx1-2 | ENSMUSG00000048528.7 |
| chr7 | 132601564 | 132602605 | 3.45285 | 4.04878 | 1.15532 | Intron (ENSMUST00000124096.2/ENSMUSG00000030849.12, intron 2 of 6) | 7 | 132594878 | 132599637 | 4760 | 2 | ENSMUST00000054562.3 | -1928 | Nkx1-2 | ENSMUSG00000048528.7 |
| chr3 | 102008237 | 102008623 | 3.764 | 4.65455 | 1.52436 | Distal Intergenic | 3 | 102010145 | 102015492 | 5348 | 1 | ENSMUST00000066187.4 | -1522 | Nhlh2 | ENSMUSG00000048540.7 |
| chr3 | 102009942 | 102010019 | 3.41497 | 3.97366 | 1.09947 | Promoter | 3 | 102010145 | 102015492 | 5348 | 1 | ENSMUST00000066187.4 | -126 | Nhlh2 | ENSMUSG00000048540.7 |
| chr3 | 102010815 | 102013909 | 3.89862 | 5.07252 | 1.85334 | Promoter | 3 | 102010145 | 102015492 | 5348 | 1 | ENSMUST00000066187.4 | 671 | Nhlh2 | ENSMUSG00000048540.7 |
| chr3 | 102016128 | 102016466 | 3.2787 | 3.74522 | 1.01733 | Distal Intergenic | 3 | 102010145 | 102015492 | 5348 | 1 | ENSMUST00000066187.4 | 5984 | Nhlh2 | ENSMUSG00000048540.7 |
| chr3 | 102016945 | 102017267 | 3.30052 | 3.78464 | 1.02326 | Distal Intergenic | 3 | 102010145 | 102015492 | 5348 | 1 | ENSMUST00000066187.4 | 6801 | Nhlh2 | ENSMUSG00000048540.7 |
| chr3 | 102017737 | 102018165 | 3.90145 | 4.92935 | 1.72008 | Distal Intergenic | 3 | 102010145 | 102015492 | 5348 | 1 | ENSMUST00000066187.4 | 7593 | Nhlh2 | ENSMUSG00000048540.7 |
| chr3 | 102021685 | 102022456 | 3.90959 | 4.97429 | 1.80454 | Distal Intergenic | 3 | 102010145 | 102015492 | 5348 | 1 | ENSMUST00000066187.4 | 11541 | Nhlh2 | ENSMUSG00000048540.7 |
| chr3 | 102023368 | 102023846 | 3.73715 | 4.55972 | 1.43421 | Distal Intergenic | 3 | 102010145 | 102015492 | 5348 | 1 | ENSMUST00000066187.4 | 13224 | Nhlh2 | ENSMUSG00000048540.7 |
| chr3 | 102024209 | 102024796 | 3.68767 | 4.5 | 1.46394 | Distal Intergenic | 3 | 102010145 | 102015492 | 5348 | 1 | ENSMUST00000066187.4 | 14065 | Nhlh2 | ENSMUSG00000048540.7 |
| chr3 | 102025197 | 102025881 | 3.44312 | 4.02738 | 1.13731 | Distal Intergenic | 3 | 102010145 | 102015492 | 5348 | 1 | ENSMUST00000066187.4 | 15053 | Nhlh2 | ENSMUSG00000048540.7 |
| chr3 | 102026614 | 102027134 | 3.64424 | 4.46031 | 1.42417 | Distal Intergenic | 3 | 102010145 | 102015492 | 5348 | 1 | ENSMUST00000066187.4 | 16470 | Nhlh2 | ENSMUSG00000048540.7 |
| chr12 | 118809994 | 118810488 | 3.37797 | 3.89648 | 1.02384 | Distal Intergenic | 12 | 118846329 | 118852578 | 6250 | 1 | ENSMUST00000063918.2 | -35841 | Sp8 | ENSMUSG00000048562.6 |
| chr12 | 118811107 | 118811287 | 3.68721 | 4.47009 | 1.39798 | Distal Intergenic | 12 | 118846329 | 118852578 | 6250 | 1 | ENSMUST00000063918.2 | -35042 | Sp8 | ENSMUSG00000048562.6 |
| chr12 | 118811664 | 118811883 | 3.77376 | 4.69317 | 1.53509 | Distal Intergenic | 12 | 118846329 | 118852578 | 6250 | 1 | ENSMUST00000063918.2 | -34446 | Sp8 | ENSMUSG00000048562.6 |
| chr12 | 118813244 | 118813561 | 3.45277 | 4.03041 | 1.09792 | Distal Intergenic | 12 | 118846329 | 118852578 | 6250 | 1 | ENSMUST00000063918.2 | -32768 | Sp8 | ENSMUSG00000048562.6 |
| chr12 | 118833094 | 118833403 | 3.42231 | 4.12892 | 1.19022 | Distal Intergenic | 12 | 118846329 | 118852578 | 6250 | 1 | ENSMUST00000063918.2 | -12926 | Sp8 | ENSMUSG00000048562.6 |
| chr12 | 118833921 | 118834166 | 3.55857 | 4.59295 | 1.49457 | Distal Intergenic | 12 | 118846329 | 118852578 | 6250 | 1 | ENSMUST00000063918.2 | -12163 | Sp8 | ENSMUSG00000048562.6 |
| chr12 | 118834564 | 118834870 | 3.76533 | 4.94475 | 1.73485 | Distal Intergenic | 12 | 118846329 | 118852578 | 6250 | 1 | ENSMUST00000063918.2 | -11459 | Sp8 | ENSMUSG00000048562.6 |
| chr12 | 118835208 | 118835477 | 3.49024 | 4.38231 | 1.37972 | Distal Intergenic | 12 | 118846329 | 118852578 | 6250 | 1 | ENSMUST00000063918.2 | -10852 | Sp8 | ENSMUSG00000048562.6 |
| chr12 | 118836002 | 118836185 | 3.23518 | 4.04961 | 1.14745 | Distal Intergenic | 12 | 118846329 | 118852578 | 6250 | 1 | ENSMUST00000063918.2 | -10144 | Sp8 | ENSMUSG00000048562.6 |
| chr12 | 118836546 | 118839944 | 3.94079 | 5.20044 | 1.97048 | Distal Intergenic | 12 | 118846329 | 118852578 | 6250 | 1 | ENSMUST00000063918.2 | -6385 | Sp8 | ENSMUSG00000048562.6 |
| chr12 | 118840275 | 118841105 | 3.9811 | 5.06422 | 1.8409 | Distal Intergenic | 12 | 118846329 | 118852578 | 6250 | 1 | ENSMUST00000063918.2 | -5224 | Sp8 | ENSMUSG00000048562.6 |
| chr12 | 118841437 | 118842037 | 3.36183 | 3.92383 | 1.08047 | Distal Intergenic | 12 | 118846329 | 118852578 | 6250 | 1 | ENSMUST00000063918.2 | -4292 | Sp8 | ENSMUSG00000048562.6 |
| chr12 | 118842652 | 118843647 | 4.22638 | 5.55421 | 2.15253 | Distal Intergenic | 12 | 118846329 | 118852578 | 6250 | 1 | ENSMUST00000063918.2 | -2682 | Sp8 | ENSMUSG00000048562.6 |
| chr12 | 118844129 | 118844297 | 3.56201 | 4.2388 | 1.24128 | Distal Intergenic | 12 | 118846329 | 118852578 | 6250 | 1 | ENSMUST00000063918.2 | -2032 | Sp8 | ENSMUSG00000048562.6 |
| chr12 | 118844790 | 118845420 | 4.01733 | 5.13133 | 1.83995 | Promoter | 12 | 118846329 | 118852578 | 6250 | 1 | ENSMUST00000063918.2 | -909 | Sp8 | ENSMUSG00000048562.6 |
| chr12 | 118846025 | 118846100 | 3.84542 | 4.83585 | 1.69824 | Promoter | 12 | 118846329 | 118852578 | 6250 | 1 | ENSMUST00000063918.2 | -229 | Sp8 | ENSMUSG00000048562.6 |
| chr12 | 118846628 | 118848124 | 4.30372 | 5.77868 | 2.37789 | Promoter | 12 | 118846329 | 118852578 | 6250 | 1 | ENSMUST00000063918.2 | 300 | Sp8 | ENSMUSG00000048562.6 |
| chr12 | 118848440 | 118850334 | 3.99863 | 5.32408 | 2.01204 | 3' UTR | 12 | 118846329 | 118852578 | 6250 | 1 | ENSMUST00000063918.2 | 2112 | Sp8 | ENSMUSG00000048562.6 |
| chr12 | 118851036 | 118851597 | 3.80173 | 4.7822 | 1.64113 | 3' UTR | 12 | 118846329 | 118852578 | 6250 | 1 | ENSMUST00000063918.2 | 4708 | Sp8 | ENSMUSG00000048562.6 |
| chr12 | 118852164 | 118852405 | 3.4003 | 3.9627 | 1.13921 | 3' UTR | 12 | 118846329 | 118852578 | 6250 | 1 | ENSMUST00000063918.2 | 5836 | Sp8 | ENSMUSG00000048562.6 |
| chr12 | 118852748 | 118853994 | 4.11103 | 5.45327 | 2.13526 | Distal Intergenic | 12 | 118846329 | 118852578 | 6250 | 1 | ENSMUST00000063918.2 | 6420 | Sp8 | ENSMUSG00000048562.6 |
| chr12 | 118854621 | 118855446 | 3.86129 | 5.06984 | 1.88855 | Distal Intergenic | 12 | 118846329 | 118852578 | 6250 | 1 | ENSMUST00000063918.2 | 8293 | Sp8 | ENSMUSG00000048562.6 |
| chr12 | 118856244 | 118857116 | 3.42604 | 4.2261 | 1.27886 | Distal Intergenic | 12 | 118846329 | 118852578 | 6250 | 1 | ENSMUST00000063918.2 | 9916 | Sp8 | ENSMUSG00000048562.6 |
| chr12 | 118857766 | 118857841 | 3.66234 | 4.45157 | 1.4595 | Distal Intergenic | 12 | 118846329 | 118852578 | 6250 | 1 | ENSMUST00000063918.2 | 11438 | Sp8 | ENSMUSG00000048562.6 |
| chr12 | 118858527 | 118859112 | 3.59023 | 4.33543 | 1.35741 | Distal Intergenic | 12 | 118846329 | 118852578 | 6250 | 1 | ENSMUST00000063918.2 | 12199 | Sp8 | ENSMUSG00000048562.6 |
| chr12 | 118859617 | 118860193 | 3.93994 | 5.19755 | 1.90049 | Distal Intergenic | 12 | 118846329 | 118852578 | 6250 | 1 | ENSMUST00000063918.2 | 13289 | Sp8 | ENSMUSG00000048562.6 |
| chr12 | 118862312 | 118862558 | 3.35198 | 3.89358 | 1.13247 | Distal Intergenic | 12 | 118846329 | 118852578 | 6250 | 1 | ENSMUST00000063918.2 | 15984 | Sp8 | ENSMUSG00000048562.6 |
| chr12 | 118865540 | 118865885 | 3.36299 | 3.87203 | 1.01733 | Downstream (1-2kb) | 12 | 118846329 | 118852578 | 6250 | 1 | ENSMUST00000063918.2 | 19212 | Sp8 | ENSMUSG00000048562.6 |
| chr7 | 142656106 | 142656530 | 3.53508 | 4.23228 | 1.22952 | Promoter | 7 | 142654352 | 142656018 | 1667 | 2 | ENSMUST00000178921.1 | -89 | Igf2 | ENSMUSG00000048583.11 |
| chr7 | 142657293 | 142657391 | 3.49347 | 4.13109 | 1.22165 | Promoter | 7 | 142650769 | 142657466 | 6698 | 2 | ENSMUST00000097936.3 | 75 | Igf2 | ENSMUSG00000048583.11 |
| chr7 | 142657844 | 142658419 | 3.71032 | 4.51225 | 1.41129 | Promoter | 7 | 142650766 | 142657496 | 6731 | 2 | ENSMUST00000000033.6 | -349 | Igf2 | ENSMUSG00000048583.11 |
| chr7 | 142658923 | 142659387 | 3.55204 | 4.36969 | 1.3452 | Promoter | 7 | 142650769 | 142659489 | 8721 | 2 | ENSMUST00000121128.2 | 102 | Igf2 | ENSMUSG00000048583.11 |
| chr7 | 142660102 | 142662006 | 3.74294 | 4.7077 | 1.60613 | Promoter | 7 | 142650769 | 142661305 | 10537 | 2 | ENSMUST00000105936.2 | 0 | Igf2 | ENSMUSG00000048583.11 |
| chr7 | 142662549 | 142662857 | 3.4579 | 4.03972 | 1.10345 | Promoter | 7 | 142654049 | 142661870 | 7822 | 2 | ENSMUST00000145896.2 | -680 | Igf2 | ENSMUSG00000048583.11 |
| chr7 | 142667600 | 142667676 | 3.63753 | 4.36905 | 1.30503 | Promoter | 7 | 142653756 | 142666816 | 13061 | 2 | ENSMUST00000105935.2 | -785 | Igf2 | ENSMUSG00000048583.11 |
| chr11 | 89297587 | 89297833 | 3.76314 | 4.66324 | 1.54311 | Downstream (2-3kb) | 11 | 89300638 | 89302332 | 1695 | 2 | ENSMUST00000061728.4 | 4499 | Nog | ENSMUSG00000048616.4 |
| chr11 | 89298435 | 89298730 | 3.81995 | 5.00412 | 1.79234 | Downstream (1-2kb) | 11 | 89300638 | 89302332 | 1695 | 2 | ENSMUST00000061728.4 | 3602 | Nog | ENSMUSG00000048616.4 |
| chr11 | 89299705 | 89300552 | 4.01451 | 5.15828 | 1.86571 | Downstream (<1kb) | 11 | 89300638 | 89302332 | 1695 | 2 | ENSMUST00000061728.4 | 1780 | Nog | ENSMUSG00000048616.4 |
| chr11 | 89301396 | 89303175 | 3.91944 | 5.03274 | 1.80921 | Promoter | 11 | 89300638 | 89302332 | 1695 | 2 | ENSMUST00000061728.4 | 0 | Nog | ENSMUSG00000048616.4 |
| chr8 | 84943340 | 84943438 | 3.34153 | 3.83966 | 1.01733 | Distal Intergenic | 8 | 84946991 | 84956603 | 9613 | 1 | ENSMUST00000067472.8 | -3553 | Rtbdn | ENSMUSG00000048617.11 |
| chr8 | 84944288 | 84944363 | 3.83665 | 4.74855 | 1.55778 | Distal Intergenic | 8 | 84946991 | 84956603 | 9613 | 1 | ENSMUST00000067472.8 | -2628 | Rtbdn | ENSMUSG00000048617.11 |
| chr8 | 84944709 | 84946203 | 4.15658 | 5.42804 | 2.07205 | Promoter | 8 | 84946991 | 84956603 | 9613 | 1 | ENSMUST00000067472.8 | -788 | Rtbdn | ENSMUSG00000048617.11 |
| chr8 | 84946534 | 84947165 | 3.50266 | 4.17646 | 1.20401 | Promoter | 8 | 84946991 | 84956603 | 9613 | 1 | ENSMUST00000067472.8 | 0 | Rtbdn | ENSMUSG00000048617.11 |
| chr8 | 84948136 | 84948740 | 3.60365 | 4.31387 | 1.29335 | Intron (ENSMUST00000067472.8/ENSMUSG00000048617.11, intron 1 of 5) | 8 | 84949860 | 84956603 | 6744 | 1 | ENSMUST00000121880.2 | -1120 | Rtbdn | ENSMUSG00000048617.11 |
| chr8 | 84950063 | 84950688 | 3.4745 | 4.16166 | 1.26689 | Promoter | 8 | 84949860 | 84956603 | 6744 | 1 | ENSMUST00000121880.2 | 204 | Rtbdn | ENSMUSG00000048617.11 |
| chr8 | 84952738 | 84953112 | 3.64895 | 4.41647 | 1.39836 | Exon (ENSMUST00000067472.8/ENSMUSG00000048617.11, exon 3 of 6) | 8 | 84951181 | 84956511 | 5331 | 1 | ENSMUST00000152378.2 | 1558 | Rtbdn | ENSMUSG00000048617.11 |
| chr3 | 146685231 | 146685486 | 3.44806 | 4.21747 | 1.27755 | Promoter | 3 | 146680450 | 146685471 | 5022 | 2 | ENSMUST00000144837.1 | 0 | Samd13 | ENSMUSG00000048652.7 |
| chr4 | 97581734 | 97582263 | 3.76034 | 4.63704 | 1.57116 | Intron (ENSMUST00000148653.2/ENSMUSG00000048747.9, intron 1 of 4) | 4 | 97568133 | 97584596 | 16464 | 2 | ENSMUST00000148653.2 | 2333 | E130114P18Rik | ENSMUSG00000048747.9 |
| chr4 | 97584435 | 97584741 | 3.73745 | 4.57026 | 1.47337 | Promoter | 4 | 97568133 | 97584596 | 16464 | 2 | ENSMUST00000148653.2 | 0 | E130114P18Rik | ENSMUSG00000048747.9 |
| chr4 | 97585259 | 97585555 | 3.46115 | 4.04906 | 1.11903 | Promoter | 4 | 97568133 | 97584636 | 16504 | 2 | ENSMUST00000147925.2 | -624 | E130114P18Rik | ENSMUSG00000048747.9 |
| chr9 | 110857763 | 110858271 | 3.43965 | 4.11491 | 1.20423 | Promoter | 9 | 110857967 | 110864628 | 6662 | 1 | ENSMUST00000051097.1 | 0 | Prss50 | ENSMUSG00000048752.3 |
| chr11 | 96330994 | 96331072 | 3.69626 | 4.54584 | 1.47286 | Intron (ENSMUST00000093944.4/ENSMUSG00000048763.6, intron 1 of 3) | 11 | 96328062 | 96344303 | 16242 | 1 | ENSMUST00000123091.2 | 2933 | Hoxb3 | ENSMUSG00000048763.6 |
| chr11 | 96332880 | 96333246 | 3.69119 | 4.79708 | 1.62502 | Intron (ENSMUST00000093944.4/ENSMUSG00000048763.6, intron 1 of 3) | 11 | 96328062 | 96344303 | 16242 | 1 | ENSMUST00000123091.2 | 4819 | Hoxb3 | ENSMUSG00000048763.6 |
| chr11 | 96340945 | 96341096 | 3.29429 | 3.76863 | 1.01733 | Promoter | 11 | 96340945 | 96341774 | 830 | 1 | ENSMUST00000143462.1 | 1 | Hoxb3 | ENSMUSG00000048763.6 |
| chr6 | 147261307 | 147261668 | 3.2709 | 3.82286 | 1.02941 | Intron (ENSMUST00000052296.6/ENSMUSG00000048776.6, intron 2 of 3) | 6 | 147252101 | 147264167 | 12067 | 2 | ENSMUST00000052296.6 | 2499 | Pthlh | ENSMUSG00000048776.6 |
| chr6 | 147262998 | 147263262 | 3.50967 | 4.47088 | 1.43026 | Promoter | 6 | 147252101 | 147264167 | 12067 | 2 | ENSMUST00000052296.6 | 905 | Pthlh | ENSMUSG00000048776.6 |
| chr6 | 147264007 | 147264489 | 3.69355 | 4.55382 | 1.50533 | Promoter | 6 | 147252101 | 147264167 | 12067 | 2 | ENSMUST00000052296.6 | 0 | Pthlh | ENSMUSG00000048776.6 |
| chr6 | 147266133 | 147266315 | 3.82475 | 4.80788 | 1.69045 | Distal Intergenic | 6 | 147252101 | 147264167 | 12067 | 2 | ENSMUST00000052296.6 | -1967 | Pthlh | ENSMUSG00000048776.6 |
| chr7 | 100970717 | 100970898 | 3.5454 | 4.21351 | 1.24213 | Distal Intergenic | 7 | 100937638 | 100964391 | 26754 | 2 | ENSMUST00000060174.4 | -6327 | P2ry6 | ENSMUSG00000048779.4 |
| chr7 | 100974005 | 100974529 | 3.91636 | 4.97475 | 1.77173 | Distal Intergenic | 7 | 100937638 | 100964391 | 26754 | 2 | ENSMUST00000060174.4 | -9615 | P2ry6 | ENSMUSG00000048779.4 |
| chr7 | 114745615 | 114745992 | 3.68061 | 4.48818 | 1.47245 | Promoter | 7 | 114745740 | 114791296 | 45557 | 1 | ENSMUST00000136645.2 | 0 | Insc | ENSMUSG00000048782.10 |
| chr6 | 90423023 | 90423098 | 3.33509 | 3.84032 | 1.05293 | Intron (ENSMUST00000062750.6/ENSMUSG00000048794.9, intron 1 of 14) | 6 | 90426484 | 90427544 | 1061 | 2 | ENSMUST00000135757.1 | 4446 | Ccdc37 | ENSMUSG00000048794.9 |
| chr6 | 90428107 | 90428402 | 3.23338 | 3.67724 | 1.01733 | Promoter | 6 | 90415755 | 90428413 | 12659 | 2 | ENSMUST00000129666.2 | 11 | Ccdc37 | ENSMUSG00000048794.9 |
| chr6 | 90428761 | 90429015 | 3.60529 | 4.32667 | 1.32778 | Promoter | 6 | 90403738 | 90428787 | 25050 | 2 | ENSMUST00000153843.2 | 0 | Ccdc37 | ENSMUSG00000048794.9 |
| chr1 | 38835629 | 38836053 | 4.46434 | 6.26901 | 2.72768 | Promoter | 1 | 38793669 | 38836090 | 42422 | 2 | ENSMUST00000147695.2 | 37 | Lonrf2 | ENSMUSG00000048814.5 |
| chr4 | 47011079 | 47011154 | 3.41324 | 3.99786 | 1.18423 | Promoter | 4 | 47008316 | 47010781 | 2466 | 2 | ENSMUST00000135777.1 | -299 | Gm568 | ENSMUSG00000048824.7 |
| chr17 | 14165607 | 14166284 | 3.64286 | 4.38886 | 1.34449 | Distal Intergenic | 17 | 14195231 | 14203831 | 8601 | 2 | ENSMUST00000053218.5 | 37547 | Dact2 | ENSMUSG00000048826.6 |
| chr17 | 14167322 | 14167567 | 3.59325 | 4.29091 | 1.26859 | Distal Intergenic | 17 | 14195231 | 14203831 | 8601 | 2 | ENSMUST00000053218.5 | 36264 | Dact2 | ENSMUSG00000048826.6 |
| chr17 | 14202650 | 14203969 | 3.9759 | 5.18637 | 1.93195 | Promoter | 17 | 14195231 | 14203831 | 8601 | 2 | ENSMUST00000053218.5 | 0 | Dact2 | ENSMUSG00000048826.6 |
| chr17 | 14204313 | 14204574 | 3.39433 | 3.92747 | 1.04619 | Promoter | 17 | 14195231 | 14203831 | 8601 | 2 | ENSMUST00000053218.5 | -483 | Dact2 | ENSMUSG00000048826.6 |
| chr17 | 14230947 | 14231418 | 3.44657 | 4.04235 | 1.13831 | Distal Intergenic | 17 | 14195231 | 14203831 | 8601 | 2 | ENSMUST00000053218.5 | -27117 | Dact2 | ENSMUSG00000048826.6 |
| chr17 | 14233393 | 14233665 | 3.68615 | 4.62202 | 1.53127 | Distal Intergenic | 17 | 14195231 | 14203831 | 8601 | 2 | ENSMUST00000053218.5 | -29563 | Dact2 | ENSMUSG00000048826.6 |
| chr11 | 16257081 | 16258008 | 3.9802 | 5.04554 | 1.79969 | Promoter | 11 | 16257724 | 16284186 | 26463 | 1 | ENSMUST00000109645.3 | 0 | Vstm2a | ENSMUSG00000048834.10 |
| chr11 | 16258905 | 16259723 | 4.01461 | 5.14362 | 1.87865 | Intron (ENSMUST00000109645.3/ENSMUSG00000048834.10, intron 1 of 4) | 11 | 16257803 | 16427310 | 169508 | 1 | ENSMUST00000109647.2 | 1103 | Vstm2a | ENSMUSG00000048834.10 |
| chr11 | 16260780 | 16261016 | 4.0013 | 5.1966 | 1.8951 | Intron (ENSMUST00000109645.3/ENSMUSG00000048834.10, intron 2 of 4) | 11 | 16257803 | 16427310 | 169508 | 1 | ENSMUST00000109647.2 | 2978 | Vstm2a | ENSMUSG00000048834.10 |
| chr4 | 119492188 | 119492385 | 3.24329 | 3.6921 | 1.01733 | Promoter | 4 | 119474549 | 119492509 | 17961 | 2 | ENSMUST00000126728.1 | 124 | Rimkla | ENSMUSG00000048899.8 |
| chr13 | 56247893 | 56249416 | 3.72656 | 4.64815 | 1.56916 | Downstream (1-2kb) | 13 | 56250505 | 56252163 | 1659 | 2 | ENSMUST00000058475.4 | 2747 | Neurog1 | ENSMUSG00000048904.4 |
| chr13 | 56249811 | 56250330 | 3.5282 | 4.19338 | 1.25731 | Downstream (<1kb) | 13 | 56250505 | 56252163 | 1659 | 2 | ENSMUST00000058475.4 | 1833 | Neurog1 | ENSMUSG00000048904.4 |
| chr13 | 56250691 | 56253669 | 3.73171 | 4.73178 | 1.57976 | Promoter | 13 | 56250505 | 56252163 | 1659 | 2 | ENSMUST00000058475.4 | 0 | Neurog1 | ENSMUSG00000048904.4 |
| chr13 | 56254285 | 56254405 | 3.72517 | 4.55302 | 1.46334 | Distal Intergenic | 13 | 56250505 | 56252163 | 1659 | 2 | ENSMUST00000058475.4 | -2123 | Neurog1 | ENSMUSG00000048904.4 |

| | | | | | | | | | | | | | | | |
|---|---|---|---|---|---|---|---|---|---|---|---|---|---|---|---|
| chr13 | 56255345 | 56255847 | 3.90662 | 5.10511 | 1.87693 | Distal Intergenic | 13 | 56250505 | 56252163 | 1659 | 2 | ENSMUST00000058475.4 | -3183 | Neurog1 | ENSMUSG00000048904.4 |
| chr13 | 56256487 | 56257290 | 4.23645 | 5.80375 | 2.4226 | Distal Intergenic | 13 | 56250505 | 56252163 | 1659 | 2 | ENSMUST00000058475.4 | -4325 | Neurog1 | ENSMUSG00000048904.4 |
| chr13 | 56257719 | 56257919 | 3.33125 | 3.82424 | 1.01733 | Distal Intergenic | 13 | 56250505 | 56252163 | 1659 | 2 | ENSMUST00000058475.4 | -5557 | Neurog1 | ENSMUSG00000048904.4 |
| chr13 | 56258304 | 56259530 | 3.71232 | 4.58196 | 1.48316 | Distal Intergenic | 13 | 56250505 | 56252163 | 1659 | 2 | ENSMUST00000058475.4 | -6142 | Neurog1 | ENSMUSG00000048904.4 |
| chr13 | 56259891 | 56260435 | 3.80675 | 4.70385 | 1.55014 | Distal Intergenic | 13 | 56250505 | 56252163 | 1659 | 2 | ENSMUST00000058475.4 | -7729 | Neurog1 | ENSMUSG00000048904.4 |
| chr13 | 56260787 | 56261399 | 3.88082 | 4.83954 | 1.63245 | Distal Intergenic | 13 | 56250505 | 56252163 | 1659 | 2 | ENSMUST00000058475.4 | -8625 | Neurog1 | ENSMUSG00000048904.4 |
| chr13 | 56262034 | 56262276 | 3.62698 | 4.62076 | 1.5049 | Distal Intergenic | 13 | 56250505 | 56252163 | 1659 | 2 | ENSMUST00000058475.4 | -9872 | Neurog1 | ENSMUSG00000048904.4 |
| chr17 | 62880494 | 62880728 | 3.54545 | 4.2939 | 1.33535 | Promoter | 17 | 62607319 | 62881144 | 273826 | 2 | ENSMUST00000078839.4 | 416 | Efna5 | ENSMUSG00000048915.7 |
| chr1 | 10993199 | 10994081 | 3.56148 | 4.2496 | 1.26658 | Promoter | 1 | 10993465 | 11303681 | 310217 | 1 | ENSMUST00000027056.6 | 0 | Prex2 | ENSMUSG00000048960.8 |
| chr1 | 11329411 | 11329486 | 3.42058 | 3.97107 | 1.06004 | Distal Intergenic | 1 | 11283993 | 11298204 | 14212 | 1 | ENSMUST00000188154.1 | 45419 | Prex2 | ENSMUSG00000048960.8 |
| chr8 | 69889012 | 69889112 | 3.40231 | 3.97133 | 1.15137 | Promoter | 8 | 69887790 | 69888185 | 396 | 2 | ENSMUST00000152456.1 | -828 | Yjefn3 | ENSMUSG00000048967.10 |
| chr5 | 139885213 | 139885371 | 3.4981 | 4.13042 | 1.19961 | Distal Intergenic | 5 | 139907943 | 139974711 | 66769 | 1 | ENSMUST00000050519.6 | -22572 | Elfn1 | ENSMUSG00000048988.7 |
| chr5 | 139888237 | 139888696 | 3.7029 | 4.53189 | 1.47225 | Distal Intergenic | 5 | 139907943 | 139974711 | 66769 | 1 | ENSMUST00000050519.6 | -19247 | Elfn1 | ENSMUSG00000048988.7 |
| chr5 | 139889045 | 139889172 | 4.05621 | 5.31326 | 1.98843 | Distal Intergenic | 5 | 139907943 | 139974711 | 66769 | 1 | ENSMUST00000050519.6 | -18771 | Elfn1 | ENSMUSG00000048988.7 |
| chr5 | 139890643 | 139891336 | 4.04698 | 5.17135 | 1.87929 | Distal Intergenic | 5 | 139907943 | 139974711 | 66769 | 1 | ENSMUST00000050519.6 | -16607 | Elfn1 | ENSMUSG00000048988.7 |
| chr5 | 139892450 | 139892750 | 3.35536 | 3.88211 | 1.09361 | Distal Intergenic | 5 | 139907943 | 139974711 | 66769 | 1 | ENSMUST00000050519.6 | -15193 | Elfn1 | ENSMUSG00000048988.7 |
| chr5 | 139893174 | 139893407 | 3.86069 | 4.84248 | 1.71634 | Distal Intergenic | 5 | 139907943 | 139974711 | 66769 | 1 | ENSMUST00000050519.6 | -14536 | Elfn1 | ENSMUSG00000048988.7 |
| chr5 | 139895353 | 139895958 | 3.69566 | 4.52492 | 1.43404 | Distal Intergenic | 5 | 139907943 | 139974711 | 66769 | 1 | ENSMUST00000050519.6 | -11985 | Elfn1 | ENSMUSG00000048988.7 |
| chr5 | 139899853 | 139900145 | 3.24853 | 3.84417 | 1.09566 | Distal Intergenic | 5 | 139907943 | 139974711 | 66769 | 1 | ENSMUST00000050519.6 | -7798 | Elfn1 | ENSMUSG00000048988.7 |
| chr5 | 139901740 | 139902032 | 3.2695 | 3.73138 | 1.01733 | Distal Intergenic | 5 | 139907943 | 139974711 | 66769 | 1 | ENSMUST00000050519.6 | -5911 | Elfn1 | ENSMUSG00000048988.7 |
| chr5 | 139904730 | 139905000 | 3.6196 | 4.64629 | 1.51644 | Distal Intergenic | 5 | 139907943 | 139974711 | 66769 | 1 | ENSMUST00000050519.6 | -2943 | Elfn1 | ENSMUSG00000048988.7 |
| chr5 | 139907018 | 139910131 | 3.8709 | 4.93829 | 1.71804 | Promoter | 5 | 139907943 | 139974711 | 66769 | 1 | ENSMUST00000050519.6 | 0 | Elfn1 | ENSMUSG00000048988.7 |
| chr5 | 139910483 | 139911782 | 3.52563 | 4.20757 | 1.23083 | Intron (ENSMUST00000050519.6/ENSMUSG00000048988.7, intron 1 of 1) | 5 | 139907943 | 139974711 | 66769 | 1 | ENSMUST00000050519.6 | 2541 | Elfn1 | ENSMUSG00000048988.7 |
| chr5 | 139912299 | 139913188 | 3.94019 | 5.03228 | 1.80659 | Intron (ENSMUST00000050519.6/ENSMUSG00000048988.7, intron 1 of 1) | 5 | 139907943 | 139974711 | 66769 | 1 | ENSMUST00000050519.6 | 4357 | Elfn1 | ENSMUSG00000048988.7 |
| chr5 | 139914200 | 139914357 | 3.37288 | 3.88694 | 1.01733 | Intron (ENSMUST00000050519.6/ENSMUSG00000048988.7, intron 1 of 1) | 5 | 139907943 | 139974711 | 66769 | 1 | ENSMUST00000050519.6 | 6258 | Elfn1 | ENSMUSG00000048988.7 |
| chr5 | 139914720 | 139915135 | 3.46872 | 4.06266 | 1.12669 | Intron (ENSMUST00000050519.6/ENSMUSG00000048988.7, intron 1 of 1) | 5 | 139907943 | 139974711 | 66769 | 1 | ENSMUST00000050519.6 | 6778 | Elfn1 | ENSMUSG00000048988.7 |
| chr5 | 139915883 | 139915987 | 3.83507 | 4.79634 | 1.68826 | Intron (ENSMUST00000050519.6/ENSMUSG00000048988.7, intron 1 of 1) | 5 | 139907943 | 139974711 | 66769 | 1 | ENSMUST00000050519.6 | 7941 | Elfn1 | ENSMUSG00000048988.7 |
| chr5 | 139930787 | 139932227 | 3.72088 | 4.54971 | 1.46494 | Intron (ENSMUST00000050519.6/ENSMUSG00000048988.7, intron 1 of 1) | 5 | 139907943 | 139974711 | 66769 | 1 | ENSMUST00000050519.6 | 22845 | Elfn1 | ENSMUSG00000048988.7 |
| chr5 | 139936372 | 139936497 | 3.43583 | 4.01578 | 1.13674 | Intron (ENSMUST00000050519.6/ENSMUSG00000048988.7, intron 1 of 1) | 5 | 139907943 | 139974711 | 66769 | 1 | ENSMUST00000050519.6 | 28430 | Elfn1 | ENSMUSG00000048988.7 |
| chr5 | 139970546 | 139970702 | 3.45388 | 4.11773 | 1.23806 | Intron (ENSMUST00000050519.6/ENSMUSG00000048988.7, intron 1 of 1) | 5 | 139907943 | 139974711 | 66769 | 1 | ENSMUST00000050519.6 | 62604 | Elfn1 | ENSMUSG00000048988.7 |
| chr5 | 139973299 | 139973430 | 3.4739 | 4.13833 | 1.24635 | Exon (ENSMUST00000050519.6/ENSMUSG00000048988.7, exon 2 of 2) | 5 | 139907943 | 139974711 | 66769 | 1 | ENSMUST00000050519.6 | 65357 | Elfn1 | ENSMUSG00000048988.7 |
| chr5 | 139977240 | 139977786 | 3.40237 | 3.93906 | 1.04403 | Distal Intergenic | 5 | 139907943 | 139974711 | 66769 | 1 | ENSMUST00000050519.6 | 69298 | Elfn1 | ENSMUSG00000048988.7 |
| chr12 | 113185515 | 113188338 | 3.70079 | 4.56564 | 1.50016 | Promoter | 12 | 113185903 | 113189520 | 3618 | 1 | ENSMUST00000058491.6 | 0 | Tmem121 | ENSMUSG00000049036.6 |
| chr2 | 71979963 | 71980191 | 3.7948 | 4.90933 | 1.74681 | Distal Intergenic | 2 | 71981240 | 72257474 | 276235 | 1 | ENSMUST00000090826.6 | -1049 | Rapgef4 | ENSMUSG00000049044.11 |
| chr2 | 71980805 | 71981430 | 3.88273 | 4.97295 | 1.81765 | Promoter | 2 | 71981240 | 72257474 | 276235 | 1 | ENSMUST00000090826.6 | 0 | Rapgef4 | ENSMUSG00000049044.11 |
| chr2 | 72092122 | 72093300 | 3.63213 | 4.4393 | 1.40409 | Intron (ENSMUST00000090826.6/ENSMUSG00000049044.11, intron 4 of 30) | 2 | 72054985 | 72141477 | 86493 | 1 | ENSMUST00000124004.1 | 37138 | Rapgef4 | ENSMUSG00000049044.11 |
| chr2 | 72094950 | 72095089 | 4.04076 | 5.19499 | 1.95826 | Intron (ENSMUST00000090826.6/ENSMUSG00000049044.11, intron 4 of 30) | 2 | 72054985 | 72141477 | 86493 | 1 | ENSMUST00000124004.1 | 39966 | Rapgef4 | ENSMUSG00000049044.11 |
| chr3 | 45378726 | 45379587 | 3.6812 | 4.54997 | 1.44312 | Promoter | 3 | 45378398 | 45434566 | 56169 | 1 | ENSMUST00000166126.1 | 329 | Pcdh10 | ENSMUSG00000049100.10 |
| chr3 | 45380355 | 45380497 | 4.00815 | 5.08094 | 1.79795 | Exon (ENSMUST00000171554.2/ENSMUSG00000049100.10, exon 1 of 5) | 3 | 45378398 | 45434566 | 56169 | 1 | ENSMUST00000166126.1 | 1958 | Pcdh10 | ENSMUSG00000049100.10 |
| chr3 | 45380851 | 45381910 | 4.26269 | 5.66885 | 2.25645 | Exon (ENSMUST00000171554.2/ENSMUSG00000049100.10, exon 1 of 5) | 3 | 45378398 | 45434566 | 56169 | 1 | ENSMUST00000166126.1 | 2454 | Pcdh10 | ENSMUSG00000049100.10 |
| chr3 | 45382822 | 45383036 | 3.63135 | 4.43562 | 1.43365 | Intron (ENSMUST00000171554.2/ENSMUSG00000049100.10, intron 1 of 4) | 3 | 45378398 | 45434566 | 56169 | 1 | ENSMUST00000166126.1 | 4425 | Pcdh10 | ENSMUSG00000049100.10 |
| chr3 | 45383862 | 45383996 | 3.34697 | 3.84786 | 1.01733 | Intron (ENSMUST00000171554.2/ENSMUSG00000049100.10, intron 2 of 4) | 3 | 45378398 | 45434566 | 56169 | 1 | ENSMUST00000166126.1 | 5465 | Pcdh10 | ENSMUSG00000049100.10 |
| chr3 | 45384363 | 45384542 | 3.80972 | 4.79189 | 1.6433 | Intron (ENSMUST00000171554.2/ENSMUSG00000049100.10, intron 3 of 4) | 3 | 45378398 | 45434566 | 56169 | 1 | ENSMUST00000166126.1 | 5966 | Pcdh10 | ENSMUSG00000049100.10 |
| chr6 | 126160012 | 126160133 | 3.64302 | 4.43365 | 1.44476 | Intron (ENSMUST00000050484.7/ENSMUSG00000049107.8, intron 1 of 1) | 6 | 126101413 | 126166744 | 65332 | 2 | ENSMUST00000050484.7 | 6611 | Ntf3 | ENSMUSG00000049107.8 |
| chr6 | 126161090 | 126161962 | 3.36837 | 4.17981 | 1.24071 | Intron (ENSMUST00000050484.7/ENSMUSG00000049107.8, intron 1 of 1) | 6 | 126101413 | 126166744 | 65332 | 2 | ENSMUST00000050484.7 | 4782 | Ntf3 | ENSMUSG00000049107.8 |
| chr6 | 126162417 | 126162583 | 3.62217 | 4.37063 | 1.33227 | Intron (ENSMUST00000050484.7/ENSMUSG00000049107.8, intron 1 of 1) | 6 | 126101413 | 126166744 | 65332 | 2 | ENSMUST00000050484.7 | 4161 | Ntf3 | ENSMUSG00000049107.8 |
| chr6 | 126163457 | 126164237 | 3.86743 | 4.8143 | 1.61808 | Intron (ENSMUST00000050484.7/ENSMUSG00000049107.8, intron 1 of 1) | 6 | 126101413 | 126166744 | 65332 | 2 | ENSMUST00000050484.7 | 2507 | Ntf3 | ENSMUSG00000049107.8 |
| chr6 | 126164875 | 126165154 | 3.37828 | 3.89509 | 1.01733 | Intron (ENSMUST00000050484.7/ENSMUSG00000049107.8, intron 1 of 1) | 6 | 126101413 | 126166744 | 65332 | 2 | ENSMUST00000050484.7 | 1590 | Ntf3 | ENSMUSG00000049107.8 |
| chr10 | 28633655 | 28633902 | 3.63187 | 4.41542 | 1.45323 | Distal Intergenic | 10 | 28668360 | 28789920 | 121561 | 1 | ENSMUST00000060409.7 | -34458 | Themis | ENSMUSG00000049109.9 |
| chr10 | 28638563 | 28639176 | 3.86694 | 4.88478 | 1.72868 | Distal Intergenic | 10 | 28668360 | 28789920 | 121561 | 1 | ENSMUST00000060409.7 | -29184 | Themis | ENSMUSG00000049109.9 |
| chr6 | 112488416 | 112488728 | 3.43355 | 4.04406 | 1.21641 | Intron (ENSMUST00000053306.6/ENSMUSG00000049112.8, intron 1 of 1) | 6 | 112473684 | 112489808 | 16125 | 2 | ENSMUST00000053306.6 | 1080 | Oxtr | ENSMUSG00000049112.8 |
| chr6 | 112489140 | 112490223 | 3.84896 | 4.99497 | 1.80121 | Promoter | 6 | 112473684 | 112489808 | 16125 | 2 | ENSMUST00000053306.6 | 0 | Oxtr | ENSMUSG00000049112.8 |
| chr13 | 30337010 | 30337341 | 3.8965 | 4.89682 | 1.71591 | Promoter | 13 | 30336441 | 30382867 | 46427 | 1 | ENSMUST00000066412.7 | 570 | Agtr1a | ENSMUSG00000049115.9 |
| chr4 | 74014791 | 74015460 | 3.90219 | 4.9997 | 1.83986 | Intron (ENSMUST00000098006.3/ENSMUSG00000049122.12, intron 1 of 14) | 4 | 74013444 | 74189928 | 176485 | 1 | ENSMUST00000084474.5 | 1348 | Frmd3 | ENSMUSG00000049122.12 |
| chr1 | 161171075 | 161171458 | 3.52102 | 4.2976 | 1.30993 | Distal Intergenic | 1 | 161156843 | 161159314 | 2472 | 2 | ENSMUST00000178511.2 | -11762 | 4930469G21Rik | ENSMUSG00000049160.10 |
| chr3 | 107976628 | 107976703 | 3.27364 | 3.7401 | 1.02584 | Distal Intergenic | 3 | 107979632 | 107979868 | 237 | 1 | ENSMUST00000138778.1 | -2929 | Gm12497 | ENSMUSG00000049231.6 |
| chr13 | 107249033 | 107249285 | 3.24788 | 3.69896 | 1.01733 | Distal Intergenic | 13 | 107413865 | 107414767 | 903 | 2 | ENSMUST00000061241.6 | 165482 | Apoo-ps | ENSMUSG00000049233.6 |
| chr13 | 107503886 | 107503996 | 3.28527 | 3.75505 | 1.01733 | Distal Intergenic | 13 | 107413865 | 107414767 | 903 | 2 | ENSMUST00000061241.6 | -89120 | Apoo-ps | ENSMUSG00000049233.6 |
| chr5 | 123878398 | 123878749 | 3.69622 | 4.66322 | 1.55752 | 3' UTR | 5 | 123876736 | 123880020 | 3285 | 2 | ENSMUST00000164267.1 | 1271 | Hcar1 | ENSMUSG00000049241.5 |
| chr2 | 42651300 | 42651434 | 3.2331 | 3.67684 | 1.01733 | Exon (ENSMUST00000142688.1/ENSMUSG00000049252.12, exon 2 of 2) | 2 | 42649878 | 42653015 | 3138 | 2 | ENSMUST00000142688.1 | 1581 | Lrp1b | ENSMUSG00000049252.12 |
| chr2 | 42652149 | 42652422 | 3.52483 | 4.27363 | 1.31485 | Promoter | 2 | 42649878 | 42653015 | 3138 | 2 | ENSMUST00000142688.1 | 593 | Lrp1b | ENSMUSG00000049252.12 |
| chr5 | 30588459 | 30590167 | 3.91547 | 4.92479 | 1.70869 | Promoter | 5 | 30588170 | 30625270 | 37101 | 1 | ENSMUST00000066295.2 | 290 | Kcnk3 | ENSMUSG00000049265.6 |
| chr5 | 30590536 | 30590819 | 3.77244 | 4.69396 | 1.60479 | Intron (ENSMUST00000066295.2/ENSMUSG00000049265.6, intron 1 of 1) | 5 | 30588170 | 30625270 | 37101 | 1 | ENSMUST00000066295.2 | 2367 | Kcnk3 | ENSMUSG00000049265.6 |
| chr5 | 30591220 | 30591363 | 3.35947 | 3.8786 | 1.05904 | Intron (ENSMUST00000066295.2/ENSMUSG00000049265.6, intron 1 of 1) | 5 | 30588170 | 30625270 | 37101 | 1 | ENSMUST00000066295.2 | 3051 | Kcnk3 | ENSMUSG00000049265.6 |
| chr5 | 30592420 | 30592496 | 3.40708 | 4.45468 | 1.40707 | Intron (ENSMUST00000066295.2/ENSMUSG00000049265.6, intron 1 of 1) | 5 | 30588170 | 30625270 | 37101 | 1 | ENSMUST00000066295.2 | 4251 | Kcnk3 | ENSMUSG00000049265.6 |
| chr9 | 40267108 | 40267360 | 3.26677 | 3.79156 | 1.07091 | Distal Intergenic | 9 | 40269216 | 40291617 | 22402 | 1 | ENSMUST00000114956.4 | -1856 | Scn3b | ENSMUSG00000049281.10 |
| chr19 | 4456564 | 4456981 | 3.886 | 4.8467 | 1.62807 | Exon (ENSMUST00000059295.4/ENSMUSG00000049303.5, exon 4 of 8) | 19 | 4447717 | 4452151 | 4435 | 2 | ENSMUST00000128114.1 | -4414 | Syt12 | ENSMUSG00000049303.5 |
| chr19 | 4476680 | 4477244 | 4.11043 | 5.39051 | 2.12339 | Promoter | 19 | 4456837 | 4477126 | 20290 | 2 | ENSMUST00000166191.1 | 0 | Syt12 | ENSMUSG00000049303.5 |
| chr9 | 14750735 | 14751011 | 3.88066 | 4.97147 | 1.75143 | Exon (ENSMUST00000061498.5/ENSMUSG00000049307.5, exon 1 of 1) | 9 | 14748459 | 14752122 | 3664 | 2 | ENSMUST00000061498.5 | 1111 | Fut4 | ENSMUSG00000049307.5 |
| chr9 | 14751343 | 14751655 | 3.51817 | 4.41532 | 1.42961 | Promoter | 9 | 14748459 | 14752122 | 3664 | 2 | ENSMUST00000061498.5 | 467 | Fut4 | ENSMUSG00000049307.5 |
| chr9 | 14752210 | 14752395 | 3.51108 | 4.19273 | 1.32173 | Promoter | 9 | 14748459 | 14752122 | 3664 | 2 | ENSMUST00000061498.5 | -89 | Fut4 | ENSMUSG00000049307.5 |
| chr9 | 41890731 | 41890806 | 3.54553 | 4.38681 | 1.39059 | Distal Intergenic | 9 | 41969739 | 41974587 | 4849 | 2 | ENSMUST00000134560.1 | 83781 | Sorl1 | ENSMUSG00000049313.8 |
| chr9 | 41935180 | 41935822 | 4.07234 | 5.36092 | 2.02549 | Distal Intergenic | 9 | 41969739 | 41974587 | 4849 | 2 | ENSMUST00000134560.1 | 38765 | Sorl1 | ENSMUSG00000049313.8 |
| chr9 | 41966518 | 41966628 | 3.83352 | 4.80326 | 1.6986 | 3' UTR | 9 | 41969739 | 41974587 | 4849 | 2 | ENSMUST00000134560.1 | 7959 | Sorl1 | ENSMUSG00000049313.8 |
| chr9 | 42122513 | 42124394 | 3.77564 | 4.71775 | 1.59736 | Promoter | 9 | 41964720 | 42124297 | 159578 | 2 | ENSMUST00000060989.8 | 0 | Sorl1 | ENSMUSG00000049313.8 |
| chr11 | 36537795 | 36538434 | 3.36561 | 3.91408 | 1.10287 | Intron (ENSMUST00000102801.2/ENSMUSG00000049336.11, intron 4 of 30) | 11 | 36163719 | 36495139 | 331421 | 2 | ENSMUST00000155147.1 | -42657 | Tenm2 | ENSMUSG00000049336.11 |
| chr11 | 36676060 | 36676135 | 3.54724 | 4.25545 | 1.33836 | Intron (ENSMUST00000102801.2/ENSMUSG00000049336.11, intron 4 of 30) | 11 | 36140752 | 36677745 | 536994 | 2 | ENSMUST00000149787.2 | 1610 | Tenm2 | ENSMUSG00000049336.11 |
| chr11 | 36676495 | 36677116 | 3.70938 | 4.58852 | 1.52008 | Promoter | 11 | 36140752 | 36677745 | 536994 | 2 | ENSMUST00000149787.2 | 629 | Tenm2 | ENSMUSG00000049336.11 |
| chr11 | 36677457 | 36677846 | 3.89485 | 4.89687 | 1.72438 | Promoter | 11 | 36140752 | 36677745 | 536994 | 2 | ENSMUST00000149787.2 | 0 | Tenm2 | ENSMUSG00000049336.11 |
| chr11 | 37231890 | 37232171 | 3.57482 | 4.44646 | 1.37218 | Intron (ENSMUST00000102801.2/ENSMUSG00000049336.11, intron 1 of 30) | 11 | 36006656 | 37235882 | 1229227 | 2 | ENSMUST00000102801.2 | 3711 | Tenm2 | ENSMUSG00000049336.11 |
| chr11 | 37233493 | 37234064 | 3.70451 | 4.5474 | 1.49824 | Intron (ENSMUST00000102801.2/ENSMUSG00000049336.11, intron 1 of 30) | 11 | 36006656 | 37235882 | 1229227 | 2 | ENSMUST00000102801.2 | 1818 | Tenm2 | ENSMUSG00000049336.11 |
| chr11 | 37234560 | 37235469 | 3.52359 | 4.19487 | 1.2401 | Promoter | 11 | 36006656 | 37235882 | 1229227 | 2 | ENSMUST00000102801.2 | 413 | Tenm2 | ENSMUSG00000049336.11 |
| chr11 | 37236134 | 37236964 | 4.19054 | 5.55479 | 2.15531 | Promoter | 11 | 36944205 | 37235964 | 291760 | 2 | ENSMUST00000157030.1 | -171 | Tenm2 | ENSMUSG00000049336.11 |
| chr3 | 138067019 | 138067305 | 3.59121 | 4.59725 | 1.52015 | Promoter | 3 | 138048761 | 138067388 | 18628 | 2 | ENSMUST00000053318.3 | 83 | Gm5105 | ENSMUSG00000049349.3 |
| chr3 | 138067704 | 138067802 | 3.22229 | 3.66064 | 1.0169 | Promoter | 3 | 138048761 | 138067388 | 18628 | 2 | ENSMUST00000053318.3 | -317 | Gm5105 | ENSMUSG00000049349.3 |
| chr7 | 127054351 | 127054663 | 3.28721 | 3.84991 | 1.07862 | Distal Intergenic | 7 | 127050156 | 127051977 | 1822 | 2 | ENSMUST00000051122.5 | -2375 | Zg16 | ENSMUSG00000049350.5 |
| chr10 | 26079093 | 26079393 | 3.59527 | 4.53851 | 1.48058 | Promoter | 10 | 25991186 | 26079052 | 87867 | 2 | ENSMUST00000066049.6 | -42 | Tmem200a | ENSMUSG00000049420.8 |
| chr11 | 53469701 | 53469868 | 3.51685 | 4.16782 | 1.23884 | Promoter | 11 | 53469749 | 53470479 | 731 | 2 | ENSMUST00000057722.2 | 611 | Gm9837 | ENSMUSG00000049427.2 |
| chr11 | 53470357 | 53470507 | 3.57479 | 4.2779 | 1.3191 | Promoter | 11 | 53469749 | 53470479 | 731 | 2 | ENSMUST00000057722.2 | 0 | Gm9837 | ENSMUSG00000049427.2 |
| chr11 | 53474652 | 53474840 | 3.37704 | 3.93542 | 1.1541 | Downstream (1-2kb) | 11 | 53469749 | 53470479 | 731 | 2 | ENSMUST00000057722.2 | -4174 | Gm9837 | ENSMUSG00000049427.2 |
| chr9 | 95843963 | 95844051 | 3.47129 | 4.28595 | 1.31208 | Intron (ENSMUST00000093800.3/ENSMUSG00000049493.8, intron 1 of 15) | 9 | 95752642 | 95845286 | 92645 | 2 | ENSMUST00000093800.3 | 1235 | Pls1 | ENSMUSG00000049493.8 |
| chr9 | 95844749 | 95845256 | 4.16624 | 5.59626 | 2.19628 | Promoter | 9 | 95752642 | 95845286 | 92645 | 2 | ENSMUST00000093800.3 | 30 | Pls1 | ENSMUSG00000049493.8 |
| chr11 | 104160489 | 104160713 | 3.38636 | 3.92101 | 1.06587 | Intron (ENSMUST00000093925.4/ENSMUSG00000018634.10, intron 3 of 12) | 11 | 104186327 | 104188551 | 2225 | 1 | ENSMUST00000107000.1 | -25614 | Sppl2c | ENSMUSG00000049506.7 |
| chr9 | 81632305 | 81633408 | 3.68177 | 4.57962 | 1.54583 | Promoter | 9 | 81631343 | 81632615 | 1273 | 2 | ENSMUST00000051005.4 | 0 | Htr1b | ENSMUSG00000049511.5 |
| chr9 | 81634214 | 81635506 | 3.63996 | 4.54138 | 1.48826 | Promoter | 9 | 81628291 | 81633828 | 5538 | 2 | ENSMUST00000183482.1 | -387 | Htr1b | ENSMUSG00000049511.5 |
| chr9 | 81636511 | 81636711 | 3.84279 | 4.83334 | 1.68701 | Intron (ENSMUST00000181893.2/ENSMUSG00000097466.3, intron 1 of 2) | 9 | 81628291 | 81633828 | 5538 | 2 | ENSMUST00000183482.1 | -2684 | Htr1b | ENSMUSG00000049511.5 |
| chr15 | 101550224 | 101550392 | 3.57275 | 4.25757 | 1.25615 | Promoter | 15 | 101541229 | 101550659 | 9431 | 2 | ENSMUST00000023713.8 | 267 | Krt82 | ENSMUSG00000049548.8 |
| chr15 | 101550964 | 101551052 | 3.45685 | 4.11621 | 1.20829 | Promoter | 15 | 101541229 | 101550659 | 9431 | 2 | ENSMUST00000023713.8 | -306 | Krt82 | ENSMUSG00000049548.8 |
| chr15 | 101552028 | 101552163 | 3.39325 | 3.96529 | 1.17358 | Downstream (<1kb) | 15 | 101541229 | 101550659 | 9431 | 2 | ENSMUST00000023713.8 | -1370 | Krt82 | ENSMUSG00000049548.8 |
| chr5 | 135250474 | 135251063 | 3.77465 | 4.84568 | 1.6849 | Promoter | 5 | 135248938 | 135251230 | 2293 | 2 | ENSMUST00000062572.2 | 167 | Fzd9 | ENSMUSG00000049551.2 |
| chr5 | 135275840 | 135276136 | 3.28597 | 4.10547 | 1.18863 | Distal Intergenic | 5 | 135248938 | 135251230 | 2293 | 2 | ENSMUST00000062572.2 | -24611 | Fzd9 | ENSMUSG00000049551.2 |
| chr9 | 56634821 | 56635265 | 3.69225 | 4.58674 | 1.50159 | Promoter | 9 | 56619088 | 56635628 | 16541 | 2 | ENSMUST00000114256.1 | 363 | Lingo1 | ENSMUSG00000049556.4 |
| chr9 | 56713101 | 56713813 | 3.34448 | 3.86912 | 1.10414 | Distal Intergenic | 9 | 56618475 | 56685253 | 66779 | 2 | ENSMUST00000053568.1 | -27849 | Lingo1 | ENSMUSG00000049556.4 |
| chr17 | 37210930 | 37211935 | 3.73943 | 4.57577 | 1.47052 | Promoter | 17 | 37210894 | 37211867 | 974 | 2 | ENSMUST00000060728.5 | 0 | Olfr95 | ENSMUSG00000049561.7 |
| chr17 | 37212271 | 37212462 | 3.46694 | 4.06819 | 1.15662 | Promoter | 17 | 37210894 | 37211867 | 974 | 2 | ENSMUST00000060728.5 | -405 | Olfr95 | ENSMUSG00000049561.7 |
| chr8 | 122341859 | 122343549 | 3.57899 | 4.35234 | 1.37578 | Intron (ENSMUST00000127664.1/ENSMUSG00000092329.1, intron 1 of 15) | 8 | 122333786 | 122335761 | 1976 | 1 | ENSMUST00000176690.1 | 8074 | Zfpm1 | ENSMUSG00000049577.9 |
| chr8 | 122344365 | 122344678 | 3.47516 | 4.09755 | 1.21755 | Intron (ENSMUST00000127664.1/ENSMUSG00000092329.1, intron 1 of 15) | 8 | 122333786 | 122335761 | 1976 | 1 | ENSMUST00000176690.1 | 10580 | Zfpm1 | ENSMUSG00000049577.9 |
| chr7 | 98360412 | 98361396 | 3.96567 | 5.06062 | 1.82935 | Promoter | 7 | 98350668 | 98361288 | 10621 | 2 | ENSMUST00000094161.5 | 0 | Tsku | ENSMUSG00000049580.7 |
| chr7 | 87583821 | 87584624 | 3.33064 | 3.9147 | 1.06675 | Distal Intergenic | 7 | 87602544 | 88130869 | 528326 | 1 | ENSMUST00000107263.3 | -17920 | Grm5 | ENSMUSG00000049583.9 |
| chr1 | 172561950 | 172562241 | 3.53864 | 4.23674 | 1.33789 | Exon (ENSMUST00000061835.4/ENSMUSG00000049598.4, exon 6 of 7) | 1 | 172559906 | 172563716 | 3811 | 1 | ENSMUST00000177086.1 | 2045 | Vsig8 | ENSMUSG00000049598.4 |
| chr11 | 96187393 | 96187688 | 3.64468 | 4.40873 | 1.39499 | Distal Intergenic | 11 | 96194316 | 96197447 | 3132 | 1 | ENSMUST00000062709.3 | -6628 | Hoxb13 | ENSMUSG00000049604.3 |
| chr11 | 96188095 | 96188508 | 3.64927 | 4.44206 | 1.4395 | Distal Intergenic | 11 | 96194316 | 96197447 | 3132 | 1 | ENSMUST00000062709.3 | -5808 | Hoxb13 | ENSMUSG00000049604.3 |
| chr11 | 96188869 | 96189477 | 3.94566 | 5.0299 | 1.81896 | Distal Intergenic | 11 | 96194316 | 96197447 | 3132 | 1 | ENSMUST00000062709.3 | -4839 | Hoxb13 | ENSMUSG00000049604.3 |
| chr11 | 96189894 | 96190442 | 3.94129 | 5.20942 | 1.91372 | Distal Intergenic | 11 | 96194316 | 96197447 | 3132 | 1 | ENSMUST00000062709.3 | -3874 | Hoxb13 | ENSMUSG00000049604.3 |
| chr11 | 96190948 | 96192999 | 3.91008 | 4.95983 | 1.77517 | Distal Intergenic | 11 | 96194316 | 96197447 | 3132 | 1 | ENSMUST00000062709.3 | -1317 | Hoxb13 | ENSMUSG00000049604.3 |
| chr11 | 96193543 | 96195124 | 4.42347 | 5.96021 | 2.45968 | Promoter | 11 | 96194316 | 96197447 | 3132 | 1 | ENSMUST00000062709.3 | 0 | Hoxb13 | ENSMUSG00000049604.3 |
| chr11 | 96195554 | 96195790 | 3.44128 | 4.01781 | 1.11493 | Intron (ENSMUST00000062709.3/ENSMUSG00000049604.3, intron 1 of 1) | 11 | 96194316 | 96197447 | 3132 | 1 | ENSMUST00000062709.3 | 1239 | Hoxb13 | ENSMUSG00000049604.3 |
| chr11 | 96196186 | 96196261 | 3.60009 | 4.3932 | 1.40492 | 3' UTR | 11 | 96194316 | 96197447 | 3132 | 1 | ENSMUST00000062709.3 | 1871 | Hoxb13 | ENSMUSG00000049604.3 |
| chr5 | 106756060 | 106756814 | 3.42897 | 4.02615 | 1.17451 | Distal Intergenic | 5 | 106695055 | 106697287 | 2233 | 2 | ENSMUST00000127932.1 | -58774 | Zfp644 | ENSMUSG00000049606.11 |
| chr5 | 106757626 | 106757946 | 3.3155 | 4.00375 | 1.14701 | Distal Intergenic | 5 | 106695055 | 106697287 | 2233 | 2 | ENSMUST00000127932.1 | -60340 | Zfp644 | ENSMUSG00000049606.11 |
| chr5 | 106774673 | 106774992 | 3.60399 | 4.42049 | 1.40864 | Distal Intergenic | 5 | 106695055 | 106697287 | 2233 | 2 | ENSMUST00000127932.1 | -77387 | Zfp644 | ENSMUSG00000049606.11 |
| chr5 | 106775535 | 106775629 | 3.39781 | 4.0594 | 1.20773 | Distal Intergenic | 5 | 106695055 | 106697287 | 2233 | 2 | ENSMUST00000127932.1 | -78249 | Zfp644 | ENSMUSG00000049606.11 |
| chr5 | 106784412 | 106784487 | 3.38649 | 3.91092 | 1.02946 | Distal Intergenic | 5 | 106695055 | 106697287 | 2233 | 2 | ENSMUST00000127932.1 | -87126 | Zfp644 | ENSMUSG00000049606.11 |
| chr5 | 106785887 | 106786217 | 3.37101 | 3.88413 | 1.01733 | Distal Intergenic | 5 | 106695055 | 106697287 | 2233 | 2 | ENSMUST00000127932.1 | -88601 | Zfp644 | ENSMUSG00000049606.11 |
| chr17 | 23834647 | 23835045 | 3.68645 | 4.57428 | 1.54186 | Promoter | 17 | 23833740 | 23835739 | 2000 | 2 | ENSMUST00000115444.2 | 694 | Prss33 | ENSMUSG00000049620.8 |
| chr13 | 56176605 | 56176698 | 4.89569 | 6.91422 | 3.14449 | 5' UTR | 13 | 56173703 | 56178885 | 5183 | 2 | ENSMUST00000169652.1 | 2187 | Tifab | ENSMUSG00000049625.5 |
| chr13 | 56198413 | 56198706 | 3.62703 | 4.48506 | 1.45066 | Distal Intergenic | 13 | 56173703 | 56178885 | 5183 | 2 | ENSMUST00000169652.1 | -19529 | Tifab | ENSMUSG00000049625.5 |
| chr13 | 56208379 | 56209145 | 3.96229 | 5.0011 | 1.7517 | Distal Intergenic | 13 | 56173703 | 56178885 | 5183 | 2 | ENSMUST00000169652.1 | -29495 | Tifab | ENSMUSG00000049625.5 |
| chr2 | 13009419 | 13009829 | 3.68094 | 4.67994 | 1.57192 | Intron (ENSMUST00000061545.5/ENSMUSG00000049630.5, intron 1 of 1) | 2 | 13003457 | 13011747 | 8291 | 2 | ENSMUST00000061545.5 | 1918 | C1ql3 | ENSMUSG00000049630.5 |
| chr2 | 13010160 | 13012003 | 4.38321 | 6.09049 | 2.67702 | Promoter | 2 | 13003457 | 13011747 | 8291 | 2 | ENSMUST00000061545.5 | 0 | C1ql3 | ENSMUSG00000049630.5 |
| chr10 | 52016964 | 52017461 | 3.44484 | 4.08863 | 1.16729 | Distal Intergenic | 10 | 52022502 | 52028471 | 5970 | 1 | ENSMUST00000163017.2 | -5041 | Vgll2 | ENSMUSG00000049641.7 |
| chr10 | 52019726 | 52019966 | 4.05278 | 5.21647 | 1.95977 | Distal Intergenic | 10 | 52022502 | 52028471 | 5970 | 1 | ENSMUST00000163017.2 | -2536 | Vgll2 | ENSMUSG00000049641.7 |
| chr10 | 52020442 | 52020911 | 3.39555 | 4.02273 | 1.19311 | Distal Intergenic | 10 | 52022502 | 52028471 | 5970 | 1 | ENSMUST00000163017.2 | -1591 | Vgll2 | ENSMUSG00000049641.7 |
| chr10 | 52021543 | 52021719 | 3.30358 | 3.78255 | 1.01733 | Promoter | 10 | 52022502 | 52028471 | 5970 | 1 | ENSMUST00000163017.2 | -783 | Vgll2 | ENSMUSG00000049641.7 |
| chr10 | 52022075 | 52022681 | 3.37859 | 4.14555 | 1.25679 | Promoter | 10 | 52022502 | 52028471 | 5970 | 1 | ENSMUST00000163017.2 | 0 | Vgll2 | ENSMUSG00000049641.7 |
| chr10 | 52025220 | 52026072 | 3.54993 | 4.34849 | 1.39823 | Exon (ENSMUST00000163017.2/ENSMUSG00000049641.7, exon 2 of 3) | 10 | 52022528 | 52028343 | 5816 | 1 | ENSMUST00000058347.4 | 2693 | Vgll2 | ENSMUSG00000049641.7 |
| chr11 | 20670312 | 20670996 | 3.42544 | 3.9852 | 1.08407 | Distal Intergenic | 11 | 20685981 | 20692802 | 6822 | 2 | ENSMUST00000134841.1 | 21806 | Aftph | ENSMUSG00000049659.10 |
| chr11 | 20675479 | 20675673 | 3.70082 | 4.569 | 1.51877 | Distal Intergenic | 11 | 20685981 | 20692802 | 6822 | 2 | ENSMUST00000134841.1 | 17129 | Aftph | ENSMUSG00000049659.10 |
| chr7 | 45363038 | 45363723 | 3.61253 | 4.50369 | 1.45197 | Intron (ENSMUST00000107779.2/ENSMUSG00000003863.11, intron 1 of 29) | 7 | 45366163 | 45367948 | 1786 | 1 | ENSMUST00000136683.1 | -2440 | Mtag2 | ENSMUSG00000049661.10 |
| chr1 | 126492650 | 126492835 | 3.43652 | 4.00742 | 1.0911 | Promoter | 1 | 125913648 | 126492742 | 579095 | 2 | ENSMUST00000162877.2 | 0 | Nckap5 | ENSMUSG00000049690.10 |
| chr1 | 126736607 | 126737108 | 3.46481 | 4.16989 | 1.23254 | Intron (ENSMUST00000094609.4/ENSMUSG00000049690.10, intron 1 of 12) | 1 | 126525039 | 126738190 | 213152 | 2 | ENSMUST00000162664.2 | 1082 | Nckap5 | ENSMUSG00000049690.10 |
| chr1 | 126738425 | 126738575 | 3.76204 | 4.8677 | 1.6755 | Promoter | 1 | 126258401 | 126738202 | 479802 | 2 | ENSMUST00000160693.1 | -224 | Nckap5 | ENSMUSG00000049690.10 |
| chr5 | 41740447 | 41740745 | 3.5167 | 4.4849 | 1.44372 | Distal Intergenic | 5 | 41761483 | 41764501 | 3019 | 2 | ENSMUST00000060820.7 | 23756 | Nkx3-2 | ENSMUSG00000049691.8 |
| chr5 | 41744001 | 41744223 | 3.51409 | 4.34082 | 1.34669 | Distal Intergenic | 5 | 41761483 | 41764501 | 3019 | 2 | ENSMUST00000060820.7 | 20278 | Nkx3-2 | ENSMUSG00000049691.8 |
| chr5 | 41747047 | 41747315 | 3.43375 | 4.09138 | 1.22821 | Distal Intergenic | 5 | 41761483 | 41764501 | 3019 | 2 | ENSMUST00000060820.7 | 17186 | Nkx3-2 | ENSMUSG00000049691.8 |
| chr5 | 41748408 | 41748826 | 3.6593 | 4.47692 | 1.48717 | Distal Intergenic | 5 | 41761483 | 41764501 | 3019 | 2 | ENSMUST00000060820.7 | 15675 | Nkx3-2 | ENSMUSG00000049691.8 |
| chr5 | 41749912 | 41750071 | 3.27899 | 3.74566 | 1.01733 | Distal Intergenic | 5 | 41761483 | 41764501 | 3019 | 2 | ENSMUST00000060820.7 | 14430 | Nkx3-2 | ENSMUSG00000049691.8 |
| chr5 | 41751522 | 41751788 | 3.62016 | 4.3775 | 1.3945 | Distal Intergenic | 5 | 41761483 | 41764501 | 3019 | 2 | ENSMUST00000060820.7 | 12713 | Nkx3-2 | ENSMUSG00000049691.8 |
| chr5 | 41752316 | 41753105 | 3.81417 | 4.72809 | 1.59993 | Distal Intergenic | 5 | 41761483 | 41764501 | 3019 | 2 | ENSMUST00000060820.7 | 11396 | Nkx3-2 | ENSMUSG00000049691.8 |
| chr5 | 41753442 | 41754329 | 3.79765 | 4.76031 | 1.62574 | Distal Intergenic | 5 | 41761483 | 41764501 | 3019 | 2 | ENSMUST00000060820.7 | 10172 | Nkx3-2 | ENSMUSG00000049691.8 |
| chr5 | 41754701 | 41755140 | 3.87696 | 4.83705 | 1.63739 | Distal Intergenic | 5 | 41761483 | 41764501 | 3019 | 2 | ENSMUST00000060820.7 | 9361 | Nkx3-2 | ENSMUSG00000049691.8 |
| chr5 | 41755505 | 41755831 | 3.63465 | 4.36625 | 1.31205 | Distal Intergenic | 5 | 41761483 | 41764501 | 3019 | 2 | ENSMUST00000060820.7 | 8670 | Nkx3-2 | ENSMUSG00000049691.8 |
| chr5 | 41756571 | 41757181 | 3.54501 | 4.58187 | 1.48069 | Distal Intergenic | 5 | 41761483 | 41764501 | 3019 | 2 | ENSMUST00000060820.7 | 7320 | Nkx3-2 | ENSMUSG00000049691.8 |
| chr5 | 41758391 | 41759154 | 4.22643 | 5.57126 | 2.15862 | Downstream (2-3kb) | 5 | 41761483 | 41764501 | 3019 | 2 | ENSMUST00000060820.7 | 5347 | Nkx3-2 | ENSMUSG00000049691.8 |
| chr5 | 41760349 | 41762305 | 4.53856 | 6.28133 | 2.74854 | 3' UTR | 5 | 41761483 | 41764501 | 3019 | 2 | ENSMUST00000060820.7 | 2196 | Nkx3-2 | ENSMUSG00000049691.8 |
| chr5 | 41762757 | 41763114 | 3.67812 | 4.45656 | 1.38742 | Intron (ENSMUST00000060820.7/ENSMUSG00000049691.8, intron 1 of 1) | 5 | 41761483 | 41764501 | 3019 | 2 | ENSMUST00000060820.7 | 1387 | Nkx3-2 | ENSMUSG00000049691.8 |
| chr5 | 41763555 | 41764460 | 4.35544 | 5.81801 | 2.34921 | Promoter | 5 | 41761483 | 41764501 | 3019 | 2 | ENSMUST00000060820.7 | 41 | Nkx3-2 | ENSMUSG00000049691.8 |
| chr5 | 41764784 | 41765258 | 4.26837 | 5.63757 | 2.22274 | Promoter | 5 | 41761483 | 41764501 | 3019 | 2 | ENSMUST00000060820.7 | -284 | Nkx3-2 | ENSMUSG00000049691.8 |
| chr5 | 41765642 | 41766918 | 4.08686 | 5.56745 | 2.19539 | Distal Intergenic | 5 | 41761483 | 41764501 | 3019 | 2 | ENSMUST00000060820.7 | -1142 | Nkx3-2 | ENSMUSG00000049691.8 |
| chr5 | 41767649 | 41768178 | 3.58188 | 4.53726 | 1.47782 | Distal Intergenic | 5 | 41761483 | 41764501 | 3019 | 2 | ENSMUST00000060820.7 | -3149 | Nkx3-2 | ENSMUSG00000049691.8 |
| chr5 | 41769347 | 41769997 | 3.67116 | 4.49423 | 1.44413 | Distal Intergenic | 5 | 41761483 | 41764501 | 3019 | 2 | ENSMUST00000060820.7 | -4847 | Nkx3-2 | ENSMUSG00000049691.8 |
| chr11 | 3983196 | 3983725 | 3.36939 | 3.92895 | 1.16571 | Promoter | 11 | 3983636 | 3999326 | 15691 | 1 | ENSMUST00000078757.2 | 0 | Gal3st1 | ENSMUSG00000049721.9 |
| chr11 | 3984654 | 3984952 | 3.67811 | 4.6321 | 1.5874 | Promoter | 11 | 3983713 | 3999315 | 15603 | 1 | ENSMUST00000063004.8 | 942 | Gal3st1 | ENSMUSG00000049721.9 |

| | | | | | | | | | | | | | | | |
|---|---|---|---|---|---|---|---|---|---|---|---|---|---|---|---|
| chr3 | 19694807 | 19695092 | 3.50454 | 4.3484 | 1.3588 | Promoter | 3 | 19693401 | 19695396 | 1996 | 2 | ENSMUST00000061294.4 | 304 | Crh | ENSMUSG00000049796.4 |
| chr11 | 20669719 | 20669940 | 3.37385 | 3.88841 | 1.01733 | Distal Intergenic | 11 | 20647155 | 20649065 | 1911 | 1 | ENSMUST00000109585.1 | 22565 | Sertad2 | ENSMUSG00000049800.8 |
| chr13 | 65153069 | 65153330 | 3.63615 | 4.43172 | 1.43475 | Promoter | 13 | 65152226 | 65153152 | 927 | 1 | ENSMUST00000058907.3 | 844 | Olfr466 | ENSMUSG00000049806.3 |
| chr11 | 99597884 | 99598083 | 3.40481 | 3.97916 | 1.15807 | Promoter | 11 | 99597347 | 99598105 | 759 | 2 | ENSMUST00000062683.2 | 22 | Krtap9-3 | ENSMUSG00000049809.2 |
| chr8 | 58912563 | 58912675 | 3.59252 | 4.88859 | 1.68482 | Promoter | 8 | 58912257 | 58913196 | 940 | 1 | ENSMUST00000160055.1 | 307 | BC030500 | ENSMUSG00000049946.7 |
| chr8 | 58913483 | 58913776 | 3.49185 | 4.32181 | 1.32799 | 3' UTR | 8 | 58912257 | 58913196 | 940 | 1 | ENSMUST00000160055.1 | 1227 | BC030500 | ENSMUSG00000049946.7 |
| chr13 | 112186646 | 112186931 | 3.37086 | 3.91176 | 1.11199 | Distal Intergenic | 13 | 112288451 | 112344641 | 56191 | 1 | ENSMUST00000056047.8 | -101520 | Ankrd55 | ENSMUSG00000049985.9 |
| chr13 | 112211871 | 112212135 | 3.22301 | 3.66172 | 1.01523 | Distal Intergenic | 13 | 112288451 | 112344641 | 56191 | 1 | ENSMUST00000056047.8 | -76316 | Ankrd55 | ENSMUSG00000049985.9 |
| chr13 | 112228492 | 112228572 | 4.46917 | 6.06775 | 2.54407 | Distal Intergenic | 13 | 112288451 | 112344641 | 56191 | 1 | ENSMUST00000056047.8 | -59879 | Ankrd55 | ENSMUSG00000049985.9 |
| chr5 | 140733950 | 140734178 | 3.30899 | 3.79069 | 1.01733 | Intron (ENSMUST00000060918.5/ENSMUSG00000050022.11, intron 2 of 7) | 5 | 140735526 | 140754691 | 19166 | 1 | ENSMUST00000120630.2 | -1348 | Amz1 | ENSMUSG00000050022.11 |
| chr5 | 140734538 | 140734817 | 3.61497 | 4.35841 | 1.37237 | Promoter | 5 | 140735526 | 140754691 | 19166 | 1 | ENSMUST00000120630.2 | -709 | Amz1 | ENSMUSG00000050022.11 |
| chr5 | 140735504 | 140735937 | 3.88927 | 4.95855 | 1.75294 | Promoter | 5 | 140735526 | 140754691 | 19166 | 1 | ENSMUST00000120630.2 | 0 | Amz1 | ENSMUSG00000050022.11 |
| chr10 | 85044355 | 85044698 | 3.41937 | 3.97324 | 1.07454 | Distal Intergenic | 10 | 85097019 | 85102495 | 5477 | 2 | ENSMUST00000059383.6 | 57797 | Fhl4 | ENSMUSG00000050035.6 |
| chr5 | 92675137 | 92675224 | 4.25022 | 5.65661 | 2.28637 | Promoter | 5 | 92608253 | 92675125 | 66873 | 2 | ENSMUST00000060930.4 | -13 | Ccdc158 | ENSMUSG00000050050.11 |
| chr1 | 174739047 | 174739344 | 3.34892 | 3.95236 | 1.12659 | Intron (ENSMUST00000030039.7/ENSMUSG00000028354.8, intron 14 of 17) | 1 | 174833785 | 174921819 | 88035 | 2 | ENSMUST00000055294.3 | 182475 | Grem2 | ENSMUSG00000050069.3 |
| chr1 | 174757767 | 174757971 | 3.41413 | 4.14048 | 1.20015 | Intron (ENSMUST00000030039.7/ENSMUSG00000028354.8, intron 15 of 17) | 1 | 174833785 | 174921819 | 88035 | 2 | ENSMUST00000055294.3 | 163848 | Grem2 | ENSMUSG00000050069.3 |
| chr1 | 174766631 | 174766837 | 3.21983 | 3.65697 | 1.01431 | Intron (ENSMUST00000030039.7/ENSMUSG00000028354.8, intron 15 of 17) | 1 | 174833785 | 174921819 | 88035 | 2 | ENSMUST00000055294.3 | 154982 | Grem2 | ENSMUSG00000050069.3 |
| chr1 | 174921644 | 174921768 | 3.49133 | 4.10989 | 1.16699 | Promoter | 1 | 174833785 | 174921819 | 88035 | 2 | ENSMUST00000055294.3 | 51 | Grem2 | ENSMUSG00000050069.3 |
| chr7 | 131547204 | 131547768 | 3.7966 | 4.75538 | 1.64965 | Intron (ENSMUST00000124096.2/ENSMUSG00000030849.12, intron 2 of 6) | 7 | 131548773 | 131556589 | 7817 | 1 | ENSMUST00000183219.2 | -1005 | Hmx2 | ENSMUSG00000050100.8 |
| chr7 | 131548238 | 131548661 | 3.79687 | 5.03872 | 1.77892 | Promoter | 7 | 131548773 | 131556589 | 7817 | 1 | ENSMUST00000183219.2 | -112 | Hmx2 | ENSMUSG00000050100.8 |
| chr7 | 131549292 | 131550520 | 4.01557 | 5.34522 | 2.08188 | Promoter | 7 | 131548773 | 131556589 | 7817 | 1 | ENSMUST00000183219.2 | 520 | Hmx2 | ENSMUSG00000050100.8 |
| chr7 | 131550850 | 131551243 | 3.70182 | 4.70759 | 1.59305 | Intron (ENSMUST00000183219.2/ENSMUSG00000050100.8, intron 1 of 2) | 7 | 131548773 | 131556589 | 7817 | 1 | ENSMUST00000183219.2 | 2078 | Hmx2 | ENSMUSG00000050100.8 |
| chr7 | 131551893 | 131552715 | 3.74593 | 4.83014 | 1.65647 | Intron (ENSMUST00000183219.2/ENSMUSG00000050100.8, intron 1 of 2) | 7 | 131554061 | 131556589 | 2529 | 1 | ENSMUST00000051997.7 | -1346 | Hmx2 | ENSMUSG00000050100.8 |
| chr7 | 131553073 | 131553620 | 3.73767 | 4.7171 | 1.6052 | Promoter | 7 | 131554061 | 131556589 | 2529 | 1 | ENSMUST00000051997.7 | -441 | Hmx2 | ENSMUSG00000050100.8 |
| chr7 | 131553961 | 131554055 | 4.05028 | 5.47461 | 2.10553 | Promoter | 7 | 131554061 | 131556589 | 2529 | 1 | ENSMUST00000051997.7 | -6 | Hmx2 | ENSMUSG00000050100.8 |
| chr7 | 131554415 | 131554813 | 4.2174 | 5.96028 | 2.45709 | Promoter | 7 | 131554061 | 131556589 | 2529 | 1 | ENSMUST00000051997.7 | 355 | Hmx2 | ENSMUSG00000050100.8 |
| chr7 | 131555335 | 131557504 | 4.05466 | 5.55065 | 2.17572 | 3' UTR | 7 | 131554061 | 131556589 | 2529 | 1 | ENSMUST00000051997.7 | 1275 | Hmx2 | ENSMUSG00000050100.8 |
| chr4 | 134249934 | 134250099 | 3.65283 | 4.53049 | 1.48134 | Downstream (1-2kb) | 4 | 134251706 | 134253023 | 1318 | 2 | ENSMUST00000138366.1 | 2924 | Grrp1 | ENSMUSG00000050105.5 |
| chr4 | 134250634 | 134251872 | 3.53201 | 4.33796 | 1.34689 | 3' UTR | 4 | 134251706 | 134253023 | 1318 | 2 | ENSMUST00000138366.1 | 1151 | Grrp1 | ENSMUSG00000050105.5 |
| chr4 | 134252178 | 134252779 | 3.43108 | 4.05916 | 1.19907 | Promoter | 4 | 134251706 | 134253023 | 1318 | 2 | ENSMUST00000138366.1 | 244 | Grrp1 | ENSMUSG00000050105.5 |
| chr4 | 134253285 | 134253482 | 3.79749 | 4.72372 | 1.63205 | Promoter | 4 | 134251706 | 134253023 | 1318 | 2 | ENSMUST00000138366.1 | -263 | Grrp1 | ENSMUSG00000050105.5 |
| chr4 | 134253891 | 134254037 | 3.39455 | 4.18166 | 1.24344 | Promoter | 4 | 134251110 | 134254106 | 2997 | 2 | ENSMUST00000060050.5 | 69 | Grrp1 | ENSMUSG00000050105.5 |
| chr4 | 134254378 | 134254705 | 3.49335 | 4.13207 | 1.21207 | Promoter | 4 | 134251110 | 134254106 | 2997 | 2 | ENSMUST00000060050.5 | -273 | Grrp1 | ENSMUSG00000050105.5 |
| chr1 | 37028635 | 37028737 | 3.5094 | 4.14319 | 1.19531 | Promoter | 1 | 37029304 | 37063649 | 34346 | 1 | ENSMUST00000124404.2 | -567 | Vwa3b | ENSMUSG00000050122.13 |
| chr11 | 78487766 | 78487841 | 3.91712 | 5.12027 | 1.86498 | Promoter | 11 | 78482834 | 78487295 | 4462 | 2 | ENSMUST00000170674.1 | -472 | Sarm1 | ENSMUSG00000050132.8 |
| chr11 | 78494554 | 78494945 | 3.65655 | 4.42962 | 1.40425 | Intron (ENSMUST00000108287.4/ENSMUSG00000050132.8, intron 1 of 8) | 11 | 78472330 | 78497458 | 25129 | 2 | ENSMUST00000108287.4 | 2513 | Sarm1 | ENSMUSG00000050132.8 |
| chr11 | 78495528 | 78495747 | 3.34193 | 3.84033 | 1.01733 | Intron (ENSMUST00000108287.4/ENSMUSG00000050132.8, intron 1 of 8) | 11 | 78472330 | 78497458 | 25129 | 2 | ENSMUST00000108287.4 | 1711 | Sarm1 | ENSMUSG00000050132.8 |
| chr11 | 78496770 | 78496875 | 4.41362 | 5.94007 | 2.45379 | Promoter | 11 | 78472330 | 78497458 | 25129 | 2 | ENSMUST00000108287.4 | 583 | Sarm1 | ENSMUSG00000050132.8 |
| chr11 | 78497247 | 78497378 | 4.09922 | 5.30688 | 2.00243 | Promoter | 11 | 78472330 | 78497458 | 25129 | 2 | ENSMUST00000108287.4 | 80 | Sarm1 | ENSMUSG00000050132.8 |
| chr17 | 87784451 | 87784586 | 3.65605 | 4.41069 | 1.35904 | Intron (ENSMUST00000055221.7/ENSMUSG00000050138.7, intron 1 of 1) | 17 | 87745837 | 87797994 | 52158 | 2 | ENSMUST00000055221.7 | 13408 | Kcnk12 | ENSMUSG00000050138.7 |
| chr17 | 87792800 | 87793174 | 3.58779 | 4.4038 | 1.40951 | Intron (ENSMUST00000055221.7/ENSMUSG00000050138.7, intron 1 of 1) | 17 | 87745837 | 87797994 | 52158 | 2 | ENSMUST00000055221.7 | 4820 | Kcnk12 | ENSMUSG00000050138.7 |
| chr17 | 87794715 | 87794898 | 3.60455 | 4.3457 | 1.38687 | Intron (ENSMUST00000055221.7/ENSMUSG00000050138.7, intron 1 of 1) | 17 | 87745837 | 87797994 | 52158 | 2 | ENSMUST00000055221.7 | 3096 | Kcnk12 | ENSMUSG00000050138.7 |
| chr17 | 87795251 | 87795710 | 3.5926 | 4.29797 | 1.29117 | Intron (ENSMUST00000055221.7/ENSMUSG00000050138.7, intron 1 of 1) | 17 | 87745837 | 87797994 | 52158 | 2 | ENSMUST00000055221.7 | 2284 | Kcnk12 | ENSMUSG00000050138.7 |
| chr17 | 87796720 | 87798900 | 4.25231 | 5.82188 | 2.39057 | Promoter | 17 | 87745837 | 87797994 | 52158 | 2 | ENSMUST00000055221.7 | 0 | Kcnk12 | ENSMUSG00000050138.7 |
| chr17 | 87799484 | 87799605 | 3.80261 | 4.69439 | 1.54922 | Distal Intergenic | 17 | 87745837 | 87797994 | 52158 | 2 | ENSMUST00000055221.7 | -1491 | Kcnk12 | ENSMUSG00000050138.7 |
| chr17 | 87799971 | 87800192 | 3.48711 | 4.09422 | 1.13906 | Distal Intergenic | 17 | 87745837 | 87797994 | 52158 | 2 | ENSMUST00000055221.7 | -1978 | Kcnk12 | ENSMUSG00000050138.7 |
| chr17 | 87826397 | 87826567 | 3.89411 | 4.86787 | 1.65802 | Distal Intergenic | 17 | 87745837 | 87797994 | 52158 | 2 | ENSMUST00000055221.7 | -28404 | Kcnk12 | ENSMUSG00000050138.7 |
| chr17 | 87827178 | 87827368 | 3.90036 | 5.22284 | 1.92667 | Distal Intergenic | 17 | 87745837 | 87797994 | 52158 | 2 | ENSMUST00000055221.7 | -29185 | Kcnk12 | ENSMUSG00000050138.7 |
| chr17 | 87836751 | 87836888 | 3.35836 | 3.86505 | 1.01733 | Distal Intergenic | 17 | 87745837 | 87797994 | 52158 | 2 | ENSMUST00000055221.7 | -38758 | Kcnk12 | ENSMUSG00000050138.7 |
| chr17 | 87838330 | 87838989 | 3.63755 | 4.36732 | 1.30236 | Distal Intergenic | 17 | 87745837 | 87797994 | 52158 | 2 | ENSMUST00000055221.7 | -40337 | Kcnk12 | ENSMUSG00000050138.7 |
| chr15 | 81234994 | 81235219 | 3.79888 | 4.85873 | 1.65914 | Promoter | 15 | 81235499 | 81238963 | 3465 | 1 | ENSMUST00000166855.1 | -280 | Mchr1 | ENSMUSG00000050164.10 |
| chr15 | 81272312 | 81272881 | 3.44722 | 4.02164 | 1.09529 | Distal Intergenic | 15 | 81235499 | 81238963 | 3465 | 1 | ENSMUST00000166855.1 | 36814 | Mchr1 | ENSMUSG00000050164.10 |
| chr15 | 81277229 | 81277541 | 3.38996 | 3.91706 | 1.03259 | Distal Intergenic | 15 | 81235499 | 81238963 | 3465 | 1 | ENSMUST00000166855.1 | 41731 | Mchr1 | ENSMUSG00000050164.10 |
| chr11 | 115306379 | 115306955 | 3.77684 | 4.66827 | 1.5238 | Promoter | 11 | 115307163 | 115332303 | 25141 | 1 | ENSMUST00000055490.3 | -208 | Otop2 | ENSMUSG00000050201.4 |
| chr11 | 115314294 | 115314408 | 3.70056 | 4.49746 | 1.41843 | Intron (ENSMUST00000055490.3/ENSMUSG00000050201.4, intron 1 of 7) | 11 | 115307163 | 115332303 | 25141 | 1 | ENSMUST00000055490.3 | 7132 | Otop2 | ENSMUSG00000050201.4 |
| chr11 | 115322627 | 115324031 | 3.75391 | 4.83307 | 1.66953 | Promoter | 11 | 115322761 | 115323504 | 744 | 1 | ENSMUST00000131050.1 | 0 | Otop2 | ENSMUSG00000050201.4 |
| chr2 | 120244730 | 120245181 | 3.53713 | 4.38097 | 1.3557 | Promoter | 2 | 120172173 | 120245082 | 72910 | 2 | ENSMUST00000127009.2 | 0 | Pla2g4e | ENSMUSG00000050211.9 |
| chr14 | 57524628 | 57524895 | 3.31498 | 3.80423 | 1.03327 | Promoter | 14 | 57524829 | 57543165 | 18337 | 1 | ENSMUST00000089494.4 | 0 | Il17d | ENSMUSG00000050222.10 |
| chr4 | 127313983 | 127314240 | 3.4794 | 4.30087 | 1.30732 | Promoter | 4 | 127311421 | 127314039 | 2619 | 2 | ENSMUST00000053753.7 | 0 | Gja4 | ENSMUSG00000050234.7 |
| chr16 | 17222334 | 17222421 | 3.81545 | 20.20032 | 15.09839 | Distal Intergenic | 16 | 17233572 | 17263430 | 29859 | 1 | ENSMUST00000090190.6 | -11151 | Hic2 | ENSMUSG00000050240.9 |
| chr8 | 36095775 | 36095983 | 3.3595 | 3.91267 | 1.14792 | Promoter | 8 | 36094861 | 36106609 | 11749 | 1 | ENSMUST00000150295.1 | 915 | D8Ertd82e | ENSMUSG00000050271.7 |
| chr8 | 36097763 | 36097983 | 3.98501 | 5.35783 | 2.00216 | Intron (ENSMUST00000110492.1/ENSMUSG00000050271.7, intron 1 of 5) | 8 | 36094861 | 36106609 | 11749 | 1 | ENSMUST00000150295.1 | 2903 | D8Ertd82e | ENSMUSG00000050271.7 |
| chr16 | 97165959 | 97166131 | 3.43681 | 4.01317 | 1.12271 | Intron (ENSMUST00000056102.7/ENSMUSG00000050272.9, intron 1 of 32) | 16 | 96592079 | 97170752 | 578674 | 2 | ENSMUST00000056102.7 | 4621 | Dscam | ENSMUSG00000050272.9 |
| chr16 | 97167344 | 97167667 | 3.64331 | 4.50984 | 1.42442 | Intron (ENSMUST00000056102.7/ENSMUSG00000050272.9, intron 1 of 32) | 16 | 96592079 | 97170752 | 578674 | 2 | ENSMUST00000056102.7 | 3085 | Dscam | ENSMUSG00000050272.9 |
| chr16 | 97168593 | 97169043 | 3.76101 | 4.60515 | 1.47187 | Intron (ENSMUST00000056102.7/ENSMUSG00000050272.9, intron 1 of 32) | 16 | 96592079 | 97170752 | 578674 | 2 | ENSMUST00000056102.7 | 1709 | Dscam | ENSMUSG00000050272.9 |
| chr16 | 97170090 | 97170333 | 3.48088 | 4.20496 | 1.26403 | Promoter | 16 | 96592079 | 97170752 | 578674 | 2 | ENSMUST00000056102.7 | 419 | Dscam | ENSMUSG00000050272.9 |
| chr18 | 86393979 | 86394200 | 3.35795 | 3.86886 | 1.03303 | Promoter | 18 | 86394952 | 86501897 | 106946 | 1 | ENSMUST00000058829.2 | -752 | Neto1 | ENSMUSG00000050321.2 |
| chr18 | 86394723 | 86395027 | 5.01473 | 7.13703 | 3.30942 | Promoter | 18 | 86394952 | 86501897 | 106946 | 1 | ENSMUST00000058829.2 | 0 | Neto1 | ENSMUSG00000050321.2 |
| chr18 | 86395620 | 86396405 | 3.5674 | 4.24105 | 1.22896 | Promoter | 18 | 86394952 | 86501897 | 106946 | 1 | ENSMUST00000058829.2 | 669 | Neto1 | ENSMUSG00000050321.2 |
| chr15 | 102930006 | 102930244 | 3.71743 | 4.78354 | 1.62133 | Distal Intergenic | 15 | 102936757 | 102938609 | 1853 | 1 | ENSMUST00000055562.2 | -6513 | Hoxc12 | ENSMUSG00000050328.2 |
| chr15 | 102931967 | 102932085 | 3.85413 | 5.00911 | 1.80111 | Distal Intergenic | 15 | 102936757 | 102938609 | 1853 | 1 | ENSMUST00000055562.2 | -4672 | Hoxc12 | ENSMUSG00000050328.2 |
| chr15 | 102933294 | 102935864 | 3.94915 | 5.09517 | 1.88931 | Promoter | 15 | 102936757 | 102938609 | 1853 | 1 | ENSMUST00000055562.2 | -893 | Hoxc12 | ENSMUSG00000050328.2 |
| chr15 | 102936200 | 102937712 | 3.85643 | 5.30048 | 1.98141 | Promoter | 15 | 102936757 | 102938609 | 1853 | 1 | ENSMUST00000055562.2 | 0 | Hoxc12 | ENSMUSG00000050328.2 |
| chr15 | 102938051 | 102938521 | 3.87917 | 4.87396 | 1.70395 | 3' UTR | 15 | 102936757 | 102938609 | 1853 | 1 | ENSMUST00000055562.2 | 1295 | Hoxc12 | ENSMUSG00000050328.2 |
| chr15 | 102938876 | 102939910 | 3.85178 | 4.8638 | 1.72049 | Distal Intergenic | 15 | 102936757 | 102938609 | 1853 | 1 | ENSMUST00000055562.2 | 2120 | Hoxc12 | ENSMUSG00000050328.2 |
| chr15 | 102940228 | 102940324 | 4.26657 | 5.95036 | 2.44503 | Distal Intergenic | 15 | 102936757 | 102938609 | 1853 | 1 | ENSMUST00000055562.2 | 3472 | Hoxc12 | ENSMUSG00000050328.2 |
| chr13 | 83709213 | 83710051 | 4.0193 | 5.36112 | 2.0007 | Distal Intergenic | 13 | 83721381 | 83741308 | 19928 | 1 | ENSMUST00000131907.3 | -11330 | C130071C03Rik | ENSMUSG00000050334.7 |
| chr13 | 83711218 | 83711712 | 3.80409 | 4.69803 | 1.54188 | Distal Intergenic | 13 | 83721381 | 83741308 | 19928 | 1 | ENSMUST00000131907.3 | -9669 | C130071C03Rik | ENSMUSG00000050334.7 |
| chr13 | 83712415 | 83713624 | 3.8317 | 4.84143 | 1.65468 | Distal Intergenic | 13 | 83721381 | 83741308 | 19928 | 1 | ENSMUST00000131907.3 | -7757 | C130071C03Rik | ENSMUSG00000050334.7 |
| chr13 | 83714026 | 83716257 | 3.84936 | 4.85351 | 1.69039 | Distal Intergenic | 13 | 83721381 | 83741308 | 19928 | 1 | ENSMUST00000131907.3 | -5124 | C130071C03Rik | ENSMUSG00000050334.7 |
| chr13 | 83716571 | 83717426 | 3.74549 | 4.8638 | 1.70907 | Distal Intergenic | 13 | 83721381 | 83741308 | 19928 | 1 | ENSMUST00000131907.3 | -3955 | C130071C03Rik | ENSMUSG00000050334.7 |
| chr13 | 83717869 | 83718804 | 3.84526 | 4.83301 | 1.72016 | Distal Intergenic | 13 | 83721381 | 83741308 | 19928 | 1 | ENSMUST00000131907.3 | -2577 | C130071C03Rik | ENSMUSG00000050334.7 |
| chr13 | 83719112 | 83719691 | 4.35187 | 5.94206 | 2.43662 | Distal Intergenic | 13 | 83721381 | 83741308 | 19928 | 1 | ENSMUST00000131907.3 | -1690 | C130071C03Rik | ENSMUSG00000050334.7 |
| chr13 | 83720274 | 83720562 | 3.5636 | 4.49338 | 1.45574 | Promoter | 13 | 83721381 | 83741308 | 19928 | 1 | ENSMUST00000131907.3 | -819 | C130071C03Rik | ENSMUSG00000050334.7 |
| chr13 | 83721039 | 83721396 | 3.80687 | 5.07424 | 1.81366 | Promoter | 13 | 83721381 | 83741308 | 19928 | 1 | ENSMUST00000131907.3 | 0 | C130071C03Rik | ENSMUSG00000050334.7 |
| chr13 | 83724240 | 83725178 | 4.15965 | 5.54802 | 2.19956 | Exon (ENSMUST00000181705.1/ENSMUSG00000097265.1, exon 2 of 2) | 13 | 83721381 | 83872680 | 151300 | 1 | ENSMUST00000185281.1 | 2860 | C130071C03Rik | ENSMUSG00000050334.7 |
| chr13 | 83725594 | 83725998 | 3.77935 | 4.79495 | 1.6443 | Intron (ENSMUST00000131907.3/ENSMUSG00000050334.7, intron 1 of 3) | 13 | 83728106 | 83735669 | 7564 | 1 | ENSMUST00000182788.1 | -2108 | C130071C03Rik | ENSMUSG00000050334.7 |
| chr13 | 83726545 | 83727960 | 4.10646 | 5.45083 | 2.12409 | Promoter | 13 | 83728106 | 83735669 | 7564 | 1 | ENSMUST00000182788.1 | -146 | C130071C03Rik | ENSMUSG00000050334.7 |
| chr13 | 83728368 | 83729110 | 4.11222 | 5.52125 | 2.13295 | Promoter | 13 | 83728106 | 83735669 | 7564 | 1 | ENSMUST00000182788.1 | 263 | C130071C03Rik | ENSMUSG00000050334.7 |
| chr13 | 83746043 | 83746249 | 3.78011 | 4.66229 | 1.54801 | Intron (ENSMUST00000185281.1/ENSMUSG00000050334.7, intron 3 of 3) | 13 | 83738881 | 83741307 | 2427 | 1 | ENSMUST00000052354.4 | 7163 | C130071C03Rik | ENSMUSG00000050334.7 |
| chr13 | 83736455 | 83743153 | 3.70421 | 4.58149 | 1.5104 | Promoter | 13 | 83738863 | 83884194 | 145332 | 1 | ENSMUST00000187663.1 | 0 | C130071C03Rik | ENSMUSG00000050334.7 |
| chr13 | 83732067 | 83734461 | 3.97689 | 5.14686 | 1.86951 | Promoter | 13 | 83732566 | 83735521 | 2956 | 1 | ENSMUST00000182701.1 | 0 | C130071C03Rik | ENSMUSG00000050334.7 |
| chr13 | 83734784 | 83734931 | 3.63477 | 4.68978 | 1.55916 | Exon (ENSMUST00000182788.1/ENSMUSG00000050334.7, exon 4 of 4) | 13 | 83732566 | 83735521 | 2956 | 1 | ENSMUST00000182701.1 | 2219 | C130071C03Rik | ENSMUSG00000050334.7 |
| chr13 | 83746719 | 83746935 | 3.30415 | 3.78342 | 1.01733 | Intron (ENSMUST00000185281.1/ENSMUSG00000050334.7, intron 3 of 3) | 13 | 83738881 | 83741307 | 2427 | 1 | ENSMUST00000052354.4 | 7839 | C130071C03Rik | ENSMUSG00000050334.7 |
| chr13 | 83744079 | 83744370 | 3.47525 | 4.0869 | 1.17424 | Intron (ENSMUST00000185281.1/ENSMUSG00000050334.7, intron 3 of 3) | 13 | 83738881 | 83741307 | 2427 | 1 | ENSMUST00000052354.4 | 5199 | C130071C03Rik | ENSMUSG00000050334.7 |
| chr13 | 83744692 | 83745554 | 3.88807 | 5.14944 | 1.88406 | Intron (ENSMUST00000185281.1/ENSMUSG00000050334.7, intron 3 of 3) | 13 | 83738881 | 83741307 | 2427 | 1 | ENSMUST00000052354.4 | 5812 | C130071C03Rik | ENSMUSG00000050334.7 |
| chr13 | 83747482 | 83748978 | 3.43211 | 4.26243 | 1.30217 | Intron (ENSMUST00000185281.1/ENSMUSG00000050334.7, intron 3 of 3) | 13 | 83738881 | 83741307 | 2427 | 1 | ENSMUST00000052354.4 | 8602 | C130071C03Rik | ENSMUSG00000050334.7 |
| chr13 | 83749605 | 83750798 | 3.47866 | 4.13596 | 1.28496 | Intron (ENSMUST00000185281.1/ENSMUSG00000050334.7, intron 3 of 3) | 13 | 83738881 | 83741307 | 2427 | 1 | ENSMUST00000052354.4 | 10725 | C130071C03Rik | ENSMUSG00000050334.7 |
| chr14 | 47365470 | 47365693 | 3.17956 | 3.73971 | 1.01733 | Distal Intergenic | 14 | 47367751 | 47380418 | 12668 | 1 | ENSMUST00000151405.2 | -2058 | Lgals3 | ENSMUSG00000050335.11 |
| chr14 | 47374380 | 47374764 | 3.99312 | 5.25541 | 1.97591 | Promoter | 14 | 47373974 | 47385794 | 11821 | 1 | ENSMUST00000146468.1 | 407 | Lgals3 | ENSMUSG00000050335.11 |
| chr2 | 74691361 | 74691546 | 3.62347 | 4.34401 | 1.29497 | Promoter | 2 | 74691924 | 74695105 | 3182 | 1 | ENSMUST00000061745.4 | -378 | Hoxd10 | ENSMUSG00000050368.4 |
| chr13 | 112578187 | 112578296 | 2.63904 | 11.9129 | 7.60136 | Intron (ENSMUST00000051756.6/ENSMUSG00000050377.7, intron 1 of 16) | 13 | 112522810 | 112580662 | 57853 | 2 | ENSMUST00000051756.6 | 2366 | Il31ra | ENSMUSG00000050377.7 |
| chr7 | 79719062 | 79719144 | 4.19209 | 5.45748 | 2.06466 | Downstream (2-3kb) | 7 | 79698098 | 79715720 | 17623 | 2 | ENSMUST00000183846.2 | -3343 | Kif7 | ENSMUSG00000050382.9 |
| chr9 | 98953912 | 98958838 | 4.13315 | 5.54445 | 2.16482 | Promoter | 9 | 98955607 | 98956847 | 1241 | 1 | ENSMUST00000051312.3 | 0 | Foxl2 | ENSMUSG00000050397.3 |
| chr9 | 98959172 | 98960255 | 4.03327 | 5.13281 | 1.83174 | Distal Intergenic | 9 | 98955607 | 98956847 | 1241 | 1 | ENSMUST00000051312.3 | 3566 | Foxl2 | ENSMUSG00000050397.3 |
| chr9 | 98961083 | 98961782 | 3.98356 | 5.16049 | 1.89693 | Distal Intergenic | 9 | 98955607 | 98956847 | 1241 | 1 | ENSMUST00000051312.3 | 5477 | Foxl2 | ENSMUSG00000050397.3 |
| chr9 | 98962313 | 98963102 | 3.57104 | 4.24977 | 1.24146 | Distal Intergenic | 9 | 98955607 | 98956847 | 1241 | 1 | ENSMUST00000051312.3 | 6707 | Foxl2 | ENSMUSG00000050397.3 |
| chr9 | 98963465 | 98964541 | 3.6056 | 4.31763 | 1.28709 | Distal Intergenic | 9 | 98955607 | 98956847 | 1241 | 1 | ENSMUST00000051312.3 | 7859 | Foxl2 | ENSMUSG00000050397.3 |
| chr9 | 98964869 | 98965344 | 3.7411 | 4.56875 | 1.45117 | Distal Intergenic | 9 | 98955607 | 98956847 | 1241 | 1 | ENSMUST00000051312.3 | 9263 | Foxl2 | ENSMUSG00000050397.3 |
| chr9 | 98967115 | 98967701 | 3.61826 | 4.39752 | 1.42241 | Distal Intergenic | 9 | 98955607 | 98956847 | 1241 | 1 | ENSMUST00000051312.3 | 11509 | Foxl2 | ENSMUSG00000050397.3 |
| chr9 | 98969482 | 98969641 | 3.61647 | 4.33574 | 1.29739 | Distal Intergenic | 9 | 98955607 | 98956847 | 1241 | 1 | ENSMUST00000051312.3 | 13876 | Foxl2 | ENSMUSG00000050397.3 |
| chr13 | 35947939 | 35948502 | 3.44821 | 4.09478 | 1.24059 | Distal Intergenic | 13 | 35958839 | 35968324 | 9486 | 1 | ENSMUST00000172766.1 | -10337 | Ppp1r3g | ENSMUSG00000050423.10 |
| chr13 | 35967904 | 35969350 | 3.64478 | 4.48554 | 1.436 | Promoter | 13 | 35967853 | 35970388 | 2536 | 1 | ENSMUST00000132661.1 | 52 | Ppp1r3g | ENSMUSG00000050423.10 |
| chr13 | 35969769 | 35970584 | 3.55861 | 4.25341 | 1.28635 | 3' UTR | 13 | 35967853 | 35970388 | 2536 | 1 | ENSMUST00000132661.1 | 1917 | Ppp1r3g | ENSMUSG00000050423.10 |
| chr15 | 80560074 | 80560446 | 3.39539 | 3.94595 | 1.10632 | Promoter | 15 | 80452240 | 80560470 | 108231 | 2 | ENSMUST00000096347.3 | 24 | Enthd1 | ENSMUSG00000050439.6 |
| chr15 | 80561202 | 80561277 | 4.08322 | 5.36928 | 2.03236 | Promoter | 15 | 80452240 | 80560470 | 108231 | 2 | ENSMUST00000096347.3 | -733 | Enthd1 | ENSMUSG00000050439.6 |
| chr15 | 80577760 | 80577949 | 3.4641 | 4.11162 | 1.25455 | Distal Intergenic | 15 | 80452240 | 80560470 | 108231 | 2 | ENSMUST00000096347.3 | -17291 | Enthd1 | ENSMUSG00000050439.6 |
| chr2 | 50067667 | 50067908 | 3.52037 | 4.20276 | 1.31643 | Intron (ENSMUST00000112712.4/ENSMUSG00000050447.10, intron 1 of 4) | 2 | 50066464 | 50193564 | 127101 | 1 | ENSMUST00000053208.8 | 1204 | Lypd6 | ENSMUSG00000050447.10 |
| chr10 | 19849460 | 19850110 | 3.74615 | 4.81935 | 1.65615 | Exon (ENSMUST00000059805.4/ENSMUSG00000050473.4, exon 2 of 2) | 10 | 19847917 | 19851459 | 3543 | 2 | ENSMUST00000059805.4 | 1349 | Slc35d3 | ENSMUSG00000050473.4 |
| chr10 | 19850464 | 19850562 | 3.78699 | 4.98286 | 1.74257 | Promoter | 10 | 19847917 | 19851459 | 3543 | 2 | ENSMUST00000059805.4 | 897 | Slc35d3 | ENSMUSG00000050473.4 |
| chr10 | 19850896 | 19853253 | 3.80329 | 4.88131 | 1.70223 | Promoter | 10 | 19847917 | 19851459 | 3543 | 2 | ENSMUST00000059805.4 | 0 | Slc35d3 | ENSMUSG00000050473.4 |
| chr14 | 88469128 | 88469649 | 3.59055 | 4.28671 | 1.26224 | Exon (ENSMUST00000061628.5/ENSMUSG00000050505.6, exon 3 of 3) | 14 | 88464747 | 88471396 | 6650 | 2 | ENSMUST00000061628.5 | 1747 | Pcdh20 | ENSMUSG00000050505.6 |
| chr4 | 132141448 | 132141555 | 3.88932 | 4.86667 | 1.67143 | Intron (ENSMUST00000056336.1/ENSMUSG00000050511.1, intron 1 of 2) | 4 | 132110726 | 132144486 | 33761 | 2 | ENSMUST00000056336.1 | 2931 | Oprd1 | ENSMUSG00000050511.1 |
| chr4 | 132143585 | 132144602 | 3.82559 | 4.89229 | 1.74188 | Promoter | 4 | 132110726 | 132144486 | 33761 | 2 | ENSMUST00000056336.1 | 0 | Oprd1 | ENSMUSG00000050511.1 |
| chr4 | 132144992 | 132145458 | 3.86192 | 4.7917 | 1.58279 | Promoter | 4 | 132110726 | 132144486 | 33761 | 2 | ENSMUST00000056336.1 | -507 | Oprd1 | ENSMUSG00000050511.1 |
| chr4 | 132146321 | 132146504 | 3.96042 | 4.98262 | 1.71132 | Downstream (1-2kb) | 4 | 132110726 | 132144486 | 33761 | 2 | ENSMUST00000056336.1 | -1836 | Oprd1 | ENSMUSG00000050511.1 |
| chr1 | 121527394 | 121528197 | 4.61929 | 6.42945 | 2.79831 | Promoter | 1 | 121509842 | 121528465 | 18624 | 2 | ENSMUST00000055884.7 | 268 | Htr5b | ENSMUSG00000050534.8 |
| chr11 | 43835255 | 43835467 | 3.23338 | 3.67724 | 1.01733 | Promoter | 11 | 43774606 | 43836332 | 61727 | 2 | ENSMUST00000124306.1 | 865 | Adra1b | ENSMUSG00000050541.9 |
| chr11 | 43836259 | 43836835 | 3.48491 | 4.21189 | 1.26263 | Promoter | 11 | 43774606 | 43836332 | 61727 | 2 | ENSMUST00000124306.1 | 0 | Adra1b | ENSMUSG00000050541.9 |
| chr2 | 167190076 | 167190765 | 3.62711 | 4.64309 | 1.56002 | Promoter | 2 | 167188422 | 167190155 | 1734 | 2 | ENSMUST00000131267.1 | 0 | Kcnb1 | ENSMUSG00000050556.8 |
| chr2 | 167191076 | 167191282 | 3.3529 | 4.15283 | 1.20809 | Promoter | 2 | 167188422 | 167190155 | 1734 | 2 | ENSMUST00000131267.1 | -922 | Kcnb1 | ENSMUSG00000050556.8 |
| chr2 | 167191802 | 167192167 | 3.92067 | 5.10852 | 1.85946 | 3' UTR | 2 | 167188422 | 167190155 | 1734 | 2 | ENSMUST00000131267.1 | -1648 | Kcnb1 | ENSMUSG00000050556.8 |
| chr2 | 132384292 | 132384481 | 3.37064 | 3.9162 | 1.11993 | Promoter | 2 | 132370329 | 132385400 | 15072 | 2 | ENSMUST00000049997.8 | 919 | Prokr2 | ENSMUSG00000050558.8 |
| chr2 | 132385239 | 132385408 | 4.26752 | 5.6542 | 2.27791 | Promoter | 2 | 132370329 | 132385400 | 15072 | 2 | ENSMUST00000049997.8 | 0 | Prokr2 | ENSMUSG00000050558.8 |
| chr2 | 132386248 | 132386373 | 3.68474 | 4.48602 | 1.45537 | Promoter | 2 | 132371165 | 132385447 | 14283 | 2 | ENSMUST00000110156.1 | -802 | Prokr2 | ENSMUSG00000050558.8 |
| chr1 | 156079003 | 156079137 | 3.65913 | 4.48328 | 1.50204 | Exon (ENSMUST00000015628.3/ENSMUSG00000015484.3, exon 5 of 5) | 1 | 156063213 | 156064842 | 1630 | 1 | ENSMUST00000125371.1 | 15791 | Tor1aip2 | ENSMUSG00000050565.11 |
| chr1 | 156104179 | 156104457 | 3.47507 | 4.07455 | 1.13479 | Intron (ENSMUST00000015628.3/ENSMUSG00000015484.3, intron 1 of 4) | 1 | 156063213 | 156064842 | 1630 | 1 | ENSMUST00000125371.1 | 40967 | Tor1aip2 | ENSMUSG00000050565.11 |
| chr1 | 156108376 | 156109178 | 3.54666 | 4.21137 | 1.23266 | Intron (ENSMUST00000015628.3/ENSMUSG00000015484.3, intron 1 of 4) | 1 | 156063213 | 156064842 | 1630 | 1 | ENSMUST00000125371.1 | 45164 | Tor1aip2 | ENSMUSG00000050565.11 |
| chr1 | 156109501 | 156110023 | 3.62968 | 4.35333 | 1.29476 | Intron (ENSMUST00000015628.3/ENSMUSG00000015484.3, intron 1 of 4) | 1 | 156063213 | 156064842 | 1630 | 1 | ENSMUST00000125371.1 | 46289 | Tor1aip2 | ENSMUSG00000050565.11 |
| chr1 | 156113090 | 156113237 | 3.3653 | 3.87551 | 1.01733 | Intron (ENSMUST00000015628.3/ENSMUSG00000015484.3, intron 1 of 4) | 1 | 156063213 | 156064842 | 1630 | 1 | ENSMUST00000125371.1 | 49878 | Tor1aip2 | ENSMUSG00000050565.11 |
| chr2 | 96319332 | 96319851 | 3.65946 | 4.51649 | 1.44891 | Intron (ENSMUST00000135431.2/ENSMUSG00000050587.9, intron 1 of 6) | 2 | 96318216 | 97630983 | 1312768 | 1 | ENSMUST00000162807.3 | 1117 | Lrrc4c | ENSMUSG00000050587.9 |
| chr2 | 97467783 | 97468199 | 3.69917 | 4.48895 | 1.39533 | Promoter | 2 | 97468089 | 97629063 | 160975 | 1 | ENSMUST00000170144.1 | 0 | Lrrc4c | ENSMUSG00000050587.9 |
| chr2 | 32082641 | 32082852 | 4.04034 | 5.68175 | 2.21852 | Promoter | 2 | 32066867 | 32083783 | 16917 | 2 | ENSMUST00000056406.6 | 931 | Fam78a | ENSMUSG00000050592.8 |
| chr2 | 32083601 | 32083740 | 4.78508 | 6.91138 | 3.16909 | Promoter | 2 | 32066867 | 32083783 | 16917 | 2 | ENSMUST00000056406.6 | 43 | Fam78a | ENSMUSG00000050592.8 |
| chr2 | 32088526 | 32089386 | 4.11241 | 5.34388 | 2.03591 | Distal Intergenic | 2 | 32083272 | 32084857 | 1586 | 2 | ENSMUST00000143795.1 | -3670 | Fam78a | ENSMUSG00000050592.8 |
| chr2 | 174672179 | 174672265 | 3.51056 | 4.18304 | 1.2904 | Intron (ENSMUST00000059452.5/ENSMUSG00000050600.5, intron 3 of 4) | 2 | 174643534 | 174710832 | 67299 | 1 | ENSMUST00000059452.5 | 28646 | Zfp831 | ENSMUSG00000050600.5 |
| chr2 | 174673238 | 174673782 | 3.70185 | 4.56725 | 1.5103 | Intron (ENSMUST00000059452.5/ENSMUSG00000050600.5, intron 3 of 4) | 2 | 174643534 | 174710832 | 67299 | 1 | ENSMUST00000059452.5 | 29705 | Zfp831 | ENSMUSG00000050600.5 |
| chr2 | 174674342 | 174674483 | 3.34344 | 4.08913 | 1.21391 | Intron (ENSMUST00000059452.5/ENSMUSG00000050600.5, intron 3 of 4) | 2 | 174643534 | 174710832 | 67299 | 1 | ENSMUST00000059452.5 | 30809 | Zfp831 | ENSMUSG00000050600.5 |
| chr2 | 174675476 | 174675716 | 3.22718 | 3.66796 | 1.01733 | Intron (ENSMUST00000059452.5/ENSMUSG00000050600.5, intron 3 of 4) | 2 | 174643534 | 174710832 | 67299 | 1 | ENSMUST00000059452.5 | 31943 | Zfp831 | ENSMUSG00000050600.5 |
| chr19 | 6924975 | 6927131 | 3.89817 | 4.92372 | 1.71312 | Promoter | 19 | 6922426 | 6925380 | 2955 | 2 | ENSMUST00000057716.4 | 0 | Tex40 | ENSMUSG00000050623.4 |
| chr19 | 6927658 | 6927930 | 4.28521 | 5.69213 | 2.29258 | Exon (ENSMUST00000025908.6/ENSMUSG00000024957.8, exon 5 of 7) | 19 | 6922426 | 6925380 | 2955 | 2 | ENSMUST00000057716.4 | -2279 | Tex40 | ENSMUSG00000050623.4 |
| chr1 | 181483896 | 181484040 | 3.71107 | 4.62964 | 1.57097 | Distal Intergenic | 1 | 181509633 | 181511451 | 1819 | 2 | ENSMUST00000058825.4 | 27411 | Ccdc121 | ENSMUSG00000050625.4 |
| chr1 | 181545434 | 181545726 | 3.79679 | 5.04027 | 1.83204 | Distal Intergenic | 1 | 181509633 | 181511451 | 1819 | 2 | ENSMUST00000058825.4 | -33984 | Ccdc121 | ENSMUSG00000050625.4 |
| chr10 | 114798863 | 114799309 | 3.61815 | 4.34108 | 1.29892 | Intron (ENSMUST00000061632.7/ENSMUSG00000050663.7, intron 1 of 18) | 10 | 114398823 | 114801364 | 402542 | 2 | ENSMUST00000061632.7 | 2055 | Trhde | ENSMUSG00000050663.7 |
| chr10 | 114800700 | 114802028 | 3.69444 | 4.59753 | 1.51565 | Promoter | 10 | 114398823 | 114801364 | 402542 | 2 | ENSMUST00000061632.7 | 0 | Trhde | ENSMUSG00000050663.7 |
| chr10 | 114802493 | 114802650 | 3.58477 | 4.28007 | 1.27667 | Promoter | 10 | 114566167 | 114802307 | 236141 | 2 | ENSMUST00000152702.1 | -187 | Trhde | ENSMUSG00000050663.7 |
| chr10 | 114803244 | 114803458 | 3.37238 | 3.8862 | 1.01733 | Promoter | 10 | 114566167 | 114802307 | 236141 | 2 | ENSMUST00000152702.1 | -938 | Trhde | ENSMUSG00000050663.7 |
| chr12 | 87299199 | 87300758 | 3.91653 | 5.03996 | 1.85769 | Promoter | 12 | 87278638 | 87299705 | 21068 | 2 | ENSMUST00000125733.2 | 0 | Ism2 | ENSMUSG00000050671.7 |
| chr2 | 160911693 | 160912614 | 3.98611 | 5.57596 | 2.18852 | Promoter | 2 | 160906437 | 160912317 | 5881 | 2 | ENSMUST00000109454.2 | 0 | Emilin3 | ENSMUSG00000050700.6 |
| chr2 | 160913130 | 160913378 | 3.32379 | 3.81529 | 1.02554 | Promoter | 2 | 160906438 | 160912328 | 5891 | 2 | ENSMUST00000057169.4 | -803 | Emilin3 | ENSMUSG00000050700.6 |
| chr2 | 160913831 | 160914090 | 3.35567 | 3.86099 | 1.01733 | Distal Intergenic | 2 | 160906438 | 160912328 | 5891 | 2 | ENSMUST00000057169.4 | -1504 | Emilin3 | ENSMUSG00000050700.6 |
| chr2 | 160914746 | 160915016 | 3.56072 | 4.26726 | 1.30407 | Distal Intergenic | 2 | 160906438 | 160912328 | 5891 | 2 | ENSMUST00000057169.4 | -2419 | Emilin3 | ENSMUSG00000050700.6 |
| chr2 | 30902878 | 30903302 | 3.72468 | 4.56584 | 1.48003 | Promoter | 2 | 30889471 | 30903297 | 13827 | 2 | ENSMUST00000102852.4 | 0 | Ptges | ENSMUSG00000050737.12 |

| | | | | | | | | | | | | | | | |
|---|---|---|---|---|---|---|---|---|---|---|---|---|---|---|---|
| chr17 | 36865736 | 36866205 | 3.53439 | 4.20885 | 1.2924 | Promoter | 17 | 36861318 | 36866746 | 5429 | 2 | ENSMUST00000173639.1 | 541 | Trim15 | ENSMUSG00000050747.9 |
| chr17 | 36866727 | 36867756 | 3.72678 | 4.62433 | 1.50039 | Promoter | 17 | 36860691 | 36867187 | 6497 | 2 | ENSMUST00000025329.7 | 0 | Trim15 | ENSMUSG00000050747.9 |
| chr8 | 124431036 | 124431364 | 3.65396 | 4.42451 | 1.40924 | Intron (ENSMUST00000052580.4/ENSMUSG00000050751.9, intron 1 of 6) | 8 | 124369049 | 124433936 | 64888 | 2 | ENSMUST00000052580.4 | 2572 | Pgbd5 | ENSMUSG00000050751.9 |
| chr8 | 124432049 | 124432264 | 3.5079 | 4.26719 | 1.32687 | Intron (ENSMUST00000052580.4/ENSMUSG00000050751.9, intron 1 of 6) | 8 | 124369049 | 124433936 | 64888 | 2 | ENSMUST00000052580.4 | 1672 | Pgbd5 | ENSMUSG00000050751.9 |
| chr8 | 124433518 | 124433925 | 4.35196 | 5.80911 | 2.35998 | Promoter | 8 | 124369049 | 124433936 | 64888 | 2 | ENSMUST00000052580.4 | 11 | Pgbd5 | ENSMUSG00000050751.9 |
| chr16 | 18608445 | 18608597 | 3.4604 | 4.05633 | 1.14949 | Exon (ENSMUST00000156827.1/ENSMUSG00000075335.3, exon 2 of 2) | 16 | 18620722 | 18621366 | 645 | 2 | ENSMUST00000051160.2 | 12769 | Gp1bb | ENSMUSG00000050761.3 |
| chr16 | 18620603 | 18622197 | 4.27396 | 5.74936 | 2.34895 | Promoter | 16 | 18620722 | 18621366 | 645 | 2 | ENSMUST00000051160.2 | 0 | Gp1bb | ENSMUSG00000050761.3 |
| chr16 | 18625969 | 18626144 | 3.85969 | 4.7966 | 1.5991 | Intron (ENSMUST00000096987.5/ENSMUSG00000072214.6, intron 2 of 11) | 16 | 18620319 | 18622403 | 2085 | 2 | ENSMUST00000167388.1 | -3567 | Gp1bb | ENSMUSG00000050761.3 |
| chr16 | 65104748 | 65104842 | 3.34412 | 3.84357 | 1.01733 | Promoter | 16 | 64924729 | 65105784 | 181056 | 2 | ENSMUST00000063076.4 | 942 | Htr1f | ENSMUSG00000050783.4 |
| chr16 | 65105196 | 65105739 | 3.78103 | 4.72879 | 1.61072 | Promoter | 16 | 64924729 | 65105784 | 181056 | 2 | ENSMUST00000063076.4 | 45 | Htr1f | ENSMUSG00000050783.4 |
| chr5 | 142692792 | 142693279 | 3.86793 | 5.05269 | 1.82072 | Distal Intergenic | 5 | 142702101 | 142722490 | 20390 | 1 | ENSMUST00000058418.7 | -8822 | Slc29a4 | ENSMUSG00000050822.7 |
| chr5 | 142699363 | 142699595 | 3.96393 | 5.08802 | 1.82985 | Distal Intergenic | 5 | 142702101 | 142722490 | 20390 | 1 | ENSMUST00000058418.7 | -2506 | Slc29a4 | ENSMUSG00000050822.7 |
| chr5 | 142700776 | 142700924 | 3.47176 | 4.083 | 1.18303 | Distal Intergenic | 5 | 142702101 | 142722490 | 20390 | 1 | ENSMUST00000058418.7 | -1177 | Slc29a4 | ENSMUSG00000050822.7 |
| chr5 | 142702163 | 142702844 | 3.64209 | 4.50876 | 1.44069 | Promoter | 5 | 142702101 | 142722490 | 20390 | 1 | ENSMUST00000058418.7 | 63 | Slc29a4 | ENSMUSG00000050822.7 |
| chr5 | 142703326 | 142703708 | 3.5563 | 4.22085 | 1.21699 | Intron (ENSMUST00000058418.7/ENSMUSG00000050822.7, intron 1 of 10) | 5 | 142702101 | 142722490 | 20390 | 1 | ENSMUST00000058418.7 | 1226 | Slc29a4 | ENSMUSG00000050822.7 |
| chr5 | 142704456 | 142704938 | 3.4364 | 4.00476 | 1.09519 | Intron (ENSMUST00000058418.7/ENSMUSG00000050822.7, intron 1 of 10) | 5 | 142702101 | 142722490 | 20390 | 1 | ENSMUST00000058418.7 | 2356 | Slc29a4 | ENSMUSG00000050822.7 |
| chr17 | 25495869 | 25495995 | 3.34303 | 3.84192 | 1.01733 | Intron (ENSMUST00000051864.4/ENSMUSG00000050824.6, intron 1 of 2) | 17 | 25489875 | 25497288 | 7414 | 2 | ENSMUST00000051864.4 | 1293 | Sstr5 | ENSMUSG00000050824.6 |
| chr17 | 25496619 | 25497164 | 3.5936 | 4.29617 | 1.27489 | Promoter | 17 | 25489875 | 25497288 | 7414 | 2 | ENSMUST00000051864.4 | 124 | Sstr5 | ENSMUSG00000050824.6 |
| chr11 | 11113846 | 11114762 | 3.87518 | 4.85089 | 1.6548 | Promoter | 11 | 11114223 | 11263526 | 149304 | 1 | ENSMUST00000109681.2 | 0 | Vwc2 | ENSMUSG00000050830.12 |
| chr11 | 11115590 | 11116314 | 3.86208 | 4.93641 | 1.71691 | Promoter | 11 | 11115851 | 11266638 | 150788 | 1 | ENSMUST00000056344.4 | 0 | Vwc2 | ENSMUSG00000050830.12 |
| chr11 | 11117571 | 11117703 | 3.46037 | 4.04737 | 1.11744 | Intron (ENSMUST00000109681.2/ENSMUSG00000050830.12, intron 2 of 3) | 11 | 11116485 | 11131183 | 14699 | 1 | ENSMUST00000154155.1 | 1087 | Vwc2 | ENSMUSG00000050830.12 |
| chr1 | 104768657 | 104769228 | 3.91473 | 5.15963 | 1.88023 | Promoter | 1 | 104768529 | 104995481 | 226953 | 1 | ENSMUST00000062528.8 | 129 | Cdh20 | ENSMUSG00000050840.8 |
| chr4 | 118547297 | 118548701 | 2.35813 | 8.1706 | 4.28827 | Distal Intergenic | 4 | 118541859 | 118544044 | 2186 | 2 | ENSMUST00000128098.2 | -3254 | Tmem125 | ENSMUSG00000050854.4 |
| chr11 | 95796148 | 95797003 | 3.84971 | 5.13969 | 1.89843 | Intron (ENSMUST00000133070.2/ENSMUSG00000075595.4, intron 5 of 6) | 11 | 95824499 | 95830701 | 6203 | 1 | ENSMUST00000150134.1 | -27496 | Phospho1 | ENSMUSG00000050860.6 |
| chr11 | 95828823 | 95829467 | 3.55795 | 4.22997 | 1.24293 | Intron (ENSMUST00000133070.2/ENSMUSG00000075595.4, intron 6 of 6) | 11 | 95830763 | 95831592 | 830 | 1 | ENSMUST00000176538.1 | -1296 | Phospho1 | ENSMUSG00000050860.6 |
| chr11 | 95830020 | 95830102 | 3.39228 | 3.9332 | 1.07666 | Promoter | 11 | 95830763 | 95831592 | 830 | 1 | ENSMUST00000176538.1 | -661 | Phospho1 | ENSMUSG00000050860.6 |
| chr11 | 95830512 | 95831552 | 3.84482 | 4.82208 | 1.64584 | Promoter | 11 | 95830763 | 95831592 | 830 | 1 | ENSMUST00000176538.1 | 0 | Phospho1 | ENSMUSG00000050860.6 |
| chr2 | 84865757 | 84867323 | 3.54299 | 4.30191 | 1.30543 | Distal Intergenic | 2 | 84880523 | 84886476 | 5954 | 2 | ENSMUST00000151759.1 | 19153 | Rtn4rl2 | ENSMUSG00000050896.7 |
| chr2 | 84871490 | 84872055 | 3.64528 | 4.59601 | 1.51318 | 3' UTR | 2 | 84880523 | 84886476 | 5954 | 2 | ENSMUST00000151759.1 | 14421 | Rtn4rl2 | ENSMUSG00000050896.7 |
| chr2 | 84872469 | 84872771 | 3.7436 | 4.82235 | 1.64641 | Exon (ENSMUST00000151799.2/ENSMUSG00000050896.7, exon 3 of 3) | 2 | 84880523 | 84886476 | 5954 | 2 | ENSMUST00000151759.1 | 13705 | Rtn4rl2 | ENSMUSG00000050896.7 |
| chr2 | 84874001 | 84874306 | 3.22885 | 3.67352 | 1.0255 | Intron (ENSMUST00000151799.2/ENSMUSG00000050896.7, intron 2 of 2) | 2 | 84880523 | 84886476 | 5954 | 2 | ENSMUST00000151759.1 | 12170 | Rtn4rl2 | ENSMUSG00000050896.7 |
| chr2 | 84880465 | 84880540 | 3.40504 | 3.95433 | 1.08407 | Exon (ENSMUST00000151799.2/ENSMUSG00000050896.7, exon 2 of 3) | 2 | 84880523 | 84886476 | 5954 | 2 | ENSMUST00000151759.1 | 5936 | Rtn4rl2 | ENSMUSG00000050896.7 |
| chr2 | 84882970 | 84883271 | 3.34658 | 3.84728 | 1.01733 | Intron (ENSMUST00000151799.2/ENSMUSG00000050896.7, intron 1 of 2) | 2 | 84880523 | 84886476 | 5954 | 2 | ENSMUST00000151759.1 | 3205 | Rtn4rl2 | ENSMUSG00000050896.7 |
| chr2 | 84885077 | 84885152 | 3.5137 | 4.14083 | 1.16295 | Intron (ENSMUST00000151799.2/ENSMUSG00000050896.7, intron 1 of 2) | 2 | 84880523 | 84886476 | 5954 | 2 | ENSMUST00000151759.1 | 1324 | Rtn4rl2 | ENSMUSG00000050896.7 |
| chr2 | 84886352 | 84886905 | 4.17334 | 5.43288 | 2.06656 | Promoter | 2 | 84871924 | 84886710 | 14787 | 2 | ENSMUST00000151799.2 | 0 | Rtn4rl2 | ENSMUSG00000050896.7 |
| chr2 | 84887550 | 84888374 | 4.18485 | 5.44367 | 2.05713 | Promoter | 2 | 84871924 | 84886710 | 14787 | 2 | ENSMUST00000151799.2 | -841 | Rtn4rl2 | ENSMUSG00000050896.7 |
| chr9 | 15874984 | 15875101 | 3.36292 | 3.89745 | 1.10783 | Promoter | 9 | 15824528 | 15874556 | 50029 | 2 | ENSMUST00000182947.1 | -429 | Mtnr1b | ENSMUSG00000050901.6 |
| chr7 | 144860862 | 144861934 | 3.99516 | 5.10478 | 1.83697 | Promoter | 7 | 144861386 | 144865243 | 3858 | 1 | ENSMUST00000060336.4 | 0 | Fgf4 | ENSMUSG00000050917.4 |
| chr7 | 144862283 | 144862417 | 3.55781 | 4.36651 | 1.34113 | Promoter | 7 | 144861386 | 144865243 | 3858 | 1 | ENSMUST00000060336.4 | 898 | Fgf4 | ENSMUSG00000050917.4 |
| chr7 | 144862826 | 144863666 | 3.91996 | 5.04012 | 1.8319 | 3' UTR | 7 | 144861386 | 144865243 | 3858 | 1 | ENSMUST00000060336.4 | 1441 | Fgf4 | ENSMUSG00000050917.4 |
| chr7 | 144864704 | 144864905 | 3.29303 | 3.9421 | 1.15443 | 3' UTR | 7 | 144861386 | 144865243 | 3858 | 1 | ENSMUST00000060336.4 | 3319 | Fgf4 | ENSMUSG00000050917.4 |
| chr11 | 77099748 | 77099978 | 3.66202 | 4.42369 | 1.37328 | Intron (ENSMUST00000108400.2/ENSMUSG00000050944.9, intron 17 of 17) | 11 | 77089944 | 77103062 | 13119 | 2 | ENSMUST00000151731.1 | 3084 | Efcab5 | ENSMUSG00000050944.9 |
| chr10 | 56378716 | 56379511 | 3.88849 | 4.95281 | 1.74622 | Intron (ENSMUST00000068581.7/ENSMUSG00000050953.9, intron 1 of 1) | 10 | 56377300 | 56390419 | 13120 | 1 | ENSMUST00000068581.7 | 1417 | Gja1 | ENSMUSG00000050953.9 |
| chr13 | 48390459 | 48390689 | 3.5476 | 4.21013 | 1.22707 | Distal Intergenic | 13 | 48489202 | 48506939 | 17738 | 2 | ENSMUST00000167682.2 | 116250 | Zfp169 | ENSMUSG00000050954.10 |
| chr13 | 48436005 | 48436472 | 3.81296 | 4.72605 | 1.59146 | Distal Intergenic | 13 | 48489202 | 48506939 | 17738 | 2 | ENSMUST00000167682.2 | 70467 | Zfp169 | ENSMUSG00000050954.10 |
| chr15 | 34837553 | 34838251 | 4.10068 | 5.62905 | 2.2471 | Promoter | 15 | 34837381 | 34842337 | 4957 | 1 | ENSMUST00000072868.3 | 173 | Kcns2 | ENSMUSG00000050963.6 |
| chr4 | 134010979 | 134011194 | 3.41448 | 4.08861 | 1.17583 | Intron (ENSMUST00000051674.2/ENSMUSG00000050966.4, intron 2 of 3) | 4 | 134006244 | 134012386 | 6143 | 2 | ENSMUST00000176113.2 | 1192 | Lin28a | ENSMUSG00000050966.4 |
| chr4 | 134013311 | 134013996 | 4.08102 | 5.32439 | 2.00452 | Promoter | 4 | 134006244 | 134012386 | 6143 | 2 | ENSMUST00000176113.2 | -926 | Lin28a | ENSMUSG00000050966.4 |
| chr4 | 134016998 | 134019067 | 4.0954 | 5.36136 | 2.06779 | Promoter | 4 | 134003330 | 134018841 | 15512 | 2 | ENSMUST00000051674.2 | 0 | Lin28a | ENSMUSG00000050966.4 |
| chr4 | 134019574 | 134020238 | 3.3399 | 3.96549 | 1.10223 | Promoter | 4 | 134003330 | 134018841 | 15512 | 2 | ENSMUST00000051674.2 | -734 | Lin28a | ENSMUSG00000050966.4 |
| chr1 | 39651396 | 39651841 | 3.72556 | 4.59089 | 1.56205 | Promoter | 1 | 39618406 | 39651182 | 32777 | 2 | ENSMUST00000053355.4 | -215 | Creg2 | ENSMUSG00000050967.4 |
| chr15 | 77469334 | 77469466 | 3.68544 | 4.62741 | 1.54405 | Distal Intergenic | 15 | 77477047 | 77490118 | 13072 | 1 | ENSMUST00000126154.2 | -7581 | Apol10a | ENSMUSG00000050982.7 |
| chr5 | 39637404 | 39637745 | 3.3748 | 3.88985 | 1.01733 | Intron (ENSMUST00000117944.1/ENSMUSG00000051022.7, intron 2 of 2) | 5 | 39615074 | 39644597 | 29524 | 2 | ENSMUST00000152057.1 | 6852 | Hs3st1 | ENSMUSG00000051022.7 |
| chr5 | 39642930 | 39643084 | 3.42238 | 4.15142 | 1.22405 | Intron (ENSMUST00000117944.1/ENSMUSG00000051022.7, intron 2 of 2) | 5 | 39615074 | 39644597 | 29524 | 2 | ENSMUST00000152057.1 | 1513 | Hs3st1 | ENSMUSG00000051022.7 |
| chr5 | 39643872 | 39644541 | 3.68139 | 4.4558 | 1.36773 | Promoter | 5 | 39615074 | 39644597 | 29524 | 2 | ENSMUST00000152057.1 | 56 | Hs3st1 | ENSMUSG00000051022.7 |
| chr5 | 39645213 | 39645531 | 3.76165 | 4.6419 | 1.54281 | Promoter | 5 | 39614769 | 39645246 | 30478 | 2 | ENSMUST00000137142.1 | 0 | Hs3st1 | ENSMUSG00000051022.7 |
| chr5 | 39645998 | 39646427 | 3.5827 | 4.47488 | 1.41922 | Promoter | 5 | 39614769 | 39645246 | 30478 | 2 | ENSMUST00000137142.1 | -753 | Hs3st1 | ENSMUSG00000051022.7 |
| chr6 | 116564614 | 116564706 | 3.47461 | 4.19818 | 1.27321 | Distal Intergenic | 6 | 116556405 | 116557412 | 1008 | 1 | ENSMUST00000060204.5 | 8210 | Olfr214 | ENSMUSG00000051046.5 |
| chr7 | 100285536 | 100285671 | 3.74487 | 4.75334 | 1.59101 | Promoter | 7 | 100285542 | 100298687 | 13146 | 1 | ENSMUST00000148620.1 | 0 | P4ha3 | ENSMUSG00000051048.11 |
| chr11 | 35796427 | 35796503 | 3.85866 | 4.78961 | 1.59046 | Downstream (<1kb) | 11 | 35797382 | 35798884 | 1503 | 2 | ENSMUST00000160726.3 | 2381 | Fbll1 | ENSMUSG00000051062.6 |
| chr11 | 35797261 | 35799081 | 3.78145 | 4.70156 | 1.56842 | Promoter | 11 | 35797382 | 35798884 | 1503 | 2 | ENSMUST00000160726.3 | 0 | Fbll1 | ENSMUSG00000051062.6 |
| chr10 | 80843455 | 80844639 | 4.06003 | 5.39484 | 2.09488 | Promoter | 10 | 80832807 | 80844039 | 11233 | 2 | ENSMUST00000053986.7 | 0 | Lingo3 | ENSMUSG00000051067.7 |
| chr10 | 80845342 | 80845509 | 3.45152 | 4.03498 | 1.11874 | Distal Intergenic | 10 | 80832807 | 80844039 | 11233 | 2 | ENSMUST00000053986.7 | -1304 | Lingo3 | ENSMUSG00000051067.7 |
| chr13 | 51702452 | 51702831 | 3.38287 | 3.90783 | 1.03787 | 3' UTR | 13 | 51685529 | 51701041 | 15513 | 2 | ENSMUST00000110042.1 | -1412 | Gm15440 | ENSMUSG00000051107.4 |
| chr13 | 51703278 | 51703510 | 3.36173 | 3.87013 | 1.01733 | Exon (ENSMUST00000110040.3/ENSMUSG00000021451.10, exon 17 of 17) | 13 | 51685529 | 51701041 | 15513 | 2 | ENSMUST00000110042.1 | -2238 | Gm15440 | ENSMUSG00000051107.4 |
| chr13 | 95989539 | 95989852 | 3.36981 | 3.88232 | 1.01733 | Exon (ENSMUST00000161263.2/ENSMUSG00000051111.10, exon 5 of 13) | 13 | 95961163 | 95984850 | 23688 | 2 | ENSMUST00000182180.1 | -4690 | Sv2c | ENSMUSG00000051111.10 |
| chr13 | 96007007 | 96007241 | 3.34153 | 3.83966 | 1.01733 | Intron (ENSMUST00000161263.2/ENSMUSG00000051111.10, intron 4 of 12) | 13 | 95961163 | 95984850 | 23688 | 2 | ENSMUST00000182180.1 | -22158 | Sv2c | ENSMUSG00000051111.10 |
| chr13 | 96128602 | 96130790 | 3.81134 | 4.73495 | 1.59154 | Intron (ENSMUST00000161263.2/ENSMUSG00000051111.10, intron 1 of 12) | 13 | 95959443 | 96132577 | 173135 | 2 | ENSMUST00000161263.2 | 1787 | Sv2c | ENSMUSG00000051111.10 |
| chr13 | 96131976 | 96133928 | 3.60036 | 4.31459 | 1.29694 | Promoter | 13 | 95959443 | 96132577 | 173135 | 2 | ENSMUST00000161263.2 | 0 | Sv2c | ENSMUSG00000051111.10 |
| chr7 | 44509829 | 44510029 | 3.93129 | 4.92583 | 1.67551 | Exon (ENSMUST00000165208.2/ENSMUSG00000038670.9, exon 17 of 28) | 7 | 44496588 | 44501134 | 4547 | 1 | ENSMUST00000107927.3 | 13242 | Fam71e1 | ENSMUSG00000051113.8 |
| chr3 | 27370153 | 27370443 | 3.39815 | 3.93967 | 1.069 | Promoter | 3 | 27371351 | 27378010 | 6660 | 1 | ENSMUST00000057186.1 | -908 | Ghsr | ENSMUSG00000051136.1 |
| chr3 | 27370905 | 27372997 | 3.79296 | 4.76871 | 1.59793 | Promoter | 3 | 27371351 | 27378010 | 6660 | 1 | ENSMUST00000057186.1 | 0 | Ghsr | ENSMUSG00000051136.1 |
| chr3 | 27373376 | 27373692 | 3.71182 | 4.53567 | 1.46095 | Intron (ENSMUST00000057186.1/ENSMUSG00000051136.1, intron 2 of 2) | 3 | 27371351 | 27378010 | 6660 | 1 | ENSMUST00000057186.1 | 2026 | Ghsr | ENSMUSG00000051136.1 |
| chr3 | 27374114 | 27374275 | 4.24036 | 5.63726 | 2.26589 | Intron (ENSMUST00000057186.1/ENSMUSG00000051136.1, intron 2 of 2) | 3 | 27371351 | 27378010 | 6660 | 1 | ENSMUST00000057186.1 | 2764 | Ghsr | ENSMUSG00000051136.1 |
| chr3 | 27374668 | 27374857 | 4.10719 | 5.39241 | 2.04649 | Exon (ENSMUST00000057186.1/ENSMUSG00000051136.1, exon 3 of 3) | 3 | 27371351 | 27378010 | 6660 | 1 | ENSMUST00000057186.1 | 3318 | Ghsr | ENSMUSG00000051136.1 |
| chr16 | 20623408 | 20623486 | 4.25949 | 5.67729 | 2.26103 | Intron (ENSMUST00000079600.5/ENSMUSG00000022842.11, intron 2 of 18) | 16 | 20619215 | 20621287 | 2073 | 2 | ENSMUST00000052939.2 | -2122 | Camk2n2 | ENSMUSG00000051146.2 |
| chr4 | 124607495 | 124607834 | 3.61403 | 4.32979 | 1.29441 | Intron (ENSMUST00000184062.1/ENSMUSG00000098760.1, intron 1 of 1) | 4 | 124609931 | 124610803 | 873 | 2 | ENSMUST00000053604.2 | 2969 | 3100002H09Rik | ENSMUSG00000051219.2 |
| chr4 | 124633155 | 124633230 | 3.32822 | 3.81962 | 1.01733 | Intron (ENSMUST00000184062.1/ENSMUSG00000098760.1, intron 1 of 1) | 4 | 124609931 | 124610803 | 873 | 2 | ENSMUST00000053604.2 | -22353 | 3100002H09Rik | ENSMUSG00000051219.2 |
| chr15 | 57994220 | 57994593 | 3.44003 | 4.12916 | 1.18797 | Intron (ENSMUST00000160942.2/ENSMUSG00000051225.4, intron 2 of 3) | 15 | 58002146 | 58010047 | 7902 | 1 | ENSMUST00000161231.1 | -7553 | Fam83a | ENSMUSG00000051225.4 |
| chr15 | 57994965 | 57995108 | 3.38114 | 3.89941 | 1.01733 | Intron (ENSMUST00000160942.2/ENSMUSG00000051225.4, intron 2 of 3) | 15 | 58002146 | 58010047 | 7902 | 1 | ENSMUST00000161231.1 | -7038 | Fam83a | ENSMUSG00000051225.4 |
| chr9 | 58191193 | 58191276 | 3.50121 | 4.12149 | 1.16164 | Distal Intergenic | 9 | 58196298 | 58201715 | 5418 | 2 | ENSMUST00000163200.1 | 10439 | Islr2 | ENSMUSG00000051243.8 |
| chr9 | 58194207 | 58194913 | 3.90175 | 4.8751 | 1.65001 | Downstream (1-2kb) | 9 | 58196298 | 58201715 | 5418 | 2 | ENSMUST00000163200.1 | 6802 | Islr2 | ENSMUSG00000051243.8 |
| chr9 | 58196020 | 58199786 | 3.83287 | 4.87497 | 1.68941 | 3' UTR | 9 | 58196298 | 58201715 | 5418 | 2 | ENSMUST00000163200.1 | 1929 | Islr2 | ENSMUSG00000051243.8 |
| chr9 | 58200715 | 58200905 | 3.43484 | 4.09741 | 1.23746 | Promoter | 9 | 58196298 | 58201715 | 5418 | 2 | ENSMUST00000163200.1 | 810 | Islr2 | ENSMUSG00000051243.8 |
| chr9 | 58201232 | 58201627 | 3.64489 | 4.48791 | 1.46399 | Promoter | 9 | 58196298 | 58201715 | 5418 | 2 | ENSMUST00000163200.1 | 88 | Islr2 | ENSMUSG00000051243.8 |
| chr9 | 58202019 | 58204311 | 3.87813 | 4.96083 | 1.78406 | Promoter | 9 | 58196298 | 58202285 | 5988 | 2 | ENSMUST00000163897.2 | 0 | Islr2 | ENSMUSG00000051243.8 |
| chr1 | 172052937 | 172053663 | 4.03058 | 5.18729 | 1.93581 | 3' UTR | 1 | 172052292 | 172057573 | 5282 | 2 | ENSMUST00000059794.3 | 3910 | Nhlh1 | ENSMUSG00000051251.3 |
| chr1 | 172054012 | 172054456 | 4.00302 | 5.07105 | 1.77898 | 5' UTR | 1 | 172052292 | 172057573 | 5282 | 2 | ENSMUST00000059794.3 | 3117 | Nhlh1 | ENSMUSG00000051251.3 |
| chr1 | 172055448 | 172055598 | 3.42676 | 4.01247 | 1.13831 | Intron (ENSMUST00000059794.3/ENSMUSG00000051251.3, intron 1 of 1) | 1 | 172052292 | 172057573 | 5282 | 2 | ENSMUST00000059794.3 | 1975 | Nhlh1 | ENSMUSG00000051251.3 |
| chr4 | 154661820 | 154662059 | 3.48096 | 4.10927 | 1.18611 | Distal Intergenic | 4 | 154666433 | 154667867 | 1435 | 2 | ENSMUST00000060062.4 | 5808 | Actrt2 | ENSMUSG00000051276.4 |
| chr4 | 154671242 | 154671616 | 3.33686 | 3.83266 | 1.01733 | Distal Intergenic | 4 | 154666433 | 154667867 | 1435 | 2 | ENSMUST00000060062.4 | -3376 | Actrt2 | ENSMUSG00000051276.4 |
| chr4 | 154675720 | 154675814 | 3.60334 | 4.31054 | 1.27846 | Distal Intergenic | 4 | 154666433 | 154667867 | 1435 | 2 | ENSMUST00000060062.4 | -7854 | Actrt2 | ENSMUSG00000051276.4 |
| chr4 | 154683063 | 154683154 | 3.73942 | 4.56122 | 1.43187 | Distal Intergenic | 4 | 154666433 | 154667867 | 1435 | 2 | ENSMUST00000060062.4 | -15197 | Actrt2 | ENSMUSG00000051276.4 |
| chr4 | 154683466 | 154683740 | 3.45435 | 4.03734 | 1.11441 | Distal Intergenic | 4 | 154666433 | 154667867 | 1435 | 2 | ENSMUST00000060062.4 | -15600 | Actrt2 | ENSMUSG00000051276.4 |
| chr4 | 154684497 | 154684668 | 3.29215 | 3.76549 | 1.01733 | Distal Intergenic | 4 | 154666433 | 154667867 | 1435 | 2 | ENSMUST00000060062.4 | -16631 | Actrt2 | ENSMUSG00000051276.4 |
| chr4 | 154696319 | 154696575 | 3.52092 | 4.15735 | 1.18303 | Distal Intergenic | 4 | 154666433 | 154667867 | 1435 | 2 | ENSMUST00000060062.4 | -28453 | Actrt2 | ENSMUSG00000051276.4 |
| chr4 | 154729200 | 154729532 | 3.48714 | 4.11817 | 1.22626 | Distal Intergenic | 4 | 154666433 | 154667867 | 1435 | 2 | ENSMUST00000060062.4 | -61334 | Actrt2 | ENSMUSG00000051276.4 |
| chr4 | 154730345 | 154730862 | 3.44021 | 4.02807 | 1.15755 | Distal Intergenic | 4 | 154666433 | 154667867 | 1435 | 2 | ENSMUST00000060062.4 | -62479 | Actrt2 | ENSMUSG00000051276.4 |
| chr4 | 154739823 | 154739916 | 3.92529 | 5.10928 | 1.89365 | Distal Intergenic | 4 | 154666433 | 154667867 | 1435 | 2 | ENSMUST00000060062.4 | -71957 | Actrt2 | ENSMUSG00000051276.4 |
| chr4 | 154741430 | 154741735 | 3.66348 | 4.42295 | 1.36281 | Distal Intergenic | 4 | 154666433 | 154667867 | 1435 | 2 | ENSMUST00000060062.4 | -73564 | Actrt2 | ENSMUSG00000051276.4 |
| chr4 | 154742109 | 154742383 | 3.82077 | 4.73722 | 1.57604 | Distal Intergenic | 4 | 154666433 | 154667867 | 1435 | 2 | ENSMUST00000060062.4 | -74243 | Actrt2 | ENSMUSG00000051276.4 |
| chr4 | 154743128 | 154743518 | 3.92364 | 4.92415 | 1.69069 | Distal Intergenic | 4 | 154666433 | 154667867 | 1435 | 2 | ENSMUST00000060062.4 | -75262 | Actrt2 | ENSMUSG00000051276.4 |
| chr4 | 154747638 | 154748099 | 3.51726 | 4.36535 | 1.38861 | Distal Intergenic | 4 | 154666433 | 154667867 | 1435 | 2 | ENSMUST00000060062.4 | -79772 | Actrt2 | ENSMUSG00000051276.4 |
| chr4 | 9842950 | 9850004 | 3.79926 | 4.802 | 1.67224 | Promoter | 4 | 9844372 | 9862345 | 17974 | 1 | ENSMUST00000057613.2 | 0 | Gdf6 | ENSMUSG00000051279.2 |
| chr4 | 9850607 | 9851223 | 3.42074 | 4.12434 | 1.23971 | Intron (ENSMUST00000057613.2/ENSMUSG00000051279.2, intron 1 of 1) | 4 | 9844372 | 9862345 | 17974 | 1 | ENSMUST00000057613.2 | 6236 | Gdf6 | ENSMUSG00000051279.2 |
| chr4 | 9855679 | 9856559 | 3.68873 | 4.52103 | 1.51631 | Intron (ENSMUST00000057613.2/ENSMUSG00000051279.2, intron 1 of 1) | 4 | 9844372 | 9862345 | 17974 | 1 | ENSMUST00000057613.2 | 11308 | Gdf6 | ENSMUSG00000051279.2 |
| chr4 | 9857209 | 9860711 | 3.77817 | 4.73367 | 1.62093 | 3' UTR | 4 | 9844372 | 9862345 | 17974 | 1 | ENSMUST00000057613.2 | 12838 | Gdf6 | ENSMUSG00000051279.2 |
| chr4 | 9861162 | 9862108 | 3.78741 | 4.83987 | 1.69881 | 3' UTR | 4 | 9844372 | 9862345 | 17974 | 1 | ENSMUST00000057613.2 | 16791 | Gdf6 | ENSMUSG00000051279.2 |
| chr16 | 92478986 | 92479288 | 3.33517 | 3.83011 | 1.01733 | Promoter | 16 | 92478742 | 92497366 | 18625 | 1 | ENSMUST00000160494.2 | 245 | 2410124H12Rik | ENSMUSG00000051297.8 |
| chr16 | 92479861 | 92479991 | 3.98333 | 5.03667 | 1.76979 | Intron (ENSMUST00000160494.2/ENSMUSG00000051297.8, intron 1 of 2) | 16 | 92478742 | 92497366 | 18625 | 1 | ENSMUST00000160494.2 | 1120 | 2410124H12Rik | ENSMUSG00000051297.8 |
| chr18 | 37645709 | 37646345 | 3.39281 | 3.98328 | 1.17925 | Distal Intergenic | 18 | 37640902 | 37644204 | 3303 | 2 | ENSMUST00000066272.4 | -1506 | Taf7 | ENSMUSG00000051316.7 |
| chr18 | 37646752 | 37646910 | 3.66679 | 4.47302 | 1.47889 | Distal Intergenic | 18 | 37640902 | 37644204 | 3303 | 2 | ENSMUST00000066272.4 | -2549 | Taf7 | ENSMUSG00000051316.7 |
| chr18 | 37647501 | 37649060 | 3.56245 | 4.25776 | 1.29678 | Distal Intergenic | 18 | 37640902 | 37644204 | 3303 | 2 | ENSMUST00000066272.4 | -3298 | Taf7 | ENSMUSG00000051316.7 |
| chr18 | 37649695 | 37649878 | 3.6702 | 4.43705 | 1.37107 | Distal Intergenic | 18 | 37640902 | 37644204 | 3303 | 2 | ENSMUST00000066272.4 | -5492 | Taf7 | ENSMUSG00000051316.7 |
| chr6 | 118592697 | 118592772 | 3.45502 | 4.04342 | 1.13233 | 3' UTR | 6 | 118592304 | 118595426 | 3123 | 2 | ENSMUST00000191196.1 | 2654 | Cacna1c | ENSMUSG00000051331.9 |
| chr6 | 118593328 | 118593627 | 3.65339 | 4.57803 | 1.50285 | 3' UTR | 6 | 118592304 | 118595426 | 3123 | 2 | ENSMUST00000191196.1 | 1799 | Cacna1c | ENSMUSG00000051331.9 |
| chr6 | 119107460 | 119108628 | 3.83528 | 4.78047 | 1.61972 | Promoter | 6 | 118592464 | 119108189 | 515726 | 2 | ENSMUST00000078320.8 | 0 | Cacna1c | ENSMUSG00000051331.9 |
| chr6 | 119112458 | 119112533 | 3.3081 | 3.79811 | 1.04784 | Intron (ENSMUST00000186889.1/ENSMUSG00000051331.9, intron 1 of 46) | 6 | 118593647 | 119108573 | 514927 | 2 | ENSMUST00000188522.1 | -3886 | Cacna1c | ENSMUSG00000051331.9 |
| chr5 | 113138854 | 113139119 | 3.49251 | 4.17839 | 1.29458 | Intron (ENSMUST00000050125.8/ENSMUSG00000051339.8, intron 1 of 7) | 5 | 113086323 | 113163351 | 77029 | 2 | ENSMUST00000050125.8 | 24232 | 2900026A02Rik | ENSMUSG00000051339.8 |
| chr5 | 113162481 | 113162643 | 3.38193 | 3.9006 | 1.01733 | Promoter | 5 | 113086323 | 113163351 | 77029 | 2 | ENSMUST00000050125.8 | 708 | 2900026A02Rik | ENSMUSG00000051339.8 |
| chr5 | 113163118 | 113163258 | 3.92033 | 4.99239 | 1.74995 | Promoter | 5 | 113086323 | 113163351 | 77029 | 2 | ENSMUST00000050125.8 | 93 | 2900026A02Rik | ENSMUSG00000051339.8 |
| chr12 | 73004096 | 73004303 | 3.33785 | 3.83961 | 1.0364 | Distal Intergenic | 12 | 73040015 | 73042815 | 2801 | 2 | ENSMUST00000176310.1 | 38512 | Six1 | ENSMUSG00000051367.8 |
| chr12 | 73021649 | 73021943 | 3.57973 | 4.40402 | 1.37785 | Distal Intergenic | 12 | 73040015 | 73042815 | 2801 | 2 | ENSMUST00000176310.1 | 20872 | Six1 | ENSMUSG00000051367.8 |
| chr12 | 73025258 | 73025422 | 3.58764 | 4.32242 | 1.30857 | Distal Intergenic | 12 | 73040015 | 73042815 | 2801 | 2 | ENSMUST00000176310.1 | 17393 | Six1 | ENSMUSG00000051367.8 |
| chr12 | 73025873 | 73025954 | 3.6046 | 4.34933 | 1.37913 | Distal Intergenic | 12 | 73040015 | 73042815 | 2801 | 2 | ENSMUST00000176310.1 | 16861 | Six1 | ENSMUSG00000051367.8 |
| chr12 | 73030277 | 73030359 | 4.05974 | 5.21722 | 1.94338 | Distal Intergenic | 12 | 73040015 | 73042815 | 2801 | 2 | ENSMUST00000176310.1 | 12456 | Six1 | ENSMUSG00000051367.8 |
| chr12 | 73031327 | 73031811 | 3.5582 | 4.35319 | 1.34357 | Distal Intergenic | 12 | 73040015 | 73042815 | 2801 | 2 | ENSMUST00000176310.1 | 11004 | Six1 | ENSMUSG00000051367.8 |
| chr12 | 73033142 | 73035494 | 4.08317 | 5.3996 | 2.11225 | Distal Intergenic | 12 | 73040015 | 73042815 | 2801 | 2 | ENSMUST00000176310.1 | 7321 | Six1 | ENSMUSG00000051367.8 |
| chr12 | 73036215 | 73036762 | 4.36788 | 5.90559 | 2.43868 | Distal Intergenic | 12 | 73040015 | 73042815 | 2801 | 2 | ENSMUST00000176310.1 | 6053 | Six1 | ENSMUSG00000051367.8 |
| chr12 | 73037095 | 73038644 | 3.74394 | 4.62905 | 1.57092 | Downstream (1-2kb) | 12 | 73040015 | 73042815 | 2801 | 2 | ENSMUST00000176310.1 | 4171 | Six1 | ENSMUSG00000051367.8 |
| chr12 | 73039147 | 73039497 | 3.70418 | 4.61233 | 1.53829 | Downstream (<1kb) | 12 | 73040015 | 73042815 | 2801 | 2 | ENSMUST00000176310.1 | 3318 | Six1 | ENSMUSG00000051367.8 |
| chr12 | 73039815 | 73040340 | 4.0526 | 5.4525 | 2.08303 | Exon (ENSMUST00000176310.1/ENSMUSG00000051367.8, exon 2 of 2) | 12 | 73040015 | 73042815 | 2801 | 2 | ENSMUST00000176310.1 | 2475 | Six1 | ENSMUSG00000051367.8 |
| chr12 | 73041118 | 73041532 | 3.83377 | 4.86507 | 1.70753 | Intron (ENSMUST00000176310.1/ENSMUSG00000051367.8, intron 1 of 1) | 12 | 73040015 | 73042815 | 2801 | 2 | ENSMUST00000176310.1 | 1283 | Six1 | ENSMUSG00000051367.8 |
| chr12 | 73041999 | 73042481 | 3.89594 | 4.94251 | 1.76585 | Promoter | 12 | 73040015 | 73042815 | 2801 | 2 | ENSMUST00000176310.1 | 334 | Six1 | ENSMUSG00000051367.8 |
| chr12 | 73042836 | 73044039 | 4.15751 | 5.4869 | 2.13213 | Promoter | 12 | 73040015 | 73042815 | 2801 | 2 | ENSMUST00000176310.1 | -22 | Six1 | ENSMUSG00000051367.8 |
| chr12 | 73044513 | 73046901 | 4.10433 | 5.55366 | 2.19218 | Promoter | 12 | 73043774 | 73045291 | 1518 | 2 | ENSMUST00000175677.1 | 0 | Six1 | ENSMUSG00000051367.8 |
| chr12 | 73047897 | 73048713 | 4.15209 | 5.61973 | 2.25931 | Promoter | 12 | 73041827 | 73047179 | 5353 | 2 | ENSMUST00000050029.7 | -719 | Six1 | ENSMUSG00000051367.8 |
| chr12 | 73049904 | 73050998 | 3.9481 | 5.09531 | 1.88132 | Intron (ENSMUST00000176091.1/ENSMUSG00000051367.8, intron 1 of 1) | 12 | 73041827 | 73047179 | 5353 | 2 | ENSMUST00000050029.7 | -2726 | Six1 | ENSMUSG00000051367.8 |
| chr12 | 73051365 | 73051630 | 3.86028 | 5.09328 | 1.83276 | Intron (ENSMUST00000176091.1/ENSMUSG00000051367.8, intron 1 of 1) | 12 | 73043531 | 73053887 | 10357 | 2 | ENSMUST00000176091.1 | 2257 | Six1 | ENSMUSG00000051367.8 |
| chr12 | 73052223 | 73053932 | 3.88972 | 5.02347 | 1.82202 | Promoter | 12 | 73043531 | 73053887 | 10357 | 2 | ENSMUST00000176091.1 | 0 | Six1 | ENSMUSG00000051367.8 |
| chr12 | 73054793 | 73055843 | 3.4433 | 4.04976 | 1.21088 | Promoter | 12 | 73043531 | 73053887 | 10357 | 2 | ENSMUST00000176091.1 | -907 | Six1 | ENSMUSG00000051367.8 |
| chr12 | 73056244 | 73056933 | 3.87742 | 4.88819 | 1.75889 | Distal Intergenic | 12 | 73043531 | 73053887 | 10357 | 2 | ENSMUST00000176091.1 | -2358 | Six1 | ENSMUSG00000051367.8 |
| chr12 | 73058466 | 73058712 | 3.25662 | 3.82364 | 1.01733 | Distal Intergenic | 12 | 73043531 | 73053887 | 10357 | 2 | ENSMUST00000176091.1 | -4580 | Six1 | ENSMUSG00000051367.8 |
| chr12 | 73063384 | 73063539 | 3.31742 | 3.95486 | 1.07632 | Distal Intergenic | 12 | 73043531 | 73053887 | 10357 | 2 | ENSMUST00000176091.1 | -9498 | Six1 | ENSMUSG00000051367.8 |
| chr12 | 73064830 | 73065178 | 3.42628 | 4.01047 | 1.16988 | Distal Intergenic | 12 | 73043531 | 73053887 | 10357 | 2 | ENSMUST00000176091.1 | -10944 | Six1 | ENSMUSG00000051367.8 |
| chr18 | 38146889 | 38147009 | 3.42518 | 3.99556 | 1.12261 | Distal Intergenic | 18 | 38188888 | 38203163 | 14276 | 2 | ENSMUST00000161701.1 | 56154 | Pcdh1 | ENSMUSG00000051375.10 |
| chr18 | 38147577 | 38147711 | 3.41285 | 3.98168 | 1.13937 | Distal Intergenic | 18 | 38188888 | 38203163 | 14276 | 2 | ENSMUST00000161701.1 | 55452 | Pcdh1 | ENSMUSG00000051375.10 |
| chr18 | 38148062 | 38148331 | 3.78467 | 4.67553 | 1.5777 | Distal Intergenic | 18 | 38188888 | 38203163 | 14276 | 2 | ENSMUST00000161701.1 | 54832 | Pcdh1 | ENSMUSG00000051375.10 |
| chr18 | 38148972 | 38149078 | 3.36327 | 3.87765 | 1.03574 | Distal Intergenic | 18 | 38188888 | 38203163 | 14276 | 2 | ENSMUST00000161701.1 | 54085 | Pcdh1 | ENSMUSG00000051375.10 |
| chr18 | 38208312 | 38208943 | 3.73855 | 4.57345 | 1.47619 | Promoter | 18 | 38196694 | 38209762 | 13069 | 2 | ENSMUST00000057185.7 | 819 | Pcdh1 | ENSMUSG00000051375.10 |
| chr18 | 38209466 | 38210552 | 3.84389 | 4.90959 | 1.73186 | Promoter | 18 | 38196694 | 38209762 | 13069 | 2 | ENSMUST00000057185.7 | 0 | Pcdh1 | ENSMUSG00000051375.10 |
| chr18 | 38211137 | 38211989 | 3.79022 | 4.85652 | 1.70985 | Promoter | 18 | 38196743 | 38211970 | 15228 | 2 | ENSMUST00000159405.1 | 0 | Pcdh1 | ENSMUSG00000051375.10 |
| chr18 | 38212352 | 38212774 | 3.9478 | 5.07036 | 1.88188 | Promoter | 18 | 38185915 | 38212053 | 26139 | 2 | ENSMUST00000160721.2 | -300 | Pcdh1 | ENSMUSG00000051375.10 |
| chr18 | 38213422 | 38213625 | 3.35822 | 3.86482 | 1.01733 | Intron (ENSMUST00000160935.1/ENSMUSG00000089983.1, intron 1 of 1) | 18 | 38185915 | 38212053 | 26139 | 2 | ENSMUST00000160721.2 | -1370 | Pcdh1 | ENSMUSG00000051375.10 |
| chr18 | 38215903 | 38215978 | 3.61512 | 4.33279 | 1.29957 | Intron (ENSMUST00000160935.1/ENSMUSG00000089983.1, intron 1 of 1) | 18 | 38185915 | 38212053 | 26139 | 2 | ENSMUST00000160721.2 | -3851 | Pcdh1 | ENSMUSG00000051375.10 |
| chr18 | 38216723 | 38217042 | 3.7956 | 4.72819 | 1.58193 | Intron (ENSMUST00000160935.1/ENSMUSG00000089983.1, intron 1 of 1) | 18 | 38185915 | 38212053 | 26139 | 2 | ENSMUST00000160721.2 | -4671 | Pcdh1 | ENSMUSG00000051375.10 |
| chr18 | 38217991 | 38218526 | 3.44167 | 4.01209 | 1.09241 | Intron (ENSMUST00000160935.1/ENSMUSG00000089983.1, intron 1 of 1) | 18 | 38185915 | 38212053 | 26139 | 2 | ENSMUST00000160721.2 | -5939 | Pcdh1 | ENSMUSG00000051375.10 |
| chr2 | 140670565 | 140670693 | 3.65759 | 4.43937 | 1.38927 | Promoter | 2 | 140659325 | 140671400 | 12076 | 2 | ENSMUST00000056760.3 | 707 | Flrt3 | ENSMUSG00000051379.7 |
| chr2 | 140671017 | 140671389 | 3.42506 | 3.98728 | 1.09361 | Promoter | 2 | 140659325 | 140671400 | 12076 | 2 | ENSMUST00000056760.3 | 11 | Flrt3 | ENSMUSG00000051379.7 |
| chr18 | 40258406 | 40258771 | 3.56468 | 4.2642 | 1.31052 | Promoter | 18 | 40258361 | 40531168 | 272808 | 1 | ENSMUST00000091927.4 | 46 | Kctd16 | ENSMUSG00000051401.6 |
| chr18 | 40259155 | 40259380 | 3.2899 | 3.78428 | 1.08024 | Promoter | 18 | 40258361 | 40531168 | 272808 | 1 | ENSMUST00000091927.4 | 795 | Kctd16 | ENSMUSG00000051401.6 |
| chr18 | 40260741 | 40260879 | 3.8001 | 4.99399 | 1.76735 | Intron (ENSMUST00000091927.4/ENSMUSG00000051401.6, intron 1 of 1) | 18 | 40258361 | 40531168 | 272808 | 1 | ENSMUST00000091927.4 | 2381 | Kctd16 | ENSMUSG00000051401.6 |
| chr18 | 40308059 | 40308388 | 1.97754 | 9.85045 | 5.69155 | Intron (ENSMUST00000091927.4/ENSMUSG00000051401.6, intron 1 of 1) | 18 | 40258361 | 40531168 | 272808 | 1 | ENSMUST00000091927.4 | 49699 | Kctd16 | ENSMUSG00000051401.6 |

| | | | | | | | | | | | | | | | |
|---|---|---|---|---|---|---|---|---|---|---|---|---|---|---|---|
| chr4 | 142005003 | 142005162 | 3.5218 | 4.32218 | 1.32148 | Intron (ENSMUST00000105780.2/ENSMUSG00000051435.6, intron 2 of 32) | 4 | 141890851 | 142011637 | 120787 | 2 | ENSMUST00000105779.3 | 6475 | Fhad1 | ENSMUSG00000051435.6 |
| chr4 | 142014782 | 142015114 | 3.44872 | 4.03113 | 1.12237 | Promoter | 4 | 141890438 | 142015067 | 124630 | 2 | ENSMUST00000105780.2 | 0 | Fhad1 | ENSMUSG00000051435.6 |
| chr4 | 142015638 | 142016540 | 3.77921 | 4.75228 | 1.60684 | Promoter | 4 | 141962875 | 142015082 | 52208 | 2 | ENSMUST00000123068.1 | -557 | Fhad1 | ENSMUSG00000051435.6 |
| chr18 | 36725381 | 36725971 | 3.77017 | 4.66957 | 1.5821 | Promoter | 18 | 36725074 | 36726736 | 1663 | 2 | ENSMUST00000061829.6 | 765 | Cd14 | ENSMUSG00000051439.6 |
| chr18 | 36727463 | 36727759 | 3.59575 | 4.52325 | 1.45883 | Promoter | 18 | 36725074 | 36726736 | 1663 | 2 | ENSMUST00000061829.6 | -728 | Cd14 | ENSMUSG00000051439.6 |
| chr13 | 113722266 | 113722615 | 3.49794 | 4.49056 | 1.41575 | Distal Intergenic | 13 | 113662896 | 113663676 | 781 | 2 | ENSMUST00000054650.4 | -58591 | Hspb3 | ENSMUSG00000051456.4 |
| chr19 | 24894487 | 24895159 | 3.54778 | 4.31425 | 1.35203 | Distal Intergenic | 19 | 24898965 | 24901309 | 2345 | 2 | ENSMUST00000058600.2 | 6150 | Foxd4 | ENSMUSG00000051490.2 |
| chr19 | 24895555 | 24896070 | 3.49189 | 4.15782 | 1.29819 | Downstream (2-3kb) | 19 | 24898965 | 24901309 | 2345 | 2 | ENSMUST00000058600.2 | 5239 | Foxd4 | ENSMUSG00000051490.2 |
| chr19 | 24896710 | 24898225 | 3.82489 | 4.91553 | 1.75005 | Downstream (<1kb) | 19 | 24898965 | 24901309 | 2345 | 2 | ENSMUST00000058600.2 | 3084 | Foxd4 | ENSMUSG00000051490.2 |
| chr19 | 24899136 | 24899641 | 4.45112 | 6.16294 | 2.68041 | 3' UTR | 19 | 24898965 | 24901309 | 2345 | 2 | ENSMUST00000058600.2 | 1668 | Foxd4 | ENSMUSG00000051490.2 |
| chr19 | 24899995 | 24900107 | 3.97824 | 5.26917 | 1.94329 | Exon (ENSMUST00000058600.2/ENSMUSG00000051490.2, exon 1 of 1) | 19 | 24898965 | 24901309 | 2345 | 2 | ENSMUST00000058600.2 | 1202 | Foxd4 | ENSMUSG00000051490.2 |
| chr19 | 24900469 | 24903185 | 3.75957 | 4.86517 | 1.69662 | Promoter | 19 | 24898965 | 24901309 | 2345 | 2 | ENSMUST00000058600.2 | 0 | Foxd4 | ENSMUSG00000051490.2 |
| chr19 | 24906643 | 24907012 | 4.14662 | 5.42804 | 2.11102 | Distal Intergenic | 19 | 24898965 | 24901309 | 2345 | 2 | ENSMUST00000058600.2 | -5335 | Foxd4 | ENSMUSG00000051490.2 |
| chr7 | 93079665 | 93081677 | 4.10836 | 5.47781 | 2.1435 | Promoter | 7 | 93079879 | 93081715 | 1837 | 1 | ENSMUST00000051179.8 | 0 | Fam181b | ENSMUSG00000051515.8 |
| chr5 | 38275003 | 38275097 | 3.82034 | 4.78065 | 1.66441 | Promoter | 5 | 38275972 | 38304209 | 28238 | 1 | ENSMUST00000063136.8 | -875 | Otop1 | ENSMUSG00000051596.8 |
| chr5 | 38276476 | 38276981 | 3.45596 | 4.30964 | 1.34795 | Promoter | 5 | 38277404 | 38304208 | 26805 | 1 | ENSMUST00000114099.3 | -423 | Otop1 | ENSMUSG00000051596.8 |
| chr5 | 38277351 | 38277639 | 4.0656 | 5.93773 | 2.43657 | Promoter | 5 | 38277404 | 38304208 | 26805 | 1 | ENSMUST00000114099.3 | 0 | Otop1 | ENSMUSG00000051596.8 |
| chr5 | 38278046 | 38278130 | 3.81224 | 4.95963 | 1.74095 | Promoter | 5 | 38277404 | 38304217 | 26814 | 1 | ENSMUST00000185357.1 | 643 | Otop1 | ENSMUSG00000051596.8 |
| chr5 | 38278447 | 38278631 | 3.40259 | 3.98678 | 1.19416 | Intron (ENSMUST00000063136.8/ENSMUSG00000051596.8, intron 2 of 7) | 5 | 38277404 | 38304217 | 26814 | 1 | ENSMUST00000185357.1 | 1044 | Otop1 | ENSMUSG00000051596.8 |
| chr18 | 37264814 | 37265042 | 3.51532 | 4.24857 | 1.28991 | Promoter | 18 | 37264998 | 37267454 | 2457 | 1 | ENSMUST00000052366.3 | 0 | Pcdhb1 | ENSMUSG00000051663.3 |
| chr1 | 75482485 | 75483004 | 3.81049 | 4.95822 | 1.74858 | Exon (ENSMUST00000113575.3/ENSMUSG00000051703.9, exon 3 of 5) | 1 | 75479704 | 75482675 | 2972 | 1 | ENSMUST00000187411.1 | 2782 | Tmem198 | ENSMUSG00000051703.9 |
| chr2 | 76872810 | 76873084 | 3.63903 | 4.47485 | 1.49061 | Exon (ENSMUST00000099981.3/ENSMUSG00000051747.9, exon 99 of 347) | 2 | 76849271 | 76870611 | 21341 | 2 | ENSMUST00000138542.1 | -2200 | Ttn | ENSMUSG00000051747.9 |
| chr2 | 76873800 | 76874127 | 3.4965 | 4.15305 | 1.28409 | Exon (ENSMUST00000099981.3/ENSMUSG00000051747.9, exon 98 of 347) | 2 | 76849271 | 76870611 | 21341 | 2 | ENSMUST00000138542.1 | -3190 | Ttn | ENSMUSG00000051747.9 |
| chr6 | 30733172 | 30733545 | 3.48431 | 4.41542 | 1.38899 | Promoter | 6 | 30733454 | 30742836 | 9383 | 1 | ENSMUST00000147400.2 | 0 | Mest | ENSMUSG00000051855.10 |
| chr6 | 30738189 | 30738851 | 3.55659 | 4.32931 | 1.36563 | Promoter | 6 | 30738055 | 30738878 | 824 | 1 | ENSMUST00000131465.1 | 135 | Mest | ENSMUSG00000051855.10 |
| chr6 | 70909539 | 70909782 | 3.71198 | 4.5118 | 1.40216 | Distal Intergenic | 6 | 70913087 | 70918927 | 5841 | 2 | ENSMUST00000063456.4 | 9145 | Tex37 | ENSMUSG00000051896.4 |
| chr6 | 70910271 | 70910412 | 3.89409 | 4.90223 | 1.69072 | Downstream (2-3kb) | 6 | 70913087 | 70918927 | 5841 | 2 | ENSMUST00000063456.4 | 8515 | Tex37 | ENSMUSG00000051896.4 |
| chr6 | 70910948 | 70911028 | 3.84494 | 4.75743 | 1.55811 | Downstream (2-3kb) | 6 | 70913087 | 70918927 | 5841 | 2 | ENSMUST00000063456.4 | 7899 | Tex37 | ENSMUSG00000051896.4 |
| chr6 | 70912054 | 70912443 | 3.69899 | 4.49159 | 1.39383 | Downstream (<1kb) | 6 | 70913087 | 70918927 | 5841 | 2 | ENSMUST00000063456.4 | 6484 | Tex37 | ENSMUSG00000051896.4 |
| chr6 | 70921286 | 70921713 | 3.90231 | 5.0494 | 1.81894 | Distal Intergenic | 6 | 70913087 | 70918927 | 5841 | 2 | ENSMUST00000063456.4 | -2360 | Tex37 | ENSMUSG00000051896.4 |
| chr7 | 120504414 | 120504739 | 3.48219 | 4.09204 | 1.15898 | Intron (ENSMUST00000056042.5/ENSMUSG00000051900.7, intron 18 of 30) | 7 | 120475422 | 120489315 | 13894 | 1 | ENSMUST00000153845.1 | 28993 | Abca16 | ENSMUSG00000051900.7 |
| chr7 | 115667015 | 115667130 | 3.64023 | 4.45123 | 1.44936 | Intron (ENSMUST00000072804.5/ENSMUSG00000051910.8, intron 5 of 16) | 7 | 115474475 | 115824707 | 350233 | 2 | ENSMUST00000169129.2 | 157577 | Sox6 | ENSMUSG00000051910.8 |
| chr7 | 115826798 | 115826883 | 4.7497 | 6.68067 | 3.0269 | Intron (ENSMUST00000072804.5/ENSMUSG00000051910.8, intron 2 of 16) | 7 | 115474475 | 115824707 | 350233 | 2 | ENSMUST00000169129.2 | -2092 | Sox6 | ENSMUSG00000051910.8 |
| chr7 | 116039235 | 116039992 | 3.73921 | 4.75414 | 1.61198 | Promoter | 7 | 115471344 | 116038744 | 567401 | 2 | ENSMUST00000166877.2 | -492 | Sox6 | ENSMUSG00000051910.8 |
| chr7 | 116036409 | 116036704 | 3.5721 | 4.35125 | 1.36787 | Intron (ENSMUST00000166877.2/ENSMUSG00000051910.8, intron 1 of 15) | 7 | 115471344 | 116038744 | 567401 | 2 | ENSMUST00000166877.2 | 2040 | Sox6 | ENSMUSG00000051910.8 |
| chr7 | 116037161 | 116037493 | 3.53963 | 4.44457 | 1.41804 | Intron (ENSMUST00000166877.2/ENSMUSG00000051910.8, intron 1 of 15) | 7 | 115471344 | 116038744 | 567401 | 2 | ENSMUST00000166877.2 | 1251 | Sox6 | ENSMUSG00000051910.8 |
| chr7 | 116038190 | 116038723 | 3.68426 | 4.59382 | 1.50995 | Promoter | 7 | 115471344 | 116038744 | 567401 | 2 | ENSMUST00000166877.2 | 21 | Sox6 | ENSMUSG00000051910.8 |
| chr15 | 43169413 | 43170017 | 3.58714 | 4.3286 | 1.39343 | Promoter | 15 | 43020811 | 43170818 | 150008 | 2 | ENSMUST00000063492.6 | 801 | Rspo2 | ENSMUSG00000051920.6 |
| chr1 | 3671213 | 3671519 | 3.25063 | 3.71281 | 1.04697 | Promoter | 1 | 3214482 | 3671498 | 457017 | 2 | ENSMUST00000070533.4 | 0 | Xkr4 | ENSMUSG00000051951.5 |
| chr16 | 36560467 | 36560754 | 3.62074 | 4.38243 | 1.40876 | Promoter | 16 | 36494467 | 36561606 | 67140 | 2 | ENSMUST00000172826.1 | 852 | Casr | ENSMUSG00000051980.8 |
| chr16 | 36561417 | 36562052 | 3.97042 | 5.32848 | 2.00098 | Promoter | 16 | 36494467 | 36561606 | 67140 | 2 | ENSMUST00000172826.1 | 0 | Casr | ENSMUSG00000051980.8 |
| chr1 | 135963848 | 135964149 | 3.3514 | 3.85627 | 1.0234 | Promoter | 1 | 135953578 | 135963140 | 9563 | 2 | ENSMUST00000140703.2 | -709 | Igfn1 | ENSMUSG00000051985.7 |
| chr2 | 147708333 | 147708679 | 3.65802 | 4.47375 | 1.49327 | Distal Intergenic | 2 | 147710693 | 147717632 | 6940 | 1 | ENSMUST00000152542.1 | -2014 | A530006G24Rik | ENSMUSG00000051986.6 |
| chr2 | 147709301 | 147709554 | 3.54129 | 4.36799 | 1.38903 | Distal Intergenic | 2 | 147710693 | 147717632 | 6940 | 1 | ENSMUST00000152542.1 | -1139 | A530006G24Rik | ENSMUSG00000051986.6 |
| chr18 | 60989451 | 60989645 | 3.26 | 3.72054 | 1.02793 | Distal Intergenic | 18 | 60995381 | 61014199 | 18819 | 2 | ENSMUST00000025520.8 | 24554 | Slc6a7 | ENSMUSG00000052026.7 |
| chr18 | 61013328 | 61014634 | 4.09774 | 5.3718 | 2.05331 | Promoter | 18 | 60995381 | 61014199 | 18819 | 2 | ENSMUST00000025520.8 | 0 | Slc6a7 | ENSMUSG00000052026.7 |
| chr18 | 61014973 | 61015125 | 3.3166 | 3.80213 | 1.01733 | Promoter | 18 | 60995381 | 61014199 | 18819 | 2 | ENSMUST00000025520.8 | -775 | Slc6a7 | ENSMUSG00000052026.7 |
| chr18 | 61020842 | 61021209 | 3.79146 | 4.66198 | 1.51003 | Intron (ENSMUST00000025521.8/ENSMUSG00000024619.8, intron 1 of 2) | 18 | 60995381 | 61014199 | 18819 | 2 | ENSMUST00000025520.8 | -6644 | Slc6a7 | ENSMUSG00000052026.7 |
| chr2 | 170484071 | 170484267 | 3.89029 | 4.98609 | 1.81939 | 3' UTR | 2 | 170496428 | 170519067 | 22640 | 1 | ENSMUST00000075087.2 | -12161 | Pfdn4 | ENSMUSG00000052033.8 |
| chr2 | 170492815 | 170493606 | 3.51008 | 4.1459 | 1.19754 | Intron (ENSMUST00000038824.5/ENSMUSG00000038567.5, intron 4 of 11) | 2 | 170496428 | 170519067 | 22640 | 1 | ENSMUST00000075087.2 | -2822 | Pfdn4 | ENSMUSG00000052033.8 |
| chr2 | 170505316 | 170505908 | 3.52756 | 4.26289 | 1.35215 | Intron (ENSMUST00000075087.2/ENSMUSG00000052033.8, intron 1 of 3) | 2 | 170511235 | 170519123 | 7889 | 1 | ENSMUST00000063682.6 | -5327 | Pfdn4 | ENSMUSG00000052033.8 |
| chr2 | 170506669 | 170506930 | 3.91441 | 4.9758 | 1.78163 | Intron (ENSMUST00000075087.2/ENSMUSG00000052033.8, intron 1 of 3) | 2 | 170511235 | 170519123 | 7889 | 1 | ENSMUST00000063682.6 | -4305 | Pfdn4 | ENSMUSG00000052033.8 |
| chr2 | 170504322 | 170504784 | 3.74909 | 4.66163 | 1.5969 | Exon (ENSMUST00000126086.1/ENSMUSG00000085495.1, exon 2 of 3) | 2 | 170511235 | 170519123 | 7889 | 1 | ENSMUST00000063682.6 | -6451 | Pfdn4 | ENSMUSG00000052033.8 |
| chr13 | 96982987 | 96983318 | 3.31062 | 3.79727 | 1.03176 | Distal Intergenic | 13 | 97021864 | 97034362 | 12499 | 2 | ENSMUST00000091377.4 | 51044 | 1700029F12Rik | ENSMUSG00000052075.6 |
| chr13 | 96984004 | 96984311 | 3.40713 | 4.14964 | 1.22129 | Distal Intergenic | 13 | 97021864 | 97034362 | 12499 | 2 | ENSMUST00000091377.4 | 50051 | 1700029F12Rik | ENSMUSG00000052075.6 |
| chr13 | 96985587 | 96985759 | 3.65545 | 4.44722 | 1.45831 | Distal Intergenic | 13 | 97021864 | 97034362 | 12499 | 2 | ENSMUST00000091377.4 | 48603 | 1700029F12Rik | ENSMUSG00000052075.6 |
| chr13 | 97007293 | 97007658 | 3.83566 | 4.76153 | 1.59598 | Distal Intergenic | 13 | 97021864 | 97034362 | 12499 | 2 | ENSMUST00000091377.4 | 26704 | 1700029F12Rik | ENSMUSG00000052075.6 |
| chr13 | 55379359 | 55380258 | 3.58926 | 4.3281 | 1.34076 | Promoter | 13 | 55379169 | 55384686 | 5518 | 1 | ENSMUST00000135851.1 | 191 | Rgs14 | ENSMUSG00000052087.8 |
| chr13 | 55380789 | 55381346 | 3.50588 | 4.19644 | 1.25639 | Exon (ENSMUST00000063771.8/ENSMUSG00000052087.8, exon 9 of 15) | 13 | 55379169 | 55384686 | 5518 | 1 | ENSMUST00000135851.1 | 1621 | Rgs14 | ENSMUSG00000052087.8 |
| chr4 | 57916395 | 57916563 | 3.54998 | 4.20698 | 1.20304 | Promoter | 4 | 57908483 | 57916297 | 7815 | 2 | ENSMUST00000063816.5 | -99 | D630039A03Rik | ENSMUSG00000052117.5 |
| chr12 | 33146312 | 33146387 | 3.79742 | 4.91958 | 1.69026 | Promoter | 12 | 33125880 | 33146909 | 21030 | 2 | ENSMUST00000151637.2 | 522 | F730043M19Rik | ENSMUSG00000052125.11 |
| chr16 | 35534534 | 35534615 | 3.49746 | 4.16302 | 1.29584 | Distal Intergenic | 16 | 35541145 | 35648140 | 106996 | 1 | ENSMUST00000133554.1 | -6530 | Sema5b | ENSMUSG00000052133.11 |
| chr16 | 35540573 | 35541607 | 3.69975 | 4.56139 | 1.51317 | Promoter | 16 | 35541145 | 35648140 | 106996 | 1 | ENSMUST00000133554.1 | 0 | Sema5b | ENSMUSG00000052133.11 |
| chr16 | 35542004 | 35543146 | 3.89356 | 5.00182 | 1.8147 | Promoter | 16 | 35541571 | 35664261 | 122691 | 1 | ENSMUST00000050625.9 | 434 | Sema5b | ENSMUSG00000052133.11 |
| chr16 | 35543617 | 35543856 | 3.41445 | 4.12922 | 1.23832 | Intron (ENSMUST00000133554.1/ENSMUSG00000052133.11, intron 1 of 5) | 16 | 35541571 | 35664261 | 122691 | 1 | ENSMUST00000050625.9 | 2047 | Sema5b | ENSMUSG00000052133.11 |
| chr16 | 35659600 | 35660662 | 4.24256 | 5.7161 | 2.35633 | Promoter | 16 | 35659435 | 35660280 | 846 | 1 | ENSMUST00000149097.1 | 166 | Sema5b | ENSMUSG00000052133.11 |
| chr16 | 35661064 | 35661536 | 3.395 | 3.95305 | 1.07913 | Promoter | 16 | 35661305 | 35663314 | 2010 | 1 | ENSMUST00000128347.1 | 0 | Sema5b | ENSMUSG00000052133.11 |
| chr16 | 35685100 | 35685175 | 3.52097 | 4.17741 | 1.25569 | Distal Intergenic | 16 | 35662471 | 35663454 | 984 | 1 | ENSMUST00000128966.1 | 22630 | Sema5b | ENSMUSG00000052133.11 |
| chr4 | 120238367 | 120238442 | 3.94268 | 4.95512 | 1.71577 | Distal Intergenic | 4 | 120267079 | 120287349 | 20271 | 2 | ENSMUST00000102656.3 | 48907 | Foxo6 | ENSMUSG00000052135.8 |
| chr4 | 120239678 | 120239840 | 3.35668 | 3.86251 | 1.01733 | Distal Intergenic | 4 | 120267079 | 120287349 | 20271 | 2 | ENSMUST00000102656.3 | 47509 | Foxo6 | ENSMUSG00000052135.8 |
| chr4 | 120241474 | 120241645 | 3.58459 | 4.27926 | 1.26305 | Distal Intergenic | 4 | 120267079 | 120287349 | 20271 | 2 | ENSMUST00000102656.3 | 45704 | Foxo6 | ENSMUSG00000052135.8 |
| chr4 | 120242278 | 120242526 | 3.55141 | 4.26352 | 1.31062 | Distal Intergenic | 4 | 120267079 | 120287349 | 20271 | 2 | ENSMUST00000102656.3 | 44823 | Foxo6 | ENSMUSG00000052135.8 |
| chr4 | 120251719 | 120251826 | 3.45973 | 4.14338 | 1.25472 | Distal Intergenic | 4 | 120267079 | 120287349 | 20271 | 2 | ENSMUST00000102656.3 | 35523 | Foxo6 | ENSMUSG00000052135.8 |
| chr4 | 120260027 | 120260232 | 3.47726 | 4.09709 | 1.20107 | Distal Intergenic | 4 | 120267079 | 120287349 | 20271 | 2 | ENSMUST00000102656.3 | 27117 | Foxo6 | ENSMUSG00000052135.8 |
| chr4 | 120262427 | 120262509 | 3.84119 | 4.86742 | 1.70761 | Distal Intergenic | 4 | 120267079 | 120287349 | 20271 | 2 | ENSMUST00000102656.3 | 24840 | Foxo6 | ENSMUSG00000052135.8 |
| chr4 | 120264596 | 120264843 | 3.80993 | 4.69107 | 1.51227 | Downstream (2-3kb) | 4 | 120267079 | 120287349 | 20271 | 2 | ENSMUST00000102656.3 | 22506 | Foxo6 | ENSMUSG00000052135.8 |
| chr4 | 120265185 | 120265260 | 3.43349 | 4.34487 | 1.34432 | Downstream (1-2kb) | 4 | 120267079 | 120287349 | 20271 | 2 | ENSMUST00000102656.3 | 22089 | Foxo6 | ENSMUSG00000052135.8 |
| chr4 | 120265588 | 120266089 | 4.03207 | 5.13605 | 1.83206 | Downstream (<1kb) | 4 | 120267079 | 120287349 | 20271 | 2 | ENSMUST00000102656.3 | 21260 | Foxo6 | ENSMUSG00000052135.8 |
| chr4 | 120267014 | 120268092 | 3.83199 | 5.15691 | 1.90521 | 3' UTR | 4 | 120267079 | 120287349 | 20271 | 2 | ENSMUST00000102656.3 | 19257 | Foxo6 | ENSMUSG00000052135.8 |
| chr4 | 120268522 | 120269126 | 3.83768 | 4.91823 | 1.74154 | Exon (ENSMUST00000102656.3/ENSMUSG00000052135.8, exon 2 of 2) | 4 | 120267079 | 120287349 | 20271 | 2 | ENSMUST00000102656.3 | 18223 | Foxo6 | ENSMUSG00000052135.8 |
| chr4 | 120269662 | 120270002 | 3.44744 | 4.2756 | 1.31063 | Intron (ENSMUST00000102656.3/ENSMUSG00000052135.8, intron 1 of 1) | 4 | 120267079 | 120287349 | 20271 | 2 | ENSMUST00000102656.3 | 17347 | Foxo6 | ENSMUSG00000052135.8 |
| chr4 | 120273686 | 120273784 | 3.62913 | 4.42106 | 1.42708 | Intron (ENSMUST00000102656.3/ENSMUSG00000052135.8, intron 1 of 1) | 4 | 120267079 | 120287349 | 20271 | 2 | ENSMUST00000102656.3 | 13565 | Foxo6 | ENSMUSG00000052135.8 |
| chr4 | 120286692 | 120287180 | 3.59705 | 4.29838 | 1.26688 | Promoter | 4 | 120267079 | 120287349 | 20271 | 2 | ENSMUST00000102656.3 | 169 | Foxo6 | ENSMUSG00000052135.8 |
| chr17 | 32395074 | 32395677 | 3.66732 | 4.43695 | 1.38416 | Promoter | 17 | 32393923 | 32395371 | 1449 | 2 | ENSMUST00000143808.1 | 0 | Rasal3 | ENSMUSG00000052142.10 |
| chr17 | 32396320 | 32397020 | 3.72484 | 4.62759 | 1.53396 | Promoter | 17 | 32395567 | 32396709 | 1143 | 2 | ENSMUST00000135968.1 | 0 | Rasal3 | ENSMUSG00000052142.10 |
| chr17 | 32397343 | 32398183 | 3.35832 | 3.87665 | 1.0584 | Promoter | 17 | 32395567 | 32396709 | 1143 | 2 | ENSMUST00000135968.1 | -635 | Rasal3 | ENSMUSG00000052142.10 |
| chr9 | 60918651 | 60918848 | 3.34153 | 3.83966 | 1.01733 | Distal Intergenic | 9 | 60821855 | 60838200 | 16346 | 2 | ENSMUST00000063858.7 | -80452 | Gm9869 | ENSMUSG00000052143.7 |
| chr9 | 60938638 | 60938746 | 3.6209 | 4.47879 | 1.43891 | Distal Intergenic | 9 | 60821855 | 60838200 | 16346 | 2 | ENSMUST00000063858.7 | -100439 | Gm9869 | ENSMUSG00000052143.7 |
| chr9 | 60940073 | 60940554 | 4.37896 | 6.19361 | 2.71633 | Distal Intergenic | 9 | 60821855 | 60838200 | 16346 | 2 | ENSMUST00000063858.7 | -101874 | Gm9869 | ENSMUSG00000052143.7 |
| chr9 | 60942140 | 60942599 | 3.86612 | 4.97268 | 1.7809 | Distal Intergenic | 9 | 60821855 | 60838200 | 16346 | 2 | ENSMUST00000063858.7 | -103941 | Gm9869 | ENSMUSG00000052143.7 |
| chr9 | 60945004 | 60945776 | 3.40193 | 3.96373 | 1.13065 | Distal Intergenic | 9 | 60821855 | 60838200 | 16346 | 2 | ENSMUST00000063858.7 | -106805 | Gm9869 | ENSMUSG00000052143.7 |
| chr9 | 60946607 | 60946956 | 3.65265 | 4.44815 | 1.45136 | Distal Intergenic | 9 | 60821855 | 60838200 | 16346 | 2 | ENSMUST00000063858.7 | -108408 | Gm9869 | ENSMUSG00000052143.7 |
| chr9 | 60947307 | 60947453 | 3.73319 | 4.61709 | 1.5572 | Distal Intergenic | 9 | 60821855 | 60838200 | 16346 | 2 | ENSMUST00000063858.7 | -109108 | Gm9869 | ENSMUSG00000052143.7 |
| chr9 | 60949954 | 60950454 | 3.52869 | 4.18146 | 1.22754 | Distal Intergenic | 9 | 60821855 | 60838200 | 16346 | 2 | ENSMUST00000063858.7 | -111755 | Gm9869 | ENSMUSG00000052143.7 |
| chr9 | 60968907 | 60969086 | 3.42456 | 3.97832 | 1.06444 | Distal Intergenic | 9 | 60821855 | 60838200 | 16346 | 2 | ENSMUST00000063858.7 | -130708 | Gm9869 | ENSMUSG00000052143.7 |
| chr9 | 61016913 | 61017032 | 3.31866 | 3.81586 | 1.05471 | Distal Intergenic | 9 | 60821855 | 60838200 | 16346 | 2 | ENSMUST00000063858.7 | -178714 | Gm9869 | ENSMUSG00000052143.7 |
| chr9 | 61045872 | 61045947 | 4.49135 | 6.02703 | 2.44366 | Distal Intergenic | 9 | 60821855 | 60838200 | 16346 | 2 | ENSMUST00000063858.7 | -207673 | Gm9869 | ENSMUSG00000052143.7 |
| chr9 | 61048705 | 61048981 | 3.5315 | 4.18596 | 1.22195 | Distal Intergenic | 9 | 60821855 | 60838200 | 16346 | 2 | ENSMUST00000063858.7 | -210506 | Gm9869 | ENSMUSG00000052143.7 |
| chr9 | 61054538 | 61054613 | 3.40518 | 3.95579 | 1.08852 | Distal Intergenic | 9 | 60821855 | 60838200 | 16346 | 2 | ENSMUST00000063858.7 | -216339 | Gm9869 | ENSMUSG00000052143.7 |
| chr9 | 61055732 | 61055807 | 3.34638 | 3.84697 | 1.01733 | Distal Intergenic | 9 | 60821855 | 60838200 | 16346 | 2 | ENSMUST00000063858.7 | -217533 | Gm9869 | ENSMUSG00000052143.7 |
| chr9 | 61057554 | 61057709 | 3.6415 | 4.41865 | 1.42462 | Distal Intergenic | 9 | 60821855 | 60838200 | 16346 | 2 | ENSMUST00000063858.7 | -219355 | Gm9869 | ENSMUSG00000052143.7 |
| chr9 | 61058124 | 61058424 | 3.33568 | 3.83089 | 1.01733 | Distal Intergenic | 9 | 60821855 | 60838200 | 16346 | 2 | ENSMUST00000063858.7 | -219925 | Gm9869 | ENSMUSG00000052143.7 |
| chr9 | 61059007 | 61059721 | 3.60003 | 4.31646 | 1.30114 | Distal Intergenic | 9 | 60821855 | 60838200 | 16346 | 2 | ENSMUST00000063858.7 | -220808 | Gm9869 | ENSMUSG00000052143.7 |
| chr9 | 61069235 | 61069596 | 3.44402 | 4.02873 | 1.13868 | Distal Intergenic | 9 | 60821855 | 60838200 | 16346 | 2 | ENSMUST00000063858.7 | -231036 | Gm9869 | ENSMUSG00000052143.7 |
| chr9 | 61070192 | 61070600 | 3.50993 | 4.14416 | 1.19385 | Distal Intergenic | 9 | 60821855 | 60838200 | 16346 | 2 | ENSMUST00000063858.7 | -231993 | Gm9869 | ENSMUSG00000052143.7 |
| chr9 | 61074422 | 61074519 | 3.3668 | 3.8808 | 1.02801 | Distal Intergenic | 9 | 60821855 | 60838200 | 16346 | 2 | ENSMUST00000063858.7 | -236223 | Gm9869 | ENSMUSG00000052143.7 |
| chr9 | 61081478 | 61081716 | 3.71427 | 4.57278 | 1.54474 | Distal Intergenic | 9 | 60821855 | 60838200 | 16346 | 2 | ENSMUST00000063858.7 | -243279 | Gm9869 | ENSMUSG00000052143.7 |
| chr9 | 61082532 | 61082651 | 3.84239 | 4.78188 | 1.618 | Distal Intergenic | 9 | 60821855 | 60838200 | 16346 | 2 | ENSMUST00000063858.7 | -244333 | Gm9869 | ENSMUSG00000052143.7 |
| chr9 | 61083545 | 61083794 | 3.74428 | 4.56123 | 1.41778 | Distal Intergenic | 9 | 60821855 | 60838200 | 16346 | 2 | ENSMUST00000063858.7 | -245346 | Gm9869 | ENSMUSG00000052143.7 |
| chr9 | 61084200 | 61084330 | 3.59187 | 4.28802 | 1.2665 | Distal Intergenic | 9 | 60821855 | 60838200 | 16346 | 2 | ENSMUST00000063858.7 | -246001 | Gm9869 | ENSMUSG00000052143.7 |
| chr9 | 61084873 | 61085201 | 3.43554 | 4.01199 | 1.12519 | Distal Intergenic | 9 | 60821855 | 60838200 | 16346 | 2 | ENSMUST00000063858.7 | -246674 | Gm9869 | ENSMUSG00000052143.7 |
| chr9 | 61085853 | 61086275 | 3.89351 | 4.87893 | 1.70603 | Distal Intergenic | 9 | 60821855 | 60838200 | 16346 | 2 | ENSMUST00000063858.7 | -247654 | Gm9869 | ENSMUSG00000052143.7 |
| chr6 | 100866335 | 100866452 | 3.52942 | 5.00004 | 1.77265 | Exon (ENSMUST00000063854.4/ENSMUSG00000052144.6, exon 9 of 9) | 6 | 100833638 | 100868717 | 35080 | 1 | ENSMUST00000063854.4 | 32698 | Ppp4r2 | ENSMUSG00000052144.6 |
| chr6 | 100880821 | 100880982 | 3.85972 | 4.91519 | 1.73364 | Distal Intergenic | 6 | 100833638 | 100868717 | 35080 | 1 | ENSMUST00000063854.4 | 47184 | Ppp4r2 | ENSMUSG00000052144.6 |
| chr6 | 100938941 | 100939027 | 3.7054 | 4.55949 | 1.54637 | Distal Intergenic | 6 | 100833638 | 100868717 | 35080 | 1 | ENSMUST00000063854.4 | 105304 | Ppp4r2 | ENSMUSG00000052144.6 |
| chr12 | 112750975 | 112751142 | 3.56621 | 4.24176 | 1.2398 | Distal Intergenic | 12 | 112760655 | 112768986 | 8332 | 1 | ENSMUST00000063888.3 | -9513 | Pld4 | ENSMUSG00000052160.6 |
| chr19 | 6530246 | 6530715 | 3.56477 | 4.26992 | 1.33673 | Exon (ENSMUST00000149574.1/ENSMUSG00000052188.6, exon 2 of 2) | 19 | 6530789 | 6531788 | 1000 | 2 | ENSMUST00000137184.1 | 1073 | Gm14964 | ENSMUSG00000052188.6 |
| chr19 | 6531186 | 6532383 | 4.69248 | 6.58531 | 3.01377 | Promoter | 19 | 6530711 | 6532011 | 1301 | 2 | ENSMUST00000149574.1 | 0 | Gm14964 | ENSMUSG00000052188.6 |
| chr16 | 35567454 | 35568137 | 3.52401 | 4.18341 | 1.26051 | Intron (ENSMUST00000133554.1/ENSMUSG00000052133.11, intron 1 of 5) | 16 | 35573321 | 35573556 | 236 | 2 | ENSMUST00000143451.1 | 5419 | Gm5963 | ENSMUSG00000052192.5 |
| chr16 | 35587627 | 35587839 | 3.36665 | 3.8855 | 1.04543 | Intron (ENSMUST00000133554.1/ENSMUSG00000052133.11, intron 1 of 5) | 16 | 35573321 | 35573556 | 236 | 2 | ENSMUST00000143451.1 | -14072 | Gm5963 | ENSMUSG00000052192.5 |
| chr16 | 35570432 | 35570599 | 3.41196 | 3.98401 | 1.15148 | Intron (ENSMUST00000133554.1/ENSMUSG00000052133.11, intron 1 of 5) | 16 | 35573321 | 35573556 | 236 | 2 | ENSMUST00000143451.1 | 2957 | Gm5963 | ENSMUSG00000052192.5 |
| chr16 | 35570978 | 35571233 | 3.57497 | 4.29703 | 1.36211 | Intron (ENSMUST00000133554.1/ENSMUSG00000052133.11, intron 1 of 5) | 16 | 35573321 | 35573556 | 236 | 2 | ENSMUST00000143451.1 | 2323 | Gm5963 | ENSMUSG00000052192.5 |
| chr16 | 35571722 | 35571931 | 3.82116 | 4.74957 | 1.63443 | Intron (ENSMUST00000133554.1/ENSMUSG00000052133.11, intron 1 of 5) | 16 | 35573321 | 35573556 | 236 | 2 | ENSMUST00000143451.1 | 1625 | Gm5963 | ENSMUSG00000052192.5 |
| chr12 | 76417383 | 76417853 | 3.49767 | 4.11386 | 1.15365 | Promoter | 12 | 76417599 | 76439473 | 21875 | 1 | ENSMUST00000063977.7 | 0 | Ppp1r36 | ENSMUSG00000052221.7 |
| chr12 | 76418273 | 76418748 | 3.83029 | 4.93648 | 1.75337 | Promoter | 12 | 76417599 | 76439473 | 21875 | 1 | ENSMUST00000063977.7 | 675 | Ppp1r36 | ENSMUSG00000052221.7 |
| chr17 | 17886189 | 17886461 | 3.43057 | 4.18166 | 1.25787 | Distal Intergenic | 17 | 17887824 | 17893952 | 6129 | 1 | ENSMUST00000149944.1 | -1363 | Fpr2 | ENSMUSG00000052270.7 |
| chr7 | 126847780 | 126849050 | 3.67149 | 4.70445 | 1.6109 | Promoter | 7 | 126847927 | 126848836 | 910 | 1 | ENSMUST00000147257.1 | 0 | Doc2a | ENSMUSG00000052301.9 |
| chr7 | 126851045 | 126851240 | 3.55035 | 4.30086 | 1.35732 | Exon (ENSMUST00000064110.8/ENSMUSG00000052301.9, exon 6 of 11) | 7 | 126848797 | 126852036 | 3240 | 1 | ENSMUST00000176771.1 | 2249 | Doc2a | ENSMUSG00000052301.9 |
| chr10 | 121243203 | 121243636 | 3.72381 | 4.59371 | 1.52914 | Distal Intergenic | 10 | 121263824 | 121311189 | 47366 | 2 | ENSMUST00000064107.5 | 67553 | Tbc1d30 | ENSMUSG00000052302.5 |
| chr10 | 121266957 | 121267462 | 3.52169 | 4.20332 | 1.29952 | Exon (ENSMUST00000064107.5/ENSMUSG00000052302.5, exon 12 of 12) | 10 | 121263824 | 121311189 | 47366 | 2 | ENSMUST00000064107.5 | 43727 | Tbc1d30 | ENSMUSG00000052302.5 |
| chr10 | 121285521 | 121285748 | 3.45284 | 4.03863 | 1.12685 | Intron (ENSMUST00000064107.5/ENSMUSG00000052302.5, intron 8 of 11) | 10 | 121263824 | 121311189 | 47366 | 2 | ENSMUST00000064107.5 | 25441 | Tbc1d30 | ENSMUSG00000052302.5 |
| chr10 | 121295204 | 121295285 | 4.25819 | 5.65255 | 2.26303 | Intron (ENSMUST00000064107.5/ENSMUSG00000052302.5, intron 6 of 11) | 10 | 121263824 | 121311189 | 47366 | 2 | ENSMUST00000064107.5 | 15904 | Tbc1d30 | ENSMUSG00000052302.5 |
| chr10 | 121296524 | 121296766 | 3.48407 | 4.09831 | 1.17158 | Exon (ENSMUST00000064107.5/ENSMUSG00000052302.5, exon 6 of 12) | 10 | 121263824 | 121311189 | 47366 | 2 | ENSMUST00000064107.5 | 14423 | Tbc1d30 | ENSMUSG00000052302.5 |
| chr10 | 121299937 | 121300225 | 3.42496 | 4.10468 | 1.20112 | Intron (ENSMUST00000064107.5/ENSMUSG00000052302.5, intron 5 of 11) | 10 | 121263824 | 121311189 | 47366 | 2 | ENSMUST00000064107.5 | 10964 | Tbc1d30 | ENSMUSG00000052302.5 |
| chr10 | 121305597 | 121305870 | 3.36205 | 4.03597 | 1.17358 | Intron (ENSMUST00000064107.5/ENSMUSG00000052302.5, intron 3 of 11) | 10 | 121263824 | 121311189 | 47366 | 2 | ENSMUST00000064107.5 | 5319 | Tbc1d30 | ENSMUSG00000052302.5 |
| chr10 | 121306420 | 121307303 | 3.55752 | 4.27614 | 1.36321 | Exon (ENSMUST00000064107.5/ENSMUSG00000052302.5, exon 2 of 12) | 10 | 121263824 | 121311189 | 47366 | 2 | ENSMUST00000064107.5 | 3886 | Tbc1d30 | ENSMUSG00000052302.5 |
| chr10 | 121308359 | 121308735 | 3.4696 | 4.11826 | 1.27944 | Intron (ENSMUST00000064107.5/ENSMUSG00000052302.5, intron 1 of 11) | 10 | 121263824 | 121311189 | 47366 | 2 | ENSMUST00000064107.5 | 2454 | Tbc1d30 | ENSMUSG00000052302.5 |
| chr10 | 121309861 | 121311308 | 3.95297 | 5.20526 | 1.95232 | Promoter | 10 | 121263824 | 121311189 | 47366 | 2 | ENSMUST00000064107.5 | 0 | Tbc1d30 | ENSMUSG00000052302.5 |
| chr10 | 121312199 | 121312438 | 3.87979 | 5.15783 | 1.87172 | Distal Intergenic | 10 | 121263824 | 121311189 | 47366 | 2 | ENSMUST00000064107.5 | -1011 | Tbc1d30 | ENSMUSG00000052302.5 |
| chr16 | 30281248 | 30281396 | 3.3097 | 3.79176 | 1.01733 | Intron (ENSMUST00000064606.7/ENSMUSG00000052316.9, intron 1 of 1) | 16 | 30269302 | 30283256 | 13955 | 2 | ENSMUST00000064606.7 | 1860 | Lrrc15 | ENSMUSG00000052316.9 |
| chr16 | 30282753 | 30283460 | 4.01512 | 5.19257 | 1.98513 | Promoter | 16 | 30269302 | 30283256 | 13955 | 2 | ENSMUST00000064606.7 | 0 | Lrrc15 | ENSMUSG00000052316.9 |
| chr16 | 30283910 | 30284007 | 4.77499 | 6.68655 | 2.99436 | Promoter | 16 | 30269302 | 30283256 | 13955 | 2 | ENSMUST00000064606.7 | -655 | Lrrc15 | ENSMUSG00000052316.9 |
| chr16 | 30284432 | 30284878 | 4.0162 | 5.48778 | 2.12081 | Distal Intergenic | 16 | 30269302 | 30283256 | 13955 | 2 | ENSMUST00000064606.7 | -1177 | Lrrc15 | ENSMUSG00000052316.9 |
| chr7 | 84084936 | 84085650 | 3.84177 | 4.99526 | 1.7752 | Promoter | 7 | 83932857 | 84086502 | 153646 | 2 | ENSMUST00000064174.6 | 852 | Cemip | ENSMUSG00000052353.8 |
| chr16 | 32503604 | 32504020 | 3.62517 | 4.35432 | 1.31558 | Intron (ENSMUST00000064192.7/ENSMUSG00000052363.7, intron 6 of 8) | 16 | 32499611 | 32561966 | 62356 | 1 | ENSMUST00000160832.1 | 3994 | Zdhhc19 | ENSMUSG00000052363.7 |
| chr13 | 12339743 | 12340827 | 3.55778 | 4.25161 | 1.30714 | Promoter | 13 | 12269426 | 12340760 | 71335 | 2 | ENSMUST00000064204.8 | 0 | Actn2 | ENSMUSG00000052374.9 |
| chr7 | 99237427 | 99237584 | 3.72345 | 4.53744 | 1.43344 | Intron (ENSMUST00000064231.7/ENSMUSG00000052396.7, intron 1 of 5) | 7 | 99226577 | 99238594 | 12018 | 2 | ENSMUST00000132343.1 | 1010 | Mogat2 | ENSMUSG00000052396.7 |
| chr7 | 99238087 | 99238731 | 3.66777 | 4.43621 | 1.35247 | Promoter | 7 | 99219084 | 99238619 | 19536 | 2 | ENSMUST00000064231.7 | 0 | Mogat2 | ENSMUSG00000052396.7 |
| chr7 | 99249126 | 99249343 | 4.21513 | 5.67561 | 2.24467 | Distal Intergenic | 7 | 99219084 | 99238619 | 19536 | 2 | ENSMUST00000064231.7 | -10508 | Mogat2 | ENSMUSG00000052396.7 |
| chr7 | 99250351 | 99250739 | 3.49741 | 4.3998 | 1.40847 | Distal Intergenic | 7 | 99219084 | 99238619 | 19536 | 2 | ENSMUST00000064231.7 | -11733 | Mogat2 | ENSMUSG00000052396.7 |
| chr2 | 25616036 | 25616643 | 3.94508 | 5.42952 | 2.10614 | Promoter | 2 | 25616007 | 25631758 | 15752 | 1 | ENSMUST00000127642.2 | 30 | Fcnaos | ENSMUSG00000052403.6 |
| chr3 | 93444363 | 93444530 | 3.46497 | 4.31515 | 1.32784 | Exon (ENSMUST00000064257.5/ENSMUSG00000052415.5, exon 2 of 2) | 3 | 93442330 | 93449077 | 6748 | 1 | ENSMUST00000064257.5 | 2034 | Tchh | ENSMUSG00000052415.5 |
| chr16 | 91156554 | 91157108 | 3.35334 | 3.90117 | 1.07175 | Distal Intergenic | 16 | 91168197 | 91171401 | 3205 | 2 | ENSMUST00000068783.2 | 14293 | Gm9881 | ENSMUSG00000052471.2 |
| chr16 | 91162957 | 91163136 | 3.52514 | 4.19819 | 1.28524 | Distal Intergenic | 16 | 91168197 | 91171401 | 3205 | 2 | ENSMUST00000068783.2 | 8265 | Gm9881 | ENSMUSG00000052471.2 |
| chr16 | 91167199 | 91167419 | 4.10409 | 5.28265 | 1.95903 | Downstream (<1kb) | 16 | 91168197 | 91171401 | 3205 | 2 | ENSMUST00000068783.2 | 3982 | Gm9881 | ENSMUSG00000052471.2 |
| chr16 | 91167897 | 91167992 | 3.81903 | 4.73084 | 1.57579 | Downstream (<1kb) | 16 | 91168197 | 91171401 | 3205 | 2 | ENSMUST00000068783.2 | 3409 | Gm9881 | ENSMUSG00000052471.2 |
| chr16 | 91169343 | 91170143 | 3.74346 | 4.68453 | 1.56162 | 3' UTR | 16 | 91168197 | 91171401 | 3205 | 2 | ENSMUST00000068783.2 | 1258 | Gm9881 | ENSMUSG00000052471.2 |
| chr16 | 91170480 | 91170687 | 4.03282 | 5.12516 | 1.82442 | Promoter | 16 | 91168197 | 91171401 | 3205 | 2 | ENSMUST00000068783.2 | 714 | Gm9881 | ENSMUSG00000052471.2 |
| chr16 | 91172101 | 91172336 | 3.84287 | 4.94635 | 1.74742 | Promoter | 16 | 91168197 | 91171401 | 3205 | 2 | ENSMUST00000068783.2 | -701 | Gm9881 | ENSMUSG00000052471.2 |
| chr16 | 91173855 | 91174242 | 3.43942 | 4.02806 | 1.15851 | Distal Intergenic | 16 | 91168197 | 91171401 | 3205 | 2 | ENSMUST00000068783.2 | -2455 | Gm9881 | ENSMUSG00000052471.2 |
| chr16 | 91178194 | 91178375 | 3.30923 | 3.93599 | 1.08125 | Distal Intergenic | 16 | 91168197 | 91171401 | 3205 | 2 | ENSMUST00000068783.2 | -6794 | Gm9881 | ENSMUSG00000052471.2 |
| chr16 | 91182858 | 91183142 | 3.42467 | 3.98249 | 1.07851 | Distal Intergenic | 16 | 91168197 | 91171401 | 3205 | 2 | ENSMUST00000068783.2 | -11458 | Gm9881 | ENSMUSG00000052471.2 |
| chr16 | 91193542 | 91193771 | 3.75582 | 4.67127 | 1.59549 | Distal Intergenic | 16 | 91168197 | 91171401 | 3205 | 2 | ENSMUST00000068783.2 | -22142 | Gm9881 | ENSMUSG00000052471.2 |

| | | | | | | | | | | | | | | | |
|---|---|---|---|---|---|---|---|---|---|---|---|---|---|---|---|
| chr8 | 99416628 | 99416887 | 3.57186 | 4.2712 | 1.30955 | Promoter | 8 | 99416642 | 99424644 | 8003 | 1 | ENSMUST00000064349.5 | 0 | A330008L17Rik | ENSMUSG00000052479.5 |
| chr13 | 98694719 | 98694868 | 3.33379 | 3.82803 | 1.01733 | Promoter | 13 | 98686235 | 98694834 | 8600 | 2 | ENSMUST00000064347.7 | 0 | Tmem171 | ENSMUSG00000052485.7 |
| chr15 | 85820395 | 85820657 | 3.33922 | 3.8362 | 1.01733 | Exon (ENSMUST00000064370.4/ENSMUSG00000052496.4, exon 1 of 1) | 15 | 85814676 | 85821733 | 7058 | 2 | ENSMUST00000064370.4 | 1076 | Pkdrej | ENSMUSG00000052496.4 |
| chr15 | 85821079 | 85821366 | 3.46058 | 4.12186 | 1.22945 | Promoter | 15 | 85814676 | 85821733 | 7058 | 2 | ENSMUST00000064370.4 | 367 | Pkdrej | ENSMUSG00000052496.4 |
| chr15 | 85821720 | 85822037 | 3.4416 | 4.02052 | 1.12294 | Promoter | 15 | 85814676 | 85821733 | 7058 | 2 | ENSMUST00000064370.4 | 0 | Pkdrej | ENSMUSG00000052496.4 |
| chr15 | 85822349 | 85822424 | 3.65949 | 4.41436 | 1.33993 | Promoter | 15 | 85814676 | 85821733 | 7058 | 2 | ENSMUST00000064370.4 | -617 | Pkdrej | ENSMUSG00000052496.4 |
| chr16 | 63857150 | 63857666 | 3.44179 | 4.07296 | 1.25434 | Intron (ENSMUST00000064405.6/ENSMUSG00000052504.6, intron 1 of 16) | 16 | 63543538 | 63864158 | 320621 | 2 | ENSMUST00000064405.6 | 6492 | Epha3 | ENSMUSG00000052504.6 |
| chr16 | 63862258 | 63864193 | 3.81835 | 4.9633 | 1.77574 | Promoter | 16 | 63543538 | 63864158 | 320621 | 2 | ENSMUST00000064405.6 | 0 | Epha3 | ENSMUSG00000052504.6 |
| chr7 | 140538836 | 140539122 | 3.24988 | 3.70196 | 1.01733 | Distal Intergenic | 7 | 140500856 | 140507318 | 6463 | 2 | ENSMUST00000064392.6 | -31519 | Olfr536 | ENSMUSG00000052508.8 |
| chr7 | 48958406 | 48959748 | 3.36601 | 4.09292 | 1.20096 | Promoter | 7 | 48959073 | 48967836 | 8764 | 1 | ENSMUST00000183531.1 | 0 | Nav2 | ENSMUSG00000052512.10 |
| chr7 | 48960559 | 48960729 | 4.09111 | 5.51849 | 2.14831 | Intron (ENSMUST00000184336.1/ENSMUSG00000052512.10, intron 4 of 4) | 7 | 48959097 | 49606821 | 647725 | 1 | ENSMUST00000183659.2 | 1463 | Nav2 | ENSMUSG00000052512.10 |
| chr7 | 48962161 | 48962573 | 3.45595 | 4.0434 | 1.12442 | Intron (ENSMUST00000184336.1/ENSMUSG00000052512.10, intron 4 of 4) | 7 | 48959097 | 49606821 | 647725 | 1 | ENSMUST00000183659.2 | 3065 | Nav2 | ENSMUSG00000052512.10 |
| chr7 | 48963202 | 48963932 | 3.65098 | 4.40246 | 1.33691 | 5' UTR | 7 | 48959097 | 49606821 | 647725 | 1 | ENSMUST00000183659.2 | 4106 | Nav2 | ENSMUSG00000052512.10 |
| chr7 | 48966325 | 48966820 | 3.79927 | 4.67549 | 1.50727 | Exon (ENSMUST00000183531.1/ENSMUSG00000052512.10, exon 2 of 2) | 7 | 48959097 | 49606821 | 647725 | 1 | ENSMUST00000183659.2 | 7229 | Nav2 | ENSMUSG00000052512.10 |
| chr7 | 48968309 | 48968601 | 3.71299 | 4.51582 | 1.41621 | Intron (ENSMUST00000183659.2/ENSMUSG00000052512.10, intron 2 of 39) | 7 | 48959097 | 49606821 | 647725 | 1 | ENSMUST00000183659.2 | 9213 | Nav2 | ENSMUSG00000052512.10 |
| chr7 | 49120280 | 49120874 | 3.41922 | 4.00108 | 1.17212 | Intron (ENSMUST00000183659.2/ENSMUSG00000052512.10, intron 2 of 39) | 7 | 49246189 | 49409643 | 163455 | 1 | ENSMUST00000184809.2 | -125315 | Nav2 | ENSMUSG00000052512.10 |
| chr7 | 49304337 | 49304412 | 4.09797 | 5.4374 | 2.08357 | Promoter | 7 | 49304254 | 49305917 | 1664 | 1 | ENSMUST00000184462.1 | 84 | Nav2 | ENSMUSG00000052512.10 |
| chr13 | 8202850 | 8203954 | 3.68882 | 4.53456 | 1.45941 | Promoter | 13 | 8202866 | 8761530 | 558665 | 1 | ENSMUST00000064473.7 | 0 | Adarb2 | ENSMUSG00000052551.10 |
| chr7 | 91090930 | 91091169 | 4.13597 | 5.67266 | 2.25605 | Promoter | 7 | 91090728 | 92447003 | 1356276 | 1 | ENSMUST00000074273.4 | 203 | Dlg2 | ENSMUSG00000052572.10 |
| chr7 | 92179982 | 92180222 | 3.58419 | 4.28318 | 1.28297 | Promoter | 7 | 92180684 | 92288339 | 107656 | 1 | ENSMUST00000129818.1 | -462 | Dlg2 | ENSMUSG00000052572.10 |
| chr7 | 92181309 | 92181731 | 4.54376 | 6.20948 | 2.64639 | Promoter | 7 | 92180684 | 92288339 | 107656 | 1 | ENSMUST00000129818.1 | 626 | Dlg2 | ENSMUSG00000052572.10 |
| chr10 | 73096243 | 73096351 | 3.43845 | 4.07844 | 1.17015 | Distal Intergenic | 10 | 73099342 | 74317264 | 1217923 | 1 | ENSMUST00000123398.3 | -2991 | Pcdh15 | ENSMUSG00000052613.11 |
| chr10 | 73096673 | 73096980 | 3.84879 | 4.8036 | 1.65895 | Distal Intergenic | 10 | 73099342 | 74317264 | 1217923 | 1 | ENSMUST00000123398.3 | -2362 | Pcdh15 | ENSMUSG00000052613.11 |
| chr10 | 73097825 | 73098415 | 3.55711 | 4.24074 | 1.25321 | Promoter | 10 | 73099342 | 74317264 | 1217923 | 1 | ENSMUST00000123398.3 | -927 | Pcdh15 | ENSMUSG00000052613.11 |
| chr10 | 73099177 | 73099350 | 4.67295 | 6.70457 | 2.98083 | Promoter | 10 | 73099342 | 74317264 | 1217923 | 1 | ENSMUST00000123398.3 | 0 | Pcdh15 | ENSMUSG00000052613.11 |
| chr10 | 73099823 | 73100252 | 3.61819 | 4.49736 | 1.4547 | Promoter | 10 | 73099453 | 73484120 | 384668 | 1 | ENSMUST00000156999.1 | 371 | Pcdh15 | ENSMUSG00000052613.11 |
| chr9 | 120675681 | 120675915 | 3.59937 | 4.32213 | 1.33841 | Distal Intergenic | 9 | 120577346 | 120578072 | 727 | 1 | ENSMUST00000064646.4 | 98336 | 5830454E08Rik | ENSMUSG00000052658.4 |
| chr9 | 120677352 | 120677607 | 3.58815 | 4.32847 | 1.36817 | Distal Intergenic | 9 | 120577346 | 120578072 | 727 | 1 | ENSMUST00000064646.4 | 100007 | 5830454E08Rik | ENSMUSG00000052658.4 |
| chr9 | 67629508 | 67629840 | 3.30344 | 3.78236 | 1.01733 | Distal Intergenic | 9 | 67217087 | 67559703 | 342617 | 2 | ENSMUST00000040025.8 | -69806 | Tln2 | ENSMUSG00000052698.9 |
| chr9 | 67631539 | 67631715 | 3.55087 | 4.2522 | 1.3431 | Distal Intergenic | 9 | 67217087 | 67559703 | 342617 | 2 | ENSMUST00000040025.8 | -71837 | Tln2 | ENSMUSG00000052698.9 |
| chr1 | 140247423 | 140247677 | 3.4983 | 4.11687 | 1.16077 | Intron (ENSMUST00000119786.2/ENSMUSG00000052726.10, intron 1 of 26) | 1 | 140246226 | 140612063 | 365838 | 1 | ENSMUST00000120709.2 | 1198 | Kcnt2 | ENSMUSG00000052726.10 |
| chr1 | 136259716 | 136261122 | 3.74832 | 4.66644 | 1.57242 | Promoter | 1 | 136258914 | 136260873 | 1960 | 2 | ENSMUST00000086395.5 | 0 | Gpr25 | ENSMUSG00000052759.6 |
| chr1 | 136261430 | 136261614 | 3.93336 | 4.93695 | 1.70376 | Promoter | 1 | 136258914 | 136260873 | 1960 | 2 | ENSMUST00000086395.5 | -558 | Gpr25 | ENSMUSG00000052759.6 |
| chr1 | 136261952 | 136262288 | 3.53234 | 4.47667 | 1.41855 | Intron (ENSMUST00000181385.1/ENSMUSG00000097433.1, intron 1 of 1) | 1 | 136258914 | 136260873 | 1960 | 2 | ENSMUST00000086395.5 | -1080 | Gpr25 | ENSMUSG00000052759.6 |
| chr1 | 136264337 | 136264630 | 3.49562 | 4.23665 | 1.3409 | Intron (ENSMUST00000181385.1/ENSMUSG00000097433.1, intron 1 of 1) | 1 | 136258914 | 136260873 | 1960 | 2 | ENSMUST00000086395.5 | -3465 | Gpr25 | ENSMUSG00000052759.6 |
| chr7 | 81988611 | 81989012 | 3.93892 | 4.96346 | 1.74396 | Intron (ENSMUST00000026096.7/ENSMUSG00000025105.8, intron 1 of 4) | 7 | 81992002 | 81992618 | 617 | 1 | ENSMUST00000064826.4 | -2990 | 4833418N17Rik | ENSMUSG00000052779.4 |
| chr7 | 81989356 | 81989733 | 3.50372 | 4.35042 | 1.35786 | Intron (ENSMUST00000026096.7/ENSMUSG00000025105.8, intron 1 of 4) | 7 | 81992002 | 81992618 | 617 | 1 | ENSMUST00000064826.4 | -2269 | 4833418N17Rik | ENSMUSG00000052779.4 |
| chr7 | 81990607 | 81990774 | 3.33237 | 3.85537 | 1.11053 | Intron (ENSMUST00000026096.7/ENSMUSG00000025105.8, intron 1 of 4) | 7 | 81992002 | 81992618 | 617 | 1 | ENSMUST00000064826.4 | -1228 | 4833418N17Rik | ENSMUSG00000052779.4 |
| chr8 | 85007685 | 85008014 | 3.50276 | 4.24003 | 1.29365 | Exon (ENSMUST00000059072.4/ENSMUSG00000052819.5, exon 10 of 10) | 8 | 85007202 | 85014408 | 7207 | 2 | ENSMUST00000059072.4 | 6394 | Best2 | ENSMUSG00000052819.5 |
| chr8 | 85009228 | 85009303 | 3.46481 | 4.06154 | 1.14446 | Exon (ENSMUST00000059072.4/ENSMUSG00000052819.5, exon 9 of 10) | 8 | 85007202 | 85014408 | 7207 | 2 | ENSMUST00000059072.4 | 5105 | Best2 | ENSMUSG00000052819.5 |
| chr8 | 85011796 | 85012024 | 3.56911 | 4.2469 | 1.24214 | Intron (ENSMUST00000059072.4/ENSMUSG00000052819.5, intron 3 of 9) | 8 | 85007202 | 85014408 | 7207 | 2 | ENSMUST00000059072.4 | 2384 | Best2 | ENSMUSG00000052819.5 |
| chr5 | 111502045 | 111502129 | 3.95749 | 4.9747 | 1.71377 | Distal Intergenic | 5 | 111581422 | 111587945 | 6524 | 1 | ENSMUST00000064930.3 | -79293 | C130026L21Rik | ENSMUSG00000052848.3 |
| chr5 | 111505872 | 111506073 | 3.34733 | 3.85742 | 1.04874 | Distal Intergenic | 5 | 111581422 | 111587945 | 6524 | 1 | ENSMUST00000064930.3 | -75349 | C130026L21Rik | ENSMUSG00000052848.3 |
| chr5 | 111507187 | 111507339 | 3.37744 | 3.89383 | 1.01733 | Distal Intergenic | 5 | 111581422 | 111587945 | 6524 | 1 | ENSMUST00000064930.3 | -74083 | C130026L21Rik | ENSMUSG00000052848.3 |
| chr5 | 111510838 | 111511117 | 3.36142 | 3.88767 | 1.07931 | Distal Intergenic | 5 | 111581422 | 111587945 | 6524 | 1 | ENSMUST00000064930.3 | -70305 | C130026L21Rik | ENSMUSG00000052848.3 |
| chr5 | 111511966 | 111512379 | 4.46835 | 6.01944 | 2.49095 | Distal Intergenic | 5 | 111581422 | 111587945 | 6524 | 1 | ENSMUST00000064930.3 | -69043 | C130026L21Rik | ENSMUSG00000052848.3 |
| chr5 | 111525432 | 111525834 | 3.81822 | 4.76507 | 1.6696 | Distal Intergenic | 5 | 111581422 | 111587945 | 6524 | 1 | ENSMUST00000064930.3 | -55588 | C130026L21Rik | ENSMUSG00000052848.3 |
| chr5 | 111529075 | 111530313 | 3.55901 | 4.23357 | 1.24202 | Distal Intergenic | 5 | 111581422 | 111587945 | 6524 | 1 | ENSMUST00000064930.3 | -51109 | C130026L21Rik | ENSMUSG00000052848.3 |
| chr5 | 111530935 | 111531576 | 3.78072 | 4.6977 | 1.59943 | Distal Intergenic | 5 | 111581422 | 111587945 | 6524 | 1 | ENSMUST00000064930.3 | -49846 | C130026L21Rik | ENSMUSG00000052848.3 |
| chr5 | 111532103 | 111532944 | 3.96566 | 4.99777 | 1.73044 | Distal Intergenic | 5 | 111581422 | 111587945 | 6524 | 1 | ENSMUST00000064930.3 | -48478 | C130026L21Rik | ENSMUSG00000052848.3 |
| chr5 | 111534493 | 111535067 | 3.59099 | 4.29056 | 1.28053 | Distal Intergenic | 5 | 111581422 | 111587945 | 6524 | 1 | ENSMUST00000064930.3 | -46355 | C130026L21Rik | ENSMUSG00000052848.3 |
| chr5 | 111535415 | 111535938 | 3.74189 | 4.57242 | 1.44927 | Distal Intergenic | 5 | 111581422 | 111587945 | 6524 | 1 | ENSMUST00000064930.3 | -45484 | C130026L21Rik | ENSMUSG00000052848.3 |
| chr5 | 111536244 | 111536325 | 3.82315 | 4.72521 | 1.54651 | Distal Intergenic | 5 | 111581422 | 111587945 | 6524 | 1 | ENSMUST00000064930.3 | -45097 | C130026L21Rik | ENSMUSG00000052848.3 |
| chr5 | 111549670 | 111549777 | 4.80195 | 6.69588 | 2.98124 | Distal Intergenic | 5 | 111581422 | 111587945 | 6524 | 1 | ENSMUST00000064930.3 | -31645 | C130026L21Rik | ENSMUSG00000052848.3 |
| chr5 | 111554605 | 111554998 | 3.36018 | 3.86779 | 1.01733 | Distal Intergenic | 5 | 111581422 | 111587945 | 6524 | 1 | ENSMUST00000064930.3 | -26424 | C130026L21Rik | ENSMUSG00000052848.3 |
| chr5 | 111556700 | 111556775 | 4.03511 | 5.18367 | 1.90312 | Distal Intergenic | 5 | 111581422 | 111587945 | 6524 | 1 | ENSMUST00000064930.3 | -24647 | C130026L21Rik | ENSMUSG00000052848.3 |
| chr5 | 111557358 | 111557653 | 3.48826 | 4.10945 | 1.18114 | Distal Intergenic | 5 | 111581422 | 111587945 | 6524 | 1 | ENSMUST00000064930.3 | -23769 | C130026L21Rik | ENSMUSG00000052848.3 |
| chr5 | 111559975 | 111560075 | 3.40543 | 3.95113 | 1.07072 | Distal Intergenic | 5 | 111581422 | 111587945 | 6524 | 1 | ENSMUST00000064930.3 | -21347 | C130026L21Rik | ENSMUSG00000052848.3 |
| chr5 | 111560771 | 111561053 | 3.56903 | 4.24715 | 1.24367 | Distal Intergenic | 5 | 111581422 | 111587945 | 6524 | 1 | ENSMUST00000064930.3 | -20369 | C130026L21Rik | ENSMUSG00000052848.3 |
| chr5 | 111569531 | 111569606 | 3.83769 | 4.93763 | 1.71517 | Distal Intergenic | 5 | 111581422 | 111587945 | 6524 | 1 | ENSMUST00000064930.3 | -11816 | C130026L21Rik | ENSMUSG00000052848.3 |
| chr5 | 111570847 | 111570942 | 4.501 | 6.07114 | 2.49408 | Distal Intergenic | 5 | 111581422 | 111587945 | 6524 | 1 | ENSMUST00000064930.3 | -10480 | C130026L21Rik | ENSMUSG00000052848.3 |
| chr5 | 111571504 | 111571579 | 3.83822 | 4.74079 | 1.5383 | Distal Intergenic | 5 | 111581422 | 111587945 | 6524 | 1 | ENSMUST00000064930.3 | -9843 | C130026L21Rik | ENSMUSG00000052848.3 |
| chr5 | 111573478 | 111573615 | 3.34153 | 3.83966 | 1.01733 | Distal Intergenic | 5 | 111581422 | 111587945 | 6524 | 1 | ENSMUST00000064930.3 | -7807 | C130026L21Rik | ENSMUSG00000052848.3 |
| chr5 | 111574103 | 111574592 | 3.47194 | 4.19439 | 1.25375 | Distal Intergenic | 5 | 111581422 | 111587945 | 6524 | 1 | ENSMUST00000064930.3 | -6830 | C130026L21Rik | ENSMUSG00000052848.3 |
| chr5 | 111576952 | 111577126 | 3.66389 | 4.6422 | 1.52117 | Distal Intergenic | 5 | 111581422 | 111587945 | 6524 | 1 | ENSMUST00000064930.3 | -4296 | C130026L21Rik | ENSMUSG00000052848.3 |
| chr5 | 111578593 | 111578696 | 3.25266 | 3.70616 | 1.01733 | Distal Intergenic | 5 | 111581422 | 111587945 | 6524 | 1 | ENSMUST00000064930.3 | -2726 | C130026L21Rik | ENSMUSG00000052848.3 |
| chr5 | 111579327 | 111579481 | 3.48044 | 4.08969 | 1.16058 | Distal Intergenic | 5 | 111581422 | 111587945 | 6524 | 1 | ENSMUST00000064930.3 | -1941 | C130026L21Rik | ENSMUSG00000052848.3 |
| chr5 | 111580135 | 111581429 | 3.57954 | 4.65731 | 1.56179 | Promoter | 5 | 111581422 | 111587945 | 6524 | 1 | ENSMUST00000064930.3 | 0 | C130026L21Rik | ENSMUSG00000052848.3 |
| chr5 | 111582627 | 111582831 | 3.62762 | 4.85742 | 1.66168 | Intron (ENSMUST00000064930.3/ENSMUSG00000052848.3, intron 3 of 3) | 5 | 111581422 | 111587945 | 6524 | 1 | ENSMUST00000064930.3 | 1206 | C130026L21Rik | ENSMUSG00000052848.3 |
| chr5 | 111584643 | 111584830 | 3.64813 | 4.46068 | 1.44699 | Intron (ENSMUST00000064930.3/ENSMUSG00000052848.3, intron 3 of 3) | 5 | 111581422 | 111587945 | 6524 | 1 | ENSMUST00000064930.3 | 3222 | C130026L21Rik | ENSMUSG00000052848.3 |
| chr5 | 111590338 | 111590504 | 3.38067 | 3.8987 | 1.01733 | Distal Intergenic | 5 | 111581422 | 111587945 | 6524 | 1 | ENSMUST00000064930.3 | 8917 | C130026L21Rik | ENSMUSG00000052848.3 |
| chr5 | 111635563 | 111635753 | 3.62255 | 4.37739 | 1.35117 | Distal Intergenic | 5 | 111581422 | 111587945 | 6524 | 1 | ENSMUST00000064930.3 | 54142 | C130026L21Rik | ENSMUSG00000052848.3 |
| chr5 | 111640086 | 111640457 | 3.70032 | 4.5258 | 1.49724 | Distal Intergenic | 5 | 111581422 | 111587945 | 6524 | 1 | ENSMUST00000064930.3 | 58665 | C130026L21Rik | ENSMUSG00000052848.3 |
| chr5 | 111640941 | 111641026 | 3.83668 | 4.78739 | 1.67402 | Distal Intergenic | 5 | 111581422 | 111587945 | 6524 | 1 | ENSMUST00000064930.3 | 59520 | C130026L21Rik | ENSMUSG00000052848.3 |
| chr5 | 111645969 | 111646187 | 3.52261 | 4.42743 | 1.40972 | Distal Intergenic | 5 | 111581422 | 111587945 | 6524 | 1 | ENSMUST00000064930.3 | 64548 | C130026L21Rik | ENSMUSG00000052848.3 |
| chr5 | 111652527 | 111652631 | 4.20016 | 5.66255 | 2.22253 | Distal Intergenic | 5 | 111581422 | 111587945 | 6524 | 1 | ENSMUST00000064930.3 | 71106 | C130026L21Rik | ENSMUSG00000052848.3 |
| chr6 | 71707681 | 71707840 | 3.35913 | 3.9011 | 1.13216 | Promoter | 6 | 71707561 | 71810710 | 103150 | 1 | ENSMUST00000121469.1 | 121 | Reep1 | ENSMUSG00000052852.7 |
| chr6 | 71708403 | 71708816 | 3.43924 | 4.08392 | 1.25515 | Promoter | 6 | 71707561 | 71810710 | 103150 | 1 | ENSMUST00000121469.1 | 843 | Reep1 | ENSMUSG00000052852.7 |
| chr7 | 122289433 | 122290497 | 3.53919 | 4.20783 | 1.22467 | Promoter | 7 | 122290028 | 122425041 | 135014 | 1 | ENSMUST00000131167.2 | 0 | Prkcb | ENSMUSG00000052889.6 |
| chr7 | 122381576 | 122381720 | 3.39221 | 3.92385 | 1.04471 | Intron (ENSMUST00000143692.2/ENSMUSG00000052889.6, intron 2 of 17) | 7 | 122416068 | 122457326 | 41259 | 1 | ENSMUST00000149583.1 | -34348 | Prkcb | ENSMUSG00000052889.6 |
| chr7 | 122404722 | 122404797 | 3.3071 | 3.8131 | 1.09682 | Intron (ENSMUST00000143692.2/ENSMUSG00000052889.6, intron 2 of 17) | 7 | 122416068 | 122457326 | 41259 | 1 | ENSMUST00000149583.1 | -11271 | Prkcb | ENSMUSG00000052889.6 |
| chr4 | 145464041 | 145464285 | 4.03438 | 5.60748 | 2.18841 | Promoter | 4 | 145464204 | 145465487 | 1284 | 1 | ENSMUST00000171805.1 | 0 | Smarca5-ps | ENSMUSG00000052912.7 |
| chr11 | 68954503 | 68954592 | 3.36704 | 4.12893 | 1.25332 | Promoter | 11 | 68943155 | 68955074 | 11920 | 2 | ENSMUST00000065040.7 | 482 | Arhgef15 | ENSMUSG00000052921.8 |
| chr11 | 68956241 | 68956349 | 3.54129 | 4.21224 | 1.26629 | Intron (ENSMUST00000108671.1/ENSMUSG00000052921.8, intron 2 of 16) | 11 | 68943157 | 68957480 | 14324 | 2 | ENSMUST00000108671.1 | 1131 | Arhgef15 | ENSMUSG00000052921.8 |
| chr19 | 28679583 | 28679678 | 3.4181 | 4.01707 | 1.2079 | Promoter | 19 | 28262364 | 28679666 | 417303 | 2 | ENSMUST00000065113.8 | 0 | Glis3 | ENSMUSG00000052942.8 |
| chr11 | 116412372 | 116412728 | 3.4576 | 4.21916 | 1.30408 | Promoter | 11 | 116336353 | 116413004 | 76652 | 2 | ENSMUST00000100202.4 | 276 | Rnf157 | ENSMUSG00000052949.9 |
| chr6 | 53943017 | 53943153 | 3.72264 | 4.62847 | 1.54226 | Intron (ENSMUST00000166545.1/ENSMUSG00000052955.3, intron 7 of 12) | 6 | 53873379 | 53978662 | 105284 | 2 | ENSMUST00000166545.1 | 35509 | Cpvl | ENSMUSG00000052955.3 |
| chr6 | 53944657 | 53945017 | 3.6381 | 4.50225 | 1.45674 | Intron (ENSMUST00000166545.1/ENSMUSG00000052955.3, intron 7 of 12) | 6 | 53873379 | 53978662 | 105284 | 2 | ENSMUST00000166545.1 | 33645 | Cpvl | ENSMUSG00000052955.3 |
| chr6 | 114397982 | 114398323 | 3.46398 | 4.14514 | 1.24848 | Promoter | 6 | 114398086 | 114479866 | 81781 | 1 | ENSMUST00000160212.1 | 0 | Hrh1 | ENSMUSG00000053004.8 |
| chr6 | 114398663 | 114398872 | 3.53098 | 4.20865 | 1.28442 | Promoter | 6 | 114398086 | 114479866 | 81781 | 1 | ENSMUST00000160212.1 | 578 | Hrh1 | ENSMUSG00000053004.8 |
| chr6 | 114407841 | 114408530 | 4.09021 | 5.25843 | 1.92407 | Intron (ENSMUST00000088987.2/ENSMUSG00000053004.8, intron 1 of 2) | 6 | 114403825 | 114482404 | 78580 | 1 | ENSMUST00000161220.1 | 4017 | Hrh1 | ENSMUSG00000053004.8 |
| chr6 | 53516653 | 53516872 | 4.25412 | 5.66763 | 2.23645 | Distal Intergenic | 6 | 53573384 | 53696007 | 122624 | 1 | ENSMUST00000047450.7 | -56512 | Creb5 | ENSMUSG00000053007.6 |
| chr6 | 53527306 | 53527403 | 3.74722 | 4.58034 | 1.45872 | Distal Intergenic | 6 | 53573384 | 53696007 | 122624 | 1 | ENSMUST00000047450.7 | -45981 | Creb5 | ENSMUSG00000053007.6 |
| chr6 | 53539341 | 53539902 | 3.49971 | 4.12035 | 1.16273 | Distal Intergenic | 6 | 53573384 | 53696007 | 122624 | 1 | ENSMUST00000047450.7 | -33482 | Creb5 | ENSMUSG00000053007.6 |
| chr6 | 53540783 | 53540900 | 3.45618 | 4.04293 | 1.12412 | Distal Intergenic | 6 | 53573384 | 53696007 | 122624 | 1 | ENSMUST00000047450.7 | -32484 | Creb5 | ENSMUSG00000053007.6 |
| chr6 | 53544902 | 53545137 | 3.3471 | 3.84805 | 1.01733 | Distal Intergenic | 6 | 53573384 | 53696007 | 122624 | 1 | ENSMUST00000047450.7 | -28247 | Creb5 | ENSMUSG00000053007.6 |
| chr6 | 53546123 | 53546425 | 3.90761 | 5.11422 | 1.83523 | Distal Intergenic | 6 | 53573384 | 53696007 | 122624 | 1 | ENSMUST00000047450.7 | -26959 | Creb5 | ENSMUSG00000053007.6 |
| chr6 | 53547216 | 53547505 | 3.52226 | 4.23762 | 1.31334 | Distal Intergenic | 6 | 53573384 | 53696007 | 122624 | 1 | ENSMUST00000047450.7 | -25879 | Creb5 | ENSMUSG00000053007.6 |
| chr6 | 53549108 | 53549183 | 3.32777 | 4.27101 | 1.29829 | Distal Intergenic | 6 | 53573384 | 53696007 | 122624 | 1 | ENSMUST00000047450.7 | -24201 | Creb5 | ENSMUSG00000053007.6 |
| chr6 | 53553232 | 53553317 | 4.13958 | 5.49642 | 2.11753 | Distal Intergenic | 6 | 53573384 | 53696007 | 122624 | 1 | ENSMUST00000047450.7 | -20067 | Creb5 | ENSMUSG00000053007.6 |
| chr6 | 53563478 | 53564256 | 3.62092 | 4.35321 | 1.33721 | Distal Intergenic | 6 | 53573384 | 53696007 | 122624 | 1 | ENSMUST00000047450.7 | -9128 | Creb5 | ENSMUSG00000053007.6 |
| chr6 | 53566499 | 53566848 | 3.36282 | 3.89726 | 1.10757 | Distal Intergenic | 6 | 53573384 | 53696007 | 122624 | 1 | ENSMUST00000047450.7 | -6536 | Creb5 | ENSMUSG00000053007.6 |
| chr6 | 53568938 | 53569088 | 3.60553 | 4.53204 | 1.46709 | Distal Intergenic | 6 | 53573384 | 53696007 | 122624 | 1 | ENSMUST00000047450.7 | -4296 | Creb5 | ENSMUSG00000053007.6 |
| chr6 | 53572439 | 53572780 | 3.70908 | 4.61829 | 1.54988 | Promoter | 6 | 53573384 | 53696007 | 122624 | 1 | ENSMUST00000047450.7 | -604 | Creb5 | ENSMUSG00000053007.6 |
| chr6 | 53573109 | 53574415 | 3.5713 | 4.40213 | 1.39233 | Promoter | 6 | 53573384 | 53696007 | 122624 | 1 | ENSMUST00000047450.7 | 0 | Creb5 | ENSMUSG00000053007.6 |
| chr6 | 53576406 | 53577029 | 3.81942 | 4.79728 | 1.69418 | Intron (ENSMUST00000047450.7/ENSMUSG00000053007.6, intron 1 of 6) | 6 | 53573384 | 53696007 | 122624 | 1 | ENSMUST00000047450.7 | 3023 | Creb5 | ENSMUSG00000053007.6 |
| chr6 | 53582207 | 53582282 | 3.5142 | 4.16512 | 1.24843 | Intron (ENSMUST00000047450.7/ENSMUSG00000053007.6, intron 1 of 6) | 6 | 53573384 | 53696007 | 122624 | 1 | ENSMUST00000047450.7 | 8824 | Creb5 | ENSMUSG00000053007.6 |
| chr6 | 53584559 | 53585228 | 3.76051 | 4.83636 | 1.67308 | Intron (ENSMUST00000047450.7/ENSMUSG00000053007.6, intron 1 of 6) | 6 | 53573384 | 53696007 | 122624 | 1 | ENSMUST00000047450.7 | 11176 | Creb5 | ENSMUSG00000053007.6 |
| chr6 | 53585687 | 53586025 | 4.12565 | 5.34148 | 2.00076 | Intron (ENSMUST00000047450.7/ENSMUSG00000053007.6, intron 1 of 6) | 6 | 53573384 | 53696007 | 122624 | 1 | ENSMUST00000047450.7 | 12304 | Creb5 | ENSMUSG00000053007.6 |
| chr6 | 53593728 | 53593955 | 3.77578 | 4.79491 | 1.62358 | Intron (ENSMUST00000047450.7/ENSMUSG00000053007.6, intron 1 of 6) | 6 | 53573384 | 53696007 | 122624 | 1 | ENSMUST00000047450.7 | 20345 | Creb5 | ENSMUSG00000053007.6 |
| chr6 | 53594403 | 53594747 | 3.50589 | 4.1684 | 1.29029 | Intron (ENSMUST00000047450.7/ENSMUSG00000053007.6, intron 1 of 6) | 6 | 53573384 | 53696007 | 122624 | 1 | ENSMUST00000047450.7 | 21020 | Creb5 | ENSMUSG00000053007.6 |
| chr6 | 53596309 | 53597358 | 3.48423 | 4.13634 | 1.27636 | Intron (ENSMUST00000047450.7/ENSMUSG00000053007.6, intron 1 of 6) | 6 | 53573384 | 53696007 | 122624 | 1 | ENSMUST00000047450.7 | 22926 | Creb5 | ENSMUSG00000053007.6 |
| chr6 | 53600429 | 53600994 | 3.46048 | 4.33708 | 1.34702 | Intron (ENSMUST00000047450.7/ENSMUSG00000053007.6, intron 1 of 6) | 6 | 53573384 | 53696007 | 122624 | 1 | ENSMUST00000047450.7 | 27046 | Creb5 | ENSMUSG00000053007.6 |
| chr6 | 53602126 | 53602538 | 3.56702 | 4.42143 | 1.40911 | Intron (ENSMUST00000047450.7/ENSMUSG00000053007.6, intron 1 of 6) | 6 | 53573384 | 53696007 | 122624 | 1 | ENSMUST00000047450.7 | 28743 | Creb5 | ENSMUSG00000053007.6 |
| chr6 | 53607380 | 53607764 | 3.38812 | 3.95228 | 1.15792 | Intron (ENSMUST00000047450.7/ENSMUSG00000053007.6, intron 2 of 6) | 6 | 53573384 | 53696007 | 122624 | 1 | ENSMUST00000047450.7 | 33997 | Creb5 | ENSMUSG00000053007.6 |
| chr6 | 53608152 | 53608602 | 3.46261 | 4.20918 | 1.28038 | Intron (ENSMUST00000047450.7/ENSMUSG00000053007.6, intron 2 of 6) | 6 | 53573384 | 53696007 | 122624 | 1 | ENSMUST00000047450.7 | 34769 | Creb5 | ENSMUSG00000053007.6 |
| chr6 | 53610230 | 53610761 | 3.76398 | 4.70022 | 1.64158 | Exon (ENSMUST00000047450.7/ENSMUSG00000053007.6, exon 3 of 7) | 6 | 53573384 | 53696007 | 122624 | 1 | ENSMUST00000047450.7 | 36847 | Creb5 | ENSMUSG00000053007.6 |
| chr6 | 53611329 | 53611723 | 3.76142 | 4.67742 | 1.63517 | Intron (ENSMUST00000047450.7/ENSMUSG00000053007.6, intron 3 of 6) | 6 | 53573384 | 53696007 | 122624 | 1 | ENSMUST00000047450.7 | 37946 | Creb5 | ENSMUSG00000053007.6 |
| chr6 | 53612119 | 53613385 | 3.9135 | 5.11183 | 1.90188 | Intron (ENSMUST00000047450.7/ENSMUSG00000053007.6, intron 3 of 6) | 6 | 53573384 | 53696007 | 122624 | 1 | ENSMUST00000047450.7 | 38736 | Creb5 | ENSMUSG00000053007.6 |
| chr6 | 53613752 | 53614278 | 3.57737 | 4.496 | 1.43075 | Intron (ENSMUST00000047450.7/ENSMUSG00000053007.6, intron 3 of 6) | 6 | 53573384 | 53696007 | 122624 | 1 | ENSMUST00000047450.7 | 40369 | Creb5 | ENSMUSG00000053007.6 |
| chr6 | 53614713 | 53614960 | 3.60855 | 4.55967 | 1.4932 | Intron (ENSMUST00000047450.7/ENSMUSG00000053007.6, intron 3 of 6) | 6 | 53573384 | 53696007 | 122624 | 1 | ENSMUST00000047450.7 | 41330 | Creb5 | ENSMUSG00000053007.6 |
| chr6 | 53617397 | 53617541 | 4.07141 | 5.43422 | 2.04981 | Intron (ENSMUST00000047450.7/ENSMUSG00000053007.6, intron 3 of 6) | 6 | 53573384 | 53696007 | 122624 | 1 | ENSMUST00000047450.7 | 44014 | Creb5 | ENSMUSG00000053007.6 |
| chr6 | 53621247 | 53621451 | 3.39391 | 4.29936 | 1.30916 | Intron (ENSMUST00000047450.7/ENSMUSG00000053007.6, intron 3 of 6) | 6 | 53573384 | 53696007 | 122624 | 1 | ENSMUST00000047450.7 | 47864 | Creb5 | ENSMUSG00000053007.6 |
| chr6 | 53622179 | 53622576 | 3.64227 | 4.5687 | 1.49823 | Intron (ENSMUST00000047450.7/ENSMUSG00000053007.6, intron 3 of 6) | 6 | 53573384 | 53696007 | 122624 | 1 | ENSMUST00000047450.7 | 48796 | Creb5 | ENSMUSG00000053007.6 |
| chr6 | 53624709 | 53625076 | 4.1032 | 5.34094 | 2.05418 | Intron (ENSMUST00000047450.7/ENSMUSG00000053007.6, intron 3 of 6) | 6 | 53573384 | 53696007 | 122624 | 1 | ENSMUST00000047450.7 | 51326 | Creb5 | ENSMUSG00000053007.6 |
| chr6 | 53628052 | 53628854 | 3.49704 | 4.14455 | 1.25359 | Intron (ENSMUST00000047450.7/ENSMUSG00000053007.6, intron 3 of 6) | 6 | 53573384 | 53696007 | 122624 | 1 | ENSMUST00000047450.7 | 54669 | Creb5 | ENSMUSG00000053007.6 |
| chr6 | 53629210 | 53629451 | 3.68515 | 4.53661 | 1.50419 | Intron (ENSMUST00000047450.7/ENSMUSG00000053007.6, intron 3 of 6) | 6 | 53573384 | 53696007 | 122624 | 1 | ENSMUST00000047450.7 | 55827 | Creb5 | ENSMUSG00000053007.6 |
| chr6 | 53631378 | 53631521 | 3.20927 | 3.64116 | 1.00681 | Intron (ENSMUST00000047450.7/ENSMUSG00000053007.6, intron 3 of 6) | 6 | 53573384 | 53696007 | 122624 | 1 | ENSMUST00000047450.7 | 57995 | Creb5 | ENSMUSG00000053007.6 |
| chr6 | 53631891 | 53632113 | 3.87892 | 5.01668 | 1.82818 | Intron (ENSMUST00000047450.7/ENSMUSG00000053007.6, intron 3 of 6) | 6 | 53573384 | 53696007 | 122624 | 1 | ENSMUST00000047450.7 | 58508 | Creb5 | ENSMUSG00000053007.6 |
| chr6 | 53637822 | 53638512 | 4.2411 | 5.76291 | 2.37202 | Intron (ENSMUST00000047450.7/ENSMUSG00000053007.6, intron 3 of 6) | 6 | 53573384 | 53696007 | 122624 | 1 | ENSMUST00000047450.7 | 64439 | Creb5 | ENSMUSG00000053007.6 |
| chr6 | 53646609 | 53647303 | 3.81001 | 4.72086 | 1.58124 | Intron (ENSMUST00000047450.7/ENSMUSG00000053007.6, intron 3 of 6) | 6 | 53573384 | 53696007 | 122624 | 1 | ENSMUST00000047450.7 | 73226 | Creb5 | ENSMUSG00000053007.6 |
| chr6 | 53694622 | 53694739 | 3.58819 | 4.33521 | 1.37391 | 3' UTR | 6 | 53573384 | 53696007 | 122624 | 1 | ENSMUST00000047450.7 | 121239 | Creb5 | ENSMUSG00000053007.6 |
| chr1 | 132541847 | 132541943 | 3.34153 | 3.83966 | 1.01733 | Promoter | 1 | 132511792 | 132542865 | 31074 | 2 | ENSMUST00000190601.1 | 922 | Cntn2 | ENSMUSG00000053024.9 |
| chr1 | 132542305 | 132542403 | 3.87195 | 4.83002 | 1.63154 | Promoter | 1 | 132511792 | 132542865 | 31074 | 2 | ENSMUST00000190601.1 | 462 | Cntn2 | ENSMUSG00000053024.9 |
| chr1 | 132543084 | 132543336 | 3.70617 | 4.72043 | 1.59259 | Promoter | 1 | 132533657 | 132543256 | 9600 | 2 | ENSMUST00000186530.1 | 0 | Cntn2 | ENSMUSG00000053024.9 |
| chr1 | 132559592 | 132559970 | 3.35944 | 3.96486 | 1.10282 | Distal Intergenic | 1 | 132533657 | 132543256 | 9600 | 2 | ENSMUST00000186530.1 | -16337 | Cntn2 | ENSMUSG00000053024.9 |
| chr7 | 75307704 | 75308565 | 3.67776 | 4.52567 | 1.51924 | Promoter | 7 | 75114897 | 75308386 | 193490 | 2 | ENSMUST00000085164.5 | 0 | Sv2b | ENSMUSG00000053025.8 |
| chr7 | 75308885 | 75309207 | 3.73185 | 4.66664 | 1.59096 | Promoter | 7 | 75114897 | 75309262 | 194366 | 2 | ENSMUST00000165175.2 | 55 | Sv2b | ENSMUSG00000053025.8 |
| chr7 | 141947963 | 141948222 | 3.44938 | 4.03029 | 1.11621 | Distal Intergenic | 7 | 141949751 | 141999005 | 49255 | 1 | ENSMUST00000078200.6 | -1529 | Brsk2 | ENSMUSG00000053046.9 |
| chr7 | 141948530 | 141949046 | 4.23234 | 5.60887 | 2.25816 | Promoter | 7 | 141949751 | 141999005 | 49255 | 1 | ENSMUST00000078200.6 | -705 | Brsk2 | ENSMUSG00000053046.9 |
| chr7 | 141950407 | 141951192 | 4.35733 | 5.92059 | 2.49975 | Promoter | 7 | 141950086 | 142004247 | 54162 | 1 | ENSMUST00000174499.2 | 322 | Brsk2 | ENSMUSG00000053046.9 |
| chr7 | 96799320 | 96799395 | 3.66554 | 4.59401 | 1.52243 | Intron (ENSMUST00000107162.2/ENSMUSG00000048078.11, intron 17 of 32) | 7 | 96791433 | 96801549 | 10117 | 2 | ENSMUST00000155101.1 | 2154 | Gm15413 | ENSMUSG00000053049.3 |
| chr16 | 84771272 | 84771352 | 4.15024 | 5.37354 | 2.04148 | Distal Intergenic | 16 | 84774123 | 84825928 | 51806 | 1 | ENSMUST00000114195.2 | -2771 | Jam2 | ENSMUSG00000053062.10 |
| chr16 | 84772406 | 84773212 | 3.59209 | 4.35552 | 1.39259 | Promoter | 16 | 84774123 | 84825928 | 51806 | 1 | ENSMUST00000114195.2 | -911 | Jam2 | ENSMUSG00000053062.10 |
| chr16 | 84774046 | 84774631 | 3.86875 | 4.92114 | 1.79294 | Promoter | 16 | 84774123 | 84825928 | 51806 | 1 | ENSMUST00000114195.2 | 0 | Jam2 | ENSMUSG00000053062.10 |
| chr16 | 84775232 | 84775540 | 3.51434 | 4.47836 | 1.42397 | Promoter | 16 | 84774386 | 84810996 | 36611 | 1 | ENSMUST00000098407.2 | 847 | Jam2 | ENSMUSG00000053062.10 |
| chr7 | 121621822 | 121622152 | 3.3473 | 3.84837 | 1.01733 | Distal Intergenic | 7 | 121707075 | 121708013 | 939 | 1 | ENSMUST00000191155.1 | -84923 | 1700069B07Rik | ENSMUSG00000053081.3 |
| chr19 | 60051455 | 60051727 | 3.30981 | 3.81997 | 1.10474 | Distal Intergenic | 19 | 60144675 | 60162591 | 17917 | 1 | ENSMUST00000065383.5 | -92948 | E330013P04Rik | ENSMUSG00000053117.5 |
| chr5 | 147155139 | 147155224 | 3.85061 | 4.77271 | 1.57334 | Distal Intergenic | 5 | 147188696 | 147190947 | 2252 | 1 | ENSMUST00000065382.5 | -33472 | Gsx1 | ENSMUSG00000053129.5 |
| chr5 | 147157625 | 147157714 | 3.621 | 4.57694 | 1.49935 | Distal Intergenic | 5 | 147188696 | 147190947 | 2252 | 1 | ENSMUST00000065382.5 | -30982 | Gsx1 | ENSMUSG00000053129.5 |
| chr5 | 147182836 | 147183898 | 3.86235 | 4.82703 | 1.6315 | Distal Intergenic | 5 | 147188696 | 147190947 | 2252 | 1 | ENSMUST00000065382.5 | -4798 | Gsx1 | ENSMUSG00000053129.5 |
| chr5 | 147185212 | 147185697 | 3.35953 | 3.8876 | 1.07117 | Distal Intergenic | 5 | 147188696 | 147190947 | 2252 | 1 | ENSMUST00000065382.5 | -2999 | Gsx1 | ENSMUSG00000053129.5 |
| chr5 | 147186046 | 147186254 | 3.6497 | 4.3953 | 1.32669 | Distal Intergenic | 5 | 147188696 | 147190947 | 2252 | 1 | ENSMUST00000065382.5 | -2442 | Gsx1 | ENSMUSG00000053129.5 |
| chr5 | 147186790 | 147187068 | 4.26076 | 5.61306 | 2.20704 | Distal Intergenic | 5 | 147188696 | 147190947 | 2252 | 1 | ENSMUST00000065382.5 | -1628 | Gsx1 | ENSMUSG00000053129.5 |
| chr5 | 147187512 | 147189319 | 3.93693 | 5.20154 | 1.92278 | Promoter | 5 | 147188696 | 147190947 | 2252 | 1 | ENSMUST00000065382.5 | 0 | Gsx1 | ENSMUSG00000053129.5 |
| chr5 | 147189827 | 147190115 | 3.9897 | 5.07544 | 1.81156 | Exon (ENSMUST00000065382.5/ENSMUSG00000053129.5, exon 2 of 2) | 5 | 147188696 | 147190947 | 2252 | 1 | ENSMUST00000065382.5 | 1132 | Gsx1 | ENSMUSG00000053129.5 |
| chr5 | 147190433 | 147191451 | 3.79153 | 4.69255 | 1.57424 | 3' UTR | 5 | 147188696 | 147190947 | 2252 | 1 | ENSMUST00000065382.5 | 1738 | Gsx1 | ENSMUSG00000053129.5 |
| chr5 | 147192940 | 147193299 | 3.37467 | 3.92875 | 1.14714 | Distal Intergenic | 5 | 147188696 | 147190947 | 2252 | 1 | ENSMUST00000065382.5 | 4245 | Gsx1 | ENSMUSG00000053129.5 |
| chr5 | 147194065 | 147194567 | 3.64035 | 4.39216 | 1.37117 | Distal Intergenic | 5 | 147188696 | 147190947 | 2252 | 1 | ENSMUST00000065382.5 | 5370 | Gsx1 | ENSMUSG00000053129.5 |
| chr2 | 162563824 | 162564155 | 3.39782 | 4.30042 | 1.29763 | Intron (ENSMUST00000109443.2/ENSMUSG00000053141.11, intron 1 of 30) | 2 | 161526357 | 162661094 | 1134738 | 2 | ENSMUST00000109442.2 | 96939 | Ptprt | ENSMUSG00000053141.11 |
| chr2 | 162658026 | 162658558 | 3.61649 | 4.33323 | 1.29542 | Intron (ENSMUST00000109443.2/ENSMUSG00000053141.11, intron 1 of 30) | 2 | 161526357 | 162661094 | 1134738 | 2 | ENSMUST00000109442.2 | 2536 | Ptprt | ENSMUSG00000053141.11 |
| chr2 | 162659259 | 162659375 | 3.43708 | 4.03862 | 1.18896 | Intron (ENSMUST00000109443.2/ENSMUSG00000053141.11, intron 1 of 30) | 2 | 161526357 | 162661094 | 1134738 | 2 | ENSMUST00000109442.2 | 1719 | Ptprt | ENSMUSG00000053141.11 |
| chr2 | 162659682 | 162661310 | 4.0808 | 5.35006 | 2.02233 | Promoter | 2 | 161521990 | 162661147 | 1139158 | 2 | ENSMUST00000109443.2 | 0 | Ptprt | ENSMUSG00000053141.11 |

| | | | | | | | | | | | | | | | |
|---|---|---|---|---|---|---|---|---|---|---|---|---|---|---|---|
| chr1 | 69826902 | 69827068 | 3.81557 | 4.74236 | 1.63245 | Promoter | 1 | 69826970 | 70267277 | 440308 | 1 | ENSMUST00000187159.1 | 0 | Spag16 | ENSMUSG00000053153.9 |
| chr2 | 165176632 | 165177045 | 3.37004 | 3.88541 | 1.02704 | Intron (ENSMUST00000065438.7/ENSMUSG00000053166.9, intron 2 of 11) | 2 | 165112327 | 165181459 | 69133 | 2 | ENSMUST00000138643.1 | 4414 | Cdh22 | ENSMUSG00000053166.9 |
| chr2 | 165177379 | 165177533 | 3.34153 | 3.83966 | 1.01733 | Intron (ENSMUST00000065438.7/ENSMUSG00000053166.9, intron 2 of 11) | 2 | 165112327 | 165181459 | 69133 | 2 | ENSMUST00000138643.1 | 3926 | Cdh22 | ENSMUSG00000053166.9 |
| chr2 | 165180976 | 165181624 | 3.76569 | 4.69758 | 1.60517 | Promoter | 2 | 165112327 | 165181459 | 69133 | 2 | ENSMUST00000138643.1 | 0 | Cdh22 | ENSMUSG00000053166.9 |
| chr2 | 165181954 | 165182183 | 3.3694 | 3.89807 | 1.07413 | Promoter | 2 | 165112327 | 165181459 | 69133 | 2 | ENSMUST00000138643.1 | -496 | Cdh22 | ENSMUSG00000053166.9 |
| chr2 | 165182731 | 165183220 | 4.03762 | 5.27025 | 1.98609 | Intron (ENSMUST00000065438.7/ENSMUSG00000053166.9, intron 1 of 11) | 2 | 165112327 | 165181459 | 69133 | 2 | ENSMUST00000138643.1 | -1273 | Cdh22 | ENSMUSG00000053166.9 |
| chr2 | 165183798 | 165183873 | 3.86055 | 4.82107 | 1.67048 | Intron (ENSMUST00000065438.7/ENSMUSG00000053166.9, intron 1 of 11) | 2 | 165112327 | 165181459 | 69133 | 2 | ENSMUST00000138643.1 | -2340 | Cdh22 | ENSMUSG00000053166.9 |
| chr2 | 165184194 | 165184320 | 3.42913 | 4.01936 | 1.18594 | Intron (ENSMUST00000065438.7/ENSMUSG00000053166.9, intron 1 of 11) | 2 | 165112327 | 165181459 | 69133 | 2 | ENSMUST00000138643.1 | -2736 | Cdh22 | ENSMUSG00000053166.9 |
| chr2 | 165185618 | 165185742 | 3.36755 | 3.87889 | 1.01733 | Intron (ENSMUST00000065438.7/ENSMUSG00000053166.9, intron 1 of 11) | 2 | 165112327 | 165181459 | 69133 | 2 | ENSMUST00000138643.1 | -4160 | Cdh22 | ENSMUSG00000053166.9 |
| chr2 | 165190058 | 165190484 | 3.44954 | 4.02784 | 1.10632 | Intron (ENSMUST00000065438.7/ENSMUSG00000053166.9, intron 1 of 11) | 2 | 165112327 | 165181459 | 69133 | 2 | ENSMUST00000138643.1 | -8600 | Cdh22 | ENSMUSG00000053166.9 |
| chr2 | 165192118 | 165192501 | 3.7757 | 4.79305 | 1.64917 | Intron (ENSMUST00000065438.7/ENSMUSG00000053166.9, intron 1 of 11) | 2 | 165112327 | 165181459 | 69133 | 2 | ENSMUST00000138643.1 | -10660 | Cdh22 | ENSMUSG00000053166.9 |
| chr2 | 165194521 | 165194923 | 3.61217 | 4.3549 | 1.34748 | Intron (ENSMUST00000065438.7/ENSMUSG00000053166.9, intron 1 of 11) | 2 | 165112327 | 165181459 | 69133 | 2 | ENSMUST00000138643.1 | -13063 | Cdh22 | ENSMUSG00000053166.9 |
| chr2 | 165195778 | 165196574 | 3.7783 | 4.76686 | 1.61548 | Intron (ENSMUST00000065438.7/ENSMUSG00000053166.9, intron 1 of 11) | 2 | 165112327 | 165181459 | 69133 | 2 | ENSMUST00000138643.1 | -14320 | Cdh22 | ENSMUSG00000053166.9 |
| chr2 | 165198738 | 165198878 | 3.34637 | 3.8569 | 1.05215 | Intron (ENSMUST00000065438.7/ENSMUSG00000053166.9, intron 1 of 11) | 2 | 165112327 | 165181459 | 69133 | 2 | ENSMUST00000138643.1 | -17280 | Cdh22 | ENSMUSG00000053166.9 |
| chr2 | 165199759 | 165200782 | 3.63563 | 4.38282 | 1.342 | Intron (ENSMUST00000065438.7/ENSMUSG00000053166.9, intron 1 of 11) | 2 | 165112327 | 165181459 | 69133 | 2 | ENSMUST00000138643.1 | -18301 | Cdh22 | ENSMUSG00000053166.9 |
| chr2 | 165201157 | 165201839 | 3.98783 | 5.17609 | 1.90648 | Intron (ENSMUST00000065438.7/ENSMUSG00000053166.9, intron 1 of 11) | 2 | 165112327 | 165181459 | 69133 | 2 | ENSMUST00000138643.1 | -19699 | Cdh22 | ENSMUSG00000053166.9 |
| chr2 | 165202343 | 165202949 | 3.81578 | 4.72417 | 1.56071 | Intron (ENSMUST00000065438.7/ENSMUSG00000053166.9, intron 1 of 11) | 2 | 165112327 | 165181459 | 69133 | 2 | ENSMUST00000138643.1 | -20885 | Cdh22 | ENSMUSG00000053166.9 |
| chr2 | 165204879 | 165205193 | 3.416 | 4.25529 | 1.26733 | Intron (ENSMUST00000065438.7/ENSMUSG00000053166.9, intron 1 of 11) | 2 | 165112327 | 165181459 | 69133 | 2 | ENSMUST00000138643.1 | -23421 | Cdh22 | ENSMUSG00000053166.9 |
| chr2 | 165209375 | 165214300 | 3.95777 | 5.18246 | 1.9623 | Intron (ENSMUST00000065438.7/ENSMUSG00000053166.9, intron 1 of 11) | 2 | 165111507 | 165234853 | 123347 | 2 | ENSMUST00000065438.7 | 20553 | Cdh22 | ENSMUSG00000053166.9 |
| chr2 | 165214681 | 165215766 | 3.75113 | 4.79542 | 1.65527 | Intron (ENSMUST00000065438.7/ENSMUSG00000053166.9, intron 1 of 11) | 2 | 165111507 | 165234853 | 123347 | 2 | ENSMUST00000065438.7 | 19087 | Cdh22 | ENSMUSG00000053166.9 |
| chr2 | 165219919 | 165220698 | 3.36477 | 3.88198 | 1.043 | Intron (ENSMUST00000065438.7/ENSMUSG00000053166.9, intron 1 of 11) | 2 | 165111507 | 165234853 | 123347 | 2 | ENSMUST00000065438.7 | 14155 | Cdh22 | ENSMUSG00000053166.9 |
| chr2 | 165222918 | 165223246 | 3.41496 | 4.1061 | 1.18734 | Intron (ENSMUST00000065438.7/ENSMUSG00000053166.9, intron 1 of 11) | 2 | 165111507 | 165234853 | 123347 | 2 | ENSMUST00000065438.7 | 11607 | Cdh22 | ENSMUSG00000053166.9 |
| chr2 | 165225145 | 165225343 | 3.49 | 4.18996 | 1.22946 | Intron (ENSMUST00000065438.7/ENSMUSG00000053166.9, intron 1 of 11) | 2 | 165111507 | 165234853 | 123347 | 2 | ENSMUST00000065438.7 | 9510 | Cdh22 | ENSMUSG00000053166.9 |
| chr2 | 165231452 | 165232145 | 3.62151 | 4.39198 | 1.4311 | Intron (ENSMUST00000065438.7/ENSMUSG00000053166.9, intron 1 of 11) | 2 | 165111507 | 165234853 | 123347 | 2 | ENSMUST00000065438.7 | 2708 | Cdh22 | ENSMUSG00000053166.9 |
| chr2 | 165233109 | 165236121 | 3.98317 | 5.26602 | 2.02088 | Promoter | 2 | 165111507 | 165234853 | 123347 | 2 | ENSMUST00000065438.7 | 0 | Cdh22 | ENSMUSG00000053166.9 |
| chr2 | 165237849 | 165238203 | 3.58121 | 4.2721 | 1.26536 | Distal Intergenic | 2 | 165111507 | 165234853 | 123347 | 2 | ENSMUST00000065438.7 | -2997 | Cdh22 | ENSMUSG00000053166.9 |
| chr2 | 165238624 | 165239709 | 3.54047 | 4.27471 | 1.32877 | Distal Intergenic | 2 | 165111507 | 165234853 | 123347 | 2 | ENSMUST00000065438.7 | -3772 | Cdh22 | ENSMUSG00000053166.9 |
| chr2 | 165240064 | 165240716 | 3.55853 | 4.31113 | 1.32362 | Distal Intergenic | 2 | 165111507 | 165234853 | 123347 | 2 | ENSMUST00000065438.7 | -5212 | Cdh22 | ENSMUSG00000053166.9 |
| chr2 | 165244137 | 165244612 | 3.44292 | 4.14115 | 1.21959 | Distal Intergenic | 2 | 165111507 | 165234853 | 123347 | 2 | ENSMUST00000065438.7 | -9285 | Cdh22 | ENSMUSG00000053166.9 |
| chr2 | 165245326 | 165245824 | 3.577 | 4.34465 | 1.38222 | Distal Intergenic | 2 | 165111507 | 165234853 | 123347 | 2 | ENSMUST00000065438.7 | -10474 | Cdh22 | ENSMUSG00000053166.9 |
| chr7 | 19822492 | 19822812 | 3.89751 | 4.87709 | 1.66285 | Promoter | 7 | 19808462 | 19822770 | 14309 | 2 | ENSMUST00000120537.2 | 0 | Bcl3 | ENSMUSG00000053175.11 |
| chr9 | 51765573 | 51766066 | 3.6471 | 4.59021 | 1.49956 | Promoter | 9 | 51765660 | 51850651 | 84992 | 1 | ENSMUST00000130405.1 | 0 | Arhgap20 | ENSMUSG00000053199.8 |
| chr10 | 22156737 | 22157507 | 3.44948 | 4.09138 | 1.21973 | Distal Intergenic | 10 | 22158569 | 22183917 | 25349 | 1 | ENSMUST00000181645.2 | -1062 | Raet1e | ENSMUSG00000053219.10 |
| chr10 | 22158390 | 22158646 | 4.67756 | 6.59584 | 2.98593 | Promoter | 10 | 22158569 | 22183917 | 25349 | 1 | ENSMUST00000181645.2 | 0 | Raet1e | ENSMUSG00000053219.10 |
| chr10 | 22169207 | 22169312 | 3.60664 | 4.6952 | 1.57489 | Intron (ENSMUST00000181645.2/ENSMUSG00000053219.10, intron 1 of 8) | 10 | 22165638 | 22180726 | 15089 | 1 | ENSMUST00000180648.2 | 3570 | Raet1e | ENSMUSG00000053219.10 |
| chr6 | 48625817 | 48626011 | 3.47319 | 4.11882 | 1.2373 | Promoter | 6 | 48626764 | 48633685 | 6922 | 1 | ENSMUST00000095938.4 | -753 | AI854703 | ENSMUSG00000053297.9 |
| chr6 | 48626777 | 48627357 | 3.44693 | 4.13194 | 1.25034 | Promoter | 6 | 48626764 | 48633685 | 6922 | 1 | ENSMUST00000095938.4 | 14 | AI854703 | ENSMUSG00000053297.9 |
| chr6 | 48628821 | 48628922 | 3.86565 | 4.87578 | 1.7188 | Promoter | 6 | 48627935 | 48633689 | 5755 | 1 | ENSMUST00000154570.1 | 887 | AI854703 | ENSMUSG00000053297.9 |
| chr6 | 48629274 | 48629661 | 3.57837 | 4.32246 | 1.4046 | Exon (ENSMUST00000095938.4/ENSMUSG00000053297.9, exon 2 of 4) | 6 | 48627935 | 48633689 | 5755 | 1 | ENSMUST00000154570.1 | 1340 | AI854703 | ENSMUSG00000053297.9 |
| chr9 | 37552220 | 37552999 | 3.72282 | 4.82085 | 1.66862 | Promoter | 9 | 37544492 | 37552904 | 8413 | 2 | ENSMUST00000065668.6 | 0 | Nrgn | ENSMUSG00000053310.6 |
| chr9 | 37567643 | 37567749 | 3.74073 | 4.72942 | 1.66274 | Distal Intergenic | 9 | 37544492 | 37552904 | 8413 | 2 | ENSMUST00000065668.6 | -14740 | Nrgn | ENSMUSG00000053310.6 |
| chr5 | 135353268 | 135353739 | 4.03255 | 5.25724 | 1.96011 | Promoter | 5 | 135353295 | 135367654 | 14360 | 1 | ENSMUST00000111180.3 | 0 | Trim50 | ENSMUSG00000053388.5 |
| chr7 | 3366161 | 3366236 | 3.46087 | 4.0574 | 1.15081 | Intron (ENSMUST00000092891.4/ENSMUSG00000069806.4, intron 5 of 5) | 7 | 3390683 | 3415648 | 24966 | 1 | ENSMUST00000182222.2 | -24447 | Cacng8 | ENSMUSG00000053395.10 |
| chr7 | 3367143 | 3368085 | 3.65233 | 4.43903 | 1.38058 | 3' UTR | 7 | 3390683 | 3415648 | 24966 | 1 | ENSMUST00000182222.2 | -22598 | Cacng8 | ENSMUSG00000053395.10 |
| chr7 | 3388867 | 3388942 | 4.4287 | 5.90342 | 2.36378 | Distal Intergenic | 7 | 3390683 | 3415648 | 24966 | 1 | ENSMUST00000182222.2 | -1741 | Cacng8 | ENSMUSG00000053395.10 |
| chr7 | 3391220 | 3391713 | 3.51535 | 4.15084 | 1.18957 | Promoter | 7 | 3390683 | 3415648 | 24966 | 1 | ENSMUST00000182222.2 | 538 | Cacng8 | ENSMUSG00000053395.10 |
| chr8 | 113846535 | 113846713 | 3.44397 | 4.01412 | 1.08735 | Intron (ENSMUST00000093113.4/ENSMUSG00000053399.8, intron 2 of 22) | 8 | 113697126 | 113848615 | 151490 | 2 | ENSMUST00000093113.4 | 1902 | Adamts18 | ENSMUSG00000053399.8 |
| chr8 | 113847976 | 113848886 | 3.76836 | 4.77895 | 1.66311 | Promoter | 8 | 113697126 | 113848615 | 151490 | 2 | ENSMUST00000093113.4 | 0 | Adamts18 | ENSMUSG00000053399.8 |
| chr8 | 113985444 | 113985650 | 3.67119 | 4.49212 | 1.42834 | Distal Intergenic | 8 | 113697126 | 113848615 | 151490 | 2 | ENSMUST00000093113.4 | -136830 | Adamts18 | ENSMUSG00000053399.8 |
| chr8 | 113990786 | 113991125 | 3.37005 | 3.88268 | 1.01733 | Distal Intergenic | 8 | 113697126 | 113848615 | 151490 | 2 | ENSMUST00000093113.4 | -142172 | Adamts18 | ENSMUSG00000053399.8 |
| chr15 | 79933807 | 79934736 | 3.47274 | 4.08785 | 1.17918 | Promoter | 15 | 79918148 | 79933899 | 15752 | 2 | ENSMUST00000146719.3 | 0 | Cbx7 | ENSMUSG00000053411.11 |
| chr15 | 79935396 | 79936474 | 3.70136 | 4.596 | 1.50183 | Intron (ENSMUST00000128931.2/ENSMUSG00000053411.11, intron 1 of 5) | 15 | 79918148 | 79933899 | 15752 | 2 | ENSMUST00000146719.3 | -1498 | Cbx7 | ENSMUSG00000053411.11 |
| chr16 | 90335621 | 90336046 | 3.76467 | 4.71938 | 1.64177 | Distal Intergenic | 16 | 90386397 | 90499553 | 113157 | 1 | ENSMUST00000065856.6 | -50351 | Hunk | ENSMUSG00000053414.7 |
| chr16 | 90385812 | 90386079 | 3.42004 | 4.10975 | 1.20932 | Promoter | 16 | 90386397 | 90499553 | 113157 | 1 | ENSMUST00000065856.6 | -318 | Hunk | ENSMUSG00000053414.7 |
| chr10 | 94249690 | 94250144 | 3.34655 | 3.85842 | 1.05667 | Distal Intergenic | 10 | 94297024 | 94298607 | 1584 | 2 | ENSMUST00000180826.1 | 48463 | Gm4792 | ENSMUSG00000053420.5 |
| chr18 | 58836027 | 58836456 | 3.75828 | 4.67757 | 1.55207 | Promoter | 18 | 58836764 | 59053678 | 216915 | 1 | ENSMUST00000052907.5 | -308 | Adamts19 | ENSMUSG00000053441.4 |
| chr18 | 58837428 | 58838145 | 3.89796 | 4.90306 | 1.68117 | Promoter | 18 | 58836764 | 59053678 | 216915 | 1 | ENSMUST00000052907.5 | 665 | Adamts19 | ENSMUSG00000053441.4 |
| chr1 | 183442534 | 183442711 | 3.23338 | 3.67724 | 1.01733 | Distal Intergenic | 1 | 183418503 | 183437053 | 18551 | 1 | ENSMUST00000065900.4 | 24032 | Hhipl2 | ENSMUSG00000053461.4 |
| chr14 | 119136895 | 119137177 | 3.49627 | 4.16971 | 1.24912 | Distal Intergenic | 14 | 119138415 | 119869594 | 731180 | 1 | ENSMUST00000065904.3 | -1238 | Hs6st3 | ENSMUSG00000053465.5 |
| chr14 | 119137586 | 119138567 | 4.43649 | 6.05442 | 2.58404 | Promoter | 14 | 119138415 | 119869594 | 731180 | 1 | ENSMUST00000065904.3 | 0 | Hs6st3 | ENSMUSG00000053465.5 |
| chr14 | 119138977 | 119139377 | 3.87791 | 5.08749 | 1.82398 | Promoter | 14 | 119138415 | 119869594 | 731180 | 1 | ENSMUST00000065904.3 | 563 | Hs6st3 | ENSMUSG00000053465.5 |
| chr14 | 119140288 | 119140395 | 3.34335 | 3.92598 | 1.14062 | Intron (ENSMUST00000065904.3/ENSMUSG00000053465.5, intron 1 of 1) | 14 | 119138415 | 119869594 | 731180 | 1 | ENSMUST00000065904.3 | 1874 | Hs6st3 | ENSMUSG00000053465.5 |
| chr14 | 119141056 | 119141176 | 3.54466 | 4.2927 | 1.3312 | Intron (ENSMUST00000065904.3/ENSMUSG00000053465.5, intron 1 of 1) | 14 | 119138415 | 119869594 | 731180 | 1 | ENSMUST00000065904.3 | 2642 | Hs6st3 | ENSMUSG00000053465.5 |
| chr2 | 52037919 | 52038023 | 4.33571 | 5.76089 | 2.2741 | Promoter | 2 | 52038009 | 52056686 | 18678 | 1 | ENSMUST00000065927.5 | 0 | Tnfaip6 | ENSMUSG00000053475.5 |
| chr2 | 52038752 | 52038990 | 3.55844 | 4.42019 | 1.34849 | Promoter | 2 | 52038009 | 52056686 | 18678 | 1 | ENSMUST00000065927.5 | 744 | Tnfaip6 | ENSMUSG00000053475.5 |
| chr2 | 52039685 | 52039808 | 3.6042 | 4.69164 | 1.55697 | Intron (ENSMUST00000065927.5/ENSMUSG00000053475.5, intron 1 of 5) | 2 | 52038009 | 52056686 | 18678 | 1 | ENSMUST00000065927.5 | 1677 | Tnfaip6 | ENSMUSG00000053475.5 |
| chr11 | 33779628 | 33779932 | 3.36084 | 4.06344 | 1.18289 | Intron (ENSMUST00000101368.3/ENSMUSG00000053519.10, intron 1 of 7) | 11 | 33629339 | 33843461 | 214123 | 2 | ENSMUST00000101368.3 | 63529 | Kcnip1 | ENSMUSG00000053519.10 |
| chr11 | 33782586 | 33782661 | 3.423 | 3.99848 | 1.14484 | Intron (ENSMUST00000101368.3/ENSMUSG00000053519.10, intron 1 of 7) | 11 | 33629339 | 33843461 | 214123 | 2 | ENSMUST00000101368.3 | 60800 | Kcnip1 | ENSMUSG00000053519.10 |
| chr11 | 33794233 | 33794391 | 3.65651 | 4.41061 | 1.35549 | Intron (ENSMUST00000101368.3/ENSMUSG00000053519.10, intron 1 of 7) | 11 | 33629339 | 33843461 | 214123 | 2 | ENSMUST00000101368.3 | 49070 | Kcnip1 | ENSMUSG00000053519.10 |
| chr11 | 33797156 | 33797539 | 3.36842 | 3.90798 | 1.11552 | Intron (ENSMUST00000101368.3/ENSMUSG00000053519.10, intron 1 of 7) | 11 | 33629339 | 33843461 | 214123 | 2 | ENSMUST00000101368.3 | 45922 | Kcnip1 | ENSMUSG00000053519.10 |
| chr11 | 33842768 | 33843798 | 3.68321 | 4.48947 | 1.4086 | Promoter | 11 | 33629339 | 33843461 | 214123 | 2 | ENSMUST00000101368.3 | 0 | Kcnip1 | ENSMUSG00000053519.10 |
| chr11 | 33849692 | 33849769 | 3.61674 | 4.37775 | 1.42249 | Intron (ENSMUST00000154760.1/ENSMUSG00000053519.10, intron 1 of 2) | 11 | 33629790 | 33843585 | 213796 | 2 | ENSMUST00000109340.3 | -6108 | Kcnip1 | ENSMUSG00000053519.10 |
| chr11 | 33992241 | 33992525 | 3.64991 | 4.62305 | 1.50293 | Promoter | 11 | 33641504 | 33992988 | 351485 | 2 | ENSMUST00000135034.1 | 463 | Kcnip1 | ENSMUSG00000053519.10 |
| chr11 | 33992983 | 33993226 | 3.28145 | 3.78579 | 1.0446 | Promoter | 11 | 33641180 | 33993152 | 351973 | 2 | ENSMUST00000154760.1 | 0 | Kcnip1 | ENSMUSG00000053519.10 |
| chr2 | 147190168 | 147190307 | 3.5894 | 4.3216 | 1.37019 | Intron (ENSMUST00000136998.2/ENSMUSG00000086509.2, intron 1 of 5) | 2 | 147187424 | 147188407 | 984 | 1 | ENSMUST00000144411.1 | 2745 | 6430503K07Rik | ENSMUSG00000053545.6 |
| chr2 | 147187463 | 147188392 | 3.50803 | 4.25884 | 1.3212 | Promoter | 2 | 147187424 | 147188407 | 984 | 1 | ENSMUST00000144411.1 | 40 | 6430503K07Rik | ENSMUSG00000053545.6 |
| chr2 | 147189262 | 147189376 | 3.47296 | 4.10451 | 1.23801 | Intron (ENSMUST00000136998.2/ENSMUSG00000086509.2, intron 1 of 5) | 2 | 147187424 | 147188407 | 984 | 1 | ENSMUST00000144411.1 | 1839 | 6430503K07Rik | ENSMUSG00000053545.6 |
| chr7 | 4829782 | 4830416 | 3.58748 | 4.328 | 1.39153 | Promoter | 7 | 4830177 | 4831127 | 951 | 2 | ENSMUST00000157068.1 | 711 | Shisa7 | ENSMUSG00000053550.8 |
| chr7 | 4843116 | 4843209 | 3.65411 | 4.54817 | 1.51144 | Intron (ENSMUST00000066041.6/ENSMUSG00000053550.8, intron 1 of 5) | 7 | 4825553 | 4844696 | 19144 | 2 | ENSMUST00000066041.6 | 1487 | Shisa7 | ENSMUSG00000053550.8 |
| chr7 | 4844155 | 4844392 | 3.45469 | 4.22018 | 1.29765 | Promoter | 7 | 4825553 | 4844696 | 19144 | 2 | ENSMUST00000066041.6 | 304 | Shisa7 | ENSMUSG00000053550.8 |
| chr2 | 130294708 | 130294929 | 3.79901 | 4.97553 | 1.74588 | Promoter | 2 | 130295169 | 130370481 | 75313 | 1 | ENSMUST00000110288.3 | -240 | Ebf4 | ENSMUSG00000053552.9 |
| chr2 | 130295275 | 130296149 | 4.38983 | 6.16512 | 2.66929 | Promoter | 2 | 130295939 | 130370481 | 74543 | 1 | ENSMUST00000110286.2 | 0 | Ebf4 | ENSMUSG00000053552.9 |
| chr5 | 139495111 | 139495429 | 3.62753 | 4.4243 | 1.43528 | Distal Intergenic | 5 | 139471211 | 139484549 | 13339 | 2 | ENSMUST00000079996.7 | -10563 | Zfand2a | ENSMUSG00000053581.8 |
| chr5 | 139496198 | 139496500 | 4.20432 | 5.87512 | 2.45375 | Distal Intergenic | 5 | 139471211 | 139484549 | 13339 | 2 | ENSMUST00000079996.7 | -11650 | Zfand2a | ENSMUSG00000053581.8 |
| chr5 | 139497039 | 139497192 | 3.38193 | 3.9006 | 1.01733 | Distal Intergenic | 5 | 139471211 | 139484549 | 13339 | 2 | ENSMUST00000079996.7 | -12491 | Zfand2a | ENSMUSG00000053581.8 |
| chr5 | 139497780 | 139498247 | 3.5484 | 4.30589 | 1.27384 | Distal Intergenic | 5 | 139471211 | 139484549 | 13339 | 2 | ENSMUST00000079996.7 | -13232 | Zfand2a | ENSMUSG00000053581.8 |
| chr5 | 139498896 | 139498980 | 3.91335 | 4.88594 | 1.64183 | Distal Intergenic | 5 | 139471211 | 139484549 | 13339 | 2 | ENSMUST00000079996.7 | -14348 | Zfand2a | ENSMUSG00000053581.8 |
| chr5 | 139510987 | 139511062 | 4.07565 | 5.43723 | 2.04649 | Distal Intergenic | 5 | 139471211 | 139484549 | 13339 | 2 | ENSMUST00000079996.7 | -26439 | Zfand2a | ENSMUSG00000053581.8 |
| chr5 | 139512111 | 139512537 | 3.75886 | 4.6132 | 1.50129 | Distal Intergenic | 5 | 139471211 | 139484549 | 13339 | 2 | ENSMUST00000079996.7 | -27563 | Zfand2a | ENSMUSG00000053581.8 |
| chr11 | 120659273 | 120660390 | 3.82986 | 5.19509 | 1.90835 | Promoter | 11 | 120659157 | 120661815 | 2659 | 1 | ENSMUST00000151998.1 | 117 | Notumos | ENSMUSG00000053613.6 |
| chr2 | 125504677 | 125504752 | 3.49154 | 4.10217 | 1.14484 | Promoter | 2 | 125505089 | 125507881 | 2793 | 1 | ENSMUST00000066157.1 | -337 | Gm9913 | ENSMUSG00000053615.1 |
| chr19 | 47443925 | 47444154 | 3.28669 | 3.75724 | 1.01733 | Intron (ENSMUST00000081619.3/ENSMUSG00000053617.6, intron 1 of 14) | 19 | 47260174 | 47464411 | 204238 | 2 | ENSMUST00000081619.3 | 20257 | Sh3pxd2a | ENSMUSG00000053617.6 |
| chr19 | 47458031 | 47458151 | 3.30531 | 3.78516 | 1.01733 | Intron (ENSMUST00000081619.3/ENSMUSG00000053617.6, intron 1 of 14) | 19 | 47260174 | 47464411 | 204238 | 2 | ENSMUST00000081619.3 | 6260 | Sh3pxd2a | ENSMUSG00000053617.6 |
| chr19 | 47461892 | 47462449 | 3.8484 | 4.85476 | 1.69045 | Intron (ENSMUST00000081619.3/ENSMUSG00000053617.6, intron 1 of 14) | 19 | 47260174 | 47464411 | 204238 | 2 | ENSMUST00000081619.3 | 1962 | Sh3pxd2a | ENSMUSG00000053617.6 |
| chr19 | 47462793 | 47463275 | 3.53523 | 4.28722 | 1.29484 | Intron (ENSMUST00000081619.3/ENSMUSG00000053617.6, intron 1 of 14) | 19 | 47260174 | 47464411 | 204238 | 2 | ENSMUST00000081619.3 | 1136 | Sh3pxd2a | ENSMUSG00000053617.6 |
| chr19 | 47463888 | 47464564 | 4.08297 | 5.42925 | 2.1132 | Promoter | 19 | 47260174 | 47464411 | 204238 | 2 | ENSMUST00000081619.3 | 0 | Sh3pxd2a | ENSMUSG00000053617.6 |
| chr19 | 47465734 | 47465911 | 3.31195 | 3.79515 | 1.01733 | Distal Intergenic | 19 | 47260174 | 47464411 | 204238 | 2 | ENSMUST00000111800.2 | -1324 | Sh3pxd2a | ENSMUSG00000053617.6 |
| chr8 | 64204104 | 64204529 | 3.36787 | 3.92525 | 1.16256 | Intron (ENSMUST00000066166.4/ENSMUSG00000053626.4, intron 1 of 20) | 8 | 64014770 | 64205993 | 191224 | 2 | ENSMUST00000066166.4 | 1464 | Tll1 | ENSMUSG00000053626.4 |
| chr8 | 64205129 | 64205898 | 3.92207 | 5.1963 | 1.91533 | Promoter | 8 | 64014770 | 64205993 | 191224 | 2 | ENSMUST00000066166.4 | 95 | Tll1 | ENSMUSG00000053626.4 |
| chr8 | 84923867 | 84924021 | 3.46984 | 4.17831 | 1.25714 | Promoter | 8 | 84911906 | 84924114 | 12209 | 2 | ENSMUST00000153000.1 | 93 | Mast1 | ENSMUSG00000053693.11 |
| chr8 | 84929247 | 84929467 | 3.41276 | 3.96169 | 1.06853 | Promoter | 8 | 84923729 | 84928775 | 5047 | 2 | ENSMUST00000130923.1 | -473 | Mast1 | ENSMUSG00000053693.11 |
| chr8 | 84930259 | 84930439 | 3.45306 | 4.21156 | 1.26197 | Intron (ENSMUST00000109741.3/ENSMUSG00000053693.11, intron 4 of 25) | 8 | 84923729 | 84928775 | 5047 | 2 | ENSMUST00000130923.1 | -1485 | Mast1 | ENSMUSG00000053693.11 |
| chr8 | 84931395 | 84932974 | 3.52803 | 4.20418 | 1.22609 | Intron (ENSMUST00000109741.3/ENSMUSG00000053693.11, intron 3 of 25) | 8 | 84923729 | 84928775 | 5047 | 2 | ENSMUST00000130923.1 | -2621 | Mast1 | ENSMUSG00000053693.11 |
| chr8 | 84934447 | 84934537 | 4.04518 | 5.2258 | 1.91264 | Intron (ENSMUST00000109741.3/ENSMUSG00000053693.11, intron 3 of 25) | 8 | 84934924 | 84937344 | 2421 | 2 | ENSMUST00000128356.1 | 2807 | Mast1 | ENSMUSG00000053693.11 |
| chr8 | 84934992 | 84935702 | 3.76762 | 4.75543 | 1.59938 | Exon (ENSMUST00000109741.3/ENSMUSG00000053693.11, exon 3 of 26) | 8 | 84934924 | 84937344 | 2421 | 2 | ENSMUST00000128356.1 | 1642 | Mast1 | ENSMUSG00000053693.11 |
| chr8 | 84936171 | 84937765 | 4.14513 | 5.41468 | 2.06193 | Promoter | 8 | 84911903 | 84937353 | 25451 | 2 | ENSMUST00000109741.3 | 0 | Mast1 | ENSMUSG00000053693.11 |
| chr8 | 84938163 | 84938715 | 3.5646 | 4.24595 | 1.25682 | Promoter | 8 | 84911908 | 84937359 | 25452 | 2 | ENSMUST00000119820.1 | -805 | Mast1 | ENSMUSG00000053693.11 |
| chr9 | 99814085 | 99814172 | 3.96157 | 5.3172 | 1.95883 | Distal Intergenic | 9 | 99874220 | 99875655 | 1436 | 2 | ENSMUST00000183065.1 | 61483 | Sox14 | ENSMUSG00000053747.9 |
| chr9 | 99862975 | 99863220 | 3.25963 | 3.71664 | 1.01733 | Distal Intergenic | 9 | 99874220 | 99875655 | 1436 | 2 | ENSMUST00000183065.1 | 12435 | Sox14 | ENSMUSG00000053747.9 |
| chr9 | 99865156 | 99865326 | 3.75868 | 4.73566 | 1.60418 | Distal Intergenic | 9 | 99874220 | 99875655 | 1436 | 2 | ENSMUST00000183065.1 | 10329 | Sox14 | ENSMUSG00000053747.9 |
| chr9 | 99866534 | 99867278 | 3.41258 | 4.09255 | 1.18417 | Distal Intergenic | 9 | 99874220 | 99875655 | 1436 | 2 | ENSMUST00000183065.1 | 8377 | Sox14 | ENSMUSG00000053747.9 |
| chr9 | 99867605 | 99867771 | 3.93686 | 5.01394 | 1.79978 | Distal Intergenic | 9 | 99874220 | 99875655 | 1436 | 2 | ENSMUST00000183065.1 | 7884 | Sox14 | ENSMUSG00000053747.9 |
| chr9 | 99868357 | 99869465 | 3.90389 | 5.16933 | 1.93825 | Distal Intergenic | 9 | 99874220 | 99875655 | 1436 | 2 | ENSMUST00000183065.1 | 6190 | Sox14 | ENSMUSG00000053747.9 |
| chr9 | 99869800 | 99871034 | 3.67507 | 4.49421 | 1.43581 | Distal Intergenic | 9 | 99874220 | 99875655 | 1436 | 2 | ENSMUST00000183065.1 | 4621 | Sox14 | ENSMUSG00000053747.9 |
| chr9 | 99871490 | 99871785 | 5.83626 | 9.01875 | 4.94623 | Downstream (2-3kb) | 9 | 99874220 | 99875655 | 1436 | 2 | ENSMUST00000183065.1 | 3870 | Sox14 | ENSMUSG00000053747.9 |
| chr9 | 99872093 | 99872445 | 3.73044 | 4.74212 | 1.60098 | Downstream (1-2kb) | 9 | 99874220 | 99875655 | 1436 | 2 | ENSMUST00000183065.1 | 3210 | Sox14 | ENSMUSG00000053747.9 |
| chr9 | 99873194 | 99873269 | 4.08828 | 5.41971 | 2.04469 | Downstream (<1kb) | 9 | 99874220 | 99875655 | 1436 | 2 | ENSMUST00000183065.1 | 2386 | Sox14 | ENSMUSG00000053747.9 |
| chr9 | 99873686 | 99874177 | 3.74883 | 4.60633 | 1.51874 | 3' UTR | 9 | 99874220 | 99875655 | 1436 | 2 | ENSMUST00000183065.1 | 1478 | Sox14 | ENSMUSG00000053747.9 |
| chr9 | 99874521 | 99874754 | 3.63869 | 4.38164 | 1.3379 | Promoter | 9 | 99874220 | 99875655 | 1436 | 2 | ENSMUST00000183065.1 | 901 | Sox14 | ENSMUSG00000053747.9 |
| chr9 | 99875164 | 99876354 | 4.60642 | 6.44227 | 2.88555 | Promoter | 9 | 99874106 | 99876170 | 2065 | 2 | ENSMUST00000054819.9 | 0 | Sox14 | ENSMUSG00000053747.9 |
| chr9 | 99876735 | 99880154 | 3.9431 | 5.03484 | 1.80941 | Promoter | 9 | 99874106 | 99876170 | 2065 | 2 | ENSMUST00000054819.9 | -566 | Sox14 | ENSMUSG00000053747.9 |
| chr6 | 32710412 | 32710517 | 3.40294 | 3.9389 | 1.04054 | Distal Intergenic | 6 | 32792654 | 32803992 | 11339 | 2 | ENSMUST00000129654.1 | 93475 | Chchd3 | ENSMUSG00000053768.8 |
| chr6 | 32710966 | 32711050 | 3.84623 | 5.07435 | 1.80108 | Distal Intergenic | 6 | 32792654 | 32803992 | 11339 | 2 | ENSMUST00000129654.1 | 92942 | Chchd3 | ENSMUSG00000053768.8 |
| chr11 | 75994576 | 75994781 | 3.53743 | 4.38025 | 1.35 | Distal Intergenic | 11 | 75999912 | 76003569 | 3658 | 1 | ENSMUST00000066408.5 | -5131 | 1700016K19Rik | ENSMUSG00000053783.5 |
| chr7 | 45832595 | 45834349 | 3.93014 | 5.25343 | 1.95709 | 3' UTR | 7 | 45825232 | 45830944 | 5713 | 2 | ENSMUST00000131384.1 | -1652 | Grwd1 | ENSMUSG00000053801.11 |
| chr10 | 106468055 | 106468339 | 3.89207 | 4.85412 | 1.62919 | Exon (ENSMUST00000181196.1/ENSMUSG00000097869.1, exon 2 of 2) | 10 | 106470310 | 106933467 | 463158 | 1 | ENSMUST00000029404.10 | -1971 | Ppfia2 | ENSMUSG00000053825.9 |
| chr10 | 106469907 | 106470290 | 3.91784 | 4.96602 | 1.79996 | Promoter | 10 | 106470310 | 106933467 | 463158 | 1 | ENSMUST00000029404.10 | -20 | Ppfia2 | ENSMUSG00000053825.9 |
| chr18 | 74959270 | 74959488 | 3.93283 | 5.03796 | 1.82015 | Intron (ENSMUST00000066532.4/ENSMUSG00000053846.4, intron 1 of 9) | 18 | 74939322 | 74961263 | 21942 | 2 | ENSMUST00000066532.4 | 1775 | Lipg | ENSMUSG00000053846.4 |
| chr18 | 74960677 | 74962749 | 3.83186 | 4.85163 | 1.66369 | Promoter | 18 | 74939322 | 74961263 | 21942 | 2 | ENSMUST00000066532.4 | 0 | Lipg | ENSMUSG00000053846.4 |
| chr18 | 74963064 | 74963918 | 3.34635 | 3.86494 | 1.08089 | Distal Intergenic | 18 | 74939322 | 74961263 | 21942 | 2 | ENSMUST00000066532.4 | -1802 | Lipg | ENSMUSG00000053846.4 |
| chr18 | 74968048 | 74968196 | 3.60084 | 4.37002 | 1.43722 | Distal Intergenic | 18 | 74939322 | 74961263 | 21942 | 2 | ENSMUST00000066532.4 | -6786 | Lipg | ENSMUSG00000053846.4 |
| chr5 | 31105668 | 31105955 | 3.25027 | 3.70258 | 1.01733 | Promoter | 5 | 31106285 | 31111984 | 5700 | 1 | ENSMUST00000190241.1 | -330 | Dnajc5g | ENSMUSG00000053856.2 |
| chr5 | 113335099 | 113335534 | 4.40871 | 6.23772 | 2.67487 | Distal Intergenic | 5 | 113357294 | 113357821 | 528 | 1 | ENSMUST00000066573.1 | -21760 | Aym1 | ENSMUSG00000053873.1 |
| chr5 | 113336136 | 113336314 | 3.44364 | 4.07029 | 1.24595 | Distal Intergenic | 5 | 113357294 | 113357821 | 528 | 1 | ENSMUST00000066573.1 | -20980 | Aym1 | ENSMUSG00000053873.1 |
| chr5 | 113336672 | 113336888 | 3.33439 | 3.86925 | 1.13503 | Distal Intergenic | 5 | 113357294 | 113357821 | 528 | 1 | ENSMUST00000066573.1 | -20406 | Aym1 | ENSMUSG00000053873.1 |
| chr5 | 113339742 | 113339817 | 3.63078 | 4.38431 | 1.38586 | Distal Intergenic | 5 | 113357294 | 113357821 | 528 | 1 | ENSMUST00000066573.1 | -17477 | Aym1 | ENSMUSG00000053873.1 |
| chr5 | 113346786 | 113347113 | 3.74166 | 4.61549 | 1.544 | Distal Intergenic | 5 | 113357294 | 113357821 | 528 | 1 | ENSMUST00000066573.1 | -10181 | Aym1 | ENSMUSG00000053873.1 |
| chr5 | 113347458 | 113347680 | 3.94732 | 4.95978 | 1.69908 | Distal Intergenic | 5 | 113357294 | 113357821 | 528 | 1 | ENSMUST00000066573.1 | -9614 | Aym1 | ENSMUSG00000053873.1 |
| chr5 | 113354241 | 113354702 | 3.58384 | 4.43353 | 1.39061 | Distal Intergenic | 5 | 113357294 | 113357821 | 528 | 1 | ENSMUST00000066573.1 | -2592 | Aym1 | ENSMUSG00000053873.1 |
| chr5 | 113364483 | 113364741 | 3.65281 | 4.4051 | 1.35617 | Distal Intergenic | 5 | 113357294 | 113357821 | 528 | 1 | ENSMUST00000066573.1 | 7190 | Aym1 | ENSMUSG00000053873.1 |
| chr5 | 113372067 | 113372821 | 3.64152 | 4.42247 | 1.36634 | Distal Intergenic | 5 | 113357294 | 113357821 | 528 | 1 | ENSMUST00000066573.1 | 14774 | Aym1 | ENSMUSG00000053873.1 |
| chr5 | 113373727 | 113374026 | 3.4715 | 4.08128 | 1.16604 | Distal Intergenic | 5 | 113357294 | 113357821 | 528 | 1 | ENSMUST00000066573.1 | 16434 | Aym1 | ENSMUSG00000053873.1 |
| chr5 | 113397736 | 113398292 | 3.45595 | 4.06018 | 1.15155 | Distal Intergenic | 5 | 113357294 | 113357821 | 528 | 1 | ENSMUST00000066573.1 | 40443 | Aym1 | ENSMUSG00000053873.1 |
| chr5 | 113403525 | 113403620 | 3.51493 | 4.16567 | 1.2157 | Distal Intergenic | 5 | 113357294 | 113357821 | 528 | 1 | ENSMUST00000066573.1 | 46232 | Aym1 | ENSMUSG00000053873.1 |
| chr3 | 135826289 | 135826511 | 4.05364 | 5.30804 | 1.98159 | Promoter | 3 | 135826090 | 135888386 | 62297 | 1 | ENSMUST00000029810.5 | 200 | Slc39a8 | ENSMUSG00000053897.10 |
| chr11 | 66508004 | 66508244 | 3.65217 | 4.47673 | 1.4791 | Intron (ENSMUST00000123454.2/ENSMUSG00000053930.8, intron 1 of 5) | 11 | 66375133 | 66502814 | 127682 | 2 | ENSMUST00000134562.1 | -5191 | Shisa6 | ENSMUSG00000053930.8 |
| chr11 | 66522554 | 66522976 | 3.47687 | 4.1327 | 1.19716 | Intron (ENSMUST00000123454.2/ENSMUSG00000053930.8, intron 1 of 5) | 11 | 66211725 | 66525795 | 314071 | 2 | ENSMUST00000123454.2 | 2819 | Shisa6 | ENSMUSG00000053930.8 |
| chr11 | 66523394 | 66523744 | 3.55259 | 4.21582 | 1.22022 | Intron (ENSMUST00000123454.2/ENSMUSG00000053930.8, intron 1 of 5) | 11 | 66211725 | 66525795 | 314071 | 2 | ENSMUST00000123454.2 | 2051 | Shisa6 | ENSMUSG00000053930.8 |
| chr11 | 66524728 | 66526751 | 3.72844 | 4.66857 | 1.57602 | Promoter | 11 | 66211725 | 66525795 | 314071 | 2 | ENSMUST00000123454.2 | 0 | Shisa6 | ENSMUSG00000053930.8 |
| chr11 | 66527693 | 66528386 | 3.92419 | 4.95442 | 1.75694 | Distal Intergenic | 11 | 66217472 | 66525964 | 308493 | 2 | ENSMUST00000066679.6 | -1730 | Shisa6 | ENSMUSG00000053930.8 |
| chr1 | 180407568 | 180408210 | 3.58763 | 4.28088 | 1.25958 | Intron (ENSMUST00000070181.6/ENSMUSG00000038855.9, intron 2 of 7) | 1 | 180432387 | 180483467 | 51081 | 2 | ENSMUST00000136521.1 | 75257 | 6330403A02Rik | ENSMUSG00000053963.6 |
| chr1 | 180408635 | 180410213 | 3.57247 | 4.30004 | 1.34314 | Intron (ENSMUST00000070181.6/ENSMUSG00000038855.9, intron 2 of 7) | 1 | 180432387 | 180483467 | 51081 | 2 | ENSMUST00000136521.1 | 73254 | 6330403A02Rik | ENSMUSG00000053963.6 |
| chr1 | 180416203 | 180416278 | 3.96572 | 4.99159 | 1.71528 | Intron (ENSMUST00000070181.6/ENSMUSG00000038855.9, intron 4 of 7) | 1 | 180432387 | 180483467 | 51081 | 2 | ENSMUST00000136521.1 | 67189 | 6330403A02Rik | ENSMUSG00000053963.6 |
| chr1 | 180417817 | 180417936 | 3.50664 | 4.13468 | 1.17942 | Intron (ENSMUST00000070181.6/ENSMUSG00000038855.9, intron 4 of 7) | 1 | 180432387 | 180483467 | 51081 | 2 | ENSMUST00000136521.1 | 65531 | 6330403A02Rik | ENSMUSG00000053963.6 |
| chr1 | 180420089 | 180420301 | 3.5365 | 4.28035 | 1.31666 | Exon (ENSMUST00000070181.6/ENSMUSG00000038855.9, exon 6 of 8) | 1 | 180432387 | 180483467 | 51081 | 2 | ENSMUST00000136521.1 | 63166 | 6330403A02Rik | ENSMUSG00000053963.6 |
| chr1 | 180421715 | 180421998 | 3.8145 | 4.79676 | 1.65454 | 3' UTR | 1 | 180432387 | 180483467 | 51081 | 2 | ENSMUST00000136521.1 | 61469 | 6330403A02Rik | ENSMUSG00000053963.6 |
| chr1 | 180431651 | 180432356 | 3.52734 | 4.1901 | 1.23152 | Downstream (<1kb) | 1 | 180432387 | 180483467 | 51081 | 2 | ENSMUST00000136521.1 | 51111 | 6330403A02Rik | ENSMUSG00000053963.6 |
| chr1 | 180433317 | 180433450 | 3.60557 | 4.31584 | 1.28612 | 3' UTR | 1 | 180432387 | 180483467 | 51081 | 2 | ENSMUST00000136521.1 | 50017 | 6330403A02Rik | ENSMUSG00000053963.6 |
| chr1 | 180434743 | 180435053 | 3.62868 | 4.38257 | 1.39317 | 3' UTR | 1 | 180432387 | 180483467 | 51081 | 2 | ENSMUST00000136521.1 | 48414 | 6330403A02Rik | ENSMUSG00000053963.6 |
| chr1 | 180482986 | 180483729 | 4.29361 | 5.67237 | 2.22315 | Promoter | 1 | 180432387 | 180483467 | 51081 | 2 | ENSMUST00000136521.1 | 0 | 6330403A02Rik | ENSMUSG00000053963.6 |
| chr15 | 98495016 | 98495189 | 3.35208 | 3.85555 | 1.01733 | Distal Intergenic | 15 | 98497474 | 98498406 | 933 | 1 | ENSMUST00000050451.5 | -2285 | Olfr279 | ENSMUSG00000054036.4 |
| chr7 | 93096446 | 93096613 | 3.32727 | 3.9634 | 1.11882 | Intron (ENSMUST00000181886.1/ENSMUSG00000097644.1, intron 3 of 3) | 7 | 93052081 | 93081027 | 28947 | 2 | ENSMUST00000098303.1 | -15420 | Gm9934 | ENSMUSG00000054061.8 |
| chr7 | 141069773 | 141070013 | 3.23391 | 3.69266 | 1.03027 | Intron (ENSMUST00000048002.5/ENSMUSG00000055629.4, intron 14 of 19) | 7 | 141078218 | 141090510 | 12293 | 1 | ENSMUST00000106039.3 | -8205 | Pkp3 | ENSMUSG00000054065.6 |

| | | | | | | | | | | | | | | | |
|---|---|---|---|---|---|---|---|---|---|---|---|---|---|---|---|
| chr7 | 141070759 | 141071362 | 3.4891 | 4.21376 | 1.293 | Exon (ENSMUST00000048002.5/ENSMUSG00000055629.4, exon 16 of 20) | 7 | 141078218 | 141090510 | 12293 | 1 | ENSMUST00000106039.3 | -6856 | Pkp3 | ENSMUSG00000054065.6 |
| chr7 | 141071993 | 141072307 | 3.81973 | 4.79215 | 1.66678 | Exon (ENSMUST00000048002.5/ENSMUSG00000055629.4, exon 20 of 20) | 7 | 141078218 | 141090510 | 12293 | 1 | ENSMUST00000106039.3 | -5911 | Pkp3 | ENSMUSG00000054065.6 |
| chr7 | 141082605 | 141083054 | 3.45214 | 4.02991 | 1.09887 | 3' UTR | 7 | 141079780 | 141082695 | 2916 | 1 | ENSMUST00000160869.1 | 2826 | Pkp3 | ENSMUSG00000054065.6 |
| chr11 | 93816762 | 93817099 | 3.49002 | 4.11726 | 1.20643 | Distal Intergenic | 11 | 93875383 | 93876143 | 761 | 2 | ENSMUST00000134807.1 | 59044 | Utp18 | ENSMUSG00000054079.7 |
| chr11 | 93817633 | 93817966 | 3.33527 | 3.87976 | 1.03415 | Distal Intergenic | 11 | 93875383 | 93876143 | 761 | 2 | ENSMUST00000134807.1 | 58177 | Utp18 | ENSMUSG00000054079.7 |
| chr15 | 88290118 | 88290386 | 3.46393 | 4.06299 | 1.15423 | Distal Intergenic | 15 | 88600302 | 88601669 | 1368 | 1 | ENSMUST00000066949.8 | -309916 | Zdhhc25 | ENSMUSG00000054117.8 |
| chr15 | 88291295 | 88291391 | 3.4448 | 4.03028 | 1.14054 | Distal Intergenic | 15 | 88600302 | 88601669 | 1368 | 1 | ENSMUST00000066949.8 | -308911 | Zdhhc25 | ENSMUSG00000054117.8 |
| chr15 | 88292038 | 88292277 | 3.50085 | 4.11798 | 1.15081 | Distal Intergenic | 15 | 88600302 | 88601669 | 1368 | 1 | ENSMUST00000066949.8 | -308025 | Zdhhc25 | ENSMUSG00000054117.8 |
| chr15 | 88560883 | 88561118 | 3.32996 | 3.82223 | 1.01733 | Distal Intergenic | 15 | 88600302 | 88601669 | 1368 | 1 | ENSMUST00000066949.8 | -39184 | Zdhhc25 | ENSMUSG00000054117.8 |
| chr17 | 36188935 | 36189353 | 3.85579 | 4.92512 | 1.74307 | Promoter | 17 | 36186576 | 36190122 | 3547 | 2 | ENSMUST00000172663.2 | 769 | H2-T3 | ENSMUSG00000054128.11 |
| chr17 | 36190292 | 36190873 | 3.46084 | 4.09921 | 1.2605 | Promoter | 17 | 36185573 | 36190287 | 4715 | 2 | ENSMUST00000102675.4 | -6 | H2-T3 | ENSMUSG00000054128.11 |
| chr17 | 36192048 | 36192406 | 3.40797 | 4.00519 | 1.20944 | Downstream (2-3kb) | 17 | 36185573 | 36190287 | 4715 | 2 | ENSMUST00000102675.4 | -1762 | H2-T3 | ENSMUSG00000054128.11 |
| chr15 | 89323226 | 89323338 | 3.60347 | 4.52765 | 1.44757 | Promoter | 15 | 89322972 | 89324728 | 1757 | 1 | ENSMUST00000066991.5 | 255 | Adm2 | ENSMUSG00000054136.5 |
| chr11 | 100136231 | 100136390 | 3.22908 | 3.91296 | 1.10994 | Promoter | 11 | 100131758 | 100135928 | 4171 | 2 | ENSMUST00000107411.2 | -304 | Krt15 | ENSMUSG00000054146.8 |
| chr11 | 100139534 | 100139618 | 3.34848 | 3.89866 | 1.14079 | Downstream (1-2kb) | 11 | 100131758 | 100135928 | 4171 | 2 | ENSMUST00000107411.2 | -3607 | Krt15 | ENSMUSG00000054146.8 |
| chr2 | 147080559 | 147080927 | 3.72344 | 4.64047 | 1.54564 | Downstream (2-3kb) | 2 | 147083316 | 147085445 | 2130 | 2 | ENSMUST00000067020.2 | 4518 | Nkx2-4 | ENSMUSG00000054160.2 |
| chr2 | 147081249 | 147082172 | 4.12819 | 5.52834 | 2.13979 | Downstream (1-2kb) | 2 | 147083316 | 147085445 | 2130 | 2 | ENSMUST00000067020.2 | 3273 | Nkx2-4 | ENSMUSG00000054160.2 |
| chr2 | 147083211 | 147083380 | 3.7227 | 4.92352 | 1.72816 | 3' UTR | 2 | 147083316 | 147085445 | 2130 | 2 | ENSMUST00000067020.2 | 2065 | Nkx2-4 | ENSMUSG00000054160.2 |
| chr2 | 147083938 | 147085532 | 4.06099 | 5.47283 | 2.12537 | Promoter | 2 | 147083316 | 147085445 | 2130 | 2 | ENSMUST00000067020.2 | 0 | Nkx2-4 | ENSMUSG00000054160.2 |
| chr2 | 147086747 | 147087052 | 3.52419 | 4.52809 | 1.46894 | Distal Intergenic | 2 | 147083316 | 147085445 | 2130 | 2 | ENSMUST00000067020.2 | -1303 | Nkx2-4 | ENSMUSG00000054160.2 |
| chr2 | 147087478 | 147087553 | 3.68509 | 4.73671 | 1.6231 | Distal Intergenic | 2 | 147083316 | 147085445 | 2130 | 2 | ENSMUST00000067020.2 | -2034 | Nkx2-4 | ENSMUSG00000054160.2 |
| chr2 | 147088084 | 147089592 | 3.8729 | 5.05161 | 1.8219 | Distal Intergenic | 2 | 147083316 | 147085445 | 2130 | 2 | ENSMUST00000067020.2 | -2640 | Nkx2-4 | ENSMUSG00000054160.2 |
| chr2 | 147089936 | 147090535 | 3.8637 | 5.01431 | 1.8028 | Distal Intergenic | 2 | 147083316 | 147085445 | 2130 | 2 | ENSMUST00000067020.2 | -4492 | Nkx2-4 | ENSMUSG00000054160.2 |
| chr2 | 147092900 | 147094108 | 3.46157 | 4.06382 | 1.14893 | Distal Intergenic | 2 | 147083316 | 147085445 | 2130 | 2 | ENSMUST00000067020.2 | -7456 | Nkx2-4 | ENSMUSG00000054160.2 |
| chr2 | 147097353 | 147098168 | 3.66459 | 4.47233 | 1.39694 | Distal Intergenic | 2 | 147083316 | 147085445 | 2130 | 2 | ENSMUST00000067020.2 | -11909 | Nkx2-4 | ENSMUSG00000054160.2 |
| chr2 | 147099531 | 147100765 | 3.9588 | 5.01341 | 1.77682 | Distal Intergenic | 2 | 147083316 | 147085445 | 2130 | 2 | ENSMUST00000067020.2 | -14087 | Nkx2-4 | ENSMUSG00000054160.2 |
| chr2 | 147102345 | 147102477 | 3.36842 | 3.88358 | 1.02919 | Distal Intergenic | 2 | 147083316 | 147085445 | 2130 | 2 | ENSMUST00000067020.2 | -16901 | Nkx2-4 | ENSMUSG00000054160.2 |
| chr2 | 147107016 | 147107251 | 3.50216 | 4.15333 | 1.2631 | Distal Intergenic | 2 | 147083316 | 147085445 | 2130 | 2 | ENSMUST00000067020.2 | -21572 | Nkx2-4 | ENSMUSG00000054160.2 |
| chr2 | 147108936 | 147109106 | 3.41199 | 3.97651 | 1.12578 | Distal Intergenic | 2 | 147083316 | 147085445 | 2130 | 2 | ENSMUST00000067020.2 | -23492 | Nkx2-4 | ENSMUSG00000054160.2 |
| chr8 | 62950593 | 62950763 | 3.35482 | 3.85972 | 1.01733 | Promoter | 8 | 62951009 | 63356852 | 405844 | 1 | ENSMUST00000117377.2 | -246 | Spock3 | ENSMUSG00000054162.10 |
| chr8 | 62951148 | 62951648 | 3.41046 | 3.96313 | 1.08622 | Promoter | 8 | 62951232 | 63356860 | 405629 | 1 | ENSMUST00000118003.2 | 0 | Spock3 | ENSMUSG00000054162.10 |
| chr15 | 39076868 | 39077637 | 3.67059 | 4.4906 | 1.46471 | Promoter | 15 | 39076932 | 39087119 | 10188 | 1 | ENSMUST00000067072.3 | 0 | Cthrc1 | ENSMUSG00000054196.6 |
| chr19 | 38096031 | 38096468 | 3.64877 | 4.40774 | 1.35148 | Promoter | 19 | 38097079 | 38114263 | 17185 | 1 | ENSMUST00000067098.6 | -611 | Ffar4 | ENSMUSG00000054200.6 |
| chr19 | 38096948 | 38097089 | 3.48489 | 4.09882 | 1.16808 | Promoter | 19 | 38097079 | 38114263 | 17185 | 1 | ENSMUST00000067098.6 | 0 | Ffar4 | ENSMUSG00000054200.6 |
| chr19 | 38097885 | 38097994 | 3.85636 | 4.94118 | 1.74066 | Promoter | 19 | 38097079 | 38114263 | 17185 | 1 | ENSMUST00000067098.6 | 807 | Ffar4 | ENSMUSG00000054200.6 |
| chr12 | 30882168 | 30882554 | 3.40779 | 3.98793 | 1.14901 | Distal Intergenic | 12 | 30884322 | 30893854 | 9533 | 1 | ENSMUST00000067087.6 | -1768 | Fam150b | ENSMUSG00000054204.6 |
| chr12 | 30882874 | 30883984 | 3.64746 | 4.4979 | 1.45971 | Promoter | 12 | 30884322 | 30893854 | 9533 | 1 | ENSMUST00000067087.6 | -338 | Fam150b | ENSMUSG00000054204.6 |
| chr12 | 30884413 | 30885005 | 4.20523 | 5.57735 | 2.1549 | Promoter | 12 | 30884322 | 30893854 | 9533 | 1 | ENSMUST00000067087.6 | 92 | Fam150b | ENSMUSG00000054204.6 |
| chr12 | 30885680 | 30885966 | 3.45459 | 4.0463 | 1.14501 | Intron (ENSMUST00000067087.6/ENSMUSG00000054204.6, intron 1 of 4) | 12 | 30884322 | 30893854 | 9533 | 1 | ENSMUST00000067087.6 | 1359 | Fam150b | ENSMUSG00000054204.6 |
| chr12 | 30887122 | 30887507 | 3.47261 | 4.08876 | 1.18846 | Exon (ENSMUST00000067087.6/ENSMUSG00000054204.6, exon 3 of 5) | 12 | 30884322 | 30893854 | 9533 | 1 | ENSMUST00000067087.6 | 2801 | Fam150b | ENSMUSG00000054204.6 |
| chr5 | 33695404 | 33695972 | 3.41977 | 4.04506 | 1.1867 | Distal Intergenic | 5 | 33721724 | 33737067 | 15344 | 1 | ENSMUST00000067150.8 | -25752 | Fgfr3 | ENSMUSG00000054252.12 |
| chr5 | 33723267 | 33723850 | 3.69113 | 4.60062 | 1.5603 | Promoter | 5 | 33723772 | 33729850 | 6079 | 1 | ENSMUST00000155022.1 | 0 | Fgfr3 | ENSMUSG00000054252.12 |
| chr5 | 115395910 | 115395985 | 3.78696 | 4.93563 | 1.729 | Intron (ENSMUST00000146546.1/ENSMUSG00000087292.1, intron 2 of 2) | 5 | 115429599 | 115455698 | 26100 | 1 | ENSMUST00000150779.2 | -33614 | Msi1 | ENSMUSG00000054256.6 |
| chr5 | 115397373 | 115397610 | 3.65137 | 4.40314 | 1.34579 | Intron (ENSMUST00000151159.1/ENSMUSG00000087292.1, intron 1 of 2) | 5 | 115429599 | 115455698 | 26100 | 1 | ENSMUST00000150779.2 | -31989 | Msi1 | ENSMUSG00000054256.6 |
| chr5 | 115398409 | 115398876 | 4.36768 | 5.86947 | 2.40492 | Intron (ENSMUST00000151159.1/ENSMUSG00000087292.1, intron 1 of 2) | 5 | 115429599 | 115455698 | 26100 | 1 | ENSMUST00000150779.2 | -30723 | Msi1 | ENSMUSG00000054256.6 |
| chr5 | 115405170 | 115405472 | 3.79833 | 4.67682 | 1.5117 | Intron (ENSMUST00000151159.1/ENSMUSG00000087292.1, intron 1 of 2) | 5 | 115429599 | 115455698 | 26100 | 1 | ENSMUST00000150779.2 | -24127 | Msi1 | ENSMUSG00000054256.6 |
| chr5 | 115415775 | 115415948 | 3.34323 | 3.84223 | 1.01733 | Intron (ENSMUST00000151159.1/ENSMUSG00000087292.1, intron 2 of 2) | 5 | 115429599 | 115455698 | 26100 | 1 | ENSMUST00000150779.2 | -13651 | Msi1 | ENSMUSG00000054256.6 |
| chr5 | 115428586 | 115431240 | 4.21202 | 5.5752 | 2.22205 | Promoter | 5 | 115429599 | 115455698 | 26100 | 1 | ENSMUST00000150779.2 | 0 | Msi1 | ENSMUSG00000054256.6 |
| chr5 | 115436458 | 115436869 | 3.83745 | 4.75937 | 1.58993 | Promoter | 5 | 115435498 | 115445694 | 10197 | 1 | ENSMUST00000130849.1 | 961 | Msi1 | ENSMUSG00000054256.6 |
| chr7 | 24377321 | 24377543 | 3.33753 | 4.08356 | 1.2074 | Exon (ENSMUST00000171904.1/ENSMUSG00000054342.7, exon 3 of 9) | 7 | 24370263 | 24385209 | 14947 | 1 | ENSMUST00000171904.1 | 7059 | Kcnn4 | ENSMUSG00000054342.7 |
| chr9 | 40854003 | 40854292 | 3.25114 | 3.70386 | 1.01733 | Distal Intergenic | 9 | 40874127 | 40877972 | 3846 | 1 | ENSMUST00000067375.3 | -19835 | Bsx | ENSMUSG00000054360.3 |
| chr9 | 40856840 | 40857022 | 3.30003 | 3.80031 | 1.07937 | Distal Intergenic | 9 | 40874127 | 40877972 | 3846 | 1 | ENSMUST00000067375.3 | -17105 | Bsx | ENSMUSG00000054360.3 |
| chr9 | 40858915 | 40859794 | 3.85687 | 4.85978 | 1.72896 | Distal Intergenic | 9 | 40874127 | 40877972 | 3846 | 1 | ENSMUST00000067375.3 | -14333 | Bsx | ENSMUSG00000054360.3 |
| chr9 | 40868564 | 40868993 | 3.63477 | 4.37699 | 1.33968 | Distal Intergenic | 9 | 40874127 | 40877972 | 3846 | 1 | ENSMUST00000067375.3 | -5134 | Bsx | ENSMUSG00000054360.3 |
| chr9 | 40869449 | 40869976 | 3.81844 | 4.73134 | 1.57489 | Distal Intergenic | 9 | 40874127 | 40877972 | 3846 | 1 | ENSMUST00000067375.3 | -4151 | Bsx | ENSMUSG00000054360.3 |
| chr9 | 40870294 | 40870660 | 3.82539 | 4.75786 | 1.59975 | Distal Intergenic | 9 | 40874127 | 40877972 | 3846 | 1 | ENSMUST00000067375.3 | -3467 | Bsx | ENSMUSG00000054360.3 |
| chr9 | 40871161 | 40872291 | 3.99076 | 5.2025 | 1.92026 | Distal Intergenic | 9 | 40874127 | 40877972 | 3846 | 1 | ENSMUST00000067375.3 | -1836 | Bsx | ENSMUSG00000054360.3 |
| chr9 | 40872631 | 40874966 | 4.2134 | 5.51415 | 2.12962 | Promoter | 9 | 40874127 | 40877972 | 3846 | 1 | ENSMUST00000067375.3 | 0 | Bsx | ENSMUSG00000054360.3 |
| chr9 | 40875360 | 40875680 | 3.43067 | 4.03775 | 1.12817 | Intron (ENSMUST00000067375.3/ENSMUSG00000054360.3, intron 1 of 2) | 9 | 40874127 | 40877972 | 3846 | 1 | ENSMUST00000067375.3 | 1234 | Bsx | ENSMUSG00000054360.3 |
| chr9 | 40876169 | 40876533 | 3.68206 | 4.58873 | 1.53832 | Exon (ENSMUST00000067375.3/ENSMUSG00000054360.3, exon 2 of 3) | 9 | 40874127 | 40877972 | 3846 | 1 | ENSMUST00000067375.3 | 2043 | Bsx | ENSMUSG00000054360.3 |
| chr9 | 40877195 | 40877372 | 3.66813 | 4.70658 | 1.56719 | Intron (ENSMUST00000067375.3/ENSMUSG00000054360.3, intron 2 of 2) | 9 | 40874127 | 40877972 | 3846 | 1 | ENSMUST00000067375.3 | 3069 | Bsx | ENSMUSG00000054360.3 |
| chr9 | 40877773 | 40879946 | 3.7045 | 4.58583 | 1.4932 | 3' UTR | 9 | 40874127 | 40877972 | 3846 | 1 | ENSMUST00000067375.3 | 3647 | Bsx | ENSMUSG00000054360.3 |
| chr9 | 40884280 | 40884545 | 3.51309 | 4.19785 | 1.30955 | Distal Intergenic | 9 | 40874127 | 40877972 | 3846 | 1 | ENSMUST00000067375.3 | 10154 | Bsx | ENSMUSG00000054360.3 |
| chr11 | 82952195 | 82952389 | 3.33669 | 4.04432 | 1.13659 | Promoter | 11 | 82952102 | 82962941 | 10840 | 1 | ENSMUST00000067443.7 | 94 | Slfn5 | ENSMUSG00000054404.7 |
| chr11 | 82954106 | 82955071 | 3.75667 | 4.7044 | 1.62936 | Promoter | 11 | 82954661 | 82961906 | 7246 | 1 | ENSMUST00000108157.1 | 0 | Slfn5 | ENSMUSG00000054404.7 |
| chr1 | 135548915 | 135549220 | 3.21977 | 3.65687 | 1.0162 | Intron (ENSMUST00000040599.9/ENSMUSG00000009418.10, intron 1 of 29) | 1 | 135584773 | 135599948 | 15176 | 1 | ENSMUST00000067468.5 | -35553 | Gm4793 | ENSMUSG00000054412.5 |
| chr1 | 135583250 | 135583336 | 3.93754 | 5.24327 | 1.96158 | Intron (ENSMUST00000040599.9/ENSMUSG00000009418.10, intron 1 of 29) | 1 | 135584773 | 135599948 | 15176 | 1 | ENSMUST00000067468.5 | -1437 | Gm4793 | ENSMUSG00000054412.5 |
| chr1 | 135583998 | 135584416 | 3.54019 | 4.35966 | 1.34011 | Promoter | 1 | 135584773 | 135599948 | 15176 | 1 | ENSMUST00000067468.5 | -357 | Gm4793 | ENSMUSG00000054412.5 |
| chr1 | 135584813 | 135585256 | 3.76615 | 4.93328 | 1.72375 | Promoter | 1 | 135584773 | 135599948 | 15176 | 1 | ENSMUST00000067468.5 | 41 | Gm4793 | ENSMUSG00000054412.5 |
| chr14 | 12821586 | 12822354 | 3.7619 | 4.88546 | 1.72484 | Promoter | 14 | 12372563 | 12823079 | 450517 | 2 | ENSMUST00000067491.7 | 725 | Cadps | ENSMUSG00000054423.8 |
| chr14 | 12822822 | 12823179 | 4.04895 | 5.43565 | 2.05903 | Promoter | 14 | 12372563 | 12823079 | 450517 | 2 | ENSMUST00000067491.7 | 0 | Cadps | ENSMUSG00000054423.8 |
| chr14 | 12823568 | 12823745 | 3.54127 | 4.22721 | 1.28697 | Promoter | 14 | 12372565 | 12823079 | 450515 | 2 | ENSMUST00000177814.1 | -490 | Cadps | ENSMUSG00000054423.8 |
| chr11 | 53478046 | 53480141 | 3.9604 | 5.23302 | 1.95745 | Promoter | 11 | 53480166 | 53481097 | 932 | 1 | ENSMUST00000067523.1 | -25 | Gm9945 | ENSMUSG00000054450.1 |
| chr12 | 11433846 | 11433921 | 3.34153 | 3.83966 | 1.01733 | Intron (ENSMUST00000072299.5/ENSMUSG00000054459.6, intron 1 of 3) | 12 | 11325247 | 11436649 | 111403 | 2 | ENSMUST00000072299.5 | 2728 | Vsnl1 | ENSMUSG00000054459.6 |
| chr12 | 11436512 | 11437667 | 3.66833 | 4.45176 | 1.38595 | Promoter | 12 | 11325247 | 11436649 | 111403 | 2 | ENSMUST00000072299.5 | 0 | Vsnl1 | ENSMUSG00000054459.6 |
| chr18 | 45268921 | 45269299 | 3.4351 | 4.04231 | 1.21956 | Promoter | 18 | 45269005 | 45685883 | 416879 | 1 | ENSMUST00000183850.2 | 0 | Kcnn2 | ENSMUSG00000054477.9 |
| chr18 | 45558487 | 45559731 | 3.55093 | 4.3652 | 1.38484 | Promoter | 18 | 45559469 | 45671249 | 111781 | 1 | ENSMUST00000184101.2 | 0 | Kcnn2 | ENSMUSG00000054477.9 |
| chr18 | 45560092 | 45560728 | 3.54923 | 4.2346 | 1.29671 | Promoter | 18 | 45560154 | 45675362 | 115209 | 1 | ENSMUST00000167895.1 | 0 | Kcnn2 | ENSMUSG00000054477.9 |
| chr1 | 14753562 | 14753891 | 3.36173 | 3.87013 | 1.01733 | Promoter | 1 | 14752937 | 14776931 | 23995 | 1 | ENSMUST00000187401.1 | 626 | Gm9947 | ENSMUSG00000054493.2 |
| chr7 | 134224101 | 134225727 | 4.11518 | 5.51382 | 2.16249 | Promoter | 7 | 133883199 | 134225097 | 341899 | 2 | ENSMUST00000067680.5 | 0 | Adam12 | ENSMUSG00000054555.6 |
| chr2 | 60452383 | 60452644 | 3.3669 | 3.87793 | 1.01733 | Exon (ENSMUST00000067708.5/ENSMUSG00000054580.9, exon 14 of 30) | 2 | 60451992 | 60455254 | 3263 | 2 | ENSMUST00000134583.1 | 2610 | Pla2r1 | ENSMUSG00000054580.9 |
| chr7 | 136803787 | 136804094 | 3.5636 | 4.25006 | 1.27558 | Distal Intergenic | 7 | 136894611 | 137128187 | 233577 | 1 | ENSMUST00000081510.2 | -90517 | Mgmt | ENSMUSG00000054612.3 |
| chr7 | 136823047 | 136823226 | 3.29539 | 3.77025 | 1.01733 | Distal Intergenic | 7 | 136894611 | 137128187 | 233577 | 1 | ENSMUST00000081510.2 | -71385 | Mgmt | ENSMUSG00000054612.3 |
| chr7 | 136823761 | 136824224 | 3.67384 | 4.55751 | 1.51381 | Distal Intergenic | 7 | 136894611 | 137128187 | 233577 | 1 | ENSMUST00000081510.2 | -70387 | Mgmt | ENSMUSG00000054612.3 |
| chr7 | 136824875 | 136824950 | 3.28195 | 3.85142 | 1.09436 | Distal Intergenic | 7 | 136894611 | 137128187 | 233577 | 1 | ENSMUST00000081510.2 | -69661 | Mgmt | ENSMUSG00000054612.3 |
| chr7 | 136826437 | 136826532 | 3.64474 | 4.58396 | 1.51291 | Distal Intergenic | 7 | 136894611 | 137128187 | 233577 | 1 | ENSMUST00000081510.2 | -68079 | Mgmt | ENSMUSG00000054612.3 |
| chr7 | 136826881 | 136826970 | 3.56068 | 4.30611 | 1.34011 | Distal Intergenic | 7 | 136894611 | 137128187 | 233577 | 1 | ENSMUST00000081510.2 | -67641 | Mgmt | ENSMUSG00000054612.3 |
| chr7 | 136871483 | 136871558 | 3.41817 | 3.96666 | 1.05737 | Distal Intergenic | 7 | 136894611 | 137128187 | 233577 | 1 | ENSMUST00000081510.2 | -23053 | Mgmt | ENSMUSG00000054612.3 |
| chr17 | 81670412 | 81670487 | 3.73442 | 4.59265 | 1.54605 | Distal Intergenic | 17 | 81388691 | 81649607 | 260917 | 2 | ENSMUST00000163680.3 | -20806 | Slc8a1 | ENSMUSG00000054640.9 |
| chr17 | 81672232 | 81672488 | 3.35208 | 3.85555 | 1.01733 | Distal Intergenic | 17 | 81388691 | 81649607 | 260917 | 2 | ENSMUST00000163680.3 | -22626 | Slc8a1 | ENSMUSG00000054640.9 |
| chr17 | 33265483 | 33265786 | 3.36841 | 3.89473 | 1.06818 | Distal Intergenic | 17 | 33268722 | 33269676 | 955 | 1 | ENSMUST00000067840.4 | -2936 | Olfr63 | ENSMUSG00000054666.4 |
| chr17 | 33268858 | 33269246 | 3.72342 | 4.53963 | 1.43636 | Promoter | 17 | 33268722 | 33269676 | 955 | 1 | ENSMUST00000067840.4 | 137 | Olfr63 | ENSMUSG00000054666.4 |
| chr17 | 33274847 | 33275074 | 3.47887 | 4.08831 | 1.16352 | Distal Intergenic | 17 | 33268722 | 33269676 | 955 | 1 | ENSMUST00000067840.4 | 6126 | Olfr63 | ENSMUSG00000054666.4 |
| chr17 | 33276182 | 33276626 | 3.97489 | 5.03984 | 1.80393 | Distal Intergenic | 17 | 33268722 | 33269676 | 955 | 1 | ENSMUST00000067840.4 | 7461 | Olfr63 | ENSMUSG00000054666.4 |
| chr5 | 113797244 | 113797567 | 3.53196 | 4.18812 | 1.23255 | Intron (ENSMUST00000067853.5/ENSMUSG00000054675.5, intron 1 of 1) | 5 | 113795638 | 113800446 | 4809 | 2 | ENSMUST00000160374.1 | 2879 | Tmem119 | ENSMUSG00000054675.5 |
| chr4 | 33209183 | 33209387 | 3.74598 | 4.74412 | 1.60703 | Promoter | 4 | 33209263 | 33231293 | 22031 | 1 | ENSMUST00000108159.1 | 0 | Srsf12 | ENSMUSG00000054679.3 |
| chr4 | 33209695 | 33209779 | 3.67804 | 4.70849 | 1.6004 | Promoter | 4 | 33209263 | 33231293 | 22031 | 1 | ENSMUST00000108159.1 | 433 | Srsf12 | ENSMUSG00000054679.3 |
| chr5 | 63651682 | 63652342 | 3.31999 | 3.81167 | 1.03259 | Intron (ENSMUST00000160066.2/ENSMUSG00000090061.4, intron 1 of 3) | 5 | 63649950 | 63650264 | 315 | 2 | ENSMUST00000067917.3 | -1419 | Gm9954 | ENSMUSG00000054714.3 |
| chr5 | 105520075 | 105520175 | 3.84606 | 4.84047 | 1.69417 | Promoter | 5 | 105519473 | 105544640 | 25168 | 1 | ENSMUST00000153955.1 | 603 | Lrrc8c | ENSMUSG00000054720.7 |
| chr5 | 105520673 | 105520761 | 3.19141 | 3.83615 | 1.03992 | Intron (ENSMUST00000067924.7/ENSMUSG00000054720.7, intron 1 of 3) | 5 | 105519473 | 105544640 | 25168 | 1 | ENSMUST00000153955.1 | 1201 | Lrrc8c | ENSMUSG00000054720.7 |
| chr5 | 105521189 | 105521358 | 3.54697 | 4.29622 | 1.32992 | Intron (ENSMUST00000067924.7/ENSMUSG00000054720.7, intron 1 of 3) | 5 | 105519473 | 105544640 | 25168 | 1 | ENSMUST00000153955.1 | 1717 | Lrrc8c | ENSMUSG00000054720.7 |
| chr13 | 42708597 | 42708958 | 3.3074 | 3.93321 | 1.07073 | Promoter | 13 | 42709543 | 43135401 | 425859 | 1 | ENSMUST00000066928.6 | -585 | Phactr1 | ENSMUSG00000054728.11 |
| chr7 | 120330825 | 120330946 | 3.38673 | 4.24212 | 1.29559 | Intron (ENSMUST00000140459.2/ENSMUSG00000054746.12, intron 1 of 14) | 7 | 120328684 | 120407687 | 79004 | 1 | ENSMUST00000121265.2 | 2142 | Abca15 | ENSMUSG00000054746.12 |
| chr7 | 120331460 | 120331787 | 3.25237 | 3.73393 | 1.03083 | Intron (ENSMUST00000140459.2/ENSMUSG00000054746.12, intron 1 of 14) | 7 | 120328684 | 120407687 | 79004 | 1 | ENSMUST00000121265.2 | 2777 | Abca15 | ENSMUSG00000054746.12 |
| chr7 | 120338234 | 120338563 | 3.36759 | 3.90592 | 1.11267 | Intron (ENSMUST00000140459.2/ENSMUSG00000054746.12, intron 5 of 14) | 7 | 120328684 | 120407687 | 79004 | 1 | ENSMUST00000121265.2 | 9551 | Abca15 | ENSMUSG00000054746.12 |
| chr14 | 63067522 | 63067732 | 3.55203 | 4.21281 | 1.21285 | Distal Intergenic | 14 | 63055073 | 63056940 | 1868 | 1 | ENSMUST00000138708.1 | 12450 | Defb42 | ENSMUSG00000054763.4 |
| chr14 | 63075631 | 63075805 | 3.54113 | 4.44219 | 1.40843 | Distal Intergenic | 14 | 63055073 | 63056940 | 1868 | 1 | ENSMUST00000138708.1 | 20559 | Defb42 | ENSMUSG00000054763.4 |
| chr8 | 45069146 | 45069227 | 3.53262 | 4.23448 | 1.33368 | Promoter | 8 | 45069137 | 45088356 | 19220 | 1 | ENSMUST00000067984.7 | 10 | Mtnr1a | ENSMUSG00000054764.8 |
| chr19 | 57543190 | 57543732 | 3.35206 | 3.85559 | 1.01733 | Distal Intergenic | 19 | 57611034 | 58133338 | 522305 | 1 | ENSMUST00000077282.5 | -67302 | Atrnl1 | ENSMUSG00000054843.8 |
| chr19 | 57544610 | 57544712 | 3.78549 | 4.65437 | 1.49924 | Distal Intergenic | 19 | 57611034 | 58133338 | 522305 | 1 | ENSMUST00000077282.5 | -66322 | Atrnl1 | ENSMUSG00000054843.8 |
| chr19 | 57557870 | 57558142 | 3.30586 | 3.78599 | 1.01733 | Distal Intergenic | 19 | 57611034 | 58133338 | 522305 | 1 | ENSMUST00000077282.5 | -52892 | Atrnl1 | ENSMUSG00000054843.8 |
| chr19 | 57647427 | 57647502 | 3.71956 | 4.57215 | 1.54559 | Intron (ENSMUST00000077282.5/ENSMUSG00000054843.8, intron 5 of 28) | 19 | 57611034 | 58133338 | 522305 | 1 | ENSMUST00000077282.5 | 36394 | Atrnl1 | ENSMUSG00000054843.8 |
| chr15 | 98674840 | 98674932 | 3.52077 | 4.15486 | 1.17477 | Intron (ENSMUST00000109149.3/ENSMUSG00000054855.8, intron 2 of 4) | 15 | 98672217 | 98677458 | 5242 | 2 | ENSMUST00000120997.1 | 2526 | Rnd1 | ENSMUSG00000054855.8 |
| chr15 | 87543237 | 87543638 | 3.48466 | 4.24131 | 1.31434 | Distal Intergenic | 15 | 87625230 | 87759336 | 134107 | 1 | ENSMUST00000068088.6 | -81592 | Fam19a5 | ENSMUSG00000054863.8 |
| chr15 | 87544049 | 87544176 | 3.39291 | 3.92062 | 1.02946 | Distal Intergenic | 15 | 87625230 | 87759336 | 134107 | 1 | ENSMUST00000068088.6 | -81054 | Fam19a5 | ENSMUSG00000054863.8 |
| chr15 | 87544741 | 87545303 | 3.52738 | 4.26096 | 1.27101 | Distal Intergenic | 15 | 87625230 | 87759336 | 134107 | 1 | ENSMUST00000068088.6 | -79927 | Fam19a5 | ENSMUSG00000054863.8 |
| chr15 | 87546144 | 87546246 | 3.94819 | 4.95275 | 1.683 | Distal Intergenic | 15 | 87625230 | 87759336 | 134107 | 1 | ENSMUST00000068088.6 | -78984 | Fam19a5 | ENSMUSG00000054863.8 |
| chr15 | 87548660 | 87548842 | 3.43084 | 3.99169 | 1.07816 | Distal Intergenic | 15 | 87625230 | 87759336 | 134107 | 1 | ENSMUST00000068088.6 | -76388 | Fam19a5 | ENSMUSG00000054863.8 |
| chr15 | 87622633 | 87623277 | 3.93525 | 4.95919 | 1.74009 | Distal Intergenic | 15 | 87625230 | 87759336 | 134107 | 1 | ENSMUST00000068088.6 | -1953 | Fam19a5 | ENSMUSG00000054863.8 |
| chr15 | 87623809 | 87623916 | 3.41397 | 4.61325 | 1.52817 | Distal Intergenic | 15 | 87625230 | 87759336 | 134107 | 1 | ENSMUST00000068088.6 | -1314 | Fam19a5 | ENSMUSG00000054863.8 |
| chr15 | 87624797 | 87625694 | 4.00402 | 5.2401 | 1.97544 | Promoter | 15 | 87625230 | 87759336 | 134107 | 1 | ENSMUST00000068088.6 | 0 | Fam19a5 | ENSMUSG00000054863.8 |
| chr9 | 123259434 | 123259649 | 4.43244 | 6.07047 | 2.52501 | 3' UTR | 9 | 123259058 | 123260789 | 1732 | 2 | ENSMUST00000068140.4 | 1140 | Tmem158 | ENSMUSG00000054871.4 |
| chr9 | 123260033 | 123260299 | 3.42956 | 4.139 | 1.20083 | Promoter | 9 | 123259058 | 123260789 | 1732 | 2 | ENSMUST00000068140.4 | 490 | Tmem158 | ENSMUSG00000054871.4 |
| chr5 | 72696758 | 72697016 | 3.38407 | 3.90794 | 1.03176 | Intron (ENSMUST00000113604.4/ENSMUSG00000054892.9, intron 14 of 14) | 5 | 72695982 | 72736589 | 40608 | 2 | ENSMUST00000169534.1 | 39573 | Txk | ENSMUSG00000054892.9 |
| chr5 | 72705666 | 72706104 | 3.78109 | 4.6945 | 1.61794 | Intron (ENSMUST00000113604.4/ENSMUSG00000054892.9, intron 11 of 14) | 5 | 72695982 | 72736589 | 40608 | 2 | ENSMUST00000169534.1 | 30485 | Txk | ENSMUSG00000054892.9 |
| chr10 | 116473802 | 116474507 | 4.08644 | 5.66474 | 2.24783 | Promoter | 10 | 116417861 | 116473878 | 56018 | 2 | ENSMUST00000068233.5 | 0 | Kcnmb4 | ENSMUSG00000054934.5 |
| chr4 | 123200183 | 123200449 | 3.35567 | 3.86099 | 1.01733 | Promoter | 4 | 123201504 | 123216275 | 14772 | 1 | ENSMUST00000068262.5 | -1055 | Nt5c1a | ENSMUSG00000054958.6 |
| chr4 | 123201297 | 123202415 | 3.74317 | 4.67249 | 1.53147 | Promoter | 4 | 123201504 | 123216275 | 14772 | 1 | ENSMUST00000068262.5 | 0 | Nt5c1a | ENSMUSG00000054958.6 |
| chr4 | 123207613 | 123208576 | 3.37564 | 4.08786 | 1.17433 | Exon (ENSMUST00000068262.5/ENSMUSG00000054958.6, exon 2 of 6) | 4 | 123201504 | 123216275 | 14772 | 1 | ENSMUST00000068262.5 | 6110 | Nt5c1a | ENSMUSG00000054958.6 |
| chr4 | 123216013 | 123216176 | 3.44226 | 4.13741 | 1.22454 | Exon (ENSMUST00000068262.5/ENSMUSG00000054958.6, exon 6 of 6) | 4 | 123201504 | 123216275 | 14772 | 1 | ENSMUST00000068262.5 | 14510 | Nt5c1a | ENSMUSG00000054958.6 |
| chr15 | 76930715 | 76930969 | 3.2207 | 3.65826 | 1.01621 | Distal Intergenic | 15 | 76910851 | 76918010 | 7160 | 2 | ENSMUST00000048854.7 | -12706 | Zfp647 | ENSMUSG00000054967.6 |
| chr9 | 65390917 | 65391521 | 3.71194 | 4.58277 | 1.55459 | Promoter | 9 | 65388684 | 65391652 | 2969 | 2 | ENSMUST00000068307.3 | 131 | Kbtbd13 | ENSMUSG00000054978.3 |
| chr11 | 24049501 | 24049829 | 3.453 | 4.44355 | 1.39275 | Exon (ENSMUST00000068360.1/ENSMUSG00000055010.1, exon 3 of 3) | 11 | 24048955 | 24075054 | 26100 | 2 | ENSMUST00000068360.1 | 25225 | A830031A19Rik | ENSMUSG00000055010.1 |
| chr11 | 24056334 | 24056438 | 3.3553 | 3.86042 | 1.01733 | Intron (ENSMUST00000068360.1/ENSMUSG00000055010.1, intron 2 of 2) | 11 | 24048955 | 24075054 | 26100 | 2 | ENSMUST00000068360.1 | 18616 | A830031A19Rik | ENSMUSG00000055010.1 |
| chr11 | 24057012 | 24057374 | 3.40261 | 3.9437 | 1.05948 | Intron (ENSMUST00000068360.1/ENSMUSG00000055010.1, intron 2 of 2) | 11 | 24048955 | 24075054 | 26100 | 2 | ENSMUST00000068360.1 | 17680 | A830031A19Rik | ENSMUSG00000055010.1 |
| chr11 | 24057734 | 24058006 | 3.32548 | 3.81568 | 1.01733 | Intron (ENSMUST00000068360.1/ENSMUSG00000055010.1, intron 2 of 2) | 11 | 24048955 | 24075054 | 26100 | 2 | ENSMUST00000068360.1 | 17048 | A830031A19Rik | ENSMUSG00000055010.1 |
| chr11 | 24058500 | 24058904 | 3.72627 | 4.66802 | 1.54061 | 5' UTR | 11 | 24048955 | 24075054 | 26100 | 2 | ENSMUST00000068360.1 | 16150 | A830031A19Rik | ENSMUSG00000055010.1 |
| chr11 | 24059356 | 24059786 | 3.8562 | 4.80066 | 1.6017 | Intron (ENSMUST00000068360.1/ENSMUSG00000055010.1, intron 1 of 2) | 11 | 24048955 | 24075054 | 26100 | 2 | ENSMUST00000068360.1 | 15268 | A830031A19Rik | ENSMUSG00000055010.1 |
| chr11 | 24060338 | 24060589 | 3.74017 | 4.59424 | 1.51039 | Intron (ENSMUST00000068360.1/ENSMUSG00000055010.1, intron 1 of 2) | 11 | 24048955 | 24075054 | 26100 | 2 | ENSMUST00000068360.1 | 14465 | A830031A19Rik | ENSMUSG00000055010.1 |
| chr11 | 24060948 | 24061297 | 4.51692 | 6.52206 | 2.8856 | Intron (ENSMUST00000068360.1/ENSMUSG00000055010.1, intron 1 of 2) | 11 | 24048955 | 24075054 | 26100 | 2 | ENSMUST00000068360.1 | 13757 | A830031A19Rik | ENSMUSG00000055010.1 |
| chr11 | 24061934 | 24063095 | 3.51247 | 4.17366 | 1.27124 | Intron (ENSMUST00000068360.1/ENSMUSG00000055010.1, intron 1 of 2) | 11 | 24048955 | 24075054 | 26100 | 2 | ENSMUST00000068360.1 | 11959 | A830031A19Rik | ENSMUSG00000055010.1 |
| chr11 | 24066094 | 24066631 | 3.36298 | 3.9235 | 1.16135 | Intron (ENSMUST00000068360.1/ENSMUSG00000055010.1, intron 1 of 2) | 11 | 24048955 | 24075054 | 26100 | 2 | ENSMUST00000068360.1 | 8423 | A830031A19Rik | ENSMUSG00000055010.1 |
| chr11 | 24069642 | 24070406 | 3.90648 | 4.95698 | 1.79406 | Intron (ENSMUST00000068360.1/ENSMUSG00000055010.1, intron 1 of 2) | 11 | 24048955 | 24075054 | 26100 | 2 | ENSMUST00000068360.1 | 4648 | A830031A19Rik | ENSMUSG00000055010.1 |
| chr11 | 24071065 | 24071900 | 4.1339 | 5.39752 | 2.07085 | Intron (ENSMUST00000068360.1/ENSMUSG00000055010.1, intron 1 of 2) | 11 | 24048955 | 24075054 | 26100 | 2 | ENSMUST00000068360.1 | 3154 | A830031A19Rik | ENSMUSG00000055010.1 |
| chr11 | 24073860 | 24074916 | 3.74017 | 4.62882 | 1.50066 | Promoter | 11 | 24048955 | 24075054 | 26100 | 2 | ENSMUST00000068360.1 | 138 | A830031A19Rik | ENSMUSG00000055010.1 |
| chr11 | 24075287 | 24075596 | 4.32762 | 5.72659 | 2.26125 | Promoter | 11 | 24048955 | 24075054 | 26100 | 2 | ENSMUST00000068360.1 | -234 | A830031A19Rik | ENSMUSG00000055010.1 |
| chr15 | 92051229 | 92052353 | 3.7469 | 4.62526 | 1.51915 | Promoter | 15 | 92051165 | 92341967 | 290803 | 1 | ENSMUST00000169825.1 | 65 | Cntn1 | ENSMUSG00000055022.9 |
| chr7 | 57386496 | 57387154 | 3.49251 | 4.1481 | 1.28171 | Promoter | 7 | 57178382 | 57386871 | 208490 | 2 | ENSMUST00000068394.6 | 0 | Gabrg3 | ENSMUSG00000055026.8 |
| chr11 | 77914157 | 77914915 | 3.72114 | 4.59938 | 1.54156 | Exon (ENSMUST00000132487.1/ENSMUSG00000055045.5, exon 3 of 3) | 11 | 77912127 | 77929288 | 17162 | 2 | ENSMUST00000132487.1 | 14373 | Gm11190 | ENSMUSG00000055045.5 |
| chr11 | 77920473 | 77920650 | 3.23338 | 3.67724 | 1.01733 | Intron (ENSMUST00000132487.1/ENSMUSG00000055045.5, intron 1 of 2) | 11 | 77912127 | 77929288 | 17162 | 2 | ENSMUST00000132487.1 | 8638 | Gm11190 | ENSMUSG00000055045.5 |
| chr11 | 77927243 | 77927544 | 3.64234 | 4.39483 | 1.3655 | Intron (ENSMUST00000132487.1/ENSMUSG00000055045.5, intron 1 of 2) | 11 | 77912127 | 77929288 | 17162 | 2 | ENSMUST00000132487.1 | 1744 | Gm11190 | ENSMUSG00000055045.5 |
| chr11 | 77929813 | 77929919 | 3.85916 | 4.88633 | 1.75347 | Promoter | 11 | 77912127 | 77929288 | 17162 | 2 | ENSMUST00000132487.1 | -526 | Gm11190 | ENSMUSG00000055045.5 |
| chr15 | 79552068 | 79552546 | 3.89251 | 4.93463 | 1.8028 | Distal Intergenic | 15 | 79527707 | 79546741 | 19035 | 2 | ENSMUST00000054014.7 | -5328 | Ddx17 | ENSMUSG00000055065.6 |
| chr1 | 179058494 | 179058987 | 3.36739 | 3.87866 | 1.01733 | Intron (ENSMUST00000128302.2/ENSMUSG00000055067.10, intron 8 of 11) | 1 | 178957250 | 178989345 | 32096 | 2 | ENSMUST00000129393.1 | -69150 | Smyd3 | ENSMUSG00000055067.10 |
| chr9 | 53705152 | 53705417 | 3.46204 | 4.21069 | 1.24614 | Promoter | 9 | 53684110 | 53706232 | 22123 | 2 | ENSMUST00000068449.3 | 815 | Rab39 | ENSMUSG00000055069.3 |
| chr7 | 57509415 | 57509942 | 3.89335 | 4.92979 | 1.78255 | Promoter | 7 | 57407690 | 57510009 | 102320 | 2 | ENSMUST00000068456.6 | 67 | Gabra5 | ENSMUSG00000055078.6 |
| chr7 | 57512841 | 57512995 | 3.79434 | 4.75059 | 1.62576 | Distal Intergenic | 7 | 57407690 | 57510009 | 102320 | 2 | ENSMUST00000068456.6 | -2833 | Gabra5 | ENSMUSG00000055078.6 |
| chr15 | 58383951 | 58384212 | 3.79249 | 4.65926 | 1.49196 | Distal Intergenic | 15 | 58341874 | 58364148 | 22275 | 2 | ENSMUST00000068515.7 | -19804 | Anxa13 | ENSMUSG00000055114.7 |
| chr9 | 67759167 | 67761208 | 3.95839 | 5.08422 | 1.88558 | Promoter | 9 | 67758712 | 67760208 | 1497 | 2 | ENSMUST00000068526.6 | 0 | M5C1000I18Rik | ENSMUSG00000055125.6 |
| chr9 | 67762296 | 67762599 | 3.38875 | 3.94832 | 1.14175 | Distal Intergenic | 9 | 67758713 | 67760230 | 1518 | 2 | ENSMUST00000186205.1 | -2067 | M5C1000I18Rik | ENSMUSG00000055125.6 |
| chr9 | 67763307 | 67763910 | 3.34669 | 3.88971 | 1.10929 | Distal Intergenic | 9 | 67758713 | 67760230 | 1518 | 2 | ENSMUST00000186205.1 | -3078 | M5C1000I18Rik | ENSMUSG00000055125.6 |
| chr10 | 118582775 | 118582907 | 3.22276 | 3.66135 | 1.01733 | Distal Intergenic | 10 | 118441047 | 118445892 | 4846 | 1 | ENSMUST00000068592.3 | 141729 | Ifng | ENSMUSG00000055170.3 |
| chr1 | 74879449 | 74879867 | 3.37143 | 3.92757 | 1.15801 | Exon (ENSMUST00000159627.1/ENSMUSG00000089964.1, exon 4 of 4) | 1 | 74882407 | 74884608 | 2202 | 2 | ENSMUST00000162938.1 | 4741 | Fev | ENSMUSG00000055197.4 |

| | | | | | | | | | | | | | | | |
|---|---|---|---|---|---|---|---|---|---|---|---|---|---|---|---|
| chr1 | 74881474 | 74882486 | 4.35085 | 5.94106 | 2.50147 | 3' UTR | 1 | 74882407 | 74884608 | 2202 | 2 | ENSMUST00000162938.1 | 2122 | Fev | ENSMUSG00000055197.4 |
| chr1 | 74883087 | 74883420 | 3.35407 | 3.85857 | 1.01733 | Intron (ENSMUST00000068631.3/ENSMUSG00000055197.4, intron 2 of 2) | 1 | 74882407 | 74884608 | 2202 | 2 | ENSMUST00000162938.1 | 1188 | Fev | ENSMUSG00000055197.4 |
| chr1 | 74884295 | 74884370 | 3.9824 | 5.01098 | 1.71243 | Promoter | 1 | 74882407 | 74884608 | 2202 | 2 | ENSMUST00000162938.1 | 238 | Fev | ENSMUSG00000055197.4 |
| chr1 | 74884908 | 74886188 | 4.17082 | 5.53215 | 2.16534 | Promoter | 1 | 74881509 | 74885419 | 3911 | 2 | ENSMUST00000068631.3 | 0 | Fev | ENSMUSG00000055197.4 |
| chr1 | 176274548 | 176275905 | 4.08578 | 5.33585 | 2.02695 | Promoter | 1 | 175962306 | 176275298 | 312993 | 2 | ENSMUST00000065967.8 | 0 | Pld5 | ENSMUSG00000055214.10 |
| chr1 | 176276392 | 176276586 | 3.54716 | 4.25394 | 1.26017 | Distal Intergenic | 1 | 175962407 | 176275312 | 312906 | 2 | ENSMUST00000125404.2 | -1081 | Pld5 | ENSMUSG00000055214.10 |
| chr13 | 58768406 | 58768481 | 4.86672 | 7.09642 | 3.33096 | Distal Intergenic | 13 | 58806596 | 58937030 | 130435 | 1 | ENSMUST00000109838.3 | -38115 | Ntrk2 | ENSMUSG00000055254.9 |
| chr13 | 58805661 | 58805745 | 3.59372 | 4.49118 | 1.4533 | Promoter | 13 | 58806596 | 58937030 | 130435 | 1 | ENSMUST00000109838.3 | -851 | Ntrk2 | ENSMUSG00000055254.9 |
| chr13 | 58806068 | 58806329 | 3.77792 | 4.63605 | 1.48819 | Promoter | 13 | 58806596 | 58937030 | 130435 | 1 | ENSMUST00000109838.3 | -267 | Ntrk2 | ENSMUSG00000055254.9 |
| chr13 | 58806690 | 58806917 | 3.63278 | 4.45752 | 1.3896 | Promoter | 13 | 58806596 | 58937030 | 130435 | 1 | ENSMUST00000109838.3 | 95 | Ntrk2 | ENSMUSG00000055254.9 |
| chr13 | 58807594 | 58809224 | 3.92388 | 4.95226 | 1.72334 | Promoter | 13 | 58807900 | 59133970 | 326071 | 1 | ENSMUST00000079828.5 | 0 | Ntrk2 | ENSMUSG00000055254.9 |
| chr13 | 58945768 | 58946108 | 3.36197 | 3.8794 | 1.04874 | Intron (ENSMUST00000079828.5/ENSMUSG00000055254.9, intron 12 of 17) | 13 | 58807900 | 59133970 | 326071 | 1 | ENSMUST00000079828.5 | 137869 | Ntrk2 | ENSMUSG00000055254.9 |
| chr13 | 59011585 | 59011798 | 3.73584 | 4.68548 | 1.59394 | Intron (ENSMUST00000079828.5/ENSMUSG00000055254.9, intron 13 of 17) | 13 | 58807900 | 59133970 | 326071 | 1 | ENSMUST00000079828.5 | 203686 | Ntrk2 | ENSMUSG00000055254.9 |
| chr13 | 59059350 | 59059430 | 4.44539 | 5.95991 | 2.43304 | Intron (ENSMUST00000079828.5/ENSMUSG00000055254.9, intron 15 of 17) | 13 | 58807900 | 59133970 | 326071 | 1 | ENSMUST00000079828.5 | 251451 | Ntrk2 | ENSMUSG00000055254.9 |
| chr13 | 59074182 | 59074329 | 3.51089 | 4.24145 | 1.30242 | Intron (ENSMUST00000079828.5/ENSMUSG00000055254.9, intron 16 of 17) | 13 | 58807900 | 59133970 | 326071 | 1 | ENSMUST00000079828.5 | 266283 | Ntrk2 | ENSMUSG00000055254.9 |
| chr1 | 74037062 | 74037182 | 3.79038 | 4.7183 | 1.63544 | Promoter | 1 | 73993163 | 74037133 | 43971 | 2 | ENSMUST00000187281.1 | 0 | Tns1 | ENSMUSG00000055322.9 |
| chr1 | 74047783 | 74048211 | 3.54576 | 4.21071 | 1.2305 | Intron (ENSMUST00000169786.2/ENSMUSG00000055322.9, intron 3 of 34) | 1 | 74002316 | 74037157 | 34842 | 2 | ENSMUST00000190599.1 | -10627 | Tns1 | ENSMUSG00000055322.9 |
| chr1 | 74051204 | 74051592 | 3.69182 | 4.48673 | 1.41455 | Intron (ENSMUST00000169786.2/ENSMUSG00000055322.9, intron 3 of 34) | 1 | 74002316 | 74037157 | 34842 | 2 | ENSMUST00000190599.1 | -14048 | Tns1 | ENSMUSG00000055322.9 |
| chr1 | 74067703 | 74068349 | 3.4462 | 4.02642 | 1.11926 | Intron (ENSMUST00000169786.2/ENSMUSG00000055322.9, intron 3 of 34) | 1 | 73910232 | 74091351 | 181120 | 2 | ENSMUST00000187584.1 | 23002 | Tns1 | ENSMUSG00000055322.9 |
| chr1 | 74090329 | 74090404 | 3.75782 | 4.60152 | 1.46562 | Promoter | 1 | 73910232 | 74091351 | 181120 | 2 | ENSMUST00000187584.1 | 947 | Tns1 | ENSMUSG00000055322.9 |
| chr1 | 74091277 | 74091582 | 3.44121 | 4.0298 | 1.15866 | Promoter | 1 | 73910232 | 74091351 | 181120 | 2 | ENSMUST00000187584.1 | 0 | Tns1 | ENSMUSG00000055322.9 |
| chr1 | 74098087 | 74098541 | 3.86839 | 5.0543 | 1.80118 | Intron (ENSMUST00000169786.2/ENSMUSG00000055322.9, intron 1 of 34) | 1 | 73997055 | 74096868 | 99814 | 2 | ENSMUST00000188208.1 | -1220 | Tns1 | ENSMUSG00000055322.9 |
| chr1 | 74098963 | 74099474 | 3.69039 | 4.51939 | 1.50751 | Intron (ENSMUST00000169786.2/ENSMUSG00000055322.9, intron 1 of 34) | 1 | 73997055 | 74096868 | 99814 | 2 | ENSMUST00000188208.1 | -2096 | Tns1 | ENSMUSG00000055322.9 |
| chr1 | 74099950 | 74100135 | 3.82715 | 4.76037 | 1.64277 | Intron (ENSMUST00000169786.2/ENSMUSG00000055322.9, intron 1 of 34) | 1 | 73997055 | 74096868 | 99814 | 2 | ENSMUST00000188208.1 | -3083 | Tns1 | ENSMUSG00000055322.9 |
| chr11 | 55349830 | 55350131 | 3.6762 | 4.44234 | 1.3618 | Distal Intergenic | 11 | 55250609 | 55336564 | 85956 | 2 | ENSMUST00000068853.7 | -13267 | Fat2 | ENSMUSG00000055333.9 |
| chr8 | 92960418 | 92960745 | 4.49582 | 6.13034 | 2.60945 | Promoter | 8 | 92961047 | 93001667 | 40621 | 1 | ENSMUST00000072939.6 | -302 | Slc6a2 | ENSMUSG00000055368.8 |
| chr4 | 25798825 | 25798900 | 3.97611 | 5.24777 | 1.97075 | Promoter | 4 | 25797703 | 25799855 | 2153 | 2 | ENSMUST00000149875.1 | 955 | Fut9 | ENSMUSG00000055373.8 |
| chr6 | 88976717 | 88977011 | 3.57163 | 4.25637 | 1.24753 | Intron (ENSMUST00000168837.3/ENSMUSG00000055403.8, intron 1 of 2) | 6 | 88950683 | 89110030 | 159348 | 1 | ENSMUST00000168837.3 | 26035 | 4933427D06Rik | ENSMUSG00000055403.8 |
| chr7 | 99266438 | 99269306 | 3.88834 | 4.99366 | 1.81864 | Promoter | 7 | 99267447 | 99337136 | 69690 | 1 | ENSMUST00000068973.5 | 0 | Map6 | ENSMUSG00000055407.8 |
| chr7 | 99270382 | 99270524 | 3.34899 | 3.85095 | 1.01733 | Intron (ENSMUST00000068973.5/ENSMUSG00000055407.8, intron 1 of 3) | 7 | 99268346 | 99337137 | 68792 | 1 | ENSMUST00000107100.2 | 2037 | Map6 | ENSMUSG00000055407.8 |
| chr7 | 99320595 | 99321280 | 3.86139 | 4.80996 | 1.62147 | Promoter | 7 | 99321200 | 99322455 | 1256 | 1 | ENSMUST00000129058.1 | 0 | Map6 | ENSMUSG00000055407.8 |
| chr7 | 99325759 | 99325885 | 3.34872 | 3.8505 | 1.01733 | Intron (ENSMUST00000068973.5/ENSMUSG00000055407.8, intron 3 of 3) | 7 | 99321200 | 99322455 | 1256 | 1 | ENSMUST00000129058.1 | 4560 | Map6 | ENSMUSG00000055407.8 |
| chr6 | 52261759 | 52262895 | 4.1551 | 5.46691 | 2.16502 | Promoter | 6 | 52262775 | 52267603 | 4829 | 1 | ENSMUST00000152875.1 | 0 | Hottip | ENSMUSG00000055408.7 |
| chr7 | 49974759 | 49976242 | 4.37854 | 6.03615 | 2.62242 | Promoter | 7 | 49974864 | 50560889 | 586026 | 1 | ENSMUST00000151721.2 | 0 | Nell1 | ENSMUSG00000055409.8 |
| chr14 | 93889328 | 93889894 | 3.69952 | 4.49352 | 1.40636 | Promoter | 14 | 93015512 | 93888732 | 873221 | 2 | ENSMUST00000068992.2 | -597 | Pcdh9 | ENSMUSG00000055421.3 |
| chr14 | 93890352 | 93890850 | 3.96196 | 5.17985 | 1.91303 | Distal Intergenic | 14 | 93015512 | 93888732 | 873221 | 2 | ENSMUST00000068992.2 | -1621 | Pcdh9 | ENSMUSG00000055421.3 |
| chr14 | 93891840 | 93892037 | 3.38193 | 3.9006 | 1.01733 | Distal Intergenic | 14 | 93015512 | 93888732 | 873221 | 2 | ENSMUST00000068992.2 | -3109 | Pcdh9 | ENSMUSG00000055421.3 |
| chr14 | 93892568 | 93893013 | 3.38037 | 3.90821 | 1.05215 | Distal Intergenic | 14 | 93015512 | 93888732 | 873221 | 2 | ENSMUST00000068992.2 | -3837 | Pcdh9 | ENSMUSG00000055421.3 |
| chr14 | 93894055 | 93894130 | 3.3862 | 3.91247 | 1.0364 | Distal Intergenic | 14 | 93015512 | 93888732 | 873221 | 2 | ENSMUST00000068992.2 | -5324 | Pcdh9 | ENSMUSG00000055421.3 |
| chr8 | 115706101 | 115706950 | 3.60173 | 4.33699 | 1.32316 | Promoter | 8 | 115682942 | 115707096 | 24155 | 2 | ENSMUST00000069009.6 | 146 | Maf | ENSMUSG00000055435.6 |
| chr17 | 72602500 | 72602766 | 3.92269 | 5.00889 | 1.8113 | Promoter | 17 | 71869442 | 72603709 | 734268 | 2 | ENSMUST00000086639.4 | 943 | Alk | ENSMUSG00000055471.6 |
| chr17 | 72604358 | 72604647 | 4.23891 | 5.83224 | 2.37708 | Promoter | 17 | 71869442 | 72603709 | 734268 | 2 | ENSMUST00000086639.4 | -650 | Alk | ENSMUSG00000055471.6 |
| chr2 | 156584293 | 156584463 | 3.41244 | 4.01329 | 1.21336 | Distal Intergenic | 2 | 156571792 | 156577457 | 5666 | 1 | ENSMUST00000144818.1 | 12502 | Gm14168 | ENSMUSG00000055494.5 |
| chr14 | 25766816 | 25766942 | 3.99448 | 5.06197 | 1.78829 | Intron (ENSMUST00000069180.7/ENSMUSG00000055538.7, intron 1 of 3) | 14 | 25711642 | 25769039 | 57398 | 2 | ENSMUST00000069180.7 | 2097 | Zcchc24 | ENSMUSG00000055538.7 |
| chr14 | 25768513 | 25768710 | 3.47572 | 4.07338 | 1.12726 | Promoter | 14 | 25711642 | 25769039 | 57398 | 2 | ENSMUST00000069180.7 | 329 | Zcchc24 | ENSMUSG00000055538.7 |
| chr14 | 25787392 | 25787669 | 3.77366 | 4.64777 | 1.5212 | Distal Intergenic | 14 | 25711642 | 25769039 | 57398 | 2 | ENSMUST00000069180.7 | -18354 | Zcchc24 | ENSMUSG00000055538.7 |
| chr16 | 60604868 | 60605177 | 4.05304 | 5.2357 | 1.9856 | Promoter | 16 | 60483597 | 60605219 | 121623 | 2 | ENSMUST00000162129.1 | 42 | Epha6 | ENSMUSG00000055540.10 |
| chr1 | 66467790 | 66468677 | 3.88774 | 4.94216 | 1.76738 | Promoter | 1 | 66468367 | 66699866 | 231500 | 1 | ENSMUST00000061620.10 | 0 | Unc80 | ENSMUSG00000055567.11 |
| chr2 | 72474878 | 72475349 | 3.44736 | 4.05903 | 1.21641 | Promoter | 2 | 72476159 | 72486893 | 10735 | 1 | ENSMUST00000102691.5 | -810 | Cdca7 | ENSMUSG00000055612.10 |
| chr2 | 72476336 | 72476583 | 3.94567 | 5.26961 | 1.96642 | Promoter | 2 | 72476390 | 72481860 | 5471 | 1 | ENSMUST00000127344.1 | 0 | Cdca7 | ENSMUSG00000055612.10 |
| chr2 | 72477165 | 72477352 | 3.26121 | 3.71894 | 1.01733 | Promoter | 2 | 72476390 | 72481860 | 5471 | 1 | ENSMUST00000127344.1 | 776 | Cdca7 | ENSMUSG00000055612.10 |
| chr2 | 72496158 | 72496405 | 3.34883 | 3.86602 | 1.0715 | Distal Intergenic | 2 | 72485107 | 72486202 | 1096 | 1 | ENSMUST00000137880.1 | 11052 | Cdca7 | ENSMUSG00000055612.10 |
| chr2 | 72499309 | 72499791 | 3.44481 | 4.02084 | 1.10694 | Distal Intergenic | 2 | 72485107 | 72486202 | 1096 | 1 | ENSMUST00000137880.1 | 14203 | Cdca7 | ENSMUSG00000055612.10 |
| chr2 | 72508120 | 72508325 | 3.46892 | 4.17782 | 1.25839 | Distal Intergenic | 2 | 72485107 | 72486202 | 1096 | 1 | ENSMUST00000137880.1 | 23014 | Cdca7 | ENSMUSG00000055612.10 |
| chr7 | 141058509 | 141058848 | 4.21194 | 5.56334 | 2.18033 | Distal Intergenic | 7 | 141061274 | 141072119 | 10846 | 1 | ENSMUST00000048002.5 | -2426 | B4galnt4 | ENSMUSG00000055629.4 |
| chr7 | 141059663 | 141060145 | 3.35955 | 4.2435 | 1.30715 | Distal Intergenic | 7 | 141061274 | 141072119 | 10846 | 1 | ENSMUST00000048002.5 | -1129 | B4galnt4 | ENSMUSG00000055629.4 |
| chr7 | 141060590 | 141061769 | 3.98191 | 5.2471 | 1.99603 | Promoter | 7 | 141061274 | 141072119 | 10846 | 1 | ENSMUST00000048002.5 | 0 | B4galnt4 | ENSMUSG00000055629.4 |
| chr7 | 141069284 | 141069359 | 3.84725 | 5.1661 | 1.91459 | Intron (ENSMUST00000048002.5/ENSMUSG00000055629.4, intron 14 of 19) | 7 | 141061274 | 141072119 | 10846 | 1 | ENSMUST00000048002.5 | 8011 | B4galnt4 | ENSMUSG00000055629.4 |
| chr2 | 31312600 | 31312800 | 3.36235 | 3.87109 | 1.01733 | Distal Intergenic | 2 | 31314415 | 31460738 | 146324 | 1 | ENSMUST00000113532.3 | -1615 | Hmcn2 | ENSMUSG00000055632.12 |
| chr2 | 31313635 | 31313757 | 3.82971 | 4.72935 | 1.53657 | Promoter | 2 | 31314415 | 31460738 | 146324 | 1 | ENSMUST00000113532.3 | -658 | Hmcn2 | ENSMUSG00000055632.12 |
| chr2 | 31314176 | 31314443 | 3.63688 | 4.38145 | 1.3245 | Promoter | 2 | 31314415 | 31460738 | 146324 | 1 | ENSMUST00000113532.3 | 0 | Hmcn2 | ENSMUSG00000055632.12 |
| chr2 | 31315092 | 31315432 | 3.54333 | 4.20371 | 1.22526 | Promoter | 2 | 31314415 | 31460738 | 146324 | 1 | ENSMUST00000113532.3 | 678 | Hmcn2 | ENSMUSG00000055632.12 |
| chr2 | 31383613 | 31383976 | 3.41754 | 4.25046 | 1.2744 | Exon (ENSMUST00000113532.3/ENSMUSG00000055632.12, exon 27 of 98) | 2 | 31400277 | 31405814 | 5538 | 1 | ENSMUST00000130375.1 | -16301 | Hmcn2 | ENSMUSG00000055632.12 |
| chr2 | 31459383 | 31459458 | 3.74845 | 4.63783 | 1.58572 | Intron (ENSMUST00000113532.3/ENSMUSG00000055632.12, intron 97 of 97) | 2 | 31454708 | 31457989 | 3282 | 1 | ENSMUST00000141294.1 | 4676 | Hmcn2 | ENSMUSG00000055632.12 |
| chr2 | 31460007 | 31460259 | 4.22883 | 5.92789 | 2.43783 | Exon (ENSMUST00000113532.3/ENSMUSG00000055632.12, exon 98 of 98) | 2 | 31454708 | 31457989 | 3282 | 1 | ENSMUST00000141294.1 | 5300 | Hmcn2 | ENSMUSG00000055632.12 |
| chr7 | 5052615 | 5053492 | 3.42998 | 4.24129 | 1.30899 | 5' UTR | 7 | 5051532 | 5053722 | 2191 | 1 | ENSMUST00000069324.5 | 1084 | Zfp580 | ENSMUSG00000055633.5 |
| chr7 | 75983803 | 75984119 | 3.45565 | 4.10831 | 1.2409 | Distal Intergenic | 7 | 75848338 | 75874130 | 25793 | 1 | ENSMUST00000171155.2 | 135466 | Klhl25 | ENSMUSG00000055652.9 |
| chr8 | 15009374 | 15010149 | 3.57925 | 4.3914 | 1.40411 | Promoter | 8 | 15011025 | 15033333 | 22309 | 1 | ENSMUST00000069399.6 | -876 | Kbtbd11 | ENSMUSG00000055675.6 |
| chr8 | 15010879 | 15011614 | 3.57525 | 4.30199 | 1.32055 | Promoter | 8 | 15011025 | 15033333 | 22309 | 1 | ENSMUST00000069399.6 | 0 | Kbtbd11 | ENSMUSG00000055675.6 |
| chr8 | 15011929 | 15012455 | 4.50515 | 6.19848 | 2.63398 | Promoter | 8 | 15011306 | 15027093 | 15788 | 1 | ENSMUST00000184598.1 | 624 | Kbtbd11 | ENSMUSG00000055675.6 |
| chr8 | 15028484 | 15028702 | 3.28414 | 3.75677 | 1.02471 | Exon (ENSMUST00000069399.6/ENSMUSG00000055675.6, exon 2 of 2) | 8 | 15027290 | 15033333 | 6044 | 1 | ENSMUST00000183471.1 | 1195 | Kbtbd11 | ENSMUSG00000055675.6 |
| chr1 | 180330567 | 180331225 | 4.31133 | 5.81309 | 2.4088 | Promoter | 1 | 180326970 | 180330550 | 3581 | 2 | ENSMUST00000050581.3 | -18 | Gm5069 | ENSMUSG00000055676.3 |
| chr1 | 180331709 | 180332713 | 3.56468 | 4.3899 | 1.37176 | 5' UTR | 1 | 180326970 | 180330550 | 3581 | 2 | ENSMUST00000050581.3 | -1160 | Gm5069 | ENSMUSG00000055676.3 |
| chr1 | 180333223 | 180333508 | 3.76328 | 5.03732 | 1.81964 | Exon (ENSMUST00000070181.6/ENSMUSG00000038855.9, exon 2 of 8) | 1 | 180326970 | 180330550 | 3581 | 2 | ENSMUST00000050581.3 | -2674 | Gm5069 | ENSMUSG00000055676.3 |
| chr1 | 180334081 | 180334406 | 3.46318 | 4.24759 | 1.27433 | Exon (ENSMUST00000070181.6/ENSMUSG00000038855.9, exon 2 of 8) | 1 | 180326970 | 180330550 | 3581 | 2 | ENSMUST00000050581.3 | -3532 | Gm5069 | ENSMUSG00000055676.3 |
| chr1 | 180354280 | 180354522 | 3.35517 | 3.92275 | 1.10905 | Intron (ENSMUST00000070181.6/ENSMUSG00000038855.9, intron 2 of 7) | 1 | 180326970 | 180330550 | 3581 | 2 | ENSMUST00000050581.3 | -23731 | Gm5069 | ENSMUSG00000055676.3 |
| chr1 | 180402953 | 180403165 | 3.46629 | 4.06311 | 1.14191 | Intron (ENSMUST00000070181.6/ENSMUSG00000038855.9, intron 2 of 7) | 1 | 180326970 | 180330550 | 3581 | 2 | ENSMUST00000050581.3 | -72404 | Gm5069 | ENSMUSG00000055676.3 |
| chr1 | 180405432 | 180405531 | 3.35769 | 3.86404 | 1.01733 | Intron (ENSMUST00000070181.6/ENSMUSG00000038855.9, intron 2 of 7) | 1 | 180326970 | 180330550 | 3581 | 2 | ENSMUST00000050581.3 | -74883 | Gm5069 | ENSMUSG00000055676.3 |
| chr1 | 180406721 | 180406833 | 3.68616 | 4.6791 | 1.54134 | Intron (ENSMUST00000070181.6/ENSMUSG00000038855.9, intron 2 of 7) | 1 | 180326970 | 180330550 | 3581 | 2 | ENSMUST00000050581.3 | -76172 | Gm5069 | ENSMUSG00000055676.3 |
| chr8 | 70329949 | 70330024 | 4.09492 | 5.40305 | 2.05494 | Promoter | 8 | 70329817 | 70331587 | 1771 | 1 | ENSMUST00000046516.10 | 133 | Gdf1 | ENSMUSG00000055694.10 |
| chr8 | 70331104 | 70331612 | 3.89904 | 4.95742 | 1.76499 | 3' UTR | 8 | 70329817 | 70331587 | 1771 | 1 | ENSMUST00000046516.10 | 1288 | Gdf1 | ENSMUSG00000055694.10 |
| chr10 | 78481877 | 78482137 | 3.32923 | 3.96625 | 1.09914 | Downstream (2-3kb) | 10 | 78484570 | 78487842 | 3273 | 2 | ENSMUST00000069431.4 | 5705 | Gm9978 | ENSMUSG00000055704.4 |
| chr14 | 103650021 | 103651036 | 3.99052 | 5.23384 | 1.9908 | Promoter | 14 | 103650228 | 103704907 | 54680 | 1 | ENSMUST00000069443.8 | 0 | Slain1 | ENSMUSG00000055717.8 |
| chr14 | 103651413 | 103651893 | 4.18289 | 5.92623 | 2.50448 | Promoter | 14 | 103650646 | 103704310 | 53665 | 1 | ENSMUST00000160805.2 | 768 | Slain1 | ENSMUSG00000055717.8 |
| chr14 | 103652349 | 103652763 | 3.77526 | 4.78666 | 1.65496 | Intron (ENSMUST00000069443.8/ENSMUSG00000055717.8, intron 1 of 6) | 14 | 103650646 | 103704310 | 53665 | 1 | ENSMUST00000160805.2 | 1704 | Slain1 | ENSMUSG00000055717.8 |
| chr14 | 103653513 | 103653672 | 3.34402 | 3.98866 | 1.10891 | Intron (ENSMUST00000069443.8/ENSMUSG00000055717.8, intron 1 of 6) | 14 | 103655771 | 103688357 | 32587 | 1 | ENSMUST00000162818.2 | -2099 | Slain1 | ENSMUSG00000055717.8 |
| chr14 | 103656390 | 103656606 | 3.62447 | 4.42394 | 1.43055 | Promoter | 14 | 103655771 | 103688357 | 32587 | 1 | ENSMUST00000162818.2 | 620 | Slain1 | ENSMUSG00000055717.8 |
| chr13 | 40606560 | 40606796 | 3.74501 | 4.65391 | 1.57455 | Distal Intergenic | 13 | 40704005 | 40707949 | 3945 | 1 | ENSMUST00000069457.1 | -97209 | Gm9979 | ENSMUSG00000055732.1 |
| chr13 | 40676593 | 40676773 | 3.68306 | 4.5365 | 1.54392 | Distal Intergenic | 13 | 40704005 | 40707949 | 3945 | 1 | ENSMUST00000069457.1 | -27232 | Gm9979 | ENSMUSG00000055732.1 |
| chr13 | 40696148 | 40696470 | 4.00433 | 5.3198 | 2.00137 | Distal Intergenic | 13 | 40704005 | 40707949 | 3945 | 1 | ENSMUST00000069457.1 | -7535 | Gm9979 | ENSMUSG00000055732.1 |
| chr13 | 40698193 | 40698356 | 4.09952 | 5.35011 | 2.05136 | Distal Intergenic | 13 | 40704005 | 40707949 | 3945 | 1 | ENSMUST00000069457.1 | -5649 | Gm9979 | ENSMUSG00000055732.1 |
| chr13 | 40701925 | 40704161 | 3.84432 | 4.85443 | 1.6723 | Promoter | 13 | 40704005 | 40707949 | 3945 | 1 | ENSMUST00000069457.1 | 0 | Gm9979 | ENSMUSG00000055732.1 |
| chr13 | 40704623 | 40705824 | 3.82437 | 4.76434 | 1.59442 | Promoter | 13 | 40704005 | 40707949 | 3945 | 1 | ENSMUST00000069457.1 | 619 | Gm9979 | ENSMUSG00000055732.1 |
| chr13 | 40706197 | 40706985 | 3.69004 | 4.58892 | 1.47643 | 3' UTR | 13 | 40704005 | 40707949 | 3945 | 1 | ENSMUST00000069457.1 | 2193 | Gm9979 | ENSMUSG00000055732.1 |
| chr13 | 40707912 | 40708135 | 3.55297 | 4.29847 | 1.30507 | 3' UTR | 13 | 40704005 | 40707949 | 3945 | 1 | ENSMUST00000069457.1 | 3908 | Gm9979 | ENSMUSG00000055732.1 |
| chr13 | 40708928 | 40709250 | 3.83655 | 4.78664 | 1.60905 | Distal Intergenic | 13 | 40704005 | 40707949 | 3945 | 1 | ENSMUST00000069457.1 | 4924 | Gm9979 | ENSMUSG00000055732.1 |
| chr13 | 40709880 | 40710461 | 3.95875 | 5.09677 | 1.87755 | Distal Intergenic | 13 | 40704005 | 40707949 | 3945 | 1 | ENSMUST00000069457.1 | 5876 | Gm9979 | ENSMUSG00000055732.1 |
| chr13 | 40711164 | 40712602 | 3.59353 | 4.39364 | 1.43155 | Distal Intergenic | 13 | 40704005 | 40707949 | 3945 | 1 | ENSMUST00000069457.1 | 7160 | Gm9979 | ENSMUSG00000055732.1 |
| chr13 | 40712937 | 40713021 | 3.58965 | 4.52516 | 1.46791 | Downstream (2-3kb) | 13 | 40704005 | 40707949 | 3945 | 1 | ENSMUST00000069457.1 | 8933 | Gm9979 | ENSMUSG00000055732.1 |
| chr13 | 40713846 | 40715263 | 3.48445 | 4.28637 | 1.32958 | Downstream (<1kb) | 13 | 40704005 | 40707949 | 3945 | 1 | ENSMUST00000069457.1 | 9842 | Gm9979 | ENSMUSG00000055732.1 |
| chr15 | 84511016 | 84511514 | 3.35843 | 3.86514 | 1.01733 | Distal Intergenic | 15 | 84553398 | 84557823 | 4426 | 2 | ENSMUST00000069476.4 | 46309 | Ldoc1l | ENSMUSG00000055745.4 |
| chr15 | 84516787 | 84516884 | 3.61539 | 4.43367 | 1.39572 | Distal Intergenic | 15 | 84553398 | 84557823 | 4426 | 2 | ENSMUST00000069476.4 | 40939 | Ldoc1l | ENSMUSG00000055745.4 |
| chr15 | 84540709 | 84540950 | 3.35766 | 3.86599 | 1.02436 | Distal Intergenic | 15 | 84553398 | 84557823 | 4426 | 2 | ENSMUST00000069476.4 | 16873 | Ldoc1l | ENSMUSG00000055745.4 |
| chr4 | 20778975 | 20779239 | 3.47915 | 4.13683 | 1.2725 | Promoter | 4 | 20243842 | 20778866 | 535025 | 2 | ENSMUST00000102998.3 | -110 | Nkain3 | ENSMUSG00000055761.9 |
| chr4 | 20779583 | 20780340 | 3.47807 | 4.12463 | 1.25335 | Promoter | 4 | 20243842 | 20778866 | 535025 | 2 | ENSMUST00000102998.3 | -718 | Nkain3 | ENSMUSG00000055761.9 |
| chr18 | 30489938 | 30490300 | 3.53888 | 4.39617 | 1.37104 | Distal Intergenic | 18 | 30495001 | 30495375 | 375 | 2 | ENSMUST00000069529.1 | 5075 | Gm7936 | ENSMUSG00000055771.1 |
| chr18 | 30491533 | 30491758 | 3.48344 | 4.27874 | 1.32006 | Distal Intergenic | 18 | 30495001 | 30495375 | 375 | 2 | ENSMUST00000069529.1 | 3617 | Gm7936 | ENSMUSG00000055771.1 |
| chr18 | 30493096 | 30493269 | 3.72098 | 4.67862 | 1.56149 | Downstream (1-2kb) | 18 | 30495001 | 30495375 | 375 | 2 | ENSMUST00000069529.1 | 2106 | Gm7936 | ENSMUSG00000055771.1 |
| chr18 | 30494965 | 30495560 | 3.76143 | 4.76547 | 1.63476 | Promoter | 18 | 30495001 | 30495375 | 375 | 2 | ENSMUST00000069529.1 | 0 | Gm7936 | ENSMUSG00000055771.1 |
| chr18 | 30507386 | 30507636 | 3.72327 | 4.56043 | 1.50554 | Distal Intergenic | 18 | 30495001 | 30495375 | 375 | 2 | ENSMUST00000069529.1 | -12012 | Gm7936 | ENSMUSG00000055771.1 |
| chr18 | 30523186 | 30523261 | 4.08382 | 5.36564 | 2.04942 | Distal Intergenic | 18 | 30495001 | 30495375 | 375 | 2 | ENSMUST00000069529.1 | -27812 | Gm7936 | ENSMUSG00000055771.1 |
| chr18 | 30528879 | 30528988 | 3.26507 | 3.86915 | 1.04724 | Distal Intergenic | 18 | 30495001 | 30495375 | 375 | 2 | ENSMUST00000069529.1 | -33505 | Gm7936 | ENSMUSG00000055771.1 |
| chr15 | 91189853 | 91189994 | 3.5218 | 4.36314 | 1.36868 | Intron (ENSMUST00000069511.6/ENSMUSG00000055782.8, intron 1 of 9) | 15 | 91145884 | 91191807 | 45924 | 2 | ENSMUST00000069511.6 | 1813 | Abcd2 | ENSMUSG00000055782.8 |
| chr15 | 91190408 | 91190676 | 4.07195 | 5.3188 | 2.02265 | Exon (ENSMUST00000069511.6/ENSMUSG00000055782.8, exon 1 of 10) | 15 | 91145884 | 91191807 | 45924 | 2 | ENSMUST00000069511.6 | 1131 | Abcd2 | ENSMUSG00000055782.8 |
| chr6 | 72828568 | 72828867 | 3.2747 | 3.74277 | 1.02842 | Distal Intergenic | 6 | 72626378 | 72789254 | 162877 | 2 | ENSMUST00000069536.6 | -39315 | Tcf7l1 | ENSMUSG00000055799.8 |
| chr6 | 72832774 | 72832880 | 3.41757 | 4.0458 | 1.17133 | Distal Intergenic | 6 | 72626378 | 72789254 | 162877 | 2 | ENSMUST00000069536.6 | -43521 | Tcf7l1 | ENSMUSG00000055799.8 |
| chr11 | 103170861 | 103171807 | 4.25953 | 5.6691 | 2.31878 | Promoter | 11 | 103171107 | 103198901 | 27795 | 1 | ENSMUST00000042286.6 | 0 | Fmnl1 | ENSMUSG00000055805.9 |
| chr11 | 103199383 | 103200184 | 3.96988 | 5.10472 | 1.8964 | 3' UTR | 11 | 103196801 | 103197876 | 1076 | 1 | ENSMUST00000129726.2 | 2583 | Fmnl1 | ENSMUSG00000055805.9 |
| chr11 | 103195333 | 103195775 | 3.95349 | 4.98924 | 1.73854 | Exon (ENSMUST00000042286.6/ENSMUSG00000055805.9, exon 18 of 26) | 11 | 103196801 | 103197876 | 1076 | 1 | ENSMUST00000129726.2 | -1026 | Fmnl1 | ENSMUSG00000055805.9 |
| chr11 | 103198203 | 103199046 | 3.45337 | 4.03729 | 1.1194 | 3' UTR | 11 | 103196801 | 103197876 | 1076 | 1 | ENSMUST00000129726.2 | 1403 | Fmnl1 | ENSMUSG00000055805.9 |
| chr16 | 88290990 | 88291129 | 3.46133 | 4.08443 | 1.2324 | Distal Intergenic | 16 | 88505807 | 88506978 | 1172 | 2 | ENSMUST00000069549.2 | 215849 | Cldn17 | ENSMUSG00000055811.2 |
| chr7 | 24332541 | 24332876 | 4.32816 | 5.75392 | 2.31005 | Downstream (<1kb) | 7 | 24333080 | 24333965 | 886 | 2 | ENSMUST00000069562.4 | 1089 | Tescl | ENSMUSG00000055826.4 |
| chr17 | 23831580 | 23832192 | 3.62804 | 4.40177 | 1.4185 | Downstream (1-2kb) | 17 | 23824740 | 23829109 | 4370 | 2 | ENSMUST00000069579.5 | -2472 | Tceb2 | ENSMUSG00000055839.5 |
| chr3 | 104776075 | 104776296 | 3.26884 | 3.88877 | 1.06396 | 5' UTR | 3 | 104767406 | 104777547 | 10142 | 2 | ENSMUST00000139783.1 | 1251 | Fam19a3 | ENSMUSG00000055865.8 |
| chr3 | 104777183 | 104777661 | 3.66116 | 4.56022 | 1.47043 | Promoter | 3 | 104767406 | 104777547 | 10142 | 2 | ENSMUST00000139783.1 | 0 | Fam19a3 | ENSMUSG00000055865.8 |
| chr3 | 104778549 | 104779655 | 3.72296 | 4.67466 | 1.59698 | Intron (ENSMUST00000162644.1/ENSMUSG00000055865.8, intron 1 of 1) | 3 | 104767406 | 104777547 | 10142 | 2 | ENSMUST00000139783.1 | -1003 | Fam19a3 | ENSMUSG00000055865.8 |
| chr3 | 104781420 | 104781990 | 3.73639 | 4.71702 | 1.60301 | Promoter | 3 | 104777361 | 104781840 | 4480 | 2 | ENSMUST00000162644.1 | 0 | Fam19a3 | ENSMUSG00000055865.8 |
| chr6 | 70948679 | 70948907 | 3.42535 | 4.00307 | 1.14876 | Distal Intergenic | 6 | 70956531 | 70961063 | 4533 | 1 | ENSMUST00000163089.1 | -7624 | Foxi3 | ENSMUSG00000055874.6 |
| chr6 | 70950285 | 70950592 | 3.44874 | 4.03996 | 1.15402 | Distal Intergenic | 6 | 70956531 | 70961063 | 4533 | 1 | ENSMUST00000163089.1 | -5939 | Foxi3 | ENSMUSG00000055874.6 |
| chr6 | 70951073 | 70951477 | 3.57583 | 4.27699 | 1.30264 | Distal Intergenic | 6 | 70956531 | 70961063 | 4533 | 1 | ENSMUST00000163089.1 | -5054 | Foxi3 | ENSMUSG00000055874.6 |
| chr6 | 70951838 | 70951940 | 3.37164 | 3.89766 | 1.06182 | Distal Intergenic | 6 | 70956531 | 70961063 | 4533 | 1 | ENSMUST00000163089.1 | -4591 | Foxi3 | ENSMUSG00000055874.6 |
| chr6 | 70952506 | 70952983 | 3.52866 | 4.18726 | 1.24267 | Distal Intergenic | 6 | 70956531 | 70961063 | 4533 | 1 | ENSMUST00000163089.1 | -3548 | Foxi3 | ENSMUSG00000055874.6 |
| chr6 | 70953289 | 70954017 | 3.91382 | 5.13592 | 1.91703 | Distal Intergenic | 6 | 70956531 | 70961063 | 4533 | 1 | ENSMUST00000163089.1 | -2514 | Foxi3 | ENSMUSG00000055874.6 |
| chr6 | 70954918 | 70955370 | 3.42726 | 4.26008 | 1.29735 | Distal Intergenic | 6 | 70956531 | 70961063 | 4533 | 1 | ENSMUST00000163089.1 | -1161 | Foxi3 | ENSMUSG00000055874.6 |
| chr6 | 70955743 | 70957294 | 3.96147 | 5.07113 | 1.85856 | Promoter | 6 | 70956531 | 70961063 | 4533 | 1 | ENSMUST00000163089.1 | 0 | Foxi3 | ENSMUSG00000055874.6 |
| chr6 | 70959582 | 70959857 | 3.42606 | 4.02439 | 1.18995 | Intron (ENSMUST00000163089.1/ENSMUSG00000055874.6, intron 1 of 1) | 6 | 70956606 | 70961066 | 4461 | 1 | ENSMUST00000069634.5 | 2977 | Foxi3 | ENSMUSG00000055874.6 |
| chr6 | 70960585 | 70960879 | 4.20416 | 5.53945 | 2.15131 | Exon (ENSMUST00000163089.1/ENSMUSG00000055874.6, exon 2 of 2) | 6 | 70956606 | 70961066 | 4461 | 1 | ENSMUST00000069634.5 | 3980 | Foxi3 | ENSMUSG00000055874.6 |
| chr6 | 70967017 | 70967197 | 3.77856 | 4.73881 | 1.62573 | Distal Intergenic | 6 | 70956606 | 70961066 | 4461 | 1 | ENSMUST00000069634.5 | 10412 | Foxi3 | ENSMUSG00000055874.6 |
| chr6 | 70967680 | 70967789 | 3.95745 | 5.03278 | 1.84273 | Distal Intergenic | 6 | 70956606 | 70961066 | 4461 | 1 | ENSMUST00000069634.5 | 11075 | Foxi3 | ENSMUSG00000055874.6 |
| chr3 | 107536910 | 107537332 | 4.14237 | 5.44329 | 2.10388 | Distal Intergenic | 3 | 107553698 | 107555073 | 1376 | 2 | ENSMUST00000052853.6 | 17741 | Ubl4b | ENSMUSG00000055891.6 |
| chr3 | 107538293 | 107538634 | 3.27579 | 3.78098 | 1.05291 | Distal Intergenic | 3 | 107553698 | 107555073 | 1376 | 2 | ENSMUST00000052853.6 | 16439 | Ubl4b | ENSMUSG00000055891.6 |
| chr3 | 107539442 | 107539659 | 3.43285 | 4.13367 | 1.24982 | Distal Intergenic | 3 | 107553698 | 107555073 | 1376 | 2 | ENSMUST00000052853.6 | 15414 | Ubl4b | ENSMUSG00000055891.6 |
| chr2 | 173586730 | 173586865 | 3.57374 | 4.32871 | 1.40203 | Exon (ENSMUST00000153162.2/ENSMUSG00000055897.8, exon 13 of 13) | 2 | 173585853 | 173590762 | 4910 | 2 | ENSMUST00000149947.2 | 3897 | Ppp4r1l-ps | ENSMUSG00000055897.8 |
| chr5 | 35349596 | 35349673 | 3.70751 | 4.74934 | 1.60819 | Distal Intergenic | 5 | 35379152 | 35388476 | 9325 | 2 | ENSMUST00000069741.3 | 38803 | E130018O15Rik | ENSMUSG00000055944.3 |
| chr5 | 35367224 | 35367528 | 3.84015 | 4.76725 | 1.60386 | Distal Intergenic | 5 | 35379152 | 35388476 | 9325 | 2 | ENSMUST00000069741.3 | 20948 | E130018O15Rik | ENSMUSG00000055944.3 |
| chr5 | 35379424 | 35379601 | 3.76822 | 4.72348 | 1.59081 | Exon (ENSMUST00000069741.3/ENSMUSG00000055944.3, exon 2 of 2) | 5 | 35379152 | 35388476 | 9325 | 2 | ENSMUST00000069741.3 | 8875 | E130018O15Rik | ENSMUSG00000055944.3 |
| chr5 | 35380123 | 35381076 | 3.51522 | 4.16172 | 1.20559 | Exon (ENSMUST00000069741.3/ENSMUSG00000055944.3, exon 2 of 2) | 5 | 35379152 | 35388476 | 9325 | 2 | ENSMUST00000069741.3 | 7400 | E130018O15Rik | ENSMUSG00000055944.3 |
| chr5 | 35382057 | 35382237 | 3.30171 | 3.77975 | 1.01733 | Exon (ENSMUST00000069741.3/ENSMUSG00000055944.3, exon 2 of 2) | 5 | 35379152 | 35388476 | 9325 | 2 | ENSMUST00000069741.3 | 6239 | E130018O15Rik | ENSMUSG00000055944.3 |
| chr5 | 35383115 | 35383190 | 3.51867 | 4.14989 | 1.16844 | Intron (ENSMUST00000069741.3/ENSMUSG00000055944.3, intron 1 of 1) | 5 | 35379152 | 35388476 | 9325 | 2 | ENSMUST00000069741.3 | 5286 | E130018O15Rik | ENSMUSG00000055944.3 |
| chr5 | 35383936 | 35384094 | 3.60931 | 4.3164 | 1.27301 | Intron (ENSMUST00000069741.3/ENSMUSG00000055944.3, intron 1 of 1) | 5 | 35379152 | 35388476 | 9325 | 2 | ENSMUST00000069741.3 | 4382 | E130018O15Rik | ENSMUSG00000055944.3 |
| chr5 | 35384598 | 35384778 | 3.44951 | 4.04187 | 1.15738 | Intron (ENSMUST00000069741.3/ENSMUSG00000055944.3, intron 1 of 1) | 5 | 35379152 | 35388476 | 9325 | 2 | ENSMUST00000069741.3 | 3698 | E130018O15Rik | ENSMUSG00000055944.3 |
| chr5 | 35385471 | 35385964 | 3.84694 | 4.95206 | 1.7821 | Intron (ENSMUST00000069741.3/ENSMUSG00000055944.3, intron 1 of 1) | 5 | 35379152 | 35388476 | 9325 | 2 | ENSMUST00000069741.3 | 2512 | E130018O15Rik | ENSMUSG00000055944.3 |
| chr5 | 35386521 | 35386812 | 4.1808 | 5.44591 | 2.08401 | Intron (ENSMUST00000069741.3/ENSMUSG00000055944.3, intron 1 of 1) | 5 | 35379152 | 35388476 | 9325 | 2 | ENSMUST00000069741.3 | 1664 | E130018O15Rik | ENSMUSG00000055944.3 |
| chr5 | 35387148 | 35387916 | 3.59496 | 4.30064 | 1.28921 | Promoter | 5 | 35379152 | 35388476 | 9325 | 2 | ENSMUST00000069741.3 | 560 | E130018O15Rik | ENSMUSG00000055944.3 |
| chr5 | 35388307 | 35390152 | 4.02486 | 5.23692 | 1.98667 | Promoter | 5 | 35379152 | 35388476 | 9325 | 2 | ENSMUST00000069741.3 | 0 | E130018O15Rik | ENSMUSG00000055944.3 |
| chr17 | 8338770 | 8338955 | 3.40513 | 3.94601 | 1.0539 | Intron (ENSMUST00000140890.2/ENSMUSG00000073468.6, intron 1 of 5) | 17 | 8340406 | 8344112 | 3707 | 1 | ENSMUST00000069742.6 | -1451 | Prr18 | ENSMUSG00000055945.7 |
| chr17 | 8340375 | 8340659 | 3.65659 | 4.61676 | 1.47898 | Promoter | 17 | 8340406 | 8344112 | 3707 | 1 | ENSMUST00000069742.6 | 0 | Prr18 | ENSMUSG00000055945.7 |
| chr17 | 8341235 | 8341604 | 3.53967 | 4.50533 | 1.44427 | Promoter | 17 | 8340739 | 8344112 | 3374 | 1 | ENSMUST00000163887.1 | 497 | Prr18 | ENSMUSG00000055945.7 |
| chr17 | 8342184 | 8342651 | 3.90125 | 4.9922 | 1.81211 | 3' UTR | 17 | 8340739 | 8344112 | 3374 | 1 | ENSMUST00000163887.1 | 1446 | Prr18 | ENSMUSG00000055945.7 |
| chr8 | 35822525 | 35823561 | 3.79521 | 4.76113 | 1.60608 | Downstream (1-2kb) | 8 | 35824709 | 35826559 | 1851 | 2 | ENSMUST00000060128.5 | 2998 | Cldn23 | ENSMUSG00000055976.5 |
| chr8 | 35824835 | 35825816 | 3.5109 | 4.14555 | 1.19531 | Promoter | 8 | 35824709 | 35826559 | 1851 | 2 | ENSMUST00000060128.5 | 743 | Cldn23 | ENSMUSG00000055976.5 |
| chr8 | 35826135 | 35826522 | 3.6969 | 4.67022 | 1.54931 | Promoter | 8 | 35824709 | 35826559 | 1851 | 2 | ENSMUST00000060128.5 | 37 | Cldn23 | ENSMUSG00000055976.5 |
| chr8 | 35827117 | 35828354 | 3.71688 | 4.55922 | 1.48773 | Promoter | 8 | 35824709 | 35826559 | 1851 | 2 | ENSMUST00000060128.5 | -559 | Cldn23 | ENSMUSG00000055976.5 |

| | | | | | | | | | | | | | | | |
|---|---|---|---|---|---|---|---|---|---|---|---|---|---|---|---|
| chr8 | 35831115 | 35831206 | 3.7651 | 4.63316 | 1.51806 | Distal Intergenic | 8 | 35824709 | 35826559 | 1851 | 2 | ENSMUST00000060128.5 | -4557 | Cldn23 | ENSMUSG00000055976.5 |
| chr8 | 35850109 | 35850384 | 3.83978 | 4.86891 | 1.67748 | Distal Intergenic | 8 | 35824709 | 35826559 | 1851 | 2 | ENSMUST00000060128.5 | -23551 | Cldn23 | ENSMUSG00000055976.5 |
| chr8 | 35865495 | 35865575 | 3.76343 | 4.6809 | 1.59911 | Distal Intergenic | 8 | 35824709 | 35826559 | 1851 | 2 | ENSMUST00000060128.5 | -38937 | Cldn23 | ENSMUSG00000055976.5 |
| chr8 | 35887595 | 35887869 | 3.68909 | 4.49761 | 1.47608 | Distal Intergenic | 8 | 35824709 | 35826559 | 1851 | 2 | ENSMUST00000060128.5 | -61037 | Cldn23 | ENSMUSG00000055976.5 |
| chr7 | 45657390 | 45657658 | 3.32528 | 3.92086 | 1.09269 | Intron (ENSMUST00000069800.4/ENSMUSG00000055978.4, intron 1 of 1) | 7 | 45648595 | 45666308 | 17714 | 2 | ENSMUST00000069800.4 | 8650 | Fut2 | ENSMUSG00000055978.4 |
| chr8 | 88647983 | 88648059 | 4.43018 | 5.94903 | 2.45437 | Promoter | 8 | 88647384 | 88665700 | 18317 | 1 | ENSMUST00000124091.1 | 600 | Nod2 | ENSMUSG00000055994.10 |
| chr8 | 88649627 | 88649887 | 3.3569 | 3.86283 | 1.01733 | Intron (ENSMUST00000054324.9/ENSMUSG00000055994.10, intron 1 of 12) | 8 | 88651341 | 88688474 | 37134 | 1 | ENSMUST00000109634.2 | -1454 | Nod2 | ENSMUSG00000055994.10 |
| chr5 | 9266301 | 9266458 | 3.90408 | 5.04017 | 1.77698 | Promoter | 5 | 9266175 | 9461486 | 195312 | 1 | ENSMUST00000154662.2 | 127 | 9330182L06Rik | ENSMUSG00000056004.11 |
| chr11 | 94701436 | 94701792 | 3.38495 | 3.91461 | 1.0507 | Intron (ENSMUST00000069852.1/ENSMUSG00000056008.1, intron 1 of 1) | 11 | 94694498 | 94704499 | 10002 | 2 | ENSMUST00000069852.1 | 2707 | Gm11541 | ENSMUSG00000056008.1 |
| chr8 | 71808874 | 71808949 | 2.42863 | 4.11444 | 1.22396 | Distal Intergenic | 8 | 71882068 | 71892565 | 10498 | 1 | ENSMUST00000188685.1 | -73119 | Zfp709 | ENSMUSG00000056019.7 |
| chr8 | 71838067 | 71838310 | 2.60703 | 3.99581 | 1.16221 | Distal Intergenic | 8 | 71882068 | 71892565 | 10498 | 1 | ENSMUST00000188685.1 | -43758 | Zfp709 | ENSMUSG00000056019.7 |
| chr8 | 71852606 | 71852948 | 3.07631 | 5.47039 | 2.10927 | Distal Intergenic | 8 | 71882068 | 71892565 | 10498 | 1 | ENSMUST00000188685.1 | -29120 | Zfp709 | ENSMUSG00000056019.7 |
| chr8 | 71853718 | 71854008 | 2.91043 | 4.51214 | 1.48808 | Distal Intergenic | 8 | 71882068 | 71892565 | 10498 | 1 | ENSMUST00000188685.1 | -28060 | Zfp709 | ENSMUSG00000056019.7 |
| chr9 | 85844144 | 85844345 | 3.40597 | 4.20988 | 1.29647 | Promoter | 9 | 85820302 | 85844323 | 24022 | 2 | ENSMUST00000185960.1 | 0 | 9330154J02Rik | ENSMUSG00000056031.8 |
| chr7 | 35583732 | 35583910 | 3.44374 | 4.16731 | 1.2812 | 3' UTR | 7 | 35578993 | 35585582 | 6590 | 2 | ENSMUST00000069912.5 | 1672 | Rgs9bp | ENSMUSG00000056043.5 |
| chr7 | 35584450 | 35584615 | 4.67029 | 6.68993 | 3.02995 | Promoter | 7 | 35578993 | 35585582 | 6590 | 2 | ENSMUST00000069912.5 | 967 | Rgs9bp | ENSMUSG00000056043.5 |
| chr10 | 49780568 | 49780687 | 3.42858 | 4.01816 | 1.18405 | Intron (ENSMUST00000105487.2/ENSMUSG00000056073.9, intron 2 of 16) | 10 | 49099843 | 49783259 | 683417 | 2 | ENSMUST00000105484.4 | 2572 | Grik2 | ENSMUSG00000056073.9 |
| chr10 | 49782757 | 49782855 | 3.53231 | 4.18474 | 1.21818 | Promoter | 10 | 49099843 | 49783259 | 683417 | 2 | ENSMUST00000105484.4 | 404 | Grik2 | ENSMUSG00000056073.9 |
| chr10 | 49783406 | 49783585 | 4.16198 | 5.40911 | 2.05221 | Promoter | 10 | 49100989 | 49783409 | 682421 | 2 | ENSMUST00000105485.2 | 0 | Grik2 | ENSMUSG00000056073.9 |
| chr10 | 49783994 | 49784748 | 3.72018 | 4.57196 | 1.4909 | Promoter | 10 | 49100989 | 49783409 | 682421 | 2 | ENSMUST00000105485.2 | -586 | Grik2 | ENSMUSG00000056073.9 |
| chr10 | 49785264 | 49785683 | 4.15211 | 5.38093 | 2.00149 | Intron (ENSMUST00000105487.2/ENSMUSG00000056073.9, intron 1 of 16) | 10 | 49100989 | 49783409 | 682421 | 2 | ENSMUST00000105485.2 | -1856 | Grik2 | ENSMUSG00000056073.9 |
| chr10 | 49786740 | 49787770 | 3.34997 | 3.90065 | 1.079 | Promoter | 10 | 49495438 | 49788754 | 293317 | 2 | ENSMUST00000105483.1 | 984 | Grik2 | ENSMUSG00000056073.9 |
| chr10 | 49788401 | 49788675 | 3.69245 | 4.56306 | 1.50797 | Promoter | 10 | 49495438 | 49788754 | 293317 | 2 | ENSMUST00000105483.1 | 79 | Grik2 | ENSMUSG00000056073.9 |
| chr15 | 25414295 | 25414607 | 3.73182 | 4.65725 | 1.50285 | Promoter | 15 | 25414192 | 25414844 | 653 | 1 | ENSMUST00000069992.5 | 104 | Gm5468 | ENSMUSG00000056089.5 |
| chr15 | 25420657 | 25420732 | 3.35061 | 3.87278 | 1.08578 | Distal Intergenic | 15 | 25414192 | 25414844 | 653 | 1 | ENSMUST00000069992.5 | 6466 | Gm5468 | ENSMUSG00000056089.5 |
| chr18 | 20744776 | 20744981 | 3.43853 | 4.01464 | 1.11548 | Intron (ENSMUST00000070080.5/ENSMUSG00000056124.5, intron 1 of 8) | 18 | 20684599 | 20746404 | 61806 | 2 | ENSMUST00000070080.5 | 1423 | B4galt6 | ENSMUSG00000056124.5 |
| chr18 | 20745978 | 20746598 | 3.86455 | 4.87853 | 1.6924 | Promoter | 18 | 20684599 | 20746404 | 61806 | 2 | ENSMUST00000070080.5 | 0 | B4galt6 | ENSMUSG00000056124.5 |
| chr18 | 20747139 | 20747502 | 3.50927 | 4.13684 | 1.17005 | Promoter | 18 | 20684599 | 20746404 | 61806 | 2 | ENSMUST00000070080.5 | -736 | B4galt6 | ENSMUSG00000056124.5 |
| chr17 | 7397084 | 7397440 | 3.37604 | 4.33366 | 1.36742 | Distal Intergenic | 17 | 7363712 | 7385305 | 21594 | 2 | ENSMUST00000070059.3 | -11780 | Gm9992 | ENSMUSG00000056133.4 |
| chr11 | 93097348 | 93097699 | 3.94724 | 5.16574 | 1.87248 | Promoter | 11 | 93098028 | 93099866 | 1839 | 1 | ENSMUST00000149611.1 | -329 | Car10 | ENSMUSG00000056158.9 |
| chr11 | 93099911 | 93100338 | 3.38222 | 3.97278 | 1.16476 | Promoter | 11 | 93099290 | 93601749 | 502460 | 1 | ENSMUST00000107863.2 | 622 | Car10 | ENSMUSG00000056158.9 |
| chr11 | 93101375 | 93101589 | 3.31289 | 3.80575 | 1.04969 | Intron (ENSMUST00000107859.2/ENSMUSG00000056158.9, intron 2 of 4) | 11 | 93099290 | 93601749 | 502460 | 1 | ENSMUST00000107863.2 | 2086 | Car10 | ENSMUSG00000056158.9 |
| chr11 | 93106329 | 93106827 | 3.47061 | 4.06619 | 1.12908 | Intron (ENSMUST00000107859.2/ENSMUSG00000056158.9, intron 2 of 4) | 11 | 93099290 | 93601749 | 502460 | 1 | ENSMUST00000107863.2 | 7040 | Car10 | ENSMUSG00000056158.9 |
| chr9 | 114645246 | 114645570 | 3.93777 | 4.95924 | 1.72865 | Distal Intergenic | 9 | 114585878 | 114640200 | 54323 | 2 | ENSMUST00000070117.6 | -5047 | Cnot10 | ENSMUSG00000056167.6 |
| chr4 | 126284616 | 126284870 | 3.62487 | 4.52762 | 1.44277 | Distal Intergenic | 4 | 126286793 | 126314330 | 27538 | 1 | ENSMUST00000070132.6 | -1923 | Col8a2 | ENSMUSG00000056174.8 |
| chr4 | 126285909 | 126287447 | 3.99423 | 5.09059 | 1.82959 | Promoter | 4 | 126286793 | 126314330 | 27538 | 1 | ENSMUST00000070132.6 | 0 | Col8a2 | ENSMUSG00000056174.8 |
| chr4 | 126287767 | 126288102 | 3.35658 | 3.86236 | 1.01733 | Promoter | 4 | 126286793 | 126314330 | 27538 | 1 | ENSMUST00000070132.6 | 975 | Col8a2 | ENSMUSG00000056174.8 |
| chr4 | 126299484 | 126300435 | 3.61535 | 4.50172 | 1.43725 | Promoter | 4 | 126300530 | 126309785 | 9256 | 1 | ENSMUST00000128435.1 | -95 | Col8a2 | ENSMUSG00000056174.8 |
| chr4 | 126300899 | 126301332 | 3.3082 | 4.24275 | 1.29486 | Promoter | 4 | 126300530 | 126309785 | 9256 | 1 | ENSMUST00000128435.1 | 370 | Col8a2 | ENSMUSG00000056174.8 |
| chr13 | 57429165 | 57429904 | 3.52755 | 4.1746 | 1.20452 | 3' UTR | 13 | 57429295 | 57907587 | 478293 | 2 | ENSMUST00000172326.1 | 477683 | Spock1 | ENSMUSG00000056222.9 |
| chr13 | 57666678 | 57666859 | 3.38428 | 4.06402 | 1.14223 | Intron (ENSMUST00000187852.1/ENSMUSG00000056222.9, intron 3 of 9) | 13 | 57429295 | 57907587 | 478293 | 2 | ENSMUST00000172326.1 | 240728 | Spock1 | ENSMUSG00000056222.9 |
| chr13 | 57741935 | 57742010 | 3.79224 | 4.66911 | 1.53328 | Intron (ENSMUST00000187852.1/ENSMUSG00000056222.9, intron 2 of 9) | 13 | 57429295 | 57907587 | 478293 | 2 | ENSMUST00000172326.1 | 165577 | Spock1 | ENSMUSG00000056222.9 |
| chr13 | 57889884 | 57890038 | 3.39039 | 3.92568 | 1.06092 | Intron (ENSMUST00000187852.1/ENSMUSG00000056222.9, intron 2 of 9) | 13 | 57429295 | 57907587 | 478293 | 2 | ENSMUST00000172326.1 | 17549 | Spock1 | ENSMUSG00000056222.9 |
| chr13 | 57906634 | 57906772 | 3.38193 | 3.9006 | 1.01733 | Promoter | 13 | 57429295 | 57907587 | 478293 | 2 | ENSMUST00000172326.1 | 815 | Spock1 | ENSMUSG00000056222.9 |
| chr13 | 57907107 | 57908216 | 3.59576 | 4.41999 | 1.38163 | Promoter | 13 | 57429295 | 57907587 | 478293 | 2 | ENSMUST00000172326.1 | 0 | Spock1 | ENSMUSG00000056222.9 |
| chr13 | 57909129 | 57909339 | 3.34744 | 3.84858 | 1.01733 | Promoter | 13 | 57421195 | 57908314 | 487120 | 2 | ENSMUST00000187852.1 | -816 | Spock1 | ENSMUSG00000056222.9 |
| chr15 | 66283462 | 66283813 | 3.97805 | 5.2421 | 1.91149 | Exon (ENSMUST00000183633.1/ENSMUSG00000056258.8, exon 1 of 1) | 15 | 66283073 | 66286051 | 2979 | 2 | ENSMUST00000183633.1 | 2238 | Kcnq3 | ENSMUSG00000056258.8 |
| chr15 | 66284450 | 66284860 | 3.58694 | 4.49663 | 1.45456 | Exon (ENSMUST00000183633.1/ENSMUSG00000056258.8, exon 1 of 1) | 15 | 66283073 | 66286051 | 2979 | 2 | ENSMUST00000183633.1 | 1191 | Kcnq3 | ENSMUSG00000056258.8 |
| chr15 | 66285250 | 66286096 | 4.11746 | 5.51114 | 2.15701 | Promoter | 15 | 66283073 | 66286051 | 2979 | 2 | ENSMUST00000183633.1 | 0 | Kcnq3 | ENSMUSG00000056258.8 |
| chr3 | 106716367 | 106716713 | 3.33918 | 3.83612 | 1.01733 | Intron (ENSMUST00000098751.4/ENSMUSG00000056260.10, intron 1 of 2) | 3 | 106721682 | 106736574 | 14893 | 1 | ENSMUST00000098750.3 | -4969 | Lrif1 | ENSMUSG00000056260.10 |
| chr14 | 13284808 | 13284946 | 3.55301 | 4.21568 | 1.21755 | Promoter | 14 | 13284780 | 13615459 | 330680 | 1 | ENSMUST00000070323.6 | 29 | Synpr | ENSMUSG00000056296.11 |
| chr3 | 54914883 | 54916033 | 3.6327 | 4.44499 | 1.40753 | Promoter | 3 | 54897068 | 54915031 | 17964 | 2 | ENSMUST00000162201.1 | 0 | Sertm1 | ENSMUSG00000056306.5 |
| chr3 | 54916872 | 54917218 | 3.47459 | 4.09511 | 1.21149 | Promoter | 3 | 54897071 | 54915887 | 18817 | 2 | ENSMUST00000070342.3 | -986 | Sertm1 | ENSMUSG00000056306.5 |
| chr10 | 29145482 | 29145864 | 3.55003 | 4.22175 | 1.25439 | Intron (ENSMUST00000092629.2/ENSMUSG00000038916.6, intron 1 of 6) | 10 | 29143282 | 29144194 | 913 | 2 | ENSMUST00000070359.2 | -1289 | Gm9996 | ENSMUSG00000056316.2 |
| chr10 | 29146240 | 29148663 | 3.8724 | 4.9045 | 1.72221 | 5' UTR | 10 | 29143282 | 29144194 | 913 | 2 | ENSMUST00000070359.2 | -2047 | Gm9996 | ENSMUSG00000056316.2 |
| chr2 | 19451836 | 19452233 | 3.64988 | 4.46212 | 1.44507 | Promoter | 2 | 19451828 | 19453151 | 1324 | 1 | ENSMUST00000070378.2 | 9 | 1810010K12Rik | ENSMUSG00000056329.3 |
| chr2 | 19452637 | 19452712 | 3.47931 | 4.08874 | 1.16295 | Promoter | 2 | 19452470 | 19454995 | 2526 | 1 | ENSMUST00000149577.1 | 168 | 1810010K12Rik | ENSMUSG00000056329.3 |
| chr2 | 19453645 | 19454415 | 3.50718 | 4.16166 | 1.24831 | Exon (ENSMUST00000149577.1/ENSMUSG00000056329.3, exon 2 of 2) | 2 | 19452470 | 19454995 | 2526 | 1 | ENSMUST00000149577.1 | 1176 | 1810010K12Rik | ENSMUSG00000056329.3 |
| chr2 | 19454839 | 19455062 | 3.61552 | 4.41263 | 1.43653 | Exon (ENSMUST00000149577.1/ENSMUSG00000056329.3, exon 2 of 2) | 2 | 19452470 | 19454995 | 2526 | 1 | ENSMUST00000149577.1 | 2370 | 1810010K12Rik | ENSMUSG00000056329.3 |
| chr2 | 19456725 | 19457356 | 3.63456 | 4.44374 | 1.42153 | Distal Intergenic | 2 | 19452470 | 19454995 | 2526 | 1 | ENSMUST00000149577.1 | 4256 | 1810010K12Rik | ENSMUSG00000056329.3 |
| chr2 | 19457705 | 19458451 | 3.36219 | 4.17278 | 1.27538 | Distal Intergenic | 2 | 19452470 | 19454995 | 2526 | 1 | ENSMUST00000149577.1 | 5236 | 1810010K12Rik | ENSMUSG00000056329.3 |
| chr12 | 73367465 | 73367773 | 3.54412 | 4.26908 | 1.34278 | Intron (ENSMUST00000070425.5/ENSMUSG00000056359.5, intron 1 of 2) | 12 | 73364075 | 73371445 | 7371 | 2 | ENSMUST00000070425.5 | 3672 | D830013O20Rik | ENSMUSG00000056359.5 |
| chr12 | 73427995 | 73428078 | 3.79617 | 4.73853 | 1.63055 | Distal Intergenic | 12 | 73364075 | 73371445 | 7371 | 2 | ENSMUST00000070425.5 | -56551 | D830013O20Rik | ENSMUSG00000056359.5 |
| chr12 | 73447703 | 73447927 | 3.6124 | 4.39559 | 1.41519 | Distal Intergenic | 12 | 73364075 | 73371445 | 7371 | 2 | ENSMUST00000070425.5 | -76259 | D830013O20Rik | ENSMUSG00000056359.5 |
| chr12 | 73449855 | 73450130 | 3.71222 | 4.60765 | 1.53862 | Distal Intergenic | 12 | 73364075 | 73371445 | 7371 | 2 | ENSMUST00000070425.5 | -78411 | D830013O20Rik | ENSMUSG00000056359.5 |
| chr5 | 139281218 | 139281441 | 3.55383 | 4.29226 | 1.29511 | Intron (ENSMUST00000151450.2/ENSMUSG00000056413.11, intron 4 of 10) | 5 | 139272930 | 139274641 | 1712 | 2 | ENSMUST00000123748.1 | -6578 | Adap1 | ENSMUSG00000056413.11 |
| chr5 | 139324570 | 139325205 | 3.855 | 4.86164 | 1.71603 | Promoter | 5 | 139271876 | 139325622 | 53747 | 2 | ENSMUST00000110865.1 | 417 | Adap1 | ENSMUSG00000056413.11 |
| chr11 | 35120363 | 35120545 | 3.32151 | 3.84098 | 1.11576 | Promoter | 11 | 35121224 | 35708507 | 587284 | 1 | ENSMUST00000069837.3 | -679 | Slit3 | ENSMUSG00000056427.5 |
| chr11 | 35120931 | 35121143 | 3.66596 | 4.51196 | 1.52586 | Promoter | 11 | 35121224 | 35708507 | 587284 | 1 | ENSMUST00000069837.3 | -81 | Slit3 | ENSMUSG00000056427.5 |
| chr11 | 35121533 | 35122062 | 3.73026 | 4.97557 | 1.75304 | Promoter | 11 | 35121224 | 35708507 | 587284 | 1 | ENSMUST00000069837.3 | 310 | Slit3 | ENSMUSG00000056427.5 |
| chr11 | 35123454 | 35123529 | 4.25222 | 5.71651 | 2.3147 | Intron (ENSMUST00000069837.3/ENSMUSG00000056427.5, intron 1 of 35) | 11 | 35121224 | 35708507 | 587284 | 1 | ENSMUST00000069837.3 | 2231 | Slit3 | ENSMUSG00000056427.5 |
| chr11 | 35123871 | 35124094 | 3.62324 | 4.39996 | 1.44565 | Intron (ENSMUST00000069837.3/ENSMUSG00000056427.5, intron 1 of 35) | 11 | 35121224 | 35708507 | 587284 | 1 | ENSMUST00000069837.3 | 2648 | Slit3 | ENSMUSG00000056427.5 |
| chr11 | 35125503 | 35125961 | 3.53784 | 4.26083 | 1.3229 | Intron (ENSMUST00000069837.3/ENSMUSG00000056427.5, intron 1 of 35) | 11 | 35121224 | 35708507 | 587284 | 1 | ENSMUST00000069837.3 | 4280 | Slit3 | ENSMUSG00000056427.5 |
| chr11 | 35126336 | 35126426 | 3.83854 | 4.84803 | 1.74106 | Intron (ENSMUST00000069837.3/ENSMUSG00000056427.5, intron 1 of 35) | 11 | 35121224 | 35708507 | 587284 | 1 | ENSMUST00000069837.3 | 5113 | Slit3 | ENSMUSG00000056427.5 |
| chr11 | 35127410 | 35127812 | 3.28784 | 3.75895 | 1.01733 | Intron (ENSMUST00000069837.3/ENSMUSG00000056427.5, intron 1 of 35) | 11 | 35121224 | 35708507 | 587284 | 1 | ENSMUST00000069837.3 | 6187 | Slit3 | ENSMUSG00000056427.5 |
| chr11 | 35128939 | 35129464 | 3.53639 | 4.21696 | 1.31322 | Intron (ENSMUST00000069837.3/ENSMUSG00000056427.5, intron 1 of 35) | 11 | 35121224 | 35708507 | 587284 | 1 | ENSMUST00000069837.3 | 7716 | Slit3 | ENSMUSG00000056427.5 |
| chr11 | 35171711 | 35171790 | 4.00176 | 5.17678 | 1.89443 | Promoter | 11 | 35171629 | 35235207 | 63579 | 1 | ENSMUST00000156571.2 | 83 | Slit3 | ENSMUSG00000056427.5 |
| chr11 | 35178291 | 35178620 | 3.41167 | 4.14674 | 1.26485 | Promoter | 11 | 35177783 | 35510128 | 332346 | 1 | ENSMUST00000153098.1 | 509 | Slit3 | ENSMUSG00000056427.5 |
| chr11 | 35511260 | 35511490 | 3.38964 | 3.95638 | 1.16541 | Intron (ENSMUST00000069837.3/ENSMUSG00000056427.5, intron 5 of 35) | 11 | 35544685 | 35612636 | 67952 | 1 | ENSMUST00000124438.1 | -33195 | Slit3 | ENSMUSG00000056427.5 |
| chr11 | 35513495 | 35513773 | 3.43368 | 4.02754 | 1.18067 | Intron (ENSMUST00000069837.3/ENSMUSG00000056427.5, intron 5 of 35) | 11 | 35544685 | 35612636 | 67952 | 1 | ENSMUST00000124438.1 | -30912 | Slit3 | ENSMUSG00000056427.5 |
| chr11 | 35514541 | 35514804 | 3.34563 | 4.14006 | 1.25231 | Intron (ENSMUST00000069837.3/ENSMUSG00000056427.5, intron 5 of 35) | 11 | 35544685 | 35612636 | 67952 | 1 | ENSMUST00000124438.1 | -29881 | Slit3 | ENSMUSG00000056427.5 |
| chr11 | 35537352 | 35537623 | 3.61743 | 4.40325 | 1.41936 | Intron (ENSMUST00000069837.3/ENSMUSG00000056427.5, intron 5 of 35) | 11 | 35544685 | 35612636 | 67952 | 1 | ENSMUST00000124438.1 | -7062 | Slit3 | ENSMUSG00000056427.5 |
| chr11 | 35541137 | 35541212 | 3.55807 | 4.30748 | 1.37545 | Intron (ENSMUST00000069837.3/ENSMUSG00000056427.5, intron 5 of 35) | 11 | 35544685 | 35612636 | 67952 | 1 | ENSMUST00000124438.1 | -3473 | Slit3 | ENSMUSG00000056427.5 |
| chr11 | 35543978 | 35544815 | 4.02968 | 5.19991 | 1.97288 | Promoter | 11 | 35544685 | 35612636 | 67952 | 1 | ENSMUST00000124438.1 | 0 | Slit3 | ENSMUSG00000056427.5 |
| chr11 | 35502972 | 35503097 | 3.30552 | 3.78557 | 1.01733 | Intron (ENSMUST00000069837.3/ENSMUSG00000056427.5, intron 4 of 35) | 11 | 35544685 | 35612636 | 67952 | 1 | ENSMUST00000124438.1 | -41588 | Slit3 | ENSMUSG00000056427.5 |
| chr11 | 35504574 | 35505005 | 3.38193 | 3.9006 | 1.01733 | Intron (ENSMUST00000069837.3/ENSMUSG00000056427.5, intron 4 of 35) | 11 | 35544685 | 35612636 | 67952 | 1 | ENSMUST00000124438.1 | -39680 | Slit3 | ENSMUSG00000056427.5 |
| chr11 | 35508106 | 35508317 | 3.31505 | 3.83358 | 1.02416 | Exon (ENSMUST00000069837.3/ENSMUSG00000056427.5, exon 5 of 36) | 11 | 35544685 | 35612636 | 67952 | 1 | ENSMUST00000124438.1 | -36368 | Slit3 | ENSMUSG00000056427.5 |
| chr11 | 35492598 | 35492673 | 3.87142 | 4.81519 | 1.60051 | Intron (ENSMUST00000069837.3/ENSMUSG00000056427.5, intron 4 of 35) | 11 | 35544685 | 35612636 | 67952 | 1 | ENSMUST00000124438.1 | -52012 | Slit3 | ENSMUSG00000056427.5 |
| chr2 | 73777446 | 73777568 | 4.02205 | 5.2288 | 1.94784 | Distal Intergenic | 2 | 73610660 | 73775346 | 164687 | 2 | ENSMUST00000180045.2 | -2101 | Chn1 | ENSMUSG00000056486.12 |
| chr3 | 84664361 | 84665097 | 3.91323 | 4.93785 | 1.73783 | Intron (ENSMUST00000154148.2/ENSMUSG00000074513.4, intron 3 of 9) | 3 | 84666192 | 84704575 | 38384 | 1 | ENSMUST00000107682.1 | -1095 | Tmem154 | ENSMUSG00000056498.8 |
| chr3 | 84682481 | 84682626 | 3.32279 | 3.81145 | 1.01733 | Intron (ENSMUST00000107682.1/ENSMUSG00000056498.8, intron 1 of 6) | 3 | 84690223 | 84692544 | 2322 | 1 | ENSMUST00000128344.1 | -7597 | Tmem154 | ENSMUSG00000056498.8 |
| chr3 | 84732614 | 84732986 | 3.78019 | 4.65478 | 1.5358 | Intron (ENSMUST00000154148.2/ENSMUSG00000074513.4, intron 3 of 9) | 3 | 84690223 | 84692544 | 2322 | 1 | ENSMUST00000128344.1 | 42392 | Tmem154 | ENSMUSG00000056498.8 |
| chr3 | 84733420 | 84733556 | 3.90668 | 4.87542 | 1.64534 | Intron (ENSMUST00000154148.2/ENSMUSG00000074513.4, intron 3 of 9) | 3 | 84690223 | 84692544 | 2322 | 1 | ENSMUST00000128344.1 | 43198 | Tmem154 | ENSMUSG00000056498.8 |
| chr3 | 84739678 | 84739815 | 3.39947 | 3.96672 | 1.15784 | Intron (ENSMUST00000154148.2/ENSMUSG00000074513.4, intron 3 of 9) | 3 | 84690223 | 84692544 | 2322 | 1 | ENSMUST00000128344.1 | 49456 | Tmem154 | ENSMUSG00000056498.8 |
| chr3 | 84745756 | 84746933 | 3.84202 | 4.78197 | 1.63099 | Intron (ENSMUST00000154148.2/ENSMUSG00000074513.4, intron 3 of 9) | 3 | 84690223 | 84692544 | 2322 | 1 | ENSMUST00000128344.1 | 55534 | Tmem154 | ENSMUSG00000056498.8 |
| chr12 | 116485809 | 116486256 | 3.83678 | 4.80066 | 1.6684 | Promoter | 12 | 116485833 | 117276793 | 790961 | 1 | ENSMUST00000070733.8 | 0 | Ptprn2 | ENSMUSG00000056553.9 |
| chr1 | 171158904 | 171158980 | 3.81851 | 4.77358 | 1.64236 | Promoter | 1 | 171159594 | 171160230 | 637 | 1 | ENSMUST00000149352.1 | -614 | Mpz | ENSMUSG00000056569.5 |
| chr4 | 133496402 | 133496620 | 3.33499 | 4.04588 | 1.18745 | Intron (ENSMUST00000125541.1/ENSMUSG00000056596.8, intron 1 of 1) | 4 | 133491118 | 133497719 | 6602 | 2 | ENSMUST00000123169.1 | 1099 | Trnp1 | ENSMUSG00000056596.8 |
| chr4 | 133497699 | 133498138 | 4.03658 | 5.3316 | 1.99635 | Promoter | 4 | 133491118 | 133497719 | 6602 | 2 | ENSMUST00000123169.1 | 0 | Trnp1 | ENSMUSG00000056596.8 |
| chr4 | 133498461 | 133498597 | 3.7062 | 4.92553 | 1.71241 | Promoter | 4 | 133491100 | 133498550 | 7451 | 2 | ENSMUST00000125541.1 | 0 | Trnp1 | ENSMUSG00000056596.8 |
| chr4 | 133499745 | 133499900 | 3.48126 | 4.40285 | 1.39454 | Distal Intergenic | 4 | 133491100 | 133498550 | 7451 | 2 | ENSMUST00000125541.1 | -1196 | Trnp1 | ENSMUSG00000056596.8 |
| chr4 | 133500476 | 133501905 | 3.78834 | 4.72561 | 1.64325 | Distal Intergenic | 4 | 133491100 | 133498550 | 7451 | 2 | ENSMUST00000125541.1 | -1927 | Trnp1 | ENSMUSG00000056596.8 |
| chr4 | 133503418 | 133503869 | 3.32046 | 3.8378 | 1.07845 | Distal Intergenic | 4 | 133491100 | 133498550 | 7451 | 2 | ENSMUST00000125541.1 | -4869 | Trnp1 | ENSMUSG00000056596.8 |
| chr4 | 133505677 | 133505769 | 3.58931 | 4.29023 | 1.27277 | Distal Intergenic | 4 | 133491100 | 133498550 | 7451 | 2 | ENSMUST00000125541.1 | -7128 | Trnp1 | ENSMUSG00000056596.8 |
| chr15 | 101773103 | 101773282 | 3.58322 | 4.27251 | 1.25206 | Downstream (2-3kb) | 15 | 101776333 | 101778693 | 2361 | 2 | ENSMUST00000127671.2 | 5411 | Krt72 | ENSMUSG00000056605.6 |
| chr15 | 101789761 | 101789953 | 4.22429 | 5.6541 | 2.22639 | Distal Intergenic | 15 | 101776172 | 101786458 | 10287 | 2 | ENSMUST00000071104.4 | -3304 | Krt72 | ENSMUSG00000056605.6 |
| chr12 | 104474180 | 104477318 | 4.11207 | 5.45453 | 2.12365 | Promoter | 12 | 104474488 | 104477489 | 3002 | 1 | ENSMUST00000070814.2 | 0 | Gm10000 | ENSMUSG00000056606.2 |
| chr12 | 104477808 | 104478567 | 4.23729 | 5.70799 | 2.35394 | Distal Intergenic | 12 | 104474488 | 104477489 | 3002 | 1 | ENSMUST00000070814.2 | 3321 | Gm10000 | ENSMUSG00000056606.2 |
| chr12 | 104479385 | 104480623 | 3.6104 | 4.37357 | 1.35387 | Distal Intergenic | 12 | 104474488 | 104477489 | 3002 | 1 | ENSMUST00000070814.2 | 4898 | Gm10000 | ENSMUSG00000056606.2 |
| chr12 | 104483621 | 104483696 | 4.12452 | 5.36389 | 2.00645 | Distal Intergenic | 12 | 104474488 | 104477489 | 3002 | 1 | ENSMUST00000070814.2 | 9134 | Gm10000 | ENSMUSG00000056606.2 |
| chr12 | 104531690 | 104532276 | 3.4624 | 4.05694 | 1.14054 | Distal Intergenic | 12 | 104474488 | 104477489 | 3002 | 1 | ENSMUST00000070814.2 | 57203 | Gm10000 | ENSMUSG00000056606.2 |
| chr12 | 104567135 | 104567271 | 3.46691 | 4.21419 | 1.25108 | Distal Intergenic | 12 | 104474488 | 104477489 | 3002 | 1 | ENSMUST00000070814.2 | 92648 | Gm10000 | ENSMUSG00000056606.2 |
| chr12 | 104567711 | 104568257 | 3.40483 | 4.09976 | 1.18191 | Distal Intergenic | 12 | 104474488 | 104477489 | 3002 | 1 | ENSMUST00000070814.2 | 93224 | Gm10000 | ENSMUSG00000056606.2 |
| chr12 | 104573651 | 104573943 | 3.73727 | 4.56767 | 1.46109 | Distal Intergenic | 12 | 104474488 | 104477489 | 3002 | 1 | ENSMUST00000070814.2 | 99164 | Gm10000 | ENSMUSG00000056606.2 |
| chr12 | 104574394 | 104574561 | 3.369 | 3.8811 | 1.01733 | Distal Intergenic | 12 | 104474488 | 104477489 | 3002 | 1 | ENSMUST00000070814.2 | 99907 | Gm10000 | ENSMUSG00000056606.2 |
| chr8 | 90669421 | 90669506 | 3.39911 | 3.93859 | 1.06004 | Distal Intergenic | 8 | 90828835 | 91054508 | 225674 | 1 | ENSMUST00000048665.6 | -159329 | Chd9 | ENSMUSG00000056608.8 |
| chr8 | 90670075 | 90670209 | 3.36622 | 3.8769 | 1.01733 | Distal Intergenic | 8 | 90828835 | 91054508 | 225674 | 1 | ENSMUST00000048665.6 | -158626 | Chd9 | ENSMUSG00000056608.8 |
| chr11 | 96277260 | 96278580 | 3.80126 | 4.77627 | 1.63112 | Distal Intergenic | 11 | 96281905 | 96285315 | 3411 | 1 | ENSMUST00000052650.3 | -3325 | Hoxb8 | ENSMUSG00000056648.5 |
| chr11 | 96279321 | 96279796 | 3.53013 | 4.27431 | 1.33269 | Distal Intergenic | 11 | 96281905 | 96285315 | 3411 | 1 | ENSMUST00000052650.3 | -2109 | Hoxb8 | ENSMUSG00000056648.5 |
| chr11 | 96281533 | 96282110 | 3.75045 | 4.7119 | 1.56886 | Promoter | 11 | 96281905 | 96285315 | 3411 | 1 | ENSMUST00000052650.3 | 0 | Hoxb8 | ENSMUSG00000056648.5 |
| chr11 | 96283193 | 96283604 | 4.55536 | 6.30847 | 2.72221 | Promoter | 11 | 96282725 | 96284693 | 1969 | 1 | ENSMUST00000168043.1 | 469 | Hoxb8 | ENSMUSG00000056648.5 |
| chr13 | 52990306 | 52990446 | 3.60492 | 4.33504 | 1.36204 | Distal Intergenic | 13 | 52967209 | 52981073 | 13865 | 2 | ENSMUST00000071065.7 | -9234 | Nfil3 | ENSMUSG00000056749.7 |
| chr13 | 53011577 | 53011895 | 3.69423 | 4.51787 | 1.47705 | Distal Intergenic | 13 | 52967209 | 52981073 | 13865 | 2 | ENSMUST00000071065.7 | -30505 | Nfil3 | ENSMUSG00000056749.7 |
| chr13 | 53053129 | 53053437 | 3.35214 | 3.8966 | 1.1243 | Distal Intergenic | 13 | 52967209 | 52981073 | 13865 | 2 | ENSMUST00000071065.7 | -72057 | Nfil3 | ENSMUSG00000056749.7 |
| chr13 | 53096047 | 53096461 | 3.42801 | 4.00069 | 1.12412 | Distal Intergenic | 13 | 52967209 | 52981073 | 13865 | 2 | ENSMUST00000071065.7 | -114975 | Nfil3 | ENSMUSG00000056749.7 |
| chr13 | 53097238 | 53097679 | 3.79332 | 4.72391 | 1.59783 | Distal Intergenic | 13 | 52967209 | 52981073 | 13865 | 2 | ENSMUST00000071065.7 | -116166 | Nfil3 | ENSMUSG00000056749.7 |
| chr13 | 53118631 | 53118754 | 3.55722 | 4.41393 | 1.40626 | Exon (ENSMUST00000130235.2/ENSMUSG00000021464.9, exon 6 of 9) | 13 | 52967209 | 52981073 | 13865 | 2 | ENSMUST00000071065.7 | -137559 | Nfil3 | ENSMUSG00000056749.7 |
| chr13 | 53120016 | 53120091 | 3.53386 | 4.24106 | 1.35333 | Intron (ENSMUST00000130235.2/ENSMUSG00000021464.9, intron 5 of 8) | 13 | 52967209 | 52981073 | 13865 | 2 | ENSMUST00000071065.7 | -138944 | Nfil3 | ENSMUSG00000056749.7 |
| chr13 | 53121308 | 53121600 | 3.62611 | 4.41613 | 1.45368 | Intron (ENSMUST00000130235.2/ENSMUSG00000021464.9, intron 5 of 8) | 13 | 52967209 | 52981073 | 13865 | 2 | ENSMUST00000071065.7 | -140236 | Nfil3 | ENSMUSG00000056749.7 |
| chr10 | 120462996 | 120463073 | 3.7843 | 4.80582 | 1.65228 | Intron (ENSMUST00000072777.8/ENSMUSG00000056758.9, intron 2 of 4) | 10 | 120470290 | 120476292 | 6003 | 2 | ENSMUST00000159310.1 | 13219 | Hmga2 | ENSMUSG00000056758.9 |
| chr10 | 120473553 | 120473957 | 3.86392 | 4.89748 | 1.72657 | Intron (ENSMUST00000072777.8/ENSMUSG00000056758.9, intron 1 of 4) | 10 | 120470290 | 120476292 | 6003 | 2 | ENSMUST00000159310.1 | 2335 | Hmga2 | ENSMUSG00000056758.9 |
| chr10 | 120474420 | 120474750 | 3.53986 | 4.20011 | 1.22369 | Intron (ENSMUST00000072777.8/ENSMUSG00000056758.9, intron 1 of 4) | 10 | 120470290 | 120476292 | 6003 | 2 | ENSMUST00000159310.1 | 1542 | Hmga2 | ENSMUSG00000056758.9 |
| chr10 | 120475059 | 120476621 | 4.38931 | 6.23673 | 2.71781 | Promoter | 10 | 120361275 | 120476469 | 115195 | 2 | ENSMUST00000072777.8 | 0 | Hmga2 | ENSMUSG00000056758.9 |
| chr10 | 120476928 | 120478078 | 3.62517 | 4.50149 | 1.49574 | Promoter | 10 | 120361275 | 120476469 | 115195 | 2 | ENSMUST00000159699.1 | -460 | Hmga2 | ENSMUSG00000056758.9 |
| chr10 | 120479235 | 120479982 | 3.86743 | 4.87958 | 1.75468 | Distal Intergenic | 10 | 120361275 | 120476469 | 115195 | 2 | ENSMUST00000159699.1 | -2767 | Hmga2 | ENSMUSG00000056758.9 |
| chr10 | 120503889 | 120504121 | 3.30329 | 3.80142 | 1.07897 | Distal Intergenic | 10 | 120361275 | 120476469 | 115195 | 2 | ENSMUST00000159699.1 | -27421 | Hmga2 | ENSMUSG00000056758.9 |
| chr8 | 88774243 | 88774516 | 3.27582 | 3.74106 | 1.01733 | Distal Intergenic | 8 | 88806391 | 88807140 | 750 | 1 | ENSMUST00000079841.2 | -31875 | Rps6-ps2 | ENSMUSG00000056772.2 |
| chr8 | 88913757 | 88914392 | 3.35601 | 3.86586 | 1.03244 | Distal Intergenic | 8 | 88806391 | 88807140 | 750 | 1 | ENSMUST00000079841.2 | 107367 | Rps6-ps2 | ENSMUSG00000056772.2 |
| chr7 | 139707260 | 139707351 | 4.41702 | 6.01008 | 2.51305 | Distal Intergenic | 7 | 139719399 | 139719881 | 483 | 1 | ENSMUST00000072532.5 | -12048 | Rpl21-ps13 | ENSMUSG00000056777.5 |
| chr18 | 64263892 | 64264116 | 3.57966 | 4.27122 | 1.26103 | Intron (ENSMUST00000025477.8/ENSMUSG00000056812.8, intron 2 of 5) | 18 | 64265996 | 64269979 | 3984 | 1 | ENSMUST00000139622.1 | -1880 | St8sia3 | ENSMUSG00000056812.8 |
| chr18 | 64265183 | 64267035 | 3.83266 | 4.82635 | 1.65526 | Promoter | 18 | 64265996 | 64269979 | 3984 | 1 | ENSMUST00000139622.1 | 0 | St8sia3 | ENSMUSG00000056812.8 |
| chr18 | 64271642 | 64271963 | 3.33275 | 3.82648 | 1.01733 | 3' UTR | 18 | 64266808 | 64269908 | 3101 | 1 | ENSMUST00000129518.1 | 4835 | St8sia3 | ENSMUSG00000056812.8 |
| chr2 | 165349738 | 165349889 | 3.36068 | 3.89187 | 1.09823 | Downstream (1-2kb) | 2 | 165353811 | 165359401 | 5591 | 2 | ENSMUST00000136482.1 | 9512 | Zfp663 | ENSMUSG00000056824.10 |
| chr2 | 165368143 | 165369398 | 3.37874 | 4.18407 | 1.25567 | Promoter | 2 | 165351297 | 165368729 | 17433 | 2 | ENSMUST00000103085.2 | 0 | Zfp663 | ENSMUSG00000056824.10 |
| chr19 | 16829188 | 16829341 | 3.50848 | 4.18208 | 1.30124 | Distal Intergenic | 19 | 16872316 | 16873830 | 1515 | 2 | ENSMUST00000072915.2 | 44489 | Foxb2 | ENSMUSG00000056829.3 |
| chr19 | 16865512 | 16866612 | 3.69328 | 4.51335 | 1.43286 | Distal Intergenic | 19 | 16872316 | 16873830 | 1515 | 2 | ENSMUST00000072915.2 | 7218 | Foxb2 | ENSMUSG00000056829.3 |
| chr19 | 16867543 | 16868138 | 3.25865 | 3.71511 | 1.01733 | Distal Intergenic | 19 | 16872316 | 16873830 | 1515 | 2 | ENSMUST00000072915.2 | 5692 | Foxb2 | ENSMUSG00000056829.3 |
| chr19 | 16868642 | 16871679 | 4.00141 | 5.24911 | 1.98875 | Downstream (<1kb) | 19 | 16872316 | 16873830 | 1515 | 2 | ENSMUST00000072915.2 | 2151 | Foxb2 | ENSMUSG00000056829.3 |
| chr19 | 16872581 | 16872965 | 4.19662 | 5.99999 | 2.49855 | Promoter | 19 | 16872316 | 16873830 | 1515 | 2 | ENSMUST00000072915.2 | 865 | Foxb2 | ENSMUSG00000056829.3 |
| chr19 | 16873408 | 16874891 | 3.90713 | 5.00418 | 1.87695 | Promoter | 19 | 16872316 | 16873830 | 1515 | 2 | ENSMUST00000072915.2 | 0 | Foxb2 | ENSMUSG00000056829.3 |
| chr19 | 16875241 | 16875405 | 3.56599 | 4.50077 | 1.43767 | Distal Intergenic | 19 | 16872316 | 16873830 | 1515 | 2 | ENSMUST00000072915.2 | -1412 | Foxb2 | ENSMUSG00000056829.3 |
| chr19 | 16876557 | 16876754 | 3.82753 | 4.81257 | 1.68164 | Distal Intergenic | 19 | 16872316 | 16873830 | 1515 | 2 | ENSMUST00000072915.2 | -2728 | Foxb2 | ENSMUSG00000056829.3 |
| chr19 | 16877904 | 16878577 | 3.74941 | 4.64407 | 1.58114 | Distal Intergenic | 19 | 16872316 | 16873830 | 1515 | 2 | ENSMUST00000072915.2 | -4075 | Foxb2 | ENSMUSG00000056829.3 |
| chr19 | 16879045 | 16879693 | 3.68837 | 4.58691 | 1.52141 | Distal Intergenic | 19 | 16872316 | 16873830 | 1515 | 2 | ENSMUST00000072915.2 | -5216 | Foxb2 | ENSMUSG00000056829.3 |
| chr19 | 16880062 | 16880443 | 4.17047 | 5.42064 | 2.07221 | Distal Intergenic | 19 | 16872316 | 16873830 | 1515 | 2 | ENSMUST00000072915.2 | -6233 | Foxb2 | ENSMUSG00000056829.3 |
| chr19 | 16880755 | 16881021 | 3.43931 | 4.13248 | 1.22454 | Distal Intergenic | 19 | 16872316 | 16873830 | 1515 | 2 | ENSMUST00000072915.2 | -6926 | Foxb2 | ENSMUSG00000056829.3 |
| chr19 | 16882538 | 16884491 | 4.06152 | 5.2279 | 1.93352 | Distal Intergenic | 19 | 16872316 | 16873830 | 1515 | 2 | ENSMUST00000072915.2 | -8709 | Foxb2 | ENSMUSG00000056829.3 |
| chr19 | 16888362 | 16888501 | 3.35887 | 3.87588 | 1.05293 | Distal Intergenic | 19 | 16872316 | 16873830 | 1515 | 2 | ENSMUST00000072915.2 | -14533 | Foxb2 | ENSMUSG00000056829.3 |
| chr19 | 16897679 | 16898023 | 3.43605 | 4.05651 | 1.23186 | Distal Intergenic | 19 | 16872316 | 16873830 | 1515 | 2 | ENSMUST00000072915.2 | -23850 | Foxb2 | ENSMUSG00000056829.3 |
| chr18 | 42418152 | 42418394 | 3.38756 | 3.92744 | 1.08177 | Distal Intergenic | 18 | 42441019 | 42441507 | 489 | 1 | ENSMUST00000080097.6 | -22625 | Gm16415 | ENSMUSG00000056849.6 |
| chr18 | 42429124 | 42429485 | 3.29117 | 3.78456 | 1.07413 | Distal Intergenic | 18 | 42441019 | 42441507 | 489 | 1 | ENSMUST00000080097.6 | -11534 | Gm16415 | ENSMUSG00000056849.6 |
| chr18 | 42469780 | 42470074 | 3.51558 | 4.17975 | 1.28926 | Distal Intergenic | 18 | 42441019 | 42441507 | 489 | 1 | ENSMUST00000080097.6 | 28762 | Gm16415 | ENSMUSG00000056849.6 |
| chr7 | 138938103 | 138938301 | 3.45764 | 4.05245 | 1.15081 | Distal Intergenic | 7 | 138940730 | 139026888 | 86159 | 1 | ENSMUST00000106111.2 | -2429 | Jakmip3 | ENSMUSG00000056856.6 |

| | | | | | | | | | | | | | | | |
|---|---|---|---|---|---|---|---|---|---|---|---|---|---|---|---|
| chr7 | 138938789 | 138938867 | 3.558 | 4.23612 | 1.26219 | Distal Intergenic | 7 | 138940730 | 139026888 | 86159 | 1 | ENSMUST00000106111.2 | -1863 | Jakmip3 | ENSMUSG00000056856.6 |
| chr7 | 138939207 | 138939373 | 3.44371 | 4.04592 | 1.18426 | Distal Intergenic | 7 | 138940730 | 139026888 | 86159 | 1 | ENSMUST00000106111.2 | -1357 | Jakmip3 | ENSMUSG00000056856.6 |
| chr7 | 138940526 | 138941672 | 3.68202 | 4.46385 | 1.39123 | Promoter | 7 | 138940730 | 139026888 | 86159 | 1 | ENSMUST00000106111.2 | 0 | Jakmip3 | ENSMUSG00000056856.6 |
| chr7 | 138943603 | 138943746 | 3.62481 | 4.34726 | 1.3 | Intron (ENSMUST00000106111.2/ENSMUSG00000056856.6, intron 1 of 9) | 7 | 138940750 | 139026882 | 86133 | 1 | ENSMUST00000166163.1 | 2854 | Jakmip3 | ENSMUSG00000056856.6 |
| chr3 | 55780335 | 55781078 | 3.87243 | 5.02659 | 1.79608 | Intron (ENSMUST00000029374.6/ENSMUSG00000027799.8, intron 41 of 57) | 3 | 55782510 | 55785285 | 2776 | 1 | ENSMUST00000075422.4 | -1432 | Mab21l1 | ENSMUSG00000056947.4 |
| chr3 | 55781776 | 55782436 | 3.52159 | 4.3305 | 1.33419 | Promoter | 3 | 55782510 | 55785285 | 2776 | 1 | ENSMUST00000075422.4 | -74 | Mab21l1 | ENSMUSG00000056947.4 |
| chr3 | 55783053 | 55783889 | 3.90558 | 5.16163 | 1.87021 | Promoter | 3 | 55782510 | 55785285 | 2776 | 1 | ENSMUST00000075422.4 | 544 | Mab21l1 | ENSMUSG00000056947.4 |
| chr3 | 55785418 | 55785498 | 3.56512 | 4.33872 | 1.35887 | Intron (ENSMUST00000029374.6/ENSMUSG00000027799.8, intron 41 of 57) | 3 | 55782510 | 55785285 | 2776 | 1 | ENSMUST00000075422.4 | 2909 | Mab21l1 | ENSMUSG00000056947.4 |
| chr3 | 55786046 | 55787876 | 3.73425 | 4.62869 | 1.58085 | Exon (ENSMUST00000029374.6/ENSMUSG00000027799.8, exon 41 of 58) | 3 | 55782510 | 55785285 | 2776 | 1 | ENSMUST00000075422.4 | 3537 | Mab21l1 | ENSMUSG00000056947.4 |
| chr3 | 55788751 | 55788857 | 4.63491 | 6.39639 | 2.78769 | Intron (ENSMUST00000029374.6/ENSMUSG00000027799.8, intron 40 of 57) | 3 | 55782510 | 55785285 | 2776 | 1 | ENSMUST00000075422.4 | 6242 | Mab21l1 | ENSMUSG00000056947.4 |
| chr3 | 55789243 | 55789502 | 3.36191 | 3.8742 | 1.03068 | Intron (ENSMUST00000029374.6/ENSMUSG00000027799.8, intron 40 of 57) | 3 | 55782510 | 55785285 | 2776 | 1 | ENSMUST00000075422.4 | 6734 | Mab21l1 | ENSMUSG00000056947.4 |
| chr7 | 62376528 | 62377278 | 3.42657 | 4.0166 | 1.17545 | Promoter | 7 | 62376979 | 62381640 | 4662 | 1 | ENSMUST00000080403.5 | 0 | Magel2 | ENSMUSG00000056972.5 |
| chr11 | 96464177 | 96465067 | 3.8774 | 4.92485 | 1.78312 | Promoter | 11 | 96464593 | 96758941 | 294349 | 1 | ENSMUST00000103154.5 | 0 | Skap1 | ENSMUSG00000057058.11 |
| chr11 | 96466583 | 96466869 | 3.76147 | 4.73604 | 1.63475 | Intron (ENSMUST00000103154.5/ENSMUSG00000057058.11, intron 1 of 12) | 11 | 96464681 | 96678371 | 213691 | 1 | ENSMUST00000107663.3 | 1903 | Skap1 | ENSMUSG00000057058.11 |
| chr11 | 96467289 | 96467692 | 4.03299 | 5.1891 | 1.9082 | Intron (ENSMUST00000103154.5/ENSMUSG00000057058.11, intron 1 of 12) | 11 | 96464681 | 96678371 | 213691 | 1 | ENSMUST00000107663.3 | 2609 | Skap1 | ENSMUSG00000057058.11 |
| chr11 | 96480064 | 96480333 | 3.53655 | 4.26368 | 1.36016 | Intron (ENSMUST00000103154.5/ENSMUSG00000057058.11, intron 1 of 12) | 11 | 96483782 | 96493718 | 9937 | 1 | ENSMUST00000149988.1 | -3449 | Skap1 | ENSMUSG00000057058.11 |
| chr11 | 96490603 | 96490723 | 3.58025 | 4.45027 | 1.392 | Promoter | 11 | 96489861 | 96704431 | 214571 | 1 | ENSMUST00000127639.1 | 743 | Skap1 | ENSMUSG00000057058.11 |
| chr11 | 96501735 | 96501960 | 3.51222 | 4.14506 | 1.18641 | Intron (ENSMUST00000103154.5/ENSMUSG00000057058.11, intron 2 of 12) | 11 | 96489861 | 96704431 | 214571 | 1 | ENSMUST00000127639.1 | 11875 | Skap1 | ENSMUSG00000057058.11 |
| chr11 | 96528073 | 96528244 | 3.30336 | 3.78233 | 1.01733 | Intron (ENSMUST00000103154.5/ENSMUSG00000057058.11, intron 3 of 12) | 11 | 96489861 | 96704431 | 214571 | 1 | ENSMUST00000127639.1 | 38213 | Skap1 | ENSMUSG00000057058.11 |
| chr11 | 96536651 | 96537240 | 3.74109 | 4.77807 | 1.65367 | Intron (ENSMUST00000103154.5/ENSMUSG00000057058.11, intron 3 of 12) | 11 | 96489861 | 96704431 | 214571 | 1 | ENSMUST00000127639.1 | 46791 | Skap1 | ENSMUSG00000057058.11 |
| chr11 | 96537553 | 96538043 | 3.56497 | 4.42689 | 1.40313 | Intron (ENSMUST00000103154.5/ENSMUSG00000057058.11, intron 3 of 12) | 11 | 96489861 | 96704431 | 214571 | 1 | ENSMUST00000127639.1 | 47693 | Skap1 | ENSMUSG00000057058.11 |
| chr11 | 96539163 | 96539481 | 3.23346 | 3.68282 | 1.02961 | Intron (ENSMUST00000103154.5/ENSMUSG00000057058.11, intron 3 of 12) | 11 | 96489861 | 96704431 | 214571 | 1 | ENSMUST00000127639.1 | 49303 | Skap1 | ENSMUSG00000057058.11 |
| chr11 | 96543640 | 96543865 | 4.49647 | 6.07459 | 2.53249 | Intron (ENSMUST00000103154.5/ENSMUSG00000057058.11, intron 4 of 12) | 11 | 96489861 | 96704431 | 214571 | 1 | ENSMUST00000127639.1 | 53780 | Skap1 | ENSMUSG00000057058.11 |
| chr11 | 96544252 | 96544548 | 3.30879 | 3.79053 | 1.01733 | Intron (ENSMUST00000103154.5/ENSMUSG00000057058.11, intron 4 of 12) | 11 | 96489861 | 96704431 | 214571 | 1 | ENSMUST00000127639.1 | 54392 | Skap1 | ENSMUSG00000057058.11 |
| chr11 | 96545614 | 96546114 | 3.49765 | 4.22406 | 1.29851 | Intron (ENSMUST00000103154.5/ENSMUSG00000057058.11, intron 4 of 12) | 11 | 96489861 | 96704431 | 214571 | 1 | ENSMUST00000127639.1 | 55754 | Skap1 | ENSMUSG00000057058.11 |
| chr11 | 96562482 | 96562759 | 3.34153 | 3.83966 | 1.01733 | Intron (ENSMUST00000103154.5/ENSMUSG00000057058.11, intron 4 of 12) | 11 | 96489861 | 96704431 | 214571 | 1 | ENSMUST00000127639.1 | 72622 | Skap1 | ENSMUSG00000057058.11 |
| chr11 | 96575488 | 96575563 | 3.50065 | 4.12998 | 1.19531 | Intron (ENSMUST00000103154.5/ENSMUSG00000057058.11, intron 4 of 12) | 11 | 96489861 | 96704431 | 214571 | 1 | ENSMUST00000127639.1 | 85628 | Skap1 | ENSMUSG00000057058.11 |
| chr11 | 96576440 | 96576543 | 3.38343 | 3.9138 | 1.05602 | Intron (ENSMUST00000103154.5/ENSMUSG00000057058.11, intron 4 of 12) | 11 | 96489861 | 96704431 | 214571 | 1 | ENSMUST00000127639.1 | 86580 | Skap1 | ENSMUSG00000057058.11 |
| chr11 | 96576893 | 96577113 | 3.63356 | 4.39821 | 1.41995 | Intron (ENSMUST00000103154.5/ENSMUSG00000057058.11, intron 4 of 12) | 11 | 96489861 | 96704431 | 214571 | 1 | ENSMUST00000127639.1 | 87033 | Skap1 | ENSMUSG00000057058.11 |
| chr11 | 96578253 | 96579400 | 3.51749 | 4.20352 | 1.31059 | Intron (ENSMUST00000103154.5/ENSMUSG00000057058.11, intron 4 of 12) | 11 | 96489861 | 96704431 | 214571 | 1 | ENSMUST00000127639.1 | 88393 | Skap1 | ENSMUSG00000057058.11 |
| chr11 | 96579745 | 96579843 | 3.30452 | 3.8079 | 1.09264 | Intron (ENSMUST00000103154.5/ENSMUSG00000057058.11, intron 4 of 12) | 11 | 96489861 | 96704431 | 214571 | 1 | ENSMUST00000127639.1 | 89885 | Skap1 | ENSMUSG00000057058.11 |
| chr11 | 96586427 | 96586734 | 3.24509 | 3.79099 | 1.01733 | Intron (ENSMUST00000103154.5/ENSMUSG00000057058.11, intron 4 of 12) | 11 | 96489861 | 96704431 | 214571 | 1 | ENSMUST00000127639.1 | 96567 | Skap1 | ENSMUSG00000057058.11 |
| chr11 | 96593977 | 96594341 | 3.4085 | 3.95489 | 1.06739 | Intron (ENSMUST00000103154.5/ENSMUSG00000057058.11, intron 4 of 12) | 11 | 96489861 | 96704431 | 214571 | 1 | ENSMUST00000127639.1 | 104117 | Skap1 | ENSMUSG00000057058.11 |
| chr11 | 96622079 | 96622905 | 3.53039 | 4.19685 | 1.26768 | Intron (ENSMUST00000103154.5/ENSMUSG00000057058.11, intron 4 of 12) | 11 | 96708235 | 96713202 | 4968 | 1 | ENSMUST00000123814.1 | -85330 | Skap1 | ENSMUSG00000057058.11 |
| chr11 | 96623886 | 96623972 | 3.36126 | 3.8694 | 1.01733 | Intron (ENSMUST00000103154.5/ENSMUSG00000057058.11, intron 4 of 12) | 11 | 96708235 | 96713202 | 4968 | 1 | ENSMUST00000123814.1 | -84263 | Skap1 | ENSMUSG00000057058.11 |
| chr11 | 96624590 | 96624776 | 3.92867 | 4.94391 | 1.73178 | Intron (ENSMUST00000103154.5/ENSMUSG00000057058.11, intron 4 of 12) | 11 | 96708235 | 96713202 | 4968 | 1 | ENSMUST00000123814.1 | -83459 | Skap1 | ENSMUSG00000057058.11 |
| chr11 | 96626838 | 96627215 | 3.48284 | 4.10582 | 1.1973 | Intron (ENSMUST00000103154.5/ENSMUSG00000057058.11, intron 4 of 12) | 11 | 96708235 | 96713202 | 4968 | 1 | ENSMUST00000123814.1 | -81020 | Skap1 | ENSMUSG00000057058.11 |
| chr11 | 96634155 | 96634304 | 3.36612 | 3.87675 | 1.01733 | Intron (ENSMUST00000103154.5/ENSMUSG00000057058.11, intron 4 of 12) | 11 | 96708235 | 96713202 | 4968 | 1 | ENSMUST00000123814.1 | -73931 | Skap1 | ENSMUSG00000057058.11 |
| chr11 | 96638157 | 96638232 | 3.85748 | 4.82605 | 1.64119 | Intron (ENSMUST00000103154.5/ENSMUSG00000057058.11, intron 4 of 12) | 11 | 96708235 | 96713202 | 4968 | 1 | ENSMUST00000123814.1 | -70003 | Skap1 | ENSMUSG00000057058.11 |
| chr11 | 96670689 | 96670790 | 3.81018 | 4.70037 | 1.53171 | Intron (ENSMUST00000103154.5/ENSMUSG00000057058.11, intron 4 of 12) | 11 | 96708235 | 96713202 | 4968 | 1 | ENSMUST00000123814.1 | -37445 | Skap1 | ENSMUSG00000057058.11 |
| chr11 | 96677734 | 96678195 | 4.22586 | 5.64036 | 2.23054 | 3' UTR | 11 | 96708235 | 96713202 | 4968 | 1 | ENSMUST00000123814.1 | -30040 | Skap1 | ENSMUSG00000057058.11 |
| chr11 | 96679496 | 96680264 | 4.00769 | 5.14686 | 1.9378 | Intron (ENSMUST00000103154.5/ENSMUSG00000057058.11, intron 4 of 12) | 11 | 96708235 | 96713202 | 4968 | 1 | ENSMUST00000123814.1 | -27971 | Skap1 | ENSMUSG00000057058.11 |
| chr11 | 96682005 | 96682105 | 3.71729 | 4.53791 | 1.46957 | Intron (ENSMUST00000103154.5/ENSMUSG00000057058.11, intron 4 of 12) | 11 | 96708235 | 96713202 | 4968 | 1 | ENSMUST00000123814.1 | -26130 | Skap1 | ENSMUSG00000057058.11 |
| chr11 | 96685467 | 96685610 | 3.37135 | 4.03 | 1.12641 | Intron (ENSMUST00000103154.5/ENSMUSG00000057058.11, intron 4 of 12) | 11 | 96708235 | 96713202 | 4968 | 1 | ENSMUST00000123814.1 | -22625 | Skap1 | ENSMUSG00000057058.11 |
| chr11 | 96695506 | 96696513 | 3.58724 | 4.29273 | 1.26873 | Intron (ENSMUST00000103154.5/ENSMUSG00000057058.11, intron 4 of 12) | 11 | 96708235 | 96713202 | 4968 | 1 | ENSMUST00000123814.1 | -11722 | Skap1 | ENSMUSG00000057058.11 |
| chr11 | 96696846 | 96697988 | 3.7257 | 4.55746 | 1.46681 | Intron (ENSMUST00000103154.5/ENSMUSG00000057058.11, intron 4 of 12) | 11 | 96708235 | 96713202 | 4968 | 1 | ENSMUST00000123814.1 | -10247 | Skap1 | ENSMUSG00000057058.11 |
| chr11 | 96699219 | 96699375 | 4.5964 | 6.26263 | 2.64096 | Intron (ENSMUST00000103154.5/ENSMUSG00000057058.11, intron 4 of 12) | 11 | 96708235 | 96713202 | 4968 | 1 | ENSMUST00000123814.1 | -8860 | Skap1 | ENSMUSG00000057058.11 |
| chr11 | 96702589 | 96702709 | 3.87059 | 4.82387 | 1.6336 | Exon (ENSMUST00000103154.5/ENSMUSG00000057058.11, exon 5 of 13) | 11 | 96708235 | 96713202 | 4968 | 1 | ENSMUST00000123814.1 | -5526 | Skap1 | ENSMUSG00000057058.11 |
| chr11 | 96705658 | 96706022 | 3.25685 | 3.71241 | 1.01733 | Intron (ENSMUST00000103154.5/ENSMUSG00000057058.11, intron 6 of 12) | 11 | 96708235 | 96713202 | 4968 | 1 | ENSMUST00000123814.1 | -2213 | Skap1 | ENSMUSG00000057058.11 |
| chr11 | 96706349 | 96706675 | 3.36262 | 4.05907 | 1.20522 | Intron (ENSMUST00000103154.5/ENSMUSG00000057058.11, intron 6 of 12) | 11 | 96708235 | 96713202 | 4968 | 1 | ENSMUST00000123814.1 | -1560 | Skap1 | ENSMUSG00000057058.11 |
| chr11 | 96707171 | 96707273 | 3.86446 | 4.85549 | 1.73942 | Promoter | 11 | 96708235 | 96713202 | 4968 | 1 | ENSMUST00000123814.1 | -962 | Skap1 | ENSMUSG00000057058.11 |
| chr11 | 96708238 | 96708884 | 3.59065 | 4.38193 | 1.40898 | Promoter | 11 | 96708235 | 96713202 | 4968 | 1 | ENSMUST00000123814.1 | 4 | Skap1 | ENSMUSG00000057058.11 |
| chr11 | 96709955 | 96710179 | 3.28367 | 3.75267 | 1.01733 | Intron (ENSMUST00000103154.5/ENSMUSG00000057058.11, intron 7 of 12) | 11 | 96708235 | 96713202 | 4968 | 1 | ENSMUST00000123814.1 | 1721 | Skap1 | ENSMUSG00000057058.11 |
| chr11 | 96711386 | 96711842 | 3.50461 | 4.13871 | 1.20504 | Intron (ENSMUST00000103154.5/ENSMUSG00000057058.11, intron 8 of 12) | 11 | 96708235 | 96713202 | 4968 | 1 | ENSMUST00000123814.1 | 3152 | Skap1 | ENSMUSG00000057058.11 |
| chr11 | 96712442 | 96712683 | 3.60372 | 4.31897 | 1.3114 | Intron (ENSMUST00000103154.5/ENSMUSG00000057058.11, intron 8 of 12) | 11 | 96708235 | 96713202 | 4968 | 1 | ENSMUST00000123814.1 | 4208 | Skap1 | ENSMUSG00000057058.11 |
| chr11 | 96713678 | 96713985 | 4.16503 | 5.42164 | 2.0581 | Exon (ENSMUST00000181758.1/ENSMUSG00000097289.1, exon 2 of 2) | 11 | 96708235 | 96713202 | 4968 | 1 | ENSMUST00000123814.1 | 5444 | Skap1 | ENSMUSG00000057058.11 |
| chr8 | 126188072 | 126188471 | 3.66272 | 4.42099 | 1.35323 | Distal Intergenic | 8 | 126298558 | 126395482 | 96925 | 1 | ENSMUST00000108759.2 | -110087 | Slc35f3 | ENSMUSG00000057060.7 |
| chr8 | 126297898 | 126298023 | 3.59538 | 4.29235 | 1.26265 | Promoter | 8 | 126298558 | 126395482 | 96925 | 1 | ENSMUST00000108759.2 | -535 | Slc35f3 | ENSMUSG00000057060.7 |
| chr8 | 126298348 | 126299758 | 3.97586 | 5.09722 | 1.8994 | Promoter | 8 | 126298558 | 126395482 | 96925 | 1 | ENSMUST00000108759.2 | 0 | Slc35f3 | ENSMUSG00000057060.7 |
| chr11 | 44610378 | 44610475 | 3.93033 | 5.0332 | 1.8076 | Distal Intergenic | 11 | 44617317 | 45004995 | 387679 | 1 | ENSMUST00000109268.1 | -6842 | Ebf1 | ENSMUSG00000057098.9 |
| chr11 | 44611426 | 44611544 | 4.0502 | 5.29111 | 1.99416 | Distal Intergenic | 11 | 44617317 | 45004995 | 387679 | 1 | ENSMUST00000109268.1 | -5773 | Ebf1 | ENSMUSG00000057098.9 |
| chr11 | 44612292 | 44612740 | 3.88726 | 4.9143 | 1.76496 | Distal Intergenic | 11 | 44617317 | 45004995 | 387679 | 1 | ENSMUST00000109268.1 | -4577 | Ebf1 | ENSMUSG00000057098.9 |
| chr11 | 44613302 | 44613467 | 3.90663 | 5.07117 | 1.81525 | Distal Intergenic | 11 | 44617317 | 45004995 | 387679 | 1 | ENSMUST00000109268.1 | -3850 | Ebf1 | ENSMUSG00000057098.9 |
| chr11 | 44613780 | 44614030 | 3.25574 | 3.7108 | 1.01733 | Distal Intergenic | 11 | 44617317 | 45004995 | 387679 | 1 | ENSMUST00000109268.1 | -3287 | Ebf1 | ENSMUSG00000057098.9 |
| chr11 | 44615826 | 44615937 | 3.93266 | 5.17233 | 1.8658 | Distal Intergenic | 11 | 44617317 | 45004995 | 387679 | 1 | ENSMUST00000109268.1 | -1380 | Ebf1 | ENSMUSG00000057098.9 |
| chr11 | 44616778 | 44618580 | 4.18928 | 5.61918 | 2.30629 | Promoter | 11 | 44617317 | 45004995 | 387679 | 1 | ENSMUST00000109268.1 | 0 | Ebf1 | ENSMUSG00000057098.9 |
| chr11 | 44619103 | 44619359 | 3.59381 | 4.45087 | 1.44809 | Promoter | 11 | 44618179 | 44643469 | 25291 | 1 | ENSMUST00000135993.2 | 925 | Ebf1 | ENSMUSG00000057098.9 |
| chr11 | 44621138 | 44621277 | 3.56759 | 4.30363 | 1.35339 | Promoter | 11 | 44621761 | 44643459 | 21699 | 1 | ENSMUST00000138452.1 | -484 | Ebf1 | ENSMUSG00000057098.9 |
| chr11 | 44917784 | 44917937 | 3.34888 | 3.85074 | 1.01733 | Intron (ENSMUST00000109268.1/ENSMUSG00000057098.9, intron 9 of 15) | 11 | 44912138 | 44972914 | 60777 | 1 | ENSMUST00000140822.1 | 5647 | Ebf1 | ENSMUSG00000057098.9 |
| chr11 | 45013984 | 45014063 | 3.5537 | 4.23744 | 1.28585 | Distal Intergenic | 11 | 44912138 | 44972914 | 60777 | 1 | ENSMUST00000140822.1 | 101847 | Ebf1 | ENSMUSG00000057098.9 |
| chr11 | 44832300 | 44833002 | 3.93515 | 4.96063 | 1.75274 | Intron (ENSMUST00000109268.1/ENSMUSG00000057098.9, intron 6 of 15) | 11 | 44912138 | 44972914 | 60777 | 1 | ENSMUST00000140822.1 | -79136 | Ebf1 | ENSMUSG00000057098.9 |
| chr11 | 44833761 | 44833942 | 3.37744 | 4.17202 | 1.27414 | Intron (ENSMUST00000109268.1/ENSMUSG00000057098.9, intron 6 of 15) | 11 | 44912138 | 44972914 | 60777 | 1 | ENSMUST00000140822.1 | -78196 | Ebf1 | ENSMUSG00000057098.9 |
| chr5 | 102481485 | 102481674 | 3.77722 | 4.6342 | 1.48022 | Promoter | 5 | 102481530 | 102667832 | 186303 | 1 | ENSMUST00000126125.1 | 0 | Arhgap24 | ENSMUSG00000057315.9 |
| chr11 | 114666767 | 114667094 | 3.41575 | 3.96544 | 1.06587 | Distal Intergenic | 11 | 114668524 | 114672331 | 3808 | 1 | ENSMUST00000106602.4 | -1430 | Rpl38 | ENSMUSG00000057322.7 |
| chr10 | 60191097 | 60191456 | 3.5116 | 4.30647 | 1.35567 | Intron (ENSMUST00000135158.3/ENSMUSG00000057337.8, intron 2 of 3) | 10 | 60181534 | 60188707 | 7174 | 2 | ENSMUST00000068690.6 | -2391 | Chst3 | ENSMUSG00000057337.8 |
| chr10 | 60192566 | 60192641 | 3.49281 | 4.32953 | 1.33446 | Intron (ENSMUST00000135158.3/ENSMUSG00000057337.8, intron 2 of 3) | 10 | 60185605 | 60194239 | 8635 | 2 | ENSMUST00000167915.1 | 1598 | Chst3 | ENSMUSG00000057337.8 |
| chr10 | 60193573 | 60194027 | 4.04414 | 5.27168 | 1.97282 | Promoter | 10 | 60185605 | 60194239 | 8635 | 2 | ENSMUST00000167915.1 | 212 | Chst3 | ENSMUSG00000057337.8 |
| chr10 | 60217918 | 60218298 | 3.85864 | 4.81691 | 1.674 | Promoter | 10 | 60181532 | 60219260 | 37729 | 2 | ENSMUST00000135158.3 | 962 | Chst3 | ENSMUSG00000057337.8 |
| chr10 | 60219363 | 60219449 | 4.45457 | 5.96182 | 2.40874 | Promoter | 10 | 60181532 | 60219260 | 37729 | 2 | ENSMUST00000135158.3 | -104 | Chst3 | ENSMUSG00000057337.8 |
| chr10 | 60220196 | 60220340 | 3.39535 | 3.92507 | 1.03216 | Promoter | 10 | 60181532 | 60219260 | 37729 | 2 | ENSMUST00000135158.3 | -937 | Chst3 | ENSMUSG00000057337.8 |
| chr2 | 113216019 | 113216572 | 4.0652 | 5.29222 | 2.02136 | Promoter | 2 | 112631380 | 113217051 | 585672 | 2 | ENSMUST00000080673.6 | 479 | Ryr3 | ENSMUSG00000057378.9 |
| chr2 | 113216906 | 113217730 | 3.56657 | 4.41179 | 1.42083 | Promoter | 2 | 112631380 | 113217051 | 585672 | 2 | ENSMUST00000080673.6 | 0 | Ryr3 | ENSMUSG00000057378.9 |
| chr2 | 113218231 | 113218317 | 3.72261 | 4.53564 | 1.42926 | Distal Intergenic | 2 | 112996535 | 113217096 | 220562 | 2 | ENSMUST00000128192.1 | -1136 | Ryr3 | ENSMUSG00000057378.9 |
| chr7 | 24636349 | 24636877 | 3.99243 | 5.18123 | 1.93271 | Promoter | 7 | 24636570 | 24641116 | 4547 | 1 | ENSMUST00000080718.4 | 0 | Lypd3 | ENSMUSG00000057454.4 |
| chr4 | 137830378 | 137830732 | 3.5273 | 4.18532 | 1.23608 | Distal Intergenic | 4 | 137862237 | 137965229 | 102993 | 1 | ENSMUST00000102518.4 | -31505 | Ece1 | ENSMUSG00000057530.9 |
| chr6 | 39873047 | 39873352 | 4.11975 | 5.89603 | 2.38323 | Promoter | 6 | 39873271 | 40255075 | 381805 | 1 | ENSMUST00000180886.1 | 0 | Tmem178b | ENSMUSG00000057716.6 |
| chr6 | 39873854 | 39873991 | 3.61434 | 4.40884 | 1.40813 | Promoter | 6 | 39873271 | 40255075 | 381805 | 1 | ENSMUST00000180886.1 | 584 | Tmem178b | ENSMUSG00000057716.6 |
| chr6 | 39874601 | 39875717 | 3.58772 | 4.34392 | 1.35505 | Intron (ENSMUST00000180886.1/ENSMUSG00000057716.6, intron 1 of 3) | 6 | 39873271 | 40255075 | 381805 | 1 | ENSMUST00000180886.1 | 1331 | Tmem178b | ENSMUSG00000057716.6 |
| chr6 | 39876350 | 39876667 | 3.71682 | 4.56854 | 1.52361 | Intron (ENSMUST00000180886.1/ENSMUSG00000057716.6, intron 1 of 3) | 6 | 39873271 | 40255075 | 381805 | 1 | ENSMUST00000180886.1 | 3080 | Tmem178b | ENSMUSG00000057716.6 |
| chr4 | 101717269 | 101717477 | 4.16768 | 5.72817 | 2.26264 | Promoter | 4 | 101717404 | 101792258 | 74855 | 1 | ENSMUST00000102777.4 | 0 | Lepr | ENSMUSG00000057722.12 |
| chr4 | 101717839 | 101717914 | 3.76148 | 4.88084 | 1.68559 | Promoter | 4 | 101717407 | 101733317 | 15911 | 1 | ENSMUST00000145024.1 | 433 | Lepr | ENSMUSG00000057722.12 |
| chr4 | 101719126 | 101719440 | 3.7276 | 4.71229 | 1.58979 | Intron (ENSMUST00000102777.4/ENSMUSG00000057722.12, intron 1 of 17) | 4 | 101717407 | 101733317 | 15911 | 1 | ENSMUST00000145024.1 | 1720 | Lepr | ENSMUSG00000057722.12 |
| chr4 | 154168636 | 154168855 | 3.44347 | 4.05481 | 1.22634 | Distal Intergenic | 4 | 154170730 | 154275713 | 104984 | 1 | ENSMUST00000030897.9 | -1875 | Megf6 | ENSMUSG00000057751.9 |
| chr4 | 154169218 | 154171541 | 3.72309 | 4.67322 | 1.61405 | Promoter | 4 | 154170730 | 154275713 | 104984 | 1 | ENSMUST00000030897.9 | 0 | Megf6 | ENSMUSG00000057751.9 |
| chr4 | 154173386 | 154173550 | 3.30538 | 3.82246 | 1.09851 | Intron (ENSMUST00000030897.9/ENSMUSG00000057751.9, intron 1 of 36) | 4 | 154170730 | 154275713 | 104984 | 1 | ENSMUST00000030897.9 | 2657 | Megf6 | ENSMUSG00000057751.9 |
| chr5 | 98292380 | 98292679 | 3.92803 | 4.93711 | 1.71229 | Distal Intergenic | 5 | 98329354 | 98801910 | 472557 | 1 | ENSMUST00000080333.3 | -36675 | 1700007G11Rik | ENSMUSG00000057816.3 |
| chr5 | 98293442 | 98294347 | 3.71238 | 4.63694 | 1.51363 | Distal Intergenic | 5 | 98329354 | 98801910 | 472557 | 1 | ENSMUST00000080333.3 | -35007 | 1700007G11Rik | ENSMUSG00000057816.3 |
| chr5 | 98327377 | 98327536 | 3.18598 | 3.74943 | 1.01733 | Distal Intergenic | 5 | 98329354 | 98801910 | 472557 | 1 | ENSMUST00000080333.3 | -1818 | 1700007G11Rik | ENSMUSG00000057816.3 |
| chr5 | 98328214 | 98328400 | 3.48471 | 4.1375 | 1.23046 | Promoter | 5 | 98329354 | 98801910 | 472557 | 1 | ENSMUST00000080333.3 | -954 | 1700007G11Rik | ENSMUSG00000057816.3 |
| chr5 | 98330184 | 98330309 | 3.61584 | 4.3843 | 1.36228 | Promoter | 5 | 98329354 | 98801910 | 472557 | 1 | ENSMUST00000080333.3 | 831 | 1700007G11Rik | ENSMUSG00000057816.3 |
| chr5 | 98330655 | 98330772 | 3.38193 | 3.9006 | 1.01733 | Intron (ENSMUST00000080333.3/ENSMUSG00000057816.3, intron 1 of 5) | 5 | 98329354 | 98801910 | 472557 | 1 | ENSMUST00000080333.3 | 1302 | 1700007G11Rik | ENSMUSG00000057816.3 |
| chr11 | 6021943 | 6022341 | 3.55396 | 4.22291 | 1.23872 | Intron (ENSMUST00000109813.3/ENSMUSG00000057897.9, intron 2 of 20) | 11 | 5990395 | 6000448 | 10054 | 2 | ENSMUST00000130427.2 | -21496 | Camk2b | ENSMUSG00000057897.9 |
| chr11 | 6037389 | 6037661 | 3.39636 | 3.9623 | 1.15725 | Intron (ENSMUST00000109813.3/ENSMUSG00000057897.9, intron 1 of 20) | 11 | 5971735 | 6065538 | 93804 | 2 | ENSMUST00000101585.4 | 27877 | Camk2b | ENSMUSG00000057897.9 |
| chr11 | 6064647 | 6066321 | 3.69508 | 4.50294 | 1.42655 | Promoter | 11 | 5969644 | 6065748 | 96105 | 2 | ENSMUST00000109813.3 | 0 | Camk2b | ENSMUSG00000057897.9 |
| chr11 | 6066742 | 6067037 | 3.42048 | 4.0141 | 1.19398 | Promoter | 11 | 5992820 | 6066362 | 73543 | 2 | ENSMUST00000123391.2 | -381 | Camk2b | ENSMUSG00000057897.9 |
| chr11 | 6067470 | 6067545 | 4.26853 | 5.6196 | 2.21183 | Distal Intergenic | 11 | 5992820 | 6066362 | 73543 | 2 | ENSMUST00000123391.2 | -1109 | Camk2b | ENSMUSG00000057897.9 |
| chr11 | 6068019 | 6068306 | 3.71759 | 4.5183 | 1.40993 | Distal Intergenic | 11 | 5992820 | 6066362 | 73543 | 2 | ENSMUST00000123391.2 | -1658 | Camk2b | ENSMUSG00000057897.9 |
| chr11 | 6068827 | 6069208 | 3.49558 | 4.11146 | 1.15601 | Distal Intergenic | 11 | 5992820 | 6066362 | 73543 | 2 | ENSMUST00000123391.2 | -2466 | Camk2b | ENSMUSG00000057897.9 |
| chr11 | 6070081 | 6070787 | 3.36776 | 3.8831 | 1.03102 | Distal Intergenic | 11 | 5992820 | 6066362 | 73543 | 2 | ENSMUST00000123391.2 | -3720 | Camk2b | ENSMUSG00000057897.9 |
| chr2 | 14604143 | 14604341 | 3.46298 | 4.18305 | 1.27791 | Promoter | 2 | 14604306 | 14987908 | 383603 | 1 | ENSMUST00000114723.3 | 0 | Cacnb2 | ENSMUSG00000057914.10 |
| chr2 | 14605087 | 14605227 | 3.48188 | 4.10866 | 1.22066 | Promoter | 2 | 14605116 | 14961042 | 355927 | 1 | ENSMUST00000156027.2 | 0 | Cacnb2 | ENSMUSG00000057914.10 |
| chr11 | 33108301 | 33108820 | 3.5478 | 4.53146 | 1.4777 | Distal Intergenic | 11 | 33117430 | 33146795 | 29366 | 2 | ENSMUST00000109363.2 | 37975 | Fgf18 | ENSMUSG00000057967.7 |
| chr11 | 33114505 | 33114917 | 3.41611 | 3.96363 | 1.05763 | Downstream (2-3kb) | 11 | 33117430 | 33146795 | 29366 | 2 | ENSMUST00000109363.2 | 31878 | Fgf18 | ENSMUSG00000057967.7 |
| chr11 | 33119490 | 33119711 | 3.35769 | 3.86404 | 1.01733 | Intron (ENSMUST00000109363.2/ENSMUSG00000057967.7, intron 3 of 3) | 11 | 33117430 | 33146795 | 29366 | 2 | ENSMUST00000109363.2 | 27084 | Fgf18 | ENSMUSG00000057967.7 |
| chr11 | 33121724 | 33122216 | 3.40626 | 3.95341 | 1.07396 | Intron (ENSMUST00000109363.2/ENSMUSG00000057967.7, intron 3 of 3) | 11 | 33117430 | 33146795 | 29366 | 2 | ENSMUST00000109363.2 | 24579 | Fgf18 | ENSMUSG00000057967.7 |
| chr11 | 33122603 | 33123364 | 3.9084 | 4.98838 | 1.74188 | Intron (ENSMUST00000109363.2/ENSMUSG00000057967.7, intron 3 of 3) | 11 | 33117430 | 33146795 | 29366 | 2 | ENSMUST00000109363.2 | 23431 | Fgf18 | ENSMUSG00000057967.7 |
| chr11 | 33124324 | 33124707 | 3.5977 | 4.29881 | 1.26552 | 5' UTR | 11 | 33117430 | 33146795 | 29366 | 2 | ENSMUST00000109363.2 | 22088 | Fgf18 | ENSMUSG00000057967.7 |
| chr11 | 33143730 | 33144309 | 3.46062 | 4.04769 | 1.11744 | Intron (ENSMUST00000109363.2/ENSMUSG00000057967.7, intron 1 of 3) | 11 | 33117430 | 33146795 | 29366 | 2 | ENSMUST00000109363.2 | 2486 | Fgf18 | ENSMUSG00000057967.7 |
| chr11 | 33145044 | 33145262 | 3.9767 | 5.03463 | 1.78661 | Intron (ENSMUST00000109363.2/ENSMUSG00000057967.7, intron 1 of 3) | 11 | 33117430 | 33146795 | 29366 | 2 | ENSMUST00000109363.2 | 1533 | Fgf18 | ENSMUSG00000057967.7 |
| chr11 | 33146356 | 33146899 | 3.8439 | 4.78073 | 1.61124 | Promoter | 11 | 33117430 | 33146795 | 29366 | 2 | ENSMUST00000109363.2 | 0 | Fgf18 | ENSMUSG00000057967.7 |
| chr11 | 33147495 | 33148190 | 3.93258 | 4.99704 | 1.78619 | Promoter | 11 | 33117430 | 33147400 | 29971 | 2 | ENSMUST00000020507.7 | -96 | Fgf18 | ENSMUSG00000057967.7 |
| chr7 | 140698399 | 140698697 | 3.35163 | 3.85489 | 1.01733 | Distal Intergenic | 7 | 140704165 | 140705267 | 1103 | 1 | ENSMUST00000080681.3 | -5468 | Olfr541 | ENSMUSG00000057997.3 |
| chr13 | 41821044 | 41821395 | 3.59311 | 4.30838 | 1.32698 | Intron (ENSMUST00000072012.4/ENSMUSG00000058022.8, intron 2 of 6) | 13 | 41777513 | 41828453 | 50941 | 2 | ENSMUST00000137905.1 | 7058 | Adtrp | ENSMUSG00000058022.8 |
| chr12 | 108353488 | 108353995 | 3.85995 | 5.06448 | 1.81495 | Intron (ENSMUST00000021684.4/ENSMUSG00000021259.4, intron 9 of 14) | 12 | 108370957 | 108508538 | 137582 | 1 | ENSMUST00000138456.2 | -16962 | Eml1 | ENSMUSG00000058070.9 |
| chr15 | 83007403 | 83007796 | 3.63961 | 4.3844 | 1.34254 | Intron (ENSMUST00000109503.4/ENSMUSG00000058099.10, intron 4 of 5) | 15 | 82997721 | 83023138 | 25418 | 2 | ENSMUST00000109503.4 | 15342 | Nfam1 | ENSMUSG00000058099.10 |
| chr15 | 83008337 | 83008619 | 3.34555 | 3.85788 | 1.05025 | Intron (ENSMUST00000109503.4/ENSMUSG00000058099.10, intron 4 of 5) | 15 | 82997721 | 83023138 | 25418 | 2 | ENSMUST00000109503.4 | 14519 | Nfam1 | ENSMUSG00000058099.10 |
| chr15 | 83009547 | 83009622 | 4.08249 | 5.22375 | 1.90155 | Intron (ENSMUST00000109503.4/ENSMUSG00000058099.10, intron 4 of 5) | 15 | 82997721 | 83023138 | 25418 | 2 | ENSMUST00000109503.4 | 13516 | Nfam1 | ENSMUSG00000058099.10 |
| chr15 | 83010043 | 83010118 | 3.23338 | 3.67724 | 1.01733 | Intron (ENSMUST00000109503.4/ENSMUSG00000058099.10, intron 4 of 5) | 15 | 82997721 | 83023138 | 25418 | 2 | ENSMUST00000109503.4 | 13020 | Nfam1 | ENSMUSG00000058099.10 |
| chr15 | 83013622 | 83014765 | 3.77506 | 4.66371 | 1.55288 | Intron (ENSMUST00000109503.4/ENSMUSG00000058099.10, intron 3 of 5) | 15 | 82997721 | 83023138 | 25418 | 2 | ENSMUST00000109503.4 | 8373 | Nfam1 | ENSMUSG00000058099.10 |
| chr15 | 83017360 | 83018141 | 3.4415 | 4.04309 | 1.13025 | Intron (ENSMUST00000109503.4/ENSMUSG00000058099.10, intron 1 of 5) | 15 | 82997721 | 83023138 | 25418 | 2 | ENSMUST00000109503.4 | 4997 | Nfam1 | ENSMUSG00000058099.10 |
| chr15 | 83019812 | 83020030 | 3.6201 | 4.33482 | 1.28053 | Intron (ENSMUST00000109503.4/ENSMUSG00000058099.10, intron 1 of 5) | 15 | 82997721 | 83023138 | 25418 | 2 | ENSMUST00000109503.4 | 3108 | Nfam1 | ENSMUSG00000058099.10 |
| chr15 | 83023387 | 83023697 | 3.66324 | 4.46401 | 1.43358 | Promoter | 15 | 82997721 | 83023138 | 25418 | 2 | ENSMUST00000109503.4 | -250 | Nfam1 | ENSMUSG00000058099.10 |
| chr15 | 83025120 | 83025614 | 3.61733 | 4.34367 | 1.32116 | Intron (ENSMUST00000023076.5/ENSMUSG00000058099.10, intron 1 of 5) | 15 | 82997721 | 83023138 | 25418 | 2 | ENSMUST00000109503.4 | -1983 | Nfam1 | ENSMUSG00000058099.10 |
| chr15 | 83026833 | 83027327 | 3.55867 | 4.27459 | 1.30602 | Intron (ENSMUST00000023076.5/ENSMUSG00000058099.10, intron 1 of 5) | 15 | 82997726 | 83029950 | 32225 | 2 | ENSMUST00000023076.5 | 2623 | Nfam1 | ENSMUSG00000058099.10 |
| chr15 | 83031384 | 83031673 | 3.90795 | 5.11112 | 1.84421 | Intron (ENSMUST00000100377.5/ENSMUSG00000058099.10, intron 1 of 5) | 15 | 82997726 | 83029950 | 32225 | 2 | ENSMUST00000023076.5 | -1435 | Nfam1 | ENSMUSG00000058099.10 |
| chr15 | 83032442 | 83032659 | 3.31228 | 3.79566 | 1.01733 | Promoter | 15 | 82997726 | 83033306 | 35581 | 2 | ENSMUST00000100377.5 | 647 | Nfam1 | ENSMUSG00000058099.10 |
| chr15 | 83032978 | 83033470 | 3.61757 | 4.33435 | 1.29184 | Promoter | 15 | 82997726 | 83033306 | 35581 | 2 | ENSMUST00000100377.5 | 0 | Nfam1 | ENSMUSG00000058099.10 |
| chr15 | 83034415 | 83034919 | 3.44548 | 4.15058 | 1.22826 | Distal Intergenic | 15 | 82997726 | 83033306 | 35581 | 2 | ENSMUST00000100377.5 | -1110 | Nfam1 | ENSMUSG00000058099.10 |
| chr15 | 83035722 | 83035851 | 3.55669 | 4.2311 | 1.25388 | Distal Intergenic | 15 | 82997726 | 83033306 | 35581 | 2 | ENSMUST00000100377.5 | -2417 | Nfam1 | ENSMUSG00000058099.10 |
| chr15 | 83053177 | 83053258 | 3.89003 | 4.85935 | 1.65554 | Distal Intergenic | 15 | 82997726 | 83033306 | 35581 | 2 | ENSMUST00000100377.5 | -19872 | Nfam1 | ENSMUSG00000058099.10 |
| chr15 | 83054428 | 83054842 | 3.47153 | 4.09019 | 1.19489 | Distal Intergenic | 15 | 82997726 | 83033306 | 35581 | 2 | ENSMUST00000100377.5 | -21123 | Nfam1 | ENSMUSG00000058099.10 |
| chr15 | 83056590 | 83056870 | 3.32085 | 3.80857 | 1.01733 | Distal Intergenic | 15 | 82997726 | 83033306 | 35581 | 2 | ENSMUST00000100377.5 | -23285 | Nfam1 | ENSMUSG00000058099.10 |
| chr15 | 83057298 | 83058174 | 4.15805 | 5.45874 | 2.10588 | Distal Intergenic | 15 | 82997726 | 83033306 | 35581 | 2 | ENSMUST00000100377.5 | -23993 | Nfam1 | ENSMUSG00000058099.10 |
| chr7 | 66840248 | 66840631 | 3.57562 | 4.66426 | 1.54925 | Promoter | 7 | 66839813 | 67152625 | 312813 | 1 | ENSMUST00000107478.2 | 436 | Adamts17 | ENSMUSG00000058145.10 |
| chr7 | 66842205 | 66842357 | 3.56912 | 4.32985 | 1.36058 | Promoter | 7 | 66842255 | 67062157 | 219903 | 1 | ENSMUST00000147880.2 | 0 | Adamts17 | ENSMUSG00000058145.10 |
| chr18 | 59175796 | 59176706 | 3.59254 | 4.45613 | 1.42392 | Promoter | 18 | 59175401 | 59410446 | 235046 | 1 | ENSMUST00000080721.4 | 396 | Chsy3 | ENSMUSG00000058152.7 |
| chr5 | 112520367 | 112520487 | 3.39352 | 4.07129 | 1.15346 | Intron (ENSMUST00000075387.6/ENSMUSG00000058153.9, intron 1 of 16) | 5 | 112419151 | 112577198 | 158048 | 2 | ENSMUST00000075387.6 | 56711 | Sez6l | ENSMUSG00000058153.9 |
| chr5 | 112532691 | 112532843 | 3.31248 | 3.79594 | 1.01733 | Intron (ENSMUST00000075387.6/ENSMUSG00000058153.9, intron 1 of 16) | 5 | 112419151 | 112577198 | 158048 | 2 | ENSMUST00000075387.6 | 44355 | Sez6l | ENSMUSG00000058153.9 |
| chr5 | 112563608 | 112563824 | 3.51137 | 4.14284 | 1.18303 | Intron (ENSMUST00000075387.6/ENSMUSG00000058153.9, intron 1 of 16) | 5 | 112419151 | 112577198 | 158048 | 2 | ENSMUST00000075387.6 | 13374 | Sez6l | ENSMUSG00000058153.9 |
| chr5 | 112565064 | 112565503 | 3.84032 | 4.76012 | 1.58852 | Intron (ENSMUST00000075387.6/ENSMUSG00000058153.9, intron 1 of 16) | 5 | 112419151 | 112577198 | 158048 | 2 | ENSMUST00000075387.6 | 11695 | Sez6l | ENSMUSG00000058153.9 |
| chr5 | 112575506 | 112575666 | 3.7063 | 4.5033 | 1.40465 | Intron (ENSMUST00000075387.6/ENSMUSG00000058153.9, intron 1 of 16) | 5 | 112419151 | 112577198 | 158048 | 2 | ENSMUST00000075387.6 | 1532 | Sez6l | ENSMUSG00000058153.9 |
| chr5 | 112576561 | 112577306 | 3.9161 | 5.07455 | 1.82645 | Promoter | 5 | 112419151 | 112577198 | 158048 | 2 | ENSMUST00000075387.6 | 0 | Sez6l | ENSMUSG00000058153.9 |
| chr5 | 112578005 | 112578099 | 3.56726 | 4.25057 | 1.25975 | Promoter | 5 | 112419151 | 112577198 | 158048 | 2 | ENSMUST00000079491.8 | -808 | Sez6l | ENSMUSG00000058153.9 |
| chr5 | 112579033 | 112579111 | 4.43193 | 5.92424 | 2.38994 | Distal Intergenic | 5 | 112419151 | 112577198 | 158048 | 2 | ENSMUST00000079491.8 | -1836 | Sez6l | ENSMUSG00000058153.9 |
| chr9 | 20569170 | 20569454 | 3.57758 | 4.27014 | 1.28006 | Distal Intergenic | 9 | 20581325 | 20595192 | 13868 | 1 | ENSMUST00000115557.3 | -11871 | Zfp846 | ENSMUSG00000058192.10 |
| chr9 | 20575873 | 20575948 | 4.21724 | 5.58257 | 2.18198 | Distal Intergenic | 9 | 20581325 | 20595192 | 13868 | 1 | ENSMUST00000115557.3 | -5377 | Zfp846 | ENSMUSG00000058192.10 |
| chr9 | 20576404 | 20576479 | 3.86636 | 4.87985 | 1.72467 | Distal Intergenic | 9 | 20581325 | 20595192 | 13868 | 1 | ENSMUST00000115557.3 | -4846 | Zfp846 | ENSMUSG00000058192.10 |
| chr1 | 192189891 | 192190669 | 3.69818 | 4.54084 | 1.53144 | Promoter | 1 | 192190784 | 192510158 | 319375 | 1 | ENSMUST00000110844.1 | -115 | Kcnh1 | ENSMUSG00000058248.7 |
| chr1 | 192191491 | 192192152 | 3.58558 | 4.46971 | 1.43869 | Promoter | 1 | 192190784 | 192510159 | 319376 | 1 | ENSMUST00000078470.6 | 708 | Kcnh1 | ENSMUSG00000058248.7 |
| chr1 | 192368425 | 192368692 | 3.56155 | 4.40149 | 1.36189 | Intron (ENSMUST00000110844.1/ENSMUSG00000058248.7, intron 7 of 10) | 1 | 192434740 | 192505645 | 70906 | 1 | ENSMUST00000151152.1 | -66048 | Kcnh1 | ENSMUSG00000058248.7 |
| chr12 | 104012743 | 104012932 | 3.27006 | 3.73913 | 1.03901 | Promoter | 12 | 103994743 | 104013652 | 18910 | 2 | ENSMUST00000058464.4 | 720 | Serpina9 | ENSMUSG00000058260.2 |
| chr11 | 74371534 | 74371769 | 3.88318 | 4.84624 | 1.6372 | Distal Intergenic | 11 | 74364671 | 74365609 | 939 | 1 | ENSMUST00000077794.1 | 6864 | Olfr412 | ENSMUSG00000058275.1 |
| chr10 | 60106375 | 60106908 | 3.93381 | 5.36788 | 2.03241 | Promoter | 10 | 60106468 | 60133908 | 27441 | 1 | ENSMUST00000165024.2 | 0 | Spock2 | ENSMUSG00000058297.11 |

| | | | | | | | | | | | | | | | |
|---|---|---|---|---|---|---|---|---|---|---|---|---|---|---|---|
| chr10 | 60107371 | 60107622 | 3.50454 | 4.39027 | 1.36848 | Promoter | 10 | 60106468 | 60133908 | 27441 | 1 | ENSMUST00000165024.2 | 904 | Spock2 | ENSMUSG00000058297.11 |
| chr15 | 101690689 | 101690816 | 3.27434 | 3.75803 | 1.07332 | Exon (ENSMUST00000023788.6/ENSMUSG00000058354.6, exon 8 of 9) | 15 | 101689932 | 101694307 | 4376 | 2 | ENSMUST00000023788.6 | 3491 | Krt6a | ENSMUSG00000058354.6 |
| chr3 | 68718696 | 68718980 | 3.30582 | 3.96892 | 1.12458 | Distal Intergenic | 3 | 68733137 | 68738077 | 4941 | 1 | ENSMUST00000080726.5 | -14157 | Gm10040 | ENSMUSG00000058360.5 |
| chr3 | 68779133 | 68779448 | 3.38885 | 4.08221 | 1.16918 | Distal Intergenic | 3 | 68733137 | 68738077 | 4941 | 1 | ENSMUST00000080726.5 | 45997 | Gm10040 | ENSMUSG00000058360.5 |
| chr3 | 36220653 | 36220909 | 3.82366 | 4.99534 | 1.79874 | Intron (ENSMUST00000091227.6/ENSMUSG00000058400.8, intron 1 of 5) | 3 | 36187805 | 36222180 | 34376 | 2 | ENSMUST00000142552.1 | 1271 | Qrfpr | ENSMUSG00000058400.8 |
| chr3 | 36222036 | 36222191 | 4.34522 | 5.76094 | 2.28114 | Promoter | 3 | 36187805 | 36222180 | 34376 | 2 | ENSMUST00000142552.1 | 0 | Qrfpr | ENSMUSG00000058400.8 |
| chr5 | 90902046 | 90902646 | 3.35164 | 3.86194 | 1.04216 | Distal Intergenic | 5 | 90903899 | 90905909 | 2011 | 1 | ENSMUST00000075433.6 | -1253 | Cxcl2 | ENSMUSG00000058427.7 |
| chr15 | 89059257 | 89059332 | 4.72154 | 6.65451 | 2.94895 | Promoter | 15 | 89059734 | 89069878 | 10145 | 1 | ENSMUST00000161372.1 | -402 | Panx2 | ENSMUSG00000058441.7 |
| chr15 | 89059724 | 89060421 | 3.72731 | 4.64601 | 1.51397 | Promoter | 15 | 89059734 | 89069878 | 10145 | 1 | ENSMUST00000161372.1 | 0 | Panx2 | ENSMUSG00000058441.7 |
| chr15 | 89063282 | 89063542 | 3.4341 | 4.02311 | 1.17306 | Intron (ENSMUST00000161372.1/ENSMUSG00000058441.7, intron 1 of 3) | 15 | 89059747 | 89070907 | 11161 | 1 | ENSMUST00000162424.1 | 3536 | Panx2 | ENSMUSG00000058441.7 |
| chr15 | 89065917 | 89066174 | 3.5135 | 4.14356 | 1.1736 | Intron (ENSMUST00000161372.1/ENSMUSG00000058441.7, intron 1 of 3) | 15 | 89069673 | 89073567 | 3895 | 1 | ENSMUST00000159960.1 | -3499 | Panx2 | ENSMUSG00000058441.7 |
| chr15 | 89067303 | 89068830 | 3.9129 | 5.09971 | 1.84982 | Promoter | 15 | 89069673 | 89073567 | 3895 | 1 | ENSMUST00000159960.1 | -843 | Panx2 | ENSMUSG00000058441.7 |
| chr5 | 150952240 | 150953610 | 4.16667 | 5.48583 | 2.11333 | Promoter | 5 | 150952607 | 150993809 | 41203 | 1 | ENSMUST00000078856.6 | 0 | Kl | ENSMUSG00000058488.6 |
| chr4 | 152311864 | 152311996 | 3.47557 | 4.4599 | 1.40908 | Intron (ENSMUST00000076183.6/ENSMUSG00000058498.7, intron 13 of 16) | 4 | 152311716 | 152313348 | 1633 | 2 | ENSMUST00000127565.1 | 1352 | Rnf207 | ENSMUSG00000058498.7 |
| chr4 | 152312379 | 152312625 | 3.38063 | 4.12312 | 1.19775 | Promoter | 4 | 152311716 | 152313348 | 1633 | 2 | ENSMUST00000127565.1 | 723 | Rnf207 | ENSMUSG00000058498.7 |
| chr4 | 152314688 | 152314796 | 3.29961 | 3.78765 | 1.05524 | Exon (ENSMUST00000134269.3/ENSMUSG00000058498.7, exon 4 of 4) | 4 | 152313403 | 152315965 | 2563 | 2 | ENSMUST00000143968.2 | 1169 | Rnf207 | ENSMUSG00000058498.7 |
| chr4 | 152315105 | 152315968 | 3.89257 | 4.91294 | 1.76439 | Promoter | 4 | 152313403 | 152315965 | 2563 | 2 | ENSMUST00000143968.2 | 0 | Rnf207 | ENSMUSG00000058498.7 |
| chr4 | 152316914 | 152317183 | 3.52047 | 4.22445 | 1.29669 | Promoter | 4 | 152315461 | 152316328 | 868 | 2 | ENSMUST00000146496.1 | -587 | Rnf207 | ENSMUSG00000058498.7 |
| chr14 | 116927606 | 116927787 | 3.87382 | 4.89419 | 1.74684 | Intron (ENSMUST00000078849.5/ENSMUSG00000058571.5, intron 1 of 8) | 14 | 116925531 | 117975406 | 1049876 | 1 | ENSMUST00000125435.1 | 2076 | Gpc6 | ENSMUSG00000058571.5 |
| chr15 | 83088943 | 83089236 | 3.28219 | 3.75045 | 1.01733 | Promoter | 15 | 83089506 | 83101767 | 12262 | 1 | ENSMUST00000142265.3 | -270 | Serhl | ENSMUSG00000058586.6 |
| chr10 | 89873248 | 89873616 | 4.60963 | 6.47075 | 2.89432 | Promoter | 10 | 89873509 | 90970219 | 1096711 | 1 | ENSMUST00000183156.2 | 0 | Anks1b | ENSMUSG00000058589.9 |
| chr10 | 90827397 | 90827851 | 3.48626 | 4.10776 | 1.18836 | Intron (ENSMUST00000183156.2/ENSMUSG00000058589.9, intron 17 of 25) | 10 | 90829111 | 90941465 | 112355 | 1 | ENSMUST00000182192.2 | -1260 | Anks1b | ENSMUSG00000058589.9 |
| chr10 | 90828346 | 90830576 | 4.04117 | 5.20184 | 1.93317 | Promoter | 10 | 90829111 | 90941465 | 112355 | 1 | ENSMUST00000182192.2 | 0 | Anks1b | ENSMUSG00000058589.9 |
| chr10 | 90831323 | 90832571 | 3.67932 | 4.45746 | 1.38947 | Intron (ENSMUST00000183156.2/ENSMUSG00000058589.9, intron 17 of 25) | 10 | 90829843 | 90921338 | 91496 | 1 | ENSMUST00000183145.1 | 1481 | Anks1b | ENSMUSG00000058589.9 |
| chr2 | 127360297 | 127360781 | 3.74981 | 4.60226 | 1.49886 | Distal Intergenic | 2 | 127363208 | 127367221 | 4014 | 1 | ENSMUST00000104934.1 | -2427 | Adra2b | ENSMUSG00000058620.5 |
| chr2 | 127362840 | 127364446 | 3.7064 | 4.63502 | 1.57303 | Promoter | 2 | 127363208 | 127367221 | 4014 | 1 | ENSMUST00000104934.1 | 0 | Adra2b | ENSMUSG00000058620.5 |
| chr2 | 127365559 | 127365831 | 4.2431 | 5.6056 | 2.22568 | 3' UTR | 2 | 127363286 | 127367221 | 3936 | 1 | ENSMUST00000071902.4 | 2274 | Adra2b | ENSMUSG00000058620.5 |
| chr19 | 21472472 | 21472782 | 3.50904 | 4.33145 | 1.30811 | Promoter | 19 | 21391307 | 21472672 | 81366 | 2 | ENSMUST00000087600.4 | 0 | Gda | ENSMUSG00000058624.7 |
| chr15 | 102051740 | 102051834 | 4.32782 | 5.75535 | 2.33101 | Distal Intergenic | 15 | 102073773 | 102097172 | 23400 | 1 | ENSMUST00000169681.1 | -21939 | Eif4b | ENSMUSG00000058655.8 |
| chr1 | 120581627 | 120581873 | 3.5174 | 4.1481 | 1.16885 | Distal Intergenic | 1 | 120602418 | 120607992 | 5575 | 1 | ENSMUST00000079721.8 | -20545 | En1 | ENSMUSG00000058665.8 |
| chr1 | 120583823 | 120583974 | 3.2957 | 3.79331 | 1.0815 | Distal Intergenic | 1 | 120602418 | 120607992 | 5575 | 1 | ENSMUST00000079721.8 | -18444 | En1 | ENSMUSG00000058665.8 |
| chr1 | 120587730 | 120588183 | 3.48916 | 4.09905 | 1.14641 | Distal Intergenic | 1 | 120602418 | 120607992 | 5575 | 1 | ENSMUST00000079721.8 | -14235 | En1 | ENSMUSG00000058665.8 |
| chr1 | 120590684 | 120591353 | 3.95991 | 5.01436 | 1.77913 | Distal Intergenic | 1 | 120602418 | 120607992 | 5575 | 1 | ENSMUST00000079721.8 | -11065 | En1 | ENSMUSG00000058665.8 |
| chr1 | 120592927 | 120594733 | 3.7858 | 4.77197 | 1.62765 | Distal Intergenic | 1 | 120602418 | 120607992 | 5575 | 1 | ENSMUST00000079721.8 | -7685 | En1 | ENSMUSG00000058665.8 |
| chr1 | 120595271 | 120596167 | 3.59779 | 4.55866 | 1.50445 | Distal Intergenic | 1 | 120602418 | 120607992 | 5575 | 1 | ENSMUST00000079721.8 | -6251 | En1 | ENSMUSG00000058665.8 |
| chr1 | 120596589 | 120597595 | 3.92834 | 5.00855 | 1.78583 | Distal Intergenic | 1 | 120602418 | 120607992 | 5575 | 1 | ENSMUST00000079721.8 | -4823 | En1 | ENSMUSG00000058665.8 |
| chr1 | 120598419 | 120599147 | 3.59931 | 4.71834 | 1.60722 | Distal Intergenic | 1 | 120602418 | 120607992 | 5575 | 1 | ENSMUST00000079721.8 | -3271 | En1 | ENSMUSG00000058665.8 |
| chr1 | 120599497 | 120603775 | 4.05916 | 5.34526 | 2.0754 | Promoter | 1 | 120602418 | 120607992 | 5575 | 1 | ENSMUST00000079721.8 | 0 | En1 | ENSMUSG00000058665.8 |
| chr1 | 120604178 | 120605596 | 4.07289 | 5.56879 | 2.1795 | Intron (ENSMUST00000079721.8/ENSMUSG00000058665.8, intron 1 of 1) | 1 | 120602418 | 120607992 | 5575 | 1 | ENSMUST00000079721.8 | 1761 | En1 | ENSMUSG00000058665.8 |
| chr1 | 120606535 | 120606643 | 3.87631 | 4.91568 | 1.74719 | Intron (ENSMUST00000079721.8/ENSMUSG00000058665.8, intron 1 of 1) | 1 | 120602418 | 120607992 | 5575 | 1 | ENSMUST00000079721.8 | 4118 | En1 | ENSMUSG00000058665.8 |
| chr1 | 120607050 | 120607419 | 3.98252 | 5.16094 | 1.91457 | 3' UTR | 1 | 120602418 | 120607992 | 5575 | 1 | ENSMUST00000079721.8 | 4633 | En1 | ENSMUSG00000058665.8 |
| chr1 | 120607888 | 120608606 | 4.12015 | 5.44297 | 2.12044 | 3' UTR | 1 | 120602418 | 120607992 | 5575 | 1 | ENSMUST00000079721.8 | 5471 | En1 | ENSMUSG00000058665.8 |
| chr1 | 120609036 | 120609664 | 3.67539 | 4.61535 | 1.54842 | Distal Intergenic | 1 | 120602418 | 120607992 | 5575 | 1 | ENSMUST00000079721.8 | 6619 | En1 | ENSMUSG00000058665.8 |
| chr1 | 120610268 | 120610992 | 3.56047 | 4.24629 | 1.2777 | Distal Intergenic | 1 | 120602418 | 120607992 | 5575 | 1 | ENSMUST00000079721.8 | 7851 | En1 | ENSMUSG00000058665.8 |
| chr12 | 56602190 | 56602332 | 3.34153 | 3.83966 | 1.01733 | Intron (ENSMUST00000182733.2/ENSMUSG00000098172.2, intron 2 of 2) | 12 | 56611389 | 56613284 | 1896 | 2 | ENSMUST00000072631.5 | 10952 | Nkx2-9 | ENSMUSG00000058669.7 |
| chr12 | 56602640 | 56602802 | 3.6099 | 4.34451 | 1.37328 | Intron (ENSMUST00000182733.2/ENSMUSG00000098172.2, intron 2 of 2) | 12 | 56611389 | 56613284 | 1896 | 2 | ENSMUST00000072631.5 | 10482 | Nkx2-9 | ENSMUSG00000058669.7 |
| chr12 | 56608122 | 56608305 | 3.39814 | 3.93545 | 1.05415 | Distal Intergenic | 12 | 56611389 | 56613284 | 1896 | 2 | ENSMUST00000072631.5 | 4979 | Nkx2-9 | ENSMUSG00000058669.7 |
| chr12 | 56611074 | 56611323 | 3.87005 | 4.84344 | 1.68253 | Downstream (<1kb) | 12 | 56611389 | 56613284 | 1896 | 2 | ENSMUST00000072631.5 | 1961 | Nkx2-9 | ENSMUSG00000058669.7 |
| chr12 | 56611790 | 56613987 | 4.0276 | 5.15728 | 1.88582 | Promoter | 12 | 56611389 | 56613284 | 1896 | 2 | ENSMUST00000072631.5 | 0 | Nkx2-9 | ENSMUSG00000058669.7 |
| chr12 | 56614463 | 56614917 | 3.88643 | 4.94199 | 1.75724 | Distal Intergenic | 12 | 56611389 | 56613284 | 1896 | 2 | ENSMUST00000072631.5 | -1180 | Nkx2-9 | ENSMUSG00000058669.7 |
| chr12 | 56618305 | 56618925 | 4.28727 | 5.87072 | 2.47529 | Distal Intergenic | 12 | 56611389 | 56613284 | 1896 | 2 | ENSMUST00000072631.5 | -5022 | Nkx2-9 | ENSMUSG00000058669.7 |
| chr12 | 56619270 | 56620501 | 3.49734 | 4.14342 | 1.21318 | Distal Intergenic | 12 | 56611389 | 56613284 | 1896 | 2 | ENSMUST00000072631.5 | -5987 | Nkx2-9 | ENSMUSG00000058669.7 |
| chr12 | 56621277 | 56621788 | 3.6012 | 4.49734 | 1.4198 | Distal Intergenic | 12 | 56611389 | 56613284 | 1896 | 2 | ENSMUST00000072631.5 | -7994 | Nkx2-9 | ENSMUSG00000058669.7 |
| chr12 | 56623003 | 56623121 | 3.78865 | 4.65176 | 1.49407 | Distal Intergenic | 12 | 56611389 | 56613284 | 1896 | 2 | ENSMUST00000072631.5 | -9720 | Nkx2-9 | ENSMUSG00000058669.7 |
| chr12 | 56625846 | 56626138 | 3.23338 | 3.67724 | 1.01733 | Distal Intergenic | 12 | 56611389 | 56613284 | 1896 | 2 | ENSMUST00000072631.5 | -12563 | Nkx2-9 | ENSMUSG00000058669.7 |
| chr12 | 56627788 | 56628122 | 3.5335 | 4.17694 | 1.18484 | Distal Intergenic | 12 | 56611389 | 56613284 | 1896 | 2 | ENSMUST00000072631.5 | -14505 | Nkx2-9 | ENSMUSG00000058669.7 |
| chr12 | 56634949 | 56635232 | 3.618 | 4.34986 | 1.32695 | Distal Intergenic | 12 | 56611389 | 56613284 | 1896 | 2 | ENSMUST00000072631.5 | -21666 | Nkx2-9 | ENSMUSG00000058669.7 |
| chr2 | 25863155 | 25863505 | 3.38103 | 3.89925 | 1.01733 | Promoter | 2 | 25863795 | 25888942 | 25148 | 1 | ENSMUST00000150788.1 | -290 | Kcnt1 | ENSMUSG00000058740.9 |
| chr2 | 25863914 | 25864087 | 3.65902 | 4.41176 | 1.33862 | Promoter | 2 | 25863854 | 25916117 | 52264 | 1 | ENSMUST00000171268.2 | 61 | Kcnt1 | ENSMUSG00000058740.9 |
| chr2 | 25864573 | 25864694 | 3.86574 | 4.80622 | 1.60879 | Promoter | 2 | 25863854 | 25916117 | 52264 | 1 | ENSMUST00000171268.2 | 720 | Kcnt1 | ENSMUSG00000058740.9 |
| chr2 | 25865363 | 25865724 | 3.97173 | 5.04434 | 1.79724 | Intron (ENSMUST00000150788.1/ENSMUSG00000058740.9, intron 1 of 2) | 2 | 25863854 | 25916117 | 52264 | 1 | ENSMUST00000171268.2 | 1510 | Kcnt1 | ENSMUSG00000058740.9 |
| chr2 | 25866415 | 25866496 | 3.57202 | 4.24716 | 1.2274 | Intron (ENSMUST00000150788.1/ENSMUSG00000058740.9, intron 1 of 2) | 2 | 25863854 | 25916117 | 52264 | 1 | ENSMUST00000171268.2 | 2562 | Kcnt1 | ENSMUSG00000058740.9 |
| chr2 | 25877585 | 25878388 | 4.68432 | 6.49699 | 2.864 | Promoter | 2 | 25877857 | 25895175 | 17319 | 1 | ENSMUST00000114170.2 | 0 | Kcnt1 | ENSMUSG00000058740.9 |
| chr7 | 45819889 | 45820172 | 3.38523 | 3.91204 | 1.04021 | 5' UTR | 7 | 45816461 | 45824756 | 8296 | 2 | ENSMUST00000071937.5 | 4584 | Kcnj14 | ENSMUSG00000058743.5 |
| chr10 | 61929500 | 61929588 | 3.87066 | 4.88145 | 1.7237 | Intron (ENSMUST00000105452.3/ENSMUSG00000058806.9, intron 2 of 39) | 10 | 61848732 | 61978121 | 129390 | 2 | ENSMUST00000051826.8 | 48533 | Col13a1 | ENSMUSG00000058806.9 |
| chr10 | 61929918 | 61930025 | 3.36308 | 3.87216 | 1.01733 | Intron (ENSMUST00000105452.3/ENSMUSG00000058806.9, intron 2 of 39) | 10 | 61848732 | 61978121 | 129390 | 2 | ENSMUST00000051826.8 | 48096 | Col13a1 | ENSMUSG00000058806.9 |
| chr10 | 61936392 | 61936974 | 3.66523 | 4.43122 | 1.37734 | Intron (ENSMUST00000105452.3/ENSMUSG00000058806.9, intron 2 of 39) | 10 | 61848732 | 61978121 | 129390 | 2 | ENSMUST00000051826.8 | 41147 | Col13a1 | ENSMUSG00000058806.9 |
| chr10 | 61977094 | 61979341 | 3.82126 | 4.77212 | 1.63657 | Promoter | 10 | 61838236 | 61979073 | 140838 | 2 | ENSMUST00000105452.3 | 0 | Col13a1 | ENSMUSG00000058806.9 |
| chr10 | 62023407 | 62023544 | 3.39157 | 3.92496 | 1.052 | Distal Intergenic | 10 | 61838236 | 61979108 | 140873 | 2 | ENSMUST00000105454.2 | -44300 | Col13a1 | ENSMUSG00000058806.9 |
| chr19 | 30504688 | 30504823 | 3.75273 | 4.60809 | 1.48215 | Distal Intergenic | 19 | 30538928 | 30539554 | 627 | 2 | ENSMUST00000078012.4 | 34731 | Ppp1r2-ps3 | ENSMUSG00000058816.4 |
| chr19 | 30539107 | 30539915 | 4.29179 | 6.415 | 2.8969 | Promoter | 19 | 30538928 | 30539554 | 627 | 2 | ENSMUST00000078012.4 | 0 | Ppp1r2-ps3 | ENSMUSG00000058816.4 |
| chr6 | 29388457 | 29388739 | 3.42516 | 4.20721 | 1.25689 | Promoter | 6 | 29379373 | 29388468 | 9096 | 2 | ENSMUST00000173653.1 | 0 | Opn1sw | ENSMUSG00000058831.8 |
| chr3 | 130181451 | 130182298 | 3.41787 | 4.00077 | 1.17372 | Promoter | 3 | 130180891 | 130599877 | 418987 | 1 | ENSMUST00000106353.1 | 561 | Col25a1 | ENSMUSG00000058897.12 |
| chr18 | 57563979 | 57564086 | 3.49762 | 4.56948 | 1.47157 | Promoter | 18 | 57563176 | 57731006 | 167831 | 1 | ENSMUST00000123872.1 | 804 | 1700011I03Rik | ENSMUSG00000058925.6 |
| chr18 | 57564401 | 57564648 | 3.49306 | 4.37548 | 1.40051 | Intron (ENSMUST00000079738.4/ENSMUSG00000058925.6, intron 3 of 6) | 18 | 57563176 | 57731006 | 167831 | 1 | ENSMUST00000123872.1 | 1226 | 1700011I03Rik | ENSMUSG00000058925.6 |
| chr18 | 57567173 | 57567442 | 3.45006 | 4.14923 | 1.23515 | Intron (ENSMUST00000079738.4/ENSMUSG00000058925.6, intron 3 of 6) | 18 | 57563176 | 57731006 | 167831 | 1 | ENSMUST00000123872.1 | 3998 | 1700011I03Rik | ENSMUSG00000058925.6 |
| chr18 | 57575617 | 57576453 | 3.26284 | 3.7313 | 1.0482 | Intron (ENSMUST00000079738.4/ENSMUSG00000058925.6, intron 3 of 6) | 18 | 57563176 | 57731006 | 167831 | 1 | ENSMUST00000123872.1 | 12442 | 1700011I03Rik | ENSMUSG00000058925.6 |
| chr18 | 57608896 | 57608971 | 3.74428 | 4.56384 | 1.42305 | Intron (ENSMUST00000079738.4/ENSMUSG00000058925.6, intron 5 of 6) | 18 | 57563176 | 57731006 | 167831 | 1 | ENSMUST00000123872.1 | 45721 | 1700011I03Rik | ENSMUSG00000058925.6 |
| chr18 | 57609399 | 57609615 | 3.355 | 3.85997 | 1.01733 | Intron (ENSMUST00000079738.4/ENSMUSG00000058925.6, intron 5 of 6) | 18 | 57563176 | 57731006 | 167831 | 1 | ENSMUST00000123872.1 | 46224 | 1700011I03Rik | ENSMUSG00000058925.6 |
| chr18 | 57610006 | 57610149 | 3.35429 | 3.8589 | 1.01733 | Intron (ENSMUST00000079738.4/ENSMUSG00000058925.6, intron 5 of 6) | 18 | 57563176 | 57731006 | 167831 | 1 | ENSMUST00000123872.1 | 46831 | 1700011I03Rik | ENSMUSG00000058925.6 |
| chr19 | 24871703 | 24871830 | 3.4295 | 4.04647 | 1.21897 | Distal Intergenic | 19 | 24875686 | 24876634 | 949 | 1 | ENSMUST00000073080.5 | -3856 | Gm10053 | ENSMUSG00000058927.5 |
| chr19 | 24875783 | 24875976 | 3.88892 | 4.9987 | 1.76707 | Promoter | 19 | 24875686 | 24876634 | 949 | 1 | ENSMUST00000073080.5 | 98 | Gm10053 | ENSMUSG00000058927.5 |
| chr19 | 24883245 | 24883437 | 3.31805 | 3.80431 | 1.01733 | Distal Intergenic | 19 | 24875686 | 24876634 | 949 | 1 | ENSMUST00000073080.5 | 7560 | Gm10053 | ENSMUSG00000058927.5 |
| chr10 | 87863005 | 87863134 | 3.83423 | 4.95617 | 1.75617 | Promoter | 10 | 87826389 | 87863381 | 36993 | 2 | ENSMUST00000146581.1 | 247 | Igf1os | ENSMUSG00000058934.8 |
| chr7 | 126823322 | 126823484 | 3.66977 | 4.43093 | 1.35063 | Promoter | 7 | 126823303 | 126830217 | 6915 | 1 | ENSMUST00000079423.5 | 20 | Fam57b | ENSMUSG00000058966.8 |
| chr7 | 46394256 | 46395136 | 3.80818 | 4.78073 | 1.63876 | Distal Intergenic | 7 | 46396497 | 46433353 | 36857 | 1 | ENSMUST00000025202.6 | -1361 | Kcnc1 | ENSMUSG00000058975.6 |
| chr7 | 46395555 | 46398268 | 4.2375 | 5.75852 | 2.34887 | Promoter | 7 | 46396497 | 46433353 | 36857 | 1 | ENSMUST00000025202.6 | 0 | Kcnc1 | ENSMUSG00000058975.6 |
| chr7 | 46398840 | 46399417 | 3.52355 | 4.43269 | 1.39686 | Intron (ENSMUST00000025202.6/ENSMUSG00000058975.6, intron 1 of 1) | 7 | 46397648 | 46438375 | 40728 | 1 | ENSMUST00000160433.1 | 1193 | Kcnc1 | ENSMUSG00000058975.6 |
| chr6 | 120578792 | 120579418 | 3.73915 | 4.5665 | 1.44469 | Distal Intergenic | 6 | 120509494 | 120531319 | 21826 | 2 | ENSMUST00000075303.6 | -47474 | Cecr5 | ENSMUSG00000058979.6 |
| chr16 | 9992114 | 9992426 | 3.3958 | 4.03334 | 1.18909 | Promoter | 16 | 9577827 | 9992533 | 414707 | 2 | ENSMUST00000032331.7 | 107 | Grin2a | ENSMUSG00000059003.7 |
| chr16 | 9992839 | 9993338 | 3.61513 | 4.33599 | 1.30477 | Promoter | 16 | 9577827 | 9992533 | 414707 | 2 | ENSMUST00000032331.7 | -307 | Grin2a | ENSMUSG00000059003.7 |
| chr16 | 9994085 | 9994461 | 4.55626 | 6.2921 | 2.76001 | Promoter | 16 | 9991655 | 9994260 | 2606 | 2 | ENSMUST00000156482.1 | 0 | Grin2a | ENSMUSG00000059003.7 |
| chr16 | 9994797 | 9995624 | 4.0049 | 5.09969 | 1.83206 | Promoter | 16 | 9577710 | 9994921 | 417212 | 2 | ENSMUST00000115835.2 | 0 | Grin2a | ENSMUSG00000059003.7 |
| chr2 | 32742456 | 32742551 | 3.37566 | 4.04375 | 1.18514 | Promoter | 2 | 32741492 | 32743676 | 2185 | 1 | ENSMUST00000139563.1 | 965 | Sh2d3c | ENSMUSG00000059013.7 |
| chr6 | 29485470 | 29485816 | 3.49927 | 4.21141 | 1.28285 | Promoter | 6 | 29483693 | 29484972 | 1280 | 2 | ENSMUST00000161655.1 | -499 | Kcp | ENSMUSG00000059022.9 |
| chr6 | 29496288 | 29496436 | 3.58368 | 4.53096 | 1.46279 | Promoter | 6 | 29482036 | 29495914 | 13879 | 2 | ENSMUST00000159479.2 | -375 | Kcp | ENSMUSG00000059022.9 |
| chr6 | 29509232 | 29509501 | 3.45132 | 4.25577 | 1.27916 | Exon (ENSMUST00000181540.1/ENSMUSG00000097137.1, exon 2 of 2) | 6 | 29502290 | 29507952 | 5663 | 2 | ENSMUST00000161237.1 | -1281 | Kcp | ENSMUSG00000059022.9 |
| chr4 | 109653473 | 109653666 | 4.01028 | 5.11115 | 1.84805 | Promoter | 4 | 109652613 | 109654131 | 1519 | 2 | ENSMUST00000127391.1 | 465 | 9630013D21Rik | ENSMUSG00000059027.9 |
| chr4 | 109653981 | 109654072 | 4.0533 | 5.16199 | 1.83221 | Promoter | 4 | 109652613 | 109654131 | 1519 | 2 | ENSMUST00000127391.1 | 59 | 9630013D21Rik | ENSMUSG00000059027.9 |
| chr4 | 109655618 | 109655945 | 3.93019 | 5.00136 | 1.79338 | Intron (ENSMUST00000125237.1/ENSMUSG00000087641.1, intron 2 of 2) | 4 | 109652613 | 109654131 | 1519 | 2 | ENSMUST00000127391.1 | -1488 | 9630013D21Rik | ENSMUSG00000059027.9 |
| chr4 | 109656343 | 109658291 | 3.80432 | 4.76397 | 1.62973 | Promoter | 4 | 109651529 | 109657827 | 6299 | 2 | ENSMUST00000156790.2 | 0 | 9630013D21Rik | ENSMUSG00000059027.9 |
| chr4 | 109659328 | 109659953 | 3.84193 | 4.90584 | 1.75684 | Downstream (<1kb) | 4 | 109652484 | 109657853 | 5370 | 2 | ENSMUST00000131216.2 | -1476 | 9630013D21Rik | ENSMUSG00000059027.9 |
| chr12 | 79530364 | 79530571 | 4.00457 | 5.28206 | 1.93697 | Distal Intergenic | 12 | 79297351 | 79508656 | 211306 | 1 | ENSMUST00000079533.6 | 233014 | Rad51b | ENSMUSG00000059060.8 |
| chr7 | 28506250 | 28506540 | 3.34118 | 4.00008 | 1.15438 | Intron (ENSMUST00000040531.8/ENSMUSG00000030602.7, intron 1 of 12) | 7 | 28508836 | 28510455 | 1620 | 2 | ENSMUST00000081684.2 | 3915 | Ifnl2 | ENSMUSG00000059128.3 |
| chr1 | 132067250 | 132067535 | 3.92104 | 5.21742 | 1.93695 | Promoter | 1 | 132026808 | 132067826 | 41019 | 2 | ENSMUST00000160656.2 | 291 | Mfsd4 | ENSMUSG00000059149.12 |
| chr1 | 132067882 | 132068350 | 3.59744 | 4.46396 | 1.4142 | Promoter | 1 | 132022806 | 132067911 | 45106 | 2 | ENSMUST00000126927.2 | 0 | Mfsd4 | ENSMUSG00000059149.12 |
| chr6 | 51844890 | 51845196 | 3.23967 | 3.8307 | 1.02941 | Distal Intergenic | 6 | 51859166 | 52012549 | 153384 | 2 | ENSMUST00000078214.5 | 167353 | Skap2 | ENSMUSG00000059182.5 |
| chr6 | 51848350 | 51848786 | 3.44733 | 4.41656 | 1.38822 | Distal Intergenic | 6 | 51859166 | 52012549 | 153384 | 2 | ENSMUST00000078214.5 | 163763 | Skap2 | ENSMUSG00000059182.5 |
| chr9 | 66805936 | 66806292 | 3.87559 | 4.96071 | 1.77058 | Promoter | 9 | 66805306 | 66805938 | 633 | 2 | ENSMUST00000112182.2 | 0 | Ppp1r2-ps4 | ENSMUSG00000059237.6 |
| chr5 | 25655083 | 25655379 | 3.76394 | 4.6415 | 1.56657 | Distal Intergenic | 5 | 25659809 | 25661275 | 1467 | 2 | ENSMUST00000181398.1 | 5896 | Gm10062 | ENSMUSG00000059244.5 |
| chr9 | 69759098 | 69761350 | 3.77859 | 4.73374 | 1.58564 | Promoter | 9 | 69757710 | 69760940 | 3231 | 2 | ENSMUST00000071281.4 | 0 | Foxb1 | ENSMUSG00000059246.4 |
| chr11 | 117199161 | 117199882 | 3.61618 | 4.46322 | 1.38994 | Promoter | 11 | 117199661 | 117362325 | 162665 | 1 | ENSMUST00000093907.5 | 0 | Sep-09 | ENSMUSG00000059248.8 |
| chr11 | 117232353 | 117232591 | 3.533 | 4.22505 | 1.3267 | Promoter | 11 | 117232285 | 117362313 | 130029 | 1 | ENSMUST00000106354.3 | 69 | Sep-09 | ENSMUSG00000059248.8 |
| chr11 | 117266027 | 117266462 | 3.49761 | 4.14864 | 1.27328 | Promoter | 11 | 117266246 | 117362313 | 96068 | 1 | ENSMUST00000019038.9 | 0 | Sep-09 | ENSMUSG00000059248.8 |
| chr11 | 117266787 | 117267459 | 3.64454 | 4.45549 | 1.44535 | Promoter | 11 | 117266246 | 117362313 | 96068 | 1 | ENSMUST00000019038.9 | 542 | Sep-09 | ENSMUSG00000059248.8 |
| chr11 | 117332555 | 117332955 | 3.84078 | 4.89099 | 1.70783 | Promoter | 11 | 117332428 | 117361629 | 29202 | 1 | ENSMUST00000155331.2 | 128 | Sep-09 | ENSMUSG00000059248.8 |
| chr11 | 117347816 | 117348032 | 3.41797 | 3.97879 | 1.10163 | Intron (ENSMUST00000093907.5/ENSMUSG00000059248.8, intron 3 of 11) | 11 | 117353180 | 117356999 | 3820 | 1 | ENSMUST00000134852.1 | -5148 | Sep-09 | ENSMUSG00000059248.8 |
| chr5 | 77095103 | 77095680 | 4.34073 | 5.82565 | 2.39779 | Promoter | 5 | 77086988 | 77095253 | 8266 | 2 | ENSMUST00000120827.3 | 0 | Hopx | ENSMUSG00000059325.9 |
| chr5 | 77096408 | 77096972 | 3.51339 | 4.30219 | 1.31186 | Intron (ENSMUST00000148568.1/ENSMUSG00000087541.1, intron 1 of 1) | 5 | 77086988 | 77095277 | 8290 | 2 | ENSMUST00000113453.3 | -1132 | Hopx | ENSMUSG00000059325.9 |
| chr5 | 77097428 | 77097894 | 3.34498 | 3.84485 | 1.01733 | Intron (ENSMUST00000148568.1/ENSMUSG00000087541.1, intron 1 of 1) | 5 | 77086988 | 77095277 | 8290 | 2 | ENSMUST00000113453.3 | -2152 | Hopx | ENSMUSG00000059325.9 |
| chr18 | 78122915 | 78123324 | 3.70425 | 4.52171 | 1.42143 | Promoter | 18 | 78101649 | 78123331 | 21683 | 2 | ENSMUST00000160292.2 | 7 | Slc14a1 | ENSMUSG00000059336.9 |
| chr18 | 78124166 | 78125391 | 3.61703 | 4.34197 | 1.30868 | Promoter | 18 | 78100091 | 78123426 | 23336 | 2 | ENSMUST00000091813.6 | -741 | Slc14a1 | ENSMUSG00000059336.9 |
| chr9 | 78821687 | 78822257 | 3.65167 | 4.43579 | 1.41091 | Promoter | 9 | 78820638 | 78821838 | 1201 | 2 | ENSMUST00000182917.1 | 0 | Gm8116 | ENSMUSG00000059422.5 |
| chr14 | 66190178 | 66190275 | 4.13641 | 5.42204 | 2.11228 | Exon (ENSMUST00000022622.8/ENSMUSG00000059456.8, exon 3 of 31) | 14 | 66153262 | 66170282 | 17021 | 2 | ENSMUST00000154865.2 | -19897 | Ptk2b | ENSMUSG00000059456.8 |
| chr14 | 66190760 | 66190934 | 3.27393 | 3.73815 | 1.01733 | Intron (ENSMUST00000022622.8/ENSMUSG00000059456.8, intron 2 of 30) | 14 | 66153262 | 66170282 | 17021 | 2 | ENSMUST00000154865.2 | -20479 | Ptk2b | ENSMUSG00000059456.8 |
| chr14 | 66213547 | 66213623 | 4.14015 | 5.31758 | 1.92384 | Promoter | 14 | 66153258 | 66213552 | 60295 | 2 | ENSMUST00000178730.2 | 0 | Ptk2b | ENSMUSG00000059456.8 |
| chr14 | 66280654 | 66281496 | 4.00116 | 5.23404 | 1.98631 | Promoter | 14 | 66153257 | 66281052 | 127796 | 2 | ENSMUST00000022622.8 | 0 | Ptk2b | ENSMUSG00000059456.8 |
| chr14 | 66281860 | 66282209 | 3.56938 | 4.31169 | 1.37068 | Promoter | 14 | 66153262 | 66281052 | 127791 | 2 | ENSMUST00000089250.3 | -809 | Ptk2b | ENSMUSG00000059456.8 |
| chr1 | 171064460 | 171065402 | 4.03234 | 12.36875 | 7.8527 | Promoter | 1 | 171057828 | 171064935 | 7108 | 2 | ENSMUST00000164179.1 | 0 | Fcgr3 | ENSMUSG00000059498.8 |
| chr1 | 171065921 | 171066376 | 4.65618 | 13.14815 | 8.67613 | Promoter | 1 | 171057828 | 171064935 | 7108 | 2 | ENSMUST00000164179.1 | -987 | Fcgr3 | ENSMUSG00000059498.8 |
| chr1 | 171066767 | 171067287 | 3.19368 | 6.01666 | 2.58026 | Distal Intergenic | 1 | 171057828 | 171064935 | 7108 | 2 | ENSMUST00000164179.1 | -1833 | Fcgr3 | ENSMUSG00000059498.8 |
| chr1 | 171067786 | 171068731 | 5.34788 | 14.65496 | 10.03011 | Distal Intergenic | 1 | 171057828 | 171064935 | 7108 | 2 | ENSMUST00000164179.1 | -2852 | Fcgr3 | ENSMUSG00000059498.8 |
| chr12 | 96134578 | 96134835 | 3.4303 | 4.00358 | 1.12412 | Distal Intergenic | 12 | 96046767 | 96047120 | 354 | 2 | ENSMUST00000073560.4 | -87459 | 1700019M22Rik | ENSMUSG00000059695.4 |
| chr12 | 96142535 | 96142871 | 3.30125 | 3.77911 | 1.01733 | Distal Intergenic | 12 | 96046767 | 96047120 | 354 | 2 | ENSMUST00000073560.4 | -95416 | 1700019M22Rik | ENSMUSG00000059695.4 |
| chr2 | 63183857 | 63184250 | 3.68841 | 4.55495 | 1.49633 | Promoter | 2 | 62786367 | 63184170 | 397804 | 2 | ENSMUST00000112452.1 | 0 | Kcnh7 | ENSMUSG00000059742.5 |
| chr2 | 63184852 | 63185522 | 3.49039 | 4.1168 | 1.20378 | Promoter | 2 | 62786281 | 63184287 | 398007 | 2 | ENSMUST00000112454.2 | -566 | Kcnh7 | ENSMUSG00000059742.5 |
| chr4 | 62674239 | 62674357 | 3.26873 | 3.73023 | 1.01733 | Promoter | 4 | 62674352 | 62690001 | 15650 | 1 | ENSMUST00000138674.1 | 0 | Rgs3 | ENSMUSG00000059810.13 |
| chr17 | 79939686 | 79939784 | 3.78915 | 4.67173 | 1.54226 | Distal Intergenic | 17 | 79848392 | 79896123 | 47732 | 2 | ENSMUST00000112437.2 | -43564 | Atl2 | ENSMUSG00000059811.7 |
| chr17 | 79958751 | 79958909 | 3.71309 | 4.56586 | 1.51195 | Distal Intergenic | 17 | 79848392 | 79896123 | 47732 | 2 | ENSMUST00000112437.2 | -62629 | Atl2 | ENSMUSG00000059811.7 |
| chr4 | 108382624 | 108384256 | 3.67183 | 4.52673 | 1.45399 | Promoter | 4 | 108367777 | 108383349 | 15573 | 2 | ENSMUST00000053157.6 | 0 | Fam159a | ENSMUSG00000059816.5 |
| chr3 | 109781996 | 109782575 | 3.35864 | 3.90763 | 1.14568 | 3' UTR | 3 | 109830958 | 109833259 | 2302 | 2 | ENSMUST00000191116.1 | 50684 | Ntng1 | ENSMUSG00000059857.10 |
| chr3 | 110140914 | 110141555 | 3.41744 | 3.96736 | 1.06304 | Intron (ENSMUST00000156177.3/ENSMUSG00000059857.10, intron 1 of 6) | 3 | 109780040 | 110143004 | 362965 | 2 | ENSMUST00000156177.3 | 1449 | Ntng1 | ENSMUSG00000059857.10 |
| chr3 | 110142288 | 110143665 | 3.62358 | 4.55921 | 1.45652 | Promoter | 3 | 109780040 | 110143004 | 362965 | 2 | ENSMUST00000156177.3 | 0 | Ntng1 | ENSMUSG00000059857.10 |
| chr3 | 110143972 | 110144316 | 3.57834 | 4.26399 | 1.25463 | Promoter | 3 | 109780046 | 110144011 | 363966 | 2 | ENSMUST00000133268.3 | 0 | Ntng1 | ENSMUSG00000059857.10 |
| chr3 | 110144787 | 110145254 | 4.22208 | 5.51597 | 2.11198 | Promoter | 3 | 110143685 | 110144011 | 327 | 2 | ENSMUST00000051253.3 | -777 | Ntng1 | ENSMUSG00000059857.10 |
| chr15 | 73720495 | 73721162 | 3.37846 | 3.93832 | 1.12887 | Distal Intergenic | 15 | 73723145 | 73758766 | 35622 | 1 | ENSMUST00000165541.2 | -1983 | Ptp4a3 | ENSMUSG00000059895.7 |
| chr15 | 73745653 | 73745879 | 3.31488 | 3.81471 | 1.06925 | Intron (ENSMUST00000165541.2/ENSMUSG00000059895.7, intron 1 of 5) | 15 | 73748026 | 73758766 | 10741 | 1 | ENSMUST00000053232.6 | -2147 | Ptp4a3 | ENSMUSG00000059895.7 |
| chr18 | 20001761 | 20002248 | 3.37099 | 4.04535 | 1.13796 | Promoter | 18 | 19960930 | 20002097 | 41168 | 2 | ENSMUST00000115848.3 | 0 | Dsc3 | ENSMUSG00000059898.8 |
| chr10 | 61273031 | 61273106 | 4.05356 | 5.5642 | 2.13409 | Promoter | 10 | 61198468 | 61273242 | 74775 | 2 | ENSMUST00000120336.1 | 136 | Adamts14 | ENSMUSG00000059901.7 |
| chr12 | 76405164 | 76405814 | 3.74601 | 4.74838 | 1.61731 | Promoter | 12 | 76404176 | 76406934 | 2759 | 1 | ENSMUST00000080449.5 | 989 | Hspa2 | ENSMUSG00000059970.6 |
| chr9 | 29411913 | 29412895 | 4.2403 | 5.59465 | 2.21534 | Promoter | 9 | 28996012 | 29412204 | 416193 | 2 | ENSMUST00000115237.2 | 0 | Ntm | ENSMUSG00000059974.5 |
| chr9 | 29803216 | 29803370 | 3.44649 | 4.03924 | 1.16295 | Intron (ENSMUST00000075069.5/ENSMUSG00000059974.5, intron 1 of 7) | 9 | 29411588 | 29963119 | 551532 | 2 | ENSMUST00000146651.1 | 159749 | Ntm | ENSMUSG00000059974.5 |
| chr9 | 29961409 | 29961691 | 3.54145 | 4.47706 | 1.42547 | Intron (ENSMUST00000075069.5/ENSMUSG00000059974.5, intron 1 of 7) | 9 | 29411588 | 29963119 | 551532 | 2 | ENSMUST00000146651.1 | 1428 | Ntm | ENSMUSG00000059974.5 |
| chr9 | 29962094 | 29962172 | 4.11349 | 5.63059 | 2.22382 | Promoter | 9 | 29411588 | 29963119 | 551532 | 2 | ENSMUST00000146651.1 | 947 | Ntm | ENSMUSG00000059974.5 |
| chr5 | 144543196 | 144543381 | 3.30707 | 3.78782 | 1.01733 | Distal Intergenic | 5 | 144545887 | 144557478 | 11592 | 1 | ENSMUST00000071782.6 | -2506 | Nptx2 | ENSMUSG00000059991.6 |
| chr5 | 144545928 | 144548162 | 3.80678 | 4.81735 | 1.67128 | Promoter | 5 | 144545887 | 144557478 | 11592 | 1 | ENSMUST00000071782.6 | 42 | Nptx2 | ENSMUSG00000059991.6 |
| chr5 | 149269390 | 149269709 | 3.75322 | 4.60166 | 1.48874 | Intron (ENSMUST00000071130.3/ENSMUSG00000060063.6, intron 1 of 4) | 5 | 149265058 | 149288045 | 22988 | 1 | ENSMUST00000071130.3 | 4333 | Alox5ap | ENSMUSG00000060063.6 |
| chr18 | 64437802 | 64438001 | 3.49263 | 4.11501 | 1.18514 | Distal Intergenic | 18 | 64448519 | 64450014 | 1496 | 2 | ENSMUST00000072796.4 | 12013 | Amd-ps3 | ENSMUSG00000060096.4 |
| chr5 | 115221848 | 115222196 | 3.42743 | 4.05191 | 1.22365 | Distal Intergenic | 5 | 115235836 | 115240991 | 5156 | 1 | ENSMUST00000081497.7 | -13640 | Pop5 | ENSMUSG00000060152.9 |
| chr10 | 117021399 | 117021546 | 3.59763 | 4.31126 | 1.31624 | Intron (ENSMUST00000020378.4/ENSMUSG00000020169.4, intron 9 of 9) | 10 | 117045341 | 117046768 | 1428 | 1 | ENSMUST00000073834.4 | -23795 | Lrrc10 | ENSMUSG00000060187.4 |
| chr10 | 117022137 | 117022613 | 3.80282 | 4.69523 | 1.55154 | Intron (ENSMUST00000020378.4/ENSMUSG00000020169.4, intron 9 of 9) | 10 | 117045341 | 117046768 | 1428 | 1 | ENSMUST00000073834.4 | -22728 | Lrrc10 | ENSMUSG00000060187.4 |
| chr10 | 117024047 | 117024385 | 3.42352 | 4.05383 | 1.20937 | Exon (ENSMUST00000020378.4/ENSMUSG00000020169.4, exon 10 of 10) | 10 | 117045341 | 117046768 | 1428 | 1 | ENSMUST00000073834.4 | -20956 | Lrrc10 | ENSMUSG00000060187.4 |
| chr5 | 72584219 | 72584575 | 3.69657 | 4.55031 | 1.46399 | Intron (ENSMUST00000031124.4/ENSMUSG00000060204.4, intron 2 of 2) | 5 | 72581639 | 72587550 | 5912 | 2 | ENSMUST00000031124.4 | 2975 | Gm5868 | ENSMUSG00000060204.4 |

| | | | | | | | | | | | | | | | |
|---|---|---|---|---|---|---|---|---|---|---|---|---|---|---|---|
| chr5 | 72586120 | 72587698 | 4.06445 | 5.33826 | 2.0563 | Promoter | 5 | 72581639 | 72587550 | 5912 | 2 | ENSMUST00000031124.4 | 0 | Gm5868 | ENSMUSG00000060204.4 |
| chr5 | 72589636 | 72589923 | 3.51252 | 4.17336 | 1.28636 | Distal Intergenic | 5 | 72581639 | 72587550 | 5912 | 2 | ENSMUST00000031124.4 | -2087 | Gm5868 | ENSMUSG00000060204.4 |
| chr8 | 125779078 | 125779153 | 2.75072 | 13.33764 | 8.73446 | Intron (ENSMUST00000047239.7/ENSMUSG00000060212.8, intron 25 of 33) | 8 | 125751511 | 125774128 | 22618 | 2 | ENSMUST00000119305.2 | -4951 | Pcnxl2 | ENSMUSG00000060212.8 |
| chr8 | 125898207 | 125898447 | 4.77654 | 6.97654 | 3.23169 | Promoter | 8 | 125751508 | 125898317 | 146810 | 2 | ENSMUST00000047239.7 | 0 | Pcnxl2 | ENSMUSG00000060212.8 |
| chr2 | 152079839 | 152079959 | 3.80865 | 4.71757 | 1.59382 | Distal Intergenic | 2 | 152081529 | 152095802 | 14274 | 1 | ENSMUST00000064061.3 | -1570 | Scrt2 | ENSMUSG00000060257.2 |
| chr2 | 152080555 | 152081337 | 3.68643 | 4.54101 | 1.43865 | Promoter | 2 | 152081529 | 152095802 | 14274 | 1 | ENSMUST00000064061.3 | -192 | Scrt2 | ENSMUSG00000060257.2 |
| chr2 | 152082535 | 152082693 | 3.77538 | 4.65277 | 1.5345 | Intron (ENSMUST00000064061.3/ENSMUSG00000060257.2, intron 1 of 1) | 2 | 152081529 | 152095802 | 14274 | 1 | ENSMUST00000064061.3 | 1007 | Scrt2 | ENSMUSG00000060257.2 |
| chr2 | 152083803 | 152083930 | 3.33498 | 3.83479 | 1.03455 | Intron (ENSMUST00000064061.3/ENSMUSG00000060257.2, intron 1 of 1) | 2 | 152081529 | 152095802 | 14274 | 1 | ENSMUST00000064061.3 | 2275 | Scrt2 | ENSMUSG00000060257.2 |
| chr2 | 152089946 | 152090160 | 3.51084 | 4.14234 | 1.18364 | Intron (ENSMUST00000064061.3/ENSMUSG00000060257.2, intron 1 of 1) | 2 | 152081529 | 152095802 | 14274 | 1 | ENSMUST00000064061.3 | 8418 | Scrt2 | ENSMUSG00000060257.2 |
| chr2 | 152091342 | 152091417 | 3.6353 | 4.53316 | 1.44506 | Intron (ENSMUST00000064061.3/ENSMUSG00000060257.2, intron 1 of 1) | 2 | 152081529 | 152095802 | 14274 | 1 | ENSMUST00000064061.3 | 9814 | Scrt2 | ENSMUSG00000060257.2 |
| chr18 | 36196667 | 36196997 | 3.37439 | 4.08299 | 1.19294 | Promoter | 18 | 36027396 | 36197232 | 169837 | 2 | ENSMUST00000115705.1 | 235 | Nrg2 | ENSMUSG00000060275.6 |
| chr18 | 36210624 | 36210799 | 3.77053 | 4.62935 | 1.50872 | Distal Intergenic | 18 | 36017707 | 36197380 | 179674 | 2 | ENSMUST00000115713.2 | -13245 | Nrg2 | ENSMUSG00000060275.6 |
| chr5 | 130447606 | 130447874 | 3.3497 | 3.87078 | 1.07695 | Promoter | 5 | 130448012 | 130840636 | 392625 | 1 | ENSMUST00000111287.2 | -138 | Caln1 | ENSMUSG00000060371.7 |
| chr5 | 130449028 | 130449615 | 3.57394 | 4.28538 | 1.27442 | Promoter | 5 | 130448801 | 130840645 | 391845 | 1 | ENSMUST00000111288.2 | 228 | Caln1 | ENSMUSG00000060371.7 |
| chr7 | 34800711 | 34800894 | 3.24478 | 3.69434 | 1.01733 | Intron (ENSMUST00000078686.6/ENSMUSG00000060402.6, intron 1 of 3) | 7 | 34674468 | 34812686 | 138219 | 2 | ENSMUST00000078686.6 | 11792 | Chst8 | ENSMUSG00000060402.6 |
| chr7 | 34809502 | 34809938 | 3.59327 | 4.303 | 1.30974 | Intron (ENSMUST00000078686.6/ENSMUSG00000060402.6, intron 1 of 3) | 7 | 34674468 | 34812686 | 138219 | 2 | ENSMUST00000078686.6 | 2748 | Chst8 | ENSMUSG00000060402.6 |
| chr7 | 34811428 | 34811729 | 3.75672 | 4.6195 | 1.517 | Promoter | 7 | 34674468 | 34812686 | 138219 | 2 | ENSMUST00000078686.6 | 957 | Chst8 | ENSMUSG00000060402.6 |
| chr5 | 129152522 | 129153130 | 3.51383 | 4.14678 | 1.18355 | Intron (ENSMUST00000056617.8/ENSMUSG00000044017.11, intron 14 of 25) | 5 | 129128077 | 129128517 | 441 | 2 | ENSMUST00000140155.1 | -24006 | Rps16-ps2 | ENSMUSG00000060419.8 |
| chr5 | 129215324 | 129215495 | 3.73892 | 4.64261 | 1.50755 | Distal Intergenic | 5 | 129128077 | 129128517 | 441 | 2 | ENSMUST00000140155.1 | -86808 | Rps16-ps2 | ENSMUSG00000060419.8 |
| chr5 | 129218410 | 129218697 | 4.1026 | 5.26694 | 1.92182 | Distal Intergenic | 5 | 129128077 | 129128517 | 441 | 2 | ENSMUST00000140155.1 | -89894 | Rps16-ps2 | ENSMUSG00000060419.8 |
| chr1 | 42539655 | 42540394 | 3.45762 | 4.06433 | 1.14277 | Distal Intergenic | 1 | 42648200 | 42651679 | 3480 | 2 | ENSMUST00000176977.2 | 111285 | Pou3f3os | ENSMUSG00000060424.9 |
| chr1 | 42541335 | 42541410 | 3.50074 | 4.12127 | 1.15684 | Distal Intergenic | 1 | 42648200 | 42651679 | 3480 | 2 | ENSMUST00000176977.2 | 110269 | Pou3f3os | ENSMUSG00000060424.9 |
| chr1 | 42678781 | 42679175 | 3.64059 | 4.49475 | 1.42755 | Intron (ENSMUST00000181660.2/ENSMUSG00000060424.9, intron 1 of 3) | 1 | 42648201 | 42667672 | 19472 | 2 | ENSMUST00000176052.2 | -11110 | Pou3f3os | ENSMUSG00000060424.9 |
| chr1 | 42680765 | 42681267 | 3.45348 | 4.03855 | 1.1201 | Intron (ENSMUST00000181660.2/ENSMUSG00000060424.9, intron 1 of 3) | 1 | 42648201 | 42693093 | 44893 | 2 | ENSMUST00000186656.1 | 11826 | Pou3f3os | ENSMUSG00000060424.9 |
| chr1 | 42681989 | 42683456 | 3.93423 | 5.03767 | 1.78415 | Intron (ENSMUST00000181660.2/ENSMUSG00000060424.9, intron 1 of 3) | 1 | 42648201 | 42693093 | 44893 | 2 | ENSMUST00000186656.1 | 9637 | Pou3f3os | ENSMUSG00000060424.9 |
| chr1 | 42684215 | 42685018 | 4.12631 | 5.34617 | 2.00273 | Intron (ENSMUST00000181660.2/ENSMUSG00000060424.9, intron 1 of 3) | 1 | 42648201 | 42693093 | 44893 | 2 | ENSMUST00000186656.1 | 8075 | Pou3f3os | ENSMUSG00000060424.9 |
| chr1 | 42686102 | 42686519 | 3.46236 | 4.04978 | 1.11552 | Intron (ENSMUST00000181660.2/ENSMUSG00000060424.9, intron 1 of 3) | 1 | 42648201 | 42693093 | 44893 | 2 | ENSMUST00000186656.1 | 6574 | Pou3f3os | ENSMUSG00000060424.9 |
| chr1 | 42686935 | 42687333 | 3.91296 | 4.90994 | 1.69799 | Intron (ENSMUST00000181660.2/ENSMUSG00000060424.9, intron 1 of 3) | 1 | 42648201 | 42693093 | 44893 | 2 | ENSMUST00000186656.1 | 5760 | Pou3f3os | ENSMUSG00000060424.9 |
| chr1 | 42688382 | 42688457 | 3.44462 | 4.20174 | 1.273 | Intron (ENSMUST00000181660.2/ENSMUSG00000060424.9, intron 1 of 3) | 1 | 42648201 | 42693093 | 44893 | 2 | ENSMUST00000186656.1 | 4636 | Pou3f3os | ENSMUSG00000060424.9 |
| chr1 | 42688946 | 42689326 | 4.00207 | 5.12348 | 1.89199 | Intron (ENSMUST00000181660.2/ENSMUSG00000060424.9, intron 1 of 3) | 1 | 42648201 | 42693093 | 44893 | 2 | ENSMUST00000186656.1 | 3767 | Pou3f3os | ENSMUSG00000060424.9 |
| chr1 | 42689696 | 42690087 | 3.97701 | 5.2781 | 1.96991 | Intron (ENSMUST00000181660.2/ENSMUSG00000060424.9, intron 1 of 3) | 1 | 42648201 | 42693093 | 44893 | 2 | ENSMUST00000186656.1 | 3006 | Pou3f3os | ENSMUSG00000060424.9 |
| chr1 | 42691638 | 42692093 | 3.91689 | 4.91135 | 1.68901 | Promoter | 1 | 42648201 | 42693093 | 44893 | 2 | ENSMUST00000186656.1 | 1000 | Pou3f3os | ENSMUSG00000060424.9 |
| chr1 | 42692462 | 42692990 | 3.71689 | 4.5601 | 1.45675 | Promoter | 1 | 42648201 | 42693093 | 44893 | 2 | ENSMUST00000186656.1 | 103 | Pou3f3os | ENSMUSG00000060424.9 |
| chr1 | 42693647 | 42695404 | 3.92399 | 4.93578 | 1.7085 | Promoter | 1 | 42590782 | 42694808 | 104027 | 2 | ENSMUST00000181660.2 | 0 | Pou3f3os | ENSMUSG00000060424.9 |
| chr15 | 55905701 | 55906006 | 3.58432 | 4.42654 | 1.40517 | Promoter | 15 | 55791938 | 55906310 | 114373 | 2 | ENSMUST00000140574.1 | 304 | Sntb1 | ENSMUSG00000060429.6 |
| chr15 | 55906359 | 55906915 | 3.88037 | 4.90946 | 1.68881 | Promoter | 15 | 55790964 | 55906722 | 115759 | 2 | ENSMUST00000110200.2 | 0 | Sntb1 | ENSMUSG00000060429.6 |
| chr15 | 55907720 | 55907991 | 3.60948 | 4.32578 | 1.30654 | Promoter | 15 | 55638843 | 55906949 | 268107 | 2 | ENSMUST00000039769.6 | -772 | Sntb1 | ENSMUSG00000060429.6 |
| chr8 | 95011029 | 95011177 | 3.36547 | 3.88035 | 1.03351 | Intron (ENSMUST00000179619.2/ENSMUSG00000031785.10, intron 12 of 13) | 8 | 95017692 | 95045247 | 27556 | 1 | ENSMUST00000051259.8 | -6515 | Gpr97 | ENSMUSG00000060470.6 |
| chr18 | 72348648 | 72348843 | 3.65862 | 4.41213 | 1.34165 | Intron (ENSMUST00000114943.4/ENSMUSG00000060534.10, intron 1 of 28) | 18 | 71258990 | 72351017 | 1092028 | 2 | ENSMUST00000073379.5 | 2174 | Dcc | ENSMUSG00000060534.10 |
| chr18 | 72349303 | 72350030 | 3.40314 | 3.94704 | 1.06845 | Promoter | 18 | 71258990 | 72351017 | 1092028 | 2 | ENSMUST00000073379.5 | 987 | Dcc | ENSMUSG00000060534.10 |
| chr18 | 72350380 | 72350626 | 3.60886 | 4.31433 | 1.26812 | Promoter | 18 | 71258990 | 72351017 | 1092028 | 2 | ENSMUST00000073379.5 | 391 | Dcc | ENSMUSG00000060534.10 |
| chr18 | 72351226 | 72351336 | 4.37921 | 5.83003 | 2.33942 | Promoter | 18 | 71258738 | 72351069 | 1092332 | 2 | ENSMUST00000114943.4 | -158 | Dcc | ENSMUSG00000060534.10 |
| chr14 | 61055980 | 61056151 | 3.61801 | 4.36594 | 1.39056 | Distal Intergenic | 14 | 60963834 | 61046855 | 83022 | 2 | ENSMUST00000111234.3 | -9126 | Tnfrsf19 | ENSMUSG00000060548.7 |
| chr8 | 47408200 | 47408764 | 3.54113 | 4.28161 | 1.3192 | Intron (ENSMUST00000181417.1/ENSMUSG00000097922.1, intron 7 of 7) | 8 | 47413129 | 47415359 | 2231 | 1 | ENSMUST00000074466.5 | -4365 | Gm10083 | ENSMUSG00000060559.5 |
| chr8 | 47409570 | 47411081 | 3.98818 | 5.0994 | 1.87139 | Intron (ENSMUST00000181417.1/ENSMUSG00000097922.1, intron 7 of 7) | 8 | 47413129 | 47415359 | 2231 | 1 | ENSMUST00000074466.5 | -2048 | Gm10083 | ENSMUSG00000060559.5 |
| chr8 | 47411421 | 47411496 | 3.93059 | 5.22388 | 1.89637 | Intron (ENSMUST00000181417.1/ENSMUSG00000097922.1, intron 7 of 7) | 8 | 47413129 | 47415359 | 2231 | 1 | ENSMUST00000074466.5 | -1633 | Gm10083 | ENSMUSG00000060559.5 |
| chr8 | 47411877 | 47412012 | 3.75107 | 4.74198 | 1.61605 | Intron (ENSMUST00000181417.1/ENSMUSG00000097922.1, intron 7 of 7) | 8 | 47413129 | 47415359 | 2231 | 1 | ENSMUST00000074466.5 | -1117 | Gm10083 | ENSMUSG00000060559.5 |
| chr8 | 47412364 | 47412880 | 3.99585 | 5.70766 | 2.32203 | Promoter | 8 | 47413129 | 47415359 | 2231 | 1 | ENSMUST00000074466.5 | -249 | Gm10083 | ENSMUSG00000060559.5 |
| chr8 | 47413324 | 47413713 | 3.65176 | 4.59194 | 1.51979 | Promoter | 8 | 47413129 | 47415359 | 2231 | 1 | ENSMUST00000074466.5 | 196 | Gm10083 | ENSMUSG00000060559.5 |
| chr4 | 141010603 | 141011135 | 3.38921 | 4.13762 | 1.23573 | Promoter | 4 | 141010637 | 141015012 | 4376 | 1 | ENSMUST00000124506.1 | 0 | Mfap2 | ENSMUSG00000060572.12 |
| chr7 | 19520285 | 19520421 | 3.8661 | 4.9281 | 1.74386 | Intron (ENSMUST00000078908.4/ENSMUSG00000060621.4, intron 2 of 3) | 7 | 19518731 | 19525050 | 6320 | 1 | ENSMUST00000078908.4 | 1555 | Nkpd1 | ENSMUSG00000060621.4 |
| chr7 | 19521766 | 19522599 | 3.64025 | 4.4136 | 1.42618 | Exon (ENSMUST00000078908.4/ENSMUSG00000060621.4, exon 3 of 4) | 7 | 19518731 | 19525050 | 6320 | 1 | ENSMUST00000078908.4 | 3036 | Nkpd1 | ENSMUSG00000060621.4 |
| chr7 | 19522907 | 19523709 | 3.95429 | 5.34412 | 2.02908 | Exon (ENSMUST00000078908.4/ENSMUSG00000060621.4, exon 4 of 4) | 7 | 19518731 | 19525050 | 6320 | 1 | ENSMUST00000078908.4 | 4177 | Nkpd1 | ENSMUSG00000060621.4 |
| chr7 | 19525008 | 19525604 | 3.38762 | 3.96159 | 1.16251 | 3' UTR | 7 | 19518731 | 19525050 | 6320 | 1 | ENSMUST00000078908.4 | 6278 | Nkpd1 | ENSMUSG00000060621.4 |
| chr10 | 71465427 | 71465526 | 3.692 | 4.57069 | 1.55204 | Distal Intergenic | 10 | 71409363 | 71433327 | 23965 | 1 | ENSMUST00000145521.1 | 56065 | Ipmk | ENSMUSG00000060733.8 |
| chr15 | 11034947 | 11035459 | 3.54272 | 4.2623 | 1.36847 | 3' UTR | 15 | 11033717 | 11037991 | 4275 | 2 | ENSMUST00000058007.6 | 2532 | Rxfp3 | ENSMUSG00000060735.4 |
| chr15 | 11036001 | 11036873 | 3.67377 | 4.55681 | 1.50775 | Exon (ENSMUST00000058007.6/ENSMUSG00000060735.4, exon 1 of 1) | 15 | 11033717 | 11037991 | 4275 | 2 | ENSMUST00000058007.6 | 1118 | Rxfp3 | ENSMUSG00000060735.4 |
| chr15 | 11037264 | 11038310 | 3.9343 | 5.0302 | 1.85765 | Promoter | 15 | 11033717 | 11037991 | 4275 | 2 | ENSMUST00000058007.6 | 0 | Rxfp3 | ENSMUSG00000060735.4 |
| chr7 | 28528320 | 28528826 | 3.44399 | 4.09279 | 1.23855 | Intron (ENSMUST00000040531.8/ENSMUSG00000030602.7, intron 1 of 12) | 7 | 28522836 | 28524322 | 1487 | 1 | ENSMUST00000078364.3 | 5485 | Ifnl3 | ENSMUSG00000060747.3 |
| chr6 | 77242733 | 77243060 | 3.64461 | 4.4254 | 1.45057 | Promoter | 6 | 77242922 | 77244112 | 1191 | 1 | ENSMUST00000161811.1 | 0 | Lrrtm1 | ENSMUSG00000060780.2 |
| chr6 | 77243574 | 77245230 | 3.67537 | 4.5159 | 1.45018 | Promoter | 6 | 77242922 | 77244112 | 1191 | 1 | ENSMUST00000161811.1 | 653 | Lrrtm1 | ENSMUSG00000060780.2 |
| chr3 | 40531395 | 40532160 | 3.71774 | 4.77741 | 1.62506 | Promoter | 3 | 40531286 | 40654617 | 123332 | 1 | ENSMUST00000061590.4 | 110 | Intu | ENSMUSG00000060798.4 |
| chr3 | 40533632 | 40534280 | 4.02193 | 5.21442 | 1.97384 | Intron (ENSMUST00000061590.4/ENSMUSG00000060798.4, intron 1 of 1) | 3 | 40531286 | 40654617 | 123332 | 1 | ENSMUST00000061590.4 | 2347 | Intu | ENSMUSG00000060798.4 |
| chr4 | 137050035 | 137050156 | 3.85468 | 4.90087 | 1.74584 | Distal Intergenic | 4 | 137014478 | 137048801 | 34324 | 2 | ENSMUST00000145505.1 | -1235 | Zbtb40 | ENSMUSG00000060862.9 |
| chr6 | 21211228 | 21211774 | 3.3169 | 3.80258 | 1.01733 | Distal Intergenic | 6 | 21215503 | 21729805 | 514303 | 1 | ENSMUST00000081542.5 | -3729 | Kcnd2 | ENSMUSG00000060882.5 |
| chr6 | 21212443 | 21212662 | 3.54417 | 4.22354 | 1.28776 | Distal Intergenic | 6 | 21215503 | 21729805 | 514303 | 1 | ENSMUST00000081542.5 | -2841 | Kcnd2 | ENSMUSG00000060882.5 |
| chr6 | 21213026 | 21213389 | 3.44724 | 4.23462 | 1.25097 | Distal Intergenic | 6 | 21215503 | 21729805 | 514303 | 1 | ENSMUST00000081542.5 | -2114 | Kcnd2 | ENSMUSG00000060882.5 |
| chr6 | 21214473 | 21216538 | 3.65897 | 4.54264 | 1.52482 | Promoter | 6 | 21215503 | 21729805 | 514303 | 1 | ENSMUST00000081542.5 | 0 | Kcnd2 | ENSMUSG00000060882.5 |
| chr8 | 17532333 | 17532933 | 3.72045 | 4.68457 | 1.54462 | Intron (ENSMUST00000082104.6/ENSMUSG00000060924.9, intron 1 of 69) | 8 | 15892537 | 17535281 | 1642745 | 2 | ENSMUST00000082104.6 | 2348 | Csmd1 | ENSMUSG00000060924.9 |
| chr8 | 17533295 | 17533517 | 3.44954 | 4.02921 | 1.11155 | Intron (ENSMUST00000082104.6/ENSMUSG00000060924.9, intron 1 of 69) | 8 | 15892537 | 17535281 | 1642745 | 2 | ENSMUST00000082104.6 | 1764 | Csmd1 | ENSMUSG00000060924.9 |
| chr8 | 17534004 | 17534415 | 3.91498 | 4.93144 | 1.7089 | Promoter | 8 | 15892537 | 17535281 | 1642745 | 2 | ENSMUST00000082104.6 | 866 | Csmd1 | ENSMUSG00000060924.9 |
| chr8 | 17535032 | 17535621 | 3.69194 | 4.48304 | 1.40014 | Promoter | 8 | 15892537 | 17535281 | 1642745 | 2 | ENSMUST00000082104.6 | 0 | Csmd1 | ENSMUSG00000060924.9 |
| chr8 | 17536775 | 17537072 | 3.47417 | 4.14756 | 1.23746 | Distal Intergenic | 8 | 16344863 | 17535586 | 1190724 | 2 | ENSMUST00000137947.1 | -1190 | Csmd1 | ENSMUSG00000060924.9 |
| chr5 | 88886611 | 88887062 | 4.08543 | 5.67961 | 2.24955 | Promoter | 5 | 88886818 | 89239653 | 352836 | 1 | ENSMUST00000148750.2 | 0 | Slc4a4 | ENSMUSG00000060961.9 |
| chr5 | 88887431 | 88888488 | 3.77881 | 4.85123 | 1.68205 | Promoter | 5 | 88887299 | 89235794 | 348496 | 1 | ENSMUST00000156238.2 | 133 | Slc4a4 | ENSMUSG00000060961.9 |
| chr5 | 88889042 | 88889562 | 4.06096 | 5.40091 | 2.04803 | Intron (ENSMUST00000148750.2/ENSMUSG00000060961.9, intron 1 of 25) | 5 | 88887299 | 89235794 | 348496 | 1 | ENSMUST00000156238.2 | 1744 | Slc4a4 | ENSMUSG00000060961.9 |
| chr5 | 89147351 | 89147774 | 3.23755 | 3.68349 | 1.01733 | Intron (ENSMUST00000148750.2/ENSMUSG00000060961.9, intron 12 of 25) | 5 | 89028092 | 89122453 | 94362 | 1 | ENSMUST00000134303.1 | 119260 | Slc4a4 | ENSMUSG00000060961.9 |
| chr13 | 71949177 | 71949380 | 3.64195 | 4.3867 | 1.33437 | Distal Intergenic | 13 | 71957921 | 71963723 | 5803 | 2 | ENSMUST00000077337.8 | 14343 | Irx1 | ENSMUSG00000060969.8 |
| chr13 | 71951120 | 71951726 | 3.51065 | 4.14404 | 1.18463 | Distal Intergenic | 13 | 71957921 | 71963723 | 5803 | 2 | ENSMUST00000077337.8 | 11997 | Irx1 | ENSMUSG00000060969.8 |
| chr13 | 71952107 | 71952895 | 3.6448 | 4.4945 | 1.47087 | Distal Intergenic | 13 | 71957921 | 71963723 | 5803 | 2 | ENSMUST00000077337.8 | 10828 | Irx1 | ENSMUSG00000060969.8 |
| chr13 | 71954887 | 71955552 | 3.43971 | 4.02287 | 1.13787 | Downstream (2-3kb) | 13 | 71957921 | 71963723 | 5803 | 2 | ENSMUST00000077337.8 | 8171 | Irx1 | ENSMUSG00000060969.8 |
| chr13 | 71955931 | 71956313 | 3.78642 | 4.66145 | 1.51099 | Downstream (1-2kb) | 13 | 71957921 | 71963723 | 5803 | 2 | ENSMUST00000077337.8 | 7410 | Irx1 | ENSMUSG00000060969.8 |
| chr13 | 71956898 | 71957044 | 3.95428 | 5.02103 | 1.81891 | Downstream (<1kb) | 13 | 71957921 | 71963723 | 5803 | 2 | ENSMUST00000077337.8 | 6679 | Irx1 | ENSMUSG00000060969.8 |
| chr13 | 71957893 | 71958387 | 4.20182 | 5.49507 | 2.12274 | 3' UTR | 13 | 71957921 | 71963723 | 5803 | 2 | ENSMUST00000077337.8 | 5336 | Irx1 | ENSMUSG00000060969.8 |
| chr13 | 71958854 | 71960094 | 3.76377 | 4.67685 | 1.56044 | Exon (ENSMUST00000077337.8/ENSMUSG00000060969.8, exon 2 of 4) | 13 | 71957921 | 71963723 | 5803 | 2 | ENSMUST00000077337.8 | 3629 | Irx1 | ENSMUSG00000060969.8 |
| chr13 | 71960641 | 71961429 | 3.57657 | 4.32921 | 1.33763 | Intron (ENSMUST00000077337.8/ENSMUSG00000060969.8, intron 1 of 3) | 13 | 71957921 | 71963723 | 5803 | 2 | ENSMUST00000077337.8 | 2294 | Irx1 | ENSMUSG00000060969.8 |
| chr13 | 71961843 | 71962480 | 3.33301 | 4.13812 | 1.23084 | Intron (ENSMUST00000077337.8/ENSMUSG00000060969.8, intron 1 of 3) | 13 | 71957921 | 71963723 | 5803 | 2 | ENSMUST00000077337.8 | 1243 | Irx1 | ENSMUSG00000060969.8 |
| chr13 | 71962817 | 71964123 | 3.67271 | 4.59691 | 1.5395 | Promoter | 13 | 71957921 | 71963723 | 5803 | 2 | ENSMUST00000077337.8 | 0 | Irx1 | ENSMUSG00000060969.8 |
| chr13 | 71966993 | 71967324 | 3.53271 | 4.2408 | 1.34701 | Distal Intergenic | 13 | 71957921 | 71963723 | 5803 | 2 | ENSMUST00000077337.8 | -3271 | Irx1 | ENSMUSG00000060969.8 |
| chr13 | 71967794 | 71967872 | 4.57406 | 6.53275 | 2.8576 | Distal Intergenic | 13 | 71957921 | 71963723 | 5803 | 2 | ENSMUST00000077337.8 | -4072 | Irx1 | ENSMUSG00000060969.8 |
| chr13 | 71969127 | 71969595 | 3.78519 | 4.7851 | 1.6649 | Distal Intergenic | 13 | 71957921 | 71963723 | 5803 | 2 | ENSMUST00000077337.8 | -5405 | Irx1 | ENSMUSG00000060969.8 |
| chr13 | 71972853 | 71973250 | 3.6828 | 4.5311 | 1.51224 | Distal Intergenic | 13 | 71957921 | 71963723 | 5803 | 2 | ENSMUST00000077337.8 | -9131 | Irx1 | ENSMUSG00000060969.8 |
| chr13 | 71974377 | 71974766 | 3.44146 | 4.08185 | 1.23882 | Distal Intergenic | 13 | 71957921 | 71963723 | 5803 | 2 | ENSMUST00000077337.8 | -10655 | Irx1 | ENSMUSG00000060969.8 |
| chr13 | 71977134 | 71977604 | 4.08742 | 5.30511 | 2.01589 | Distal Intergenic | 13 | 71957921 | 71963723 | 5803 | 2 | ENSMUST00000077337.8 | -13412 | Irx1 | ENSMUSG00000060969.8 |
| chr13 | 71983152 | 71983264 | 3.58284 | 4.3215 | 1.33367 | Distal Intergenic | 13 | 71957921 | 71963723 | 5803 | 2 | ENSMUST00000077337.8 | -19430 | Irx1 | ENSMUSG00000060969.8 |
| chr13 | 72027770 | 72029501 | 3.68942 | 4.55548 | 1.47656 | Distal Intergenic | 13 | 71957921 | 71963723 | 5803 | 2 | ENSMUST00000077337.8 | -64048 | Irx1 | ENSMUSG00000060969.8 |
| chr13 | 72030406 | 72030638 | 3.52038 | 4.21571 | 1.26765 | Distal Intergenic | 13 | 71957921 | 71963723 | 5803 | 2 | ENSMUST00000077337.8 | -66684 | Irx1 | ENSMUSG00000060969.8 |
| chr4 | 12241597 | 12241775 | 3.26486 | 3.72443 | 1.01733 | Distal Intergenic | 4 | 12232833 | 12233951 | 1119 | 1 | ENSMUST00000125165.1 | 8765 | Gm11847 | ENSMUSG00000060989.8 |
| chr18 | 7000578 | 7000679 | 3.58342 | 4.46298 | 1.41854 | Exon (ENSMUST00000079788.5/ENSMUSG00000061013.5, exon 3 of 7) | 18 | 7000636 | 7003782 | 3147 | 2 | ENSMUST00000176608.1 | 3103 | Mkx | ENSMUSG00000061013.5 |
| chr18 | 7001081 | 7001386 | 3.71455 | 4.6053 | 1.5391 | Intron (ENSMUST00000079788.5/ENSMUSG00000061013.5, intron 2 of 6) | 18 | 7000636 | 7003782 | 3147 | 2 | ENSMUST00000176608.1 | 2396 | Mkx | ENSMUSG00000061013.5 |
| chr18 | 7001759 | 7001910 | 3.81572 | 5.07065 | 1.82329 | Intron (ENSMUST00000079788.5/ENSMUSG00000061013.5, intron 2 of 6) | 18 | 7000636 | 7003782 | 3147 | 2 | ENSMUST00000176608.1 | 1872 | Mkx | ENSMUSG00000061013.5 |
| chr18 | 7003400 | 7003523 | 3.46204 | 4.08927 | 1.2434 | Promoter | 18 | 7000636 | 7003782 | 3147 | 2 | ENSMUST00000176608.1 | 259 | Mkx | ENSMUSG00000061013.5 |
| chr18 | 7003874 | 7003949 | 3.91828 | 5.59845 | 2.20897 | Promoter | 18 | 7000636 | 7003782 | 3147 | 2 | ENSMUST00000176608.1 | -93 | Mkx | ENSMUSG00000061013.5 |
| chr18 | 7004277 | 7005582 | 3.82634 | 4.95453 | 1.75849 | Promoter | 18 | 6934966 | 7004780 | 69815 | 2 | ENSMUST00000079788.5 | 0 | Mkx | ENSMUSG00000061013.5 |
| chr18 | 7005967 | 7006323 | 4.11 | 5.31311 | 2.00967 | Distal Intergenic | 18 | 6934966 | 7004780 | 69815 | 2 | ENSMUST00000079788.5 | -1188 | Mkx | ENSMUSG00000061013.5 |
| chr18 | 7006762 | 7007027 | 4.03047 | 5.15661 | 1.90556 | Distal Intergenic | 18 | 6934966 | 7004780 | 69815 | 2 | ENSMUST00000079788.5 | -1983 | Mkx | ENSMUSG00000061013.5 |
| chr8 | 106506837 | 106508687 | 3.78997 | 4.70178 | 1.56453 | Exon (ENSMUST00000066875.7/ENSMUSG00000054069.7, exon 3 of 3) | 8 | 106510913 | 106556908 | 45996 | 1 | ENSMUST00000080797.6 | -2226 | Cdh3 | ENSMUSG00000061048.7 |
| chr8 | 106509236 | 106511461 | 4.16678 | 5.46289 | 2.09582 | Promoter | 8 | 106510913 | 106556908 | 45996 | 1 | ENSMUST00000080797.6 | 0 | Cdh3 | ENSMUSG00000061048.7 |
| chr8 | 106511803 | 106512581 | 3.73482 | 4.60197 | 1.50944 | Promoter | 8 | 106510913 | 106556908 | 45996 | 1 | ENSMUST00000080797.6 | 891 | Cdh3 | ENSMUSG00000061048.7 |
| chr16 | 39985235 | 39985502 | 3.38552 | 3.98225 | 1.16373 | Promoter | 16 | 39984595 | 40657886 | 673292 | 1 | ENSMUST00000189829.1 | 641 | Lsamp | ENSMUSG00000061080.6 |
| chr16 | 41532719 | 41533016 | 3.42825 | 4.02798 | 1.2069 | Promoter | 16 | 41532858 | 42146213 | 613356 | 1 | ENSMUST00000078873.5 | 0 | Lsamp | ENSMUSG00000061080.6 |
| chr16 | 41534107 | 41534577 | 3.40026 | 3.97743 | 1.17964 | Promoter | 16 | 41533419 | 41539253 | 5835 | 1 | ENSMUST00000187185.1 | 689 | Lsamp | ENSMUSG00000061080.6 |
| chr17 | 32453342 | 32453748 | 3.64644 | 4.63044 | 1.54382 | Distal Intergenic | 17 | 32468462 | 32492479 | 24018 | 1 | ENSMUST00000003413.6 | -14714 | Cyp4f39 | ENSMUSG00000061126.4 |
| chr19 | 40993983 | 40994272 | 3.32584 | 3.81617 | 1.01733 | Promoter | 19 | 40928927 | 40994148 | 65222 | 2 | ENSMUST00000117695.2 | 0 | Blnk | ENSMUSG00000061132.8 |
| chr2 | 10297788 | 10298019 | 4.25226 | 5.7051 | 2.26502 | Distal Intergenic | 2 | 10370510 | 10439137 | 68628 | 1 | ENSMUST00000114862.2 | -72491 | Sfmbt2 | ENSMUSG00000061186.10 |
| chr2 | 10367953 | 10368601 | 3.74541 | 4.61283 | 1.53285 | Intron (ENSMUST00000137351.1/ENSMUSG00000086748.1, intron 1 of 3) | 2 | 10370510 | 10439137 | 68628 | 1 | ENSMUST00000114862.2 | -1909 | Sfmbt2 | ENSMUSG00000061186.10 |
| chr2 | 10371575 | 10371920 | 3.67066 | 4.44566 | 1.38617 | Promoter | 2 | 10372436 | 10439137 | 66702 | 1 | ENSMUST00000114864.3 | -516 | Sfmbt2 | ENSMUSG00000061186.10 |
| chr4 | 41629192 | 41629631 | 3.6703 | 4.44658 | 1.40161 | Intron (ENSMUST00000102963.4/ENSMUSG00000061322.10, intron 13 of 19) | 4 | 41631644 | 41638158 | 6515 | 1 | ENSMUST00000143198.1 | -2013 | Dnaic1 | ENSMUSG00000061322.10 |
| chr4 | 41631353 | 41631865 | 3.87955 | 4.88813 | 1.69811 | Promoter | 4 | 41631644 | 41638158 | 6515 | 1 | ENSMUST00000143198.1 | 0 | Dnaic1 | ENSMUSG00000061322.10 |
| chr10 | 81991828 | 81992277 | 3.70863 | 4.58731 | 1.49744 | Distal Intergenic | 10 | 82058417 | 82061329 | 2913 | 1 | ENSMUST00000105313.1 | -66140 | Zfp873 | ENSMUSG00000061371.6 |
| chr6 | 127541640 | 127541722 | 4.07265 | 5.22409 | 1.92306 | Distal Intergenic | 6 | 127577975 | 127629938 | 51964 | 1 | ENSMUST00000071563.4 | -36253 | Efcab4b | ENSMUSG00000061414.4 |
| chr19 | 5084527 | 5084884 | 3.62281 | 4.38402 | 1.41838 | Promoter | 19 | 5082902 | 5084881 | 1980 | 2 | ENSMUST00000136372.1 | 0 | Tmem151a | ENSMUSG00000061451.8 |
| chr5 | 53256624 | 53256879 | 4.1242 | 5.41517 | 2.07671 | Distal Intergenic | 5 | 53267106 | 53278487 | 11382 | 1 | ENSMUST00000136866.2 | -10227 | Smim20 | ENSMUSG00000061461.7 |
| chr5 | 53257317 | 53257613 | 4.34622 | 5.80964 | 2.36899 | Distal Intergenic | 5 | 53267106 | 53278487 | 11382 | 1 | ENSMUST00000136866.2 | -9493 | Smim20 | ENSMUSG00000061461.7 |
| chr5 | 53258020 | 53258144 | 3.81745 | 4.8266 | 1.71611 | Distal Intergenic | 5 | 53267106 | 53278487 | 11382 | 1 | ENSMUST00000136866.2 | -8962 | Smim20 | ENSMUSG00000061461.7 |
| chr11 | 59134441 | 59134721 | 4.11028 | 5.2885 | 1.94337 | Exon (ENSMUST00000047441.8/ENSMUSG00000061462.10, exon 2 of 104) | 11 | 58994256 | 59136375 | 142120 | 2 | ENSMUST00000047441.8 | 1654 | Obscn | ENSMUSG00000061462.10 |
| chr11 | 59135611 | 59136077 | 3.53127 | 4.1835 | 1.21109 | Promoter | 11 | 58994256 | 59136375 | 142120 | 2 | ENSMUST00000047441.8 | 298 | Obscn | ENSMUSG00000061462.10 |
| chr11 | 59138607 | 59138730 | 3.72076 | 4.54214 | 1.45821 | Distal Intergenic | 11 | 59012141 | 59136402 | 124262 | 2 | ENSMUST00000020732.5 | -2206 | Obscn | ENSMUSG00000061462.10 |
| chr11 | 59139151 | 59139401 | 3.2513 | 3.8483 | 1.02941 | Distal Intergenic | 11 | 59012141 | 59136402 | 124262 | 2 | ENSMUST00000020732.5 | -2750 | Obscn | ENSMUSG00000061462.10 |
| chr9 | 101600487 | 101600749 | 3.34442 | 3.84408 | 1.01733 | Distal Intergenic | 9 | 101448501 | 101449353 | 853 | 2 | ENSMUST00000075243.7 | -151135 | Gm5161 | ENSMUSG00000061486.7 |
| chr9 | 101617626 | 101617762 | 3.55072 | 4.26257 | 1.34948 | Distal Intergenic | 9 | 101448501 | 101449353 | 853 | 2 | ENSMUST00000075243.7 | -168274 | Gm5161 | ENSMUSG00000061486.7 |
| chr14 | 118233316 | 118238775 | 4.18675 | 5.62472 | 2.23818 | Promoter | 14 | 118233232 | 118237030 | 3799 | 2 | ENSMUST00000170662.1 | 0 | Sox21 | ENSMUSG00000061517.8 |
| chr14 | 118239389 | 118241073 | 3.68311 | 4.48625 | 1.44865 | Intron (ENSMUST00000163396.1/ENSMUSG00000090534.1, intron 1 of 2) | 14 | 118233232 | 118237030 | 3799 | 2 | ENSMUST00000170662.1 | -2360 | Sox21 | ENSMUSG00000061517.8 |
| chr14 | 118242115 | 118242380 | 3.35234 | 3.85596 | 1.01733 | Intron (ENSMUST00000163396.1/ENSMUSG00000090534.1, intron 1 of 2) | 14 | 118233232 | 118237030 | 3799 | 2 | ENSMUST00000170662.1 | -5086 | Sox21 | ENSMUSG00000061517.8 |
| chr14 | 118242818 | 118242897 | 4.11024 | 5.28859 | 1.95616 | Intron (ENSMUST00000163396.1/ENSMUSG00000090534.1, intron 1 of 2) | 14 | 118233232 | 118237030 | 3799 | 2 | ENSMUST00000170662.1 | -5789 | Sox21 | ENSMUSG00000061517.8 |
| chr14 | 118243641 | 118243804 | 3.26759 | 3.7398 | 1.05266 | Intron (ENSMUST00000163396.1/ENSMUSG00000090534.1, intron 1 of 2) | 14 | 118233232 | 118237030 | 3799 | 2 | ENSMUST00000170662.1 | -6612 | Sox21 | ENSMUSG00000061517.8 |
| chr14 | 122474856 | 122476770 | 4.25756 | 5.73231 | 2.34859 | Promoter | 14 | 122475435 | 122479852 | 4418 | 1 | ENSMUST00000075888.5 | 0 | Zic2 | ENSMUSG00000061524.8 |
| chr14 | 122477157 | 122477587 | 3.54288 | 4.50426 | 1.44377 | Promoter | 14 | 122478100 | 122479050 | 951 | 1 | ENSMUST00000137059.1 | -513 | Zic2 | ENSMUSG00000061524.8 |
| chr14 | 122478021 | 122479244 | 3.73218 | 4.74907 | 1.60841 | Promoter | 14 | 122478100 | 122479050 | 951 | 1 | ENSMUST00000137059.1 | 0 | Zic2 | ENSMUSG00000061524.8 |
| chr14 | 122479556 | 122480579 | 3.58839 | 4.37706 | 1.35221 | 3' UTR | 14 | 122478100 | 122479050 | 951 | 1 | ENSMUST00000137059.1 | 1457 | Zic2 | ENSMUSG00000061524.8 |
| chr14 | 122480885 | 122483654 | 3.93437 | 5.16977 | 1.89295 | Distal Intergenic | 14 | 122478100 | 122479050 | 951 | 1 | ENSMUST00000137059.1 | 2786 | Zic2 | ENSMUSG00000061524.8 |
| chr5 | 26817396 | 26817798 | 3.51295 | 4.16415 | 1.24902 | Promoter | 5 | 26817434 | 27452794 | 635361 | 1 | ENSMUST00000136574.1 | 0 | Dpp6 | ENSMUSG00000061576.10 |
| chr5 | 27049350 | 27049815 | 3.57285 | 4.2889 | 1.35162 | Promoter | 5 | 27049199 | 27727501 | 678303 | 1 | ENSMUST00000122171.2 | 152 | Dpp6 | ENSMUSG00000061576.10 |
| chr8 | 94942485 | 94943179 | 3.94941 | 4.96385 | 1.70778 | 3' UTR | 8 | 94934167 | 94935084 | 918 | 1 | ENSMUST00000142991.1 | 8319 | Gpr114 | ENSMUSG00000061577.6 |
| chr8 | 94949710 | 94949785 | 3.43683 | 4.00069 | 1.078 | Distal Intergenic | 8 | 94934167 | 94935084 | 918 | 1 | ENSMUST00000142991.1 | 15544 | Gpr114 | ENSMUSG00000061577.6 |
| chr5 | 117413969 | 117414107 | 3.37261 | 3.88654 | 1.01733 | Promoter | 5 | 117414000 | 117555942 | 141943 | 1 | ENSMUST00000073347.4 | 0 | Ksr2 | ENSMUSG00000061578.8 |
| chr5 | 117416821 | 117416896 | 3.61017 | 4.38038 | 1.40324 | Intron (ENSMUST00000073347.4/ENSMUSG00000061578.8, intron 1 of 3) | 5 | 117414000 | 117775003 | 361004 | 1 | ENSMUST00000180430.1 | 2822 | Ksr2 | ENSMUSG00000061578.8 |
| chr2 | 156612254 | 156615211 | 4.06643 | 5.23489 | 1.91581 | Promoter | 2 | 156613705 | 156764361 | 150657 | 1 | ENSMUST00000109567.4 | 0 | Dlgap4 | ENSMUSG00000061689.10 |
| chr2 | 156686611 | 156686985 | 3.40516 | 4.31222 | 1.31319 | Intron (ENSMUST00000109567.4/ENSMUSG00000061689.10, intron 2 of 12) | 2 | 156700622 | 156763174 | 62553 | 1 | ENSMUST00000109568.4 | -13637 | Dlgap4 | ENSMUSG00000061689.10 |
| chr2 | 156683936 | 156684547 | 4.05786 | 5.18623 | 1.87407 | Intron (ENSMUST00000109567.4/ENSMUSG00000061689.10, intron 2 of 12) | 2 | 156700622 | 156763174 | 62553 | 1 | ENSMUST00000109568.4 | -16075 | Dlgap4 | ENSMUSG00000061689.10 |
| chr7 | 25675138 | 25675682 | 4.01386 | 5.50184 | 2.12902 | Promoter | 7 | 25669139 | 25675166 | 6028 | 2 | ENSMUST00000079439.4 | 0 | Tmem91 | ENSMUSG00000061702.5 |
| chr11 | 98339386 | 98340087 | 3.95696 | 5.10137 | 1.85975 | Distal Intergenic | 11 | 98348404 | 98357796 | 9393 | 1 | ENSMUST00000078694.7 | -8317 | Ppp1r1b | ENSMUSG00000061718.7 |
| chr11 | 98341690 | 98342439 | 3.78689 | 4.66477 | 1.51668 | Distal Intergenic | 11 | 98348404 | 98357796 | 9393 | 1 | ENSMUST00000078694.7 | -5965 | Ppp1r1b | ENSMUSG00000061718.7 |
| chr11 | 98346030 | 98347115 | 3.66971 | 4.44923 | 1.3942 | Distal Intergenic | 11 | 98348404 | 98357796 | 9393 | 1 | ENSMUST00000078694.7 | -1289 | Ppp1r1b | ENSMUSG00000061718.7 |
| chr11 | 98348269 | 98349174 | 3.92221 | 5.14321 | 1.88204 | Promoter | 11 | 98348404 | 98357796 | 9393 | 1 | ENSMUST00000078694.7 | 0 | Ppp1r1b | ENSMUSG00000061718.7 |
| chr11 | 98349725 | 98350388 | 3.71075 | 4.52689 | 1.43028 | Promoter | 11 | 98349996 | 98357794 | 7799 | 1 | ENSMUST00000137634.1 | 0 | Ppp1r1b | ENSMUSG00000061718.7 |
| chr11 | 98351401 | 98351488 | 3.52812 | 4.17172 | 1.18904 | Promoter | 11 | 98351489 | 98357286 | 5798 | 1 | ENSMUST00000133700.2 | -1 | Ppp1r1b | ENSMUSG00000061718.7 |
| chr19 | 6538502 | 6538763 | 3.66629 | 4.43131 | 1.37742 | Exon (ENSMUST00000113451.2/ENSMUSG00000061742.6, exon 5 of 10) | 19 | 6536065 | 6541549 | 5485 | 2 | ENSMUST00000129698.1 | 2786 | Slc22a12 | ENSMUSG00000061742.6 |
| chr16 | 34057518 | 34057782 | 3.4134 | 4.21853 | 1.27533 | Intron (ENSMUST00000114973.2/ENSMUSG00000061751.10, intron 1 of 26) | 16 | 33974258 | 34095987 | 121730 | 2 | ENSMUST00000114973.2 | 38205 | Kalrn | ENSMUSG00000061751.10 |
| chr16 | 34572615 | 34573067 | 3.60046 | 4.34545 | 1.39031 | Promoter | 16 | 34049689 | 34573532 | 523844 | 2 | ENSMUST00000114961.2 | 465 | Kalrn | ENSMUSG00000061751.10 |
| chr16 | 34603247 | 34603402 | 3.56043 | 4.26467 | 1.33709 | Distal Intergenic | 16 | 34049689 | 34573532 | 523844 | 2 | ENSMUST00000114961.2 | -29716 | Kalrn | ENSMUSG00000061751.10 |
| chr6 | 7555048 | 7555496 | 3.74162 | 4.76517 | 1.62729 | Promoter | 6 | 7555074 | 7562976 | 7903 | 1 | ENSMUST00000090679.3 | 0 | Tac1 | ENSMUSG00000061762.7 |
| chr6 | 7555887 | 7556011 | 3.21193 | 3.7887 | 1.02768 | Promoter | 6 | 7556091 | 7559114 | 3024 | 1 | ENSMUST00000185130.1 | -80 | Tac1 | ENSMUSG00000061762.7 |

| | | | | | | | | | | | | | | | |
|---|---|---|---|---|---|---|---|---|---|---|---|---|---|---|---|
| chr8 | 58910955 | 58911304 | 3.4337 | 4.01106 | 1.13174 | Promoter | 8 | 58427580 | 58911576 | 483997 | 2 | ENSMUST00000146513.2 | 272 | Galntl6 | ENSMUSG00000061864.7 |
| chr7 | 81497299 | 81497427 | 3.58128 | 4.30424 | 1.29711 | Promoter | 7 | 81494275 | 81498281 | 4007 | 2 | ENSMUST00000073406.6 | 854 | BC048679 | ENSMUSG00000061877.7 |
| chr7 | 81497894 | 81498242 | 3.3635 | 3.87737 | 1.03351 | Promoter | 7 | 81494275 | 81498281 | 4007 | 2 | ENSMUST00000073406.6 | 39 | BC048679 | ENSMUSG00000061877.7 |
| chr11 | 116531419 | 116533129 | 3.6957 | 4.58495 | 1.53839 | Promoter | 11 | 116531744 | 116536657 | 4914 | 1 | ENSMUST00000106387.3 | 0 | Sphk1 | ENSMUSG00000061878.10 |
| chr11 | 116533782 | 116534148 | 3.60197 | 4.53834 | 1.46798 | Promoter | 11 | 116534160 | 116535288 | 1129 | 1 | ENSMUST00000148185.1 | -12 | Sphk1 | ENSMUSG00000061878.10 |
| chr2 | 179109476 | 179109687 | 3.29883 | 3.7755 | 1.01733 | Distal Intergenic | 2 | 179177425 | 179177880 | 456 | 1 | ENSMUST00000118761.1 | -67738 | Gm14292 | ENSMUSG00000061897.5 |
| chr5 | 143157519 | 143157804 | 3.27746 | 3.7434 | 1.01733 | Distal Intergenic | 5 | 143172187 | 143176734 | 4548 | 2 | ENSMUST00000166382.2 | 18930 | Rbak | ENSMUSG00000061898.5 |
| chr5 | 143158615 | 143158761 | 3.60982 | 4.44187 | 1.41422 | Distal Intergenic | 5 | 143172187 | 143176734 | 4548 | 2 | ENSMUST00000166382.2 | 17973 | Rbak | ENSMUSG00000061898.5 |
| chr12 | 29527033 | 29527770 | 3.58472 | 4.33229 | 1.33758 | Promoter | 12 | 29528384 | 29923209 | 394826 | 1 | ENSMUST00000049784.10 | -614 | Myt1l | ENSMUSG00000061911.9 |
| chr12 | 29528408 | 29528533 | 4.19245 | 5.45616 | 2.09123 | Promoter | 12 | 29528384 | 29923209 | 394826 | 1 | ENSMUST00000049784.10 | 25 | Myt1l | ENSMUSG00000061911.9 |
| chr12 | 29533315 | 29533420 | 3.53692 | 4.79104 | 1.63589 | Intron (ENSMUST00000049784.10/ENSMUSG00000061911.9, intron 1 of 24) | 12 | 29534905 | 29923209 | 388305 | 1 | ENSMUST00000021009.8 | -1485 | Myt1l | ENSMUSG00000061911.9 |
| chr12 | 29534614 | 29535335 | 3.69711 | 4.55922 | 1.52526 | Promoter | 12 | 29534905 | 29923209 | 388305 | 1 | ENSMUST00000021009.8 | 0 | Myt1l | ENSMUSG00000061911.9 |
| chr12 | 29535812 | 29536038 | 3.4106 | 3.98524 | 1.16462 | Promoter | 12 | 29534905 | 29923209 | 388305 | 1 | ENSMUST00000021009.8 | 908 | Myt1l | ENSMUSG00000061911.9 |
| chr3 | 130737782 | 130738488 | 3.52267 | 4.16771 | 1.21088 | Distal Intergenic | 3 | 130726839 | 130730398 | 3560 | 2 | ENSMUST00000079085.6 | -7385 | Rpl34 | ENSMUSG00000062006.7 |
| chr7 | 45779200 | 45779294 | 3.35599 | 3.86146 | 1.01733 | Distal Intergenic | 7 | 45783738 | 45804144 | 20407 | 1 | ENSMUST00000118564.2 | -4444 | Lmtk3 | ENSMUSG00000062044.8 |
| chr7 | 45782068 | 45782253 | 3.31659 | 3.80217 | 1.01733 | Distal Intergenic | 7 | 45783738 | 45804144 | 20407 | 1 | ENSMUST00000118564.2 | -1485 | Lmtk3 | ENSMUSG00000062044.8 |
| chr7 | 45783971 | 45784191 | 3.51825 | 4.15115 | 1.17508 | Promoter | 7 | 45783947 | 45804109 | 20163 | 1 | ENSMUST00000072580.6 | 25 | Lmtk3 | ENSMUSG00000062044.8 |
| chr7 | 45784931 | 45785087 | 3.45859 | 4.04279 | 1.11056 | Promoter | 7 | 45785380 | 45804144 | 18765 | 1 | ENSMUST00000120005.2 | -293 | Lmtk3 | ENSMUSG00000062044.8 |
| chr7 | 45785406 | 45786337 | 4.12 | 5.33826 | 1.98496 | Promoter | 7 | 45785510 | 45787758 | 2249 | 1 | ENSMUST00000134347.1 | 0 | Lmtk3 | ENSMUSG00000062044.8 |
| chr7 | 45798221 | 45798708 | 3.66736 | 4.55811 | 1.50656 | Exon (ENSMUST00000118564.2/ENSMUSG00000062044.8, exon 13 of 16) | 7 | 45787904 | 45792586 | 4683 | 1 | ENSMUST00000146738.1 | 10318 | Lmtk3 | ENSMUSG00000062044.8 |
| chr5 | 31115586 | 31115843 | 3.37542 | 3.89077 | 1.01733 | Promoter | 5 | 31116712 | 31137628 | 20917 | 1 | ENSMUST00000013771.9 | -869 | Trim54 | ENSMUSG00000062077.9 |
| chr5 | 31116712 | 31116928 | 3.47718 | 4.0838 | 1.15717 | Promoter | 5 | 31116734 | 31137630 | 20897 | 1 | ENSMUST00000114637.1 | 0 | Trim54 | ENSMUSG00000062077.9 |
| chr3 | 105059839 | 105060079 | 3.25637 | 3.71169 | 1.01733 | Distal Intergenic | 3 | 105001915 | 105053146 | 51232 | 2 | ENSMUST00000077548.6 | -6694 | Cttnbp2nl | ENSMUSG00000062127.6 |
| chr9 | 73968861 | 73969097 | 3.69195 | 4.55138 | 1.51111 | Promoter | 9 | 73479422 | 73968966 | 489545 | 2 | ENSMUST00000184666.2 | 0 | Unc13c | ENSMUSG00000062151.8 |
| chr15 | 78398727 | 78399195 | 3.34974 | 3.89317 | 1.1367 | Downstream (<1kb) | 15 | 78378400 | 78395912 | 17513 | 2 | ENSMUST00000165170.2 | -2816 | Tex33 | ENSMUSG00000062154.8 |
| chr9 | 27790719 | 27791524 | 3.42072 | 3.98379 | 1.10049 | Promoter | 9 | 27790775 | 27791518 | 744 | 1 | ENSMUST00000134002.1 | 0 | Opcml | ENSMUSG00000062257.7 |
| chr9 | 27791888 | 27792388 | 3.6772 | 4.45237 | 1.37316 | Promoter | 9 | 27791302 | 28030026 | 238725 | 1 | ENSMUST00000123779.1 | 587 | Opcml | ENSMUSG00000062257.7 |
| chr9 | 27793516 | 27793819 | 3.3074 | 3.78831 | 1.01733 | Intron (ENSMUST00000115243.3/ENSMUSG00000062257.7, intron 1 of 7) | 9 | 27791302 | 28030026 | 238725 | 1 | ENSMUST00000123779.1 | 2215 | Opcml | ENSMUSG00000062257.7 |
| chr9 | 28404179 | 28404359 | 3.83882 | 4.87413 | 1.70652 | Promoter | 9 | 28404169 | 28650288 | 246120 | 1 | ENSMUST00000150098.1 | 11 | Opcml | ENSMUSG00000062257.7 |
| chr13 | 46201877 | 46202033 | 3.33349 | 3.82756 | 1.01733 | Distal Intergenic | 13 | 46175428 | 46191048 | 15621 | 2 | ENSMUST00000072437.3 | -10830 | Gm10113 | ENSMUSG00000062282.3 |
| chr13 | 46217343 | 46217601 | 3.59045 | 4.40319 | 1.39721 | Distal Intergenic | 13 | 46175428 | 46191048 | 15621 | 2 | ENSMUST00000072437.3 | -26296 | Gm10113 | ENSMUSG00000062282.3 |
| chr13 | 46218472 | 46218862 | 3.30266 | 3.80774 | 1.09504 | Distal Intergenic | 13 | 46175428 | 46191048 | 15621 | 2 | ENSMUST00000072437.3 | -27425 | Gm10113 | ENSMUSG00000062282.3 |
| chr13 | 46219219 | 46219365 | 3.30837 | 4.02633 | 1.17021 | Distal Intergenic | 13 | 46175428 | 46191048 | 15621 | 2 | ENSMUST00000072437.3 | -28172 | Gm10113 | ENSMUSG00000062282.3 |
| chr9 | 111310715 | 111313073 | 3.77703 | 4.8707 | 1.68784 | Promoter | 9 | 111311739 | 111395774 | 84036 | 1 | ENSMUST00000078626.3 | 0 | Trank1 | ENSMUSG00000062296.4 |
| chr9 | 111313853 | 111314233 | 3.72866 | 4.55414 | 1.45904 | Intron (ENSMUST00000078626.3/ENSMUSG00000062296.4, intron 1 of 21) | 9 | 111311739 | 111395774 | 84036 | 1 | ENSMUST00000078626.3 | 2115 | Trank1 | ENSMUSG00000062296.4 |
| chr9 | 111314635 | 111314761 | 3.26623 | 3.72651 | 1.01733 | Intron (ENSMUST00000078626.3/ENSMUSG00000062296.4, intron 1 of 21) | 9 | 111311739 | 111395774 | 84036 | 1 | ENSMUST00000078626.3 | 2897 | Trank1 | ENSMUSG00000062296.4 |
| chr1 | 88504840 | 88505034 | 3.24916 | 3.70802 | 1.03816 | Intron (ENSMUST00000067625.8/ENSMUSG00000062310.7, intron 1 of 2) | 1 | 88499871 | 88510066 | 10196 | 2 | ENSMUST00000067625.8 | 5032 | Glrp1 | ENSMUSG00000062310.7 |
| chr17 | 8427789 | 8428297 | 3.47022 | 4.08387 | 1.18771 | Distal Intergenic | 17 | 8434423 | 8442496 | 8074 | 1 | ENSMUST00000074667.3 | -6126 | T | ENSMUSG00000062327.5 |
| chr17 | 8428607 | 8430130 | 3.67164 | 4.48537 | 1.43001 | Distal Intergenic | 17 | 8434423 | 8442496 | 8074 | 1 | ENSMUST00000074667.3 | -4293 | T | ENSMUSG00000062327.5 |
| chr17 | 8430673 | 8431437 | 3.6852 | 4.49443 | 1.438 | Distal Intergenic | 17 | 8434423 | 8442496 | 8074 | 1 | ENSMUST00000074667.3 | -2986 | T | ENSMUSG00000062327.5 |
| chr17 | 8432810 | 8432885 | 3.43876 | 4.01405 | 1.11522 | Distal Intergenic | 17 | 8434423 | 8442496 | 8074 | 1 | ENSMUST00000074667.3 | -1538 | T | ENSMUSG00000062327.5 |
| chr17 | 8433825 | 8434642 | 3.96281 | 5.01155 | 1.76269 | Promoter | 17 | 8434423 | 8442496 | 8074 | 1 | ENSMUST00000074667.3 | 0 | T | ENSMUSG00000062327.5 |
| chr17 | 8434994 | 8435492 | 3.82061 | 4.91787 | 1.73486 | Promoter | 17 | 8435411 | 8441742 | 6332 | 1 | ENSMUST00000177118.1 | 0 | T | ENSMUSG00000062327.5 |
| chr17 | 8437228 | 8437473 | 3.49603 | 4.115 | 1.16564 | Intron (ENSMUST00000074667.3/ENSMUSG00000062327.5, intron 5 of 7) | 17 | 8435411 | 8441742 | 6332 | 1 | ENSMUST00000177118.1 | 1818 | T | ENSMUSG00000062327.5 |
| chr17 | 8447734 | 8447929 | 3.58967 | 4.33791 | 1.40599 | Distal Intergenic | 17 | 8435411 | 8441742 | 6332 | 1 | ENSMUST00000177118.1 | 12324 | T | ENSMUSG00000062327.5 |
| chr17 | 8456336 | 8456465 | 3.23866 | 3.68516 | 1.01733 | Distal Intergenic | 17 | 8435411 | 8441742 | 6332 | 1 | ENSMUST00000177118.1 | 20926 | T | ENSMUSG00000062327.5 |
| chr18 | 74981786 | 74981998 | 3.39031 | 3.96489 | 1.1818 | Distal Intergenic | 18 | 75000477 | 75003381 | 2905 | 1 | ENSMUST00000079716.5 | -18479 | Rpl17 | ENSMUSG00000062328.7 |
| chr18 | 74990272 | 74990504 | 3.33241 | 3.82615 | 1.01733 | Distal Intergenic | 18 | 75000477 | 75003381 | 2905 | 1 | ENSMUST00000079716.5 | -9973 | Rpl17 | ENSMUSG00000062328.7 |
| chr5 | 37734649 | 37734903 | 3.34153 | 3.83966 | 1.01733 | Promoter | 5 | 37735519 | 37739820 | 4302 | 1 | ENSMUST00000073554.3 | -616 | Cytl1 | ENSMUSG00000062329.3 |
| chr6 | 127251755 | 127251904 | 3.41163 | 4.1533 | 1.21943 | Distal Intergenic | 6 | 127233762 | 127234619 | 858 | 1 | ENSMUST00000071458.4 | 17994 | Gm4968 | ENSMUSG00000062365.4 |
| chr8 | 123409291 | 123410300 | 3.69608 | 4.57523 | 1.49775 | 3' UTR | 8 | 123411424 | 123422015 | 10592 | 1 | ENSMUST00000071134.3 | -1124 | Tubb3 | ENSMUSG00000062380.3 |
| chr8 | 123410890 | 123412488 | 3.64764 | 4.40022 | 1.3522 | Promoter | 8 | 123411424 | 123422015 | 10592 | 1 | ENSMUST00000071134.3 | 0 | Tubb3 | ENSMUSG00000062380.3 |
| chr8 | 123413905 | 123414824 | 3.85212 | 4.88221 | 1.68772 | Intron (ENSMUST00000127664.1/ENSMUSG00000092329.1, intron 1 of 15) | 8 | 123411424 | 123422015 | 10592 | 1 | ENSMUST00000071134.3 | 2482 | Tubb3 | ENSMUSG00000062380.3 |
| chr8 | 123421515 | 123421669 | 3.45442 | 4.08995 | 1.22007 | Exon (ENSMUST00000071134.3/ENSMUSG00000062380.3, exon 4 of 4) | 8 | 123411424 | 123422015 | 10592 | 1 | ENSMUST00000071134.3 | 10092 | Tubb3 | ENSMUSG00000062380.3 |
| chr11 | 114423455 | 114423913 | 3.34124 | 3.84725 | 1.04543 | Promoter | 11 | 114413643 | 114423887 | 10245 | 2 | ENSMUST00000141088.1 | 0 | 4932435O22Rik | ENSMUSG00000062391.3 |
| chr11 | 114466091 | 114466512 | 3.53264 | 4.19997 | 1.21634 | Distal Intergenic | 11 | 114413401 | 114448860 | 35460 | 2 | ENSMUST00000148405.1 | -17232 | 4932435O22Rik | ENSMUSG00000062391.3 |
| chr11 | 114467640 | 114468169 | 3.5563 | 4.4455 | 1.3911 | Distal Intergenic | 11 | 114413401 | 114448860 | 35460 | 2 | ENSMUST00000148405.1 | -18781 | 4932435O22Rik | ENSMUSG00000062391.3 |
| chr11 | 114485572 | 114485657 | 3.56074 | 4.24431 | 1.27351 | Distal Intergenic | 11 | 114413401 | 114448860 | 35460 | 2 | ENSMUST00000148405.1 | -36713 | 4932435O22Rik | ENSMUSG00000062391.3 |
| chr11 | 114493285 | 114493494 | 3.38193 | 3.9006 | 1.01733 | Distal Intergenic | 11 | 114413401 | 114448860 | 35460 | 2 | ENSMUST00000148405.1 | -44426 | 4932435O22Rik | ENSMUSG00000062391.3 |
| chr19 | 37684247 | 37684719 | 3.62323 | 4.54588 | 1.48923 | Promoter | 19 | 37685581 | 37693398 | 7818 | 1 | ENSMUST00000073391.4 | -862 | Cyp26c1 | ENSMUSG00000062432.4 |
| chr19 | 37687120 | 37687365 | 3.92823 | 5.02966 | 1.8202 | Exon (ENSMUST00000073391.4/ENSMUSG00000062432.4, exon 3 of 6) | 19 | 37685581 | 37693398 | 7818 | 1 | ENSMUST00000073391.4 | 1540 | Cyp26c1 | ENSMUSG00000062432.4 |
| chr19 | 37690629 | 37691045 | 3.50205 | 4.16944 | 1.3007 | Exon (ENSMUST00000073391.4/ENSMUSG00000062432.4, exon 5 of 6) | 19 | 37685581 | 37693398 | 7818 | 1 | ENSMUST00000073391.4 | 5049 | Cyp26c1 | ENSMUSG00000062432.4 |
| chr7 | 81461754 | 81461880 | 3.33963 | 4.00232 | 1.12372 | Intron (ENSMUST00000147624.1/ENSMUSG00000062444.9, intron 7 of 8) | 7 | 81460399 | 81465295 | 4897 | 2 | ENSMUST00000147624.1 | 3415 | Ap3b2 | ENSMUSG00000062444.9 |
| chr7 | 81476956 | 81477094 | 3.6091 | 4.32952 | 1.31499 | Exon (ENSMUST00000082090.8/ENSMUSG00000062444.9, exon 8 of 26) | 7 | 81477123 | 81478410 | 1288 | 2 | ENSMUST00000125634.1 | 1316 | Ap3b2 | ENSMUSG00000062444.9 |
| chr7 | 81493637 | 81493744 | 3.52504 | 4.27167 | 1.3198 | Promoter | 7 | 81474380 | 81493706 | 19327 | 2 | ENSMUST00000152355.2 | 0 | Ap3b2 | ENSMUSG00000062444.9 |
| chr7 | 81495722 | 81496122 | 3.80933 | 4.70923 | 1.56205 | Exon (ENSMUST00000073406.6/ENSMUSG00000061877.7, exon 4 of 6) | 7 | 81460401 | 81493925 | 33525 | 2 | ENSMUST00000082090.8 | -1798 | Ap3b2 | ENSMUSG00000062444.9 |
| chr17 | 32634867 | 32635024 | 3.48898 | 4.15377 | 1.28955 | Exon (ENSMUST00000077639.5/ENSMUSG00000062464.5, exon 13 of 13) | 17 | 32621319 | 32636184 | 14866 | 1 | ENSMUST00000077639.5 | 13549 | Cyp4f37 | ENSMUSG00000062464.5 |
| chr16 | 95258251 | 95258482 | 3.24227 | 3.69056 | 1.01733 | Promoter | 16 | 95258252 | 95300211 | 41960 | 1 | ENSMUST00000113858.1 | 0 | Kcnj15 | ENSMUSG00000062609.8 |
| chr16 | 95259347 | 95259729 | 4.00159 | 5.15874 | 1.91648 | Intron (ENSMUST00000113861.2/ENSMUSG00000062609.8, intron 3 of 4) | 16 | 95258252 | 95300211 | 41960 | 1 | ENSMUST00000113858.1 | 1096 | Kcnj15 | ENSMUSG00000062609.8 |
| chr2 | 31243666 | 31243835 | 3.33705 | 4.02539 | 1.18811 | Distal Intergenic | 2 | 31245823 | 31295989 | 50167 | 1 | ENSMUST00000000199.7 | -1988 | Ncs1 | ENSMUSG00000062661.6 |
| chr2 | 31246122 | 31246462 | 3.34583 | 3.94604 | 1.10777 | Promoter | 2 | 31246166 | 31292319 | 46154 | 1 | ENSMUST00000150670.1 | 0 | Ncs1 | ENSMUSG00000062661.6 |
| chr16 | 94073381 | 94073587 | 3.42214 | 4.06584 | 1.21546 | Distal Intergenic | 16 | 94085260 | 94127032 | 41773 | 1 | ENSMUST00000072182.7 | -11673 | Sim2 | ENSMUSG00000062713.7 |
| chr16 | 94074502 | 94074846 | 3.42104 | 4.00898 | 1.18976 | Distal Intergenic | 16 | 94085260 | 94127032 | 41773 | 1 | ENSMUST00000072182.7 | -10414 | Sim2 | ENSMUSG00000062713.7 |
| chr16 | 94075417 | 94075999 | 3.77546 | 4.81539 | 1.64699 | Distal Intergenic | 16 | 94085260 | 94127032 | 41773 | 1 | ENSMUST00000072182.7 | -9261 | Sim2 | ENSMUSG00000062713.7 |
| chr16 | 94076467 | 94076803 | 4.21289 | 5.60963 | 2.25501 | Distal Intergenic | 16 | 94085260 | 94127032 | 41773 | 1 | ENSMUST00000072182.7 | -8457 | Sim2 | ENSMUSG00000062713.7 |
| chr16 | 94077116 | 94077318 | 3.43694 | 4.00489 | 1.09241 | Distal Intergenic | 16 | 94085260 | 94127032 | 41773 | 1 | ENSMUST00000072182.7 | -7942 | Sim2 | ENSMUSG00000062713.7 |
| chr16 | 94077930 | 94078601 | 3.66915 | 4.44246 | 1.38444 | Distal Intergenic | 16 | 94085260 | 94127032 | 41773 | 1 | ENSMUST00000072182.7 | -6659 | Sim2 | ENSMUSG00000062713.7 |
| chr16 | 94078929 | 94079868 | 4.0878 | 5.30214 | 1.95858 | Distal Intergenic | 16 | 94085260 | 94127032 | 41773 | 1 | ENSMUST00000072182.7 | -5392 | Sim2 | ENSMUSG00000062713.7 |
| chr16 | 94080261 | 94080995 | 4.07611 | 5.44734 | 2.11755 | Distal Intergenic | 16 | 94085260 | 94127032 | 41773 | 1 | ENSMUST00000072182.7 | -4265 | Sim2 | ENSMUSG00000062713.7 |
| chr16 | 94081600 | 94084312 | 4.03466 | 5.19491 | 1.9031 | Promoter | 16 | 94085260 | 94127032 | 41773 | 1 | ENSMUST00000072182.7 | -948 | Sim2 | ENSMUSG00000062713.7 |
| chr16 | 94085026 | 94086137 | 4.17655 | 5.71772 | 2.28377 | Promoter | 16 | 94085260 | 94127032 | 41773 | 1 | ENSMUST00000072182.7 | 0 | Sim2 | ENSMUSG00000062713.7 |
| chr16 | 94086537 | 94087571 | 3.68085 | 4.61797 | 1.50836 | Intron (ENSMUST00000072182.7/ENSMUSG00000062713.7, intron 1 of 10) | 16 | 94085260 | 94127032 | 41773 | 1 | ENSMUST00000072182.7 | 1278 | Sim2 | ENSMUSG00000062713.7 |
| chr16 | 94088104 | 94088617 | 3.50515 | 4.22948 | 1.3208 | Intron (ENSMUST00000072182.7/ENSMUSG00000062713.7, intron 1 of 10) | 16 | 94085260 | 94127032 | 41773 | 1 | ENSMUST00000072182.7 | 2845 | Sim2 | ENSMUSG00000062713.7 |
| chr16 | 94089975 | 94090512 | 3.62796 | 4.45631 | 1.46831 | Intron (ENSMUST00000072182.7/ENSMUSG00000062713.7, intron 1 of 10) | 16 | 94085260 | 94127032 | 41773 | 1 | ENSMUST00000072182.7 | 4716 | Sim2 | ENSMUSG00000062713.7 |
| chr16 | 94091906 | 94092165 | 3.47715 | 4.19043 | 1.24838 | Intron (ENSMUST00000072182.7/ENSMUSG00000062713.7, intron 1 of 10) | 16 | 94085260 | 94127032 | 41773 | 1 | ENSMUST00000072182.7 | 6647 | Sim2 | ENSMUSG00000062713.7 |
| chr16 | 94092941 | 94093560 | 3.8876 | 5.01636 | 1.80791 | Intron (ENSMUST00000072182.7/ENSMUSG00000062713.7, intron 1 of 10) | 16 | 94085260 | 94127032 | 41773 | 1 | ENSMUST00000072182.7 | 7682 | Sim2 | ENSMUSG00000062713.7 |
| chr16 | 94094261 | 94096834 | 3.78266 | 4.7319 | 1.6252 | Exon (ENSMUST00000072182.7/ENSMUSG00000062713.7, exon 2 of 11) | 16 | 94085260 | 94127032 | 41773 | 1 | ENSMUST00000072182.7 | 9002 | Sim2 | ENSMUSG00000062713.7 |
| chr16 | 94097228 | 94098421 | 3.60455 | 4.45866 | 1.44424 | Exon (ENSMUST00000072182.7/ENSMUSG00000062713.7, exon 3 of 11) | 16 | 94085260 | 94127032 | 41773 | 1 | ENSMUST00000072182.7 | 11969 | Sim2 | ENSMUSG00000062713.7 |
| chr16 | 94098967 | 94099176 | 3.36214 | 3.87075 | 1.01733 | Intron (ENSMUST00000072182.7/ENSMUSG00000062713.7, intron 3 of 10) | 16 | 94085260 | 94127032 | 41773 | 1 | ENSMUST00000072182.7 | 13708 | Sim2 | ENSMUSG00000062713.7 |
| chr16 | 94099781 | 94100131 | 3.48244 | 4.12833 | 1.24267 | Intron (ENSMUST00000072182.7/ENSMUSG00000062713.7, intron 3 of 10) | 16 | 94085260 | 94127032 | 41773 | 1 | ENSMUST00000072182.7 | 14522 | Sim2 | ENSMUSG00000062713.7 |
| chr16 | 94100993 | 94101136 | 3.58043 | 4.26822 | 1.24955 | Intron (ENSMUST00000072182.7/ENSMUSG00000062713.7, intron 3 of 10) | 16 | 94085260 | 94127032 | 41773 | 1 | ENSMUST00000072182.7 | 15734 | Sim2 | ENSMUSG00000062713.7 |
| chr16 | 94101537 | 94101806 | 3.68926 | 4.49801 | 1.44282 | Intron (ENSMUST00000072182.7/ENSMUSG00000062713.7, intron 3 of 10) | 16 | 94085260 | 94127032 | 41773 | 1 | ENSMUST00000072182.7 | 16278 | Sim2 | ENSMUSG00000062713.7 |
| chr16 | 94103050 | 94103351 | 4.00558 | 5.23074 | 1.91456 | Intron (ENSMUST00000072182.7/ENSMUSG00000062713.7, intron 3 of 10) | 16 | 94085260 | 94127032 | 41773 | 1 | ENSMUST00000072182.7 | 17791 | Sim2 | ENSMUSG00000062713.7 |
| chr16 | 94104904 | 94104979 | 3.65488 | 4.40341 | 1.33769 | Intron (ENSMUST00000072182.7/ENSMUSG00000062713.7, intron 4 of 10) | 16 | 94085260 | 94127032 | 41773 | 1 | ENSMUST00000072182.7 | 19645 | Sim2 | ENSMUSG00000062713.7 |
| chr16 | 94109384 | 94109491 | 3.91353 | 4.91918 | 1.68834 | Exon (ENSMUST00000072182.7/ENSMUSG00000062713.7, exon 6 of 11) | 16 | 94085260 | 94127032 | 41773 | 1 | ENSMUST00000072182.7 | 24125 | Sim2 | ENSMUSG00000062713.7 |
| chr16 | 94114916 | 94115409 | 3.58021 | 4.26724 | 1.25145 | Exon (ENSMUST00000072182.7/ENSMUSG00000062713.7, exon 7 of 11) | 16 | 94085260 | 94127032 | 41773 | 1 | ENSMUST00000072182.7 | 29657 | Sim2 | ENSMUSG00000062713.7 |
| chr16 | 94123292 | 94123888 | 3.82669 | 5.02326 | 1.83989 | Exon (ENSMUST00000072182.7/ENSMUSG00000062713.7, exon 10 of 11) | 16 | 94085260 | 94127032 | 41773 | 1 | ENSMUST00000072182.7 | 38033 | Sim2 | ENSMUSG00000062713.7 |
| chr16 | 94125458 | 94126133 | 4.12926 | 5.34207 | 2.00757 | 3' UTR | 16 | 94085260 | 94127032 | 41773 | 1 | ENSMUST00000072182.7 | 40199 | Sim2 | ENSMUSG00000062713.7 |
| chr13 | 22036191 | 22037323 | 3.95779 | 4.99127 | 1.73348 | Promoter | 13 | 22035870 | 22036345 | 476 | 1 | ENSMUST00000110455.3 | 322 | Hist1h2bk | ENSMUSG00000062727.4 |
| chr7 | 131598098 | 131598398 | 3.65249 | 4.41203 | 1.37351 | Intron (ENSMUST00000124096.2/ENSMUSG00000030849.12, intron 2 of 6) | 7 | 131614188 | 131614988 | 801 | 2 | ENSMUST00000085365.5 | 16590 | Gm16477 | ENSMUSG00000062758.6 |
| chr7 | 131609702 | 131609790 | 3.34153 | 3.83966 | 1.01733 | Intron (ENSMUST00000124096.2/ENSMUSG00000030849.12, intron 2 of 6) | 7 | 131614188 | 131614988 | 801 | 2 | ENSMUST00000085365.5 | 5198 | Gm16477 | ENSMUSG00000062758.6 |
| chr7 | 131644892 | 131645027 | 3.6025 | 4.44036 | 1.37684 | Intron (ENSMUST00000124096.2/ENSMUSG00000030849.12, intron 2 of 6) | 7 | 131614188 | 131614988 | 801 | 2 | ENSMUST00000085365.5 | -29905 | Gm16477 | ENSMUSG00000062758.6 |
| chr7 | 131645372 | 131645760 | 3.43191 | 4.04267 | 1.13656 | Intron (ENSMUST00000124096.2/ENSMUSG00000030849.12, intron 2 of 6) | 7 | 131614188 | 131614988 | 801 | 2 | ENSMUST00000085365.5 | -30385 | Gm16477 | ENSMUSG00000062758.6 |
| chr7 | 131721995 | 131722121 | 3.44543 | 4.02343 | 1.11288 | Intron (ENSMUST00000124096.2/ENSMUSG00000030849.12, intron 2 of 6) | 7 | 131614188 | 131614988 | 801 | 2 | ENSMUST00000085365.5 | -107008 | Gm16477 | ENSMUSG00000062758.6 |
| chr7 | 131722855 | 131723218 | 3.48198 | 4.08586 | 1.13835 | Intron (ENSMUST00000124096.2/ENSMUSG00000030849.12, intron 2 of 6) | 7 | 131614188 | 131614988 | 801 | 2 | ENSMUST00000085365.5 | -107868 | Gm16477 | ENSMUSG00000062758.6 |
| chr7 | 131744396 | 131744473 | 3.50631 | 4.13365 | 1.17751 | Intron (ENSMUST00000124096.2/ENSMUSG00000030849.12, intron 2 of 6) | 7 | 131614188 | 131614988 | 801 | 2 | ENSMUST00000085365.5 | -129409 | Gm16477 | ENSMUSG00000062758.6 |
| chr7 | 131769533 | 131769651 | 3.79152 | 4.75952 | 1.63968 | Intron (ENSMUST00000124096.2/ENSMUSG00000030849.12, intron 2 of 6) | 7 | 131614188 | 131614988 | 801 | 2 | ENSMUST00000085365.5 | -154546 | Gm16477 | ENSMUSG00000062758.6 |
| chr7 | 131774626 | 131775335 | 3.51932 | 4.16808 | 1.22405 | Intron (ENSMUST00000124096.2/ENSMUSG00000030849.12, intron 2 of 6) | 7 | 131614188 | 131614988 | 801 | 2 | ENSMUST00000085365.5 | -159639 | Gm16477 | ENSMUSG00000062758.6 |
| chr15 | 84420228 | 84420495 | 3.34153 | 3.83966 | 1.01733 | 5' UTR | 15 | 84417253 | 84443697 | 26445 | 2 | ENSMUST00000186527.1 | 23202 | 1810041L15Rik | ENSMUSG00000062760.5 |
| chr15 | 84445715 | 84446291 | 3.76949 | 4.62682 | 1.49437 | Promoter | 15 | 84380047 | 84446545 | 66499 | 2 | ENSMUST00000189248.1 | 254 | 1810041L15Rik | ENSMUSG00000062760.5 |
| chr15 | 84449039 | 84449169 | 3.68875 | 4.48262 | 1.42955 | Distal Intergenic | 15 | 84379203 | 84447097 | 67895 | 2 | ENSMUST00000080751.3 | -1943 | 1810041L15Rik | ENSMUSG00000062760.5 |
| chr7 | 44590622 | 44590740 | 3.43663 | 4.00032 | 1.07777 | Promoter | 7 | 44590664 | 44602141 | 11478 | 1 | ENSMUST00000071062.5 | 0 | Kcnc3 | ENSMUSG00000062785.8 |
| chr7 | 44591282 | 44591867 | 4.2336 | 5.73704 | 2.29279 | Promoter | 7 | 44590886 | 44604754 | 13869 | 1 | ENSMUST00000107906.3 | 397 | Kcnc3 | ENSMUSG00000062785.8 |
| chr7 | 44592399 | 44593982 | 3.71172 | 4.61534 | 1.49158 | Intron (ENSMUST00000071062.5/ENSMUSG00000062785.8, intron 1 of 4) | 7 | 44590886 | 44604754 | 13869 | 1 | ENSMUST00000107906.3 | 1514 | Kcnc3 | ENSMUSG00000062785.8 |
| chr7 | 44601277 | 44601545 | 3.68025 | 4.45279 | 1.36952 | Intron (ENSMUST00000071062.5/ENSMUSG00000062785.8, intron 4 of 4) | 7 | 44590886 | 44604754 | 13869 | 1 | ENSMUST00000107906.3 | 10392 | Kcnc3 | ENSMUSG00000062785.8 |
| chr7 | 44601919 | 44602184 | 4.04253 | 5.14982 | 1.83422 | 3' UTR | 7 | 44590886 | 44604754 | 13869 | 1 | ENSMUST00000107906.3 | 11034 | Kcnc3 | ENSMUSG00000062785.8 |
| chr17 | 28071242 | 28071791 | 3.47813 | 4.08343 | 1.15081 | Exon (ENSMUST00000042692.6/ENSMUSG00000062859.9, exon 5 of 10) | 17 | 28066797 | 28068476 | 1680 | 2 | ENSMUST00000123628.1 | -2767 | Tcp11 | ENSMUSG00000062859.9 |
| chr17 | 28080038 | 28080416 | 3.52862 | 4.23481 | 1.35516 | Promoter | 17 | 28066751 | 28080567 | 13817 | 2 | ENSMUST00000114836.2 | 151 | Tcp11 | ENSMUSG00000062859.9 |
| chr17 | 28106241 | 28106371 | 3.78672 | 4.65203 | 1.49177 | Distal Intergenic | 17 | 28069635 | 28080639 | 11005 | 2 | ENSMUST00000129046.2 | -25603 | Tcp11 | ENSMUSG00000062859.9 |
| chr10 | 13473588 | 13474199 | 3.30239 | 3.90192 | 1.1373 | Promoter | 10 | 13213395 | 13474396 | 261002 | 2 | ENSMUST00000105547.2 | 197 | Phactr2 | ENSMUSG00000062866.10 |
| chr5 | 75976000 | 75976378 | 3.34319 | 3.87805 | 1.13089 | Intron (ENSMUST00000113516.1/ENSMUSG00000062960.7, intron 1 of 29) | 5 | 75932827 | 75978458 | 45632 | 2 | ENSMUST00000113516.1 | 2080 | Kdr | ENSMUSG00000062960.7 |
| chr5 | 75977757 | 75978466 | 3.73082 | 4.71803 | 1.64043 | Promoter | 5 | 75932827 | 75978458 | 45632 | 2 | ENSMUST00000113516.1 | 0 | Kdr | ENSMUSG00000062960.7 |
| chr12 | 80758508 | 80760487 | 4.29625 | 5.7412 | 2.31254 | Promoter | 12 | 80754046 | 80760541 | 6496 | 2 | ENSMUST00000073251.6 | 54 | Ccdc177 | ENSMUSG00000062961.6 |
| chr12 | 80760888 | 80761175 | 3.92537 | 5.21124 | 1.90942 | Promoter | 12 | 80754046 | 80760541 | 6496 | 2 | ENSMUST00000073251.6 | -348 | Ccdc177 | ENSMUSG00000062961.6 |
| chr12 | 80761598 | 80761799 | 3.57807 | 4.26479 | 1.25831 | Distal Intergenic | 12 | 80754046 | 80760541 | 6496 | 2 | ENSMUST00000073251.6 | -1058 | Ccdc177 | ENSMUSG00000062961.6 |
| chr12 | 80762713 | 80763155 | 3.53663 | 4.19011 | 1.20831 | Distal Intergenic | 12 | 80754046 | 80760541 | 6496 | 2 | ENSMUST00000073251.6 | -2173 | Ccdc177 | ENSMUSG00000062961.6 |
| chr8 | 31917374 | 31918948 | 3.7956 | 4.77961 | 1.6759 | Promoter | 8 | 31818028 | 31918203 | 100176 | 2 | ENSMUST00000073884.4 | 0 | Nrg1 | ENSMUSG00000062991.6 |
| chr6 | 8778135 | 8778784 | 3.58873 | 4.28425 | 1.26495 | Promoter | 6 | 8630527 | 8778154 | 147628 | 2 | ENSMUST00000115519.2 | 0 | Ica1 | ENSMUSG00000062995.7 |
| chr14 | 63938631 | 63938706 | 3.39083 | 3.95752 | 1.14688 | Distal Intergenic | 14 | 63943674 | 63950732 | 7059 | 1 | ENSMUST00000079652.5 | -4968 | Sox7 | ENSMUSG00000063060.5 |
| chr14 | 63942959 | 63943270 | 3.66605 | 4.67569 | 1.56217 | Promoter | 14 | 63943674 | 63950732 | 7059 | 1 | ENSMUST00000079652.5 | -404 | Sox7 | ENSMUSG00000063060.5 |
| chr14 | 63943585 | 63944388 | 4.06145 | 5.2753 | 1.9926 | Promoter | 14 | 63943674 | 63950732 | 7059 | 1 | ENSMUST00000079652.5 | 0 | Sox7 | ENSMUSG00000063060.5 |
| chr14 | 63945551 | 63945831 | 3.66735 | 4.53559 | 1.48822 | Intron (ENSMUST00000079652.5/ENSMUSG00000063060.5, intron 1 of 1) | 14 | 63943674 | 63950732 | 7059 | 1 | ENSMUST00000079652.5 | 1878 | Sox7 | ENSMUSG00000063060.5 |
| chr6 | 77979482 | 77979770 | 3.58596 | 4.76157 | 1.63 | Promoter | 6 | 76881637 | 77979699 | 1098063 | 2 | ENSMUST00000160894.2 | 0 | Ctnna2 | ENSMUSG00000063063.7 |
| chr12 | 55536226 | 55536591 | 3.73327 | 4.57405 | 1.48409 | Distal Intergenic | 12 | 55565239 | 55566896 | 1658 | 1 | ENSMUST00000080123.1 | -28648 | Aldoart2 | ENSMUSG00000063129.1 |
| chr12 | 55540519 | 55540679 | 3.43151 | 4.12112 | 1.20877 | Distal Intergenic | 12 | 55565239 | 55566896 | 1658 | 1 | ENSMUST00000080123.1 | -24560 | Aldoart2 | ENSMUSG00000063129.1 |
| chr14 | 24002723 | 24002910 | 3.40778 | 4.27032 | 1.32315 | 3' UTR | 14 | 23299891 | 24003977 | 704087 | 2 | ENSMUST00000190985.1 | 1067 | Kcnma1 | ENSMUSG00000063142.8 |
| chr14 | 24003338 | 24003783 | 4.05632 | 5.3128 | 2.0107 | Promoter | 14 | 23299891 | 24003977 | 704087 | 2 | ENSMUST00000190985.1 | 194 | Kcnma1 | ENSMUSG00000063142.8 |
| chr14 | 24004104 | 24005081 | 3.62909 | 4.80907 | 1.65256 | Promoter | 14 | 23299381 | 24004205 | 704825 | 2 | ENSMUST00000188285.1 | 0 | Kcnma1 | ENSMUSG00000063142.8 |
| chr14 | 24005557 | 24005813 | 3.6832 | 4.60893 | 1.51589 | Promoter | 14 | 23299395 | 24004859 | 705465 | 2 | ENSMUST00000188210.1 | -699 | Kcnma1 | ENSMUSG00000063142.8 |
| chr2 | 69629844 | 69630091 | 3.34692 | 3.84779 | 1.01733 | Distal Intergenic | 2 | 69647171 | 69667318 | 20148 | 1 | ENSMUST00000134659.2 | -17080 | Bbs5 | ENSMUSG00000063145.5 |
| chr13 | 53507065 | 53507267 | 3.47353 | 4.07233 | 1.13549 | Distal Intergenic | 13 | 53525703 | 53526059 | 357 | 1 | ENSMUST00000081132.4 | -18436 | Gm5449 | ENSMUSG00000063166.4 |
| chr13 | 53508539 | 53508847 | 4.07211 | 5.30309 | 2.00608 | Distal Intergenic | 13 | 53525703 | 53526059 | 357 | 1 | ENSMUST00000081132.4 | -16856 | Gm5449 | ENSMUSG00000063166.4 |
| chr13 | 53607984 | 53608126 | 3.59153 | 4.34723 | 1.39184 | Distal Intergenic | 13 | 53525703 | 53526059 | 357 | 1 | ENSMUST00000081132.4 | 82282 | Gm5449 | ENSMUSG00000063166.4 |
| chr13 | 53681831 | 53682072 | 3.49936 | 4.4158 | 1.39888 | Distal Intergenic | 13 | 53525703 | 53526059 | 357 | 1 | ENSMUST00000081132.4 | 156129 | Gm5449 | ENSMUSG00000063166.4 |
| chr17 | 27428559 | 27428768 | 3.542 | 4.20088 | 1.22547 | Intron (ENSMUST00000118489.2/ENSMUSG00000063239.11, intron 8 of 8) | 17 | 27451629 | 27473067 | 21439 | 2 | ENSMUST00000147865.1 | 44299 | Grm4 | ENSMUSG00000063239.11 |
| chr17 | 27434692 | 27434901 | 4.13129 | 5.36647 | 2.05003 | Exon (ENSMUST00000118489.2/ENSMUSG00000063239.11, exon 8 of 9) | 17 | 27451629 | 27473067 | 21439 | 2 | ENSMUST00000147865.1 | 38166 | Grm4 | ENSMUSG00000063239.11 |
| chr17 | 27501273 | 27501405 | 3.57491 | 4.26547 | 1.27836 | Intron (ENSMUST00000118489.2/ENSMUSG00000063239.11, intron 1 of 8) | 17 | 27422387 | 27503304 | 80918 | 2 | ENSMUST00000118489.2 | 1899 | Grm4 | ENSMUSG00000063239.11 |
| chr17 | 27505436 | 27505680 | 3.73406 | 4.5593 | 1.44073 | Intron (ENSMUST00000118161.1/ENSMUSG00000063239.11, intron 1 of 10) | 17 | 27422387 | 27503304 | 80918 | 2 | ENSMUST00000118489.2 | -2133 | Grm4 | ENSMUSG00000063239.11 |
| chr17 | 27506025 | 27506262 | 3.36846 | 3.88029 | 1.01733 | Intron (ENSMUST00000118161.1/ENSMUSG00000063239.11, intron 1 of 10) | 17 | 27422387 | 27503304 | 80918 | 2 | ENSMUST00000118489.2 | -2722 | Grm4 | ENSMUSG00000063239.11 |
| chr17 | 27506648 | 27506947 | 4.19685 | 5.50545 | 2.11166 | Intron (ENSMUST00000118161.1/ENSMUSG00000063239.11, intron 1 of 10) | 17 | 27422387 | 27503304 | 80918 | 2 | ENSMUST00000118489.2 | -3345 | Grm4 | ENSMUSG00000063239.11 |
| chr17 | 27510683 | 27512793 | 3.94548 | 5.06168 | 1.82149 | Promoter | 17 | 27425904 | 27513341 | 87438 | 2 | ENSMUST00000118161.1 | 548 | Grm4 | ENSMUSG00000063239.11 |
| chr17 | 27513107 | 27514303 | 3.75234 | 4.60905 | 1.49488 | Promoter | 17 | 27425904 | 27513341 | 87438 | 2 | ENSMUST00000118161.1 | 0 | Grm4 | ENSMUSG00000063239.11 |
| chr17 | 27515329 | 27515675 | 3.40649 | 3.99966 | 1.19559 | Distal Intergenic | 17 | 27425904 | 27513341 | 87438 | 2 | ENSMUST00000118161.1 | -1989 | Grm4 | ENSMUSG00000063239.11 |
| chr4 | 146718600 | 146718817 | 2.93222 | 5.32581 | 1.9877 | Intron (ENSMUST00000181199.1/ENSMUSG00000096958.1, intron 2 of 3) | 4 | 146654927 | 146657607 | 2681 | 1 | ENSMUST00000070932.3 | 63674 | Gm13248 | ENSMUSG00000063245.6 |
| chr15 | 89841625 | 89841951 | 3.42087 | 4.00781 | 1.108 | Promoter | 15 | 89782393 | 89841860 | 59468 | 2 | ENSMUST00000029441.3 | 0 | Syt10 | ENSMUSG00000063260.2 |
| chr15 | 89842599 | 89842721 | 3.37969 | 3.89721 | 1.01733 | Promoter | 15 | 89782393 | 89841860 | 59468 | 2 | ENSMUST00000029441.3 | -740 | Syt10 | ENSMUSG00000063260.2 |
| chr14 | 7657990 | 7659179 | 4.33074 | 6.56397 | 2.98182 | Distal Intergenic | 14 | 7640895 | 7643972 | 3078 | 2 | ENSMUST00000171360.2 | -14019 | Gm10128 | ENSMUSG00000063277.7 |
| chr15 | 94629333 | 94629463 | 3.93747 | 4.98749 | 1.7912 | Promoter | 15 | 94629185 | 95096097 | 466913 | 1 | ENSMUST00000080141.4 | 149 | Tmem117 | ENSMUSG00000063296.4 |
| chr15 | 94651486 | 94651608 | 3.53933 | 4.23575 | 1.31929 | Intron (ENSMUST00000080141.4/ENSMUSG00000063296.4, intron 2 of 7) | 15 | 94629185 | 95096097 | 466913 | 1 | ENSMUST00000080141.4 | 22302 | Tmem117 | ENSMUSG00000063296.4 |
| chr7 | 54835562 | 54835637 | 3.30985 | 3.82132 | 1.10568 | Promoter | 7 | 54835959 | 55265366 | 429408 | 1 | ENSMUST00000082373.6 | -322 | Luzp2 | ENSMUSG00000063297.6 |
| chr3 | 130104187 | 130104614 | 3.51428 | 4.21167 | 1.31701 | Distal Intergenic | 3 | 130068412 | 130068978 | 567 | 1 | ENSMUST00000076703.6 | 35776 | Gm9396 | ENSMUSG00000063328.6 |
| chr5 | 123489867 | 123490193 | 3.33249 | 3.84132 | 1.06985 | Promoter | 5 | 123489327 | 123508198 | 18872 | 1 | ENSMUST00000121444.1 | 541 | Lrrc43 | ENSMUSG00000063409.7 |
| chr5 | 123497360 | 123497435 | 3.5402 | 4.19343 | 1.20088 | Intron (ENSMUST00000094327.4/ENSMUSG00000063409.7, intron 4 of 11) | 5 | 123489327 | 123508198 | 18872 | 1 | ENSMUST00000121444.1 | 8034 | Lrrc43 | ENSMUSG00000063409.7 |

| | | | | | | | | | | | | | | | |
|---|---|---|---|---|---|---|---|---|---|---|---|---|---|---|---|
| chr6 | 84406723 | 84406866 | 3.50753 | 4.32067 | 1.3251 | Distal Intergenic | 6 | 84571414 | 84593810 | 22397 | 2 | ENSMUST00000077705.3 | 186944 | Cyp26b1 | ENSMUSG00000063415.7 |
| chr6 | 84477129 | 84477352 | 3.39552 | 4.12214 | 1.18995 | Distal Intergenic | 6 | 84571414 | 84593810 | 22397 | 2 | ENSMUST00000077705.3 | 116458 | Cyp26b1 | ENSMUSG00000063415.7 |
| chr6 | 84552911 | 84553073 | 3.93838 | 5.03537 | 1.87305 | Distal Intergenic | 6 | 84571414 | 84593810 | 22397 | 2 | ENSMUST00000077705.3 | 40737 | Cyp26b1 | ENSMUSG00000063415.7 |
| chr6 | 84553865 | 84554000 | 3.44386 | 4.05201 | 1.21829 | Distal Intergenic | 6 | 84571414 | 84593810 | 22397 | 2 | ENSMUST00000077705.3 | 39810 | Cyp26b1 | ENSMUSG00000063415.7 |
| chr6 | 84573448 | 84573745 | 3.36562 | 3.88833 | 1.06062 | 3' UTR | 6 | 84571414 | 84593810 | 22397 | 2 | ENSMUST00000077705.3 | 20065 | Cyp26b1 | ENSMUSG00000063415.7 |
| chr6 | 84574685 | 84575978 | 3.37826 | 3.89506 | 1.01733 | Exon (ENSMUST00000077705.3/ENSMUSG00000063415.7, exon 6 of 7) | 6 | 84571414 | 84593810 | 22397 | 2 | ENSMUST00000077705.3 | 17832 | Cyp26b1 | ENSMUSG00000063415.7 |
| chr6 | 84583234 | 84584356 | 3.78157 | 4.66013 | 1.52965 | Exon (ENSMUST00000077705.3/ENSMUSG00000063415.7, exon 3 of 7) | 6 | 84571414 | 84593810 | 22397 | 2 | ENSMUST00000077705.3 | 9454 | Cyp26b1 | ENSMUSG00000063415.7 |
| chr6 | 84584679 | 84585648 | 3.80007 | 4.89232 | 1.68787 | Intron (ENSMUST00000077705.3/ENSMUSG00000063415.7, intron 2 of 6) | 6 | 84571414 | 84593810 | 22397 | 2 | ENSMUST00000077705.3 | 8162 | Cyp26b1 | ENSMUSG00000063415.7 |
| chr6 | 84586210 | 84586764 | 4.00698 | 5.16375 | 1.90034 | Intron (ENSMUST00000077705.3/ENSMUSG00000063415.7, intron 2 of 6) | 6 | 84571414 | 84593810 | 22397 | 2 | ENSMUST00000077705.3 | 7046 | Cyp26b1 | ENSMUSG00000063415.7 |
| chr6 | 84587099 | 84588058 | 3.70538 | 4.52701 | 1.46087 | 5' UTR | 6 | 84571414 | 84593810 | 22397 | 2 | ENSMUST00000077705.3 | 5752 | Cyp26b1 | ENSMUSG00000063415.7 |
| chr6 | 84588365 | 84589416 | 4.03523 | 5.30587 | 2.02893 | Intron (ENSMUST00000077705.3/ENSMUSG00000063415.7, intron 1 of 6) | 6 | 84571414 | 84593810 | 22397 | 2 | ENSMUST00000077705.3 | 4394 | Cyp26b1 | ENSMUSG00000063415.7 |
| chr6 | 84591302 | 84591395 | 3.82692 | 4.73198 | 1.54892 | Intron (ENSMUST00000077705.3/ENSMUSG00000063415.7, intron 1 of 6) | 6 | 84571414 | 84593810 | 22397 | 2 | ENSMUST00000077705.3 | 2415 | Cyp26b1 | ENSMUSG00000063415.7 |
| chr5 | 113461547 | 113462174 | 3.60205 | 4.34955 | 1.32804 | Distal Intergenic | 5 | 113490333 | 113493141 | 2809 | 1 | ENSMUST00000138757.2 | -28159 | Wscd2 | ENSMUSG00000063430.8 |
| chr5 | 113463791 | 113463945 | 4.09261 | 5.26077 | 1.93883 | Distal Intergenic | 5 | 113490333 | 113493141 | 2809 | 1 | ENSMUST00000138757.2 | -26388 | Wscd2 | ENSMUSG00000063430.8 |
| chr5 | 113465909 | 113466175 | 3.77241 | 4.65887 | 1.53627 | Distal Intergenic | 5 | 113490333 | 113493141 | 2809 | 1 | ENSMUST00000138757.2 | -24158 | Wscd2 | ENSMUSG00000063430.8 |
| chr5 | 113488501 | 113488616 | 3.35178 | 3.85511 | 1.01733 | Distal Intergenic | 5 | 113490333 | 113493141 | 2809 | 1 | ENSMUST00000138757.2 | -1717 | Wscd2 | ENSMUSG00000063430.8 |
| chr5 | 113490422 | 113491235 | 3.65145 | 4.45009 | 1.38702 | Promoter | 5 | 113490447 | 113589725 | 99279 | 1 | ENSMUST00000094452.3 | 0 | Wscd2 | ENSMUSG00000063430.8 |
| chr5 | 113503539 | 113503635 | 3.43472 | 4.00161 | 1.09241 | Intron (ENSMUST00000094452.3/ENSMUSG00000063430.8, intron 1 of 8) | 5 | 113490752 | 113554584 | 63833 | 1 | ENSMUST00000181650.1 | 12788 | Wscd2 | ENSMUSG00000063430.8 |
| chr5 | 113536975 | 113537607 | 3.62769 | 4.36187 | 1.32376 | Intron (ENSMUST00000094452.3/ENSMUSG00000063430.8, intron 1 of 8) | 5 | 113549985 | 113556509 | 6525 | 1 | ENSMUST00000180565.1 | -12378 | Wscd2 | ENSMUSG00000063430.8 |
| chr5 | 113551115 | 113551190 | 3.60827 | 4.33527 | 1.34734 | Exon (ENSMUST00000094452.3/ENSMUSG00000063430.8, exon 2 of 9) | 5 | 113549985 | 113556509 | 6525 | 1 | ENSMUST00000180565.1 | 1131 | Wscd2 | ENSMUSG00000063430.8 |
| chr5 | 113552011 | 113552325 | 3.29508 | 3.76979 | 1.01733 | Exon (ENSMUST00000181650.1/ENSMUSG00000063430.8, exon 2 of 2) | 5 | 113549985 | 113556509 | 6525 | 1 | ENSMUST00000180565.1 | 2027 | Wscd2 | ENSMUSG00000063430.8 |
| chr19 | 48204984 | 48205194 | 3.43733 | 4.01729 | 1.13453 | Promoter | 19 | 48206025 | 48805505 | 599481 | 1 | ENSMUST00000078880.5 | -831 | Sorcs3 | ENSMUSG00000063434.5 |
| chr19 | 48206201 | 48206811 | 4.25742 | 5.72876 | 2.34965 | Promoter | 19 | 48206025 | 48805505 | 599481 | 1 | ENSMUST00000078880.5 | 177 | Sorcs3 | ENSMUSG00000063434.5 |
| chr4 | 49058749 | 49059125 | 3.52373 | 4.18008 | 1.25007 | Promoter | 4 | 49059273 | 49208600 | 149328 | 1 | ENSMUST00000137884.1 | -148 | E130309F12Rik | ENSMUSG00000063446.4 |
| chr4 | 49059681 | 49060470 | 3.77946 | 4.76286 | 1.61591 | Promoter | 4 | 49059456 | 49302269 | 242814 | 1 | ENSMUST00000151692.1 | 226 | E130309F12Rik | ENSMUSG00000063446.4 |
| chr14 | 33214173 | 33214422 | 4.72112 | 6.85238 | 3.21409 | Promoter | 14 | 33214026 | 33217006 | 2981 | 1 | ENSMUST00000165792.1 | 148 | Arhgap22 | ENSMUSG00000063506.8 |
| chr14 | 33250288 | 33250594 | 3.83046 | 4.9967 | 1.76532 | Promoter | 14 | 33250390 | 33271983 | 21594 | 1 | ENSMUST00000131086.1 | 0 | Arhgap22 | ENSMUSG00000063506.8 |
| chr14 | 33251334 | 33251714 | 3.87572 | 4.92981 | 1.73181 | Promoter | 14 | 33250390 | 33271983 | 21594 | 1 | ENSMUST00000131086.1 | 945 | Arhgap22 | ENSMUSG00000063506.8 |
| chr14 | 33252512 | 33252760 | 3.65734 | 4.42107 | 1.37879 | Intron (ENSMUST00000111956.3/ENSMUSG00000063506.8, intron 2 of 10) | 14 | 33250390 | 33271983 | 21594 | 1 | ENSMUST00000131086.1 | 2123 | Arhgap22 | ENSMUSG00000063506.8 |
| chr14 | 33290251 | 33290532 | 3.52099 | 4.15996 | 1.19224 | Intron (ENSMUST00000111956.3/ENSMUSG00000063506.8, intron 3 of 10) | 14 | 33306118 | 33369932 | 63815 | 1 | ENSMUST00000137590.2 | -15586 | Arhgap22 | ENSMUSG00000063506.8 |
| chr14 | 33351495 | 33351695 | 4.07306 | 5.53296 | 2.18206 | Promoter | 14 | 33351312 | 33362575 | 11264 | 1 | ENSMUST00000140711.1 | 184 | Arhgap22 | ENSMUSG00000063506.8 |
| chr14 | 33352705 | 33352799 | 3.46444 | 4.08405 | 1.22065 | Intron (ENSMUST00000111956.3/ENSMUSG00000063506.8, intron 5 of 10) | 14 | 33351312 | 33362575 | 11264 | 1 | ENSMUST00000140711.1 | 1394 | Arhgap22 | ENSMUSG00000063506.8 |
| chr14 | 33362389 | 33362718 | 3.75698 | 4.80347 | 1.63456 | Promoter | 14 | 33362376 | 33363325 | 950 | 1 | ENSMUST00000132659.1 | 14 | Arhgap22 | ENSMUSG00000063506.8 |
| chr15 | 85521196 | 85521287 | 3.3637 | 3.9069 | 1.13198 | Distal Intergenic | 15 | 85510812 | 85511219 | 408 | 1 | ENSMUST00000079690.3 | 10385 | Gm4825 | ENSMUSG00000063548.3 |
| chr11 | 51389515 | 51390043 | 3.65793 | 4.41478 | 1.34887 | Intron (ENSMUST00000102765.3/ENSMUSG00000063564.8, intron 2 of 28) | 11 | 51289920 | 51583918 | 293999 | 1 | ENSMUST00000102765.3 | 99596 | Col23a1 | ENSMUSG00000063564.8 |
| chr11 | 51395029 | 51395207 | 3.3732 | 3.9153 | 1.1159 | Intron (ENSMUST00000102765.3/ENSMUSG00000063564.8, intron 2 of 28) | 11 | 51289920 | 51583918 | 293999 | 1 | ENSMUST00000102765.3 | 105110 | Col23a1 | ENSMUSG00000063564.8 |
| chr11 | 51429658 | 51429862 | 3.31791 | 3.81466 | 1.05415 | Intron (ENSMUST00000102765.3/ENSMUSG00000063564.8, intron 2 of 28) | 11 | 51289920 | 51583918 | 293999 | 1 | ENSMUST00000102765.3 | 139739 | Col23a1 | ENSMUSG00000063564.8 |
| chr11 | 51433071 | 51433146 | 3.85884 | 4.78731 | 1.58148 | Intron (ENSMUST00000102765.3/ENSMUSG00000063564.8, intron 2 of 28) | 11 | 51574432 | 51580299 | 5868 | 1 | ENSMUST00000151098.1 | -141286 | Col23a1 | ENSMUSG00000063564.8 |
| chr11 | 51469075 | 51469896 | 3.57768 | 4.26185 | 1.2442 | Intron (ENSMUST00000102765.3/ENSMUSG00000063564.8, intron 2 of 28) | 11 | 51574432 | 51580299 | 5868 | 1 | ENSMUST00000151098.1 | -104536 | Col23a1 | ENSMUSG00000063564.8 |
| chr11 | 51470998 | 51471346 | 3.56427 | 4.33449 | 1.36974 | Intron (ENSMUST00000102765.3/ENSMUSG00000063564.8, intron 2 of 28) | 11 | 51574432 | 51580299 | 5868 | 1 | ENSMUST00000151098.1 | -103086 | Col23a1 | ENSMUSG00000063564.8 |
| chr6 | 53231158 | 53231288 | 3.39595 | 4.0673 | 1.16761 | Distal Intergenic | 6 | 52768797 | 53068631 | 299835 | 2 | ENSMUST00000074541.5 | -162528 | Jazf1 | ENSMUSG00000063568.6 |
| chr1 | 92610430 | 92610809 | 3.68758 | 4.70272 | 1.56939 | Distal Intergenic | 1 | 92607839 | 92608847 | 1009 | 1 | ENSMUST00000079790.4 | 2592 | Olfr1410 | ENSMUSG00000063583.5 |
| chr19 | 14892303 | 14892403 | 4.69604 | 6.5579 | 2.9523 | Distal Intergenic | 19 | 14959549 | 14960766 | 1218 | 2 | ENSMUST00000072982.1 | 68363 | Gm5513 | ENSMUSG00000063586.1 |
| chr19 | 14892729 | 14893036 | 3.2728 | 3.77803 | 1.0485 | Distal Intergenic | 19 | 14959549 | 14960766 | 1218 | 2 | ENSMUST00000072982.1 | 67730 | Gm5513 | ENSMUSG00000063586.1 |
| chr7 | 16885393 | 16886276 | 4.10632 | 5.42533 | 2.15167 | Exon (ENSMUST00000108493.1/ENSMUSG00000078794.3, exon 4 of 4) | 7 | 16891786 | 16895435 | 3650 | 1 | ENSMUST00000078182.4 | -5510 | Gng8 | ENSMUSG00000063594.4 |
| chr3 | 29082124 | 29083060 | 3.96343 | 5.05948 | 1.80977 | Promoter | 3 | 29082300 | 29582912 | 500613 | 1 | ENSMUST00000146943.2 | 0 | Egfem1 | ENSMUSG00000063600.9 |
| chr3 | 29083608 | 29083683 | 3.69396 | 4.47465 | 1.3726 | Promoter | 3 | 29082771 | 29690510 | 607740 | 1 | ENSMUST00000118531.2 | 838 | Egfem1 | ENSMUSG00000063600.9 |
| chr3 | 29084177 | 29084277 | 3.7611 | 4.61051 | 1.47305 | Intron (ENSMUST00000124809.2/ENSMUSG00000063600.9, intron 1 of 3) | 3 | 29082771 | 29690510 | 607740 | 1 | ENSMUST00000118531.2 | 1407 | Egfem1 | ENSMUSG00000063600.9 |
| chr8 | 94899257 | 94899332 | 4.26861 | 5.87891 | 2.37684 | Distal Intergenic | 8 | 94913565 | 94917521 | 3957 | 2 | ENSMUST00000162020.1 | 18189 | Ccdc102a | ENSMUSG00000063605.5 |
| chr8 | 94913046 | 94913463 | 3.56559 | 4.37025 | 1.38509 | Exon (ENSMUST00000077955.5/ENSMUSG00000063605.5, exon 2 of 9) | 8 | 94913565 | 94917521 | 3957 | 2 | ENSMUST00000162020.1 | 4058 | Ccdc102a | ENSMUSG00000063605.5 |
| chr8 | 94917969 | 94918173 | 3.29439 | 3.95722 | 1.08849 | Promoter | 8 | 94902869 | 94918098 | 15230 | 2 | ENSMUST00000077955.5 | 0 | Ccdc102a | ENSMUSG00000063605.5 |
| chr8 | 29217332 | 29217952 | 3.3157 | 3.85369 | 1.08609 | Intron (ENSMUST00000168630.2/ENSMUSG00000063626.5, intron 1 of 17) | 8 | 28646717 | 29219338 | 572622 | 2 | ENSMUST00000168630.2 | 1386 | Unc5d | ENSMUSG00000063626.5 |
| chr8 | 29218643 | 29219735 | 3.75412 | 4.64221 | 1.56449 | Promoter | 8 | 28646717 | 29219338 | 572622 | 2 | ENSMUST00000168630.2 | 0 | Unc5d | ENSMUSG00000063626.5 |
| chr8 | 29220295 | 29220472 | 3.25346 | 3.70736 | 1.01733 | Promoter | 8 | 28646717 | 29219338 | 572622 | 2 | ENSMUST00000168630.2 | -958 | Unc5d | ENSMUSG00000063626.5 |
| chr12 | 27338565 | 27339555 | 4.02035 | 5.2809 | 1.99886 | 3' UTR | 12 | 27334264 | 27342574 | 8311 | 2 | ENSMUST00000079063.6 | 3019 | Sox11 | ENSMUSG00000063632.6 |
| chr12 | 27340293 | 27343515 | 6.05343 | 11.15454 | 6.99148 | Promoter | 12 | 27334264 | 27342574 | 8311 | 2 | ENSMUST00000079063.6 | 0 | Sox11 | ENSMUSG00000063632.6 |
| chr5 | 37015729 | 37016043 | 3.51595 | 4.50815 | 1.43454 | Distal Intergenic | 5 | 37028335 | 37091447 | 63113 | 1 | ENSMUST00000173836.2 | -12292 | Jakmip1 | ENSMUSG00000063646.8 |
| chr5 | 37019952 | 37020138 | 3.66477 | 4.42916 | 1.36014 | Distal Intergenic | 5 | 37028335 | 37091447 | 63113 | 1 | ENSMUST00000173836.2 | -8197 | Jakmip1 | ENSMUSG00000063646.8 |
| chr5 | 37023814 | 37024504 | 3.63429 | 4.36381 | 1.30623 | Distal Intergenic | 5 | 37028335 | 37091447 | 63113 | 1 | ENSMUST00000173836.2 | -3831 | Jakmip1 | ENSMUSG00000063646.8 |
| chr5 | 37025758 | 37025845 | 4.04468 | 5.1914 | 1.93896 | Distal Intergenic | 5 | 37028335 | 37091447 | 63113 | 1 | ENSMUST00000173836.2 | -2490 | Jakmip1 | ENSMUSG00000063646.8 |
| chr5 | 37026208 | 37026350 | 3.51443 | 4.16516 | 1.24712 | Distal Intergenic | 5 | 37028335 | 37091447 | 63113 | 1 | ENSMUST00000173836.2 | -1985 | Jakmip1 | ENSMUSG00000063646.8 |
| chr5 | 37027792 | 37029720 | 3.83681 | 4.86461 | 1.67524 | Promoter | 5 | 37028335 | 37091447 | 63113 | 1 | ENSMUST00000173836.2 | 0 | Jakmip1 | ENSMUSG00000063646.8 |
| chr5 | 37030845 | 37031423 | 4.3259 | 4.0715 | 1.19256 | Promoter | 5 | 37029887 | 37091242 | 61356 | 1 | ENSMUST00000174097.2 | 959 | Jakmip1 | ENSMUSG00000063646.8 |
| chr5 | 37076544 | 37076850 | 3.70756 | 4.50418 | 1.41197 | Intron (ENSMUST00000173836.2/ENSMUSG00000063646.8, intron 1 of 2) | 5 | 37085370 | 37148001 | 62632 | 1 | ENSMUST00000121010.3 | -8520 | Jakmip1 | ENSMUSG00000063646.8 |
| chr5 | 37077911 | 37078035 | 3.37057 | 3.88346 | 1.01733 | Intron (ENSMUST00000173836.2/ENSMUSG00000063646.8, intron 1 of 2) | 5 | 37085370 | 37148001 | 62632 | 1 | ENSMUST00000121010.3 | -7335 | Jakmip1 | ENSMUSG00000063646.8 |
| chr7 | 25368367 | 25368949 | 3.67187 | 4.43221 | 1.34561 | Promoter | 7 | 25367616 | 25369724 | 2109 | 2 | ENSMUST00000076276.3 | 775 | Cnfn | ENSMUSG00000063651.6 |
| chr7 | 25369863 | 25370211 | 4.2192 | 5.59899 | 2.18748 | Promoter | 7 | 25367616 | 25369724 | 2109 | 2 | ENSMUST00000076276.3 | -140 | Cnfn | ENSMUSG00000063651.6 |
| chr8 | 23157593 | 23158284 | 3.42417 | 4.03379 | 1.18968 | 3' UTR | 8 | 23153269 | 23158948 | 5680 | 1 | ENSMUST00000071588.7 | 4325 | Nkx6-3 | ENSMUSG00000063672.7 |
| chr8 | 23158635 | 23158710 | 3.83142 | 4.76021 | 1.60561 | 3' UTR | 8 | 23153269 | 23158948 | 5680 | 1 | ENSMUST00000071588.7 | 5367 | Nkx6-3 | ENSMUSG00000063672.7 |
| chr8 | 23159031 | 23159160 | 3.40403 | 3.95861 | 1.1037 | Distal Intergenic | 8 | 23153269 | 23158948 | 5680 | 1 | ENSMUST00000071588.7 | 5763 | Nkx6-3 | ENSMUSG00000063672.7 |
| chr1 | 139379392 | 139379467 | 3.37409 | 3.88875 | 1.01733 | Distal Intergenic | 1 | 139231216 | 139377100 | 145885 | 2 | ENSMUST00000145244.2 | -2293 | Crb1 | ENSMUSG00000063681.9 |
| chr15 | 75993030 | 75993251 | 3.53783 | 4.35241 | 1.37314 | Promoter | 15 | 75993769 | 75995824 | 2056 | 1 | ENSMUST00000161961.1 | -518 | Mapk15 | ENSMUSG00000063704.7 |
| chr15 | 75993873 | 75994062 | 3.36424 | 4.22741 | 1.27732 | Promoter | 15 | 75993769 | 75999152 | 5384 | 1 | ENSMUST00000089669.4 | 105 | Mapk15 | ENSMUSG00000063704.7 |
| chr15 | 75996332 | 75997056 | 3.49484 | 4.25488 | 1.33358 | Promoter | 15 | 75996713 | 75999152 | 2440 | 1 | ENSMUST00000160637.1 | 0 | Mapk15 | ENSMUSG00000063704.7 |
| chr15 | 75998263 | 76000004 | 3.8552 | 4.96307 | 1.78023 | 3' UTR | 15 | 75996713 | 75999152 | 2440 | 1 | ENSMUST00000160637.1 | 1551 | Mapk15 | ENSMUSG00000063704.7 |
| chr15 | 76000476 | 76000715 | 3.31871 | 3.95033 | 1.07565 | Downstream (<1kb) | 15 | 75996713 | 75999152 | 2440 | 1 | ENSMUST00000160637.1 | 3764 | Mapk15 | ENSMUSG00000063704.7 |
| chr15 | 76001066 | 76001552 | 3.92719 | 5.26447 | 1.95263 | 3' UTR | 15 | 75996713 | 75999152 | 2440 | 1 | ENSMUST00000160637.1 | 4354 | Mapk15 | ENSMUSG00000063704.7 |
| chr15 | 76002475 | 76003704 | 3.8763 | 4.88713 | 1.73796 | Exon (ENSMUST00000170153.1/ENSMUSG00000046761.7, exon 5 of 5) | 15 | 75996713 | 75999152 | 2440 | 1 | ENSMUST00000160637.1 | 5763 | Mapk15 | ENSMUSG00000063704.7 |
| chr15 | 81691542 | 81692276 | 3.64625 | 4.40196 | 1.36875 | 3' UTR | 15 | 81678296 | 81694028 | 15733 | 2 | ENSMUST00000173598.2 | 1752 | Chadl | ENSMUSG00000063765.6 |
| chr15 | 81692928 | 81693731 | 3.75274 | 4.62731 | 1.50904 | Promoter | 15 | 81678296 | 81694028 | 15733 | 2 | ENSMUST00000173598.2 | 297 | Chadl | ENSMUSG00000063765.6 |
| chr15 | 81694827 | 81695253 | 3.79707 | 4.89248 | 1.74351 | Promoter | 15 | 81686181 | 81694207 | 8027 | 2 | ENSMUST00000173898.1 | -621 | Chadl | ENSMUSG00000063765.6 |
| chr15 | 98424081 | 98424559 | 3.48991 | 4.15799 | 1.30611 | Distal Intergenic | 15 | 98437470 | 98438397 | 928 | 1 | ENSMUST00000075851.3 | -12911 | Olfr282 | ENSMUSG00000063780.4 |
| chr10 | 45464995 | 45465678 | 3.43343 | 4.10363 | 1.24023 | Intron (ENSMUST00000079390.6/ENSMUSG00000063804.7, intron 2 of 4) | 10 | 45376620 | 45470201 | 93582 | 2 | ENSMUST00000079390.6 | 4523 | Lin28b | ENSMUSG00000063804.7 |
| chr10 | 45466348 | 45466863 | 3.44029 | 4.01288 | 1.10276 | Intron (ENSMUST00000079390.6/ENSMUSG00000063804.7, intron 2 of 4) | 10 | 45376620 | 45470201 | 93582 | 2 | ENSMUST00000079390.6 | 3338 | Lin28b | ENSMUSG00000063804.7 |
| chr10 | 45469026 | 45470241 | 3.93183 | 5.01334 | 1.80881 | Promoter | 10 | 45376620 | 45470201 | 93582 | 2 | ENSMUST00000079390.6 | 0 | Lin28b | ENSMUSG00000063804.7 |
| chr10 | 45470549 | 45470825 | 3.31104 | 3.79777 | 1.03138 | Promoter | 10 | 45376620 | 45470201 | 93582 | 2 | ENSMUST00000079390.6 | -349 | Lin28b | ENSMUSG00000063804.7 |
| chr10 | 45471164 | 45471721 | 3.71951 | 4.54684 | 1.47894 | Promoter | 10 | 45376620 | 45470201 | 93582 | 2 | ENSMUST00000079390.6 | -964 | Lin28b | ENSMUSG00000063804.7 |
| chr10 | 45472156 | 45473997 | 3.84748 | 5.00105 | 1.81306 | Distal Intergenic | 10 | 45376620 | 45470201 | 93582 | 2 | ENSMUST00000079390.6 | -1956 | Lin28b | ENSMUSG00000063804.7 |
| chr10 | 45474360 | 45474152 | 3.77029 | 4.9522 | 1.7759 | Distal Intergenic | 10 | 45376620 | 45470201 | 93582 | 2 | ENSMUST00000079390.6 | -4160 | Lin28b | ENSMUSG00000063804.7 |
| chr10 | 45475507 | 45475582 | 3.99338 | 5.10229 | 1.85877 | Distal Intergenic | 10 | 45376620 | 45470201 | 93582 | 2 | ENSMUST00000079390.6 | -5307 | Lin28b | ENSMUSG00000063804.7 |
| chr10 | 45475975 | 45476324 | 3.49667 | 4.14001 | 1.22821 | Distal Intergenic | 10 | 45376620 | 45470201 | 93582 | 2 | ENSMUST00000079390.6 | -5775 | Lin28b | ENSMUSG00000063804.7 |
| chr10 | 45478631 | 45478826 | 3.29099 | 3.80595 | 1.0279 | Distal Intergenic | 10 | 45376620 | 45470201 | 93582 | 2 | ENSMUST00000079390.6 | -8431 | Lin28b | ENSMUSG00000063804.7 |
| chr10 | 45479580 | 45480608 | 3.44832 | 4.05184 | 1.18617 | Distal Intergenic | 10 | 45376620 | 45470201 | 93582 | 2 | ENSMUST00000079390.6 | -9380 | Lin28b | ENSMUSG00000063804.7 |
| chr10 | 45481162 | 45481805 | 3.67902 | 4.48202 | 1.47449 | Distal Intergenic | 10 | 45376620 | 45470201 | 93582 | 2 | ENSMUST00000079390.6 | -10962 | Lin28b | ENSMUSG00000063804.7 |
| chr10 | 45484642 | 45487430 | 3.81618 | 5.01148 | 1.79513 | Distal Intergenic | 10 | 45376620 | 45470201 | 93582 | 2 | ENSMUST00000079390.6 | -14442 | Lin28b | ENSMUSG00000063804.7 |
| chr10 | 45489034 | 45489198 | 3.68959 | 4.81638 | 1.64652 | Distal Intergenic | 10 | 45376620 | 45470201 | 93582 | 2 | ENSMUST00000079390.6 | -18834 | Lin28b | ENSMUSG00000063804.7 |
| chr10 | 45489747 | 45490173 | 3.96019 | 5.08433 | 1.86219 | Distal Intergenic | 10 | 45376620 | 45470201 | 93582 | 2 | ENSMUST00000079390.6 | -19547 | Lin28b | ENSMUSG00000063804.7 |
| chr14 | 21686220 | 21687056 | 3.62238 | 4.40914 | 1.43587 | Exon (ENSMUST00000073870.5/ENSMUSG00000063821.5, exon 3 of 4) | 14 | 21676582 | 21714576 | 37995 | 2 | ENSMUST00000073870.5 | 27520 | Dupd1 | ENSMUSG00000063821.5 |
| chr2 | 145168564 | 145168965 | 3.46732 | 4.14932 | 1.26931 | Promoter | 2 | 145167919 | 145518947 | 351029 | 1 | ENSMUST00000153249.2 | 646 | Slc24a3 | ENSMUSG00000063873.5 |
| chr2 | 145169290 | 145170046 | 4.17095 | 5.50556 | 2.22051 | Intron (ENSMUST00000110007.2/ENSMUSG00000063873.5, intron 1 of 16) | 2 | 145167919 | 145518947 | 351029 | 1 | ENSMUST00000153249.2 | 1372 | Slc24a3 | ENSMUSG00000063873.5 |
| chr3 | 26151884 | 26151959 | 4.27482 | 5.69936 | 2.32961 | Distal Intergenic | 3 | 25431811 | 26133734 | 701924 | 2 | ENSMUST00000108308.4 | -18151 | Nlgn1 | ENSMUSG00000063887.8 |
| chr3 | 26152482 | 26153079 | 3.51614 | 4.19666 | 1.31649 | Distal Intergenic | 3 | 25431811 | 26133734 | 701924 | 2 | ENSMUST00000108308.4 | -18749 | Nlgn1 | ENSMUSG00000063887.8 |
| chr3 | 26153522 | 26153608 | 4.12078 | 5.38829 | 2.07086 | Distal Intergenic | 3 | 25431811 | 26133734 | 701924 | 2 | ENSMUST00000108308.4 | -19789 | Nlgn1 | ENSMUSG00000063887.8 |
| chr5 | 116569775 | 116569962 | 3.37789 | 3.89451 | 1.01733 | Intron (ENSMUST00000076124.5/ENSMUSG00000063919.7, intron 1 of 12) | 5 | 116439272 | 116591817 | 152546 | 2 | ENSMUST00000076124.5 | 21855 | Srrm4 | ENSMUSG00000063919.7 |
| chr5 | 116570408 | 116570483 | 3.86594 | 4.81251 | 1.6219 | Intron (ENSMUST00000076124.5/ENSMUSG00000063919.7, intron 1 of 12) | 5 | 116439272 | 116591817 | 152546 | 2 | ENSMUST00000076124.5 | 21334 | Srrm4 | ENSMUSG00000063919.7 |
| chr5 | 116571013 | 116571840 | 3.5785 | 4.32785 | 1.31563 | Intron (ENSMUST00000076124.5/ENSMUSG00000063919.7, intron 1 of 12) | 5 | 116439272 | 116591817 | 152546 | 2 | ENSMUST00000076124.5 | 19977 | Srrm4 | ENSMUSG00000063919.7 |
| chr5 | 116587955 | 116588179 | 3.69383 | 4.47176 | 1.36709 | Intron (ENSMUST00000076124.5/ENSMUSG00000063919.7, intron 1 of 12) | 5 | 116439272 | 116591817 | 152546 | 2 | ENSMUST00000076124.5 | 3638 | Srrm4 | ENSMUSG00000063919.7 |
| chr5 | 116588587 | 116589021 | 3.55116 | 4.24378 | 1.27757 | Intron (ENSMUST00000076124.5/ENSMUSG00000063919.7, intron 1 of 12) | 5 | 116439272 | 116591817 | 152546 | 2 | ENSMUST00000076124.5 | 2796 | Srrm4 | ENSMUSG00000063919.7 |
| chr5 | 116589799 | 116590002 | 3.67666 | 4.44188 | 1.35407 | Intron (ENSMUST00000076124.5/ENSMUSG00000063919.7, intron 1 of 12) | 5 | 116439272 | 116591817 | 152546 | 2 | ENSMUST00000076124.5 | 1815 | Srrm4 | ENSMUSG00000063919.7 |
| chr5 | 116591273 | 116592453 | 4.24891 | 5.66191 | 2.24322 | Promoter | 5 | 116439272 | 116591817 | 152546 | 2 | ENSMUST00000076124.5 | 0 | Srrm4 | ENSMUSG00000063919.7 |
| chr5 | 116593838 | 116594132 | 4.26314 | 5.70644 | 2.24103 | Distal Intergenic | 5 | 116439272 | 116591817 | 152546 | 2 | ENSMUST00000076124.5 | -2022 | Srrm4 | ENSMUSG00000063919.7 |
| chr5 | 116596720 | 116596859 | 3.37744 | 3.89383 | 1.01733 | Distal Intergenic | 5 | 116439272 | 116591817 | 152546 | 2 | ENSMUST00000076124.5 | -4904 | Srrm4 | ENSMUSG00000063919.7 |
| chr5 | 72578424 | 72579686 | 3.7363 | 4.61176 | 1.50972 | Exon (ENSMUST00000073528.2/ENSMUSG00000063935.2, exon 2 of 4) | 5 | 72577114 | 72581084 | 3971 | 2 | ENSMUST00000073528.2 | 1398 | Zar1 | ENSMUSG00000063935.2 |
| chr5 | 72580018 | 72581144 | 4.66794 | 7.01205 | 3.3956 | Promoter | 5 | 72577114 | 72581084 | 3971 | 2 | ENSMUST00000073528.2 | 0 | Zar1 | ENSMUSG00000063935.2 |
| chr7 | 43796313 | 43796530 | 3.37977 | 3.89734 | 1.01733 | 3' UTR | 7 | 43797577 | 43803822 | 6246 | 1 | ENSMUST00000085461.2 | -1047 | Klk8 | ENSMUSG00000064023.3 |
| chr18 | 73865251 | 73865404 | 4.50268 | 6.15952 | 2.64298 | Intron (ENSMUST00000120033.2/ENSMUSG00000064036.10, intron 3 of 8) | 18 | 73863672 | 73871980 | 8309 | 1 | ENSMUST00000134847.1 | 1580 | Mro | ENSMUSG00000064036.10 |
| chr14 | 122426487 | 122426667 | 3.60798 | 4.33632 | 1.33907 | Distal Intergenic | 14 | 122434854 | 122451115 | 16262 | 2 | ENSMUST00000081580.2 | 24448 | Gm5089 | ENSMUSG00000064052.2 |
| chr14 | 122427062 | 122427252 | 3.43586 | 4.01171 | 1.12261 | Distal Intergenic | 14 | 122434854 | 122451115 | 16262 | 2 | ENSMUST00000081580.2 | 23863 | Gm5089 | ENSMUSG00000064052.2 |
| chr14 | 122429207 | 122429358 | 3.9242 | 4.92539 | 1.71302 | Distal Intergenic | 14 | 122434854 | 122451115 | 16262 | 2 | ENSMUST00000081580.2 | 21757 | Gm5089 | ENSMUSG00000064052.2 |
| chr14 | 122434279 | 122434354 | 3.34676 | 3.84754 | 1.01733 | Downstream (<1kb) | 14 | 122434854 | 122451115 | 16262 | 2 | ENSMUST00000081580.2 | 16761 | Gm5089 | ENSMUSG00000064052.2 |
| chr14 | 122450063 | 122450274 | 4.20022 | 5.55319 | 2.20193 | Promoter | 14 | 122434854 | 122451115 | 16262 | 2 | ENSMUST00000081580.2 | 841 | Gm5089 | ENSMUSG00000064052.2 |
| chr14 | 122450584 | 122451347 | 4.09165 | 5.35432 | 2.0812 | Promoter | 14 | 122434854 | 122451115 | 16262 | 2 | ENSMUST00000081580.2 | 0 | Gm5089 | ENSMUSG00000064052.2 |
| chr14 | 122452152 | 122452280 | 3.29121 | 3.76397 | 1.01733 | Distal Intergenic | 14 | 122434854 | 122451115 | 16262 | 2 | ENSMUST00000081580.2 | -1038 | Gm5089 | ENSMUSG00000064052.2 |
| chr14 | 122453089 | 122454063 | 3.79342 | 5.08433 | 1.83052 | Downstream (2-3kb) | 14 | 122434854 | 122451115 | 16262 | 2 | ENSMUST00000081580.2 | -1975 | Gm5089 | ENSMUSG00000064052.2 |
| chr14 | 122454416 | 122454743 | 3.69876 | 4.68689 | 1.57962 | Downstream (2-3kb) | 14 | 122434854 | 122451115 | 16262 | 2 | ENSMUST00000081580.2 | -3302 | Gm5089 | ENSMUSG00000064052.2 |
| chr14 | 122455151 | 122455613 | 4.58009 | 6.4801 | 2.89488 | Downstream (1-2kb) | 14 | 122434854 | 122451115 | 16262 | 2 | ENSMUST00000081580.2 | -4037 | Gm5089 | ENSMUSG00000064052.2 |
| chr6 | 91213255 | 91213434 | 3.36505 | 3.99245 | 1.17689 | Promoter | 6 | 91212831 | 91233304 | 20474 | 1 | ENSMUST00000134974.1 | 425 | Fbln2 | ENSMUSG00000064080.7 |
| chr6 | 91214125 | 91214451 | 3.49203 | 4.1061 | 1.15576 | Intron (ENSMUST00000113498.3/ENSMUSG00000064080.7, intron 1 of 16) | 6 | 91212831 | 91233304 | 20474 | 1 | ENSMUST00000134974.1 | 1295 | Fbln2 | ENSMUSG00000064080.7 |
| chr6 | 91284867 | 91285416 | 3.8489 | 4.79652 | 1.62315 | Distal Intergenic | 6 | 91265879 | 91272431 | 6553 | 1 | ENSMUST00000137029.1 | 18989 | Fbln2 | ENSMUSG00000064080.7 |
| chr6 | 91285772 | 91286944 | 3.67409 | 4.57206 | 1.48579 | Distal Intergenic | 6 | 91265879 | 91272431 | 6553 | 1 | ENSMUST00000137029.1 | 19894 | Fbln2 | ENSMUSG00000064080.7 |
| chr16 | 67613931 | 67614251 | 3.78595 | 4.7017 | 1.57607 | Intron (ENSMUST00000114292.2/ENSMUSG00000064115.8, intron 1 of 9) | 16 | 66784789 | 67620493 | 835705 | 2 | ENSMUST00000123266.1 | 6242 | Cadm2 | ENSMUSG00000064115.8 |
| chr16 | 67615684 | 67616233 | 3.45275 | 4.41769 | 1.3934 | Intron (ENSMUST00000114292.2/ENSMUSG00000064115.8, intron 1 of 9) | 16 | 66784789 | 67620493 | 835705 | 2 | ENSMUST00000123266.1 | 4260 | Cadm2 | ENSMUSG00000064115.8 |
| chr16 | 67616749 | 67617209 | 3.69716 | 4.51217 | 1.46969 | Intron (ENSMUST00000114292.2/ENSMUSG00000064115.8, intron 1 of 9) | 16 | 66784789 | 67620493 | 835705 | 2 | ENSMUST00000123266.1 | 3284 | Cadm2 | ENSMUSG00000064115.8 |
| chr16 | 67617868 | 67618529 | 3.46275 | 4.0566 | 1.13024 | Intron (ENSMUST00000114292.2/ENSMUSG00000064115.8, intron 1 of 9) | 16 | 66784789 | 67620493 | 835705 | 2 | ENSMUST00000123266.1 | 1964 | Cadm2 | ENSMUSG00000064115.8 |
| chr16 | 67618842 | 67619289 | 4.57437 | 6.36164 | 2.74721 | Intron (ENSMUST00000114292.2/ENSMUSG00000064115.8, intron 1 of 9) | 16 | 66784789 | 67620493 | 835705 | 2 | ENSMUST00000123266.1 | 1204 | Cadm2 | ENSMUSG00000064115.8 |
| chr16 | 67619762 | 67620756 | 3.46436 | 4.08343 | 1.17636 | Promoter | 16 | 66784789 | 67620493 | 835705 | 2 | ENSMUST00000123266.1 | 0 | Cadm2 | ENSMUSG00000064115.8 |
| chr8 | 4215137 | 4215399 | 3.47719 | 4.12884 | 1.28514 | Exon (ENSMUST00000168386.3/ENSMUSG00000064125.9, exon 5 of 7) | 8 | 4215049 | 4216912 | 1864 | 2 | ENSMUST00000177491.2 | 1513 | BC068157 | ENSMUSG00000064125.9 |
| chr8 | 4215879 | 4216773 | 3.7849 | 4.66432 | 1.53239 | Promoter | 8 | 4215049 | 4216912 | 1864 | 2 | ENSMUST00000177491.2 | 139 | BC068157 | ENSMUSG00000064125.9 |
| chr8 | 4217151 | 4217690 | 4.14129 | 5.36469 | 2.00991 | Promoter | 8 | 4209543 | 4217312 | 7770 | 2 | ENSMUST00000168386.3 | 0 | BC068157 | ENSMUSG00000064125.9 |
| chr7 | 45623674 | 45623757 | 5.23906 | 8.09827 | 4.13976 | Promoter | 7 | 45624393 | 45628823 | 4431 | 1 | ENSMUST00000130327.1 | -636 | Izumo1 | ENSMUSG00000064158.4 |
| chr15 | 101812692 | 101812967 | 3.31092 | 3.79377 | 1.01733 | Intron (ENSMUST00000023712.7/ENSMUSG00000064201.8, intron 7 of 8) | 15 | 101810689 | 101818169 | 7481 | 2 | ENSMUST00000023712.7 | 5202 | Krt2 | ENSMUSG00000064201.8 |
| chr9 | 95549409 | 95549690 | 3.43937 | 4.03833 | 1.19158 | Distal Intergenic | 9 | 95559657 | 95568023 | 8367 | 1 | ENSMUST00000079597.6 | -9967 | Paqr9 | ENSMUSG00000064225.6 |
| chr9 | 95558256 | 95560453 | 3.84062 | 4.86008 | 1.67547 | Promoter | 9 | 95559657 | 95568023 | 8367 | 1 | ENSMUST00000079597.6 | 0 | Paqr9 | ENSMUSG00000064225.6 |
| chr9 | 95560784 | 95561397 | 3.73408 | 4.55495 | 1.43624 | 3' UTR | 9 | 95559657 | 95568023 | 8367 | 1 | ENSMUST00000079597.6 | 1128 | Paqr9 | ENSMUSG00000064225.6 |
| chr9 | 95562979 | 95563285 | 3.94211 | 4.96171 | 1.72243 | 3' UTR | 9 | 95559657 | 95568023 | 8367 | 1 | ENSMUST00000079597.6 | 3323 | Paqr9 | ENSMUSG00000064225.6 |
| chr9 | 95565476 | 95565633 | 4.21212 | 5.54522 | 2.11708 | 3' UTR | 9 | 95559657 | 95568023 | 8367 | 1 | ENSMUST00000079597.6 | 5820 | Paqr9 | ENSMUSG00000064225.6 |
| chr9 | 95581026 | 95581282 | 3.62097 | 4.418 | 1.43405 | Distal Intergenic | 9 | 95559657 | 95568023 | 8367 | 1 | ENSMUST00000079597.6 | 21370 | Paqr9 | ENSMUSG00000064225.6 |
| chr7 | 24585953 | 24586088 | 3.44781 | 4.08203 | 1.2274 | Exon (ENSMUST00000094705.2/ENSMUSG00000066721.3, exon 4 of 4) | 7 | 24587543 | 24608925 | 21383 | 1 | ENSMUST00000077191.6 | -1455 | Ethe1 | ENSMUSG00000064254.6 |
| chr2 | 71652176 | 71652404 | 3.3615 | 3.89135 | 1.09361 | Distal Intergenic | 2 | 71719417 | 71730966 | 11550 | 1 | ENSMUST00000079720.5 | -67013 | Gm1631 | ENSMUSG00000064263.6 |
| chr2 | 71656787 | 71656882 | 4.02168 | 5.4752 | 2.12651 | Distal Intergenic | 2 | 71719417 | 71730966 | 11550 | 1 | ENSMUST00000079720.5 | -62535 | Gm1631 | ENSMUSG00000064263.6 |
| chr2 | 71675201 | 71675403 | 3.22942 | 3.6758 | 1.02637 | Distal Intergenic | 2 | 71719417 | 71730966 | 11550 | 1 | ENSMUST00000079720.5 | -44014 | Gm1631 | ENSMUSG00000064263.6 |
| chr6 | 105679569 | 105679669 | 4.94704 | 7.04926 | 3.29219 | Intron (ENSMUST00000132044.1/ENSMUSG00000064293.9, intron 1 of 2) | 6 | 105677775 | 106699310 | 1021536 | 1 | ENSMUST00000089208.3 | 1795 | Cntn4 | ENSMUSG00000064293.9 |
| chr8 | 80044059 | 80044254 | 3.29457 | 3.78281 | 1.06069 | Intron (ENSMUST00000079038.2/ENSMUSG00000064325.3, exon 4 of 13) | 8 | 79970345 | 80058006 | 87662 | 2 | ENSMUST00000079038.2 | 13752 | Hhip | ENSMUSG00000064325.3 |
| chr8 | 80056201 | 80056513 | 3.82665 | 5.04726 | 1.83157 | Intron (ENSMUST00000079038.2/ENSMUSG00000064325.3, intron 1 of 12) | 8 | 79970345 | 80058006 | 87662 | 2 | ENSMUST00000079038.2 | 1493 | Hhip | ENSMUSG00000064325.3 |
| chr8 | 80057366 | 80057552 | 3.31563 | 4.07888 | 1.16385 | Promoter | 8 | 79970345 | 80058006 | 87662 | 2 | ENSMUST00000079038.2 | 454 | Hhip | ENSMUSG00000064325.3 |
| chr2 | 66439620 | 66440344 | 3.63071 | 4.4229 | 1.42669 | Promoter | 2 | 66348364 | 66440814 | 92451 | 2 | ENSMUST00000129508.1 | 470 | Scn1a | ENSMUSG00000064329.8 |
| chr8 | 84338363 | 84338655 | 3.59127 | 4.43027 | 1.42162 | Distal Intergenic | 8 | 84380449 | 84380581 | 133 | 2 | ENSMUST00000082458.1 | 41926 | Gm23626 | ENSMUSG00000064392.1 |
| chr8 | 84350580 | 84350856 | 3.37688 | 3.89298 | 1.01733 | Distal Intergenic | 8 | 84380449 | 84380581 | 133 | 2 | ENSMUST00000082458.1 | 29725 | Gm23626 | ENSMUSG00000064392.1 |
| chr8 | 84361332 | 84361422 | 3.74773 | 4.58385 | 1.45453 | Distal Intergenic | 8 | 84380449 | 84380581 | 133 | 2 | ENSMUST00000082458.1 | 19159 | Gm23626 | ENSMUSG00000064392.1 |
| chr8 | 84362606 | 84362681 | 3.54626 | 4.20259 | 1.20737 | Distal Intergenic | 8 | 84380449 | 84380581 | 133 | 2 | ENSMUST00000082458.1 | 17900 | Gm23626 | ENSMUSG00000064392.1 |
| chr9 | 120692186 | 120692299 | 3.80216 | 4.85435 | 1.70199 | Distal Intergenic | 9 | 120791465 | 120791785 | 321 | 2 | ENSMUST00000082463.1 | 99486 | Gm23624 | ENSMUSG00000064397.1 |
| chr16 | 90580659 | 90581248 | 3.56324 | 4.28009 | 1.34954 | Distal Intergenic | 16 | 90602496 | 90602617 | 122 | 1 | ENSMUST00000082537.1 | -21248 | Gm23406 | ENSMUSG00000064471.1 |
| chr16 | 90629576 | 90629861 | 3.8608 | 4.79751 | 1.5961 | Distal Intergenic | 16 | 90602496 | 90602617 | 122 | 1 | ENSMUST00000082537.1 | 27081 | Gm23406 | ENSMUSG00000064471.1 |

| | | | | | | | | | | | | | | | |
|---|---|---|---|---|---|---|---|---|---|---|---|---|---|---|---|
| chr9 | 55549978 | 55550472 | 3.73902 | 4.75358 | 1.6485 | 3' UTR | 9 | 55555622 | 55555712 | 91 | 2 | ENSMUST00000082538.1 | 5240 | Gm23407 | ENSMUSG00000064472.1 |
| chr14 | 72294827 | 72296061 | 3.78658 | 4.67157 | 1.53967 | Distal Intergenic | 14 | 72131088 | 72131221 | 134 | 1 | ENSMUST00000082551.1 | 163740 | Gm23735 | ENSMUSG00000064485.1 |
| chr11 | 79593398 | 79593723 | 3.27832 | 4.0249 | 1.11807 | Intron (ENSMUST00000017783.7/ENSMUSG00000017639.8, intron 1 of 14) | 11 | 79595165 | 79595268 | 104 | 1 | ENSMUST00000082552.1 | -1442 | Gm23293 | ENSMUSG00000064486.1 |
| chr11 | 86436372 | 86436747 | 3.78964 | 4.77455 | 1.60222 | Promoter | 11 | 86437310 | 86437415 | 106 | 2 | ENSMUST00000082592.1 | 668 | Gm26439 | ENSMUSG00000064526.1 |
| chr11 | 86437259 | 86437592 | 3.4962 | 4.21525 | 1.24285 | Promoter | 11 | 86437310 | 86437415 | 106 | 2 | ENSMUST00000082592.1 | 0 | Gm26439 | ENSMUSG00000064526.1 |
| chr11 | 86438009 | 86438399 | 3.3964 | 3.93831 | 1.07353 | Promoter | 11 | 86437310 | 86437415 | 106 | 2 | ENSMUST00000082592.1 | -595 | Gm26439 | ENSMUSG00000064526.1 |
| chr17 | 62384313 | 62384980 | 3.60831 | 4.32776 | 1.30762 | Distal Intergenic | 17 | 62456987 | 62457120 | 134 | 2 | ENSMUST00000082636.1 | 72140 | Gm25800 | ENSMUSG00000064570.1 |
| chr11 | 35008341 | 35008430 | 3.31511 | 3.80833 | 1.04699 | Distal Intergenic | 11 | 34899017 | 34899123 | 107 | 1 | ENSMUST00000082643.1 | 109325 | Gm25799 | ENSMUSG00000064577.1 |
| chr3 | 45681197 | 45681407 | 3.37428 | 4.18492 | 1.23883 | Distal Intergenic | 3 | 45817736 | 45817898 | 163 | 1 | ENSMUST00000082645.1 | -136329 | Gm25796 | ENSMUSG00000064579.1 |
| chr14 | 99367725 | 99367919 | 3.5307 | 4.25094 | 1.30687 | Distal Intergenic | 14 | 99360672 | 99360827 | 156 | 2 | ENSMUST00000082650.1 | -6899 | Gm25130 | ENSMUSG00000064584.1 |
| chr7 | 37832942 | 37833198 | 3.57516 | 4.29173 | 1.34127 | Distal Intergenic | 7 | 37824242 | 37824347 | 106 | 2 | ENSMUST00000082657.1 | -8596 | Gm22302 | ENSMUSG00000064591.1 |
| chr7 | 37834711 | 37834811 | 3.79295 | 4.87133 | 1.72866 | Distal Intergenic | 7 | 37824242 | 37824347 | 106 | 2 | ENSMUST00000082657.1 | -10365 | Gm22302 | ENSMUSG00000064591.1 |
| chr5 | 119958060 | 119958261 | 3.40577 | 3.99006 | 1.19146 | Distal Intergenic | 5 | 120064335 | 120064443 | 109 | 1 | ENSMUST00000082683.1 | -106074 | n-R5s176 | ENSMUSG00000064617.1 |
| chr5 | 119962749 | 119963405 | 3.96653 | 5.05642 | 1.79439 | Distal Intergenic | 5 | 120064335 | 120064443 | 109 | 1 | ENSMUST00000082683.1 | -100930 | n-R5s176 | ENSMUSG00000064617.1 |
| chr5 | 119964596 | 119964810 | 3.34658 | 3.84728 | 1.01733 | Distal Intergenic | 5 | 120064335 | 120064443 | 109 | 1 | ENSMUST00000082683.1 | -99525 | n-R5s176 | ENSMUSG00000064617.1 |
| chr5 | 119967385 | 119967462 | 3.63196 | 4.41972 | 1.4523 | Distal Intergenic | 5 | 120064335 | 120064443 | 109 | 1 | ENSMUST00000082683.1 | -96873 | n-R5s176 | ENSMUSG00000064617.1 |
| chr5 | 120088611 | 120088686 | 3.52052 | 4.15497 | 1.17453 | Distal Intergenic | 5 | 120064335 | 120064443 | 109 | 1 | ENSMUST00000082683.1 | 24277 | n-R5s176 | ENSMUSG00000064617.1 |
| chr5 | 120089322 | 120089446 | 3.37789 | 3.89451 | 1.01733 | Distal Intergenic | 5 | 120064335 | 120064443 | 109 | 1 | ENSMUST00000082683.1 | 24988 | n-R5s176 | ENSMUSG00000064617.1 |
| chr8 | 9459636 | 9459758 | 3.40204 | 4.07651 | 1.16834 | Intron (ENSMUST00000110969.3/ENSMUSG00000079157.3, intron 1 of 2) | 8 | 9568132 | 9568249 | 118 | 2 | ENSMUST00000082684.1 | 108491 | n-R5s95 | ENSMUSG00000064618.1 |
| chr15 | 87214948 | 87215023 | 4.84213 | 6.85971 | 3.11072 | Distal Intergenic | 15 | 87169113 | 87169216 | 104 | 1 | ENSMUST00000082697.1 | 45836 | n-R5s42 | ENSMUSG00000064631.1 |
| chr14 | 118493868 | 118493943 | 3.51243 | 4.20152 | 1.3331 | Intron (ENSMUST00000036554.7/ENSMUSG00000032849.8, intron 29 of 30) | 14 | 118585039 | 118585138 | 100 | 1 | ENSMUST00000082710.1 | -91096 | Gm23302 | ENSMUSG00000064644.1 |
| chr14 | 118533643 | 118533718 | 3.41805 | 3.98928 | 1.13868 | Intron (ENSMUST00000036554.7/ENSMUSG00000032849.8, intron 21 of 30) | 14 | 118585039 | 118585138 | 100 | 1 | ENSMUST00000082710.1 | -51321 | Gm23302 | ENSMUSG00000064644.1 |
| chr14 | 118550709 | 118551335 | 3.64494 | 4.39614 | 1.35772 | Intron (ENSMUST00000036554.7/ENSMUSG00000032849.8, intron 20 of 30) | 14 | 118585039 | 118585138 | 100 | 1 | ENSMUST00000082710.1 | -33704 | Gm23302 | ENSMUSG00000064644.1 |
| chr14 | 118551924 | 118552336 | 3.7423 | 4.59626 | 1.4998 | Intron (ENSMUST00000036554.7/ENSMUSG00000032849.8, intron 20 of 30) | 14 | 118585039 | 118585138 | 100 | 1 | ENSMUST00000082710.1 | -32703 | Gm23302 | ENSMUSG00000064644.1 |
| chr4 | 148401865 | 148402176 | 3.55846 | 4.22244 | 1.21242 | Distal Intergenic | 4 | 148443334 | 148443485 | 152 | 2 | ENSMUST00000082714.2 | 41309 | Gm23303 | ENSMUSG00000064648.2 |
| chr4 | 148402561 | 148402984 | 3.55851 | 4.23556 | 1.26003 | Distal Intergenic | 4 | 148443334 | 148443485 | 152 | 2 | ENSMUST00000082714.2 | 40501 | Gm23303 | ENSMUSG00000064648.2 |
| chr4 | 148403695 | 148404152 | 4.07693 | 5.44361 | 2.0871 | Distal Intergenic | 4 | 148443334 | 148443485 | 152 | 2 | ENSMUST00000082714.2 | 39333 | Gm23303 | ENSMUSG00000064648.2 |
| chr4 | 148404803 | 148405159 | 3.37517 | 3.89644 | 1.03869 | Distal Intergenic | 4 | 148443334 | 148443485 | 152 | 2 | ENSMUST00000082714.2 | 38326 | Gm23303 | ENSMUSG00000064648.2 |
| chr4 | 148415337 | 148416227 | 3.80417 | 4.78106 | 1.66725 | Distal Intergenic | 4 | 148443334 | 148443485 | 152 | 2 | ENSMUST00000082714.2 | 27258 | Gm23303 | ENSMUSG00000064648.2 |
| chr9 | 46672185 | 46672364 | 3.35854 | 3.86532 | 1.01733 | Distal Intergenic | 9 | 46630113 | 46630219 | 107 | 2 | ENSMUST00000082739.1 | -41967 | Gm22805 | ENSMUSG00000064673.1 |
| chr9 | 46702573 | 46702665 | 3.81727 | 4.73013 | 1.59771 | Distal Intergenic | 9 | 46630113 | 46630219 | 107 | 2 | ENSMUST00000082739.1 | -72355 | Gm22805 | ENSMUSG00000064673.1 |
| chr11 | 109068466 | 109068789 | 3.32764 | 3.81877 | 1.01733 | Distal Intergenic | 11 | 109066984 | 109067090 | 107 | 2 | ENSMUST00000082757.1 | -1377 | Gm24149 | ENSMUSG00000064691.1 |
| chr11 | 109069114 | 109069407 | 3.65447 | 4.41113 | 1.36154 | Distal Intergenic | 11 | 109066984 | 109067090 | 107 | 2 | ENSMUST00000082757.1 | -2025 | Gm24149 | ENSMUSG00000064691.1 |
| chr11 | 109076822 | 109076906 | 3.84866 | 4.77961 | 1.59711 | Distal Intergenic | 11 | 109066984 | 109067090 | 107 | 2 | ENSMUST00000082757.1 | -9733 | Gm24149 | ENSMUSG00000064691.1 |
| chr5 | 139839173 | 139839717 | 3.38025 | 3.89806 | 1.01733 | Distal Intergenic | 5 | 139844930 | 139845250 | 321 | 1 | ENSMUST00000082760.1 | -5213 | Gm24146 | ENSMUSG00000064694.1 |
| chr5 | 139847445 | 139847556 | 3.37496 | 3.99282 | 1.10869 | Distal Intergenic | 5 | 139844930 | 139845250 | 321 | 1 | ENSMUST00000082760.1 | 2516 | Gm24146 | ENSMUSG00000064694.1 |
| chr5 | 139853501 | 139853697 | 3.35707 | 3.8631 | 1.01733 | Distal Intergenic | 5 | 139844930 | 139845250 | 321 | 1 | ENSMUST00000082760.1 | 8572 | Gm24146 | ENSMUSG00000064694.1 |
| chr5 | 139871347 | 139871629 | 4.39182 | 5.84864 | 2.34856 | Distal Intergenic | 5 | 139844930 | 139845250 | 321 | 1 | ENSMUST00000082760.1 | 26418 | Gm24146 | ENSMUSG00000064694.1 |
| chr5 | 149287088 | 149287163 | 3.47929 | 4.11444 | 1.24261 | Intron (ENSMUST00000071130.3/ENSMUSG00000060063.6, intron 4 of 4) | 5 | 149281118 | 149281223 | 106 | 2 | ENSMUST00000082770.1 | -5866 | n-R5s180 | ENSMUSG00000064704.1 |
| chr5 | 149287948 | 149288140 | 3.49791 | 4.14099 | 1.25007 | 3' UTR | 5 | 149281118 | 149281223 | 106 | 2 | ENSMUST00000082770.1 | -6726 | n-R5s180 | ENSMUSG00000064704.1 |
| chr12 | 82060007 | 82060779 | 3.52304 | 4.22785 | 1.3078 | Distal Intergenic | 12 | 82150468 | 82150563 | 96 | 1 | ENSMUST00000082780.1 | -89689 | n-R5s62 | ENSMUSG00000064714.1 |
| chr17 | 47852328 | 47852648 | 3.35347 | 3.85766 | 1.01733 | Distal Intergenic | 17 | 47848834 | 47848952 | 119 | 1 | ENSMUST00000082814.1 | 3495 | n-R5s28 | ENSMUSG00000064748.1 |
| chr17 | 47862327 | 47862408 | 3.68242 | 4.45852 | 1.38511 | Distal Intergenic | 17 | 47848834 | 47848952 | 119 | 1 | ENSMUST00000082814.1 | 13494 | n-R5s28 | ENSMUSG00000064748.1 |
| chr8 | 89457327 | 89457965 | 3.40296 | 4.20208 | 1.25113 | Distal Intergenic | 8 | 89589435 | 89589541 | 107 | 2 | ENSMUST00000082824.1 | 131576 | Gm26331 | ENSMUSG00000064758.1 |
| chr8 | 89477418 | 89477616 | 3.29926 | 3.77607 | 1.01733 | Distal Intergenic | 8 | 89589435 | 89589541 | 107 | 2 | ENSMUST00000082824.1 | 111925 | Gm26331 | ENSMUSG00000064758.1 |
| chr8 | 89568698 | 89568985 | 3.39052 | 3.97127 | 1.18979 | Distal Intergenic | 8 | 89589435 | 89589541 | 107 | 2 | ENSMUST00000082824.1 | 20556 | Gm26331 | ENSMUSG00000064758.1 |
| chr3 | 96414298 | 96414842 | 4.06344 | 5.40434 | 2.03454 | Promoter | 3 | 96414437 | 96414859 | 423 | 2 | ENSMUST00000082862.1 | 17 | Terc | ENSMUSG00000064796.1 |
| chr3 | 96418297 | 96418528 | 4.0275 | 5.25265 | 1.96054 | Distal Intergenic | 3 | 96414437 | 96414859 | 423 | 2 | ENSMUST00000082862.1 | -3439 | Terc | ENSMUSG00000064796.1 |
| chr12 | 46399288 | 46399624 | 3.36131 | 3.91695 | 1.16266 | Distal Intergenic | 12 | 45984343 | 45984549 | 207 | 1 | ENSMUST00000082866.1 | 414946 | Gm24948 | ENSMUSG00000064800.1 |
| chr1 | 133595081 | 133595469 | 2.49512 | 5.24265 | 1.95954 | Distal Intergenic | 1 | 133601223 | 133601355 | 133 | 1 | ENSMUST00000082880.1 | -5754 | Gm23939 | ENSMUSG00000064814.1 |
| chr16 | 73279170 | 73279394 | 3.36135 | 3.96109 | 1.09865 | Distal Intergenic | 16 | 73422793 | 73422959 | 167 | 2 | ENSMUST00000082896.1 | 143565 | Gm25699 | ENSMUSG00000064830.1 |
| chr12 | 77517814 | 77518150 | 3.3437 | 3.84295 | 1.01733 | Distal Intergenic | 12 | 77418417 | 77418576 | 160 | 1 | ENSMUST00000082899.1 | 99398 | Gm25926 | ENSMUSG00000064833.1 |
| chr3 | 35649143 | 35649359 | 3.31466 | 3.83636 | 1.09851 | Distal Intergenic | 3 | 35735373 | 35735477 | 105 | 1 | ENSMUST00000082900.1 | -86014 | Gm25696 | ENSMUSG00000064834.1 |
| chr19 | 25461477 | 25461875 | 3.3452 | 3.85955 | 1.0677 | Distal Intergenic | 19 | 25453960 | 25454063 | 104 | 2 | ENSMUST00000082909.1 | -7415 | Gm26207 | ENSMUSG00000064843.1 |
| chr19 | 25462461 | 25462964 | 3.43949 | 4.01663 | 1.12067 | Distal Intergenic | 19 | 25453960 | 25454063 | 104 | 2 | ENSMUST00000082909.1 | -8399 | Gm26207 | ENSMUSG00000064843.1 |
| chr3 | 55534415 | 55534856 | 3.81108 | 4.71069 | 1.55068 | 3' UTR | 3 | 55548446 | 55548552 | 107 | 1 | ENSMUST00000082970.1 | -13590 | Gm25404 | ENSMUSG00000064904.1 |
| chr15 | 96658215 | 96658380 | 3.24394 | 3.69308 | 1.01733 | Distal Intergenic | 15 | 96671793 | 96671900 | 108 | 1 | ENSMUST00000082994.1 | -13413 | Gm22045 | ENSMUSG00000064928.1 |
| chr2 | 168341252 | 168341538 | 3.63266 | 4.37126 | 1.34201 | Distal Intergenic | 2 | 168335700 | 168335836 | 137 | 1 | ENSMUST00000083027.1 | 5553 | Gm22704 | ENSMUSG00000064961.1 |
| chr1 | 178684460 | 178684535 | 3.91572 | 5.04774 | 1.82805 | Intron (ENSMUST00000161017.2/ENSMUSG00000026494.7, intron 3 of 14) | 1 | 178636905 | 178637071 | 167 | 1 | ENSMUST00000083033.1 | 47556 | Gm24405 | ENSMUSG00000064967.1 |
| chr8 | 117962552 | 117962651 | 3.35102 | 3.85719 | 1.02869 | Distal Intergenic | 8 | 118120954 | 118121083 | 130 | 1 | ENSMUST00000083094.1 | -158303 | Gm25200 | ENSMUSG00000065028.1 |
| chr8 | 117963029 | 117963292 | 3.45665 | 4.13871 | 1.25231 | Distal Intergenic | 8 | 118120954 | 118121083 | 130 | 1 | ENSMUST00000083094.1 | -157662 | Gm25200 | ENSMUSG00000065028.1 |
| chr8 | 118191292 | 118191423 | 3.34281 | 3.84159 | 1.01733 | Distal Intergenic | 8 | 118120954 | 118121083 | 130 | 1 | ENSMUST00000083094.1 | 70339 | Gm25200 | ENSMUSG00000065028.1 |
| chr6 | 131377530 | 131377618 | 6.95852 | 13.58277 | 8.95748 | Intron (ENSMUST00000032309.7/ENSMUSG00000030189.10, intron 5 of 9) | 6 | 131380273 | 131380406 | 134 | 2 | ENSMUST00000083102.1 | 2788 | Gm22362 | ENSMUSG00000065036.1 |
| chr10 | 116364013 | 116364141 | 3.50851 | 4.14073 | 1.19068 | Exon (ENSMUST00000092167.5/ENSMUSG00000020154.9, exon 26 of 32) | 10 | 116340583 | 116340719 | 137 | 1 | ENSMUST00000083152.1 | 23431 | Gm26495 | ENSMUSG00000065086.1 |
| chr11 | 65576429 | 65576622 | 3.24426 | 3.91955 | 1.11063 | Distal Intergenic | 11 | 65626394 | 65626503 | 110 | 1 | ENSMUST00000083170.1 | -49772 | Gm26128 | ENSMUSG00000065104.1 |
| chr12 | 31248898 | 31249051 | 3.4528 | 4.06944 | 1.18644 | Distal Intergenic | 12 | 31258933 | 31259062 | 130 | 2 | ENSMUST00000083266.1 | 10011 | Gm24613 | ENSMUSG00000065200.1 |
| chr11 | 82508283 | 82508545 | 3.5708 | 4.25171 | 1.24064 | Distal Intergenic | 11 | 82596073 | 82596179 | 107 | 2 | ENSMUST00000083267.1 | 87634 | Gm24612 | ENSMUSG00000065201.1 |
| chr11 | 82564281 | 82564776 | 3.46783 | 4.07215 | 1.16602 | Distal Intergenic | 11 | 82596073 | 82596179 | 107 | 2 | ENSMUST00000083267.1 | 31403 | Gm24612 | ENSMUSG00000065201.1 |
| chr11 | 82600268 | 82600473 | 3.51182 | 4.13939 | 1.16809 | Distal Intergenic | 11 | 82596073 | 82596179 | 107 | 2 | ENSMUST00000083267.1 | -4090 | Gm24612 | ENSMUSG00000065201.1 |
| chr3 | 8245786 | 8246562 | 2.30576 | 9.48815 | 5.40443 | Distal Intergenic | 3 | 8429158 | 8429264 | 107 | 1 | ENSMUST00000083269.1 | -182596 | Gm24614 | ENSMUSG00000065203.1 |
| chr1 | 77929718 | 77930342 | 3.33764 | 4.0146 | 1.15765 | Distal Intergenic | 1 | 77898283 | 77898446 | 164 | 2 | ENSMUST00000083281.1 | -31273 | Gm26263 | ENSMUSG00000065215.1 |
| chr2 | 74622322 | 74622504 | 3.633 | 4.42692 | 1.43055 | Downstream (2-3kb) | 2 | 74624642 | 74624748 | 107 | 1 | ENSMUST00000083303.1 | -2138 | Gm22975 | ENSMUSG00000065237.1 |
| chr2 | 74622964 | 74623301 | 3.69315 | 4.55282 | 1.51911 | Downstream (1-2kb) | 2 | 74624642 | 74624748 | 107 | 1 | ENSMUST00000083303.1 | -1341 | Gm22975 | ENSMUSG00000065237.1 |
| chr3 | 128344786 | 128345117 | 3.61915 | 4.44612 | 1.46075 | Distal Intergenic | 3 | 128540372 | 128540480 | 109 | 1 | ENSMUST00000083314.1 | -195255 | Gm22293 | ENSMUSG00000065248.1 |
| chr13 | 39823015 | 39823200 | 3.48007 | 4.08107 | 1.12892 | Distal Intergenic | 13 | 39858863 | 39859027 | 165 | 2 | ENSMUST00000083322.1 | 35827 | Gm23972 | ENSMUSG00000065256.1 |
| chr13 | 39823772 | 39824644 | 3.61195 | 4.40008 | 1.38711 | Distal Intergenic | 13 | 39858863 | 39859027 | 165 | 2 | ENSMUST00000083322.1 | 34383 | Gm23972 | ENSMUSG00000065256.1 |
| chr13 | 39855869 | 39856316 | 3.3705 | 3.89296 | 1.05123 | Downstream (2-3kb) | 13 | 39858863 | 39859027 | 165 | 2 | ENSMUST00000083322.1 | 2711 | Gm23972 | ENSMUSG00000065256.1 |
| chr13 | 39877511 | 39877658 | 3.4914 | 4.147 | 1.27748 | Distal Intergenic | 13 | 39858863 | 39859027 | 165 | 2 | ENSMUST00000083322.1 | -18485 | Gm23972 | ENSMUSG00000065256.1 |
| chr3 | 149088448 | 149088953 | 3.62327 | 4.36345 | 1.33539 | Distal Intergenic | 3 | 149028932 | 149029038 | 107 | 1 | ENSMUST00000083336.1 | 59517 | Gm25127 | ENSMUSG00000065270.1 |
| chr1 | 138913079 | 138913461 | 3.78291 | 4.71171 | 1.64274 | Distal Intergenic | 1 | 138878262 | 138878377 | 116 | 2 | ENSMUST00000083341.1 | -34703 | n-R5s217 | ENSMUSG00000065275.1 |
| chr12 | 118456296 | 118456755 | 3.69797 | 4.55098 | 1.48747 | Distal Intergenic | 12 | 118504928 | 118505034 | 107 | 2 | ENSMUST00000083414.1 | 48279 | Gm25577 | ENSMUSG00000065348.1 |
| chr12 | 118457676 | 118458923 | 3.96429 | 5.1588 | 1.90552 | Distal Intergenic | 12 | 118504928 | 118505034 | 107 | 2 | ENSMUST00000083414.1 | 46111 | Gm25577 | ENSMUSG00000065348.1 |
| chr12 | 118460928 | 118461856 | 3.53321 | 4.20718 | 1.27794 | Distal Intergenic | 12 | 118504928 | 118505034 | 107 | 2 | ENSMUST00000083414.1 | 43178 | Gm25577 | ENSMUSG00000065348.1 |
| chr12 | 118510329 | 118510663 | 3.38506 | 3.93506 | 1.11905 | Distal Intergenic | 12 | 118504928 | 118505034 | 107 | 2 | ENSMUST00000083414.1 | -5296 | Gm25577 | ENSMUSG00000065348.1 |
| chr12 | 118528980 | 118529152 | 3.29437 | 3.76872 | 1.01733 | Distal Intergenic | 12 | 118504928 | 118505034 | 107 | 2 | ENSMUST00000083414.1 | -23947 | Gm25577 | ENSMUSG00000065348.1 |
| chr12 | 118529461 | 118529667 | 3.68084 | 4.51094 | 1.48749 | Distal Intergenic | 12 | 118504928 | 118505034 | 107 | 2 | ENSMUST00000083414.1 | -24428 | Gm25577 | ENSMUSG00000065348.1 |
| chr12 | 118530143 | 118530316 | 3.58668 | 4.54274 | 1.48388 | Distal Intergenic | 12 | 118504928 | 118505034 | 107 | 2 | ENSMUST00000083414.1 | -25110 | Gm25577 | ENSMUSG00000065348.1 |
| chr12 | 118530738 | 118530905 | 3.51036 | 4.16933 | 1.27661 | Distal Intergenic | 12 | 118504928 | 118505034 | 107 | 2 | ENSMUST00000083414.1 | -25705 | Gm25577 | ENSMUSG00000065348.1 |
| chr7 | 19469540 | 19469878 | 3.3117 | 3.97893 | 1.13495 | Distal Intergenic | 7 | 19460797 | 19460935 | 139 | 2 | ENSMUST00000083454.1 | -8606 | Gm23099 | ENSMUSG00000065388.1 |
| chr7 | 19474261 | 19474742 | 3.42248 | 4.17558 | 1.2418 | Distal Intergenic | 7 | 19460797 | 19460935 | 139 | 2 | ENSMUST00000083454.1 | -13327 | Gm23099 | ENSMUSG00000065388.1 |
| chr10 | 122964290 | 122964587 | 3.46609 | 4.13341 | 1.21049 | Distal Intergenic | 10 | 122985634 | 122985723 | 90 | 2 | ENSMUST00000083472.2 | 21136 | Mirlet7i | ENSMUSG00000065406.2 |
| chr9 | 106151411 | 106151808 | 3.36494 | 3.87496 | 1.01733 | Intron (ENSMUST00000143956.1/ENSMUSG00000086296.2, intron 2 of 6) | 9 | 106154124 | 106154213 | 90 | 1 | ENSMUST00000083473.1 | -2316 | Mir135a-1 | ENSMUSG00000065407.1 |
| chr4 | 88931238 | 88931503 | 3.98719 | 5.24993 | 1.92931 | Distal Intergenic | 4 | 88910557 | 88910662 | 106 | 2 | ENSMUST00000083474.1 | -20577 | Mir31 | ENSMUSG00000065408.1 |
| chr4 | 88932611 | 88932688 | 3.42355 | 3.98858 | 1.10632 | Distal Intergenic | 4 | 88910557 | 88910662 | 106 | 2 | ENSMUST00000083474.1 | -21950 | Mir31 | ENSMUSG00000065408.1 |
| chr4 | 88935340 | 88935416 | 3.91123 | 4.87995 | 1.6389 | Distal Intergenic | 4 | 88910557 | 88910662 | 106 | 2 | ENSMUST00000083474.1 | -24679 | Mir31 | ENSMUSG00000065408.1 |
| chr4 | 88937458 | 88937811 | 3.63 | 4.36546 | 1.32999 | Distal Intergenic | 4 | 88910557 | 88910662 | 106 | 2 | ENSMUST00000083474.1 | -26797 | Mir31 | ENSMUSG00000065408.1 |
| chr4 | 88938157 | 88939881 | 3.77551 | 4.74802 | 1.60904 | Distal Intergenic | 4 | 88910557 | 88910662 | 106 | 2 | ENSMUST00000083474.1 | -27496 | Mir31 | ENSMUSG00000065408.1 |
| chr4 | 88940428 | 88940896 | 3.5686 | 4.33474 | 1.37125 | Distal Intergenic | 4 | 88910557 | 88910662 | 106 | 2 | ENSMUST00000083474.1 | -29767 | Mir31 | ENSMUSG00000065408.1 |
| chr4 | 88941666 | 88942388 | 3.52377 | 4.21141 | 1.28726 | Distal Intergenic | 4 | 88910557 | 88910662 | 106 | 2 | ENSMUST00000083474.1 | -31005 | Mir31 | ENSMUSG00000065408.1 |
| chr4 | 88956419 | 88957273 | 3.44278 | 4.0197 | 1.11353 | Distal Intergenic | 4 | 88910557 | 88910662 | 106 | 2 | ENSMUST00000083474.1 | -45758 | Mir31 | ENSMUSG00000065408.1 |
| chr9 | 122629105 | 122629381 | 3.34435 | 3.84393 | 1.01733 | Intron (ENSMUST00000181682.2/ENSMUSG00000097736.2, intron 2 of 3) | 9 | 122682876 | 122682974 | 99 | 1 | ENSMUST00000083480.1 | -53495 | Mir138-1 | ENSMUSG00000065414.1 |
| chr13 | 83735518 | 83736050 | 4.20656 | 5.5911 | 2.1733 | Exon (ENSMUST00000182788.1/ENSMUSG00000050334.7, exon 4 of 4) | 13 | 83738806 | 83738892 | 87 | 1 | ENSMUST00000083490.1 | -2756 | Mir9-2 | ENSMUSG00000065424.1 |
| chr1 | 132197751 | 132198182 | 3.72789 | 4.54895 | 1.44523 | Promoter | 1 | 132198088 | 132198184 | 97 | 1 | ENSMUST00000083501.1 | 0 | Mir135b | ENSMUSG00000065435.1 |
| chr1 | 132199129 | 132199411 | 3.50241 | 4.26683 | 1.30818 | Intron (ENSMUST00000188169.1/ENSMUSG00000079330.3, intron 1 of 4) | 1 | 132198088 | 132198184 | 97 | 1 | ENSMUST00000083501.1 | 1042 | Mir135b | ENSMUSG00000065435.1 |
| chr7 | 101453002 | 101453311 | 3.4344 | 4.06923 | 1.18412 | Intron (ENSMUST00000084894.8/ENSMUSG00000030653.11, intron 2 of 30) | 7 | 101475376 | 101475443 | 68 | 1 | ENSMUST00000083512.1 | -22065 | Mir139 | ENSMUSG00000065446.1 |
| chr7 | 101482212 | 101482448 | 3.30718 | 3.78796 | 1.01733 | Intron (ENSMUST00000084894.8/ENSMUSG00000030653.11, intron 3 of 30) | 7 | 101475376 | 101475443 | 68 | 1 | ENSMUST00000083512.1 | 6837 | Mir139 | ENSMUSG00000065446.1 |
| chr7 | 101500999 | 101501309 | 3.73557 | 4.61406 | 1.57852 | Exon (ENSMUST00000084894.8/ENSMUSG00000030653.11, exon 8 of 31) | 7 | 101475376 | 101475443 | 68 | 1 | ENSMUST00000083512.1 | 25624 | Mir139 | ENSMUSG00000065446.1 |
| chr7 | 101503785 | 101504543 | 3.44092 | 4.31185 | 1.32924 | Exon (ENSMUST00000084894.8/ENSMUSG00000030653.11, exon 15 of 31) | 7 | 101475376 | 101475443 | 68 | 1 | ENSMUST00000083512.1 | 28410 | Mir139 | ENSMUSG00000065446.1 |
| chr7 | 101504981 | 101505132 | 3.74965 | 4.76188 | 1.63453 | Exon (ENSMUST00000084894.8/ENSMUSG00000030653.11, exon 19 of 31) | 7 | 101475376 | 101475443 | 68 | 1 | ENSMUST00000083512.1 | 29606 | Mir139 | ENSMUSG00000065446.1 |
| chr2 | 180891770 | 180892463 | 3.86755 | 4.82633 | 1.64236 | Downstream (2-3kb) | 2 | 180894031 | 180894117 | 87 | 1 | ENSMUST00000083520.1 | -1568 | Mir124a-3 | ENSMUSG00000065454.1 |
| chr2 | 180893037 | 180894880 | 3.85634 | 4.94852 | 1.73142 | Promoter | 2 | 180894031 | 180894117 | 87 | 1 | ENSMUST00000083520.1 | 0 | Mir124a-3 | ENSMUSG00000065454.1 |
| chr2 | 180422636 | 180422868 | 3.3752 | 3.89044 | 1.01733 | Distal Intergenic | 2 | 180398379 | 180398482 | 104 | 1 | ENSMUST00000083526.1 | 24258 | Mir133a-2 | ENSMUSG00000065460.1 |
| chr6 | 29016691 | 29017119 | 3.36896 | 3.92129 | 1.15123 | Intron (ENSMUST00000164104.1/ENSMUSG00000092090.1, intron 1 of 1) | 6 | 29022619 | 29022691 | 73 | 1 | ENSMUST00000083535.1 | -5500 | Mir129-1 | ENSMUSG00000065469.1 |
| chr2 | 29843510 | 29843893 | 3.49376 | 4.13162 | 1.2295 | 3' UTR | 2 | 29845631 | 29845727 | 97 | 2 | ENSMUST00000083551.1 | 1834 | Mir219a-2 | ENSMUSG00000065485.1 |
| chr6 | 51145874 | 51145949 | 3.31438 | 3.81899 | 1.08328 | Distal Intergenic | 6 | 51269812 | 51269910 | 99 | 2 | ENSMUST00000083571.1 | 123961 | Mir148a | ENSMUSG00000065505.1 |
| chr6 | 51163445 | 51163537 | 3.6385 | 4.39078 | 1.36371 | Distal Intergenic | 6 | 51269812 | 51269910 | 99 | 2 | ENSMUST00000083571.1 | 106373 | Mir148a | ENSMUSG00000065505.1 |
| chr6 | 51165627 | 51165747 | 3.97097 | 5.05999 | 1.86128 | Distal Intergenic | 6 | 51269812 | 51269910 | 99 | 2 | ENSMUST00000083571.1 | 104163 | Mir148a | ENSMUSG00000065505.1 |
| chr6 | 51166478 | 51166770 | 3.86963 | 4.88018 | 1.74023 | Distal Intergenic | 6 | 51269812 | 51269910 | 99 | 2 | ENSMUST00000083571.1 | 103140 | Mir148a | ENSMUSG00000065505.1 |
| chr6 | 51167487 | 51167755 | 3.4192 | 4.37616 | 1.33828 | Distal Intergenic | 6 | 51269812 | 51269910 | 99 | 2 | ENSMUST00000083571.1 | 102155 | Mir148a | ENSMUSG00000065505.1 |
| chr6 | 51168788 | 51169148 | 4.53086 | 6.38238 | 2.82289 | Distal Intergenic | 6 | 51269812 | 51269910 | 99 | 2 | ENSMUST00000083571.1 | 100762 | Mir148a | ENSMUSG00000065505.1 |
| chr6 | 51169613 | 51169741 | 3.80811 | 4.81362 | 1.65219 | Distal Intergenic | 6 | 51269812 | 51269910 | 99 | 2 | ENSMUST00000083571.1 | 100169 | Mir148a | ENSMUSG00000065505.1 |
| chr6 | 51170167 | 51170480 | 3.52582 | 4.28504 | 1.31981 | Distal Intergenic | 6 | 51269812 | 51269910 | 99 | 2 | ENSMUST00000083571.1 | 99430 | Mir148a | ENSMUSG00000065505.1 |
| chr6 | 51170843 | 51172027 | 3.67577 | 4.63744 | 1.52493 | Distal Intergenic | 6 | 51269812 | 51269910 | 99 | 2 | ENSMUST00000083571.1 | 97883 | Mir148a | ENSMUSG00000065505.1 |
| chr6 | 51173083 | 51173242 | 4.15355 | 5.51581 | 2.1374 | Distal Intergenic | 6 | 51269812 | 51269910 | 99 | 2 | ENSMUST00000083571.1 | 96668 | Mir148a | ENSMUSG00000065505.1 |
| chr6 | 51175618 | 51176481 | 3.39041 | 4.13641 | 1.23658 | Distal Intergenic | 6 | 51269812 | 51269910 | 99 | 2 | ENSMUST00000083571.1 | 93429 | Mir148a | ENSMUSG00000065505.1 |
| chr6 | 51176895 | 51178802 | 4.0631 | 5.35263 | 2.05164 | Distal Intergenic | 6 | 51269812 | 51269910 | 99 | 2 | ENSMUST00000083571.1 | 91108 | Mir148a | ENSMUSG00000065505.1 |
| chr6 | 51179120 | 51179248 | 3.52191 | 4.44168 | 1.43523 | Distal Intergenic | 6 | 51269812 | 51269910 | 99 | 2 | ENSMUST00000083571.1 | 90662 | Mir148a | ENSMUSG00000065505.1 |
| chr6 | 51179687 | 51181493 | 4.03027 | 5.16618 | 1.86715 | Distal Intergenic | 6 | 51269812 | 51269910 | 99 | 2 | ENSMUST00000083571.1 | 88417 | Mir148a | ENSMUSG00000065505.1 |
| chr6 | 51181875 | 51184759 | 4.04108 | 5.22104 | 1.92131 | Distal Intergenic | 6 | 51269812 | 51269910 | 99 | 2 | ENSMUST00000083571.1 | 85151 | Mir148a | ENSMUSG00000065505.1 |
| chr6 | 51185599 | 51185830 | 3.58743 | 4.3357 | 1.38878 | Distal Intergenic | 6 | 51269812 | 51269910 | 99 | 2 | ENSMUST00000083571.1 | 84080 | Mir148a | ENSMUSG00000065505.1 |
| chr6 | 51186256 | 51186402 | 3.50449 | 4.17821 | 1.25035 | Distal Intergenic | 6 | 51269812 | 51269910 | 99 | 2 | ENSMUST00000083571.1 | 83508 | Mir148a | ENSMUSG00000065505.1 |
| chr6 | 51187182 | 51187525 | 3.37492 | 4.16846 | 1.23498 | Distal Intergenic | 6 | 51269812 | 51269910 | 99 | 2 | ENSMUST00000083571.1 | 82385 | Mir148a | ENSMUSG00000065505.1 |
| chr6 | 51188148 | 51189328 | 3.85737 | 4.83267 | 1.66615 | Distal Intergenic | 6 | 51269812 | 51269910 | 99 | 2 | ENSMUST00000083571.1 | 80582 | Mir148a | ENSMUSG00000065505.1 |
| chr6 | 51189838 | 51189981 | 3.3229 | 3.82488 | 1.06376 | Distal Intergenic | 6 | 51269812 | 51269910 | 99 | 2 | ENSMUST00000083571.1 | 79929 | Mir148a | ENSMUSG00000065505.1 |
| chr6 | 51190906 | 51191129 | 3.87865 | 4.84071 | 1.64218 | Distal Intergenic | 6 | 51269812 | 51269910 | 99 | 2 | ENSMUST00000083571.1 | 78781 | Mir148a | ENSMUSG00000065505.1 |
| chr6 | 51191587 | 51191691 | 4.54901 | 6.17933 | 2.599 | Distal Intergenic | 6 | 51269812 | 51269910 | 99 | 2 | ENSMUST00000083571.1 | 78219 | Mir148a | ENSMUSG00000065505.1 |
| chr6 | 51192189 | 51192698 | 3.38207 | 4.13783 | 1.23239 | Distal Intergenic | 6 | 51269812 | 51269910 | 99 | 2 | ENSMUST00000083571.1 | 77212 | Mir148a | ENSMUSG00000065505.1 |
| chr6 | 51193024 | 51193615 | 4.17404 | 5.47815 | 2.0894 | Distal Intergenic | 6 | 51269812 | 51269910 | 99 | 2 | ENSMUST00000083571.1 | 76295 | Mir148a | ENSMUSG00000065505.1 |
| chr6 | 51194063 | 51194509 | 3.72951 | 4.69303 | 1.56636 | Distal Intergenic | 6 | 51269812 | 51269910 | 99 | 2 | ENSMUST00000083571.1 | 75401 | Mir148a | ENSMUSG00000065505.1 |
| chr6 | 51194821 | 51195224 | 3.59887 | 4.49509 | 1.42361 | Distal Intergenic | 6 | 51269812 | 51269910 | 99 | 2 | ENSMUST00000083571.1 | 74686 | Mir148a | ENSMUSG00000065505.1 |
| chr6 | 51196076 | 51196272 | 3.73188 | 4.56235 | 1.4714 | Distal Intergenic | 6 | 51269812 | 51269910 | 99 | 2 | ENSMUST00000083571.1 | 73638 | Mir148a | ENSMUSG00000065505.1 |
| chr6 | 51197088 | 51197232 | 3.60701 | 4.32458 | 1.31597 | Distal Intergenic | 6 | 51269812 | 51269910 | 99 | 2 | ENSMUST00000083571.1 | 72678 | Mir148a | ENSMUSG00000065505.1 |
| chr6 | 51197633 | 51197806 | 3.53493 | 4.18628 | 1.21014 | Distal Intergenic | 6 | 51269812 | 51269910 | 99 | 2 | ENSMUST00000083571.1 | 72104 | Mir148a | ENSMUSG00000065505.1 |
| chr6 | 51206982 | 51208439 | 3.79322 | 4.683 | 1.54189 | Distal Intergenic | 6 | 51269812 | 51269910 | 99 | 2 | ENSMUST00000083571.1 | 61471 | Mir148a | ENSMUSG00000065505.1 |
| chr6 | 51210667 | 51210767 | 3.40522 | 3.96102 | 1.07973 | Distal Intergenic | 6 | 51269812 | 51269910 | 99 | 2 | ENSMUST00000083571.1 | 59143 | Mir148a | ENSMUSG00000065505.1 |
| chr6 | 51215526 | 51215970 | 3.60518 | 4.34315 | 1.35111 | Distal Intergenic | 6 | 51269812 | 51269910 | 99 | 2 | ENSMUST00000083571.1 | 53940 | Mir148a | ENSMUSG00000065505.1 |
| chr6 | 51270500 | 51270731 | 3.46241 | 4.05156 | 1.12115 | Promoter | 6 | 51269812 | 51269910 | 99 | 2 | ENSMUST00000083571.1 | -591 | Mir148a | ENSMUSG00000065505.1 |
| chr2 | 94225884 | 94226071 | 3.35322 | 3.85731 | 1.01733 | Distal Intergenic | 2 | 94241364 | 94241453 | 90 | 2 | ENSMUST00000083577.1 | 15382 | Mir129-2 | ENSMUSG00000065511.1 |
| chr2 | 94227971 | 94228283 | 3.47825 | 4.08614 | 1.15971 | Distal Intergenic | 2 | 94241364 | 94241453 | 90 | 2 | ENSMUST00000083577.1 | 13170 | Mir129-2 | ENSMUSG00000065511.1 |
| chr2 | 94231714 | 94232003 | 3.73579 | 4.61805 | 1.54207 | Distal Intergenic | 2 | 94241364 | 94241453 | 90 | 2 | ENSMUST00000083577.1 | 9450 | Mir129-2 | ENSMUSG00000065511.1 |
| chr2 | 94234047 | 94234337 | 3.47383 | 4.0945 | 1.21404 | Distal Intergenic | 2 | 94241364 | 94241453 | 90 | 2 | ENSMUST00000083577.1 | 7116 | Mir129-2 | ENSMUSG00000065511.1 |
| chr2 | 94236438 | 94236579 | 4.01796 | 5.24961 | 1.93574 | Distal Intergenic | 2 | 94241364 | 94241453 | 90 | 2 | ENSMUST00000083577.1 | 4874 | Mir129-2 | ENSMUSG00000065511.1 |
| chr2 | 94237740 | 94238321 | 3.93437 | 5.05851 | 1.84245 | Distal Intergenic | 2 | 94241364 | 94241453 | 90 | 2 | ENSMUST00000083577.1 | 3132 | Mir129-2 | ENSMUSG00000065511.1 |
| chr2 | 94239583 | 94239998 | 4.1142 | 5.54851 | 2.19022 | Downstream (1-2kb) | 2 | 94241364 | 94241453 | 90 | 2 | ENSMUST00000083577.1 | 1455 | Mir129-2 | ENSMUSG00000065511.1 |
| chr2 | 94240395 | 94240746 | 3.80414 | 4.88058 | 1.67463 | Promoter | 2 | 94241364 | 94241453 | 90 | 2 | ENSMUST00000083577.1 | 707 | Mir129-2 | ENSMUSG00000065511.1 |
| chr2 | 94241091 | 94241498 | 4.68719 | 6.56683 | 2.98372 | Promoter | 2 | 94241364 | 94241453 | 90 | 2 | ENSMUST00000083577.1 | 0 | Mir129-2 | ENSMUSG00000065511.1 |
| chr2 | 94243303 | 94244370 | 3.68912 | 4.48892 | 1.43655 | Distal Intergenic | 2 | 94241364 | 94241453 | 90 | 2 | ENSMUST00000083577.1 | -1851 | Mir129-2 | ENSMUSG00000065511.1 |
| chr2 | 94245067 | 94245447 | 3.70061 | 4.51491 | 1.45587 | Distal Intergenic | 2 | 94241364 | 94241453 | 90 | 2 | ENSMUST00000083577.1 | -3615 | Mir129-2 | ENSMUSG00000065511.1 |
| chr2 | 94245999 | 94246481 | 3.54921 | 4.27038 | 1.27677 | Distal Intergenic | 2 | 94241364 | 94241453 | 90 | 2 | ENSMUST00000083577.1 | -4547 | Mir129-2 | ENSMUSG00000065511.1 |
| chr2 | 94247001 | 94247200 | 3.68926 | 4.50946 | 1.42867 | Distal Intergenic | 2 | 94241364 | 94241453 | 90 | 2 | ENSMUST00000083577.1 | -5549 | Mir129-2 | ENSMUSG00000065511.1 |
| chr2 | 94248277 | 94248663 | 3.82122 | 4.97825 | 1.76889 | Distal Intergenic | 2 | 94241364 | 94241453 | 90 | 2 | ENSMUST00000083577.1 | -6825 | Mir129-2 | ENSMUSG00000065511.1 |
| chr12 | 117246541 | 117246616 | 3.63955 | 4.41221 | 1.43725 | Intron (ENSMUST00000190247.1/ENSMUSG00000056553.9, intron 17 of 22) | 12 | 117250817 | 117250885 | 69 | 1 | ENSMUST00000083604.1 | -4201 | Mir153 | ENSMUSG00000065538.1 |
| chr3 | 118433437 | 118434003 | 3.72227 | 4.59387 | 1.54864 | Promoter | 3 | 118433843 | 118433944 | 102 | 1 | ENSMUST00000083635.1 | 0 | Mir137 | ENSMUSG00000065569.1 |

| | | | | | | | | | | | | | | | |
|---|---|---|---|---|---|---|---|---|---|---|---|---|---|---|---|
| chr12 | 112130493 | 112130821 | 3.48116 | 4.14077 | 1.27635 | Promoter | 12 | 112130880 | 112130955 | 76 | 1 | ENSMUST00000083640.1 | -59 | Mir203 | ENSMUSG00000065574.1 |
| chr12 | 112131187 | 112131262 | 3.58348 | 4.55505 | 1.49967 | Promoter | 12 | 112130880 | 112130955 | 76 | 1 | ENSMUST00000083640.1 | 308 | Mir203 | ENSMUSG00000065574.1 |
| chr12 | 112134856 | 112134934 | 3.88964 | 4.8833 | 1.73279 | Distal Intergenic | 12 | 112130880 | 112130955 | 76 | 1 | ENSMUST00000083640.1 | 3977 | Mir203 | ENSMUSG00000065574.1 |
| chr11 | 35591490 | 35591646 | 3.38462 | 4.11822 | 1.20708 | Intron (ENSMUST00000069837.3/ENSMUSG00000056427.5, intron 10 of 35) | 11 | 35616816 | 35616925 | 110 | 1 | ENSMUST00000083649.1 | -25170 | Mir218-2 | ENSMUSG00000065583.1 |
| chr11 | 35593815 | 35594562 | 3.39742 | 3.95507 | 1.12768 | Intron (ENSMUST00000069837.3/ENSMUSG00000056427.5, intron 10 of 35) | 11 | 35616816 | 35616925 | 110 | 1 | ENSMUST00000083649.1 | -22254 | Mir218-2 | ENSMUSG00000065583.1 |
| chr6 | 30168886 | 30169319 | 3.76006 | 4.71045 | 1.63607 | Promoter | 6 | 30169446 | 30169551 | 106 | 2 | ENSMUST00000083652.1 | 232 | Mir96 | ENSMUSG00000065586.1 |
| chr14 | 64590296 | 64590835 | 4.01697 | 5.18793 | 1.94616 | Promoter | 14 | 64590657 | 64590741 | 85 | 1 | ENSMUST00000083663.1 | 0 | Mir124a-1 | ENSMUSG00000065597.1 |
| chr7 | 78882494 | 78883260 | 3.67074 | 4.44667 | 1.38504 | Distal Intergenic | 7 | 78888277 | 78888373 | 97 | 1 | ENSMUST00000083675.1 | -5017 | Mir7-2 | ENSMUSG00000065609.1 |
| chr7 | 78884300 | 78884508 | 4.20034 | 5.46353 | 2.067 | Distal Intergenic | 7 | 78888277 | 78888373 | 97 | 1 | ENSMUST00000083675.1 | -3769 | Mir7-2 | ENSMUSG00000065609.1 |
| chr7 | 78884972 | 78885275 | 3.60167 | 4.32307 | 1.32673 | Distal Intergenic | 7 | 78888277 | 78888373 | 97 | 1 | ENSMUST00000083675.1 | -3002 | Mir7-2 | ENSMUSG00000065609.1 |
| chr7 | 78885778 | 78885909 | 3.48812 | 4.09419 | 1.13468 | Downstream (2-3kb) | 7 | 78888277 | 78888373 | 97 | 1 | ENSMUST00000083675.1 | -2368 | Mir7-2 | ENSMUSG00000065609.1 |
| chr7 | 78888702 | 78888813 | 3.43908 | 4.01081 | 1.10163 | Promoter | 7 | 78888277 | 78888373 | 97 | 1 | ENSMUST00000083675.1 | 426 | Mir7-2 | ENSMUSG00000065609.1 |
| chr7 | 78890101 | 78890244 | 3.37236 | 3.88617 | 1.01733 | Distal Intergenic | 7 | 78888277 | 78888373 | 97 | 1 | ENSMUST00000083675.1 | 1825 | Mir7-2 | ENSMUSG00000065609.1 |
| chr6 | 31017189 | 31017265 | 3.63154 | 4.38078 | 1.37795 | Distal Intergenic | 6 | 31062660 | 31062747 | 88 | 2 | ENSMUST00000083676.1 | 45482 | Mir29a | ENSMUSG00000065610.1 |
| chr6 | 31017668 | 31017743 | 4.0135 | 5.12584 | 1.88857 | Distal Intergenic | 6 | 31062660 | 31062747 | 88 | 2 | ENSMUST00000083676.1 | 45004 | Mir29a | ENSMUSG00000065610.1 |
| chr1 | 74897338 | 74898223 | 3.43376 | 4.00152 | 1.09742 | Downstream (2-3kb) | 1 | 74900652 | 74900726 | 75 | 2 | ENSMUST00000083682.2 | 2503 | Mir375 | ENSMUSG00000065616.2 |
| chr1 | 74900472 | 74901447 | 3.98253 | 5.08044 | 1.81631 | Promoter | 1 | 74900652 | 74900726 | 75 | 2 | ENSMUST00000083682.2 | 0 | Mir375 | ENSMUSG00000065616.2 |
| chr6 | 30169935 | 30170461 | 3.93193 | 5.10543 | 1.87237 | Promoter | 6 | 30169668 | 30169737 | 70 | 2 | ENSMUST00000083685.2 | -199 | Mir183 | ENSMUSG00000065619.2 |
| chr6 | 30171308 | 30172314 | 3.77176 | 4.74865 | 1.63743 | Distal Intergenic | 6 | 30169668 | 30169737 | 70 | 2 | ENSMUST00000083685.2 | -1572 | Mir183 | ENSMUSG00000065619.2 |
| chr6 | 30172709 | 30174847 | 3.77739 | 5.01226 | 1.79886 | Distal Intergenic | 6 | 30169668 | 30169737 | 70 | 2 | ENSMUST00000083685.2 | -2973 | Mir183 | ENSMUSG00000065619.2 |
| chr6 | 30175510 | 30176171 | 4.391 | 6.09995 | 2.58474 | Distal Intergenic | 6 | 30169668 | 30169737 | 70 | 2 | ENSMUST00000083685.2 | -5774 | Mir183 | ENSMUSG00000065619.2 |
| chr6 | 30176576 | 30177700 | 3.84028 | 5.08976 | 1.83582 | Distal Intergenic | 6 | 30169668 | 30169737 | 70 | 2 | ENSMUST00000083685.2 | -6840 | Mir183 | ENSMUSG00000065619.2 |
| chr6 | 30178009 | 30180203 | 3.69385 | 4.60644 | 1.56323 | Distal Intergenic | 6 | 30169668 | 30169737 | 70 | 2 | ENSMUST00000083685.2 | -8273 | Mir183 | ENSMUSG00000065619.2 |
| chr6 | 30183114 | 30183354 | 3.43012 | 4.11908 | 1.19889 | Distal Intergenic | 6 | 30169668 | 30169737 | 70 | 2 | ENSMUST00000083685.2 | -13378 | Mir183 | ENSMUSG00000065619.2 |
| chr6 | 30205029 | 30205104 | 3.36939 | 3.88169 | 1.01733 | Distal Intergenic | 6 | 30169668 | 30169737 | 70 | 2 | ENSMUST00000083685.2 | -35293 | Mir183 | ENSMUSG00000065619.2 |
| chr8 | 92036955 | 92037074 | 3.34261 | 3.84128 | 1.01733 | Intron (ENSMUST00000190257.1/ENSMUSG00000101613.1, intron 3 of 3) | 8 | 92039917 | 92040020 | 104 | 1 | ENSMUST00000083702.1 | -2843 | Gm24254 | ENSMUSG00000065636.1 |
| chr8 | 92062158 | 92062263 | 3.62279 | 4.35228 | 1.32821 | Distal Intergenic | 8 | 92039917 | 92040020 | 104 | 1 | ENSMUST00000083702.1 | 22242 | Gm24254 | ENSMUSG00000065636.1 |
| chr8 | 92096175 | 92096346 | 3.70677 | 4.52062 | 1.45578 | Distal Intergenic | 8 | 92039917 | 92040020 | 104 | 1 | ENSMUST00000083702.1 | 56259 | Gm24254 | ENSMUSG00000065636.1 |
| chr8 | 92113506 | 92113693 | 3.77363 | 4.70405 | 1.64135 | Distal Intergenic | 8 | 92039917 | 92040020 | 104 | 1 | ENSMUST00000083702.1 | 73590 | Gm24254 | ENSMUSG00000065636.1 |
| chr8 | 92114858 | 92115178 | 3.65892 | 4.47436 | 1.43519 | Distal Intergenic | 8 | 92039917 | 92040020 | 104 | 1 | ENSMUST00000083702.1 | 74942 | Gm24254 | ENSMUSG00000065636.1 |
| chr8 | 92134030 | 92134259 | 3.58021 | 4.27515 | 1.28429 | Distal Intergenic | 8 | 92039917 | 92040020 | 104 | 1 | ENSMUST00000083702.1 | 94114 | Gm24254 | ENSMUSG00000065636.1 |
| chr8 | 92135126 | 92136307 | 3.78446 | 4.68656 | 1.55275 | Distal Intergenic | 8 | 92039917 | 92040020 | 104 | 1 | ENSMUST00000083702.1 | 95210 | Gm24254 | ENSMUSG00000065636.1 |
| chr8 | 92136617 | 92136717 | 3.35971 | 3.86708 | 1.01733 | Distal Intergenic | 8 | 92039917 | 92040020 | 104 | 1 | ENSMUST00000083702.1 | 96701 | Gm24254 | ENSMUSG00000065636.1 |
| chr18 | 84768695 | 84768814 | 3.42006 | 4.01073 | 1.19329 | Distal Intergenic | 18 | 84742162 | 84742317 | 156 | 2 | ENSMUST00000083812.1 | -26379 | Gm25005 | ENSMUSG00000065746.1 |
| chr3 | 96456385 | 96459625 | 3.84651 | 6.34009 | 2.78519 | Promoter | 3 | 96460024 | 96460189 | 166 | 2 | ENSMUST00000083839.1 | 564 | Rnu1b6 | ENSMUSG00000065773.1 |
| chr3 | 96464001 | 96464352 | 3.3901 | 3.91854 | 1.0371 | Distal Intergenic | 3 | 96460024 | 96460189 | 166 | 2 | ENSMUST00000083839.1 | -3813 | Rnu1b6 | ENSMUSG00000065773.1 |
| chr3 | 96465100 | 96465531 | 3.84447 | 4.86495 | 1.71117 | Distal Intergenic | 3 | 96460024 | 96460189 | 166 | 2 | ENSMUST00000083839.1 | -4912 | Rnu1b6 | ENSMUSG00000065773.1 |
| chr3 | 96471136 | 96471277 | 3.69406 | 4.55925 | 1.51354 | Distal Intergenic | 3 | 96460024 | 96460189 | 166 | 2 | ENSMUST00000083839.1 | -10948 | Rnu1b6 | ENSMUSG00000065773.1 |
| chr3 | 96471631 | 96472052 | 4.28696 | 5.80073 | 2.39224 | Distal Intergenic | 3 | 96460024 | 96460189 | 166 | 2 | ENSMUST00000083839.1 | -11443 | Rnu1b6 | ENSMUSG00000065773.1 |
| chr3 | 103421482 | 103421718 | 3.28699 | 3.84477 | 1.1042 | Distal Intergenic | 3 | 103427437 | 103427543 | 107 | 1 | ENSMUST00000083855.1 | -5719 | Gm25869 | ENSMUSG00000065789.1 |
| chr2 | 147800314 | 147800566 | 3.68554 | 4.55642 | 1.51517 | Distal Intergenic | 2 | 147874772 | 147874878 | 107 | 2 | ENSMUST00000083857.1 | 74312 | Gm25516 | ENSMUSG00000065791.1 |
| chr2 | 147807268 | 147807418 | 3.41123 | 4.23022 | 1.28332 | Distal Intergenic | 2 | 147874772 | 147874878 | 107 | 2 | ENSMUST00000083857.1 | 67460 | Gm25516 | ENSMUSG00000065791.1 |
| chr2 | 147875603 | 147876304 | 3.56191 | 4.28693 | 1.37583 | Promoter | 2 | 147874772 | 147874878 | 107 | 2 | ENSMUST00000083857.1 | -726 | Gm25516 | ENSMUSG00000065791.1 |
| chr2 | 147877030 | 147877730 | 3.26281 | 3.73217 | 1.03908 | Distal Intergenic | 2 | 147874772 | 147874878 | 107 | 2 | ENSMUST00000083857.1 | -2153 | Gm25516 | ENSMUSG00000065791.1 |
| chr2 | 147878654 | 147878942 | 3.61022 | 4.38974 | 1.40274 | Distal Intergenic | 2 | 147874772 | 147874878 | 107 | 2 | ENSMUST00000083857.1 | -3777 | Gm25516 | ENSMUSG00000065791.1 |
| chr2 | 147905334 | 147905847 | 3.26862 | 3.88529 | 1.06724 | Distal Intergenic | 2 | 147874772 | 147874878 | 107 | 2 | ENSMUST00000083857.1 | -30457 | Gm25516 | ENSMUSG00000065791.1 |
| chr16 | 91350829 | 91351247 | 3.5906 | 4.28651 | 1.26442 | Distal Intergenic | 16 | 91349702 | 91349784 | 83 | 2 | ENSMUST00000083877.1 | -1046 | Gm24695 | ENSMUSG00000065811.1 |
| chr16 | 91353357 | 91353604 | 3.55324 | 4.30593 | 1.33623 | Distal Intergenic | 16 | 91349702 | 91349784 | 83 | 2 | ENSMUST00000083877.1 | -3574 | Gm24695 | ENSMUSG00000065811.1 |
| chr15 | 61158159 | 61158810 | 3.31566 | 3.87161 | 1.11399 | Distal Intergenic | 15 | 61357871 | 61358004 | 134 | 2 | ENSMUST00000083878.1 | 199194 | Gm24696 | ENSMUSG00000065812.1 |
| chr15 | 61584095 | 61584434 | 3.49343 | 4.17748 | 1.29172 | Distal Intergenic | 15 | 61357871 | 61358004 | 134 | 2 | ENSMUST00000083878.1 | -226092 | Gm24696 | ENSMUSG00000065812.1 |
| chr15 | 61587171 | 61587378 | 3.40862 | 3.99627 | 1.19398 | Distal Intergenic | 15 | 61357871 | 61358004 | 134 | 2 | ENSMUST00000083878.1 | -229168 | Gm24696 | ENSMUSG00000065812.1 |
| chr15 | 61593449 | 61593827 | 3.5634 | 4.27382 | 1.29139 | Distal Intergenic | 15 | 61357871 | 61358004 | 134 | 2 | ENSMUST00000083878.1 | -235446 | Gm24696 | ENSMUSG00000065812.1 |
| chr7 | 16471693 | 16472104 | 3.53327 | 4.27091 | 1.32494 | Promoter | 7 | 16471813 | 16471931 | 119 | 1 | ENSMUST00000083959.1 | 0 | n-R5s151 | ENSMUSG00000065893.1 |
| chr7 | 16473910 | 16474202 | 3.70592 | 4.53755 | 1.51103 | Intron (ENSMUST00000002053.8/ENSMUSG00000001988.8, intron 3 of 11) | 7 | 16471813 | 16471931 | 119 | 1 | ENSMUST00000083959.1 | 2098 | n-R5s151 | ENSMUSG00000065893.1 |
| chr1 | 171024364 | 171024720 | 3.29433 | 3.76865 | 1.01733 | Intron (ENSMUST00000078825.4/ENSMUSG00000059089.4, intron 3 of 4) | 1 | 171027965 | 171028129 | 165 | 1 | ENSMUST00000083971.1 | -3245 | Gm26110 | ENSMUSG00000065905.1 |
| chr1 | 171029759 | 171030213 | 3.30393 | 3.80366 | 1.08358 | 3' UTR | 1 | 171027965 | 171028129 | 165 | 1 | ENSMUST00000083971.1 | 1795 | Gm26110 | ENSMUSG00000065905.1 |
| chr15 | 93234401 | 93234478 | 3.93557 | 5.526 | 2.119 | Promoter | 15 | 93234363 | 93234700 | 338 | 1 | ENSMUST00000083977.1 | 39 | Gm24447 | ENSMUSG00000065911.1 |
| chr11 | 40928257 | 40928519 | 3.44703 | 4.03647 | 1.15081 | Distal Intergenic | 11 | 41034968 | 41035123 | 156 | 2 | ENSMUST00000083982.1 | 106604 | Gm24446 | ENSMUSG00000065916.1 |
| chr18 | 73533423 | 73533498 | 2.82802 | 8.46323 | 4.48133 | Promoter | 18 | 73533403 | 73533550 | 148 | 2 | ENSMUST00000083988.1 | 52 | n-R5-8s1 | ENSMUSG00000065922.1 |
| chr18 | 73544023 | 73544289 | 3.73697 | 4.62959 | 1.57362 | Distal Intergenic | 18 | 73533403 | 73533550 | 148 | 2 | ENSMUST00000083988.1 | -10474 | n-R5-8s1 | ENSMUSG00000065922.1 |
| chr16 | 72029406 | 72029580 | 3.33595 | 3.85822 | 1.10566 | Distal Intergenic | 16 | 71663788 | 71663918 | 131 | 1 | ENSMUST00000083990.1 | 365619 | Gm22797 | ENSMUSG00000065924.1 |
| chr16 | 13986708 | 13986843 | 3.51315 | 4.17895 | 1.29277 | Promoter | 16 | 13981702 | 13986855 | 5154 | 2 | ENSMUST00000117803.1 | 12 | Ifitm7 | ENSMUSG00000065968.7 |
| chr4 | 119071730 | 119072031 | 3.58629 | 4.39765 | 1.39389 | Distal Intergenic | 4 | 119054790 | 119069111 | 14322 | 2 | ENSMUST00000128998.1 | -2620 | Gm12866 | ENSMUSG00000066060.5 |
| chr4 | 119072782 | 119073115 | 4.38507 | 5.97735 | 2.4981 | Distal Intergenic | 4 | 119054790 | 119069111 | 14322 | 2 | ENSMUST00000128998.1 | -3672 | Gm12866 | ENSMUSG00000066060.5 |
| chr7 | 140687106 | 140687407 | 3.5729 | 4.25734 | 1.25911 | Distal Intergenic | 7 | 140690884 | 140691918 | 1035 | 1 | ENSMUST00000084454.3 | -3477 | Olfr45 | ENSMUSG00000066122.3 |
| chr7 | 139893155 | 139893636 | 3.42009 | 3.97375 | 1.07209 | Distal Intergenic | 7 | 139894696 | 139923776 | 29081 | 1 | ENSMUST00000151421.2 | -1060 | Kndc1 | ENSMUSG00000066129.9 |
| chr7 | 139894384 | 139895348 | 3.69891 | 4.50872 | 1.41905 | Promoter | 7 | 139894696 | 139923776 | 29081 | 1 | ENSMUST00000151421.2 | 0 | Kndc1 | ENSMUSG00000066129.9 |
| chr7 | 139914332 | 139914520 | 3.44611 | 4.02391 | 1.11064 | 3' UTR | 7 | 139923552 | 139941537 | 17986 | 1 | ENSMUST00000156941.1 | -9032 | Kndc1 | ENSMUSG00000066129.9 |
| chr7 | 139915735 | 139916844 | 3.4693 | 4.12136 | 1.19723 | Intron (ENSMUST00000151421.2/ENSMUSG00000066129.9, intron 9 of 16) | 7 | 139923552 | 139941537 | 17986 | 1 | ENSMUST00000156941.1 | -6708 | Kndc1 | ENSMUSG00000066129.9 |
| chr7 | 122670418 | 122670717 | 3.27595 | 3.74119 | 1.01733 | Promoter | 7 | 122670492 | 122769393 | 98902 | 1 | ENSMUST00000084615.4 | 0 | Cacng3 | ENSMUSG00000066189.4 |
| chr7 | 122671926 | 122672333 | 4.2615 | 5.93055 | 2.46772 | Promoter | 7 | 122671418 | 122768540 | 97123 | 1 | ENSMUST00000182095.1 | 509 | Cacng3 | ENSMUSG00000066189.4 |
| chr7 | 122713035 | 122713279 | 3.78298 | 4.72585 | 1.62089 | Intron (ENSMUST00000084615.4/ENSMUSG00000066189.4, intron 1 of 3) | 7 | 122671418 | 122768540 | 97123 | 1 | ENSMUST00000182095.1 | 41618 | Cacng3 | ENSMUSG00000066189.4 |
| chr7 | 119184067 | 119184307 | 3.70606 | 4.49594 | 1.38442 | Promoter | 7 | 119144323 | 119184374 | 40052 | 2 | ENSMUST00000084650.4 | 67 | Gpr139 | ENSMUSG00000066197.4 |
| chr7 | 119184975 | 119185502 | 4.33849 | 5.89563 | 2.35689 | Promoter | 7 | 119144323 | 119184374 | 40052 | 2 | ENSMUST00000084650.4 | -602 | Gpr139 | ENSMUSG00000066197.4 |
| chr4 | 41726679 | 41726836 | 3.66364 | 4.55682 | 1.47581 | Intron (ENSMUST00000108041.2/ENSMUSG00000073889.5, intron 2 of 13) | 4 | 41724139 | 41730286 | 6148 | 2 | ENSMUST00000084698.6 | 3450 | Arid3c | ENSMUSG00000066224.8 |
| chr4 | 41728206 | 41728388 | 3.67183 | 4.50241 | 1.48266 | Intron (ENSMUST00000108041.2/ENSMUSG00000073889.5, intron 2 of 13) | 4 | 41724139 | 41730286 | 6148 | 2 | ENSMUST00000084698.6 | 1898 | Arid3c | ENSMUSG00000066224.8 |
| chr4 | 41730068 | 41731546 | 4.09136 | 5.43922 | 2.09423 | Promoter | 4 | 41723836 | 41731142 | 7307 | 2 | ENSMUST00000171251.2 | 0 | Arid3c | ENSMUSG00000066224.8 |
| chr7 | 102713407 | 102714054 | 3.34882 | 3.88552 | 1.13116 | Promoter | 7 | 102713455 | 102714411 | 957 | 2 | ENSMUST00000084817.2 | 357 | Olfr33 | ENSMUSG00000066273.5 |
| chr8 | 87038837 | 87039240 | 3.50204 | 4.14261 | 1.2273 | Distal Intergenic | 8 | 87047689 | 87048226 | 538 | 1 | ENSMUST00000183478.1 | -8449 | Rps13-ps1 | ENSMUSG00000066362.5 |
| chr12 | 89810821 | 89811078 | 3.45856 | 4.0595 | 1.17135 | Intron (ENSMUST00000190626.1/ENSMUSG00000066392.6, intron 13 of 18) | 12 | 89812483 | 90332899 | 520417 | 1 | ENSMUST00000110133.3 | -1405 | Nrxn3 | ENSMUSG00000066392.6 |
| chr12 | 89811414 | 89811863 | 3.94871 | 4.98329 | 1.76228 | Promoter | 12 | 89812483 | 90332899 | 520417 | 1 | ENSMUST00000110133.3 | -620 | Nrxn3 | ENSMUSG00000066392.6 |
| chr13 | 73309895 | 73310132 | 3.61692 | 4.44588 | 1.42704 | Distal Intergenic | 13 | 73317844 | 73318143 | 300 | 1 | ENSMUST00000085163.3 | -7712 | Gm10263 | ENSMUSG00000066407.3 |
| chr12 | 80693724 | 80693926 | 3.62274 | 4.52078 | 1.46896 | Intron (ENSMUST00000140770.1/ENSMUSG00000066438.6, intron 1 of 12) | 12 | 80692591 | 80724214 | 31624 | 1 | ENSMUST00000140770.1 | 1134 | Plekhd1 | ENSMUSG00000066438.6 |
| chr9 | 74842948 | 74843052 | 3.59973 | 4.41989 | 1.40201 | Distal Intergenic | 9 | 74847983 | 74852873 | 4891 | 1 | ENSMUST00000160979.3 | -4931 | Gm16551 | ENSMUSG00000066477.8 |
| chr9 | 74845965 | 74846268 | 3.67225 | 4.55179 | 1.51073 | Distal Intergenic | 9 | 74847983 | 74852873 | 4891 | 1 | ENSMUST00000160979.3 | -1715 | Gm16551 | ENSMUSG00000066477.8 |
| chr9 | 74846873 | 74847674 | 3.89007 | 4.99373 | 1.76654 | Promoter | 9 | 74847983 | 74852873 | 4891 | 1 | ENSMUST00000160979.3 | -309 | Gm16551 | ENSMUSG00000066477.8 |
| chr9 | 74848195 | 74849365 | 4.29254 | 5.82232 | 2.44531 | Promoter | 9 | 74848437 | 74860134 | 11698 | 1 | ENSMUST00000161862.2 | 0 | Gm16551 | ENSMUSG00000066477.8 |
| chr9 | 74849816 | 74851934 | 3.91244 | 4.97122 | 1.7722 | Promoter | 9 | 74848995 | 74851204 | 2210 | 1 | ENSMUST00000160842.2 | 822 | Gm16551 | ENSMUSG00000066477.8 |
| chr9 | 73163617 | 73163729 | 3.8448 | 4.78744 | 1.64515 | Distal Intergenic | 9 | 73175429 | 73175741 | 313 | 2 | ENSMUST00000183748.1 | 12012 | Gm5745 | ENSMUSG00000066478.3 |
| chr9 | 65520162 | 65520649 | 4.68436 | 6.45842 | 2.7976 | Distal Intergenic | 9 | 65501453 | 65516999 | 15547 | 2 | ENSMUST00000061766.5 | -3164 | Ankdd1a | ENSMUSG00000066510.4 |
| chr9 | 65521361 | 65522222 | 4.085 | 5.55731 | 2.21873 | Distal Intergenic | 9 | 65501453 | 65516999 | 15547 | 2 | ENSMUST00000061766.5 | -4363 | Ankdd1a | ENSMUSG00000066510.4 |
| chr7 | 36157039 | 36157777 | 3.52427 | 4.19655 | 1.2673 | Distal Intergenic | 7 | 36118274 | 36118828 | 555 | 1 | ENSMUST00000119541.1 | 38766 | Rpl17-ps9 | ENSMUSG00000066543.8 |
| chr9 | 58479669 | 58479892 | 3.44466 | 4.07543 | 1.23276 | Distal Intergenic | 9 | 58488603 | 58499742 | 11140 | 1 | ENSMUST00000085658.4 | -8711 | 6030419C18Rik | ENSMUSG00000066607.4 |
| chr9 | 58489088 | 58489274 | 3.53947 | 4.19534 | 1.2165 | Promoter | 9 | 58488603 | 58499742 | 11140 | 1 | ENSMUST00000085658.4 | 486 | 6030419C18Rik | ENSMUSG00000066607.4 |
| chr1 | 180893122 | 180893236 | 3.36941 | 3.88171 | 1.01733 | Promoter | 1 | 180893108 | 180899103 | 5996 | 1 | ENSMUST00000085797.5 | 15 | Lefty2 | ENSMUSG00000066652.5 |
| chr1 | 180893802 | 180894159 | 3.66985 | 4.4483 | 1.40016 | Promoter | 1 | 180894458 | 180897487 | 3030 | 1 | ENSMUST00000155546.1 | -299 | Lefty2 | ENSMUSG00000066652.5 |
| chr1 | 180894557 | 180895017 | 4.11857 | 5.36081 | 2.04778 | Promoter | 1 | 180894458 | 180897487 | 3030 | 1 | ENSMUST00000155546.1 | 100 | Lefty2 | ENSMUSG00000066652.5 |
| chr9 | 48834813 | 48835471 | 3.55182 | 4.30221 | 1.34804 | Promoter | 9 | 48654311 | 48835945 | 181635 | 2 | ENSMUST00000093852.3 | 474 | Zbtb16 | ENSMUSG00000066687.4 |
| chr9 | 48854520 | 48854607 | 4.39163 | 6.36391 | 2.7505 | Distal Intergenic | 9 | 48654311 | 48835945 | 181635 | 2 | ENSMUST00000093852.3 | -18576 | Zbtb16 | ENSMUSG00000066687.4 |
| chr17 | 23683300 | 23683422 | 3.44443 | 4.08448 | 1.24209 | Promoter | 17 | 23682584 | 23684018 | 1435 | 2 | ENSMUST00000085989.6 | 596 | Cldn9 | ENSMUSG00000066720.6 |
| chr7 | 17072245 | 17072360 | 3.629 | 4.39463 | 1.42648 | Distal Intergenic | 7 | 17074040 | 17133450 | 59411 | 1 | ENSMUST00000071399.7 | -1680 | Psg16 | ENSMUSG00000066760.7 |
| chr9 | 35305217 | 35305587 | 1.87461 | 11.32151 | 6.85667 | Distal Intergenic | 9 | 35281518 | 35281835 | 318 | 2 | ENSMUST00000091097.2 | -23383 | Gm5614 | ENSMUSG00000066771.2 |
| chr1 | 150993298 | 150993665 | 3.45355 | 4.69801 | 1.5672 | Promoter | 1 | 150873748 | 150993435 | 119688 | 2 | ENSMUST00000155715.1 | 0 | Hmcn1 | ENSMUSG00000066842.13 |
| chr1 | 150994911 | 150995304 | 3.81142 | 5.24591 | 1.94177 | Distal Intergenic | 1 | 150873748 | 150993435 | 119688 | 2 | ENSMUST00000155715.1 | -1477 | Hmcn1 | ENSMUSG00000066842.13 |
| chr5 | 114309599 | 114309963 | 3.45472 | 4.05826 | 1.15862 | Distal Intergenic | 5 | 114314941 | 114362208 | 47268 | 1 | ENSMUST00000124316.2 | -4978 | Myo1h | ENSMUSG00000066952.6 |
| chr5 | 114315476 | 114315564 | 3.84891 | 4.78165 | 1.60914 | Promoter | 5 | 114314941 | 114364576 | 49636 | 1 | ENSMUST00000169347.1 | 536 | Myo1h | ENSMUSG00000066952.6 |
| chr5 | 113220417 | 113221431 | 3.83244 | 4.82027 | 1.66675 | Distal Intergenic | 5 | 113226909 | 113239263 | 12355 | 1 | ENSMUST00000086615.2 | -5478 | Tmem211 | ENSMUSG00000066964.2 |
| chr7 | 131557832 | 131558679 | 3.63799 | 4.52418 | 1.49017 | Intron (ENSMUST00000124096.2/ENSMUSG00000030849.12, intron 2 of 6) | 7 | 131560391 | 131571898 | 11508 | 1 | ENSMUST00000084502.5 | -1712 | Bub3 | ENSMUSG00000066979.5 |
| chr4 | 97109564 | 97109842 | 3.35039 | 3.85301 | 1.01733 | Distal Intergenic | 4 | 97182840 | 97183166 | 327 | 2 | ENSMUST00000086672.2 | 73324 | Gm10192 | ENSMUSG00000066990.2 |
| chr1 | 89991657 | 89992583 | 3.89986 | 4.89055 | 1.68273 | Intron (ENSMUST00000086882.2/ENSMUSG00000067081.7, intron 3 of 5) | 1 | 89950220 | 90014577 | 64358 | 2 | ENSMUST00000086882.2 | 21994 | Asb18 | ENSMUSG00000067081.7 |
| chr1 | 89992960 | 89993366 | 3.35429 | 3.8589 | 1.01733 | Exon (ENSMUST00000086882.2/ENSMUSG00000067081.7, exon 3 of 6) | 1 | 89950220 | 90014577 | 64358 | 2 | ENSMUST00000086882.2 | 21211 | Asb18 | ENSMUSG00000067081.7 |
| chr17 | 33977662 | 33977792 | 3.29859 | 4.09067 | 1.18241 | Promoter | 17 | 33974831 | 33978790 | 3960 | 2 | ENSMUST00000174250.2 | 998 | H2-K2 | ENSMUSG00000067203.7 |
| chr5 | 72645919 | 72646826 | 4.44818 | 6.12159 | 2.62943 | Promoter | 5 | 72647796 | 72671078 | 23283 | 1 | ENSMUST00000087212.3 | -970 | Nipal1 | ENSMUSG00000067219.5 |
| chr5 | 72647178 | 72648311 | 3.61147 | 4.42033 | 1.42268 | Promoter | 5 | 72647796 | 72671078 | 23283 | 1 | ENSMUST00000087212.3 | 0 | Nipal1 | ENSMUSG00000067219.5 |
| chr17 | 35470125 | 35471514 | 3.76969 | 4.85976 | 1.77355 | Promoter | 17 | 35470097 | 35474551 | 4455 | 1 | ENSMUST00000174589.1 | 29 | H2-Q10 | ENSMUSG00000067235.9 |
| chr17 | 35473163 | 35473509 | 4.12059 | 5.37239 | 2.05264 | Exon (ENSMUST00000068291.6/ENSMUSG00000067235.9, exon 4 of 7) | 17 | 35470097 | 35474551 | 4455 | 1 | ENSMUST00000174589.1 | 3067 | H2-Q10 | ENSMUSG00000067235.9 |
| chr19 | 38264577 | 38265097 | 3.68558 | 4.65974 | 1.51433 | Promoter | 19 | 38264782 | 38269046 | 4265 | 1 | ENSMUST00000134832.1 | 0 | Lgi1 | ENSMUSG00000067242.6 |
| chr19 | 38265973 | 38266222 | 3.3705 | 3.89855 | 1.06919 | Promoter | 19 | 38265410 | 38301269 | 35860 | 1 | ENSMUST00000130039.1 | 564 | Lgi1 | ENSMUSG00000067242.6 |
| chr19 | 38280698 | 38280773 | 3.41951 | 3.9769 | 1.08622 | Intron (ENSMUST00000087252.6/ENSMUSG00000067242.6, intron 2 of 7) | 19 | 38265410 | 38301269 | 35860 | 1 | ENSMUST00000130039.1 | 15289 | Lgi1 | ENSMUSG00000067242.6 |
| chr4 | 99654527 | 99660023 | 4.25913 | 5.70714 | 2.27798 | Promoter | 4 | 99656299 | 99658622 | 2324 | 1 | ENSMUST00000087285.4 | 0 | Foxd3 | ENSMUSG00000067261.4 |
| chr5 | 46445157 | 46445571 | 3.38122 | 4.07884 | 1.17504 | Distal Intergenic | 5 | 46414344 | 46414616 | 273 | 1 | ENSMUST00000121537.1 | 30814 | Gm7931 | ENSMUSG00000067321.4 |
| chr19 | 28721722 | 28722003 | 3.38193 | 3.9006 | 1.01733 | Distal Intergenic | 19 | 28742159 | 28742551 | 393 | 2 | ENSMUST00000087473.1 | 20548 | Rps15a-ps2 | ENSMUSG00000067351.1 |
| chr5 | 35391112 | 35392737 | 3.80862 | 4.74752 | 1.59129 | 3' UTR | 5 | 35389117 | 35392861 | 3745 | 1 | ENSMUST00000087674.4 | 1996 | Hmx1 | ENSMUSG00000067438.4 |
| chr5 | 35393190 | 35393676 | 4.22695 | 5.59295 | 2.21202 | Intron (ENSMUST00000172923.1/ENSMUSG00000067438.4, intron 1 of 1) | 5 | 35389117 | 35392861 | 3745 | 1 | ENSMUST00000087674.4 | 4074 | Hmx1 | ENSMUSG00000067438.4 |
| chr5 | 35394080 | 35396053 | 3.65402 | 4.62967 | 1.531 | Intron (ENSMUST00000172923.1/ENSMUSG00000067438.4, intron 1 of 1) | 5 | 35389117 | 35392861 | 3745 | 1 | ENSMUST00000087674.4 | 4964 | Hmx1 | ENSMUSG00000067438.4 |
| chr5 | 35396947 | 35398641 | 3.83307 | 4.84068 | 1.66346 | Intron (ENSMUST00000172923.1/ENSMUSG00000067438.4, intron 1 of 1) | 5 | 35389117 | 35392861 | 3745 | 1 | ENSMUST00000087674.4 | 7831 | Hmx1 | ENSMUSG00000067438.4 |
| chr5 | 35399106 | 35401597 | 3.98762 | 5.06891 | 1.80474 | Exon (ENSMUST00000172923.1/ENSMUSG00000067438.4, exon 2 of 2) | 5 | 35389117 | 35392861 | 3745 | 1 | ENSMUST00000087674.4 | 9990 | Hmx1 | ENSMUSG00000067438.4 |
| chr5 | 35403215 | 35403484 | 3.3607 | 3.86856 | 1.01733 | Distal Intergenic | 5 | 35389117 | 35392861 | 3745 | 1 | ENSMUST00000087674.4 | 14099 | Hmx1 | ENSMUSG00000067438.4 |
| chr5 | 35404318 | 35404585 | 3.4962 | 4.11368 | 1.16045 | Distal Intergenic | 5 | 35389117 | 35392861 | 3745 | 1 | ENSMUST00000087674.4 | 15202 | Hmx1 | ENSMUSG00000067438.4 |
| chr5 | 35405980 | 35406346 | 3.38193 | 3.9006 | 1.01733 | Distal Intergenic | 5 | 35389117 | 35392861 | 3745 | 1 | ENSMUST00000087674.4 | 16864 | Hmx1 | ENSMUSG00000067438.4 |
| chr5 | 35408654 | 35408881 | 4.20263 | 5.47806 | 2.08838 | Distal Intergenic | 5 | 35389117 | 35392861 | 3745 | 1 | ENSMUST00000087674.4 | 19538 | Hmx1 | ENSMUSG00000067438.4 |
| chr5 | 35412637 | 35412733 | 3.53482 | 4.18536 | 1.20695 | Distal Intergenic | 5 | 35389117 | 35392861 | 3745 | 1 | ENSMUST00000087674.4 | 23521 | Hmx1 | ENSMUSG00000067438.4 |
| chr5 | 35454465 | 35454763 | 3.37405 | 3.88871 | 1.01733 | Distal Intergenic | 5 | 35389117 | 35392861 | 3745 | 1 | ENSMUST00000087674.4 | 65349 | Hmx1 | ENSMUSG00000067438.4 |
| chr2 | 172040988 | 172041246 | 3.48298 | 4.11619 | 1.23495 | Intron (ENSMUST00000087950.3/ENSMUSG00000067578.3, intron 1 of 2) | 2 | 172036233 | 172043466 | 7234 | 2 | ENSMUST00000087950.3 | 2220 | Cbln4 | ENSMUSG00000067578.3 |
| chr2 | 172041753 | 172042263 | 3.65382 | 4.4753 | 1.49014 | Exon (ENSMUST00000087950.3/ENSMUSG00000067578.3, exon 1 of 3) | 2 | 172036233 | 172043466 | 7234 | 2 | ENSMUST00000087950.3 | 1203 | Cbln4 | ENSMUSG00000067578.3 |
| chr2 | 172042883 | 172043517 | 4.52483 | 6.33226 | 2.799 | Promoter | 2 | 172036233 | 172043466 | 7234 | 2 | ENSMUST00000087950.3 | 0 | Cbln4 | ENSMUSG00000067578.3 |
| chr13 | 51464357 | 51464854 | 3.54161 | 4.21378 | 1.23967 | Intron (ENSMUST00000021898.5/ENSMUSG00000021448.6, intron 6 of 11) | 13 | 51408618 | 51422797 | 14180 | 1 | ENSMUST00000087978.3 | 55740 | S1pr3 | ENSMUSG00000067586.3 |
| chr15 | 101861014 | 101861574 | 3.60649 | 4.50774 | 1.47013 | Exon (ENSMUST00000087996.5/ENSMUSG00000067594.5, exon 6 of 9) | 15 | 101858732 | 101869705 | 10974 | 2 | ENSMUST00000087996.5 | 8131 | Krt77 | ENSMUSG00000067594.5 |
| chr15 | 101869707 | 101870007 | 3.51634 | 4.19759 | 1.2829 | Promoter | 15 | 101858732 | 101869705 | 10974 | 2 | ENSMUST00000087996.5 | -3 | Krt77 | ENSMUSG00000067594.5 |
| chr15 | 101770839 | 101771283 | 3.6218 | 4.37004 | 1.35667 | Distal Intergenic | 15 | 101754259 | 101763504 | 9246 | 2 | ENSMUST00000088018.3 | -7336 | Krt74 | ENSMUSG00000067596.3 |
| chr15 | 101478485 | 101478666 | 3.23338 | 3.67724 | 1.01733 | Intron (ENSMUST00000088049.3/ENSMUSG00000067614.4, intron 7 of 8) | 15 | 101473478 | 101479983 | 6506 | 1 | ENSMUST00000088049.3 | 5008 | Krt86 | ENSMUSG00000067614.4 |
| chr1 | 16092545 | 16093461 | 3.43819 | 4.00835 | 1.09787 | Promoter | 1 | 16065979 | 16093325 | 27347 | 2 | ENSMUST00000145070.2 | 0 | 4930444P10Rik | ENSMUSG00000067795.8 |
| chr17 | 23535181 | 23536270 | 3.26226 | 5.01915 | 1.8154 | Distal Intergenic | 17 | 23551073 | 23553488 | 2416 | 2 | ENSMUST00000088673.4 | 17218 | Casp16 | ENSMUSG00000067882.8 |
| chr17 | 23536645 | 23536895 | 3.27629 | 4.87045 | 1.71083 | Distal Intergenic | 17 | 23551073 | 23553488 | 2416 | 2 | ENSMUST00000088673.4 | 16593 | Casp16 | ENSMUSG00000067882.8 |
| chr17 | 23537873 | 23538079 | 3.48157 | 4.46735 | 1.42887 | Distal Intergenic | 17 | 23551073 | 23553488 | 2416 | 2 | ENSMUST00000088673.4 | 15409 | Casp16 | ENSMUSG00000067882.8 |
| chr17 | 23538848 | 23539070 | 3.29264 | 4.08147 | 1.17676 | Distal Intergenic | 17 | 23551073 | 23553488 | 2416 | 2 | ENSMUST00000088673.4 | 14418 | Casp16 | ENSMUSG00000067882.8 |
| chr19 | 4708564 | 4708986 | 3.62218 | 4.34836 | 1.31568 | Distal Intergenic | 19 | 4711208 | 4752353 | 41146 | 1 | ENSMUST00000008991.6 | -2222 | Sptbn2 | ENSMUSG00000067889.4 |
| chr19 | 4709652 | 4709905 | 3.30439 | 3.78381 | 1.01733 | Distal Intergenic | 19 | 4711208 | 4752353 | 41146 | 1 | ENSMUST00000008991.6 | -1303 | Sptbn2 | ENSMUSG00000067889.4 |
| chr19 | 4710287 | 4710402 | 3.55114 | 4.35679 | 1.36471 | Promoter | 19 | 4711208 | 4752353 | 41146 | 1 | ENSMUST00000008991.6 | -806 | Sptbn2 | ENSMUSG00000067889.4 |
| chr19 | 4711878 | 4713000 | 3.39641 | 4.11806 | 1.20418 | Promoter | 19 | 4711208 | 4752353 | 41146 | 1 | ENSMUST00000008991.6 | 671 | Sptbn2 | ENSMUSG00000067889.4 |
| chr2 | 152375530 | 152375844 | 3.57663 | 4.33526 | 1.31267 | Promoter | 2 | 152374748 | 152375149 | 402 | 1 | ENSMUST00000089083.2 | 783 | Gm14164 | ENSMUSG00000068062.3 |
| chr2 | 152131852 | 152132099 | 3.52808 | 4.43116 | 1.39384 | Distal Intergenic | 2 | 152143561 | 152149097 | 5537 | 1 | ENSMUST00000089112.5 | -11462 | Tcf15 | ENSMUSG00000068079.5 |
| chr2 | 152142270 | 152142628 | 3.36173 | 3.87013 | 1.01733 | Promoter | 2 | 152143561 | 152149097 | 5537 | 1 | ENSMUST00000089112.5 | -933 | Tcf15 | ENSMUSG00000068079.5 |
| chr2 | 152143448 | 152144197 | 4.58245 | 6.34409 | 2.74907 | Promoter | 2 | 152143561 | 152149097 | 5537 | 1 | ENSMUST00000089112.5 | 0 | Tcf15 | ENSMUSG00000068079.5 |
| chr2 | 152144611 | 152144996 | 3.61895 | 4.34538 | 1.31486 | Intron (ENSMUST00000089112.5/ENSMUSG00000068079.5, intron 1 of 1) | 2 | 152143561 | 152149097 | 5537 | 1 | ENSMUST00000089112.5 | 1051 | Tcf15 | ENSMUSG00000068079.5 |
| chr2 | 152145530 | 152145605 | 3.66531 | 4.4409 | 1.39977 | Intron (ENSMUST00000089112.5/ENSMUSG00000068079.5, intron 1 of 1) | 2 | 152143561 | 152149097 | 5537 | 1 | ENSMUST00000089112.5 | 1970 | Tcf15 | ENSMUSG00000068079.5 |
| chr2 | 152147265 | 152147341 | 4.31748 | 5.7266 | 2.30889 | Intron (ENSMUST00000089112.5/ENSMUSG00000068079.5, intron 1 of 1) | 2 | 152143561 | 152149097 | 5537 | 1 | ENSMUST00000089112.5 | 3705 | Tcf15 | ENSMUSG00000068079.5 |
| chr15 | 82259584 | 82259839 | 3.35584 | 3.9323 | 1.12478 | 3' UTR | 15 | 82256023 | 82260753 | 4731 | 1 | ENSMUST00000143238.1 | 3562 | 1500009C09Rik | ENSMUSG00000068099.4 |
| chr15 | 82225184 | 82225431 | 3.94738 | 5.2287 | 1.92764 | Promoter | 15 | 82221745 | 82224339 | 2595 | 2 | ENSMUST00000109535.2 | -846 | Tnfrsf13c | ENSMUSG00000068105.5 |
| chr15 | 82064328 | 82065069 | 3.57865 | 4.30312 | 1.36193 | Exon (ENSMUST00000100396.2/ENSMUSG00000075524.2, exon 2 of 2) | 15 | 82069997 | 82126659 | 56663 | 1 | ENSMUST00000089178.4 | -4928 | Mei1 | ENSMUSG00000068117.4 |
| chr15 | 82069622 | 82069751 | 3.5271 | 4.20281 | 1.30271 | Promoter | 15 | 82069997 | 82126659 | 56663 | 1 | ENSMUST00000089178.4 | -246 | Mei1 | ENSMUSG00000068117.4 |
| chr15 | 82070569 | 82070882 | 3.54256 | 4.222 | 1.29929 | Promoter | 15 | 82069997 | 82126659 | 56663 | 1 | ENSMUST00000089178.4 | 573 | Mei1 | ENSMUSG00000068117.4 |
| chr2 | 146218199 | 146219842 | 4.48852 | 6.24253 | 2.69749 | Distal Intergenic | 2 | 146221921 | 146225020 | 3100 | 1 | ENSMUST00000089257.5 | -2079 | Insm1 | ENSMUSG00000068154.5 |

| | | | | | | | | | | | | | | | |
|---|---|---|---|---|---|---|---|---|---|---|---|---|---|---|---|
| chr2 | 146220715 | 146224465 | 4.08904 | 5.3232 | 1.99554 | Promoter | 2 | 146221921 | 146225020 | 3100 | 1 | ENSMUST00000089257.5 | 0 | Insm1 | ENSMUSG00000068154.5 |
| chr2 | 146224902 | 146225920 | 3.63966 | 4.38005 | 1.33046 | 3' UTR | 2 | 146221921 | 146225020 | 3100 | 1 | ENSMUST00000089257.5 | 2982 | Insm1 | ENSMUSG00000068154.5 |
| chr2 | 146226271 | 146226730 | 3.65136 | 4.39473 | 1.32237 | Distal Intergenic | 2 | 146221921 | 146225020 | 3100 | 1 | ENSMUST00000089257.5 | 4351 | Insm1 | ENSMUSG00000068154.5 |
| chr2 | 146227281 | 146227790 | 3.72077 | 4.62914 | 1.52402 | Distal Intergenic | 2 | 146221921 | 146225020 | 3100 | 1 | ENSMUST00000089257.5 | 5361 | Insm1 | ENSMUSG00000068154.5 |
| chr2 | 146233797 | 146233955 | 3.65019 | 4.67814 | 1.54443 | Distal Intergenic | 2 | 146221921 | 146225020 | 3100 | 1 | ENSMUST00000089257.5 | 11877 | Insm1 | ENSMUSG00000068154.5 |
| chr16 | 57754462 | 57754991 | 3.25796 | 3.7174 | 1.02808 | Promoter | 16 | 57624258 | 57754737 | 130480 | 2 | ENSMUST00000089332.4 | 0 | Col8a1 | ENSMUSG00000068196.4 |
| chr16 | 57756063 | 57756427 | 3.53396 | 4.53325 | 1.4797 | Distal Intergenic | 16 | 57624258 | 57754737 | 130480 | 2 | ENSMUST00000089332.4 | -1327 | Col8a1 | ENSMUSG00000068196.4 |
| chr15 | 79223776 | 79223973 | 4.06695 | 5.19809 | 1.86806 | Intron (ENSMUST00000144651.2/ENSMUSG00000075555.6, intron 3 of 3) | 15 | 79229173 | 79249484 | 20312 | 1 | ENSMUST00000163571.2 | -5200 | Pick1 | ENSMUSG00000068206.7 |
| chr15 | 78507744 | 78507849 | 3.50754 | 4.1416 | 1.19965 | Distal Intergenic | 15 | 78480553 | 78495066 | 14514 | 2 | ENSMUST00000089398.2 | -12679 | Il2rb | ENSMUSG00000068227.3 |
| chr15 | 77722725 | 77723332 | 3.46485 | 4.05818 | 1.13174 | Distal Intergenic | 15 | 77729121 | 77736381 | 7261 | 1 | ENSMUST00000109775.2 | -5789 | Apol9b | ENSMUSG00000068246.5 |
| chr6 | 87729793 | 87730043 | 3.53171 | 4.18483 | 1.22148 | Promoter | 6 | 87730869 | 87734017 | 3149 | 1 | ENSMUST00000160743.1 | -826 | Efcc1 | ENSMUSG00000068263.6 |
| chr6 | 87730829 | 87731552 | 4.08861 | 5.27865 | 1.97285 | Promoter | 6 | 87730869 | 87734017 | 3149 | 1 | ENSMUST00000160743.1 | 0 | Efcc1 | ENSMUSG00000068263.6 |
| chr6 | 87732682 | 87732857 | 3.93456 | 4.99615 | 1.78953 | Promoter | 6 | 87732809 | 87751896 | 19088 | 1 | ENSMUST00000159956.1 | 0 | Efcc1 | ENSMUSG00000068263.6 |
| chr6 | 87733178 | 87734221 | 3.81233 | 4.8045 | 1.66712 | Promoter | 6 | 87732809 | 87751896 | 19088 | 1 | ENSMUST00000159956.1 | 370 | Efcc1 | ENSMUSG00000068263.6 |
| chr6 | 85408255 | 85408577 | 3.74691 | 4.58863 | 1.49117 | Distal Intergenic | 6 | 85423886 | 85428877 | 4992 | 1 | ENSMUST00000089578.8 | -15309 | Noto | ENSMUSG00000068302.8 |
| chr6 | 85408891 | 85409626 | 3.64332 | 4.60971 | 1.53398 | Distal Intergenic | 6 | 85423886 | 85428877 | 4992 | 1 | ENSMUST00000089578.8 | -14260 | Noto | ENSMUSG00000068302.8 |
| chr6 | 85410630 | 85410923 | 3.26759 | 3.72852 | 1.01733 | Distal Intergenic | 6 | 85423886 | 85428877 | 4992 | 1 | ENSMUST00000089578.8 | -12963 | Noto | ENSMUSG00000068302.8 |
| chr6 | 85154988 | 85155388 | 3.53399 | 4.2285 | 1.32737 | Promoter | 6 | 85155005 | 85155968 | 964 | 1 | ENSMUST00000089584.6 | 0 | Spr-ps1 | ENSMUSG00000068303.6 |
| chr6 | 83064463 | 83064863 | 3.5846 | 4.51391 | 1.46026 | Exon (ENSMUST00000077502.2/ENSMUSG00000009145.4, exon 9 of 12) | 6 | 83068591 | 83070143 | 1553 | 2 | ENSMUST00000089641.4 | 5280 | Tlx2 | ENSMUSG00000068327.4 |
| chr6 | 83066199 | 83067623 | 3.55758 | 4.30372 | 1.28821 | 3' UTR | 6 | 83068591 | 83070143 | 1553 | 2 | ENSMUST00000089641.4 | 2520 | Tlx2 | ENSMUSG00000068327.4 |
| chr6 | 83068307 | 83071828 | 4.25987 | 5.83407 | 2.38909 | Promoter | 6 | 83068324 | 83070225 | 1902 | 2 | ENSMUST00000174674.2 | 0 | Tlx2 | ENSMUSG00000068327.4 |
| chr6 | 83072248 | 83072902 | 3.97463 | 5.1766 | 1.9383 | Distal Intergenic | 6 | 83068324 | 83070225 | 1902 | 2 | ENSMUST00000174674.2 | -2024 | Tlx2 | ENSMUSG00000068327.4 |
| chr2 | 104379204 | 104379441 | 3.37655 | 3.92127 | 1.11886 | Intron (ENSMUST00000089726.4/ENSMUSG00000068373.9, intron 1 of 20) | 2 | 104255123 | 104409992 | 154870 | 2 | ENSMUST00000149466.1 | 30551 | D430041D05Rik | ENSMUSG00000068373.9 |
| chr2 | 104409809 | 104411613 | 4.22914 | 5.72769 | 2.33483 | Promoter | 2 | 104143073 | 104410334 | 267262 | 2 | ENSMUST00000089726.4 | 0 | D430041D05Rik | ENSMUSG00000068373.9 |
| chr14 | 51357943 | 51358276 | 3.37265 | 3.89094 | 1.03258 | Distal Intergenic | 14 | 51361756 | 51569982 | 208227 | 1 | ENSMUST00000162998.1 | -3480 | Gm7247 | ENSMUSG00000068399.5 |
| chr14 | 51358582 | 51358739 | 4.06574 | 5.27294 | 1.98496 | Downstream (2-3kb) | 14 | 51361756 | 51569982 | 208227 | 1 | ENSMUST00000162998.1 | -3017 | Gm7247 | ENSMUSG00000068399.5 |
| chr16 | 26968273 | 26968406 | 3.63422 | 4.43256 | 1.46486 | Intron (ENSMUST00000089832.4/ENSMUSG00000068428.6, intron 1 of 4) | 16 | 26957236 | 26989974 | 32739 | 2 | ENSMUST00000089832.4 | 21568 | Gmnc | ENSMUSG00000068428.6 |
| chr2 | 122294311 | 122295282 | 3.45901 | 4.15191 | 1.20301 | Exon (ENSMUST00000053734.5/ENSMUSG00000068452.7, exon 8 of 32) | 2 | 122279247 | 122298165 | 18919 | 2 | ENSMUST00000053734.5 | 2883 | Duox2 | ENSMUSG00000068452.7 |
| chr2 | 122295616 | 122295869 | 3.46725 | 4.0597 | 1.12412 | Intron (ENSMUST00000053734.5/ENSMUSG00000068452.7, intron 6 of 31) | 2 | 122279247 | 122298165 | 18919 | 2 | ENSMUST00000053734.5 | 2296 | Duox2 | ENSMUSG00000068452.7 |
| chr2 | 122296178 | 122296564 | 3.60108 | 4.34921 | 1.38663 | Exon (ENSMUST00000053734.5/ENSMUSG00000068452.7, exon 5 of 32) | 2 | 122279247 | 122298165 | 18919 | 2 | ENSMUST00000053734.5 | 1601 | Duox2 | ENSMUSG00000068452.7 |
| chr2 | 122297110 | 122298877 | 3.84384 | 4.83774 | 1.71013 | Promoter | 2 | 122279247 | 122298165 | 18919 | 2 | ENSMUST00000053734.5 | 0 | Duox2 | ENSMUSG00000068452.7 |
| chr15 | 52040223 | 52040336 | 3.60903 | 4.33143 | 1.32879 | Promoter | 15 | 52040107 | 52045722 | 5616 | 1 | ENSMUST00000090025.4 | 117 | Aard | ENSMUSG00000068522.4 |
| chr3 | 116251200 | 116252736 | 4.55356 | 6.32089 | 2.79528 | Promoter | 3 | 116249654 | 116253484 | 3831 | 2 | ENSMUST00000090473.5 | 748 | Gpr88 | ENSMUSG00000068696.5 |
| chr3 | 116253051 | 116253691 | 3.65314 | 4.43571 | 1.43903 | Promoter | 3 | 116249654 | 116253484 | 3831 | 2 | ENSMUST00000090473.5 | 0 | Gpr88 | ENSMUSG00000068696.5 |
| chr3 | 116255026 | 116255353 | 3.56235 | 4.35215 | 1.42156 | Distal Intergenic | 3 | 116249654 | 116253484 | 3831 | 2 | ENSMUST00000090473.5 | -1543 | Gpr88 | ENSMUSG00000068696.5 |
| chr3 | 116256364 | 116256652 | 3.28139 | 3.79104 | 1.06026 | Distal Intergenic | 3 | 116249654 | 116253484 | 3831 | 2 | ENSMUST00000090473.5 | -2881 | Gpr88 | ENSMUSG00000068696.5 |
| chr6 | 29432845 | 29434660 | 3.9768 | 5.07324 | 1.82256 | Promoter | 6 | 29433153 | 29461888 | 28736 | 1 | ENSMUST00000090474.5 | 0 | Flnc | ENSMUSG00000068699.7 |
| chr6 | 29435449 | 29435592 | 3.80147 | 4.68074 | 1.51101 | Intron (ENSMUST00000090474.5/ENSMUSG00000068699.7, intron 1 of 47) | 6 | 29433276 | 29461849 | 28574 | 1 | ENSMUST00000101617.3 | 2174 | Flnc | ENSMUSG00000068699.7 |
| chr2 | 93177707 | 93177931 | 3.36612 | 3.87675 | 1.01733 | Distal Intergenic | 2 | 93187548 | 93198919 | 11372 | 1 | ENSMUST00000150462.2 | -9617 | Trp53i11 | ENSMUSG00000068735.9 |
| chr2 | 93178549 | 93179452 | 3.95562 | 5.01367 | 1.79704 | Distal Intergenic | 2 | 93187548 | 93198919 | 11372 | 1 | ENSMUST00000150462.2 | -8096 | Trp53i11 | ENSMUSG00000068735.9 |
| chr2 | 93180314 | 93180690 | 3.39223 | 3.92198 | 1.03787 | Distal Intergenic | 2 | 93187548 | 93198919 | 11372 | 1 | ENSMUST00000150462.2 | -6858 | Trp53i11 | ENSMUSG00000068735.9 |
| chr6 | 22874071 | 22874681 | 3.46733 | 4.06978 | 1.15971 | Promoter | 6 | 22875502 | 23052916 | 177415 | 1 | ENSMUST00000090568.3 | -821 | Ptprz1 | ENSMUSG00000068748.4 |
| chr6 | 22875177 | 22875860 | 3.51739 | 4.21149 | 1.24778 | Promoter | 6 | 22875502 | 23052916 | 177415 | 1 | ENSMUST00000090568.3 | 0 | Ptprz1 | ENSMUSG00000068748.4 |
| chr2 | 73259360 | 73259899 | 3.79503 | 4.68897 | 1.54698 | Distal Intergenic | 2 | 73271925 | 73277471 | 5547 | 1 | ENSMUST00000090813.5 | -12026 | Sp9 | ENSMUSG00000068859.5 |
| chr2 | 73260295 | 73260436 | 3.76105 | 4.63673 | 1.50086 | Distal Intergenic | 2 | 73271925 | 73277471 | 5547 | 1 | ENSMUST00000090813.5 | -11489 | Sp9 | ENSMUSG00000068859.5 |
| chr2 | 73261123 | 73261290 | 3.80748 | 4.72147 | 1.56694 | Distal Intergenic | 2 | 73271925 | 73277471 | 5547 | 1 | ENSMUST00000090813.5 | -10635 | Sp9 | ENSMUSG00000068859.5 |
| chr2 | 73261756 | 73262362 | 3.71677 | 4.52628 | 1.42008 | Distal Intergenic | 2 | 73271925 | 73277471 | 5547 | 1 | ENSMUST00000090813.5 | -9563 | Sp9 | ENSMUSG00000068859.5 |
| chr2 | 73262685 | 73263295 | 3.84076 | 4.79344 | 1.61376 | Distal Intergenic | 2 | 73271925 | 73277471 | 5547 | 1 | ENSMUST00000090813.5 | -8630 | Sp9 | ENSMUSG00000068859.5 |
| chr2 | 73263746 | 73265415 | 3.59142 | 4.43596 | 1.39454 | Distal Intergenic | 2 | 73271925 | 73277471 | 5547 | 1 | ENSMUST00000090813.5 | -6510 | Sp9 | ENSMUSG00000068859.5 |
| chr2 | 73265993 | 73266484 | 3.73069 | 4.54923 | 1.43869 | Distal Intergenic | 2 | 73271925 | 73277471 | 5547 | 1 | ENSMUST00000090813.5 | -5441 | Sp9 | ENSMUSG00000068859.5 |
| chr2 | 73266971 | 73267103 | 3.61559 | 4.33837 | 1.30567 | Distal Intergenic | 2 | 73271925 | 73277471 | 5547 | 1 | ENSMUST00000090813.5 | -4822 | Sp9 | ENSMUSG00000068859.5 |
| chr2 | 73267422 | 73267624 | 3.55893 | 4.23143 | 1.24214 | Distal Intergenic | 2 | 73271925 | 73277471 | 5547 | 1 | ENSMUST00000090813.5 | -4301 | Sp9 | ENSMUSG00000068859.5 |
| chr2 | 73267958 | 73269282 | 4.02675 | 5.20322 | 1.92095 | Distal Intergenic | 2 | 73271925 | 73277471 | 5547 | 1 | ENSMUST00000090813.5 | -2643 | Sp9 | ENSMUSG00000068859.5 |
| chr2 | 73269660 | 73273571 | 3.88868 | 4.97049 | 1.75555 | Promoter | 2 | 73271925 | 73277471 | 5547 | 1 | ENSMUST00000090813.5 | 0 | Sp9 | ENSMUSG00000068859.5 |
| chr2 | 73273884 | 73274746 | 4.31926 | 5.89198 | 2.45293 | 3' UTR | 2 | 73271966 | 73284706 | 12741 | 1 | ENSMUST00000147133.1 | 1919 | Sp9 | ENSMUSG00000068859.5 |
| chr2 | 73275096 | 73276528 | 3.86541 | 4.97149 | 1.80848 | 3' UTR | 2 | 73271966 | 73284706 | 12741 | 1 | ENSMUST00000147133.1 | 3131 | Sp9 | ENSMUSG00000068859.5 |
| chr2 | 73277055 | 73278292 | 4.23072 | 5.78689 | 2.395 | 3' UTR | 2 | 73271966 | 73284706 | 12741 | 1 | ENSMUST00000147133.1 | 5090 | Sp9 | ENSMUSG00000068859.5 |
| chr2 | 73279027 | 73279373 | 4.02402 | 5.27792 | 1.95192 | Intron (ENSMUST00000147133.1/ENSMUSG00000068859.5, intron 1 of 2) | 2 | 73271966 | 73284706 | 12741 | 1 | ENSMUST00000147133.1 | 7062 | Sp9 | ENSMUSG00000068859.5 |
| chr2 | 73279739 | 73282388 | 3.77831 | 4.90083 | 1.72443 | Exon (ENSMUST00000147133.1/ENSMUSG00000068859.5, exon 2 of 3) | 2 | 73271966 | 73284706 | 12741 | 1 | ENSMUST00000147133.1 | 7774 | Sp9 | ENSMUSG00000068859.5 |
| chr3 | 92239768 | 92240255 | 2.64342 | 4.23093 | 1.26648 | Intron (ENSMUST00000090871.2/ENSMUSG00000068893.4, intron 1 of 1) | 3 | 92250647 | 92257298 | 6652 | 1 | ENSMUST00000071805.2 | -10392 | Sprr2a2 | ENSMUSG00000068893.4 |
| chr3 | 58129821 | 58129931 | 3.8187 | 4.7683 | 1.65196 | Distal Intergenic | 3 | 58102609 | 58104269 | 1661 | 2 | ENSMUST00000091117.2 | -25553 | Gm5537 | ENSMUSG00000069008.2 |
| chr3 | 58131183 | 58131601 | 3.62351 | 4.40627 | 1.41007 | Distal Intergenic | 3 | 58102609 | 58104269 | 1661 | 2 | ENSMUST00000091117.2 | -26915 | Gm5537 | ENSMUSG00000069008.2 |
| chr3 | 31208774 | 31208991 | 3.29886 | 3.92047 | 1.11039 | 3' UTR | 3 | 31222720 | 31258394 | 35675 | 2 | ENSMUST00000108245.1 | 49403 | Slc7a14 | ENSMUSG00000069072.4 |
| chr3 | 31308664 | 31308927 | 3.66164 | 4.50949 | 1.47054 | Intron (ENSMUST00000129578.1/ENSMUSG00000085953.1, intron 1 of 3) | 3 | 31202910 | 31310378 | 107469 | 2 | ENSMUST00000091259.3 | 1451 | Slc7a14 | ENSMUSG00000069072.4 |
| chr3 | 31309722 | 31310141 | 3.61229 | 4.37437 | 1.41581 | Promoter | 3 | 31202910 | 31310378 | 107469 | 2 | ENSMUST00000091259.3 | 237 | Slc7a14 | ENSMUSG00000069072.4 |
| chr3 | 30370953 | 30371055 | 3.54924 | 4.46387 | 1.42641 | Intron (ENSMUST00000173495.2/ENSMUSG00000027684.11, intron 1 of 15) | 3 | 30267825 | 30269947 | 2123 | 2 | ENSMUST00000091270.2 | -101007 | Gm10258 | ENSMUSG00000069074.2 |
| chr3 | 25143655 | 25143941 | 3.71496 | 4.67347 | 1.56117 | Distal Intergenic | 3 | 25211016 | 25212605 | 1590 | 2 | ENSMUST00000091289.4 | 68664 | Gm10259 | ENSMUSG00000069083.4 |
| chr3 | 25144468 | 25144569 | 3.34141 | 3.83948 | 1.01733 | Distal Intergenic | 3 | 25211016 | 25212605 | 1590 | 2 | ENSMUST00000091289.4 | 68036 | Gm10259 | ENSMUSG00000069083.4 |
| chr13 | 97827199 | 97827387 | 3.50925 | 4.20741 | 1.3342 | Distal Intergenic | 13 | 97760130 | 97760588 | 459 | 2 | ENSMUST00000074072.3 | -66612 | Gm10260 | ENSMUSG00000069117.3 |
| chr2 | 23320842 | 23321869 | 3.63741 | 4.39166 | 1.344 | Promoter | 2 | 23321246 | 23401973 | 80728 | 1 | ENSMUST00000102945.1 | 0 | Nxph2 | ENSMUSG00000069132.3 |
| chr13 | 81631618 | 81631897 | 3.94083 | 5.02175 | 1.82383 | Intron (ENSMUST00000095585.5/ENSMUSG00000069170.8, intron 1 of 89) | 13 | 81560572 | 81633139 | 72568 | 2 | ENSMUST00000126444.2 | 1242 | Gpr98 | ENSMUSG00000069170.8 |
| chr13 | 81632279 | 81632421 | 3.94034 | 5.0089 | 1.82613 | Promoter | 13 | 81560572 | 81633139 | 72568 | 2 | ENSMUST00000126444.2 | 718 | Gpr98 | ENSMUSG00000069170.8 |
| chr13 | 78169593 | 78169886 | 3.60286 | 4.41494 | 1.40044 | Distal Intergenic | 13 | 78189872 | 78190443 | 572 | 2 | ENSMUST00000135284.1 | 20557 | Nr2f1 | ENSMUSG00000069171.8 |
| chr13 | 78170703 | 78171062 | 3.53138 | 4.39562 | 1.37681 | Distal Intergenic | 13 | 78189872 | 78190443 | 572 | 2 | ENSMUST00000135284.1 | 19381 | Nr2f1 | ENSMUSG00000069171.8 |
| chr13 | 78171604 | 78171834 | 3.97538 | 5.11161 | 1.91942 | Distal Intergenic | 13 | 78189872 | 78190443 | 572 | 2 | ENSMUST00000135284.1 | 18609 | Nr2f1 | ENSMUSG00000069171.8 |
| chr13 | 78172326 | 78172456 | 3.69929 | 4.8264 | 1.68807 | Distal Intergenic | 13 | 78189872 | 78190443 | 572 | 2 | ENSMUST00000135284.1 | 17987 | Nr2f1 | ENSMUSG00000069171.8 |
| chr13 | 78172868 | 78174567 | 3.51615 | 4.35907 | 1.38904 | Distal Intergenic | 13 | 78189872 | 78190443 | 572 | 2 | ENSMUST00000135284.1 | 15876 | Nr2f1 | ENSMUSG00000069171.8 |
| chr13 | 78175104 | 78176792 | 3.7011 | 4.56722 | 1.53553 | Distal Intergenic | 13 | 78189872 | 78190443 | 572 | 2 | ENSMUST00000135284.1 | 13651 | Nr2f1 | ENSMUSG00000069171.8 |
| chr13 | 78178112 | 78178892 | 3.88747 | 5.03557 | 1.80671 | Distal Intergenic | 13 | 78189872 | 78190443 | 572 | 2 | ENSMUST00000135284.1 | 11551 | Nr2f1 | ENSMUSG00000069171.8 |
| chr13 | 78179203 | 78179836 | 3.53218 | 4.38295 | 1.38124 | Distal Intergenic | 13 | 78189872 | 78190443 | 572 | 2 | ENSMUST00000135284.1 | 10607 | Nr2f1 | ENSMUSG00000069171.8 |
| chr13 | 78180219 | 78180355 | 3.41954 | 3.98808 | 1.09677 | Distal Intergenic | 13 | 78189872 | 78190443 | 572 | 2 | ENSMUST00000135284.1 | 10088 | Nr2f1 | ENSMUSG00000069171.8 |
| chr13 | 78180785 | 78182427 | 4.44599 | 6.06497 | 2.57952 | Distal Intergenic | 13 | 78189872 | 78190443 | 572 | 2 | ENSMUST00000135284.1 | 8016 | Nr2f1 | ENSMUSG00000069171.8 |
| chr13 | 78182806 | 78183155 | 3.72448 | 4.95982 | 1.74461 | Distal Intergenic | 13 | 78189872 | 78190443 | 572 | 2 | ENSMUST00000135284.1 | 7288 | Nr2f1 | ENSMUSG00000069171.8 |
| chr13 | 78183741 | 78184368 | 3.89007 | 4.86524 | 1.66336 | Distal Intergenic | 13 | 78189872 | 78190443 | 572 | 2 | ENSMUST00000135284.1 | 6075 | Nr2f1 | ENSMUSG00000069171.8 |
| chr13 | 78184787 | 78184932 | 3.62828 | 4.37107 | 1.34906 | Distal Intergenic | 13 | 78189872 | 78190443 | 572 | 2 | ENSMUST00000135284.1 | 5511 | Nr2f1 | ENSMUSG00000069171.8 |
| chr13 | 78186978 | 78187253 | 3.36385 | 3.90487 | 1.12538 | Downstream (1-2kb) | 13 | 78189872 | 78190443 | 572 | 2 | ENSMUST00000135284.1 | 3190 | Nr2f1 | ENSMUSG00000069171.8 |
| chr13 | 78194914 | 78195515 | 3.79602 | 4.80849 | 1.67677 | Promoter | 13 | 78189733 | 78196373 | 6641 | 2 | ENSMUST00000125176.2 | 858 | Nr2f1 | ENSMUSG00000069171.8 |
| chr13 | 78197286 | 78199903 | 4.10509 | 5.41656 | 2.0815 | Promoter | 13 | 78188973 | 78199757 | 10785 | 2 | ENSMUST00000091458.7 | 0 | Nr2f1 | ENSMUSG00000069171.8 |
| chr13 | 54736975 | 54737273 | 3.67715 | 4.45105 | 1.37608 | 3' UTR | 13 | 54736673 | 54749669 | 12997 | 2 | ENSMUST00000099506.1 | 12396 | Gprin1 | ENSMUSG00000069227.4 |
| chr13 | 54737862 | 54738440 | 3.66588 | 4.45523 | 1.37918 | Exon (ENSMUST00000135343.1/ENSMUSG00000069227.4, exon 2 of 2) | 13 | 54736673 | 54749669 | 12997 | 2 | ENSMUST00000099506.1 | 11229 | Gprin1 | ENSMUSG00000069227.4 |
| chr13 | 54738881 | 54739024 | 3.97291 | 5.02543 | 1.77593 | Exon (ENSMUST00000135343.1/ENSMUSG00000069227.4, exon 2 of 2) | 13 | 54736673 | 54749669 | 12997 | 2 | ENSMUST00000099506.1 | 10645 | Gprin1 | ENSMUSG00000069227.4 |
| chr13 | 54749244 | 54750207 | 4.57873 | 6.42974 | 2.87263 | Promoter | 13 | 54736671 | 54749874 | 13204 | 2 | ENSMUST00000135343.1 | 0 | Gprin1 | ENSMUSG00000069227.4 |
| chr13 | 54755418 | 54755782 | 3.71851 | 4.58915 | 1.48755 | Distal Intergenic | 13 | 54736671 | 54749874 | 13204 | 2 | ENSMUST00000135343.1 | -5545 | Gprin1 | ENSMUSG00000069227.4 |
| chr13 | 22049818 | 22050277 | 3.82011 | 4.82612 | 1.69875 | Distal Intergenic | 13 | 22043214 | 22043676 | 463 | 1 | ENSMUST00000110452.1 | 6605 | Hist1h2bj | ENSMUSG00000069300.3 |
| chr13 | 22031059 | 22031903 | 3.94136 | 4.95217 | 1.70249 | Exon (ENSMUST00000146121.1/ENSMUSG00000043145.6, exon 2 of 2) | 13 | 22035164 | 22035568 | 405 | 2 | ENSMUST00000091742.5 | 3665 | Hist1h2ah | ENSMUSG00000069302.5 |
| chr13 | 22032593 | 22032711 | 3.34153 | 3.83966 | 1.01733 | Downstream (2-3kb) | 13 | 22035164 | 22035568 | 405 | 2 | ENSMUST00000091742.5 | 2857 | Hist1h2ah | ENSMUSG00000069302.5 |
| chr13 | 22033373 | 22033707 | 3.51821 | 4.41058 | 1.37353 | Downstream (1-2kb) | 13 | 22035164 | 22035568 | 405 | 2 | ENSMUST00000091742.5 | 1861 | Hist1h2ah | ENSMUSG00000069302.5 |
| chr13 | 22034259 | 22034519 | 3.64819 | 4.38737 | 1.31633 | Downstream (<1kb) | 13 | 22035164 | 22035568 | 405 | 2 | ENSMUST00000091742.5 | 1049 | Hist1h2ah | ENSMUSG00000069302.5 |
| chr13 | 21791909 | 21792133 | 3.5005 | 4.27654 | 1.31784 | Distal Intergenic | 13 | 21787508 | 21787976 | 469 | 1 | ENSMUST00000102982.1 | 4402 | Hist1h2bp | ENSMUSG00000069308.7 |
| chr18 | 57465532 | 57465607 | 3.2586 | 3.71503 | 1.01733 | Intron (ENSMUST00000130696.2/ENSMUSG00000086607.2, intron 2 of 3) | 18 | 57468486 | 57478134 | 9649 | 1 | ENSMUST00000091892.2 | -2879 | Ctxn3 | ENSMUSG00000069372.2 |
| chr18 | 57469592 | 57469865 | 3.32729 | 3.81823 | 1.01733 | Intron (ENSMUST00000091892.2/ENSMUSG00000069372.2, intron 1 of 1) | 18 | 57468486 | 57478134 | 9649 | 1 | ENSMUST00000091892.2 | 1107 | Ctxn3 | ENSMUSG00000069372.2 |
| chr18 | 57482412 | 57482714 | 3.27961 | 3.74655 | 1.01733 | Intron (ENSMUST00000130696.2/ENSMUSG00000086607.2, intron 2 of 3) | 18 | 57468486 | 57478134 | 9649 | 1 | ENSMUST00000091892.2 | 13927 | Ctxn3 | ENSMUSG00000069372.2 |
| chr18 | 53446574 | 53447026 | 3.91839 | 5.00616 | 1.81199 | Distal Intergenic | 18 | 53464546 | 53575857 | 111312 | 1 | ENSMUST00000091900.4 | -17520 | Prdm6 | ENSMUSG00000069378.8 |
| chr18 | 53449614 | 53449689 | 3.67871 | 4.50032 | 1.49954 | Distal Intergenic | 18 | 53464546 | 53575857 | 111312 | 1 | ENSMUST00000091900.4 | -14857 | Prdm6 | ENSMUSG00000069378.8 |
| chr18 | 53452163 | 53452529 | 3.41693 | 4.02603 | 1.20478 | Distal Intergenic | 18 | 53464546 | 53575857 | 111312 | 1 | ENSMUST00000091900.4 | -12017 | Prdm6 | ENSMUSG00000069378.8 |
| chr18 | 53460103 | 53463056 | 3.77339 | 4.77124 | 1.63572 | Distal Intergenic | 18 | 53464546 | 53575857 | 111312 | 1 | ENSMUST00000091900.4 | -1490 | Prdm6 | ENSMUSG00000069378.8 |
| chr18 | 53464027 | 53466683 | 4.22314 | 5.5929 | 2.18754 | Promoter | 18 | 53464546 | 53575857 | 111312 | 1 | ENSMUST00000091900.4 | 0 | Prdm6 | ENSMUSG00000069378.8 |
| chr18 | 53467049 | 53467391 | 3.45587 | 4.04958 | 1.15081 | Intron (ENSMUST00000091900.4/ENSMUSG00000069378.8, intron 1 of 6) | 18 | 53465051 | 53575857 | 110807 | 1 | ENSMUST00000154557.2 | 1999 | Prdm6 | ENSMUSG00000069378.8 |
| chr18 | 53467854 | 53468274 | 4.54164 | 6.93145 | 3.32727 | Intron (ENSMUST00000091900.4/ENSMUSG00000069378.8, intron 1 of 6) | 18 | 53465051 | 53575857 | 110807 | 1 | ENSMUST00000154557.2 | 2804 | Prdm6 | ENSMUSG00000069378.8 |
| chr18 | 53468849 | 53471197 | 3.93272 | 5.00116 | 1.80975 | Intron (ENSMUST00000091900.4/ENSMUSG00000069378.8, intron 1 of 6) | 18 | 53472981 | 53575907 | 102927 | 1 | ENSMUST00000115399.2 | -1784 | Prdm6 | ENSMUSG00000069378.8 |
| chr18 | 53472073 | 53472556 | 3.66542 | 4.5877 | 1.52412 | Promoter | 18 | 53472981 | 53575907 | 102927 | 1 | ENSMUST00000115399.2 | -425 | Prdm6 | ENSMUSG00000069378.8 |
| chr18 | 53473569 | 53473749 | 3.50223 | 4.7631 | 1.63064 | Promoter | 18 | 53472981 | 53575907 | 102927 | 1 | ENSMUST00000115399.2 | 589 | Prdm6 | ENSMUSG00000069378.8 |
| chr18 | 53475003 | 53475160 | 3.43929 | 4.19676 | 1.24161 | Intron (ENSMUST00000091900.4/ENSMUSG00000069378.8, intron 2 of 6) | 18 | 53472981 | 53575907 | 102927 | 1 | ENSMUST00000115399.2 | 2023 | Prdm6 | ENSMUSG00000069378.8 |
| chr18 | 53476936 | 53477375 | 3.561 | 4.35181 | 1.3685 | Intron (ENSMUST00000091900.4/ENSMUSG00000069378.8, intron 2 of 6) | 18 | 53472981 | 53575907 | 102927 | 1 | ENSMUST00000115399.2 | 3956 | Prdm6 | ENSMUSG00000069378.8 |
| chr18 | 53479520 | 53479884 | 3.36342 | 4.33939 | 1.37979 | Intron (ENSMUST00000091900.4/ENSMUSG00000069378.8, intron 2 of 6) | 18 | 53472981 | 53575907 | 102927 | 1 | ENSMUST00000115399.2 | 6540 | Prdm6 | ENSMUSG00000069378.8 |
| chr18 | 53480313 | 53481376 | 4.13331 | 5.6501 | 2.24558 | Intron (ENSMUST00000091900.4/ENSMUSG00000069378.8, intron 2 of 6) | 18 | 53472981 | 53575907 | 102927 | 1 | ENSMUST00000115399.2 | 7333 | Prdm6 | ENSMUSG00000069378.8 |
| chr18 | 53482543 | 53482783 | 3.4625 | 4.22188 | 1.29048 | Intron (ENSMUST00000091900.4/ENSMUSG00000069378.8, intron 2 of 6) | 18 | 53472981 | 53575907 | 102927 | 1 | ENSMUST00000115399.2 | 9563 | Prdm6 | ENSMUSG00000069378.8 |
| chr18 | 53483104 | 53483233 | 3.41882 | 4.34177 | 1.35231 | Intron (ENSMUST00000091900.4/ENSMUSG00000069378.8, intron 2 of 6) | 18 | 53472981 | 53575907 | 102927 | 1 | ENSMUST00000115399.2 | 10124 | Prdm6 | ENSMUSG00000069378.8 |
| chr18 | 53552081 | 53553088 | 3.82559 | 4.83504 | 1.6519 | Promoter | 18 | 53551594 | 53575857 | 24264 | 1 | ENSMUST00000115398.1 | 488 | Prdm6 | ENSMUSG00000069378.8 |
| chr18 | 53554321 | 53554727 | 3.41821 | 4.16737 | 1.22888 | Intron (ENSMUST00000091900.4/ENSMUSG00000069378.8, intron 5 of 6) | 18 | 53551594 | 53575857 | 24264 | 1 | ENSMUST00000115398.1 | 2728 | Prdm6 | ENSMUSG00000069378.8 |
| chr18 | 53568161 | 53568302 | 4.07761 | 5.31246 | 2.01808 | Exon (ENSMUST00000091900.4/ENSMUSG00000069378.8, exon 6 of 7) | 18 | 53551594 | 53575857 | 24264 | 1 | ENSMUST00000115398.1 | 16568 | Prdm6 | ENSMUSG00000069378.8 |
| chr19 | 61199723 | 61199821 | 2.24735 | 13.2265 | 8.6901 | Distal Intergenic | 19 | 61183890 | 61183955 | 66 | 1 | ENSMUST00000092094.1 | 15834 | Gm6020 | ENSMUSG00000069475.1 |
| chr2 | 49571643 | 49571865 | 3.63645 | 4.43219 | 1.43871 | Distal Intergenic | 2 | 49528875 | 49537081 | 8207 | 1 | ENSMUST00000152485.1 | 42769 | Epc2 | ENSMUSG00000069495.7 |
| chr10 | 69532420 | 69532955 | 3.54442 | 4.22099 | 1.28342 | Promoter | 10 | 69533772 | 69720860 | 187089 | 1 | ENSMUST00000182379.2 | -817 | Ank3 | ENSMUSG00000069601.8 |
| chr10 | 69533346 | 69534486 | 3.48965 | 4.13353 | 1.21955 | Promoter | 10 | 69533772 | 69720860 | 187089 | 1 | ENSMUST00000182379.2 | 0 | Ank3 | ENSMUSG00000069601.8 |
| chr10 | 69534891 | 69535815 | 3.71377 | 4.60754 | 1.55759 | Promoter | 10 | 69534222 | 70015633 | 481412 | 1 | ENSMUST00000092434.6 | 670 | Ank3 | ENSMUSG00000069601.8 |
| chr10 | 69572791 | 69573047 | 3.60368 | 4.36645 | 1.38234 | Intron (ENSMUST00000182379.2/ENSMUSG00000069601.8, intron 1 of 2) | 10 | 69534222 | 70015633 | 481412 | 1 | ENSMUST00000092434.6 | 38570 | Ank3 | ENSMUSG00000069601.8 |
| chr1 | 13052336 | 13052411 | 3.94181 | 4.98603 | 1.77067 | Distal Intergenic | 1 | 13062025 | 13062345 | 321 | 1 | ENSMUST00000059065.9 | -9614 | Gm5250 | ENSMUSG00000069620.6 |
| chr10 | 37137879 | 37137996 | 3.21646 | 3.65192 | 1.01414 | Promoter | 10 | 37133375 | 37138920 | 5546 | 2 | ENSMUST00000092584.5 | 924 | Marcks | ENSMUSG00000069662.5 |
| chr10 | 37138343 | 37138796 | 3.63737 | 4.43358 | 1.43801 | Promoter | 10 | 37133375 | 37138920 | 5546 | 2 | ENSMUST00000092584.5 | 124 | Marcks | ENSMUSG00000069662.5 |
| chr10 | 32887490 | 32887612 | 3.63126 | 4.51139 | 1.43412 | Intron (ENSMUST00000191234.1/ENSMUSG00000069670.2, intron 1 of 6) | 10 | 31689310 | 32890461 | 1201152 | 2 | ENSMUST00000191234.1 | 2849 | Nkain2 | ENSMUSG00000069670.2 |
| chr10 | 32889634 | 32890189 | 3.57333 | 4.54226 | 1.47337 | Promoter | 10 | 31689310 | 32890461 | 1201152 | 2 | ENSMUST00000191234.1 | 272 | Nkain2 | ENSMUSG00000069670.2 |
| chr5 | 15661703 | 15661906 | 2.30633 | 4.415 | 1.39218 | Distal Intergenic | 5 | 15652286 | 15657059 | 4774 | 2 | ENSMUST00000092696.4 | -4645 | 4930572O03Rik | ENSMUSG00000069720.4 |
| chr11 | 90026183 | 90026747 | 4.32919 | 5.76239 | 2.32089 | Distal Intergenic | 11 | 90030348 | 90036508 | 6161 | 1 | ENSMUST00000092788.3 | -3601 | Tmem100 | ENSMUSG00000069763.3 |
| chr11 | 90029723 | 90029983 | 3.42228 | 3.98458 | 1.0991 | Promoter | 11 | 90030348 | 90036508 | 6161 | 1 | ENSMUST00000092788.3 | -365 | Tmem100 | ENSMUSG00000069763.3 |
| chr11 | 85841226 | 85842286 | 4.39793 | 6.24864 | 2.75536 | 3' UTR | 11 | 85846790 | 85850333 | 3544 | 2 | ENSMUST00000146433.1 | 8047 | Gm11444 | ENSMUSG00000069785.4 |
| chr11 | 85842660 | 85845207 | 3.73443 | 4.76051 | 1.64682 | Downstream (1-2kb) | 11 | 85846790 | 85850333 | 3544 | 2 | ENSMUST00000146433.1 | 5126 | Gm11444 | ENSMUSG00000069785.4 |
| chr11 | 85846277 | 85847457 | 3.42255 | 4.00615 | 1.16443 | 3' UTR | 11 | 85846790 | 85850333 | 3544 | 2 | ENSMUST00000146433.1 | 2876 | Gm11444 | ENSMUSG00000069785.4 |
| chr11 | 85850002 | 85850227 | 3.36919 | 3.88572 | 1.03258 | Promoter | 11 | 85846790 | 85850333 | 3544 | 2 | ENSMUST00000146433.1 | 106 | Gm11444 | ENSMUSG00000069785.4 |
| chr11 | 85850567 | 85850966 | 3.79856 | 4.83211 | 1.66966 | Promoter | 11 | 85846790 | 85850333 | 3544 | 2 | ENSMUST00000146433.1 | -235 | Gm11444 | ENSMUSG00000069785.4 |
| chr11 | 85863716 | 85863991 | 3.50495 | 4.17137 | 1.29495 | Distal Intergenic | 11 | 85846790 | 85850333 | 3544 | 2 | ENSMUST00000146433.1 | -13384 | Gm11444 | ENSMUSG00000069785.4 |
| chr11 | 85865758 | 85866105 | 3.66864 | 4.58088 | 1.48308 | Distal Intergenic | 11 | 85846790 | 85850333 | 3544 | 2 | ENSMUST00000146433.1 | -15426 | Gm11444 | ENSMUSG00000069785.4 |
| chr11 | 82991225 | 82992035 | 4.3497 | 6.00857 | 2.55396 | Promoter | 11 | 82979629 | 82991830 | 12202 | 2 | ENSMUST00000038211.6 | 0 | Slfn9 | ENSMUSG00000069793.6 |
| chr11 | 82616541 | 82616666 | 3.72596 | 4.54557 | 1.4549 | Distal Intergenic | 11 | 82633353 | 82636309 | 2957 | 1 | ENSMUST00000125600.1 | -16687 | Gm11426 | ENSMUSG00000069796.3 |
| chr11 | 82617729 | 82617962 | 3.69411 | 4.53077 | 1.41443 | Distal Intergenic | 11 | 82633353 | 82636309 | 2957 | 1 | ENSMUST00000125600.1 | -15391 | Gm11426 | ENSMUSG00000069796.3 |
| chr11 | 82618318 | 82618717 | 3.44638 | 4.035 | 1.14876 | Distal Intergenic | 11 | 82633353 | 82636309 | 2957 | 1 | ENSMUST00000125600.1 | -14636 | Gm11426 | ENSMUSG00000069796.3 |
| chr11 | 82619378 | 82619453 | 3.58726 | 4.27496 | 1.24425 | Distal Intergenic | 11 | 82633353 | 82636309 | 2957 | 1 | ENSMUST00000125600.1 | -13900 | Gm11426 | ENSMUSG00000069796.3 |
| chr7 | 3331918 | 3332178 | 3.80446 | 4.83792 | 1.68129 | Promoter | 7 | 3332955 | 3368216 | 35262 | 1 | ENSMUST00000092891.4 | -777 | Cacng7 | ENSMUSG00000069806.4 |
| chr7 | 3333001 | 3333077 | 4.04642 | 5.30563 | 1.96436 | Promoter | 7 | 3332955 | 3368216 | 35262 | 1 | ENSMUST00000092891.4 | 47 | Cacng7 | ENSMUSG00000069806.4 |
| chr7 | 3333962 | 3334243 | 3.50183 | 4.21717 | 1.30419 | Intron (ENSMUST00000092891.4/ENSMUSG00000069806.4, intron 1 of 5) | 7 | 3332955 | 3368216 | 35262 | 1 | ENSMUST00000092891.4 | 1008 | Cacng7 | ENSMUSG00000069806.4 |
| chr11 | 74619393 | 74620874 | 3.75518 | 4.6926 | 1.63214 | Promoter | 11 | 74619605 | 74641516 | 21912 | 1 | ENSMUST00000100866.2 | 0 | E130309D14Rik | ENSMUSG00000069814.6 |
| chr11 | 74621349 | 74621595 | 4.02323 | 5.42719 | 2.05158 | Intron (ENSMUST00000100866.2/ENSMUSG00000069814.6, intron 1 of 3) | 11 | 74619605 | 74641516 | 21912 | 1 | ENSMUST00000100866.2 | 1745 | E130309D14Rik | ENSMUSG00000069814.6 |
| chr11 | 61342528 | 61342670 | 3.42228 | 4.28105 | 1.33274 | Promoter | 11 | 61336901 | 61342584 | 5684 | 2 | ENSMUST00000147450.1 | 0 | Slc47a2 | ENSMUSG00000069855.4 |
| chr11 | 61343004 | 61343548 | 3.50411 | 4.32808 | 1.33982 | Promoter | 11 | 61301631 | 61342860 | 41230 | 2 | ENSMUST00000093029.3 | -145 | Slc47a2 | ENSMUSG00000069855.4 |
| chr11 | 61343899 | 61344039 | 3.31329 | 3.82319 | 1.08746 | 3' UTR | 11 | 61301631 | 61342860 | 41230 | 2 | ENSMUST00000093029.3 | -1040 | Slc47a2 | ENSMUSG00000069855.4 |
| chr11 | 61344864 | 61345042 | 3.73018 | 4.59063 | 1.54922 | Intron (ENSMUST00000010267.4/ENSMUSG00000010122.9, intron 16 of 16) | 11 | 61301631 | 61342860 | 41230 | 2 | ENSMUST00000093029.3 | -2005 | Slc47a2 | ENSMUSG00000069855.4 |
| chr11 | 49589167 | 49589259 | 4.61749 | 7.24411 | 3.49623 | Promoter | 11 | 49586641 | 49588462 | 1822 | 2 | ENSMUST00000093146.3 | -706 | 1700024J04Rik | ENSMUSG00000069890.3 |
| chr11 | 34314935 | 34316445 | 3.74542 | 4.73708 | 1.62942 | Promoter | 11 | 34314822 | 34422640 | 107819 | 1 | ENSMUST00000165963.2 | 114 | Fam196b | ENSMUSG00000069911.5 |
| chr11 | 32284102 | 32284184 | 4.39769 | 6.97819 | 3.21709 | Promoter | 11 | 32283811 | 32284438 | 628 | 1 | ENSMUST00000142555.1 | 292 | Hba-a1 | ENSMUSG00000069919.7 |
| chr8 | 89184212 | 89184551 | 3.81968 | 4.73517 | 1.5887 | Downstream (2-3kb) | 8 | 89186946 | 89187560 | 615 | 2 | ENSMUST00000093326.3 | 3009 | Gm5356 | ENSMUSG00000069962.3 |
| chr8 | 89271447 | 89272127 | 3.99177 | 5.08388 | 1.82973 | Distal Intergenic | 8 | 89186946 | 89187560 | 615 | 2 | ENSMUST00000093326.3 | -83888 | Gm5356 | ENSMUSG00000069962.3 |
| chr8 | 89289033 | 89289289 | 3.38193 | 3.9006 | 1.01733 | Distal Intergenic | 8 | 89186946 | 89187560 | 615 | 2 | ENSMUST00000093326.3 | -101474 | Gm5356 | ENSMUSG00000069962.3 |
| chr8 | 89303391 | 89303466 | 3.74267 | 4.5689 | 1.43738 | Distal Intergenic | 8 | 89186946 | 89187560 | 615 | 2 | ENSMUST00000093326.3 | -115832 | Gm5356 | ENSMUSG00000069962.3 |

| | | | | | | | | | | | | | | | |
|---|---|---|---|---|---|---|---|---|---|---|---|---|---|---|---|
| chr8 | 87535196 | 87535288 | 3.94889 | 4.96695 | 1.73546 | Distal Intergenic | 8 | 87563853 | 87589424 | 25572 | 1 | ENSMUST00000093342.5 | -28565 | 4933402J07Rik | ENSMUSG00000069971.5 |
| chr8 | 71707135 | 71707419 | 3.81288 | 4.78281 | 1.65626 | Downstream (<1kb) | 8 | 71709637 | 71711106 | 1470 | 2 | ENSMUST00000149363.2 | 3687 | Fcho1 | ENSMUSG00000070000.8 |
| chr8 | 71708649 | 71709366 | 3.88217 | 4.9523 | 1.77156 | 3' UTR | 8 | 71709637 | 71711106 | 1470 | 2 | ENSMUST00000149363.2 | 1740 | Fcho1 | ENSMUSG00000070000.8 |
| chr8 | 71709765 | 71710289 | 3.59087 | 4.36468 | 1.37845 | Promoter | 8 | 71709637 | 71711106 | 1470 | 2 | ENSMUST00000149363.2 | 817 | Fcho1 | ENSMUSG00000070000.8 |
| chr8 | 45358406 | 45358619 | 3.6728 | 4.59575 | 1.47972 | Promoter | 8 | 45338759 | 45358762 | 20004 | 2 | ENSMUST00000155230.1 | 143 | Fam149a | ENSMUSG00000070044.10 |
| chr8 | 45358930 | 45359547 | 3.67464 | 4.51243 | 1.48597 | Promoter | 8 | 45336717 | 45358865 | 22149 | 2 | ENSMUST00000135912.2 | -66 | Fam149a | ENSMUSG00000070044.10 |
| chr8 | 45381078 | 45381944 | 3.50499 | 4.18742 | 1.30981 | Promoter | 8 | 45336725 | 45382291 | 45567 | 2 | ENSMUST00000093526.7 | 347 | Fam149a | ENSMUSG00000070044.10 |
| chr8 | 45382293 | 45382419 | 3.74531 | 4.86965 | 1.67401 | Promoter | 8 | 45336725 | 45382291 | 45567 | 2 | ENSMUST00000093526.7 | -3 | Fam149a | ENSMUSG00000070044.10 |
| chr12 | 109607848 | 109607966 | 3.76555 | 4.75561 | 1.59323 | Intron (ENSMUST00000183215.2/ENSMUSG00000097451.4, intron 2 of 7) | 12 | 109611500 | 109611595 | 96 | 1 | ENSMUST00000093593.1 | -3534 | Mir341 | ENSMUSG00000070101.1 |
| chr7 | 142654372 | 142655155 | 3.91253 | 4.91813 | 1.70129 | Promoter | 7 | 142654924 | 142654996 | 73 | 2 | ENSMUST00000093631.1 | 0 | Mir483 | ENSMUSG00000070140.1 |
| chr2 | 180391008 | 180391184 | 3.27876 | 3.74536 | 1.01733 | Distal Intergenic | 2 | 180389048 | 180389124 | 77 | 1 | ENSMUST00000093635.1 | 1961 | Mir1a-1 | ENSMUSG00000070144.1 |
| chr2 | 180392846 | 180393267 | 3.50831 | 4.14041 | 1.19117 | Distal Intergenic | 2 | 180389048 | 180389124 | 77 | 1 | ENSMUST00000093635.1 | 3799 | Mir1a-1 | ENSMUSG00000070144.1 |
| chr1 | 20477993 | 20478494 | 3.77538 | 4.64961 | 1.52987 | Intron (ENSMUST00000088448.6/ENSMUSG00000043760.11, intron 35 of 66) | 1 | 20444811 | 20444912 | 102 | 2 | ENSMUST00000093665.1 | -33082 | Gm24162 | ENSMUSG00000070175.1 |
| chr17 | 44669742 | 44670002 | 3.3229 | 3.81166 | 1.01733 | Intron (ENSMUST00000162816.2/ENSMUSG00000039153.11, intron 3 of 5) | 17 | 44641025 | 44641143 | 119 | 2 | ENSMUST00000093699.1 | -28600 | n-R5s27 | ENSMUSG00000070209.1 |
| chr15 | 78367978 | 78368056 | 3.55887 | 4.34028 | 1.35657 | Downstream (1-2kb) | 15 | 78369248 | 78369357 | 110 | 1 | ENSMUST00000093723.1 | -1192 | Gm26329 | ENSMUSG00000070235.1 |
| chr15 | 78369582 | 78369780 | 3.48679 | 4.2573 | 1.30257 | Promoter | 15 | 78369248 | 78369357 | 110 | 1 | ENSMUST00000093723.1 | 335 | Gm26329 | ENSMUSG00000070235.1 |
| chr15 | 78370113 | 78370358 | 3.41859 | 4.01692 | 1.21188 | Promoter | 15 | 78369248 | 78369357 | 110 | 1 | ENSMUST00000093723.1 | 866 | Gm26329 | ENSMUSG00000070235.1 |
| chr15 | 78372073 | 78372193 | 3.34409 | 3.94786 | 1.11695 | Distal Intergenic | 15 | 78369248 | 78369357 | 110 | 1 | ENSMUST00000093723.1 | 2826 | Gm26329 | ENSMUSG00000070235.1 |
| chr9 | 78392459 | 78392837 | 4.01569 | 5.17565 | 1.92225 | Downstream (2-3kb) | 9 | 78395777 | 78423587 | 27811 | 1 | ENSMUST00000113367.1 | -2940 | Ddx43 | ENSMUSG00000070291.4 |
| chr9 | 78393173 | 78393459 | 3.93269 | 4.99067 | 1.79547 | Downstream (1-2kb) | 9 | 78395777 | 78423587 | 27811 | 1 | ENSMUST00000113367.1 | -2318 | Ddx43 | ENSMUSG00000070291.4 |
| chr9 | 78393905 | 78394436 | 3.46561 | 4.08578 | 1.2012 | Downstream (<1kb) | 9 | 78395777 | 78423587 | 27811 | 1 | ENSMUST00000113367.1 | -1341 | Ddx43 | ENSMUSG00000070291.4 |
| chr9 | 78394790 | 78394920 | 3.46867 | 4.09481 | 1.24095 | Promoter | 9 | 78395777 | 78423587 | 27811 | 1 | ENSMUST00000113367.1 | -857 | Ddx43 | ENSMUSG00000070291.4 |
| chr9 | 45051940 | 45052204 | 3.48896 | 4.12402 | 1.23718 | 3' UTR | 9 | 45055186 | 45077436 | 22251 | 1 | ENSMUST00000114664.2 | -2982 | Mpzl3 | ENSMUSG00000070305.5 |
| chr11 | 117859540 | 117859836 | 3.58439 | 4.27161 | 1.24489 | Promoter | 11 | 117860752 | 117865543 | 4792 | 1 | ENSMUST00000093905.2 | -916 | Tmem235 | ENSMUSG00000070330.2 |
| chr11 | 117860190 | 117860265 | 3.53527 | 4.1811 | 1.19007 | Promoter | 11 | 117860752 | 117865543 | 4792 | 1 | ENSMUST00000093905.2 | -487 | Tmem235 | ENSMUSG00000070330.2 |
| chr11 | 116446555 | 116446929 | 3.93013 | 4.99484 | 1.78904 | Promoter | 11 | 116444541 | 116446991 | 2451 | 2 | ENSMUST00000140697.1 | 62 | Qrich2 | ENSMUSG00000070331.8 |
| chr11 | 115435589 | 115435933 | 3.27289 | 3.76133 | 1.04973 | Distal Intergenic | 11 | 115440545 | 115448270 | 7726 | 1 | ENSMUST00000093914.4 | -4612 | 4933422H20Rik | ENSMUSG00000070332.4 |
| chr11 | 97343608 | 97344050 | 3.40864 | 3.98657 | 1.17303 | Exon (ENSMUST00000093942.4/ENSMUSG00000070337.4, exon 7 of 11) | 11 | 97332109 | 97352016 | 19908 | 2 | ENSMUST00000093942.4 | 7966 | Gpr179 | ENSMUSG00000070337.4 |
| chr7 | 144979146 | 144979283 | 3.46873 | 4.06925 | 1.15081 | Distal Intergenic | 7 | 144929931 | 144939869 | 9939 | 2 | ENSMUST00000093962.4 | -39278 | Ccnd1 | ENSMUSG00000070348.4 |
| chr7 | 144985557 | 144985866 | 3.42139 | 4.01952 | 1.17961 | Distal Intergenic | 7 | 144929931 | 144939869 | 9939 | 2 | ENSMUST00000093962.4 | -45689 | Ccnd1 | ENSMUSG00000070348.4 |
| chr7 | 144986907 | 144987118 | 3.36963 | 3.88206 | 1.01733 | Distal Intergenic | 7 | 144929931 | 144939869 | 9939 | 2 | ENSMUST00000093962.4 | -47039 | Ccnd1 | ENSMUSG00000070348.4 |
| chr7 | 144987558 | 144987700 | 3.67573 | 4.49843 | 1.44504 | Distal Intergenic | 7 | 144929931 | 144939869 | 9939 | 2 | ENSMUST00000093962.4 | -47690 | Ccnd1 | ENSMUSG00000070348.4 |
| chr7 | 144988135 | 144988494 | 3.97604 | 5.04801 | 1.83114 | Distal Intergenic | 7 | 144929931 | 144939869 | 9939 | 2 | ENSMUST00000093962.4 | -48267 | Ccnd1 | ENSMUSG00000070348.4 |
| chr7 | 144989694 | 144989804 | 3.3063 | 3.791 | 1.03259 | Distal Intergenic | 7 | 144929931 | 144939869 | 9939 | 2 | ENSMUST00000093962.4 | -49826 | Ccnd1 | ENSMUSG00000070348.4 |
| chr7 | 144990321 | 144990422 | 3.46355 | 4.20073 | 1.29 | Distal Intergenic | 7 | 144929931 | 144939869 | 9939 | 2 | ENSMUST00000093962.4 | -50453 | Ccnd1 | ENSMUSG00000070348.4 |
| chr7 | 144992392 | 144992523 | 3.41742 | 4.15046 | 1.26376 | Distal Intergenic | 7 | 144929931 | 144939869 | 9939 | 2 | ENSMUST00000093962.4 | -52524 | Ccnd1 | ENSMUSG00000070348.4 |
| chr7 | 129255845 | 129256274 | 3.87226 | 4.99541 | 1.77231 | Promoter | 7 | 129257094 | 129391307 | 134214 | 1 | ENSMUST00000094018.4 | -820 | Ppapdc1a | ENSMUSG00000070366.4 |
| chr7 | 129256646 | 129257530 | 3.81257 | 4.7555 | 1.60321 | Promoter | 7 | 129257094 | 129391307 | 134214 | 1 | ENSMUST00000094018.4 | 0 | Ppapdc1a | ENSMUSG00000070366.4 |
| chr7 | 127944566 | 127945306 | 3.66286 | 4.5769 | 1.54288 | Exon (ENSMUST00000094026.4/ENSMUSG00000070371.6, exon 4 of 15) | 7 | 127932638 | 127946725 | 14088 | 2 | ENSMUST00000094026.4 | 1419 | Prss36 | ENSMUSG00000070371.6 |
| chr11 | 72314315 | 72314586 | 3.74534 | 4.79675 | 1.62287 | Promoter | 11 | 72314444 | 72319416 | 4973 | 1 | ENSMUST00000108504.1 | 0 | Fbxo39 | ENSMUSG00000070388.3 |
| chr7 | 102106808 | 102107045 | 4.71386 | 6.50103 | 2.82349 | Exon (ENSMUST00000033300.2/ENSMUSG00000030996.7, exon 3 of 5) | 7 | 102096879 | 102102845 | 5967 | 2 | ENSMUST00000094128.3 | -3964 | Art5 | ENSMUSG00000070424.6 |
| chr7 | 82335333 | 82335580 | 3.77973 | 4.6919 | 1.56007 | Promoter | 7 | 82335694 | 82340024 | 4331 | 1 | ENSMUST00000172490.1 | -114 | Adamtsl3 | ENSMUSG00000070469.6 |
| chr7 | 82336991 | 82337119 | 3.38193 | 3.9006 | 1.01733 | Promoter | 7 | 82337218 | 82612430 | 275213 | 1 | ENSMUST00000173828.1 | -99 | Adamtsl3 | ENSMUSG00000070469.6 |
| chr7 | 82530758 | 82530833 | 3.70153 | 4.64922 | 1.56514 | Intron (ENSMUST00000173287.2/ENSMUSG00000070469.6, intron 15 of 29) | 7 | 82611777 | 82612774 | 998 | 1 | ENSMUST00000172784.1 | -80944 | Adamtsl3 | ENSMUSG00000070469.6 |
| chr7 | 82531231 | 82531915 | 3.32641 | 3.81876 | 1.02369 | Intron (ENSMUST00000173287.2/ENSMUSG00000070469.6, intron 15 of 29) | 7 | 82611777 | 82612774 | 998 | 1 | ENSMUST00000172784.1 | -79862 | Adamtsl3 | ENSMUSG00000070469.6 |
| chr5 | 134984676 | 134984929 | 3.40616 | 4.2575 | 1.30196 | Downstream (2-3kb) | 5 | 134986214 | 134987472 | 1259 | 1 | ENSMUST00000094245.3 | -1285 | Cldn3 | ENSMUSG00000070473.4 |
| chr5 | 134986129 | 134987111 | 3.925 | 5.10821 | 1.85092 | Promoter | 5 | 134986214 | 134987472 | 1259 | 1 | ENSMUST00000094245.3 | 0 | Cldn3 | ENSMUSG00000070473.4 |
| chr5 | 134987524 | 134988269 | 3.70419 | 4.60994 | 1.5498 | Exon (ENSMUST00000148729.2/ENSMUSG00000054909.6, exon 5 of 5) | 5 | 134986214 | 134987472 | 1259 | 1 | ENSMUST00000094245.3 | 1311 | Cldn3 | ENSMUSG00000070473.4 |
| chr5 | 134988655 | 134988900 | 3.38338 | 3.95091 | 1.17115 | Intron (ENSMUST00000148729.2/ENSMUSG00000054909.6, intron 4 of 4) | 5 | 134986214 | 134987472 | 1259 | 1 | ENSMUST00000094245.3 | 2442 | Cldn3 | ENSMUSG00000070473.4 |
| chr2 | 178415719 | 178415978 | 3.64854 | 4.39708 | 1.35104 | Intron (ENSMUST00000094251.5/ENSMUSG00000070476.5, intron 1 of 3) | 2 | 178414582 | 178420290 | 5709 | 1 | ENSMUST00000129003.1 | 1138 | Fam217b | ENSMUSG00000070476.5 |
| chr5 | 125530756 | 125530991 | 3.66627 | 4.57431 | 1.51977 | Promoter | 5 | 125531774 | 125622780 | 91007 | 1 | ENSMUST00000185104.1 | -783 | Tmem132b | ENSMUSG00000070498.3 |
| chr5 | 125531304 | 125531505 | 3.61096 | 4.4769 | 1.46073 | Promoter | 5 | 125531774 | 125622780 | 91007 | 1 | ENSMUST00000185104.1 | -269 | Tmem132b | ENSMUSG00000070498.3 |
| chr5 | 125531868 | 125531996 | 3.63611 | 4.39357 | 1.39327 | Promoter | 5 | 125531774 | 125622780 | 91007 | 1 | ENSMUST00000185104.1 | 95 | Tmem132b | ENSMUSG00000070498.3 |
| chr5 | 125532375 | 125532617 | 3.77462 | 4.98656 | 1.78481 | Promoter | 5 | 125532387 | 125792583 | 260197 | 1 | ENSMUST00000031446.6 | 0 | Tmem132b | ENSMUSG00000070498.3 |
| chr5 | 125532935 | 125533204 | 3.99705 | 5.24783 | 1.95497 | Promoter | 5 | 125532387 | 125792583 | 260197 | 1 | ENSMUST00000031446.6 | 549 | Tmem132b | ENSMUSG00000070498.3 |
| chr5 | 125533577 | 125535451 | 3.6704 | 4.50958 | 1.4606 | Intron (ENSMUST00000185104.1/ENSMUSG00000070498.3, intron 1 of 1) | 5 | 125532387 | 125792583 | 260197 | 1 | ENSMUST00000031446.6 | 1191 | Tmem132b | ENSMUSG00000070498.3 |
| chr5 | 125535936 | 125536011 | 3.8571 | 4.79675 | 1.61939 | Intron (ENSMUST00000185104.1/ENSMUSG00000070498.3, intron 1 of 1) | 5 | 125532387 | 125792583 | 260197 | 1 | ENSMUST00000031446.6 | 3550 | Tmem132b | ENSMUSG00000070498.3 |
| chr1 | 172615539 | 172615753 | 3.66254 | 4.44124 | 1.401 | Distal Intergenic | 1 | 172596640 | 172602551 | 5912 | 2 | ENSMUST00000094303.4 | -12989 | Fcrl6 | ENSMUSG00000070504.4 |
| chr1 | 172616189 | 172616496 | 3.82492 | 4.79228 | 1.65977 | Distal Intergenic | 1 | 172596640 | 172602551 | 5912 | 2 | ENSMUST00000094303.4 | -13639 | Fcrl6 | ENSMUSG00000070504.4 |
| chr1 | 172616876 | 172617546 | 4.27689 | 5.95088 | 2.46834 | Distal Intergenic | 1 | 172596640 | 172602551 | 5912 | 2 | ENSMUST00000094303.4 | -14326 | Fcrl6 | ENSMUSG00000070504.4 |
| chr7 | 73397808 | 73397883 | 3.50072 | 4.18105 | 1.32383 | Intron (ENSMUST00000094312.6/ENSMUSG00000070509.10, intron 2 of 3) | 7 | 73391227 | 73419899 | 28673 | 1 | ENSMUST00000139780.1 | 6582 | Rgma | ENSMUSG00000070509.10 |
| chr7 | 73400190 | 73400356 | 3.46271 | 4.12041 | 1.2338 | Intron (ENSMUST00000094312.6/ENSMUSG00000070509.10, intron 2 of 3) | 7 | 73391227 | 73419899 | 28673 | 1 | ENSMUST00000139780.1 | 8964 | Rgma | ENSMUSG00000070509.10 |
| chr1 | 170906521 | 170908968 | 3.62819 | 4.41899 | 1.40622 | Exon (ENSMUST00000094337.1/ENSMUSG00000070524.1, exon 4 of 6) | 1 | 170907273 | 170912941 | 5669 | 2 | ENSMUST00000094337.1 | 3973 | Fcrlb | ENSMUSG00000070524.1 |
| chr2 | 164608616 | 164609000 | 3.46861 | 4.06333 | 1.12974 | 5' UTR | 2 | 164596466 | 164613386 | 16921 | 2 | ENSMUST00000109339.2 | 4386 | Wfdc8 | ENSMUSG00000070533.6 |
| chr7 | 45163486 | 45164519 | 4.18257 | 5.88036 | 2.47705 | Promoter | 7 | 45163921 | 45176138 | 12218 | 1 | ENSMUST00000085374.5 | 0 | Slc17a7 | ENSMUSG00000070570.4 |
| chr7 | 45166532 | 45168904 | 3.64481 | 4.5235 | 1.50156 | Exon (ENSMUST00000085374.5/ENSMUSG00000070570.4, exon 2 of 12) | 7 | 45163921 | 45176138 | 12218 | 1 | ENSMUST00000085374.5 | 2612 | Slc17a7 | ENSMUSG00000070570.4 |
| chr5 | 111413681 | 111413768 | 4.42553 | 5.92505 | 2.40772 | Distal Intergenic | 5 | 111417362 | 111418073 | 712 | 1 | ENSMUST00000138784.1 | -3594 | Mn1 | ENSMUSG00000070576.4 |
| chr5 | 111414231 | 111414583 | 3.49991 | 4.1176 | 1.15494 | Distal Intergenic | 5 | 111417362 | 111418073 | 712 | 1 | ENSMUST00000138784.1 | -2779 | Mn1 | ENSMUSG00000070576.4 |
| chr5 | 111415125 | 111415411 | 3.34153 | 3.83966 | 1.01733 | Downstream (2-3kb) | 5 | 111417362 | 111418073 | 712 | 1 | ENSMUST00000138784.1 | -1951 | Mn1 | ENSMUSG00000070576.4 |
| chr5 | 111416387 | 111419467 | 4.08543 | 5.33914 | 2.01588 | Promoter | 5 | 111417362 | 111418073 | 712 | 1 | ENSMUST00000138784.1 | 0 | Mn1 | ENSMUSG00000070576.4 |
| chr5 | 111419824 | 111421318 | 3.90976 | 5.00995 | 1.81032 | Exon (ENSMUST00000094463.4/ENSMUSG00000070576.4, exon 1 of 2) | 5 | 111417442 | 111457033 | 39592 | 1 | ENSMUST00000094463.4 | 2383 | Mn1 | ENSMUSG00000070576.4 |
| chr5 | 111422991 | 111423135 | 3.221 | 3.65871 | 1.01733 | Intron (ENSMUST00000094463.4/ENSMUSG00000070576.4, intron 1 of 1) | 5 | 111417442 | 111457033 | 39592 | 1 | ENSMUST00000094463.4 | 5550 | Mn1 | ENSMUSG00000070576.4 |
| chr5 | 111424008 | 111424277 | 3.25178 | 3.71995 | 1.04945 | Intron (ENSMUST00000094463.4/ENSMUSG00000070576.4, intron 1 of 1) | 5 | 111417442 | 111457033 | 39592 | 1 | ENSMUST00000094463.4 | 6567 | Mn1 | ENSMUSG00000070576.4 |
| chr5 | 111458934 | 111459538 | 3.40474 | 4.16564 | 1.25486 | Distal Intergenic | 5 | 111417442 | 111457033 | 39592 | 1 | ENSMUST00000094463.4 | 41493 | Mn1 | ENSMUSG00000070576.4 |
| chr5 | 111472763 | 111473073 | 3.54265 | 4.20639 | 1.23637 | Distal Intergenic | 5 | 111417442 | 111457033 | 39592 | 1 | ENSMUST00000094463.4 | 55322 | Mn1 | ENSMUSG00000070576.4 |
| chr5 | 111473880 | 111474045 | 3.54515 | 4.22142 | 1.27705 | Distal Intergenic | 5 | 111417442 | 111457033 | 39592 | 1 | ENSMUST00000094463.4 | 56439 | Mn1 | ENSMUSG00000070576.4 |
| chr5 | 111480506 | 111480705 | 3.36029 | 3.86795 | 1.01733 | Distal Intergenic | 5 | 111417442 | 111457033 | 39592 | 1 | ENSMUST00000094463.4 | 63065 | Mn1 | ENSMUSG00000070576.4 |
| chr5 | 111482583 | 111482915 | 3.62491 | 4.39727 | 1.43205 | Distal Intergenic | 5 | 111417442 | 111457033 | 39592 | 1 | ENSMUST00000094463.4 | 65142 | Mn1 | ENSMUSG00000070576.4 |
| chr7 | 43475848 | 43476552 | 3.53916 | 4.28462 | 1.29943 | Exon (ENSMUST00000107974.1/ENSMUSG00000013367.4, exon 6 of 8) | 7 | 43463151 | 43471913 | 8763 | 1 | ENSMUST00000107977.1 | 12698 | Vsig10l | ENSMUSG00000070604.3 |
| chr4 | 136636174 | 136636397 | 3.70882 | 4.53399 | 1.48478 | Intron (ENSMUST00000178843.2/ENSMUSG00000070683.7, intron 7 of 7) | 4 | 136622621 | 136639110 | 16490 | 1 | ENSMUST00000178843.2 | 13554 | Lactbl1 | ENSMUSG00000070683.7 |
| chr4 | 136637106 | 136637843 | 4.52167 | 6.4968 | 2.97105 | Exon (ENSMUST00000178843.2/ENSMUSG00000070683.7, exon 8 of 8) | 4 | 136622621 | 136639110 | 16490 | 1 | ENSMUST00000178843.2 | 14486 | Lactbl1 | ENSMUSG00000070683.7 |
| chr4 | 136638371 | 136638452 | 3.78193 | 5.06792 | 1.82751 | 3' UTR | 4 | 136622621 | 136639110 | 16490 | 1 | ENSMUST00000178843.2 | 15751 | Lactbl1 | ENSMUSG00000070683.7 |
| chr4 | 136416908 | 136417254 | 3.51363 | 4.15229 | 1.19813 | Distal Intergenic | 4 | 136423524 | 136444398 | 20875 | 1 | ENSMUST00000088677.4 | -6270 | Htr1d | ENSMUSG00000070687.6 |
| chr4 | 136419437 | 136419699 | 3.58636 | 4.30783 | 1.31532 | Distal Intergenic | 4 | 136423524 | 136444398 | 20875 | 1 | ENSMUST00000088677.4 | -3825 | Htr1d | ENSMUSG00000070687.6 |
| chr4 | 136423187 | 136423588 | 3.49035 | 4.10173 | 1.14487 | Promoter | 4 | 136423524 | 136444398 | 20875 | 1 | ENSMUST00000088677.4 | 0 | Htr1d | ENSMUSG00000070687.6 |
| chr4 | 136424238 | 136425158 | 3.74235 | 4.65372 | 1.57777 | Promoter | 4 | 136424612 | 136441970 | 17359 | 1 | ENSMUST00000133818.1 | 0 | Htr1d | ENSMUSG00000070687.6 |
| chr4 | 135128937 | 135129539 | 3.46216 | 4.054 | 1.12963 | Intron (ENSMUST00000056977.8/ENSMUSG00000070691.5, intron 1 of 5) | 4 | 135120652 | 135177990 | 57339 | 1 | ENSMUST00000056977.8 | 8286 | Runx3 | ENSMUSG00000070691.5 |
| chr4 | 135129969 | 135130050 | 3.80418 | 4.68014 | 1.50448 | Intron (ENSMUST00000056977.8/ENSMUSG00000070691.5, intron 1 of 5) | 4 | 135120652 | 135177990 | 57339 | 1 | ENSMUST00000056977.8 | 9318 | Runx3 | ENSMUSG00000070691.5 |
| chr4 | 135158452 | 135158939 | 3.94956 | 4.97126 | 1.72722 | Intron (ENSMUST00000056977.8/ENSMUSG00000070691.5, intron 3 of 5) | 4 | 135154436 | 135155342 | 907 | 1 | ENSMUST00000127109.1 | 4017 | Runx3 | ENSMUSG00000070691.5 |
| chr4 | 135160637 | 135160984 | 3.71802 | 4.52146 | 1.4051 | Intron (ENSMUST00000056977.8/ENSMUSG00000070691.5, intron 3 of 5) | 4 | 135154436 | 135155342 | 907 | 1 | ENSMUST00000127109.1 | 6202 | Runx3 | ENSMUSG00000070691.5 |
| chr4 | 135164346 | 135164421 | 3.43025 | 3.98868 | 1.07072 | Intron (ENSMUST00000056977.8/ENSMUSG00000070691.5, intron 4 of 5) | 4 | 135154436 | 135155342 | 907 | 1 | ENSMUST00000127109.1 | 9911 | Runx3 | ENSMUSG00000070691.5 |
| chr4 | 135165393 | 135165759 | 3.63941 | 4.37926 | 1.3237 | Intron (ENSMUST00000056977.8/ENSMUSG00000070691.5, intron 4 of 5) | 4 | 135154436 | 135155342 | 907 | 1 | ENSMUST00000127109.1 | 10958 | Runx3 | ENSMUSG00000070691.5 |
| chr4 | 135166432 | 135167293 | 3.39028 | 3.94331 | 1.05384 | Intron (ENSMUST00000056977.8/ENSMUSG00000070691.5, intron 4 of 5) | 4 | 135154436 | 135155342 | 907 | 1 | ENSMUST00000127109.1 | 11997 | Runx3 | ENSMUSG00000070691.5 |
| chr4 | 135168396 | 135168933 | 3.74184 | 4.58073 | 1.48471 | Intron (ENSMUST00000056977.8/ENSMUSG00000070691.5, intron 4 of 5) | 4 | 135154436 | 135155342 | 907 | 1 | ENSMUST00000127109.1 | 13961 | Runx3 | ENSMUSG00000070691.5 |
| chr4 | 135170078 | 135170338 | 3.39311 | 3.97584 | 1.0935 | Intron (ENSMUST00000056977.8/ENSMUSG00000070691.5, intron 4 of 5) | 4 | 135154436 | 135155342 | 907 | 1 | ENSMUST00000127109.1 | 15643 | Runx3 | ENSMUSG00000070691.5 |
| chr4 | 135148830 | 135149069 | 4.23829 | 5.85177 | 2.43219 | Intron (ENSMUST00000056977.8/ENSMUSG00000070691.5, intron 1 of 5) | 4 | 135152496 | 135176181 | 23686 | 1 | ENSMUST00000119564.1 | -3427 | Runx3 | ENSMUSG00000070691.5 |
| chr4 | 135149422 | 135149649 | 3.69682 | 4.63646 | 1.52173 | Intron (ENSMUST00000056977.8/ENSMUSG00000070691.5, intron 1 of 5) | 4 | 135152496 | 135176181 | 23686 | 1 | ENSMUST00000119564.1 | -2847 | Runx3 | ENSMUSG00000070691.5 |
| chr4 | 135150094 | 135153783 | 4.29997 | 5.86364 | 2.44031 | Promoter | 4 | 135152496 | 135176181 | 23686 | 1 | ENSMUST00000119564.1 | 0 | Runx3 | ENSMUSG00000070691.5 |
| chr4 | 135154119 | 135154228 | 3.50226 | 4.13392 | 1.19948 | Promoter | 4 | 135154436 | 135155342 | 907 | 1 | ENSMUST00000127109.1 | -208 | Runx3 | ENSMUSG00000070691.5 |
| chr4 | 135154555 | 135154712 | 3.55666 | 4.22902 | 1.24616 | Promoter | 4 | 135154436 | 135155342 | 907 | 1 | ENSMUST00000127109.1 | 120 | Runx3 | ENSMUSG00000070691.5 |
| chr4 | 135156231 | 135158049 | 4.24924 | 5.64977 | 2.26461 | Intron (ENSMUST00000056977.8/ENSMUSG00000070691.5, intron 3 of 5) | 4 | 135154436 | 135155342 | 907 | 1 | ENSMUST00000127109.1 | 1796 | Runx3 | ENSMUSG00000070691.5 |
| chr4 | 135203172 | 135204603 | 3.54459 | 4.38887 | 1.40484 | Distal Intergenic | 4 | 135154436 | 135155342 | 907 | 1 | ENSMUST00000127109.1 | 48737 | Runx3 | ENSMUSG00000070691.5 |
| chr4 | 135205323 | 135206592 | 3.84345 | 4.97954 | 1.82179 | Distal Intergenic | 4 | 135154436 | 135155342 | 907 | 1 | ENSMUST00000127109.1 | 50888 | Runx3 | ENSMUSG00000070691.5 |
| chr4 | 135178202 | 135178447 | 3.41725 | 3.975 | 1.09208 | Distal Intergenic | 4 | 135154436 | 135155342 | 907 | 1 | ENSMUST00000127109.1 | 23767 | Runx3 | ENSMUSG00000070691.5 |
| chr4 | 135173711 | 135173799 | 4.09642 | 5.31427 | 2.0056 | Intron (ENSMUST00000056977.8/ENSMUSG00000070691.5, intron 5 of 5) | 4 | 135154436 | 135155342 | 907 | 1 | ENSMUST00000127109.1 | 19276 | Runx3 | ENSMUSG00000070691.5 |
| chr4 | 135174981 | 135175896 | 4.00871 | 5.12025 | 1.84696 | 3' UTR | 4 | 135154436 | 135155342 | 907 | 1 | ENSMUST00000127109.1 | 20546 | Runx3 | ENSMUSG00000070691.5 |
| chr4 | 135177252 | 135177372 | 3.69855 | 4.49301 | 1.40682 | 3' UTR | 4 | 135154436 | 135155342 | 907 | 1 | ENSMUST00000127109.1 | 22817 | Runx3 | ENSMUSG00000070691.5 |
| chr4 | 135198632 | 135198717 | 3.6457 | 4.43325 | 1.44766 | Distal Intergenic | 4 | 135154436 | 135155342 | 907 | 1 | ENSMUST00000127109.1 | 44197 | Runx3 | ENSMUSG00000070691.5 |
| chr4 | 135202055 | 135202213 | 3.24652 | 4.09181 | 1.17492 | Distal Intergenic | 4 | 135154436 | 135155342 | 907 | 1 | ENSMUST00000127109.1 | 47620 | Runx3 | ENSMUSG00000070691.5 |
| chr4 | 131920656 | 131922309 | 3.84939 | 5.02662 | 1.80434 | Promoter | 4 | 131921771 | 131923140 | 1370 | 1 | ENSMUST00000094666.3 | 0 | Tmem200b | ENSMUSG00000070720.3 |
| chr4 | 131922856 | 131923282 | 3.93329 | 5.28554 | 1.99109 | 3' UTR | 4 | 131921771 | 131923140 | 1370 | 1 | ENSMUST00000094666.3 | 1086 | Tmem200b | ENSMUSG00000070720.3 |
| chr5 | 66062072 | 66062321 | 3.49647 | 4.11138 | 1.15081 | Intron (ENSMUST00000113726.2/ENSMUSG00000070780.6, intron 4 of 8) | 5 | 66026785 | 66054550 | 27766 | 2 | ENSMUST00000145625.2 | -7523 | Rbm47 | ENSMUSG00000070780.6 |
| chr5 | 66077682 | 66077993 | 3.2429 | 3.93416 | 1.13416 | Intron (ENSMUST00000113726.2/ENSMUSG00000070780.6, intron 4 of 8) | 5 | 66027220 | 66080984 | 53765 | 2 | ENSMUST00000127275.2 | 2991 | Rbm47 | ENSMUSG00000070780.6 |
| chr5 | 66147430 | 66147634 | 3.949 | 5.08202 | 1.81521 | Intron (ENSMUST00000113726.2/ENSMUSG00000070780.6, intron 3 of 8) | 5 | 66018602 | 66150898 | 132297 | 2 | ENSMUST00000113725.2 | 3264 | Rbm47 | ENSMUSG00000070780.6 |
| chr5 | 66150315 | 66150669 | 3.32668 | 3.9905 | 1.15194 | Promoter | 5 | 66018602 | 66150898 | 132297 | 2 | ENSMUST00000113725.2 | 229 | Rbm47 | ENSMUSG00000070780.6 |
| chr5 | 66150980 | 66151467 | 3.48027 | 4.47478 | 1.42731 | Promoter | 5 | 66027240 | 66151323 | 124084 | 2 | ENSMUST00000131838.2 | 0 | Rbm47 | ENSMUSG00000070780.6 |
| chr5 | 66151847 | 66152002 | 3.74601 | 4.65456 | 1.60362 | Promoter | 5 | 66016557 | 66151924 | 135368 | 2 | ENSMUST00000167950.2 | 0 | Rbm47 | ENSMUSG00000070780.6 |
| chr5 | 66152362 | 66152545 | 3.53499 | 4.23572 | 1.34452 | Promoter | 5 | 66016557 | 66151924 | 135368 | 2 | ENSMUST00000167950.2 | -439 | Rbm47 | ENSMUSG00000070780.6 |
| chr4 | 120647364 | 120648165 | 3.34842 | 3.94505 | 1.14726 | Distal Intergenic | 4 | 120666572 | 120667820 | 1249 | 1 | ENSMUST00000094814.5 | -18407 | Cited4 | ENSMUSG00000070803.6 |
| chr4 | 120648616 | 120648939 | 3.51681 | 4.20276 | 1.31104 | Distal Intergenic | 4 | 120666572 | 120667820 | 1249 | 1 | ENSMUST00000094814.5 | -17633 | Cited4 | ENSMUSG00000070803.6 |
| chr4 | 120649735 | 120650116 | 3.76431 | 4.88658 | 1.69274 | Distal Intergenic | 4 | 120666572 | 120667820 | 1249 | 1 | ENSMUST00000094814.5 | -16456 | Cited4 | ENSMUSG00000070803.6 |
| chr4 | 120651111 | 120651212 | 3.4118 | 4.0753 | 1.19056 | Distal Intergenic | 4 | 120666572 | 120667820 | 1249 | 1 | ENSMUST00000094814.5 | -15360 | Cited4 | ENSMUSG00000070803.6 |
| chr4 | 120652550 | 120652784 | 3.67252 | 4.51623 | 1.48541 | Distal Intergenic | 4 | 120666572 | 120667820 | 1249 | 1 | ENSMUST00000094814.5 | -13788 | Cited4 | ENSMUSG00000070803.6 |
| chr4 | 120698464 | 120698584 | 3.41004 | 3.96186 | 1.08407 | Exon (ENSMUST00000030376.7/ENSMUSG00000028631.7, exon 13 of 14) | 4 | 120666572 | 120667820 | 1249 | 1 | ENSMUST00000094814.5 | 31893 | Cited4 | ENSMUSG00000070803.6 |
| chr7 | 13124129 | 13124660 | 3.48058 | 4.10949 | 1.22364 | Distal Intergenic | 7 | 13131471 | 13132358 | 888 | 2 | ENSMUST00000094827.1 | 7698 | Vmn1r87 | ENSMUSG00000070815.1 |
| chr4 | 118741038 | 118741315 | 3.37203 | 3.88831 | 1.0267 | Distal Intergenic | 4 | 118734531 | 118735478 | 948 | 1 | ENSMUST00000094832.1 | 6508 | Olfr1339 | ENSMUSG00000070820.1 |
| chr5 | 37162529 | 37163410 | 4.16559 | 5.44848 | 2.07446 | Distal Intergenic | 5 | 37185897 | 37193023 | 7127 | 1 | ENSMUST00000094840.3 | -22487 | Gm1043 | ENSMUSG00000070823.3 |
| chr5 | 37164287 | 37164413 | 3.37718 | 3.89343 | 1.01733 | Distal Intergenic | 5 | 37185897 | 37193023 | 7127 | 1 | ENSMUST00000094840.3 | -21484 | Gm1043 | ENSMUSG00000070823.3 |
| chr2 | 82052591 | 82053809 | 3.59815 | 4.33306 | 1.35765 | Promoter | 2 | 82053222 | 82259879 | 206658 | 1 | ENSMUST00000047527.7 | 0 | Zfp804a | ENSMUSG00000070866.4 |
| chr2 | 82054259 | 82054644 | 4.126 | 5.65329 | 2.23134 | Intron (ENSMUST00000047527.7/ENSMUSG00000070866.4, intron 1 of 3) | 2 | 82053222 | 82259879 | 206658 | 1 | ENSMUST00000047527.7 | 1038 | Zfp804a | ENSMUSG00000070866.4 |
| chr2 | 82055538 | 82055729 | 3.23316 | 3.67692 | 1.01733 | Intron (ENSMUST00000047527.7/ENSMUSG00000070866.4, intron 1 of 3) | 2 | 82053222 | 82259879 | 206658 | 1 | ENSMUST00000047527.7 | 2317 | Zfp804a | ENSMUSG00000070866.4 |
| chr2 | 70557985 | 70558152 | 3.36816 | 3.87982 | 1.01733 | Intron (ENSMUST00000130618.2/ENSMUSG00000070880.5, intron 1 of 3) | 2 | 70562042 | 70587136 | 25095 | 1 | ENSMUST00000130604.2 | -3890 | Gad1 | ENSMUSG00000070880.5 |
| chr2 | 70558722 | 70559726 | 3.65725 | 4.48905 | 1.42814 | Intron (ENSMUST00000130618.2/ENSMUSG00000070880.5, intron 1 of 3) | 2 | 70562042 | 70587136 | 25095 | 1 | ENSMUST00000130604.2 | -2316 | Gad1 | ENSMUSG00000070880.5 |
| chr2 | 70560161 | 70560362 | 4.16681 | 5.42181 | 2.04431 | Intron (ENSMUST00000130618.2/ENSMUSG00000070880.5, intron 1 of 3) | 2 | 70562042 | 70587136 | 25095 | 1 | ENSMUST00000130604.2 | -1680 | Gad1 | ENSMUSG00000070880.5 |
| chr2 | 70561481 | 70562337 | 3.86666 | 4.81528 | 1.61492 | Promoter | 2 | 70562042 | 70587136 | 25095 | 1 | ENSMUST00000130604.2 | 0 | Gad1 | ENSMUSG00000070880.5 |
| chr2 | 70562764 | 70563051 | 3.61788 | 4.40088 | 1.42274 | Promoter | 2 | 70562899 | 70585121 | 22223 | 1 | ENSMUST00000130998.2 | 0 | Gad1 | ENSMUSG00000070880.5 |
| chr2 | 70565163 | 70566068 | 4.31364 | 5.7705 | 2.37628 | Promoter | 2 | 70566377 | 70602012 | 35636 | 1 | ENSMUST00000140478.1 | -309 | Gad1 | ENSMUSG00000070880.5 |
| chr2 | 70566445 | 70567457 | 3.77705 | 4.77196 | 1.64923 | Promoter | 2 | 70566377 | 70602012 | 35636 | 1 | ENSMUST00000140478.1 | 69 | Gad1 | ENSMUSG00000070880.5 |
| chr2 | 70568389 | 70568919 | 3.98514 | 5.10239 | 1.8533 | Intron (ENSMUST00000130604.2/ENSMUSG00000070880.5, intron 3 of 8) | 2 | 70566377 | 70602012 | 35636 | 1 | ENSMUST00000140478.1 | 2013 | Gad1 | ENSMUSG00000070880.5 |
| chr2 | 70569489 | 70569643 | 3.34153 | 3.83966 | 1.01733 | Intron (ENSMUST00000130604.2/ENSMUSG00000070880.5, intron 3 of 8) | 2 | 70566377 | 70602012 | 35636 | 1 | ENSMUST00000140478.1 | 3113 | Gad1 | ENSMUSG00000070880.5 |
| chr4 | 99333186 | 99333546 | 3.53273 | 4.17875 | 1.19415 | Distal Intergenic | 4 | 99295900 | 99297258 | 1359 | 1 | ENSMUST00000094955.2 | 37287 | Gm12689 | ENSMUSG00000070891.2 |
| chr1 | 40324846 | 40325024 | 3.33237 | 4.03811 | 1.20234 | Promoter | 1 | 40324627 | 40365471 | 40845 | 1 | ENSMUST00000095020.3 | 220 | Il1rl2 | ENSMUSG00000070942.3 |
| chr1 | 40325389 | 40325464 | 3.60693 | 4.52779 | 1.43716 | Promoter | 1 | 40324627 | 40365471 | 40845 | 1 | ENSMUST00000095020.3 | 763 | Il1rl2 | ENSMUSG00000070942.3 |
| chr4 | 46341891 | 46345602 | 4.4718 | 6.27613 | 2.77238 | Promoter | 4 | 46343609 | 46346412 | 2804 | 1 | ENSMUST00000095097.2 | 0 | Foxe1 | ENSMUSG00000070990.2 |
| chr8 | 46257993 | 46258585 | 3.37944 | 3.93935 | 1.10013 | Distal Intergenic | 8 | 46254677 | 46255678 | 1002 | 2 | ENSMUST00000095320.1 | -2316 | Gm10313 | ENSMUSG00000071102.1 |
| chr6 | 142412498 | 142412617 | 3.69223 | 4.53914 | 1.46887 | Promoter | 6 | 142413441 | 142419730 | 6290 | 1 | ENSMUST00000088263.5 | -824 | B230216G23Rik | ENSMUSG00000071112.6 |
| chr6 | 142413579 | 142414931 | 4.08432 | 5.24869 | 1.92835 | Promoter | 6 | 142413833 | 142418571 | 4739 | 1 | ENSMUST00000126521.3 | 0 | B230216G23Rik | ENSMUSG00000071112.6 |
| chr6 | 142415642 | 142415998 | 3.71844 | 4.53464 | 1.45744 | Intron (ENSMUST00000088263.5/ENSMUSG00000071112.6, intron 4 of 5) | 6 | 142414521 | 142419730 | 5210 | 1 | ENSMUST00000123209.1 | 1122 | B230216G23Rik | ENSMUSG00000071112.6 |
| chr13 | 104938112 | 104938227 | 3.38975 | 3.91872 | 1.03958 | Distal Intergenic | 13 | 104918779 | 104919186 | 408 | 1 | ENSMUST00000171334.1 | 19334 | Gm4938 | ENSMUSG00000071193.4 |
| chr17 | 24657981 | 24658178 | 3.47639 | 4.15896 | 1.22609 | Promoter | 17 | 24657330 | 24658457 | 1128 | 2 | ENSMUST00000095544.5 | 279 | Npw | ENSMUSG00000071230.5 |
| chr12 | 84680573 | 84681232 | 3.67976 | 4.4602 | 1.39126 | 5' UTR | 12 | 84677278 | 84698807 | 21530 | 2 | ENSMUST00000095550.2 | 17575 | Syndig1l | ENSMUSG00000071234.2 |
| chr12 | 84695090 | 84695201 | 3.83041 | 4.75369 | 1.58956 | Intron (ENSMUST00000095550.2/ENSMUSG00000071234.2, intron 1 of 3) | 12 | 84677278 | 84698807 | 21530 | 2 | ENSMUST00000095550.2 | 3606 | Syndig1l | ENSMUSG00000071234.2 |
| chr12 | 84695681 | 84695908 | 3.90598 | 4.9238 | 1.71989 | Intron (ENSMUST00000095550.2/ENSMUSG00000071234.2, intron 1 of 3) | 12 | 84677278 | 84698807 | 21530 | 2 | ENSMUST00000095550.2 | 2899 | Syndig1l | ENSMUSG00000071234.2 |
| chr12 | 84696914 | 84697499 | 3.94548 | 4.98695 | 1.77329 | Intron (ENSMUST00000095550.2/ENSMUSG00000071234.2, intron 1 of 3) | 12 | 84677278 | 84698807 | 21530 | 2 | ENSMUST00000095550.2 | 1308 | Syndig1l | ENSMUSG00000071234.2 |
| chr12 | 84698058 | 84698988 | 4.08171 | 5.33212 | 2.02757 | Promoter | 12 | 84677278 | 84698807 | 21530 | 2 | ENSMUST00000095550.2 | 0 | Syndig1l | ENSMUSG00000071234.2 |
| chr12 | 84699491 | 84699615 | 3.50722 | 4.13586 | 1.17983 | Promoter | 12 | 84677278 | 84698807 | 21530 | 2 | ENSMUST00000095550.2 | -685 | Syndig1l | ENSMUSG00000071234.2 |
| chr12 | 84702113 | 84702240 | 4.13102 | 5.32568 | 1.95767 | Distal Intergenic | 12 | 84677278 | 84698807 | 21530 | 2 | ENSMUST00000095550.2 | -3307 | Syndig1l | ENSMUSG00000071234.2 |
| chr12 | 84711098 | 84711399 | 3.24456 | 3.69401 | 1.01733 | Distal Intergenic | 12 | 84677278 | 84698807 | 21530 | 2 | ENSMUST00000095550.2 | -12292 | Syndig1l | ENSMUSG00000071234.2 |
| chr12 | 84716631 | 84716961 | 3.41094 | 3.9787 | 1.13789 | Distal Intergenic | 12 | 84677278 | 84698807 | 21530 | 2 | ENSMUST00000095550.2 | -17825 | Syndig1l | ENSMUSG00000071234.2 |
| chr12 | 84631277 | 84631414 | 3.3962 | 3.9436 | 1.09361 | Distal Intergenic | 12 | 84641019 | 84651455 | 10437 | 1 | ENSMUST00000166772.3 | -9605 | Vrtn | ENSMUSG00000071235.6 |
| chr12 | 84631987 | 84632210 | 3.83633 | 4.7706 | 1.62575 | Distal Intergenic | 12 | 84641019 | 84651455 | 10437 | 1 | ENSMUST00000166772.3 | -8809 | Vrtn | ENSMUSG00000071235.6 |
| chr12 | 84640279 | 84640488 | 3.63318 | 4.38011 | 1.36002 | Promoter | 12 | 84641019 | 84651455 | 10437 | 1 | ENSMUST00000166772.3 | -531 | Vrtn | ENSMUSG00000071235.6 |
| chr12 | 84640998 | 84641656 | 3.52789 | 4.19789 | 1.27939 | Promoter | 12 | 84641019 | 84651455 | 10437 | 1 | ENSMUST00000166772.3 | 0 | Vrtn | ENSMUSG00000071235.6 |

| | | | | | | | | | | | | | | | |
|---|---|---|---|---|---|---|---|---|---|---|---|---|---|---|---|
| chr12 | 84642021 | 84642178 | 3.59553 | 4.41028 | 1.4207 | Promoter | 12 | 84642555 | 84651455 | 8901 | 1 | ENSMUST00000167227.2 | -377 | Vrtn | ENSMUSG00000071235.6 |
| chr12 | 84642590 | 84644188 | 3.57072 | 4.31345 | 1.32649 | Promoter | 12 | 84642896 | 84651455 | 8560 | 1 | ENSMUST00000095551.4 | 0 | Vrtn | ENSMUSG00000071235.6 |
| chr12 | 74283196 | 74284519 | 4.00765 | 5.11067 | 1.84243 | Promoter | 12 | 74284281 | 74295931 | 11651 | 1 | ENSMUST00000186463.1 | 0 | 1700086L19Rik | ENSMUSG00000071265.4 |
| chr12 | 74284899 | 74285016 | 4.49361 | 6.05907 | 2.4924 | Promoter | 12 | 74284289 | 74295945 | 11657 | 1 | ENSMUST00000185822.1 | 611 | 1700086L19Rik | ENSMUSG00000071265.4 |
| chr10 | 45335268 | 45336765 | 4.15718 | 5.53161 | 2.23518 | Promoter | 10 | 45335762 | 45369708 | 33947 | 1 | ENSMUST00000095715.3 | 0 | Bves | ENSMUSG00000071317.3 |
| chr6 | 85511608 | 85512356 | 4.12787 | 5.44653 | 2.09723 | 3' UTR | 6 | 85511131 | 85513589 | 2459 | 2 | ENSMUST00000095759.3 | 1233 | Egr4 | ENSMUSG00000071341.3 |
| chr6 | 85512711 | 85514378 | 4.16642 | 5.45038 | 2.08606 | Promoter | 6 | 85511131 | 85513589 | 2459 | 2 | ENSMUST00000095759.3 | 0 | Egr4 | ENSMUSG00000071341.3 |
| chr6 | 85514713 | 85515532 | 4.05157 | 5.27619 | 1.98565 | Distal Intergenic | 6 | 85511131 | 85513589 | 2459 | 2 | ENSMUST00000095759.3 | -1125 | Egr4 | ENSMUSG00000071341.3 |
| chr6 | 63253656 | 63254176 | 3.24456 | 3.69787 | 1.02737 | Distal Intergenic | 6 | 63255876 | 63314071 | 58196 | 1 | ENSMUST00000159319.1 | -1700 | Grid2 | ENSMUSG00000071424.4 |
| chr6 | 63254775 | 63254850 | 3.70528 | 4.5779 | 1.57264 | Distal Intergenic | 6 | 63255876 | 63314071 | 58196 | 1 | ENSMUST00000159319.1 | -1026 | Grid2 | ENSMUSG00000071424.4 |
| chr6 | 63255221 | 63255693 | 3.24216 | 3.71552 | 1.03759 | Promoter | 6 | 63255876 | 63314071 | 58196 | 1 | ENSMUST00000159319.1 | -183 | Grid2 | ENSMUSG00000071424.4 |
| chr6 | 63256352 | 63256736 | 3.52882 | 4.52885 | 1.49167 | Promoter | 6 | 63256826 | 64278792 | 1021967 | 1 | ENSMUST00000161105.1 | -90 | Grid2 | ENSMUSG00000071424.4 |
| chr6 | 63257175 | 63257959 | 3.67791 | 4.62458 | 1.55181 | Promoter | 6 | 63256826 | 64278792 | 1021967 | 1 | ENSMUST00000161105.1 | 350 | Grid2 | ENSMUSG00000071424.4 |
| chr6 | 38635645 | 38635920 | 3.51202 | 4.23877 | 1.30687 | Intron (ENSMUST00000096030.3/ENSMUSG00000071537.3, intron 1 of 4) | 6 | 38626660 | 38637239 | 10580 | 2 | ENSMUST00000096030.3 | 1319 | Klrg2 | ENSMUSG00000071537.3 |
| chr6 | 38636808 | 38636883 | 3.81202 | 5.41642 | 2.04867 | Promoter | 6 | 38626660 | 38637239 | 10580 | 2 | ENSMUST00000096030.3 | 356 | Klrg2 | ENSMUSG00000071537.3 |
| chr6 | 38637855 | 38638459 | 3.50464 | 4.16607 | 1.28446 | Promoter | 6 | 38626660 | 38637239 | 10580 | 2 | ENSMUST00000096030.3 | -617 | Klrg2 | ENSMUSG00000071537.3 |
| chr6 | 38638967 | 38639144 | 3.72784 | 4.55239 | 1.46772 | Distal Intergenic | 6 | 38626660 | 38637239 | 10580 | 2 | ENSMUST00000096030.3 | -1729 | Klrg2 | ENSMUSG00000071537.3 |
| chr14 | 32682505 | 32682969 | 3.50998 | 4.42417 | 1.3859 | Intron (ENSMUST00000096038.3/ENSMUSG00000071540.3, intron 1 of 3) | 14 | 32659119 | 32685272 | 26154 | 2 | ENSMUST00000096038.3 | 2303 | 3425401B19Rik | ENSMUSG00000071540.3 |
| chr6 | 30546887 | 30547469 | 3.47473 | 4.07096 | 1.12412 | Promoter | 6 | 30546642 | 30547155 | 514 | 1 | ENSMUST00000123588.1 | 246 | Cpa2 | ENSMUSG00000071553.5 |
| chr6 | 30551082 | 30551511 | 3.38193 | 3.9006 | 1.01733 | Promoter | 6 | 30551903 | 30564248 | 12346 | 1 | ENSMUST00000144282.1 | -392 | Cpa2 | ENSMUSG00000071553.5 |
| chr19 | 33224389 | 33224546 | 4.01866 | 5.43166 | 2.06891 | Intron (ENSMUST00000096114.6/ENSMUSG00000071573.8, intron 4 of 6) | 19 | 33137747 | 33392265 | 254519 | 2 | ENSMUST00000096114.6 | 167719 | Rnls | ENSMUSG00000071573.8 |
| chr19 | 24032417 | 24032492 | 3.54298 | 4.19421 | 1.19531 | Distal Intergenic | 19 | 23972751 | 24031019 | 58269 | 2 | ENSMUST00000096164.4 | -1399 | Fam189a2 | ENSMUSG00000071604.4 |
| chr16 | 17211062 | 17211137 | 3.213 | 3.64675 | 1.01109 | Exon (ENSMUST00000169803.2/ENSMUSG00000071636.6, exon 1 of 1) | 16 | 17208135 | 17213921 | 5787 | 1 | ENSMUST00000169803.2 | 2928 | Rimbp3 | ENSMUSG00000071636.6 |
| chr19 | 4681877 | 4682021 | 3.24206 | 3.69025 | 1.01733 | Intron (ENSMUST00000177696.2/ENSMUSG00000071691.5, intron 3 of 14) | 19 | 4648317 | 4698315 | 49999 | 2 | ENSMUST00000096325.3 | 16294 | Gm960 | ENSMUSG00000071691.5 |
| chr19 | 4698282 | 4698748 | 4.5603 | 6.47071 | 2.83652 | Promoter | 19 | 4625841 | 4698668 | 72828 | 2 | ENSMUST00000177696.2 | 0 | Gm960 | ENSMUSG00000071691.5 |
| chr19 | 4699601 | 4700181 | 3.7083 | 4.56988 | 1.55742 | Promoter | 19 | 4625841 | 4698668 | 72828 | 2 | ENSMUST00000177696.2 | -934 | Gm960 | ENSMUSG00000071691.5 |
| chr19 | 4700532 | 4701111 | 3.38115 | 3.94942 | 1.17622 | Distal Intergenic | 19 | 4625841 | 4698668 | 72828 | 2 | ENSMUST00000177696.2 | -1865 | Gm960 | ENSMUSG00000071691.5 |
| chr15 | 78410269 | 78410370 | 3.49218 | 4.10507 | 1.15023 | Exon (ENSMUST00000167140.2/ENSMUSG00000071711.6, exon 2 of 3) | 15 | 78407104 | 78414002 | 6899 | 1 | ENSMUST00000043865.6 | 3166 | Mpst | ENSMUSG00000071711.6 |
| chr15 | 78411910 | 78412045 | 3.6102 | 4.33394 | 1.32339 | Intron (ENSMUST00000167140.2/ENSMUSG00000071711.6, intron 2 of 2) | 15 | 78407104 | 78414002 | 6899 | 1 | ENSMUST00000043865.6 | 4807 | Mpst | ENSMUSG00000071711.6 |
| chr15 | 78254100 | 78254405 | 3.41188 | 3.95614 | 1.05373 | Promoter | 15 | 78253246 | 78260769 | 7524 | 1 | ENSMUST00000126028.2 | 855 | Ncf4 | ENSMUSG00000071715.6 |
| chr18 | 62939760 | 62940146 | 3.63398 | 4.47545 | 1.49162 | Intron (ENSMUST00000096554.5/ENSMUSG00000071847.7, intron 3 of 4) | 18 | 62922329 | 62952464 | 30136 | 1 | ENSMUST00000163716.2 | 17432 | Apcdd1 | ENSMUSG00000071847.7 |
| chr18 | 44811210 | 44812215 | 3.5517 | 4.25524 | 1.28721 | Promoter | 18 | 44425061 | 44812182 | 387122 | 2 | ENSMUST00000089874.5 | 0 | Mcc | ENSMUSG00000071856.5 |
| chr18 | 44813717 | 44814014 | 3.34826 | 3.84982 | 1.01733 | Distal Intergenic | 18 | 44425061 | 44812182 | 387122 | 2 | ENSMUST00000089874.5 | -1536 | Mcc | ENSMUSG00000071856.5 |
| chr18 | 40256042 | 40256363 | 4.02233 | 5.18821 | 1.90887 | Promoter | 18 | 40256962 | 40257684 | 723 | 1 | ENSMUST00000096572.1 | -599 | 2900055J20Rik | ENSMUSG00000071860.1 |
| chr18 | 40256773 | 40257264 | 4.09766 | 5.46718 | 2.10761 | Promoter | 18 | 40256962 | 40257684 | 723 | 1 | ENSMUST00000096572.1 | 0 | 2900055J20Rik | ENSMUSG00000071860.1 |
| chr17 | 7825278 | 7825625 | 3.40298 | 3.98877 | 1.19306 | Intron (ENSMUST00000167580.2/ENSMUSG00000071984.5, intron 1 of 3) | 17 | 7782444 | 7827302 | 44859 | 2 | ENSMUST00000167580.2 | 1677 | Fndc1 | ENSMUSG00000071984.5 |
| chr17 | 7826291 | 7826420 | 4.0149 | 5.29393 | 1.96175 | Promoter | 17 | 7782444 | 7827302 | 44859 | 2 | ENSMUST00000167580.2 | 882 | Fndc1 | ENSMUSG00000071984.5 |
| chr17 | 7826823 | 7827230 | 3.70605 | 4.56923 | 1.55106 | Promoter | 17 | 7782444 | 7827302 | 44859 | 2 | ENSMUST00000167580.2 | 72 | Fndc1 | ENSMUSG00000071984.5 |
| chr3 | 101300396 | 101300678 | 3.46852 | 4.28575 | 1.33762 | Distal Intergenic | 3 | 101306561 | 101307079 | 519 | 2 | ENSMUST00000097146.2 | 6401 | Gm10355 | ENSMUSG00000072381.2 |
| chr14 | 32379591 | 32379666 | 3.48481 | 4.11245 | 1.21865 | Intron (ENSMUST00000100723.2/ENSMUSG00000072473.3, intron 1 of 2) | 14 | 32376502 | 32388373 | 11872 | 2 | ENSMUST00000100723.2 | 8707 | 1700024G13Rik | ENSMUSG00000072473.3 |
| chr14 | 32418848 | 32419072 | 3.86345 | 4.83111 | 1.69134 | Exon (ENSMUST00000070125.4/ENSMUSG00000021919.7, exon 13 of 16) | 14 | 32376502 | 32388373 | 11872 | 2 | ENSMUST00000100723.2 | -30476 | 1700024G13Rik | ENSMUSG00000072473.3 |
| chr15 | 73838890 | 73839553 | 3.60764 | 4.35363 | 1.33771 | Promoter | 15 | 73786412 | 73839671 | 53260 | 2 | ENSMUST00000071419.6 | 118 | Mroh5 | ENSMUSG00000072487.5 |
| chr15 | 73839885 | 73840424 | 3.54747 | 4.23631 | 1.29264 | Promoter | 15 | 73796736 | 73839699 | 42964 | 2 | ENSMUST00000136698.1 | -187 | Mroh5 | ENSMUSG00000072487.5 |
| chr15 | 73870201 | 73870292 | 3.40733 | 3.95245 | 1.06479 | Distal Intergenic | 15 | 73796736 | 73839699 | 42964 | 2 | ENSMUST00000136698.1 | -30503 | Mroh5 | ENSMUSG00000072487.5 |
| chr15 | 73951824 | 73952213 | 3.4258 | 3.99008 | 1.09947 | Distal Intergenic | 15 | 73796736 | 73839699 | 42964 | 2 | ENSMUST00000136698.1 | -112126 | Mroh5 | ENSMUSG00000072487.5 |
| chr5 | 128550888 | 128551354 | 3.55767 | 4.39071 | 1.38105 | Distal Intergenic | 5 | 128579106 | 128585378 | 6273 | 2 | ENSMUST00000140028.2 | 34024 | 5930412G12Rik | ENSMUSG00000072591.5 |
| chr5 | 128568721 | 128568940 | 3.44516 | 4.14185 | 1.22116 | Distal Intergenic | 5 | 128579106 | 128585378 | 6273 | 2 | ENSMUST00000140028.2 | 16438 | 5930412G12Rik | ENSMUSG00000072591.5 |
| chr5 | 128596351 | 128597214 | 4.05641 | 5.28718 | 2.00984 | Intron (ENSMUST00000154305.2/ENSMUSG00000072591.5, intron 1 of 5) | 5 | 128584180 | 128600654 | 16475 | 2 | ENSMUST00000126472.1 | 3440 | 5930412G12Rik | ENSMUSG00000072591.5 |
| chr5 | 128597644 | 128597752 | 3.36484 | 3.87482 | 1.01733 | Intron (ENSMUST00000154305.2/ENSMUSG00000072591.5, intron 1 of 5) | 5 | 128584180 | 128600654 | 16475 | 2 | ENSMUST00000126472.1 | 2902 | 5930412G12Rik | ENSMUSG00000072591.5 |
| chr5 | 128598418 | 128598561 | 3.51417 | 4.15779 | 1.21378 | Intron (ENSMUST00000154305.2/ENSMUSG00000072591.5, intron 1 of 5) | 5 | 128584180 | 128600654 | 16475 | 2 | ENSMUST00000126472.1 | 2093 | 5930412G12Rik | ENSMUSG00000072591.5 |
| chr5 | 128599280 | 128600248 | 4.31307 | 5.72675 | 2.26947 | Promoter | 5 | 128584180 | 128600654 | 16475 | 2 | ENSMUST00000126472.1 | 406 | 5930412G12Rik | ENSMUSG00000072591.5 |
| chr5 | 128600561 | 128601478 | 3.75794 | 4.65935 | 1.53078 | Promoter | 5 | 128579106 | 128600687 | 21582 | 2 | ENSMUST00000154305.2 | 0 | 5930412G12Rik | ENSMUSG00000072591.5 |
| chr6 | 147082636 | 147083151 | 3.42659 | 3.98223 | 1.06739 | Intron (ENSMUST00000100780.2/ENSMUSG00000072662.2, intron 1 of 2) | 6 | 147081433 | 147087001 | 5569 | 2 | ENSMUST00000123367.1 | 3850 | Mansc4 | ENSMUSG00000072662.2 |
| chr6 | 147083594 | 147083686 | 3.29692 | 3.78579 | 1.06336 | Intron (ENSMUST00000100780.2/ENSMUSG00000072662.2, intron 1 of 2) | 6 | 147081433 | 147087001 | 5569 | 2 | ENSMUST00000123367.1 | 3315 | Mansc4 | ENSMUSG00000072662.2 |
| chr6 | 147085676 | 147085953 | 3.728 | 4.69102 | 1.59991 | Intron (ENSMUST00000100780.2/ENSMUSG00000072662.2, intron 1 of 2) | 6 | 147081433 | 147087001 | 5569 | 2 | ENSMUST00000123367.1 | 1048 | Mansc4 | ENSMUSG00000072662.2 |
| chr6 | 147086626 | 147087396 | 4.06057 | 5.34841 | 2.05548 | Promoter | 6 | 147075060 | 147087032 | 11973 | 2 | ENSMUST00000100780.2 | 0 | Mansc4 | ENSMUSG00000072662.2 |
| chr6 | 147087865 | 147087940 | 3.71022 | 4.69735 | 1.58893 | Promoter | 6 | 147075060 | 147087032 | 11973 | 2 | ENSMUST00000100780.2 | -834 | Mansc4 | ENSMUSG00000072662.2 |
| chr6 | 147088298 | 147088440 | 3.31065 | 3.9875 | 1.16769 | Distal Intergenic | 6 | 147075060 | 147087032 | 11973 | 2 | ENSMUST00000100780.2 | -1267 | Mansc4 | ENSMUSG00000072662.2 |
| chr6 | 143224237 | 143224312 | 3.70356 | 4.71287 | 1.61307 | Distal Intergenic | 6 | 143245888 | 143249112 | 3225 | 1 | ENSMUST00000159793.2 | -21576 | D6Ertd474e | ENSMUSG00000072679.5 |
| chr6 | 143225571 | 143226074 | 3.49427 | 4.33145 | 1.33357 | Distal Intergenic | 6 | 143245888 | 143249112 | 3225 | 1 | ENSMUST00000159793.2 | -19814 | D6Ertd474e | ENSMUSG00000072679.5 |
| chr6 | 143259444 | 143259533 | 3.45575 | 4.25463 | 1.28556 | Intron (ENSMUST00000160188.2/ENSMUSG00000072679.5, intron 4 of 4) | 6 | 143245893 | 143297272 | 51380 | 1 | ENSMUST00000100817.2 | 13552 | D6Ertd474e | ENSMUSG00000072679.5 |
| chr6 | 142813841 | 142814042 | 3.33593 | 3.83123 | 1.01733 | Promoter | 6 | 142814231 | 142833494 | 19264 | 1 | ENSMUST00000100821.2 | -189 | Gm7457 | ENSMUSG00000072683.3 |
| chr14 | 25042683 | 25042780 | 3.33502 | 3.85433 | 1.06302 | Distal Intergenic | 14 | 24909414 | 24913440 | 4027 | 1 | ENSMUST00000100826.2 | 133270 | Gm10398 | ENSMUSG00000072686.2 |
| chr14 | 25108336 | 25109156 | 3.4779 | 4.14673 | 1.2861 | Distal Intergenic | 14 | 24909414 | 24913440 | 4027 | 1 | ENSMUST00000100826.2 | 198923 | Gm10398 | ENSMUSG00000072686.2 |
| chr5 | 112748686 | 112749014 | 3.40113 | 3.95105 | 1.09278 | Intron (ENSMUST00000086617.5/ENSMUSG00000072720.4, intron 39 of 43) | 5 | 112688878 | 112765931 | 77054 | 2 | ENSMUST00000182189.1 | 16917 | Myo18b | ENSMUSG00000072720.4 |
| chr5 | 112769351 | 112769441 | 3.52431 | 4.21422 | 1.30267 | Intron (ENSMUST00000086617.5/ENSMUSG00000072720.4, intron 34 of 43) | 5 | 112688878 | 112765931 | 77054 | 2 | ENSMUST00000182189.1 | -3421 | Myo18b | ENSMUSG00000072720.4 |
| chr5 | 112847923 | 112848229 | 3.35241 | 3.85607 | 1.01733 | Intron (ENSMUST00000086617.5/ENSMUSG00000072720.4, intron 12 of 43) | 5 | 112772886 | 112871586 | 98701 | 2 | ENSMUST00000183029.2 | 23357 | Myo18b | ENSMUSG00000072720.4 |
| chr5 | 112848665 | 112848968 | 4.15898 | 5.3772 | 1.99388 | Intron (ENSMUST00000086617.5/ENSMUSG00000072720.4, intron 12 of 43) | 5 | 112772886 | 112871586 | 98701 | 2 | ENSMUST00000183029.2 | 22618 | Myo18b | ENSMUSG00000072720.4 |
| chr5 | 112853289 | 112853798 | 3.6161 | 4.36707 | 1.36052 | Intron (ENSMUST00000086617.5/ENSMUSG00000072720.4, intron 12 of 43) | 5 | 112772886 | 112871586 | 98701 | 2 | ENSMUST00000183029.2 | 17788 | Myo18b | ENSMUSG00000072720.4 |
| chr5 | 112429023 | 112429317 | 3.52943 | 4.19503 | 1.26718 | Intron (ENSMUST00000075387.6/ENSMUSG00000058153.9, intron 12 of 16) | 5 | 112449426 | 112451738 | 2313 | 1 | ENSMUST00000100882.2 | -20109 | Gm6588 | ENSMUSG00000072722.2 |
| chr5 | 112430154 | 112430400 | 3.28538 | 3.75531 | 1.01733 | Intron (ENSMUST00000075387.6/ENSMUSG00000058153.9, intron 12 of 16) | 5 | 112449426 | 112451738 | 2313 | 1 | ENSMUST00000100882.2 | -19026 | Gm6588 | ENSMUSG00000072722.2 |
| chr5 | 112433015 | 112433090 | 3.59842 | 4.30485 | 1.29132 | Intron (ENSMUST00000075387.6/ENSMUSG00000058153.9, intron 12 of 16) | 5 | 112449426 | 112451738 | 2313 | 1 | ENSMUST00000100882.2 | -16336 | Gm6588 | ENSMUSG00000072722.2 |
| chr5 | 112475406 | 112475511 | 3.94226 | 5.06398 | 1.78958 | Exon (ENSMUST00000075387.6/ENSMUSG00000058153.9, exon 2 of 17) | 5 | 112449426 | 112451738 | 2313 | 1 | ENSMUST00000100882.2 | 25981 | Gm6588 | ENSMUSG00000072722.2 |
| chr14 | 7789353 | 7789456 | 3.46426 | 4.07604 | 1.19116 | Intron (ENSMUST00000081331.5/ENSMUSG00000072723.5, intron 2 of 8) | 14 | 7771186 | 7791553 | 20368 | 2 | ENSMUST00000170177.1 | 2097 | Gm10044 | ENSMUSG00000072723.5 |
| chr14 | 7791388 | 7791610 | 3.7667 | 4.80637 | 1.64272 | Promoter | 14 | 7756361 | 7791575 | 35215 | 2 | ENSMUST00000166813.2 | 0 | Gm10044 | ENSMUSG00000072723.5 |
| chr5 | 108364663 | 108364930 | 3.50332 | 4.41691 | 1.39867 | Distal Intergenic | 5 | 108369858 | 108378239 | 8382 | 1 | ENSMUST00000100944.2 | -4928 | Gm10419 | ENSMUSG00000072769.2 |
| chr5 | 108365335 | 108365512 | 3.83259 | 5.07234 | 1.84772 | Distal Intergenic | 5 | 108369858 | 108378239 | 8382 | 1 | ENSMUST00000100944.2 | -4346 | Gm10419 | ENSMUSG00000072769.2 |
| chr5 | 108365829 | 108368360 | 3.91754 | 4.98243 | 1.79704 | Distal Intergenic | 5 | 108369858 | 108378239 | 8382 | 1 | ENSMUST00000100944.2 | -1498 | Gm10419 | ENSMUSG00000072769.2 |
| chr5 | 108368997 | 108369484 | 3.72511 | 4.84109 | 1.6867 | Promoter | 5 | 108369858 | 108378239 | 8382 | 1 | ENSMUST00000100944.2 | -374 | Gm10419 | ENSMUSG00000072769.2 |
| chr5 | 108369979 | 108370303 | 3.49228 | 4.27464 | 1.31407 | Promoter | 5 | 108369858 | 108378239 | 8382 | 1 | ENSMUST00000100944.2 | 122 | Gm10419 | ENSMUSG00000072769.2 |
| chr5 | 108371041 | 108371137 | 3.88389 | 4.84867 | 1.63623 | Intron (ENSMUST00000100944.2/ENSMUSG00000072769.2, intron 2 of 4) | 5 | 108369858 | 108378239 | 8382 | 1 | ENSMUST00000100944.2 | 1184 | Gm10419 | ENSMUSG00000072769.2 |
| chr5 | 108372549 | 108372823 | 3.54297 | 4.23563 | 1.31395 | 3' UTR | 5 | 108369858 | 108378239 | 8382 | 1 | ENSMUST00000100944.2 | 2692 | Gm10419 | ENSMUSG00000072769.2 |
| chr12 | 112800942 | 112801050 | 3.30089 | 3.84094 | 1.02996 | Intron (ENSMUST00000124526.1/ENSMUSG00000072812.4, intron 1 of 6) | 12 | 112781010 | 112802657 | 21648 | 2 | ENSMUST00000124526.1 | 1607 | Ahnak2 | ENSMUSG00000072812.4 |
| chr12 | 112802124 | 112802739 | 3.61691 | 4.33732 | 1.30416 | Promoter | 12 | 112781010 | 112802657 | 21648 | 2 | ENSMUST00000124526.1 | 0 | Ahnak2 | ENSMUSG00000072812.4 |
| chr6 | 100204354 | 100204676 | 3.30031 | 3.77765 | 1.01733 | Distal Intergenic | 6 | 100228566 | 100287441 | 58876 | 2 | ENSMUST00000101118.2 | 82765 | Rybp | ENSMUSG00000072872.2 |
| chr6 | 100206735 | 100207245 | 3.34603 | 3.89043 | 1.08629 | Distal Intergenic | 6 | 100228566 | 100287441 | 58876 | 2 | ENSMUST00000101118.2 | 80196 | Rybp | ENSMUSG00000072872.2 |
| chr6 | 100301639 | 100301897 | 3.30059 | 3.77806 | 1.01733 | Distal Intergenic | 6 | 100228566 | 100287441 | 58876 | 2 | ENSMUST00000101118.2 | -14199 | Rybp | ENSMUSG00000072872.2 |
| chr6 | 100336653 | 100336979 | 3.35263 | 3.85991 | 1.02975 | Distal Intergenic | 6 | 100228566 | 100287441 | 58876 | 2 | ENSMUST00000101118.2 | -49213 | Rybp | ENSMUSG00000072872.2 |
| chr6 | 100385497 | 100386051 | 3.91343 | 4.90384 | 1.68491 | Distal Intergenic | 6 | 100228566 | 100287441 | 58876 | 2 | ENSMUST00000101118.2 | -98057 | Rybp | ENSMUSG00000072872.2 |
| chr5 | 74931873 | 74932151 | 3.35065 | 3.85342 | 1.01733 | Distal Intergenic | 5 | 74949194 | 74966368 | 17175 | 1 | ENSMUST00000101121.1 | -17043 | Gm6116 | ENSMUSG00000072874.3 |
| chr5 | 74941267 | 74941588 | 4.08025 | 5.30151 | 1.98762 | Distal Intergenic | 5 | 74949194 | 74966368 | 17175 | 1 | ENSMUST00000101121.1 | -7606 | Gm6116 | ENSMUSG00000072874.3 |
| chr5 | 74949453 | 74949544 | 3.51298 | 4.15804 | 1.22442 | Promoter | 5 | 74949194 | 74966368 | 17175 | 1 | ENSMUST00000101121.1 | 260 | Gm6116 | ENSMUSG00000072874.3 |
| chr5 | 74955510 | 74955853 | 3.62605 | 4.40609 | 1.37838 | Intron (ENSMUST00000101121.1/ENSMUSG00000072874.3, intron 1 of 4) | 5 | 74949194 | 74966368 | 17175 | 1 | ENSMUST00000101121.1 | 6317 | Gm6116 | ENSMUSG00000072874.3 |
| chr5 | 74956398 | 74957500 | 3.88326 | 4.86876 | 1.67656 | Exon (ENSMUST00000101121.1/ENSMUSG00000072874.3, exon 2 of 5) | 5 | 74949194 | 74966368 | 17175 | 1 | ENSMUST00000101121.1 | 7205 | Gm6116 | ENSMUSG00000072874.3 |
| chr5 | 74960331 | 74960409 | 3.71989 | 4.52658 | 1.41169 | Intron (ENSMUST00000101121.1/ENSMUSG00000072874.3, intron 2 of 4) | 5 | 74949194 | 74966368 | 17175 | 1 | ENSMUST00000101121.1 | 11138 | Gm6116 | ENSMUSG00000072874.3 |
| chr5 | 74963676 | 74963751 | 3.57675 | 4.25693 | 1.23662 | Intron (ENSMUST00000101121.1/ENSMUSG00000072874.3, intron 4 of 4) | 5 | 74949194 | 74966368 | 17175 | 1 | ENSMUST00000101121.1 | 14483 | Gm6116 | ENSMUSG00000072874.3 |
| chr5 | 74979847 | 74980260 | 3.51695 | 4.15457 | 1.19531 | Distal Intergenic | 5 | 74949194 | 74966368 | 17175 | 1 | ENSMUST00000101121.1 | 30654 | Gm6116 | ENSMUSG00000072874.3 |
| chr6 | 99689200 | 99689494 | 3.51801 | 4.15635 | 1.19531 | Distal Intergenic | 6 | 99692679 | 99693818 | 1140 | 1 | ENSMUST00000101122.1 | -3185 | Gpr27 | ENSMUSG00000072875.1 |
| chr6 | 99691950 | 99694738 | 3.82276 | 4.84196 | 1.69215 | Promoter | 6 | 99692679 | 99693818 | 1140 | 1 | ENSMUST00000101122.1 | 0 | Gpr27 | ENSMUSG00000072875.1 |
| chr6 | 85125846 | 85125968 | 3.51378 | 4.14973 | 1.18856 | Promoter | 6 | 85111416 | 85126125 | 14710 | 2 | ENSMUST00000113802.3 | 157 | Gm5878 | ENSMUSG00000072952.5 |
| chr6 | 85126941 | 85127244 | 3.45613 | 4.03736 | 1.10461 | Promoter | 6 | 85111416 | 85126125 | 14710 | 2 | ENSMUST00000113802.3 | -817 | Gm5878 | ENSMUSG00000072952.5 |
| chr5 | 44100038 | 44101538 | 3.77939 | 4.73834 | 1.60271 | Promoter | 5 | 44100442 | 44102513 | 2072 | 1 | ENSMUST00000072800.4 | 0 | Gm16401 | ENSMUSG00000072962.2 |
| chr11 | 51085242 | 51085527 | 4.64634 | 6.42149 | 2.82053 | Promoter | 11 | 51085087 | 51091833 | 6747 | 1 | ENSMUST00000101255.1 | 156 | 9230009I02Rik | ENSMUSG00000072982.1 |
| chr11 | 51094519 | 51094675 | 3.36897 | 3.88105 | 1.01733 | Distal Intergenic | 11 | 51085087 | 51091833 | 6747 | 1 | ENSMUST00000101255.1 | 9433 | 9230009I02Rik | ENSMUSG00000072982.1 |
| chr12 | 76435131 | 76435392 | 3.66965 | 4.62781 | 1.52827 | Intron (ENSMUST00000063977.7/ENSMUSG00000052221.7, intron 6 of 10) | 12 | 76444560 | 76451336 | 6777 | 1 | ENSMUST00000101281.2 | -9168 | Gm10451 | ENSMUSG00000073000.2 |
| chr6 | 64688080 | 64688744 | 3.46656 | 4.08642 | 1.22373 | Distal Intergenic | 6 | 64729146 | 64731233 | 2088 | 1 | ENSMUST00000101351.4 | -40402 | Atoh1 | ENSMUSG00000073043.4 |
| chr6 | 64728099 | 64728312 | 3.46264 | 4.20797 | 1.26841 | Promoter | 6 | 64729146 | 64731233 | 2088 | 1 | ENSMUST00000101351.4 | -834 | Atoh1 | ENSMUSG00000073043.4 |
| chr6 | 64728656 | 64729046 | 3.78307 | 4.76369 | 1.63419 | Promoter | 6 | 64729146 | 64731233 | 2088 | 1 | ENSMUST00000101351.4 | -100 | Atoh1 | ENSMUSG00000073043.4 |
| chr6 | 64729396 | 64734018 | 3.80995 | 4.80879 | 1.69541 | Promoter | 6 | 64729146 | 64731233 | 2088 | 1 | ENSMUST00000101351.4 | 251 | Atoh1 | ENSMUSG00000073043.4 |
| chr6 | 64734479 | 64735224 | 4.05322 | 5.27831 | 1.96961 | Distal Intergenic | 6 | 64729146 | 64731233 | 2088 | 1 | ENSMUST00000101351.4 | 5334 | Atoh1 | ENSMUSG00000073043.4 |
| chr6 | 50809574 | 50810123 | 3.78854 | 4.69273 | 1.60138 | Distal Intergenic | 6 | 50846713 | 50847770 | 1058 | 2 | ENSMUST00000101402.4 | 37647 | G930045G22Rik | ENSMUSG00000073072.5 |
| chr6 | 50810437 | 50810848 | 3.82136 | 4.79407 | 1.65922 | Distal Intergenic | 6 | 50846713 | 50847770 | 1058 | 2 | ENSMUST00000101402.4 | 36922 | G930045G22Rik | ENSMUSG00000073072.5 |
| chr6 | 50821870 | 50821945 | 3.24273 | 3.69125 | 1.01733 | Distal Intergenic | 6 | 50846713 | 50847770 | 1058 | 2 | ENSMUST00000101402.4 | 25825 | G930045G22Rik | ENSMUSG00000073072.5 |
| chr6 | 50822359 | 50822617 | 3.21414 | 3.64845 | 1.00889 | Distal Intergenic | 6 | 50846713 | 50847770 | 1058 | 2 | ENSMUST00000101402.4 | 25153 | G930045G22Rik | ENSMUSG00000073072.5 |
| chr6 | 50924768 | 50924910 | 3.67963 | 4.59555 | 1.52099 | Distal Intergenic | 6 | 50846524 | 50878079 | 31556 | 2 | ENSMUST00000161723.1 | -46690 | G930045G22Rik | ENSMUSG00000073072.5 |
| chr6 | 50929038 | 50929344 | 3.7854 | 4.72697 | 1.63692 | Distal Intergenic | 6 | 50846524 | 50878079 | 31556 | 2 | ENSMUST00000161723.1 | -50960 | G930045G22Rik | ENSMUSG00000073072.5 |
| chr6 | 48852325 | 48853060 | 3.55165 | 4.23067 | 1.26559 | Distal Intergenic | 6 | 48860356 | 48864297 | 3942 | 1 | ENSMUST00000101425.3 | -7296 | Gm7932 | ENSMUSG00000073090.3 |
| chr5 | 30342942 | 30343088 | 3.29673 | 3.77226 | 1.01733 | Intron (ENSMUST00000101448.3/ENSMUSG00000073102.3, intron 3 of 16) | 5 | 30281388 | 30366708 | 85321 | 1 | ENSMUST00000101448.3 | 61555 | Drc1 | ENSMUSG00000073102.3 |
| chr6 | 36384036 | 36384416 | 3.55775 | 4.35719 | 1.39431 | Intron (ENSMUST00000101535.3/ENSMUSG00000073154.5, intron 1 of 4) | 6 | 36209620 | 36387851 | 178232 | 2 | ENSMUST00000101535.3 | 3435 | 9330158H04Rik | ENSMUSG00000073154.5 |
| chr6 | 36385428 | 36385556 | 3.30681 | 3.78742 | 1.01733 | Intron (ENSMUST00000101535.3/ENSMUSG00000073154.5, intron 1 of 4) | 6 | 36209620 | 36387851 | 178232 | 2 | ENSMUST00000101535.3 | 2295 | 9330158H04Rik | ENSMUSG00000073154.5 |
| chr6 | 36386106 | 36386308 | 3.30741 | 3.78832 | 1.01733 | Intron (ENSMUST00000101535.3/ENSMUSG00000073154.5, intron 1 of 4) | 6 | 36209620 | 36387851 | 178232 | 2 | ENSMUST00000101535.3 | 1543 | 9330158H04Rik | ENSMUSG00000073154.5 |
| chr6 | 36386784 | 36387670 | 3.59716 | 4.32412 | 1.34621 | Promoter | 6 | 36209620 | 36387851 | 178232 | 2 | ENSMUST00000101535.3 | 181 | 9330158H04Rik | ENSMUSG00000073154.5 |
| chr6 | 36388533 | 36389431 | 3.72931 | 4.55064 | 1.43606 | Promoter | 6 | 36387388 | 36388917 | 1530 | 2 | ENSMUST00000159032.1 | 0 | 9330158H04Rik | ENSMUSG00000073154.5 |
| chr6 | 36390333 | 36390516 | 3.58539 | 4.27819 | 1.2665 | Promoter | 6 | 36387381 | 36390191 | 2811 | 2 | ENSMUST00000160156.1 | -143 | 9330158H04Rik | ENSMUSG00000073154.5 |
| chr12 | 18514338 | 18514653 | 3.37851 | 3.90622 | 1.05547 | Promoter | 12 | 18514738 | 18534191 | 19454 | 1 | ENSMUST00000063216.4 | -85 | 5730507C01Rik | ENSMUSG00000073197.3 |
| chr6 | 30945910 | 30946200 | 3.82489 | 4.81563 | 1.6725 | Distal Intergenic | 6 | 30956023 | 30959078 | 3056 | 2 | ENSMUST00000101589.4 | 12878 | Klf14 | ENSMUSG00000073209.4 |
| chr6 | 30955376 | 30955949 | 3.24793 | 3.7523 | 1.04893 | Downstream (<1kb) | 6 | 30956023 | 30959078 | 3056 | 2 | ENSMUST00000101589.4 | 3129 | Klf14 | ENSMUSG00000073209.4 |
| chr6 | 30956440 | 30956679 | 3.59515 | 4.36938 | 1.39313 | 3' UTR | 6 | 30956023 | 30959078 | 3056 | 2 | ENSMUST00000101589.4 | 2399 | Klf14 | ENSMUSG00000073209.4 |
| chr6 | 30957191 | 30959106 | 4.18174 | 5.81268 | 2.38633 | Promoter | 6 | 30956023 | 30959078 | 3056 | 2 | ENSMUST00000101589.4 | 0 | Klf14 | ENSMUSG00000073209.4 |
| chr6 | 30959891 | 30960374 | 3.91199 | 5.00826 | 1.85025 | Promoter | 6 | 30956023 | 30959078 | 3056 | 2 | ENSMUST00000101589.4 | -814 | Klf14 | ENSMUSG00000073209.4 |
| chr6 | 30960702 | 30962006 | 3.7339 | 4.82518 | 1.66284 | Distal Intergenic | 6 | 30956023 | 30959078 | 3056 | 2 | ENSMUST00000101589.4 | -1625 | Klf14 | ENSMUSG00000073209.4 |
| chr6 | 30962383 | 30962614 | 3.51718 | 4.3459 | 1.36734 | Distal Intergenic | 6 | 30956023 | 30959078 | 3056 | 2 | ENSMUST00000101589.4 | -3306 | Klf14 | ENSMUSG00000073209.4 |
| chr6 | 30963571 | 30963910 | 3.62502 | 4.43693 | 1.43543 | Distal Intergenic | 6 | 30956023 | 30959078 | 3056 | 2 | ENSMUST00000101589.4 | -4494 | Klf14 | ENSMUSG00000073209.4 |
| chr6 | 30965317 | 30965778 | 3.44149 | 4.19963 | 1.25086 | Distal Intergenic | 6 | 30956023 | 30959078 | 3056 | 2 | ENSMUST00000101589.4 | -6240 | Klf14 | ENSMUSG00000073209.4 |
| chr6 | 30966800 | 30967314 | 3.52389 | 4.23563 | 1.34564 | Distal Intergenic | 6 | 30956023 | 30959078 | 3056 | 2 | ENSMUST00000101589.4 | -7723 | Klf14 | ENSMUSG00000073209.4 |
| chr6 | 30968268 | 30968518 | 3.35274 | 3.88979 | 1.11513 | Distal Intergenic | 6 | 30956023 | 30959078 | 3056 | 2 | ENSMUST00000101589.4 | -9191 | Klf14 | ENSMUSG00000073209.4 |
| chr6 | 30969153 | 30969353 | 3.70757 | 4.579 | 1.5691 | Distal Intergenic | 6 | 30956023 | 30959078 | 3056 | 2 | ENSMUST00000101589.4 | -10076 | Klf14 | ENSMUSG00000073209.4 |
| chr6 | 30970029 | 30970351 | 4.09616 | 5.45877 | 2.07353 | Distal Intergenic | 6 | 30956023 | 30959078 | 3056 | 2 | ENSMUST00000101589.4 | -10952 | Klf14 | ENSMUSG00000073209.4 |
| chr5 | 5575340 | 5575525 | 3.77154 | 4.69945 | 1.62037 | Exon (ENSMUST00000101627.2/ENSMUSG00000073234.2, exon 2 of 2) | 5 | 5573952 | 5576203 | 2252 | 1 | ENSMUST00000101627.2 | 1389 | Gm8773 | ENSMUSG00000073234.2 |
| chr17 | 90454383 | 90455221 | 3.95604 | 5.11471 | 1.87429 | Promoter | 17 | 90454889 | 90455501 | 613 | 1 | ENSMUST00000097270.4 | 0 | Gm10493 | ENSMUSG00000073367.4 |
| chr17 | 67629197 | 67629439 | 3.36161 | 3.86993 | 1.01733 | Downstream (1-2kb) | 17 | 67630965 | 67632723 | 1759 | 2 | ENSMUST00000097290.2 | 3284 | Lrrc30 | ENSMUSG00000073375.2 |
| chr17 | 36166340 | 36167523 | 3.82875 | 4.83404 | 1.72325 | Promoter | 17 | 36165368 | 36167844 | 2477 | 2 | ENSMUST00000174693.2 | 321 | Gm8909 | ENSMUSG00000073402.6 |
| chr17 | 36168000 | 36169811 | 3.91324 | 5.17653 | 1.92891 | Promoter | 17 | 36164443 | 36168523 | 4081 | 2 | ENSMUST00000173353.2 | 0 | Gm8909 | ENSMUSG00000073402.6 |
| chr17 | 36170126 | 36170201 | 3.85368 | 4.85805 | 1.70525 | Intron (ENSMUST00000173648.1/ENSMUSG00000092474.1, intron 2 of 2) | 17 | 36164444 | 36168537 | 4094 | 2 | ENSMUST00000040467.9 | -1590 | Gm8909 | ENSMUSG00000073402.6 |
| chr17 | 36145659 | 36145868 | 3.10249 | 4.21888 | 1.29649 | Promoter | 17 | 36141758 | 36145744 | 3987 | 2 | ENSMUST00000174778.2 | 0 | Gm10499 | ENSMUSG00000073403.6 |
| chr17 | 36146176 | 36146784 | 3.50219 | 4.17645 | 1.23964 | Promoter | 17 | 36141758 | 36145923 | 4166 | 2 | ENSMUST00000174094.1 | -254 | Gm10499 | ENSMUSG00000073403.6 |
| chr17 | 36149820 | 36150439 | 3.73317 | 4.63141 | 1.5691 | Intron (ENSMUST00000056774.10/ENSMUSG00000038311.8, intron 1 of 4) | 17 | 36141758 | 36145923 | 4166 | 2 | ENSMUST00000174094.1 | -3898 | Gm10499 | ENSMUSG00000073403.6 |
| chr17 | 36113507 | 36113730 | 3.80596 | 4.80707 | 1.66033 | Intron (ENSMUST00000174728.2/ENSMUSG00000092365.2, intron 1 of 1) | 17 | 36109234 | 36111658 | 2425 | 2 | ENSMUST00000172538.1 | -1850 | C920025E04Rik | ENSMUSG00000073405.6 |
| chr17 | 36081969 | 36082242 | 3.29116 | 3.77045 | 1.04005 | Promoter | 17 | 36080115 | 36081987 | 1873 | 2 | ENSMUST00000184850.2 | 0 | H2-Bl | ENSMUSG00000073406.5 |
| chr17 | 36083804 | 36084433 | 3.65632 | 4.52748 | 1.48148 | Promoter | 17 | 36080189 | 36084223 | 4035 | 2 | ENSMUST00000183560.2 | 0 | H2-Bl | ENSMUSG00000073406.5 |
| chr17 | 35643987 | 35644166 | 3.28041 | 3.74775 | 1.01733 | Promoter | 17 | 35635752 | 35643695 | 7944 | 2 | ENSMUST00000095467.3 | -293 | Dpcr1 | ENSMUSG00000073408.2 |
| chr17 | 34256335 | 34256410 | 3.93648 | 5.22542 | 1.91547 | Distal Intergenic | 17 | 34263209 | 34269418 | 6210 | 1 | ENSMUST00000040828.5 | -6799 | H2-Ab1 | ENSMUSG00000073421.5 |
| chr17 | 34264177 | 34264252 | 3.74199 | 4.57043 | 1.45643 | Promoter | 17 | 34263209 | 34269418 | 6210 | 1 | ENSMUST00000040828.5 | 969 | H2-Ab1 | ENSMUSG00000073421.5 |
| chr17 | 34264583 | 34265588 | 4.16937 | 5.49361 | 2.2009 | Exon (ENSMUST00000040828.5/ENSMUSG00000073421.5, exon 2 of 6) | 17 | 34263209 | 34269418 | 6210 | 1 | ENSMUST00000040828.5 | 1375 | H2-Ab1 | ENSMUSG00000073421.5 |
| chr17 | 32674217 | 32674292 | 4.56659 | 6.20913 | 2.63488 | Exon (ENSMUST00000165061.1/ENSMUSG00000090700.1, exon 8 of 12) | 17 | 32685627 | 32703352 | 17726 | 1 | ENSMUST00000168171.2 | -11335 | Cyp4f15 | ENSMUSG00000073424.4 |
| chr17 | 32675845 | 32676069 | 3.50641 | 4.14961 | 1.23424 | Exon (ENSMUST00000165061.1/ENSMUSG00000090700.1, exon 11 of 12) | 17 | 32685627 | 32703352 | 17726 | 1 | ENSMUST00000168171.2 | -9558 | Cyp4f15 | ENSMUSG00000073424.4 |
| chr17 | 26200347 | 26201352 | 3.96336 | 5.05203 | 1.80966 | Promoter | 17 | 26196226 | 26201143 | 4918 | 2 | ENSMUST00000176961.2 | 0 | Arhgdig | ENSMUSG00000073433.5 |
| chr18 | 89768776 | 89770012 | 4.14152 | 5.36801 | 2.01837 | Promoter | 18 | 89301082 | 89769528 | 468447 | 2 | ENSMUST00000097495.3 | 0 | Dok6 | ENSMUSG00000073514.3 |
| chr1 | 159523922 | 159524132 | 3.39684 | 4.00553 | 1.15619 | Distal Intergenic | 1 | 159614749 | 159684875 | 70127 | 1 | ENSMUST00000097514.2 | -90617 | Gm10530 | ENSMUSG00000073528.2 |
| chr1 | 158959056 | 158959215 | 3.32248 | 3.81102 | 1.01733 | Promoter | 1 | 158711727 | 158958367 | 246641 | 2 | ENSMUST00000159861.3 | -690 | Pappa2 | ENSMUSG00000073530.6 |
| chr18 | 51117600 | 51117931 | 3.82599 | 4.89208 | 1.71771 | Promoter | 18 | 51117898 | 51304641 | 186744 | 1 | ENSMUST00000116639.2 | 0 | Prr16 | ENSMUSG00000073565.4 |
| chr18 | 51119169 | 51119255 | 3.25198 | 3.71204 | 1.03901 | Intron (ENSMUST00000116639.2/ENSMUSG00000073565.4, intron 1 of 1) | 18 | 51117898 | 51304641 | 186744 | 1 | ENSMUST00000116639.2 | 1272 | Prr16 | ENSMUSG00000073565.4 |
| chr18 | 35653014 | 35653326 | 3.99117 | 5.25519 | 1.96346 | Exon (ENSMUST00000190196.1/ENSMUSG00000073600.2, exon 1 of 1) | 18 | 35650355 | 35652006 | 1652 | 2 | ENSMUST00000186951.1 | -1009 | Prob1 | ENSMUSG00000073600.2 |

| | | | | | | | | | | | | | | | |
|---|---|---|---|---|---|---|---|---|---|---|---|---|---|---|---|
| chr18 | 35653690 | 35655230 | 4.12985 | 5.56175 | 2.20311 | Promoter | 18 | 35652167 | 35655187 | 3021 | 2 | ENSMUST00000097619.1 | 0 | Prob1 | ENSMUSG00000073600.2 |
| chr1 | 71652930 | 71653069 | 3.43846 | 4.02188 | 1.14446 | Promoter | 1 | 71652888 | 71662053 | 9166 | 1 | ENSMUST00000127218.2 | 43 | Apol7d | ENSMUSG00000073652.5 |
| chr1 | 71653446 | 71654378 | 3.69562 | 4.53178 | 1.45422 | Promoter | 1 | 71653339 | 71657245 | 3907 | 1 | ENSMUST00000125934.1 | 108 | Apol7d | ENSMUSG00000073652.5 |
| chr4 | 156130030 | 156130140 | 3.92101 | 4.92266 | 1.68861 | Promoter | 4 | 156130513 | 156133415 | 2903 | 1 | ENSMUST00000097730.4 | -373 | Rnf223 | ENSMUSG00000073672.4 |
| chr4 | 155033662 | 155033947 | 3.61366 | 4.34193 | 1.33006 | Intron (ENSMUST00000135665.3/ENSMUSG00000029055.12, intron 1 of 21) | 4 | 155042046 | 155045422 | 3377 | 1 | ENSMUST00000097750.2 | -8099 | Gm10564 | ENSMUSG00000073686.2 |
| chr4 | 155039666 | 155039853 | 3.84154 | 4.75396 | 1.55728 | Intron (ENSMUST00000135665.3/ENSMUSG00000029055.12, intron 1 of 21) | 4 | 155042046 | 155045422 | 3377 | 1 | ENSMUST00000097750.2 | -2193 | Gm10564 | ENSMUSG00000073686.2 |
| chr4 | 152698071 | 152698165 | 4.25722 | 5.58747 | 2.15989 | Promoter | 4 | 152697540 | 152700515 | 2976 | 1 | ENSMUST00000143133.1 | 532 | Gm833 | ENSMUSG00000073699.3 |
| chr4 | 152698991 | 152699203 | 3.36248 | 3.87338 | 1.02485 | Exon (ENSMUST00000143133.1/ENSMUSG00000073699.3, exon 2 of 3) | 4 | 152697540 | 152700515 | 2976 | 1 | ENSMUST00000143133.1 | 1452 | Gm833 | ENSMUSG00000073699.3 |
| chr4 | 142070944 | 142071200 | 3.58713 | 4.30357 | 1.34573 | Intron (ENSMUST00000036572.3/ENSMUSG00000040616.3, intron 1 of 2) | 4 | 142083972 | 142088059 | 4088 | 1 | ENSMUST00000153023.1 | -12772 | Tmem51os1 | ENSMUSG00000073728.5 |
| chr4 | 142080287 | 142080521 | 3.32318 | 3.88531 | 1.11099 | Intron (ENSMUST00000036572.3/ENSMUSG00000040616.3, intron 1 of 2) | 4 | 142083972 | 142088059 | 4088 | 1 | ENSMUST00000153023.1 | -3451 | Tmem51os1 | ENSMUSG00000073728.5 |
| chr4 | 142094962 | 142095166 | 3.59518 | 4.30152 | 1.29172 | Distal Intergenic | 4 | 142084856 | 142088077 | 3222 | 1 | ENSMUST00000139653.1 | 10107 | Tmem51os1 | ENSMUSG00000073728.5 |
| chr4 | 130308610 | 130309153 | 3.97393 | 5.20542 | 1.92914 | Promoter | 4 | 130308162 | 130308674 | 513 | 2 | ENSMUST00000097865.1 | 0 | Gm10570 | ENSMUSG00000073752.1 |
| chr4 | 126618803 | 126618980 | 3.53354 | 4.21099 | 1.27775 | Distal Intergenic | 4 | 126609854 | 126614371 | 4518 | 1 | ENSMUST00000164362.1 | 8950 | 5730409E04Rik | ENSMUSG00000073755.4 |
| chr4 | 126619969 | 126621013 | 3.64034 | 4.38642 | 1.33239 | Distal Intergenic | 4 | 126609854 | 126614371 | 4518 | 1 | ENSMUST00000164362.1 | 10116 | 5730409E04Rik | ENSMUSG00000073755.4 |
| chr4 | 126621725 | 126621959 | 3.51799 | 4.30544 | 1.33655 | Distal Intergenic | 4 | 126609854 | 126614371 | 4518 | 1 | ENSMUST00000164362.1 | 11872 | 5730409E04Rik | ENSMUSG00000073755.4 |
| chr4 | 126635520 | 126635712 | 3.40665 | 3.95767 | 1.08697 | Distal Intergenic | 4 | 126609854 | 126614371 | 4518 | 1 | ENSMUST00000164362.1 | 25667 | 5730409E04Rik | ENSMUSG00000073755.4 |
| chr4 | 126636368 | 126637026 | 3.56588 | 4.2976 | 1.29527 | Distal Intergenic | 4 | 126609854 | 126614371 | 4518 | 1 | ENSMUST00000164362.1 | 26515 | 5730409E04Rik | ENSMUSG00000073755.4 |
| chr4 | 107811739 | 107811870 | 3.5086 | 4.13429 | 1.16403 | Promoter | 4 | 107806033 | 107810948 | 4916 | 2 | ENSMUST00000097930.1 | -792 | Lrp8os2 | ENSMUSG00000073779.2 |
| chr4 | 98819433 | 98819785 | 3.39105 | 3.97743 | 1.15349 | Promoter | 4 | 98819815 | 98820324 | 510 | 2 | ENSMUST00000168045.1 | 539 | I0C0044D17Rik | ENSMUSG00000073794.3 |
| chr4 | 89303036 | 89303868 | 3.78124 | 5.09194 | 1.85558 | Downstream (2-3kb) | 4 | 89306289 | 89311032 | 4744 | 2 | ENSMUST00000097981.4 | 7164 | Cdkn2b | ENSMUSG00000073802.4 |
| chr4 | 89305745 | 89307585 | 3.9457 | 5.23725 | 1.95594 | 3' UTR | 4 | 89306289 | 89311032 | 4744 | 2 | ENSMUST00000097981.4 | 3447 | Cdkn2b | ENSMUSG00000073802.4 |
| chr4 | 89307995 | 89308865 | 4.28485 | 5.7731 | 2.45679 | Intron (ENSMUST00000097981.4/ENSMUSG00000073802.4, intron 1 of 1) | 4 | 89306289 | 89311032 | 4744 | 2 | ENSMUST00000097981.4 | 2167 | Cdkn2b | ENSMUSG00000073802.4 |
| chr4 | 89309679 | 89310757 | 3.97038 | 5.05735 | 1.8287 | Promoter | 4 | 89306289 | 89311032 | 4744 | 2 | ENSMUST00000097981.4 | 275 | Cdkn2b | ENSMUSG00000073802.4 |
| chr4 | 89311825 | 89312063 | 3.37647 | 3.89236 | 1.01733 | Promoter | 4 | 89306289 | 89311032 | 4744 | 2 | ENSMUST00000097981.4 | -794 | Cdkn2b | ENSMUSG00000073802.4 |
| chr4 | 89312948 | 89313873 | 3.59916 | 4.38874 | 1.37853 | Distal Intergenic | 4 | 89306289 | 89311032 | 4744 | 2 | ENSMUST00000097981.4 | -1917 | Cdkn2b | ENSMUSG00000073802.4 |
| chr7 | 135354912 | 135355114 | 3.30172 | 3.77976 | 1.01733 | Distal Intergenic | 7 | 135396342 | 135399544 | 3203 | 1 | ENSMUST00000097982.2 | -41228 | 4930544L04Rik | ENSMUSG00000073803.2 |
| chr7 | 135398393 | 135398654 | 3.31205 | 3.79529 | 1.01733 | 3' UTR | 7 | 135396342 | 135399544 | 3203 | 1 | ENSMUST00000097982.2 | 2052 | 4930544L04Rik | ENSMUSG00000073803.2 |
| chr7 | 134937632 | 134937783 | 3.83769 | 4.89714 | 1.76116 | Promoter | 7 | 134881926 | 134938430 | 56505 | 2 | ENSMUST00000171394.1 | 647 | Fam196a | ENSMUSG00000073805.4 |
| chr4 | 45671800 | 45672035 | 3.59242 | 4.28828 | 1.26175 | Distal Intergenic | 4 | 45683589 | 45692938 | 9350 | 1 | ENSMUST00000123741.1 | -11554 | Gm829 | ENSMUSG00000073860.7 |
| chr7 | 113703721 | 113703796 | 5.53771 | 9.98333 | 5.78672 | Distal Intergenic | 7 | 113707715 | 113708200 | 486 | 2 | ENSMUST00000098100.3 | 4404 | Gm5600 | ENSMUSG00000073862.3 |
| chr4 | 41782372 | 41782615 | 3.76508 | 4.72334 | 1.61702 | Distal Intergenic | 4 | 41773188 | 41774247 | 1060 | 2 | ENSMUST00000155240.1 | -8126 | Ccl27a | ENSMUSG00000073888.7 |
| chr4 | 41698487 | 41699084 | 3.84502 | 4.95104 | 1.75056 | Promoter | 4 | 41699989 | 41769474 | 69486 | 1 | ENSMUST00000108041.2 | -905 | Il11ra1 | ENSMUSG00000073889.5 |
| chr4 | 41700237 | 41700354 | 3.52359 | 4.66966 | 1.54549 | Promoter | 4 | 41699989 | 41769474 | 69486 | 1 | ENSMUST00000108041.2 | 249 | Il11ra1 | ENSMUSG00000073889.5 |
| chr7 | 107208005 | 107208129 | 3.64701 | 4.43359 | 1.44386 | Downstream (2-3kb) | 7 | 107209445 | 107210916 | 1472 | 1 | ENSMUST00000098135.1 | -1316 | Rbmxl2 | ENSMUSG00000073894.1 |
| chr7 | 107208472 | 107208547 | 4.54585 | 6.16864 | 2.59994 | Promoter | 7 | 107209445 | 107210916 | 1472 | 1 | ENSMUST00000098135.1 | -898 | Rbmxl2 | ENSMUSG00000073894.1 |
| chr7 | 107209237 | 107210687 | 4.18728 | 5.65192 | 2.3127 | Promoter | 7 | 107209445 | 107210916 | 1472 | 1 | ENSMUST00000098135.1 | 0 | Rbmxl2 | ENSMUSG00000073894.1 |
| chr7 | 107212925 | 107213127 | 3.51568 | 4.24875 | 1.31144 | Distal Intergenic | 7 | 107209445 | 107210916 | 1472 | 1 | ENSMUST00000098135.1 | 3481 | Rbmxl2 | ENSMUSG00000073894.1 |
| chr7 | 107213585 | 107213750 | 3.80733 | 4.8083 | 1.66455 | Distal Intergenic | 7 | 107209445 | 107210916 | 1472 | 1 | ENSMUST00000098135.1 | 4141 | Rbmxl2 | ENSMUSG00000073894.1 |
| chr4 | 35156304 | 35156815 | 3.53788 | 4.19557 | 1.22299 | Promoter | 4 | 34949074 | 35157484 | 208411 | 2 | ENSMUST00000102975.4 | 669 | Mob3b | ENSMUSG00000073910.5 |
| chr4 | 20008042 | 20008842 | 4.04553 | 5.20943 | 1.91796 | Promoter | 4 | 20008357 | 20030785 | 22429 | 1 | ENSMUST00000117632.2 | 0 | Ttpa | ENSMUSG00000073988.8 |
| chr9 | 121778496 | 121778662 | 3.49081 | 4.0991 | 1.13765 | Promoter | 9 | 121777607 | 121783819 | 6213 | 1 | ENSMUST00000098272.2 | 890 | Klhl40 | ENSMUSG00000074001.2 |
| chr9 | 121779963 | 121780286 | 3.37547 | 3.89085 | 1.01733 | Exon (ENSMUST00000098272.2/ENSMUSG00000074001.2, exon 2 of 6) | 9 | 121777607 | 121783819 | 6213 | 1 | ENSMUST00000098272.2 | 2357 | Klhl40 | ENSMUSG00000074001.2 |
| chr7 | 98192105 | 98192351 | 3.75815 | 4.69414 | 1.61846 | Downstream (<1kb) | 7 | 98192420 | 98199481 | 7062 | 2 | ENSMUST00000098278.2 | 7130 | B3gnt6 | ENSMUSG00000074004.2 |
| chr7 | 98192762 | 98193810 | 3.70604 | 4.61153 | 1.55752 | 3' UTR | 7 | 98192420 | 98199481 | 7062 | 2 | ENSMUST00000098278.2 | 5671 | B3gnt6 | ENSMUSG00000074004.2 |
| chr7 | 98194214 | 98194520 | 3.63382 | 4.49882 | 1.45313 | Exon (ENSMUST00000098278.2/ENSMUSG00000074004.2, exon 2 of 2) | 7 | 98192420 | 98199481 | 7062 | 2 | ENSMUST00000098278.2 | 4961 | B3gnt6 | ENSMUSG00000074004.2 |
| chr7 | 98141322 | 98141607 | 3.35779 | 3.90958 | 1.15922 | Intron (ENSMUST00000040971.8/ENSMUSG00000035547.9, intron 3 of 12) | 7 | 98143360 | 98145502 | 2143 | 2 | ENSMUST00000098281.2 | 3895 | Omp | ENSMUSG00000074006.2 |
| chr7 | 98142706 | 98142983 | 3.47969 | 4.12775 | 1.2762 | Intron (ENSMUST00000040971.8/ENSMUSG00000035547.9, intron 3 of 12) | 7 | 98143360 | 98145502 | 2143 | 2 | ENSMUST00000098281.2 | 2519 | Omp | ENSMUSG00000074006.2 |
| chr7 | 98145538 | 98145687 | 3.64361 | 4.39565 | 1.37328 | Promoter | 7 | 98143360 | 98145502 | 2143 | 2 | ENSMUST00000098281.2 | -37 | Omp | ENSMUSG00000074006.2 |
| chr7 | 98146651 | 98147067 | 3.3592 | 3.96611 | 1.1337 | Intron (ENSMUST00000040971.8/ENSMUSG00000035547.9, intron 3 of 12) | 7 | 98143360 | 98145502 | 2143 | 2 | ENSMUST00000098281.2 | -1150 | Omp | ENSMUSG00000074006.2 |
| chr7 | 98148189 | 98148823 | 3.63853 | 4.3912 | 1.38441 | Intron (ENSMUST00000040971.8/ENSMUSG00000035547.9, intron 3 of 12) | 7 | 98143360 | 98145502 | 2143 | 2 | ENSMUST00000098281.2 | -2688 | Omp | ENSMUSG00000074006.2 |
| chr8 | 121909041 | 121909735 | 3.62116 | 4.41982 | 1.4449 | Exon (ENSMUST00000181944.1/ENSMUSG00000074052.6, exon 1 of 1) | 8 | 121907833 | 121918428 | 10596 | 1 | ENSMUST00000140928.1 | 1209 | BC048644 | ENSMUSG00000074052.6 |
| chr8 | 121912498 | 121912769 | 3.29796 | 3.78422 | 1.04913 | Intron (ENSMUST00000127664.1/ENSMUSG00000092329.1, intron 1 of 15) | 8 | 121907833 | 121918428 | 10596 | 1 | ENSMUST00000140928.1 | 4666 | BC048644 | ENSMUSG00000074052.6 |
| chr7 | 45415602 | 45415998 | 3.57244 | 4.25008 | 1.23541 | Exon (ENSMUST00000058879.6/ENSMUSG00000074121.2, exon 3 of 3) | 7 | 45413695 | 45417178 | 3484 | 1 | ENSMUST00000058879.6 | 1908 | Ntf5 | ENSMUSG00000074121.2 |
| chr8 | 84010431 | 84010776 | 3.66276 | 4.47275 | 1.47941 | Promoter | 8 | 84010239 | 84011411 | 1173 | 2 | ENSMUST00000056686.6 | 635 | 2210011C24Rik | ENSMUSG00000074217.3 |
| chr8 | 84012488 | 84012673 | 3.38858 | 4.14123 | 1.22674 | Promoter | 8 | 84011444 | 84012171 | 728 | 2 | ENSMUST00000185457.1 | -318 | 2210011C24Rik | ENSMUSG00000074217.3 |
| chr3 | 128003181 | 128003602 | 3.92246 | 4.96595 | 1.76718 | Distal Intergenic | 3 | 128038694 | 128040661 | 1968 | 1 | ENSMUST00000098625.2 | -35092 | Gm10650 | ENSMUSG00000074237.2 |
| chr3 | 128086769 | 128087497 | 3.6057 | 4.37827 | 1.36264 | Distal Intergenic | 3 | 128038694 | 128040661 | 1968 | 1 | ENSMUST00000098625.2 | 48076 | Gm10650 | ENSMUSG00000074237.2 |
| chr9 | 61116929 | 61117184 | 3.51638 | 4.28973 | 1.31481 | Distal Intergenic | 9 | 61370878 | 61371627 | 750 | 2 | ENSMUST00000187131.1 | 254443 | Gm10655 | ENSMUSG00000074256.3 |
| chr9 | 61122194 | 61122413 | 3.50597 | 4.17718 | 1.26214 | Distal Intergenic | 9 | 61370878 | 61371627 | 750 | 2 | ENSMUST00000187131.1 | 249214 | Gm10655 | ENSMUSG00000074256.3 |
| chr9 | 61130277 | 61130494 | 3.3535 | 3.85772 | 1.01733 | Distal Intergenic | 9 | 61370878 | 61371627 | 750 | 2 | ENSMUST00000187131.1 | 241133 | Gm10655 | ENSMUSG00000074256.3 |
| chr9 | 61145827 | 61145965 | 3.32175 | 3.81208 | 1.02454 | Distal Intergenic | 9 | 61370878 | 61371627 | 750 | 2 | ENSMUST00000187131.1 | 225662 | Gm10655 | ENSMUSG00000074256.3 |
| chr9 | 61156464 | 61156989 | 3.55114 | 4.26014 | 1.35546 | Distal Intergenic | 9 | 61370878 | 61371627 | 750 | 2 | ENSMUST00000187131.1 | 214638 | Gm10655 | ENSMUSG00000074256.3 |
| chr9 | 61157914 | 61158181 | 3.47294 | 4.12364 | 1.25797 | Distal Intergenic | 9 | 61370878 | 61371627 | 750 | 2 | ENSMUST00000187131.1 | 213446 | Gm10655 | ENSMUSG00000074256.3 |
| chr9 | 61158710 | 61159374 | 3.67832 | 4.64392 | 1.53361 | Distal Intergenic | 9 | 61370878 | 61371627 | 750 | 2 | ENSMUST00000187131.1 | 212253 | Gm10655 | ENSMUSG00000074256.3 |
| chr9 | 61160347 | 61161115 | 3.65322 | 4.41538 | 1.36643 | Distal Intergenic | 9 | 61370878 | 61371627 | 750 | 2 | ENSMUST00000187131.1 | 210512 | Gm10655 | ENSMUSG00000074256.3 |
| chr9 | 61161429 | 61161504 | 3.68837 | 4.48301 | 1.43132 | Distal Intergenic | 9 | 61370878 | 61371627 | 750 | 2 | ENSMUST00000187131.1 | 210123 | Gm10655 | ENSMUSG00000074256.3 |
| chr9 | 61163392 | 61163774 | 3.50514 | 4.13139 | 1.17553 | Distal Intergenic | 9 | 61370878 | 61371627 | 750 | 2 | ENSMUST00000187131.1 | 207853 | Gm10655 | ENSMUSG00000074256.3 |
| chr9 | 61164514 | 61164883 | 4.1422 | 5.3535 | 1.99578 | Distal Intergenic | 9 | 61370878 | 61371627 | 750 | 2 | ENSMUST00000187131.1 | 206744 | Gm10655 | ENSMUSG00000074256.3 |
| chr9 | 61165667 | 61166133 | 3.68555 | 4.5877 | 1.50278 | Distal Intergenic | 9 | 61370878 | 61371627 | 750 | 2 | ENSMUST00000187131.1 | 205494 | Gm10655 | ENSMUSG00000074256.3 |
| chr9 | 61179994 | 61180362 | 3.45545 | 4.07138 | 1.22552 | Distal Intergenic | 9 | 61370878 | 61371627 | 750 | 2 | ENSMUST00000187131.1 | 191265 | Gm10655 | ENSMUSG00000074256.3 |
| chr9 | 61193948 | 61194060 | 4.3054 | 5.6825 | 2.22361 | Distal Intergenic | 9 | 61370878 | 61371627 | 750 | 2 | ENSMUST00000187131.1 | 177567 | Gm10655 | ENSMUSG00000074256.3 |
| chr9 | 61208475 | 61208659 | 3.37436 | 3.88917 | 1.01733 | Distal Intergenic | 9 | 61370878 | 61371627 | 750 | 2 | ENSMUST00000187131.1 | 162968 | Gm10655 | ENSMUSG00000074256.3 |
| chr9 | 61210055 | 61210139 | 4.46229 | 5.98846 | 2.43909 | Distal Intergenic | 9 | 61370878 | 61371627 | 750 | 2 | ENSMUST00000187131.1 | 161488 | Gm10655 | ENSMUSG00000074256.3 |
| chr9 | 61223291 | 61223467 | 3.93834 | 5.00204 | 1.77465 | Distal Intergenic | 9 | 61370878 | 61371627 | 750 | 2 | ENSMUST00000187131.1 | 148160 | Gm10655 | ENSMUSG00000074256.3 |
| chr9 | 61227982 | 61229067 | 3.41696 | 4.16852 | 1.23506 | Distal Intergenic | 9 | 61370878 | 61371627 | 750 | 2 | ENSMUST00000187131.1 | 142560 | Gm10655 | ENSMUSG00000074256.3 |
| chr9 | 61238333 | 61238492 | 3.23338 | 3.67724 | 1.01733 | Distal Intergenic | 9 | 61370878 | 61371627 | 750 | 2 | ENSMUST00000187131.1 | 133135 | Gm10655 | ENSMUSG00000074256.3 |
| chr9 | 61241625 | 61242288 | 3.6771 | 4.50376 | 1.42483 | Distal Intergenic | 9 | 61370878 | 61371627 | 750 | 2 | ENSMUST00000187131.1 | 129339 | Gm10655 | ENSMUSG00000074256.3 |
| chr9 | 61258235 | 61258470 | 3.5285 | 4.20319 | 1.29731 | Distal Intergenic | 9 | 61370878 | 61371627 | 750 | 2 | ENSMUST00000187131.1 | 113157 | Gm10655 | ENSMUSG00000074256.3 |
| chr9 | 61260595 | 61260745 | 3.21625 | 3.65161 | 1.01259 | Distal Intergenic | 9 | 61370878 | 61371627 | 750 | 2 | ENSMUST00000187131.1 | 110882 | Gm10655 | ENSMUSG00000074256.3 |
| chr9 | 61281155 | 61281296 | 3.63441 | 4.39018 | 1.32539 | Distal Intergenic | 9 | 61370878 | 61371627 | 750 | 2 | ENSMUST00000187131.1 | 90331 | Gm10655 | ENSMUSG00000074256.3 |
| chr9 | 61281987 | 61282101 | 3.34153 | 3.83966 | 1.01733 | Distal Intergenic | 9 | 61370878 | 61371627 | 750 | 2 | ENSMUST00000187131.1 | 89526 | Gm10655 | ENSMUSG00000074256.3 |
| chr9 | 61300713 | 61301038 | 3.58407 | 4.28288 | 1.28133 | Distal Intergenic | 9 | 61370878 | 61371627 | 750 | 2 | ENSMUST00000187131.1 | 70589 | Gm10655 | ENSMUSG00000074256.3 |
| chr9 | 61301493 | 61301946 | 3.33098 | 3.82378 | 1.01733 | Distal Intergenic | 9 | 61370878 | 61371627 | 750 | 2 | ENSMUST00000187131.1 | 69681 | Gm10655 | ENSMUSG00000074256.3 |
| chr9 | 61315357 | 61315562 | 3.55214 | 4.25678 | 1.34598 | Distal Intergenic | 9 | 61370878 | 61371627 | 750 | 2 | ENSMUST00000187131.1 | 56065 | Gm10655 | ENSMUSG00000074256.3 |
| chr9 | 61321820 | 61322192 | 3.46276 | 4.09678 | 1.24142 | Distal Intergenic | 9 | 61370878 | 61371627 | 750 | 2 | ENSMUST00000187131.1 | 49435 | Gm10655 | ENSMUSG00000074256.3 |
| chr9 | 61324458 | 61324615 | 3.5966 | 4.50955 | 1.46986 | Distal Intergenic | 9 | 61370878 | 61371627 | 750 | 2 | ENSMUST00000187131.1 | 47012 | Gm10655 | ENSMUSG00000074256.3 |
| chr9 | 61326968 | 61327102 | 3.27973 | 3.74685 | 1.01733 | Distal Intergenic | 9 | 61370878 | 61371627 | 750 | 2 | ENSMUST00000187131.1 | 44525 | Gm10655 | ENSMUSG00000074256.3 |
| chr9 | 61328573 | 61328871 | 3.83271 | 4.75678 | 1.61024 | Distal Intergenic | 9 | 61370878 | 61371627 | 750 | 2 | ENSMUST00000187131.1 | 42756 | Gm10655 | ENSMUSG00000074256.3 |
| chr9 | 61338588 | 61338712 | 3.49712 | 4.19877 | 1.28346 | Distal Intergenic | 9 | 61370878 | 61371627 | 750 | 2 | ENSMUST00000187131.1 | 32915 | Gm10655 | ENSMUSG00000074256.3 |
| chr9 | 61341868 | 61342024 | 3.38193 | 3.9006 | 1.01733 | Distal Intergenic | 9 | 61370878 | 61371627 | 750 | 2 | ENSMUST00000187131.1 | 29603 | Gm10655 | ENSMUSG00000074256.3 |
| chr9 | 61343267 | 61343342 | 3.34424 | 3.84742 | 1.02896 | Distal Intergenic | 9 | 61370878 | 61371627 | 750 | 2 | ENSMUST00000187131.1 | 28285 | Gm10655 | ENSMUSG00000074256.3 |
| chr9 | 61345010 | 61345291 | 3.42535 | 3.98209 | 1.07353 | Distal Intergenic | 9 | 61370878 | 61371627 | 750 | 2 | ENSMUST00000187131.1 | 26336 | Gm10655 | ENSMUSG00000074256.3 |
| chr9 | 61347571 | 61347703 | 3.61367 | 4.32706 | 1.28355 | Distal Intergenic | 9 | 61370878 | 61371627 | 750 | 2 | ENSMUST00000187131.1 | 23924 | Gm10655 | ENSMUSG00000074256.3 |
| chr9 | 61352401 | 61352629 | 3.36276 | 3.8764 | 1.03402 | Distal Intergenic | 9 | 61370878 | 61371627 | 750 | 2 | ENSMUST00000187131.1 | 18998 | Gm10655 | ENSMUSG00000074256.3 |
| chr9 | 61354327 | 61354495 | 3.41182 | 3.95978 | 1.067 | Distal Intergenic | 9 | 61370878 | 61371627 | 750 | 2 | ENSMUST00000187131.1 | 17132 | Gm10655 | ENSMUSG00000074256.3 |
| chr9 | 59680333 | 59680485 | 3.35688 | 4.16481 | 1.23985 | Promoter | 9 | 59680144 | 59713923 | 33780 | 1 | ENSMUST00000123914.2 | 190 | Gramd2 | ENSMUSG00000074259.5 |
| chr7 | 25149908 | 25149983 | 3.65625 | 4.4076 | 1.33856 | Intron (ENSMUST00000175754.2/ENSMUSG00000008496.13, intron 1 of 3) | 7 | 25152457 | 25156627 | 4171 | 1 | ENSMUST00000098678.2 | -2474 | D930028M14Rik | ENSMUSG00000074274.2 |
| chr7 | 25151059 | 25151579 | 4.1246 | 5.31966 | 1.96683 | Promoter | 7 | 25152457 | 25156627 | 4171 | 1 | ENSMUST00000098678.2 | -878 | D930028M14Rik | ENSMUSG00000074274.2 |
| chr9 | 54066618 | 54066694 | 3.64491 | 4.39248 | 1.35394 | Intron (ENSMUST00000098760.3/ENSMUSG00000074345.3, intron 1 of 1) | 9 | 54025606 | 54068411 | 42806 | 2 | ENSMUST00000098760.3 | 1717 | Tnfaip8l3 | ENSMUSG00000074345.3 |
| chr9 | 54067206 | 54068358 | 3.54995 | 4.25924 | 1.285 | Promoter | 9 | 54025606 | 54068411 | 42806 | 2 | ENSMUST00000098760.3 | 53 | Tnfaip8l3 | ENSMUSG00000074345.3 |
| chr9 | 54068921 | 54069045 | 3.40569 | 3.97472 | 1.15307 | Promoter | 9 | 54025606 | 54068411 | 42806 | 2 | ENSMUST00000098760.3 | -511 | Tnfaip8l3 | ENSMUSG00000074345.3 |
| chr3 | 105448389 | 105448560 | 3.42705 | 4.0169 | 1.18895 | Exon (ENSMUST00000130683.1/ENSMUSG00000074346.6, exon 4 of 4) | 3 | 105441595 | 105451727 | 10133 | 2 | ENSMUST00000143893.1 | 3167 | Kcnd3os | ENSMUSG00000074346.6 |
| chr3 | 105450298 | 105450373 | 4.03636 | 5.41603 | 2.05181 | Intron (ENSMUST00000143893.1/ENSMUSG00000074346.6, intron 1 of 1) | 3 | 105441595 | 105451727 | 10133 | 2 | ENSMUST00000143893.1 | 1354 | Kcnd3os | ENSMUSG00000074346.6 |
| chr9 | 51874625 | 51875000 | 3.81132 | 4.81578 | 1.66264 | Promoter | 9 | 51840372 | 51875788 | 35417 | 2 | ENSMUST00000130086.1 | 788 | Arhgap20os | ENSMUSG00000074354.6 |
| chr9 | 51876602 | 51876792 | 3.77908 | 4.73659 | 1.61742 | Promoter | 9 | 51832814 | 51875886 | 43073 | 2 | ENSMUST00000152203.2 | -717 | Arhgap20os | ENSMUSG00000074354.6 |
| chr9 | 45134405 | 45134703 | 3.61632 | 4.32991 | 1.28034 | Promoter | 9 | 45107176 | 45135606 | 28431 | 2 | ENSMUST00000098827.2 | 903 | Gm10684 | ENSMUSG00000074385.2 |
| chr9 | 45135610 | 45136063 | 3.3659 | 3.908 | 1.08089 | Promoter | 9 | 45107176 | 45135606 | 28431 | 2 | ENSMUST00000098827.2 | -5 | Gm10684 | ENSMUSG00000074385.2 |
| chr9 | 41376685 | 41377167 | 3.71311 | 4.57409 | 1.52127 | Promoter | 9 | 41376561 | 41432079 | 55519 | 1 | ENSMUST00000190109.1 | 125 | 2610203C20Rik | ENSMUSG00000074415.5 |
| chr9 | 41377763 | 41378009 | 3.47353 | 4.08475 | 1.17505 | Intron (ENSMUST00000188945.1/ENSMUSG00000074415.5, intron 1 of 4) | 9 | 41376561 | 41432079 | 55519 | 1 | ENSMUST00000190109.1 | 1203 | 2610203C20Rik | ENSMUSG00000074415.5 |
| chr9 | 41579447 | 41579699 | 3.53652 | 4.19262 | 1.22547 | Promoter | 9 | 41579494 | 41592826 | 13333 | 1 | ENSMUST00000188774.1 | 0 | 2610203C20Rik | ENSMUSG00000074415.5 |
| chr9 | 41580786 | 41581039 | 3.61383 | 4.35807 | 1.32473 | Promoter | 9 | 41580741 | 41592827 | 12087 | 1 | ENSMUST00000098868.3 | 46 | 2610203C20Rik | ENSMUSG00000074415.5 |
| chr9 | 22188529 | 22189134 | 3.43016 | 4.04519 | 1.1718 | Promoter | 9 | 22188168 | 22202123 | 13956 | 1 | ENSMUST00000178901.2 | 362 | Zfp872 | ENSMUSG00000074472.4 |
| chr8 | 12060324 | 12060667 | 4.07104 | 5.24777 | 1.97101 | Distal Intergenic | 8 | 12278819 | 12280657 | 1839 | 1 | ENSMUST00000098935.1 | -218152 | A230072I06Rik | ENSMUSG00000074473.1 |
| chr8 | 12126788 | 12127172 | 3.37872 | 4.31136 | 1.31015 | Distal Intergenic | 8 | 12278819 | 12280657 | 1839 | 1 | ENSMUST00000098935.1 | -151647 | A230072I06Rik | ENSMUSG00000074473.1 |
| chr8 | 12152366 | 12152559 | 3.36743 | 3.87872 | 1.01733 | Distal Intergenic | 8 | 12278819 | 12280657 | 1839 | 1 | ENSMUST00000098935.1 | -126260 | A230072I06Rik | ENSMUSG00000074473.1 |
| chr8 | 12162391 | 12162599 | 4.17986 | 5.55084 | 2.15882 | Distal Intergenic | 8 | 12278819 | 12280657 | 1839 | 1 | ENSMUST00000098935.1 | -116220 | A230072I06Rik | ENSMUSG00000074473.1 |
| chr8 | 12163608 | 12163836 | 3.60195 | 4.51313 | 1.44342 | Distal Intergenic | 8 | 12278819 | 12280657 | 1839 | 1 | ENSMUST00000098935.1 | -114983 | A230072I06Rik | ENSMUSG00000074473.1 |
| chr8 | 12266647 | 12266932 | 3.58371 | 4.37363 | 1.36393 | Distal Intergenic | 8 | 12278819 | 12280657 | 1839 | 1 | ENSMUST00000098935.1 | -11887 | A230072I06Rik | ENSMUSG00000074473.1 |
| chr9 | 16497851 | 16498005 | 3.58878 | 4.35136 | 1.38846 | Distal Intergenic | 9 | 15910205 | 16378231 | 468027 | 2 | ENSMUST00000082170.5 | -119621 | Fat3 | ENSMUSG00000074505.4 |
| chr9 | 16499670 | 16502185 | 3.95511 | 5.20897 | 1.93811 | Distal Intergenic | 9 | 15910205 | 16378231 | 468027 | 2 | ENSMUST00000082170.5 | -121440 | Fat3 | ENSMUSG00000074505.4 |
| chr9 | 16503950 | 16504177 | 3.5066 | 4.2196 | 1.27081 | Distal Intergenic | 9 | 15910205 | 16378231 | 468027 | 2 | ENSMUST00000082170.5 | -125720 | Fat3 | ENSMUSG00000074505.4 |
| chr3 | 83112225 | 83112442 | 3.52254 | 4.1889 | 1.2538 | Distal Intergenic | 3 | 83125276 | 83129246 | 3971 | 2 | ENSMUST00000047876.6 | 16804 | Gm10710 | ENSMUSG00000074517.2 |
| chr3 | 83121381 | 83121582 | 3.45937 | 4.1181 | 1.24363 | Distal Intergenic | 3 | 83125276 | 83129246 | 3971 | 2 | ENSMUST00000047876.6 | 7664 | Gm10710 | ENSMUSG00000074517.2 |
| chr3 | 83126471 | 83126863 | 3.77811 | 4.7058 | 1.64525 | Exon (ENSMUST00000047876.6/ENSMUSG00000074517.2, exon 1 of 1) | 3 | 83125276 | 83129246 | 3971 | 2 | ENSMUST00000047876.6 | 2383 | Gm10710 | ENSMUSG00000074517.2 |
| chr3 | 83127306 | 83129938 | 3.90103 | 5.0044 | 1.82448 | Promoter | 3 | 83125276 | 83129246 | 3971 | 2 | ENSMUST00000047876.6 | 0 | Gm10710 | ENSMUSG00000074517.2 |
| chr2 | 174207412 | 174207529 | 3.30816 | 3.78944 | 1.01733 | Distal Intergenic | 2 | 174125038 | 174183918 | 58881 | 2 | ENSMUST00000131346.1 | -23495 | Gm10714 | ENSMUSG00000074555.5 |
| chr9 | 3015199 | 3016480 | 1.95532 | 11.10973 | 6.77005 | Promoter | 9 | 3015654 | 3017210 | 1557 | 1 | ENSMUST00000099050.3 | 0 | Gm10720 | ENSMUSG00000074564.3 |
| chr2 | 168272153 | 168272345 | 3.38045 | 3.92221 | 1.10163 | Intron (ENSMUST00000109191.1/ENSMUSG00000074575.4, intron 1 of 2) | 2 | 168261698 | 168269331 | 7634 | 2 | ENSMUST00000099069.2 | -2823 | Kcng1 | ENSMUSG00000074575.4 |
| chr2 | 168275163 | 168275328 | 3.61991 | 4.35357 | 1.32258 | Intron (ENSMUST00000109191.1/ENSMUSG00000074575.4, intron 1 of 2) | 2 | 168260117 | 168281132 | 21016 | 2 | ENSMUST00000109191.1 | 5804 | Kcng1 | ENSMUSG00000074575.4 |
| chr2 | 168278985 | 168279553 | 4.20897 | 5.48343 | 2.07317 | Intron (ENSMUST00000109191.1/ENSMUSG00000074575.4, intron 1 of 2) | 2 | 168260117 | 168281132 | 21016 | 2 | ENSMUST00000109191.1 | 1579 | Kcng1 | ENSMUSG00000074575.4 |
| chr2 | 168279939 | 168281382 | 3.73909 | 4.61149 | 1.48778 | Promoter | 2 | 168260117 | 168281132 | 21016 | 2 | ENSMUST00000109191.1 | 0 | Kcng1 | ENSMUSG00000074575.4 |
| chr2 | 168281707 | 168281871 | 3.58296 | 4.4419 | 1.39777 | Promoter | 2 | 168269099 | 168281736 | 12638 | 2 | ENSMUST00000131749.1 | 0 | Kcng1 | ENSMUSG00000074575.4 |
| chr2 | 163190035 | 163190305 | 3.31985 | 3.80712 | 1.01733 | Distal Intergenic | 2 | 163203125 | 163324170 | 121046 | 1 | ENSMUST00000109428.3 | -12820 | Tox2 | ENSMUSG00000074607.6 |
| chr2 | 163203713 | 163204746 | 3.49149 | 4.28376 | 1.30082 | Promoter | 2 | 163204553 | 163276098 | 71546 | 1 | ENSMUST00000128999.2 | 0 | Tox2 | ENSMUSG00000074607.6 |
| chr2 | 163206327 | 163206935 | 3.93224 | 5.32652 | 1.99764 | Intron (ENSMUST00000109428.3/ENSMUSG00000074607.6, intron 1 of 8) | 2 | 163204553 | 163276098 | 71546 | 1 | ENSMUST00000128999.2 | 1775 | Tox2 | ENSMUSG00000074607.6 |
| chr2 | 163207625 | 163207959 | 3.95546 | 5.25703 | 1.949 | Intron (ENSMUST00000109428.3/ENSMUSG00000074607.6, intron 1 of 8) | 2 | 163204553 | 163276098 | 71546 | 1 | ENSMUST00000128999.2 | 3073 | Tox2 | ENSMUSG00000074607.6 |
| chr2 | 163253159 | 163253414 | 3.53915 | 4.41575 | 1.40037 | Intron (ENSMUST00000109428.3/ENSMUSG00000074607.6, intron 2 of 8) | 2 | 163225454 | 163323102 | 97649 | 1 | ENSMUST00000165937.2 | 27706 | Tox2 | ENSMUSG00000074607.6 |
| chr2 | 163285906 | 163287088 | 3.51434 | 4.21169 | 1.31462 | Intron (ENSMUST00000109428.3/ENSMUSG00000074607.6, intron 3 of 8) | 2 | 163320378 | 163323102 | 2725 | 1 | ENSMUST00000148599.1 | -33290 | Tox2 | ENSMUSG00000074607.6 |
| chr2 | 163289414 | 163290072 | 3.55638 | 4.44548 | 1.41127 | Intron (ENSMUST00000109428.3/ENSMUSG00000074607.6, intron 3 of 8) | 2 | 163320378 | 163323102 | 2725 | 1 | ENSMUST00000148599.1 | -30306 | Tox2 | ENSMUSG00000074607.6 |
| chr2 | 163299463 | 163299749 | 3.49513 | 4.15994 | 1.29543 | Intron (ENSMUST00000109428.3/ENSMUSG00000074607.6, intron 3 of 8) | 2 | 163320378 | 163323102 | 2725 | 1 | ENSMUST00000148599.1 | -20629 | Tox2 | ENSMUSG00000074607.6 |
| chr2 | 163339671 | 163340076 | 3.59149 | 4.44467 | 1.3937 | Exon (ENSMUST00000017961.5/ENSMUSG00000017817.6, exon 4 of 6) | 2 | 163320378 | 163323102 | 2725 | 1 | ENSMUST00000148599.1 | 19294 | Tox2 | ENSMUSG00000074607.6 |
| chr2 | 160389861 | 160390164 | 3.54673 | 4.42721 | 1.3882 | Distal Intergenic | 2 | 160363703 | 160367065 | 3363 | 2 | ENSMUST00000099126.4 | -22797 | Mafb | ENSMUSG00000074622.4 |
| chr2 | 158553902 | 158554086 | 3.53372 | 4.18152 | 1.20025 | Distal Intergenic | 2 | 158541737 | 158550762 | 9026 | 1 | ENSMUST00000135838.1 | 12166 | Arhgap40 | ENSMUSG00000074625.5 |
| chr2 | 158555395 | 158555578 | 3.41083 | 3.95394 | 1.05156 | Distal Intergenic | 2 | 158541737 | 158550762 | 9026 | 1 | ENSMUST00000135838.1 | 13659 | Arhgap40 | ENSMUSG00000074625.5 |
| chr2 | 158555942 | 158556017 | 3.71326 | 4.78691 | 1.61707 | Distal Intergenic | 2 | 158541737 | 158550762 | 9026 | 1 | ENSMUST00000135838.1 | 14206 | Arhgap40 | ENSMUSG00000074625.5 |
| chr2 | 158567439 | 158567747 | 3.32581 | 3.83284 | 1.07666 | Distal Intergenic | 2 | 158541737 | 158550762 | 9026 | 1 | ENSMUST00000135838.1 | 25703 | Arhgap40 | ENSMUSG00000074625.5 |
| chr2 | 157092800 | 157093174 | 3.32936 | 3.88412 | 1.11812 | Intron (ENSMUST00000166140.2/ENSMUSG00000074628.4, intron 4 of 5) | 2 | 157088453 | 157096482 | 8030 | 1 | ENSMUST00000099140.2 | 4348 | Tldc2 | ENSMUSG00000074628.4 |
| chr3 | 34648821 | 34650577 | 3.76569 | 4.69934 | 1.5731 | Promoter | 3 | 34650405 | 34652461 | 2057 | 1 | ENSMUST00000099151.3 | 0 | Sox2 | ENSMUSG00000074637.5 |
| chr3 | 34645064 | 34645719 | 3.65585 | 4.72396 | 1.60093 | Intron (ENSMUST00000173845.2/ENSMUSG00000090828.3, intron 3 of 3) | 3 | 34650405 | 34652461 | 2057 | 1 | ENSMUST00000099151.3 | -4686 | Sox2 | ENSMUSG00000074637.5 |
| chr3 | 34646045 | 34648342 | 3.95204 | 5.05681 | 1.80331 | Intron (ENSMUST00000173845.2/ENSMUSG00000090828.3, intron 3 of 3) | 3 | 34650405 | 34652461 | 2057 | 1 | ENSMUST00000099151.3 | -2063 | Sox2 | ENSMUSG00000074637.5 |
| chr3 | 34650960 | 34651523 | 4.03843 | 5.85796 | 2.39554 | Promoter | 3 | 34650405 | 34652461 | 2057 | 1 | ENSMUST00000099151.3 | 556 | Sox2 | ENSMUSG00000074637.5 |
| chr13 | 112991452 | 112991806 | 4.01963 | 5.39767 | 2.05117 | Distal Intergenic | 13 | 112993868 | 113000394 | 6527 | 1 | ENSMUST00000092089.4 | -2062 | Mcidas | ENSMUSG00000074651.2 |
| chr13 | 112992579 | 112993218 | 4.4696 | 6.33608 | 2.81119 | Promoter | 13 | 112993868 | 113000394 | 6527 | 1 | ENSMUST00000092089.4 | -650 | Mcidas | ENSMUSG00000074651.2 |
| chr13 | 112993526 | 112994271 | 4.74209 | 6.86916 | 3.23371 | Promoter | 13 | 112993868 | 113000394 | 6527 | 1 | ENSMUST00000092089.4 | 0 | Mcidas | ENSMUSG00000074651.2 |
| chr13 | 112994644 | 112994728 | 3.62198 | 5.07545 | 1.82348 | Promoter | 13 | 112993868 | 113000394 | 6527 | 1 | ENSMUST00000092089.4 | 777 | Mcidas | ENSMUSG00000074651.2 |
| chr13 | 112995035 | 112995340 | 3.86279 | 4.87563 | 1.73326 | Intron (ENSMUST00000092089.4/ENSMUSG00000074651.2, intron 3 of 6) | 13 | 112993868 | 113000394 | 6527 | 1 | ENSMUST00000092089.4 | 1168 | Mcidas | ENSMUSG00000074651.2 |
| chr13 | 112996271 | 112996568 | 3.73475 | 4.59367 | 1.54535 | Intron (ENSMUST00000092089.4/ENSMUSG00000074651.2, intron 3 of 6) | 13 | 112993868 | 113000394 | 6527 | 1 | ENSMUST00000092089.4 | 2404 | Mcidas | ENSMUSG00000074651.2 |
| chr13 | 112996895 | 112997036 | 4.10257 | 5.59964 | 2.21524 | Exon (ENSMUST00000092089.4/ENSMUSG00000074651.2, exon 4 of 7) | 13 | 112993868 | 113000394 | 6527 | 1 | ENSMUST00000092089.4 | 3028 | Mcidas | ENSMUSG00000074651.2 |
| chr13 | 112997358 | 112997455 | 4.04456 | 5.74817 | 2.28724 | Exon (ENSMUST00000092089.4/ENSMUSG00000074651.2, exon 5 of 7) | 13 | 112993868 | 113000394 | 6527 | 1 | ENSMUST00000092089.4 | 3491 | Mcidas | ENSMUSG00000074651.2 |
| chr13 | 112998837 | 112999015 | 3.89841 | 5.05742 | 1.78951 | Exon (ENSMUST00000092089.4/ENSMUSG00000074651.2, exon 7 of 7) | 13 | 112993868 | 113000394 | 6527 | 1 | ENSMUST00000092089.4 | 4970 | Mcidas | ENSMUSG00000074651.2 |
| chr13 | 112999642 | 112999988 | 4.14049 | 5.62343 | 2.21028 | 3' UTR | 13 | 112993868 | 113000394 | 6527 | 1 | ENSMUST00000092089.4 | 5775 | Mcidas | ENSMUSG00000074651.2 |
| chr13 | 113000695 | 113000912 | 3.60401 | 4.39546 | 1.40015 | Distal Intergenic | 13 | 112993868 | 113000394 | 6527 | 1 | ENSMUST00000092089.4 | 6828 | Mcidas | ENSMUSG00000074651.2 |
| chr2 | 155614466 | 155614751 | 3.37261 | 3.88654 | 1.01733 | Exon (ENSMUST00000092995.5/ENSMUSG00000074652.3, exon 6 of 42) | 2 | 155611212 | 155634307 | 23096 | 1 | ENSMUST00000092995.5 | 3255 | Myh7b | ENSMUSG00000074652.3 |

| | | | | | | | | | | | | | | | |
|---|---|---|---|---|---|---|---|---|---|---|---|---|---|---|---|
| chr10 | 127237635 | 127237743 | 3.79471 | 4.74543 | 1.65457 | Exon (ENSMUST00000099172.3/ENSMUSG00000074657.4, exon 16 of 29) | 10 | 127225696 | 127263363 | 37668 | 2 | ENSMUST00000099172.3 | 25620 | Kif5a | ENSMUSG00000074657.4 |
| chr10 | 127262948 | 127263170 | 3.60052 | 4.41625 | 1.40554 | Promoter | 10 | 127225696 | 127263363 | 37668 | 2 | ENSMUST00000099172.3 | 193 | Kif5a | ENSMUSG00000074657.4 |
| chr3 | 26331261 | 26332097 | 3.66009 | 4.57456 | 1.51703 | Promoter | 3 | 26331150 | 26332378 | 1229 | 1 | ENSMUST00000099182.2 | 112 | A830092H15Rik | ENSMUSG00000074664.2 |
| chr3 | 26332507 | 26333210 | 3.59998 | 4.36269 | 1.41503 | Distal Intergenic | 3 | 26331150 | 26332378 | 1229 | 1 | ENSMUST00000099182.2 | 1358 | A830092H15Rik | ENSMUSG00000074664.2 |
| chr3 | 26333635 | 26334133 | 3.22167 | 3.65971 | 1.01575 | Distal Intergenic | 3 | 26331150 | 26332378 | 1229 | 1 | ENSMUST00000099182.2 | 2486 | A830092H15Rik | ENSMUSG00000074664.2 |
| chr3 | 17784619 | 17784694 | 3.26475 | 3.72425 | 1.01733 | 3' UTR | 3 | 17783514 | 17786898 | 3385 | 2 | ENSMUST00000099198.2 | 2204 | Gm10742 | ENSMUSG00000074674.2 |
| chr3 | 17785836 | 17787035 | 4.28281 | 5.84482 | 2.38201 | Promoter | 3 | 17783514 | 17786898 | 3385 | 2 | ENSMUST00000099198.2 | 0 | Gm10742 | ENSMUSG00000074674.2 |
| chr3 | 17787494 | 17787909 | 3.55582 | 4.23021 | 1.2528 | Promoter | 3 | 17783514 | 17786898 | 3385 | 2 | ENSMUST00000099198.2 | -597 | Gm10742 | ENSMUSG00000074674.2 |
| chr2 | 151663323 | 151663431 | 3.24275 | 3.69128 | 1.01733 | Intron (ENSMUST00000096439.3/ENSMUSG00000074704.4, intron 4 of 12) | 2 | 151645404 | 151668533 | 23130 | 2 | ENSMUST00000096439.3 | 5102 | Rad21l | ENSMUSG00000074704.4 |
| chr2 | 151665506 | 151666090 | 3.73755 | 4.75141 | 1.59862 | Intron (ENSMUST00000096439.3/ENSMUSG00000074704.4, intron 3 of 12) | 2 | 151645404 | 151668533 | 23130 | 2 | ENSMUST00000096439.3 | 2443 | Rad21l | ENSMUSG00000074704.4 |
| chr2 | 151666531 | 151666867 | 3.64137 | 4.38987 | 1.36438 | Intron (ENSMUST00000096439.3/ENSMUSG00000074704.4, intron 2 of 12) | 2 | 151645404 | 151668533 | 23130 | 2 | ENSMUST00000096439.3 | 1666 | Rad21l | ENSMUSG00000074704.4 |
| chr2 | 151669652 | 151669742 | 3.89818 | 5.76634 | 2.28706 | Distal Intergenic | 2 | 151645404 | 151668533 | 23130 | 2 | ENSMUST00000180195.2 | -1120 | Rad21l | ENSMUSG00000074704.4 |
| chr2 | 151670172 | 151670568 | 3.62636 | 4.42317 | 1.47359 | Distal Intergenic | 2 | 151645404 | 151668533 | 23130 | 2 | ENSMUST00000180195.2 | -1640 | Rad21l | ENSMUSG00000074704.4 |
| chr2 | 149830714 | 149831030 | 3.91597 | 4.93776 | 1.71396 | Promoter | 2 | 149830788 | 150004392 | 173605 | 1 | ENSMUST00000109935.2 | 0 | Syndig1 | ENSMUSG00000074736.5 |
| chr2 | 149831427 | 149832402 | 3.45266 | 4.03189 | 1.1037 | Promoter | 2 | 149830524 | 149831213 | 690 | 2 | ENSMUST00000099264.1 | -215 | C530025M09Rik | ENSMUSG00000074737.1 |
| chr2 | 149832730 | 149833126 | 3.69522 | 4.48878 | 1.4049 | Intron (ENSMUST00000144179.1/ENSMUSG00000074736.5, intron 1 of 1) | 2 | 149830524 | 149831213 | 690 | 2 | ENSMUST00000099264.1 | -1518 | C530025M09Rik | ENSMUSG00000074737.1 |
| chr19 | 59399540 | 59399684 | 3.37718 | 3.89343 | 1.01733 | Distal Intergenic | 19 | 59296084 | 59345780 | 49697 | 2 | ENSMUST00000099274.2 | -53761 | Pdzd8 | ENSMUSG00000074746.2 |
| chr19 | 59400159 | 59401018 | 3.76375 | 4.65928 | 1.52827 | Distal Intergenic | 19 | 59296084 | 59345780 | 49697 | 2 | ENSMUST00000099274.2 | -54380 | Pdzd8 | ENSMUSG00000074746.2 |
| chr2 | 139678653 | 139679196 | 3.45149 | 4.1527 | 1.28144 | Promoter | 2 | 139678178 | 139758581 | 80404 | 1 | ENSMUST00000099307.3 | 476 | Ism1 | ENSMUSG00000074766.5 |
| chr10 | 94904004 | 94904380 | 3.36023 | 3.88612 | 1.0814 | Intron (ENSMUST00000099337.3/ENSMUSG00000074785.4, intron 4 of 30) | 10 | 94870830 | 94922657 | 51828 | 2 | ENSMUST00000180573.1 | 18277 | Plxnc1 | ENSMUSG00000074785.4 |
| chr10 | 94905820 | 94906077 | 3.61062 | 4.46271 | 1.43824 | Intron (ENSMUST00000099337.3/ENSMUSG00000074785.4, intron 4 of 30) | 10 | 94870830 | 94922657 | 51828 | 2 | ENSMUST00000180573.1 | 16580 | Plxnc1 | ENSMUSG00000074785.4 |
| chr10 | 94942434 | 94943014 | 3.97536 | 5.0716 | 1.8429 | Intron (ENSMUST00000099337.3/ENSMUSG00000074785.4, intron 1 of 30) | 10 | 94790866 | 94944578 | 153713 | 2 | ENSMUST00000099337.3 | 1564 | Plxnc1 | ENSMUSG00000074785.4 |
| chr10 | 94943707 | 94945626 | 3.83717 | 4.99442 | 1.76597 | Promoter | 10 | 94790866 | 94944578 | 153713 | 2 | ENSMUST00000099337.3 | 0 | Plxnc1 | ENSMUSG00000074785.4 |
| chr2 | 130696196 | 130696622 | 3.28015 | 3.76115 | 1.04402 | Promoter | 2 | 130686577 | 130697469 | 10893 | 2 | ENSMUST00000134647.2 | 847 | Slc4a11 | ENSMUSG00000074796.5 |
| chr5 | 142529735 | 142530044 | 4.13426 | 5.33045 | 1.96744 | Promoter | 5 | 142525838 | 142530076 | 4239 | 2 | ENSMUST00000099400.2 | 32 | Papolb | ENSMUSG00000074817.2 |
| chr5 | 142530470 | 142530715 | 3.37756 | 3.89401 | 1.01733 | Promoter | 5 | 142525838 | 142530076 | 4239 | 2 | ENSMUST00000099400.2 | -395 | Papolb | ENSMUSG00000074817.2 |
| chr5 | 142531053 | 142531224 | 3.35163 | 3.85489 | 1.01733 | Promoter | 5 | 142525838 | 142530076 | 4239 | 2 | ENSMUST00000099400.2 | -978 | Papolb | ENSMUSG00000074817.2 |
| chr19 | 45039717 | 45040182 | 3.41066 | 3.9941 | 1.18716 | Exon (ENSMUST00000169459.2/ENSMUSG00000074818.7, exon 3 of 16) | 19 | 45026907 | 45045772 | 18866 | 2 | ENSMUST00000169459.2 | 5590 | Pdzd7 | ENSMUSG00000074818.7 |
| chr19 | 45044522 | 45045818 | 3.7417 | 4.64878 | 1.59819 | Promoter | 19 | 45026907 | 45045772 | 18866 | 2 | ENSMUST00000169459.2 | 0 | Pdzd7 | ENSMUSG00000074818.7 |
| chr19 | 43384739 | 43384964 | 3.89075 | 4.91205 | 1.70743 | Exon (ENSMUST00000099428.3/ENSMUSG00000074852.3, exon 2 of 12) | 19 | 42788947 | 43388311 | 599365 | 2 | ENSMUST00000099428.3 | 3347 | Hpse2 | ENSMUSG00000074852.3 |
| chr19 | 43385333 | 43385933 | 3.36579 | 3.89451 | 1.08205 | Intron (ENSMUST00000099428.3/ENSMUSG00000074852.3, intron 1 of 11) | 19 | 42788947 | 43388311 | 599365 | 2 | ENSMUST00000099428.3 | 2378 | Hpse2 | ENSMUSG00000074852.3 |
| chr19 | 43386322 | 43387890 | 3.73905 | 4.69037 | 1.61069 | Promoter | 19 | 42788947 | 43388311 | 599365 | 2 | ENSMUST00000099428.3 | 421 | Hpse2 | ENSMUSG00000074852.3 |
| chr19 | 43388213 | 43389298 | 3.63747 | 4.39924 | 1.3957 | Promoter | 19 | 42788947 | 43388311 | 599365 | 2 | ENSMUST00000099428.3 | 0 | Hpse2 | ENSMUSG00000074852.3 |
| chr19 | 43391390 | 43391555 | 3.31118 | 3.79403 | 1.01733 | Distal Intergenic | 19 | 42788947 | 43388311 | 599365 | 2 | ENSMUST00000099428.3 | -3080 | Hpse2 | ENSMUSG00000074852.3 |
| chr13 | 61665872 | 61666212 | 3.38092 | 3.95506 | 1.18512 | Distal Intergenic | 13 | 61564631 | 61570127 | 5497 | 2 | ENSMUST00000054702.7 | -95746 | Cts3 | ENSMUSG00000074870.3 |
| chr13 | 56373047 | 56373153 | 3.75698 | 4.60481 | 1.48656 | Distal Intergenic | 13 | 56362901 | 56365769 | 2869 | 1 | ENSMUST00000099476.3 | 10147 | Gm10782 | ENSMUSG00000074885.3 |
| chr13 | 56381617 | 56381984 | 3.64729 | 4.53797 | 1.51332 | Distal Intergenic | 13 | 56362901 | 56365769 | 2869 | 1 | ENSMUST00000099476.3 | 18717 | Gm10782 | ENSMUSG00000074885.3 |
| chr13 | 56386191 | 56386378 | 3.48839 | 4.16978 | 1.28094 | Distal Intergenic | 13 | 56362901 | 56365769 | 2869 | 1 | ENSMUST00000099476.3 | 23291 | Gm10782 | ENSMUSG00000074885.3 |
| chr16 | 96280892 | 96281110 | 3.81318 | 4.90265 | 1.68987 | Promoter | 16 | 96280798 | 96319857 | 39060 | 1 | ENSMUST00000099497.3 | 95 | B3galt5 | ENSMUSG00000074892.4 |
| chr16 | 96281916 | 96282135 | 3.32003 | 3.80753 | 1.01733 | Intron (ENSMUST00000113800.3/ENSMUSG00000074892.4, intron 2 of 3) | 16 | 96280798 | 96319857 | 39060 | 1 | ENSMUST00000099497.3 | 1119 | B3galt5 | ENSMUSG00000074892.4 |
| chr2 | 120076922 | 120077665 | 3.75182 | 4.6797 | 1.62261 | Exon (ENSMUST00000134935.1/ENSMUSG00000074899.4, exon 7 of 11) | 2 | 120061586 | 120072523 | 10938 | 2 | ENSMUST00000110756.1 | -4400 | Sptbn5 | ENSMUSG00000074899.4 |
| chr2 | 119322276 | 119322970 | 3.71831 | 4.78565 | 1.64861 | Intron (ENSMUST00000147929.2/ENSMUSG00000074912.5, intron 3 of 3) | 2 | 119321187 | 119325543 | 4357 | 2 | ENSMUST00000147929.2 | 2573 | Gm14207 | ENSMUSG00000074912.5 |
| chr2 | 119323360 | 119323489 | 3.58801 | 4.51924 | 1.46589 | Intron (ENSMUST00000147929.2/ENSMUSG00000074912.5, intron 3 of 3) | 2 | 119321187 | 119325543 | 4357 | 2 | ENSMUST00000147929.2 | 2054 | Gm14207 | ENSMUSG00000074912.5 |
| chr2 | 119324011 | 119324389 | 3.9469 | 5.17787 | 1.92675 | Intron (ENSMUST00000147929.2/ENSMUSG00000074912.5, intron 3 of 3) | 2 | 119321187 | 119325543 | 4357 | 2 | ENSMUST00000147929.2 | 1154 | Gm14207 | ENSMUSG00000074912.5 |
| chr2 | 119327261 | 119327341 | 3.94248 | 5.06353 | 1.81545 | Intron (ENSMUST00000102517.3/ENSMUSG00000027314.6, intron 3 of 10) | 2 | 119321187 | 119326194 | 5008 | 2 | ENSMUST00000156538.2 | -1068 | Gm14207 | ENSMUSG00000074912.5 |
| chr2 | 118663493 | 118664037 | 3.55493 | 4.26793 | 1.3492 | Promoter | 2 | 118663762 | 118664956 | 1195 | 1 | ENSMUST00000152550.1 | 0 | Pak6 | ENSMUSG00000074923.5 |
| chr2 | 118672744 | 118672957 | 3.49585 | 4.11529 | 1.16874 | Intron (ENSMUST00000099557.4/ENSMUSG00000074923.5, intron 2 of 10) | 2 | 118677229 | 118698018 | 20790 | 1 | ENSMUST00000110853.2 | -4272 | Pak6 | ENSMUSG00000074923.5 |
| chr2 | 118674678 | 118674874 | 4.27568 | 5.64663 | 2.20571 | Intron (ENSMUST00000099557.4/ENSMUSG00000074923.5, intron 2 of 10) | 2 | 118677229 | 118698018 | 20790 | 1 | ENSMUST00000110853.2 | -2355 | Pak6 | ENSMUSG00000074923.5 |
| chr2 | 118675867 | 118677887 | 4.12007 | 5.49653 | 2.12451 | Promoter | 2 | 118677229 | 118698018 | 20790 | 1 | ENSMUST00000110853.2 | 0 | Pak6 | ENSMUSG00000074923.5 |
| chr2 | 118685998 | 118686144 | 3.54116 | 4.2744 | 1.3268 | Intron (ENSMUST00000099557.4/ENSMUSG00000074923.5, intron 3 of 10) | 2 | 118677263 | 118697236 | 19974 | 1 | ENSMUST00000132577.1 | 8736 | Pak6 | ENSMUSG00000074923.5 |
| chr2 | 113755124 | 113755838 | 3.49461 | 4.11474 | 1.16609 | Intron (ENSMUST00000099575.3/ENSMUSG00000074934.3, intron 1 of 1) | 2 | 113746164 | 113758646 | 12483 | 2 | ENSMUST00000099575.3 | 2808 | Grem1 | ENSMUSG00000074934.3 |
| chr2 | 113756304 | 113756540 | 3.41873 | 4.15832 | 1.22145 | Intron (ENSMUST00000099575.3/ENSMUSG00000074934.3, intron 1 of 1) | 2 | 113746164 | 113758646 | 12483 | 2 | ENSMUST00000099575.3 | 2106 | Grem1 | ENSMUSG00000074934.3 |
| chr2 | 113757076 | 113758587 | 3.59463 | 4.29905 | 1.27837 | Promoter | 2 | 113746164 | 113758646 | 12483 | 2 | ENSMUST00000099575.3 | 59 | Grem1 | ENSMUSG00000074934.3 |
| chr2 | 113758990 | 113759378 | 3.78064 | 4.67795 | 1.5518 | Promoter | 2 | 113746164 | 113758646 | 12483 | 2 | ENSMUST00000099575.3 | -345 | Grem1 | ENSMUSG00000074934.3 |
| chr2 | 106180996 | 106181276 | 3.58035 | 4.56109 | 1.47509 | Intron (ENSMUST00000122388.3/ENSMUSG00000074981.7, intron 8 of 33) | 2 | 106222057 | 106385623 | 163567 | 1 | ENSMUST00000152776.2 | -40781 | Dcdc5 | ENSMUSG00000074981.7 |
| chr2 | 106194810 | 106195036 | 3.54167 | 4.28819 | 1.3315 | Exon (ENSMUST00000122388.3/ENSMUSG00000074981.7, exon 10 of 34) | 2 | 106222057 | 106385623 | 163567 | 1 | ENSMUST00000152776.2 | -27021 | Dcdc5 | ENSMUSG00000074981.7 |
| chr2 | 106428797 | 106429131 | 3.55147 | 4.51799 | 1.45152 | Distal Intergenic | 2 | 106383687 | 106406151 | 22465 | 1 | ENSMUST00000037284.7 | 45111 | Dcdc5 | ENSMUSG00000074981.7 |
| chr10 | 67637465 | 67637567 | 3.75502 | 4.59114 | 1.45355 | Distal Intergenic | 10 | 67572042 | 67574263 | 2222 | 1 | ENSMUST00000099661.2 | 65424 | Gm10797 | ENSMUSG00000074999.2 |
| chr2 | 102449272 | 102449541 | 3.53957 | 4.46926 | 1.4232 | 3' UTR | 2 | 102449366 | 102452499 | 3134 | 2 | ENSMUST00000099678.4 | 2958 | Fjx1 | ENSMUSG00000075012.4 |
| chr2 | 102450658 | 102451824 | 4.40395 | 6.19356 | 2.67704 | Promoter | 2 | 102449366 | 102452499 | 3134 | 2 | ENSMUST00000099678.4 | 675 | Fjx1 | ENSMUSG00000075012.4 |
| chr2 | 98664900 | 98665203 | 1.85969 | 9.70837 | 5.50862 | Downstream (1-2kb) | 2 | 98666547 | 98667301 | 755 | 2 | ENSMUST00000099683.1 | 2098 | Gm10800 | ENSMUSG00000075014.1 |
| chr2 | 98666246 | 98667313 | 1.86956 | 11.41176 | 6.97715 | Promoter | 2 | 98666547 | 98667301 | 755 | 2 | ENSMUST00000099683.1 | 0 | Gm10800 | ENSMUSG00000075014.1 |
| chr2 | 98662242 | 98662975 | 1.87534 | 11.49978 | 7.06138 | Promoter | 2 | 98662237 | 98664083 | 1847 | 1 | ENSMUST00000099684.3 | 6 | Gm10801 | ENSMUSG00000075015.3 |
| chr2 | 98663665 | 98664078 | 1.8348 | 10.47253 | 6.19219 | Exon (ENSMUST00000099684.3/ENSMUSG00000075015.3, exon 3 of 4) | 2 | 98662237 | 98664083 | 1847 | 1 | ENSMUST00000099684.3 | 1429 | Gm10801 | ENSMUSG00000075015.3 |
| chr2 | 94264437 | 94264563 | 4.12086 | 5.42616 | 2.09043 | Promoter | 2 | 94251879 | 94264739 | 12861 | 2 | ENSMUST00000129661.1 | 176 | E530001K10Rik | ENSMUSG00000075020.5 |
| chr2 | 93522097 | 93522412 | 3.69138 | 4.60468 | 1.52699 | Distal Intergenic | 2 | 93560568 | 93564361 | 3794 | 1 | ENSMUST00000099692.2 | -38156 | Gm10803 | ENSMUSG00000075024.2 |
| chr2 | 93480826 | 93480908 | 3.89783 | 4.89725 | 1.74194 | Distal Intergenic | 2 | 93468439 | 93468675 | 237 | 1 | ENSMUST00000167654.1 | 12388 | Gm10804 | ENSMUSG00000075025.5 |
| chr2 | 93489160 | 93489434 | 3.49598 | 4.11439 | 1.16125 | Distal Intergenic | 2 | 93468439 | 93468675 | 237 | 1 | ENSMUST00000167654.1 | 20722 | Gm10804 | ENSMUSG00000075025.5 |
| chr2 | 91921624 | 91922492 | 3.68851 | 4.73694 | 1.61979 | Promoter | 2 | 91920528 | 91920928 | 401 | 2 | ENSMUST00000099710.1 | -697 | Gm10805 | ENSMUSG00000075036.1 |
| chr2 | 91922808 | 91924012 | 3.9629 | 5.2 | 1.96349 | Distal Intergenic | 2 | 91920528 | 91920928 | 401 | 2 | ENSMUST00000099710.1 | -1881 | Gm10805 | ENSMUSG00000075036.1 |
| chr10 | 44003583 | 44003720 | 3.44799 | 4.58802 | 1.49392 | Promoter | 10 | 44004154 | 44004701 | 548 | 1 | ENSMUST00000099864.3 | -434 | Gm10812 | ENSMUSG00000075162.3 |
| chr2 | 79427843 | 79429243 | 3.98097 | 5.15598 | 1.92545 | Promoter | 2 | 79331752 | 79428954 | 97203 | 2 | ENSMUST00000099974.4 | 0 | Cerkl | ENSMUSG00000075256.4 |
| chr2 | 79463591 | 79463917 | 3.65813 | 4.49885 | 1.47538 | Distal Intergenic | 2 | 79338569 | 79456785 | 118217 | 2 | ENSMUST00000153602.2 | -6807 | Cerkl | ENSMUSG00000075256.4 |
| chr2 | 79458665 | 79459726 | 3.78718 | 5.03633 | 1.84304 | Exon (ENSMUST00000180703.1/ENSMUSG00000097843.1, exon 2 of 2) | 2 | 79338569 | 79456785 | 118217 | 2 | ENSMUST00000153602.2 | -1881 | Cerkl | ENSMUSG00000075256.4 |
| chr2 | 79460165 | 79460417 | 3.93895 | 5.01477 | 1.8495 | Exon (ENSMUST00000180703.1/ENSMUSG00000097843.1, exon 2 of 2) | 2 | 79338569 | 79456785 | 118217 | 2 | ENSMUST00000153602.2 | -3381 | Cerkl | ENSMUSG00000075256.4 |
| chr2 | 79462600 | 79462814 | 3.37523 | 3.94506 | 1.18163 | Distal Intergenic | 2 | 79338569 | 79456785 | 118217 | 2 | ENSMUST00000153602.2 | -5816 | Cerkl | ENSMUSG00000075256.4 |
| chr2 | 76028391 | 76028693 | 3.5048 | 4.26921 | 1.32076 | Intron (ENSMUST00000099992.4/ENSMUSG00000075270.6, intron 17 of 19) | 2 | 75989141 | 76006043 | 16903 | 2 | ENSMUST00000124825.1 | -22349 | Pde11a | ENSMUSG00000075270.6 |
| chr2 | 76067772 | 76067947 | 3.35031 | 3.85291 | 1.01733 | Intron (ENSMUST00000099992.4/ENSMUSG00000075270.6, intron 12 of 19) | 2 | 75989141 | 76006043 | 16903 | 2 | ENSMUST00000124825.1 | -61730 | Pde11a | ENSMUSG00000075270.6 |
| chr2 | 76068793 | 76069478 | 4.04922 | 5.32049 | 2.01946 | Intron (ENSMUST00000099992.4/ENSMUSG00000075270.6, intron 12 of 19) | 2 | 75989141 | 76006043 | 16903 | 2 | ENSMUST00000124825.1 | -62751 | Pde11a | ENSMUSG00000075270.6 |
| chr2 | 76069880 | 76070508 | 4.23252 | 5.65399 | 2.23732 | Intron (ENSMUST00000099992.4/ENSMUSG00000075270.6, intron 12 of 19) | 2 | 75989141 | 76006043 | 16903 | 2 | ENSMUST00000124825.1 | -63838 | Pde11a | ENSMUSG00000075270.6 |
| chr2 | 76323191 | 76323505 | 3.2462 | 3.69646 | 1.01733 | Intron (ENSMUST00000099992.4/ENSMUSG00000075270.6, intron 1 of 19) | 2 | 75989912 | 76338774 | 348863 | 2 | ENSMUST00000099992.4 | 15269 | Pde11a | ENSMUSG00000075270.6 |
| chr2 | 76337833 | 76339618 | 3.65992 | 4.47139 | 1.4614 | Promoter | 2 | 75989912 | 76338774 | 348863 | 2 | ENSMUST00000099992.4 | 0 | Pde11a | ENSMUSG00000075270.6 |
| chr2 | 75210909 | 75211062 | 3.5685 | 4.32889 | 1.36084 | Intron (ENSMUST00000186411.1/ENSMUSG00000100303.1, intron 2 of 10) | 2 | 75192277 | 75192759 | 483 | 1 | ENSMUST00000122260.1 | 18633 | Gm13653 | ENSMUSG00000075276.5 |
| chr2 | 75312784 | 75313192 | 3.31003 | 3.79232 | 1.01733 | Exon (ENSMUST00000186411.1/ENSMUSG00000100303.1, exon 10 of 11) | 2 | 75192277 | 75192759 | 483 | 1 | ENSMUST00000122260.1 | 120508 | Gm13653 | ENSMUSG00000075276.5 |
| chr2 | 73528653 | 73528829 | 3.37442 | 4.03467 | 1.13333 | Promoter | 2 | 73447932 | 73529382 | 81451 | 2 | ENSMUST00000129594.1 | 553 | Wipf1 | ENSMUSG00000075284.5 |
| chr16 | 29964588 | 29964799 | 3.55875 | 4.25513 | 1.3293 | Distal Intergenic | 16 | 29958493 | 29962307 | 3815 | 2 | ENSMUST00000100016.3 | -2282 | Gm1968 | ENSMUSG00000075286.3 |
| chr16 | 29966893 | 29967304 | 3.96036 | 5.08059 | 1.89869 | Distal Intergenic | 16 | 29958493 | 29962307 | 3815 | 2 | ENSMUST00000100016.3 | -4587 | Gm1968 | ENSMUSG00000075286.3 |
| chr16 | 29967943 | 29968375 | 3.34351 | 3.87286 | 1.11803 | Distal Intergenic | 16 | 29958493 | 29962307 | 3815 | 2 | ENSMUST00000100016.3 | -5637 | Gm1968 | ENSMUSG00000075286.3 |
| chr10 | 22159795 | 22160205 | 3.64841 | 4.53216 | 1.47895 | Intron (ENSMUST00000181645.2/ENSMUSG00000053219.10, intron 1 of 8) | 10 | 22158586 | 22288849 | 130264 | 1 | ENSMUST00000105522.3 | 1210 | H60b | ENSMUSG00000075297.5 |
| chr2 | 70506703 | 70508549 | 3.86468 | 4.88129 | 1.73738 | Promoter | 2 | 70508819 | 70540884 | 32066 | 1 | ENSMUST00000100041.3 | -270 | Erich2 | ENSMUSG00000075302.5 |
| chr2 | 70469675 | 70469760 | 3.79216 | 4.73022 | 1.62593 | Distal Intergenic | 2 | 70474923 | 70477729 | 2807 | 1 | ENSMUST00000100043.2 | -5163 | Sp5 | ENSMUSG00000075304.2 |
| chr2 | 70472119 | 70472381 | 3.54986 | 4.5142 | 1.45261 | Distal Intergenic | 2 | 70474923 | 70477729 | 2807 | 1 | ENSMUST00000100043.2 | -2542 | Sp5 | ENSMUSG00000075304.2 |
| chr2 | 70473537 | 70475413 | 4.28348 | 5.74987 | 2.34129 | Promoter | 2 | 70474923 | 70477729 | 2807 | 1 | ENSMUST00000100043.2 | 0 | Sp5 | ENSMUSG00000075304.2 |
| chr2 | 70475786 | 70477697 | 4.0233 | 5.15253 | 1.87931 | Promoter | 2 | 70474923 | 70477729 | 2807 | 1 | ENSMUST00000100043.2 | 864 | Sp5 | ENSMUSG00000075304.2 |
| chr2 | 70480835 | 70481536 | 3.51043 | 4.14586 | 1.19887 | Distal Intergenic | 2 | 70474923 | 70477729 | 2807 | 1 | ENSMUST00000100043.2 | 5913 | Sp5 | ENSMUSG00000075304.2 |
| chr2 | 66634066 | 66635306 | 3.95517 | 5.05224 | 1.85802 | Promoter | 2 | 66480080 | 66634952 | 154873 | 2 | ENSMUST00000100063.3 | 0 | Scn9a | ENSMUSG00000075316.6 |
| chr2 | 54081746 | 54081991 | 3.36082 | 3.86875 | 1.01733 | Downstream (2-3kb) | 2 | 54084093 | 54085552 | 1460 | 2 | ENSMUST00000100089.2 | 3561 | Rprm | ENSMUSG00000075334.2 |
| chr2 | 54083744 | 54084846 | 3.51195 | 4.24204 | 1.24574 | Promoter | 2 | 54084093 | 54085552 | 1460 | 2 | ENSMUST00000100089.2 | 706 | Rprm | ENSMUSG00000075334.2 |
| chr16 | 18589964 | 18590193 | 3.51233 | 4.50705 | 1.45872 | Distal Intergenic | 16 | 18591992 | 18609751 | 17760 | 1 | ENSMUST00000156827.1 | -1799 | 4930588K23Rik | ENSMUSG00000075335.3 |
| chr16 | 18590598 | 18590702 | 3.73481 | 4.61924 | 1.56311 | Distal Intergenic | 16 | 18591992 | 18609751 | 17760 | 1 | ENSMUST00000156827.1 | -1290 | 4930588K23Rik | ENSMUSG00000075335.3 |
| chr16 | 18591143 | 18592290 | 3.83575 | 4.85876 | 1.68596 | Promoter | 16 | 18591992 | 18609751 | 17760 | 1 | ENSMUST00000156827.1 | 0 | 4930588K23Rik | ENSMUSG00000075335.3 |
| chr16 | 18596566 | 18596720 | 3.43511 | 4.00744 | 1.11155 | Intron (ENSMUST00000156827.1/ENSMUSG00000075335.3, intron 1 of 1) | 16 | 18591992 | 18609751 | 17760 | 1 | ENSMUST00000156827.1 | 4575 | 4930588K23Rik | ENSMUSG00000075335.3 |
| chr4 | 116839767 | 116840201 | 3.40931 | 3.99935 | 1.20111 | Promoter | 4 | 116839907 | 116840221 | 315 | 2 | ENSMUST00000121187.1 | 20 | Rpl36-ps8 | ENSMUSG00000075390.5 |
| chr4 | 116840552 | 116840686 | 3.23445 | 3.67885 | 1.01733 | Promoter | 4 | 116839907 | 116840221 | 315 | 2 | ENSMUST00000121187.1 | -332 | Rpl36-ps8 | ENSMUSG00000075390.5 |
| chr11 | 114281374 | 114281553 | 3.31646 | 4.00412 | 1.11915 | Distal Intergenic | 11 | 114251843 | 114253127 | 1285 | 1 | ENSMUST00000130963.1 | 29532 | Gm11691 | ENSMUSG00000075431.6 |
| chr11 | 114293826 | 114293901 | 3.8298 | 4.74306 | 1.58528 | Distal Intergenic | 11 | 114251843 | 114253127 | 1285 | 1 | ENSMUST00000130963.1 | 41984 | Gm11691 | ENSMUSG00000075431.6 |
| chr11 | 112796202 | 112796380 | 3.30367 | 3.7828 | 1.01733 | Intron (ENSMUST00000127337.1/ENSMUSG00000075437.4, intron 2 of 2) | 11 | 112792731 | 112799008 | 6278 | 2 | ENSMUST00000127337.1 | 2628 | Gm11681 | ENSMUSG00000075437.4 |
| chr11 | 112810509 | 112810937 | 3.68138 | 4.4636 | 1.38333 | Distal Intergenic | 11 | 112792731 | 112799008 | 6278 | 2 | ENSMUST00000127337.1 | -11502 | Gm11681 | ENSMUSG00000075437.4 |
| chr14 | 122484857 | 122485022 | 3.47936 | 4.39134 | 1.37095 | Distal Intergenic | 14 | 122490586 | 122492121 | 1536 | 1 | ENSMUST00000100294.2 | -5564 | Gm10837 | ENSMUSG00000075465.2 |
| chr14 | 122485404 | 122486924 | 3.83586 | 4.86883 | 1.69102 | Distal Intergenic | 14 | 122490586 | 122492121 | 1536 | 1 | ENSMUST00000100294.2 | -3662 | Gm10837 | ENSMUSG00000075465.2 |
| chr14 | 122487497 | 122488524 | 3.94074 | 5.23627 | 1.9583 | Distal Intergenic | 14 | 122490586 | 122492121 | 1536 | 1 | ENSMUST00000100294.2 | -2062 | Gm10837 | ENSMUSG00000075465.2 |
| chr14 | 122488996 | 122490239 | 3.58097 | 4.4345 | 1.42081 | Promoter | 14 | 122490586 | 122492121 | 1536 | 1 | ENSMUST00000100294.2 | -347 | Gm10837 | ENSMUSG00000075465.2 |
| chr14 | 122490580 | 122491230 | 4.00908 | 5.33999 | 2.02785 | Promoter | 14 | 122490586 | 122492121 | 1536 | 1 | ENSMUST00000100294.2 | 0 | Gm10837 | ENSMUSG00000075465.2 |
| chr14 | 108911175 | 108911636 | 3.78146 | 5.10716 | 1.85832 | 3' UTR | 14 | 108909991 | 108914239 | 4249 | 2 | ENSMUST00000100322.2 | 2603 | Slitrk1 | ENSMUSG00000075478.2 |
| chr14 | 108912164 | 108913092 | 3.45509 | 4.10171 | 1.27108 | Exon (ENSMUST00000100322.2/ENSMUSG00000075478.2, exon 1 of 1) | 14 | 108909991 | 108914239 | 4249 | 2 | ENSMUST00000100322.2 | 1147 | Slitrk1 | ENSMUSG00000075478.2 |
| chr11 | 102887982 | 102888121 | 3.49792 | 4.12091 | 1.17751 | 3' UTR | 11 | 102885169 | 102886731 | 1563 | 1 | ENSMUST00000100369.3 | 2814 | Fam187a | ENSMUSG00000075510.3 |
| chr15 | 82061912 | 82062217 | 3.45062 | 4.1501 | 1.23322 | Exon (ENSMUST00000100396.2/ENSMUSG00000075524.2, exon 2 of 2) | 15 | 82059151 | 82066538 | 7388 | 1 | ENSMUST00000100396.2 | 2762 | 4930407I10Rik | ENSMUSG00000075524.2 |
| chr15 | 82063621 | 82063768 | 3.72764 | 4.68003 | 1.58716 | Exon (ENSMUST00000100396.2/ENSMUSG00000075524.2, exon 2 of 2) | 15 | 82059151 | 82066538 | 7388 | 1 | ENSMUST00000100396.2 | 4471 | 4930407I10Rik | ENSMUSG00000075524.2 |
| chr2 | 9898057 | 9898757 | 3.63363 | 4.39602 | 1.40656 | Distal Intergenic | 2 | 9906178 | 9914678 | 8501 | 1 | ENSMUST00000149014.1 | -7421 | Gm13262 | ENSMUSG00000075534.5 |
| chr2 | 9903944 | 9904104 | 4.0982 | 5.30992 | 2.03388 | Distal Intergenic | 2 | 9906178 | 9914678 | 8501 | 1 | ENSMUST00000149014.1 | -2074 | Gm13262 | ENSMUSG00000075534.5 |
| chr2 | 9905343 | 9906072 | 3.89912 | 4.9478 | 1.76098 | Promoter | 2 | 9906178 | 9914678 | 8501 | 1 | ENSMUST00000149014.1 | -106 | Gm13262 | ENSMUSG00000075534.5 |
| chr5 | 147314871 | 147316415 | 4.17738 | 5.49989 | 2.12745 | 3' UTR | 5 | 147314984 | 147322440 | 7457 | 2 | ENSMUST00000100433.4 | 6025 | Urad | ENSMUSG00000075543.4 |
| chr5 | 147316745 | 147317423 | 3.76454 | 4.74218 | 1.64779 | Intron (ENSMUST00000100433.4/ENSMUSG00000075543.4, intron 1 of 1) | 5 | 147314984 | 147322440 | 7457 | 2 | ENSMUST00000100433.4 | 5017 | Urad | ENSMUSG00000075543.4 |
| chr5 | 147317907 | 147318324 | 3.67597 | 4.49328 | 1.46947 | Intron (ENSMUST00000100433.4/ENSMUSG00000075543.4, intron 1 of 1) | 5 | 147314984 | 147322440 | 7457 | 2 | ENSMUST00000100433.4 | 4116 | Urad | ENSMUSG00000075543.4 |
| chr5 | 147322459 | 147322891 | 3.99419 | 5.06007 | 1.78524 | Promoter | 5 | 147314984 | 147322440 | 7457 | 2 | ENSMUST00000100433.4 | -20 | Urad | ENSMUSG00000075543.4 |
| chr5 | 147326161 | 147326361 | 3.36156 | 3.87661 | 1.04106 | Distal Intergenic | 5 | 147314984 | 147322440 | 7457 | 2 | ENSMUST00000100433.4 | -3722 | Urad | ENSMUSG00000075543.4 |
| chr14 | 67295830 | 67295926 | 3.25288 | 3.70646 | 1.01733 | Intron (ENSMUST00000022637.8/ENSMUSG00000022053.8, intron 7 of 16) | 14 | 67304888 | 67314711 | 9824 | 2 | ENSMUST00000100448.2 | 18785 | Gm6878 | ENSMUSG00000075549.2 |
| chr14 | 67308232 | 67308670 | 3.75557 | 4.59821 | 1.46595 | Intron (ENSMUST00000022637.8/ENSMUSG00000022053.8, intron 7 of 16) | 14 | 67304888 | 67314711 | 9824 | 2 | ENSMUST00000100448.2 | 6041 | Gm6878 | ENSMUSG00000075549.2 |
| chr15 | 79167173 | 79167248 | 3.8279 | 4.73157 | 1.54469 | Intron (ENSMUST00000131710.2/ENSMUSG00000075555.6, intron 1 of 2) | 15 | 79166066 | 79227524 | 61459 | 1 | ENSMUST00000144651.2 | 1108 | Gm10863 | ENSMUSG00000075555.6 |
| chr15 | 79201980 | 79202055 | 4.13915 | 5.80913 | 2.3115 | Intron (ENSMUST00000131710.2/ENSMUSG00000075555.6, intron 1 of 2) | 15 | 79209311 | 79217383 | 8073 | 1 | ENSMUST00000148028.1 | -7256 | Gm10863 | ENSMUSG00000075555.6 |
| chr15 | 79203503 | 79203901 | 3.35484 | 3.85973 | 1.01733 | Intron (ENSMUST00000131710.2/ENSMUSG00000075555.6, intron 1 of 2) | 15 | 79209311 | 79217383 | 8073 | 1 | ENSMUST00000148028.1 | -5410 | Gm10863 | ENSMUSG00000075555.6 |
| chr11 | 96447817 | 96448146 | 3.28464 | 3.75411 | 1.01733 | Downstream (<1kb) | 11 | 96448323 | 96464547 | 16225 | 2 | ENSMUST00000153012.1 | 16401 | Gm11529 | ENSMUSG00000075586.4 |
| chr11 | 96458043 | 96458331 | 3.33731 | 3.8468 | 1.06479 | Intron (ENSMUST00000153012.1/ENSMUSG00000075586.4, intron 2 of 2) | 11 | 96448323 | 96464547 | 16225 | 2 | ENSMUST00000153012.1 | 6216 | Gm11529 | ENSMUSG00000075586.4 |
| chr11 | 96463061 | 96463357 | 3.69061 | 4.49012 | 1.43493 | Intron (ENSMUST00000153012.1/ENSMUSG00000075586.4, intron 2 of 2) | 11 | 96448323 | 96464547 | 16225 | 2 | ENSMUST00000153012.1 | 1190 | Gm11529 | ENSMUSG00000075586.4 |
| chr11 | 96394980 | 96395288 | 3.51206 | 4.1758 | 1.28636 | Distal Intergenic | 11 | 96392112 | 96392666 | 555 | 1 | ENSMUST00000117614.1 | 2869 | Gm11531 | ENSMUSG00000075587.5 |
| chr11 | 96398817 | 96399125 | 3.77322 | 4.62991 | 1.48247 | Distal Intergenic | 11 | 96392112 | 96392666 | 555 | 1 | ENSMUST00000117614.1 | 6706 | Gm11531 | ENSMUSG00000075587.5 |
| chr11 | 96399915 | 96400239 | 3.40351 | 3.96518 | 1.09589 | Distal Intergenic | 11 | 96392112 | 96392666 | 555 | 1 | ENSMUST00000117614.1 | 7804 | Gm11531 | ENSMUSG00000075587.5 |
| chr11 | 96402272 | 96402604 | 3.86173 | 4.88479 | 1.70615 | Distal Intergenic | 11 | 96392112 | 96392666 | 555 | 1 | ENSMUST00000117614.1 | 10161 | Gm11531 | ENSMUSG00000075587.5 |
| chr11 | 96404871 | 96404965 | 3.52512 | 4.16509 | 1.18657 | Distal Intergenic | 11 | 96392112 | 96392666 | 555 | 1 | ENSMUST00000117614.1 | 12760 | Gm11531 | ENSMUSG00000075587.5 |
| chr11 | 96406308 | 96406536 | 3.38496 | 3.95724 | 1.16216 | Distal Intergenic | 11 | 96392112 | 96392666 | 555 | 1 | ENSMUST00000117614.1 | 14197 | Gm11531 | ENSMUSG00000075587.5 |
| chr11 | 96407109 | 96408138 | 4.03917 | 5.22867 | 1.97986 | Distal Intergenic | 11 | 96392112 | 96392666 | 555 | 1 | ENSMUST00000117614.1 | 14998 | Gm11531 | ENSMUSG00000075587.5 |
| chr11 | 96408778 | 96409254 | 3.66387 | 4.41955 | 1.3466 | Distal Intergenic | 11 | 96392112 | 96392666 | 555 | 1 | ENSMUST00000117614.1 | 16667 | Gm11531 | ENSMUSG00000075587.5 |
| chr11 | 96410784 | 96411478 | 3.45118 | 4.02735 | 1.09564 | Distal Intergenic | 11 | 96392112 | 96392666 | 555 | 1 | ENSMUST00000117614.1 | 18673 | Gm11531 | ENSMUSG00000075587.5 |
| chr11 | 96414217 | 96414377 | 3.60781 | 4.31436 | 1.27086 | Distal Intergenic | 11 | 96392112 | 96392666 | 555 | 1 | ENSMUST00000117614.1 | 22106 | Gm11531 | ENSMUSG00000075587.5 |
| chr11 | 96416091 | 96416547 | 4.12518 | 5.43476 | 2.13346 | Distal Intergenic | 11 | 96392112 | 96392666 | 555 | 1 | ENSMUST00000117614.1 | 23980 | Gm11531 | ENSMUSG00000075587.5 |
| chr11 | 96419202 | 96419733 | 3.71183 | 4.58723 | 1.51105 | Distal Intergenic | 11 | 96392112 | 96392666 | 555 | 1 | ENSMUST00000117614.1 | 27091 | Gm11531 | ENSMUSG00000075587.5 |
| chr11 | 96423007 | 96423082 | 3.49198 | 4.12838 | 1.23718 | Distal Intergenic | 11 | 96392112 | 96392666 | 555 | 1 | ENSMUST00000117614.1 | 30896 | Gm11531 | ENSMUSG00000075587.5 |
| chr11 | 96424904 | 96425739 | 3.63602 | 4.37599 | 1.33297 | Distal Intergenic | 11 | 96392112 | 96392666 | 555 | 1 | ENSMUST00000117614.1 | 32793 | Gm11531 | ENSMUSG00000075587.5 |
| chr11 | 96351378 | 96351579 | 3.41382 | 3.96018 | 1.05763 | Promoter | 11 | 96351631 | 96354012 | 2382 | 1 | ENSMUST00000100523.6 | -52 | Hoxb2 | ENSMUSG00000075588.6 |
| chr12 | 110278668 | 110279394 | 3.66989 | 4.49195 | 1.42421 | Promoter | 12 | 110279068 | 110281097 | 2030 | 1 | ENSMUST00000173014.1 | 0 | Dio3 | ENSMUSG00000075707.5 |
| chr12 | 110279905 | 110280434 | 3.55055 | 4.33226 | 1.35509 | Promoter | 12 | 110279260 | 110280708 | 1449 | 1 | ENSMUST00000097228.4 | 646 | Dio3 | ENSMUSG00000075707.5 |
| chr12 | 110280805 | 110280926 | 3.62402 | 4.35673 | 1.32664 | 3' UTR | 12 | 110279260 | 110280708 | 1449 | 1 | ENSMUST00000097228.4 | 1546 | Dio3 | ENSMUSG00000075707.5 |
| chr12 | 110281753 | 110281894 | 3.54701 | 4.32452 | 1.31484 | Distal Intergenic | 12 | 110279260 | 110280708 | 1449 | 1 | ENSMUST00000097228.4 | 2494 | Dio3 | ENSMUSG00000075707.5 |
| chr12 | 110282545 | 110283019 | 3.98209 | 5.05226 | 1.79289 | Distal Intergenic | 12 | 110279260 | 110280708 | 1449 | 1 | ENSMUST00000097228.4 | 3286 | Dio3 | ENSMUSG00000075707.5 |
| chr12 | 110372875 | 110372950 | 3.60176 | 4.31264 | 1.29691 | Distal Intergenic | 12 | 110279260 | 110280708 | 1449 | 1 | ENSMUST00000097228.4 | 93616 | Dio3 | ENSMUSG00000075707.5 |
| chr12 | 110374499 | 110375304 | 3.7188 | 4.53407 | 1.44083 | Distal Intergenic | 12 | 110279260 | 110280708 | 1449 | 1 | ENSMUST00000097228.4 | 95240 | Dio3 | ENSMUSG00000075707.5 |
| chr12 | 110375912 | 110376000 | 3.74931 | 4.58552 | 1.47585 | Distal Intergenic | 12 | 110279260 | 110280708 | 1449 | 1 | ENSMUST00000097228.4 | 96653 | Dio3 | ENSMUSG00000075707.5 |
| chr12 | 12693079 | 12693415 | 3.52346 | 4.1868 | 1.25471 | Distal Intergenic | 12 | 12700039 | 12700145 | 107 | 1 | ENSMUST00000101943.1 | -6624 | Gm25538 | ENSMUSG00000075893.1 |
| chr12 | 12694709 | 12695089 | 3.67116 | 4.57182 | 1.48242 | Distal Intergenic | 12 | 12700039 | 12700145 | 107 | 1 | ENSMUST00000101943.1 | -4950 | Gm25538 | ENSMUSG00000075893.1 |
| chr4 | 9118367 | 9118590 | 3.26034 | 3.86198 | 1.0194 | Distal Intergenic | 4 | 9210003 | 9210108 | 106 | 1 | ENSMUST00000082782.1 | -91413 | Gm23423 | ENSMUSG00000075958.1 |
| chr4 | 9135734 | 9135911 | 3.6957 | 4.7529 | 1.60801 | Distal Intergenic | 4 | 9210003 | 9210108 | 106 | 1 | ENSMUST00000082782.1 | -74092 | Gm23423 | ENSMUSG00000075958.1 |
| chr4 | 9136892 | 9137318 | 3.5329 | 4.21955 | 1.31964 | Distal Intergenic | 4 | 9210003 | 9210108 | 106 | 1 | ENSMUST00000082782.1 | -72685 | Gm23423 | ENSMUSG00000075958.1 |
| chr4 | 9144635 | 9144978 | 3.61082 | 4.32596 | 1.29941 | Distal Intergenic | 4 | 9210003 | 9210108 | 106 | 1 | ENSMUST00000082782.1 | -65025 | Gm23423 | ENSMUSG00000075958.1 |

| | | | | | | | | | | | | | | | |
|---|---|---|---|---|---|---|---|---|---|---|---|---|---|---|---|
| chr4 | 9167900 | 9168025 | 3.6658 | 4.54877 | 1.45188 | Distal Intergenic | 4 | 9210003 | 9210108 | 106 | 1 | ENSMUST00000082782.1 | -41978 | Gm23423 | ENSMUSG00000075958.1 |
| chr4 | 9174369 | 9174451 | 4.11626 | 5.34026 | 2.04605 | Distal Intergenic | 4 | 9210003 | 9210108 | 106 | 1 | ENSMUST00000082782.1 | -35552 | Gm23423 | ENSMUSG00000075958.1 |
| chr4 | 9175407 | 9175935 | 3.8434 | 4.83684 | 1.70955 | Distal Intergenic | 4 | 9210003 | 9210108 | 106 | 1 | ENSMUST00000082782.1 | -34068 | Gm23423 | ENSMUSG00000075958.1 |
| chr4 | 9180496 | 9180710 | 3.59191 | 4.34386 | 1.41046 | Distal Intergenic | 4 | 9210003 | 9210108 | 106 | 1 | ENSMUST00000082782.1 | -29293 | Gm23423 | ENSMUSG00000075958.1 |
| chr4 | 9181084 | 9181347 | 3.54016 | 4.25334 | 1.36516 | Distal Intergenic | 4 | 9210003 | 9210108 | 106 | 1 | ENSMUST00000082782.1 | -28656 | Gm23423 | ENSMUSG00000075958.1 |
| chr4 | 9181659 | 9181734 | 4.02065 | 5.1791 | 1.92002 | Distal Intergenic | 4 | 9210003 | 9210108 | 106 | 1 | ENSMUST00000082782.1 | -28269 | Gm23423 | ENSMUSG00000075958.1 |
| chr4 | 9187536 | 9187711 | 3.52584 | 4.39546 | 1.38724 | Distal Intergenic | 4 | 9210003 | 9210108 | 106 | 1 | ENSMUST00000082782.1 | -22292 | Gm23423 | ENSMUSG00000075958.1 |
| chr4 | 9206983 | 9207263 | 3.35839 | 3.88352 | 1.08227 | Distal Intergenic | 4 | 9210003 | 9210108 | 106 | 1 | ENSMUST00000082782.1 | -2740 | Gm23423 | ENSMUSG00000075958.1 |
| chr12 | 85721045 | 85721223 | 3.73385 | 4.56812 | 1.47299 | Distal Intergenic | 12 | 85718477 | 85718595 | 119 | 2 | ENSMUST00000102019.1 | -2451 | n-R5s64 | ENSMUSG00000075973.1 |
| chr14 | 63743525 | 63743871 | 3.8193 | 4.93561 | 1.73833 | Intron (ENSMUST00000119973.2/ENSMUSG00000035067.9, intron 1 of 2) | 14 | 63727189 | 63727268 | 80 | 1 | ENSMUST00000102094.3 | 16337 | Mir598 | ENSMUSG00000076049.3 |
| chr16 | 28562998 | 28563330 | 3.70486 | 4.57767 | 1.52698 | Distal Intergenic | 16 | 28599935 | 28600043 | 109 | 2 | ENSMUST00000102110.1 | 36713 | Mir690 | ENSMUSG00000076064.1 |
| chr16 | 28564246 | 28565102 | 4.0828 | 5.48903 | 2.11325 | Distal Intergenic | 16 | 28599935 | 28600043 | 109 | 2 | ENSMUST00000102110.1 | 34941 | Mir690 | ENSMUSG00000076064.1 |
| chr16 | 28752428 | 28752866 | 3.77048 | 4.65381 | 1.56422 | Distal Intergenic | 16 | 28599935 | 28600043 | 109 | 2 | ENSMUST00000102110.1 | -152386 | Mir690 | ENSMUSG00000076064.1 |
| chr16 | 28755265 | 28755690 | 3.3528 | 3.85667 | 1.01733 | Distal Intergenic | 16 | 28599935 | 28600043 | 109 | 2 | ENSMUST00000102110.1 | -155223 | Mir690 | ENSMUSG00000076064.1 |
| chr3 | 89227100 | 89227301 | 3.55084 | 4.30214 | 1.34496 | Promoter | 3 | 89227116 | 89227198 | 83 | 2 | ENSMUST00000102300.2 | 0 | Mir92b | ENSMUSG00000076255.2 |
| chr16 | 11143958 | 11144300 | 2.81895 | 19.40207 | 14.37329 | Promoter | 16 | 11144125 | 11144181 | 57 | 1 | ENSMUST00000102303.1 | 0 | Gm23935 | ENSMUSG00000076258.1 |
| chr9 | 56223613 | 56223950 | 3.92552 | 19.34602 | 14.28212 | Promoter | 9 | 56223715 | 56223771 | 57 | 2 | ENSMUST00000102326.1 | 0 | Gm24270 | ENSMUSG00000076281.1 |
| chr19 | 18056460 | 18056651 | 3.73174 | 4.64355 | 1.54177 | Distal Intergenic | 19 | 17928111 | 17928226 | 116 | 1 | ENSMUST00000102348.1 | 128350 | Gm23061 | ENSMUSG00000076303.1 |
| chr16 | 94154623 | 94155027 | 3.6584 | 4.52632 | 1.49388 | Intron (ENSMUST00000163193.2/ENSMUSG00000040820.10, intron 6 of 10) | 16 | 94169302 | 94169411 | 110 | 2 | ENSMUST00000102349.1 | 14384 | Gm23062 | ENSMUSG00000076304.1 |
| chr16 | 94179776 | 94179865 | 3.68879 | 4.48753 | 1.43193 | Intron (ENSMUST00000163193.2/ENSMUSG00000040820.10, intron 6 of 10) | 16 | 94169302 | 94169411 | 110 | 2 | ENSMUST00000102349.1 | -10366 | Gm23062 | ENSMUSG00000076304.1 |
| chr2 | 122666054 | 122666134 | 3.98734 | 5.0764 | 1.85694 | Distal Intergenic | 2 | 122640803 | 122640881 | 79 | 1 | ENSMUST00000102354.2 | 25252 | Mir147 | ENSMUSG00000076309.2 |
| chr9 | 42885357 | 42885624 | 3.85738 | 5.09329 | 1.83256 | Intron (ENSMUST00000114865.2/ENSMUSG00000032017.9, intron 1 of 20) | 9 | 42862051 | 42862129 | 79 | 2 | ENSMUST00000102400.2 | -23229 | Gm27509 | ENSMUSG00000076355.2 |
| chr9 | 42886065 | 42886662 | 3.85976 | 4.88136 | 1.76575 | Intron (ENSMUST00000114865.2/ENSMUSG00000032017.9, intron 1 of 20) | 9 | 42862051 | 42862129 | 79 | 2 | ENSMUST00000102400.2 | -23937 | Gm27509 | ENSMUSG00000076355.2 |
| chr6 | 30165423 | 30165578 | 3.59655 | 4.31582 | 1.3429 | Promoter | 6 | 30165918 | 30165992 | 75 | 2 | ENSMUST00000102406.1 | 414 | Mir182 | ENSMUSG00000076361.1 |
| chr8 | 56617466 | 56617740 | 3.30636 | 3.81911 | 1.10578 | Distal Intergenic | 8 | 56561301 | 56561409 | 109 | 1 | ENSMUST00000102437.1 | 56166 | Gm25992 | ENSMUSG00000076392.1 |
| chr4 | 123322280 | 123323693 | 4.09386 | 5.51876 | 2.17831 | Promoter | 4 | 123321875 | 123323634 | 1760 | 2 | ENSMUST00000102640.1 | 0 | Oxct2a | ENSMUSG00000076436.1 |
| chr4 | 123115966 | 123116985 | 3.59836 | 4.35117 | 1.32541 | Promoter | 4 | 123116266 | 123118000 | 1735 | 1 | ENSMUST00000102648.5 | 0 | Oxct2b | ENSMUSG00000076438.5 |
| chr4 | 123117673 | 123117866 | 3.35429 | 3.8589 | 1.01733 | 3' UTR | 4 | 123116266 | 123118000 | 1735 | 1 | ENSMUST00000102648.5 | 1408 | Oxct2b | ENSMUSG00000076438.5 |
| chr2 | 31469408 | 31469638 | 3.77272 | 4.65203 | 1.57167 | Promoter | 2 | 31470207 | 31520672 | 50466 | 1 | ENSMUST00000102840.4 | -569 | Ass1 | ENSMUSG00000076441.4 |
| chr2 | 31469952 | 31470092 | 3.28676 | 3.75736 | 1.01733 | Promoter | 2 | 31470207 | 31520672 | 50466 | 1 | ENSMUST00000102840.4 | -115 | Ass1 | ENSMUSG00000076441.4 |
| chr2 | 31470711 | 31471108 | 4.38475 | 5.90613 | 2.44245 | Promoter | 2 | 31470207 | 31520672 | 50466 | 1 | ENSMUST00000102840.4 | 505 | Ass1 | ENSMUSG00000076441.4 |
| chr2 | 31471595 | 31472409 | 3.4277 | 3.9975 | 1.11035 | Intron (ENSMUST00000102840.4/ENSMUSG00000076441.4, intron 1 of 15) | 2 | 31470207 | 31520672 | 50466 | 1 | ENSMUST00000102840.4 | 1389 | Ass1 | ENSMUSG00000076441.4 |
| chr10 | 120420926 | 120421556 | 3.85601 | 4.86219 | 1.69938 | Intron (ENSMUST00000072777.8/ENSMUSG00000056758.9, intron 3 of 4) | 10 | 120447991 | 120448110 | 120 | 2 | ENSMUST00000103249.1 | 26554 | Mir763 | ENSMUSG00000076448.1 |
| chr10 | 120426814 | 120427072 | 3.55611 | 4.31022 | 1.34889 | Intron (ENSMUST00000072777.8/ENSMUSG00000056758.9, intron 3 of 4) | 10 | 120447991 | 120448110 | 120 | 2 | ENSMUST00000103249.1 | 21038 | Mir763 | ENSMUSG00000076448.1 |
| chr10 | 120429005 | 120429892 | 3.55395 | 4.33663 | 1.36664 | Intron (ENSMUST00000072777.8/ENSMUSG00000056758.9, intron 3 of 4) | 10 | 120447991 | 120448110 | 120 | 2 | ENSMUST00000103249.1 | 18218 | Mir763 | ENSMUSG00000076448.1 |
| chr10 | 120430466 | 120430800 | 4.1208 | 5.64494 | 2.25312 | Intron (ENSMUST00000072777.8/ENSMUSG00000056758.9, intron 3 of 4) | 10 | 120447991 | 120448110 | 120 | 2 | ENSMUST00000103249.1 | 17310 | Mir763 | ENSMUSG00000076448.1 |
| chr10 | 120431517 | 120432062 | 3.77967 | 4.78928 | 1.6714 | Intron (ENSMUST00000072777.8/ENSMUSG00000056758.9, intron 3 of 4) | 10 | 120447991 | 120448110 | 120 | 2 | ENSMUST00000103249.1 | 16048 | Mir763 | ENSMUSG00000076448.1 |
| chr10 | 120432542 | 120433181 | 3.42311 | 4.05466 | 1.2231 | Intron (ENSMUST00000072777.8/ENSMUSG00000056758.9, intron 3 of 4) | 10 | 120447991 | 120448110 | 120 | 2 | ENSMUST00000103249.1 | 14929 | Mir763 | ENSMUSG00000076448.1 |
| chr10 | 120442682 | 120442797 | 3.22844 | 3.66985 | 1.01733 | Intron (ENSMUST00000072777.8/ENSMUSG00000056758.9, intron 3 of 4) | 10 | 120447991 | 120448110 | 120 | 2 | ENSMUST00000103249.1 | 5313 | Mir763 | ENSMUSG00000076448.1 |
| chr10 | 120446823 | 120447443 | 3.48854 | 4.10043 | 1.15486 | Promoter | 10 | 120447991 | 120448110 | 120 | 2 | ENSMUST00000103249.1 | 667 | Mir763 | ENSMUSG00000076448.1 |
| chr6 | 70722542 | 70722648 | 3.29033 | 3.92167 | 1.0872 | Promoter | 6 | 70722562 | 70722599 | 38 | 1 | ENSMUST00000103405.1 | 0 | Igkj1 | ENSMUSG00000076604.1 |
| chr6 | 70732715 | 70733081 | 3.79467 | 4.71952 | 1.63853 | Distal Intergenic | 6 | 70726435 | 70726757 | 323 | 1 | ENSMUST00000103410.1 | 6281 | Igkc | ENSMUSG00000076609.1 |
| chr14 | 53632874 | 53633193 | 3.44179 | 4.12865 | 1.22975 | Downstream (2-3kb) | 14 | 53634829 | 53635398 | 570 | 1 | ENSMUST00000103658.2 | -1636 | Trav13-2 | ENSMUSG00000076846.2 |
| chr14 | 54225757 | 54225832 | 3.64062 | 4.38532 | 1.34485 | Distal Intergenic | 14 | 54220521 | 54223097 | 2577 | 1 | ENSMUST00000103740.1 | 5237 | Trac | ENSMUSG00000076928.1 |
| chr14 | 54229094 | 54229315 | 3.75371 | 4.64848 | 1.59876 | Distal Intergenic | 14 | 54220521 | 54223097 | 2577 | 1 | ENSMUST00000103740.1 | 8574 | Trac | ENSMUSG00000076928.1 |
| chr18 | 73625243 | 73625573 | 3.5621 | 4.43587 | 1.41126 | Promoter | 18 | 73624928 | 73625037 | 110 | 1 | ENSMUST00000103753.1 | 316 | Gm22065 | ENSMUSG00000076941.1 |
| chr14 | 12335028 | 12335333 | 3.89265 | 5.14899 | 1.8638 | Distal Intergenic | 14 | 12333776 | 12333904 | 129 | 2 | ENSMUST00000103970.1 | -1125 | Gm24578 | ENSMUSG00000077158.1 |
| chr14 | 12335683 | 12336279 | 3.58555 | 4.45119 | 1.42272 | Distal Intergenic | 14 | 12333776 | 12333904 | 129 | 2 | ENSMUST00000103970.1 | -1780 | Gm24578 | ENSMUSG00000077158.1 |
| chr14 | 12338144 | 12338597 | 3.71541 | 4.58621 | 1.55018 | Distal Intergenic | 14 | 12333776 | 12333904 | 129 | 2 | ENSMUST00000103970.1 | -4241 | Gm24578 | ENSMUSG00000077158.1 |
| chr7 | 69992448 | 69992665 | 3.72858 | 4.63014 | 1.5555 | Distal Intergenic | 7 | 69957776 | 69957904 | 129 | 2 | ENSMUST00000103978.1 | -34545 | Gm24120 | ENSMUSG00000077166.1 |
| chr12 | 56544726 | 56544802 | 4.44876 | 6.01285 | 2.4882 | Intron (ENSMUST00000182733.2/ENSMUSG00000098172.2, intron 2 of 2) | 12 | 56551459 | 56551590 | 132 | 1 | ENSMUST00000103989.1 | -6657 | Gm25760 | ENSMUSG00000077177.1 |
| chr12 | 56547468 | 56547738 | 3.36001 | 3.95171 | 1.12421 | Intron (ENSMUST00000182733.2/ENSMUSG00000098172.2, intron 2 of 2) | 12 | 56551459 | 56551590 | 132 | 1 | ENSMUST00000103989.1 | -3721 | Gm25760 | ENSMUSG00000077177.1 |
| chr12 | 56564232 | 56564513 | 3.28652 | 3.75693 | 1.01733 | Intron (ENSMUST00000182733.2/ENSMUSG00000098172.2, intron 2 of 2) | 12 | 56551459 | 56551590 | 132 | 1 | ENSMUST00000103989.1 | 12774 | Gm25760 | ENSMUSG00000077177.1 |
| chr1 | 137221533 | 137222594 | 3.58529 | 4.29195 | 1.26415 | Distal Intergenic | 1 | 137215544 | 137215647 | 104 | 2 | ENSMUST00000104019.1 | -5887 | Gm25609 | ENSMUSG00000077207.1 |
| chr9 | 99798832 | 99799092 | 3.57707 | 4.31696 | 1.38719 | Distal Intergenic | 9 | 99750391 | 99750517 | 127 | 2 | ENSMUST00000104027.1 | -48316 | Gm23949 | ENSMUSG00000077215.1 |
| chr9 | 99799568 | 99800340 | 3.86898 | 4.99428 | 1.79181 | Distal Intergenic | 9 | 99750391 | 99750517 | 127 | 2 | ENSMUST00000104027.1 | -49052 | Gm23949 | ENSMUSG00000077215.1 |
| chr9 | 99802388 | 99802572 | 3.48139 | 4.19672 | 1.27493 | Distal Intergenic | 9 | 99750391 | 99750517 | 127 | 2 | ENSMUST00000104027.1 | -51872 | Gm23949 | ENSMUSG00000077215.1 |
| chr19 | 56134204 | 56134775 | 3.51609 | 4.15966 | 1.21755 | Distal Intergenic | 19 | 56277518 | 56277645 | 128 | 2 | ENSMUST00000104035.1 | 142870 | Gm22271 | ENSMUSG00000077223.1 |
| chr14 | 107075299 | 107075374 | 5.39353 | 9.34496 | 5.16041 | Distal Intergenic | 14 | 106766030 | 106766156 | 127 | 1 | ENSMUST00000104036.1 | 309270 | Gm23437 | ENSMUSG00000077224.1 |
| chr6 | 143446452 | 143446571 | 3.87738 | 4.91547 | 1.74875 | Distal Intergenic | 6 | 143465269 | 143465397 | 129 | 2 | ENSMUST00000104053.1 | 18826 | Gm23272 | ENSMUSG00000077241.1 |
| chr12 | 34904712 | 34905280 | 4.27838 | 6.00206 | 2.53537 | Distal Intergenic | 12 | 34895024 | 34895155 | 132 | 1 | ENSMUST00000104076.1 | 9689 | Gm24429 | ENSMUSG00000077264.1 |
| chr12 | 34906583 | 34907361 | 3.95149 | 5.0452 | 1.77663 | Distal Intergenic | 12 | 34895024 | 34895155 | 132 | 1 | ENSMUST00000104076.1 | 11560 | Gm24429 | ENSMUSG00000077264.1 |
| chr12 | 34907739 | 34908370 | 3.53398 | 4.18681 | 1.21842 | Distal Intergenic | 12 | 34895024 | 34895155 | 132 | 1 | ENSMUST00000104076.1 | 12716 | Gm24429 | ENSMUSG00000077264.1 |
| chr13 | 36582961 | 36583036 | 3.54458 | 4.29359 | 1.36845 | Distal Intergenic | 13 | 36654405 | 36654536 | 132 | 1 | ENSMUST00000104094.1 | -71369 | Gm26395 | ENSMUSG00000077282.1 |
| chr13 | 36583460 | 36583664 | 3.42176 | 4.02875 | 1.22609 | Distal Intergenic | 13 | 36654405 | 36654536 | 132 | 1 | ENSMUST00000104094.1 | -70741 | Gm26395 | ENSMUSG00000077282.1 |
| chr2 | 157691664 | 157692521 | 3.54971 | 4.22304 | 1.23019 | Distal Intergenic | 2 | 157713668 | 157713801 | 134 | 2 | ENSMUST00000104102.1 | 21280 | Gm25407 | ENSMUSG00000077290.1 |
| chr2 | 157692952 | 157693144 | 3.35019 | 3.85272 | 1.01733 | Distal Intergenic | 2 | 157713668 | 157713801 | 134 | 2 | ENSMUST00000104102.1 | 20657 | Gm25407 | ENSMUSG00000077290.1 |
| chr2 | 157693769 | 157693943 | 3.34044 | 3.87328 | 1.09243 | Distal Intergenic | 2 | 157713668 | 157713801 | 134 | 2 | ENSMUST00000104102.1 | 19858 | Gm25407 | ENSMUSG00000077290.1 |
| chr5 | 53104425 | 53104713 | 3.47218 | 4.07449 | 1.15081 | Downstream (2-3kb) | 5 | 53143160 | 53143238 | 79 | 1 | ENSMUST00000104128.1 | -38447 | n-R5s171 | ENSMUSG00000077316.1 |
| chr17 | 13678566 | 13678748 | 3.54985 | 4.24101 | 1.30335 | Intron (ENSMUST00000135850.2/ENSMUSG00000038347.9, intron 1 of 4) | 17 | 13675376 | 13675482 | 107 | 1 | ENSMUST00000104161.1 | 3191 | Gm24867 | ENSMUSG00000077349.1 |
| chr6 | 50503794 | 50503988 | 3.27548 | 3.74041 | 1.01733 | Distal Intergenic | 6 | 50539904 | 50540034 | 131 | 2 | ENSMUST00000104169.1 | 36046 | Gm26289 | ENSMUSG00000077357.1 |
| chr6 | 50510513 | 50510679 | 3.41089 | 3.95644 | 1.06004 | Distal Intergenic | 6 | 50539904 | 50540034 | 131 | 2 | ENSMUST00000104169.1 | 29355 | Gm26289 | ENSMUSG00000077357.1 |
| chr6 | 50530924 | 50531020 | 3.70027 | 4.65796 | 1.52964 | Distal Intergenic | 6 | 50539904 | 50540034 | 131 | 2 | ENSMUST00000104169.1 | 9014 | Gm26289 | ENSMUSG00000077357.1 |
| chr6 | 71013202 | 71013624 | 3.98889 | 5.19337 | 1.90895 | Distal Intergenic | 6 | 71039579 | 71039733 | 155 | 2 | ENSMUST00000104181.1 | 26109 | Gm23485 | ENSMUSG00000077369.1 |
| chr6 | 71021504 | 71021846 | 3.52357 | 4.16318 | 1.18974 | Distal Intergenic | 6 | 71039579 | 71039733 | 155 | 2 | ENSMUST00000104181.1 | 17887 | Gm23485 | ENSMUSG00000077369.1 |
| chr6 | 71048545 | 71048624 | 3.55572 | 4.35537 | 1.34736 | Distal Intergenic | 6 | 71039579 | 71039733 | 155 | 2 | ENSMUST00000104181.1 | -8813 | Gm23485 | ENSMUSG00000077369.1 |
| chr6 | 71049491 | 71050106 | 3.42648 | 4.13848 | 1.21978 | Distal Intergenic | 6 | 71039579 | 71039733 | 155 | 2 | ENSMUST00000104181.1 | -9759 | Gm23485 | ENSMUSG00000077369.1 |
| chr4 | 130861171 | 130862147 | 3.53753 | 4.25168 | 1.25351 | Distal Intergenic | 4 | 130857751 | 130857880 | 130 | 2 | ENSMUST00000104193.1 | -3292 | Gm22660 | ENSMUSG00000077381.1 |
| chr4 | 130862597 | 130862963 | 3.93795 | 4.95138 | 1.71045 | Distal Intergenic | 4 | 130857751 | 130857880 | 130 | 2 | ENSMUST00000104193.1 | -4718 | Gm22660 | ENSMUSG00000077381.1 |
| chr4 | 130865508 | 130865730 | 4.09389 | 5.25655 | 1.93279 | Distal Intergenic | 4 | 130857751 | 130857880 | 130 | 2 | ENSMUST00000104193.1 | -7629 | Gm22660 | ENSMUSG00000077381.1 |
| chr4 | 130866510 | 130866851 | 3.61551 | 4.34746 | 1.34359 | Distal Intergenic | 4 | 130857751 | 130857880 | 130 | 2 | ENSMUST00000104193.1 | -8631 | Gm22660 | ENSMUSG00000077381.1 |
| chr4 | 130876122 | 130876765 | 3.55057 | 4.21234 | 1.21687 | Distal Intergenic | 4 | 130857751 | 130857880 | 130 | 2 | ENSMUST00000104193.1 | -18243 | Gm22660 | ENSMUSG00000077381.1 |
| chr9 | 46495249 | 46495448 | 3.87355 | 4.86986 | 1.7302 | Distal Intergenic | 9 | 46562974 | 46563103 | 130 | 1 | ENSMUST00000104195.1 | -67526 | Gm22659 | ENSMUSG00000077383.1 |
| chr12 | 104425171 | 104425418 | 3.85946 | 4.92137 | 1.7492 | Distal Intergenic | 12 | 104419273 | 104419400 | 128 | 2 | ENSMUST00000104210.1 | -5772 | Gm24334 | ENSMUSG00000077398.1 |
| chr12 | 104440570 | 104440779 | 3.54842 | 4.2406 | 1.30597 | Distal Intergenic | 12 | 104419273 | 104419400 | 128 | 2 | ENSMUST00000104210.1 | -21171 | Gm24334 | ENSMUSG00000077398.1 |
| chr14 | 105022477 | 105022628 | 3.6013 | 4.31173 | 1.29873 | Distal Intergenic | 14 | 104971427 | 104971560 | 134 | 1 | ENSMUST00000104237.1 | 51051 | Gm24280 | ENSMUSG00000077425.1 |
| chr3 | 147991391 | 147991777 | 3.31921 | 3.8727 | 1.07854 | Distal Intergenic | 3 | 148354087 | 148354218 | 132 | 2 | ENSMUST00000104253.1 | 362441 | Gm23131 | ENSMUSG00000077441.1 |
| chr3 | 148336777 | 148337123 | 3.45925 | 4.09733 | 1.26762 | Distal Intergenic | 3 | 148354087 | 148354218 | 132 | 2 | ENSMUST00000104253.1 | 17095 | Gm23131 | ENSMUSG00000077441.1 |
| chr1 | 24560496 | 24560713 | 3.98083 | 5.20353 | 1.91266 | Intron (ENSMUST00000115244.3/ENSMUSG00000026141.8, intron 3 of 50) | 1 | 24548841 | 24548977 | 137 | 2 | ENSMUST00000104275.1 | -11520 | Gm26418 | ENSMUSG00000077463.1 |
| chr1 | 24564612 | 24564944 | 3.42377 | 3.9944 | 1.12593 | Intron (ENSMUST00000115244.3/ENSMUSG00000026141.8, intron 2 of 50) | 1 | 24548841 | 24548977 | 137 | 2 | ENSMUST00000104275.1 | -15636 | Gm26418 | ENSMUSG00000077463.1 |
| chr4 | 153758615 | 153758802 | 3.40615 | 4.08245 | 1.23003 | Distal Intergenic | 4 | 153876384 | 153876515 | 132 | 1 | ENSMUST00000104331.1 | -117582 | Gm25880 | ENSMUSG00000077519.1 |
| chr4 | 153791519 | 153791785 | 3.35971 | 3.86708 | 1.01733 | Distal Intergenic | 4 | 153876384 | 153876515 | 132 | 1 | ENSMUST00000104331.1 | -84599 | Gm25880 | ENSMUSG00000077519.1 |
| chr4 | 153792716 | 153793125 | 3.36732 | 3.87856 | 1.01733 | Distal Intergenic | 4 | 153876384 | 153876515 | 132 | 1 | ENSMUST00000104331.1 | -83259 | Gm25880 | ENSMUSG00000077519.1 |
| chr4 | 153806820 | 153807384 | 3.36479 | 3.92453 | 1.07882 | Distal Intergenic | 4 | 153876384 | 153876515 | 132 | 1 | ENSMUST00000104331.1 | -69000 | Gm25880 | ENSMUSG00000077519.1 |
| chr4 | 153818946 | 153819035 | 3.32159 | 4.14382 | 1.23585 | Distal Intergenic | 4 | 153876384 | 153876515 | 132 | 1 | ENSMUST00000104331.1 | -57349 | Gm25880 | ENSMUSG00000077519.1 |
| chr4 | 153844952 | 153845443 | 3.36528 | 3.93265 | 1.13586 | Distal Intergenic | 4 | 153876384 | 153876515 | 132 | 1 | ENSMUST00000104331.1 | -30941 | Gm25880 | ENSMUSG00000077519.1 |
| chr4 | 153846272 | 153846484 | 3.32919 | 3.87708 | 1.08459 | Distal Intergenic | 4 | 153876384 | 153876515 | 132 | 1 | ENSMUST00000104331.1 | -29900 | Gm25880 | ENSMUSG00000077519.1 |
| chr4 | 153862806 | 153862984 | 3.34826 | 3.84982 | 1.01733 | Distal Intergenic | 4 | 153876384 | 153876515 | 132 | 1 | ENSMUST00000104331.1 | -13400 | Gm25880 | ENSMUSG00000077519.1 |
| chr4 | 153871939 | 153872016 | 4.14268 | 5.41141 | 2.05912 | Distal Intergenic | 4 | 153876384 | 153876515 | 132 | 1 | ENSMUST00000104331.1 | -4368 | Gm25880 | ENSMUSG00000077519.1 |
| chr11 | 34845389 | 34845736 | 3.87777 | 4.89976 | 1.74325 | Distal Intergenic | 11 | 34842228 | 34842354 | 127 | 2 | ENSMUST00000104355.1 | -3036 | Gm22022 | ENSMUSG00000077543.1 |
| chr5 | 60904276 | 60904352 | 5.5066 | 8.19356 | 4.14478 | Distal Intergenic | 5 | 60717317 | 60717439 | 123 | 2 | ENSMUST00000104367.1 | -186838 | Gm23708 | ENSMUSG00000077555.1 |
| chr3 | 33904262 | 33904429 | 3.27402 | 3.75505 | 1.06927 | Distal Intergenic | 3 | 33959069 | 33959200 | 132 | 2 | ENSMUST00000104371.1 | 54771 | Gm23175 | ENSMUSG00000077559.1 |
| chr3 | 33954250 | 33954380 | 3.24929 | 3.70108 | 1.01733 | Distal Intergenic | 3 | 33959069 | 33959200 | 132 | 2 | ENSMUST00000104371.1 | 4820 | Gm23175 | ENSMUSG00000077559.1 |
| chr3 | 33955483 | 33956165 | 3.70178 | 4.56833 | 1.51537 | Downstream (2-3kb) | 3 | 33959069 | 33959200 | 132 | 2 | ENSMUST00000104371.1 | 3035 | Gm23175 | ENSMUSG00000077559.1 |
| chr11 | 79645207 | 79645520 | 3.33899 | 3.8633 | 1.10405 | Intron (ENSMUST00000017783.7/ENSMUSG00000017639.8, intron 3 of 14) | 11 | 79639021 | 79639128 | 108 | 2 | ENSMUST00000104386.1 | -6080 | Gm24887 | ENSMUSG00000077574.1 |
| chr3 | 157200766 | 157200906 | 3.30785 | 3.83821 | 1.10596 | Intron (ENSMUST00000074015.5/ENSMUSG00000040037.8, intron 6 of 6) | 3 | 157210929 | 157211060 | 132 | 1 | ENSMUST00000104393.1 | -10023 | Gm24373 | ENSMUSG00000077581.1 |
| chr3 | 157209120 | 157209196 | 3.85526 | 4.85548 | 1.69849 | Intron (ENSMUST00000074015.5/ENSMUSG00000040037.8, intron 6 of 6) | 3 | 157210929 | 157211060 | 132 | 1 | ENSMUST00000104393.1 | -1733 | Gm24373 | ENSMUSG00000077581.1 |
| chr3 | 157217556 | 157217631 | 3.39988 | 3.94653 | 1.08407 | Intron (ENSMUST00000074015.5/ENSMUSG00000040037.8, intron 6 of 6) | 3 | 157210929 | 157211060 | 132 | 1 | ENSMUST00000104393.1 | 6628 | Gm24373 | ENSMUSG00000077581.1 |
| chr4 | 44422830 | 44423030 | 3.73086 | 4.5386 | 1.40666 | Distal Intergenic | 4 | 44412469 | 44412595 | 127 | 2 | ENSMUST00000104399.1 | -10236 | Gm24376 | ENSMUSG00000077587.1 |
| chr4 | 44424084 | 44424533 | 3.45843 | 4.29679 | 1.3126 | Distal Intergenic | 4 | 44412469 | 44412595 | 127 | 2 | ENSMUST00000104399.1 | -11490 | Gm24376 | ENSMUSG00000077587.1 |
| chr4 | 44425068 | 44426369 | 3.7817 | 4.90278 | 1.7323 | Distal Intergenic | 4 | 44412469 | 44412595 | 127 | 2 | ENSMUST00000104399.1 | -12474 | Gm24376 | ENSMUSG00000077587.1 |
| chr4 | 44427554 | 44427832 | 3.60928 | 4.32524 | 1.29996 | Distal Intergenic | 4 | 44412469 | 44412595 | 127 | 2 | ENSMUST00000104399.1 | -14960 | Gm24376 | ENSMUSG00000077587.1 |
| chr18 | 81645270 | 81645433 | 3.5445 | 4.20498 | 1.22471 | Distal Intergenic | 18 | 81981069 | 81981196 | 128 | 1 | ENSMUST00000104411.1 | -335636 | Gm26012 | ENSMUSG00000077599.1 |
| chr18 | 81646199 | 81647098 | 3.77146 | 4.65457 | 1.54256 | Distal Intergenic | 18 | 81981069 | 81981196 | 128 | 1 | ENSMUST00000104411.1 | -333971 | Gm26012 | ENSMUSG00000077599.1 |
| chr18 | 81647784 | 81648133 | 3.32287 | 3.81167 | 1.01733 | Distal Intergenic | 18 | 81981069 | 81981196 | 128 | 1 | ENSMUST00000104411.1 | -332936 | Gm26012 | ENSMUSG00000077599.1 |
| chr18 | 81648867 | 81649050 | 3.62841 | 4.71711 | 1.60521 | Distal Intergenic | 18 | 81981069 | 81981196 | 128 | 1 | ENSMUST00000104411.1 | -332019 | Gm26012 | ENSMUSG00000077599.1 |
| chr18 | 81651560 | 81652013 | 3.4261 | 4.02335 | 1.17912 | Distal Intergenic | 18 | 81981069 | 81981196 | 128 | 1 | ENSMUST00000104411.1 | -329056 | Gm26012 | ENSMUSG00000077599.1 |
| chr18 | 81703550 | 81703729 | 3.31444 | 3.80899 | 1.05194 | Distal Intergenic | 18 | 81981069 | 81981196 | 128 | 1 | ENSMUST00000104411.1 | -277340 | Gm26012 | ENSMUSG00000077599.1 |
| chr18 | 81898170 | 81899244 | 3.42089 | 3.97776 | 1.08216 | Distal Intergenic | 18 | 81981069 | 81981196 | 128 | 1 | ENSMUST00000104411.1 | -81825 | Gm26012 | ENSMUSG00000077599.1 |
| chr18 | 81903029 | 81903110 | 3.83991 | 4.77004 | 1.61754 | Distal Intergenic | 18 | 81981069 | 81981196 | 128 | 1 | ENSMUST00000104411.1 | -77959 | Gm26012 | ENSMUSG00000077599.1 |
| chr18 | 81916395 | 81916988 | 3.79927 | 4.95208 | 1.75527 | Distal Intergenic | 18 | 81981069 | 81981196 | 128 | 1 | ENSMUST00000104411.1 | -64081 | Gm26012 | ENSMUSG00000077599.1 |
| chr18 | 81917626 | 81917937 | 3.89646 | 5.04152 | 1.78806 | Distal Intergenic | 18 | 81981069 | 81981196 | 128 | 1 | ENSMUST00000104411.1 | -63132 | Gm26012 | ENSMUSG00000077599.1 |
| chr11 | 77036382 | 77036611 | 3.43213 | 3.99212 | 1.0728 | Distal Intergenic | 11 | 77052947 | 77053067 | 121 | 1 | ENSMUST00000104451.1 | -16336 | Gm22772 | ENSMUSG00000077639.1 |
| chr11 | 77037030 | 77037532 | 3.37398 | 3.88861 | 1.01733 | Distal Intergenic | 11 | 77052947 | 77053067 | 121 | 1 | ENSMUST00000104451.1 | -15415 | Gm22772 | ENSMUSG00000077639.1 |
| chr12 | 106172182 | 106172531 | 3.63983 | 4.37306 | 1.31047 | Distal Intergenic | 12 | 106221075 | 106221206 | 132 | 1 | ENSMUST00000104452.1 | -48544 | Gm24986 | ENSMUSG00000077640.1 |
| chr12 | 106172945 | 106173333 | 4.62892 | 6.40587 | 2.82623 | Distal Intergenic | 12 | 106221075 | 106221206 | 132 | 1 | ENSMUST00000104452.1 | -47742 | Gm24986 | ENSMUSG00000077640.1 |
| chr12 | 106174173 | 106174669 | 3.60224 | 4.44166 | 1.41027 | Distal Intergenic | 12 | 106221075 | 106221206 | 132 | 1 | ENSMUST00000104452.1 | -46406 | Gm24986 | ENSMUSG00000077640.1 |
| chr12 | 106175162 | 106175370 | 3.40432 | 4.38565 | 1.34933 | Distal Intergenic | 12 | 106221075 | 106221206 | 132 | 1 | ENSMUST00000104452.1 | -45705 | Gm24986 | ENSMUSG00000077640.1 |
| chr12 | 106227336 | 106227754 | 4.07133 | 5.336 | 1.99804 | Distal Intergenic | 12 | 106221075 | 106221206 | 132 | 1 | ENSMUST00000104452.1 | 6262 | Gm24986 | ENSMUSG00000077640.1 |
| chr14 | 67570949 | 67571191 | 3.5824 | 4.51727 | 1.46452 | Distal Intergenic | 14 | 67566850 | 67566981 | 132 | 1 | ENSMUST00000104459.1 | 4100 | Gm24981 | ENSMUSG00000077647.1 |
| chr14 | 67606641 | 67606990 | 3.41621 | 4.0058 | 1.18139 | Distal Intergenic | 14 | 67566850 | 67566981 | 132 | 1 | ENSMUST00000104459.1 | 39792 | Gm24981 | ENSMUSG00000077647.1 |
| chr14 | 67611232 | 67611442 | 3.62645 | 4.38865 | 1.38322 | Distal Intergenic | 14 | 67566850 | 67566981 | 132 | 1 | ENSMUST00000104459.1 | 44383 | Gm24981 | ENSMUSG00000077647.1 |
| chr14 | 67611913 | 67612021 | 3.75535 | 4.81446 | 1.62452 | Distal Intergenic | 14 | 67566850 | 67566981 | 132 | 1 | ENSMUST00000104459.1 | 45064 | Gm24981 | ENSMUSG00000077647.1 |
| chr14 | 67613276 | 67613594 | 3.82882 | 4.73046 | 1.54893 | Distal Intergenic | 14 | 67566850 | 67566981 | 132 | 1 | ENSMUST00000104459.1 | 46427 | Gm24981 | ENSMUSG00000077647.1 |
| chr14 | 67614682 | 67614974 | 4.31406 | 5.74713 | 2.31311 | Distal Intergenic | 14 | 67566850 | 67566981 | 132 | 1 | ENSMUST00000104459.1 | 47833 | Gm24981 | ENSMUSG00000077647.1 |
| chr14 | 67615898 | 67616102 | 3.35395 | 3.85838 | 1.01733 | Distal Intergenic | 14 | 67566850 | 67566981 | 132 | 1 | ENSMUST00000104459.1 | 49049 | Gm24981 | ENSMUSG00000077647.1 |
| chr14 | 118840168 | 118840263 | 3.57733 | 4.28899 | 1.33125 | Intron (ENSMUST00000047761.7/ENSMUSG00000022132.10, intron 1 of 4) | 14 | 118826909 | 118827031 | 123 | 2 | ENSMUST00000104473.1 | -13138 | n-R5s51 | ENSMUSG00000077661.1 |
| chr4 | 56707588 | 56707942 | 3.40068 | 3.95161 | 1.09792 | Distal Intergenic | 4 | 56678904 | 56679035 | 132 | 2 | ENSMUST00000104475.1 | -28554 | Gm26144 | ENSMUSG00000077663.1 |
| chr6 | 56652069 | 56652580 | 3.75848 | 4.68539 | 1.62436 | Distal Intergenic | 6 | 56600818 | 56600949 | 132 | 2 | ENSMUST00000104494.1 | -51121 | Gm22493 | ENSMUSG00000077682.1 |
| chr5 | 53319969 | 53320122 | 3.21485 | 3.6495 | 1.01281 | Distal Intergenic | 5 | 53346259 | 53346372 | 114 | 2 | ENSMUST00000104508.1 | 26250 | n-R5s172 | ENSMUSG00000077696.1 |
| chr17 | 48638915 | 48639268 | 3.56265 | 4.26915 | 1.3466 | Distal Intergenic | 17 | 48617823 | 48617954 | 132 | 1 | ENSMUST00000104520.1 | 21093 | Gm24071 | ENSMUSG00000077708.1 |
| chr17 | 48647546 | 48647740 | 3.36927 | 3.89907 | 1.07962 | Distal Intergenic | 17 | 48617823 | 48617954 | 132 | 1 | ENSMUST00000104520.1 | 29724 | Gm24071 | ENSMUSG00000077708.1 |
| chr14 | 104711940 | 104712188 | 4.05928 | 8.20009 | 4.2587 | Distal Intergenic | 14 | 104628944 | 104629060 | 117 | 2 | ENSMUST00000104531.1 | -82881 | Gm25724 | ENSMUSG00000077719.1 |
| chr5 | 147195604 | 147195850 | 3.35578 | 3.91942 | 1.1468 | Distal Intergenic | 5 | 147200124 | 147200253 | 130 | 2 | ENSMUST00000104547.1 | 4403 | Gm24556 | ENSMUSG00000077735.1 |
| chr5 | 147198224 | 147198625 | 3.6477 | 4.43241 | 1.39981 | Downstream (1-2kb) | 5 | 147200124 | 147200253 | 130 | 2 | ENSMUST00000104547.1 | 1628 | Gm24556 | ENSMUSG00000077735.1 |
| chr5 | 147222007 | 147222088 | 4.45038 | 5.98465 | 2.46217 | Distal Intergenic | 5 | 147200124 | 147200253 | 130 | 2 | ENSMUST00000104547.1 | -21755 | Gm24556 | ENSMUSG00000077735.1 |
| chr5 | 147223581 | 147224110 | 3.91742 | 4.93073 | 1.71149 | Distal Intergenic | 5 | 147200124 | 147200253 | 130 | 2 | ENSMUST00000104547.1 | -23329 | Gm24556 | ENSMUSG00000077735.1 |
| chr5 | 147226219 | 147226643 | 3.69202 | 4.47774 | 1.38999 | Downstream (1-2kb) | 5 | 147200124 | 147200253 | 130 | 2 | ENSMUST00000104547.1 | -25967 | Gm24556 | ENSMUSG00000077735.1 |
| chr5 | 147227132 | 147227424 | 3.49741 | 4.21243 | 1.31253 | Downstream (1-2kb) | 5 | 147200124 | 147200253 | 130 | 2 | ENSMUST00000104547.1 | -26880 | Gm24556 | ENSMUSG00000077735.1 |
| chr12 | 33594110 | 33594324 | 3.44829 | 4.03712 | 1.14651 | Distal Intergenic | 12 | 33452712 | 33452843 | 132 | 2 | ENSMUST00000104562.1 | -141268 | Gm23529 | ENSMUSG00000077750.1 |
| chr2 | 168831097 | 168831597 | 3.62909 | 4.36751 | 1.33453 | Distal Intergenic | 2 | 168882040 | 168882168 | 129 | 2 | ENSMUST00000104573.1 | 50571 | Gm25214 | ENSMUSG00000077761.1 |
| chr2 | 168837253 | 168837526 | 3.49395 | 4.13053 | 1.22156 | Distal Intergenic | 2 | 168882040 | 168882168 | 129 | 2 | ENSMUST00000104573.1 | 44642 | Gm25214 | ENSMUSG00000077761.1 |
| chr2 | 168837850 | 168838754 | 3.85491 | 4.78748 | 1.59206 | Distal Intergenic | 2 | 168882040 | 168882168 | 129 | 2 | ENSMUST00000104573.1 | 43414 | Gm25214 | ENSMUSG00000077761.1 |
| chr2 | 168839308 | 168840526 | 3.80958 | 4.74815 | 1.58458 | Distal Intergenic | 2 | 168882040 | 168882168 | 129 | 2 | ENSMUST00000104573.1 | 41642 | Gm25214 | ENSMUSG00000077761.1 |
| chr2 | 168840901 | 168843072 | 4.03678 | 5.27002 | 1.99662 | Distal Intergenic | 2 | 168882040 | 168882168 | 129 | 2 | ENSMUST00000104573.1 | 39096 | Gm25214 | ENSMUSG00000077761.1 |
| chr2 | 168843540 | 168843676 | 3.65641 | 4.83333 | 1.64704 | Distal Intergenic | 2 | 168882040 | 168882168 | 129 | 2 | ENSMUST00000104573.1 | 38492 | Gm25214 | ENSMUSG00000077761.1 |
| chr2 | 168844787 | 168845072 | 3.30085 | 3.77849 | 1.01733 | Distal Intergenic | 2 | 168882040 | 168882168 | 129 | 2 | ENSMUST00000104573.1 | 37096 | Gm25214 | ENSMUSG00000077761.1 |
| chr2 | 168845519 | 168845724 | 3.39139 | 3.95202 | 1.12587 | Distal Intergenic | 2 | 168882040 | 168882168 | 129 | 2 | ENSMUST00000104573.1 | 36444 | Gm25214 | ENSMUSG00000077761.1 |
| chr2 | 168847815 | 168848342 | 3.34288 | 3.84169 | 1.01733 | Distal Intergenic | 2 | 168882040 | 168882168 | 129 | 2 | ENSMUST00000104573.1 | 33826 | Gm25214 | ENSMUSG00000077761.1 |
| chr9 | 52286040 | 52286232 | 3.57465 | 4.25493 | 1.241 | Distal Intergenic | 9 | 52291199 | 52291331 | 133 | 1 | ENSMUST00000104608.1 | -4967 | Gm25562 | ENSMUSG00000077796.1 |
| chr13 | 56154202 | 56154397 | 3.62407 | 4.37981 | 1.40012 | Distal Intergenic | 13 | 56167600 | 56167722 | 123 | 1 | ENSMUST00000104611.1 | -13203 | Gm25148 | ENSMUSG00000077799.1 |
| chr13 | 56156489 | 56156775 | 3.4404 | 4.03472 | 1.16847 | Distal Intergenic | 13 | 56167600 | 56167722 | 123 | 1 | ENSMUST00000104611.1 | -10825 | Gm25148 | ENSMUSG00000077799.1 |
| chr13 | 56159946 | 56160662 | 3.45229 | 4.06833 | 1.22126 | Distal Intergenic | 13 | 56167600 | 56167722 | 123 | 1 | ENSMUST00000104611.1 | -6938 | Gm25148 | ENSMUSG00000077799.1 |

| | | | | | | | | | | | | | | | |
|---|---|---|---|---|---|---|---|---|---|---|---|---|---|---|---|
| chr13 | 56161056 | 56161160 | 3.48009 | 4.09759 | 1.18866 | Distal Intergenic | 13 | 56167600 | 56167722 | 123 | 1 | ENSMUST00000104611.1 | -6440 | Gm25148 | ENSMUSG00000077799.1 |
| chr5 | 128445056 | 128445200 | 3.37062 | 3.88354 | 1.01733 | Distal Intergenic | 5 | 128435255 | 128435347 | 93 | 1 | ENSMUST00000104628.2 | 9802 | Gm22735 | ENSMUSG00000077816.2 |
| chr5 | 128470072 | 128470268 | 3.38193 | 3.9006 | 1.01733 | Distal Intergenic | 5 | 128435255 | 128435347 | 93 | 1 | ENSMUST00000104628.2 | 34818 | Gm22735 | ENSMUSG00000077816.2 |
| chr5 | 128482903 | 128483651 | 3.44151 | 4.02338 | 1.12843 | Distal Intergenic | 5 | 128435255 | 128435347 | 93 | 1 | ENSMUST00000104628.2 | 47649 | Gm22735 | ENSMUSG00000077816.2 |
| chr5 | 128483970 | 128484559 | 3.53977 | 4.19829 | 1.228 | Distal Intergenic | 5 | 128435255 | 128435347 | 93 | 1 | ENSMUST00000104628.2 | 48716 | Gm22735 | ENSMUSG00000077816.2 |
| chr16 | 31545352 | 31545623 | 3.34947 | 3.9094 | 1.13681 | Distal Intergenic | 16 | 31573894 | 31573965 | 72 | 2 | ENSMUST00000104649.2 | 28342 | Gm27939 | ENSMUSG00000077839.2 |
| chr16 | 31546020 | 31546286 | 3.38283 | 3.91557 | 1.06328 | Distal Intergenic | 16 | 31573894 | 31573965 | 72 | 2 | ENSMUST00000104649.2 | 27679 | Gm27939 | ENSMUSG00000077839.2 |
| chr12 | 17993511 | 17993961 | 3.5839 | 4.31134 | 1.36995 | Distal Intergenic | 12 | 17949458 | 17949548 | 91 | 2 | ENSMUST00000104684.2 | -43964 | Gm23913 | ENSMUSG00000077876.2 |
| chr12 | 109615070 | 109615508 | 3.31626 | 3.84854 | 1.10456 | Intron (ENSMUST00000183215.2/ENSMUSG00000097451.4, intron 2 of 7) | 12 | 109618114 | 109618200 | 87 | 1 | ENSMUST00000104687.1 | -2606 | Gm23912 | ENSMUSG00000077879.1 |
| chr13 | 57985271 | 57985418 | 3.42033 | 4.01198 | 1.19514 | Distal Intergenic | 13 | 58023125 | 58023200 | 76 | 2 | ENSMUST00000104769.1 | 37782 | Mir874 | ENSMUSG00000077962.1 |
| chr13 | 57997132 | 57997256 | 3.33272 | 4.19388 | 1.23365 | Distal Intergenic | 13 | 58023125 | 58023200 | 76 | 2 | ENSMUST00000104769.1 | 25944 | Mir874 | ENSMUSG00000077962.1 |
| chr18 | 83012044 | 83012363 | 3.29794 | 3.79608 | 1.04605 | Distal Intergenic | 18 | 82976323 | 82976414 | 92 | 1 | ENSMUST00000104783.1 | 35722 | Gm26258 | ENSMUSG00000077976.1 |
| chr13 | 113037331 | 113037406 | 3.75019 | 4.61171 | 1.48858 | Promoter | 13 | 113037419 | 113037498 | 80 | 1 | ENSMUST00000104833.1 | -13 | Mir449b | ENSMUSG00000078027.1 |
| chr2 | 118700887 | 118701366 | 4.03168 | 5.25513 | 1.9513 | 3' UTR | 2 | 118699103 | 118703963 | 4861 | 2 | ENSMUST00000104937.1 | 2597 | Ankrd63 | ENSMUSG00000078137.1 |
| chr2 | 118702192 | 118705040 | 3.73388 | 4.98252 | 1.78027 | Promoter | 2 | 118699103 | 118703963 | 4861 | 2 | ENSMUST00000104937.1 | 0 | Ankrd63 | ENSMUSG00000078137.1 |
| chr3 | 143100048 | 143101539 | 4.10446 | 5.45959 | 2.12562 | Distal Intergenic | 3 | 142962561 | 142963792 | 1232 | 1 | ENSMUST00000181051.1 | 137488 | Gm2574 | ENSMUSG00000078162.5 |
| chr6 | 53697167 | 53697252 | 3.2565 | 3.71188 | 1.01733 | Distal Intergenic | 6 | 53819975 | 53820040 | 66 | 1 | ENSMUST00000104970.2 | -122723 | Gm16499 | ENSMUSG00000078169.2 |
| chr6 | 53697961 | 53698166 | 3.77221 | 4.66843 | 1.58666 | Distal Intergenic | 6 | 53819975 | 53820040 | 66 | 1 | ENSMUST00000104970.2 | -121809 | Gm16499 | ENSMUSG00000078169.2 |
| chr6 | 53698699 | 53698910 | 3.33731 | 3.8374 | 1.03176 | Distal Intergenic | 6 | 53819975 | 53820040 | 66 | 1 | ENSMUST00000104970.2 | -121065 | Gm16499 | ENSMUSG00000078169.2 |
| chr6 | 53702975 | 53703330 | 3.63568 | 4.37295 | 1.32741 | Distal Intergenic | 6 | 53819975 | 53820040 | 66 | 1 | ENSMUST00000104970.2 | -116645 | Gm16499 | ENSMUSG00000078169.2 |
| chr6 | 53721465 | 53721733 | 3.71888 | 4.61591 | 1.56588 | Distal Intergenic | 6 | 53819975 | 53820040 | 66 | 1 | ENSMUST00000104970.2 | -98242 | Gm16499 | ENSMUSG00000078169.2 |
| chr6 | 53722173 | 53722961 | 3.58083 | 4.4178 | 1.42037 | Distal Intergenic | 6 | 53819975 | 53820040 | 66 | 1 | ENSMUST00000104970.2 | -97014 | Gm16499 | ENSMUSG00000078169.2 |
| chr6 | 53759946 | 53760088 | 3.61414 | 4.50329 | 1.45415 | Distal Intergenic | 6 | 53819975 | 53820040 | 66 | 1 | ENSMUST00000104970.2 | -59887 | Gm16499 | ENSMUSG00000078169.2 |
| chr6 | 53779822 | 53780011 | 3.87894 | 4.98095 | 1.75003 | Distal Intergenic | 6 | 53819975 | 53820040 | 66 | 1 | ENSMUST00000104970.2 | -39964 | Gm16499 | ENSMUSG00000078169.2 |
| chr6 | 53795328 | 53795452 | 4.44733 | 6.4235 | 2.78894 | Distal Intergenic | 6 | 53819975 | 53820040 | 66 | 1 | ENSMUST00000104970.2 | -24523 | Gm16499 | ENSMUSG00000078169.2 |
| chr6 | 53806219 | 53806465 | 3.64835 | 4.44627 | 1.41774 | Distal Intergenic | 6 | 53819975 | 53820040 | 66 | 1 | ENSMUST00000104970.2 | -13510 | Gm16499 | ENSMUSG00000078169.2 |
| chr6 | 53806814 | 53806967 | 3.64766 | 4.41011 | 1.38888 | Distal Intergenic | 6 | 53819975 | 53820040 | 66 | 1 | ENSMUST00000104970.2 | -13008 | Gm16499 | ENSMUSG00000078169.2 |
| chr6 | 53818053 | 53819271 | 3.79466 | 5.11769 | 1.85708 | Promoter | 6 | 53819975 | 53820040 | 66 | 1 | ENSMUST00000104970.2 | -704 | Gm16499 | ENSMUSG00000078169.2 |
| chr2 | 25163362 | 25163488 | 3.30183 | 3.78011 | 1.01733 | Distal Intergenic | 2 | 25180758 | 25183339 | 2582 | 1 | ENSMUST00000104999.3 | -17270 | Nrarp | ENSMUSG00000078202.3 |
| chr2 | 25179785 | 25179941 | 3.36986 | 3.88238 | 1.01733 | Promoter | 2 | 25180758 | 25183339 | 2582 | 1 | ENSMUST00000104999.3 | -817 | Nrarp | ENSMUSG00000078202.3 |
| chr2 | 25180260 | 25180624 | 3.63487 | 4.37189 | 1.32467 | Promoter | 2 | 25180758 | 25183339 | 2582 | 1 | ENSMUST00000104999.3 | -134 | Nrarp | ENSMUSG00000078202.3 |
| chr2 | 25180947 | 25182159 | 3.86756 | 4.88559 | 1.7067 | Promoter | 2 | 25180758 | 25183339 | 2582 | 1 | ENSMUST00000104999.3 | 190 | Nrarp | ENSMUSG00000078202.3 |
| chr4 | 139978875 | 139979079 | 3.27691 | 3.7425 | 1.01733 | Distal Intergenic | 4 | 139960220 | 139968026 | 7807 | 2 | ENSMUST00000105031.3 | -10850 | Klhdc7a | ENSMUSG00000078234.6 |
| chr4 | 139993682 | 139993778 | 4.01344 | 5.10599 | 1.83542 | Distal Intergenic | 4 | 139960220 | 139968026 | 7807 | 2 | ENSMUST00000105031.3 | -25657 | Klhdc7a | ENSMUSG00000078234.6 |
| chr4 | 140003832 | 140003961 | 3.76461 | 4.60562 | 1.45486 | Distal Intergenic | 4 | 139960220 | 139968026 | 7807 | 2 | ENSMUST00000105031.3 | -35807 | Klhdc7a | ENSMUSG00000078234.6 |
| chr4 | 140004294 | 140004910 | 3.67092 | 4.46133 | 1.39982 | Distal Intergenic | 4 | 139960220 | 139968026 | 7807 | 2 | ENSMUST00000105031.3 | -36269 | Klhdc7a | ENSMUSG00000078234.6 |
| chr4 | 140036911 | 140037127 | 3.85816 | 4.79976 | 1.60256 | Intron (ENSMUST00000039331.8/ENSMUSG00000040972.8, intron 5 of 9) | 4 | 139960220 | 139968026 | 7807 | 2 | ENSMUST00000105031.3 | -68886 | Klhdc7a | ENSMUSG00000078234.6 |
| chr4 | 140037653 | 140038144 | 3.43017 | 4.05699 | 1.16162 | Intron (ENSMUST00000039331.8/ENSMUSG00000040972.8, intron 4 of 9) | 4 | 139960220 | 139968026 | 7807 | 2 | ENSMUST00000105031.3 | -69628 | Klhdc7a | ENSMUSG00000078234.6 |
| chr4 | 140038844 | 140039158 | 3.55553 | 4.21823 | 1.21327 | Intron (ENSMUST00000039331.8/ENSMUSG00000040972.8, intron 4 of 9) | 4 | 139960220 | 139968026 | 7807 | 2 | ENSMUST00000105031.3 | -70819 | Klhdc7a | ENSMUSG00000078234.6 |
| chr4 | 140039708 | 140040104 | 3.90948 | 4.95846 | 1.73572 | Intron (ENSMUST00000039331.8/ENSMUSG00000040972.8, intron 4 of 9) | 4 | 139960220 | 139968026 | 7807 | 2 | ENSMUST00000105031.3 | -71683 | Klhdc7a | ENSMUSG00000078234.6 |
| chr4 | 140040540 | 140040863 | 3.43246 | 4.00556 | 1.11903 | Intron (ENSMUST00000039331.8/ENSMUSG00000040972.8, intron 4 of 9) | 4 | 139960220 | 139968026 | 7807 | 2 | ENSMUST00000105031.3 | -72515 | Klhdc7a | ENSMUSG00000078234.6 |
| chr4 | 138396706 | 138396851 | 3.67642 | 4.56922 | 1.50093 | Promoter | 4 | 138394092 | 138396528 | 2437 | 2 | ENSMUST00000105032.3 | -179 | Fam43b | ENSMUSG00000078235.3 |
| chr4 | 138397506 | 138398026 | 3.92152 | 5.06076 | 1.83937 | Promoter | 4 | 138394092 | 138396528 | 2437 | 2 | ENSMUST00000105032.3 | -979 | Fam43b | ENSMUSG00000078235.3 |
| chr4 | 138398337 | 138398472 | 3.91815 | 4.90982 | 1.69102 | Distal Intergenic | 4 | 138394092 | 138396528 | 2437 | 2 | ENSMUST00000105032.3 | -1810 | Fam43b | ENSMUSG00000078235.3 |
| chr11 | 99993817 | 99994008 | 3.45589 | 4.12483 | 1.25738 | Promoter | 11 | 99993234 | 99993994 | 761 | 2 | ENSMUST00000105049.1 | 0 | Krtap17-1 | ENSMUSG00000078252.1 |
| chr11 | 99804128 | 99804350 | 3.49562 | 4.69588 | 1.58087 | Promoter | 11 | 99803557 | 99804446 | 890 | 2 | ENSMUST00000105056.1 | 96 | Gm11554 | ENSMUSG00000078259.3 |
| chr11 | 99798271 | 99798439 | 4.25647 | 6.15731 | 2.55102 | Promoter | 11 | 99798064 | 99798932 | 869 | 2 | ENSMUST00000105057.1 | 493 | Gm11569 | ENSMUSG00000078260.1 |
| chr9 | 52676158 | 52676278 | 3.40636 | 3.97182 | 1.13826 | 3' UTR | 9 | 52673044 | 52679429 | 6386 | 2 | ENSMUST00000098768.2 | 3151 | AI593442 | ENSMUSG00000078307.2 |
| chr9 | 52678443 | 52678749 | 3.48966 | 4.10195 | 1.15407 | Promoter | 9 | 52673044 | 52679429 | 6386 | 2 | ENSMUST00000098768.2 | 680 | AI593442 | ENSMUSG00000078307.2 |
| chr9 | 52679418 | 52679760 | 4.68358 | 6.52834 | 2.93017 | Promoter | 9 | 52673044 | 52679429 | 6386 | 2 | ENSMUST00000098768.2 | 0 | AI593442 | ENSMUSG00000078307.2 |
| chr4 | 3870837 | 3870980 | 3.70855 | 4.53124 | 1.458 | Exon (ENSMUST00000105158.1/ENSMUSG00000078365.5, exon 1 of 1) | 4 | 3870657 | 3872105 | 1449 | 2 | ENSMUST00000105158.1 | 1125 | Mos | ENSMUSG00000078365.5 |
| chr10 | 78749139 | 78749245 | 3.42411 | 4.28526 | 1.32509 | Exon (ENSMUST00000105383.2/ENSMUSG00000078442.2, exon 3 of 7) | 10 | 78746924 | 78753067 | 6144 | 2 | ENSMUST00000105383.2 | 3822 | Ccdc105 | ENSMUSG00000078442.2 |
| chr10 | 78751336 | 78751520 | 3.39044 | 3.95356 | 1.11224 | Intron (ENSMUST00000105383.2/ENSMUSG00000078442.2, intron 1 of 6) | 10 | 78746924 | 78753067 | 6144 | 2 | ENSMUST00000105383.2 | 1547 | Ccdc105 | ENSMUSG00000078442.2 |
| chr10 | 78751850 | 78753208 | 3.96283 | 5.0858 | 1.91892 | Promoter | 10 | 78746924 | 78753067 | 6144 | 2 | ENSMUST00000105383.2 | 0 | Ccdc105 | ENSMUSG00000078442.2 |
| chr10 | 22381399 | 22381817 | 3.51666 | 4.22187 | 1.26244 | Distal Intergenic | 10 | 22361894 | 22374139 | 12246 | 1 | ENSMUST00000095795.4 | 19506 | Raet1d | ENSMUSG00000078452.5 |
| chr10 | 10689219 | 10689508 | 3.31777 | 3.82458 | 1.08689 | 3' UTR | 10 | 10681348 | 10681858 | 511 | 1 | ENSMUST00000105564.1 | 7872 | Gm10944 | ENSMUSG00000078481.1 |
| chr4 | 155409010 | 155409176 | 3.34793 | 3.87941 | 1.11272 | Promoter | 4 | 155409190 | 155415987 | 6798 | 1 | ENSMUST00000129481.1 | -14 | C030017K20Rik | ENSMUSG00000078490.4 |
| chr4 | 131487094 | 131487296 | 3.76297 | 4.72487 | 1.59856 | Promoter | 4 | 131487197 | 131505527 | 18331 | 1 | ENSMUST00000149942.1 | 0 | Gm831 | ENSMUSG00000078531.3 |
| chr4 | 131550715 | 131550855 | 3.38724 | 3.91028 | 1.0232 | Distal Intergenic | 4 | 131487197 | 131505527 | 18331 | 1 | ENSMUST00000149942.1 | 63519 | Gm831 | ENSMUSG00000078531.3 |
| chr4 | 131569920 | 131570206 | 4.08721 | 5.25711 | 1.94686 | Distal Intergenic | 4 | 131487197 | 131505527 | 18331 | 1 | ENSMUST00000149942.1 | 82724 | Gm831 | ENSMUSG00000078531.3 |
| chr4 | 130557921 | 130558124 | 3.34153 | 3.83966 | 1.01733 | Promoter | 4 | 130539063 | 130557988 | 18926 | 2 | ENSMUST00000180498.1 | 0 | Nkain1 | ENSMUSG00000078532.4 |
| chr4 | 130559269 | 130559519 | 3.49223 | 4.13028 | 1.18617 | Intron (ENSMUST00000180884.2/ENSMUSG00000078532.4, intron 1 of 6) | 4 | 130539063 | 130557988 | 18926 | 2 | ENSMUST00000180498.1 | -1282 | Nkain1 | ENSMUSG00000078532.4 |
| chr4 | 130567951 | 130568564 | 3.33244 | 3.84033 | 1.06747 | Intron (ENSMUST00000180884.2/ENSMUSG00000078532.4, intron 1 of 6) | 4 | 130531602 | 130573966 | 42365 | 2 | ENSMUST00000180884.2 | 5402 | Nkain1 | ENSMUSG00000078532.4 |
| chr4 | 130569063 | 130569200 | 3.31871 | 3.80531 | 1.01733 | Intron (ENSMUST00000180884.2/ENSMUSG00000078532.4, intron 1 of 6) | 4 | 130531602 | 130573966 | 42365 | 2 | ENSMUST00000180884.2 | 4766 | Nkain1 | ENSMUSG00000078532.4 |
| chr4 | 130570223 | 130570446 | 3.46499 | 4.07921 | 1.20679 | Intron (ENSMUST00000180884.2/ENSMUSG00000078532.4, intron 1 of 6) | 4 | 130531602 | 130573966 | 42365 | 2 | ENSMUST00000180884.2 | 3520 | Nkain1 | ENSMUSG00000078532.4 |
| chr4 | 130571114 | 130571395 | 3.92244 | 5.04354 | 1.79137 | Intron (ENSMUST00000180884.2/ENSMUSG00000078532.4, intron 1 of 6) | 4 | 130531602 | 130573966 | 42365 | 2 | ENSMUST00000180884.2 | 2571 | Nkain1 | ENSMUSG00000078532.4 |
| chr4 | 130571757 | 130572161 | 3.67352 | 4.45483 | 1.41413 | Intron (ENSMUST00000180884.2/ENSMUSG00000078532.4, intron 1 of 6) | 4 | 130531602 | 130573966 | 42365 | 2 | ENSMUST00000180884.2 | 1805 | Nkain1 | ENSMUSG00000078532.4 |
| chr4 | 130572549 | 130574126 | 3.75224 | 4.68061 | 1.55494 | Promoter | 4 | 130531602 | 130573966 | 42365 | 2 | ENSMUST00000180884.2 | 0 | Nkain1 | ENSMUSG00000078532.4 |
| chr13 | 119954803 | 119954878 | 3.42378 | 3.99369 | 1.12412 | Distal Intergenic | 13 | 119949357 | 119950806 | 1450 | 2 | ENSMUST00000180768.1 | -3998 | B020031M17Rik | ENSMUSG00000078537.3 |
| chr4 | 129491779 | 129491895 | 3.2488 | 3.70034 | 1.01733 | Promoter | 4 | 129491240 | 129491956 | 717 | 1 | ENSMUST00000135698.1 | 540 | Fam229a | ENSMUSG00000078554.2 |
| chr11 | 116725487 | 116725655 | 3.4748 | 4.07107 | 1.12412 | Distal Intergenic | 11 | 116739062 | 116749833 | 10772 | 2 | ENSMUST00000106377.2 | 24178 | Gm11735 | ENSMUSG00000078586.3 |
| chr7 | 123979671 | 123980040 | 3.32059 | 3.8082 | 1.01733 | Intron (ENSMUST00000181984.1/ENSMUSG00000098129.1, intron 1 of 3) | 7 | 123982869 | 124398989 | 416121 | 1 | ENSMUST00000106437.1 | -2829 | Hs3st4 | ENSMUSG00000078591.1 |
| chr7 | 123981950 | 123982412 | 3.9953 | 5.12877 | 1.9067 | Promoter | 7 | 123982869 | 124398989 | 416121 | 1 | ENSMUST00000106437.1 | -457 | Hs3st4 | ENSMUSG00000078591.1 |
| chr7 | 123982780 | 123983723 | 3.73742 | 4.66472 | 1.61932 | Promoter | 7 | 123982869 | 124398989 | 416121 | 1 | ENSMUST00000106437.1 | 0 | Hs3st4 | ENSMUSG00000078591.1 |
| chr11 | 114215067 | 114215484 | 3.37079 | 3.88379 | 1.01733 | Distal Intergenic | 11 | 114198258 | 114199199 | 942 | 1 | ENSMUST00000106604.1 | 16810 | 1700092K14Rik | ENSMUSG00000078600.1 |
| chr11 | 114217361 | 114217623 | 3.51292 | 4.14733 | 1.19117 | Downstream (1-2kb) | 11 | 114198258 | 114199199 | 942 | 1 | ENSMUST00000106604.1 | 19104 | 1700092K14Rik | ENSMUSG00000078600.1 |
| chr11 | 114218145 | 114218441 | 3.35805 | 3.86456 | 1.01733 | Downstream (<1kb) | 11 | 114198258 | 114199199 | 942 | 1 | ENSMUST00000106604.1 | 19888 | 1700092K14Rik | ENSMUSG00000078600.1 |
| chr11 | 109106342 | 109106695 | 3.36286 | 3.90684 | 1.13484 | Distal Intergenic | 11 | 109139291 | 109144369 | 5079 | 2 | ENSMUST00000106710.2 | 37674 | E030025P04Rik | ENSMUSG00000078605.2 |
| chr4 | 104913219 | 104913582 | 3.40676 | 3.98223 | 1.16704 | Promoter | 4 | 104913456 | 105016863 | 103408 | 1 | ENSMUST00000106803.3 | 0 | 1700024P16Rik | ENSMUSG00000078612.4 |
| chr4 | 101997505 | 101997766 | 3.18802 | 4.0502 | 1.14359 | Distal Intergenic | 4 | 101988116 | 101990014 | 1899 | 1 | ENSMUST00000106915.3 | 9390 | Gm12789 | ENSMUSG00000078625.5 |
| chr4 | 101999474 | 101999706 | 3.28761 | 3.75856 | 1.01733 | Distal Intergenic | 4 | 101988116 | 101990014 | 1899 | 1 | ENSMUST00000106915.3 | 11359 | Gm12789 | ENSMUSG00000078625.5 |
| chr4 | 102009916 | 102010314 | 3.6147 | 4.49977 | 1.43852 | Distal Intergenic | 4 | 101988116 | 101990014 | 1899 | 1 | ENSMUST00000106915.3 | 21801 | Gm12789 | ENSMUSG00000078625.5 |
| chr4 | 102014956 | 102015230 | 3.72316 | 4.56784 | 1.51175 | Distal Intergenic | 4 | 101988116 | 101990014 | 1899 | 1 | ENSMUST00000106915.3 | 26841 | Gm12789 | ENSMUSG00000078625.5 |
| chr4 | 102015844 | 102016155 | 3.71784 | 4.54211 | 1.45722 | Distal Intergenic | 4 | 101988116 | 101990014 | 1899 | 1 | ENSMUST00000106915.3 | 27729 | Gm12789 | ENSMUSG00000078625.5 |
| chr4 | 102016607 | 102016940 | 3.37577 | 4.03548 | 1.17275 | Distal Intergenic | 4 | 101988116 | 101990014 | 1899 | 1 | ENSMUST00000106915.3 | 28492 | Gm12789 | ENSMUSG00000078625.5 |
| chr11 | 105409529 | 105409645 | 3.32731 | 3.81832 | 1.01733 | Promoter | 11 | 105396695 | 105408707 | 12013 | 2 | ENSMUST00000151305.1 | -823 | Mar-10 | ENSMUSG00000078627.4 |
| chr11 | 105410171 | 105410476 | 3.7505 | 4.58502 | 1.45384 | Intron (ENSMUST00000138977.2/ENSMUSG00000078627.4, intron 4 of 11) | 11 | 105396695 | 105408707 | 12013 | 2 | ENSMUST00000151305.1 | -1465 | Mar-10 | ENSMUSG00000078627.4 |
| chr11 | 105456634 | 105456908 | 4.03173 | 5.24792 | 1.94276 | Promoter | 11 | 105360798 | 105456735 | 95938 | 2 | ENSMUST00000049995.4 | 0 | Mar-10 | ENSMUSG00000078627.4 |
| chr11 | 103482062 | 103482137 | 3.54175 | 4.24896 | 1.35126 | Intron (ENSMUST00000153273.1/ENSMUSG00000078632.3, intron 5 of 10) | 11 | 103451955 | 103504597 | 52643 | 2 | ENSMUST00000153273.1 | 22460 | Lrrc37a | ENSMUSG00000078632.3 |
| chr7 | 51749163 | 51749691 | 3.59156 | 4.29124 | 1.2761 | Downstream (1-2kb) | 7 | 51746697 | 51747698 | 1002 | 1 | ENSMUST00000107041.3 | 2467 | Gm7336 | ENSMUSG00000078636.3 |
| chr11 | 99772455 | 99772808 | 3.84073 | 4.81074 | 1.65181 | Promoter | 11 | 99771714 | 99772913 | 1200 | 2 | ENSMUST00000107440.1 | 105 | Gm11595 | ENSMUSG00000078668.3 |
| chr11 | 96250162 | 96252084 | 3.76883 | 4.66426 | 1.52905 | Promoter | 11 | 96251105 | 96264484 | 13380 | 1 | ENSMUST00000129907.2 | 0 | Gm53 | ENSMUSG00000078706.4 |
| chr11 | 96252907 | 96253049 | 3.72816 | 4.54462 | 1.4351 | Promoter | 11 | 96252532 | 96261693 | 9162 | 1 | ENSMUST00000139381.1 | 376 | Gm53 | ENSMUSG00000078706.4 |
| chr11 | 96254553 | 96256183 | 4.15569 | 5.49515 | 2.15105 | Intron (ENSMUST00000129907.2/ENSMUSG00000078706.4, intron 1 of 2) | 11 | 96252532 | 96261693 | 9162 | 1 | ENSMUST00000139381.1 | 2022 | Gm53 | ENSMUSG00000078706.4 |
| chr11 | 96256523 | 96256791 | 3.62609 | 4.36065 | 1.33331 | Intron (ENSMUST00000129907.2/ENSMUSG00000078706.4, intron 1 of 2) | 11 | 96252532 | 96261693 | 9162 | 1 | ENSMUST00000139381.1 | 3992 | Gm53 | ENSMUSG00000078706.4 |
| chr11 | 96260758 | 96261373 | 3.53316 | 4.20573 | 1.24042 | Promoter | 11 | 96261610 | 96264868 | 3259 | 1 | ENSMUST00000143887.1 | -237 | Gm53 | ENSMUSG00000078706.4 |
| chr7 | 28129713 | 28130053 | 3.42171 | 3.97598 | 1.07132 | Promoter | 7 | 28129816 | 28141299 | 11484 | 1 | ENSMUST00000130693.2 | 0 | 9530053A07Rik | ENSMUSG00000078776.4 |
| chr7 | 16657412 | 16657495 | 4.26258 | 5.56877 | 2.11139 | Distal Intergenic | 7 | 16671331 | 16675705 | 4375 | 2 | ENSMUST00000108498.2 | 18210 | Ceacam15 | ENSMUSG00000078795.2 |
| chr7 | 16061712 | 16061945 | 3.62662 | 4.55649 | 1.47079 | Distal Intergenic | 7 | 16071942 | 16096328 | 24387 | 1 | ENSMUST00000108509.1 | -9997 | Zfp541 | ENSMUSG00000078796.3 |
| chr7 | 3420276 | 3420419 | 3.33768 | 3.83386 | 1.01733 | Distal Intergenic | 7 | 3424195 | 3435667 | 11473 | 1 | ENSMUST00000183200.2 | -3776 | Cacng6 | ENSMUSG00000078815.3 |
| chr7 | 3423471 | 3423893 | 3.60545 | 4.37219 | 1.40292 | Promoter | 7 | 3424195 | 3435667 | 11473 | 1 | ENSMUST00000183200.2 | -302 | Cacng6 | ENSMUSG00000078815.3 |
| chr7 | 3424591 | 3425496 | 3.53516 | 4.18617 | 1.20521 | Promoter | 7 | 3424662 | 3435163 | 10502 | 1 | ENSMUST00000108647.2 | 0 | Cacng6 | ENSMUSG00000078815.3 |
| chr7 | 3303623 | 3304588 | 3.65051 | 4.5829 | 1.51277 | Promoter | 7 | 3303645 | 3330367 | 26723 | 1 | ENSMUST00000172109.1 | 0 | Prkcg | ENSMUSG00000078816.4 |
| chr13 | 45078797 | 45079121 | 3.2914 | 4.16752 | 1.23723 | Promoter | 13 | 45078498 | 45079127 | 630 | 2 | ENSMUST00000109187.1 | 6 | Hsp25-ps1 | ENSMUSG00000078915.1 |
| chr13 | 113028960 | 113029349 | 3.90833 | 4.87886 | 1.63648 | Distal Intergenic | 13 | 113035111 | 113038324 | 3214 | 1 | ENSMUST00000180543.2 | -5762 | Cdc20b | ENSMUSG00000078926.3 |
| chr13 | 113032220 | 113032465 | 3.98746 | 5.03691 | 1.75654 | Distal Intergenic | 13 | 113035111 | 113038324 | 3214 | 1 | ENSMUST00000180543.2 | -2646 | Cdc20b | ENSMUSG00000078926.3 |
| chr13 | 113033333 | 113033677 | 3.67447 | 4.4511 | 1.38199 | Distal Intergenic | 13 | 113035111 | 113038324 | 3214 | 1 | ENSMUST00000180543.2 | -1434 | Cdc20b | ENSMUSG00000078926.3 |
| chr13 | 113034420 | 113035673 | 3.67582 | 4.56807 | 1.47825 | Promoter | 13 | 113035111 | 113038324 | 3214 | 1 | ENSMUST00000180543.2 | 0 | Cdc20b | ENSMUSG00000078926.3 |
| chr2 | 165076990 | 165077900 | 3.55183 | 4.23127 | 1.27319 | Downstream (<1kb) | 2 | 165078693 | 165090750 | 12058 | 2 | ENSMUST00000109304.1 | 12850 | 1700025C18Rik | ENSMUSG00000078935.1 |
| chr2 | 165078414 | 165078648 | 3.31166 | 3.79473 | 1.01733 | Downstream (<1kb) | 2 | 165078693 | 165090750 | 12058 | 2 | ENSMUST00000109304.1 | 12102 | 1700025C18Rik | ENSMUSG00000078935.1 |
| chr2 | 165079028 | 165079305 | 3.56046 | 4.31232 | 1.34677 | Exon (ENSMUST00000109304.1/ENSMUSG00000078935.1, exon 3 of 3) | 2 | 165078693 | 165090750 | 12058 | 2 | ENSMUST00000109304.1 | 11445 | 1700025C18Rik | ENSMUSG00000078935.1 |
| chr2 | 165080885 | 165081559 | 3.51436 | 4.17912 | 1.29296 | Intron (ENSMUST00000109304.1/ENSMUSG00000078935.1, intron 2 of 2) | 2 | 165078693 | 165090750 | 12058 | 2 | ENSMUST00000109304.1 | 9191 | 1700025C18Rik | ENSMUSG00000078935.1 |
| chr2 | 165097594 | 165097799 | 3.31843 | 3.80489 | 1.01733 | Distal Intergenic | 2 | 165078693 | 165090750 | 12058 | 2 | ENSMUST00000109304.1 | -6845 | 1700025C18Rik | ENSMUSG00000078935.1 |
| chr2 | 165103220 | 165103555 | 3.60385 | 4.31234 | 1.28134 | Distal Intergenic | 2 | 165078693 | 165090750 | 12058 | 2 | ENSMUST00000109304.1 | -12471 | 1700025C18Rik | ENSMUSG00000078935.1 |
| chr2 | 165103909 | 165104269 | 3.45946 | 4.05333 | 1.14234 | Distal Intergenic | 2 | 165078693 | 165090750 | 12058 | 2 | ENSMUST00000109304.1 | -13160 | 1700025C18Rik | ENSMUSG00000078935.1 |
| chr2 | 165105459 | 165106019 | 4.04419 | 5.34237 | 2.05653 | Distal Intergenic | 2 | 165078693 | 165090750 | 12058 | 2 | ENSMUST00000109304.1 | -14710 | 1700025C18Rik | ENSMUSG00000078935.1 |
| chr2 | 165106615 | 165106967 | 3.47813 | 4.08394 | 1.14431 | Distal Intergenic | 2 | 165078693 | 165090750 | 12058 | 2 | ENSMUST00000109304.1 | -15866 | 1700025C18Rik | ENSMUSG00000078935.1 |
| chr2 | 165108236 | 165108772 | 3.67317 | 4.54217 | 1.46275 | Downstream (2-3kb) | 2 | 165078693 | 165090750 | 12058 | 2 | ENSMUST00000109304.1 | -17487 | 1700025C18Rik | ENSMUSG00000078935.1 |
| chr2 | 165110305 | 165110439 | 3.48685 | 4.09438 | 1.14214 | Downstream (1-2kb) | 2 | 165078693 | 165090750 | 12058 | 2 | ENSMUST00000109304.1 | -19556 | 1700025C18Rik | ENSMUSG00000078935.1 |
| chr2 | 165110829 | 165113080 | 3.80535 | 4.74325 | 1.59057 | 3' UTR | 2 | 165078693 | 165090750 | 12058 | 2 | ENSMUST00000109304.1 | -20080 | 1700025C18Rik | ENSMUSG00000078935.1 |
| chr2 | 165113593 | 165114653 | 3.57675 | 4.27988 | 1.29622 | Intron (ENSMUST00000065438.7/ENSMUSG00000053166.9, intron 11 of 11) | 2 | 165078693 | 165090750 | 12058 | 2 | ENSMUST00000109304.1 | -22844 | 1700025C18Rik | ENSMUSG00000078935.1 |
| chr2 | 165118286 | 165118630 | 3.40535 | 3.94421 | 1.04645 | Intron (ENSMUST00000065438.7/ENSMUSG00000053166.9, intron 10 of 11) | 2 | 165078693 | 165090750 | 12058 | 2 | ENSMUST00000109304.1 | -27537 | 1700025C18Rik | ENSMUSG00000078935.1 |
| chr2 | 165119669 | 165120480 | 3.60774 | 4.32079 | 1.28444 | Intron (ENSMUST00000065438.7/ENSMUSG00000053166.9, intron 10 of 11) | 2 | 165078693 | 165090750 | 12058 | 2 | ENSMUST00000109304.1 | -28920 | 1700025C18Rik | ENSMUSG00000078935.1 |
| chr2 | 165120938 | 165121480 | 3.51032 | 4.13481 | 1.15971 | Intron (ENSMUST00000065438.7/ENSMUSG00000053166.9, intron 10 of 11) | 2 | 165078693 | 165090750 | 12058 | 2 | ENSMUST00000109304.1 | -30189 | 1700025C18Rik | ENSMUSG00000078935.1 |
| chr13 | 97365636 | 97365789 | 3.5855 | 4.34032 | 1.41467 | Distal Intergenic | 13 | 97434970 | 97497007 | 62038 | 1 | ENSMUST00000109431.4 | -69181 | Lincenc1 | ENSMUSG00000078952.4 |
| chr13 | 97366348 | 97366566 | 3.51503 | 4.19294 | 1.31152 | Distal Intergenic | 13 | 97434970 | 97497007 | 62038 | 1 | ENSMUST00000109431.4 | -68404 | Lincenc1 | ENSMUSG00000078952.4 |
| chr13 | 97423819 | 97424113 | 3.65802 | 4.44528 | 1.44322 | Distal Intergenic | 13 | 97434970 | 97497007 | 62038 | 1 | ENSMUST00000109431.4 | -10857 | Lincenc1 | ENSMUSG00000078952.4 |
| chr13 | 97436089 | 97436233 | 3.45282 | 4.15343 | 1.23763 | Intron (ENSMUST00000109431.4/ENSMUSG00000078952.4, intron 1 of 1) | 13 | 97434970 | 97497007 | 62038 | 1 | ENSMUST00000109431.4 | 1120 | Lincenc1 | ENSMUSG00000078952.4 |
| chr13 | 97436640 | 97436774 | 3.54041 | 4.35605 | 1.33716 | Intron (ENSMUST00000109431.4/ENSMUSG00000078952.4, intron 1 of 1) | 13 | 97434970 | 97497007 | 62038 | 1 | ENSMUST00000109431.4 | 1671 | Lincenc1 | ENSMUSG00000078952.4 |
| chr13 | 97437685 | 97437912 | 3.29565 | 3.77071 | 1.01733 | Intron (ENSMUST00000109431.4/ENSMUSG00000078952.4, intron 1 of 1) | 13 | 97434970 | 97497007 | 62038 | 1 | ENSMUST00000109431.4 | 2716 | Lincenc1 | ENSMUSG00000078952.4 |
| chr13 | 97442395 | 97442656 | 3.55132 | 4.23362 | 1.28155 | Intron (ENSMUST00000109431.4/ENSMUSG00000078952.4, intron 1 of 1) | 13 | 97434970 | 97497007 | 62038 | 1 | ENSMUST00000109431.4 | 7426 | Lincenc1 | ENSMUSG00000078952.4 |
| chr13 | 97514530 | 97514605 | 3.70194 | 4.52176 | 1.49602 | Distal Intergenic | 13 | 97482955 | 97489700 | 6746 | 1 | ENSMUST00000180371.1 | 31576 | Lincenc1 | ENSMUSG00000078952.4 |
| chr13 | 97599304 | 97599379 | 3.42646 | 4.22388 | 1.29627 | Distal Intergenic | 13 | 97482955 | 97489700 | 6746 | 1 | ENSMUST00000180371.1 | 116350 | Lincenc1 | ENSMUSG00000078952.4 |
| chr2 | 158189023 | 158189176 | 3.38193 | 3.9006 | 1.01733 | Intron (ENSMUST00000103121.4/ENSMUSG00000037813.8, intron 14 of 15) | 2 | 158192008 | 158195749 | 3742 | 1 | ENSMUST00000109506.2 | -2832 | 1700060C20Rik | ENSMUSG00000078957.3 |
| chr2 | 158189696 | 158189817 | 3.60929 | 4.33714 | 1.34978 | Intron (ENSMUST00000103121.4/ENSMUSG00000037813.8, intron 14 of 15) | 2 | 158192008 | 158195749 | 3742 | 1 | ENSMUST00000109506.2 | -2191 | 1700060C20Rik | ENSMUSG00000078957.3 |
| chr8 | 92719675 | 92719919 | 3.42888 | 3.98831 | 1.07666 | Distal Intergenic | 8 | 92715747 | 92716975 | 1229 | 2 | ENSMUST00000181181.1 | -2701 | Gm3272 | ENSMUSG00000078967.5 |
| chr8 | 92722938 | 92723125 | 3.37429 | 3.94022 | 1.14555 | Distal Intergenic | 8 | 92715747 | 92716975 | 1229 | 2 | ENSMUST00000181181.1 | -5964 | Gm3272 | ENSMUSG00000078967.5 |
| chr8 | 92761714 | 92763111 | 3.68867 | 4.54399 | 1.51156 | Distal Intergenic | 8 | 92715747 | 92716975 | 1229 | 2 | ENSMUST00000181181.1 | -44740 | Gm3272 | ENSMUSG00000078967.5 |
| chr8 | 71684020 | 71684363 | 3.67262 | 4.45085 | 1.38864 | Exon (ENSMUST00000110013.4/ENSMUSG00000031805.12, exon 15 of 24) | 8 | 71689214 | 71690575 | 1362 | 1 | ENSMUST00000034261.7 | -4851 | Insl3 | ENSMUSG00000079019.2 |
| chr8 | 71685142 | 71685396 | 3.58752 | 4.27965 | 1.25307 | Exon (ENSMUST00000110013.4/ENSMUSG00000031805.12, exon 18 of 24) | 8 | 71689214 | 71690575 | 1362 | 1 | ENSMUST00000034261.7 | -3818 | Insl3 | ENSMUSG00000079019.2 |
| chr8 | 71685784 | 71686054 | 3.6993 | 4.52962 | 1.43569 | Exon (ENSMUST00000110013.4/ENSMUSG00000031805.12, exon 20 of 24) | 8 | 71689214 | 71690575 | 1362 | 1 | ENSMUST00000034261.7 | -3160 | Insl3 | ENSMUSG00000079019.2 |
| chr8 | 71688029 | 71688277 | 3.82221 | 4.82417 | 1.71521 | Promoter | 8 | 71689214 | 71690575 | 1362 | 1 | ENSMUST00000034261.7 | -937 | Insl3 | ENSMUSG00000079019.2 |
| chr8 | 71688765 | 71688916 | 3.80555 | 4.75396 | 1.64288 | Promoter | 8 | 71689214 | 71690575 | 1362 | 1 | ENSMUST00000034261.7 | -298 | Insl3 | ENSMUSG00000079019.2 |
| chr8 | 71689270 | 71690022 | 3.76467 | 4.71751 | 1.60024 | Promoter | 8 | 71689214 | 71690575 | 1362 | 1 | ENSMUST00000034261.7 | 57 | Insl3 | ENSMUSG00000079019.2 |
| chr8 | 71692642 | 71693156 | 3.58044 | 4.45586 | 1.43053 | Exon (ENSMUST00000034260.7/ENSMUSG00000031803.7, exon 3 of 3) | 8 | 71689214 | 71690575 | 1362 | 1 | ENSMUST00000034261.7 | 3429 | Insl3 | ENSMUSG00000079019.2 |
| chr8 | 71693519 | 71693595 | 4.11539 | 5.68039 | 2.22206 | Exon (ENSMUST00000034260.7/ENSMUSG00000031803.7, exon 2 of 3) | 8 | 71689214 | 71690575 | 1362 | 1 | ENSMUST00000034261.7 | 4306 | Insl3 | ENSMUSG00000079019.2 |
| chr2 | 129037263 | 129037346 | 3.53191 | 4.23614 | 1.32718 | Downstream (<1kb) | 2 | 129038145 | 129048172 | 10028 | 2 | ENSMUST00000145798.2 | 10826 | Gm14025 | ENSMUSG00000079051.4 |
| chr2 | 129038247 | 129038528 | 3.27907 | 4.20132 | 1.23902 | Exon (ENSMUST00000145798.2/ENSMUSG00000079051.4, exon 3 of 3) | 2 | 129038145 | 129048172 | 10028 | 2 | ENSMUST00000145798.2 | 9644 | Gm14025 | ENSMUSG00000079051.4 |
| chr2 | 129039401 | 129039816 | 3.87885 | 5.35047 | 2.03499 | Exon (ENSMUST00000145798.2/ENSMUSG00000079051.4, exon 3 of 3) | 2 | 129038145 | 129048172 | 10028 | 2 | ENSMUST00000145798.2 | 8356 | Gm14025 | ENSMUSG00000079051.4 |
| chr2 | 128331737 | 128331979 | 3.37128 | 3.89342 | 1.04874 | Intron (ENSMUST00000143065.2/ENSMUSG00000074813.7, intron 1 of 3) | 2 | 128423654 | 128424565 | 912 | 1 | ENSMUST00000131961.1 | -91675 | Gm14010 | ENSMUSG00000079053.3 |
| chr2 | 128393852 | 128394104 | 3.60254 | 4.3963 | 1.3964 | Intron (ENSMUST00000143065.2/ENSMUSG00000074813.7, intron 1 of 3) | 2 | 128423654 | 128424565 | 912 | 1 | ENSMUST00000131961.1 | -29550 | Gm14010 | ENSMUSG00000079053.3 |
| chr2 | 128395464 | 128395698 | 3.45411 | 4.08563 | 1.25656 | Intron (ENSMUST00000143065.2/ENSMUSG00000074813.7, intron 1 of 3) | 2 | 128423654 | 128424565 | 912 | 1 | ENSMUST00000131961.1 | -27956 | Gm14010 | ENSMUSG00000079053.3 |
| chr2 | 128401968 | 128402331 | 3.48983 | 4.25438 | 1.27335 | Intron (ENSMUST00000143065.2/ENSMUSG00000074813.7, intron 1 of 3) | 2 | 128423654 | 128424565 | 912 | 1 | ENSMUST00000131961.1 | -21323 | Gm14010 | ENSMUSG00000079053.3 |
| chr12 | 81278689 | 81278821 | 2.57712 | 5.45832 | 2.13713 | Intron (ENSMUST00000085238.7/ENSMUSG00000079055.5, intron 2 of 6) | 12 | 81202397 | 81294674 | 92278 | 2 | ENSMUST00000182366.1 | 15853 | Slc8a3 | ENSMUSG00000079055.5 |
| chr12 | 81331167 | 81332009 | 3.76503 | 4.61785 | 1.47918 | Promoter | 12 | 81198276 | 81332959 | 134684 | 2 | ENSMUST00000064594.8 | 950 | Slc8a3 | ENSMUSG00000079055.5 |
| chr12 | 81332517 | 81333193 | 3.67126 | 4.55955 | 1.4976 | Promoter | 12 | 81197915 | 81333164 | 135250 | 2 | ENSMUST00000085238.7 | 0 | Slc8a3 | ENSMUSG00000079055.5 |
| chr12 | 81333838 | 81334171 | 4.19409 | 5.43593 | 2.0211 | Promoter | 12 | 81214024 | 81333180 | 119157 | 2 | ENSMUST00000183102.1 | -659 | Slc8a3 | ENSMUSG00000079055.5 |
| chr12 | 81335202 | 81335350 | 3.65801 | 4.41189 | 1.33814 | Distal Intergenic | 12 | 81214024 | 81333180 | 119157 | 2 | ENSMUST00000183102.1 | -2023 | Slc8a3 | ENSMUSG00000079055.5 |
| chr12 | 81335755 | 81336091 | 3.48868 | 4.10039 | 1.15164 | Distal Intergenic | 12 | 81214024 | 81333180 | 119157 | 2 | ENSMUST00000183102.1 | -2576 | Slc8a3 | ENSMUSG00000079055.5 |
| chr2 | 127482487 | 127482596 | 3.36966 | 4.02748 | 1.13422 | Promoter | 2 | 127458130 | 127482499 | 24370 | 2 | ENSMUST00000088538.5 | 0 | Kcnip3 | ENSMUSG00000079056.7 |
| chr2 | 127484718 | 127484921 | 3.38936 | 4.23148 | 1.29567 | Intron (ENSMUST00000028850.9/ENSMUSG00000079056.7, intron 2 of 8) | 2 | 127458130 | 127482499 | 24370 | 2 | ENSMUST00000088538.5 | -2220 | Kcnip3 | ENSMUSG00000079056.7 |
| chr8 | 32883506 | 32885216 | 3.5856 | 4.35419 | 1.39267 | Distal Intergenic | 8 | 32890475 | 32942482 | 52008 | 2 | ENSMUST00000180220.2 | 57266 | Gm3985 | ENSMUSG00000079070.5 |
| chr8 | 9769405 | 9769780 | 3.49756 | 4.1477 | 1.26801 | Intron (ENSMUST00000110969.3/ENSMUSG00000079157.3, intron 1 of 2) | 8 | 9206001 | 9771018 | 565018 | 2 | ENSMUST00000110969.3 | 1238 | Fam155a | ENSMUSG00000079157.3 |

| | | | | | | | | | | | | | | | |
|---|---|---|---|---|---|---|---|---|---|---|---|---|---|---|---|
| chr8 | 9770772 | 9771093 | 3.64294 | 4.39948 | 1.33889 | Promoter | 8 | 9206001 | 9771018 | 565018 | 2 | ENSMUST00000110969.3 | 0 | Fam155a | ENSMUSG00000079157.3 |
| chr8 | 9771578 | 9772055 | 3.77631 | 4.6506 | 1.52007 | Promoter | 8 | 9206001 | 9771018 | 565018 | 2 | ENSMUST00000110969.3 | -561 | Fam155a | ENSMUSG00000079157.3 |
| chr8 | 9772437 | 9772978 | 3.72715 | 4.61553 | 1.53233 | Distal Intergenic | 8 | 9206001 | 9771018 | 565018 | 2 | ENSMUST00000110969.3 | -1420 | Fam155a | ENSMUSG00000079157.3 |
| chr8 | 9774682 | 9775334 | 3.44754 | 4.08531 | 1.15656 | Distal Intergenic | 8 | 9206001 | 9771018 | 565018 | 2 | ENSMUST00000110969.3 | -3665 | Fam155a | ENSMUSG00000079157.3 |
| chr1 | 182955191 | 182955541 | 3.45732 | 4.05809 | 1.17189 | Promoter | 1 | 182954788 | 182976044 | 21257 | 1 | ENSMUST00000110997.1 | 404 | Tlr5 | ENSMUSG00000079164.3 |
| chr5 | 137425793 | 137426137 | 3.39012 | 3.91553 | 1.02638 | Intron (ENSMUST00000164178.1/ENSMUSG00000079173.6, intron 38 of 86) | 5 | 137409581 | 137411581 | 2001 | 2 | ENSMUST00000150470.1 | -14213 | Zan | ENSMUSG00000079173.6 |
| chr5 | 137463509 | 137463862 | 3.55864 | 4.24962 | 1.31047 | Exon (ENSMUST00000164178.1/ENSMUSG00000079173.6, exon 13 of 87) | 5 | 137378637 | 137476595 | 97959 | 2 | ENSMUST00000164178.1 | 12733 | Zan | ENSMUSG00000079173.6 |
| chr5 | 137464422 | 137464562 | 3.14139 | 4.03755 | 1.14841 | Exon (ENSMUST00000164178.1/ENSMUSG00000079173.6, exon 13 of 87) | 5 | 137378637 | 137476595 | 97959 | 2 | ENSMUST00000164178.1 | 12033 | Zan | ENSMUSG00000079173.6 |
| chr5 | 137172545 | 137172674 | 3.43477 | 4.00125 | 1.09149 | Distal Intergenic | 5 | 137154029 | 137166070 | 12042 | 2 | ENSMUST00000111075.3 | -6476 | Gm3054 | ENSMUSG00000079174.3 |
| chr5 | 137186192 | 137186425 | 3.34195 | 4.29162 | 1.30047 | Distal Intergenic | 5 | 137154029 | 137166070 | 12042 | 2 | ENSMUST00000111075.3 | -20123 | Gm3054 | ENSMUSG00000079174.3 |
| chr9 | 114547709 | 114547890 | 3.51049 | 4.14487 | 1.19531 | Intron (ENSMUST00000111816.2/ENSMUSG00000079259.2, intron 1 of 3) | 9 | 114507133 | 114564369 | 57237 | 2 | ENSMUST00000111816.2 | 16479 | Trim71 | ENSMUSG00000079259.2 |
| chr9 | 114559138 | 114559318 | 3.23307 | 3.67678 | 1.01733 | Intron (ENSMUST00000111816.2/ENSMUSG00000079259.2, intron 1 of 3) | 9 | 114507133 | 114564369 | 57237 | 2 | ENSMUST00000111816.2 | 5051 | Trim71 | ENSMUSG00000079259.2 |
| chr9 | 114559836 | 114560152 | 3.225 | 3.66469 | 1.01733 | Intron (ENSMUST00000111816.2/ENSMUSG00000079259.2, intron 1 of 3) | 9 | 114507133 | 114564369 | 57237 | 2 | ENSMUST00000111816.2 | 4217 | Trim71 | ENSMUSG00000079259.2 |
| chr9 | 114561025 | 114561406 | 3.89808 | 5.10982 | 1.91467 | Intron (ENSMUST00000111816.2/ENSMUSG00000079259.2, intron 1 of 3) | 9 | 114507133 | 114564369 | 57237 | 2 | ENSMUST00000111816.2 | 2963 | Trim71 | ENSMUSG00000079259.2 |
| chr9 | 114562109 | 114564664 | 3.96902 | 5.20858 | 1.93801 | Promoter | 9 | 114507133 | 114564369 | 57237 | 2 | ENSMUST00000111816.2 | 0 | Trim71 | ENSMUSG00000079259.2 |
| chr14 | 46379603 | 46379877 | 3.30328 | 3.78212 | 1.01733 | Promoter | 14 | 46379467 | 46383608 | 4142 | 1 | ENSMUST00000111826.2 | 137 | Gm15217 | ENSMUSG00000079261.2 |
| chr14 | 46380305 | 46380708 | 3.84833 | 4.81887 | 1.67424 | Promoter | 14 | 46381127 | 46383368 | 2242 | 1 | ENSMUST00000147092.1 | -419 | Gm15217 | ENSMUSG00000079261.2 |
| chr5 | 116081827 | 116081970 | 3.73154 | 4.63359 | 1.55137 | Intron (ENSMUST00000111997.1/ENSMUSG00000079278.1, intron 1 of 2) | 5 | 116038755 | 116083244 | 44490 | 2 | ENSMUST00000111997.1 | 1274 | Tmem233 | ENSMUSG00000079278.1 |
| chr5 | 116082573 | 116083323 | 3.78012 | 4.66693 | 1.53924 | Promoter | 5 | 116038755 | 116083244 | 44490 | 2 | ENSMUST00000111997.1 | 0 | Tmem233 | ENSMUSG00000079278.1 |
| chr1 | 132183645 | 132183971 | 3.43735 | 4.16291 | 1.23781 | Distal Intergenic | 1 | 132191431 | 132259227 | 67797 | 1 | ENSMUST00000188169.1 | -7460 | Lemd1 | ENSMUSG00000079330.3 |
| chr1 | 132187547 | 132187623 | 4.505 | 6.10617 | 2.57265 | Distal Intergenic | 1 | 132191431 | 132259227 | 67797 | 1 | ENSMUST00000188169.1 | -3808 | Lemd1 | ENSMUSG00000079330.3 |
| chr1 | 132188108 | 132189039 | 3.71899 | 4.53824 | 1.45167 | Distal Intergenic | 1 | 132191431 | 132259227 | 67797 | 1 | ENSMUST00000188169.1 | -2392 | Lemd1 | ENSMUSG00000079330.3 |
| chr1 | 132189531 | 132191002 | 3.90091 | 5.28539 | 1.97768 | Promoter | 1 | 132191431 | 132259227 | 67797 | 1 | ENSMUST00000188169.1 | -429 | Lemd1 | ENSMUSG00000079330.3 |
| chr1 | 132191378 | 132193820 | 3.95936 | 5.06834 | 1.82021 | Promoter | 1 | 132191431 | 132259227 | 67797 | 1 | ENSMUST00000188169.1 | 0 | Lemd1 | ENSMUSG00000079330.3 |
| chr1 | 132211361 | 132211496 | 3.37449 | 3.88937 | 1.01733 | Intron (ENSMUST00000188169.1/ENSMUSG00000079330.3, intron 1 of 4) | 1 | 132203852 | 132204509 | 658 | 1 | ENSMUST00000185223.1 | 7510 | Lemd1 | ENSMUSG00000079330.3 |
| chr1 | 132222932 | 132223188 | 3.30587 | 3.78602 | 1.01733 | Intron (ENSMUST00000188169.1/ENSMUSG00000079330.3, intron 1 of 4) | 1 | 132226314 | 132257189 | 30876 | 1 | ENSMUST00000185436.1 | -3126 | Lemd1 | ENSMUSG00000079330.3 |
| chr1 | 132208611 | 132209836 | 3.76112 | 4.62025 | 1.49024 | Exon (ENSMUST00000188175.1/ENSMUSG00000079330.3, exon 2 of 2) | 1 | 132203852 | 132204509 | 658 | 1 | ENSMUST00000185223.1 | 4760 | Lemd1 | ENSMUSG00000079330.3 |
| chr1 | 132210472 | 132210996 | 3.94925 | 5.14476 | 1.87628 | Intron (ENSMUST00000188169.1/ENSMUSG00000079330.3, intron 1 of 4) | 1 | 132203852 | 132204509 | 658 | 1 | ENSMUST00000185223.1 | 6621 | Lemd1 | ENSMUSG00000079330.3 |
| chr14 | 7567964 | 7568308 | 3.95478 | 5.02933 | 1.76198 | Promoter | 14 | 7545151 | 7568566 | 23416 | 2 | ENSMUST00000163790.1 | 258 | Gm3558 | ENSMUSG00000079364.3 |
| chr14 | 7568736 | 7568977 | 3.58321 | 4.49701 | 1.4316 | Promoter | 14 | 7545151 | 7568566 | 23416 | 2 | ENSMUST00000163790.1 | -171 | Gm3558 | ENSMUSG00000079364.3 |
| chr19 | 21571709 | 21571799 | 3.15444 | 3.70169 | 1.01733 | Distal Intergenic | 19 | 21555674 | 21557657 | 1984 | 1 | ENSMUST00000112772.2 | 16036 | Gm3443 | ENSMUSG00000079382.2 |
| chr14 | 4499371 | 4499683 | 3.62281 | 4.35447 | 1.33471 | Intron (ENSMUST00000166848.2/ENSMUSG00000079386.3, intron 1 of 3) | 14 | 4514758 | 4519452 | 4695 | 1 | ENSMUST00000112776.2 | -15075 | Gm3173 | ENSMUSG00000079386.3 |
| chr14 | 4334080 | 4335228 | 3.97755 | 5.06215 | 1.78716 | Promoter | 14 | 4334763 | 4354688 | 19926 | 1 | ENSMUST00000165466.2 | 0 | 2610042L04Rik | ENSMUSG00000079388.4 |
| chr14 | 4182951 | 4183157 | 3.33854 | 3.83517 | 1.01733 | Promoter | 14 | 4182576 | 4204895 | 22320 | 1 | ENSMUST00000164512.2 | 376 | Gm2974 | ENSMUSG00000079391.5 |
| chr14 | 6286941 | 6287469 | 3.57547 | 4.49255 | 1.43828 | Promoter | 14 | 6259747 | 6287250 | 27504 | 2 | ENSMUST00000170104.2 | 0 | Gm3411 | ENSMUSG00000079396.4 |
| chr14 | 3652473 | 3652633 | 3.56803 | 4.24819 | 1.24901 | Promoter | 14 | 3652030 | 3673215 | 21186 | 1 | ENSMUST00000167430.2 | 444 | Gm3020 | ENSMUSG00000079402.4 |
| chr14 | 3049225 | 3049903 | 3.42763 | 5.57149 | 2.25574 | Promoter | 14 | 3049285 | 3076840 | 27556 | 1 | ENSMUST00000166494.1 | 0 | Gm2897 | ENSMUSG00000079410.4 |
| chr1 | 86303121 | 86303723 | 3.69046 | 4.54637 | 1.51711 | Promoter | 1 | 86303234 | 86307305 | 4072 | 1 | ENSMUST00000113306.3 | 0 | B3gnt7 | ENSMUSG00000079445.3 |
| chr2 | 31633585 | 31638381 | 3.73785 | 4.78932 | 1.64741 | Exon (ENSMUST00000145389.1/ENSMUSG00000085224.1, exon 3 of 3) | 2 | 31640037 | 31655795 | 15759 | 1 | ENSMUST00000113470.2 | -1656 | Prdm12 | ENSMUSG00000079466.2 |
| chr2 | 31638729 | 31643188 | 3.93558 | 5.14935 | 1.93375 | Promoter | 2 | 31640037 | 31655795 | 15759 | 1 | ENSMUST00000113470.2 | 0 | Prdm12 | ENSMUSG00000079466.2 |
| chr2 | 29746397 | 29746746 | 4.03578 | 5.21096 | 1.91873 | Promoter | 2 | 29744919 | 29747250 | 2332 | 2 | ENSMUST00000129574.1 | 504 | Gm13420 | ENSMUSG00000079497.3 |
| chr2 | 29747119 | 29748115 | 3.96811 | 5.08841 | 1.82648 | Promoter | 2 | 29744919 | 29747250 | 2332 | 2 | ENSMUST00000129574.1 | 0 | Gm13420 | ENSMUSG00000079497.3 |
| chr2 | 29750381 | 29750515 | 3.56984 | 4.38208 | 1.38386 | Distal Intergenic | 2 | 29744919 | 29747250 | 2332 | 2 | ENSMUST00000129574.1 | -3132 | Gm13420 | ENSMUSG00000079497.3 |
| chr2 | 29751119 | 29752506 | 3.61174 | 4.33207 | 1.30374 | Distal Intergenic | 2 | 29744919 | 29747250 | 2332 | 2 | ENSMUST00000129574.1 | -3870 | Gm13420 | ENSMUSG00000079497.3 |
| chr2 | 28992732 | 28993279 | 3.88776 | 5.01141 | 1.81152 | Intron (ENSMUST00000189027.1/ENSMUSG00000079502.3, intron 1 of 6) | 2 | 28927834 | 29039360 | 111527 | 2 | ENSMUST00000190049.1 | 46081 | 1700101E01Rik | ENSMUSG00000079502.3 |
| chr2 | 28997336 | 28997462 | 3.58696 | 4.38666 | 1.36164 | Intron (ENSMUST00000189027.1/ENSMUSG00000079502.3, intron 1 of 6) | 2 | 28927834 | 29039360 | 111527 | 2 | ENSMUST00000190049.1 | 41898 | 1700101E01Rik | ENSMUSG00000079502.3 |
| chr2 | 29009834 | 29010149 | 3.25545 | 4.16014 | 1.22125 | Intron (ENSMUST00000189027.1/ENSMUSG00000079502.3, intron 1 of 6) | 2 | 28927834 | 29039360 | 111527 | 2 | ENSMUST00000190049.1 | 29211 | 1700101E01Rik | ENSMUSG00000079502.3 |
| chr2 | 29010575 | 29011035 | 3.44289 | 4.04448 | 1.20121 | Intron (ENSMUST00000189027.1/ENSMUSG00000079502.3, intron 1 of 6) | 2 | 28927834 | 29039360 | 111527 | 2 | ENSMUST00000190049.1 | 28325 | 1700101E01Rik | ENSMUSG00000079502.3 |
| chr2 | 29011587 | 29011931 | 3.4245 | 3.99458 | 1.12271 | Intron (ENSMUST00000189027.1/ENSMUSG00000079502.3, intron 1 of 6) | 2 | 28927834 | 29039360 | 111527 | 2 | ENSMUST00000190049.1 | 27429 | 1700101E01Rik | ENSMUSG00000079502.3 |
| chr2 | 29014596 | 29015086 | 3.53883 | 4.20083 | 1.23829 | Intron (ENSMUST00000189027.1/ENSMUSG00000079502.3, intron 1 of 6) | 2 | 28927834 | 29039360 | 111527 | 2 | ENSMUST00000190049.1 | 24274 | 1700101E01Rik | ENSMUSG00000079502.3 |
| chr2 | 29015704 | 29016222 | 3.66613 | 4.42333 | 1.34067 | Intron (ENSMUST00000189027.1/ENSMUSG00000079502.3, intron 1 of 6) | 2 | 28927834 | 29039360 | 111527 | 2 | ENSMUST00000190049.1 | 23138 | 1700101E01Rik | ENSMUSG00000079502.3 |
| chr2 | 29016793 | 29017376 | 3.66587 | 4.48643 | 1.4276 | Intron (ENSMUST00000189027.1/ENSMUSG00000079502.3, intron 1 of 6) | 2 | 28927834 | 29039360 | 111527 | 2 | ENSMUST00000190049.1 | 21984 | 1700101E01Rik | ENSMUSG00000079502.3 |
| chr2 | 29018414 | 29018714 | 3.51787 | 4.15468 | 1.19029 | Intron (ENSMUST00000189027.1/ENSMUSG00000079502.3, intron 1 of 6) | 2 | 28927834 | 29039360 | 111527 | 2 | ENSMUST00000190049.1 | 20646 | 1700101E01Rik | ENSMUSG00000079502.3 |
| chr2 | 29020997 | 29021650 | 3.61139 | 4.33229 | 1.30516 | Intron (ENSMUST00000189027.1/ENSMUSG00000079502.3, intron 1 of 6) | 2 | 28927834 | 29039360 | 111527 | 2 | ENSMUST00000190049.1 | 17710 | 1700101E01Rik | ENSMUSG00000079502.3 |
| chr2 | 29023237 | 29023395 | 3.37763 | 3.89411 | 1.01733 | Intron (ENSMUST00000189027.1/ENSMUSG00000079502.3, intron 1 of 6) | 2 | 28927834 | 29039360 | 111527 | 2 | ENSMUST00000190049.1 | 15965 | 1700101E01Rik | ENSMUSG00000079502.3 |
| chr2 | 29030161 | 29030430 | 3.47036 | 4.23015 | 1.28958 | Intron (ENSMUST00000189027.1/ENSMUSG00000079502.3, intron 1 of 6) | 2 | 28927834 | 29039360 | 111527 | 2 | ENSMUST00000190049.1 | 8930 | 1700101E01Rik | ENSMUSG00000079502.3 |
| chr2 | 29038368 | 29039179 | 4.27428 | 5.6831 | 2.26566 | Promoter | 2 | 28927834 | 29039360 | 111527 | 2 | ENSMUST00000190049.1 | 181 | 1700101E01Rik | ENSMUSG00000079502.3 |
| chr2 | 29041763 | 29042280 | 3.68271 | 4.52342 | 1.44801 | Intron (ENSMUST00000189027.1/ENSMUSG00000079502.3, intron 1 of 6) | 2 | 28927834 | 29039360 | 111527 | 2 | ENSMUST00000190049.1 | -2404 | 1700101E01Rik | ENSMUSG00000079502.3 |
| chr2 | 29042646 | 29042895 | 3.31085 | 3.80958 | 1.07253 | Intron (ENSMUST00000189027.1/ENSMUSG00000079502.3, intron 1 of 6) | 2 | 28927834 | 29039360 | 111527 | 2 | ENSMUST00000190049.1 | -3287 | 1700101E01Rik | ENSMUSG00000079502.3 |
| chr2 | 29045715 | 29046080 | 3.51344 | 4.14541 | 1.18115 | Intron (ENSMUST00000189027.1/ENSMUSG00000079502.3, intron 1 of 6) | 2 | 28927834 | 29039360 | 111527 | 2 | ENSMUST00000190049.1 | -6356 | 1700101E01Rik | ENSMUSG00000079502.3 |
| chr2 | 29047159 | 29047781 | 3.41398 | 4.09477 | 1.19036 | Intron (ENSMUST00000189027.1/ENSMUSG00000079502.3, intron 1 of 6) | 2 | 28926949 | 29055024 | 128076 | 2 | ENSMUST00000189027.1 | 7243 | 1700101E01Rik | ENSMUSG00000079502.3 |
| chr2 | 29050542 | 29050972 | 3.61591 | 4.38203 | 1.36392 | Intron (ENSMUST00000189027.1/ENSMUSG00000079502.3, intron 1 of 6) | 2 | 28926949 | 29055024 | 128076 | 2 | ENSMUST00000189027.1 | 4052 | 1700101E01Rik | ENSMUSG00000079502.3 |
| chr17 | 35320733 | 35321335 | 3.6954 | 4.56836 | 1.51429 | Promoter | 17 | 35320558 | 35325099 | 4542 | 1 | ENSMUST00000073208.5 | 176 | H2-Q1 | ENSMUSG00000079507.5 |
| chr2 | 9883424 | 9884174 | 3.52118 | 4.2514 | 1.2992 | Promoter | 2 | 9883042 | 9883993 | 952 | 2 | ENSMUST00000114915.2 | 0 | 9230102O04Rik | ENSMUSG00000079602.2 |
| chr2 | 9884761 | 9885663 | 4.46232 | 6.07394 | 2.52999 | Promoter | 2 | 9883042 | 9883993 | 952 | 2 | ENSMUST00000114915.2 | -769 | 9230102O04Rik | ENSMUSG00000079602.2 |
| chr6 | 29172763 | 29172911 | 3.70558 | 4.61566 | 1.53224 | Intron (ENSMUST00000159200.1/ENSMUSG00000079654.3, intron 4 of 4) | 6 | 29169232 | 29179584 | 10353 | 2 | ENSMUST00000159200.1 | 6673 | Prrt4 | ENSMUSG00000079654.3 |
| chr6 | 29177220 | 29177785 | 3.3922 | 4.0652 | 1.20963 | 5' UTR | 6 | 29169232 | 29179584 | 10353 | 2 | ENSMUST00000159200.1 | 1799 | Prrt4 | ENSMUSG00000079654.3 |
| chr6 | 29178493 | 29179571 | 3.58039 | 4.28254 | 1.30026 | Promoter | 6 | 29169232 | 29179584 | 10353 | 2 | ENSMUST00000159200.1 | 13 | Prrt4 | ENSMUSG00000079654.3 |
| chr17 | 24209189 | 24209951 | 3.74851 | 4.67999 | 1.59249 | Promoter | 17 | 24203848 | 24209387 | 5540 | 2 | ENSMUST00000024931.4 | 0 | Ntn3 | ENSMUSG00000079662.1 |
| chr10 | 7390816 | 7391594 | 3.51568 | 4.46689 | 1.42778 | Distal Intergenic | 10 | 7445391 | 7473361 | 27971 | 2 | ENSMUST00000169796.2 | 81767 | Ulbp1 | ENSMUSG00000079685.4 |
| chr10 | 7392040 | 7392388 | 3.43236 | 4.25937 | 1.30174 | Distal Intergenic | 10 | 7445391 | 7473361 | 27971 | 2 | ENSMUST00000169796.2 | 80973 | Ulbp1 | ENSMUSG00000079685.4 |
| chr10 | 7392728 | 7393354 | 3.71665 | 4.6534 | 1.56936 | Distal Intergenic | 10 | 7445391 | 7473361 | 27971 | 2 | ENSMUST00000169796.2 | 80007 | Ulbp1 | ENSMUSG00000079685.4 |
| chr10 | 7393694 | 7393923 | 3.22618 | 4.1124 | 1.19782 | Distal Intergenic | 10 | 7445391 | 7473361 | 27971 | 2 | ENSMUST00000169796.2 | 79438 | Ulbp1 | ENSMUSG00000079685.4 |
| chr11 | 31243582 | 31243983 | 3.53085 | 4.39481 | 1.3831 | Distal Intergenic | 11 | 31177818 | 31184374 | 6557 | 2 | ENSMUST00000116006.2 | -59209 | Gm12106 | ENSMUSG00000079890.2 |
| chr11 | 31247719 | 31248079 | 3.46169 | 4.09354 | 1.25389 | Distal Intergenic | 11 | 31177818 | 31184374 | 6557 | 2 | ENSMUST00000116006.2 | -63346 | Gm12106 | ENSMUSG00000079890.2 |
| chr4 | 127391536 | 127391771 | 3.25257 | 3.706 | 1.01733 | Distal Intergenic | 4 | 127408379 | 127408798 | 420 | 2 | ENSMUST00000116114.2 | 17027 | Gm12947 | ENSMUSG00000079998.2 |
| chr4 | 127419253 | 127419938 | 3.91904 | 4.94002 | 1.7276 | Distal Intergenic | 4 | 127408379 | 127408798 | 420 | 2 | ENSMUST00000116114.2 | -10456 | Gm12947 | ENSMUSG00000079998.2 |
| chr12 | 110277183 | 110278188 | 3.66107 | 4.44626 | 1.37727 | Promoter | 12 | 110278048 | 110278129 | 82 | 2 | ENSMUST00000116706.1 | 0 | Mir1247 | ENSMUSG00000080356.1 |
| chr4 | 152907503 | 152907819 | 3.62968 | 4.35591 | 1.30432 | Distal Intergenic | 4 | 153013490 | 153013584 | 95 | 2 | ENSMUST00000116710.1 | 105765 | Gm25779 | ENSMUSG00000080360.1 |
| chr4 | 152953209 | 152953284 | 3.39597 | 4.15135 | 1.20765 | Distal Intergenic | 4 | 153013490 | 153013584 | 95 | 2 | ENSMUST00000116710.1 | 60300 | Gm25779 | ENSMUSG00000080360.1 |
| chr4 | 153019322 | 153019478 | 3.58889 | 4.4456 | 1.41451 | Distal Intergenic | 4 | 153013490 | 153013584 | 95 | 2 | ENSMUST00000116710.1 | -5739 | Gm25779 | ENSMUSG00000080360.1 |
| chr7 | 128814521 | 128814766 | 3.35112 | 3.87341 | 1.08549 | Distal Intergenic | 7 | 128826347 | 128826467 | 121 | 1 | ENSMUST00000116713.1 | -11581 | Gm25778 | ENSMUSG00000080363.1 |
| chr9 | 48538444 | 48538778 | 3.82278 | 4.89687 | 1.7347 | Distal Intergenic | 9 | 48506964 | 48507060 | 97 | 2 | ENSMUST00000116751.1 | -31385 | Gm23653 | ENSMUSG00000080401.1 |
| chr12 | 109612260 | 109612736 | 3.65589 | 4.41178 | 1.35537 | Promoter | 12 | 109611822 | 109611941 | 120 | 1 | ENSMUST00000116756.1 | 439 | Mir1188 | ENSMUSG00000080406.1 |
| chr5 | 120223010 | 120223160 | 3.3372 | 3.83314 | 1.01733 | Distal Intergenic | 5 | 120287726 | 120287847 | 122 | 1 | ENSMUST00000116762.1 | -64566 | Gm26474 | ENSMUSG00000080412.1 |
| chr5 | 120223740 | 120224007 | 3.63546 | 4.42978 | 1.43434 | Distal Intergenic | 5 | 120287726 | 120287847 | 122 | 1 | ENSMUST00000116762.1 | -63719 | Gm26474 | ENSMUSG00000080412.1 |
| chr5 | 120230761 | 120231039 | 3.38193 | 3.9006 | 1.01733 | Distal Intergenic | 5 | 120287726 | 120287847 | 122 | 1 | ENSMUST00000116762.1 | -56687 | Gm26474 | ENSMUSG00000080412.1 |
| chr5 | 120231514 | 120232631 | 3.59941 | 4.30099 | 1.26672 | Distal Intergenic | 5 | 120287726 | 120287847 | 122 | 1 | ENSMUST00000116762.1 | -55095 | Gm26474 | ENSMUSG00000080412.1 |
| chr5 | 120250385 | 120250623 | 3.54257 | 4.19979 | 1.21852 | Distal Intergenic | 5 | 120287726 | 120287847 | 122 | 1 | ENSMUST00000116762.1 | -37103 | Gm26474 | ENSMUSG00000080412.1 |
| chr5 | 120251180 | 120251660 | 3.43782 | 4.06441 | 1.19439 | Distal Intergenic | 5 | 120287726 | 120287847 | 122 | 1 | ENSMUST00000116762.1 | -36066 | Gm26474 | ENSMUSG00000080412.1 |
| chr5 | 120253083 | 120253364 | 3.91279 | 4.90245 | 1.68011 | Distal Intergenic | 5 | 120287726 | 120287847 | 122 | 1 | ENSMUST00000116762.1 | -34362 | Gm26474 | ENSMUSG00000080412.1 |
| chr5 | 120263530 | 120263736 | 3.27909 | 3.74597 | 1.01733 | Distal Intergenic | 5 | 120287726 | 120287847 | 122 | 1 | ENSMUST00000116762.1 | -23990 | Gm26474 | ENSMUSG00000080412.1 |
| chr5 | 120269601 | 120269980 | 3.682 | 4.50225 | 1.42307 | Distal Intergenic | 5 | 120287726 | 120287847 | 122 | 1 | ENSMUST00000116762.1 | -17746 | Gm26474 | ENSMUSG00000080412.1 |
| chr5 | 120271040 | 120271329 | 3.36453 | 4.00299 | 1.15403 | Distal Intergenic | 5 | 120287726 | 120287847 | 122 | 1 | ENSMUST00000116762.1 | -16397 | Gm26474 | ENSMUSG00000080412.1 |
| chr5 | 120281171 | 120281967 | 3.97169 | 5.17445 | 1.91522 | Distal Intergenic | 5 | 120287726 | 120287847 | 122 | 1 | ENSMUST00000116762.1 | -5759 | Gm26474 | ENSMUSG00000080412.1 |
| chr5 | 120287413 | 120287532 | 3.37929 | 4.02168 | 1.13571 | Promoter | 5 | 120287726 | 120287847 | 122 | 1 | ENSMUST00000116762.1 | -194 | Gm26474 | ENSMUSG00000080412.1 |
| chr5 | 120296931 | 120297287 | 3.60605 | 4.3232 | 1.3133 | Distal Intergenic | 5 | 120287726 | 120287847 | 122 | 1 | ENSMUST00000116762.1 | 9206 | Gm26474 | ENSMUSG00000080412.1 |
| chr5 | 120298399 | 120298803 | 3.51481 | 4.20199 | 1.2361 | Distal Intergenic | 5 | 120287726 | 120287847 | 122 | 1 | ENSMUST00000116762.1 | 10674 | Gm26474 | ENSMUSG00000080412.1 |
| chr5 | 120301654 | 120302016 | 3.51976 | 4.17241 | 1.23136 | Distal Intergenic | 5 | 120287726 | 120287847 | 122 | 1 | ENSMUST00000116762.1 | 13929 | Gm26474 | ENSMUSG00000080412.1 |
| chr5 | 120302358 | 120302684 | 3.65461 | 4.41273 | 1.36031 | Distal Intergenic | 5 | 120287726 | 120287847 | 122 | 1 | ENSMUST00000116762.1 | 14633 | Gm26474 | ENSMUSG00000080412.1 |
| chr5 | 120303990 | 120304443 | 3.63755 | 4.4688 | 1.40317 | Distal Intergenic | 5 | 120287726 | 120287847 | 122 | 1 | ENSMUST00000116762.1 | 16265 | Gm26474 | ENSMUSG00000080412.1 |
| chr5 | 120321791 | 120322705 | 3.68022 | 4.45489 | 1.37011 | Distal Intergenic | 5 | 120287726 | 120287847 | 122 | 1 | ENSMUST00000116762.1 | 34066 | Gm26474 | ENSMUSG00000080412.1 |
| chr5 | 120358137 | 120358400 | 3.36073 | 4.16417 | 1.23025 | Distal Intergenic | 5 | 120287726 | 120287847 | 122 | 1 | ENSMUST00000116762.1 | 70412 | Gm26474 | ENSMUSG00000080412.1 |
| chr14 | 67257241 | 67257563 | 3.3379 | 3.83419 | 1.01733 | Intron (ENSMUST00000022637.8/ENSMUSG00000022053.8, intron 7 of 16) | 14 | 67274869 | 67274963 | 95 | 2 | ENSMUST00000116782.1 | 17400 | Gm23178 | ENSMUSG00000080432.1 |
| chr14 | 67261270 | 67261925 | 3.51156 | 4.18715 | 1.29869 | Intron (ENSMUST00000022637.8/ENSMUSG00000022053.8, intron 7 of 16) | 14 | 67274869 | 67274963 | 95 | 2 | ENSMUST00000116782.1 | 13038 | Gm23178 | ENSMUSG00000080432.1 |
| chr14 | 67262517 | 67263122 | 3.77571 | 4.84831 | 1.66105 | Intron (ENSMUST00000022637.8/ENSMUSG00000022053.8, intron 7 of 16) | 14 | 67274869 | 67274963 | 95 | 2 | ENSMUST00000116782.1 | 11841 | Gm23178 | ENSMUSG00000080432.1 |
| chr14 | 67263973 | 67264290 | 3.54103 | 4.33143 | 1.33608 | Intron (ENSMUST00000022637.8/ENSMUSG00000022053.8, intron 7 of 16) | 14 | 67274869 | 67274963 | 95 | 2 | ENSMUST00000116782.1 | 10673 | Gm23178 | ENSMUSG00000080432.1 |
| chr14 | 67271409 | 67272255 | 3.45751 | 4.04426 | 1.12179 | Intron (ENSMUST00000022637.8/ENSMUSG00000022053.8, intron 7 of 16) | 14 | 67274869 | 67274963 | 95 | 2 | ENSMUST00000116782.1 | 2708 | Gm23178 | ENSMUSG00000080432.1 |
| chr14 | 67272586 | 67273817 | 3.90023 | 4.9362 | 1.77895 | Intron (ENSMUST00000022637.8/ENSMUSG00000022053.8, intron 7 of 16) | 14 | 67274869 | 67274963 | 95 | 2 | ENSMUST00000116782.1 | 1146 | Gm23178 | ENSMUSG00000080432.1 |
| chr14 | 67274374 | 67274670 | 3.25617 | 3.71147 | 1.01733 | Promoter | 14 | 67274869 | 67274963 | 95 | 2 | ENSMUST00000116782.1 | 293 | Gm23178 | ENSMUSG00000080432.1 |
| chr14 | 67279027 | 67279269 | 3.6347 | 4.42365 | 1.43805 | Intron (ENSMUST00000022637.8/ENSMUSG00000022053.8, intron 7 of 16) | 14 | 67274869 | 67274963 | 95 | 2 | ENSMUST00000116782.1 | -4065 | Gm23178 | ENSMUSG00000080432.1 |
| chr14 | 67288741 | 67289037 | 3.47092 | 4.08782 | 1.20578 | Intron (ENSMUST00000022637.8/ENSMUSG00000022053.8, intron 7 of 16) | 14 | 67274869 | 67274963 | 95 | 2 | ENSMUST00000116782.1 | -13779 | Gm23178 | ENSMUSG00000080432.1 |
| chr4 | 135509245 | 135509406 | 3.4493 | 4.02395 | 1.09361 | Intron (ENSMUST00000153891.2/ENSMUSG00000028801.9, intron 2 of 3) | 4 | 135512351 | 135512479 | 129 | 2 | ENSMUST00000116824.1 | 3073 | Gm25317 | ENSMUSG00000080474.1 |
| chr4 | 135524824 | 135526478 | 3.90927 | 4.98913 | 1.82023 | Exon (ENSMUST00000123221.1/ENSMUSG00000028801.9, exon 7 of 9) | 4 | 135512351 | 135512479 | 129 | 2 | ENSMUST00000116824.1 | -12346 | Gm25317 | ENSMUSG00000080474.1 |
| chr4 | 135530341 | 135530709 | 3.56135 | 4.30849 | 1.35788 | Intron (ENSMUST00000123221.1/ENSMUSG00000028801.9, intron 8 of 8) | 4 | 135512351 | 135512479 | 129 | 2 | ENSMUST00000116824.1 | -17863 | Gm25317 | ENSMUSG00000080474.1 |
| chr10 | 60957726 | 60958156 | 3.65112 | 4.40077 | 1.33399 | Distal Intergenic | 10 | 60960723 | 60960844 | 122 | 1 | ENSMUST00000116876.1 | -2567 | Mir466j | ENSMUSG00000080526.1 |
| chr10 | 60959011 | 60959186 | 3.35989 | 3.86736 | 1.01733 | Downstream (1-2kb) | 10 | 60960723 | 60960844 | 122 | 1 | ENSMUST00000116876.1 | -1537 | Mir466j | ENSMUSG00000080526.1 |
| chr10 | 60970686 | 60970766 | 3.89903 | 4.91766 | 1.75359 | Distal Intergenic | 10 | 60960723 | 60960844 | 122 | 1 | ENSMUST00000116876.1 | 9964 | Mir466j | ENSMUSG00000080526.1 |
| chr10 | 60979332 | 60979634 | 3.58898 | 4.31038 | 1.36227 | Distal Intergenic | 10 | 60960723 | 60960844 | 122 | 1 | ENSMUST00000116876.1 | 18610 | Mir466j | ENSMUSG00000080526.1 |
| chr18 | 35919549 | 35919719 | 3.23963 | 3.6866 | 1.01733 | Distal Intergenic | 18 | 35952824 | 35952907 | 84 | 1 | ENSMUST00000116879.1 | -33105 | Gm23882 | ENSMUSG00000080529.1 |
| chr18 | 35920674 | 35921073 | 3.73922 | 4.58286 | 1.50173 | Distal Intergenic | 18 | 35952824 | 35952907 | 84 | 1 | ENSMUST00000116879.1 | -31751 | Gm23882 | ENSMUSG00000080529.1 |
| chr18 | 35930716 | 35930844 | 3.45403 | 4.2414 | 1.29259 | Distal Intergenic | 18 | 35952824 | 35952907 | 84 | 1 | ENSMUST00000116879.1 | -21980 | Gm23882 | ENSMUSG00000080529.1 |
| chr18 | 35938413 | 35938690 | 3.50358 | 4.17309 | 1.28287 | Distal Intergenic | 18 | 35952824 | 35952907 | 84 | 1 | ENSMUST00000116879.1 | -14134 | Gm23882 | ENSMUSG00000080529.1 |
| chr6 | 148957316 | 148957559 | 3.62551 | 4.36791 | 1.33433 | Distal Intergenic | 6 | 148967316 | 148967437 | 122 | 1 | ENSMUST00000116886.1 | -9757 | Gm25539 | ENSMUSG00000080536.1 |
| chr7 | 27362451 | 27362682 | 3.92906 | 4.94774 | 1.73522 | Promoter | 7 | 27363314 | 27363401 | 88 | 1 | ENSMUST00000116912.1 | -632 | Gm26043 | ENSMUSG00000080562.1 |
| chr7 | 27363739 | 27364520 | 3.80487 | 5.12024 | 1.86055 | Promoter | 7 | 27363314 | 27363401 | 88 | 1 | ENSMUST00000116912.1 | 426 | Gm26043 | ENSMUSG00000080562.1 |
| chr7 | 27365588 | 27365815 | 4.42925 | 5.97462 | 2.50273 | Exon (ENSMUST00000108362.2/ENSMUSG00000011751.11, exon 12 of 18) | 7 | 27363314 | 27363401 | 88 | 1 | ENSMUST00000116912.1 | 2275 | Gm26043 | ENSMUSG00000080562.1 |
| chr7 | 27367488 | 27368231 | 3.63193 | 4.598 | 1.51562 | Exon (ENSMUST00000108362.2/ENSMUSG00000011751.11, exon 9 of 18) | 7 | 27363314 | 27363401 | 88 | 1 | ENSMUST00000116912.1 | 4175 | Gm26043 | ENSMUSG00000080562.1 |
| chr7 | 27369450 | 27369965 | 3.43746 | 4.05912 | 1.23736 | Exon (ENSMUST00000108362.2/ENSMUSG00000011751.11, exon 8 of 18) | 7 | 27363314 | 27363401 | 88 | 1 | ENSMUST00000116912.1 | 6137 | Gm26043 | ENSMUSG00000080562.1 |
| chr7 | 27370394 | 27370918 | 3.67708 | 4.70261 | 1.59033 | Exon (ENSMUST00000108362.2/ENSMUSG00000011751.11, exon 7 of 18) | 7 | 27363314 | 27363401 | 88 | 1 | ENSMUST00000116912.1 | 7081 | Gm26043 | ENSMUSG00000080562.1 |
| chr7 | 27371314 | 27371464 | 3.95679 | 5.23667 | 1.92514 | Intron (ENSMUST00000108362.2/ENSMUSG00000011751.11, intron 6 of 17) | 7 | 27363314 | 27363401 | 88 | 1 | ENSMUST00000116912.1 | 8001 | Gm26043 | ENSMUSG00000080562.1 |
| chr7 | 27372049 | 27372311 | 3.67643 | 4.7923 | 1.65216 | Exon (ENSMUST00000108362.2/ENSMUSG00000011751.11, exon 5 of 18) | 7 | 27363314 | 27363401 | 88 | 1 | ENSMUST00000116912.1 | 8736 | Gm26043 | ENSMUSG00000080562.1 |
| chr7 | 27373430 | 27374847 | 3.57377 | 4.417 | 1.40729 | Exon (ENSMUST00000108362.2/ENSMUSG00000011751.11, exon 4 of 18) | 7 | 27363314 | 27363401 | 88 | 1 | ENSMUST00000116912.1 | 10117 | Gm26043 | ENSMUSG00000080562.1 |
| chr7 | 27375178 | 27375593 | 3.83491 | 4.75227 | 1.57316 | Exon (ENSMUST00000108362.2/ENSMUSG00000011751.11, exon 3 of 18) | 7 | 27363314 | 27363401 | 88 | 1 | ENSMUST00000116912.1 | 11865 | Gm26043 | ENSMUSG00000080562.1 |
| chr7 | 27376895 | 27377114 | 3.97606 | 5.03127 | 1.76436 | Intron (ENSMUST00000108362.2/ENSMUSG00000011751.11, intron 2 of 17) | 7 | 27363314 | 27363401 | 88 | 1 | ENSMUST00000116912.1 | 13582 | Gm26043 | ENSMUSG00000080562.1 |
| chr7 | 27378452 | 27378603 | 3.4948 | 4.21709 | 1.27832 | Intron (ENSMUST00000108362.2/ENSMUSG00000011751.11, intron 2 of 17) | 7 | 27363314 | 27363401 | 88 | 1 | ENSMUST00000116912.1 | 15139 | Gm26043 | ENSMUSG00000080562.1 |
| chr12 | 101011717 | 101011878 | 3.27799 | 3.74412 | 1.01733 | Intron (ENSMUST00000068411.3/ENSMUSG00000021182.9, intron 3 of 29) | 12 | 101021673 | 101021768 | 96 | 2 | ENSMUST00000116925.2 | 9890 | Mir1190 | ENSMUSG00000080575.2 |
| chr11 | 71986833 | 71987510 | 3.62616 | 4.39852 | 1.37938 | Distal Intergenic | 11 | 72022755 | 72022857 | 103 | 1 | ENSMUST00000116927.1 | -35245 | Gm23226 | ENSMUSG00000080577.1 |
| chr11 | 72002727 | 72002998 | 3.17145 | 3.72743 | 1.02772 | Distal Intergenic | 11 | 72022755 | 72022857 | 103 | 1 | ENSMUST00000116927.1 | -19757 | Gm23226 | ENSMUSG00000080577.1 |
| chr11 | 72028146 | 72028451 | 3.48038 | 4.15034 | 1.23979 | 3' UTR | 11 | 72022755 | 72022857 | 103 | 1 | ENSMUST00000116927.1 | 5392 | Gm23226 | ENSMUSG00000080577.1 |
| chr13 | 71097722 | 71097930 | 4.02362 | 5.14719 | 1.9078 | Distal Intergenic | 13 | 71107089 | 71107209 | 121 | 1 | ENSMUST00000116930.1 | -9159 | Mir466f-4 | ENSMUSG00000080580.1 |
| chr13 | 71105677 | 71105937 | 3.79696 | 4.68269 | 1.54574 | Downstream (1-2kb) | 13 | 71107089 | 71107209 | 121 | 1 | ENSMUST00000116930.1 | -1152 | Mir466f-4 | ENSMUSG00000080580.1 |
| chr13 | 71148017 | 71148417 | 3.99189 | 5.05231 | 1.77946 | Distal Intergenic | 13 | 71107089 | 71107209 | 121 | 1 | ENSMUST00000116930.1 | 40929 | Mir466f-4 | ENSMUSG00000080580.1 |
| chr13 | 71157429 | 71157725 | 3.33992 | 3.87009 | 1.08949 | Distal Intergenic | 13 | 71107089 | 71107209 | 121 | 1 | ENSMUST00000116930.1 | 50341 | Mir466f-4 | ENSMUSG00000080580.1 |
| chr13 | 71459164 | 71459239 | 3.62015 | 4.75236 | 1.59386 | Distal Intergenic | 13 | 71107089 | 71107209 | 121 | 1 | ENSMUST00000116930.1 | 352076 | Mir466f-4 | ENSMUSG00000080580.1 |
| chr13 | 71461816 | 71461958 | 3.27259 | 3.90384 | 1.07467 | Distal Intergenic | 13 | 71107089 | 71107209 | 121 | 1 | ENSMUST00000116930.1 | 354728 | Mir466f-4 | ENSMUSG00000080580.1 |
| chr13 | 71464645 | 71465022 | 3.58061 | 4.40853 | 1.40279 | Distal Intergenic | 13 | 71107089 | 71107209 | 121 | 1 | ENSMUST00000116930.1 | 357557 | Mir466f-4 | ENSMUSG00000080580.1 |
| chr5 | 67052900 | 67053000 | 4.03528 | 5.17778 | 1.90773 | Promoter | 5 | 67053679 | 67053792 | 114 | 2 | ENSMUST00000116951.1 | 792 | Gm23841 | ENSMUSG00000080601.1 |
| chr5 | 67057853 | 67058196 | 3.43233 | 4.04555 | 1.21062 | Distal Intergenic | 5 | 67053679 | 67053792 | 114 | 2 | ENSMUST00000116951.1 | -4062 | Gm23841 | ENSMUSG00000080601.1 |
| chr5 | 67060193 | 67060463 | 3.59678 | 4.34574 | 1.39834 | Distal Intergenic | 5 | 67053679 | 67053792 | 114 | 2 | ENSMUST00000116951.1 | -6402 | Gm23841 | ENSMUSG00000080601.1 |
| chr7 | 67775974 | 67776200 | 3.32998 | 3.97317 | 1.16291 | Downstream (2-3kb) | 7 | 67778410 | 67778530 | 121 | 2 | ENSMUST00000116959.1 | 2330 | Gm23844 | ENSMUSG00000080609.1 |
| chr9 | 115392213 | 115392299 | 3.56521 | 4.23553 | 1.22269 | Distal Intergenic | 9 | 115382099 | 115382221 | 123 | 1 | ENSMUST00000116986.1 | 10115 | Gm25510 | ENSMUSG00000080636.1 |
| chr13 | 23290369 | 23290597 | 3.47727 | 4.19049 | 1.26153 | Intron (ENSMUST00000155432.1/ENSMUSG00000085488.1, intron 1 of 1) | 13 | 23298049 | 23298196 | 148 | 1 | ENSMUST00000119479.1 | -7452 | Gm11333 | ENSMUSG00000080698.1 |
| chr13 | 23292584 | 23292696 | 3.86273 | 4.90812 | 1.73488 | Intron (ENSMUST00000155432.1/ENSMUSG00000085488.1, intron 1 of 1) | 13 | 23298049 | 23298196 | 148 | 1 | ENSMUST00000119479.1 | -5353 | Gm11333 | ENSMUSG00000080698.1 |
| chr13 | 23296687 | 23298226 | 3.69705 | 4.66544 | 1.56822 | Promoter | 13 | 23298049 | 23298196 | 148 | 1 | ENSMUST00000119479.1 | 0 | Gm11333 | ENSMUSG00000080698.1 |
| chr13 | 23299910 | 23300780 | 3.68186 | 4.58026 | 1.54416 | Intron (ENSMUST00000155432.1/ENSMUSG00000085488.1, intron 1 of 1) | 13 | 23298049 | 23298196 | 148 | 1 | ENSMUST00000119479.1 | 1862 | Gm11333 | ENSMUSG00000080698.1 |
| chr13 | 23301595 | 23302047 | 3.8348 | 4.85224 | 1.71691 | Intron (ENSMUST00000155432.1/ENSMUSG00000085488.1, intron 1 of 1) | 13 | 23298049 | 23298196 | 148 | 1 | ENSMUST00000119479.1 | 3547 | Gm11333 | ENSMUSG00000080698.1 |
| chr13 | 23302450 | 23303267 | 4.21037 | 5.62557 | 2.29067 | Intron (ENSMUST00000155432.1/ENSMUSG00000085488.1, intron 1 of 1) | 13 | 23298049 | 23298196 | 148 | 1 | ENSMUST00000119479.1 | 4402 | Gm11333 | ENSMUSG00000080698.1 |
| chr13 | 23304097 | 23304340 | 3.68144 | 4.60588 | 1.53531 | Intron (ENSMUST00000155432.1/ENSMUSG00000085488.1, intron 1 of 1) | 13 | 23298049 | 23298196 | 148 | 1 | ENSMUST00000119479.1 | 6049 | Gm11333 | ENSMUSG00000080698.1 |
| chr6 | 18336451 | 18336688 | 3.40212 | 4.40519 | 1.37486 | Distal Intergenic | 6 | 18294271 | 18295231 | 961 | 1 | ENSMUST00000120840.1 | 42181 | Gm15595 | ENSMUSG00000080767.1 |
| chr13 | 21881339 | 21882421 | 3.76403 | 4.87325 | 1.68745 | Distal Intergenic | 13 | 21875233 | 21876229 | 997 | 2 | ENSMUST00000121002.1 | -5111 | Gm11278 | ENSMUSG00000080770.1 |
| chr2 | 5379139 | 5379364 | 2.81585 | 10.7187 | 6.71784 | Intron (ENSMUST00000044009.8/ENSMUSG00000039145.11, intron 3 of 10) | 2 | 5283650 | 5283874 | 225 | 1 | ENSMUST00000117204.1 | 95490 | Gm13197 | ENSMUSG00000080820.1 |
| chr11 | 40600311 | 40600670 | 3.27579 | 3.74082 | 1.01733 | Distal Intergenic | 11 | 40612718 | 40613063 | 346 | 1 | ENSMUST00000119565.1 | -12048 | Gm12137 | ENSMUSG00000080835.1 |

| | | | | | | | | | | | | | | | |
|---|---|---|---|---|---|---|---|---|---|---|---|---|---|---|---|
| chr11 | 101479140 | 101479380 | 3.5167 | 4.22384 | 1.28091 | Distal Intergenic | 11 | 101476226 | 101476521 | 296 | 1 | ENSMUST00000119535.1 | 2915 | Gm11625 | ENSMUSG00000080839.1 |
| chr2 | 105611091 | 105611172 | 3.50075 | 4.58946 | 1.48262 | Intron (ENSMUST00000149133.2/ENSMUSG00000086029.2, intron 1 of 3) | 2 | 105554201 | 105554845 | 645 | 2 | ENSMUST00000119438.1 | -56247 | Rpl10-ps1 | ENSMUSG00000080859.1 |
| chr12 | 87057135 | 87057270 | 3.43151 | 4.12112 | 1.20877 | Exon (ENSMUST00000154801.2/ENSMUSG00000034145.9, exon 5 of 26) | 12 | 87067477 | 87067591 | 115 | 1 | ENSMUST00000119807.1 | -10207 | Gm15745 | ENSMUSG00000080868.1 |
| chr2 | 146720069 | 146720383 | 3.32733 | 3.81828 | 1.01733 | Distal Intergenic | 2 | 146754707 | 146755705 | 999 | 2 | ENSMUST00000119851.1 | 35322 | Gm14111 | ENSMUSG00000080870.1 |
| chr11 | 93763208 | 93763501 | 3.34964 | 3.86179 | 1.05215 | Distal Intergenic | 11 | 93695082 | 93695551 | 470 | 1 | ENSMUST00000122414.1 | 68127 | Gm11506 | ENSMUSG00000080872.1 |
| chr11 | 52437565 | 52437641 | 4.31841 | 5.79455 | 2.34867 | Distal Intergenic | 11 | 52405316 | 52405478 | 163 | 1 | ENSMUST00000122004.1 | 32250 | Gm12208 | ENSMUSG00000080951.1 |
| chr5 | 146261008 | 146261348 | 2.74638 | 18.06478 | 13.1567 | Intron (ENSMUST00000162494.2/ENSMUSG00000029635.10, intron 2 of 11) | 5 | 146248551 | 146249422 | 872 | 1 | ENSMUST00000118049.1 | 12458 | Gm15739 | ENSMUSG00000080957.1 |
| chr11 | 81534319 | 81534394 | 3.85383 | 4.84946 | 1.7141 | Intron (ENSMUST00000066197.6/ENSMUSG00000020704.8, intron 1 of 9) | 11 | 81572655 | 81573473 | 819 | 2 | ENSMUST00000117522.1 | 39079 | 1700071K01Rik | ENSMUSG00000080980.1 |
| chr6 | 147268237 | 147269181 | 3.29057 | 3.77705 | 1.06404 | Distal Intergenic | 6 | 147271837 | 147272649 | 813 | 1 | ENSMUST00000118025.1 | -2656 | Gm15766 | ENSMUSG00000081023.1 |
| chr4 | 13565566 | 13565726 | 3.37267 | 3.92114 | 1.13738 | Distal Intergenic | 4 | 13606034 | 13606769 | 736 | 1 | ENSMUST00000120710.1 | -40308 | Gm11825 | ENSMUSG00000081063.1 |
| chr11 | 33013296 | 33013607 | 3.50313 | 4.14715 | 1.24385 | Distal Intergenic | 11 | 33038816 | 33039703 | 888 | 2 | ENSMUST00000121691.1 | 26096 | Rpsa-ps4 | ENSMUSG00000081076.3 |
| chr11 | 33048656 | 33048738 | 3.96402 | 4.98041 | 1.69806 | Distal Intergenic | 11 | 33038816 | 33039703 | 888 | 2 | ENSMUST00000121691.1 | -8954 | Rpsa-ps4 | ENSMUSG00000081076.3 |
| chr11 | 111967679 | 111967943 | 3.35682 | 3.89934 | 1.12523 | Distal Intergenic | 11 | 112042800 | 112043578 | 779 | 2 | ENSMUST00000121830.1 | 75635 | Gm11679 | ENSMUSG00000081088.1 |
| chr2 | 150526075 | 150526244 | 3.73499 | 4.90527 | 1.72814 | Distal Intergenic | 2 | 150521093 | 150522300 | 1208 | 1 | ENSMUST00000119514.1 | 4983 | Gm14128 | ENSMUSG00000081092.1 |
| chr6 | 127266509 | 127266840 | 3.7393 | 4.58782 | 1.50526 | Distal Intergenic | 6 | 127282063 | 127282509 | 447 | 2 | ENSMUST00000121087.1 | 15669 | Gm7308 | ENSMUSG00000081113.1 |
| chr6 | 127336227 | 127336479 | 3.47012 | 4.44998 | 1.39565 | Distal Intergenic | 6 | 127282063 | 127282509 | 447 | 2 | ENSMUST00000121087.1 | -53719 | Gm7308 | ENSMUSG00000081113.1 |
| chr6 | 127363387 | 127363767 | 3.2743 | 3.7386 | 1.01733 | Distal Intergenic | 6 | 127282063 | 127282509 | 447 | 2 | ENSMUST00000121087.1 | -80879 | Gm7308 | ENSMUSG00000081113.1 |
| chr2 | 18731611 | 18731922 | 3.49564 | 4.13988 | 1.24017 | Intron (ENSMUST00000095132.4/ENSMUSG00000037708.11, intron 4 of 10) | 2 | 18758984 | 18759764 | 781 | 1 | ENSMUST00000119424.1 | -27062 | Gm13333 | ENSMUSG00000081139.1 |
| chr4 | 99459383 | 99459791 | 3.78303 | 4.67552 | 1.58027 | Distal Intergenic | 4 | 99430256 | 99431254 | 999 | 2 | ENSMUST00000118675.1 | -28130 | Gm12684 | ENSMUSG00000081150.1 |
| chr16 | 91107297 | 91107447 | 3.54497 | 4.20387 | 1.21881 | Distal Intergenic | 16 | 91114189 | 91114599 | 411 | 2 | ENSMUST00000119544.1 | 7152 | H3f3a-ps2 | ENSMUSG00000081176.1 |
| chr16 | 91110143 | 91110265 | 3.4969 | 4.11622 | 1.16564 | Distal Intergenic | 16 | 91114189 | 91114599 | 411 | 2 | ENSMUST00000119544.1 | 4334 | H3f3a-ps2 | ENSMUSG00000081176.1 |
| chr16 | 91118729 | 91119295 | 3.46756 | 4.08895 | 1.20409 | Distal Intergenic | 16 | 91114189 | 91114599 | 411 | 2 | ENSMUST00000119544.1 | -4131 | H3f3a-ps2 | ENSMUSG00000081176.1 |
| chr16 | 91139527 | 91139827 | 3.99592 | 5.24513 | 1.98418 | Distal Intergenic | 16 | 91114189 | 91114599 | 411 | 2 | ENSMUST00000119544.1 | -24929 | H3f3a-ps2 | ENSMUSG00000081176.1 |
| chr16 | 91140142 | 91140575 | 3.42224 | 4.04852 | 1.21111 | Distal Intergenic | 16 | 91114189 | 91114599 | 411 | 2 | ENSMUST00000119544.1 | -25544 | H3f3a-ps2 | ENSMUSG00000081176.1 |
| chr16 | 91141238 | 91141481 | 3.51241 | 4.38774 | 1.35834 | Distal Intergenic | 16 | 91114189 | 91114599 | 411 | 2 | ENSMUST00000119544.1 | -26640 | H3f3a-ps2 | ENSMUSG00000081176.1 |
| chr4 | 147738630 | 147739103 | 3.1833 | 4.31983 | 1.33518 | Promoter | 4 | 147735477 | 147738426 | 2950 | 2 | ENSMUST00000121031.1 | -205 | Gm13136 | ENSMUSG00000081179.1 |
| chr2 | 18585122 | 18585357 | 3.55393 | 4.22949 | 1.24359 | Distal Intergenic | 2 | 18568126 | 18568198 | 73 | 1 | ENSMUST00000120891.1 | 16997 | Gm13353 | ENSMUSG00000081181.1 |
| chr2 | 18585775 | 18587809 | 3.85759 | 4.88119 | 1.69768 | Distal Intergenic | 2 | 18568126 | 18568198 | 73 | 1 | ENSMUST00000120891.1 | 17650 | Gm13353 | ENSMUSG00000081181.1 |
| chr2 | 18588183 | 18588361 | 4.10404 | 5.31613 | 1.95768 | Distal Intergenic | 2 | 18568126 | 18568198 | 73 | 1 | ENSMUST00000120891.1 | 20058 | Gm13353 | ENSMUSG00000081181.1 |
| chr2 | 18589409 | 18589613 | 3.38972 | 3.92029 | 1.04543 | Distal Intergenic | 2 | 18568126 | 18568198 | 73 | 1 | ENSMUST00000120891.1 | 21284 | Gm13353 | ENSMUSG00000081181.1 |
| chr2 | 18590158 | 18590608 | 3.66939 | 4.52915 | 1.49392 | Distal Intergenic | 2 | 18568126 | 18568198 | 73 | 1 | ENSMUST00000120891.1 | 22033 | Gm13353 | ENSMUSG00000081181.1 |
| chr2 | 18591028 | 18591258 | 3.34153 | 3.83966 | 1.01733 | Distal Intergenic | 2 | 18568126 | 18568198 | 73 | 1 | ENSMUST00000120891.1 | 22903 | Gm13353 | ENSMUSG00000081181.1 |
| chr2 | 18603034 | 18603702 | 3.59158 | 4.28982 | 1.27245 | Distal Intergenic | 2 | 18568126 | 18568198 | 73 | 1 | ENSMUST00000120891.1 | 34909 | Gm13353 | ENSMUSG00000081181.1 |
| chr2 | 18604121 | 18604675 | 3.34966 | 3.86584 | 1.06535 | Distal Intergenic | 2 | 18568126 | 18568198 | 73 | 1 | ENSMUST00000120891.1 | 35996 | Gm13353 | ENSMUSG00000081181.1 |
| chr4 | 126656664 | 126656739 | 3.87197 | 4.80893 | 1.58896 | Distal Intergenic | 4 | 126665254 | 126666125 | 872 | 1 | ENSMUST00000120203.1 | -8515 | Gm12933 | ENSMUSG00000081210.1 |
| chr4 | 126660743 | 126660889 | 3.45111 | 4.03582 | 1.12654 | Distal Intergenic | 4 | 126665254 | 126666125 | 872 | 1 | ENSMUST00000120203.1 | -4365 | Gm12933 | ENSMUSG00000081210.1 |
| chr11 | 58097703 | 58097821 | 3.40043 | 3.97847 | 1.18208 | Promoter | 11 | 58097104 | 58097773 | 670 | 2 | ENSMUST00000117585.1 | 0 | Gm12247 | ENSMUSG00000081223.1 |
| chr4 | 118517111 | 118517448 | 3.61674 | 4.42873 | 1.36848 | Distal Intergenic | 4 | 118522018 | 118522609 | 592 | 1 | ENSMUST00000117678.1 | -4570 | Gm12857 | ENSMUSG00000081233.1 |
| chr13 | 33217706 | 33217878 | 3.27853 | 3.74493 | 1.01733 | Promoter | 13 | 33217268 | 33218151 | 884 | 2 | ENSMUST00000121513.1 | 273 | Gm11385 | ENSMUSG00000081237.3 |
| chr11 | 32900652 | 32901127 | 3.98434 | 5.04352 | 1.78672 | Downstream (2-3kb) | 11 | 32910373 | 32910603 | 231 | 1 | ENSMUST00000117772.1 | -9246 | Gm12112 | ENSMUSG00000081254.1 |
| chr11 | 32908320 | 32908519 | 3.47539 | 4.07591 | 1.13789 | Intron (ENSMUST00000150073.1/ENSMUSG00000085402.1, intron 2 of 2) | 11 | 32910373 | 32910603 | 231 | 1 | ENSMUST00000117772.1 | -1854 | Gm12112 | ENSMUSG00000081254.1 |
| chr11 | 59575446 | 59575855 | 3.4348 | 4.05221 | 1.23565 | Downstream (2-3kb) | 11 | 59577992 | 59578636 | 645 | 2 | ENSMUST00000117499.1 | 2781 | Gm12261 | ENSMUSG00000081262.1 |
| chr11 | 59578806 | 59579149 | 3.33499 | 4.00995 | 1.15083 | Promoter | 11 | 59577992 | 59578636 | 645 | 2 | ENSMUST00000117499.1 | -171 | Gm12261 | ENSMUSG00000081262.1 |
| chr4 | 154207446 | 154207672 | 3.40783 | 3.96063 | 1.09149 | Intron (ENSMUST00000030897.9/ENSMUSG00000057751.9, intron 3 of 36) | 4 | 154220541 | 154221012 | 472 | 1 | ENSMUST00000121194.1 | -12869 | Gm13132 | ENSMUSG00000081276.1 |
| chr13 | 17859574 | 17859801 | 3.28149 | 3.76003 | 1.04052 | Distal Intergenic | 13 | 17820946 | 17821770 | 825 | 1 | ENSMUST00000118792.2 | 38629 | Gm18859 | ENSMUSG00000081285.2 |
| chr11 | 52461485 | 52461979 | 3.60011 | 4.30161 | 1.26681 | Distal Intergenic | 11 | 52478508 | 52479103 | 596 | 2 | ENSMUST00000120089.1 | 17124 | Gm12209 | ENSMUSG00000081297.1 |
| chr11 | 52479250 | 52479751 | 3.88034 | 5.10524 | 1.82684 | Promoter | 11 | 52478508 | 52479103 | 596 | 2 | ENSMUST00000120089.1 | -148 | Gm12209 | ENSMUSG00000081297.1 |
| chr11 | 52480913 | 52481119 | 3.83564 | 4.75308 | 1.57037 | Distal Intergenic | 11 | 52478508 | 52479103 | 596 | 2 | ENSMUST00000120089.1 | -1811 | Gm12209 | ENSMUSG00000081297.1 |
| chr11 | 50457742 | 50457875 | 3.28227 | 3.75473 | 1.03179 | Distal Intergenic | 11 | 50446577 | 50447023 | 447 | 1 | ENSMUST00000118203.1 | 11166 | Gm12197 | ENSMUSG00000081311.1 |
| chr11 | 50458231 | 50458706 | 3.49261 | 4.18973 | 1.27301 | Distal Intergenic | 11 | 50446577 | 50447023 | 447 | 1 | ENSMUST00000118203.1 | 11655 | Gm12197 | ENSMUSG00000081311.1 |
| chr11 | 50459203 | 50459278 | 3.2537 | 3.70769 | 1.01733 | Distal Intergenic | 11 | 50446577 | 50447023 | 447 | 1 | ENSMUST00000118203.1 | 12627 | Gm12197 | ENSMUSG00000081311.1 |
| chr11 | 83817106 | 83817443 | 3.91662 | 4.90654 | 1.67689 | Downstream (<1kb) | 11 | 83800464 | 83801092 | 629 | 1 | ENSMUST00000122379.1 | 16643 | Gm11434 | ENSMUSG00000081342.1 |
| chr11 | 83817785 | 83818163 | 3.57821 | 4.26258 | 1.24259 | Exon (ENSMUST00000135268.1/ENSMUSG00000085235.1, exon 2 of 2) | 11 | 83800464 | 83801092 | 629 | 1 | ENSMUST00000122379.1 | 17322 | Gm11434 | ENSMUSG00000081342.1 |
| chr4 | 81563478 | 81563875 | 3.90386 | 4.96536 | 1.78723 | Distal Intergenic | 4 | 81455943 | 81461732 | 5790 | 2 | ENSMUST00000121173.1 | -101747 | Gm11765 | ENSMUSG00000081391.1 |
| chr4 | 81566469 | 81566670 | 3.67013 | 4.43152 | 1.35606 | Distal Intergenic | 4 | 81455943 | 81461732 | 5790 | 2 | ENSMUST00000121173.1 | -104738 | Gm11765 | ENSMUSG00000081391.1 |
| chr4 | 81568066 | 81568207 | 3.41125 | 3.97555 | 1.12609 | Distal Intergenic | 4 | 81455943 | 81461732 | 5790 | 2 | ENSMUST00000121173.1 | -106335 | Gm11765 | ENSMUSG00000081391.1 |
| chr11 | 109011652 | 109012088 | 2.40907 | 16.32823 | 11.52381 | Distal Intergenic | 11 | 109007651 | 109008781 | 1131 | 1 | ENSMUST00000119655.1 | 4002 | Gm11668 | ENSMUSG00000081392.1 |
| chr4 | 147100733 | 147100808 | 3.21105 | 4.57903 | 1.49609 | Intron (ENSMUST00000181084.1/ENSMUSG00000097168.1, intron 3 of 3) | 4 | 147102327 | 147103981 | 1655 | 1 | ENSMUST00000117213.1 | -1519 | Gm13142 | ENSMUSG00000081405.1 |
| chr4 | 146263509 | 146264019 | 3.32135 | 5.649 | 2.23798 | Promoter | 4 | 146263559 | 146264190 | 632 | 1 | ENSMUST00000121771.1 | 0 | Gm13168 | ENSMUSG00000081460.1 |
| chr2 | 167588826 | 167589607 | 3.85489 | 4.83443 | 1.702 | Intron (ENSMUST00000127006.1/ENSMUSG00000085596.1, intron 2 of 3) | 2 | 167591395 | 167591870 | 476 | 1 | ENSMUST00000118482.1 | -1788 | Gm11475 | ENSMUSG00000081511.1 |
| chr2 | 167597452 | 167597666 | 3.67065 | 4.46746 | 1.45996 | Intron (ENSMUST00000127006.1/ENSMUSG00000085596.1, intron 2 of 3) | 2 | 167591395 | 167591870 | 476 | 1 | ENSMUST00000118482.1 | 6058 | Gm11475 | ENSMUSG00000081511.1 |
| chr11 | 109192688 | 109193251 | 3.75924 | 4.65233 | 1.57454 | Distal Intergenic | 11 | 109159187 | 109160194 | 1008 | 2 | ENSMUST00000120913.1 | -32495 | Gm11658 | ENSMUSG00000081524.3 |
| chr11 | 114312460 | 114312639 | 3.38407 | 3.90654 | 1.02686 | Distal Intergenic | 11 | 114357129 | 114358261 | 1133 | 1 | ENSMUST00000122121.1 | -44490 | Gm11692 | ENSMUSG00000081548.1 |
| chr11 | 114314088 | 114314364 | 3.89459 | 5.02532 | 1.80343 | Distal Intergenic | 11 | 114357129 | 114358261 | 1133 | 1 | ENSMUST00000122121.1 | -42765 | Gm11692 | ENSMUSG00000081548.1 |
| chr11 | 114315338 | 114315616 | 3.33244 | 3.89918 | 1.11305 | Distal Intergenic | 11 | 114357129 | 114358261 | 1133 | 1 | ENSMUST00000122121.1 | -41513 | Gm11692 | ENSMUSG00000081548.1 |
| chr11 | 114316016 | 114316249 | 3.70653 | 4.55935 | 1.4693 | Distal Intergenic | 11 | 114357129 | 114358261 | 1133 | 1 | ENSMUST00000122121.1 | -40880 | Gm11692 | ENSMUSG00000081548.1 |
| chr11 | 114376116 | 114376235 | 3.37275 | 3.88675 | 1.01733 | Distal Intergenic | 11 | 114357129 | 114358261 | 1133 | 1 | ENSMUST00000122121.1 | 18988 | Gm11692 | ENSMUSG00000081548.1 |
| chr4 | 45590645 | 45591691 | 3.62682 | 4.38572 | 1.3292 | Distal Intergenic | 4 | 45550017 | 45550499 | 483 | 1 | ENSMUST00000119140.1 | 40629 | Gm12411 | ENSMUSG00000081559.3 |
| chr13 | 21908914 | 21909113 | 3.40954 | 3.95093 | 1.04784 | Distal Intergenic | 13 | 21903156 | 21904784 | 1629 | 1 | ENSMUST00000121524.1 | 5759 | Gm11279 | ENSMUSG00000081576.1 |
| chr13 | 21910967 | 21911117 | 3.38675 | 3.91626 | 1.04699 | Distal Intergenic | 13 | 21903156 | 21904784 | 1629 | 1 | ENSMUST00000121524.1 | 7812 | Gm11279 | ENSMUSG00000081576.1 |
| chr13 | 21911509 | 21911654 | 3.355 | 3.85997 | 1.01733 | Distal Intergenic | 13 | 21903156 | 21904784 | 1629 | 1 | ENSMUST00000121524.1 | 8354 | Gm11279 | ENSMUSG00000081576.1 |
| chr13 | 21912524 | 21912872 | 3.89345 | 4.87228 | 1.6666 | Distal Intergenic | 13 | 21903156 | 21904784 | 1629 | 1 | ENSMUST00000121524.1 | 9369 | Gm11279 | ENSMUSG00000081576.1 |
| chr13 | 21914397 | 21914838 | 4.11656 | 5.37532 | 2.02935 | Distal Intergenic | 13 | 21903156 | 21904784 | 1629 | 1 | ENSMUST00000121524.1 | 11242 | Gm11279 | ENSMUSG00000081576.1 |
| chr13 | 21915144 | 21917760 | 3.87352 | 4.99111 | 1.80692 | Distal Intergenic | 13 | 21903156 | 21904784 | 1629 | 1 | ENSMUST00000121524.1 | 11989 | Gm11279 | ENSMUSG00000081576.1 |
| chr13 | 21918214 | 21918390 | 3.82598 | 4.77504 | 1.66706 | Distal Intergenic | 13 | 21903156 | 21904784 | 1629 | 1 | ENSMUST00000121524.1 | 15059 | Gm11279 | ENSMUSG00000081576.1 |
| chr13 | 21918845 | 21920533 | 3.87207 | 4.94226 | 1.76538 | Distal Intergenic | 13 | 21903156 | 21904784 | 1629 | 1 | ENSMUST00000121524.1 | 15690 | Gm11279 | ENSMUSG00000081576.1 |
| chr13 | 21921446 | 21921603 | 3.54839 | 4.22344 | 1.26939 | Distal Intergenic | 13 | 21903156 | 21904784 | 1629 | 1 | ENSMUST00000121524.1 | 18291 | Gm11279 | ENSMUSG00000081576.1 |
| chr13 | 21923336 | 21923428 | 3.44816 | 4.03687 | 1.14621 | Distal Intergenic | 13 | 21903156 | 21904784 | 1629 | 1 | ENSMUST00000121524.1 | 20181 | Gm11279 | ENSMUSG00000081576.1 |
| chr4 | 57447220 | 57447298 | 4.50367 | 6.05301 | 2.47654 | Intron (ENSMUST00000102905.2/ENSMUSG00000090053.4, intron 1 of 6) | 4 | 57455561 | 57457549 | 1989 | 2 | ENSMUST00000121549.1 | 10251 | Gm12539 | ENSMUSG00000081615.1 |
| chr11 | 85923932 | 85924239 | 3.2266 | 3.66709 | 1.01635 | Distal Intergenic | 11 | 85950495 | 85950973 | 479 | 1 | ENSMUST00000118790.1 | -26256 | Gm11443 | ENSMUSG00000081632.1 |
| chr11 | 85925128 | 85925422 | 3.93741 | 5.04241 | 1.83614 | Distal Intergenic | 11 | 85950495 | 85950973 | 479 | 1 | ENSMUST00000118790.1 | -25073 | Gm11443 | ENSMUSG00000081632.1 |
| chr11 | 85925786 | 85926735 | 3.78794 | 4.83596 | 1.67454 | Distal Intergenic | 11 | 85950495 | 85950973 | 479 | 1 | ENSMUST00000118790.1 | -23760 | Gm11443 | ENSMUSG00000081632.1 |
| chr11 | 85932779 | 85933080 | 3.66174 | 4.5159 | 1.47629 | Distal Intergenic | 11 | 85950495 | 85950973 | 479 | 1 | ENSMUST00000118790.1 | -17415 | Gm11443 | ENSMUSG00000081632.1 |
| chr11 | 85937825 | 85938046 | 3.36667 | 3.87758 | 1.01733 | Distal Intergenic | 11 | 85950495 | 85950973 | 479 | 1 | ENSMUST00000118790.1 | -12449 | Gm11443 | ENSMUSG00000081632.1 |
| chr11 | 85942256 | 85942628 | 3.47357 | 4.1755 | 1.26638 | Distal Intergenic | 11 | 85950495 | 85950973 | 479 | 1 | ENSMUST00000118790.1 | -7867 | Gm11443 | ENSMUSG00000081632.1 |
| chr11 | 85955707 | 85956149 | 3.29213 | 3.91805 | 1.09108 | Distal Intergenic | 11 | 85950495 | 85950973 | 479 | 1 | ENSMUST00000118790.1 | 5213 | Gm11443 | ENSMUSG00000081632.1 |
| chr11 | 85961461 | 85962164 | 3.26062 | 3.72036 | 1.02495 | Distal Intergenic | 11 | 85950495 | 85950973 | 479 | 1 | ENSMUST00000118790.1 | 10967 | Gm11443 | ENSMUSG00000081632.1 |
| chr11 | 85969535 | 85969679 | 3.95801 | 5.0685 | 1.84096 | Distal Intergenic | 11 | 85950495 | 85950973 | 479 | 1 | ENSMUST00000118790.1 | 19041 | Gm11443 | ENSMUSG00000081632.1 |
| chr11 | 85976471 | 85976853 | 3.87379 | 4.90265 | 1.76783 | Distal Intergenic | 11 | 85950495 | 85950973 | 479 | 1 | ENSMUST00000118790.1 | 25977 | Gm11443 | ENSMUSG00000081632.1 |
| chr11 | 85984442 | 85984757 | 3.72412 | 4.57254 | 1.52573 | Distal Intergenic | 11 | 85950495 | 85950973 | 479 | 1 | ENSMUST00000118790.1 | 33948 | Gm11443 | ENSMUSG00000081632.1 |
| chr11 | 85986583 | 85986884 | 3.60887 | 4.40763 | 1.40954 | Distal Intergenic | 11 | 85950495 | 85950973 | 479 | 1 | ENSMUST00000118790.1 | 36089 | Gm11443 | ENSMUSG00000081632.1 |
| chr11 | 85988094 | 85989184 | 3.65488 | 4.47801 | 1.46145 | Distal Intergenic | 11 | 85950495 | 85950973 | 479 | 1 | ENSMUST00000118790.1 | 37600 | Gm11443 | ENSMUSG00000081632.1 |
| chr11 | 85989899 | 85990352 | 3.49761 | 4.15946 | 1.29147 | Distal Intergenic | 11 | 85950495 | 85950973 | 479 | 1 | ENSMUST00000118790.1 | 39405 | Gm11443 | ENSMUSG00000081632.1 |
| chr11 | 85992121 | 85992252 | 3.82168 | 4.79781 | 1.66578 | Distal Intergenic | 11 | 85950495 | 85950973 | 479 | 1 | ENSMUST00000118790.1 | 41627 | Gm11443 | ENSMUSG00000081632.1 |
| chr11 | 85993791 | 85994229 | 3.87935 | 4.92894 | 1.77649 | Distal Intergenic | 11 | 85950495 | 85950973 | 479 | 1 | ENSMUST00000118790.1 | 43297 | Gm11443 | ENSMUSG00000081632.1 |
| chr11 | 86014095 | 86014508 | 3.58122 | 4.35981 | 1.42596 | Distal Intergenic | 11 | 85950495 | 85950973 | 479 | 1 | ENSMUST00000118790.1 | 63601 | Gm11443 | ENSMUSG00000081632.1 |
| chr11 | 86015112 | 86015457 | 3.57492 | 4.44636 | 1.41028 | Distal Intergenic | 11 | 85950495 | 85950973 | 479 | 1 | ENSMUST00000118790.1 | 64618 | Gm11443 | ENSMUSG00000081632.1 |
| chr11 | 86016025 | 86016804 | 3.99497 | 5.34538 | 2.05109 | Distal Intergenic | 11 | 85950495 | 85950973 | 479 | 1 | ENSMUST00000118790.1 | 65531 | Gm11443 | ENSMUSG00000081632.1 |
| chr11 | 86028337 | 86028519 | 4.03892 | 5.18884 | 1.94408 | Distal Intergenic | 11 | 85950495 | 85950973 | 479 | 1 | ENSMUST00000118790.1 | 77843 | Gm11443 | ENSMUSG00000081632.1 |
| chr11 | 111746242 | 111746531 | 3.79219 | 4.68892 | 1.56726 | Distal Intergenic | 11 | 111724648 | 111725624 | 977 | 2 | ENSMUST00000121349.1 | -20619 | Gm11677 | ENSMUSG00000081666.1 |
| chr6 | 85097252 | 85097327 | 4.58187 | 7.1453 | 3.43225 | Distal Intergenic | 6 | 85076192 | 85076719 | 528 | 1 | ENSMUST00000119728.1 | 21061 | Npm3-ps1 | ENSMUSG00000081669.1 |
| chr5 | 128601879 | 128602241 | 3.69759 | 4.49158 | 1.40782 | Exon (ENSMUST00000117102.3/ENSMUSG00000081683.5, exon 1 of 1) | 5 | 128600844 | 128604093 | 3250 | 1 | ENSMUST00000117102.3 | 1036 | Fzd10 | ENSMUSG00000081683.5 |
| chr5 | 128602623 | 128602759 | 3.6091 | 4.38704 | 1.40266 | Exon (ENSMUST00000117102.3/ENSMUSG00000081683.5, exon 1 of 1) | 5 | 128600844 | 128604093 | 3250 | 1 | ENSMUST00000117102.3 | 1780 | Fzd10 | ENSMUSG00000081683.5 |
| chr5 | 128605935 | 128606014 | 4.47502 | 5.98967 | 2.40834 | Distal Intergenic | 5 | 128600844 | 128604093 | 3250 | 1 | ENSMUST00000117102.3 | 5092 | Fzd10 | ENSMUSG00000081683.5 |
| chr5 | 128606385 | 128606627 | 3.35331 | 3.89325 | 1.10901 | Distal Intergenic | 5 | 128600844 | 128604093 | 3250 | 1 | ENSMUST00000117102.3 | 5542 | Fzd10 | ENSMUSG00000081683.5 |
| chr5 | 128617124 | 128617860 | 3.33076 | 3.82634 | 1.02704 | Distal Intergenic | 5 | 128600844 | 128604093 | 3250 | 1 | ENSMUST00000117102.3 | 16281 | Fzd10 | ENSMUSG00000081683.5 |
| chr5 | 128618407 | 128618690 | 3.38193 | 3.9006 | 1.01733 | Distal Intergenic | 5 | 128600844 | 128604093 | 3250 | 1 | ENSMUST00000117102.3 | 17564 | Fzd10 | ENSMUSG00000081683.5 |
| chr4 | 40676861 | 40677242 | 3.54728 | 4.58859 | 1.49337 | Downstream (1-2kb) | 4 | 40678332 | 40678778 | 447 | 1 | ENSMUST00000117626.1 | -1090 | Gm12391 | ENSMUSG00000081720.1 |
| chr2 | 104318245 | 104318423 | 3.35753 | 3.86872 | 1.03455 | Intron (ENSMUST00000089726.4/ENSMUSG00000068373.9, intron 1 of 20) | 2 | 104291574 | 104292575 | 1002 | 1 | ENSMUST00000120886.1 | 26672 | Gm13882 | ENSMUSG00000081721.3 |
| chr2 | 157507531 | 157507638 | 3.66018 | 4.42495 | 1.37403 | Distal Intergenic | 2 | 157521697 | 157522028 | 332 | 2 | ENSMUST00000117142.1 | 14390 | Gm14279 | ENSMUSG00000081740.3 |
| chr2 | 157511869 | 157512133 | 3.37105 | 3.88419 | 1.01733 | Distal Intergenic | 2 | 157521697 | 157522028 | 332 | 2 | ENSMUST00000117142.1 | 9895 | Gm14279 | ENSMUSG00000081740.3 |
| chr2 | 157513198 | 157513453 | 3.46023 | 4.04748 | 1.11834 | Distal Intergenic | 2 | 157521697 | 157522028 | 332 | 2 | ENSMUST00000117142.1 | 8575 | Gm14279 | ENSMUSG00000081740.3 |
| chr2 | 157513772 | 157514204 | 3.78176 | 4.63815 | 1.48543 | Distal Intergenic | 2 | 157521697 | 157522028 | 332 | 2 | ENSMUST00000117142.1 | 7824 | Gm14279 | ENSMUSG00000081740.3 |
| chr11 | 64800873 | 64801632 | 3.4777 | 4.19537 | 1.22603 | Distal Intergenic | 11 | 64759717 | 64760027 | 311 | 2 | ENSMUST00000121627.1 | -40847 | Gm12291 | ENSMUSG00000081756.1 |
| chr2 | 102205555 | 102206103 | 3.49381 | 4.23993 | 1.30871 | Distal Intergenic | 2 | 102209412 | 102210379 | 968 | 2 | ENSMUST00000118901.1 | 4276 | Gm13866 | ENSMUSG00000081774.1 |
| chr2 | 102258367 | 102259552 | 3.64253 | 4.45299 | 1.4717 | Distal Intergenic | 2 | 102209412 | 102210379 | 968 | 2 | ENSMUST00000118901.1 | -47989 | Gm13866 | ENSMUSG00000081774.1 |
| chr4 | 143378314 | 143378522 | 3.55122 | 4.21055 | 1.209 | Distal Intergenic | 4 | 143389659 | 143390416 | 758 | 1 | ENSMUST00000122215.1 | -11137 | Anp32b-ps1 | ENSMUSG00000081792.4 |
| chr4 | 143379869 | 143380163 | 3.96857 | 5.32623 | 1.9898 | Distal Intergenic | 4 | 143389659 | 143390416 | 758 | 1 | ENSMUST00000122215.1 | -9496 | Anp32b-ps1 | ENSMUSG00000081792.4 |
| chr4 | 22504027 | 22504173 | 3.31646 | 3.81862 | 1.07456 | Distal Intergenic | 4 | 22519628 | 22520027 | 400 | 1 | ENSMUST00000117908.1 | -15455 | Gm11878 | ENSMUSG00000081810.1 |
| chr4 | 22507543 | 22507618 | 4.00762 | 5.11547 | 1.88981 | Distal Intergenic | 4 | 22519628 | 22520027 | 400 | 1 | ENSMUST00000117908.1 | -12010 | Gm11878 | ENSMUSG00000081810.1 |
| chr3 | 137889519 | 137889696 | 3.42707 | 4.02002 | 1.19583 | Intron (ENSMUST00000090178.5/ENSMUSG00000074212.3, intron 2 of 7) | 3 | 137905009 | 137905399 | 391 | 1 | ENSMUST00000118147.1 | -15313 | Gm15460 | ENSMUSG00000081829.1 |
| chr4 | 142907322 | 142907605 | 3.2903 | 3.7626 | 1.01733 | Distal Intergenic | 4 | 143152572 | 143153030 | 459 | 1 | ENSMUST00000119886.1 | -244967 | Gm13038 | ENSMUSG00000081838.1 |
| chr4 | 142908837 | 142908995 | 3.3672 | 4.04932 | 1.16741 | Distal Intergenic | 4 | 143152572 | 143153030 | 459 | 1 | ENSMUST00000119886.1 | -243577 | Gm13038 | ENSMUSG00000081838.1 |
| chr4 | 142909669 | 142909885 | 4.14075 | 5.37277 | 2.04045 | Distal Intergenic | 4 | 143152572 | 143153030 | 459 | 1 | ENSMUST00000119886.1 | -242687 | Gm13038 | ENSMUSG00000081838.1 |
| chr4 | 142982927 | 142983002 | 3.78646 | 4.66484 | 1.52621 | Distal Intergenic | 4 | 143152572 | 143153030 | 459 | 1 | ENSMUST00000119886.1 | -169570 | Gm13038 | ENSMUSG00000081838.1 |
| chr4 | 143014530 | 143014701 | 3.38193 | 3.9006 | 1.01733 | Distal Intergenic | 4 | 143152572 | 143153030 | 459 | 1 | ENSMUST00000119886.1 | -137871 | Gm13038 | ENSMUSG00000081838.1 |
| chr4 | 143074028 | 143074399 | 3.80811 | 4.82012 | 1.67721 | Distal Intergenic | 4 | 143152572 | 143153030 | 459 | 1 | ENSMUST00000119886.1 | -78173 | Gm13038 | ENSMUSG00000081838.1 |
| chr5 | 73010612 | 73010994 | 3.59394 | 4.45753 | 1.44281 | Intron (ENSMUST00000031127.7/ENSMUSG00000029219.7, intron 2 of 2) | 5 | 73013499 | 73017376 | 3878 | 1 | ENSMUST00000121417.1 | -2505 | Gm15847 | ENSMUSG00000081849.1 |
| chr5 | 73011935 | 73012209 | 3.45289 | 4.39366 | 1.36694 | Exon (ENSMUST00000031127.7/ENSMUSG00000029219.7, exon 3 of 3) | 5 | 73013499 | 73017376 | 3878 | 1 | ENSMUST00000121417.1 | -1290 | Gm15847 | ENSMUSG00000081849.1 |
| chr5 | 73012560 | 73013057 | 3.66072 | 4.48757 | 1.46405 | Promoter | 5 | 73013499 | 73017376 | 3878 | 1 | ENSMUST00000121417.1 | -442 | Gm15847 | ENSMUSG00000081849.1 |
| chr5 | 73013378 | 73014286 | 3.78287 | 4.84506 | 1.72851 | Promoter | 5 | 73013499 | 73017376 | 3878 | 1 | ENSMUST00000121417.1 | 0 | Gm15847 | ENSMUSG00000081849.1 |
| chr5 | 73014628 | 73015117 | 3.93009 | 5.06361 | 1.83165 | 3' UTR | 5 | 73013499 | 73017376 | 3878 | 1 | ENSMUST00000121417.1 | 1130 | Gm15847 | ENSMUSG00000081849.1 |
| chr5 | 73015942 | 73016625 | 3.819 | 5.11656 | 1.89742 | Exon (ENSMUST00000121417.1/ENSMUSG00000081849.1, exon 2 of 4) | 5 | 73013499 | 73017376 | 3878 | 1 | ENSMUST00000121417.1 | 2444 | Gm15847 | ENSMUSG00000081849.1 |
| chr6 | 52837781 | 52837960 | 3.35475 | 4.00487 | 1.1056 | Intron (ENSMUST00000136250.2/ENSMUSG00000063568.6, intron 2 of 4) | 6 | 52843109 | 52843701 | 593 | 2 | ENSMUST00000119248.1 | 5741 | Gm15572 | ENSMUSG00000081872.1 |
| chr6 | 52839710 | 52839930 | 3.56726 | 4.28221 | 1.35722 | Intron (ENSMUST00000136250.2/ENSMUSG00000063568.6, intron 2 of 4) | 6 | 52843109 | 52843701 | 593 | 2 | ENSMUST00000119248.1 | 3771 | Gm15572 | ENSMUSG00000081872.1 |
| chr6 | 52840682 | 52840950 | 3.9469 | 5.08132 | 1.84742 | Intron (ENSMUST00000136250.2/ENSMUSG00000063568.6, intron 2 of 4) | 6 | 52843109 | 52843701 | 593 | 2 | ENSMUST00000119248.1 | 2751 | Gm15572 | ENSMUSG00000081872.1 |
| chr6 | 52843224 | 52843458 | 3.22443 | 3.66385 | 1.01733 | Promoter | 6 | 52843109 | 52843701 | 593 | 2 | ENSMUST00000119248.1 | 243 | Gm15572 | ENSMUSG00000081872.1 |
| chr4 | 11395885 | 11395960 | 3.4396 | 4.26964 | 1.29205 | Distal Intergenic | 4 | 11400346 | 11401324 | 979 | 1 | ENSMUST00000117335.1 | -4386 | Gm11821 | ENSMUSG00000081884.1 |
| chr4 | 145640804 | 145641334 | 3.42156 | 5.04948 | 1.83921 | Promoter | 4 | 145641357 | 145644399 | 3043 | 1 | ENSMUST00000119218.1 | -23 | Gm13231 | ENSMUSG00000081885.1 |
| chr3 | 103682531 | 103682867 | 3.60629 | 4.40571 | 1.37699 | Distal Intergenic | 3 | 103694798 | 103695197 | 400 | 1 | ENSMUST00000122153.1 | -11931 | Gm15986 | ENSMUSG00000081910.1 |
| chr3 | 103683217 | 103683523 | 4.00483 | 5.06237 | 1.76029 | Distal Intergenic | 3 | 103694798 | 103695197 | 400 | 1 | ENSMUST00000122153.1 | -11275 | Gm15986 | ENSMUSG00000081910.1 |
| chr11 | 73562483 | 73562957 | 3.82566 | 4.87056 | 1.65215 | Distal Intergenic | 11 | 73540419 | 73541353 | 935 | 2 | ENSMUST00000118895.1 | -21131 | Olfr383-ps1 | ENSMUSG00000081961.1 |
| chr2 | 77379628 | 77379839 | 3.355 | 3.85997 | 1.01733 | Distal Intergenic | 2 | 77335359 | 77335964 | 606 | 1 | ENSMUST00000122325.1 | 44270 | Gm13649 | ENSMUSG00000082006.1 |
| chr1 | 75681299 | 75681478 | 3.46381 | 4.27576 | 1.29814 | Distal Intergenic | 1 | 75636390 | 75637179 | 790 | 1 | ENSMUST00000121823.1 | 44910 | Gm5257 | ENSMUSG00000082012.1 |
| chr13 | 31077383 | 31077710 | 3.26472 | 3.86864 | 1.09372 | Distal Intergenic | 13 | 31009143 | 31013974 | 4832 | 1 | ENSMUST00000120294.1 | 68241 | Gm11371 | ENSMUSG00000082014.1 |
| chr13 | 31079754 | 31080401 | 3.62574 | 4.53957 | 1.50913 | Distal Intergenic | 13 | 31009143 | 31013974 | 4832 | 1 | ENSMUST00000120294.1 | 70612 | Gm11371 | ENSMUSG00000082014.1 |
| chr13 | 31081619 | 31081786 | 3.45058 | 4.09188 | 1.26936 | Distal Intergenic | 13 | 31009143 | 31013974 | 4832 | 1 | ENSMUST00000120294.1 | 72477 | Gm11371 | ENSMUSG00000082014.1 |
| chr13 | 31083267 | 31083355 | 4.24643 | 5.69766 | 2.33219 | Distal Intergenic | 13 | 31009143 | 31013974 | 4832 | 1 | ENSMUST00000120294.1 | 74125 | Gm11371 | ENSMUSG00000082014.1 |
| chr13 | 31088762 | 31088884 | 3.81903 | 4.78138 | 1.68233 | Distal Intergenic | 13 | 31009143 | 31013974 | 4832 | 1 | ENSMUST00000120294.1 | 79620 | Gm11371 | ENSMUSG00000082014.1 |
| chr4 | 115657254 | 115657550 | 3.3364 | 3.8532 | 1.09099 | Distal Intergenic | 4 | 115652389 | 115653480 | 1092 | 1 | ENSMUST00000119777.1 | 4866 | Gm12848 | ENSMUSG00000082043.1 |
| chr11 | 102130069 | 102130284 | 3.34249 | 3.86349 | 1.03719 | Distal Intergenic | 11 | 102124072 | 102124640 | 569 | 1 | ENSMUST00000120901.1 | 5998 | Gm11585 | ENSMUSG00000082045.4 |
| chr4 | 116874547 | 116874862 | 4.08303 | 5.33647 | 2.0827 | Promoter | 4 | 116874588 | 116875465 | 878 | 2 | ENSMUST00000119900.1 | 603 | Gm12993 | ENSMUSG00000082063.1 |
| chr2 | 154707388 | 154707717 | 3.37882 | 3.89591 | 1.01733 | Distal Intergenic | 2 | 154703809 | 154704699 | 891 | 1 | ENSMUST00000119057.1 | 3580 | Rpl5-ps2 | ENSMUSG00000082064.3 |
| chr19 | 24289646 | 24289783 | 3.29265 | 3.76613 | 1.01733 | Promoter | 19 | 24289716 | 24290826 | 1111 | 1 | ENSMUST00000118970.1 | 0 | Gm16216 | ENSMUSG00000082107.1 |
| chr4 | 137225830 | 137225905 | 3.51306 | 4.15434 | 1.2154 | Distal Intergenic | 4 | 137222641 | 137223322 | 682 | 2 | ENSMUST00000119286.1 | -2509 | Gm13002 | ENSMUSG00000082149.1 |
| chr2 | 75586077 | 75586356 | 3.89845 | 5.00646 | 1.80139 | Distal Intergenic | 2 | 75632638 | 75633382 | 745 | 2 | ENSMUST00000119695.1 | 47026 | Gm13655 | ENSMUSG00000082152.1 |
| chr2 | 75591933 | 75592421 | 3.41162 | 3.96982 | 1.10425 | Distal Intergenic | 2 | 75632638 | 75633382 | 745 | 2 | ENSMUST00000119695.1 | 40961 | Gm13655 | ENSMUSG00000082152.1 |
| chr7 | 104501978 | 104502559 | 4.27893 | 5.71283 | 2.34035 | Promoter | 7 | 104501055 | 104502149 | 1095 | 2 | ENSMUST00000117815.1 | 0 | Gm16464 | ENSMUSG00000082154.1 |
| chr2 | 147126239 | 147126479 | 3.22824 | 3.66956 | 1.01671 | Distal Intergenic | 2 | 147146408 | 147147212 | 805 | 2 | ENSMUST00000117733.1 | 20733 | Gm14116 | ENSMUSG00000082158.1 |
| chr2 | 147152484 | 147152830 | 3.83582 | 4.89123 | 1.68995 | Distal Intergenic | 2 | 147146408 | 147147212 | 805 | 2 | ENSMUST00000117733.1 | -5273 | Gm14116 | ENSMUSG00000082158.1 |
| chr11 | 101924903 | 101925072 | 3.33227 | 3.85593 | 1.1141 | Distal Intergenic | 11 | 101939501 | 101939899 | 399 | 1 | ENSMUST00000120378.1 | -14429 | Rpl27-ps2 | ENSMUSG00000082171.1 |
| chr11 | 101936342 | 101936646 | 3.36046 | 4.05392 | 1.14932 | Distal Intergenic | 11 | 101939501 | 101939899 | 399 | 1 | ENSMUST00000120378.1 | -2855 | Rpl27-ps2 | ENSMUSG00000082171.1 |
| chr11 | 101937837 | 101938111 | 3.6758 | 4.5417 | 1.51774 | Downstream (2-3kb) | 11 | 101939501 | 101939899 | 399 | 1 | ENSMUST00000120378.1 | -1390 | Rpl27-ps2 | ENSMUSG00000082171.1 |

| | | | | | | | | | | | | | | | |
|---|---|---|---|---|---|---|---|---|---|---|---|---|---|---|---|
| chr4 | 146067349 | 146068020 | 3.46139 | 4.78388 | 1.62866 | Promoter | 4 | 146067752 | 146070928 | 3177 | 1 | ENSMUST00000121342.1 | 0 | Gm13034 | ENSMUSG00000082195.3 |
| chr4 | 120893042 | 120893220 | 3.36788 | 3.8985 | 1.08407 | 3' UTR | 4 | 120901223 | 120901660 | 438 | 1 | ENSMUST00000118758.1 | -8003 | Gm12871 | ENSMUSG00000082200.1 |
| chr4 | 120894038 | 120894530 | 3.4453 | 4.2452 | 1.28449 | 3' UTR | 4 | 120901223 | 120901660 | 438 | 1 | ENSMUST00000118758.1 | -6693 | Gm12871 | ENSMUSG00000082200.1 |
| chr4 | 120895077 | 120895534 | 3.7478 | 4.76493 | 1.59929 | 3' UTR | 4 | 120901223 | 120901660 | 438 | 1 | ENSMUST00000118758.1 | -5689 | Gm12871 | ENSMUSG00000082200.1 |
| chr2 | 178785693 | 178785768 | 3.822 | 4.7253 | 1.55717 | Distal Intergenic | 2 | 179020119 | 179021133 | 1015 | 2 | ENSMUST00000118225.1 | 235365 | Gm14314 | ENSMUSG00000082215.3 |
| chr6 | 33173076 | 33173332 | 3.58115 | 4.33292 | 1.37841 | Distal Intergenic | 6 | 33247328 | 33248099 | 772 | 2 | ENSMUST00000122048.1 | 74767 | Gm1401 | ENSMUSG00000082217.1 |
| chr6 | 33174511 | 33174690 | 3.35589 | 3.87572 | 1.06818 | Distal Intergenic | 6 | 33247328 | 33248099 | 772 | 2 | ENSMUST00000122048.1 | 73409 | Gm1401 | ENSMUSG00000082217.1 |
| chr2 | 168440902 | 168441330 | 3.3143 | 3.83886 | 1.10165 | Distal Intergenic | 2 | 168417730 | 168418574 | 845 | 2 | ENSMUST00000119539.1 | -22329 | Gm14234 | ENSMUSG00000082239.1 |
| chr2 | 168444234 | 168444341 | 3.45964 | 4.1005 | 1.16156 | Distal Intergenic | 2 | 168417730 | 168418574 | 845 | 2 | ENSMUST00000119539.1 | -25661 | Gm14234 | ENSMUSG00000082239.1 |
| chr2 | 168445225 | 168445489 | 3.58664 | 4.28607 | 1.27377 | Distal Intergenic | 2 | 168417730 | 168418574 | 845 | 2 | ENSMUST00000119539.1 | -26652 | Gm14234 | ENSMUSG00000082239.1 |
| chr2 | 168447769 | 168448111 | 3.42114 | 4.01519 | 1.19397 | Distal Intergenic | 2 | 168417730 | 168418574 | 845 | 2 | ENSMUST00000119539.1 | -29196 | Gm14234 | ENSMUSG00000082239.1 |
| chr2 | 168451522 | 168451597 | 3.46856 | 4.26806 | 1.28506 | Distal Intergenic | 2 | 168417730 | 168418574 | 845 | 2 | ENSMUST00000119539.1 | -32949 | Gm14234 | ENSMUSG00000082239.1 |
| chr2 | 168471706 | 168472009 | 3.68573 | 4.52518 | 1.51384 | Distal Intergenic | 2 | 168417730 | 168418574 | 845 | 2 | ENSMUST00000119539.1 | -53133 | Gm14234 | ENSMUSG00000082239.1 |
| chr11 | 63669021 | 63669373 | 3.56991 | 4.26982 | 1.30235 | Distal Intergenic | 11 | 63599632 | 63600923 | 1292 | 2 | ENSMUST00000119770.1 | -68099 | Gm12287 | ENSMUSG00000082256.1 |
| chr11 | 55387158 | 55387909 | 3.70331 | 4.55777 | 1.51387 | Promoter | 11 | 55387587 | 55388149 | 563 | 2 | ENSMUST00000120459.1 | 240 | Gm12234 | ENSMUSG00000082257.1 |
| chr11 | 3100439 | 3100533 | 1.92936 | 8.86163 | 4.76 | Distal Intergenic | 11 | 3125904 | 3129396 | 3493 | 1 | ENSMUST00000145164.2 | -25371 | Pisd-ps1 | ENSMUSG00000082286.5 |
| chr11 | 3103784 | 3104035 | 2.24737 | 10.27283 | 6.1195 | Distal Intergenic | 11 | 3125904 | 3129396 | 3493 | 1 | ENSMUST00000145164.2 | -21869 | Pisd-ps1 | ENSMUSG00000082286.5 |
| chr11 | 35275915 | 35276101 | 3.35648 | 4.12404 | 1.20708 | Intron (ENSMUST00000069837.3/ENSMUSG00000056427.5, intron 4 of 35) | 11 | 35323199 | 35323670 | 472 | 2 | ENSMUST00000120348.1 | 47569 | Gm12123 | ENSMUSG00000082309.3 |
| chr11 | 35278003 | 35278948 | 3.79587 | 4.78262 | 1.66908 | Intron (ENSMUST00000069837.3/ENSMUSG00000056427.5, intron 4 of 35) | 11 | 35323199 | 35323670 | 472 | 2 | ENSMUST00000120348.1 | 44722 | Gm12123 | ENSMUSG00000082309.3 |
| chr11 | 35287337 | 35287463 | 3.28274 | 3.84303 | 1.09865 | Intron (ENSMUST00000069837.3/ENSMUSG00000056427.5, intron 4 of 35) | 11 | 35323199 | 35323670 | 472 | 2 | ENSMUST00000120348.1 | 36207 | Gm12123 | ENSMUSG00000082309.3 |
| chr11 | 35286749 | 35286859 | 3.5345 | 4.1911 | 1.2309 | Intron (ENSMUST00000069837.3/ENSMUSG00000056427.5, intron 4 of 35) | 11 | 35323199 | 35323670 | 472 | 2 | ENSMUST00000120348.1 | 36811 | Gm12123 | ENSMUSG00000082309.3 |
| chr2 | 6132832 | 6136067 | 4.03581 | 5.30496 | 2.00385 | Promoter | 2 | 6132208 | 6133956 | 1749 | 2 | ENSMUST00000117166.2 | 0 | Gm13384 | ENSMUSG00000082332.2 |
| chr2 | 6136406 | 6137492 | 3.63471 | 4.37364 | 1.33114 | Distal Intergenic | 2 | 6132208 | 6133956 | 1749 | 2 | ENSMUST00000117166.2 | -2451 | Gm13384 | ENSMUSG00000082332.2 |
| chr2 | 6138642 | 6138723 | 3.97871 | 5.0311 | 1.77822 | Distal Intergenic | 2 | 6132208 | 6133956 | 1749 | 2 | ENSMUST00000117166.2 | -4687 | Gm13384 | ENSMUSG00000082332.2 |
| chr2 | 6139384 | 6139639 | 3.46857 | 4.10367 | 1.24593 | Distal Intergenic | 2 | 6132208 | 6133956 | 1749 | 2 | ENSMUST00000117166.2 | -5429 | Gm13384 | ENSMUSG00000082332.2 |
| chr2 | 6140688 | 6140853 | 3.64263 | 4.53306 | 1.46277 | Distal Intergenic | 2 | 6132208 | 6133956 | 1749 | 2 | ENSMUST00000117166.2 | -6733 | Gm13384 | ENSMUSG00000082332.2 |
| chr2 | 6142826 | 6143122 | 3.56208 | 4.38421 | 1.38696 | Distal Intergenic | 2 | 6132208 | 6133956 | 1749 | 2 | ENSMUST00000117166.2 | -8871 | Gm13384 | ENSMUSG00000082332.2 |
| chr2 | 6144973 | 6145989 | 4.00188 | 5.11768 | 1.84559 | Distal Intergenic | 2 | 6132208 | 6133956 | 1749 | 2 | ENSMUST00000117166.2 | -11018 | Gm13384 | ENSMUSG00000082332.2 |
| chr2 | 6147125 | 6147423 | 3.65096 | 4.40654 | 1.3535 | Distal Intergenic | 2 | 6132208 | 6133956 | 1749 | 2 | ENSMUST00000117166.2 | -13170 | Gm13384 | ENSMUSG00000082332.2 |
| chr2 | 6148054 | 6148433 | 3.35293 | 3.88211 | 1.09343 | Distal Intergenic | 2 | 6132208 | 6133956 | 1749 | 2 | ENSMUST00000117166.2 | -14099 | Gm13384 | ENSMUSG00000082332.2 |
| chr2 | 6148873 | 6149895 | 3.73263 | 4.58227 | 1.48678 | Distal Intergenic | 2 | 6132208 | 6133956 | 1749 | 2 | ENSMUST00000117166.2 | -14918 | Gm13384 | ENSMUSG00000082332.2 |
| chr2 | 6151036 | 6151607 | 3.83138 | 4.85197 | 1.66614 | Distal Intergenic | 2 | 6132208 | 6133956 | 1749 | 2 | ENSMUST00000117166.2 | -17081 | Gm13384 | ENSMUSG00000082332.2 |
| chr2 | 6151938 | 6152458 | 3.82334 | 4.80039 | 1.6377 | Distal Intergenic | 2 | 6132208 | 6133956 | 1749 | 2 | ENSMUST00000117166.2 | -17983 | Gm13384 | ENSMUSG00000082332.2 |
| chr2 | 6153766 | 6155305 | 4.08877 | 5.48914 | 2.17996 | Distal Intergenic | 2 | 6132208 | 6133956 | 1749 | 2 | ENSMUST00000117166.2 | -19811 | Gm13384 | ENSMUSG00000082332.2 |
| chr2 | 6156976 | 6157811 | 3.61169 | 4.42786 | 1.41985 | Distal Intergenic | 2 | 6132208 | 6133956 | 1749 | 2 | ENSMUST00000117166.2 | -23021 | Gm13384 | ENSMUSG00000082332.2 |
| chr2 | 6158224 | 6158300 | 3.37924 | 3.94394 | 1.16795 | Distal Intergenic | 2 | 6132208 | 6133956 | 1749 | 2 | ENSMUST00000117166.2 | -24269 | Gm13384 | ENSMUSG00000082332.2 |
| chr2 | 6158679 | 6158973 | 3.51335 | 4.33531 | 1.3529 | Distal Intergenic | 2 | 6132208 | 6133956 | 1749 | 2 | ENSMUST00000117166.2 | -24724 | Gm13384 | ENSMUSG00000082332.2 |
| chr2 | 6159364 | 6159894 | 3.55735 | 4.44434 | 1.43238 | Distal Intergenic | 2 | 6132208 | 6133956 | 1749 | 2 | ENSMUST00000117166.2 | -25409 | Gm13384 | ENSMUSG00000082332.2 |
| chr2 | 6160480 | 6160948 | 3.91863 | 5.0022 | 1.82183 | Distal Intergenic | 2 | 6132208 | 6133956 | 1749 | 2 | ENSMUST00000117166.2 | -26525 | Gm13384 | ENSMUSG00000082332.2 |
| chr2 | 6162222 | 6163367 | 3.68981 | 4.55264 | 1.5018 | Distal Intergenic | 2 | 6132208 | 6133956 | 1749 | 2 | ENSMUST00000117166.2 | -28267 | Gm13384 | ENSMUSG00000082332.2 |
| chr4 | 145744525 | 145745381 | 4.04122 | 7.23514 | 3.56878 | Promoter | 4 | 145745746 | 145747401 | 1656 | 1 | ENSMUST00000118855.1 | -365 | Gm13244 | ENSMUSG00000082334.1 |
| chr5 | 118465647 | 118465722 | 4.01733 | 5.10887 | 1.83567 | Distal Intergenic | 5 | 118502840 | 118503317 | 478 | 1 | ENSMUST00000119191.1 | -37118 | Gm15754 | ENSMUSG00000082365.1 |
| chr5 | 118466917 | 118467635 | 3.88112 | 4.94507 | 1.73951 | Distal Intergenic | 5 | 118502840 | 118503317 | 478 | 1 | ENSMUST00000119191.1 | -35205 | Gm15754 | ENSMUSG00000082365.1 |
| chr5 | 118470237 | 118470312 | 3.38193 | 3.9006 | 1.01733 | Distal Intergenic | 5 | 118502840 | 118503317 | 478 | 1 | ENSMUST00000119191.1 | -32528 | Gm15754 | ENSMUSG00000082365.1 |
| chr5 | 118477105 | 118477603 | 3.32442 | 3.82479 | 1.04996 | Distal Intergenic | 5 | 118502840 | 118503317 | 478 | 1 | ENSMUST00000119191.1 | -25237 | Gm15754 | ENSMUSG00000082365.1 |
| chr5 | 118483722 | 118483842 | 3.36482 | 4.05789 | 1.17997 | Distal Intergenic | 5 | 118502840 | 118503317 | 478 | 1 | ENSMUST00000119191.1 | -18998 | Gm15754 | ENSMUSG00000082365.1 |
| chr5 | 118513908 | 118514931 | 3.87808 | 4.87554 | 1.71889 | Distal Intergenic | 5 | 118502840 | 118503317 | 478 | 1 | ENSMUST00000119191.1 | 11069 | Gm15754 | ENSMUSG00000082365.1 |
| chr11 | 70426374 | 70426528 | 3.42544 | 3.99249 | 1.11019 | Promoter | 11 | 70426553 | 70426973 | 421 | 1 | ENSMUST00000121347.1 | -25 | Gm12313 | ENSMUSG00000082370.1 |
| chr4 | 32074757 | 32075195 | 3.74306 | 4.58495 | 1.49145 | Distal Intergenic | 4 | 32051738 | 32052914 | 1177 | 1 | ENSMUST00000118472.1 | 23020 | Gm11927 | ENSMUSG00000082398.1 |
| chr3 | 101161241 | 101161421 | 3.65424 | 4.59449 | 1.47568 | Distal Intergenic | 3 | 101206859 | 101207399 | 541 | 2 | ENSMUST00000120887.1 | 45978 | Gm12486 | ENSMUSG00000082415.1 |
| chr4 | 87648343 | 87648552 | 3.64107 | 4.43921 | 1.44128 | Distal Intergenic | 4 | 87580789 | 87581560 | 772 | 1 | ENSMUST00000118382.1 | 67555 | Gm12604 | ENSMUSG00000082443.1 |
| chr4 | 87652886 | 87652993 | 3.38592 | 3.97249 | 1.1791 | Distal Intergenic | 4 | 87580789 | 87581560 | 772 | 1 | ENSMUST00000118382.1 | 72098 | Gm12604 | ENSMUSG00000082443.1 |
| chr11 | 111484650 | 111484761 | 3.33723 | 3.83933 | 1.03898 | Distal Intergenic | 11 | 111612817 | 111613306 | 490 | 2 | ENSMUST00000120443.1 | 128545 | Gm11676 | ENSMUSG00000082448.1 |
| chr11 | 111495257 | 111495393 | 3.33703 | 3.83896 | 1.03869 | Distal Intergenic | 11 | 111612817 | 111613306 | 490 | 2 | ENSMUST00000120443.1 | 117913 | Gm11676 | ENSMUSG00000082448.1 |
| chr11 | 111496382 | 111496481 | 4.33092 | 5.75507 | 2.32551 | Distal Intergenic | 11 | 111612817 | 111613306 | 490 | 2 | ENSMUST00000120443.1 | 116825 | Gm11676 | ENSMUSG00000082448.1 |
| chr11 | 111499113 | 111499465 | 3.87465 | 5.01338 | 1.79569 | Distal Intergenic | 11 | 111612817 | 111613306 | 490 | 2 | ENSMUST00000120443.1 | 113841 | Gm11676 | ENSMUSG00000082448.1 |
| chr11 | 111499966 | 111500133 | 3.46739 | 4.08712 | 1.2162 | Distal Intergenic | 11 | 111612817 | 111613306 | 490 | 2 | ENSMUST00000120443.1 | 113173 | Gm11676 | ENSMUSG00000082448.1 |
| chr11 | 111500677 | 111501013 | 4.33476 | 5.96021 | 2.5198 | Distal Intergenic | 11 | 111612817 | 111613306 | 490 | 2 | ENSMUST00000120443.1 | 112293 | Gm11676 | ENSMUSG00000082448.1 |
| chr11 | 111502768 | 111502981 | 3.55653 | 4.30381 | 1.38251 | Distal Intergenic | 11 | 111612817 | 111613306 | 490 | 2 | ENSMUST00000120443.1 | 110325 | Gm11676 | ENSMUSG00000082448.1 |
| chr11 | 111503965 | 111504332 | 3.97341 | 5.12163 | 1.89245 | Distal Intergenic | 11 | 111612817 | 111613306 | 490 | 2 | ENSMUST00000120443.1 | 108974 | Gm11676 | ENSMUSG00000082448.1 |
| chr11 | 111505156 | 111505236 | 4.49796 | 6.10247 | 2.5593 | Distal Intergenic | 11 | 111612817 | 111613306 | 490 | 2 | ENSMUST00000120443.1 | 108070 | Gm11676 | ENSMUSG00000082448.1 |
| chr11 | 111506305 | 111506474 | 3.7155 | 4.6647 | 1.55492 | Distal Intergenic | 11 | 111612817 | 111613306 | 490 | 2 | ENSMUST00000120443.1 | 106832 | Gm11676 | ENSMUSG00000082448.1 |
| chr11 | 111509425 | 111510029 | 3.83913 | 4.86393 | 1.72958 | Distal Intergenic | 11 | 111612817 | 111613306 | 490 | 2 | ENSMUST00000120443.1 | 103277 | Gm11676 | ENSMUSG00000082448.1 |
| chr11 | 111510441 | 111510602 | 3.34731 | 4.02169 | 1.12883 | Distal Intergenic | 11 | 111612817 | 111613306 | 490 | 2 | ENSMUST00000120443.1 | 102704 | Gm11676 | ENSMUSG00000082448.1 |
| chr11 | 111510966 | 111511158 | 3.45878 | 4.2578 | 1.31973 | Distal Intergenic | 11 | 111612817 | 111613306 | 490 | 2 | ENSMUST00000120443.1 | 102148 | Gm11676 | ENSMUSG00000082448.1 |
| chr11 | 111511629 | 111512456 | 3.67184 | 4.47596 | 1.46674 | Distal Intergenic | 11 | 111612817 | 111613306 | 490 | 2 | ENSMUST00000120443.1 | 100850 | Gm11676 | ENSMUSG00000082448.1 |
| chr11 | 111512783 | 111513533 | 3.56242 | 4.28515 | 1.29791 | Distal Intergenic | 11 | 111612817 | 111613306 | 490 | 2 | ENSMUST00000120443.1 | 99773 | Gm11676 | ENSMUSG00000082448.1 |
| chr11 | 111515859 | 111516138 | 3.27442 | 3.90223 | 1.07654 | Distal Intergenic | 11 | 111612817 | 111613306 | 490 | 2 | ENSMUST00000120443.1 | 97168 | Gm11676 | ENSMUSG00000082448.1 |
| chr11 | 111517384 | 111517592 | 3.29137 | 3.77843 | 1.06232 | Distal Intergenic | 11 | 111612817 | 111613306 | 490 | 2 | ENSMUST00000120443.1 | 95714 | Gm11676 | ENSMUSG00000082448.1 |
| chr11 | 111521222 | 111521717 | 3.7809 | 4.81324 | 1.64504 | Distal Intergenic | 11 | 111612817 | 111613306 | 490 | 2 | ENSMUST00000120443.1 | 91589 | Gm11676 | ENSMUSG00000082448.1 |
| chr11 | 111522339 | 111522414 | 3.49634 | 4.11357 | 1.15971 | Distal Intergenic | 11 | 111612817 | 111613306 | 490 | 2 | ENSMUST00000120443.1 | 90892 | Gm11676 | ENSMUSG00000082448.1 |
| chr11 | 111524164 | 111525478 | 3.72356 | 4.65485 | 1.55754 | Distal Intergenic | 11 | 111612817 | 111613306 | 490 | 2 | ENSMUST00000120443.1 | 87828 | Gm11676 | ENSMUSG00000082448.1 |
| chr11 | 111525956 | 111526259 | 3.51307 | 4.13968 | 1.16225 | Distal Intergenic | 11 | 111612817 | 111613306 | 490 | 2 | ENSMUST00000120443.1 | 87047 | Gm11676 | ENSMUSG00000082448.1 |
| chr11 | 111526803 | 111527067 | 3.73436 | 4.55291 | 1.42858 | Distal Intergenic | 11 | 111612817 | 111613306 | 490 | 2 | ENSMUST00000120443.1 | 86239 | Gm11676 | ENSMUSG00000082448.1 |
| chr11 | 111530352 | 111531025 | 3.66336 | 4.44204 | 1.40261 | Distal Intergenic | 11 | 111612817 | 111613306 | 490 | 2 | ENSMUST00000120443.1 | 82281 | Gm11676 | ENSMUSG00000082448.1 |
| chr11 | 111533883 | 111533962 | 3.95431 | 5.08649 | 1.82498 | Distal Intergenic | 11 | 111612817 | 111613306 | 490 | 2 | ENSMUST00000120443.1 | 79344 | Gm11676 | ENSMUSG00000082448.1 |
| chr11 | 111534322 | 111534513 | 3.35296 | 3.86116 | 1.03237 | Distal Intergenic | 11 | 111612817 | 111613306 | 490 | 2 | ENSMUST00000120443.1 | 78793 | Gm11676 | ENSMUSG00000082448.1 |
| chr11 | 111535717 | 111535843 | 3.49847 | 4.27986 | 1.31592 | Distal Intergenic | 11 | 111612817 | 111613306 | 490 | 2 | ENSMUST00000120443.1 | 77463 | Gm11676 | ENSMUSG00000082448.1 |
| chr11 | 111542006 | 111542236 | 3.38032 | 4.13699 | 1.22822 | Distal Intergenic | 11 | 111612817 | 111613306 | 490 | 2 | ENSMUST00000120443.1 | 71070 | Gm11676 | ENSMUSG00000082448.1 |
| chr11 | 111543017 | 111543149 | 3.68468 | 4.51973 | 1.4413 | Distal Intergenic | 11 | 111612817 | 111613306 | 490 | 2 | ENSMUST00000120443.1 | 70157 | Gm11676 | ENSMUSG00000082448.1 |
| chr11 | 57474555 | 57474733 | 3.52272 | 4.16856 | 1.21404 | Distal Intergenic | 11 | 57449107 | 57449844 | 738 | 1 | ENSMUST00000118824.1 | 25449 | Gm12242 | ENSMUSG00000082476.1 |
| chr4 | 22152087 | 22152398 | 3.44566 | 4.20812 | 1.27477 | Distal Intergenic | 4 | 22117134 | 22117532 | 399 | 2 | ENSMUST00000117612.1 | -34556 | Gm11880 | ENSMUSG00000082490.1 |
| chr4 | 22152989 | 22153749 | 3.82879 | 4.78246 | 1.678 | Distal Intergenic | 4 | 22117134 | 22117532 | 399 | 2 | ENSMUST00000117612.1 | -35458 | Gm11880 | ENSMUSG00000082490.1 |
| chr2 | 162908018 | 162908098 | 3.60822 | 4.33741 | 1.32923 | Promoter | 2 | 162907924 | 162908613 | 690 | 1 | ENSMUST00000122376.1 | 95 | Gm11452 | ENSMUSG00000082514.1 |
| chr4 | 123950460 | 123950731 | 3.33696 | 3.87556 | 1.05528 | Distal Intergenic | 4 | 123926234 | 123926555 | 322 | 1 | ENSMUST00000118389.1 | 24227 | Gm12902 | ENSMUSG00000082515.1 |
| chr4 | 123955884 | 123957313 | 3.85396 | 4.84648 | 1.65988 | Distal Intergenic | 4 | 123926234 | 123926555 | 322 | 1 | ENSMUST00000118389.1 | 29651 | Gm12902 | ENSMUSG00000082515.1 |
| chr4 | 123957751 | 123958026 | 3.6812 | 4.45833 | 1.38104 | Distal Intergenic | 4 | 123926234 | 123926555 | 322 | 1 | ENSMUST00000118389.1 | 31518 | Gm12902 | ENSMUSG00000082515.1 |
| chr4 | 123960210 | 123960328 | 3.59079 | 4.28613 | 1.26157 | Distal Intergenic | 4 | 123926234 | 123926555 | 322 | 1 | ENSMUST00000118389.1 | 33977 | Gm12902 | ENSMUSG00000082515.1 |
| chr4 | 123966969 | 123967130 | 3.35836 | 3.86505 | 1.01733 | Distal Intergenic | 4 | 123926234 | 123926555 | 322 | 1 | ENSMUST00000118389.1 | 40736 | Gm12902 | ENSMUSG00000082515.1 |
| chr4 | 123971442 | 123971517 | 3.60988 | 4.32237 | 1.28865 | Distal Intergenic | 4 | 123926234 | 123926555 | 322 | 1 | ENSMUST00000118389.1 | 45209 | Gm12902 | ENSMUSG00000082515.1 |
| chr4 | 123972517 | 123973698 | 3.51131 | 4.23359 | 1.26559 | Distal Intergenic | 4 | 123926234 | 123926555 | 322 | 1 | ENSMUST00000118389.1 | 46284 | Gm12902 | ENSMUSG00000082515.1 |
| chr4 | 123986677 | 123986933 | 3.80156 | 4.68499 | 1.5346 | Distal Intergenic | 4 | 123926234 | 123926555 | 322 | 1 | ENSMUST00000118389.1 | 60444 | Gm12902 | ENSMUSG00000082515.1 |
| chr4 | 123987897 | 123988107 | 3.51022 | 4.14467 | 1.19531 | Distal Intergenic | 4 | 123926234 | 123926555 | 322 | 1 | ENSMUST00000118389.1 | 61664 | Gm12902 | ENSMUSG00000082515.1 |
| chr4 | 123990302 | 123990719 | 3.59486 | 4.30964 | 1.31351 | Distal Intergenic | 4 | 123926234 | 123926555 | 322 | 1 | ENSMUST00000118389.1 | 64069 | Gm12902 | ENSMUSG00000082515.1 |
| chr4 | 123991265 | 123991482 | 3.38061 | 3.8986 | 1.01733 | Distal Intergenic | 4 | 123926234 | 123926555 | 322 | 1 | ENSMUST00000118389.1 | 65032 | Gm12902 | ENSMUSG00000082515.1 |
| chr4 | 124102314 | 124102681 | 3.70267 | 4.49573 | 1.39333 | Distal Intergenic | 4 | 123926234 | 123926555 | 322 | 1 | ENSMUST00000118389.1 | 176081 | Gm12902 | ENSMUSG00000082515.1 |
| chr1 | 75598935 | 75599144 | 3.73374 | 4.56403 | 1.45791 | Distal Intergenic | 1 | 75624654 | 75626329 | 1676 | 1 | ENSMUST00000120832.1 | -25510 | Gm15182 | ENSMUSG00000082528.1 |
| chr16 | 91534630 | 91534706 | 4.14989 | 11.35667 | 6.8834 | Distal Intergenic | 16 | 91539182 | 91540060 | 879 | 2 | ENSMUST00000121190.1 | 5354 | Gm7860 | ENSMUSG00000082535.1 |
| chr11 | 117930811 | 117931078 | 3.29117 | 3.76398 | 1.01733 | Distal Intergenic | 11 | 117921905 | 117922328 | 424 | 1 | ENSMUST00000121028.1 | 8907 | Gm11726 | ENSMUSG00000082562.1 |
| chr2 | 57096000 | 57096085 | 4.07707 | 5.27836 | 1.99015 | Distal Intergenic | 2 | 57079748 | 57080438 | 691 | 1 | ENSMUST00000117926.1 | 16253 | Gm13527 | ENSMUSG00000082578.1 |
| chr15 | 74146364 | 74146514 | 3.42515 | 3.9848 | 1.08407 | Distal Intergenic | 15 | 74093686 | 74094333 | 648 | 2 | ENSMUST00000120421.1 | -52032 | Gm15387 | ENSMUSG00000082585.3 |
| chr15 | 74156362 | 74156437 | 3.59678 | 4.30046 | 1.28429 | Distal Intergenic | 15 | 74093686 | 74094333 | 648 | 2 | ENSMUST00000120421.1 | -62030 | Gm15387 | ENSMUSG00000082585.3 |
| chr15 | 74174644 | 74175337 | 3.4107 | 3.95506 | 1.05616 | Distal Intergenic | 15 | 74093686 | 74094333 | 648 | 2 | ENSMUST00000120421.1 | -80312 | Gm15387 | ENSMUSG00000082585.3 |
| chr15 | 74176385 | 74176460 | 3.45923 | 4.04153 | 1.10276 | Distal Intergenic | 15 | 74093686 | 74094333 | 648 | 2 | ENSMUST00000120421.1 | -82053 | Gm15387 | ENSMUSG00000082585.3 |
| chr15 | 74176794 | 74176998 | 3.36767 | 3.87909 | 1.01733 | Distal Intergenic | 15 | 74093686 | 74094333 | 648 | 2 | ENSMUST00000120421.1 | -82462 | Gm15387 | ENSMUSG00000082585.3 |
| chr15 | 74183134 | 74183259 | 3.32385 | 3.81306 | 1.01733 | Distal Intergenic | 15 | 74093686 | 74094333 | 648 | 2 | ENSMUST00000120421.1 | -88802 | Gm15387 | ENSMUSG00000082585.3 |
| chr15 | 74193586 | 74194057 | 3.56151 | 4.24603 | 1.25989 | Distal Intergenic | 15 | 74093686 | 74094333 | 648 | 2 | ENSMUST00000120421.1 | -99254 | Gm15387 | ENSMUSG00000082585.3 |
| chr15 | 74196792 | 74197002 | 3.57521 | 4.43861 | 1.39049 | Distal Intergenic | 15 | 74093686 | 74094333 | 648 | 2 | ENSMUST00000120421.1 | -102460 | Gm15387 | ENSMUSG00000082585.3 |
| chr15 | 74294632 | 74294854 | 3.38373 | 3.9399 | 1.1457 | Distal Intergenic | 15 | 74093686 | 74094333 | 648 | 2 | ENSMUST00000120421.1 | -200300 | Gm15387 | ENSMUSG00000082585.3 |
| chr11 | 61194212 | 61194608 | 3.38297 | 3.91395 | 1.05781 | Distal Intergenic | 11 | 61205720 | 61206175 | 456 | 2 | ENSMUST00000117939.1 | 11567 | Rps13-ps5 | ENSMUSG00000082626.1 |
| chr1 | 61135069 | 61135172 | 3.55885 | 4.24507 | 1.29291 | Distal Intergenic | 1 | 61121378 | 61124004 | 2627 | 2 | ENSMUST00000118999.1 | -11066 | Gm11590 | ENSMUSG00000082661.1 |
| chr1 | 61136070 | 61136147 | 3.71305 | 4.57739 | 1.55574 | Distal Intergenic | 1 | 61121378 | 61124004 | 2627 | 2 | ENSMUST00000118999.1 | -12067 | Gm11590 | ENSMUSG00000082661.1 |
| chr1 | 61137620 | 61139974 | 4.11927 | 5.52914 | 2.19879 | Distal Intergenic | 1 | 61121378 | 61124004 | 2627 | 2 | ENSMUST00000118999.1 | -13617 | Gm11590 | ENSMUSG00000082661.1 |
| chr1 | 61140384 | 61141159 | 3.45397 | 4.04109 | 1.11746 | Distal Intergenic | 1 | 61121378 | 61124004 | 2627 | 2 | ENSMUST00000118999.1 | -16381 | Gm11590 | ENSMUSG00000082661.1 |
| chr1 | 61142432 | 61142714 | 3.35949 | 3.86674 | 1.01733 | Distal Intergenic | 1 | 61121378 | 61124004 | 2627 | 2 | ENSMUST00000118999.1 | -18429 | Gm11590 | ENSMUSG00000082661.1 |
| chr1 | 61145092 | 61146128 | 3.44253 | 4.02511 | 1.11724 | Distal Intergenic | 1 | 61121378 | 61124004 | 2627 | 2 | ENSMUST00000118999.1 | -21089 | Gm11590 | ENSMUSG00000082661.1 |
| chr4 | 21670078 | 21671153 | 3.6954 | 4.5414 | 1.49305 | Promoter | 4 | 21671040 | 21671308 | 269 | 1 | ENSMUST00000120002.1 | 0 | Gm11876 | ENSMUSG00000082672.1 |
| chr4 | 21672847 | 21673000 | 3.38193 | 3.9006 | 1.01733 | Distal Intergenic | 4 | 21671040 | 21671308 | 269 | 1 | ENSMUST00000120002.1 | 1808 | Gm11876 | ENSMUSG00000082672.1 |
| chr4 | 21673666 | 21673764 | 3.65789 | 4.41519 | 1.35389 | Distal Intergenic | 4 | 21671040 | 21671308 | 269 | 1 | ENSMUST00000120002.1 | 2627 | Gm11876 | ENSMUSG00000082672.1 |
| chr4 | 21674132 | 21674924 | 4.35486 | 5.8746 | 2.42409 | Downstream (2-3kb) | 4 | 21671040 | 21671308 | 269 | 1 | ENSMUST00000120002.1 | 3093 | Gm11876 | ENSMUSG00000082672.1 |
| chr4 | 21675576 | 21676348 | 4.0788 | 5.25339 | 1.92369 | Downstream (1-2kb) | 4 | 21671040 | 21671308 | 269 | 1 | ENSMUST00000120002.1 | 4537 | Gm11876 | ENSMUSG00000082672.1 |
| chr4 | 21676668 | 21677781 | 3.62789 | 4.36889 | 1.34006 | 3' UTR | 4 | 21671040 | 21671308 | 269 | 1 | ENSMUST00000120002.1 | 5629 | Gm11876 | ENSMUSG00000082672.1 |
| chr4 | 112979411 | 112979873 | 3.47321 | 4.27934 | 1.29057 | Promoter | 4 | 112978465 | 112979813 | 1349 | 2 | ENSMUST00000117562.1 | 0 | Gm12818 | ENSMUSG00000082678.1 |
| chr11 | 104571556 | 104571640 | 3.51183 | 4.44742 | 1.37547 | Promoter | 11 | 104571268 | 104571557 | 290 | 2 | ENSMUST00000117888.1 | 0 | Gm11663 | ENSMUSG00000082706.1 |
| chr11 | 97967481 | 97967617 | 3.35212 | 3.85565 | 1.01733 | Intron (ENSMUST00000017561.9/ENSMUSG00000017417.9, intron 2 of 13) | 11 | 97968792 | 97969235 | 444 | 1 | ENSMUST00000118176.1 | -1175 | Gm11633 | ENSMUSG00000082715.1 |
| chr11 | 82970874 | 82971222 | 3.52359 | 4.22474 | 1.32446 | Promoter | 11 | 82971092 | 82971920 | 829 | 1 | ENSMUST00000121929.1 | 0 | Gm11424 | ENSMUSG00000082726.1 |
| chr4 | 135743745 | 135743887 | 3.36195 | 3.87044 | 1.01733 | Intron (ENSMUST00000102546.3/ENSMUSG00000037157.8, intron 4 of 6) | 4 | 135756218 | 135756695 | 478 | 2 | ENSMUST00000119054.1 | 12808 | Gm12988 | ENSMUSG00000082743.1 |
| chr4 | 135745566 | 135746160 | 4.20142 | 5.5923 | 2.21978 | Intron (ENSMUST00000102546.3/ENSMUSG00000037157.8, intron 5 of 6) | 4 | 135756218 | 135756695 | 478 | 2 | ENSMUST00000119054.1 | 10535 | Gm12988 | ENSMUSG00000082743.1 |
| chr4 | 147124747 | 147125058 | 3.0157 | 6.3022 | 2.74587 | Promoter | 4 | 147125256 | 147125816 | 561 | 1 | ENSMUST00000121481.1 | -198 | Gm13162 | ENSMUSG00000082779.1 |
| chr18 | 77226551 | 77226735 | 3.57729 | 4.55738 | 1.45498 | Intron (ENSMUST00000079618.5/ENSMUSG00000025425.12, intron 1 of 6) | 18 | 77218354 | 77219327 | 974 | 2 | ENSMUST00000118893.1 | -7225 | Gm15957 | ENSMUSG00000082784.1 |
| chr6 | 118010878 | 118011948 | 3.90148 | 4.89123 | 1.67669 | Distal Intergenic | 6 | 117997207 | 117997992 | 786 | 1 | ENSMUST00000118257.1 | 13672 | Gm4875 | ENSMUSG00000082791.3 |
| chr2 | 28780488 | 28780622 | 3.62345 | 4.36219 | 1.36281 | Intron (ENSMUST00000074156.6/ENSMUSG00000026807.8, intron 11 of 12) | 2 | 28793350 | 28793569 | 220 | 2 | ENSMUST00000121089.1 | 12947 | Gm13385 | ENSMUSG00000082796.1 |
| chr1 | 61057662 | 61057954 | 3.31016 | 3.79478 | 1.02554 | Distal Intergenic | 1 | 61075674 | 61076225 | 552 | 1 | ENSMUST00000119156.1 | -17720 | Rpl17-ps1 | ENSMUSG00000082804.3 |
| chr1 | 61060675 | 61060922 | 3.43352 | 3.99556 | 1.07749 | Distal Intergenic | 1 | 61075674 | 61076225 | 552 | 1 | ENSMUST00000119156.1 | -14752 | Rpl17-ps1 | ENSMUSG00000082804.3 |
| chr11 | 3945339 | 3945554 | 3.91236 | 4.89566 | 1.66609 | Intron (ENSMUST00000042344.1/ENSMUSG00000034493.1, intron 1 of 1) | 11 | 3951714 | 3952716 | 1003 | 1 | ENSMUST00000122352.1 | -6160 | Gm11953 | ENSMUSG00000082816.1 |
| chr4 | 136070970 | 136071164 | 3.5133 | 4.25797 | 1.28988 | Distal Intergenic | 4 | 136074513 | 136074972 | 460 | 2 | ENSMUST00000121307.1 | 3808 | Gm13009 | ENSMUSG00000082877.1 |
| chr4 | 136081669 | 136082459 | 4.12698 | 5.42874 | 2.12478 | Distal Intergenic | 4 | 136074513 | 136074972 | 460 | 2 | ENSMUST00000121307.1 | -6698 | Gm13009 | ENSMUSG00000082877.1 |
| chr4 | 136105063 | 136105266 | 3.55494 | 4.26954 | 1.36155 | Distal Intergenic | 4 | 136074513 | 136074972 | 460 | 2 | ENSMUST00000121307.1 | -30092 | Gm13009 | ENSMUSG00000082877.1 |
| chr4 | 33555263 | 33556580 | 3.74433 | 4.619 | 1.52497 | Distal Intergenic | 4 | 33614832 | 33615138 | 307 | 2 | ENSMUST00000121474.1 | 58558 | Gm12398 | ENSMUSG00000082909.1 |
| chr4 | 33581437 | 33581619 | 3.40089 | 4.20291 | 1.26064 | Distal Intergenic | 4 | 33614832 | 33615138 | 307 | 2 | ENSMUST00000121474.1 | 33519 | Gm12398 | ENSMUSG00000082909.1 |
| chr2 | 165954605 | 165954912 | 3.2577 | 3.71369 | 1.01733 | Distal Intergenic | 2 | 165959789 | 165960350 | 562 | 1 | ENSMUST00000117749.1 | -4877 | Gm11462 | ENSMUSG00000082924.1 |
| chr4 | 147275292 | 147275468 | 2.9536 | 4.33133 | 1.35714 | Promoter | 4 | 147275976 | 147279096 | 3121 | 1 | ENSMUST00000117628.1 | -508 | Gm13135 | ENSMUSG00000082925.1 |
| chr4 | 147275848 | 147276118 | 3.11846 | 5.19818 | 1.96992 | Promoter | 4 | 147275976 | 147279096 | 3121 | 1 | ENSMUST00000117628.1 | 0 | Gm13135 | ENSMUSG00000082925.1 |
| chr2 | 18605860 | 18606081 | 3.46401 | 4.19249 | 1.23715 | Distal Intergenic | 2 | 18641786 | 18642102 | 317 | 2 | ENSMUST00000117996.1 | 36021 | Gm13355 | ENSMUSG00000082943.1 |
| chr2 | 18606660 | 18608148 | 3.56557 | 4.26195 | 1.28925 | Distal Intergenic | 2 | 18641786 | 18642102 | 317 | 2 | ENSMUST00000117996.1 | 33954 | Gm13355 | ENSMUSG00000082943.1 |
| chr2 | 18608624 | 18609430 | 3.86894 | 4.85905 | 1.68503 | Distal Intergenic | 2 | 18641786 | 18642102 | 317 | 2 | ENSMUST00000117996.1 | 32672 | Gm13355 | ENSMUSG00000082943.1 |
| chr2 | 18609738 | 18610037 | 4.16813 | 5.40101 | 2.01877 | Distal Intergenic | 2 | 18641786 | 18642102 | 317 | 2 | ENSMUST00000117996.1 | 32065 | Gm13355 | ENSMUSG00000082943.1 |
| chr2 | 18611883 | 18612439 | 3.38124 | 3.89955 | 1.01733 | Distal Intergenic | 2 | 18641786 | 18642102 | 317 | 2 | ENSMUST00000117996.1 | 29663 | Gm13355 | ENSMUSG00000082943.1 |
| chr2 | 18618120 | 18618320 | 3.63963 | 4.43701 | 1.43938 | Distal Intergenic | 2 | 18641786 | 18642102 | 317 | 2 | ENSMUST00000117996.1 | 23782 | Gm13355 | ENSMUSG00000082943.1 |
| chr2 | 74642407 | 74643339 | 3.60755 | 4.41207 | 1.38902 | Distal Intergenic | 2 | 74631840 | 74632298 | 459 | 1 | ENSMUST00000117554.1 | 10568 | Gm13672 | ENSMUSG00000082973.1 |
| chr4 | 28176930 | 28177014 | 2.22143 | 11.76623 | 7.41507 | Distal Intergenic | 4 | 28067177 | 28068199 | 1023 | 1 | ENSMUST00000117938.1 | 109754 | Gm11911 | ENSMUSG00000082977.1 |
| chr7 | 130362834 | 130362936 | 12.70912 | 28.455 | 22.98423 | Intron (ENSMUST00000124096.2/ENSMUSG00000030849.12, intron 2 of 6) | 7 | 130352038 | 130354362 | 2325 | 2 | ENSMUST00000121836.1 | -8473 | Gm5903 | ENSMUSG00000082988.3 |
| chr7 | 130369155 | 130369397 | 3.34153 | 3.83966 | 1.01733 | Intron (ENSMUST00000124096.2/ENSMUSG00000030849.12, intron 2 of 6) | 7 | 130352038 | 130354362 | 2325 | 2 | ENSMUST00000121836.1 | -14794 | Gm5903 | ENSMUSG00000082988.3 |
| chr11 | 3705121 | 3705196 | 3.98258 | 5.03109 | 1.74951 | Exon (ENSMUST00000102950.4/ENSMUSG00000020435.12, exon 12 of 12) | 11 | 3697430 | 3698326 | 897 | 2 | ENSMUST00000117960.1 | -6796 | Gm16518 | ENSMUSG00000083015.1 |
| chr9 | 78389731 | 78390155 | 3.49248 | 4.1335 | 1.25299 | Promoter | 9 | 78388901 | 78389467 | 567 | 2 | ENSMUST00000119091.1 | -265 | Gm8087 | ENSMUSG00000083032.1 |
| chr4 | 146580266 | 146580653 | 4.22774 | 7.21118 | 3.48697 | Promoter | 4 | 146580629 | 146582312 | 1684 | 1 | ENSMUST00000118760.1 | 0 | Gm13233 | ENSMUSG00000083038.1 |
| chr4 | 84076415 | 84076699 | 3.72627 | 4.81527 | 1.68047 | Distal Intergenic | 4 | 84104895 | 84105902 | 1008 | 1 | ENSMUST00000120911.1 | -28196 | Gm12416 | ENSMUSG00000083044.1 |
| chr4 | 84081471 | 84081757 | 3.82113 | 4.8074 | 1.67182 | Distal Intergenic | 4 | 84104895 | 84105902 | 1008 | 1 | ENSMUST00000120911.1 | -23138 | Gm12416 | ENSMUSG00000083044.1 |
| chr4 | 84088973 | 84089088 | 3.36847 | 3.88029 | 1.01733 | Distal Intergenic | 4 | 84104895 | 84105902 | 1008 | 1 | ENSMUST00000120911.1 | -15807 | Gm12416 | ENSMUSG00000083044.1 |
| chr4 | 84089811 | 84089890 | 3.57613 | 4.33409 | 1.37114 | Distal Intergenic | 4 | 84104895 | 84105902 | 1008 | 1 | ENSMUST00000120911.1 | -15005 | Gm12416 | ENSMUSG00000083044.1 |
| chr4 | 84092725 | 84093133 | 3.90451 | 5.09091 | 1.8397 | Distal Intergenic | 4 | 84104895 | 84105902 | 1008 | 1 | ENSMUST00000120911.1 | -11762 | Gm12416 | ENSMUSG00000083044.1 |
| chr4 | 84112673 | 84112794 | 3.65878 | 4.57963 | 1.46619 | Distal Intergenic | 4 | 84104895 | 84105902 | 1008 | 1 | ENSMUST00000120911.1 | 7779 | Gm12416 | ENSMUSG00000083044.1 |
| chr1 | 63668009 | 63668084 | 3.73648 | 4.56999 | 1.4884 | Intron (ENSMUST00000090313.4/ENSMUSG00000069085.5, intron 6 of 14) | 1 | 63671468 | 63671971 | 504 | 1 | ENSMUST00000120763.1 | -3384 | Gm12748 | ENSMUSG00000083066.1 |

| | | | | | | | | | | | | | | | |
|---|---|---|---|---|---|---|---|---|---|---|---|---|---|---|---|
| chr2 | 73978460 | 73978755 | 3.41179 | 4.00062 | 1.18 | Distal Intergenic | 2 | 73959669 | 73960669 | 1001 | 2 | ENSMUST00000119237.2 | -17792 | Gm13668 | ENSMUSG00000083072.2 |
| chr2 | 73980590 | 73981306 | 3.2916 | 3.77469 | 1.05022 | Distal Intergenic | 2 | 73959669 | 73960669 | 1001 | 2 | ENSMUST00000119237.2 | -19922 | Gm13668 | ENSMUSG00000083072.2 |
| chr2 | 74030168 | 74030422 | 3.30038 | 3.80983 | 1.04579 | Distal Intergenic | 2 | 73959669 | 73960669 | 1001 | 2 | ENSMUST00000119237.2 | -69500 | Gm13668 | ENSMUSG00000083072.2 |
| chr4 | 152599367 | 152599644 | 3.50957 | 4.20456 | 1.30985 | Distal Intergenic | 4 | 152677748 | 152678003 | 256 | 2 | ENSMUST00000118685.1 | 78359 | Gm13172 | ENSMUSG00000083078.1 |
| chr4 | 152625354 | 152625859 | 3.61156 | 4.4437 | 1.37918 | Distal Intergenic | 4 | 152677748 | 152678003 | 256 | 2 | ENSMUST00000118685.1 | 52144 | Gm13172 | ENSMUSG00000083078.1 |
| chr2 | 147475626 | 147475815 | 3.5224 | 4.22747 | 1.32066 | Distal Intergenic | 2 | 147402954 | 147403262 | 309 | 2 | ENSMUST00000118814.1 | -72365 | Gm14109 | ENSMUSG00000083126.1 |
| chr2 | 147481130 | 147481234 | 3.80166 | 4.72198 | 1.63217 | Distal Intergenic | 2 | 147402954 | 147403262 | 309 | 2 | ENSMUST00000118814.1 | -77869 | Gm14109 | ENSMUSG00000083126.1 |
| chr11 | 52528059 | 52528412 | 3.64705 | 4.43655 | 1.43429 | Distal Intergenic | 11 | 52487158 | 52487310 | 153 | 2 | ENSMUST00000118245.1 | -40750 | Gm12210 | ENSMUSG00000083129.1 |
| chr2 | 31650605 | 31651197 | 3.59584 | 4.41712 | 1.3819 | Intron (ENSMUST00000113470.2/ENSMUSG00000079466.2, intron 3 of 4) | 2 | 31660470 | 31661427 | 958 | 1 | ENSMUST00000122301.1 | -9273 | Gm13424 | ENSMUSG00000083143.1 |
| chr2 | 31652159 | 31655267 | 4.20277 | 5.54804 | 2.19532 | 3' UTR | 2 | 31660470 | 31661427 | 958 | 1 | ENSMUST00000122301.1 | -5203 | Gm13424 | ENSMUSG00000083143.1 |
| chr2 | 31655665 | 31656240 | 3.50603 | 4.22505 | 1.25296 | 3' UTR | 2 | 31660470 | 31661427 | 958 | 1 | ENSMUST00000122301.1 | -4230 | Gm13424 | ENSMUSG00000083143.1 |
| chr2 | 31664106 | 31664326 | 3.29096 | 3.77742 | 1.06069 | Distal Intergenic | 2 | 31660470 | 31661427 | 958 | 1 | ENSMUST00000122301.1 | 3637 | Gm13424 | ENSMUSG00000083143.1 |
| chr11 | 52188116 | 52188538 | 4.07635 | 5.30901 | 2.02825 | Promoter | 11 | 52188953 | 52189553 | 601 | 1 | ENSMUST00000119795.1 | -415 | Gm12206 | ENSMUSG00000083175.1 |
| chr5 | 115203307 | 115203426 | 4.08669 | 5.22951 | 1.88126 | Distal Intergenic | 5 | 115198343 | 115198705 | 363 | 1 | ENSMUST00000122278.1 | 4965 | Gm13828 | ENSMUSG00000083182.1 |
| chr5 | 115203752 | 115203942 | 3.41136 | 4.20939 | 1.28479 | Distal Intergenic | 5 | 115198343 | 115198705 | 363 | 1 | ENSMUST00000122278.1 | 5410 | Gm13828 | ENSMUSG00000083182.1 |
| chr5 | 115204903 | 115205116 | 3.59212 | 4.29784 | 1.29657 | Distal Intergenic | 5 | 115198343 | 115198705 | 363 | 1 | ENSMUST00000122278.1 | 6561 | Gm13828 | ENSMUSG00000083182.1 |
| chr5 | 115205815 | 115205962 | 4.09603 | 5.4166 | 2.0747 | Distal Intergenic | 5 | 115198343 | 115198705 | 363 | 1 | ENSMUST00000122278.1 | 7473 | Gm13828 | ENSMUSG00000083182.1 |
| chr5 | 115214607 | 115215026 | 3.67744 | 4.52283 | 1.44541 | Distal Intergenic | 5 | 115198343 | 115198705 | 363 | 1 | ENSMUST00000122278.1 | 16265 | Gm13828 | ENSMUSG00000083182.1 |
| chr4 | 115364340 | 115364545 | 3.17664 | 3.7353 | 1.01733 | Distal Intergenic | 4 | 115343788 | 115363115 | 19328 | 1 | ENSMUST00000120813.1 | 20553 | Cyp4a30-ps | ENSMUSG00000083190.1 |
| chr11 | 82682659 | 82683134 | 3.46365 | 4.05152 | 1.11441 | Distal Intergenic | 11 | 82725143 | 82725436 | 294 | 1 | ENSMUST00000122124.1 | -42009 | Gm11421 | ENSMUSG00000083203.1 |
| chr6 | 30714480 | 30714686 | 3.48465 | 4.1658 | 1.2445 | Distal Intergenic | 6 | 30719873 | 30720103 | 231 | 1 | ENSMUST00000120615.1 | -5187 | Gm14528 | ENSMUSG00000083221.1 |
| chr16 | 90891247 | 90891329 | 3.82051 | 4.84961 | 1.67161 | Intron (ENSMUST00000099548.3/ENSMUSG00000039903.11, intron 7 of 9) | 16 | 90915180 | 90915731 | 552 | 1 | ENSMUST00000117990.1 | -23851 | Gm7831 | ENSMUSG00000083226.1 |
| chr16 | 90891746 | 90891897 | 3.59994 | 4.35613 | 1.37232 | Intron (ENSMUST00000099548.3/ENSMUSG00000039903.11, intron 7 of 9) | 16 | 90915180 | 90915731 | 552 | 1 | ENSMUST00000117990.1 | -23283 | Gm7831 | ENSMUSG00000083226.1 |
| chr16 | 90904262 | 90904337 | 3.35104 | 3.854 | 1.01733 | 3' UTR | 16 | 90915180 | 90915731 | 552 | 1 | ENSMUST00000117990.1 | -10843 | Gm7831 | ENSMUSG00000083226.1 |
| chr11 | 114912679 | 114912928 | 3.31493 | 3.91139 | 1.07042 | Distal Intergenic | 11 | 114908731 | 114909185 | 455 | 1 | ENSMUST00000117948.1 | 3949 | Gm11698 | ENSMUSG00000083237.1 |
| chr4 | 60081781 | 60081862 | 6.00729 | 15.6445 | 10.87512 | Distal Intergenic | 4 | 60085840 | 60088577 | 2738 | 1 | ENSMUST00000120720.1 | -3978 | Mup-ps1 | ENSMUSG00000083239.1 |
| chr1 | 75563200 | 75563524 | 3.41461 | 4.04455 | 1.21808 | Distal Intergenic | 1 | 75568083 | 75568574 | 492 | 1 | ENSMUST00000117745.1 | -4559 | Gm15180 | ENSMUSG00000083244.1 |
| chr10 | 10291491 | 10291624 | 3.24263 | 3.6911 | 1.01733 | Distal Intergenic | 10 | 10343115 | 10344327 | 1213 | 1 | ENSMUST00000118589.1 | -51491 | Gm15568 | ENSMUSG00000083257.1 |
| chr10 | 10295027 | 10295198 | 3.21408 | 3.64835 | 1.01232 | Distal Intergenic | 10 | 10343115 | 10344327 | 1213 | 1 | ENSMUST00000118589.1 | -47917 | Gm15568 | ENSMUSG00000083257.1 |
| chr2 | 72619468 | 72619560 | 3.49362 | 4.13765 | 1.25998 | Distal Intergenic | 2 | 72632270 | 72633426 | 1157 | 1 | ENSMUST00000118184.1 | -12710 | Pex13-ps | ENSMUSG00000083279.1 |
| chr2 | 72630243 | 72630318 | 3.5263 | 4.16963 | 1.19781 | Distal Intergenic | 2 | 72632270 | 72633426 | 1157 | 1 | ENSMUST00000118184.1 | -1952 | Pex13-ps | ENSMUSG00000083279.1 |
| chr4 | 138411593 | 138411679 | 3.71457 | 4.53763 | 1.46928 | Distal Intergenic | 4 | 138425788 | 138426002 | 215 | 1 | ENSMUST00000117723.1 | -14109 | Rpl38-ps1 | ENSMUSG00000083326.4 |
| chr4 | 138418490 | 138418565 | 3.46976 | 4.10154 | 1.18082 | Distal Intergenic | 4 | 138425788 | 138426002 | 215 | 1 | ENSMUST00000117723.1 | -7223 | Rpl38-ps1 | ENSMUSG00000083326.4 |
| chr4 | 138419379 | 138419592 | 3.35931 | 3.86647 | 1.01733 | Distal Intergenic | 4 | 138425788 | 138426002 | 215 | 1 | ENSMUST00000117723.1 | -6196 | Rpl38-ps1 | ENSMUSG00000083326.4 |
| chr11 | 96233725 | 96234050 | 3.7309 | 4.90821 | 1.71138 | Distal Intergenic | 11 | 96248633 | 96249189 | 557 | 1 | ENSMUST00000119837.1 | -14583 | Gm11539 | ENSMUSG00000083337.1 |
| chr11 | 96235005 | 96235900 | 3.47161 | 4.12658 | 1.24954 | Distal Intergenic | 11 | 96248633 | 96249189 | 557 | 1 | ENSMUST00000119837.1 | -12733 | Gm11539 | ENSMUSG00000083337.1 |
| chr11 | 96248877 | 96249184 | 3.64545 | 4.39262 | 1.34953 | Promoter | 11 | 96248633 | 96249189 | 557 | 1 | ENSMUST00000119837.1 | 245 | Gm11539 | ENSMUSG00000083337.1 |
| chr4 | 122455263 | 122455342 | 4.71438 | 14.18006 | 9.45767 | Promoter | 4 | 122455210 | 122455435 | 226 | 1 | ENSMUST00000122214.1 | 54 | Gm12895 | ENSMUSG00000083365.1 |
| chr4 | 132113102 | 132113712 | 3.40248 | 4.00254 | 1.19256 | 3' UTR | 4 | 132093891 | 132094298 | 408 | 1 | ENSMUST00000122137.1 | 19212 | Gm13214 | ENSMUSG00000083368.1 |
| chr15 | 73801579 | 73801822 | 3.29673 | 3.77226 | 1.01733 | Intron (ENSMUST00000151999.2/ENSMUSG00000072487.5, intron 4 of 24) | 15 | 73809507 | 73809896 | 390 | 2 | ENSMUST00000120938.1 | 8074 | Gm3244 | ENSMUSG00000083380.1 |
| chr3 | 36531304 | 36531594 | 3.60175 | 4.50106 | 1.44691 | Distal Intergenic | 3 | 36539383 | 36540081 | 699 | 2 | ENSMUST00000118210.1 | 8487 | Gm11550 | ENSMUSG00000083388.1 |
| chr3 | 36532296 | 36532371 | 3.33152 | 3.85809 | 1.12281 | Distal Intergenic | 3 | 36539383 | 36540081 | 699 | 2 | ENSMUST00000118210.1 | 7710 | Gm11550 | ENSMUSG00000083388.1 |
| chr2 | 181917432 | 181918069 | 2.34349 | 11.76681 | 7.3856 | Distal Intergenic | 2 | 181896823 | 181898712 | 1890 | 1 | ENSMUST00000119395.1 | 20610 | Gm5466 | ENSMUSG00000083397.1 |
| chr2 | 181918563 | 181918713 | 2.18636 | 5.9417 | 2.51247 | Distal Intergenic | 2 | 181896823 | 181898712 | 1890 | 1 | ENSMUST00000119395.1 | 21741 | Gm5466 | ENSMUSG00000083397.1 |
| chr2 | 181929050 | 181929363 | 2.5811 | 11.4964 | 7.11072 | Distal Intergenic | 2 | 181896823 | 181898712 | 1890 | 1 | ENSMUST00000119395.1 | 32228 | Gm5466 | ENSMUSG00000083397.1 |
| chr2 | 181930902 | 181931635 | 3.00227 | 11.80567 | 7.46841 | Distal Intergenic | 2 | 181896823 | 181898712 | 1890 | 1 | ENSMUST00000119395.1 | 34080 | Gm5466 | ENSMUSG00000083397.1 |
| chr2 | 4817071 | 4817363 | 3.35208 | 3.85555 | 1.01733 | Distal Intergenic | 2 | 4834136 | 4834612 | 477 | 2 | ENSMUST00000117910.1 | 17249 | Gm13176 | ENSMUSG00000083407.1 |
| chr13 | 24295861 | 24296591 | 3.47112 | 4.07557 | 1.15933 | Distal Intergenic | 13 | 24291394 | 24291805 | 412 | 2 | ENSMUST00000119226.1 | -4057 | Gm11340 | ENSMUSG00000083409.1 |
| chr2 | 179661827 | 179661976 | 3.37436 | 3.89806 | 1.04874 | Intron (ENSMUST00000000314.7/ENSMUSG00000000305.7, intron 2 of 15) | 2 | 179612622 | 179613567 | 946 | 2 | ENSMUST00000120800.1 | -48261 | Gm14300 | ENSMUSG00000083410.1 |
| chr11 | 52701269 | 52701377 | 4.09887 | 5.2767 | 1.9437 | Distal Intergenic | 11 | 52682779 | 52683224 | 446 | 2 | ENSMUST00000120769.1 | -18046 | Gm11187 | ENSMUSG00000083442.1 |
| chr11 | 52702645 | 52702793 | 3.71944 | 4.52315 | 1.4145 | Distal Intergenic | 11 | 52682779 | 52683224 | 446 | 2 | ENSMUST00000120769.1 | -19422 | Gm11187 | ENSMUSG00000083442.1 |
| chr4 | 117509029 | 117509194 | 3.68062 | 4.5097 | 1.47449 | Distal Intergenic | 4 | 117513823 | 117514201 | 379 | 2 | ENSMUST00000121841.1 | 5007 | Gm12826 | ENSMUSG00000083446.1 |
| chr2 | 130360556 | 130360827 | 3.32144 | 3.82221 | 1.05992 | Intron (ENSMUST00000110288.3/ENSMUSG00000053552.9, intron 11 of 17) | 2 | 130384883 | 130385329 | 447 | 2 | ENSMUST00000117531.1 | 24502 | Gm14044 | ENSMUSG00000083483.3 |
| chr2 | 130361480 | 130362628 | 4.23848 | 5.56476 | 2.15844 | Exon (ENSMUST00000110288.3/ENSMUSG00000053552.9, exon 12 of 18) | 2 | 130384883 | 130385329 | 447 | 2 | ENSMUST00000117531.1 | 22701 | Gm14044 | ENSMUSG00000083483.3 |
| chr2 | 130365268 | 130365646 | 3.58754 | 4.30509 | 1.30291 | Exon (ENSMUST00000110288.3/ENSMUSG00000053552.9, exon 16 of 18) | 2 | 130384883 | 130385329 | 447 | 2 | ENSMUST00000117531.1 | 19683 | Gm14044 | ENSMUSG00000083483.3 |
| chr4 | 122802673 | 122802769 | 6.26294 | 10.00502 | 5.75591 | Distal Intergenic | 4 | 122778154 | 122793749 | 15596 | 2 | ENSMUST00000119988.1 | -8925 | Gm12873 | ENSMUSG00000083494.1 |
| chr4 | 122806792 | 122807179 | 3.57905 | 4.3208 | 1.36942 | Distal Intergenic | 4 | 122778154 | 122793749 | 15596 | 2 | ENSMUST00000119988.1 | -13044 | Gm12873 | ENSMUSG00000083494.1 |
| chr2 | 156646578 | 156646738 | 3.57474 | 4.25837 | 1.25366 | Intron (ENSMUST00000109567.4/ENSMUSG00000061689.10, intron 1 of 12) | 2 | 156632825 | 156633468 | 644 | 2 | ENSMUST00000118255.1 | -13111 | Gm14172 | ENSMUSG00000083498.1 |
| chr17 | 34392343 | 34392480 | 3.47768 | 4.08622 | 1.16295 | Promoter | 17 | 34392403 | 34392690 | 288 | 1 | ENSMUST00000121290.1 | 0 | Gm15321 | ENSMUSG00000083522.1 |
| chr3 | 101236694 | 101236994 | 3.74332 | 4.62762 | 1.58399 | Distal Intergenic | 3 | 101233460 | 101233895 | 436 | 1 | ENSMUST00000120257.1 | 3235 | Tpt1-ps1 | ENSMUSG00000083546.1 |
| chr11 | 95185811 | 95185979 | 3.42199 | 4.10602 | 1.20197 | Distal Intergenic | 11 | 95176813 | 95177244 | 432 | 1 | ENSMUST00000117670.1 | 8999 | Gm11514 | ENSMUSG00000083548.1 |
| chr11 | 95200279 | 95200384 | 3.4081 | 3.95555 | 1.07209 | Distal Intergenic | 11 | 95176813 | 95177244 | 432 | 1 | ENSMUST00000117670.1 | 23467 | Gm11514 | ENSMUSG00000083548.1 |
| chr11 | 95201057 | 95201252 | 3.49739 | 4.11836 | 1.17158 | Distal Intergenic | 11 | 95176813 | 95177244 | 432 | 1 | ENSMUST00000117670.1 | 24245 | Gm11514 | ENSMUSG00000083548.1 |
| chr4 | 122122907 | 122122991 | 3.79366 | 11.24354 | 6.87868 | Promoter | 4 | 122122847 | 122123072 | 226 | 1 | ENSMUST00000118430.1 | 61 | Gm12896 | ENSMUSG00000083558.1 |
| chr11 | 19746133 | 19746312 | 3.72411 | 4.53415 | 1.43159 | Distal Intergenic | 11 | 19749419 | 19750066 | 648 | 2 | ENSMUST00000118065.1 | 3754 | Gm12028 | ENSMUSG00000083601.1 |
| chr11 | 19760397 | 19760975 | 3.39286 | 4.10129 | 1.21482 | Distal Intergenic | 11 | 19749419 | 19750066 | 648 | 2 | ENSMUST00000118065.1 | -10332 | Gm12028 | ENSMUSG00000083601.1 |
| chr11 | 19761713 | 19761861 | 3.21794 | 3.65414 | 1.01175 | Distal Intergenic | 11 | 19749419 | 19750066 | 648 | 2 | ENSMUST00000118065.1 | -11648 | Gm12028 | ENSMUSG00000083601.1 |
| chr7 | 27284919 | 27285029 | 3.7586 | 4.67212 | 1.62574 | Distal Intergenic | 7 | 27271768 | 27272392 | 625 | 2 | ENSMUST00000122093.1 | -12528 | Gm15570 | ENSMUSG00000083622.1 |
| chr2 | 127559322 | 127559397 | 3.56192 | 4.22875 | 1.21624 | Distal Intergenic | 2 | 127568577 | 127569426 | 850 | 2 | ENSMUST00000121178.1 | 10029 | Gm14229 | ENSMUSG00000083648.1 |
| chr2 | 127561648 | 127561784 | 3.3534 | 3.85755 | 1.01733 | Distal Intergenic | 2 | 127568577 | 127569426 | 850 | 2 | ENSMUST00000121178.1 | 7642 | Gm14229 | ENSMUSG00000083648.1 |
| chr13 | 23395095 | 23395547 | 4.12229 | 5.49887 | 2.12787 | Distal Intergenic | 13 | 23382505 | 23382660 | 156 | 1 | ENSMUST00000121969.1 | 12591 | Gm11334 | ENSMUSG00000083667.1 |
| chr13 | 23396116 | 23396805 | 3.70185 | 4.50458 | 1.43539 | Distal Intergenic | 13 | 23382505 | 23382660 | 156 | 1 | ENSMUST00000121969.1 | 13612 | Gm11334 | ENSMUSG00000083667.1 |
| chr13 | 23397119 | 23397524 | 3.92052 | 5.29613 | 2.00641 | Distal Intergenic | 13 | 23382505 | 23382660 | 156 | 1 | ENSMUST00000121969.1 | 14615 | Gm11334 | ENSMUSG00000083667.1 |
| chr13 | 23398001 | 23398305 | 3.69802 | 4.49224 | 1.42413 | Distal Intergenic | 13 | 23382505 | 23382660 | 156 | 1 | ENSMUST00000121969.1 | 15497 | Gm11334 | ENSMUSG00000083667.1 |
| chr13 | 23399867 | 23400292 | 3.28723 | 3.76159 | 1.02975 | Distal Intergenic | 13 | 23382505 | 23382660 | 156 | 1 | ENSMUST00000121969.1 | 17363 | Gm11334 | ENSMUSG00000083667.1 |
| chr13 | 23401186 | 23401604 | 3.34402 | 3.85317 | 1.0246 | Distal Intergenic | 13 | 23382505 | 23382660 | 156 | 1 | ENSMUST00000121969.1 | 18682 | Gm11334 | ENSMUSG00000083667.1 |
| chr4 | 99624887 | 99625179 | 4.99363 | 7.17535 | 3.42986 | Distal Intergenic | 4 | 99615817 | 99616522 | 706 | 2 | ENSMUST00000119752.1 | -8366 | Gm12687 | ENSMUSG00000083698.1 |
| chr4 | 99625685 | 99626162 | 3.61683 | 4.34934 | 1.30773 | Distal Intergenic | 4 | 99615817 | 99616522 | 706 | 2 | ENSMUST00000119752.1 | -9164 | Gm12687 | ENSMUSG00000083698.1 |
| chr11 | 90844722 | 90844973 | 3.36563 | 3.87601 | 1.01733 | Intron (ENSMUST00000142521.1/ENSMUSG00000085940.1, intron 2 of 2) | 11 | 90859767 | 90861348 | 1582 | 2 | ENSMUST00000118450.1 | 16375 | Gm11516 | ENSMUSG00000083764.1 |
| chr11 | 114567049 | 114567426 | 3.45733 | 4.04559 | 1.12742 | Distal Intergenic | 11 | 114664608 | 114664909 | 302 | 1 | ENSMUST00000120766.1 | -97182 | Gm11689 | ENSMUSG00000083769.1 |
| chr11 | 114571224 | 114571508 | 3.54398 | 4.20113 | 1.21508 | Distal Intergenic | 11 | 114664608 | 114664909 | 302 | 1 | ENSMUST00000120766.1 | -93100 | Gm11689 | ENSMUSG00000083769.1 |
| chr11 | 114597219 | 114597941 | 3.51319 | 4.14438 | 1.17708 | Distal Intergenic | 11 | 114664608 | 114664909 | 302 | 1 | ENSMUST00000120766.1 | -66667 | Gm11689 | ENSMUSG00000083769.1 |
| chr11 | 114637197 | 114637580 | 4.13548 | 5.55378 | 2.17722 | Distal Intergenic | 11 | 114664608 | 114664909 | 302 | 1 | ENSMUST00000120766.1 | -27028 | Gm11689 | ENSMUSG00000083769.1 |
| chr11 | 114638046 | 114638368 | 4.20399 | 5.47438 | 2.06884 | Distal Intergenic | 11 | 114664608 | 114664909 | 302 | 1 | ENSMUST00000120766.1 | -26240 | Gm11689 | ENSMUSG00000083769.1 |
| chr11 | 114638918 | 114640011 | 3.87909 | 4.98248 | 1.783 | Distal Intergenic | 11 | 114664608 | 114664909 | 302 | 1 | ENSMUST00000120766.1 | -24597 | Gm11689 | ENSMUSG00000083769.1 |
| chr11 | 114652534 | 114653165 | 3.71429 | 4.5247 | 1.42651 | Distal Intergenic | 11 | 114664608 | 114664909 | 302 | 1 | ENSMUST00000120766.1 | -11443 | Gm11689 | ENSMUSG00000083769.1 |
| chr11 | 114656701 | 114656819 | 3.57814 | 4.65037 | 1.52742 | Distal Intergenic | 11 | 114664608 | 114664909 | 302 | 1 | ENSMUST00000120766.1 | -7789 | Gm11689 | ENSMUSG00000083769.1 |
| chr11 | 114657311 | 114657635 | 3.3752 | 3.89044 | 1.01733 | Distal Intergenic | 11 | 114664608 | 114664909 | 302 | 1 | ENSMUST00000120766.1 | -6973 | Gm11689 | ENSMUSG00000083769.1 |
| chr11 | 114657944 | 114658019 | 3.40356 | 3.94004 | 1.04124 | Distal Intergenic | 11 | 114664608 | 114664909 | 302 | 1 | ENSMUST00000120766.1 | -6589 | Gm11689 | ENSMUSG00000083769.1 |
| chr2 | 28932096 | 28932436 | 3.451 | 4.21051 | 1.24712 | Intron (ENSMUST00000189027.1/ENSMUSG00000079502.3, intron 6 of 6) | 2 | 28933610 | 28934609 | 1000 | 2 | ENSMUST00000121658.1 | 2173 | Gm13394 | ENSMUSG00000083773.4 |
| chr2 | 28936007 | 28936122 | 3.76799 | 4.68433 | 1.58932 | Intron (ENSMUST00000189027.1/ENSMUSG00000079502.3, intron 6 of 6) | 2 | 28933610 | 28934609 | 1000 | 2 | ENSMUST00000121658.1 | -1399 | Gm13394 | ENSMUSG00000083773.4 |
| chr2 | 28937846 | 28938738 | 3.61623 | 4.45317 | 1.39771 | Intron (ENSMUST00000189027.1/ENSMUSG00000079502.3, intron 6 of 6) | 2 | 28933610 | 28934609 | 1000 | 2 | ENSMUST00000121658.1 | -3238 | Gm13394 | ENSMUSG00000083773.4 |
| chr2 | 28939158 | 28939360 | 3.46227 | 4.07088 | 1.19207 | Intron (ENSMUST00000189027.1/ENSMUSG00000079502.3, intron 6 of 6) | 2 | 28933610 | 28934609 | 1000 | 2 | ENSMUST00000121658.1 | -4550 | Gm13394 | ENSMUSG00000083773.4 |
| chr2 | 28944142 | 28944357 | 3.55026 | 4.20982 | 1.21206 | Intron (ENSMUST00000189027.1/ENSMUSG00000079502.3, intron 6 of 6) | 2 | 28933610 | 28934609 | 1000 | 2 | ENSMUST00000121658.1 | -9534 | Gm13394 | ENSMUSG00000083773.4 |
| chr2 | 28944705 | 28944863 | 3.72748 | 4.6016 | 1.52789 | Intron (ENSMUST00000189027.1/ENSMUSG00000079502.3, intron 6 of 6) | 2 | 28933610 | 28934609 | 1000 | 2 | ENSMUST00000121658.1 | -10097 | Gm13394 | ENSMUSG00000083773.4 |
| chr2 | 28945993 | 28946151 | 3.37039 | 3.88319 | 1.01733 | Intron (ENSMUST00000189027.1/ENSMUSG00000079502.3, intron 6 of 6) | 2 | 28933610 | 28934609 | 1000 | 2 | ENSMUST00000121658.1 | -11385 | Gm13394 | ENSMUSG00000083773.4 |
| chr2 | 28947341 | 28947724 | 3.61432 | 4.52053 | 1.4641 | Intron (ENSMUST00000189027.1/ENSMUSG00000079502.3, intron 6 of 6) | 2 | 28933610 | 28934609 | 1000 | 2 | ENSMUST00000121658.1 | -12733 | Gm13394 | ENSMUSG00000083773.4 |
| chr2 | 28948558 | 28948852 | 4.02248 | 5.11189 | 1.81105 | Intron (ENSMUST00000189027.1/ENSMUSG00000079502.3, intron 6 of 6) | 2 | 28933610 | 28934609 | 1000 | 2 | ENSMUST00000121658.1 | -13950 | Gm13394 | ENSMUSG00000083773.4 |
| chr2 | 28949924 | 28950032 | 3.35085 | 3.85372 | 1.01733 | Intron (ENSMUST00000189027.1/ENSMUSG00000079502.3, intron 6 of 6) | 2 | 28933610 | 28934609 | 1000 | 2 | ENSMUST00000121658.1 | -15316 | Gm13394 | ENSMUSG00000083773.4 |
| chr2 | 28954590 | 28954722 | 3.29258 | 3.76604 | 1.01733 | Intron (ENSMUST00000189027.1/ENSMUSG00000079502.3, intron 6 of 6) | 2 | 28933610 | 28934609 | 1000 | 2 | ENSMUST00000121658.1 | -19982 | Gm13394 | ENSMUSG00000083773.4 |
| chr2 | 28956068 | 28956398 | 3.78004 | 4.67236 | 1.57428 | 3' UTR | 2 | 28933610 | 28934609 | 1000 | 2 | ENSMUST00000121658.1 | -21460 | Gm13394 | ENSMUSG00000083773.4 |
| chr2 | 28959656 | 28959867 | 3.66773 | 4.47693 | 1.4787 | Intron (ENSMUST00000189027.1/ENSMUSG00000079502.3, intron 5 of 6) | 2 | 28933610 | 28934609 | 1000 | 2 | ENSMUST00000121658.1 | -25048 | Gm13394 | ENSMUSG00000083773.4 |
| chr2 | 28961139 | 28961486 | 3.32864 | 4.03824 | 1.16834 | Intron (ENSMUST00000189027.1/ENSMUSG00000079502.3, intron 5 of 6) | 2 | 28933610 | 28934609 | 1000 | 2 | ENSMUST00000121658.1 | -26531 | Gm13394 | ENSMUSG00000083773.4 |
| chr2 | 28964509 | 28964641 | 3.76194 | 4.68378 | 1.62884 | Intron (ENSMUST00000189027.1/ENSMUSG00000079502.3, intron 4 of 6) | 2 | 28933610 | 28934609 | 1000 | 2 | ENSMUST00000121658.1 | -29901 | Gm13394 | ENSMUSG00000083773.4 |
| chr2 | 28965893 | 28966148 | 3.55989 | 4.2868 | 1.36121 | Intron (ENSMUST00000189027.1/ENSMUSG00000079502.3, intron 4 of 6) | 2 | 28933610 | 28934609 | 1000 | 2 | ENSMUST00000121658.1 | -31285 | Gm13394 | ENSMUSG00000083773.4 |
| chr2 | 28966486 | 28966578 | 3.90554 | 4.96413 | 1.82 | Intron (ENSMUST00000189027.1/ENSMUSG00000079502.3, intron 4 of 6) | 2 | 28933610 | 28934609 | 1000 | 2 | ENSMUST00000121658.1 | -31878 | Gm13394 | ENSMUSG00000083773.4 |
| chr2 | 28969050 | 28969646 | 3.44988 | 4.07395 | 1.2458 | Intron (ENSMUST00000189027.1/ENSMUSG00000079502.3, intron 4 of 6) | 2 | 28933610 | 28934609 | 1000 | 2 | ENSMUST00000121658.1 | -34442 | Gm13394 | ENSMUSG00000083773.4 |
| chr2 | 28976383 | 28976458 | 4.75507 | 6.69049 | 3.01133 | Intron (ENSMUST00000189027.1/ENSMUSG00000079502.3, intron 4 of 6) | 2 | 28933610 | 28934609 | 1000 | 2 | ENSMUST00000121658.1 | -41775 | Gm13394 | ENSMUSG00000083773.4 |
| chr2 | 28980763 | 28981013 | 3.67935 | 4.44646 | 1.35959 | Intron (ENSMUST00000189027.1/ENSMUSG00000079502.3, intron 4 of 6) | 2 | 28933610 | 28934609 | 1000 | 2 | ENSMUST00000121658.1 | -46155 | Gm13394 | ENSMUSG00000083773.4 |
| chr2 | 28981393 | 28981663 | 3.46832 | 4.09055 | 1.21096 | Intron (ENSMUST00000189027.1/ENSMUSG00000079502.3, intron 4 of 6) | 2 | 28933610 | 28934609 | 1000 | 2 | ENSMUST00000121658.1 | -46785 | Gm13394 | ENSMUSG00000083773.4 |
| chr4 | 8909388 | 8909496 | 4.49453 | 6.26045 | 2.67863 | Distal Intergenic | 4 | 8941259 | 8941656 | 398 | 1 | ENSMUST00000119493.1 | -31763 | Rps18-ps2 | ENSMUSG00000083786.1 |
| chr4 | 8910156 | 8910343 | 3.33032 | 3.8979 | 1.10894 | Distal Intergenic | 4 | 8941259 | 8941656 | 398 | 1 | ENSMUST00000119493.1 | -30916 | Rps18-ps2 | ENSMUSG00000083786.1 |
| chr4 | 8910863 | 8911023 | 3.3504 | 3.87515 | 1.09547 | Distal Intergenic | 4 | 8941259 | 8941656 | 398 | 1 | ENSMUST00000119493.1 | -30236 | Rps18-ps2 | ENSMUSG00000083786.1 |
| chr4 | 8911460 | 8911709 | 4.68113 | 6.51964 | 2.86334 | Distal Intergenic | 4 | 8941259 | 8941656 | 398 | 1 | ENSMUST00000119493.1 | -29550 | Rps18-ps2 | ENSMUSG00000083786.1 |
| chr4 | 8912687 | 8913095 | 3.42767 | 4.04887 | 1.19519 | Distal Intergenic | 4 | 8941259 | 8941656 | 398 | 1 | ENSMUST00000119493.1 | -28164 | Rps18-ps2 | ENSMUSG00000083786.1 |
| chr4 | 8921233 | 8921501 | 3.54912 | 4.21834 | 1.2451 | Distal Intergenic | 4 | 8941259 | 8941656 | 398 | 1 | ENSMUST00000119493.1 | -19758 | Rps18-ps2 | ENSMUSG00000083786.1 |
| chr4 | 8987556 | 8987887 | 3.39652 | 3.9535 | 1.12685 | Distal Intergenic | 4 | 8941259 | 8941656 | 398 | 1 | ENSMUST00000119493.1 | 46298 | Rps18-ps2 | ENSMUSG00000083786.1 |
| chr4 | 8988227 | 8988356 | 3.25999 | 3.71712 | 1.01733 | Distal Intergenic | 4 | 8941259 | 8941656 | 398 | 1 | ENSMUST00000119493.1 | 46969 | Rps18-ps2 | ENSMUSG00000083786.1 |
| chr4 | 9012784 | 9013019 | 3.54823 | 4.23302 | 1.30144 | Distal Intergenic | 4 | 8941259 | 8941656 | 398 | 1 | ENSMUST00000119493.1 | 71526 | Rps18-ps2 | ENSMUSG00000083786.1 |
| chr4 | 9019448 | 9019523 | 3.78672 | 4.81402 | 1.6501 | Distal Intergenic | 4 | 8941259 | 8941656 | 398 | 1 | ENSMUST00000119493.1 | 78190 | Rps18-ps2 | ENSMUSG00000083786.1 |
| chr4 | 9022888 | 9023083 | 3.33535 | 3.83703 | 1.04054 | Distal Intergenic | 4 | 8941259 | 8941656 | 398 | 1 | ENSMUST00000119493.1 | 81630 | Rps18-ps2 | ENSMUSG00000083786.1 |
| chr4 | 9028258 | 9029159 | 3.3317 | 3.84058 | 1.07227 | Distal Intergenic | 4 | 8941259 | 8941656 | 398 | 1 | ENSMUST00000119493.1 | 87000 | Rps18-ps2 | ENSMUSG00000083786.1 |
| chr13 | 22050835 | 22051412 | 3.35676 | 3.87955 | 1.07726 | Distal Intergenic | 13 | 22057346 | 22057862 | 517 | 2 | ENSMUST00000120775.1 | 6450 | Vmn1r-ps92 | ENSMUSG00000083873.1 |
| chr16 | 31905961 | 31906592 | 3.59078 | 4.32123 | 1.32514 | Distal Intergenic | 16 | 31924989 | 31925868 | 880 | 1 | ENSMUST00000120250.1 | -18397 | Gm15703 | ENSMUSG00000083890.1 |
| chr11 | 16412698 | 16412773 | 3.51166 | 4.17432 | 1.29156 | Intron (ENSMUST00000109647.2/ENSMUSG00000048834.10, intron 6 of 6) | 11 | 16439254 | 16440319 | 1066 | 1 | ENSMUST00000118857.1 | -26481 | Gm12012 | ENSMUSG00000083925.1 |
| chr6 | 29970675 | 29971016 | 3.52911 | 4.28753 | 1.32378 | Distal Intergenic | 6 | 29963644 | 29963850 | 207 | 1 | ENSMUST00000122139.1 | 7032 | Gm13725 | ENSMUSG00000083961.1 |
| chr6 | 29972829 | 29973304 | 3.67143 | 4.4653 | 1.41612 | Distal Intergenic | 6 | 29963644 | 29963850 | 207 | 1 | ENSMUST00000122139.1 | 9186 | Gm13725 | ENSMUSG00000083961.1 |
| chr2 | 24678987 | 24679447 | 3.82717 | 4.83472 | 1.71694 | Exon (ENSMUST00000070864.8/ENSMUSG00000004113.13, exon 19 of 46) | 2 | 24674701 | 24675346 | 646 | 1 | ENSMUST00000121558.1 | 4287 | Gm13459 | ENSMUSG00000083973.1 |
| chr5 | 147133626 | 147133869 | 3.48388 | 4.2005 | 1.27824 | Distal Intergenic | 5 | 147100562 | 147101053 | 492 | 2 | ENSMUST00000120308.1 | -32574 | Rps10-ps4 | ENSMUSG00000083988.1 |
| chr11 | 30754784 | 30755407 | 3.79807 | 4.84635 | 1.70185 | Promoter | 11 | 30754572 | 30754961 | 390 | 2 | ENSMUST00000121590.1 | 0 | Gm12101 | ENSMUSG00000083990.1 |
| chr3 | 36835877 | 36836000 | 3.62837 | 4.38793 | 1.42034 | Distal Intergenic | 3 | 36824352 | 36824958 | 607 | 2 | ENSMUST00000120538.1 | -10920 | Gm12376 | ENSMUSG00000084027.1 |
| chr2 | 118511978 | 118512472 | 3.54435 | 4.20125 | 1.2131 | Distal Intergenic | 2 | 118483313 | 118484177 | 865 | 2 | ENSMUST00000118018.1 | -27802 | Gm13983 | ENSMUSG00000084028.1 |
| chr4 | 135380991 | 135381101 | 3.29416 | 3.7684 | 1.01733 | Intron (ENSMUST00000105858.2/ENSMUSG00000043924.11, intron 1 of 4) | 4 | 135377158 | 135377235 | 78 | 1 | ENSMUST00000120561.1 | 3834 | Gm12991 | ENSMUSG00000084032.1 |
| chr2 | 170447589 | 170447769 | 3.28433 | 3.75363 | 1.01733 | Distal Intergenic | 2 | 170463514 | 170463819 | 306 | 2 | ENSMUST00000120455.1 | 16050 | Gm14269 | ENSMUSG00000084067.3 |
| chr2 | 170455230 | 170455427 | 3.50637 | 4.13626 | 1.18544 | Distal Intergenic | 2 | 170463514 | 170463819 | 306 | 2 | ENSMUST00000120455.1 | 8392 | Gm14269 | ENSMUSG00000084067.3 |
| chr2 | 170458053 | 170458516 | 3.41261 | 3.99716 | 1.17942 | Distal Intergenic | 2 | 170463514 | 170463819 | 306 | 2 | ENSMUST00000120455.1 | 5303 | Gm14269 | ENSMUSG00000084067.3 |
| chr2 | 170461463 | 170462034 | 3.38765 | 3.95127 | 1.12986 | Downstream (1-2kb) | 2 | 170463514 | 170463819 | 306 | 2 | ENSMUST00000120455.1 | 1785 | Gm14269 | ENSMUSG00000084067.3 |
| chr2 | 170463034 | 170463119 | 4.20184 | 5.49786 | 2.13294 | Promoter | 2 | 170463514 | 170463819 | 306 | 2 | ENSMUST00000120455.1 | 700 | Gm14269 | ENSMUSG00000084067.3 |
| chr2 | 170469655 | 170469743 | 3.27089 | 3.75068 | 1.0548 | Distal Intergenic | 2 | 170463514 | 170463819 | 306 | 2 | ENSMUST00000120455.1 | -5837 | Gm14269 | ENSMUSG00000084067.3 |
| chr2 | 170470353 | 170470593 | 3.37562 | 3.92033 | 1.12067 | Distal Intergenic | 2 | 170463514 | 170463819 | 306 | 2 | ENSMUST00000120455.1 | -6535 | Gm14269 | ENSMUSG00000084067.3 |
| chr11 | 97084880 | 97084977 | 3.67857 | 4.50923 | 1.48766 | Distal Intergenic | 11 | 97096933 | 97097326 | 394 | 1 | ENSMUST00000119956.1 | -11956 | Gm11573 | ENSMUSG00000084068.1 |
| chr11 | 97091880 | 97092160 | 3.67362 | 4.44843 | 1.38713 | Distal Intergenic | 11 | 97096933 | 97097326 | 394 | 1 | ENSMUST00000119956.1 | -4773 | Gm11573 | ENSMUSG00000084068.1 |
| chr11 | 97098695 | 97098991 | 3.99473 | 5.06709 | 1.79828 | 3' UTR | 11 | 97096933 | 97097326 | 394 | 1 | ENSMUST00000119956.1 | 1763 | Gm11573 | ENSMUSG00000084068.1 |
| chr11 | 97102027 | 97102256 | 3.39671 | 4.0225 | 1.17598 | Exon (ENSMUST00000001484.2/ENSMUSG00000001444.2, exon 2 of 6) | 11 | 97096933 | 97097326 | 394 | 1 | ENSMUST00000119956.1 | 5095 | Gm11573 | ENSMUSG00000084068.1 |
| chr13 | 23269831 | 23269989 | 3.62581 | 4.4091 | 1.42153 | Promoter | 13 | 23268880 | 23269474 | 595 | 1 | ENSMUST00000120909.1 | 952 | Vmn1r-ps135 | ENSMUSG00000084096.1 |
| chr13 | 23270523 | 23271021 | 4.19878 | 5.57828 | 2.22431 | Distal Intergenic | 13 | 23268880 | 23269474 | 595 | 1 | ENSMUST00000120909.1 | 1644 | Vmn1r-ps135 | ENSMUSG00000084096.1 |
| chr2 | 23233344 | 23233685 | 3.41228 | 3.99278 | 1.08737 | Intron (ENSMUST00000028113.4/ENSMUSG00000026774.6, intron 5 of 10) | 2 | 23235236 | 23235739 | 504 | 1 | ENSMUST00000118834.1 | -1551 | Gm13422 | ENSMUSG00000084098.1 |
| chr2 | 23236391 | 23237073 | 3.37861 | 3.90547 | 1.05226 | Intron (ENSMUST00000028113.4/ENSMUSG00000026774.6, intron 6 of 10) | 2 | 23235236 | 23235739 | 504 | 1 | ENSMUST00000118834.1 | 1156 | Gm13422 | ENSMUSG00000084098.1 |
| chr4 | 146239042 | 146240120 | 3.91081 | 6.60479 | 3.06191 | Promoter | 4 | 146239954 | 146241633 | 1680 | 1 | ENSMUST00000122090.1 | 0 | Gm13169 | ENSMUSG00000084127.1 |
| chr8 | 106135905 | 106137090 | 3.6243 | 4.48155 | 1.40285 | Promoter | 8 | 106130319 | 106136974 | 6656 | 2 | ENSMUST00000115979.3 | 0 | Esrp2 | ENSMUSG00000084128.5 |
| chr2 | 19519561 | 19519817 | 3.70352 | 4.49375 | 1.38768 | Intron (ENSMUST00000062060.4/ENSMUSG00000026734.9, intron 5 of 16) | 2 | 19509090 | 19509317 | 228 | 2 | ENSMUST00000121928.1 | -10245 | Gm13345 | ENSMUSG00000084160.1 |
| chr11 | 3998630 | 3998887 | 3.86587 | 5.01142 | 1.78601 | Exon (ENSMUST00000078757.2/ENSMUSG00000049721.9, exon 3 of 3) | 11 | 4002457 | 4002805 | 349 | 1 | ENSMUST00000121403.1 | -3570 | Gm11954 | ENSMUSG00000084170.1 |
| chr11 | 3999553 | 3999865 | 3.75074 | 4.65119 | 1.60877 | Distal Intergenic | 11 | 4002457 | 4002805 | 349 | 1 | ENSMUST00000121403.1 | -2592 | Gm11954 | ENSMUSG00000084170.1 |
| chr13 | 31204937 | 31205053 | 3.43624 | 4.01272 | 1.12412 | Distal Intergenic | 13 | 31199086 | 31199655 | 570 | 1 | ENSMUST00000117348.1 | 5852 | Gm11372 | ENSMUSG00000084175.1 |
| chr11 | 23954791 | 23955057 | 4.03859 | 5.13266 | 1.8253 | Distal Intergenic | 11 | 23928596 | 23928854 | 259 | 2 | ENSMUST00000117533.1 | -25938 | Gm12062 | ENSMUSG00000084206.1 |
| chr19 | 6595155 | 6595431 | 3.36066 | 3.86851 | 1.01733 | Distal Intergenic | 19 | 6592602 | 6593236 | 635 | 2 | ENSMUST00000118281.1 | -1920 | Hmgb1-ps4 | ENSMUSG00000084221.1 |
| chr19 | 6601352 | 6602064 | 3.63355 | 4.3726 | 1.32559 | Distal Intergenic | 19 | 6592602 | 6593236 | 635 | 2 | ENSMUST00000118281.1 | -8117 | Hmgb1-ps4 | ENSMUSG00000084221.1 |
| chr11 | 57883069 | 57883331 | 3.36501 | 3.87508 | 1.01733 | Distal Intergenic | 11 | 57880896 | 57881186 | 291 | 1 | ENSMUST00000119500.1 | 2174 | Gm12244 | ENSMUSG00000084223.1 |
| chr11 | 57886369 | 57886589 | 3.37882 | 3.89591 | 1.01733 | Distal Intergenic | 11 | 57880896 | 57881186 | 291 | 1 | ENSMUST00000119500.1 | 5474 | Gm12244 | ENSMUSG00000084223.1 |
| chr6 | 52443218 | 52443842 | 3.86949 | 4.85487 | 1.71077 | Distal Intergenic | 6 | 52357544 | 52357824 | 281 | 2 | ENSMUST00000117393.1 | -85395 | Gm15054 | ENSMUSG00000084282.1 |
| chr6 | 52444188 | 52444324 | 3.65305 | 4.52324 | 1.46773 | Distal Intergenic | 6 | 52357544 | 52357824 | 281 | 2 | ENSMUST00000117393.1 | -86365 | Gm15054 | ENSMUSG00000084282.1 |
| chr17 | 47954106 | 47954306 | 3.40781 | 3.94779 | 1.04593 | Distal Intergenic | 17 | 47936090 | 47936446 | 357 | 1 | ENSMUST00000117186.1 | 18017 | Gm5228 | ENSMUSG00000084288.1 |
| chr11 | 3628731 | 3629669 | 4.18414 | 5.6775 | 2.27977 | Promoter | 11 | 3628824 | 3629661 | 838 | 1 | ENSMUST00000121615.1 | 0 | Gm11950 | ENSMUSG00000084310.1 |

| | | | | | | | | | | | | | | | |
|---|---|---|---|---|---|---|---|---|---|---|---|---|---|---|---|
| chr11 | 3630105 | 3630670 | 3.89393 | 4.9629 | 1.77517 | Downstream (2-3kb) | 11 | 3628824 | 3629661 | 838 | 1 | ENSMUST00000121615.1 | 1282 | Gm11950 | ENSMUSG00000084310.1 |
| chr4 | 82617313 | 82617783 | 3.81219 | 4.76116 | 1.61551 | Intron (ENSMUST00000155821.1/ENSMUSG00000008575.12, intron 1 of 1) | 4 | 82642400 | 82642666 | 267 | 1 | ENSMUST00000118347.1 | -24617 | Gm11268 | ENSMUSG00000084357.1 |
| chr4 | 82618814 | 82619641 | 3.54749 | 4.22578 | 1.27493 | Intron (ENSMUST00000155821.1/ENSMUSG00000008575.12, intron 1 of 1) | 4 | 82642400 | 82642666 | 267 | 1 | ENSMUST00000118347.1 | -22759 | Gm11268 | ENSMUSG00000084357.1 |
| chr2 | 84025228 | 84025636 | 3.71729 | 4.60247 | 1.53233 | Distal Intergenic | 2 | 84034243 | 84034613 | 371 | 2 | ENSMUST00000120774.1 | 8977 | Gm13692 | ENSMUSG00000084359.1 |
| chr2 | 84026103 | 84027033 | 3.69196 | 4.58999 | 1.53942 | Distal Intergenic | 2 | 84034243 | 84034613 | 371 | 2 | ENSMUST00000120774.1 | 7580 | Gm13692 | ENSMUSG00000084359.1 |
| chr2 | 84027479 | 84028032 | 3.74544 | 4.64214 | 1.56833 | Distal Intergenic | 2 | 84034243 | 84034613 | 371 | 2 | ENSMUST00000120774.1 | 6581 | Gm13692 | ENSMUSG00000084359.1 |
| chr11 | 97117893 | 97118078 | 3.38193 | 3.9006 | 1.01733 | Promoter | 11 | 97116796 | 97116915 | 120 | 2 | ENSMUST00000119164.1 | -979 | Gm11572 | ENSMUSG00000084368.1 |
| chr11 | 97131394 | 97131568 | 3.30669 | 3.78723 | 1.01733 | Distal Intergenic | 11 | 97116796 | 97116915 | 120 | 2 | ENSMUST00000119164.1 | -14480 | Gm11572 | ENSMUSG00000084368.1 |
| chr11 | 18011223 | 18011559 | 3.34298 | 3.99326 | 1.18051 | Distal Intergenic | 11 | 17973788 | 17974790 | 1003 | 2 | ENSMUST00000122453.1 | -36434 | Gm12017 | ENSMUSG00000084413.1 |
| chr11 | 18014777 | 18015020 | 3.77716 | 4.7595 | 1.61708 | Distal Intergenic | 11 | 17973788 | 17974790 | 1003 | 2 | ENSMUST00000122453.1 | -39988 | Gm12017 | ENSMUSG00000084413.1 |
| chr13 | 94594965 | 94596481 | 4.18003 | 5.5396 | 2.21045 | Distal Intergenic | 13 | 94517162 | 94517275 | 114 | 1 | ENSMUST00000122474.1 | 77804 | Gm25109 | ENSMUSG00000084423.1 |
| chr13 | 94597027 | 94597462 | 3.76791 | 4.86246 | 1.67572 | Distal Intergenic | 13 | 94517162 | 94517275 | 114 | 1 | ENSMUST00000122474.1 | 79866 | Gm25109 | ENSMUSG00000084423.1 |
| chr5 | 99821001 | 99821353 | 3.33543 | 3.83568 | 1.03323 | Distal Intergenic | 5 | 99870248 | 99870323 | 76 | 1 | ENSMUST00000122496.1 | -48895 | Gm26252 | ENSMUSG00000084445.1 |
| chr5 | 99857173 | 99857461 | 3.70482 | 4.67397 | 1.587 | Distal Intergenic | 5 | 99870248 | 99870323 | 76 | 1 | ENSMUST00000122496.1 | -12787 | Gm26252 | ENSMUSG00000084445.1 |
| chr12 | 71577271 | 71577751 | 4.1059 | 5.34291 | 2.06463 | Distal Intergenic | 12 | 71724652 | 71724954 | 303 | 1 | ENSMUST00000122514.1 | -146901 | Gm22959 | ENSMUSG00000084463.1 |
| chr12 | 71578409 | 71579198 | 3.30194 | 3.78009 | 1.01733 | Distal Intergenic | 12 | 71724652 | 71724954 | 303 | 1 | ENSMUST00000122514.1 | -145454 | Gm22959 | ENSMUSG00000084463.1 |
| chr14 | 120804143 | 120804405 | 3.72788 | 4.57383 | 1.47566 | Distal Intergenic | 14 | 120798853 | 120798936 | 84 | 1 | ENSMUST00000122527.1 | 5291 | Gm25781 | ENSMUSG00000084476.1 |
| chr4 | 49479218 | 49479526 | 3.58364 | 4.35702 | 1.37839 | Distal Intergenic | 4 | 49474307 | 49474410 | 104 | 1 | ENSMUST00000122530.1 | 4912 | Gm26424 | ENSMUSG00000084479.1 |
| chr4 | 49480732 | 49480847 | 3.20468 | 3.77774 | 1.01733 | Distal Intergenic | 4 | 49474307 | 49474410 | 104 | 1 | ENSMUST00000122530.1 | 6426 | Gm26424 | ENSMUSG00000084479.1 |
| chr4 | 49482097 | 49482334 | 3.63873 | 4.55272 | 1.48795 | Distal Intergenic | 4 | 49474307 | 49474410 | 104 | 1 | ENSMUST00000122530.1 | 7791 | Gm26424 | ENSMUSG00000084479.1 |
| chr7 | 90301889 | 90302202 | 4.03944 | 5.16383 | 1.85995 | Distal Intergenic | 7 | 90293734 | 90293846 | 113 | 1 | ENSMUST00000122531.1 | 8156 | n-R5s155 | ENSMUSG00000084480.1 |
| chr7 | 90304005 | 90304166 | 3.59543 | 4.29802 | 1.27645 | Distal Intergenic | 7 | 90293734 | 90293846 | 113 | 1 | ENSMUST00000122531.1 | 10272 | n-R5s155 | ENSMUSG00000084480.1 |
| chr16 | 22572904 | 22573084 | 3.32826 | 3.81971 | 1.01733 | Intron (ENSMUST00000089925.4/ENSMUSG00000022861.11, intron 11 of 24) | 16 | 22595029 | 22595220 | 192 | 1 | ENSMUST00000122562.1 | -21945 | Gm22481 | ENSMUSG00000084511.1 |
| chr19 | 32955619 | 32955784 | 3.63982 | 4.52898 | 1.46778 | Distal Intergenic | 19 | 32969950 | 32970308 | 359 | 2 | ENSMUST00000122563.1 | 14524 | Gm22482 | ENSMUSG00000084512.1 |
| chr19 | 33035939 | 33036159 | 3.40408 | 4.04798 | 1.13763 | Distal Intergenic | 19 | 32969950 | 32970308 | 359 | 2 | ENSMUST00000122563.1 | -65632 | Gm22482 | ENSMUSG00000084512.1 |
| chr19 | 33057807 | 33057924 | 3.34153 | 3.83966 | 1.01733 | Distal Intergenic | 19 | 32969950 | 32970308 | 359 | 2 | ENSMUST00000122563.1 | -87500 | Gm22482 | ENSMUSG00000084512.1 |
| chr19 | 33116129 | 33116235 | 3.35645 | 3.86688 | 1.03402 | Distal Intergenic | 19 | 32969950 | 32970308 | 359 | 2 | ENSMUST00000122563.1 | -145822 | Gm22482 | ENSMUSG00000084512.1 |
| chr19 | 33119811 | 33120741 | 3.44035 | 4.05107 | 1.12418 | Distal Intergenic | 19 | 32969950 | 32970308 | 359 | 2 | ENSMUST00000122563.1 | -149504 | Gm22482 | ENSMUSG00000084512.1 |
| chr19 | 33121307 | 33121749 | 4.58721 | 6.51416 | 2.88909 | Distal Intergenic | 19 | 32969950 | 32970308 | 359 | 2 | ENSMUST00000122563.1 | -151000 | Gm22482 | ENSMUSG00000084512.1 |
| chr19 | 33122088 | 33122402 | 3.40223 | 3.98397 | 1.08945 | Distal Intergenic | 19 | 32969950 | 32970308 | 359 | 2 | ENSMUST00000122563.1 | -151781 | Gm22482 | ENSMUSG00000084512.1 |
| chr19 | 33122809 | 33123212 | 3.88725 | 4.93027 | 1.71017 | Distal Intergenic | 19 | 32969950 | 32970308 | 359 | 2 | ENSMUST00000122563.1 | -152502 | Gm22482 | ENSMUSG00000084512.1 |
| chr19 | 33123524 | 33123890 | 3.49225 | 4.10938 | 1.16703 | Distal Intergenic | 19 | 32969950 | 32970308 | 359 | 2 | ENSMUST00000122563.1 | -153217 | Gm22482 | ENSMUSG00000084512.1 |
| chr19 | 33124358 | 33125079 | 3.91249 | 4.90666 | 1.68288 | Distal Intergenic | 19 | 32969950 | 32970308 | 359 | 2 | ENSMUST00000122563.1 | -154051 | Gm22482 | ENSMUSG00000084512.1 |
| chr19 | 33126260 | 33126335 | 3.8041 | 4.67863 | 1.50247 | Distal Intergenic | 19 | 32969950 | 32970308 | 359 | 2 | ENSMUST00000122563.1 | -155953 | Gm22482 | ENSMUSG00000084512.1 |
| chr19 | 33127065 | 33127527 | 3.79618 | 4.73561 | 1.60939 | Distal Intergenic | 19 | 32969950 | 32970308 | 359 | 2 | ENSMUST00000122563.1 | -156758 | Gm22482 | ENSMUSG00000084512.1 |
| chr19 | 33130783 | 33131020 | 3.69892 | 4.5913 | 1.53887 | Distal Intergenic | 19 | 32969950 | 32970308 | 359 | 2 | ENSMUST00000122563.1 | -160476 | Gm22482 | ENSMUSG00000084512.1 |
| chr19 | 33131565 | 33131871 | 3.82817 | 4.80863 | 1.67921 | Distal Intergenic | 19 | 32969950 | 32970308 | 359 | 2 | ENSMUST00000122563.1 | -161258 | Gm22482 | ENSMUSG00000084512.1 |
| chr19 | 33136737 | 33136927 | 3.56864 | 4.32911 | 1.36008 | Downstream (<1kb) | 19 | 32969950 | 32970308 | 359 | 2 | ENSMUST00000122563.1 | -166430 | Gm22482 | ENSMUSG00000084512.1 |
| chr19 | 33143329 | 33143671 | 3.66455 | 4.7502 | 1.61479 | Intron (ENSMUST00000096114.6/ENSMUSG00000071573.8, intron 6 of 6) | 19 | 32969950 | 32970308 | 359 | 2 | ENSMUST00000122563.1 | -173022 | Gm22482 | ENSMUSG00000084512.1 |
| chr19 | 33168048 | 33168459 | 3.40847 | 3.95034 | 1.05141 | Exon (ENSMUST00000096114.6/ENSMUSG00000071573.8, exon 6 of 7) | 19 | 32969950 | 32970308 | 359 | 2 | ENSMUST00000122563.1 | -197741 | Gm22482 | ENSMUSG00000084512.1 |
| chr19 | 33169241 | 33169404 | 3.6271 | 4.36642 | 1.34501 | Intron (ENSMUST00000096114.6/ENSMUSG00000071573.8, intron 5 of 6) | 19 | 32969950 | 32970308 | 359 | 2 | ENSMUST00000122563.1 | -198934 | Gm22482 | ENSMUSG00000084512.1 |
| chr5 | 148757277 | 148757505 | 4.15619 | 5.40507 | 2.08637 | Distal Intergenic | 5 | 148753323 | 148753440 | 118 | 2 | ENSMUST00000122564.1 | -3838 | n-R5s179 | ENSMUSG00000084513.1 |
| chr5 | 148759048 | 148759179 | 3.29865 | 3.77515 | 1.01733 | Distal Intergenic | 5 | 148753323 | 148753440 | 118 | 2 | ENSMUST00000122564.1 | -5609 | n-R5s179 | ENSMUSG00000084513.1 |
| chr9 | 118453803 | 118454384 | 3.56965 | 4.25111 | 1.24516 | Distal Intergenic | 9 | 118466984 | 118467064 | 81 | 2 | ENSMUST00000122566.1 | 12680 | Gm22479 | ENSMUSG00000084515.1 |
| chr9 | 118455536 | 118455707 | 3.59558 | 4.30384 | 1.30364 | Distal Intergenic | 9 | 118466984 | 118467064 | 81 | 2 | ENSMUST00000122566.1 | 11357 | Gm22479 | ENSMUSG00000084515.1 |
| chr9 | 118457387 | 118457539 | 3.73459 | 4.55928 | 1.44934 | Distal Intergenic | 9 | 118466984 | 118467064 | 81 | 2 | ENSMUST00000122566.1 | 9525 | Gm22479 | ENSMUSG00000084515.1 |
| chr9 | 118465742 | 118465961 | 3.40493 | 4.16465 | 1.25072 | Downstream (1-2kb) | 9 | 118466984 | 118467064 | 81 | 2 | ENSMUST00000122566.1 | 1103 | Gm22479 | ENSMUSG00000084515.1 |
| chr9 | 118467766 | 118469870 | 4.43306 | 5.99388 | 2.52016 | Promoter | 9 | 118466984 | 118467064 | 81 | 2 | ENSMUST00000122566.1 | -703 | Gm22479 | ENSMUSG00000084515.1 |
| chr9 | 118471126 | 118471503 | 4.02171 | 5.10056 | 1.80112 | Distal Intergenic | 9 | 118466984 | 118467064 | 81 | 2 | ENSMUST00000122566.1 | -4063 | Gm22479 | ENSMUSG00000084515.1 |
| chr4 | 11914269 | 11914799 | 3.35258 | 3.91832 | 1.1383 | Distal Intergenic | 4 | 11928424 | 11928739 | 316 | 1 | ENSMUST00000122569.1 | -13625 | Gm22477 | ENSMUSG00000084518.1 |
| chr18 | 25507287 | 25507553 | 3.69488 | 4.56108 | 1.49786 | Intron (ENSMUST00000025117.7/ENSMUSG00000024268.10, intron 3 of 12) | 18 | 25598145 | 25598454 | 310 | 2 | ENSMUST00000122588.1 | 90901 | Gm25845 | ENSMUSG00000084537.1 |
| chr18 | 25507899 | 25508221 | 3.27223 | 3.73553 | 1.01733 | Intron (ENSMUST00000025117.7/ENSMUSG00000024268.10, intron 3 of 12) | 18 | 25598145 | 25598454 | 310 | 2 | ENSMUST00000122588.1 | 90233 | Gm25845 | ENSMUSG00000084537.1 |
| chr18 | 25508867 | 25509031 | 3.39379 | 3.98404 | 1.18037 | Intron (ENSMUST00000025117.7/ENSMUSG00000024268.10, intron 3 of 12) | 18 | 25598145 | 25598454 | 310 | 2 | ENSMUST00000122588.1 | 89423 | Gm25845 | ENSMUSG00000084537.1 |
| chr18 | 25510716 | 25511043 | 3.34685 | 3.88979 | 1.12596 | Intron (ENSMUST00000025117.7/ENSMUSG00000024268.10, intron 3 of 12) | 18 | 25598145 | 25598454 | 310 | 2 | ENSMUST00000122588.1 | 87411 | Gm25845 | ENSMUSG00000084537.1 |
| chr18 | 25537132 | 25537245 | 3.37132 | 3.88465 | 1.01733 | Intron (ENSMUST00000025117.7/ENSMUSG00000024268.10, intron 3 of 12) | 18 | 25598145 | 25598454 | 310 | 2 | ENSMUST00000122588.1 | 61209 | Gm25845 | ENSMUSG00000084537.1 |
| chr18 | 25538141 | 25538258 | 3.45286 | 4.03992 | 1.13174 | Intron (ENSMUST00000025117.7/ENSMUSG00000024268.10, intron 2 of 12) | 18 | 25598145 | 25598454 | 310 | 2 | ENSMUST00000122588.1 | 60196 | Gm25845 | ENSMUSG00000084537.1 |
| chr18 | 25566532 | 25566917 | 3.53906 | 4.21239 | 1.27832 | Intron (ENSMUST00000025117.7/ENSMUSG00000024268.10, intron 2 of 12) | 18 | 25598145 | 25598454 | 310 | 2 | ENSMUST00000122588.1 | 31537 | Gm25845 | ENSMUSG00000084537.1 |
| chr18 | 25567829 | 25568092 | 3.62372 | 4.388 | 1.39627 | Intron (ENSMUST00000025117.7/ENSMUSG00000024268.10, intron 2 of 12) | 18 | 25598145 | 25598454 | 310 | 2 | ENSMUST00000122588.1 | 30362 | Gm25845 | ENSMUSG00000084537.1 |
| chr18 | 25568437 | 25568564 | 3.36578 | 3.87622 | 1.01733 | Intron (ENSMUST00000025117.7/ENSMUSG00000024268.10, intron 2 of 12) | 18 | 25598145 | 25598454 | 310 | 2 | ENSMUST00000122588.1 | 29890 | Gm25845 | ENSMUSG00000084537.1 |
| chr18 | 25577036 | 25577465 | 3.60826 | 4.34901 | 1.37859 | Intron (ENSMUST00000025117.7/ENSMUSG00000024268.10, intron 2 of 12) | 18 | 25598145 | 25598454 | 310 | 2 | ENSMUST00000122588.1 | 20989 | Gm25845 | ENSMUSG00000084537.1 |
| chr18 | 25578258 | 25578865 | 3.47372 | 4.12836 | 1.19779 | Intron (ENSMUST00000025117.7/ENSMUSG00000024268.10, intron 2 of 12) | 18 | 25598145 | 25598454 | 310 | 2 | ENSMUST00000122588.1 | 19589 | Gm25845 | ENSMUSG00000084537.1 |
| chr18 | 25590514 | 25590589 | 4.00386 | 5.07477 | 1.78212 | Intron (ENSMUST00000025117.7/ENSMUSG00000024268.10, intron 2 of 12) | 18 | 25598145 | 25598454 | 310 | 2 | ENSMUST00000122588.1 | 7865 | Gm25845 | ENSMUSG00000084537.1 |
| chr18 | 25621797 | 25622479 | 3.65107 | 4.48607 | 1.43651 | Intron (ENSMUST00000025117.7/ENSMUSG00000024268.10, intron 2 of 12) | 18 | 25598145 | 25598454 | 310 | 2 | ENSMUST00000122588.1 | -23344 | Gm25845 | ENSMUSG00000084537.1 |
| chr18 | 25675515 | 25675590 | 3.43862 | 4.01629 | 1.12412 | Intron (ENSMUST00000025117.7/ENSMUSG00000024268.10, intron 2 of 12) | 18 | 25598145 | 25598454 | 310 | 2 | ENSMUST00000122588.1 | -77062 | Gm25845 | ENSMUSG00000084537.1 |
| chr5 | 119271037 | 119271420 | 3.47246 | 4.07136 | 1.13746 | Distal Intergenic | 5 | 119297245 | 119297361 | 117 | 1 | ENSMUST00000122599.1 | -25825 | n-R5s175 | ENSMUSG00000084548.1 |
| chr5 | 119371226 | 119371496 | 3.35019 | 3.85272 | 1.01733 | Distal Intergenic | 5 | 119297245 | 119297361 | 117 | 1 | ENSMUST00000122599.1 | 73982 | n-R5s175 | ENSMUSG00000084548.1 |
| chr5 | 119381980 | 119382127 | 3.35506 | 3.86007 | 1.01733 | Distal Intergenic | 5 | 119297245 | 119297361 | 117 | 1 | ENSMUST00000122599.1 | 84736 | n-R5s175 | ENSMUSG00000084548.1 |
| chr5 | 119410778 | 119411301 | 3.52102 | 4.15417 | 1.17104 | Distal Intergenic | 5 | 119297245 | 119297361 | 117 | 1 | ENSMUST00000122599.1 | 113534 | n-R5s175 | ENSMUSG00000084548.1 |
| chr1 | 188711014 | 188711230 | 3.42762 | 4.01863 | 1.19162 | Intron (ENSMUST00000060479.8/ENSMUSG00000026609.10, intron 37 of 70) | 1 | 188760476 | 188760579 | 104 | 1 | ENSMUST00000122601.1 | -49246 | Gm25269 | ENSMUSG00000084550.1 |
| chr16 | 5307117 | 5307654 | 3.60284 | 4.34995 | 1.39771 | Distal Intergenic | 16 | 5324036 | 5324157 | 122 | 2 | ENSMUST00000122624.1 | 16503 | n-R5s30 | ENSMUSG00000084573.1 |
| chr2 | 74965685 | 74965800 | 3.75037 | 4.60719 | 1.52063 | Distal Intergenic | 2 | 74935415 | 74935514 | 100 | 2 | ENSMUST00000122625.1 | -30172 | n-R5s198 | ENSMUSG00000084574.1 |
| chr2 | 74966119 | 74966236 | 3.37992 | 3.91225 | 1.06924 | Distal Intergenic | 2 | 74935415 | 74935514 | 100 | 2 | ENSMUST00000122625.1 | -30606 | n-R5s198 | ENSMUSG00000084574.1 |
| chr2 | 75002221 | 75002476 | 3.42391 | 3.997 | 1.13103 | Distal Intergenic | 2 | 74935415 | 74935514 | 100 | 2 | ENSMUST00000122625.1 | -66708 | n-R5s198 | ENSMUSG00000084574.1 |
| chr2 | 75003290 | 75003516 | 3.58857 | 4.31602 | 1.28041 | Distal Intergenic | 2 | 74935415 | 74935514 | 100 | 2 | ENSMUST00000122625.1 | -67777 | n-R5s198 | ENSMUSG00000084574.1 |
| chr4 | 82303634 | 82304331 | 3.2691 | 3.73078 | 1.01733 | Intron (ENSMUST00000050872.9/ENSMUSG00000008575.12, intron 11 of 11) | 4 | 82439410 | 82439514 | 105 | 1 | ENSMUST00000122646.1 | -135079 | n-R5s188 | ENSMUSG00000084595.1 |
| chr4 | 82305615 | 82306056 | 3.41947 | 3.99544 | 1.15293 | Intron (ENSMUST00000050872.9/ENSMUSG00000008575.12, intron 11 of 11) | 4 | 82439410 | 82439514 | 105 | 1 | ENSMUST00000122646.1 | -133354 | n-R5s188 | ENSMUSG00000084595.1 |
| chr4 | 82311701 | 82311804 | 3.29767 | 3.77367 | 1.01733 | Intron (ENSMUST00000050872.9/ENSMUSG00000008575.12, intron 10 of 11) | 4 | 82439410 | 82439514 | 105 | 1 | ENSMUST00000122646.1 | -127606 | n-R5s188 | ENSMUSG00000084595.1 |
| chr4 | 82407436 | 82407821 | 3.3545 | 3.91066 | 1.13656 | Intron (ENSMUST00000050872.9/ENSMUSG00000008575.12, intron 2 of 11) | 4 | 82439410 | 82439514 | 105 | 1 | ENSMUST00000122646.1 | -31589 | n-R5s188 | ENSMUSG00000084595.1 |
| chr4 | 82408784 | 82408909 | 3.47827 | 4.19199 | 1.2708 | Intron (ENSMUST00000050872.9/ENSMUSG00000008575.12, intron 2 of 11) | 4 | 82439410 | 82439514 | 105 | 1 | ENSMUST00000122646.1 | -30501 | n-R5s188 | ENSMUSG00000084595.1 |
| chr4 | 82410908 | 82411266 | 3.34521 | 3.91154 | 1.13803 | Intron (ENSMUST00000050872.9/ENSMUSG00000008575.12, intron 2 of 11) | 4 | 82439410 | 82439514 | 105 | 1 | ENSMUST00000122646.1 | -28144 | n-R5s188 | ENSMUSG00000084595.1 |
| chr4 | 82432800 | 82432959 | 3.43232 | 4.02856 | 1.19408 | Intron (ENSMUST00000050872.9/ENSMUSG00000008575.12, intron 2 of 11) | 4 | 82439410 | 82439514 | 105 | 1 | ENSMUST00000122646.1 | -6451 | n-R5s188 | ENSMUSG00000084595.1 |
| chr4 | 82436841 | 82437356 | 4.28206 | 5.64503 | 2.21115 | Intron (ENSMUST00000050872.9/ENSMUSG00000008575.12, intron 2 of 11) | 4 | 82439410 | 82439514 | 105 | 1 | ENSMUST00000122646.1 | -2054 | n-R5s188 | ENSMUSG00000084595.1 |
| chr4 | 82438071 | 82438154 | 4.30783 | 5.74862 | 2.30102 | Intron (ENSMUST00000050872.9/ENSMUSG00000008575.12, intron 2 of 11) | 4 | 82439410 | 82439514 | 105 | 1 | ENSMUST00000122646.1 | -1256 | n-R5s188 | ENSMUSG00000084595.1 |
| chr4 | 82438685 | 82438879 | 3.9834 | 5.14868 | 1.8937 | Promoter | 4 | 82439410 | 82439514 | 105 | 1 | ENSMUST00000122646.1 | -531 | n-R5s188 | ENSMUSG00000084595.1 |
| chr4 | 82439337 | 82440016 | 3.56222 | 4.30376 | 1.35599 | Promoter | 4 | 82439410 | 82439514 | 105 | 1 | ENSMUST00000122646.1 | 0 | n-R5s188 | ENSMUSG00000084595.1 |
| chr4 | 82441263 | 82441604 | 4.09102 | 5.3291 | 2.04894 | Intron (ENSMUST00000050872.9/ENSMUSG00000008575.12, intron 2 of 11) | 4 | 82439410 | 82439514 | 105 | 1 | ENSMUST00000122646.1 | 1854 | n-R5s188 | ENSMUSG00000084595.1 |
| chr4 | 82443753 | 82443836 | 3.57013 | 4.6553 | 1.53876 | Intron (ENSMUST00000050872.9/ENSMUSG00000008575.12, intron 2 of 11) | 4 | 82439410 | 82439514 | 105 | 1 | ENSMUST00000122646.1 | 4344 | n-R5s188 | ENSMUSG00000084595.1 |
| chr4 | 82444238 | 82445114 | 3.47927 | 4.14038 | 1.28731 | Intron (ENSMUST00000050872.9/ENSMUSG00000008575.12, intron 2 of 11) | 4 | 82439410 | 82439514 | 105 | 1 | ENSMUST00000122646.1 | 4829 | n-R5s188 | ENSMUSG00000084595.1 |
| chr4 | 82446244 | 82446353 | 3.53427 | 4.27694 | 1.32307 | Intron (ENSMUST00000050872.9/ENSMUSG00000008575.12, intron 2 of 11) | 4 | 82439410 | 82439514 | 105 | 1 | ENSMUST00000122646.1 | 6835 | n-R5s188 | ENSMUSG00000084595.1 |
| chr4 | 82449191 | 82449461 | 3.60544 | 4.69252 | 1.56596 | Intron (ENSMUST00000050872.9/ENSMUSG00000008575.12, intron 2 of 11) | 4 | 82439410 | 82439514 | 105 | 1 | ENSMUST00000122646.1 | 9782 | n-R5s188 | ENSMUSG00000084595.1 |
| chr4 | 82449777 | 82449852 | 3.79196 | 4.7263 | 1.63558 | Intron (ENSMUST00000050872.9/ENSMUSG00000008575.12, intron 2 of 11) | 4 | 82439410 | 82439514 | 105 | 1 | ENSMUST00000122646.1 | 10368 | n-R5s188 | ENSMUSG00000084595.1 |
| chr4 | 82450967 | 82451049 | 3.72273 | 4.59888 | 1.54029 | Intron (ENSMUST00000050872.9/ENSMUSG00000008575.12, intron 2 of 11) | 4 | 82439410 | 82439514 | 105 | 1 | ENSMUST00000122646.1 | 11558 | n-R5s188 | ENSMUSG00000084595.1 |
| chr4 | 82451735 | 82452036 | 3.1925 | 3.75931 | 1.01733 | Intron (ENSMUST00000050872.9/ENSMUSG00000008575.12, intron 2 of 11) | 4 | 82439410 | 82439514 | 105 | 1 | ENSMUST00000122646.1 | 12326 | n-R5s188 | ENSMUSG00000084595.1 |
| chr4 | 82454279 | 82454475 | 3.61132 | 4.43937 | 1.47281 | Intron (ENSMUST00000050872.9/ENSMUSG00000008575.12, intron 2 of 11) | 4 | 82439410 | 82439514 | 105 | 1 | ENSMUST00000122646.1 | 14870 | n-R5s188 | ENSMUSG00000084595.1 |
| chr4 | 82455759 | 82456004 | 4.41173 | 5.9877 | 2.49067 | Intron (ENSMUST00000050872.9/ENSMUSG00000008575.12, intron 2 of 11) | 4 | 82439410 | 82439514 | 105 | 1 | ENSMUST00000122646.1 | 16350 | n-R5s188 | ENSMUSG00000084595.1 |
| chr4 | 82459542 | 82459731 | 3.49082 | 4.12597 | 1.22853 | Intron (ENSMUST00000050872.9/ENSMUSG00000008575.12, intron 2 of 11) | 4 | 82439410 | 82439514 | 105 | 1 | ENSMUST00000122646.1 | 20133 | n-R5s188 | ENSMUSG00000084595.1 |
| chr4 | 82460067 | 82460396 | 3.30191 | 3.80212 | 1.08799 | Intron (ENSMUST00000050872.9/ENSMUSG00000008575.12, intron 2 of 11) | 4 | 82439410 | 82439514 | 105 | 1 | ENSMUST00000122646.1 | 20658 | n-R5s188 | ENSMUSG00000084595.1 |
| chr4 | 82461239 | 82461446 | 3.89207 | 4.95047 | 1.80315 | Intron (ENSMUST00000050872.9/ENSMUSG00000008575.12, intron 2 of 11) | 4 | 82439410 | 82439514 | 105 | 1 | ENSMUST00000122646.1 | 21830 | n-R5s188 | ENSMUSG00000084595.1 |
| chr4 | 82462341 | 82462537 | 3.55929 | 4.31843 | 1.36736 | Intron (ENSMUST00000050872.9/ENSMUSG00000008575.12, intron 2 of 11) | 4 | 82439410 | 82439514 | 105 | 1 | ENSMUST00000122646.1 | 22932 | n-R5s188 | ENSMUSG00000084595.1 |
| chr4 | 82463455 | 82463557 | 3.5091 | 4.18529 | 1.31037 | Intron (ENSMUST00000050872.9/ENSMUSG00000008575.12, intron 2 of 11) | 4 | 82439410 | 82439514 | 105 | 1 | ENSMUST00000122646.1 | 24046 | n-R5s188 | ENSMUSG00000084595.1 |
| chr4 | 82463925 | 82464068 | 3.32365 | 3.82791 | 1.07072 | Intron (ENSMUST00000050872.9/ENSMUSG00000008575.12, intron 2 of 11) | 4 | 82439410 | 82439514 | 105 | 1 | ENSMUST00000122646.1 | 24516 | n-R5s188 | ENSMUSG00000084595.1 |
| chr4 | 82466883 | 82467063 | 4.1146 | 5.37444 | 2.08824 | Intron (ENSMUST00000050872.9/ENSMUSG00000008575.12, intron 2 of 11) | 4 | 82439410 | 82439514 | 105 | 1 | ENSMUST00000122646.1 | 27474 | n-R5s188 | ENSMUSG00000084595.1 |
| chr4 | 82467494 | 82468229 | 3.34412 | 3.87195 | 1.11494 | Intron (ENSMUST00000050872.9/ENSMUSG00000008575.12, intron 2 of 11) | 4 | 82439410 | 82439514 | 105 | 1 | ENSMUST00000122646.1 | 28085 | n-R5s188 | ENSMUSG00000084595.1 |
| chr9 | 118227874 | 118228167 | 3.47363 | 4.09963 | 1.23202 | Distal Intergenic | 9 | 118287786 | 118288092 | 307 | 1 | ENSMUST00000122659.1 | -59619 | Gm22198 | ENSMUSG00000084608.1 |
| chr9 | 118228853 | 118229136 | 3.59173 | 4.31077 | 1.33692 | Distal Intergenic | 9 | 118287786 | 118288092 | 307 | 1 | ENSMUST00000122659.1 | -58650 | Gm22198 | ENSMUSG00000084608.1 |
| chr9 | 118249756 | 118249991 | 3.35659 | 3.87637 | 1.06677 | Distal Intergenic | 9 | 118287786 | 118288092 | 307 | 1 | ENSMUST00000122659.1 | -37795 | Gm22198 | ENSMUSG00000084608.1 |
| chr9 | 118269911 | 118269988 | 3.53182 | 4.19839 | 1.27021 | Distal Intergenic | 9 | 118287786 | 118288092 | 307 | 1 | ENSMUST00000122659.1 | -17798 | Gm22198 | ENSMUSG00000084608.1 |
| chr10 | 75438154 | 75438248 | 3.40524 | 3.98692 | 1.18875 | Exon (ENSMUST00000039925.7/ENSMUSG00000033427.8, exon 9 of 10) | 10 | 75442620 | 75442942 | 323 | 1 | ENSMUST00000122673.1 | -4372 | Gm25545 | ENSMUSG00000084622.1 |
| chr10 | 75447228 | 75447313 | 3.67801 | 4.45621 | 1.38498 | Intron (ENSMUST00000145890.2/ENSMUSG00000033416.9, intron 5 of 8) | 10 | 75442620 | 75442942 | 323 | 1 | ENSMUST00000122673.1 | 4609 | Gm25545 | ENSMUSG00000084622.1 |
| chr7 | 132026337 | 132026474 | 3.51777 | 4.1489 | 1.16988 | Intron (ENSMUST00000124096.2/ENSMUSG00000030849.12, intron 2 of 6) | 7 | 132058641 | 132058760 | 120 | 1 | ENSMUST00000122681.1 | -32167 | n-R5s159 | ENSMUSG00000084630.1 |
| chr7 | 132043590 | 132044342 | 3.38173 | 3.90443 | 1.03196 | 3' UTR | 7 | 132058641 | 132058760 | 120 | 1 | ENSMUST00000122681.1 | -14299 | n-R5s159 | ENSMUSG00000084630.1 |
| chr3 | 107374828 | 107375208 | 3.38611 | 3.91319 | 1.03958 | Distal Intergenic | 3 | 107415863 | 107416195 | 333 | 2 | ENSMUST00000122691.1 | 40987 | Gm24383 | ENSMUSG00000084640.1 |
| chr8 | 83747697 | 83747912 | 4.36277 | 6.07601 | 2.60411 | Distal Intergenic | 8 | 83752918 | 83753204 | 287 | 1 | ENSMUST00000122733.1 | -5006 | Gm22024 | ENSMUSG00000084682.1 |
| chr9 | 121221409 | 121221866 | 3.77399 | 4.65113 | 1.52138 | Intron (ENSMUST00000171923.2/ENSMUSG00000040936.9, intron 17 of 36) | 9 | 121242342 | 121242587 | 246 | 1 | ENSMUST00000122757.1 | -20476 | Gm22987 | ENSMUSG00000084706.1 |
| chr9 | 121225175 | 121225372 | 3.78247 | 4.65287 | 1.51575 | Intron (ENSMUST00000171923.2/ENSMUSG00000040936.9, intron 17 of 36) | 9 | 121242342 | 121242587 | 246 | 1 | ENSMUST00000122757.1 | -16970 | Gm22987 | ENSMUSG00000084706.1 |
| chr9 | 121226421 | 121226775 | 3.98941 | 5.06369 | 1.77511 | Intron (ENSMUST00000171923.2/ENSMUSG00000040936.9, intron 17 of 36) | 9 | 121242342 | 121242587 | 246 | 1 | ENSMUST00000122757.1 | -15567 | Gm22987 | ENSMUSG00000084706.1 |
| chr9 | 121229800 | 121229888 | 7.51059 | 13.10292 | 8.53237 | Intron (ENSMUST00000171923.2/ENSMUSG00000040936.9, intron 17 of 36) | 9 | 121242342 | 121242587 | 246 | 1 | ENSMUST00000122757.1 | -12454 | Gm22987 | ENSMUSG00000084706.1 |
| chr9 | 121231350 | 121231425 | 4.03416 | 5.12775 | 1.81311 | Intron (ENSMUST00000171923.2/ENSMUSG00000040936.9, intron 17 of 36) | 9 | 121242342 | 121242587 | 246 | 1 | ENSMUST00000122757.1 | -10917 | Gm22987 | ENSMUSG00000084706.1 |
| chr9 | 121234680 | 121234803 | 4.05395 | 5.69398 | 2.25965 | Intron (ENSMUST00000171923.2/ENSMUSG00000040936.9, intron 17 of 36) | 9 | 121242342 | 121242587 | 246 | 1 | ENSMUST00000122757.1 | -7539 | Gm22987 | ENSMUSG00000084706.1 |
| chr9 | 121237657 | 121237888 | 4.12429 | 5.29782 | 1.92619 | Intron (ENSMUST00000171923.2/ENSMUSG00000040936.9, intron 17 of 36) | 9 | 121242342 | 121242587 | 246 | 1 | ENSMUST00000122757.1 | -4454 | Gm22987 | ENSMUSG00000084706.1 |
| chr9 | 121238334 | 121238607 | 3.54853 | 4.20434 | 1.20144 | Intron (ENSMUST00000171923.2/ENSMUSG00000040936.9, intron 17 of 36) | 9 | 121242342 | 121242587 | 246 | 1 | ENSMUST00000122757.1 | -3735 | Gm22987 | ENSMUSG00000084706.1 |
| chr9 | 121239305 | 121239511 | 4.05326 | 5.68295 | 2.2723 | Intron (ENSMUST00000171923.2/ENSMUSG00000040936.9, intron 17 of 36) | 9 | 121242342 | 121242587 | 246 | 1 | ENSMUST00000122757.1 | -2831 | Gm22987 | ENSMUSG00000084706.1 |
| chr9 | 106004060 | 106004807 | 3.69309 | 4.5595 | 1.5205 | Intron (ENSMUST00000121963.1/ENSMUSG00000032572.7, intron 33 of 37) | 9 | 106001718 | 106001798 | 81 | 1 | ENSMUST00000122765.1 | 2343 | Gm24631 | ENSMUSG00000084714.1 |
| chr9 | 106005386 | 106005466 | 3.92275 | 5.37131 | 2.00914 | Intron (ENSMUST00000121963.1/ENSMUSG00000032572.7, intron 33 of 37) | 9 | 106001718 | 106001798 | 81 | 1 | ENSMUST00000122765.1 | 3669 | Gm24631 | ENSMUSG00000084714.1 |
| chr9 | 106009209 | 106009450 | 3.65886 | 4.47813 | 1.46393 | Intron (ENSMUST00000121963.1/ENSMUSG00000032572.7, intron 33 of 37) | 9 | 106001718 | 106001798 | 81 | 1 | ENSMUST00000122765.1 | 7492 | Gm24631 | ENSMUSG00000084714.1 |
| chr9 | 106000670 | 106000931 | 3.43146 | 4.08647 | 1.23604 | Promoter | 9 | 106001718 | 106001798 | 81 | 1 | ENSMUST00000122765.1 | -787 | Gm24631 | ENSMUSG00000084714.1 |
| chr17 | 36242358 | 36242515 | 3.53684 | 4.38042 | 1.3738 | Promoter | 17 | 36241789 | 36242106 | 318 | 2 | ENSMUST00000122801.1 | -253 | Gm22453 | ENSMUSG00000084750.1 |
| chr4 | 124441635 | 124441733 | 3.52492 | 4.17516 | 1.22378 | Intron (ENSMUST00000127913.2/ENSMUSG00000084757.2, intron 2 of 3) | 4 | 124450235 | 124451285 | 1051 | 2 | ENSMUST00000156566.1 | 9552 | 1700057H15Rik | ENSMUSG00000084757.2 |
| chr4 | 124442869 | 124443111 | 4.20105 | 5.47087 | 2.07493 | Intron (ENSMUST00000127913.2/ENSMUSG00000084757.2, intron 2 of 3) | 4 | 124450235 | 124451285 | 1051 | 2 | ENSMUST00000156566.1 | 8174 | 1700057H15Rik | ENSMUSG00000084757.2 |
| chr4 | 124444075 | 124444266 | 3.37029 | 3.88304 | 1.01733 | Intron (ENSMUST00000127913.2/ENSMUSG00000084757.2, intron 2 of 3) | 4 | 124450235 | 124451285 | 1051 | 2 | ENSMUST00000156566.1 | 7019 | 1700057H15Rik | ENSMUSG00000084757.2 |
| chr4 | 124444589 | 124444664 | 3.90762 | 4.8896 | 1.66162 | Intron (ENSMUST00000127913.2/ENSMUSG00000084757.2, intron 2 of 3) | 4 | 124450235 | 124451285 | 1051 | 2 | ENSMUST00000156566.1 | 6621 | 1700057H15Rik | ENSMUSG00000084757.2 |
| chr4 | 124456355 | 124456603 | 3.33237 | 3.82594 | 1.01733 | Intron (ENSMUST00000143859.2/ENSMUSG00000084757.2, intron 2 of 3) | 4 | 124414416 | 124451295 | 36880 | 2 | ENSMUST00000127913.2 | -5061 | 1700057H15Rik | ENSMUSG00000084757.2 |
| chr4 | 124472646 | 124472731 | 5.17207 | 7.45428 | 3.53527 | Intron (ENSMUST00000143859.2/ENSMUSG00000084757.2, intron 2 of 3) | 4 | 124449902 | 124485959 | 36058 | 2 | ENSMUST00000143859.2 | 13228 | 1700057H15Rik | ENSMUSG00000084757.2 |
| chr4 | 124476432 | 124478512 | 3.65612 | 4.463 | 1.43596 | Intron (ENSMUST00000143859.2/ENSMUSG00000084757.2, intron 2 of 3) | 4 | 124449902 | 124485959 | 36058 | 2 | ENSMUST00000143859.2 | 7447 | 1700057H15Rik | ENSMUSG00000084757.2 |
| chr4 | 124483290 | 124483458 | 3.38001 | 4.00178 | 1.1556 | Intron (ENSMUST00000143859.2/ENSMUSG00000084757.2, intron 1 of 3) | 4 | 124449902 | 124485959 | 36058 | 2 | ENSMUST00000143859.2 | 2501 | 1700057H15Rik | ENSMUSG00000084757.2 |
| chr4 | 124483849 | 124484073 | 3.36462 | 3.90157 | 1.11316 | Intron (ENSMUST00000143859.2/ENSMUSG00000084757.2, intron 1 of 3) | 4 | 124449902 | 124485959 | 36058 | 2 | ENSMUST00000143859.2 | 1886 | 1700057H15Rik | ENSMUSG00000084757.2 |
| chr4 | 124485098 | 124485173 | 3.36539 | 3.9174 | 1.13176 | Promoter | 4 | 124449902 | 124485959 | 36058 | 2 | ENSMUST00000143859.2 | 786 | 1700057H15Rik | ENSMUSG00000084757.2 |
| chr4 | 41636797 | 41637064 | 3.28814 | 3.90408 | 1.05015 | Intron (ENSMUST00000102963.4/ENSMUSG00000061322.10, intron 19 of 19) | 4 | 41639378 | 41641416 | 2039 | 1 | ENSMUST00000155382.1 | -2314 | Gm12406 | ENSMUSG00000084761.1 |
| chr6 | 16995304 | 16995913 | 3.75688 | 4.70644 | 1.63501 | Distal Intergenic | 6 | 17058806 | 17064996 | 6191 | 2 | ENSMUST00000140746.1 | 69083 | Gm15473 | ENSMUSG00000084772.1 |
| chr6 | 16999723 | 16999798 | 3.49613 | 4.16561 | 1.30693 | Distal Intergenic | 6 | 17058806 | 17064996 | 6191 | 2 | ENSMUST00000140746.1 | 65198 | Gm15473 | ENSMUSG00000084772.1 |
| chr11 | 45025560 | 45026536 | 3.53021 | 4.26538 | 1.30351 | Distal Intergenic | 11 | 45102866 | 45116928 | 14063 | 2 | ENSMUST00000143869.1 | 90392 | Gm12159 | ENSMUSG00000084773.1 |
| chr11 | 45026901 | 45027490 | 3.3331 | 3.82697 | 1.01733 | Distal Intergenic | 11 | 45102866 | 45116928 | 14063 | 2 | ENSMUST00000143869.1 | 89438 | Gm12159 | ENSMUSG00000084773.1 |
| chr2 | 147321166 | 147321409 | 3.39891 | 4.02976 | 1.13592 | Intron (ENSMUST00000136998.2/ENSMUSG00000086509.2, intron 5 of 5) | 2 | 147323562 | 147325087 | 1526 | 2 | ENSMUST00000127688.1 | 3678 | Gm14110 | ENSMUSG00000084774.1 |
| chr2 | 147323166 | 147323494 | 3.37591 | 3.89152 | 1.01733 | Intron (ENSMUST00000136998.2/ENSMUSG00000086509.2, intron 5 of 5) | 2 | 147323562 | 147325087 | 1526 | 2 | ENSMUST00000127688.1 | 1593 | Gm14110 | ENSMUSG00000084774.1 |
| chr2 | 74199817 | 74200285 | 3.556 | 4.25452 | 1.32408 | Distal Intergenic | 2 | 74135757 | 74150834 | 15078 | 1 | ENSMUST00000132332.1 | 64061 | Gm13667 | ENSMUSG00000084791.1 |
| chr12 | 57520754 | 57520856 | 3.34205 | 3.84045 | 1.01733 | Promoter | 12 | 57519043 | 57520624 | 1582 | 2 | ENSMUST00000146126.1 | -131 | Gm2568 | ENSMUSG00000084793.1 |
| chr12 | 57526004 | 57526109 | 3.34734 | 3.84842 | 1.01733 | Distal Intergenic | 12 | 57519043 | 57520624 | 1582 | 2 | ENSMUST00000146126.1 | -5381 | Gm2568 | ENSMUSG00000084793.1 |
| chr12 | 57527905 | 57527995 | 4.44357 | 6.00934 | 2.53521 | Distal Intergenic | 12 | 57519043 | 57520624 | 1582 | 2 | ENSMUST00000146126.1 | -7282 | Gm2568 | ENSMUSG00000084793.1 |
| chr12 | 57528510 | 57528690 | 3.73804 | 4.61955 | 1.57655 | Distal Intergenic | 12 | 57519043 | 57520624 | 1582 | 2 | ENSMUST00000146126.1 | -7887 | Gm2568 | ENSMUSG00000084793.1 |
| chr12 | 57529298 | 57529755 | 3.67141 | 4.4783 | 1.48259 | Distal Intergenic | 12 | 57519043 | 57520624 | 1582 | 2 | ENSMUST00000146126.1 | -8675 | Gm2568 | ENSMUSG00000084793.1 |
| chr12 | 57530254 | 57530612 | 3.73025 | 4.65364 | 1.54297 | Distal Intergenic | 12 | 57519043 | 57520624 | 1582 | 2 | ENSMUST00000146126.1 | -9631 | Gm2568 | ENSMUSG00000084793.1 |
| chr2 | 118975873 | 118976071 | 3.82151 | 4.82546 | 1.66668 | Distal Intergenic | 2 | 118997068 | 119000684 | 3617 | 1 | ENSMUST00000130025.1 | -20997 | Gm14089 | ENSMUSG00000084802.1 |
| chr2 | 118976904 | 118977726 | 3.45107 | 4.32458 | 1.32436 | Distal Intergenic | 2 | 118997068 | 119000684 | 3617 | 1 | ENSMUST00000130025.1 | -19342 | Gm14089 | ENSMUSG00000084802.1 |
| chr2 | 118997332 | 118997618 | 3.29018 | 3.76245 | 1.01733 | Promoter | 2 | 118997068 | 119000684 | 3617 | 1 | ENSMUST00000130025.1 | 265 | Gm14089 | ENSMUSG00000084802.1 |
| chr19 | 6663645 | 6663885 | 3.23338 | 3.67724 | 1.01733 | Intron (ENSMUST00000137405.1/ENSMUSG00000084825.1, intron 1 of 1) | 19 | 6617671 | 6674548 | 56878 | 2 | ENSMUST00000137405.1 | 10663 | 2410152P15Rik | ENSMUSG00000084825.1 |
| chr2 | 166272762 | 166273074 | 3.65067 | 4.4122 | 1.37861 | Promoter | 2 | 166273908 | 166294831 | 20924 | 1 | ENSMUST00000152987.1 | -834 | Gm11468 | ENSMUSG00000084840.1 |
| chr2 | 166273586 | 166273948 | 3.49673 | 4.13092 | 1.22073 | Promoter | 2 | 166273908 | 166294831 | 20924 | 1 | ENSMUST00000152987.1 | 0 | Gm11468 | ENSMUSG00000084840.1 |
| chr2 | 166291024 | 166291129 | 3.51278 | 4.13916 | 1.16194 | Intron (ENSMUST00000152987.1/ENSMUSG00000084840.1, intron 2 of 2) | 2 | 166273908 | 166294831 | 20924 | 1 | ENSMUST00000152987.1 | 17117 | Gm11468 | ENSMUSG00000084840.1 |
| chr2 | 166294821 | 166295139 | 3.5327 | 4.24787 | 1.34867 | Exon (ENSMUST00000152987.1/ENSMUSG00000084840.1, exon 3 of 3) | 2 | 166273908 | 166294831 | 20924 | 1 | ENSMUST00000152987.1 | 20914 | Gm11468 | ENSMUSG00000084840.1 |
| chr2 | 166295803 | 166295938 | 3.3468 | 3.84761 | 1.01733 | Distal Intergenic | 2 | 166273908 | 166294831 | 20924 | 1 | ENSMUST00000152987.1 | 21896 | Gm11468 | ENSMUSG00000084840.1 |
| chr2 | 166330693 | 166330916 | 3.49116 | 4.10654 | 1.16235 | Distal Intergenic | 2 | 166273908 | 166294831 | 20924 | 1 | ENSMUST00000152987.1 | 56786 | Gm11468 | ENSMUSG00000084840.1 |
| chr2 | 180384618 | 180384887 | 3.6199 | 4.35739 | 1.34646 | Promoter | 2 | 180370491 | 180385885 | 15395 | 2 | ENSMUST00000145549.1 | 998 | B230312C02Rik | ENSMUSG00000084843.1 |
| chr2 | 180385420 | 180385786 | 3.51922 | 4.15769 | 1.19358 | Promoter | 2 | 180370491 | 180385885 | 15395 | 2 | ENSMUST00000145549.1 | 99 | B230312C02Rik | ENSMUSG00000084843.1 |

| | | | | | | | | | | | | | | | |
|---|---|---|---|---|---|---|---|---|---|---|---|---|---|---|---|
| chr11 | 96345567 | 96346195 | 3.49379 | 4.1273 | 1.1854 | Exon (ENSMUST00000093944.4/ENSMUSG00000048763.6, exon 4 of 4) | 11 | 96345481 | 96347923 | 2443 | 2 | ENSMUST00000131275.2 | 1728 | Hoxb3os | ENSMUSG00000084844.3 |
| chr11 | 96356372 | 96356667 | 3.52414 | 4.23787 | 1.31243 | Distal Intergenic | 11 | 96343072 | 96354691 | 11620 | 2 | ENSMUST00000147410.3 | -1682 | Hoxb3os | ENSMUSG00000084844.3 |
| chr11 | 96353213 | 96353460 | 3.34717 | 3.84816 | 1.01733 | Exon (ENSMUST00000182030.1/ENSMUSG00000075588.6, exon 1 of 1) | 11 | 96343072 | 96354691 | 11620 | 2 | ENSMUST00000147410.3 | 1231 | Hoxb3os | ENSMUSG00000084844.3 |
| chr11 | 96353945 | 96354161 | 3.40452 | 3.99266 | 1.11346 | Promoter | 11 | 96343072 | 96354691 | 11620 | 2 | ENSMUST00000147410.3 | 530 | Hoxb3os | ENSMUSG00000084844.3 |
| chr11 | 96354497 | 96354808 | 3.53508 | 4.23938 | 1.29215 | Promoter | 11 | 96343072 | 96354691 | 11620 | 2 | ENSMUST00000147410.3 | 0 | Hoxb3os | ENSMUSG00000084844.3 |
| chr10 | 81451472 | 81451577 | 3.68923 | 4.76037 | 1.61548 | Distal Intergenic | 10 | 81461159 | 81462972 | 1814 | 1 | ENSMUST00000156926.1 | -9582 | Gm16105 | ENSMUSG00000084849.1 |
| chr10 | 81452421 | 81452688 | 3.88135 | 4.91001 | 1.74457 | Distal Intergenic | 10 | 81461159 | 81462972 | 1814 | 1 | ENSMUST00000156926.1 | -8471 | Gm16105 | ENSMUSG00000084849.1 |
| chr10 | 81453415 | 81453988 | 3.45386 | 4.06156 | 1.16948 | Distal Intergenic | 10 | 81461159 | 81462972 | 1814 | 1 | ENSMUST00000156926.1 | -7171 | Gm16105 | ENSMUSG00000084849.1 |
| chr10 | 81456507 | 81456659 | 3.42682 | 3.98846 | 1.08852 | Downstream (2-3kb) | 10 | 81461159 | 81462972 | 1814 | 1 | ENSMUST00000156926.1 | -4500 | Gm16105 | ENSMUSG00000084849.1 |
| chr10 | 81458399 | 81458550 | 3.31734 | 3.94683 | 1.14466 | Downstream (<1kb) | 10 | 81461159 | 81462972 | 1814 | 1 | ENSMUST00000156926.1 | -2609 | Gm16105 | ENSMUSG00000084849.1 |
| chr10 | 81459173 | 81460560 | 3.98379 | 5.08058 | 1.82084 | Promoter | 10 | 81461159 | 81462972 | 1814 | 1 | ENSMUST00000156926.1 | -599 | Gm16105 | ENSMUSG00000084849.1 |
| chr10 | 81461705 | 81462194 | 3.73771 | 4.66357 | 1.56612 | Promoter | 10 | 81461159 | 81462972 | 1814 | 1 | ENSMUST00000156926.1 | 547 | Gm16105 | ENSMUSG00000084849.1 |
| chr10 | 81463638 | 81464930 | 4.22668 | 5.54955 | 2.14784 | Exon (ENSMUST00000118763.2/ENSMUSG00000034818.11, exon 10 of 12) | 10 | 81461159 | 81462972 | 1814 | 1 | ENSMUST00000156926.1 | 2480 | Gm16105 | ENSMUSG00000084849.1 |
| chr10 | 81465369 | 81465540 | 3.42676 | 4.47178 | 1.41951 | Intron (ENSMUST00000118763.2/ENSMUSG00000034818.11, intron 9 of 11) | 10 | 81461159 | 81462972 | 1814 | 1 | ENSMUST00000156926.1 | 4211 | Gm16105 | ENSMUSG00000084849.1 |
| chr4 | 82604126 | 82604401 | 3.43105 | 4.05029 | 1.24114 | Intron (ENSMUST00000155821.1/ENSMUSG00000008575.12, intron 1 of 1) | 4 | 82585048 | 82587319 | 2272 | 2 | ENSMUST00000137477.1 | -16808 | Gm11267 | ENSMUSG00000084851.1 |
| chr4 | 82604770 | 82604917 | 3.66859 | 4.65727 | 1.59695 | Intron (ENSMUST00000155821.1/ENSMUSG00000008575.12, intron 1 of 1) | 4 | 82585048 | 82587319 | 2272 | 2 | ENSMUST00000137477.1 | -17452 | Gm11267 | ENSMUSG00000084851.1 |
| chr4 | 82610395 | 82610470 | 3.62292 | 4.76907 | 1.6141 | Intron (ENSMUST00000155821.1/ENSMUSG00000008575.12, intron 1 of 1) | 4 | 82585048 | 82587319 | 2272 | 2 | ENSMUST00000137477.1 | -23077 | Gm11267 | ENSMUSG00000084851.1 |
| chr11 | 95610279 | 95610657 | 3.36375 | 3.87318 | 1.01733 | Distal Intergenic | 11 | 95622660 | 95629836 | 7177 | 1 | ENSMUST00000156822.1 | -12003 | Gm11528 | ENSMUSG00000084856.1 |
| chr11 | 95617525 | 95617681 | 3.37941 | 3.89679 | 1.01733 | Distal Intergenic | 11 | 95622660 | 95629836 | 7177 | 1 | ENSMUST00000156822.1 | -4979 | Gm11528 | ENSMUSG00000084856.1 |
| chr11 | 95644025 | 95644540 | 3.76222 | 4.75745 | 1.6086 | Distal Intergenic | 11 | 95622660 | 95629836 | 7177 | 1 | ENSMUST00000156822.1 | 21366 | Gm11528 | ENSMUSG00000084856.1 |
| chr19 | 6417359 | 6417675 | 3.3116 | 3.79472 | 1.01733 | Promoter | 19 | 6409400 | 6418606 | 9207 | 2 | ENSMUST00000124118.2 | 931 | Gm14965 | ENSMUSG00000084876.2 |
| chr19 | 6418131 | 6418322 | 3.36635 | 4.02244 | 1.12109 | Promoter | 19 | 6409400 | 6418606 | 9207 | 2 | ENSMUST00000124118.2 | 284 | Gm14965 | ENSMUSG00000084876.2 |
| chr11 | 118873206 | 118873281 | 3.77075 | 4.61545 | 1.45985 | Intron (ENSMUST00000117731.2/ENSMUSG00000025576.12, intron 1 of 13) | 11 | 118866257 | 118867220 | 964 | 2 | ENSMUST00000127327.1 | -5987 | Gm11749 | ENSMUSG00000084887.1 |
| chr11 | 99403652 | 99403946 | 3.40204 | 3.93938 | 1.04699 | Distal Intergenic | 11 | 99416956 | 99417752 | 797 | 1 | ENSMUST00000126470.1 | -13010 | A830036E02Rik | ENSMUSG00000084890.1 |
| chr4 | 36952419 | 36952643 | 3.4136 | 4.0774 | 1.22041 | Promoter | 4 | 36952966 | 36988032 | 35067 | 1 | ENSMUST00000144029.1 | -323 | Gm12371 | ENSMUSG00000084898.1 |
| chr3 | 122055324 | 122055610 | 3.33067 | 3.82331 | 1.01733 | Promoter | 3 | 122054750 | 122055548 | 799 | 2 | ENSMUST00000145966.1 | 0 | Gm16078 | ENSMUSG00000084912.1 |
| chr4 | 120302618 | 120302845 | 4.26384 | 5.59628 | 2.15581 | Exon (ENSMUST00000152384.1/ENSMUSG00000084929.1, exon 2 of 2) | 4 | 120291350 | 120303892 | 12543 | 1 | ENSMUST00000152384.1 | 11269 | Foxo6os | ENSMUSG00000084929.1 |
| chr2 | 71521794 | 71522229 | 3.60185 | 4.35413 | 1.35493 | Intron (ENSMUST00000037210.8/ENSMUSG00000041921.11, intron 6 of 9) | 2 | 71516454 | 71527919 | 11466 | 2 | ENSMUST00000140271.1 | 5690 | Dlx1as | ENSMUSG00000084946.2 |
| chr2 | 71522557 | 71523170 | 3.39614 | 3.97434 | 1.1822 | Exon (ENSMUST00000037210.8/ENSMUSG00000041921.11, exon 7 of 10) | 2 | 71516454 | 71527919 | 11466 | 2 | ENSMUST00000140271.1 | 4749 | Dlx1as | ENSMUSG00000084946.2 |
| chr2 | 71525037 | 71525402 | 3.86179 | 4.80986 | 1.63052 | 3' UTR | 2 | 71516454 | 71527919 | 11466 | 2 | ENSMUST00000140271.1 | 2517 | Dlx1as | ENSMUSG00000084946.2 |
| chr2 | 71525938 | 71526214 | 4.3668 | 5.85786 | 2.39395 | Intron (ENSMUST00000140271.1/ENSMUSG00000084946.2, intron 1 of 2) | 2 | 71516454 | 71527919 | 11466 | 2 | ENSMUST00000140271.1 | 1705 | Dlx1as | ENSMUSG00000084946.2 |
| chr2 | 71526544 | 71526666 | 3.65201 | 4.50684 | 1.422 | Intron (ENSMUST00000140271.1/ENSMUSG00000084946.2, intron 1 of 2) | 2 | 71516454 | 71527919 | 11466 | 2 | ENSMUST00000140271.1 | 1253 | Dlx1as | ENSMUSG00000084946.2 |
| chr2 | 71527207 | 71527740 | 4.35551 | 5.89501 | 2.4599 | Promoter | 2 | 71516454 | 71527919 | 11466 | 2 | ENSMUST00000140271.1 | 179 | Dlx1as | ENSMUSG00000084946.2 |
| chr2 | 71536021 | 71537225 | 3.79083 | 4.71345 | 1.62577 | Promoter | 2 | 71527839 | 71537810 | 9972 | 2 | ENSMUST00000143767.1 | 585 | Dlx1as | ENSMUSG00000084946.2 |
| chr2 | 71537550 | 71538578 | 3.8585 | 4.85067 | 1.67676 | Promoter | 2 | 71527839 | 71537810 | 9972 | 2 | ENSMUST00000143767.1 | 0 | Dlx1as | ENSMUSG00000084946.2 |
| chr2 | 71534058 | 71534647 | 4.24678 | 5.64002 | 2.29714 | Intron (ENSMUST00000143767.1/ENSMUSG00000084946.2, intron 1 of 1) | 2 | 71527839 | 71537810 | 9972 | 2 | ENSMUST00000143767.1 | 3163 | Dlx1as | ENSMUSG00000084946.2 |
| chr2 | 71535256 | 71535507 | 3.88502 | 5.31003 | 1.96972 | Intron (ENSMUST00000143767.1/ENSMUSG00000084946.2, intron 1 of 1) | 2 | 71527839 | 71537810 | 9972 | 2 | ENSMUST00000143767.1 | 2303 | Dlx1as | ENSMUSG00000084946.2 |
| chr2 | 71539024 | 71540228 | 3.73353 | 4.57844 | 1.48806 | Distal Intergenic | 2 | 71532784 | 71537891 | 5108 | 2 | ENSMUST00000137251.2 | -1134 | Dlx1as | ENSMUSG00000084946.2 |
| chr2 | 71540641 | 71541893 | 4.05066 | 5.2106 | 1.89375 | Downstream (1-2kb) | 2 | 71532784 | 71537891 | 5108 | 2 | ENSMUST00000137251.2 | -2751 | Dlx1as | ENSMUSG00000084946.2 |
| chr7 | 128414462 | 128414961 | 3.87078 | 4.82758 | 1.65527 | Intron (ENSMUST00000033133.6/ENSMUSG00000030844.6, intron 1 of 4) | 7 | 128410420 | 128412341 | 1922 | 2 | ENSMUST00000141541.1 | -2122 | Gm15503 | ENSMUSG00000084964.1 |
| chr11 | 120531800 | 120532657 | 3.74533 | 4.73552 | 1.61526 | Promoter | 11 | 120526907 | 120531784 | 4878 | 2 | ENSMUST00000148579.1 | -17 | Gm11789 | ENSMUSG00000084983.2 |
| chr4 | 135144426 | 135144652 | 3.88126 | 4.9197 | 1.74908 | Promoter | 4 | 135143421 | 135144285 | 865 | 2 | ENSMUST00000127711.1 | -142 | Gm16224 | ENSMUSG00000084986.1 |
| chr4 | 135147741 | 135147881 | 3.71979 | 4.54246 | 1.46634 | Intron (ENSMUST00000056977.8/ENSMUSG00000070691.5, intron 1 of 5) | 4 | 135143421 | 135144285 | 865 | 2 | ENSMUST00000127711.1 | -3457 | Gm16224 | ENSMUSG00000084986.1 |
| chr4 | 135148231 | 135148505 | 3.54988 | 4.33761 | 1.34742 | Intron (ENSMUST00000056977.8/ENSMUSG00000070691.5, intron 1 of 5) | 4 | 135143421 | 135144285 | 865 | 2 | ENSMUST00000127711.1 | -3947 | Gm16224 | ENSMUSG00000084986.1 |
| chr4 | 135145038 | 135145392 | 4.05819 | 5.23153 | 1.95497 | Promoter | 4 | 135143421 | 135144285 | 865 | 2 | ENSMUST00000127711.1 | -754 | Gm16224 | ENSMUSG00000084986.1 |
| chr4 | 135145839 | 135147337 | 3.7069 | 4.55557 | 1.51081 | Intron (ENSMUST00000056977.8/ENSMUSG00000070691.5, intron 1 of 5) | 4 | 135143421 | 135144285 | 865 | 2 | ENSMUST00000127711.1 | -1555 | Gm16224 | ENSMUSG00000084986.1 |
| chr4 | 135138618 | 135139080 | 3.79062 | 4.66716 | 1.5181 | Intron (ENSMUST00000056977.8/ENSMUSG00000070691.5, intron 1 of 5) | 4 | 135143421 | 135144285 | 865 | 2 | ENSMUST00000127711.1 | 5205 | Gm16224 | ENSMUSG00000084986.1 |
| chr4 | 135140724 | 135141646 | 3.45087 | 4.19176 | 1.25143 | Intron (ENSMUST00000056977.8/ENSMUSG00000070691.5, intron 1 of 5) | 4 | 135143421 | 135144285 | 865 | 2 | ENSMUST00000127711.1 | 2639 | Gm16224 | ENSMUSG00000084986.1 |
| chr4 | 154599865 | 154600045 | 3.42137 | 3.9725 | 1.06092 | Intron (ENSMUST00000030902.7/ENSMUSG00000039410.11, intron 1 of 16) | 4 | 154601197 | 154604202 | 3006 | 1 | ENSMUST00000144347.1 | -1152 | Gm13134 | ENSMUSG00000084987.1 |
| chr4 | 154604926 | 154605184 | 3.47362 | 4.0894 | 1.19531 | Intron (ENSMUST00000030902.7/ENSMUSG00000039410.11, intron 1 of 16) | 4 | 154601315 | 154602704 | 1390 | 1 | ENSMUST00000132106.1 | 3612 | Gm13134 | ENSMUSG00000084987.1 |
| chr4 | 154592255 | 154592417 | 3.33626 | 3.83178 | 1.01733 | Intron (ENSMUST00000030902.7/ENSMUSG00000039410.11, intron 1 of 16) | 4 | 154601197 | 154604202 | 3006 | 1 | ENSMUST00000144347.1 | -8780 | Gm13134 | ENSMUSG00000084987.1 |
| chr4 | 154594898 | 154594973 | 3.90666 | 4.87801 | 1.64243 | Intron (ENSMUST00000030902.7/ENSMUSG00000039410.11, intron 1 of 16) | 4 | 154601197 | 154604202 | 3006 | 1 | ENSMUST00000144347.1 | -6224 | Gm13134 | ENSMUSG00000084987.1 |
| chr1 | 93177817 | 93178077 | 3.36569 | 3.87609 | 1.01733 | Distal Intergenic | 1 | 93194049 | 93198641 | 4593 | 1 | ENSMUST00000188817.1 | -15972 | E030010N08Rik | ENSMUSG00000084989.2 |
| chr1 | 93180802 | 93181117 | 3.5155 | 4.17005 | 1.25927 | Distal Intergenic | 1 | 93194049 | 93198641 | 4593 | 1 | ENSMUST00000188817.1 | -12932 | E030010N08Rik | ENSMUSG00000084989.2 |
| chr1 | 93184314 | 93184597 | 3.60146 | 4.3069 | 1.27544 | Distal Intergenic | 1 | 93194049 | 93198641 | 4593 | 1 | ENSMUST00000188817.1 | -9452 | E030010N08Rik | ENSMUSG00000084989.2 |
| chr1 | 93189516 | 93189660 | 3.37163 | 3.88507 | 1.01733 | Distal Intergenic | 1 | 93194049 | 93198641 | 4593 | 1 | ENSMUST00000188817.1 | -4389 | E030010N08Rik | ENSMUSG00000084989.2 |
| chr14 | 100285437 | 100286049 | 3.51306 | 4.20753 | 1.28511 | Distal Intergenic | 14 | 100219486 | 100219789 | 304 | 1 | ENSMUST00000155141.1 | 65952 | Gm16260 | ENSMUSG00000084991.1 |
| chr3 | 36516425 | 36516591 | 3.24838 | 3.69971 | 1.01733 | Exon (ENSMUST00000124606.1/ENSMUSG00000085007.1, exon 3 of 3) | 3 | 36515057 | 36521813 | 6757 | 2 | ENSMUST00000124606.1 | 5222 | Gm11549 | ENSMUSG00000085007.1 |
| chr3 | 36521168 | 36521588 | 3.78497 | 5.01653 | 1.77467 | Promoter | 3 | 36515057 | 36521813 | 6757 | 2 | ENSMUST00000124606.1 | 225 | Gm11549 | ENSMUSG00000085007.1 |
| chr4 | 82063538 | 82063738 | 3.54145 | 4.32083 | 1.36444 | Intron (ENSMUST00000143755.1/ENSMUSG00000087110.1, intron 1 of 5) | 4 | 82065431 | 82102808 | 37378 | 1 | ENSMUST00000137810.2 | -1693 | Gm5860 | ENSMUSG00000085022.2 |
| chr4 | 82196330 | 82196531 | 3.83908 | 4.86723 | 1.67079 | Distal Intergenic | 4 | 82065957 | 82102673 | 36717 | 1 | ENSMUST00000161588.1 | 130374 | Gm5860 | ENSMUSG00000085022.2 |
| chr11 | 84639868 | 84639951 | 3.39122 | 3.91754 | 1.0276 | Distal Intergenic | 11 | 84716976 | 84720432 | 3457 | 2 | ENSMUST00000150735.1 | 80481 | 1700109G15Rik | ENSMUSG00000085044.1 |
| chr11 | 84641629 | 84642120 | 3.44965 | 4.02949 | 1.11195 | Distal Intergenic | 11 | 84716976 | 84720432 | 3457 | 2 | ENSMUST00000150735.1 | 78312 | 1700109G15Rik | ENSMUSG00000085044.1 |
| chr11 | 84642510 | 84642994 | 3.55324 | 4.22274 | 1.23825 | Distal Intergenic | 11 | 84716976 | 84720432 | 3457 | 2 | ENSMUST00000150735.1 | 77438 | 1700109G15Rik | ENSMUSG00000085044.1 |
| chr11 | 84659418 | 84659574 | 3.85521 | 4.79328 | 1.59841 | Distal Intergenic | 11 | 84716976 | 84720432 | 3457 | 2 | ENSMUST00000150735.1 | 60858 | 1700109G15Rik | ENSMUSG00000085044.1 |
| chr11 | 84701277 | 84701583 | 3.69528 | 4.56986 | 1.45356 | Distal Intergenic | 11 | 84716976 | 84720432 | 3457 | 2 | ENSMUST00000150735.1 | 18849 | 1700109G15Rik | ENSMUSG00000085044.1 |
| chr11 | 84724007 | 84724094 | 3.8495 | 5.10501 | 1.85413 | Distal Intergenic | 11 | 84716976 | 84720432 | 3457 | 2 | ENSMUST00000150735.1 | -3576 | 1700109G15Rik | ENSMUSG00000085044.1 |
| chr11 | 84745129 | 84745242 | 3.31671 | 3.80242 | 1.01733 | Distal Intergenic | 11 | 84716976 | 84720432 | 3457 | 2 | ENSMUST00000150735.1 | -24698 | 1700109G15Rik | ENSMUSG00000085044.1 |
| chr11 | 62757986 | 62758146 | 3.56678 | 4.33258 | 1.37081 | Exon (ENSMUST00000138105.1/ENSMUSG00000085045.1, exon 2 of 2) | 11 | 62757178 | 62760558 | 3381 | 2 | ENSMUST00000138105.1 | 2412 | Zfp286it | ENSMUSG00000085045.1 |
| chr2 | 166215635 | 166215738 | 3.36084 | 3.88619 | 1.07894 | Distal Intergenic | 2 | 166207230 | 166208627 | 1398 | 2 | ENSMUST00000129407.1 | -7009 | Gm11467 | ENSMUSG00000085046.1 |
| chr4 | 140435606 | 140435789 | 3.79746 | 4.67715 | 1.5235 | Distal Intergenic | 4 | 140437963 | 140461291 | 23329 | 1 | ENSMUST00000140113.1 | -2174 | Gm13026 | ENSMUSG00000085047.1 |
| chr4 | 140458713 | 140459013 | 3.46127 | 4.0909 | 1.23975 | Intron (ENSMUST00000140113.1/ENSMUSG00000085047.1, intron 1 of 1) | 4 | 140437963 | 140461291 | 23329 | 1 | ENSMUST00000140113.1 | 20751 | Gm13026 | ENSMUSG00000085047.1 |
| chr4 | 140461238 | 140461367 | 3.62142 | 4.40932 | 1.41569 | Exon (ENSMUST00000140113.1/ENSMUSG00000085047.1, exon 2 of 2) | 4 | 140437963 | 140461291 | 23329 | 1 | ENSMUST00000140113.1 | 23276 | Gm13026 | ENSMUSG00000085047.1 |
| chr4 | 140461698 | 140462040 | 3.80073 | 4.7456 | 1.63675 | Distal Intergenic | 4 | 140437963 | 140461291 | 23329 | 1 | ENSMUST00000140113.1 | 23736 | Gm13026 | ENSMUSG00000085047.1 |
| chr4 | 140463456 | 140464020 | 3.75355 | 4.72836 | 1.60286 | Distal Intergenic | 4 | 140437963 | 140461291 | 23329 | 1 | ENSMUST00000140113.1 | 25494 | Gm13026 | ENSMUSG00000085047.1 |
| chr4 | 140466402 | 140466994 | 3.43818 | 4.04084 | 1.16735 | Distal Intergenic | 4 | 140437963 | 140461291 | 23329 | 1 | ENSMUST00000140113.1 | 28440 | Gm13026 | ENSMUSG00000085047.1 |
| chr4 | 140467367 | 140467620 | 3.26098 | 3.7186 | 1.01733 | Distal Intergenic | 4 | 140437963 | 140461291 | 23329 | 1 | ENSMUST00000140113.1 | 29405 | Gm13026 | ENSMUSG00000085047.1 |
| chr4 | 140470169 | 140470307 | 3.43353 | 4.03075 | 1.19874 | Distal Intergenic | 4 | 140437963 | 140461291 | 23329 | 1 | ENSMUST00000140113.1 | 32207 | Gm13026 | ENSMUSG00000085047.1 |
| chr4 | 140476202 | 140476429 | 3.38075 | 3.91214 | 1.06444 | Distal Intergenic | 4 | 140437963 | 140461291 | 23329 | 1 | ENSMUST00000140113.1 | 38240 | Gm13026 | ENSMUSG00000085047.1 |
| chr4 | 140481870 | 140482194 | 3.3867 | 3.99517 | 1.09496 | Distal Intergenic | 4 | 140437963 | 140461291 | 23329 | 1 | ENSMUST00000140113.1 | 43908 | Gm13026 | ENSMUSG00000085047.1 |
| chr4 | 140492797 | 140492983 | 3.722 | 4.53635 | 1.43335 | Distal Intergenic | 4 | 140437963 | 140461291 | 23329 | 1 | ENSMUST00000140113.1 | 54835 | Gm13026 | ENSMUSG00000085047.1 |
| chr4 | 140493408 | 140493927 | 3.54206 | 4.2541 | 1.25665 | Distal Intergenic | 4 | 140437963 | 140461291 | 23329 | 1 | ENSMUST00000140113.1 | 55446 | Gm13026 | ENSMUSG00000085047.1 |
| chr4 | 140496754 | 140496876 | 3.46306 | 4.11566 | 1.26845 | Distal Intergenic | 4 | 140437963 | 140461291 | 23329 | 1 | ENSMUST00000140113.1 | 58792 | Gm13026 | ENSMUSG00000085047.1 |
| chr4 | 140497786 | 140498162 | 3.24907 | 3.70076 | 1.01733 | Distal Intergenic | 4 | 140437963 | 140461291 | 23329 | 1 | ENSMUST00000140113.1 | 59824 | Gm13026 | ENSMUSG00000085047.1 |
| chr4 | 140501815 | 140502039 | 4.02852 | 5.22544 | 1.97008 | Distal Intergenic | 4 | 140437963 | 140461291 | 23329 | 1 | ENSMUST00000140113.1 | 63853 | Gm13026 | ENSMUSG00000085047.1 |
| chr4 | 140502525 | 140502709 | 3.43402 | 4.18902 | 1.26165 | Distal Intergenic | 4 | 140437963 | 140461291 | 23329 | 1 | ENSMUST00000140113.1 | 64563 | Gm13026 | ENSMUSG00000085047.1 |
| chr11 | 118808049 | 118808124 | 3.65933 | 4.57357 | 1.49025 | Intron (ENSMUST00000117731.2/ENSMUSG00000025576.12, intron 2 of 13) | 11 | 118791372 | 118793828 | 2457 | 1 | ENSMUST00000146016.1 | 16678 | Gm11750 | ENSMUSG00000085059.1 |
| chr6 | 105677414 | 105677645 | 3.57712 | 4.29617 | 1.36274 | Promoter | 6 | 105674755 | 105677581 | 2827 | 2 | ENSMUST00000142962.1 | 0 | Gm15631 | ENSMUSG00000085067.1 |
| chr4 | 154637382 | 154638022 | 4.26567 | 5.68976 | 2.31153 | Promoter | 4 | 154637576 | 154644724 | 7149 | 1 | ENSMUST00000136217.2 | 0 | Gm13111 | ENSMUSG00000085069.2 |
| chr4 | 154638537 | 154638745 | 4.08617 | 5.27502 | 1.98888 | Promoter | 4 | 154637576 | 154644724 | 7149 | 1 | ENSMUST00000136217.2 | 962 | Gm13111 | ENSMUSG00000085069.2 |
| chr4 | 154639236 | 154639657 | 3.72405 | 4.62711 | 1.57393 | Intron (ENSMUST00000136217.2/ENSMUSG00000085069.2, intron 1 of 1) | 4 | 154637576 | 154644724 | 7149 | 1 | ENSMUST00000136217.2 | 1661 | Gm13111 | ENSMUSG00000085069.2 |
| chr4 | 154640195 | 154640474 | 3.75771 | 4.63274 | 1.54556 | Intron (ENSMUST00000136217.2/ENSMUSG00000085069.2, intron 1 of 1) | 4 | 154637576 | 154644724 | 7149 | 1 | ENSMUST00000136217.2 | 2620 | Gm13111 | ENSMUSG00000085069.2 |
| chr4 | 154641341 | 154641607 | 4.03104 | 5.73015 | 2.26101 | Exon (ENSMUST00000136217.2/ENSMUSG00000085069.2, exon 2 of 2) | 4 | 154637576 | 154644724 | 7149 | 1 | ENSMUST00000136217.2 | 3766 | Gm13111 | ENSMUSG00000085069.2 |
| chr2 | 139633496 | 139634015 | 3.59445 | 4.31648 | 1.31693 | Intron (ENSMUST00000110083.2/ENSMUSG00000039092.4, intron 11 of 11) | 2 | 139256876 | 139677714 | 420839 | 2 | ENSMUST00000134801.1 | 43699 | Gm14066 | ENSMUSG00000085071.1 |
| chr11 | 90117267 | 90117690 | 3.67975 | 4.65356 | 1.5625 | Distal Intergenic | 11 | 90134842 | 90143624 | 8783 | 2 | ENSMUST00000142279.1 | 25934 | Gm11494 | ENSMUSG00000085074.1 |
| chr7 | 99658902 | 99659172 | 3.3792 | 3.91968 | 1.09947 | 3' UTR | 7 | 99661439 | 99667021 | 5583 | 1 | ENSMUST00000131160.1 | -2267 | Gm15635 | ENSMUSG00000085095.1 |
| chr7 | 99666667 | 99667367 | 3.63245 | 4.37139 | 1.31778 | Exon (ENSMUST00000131160.1/ENSMUSG00000085095.1, exon 2 of 2) | 7 | 99665153 | 99667394 | 2242 | 1 | ENSMUST00000148972.1 | 1515 | Gm15635 | ENSMUSG00000085095.1 |
| chr7 | 99667823 | 99668015 | 3.35067 | 3.86188 | 1.02482 | Intron (ENSMUST00000032985.5/ENSMUSG00000030737.11, intron 8 of 13) | 7 | 99665153 | 99667394 | 2242 | 1 | ENSMUST00000148972.1 | 2671 | Gm15635 | ENSMUSG00000085095.1 |
| chr8 | 66449560 | 66449761 | 3.70543 | 4.51204 | 1.43986 | Intron (ENSMUST00000152320.2/ENSMUSG00000036469.11, intron 7 of 8) | 8 | 66439455 | 66442809 | 3355 | 2 | ENSMUST00000140669.1 | -6752 | Gm15356 | ENSMUSG00000085109.1 |
| chr10 | 85930011 | 85930466 | 3.8019 | 4.76032 | 1.65837 | Intron (ENSMUST00000156605.1/ENSMUSG00000085111.3, intron 1 of 2) | 10 | 85928774 | 85946206 | 17433 | 1 | ENSMUST00000156605.1 | 1238 | Ascl4 | ENSMUSG00000085111.3 |
| chr12 | 107111282 | 107111518 | 4.1924 | 5.43308 | 2.02309 | Distal Intergenic | 12 | 107079360 | 107099673 | 20314 | 2 | ENSMUST00000143652.1 | -11610 | Gm16086 | ENSMUSG00000085120.1 |
| chr13 | 23285195 | 23286805 | 3.52872 | 4.5741 | 1.50436 | Intron (ENSMUST00000155432.1/ENSMUSG00000085488.1, intron 1 of 1) | 13 | 23278350 | 23282267 | 3918 | 1 | ENSMUST00000156977.1 | 6846 | 4933404K08Rik | ENSMUSG00000085122.1 |
| chr13 | 23287686 | 23288040 | 3.93487 | 5.16328 | 1.87864 | Intron (ENSMUST00000155432.1/ENSMUSG00000085488.1, intron 1 of 1) | 13 | 23278350 | 23282267 | 3918 | 1 | ENSMUST00000156977.1 | 9337 | 4933404K08Rik | ENSMUSG00000085122.1 |
| chr14 | 46593941 | 46594514 | 3.89848 | 4.95289 | 1.79297 | Distal Intergenic | 14 | 46570213 | 46575847 | 5635 | 2 | ENSMUST00000151828.1 | -18095 | Gm15219 | ENSMUSG00000085123.1 |
| chr2 | 36198211 | 36198331 | 4.20446 | 5.48557 | 2.08668 | Distal Intergenic | 2 | 36202314 | 36204259 | 1946 | 1 | ENSMUST00000123224.1 | -3983 | Gm13429 | ENSMUSG00000085141.1 |
| chr2 | 36204266 | 36204804 | 3.46886 | 4.11818 | 1.2563 | Intron (ENSMUST00000138755.1/ENSMUSG00000085141.1, intron 1 of 1) | 2 | 36202822 | 36212903 | 10082 | 1 | ENSMUST00000138755.1 | 1445 | Gm13429 | ENSMUSG00000085141.1 |
| chr2 | 36206819 | 36207033 | 3.3466 | 3.93088 | 1.12672 | Intron (ENSMUST00000138755.1/ENSMUSG00000085141.1, intron 1 of 1) | 2 | 36210772 | 36212902 | 2131 | 1 | ENSMUST00000128242.1 | -3739 | Gm13429 | ENSMUSG00000085141.1 |
| chr2 | 36210659 | 36210924 | 3.89603 | 4.90023 | 1.74183 | Promoter | 2 | 36210772 | 36212902 | 2131 | 1 | ENSMUST00000128242.1 | 0 | Gm13429 | ENSMUSG00000085141.1 |
| chr2 | 36211894 | 36211969 | 3.80377 | 5.06384 | 1.79943 | Intron (ENSMUST00000138755.1/ENSMUSG00000085141.1, intron 1 of 1) | 2 | 36210772 | 36212902 | 2131 | 1 | ENSMUST00000128242.1 | 1123 | Gm13429 | ENSMUSG00000085141.1 |
| chr4 | 100035356 | 100035637 | 3.3201 | 3.83377 | 1.02392 | Distal Intergenic | 4 | 100052520 | 100065984 | 13465 | 1 | ENSMUST00000123258.1 | -16883 | Gm12702 | ENSMUSG00000085142.1 |
| chr4 | 89351074 | 89351509 | 3.38358 | 3.94046 | 1.133 | Distal Intergenic | 4 | 89381957 | 89399880 | 17924 | 1 | ENSMUST00000129908.1 | -30448 | Gm12609 | ENSMUSG00000085147.1 |
| chr2 | 167189304 | 167189742 | 3.5624 | 4.24743 | 1.25954 | Promoter | 2 | 167189661 | 167192808 | 3148 | 1 | ENSMUST00000147446.1 | 0 | 1110018N20Rik | ENSMUSG00000085151.2 |
| chr11 | 29520543 | 29520634 | 3.66217 | 4.57135 | 1.48689 | Distal Intergenic | 11 | 29515086 | 29517355 | 2270 | 1 | ENSMUST00000138402.1 | 5458 | Gm12089 | ENSMUSG00000085165.1 |
| chr11 | 24075912 | 24077530 | 3.73526 | 4.79654 | 1.64106 | Promoter | 11 | 24076559 | 24077367 | 809 | 1 | ENSMUST00000148087.1 | 0 | Gm12063 | ENSMUSG00000085167.1 |
| chr4 | 152351165 | 152351278 | 3.44329 | 4.03301 | 1.15866 | Intron (ENSMUST00000030775.6/ENSMUSG00000005045.11, intron 2 of 41) | 4 | 152352198 | 152353641 | 1444 | 2 | ENSMUST00000145163.1 | 2363 | Gm13645 | ENSMUSG00000085179.1 |
| chr4 | 152351708 | 152352156 | 3.53942 | 4.22178 | 1.30587 | Intron (ENSMUST00000030775.6/ENSMUSG00000005045.11, intron 2 of 41) | 4 | 152352198 | 152353641 | 1444 | 2 | ENSMUST00000145163.1 | 1485 | Gm13645 | ENSMUSG00000085179.1 |
| chr4 | 152360425 | 152360811 | 3.54989 | 4.24782 | 1.32619 | Exon (ENSMUST00000030775.6/ENSMUSG00000005045.11, exon 8 of 42) | 4 | 152352198 | 152353641 | 1444 | 2 | ENSMUST00000145163.1 | -6785 | Gm13645 | ENSMUSG00000085179.1 |
| chr17 | 24211840 | 24212252 | 3.60615 | 4.37446 | 1.42915 | Promoter | 17 | 24212515 | 24221730 | 9216 | 1 | ENSMUST00000137883.1 | -263 | BC028777 | ENSMUSG00000085185.4 |
| chr4 | 84017589 | 84017664 | 3.99653 | 5.22888 | 1.93629 | Distal Intergenic | 4 | 84042039 | 84044879 | 2841 | 2 | ENSMUST00000125884.1 | 27215 | 6030471H07Rik | ENSMUSG00000085217.1 |
| chr4 | 84037213 | 84037302 | 4.15054 | 5.54596 | 2.16169 | Distal Intergenic | 4 | 84042039 | 84044879 | 2841 | 2 | ENSMUST00000125884.1 | 7577 | 6030471H07Rik | ENSMUSG00000085217.1 |
| chr4 | 84037704 | 84038038 | 3.79039 | 4.66547 | 1.50789 | Distal Intergenic | 4 | 84042039 | 84044879 | 2841 | 2 | ENSMUST00000125884.1 | 6841 | 6030471H07Rik | ENSMUSG00000085217.1 |
| chr4 | 84065203 | 84065588 | 3.44642 | 4.27853 | 1.30002 | Distal Intergenic | 4 | 84042039 | 84044879 | 2841 | 2 | ENSMUST00000125884.1 | -20325 | 6030471H07Rik | ENSMUSG00000085217.1 |
| chr4 | 84066300 | 84066518 | 3.97555 | 5.11413 | 1.87273 | Distal Intergenic | 4 | 84042039 | 84044879 | 2841 | 2 | ENSMUST00000125884.1 | -21422 | 6030471H07Rik | ENSMUSG00000085217.1 |
| chr4 | 84069796 | 84070041 | 3.52058 | 4.21059 | 1.3286 | Distal Intergenic | 4 | 84042039 | 84044879 | 2841 | 2 | ENSMUST00000125884.1 | -24918 | 6030471H07Rik | ENSMUSG00000085217.1 |
| chr4 | 84070516 | 84070883 | 3.65985 | 4.49139 | 1.46884 | Distal Intergenic | 4 | 84042039 | 84044879 | 2841 | 2 | ENSMUST00000125884.1 | -25638 | 6030471H07Rik | ENSMUSG00000085217.1 |
| chr2 | 31644424 | 31645110 | 3.33691 | 3.85004 | 1.04728 | Intron (ENSMUST00000113470.2/ENSMUSG00000079466.2, intron 3 of 4) | 2 | 31638170 | 31640589 | 2420 | 2 | ENSMUST00000145389.1 | -3836 | Gm13425 | ENSMUSG00000085224.1 |
| chr2 | 31645880 | 31646162 | 3.53547 | 4.52644 | 1.44012 | Intron (ENSMUST00000113470.2/ENSMUSG00000079466.2, intron 3 of 4) | 2 | 31638170 | 31640589 | 2420 | 2 | ENSMUST00000145389.1 | -5292 | Gm13425 | ENSMUSG00000085224.1 |
| chr2 | 31647459 | 31648310 | 3.78887 | 4.74102 | 1.61991 | Intron (ENSMUST00000113470.2/ENSMUSG00000079466.2, intron 3 of 4) | 2 | 31638170 | 31640589 | 2420 | 2 | ENSMUST00000145389.1 | -6871 | Gm13425 | ENSMUSG00000085224.1 |
| chr2 | 31648868 | 31648943 | 4.11233 | 5.27196 | 1.91444 | Intron (ENSMUST00000113470.2/ENSMUSG00000079466.2, intron 3 of 4) | 2 | 31638170 | 31640589 | 2420 | 2 | ENSMUST00000145389.1 | -8280 | Gm13425 | ENSMUSG00000085224.1 |
| chr2 | 170586128 | 170586439 | 3.41862 | 4.01139 | 1.19892 | Intron (ENSMUST00000062355.1/ENSMUSG00000043583.1, intron 1 of 1) | 2 | 170576054 | 170578424 | 2371 | 1 | ENSMUST00000156516.1 | 10075 | Gm14637 | ENSMUSG00000085225.1 |
| chr11 | 71439547 | 71440086 | 3.48921 | 4.10376 | 1.16295 | Distal Intergenic | 11 | 71429828 | 71436180 | 6353 | 2 | ENSMUST00000152249.1 | -3368 | Gm16013 | ENSMUSG00000085230.1 |
| chr11 | 33676442 | 33676651 | 3.26718 | 3.728 | 1.01733 | Intron (ENSMUST00000101368.3/ENSMUSG00000053519.10, intron 1 of 7) | 11 | 33695894 | 33697908 | 2015 | 1 | ENSMUST00000136083.1 | -19243 | Gm12119 | ENSMUSG00000085240.1 |
| chr11 | 106066780 | 106066856 | 3.29102 | 15.539 | 10.71122 | Promoter | 11 | 106065077 | 106066305 | 1229 | 2 | ENSMUST00000140736.1 | -476 | Taco1os | ENSMUSG00000085255.1 |
| chr4 | 109969942 | 109970452 | 3.24559 | 3.69554 | 1.01665 | Distal Intergenic | 4 | 109976747 | 109977491 | 745 | 2 | ENSMUST00000124754.1 | 7039 | Dmrta2os | ENSMUSG00000085256.1 |
| chr4 | 109971120 | 109972071 | 3.56404 | 4.35313 | 1.38862 | Distal Intergenic | 4 | 109976747 | 109977491 | 745 | 2 | ENSMUST00000124754.1 | 5420 | Dmrta2os | ENSMUSG00000085256.1 |
| chr4 | 109972735 | 109975207 | 3.90486 | 5.00942 | 1.83563 | Downstream (1-2kb) | 4 | 109976747 | 109977491 | 745 | 2 | ENSMUST00000124754.1 | 2284 | Dmrta2os | ENSMUSG00000085256.1 |
| chr4 | 109975527 | 109976582 | 3.50859 | 4.32442 | 1.34651 | Promoter | 4 | 109976747 | 109977491 | 745 | 2 | ENSMUST00000124754.1 | 909 | Dmrta2os | ENSMUSG00000085256.1 |
| chr4 | 109976944 | 109977203 | 3.66313 | 4.5562 | 1.51382 | Promoter | 4 | 109976747 | 109977491 | 745 | 2 | ENSMUST00000124754.1 | 288 | Dmrta2os | ENSMUSG00000085256.1 |
| chr4 | 124892541 | 124892820 | 3.46244 | 4.04849 | 1.11019 | Intron (ENSMUST00000149146.1/ENSMUSG00000028876.7, intron 1 of 14) | 4 | 124890888 | 124894782 | 3895 | 2 | ENSMUST00000154947.1 | 1962 | 1700125G02Rik | ENSMUSG00000085263.1 |
| chr4 | 124894106 | 124895608 | 4.14575 | 5.41355 | 2.04626 | Promoter | 4 | 124890888 | 124894782 | 3895 | 2 | ENSMUST00000154947.1 | 0 | 1700125G02Rik | ENSMUSG00000085263.1 |
| chr5 | 111771350 | 111771483 | 3.32034 | 3.80775 | 1.01733 | Distal Intergenic | 5 | 111734303 | 111761725 | 27423 | 2 | ENSMUST00000129065.2 | -9626 | E130006D01Rik | ENSMUSG00000085271.2 |
| chr5 | 111836650 | 111837487 | 4.2103 | 6.01905 | 2.55171 | Distal Intergenic | 5 | 111734303 | 111761725 | 27423 | 2 | ENSMUST00000129065.2 | -74926 | E130006D01Rik | ENSMUSG00000085271.2 |
| chr5 | 111842079 | 111842644 | 3.54138 | 4.22482 | 1.30782 | Distal Intergenic | 5 | 111734303 | 111761725 | 27423 | 2 | ENSMUST00000129065.2 | -80355 | E130006D01Rik | ENSMUSG00000085271.2 |
| chr5 | 111857848 | 111858072 | 3.33224 | 3.83778 | 1.05518 | Distal Intergenic | 5 | 111734303 | 111761725 | 27423 | 2 | ENSMUST00000129065.2 | -96124 | E130006D01Rik | ENSMUSG00000085271.2 |
| chr5 | 111886118 | 111886369 | 3.38158 | 3.90538 | 1.03606 | Distal Intergenic | 5 | 111734303 | 111761725 | 27423 | 2 | ENSMUST00000129065.2 | -124394 | E130006D01Rik | ENSMUSG00000085271.2 |
| chr5 | 111894743 | 111895478 | 3.62157 | 4.35054 | 1.32007 | Distal Intergenic | 5 | 111734303 | 111761725 | 27423 | 2 | ENSMUST00000129065.2 | -133019 | E130006D01Rik | ENSMUSG00000085271.2 |
| chr5 | 111898402 | 111898478 | 3.90107 | 4.87096 | 1.64874 | Distal Intergenic | 5 | 111734303 | 111761725 | 27423 | 2 | ENSMUST00000129065.2 | -136678 | E130006D01Rik | ENSMUSG00000085271.2 |
| chr5 | 111900952 | 111901516 | 3.4252 | 3.97948 | 1.06514 | Distal Intergenic | 5 | 111734303 | 111761725 | 27423 | 2 | ENSMUST00000129065.2 | -139228 | E130006D01Rik | ENSMUSG00000085271.2 |
| chr5 | 111909786 | 111910409 | 3.27459 | 3.75646 | 1.06896 | Distal Intergenic | 5 | 111734303 | 111761725 | 27423 | 2 | ENSMUST00000129065.2 | -148062 | E130006D01Rik | ENSMUSG00000085271.2 |
| chr5 | 111918470 | 111918639 | 4.18166 | 5.43082 | 2.03675 | Distal Intergenic | 5 | 111734303 | 111761725 | 27423 | 2 | ENSMUST00000129065.2 | -156746 | E130006D01Rik | ENSMUSG00000085271.2 |
| chr5 | 111921055 | 111921451 | 3.80362 | 4.69993 | 1.56315 | Distal Intergenic | 5 | 111734303 | 111761725 | 27423 | 2 | ENSMUST00000129065.2 | -159331 | E130006D01Rik | ENSMUSG00000085271.2 |
| chr5 | 111945100 | 111945302 | 3.47744 | 4.07472 | 1.12287 | Distal Intergenic | 5 | 111734303 | 111761725 | 27423 | 2 | ENSMUST00000129065.2 | -183376 | E130006D01Rik | ENSMUSG00000085271.2 |
| chr2 | 77317084 | 77317439 | 3.68626 | 4.60113 | 1.51829 | Distal Intergenic | 2 | 77314751 | 77317035 | 2285 | 1 | ENSMUST00000144043.1 | 2334 | Gm13943 | ENSMUSG00000085277.1 |
| chr3 | 108165971 | 108166112 | 3.35191 | 3.89538 | 1.1396 | Promoter | 3 | 108166443 | 108185907 | 19465 | 1 | ENSMUST00000143109.1 | -331 | Gm12524 | ENSMUSG00000085318.1 |
| chr3 | 108174954 | 108175299 | 3.39318 | 3.94077 | 1.09947 | Intron (ENSMUST00000143109.1/ENSMUSG00000085318.1, intron 1 of 1) | 3 | 108166443 | 108185907 | 19465 | 1 | ENSMUST00000143109.1 | 8512 | Gm12524 | ENSMUSG00000085318.1 |
| chr11 | 112137633 | 112137825 | 3.71188 | 4.52121 | 1.41763 | Distal Intergenic | 11 | 112107438 | 112132196 | 24759 | 1 | ENSMUST00000146550.1 | 30196 | Gm11680 | ENSMUSG00000085324.1 |
| chr11 | 112198300 | 112198375 | 3.95262 | 5.15459 | 1.86421 | Distal Intergenic | 11 | 112107438 | 112132196 | 24759 | 1 | ENSMUST00000146550.1 | 90863 | Gm11680 | ENSMUSG00000085324.1 |
| chr17 | 28908282 | 28908870 | 3.51173 | 4.14923 | 1.19992 | Exon (ENSMUST00000062357.4/ENSMUSG00000048905.4, exon 3 of 12) | 17 | 28905265 | 28906771 | 1507 | 1 | ENSMUST00000125317.1 | 3018 | 1700030A11Rik | ENSMUSG00000085333.1 |
| chr17 | 28909183 | 28909867 | 4.03415 | 5.1963 | 1.93189 | Exon (ENSMUST00000062357.4/ENSMUSG00000048905.4, exon 2 of 12) | 17 | 28905265 | 28906771 | 1507 | 1 | ENSMUST00000125317.1 | 3919 | 1700030A11Rik | ENSMUSG00000085333.1 |
| chr11 | 102962065 | 102962372 | 3.6972 | 4.63359 | 1.51836 | Intron (ENSMUST00000181125.1/ENSMUSG00000097410.1, intron 1 of 1) | 11 | 102958300 | 102964387 | 6088 | 2 | ENSMUST00000138424.1 | 2015 | 2410004I01Rik | ENSMUSG00000085338.1 |
| chr11 | 102962943 | 102963135 | 4.2338 | 5.63171 | 2.27562 | Intron (ENSMUST00000181125.1/ENSMUSG00000097410.1, intron 1 of 1) | 11 | 102958300 | 102964387 | 6088 | 2 | ENSMUST00000138424.1 | 1252 | 2410004I01Rik | ENSMUSG00000085338.1 |
| chr11 | 102963869 | 102964070 | 3.51257 | 4.33803 | 1.30962 | Promoter | 11 | 102958300 | 102964387 | 6088 | 2 | ENSMUST00000138424.1 | 317 | 2410004I01Rik | ENSMUSG00000085338.1 |
| chr11 | 102972541 | 102972705 | 3.4283 | 3.98707 | 1.07537 | Intron (ENSMUST00000181125.1/ENSMUSG00000097410.1, intron 1 of 1) | 11 | 102958300 | 102964387 | 6088 | 2 | ENSMUST00000138424.1 | -8155 | 2410004I01Rik | ENSMUSG00000085338.1 |

| | | | | | | | | | | | | | | | |
|---|---|---|---|---|---|---|---|---|---|---|---|---|---|---|---|
| chr5 | 119641153 | 119641694 | 3.42566 | 4.01915 | 1.10393 | Intron (ENSMUST00000129108.1/ENSMUSG00000087516.2, intron 1 of 2) | 5 | 119646134 | 119659262 | 13129 | 1 | ENSMUST00000154997.1 | -4440 | Gm16064 | ENSMUSG00000085345.1 |
| chr5 | 119647851 | 119648010 | 3.59383 | 4.28693 | 1.25151 | Intron (ENSMUST00000154997.1/ENSMUSG00000085345.1, intron 1 of 1) | 5 | 119646134 | 119659262 | 13129 | 1 | ENSMUST00000154997.1 | 1718 | Gm16064 | ENSMUSG00000085345.1 |
| chr5 | 119649866 | 119650168 | 3.86438 | 4.82778 | 1.64641 | Intron (ENSMUST00000154997.1/ENSMUSG00000085345.1, intron 1 of 1) | 5 | 119646134 | 119659262 | 13129 | 1 | ENSMUST00000154997.1 | 3733 | Gm16064 | ENSMUSG00000085345.1 |
| chr5 | 119655311 | 119655696 | 3.49817 | 4.1198 | 1.17189 | Intron (ENSMUST00000154997.1/ENSMUSG00000085345.1, intron 1 of 1) | 5 | 119646134 | 119659262 | 13129 | 1 | ENSMUST00000154997.1 | 9178 | Gm16064 | ENSMUSG00000085345.1 |
| chr5 | 119656764 | 119657047 | 3.41854 | 3.97774 | 1.09471 | Intron (ENSMUST00000154997.1/ENSMUSG00000085345.1, intron 1 of 1) | 5 | 119646134 | 119659262 | 13129 | 1 | ENSMUST00000154997.1 | 10631 | Gm16064 | ENSMUSG00000085345.1 |
| chr5 | 119657902 | 119658088 | 3.39905 | 3.93486 | 1.04699 | Intron (ENSMUST00000154997.1/ENSMUSG00000085345.1, intron 1 of 1) | 5 | 119646134 | 119659262 | 13129 | 1 | ENSMUST00000154997.1 | 11769 | Gm16064 | ENSMUSG00000085345.1 |
| chr5 | 119650999 | 119652140 | 3.88759 | 4.85579 | 1.64241 | Intron (ENSMUST00000154997.1/ENSMUSG00000085345.1, intron 1 of 1) | 5 | 119646134 | 119659262 | 13129 | 1 | ENSMUST00000154997.1 | 4866 | Gm16064 | ENSMUSG00000085345.1 |
| chr2 | 22744825 | 22744941 | 2.58436 | 15.81988 | 11.09709 | Distal Intergenic | 2 | 22755984 | 22762273 | 6290 | 1 | ENSMUST00000135792.1 | -11043 | 1700092C17Rik | ENSMUSG00000085353.2 |
| chr4 | 55648577 | 55648952 | 3.50392 | 4.13603 | 1.19895 | Distal Intergenic | 4 | 55605149 | 55636901 | 31753 | 1 | ENSMUST00000123620.1 | 43429 | Gm12506 | ENSMUSG00000085375.1 |
| chr4 | 55649339 | 55649947 | 3.58608 | 4.31705 | 1.36666 | Distal Intergenic | 4 | 55605149 | 55636901 | 31753 | 1 | ENSMUST00000123620.1 | 44191 | Gm12506 | ENSMUSG00000085375.1 |
| chr4 | 55650891 | 55651681 | 3.97093 | 5.0428 | 1.7861 | Distal Intergenic | 4 | 55605149 | 55636901 | 31753 | 1 | ENSMUST00000123620.1 | 45743 | Gm12506 | ENSMUSG00000085375.1 |
| chr4 | 55757224 | 55757456 | 3.39716 | 3.98482 | 1.19792 | Distal Intergenic | 4 | 55605149 | 55636901 | 31753 | 1 | ENSMUST00000123620.1 | 152076 | Gm12506 | ENSMUSG00000085375.1 |
| chr2 | 74054803 | 74055073 | 3.31942 | 3.82304 | 1.07454 | Distal Intergenic | 2 | 74100924 | 74102342 | 1419 | 2 | ENSMUST00000151443.1 | 47269 | A630050E04Rik | ENSMUSG00000085387.1 |
| chr2 | 74065250 | 74065546 | 3.68962 | 4.58141 | 1.50767 | Distal Intergenic | 2 | 74100924 | 74102342 | 1419 | 2 | ENSMUST00000151443.1 | 36796 | A630050E04Rik | ENSMUSG00000085387.1 |
| chr2 | 74066772 | 74067106 | 3.33123 | 4.02634 | 1.15898 | Distal Intergenic | 2 | 74100924 | 74102342 | 1419 | 2 | ENSMUST00000151443.1 | 35236 | A630050E04Rik | ENSMUSG00000085387.1 |
| chr2 | 74067802 | 74067892 | 3.45355 | 4.72903 | 1.60201 | Distal Intergenic | 2 | 74100924 | 74102342 | 1419 | 2 | ENSMUST00000151443.1 | 34450 | A630050E04Rik | ENSMUSG00000085387.1 |
| chr2 | 19447693 | 19448517 | 3.64001 | 4.51056 | 1.4473 | Distal Intergenic | 2 | 19441508 | 19446674 | 5167 | 2 | ENSMUST00000122978.1 | -1020 | Ptf1aos | ENSMUSG00000085388.1 |
| chr4 | 129957032 | 129957256 | 3.5513 | 4.21673 | 1.2309 | 3' UTR | 4 | 129960374 | 129963520 | 3147 | 1 | ENSMUST00000147634.2 | -3118 | 1700003M07Rik | ENSMUSG00000085389.2 |
| chr4 | 129967471 | 129967780 | 3.38529 | 3.94103 | 1.13029 | Distal Intergenic | 4 | 129960808 | 129963657 | 2850 | 1 | ENSMUST00000139884.1 | 6664 | 1700003M07Rik | ENSMUSG00000085389.2 |
| chr4 | 129971952 | 129972080 | 3.37703 | 3.90159 | 1.04699 | Distal Intergenic | 4 | 129960808 | 129963657 | 2850 | 1 | ENSMUST00000139884.1 | 11145 | 1700003M07Rik | ENSMUSG00000085389.2 |
| chr4 | 153924875 | 153924950 | 3.38193 | 3.9006 | 1.01733 | Distal Intergenic | 4 | 153907386 | 153908666 | 1281 | 1 | ENSMUST00000156438.1 | 17490 | Gm13115 | ENSMUSG00000085390.1 |
| chr11 | 32925013 | 32925213 | 3.92006 | 4.91136 | 1.67614 | Promoter | 11 | 32903137 | 32926160 | 23024 | 2 | ENSMUST00000150073.1 | 947 | Gm12111 | ENSMUSG00000085402.1 |
| chr11 | 32925587 | 32925757 | 3.88438 | 4.96645 | 1.74962 | Promoter | 11 | 32903137 | 32926160 | 23024 | 2 | ENSMUST00000150073.1 | 403 | Gm12111 | ENSMUSG00000085402.1 |
| chr11 | 32930858 | 32931174 | 3.2537 | 3.70951 | 1.02332 | Distal Intergenic | 11 | 32903137 | 32926160 | 23024 | 2 | ENSMUST00000150073.1 | -4699 | Gm12111 | ENSMUSG00000085402.1 |
| chr11 | 32932623 | 32932793 | 3.30318 | 4.01964 | 1.17777 | Distal Intergenic | 11 | 32903137 | 32926160 | 23024 | 2 | ENSMUST00000150073.1 | -6464 | Gm12111 | ENSMUSG00000085402.1 |
| chr2 | 74257843 | 74258181 | 3.66599 | 4.44225 | 1.39809 | Distal Intergenic | 2 | 74298940 | 74307717 | 8778 | 2 | ENSMUST00000153743.1 | 49536 | 4930441J16Rik | ENSMUSG00000085405.1 |
| chr2 | 74296332 | 74297026 | 3.5528 | 4.38048 | 1.38069 | Downstream (1-2kb) | 2 | 74298940 | 74307717 | 8778 | 2 | ENSMUST00000153743.1 | 10691 | 4930441J16Rik | ENSMUSG00000085405.1 |
| chr2 | 74297850 | 74298004 | 3.42962 | 4.16542 | 1.23025 | Downstream (<1kb) | 2 | 74298940 | 74307717 | 8778 | 2 | ENSMUST00000153743.1 | 9713 | 4930441J16Rik | ENSMUSG00000085405.1 |
| chr2 | 74298322 | 74298533 | 3.82489 | 4.80024 | 1.69612 | Downstream (<1kb) | 2 | 74298940 | 74307717 | 8778 | 2 | ENSMUST00000153743.1 | 9184 | 4930441J16Rik | ENSMUSG00000085405.1 |
| chr2 | 74299214 | 74300139 | 3.6176 | 4.4395 | 1.42018 | Intron (ENSMUST00000153743.1/ENSMUSG00000085405.1, intron 4 of 4) | 2 | 74298940 | 74307717 | 8778 | 2 | ENSMUST00000153743.1 | 7578 | 4930441J16Rik | ENSMUSG00000085405.1 |
| chr2 | 74302763 | 74302895 | 3.30621 | 3.7865 | 1.01733 | Intron (ENSMUST00000153743.1/ENSMUSG00000085405.1, intron 3 of 4) | 2 | 74298940 | 74307717 | 8778 | 2 | ENSMUST00000153743.1 | 4822 | 4930441J16Rik | ENSMUSG00000085405.1 |
| chr2 | 74304475 | 74304821 | 3.53827 | 4.19884 | 1.23827 | Intron (ENSMUST00000153743.1/ENSMUSG00000085405.1, intron 2 of 4) | 2 | 74298940 | 74307717 | 8778 | 2 | ENSMUST00000153743.1 | 2896 | 4930441J16Rik | ENSMUSG00000085405.1 |
| chr6 | 6006029 | 6006327 | 3.66884 | 4.54263 | 1.51845 | Exon (ENSMUST00000115556.2/ENSMUSG00000029757.11, exon 15 of 17) | 6 | 5963909 | 5977393 | 13485 | 2 | ENSMUST00000130004.1 | -28637 | 1700019G24Rik | ENSMUSG00000085416.1 |
| chr11 | 113223995 | 113224075 | 3.30868 | 3.79024 | 1.01733 | Distal Intergenic | 11 | 113203315 | 113209669 | 6355 | 2 | ENSMUST00000130373.1 | -14327 | 4732490B19Rik | ENSMUSG00000085421.1 |
| chr2 | 72810298 | 72810808 | 3.32415 | 3.81356 | 1.01733 | Intron (ENSMUST00000136807.2/ENSMUSG00000085427.2, intron 1 of 5) | 2 | 72745846 | 72813672 | 67827 | 2 | ENSMUST00000136807.2 | 2864 | 6430710C18Rik | ENSMUSG00000085427.2 |
| chr2 | 72811954 | 72812075 | 4.09536 | 5.28028 | 1.92987 | Intron (ENSMUST00000136807.2/ENSMUSG00000085427.2, intron 1 of 5) | 2 | 72745846 | 72813672 | 67827 | 2 | ENSMUST00000136807.2 | 1597 | 6430710C18Rik | ENSMUSG00000085427.2 |
| chr2 | 72812513 | 72813365 | 4.03963 | 5.17911 | 1.89914 | Promoter | 2 | 72745846 | 72813672 | 67827 | 2 | ENSMUST00000136807.2 | 307 | 6430710C18Rik | ENSMUSG00000085427.2 |
| chr10 | 119455699 | 119456314 | 3.71169 | 4.52061 | 1.42538 | Promoter | 10 | 119438800 | 119455379 | 16580 | 2 | ENSMUST00000146294.1 | -321 | Grip1os3 | ENSMUSG00000085454.1 |
| chr2 | 33700689 | 33700764 | 3.41574 | 4.16623 | 1.21631 | Distal Intergenic | 2 | 33716361 | 33718899 | 2539 | 2 | ENSMUST00000130532.1 | 18135 | 9430024E24Rik | ENSMUSG00000085463.1 |
| chr2 | 33707547 | 33707768 | 3.3705 | 3.88337 | 1.01733 | Distal Intergenic | 2 | 33716361 | 33718899 | 2539 | 2 | ENSMUST00000130532.1 | 11131 | 9430024E24Rik | ENSMUSG00000085463.1 |
| chr2 | 33715077 | 33715236 | 3.5186 | 4.31202 | 1.31579 | Downstream (1-2kb) | 2 | 33716361 | 33718899 | 2539 | 2 | ENSMUST00000130532.1 | 3663 | 9430024E24Rik | ENSMUSG00000085463.1 |
| chr2 | 33716927 | 33717163 | 3.6732 | 4.596 | 1.49458 | Intron (ENSMUST00000130532.1/ENSMUSG00000085463.1, intron 1 of 1) | 2 | 33716361 | 33718899 | 2539 | 2 | ENSMUST00000130532.1 | 1736 | 9430024E24Rik | ENSMUSG00000085463.1 |
| chr2 | 33717815 | 33718070 | 3.66366 | 4.44771 | 1.38787 | Promoter | 2 | 33716361 | 33718899 | 2539 | 2 | ENSMUST00000130532.1 | 829 | 9430024E24Rik | ENSMUSG00000085463.1 |
| chr2 | 33718447 | 33719127 | 3.83463 | 4.74414 | 1.56048 | Promoter | 2 | 33716361 | 33718899 | 2539 | 2 | ENSMUST00000130532.1 | 0 | 9430024E24Rik | ENSMUSG00000085463.1 |
| chr11 | 45550257 | 45550671 | 3.49429 | 4.11488 | 1.17243 | Distal Intergenic | 11 | 45452399 | 45468504 | 16106 | 2 | ENSMUST00000123422.1 | -81754 | Gm12162 | ENSMUSG00000085476.1 |
| chr11 | 101597780 | 101597855 | 3.64457 | 4.47327 | 1.44511 | Intron (ENSMUST00000154147.1/ENSMUSG00000085486.1, intron 1 of 2) | 11 | 101593743 | 101599256 | 5514 | 2 | ENSMUST00000154147.1 | 1401 | Gm11634 | ENSMUSG00000085486.1 |
| chr13 | 23306916 | 23307294 | 4.06163 | 5.63208 | 2.19185 | Intron (ENSMUST00000155432.1/ENSMUSG00000085488.1, intron 1 of 1) | 13 | 23276204 | 23313400 | 37197 | 2 | ENSMUST00000155432.1 | 6106 | 4930557F10Rik | ENSMUSG00000085488.1 |
| chr13 | 23307668 | 23309888 | 4.11799 | 5.6661 | 2.28982 | Intron (ENSMUST00000155432.1/ENSMUSG00000085488.1, intron 1 of 1) | 13 | 23276204 | 23313400 | 37197 | 2 | ENSMUST00000155432.1 | 3512 | 4930557F10Rik | ENSMUSG00000085488.1 |
| chr13 | 23310214 | 23310411 | 4.39272 | 6.17894 | 2.62615 | Intron (ENSMUST00000155432.1/ENSMUSG00000085488.1, intron 1 of 1) | 13 | 23276204 | 23313400 | 37197 | 2 | ENSMUST00000155432.1 | 2989 | 4930557F10Rik | ENSMUSG00000085488.1 |
| chr13 | 23311593 | 23312559 | 3.78106 | 4.82052 | 1.6612 | Promoter | 13 | 23276204 | 23313400 | 37197 | 2 | ENSMUST00000155432.1 | 841 | 4930557F10Rik | ENSMUSG00000085488.1 |
| chr13 | 23312950 | 23313080 | 3.94072 | 5.08534 | 1.82769 | Promoter | 13 | 23276204 | 23313400 | 37197 | 2 | ENSMUST00000155432.1 | 320 | 4930557F10Rik | ENSMUSG00000085488.1 |
| chr5 | 31135776 | 31136231 | 3.96891 | 5.04545 | 1.7913 | Exon (ENSMUST00000013771.9/ENSMUSG00000062077.9, exon 5 of 9) | 5 | 31135753 | 31137344 | 1592 | 2 | ENSMUST00000136014.1 | 1113 | Gm15469 | ENSMUSG00000085493.1 |
| chr5 | 31136736 | 31137403 | 3.42229 | 4.0015 | 1.09884 | Promoter | 5 | 31135753 | 31137344 | 1592 | 2 | ENSMUST00000136014.1 | 0 | Gm15469 | ENSMUSG00000085493.1 |
| chr5 | 74722166 | 74722763 | 3.55256 | 4.3976 | 1.36821 | Intron (ENSMUST00000156466.2/ENSMUSG00000085657.2, intron 4 of 6) | 5 | 74731517 | 74734450 | 2934 | 2 | ENSMUST00000137731.1 | 11687 | Gm15985 | ENSMUSG00000085497.1 |
| chr5 | 74734394 | 74734571 | 3.84146 | 5.04204 | 1.79986 | Promoter | 5 | 74731517 | 74734450 | 2934 | 2 | ENSMUST00000137731.1 | 0 | Gm15985 | ENSMUSG00000085497.1 |
| chr5 | 74739295 | 74739370 | 3.68853 | 4.53638 | 1.49748 | Distal Intergenic | 5 | 74731517 | 74734450 | 2934 | 2 | ENSMUST00000137731.1 | -4846 | Gm15985 | ENSMUSG00000085497.1 |
| chr5 | 74839271 | 74839563 | 3.29499 | 3.79491 | 1.09399 | Distal Intergenic | 5 | 74731517 | 74734450 | 2934 | 2 | ENSMUST00000137731.1 | -104822 | Gm15985 | ENSMUSG00000085497.1 |
| chr14 | 44858841 | 44859126 | 3.40796 | 3.95299 | 1.06347 | Promoter | 14 | 44858954 | 44974944 | 115991 | 1 | ENSMUST00000127408.1 | 0 | Gm16976 | ENSMUSG00000085500.3 |
| chr11 | 71967679 | 71967754 | 3.8248 | 4.71839 | 1.52765 | Distal Intergenic | 11 | 71928471 | 71940122 | 11652 | 2 | ENSMUST00000128489.1 | -27558 | 4930401O10Rik | ENSMUSG00000085528.1 |
| chr4 | 137115990 | 137116099 | 3.47665 | 4.2657 | 1.31601 | Downstream (<1kb) | 4 | 137116709 | 137117582 | 874 | 2 | ENSMUST00000132824.1 | 1483 | 1700037C06Rik | ENSMUSG00000085542.1 |
| chr4 | 137116941 | 137117040 | 3.12524 | 3.96537 | 1.10745 | Promoter | 4 | 137116709 | 137117582 | 874 | 2 | ENSMUST00000132824.1 | 542 | 1700037C06Rik | ENSMUSG00000085542.1 |
| chr2 | 26191601 | 26191875 | 3.67491 | 4.43773 | 1.35053 | Promoter | 2 | 26186729 | 26191678 | 4950 | 2 | ENSMUST00000133025.1 | 0 | Gm13553 | ENSMUSG00000085545.1 |
| chr2 | 26193361 | 26193762 | 3.75862 | 4.62291 | 1.51503 | Distal Intergenic | 2 | 26186729 | 26191678 | 4950 | 2 | ENSMUST00000133025.1 | -1684 | Gm13553 | ENSMUSG00000085545.1 |
| chr2 | 26195928 | 26196088 | 3.73398 | 4.57991 | 1.5055 | Distal Intergenic | 2 | 26186729 | 26191678 | 4950 | 2 | ENSMUST00000133025.1 | -4251 | Gm13553 | ENSMUSG00000085545.1 |
| chr2 | 26196676 | 26197112 | 3.36889 | 4.25902 | 1.28781 | Distal Intergenic | 2 | 26186729 | 26191678 | 4950 | 2 | ENSMUST00000133025.1 | -4999 | Gm13553 | ENSMUSG00000085545.1 |
| chr4 | 107731439 | 107731584 | 3.47224 | 4.1697 | 1.25825 | Intron (ENSMUST00000143064.2/ENSMUSG00000085549.2, intron 3 of 3) | 4 | 107707057 | 107708644 | 1588 | 1 | ENSMUST00000147654.1 | 24383 | 1700047F07Rik | ENSMUSG00000085549.2 |
| chr14 | 122470453 | 122472432 | 3.9233 | 5.15406 | 1.87357 | Exon (ENSMUST00000135485.2/ENSMUSG00000085555.4, exon 3 of 3) | 14 | 122470284 | 122473579 | 3296 | 2 | ENSMUST00000135485.2 | 1147 | 2610035F20Rik | ENSMUSG00000085555.4 |
| chr14 | 122472754 | 122474497 | 3.8129 | 4.77491 | 1.67053 | Promoter | 14 | 122470284 | 122473579 | 3296 | 2 | ENSMUST00000135485.2 | 0 | 2610035F20Rik | ENSMUSG00000085555.4 |
| chr4 | 125718566 | 125718647 | 4.72528 | 6.50757 | 2.79668 | Distal Intergenic | 4 | 125887851 | 125898982 | 11132 | 2 | ENSMUST00000123368.2 | 180335 | 2610028E06Rik | ENSMUSG00000085562.2 |
| chr4 | 125726791 | 125727123 | 3.80926 | 4.72359 | 1.58617 | Distal Intergenic | 4 | 125887851 | 125898982 | 11132 | 2 | ENSMUST00000123368.2 | 171859 | 2610028E06Rik | ENSMUSG00000085562.2 |
| chr4 | 125727982 | 125728102 | 3.94401 | 5.21379 | 1.92603 | Distal Intergenic | 4 | 125887851 | 125898982 | 11132 | 2 | ENSMUST00000123368.2 | 170880 | 2610028E06Rik | ENSMUSG00000085562.2 |
| chr4 | 125729771 | 125730340 | 4.05726 | 5.21171 | 1.88285 | Distal Intergenic | 4 | 125887851 | 125898982 | 11132 | 2 | ENSMUST00000123368.2 | 168642 | 2610028E06Rik | ENSMUSG00000085562.2 |
| chr4 | 125833228 | 125833308 | 4.38038 | 5.86813 | 2.41949 | Distal Intergenic | 4 | 125887851 | 125898982 | 11132 | 2 | ENSMUST00000123368.2 | 65674 | 2610028E06Rik | ENSMUSG00000085562.2 |
| chr4 | 125837824 | 125838003 | 3.58113 | 4.28892 | 1.31824 | Distal Intergenic | 4 | 125887851 | 125898982 | 11132 | 2 | ENSMUST00000123368.2 | 60979 | 2610028E06Rik | ENSMUSG00000085562.2 |
| chr4 | 125864950 | 125865237 | 3.63365 | 4.42792 | 1.43143 | Distal Intergenic | 4 | 125887851 | 125898982 | 11132 | 2 | ENSMUST00000123368.2 | 33745 | 2610028E06Rik | ENSMUSG00000085562.2 |
| chr4 | 125910806 | 125911318 | 3.59378 | 4.40138 | 1.40751 | Intron (ENSMUST00000148534.2/ENSMUSG00000085562.2, intron 2 of 5) | 4 | 125890415 | 125917171 | 26757 | 2 | ENSMUST00000153589.1 | 5853 | 2610028E06Rik | ENSMUSG00000085562.2 |
| chr4 | 125911654 | 125912034 | 3.64419 | 4.42843 | 1.42054 | Intron (ENSMUST00000148534.2/ENSMUSG00000085562.2, intron 2 of 5) | 4 | 125890415 | 125917171 | 26757 | 2 | ENSMUST00000153589.1 | 5137 | 2610028E06Rik | ENSMUSG00000085562.2 |
| chr4 | 125912849 | 125913160 | 3.79552 | 4.6855 | 1.56272 | Intron (ENSMUST00000148534.2/ENSMUSG00000085562.2, intron 2 of 5) | 4 | 125890415 | 125917171 | 26757 | 2 | ENSMUST00000153589.1 | 4011 | 2610028E06Rik | ENSMUSG00000085562.2 |
| chr4 | 125927891 | 125928044 | 3.59335 | 4.3859 | 1.41131 | Distal Intergenic | 4 | 125918575 | 125922417 | 3843 | 2 | ENSMUST00000144539.1 | -5475 | 2610028E06Rik | ENSMUSG00000085562.2 |
| chr4 | 125930382 | 125930568 | 3.39718 | 4.02893 | 1.13234 | Distal Intergenic | 4 | 125918575 | 125922417 | 3843 | 2 | ENSMUST00000144539.1 | -7966 | 2610028E06Rik | ENSMUSG00000085562.2 |
| chr4 | 125931803 | 125932577 | 3.58205 | 4.3444 | 1.38027 | Distal Intergenic | 4 | 125918575 | 125922417 | 3843 | 2 | ENSMUST00000144539.1 | -9387 | 2610028E06Rik | ENSMUSG00000085562.2 |
| chr4 | 125933304 | 125933379 | 3.7013 | 4.49172 | 1.39243 | Distal Intergenic | 4 | 125918575 | 125922417 | 3843 | 2 | ENSMUST00000144539.1 | -10888 | 2610028E06Rik | ENSMUSG00000085562.2 |
| chr11 | 50602883 | 50603852 | 3.83906 | 4.81668 | 1.71587 | Promoter | 11 | 50599298 | 50602448 | 3151 | 2 | ENSMUST00000154068.1 | -436 | Gm12198 | ENSMUSG00000085564.1 |
| chr11 | 50605501 | 50605685 | 3.59603 | 4.31644 | 1.34786 | Intron (ENSMUST00000040523.8/ENSMUSG00000036545.8, intron 2 of 21) | 11 | 50599298 | 50602448 | 3151 | 2 | ENSMUST00000154068.1 | -3054 | Gm12198 | ENSMUSG00000085564.1 |
| chr5 | 147955322 | 147956240 | 3.49756 | 4.36539 | 1.35224 | Promoter | 5 | 147949882 | 147957166 | 7285 | 2 | ENSMUST00000138406.1 | 926 | Gm15721 | ENSMUSG00000085565.1 |
| chr8 | 11182577 | 11182982 | 3.53984 | 4.19714 | 1.21773 | Distal Intergenic | 8 | 11187755 | 11204503 | 16749 | 1 | ENSMUST00000151014.1 | -4773 | Gm15418 | ENSMUSG00000085573.1 |
| chr8 | 11184485 | 11184808 | 3.3639 | 3.87339 | 1.01733 | Distal Intergenic | 8 | 11187755 | 11204503 | 16749 | 1 | ENSMUST00000151014.1 | -2947 | Gm15418 | ENSMUSG00000085573.1 |
| chr8 | 11187853 | 11188965 | 3.7534 | 4.73772 | 1.65625 | Promoter | 8 | 11187755 | 11204503 | 16749 | 1 | ENSMUST00000151014.1 | 99 | Gm15418 | ENSMUSG00000085573.1 |
| chr8 | 11193445 | 11193520 | 3.61102 | 4.32488 | 1.29456 | Intron (ENSMUST00000151014.1/ENSMUSG00000085573.1, intron 1 of 2) | 8 | 11187755 | 11204503 | 16749 | 1 | ENSMUST00000151014.1 | 5691 | Gm15418 | ENSMUSG00000085573.1 |
| chr8 | 11193905 | 11193980 | 3.82535 | 4.73391 | 1.5715 | Intron (ENSMUST00000151014.1/ENSMUSG00000085573.1, intron 1 of 2) | 8 | 11187755 | 11204503 | 16749 | 1 | ENSMUST00000151014.1 | 6151 | Gm15418 | ENSMUSG00000085573.1 |
| chr8 | 11201869 | 11202099 | 3.71121 | 4.6839 | 1.5457 | Intron (ENSMUST00000151014.1/ENSMUSG00000085573.1, intron 2 of 2) | 8 | 11187755 | 11204503 | 16749 | 1 | ENSMUST00000151014.1 | 14115 | Gm15418 | ENSMUSG00000085573.1 |
| chr4 | 142042215 | 142042441 | 3.9221 | 5.25916 | 1.93175 | Intron (ENSMUST00000036572.3/ENSMUSG00000040616.3, intron 1 of 2) | 4 | 142042912 | 142048578 | 5667 | 2 | ENSMUST00000128385.1 | 6137 | Gm13053 | ENSMUSG00000085574.1 |
| chr9 | 24695798 | 24696976 | 3.46827 | 4.10498 | 1.19033 | Promoter | 9 | 24557048 | 24696293 | 139246 | 2 | ENSMUST00000133010.2 | 0 | Dpy19l2 | ENSMUSG00000085576.2 |
| chr4 | 151576422 | 151576668 | 3.29256 | 3.91797 | 1.05816 | Intron (ENSMUST00000169423.3/ENSMUSG00000014592.14, intron 4 of 24) | 4 | 151376470 | 151379582 | 3113 | 1 | ENSMUST00000129097.1 | 199953 | 4930589P08Rik | ENSMUSG00000085593.1 |
| chr4 | 151587161 | 151587421 | 3.42616 | 4.18284 | 1.26688 | Intron (ENSMUST00000169423.3/ENSMUSG00000014592.14, intron 3 of 24) | 4 | 151376470 | 151379582 | 3113 | 1 | ENSMUST00000129097.1 | 210692 | 4930589P08Rik | ENSMUSG00000085593.1 |
| chr4 | 151467068 | 151467410 | 3.45095 | 4.02643 | 1.09361 | Intron (ENSMUST00000169423.3/ENSMUSG00000014592.14, intron 4 of 24) | 4 | 151376470 | 151379582 | 3113 | 1 | ENSMUST00000129097.1 | 90599 | 4930589P08Rik | ENSMUSG00000085593.1 |
| chr4 | 151474441 | 151475750 | 3.57759 | 4.36786 | 1.37783 | Intron (ENSMUST00000169423.3/ENSMUSG00000014592.14, intron 4 of 24) | 4 | 151376470 | 151379582 | 3113 | 1 | ENSMUST00000129097.1 | 97972 | 4930589P08Rik | ENSMUSG00000085593.1 |
| chr4 | 151538564 | 151538828 | 3.46821 | 4.06525 | 1.13868 | Intron (ENSMUST00000169423.3/ENSMUSG00000014592.14, intron 4 of 24) | 4 | 151376470 | 151379582 | 3113 | 1 | ENSMUST00000129097.1 | 162095 | 4930589P08Rik | ENSMUSG00000085593.1 |
| chr4 | 151550534 | 151550634 | 3.77675 | 4.71704 | 1.62356 | Intron (ENSMUST00000169423.3/ENSMUSG00000014592.14, intron 4 of 24) | 4 | 151376470 | 151379582 | 3113 | 1 | ENSMUST00000129097.1 | 174065 | 4930589P08Rik | ENSMUSG00000085593.1 |
| chr4 | 151553580 | 151553724 | 3.70031 | 4.67334 | 1.55652 | Intron (ENSMUST00000169423.3/ENSMUSG00000014592.14, intron 4 of 24) | 4 | 151376470 | 151379582 | 3113 | 1 | ENSMUST00000129097.1 | 177111 | 4930589P08Rik | ENSMUSG00000085593.1 |
| chr11 | 101870740 | 101870850 | 3.37844 | 3.9002 | 1.03455 | Distal Intergenic | 11 | 101886237 | 101891206 | 4970 | 1 | ENSMUST00000147438.1 | -15387 | Gm11551 | ENSMUSG00000085594.1 |
| chr11 | 101881580 | 101882279 | 3.54787 | 4.24695 | 1.33812 | Intron (ENSMUST00000057054.7/ENSMUSG00000001493.9, intron 1 of 2) | 11 | 101886237 | 101891206 | 4970 | 1 | ENSMUST00000147438.1 | -3958 | Gm11551 | ENSMUSG00000085594.1 |
| chr11 | 101888816 | 101888989 | 3.48176 | 4.18551 | 1.2512 | Intron (ENSMUST00000147438.1/ENSMUSG00000085594.1, intron 1 of 1) | 11 | 101886237 | 101891206 | 4970 | 1 | ENSMUST00000147438.1 | 2580 | Gm11551 | ENSMUSG00000085594.1 |
| chr11 | 101889982 | 101890258 | 4.05642 | 5.17339 | 1.8531 | Intron (ENSMUST00000147438.1/ENSMUSG00000085594.1, intron 1 of 1) | 11 | 101886237 | 101891206 | 4970 | 1 | ENSMUST00000147438.1 | 3746 | Gm11551 | ENSMUSG00000085594.1 |
| chr9 | 29594948 | 29595201 | 4.09973 | 5.55812 | 2.16828 | Intron (ENSMUST00000075069.5/ENSMUSG00000059974.5, intron 1 of 7) | 9 | 29590484 | 29592510 | 2027 | 2 | ENSMUST00000141752.1 | -2439 | Gm15521 | ENSMUSG00000085638.1 |
| chr12 | 107684754 | 107685018 | 3.24243 | 3.69081 | 1.01733 | Distal Intergenic | 12 | 107723740 | 107735488 | 11749 | 2 | ENSMUST00000130564.1 | 50470 | 4930465M20Rik | ENSMUSG00000085641.1 |
| chr2 | 164175880 | 164176656 | 3.72966 | 4.56724 | 1.47616 | Promoter | 2 | 164177584 | 164196564 | 18981 | 1 | ENSMUST00000143202.2 | -928 | A730032A03Rik | ENSMUSG00000085649.2 |
| chr2 | 164177193 | 164177268 | 3.25561 | 3.71058 | 1.01733 | Promoter | 2 | 164177584 | 164196564 | 18981 | 1 | ENSMUST00000143202.2 | -316 | A730032A03Rik | ENSMUSG00000085649.2 |
| chr2 | 164178494 | 164178789 | 3.68564 | 4.52153 | 1.46393 | Promoter | 2 | 164177584 | 164196564 | 18981 | 1 | ENSMUST00000143202.2 | 911 | A730032A03Rik | ENSMUSG00000085649.2 |
| chr4 | 131913653 | 131913778 | 4.23664 | 5.56209 | 2.14959 | Exon (ENSMUST00000132111.1/ENSMUSG00000085667.1, exon 2 of 2) | 4 | 131912416 | 131915516 | 3101 | 2 | ENSMUST00000132111.1 | 1738 | Gm12992 | ENSMUSG00000085667.1 |
| chr4 | 131914557 | 131915680 | 3.65644 | 4.51193 | 1.4491 | Promoter | 4 | 131912416 | 131915516 | 3101 | 2 | ENSMUST00000132111.1 | 0 | Gm12992 | ENSMUSG00000085667.1 |
| chr4 | 131916272 | 131917314 | 3.88685 | 4.90245 | 1.71934 | Promoter | 4 | 131912416 | 131915516 | 3101 | 2 | ENSMUST00000132111.1 | -757 | Gm12992 | ENSMUSG00000085667.1 |
| chr2 | 35513956 | 35514187 | 3.29315 | 3.767 | 1.01733 | Distal Intergenic | 2 | 35503531 | 35506090 | 2560 | 2 | ENSMUST00000130083.1 | -7867 | Gm13447 | ENSMUSG00000085668.1 |
| chr2 | 21368455 | 21369098 | 3.46551 | 4.33103 | 1.36182 | Promoter | 2 | 21341925 | 21369111 | 27187 | 2 | ENSMUST00000133391.1 | 13 | Gm13327 | ENSMUSG00000085680.1 |
| chr2 | 166518109 | 166518293 | 3.24393 | 3.69305 | 1.01733 | Distal Intergenic | 2 | 166502101 | 166505584 | 3484 | 2 | ENSMUST00000123480.1 | -12526 | Gm14267 | ENSMUSG00000085682.1 |
| chr4 | 90045968 | 90046079 | 9.63489 | 20.73017 | 15.68358 | Distal Intergenic | 4 | 90185527 | 90190009 | 4483 | 1 | ENSMUST00000145751.1 | -139448 | Gm12629 | ENSMUSG00000085690.1 |
| chr2 | 154735081 | 154735432 | 3.4861 | 4.0989 | 1.16295 | Intron (ENSMUST00000155334.1/ENSMUSG00000085691.1, intron 1 of 2) | 2 | 154722370 | 154738451 | 16082 | 2 | ENSMUST00000155334.1 | 3019 | Gm14216 | ENSMUSG00000085691.1 |
| chr3 | 30011980 | 30012533 | 3.62819 | 4.56724 | 1.48688 | Promoter | 3 | 30012379 | 30053185 | 40807 | 1 | ENSMUST00000125611.2 | 0 | Mecomos | ENSMUSG00000085702.3 |
| chr11 | 53462566 | 53462709 | 3.56091 | 4.3175 | 1.34999 | Promoter | 11 | 53462379 | 53463613 | 1235 | 2 | ENSMUST00000141032.1 | 904 | Gm12212 | ENSMUSG00000085707.1 |
| chr11 | 53463385 | 53464474 | 3.72427 | 4.65857 | 1.57338 | Promoter | 11 | 53462379 | 53463613 | 1235 | 2 | ENSMUST00000141032.1 | 0 | Gm12212 | ENSMUSG00000085707.1 |
| chr5 | 119518989 | 119519064 | 3.58064 | 4.26505 | 1.24453 | Distal Intergenic | 5 | 119625942 | 119634464 | 8523 | 1 | ENSMUST00000156235.1 | -106878 | Gm16063 | ENSMUSG00000085708.1 |
| chr5 | 119562684 | 119562767 | 4.07212 | 5.19858 | 1.858 | Distal Intergenic | 5 | 119625942 | 119634464 | 8523 | 1 | ENSMUST00000156235.1 | -63175 | Gm16063 | ENSMUSG00000085708.1 |
| chr5 | 119568151 | 119568444 | 3.34317 | 3.8423 | 1.01733 | Distal Intergenic | 5 | 119625942 | 119634464 | 8523 | 1 | ENSMUST00000156235.1 | -57498 | Gm16063 | ENSMUSG00000085708.1 |
| chr5 | 119569178 | 119569413 | 3.68238 | 4.45116 | 1.35862 | Distal Intergenic | 5 | 119625942 | 119634464 | 8523 | 1 | ENSMUST00000156235.1 | -56529 | Gm16063 | ENSMUSG00000085708.1 |
| chr5 | 119587436 | 119587705 | 3.54717 | 4.21335 | 1.24124 | Distal Intergenic | 5 | 119625942 | 119634464 | 8523 | 1 | ENSMUST00000156235.1 | -38237 | Gm16063 | ENSMUSG00000085708.1 |
| chr2 | 180854645 | 180855106 | 3.65883 | 4.46351 | 1.44793 | Intron (ENSMUST00000137629.1/ENSMUSG00000086166.1, intron 1 of 3) | 2 | 180847652 | 180849046 | 1395 | 1 | ENSMUST00000146224.1 | 6994 | Gm14339 | ENSMUSG00000085728.1 |
| chr2 | 180855569 | 180855720 | 3.36932 | 3.88158 | 1.01733 | Intron (ENSMUST00000137629.1/ENSMUSG00000086166.1, intron 1 of 3) | 2 | 180847652 | 180849046 | 1395 | 1 | ENSMUST00000146224.1 | 7918 | Gm14339 | ENSMUSG00000085728.1 |
| chr11 | 18868365 | 18869202 | 3.38128 | 4.02751 | 1.15091 | Exon (ENSMUST00000133948.1/ENSMUSG00000085743.1, exon 2 of 3) | 11 | 18845851 | 18872155 | 26305 | 2 | ENSMUST00000128654.1 | 2953 | 8430419K02Rik | ENSMUSG00000085743.1 |
| chr11 | 18869714 | 18870292 | 3.69321 | 4.84935 | 1.71847 | Intron (ENSMUST00000128654.1/ENSMUSG00000085743.1, intron 1 of 2) | 11 | 18845851 | 18872155 | 26305 | 2 | ENSMUST00000128654.1 | 1863 | 8430419K02Rik | ENSMUSG00000085743.1 |
| chr11 | 18871260 | 18872110 | 3.9641 | 5.03669 | 1.80871 | Promoter | 11 | 18845851 | 18872155 | 26305 | 2 | ENSMUST00000128654.1 | 45 | 8430419K02Rik | ENSMUSG00000085743.1 |
| chr11 | 18872854 | 18873745 | 3.88007 | 5.00024 | 1.7886 | Promoter | 11 | 18854043 | 18874309 | 20267 | 2 | ENSMUST00000133948.1 | 564 | 8430419K02Rik | ENSMUSG00000085743.1 |
| chr11 | 18874231 | 18874935 | 3.53451 | 4.44001 | 1.39351 | Promoter | 11 | 18854043 | 18874309 | 20267 | 2 | ENSMUST00000133948.1 | 0 | 8430419K02Rik | ENSMUSG00000085743.1 |
| chr11 | 18875435 | 18875913 | 3.50071 | 4.12967 | 1.17679 | Distal Intergenic | 11 | 18854043 | 18874309 | 20267 | 2 | ENSMUST00000133948.1 | -1127 | 8430419K02Rik | ENSMUSG00000085743.1 |
| chr11 | 18876283 | 18876796 | 3.73954 | 4.57547 | 1.46323 | Distal Intergenic | 11 | 18854043 | 18874309 | 20267 | 2 | ENSMUST00000133948.1 | -1975 | 8430419K02Rik | ENSMUSG00000085743.1 |
| chr11 | 18877280 | 18877515 | 4.12382 | 5.37361 | 2.04387 | Downstream (2-3kb) | 11 | 18854043 | 18874309 | 20267 | 2 | ENSMUST00000133948.1 | -2972 | 8430419K02Rik | ENSMUSG00000085743.1 |
| chr11 | 78398562 | 78398886 | 3.21311 | 3.64691 | 1.00882 | Intron (ENSMUST00000149865.1/ENSMUSG00000085747.1, intron 4 of 4) | 11 | 78394485 | 78405657 | 11173 | 1 | ENSMUST00000149865.1 | 4078 | Slc13a2os | ENSMUSG00000085747.1 |
| chr2 | 128179337 | 128179555 | 3.79917 | 4.80367 | 1.64115 | Distal Intergenic | 2 | 128189987 | 128204634 | 14648 | 1 | ENSMUST00000152439.1 | -10432 | Gm14009 | ENSMUSG00000085750.1 |
| chr4 | 142017237 | 142017590 | 4.21268 | 5.63965 | 2.22939 | Promoter | 4 | 142017898 | 142028996 | 11099 | 1 | ENSMUST00000126231.1 | -308 | 4930455G09Rik | ENSMUSG00000085761.1 |
| chr4 | 142017946 | 142018411 | 3.62339 | 4.35064 | 1.31221 | Promoter | 4 | 142017898 | 142028996 | 11099 | 1 | ENSMUST00000126231.1 | 49 | 4930455G09Rik | ENSMUSG00000085761.1 |
| chr4 | 142020380 | 142020481 | 4.41896 | 5.98947 | 2.4773 | Intron (ENSMUST00000126231.1/ENSMUSG00000085761.1, intron 1 of 1) | 4 | 142017898 | 142028996 | 11099 | 1 | ENSMUST00000126231.1 | 2483 | 4930455G09Rik | ENSMUSG00000085761.1 |
| chr2 | 27887625 | 27888095 | 3.49279 | 4.18301 | 1.29379 | Promoter | 2 | 27882925 | 27887052 | 4128 | 2 | ENSMUST00000154246.1 | -574 | 2810430I11Rik | ENSMUSG00000085766.1 |
| chr2 | 27888482 | 27888717 | 3.47007 | 4.11826 | 1.23651 | Intron (ENSMUST00000028280.8/ENSMUSG00000026837.10, intron 1 of 65) | 2 | 27882925 | 27887052 | 4128 | 2 | ENSMUST00000154246.1 | -1431 | 2810430I11Rik | ENSMUSG00000085766.1 |
| chr2 | 27891259 | 27891514 | 3.43986 | 4.05515 | 1.2162 | Intron (ENSMUST00000028280.8/ENSMUSG00000026837.10, intron 1 of 65) | 2 | 27882925 | 27887052 | 4128 | 2 | ENSMUST00000154246.1 | -4208 | 2810430I11Rik | ENSMUSG00000085766.1 |
| chr10 | 81194697 | 81194863 | 3.44258 | 4.02133 | 1.12067 | Promoter | 10 | 81194691 | 81210877 | 16187 | 1 | ENSMUST00000150782.1 | 7 | Atcayos | ENSMUSG00000085779.1 |
| chr11 | 44665805 | 44666122 | 3.30623 | 3.82417 | 1.01733 | Intron (ENSMUST00000109268.1/ENSMUSG00000057098.9, intron 6 of 15) | 11 | 44659522 | 44662676 | 3155 | 2 | ENSMUST00000130493.1 | -3130 | Gm12158 | ENSMUSG00000085784.1 |
| chr11 | 44761031 | 44761264 | 3.63877 | 4.39892 | 1.40401 | Intron (ENSMUST00000109268.1/ENSMUSG00000057098.9, intron 6 of 15) | 11 | 44659522 | 44662676 | 3155 | 2 | ENSMUST00000130493.1 | -98356 | Gm12158 | ENSMUSG00000085784.1 |
| chr6 | 83702386 | 83703767 | 3.89664 | 4.9127 | 1.7311 | Promoter | 6 | 83702130 | 83703543 | 1414 | 2 | ENSMUST00000150233.2 | 0 | Vax2os | ENSMUSG00000085794.5 |
| chr6 | 83704117 | 83705882 | 3.83918 | 4.89346 | 1.69618 | Promoter | 6 | 83702480 | 83703622 | 1143 | 2 | ENSMUST00000173883.1 | -496 | Vax2os | ENSMUSG00000085794.5 |
| chr6 | 83706251 | 83706676 | 3.52033 | 4.17424 | 1.22863 | Intron (ENSMUST00000123402.1/ENSMUSG00000085794.5, intron 2 of 5) | 6 | 83702480 | 83703622 | 1143 | 2 | ENSMUST00000173883.1 | -2630 | Vax2os | ENSMUSG00000085794.5 |
| chr6 | 83711847 | 83716233 | 4.17502 | 5.63287 | 2.28482 | Promoter | 6 | 83692806 | 83712201 | 19396 | 2 | ENSMUST00000123402.1 | 0 | Vax2os | ENSMUSG00000085794.5 |
| chr2 | 146646782 | 146647102 | 3.70077 | 4.51013 | 1.43669 | Distal Intergenic | 2 | 146542931 | 146546108 | 3178 | 1 | ENSMUST00000147212.1 | 103852 | 4933406D12Rik | ENSMUSG00000085796.1 |
| chr11 | 89312413 | 89312640 | 3.51732 | 4.14876 | 1.17031 | Distal Intergenic | 11 | 89302723 | 89309428 | 6706 | 1 | ENSMUST00000140754.1 | 9691 | Gm2018 | ENSMUSG00000085798.1 |
| chr12 | 106829465 | 106829629 | 3.34791 | 3.84928 | 1.01733 | Distal Intergenic | 12 | 106837982 | 106878841 | 40860 | 1 | ENSMUST00000156376.1 | -8353 | 1700013N06Rik | ENSMUSG00000085801.1 |
| chr12 | 106840598 | 106840684 | 3.83634 | 4.74204 | 1.55129 | Intron (ENSMUST00000156376.1/ENSMUSG00000085801.1, intron 2 of 4) | 12 | 106837982 | 106878841 | 40860 | 1 | ENSMUST00000156376.1 | 2617 | 1700013N06Rik | ENSMUSG00000085801.1 |
| chr4 | 130985533 | 130985908 | 3.38079 | 3.91465 | 1.07311 | Intron (ENSMUST00000143967.1/ENSMUSG00000085804.1, intron 2 of 3) | 4 | 130974790 | 131001339 | 26550 | 1 | ENSMUST00000143967.1 | 10744 | Gm12973 | ENSMUSG00000085804.1 |
| chr3 | 146628928 | 146629150 | 3.26412 | 3.72331 | 1.01733 | Intron (ENSMUST00000029837.8/ENSMUSG00000028186.9, intron 7 of 7) | 3 | 146641932 | 146651715 | 9784 | 1 | ENSMUST00000141199.2 | -12782 | Gm16325 | ENSMUSG00000085810.2 |
| chr4 | 142304330 | 142304579 | 3.58536 | 4.27584 | 1.25094 | Distal Intergenic | 4 | 142307765 | 142346475 | 38711 | 2 | ENSMUST00000125582.1 | 41896 | Gm13052 | ENSMUSG00000085816.1 |
| chr4 | 142306325 | 142307209 | 3.74381 | 4.5875 | 1.47402 | Downstream (<1kb) | 4 | 142307765 | 142346475 | 38711 | 2 | ENSMUST00000125582.1 | 39266 | Gm13052 | ENSMUSG00000085816.1 |
| chr4 | 142307659 | 142308028 | 3.96908 | 5.00879 | 1.75357 | Exon (ENSMUST00000125582.1/ENSMUSG00000085816.1, exon 4 of 4) | 4 | 142307765 | 142346475 | 38711 | 2 | ENSMUST00000125582.1 | 38447 | Gm13052 | ENSMUSG00000085816.1 |
| chr4 | 142308400 | 142308620 | 3.50302 | 4.2333 | 1.29255 | Intron (ENSMUST00000125582.1/ENSMUSG00000085816.1, intron 3 of 3) | 4 | 142307765 | 142346475 | 38711 | 2 | ENSMUST00000125582.1 | 37855 | Gm13052 | ENSMUSG00000085816.1 |
| chr4 | 142315424 | 142315670 | 3.38041 | 3.98642 | 1.17805 | Intron (ENSMUST00000125582.1/ENSMUSG00000085816.1, intron 3 of 3) | 4 | 142307765 | 142346475 | 38711 | 2 | ENSMUST00000125582.1 | 30805 | Gm13052 | ENSMUSG00000085816.1 |
| chr4 | 142316991 | 142317469 | 3.56761 | 4.3921 | 1.39165 | Intron (ENSMUST00000125582.1/ENSMUSG00000085816.1, intron 3 of 3) | 4 | 142307765 | 142346475 | 38711 | 2 | ENSMUST00000125582.1 | 29006 | Gm13052 | ENSMUSG00000085816.1 |
| chr4 | 142321948 | 142322088 | 3.2459 | 3.70344 | 1.0406 | Intron (ENSMUST00000125582.1/ENSMUSG00000085816.1, intron 3 of 3) | 4 | 142307765 | 142346475 | 38711 | 2 | ENSMUST00000125582.1 | 24387 | Gm13052 | ENSMUSG00000085816.1 |

| | | | | | | | | | | | | | | | |
|---|---|---|---|---|---|---|---|---|---|---|---|---|---|---|---|
| chr12 | 106665086 | 106665434 | 3.50088 | 4.14248 | 1.2356 | Distal Intergenic | 12 | 106695716 | 106716324 | 20609 | 2 | ENSMUST00000124363.2 | 50890 | 4933406K04Rik | ENSMUSG00000085820.2 |
| chr13 | 22012491 | 22013493 | 3.48466 | 4.18677 | 1.28752 | Promoter | 13 | 22014468 | 22015348 | 881 | 1 | ENSMUST00000133204.2 | -975 | Gm11290 | ENSMUSG00000085827.2 |
| chr13 | 22013814 | 22015988 | 3.88102 | 5.13935 | 1.90677 | Promoter | 13 | 22014468 | 22015348 | 881 | 1 | ENSMUST00000133204.2 | 0 | Gm11290 | ENSMUSG00000085827.2 |
| chr2 | 25291376 | 25292245 | 3.58309 | 4.34273 | 1.37418 | Promoter | 2 | 25291220 | 25298925 | 7706 | 1 | ENSMUST00000129723.1 | 157 | Grin1os | ENSMUSG00000085830.1 |
| chr4 | 137142375 | 137142730 | 3.61877 | 4.33825 | 1.29741 | Distal Intergenic | 4 | 137159082 | 137165066 | 5985 | 2 | ENSMUST00000151040.2 | 22336 | Gm13003 | ENSMUSG00000085833.2 |
| chr4 | 137156145 | 137156231 | 3.94651 | 5.03338 | 1.84524 | Downstream (2-3kb) | 4 | 137159082 | 137165066 | 5985 | 2 | ENSMUST00000151040.2 | 8835 | Gm13003 | ENSMUSG00000085833.2 |
| chr4 | 137164161 | 137164677 | 3.48611 | 4.17278 | 1.30466 | Promoter | 4 | 137159082 | 137165066 | 5985 | 2 | ENSMUST00000151040.2 | 389 | Gm13003 | ENSMUSG00000085833.2 |
| chr10 | 116472967 | 116473348 | 4.21096 | 5.72292 | 2.2884 | Promoter | 10 | 116473163 | 116474982 | 1820 | 1 | ENSMUST00000080943.8 | 0 | Kcnmb4os2 | ENSMUSG00000085837.3 |
| chr10 | 116475610 | 116475996 | 3.25128 | 3.70868 | 1.03178 | Distal Intergenic | 10 | 116474345 | 116474987 | 643 | 1 | ENSMUST00000123348.1 | 1266 | Kcnmb4os2 | ENSMUSG00000085837.3 |
| chr10 | 116476546 | 116477190 | 3.62265 | 4.45789 | 1.44449 | Distal Intergenic | 10 | 116474345 | 116474987 | 643 | 1 | ENSMUST00000123348.1 | 2202 | Kcnmb4os2 | ENSMUSG00000085837.3 |
| chr4 | 98795232 | 98795424 | 3.80464 | 4.83168 | 1.6555 | Intron (ENSMUST00000133455.1/ENSMUSG00000085843.1, intron 1 of 2) | 4 | 98793768 | 98798900 | 5133 | 1 | ENSMUST00000133455.1 | 1465 | Kank4os | ENSMUSG00000085843.1 |
| chr4 | 98796537 | 98796650 | 3.49755 | 4.35178 | 1.33589 | Intron (ENSMUST00000133455.1/ENSMUSG00000085843.1, intron 2 of 2) | 4 | 98793768 | 98798900 | 5133 | 1 | ENSMUST00000133455.1 | 2770 | Kank4os | ENSMUSG00000085843.1 |
| chr4 | 98788422 | 98788799 | 3.56897 | 4.30121 | 1.35918 | Intron (ENSMUST00000102790.3/ENSMUSG00000035407.8, intron 1 of 9) | 4 | 98793768 | 98798900 | 5133 | 1 | ENSMUST00000133455.1 | -4969 | Kank4os | ENSMUSG00000085843.1 |
| chr4 | 98793600 | 98794136 | 3.507 | 4.41578 | 1.38378 | Promoter | 4 | 98793768 | 98798900 | 5133 | 1 | ENSMUST00000133455.1 | 0 | Kank4os | ENSMUSG00000085843.1 |
| chr4 | 98804161 | 98804263 | 4.11017 | 5.29018 | 1.96528 | Intron (ENSMUST00000102790.3/ENSMUSG00000035407.8, intron 1 of 9) | 4 | 98793768 | 98798900 | 5133 | 1 | ENSMUST00000133455.1 | 10394 | Kank4os | ENSMUSG00000085843.1 |
| chr4 | 98805276 | 98805477 | 3.36963 | 3.92071 | 1.09982 | Intron (ENSMUST00000102790.3/ENSMUSG00000035407.8, intron 1 of 9) | 4 | 98793768 | 98798900 | 5133 | 1 | ENSMUST00000133455.1 | 11509 | Kank4os | ENSMUSG00000085843.1 |
| chr4 | 98797181 | 98797383 | 3.74023 | 4.81624 | 1.63568 | Exon (ENSMUST00000133455.1/ENSMUSG00000085843.1, exon 3 of 3) | 4 | 98793768 | 98798900 | 5133 | 1 | ENSMUST00000133455.1 | 3414 | Kank4os | ENSMUSG00000085843.1 |
| chr4 | 98803107 | 98803754 | 3.39627 | 3.96216 | 1.15774 | Intron (ENSMUST00000102790.3/ENSMUSG00000035407.8, intron 1 of 9) | 4 | 98793768 | 98798900 | 5133 | 1 | ENSMUST00000133455.1 | 9340 | Kank4os | ENSMUSG00000085843.1 |
| chr5 | 77158238 | 77158544 | 3.58352 | 4.28501 | 1.29738 | Distal Intergenic | 5 | 77116810 | 77122348 | 5539 | 2 | ENSMUST00000156195.1 | -35891 | Gm15831 | ENSMUSG00000085858.1 |
| chr11 | 97612852 | 97613070 | 3.54398 | 4.20373 | 1.22066 | Intron (ENSMUST00000131687.1/ENSMUSG00000085860.1, intron 1 of 3) | 11 | 97614982 | 97619031 | 4050 | 1 | ENSMUST00000145262.1 | -1912 | 2410003L11Rik | ENSMUSG00000085860.1 |
| chr11 | 97617952 | 97618182 | 3.37544 | 3.92843 | 1.14631 | Intron (ENSMUST00000131687.1/ENSMUSG00000085860.1, intron 2 of 3) | 11 | 97614982 | 97619031 | 4050 | 1 | ENSMUST00000145262.1 | 2971 | 2410003L11Rik | ENSMUSG00000085860.1 |
| chr11 | 107510789 | 107510864 | 3.27196 | 3.73507 | 1.01733 | Downstream (<1kb) | 11 | 107511403 | 107527220 | 15818 | 2 | ENSMUST00000149428.1 | 16356 | C330019F10Rik | ENSMUSG00000085864.1 |
| chr16 | 91305716 | 91305845 | 3.34765 | 3.84892 | 1.01733 | Distal Intergenic | 16 | 91320677 | 91325740 | 5064 | 2 | ENSMUST00000146994.1 | 19895 | Gm15966 | ENSMUSG00000085865.1 |
| chr16 | 91308374 | 91308697 | 3.46286 | 4.0524 | 1.11926 | Distal Intergenic | 16 | 91320677 | 91325740 | 5064 | 2 | ENSMUST00000146994.1 | 17043 | Gm15966 | ENSMUSG00000085865.1 |
| chr16 | 91309656 | 91309844 | 3.37076 | 3.88374 | 1.01733 | Distal Intergenic | 16 | 91320677 | 91325740 | 5064 | 2 | ENSMUST00000146994.1 | 15896 | Gm15966 | ENSMUSG00000085865.1 |
| chr16 | 91310670 | 91310956 | 3.91975 | 4.92576 | 1.70498 | Distal Intergenic | 16 | 91320677 | 91325740 | 5064 | 2 | ENSMUST00000146994.1 | 14784 | Gm15966 | ENSMUSG00000085865.1 |
| chr16 | 91312183 | 91312667 | 3.38193 | 3.9006 | 1.01733 | Distal Intergenic | 16 | 91320677 | 91325740 | 5064 | 2 | ENSMUST00000146994.1 | 13073 | Gm15966 | ENSMUSG00000085865.1 |
| chr16 | 91318297 | 91318670 | 3.54415 | 4.21423 | 1.25661 | Downstream (1-2kb) | 16 | 91320677 | 91325740 | 5064 | 2 | ENSMUST00000146994.1 | 7070 | Gm15966 | ENSMUSG00000085865.1 |
| chr16 | 91320486 | 91320678 | 3.9992 | 5.13467 | 1.86356 | Exon (ENSMUST00000136853.1/ENSMUSG00000085865.1, exon 2 of 2) | 16 | 91320677 | 91325740 | 5064 | 2 | ENSMUST00000146994.1 | 5062 | Gm15966 | ENSMUSG00000085865.1 |
| chr16 | 91321463 | 91321716 | 3.3695 | 3.88185 | 1.01733 | Intron (ENSMUST00000136853.1/ENSMUSG00000085865.1, intron 1 of 1) | 16 | 91320677 | 91325740 | 5064 | 2 | ENSMUST00000146994.1 | 4024 | Gm15966 | ENSMUSG00000085865.1 |
| chr16 | 91325022 | 91325097 | 3.49649 | 4.12596 | 1.20233 | Promoter | 16 | 91320677 | 91325740 | 5064 | 2 | ENSMUST00000146994.1 | 643 | Gm15966 | ENSMUSG00000085865.1 |
| chr16 | 91332895 | 91332989 | 3.56448 | 4.24279 | 1.25378 | Distal Intergenic | 16 | 91320452 | 91328995 | 8544 | 2 | ENSMUST00000136853.1 | -3901 | Gm15966 | ENSMUSG00000085865.1 |
| chr16 | 91334351 | 91334731 | 3.91729 | 4.91004 | 1.67681 | Distal Intergenic | 16 | 91320452 | 91328995 | 8544 | 2 | ENSMUST00000136853.1 | -5357 | Gm15966 | ENSMUSG00000085865.1 |
| chr2 | 70888548 | 70888807 | 3.60308 | 4.37741 | 1.3366 | Distal Intergenic | 2 | 70877829 | 70884408 | 6580 | 1 | ENSMUST00000150119.1 | 10720 | Gm13632 | ENSMUSG00000085868.1 |
| chr4 | 45770458 | 45770633 | 3.46178 | 4.17945 | 1.26834 | Distal Intergenic | 4 | 45762377 | 45764223 | 1847 | 2 | ENSMUST00000147231.1 | -6236 | Gm12409 | ENSMUSG00000085876.1 |
| chr13 | 54911046 | 54911312 | 3.30713 | 3.8076 | 1.06897 | Distal Intergenic | 13 | 54913235 | 54914479 | 1245 | 1 | ENSMUST00000154615.2 | -1923 | 5330429C05Rik | ENSMUSG00000085896.2 |
| chr2 | 172328981 | 172329168 | 3.98534 | 5.08805 | 1.87511 | Intron (ENSMUST00000136315.1/ENSMUSG00000085905.1, intron 1 of 4) | 2 | 172324334 | 172338663 | 14330 | 1 | ENSMUST00000136315.1 | 4648 | Gm14273 | ENSMUSG00000085905.1 |
| chr17 | 29078437 | 29078622 | 3.32736 | 3.96342 | 1.0718 | Promoter | 17 | 29057474 | 29078961 | 21488 | 2 | ENSMUST00000133221.1 | 339 | Trp53cor1 | ENSMUSG00000085912.1 |
| chr5 | 100328323 | 100328765 | 3.38637 | 3.93328 | 1.10514 | Intron (ENSMUST00000150506.1/ENSMUSG00000085915.1, intron 1 of 4) | 5 | 100220440 | 100355119 | 134680 | 2 | ENSMUST00000150506.1 | 26354 | Gm8091 | ENSMUSG00000085915.1 |
| chr8 | 122686162 | 122686274 | 3.30434 | 3.81752 | 1.04729 | Intron (ENSMUST00000127664.1/ENSMUSG00000092329.1, intron 1 of 15) | 8 | 122683602 | 122683839 | 238 | 2 | ENSMUST00000128586.1 | -2324 | Gm15899 | ENSMUSG00000085917.1 |
| chr5 | 111743026 | 111743383 | 3.4234 | 3.97972 | 1.07558 | Promoter | 5 | 111742849 | 111748176 | 5328 | 1 | ENSMUST00000149538.1 | 178 | Gm16019 | ENSMUSG00000085919.1 |
| chr12 | 109598016 | 109598389 | 3.46811 | 4.0592 | 1.11793 | Intron (ENSMUST00000149046.2/ENSMUSG00000085925.2, intron 1 of 1) | 12 | 109589194 | 109600330 | 11137 | 2 | ENSMUST00000149046.2 | 1941 | Rtl1 | ENSMUSG00000085925.2 |
| chr12 | 109598972 | 109599096 | 3.34759 | 3.8488 | 1.01733 | Intron (ENSMUST00000149046.2/ENSMUSG00000085925.2, intron 1 of 1) | 12 | 109589194 | 109600330 | 11137 | 2 | ENSMUST00000149046.2 | 1234 | Rtl1 | ENSMUSG00000085925.2 |
| chr4 | 139933285 | 139933657 | 3.91327 | 5.15013 | 1.8955 | Promoter | 4 | 139926683 | 139933074 | 6392 | 2 | ENSMUST00000138672.1 | -212 | 4933427I22Rik | ENSMUSG00000085928.1 |
| chr4 | 139934604 | 139934886 | 3.43595 | 4.00314 | 1.09126 | Intron (ENSMUST00000153921.1/ENSMUSG00000085928.1, intron 2 of 2) | 4 | 139926683 | 139933074 | 6392 | 2 | ENSMUST00000138672.1 | -1531 | 4933427I22Rik | ENSMUSG00000085928.1 |
| chr4 | 139948679 | 139948988 | 3.41737 | 3.96952 | 1.07163 | Distal Intergenic | 4 | 139926111 | 139943519 | 17409 | 2 | ENSMUST00000153921.1 | -5161 | 4933427I22Rik | ENSMUSG00000085928.1 |
| chr2 | 27559356 | 27559610 | 3.48754 | 4.1025 | 1.16711 | Distal Intergenic | 2 | 27540429 | 27543185 | 2757 | 1 | ENSMUST00000132898.1 | 18928 | Gm13421 | ENSMUSG00000085929.1 |
| chr2 | 27560759 | 27560888 | 3.64282 | 4.37625 | 1.30565 | Distal Intergenic | 2 | 27540429 | 27543185 | 2757 | 1 | ENSMUST00000132898.1 | 20331 | Gm13421 | ENSMUSG00000085929.1 |
| chr2 | 27562350 | 27562775 | 3.82411 | 4.77369 | 1.6155 | Distal Intergenic | 2 | 27540429 | 27543185 | 2757 | 1 | ENSMUST00000132898.1 | 21922 | Gm13421 | ENSMUSG00000085929.1 |
| chr2 | 27565261 | 27565763 | 3.57701 | 4.26526 | 1.25566 | Distal Intergenic | 2 | 27540429 | 27543185 | 2757 | 1 | ENSMUST00000132898.1 | 24833 | Gm13421 | ENSMUSG00000085929.1 |
| chr2 | 27573386 | 27573774 | 3.64064 | 4.57919 | 1.50814 | Distal Intergenic | 2 | 27540429 | 27543185 | 2757 | 1 | ENSMUST00000132898.1 | 32958 | Gm13421 | ENSMUSG00000085929.1 |
| chr2 | 27578936 | 27579043 | 3.66483 | 4.43258 | 1.37069 | Distal Intergenic | 2 | 27540429 | 27543185 | 2757 | 1 | ENSMUST00000132898.1 | 38508 | Gm13421 | ENSMUSG00000085929.1 |
| chr13 | 28885717 | 28886688 | 3.68617 | 4.54653 | 1.52111 | Promoter | 13 | 28460778 | 28885620 | 424843 | 2 | ENSMUST00000148885.2 | -98 | 2610307P16Rik | ENSMUSG00000085936.2 |
| chr2 | 152046785 | 152047348 | 3.71956 | 4.5306 | 1.42009 | Distal Intergenic | 2 | 152074855 | 152076808 | 1954 | 2 | ENSMUST00000147463.1 | 29460 | AA387200 | ENSMUSG00000085946.1 |
| chr2 | 152048255 | 152050842 | 3.93648 | 4.99312 | 1.75144 | Distal Intergenic | 2 | 152074855 | 152076808 | 1954 | 2 | ENSMUST00000147463.1 | 25966 | AA387200 | ENSMUSG00000085946.1 |
| chr2 | 152051705 | 152052186 | 3.64886 | 4.39506 | 1.3337 | Distal Intergenic | 2 | 152074855 | 152076808 | 1954 | 2 | ENSMUST00000147463.1 | 24622 | AA387200 | ENSMUSG00000085946.1 |
| chr2 | 38413116 | 38413259 | 3.43148 | 3.99094 | 1.07209 | Distal Intergenic | 2 | 38424002 | 38455851 | 31850 | 1 | ENSMUST00000146404.1 | -10743 | Gm13589 | ENSMUSG00000085950.1 |
| chr2 | 38423952 | 38424259 | 3.75014 | 4.62485 | 1.57035 | Promoter | 2 | 38424002 | 38455851 | 31850 | 1 | ENSMUST00000146404.1 | 0 | Gm13589 | ENSMUSG00000085950.1 |
| chr4 | 140058050 | 140058333 | 4.06148 | 5.24364 | 1.90417 | Intron (ENSMUST00000039331.8/ENSMUSG00000040972.8, intron 4 of 9) | 4 | 140131877 | 140133351 | 1475 | 1 | ENSMUST00000135858.1 | -73544 | Gm13027 | ENSMUSG00000085968.1 |
| chr4 | 140068645 | 140069676 | 3.45461 | 4.04082 | 1.12535 | Intron (ENSMUST00000039331.8/ENSMUSG00000040972.8, intron 3 of 9) | 4 | 140131877 | 140133351 | 1475 | 1 | ENSMUST00000135858.1 | -62201 | Gm13027 | ENSMUSG00000085968.1 |
| chr4 | 140103644 | 140103924 | 3.65108 | 4.52814 | 1.46566 | Intron (ENSMUST00000039331.8/ENSMUSG00000040972.8, intron 3 of 9) | 4 | 140131877 | 140133351 | 1475 | 1 | ENSMUST00000135858.1 | -27953 | Gm13027 | ENSMUSG00000085968.1 |
| chr4 | 140107978 | 140108196 | 3.71356 | 4.51935 | 1.42489 | Intron (ENSMUST00000039331.8/ENSMUSG00000040972.8, intron 2 of 9) | 4 | 140131877 | 140133351 | 1475 | 1 | ENSMUST00000135858.1 | -23681 | Gm13027 | ENSMUSG00000085968.1 |
| chr4 | 140108708 | 140109074 | 3.37095 | 3.89308 | 1.04911 | Intron (ENSMUST00000039331.8/ENSMUSG00000040972.8, intron 2 of 9) | 4 | 140131877 | 140133351 | 1475 | 1 | ENSMUST00000135858.1 | -22803 | Gm13027 | ENSMUSG00000085968.1 |
| chr4 | 140109744 | 140110250 | 3.55419 | 4.44423 | 1.42206 | Intron (ENSMUST00000039331.8/ENSMUSG00000040972.8, intron 2 of 9) | 4 | 140131877 | 140133351 | 1475 | 1 | ENSMUST00000135858.1 | -21627 | Gm13027 | ENSMUSG00000085968.1 |
| chr11 | 87710449 | 87710831 | 3.70212 | 4.50847 | 1.43136 | Intron (ENSMUST00000134684.2/ENSMUSG00000034177.10, intron 1 of 5) | 11 | 87702123 | 87704989 | 2867 | 1 | ENSMUST00000142895.1 | 8327 | 1110028F11Rik | ENSMUSG00000085972.1 |
| chr4 | 45613729 | 45613963 | 3.26364 | 3.72259 | 1.01733 | Distal Intergenic | 4 | 45639936 | 45642304 | 2369 | 1 | ENSMUST00000151883.1 | -25973 | Gm12408 | ENSMUSG00000085980.1 |
| chr4 | 45654653 | 45654861 | 3.22133 | 3.84527 | 1.03772 | Distal Intergenic | 4 | 45639936 | 45642304 | 2369 | 1 | ENSMUST00000151883.1 | 14718 | Gm12408 | ENSMUSG00000085980.1 |
| chr11 | 110605400 | 110605576 | 3.71467 | 4.68083 | 1.5966 | Distal Intergenic | 11 | 110544507 | 110546399 | 1893 | 1 | ENSMUST00000144689.1 | 60894 | Gm11682 | ENSMUSG00000085985.1 |
| chr2 | 33922671 | 33922998 | 3.62409 | 4.40193 | 1.42478 | Distal Intergenic | 2 | 33938758 | 33942111 | 3354 | 2 | ENSMUST00000130988.1 | 19113 | Gm13403 | ENSMUSG00000085987.1 |
| chr11 | 95201724 | 95202742 | 4.04535 | 5.19367 | 1.90814 | Distal Intergenic | 11 | 95223290 | 95225632 | 2343 | 2 | ENSMUST00000150818.1 | 22890 | Gm11515 | ENSMUSG00000085992.1 |
| chr11 | 95210314 | 95210680 | 3.57821 | 4.26494 | 1.25468 | Distal Intergenic | 11 | 95223290 | 95225632 | 2343 | 2 | ENSMUST00000150818.1 | 14952 | Gm11515 | ENSMUSG00000085992.1 |
| chr11 | 95211383 | 95211745 | 3.34901 | 3.85094 | 1.01733 | Distal Intergenic | 11 | 95223290 | 95225632 | 2343 | 2 | ENSMUST00000150818.1 | 13887 | Gm11515 | ENSMUSG00000085992.1 |
| chr11 | 95214051 | 95214609 | 3.37519 | 3.89268 | 1.0253 | Distal Intergenic | 11 | 95223290 | 95225632 | 2343 | 2 | ENSMUST00000150818.1 | 11023 | Gm11515 | ENSMUSG00000085992.1 |
| chr11 | 95219441 | 95219815 | 3.88742 | 4.86489 | 1.69083 | Distal Intergenic | 11 | 95223290 | 95225632 | 2343 | 2 | ENSMUST00000150818.1 | 5817 | Gm11515 | ENSMUSG00000085992.1 |
| chr11 | 95222796 | 95223748 | 4.10402 | 5.4403 | 2.10181 | Exon (ENSMUST00000150818.1/ENSMUSG00000085992.1, exon 2 of 2) | 11 | 95223290 | 95225632 | 2343 | 2 | ENSMUST00000150818.1 | 1884 | Gm11515 | ENSMUSG00000085992.1 |
| chr4 | 4136354 | 4136826 | 3.88254 | 4.84262 | 1.63316 | Intron (ENSMUST00000070375.7/ENSMUSG00000045573.9, intron 1 of 1) | 4 | 4138321 | 4188703 | 50383 | 1 | ENSMUST00000131498.1 | -1495 | A830012C17Rik | ENSMUSG00000085996.1 |
| chr12 | 11455589 | 11456690 | 3.87983 | 4.97085 | 1.78638 | Promoter | 12 | 11456079 | 11462928 | 6850 | 1 | ENSMUST00000124065.1 | 0 | Rad51ap2 | ENSMUSG00000086022.1 |
| chr11 | 19050388 | 19050477 | 3.40435 | 3.98438 | 1.18381 | Distal Intergenic | 11 | 19065276 | 19075740 | 10465 | 1 | ENSMUST00000144854.1 | -14799 | 4933406G16Rik | ENSMUSG00000086025.1 |
| chr2 | 163178350 | 163178594 | 3.5809 | 4.29935 | 1.34561 | Distal Intergenic | 2 | 163152895 | 163156106 | 3212 | 2 | ENSMUST00000136371.1 | -22245 | Gm14254 | ENSMUSG00000086036.1 |
| chr4 | 146537491 | 146537917 | 2.92058 | 5.5673 | 2.17739 | Exon (ENSMUST00000105735.3/ENSMUSG00000056300.9, exon 5 of 5) | 4 | 146519039 | 146546873 | 27835 | 2 | ENSMUST00000145269.1 | 8956 | 1700095A21Rik | ENSMUSG00000086046.1 |
| chr5 | 117046989 | 117047278 | 3.63327 | 4.41552 | 1.4313 | Distal Intergenic | 5 | 117115218 | 117118723 | 3506 | 1 | ENSMUST00000149413.1 | -67940 | 9530046B11Rik | ENSMUSG00000086047.1 |
| chr5 | 117088186 | 117088778 | 3.45468 | 4.10903 | 1.23645 | Downstream (2-3kb) | 5 | 117115218 | 117118723 | 3506 | 1 | ENSMUST00000149413.1 | -26440 | 9530046B11Rik | ENSMUSG00000086047.1 |
| chr5 | 114968252 | 114968543 | 3.46024 | 4.07179 | 1.20616 | Promoter | 5 | 114968784 | 114997849 | 29066 | 1 | ENSMUST00000129609.1 | -241 | Hnf1aos1 | ENSMUSG00000086054.2 |
| chr11 | 82930204 | 82933144 | 4.46254 | 6.17926 | 2.70536 | Promoter | 11 | 82924903 | 82933382 | 8480 | 2 | ENSMUST00000156395.1 | 238 | Unc45bos | ENSMUSG00000086058.1 |
| chr8 | 84617034 | 84617412 | 3.5206 | 4.15757 | 1.18594 | Promoter | 8 | 84611202 | 84617184 | 5983 | 2 | ENSMUST00000127527.1 | 0 | Gm16183 | ENSMUSG00000086067.1 |
| chr4 | 124143170 | 124143414 | 3.47562 | 4.11826 | 1.19561 | Distal Intergenic | 4 | 124293693 | 124299848 | 6156 | 2 | ENSMUST00000139012.1 | 156434 | Gm12916 | ENSMUSG00000086087.1 |
| chr4 | 124144841 | 124145165 | 3.55353 | 4.3062 | 1.34494 | Distal Intergenic | 4 | 124293693 | 124299848 | 6156 | 2 | ENSMUST00000139012.1 | 154683 | Gm12916 | ENSMUSG00000086087.1 |
| chr4 | 124145788 | 124146690 | 3.53442 | 4.24439 | 1.3231 | Distal Intergenic | 4 | 124293693 | 124299848 | 6156 | 2 | ENSMUST00000139012.1 | 153158 | Gm12916 | ENSMUSG00000086087.1 |
| chr4 | 124147514 | 124147663 | 3.6986 | 4.50441 | 1.44125 | Distal Intergenic | 4 | 124293693 | 124299848 | 6156 | 2 | ENSMUST00000139012.1 | 152185 | Gm12916 | ENSMUSG00000086087.1 |
| chr4 | 124149986 | 124150243 | 3.60425 | 4.32206 | 1.31602 | Distal Intergenic | 4 | 124293693 | 124299848 | 6156 | 2 | ENSMUST00000139012.1 | 149605 | Gm12916 | ENSMUSG00000086087.1 |
| chr4 | 124176073 | 124176148 | 4.08648 | 5.23153 | 1.89841 | Distal Intergenic | 4 | 124293693 | 124299848 | 6156 | 2 | ENSMUST00000139012.1 | 123700 | Gm12916 | ENSMUSG00000086087.1 |
| chr4 | 124177196 | 124177354 | 3.91738 | 4.97963 | 1.75167 | Distal Intergenic | 4 | 124293693 | 124299848 | 6156 | 2 | ENSMUST00000139012.1 | 122494 | Gm12916 | ENSMUSG00000086087.1 |
| chr4 | 124177950 | 124178025 | 3.35685 | 3.86277 | 1.01733 | Distal Intergenic | 4 | 124293693 | 124299848 | 6156 | 2 | ENSMUST00000139012.1 | 121823 | Gm12916 | ENSMUSG00000086087.1 |
| chr4 | 124178460 | 124179682 | 3.6363 | 4.55798 | 1.46481 | Distal Intergenic | 4 | 124293693 | 124299848 | 6156 | 2 | ENSMUST00000139012.1 | 120166 | Gm12916 | ENSMUSG00000086087.1 |
| chr4 | 124182105 | 124182240 | 3.36825 | 3.98431 | 1.13727 | Distal Intergenic | 4 | 124293693 | 124299848 | 6156 | 2 | ENSMUST00000139012.1 | 117608 | Gm12916 | ENSMUSG00000086087.1 |
| chr4 | 124188911 | 124189521 | 4.0088 | 5.10376 | 1.83569 | Distal Intergenic | 4 | 124293693 | 124299848 | 6156 | 2 | ENSMUST00000139012.1 | 110327 | Gm12916 | ENSMUSG00000086087.1 |
| chr4 | 124220864 | 124221285 | 3.71293 | 4.69584 | 1.56441 | Distal Intergenic | 4 | 124293693 | 124299848 | 6156 | 2 | ENSMUST00000139012.1 | 78563 | Gm12916 | ENSMUSG00000086087.1 |
| chr4 | 124221896 | 124222111 | 3.31858 | 3.80511 | 1.01733 | Distal Intergenic | 4 | 124293693 | 124299848 | 6156 | 2 | ENSMUST00000139012.1 | 77737 | Gm12916 | ENSMUSG00000086087.1 |
| chr4 | 124241506 | 124241675 | 3.66297 | 4.52621 | 1.48238 | Distal Intergenic | 4 | 124293693 | 124299848 | 6156 | 2 | ENSMUST00000139012.1 | 58173 | Gm12916 | ENSMUSG00000086087.1 |
| chr4 | 124253062 | 124253260 | 3.40875 | 4.10607 | 1.1922 | Distal Intergenic | 4 | 124293693 | 124299848 | 6156 | 2 | ENSMUST00000139012.1 | 46588 | Gm12916 | ENSMUSG00000086087.1 |
| chr4 | 124270322 | 124270510 | 3.72182 | 4.54293 | 1.46099 | Distal Intergenic | 4 | 124293693 | 124299848 | 6156 | 2 | ENSMUST00000139012.1 | 29338 | Gm12916 | ENSMUSG00000086087.1 |
| chr4 | 124271276 | 124271406 | 3.35167 | 3.86102 | 1.03869 | Distal Intergenic | 4 | 124293693 | 124299848 | 6156 | 2 | ENSMUST00000139012.1 | 28442 | Gm12916 | ENSMUSG00000086087.1 |
| chr4 | 124276817 | 124277152 | 3.42527 | 4.14263 | 1.19254 | Distal Intergenic | 4 | 124293693 | 124299848 | 6156 | 2 | ENSMUST00000139012.1 | 22696 | Gm12916 | ENSMUSG00000086087.1 |
| chr4 | 124280277 | 124280352 | 3.84376 | 4.77863 | 1.62252 | Distal Intergenic | 4 | 124293693 | 124299848 | 6156 | 2 | ENSMUST00000139012.1 | 19496 | Gm12916 | ENSMUSG00000086087.1 |
| chr4 | 124301339 | 124301656 | 3.69126 | 4.70864 | 1.5549 | Downstream (1-2kb) | 4 | 124293693 | 124299848 | 6156 | 2 | ENSMUST00000139012.1 | -1492 | Gm12916 | ENSMUSG00000086087.1 |
| chr4 | 99636568 | 99636780 | 3.53314 | 4.19175 | 1.24046 | Distal Intergenic | 4 | 99653607 | 99654983 | 1377 | 2 | ENSMUST00000136525.1 | 18203 | Gm12688 | ENSMUSG00000086096.1 |
| chr4 | 99640785 | 99640938 | 3.94535 | 5.04679 | 1.83867 | Distal Intergenic | 4 | 99653607 | 99654983 | 1377 | 2 | ENSMUST00000136525.1 | 14045 | Gm12688 | ENSMUSG00000086096.1 |
| chr4 | 99641286 | 99641695 | 3.57112 | 4.37717 | 1.39844 | Distal Intergenic | 4 | 99653607 | 99654983 | 1377 | 2 | ENSMUST00000136525.1 | 13288 | Gm12688 | ENSMUSG00000086096.1 |
| chr4 | 99643520 | 99643950 | 3.34205 | 3.8502 | 1.04386 | Distal Intergenic | 4 | 99653607 | 99654983 | 1377 | 2 | ENSMUST00000136525.1 | 11033 | Gm12688 | ENSMUSG00000086096.1 |
| chr4 | 99646058 | 99646394 | 3.44733 | 4.03354 | 1.1274 | Distal Intergenic | 4 | 99653607 | 99654983 | 1377 | 2 | ENSMUST00000136525.1 | 8589 | Gm12688 | ENSMUSG00000086096.1 |
| chr4 | 99648144 | 99652922 | 4.35248 | 5.86073 | 2.42769 | Downstream (<1kb) | 4 | 99653607 | 99654983 | 1377 | 2 | ENSMUST00000136525.1 | 2061 | Gm12688 | ENSMUSG00000086096.1 |
| chr4 | 99653312 | 99654199 | 4.22809 | 5.83226 | 2.37161 | Promoter | 4 | 99653607 | 99654983 | 1377 | 2 | ENSMUST00000136525.1 | 784 | Gm12688 | ENSMUSG00000086096.1 |
| chr13 | 54838742 | 54839244 | 3.91721 | 4.96456 | 1.77194 | Downstream (1-2kb) | 13 | 54840649 | 54840904 | 256 | 2 | ENSMUST00000148974.1 | 1660 | Gm16250 | ENSMUSG00000086097.1 |
| chr13 | 54839960 | 54840055 | 3.74075 | 4.58001 | 1.48789 | Promoter | 13 | 54840649 | 54840904 | 256 | 2 | ENSMUST00000148974.1 | 849 | Gm16250 | ENSMUSG00000086097.1 |
| chr2 | 6505159 | 6505434 | 3.45047 | 4.04547 | 1.13075 | Distal Intergenic | 2 | 6472272 | 6472903 | 632 | 1 | ENSMUST00000128589.1 | 32888 | Gm13391 | ENSMUSG00000086109.1 |
| chr11 | 24282165 | 24283008 | 3.66322 | 4.43915 | 1.40051 | Distal Intergenic | 11 | 24198737 | 24202624 | 3888 | 2 | ENSMUST00000149592.1 | -79542 | Gm12065 | ENSMUSG00000086117.1 |
| chr11 | 24285404 | 24285500 | 3.7886 | 4.67411 | 1.54118 | Distal Intergenic | 11 | 24198737 | 24202624 | 3888 | 2 | ENSMUST00000149592.1 | -82781 | Gm12065 | ENSMUSG00000086117.1 |
| chr11 | 24287568 | 24287877 | 3.54396 | 4.19903 | 1.20662 | Distal Intergenic | 11 | 24198737 | 24202624 | 3888 | 2 | ENSMUST00000149592.1 | -84945 | Gm12065 | ENSMUSG00000086117.1 |
| chr11 | 24306001 | 24306229 | 3.64438 | 4.42747 | 1.42001 | Distal Intergenic | 11 | 24198737 | 24202624 | 3888 | 2 | ENSMUST00000149592.1 | -103378 | Gm12065 | ENSMUSG00000086117.1 |
| chr5 | 116193339 | 116193657 | 3.33109 | 3.83813 | 1.06739 | Intron (ENSMUST00000050178.7/ENSMUSG00000043913.9, intron 2 of 13) | 5 | 116243706 | 116254953 | 11248 | 1 | ENSMUST00000156169.1 | -50049 | Gm14507 | ENSMUSG00000086122.1 |
| chr5 | 116241156 | 116241445 | 3.68197 | 4.45722 | 1.37712 | Intron (ENSMUST00000050178.7/ENSMUSG00000043913.9, intron 1 of 13) | 5 | 116243706 | 116254953 | 11248 | 1 | ENSMUST00000156169.1 | -2261 | Gm14507 | ENSMUSG00000086122.1 |
| chr6 | 52314697 | 52317604 | 3.93089 | 5.11652 | 1.90919 | Promoter | 6 | 52308389 | 52314832 | 6444 | 2 | ENSMUST00000125305.1 | 0 | 5730457N03Rik | ENSMUSG00000086126.1 |
| chr6 | 52318013 | 52318139 | 3.98007 | 5.05593 | 1.80105 | 3' UTR | 6 | 52308389 | 52314832 | 6444 | 2 | ENSMUST00000125305.1 | -3182 | 5730457N03Rik | ENSMUSG00000086126.1 |
| chr6 | 52318568 | 52319382 | 3.50936 | 4.21458 | 1.24971 | Distal Intergenic | 6 | 52308389 | 52314832 | 6444 | 2 | ENSMUST00000125305.1 | -3737 | 5730457N03Rik | ENSMUSG00000086126.1 |
| chr6 | 52319802 | 52323910 | 4.06137 | 5.29171 | 1.99724 | Distal Intergenic | 6 | 52308389 | 52314832 | 6444 | 2 | ENSMUST00000125305.1 | -4971 | 5730457N03Rik | ENSMUSG00000086126.1 |
| chr6 | 52324883 | 52325412 | 3.61857 | 4.33862 | 1.29156 | Distal Intergenic | 6 | 52308389 | 52314832 | 6444 | 2 | ENSMUST00000125305.1 | -10052 | 5730457N03Rik | ENSMUSG00000086126.1 |
| chr6 | 52326081 | 52326311 | 3.50398 | 4.12837 | 1.17135 | Distal Intergenic | 6 | 52308389 | 52314832 | 6444 | 2 | ENSMUST00000125305.1 | -11250 | 5730457N03Rik | ENSMUSG00000086126.1 |
| chr6 | 52328892 | 52329038 | 3.37261 | 3.88654 | 1.01733 | Distal Intergenic | 6 | 52308389 | 52314832 | 6444 | 2 | ENSMUST00000125305.1 | -14061 | 5730457N03Rik | ENSMUSG00000086126.1 |
| chr6 | 52329701 | 52329814 | 4.10895 | 5.29381 | 1.95842 | Distal Intergenic | 6 | 52308389 | 52314832 | 6444 | 2 | ENSMUST00000125305.1 | -14870 | 5730457N03Rik | ENSMUSG00000086126.1 |
| chr11 | 114167713 | 114167913 | 3.52454 | 4.17307 | 1.22022 | Distal Intergenic | 11 | 114185973 | 114186674 | 702 | 2 | ENSMUST00000126577.1 | 18761 | 1700001J04Rik | ENSMUSG00000086135.1 |
| chr13 | 54386367 | 54386639 | 3.73136 | 4.63375 | 1.51685 | Distal Intergenic | 13 | 54399092 | 54399334 | 243 | 2 | ENSMUST00000124238.1 | 12695 | Gm16248 | ENSMUSG00000086137.1 |
| chr5 | 114968911 | 114969611 | 3.51768 | 4.16366 | 1.22256 | Promoter | 5 | 114968823 | 114971400 | 2578 | 1 | ENSMUST00000138108.1 | 89 | Hnf1aos2 | ENSMUSG00000086140.1 |
| chr2 | 147968512 | 147968964 | 3.63609 | 4.38023 | 1.35444 | Intron (ENSMUST00000129082.1/ENSMUSG00000086141.2, intron 2 of 6) | 2 | 147944740 | 147976185 | 31446 | 2 | ENSMUST00000129082.1 | 7221 | 9030622O22Rik | ENSMUSG00000086141.2 |
| chr2 | 147989356 | 147989510 | 3.52353 | 4.34873 | 1.37933 | Intron (ENSMUST00000136555.2/ENSMUSG00000086141.2, intron 1 of 2) | 2 | 147944740 | 147976185 | 31446 | 2 | ENSMUST00000129082.1 | -13172 | 9030622O22Rik | ENSMUSG00000086141.2 |
| chr2 | 148017596 | 148018416 | 3.59728 | 4.38712 | 1.41994 | Intron (ENSMUST00000136555.2/ENSMUSG00000086141.2, intron 1 of 2) | 2 | 147972830 | 148038270 | 65441 | 2 | ENSMUST00000132070.1 | 19854 | 9030622O22Rik | ENSMUSG00000086141.2 |
| chr2 | 148018861 | 148018936 | 3.55492 | 4.30828 | 1.33922 | Intron (ENSMUST00000136555.2/ENSMUSG00000086141.2, intron 1 of 2) | 2 | 147972830 | 148038270 | 65441 | 2 | ENSMUST00000132070.1 | 19334 | 9030622O22Rik | ENSMUSG00000086141.2 |
| chr2 | 148019770 | 148020306 | 3.78915 | 4.79166 | 1.64587 | Intron (ENSMUST00000136555.2/ENSMUSG00000086141.2, intron 1 of 2) | 2 | 147972830 | 148038270 | 65441 | 2 | ENSMUST00000132070.1 | 17964 | 9030622O22Rik | ENSMUSG00000086141.2 |
| chr2 | 148021099 | 148021633 | 3.51538 | 4.20959 | 1.33456 | Intron (ENSMUST00000136555.2/ENSMUSG00000086141.2, intron 1 of 2) | 2 | 147972830 | 148038270 | 65441 | 2 | ENSMUST00000132070.1 | 16637 | 9030622O22Rik | ENSMUSG00000086141.2 |
| chr2 | 148022427 | 148023021 | 3.32456 | 3.99682 | 1.13902 | Intron (ENSMUST00000136555.2/ENSMUSG00000086141.2, intron 1 of 2) | 2 | 147972830 | 148038270 | 65441 | 2 | ENSMUST00000132070.1 | 15249 | 9030622O22Rik | ENSMUSG00000086141.2 |
| chr2 | 148023898 | 148024284 | 3.44835 | 4.1467 | 1.22745 | Intron (ENSMUST00000136555.2/ENSMUSG00000086141.2, intron 1 of 2) | 2 | 147972830 | 148038270 | 65441 | 2 | ENSMUST00000132070.1 | 13986 | 9030622O22Rik | ENSMUSG00000086141.2 |
| chr2 | 148024685 | 148024760 | 3.42781 | 4.03186 | 1.20654 | Intron (ENSMUST00000136555.2/ENSMUSG00000086141.2, intron 1 of 2) | 2 | 147972830 | 148038270 | 65441 | 2 | ENSMUST00000132070.1 | 13510 | 9030622O22Rik | ENSMUSG00000086141.2 |
| chr2 | 148025840 | 148026068 | 3.23007 | 3.8915 | 1.09558 | Intron (ENSMUST00000136555.2/ENSMUSG00000086141.2, intron 1 of 2) | 2 | 147972830 | 148038270 | 65441 | 2 | ENSMUST00000132070.1 | 12202 | 9030622O22Rik | ENSMUSG00000086141.2 |
| chr2 | 148026401 | 148026705 | 4.05906 | 5.24258 | 1.9896 | Intron (ENSMUST00000136555.2/ENSMUSG00000086141.2, intron 1 of 2) | 2 | 147972830 | 148038270 | 65441 | 2 | ENSMUST00000132070.1 | 11565 | 9030622O22Rik | ENSMUSG00000086141.2 |
| chr2 | 148027390 | 148028328 | 3.57965 | 4.31194 | 1.34128 | Intron (ENSMUST00000136555.2/ENSMUSG00000086141.2, intron 1 of 2) | 2 | 147972830 | 148038270 | 65441 | 2 | ENSMUST00000132070.1 | 9942 | 9030622O22Rik | ENSMUSG00000086141.2 |
| chr2 | 148028676 | 148028896 | 3.44057 | 4.18105 | 1.2361 | Intron (ENSMUST00000136555.2/ENSMUSG00000086141.2, intron 1 of 2) | 2 | 147972830 | 148038270 | 65441 | 2 | ENSMUST00000132070.1 | 9374 | 9030622O22Rik | ENSMUSG00000086141.2 |
| chr2 | 148029751 | 148030146 | 3.43528 | 4.28932 | 1.32253 | Intron (ENSMUST00000136555.2/ENSMUSG00000086141.2, intron 1 of 2) | 2 | 147972830 | 148038270 | 65441 | 2 | ENSMUST00000132070.1 | 8124 | 9030622O22Rik | ENSMUSG00000086141.2 |
| chr2 | 148030575 | 148031068 | 4.1316 | 5.38077 | 2.05752 | Intron (ENSMUST00000136555.2/ENSMUSG00000086141.2, intron 1 of 2) | 2 | 147972830 | 148038270 | 65441 | 2 | ENSMUST00000132070.1 | 7202 | 9030622O22Rik | ENSMUSG00000086141.2 |
| chr2 | 148031510 | 148031849 | 3.90475 | 5.23687 | 1.95289 | Intron (ENSMUST00000136555.2/ENSMUSG00000086141.2, intron 1 of 2) | 2 | 147972830 | 148038270 | 65441 | 2 | ENSMUST00000132070.1 | 6421 | 9030622O22Rik | ENSMUSG00000086141.2 |
| chr2 | 148032287 | 148032752 | 3.39846 | 3.96386 | 1.15251 | Intron (ENSMUST00000136555.2/ENSMUSG00000086141.2, intron 1 of 2) | 2 | 147972830 | 148038270 | 65441 | 2 | ENSMUST00000132070.1 | 5518 | 9030622O22Rik | ENSMUSG00000086141.2 |
| chr2 | 148033120 | 148033526 | 3.58856 | 4.32849 | 1.38276 | Intron (ENSMUST00000136555.2/ENSMUSG00000086141.2, intron 1 of 2) | 2 | 147972830 | 148038270 | 65441 | 2 | ENSMUST00000132070.1 | 4744 | 9030622O22Rik | ENSMUSG00000086141.2 |
| chr2 | 148034276 | 148035483 | 3.6495 | 4.49924 | 1.4501 | Intron (ENSMUST00000136555.2/ENSMUSG00000086141.2, intron 1 of 2) | 2 | 147972830 | 148038270 | 65441 | 2 | ENSMUST00000132070.1 | 2787 | 9030622O22Rik | ENSMUSG00000086141.2 |
| chr2 | 148036787 | 148038107 | 3.66306 | 4.52471 | 1.4896 | Promoter | 2 | 147972830 | 148038270 | 65441 | 2 | ENSMUST00000132070.1 | 163 | 9030622O22Rik | ENSMUSG00000086141.2 |
| chr2 | 148038484 | 148040913 | 4.2097 | 5.66434 | 2.31741 | Promoter | 2 | 147972676 | 148040203 | 67528 | 2 | ENSMUST00000136555.2 | 0 | 9030622O22Rik | ENSMUSG00000086141.2 |
| chr2 | 148041973 | 148042481 | 3.92724 | 5.0117 | 1.81641 | Downstream (<1kb) | 2 | 147972676 | 148040203 | 67528 | 2 | ENSMUST00000136555.2 | -1771 | 9030622O22Rik | ENSMUSG00000086141.2 |
| chr2 | 163119292 | 163119683 | 3.36606 | 3.9161 | 1.14486 | Distal Intergenic | 2 | 163122606 | 163143959 | 21354 | 1 | ENSMUST00000144092.1 | -2923 | Gm11454 | ENSMUSG00000086152.1 |
| chr11 | 102739556 | 102739749 | 3.3398 | 4.03219 | 1.12453 | Intron (ENSMUST00000124025.1/ENSMUSG00000086162.1, intron 4 of 10) | 11 | 102732405 | 102752861 | 20457 | 1 | ENSMUST00000124025.1 | 7152 | Gm16342 | ENSMUSG00000086162.1 |
| chr11 | 102741166 | 102741629 | 3.4569 | 4.03896 | 1.10632 | Exon (ENSMUST00000124025.1/ENSMUSG00000086162.1, exon 5 of 11) | 11 | 102732405 | 102752861 | 20457 | 1 | ENSMUST00000124025.1 | 8762 | Gm16342 | ENSMUSG00000086162.1 |
| chr4 | 140138932 | 140139101 | 3.30393 | 3.78309 | 1.01733 | Intron (ENSMUST00000039331.8/ENSMUSG00000040972.8, intron 2 of 9) | 4 | 140142873 | 140144633 | 1761 | 1 | ENSMUST00000137283.1 | -3772 | Gm13029 | ENSMUSG00000086164.1 |
| chr4 | 140143453 | 140144100 | 3.76613 | 4.70763 | 1.57877 | Promoter | 4 | 140142873 | 140144633 | 1761 | 1 | ENSMUST00000137283.1 | 581 | Gm13029 | ENSMUSG00000086164.1 |
| chr4 | 140144749 | 140145075 | 3.79547 | 4.704 | 1.59694 | Intron (ENSMUST00000039331.8/ENSMUSG00000040972.8, intron 2 of 9) | 4 | 140142873 | 140144633 | 1761 | 1 | ENSMUST00000137283.1 | 1877 | Gm13029 | ENSMUSG00000086164.1 |
| chr2 | 180888419 | 180888898 | 3.73673 | 4.60571 | 1.48193 | Promoter | 2 | 180829490 | 180889660 | 60171 | 2 | ENSMUST00000137629.1 | 762 | Gm14342 | ENSMUSG00000086166.1 |
| chr2 | 180889357 | 180889712 | 3.515 | 4.14709 | 1.17836 | Promoter | 2 | 180829490 | 180889660 | 60171 | 2 | ENSMUST00000137629.1 | 0 | Gm14342 | ENSMUSG00000086166.1 |
| chr2 | 180890301 | 180890437 | 3.50646 | 4.13372 | 1.1728 | Promoter | 2 | 180829490 | 180889660 | 60171 | 2 | ENSMUST00000137629.1 | -642 | Gm14342 | ENSMUSG00000086166.1 |

| | | | | | | | | | | | | | | | |
|---|---|---|---|---|---|---|---|---|---|---|---|---|---|---|---|
| chr2 | 180890882 | 180891249 | 3.46128 | 4.12001 | 1.24327 | Distal Intergenic | 2 | 180829490 | 180889660 | 60171 | 2 | ENSMUST00000137629.1 | -1223 | Gm14342 | ENSMUSG00000086166.1 |
| chr2 | 143544539 | 143544752 | 3.33795 | 3.83426 | 1.01733 | Intron (ENSMUST00000126735.2/ENSMUSG00000086171.2, intron 1 of 2) | 2 | 143537970 | 143545977 | 8008 | 2 | ENSMUST00000156780.1 | 1225 | Pcsk2os1 | ENSMUSG00000086171.2 |
| chr2 | 143547398 | 143547909 | 4.02534 | 5.24298 | 1.94927 | Promoter | 2 | 143535476 | 143547837 | 12362 | 2 | ENSMUST00000126735.2 | 0 | Pcsk2os1 | ENSMUSG00000086171.2 |
| chr2 | 143548755 | 143548865 | 3.53329 | 4.49838 | 1.4618 | Promoter | 2 | 143535476 | 143547837 | 12362 | 2 | ENSMUST00000126735.2 | -919 | Pcsk2os1 | ENSMUSG00000086171.2 |
| chr16 | 38856241 | 38856570 | 3.91436 | 4.97321 | 1.7865 | Distal Intergenic | 16 | 38828908 | 38829244 | 337 | 1 | ENSMUST00000148657.1 | 27334 | Gm15802 | ENSMUSG00000086175.1 |
| chr2 | 164550710 | 164551160 | 3.64107 | 4.42856 | 1.41364 | Promoter | 2 | 164550530 | 164559730 | 9201 | 1 | ENSMUST00000151754.1 | 181 | Gm14317 | ENSMUSG00000086179.1 |
| chr4 | 44636456 | 44636668 | 3.34376 | 3.84302 | 1.01733 | Intron (ENSMUST00000102932.4/ENSMUSG00000014030.10, intron 6 of 8) | 4 | 44664415 | 44685347 | 20933 | 1 | ENSMUST00000140826.1 | -27747 | Gm12462 | ENSMUSG00000086211.1 |
| chr4 | 44660830 | 44660935 | 3.44268 | 4.05478 | 1.22185 | Intron (ENSMUST00000102932.4/ENSMUSG00000014030.10, intron 5 of 8) | 4 | 44664415 | 44685347 | 20933 | 1 | ENSMUST00000140826.1 | -3480 | Gm12462 | ENSMUSG00000086211.1 |
| chr4 | 44661519 | 44663095 | 3.46195 | 4.11431 | 1.2464 | Intron (ENSMUST00000102932.4/ENSMUSG00000014030.10, intron 5 of 8) | 4 | 44664415 | 44685347 | 20933 | 1 | ENSMUST00000140826.1 | -1320 | Gm12462 | ENSMUSG00000086211.1 |
| chr4 | 44663635 | 44664752 | 3.99832 | 5.23602 | 1.94716 | Promoter | 4 | 44664415 | 44685347 | 20933 | 1 | ENSMUST00000140826.1 | 0 | Gm12462 | ENSMUSG00000086211.1 |
| chr4 | 44665348 | 44665742 | 3.50545 | 4.12721 | 1.1588 | Promoter | 4 | 44664415 | 44685347 | 20933 | 1 | ENSMUST00000140826.1 | 934 | Gm12462 | ENSMUSG00000086211.1 |
| chr3 | 5219310 | 5220101 | 4.67119 | 6.74614 | 3.18411 | Promoter | 3 | 5177586 | 5220823 | 43238 | 2 | ENSMUST00000128815.3 | 722 | 2700069I18Rik | ENSMUSG00000086224.3 |
| chr11 | 50852530 | 50853351 | 3.6373 | 4.41121 | 1.43626 | Promoter | 11 | 50851332 | 50852582 | 1251 | 2 | ENSMUST00000123764.1 | 0 | Gm12660 | ENSMUSG00000086226.1 |
| chr4 | 9296969 | 9297047 | 3.84785 | 4.89637 | 1.69796 | Intron (ENSMUST00000128117.2/ENSMUSG00000041216.10, intron 2 of 2) | 4 | 9283972 | 9287391 | 3420 | 2 | ENSMUST00000149848.1 | -9579 | Gm11816 | ENSMUSG00000086233.1 |
| chr2 | 169133490 | 169133661 | 3.49102 | 4.3693 | 1.38594 | Distal Intergenic | 2 | 169106453 | 169110415 | 3963 | 1 | ENSMUST00000144607.1 | 27038 | Gm14258 | ENSMUSG00000086238.1 |
| chr2 | 169134950 | 169135251 | 3.34989 | 3.85227 | 1.01733 | Distal Intergenic | 2 | 169106453 | 169110415 | 3963 | 1 | ENSMUST00000144607.1 | 28498 | Gm14258 | ENSMUSG00000086238.1 |
| chr2 | 169143967 | 169144309 | 3.43745 | 4.0064 | 1.09519 | Distal Intergenic | 2 | 169106453 | 169110415 | 3963 | 1 | ENSMUST00000144607.1 | 37515 | Gm14258 | ENSMUSG00000086238.1 |
| chr2 | 169155063 | 169155565 | 3.49428 | 4.1122 | 1.16535 | Distal Intergenic | 2 | 169106453 | 169110415 | 3963 | 1 | ENSMUST00000144607.1 | 48611 | Gm14258 | ENSMUSG00000086238.1 |
| chr2 | 169156223 | 169156551 | 3.492 | 4.1568 | 1.26359 | Distal Intergenic | 2 | 169106453 | 169110415 | 3963 | 1 | ENSMUST00000144607.1 | 49771 | Gm14258 | ENSMUSG00000086238.1 |
| chr2 | 169158452 | 169158676 | 3.45882 | 4.07294 | 1.20895 | Distal Intergenic | 2 | 169106453 | 169110415 | 3963 | 1 | ENSMUST00000144607.1 | 52000 | Gm14258 | ENSMUSG00000086238.1 |
| chr9 | 95639957 | 95640144 | 3.33835 | 3.84515 | 1.05333 | Intron (ENSMUST00000015498.8/ENSMUSG00000015354.8, intron 2 of 8) | 9 | 95636972 | 95638265 | 1294 | 2 | ENSMUST00000131619.2 | -1693 | Gm16127 | ENSMUSG00000086248.2 |
| chr4 | 131100956 | 131101068 | 3.72403 | 4.54104 | 1.43344 | Distal Intergenic | 4 | 131000832 | 131010474 | 9643 | 2 | ENSMUST00000153810.1 | -90483 | A930031H19Rik | ENSMUSG00000086262.2 |
| chr4 | 131108272 | 131108608 | 3.57873 | 4.30912 | 1.37716 | Distal Intergenic | 4 | 131000832 | 131010474 | 9643 | 2 | ENSMUST00000153810.1 | -97799 | A930031H19Rik | ENSMUSG00000086262.2 |
| chr7 | 142663417 | 142664497 | 3.6621 | 4.53298 | 1.45555 | Promoter | 7 | 142665138 | 142666092 | 955 | 1 | ENSMUST00000124393.1 | -641 | Igf2os | ENSMUSG00000086266.1 |
| chr7 | 142665253 | 142666238 | 3.67032 | 4.44429 | 1.38008 | Promoter | 7 | 142665138 | 142666092 | 955 | 1 | ENSMUST00000124393.1 | 116 | Igf2os | ENSMUSG00000086266.1 |
| chr17 | 13749727 | 13750103 | 3.83094 | 4.90189 | 1.71629 | Intron (ENSMUST00000135850.2/ENSMUSG00000038347.9, intron 1 of 4) | 17 | 13759370 | 13759985 | 616 | 2 | ENSMUST00000129681.1 | 9882 | Gm16052 | ENSMUSG00000086292.1 |
| chr17 | 13758633 | 13758708 | 4.1729 | 5.83191 | 2.35465 | Intron (ENSMUST00000135850.2/ENSMUSG00000038347.9, intron 1 of 4) | 17 | 13759370 | 13759985 | 616 | 2 | ENSMUST00000129681.1 | 1277 | Gm16052 | ENSMUSG00000086292.1 |
| chr17 | 13753648 | 13753754 | 3.75446 | 4.65838 | 1.57945 | Intron (ENSMUST00000135850.2/ENSMUSG00000038347.9, intron 1 of 4) | 17 | 13759370 | 13759985 | 616 | 2 | ENSMUST00000129681.1 | 6231 | Gm16052 | ENSMUSG00000086292.1 |
| chr17 | 13755000 | 13755111 | 3.5362 | 4.27995 | 1.32693 | Intron (ENSMUST00000135850.2/ENSMUSG00000038347.9, intron 1 of 4) | 17 | 13759370 | 13759985 | 616 | 2 | ENSMUST00000129681.1 | 4874 | Gm16052 | ENSMUSG00000086292.1 |
| chr17 | 13756195 | 13756270 | 3.47715 | 4.20412 | 1.28007 | Intron (ENSMUST00000135850.2/ENSMUSG00000038347.9, intron 1 of 4) | 17 | 13759370 | 13759985 | 616 | 2 | ENSMUST00000129681.1 | 3715 | Gm16052 | ENSMUSG00000086292.1 |
| chr9 | 106147231 | 106147740 | 3.46638 | 4.22342 | 1.2566 | Promoter | 9 | 106148687 | 106159705 | 11019 | 1 | ENSMUST00000143956.1 | -947 | D030055H07Rik | ENSMUSG00000086296.2 |
| chr9 | 106148952 | 106149773 | 3.64664 | 4.39605 | 1.34501 | Promoter | 9 | 106148687 | 106159705 | 11019 | 1 | ENSMUST00000143956.1 | 266 | D030055H07Rik | ENSMUSG00000086296.2 |
| chr9 | 106150385 | 106150538 | 3.68159 | 4.60641 | 1.52847 | Intron (ENSMUST00000143956.1/ENSMUSG00000086296.2, intron 1 of 6) | 9 | 106148687 | 106159705 | 11019 | 1 | ENSMUST00000143956.1 | 1699 | D030055H07Rik | ENSMUSG00000086296.2 |
| chr5 | 114512585 | 114512699 | 3.38018 | 3.89795 | 1.01733 | Distal Intergenic | 5 | 114551289 | 114554016 | 2728 | 1 | ENSMUST00000156209.1 | -38590 | Gm13790 | ENSMUSG00000086302.1 |
| chr5 | 114536960 | 114538499 | 3.55145 | 4.24842 | 1.28266 | Distal Intergenic | 5 | 114551289 | 114554016 | 2728 | 1 | ENSMUST00000156209.1 | -12790 | Gm13790 | ENSMUSG00000086302.1 |
| chr11 | 119119296 | 119119509 | 3.53818 | 4.21478 | 1.26008 | Distal Intergenic | 11 | 119106328 | 119111425 | 5098 | 2 | ENSMUST00000156576.1 | -7872 | Gm11754 | ENSMUSG00000086306.1 |
| chr2 | 116072061 | 116072136 | 4.27972 | 5.71116 | 2.29185 | Intron (ENSMUST00000152412.1/ENSMUSG00000086308.1, intron 2 of 4) | 2 | 116067968 | 116085910 | 17943 | 1 | ENSMUST00000156095.1 | 4094 | G630016G05Rik | ENSMUSG00000086308.1 |
| chr2 | 116074682 | 116075104 | 3.53311 | 4.36585 | 1.37043 | Intron (ENSMUST00000152412.1/ENSMUSG00000086308.1, intron 2 of 4) | 2 | 116067968 | 116085910 | 17943 | 1 | ENSMUST00000156095.1 | 6715 | G630016G05Rik | ENSMUSG00000086308.1 |
| chr2 | 116075889 | 116076089 | 4.09534 | 5.66384 | 2.23905 | Intron (ENSMUST00000152412.1/ENSMUSG00000086308.1, intron 2 of 4) | 2 | 116067968 | 116085910 | 17943 | 1 | ENSMUST00000156095.1 | 7922 | G630016G05Rik | ENSMUSG00000086308.1 |
| chr11 | 35387841 | 35387923 | 3.91273 | 4.99948 | 1.7662 | Intron (ENSMUST00000069837.3/ENSMUSG00000056427.5, intron 4 of 35) | 11 | 35350069 | 35350785 | 717 | 1 | ENSMUST00000124503.1 | 37773 | Gm12122 | ENSMUSG00000086311.1 |
| chr11 | 35431883 | 35432119 | 4.06315 | 5.21442 | 1.91491 | Intron (ENSMUST00000069837.3/ENSMUSG00000056427.5, intron 4 of 35) | 11 | 35350069 | 35350785 | 717 | 1 | ENSMUST00000124503.1 | 81815 | Gm12122 | ENSMUSG00000086311.1 |
| chr11 | 35337556 | 35337855 | 3.71754 | 4.62552 | 1.54369 | Intron (ENSMUST00000069837.3/ENSMUSG00000056427.5, intron 4 of 35) | 11 | 35350069 | 35350785 | 717 | 1 | ENSMUST00000124503.1 | -12214 | Gm12122 | ENSMUSG00000086311.1 |
| chr16 | 35981698 | 35981792 | 2.84849 | 18.07687 | 13.17811 | Promoter | 16 | 35966775 | 35981141 | 14367 | 2 | ENSMUST00000144134.2 | -558 | Gm15564 | ENSMUSG00000086324.3 |
| chr4 | 148389692 | 148390072 | 3.49376 | 4.11044 | 1.16295 | Distal Intergenic | 4 | 148356379 | 148359460 | 3082 | 2 | ENSMUST00000138802.1 | -30233 | Gm13200 | ENSMUSG00000086326.1 |
| chr4 | 148400201 | 148400417 | 3.56294 | 4.38077 | 1.36491 | Distal Intergenic | 4 | 148356379 | 148359460 | 3082 | 2 | ENSMUST00000138802.1 | -40742 | Gm13200 | ENSMUSG00000086326.1 |
| chr11 | 96156423 | 96157099 | 3.61029 | 4.38419 | 1.44027 | Exon (ENSMUST00000167258.2/ENSMUSG00000038756.8, exon 13 of 15) | 11 | 96177551 | 96178800 | 1250 | 1 | ENSMUST00000145517.1 | -20452 | Gm11535 | ENSMUSG00000086337.1 |
| chr11 | 96160700 | 96161618 | 3.63512 | 4.46407 | 1.46974 | Intron (ENSMUST00000167258.2/ENSMUSG00000038756.8, intron 14 of 14) | 11 | 96177551 | 96178800 | 1250 | 1 | ENSMUST00000145517.1 | -15933 | Gm11535 | ENSMUSG00000086337.1 |
| chr11 | 96162377 | 96162622 | 3.3851 | 3.95987 | 1.18736 | Intron (ENSMUST00000167258.2/ENSMUSG00000038756.8, intron 14 of 14) | 11 | 96177551 | 96178800 | 1250 | 1 | ENSMUST00000145517.1 | -14929 | Gm11535 | ENSMUSG00000086337.1 |
| chr11 | 96164290 | 96164667 | 3.5998 | 4.33822 | 1.34948 | Intron (ENSMUST00000167258.2/ENSMUSG00000038756.8, intron 14 of 14) | 11 | 96177551 | 96178800 | 1250 | 1 | ENSMUST00000145517.1 | -12884 | Gm11535 | ENSMUSG00000086337.1 |
| chr11 | 96164982 | 96165164 | 4.71149 | 6.58637 | 2.94553 | 3' UTR | 11 | 96177551 | 96178800 | 1250 | 1 | ENSMUST00000145517.1 | -12387 | Gm11535 | ENSMUSG00000086337.1 |
| chr11 | 96167239 | 96167645 | 3.4648 | 4.09739 | 1.24178 | Distal Intergenic | 11 | 96177551 | 96178800 | 1250 | 1 | ENSMUST00000145517.1 | -9906 | Gm11535 | ENSMUSG00000086337.1 |
| chr11 | 96168843 | 96169504 | 3.57995 | 4.41716 | 1.38279 | Distal Intergenic | 11 | 96177551 | 96178800 | 1250 | 1 | ENSMUST00000145517.1 | -8047 | Gm11535 | ENSMUSG00000086337.1 |
| chr11 | 96169902 | 96174502 | 3.8786 | 4.95342 | 1.7922 | Distal Intergenic | 11 | 96177551 | 96178800 | 1250 | 1 | ENSMUST00000145517.1 | -3049 | Gm11535 | ENSMUSG00000086337.1 |
| chr11 | 96174963 | 96175417 | 3.96206 | 5.44689 | 2.06829 | Distal Intergenic | 11 | 96177551 | 96178800 | 1250 | 1 | ENSMUST00000145517.1 | -2134 | Gm11535 | ENSMUSG00000086337.1 |
| chr11 | 96175962 | 96176500 | 3.58684 | 4.5838 | 1.49502 | Downstream (2-3kb) | 11 | 96177551 | 96178800 | 1250 | 1 | ENSMUST00000145517.1 | -1051 | Gm11535 | ENSMUSG00000086337.1 |
| chr11 | 96177258 | 96178057 | 3.96191 | 5.16619 | 1.88771 | Promoter | 11 | 96177551 | 96178800 | 1250 | 1 | ENSMUST00000145517.1 | 0 | Gm11535 | ENSMUSG00000086337.1 |
| chr11 | 96179323 | 96179504 | 3.60697 | 4.6714 | 1.56026 | Distal Intergenic | 11 | 96177551 | 96178800 | 1250 | 1 | ENSMUST00000145517.1 | 1773 | Gm11535 | ENSMUSG00000086337.1 |
| chr11 | 96180974 | 96181514 | 3.88924 | 5.0584 | 1.81142 | Distal Intergenic | 11 | 96177551 | 96178800 | 1250 | 1 | ENSMUST00000145517.1 | 3424 | Gm11535 | ENSMUSG00000086337.1 |
| chr11 | 96182455 | 96184367 | 4.09627 | 5.4293 | 2.10909 | Distal Intergenic | 11 | 96177551 | 96178800 | 1250 | 1 | ENSMUST00000145517.1 | 4905 | Gm11535 | ENSMUSG00000086337.1 |
| chr11 | 96185500 | 96186306 | 3.47868 | 4.36605 | 1.37878 | Distal Intergenic | 11 | 96177551 | 96178800 | 1250 | 1 | ENSMUST00000145517.1 | 7950 | Gm11535 | ENSMUSG00000086337.1 |
| chr5 | 115394159 | 115395422 | 3.66181 | 4.44347 | 1.38622 | Intron (ENSMUST00000146546.1/ENSMUSG00000087292.1, intron 2 of 2) | 5 | 115358700 | 115360123 | 1424 | 2 | ENSMUST00000139500.1 | -34037 | 4930401G09Rik | ENSMUSG00000086352.1 |
| chr9 | 42055105 | 42055606 | 3.49927 | 4.19891 | 1.23309 | Intron (ENSMUST00000060989.8/ENSMUSG00000049313.8, intron 12 of 47) | 9 | 42022615 | 42034113 | 11499 | 1 | ENSMUST00000143710.1 | 32491 | Gm16214 | ENSMUSG00000086360.1 |
| chr13 | 29044375 | 29044760 | 3.89828 | 4.87901 | 1.65787 | Distal Intergenic | 13 | 29016708 | 29040336 | 23629 | 1 | ENSMUST00000145801.1 | 27668 | A330102I10Rik | ENSMUSG00000086363.2 |
| chr11 | 32137999 | 32138292 | 3.60682 | 4.35998 | 1.33903 | Distal Intergenic | 11 | 32095261 | 32101228 | 5968 | 2 | ENSMUST00000145338.1 | -36772 | Gm12108 | ENSMUSG00000086386.1 |
| chr3 | 89315953 | 89316121 | 4.23896 | 5.54638 | 2.12258 | Promoter | 3 | 89315439 | 89318907 | 3469 | 1 | ENSMUST00000135144.1 | 515 | Gm15998 | ENSMUSG00000086389.1 |
| chr3 | 89316481 | 89316646 | 3.70009 | 4.50092 | 1.41721 | Exon (ENSMUST00000135144.1/ENSMUSG00000086389.1, exon 2 of 3) | 3 | 89315439 | 89318907 | 3469 | 1 | ENSMUST00000135144.1 | 1043 | Gm15998 | ENSMUSG00000086389.1 |
| chr3 | 89317369 | 89317669 | 3.66065 | 4.6058 | 1.50039 | Intron (ENSMUST00000135144.1/ENSMUSG00000086389.1, intron 2 of 2) | 3 | 89315439 | 89318907 | 3469 | 1 | ENSMUST00000135144.1 | 1931 | Gm15998 | ENSMUSG00000086389.1 |
| chr3 | 89318008 | 89318595 | 4.04315 | 5.19548 | 1.87628 | Exon (ENSMUST00000135144.1/ENSMUSG00000086389.1, exon 3 of 3) | 3 | 89315439 | 89318907 | 3469 | 1 | ENSMUST00000135144.1 | 2570 | Gm15998 | ENSMUSG00000086389.1 |
| chr11 | 97545032 | 97545436 | 3.51598 | 4.17699 | 1.22271 | Intron (ENSMUST00000107596.3/ENSMUSG00000038453.10, intron 3 of 19) | 11 | 97566711 | 97576186 | 9476 | 1 | ENSMUST00000145845.1 | -21275 | Gm11612 | ENSMUSG00000086394.1 |
| chr11 | 97546287 | 97547565 | 4.06485 | 5.26623 | 1.96508 | Intron (ENSMUST00000107596.3/ENSMUSG00000038453.10, intron 3 of 19) | 11 | 97566711 | 97576186 | 9476 | 1 | ENSMUST00000145845.1 | -19146 | Gm11612 | ENSMUSG00000086394.1 |
| chr11 | 97548432 | 97549205 | 3.68667 | 4.54915 | 1.47397 | Exon (ENSMUST00000107596.3/ENSMUSG00000038453.10, exon 3 of 20) | 11 | 97566711 | 97576186 | 9476 | 1 | ENSMUST00000145845.1 | -17506 | Gm11612 | ENSMUSG00000086394.1 |
| chr11 | 97549850 | 97550390 | 4.12323 | 5.32056 | 1.97201 | Intron (ENSMUST00000107596.3/ENSMUSG00000038453.10, intron 2 of 19) | 11 | 97566711 | 97576186 | 9476 | 1 | ENSMUST00000145845.1 | -16321 | Gm11612 | ENSMUSG00000086394.1 |
| chr11 | 97550750 | 97551072 | 3.74241 | 4.82647 | 1.64285 | Intron (ENSMUST00000107596.3/ENSMUSG00000038453.10, intron 2 of 19) | 11 | 97566711 | 97576186 | 9476 | 1 | ENSMUST00000145845.1 | -15639 | Gm11612 | ENSMUSG00000086394.1 |
| chr11 | 97551784 | 97552018 | 3.19744 | 3.9558 | 1.11403 | Exon (ENSMUST00000107596.3/ENSMUSG00000038453.10, exon 2 of 20) | 11 | 97566711 | 97576186 | 9476 | 1 | ENSMUST00000145845.1 | -14693 | Gm11612 | ENSMUSG00000086394.1 |
| chr11 | 97552397 | 97552627 | 3.75265 | 4.61957 | 1.47893 | Intron (ENSMUST00000107596.3/ENSMUSG00000038453.10, intron 1 of 19) | 11 | 97566711 | 97576186 | 9476 | 1 | ENSMUST00000145845.1 | -14084 | Gm11612 | ENSMUSG00000086394.1 |
| chr11 | 97555801 | 97555975 | 3.36961 | 3.89089 | 1.04874 | Intron (ENSMUST00000107596.3/ENSMUSG00000038453.10, intron 1 of 19) | 11 | 97566711 | 97576186 | 9476 | 1 | ENSMUST00000145845.1 | -10736 | Gm11612 | ENSMUSG00000086394.1 |
| chr11 | 97560056 | 97560363 | 3.53287 | 4.33115 | 1.31736 | Intron (ENSMUST00000107596.3/ENSMUSG00000038453.10, intron 1 of 19) | 11 | 97566711 | 97576186 | 9476 | 1 | ENSMUST00000145845.1 | -6348 | Gm11612 | ENSMUSG00000086394.1 |
| chr11 | 97564055 | 97564278 | 3.38193 | 3.9006 | 1.01733 | Intron (ENSMUST00000107596.3/ENSMUSG00000038453.10, intron 1 of 19) | 11 | 97566711 | 97576186 | 9476 | 1 | ENSMUST00000145845.1 | -2433 | Gm11612 | ENSMUSG00000086394.1 |
| chr11 | 97565708 | 97565789 | 3.48327 | 4.11792 | 1.23788 | Promoter | 11 | 97566711 | 97576186 | 9476 | 1 | ENSMUST00000145845.1 | -922 | Gm11612 | ENSMUSG00000086394.1 |
| chr11 | 97566941 | 97567088 | 3.39134 | 3.92374 | 1.04874 | Promoter | 11 | 97566711 | 97576186 | 9476 | 1 | ENSMUST00000145845.1 | 231 | Gm11612 | ENSMUSG00000086394.1 |
| chr11 | 97567474 | 97568248 | 3.8028 | 4.75659 | 1.57478 | Promoter | 11 | 97566711 | 97576186 | 9476 | 1 | ENSMUST00000145845.1 | 764 | Gm11612 | ENSMUSG00000086394.1 |
| chr11 | 97569289 | 97569680 | 3.4317 | 3.99632 | 1.08918 | Intron (ENSMUST00000145845.1/ENSMUSG00000086394.1, intron 1 of 1) | 11 | 97566711 | 97576186 | 9476 | 1 | ENSMUST00000145845.1 | 2579 | Gm11612 | ENSMUSG00000086394.1 |
| chr7 | 68409888 | 68410112 | 3.22895 | 3.67061 | 1.01733 | Distal Intergenic | 7 | 68338668 | 68353201 | 14534 | 2 | ENSMUST00000129965.1 | -56688 | Gm16158 | ENSMUSG00000086410.2 |
| chr7 | 68410596 | 68410885 | 3.30396 | 3.78313 | 1.01733 | Distal Intergenic | 7 | 68338668 | 68353201 | 14534 | 2 | ENSMUST00000129965.1 | -57396 | Gm16158 | ENSMUSG00000086410.2 |
| chr7 | 68480562 | 68480838 | 3.67462 | 4.61662 | 1.525 | Distal Intergenic | 7 | 68338668 | 68353201 | 14534 | 2 | ENSMUST00000129965.1 | -127362 | Gm16158 | ENSMUSG00000086410.2 |
| chr13 | 29233011 | 29233095 | 4.09081 | 5.57476 | 2.16501 | Intron (ENSMUST00000140278.2/ENSMUSG00000006191.12, intron 15 of 16) | 13 | 29235777 | 29236620 | 844 | 2 | ENSMUST00000137593.1 | 3525 | Gm11364 | ENSMUSG00000086423.1 |
| chr13 | 29285457 | 29285691 | 3.44992 | 4.03723 | 1.13789 | Intron (ENSMUST00000140278.2/ENSMUSG00000006191.12, intron 15 of 16) | 13 | 29235777 | 29236620 | 844 | 2 | ENSMUST00000137593.1 | -48838 | Gm11364 | ENSMUSG00000086423.1 |
| chr2 | 28098960 | 28099047 | 4.09459 | 5.41144 | 2.06245 | Intron (ENSMUST00000126140.2/ENSMUSG00000086425.2, intron 1 of 7) | 2 | 28095483 | 28113851 | 18369 | 1 | ENSMUST00000156508.1 | 3478 | F730016J06Rik | ENSMUSG00000086425.2 |
| chr2 | 28099841 | 28099981 | 3.78366 | 4.65613 | 1.51364 | Exon (ENSMUST00000126140.2/ENSMUSG00000086425.2, exon 2 of 8) | 2 | 28095483 | 28113851 | 18369 | 1 | ENSMUST00000156508.1 | 4359 | F730016J06Rik | ENSMUSG00000086425.2 |
| chr2 | 28150262 | 28150540 | 3.48047 | 4.11578 | 1.17542 | Distal Intergenic | 2 | 28118207 | 28125203 | 6997 | 1 | ENSMUST00000128888.1 | 32056 | F730016J06Rik | ENSMUSG00000086425.2 |
| chr2 | 28150966 | 28151120 | 3.38193 | 3.9006 | 1.01733 | Distal Intergenic | 2 | 28118207 | 28125203 | 6997 | 1 | ENSMUST00000128888.1 | 32760 | F730016J06Rik | ENSMUSG00000086425.2 |
| chr6 | 52242952 | 52244180 | 4.00934 | 5.10828 | 1.85141 | Promoter | 6 | 52244393 | 52249791 | 5399 | 1 | ENSMUST00000156515.3 | -213 | Hoxa11os | ENSMUSG00000086427.3 |
| chr6 | 52248672 | 52248903 | 3.49238 | 4.17987 | 1.28723 | Promoter | 6 | 52248422 | 52249799 | 1378 | 1 | ENSMUST00000154027.2 | 251 | Hoxa11os | ENSMUSG00000086427.3 |
| chr10 | 42714679 | 42714896 | 3.63643 | 4.39188 | 1.38217 | Distal Intergenic | 10 | 42698036 | 42698518 | 483 | 2 | ENSMUST00000149950.1 | -16162 | Gm15200 | ENSMUSG00000086434.1 |
| chr4 | 140420112 | 140420322 | 3.40466 | 3.94967 | 1.06942 | Distal Intergenic | 4 | 140424644 | 140431931 | 7288 | 1 | ENSMUST00000135716.1 | -4322 | Gm13021 | ENSMUSG00000086437.1 |
| chr4 | 140423375 | 140423626 | 3.44995 | 4.02844 | 1.10632 | Distal Intergenic | 4 | 140424644 | 140431931 | 7288 | 1 | ENSMUST00000135716.1 | -1018 | Gm13021 | ENSMUSG00000086437.1 |
| chr4 | 140425236 | 140425559 | 3.61924 | 4.4237 | 1.3781 | Promoter | 4 | 140424644 | 140431931 | 7288 | 1 | ENSMUST00000135716.1 | 593 | Gm13021 | ENSMUSG00000086437.1 |
| chr11 | 105362148 | 105362365 | 3.42794 | 4.14495 | 1.22687 | Promoter | 11 | 105363252 | 105390642 | 27391 | 1 | ENSMUST00000153672.1 | -887 | Gm11638 | ENSMUSG00000086487.1 |
| chr2 | 158595449 | 158596015 | 3.96841 | 5.08745 | 1.87004 | Exon (ENSMUST00000145773.1/ENSMUSG00000086496.1, exon 3 of 3) | 2 | 158603188 | 158610590 | 7403 | 2 | ENSMUST00000155319.1 | 14575 | Gm14204 | ENSMUSG00000086496.1 |
| chr2 | 158596328 | 158596524 | 3.5031 | 4.35364 | 1.38464 | Exon (ENSMUST00000145773.1/ENSMUSG00000086496.1, exon 3 of 3) | 2 | 158603188 | 158610590 | 7403 | 2 | ENSMUST00000155319.1 | 14066 | Gm14204 | ENSMUSG00000086496.1 |
| chr2 | 158598005 | 158598081 | 3.66628 | 4.60076 | 1.53125 | Intron (ENSMUST00000145773.1/ENSMUSG00000086496.1, intron 1 of 2) | 2 | 158603188 | 158610590 | 7403 | 2 | ENSMUST00000155319.1 | 12509 | Gm14204 | ENSMUSG00000086496.1 |
| chr2 | 158604140 | 158604217 | 4.14459 | 5.69202 | 2.23901 | Exon (ENSMUST00000123944.1/ENSMUSG00000086496.1, exon 2 of 3) | 2 | 158603188 | 158610590 | 7403 | 2 | ENSMUST00000155319.1 | 6373 | Gm14204 | ENSMUSG00000086496.1 |
| chr2 | 158605643 | 158605718 | 3.96092 | 4.99971 | 1.74943 | Intron (ENSMUST00000145773.1/ENSMUSG00000086496.1, intron 1 of 2) | 2 | 158603188 | 158610590 | 7403 | 2 | ENSMUST00000155319.1 | 4872 | Gm14204 | ENSMUSG00000086496.1 |
| chr2 | 158608718 | 158609203 | 3.29348 | 3.81774 | 1.06151 | Intron (ENSMUST00000145773.1/ENSMUSG00000086496.1, intron 1 of 2) | 2 | 158603188 | 158610590 | 7403 | 2 | ENSMUST00000155319.1 | 1387 | Gm14204 | ENSMUSG00000086496.1 |
| chr13 | 22016376 | 22018165 | 3.63824 | 4.42087 | 1.40958 | Promoter | 13 | 22016690 | 22018249 | 1560 | 1 | ENSMUST00000147811.1 | 0 | Gm11291 | ENSMUSG00000086508.1 |
| chr13 | 22018790 | 22018964 | 3.54254 | 4.26257 | 1.3384 | Distal Intergenic | 13 | 22016690 | 22018249 | 1560 | 1 | ENSMUST00000147811.1 | 2101 | Gm11291 | ENSMUSG00000086508.1 |
| chr13 | 22019504 | 22022798 | 4.02674 | 5.19526 | 1.90982 | Distal Intergenic | 13 | 22016690 | 22018249 | 1560 | 1 | ENSMUST00000147811.1 | 2815 | Gm11291 | ENSMUSG00000086508.1 |
| chr2 | 147170656 | 147170871 | 4.00327 | 5.09473 | 1.79786 | Distal Intergenic | 2 | 147184083 | 147331681 | 147599 | 1 | ENSMUST00000136998.2 | -13212 | Nkx2-2os | ENSMUSG00000086509.2 |
| chr2 | 147173157 | 147173498 | 3.70999 | 4.53352 | 1.48054 | Distal Intergenic | 2 | 147184083 | 147331681 | 147599 | 1 | ENSMUST00000136998.2 | -10585 | Nkx2-2os | ENSMUSG00000086509.2 |
| chr2 | 147174528 | 147174967 | 3.72534 | 4.57152 | 1.51126 | Downstream (2-3kb) | 2 | 147184083 | 147331681 | 147599 | 1 | ENSMUST00000136998.2 | -9116 | Nkx2-2os | ENSMUSG00000086509.2 |
| chr2 | 147176790 | 147177322 | 3.53216 | 4.40263 | 1.39922 | Downstream (<1kb) | 2 | 147184083 | 147331681 | 147599 | 1 | ENSMUST00000136998.2 | -6761 | Nkx2-2os | ENSMUSG00000086509.2 |
| chr2 | 147178266 | 147179901 | 3.80748 | 4.78174 | 1.61707 | Intron (ENSMUST00000109970.4/ENSMUSG00000027434.11, intron 2 of 2) | 2 | 147184083 | 147331681 | 147599 | 1 | ENSMUST00000136998.2 | -4182 | Nkx2-2os | ENSMUSG00000086509.2 |
| chr2 | 147180894 | 147181745 | 3.62053 | 4.39674 | 1.3498 | Intron (ENSMUST00000109970.4/ENSMUSG00000027434.11, intron 2 of 2) | 2 | 147184083 | 147331681 | 147599 | 1 | ENSMUST00000136998.2 | -2338 | Nkx2-2os | ENSMUSG00000086509.2 |
| chr2 | 147182065 | 147183462 | 3.75213 | 4.59353 | 1.46872 | Promoter | 2 | 147184083 | 147331681 | 147599 | 1 | ENSMUST00000136998.2 | -621 | Nkx2-2os | ENSMUSG00000086509.2 |
| chr2 | 147183818 | 147184959 | 4.15184 | 5.42356 | 2.05021 | Promoter | 2 | 147184083 | 147331681 | 147599 | 1 | ENSMUST00000136998.2 | 0 | Nkx2-2os | ENSMUSG00000086509.2 |
| chr2 | 70508861 | 70509411 | 3.38209 | 3.91755 | 1.07611 | Promoter | 2 | 70503433 | 70509199 | 5767 | 2 | ENSMUST00000150702.1 | 0 | Erich2os | ENSMUSG00000086515.1 |
| chr17 | 29891112 | 29891359 | 3.49841 | 4.12509 | 1.18983 | Exon (ENSMUST00000149552.2/ENSMUSG00000086534.2, exon 3 of 6) | 17 | 29890084 | 29897067 | 6984 | 1 | ENSMUST00000149552.2 | 1029 | Gm16758 | ENSMUSG00000086534.2 |
| chr17 | 29904374 | 29904822 | 3.37141 | 3.90409 | 1.08151 | Intron (ENSMUST00000167190.2/ENSMUSG00000043557.10, intron 4 of 12) | 17 | 29890084 | 29897067 | 6984 | 1 | ENSMUST00000149552.2 | 14291 | Gm16758 | ENSMUSG00000086534.2 |
| chr2 | 174295307 | 174296405 | 3.62196 | 4.37662 | 1.32604 | Promoter | 2 | 174281237 | 174295436 | 14200 | 2 | ENSMUST00000151472.2 | 0 | Nespas | ENSMUSG00000086537.2 |
| chr15 | 56692158 | 56693774 | 3.79896 | 4.75204 | 1.64983 | Promoter | 15 | 56693552 | 56778399 | 84848 | 1 | ENSMUST00000166290.1 | 0 | Has2os | ENSMUSG00000086541.3 |
| chr11 | 98598764 | 98599134 | 3.30967 | 3.79186 | 1.01733 | Promoter | 11 | 98598291 | 98599073 | 783 | 1 | ENSMUST00000142268.1 | 474 | Gm12 | ENSMUSG00000086545.3 |
| chr11 | 98599637 | 98599947 | 3.70313 | 4.48758 | 1.37715 | Distal Intergenic | 11 | 98598291 | 98599073 | 783 | 1 | ENSMUST00000142268.1 | 1347 | Gm12 | ENSMUSG00000086545.3 |
| chr11 | 119135470 | 119135584 | 3.444 | 4.14007 | 1.22475 | Intron (ENSMUST00000144096.1/ENSMUSG00000086547.1, intron 2 of 5) | 11 | 119123011 | 119138421 | 15411 | 2 | ENSMUST00000144096.1 | 2837 | Gm11755 | ENSMUSG00000086547.1 |
| chr4 | 156078068 | 156078326 | 3.8792 | 4.86556 | 1.69705 | Distal Intergenic | 4 | 156052715 | 156059802 | 7088 | 2 | ENSMUST00000138361.1 | -18267 | Gm13648 | ENSMUSG00000086549.1 |
| chr11 | 95133417 | 95133695 | 3.34026 | 3.83794 | 1.01733 | Distal Intergenic | 11 | 95145052 | 95159047 | 13996 | 1 | ENSMUST00000156477.3 | -11357 | A730090H04Rik | ENSMUSG00000086552.3 |
| chr11 | 95138186 | 95138415 | 3.54532 | 4.23729 | 1.29043 | Downstream (2-3kb) | 11 | 95145052 | 95159047 | 13996 | 1 | ENSMUST00000156477.3 | -6637 | A730090H04Rik | ENSMUSG00000086552.3 |
| chr11 | 95139158 | 95139936 | 3.72244 | 4.58161 | 1.52474 | Downstream (<1kb) | 11 | 95145052 | 95159047 | 13996 | 1 | ENSMUST00000156477.3 | -5116 | A730090H04Rik | ENSMUSG00000086552.3 |
| chr11 | 95141462 | 95141985 | 3.49995 | 4.2981 | 1.35127 | Exon (ENSMUST00000021241.7/ENSMUSG00000020871.8, exon 2 of 3) | 11 | 95145052 | 95159047 | 13996 | 1 | ENSMUST00000156477.3 | -3067 | A730090H04Rik | ENSMUSG00000086552.3 |
| chr11 | 95142582 | 95143506 | 4.04564 | 5.3902 | 2.06256 | Intron (ENSMUST00000021241.7/ENSMUSG00000020871.8, intron 1 of 2) | 11 | 95145052 | 95159047 | 13996 | 1 | ENSMUST00000156477.3 | -1546 | A730090H04Rik | ENSMUSG00000086552.3 |
| chr11 | 95144305 | 95145564 | 3.79862 | 4.72537 | 1.62767 | Promoter | 11 | 95145052 | 95159047 | 13996 | 1 | ENSMUST00000156477.3 | 0 | A730090H04Rik | ENSMUSG00000086552.3 |
| chr4 | 154543321 | 154543812 | 3.48043 | 4.20355 | 1.25207 | Intron (ENSMUST00000030902.7/ENSMUSG00000039410.11, intron 1 of 16) | 4 | 154550282 | 154557645 | 7364 | 1 | ENSMUST00000143888.1 | -6470 | Gm13133 | ENSMUSG00000086565.1 |
| chr4 | 154565763 | 154566135 | 3.42139 | 4.01189 | 1.15423 | Intron (ENSMUST00000030902.7/ENSMUSG00000039410.11, intron 1 of 16) | 4 | 154550282 | 154557645 | 7364 | 1 | ENSMUST00000143888.1 | 15482 | Gm13133 | ENSMUSG00000086565.1 |
| chr4 | 154568789 | 154570158 | 3.61442 | 4.45747 | 1.4376 | Intron (ENSMUST00000030902.7/ENSMUSG00000039410.11, intron 1 of 16) | 4 | 154550282 | 154557645 | 7364 | 1 | ENSMUST00000143888.1 | 18508 | Gm13133 | ENSMUSG00000086565.1 |
| chr4 | 154573884 | 154574125 | 4.14895 | 5.33541 | 1.938 | Intron (ENSMUST00000030902.7/ENSMUSG00000039410.11, intron 1 of 16) | 4 | 154550282 | 154557645 | 7364 | 1 | ENSMUST00000143888.1 | 23603 | Gm13133 | ENSMUSG00000086565.1 |
| chr4 | 154544125 | 154544200 | 3.37882 | 3.89591 | 1.01733 | Intron (ENSMUST00000030902.7/ENSMUSG00000039410.11, intron 1 of 16) | 4 | 154550282 | 154557645 | 7364 | 1 | ENSMUST00000143888.1 | -6082 | Gm13133 | ENSMUSG00000086565.1 |
| chr4 | 154545020 | 154545109 | 4.16851 | 5.41485 | 2.04366 | Intron (ENSMUST00000030902.7/ENSMUSG00000039410.11, intron 1 of 16) | 4 | 154550282 | 154557645 | 7364 | 1 | ENSMUST00000143888.1 | -5173 | Gm13133 | ENSMUSG00000086565.1 |
| chr4 | 154545989 | 154546801 | 3.4477 | 4.0207 | 1.09069 | Intron (ENSMUST00000030902.7/ENSMUSG00000039410.11, intron 1 of 16) | 4 | 154550282 | 154557645 | 7364 | 1 | ENSMUST00000143888.1 | -3481 | Gm13133 | ENSMUSG00000086565.1 |
| chr4 | 154549709 | 154549812 | 3.98895 | 5.11055 | 1.88591 | Promoter | 4 | 154550282 | 154557645 | 7364 | 1 | ENSMUST00000143888.1 | -470 | Gm13133 | ENSMUSG00000086565.1 |
| chr4 | 154558328 | 154558589 | 3.46196 | 4.09304 | 1.17047 | Intron (ENSMUST00000030902.7/ENSMUSG00000039410.11, intron 1 of 16) | 4 | 154550282 | 154557645 | 7364 | 1 | ENSMUST00000143888.1 | 8047 | Gm13133 | ENSMUSG00000086565.1 |
| chr4 | 154564537 | 154564844 | 3.54483 | 4.20328 | 1.21755 | Intron (ENSMUST00000030902.7/ENSMUSG00000039410.11, intron 1 of 16) | 4 | 154550282 | 154557645 | 7364 | 1 | ENSMUST00000143888.1 | 14256 | Gm13133 | ENSMUSG00000086565.1 |
| chr2 | 75484010 | 75484327 | 3.25182 | 3.7129 | 1.04243 | Distal Intergenic | 2 | 75458966 | 75464788 | 5823 | 2 | ENSMUST00000142828.1 | -19223 | Gm13660 | ENSMUSG00000086576.1 |
| chr2 | 75493288 | 75493401 | 3.57014 | 4.26726 | 1.27489 | Distal Intergenic | 2 | 75458966 | 75464788 | 5823 | 2 | ENSMUST00000142828.1 | -28501 | Gm13660 | ENSMUSG00000086576.1 |
| chr2 | 75538245 | 75538407 | 3.49742 | 4.3195 | 1.33995 | Distal Intergenic | 2 | 75458966 | 75464788 | 5823 | 2 | ENSMUST00000142828.1 | -73458 | Gm13660 | ENSMUSG00000086576.1 |
| chr2 | 60552457 | 60552537 | 3.57985 | 4.30691 | 1.37675 | Promoter | 2 | 60552355 | 60553274 | 920 | 1 | ENSMUST00000126361.1 | 103 | Gm13583 | ENSMUSG00000086578.2 |
| chr9 | 114057172 | 114058242 | 4.38567 | 5.84144 | 2.34158 | Promoter | 9 | 114057354 | 114098733 | 41380 | 1 | ENSMUST00000135338.1 | 0 | Susd5 | ENSMUSG00000086596.1 |
| chr15 | 89522849 | 89523649 | 3.47591 | 4.08598 | 1.17149 | Promoter | 15 | 89520096 | 89523677 | 3582 | 2 | ENSMUST00000155610.1 | 28 | Gm15609 | ENSMUSG00000086602.1 |
| chr2 | 167145125 | 167145200 | 3.61629 | 4.37845 | 1.40755 | Intron (ENSMUST00000059826.8/ENSMUSG00000050556.8, intron 1 of 1) | 2 | 167098794 | 167101826 | 3033 | 2 | ENSMUST00000144351.1 | -43300 | Gm14290 | ENSMUSG00000086605.1 |
| chr11 | 24098730 | 24098991 | 3.44034 | 4.2405 | 1.29451 | Intron (ENSMUST00000000881.7/ENSMUSG00000000861.10, intron 2 of 4) | 11 | 24099688 | 24115405 | 15718 | 2 | ENSMUST00000133819.1 | 16414 | Gm12064 | ENSMUSG00000086612.1 |
| chr6 | 135907236 | 135907379 | 3.84533 | 4.80696 | 1.66982 | Intron (ENSMUST00000053880.7/ENSMUSG00000030209.9, intron 3 of 12) | 6 | 135875505 | 135884792 | 9288 | 1 | ENSMUST00000155455.2 | 31732 | Gm14330 | ENSMUSG00000086614.2 |
| chr2 | 9876724 | 9877684 | 3.76611 | 4.79375 | 1.64048 | Promoter | 2 | 9877256 | 9878869 | 1614 | 1 | ENSMUST00000131297.1 | 0 | Gm13256 | ENSMUSG00000086618.1 |
| chr2 | 165130935 | 165131541 | 3.43804 | 4.16565 | 1.2414 | Intron (ENSMUST00000065438.7/ENSMUSG00000053166.9, intron 8 of 11) | 2 | 165159412 | 165163969 | 4558 | 1 | ENSMUST00000124728.1 | -27871 | Gm14397 | ENSMUSG00000086630.1 |
| chr2 | 165133363 | 165134241 | 3.6686 | 4.4393 | 1.37112 | Intron (ENSMUST00000065438.7/ENSMUSG00000053166.9, intron 8 of 11) | 2 | 165159412 | 165163969 | 4558 | 1 | ENSMUST00000124728.1 | -25171 | Gm14397 | ENSMUSG00000086630.1 |
| chr2 | 165135691 | 165135989 | 3.5402 | 4.18915 | 1.19224 | Intron (ENSMUST00000065438.7/ENSMUSG00000053166.9, intron 7 of 11) | 2 | 165159412 | 165163969 | 4558 | 1 | ENSMUST00000124728.1 | -23423 | Gm14397 | ENSMUSG00000086630.1 |
| chr2 | 165137121 | 165137429 | 3.36398 | 3.87352 | 1.01733 | Intron (ENSMUST00000065438.7/ENSMUSG00000053166.9, intron 7 of 11) | 2 | 165159412 | 165163969 | 4558 | 1 | ENSMUST00000124728.1 | -21983 | Gm14397 | ENSMUSG00000086630.1 |
| chr2 | 165140290 | 165140405 | 4.27911 | 5.70634 | 2.26538 | Intron (ENSMUST00000065438.7/ENSMUSG00000053166.9, intron 7 of 11) | 2 | 165159412 | 165163969 | 4558 | 1 | ENSMUST00000124728.1 | -19007 | Gm14397 | ENSMUSG00000086630.1 |
| chr2 | 165142217 | 165142413 | 4.40968 | 5.92919 | 2.44545 | Exon (ENSMUST00000065438.7/ENSMUSG00000053166.9, exon 7 of 12) | 2 | 165159412 | 165163969 | 4558 | 1 | ENSMUST00000124728.1 | -16999 | Gm14397 | ENSMUSG00000086630.1 |
| chr2 | 165143576 | 165143822 | 3.33221 | 4.18233 | 1.22592 | Exon (ENSMUST00000065438.7/ENSMUSG00000053166.9, exon 6 of 12) | 2 | 165159412 | 165163969 | 4558 | 1 | ENSMUST00000124728.1 | -15590 | Gm14397 | ENSMUSG00000086630.1 |
| chr2 | 165144789 | 165144930 | 3.62951 | 4.36102 | 1.32243 | Intron (ENSMUST00000065438.7/ENSMUSG00000053166.9, intron 5 of 11) | 2 | 165159412 | 165163969 | 4558 | 1 | ENSMUST00000124728.1 | -14482 | Gm14397 | ENSMUSG00000086630.1 |
| chr2 | 165159611 | 165159809 | 3.74155 | 4.70384 | 1.56658 | Promoter | 2 | 165159412 | 165163969 | 4558 | 1 | ENSMUST00000124728.1 | 200 | Gm14397 | ENSMUSG00000086630.1 |
| chr2 | 165165747 | 165166224 | 3.36935 | 3.88162 | 1.01733 | Intron (ENSMUST00000065438.7/ENSMUSG00000053166.9, intron 3 of 11) | 2 | 165159412 | 165163969 | 4558 | 1 | ENSMUST00000124728.1 | 6336 | Gm14397 | ENSMUSG00000086630.1 |
| chr2 | 165167450 | 165167891 | 3.36981 | 3.88232 | 1.01733 | Intron (ENSMUST00000065438.7/ENSMUSG00000053166.9, intron 3 of 11) | 2 | 165159412 | 165163969 | 4558 | 1 | ENSMUST00000124728.1 | 8039 | Gm14397 | ENSMUSG00000086630.1 |
| chr1 | 130300480 | 130300748 | 3.27298 | 3.74611 | 1.04714 | Distal Intergenic | 1 | 130269950 | 130281258 | 11309 | 2 | ENSMUST00000155992.1 | -19223 | Gm16081 | ENSMUSG00000086633.1 |
| chr16 | 31426277 | 31426352 | 3.34153 | 3.83966 | 1.01733 | Intron (ENSMUST00000089759.3/ENSMUSG00000046598.9, intron 1 of 6) | 16 | 31427876 | 31428291 | 416 | 2 | ENSMUST00000129645.1 | 1939 | Gm15743 | ENSMUSG00000086645.2 |

| | | | | | | | | | | | | | | | |
|---|---|---|---|---|---|---|---|---|---|---|---|---|---|---|---|
| chr16 | 31427231 | 31427552 | 3.64877 | 4.46144 | 1.44451 | Promoter | 16 | 31427876 | 31428291 | 416 | 2 | ENSMUST00000129645.1 | 739 | Gm15743 | ENSMUSG00000086645.2 |
| chr5 | 116158365 | 116159005 | 3.64631 | 4.47271 | 1.42935 | Intron (ENSMUST00000050178.7/ENSMUSG00000043913.9, intron 6 of 13) | 5 | 116123614 | 116129110 | 5497 | 1 | ENSMUST00000127792.1 | 34752 | C330018A13Rik | ENSMUSG00000086655.1 |
| chr7 | 65527254 | 65527329 | 3.99364 | 5.07317 | 1.84039 | Distal Intergenic | 7 | 65582031 | 65618869 | 36839 | 1 | ENSMUST00000134646.2 | -54702 | BC046251 | ENSMUSG00000086662.2 |
| chr7 | 65527707 | 65527786 | 4.0853 | 5.29868 | 2.0241 | Distal Intergenic | 7 | 65582031 | 65618869 | 36839 | 1 | ENSMUST00000134646.2 | -54245 | BC046251 | ENSMUSG00000086662.2 |
| chr7 | 65529134 | 65529209 | 3.33355 | 3.97278 | 1.08049 | Distal Intergenic | 7 | 65582031 | 65618869 | 36839 | 1 | ENSMUST00000134646.2 | -52822 | BC046251 | ENSMUSG00000086662.2 |
| chr4 | 150033163 | 150034150 | 3.36811 | 3.90362 | 1.10108 | Distal Intergenic | 4 | 150049013 | 150056976 | 7964 | 1 | ENSMUST00000134236.1 | -14863 | Gm13067 | ENSMUSG00000086665.1 |
| chr6 | 32377805 | 32377881 | 3.82096 | 4.76659 | 1.66834 | Intron (ENSMUST00000115096.3/ENSMUSG00000029765.11, intron 3 of 31) | 6 | 32451170 | 32454750 | 3581 | 1 | ENSMUST00000152538.1 | -73289 | Plxna4os2 | ENSMUSG00000086675.1 |
| chr6 | 32394684 | 32394875 | 3.74777 | 4.64897 | 1.57017 | Intron (ENSMUST00000115096.3/ENSMUSG00000029765.11, intron 3 of 31) | 6 | 32451170 | 32454750 | 3581 | 1 | ENSMUST00000152538.1 | -56295 | Plxna4os2 | ENSMUSG00000086675.1 |
| chr11 | 45810006 | 45810227 | 3.75197 | 4.76289 | 1.62722 | Intron (ENSMUST00000137002.1/ENSMUSG00000086686.1, intron 1 of 3) | 11 | 45808087 | 45841171 | 33085 | 1 | ENSMUST00000137002.1 | 1920 | F630206G17Rik | ENSMUSG00000086686.1 |
| chr5 | 23830959 | 23831108 | 3.23467 | 3.67918 | 1.01733 | Promoter | 5 | 23831080 | 23831623 | 544 | 2 | ENSMUST00000126393.1 | 515 | 4933427G23Rik | ENSMUSG00000086697.1 |
| chr5 | 23831415 | 23831731 | 3.65807 | 4.6548 | 1.54725 | Promoter | 5 | 23831080 | 23831623 | 544 | 2 | ENSMUST00000126393.1 | 0 | 4933427G23Rik | ENSMUSG00000086697.1 |
| chr13 | 118715727 | 118716432 | 3.46877 | 4.09121 | 1.21905 | Promoter | 13 | 118695292 | 118715396 | 20105 | 2 | ENSMUST00000144680.1 | -332 | Gm16263 | ENSMUSG00000086709.1 |
| chr5 | 116322611 | 116323118 | 3.65567 | 4.42726 | 1.39321 | Distal Intergenic | 5 | 116312360 | 116318281 | 5922 | 1 | ENSMUST00000128473.1 | 10252 | B230112J18Rik | ENSMUSG00000086715.1 |
| chr5 | 116323839 | 116324396 | 3.929 | 5.09581 | 1.82518 | Distal Intergenic | 5 | 116312360 | 116318281 | 5922 | 1 | ENSMUST00000128473.1 | 11480 | B230112J18Rik | ENSMUSG00000086715.1 |
| chr6 | 32327323 | 32327631 | 4.24586 | 5.57588 | 2.16424 | Intron (ENSMUST00000115096.3/ENSMUSG00000029765.11, intron 3 of 31) | 6 | 32303913 | 32316606 | 12694 | 1 | ENSMUST00000143328.1 | 23411 | Plxna4os3 | ENSMUSG00000086720.1 |
| chr6 | 32358920 | 32359096 | 3.24724 | 3.69802 | 1.01733 | Intron (ENSMUST00000115096.3/ENSMUSG00000029765.11, intron 3 of 31) | 6 | 32303913 | 32316606 | 12694 | 1 | ENSMUST00000143328.1 | 55008 | Plxna4os3 | ENSMUSG00000086720.1 |
| chr6 | 32376740 | 32377254 | 3.55503 | 4.26719 | 1.35304 | Intron (ENSMUST00000115096.3/ENSMUSG00000029765.11, intron 3 of 31) | 6 | 32303913 | 32316606 | 12694 | 1 | ENSMUST00000143328.1 | 72828 | Plxna4os3 | ENSMUSG00000086720.1 |
| chr8 | 23149613 | 23150157 | 3.89008 | 4.99347 | 1.76095 | Promoter | 8 | 23138794 | 23149585 | 10792 | 2 | ENSMUST00000141930.1 | -29 | Gm15816 | ENSMUSG00000086741.1 |
| chr8 | 23150845 | 23150953 | 4.5718 | 6.23463 | 2.65055 | Distal Intergenic | 8 | 23138794 | 23149585 | 10792 | 2 | ENSMUST00000141930.1 | -1261 | Gm15816 | ENSMUSG00000086741.1 |
| chr14 | 46385886 | 46386017 | 3.76733 | 4.68214 | 1.53654 | Exon (ENSMUST00000074077.6/ENSMUSG00000021835.9, exon 3 of 4) | 14 | 46387519 | 46389282 | 1764 | 1 | ENSMUST00000127359.1 | -1502 | Gm15222 | ENSMUSG00000086746.1 |
| chr14 | 46387032 | 46387150 | 4.31344 | 5.6913 | 2.22685 | Promoter | 14 | 46387519 | 46389282 | 1764 | 1 | ENSMUST00000127359.1 | -369 | Gm15222 | ENSMUSG00000086746.1 |
| chr14 | 46388718 | 46388978 | 3.52053 | 4.1588 | 1.1882 | Intron (ENSMUST00000127359.1/ENSMUSG00000086746.1, intron 1 of 1) | 14 | 46387519 | 46389282 | 1764 | 1 | ENSMUST00000127359.1 | 1200 | Gm15222 | ENSMUSG00000086746.1 |
| chr2 | 10373391 | 10373686 | 3.39579 | 3.93232 | 1.05547 | Promoter | 2 | 10339283 | 10374041 | 34759 | 2 | ENSMUST00000137351.1 | 355 | Gm13261 | ENSMUSG00000086748.1 |
| chr2 | 10374001 | 10374632 | 3.50552 | 4.40563 | 1.40468 | Promoter | 2 | 10339283 | 10374041 | 34759 | 2 | ENSMUST00000137351.1 | 0 | Gm13261 | ENSMUSG00000086748.1 |
| chr2 | 10375008 | 10375796 | 3.69816 | 4.55081 | 1.50652 | Promoter | 2 | 10339283 | 10374041 | 34759 | 2 | ENSMUST00000137351.1 | -968 | Gm13261 | ENSMUSG00000086748.1 |
| chr2 | 10376465 | 10376597 | 3.44464 | 4.01979 | 1.1037 | Intron (ENSMUST00000114862.2/ENSMUSG00000061186.10, intron 1 of 5) | 2 | 10339283 | 10374041 | 34759 | 2 | ENSMUST00000137351.1 | -2425 | Gm13261 | ENSMUSG00000086748.1 |
| chr5 | 123506876 | 123507108 | 3.73331 | 4.55181 | 1.42922 | Intron (ENSMUST00000094327.4/ENSMUSG00000063409.7, intron 11 of 11) | 5 | 123508395 | 123514011 | 5617 | 1 | ENSMUST00000134552.1 | -1287 | Gm15751 | ENSMUSG00000086753.1 |
| chr5 | 123507421 | 123507833 | 3.5913 | 4.30375 | 1.32395 | Promoter | 5 | 123508395 | 123514011 | 5617 | 1 | ENSMUST00000134552.1 | -562 | Gm15751 | ENSMUSG00000086753.1 |
| chr11 | 114091408 | 114091607 | 3.31306 | 3.96349 | 1.10464 | Promoter | 11 | 114091406 | 114095336 | 3931 | 1 | ENSMUST00000129365.1 | 3 | 1700025D23Rik | ENSMUSG00000086760.1 |
| chr11 | 114114444 | 114114835 | 3.59913 | 4.30589 | 1.27966 | Distal Intergenic | 11 | 114091406 | 114095336 | 3931 | 1 | ENSMUST00000129365.1 | 23039 | 1700025D23Rik | ENSMUSG00000086760.1 |
| chr11 | 114119266 | 114119776 | 3.75568 | 4.60512 | 1.47586 | Distal Intergenic | 11 | 114091406 | 114095336 | 3931 | 1 | ENSMUST00000129365.1 | 27861 | 1700025D23Rik | ENSMUSG00000086760.1 |
| chr11 | 114129027 | 114129367 | 3.37283 | 3.88997 | 1.02823 | Distal Intergenic | 11 | 114091406 | 114095336 | 3931 | 1 | ENSMUST00000129365.1 | 37622 | 1700025D23Rik | ENSMUSG00000086760.1 |
| chr11 | 114129710 | 114130340 | 3.49841 | 4.14257 | 1.2288 | Distal Intergenic | 11 | 114091406 | 114095336 | 3931 | 1 | ENSMUST00000129365.1 | 38305 | 1700025D23Rik | ENSMUSG00000086760.1 |
| chr6 | 34252369 | 34252859 | 4.0084 | 5.17464 | 1.9274 | Distal Intergenic | 6 | 34290265 | 34324249 | 33985 | 1 | ENSMUST00000138275.1 | -37406 | Gm14546 | ENSMUSG00000086762.1 |
| chr4 | 135618736 | 135618811 | 3.49789 | 4.13326 | 1.22269 | Distal Intergenic | 4 | 135626655 | 135630198 | 3544 | 1 | ENSMUST00000153347.1 | -7844 | 1700029M20Rik | ENSMUSG00000086788.2 |
| chr4 | 135626767 | 135626973 | 3.26294 | 3.72158 | 1.01733 | Promoter | 4 | 135626774 | 135628485 | 1712 | 1 | ENSMUST00000134416.1 | 0 | 1700029M20Rik | ENSMUSG00000086788.2 |
| chr4 | 135644805 | 135645145 | 3.34121 | 3.87925 | 1.05291 | Distal Intergenic | 4 | 135627253 | 135627985 | 733 | 1 | ENSMUST00000148529.1 | 17553 | 1700029M20Rik | ENSMUSG00000086788.2 |
| chr4 | 135645992 | 135646069 | 4.0323 | 5.12769 | 1.81487 | Distal Intergenic | 4 | 135627253 | 135627985 | 733 | 1 | ENSMUST00000148529.1 | 18740 | 1700029M20Rik | ENSMUSG00000086788.2 |
| chr4 | 34961929 | 34962128 | 3.53634 | 4.24436 | 1.35449 | Promoter | 4 | 34962220 | 34966965 | 4746 | 1 | ENSMUST00000144419.1 | -92 | Gm12364 | ENSMUSG00000086792.1 |
| chr4 | 34969404 | 34969637 | 3.244 | 3.69316 | 1.01733 | Intron (ENSMUST00000102975.4/ENSMUSG00000073910.5, intron 3 of 3) | 4 | 34962220 | 34966965 | 4746 | 1 | ENSMUST00000144419.1 | 7185 | Gm12364 | ENSMUSG00000086792.1 |
| chr4 | 154651213 | 154651288 | 3.54161 | 4.27818 | 1.29199 | Intron (ENSMUST00000138077.1/ENSMUSG00000086810.1, intron 1 of 1) | 4 | 154645754 | 154655318 | 9565 | 2 | ENSMUST00000138077.1 | 4030 | Gm13110 | ENSMUSG00000086810.1 |
| chr4 | 154652666 | 154653203 | 3.92389 | 5.18513 | 1.90501 | Intron (ENSMUST00000138077.1/ENSMUSG00000086810.1, intron 1 of 1) | 4 | 154645754 | 154655318 | 9565 | 2 | ENSMUST00000138077.1 | 2115 | Gm13110 | ENSMUSG00000086810.1 |
| chr4 | 154659522 | 154659997 | 3.26082 | 3.71836 | 1.01733 | Distal Intergenic | 4 | 154645754 | 154655318 | 9565 | 2 | ENSMUST00000138077.1 | -4205 | Gm13110 | ENSMUSG00000086810.1 |
| chr7 | 132151698 | 132151798 | 3.76317 | 4.60682 | 1.46432 | Intron (ENSMUST00000124096.2/ENSMUSG00000030849.12, intron 2 of 6) | 7 | 132154171 | 132155301 | 1131 | 1 | ENSMUST00000152302.1 | -2373 | Gm15677 | ENSMUSG00000086814.1 |
| chr5 | 30106566 | 30106815 | 3.70454 | 4.55883 | 1.53222 | Intron (ENSMUST00000058045.4/ENSMUSG00000044576.6, intron 1 of 5) | 5 | 30103889 | 30105359 | 1471 | 2 | ENSMUST00000127749.3 | -1208 | 3110082J24Rik | ENSMUSG00000086815.3 |
| chr5 | 30107444 | 30107929 | 3.91497 | 4.98841 | 1.83414 | Intron (ENSMUST00000058045.4/ENSMUSG00000044576.6, intron 1 of 5) | 5 | 30103889 | 30105359 | 1471 | 2 | ENSMUST00000127749.3 | -2086 | 3110082J24Rik | ENSMUSG00000086815.3 |
| chr2 | 21141811 | 21142184 | 3.47462 | 4.12374 | 1.2773 | Promoter | 2 | 21044808 | 21141954 | 97147 | 2 | ENSMUST00000137984.1 | 0 | Gm13377 | ENSMUSG00000086821.3 |
| chr2 | 21142560 | 21143013 | 3.64811 | 4.50684 | 1.50332 | Promoter | 2 | 21044808 | 21141954 | 97147 | 2 | ENSMUST00000137984.1 | -607 | Gm13377 | ENSMUSG00000086821.3 |
| chr5 | 119685993 | 119686370 | 3.46142 | 4.05088 | 1.12412 | Exon (ENSMUST00000135385.1/ENSMUSG00000086847.1, exon 3 of 3) | 5 | 119685095 | 119691218 | 6124 | 2 | ENSMUST00000135385.1 | 4848 | Tbx3os2 | ENSMUSG00000086847.1 |
| chr5 | 119686955 | 119687387 | 3.91849 | 5.11003 | 1.87148 | Intron (ENSMUST00000135385.1/ENSMUSG00000086847.1, intron 2 of 2) | 5 | 119685095 | 119691218 | 6124 | 2 | ENSMUST00000135385.1 | 3831 | Tbx3os2 | ENSMUSG00000086847.1 |
| chr5 | 119687773 | 119688001 | 3.47173 | 4.0653 | 1.12001 | Intron (ENSMUST00000135385.1/ENSMUSG00000086847.1, intron 2 of 2) | 5 | 119685095 | 119691218 | 6124 | 2 | ENSMUST00000135385.1 | 3217 | Tbx3os2 | ENSMUSG00000086847.1 |
| chr5 | 119688963 | 119689189 | 3.79354 | 4.67986 | 1.53794 | Intron (ENSMUST00000135385.1/ENSMUSG00000086847.1, intron 1 of 2) | 5 | 119685095 | 119691218 | 6124 | 2 | ENSMUST00000135385.1 | 2029 | Tbx3os2 | ENSMUSG00000086847.1 |
| chr5 | 119691133 | 119691608 | 3.77778 | 4.72347 | 1.61884 | Promoter | 5 | 119685095 | 119691218 | 6124 | 2 | ENSMUST00000135385.1 | 0 | Tbx3os2 | ENSMUSG00000086847.1 |
| chr5 | 119692530 | 119692940 | 3.53596 | 4.5227 | 1.47168 | Distal Intergenic | 5 | 119685095 | 119691218 | 6124 | 2 | ENSMUST00000135385.1 | -1313 | Tbx3os2 | ENSMUSG00000086847.1 |
| chr5 | 119701617 | 119701967 | 3.27169 | 3.73469 | 1.01733 | Distal Intergenic | 5 | 119685095 | 119691218 | 6124 | 2 | ENSMUST00000135385.1 | -10400 | Tbx3os2 | ENSMUSG00000086847.1 |
| chr5 | 119703930 | 119704016 | 3.50707 | 4.1425 | 1.20178 | Distal Intergenic | 5 | 119685095 | 119691218 | 6124 | 2 | ENSMUST00000135385.1 | -12713 | Tbx3os2 | ENSMUSG00000086847.1 |
| chr5 | 119705698 | 119706724 | 3.64062 | 4.46926 | 1.42302 | Distal Intergenic | 5 | 119685095 | 119691218 | 6124 | 2 | ENSMUST00000135385.1 | -14481 | Tbx3os2 | ENSMUSG00000086847.1 |
| chr5 | 119708275 | 119708839 | 3.62365 | 4.54073 | 1.49998 | Distal Intergenic | 5 | 119685095 | 119691218 | 6124 | 2 | ENSMUST00000135385.1 | -17058 | Tbx3os2 | ENSMUSG00000086847.1 |
| chr5 | 119710103 | 119710362 | 3.83149 | 4.74866 | 1.59281 | Distal Intergenic | 5 | 119685095 | 119691218 | 6124 | 2 | ENSMUST00000135385.1 | -18886 | Tbx3os2 | ENSMUSG00000086847.1 |
| chr5 | 119710995 | 119711367 | 3.45791 | 4.09461 | 1.1858 | Distal Intergenic | 5 | 119685095 | 119691218 | 6124 | 2 | ENSMUST00000135385.1 | -19778 | Tbx3os2 | ENSMUSG00000086847.1 |
| chr5 | 119739560 | 119739687 | 3.88579 | 4.98981 | 1.75627 | Distal Intergenic | 5 | 119685095 | 119691218 | 6124 | 2 | ENSMUST00000135385.1 | -48343 | Tbx3os2 | ENSMUSG00000086847.1 |
| chr5 | 119740969 | 119741132 | 3.36704 | 3.89441 | 1.07491 | Distal Intergenic | 5 | 119685095 | 119691218 | 6124 | 2 | ENSMUST00000135385.1 | -49752 | Tbx3os2 | ENSMUSG00000086847.1 |
| chr5 | 119742281 | 119742693 | 3.6158 | 4.51184 | 1.50604 | Distal Intergenic | 5 | 119685095 | 119691218 | 6124 | 2 | ENSMUST00000135385.1 | -51064 | Tbx3os2 | ENSMUSG00000086847.1 |
| chr5 | 119748564 | 119748857 | 3.9701 | 5.10641 | 1.85024 | Distal Intergenic | 5 | 119685095 | 119691218 | 6124 | 2 | ENSMUST00000135385.1 | -57347 | Tbx3os2 | ENSMUSG00000086847.1 |
| chr5 | 119749301 | 119749755 | 3.45789 | 4.05615 | 1.16248 | Distal Intergenic | 5 | 119685095 | 119691218 | 6124 | 2 | ENSMUST00000135385.1 | -58084 | Tbx3os2 | ENSMUSG00000086847.1 |
| chr5 | 119750571 | 119751056 | 3.80005 | 4.70566 | 1.56776 | Distal Intergenic | 5 | 119685095 | 119691218 | 6124 | 2 | ENSMUST00000135385.1 | -59354 | Tbx3os2 | ENSMUSG00000086847.1 |
| chr5 | 119752807 | 119753050 | 3.78791 | 4.7777 | 1.61064 | Distal Intergenic | 5 | 119685095 | 119691218 | 6124 | 2 | ENSMUST00000135385.1 | -61590 | Tbx3os2 | ENSMUSG00000086847.1 |
| chr5 | 119755412 | 119755726 | 3.53717 | 4.22428 | 1.30909 | Distal Intergenic | 5 | 119685095 | 119691218 | 6124 | 2 | ENSMUST00000135385.1 | -64195 | Tbx3os2 | ENSMUSG00000086847.1 |
| chr5 | 119756353 | 119756566 | 3.47067 | 4.07856 | 1.17331 | Distal Intergenic | 5 | 119685095 | 119691218 | 6124 | 2 | ENSMUST00000135385.1 | -65136 | Tbx3os2 | ENSMUSG00000086847.1 |
| chr6 | 145180040 | 145180192 | 3.35921 | 3.86638 | 1.01733 | Intron (ENSMUST00000111727.2/ENSMUSG00000043541.11, intron 10 of 11) | 6 | 145182275 | 145187630 | 5356 | 1 | ENSMUST00000141307.1 | -2083 | Gm15543 | ENSMUSG00000086863.1 |
| chr4 | 117430182 | 117430393 | 3.37055 | 3.98231 | 1.15413 | Intron (ENSMUST00000030439.9/ENSMUSG00000028677.13, intron 2 of 14) | 4 | 117445950 | 117455040 | 9091 | 1 | ENSMUST00000136115.1 | -15557 | Gm12828 | ENSMUSG00000086876.1 |
| chr4 | 117415885 | 117416230 | 3.86486 | 4.80835 | 1.61504 | Intron (ENSMUST00000030439.9/ENSMUSG00000028677.13, intron 2 of 14) | 4 | 117445950 | 117455040 | 9091 | 1 | ENSMUST00000136115.1 | -29720 | Gm12828 | ENSMUSG00000086876.1 |
| chr4 | 117465595 | 117466033 | 3.85752 | 4.8674 | 1.68744 | Intron (ENSMUST00000030439.9/ENSMUSG00000028677.13, intron 2 of 14) | 4 | 117445950 | 117455040 | 9091 | 1 | ENSMUST00000136115.1 | 19646 | Gm12828 | ENSMUSG00000086876.1 |
| chr4 | 117441421 | 117441737 | 3.38193 | 3.9006 | 1.01733 | Intron (ENSMUST00000030439.9/ENSMUSG00000028677.13, intron 2 of 14) | 4 | 117445950 | 117455040 | 9091 | 1 | ENSMUST00000136115.1 | -4213 | Gm12828 | ENSMUSG00000086876.1 |
| chr4 | 117442253 | 117442515 | 3.96211 | 4.98731 | 1.72198 | Intron (ENSMUST00000030439.9/ENSMUSG00000028677.13, intron 2 of 14) | 4 | 117445950 | 117455040 | 9091 | 1 | ENSMUST00000136115.1 | -3435 | Gm12828 | ENSMUSG00000086876.1 |
| chr4 | 117450957 | 117451307 | 3.64859 | 4.46044 | 1.4576 | Intron (ENSMUST00000136115.1/ENSMUSG00000086876.1, intron 2 of 2) | 4 | 117445950 | 117455040 | 9091 | 1 | ENSMUST00000136115.1 | 5008 | Gm12828 | ENSMUSG00000086876.1 |
| chr4 | 135026579 | 135026727 | 3.5463 | 4.25689 | 1.30471 | Distal Intergenic | 4 | 135118221 | 135120447 | 2227 | 2 | ENSMUST00000146152.1 | 93720 | Gm16225 | ENSMUSG00000086884.1 |
| chr4 | 135027183 | 135027422 | 3.30725 | 3.8037 | 1.07146 | Distal Intergenic | 4 | 135118221 | 135120447 | 2227 | 2 | ENSMUST00000146152.1 | 93025 | Gm16225 | ENSMUSG00000086884.1 |
| chr11 | 117006815 | 117006983 | 3.32835 | 3.81984 | 1.01733 | Distal Intergenic | 11 | 117038289 | 117040188 | 1900 | 2 | ENSMUST00000152401.1 | 33205 | Gm11728 | ENSMUSG00000086902.2 |
| chr2 | 33965887 | 33967483 | 3.73667 | 4.65664 | 1.59135 | Distal Intergenic | 2 | 33978246 | 33980324 | 2079 | 1 | ENSMUST00000156633.1 | -10763 | Gm13404 | ENSMUSG00000086904.1 |
| chr3 | 100438490 | 100438803 | 3.38043 | 3.91166 | 1.04685 | Distal Intergenic | 3 | 100426439 | 100437754 | 11316 | 1 | ENSMUST00000152827.1 | 12052 | 4930406D18Rik | ENSMUSG00000086923.1 |
| chr11 | 119934368 | 119934649 | 3.52582 | 4.19258 | 1.28429 | Downstream (1-2kb) | 11 | 119938511 | 119942074 | 3564 | 1 | ENSMUST00000143428.1 | -3862 | Gm11766 | ENSMUSG00000086924.1 |
| chr2 | 33543236 | 33544854 | 4.07175 | 5.3436 | 2.07065 | Distal Intergenic | 2 | 33489782 | 33503632 | 13851 | 1 | ENSMUST00000134014.1 | 53455 | Gm13530 | ENSMUSG00000086939.1 |
| chr2 | 33546153 | 33546431 | 3.46868 | 4.1376 | 1.26332 | Distal Intergenic | 2 | 33489782 | 33503632 | 13851 | 1 | ENSMUST00000134014.1 | 56372 | Gm13530 | ENSMUSG00000086939.1 |
| chr11 | 72265567 | 72265840 | 3.38729 | 4.34432 | 1.33193 | Promoter | 11 | 72266421 | 72268575 | 2155 | 1 | ENSMUST00000147937.1 | -581 | 1700051A21Rik | ENSMUSG00000086951.1 |
| chr11 | 72266320 | 72266501 | 3.56971 | 4.43721 | 1.38173 | Promoter | 11 | 72266421 | 72268575 | 2155 | 1 | ENSMUST00000147937.1 | 0 | 1700051A21Rik | ENSMUSG00000086951.1 |
| chr4 | 124309393 | 124309741 | 3.21 | 3.64226 | 1.00765 | Intron (ENSMUST00000153719.1/ENSMUSG00000086955.1, intron 1 of 1) | 4 | 124302837 | 124311461 | 8625 | 2 | ENSMUST00000153719.1 | 1720 | 1700021L23Rik | ENSMUSG00000086955.1 |
| chr4 | 124310690 | 124310871 | 3.26565 | 3.73246 | 1.04045 | Promoter | 4 | 124302837 | 124311461 | 8625 | 2 | ENSMUST00000153719.1 | 590 | 1700021L23Rik | ENSMUSG00000086955.1 |
| chr4 | 124317127 | 124317202 | 3.35467 | 4.02319 | 1.10333 | Distal Intergenic | 4 | 124302837 | 124311461 | 8625 | 2 | ENSMUST00000153719.1 | -5667 | 1700021L23Rik | ENSMUSG00000086955.1 |
| chr4 | 124317869 | 124318240 | 3.22958 | 3.67155 | 1.01623 | Distal Intergenic | 4 | 124302837 | 124311461 | 8625 | 2 | ENSMUST00000153719.1 | -6409 | 1700021L23Rik | ENSMUSG00000086955.1 |
| chr4 | 140386655 | 140386924 | 3.377 | 3.91393 | 1.09098 | Distal Intergenic | 4 | 140391148 | 140396604 | 5457 | 1 | ENSMUST00000156409.1 | -4224 | Gm13017 | ENSMUSG00000086960.1 |
| chr2 | 170400523 | 170400866 | 3.69715 | 4.50123 | 1.39985 | Intron (ENSMUST00000154650.2/ENSMUSG00000013523.8, intron 1 of 6) | 2 | 170405181 | 170407175 | 1995 | 1 | ENSMUST00000133586.1 | -4315 | Bcas1os1 | ENSMUSG00000086970.1 |
| chr2 | 170401372 | 170401477 | 3.46818 | 4.34042 | 1.35191 | Intron (ENSMUST00000154650.2/ENSMUSG00000013523.8, intron 1 of 6) | 2 | 170405181 | 170407175 | 1995 | 1 | ENSMUST00000133586.1 | -3704 | Bcas1os1 | ENSMUSG00000086970.1 |
| chr2 | 170399914 | 170400067 | 3.47191 | 4.22972 | 1.27488 | Intron (ENSMUST00000154650.2/ENSMUSG00000013523.8, intron 1 of 6) | 2 | 170405181 | 170407175 | 1995 | 1 | ENSMUST00000133586.1 | -5114 | Bcas1os1 | ENSMUSG00000086970.1 |
| chr9 | 35574029 | 35574793 | 4.04449 | 5.34792 | 2.038 | Promoter | 9 | 35574112 | 35574400 | 289 | 2 | ENSMUST00000121769.1 | 0 | Gm6762 | ENSMUSG00000086984.4 |
| chr6 | 84106855 | 84107030 | 3.3861 | 3.92572 | 1.08407 | Promoter | 6 | 84101206 | 84107002 | 5797 | 2 | ENSMUST00000137673.1 | 0 | Gm15475 | ENSMUSG00000087001.1 |
| chr19 | 58140331 | 58140669 | 3.3614 | 4.04749 | 1.14686 | Distal Intergenic | 19 | 58204139 | 58236047 | 31909 | 1 | ENSMUST00000143595.1 | -63470 | Gm16277 | ENSMUSG00000087002.1 |
| chr19 | 58219746 | 58219937 | 3.26614 | 3.72634 | 1.01733 | Intron (ENSMUST00000143595.1/ENSMUSG00000087002.1, intron 1 of 1) | 19 | 58204139 | 58236047 | 31909 | 1 | ENSMUST00000143595.1 | 15608 | Gm16277 | ENSMUSG00000087002.1 |
| chr19 | 58221140 | 58221475 | 3.2914 | 3.76426 | 1.01733 | Intron (ENSMUST00000143595.1/ENSMUSG00000087002.1, intron 1 of 1) | 19 | 58204139 | 58236047 | 31909 | 1 | ENSMUST00000143595.1 | 17002 | Gm16277 | ENSMUSG00000087002.1 |
| chr19 | 58228632 | 58229741 | 3.5017 | 4.12279 | 1.16224 | Intron (ENSMUST00000143595.1/ENSMUSG00000087002.1, intron 1 of 1) | 19 | 58204139 | 58236047 | 31909 | 1 | ENSMUST00000143595.1 | 24494 | Gm16277 | ENSMUSG00000087002.1 |
| chr2 | 93954198 | 93954666 | 3.41259 | 3.95827 | 1.05778 | Downstream (1-2kb) | 2 | 93955810 | 93957049 | 1240 | 2 | ENSMUST00000148314.2 | 2383 | Gm13889 | ENSMUSG00000087006.3 |
| chr2 | 93955010 | 93957252 | 3.75915 | 4.68536 | 1.56346 | Promoter | 2 | 93955810 | 93957049 | 1240 | 2 | ENSMUST00000148314.2 | 0 | Gm13889 | ENSMUSG00000087006.3 |
| chr4 | 125191095 | 125191656 | 3.64488 | 4.62885 | 1.53708 | Distal Intergenic | 4 | 125228597 | 125242052 | 13456 | 1 | ENSMUST00000130055.1 | -36941 | 1700041M05Rik | ENSMUSG00000087010.1 |
| chr4 | 125192202 | 125192422 | 3.31465 | 3.79922 | 1.01733 | Distal Intergenic | 4 | 125228597 | 125242052 | 13456 | 1 | ENSMUST00000130055.1 | -36175 | 1700041M05Rik | ENSMUSG00000087010.1 |
| chr4 | 125193784 | 125193876 | 3.36629 | 3.87701 | 1.01733 | Distal Intergenic | 4 | 125228597 | 125242052 | 13456 | 1 | ENSMUST00000130055.1 | -34721 | 1700041M05Rik | ENSMUSG00000087010.1 |
| chr4 | 125195065 | 125195512 | 4.26113 | 5.69508 | 2.25518 | Distal Intergenic | 4 | 125228597 | 125242052 | 13456 | 1 | ENSMUST00000130055.1 | -33085 | 1700041M05Rik | ENSMUSG00000087010.1 |
| chr4 | 125219790 | 125219865 | 3.41456 | 3.98035 | 1.12412 | Distal Intergenic | 4 | 125228597 | 125242052 | 13456 | 1 | ENSMUST00000130055.1 | -8732 | 1700041M05Rik | ENSMUSG00000087010.1 |
| chr4 | 125223148 | 125223370 | 3.5287 | 4.38216 | 1.39852 | Distal Intergenic | 4 | 125228597 | 125242052 | 13456 | 1 | ENSMUST00000130055.1 | -5227 | 1700041M05Rik | ENSMUSG00000087010.1 |
| chr4 | 125236721 | 125236965 | 3.81362 | 4.71327 | 1.5643 | Intron (ENSMUST00000130055.1/ENSMUSG00000087010.1, intron 1 of 1) | 4 | 125228597 | 125242052 | 13456 | 1 | ENSMUST00000130055.1 | 8125 | 1700041M05Rik | ENSMUSG00000087010.1 |
| chr4 | 125246930 | 125247046 | 3.30097 | 3.77875 | 1.01733 | Distal Intergenic | 4 | 125228597 | 125242052 | 13456 | 1 | ENSMUST00000130055.1 | 18334 | 1700041M05Rik | ENSMUSG00000087010.1 |
| chr4 | 125264331 | 125264594 | 3.66314 | 4.41463 | 1.33283 | Distal Intergenic | 4 | 125228597 | 125242052 | 13456 | 1 | ENSMUST00000130055.1 | 35735 | 1700041M05Rik | ENSMUSG00000087010.1 |
| chr4 | 125269147 | 125269330 | 3.24622 | 3.84061 | 1.02488 | Distal Intergenic | 4 | 125228597 | 125242052 | 13456 | 1 | ENSMUST00000130055.1 | 40551 | 1700041M05Rik | ENSMUSG00000087010.1 |
| chr4 | 125275168 | 125275248 | 3.6332 | 4.58465 | 1.50619 | Distal Intergenic | 4 | 125228597 | 125242052 | 13456 | 1 | ENSMUST00000130055.1 | 46572 | 1700041M05Rik | ENSMUSG00000087010.1 |
| chr4 | 125276340 | 125276863 | 3.49767 | 4.12486 | 1.19305 | Distal Intergenic | 4 | 125228597 | 125242052 | 13456 | 1 | ENSMUST00000130055.1 | 47744 | 1700041M05Rik | ENSMUSG00000087010.1 |
| chr4 | 125336817 | 125337034 | 3.30271 | 3.78925 | 1.04521 | Distal Intergenic | 4 | 125228597 | 125242052 | 13456 | 1 | ENSMUST00000130055.1 | 108221 | 1700041M05Rik | ENSMUSG00000087010.1 |
| chr4 | 124987117 | 124987272 | 4.56351 | 6.36722 | 2.78366 | Promoter | 4 | 124957647 | 124987009 | 29363 | 2 | ENSMUST00000146679.1 | -109 | Gm12930 | ENSMUSG00000087011.1 |
| chr4 | 124987964 | 124988134 | 3.33802 | 3.83437 | 1.01733 | Promoter | 4 | 124957647 | 124987009 | 29363 | 2 | ENSMUST00000146679.1 | -956 | Gm12930 | ENSMUSG00000087011.1 |
| chr4 | 124988768 | 124989059 | 3.46112 | 4.05852 | 1.1525 | Intron (ENSMUST00000030687.7/ENSMUSG00000028871.7, intron 1 of 5) | 4 | 124957647 | 124987009 | 29363 | 2 | ENSMUST00000146679.1 | -1760 | Gm12930 | ENSMUSG00000087011.1 |
| chr4 | 124990374 | 124990562 | 3.28798 | 4.19246 | 1.24325 | Intron (ENSMUST00000030687.7/ENSMUSG00000028871.7, intron 1 of 5) | 4 | 124957647 | 124987009 | 29363 | 2 | ENSMUST00000146679.1 | -3366 | Gm12930 | ENSMUSG00000087011.1 |
| chr4 | 154771995 | 154772240 | 3.43862 | 4.03632 | 1.15444 | Distal Intergenic | 4 | 154837459 | 154839275 | 1817 | 1 | ENSMUST00000130812.1 | -65219 | Gm13113 | ENSMUSG00000087012.1 |
| chr4 | 154780097 | 154780445 | 3.71215 | 4.61802 | 1.54876 | Distal Intergenic | 4 | 154837459 | 154839275 | 1817 | 1 | ENSMUST00000130812.1 | -57014 | Gm13113 | ENSMUSG00000087012.1 |
| chr4 | 154786131 | 154786441 | 3.389 | 3.9222 | 1.05602 | Distal Intergenic | 4 | 154837459 | 154839275 | 1817 | 1 | ENSMUST00000130812.1 | -51018 | Gm13113 | ENSMUSG00000087012.1 |
| chr4 | 154786946 | 154787637 | 3.58227 | 4.27118 | 1.25527 | Distal Intergenic | 4 | 154837459 | 154839275 | 1817 | 1 | ENSMUST00000130812.1 | -49822 | Gm13113 | ENSMUSG00000087012.1 |
| chr4 | 154788251 | 154788734 | 3.33975 | 3.98878 | 1.10047 | Distal Intergenic | 4 | 154837459 | 154839275 | 1817 | 1 | ENSMUST00000130812.1 | -48725 | Gm13113 | ENSMUSG00000087012.1 |
| chr4 | 154802452 | 154802918 | 4.18642 | 5.4719 | 2.07484 | Distal Intergenic | 4 | 154837459 | 154839275 | 1817 | 1 | ENSMUST00000130812.1 | -34541 | Gm13113 | ENSMUSG00000087012.1 |
| chr4 | 154812211 | 154812675 | 3.60289 | 4.31345 | 1.28866 | Distal Intergenic | 4 | 154837459 | 154839275 | 1817 | 1 | ENSMUST00000130812.1 | -24784 | Gm13113 | ENSMUSG00000087012.1 |
| chr4 | 154839300 | 154839526 | 3.33529 | 4.00579 | 1.13289 | Distal Intergenic | 4 | 154837459 | 154839275 | 1817 | 1 | ENSMUST00000130812.1 | 1842 | Gm13113 | ENSMUSG00000087012.1 |
| chr11 | 85828627 | 85828717 | 3.30299 | 3.78167 | 1.01733 | Intron (ENSMUST00000140673.2/ENSMUSG00000087013.2, intron 1 of 4) | 11 | 85791661 | 85832209 | 40549 | 2 | ENSMUST00000140673.2 | 3492 | 2610027K06Rik | ENSMUSG00000087013.2 |
| chr11 | 85829091 | 85831183 | 3.9633 | 5.17891 | 1.94347 | Intron (ENSMUST00000140673.2/ENSMUSG00000087013.2, intron 1 of 4) | 11 | 85791661 | 85832209 | 40549 | 2 | ENSMUST00000140673.2 | 1026 | 2610027K06Rik | ENSMUSG00000087013.2 |
| chr2 | 129573336 | 129573677 | 3.54351 | 4.20014 | 1.21347 | Distal Intergenic | 2 | 129565047 | 129566068 | 1022 | 2 | ENSMUST00000150181.1 | -7269 | Gm14041 | ENSMUSG00000087015.1 |
| chr2 | 129574516 | 129574746 | 3.95192 | 5.09093 | 1.87946 | Distal Intergenic | 2 | 129565047 | 129566068 | 1022 | 2 | ENSMUST00000150181.1 | -8449 | Gm14041 | ENSMUSG00000087015.1 |
| chr4 | 148305670 | 148306169 | 3.60203 | 4.43112 | 1.41022 | Promoter | 4 | 148306853 | 148316569 | 9717 | 1 | ENSMUST00000129579.2 | -684 | Gm13206 | ENSMUSG00000087027.2 |
| chr4 | 148318120 | 148318357 | 3.55832 | 4.2314 | 1.23966 | Distal Intergenic | 4 | 148306853 | 148316569 | 9717 | 1 | ENSMUST00000129579.2 | 11268 | Gm13206 | ENSMUSG00000087027.2 |
| chr4 | 148319516 | 148319617 | 4.19395 | 5.45905 | 2.06682 | Distal Intergenic | 4 | 148306853 | 148316569 | 9717 | 1 | ENSMUST00000129579.2 | 12664 | Gm13206 | ENSMUSG00000087027.2 |
| chr4 | 148321557 | 148322614 | 3.42873 | 4.00502 | 1.13163 | Distal Intergenic | 4 | 148306853 | 148316569 | 9717 | 1 | ENSMUST00000129579.2 | 14705 | Gm13206 | ENSMUSG00000087027.2 |
| chr11 | 97532281 | 97532846 | 4.00263 | 5.16083 | 1.90735 | Intron (ENSMUST00000107596.3/ENSMUSG00000038453.10, intron 11 of 19) | 11 | 97519612 | 97521895 | 2284 | 2 | ENSMUST00000134289.1 | -10387 | Gm11611 | ENSMUSG00000087042.1 |
| chr11 | 97533236 | 97533895 | 3.95741 | 5.0183 | 1.78396 | Exon (ENSMUST00000107596.3/ENSMUSG00000038453.10, exon 9 of 20) | 11 | 97519612 | 97521895 | 2284 | 2 | ENSMUST00000134289.1 | -11342 | Gm11611 | ENSMUSG00000087042.1 |
| chr11 | 97534462 | 97535440 | 4.26785 | 5.75182 | 2.34467 | Exon (ENSMUST00000107596.3/ENSMUSG00000038453.10, exon 8 of 20) | 11 | 97519612 | 97521895 | 2284 | 2 | ENSMUST00000134289.1 | -12568 | Gm11611 | ENSMUSG00000087042.1 |
| chr11 | 97536204 | 97537262 | 4.19202 | 5.49621 | 2.13574 | Exon (ENSMUST00000107596.3/ENSMUSG00000038453.10, exon 5 of 20) | 11 | 97519612 | 97521895 | 2284 | 2 | ENSMUST00000134289.1 | -14310 | Gm11611 | ENSMUSG00000087042.1 |
| chr11 | 97537764 | 97538975 | 3.86894 | 4.93939 | 1.73566 | Intron (ENSMUST00000107596.3/ENSMUSG00000038453.10, intron 4 of 19) | 11 | 97519612 | 97521895 | 2284 | 2 | ENSMUST00000134289.1 | -15870 | Gm11611 | ENSMUSG00000087042.1 |
| chr11 | 97539625 | 97539700 | 3.62061 | 4.34282 | 1.30718 | Intron (ENSMUST00000107596.3/ENSMUSG00000038453.10, intron 4 of 19) | 11 | 97519612 | 97521895 | 2284 | 2 | ENSMUST00000134289.1 | -17731 | Gm11611 | ENSMUSG00000087042.1 |
| chr11 | 97541304 | 97541394 | 3.44418 | 4.32382 | 1.32576 | Intron (ENSMUST00000107596.3/ENSMUSG00000038453.10, intron 4 of 19) | 11 | 97519612 | 97521895 | 2284 | 2 | ENSMUST00000134289.1 | -19410 | Gm11611 | ENSMUSG00000087042.1 |
| chr11 | 97541722 | 97542169 | 3.54034 | 4.20222 | 1.23896 | Intron (ENSMUST00000107596.3/ENSMUSG00000038453.10, intron 4 of 19) | 11 | 97519612 | 97521895 | 2284 | 2 | ENSMUST00000134289.1 | -19828 | Gm11611 | ENSMUSG00000087042.1 |
| chr4 | 57339206 | 57339362 | 3.42118 | 4.00015 | 1.16021 | Distal Intergenic | 4 | 57361294 | 57364252 | 2959 | 2 | ENSMUST00000143574.1 | 24890 | 1700042G15Rik | ENSMUSG00000087044.1 |
| chr11 | 75794402 | 75794488 | 3.41917 | 3.97696 | 1.08852 | Promoter | 11 | 75794768 | 75825223 | 30456 | 1 | ENSMUST00000127424.2 | -280 | Gm12339 | ENSMUSG00000087059.2 |
| chr5 | 123324396 | 123324709 | 3.51543 | 4.31782 | 1.29769 | 3' UTR | 5 | 123293638 | 123310324 | 16687 | 2 | ENSMUST00000133476.1 | -14073 | Gm15857 | ENSMUSG00000087063.1 |
| chr12 | 111401894 | 111402000 | 3.62456 | 4.35076 | 1.31396 | Distal Intergenic | 12 | 111406809 | 111412077 | 5269 | 1 | ENSMUST00000150384.1 | -4809 | A230065H16Rik | ENSMUSG00000087075.1 |
| chr11 | 50673383 | 50673843 | 3.32656 | 3.83099 | 1.06556 | Intron (ENSMUST00000040523.8/ENSMUSG00000036545.8, intron 3 of 21) | 11 | 50717670 | 50718915 | 1246 | 2 | ENSMUST00000137567.1 | 45072 | Gm12199 | ENSMUSG00000087080.1 |
| chr4 | 140371712 | 140371814 | 3.37387 | 3.90868 | 1.08905 | Distal Intergenic | 4 | 140380630 | 140385574 | 4945 | 1 | ENSMUST00000126252.1 | -8816 | Gm13016 | ENSMUSG00000087084.1 |
| chr4 | 140372670 | 140372858 | 3.52237 | 4.16097 | 1.18773 | Distal Intergenic | 4 | 140380630 | 140385574 | 4945 | 1 | ENSMUST00000126252.1 | -7772 | Gm13016 | ENSMUSG00000087084.1 |
| chr13 | 23313439 | 23313709 | 3.70619 | 4.62453 | 1.54348 | Promoter | 13 | 23313525 | 23385799 | 72275 | 1 | ENSMUST00000122940.1 | 0 | 4930586N03Rik | ENSMUSG00000087097.1 |
| chr13 | 23314618 | 23314742 | 4.13048 | 5.61637 | 2.19321 | Intron (ENSMUST00000122940.1/ENSMUSG00000087097.1, intron 2 of 3) | 13 | 23313525 | 23385799 | 72275 | 1 | ENSMUST00000122940.1 | 1094 | 4930586N03Rik | ENSMUSG00000087097.1 |
| chr13 | 23317493 | 23318458 | 3.51907 | 4.19596 | 1.30927 | Intron (ENSMUST00000122940.1/ENSMUSG00000087097.1, intron 2 of 3) | 13 | 23313525 | 23385799 | 72275 | 1 | ENSMUST00000122940.1 | 3969 | 4930586N03Rik | ENSMUSG00000087097.1 |
| chr13 | 23318775 | 23319097 | 3.4286 | 4.0365 | 1.22227 | Intron (ENSMUST00000122940.1/ENSMUSG00000087097.1, intron 2 of 3) | 13 | 23313525 | 23385799 | 72275 | 1 | ENSMUST00000122940.1 | 5251 | 4930586N03Rik | ENSMUSG00000087097.1 |
| chr2 | 171996515 | 171996820 | 3.45605 | 4.079 | 1.22241 | Intron (ENSMUST00000136486.1/ENSMUSG00000087100.1, intron 1 of 1) | 2 | 171995941 | 172003995 | 8055 | 2 | ENSMUST00000136486.1 | 7175 | Gm14641 | ENSMUSG00000087100.1 |
| chr2 | 19422173 | 19422486 | 3.45682 | 4.04492 | 1.12543 | Distal Intergenic | 2 | 19429104 | 19431983 | 2880 | 2 | ENSMUST00000137714.1 | 9497 | Gm13344 | ENSMUSG00000087103.1 |
| chr2 | 19422806 | 19423387 | 3.62489 | 4.37699 | 1.35687 | Distal Intergenic | 2 | 19429104 | 19431983 | 2880 | 2 | ENSMUST00000137714.1 | 8596 | Gm13344 | ENSMUSG00000087103.1 |
| chr2 | 19429387 | 19429508 | 4.17983 | 5.68409 | 2.27442 | Exon (ENSMUST00000137714.1/ENSMUSG00000087103.1, exon 2 of 2) | 2 | 19429104 | 19431983 | 2880 | 2 | ENSMUST00000137714.1 | 2475 | Gm13344 | ENSMUSG00000087103.1 |
| chr2 | 19431556 | 19431958 | 3.55988 | 4.46229 | 1.46174 | Promoter | 2 | 19429104 | 19431983 | 2880 | 2 | ENSMUST00000137714.1 | 25 | Gm13344 | ENSMUSG00000087103.1 |
| chr11 | 83219896 | 83219980 | 3.7397 | 4.67237 | 1.56898 | Distal Intergenic | 11 | 83223576 | 83225869 | 2294 | 1 | ENSMUST00000145461.2 | -3596 | AI662270 | ENSMUSG00000087107.2 |
| chr12 | 71339564 | 71339697 | 3.37343 | 3.89336 | 1.0371 | Distal Intergenic | 12 | 71317621 | 71321660 | 4040 | 2 | ENSMUST00000132822.1 | -17905 | 4930474H06Rik | ENSMUSG00000087109.1 |
| chr10 | 81022948 | 81023346 | 3.64795 | 4.39955 | 1.35266 | Intron (ENSMUST00000055125.4/ENSMUSG00000043670.4, intron 1 of 1) | 10 | 81024811 | 81026273 | 1463 | 1 | ENSMUST00000151303.1 | -1465 | Gm16099 | ENSMUSG00000087114.1 |
| chr11 | 96875578 | 96877410 | 4.02701 | 5.16121 | 1.88699 | Distal Intergenic | 11 | 96863290 | 96873954 | 10665 | 2 | ENSMUST00000184482.2 | -1625 | Gm11523 | ENSMUSG00000087117.3 |

| | | | | | | | | | | | | | | | |
|---|---|---|---|---|---|---|---|---|---|---|---|---|---|---|---|
| chr2 | 6163739 | 6164308 | 4.0626 | 5.40335 | 2.05818 | Distal Intergenic | 2 | 6193251 | 6321611 | 128361 | 1 | ENSMUST00000151782.1 | -28943 | A230108P19Rik | ENSMUSG00000087125.1 |
| chr2 | 6165804 | 6166368 | 3.33753 | 3.90109 | 1.09832 | Distal Intergenic | 2 | 6193251 | 6321611 | 128361 | 1 | ENSMUST00000151782.1 | -26883 | A230108P19Rik | ENSMUSG00000087125.1 |
| chr2 | 6168506 | 6169285 | 3.55336 | 4.28123 | 1.36892 | Distal Intergenic | 2 | 6193251 | 6321611 | 128361 | 1 | ENSMUST00000151782.1 | -23966 | A230108P19Rik | ENSMUSG00000087125.1 |
| chr2 | 6169696 | 6170216 | 3.53524 | 4.43116 | 1.38459 | Distal Intergenic | 2 | 6193251 | 6321611 | 128361 | 1 | ENSMUST00000151782.1 | -23035 | A230108P19Rik | ENSMUSG00000087125.1 |
| chr2 | 6170588 | 6172131 | 4.05776 | 5.44906 | 2.09936 | Distal Intergenic | 2 | 6193251 | 6321611 | 128361 | 1 | ENSMUST00000151782.1 | -21120 | A230108P19Rik | ENSMUSG00000087125.1 |
| chr2 | 6172667 | 6173043 | 3.21968 | 3.65674 | 1.01309 | Distal Intergenic | 2 | 6193251 | 6321611 | 128361 | 1 | ENSMUST00000151782.1 | -20208 | A230108P19Rik | ENSMUSG00000087125.1 |
| chr2 | 6173713 | 6173998 | 3.56002 | 4.31608 | 1.35641 | Distal Intergenic | 2 | 6193251 | 6321611 | 128361 | 1 | ENSMUST00000151782.1 | -19253 | A230108P19Rik | ENSMUSG00000087125.1 |
| chr2 | 6174363 | 6174763 | 3.91725 | 5.2065 | 1.90349 | Distal Intergenic | 2 | 6193251 | 6321611 | 128361 | 1 | ENSMUST00000151782.1 | -18488 | A230108P19Rik | ENSMUSG00000087125.1 |
| chr2 | 6175924 | 6176339 | 3.91526 | 5.03099 | 1.85293 | Distal Intergenic | 2 | 6193251 | 6321611 | 128361 | 1 | ENSMUST00000151782.1 | -16912 | A230108P19Rik | ENSMUSG00000087125.1 |
| chr2 | 6177062 | 6177834 | 3.87098 | 4.924 | 1.77135 | Distal Intergenic | 2 | 6193251 | 6321611 | 128361 | 1 | ENSMUST00000151782.1 | -15417 | A230108P19Rik | ENSMUSG00000087125.1 |
| chr2 | 6178341 | 6179409 | 3.70948 | 4.74637 | 1.61075 | Distal Intergenic | 2 | 6193251 | 6321611 | 128361 | 1 | ENSMUST00000151782.1 | -13842 | A230108P19Rik | ENSMUSG00000087125.1 |
| chr2 | 6179762 | 6180509 | 3.60629 | 4.34056 | 1.30019 | Distal Intergenic | 2 | 6193251 | 6321611 | 128361 | 1 | ENSMUST00000151782.1 | -12742 | A230108P19Rik | ENSMUSG00000087125.1 |
| chr2 | 6181061 | 6181502 | 4.16862 | 5.41331 | 2.04034 | Distal Intergenic | 2 | 6193251 | 6321611 | 128361 | 1 | ENSMUST00000151782.1 | -11749 | A230108P19Rik | ENSMUSG00000087125.1 |
| chr2 | 6183799 | 6184334 | 3.88733 | 4.85161 | 1.64625 | Distal Intergenic | 2 | 6193251 | 6321611 | 128361 | 1 | ENSMUST00000151782.1 | -8917 | A230108P19Rik | ENSMUSG00000087125.1 |
| chr2 | 6184669 | 6185166 | 4.04095 | 5.17492 | 1.87597 | Distal Intergenic | 2 | 6193251 | 6321611 | 128361 | 1 | ENSMUST00000151782.1 | -8085 | A230108P19Rik | ENSMUSG00000087125.1 |
| chr2 | 6186072 | 6186374 | 3.44219 | 4.02256 | 1.12726 | Downstream (2-3kb) | 2 | 6193251 | 6321611 | 128361 | 1 | ENSMUST00000151782.1 | -6877 | A230108P19Rik | ENSMUSG00000087125.1 |
| chr2 | 6187115 | 6187474 | 3.96711 | 4.99267 | 1.71419 | Downstream (<1kb) | 2 | 6193251 | 6321611 | 128361 | 1 | ENSMUST00000151782.1 | -5777 | A230108P19Rik | ENSMUSG00000087125.1 |
| chr2 | 6188687 | 6188978 | 3.57595 | 4.34636 | 1.37767 | 3' UTR | 2 | 6193251 | 6321611 | 128361 | 1 | ENSMUST00000151782.1 | -4273 | A230108P19Rik | ENSMUSG00000087125.1 |
| chr2 | 6191875 | 6192146 | 3.62076 | 4.46158 | 1.43569 | Intron (ENSMUST00000042658.3/ENSMUSG00000039063.4, intron 4 of 4) | 2 | 6193251 | 6321611 | 128361 | 1 | ENSMUST00000151782.1 | -1105 | A230108P19Rik | ENSMUSG00000087125.1 |
| chr2 | 6193502 | 6193700 | 3.39945 | 4.13932 | 1.19699 | Promoter | 2 | 6193251 | 6321611 | 128361 | 1 | ENSMUST00000151782.1 | 252 | A230108P19Rik | ENSMUSG00000087125.1 |
| chr7 | 34675178 | 34675775 | 4.07476 | 5.21911 | 1.90422 | 3' UTR | 7 | 34665413 | 34673999 | 8587 | 2 | ENSMUST00000126294.1 | -1180 | Gm12756 | ENSMUSG00000087127.1 |
| chr7 | 34676105 | 34676180 | 3.60064 | 4.33558 | 1.36234 | 3' UTR | 7 | 34665413 | 34673999 | 8587 | 2 | ENSMUST00000126294.1 | -2107 | Gm12756 | ENSMUSG00000087127.1 |
| chr6 | 82131109 | 82131471 | 3.55061 | 4.58716 | 1.52122 | Distal Intergenic | 6 | 82050453 | 82052581 | 2129 | 2 | ENSMUST00000145950.1 | -78529 | Gm15864 | ENSMUSG00000087136.1 |
| chr6 | 82131782 | 82132199 | 3.69937 | 4.75838 | 1.60543 | Distal Intergenic | 6 | 82050453 | 82052581 | 2129 | 2 | ENSMUST00000145950.1 | -79202 | Gm15864 | ENSMUSG00000087136.1 |
| chr6 | 82134300 | 82134449 | 3.88029 | 5.0299 | 1.84752 | Distal Intergenic | 6 | 82050453 | 82052581 | 2129 | 2 | ENSMUST00000145950.1 | -81720 | Gm15864 | ENSMUSG00000087136.1 |
| chr6 | 82141735 | 82141994 | 3.46511 | 4.11199 | 1.25555 | Distal Intergenic | 6 | 82050453 | 82052581 | 2129 | 2 | ENSMUST00000145950.1 | -89155 | Gm15864 | ENSMUSG00000087136.1 |
| chr13 | 78202191 | 78202290 | 3.52939 | 4.19106 | 1.25812 | Promoter | 13 | 78202374 | 78236564 | 34191 | 1 | ENSMUST00000123998.1 | -84 | A830082K12Rik | ENSMUSG00000087143.4 |
| chr13 | 78203049 | 78203233 | 3.64872 | 4.38773 | 1.3148 | Promoter | 13 | 78202787 | 78279391 | 76605 | 1 | ENSMUST00000183171.1 | 263 | A830082K12Rik | ENSMUSG00000087143.4 |
| chr13 | 78203796 | 78203871 | 3.76131 | 4.61199 | 1.48939 | Intron (ENSMUST00000145613.2/ENSMUSG00000087143.4, intron 1 of 2) | 13 | 78202787 | 78279391 | 76605 | 1 | ENSMUST00000183171.1 | 1010 | A830082K12Rik | ENSMUSG00000087143.4 |
| chr13 | 78204200 | 78204703 | 4.01739 | 5.14186 | 1.89457 | Intron (ENSMUST00000145613.2/ENSMUSG00000087143.4, intron 1 of 2) | 13 | 78202787 | 78279391 | 76605 | 1 | ENSMUST00000183171.1 | 1414 | A830082K12Rik | ENSMUSG00000087143.4 |
| chr13 | 78208814 | 78209479 | 3.79075 | 4.85039 | 1.66698 | Intron (ENSMUST00000145613.2/ENSMUSG00000087143.4, intron 1 of 2) | 13 | 78202787 | 78279391 | 76605 | 1 | ENSMUST00000183171.1 | 6028 | A830082K12Rik | ENSMUSG00000087143.4 |
| chr13 | 78209825 | 78210545 | 3.81863 | 5.02766 | 1.80549 | Intron (ENSMUST00000145613.2/ENSMUSG00000087143.4, intron 1 of 2) | 13 | 78202787 | 78279391 | 76605 | 1 | ENSMUST00000183171.1 | 7039 | A830082K12Rik | ENSMUSG00000087143.4 |
| chr13 | 78211268 | 78212156 | 3.41514 | 4.0603 | 1.21781 | Intron (ENSMUST00000145613.2/ENSMUSG00000087143.4, intron 1 of 2) | 13 | 78202787 | 78279391 | 76605 | 1 | ENSMUST00000183171.1 | 8482 | A830082K12Rik | ENSMUSG00000087143.4 |
| chr2 | 38713496 | 38713635 | 3.43682 | 4.00183 | 1.08205 | Promoter | 2 | 38712676 | 38720638 | 7963 | 1 | ENSMUST00000146954.1 | 821 | Nr5a1os | ENSMUSG00000087164.1 |
| chr4 | 98722123 | 98722345 | 3.31107 | 3.79382 | 1.01733 | Distal Intergenic | 4 | 98726734 | 98733644 | 6911 | 1 | ENSMUST00000152889.3 | -4389 | L1td1 | ENSMUSG00000087166.4 |
| chr4 | 98726881 | 98726964 | 3.94483 | 5.05558 | 1.82258 | Promoter | 4 | 98726793 | 98733560 | 6768 | 1 | ENSMUST00000171708.1 | 89 | L1td1 | ENSMUSG00000087166.4 |
| chr4 | 98727405 | 98727480 | 3.71602 | 4.63528 | 1.56445 | Promoter | 4 | 98726793 | 98733560 | 6768 | 1 | ENSMUST00000171708.1 | 613 | L1td1 | ENSMUSG00000087166.4 |
| chr4 | 98729732 | 98729882 | 3.59949 | 4.37598 | 1.39946 | Intron (ENSMUST00000152889.3/ENSMUSG00000087166.4, intron 2 of 2) | 4 | 98726793 | 98733560 | 6768 | 1 | ENSMUST00000171708.1 | 2940 | L1td1 | ENSMUSG00000087166.4 |
| chr4 | 131631090 | 131631647 | 3.3126 | 3.82364 | 1.10397 | Distal Intergenic | 4 | 131647048 | 131672141 | 25094 | 2 | ENSMUST00000144212.1 | 40494 | Gm12962 | ENSMUSG00000087171.1 |
| chr4 | 131640425 | 131640650 | 3.31829 | 3.86747 | 1.09937 | Distal Intergenic | 4 | 131647048 | 131672141 | 25094 | 2 | ENSMUST00000144212.1 | 31491 | Gm12962 | ENSMUSG00000087171.1 |
| chr4 | 131678871 | 131679197 | 3.38777 | 3.97866 | 1.16881 | Distal Intergenic | 4 | 131647376 | 131677845 | 30470 | 2 | ENSMUST00000138697.1 | -1027 | Gm12962 | ENSMUSG00000087171.1 |
| chr2 | 18687219 | 18690821 | 4.21074 | 5.73778 | 2.32617 | Distal Intergenic | 2 | 18694032 | 18698884 | 4853 | 1 | ENSMUST00000137796.2 | -3211 | Gm13334 | ENSMUSG00000087182.2 |
| chr2 | 18691160 | 18691433 | 3.33778 | 4.08853 | 1.22838 | Distal Intergenic | 2 | 18694032 | 18698884 | 4853 | 1 | ENSMUST00000137796.2 | -2599 | Gm13334 | ENSMUSG00000087182.2 |
| chr2 | 18692057 | 18693185 | 4.12182 | 5.55007 | 2.13442 | Promoter | 2 | 18694032 | 18698884 | 4853 | 1 | ENSMUST00000137796.2 | -847 | Gm13334 | ENSMUSG00000087182.2 |
| chr2 | 18694353 | 18694770 | 4.27208 | 5.7554 | 2.37964 | Promoter | 2 | 18694052 | 18699037 | 4986 | 1 | ENSMUST00000138017.1 | 302 | Gm13334 | ENSMUSG00000087182.2 |
| chr2 | 18695649 | 18695792 | 3.67979 | 4.5104 | 1.51344 | Intron (ENSMUST00000137796.2/ENSMUSG00000087182.2, intron 1 of 2) | 2 | 18694052 | 18699037 | 4986 | 1 | ENSMUST00000138017.1 | 1598 | Gm13334 | ENSMUSG00000087182.2 |
| chr11 | 107792104 | 107792513 | 3.8635 | 4.88062 | 1.70388 | Intron (ENSMUST00000021066.3/ENSMUSG00000020723.3, intron 1 of 3) | 11 | 107793866 | 107795982 | 2117 | 1 | ENSMUST00000141271.1 | -1353 | Gm11650 | ENSMUSG00000087184.1 |
| chr11 | 107792819 | 107794365 | 4.2237 | 5.6953 | 2.29836 | Promoter | 11 | 107793866 | 107795982 | 2117 | 1 | ENSMUST00000141271.1 | 0 | Gm11650 | ENSMUSG00000087184.1 |
| chr11 | 84526795 | 84527099 | 4.21275 | 5.53315 | 2.17937 | Intron (ENSMUST00000134800.2/ENSMUSG00000087211.2, intron 1 of 3) | 11 | 84525684 | 84535830 | 10147 | 1 | ENSMUST00000128121.1 | 1112 | Lhx1os | ENSMUSG00000087211.2 |
| chr11 | 84528173 | 84528837 | 3.83601 | 4.87045 | 1.66921 | Intron (ENSMUST00000134800.2/ENSMUSG00000087211.2, intron 1 of 3) | 11 | 84525684 | 84535830 | 10147 | 1 | ENSMUST00000128121.1 | 2490 | Lhx1os | ENSMUSG00000087211.2 |
| chr11 | 84529250 | 84529432 | 3.67993 | 4.49632 | 1.43798 | Intron (ENSMUST00000134800.2/ENSMUSG00000087211.2, intron 1 of 3) | 11 | 84525684 | 84535830 | 10147 | 1 | ENSMUST00000128121.1 | 3567 | Lhx1os | ENSMUSG00000087211.2 |
| chr11 | 84531701 | 84532478 | 3.38193 | 3.9006 | 1.01733 | Intron (ENSMUST00000134800.2/ENSMUSG00000087211.2, intron 1 of 3) | 11 | 84525684 | 84535830 | 10147 | 1 | ENSMUST00000128121.1 | 6018 | Lhx1os | ENSMUSG00000087211.2 |
| chr11 | 84538395 | 84538567 | 3.53374 | 4.45977 | 1.42344 | Distal Intergenic | 11 | 84525684 | 84535830 | 10147 | 1 | ENSMUST00000128121.1 | 12712 | Lhx1os | ENSMUSG00000087211.2 |
| chr11 | 84551512 | 84551673 | 3.62641 | 4.34904 | 1.29646 | Distal Intergenic | 11 | 84525684 | 84535830 | 10147 | 1 | ENSMUST00000128121.1 | 25829 | Lhx1os | ENSMUSG00000087211.2 |
| chr11 | 84554376 | 84554524 | 3.34763 | 3.84885 | 1.01733 | Distal Intergenic | 11 | 84525684 | 84535830 | 10147 | 1 | ENSMUST00000128121.1 | 28693 | Lhx1os | ENSMUSG00000087211.2 |
| chr11 | 84563139 | 84563503 | 3.50447 | 4.13457 | 1.19117 | Distal Intergenic | 11 | 84525684 | 84535830 | 10147 | 1 | ENSMUST00000128121.1 | 37456 | Lhx1os | ENSMUSG00000087211.2 |
| chr11 | 84570276 | 84570688 | 3.95455 | 5.01799 | 1.7556 | Distal Intergenic | 11 | 84525684 | 84535830 | 10147 | 1 | ENSMUST00000128121.1 | 44593 | Lhx1os | ENSMUSG00000087211.2 |
| chr13 | 31562521 | 31563035 | 4.02471 | 5.14416 | 1.86449 | Distal Intergenic | 13 | 31564749 | 31573562 | 8814 | 1 | ENSMUST00000131539.1 | -1714 | Gm11377 | ENSMUSG00000087220.1 |
| chr13 | 31563617 | 31564254 | 3.93501 | 4.94019 | 1.69225 | Promoter | 13 | 31564749 | 31573562 | 8814 | 1 | ENSMUST00000131539.1 | -495 | Gm11377 | ENSMUSG00000087220.1 |
| chr13 | 31565133 | 31566478 | 3.68244 | 4.60469 | 1.50213 | Promoter | 13 | 31564749 | 31573562 | 8814 | 1 | ENSMUST00000131539.1 | 385 | Gm11377 | ENSMUSG00000087220.1 |
| chr1 | 6358732 | 6359455 | 3.58661 | 4.38961 | 1.40913 | Promoter | 1 | 6359331 | 6394731 | 35401 | 1 | ENSMUST00000133144.2 | 0 | Fam150a | ENSMUSG00000087247.2 |
| chr1 | 6382888 | 6383110 | 3.16829 | 3.78339 | 1.03481 | Intron (ENSMUST00000133144.2/ENSMUSG00000087247.2, intron 1 of 4) | 1 | 6359331 | 6394731 | 35401 | 1 | ENSMUST00000133144.2 | 23558 | Fam150a | ENSMUSG00000087247.2 |
| chr11 | 113192290 | 113192379 | 4.13709 | 5.40138 | 2.02563 | Intron (ENSMUST00000145206.1/ENSMUSG00000087259.2, intron 1 of 1) | 11 | 113043895 | 113201822 | 157928 | 2 | ENSMUST00000145206.1 | 9443 | 2610035D17Rik | ENSMUSG00000087259.2 |
| chr11 | 113200677 | 113201064 | 3.58666 | 4.38249 | 1.39559 | Promoter | 11 | 113043895 | 113201822 | 157928 | 2 | ENSMUST00000145206.1 | 758 | 2610035D17Rik | ENSMUSG00000087259.2 |
| chr2 | 70489166 | 70489328 | 3.58875 | 4.28382 | 1.26859 | Downstream (<1kb) | 2 | 70490244 | 70491468 | 1225 | 2 | ENSMUST00000154284.1 | 2140 | Gad1os | ENSMUSG00000087264.1 |
| chr2 | 70489660 | 70490025 | 3.45468 | 4.10461 | 1.24708 | Exon (ENSMUST00000151851.1/ENSMUSG00000087264.1, exon 3 of 3) | 2 | 70490244 | 70491468 | 1225 | 2 | ENSMUST00000154284.1 | 1443 | Gad1os | ENSMUSG00000087264.1 |
| chr2 | 70490518 | 70490698 | 3.2566 | 3.7227 | 1.04052 | Promoter | 2 | 70490244 | 70491468 | 1225 | 2 | ENSMUST00000154284.1 | 770 | Gad1os | ENSMUSG00000087264.1 |
| chr2 | 70491036 | 70491123 | 3.38879 | 3.93255 | 1.08954 | Promoter | 2 | 70490244 | 70491468 | 1225 | 2 | ENSMUST00000154284.1 | 345 | Gad1os | ENSMUSG00000087264.1 |
| chr2 | 70495656 | 70495901 | 3.22732 | 3.66817 | 1.01707 | Intron (ENSMUST00000151851.1/ENSMUSG00000087264.1, intron 1 of 2) | 2 | 70490244 | 70491468 | 1225 | 2 | ENSMUST00000154284.1 | -4189 | Gad1os | ENSMUSG00000087264.1 |
| chr2 | 30677029 | 30677639 | 3.69633 | 4.86376 | 1.70481 | Promoter | 2 | 30658613 | 30678011 | 19399 | 2 | ENSMUST00000144704.1 | 372 | Gm14486 | ENSMUSG00000087268.1 |
| chr2 | 30687265 | 30687592 | 3.91332 | 4.93842 | 1.77727 | Intron (ENSMUST00000146702.1/ENSMUSG00000086405.1, intron 2 of 3) | 2 | 30658613 | 30678011 | 19399 | 2 | ENSMUST00000144704.1 | -9255 | Gm14486 | ENSMUSG00000087268.1 |
| chr4 | 135291827 | 135291985 | 3.37934 | 3.90486 | 1.04619 | Distal Intergenic | 4 | 135290366 | 135290675 | 310 | 2 | ENSMUST00000120974.1 | -1153 | Gm12983 | ENSMUSG00000087270.2 |
| chr4 | 148686338 | 148686859 | 3.3209 | 3.81971 | 1.05616 | Distal Intergenic | 4 | 148713703 | 148716581 | 2879 | 1 | ENSMUST00000139895.1 | -26844 | Gm13203 | ENSMUSG00000087273.1 |
| chr4 | 148724377 | 148725499 | 3.53467 | 4.22498 | 1.28481 | Distal Intergenic | 4 | 148713703 | 148716581 | 2879 | 1 | ENSMUST00000139895.1 | 10675 | Gm13203 | ENSMUSG00000087273.1 |
| chr4 | 148726270 | 148726452 | 3.67408 | 4.44216 | 1.37328 | Distal Intergenic | 4 | 148713703 | 148716581 | 2879 | 1 | ENSMUST00000139895.1 | 12568 | Gm13203 | ENSMUSG00000087273.1 |
| chr4 | 148732336 | 148732783 | 3.42339 | 3.99762 | 1.12826 | Distal Intergenic | 4 | 148713703 | 148716581 | 2879 | 1 | ENSMUST00000139895.1 | 18634 | Gm13203 | ENSMUSG00000087273.1 |
| chr13 | 31606292 | 31606417 | 3.84038 | 4.77733 | 1.62402 | Distal Intergenic | 13 | 31619168 | 31620369 | 1202 | 1 | ENSMUST00000130966.1 | -12751 | Gm11378 | ENSMUSG00000087276.1 |
| chr13 | 31607795 | 31607870 | 4.10103 | 5.2985 | 1.94724 | Distal Intergenic | 13 | 31619168 | 31620369 | 1202 | 1 | ENSMUST00000130966.1 | -11298 | Gm11378 | ENSMUSG00000087276.1 |
| chr13 | 31610658 | 31611025 | 3.49963 | 4.12049 | 1.16665 | Distal Intergenic | 13 | 31619168 | 31620369 | 1202 | 1 | ENSMUST00000130966.1 | -8143 | Gm11378 | ENSMUSG00000087276.1 |
| chr13 | 31611963 | 31612193 | 4.18284 | 5.46083 | 2.08945 | Distal Intergenic | 13 | 31619168 | 31620369 | 1202 | 1 | ENSMUST00000130966.1 | -6975 | Gm11378 | ENSMUSG00000087276.1 |
| chr13 | 31613290 | 31613718 | 3.97364 | 5.07274 | 1.84756 | Distal Intergenic | 13 | 31619168 | 31620369 | 1202 | 1 | ENSMUST00000130966.1 | -5450 | Gm11378 | ENSMUSG00000087276.1 |
| chr13 | 31614718 | 31617198 | 3.81953 | 4.85872 | 1.70911 | Distal Intergenic | 13 | 31619168 | 31620369 | 1202 | 1 | ENSMUST00000130966.1 | -1970 | Gm11378 | ENSMUSG00000087276.1 |
| chr13 | 31617693 | 31618688 | 3.29171 | 3.85016 | 1.08368 | Promoter | 13 | 31619168 | 31620369 | 1202 | 1 | ENSMUST00000130966.1 | -480 | Gm11378 | ENSMUSG00000087276.1 |
| chr13 | 31619094 | 31619484 | 4.14265 | 5.43562 | 2.06808 | Promoter | 13 | 31619168 | 31620369 | 1202 | 1 | ENSMUST00000130966.1 | 0 | Gm11378 | ENSMUSG00000087276.1 |
| chr13 | 31620073 | 31620480 | 3.85456 | 4.83765 | 1.6479 | Promoter | 13 | 31619168 | 31620369 | 1202 | 1 | ENSMUST00000130966.1 | 906 | Gm11378 | ENSMUSG00000087276.1 |
| chr2 | 103349066 | 103349144 | 3.44575 | 4.02808 | 1.1278 | Intron (ENSMUST00000139065.1/ENSMUSG00000087278.1, intron 1 of 2) | 2 | 103338601 | 103372764 | 34164 | 2 | ENSMUST00000139065.1 | 23620 | A930006I01Rik | ENSMUSG00000087278.1 |
| chr2 | 60402746 | 60402841 | 9.78054 | 29.34007 | 23.83413 | Distal Intergenic | 2 | 60411525 | 60412579 | 1055 | 1 | ENSMUST00000131457.1 | -8684 | Gm13580 | ENSMUSG00000087283.1 |
| chr14 | 59843391 | 59843466 | 3.49266 | 4.12902 | 1.23576 | Intron (ENSMUST00000080368.6/ENSMUSG00000021983.10, intron 26 of 35) | 14 | 60039610 | 60040491 | 882 | 2 | ENSMUST00000132939.1 | 197025 | Gm9013 | ENSMUSG00000087286.1 |
| chr14 | 59847378 | 59847898 | 3.67784 | 4.45817 | 1.39621 | Intron (ENSMUST00000080368.6/ENSMUSG00000021983.10, intron 26 of 35) | 14 | 60039610 | 60040491 | 882 | 2 | ENSMUST00000132939.1 | 192593 | Gm9013 | ENSMUSG00000087286.1 |
| chr14 | 59890800 | 59890948 | 3.26084 | 3.71839 | 1.01733 | Intron (ENSMUST00000080368.6/ENSMUSG00000021983.10, intron 24 of 35) | 14 | 60039610 | 60040491 | 882 | 2 | ENSMUST00000132939.1 | 149543 | Gm9013 | ENSMUSG00000087286.1 |
| chr14 | 59960381 | 59960702 | 3.30681 | 3.78741 | 1.01733 | Intron (ENSMUST00000080368.6/ENSMUSG00000021983.10, intron 23 of 35) | 14 | 60039610 | 60040491 | 882 | 2 | ENSMUST00000132939.1 | 79789 | Gm9013 | ENSMUSG00000087286.1 |
| chr14 | 59961601 | 59961705 | 3.6188 | 4.38233 | 1.42247 | Intron (ENSMUST00000080368.6/ENSMUSG00000021983.10, intron 23 of 35) | 14 | 60039610 | 60040491 | 882 | 2 | ENSMUST00000132939.1 | 78786 | Gm9013 | ENSMUSG00000087286.1 |
| chr4 | 134015568 | 134015751 | 3.28403 | 3.76279 | 1.04307 | Intron (ENSMUST00000051674.2/ENSMUSG00000050966.4, intron 2 of 3) | 4 | 134018147 | 134019869 | 1723 | 1 | ENSMUST00000137962.1 | -2396 | Gm13061 | ENSMUSG00000087288.1 |
| chr4 | 134016400 | 134016658 | 4.1287 | 5.3577 | 2.00285 | Intron (ENSMUST00000051674.2/ENSMUSG00000050966.4, intron 2 of 3) | 4 | 134018147 | 134019869 | 1723 | 1 | ENSMUST00000137962.1 | -1489 | Gm13061 | ENSMUSG00000087288.1 |
| chr11 | 33075672 | 33075999 | 3.8852 | 4.8592 | 1.66884 | Intron (ENSMUST00000147896.1/ENSMUSG00000087318.1, intron 1 of 3) | 11 | 33068551 | 33080521 | 11971 | 1 | ENSMUST00000147896.1 | 7122 | Gm12114 | ENSMUSG00000087318.1 |
| chr11 | 33077689 | 33077905 | 3.36878 | 3.88076 | 1.01733 | Exon (ENSMUST00000147896.1/ENSMUSG00000087318.1, exon 3 of 4) | 11 | 33068551 | 33080521 | 11971 | 1 | ENSMUST00000147896.1 | 9139 | Gm12114 | ENSMUSG00000087318.1 |
| chr11 | 33097846 | 33098063 | 3.26116 | 3.71887 | 1.01733 | Distal Intergenic | 11 | 33068551 | 33080521 | 11971 | 1 | ENSMUST00000147896.1 | 29296 | Gm12114 | ENSMUSG00000087318.1 |
| chr11 | 33098463 | 33098685 | 3.26594 | 3.72605 | 1.01733 | Distal Intergenic | 11 | 33068551 | 33080521 | 11971 | 1 | ENSMUST00000147896.1 | 29913 | Gm12114 | ENSMUSG00000087318.1 |
| chr11 | 33101595 | 33102043 | 3.30866 | 3.7902 | 1.01733 | Distal Intergenic | 11 | 33068551 | 33080521 | 11971 | 1 | ENSMUST00000147896.1 | 33045 | Gm12114 | ENSMUSG00000087318.1 |
| chr11 | 33103610 | 33103753 | 3.47865 | 4.10849 | 1.23827 | Distal Intergenic | 11 | 33068551 | 33080521 | 11971 | 1 | ENSMUST00000147896.1 | 35060 | Gm12114 | ENSMUSG00000087318.1 |
| chr11 | 33104136 | 33104256 | 3.96769 | 5.06682 | 1.87251 | Distal Intergenic | 11 | 33068551 | 33080521 | 11971 | 1 | ENSMUST00000147896.1 | 35586 | Gm12114 | ENSMUSG00000087318.1 |
| chr11 | 33106312 | 33106542 | 3.47843 | 4.1113 | 1.17877 | Distal Intergenic | 11 | 33068551 | 33080521 | 11971 | 1 | ENSMUST00000147896.1 | 37762 | Gm12114 | ENSMUSG00000087318.1 |
| chr11 | 33107236 | 33107452 | 3.48855 | 4.15459 | 1.27798 | Distal Intergenic | 11 | 33068551 | 33080521 | 11971 | 1 | ENSMUST00000147896.1 | 38686 | Gm12114 | ENSMUSG00000087318.1 |
| chr13 | 94144850 | 94144925 | 3.47506 | 4.0752 | 1.13765 | Intron (ENSMUST00000054274.2/ENSMUSG00000045312.6, intron 2 of 3) | 13 | 94137447 | 94140117 | 2671 | 2 | ENSMUST00000134284.1 | -4734 | Gm15907 | ENSMUSG00000087319.1 |
| chr13 | 94145521 | 94145891 | 3.55127 | 4.21176 | 1.21333 | Intron (ENSMUST00000054274.2/ENSMUSG00000045312.6, intron 2 of 3) | 13 | 94137447 | 94140117 | 2671 | 2 | ENSMUST00000134284.1 | -5405 | Gm15907 | ENSMUSG00000087319.1 |
| chr13 | 94142562 | 94143198 | 3.67373 | 4.46508 | 1.39294 | Intron (ENSMUST00000054274.2/ENSMUSG00000045312.6, intron 2 of 3) | 13 | 94137447 | 94140117 | 2671 | 2 | ENSMUST00000134284.1 | -2446 | Gm15907 | ENSMUSG00000087319.1 |
| chr11 | 75866729 | 75866997 | 3.34183 | 3.8659 | 1.10716 | Intron (ENSMUST00000121287.2/ENSMUSG00000020847.10, intron 5 of 8) | 11 | 75860969 | 75861494 | 526 | 1 | ENSMUST00000135873.1 | 5761 | Gm16075 | ENSMUSG00000087322.1 |
| chr11 | 75873659 | 75873992 | 3.72396 | 4.56297 | 1.4961 | Intron (ENSMUST00000121287.2/ENSMUSG00000020847.10, intron 5 of 8) | 11 | 75860969 | 75861494 | 526 | 1 | ENSMUST00000135873.1 | 12691 | Gm16075 | ENSMUSG00000087322.1 |
| chr11 | 75875759 | 75876063 | 3.5134 | 4.15044 | 1.19965 | Intron (ENSMUST00000121287.2/ENSMUSG00000020847.10, intron 5 of 8) | 11 | 75860969 | 75861494 | 526 | 1 | ENSMUST00000135873.1 | 14791 | Gm16075 | ENSMUSG00000087322.1 |
| chr11 | 75878474 | 75878692 | 3.292 | 3.77481 | 1.04975 | Intron (ENSMUST00000121287.2/ENSMUSG00000020847.10, intron 5 of 8) | 11 | 75860969 | 75861494 | 526 | 1 | ENSMUST00000135873.1 | 17506 | Gm16075 | ENSMUSG00000087322.1 |
| chr4 | 44103488 | 44103605 | 3.74825 | 4.61941 | 1.56338 | Intron (ENSMUST00000146576.1/ENSMUSG00000087326.1, intron 1 of 1) | 4 | 44092632 | 44109311 | 16680 | 1 | ENSMUST00000146576.1 | 10857 | Gm12503 | ENSMUSG00000087326.1 |
| chr4 | 44105825 | 44105977 | 3.27363 | 3.73762 | 1.01733 | Intron (ENSMUST00000146576.1/ENSMUSG00000087326.1, intron 1 of 1) | 4 | 44092632 | 44109311 | 16680 | 1 | ENSMUST00000146576.1 | 13194 | Gm12503 | ENSMUSG00000087326.1 |
| chr11 | 33734248 | 33734617 | 3.57432 | 4.38428 | 1.36056 | Intron (ENSMUST00000101368.3/ENSMUSG00000053519.10, intron 1 of 7) | 11 | 33705550 | 33711435 | 5886 | 1 | ENSMUST00000125068.1 | 28699 | Gm12120 | ENSMUSG00000087328.1 |
| chr11 | 33754947 | 33755029 | 3.25006 | 17.05679 | 12.14569 | Intron (ENSMUST00000101368.3/ENSMUSG00000053519.10, intron 1 of 7) | 11 | 33705550 | 33711435 | 5886 | 1 | ENSMUST00000125068.1 | 49398 | Gm12120 | ENSMUSG00000087328.1 |
| chr11 | 33770476 | 33770573 | 3.40469 | 3.99687 | 1.15466 | Intron (ENSMUST00000101368.3/ENSMUSG00000053519.10, intron 1 of 7) | 11 | 33705550 | 33711435 | 5886 | 1 | ENSMUST00000125068.1 | 64927 | Gm12120 | ENSMUSG00000087328.1 |
| chr4 | 99560703 | 99560852 | 3.34153 | 3.83966 | 1.01733 | Distal Intergenic | 4 | 99569500 | 99573011 | 3512 | 1 | ENSMUST00000132108.1 | -8648 | Gm12690 | ENSMUSG00000087332.1 |
| chr4 | 99572737 | 99573315 | 3.52337 | 4.20149 | 1.27145 | Exon (ENSMUST00000132108.1/ENSMUSG00000087332.1, exon 2 of 2) | 4 | 99569500 | 99573011 | 3512 | 1 | ENSMUST00000132108.1 | 3238 | Gm12690 | ENSMUSG00000087332.1 |
| chr2 | 75064229 | 75064822 | 3.66409 | 4.43955 | 1.38897 | Distal Intergenic | 2 | 75124871 | 75137798 | 12928 | 2 | ENSMUST00000143898.1 | 72976 | Gm13652 | ENSMUSG00000087333.1 |
| chr2 | 75086201 | 75086537 | 4.11045 | 5.31019 | 1.96532 | Distal Intergenic | 2 | 75124871 | 75137798 | 12928 | 2 | ENSMUST00000143898.1 | 51261 | Gm13652 | ENSMUSG00000087333.1 |
| chr2 | 75086971 | 75087227 | 3.4882 | 4.50034 | 1.44307 | Distal Intergenic | 2 | 75124871 | 75137798 | 12928 | 2 | ENSMUST00000143898.1 | 50571 | Gm13652 | ENSMUSG00000087333.1 |
| chr2 | 75089160 | 75089545 | 3.65096 | 4.41752 | 1.35617 | Distal Intergenic | 2 | 75124871 | 75137798 | 12928 | 2 | ENSMUST00000143898.1 | 48253 | Gm13652 | ENSMUSG00000087333.1 |
| chr2 | 75090683 | 75090978 | 3.79145 | 4.67291 | 1.55928 | Distal Intergenic | 2 | 75124871 | 75137798 | 12928 | 2 | ENSMUST00000143898.1 | 46820 | Gm13652 | ENSMUSG00000087333.1 |
| chr2 | 75100210 | 75100480 | 3.69962 | 4.50626 | 1.44061 | Distal Intergenic | 2 | 75124871 | 75137798 | 12928 | 2 | ENSMUST00000143898.1 | 37318 | Gm13652 | ENSMUSG00000087333.1 |
| chr2 | 75136748 | 75137782 | 4.27208 | 5.73084 | 2.29971 | Promoter | 2 | 75124871 | 75137798 | 12928 | 2 | ENSMUST00000143898.1 | 16 | Gm13652 | ENSMUSG00000087333.1 |
| chr13 | 54935092 | 54935236 | 3.23584 | 3.68093 | 1.01733 | Distal Intergenic | 13 | 54926763 | 54930256 | 3494 | 2 | ENSMUST00000144718.2 | -4837 | 4930526F13Rik | ENSMUSG00000087335.2 |
| chr13 | 54938202 | 54938420 | 3.60563 | 4.47858 | 1.42642 | Distal Intergenic | 13 | 54926763 | 54930256 | 3494 | 2 | ENSMUST00000144718.2 | -7947 | 4930526F13Rik | ENSMUSG00000087335.2 |
| chr6 | 108709718 | 108710070 | 3.48006 | 4.21764 | 1.30723 | Distal Intergenic | 6 | 108577039 | 108660934 | 83896 | 2 | ENSMUST00000137478.1 | -48785 | 0610040F04Rik | ENSMUSG00000087341.2 |
| chr4 | 85514378 | 85515317 | 3.38867 | 3.92514 | 1.06799 | Promoter | 4 | 85514327 | 85783127 | 268801 | 1 | ENSMUST00000147066.1 | 52 | Gm12680 | ENSMUSG00000087349.1 |
| chr2 | 4301513 | 4301659 | 3.47649 | 4.13913 | 1.28097 | Promoter | 2 | 4300511 | 4301167 | 657 | 2 | ENSMUST00000156609.1 | -347 | Gm13187 | ENSMUSG00000087355.2 |
| chr2 | 4303126 | 4303281 | 3.33512 | 3.83001 | 1.01733 | Intron (ENSMUST00000177457.2/ENSMUSG00000026657.11, intron 2 of 24) | 2 | 4300511 | 4301167 | 657 | 2 | ENSMUST00000156609.1 | -1960 | Gm13187 | ENSMUSG00000087355.2 |
| chr7 | 44263138 | 44263638 | 4.02555 | 5.17541 | 1.86923 | Distal Intergenic | 7 | 44261561 | 44261889 | 329 | 1 | ENSMUST00000147088.1 | 1578 | Gm15517 | ENSMUSG00000087376.2 |
| chr4 | 84120535 | 84120831 | 3.52484 | 4.25458 | 1.26861 | Distal Intergenic | 4 | 84126350 | 84130289 | 3940 | 2 | ENSMUST00000150794.1 | 9458 | Gm12414 | ENSMUSG00000087378.1 |
| chr4 | 84122233 | 84122759 | 3.56398 | 4.32877 | 1.36777 | Distal Intergenic | 4 | 84126350 | 84130289 | 3940 | 2 | ENSMUST00000150794.1 | 7530 | Gm12414 | ENSMUSG00000087378.1 |
| chr4 | 84124647 | 84125067 | 3.51177 | 4.14688 | 1.19531 | Downstream (1-2kb) | 4 | 84126350 | 84130289 | 3940 | 2 | ENSMUST00000150794.1 | 5222 | Gm12414 | ENSMUSG00000087378.1 |
| chr4 | 84137767 | 84137856 | 4.52596 | 6.28396 | 2.70114 | Distal Intergenic | 4 | 84126350 | 84130289 | 3940 | 2 | ENSMUST00000150794.1 | -7479 | Gm12414 | ENSMUSG00000087378.1 |
| chr4 | 84139270 | 84139513 | 3.33249 | 3.86724 | 1.10752 | Distal Intergenic | 4 | 84126350 | 84130289 | 3940 | 2 | ENSMUST00000150794.1 | -8982 | Gm12414 | ENSMUSG00000087378.1 |
| chr4 | 84162308 | 84162623 | 3.51206 | 4.14175 | 1.17488 | Distal Intergenic | 4 | 84126350 | 84130289 | 3940 | 2 | ENSMUST00000150794.1 | -32020 | Gm12414 | ENSMUSG00000087378.1 |
| chr6 | 31861362 | 31861466 | 3.29079 | 3.76341 | 1.01733 | Intron (ENSMUST00000153328.1/ENSMUSG00000087379.1, intron 1 of 2) | 6 | 31786675 | 31922270 | 135596 | 1 | ENSMUST00000153328.1 | 74688 | Gm13849 | ENSMUSG00000087379.1 |
| chr4 | 156034841 | 156035161 | 3.36041 | 4.05259 | 1.14209 | 3' UTR | 4 | 156023727 | 156031643 | 7917 | 2 | ENSMUST00000151836.2 | -3199 | Gm16008 | ENSMUSG00000087381.3 |
| chr2 | 173107172 | 173107342 | 3.82135 | 4.72147 | 1.55032 | Intron (ENSMUST00000094287.4/ENSMUSG00000070495.6, intron 7 of 10) | 2 | 173110818 | 173114709 | 3892 | 1 | ENSMUST00000144256.1 | -3476 | Ctcflos | ENSMUSG00000087382.2 |
| chr2 | 173129302 | 173129581 | 3.38193 | 3.9006 | 1.01733 | Exon (ENSMUST00000142207.2/ENSMUSG00000087382.2, exon 2 of 4) | 2 | 173124749 | 173133204 | 8456 | 1 | ENSMUST00000153523.1 | 4554 | Ctcflos | ENSMUSG00000087382.2 |
| chr2 | 173133004 | 173133086 | 3.80648 | 4.69166 | 1.52594 | Exon (ENSMUST00000153523.1/ENSMUSG00000087382.2, exon 3 of 3) | 2 | 173124749 | 173133204 | 8456 | 1 | ENSMUST00000153523.1 | 8256 | Ctcflos | ENSMUSG00000087382.2 |
| chr4 | 46336176 | 46336483 | 3.66812 | 4.66888 | 1.55757 | Intron (ENSMUST00000123933.2/ENSMUSG00000087383.2, intron 1 of 4) | 4 | 46300007 | 46343431 | 43425 | 2 | ENSMUST00000123933.2 | 6948 | Gm12446 | ENSMUSG00000087383.2 |
| chr4 | 46336908 | 46337261 | 3.75526 | 4.63381 | 1.56997 | Intron (ENSMUST00000123933.2/ENSMUSG00000087383.2, intron 1 of 4) | 4 | 46300007 | 46343431 | 43425 | 2 | ENSMUST00000123933.2 | 6170 | Gm12446 | ENSMUSG00000087383.2 |
| chr4 | 46337927 | 46338236 | 3.94655 | 5.28938 | 2.01516 | Intron (ENSMUST00000123933.2/ENSMUSG00000087383.2, intron 1 of 4) | 4 | 46300007 | 46343431 | 43425 | 2 | ENSMUST00000123933.2 | 5195 | Gm12446 | ENSMUSG00000087383.2 |
| chr4 | 46339006 | 46339486 | 3.86898 | 4.82702 | 1.65863 | Intron (ENSMUST00000123933.2/ENSMUSG00000087383.2, intron 1 of 4) | 4 | 46300007 | 46343431 | 43425 | 2 | ENSMUST00000123933.2 | 3945 | Gm12446 | ENSMUSG00000087383.2 |
| chr4 | 46339820 | 46339895 | 4.44037 | 5.96004 | 2.45894 | Intron (ENSMUST00000123933.2/ENSMUSG00000087383.2, intron 1 of 4) | 4 | 46300007 | 46343431 | 43425 | 2 | ENSMUST00000123933.2 | 3536 | Gm12446 | ENSMUSG00000087383.2 |
| chr4 | 46340316 | 46341311 | 3.83607 | 4.77467 | 1.62615 | Intron (ENSMUST00000123933.2/ENSMUSG00000087383.2, intron 1 of 4) | 4 | 46300007 | 46343431 | 43425 | 2 | ENSMUST00000123933.2 | 2120 | Gm12446 | ENSMUSG00000087383.2 |
| chr4 | 46352127 | 46352483 | 3.76559 | 4.97602 | 1.76136 | Distal Intergenic | 4 | 46328464 | 46345515 | 17052 | 2 | ENSMUST00000152069.1 | -6613 | Gm12446 | ENSMUSG00000087383.2 |
| chr4 | 46345940 | 46347275 | 4.0229 | 5.32897 | 2.0648 | Promoter | 4 | 46328464 | 46345515 | 17052 | 2 | ENSMUST00000152069.1 | -426 | Gm12446 | ENSMUSG00000087383.2 |
| chr4 | 46347717 | 46348277 | 3.785 | 4.85742 | 1.68359 | Distal Intergenic | 4 | 46328464 | 46345515 | 17052 | 2 | ENSMUST00000152069.1 | -2203 | Gm12446 | ENSMUSG00000087383.2 |
| chr4 | 46348646 | 46349969 | 3.70973 | 4.62982 | 1.55296 | Distal Intergenic | 4 | 46328464 | 46345515 | 17052 | 2 | ENSMUST00000152069.1 | -3132 | Gm12446 | ENSMUSG00000087383.2 |
| chr4 | 46352911 | 46354277 | 3.45261 | 4.05368 | 1.17042 | Distal Intergenic | 4 | 46328464 | 46345515 | 17052 | 2 | ENSMUST00000152069.1 | -7397 | Gm12446 | ENSMUSG00000087383.2 |
| chr11 | 109847208 | 109847351 | 3.30659 | 3.93875 | 1.0691 | Intron (ENSMUST00000143247.1/ENSMUSG00000087393.1, intron 1 of 2) | 11 | 109845878 | 109848476 | 2599 | 1 | ENSMUST00000143247.1 | 1331 | 1700023C21Rik | ENSMUSG00000087393.1 |
| chr11 | 109849689 | 109849764 | 4.21406 | 5.56937 | 2.19641 | Distal Intergenic | 11 | 109845878 | 109848476 | 2599 | 1 | ENSMUST00000143247.1 | 3812 | 1700023C21Rik | ENSMUSG00000087393.1 |
| chr11 | 109850241 | 109850514 | 3.47327 | 4.12313 | 1.2752 | Distal Intergenic | 11 | 109845878 | 109848476 | 2599 | 1 | ENSMUST00000143247.1 | 4364 | 1700023C21Rik | ENSMUSG00000087393.1 |
| chr11 | 109859103 | 109859421 | 3.26197 | 3.72009 | 1.01733 | Distal Intergenic | 11 | 109845878 | 109848476 | 2599 | 1 | ENSMUST00000143247.1 | 13226 | 1700023C21Rik | ENSMUSG00000087393.1 |
| chr11 | 109860259 | 109860334 | 3.61757 | 4.37732 | 1.41875 | Distal Intergenic | 11 | 109845878 | 109848476 | 2599 | 1 | ENSMUST00000143247.1 | 14382 | 1700023C21Rik | ENSMUSG00000087393.1 |
| chr11 | 84764546 | 84765204 | 3.63469 | 4.39651 | 1.36888 | Intron (ENSMUST00000155046.1/ENSMUSG00000087407.1, intron 2 of 2) | 11 | 84763855 | 84777829 | 13975 | 2 | ENSMUST00000155046.1 | 12625 | Gm11431 | ENSMUSG00000087407.1 |
| chr11 | 84780150 | 84780805 | 3.56703 | 4.34966 | 1.37647 | Distal Intergenic | 11 | 84763855 | 84777829 | 13975 | 2 | ENSMUST00000155046.1 | -2322 | Gm11431 | ENSMUSG00000087407.1 |
| chr11 | 84784181 | 84784572 | 3.74044 | 4.6859 | 1.59346 | Distal Intergenic | 11 | 84763855 | 84777829 | 13975 | 2 | ENSMUST00000155046.1 | -6353 | Gm11431 | ENSMUSG00000087407.1 |
| chr8 | 70315628 | 70315982 | 3.45026 | 4.06363 | 1.22768 | Promoter | 8 | 70315775 | 70331592 | 15818 | 1 | ENSMUST00000165819.3 | 0 | Cers1 | ENSMUSG00000087408.4 |
| chr4 | 82507188 | 82508088 | 3.93161 | 4.96702 | 1.75062 | Promoter | 4 | 82508016 | 82547292 | 39277 | 1 | ENSMUST00000139835.1 | 0 | Gm11266 | ENSMUSG00000087413.1 |

| | | | | | | | | | | | | | | | |
|---|---|---|---|---|---|---|---|---|---|---|---|---|---|---|---|
| chr4 | 82509801 | 82510341 | 3.78469 | 4.65569 | 1.51943 | Intron (ENSMUST00000139835.1/ENSMUSG00000087413.1, intron 1 of 1) | 4 | 82508016 | 82547292 | 39277 | 1 | ENSMUST00000139835.1 | 1786 | Gm11266 | ENSMUSG00000087413.1 |
| chr4 | 82510815 | 82511961 | 3.58966 | 4.42398 | 1.37436 | Intron (ENSMUST00000139835.1/ENSMUSG00000087413.1, intron 1 of 1) | 4 | 82508016 | 82547292 | 39277 | 1 | ENSMUST00000139835.1 | 2800 | Gm11266 | ENSMUSG00000087413.1 |
| chr4 | 82513167 | 82514118 | 3.72218 | 4.54664 | 1.43995 | Intron (ENSMUST00000139835.1/ENSMUSG00000087413.1, intron 1 of 1) | 4 | 82508016 | 82547292 | 39277 | 1 | ENSMUST00000139835.1 | 5152 | Gm11266 | ENSMUSG00000087413.1 |
| chr4 | 82514648 | 82515301 | 3.53532 | 4.3946 | 1.37971 | Intron (ENSMUST00000139835.1/ENSMUSG00000087413.1, intron 1 of 1) | 4 | 82508016 | 82547292 | 39277 | 1 | ENSMUST00000139835.1 | 6633 | Gm11266 | ENSMUSG00000087413.1 |
| chr4 | 82515610 | 82515778 | 3.51416 | 4.14167 | 1.16346 | Intron (ENSMUST00000139835.1/ENSMUSG00000087413.1, intron 1 of 1) | 4 | 82508016 | 82547292 | 39277 | 1 | ENSMUST00000139835.1 | 7595 | Gm11266 | ENSMUSG00000087413.1 |
| chr4 | 82516115 | 82516224 | 3.34856 | 3.85026 | 1.01733 | Intron (ENSMUST00000139835.1/ENSMUSG00000087413.1, intron 1 of 1) | 4 | 82508016 | 82547292 | 39277 | 1 | ENSMUST00000139835.1 | 8100 | Gm11266 | ENSMUSG00000087413.1 |
| chr4 | 82517163 | 82517605 | 3.39894 | 3.93678 | 1.05458 | Intron (ENSMUST00000139835.1/ENSMUSG00000087413.1, intron 1 of 1) | 4 | 82508016 | 82547292 | 39277 | 1 | ENSMUST00000139835.1 | 9148 | Gm11266 | ENSMUSG00000087413.1 |
| chr4 | 82519160 | 82519235 | 3.50945 | 4.18985 | 1.29385 | Intron (ENSMUST00000139835.1/ENSMUSG00000087413.1, intron 1 of 1) | 4 | 82508016 | 82547292 | 39277 | 1 | ENSMUST00000139835.1 | 11145 | Gm11266 | ENSMUSG00000087413.1 |
| chr4 | 82524037 | 82524538 | 3.3198 | 3.83288 | 1.09909 | Intron (ENSMUST00000139835.1/ENSMUSG00000087413.1, intron 1 of 1) | 4 | 82508016 | 82547292 | 39277 | 1 | ENSMUST00000139835.1 | 16022 | Gm11266 | ENSMUSG00000087413.1 |
| chr4 | 82527926 | 82528200 | 3.6954 | 4.47621 | 1.37902 | Intron (ENSMUST00000139835.1/ENSMUSG00000087413.1, intron 1 of 1) | 4 | 82508016 | 82547292 | 39277 | 1 | ENSMUST00000139835.1 | 19911 | Gm11266 | ENSMUSG00000087413.1 |
| chr4 | 82533019 | 82533137 | 3.40704 | 3.95476 | 1.07483 | Intron (ENSMUST00000139835.1/ENSMUSG00000087413.1, intron 1 of 1) | 4 | 82508016 | 82547292 | 39277 | 1 | ENSMUST00000139835.1 | 25004 | Gm11266 | ENSMUSG00000087413.1 |
| chr4 | 82533507 | 82533619 | 3.35855 | 3.87328 | 1.04543 | Intron (ENSMUST00000139835.1/ENSMUSG00000087413.1, intron 1 of 1) | 4 | 82508016 | 82547292 | 39277 | 1 | ENSMUST00000139835.1 | 25492 | Gm11266 | ENSMUSG00000087413.1 |
| chr4 | 82533980 | 82534238 | 3.46495 | 4.18077 | 1.26764 | Intron (ENSMUST00000139835.1/ENSMUSG00000087413.1, intron 1 of 1) | 4 | 82508016 | 82547292 | 39277 | 1 | ENSMUST00000139835.1 | 25965 | Gm11266 | ENSMUSG00000087413.1 |
| chr4 | 82535283 | 82535886 | 3.69707 | 4.59109 | 1.50505 | Intron (ENSMUST00000139835.1/ENSMUSG00000087413.1, intron 1 of 1) | 4 | 82508016 | 82547292 | 39277 | 1 | ENSMUST00000139835.1 | 27268 | Gm11266 | ENSMUSG00000087413.1 |
| chr4 | 82537275 | 82537377 | 3.18033 | 4.13295 | 1.21014 | Intron (ENSMUST00000139835.1/ENSMUSG00000087413.1, intron 1 of 1) | 4 | 82508016 | 82547292 | 39277 | 1 | ENSMUST00000139835.1 | 29260 | Gm11266 | ENSMUSG00000087413.1 |
| chr4 | 82537836 | 82538023 | 3.76141 | 4.77449 | 1.62162 | Intron (ENSMUST00000139835.1/ENSMUSG00000087413.1, intron 1 of 1) | 4 | 82508016 | 82547292 | 39277 | 1 | ENSMUST00000139835.1 | 29821 | Gm11266 | ENSMUSG00000087413.1 |
| chr4 | 82538759 | 82539046 | 3.7005 | 4.63179 | 1.58125 | Intron (ENSMUST00000139835.1/ENSMUSG00000087413.1, intron 1 of 1) | 4 | 82508016 | 82547292 | 39277 | 1 | ENSMUST00000139835.1 | 30744 | Gm11266 | ENSMUSG00000087413.1 |
| chr4 | 82541350 | 82541645 | 3.52561 | 4.24997 | 1.32404 | Intron (ENSMUST00000139835.1/ENSMUSG00000087413.1, intron 1 of 1) | 4 | 82508016 | 82547292 | 39277 | 1 | ENSMUST00000139835.1 | 33335 | Gm11266 | ENSMUSG00000087413.1 |
| chr4 | 82542104 | 82542391 | 3.76706 | 4.87929 | 1.67658 | Intron (ENSMUST00000139835.1/ENSMUSG00000087413.1, intron 1 of 1) | 4 | 82508016 | 82547292 | 39277 | 1 | ENSMUST00000139835.1 | 34089 | Gm11266 | ENSMUSG00000087413.1 |
| chr4 | 137118874 | 137119441 | 3.47478 | 4.12378 | 1.27889 | Promoter | 4 | 137117143 | 137118143 | 1001 | 2 | ENSMUST00000154285.1 | -732 | Gm13001 | ENSMUSG00000087417.1 |
| chr4 | 137124335 | 137124555 | 3.57305 | 4.30843 | 1.37009 | Distal Intergenic | 4 | 137117143 | 137118143 | 1001 | 2 | ENSMUST00000154285.1 | -6193 | Gm13001 | ENSMUSG00000087417.1 |
| chr4 | 137127761 | 137128397 | 3.55493 | 4.28194 | 1.35415 | Distal Intergenic | 4 | 137117143 | 137118143 | 1001 | 2 | ENSMUST00000154285.1 | -9619 | Gm13001 | ENSMUSG00000087417.1 |
| chr2 | 34105792 | 34105935 | 3.58462 | 4.34018 | 1.41708 | Intron (ENSMUST00000134921.1/ENSMUSG00000087461.1, intron 1 of 3) | 2 | 34018289 | 34107729 | 89441 | 2 | ENSMUST00000134921.1 | 1794 | C230014O12Rik | ENSMUSG00000087461.1 |
| chr2 | 34106373 | 34106915 | 3.84476 | 4.93641 | 1.7666 | Promoter | 2 | 34018289 | 34107729 | 89441 | 2 | ENSMUST00000134921.1 | 814 | C230014O12Rik | ENSMUSG00000087461.1 |
| chr2 | 34107278 | 34107581 | 3.49929 | 4.17788 | 1.32301 | Promoter | 2 | 34018289 | 34107729 | 89441 | 2 | ENSMUST00000134921.1 | 148 | C230014O12Rik | ENSMUSG00000087461.1 |
| chr4 | 119028360 | 119028689 | 3.83416 | 4.80494 | 1.67075 | Distal Intergenic | 4 | 118999660 | 119000485 | 826 | 2 | ENSMUST00000150754.1 | -27876 | Gm12862 | ENSMUSG00000087464.1 |
| chr9 | 115978045 | 115978204 | 3.70499 | 4.49376 | 1.38414 | Intron (ENSMUST00000119291.2/ENSMUSG00000056880.7, intron 11 of 14) | 9 | 115990956 | 116001788 | 10833 | 2 | ENSMUST00000130584.1 | 23584 | Gm16142 | ENSMUSG00000087469.1 |
| chr2 | 6267260 | 6267354 | 3.40887 | 3.97678 | 1.14353 | Intron (ENSMUST00000151782.1/ENSMUSG00000087125.1, intron 3 of 6) | 2 | 6255946 | 6257266 | 1321 | 2 | ENSMUST00000131190.1 | -9995 | Gm13383 | ENSMUSG00000087485.1 |
| chr5 | 36872841 | 36873258 | 3.88319 | 4.88942 | 1.75192 | Intron (ENSMUST00000031003.7/ENSMUSG00000029120.7, intron 1 of 8) | 5 | 36873595 | 36876879 | 3285 | 2 | ENSMUST00000152862.1 | 3621 | Ppp2r2cos | ENSMUSG00000087487.1 |
| chr5 | 36887646 | 36887875 | 3.57855 | 4.27449 | 1.28776 | Intron (ENSMUST00000031003.7/ENSMUSG00000029120.7, intron 1 of 8) | 5 | 36873595 | 36876879 | 3285 | 2 | ENSMUST00000152862.1 | -10768 | Ppp2r2cos | ENSMUSG00000087487.1 |
| chr11 | 33204956 | 33206470 | 3.79403 | 4.70801 | 1.59969 | Promoter | 11 | 33204055 | 33221477 | 17423 | 1 | ENSMUST00000131851.1 | 902 | Gm12116 | ENSMUSG00000087501.1 |
| chr11 | 33207316 | 33208095 | 3.61465 | 4.4706 | 1.46806 | Intron (ENSMUST00000131851.1/ENSMUSG00000087501.1, intron 1 of 3) | 11 | 33204055 | 33221477 | 17423 | 1 | ENSMUST00000131851.1 | 3262 | Gm12116 | ENSMUSG00000087501.1 |
| chr2 | 75315396 | 75315549 | 3.37783 | 3.94358 | 1.16713 | Exon (ENSMUST00000186411.1/ENSMUSG00000100303.1, exon 11 of 11) | 2 | 75436057 | 75444179 | 8123 | 1 | ENSMUST00000135329.1 | -120508 | Gm13658 | ENSMUSG00000087504.1 |
| chr2 | 75324735 | 75324977 | 3.56777 | 4.32777 | 1.35524 | Distal Intergenic | 2 | 75436057 | 75444179 | 8123 | 1 | ENSMUST00000135329.1 | -111080 | Gm13658 | ENSMUSG00000087504.1 |
| chr2 | 75371118 | 75371467 | 3.4168 | 4.00321 | 1.18358 | Distal Intergenic | 2 | 75436057 | 75444179 | 8123 | 1 | ENSMUST00000135329.1 | -64590 | Gm13658 | ENSMUSG00000087504.1 |
| chr2 | 75423053 | 75423171 | 3.4885 | 4.15348 | 1.27996 | Distal Intergenic | 2 | 75436057 | 75444179 | 8123 | 1 | ENSMUST00000135329.1 | -12886 | Gm13658 | ENSMUSG00000087504.1 |
| chr2 | 75424051 | 75424127 | 4.61519 | 6.7346 | 2.9971 | Distal Intergenic | 2 | 75436057 | 75444179 | 8123 | 1 | ENSMUST00000135329.1 | -11930 | Gm13658 | ENSMUSG00000087504.1 |
| chr2 | 75425124 | 75425216 | 3.76466 | 4.69601 | 1.62428 | Distal Intergenic | 2 | 75436057 | 75444179 | 8123 | 1 | ENSMUST00000135329.1 | -10841 | Gm13658 | ENSMUSG00000087504.1 |
| chr2 | 156617551 | 156617872 | 3.35717 | 3.86325 | 1.01733 | Intron (ENSMUST00000109567.4/ENSMUSG00000061689.10, intron 1 of 12) | 2 | 156614447 | 156615448 | 1002 | 2 | ENSMUST00000153158.1 | -2104 | Gm14174 | ENSMUSG00000087508.1 |
| chr2 | 156616955 | 156617148 | 3.34557 | 3.84575 | 1.01733 | Intron (ENSMUST00000109567.4/ENSMUSG00000061689.10, intron 1 of 12) | 2 | 156614447 | 156615448 | 1002 | 2 | ENSMUST00000153158.1 | -1508 | Gm14174 | ENSMUSG00000087508.1 |
| chr5 | 119626658 | 119626920 | 3.53838 | 4.41949 | 1.37525 | Promoter | 5 | 119611263 | 119627382 | 16120 | 2 | ENSMUST00000134493.2 | 462 | Tbx3os1 | ENSMUSG00000087516.2 |
| chr5 | 119628062 | 119628365 | 3.71192 | 4.56359 | 1.54234 | Promoter | 5 | 119611900 | 119627593 | 15694 | 2 | ENSMUST00000140101.1 | -470 | Tbx3os1 | ENSMUSG00000087516.2 |
| chr5 | 119630132 | 119630447 | 3.25795 | 3.71409 | 1.01733 | Intron (ENSMUST00000156235.1/ENSMUSG00000085708.1, intron 1 of 3) | 5 | 119611900 | 119627593 | 15694 | 2 | ENSMUST00000140101.1 | -2540 | Tbx3os1 | ENSMUSG00000087516.2 |
| chr5 | 119671249 | 119672727 | 4.07891 | 5.47183 | 2.15509 | Promoter | 5 | 119617901 | 119671365 | 53465 | 2 | ENSMUST00000129108.1 | 0 | Tbx3os1 | ENSMUSG00000087516.2 |
| chr2 | 147363310 | 147363610 | 4.20581 | 5.77288 | 2.30908 | Promoter | 2 | 147361541 | 147362409 | 869 | 2 | ENSMUST00000140987.1 | -902 | AI646519 | ENSMUSG00000087521.1 |
| chr2 | 147362359 | 147362949 | 4.20367 | 5.69075 | 2.2762 | Promoter | 2 | 147361541 | 147362409 | 869 | 2 | ENSMUST00000140987.1 | 0 | AI646519 | ENSMUSG00000087521.1 |
| chr2 | 131546244 | 131546766 | 4.00768 | 5.19442 | 1.91533 | Exon (ENSMUST00000103184.3/ENSMUSG00000027335.9, exon 2 of 2) | 2 | 131561614 | 131563179 | 1566 | 1 | ENSMUST00000146049.1 | -14848 | Gm14285 | ENSMUSG00000087524.1 |
| chr2 | 131548544 | 131549041 | 3.40073 | 3.97281 | 1.17111 | Intron (ENSMUST00000103184.3/ENSMUSG00000027335.9, intron 1 of 1) | 2 | 131561614 | 131563179 | 1566 | 1 | ENSMUST00000146049.1 | -12573 | Gm14285 | ENSMUSG00000087524.1 |
| chr2 | 131559749 | 131559967 | 3.43522 | 4.03961 | 1.20403 | Intron (ENSMUST00000103184.3/ENSMUSG00000027335.9, intron 1 of 1) | 2 | 131561614 | 131563179 | 1566 | 1 | ENSMUST00000146049.1 | -1647 | Gm14285 | ENSMUSG00000087524.1 |
| chr2 | 131561106 | 131561198 | 3.70787 | 4.93071 | 1.73407 | Promoter | 2 | 131561614 | 131563179 | 1566 | 1 | ENSMUST00000146049.1 | -416 | Gm14285 | ENSMUSG00000087524.1 |
| chr7 | 128010038 | 128010275 | 3.4881 | 4.14709 | 1.28001 | Exon (ENSMUST00000106248.2/ENSMUSG00000042828.7, exon 7 of 7) | 7 | 128002849 | 128005190 | 2342 | 2 | ENSMUST00000126205.1 | -4849 | Gm15533 | ENSMUSG00000087530.1 |
| chr11 | 81614222 | 81614409 | 3.66553 | 4.50486 | 1.47802 | Intron (ENSMUST00000066197.6/ENSMUSG00000020704.8, intron 1 of 9) | 11 | 81587361 | 81593478 | 6118 | 1 | ENSMUST00000156193.1 | 26862 | Gm11418 | ENSMUSG00000087534.1 |
| chr11 | 81648219 | 81648428 | 3.46413 | 4.05739 | 1.13249 | Intron (ENSMUST00000066197.6/ENSMUSG00000020704.8, intron 1 of 9) | 11 | 81587361 | 81593478 | 6118 | 1 | ENSMUST00000156193.1 | 60859 | Gm11418 | ENSMUSG00000087534.1 |
| chr2 | 162682880 | 162683120 | 3.27524 | 3.74274 | 1.02686 | Distal Intergenic | 2 | 162679455 | 162680529 | 1075 | 1 | ENSMUST00000155173.1 | 3426 | Gm14246 | ENSMUSG00000087536.1 |
| chr2 | 38358731 | 38359705 | 3.74129 | 4.68196 | 1.59894 | Exon (ENSMUST00000137134.1/ENSMUSG00000087539.1, exon 3 of 3) | 2 | 38359580 | 38363980 | 4401 | 2 | ENSMUST00000137134.1 | 4275 | Gm13584 | ENSMUSG00000087539.1 |
| chr2 | 38360106 | 38360333 | 3.60703 | 4.36593 | 1.39969 | Intron (ENSMUST00000143783.3/ENSMUSG00000000247.6, intron 4 of 4) | 2 | 38359580 | 38363980 | 4401 | 2 | ENSMUST00000137134.1 | 3647 | Gm13584 | ENSMUSG00000087539.1 |
| chr2 | 38360903 | 38361249 | 3.50636 | 4.15933 | 1.23517 | Exon (ENSMUST00000137134.1/ENSMUSG00000087539.1, exon 2 of 3) | 2 | 38359580 | 38363980 | 4401 | 2 | ENSMUST00000137134.1 | 2731 | Gm13584 | ENSMUSG00000087539.1 |
| chr2 | 38361647 | 38361963 | 3.31841 | 3.82958 | 1.07359 | Intron (ENSMUST00000143783.3/ENSMUSG00000000247.6, intron 4 of 4) | 2 | 38359580 | 38363980 | 4401 | 2 | ENSMUST00000137134.1 | 2017 | Gm13584 | ENSMUSG00000087539.1 |
| chr2 | 38363626 | 38363755 | 3.38883 | 3.92015 | 1.04969 | Promoter | 2 | 38359580 | 38363980 | 4401 | 2 | ENSMUST00000137134.1 | 225 | Gm13584 | ENSMUSG00000087539.1 |
| chr2 | 38364097 | 38365056 | 3.60812 | 4.31969 | 1.2884 | Promoter | 2 | 38359580 | 38363980 | 4401 | 2 | ENSMUST00000137134.1 | -118 | Gm13584 | ENSMUSG00000087539.1 |
| chr2 | 38365609 | 38366340 | 3.41752 | 3.99827 | 1.15303 | Intron (ENSMUST00000143783.3/ENSMUSG00000000247.6, intron 4 of 4) | 2 | 38359580 | 38363980 | 4401 | 2 | ENSMUST00000137134.1 | -1630 | Gm13584 | ENSMUSG00000087539.1 |
| chr2 | 38366699 | 38367360 | 3.75453 | 4.71434 | 1.57508 | Intron (ENSMUST00000143783.3/ENSMUSG00000000247.6, intron 4 of 4) | 2 | 38359580 | 38363980 | 4401 | 2 | ENSMUST00000137134.1 | -2720 | Gm13584 | ENSMUSG00000087539.1 |
| chr2 | 38368589 | 38369395 | 4.1644 | 5.51001 | 2.11618 | 3' UTR | 2 | 38359580 | 38363980 | 4401 | 2 | ENSMUST00000137134.1 | -4610 | Gm13584 | ENSMUSG00000087539.1 |
| chr2 | 38369738 | 38370683 | 3.81699 | 4.72752 | 1.56615 | Distal Intergenic | 2 | 38359580 | 38363980 | 4401 | 2 | ENSMUST00000137134.1 | -5759 | Gm13584 | ENSMUSG00000087539.1 |
| chr2 | 38370997 | 38371555 | 4.06156 | 5.21671 | 1.92019 | Distal Intergenic | 2 | 38359580 | 38363980 | 4401 | 2 | ENSMUST00000137134.1 | -7018 | Gm13584 | ENSMUSG00000087539.1 |
| chr2 | 38378634 | 38379208 | 3.5133 | 4.24634 | 1.32839 | Distal Intergenic | 2 | 38359580 | 38363980 | 4401 | 2 | ENSMUST00000137134.1 | -14655 | Gm13584 | ENSMUSG00000087539.1 |
| chr2 | 38379581 | 38380279 | 4.21746 | 5.57365 | 2.22582 | Distal Intergenic | 2 | 38359580 | 38363980 | 4401 | 2 | ENSMUST00000137134.1 | -15602 | Gm13584 | ENSMUSG00000087539.1 |
| chr2 | 38380613 | 38381227 | 3.37193 | 3.89288 | 1.04338 | Distal Intergenic | 2 | 38359580 | 38363980 | 4401 | 2 | ENSMUST00000137134.1 | -16634 | Gm13584 | ENSMUSG00000087539.1 |
| chr2 | 38381533 | 38381971 | 3.40733 | 3.95048 | 1.05795 | Distal Intergenic | 2 | 38359580 | 38363980 | 4401 | 2 | ENSMUST00000137134.1 | -17554 | Gm13584 | ENSMUSG00000087539.1 |
| chr5 | 77093802 | 77094041 | 3.35384 | 3.85822 | 1.01733 | Promoter | 5 | 77094532 | 77102303 | 7772 | 1 | ENSMUST00000148568.1 | -491 | Hopxos | ENSMUSG00000087541.1 |
| chr2 | 157605034 | 157605141 | 4.37221 | 5.90916 | 2.43935 | Distal Intergenic | 2 | 157595583 | 157597320 | 1738 | 2 | ENSMUST00000131365.1 | -7715 | 2410019O14Rik | ENSMUSG00000087547.1 |
| chr2 | 180773371 | 180773972 | 3.55521 | 4.3456 | 1.37282 | Intron (ENSMUST00000154524.1/ENSMUSG00000087550.1, intron 1 of 2) | 2 | 180766214 | 180775403 | 9190 | 2 | ENSMUST00000154524.1 | 1431 | Gm14343 | ENSMUSG00000087550.1 |
| chr2 | 120246497 | 120246656 | 3.35304 | 3.87595 | 1.08407 | Promoter | 2 | 120244921 | 120245598 | 678 | 2 | ENSMUST00000155641.1 | -900 | Gm13997 | ENSMUSG00000087555.1 |
| chr2 | 93085902 | 93086170 | 3.35436 | 3.85901 | 1.01733 | Exon (ENSMUST00000150159.1/ENSMUSG00000087559.1, exon 2 of 2) | 2 | 93085942 | 93098244 | 12303 | 2 | ENSMUST00000150159.1 | 12074 | Gm13802 | ENSMUSG00000087559.1 |
| chr2 | 93089743 | 93089960 | 3.29768 | 3.78016 | 1.03763 | Intron (ENSMUST00000150159.1/ENSMUSG00000087559.1, intron 1 of 1) | 2 | 93085942 | 93098244 | 12303 | 2 | ENSMUST00000150159.1 | 8284 | Gm13802 | ENSMUSG00000087559.1 |
| chr7 | 44582493 | 44582782 | 3.44842 | 4.14002 | 1.26897 | Exon (ENSMUST00000153124.1/ENSMUSG00000002204.8, exon 5 of 5) | 7 | 44579968 | 44590624 | 10657 | 2 | ENSMUST00000148180.1 | 7842 | Gm15396 | ENSMUSG00000087560.1 |
| chr7 | 44585113 | 44585873 | 3.58675 | 4.46266 | 1.42038 | Exon (ENSMUST00000002274.8/ENSMUSG00000002204.8, exon 6 of 9) | 7 | 44579968 | 44590624 | 10657 | 2 | ENSMUST00000148180.1 | 4751 | Gm15396 | ENSMUSG00000087560.1 |
| chr7 | 44586226 | 44587630 | 3.84453 | 4.92784 | 1.76271 | 3' UTR | 7 | 44579968 | 44590624 | 10657 | 2 | ENSMUST00000148180.1 | 2994 | Gm15396 | ENSMUSG00000087560.1 |
| chr7 | 44588101 | 44588701 | 4.05737 | 5.29125 | 2.00158 | Intron (ENSMUST00000148180.1/ENSMUSG00000087560.1, intron 1 of 2) | 7 | 44579968 | 44590624 | 10657 | 2 | ENSMUST00000148180.1 | 1923 | Gm15396 | ENSMUSG00000087560.1 |
| chr7 | 44589163 | 44589351 | 3.47038 | 4.3007 | 1.32445 | Intron (ENSMUST00000148180.1/ENSMUSG00000087560.1, intron 1 of 2) | 7 | 44579968 | 44590624 | 10657 | 2 | ENSMUST00000148180.1 | 1273 | Gm15396 | ENSMUSG00000087560.1 |
| chr1 | 63244416 | 63245052 | 4.4078 | 5.89884 | 2.39932 | Distal Intergenic | 1 | 63266848 | 63273115 | 6268 | 2 | ENSMUST00000153049.2 | 28063 | Gm11608 | ENSMUSG00000087561.2 |
| chr1 | 63245362 | 63245796 | 3.87843 | 4.89561 | 1.74733 | Distal Intergenic | 1 | 63266848 | 63273115 | 6268 | 2 | ENSMUST00000153049.2 | 27319 | Gm11608 | ENSMUSG00000087561.2 |
| chr1 | 63246586 | 63248140 | 3.82925 | 4.80238 | 1.701 | Distal Intergenic | 1 | 63266848 | 63273115 | 6268 | 2 | ENSMUST00000153049.2 | 24975 | Gm11608 | ENSMUSG00000087561.2 |
| chr1 | 63248469 | 63249916 | 3.57735 | 4.5052 | 1.44666 | Distal Intergenic | 1 | 63266848 | 63273115 | 6268 | 2 | ENSMUST00000153049.2 | 23199 | Gm11608 | ENSMUSG00000087561.2 |
| chr1 | 63251750 | 63251953 | 4.01965 | 5.78835 | 2.33592 | Distal Intergenic | 1 | 63266848 | 63273115 | 6268 | 2 | ENSMUST00000153049.2 | 21162 | Gm11608 | ENSMUSG00000087561.2 |
| chr1 | 63252839 | 63253060 | 3.37847 | 4.36136 | 1.3445 | Distal Intergenic | 1 | 63266848 | 63273115 | 6268 | 2 | ENSMUST00000153049.2 | 20055 | Gm11608 | ENSMUSG00000087561.2 |
| chr1 | 63253530 | 63254006 | 3.74404 | 4.63858 | 1.5959 | Distal Intergenic | 1 | 63266848 | 63273115 | 6268 | 2 | ENSMUST00000153049.2 | 19109 | Gm11608 | ENSMUSG00000087561.2 |
| chr1 | 63254523 | 63254680 | 3.67916 | 4.48886 | 1.48585 | Distal Intergenic | 1 | 63266848 | 63273115 | 6268 | 2 | ENSMUST00000153049.2 | 18435 | Gm11608 | ENSMUSG00000087561.2 |
| chr1 | 63256436 | 63257880 | 3.87187 | 5.01812 | 1.85939 | Distal Intergenic | 1 | 63266848 | 63273115 | 6268 | 2 | ENSMUST00000153049.2 | 15235 | Gm11608 | ENSMUSG00000087561.2 |
| chr1 | 63258562 | 63258647 | 3.93406 | 5.12305 | 1.87139 | Distal Intergenic | 1 | 63266848 | 63273115 | 6268 | 2 | ENSMUST00000153049.2 | 14468 | Gm11608 | ENSMUSG00000087561.2 |
| chr1 | 63259095 | 63259388 | 3.49897 | 4.23637 | 1.29716 | Distal Intergenic | 1 | 63266848 | 63273115 | 6268 | 2 | ENSMUST00000153049.2 | 13727 | Gm11608 | ENSMUSG00000087561.2 |
| chr1 | 63259695 | 63260481 | 3.78143 | 4.84197 | 1.70739 | Distal Intergenic | 1 | 63266848 | 63273115 | 6268 | 2 | ENSMUST00000153049.2 | 12634 | Gm11608 | ENSMUSG00000087561.2 |
| chr1 | 63261756 | 63262088 | 4.17411 | 5.5477 | 2.16933 | Distal Intergenic | 1 | 63266848 | 63273115 | 6268 | 2 | ENSMUST00000153049.2 | 11027 | Gm11608 | ENSMUSG00000087561.2 |
| chr1 | 63262504 | 63262841 | 3.47774 | 4.13541 | 1.28261 | Distal Intergenic | 1 | 63266848 | 63273115 | 6268 | 2 | ENSMUST00000153049.2 | 10274 | Gm11608 | ENSMUSG00000087561.2 |
| chr1 | 63263585 | 63263974 | 3.63834 | 4.44184 | 1.4479 | Downstream (2-3kb) | 1 | 63266848 | 63273115 | 6268 | 2 | ENSMUST00000153049.2 | 9141 | Gm11608 | ENSMUSG00000087561.2 |
| chr1 | 63265307 | 63265490 | 3.73173 | 4.60979 | 1.56005 | Downstream (1-2kb) | 1 | 63266848 | 63273115 | 6268 | 2 | ENSMUST00000153049.2 | 7625 | Gm11608 | ENSMUSG00000087561.2 |
| chr1 | 63267934 | 63268614 | 3.29607 | 3.80795 | 1.04986 | Intron (ENSMUST00000153049.2/ENSMUSG00000087561.2, intron 4 of 4) | 1 | 63266848 | 63273115 | 6268 | 2 | ENSMUST00000153049.2 | 4501 | Gm11608 | ENSMUSG00000087561.2 |
| chr2 | 158612055 | 158614886 | 3.90436 | 4.98861 | 1.76523 | Promoter | 2 | 158610910 | 158611820 | 911 | 2 | ENSMUST00000138427.1 | -236 | Gm14205 | ENSMUSG00000087563.1 |
| chr2 | 158615340 | 158615549 | 3.43908 | 4.01081 | 1.10163 | 3' UTR | 2 | 158610910 | 158611820 | 911 | 2 | ENSMUST00000138427.1 | -3521 | Gm14205 | ENSMUSG00000087563.1 |
| chr2 | 158616733 | 158616970 | 3.81475 | 4.7746 | 1.67355 | Distal Intergenic | 2 | 158610910 | 158611820 | 911 | 2 | ENSMUST00000138427.1 | -4914 | Gm14205 | ENSMUSG00000087563.1 |
| chr2 | 158617680 | 158618218 | 3.65108 | 4.44513 | 1.39715 | Distal Intergenic | 2 | 158610910 | 158611820 | 911 | 2 | ENSMUST00000138427.1 | -5861 | Gm14205 | ENSMUSG00000087563.1 |
| chr18 | 33776976 | 33777288 | 3.54724 | 4.25311 | 1.30847 | Distal Intergenic | 18 | 33794892 | 33795986 | 1095 | 1 | ENSMUST00000146010.2 | -17604 | 2410004N09Rik | ENSMUSG00000087590.2 |
| chr5 | 66674906 | 66675554 | 3.53061 | 4.23317 | 1.34862 | Promoter | 5 | 66655792 | 66676033 | 20242 | 2 | ENSMUST00000177319.1 | 479 | Uchl1os | ENSMUSG00000087601.2 |
| chr11 | 63004944 | 63005052 | 3.38627 | 4.12046 | 1.19035 | Promoter | 11 | 62999115 | 63004134 | 5020 | 2 | ENSMUST00000130177.1 | -811 | Cdrt4os1 | ENSMUSG00000087604.2 |
| chr5 | 100943225 | 100943478 | 3.53005 | 4.18098 | 1.21881 | Distal Intergenic | 5 | 100840797 | 100847167 | 6371 | 2 | ENSMUST00000142871.1 | -96059 | 4930458D05Rik | ENSMUSG00000087611.1 |
| chr5 | 100944080 | 100944513 | 3.60783 | 4.41947 | 1.40737 | Distal Intergenic | 5 | 100840797 | 100847167 | 6371 | 2 | ENSMUST00000142871.1 | -96914 | 4930458D05Rik | ENSMUSG00000087611.1 |
| chr5 | 100952053 | 100952133 | 3.74603 | 4.59144 | 1.49626 | Distal Intergenic | 5 | 100840797 | 100847167 | 6371 | 2 | ENSMUST00000142871.1 | -104887 | 4930458D05Rik | ENSMUSG00000087611.1 |
| chr5 | 100988414 | 100988677 | 3.40731 | 3.95229 | 1.06479 | Distal Intergenic | 5 | 100840797 | 100847167 | 6371 | 2 | ENSMUST00000142871.1 | -141248 | 4930458D05Rik | ENSMUSG00000087611.1 |
| chr2 | 166339129 | 166339581 | 3.98305 | 5.23544 | 1.93833 | Distal Intergenic | 2 | 166380828 | 166398157 | 17330 | 2 | ENSMUST00000151070.1 | 58576 | Gm14268 | ENSMUSG00000087617.1 |
| chr2 | 166344695 | 166344974 | 3.5111 | 4.16105 | 1.25092 | Distal Intergenic | 2 | 166380828 | 166398157 | 17330 | 2 | ENSMUST00000151070.1 | 53183 | Gm14268 | ENSMUSG00000087617.1 |
| chr2 | 166346367 | 166346822 | 3.38346 | 3.93616 | 1.13516 | Distal Intergenic | 2 | 166380828 | 166398157 | 17330 | 2 | ENSMUST00000151070.1 | 51335 | Gm14268 | ENSMUSG00000087617.1 |
| chr2 | 166348208 | 166348283 | 3.60632 | 4.37566 | 1.43521 | Distal Intergenic | 2 | 166380828 | 166398157 | 17330 | 2 | ENSMUST00000151070.1 | 49874 | Gm14268 | ENSMUSG00000087617.1 |
| chr2 | 166349316 | 166349654 | 3.45492 | 4.15791 | 1.24502 | Distal Intergenic | 2 | 166380828 | 166398157 | 17330 | 2 | ENSMUST00000151070.1 | 48503 | Gm14268 | ENSMUSG00000087617.1 |
| chr2 | 166355442 | 166356053 | 3.50013 | 4.14611 | 1.25255 | Distal Intergenic | 2 | 166380828 | 166398157 | 17330 | 2 | ENSMUST00000151070.1 | 42104 | Gm14268 | ENSMUSG00000087617.1 |
| chr2 | 166356466 | 166356568 | 4.1964 | 5.44975 | 2.04292 | Distal Intergenic | 2 | 166380828 | 166398157 | 17330 | 2 | ENSMUST00000151070.1 | 41589 | Gm14268 | ENSMUSG00000087617.1 |
| chr2 | 166406287 | 166406560 | 3.91765 | 4.91602 | 1.70172 | Distal Intergenic | 2 | 166381114 | 166398187 | 17074 | 2 | ENSMUST00000132465.1 | -8101 | Gm14268 | ENSMUSG00000087617.1 |
| chr2 | 166407770 | 166408084 | 3.54551 | 4.20307 | 1.21347 | Distal Intergenic | 2 | 166381114 | 166398187 | 17074 | 2 | ENSMUST00000132465.1 | -9584 | Gm14268 | ENSMUSG00000087617.1 |
| chr2 | 166411056 | 166411235 | 3.47388 | 4.07462 | 1.14229 | Distal Intergenic | 2 | 166381114 | 166398187 | 17074 | 2 | ENSMUST00000132465.1 | -12870 | Gm14268 | ENSMUSG00000087617.1 |
| chr2 | 166412221 | 166412797 | 3.60294 | 4.40192 | 1.38329 | Distal Intergenic | 2 | 166381114 | 166398187 | 17074 | 2 | ENSMUST00000132465.1 | -14035 | Gm14268 | ENSMUSG00000087617.1 |
| chr2 | 146823773 | 146824074 | 4.76387 | 6.75263 | 3.07718 | Distal Intergenic | 2 | 146828327 | 146839732 | 11406 | 2 | ENSMUST00000139345.1 | 15658 | Gm14114 | ENSMUSG00000087618.1 |
| chr2 | 146826272 | 146826797 | 3.39498 | 3.95476 | 1.13846 | Downstream (1-2kb) | 2 | 146828327 | 146839732 | 11406 | 2 | ENSMUST00000139345.1 | 12935 | Gm14114 | ENSMUSG00000087618.1 |
| chr2 | 146829024 | 146829110 | 3.53188 | 4.45018 | 1.44281 | Intron (ENSMUST00000139345.1/ENSMUSG00000087618.1, intron 1 of 1) | 2 | 146828327 | 146839732 | 11406 | 2 | ENSMUST00000139345.1 | 10622 | Gm14114 | ENSMUSG00000087618.1 |
| chr2 | 146833189 | 146835349 | 3.67125 | 4.67008 | 1.55328 | Intron (ENSMUST00000139345.1/ENSMUSG00000087618.1, intron 1 of 1) | 2 | 146828327 | 146839732 | 11406 | 2 | ENSMUST00000139345.1 | 4383 | Gm14114 | ENSMUSG00000087618.1 |
| chr2 | 146835713 | 146837088 | 3.68067 | 4.57328 | 1.53512 | Intron (ENSMUST00000139345.1/ENSMUSG00000087618.1, intron 1 of 1) | 2 | 146828327 | 146839732 | 11406 | 2 | ENSMUST00000139345.1 | 2644 | Gm14114 | ENSMUSG00000087618.1 |
| chr2 | 146838629 | 146839379 | 4.52383 | 6.52263 | 2.94035 | Promoter | 2 | 146828327 | 146839732 | 11406 | 2 | ENSMUST00000139345.1 | 353 | Gm14114 | ENSMUSG00000087618.1 |
| chr2 | 146839712 | 146840879 | 3.86376 | 4.88585 | 1.73895 | Promoter | 2 | 146828327 | 146839732 | 11406 | 2 | ENSMUST00000139345.1 | 0 | Gm14114 | ENSMUSG00000087618.1 |
| chr2 | 146841237 | 146842363 | 3.87787 | 4.85623 | 1.68659 | Distal Intergenic | 2 | 146828327 | 146839732 | 11406 | 2 | ENSMUST00000139345.1 | -1506 | Gm14114 | ENSMUSG00000087618.1 |
| chr4 | 139925311 | 139925743 | 4.01107 | 5.30724 | 2.01135 | Intron (ENSMUST00000128563.1/ENSMUSG00000087628.1, intron 1 of 1) | 4 | 139923349 | 139933170 | 9822 | 1 | ENSMUST00000128563.1 | 1963 | Gm13028 | ENSMUSG00000087628.1 |
| chr4 | 139926877 | 139927203 | 3.36577 | 3.87622 | 1.01733 | Exon (ENSMUST00000153921.1/ENSMUSG00000085928.1, exon 3 of 3) | 4 | 139923349 | 139933170 | 9822 | 1 | ENSMUST00000128563.1 | 3529 | Gm13028 | ENSMUSG00000087628.1 |
| chr14 | 33367141 | 33367445 | 3.71226 | 4.6278 | 1.54712 | Exon (ENSMUST00000111956.3/ENSMUSG00000063506.8, exon 10 of 11) | 14 | 33365951 | 33370143 | 4193 | 2 | ENSMUST00000137359.1 | 2698 | Gm15512 | ENSMUSG00000087639.1 |
| chr14 | 33369845 | 33369999 | 3.4509 | 4.07873 | 1.22016 | Promoter | 14 | 33365951 | 33370143 | 4193 | 2 | ENSMUST00000137359.1 | 144 | Gm15512 | ENSMUSG00000087639.1 |
| chr4 | 139905699 | 139906010 | 3.34153 | 3.83966 | 1.01733 | Distal Intergenic | 4 | 139910488 | 139912133 | 1646 | 1 | ENSMUST00000151651.1 | -4478 | Gm1667 | ENSMUSG00000087646.1 |
| chr4 | 139911588 | 139912064 | 3.5457 | 4.22435 | 1.28083 | Exon (ENSMUST00000151651.1/ENSMUSG00000087646.1, exon 3 of 3) | 4 | 139910488 | 139912133 | 1646 | 1 | ENSMUST00000151651.1 | 1101 | Gm1667 | ENSMUSG00000087646.1 |
| chr5 | 148821521 | 148821606 | 3.65818 | 4.41177 | 1.34608 | Promoter | 5 | 148820767 | 148829408 | 8642 | 1 | ENSMUST00000136919.1 | 755 | Gm15407 | ENSMUSG00000087647.1 |
| chr5 | 148831495 | 148832343 | 3.68196 | 4.53571 | 1.47065 | Distal Intergenic | 5 | 148820767 | 148829408 | 8642 | 1 | ENSMUST00000136919.1 | 10729 | Gm15407 | ENSMUSG00000087647.1 |
| chr10 | 83722761 | 83723045 | 4.00136 | 5.36141 | 2.0096 | Promoter | 10 | 83722865 | 83762761 | 39897 | 1 | ENSMUST00000150459.1 | 0 | 1500009L16Rik | ENSMUSG00000087651.2 |
| chr10 | 83725064 | 83725142 | 3.55936 | 4.2931 | 1.36534 | Intron (ENSMUST00000150459.1/ENSMUSG00000087651.2, intron 1 of 5) | 10 | 83722865 | 83762761 | 39897 | 1 | ENSMUST00000150459.1 | 2200 | 1500009L16Rik | ENSMUSG00000087651.2 |
| chr10 | 83954493 | 83954665 | 8.66146 | 18.32876 | 13.51614 | Distal Intergenic | 10 | 83722865 | 83762761 | 39897 | 1 | ENSMUST00000150459.1 | 231629 | 1500009L16Rik | ENSMUSG00000087651.2 |
| chr7 | 132595614 | 132595721 | 3.36764 | 3.9117 | 1.13277 | 3' UTR | 7 | 132597387 | 132599637 | 2251 | 1 | ENSMUST00000138891.2 | -1666 | Gm16764 | ENSMUSG00000087657.3 |
| chr7 | 132596419 | 132597959 | 3.79822 | 5.09408 | 1.86086 | Promoter | 7 | 132597387 | 132599637 | 2251 | 1 | ENSMUST00000138891.2 | 0 | Gm16764 | ENSMUSG00000087657.3 |
| chr6 | 52150765 | 52151096 | 3.73673 | 4.58835 | 1.51765 | Distal Intergenic | 6 | 52156902 | 52158528 | 1627 | 1 | ENSMUST00000129546.2 | -5806 | Hotairm1 | ENSMUSG00000087658.2 |
| chr4 | 89253601 | 89253884 | 3.99747 | 5.09354 | 1.84232 | Exon (ENSMUST00000129527.2/ENSMUSG00000087659.2, exon 2 of 4) | 4 | 89235699 | 89272587 | 36889 | 2 | ENSMUST00000129527.2 | 18703 | Gm12606 | ENSMUSG00000087659.2 |
| chr4 | 89266550 | 89266845 | 3.77409 | 4.73312 | 1.5873 | Intron (ENSMUST00000129527.2/ENSMUSG00000087659.2, intron 1 of 3) | 4 | 89235699 | 89272587 | 36889 | 2 | ENSMUST00000129527.2 | 5742 | Gm12606 | ENSMUSG00000087659.2 |
| chr4 | 89269541 | 89270023 | 3.45865 | 4.1518 | 1.26301 | Intron (ENSMUST00000129527.2/ENSMUSG00000087659.2, intron 1 of 3) | 4 | 89235699 | 89272587 | 36889 | 2 | ENSMUST00000129527.2 | 2564 | Gm12606 | ENSMUSG00000087659.2 |
| chr4 | 89270601 | 89271070 | 3.46735 | 4.06123 | 1.12908 | Intron (ENSMUST00000129527.2/ENSMUSG00000087659.2, intron 1 of 3) | 4 | 89235699 | 89272587 | 36889 | 2 | ENSMUST00000129527.2 | 1517 | Gm12606 | ENSMUSG00000087659.2 |
| chr4 | 89271946 | 89272604 | 4.34505 | 6.12648 | 2.6666 | Promoter | 4 | 89235699 | 89272587 | 36889 | 2 | ENSMUST00000129527.2 | 0 | Gm12606 | ENSMUSG00000087659.2 |
| chr4 | 89272995 | 89273840 | 4.01813 | 5.15136 | 1.87578 | Promoter | 4 | 89235717 | 89273403 | 37687 | 2 | ENSMUST00000127174.1 | 0 | Gm12606 | ENSMUSG00000087659.2 |
| chr4 | 89274719 | 89277278 | 3.82719 | 4.81699 | 1.64027 | 3' UTR | 4 | 89235717 | 89273403 | 37687 | 2 | ENSMUST00000127174.1 | -1317 | Gm12606 | ENSMUSG00000087659.2 |
| chr11 | 53002243 | 53002500 | 3.50052 | 4.17111 | 1.21386 | Intron (ENSMUST00000036796.7/ENSMUSG00000036264.9, intron 4 of 15) | 11 | 53111451 | 53120056 | 8606 | 2 | ENSMUST00000137096.1 | 117556 | Gm11186 | ENSMUSG00000087668.1 |
| chr11 | 53057334 | 53057768 | 3.36244 | 3.92005 | 1.10871 | Intron (ENSMUST00000036796.7/ENSMUSG00000036264.9, intron 4 of 15) | 11 | 53111451 | 53120056 | 8606 | 2 | ENSMUST00000137096.1 | 62288 | Gm11186 | ENSMUSG00000087668.1 |
| chr11 | 119548495 | 119548848 | 3.88285 | 5.13083 | 1.8594 | Promoter | 11 | 119548630 | 119569046 | 20417 | 1 | ENSMUST00000135630.1 | 0 | Gm11762 | ENSMUSG00000087675.1 |
| chr11 | 119549186 | 119549956 | 3.77924 | 4.95134 | 1.78636 | Promoter | 11 | 119548630 | 119569046 | 20417 | 1 | ENSMUST00000135630.1 | 557 | Gm11762 | ENSMUSG00000087675.1 |
| chr11 | 119551152 | 119551302 | 3.43151 | 4.12112 | 1.20877 | Intron (ENSMUST00000135630.1/ENSMUSG00000087675.1, intron 1 of 3) | 11 | 119548630 | 119569046 | 20417 | 1 | ENSMUST00000135630.1 | 2523 | Gm11762 | ENSMUSG00000087675.1 |
| chr11 | 119563234 | 119563400 | 4.44533 | 6.23351 | 2.66376 | Intron (ENSMUST00000135630.1/ENSMUSG00000087675.1, intron 1 of 3) | 11 | 119548630 | 119569046 | 20417 | 1 | ENSMUST00000135630.1 | 14605 | Gm11762 | ENSMUSG00000087675.1 |
| chr15 | 101131464 | 101131759 | 3.69165 | 4.525 | 1.5114 | Promoter | 15 | 101129916 | 101131592 | 1677 | 2 | ENSMUST00000153253.1 | 0 | Gm16031 | ENSMUSG00000087690.1 |
| chr2 | 114014650 | 114014884 | 3.23758 | 3.68354 | 1.01733 | Promoter | 2 | 114013745 | 114031451 | 17707 | 1 | ENSMUST00000134129.1 | 906 | A530058N18Rik | ENSMUSG00000087694.3 |
| chr2 | 114018418 | 114018493 | 3.8249 | 4.88169 | 1.69235 | Exon (ENSMUST00000147411.1/ENSMUSG00000087694.3, exon 2 of 2) | 2 | 114013745 | 114031451 | 17707 | 1 | ENSMUST00000134129.1 | 4674 | A530058N18Rik | ENSMUSG00000087694.3 |
| chr9 | 20793702 | 20794349 | 3.72496 | 4.58502 | 1.49442 | Exon (ENSMUST00000004201.7/ENSMUSG00000004098.7, exon 28 of 67) | 9 | 20807034 | 20807354 | 321 | 1 | ENSMUST00000157103.1 | -12685 | Gm26274 | ENSMUSG00000087728.1 |
| chr14 | 48888725 | 48889206 | 3.39124 | 3.98723 | 1.17969 | Distal Intergenic | 14 | 49001717 | 49001836 | 120 | 2 | ENSMUST00000157129.1 | 112630 | Gm23470 | ENSMUSG00000087754.1 |
| chr14 | 48985724 | 48985832 | 3.38843 | 3.92394 | 1.06525 | Distal Intergenic | 14 | 49001717 | 49001836 | 120 | 2 | ENSMUST00000157129.1 | 16004 | Gm23470 | ENSMUSG00000087754.1 |
| chr14 | 48990324 | 48990960 | 3.42023 | 3.98467 | 1.10741 | Distal Intergenic | 14 | 49001717 | 49001836 | 120 | 2 | ENSMUST00000157129.1 | 10876 | Gm23470 | ENSMUSG00000087754.1 |
| chr14 | 48565653 | 48565728 | 3.41963 | 4.00603 | 1.18545 | Distal Intergenic | 14 | 48652499 | 48652602 | 104 | 1 | ENSMUST00000157131.1 | -86771 | Gm23469 | ENSMUSG00000087756.1 |
| chr14 | 48566241 | 48566437 | 3.37857 | 3.92156 | 1.10939 | Distal Intergenic | 14 | 48652499 | 48652602 | 104 | 1 | ENSMUST00000157131.1 | -86062 | Gm23469 | ENSMUSG00000087756.1 |

| | | | | | | | | | | | | | | | |
|---|---|---|---|---|---|---|---|---|---|---|---|---|---|---|---|
| chr14 | 48566760 | 48566893 | 3.88494 | 5.1183 | 1.89588 | Distal Intergenic | 14 | 48652499 | 48652602 | 104 | 1 | ENSMUST00000157131.1 | -85606 | Gm23469 | ENSMUSG00000087756.1 |
| chr14 | 48580936 | 48581069 | 3.3938 | 3.94141 | 1.09823 | Distal Intergenic | 14 | 48652499 | 48652602 | 104 | 1 | ENSMUST00000157131.1 | -71430 | Gm23469 | ENSMUSG00000087756.1 |
| chr14 | 48649952 | 48650223 | 3.51539 | 4.15344 | 1.19935 | Downstream (2-3kb) | 14 | 48652499 | 48652602 | 104 | 1 | ENSMUST00000157131.1 | -2276 | Gm23469 | ENSMUSG00000087756.1 |
| chr14 | 48650767 | 48651132 | 3.81273 | 4.90135 | 1.721 | Downstream (1-2kb) | 14 | 48652499 | 48652602 | 104 | 1 | ENSMUST00000157131.1 | -1367 | Gm23469 | ENSMUSG00000087756.1 |
| chr14 | 48651515 | 48651728 | 4.05038 | 5.40168 | 2.03508 | Promoter | 14 | 48652499 | 48652602 | 104 | 1 | ENSMUST00000157131.1 | -771 | Gm23469 | ENSMUSG00000087756.1 |
| chr14 | 48652068 | 48653871 | 3.8625 | 4.92291 | 1.76491 | Promoter | 14 | 48652499 | 48652602 | 104 | 1 | ENSMUST00000157131.1 | 0 | Gm23469 | ENSMUSG00000087756.1 |
| chr14 | 48654276 | 48657012 | 4.06967 | 5.2539 | 1.94856 | Downstream (<1kb) | 14 | 48652499 | 48652602 | 104 | 1 | ENSMUST00000157131.1 | 1778 | Gm23469 | ENSMUSG00000087756.1 |
| chr8 | 50913888 | 50914018 | 3.46882 | 4.07547 | 1.169 | Distal Intergenic | 8 | 50563241 | 50563303 | 63 | 1 | ENSMUST00000157144.1 | 350648 | Gm23986 | ENSMUSG00000087769.1 |
| chr8 | 50914768 | 50916060 | 3.57886 | 4.27276 | 1.27015 | Distal Intergenic | 8 | 50563241 | 50563303 | 63 | 1 | ENSMUST00000157144.1 | 351528 | Gm23986 | ENSMUSG00000087769.1 |
| chr8 | 50917205 | 50917283 | 4.2678 | 5.68133 | 2.26186 | Distal Intergenic | 8 | 50563241 | 50563303 | 63 | 1 | ENSMUST00000157144.1 | 353965 | Gm23986 | ENSMUSG00000087769.1 |
| chr2 | 165253690 | 165254057 | 3.77207 | 4.80491 | 1.65335 | Distal Intergenic | 2 | 165270046 | 165270176 | 131 | 2 | ENSMUST00000157175.1 | 16119 | Gm25569 | ENSMUSG00000087800.1 |
| chr2 | 165256506 | 165256800 | 3.65762 | 4.52301 | 1.48974 | Distal Intergenic | 2 | 165270046 | 165270176 | 131 | 2 | ENSMUST00000157175.1 | 13376 | Gm25569 | ENSMUSG00000087800.1 |
| chr2 | 165262261 | 165262463 | 3.28089 | 3.74851 | 1.01733 | Distal Intergenic | 2 | 165270046 | 165270176 | 131 | 2 | ENSMUST00000157175.1 | 7713 | Gm25569 | ENSMUSG00000087800.1 |
| chr1 | 135340144 | 135340252 | 3.2426 | 3.69105 | 1.01733 | Intron (ENSMUST00000059352.2/ENSMUSG00000048096.7, intron 1 of 2) | 1 | 135353717 | 135353822 | 106 | 2 | ENSMUST00000157234.1 | 13570 | Gm22796 | ENSMUSG00000087859.1 |
| chr7 | 136235213 | 136235415 | 3.71253 | 4.53876 | 1.43617 | Distal Intergenic | 7 | 136509026 | 136509333 | 308 | 2 | ENSMUST00000157244.1 | 273918 | Gm24444 | ENSMUSG00000087869.1 |
| chr7 | 136404950 | 136405147 | 3.57108 | 4.25244 | 1.24397 | Distal Intergenic | 7 | 136509026 | 136509333 | 308 | 2 | ENSMUST00000157244.1 | 104186 | Gm24444 | ENSMUSG00000087869.1 |
| chr7 | 136406337 | 136406578 | 4.23075 | 5.90385 | 2.39693 | Distal Intergenic | 7 | 136509026 | 136509333 | 308 | 2 | ENSMUST00000157244.1 | 102755 | Gm24444 | ENSMUSG00000087869.1 |
| chr7 | 136406885 | 136407134 | 3.5219 | 4.16127 | 1.19188 | Distal Intergenic | 7 | 136509026 | 136509333 | 308 | 2 | ENSMUST00000157244.1 | 102199 | Gm24444 | ENSMUSG00000087869.1 |
| chr7 | 136430540 | 136430923 | 3.85243 | 4.77922 | 1.58535 | Distal Intergenic | 7 | 136509026 | 136509333 | 308 | 2 | ENSMUST00000157244.1 | 78410 | Gm24444 | ENSMUSG00000087869.1 |
| chr7 | 136431472 | 136431678 | 3.48614 | 4.09754 | 1.15738 | Distal Intergenic | 7 | 136509026 | 136509333 | 308 | 2 | ENSMUST00000157244.1 | 77655 | Gm24444 | ENSMUSG00000087869.1 |
| chr7 | 136479589 | 136479833 | 3.4136 | 4.21358 | 1.27884 | Distal Intergenic | 7 | 136509026 | 136509333 | 308 | 2 | ENSMUST00000157244.1 | 29500 | Gm24444 | ENSMUSG00000087869.1 |
| chr7 | 136489663 | 136489979 | 3.63864 | 4.50763 | 1.47707 | Distal Intergenic | 7 | 136509026 | 136509333 | 308 | 2 | ENSMUST00000157244.1 | 19354 | Gm24444 | ENSMUSG00000087869.1 |
| chr7 | 136491254 | 136491510 | 3.59772 | 4.41554 | 1.41434 | Distal Intergenic | 7 | 136509026 | 136509333 | 308 | 2 | ENSMUST00000157244.1 | 17823 | Gm24444 | ENSMUSG00000087869.1 |
| chr7 | 136495549 | 136495798 | 3.59118 | 4.2933 | 1.27931 | Distal Intergenic | 7 | 136509026 | 136509333 | 308 | 2 | ENSMUST00000157244.1 | 13535 | Gm24444 | ENSMUSG00000087869.1 |
| chr7 | 136496513 | 136496651 | 4.41895 | 5.90685 | 2.39475 | Distal Intergenic | 7 | 136509026 | 136509333 | 308 | 2 | ENSMUST00000157244.1 | 12682 | Gm24444 | ENSMUSG00000087869.1 |
| chr7 | 136497285 | 136497455 | 3.34153 | 3.83966 | 1.01733 | Distal Intergenic | 7 | 136509026 | 136509333 | 308 | 2 | ENSMUST00000157244.1 | 11878 | Gm24444 | ENSMUSG00000087869.1 |
| chr7 | 136501355 | 136501583 | 3.82952 | 4.73641 | 1.56067 | Distal Intergenic | 7 | 136509026 | 136509333 | 308 | 2 | ENSMUST00000157244.1 | 7750 | Gm24444 | ENSMUSG00000087869.1 |
| chr7 | 136502081 | 136503012 | 3.54984 | 4.24479 | 1.26363 | Distal Intergenic | 7 | 136509026 | 136509333 | 308 | 2 | ENSMUST00000157244.1 | 6321 | Gm24444 | ENSMUSG00000087869.1 |
| chr7 | 136582686 | 136583093 | 3.58132 | 4.31461 | 1.37833 | Distal Intergenic | 7 | 136509026 | 136509333 | 308 | 2 | ENSMUST00000157244.1 | -73354 | Gm24444 | ENSMUSG00000087869.1 |
| chr7 | 136676355 | 136676706 | 3.65664 | 4.464 | 1.48439 | Distal Intergenic | 7 | 136509026 | 136509333 | 308 | 2 | ENSMUST00000157244.1 | -167023 | Gm24444 | ENSMUSG00000087869.1 |
| chr7 | 136677989 | 136678314 | 3.70826 | 4.59547 | 1.50194 | Distal Intergenic | 7 | 136509026 | 136509333 | 308 | 2 | ENSMUST00000157244.1 | -168657 | Gm24444 | ENSMUSG00000087869.1 |
| chr3 | 17809588 | 17810055 | 3.36451 | 3.87432 | 1.01733 | Distal Intergenic | 3 | 17803621 | 17803747 | 127 | 2 | ENSMUST00000157268.1 | -5842 | Gm23441 | ENSMUSG00000087893.1 |
| chr4 | 138826394 | 138826820 | 3.34172 | 3.84643 | 1.04005 | Promoter | 4 | 138827376 | 138827476 | 101 | 2 | ENSMUST00000157269.1 | 656 | Gm25280 | ENSMUSG00000087894.1 |
| chr10 | 85349141 | 85349385 | 3.47123 | 4.18766 | 1.31065 | Distal Intergenic | 10 | 85333901 | 85334004 | 104 | 1 | ENSMUST00000157270.1 | 15241 | Gm25281 | ENSMUSG00000087895.1 |
| chr6 | 47654430 | 47654853 | 3.98085 | 8.14035 | 4.26334 | Distal Intergenic | 6 | 47637143 | 47637219 | 77 | 1 | ENSMUST00000157272.1 | 17288 | Gm25283 | ENSMUSG00000087897.1 |
| chr6 | 47655733 | 47655834 | 3.01657 | 7.92314 | 4.05735 | Distal Intergenic | 6 | 47637143 | 47637219 | 77 | 1 | ENSMUST00000157272.1 | 18591 | Gm25283 | ENSMUSG00000087897.1 |
| chr6 | 47669024 | 47669161 | 4.25486 | 6.94726 | 3.28169 | Distal Intergenic | 6 | 47637143 | 47637219 | 77 | 1 | ENSMUST00000157272.1 | 31882 | Gm25283 | ENSMUSG00000087897.1 |
| chr6 | 47672994 | 47673127 | 2.51832 | 7.65824 | 3.7378 | Distal Intergenic | 6 | 47637143 | 47637219 | 77 | 1 | ENSMUST00000157272.1 | 35852 | Gm25283 | ENSMUSG00000087897.1 |
| chr4 | 132946060 | 132946349 | 3.49681 | 4.11038 | 1.14547 | Distal Intergenic | 4 | 132931211 | 132931317 | 107 | 2 | ENSMUST00000157286.1 | -14744 | Gm24913 | ENSMUSG00000087911.1 |
| chr13 | 46570770 | 46571144 | 3.49935 | 4.12432 | 1.18212 | Intron (ENSMUST00000021802.9/ENSMUSG00000021373.10, intron 4 of 12) | 13 | 46518684 | 46518761 | 78 | 1 | ENSMUST00000157288.1 | 52087 | Gm24915 | ENSMUSG00000087913.1 |
| chr14 | 20230548 | 20230623 | 4.47196 | 6.08115 | 2.55259 | Distal Intergenic | 14 | 20199996 | 20200112 | 117 | 2 | ENSMUST00000157291.1 | -30437 | Gm24912 | ENSMUSG00000087916.1 |
| chr6 | 100433046 | 100433558 | 3.29711 | 4.08817 | 1.1688 | Distal Intergenic | 6 | 100494681 | 100494814 | 134 | 2 | ENSMUST00000157304.1 | 61256 | Gm23234 | ENSMUSG00000087929.1 |
| chr6 | 100438124 | 100438293 | 3.81944 | 4.72638 | 1.57119 | Distal Intergenic | 6 | 100494681 | 100494814 | 134 | 2 | ENSMUST00000157304.1 | 56521 | Gm23234 | ENSMUSG00000087929.1 |
| chr6 | 100438631 | 100438725 | 4.03342 | 5.12788 | 1.82361 | Distal Intergenic | 6 | 100494681 | 100494814 | 134 | 2 | ENSMUST00000157304.1 | 56089 | Gm23234 | ENSMUSG00000087929.1 |
| chr6 | 100498723 | 100498798 | 3.97946 | 5.02742 | 1.7668 | Distal Intergenic | 6 | 100494681 | 100494814 | 134 | 2 | ENSMUST00000157304.1 | -3910 | Gm23234 | ENSMUSG00000087929.1 |
| chr3 | 53572071 | 53572335 | 3.75182 | 4.66889 | 1.58402 | Exon (ENSMUST00000091137.4/ENSMUSG00000037016.7, exon 6 of 24) | 3 | 53512002 | 53512126 | 125 | 1 | ENSMUST00000157311.1 | 60070 | Gm23675 | ENSMUSG00000087936.1 |
| chr6 | 99887837 | 99888260 | 3.54902 | 4.22497 | 1.2656 | Distal Intergenic | 6 | 99893226 | 99893547 | 322 | 2 | ENSMUST00000157321.1 | 5287 | Gm24248 | ENSMUSG00000087946.1 |
| chr6 | 99892188 | 99892288 | 3.46993 | 4.08895 | 1.21508 | Downstream (<1kb) | 6 | 99893226 | 99893547 | 322 | 2 | ENSMUST00000157321.1 | 1259 | Gm24248 | ENSMUSG00000087946.1 |
| chr6 | 99909482 | 99909811 | 3.63026 | 4.42868 | 1.43563 | Distal Intergenic | 6 | 99893226 | 99893547 | 322 | 2 | ENSMUST00000157321.1 | -15936 | Gm24248 | ENSMUSG00000087946.1 |
| chr7 | 88128300 | 88128458 | 3.37789 | 3.89451 | 1.01733 | Intron (ENSMUST00000107263.3/ENSMUSG00000049583.9, intron 7 of 7) | 7 | 88087188 | 88087251 | 64 | 2 | ENSMUST00000157322.1 | -41050 | Gm24249 | ENSMUSG00000087947.1 |
| chr7 | 88130118 | 88131136 | 4.00546 | 5.21836 | 1.93758 | 3' UTR | 7 | 88087188 | 88087251 | 64 | 2 | ENSMUST00000157322.1 | -42868 | Gm24249 | ENSMUSG00000087947.1 |
| chr5 | 64149560 | 64149855 | 3.39322 | 4.06317 | 1.15765 | Distal Intergenic | 5 | 64144248 | 64144381 | 134 | 1 | ENSMUST00000157329.1 | 5313 | Gm22559 | ENSMUSG00000087954.1 |
| chr2 | 57698072 | 57698365 | 3.94249 | 5.03775 | 1.84798 | Distal Intergenic | 2 | 57653672 | 57653822 | 151 | 2 | ENSMUST00000157339.1 | -44251 | Gm25388 | ENSMUSG00000087964.1 |
| chr8 | 26308135 | 26308951 | 3.57441 | 4.31519 | 1.37982 | Distal Intergenic | 8 | 26240150 | 26240408 | 259 | 2 | ENSMUST00000157356.1 | -67728 | Gm22227 | ENSMUSG00000087981.1 |
| chr8 | 26354724 | 26355008 | 3.35721 | 3.91275 | 1.15541 | Distal Intergenic | 8 | 26240150 | 26240408 | 259 | 2 | ENSMUST00000157356.1 | -114317 | Gm22227 | ENSMUSG00000087981.1 |
| chr8 | 26381718 | 26382113 | 3.58839 | 4.30161 | 1.3251 | Distal Intergenic | 8 | 26240150 | 26240408 | 259 | 2 | ENSMUST00000157356.1 | -141311 | Gm22227 | ENSMUSG00000087981.1 |
| chr8 | 26382469 | 26382561 | 3.41349 | 4.18692 | 1.25258 | Distal Intergenic | 8 | 26240150 | 26240408 | 259 | 2 | ENSMUST00000157356.1 | -142062 | Gm22227 | ENSMUSG00000087981.1 |
| chr10 | 80834579 | 80834927 | 3.49316 | 4.1862 | 1.24568 | Exon (ENSMUST00000053986.7/ENSMUSG00000051067.7, exon 2 of 2) | 10 | 80828797 | 80828870 | 74 | 1 | ENSMUST00000157368.1 | 5783 | Mir1982 | ENSMUSG00000087993.1 |
| chr10 | 80835301 | 80835571 | 3.67968 | 4.71011 | 1.5629 | Exon (ENSMUST00000053986.7/ENSMUSG00000051067.7, exon 2 of 2) | 10 | 80828797 | 80828870 | 74 | 1 | ENSMUST00000157368.1 | 6505 | Mir1982 | ENSMUSG00000087993.1 |
| chr6 | 83716552 | 83719445 | 4.05099 | 5.22631 | 1.92893 | Intron (ENSMUST00000037807.2/ENSMUSG00000034777.2, intron 1 of 2) | 6 | 83721296 | 83721392 | 97 | 1 | ENSMUST00000157370.1 | -1851 | Gm25054 | ENSMUSG00000087995.1 |
| chr6 | 83719831 | 83721110 | 4.03459 | 5.31786 | 2.03007 | Promoter | 6 | 83721296 | 83721392 | 97 | 1 | ENSMUST00000157370.1 | -186 | Gm25054 | ENSMUSG00000087995.1 |
| chr6 | 83721463 | 83722609 | 3.79389 | 4.67789 | 1.53658 | Promoter | 6 | 83721296 | 83721392 | 97 | 1 | ENSMUST00000157370.1 | 168 | Gm25054 | ENSMUSG00000087995.1 |
| chr6 | 83722950 | 83723050 | 3.43416 | 4.00111 | 1.09361 | Intron (ENSMUST00000037807.2/ENSMUSG00000034777.2, intron 1 of 2) | 6 | 83721296 | 83721392 | 97 | 1 | ENSMUST00000157370.1 | 1655 | Gm25054 | ENSMUSG00000087995.1 |
| chr6 | 83723490 | 83724018 | 4.0989 | 5.3944 | 2.05533 | Intron (ENSMUST00000037807.2/ENSMUSG00000034777.2, intron 1 of 2) | 6 | 83721296 | 83721392 | 97 | 1 | ENSMUST00000157370.1 | 2195 | Gm25054 | ENSMUSG00000087995.1 |
| chr6 | 83724575 | 83725751 | 3.78439 | 4.67282 | 1.55941 | Intron (ENSMUST00000037807.2/ENSMUSG00000034777.2, intron 1 of 2) | 6 | 83721296 | 83721392 | 97 | 1 | ENSMUST00000157370.1 | 3280 | Gm25054 | ENSMUSG00000087995.1 |
| chr6 | 83727148 | 83727286 | 3.73854 | 4.80708 | 1.64445 | Intron (ENSMUST00000037807.2/ENSMUSG00000034777.2, intron 1 of 2) | 6 | 83721296 | 83721392 | 97 | 1 | ENSMUST00000157370.1 | 5853 | Gm25054 | ENSMUSG00000087995.1 |
| chr6 | 83727639 | 83727779 | 3.29823 | 3.77451 | 1.01733 | Intron (ENSMUST00000037807.2/ENSMUSG00000034777.2, intron 1 of 2) | 6 | 83721296 | 83721392 | 97 | 1 | ENSMUST00000157370.1 | 6344 | Gm25054 | ENSMUSG00000087995.1 |
| chr6 | 83730524 | 83730753 | 3.43873 | 4.07923 | 1.2224 | Intron (ENSMUST00000037807.2/ENSMUSG00000034777.2, intron 1 of 2) | 6 | 83721296 | 83721392 | 97 | 1 | ENSMUST00000157370.1 | 9229 | Gm25054 | ENSMUSG00000087995.1 |
| chr4 | 131193882 | 131194405 | 4.26452 | 5.62041 | 2.2095 | Distal Intergenic | 4 | 131239277 | 131239410 | 134 | 1 | ENSMUST00000157416.1 | -44872 | Gm25261 | ENSMUSG00000088041.1 |
| chr4 | 131253011 | 131253158 | 3.67904 | 4.87499 | 1.69145 | Distal Intergenic | 4 | 131239277 | 131239410 | 134 | 1 | ENSMUST00000157416.1 | 13735 | Gm25261 | ENSMUSG00000088041.1 |
| chr9 | 43869827 | 43870076 | 3.36522 | 3.87539 | 1.01733 | Distal Intergenic | 9 | 43905898 | 43906002 | 105 | 2 | ENSMUST00000157419.1 | 35926 | Gm23326 | ENSMUSG00000088044.1 |
| chr9 | 43876402 | 43877743 | 3.49087 | 4.1309 | 1.21953 | Distal Intergenic | 9 | 43905898 | 43906002 | 105 | 2 | ENSMUST00000157419.1 | 28259 | Gm23326 | ENSMUSG00000088044.1 |
| chr9 | 43878708 | 43878822 | 4.32628 | 5.90644 | 2.40516 | Distal Intergenic | 9 | 43905898 | 43906002 | 105 | 2 | ENSMUST00000157419.1 | 27180 | Gm23326 | ENSMUSG00000088044.1 |
| chr9 | 43879268 | 43881064 | 3.88262 | 4.95397 | 1.74017 | Distal Intergenic | 9 | 43905898 | 43906002 | 105 | 2 | ENSMUST00000157419.1 | 24938 | Gm23326 | ENSMUSG00000088044.1 |
| chr9 | 43887211 | 43887757 | 3.45434 | 4.04265 | 1.1312 | Distal Intergenic | 9 | 43905898 | 43906002 | 105 | 2 | ENSMUST00000157419.1 | 18245 | Gm23326 | ENSMUSG00000088044.1 |
| chr9 | 43917406 | 43918097 | 3.61749 | 4.38642 | 1.38822 | Distal Intergenic | 9 | 43905898 | 43906002 | 105 | 2 | ENSMUST00000157419.1 | -11405 | Gm23326 | ENSMUSG00000088044.1 |
| chr9 | 43919033 | 43919156 | 3.25557 | 3.85477 | 1.02216 | Distal Intergenic | 9 | 43905898 | 43906002 | 105 | 2 | ENSMUST00000157419.1 | -13032 | Gm23326 | ENSMUSG00000088044.1 |
| chr9 | 43919828 | 43920874 | 3.49441 | 4.12889 | 1.21775 | Distal Intergenic | 9 | 43905898 | 43906002 | 105 | 2 | ENSMUST00000157419.1 | -13827 | Gm23326 | ENSMUSG00000088044.1 |
| chr9 | 43921912 | 43922274 | 3.66096 | 4.4356 | 1.41126 | Distal Intergenic | 9 | 43905898 | 43906002 | 105 | 2 | ENSMUST00000157419.1 | -15911 | Gm23326 | ENSMUSG00000088044.1 |
| chr9 | 43927348 | 43927489 | 3.63616 | 4.36709 | 1.30948 | Distal Intergenic | 9 | 43905898 | 43906002 | 105 | 2 | ENSMUST00000157419.1 | -21347 | Gm23326 | ENSMUSG00000088044.1 |
| chr9 | 43929541 | 43929666 | 3.39573 | 3.93031 | 1.04874 | Distal Intergenic | 9 | 43905898 | 43906002 | 105 | 2 | ENSMUST00000157419.1 | -23540 | Gm23326 | ENSMUSG00000088044.1 |
| chr9 | 43935518 | 43935610 | 3.53639 | 4.20959 | 1.27816 | Distal Intergenic | 9 | 43905898 | 43906002 | 105 | 2 | ENSMUST00000157419.1 | -29517 | Gm23326 | ENSMUSG00000088044.1 |
| chr9 | 43939470 | 43939643 | 3.66671 | 4.43308 | 1.3705 | Distal Intergenic | 9 | 43905898 | 43906002 | 105 | 2 | ENSMUST00000157419.1 | -33469 | Gm23326 | ENSMUSG00000088044.1 |
| chr9 | 43957533 | 43957761 | 3.33612 | 3.83154 | 1.01733 | Distal Intergenic | 9 | 43905898 | 43906002 | 105 | 2 | ENSMUST00000157419.1 | -51532 | Gm23326 | ENSMUSG00000088044.1 |
| chr7 | 81626517 | 81626662 | 3.68487 | 4.50515 | 1.4946 | Intron (ENSMUST00000098326.2/ENSMUSG00000025813.8, intron 3 of 8) | 7 | 81669728 | 81669835 | 108 | 1 | ENSMUST00000157432.1 | -43066 | Gm26149 | ENSMUSG00000088057.1 |
| chr2 | 166758128 | 166758403 | 3.43934 | 4.01464 | 1.11441 | Distal Intergenic | 2 | 166768745 | 166768847 | 103 | 1 | ENSMUST00000157438.1 | -10342 | Gm24477 | ENSMUSG00000088063.1 |
| chr18 | 84798541 | 84798829 | 3.9935 | 5.0992 | 1.86818 | Distal Intergenic | 18 | 84822002 | 84822304 | 303 | 1 | ENSMUST00000157445.1 | -23173 | Gm22819 | ENSMUSG00000088070.1 |
| chr2 | 106797169 | 106797363 | 3.61259 | 4.3238 | 1.27991 | Intron (ENSMUST00000111063.2/ENSMUSG00000016386.10, intron 4 of 6) | 2 | 106809336 | 106809458 | 123 | 1 | ENSMUST00000157453.2 | -11973 | Gm22813 | ENSMUSG00000088078.2 |
| chr2 | 106855095 | 106855470 | 3.47237 | 4.09947 | 1.23129 | Intron (ENSMUST00000111063.2/ENSMUSG00000016386.10, intron 4 of 6) | 2 | 106809336 | 106809458 | 123 | 1 | ENSMUST00000157453.2 | 45760 | Gm22813 | ENSMUSG00000088078.2 |
| chr2 | 106856152 | 106856231 | 3.38731 | 3.93658 | 1.11621 | Intron (ENSMUST00000111063.2/ENSMUSG00000016386.10, intron 4 of 6) | 2 | 106809336 | 106809458 | 123 | 1 | ENSMUST00000157453.2 | 46817 | Gm22813 | ENSMUSG00000088078.2 |
| chr14 | 118322881 | 118323816 | 3.53761 | 4.18992 | 1.20925 | Distal Intergenic | 14 | 118299990 | 118300117 | 128 | 1 | ENSMUST00000157470.1 | 22892 | Gm23669 | ENSMUSG00000088095.1 |
| chr11 | 109494326 | 109494643 | 3.56069 | 4.23428 | 1.24385 | Intron (ENSMUST00000020928.7/ENSMUSG00000020604.8, intron 4 of 13) | 11 | 109499714 | 109499824 | 111 | 2 | ENSMUST00000157481.1 | 5181 | Gm22378 | ENSMUSG00000088106.1 |
| chr11 | 109496141 | 109496307 | 3.36532 | 4.0014 | 1.12915 | Intron (ENSMUST00000020928.7/ENSMUSG00000020604.8, intron 4 of 13) | 11 | 109499714 | 109499824 | 111 | 2 | ENSMUST00000157481.1 | 3517 | Gm22378 | ENSMUSG00000088106.1 |
| chr11 | 35262681 | 35262997 | 3.60446 | 4.37289 | 1.43361 | Intron (ENSMUST00000069837.3/ENSMUSG00000056427.5, intron 4 of 35) | 11 | 35214401 | 35214537 | 137 | 1 | ENSMUST00000157500.1 | 48281 | Gm26070 | ENSMUSG00000088125.1 |
| chr11 | 35204380 | 35204919 | 3.39649 | 3.97223 | 1.18017 | Intron (ENSMUST00000069837.3/ENSMUSG00000056427.5, intron 3 of 35) | 11 | 35214401 | 35214537 | 137 | 1 | ENSMUST00000157500.1 | -9482 | Gm26070 | ENSMUSG00000088125.1 |
| chr18 | 30472726 | 30472852 | 3.81373 | 4.81763 | 1.68273 | Distal Intergenic | 18 | 30476407 | 30476518 | 112 | 1 | ENSMUST00000157501.1 | -3555 | Gm22870 | ENSMUSG00000088126.1 |
| chr18 | 30473238 | 30473367 | 3.19975 | 3.91644 | 1.07608 | Distal Intergenic | 18 | 30476407 | 30476518 | 112 | 1 | ENSMUST00000157501.1 | -3040 | Gm22870 | ENSMUSG00000088126.1 |
| chr18 | 30479641 | 30479967 | 3.58316 | 4.3979 | 1.39064 | Distal Intergenic | 18 | 30476407 | 30476518 | 112 | 1 | ENSMUST00000157501.1 | 3235 | Gm22870 | ENSMUSG00000088126.1 |
| chr18 | 30482551 | 30483197 | 3.6832 | 4.50205 | 1.47397 | Distal Intergenic | 18 | 30476407 | 30476518 | 112 | 1 | ENSMUST00000157501.1 | 6145 | Gm22870 | ENSMUSG00000088126.1 |
| chr18 | 30484539 | 30486438 | 3.63676 | 4.53726 | 1.50509 | Distal Intergenic | 18 | 30476407 | 30476518 | 112 | 1 | ENSMUST00000157501.1 | 8133 | Gm22870 | ENSMUSG00000088126.1 |
| chr18 | 83282532 | 83283178 | 3.3518 | 3.85666 | 1.02267 | Distal Intergenic | 18 | 83355199 | 83355364 | 166 | 2 | ENSMUST00000157518.1 | 72186 | Gm24720 | ENSMUSG00000088143.1 |
| chr18 | 83285737 | 83286071 | 3.57645 | 4.45039 | 1.38194 | Distal Intergenic | 18 | 83355199 | 83355364 | 166 | 2 | ENSMUST00000157518.1 | 69293 | Gm24720 | ENSMUSG00000088143.1 |
| chr18 | 83335262 | 83335464 | 3.53154 | 4.51008 | 1.46353 | Distal Intergenic | 18 | 83355199 | 83355364 | 166 | 2 | ENSMUST00000157518.1 | 19900 | Gm24720 | ENSMUSG00000088143.1 |
| chr18 | 83635116 | 83635278 | 3.32627 | 3.85909 | 1.1101 | Distal Intergenic | 18 | 83355199 | 83355364 | 166 | 2 | ENSMUST00000157518.1 | -279753 | Gm24720 | ENSMUSG00000088143.1 |
| chr13 | 21897360 | 21897456 | 3.72492 | 4.56901 | 1.5175 | Promoter | 13 | 21896918 | 21897049 | 132 | 2 | ENSMUST00000157523.1 | -312 | Mir1983 | ENSMUSG00000088148.1 |
| chr18 | 69197834 | 69197952 | 3.25229 | 3.70558 | 1.01733 | Distal Intergenic | 18 | 69173987 | 69174120 | 134 | 2 | ENSMUST00000157541.1 | -23715 | Gm23536 | ENSMUSG00000088166.1 |
| chr18 | 69198275 | 69198350 | 3.82459 | 4.78617 | 1.68545 | Distal Intergenic | 18 | 69173987 | 69174120 | 134 | 2 | ENSMUST00000157541.1 | -24156 | Gm23536 | ENSMUSG00000088166.1 |
| chr18 | 69199495 | 69199736 | 3.79946 | 4.73929 | 1.64907 | Distal Intergenic | 18 | 69173987 | 69174120 | 134 | 2 | ENSMUST00000157541.1 | -25376 | Gm23536 | ENSMUSG00000088166.1 |
| chr14 | 56865061 | 56865353 | 4.44113 | 6.19162 | 2.67372 | Distal Intergenic | 14 | 56852830 | 56852888 | 59 | 2 | ENSMUST00000157555.1 | -12174 | Gm22218 | ENSMUSG00000088180.1 |
| chr14 | 56865802 | 56866254 | 3.71567 | 4.61041 | 1.55748 | Distal Intergenic | 14 | 56852830 | 56852888 | 59 | 2 | ENSMUST00000157555.1 | -12915 | Gm22218 | ENSMUSG00000088180.1 |
| chr8 | 35765811 | 35765965 | 3.62907 | 4.56068 | 1.46918 | Distal Intergenic | 8 | 35747449 | 35747562 | 114 | 1 | ENSMUST00000157557.1 | 18363 | Gm22216 | ENSMUSG00000088182.1 |
| chr17 | 79979729 | 79979914 | 3.45372 | 4.04023 | 1.1278 | Distal Intergenic | 17 | 80033514 | 80033609 | 96 | 1 | ENSMUST00000157558.1 | -53600 | Gm22215 | ENSMUSG00000088183.1 |
| chr19 | 41744429 | 41744795 | 3.349 | 3.85586 | 1.03474 | Promoter | 19 | 41744804 | 41745053 | 250 | 1 | ENSMUST00000157595.1 | -9 | Gm22601 | ENSMUSG00000088220.1 |
| chr1 | 167340302 | 167340384 | 5.58253 | 18.89637 | 13.94903 | Promoter | 1 | 167340369 | 167340424 | 56 | 2 | ENSMUST00000157621.1 | 40 | Gm25911 | ENSMUSG00000088246.1 |
| chr6 | 103649072 | 103649274 | 1.93267 | 11.96109 | 7.47325 | Intron (ENSMUST00000066905.6/ENSMUSG00000030077.9, intron 4 of 26) | 6 | 103588704 | 103588765 | 62 | 1 | ENSMUST00000157640.1 | 60369 | Gm24784 | ENSMUSG00000088265.1 |
| chr11 | 107816990 | 107817158 | 3.51131 | 4.15927 | 1.23392 | Distal Intergenic | 11 | 107827374 | 107827660 | 287 | 1 | ENSMUST00000157654.1 | -10216 | Gm23120 | ENSMUSG00000088279.1 |
| chr15 | 102370880 | 102371076 | 3.37651 | 4.32281 | 1.32596 | Distal Intergenic | 15 | 102373880 | 102373983 | 104 | 1 | ENSMUST00000157661.1 | -2804 | Gm23590 | ENSMUSG00000088286.1 |
| chr9 | 102544965 | 102546153 | 3.72299 | 4.87176 | 1.68557 | 3' UTR | 9 | 102547766 | 102547877 | 112 | 1 | ENSMUST00000157699.1 | -1613 | Gm25669 | ENSMUSG00000088324.1 |
| chr9 | 102546584 | 102547110 | 3.49941 | 4.17506 | 1.31477 | Promoter | 9 | 102547766 | 102547877 | 112 | 1 | ENSMUST00000157699.1 | -656 | Gm25669 | ENSMUSG00000088324.1 |
| chr5 | 30528503 | 30528610 | 3.69003 | 4.62125 | 1.50218 | Intron (ENSMUST00000065486.2/ENSMUSG00000053194.2, intron 3 of 6) | 5 | 30514578 | 30514684 | 107 | 2 | ENSMUST00000157702.1 | -13820 | Gm25666 | ENSMUSG00000088327.1 |
| chr8 | 94697832 | 94698236 | 3.65715 | 4.62351 | 1.54741 | Promoter | 8 | 94698016 | 94698285 | 270 | 2 | ENSMUST00000157706.1 | 49 | Gm22846 | ENSMUSG00000088331.1 |
| chr12 | 9904737 | 9904925 | 3.64571 | 4.39573 | 1.36142 | Distal Intergenic | 12 | 9865347 | 9865450 | 104 | 2 | ENSMUST00000157712.1 | -39288 | Gm22845 | ENSMUSG00000088337.1 |
| chr12 | 9918431 | 9919082 | 3.47522 | 4.09724 | 1.21624 | Distal Intergenic | 12 | 9865347 | 9865450 | 104 | 2 | ENSMUST00000157712.1 | -52982 | Gm22845 | ENSMUSG00000088337.1 |
| chr7 | 24335237 | 24335418 | 3.57022 | 4.36657 | 1.3834 | Downstream (<1kb) | 7 | 24336340 | 24336465 | 126 | 2 | ENSMUST00000157729.2 | 1047 | Gm27592 | ENSMUSG00000088354.2 |
| chr12 | 56297740 | 56297825 | 4.06386 | 5.19449 | 1.87579 | Distal Intergenic | 12 | 56266920 | 56267204 | 285 | 2 | ENSMUST00000157764.1 | -30537 | Gm24685 | ENSMUSG00000088389.1 |
| chr12 | 56298207 | 56298806 | 3.59112 | 4.339 | 1.30932 | Distal Intergenic | 12 | 56266920 | 56267204 | 285 | 2 | ENSMUST00000157764.1 | -31004 | Gm24685 | ENSMUSG00000088389.1 |
| chr8 | 120695655 | 120696258 | 3.35949 | 3.90807 | 1.11031 | Intron (ENSMUST00000127664.1/ENSMUSG00000092329.1, intron 1 of 15) | 8 | 120717500 | 120717754 | 255 | 2 | ENSMUST00000157791.1 | 21496 | Gm24239 | ENSMUSG00000088416.1 |
| chr8 | 114958893 | 114959039 | 3.27998 | 3.84705 | 1.10588 | Intron (ENSMUST00000004756.8/ENSMUSG00000004637.10, intron 8 of 8) | 8 | 115052814 | 115052913 | 100 | 2 | ENSMUST00000157803.1 | 93874 | Gm22556 | ENSMUSG00000088428.1 |
| chr8 | 115105626 | 115106186 | 3.54472 | 4.20233 | 1.21025 | Intron (ENSMUST00000004756.8/ENSMUSG00000004637.10, intron 8 of 8) | 8 | 115052814 | 115052913 | 100 | 2 | ENSMUST00000157803.1 | -52714 | Gm22556 | ENSMUSG00000088428.1 |
| chr8 | 115150552 | 115150837 | 3.42127 | 3.9815 | 1.09361 | Intron (ENSMUST00000004756.8/ENSMUSG00000004637.10, intron 8 of 8) | 8 | 115052814 | 115052913 | 100 | 2 | ENSMUST00000157803.1 | -97640 | Gm22556 | ENSMUSG00000088428.1 |
| chr8 | 115257333 | 115257578 | 3.37636 | 3.89219 | 1.01733 | Intron (ENSMUST00000004756.8/ENSMUSG00000004637.10, intron 8 of 8) | 8 | 115052814 | 115052913 | 100 | 2 | ENSMUST00000157803.1 | -204421 | Gm22556 | ENSMUSG00000088428.1 |
| chr13 | 77438941 | 77439050 | 2.10602 | 12.28118 | 7.81703 | Intron (ENSMUST00000168779.1/ENSMUSG00000071252.5, intron 19 of 19) | 13 | 77676821 | 77676951 | 131 | 1 | ENSMUST00000157896.1 | -237771 | Gm26088 | ENSMUSG00000088521.1 |
| chr3 | 130924026 | 130924274 | 3.62607 | 4.39113 | 1.38286 | Distal Intergenic | 3 | 130870949 | 130871076 | 128 | 2 | ENSMUST00000157897.1 | -52951 | Gm26090 | ENSMUSG00000088522.1 |
| chr3 | 130944264 | 130944566 | 3.31152 | 3.83875 | 1.09957 | Distal Intergenic | 3 | 130870949 | 130871076 | 128 | 2 | ENSMUST00000157897.1 | -73189 | Gm26090 | ENSMUSG00000088522.1 |
| chr19 | 47703270 | 47703448 | 3.38836 | 3.9157 | 1.0364 | Distal Intergenic | 19 | 47710866 | 47710981 | 116 | 1 | ENSMUST00000157900.1 | -7418 | Gm26085 | ENSMUSG00000088525.1 |
| chr9 | 60403852 | 60403931 | 4.30362 | 5.73961 | 2.33827 | Intron (ENSMUST00000171654.2/ENSMUSG00000032289.10, intron 3 of 16) | 9 | 60419863 | 60419985 | 123 | 1 | ENSMUST00000157925.1 | -15932 | Gm22099 | ENSMUSG00000088550.1 |
| chr9 | 60410408 | 60410718 | 3.6212 | 4.40979 | 1.42765 | Intron (ENSMUST00000171654.2/ENSMUSG00000032289.10, intron 3 of 16) | 9 | 60419863 | 60419985 | 123 | 1 | ENSMUST00000157925.1 | -9145 | Gm22099 | ENSMUSG00000088550.1 |
| chr13 | 34249269 | 34249399 | 3.65984 | 4.43169 | 1.40835 | Intron (ENSMUST00000040336.6/ENSMUSG00000038267.9, intron 3 of 10) | 13 | 34276965 | 34277252 | 288 | 2 | ENSMUST00000157948.1 | 27853 | Gm24530 | ENSMUSG00000088573.1 |
| chr5 | 122509272 | 122509662 | 3.65092 | 4.40295 | 1.34468 | Distal Intergenic | 5 | 122506789 | 122506889 | 101 | 1 | ENSMUST00000157958.1 | 2484 | Gm22965 | ENSMUSG00000088583.1 |
| chr4 | 136135080 | 136135734 | 3.54332 | 4.31262 | 1.36735 | Downstream (2-3kb) | 4 | 136137496 | 136137598 | 103 | 1 | ENSMUST00000157989.1 | -1762 | Gm26001 | ENSMUSG00000088614.1 |
| chr4 | 136136184 | 136136508 | 4.07387 | 5.25292 | 1.99039 | Promoter | 4 | 136137496 | 136137598 | 103 | 1 | ENSMUST00000157989.1 | -988 | Gm26001 | ENSMUSG00000088614.1 |
| chr6 | 93716594 | 93716923 | 3.54315 | 4.23328 | 1.28673 | Intron (ENSMUST00000093769.5/ENSMUSG00000045095.11, intron 12 of 22) | 6 | 93793772 | 93793877 | 106 | 1 | ENSMUST00000158002.1 | -76849 | Gm23035 | ENSMUSG00000088627.1 |
| chr6 | 93717798 | 93718199 | 3.44415 | 4.05785 | 1.19691 | Intron (ENSMUST00000093769.5/ENSMUSG00000045095.11, intron 11 of 22) | 6 | 93793772 | 93793877 | 106 | 1 | ENSMUST00000158002.1 | -75573 | Gm23035 | ENSMUSG00000088627.1 |
| chr14 | 28797778 | 28797955 | 3.4742 | 4.0702 | 1.12412 | Distal Intergenic | 14 | 29009696 | 29009799 | 104 | 2 | ENSMUST00000158057.1 | 211844 | Gm23999 | ENSMUSG00000088682.1 |
| chr5 | 37091342 | 37091549 | 3.33439 | 3.85967 | 1.11563 | Promoter | 5 | 37092258 | 37092360 | 103 | 2 | ENSMUST00000158082.1 | 811 | Gm25551 | ENSMUSG00000088707.1 |
| chr9 | 24952969 | 24953079 | 3.91282 | 4.90202 | 1.67831 | Distal Intergenic | 9 | 24964536 | 24964669 | 134 | 1 | ENSMUST00000158106.1 | -11457 | Gm23893 | ENSMUSG00000088731.1 |
| chr15 | 80452334 | 80452417 | 3.82109 | 4.77775 | 1.68543 | 3' UTR | 15 | 80458038 | 80458174 | 137 | 2 | ENSMUST00000158115.1 | 5757 | Gm23220 | ENSMUSG00000088740.1 |
| chr15 | 80459776 | 80460207 | 3.35339 | 3.8998 | 1.14691 | Intron (ENSMUST00000096347.3/ENSMUSG00000050439.6, intron 5 of 5) | 15 | 80458038 | 80458174 | 137 | 2 | ENSMUST00000158115.1 | -1603 | Gm23220 | ENSMUSG00000088740.1 |
| chr15 | 80461738 | 80462095 | 3.35289 | 3.87362 | 1.07666 | Intron (ENSMUST00000096347.3/ENSMUSG00000050439.6, intron 5 of 5) | 15 | 80458038 | 80458174 | 137 | 2 | ENSMUST00000158115.1 | -3565 | Gm23220 | ENSMUSG00000088740.1 |
| chr3 | 153859106 | 153859397 | 3.32562 | 3.81571 | 1.01733 | Intron (ENSMUST00000005630.5/ENSMUSG00000005493.10, intron 19 of 19) | 3 | 153864807 | 153864931 | 125 | 1 | ENSMUST00000158139.1 | -5410 | Gm24395 | ENSMUSG00000088764.1 |
| chr4 | 154136735 | 154137344 | 3.37016 | 4.02773 | 1.20277 | Intron (ENSMUST00000105644.3/ENSMUSG00000029026.11, intron 1 of 13) | 4 | 154134030 | 154134173 | 144 | 1 | ENSMUST00000158149.1 | 2706 | Gm26036 | ENSMUSG00000088774.1 |
| chr7 | 142134262 | 142134707 | 3.78035 | 4.84271 | 1.68876 | Distal Intergenic | 7 | 142153864 | 142153957 | 94 | 1 | ENSMUST00000158159.1 | -19157 | Gm25370 | ENSMUSG00000088784.1 |
| chr7 | 142144983 | 142145177 | 3.47299 | 4.08456 | 1.14979 | Distal Intergenic | 7 | 142153864 | 142153957 | 94 | 1 | ENSMUST00000158159.1 | -8687 | Gm25370 | ENSMUSG00000088784.1 |
| chr12 | 118658614 | 118658987 | 3.60421 | 4.37911 | 1.39833 | Distal Intergenic | 12 | 118753554 | 118753752 | 199 | 1 | ENSMUST00000158172.1 | -94567 | Gm22537 | ENSMUSG00000088797.1 |
| chr12 | 118659511 | 118659590 | 5.39347 | 8.30957 | 4.27338 | Distal Intergenic | 12 | 118753554 | 118753752 | 199 | 1 | ENSMUST00000158172.1 | -93964 | Gm22537 | ENSMUSG00000088797.1 |
| chr12 | 118660046 | 118660464 | 3.43634 | 4.05327 | 1.23232 | Distal Intergenic | 12 | 118753554 | 118753752 | 199 | 1 | ENSMUST00000158172.1 | -93090 | Gm22537 | ENSMUSG00000088797.1 |
| chr12 | 118661356 | 118662191 | 3.44781 | 4.19405 | 1.25626 | Distal Intergenic | 12 | 118753554 | 118753752 | 199 | 1 | ENSMUST00000158172.1 | -91363 | Gm22537 | ENSMUSG00000088797.1 |
| chr14 | 114855377 | 114855717 | 3.4992 | 4.33694 | 1.32654 | Distal Intergenic | 14 | 114980428 | 114980533 | 106 | 2 | ENSMUST00000158177.1 | 124816 | Gm24073 | ENSMUSG00000088802.1 |
| chr3 | 143597111 | 143597311 | 3.73217 | 4.61599 | 1.57036 | Distal Intergenic | 3 | 143674943 | 143675067 | 125 | 1 | ENSMUST00000158226.1 | -77632 | Gm24728 | ENSMUSG00000088851.1 |
| chr11 | 95918386 | 95919460 | 3.71349 | 4.59581 | 1.52181 | Downstream (2-3kb) | 11 | 95921102 | 95921226 | 125 | 1 | ENSMUST00000158229.1 | -1642 | Gm24725 | ENSMUSG00000088854.1 |
| chr11 | 95919932 | 95920384 | 3.88758 | 4.90057 | 1.721 | Promoter | 11 | 95921102 | 95921226 | 125 | 1 | ENSMUST00000158229.1 | -718 | Gm24725 | ENSMUSG00000088854.1 |
| chr11 | 95920804 | 95922490 | 3.62132 | 4.40358 | 1.43667 | Promoter | 11 | 95921102 | 95921226 | 125 | 1 | ENSMUST00000158229.1 | 0 | Gm24725 | ENSMUSG00000088854.1 |
| chr11 | 95922836 | 95923277 | 3.60242 | 4.37108 | 1.4198 | Distal Intergenic | 11 | 95921102 | 95921226 | 125 | 1 | ENSMUST00000158229.1 | 1735 | Gm24725 | ENSMUSG00000088854.1 |

| | | | | | | | | | | | | | | | |
|---|---|---|---|---|---|---|---|---|---|---|---|---|---|---|---|
| chr11 | 95926287 | 95926782 | 3.66877 | 4.90394 | 1.71068 | Distal Intergenic | 11 | 95921102 | 95921226 | 125 | 1 | ENSMUST00000158229.1 | 5186 | Gm24725 | ENSMUSG00000088854.1 |
| chr10 | 7866908 | 7867644 | 4.48184 | 6.15021 | 2.66522 | Exon (ENSMUST00000039484.4/ENSMUSG00000039981.5, exon 6 of 6) | 10 | 7848361 | 7848481 | 121 | 2 | ENSMUST00000158232.1 | -18428 | Gm24726 | ENSMUSG00000088857.1 |
| chr10 | 7868032 | 7868241 | 3.49837 | 4.15867 | 1.2282 | 3' UTR | 10 | 7848361 | 7848481 | 121 | 2 | ENSMUST00000158232.1 | -19552 | Gm24726 | ENSMUSG00000088857.1 |
| chr8 | 89655948 | 89656118 | 3.39194 | 3.96885 | 1.18573 | Distal Intergenic | 8 | 89667173 | 89667304 | 132 | 2 | ENSMUST00000158258.1 | 11186 | Gm24212 | ENSMUSG00000088883.1 |
| chr8 | 89657487 | 89657933 | 3.31911 | 3.82504 | 1.0833 | Distal Intergenic | 8 | 89667173 | 89667304 | 132 | 2 | ENSMUST00000158258.1 | 9371 | Gm24212 | ENSMUSG00000088883.1 |
| chr8 | 89659654 | 89659729 | 3.8213 | 4.8933 | 1.69357 | Distal Intergenic | 8 | 89667173 | 89667304 | 132 | 2 | ENSMUST00000158258.1 | 7575 | Gm24212 | ENSMUSG00000088883.1 |
| chr13 | 81945862 | 81946372 | 3.74982 | 4.65366 | 1.60852 | Distal Intergenic | 13 | 82050751 | 82050881 | 131 | 1 | ENSMUST00000158298.1 | -104379 | Gm24295 | ENSMUSG00000088923.1 |
| chr10 | 120550235 | 120550464 | 3.90057 | 5.12295 | 1.87754 | Distal Intergenic | 10 | 120605537 | 120605596 | 60 | 1 | ENSMUST00000158299.1 | -55073 | Gm24298 | ENSMUSG00000088924.1 |
| chr10 | 120550797 | 120551485 | 3.80957 | 4.96937 | 1.76295 | Distal Intergenic | 10 | 120605537 | 120605596 | 60 | 1 | ENSMUST00000158299.1 | -54052 | Gm24298 | ENSMUSG00000088924.1 |
| chr10 | 120551859 | 120554020 | 4.15732 | 5.71226 | 2.31024 | Distal Intergenic | 10 | 120605537 | 120605596 | 60 | 1 | ENSMUST00000158299.1 | -51517 | Gm24298 | ENSMUSG00000088924.1 |
| chr10 | 120570597 | 120570755 | 3.48197 | 4.08761 | 1.14446 | Distal Intergenic | 10 | 120605537 | 120605596 | 60 | 1 | ENSMUST00000158299.1 | -34782 | Gm24298 | ENSMUSG00000088924.1 |
| chr10 | 120572494 | 120572580 | 4.19682 | 5.60072 | 2.18719 | Distal Intergenic | 10 | 120605537 | 120605596 | 60 | 1 | ENSMUST00000158299.1 | -32957 | Gm24298 | ENSMUSG00000088924.1 |
| chr10 | 120582203 | 120582937 | 3.64892 | 4.60646 | 1.52009 | Distal Intergenic | 10 | 120605537 | 120605596 | 60 | 1 | ENSMUST00000158299.1 | -22600 | Gm24298 | ENSMUSG00000088924.1 |
| chr2 | 166176056 | 166176279 | 3.65405 | 4.47632 | 1.50587 | Distal Intergenic | 2 | 166189697 | 166189801 | 105 | 1 | ENSMUST00000158301.1 | -13418 | Gm22687 | ENSMUSG00000088926.1 |
| chr4 | 124358705 | 124358832 | 3.35858 | 3.87076 | 1.0364 | Distal Intergenic | 4 | 124335578 | 124335681 | 104 | 2 | ENSMUST00000158308.1 | -23025 | Gm22983 | ENSMUSG00000088933.1 |
| chr4 | 124359831 | 124359944 | 3.69805 | 4.50015 | 1.43755 | Distal Intergenic | 4 | 124335578 | 124335681 | 104 | 2 | ENSMUST00000158308.1 | -24151 | Gm22983 | ENSMUSG00000088933.1 |
| chr1 | 163358024 | 163358193 | 3.48393 | 4.10466 | 1.19531 | Distal Intergenic | 1 | 163402283 | 163402390 | 108 | 1 | ENSMUST00000158326.1 | -44090 | Gm24940 | ENSMUSG00000088951.1 |
| chr6 | 125876899 | 125877189 | 17.49243 | 46.88733 | 41.05278 | Intron (ENSMUST00000159111.2/ENSMUSG00000038115.11, intron 13 of 15) | 6 | 125884415 | 125884524 | 110 | 1 | ENSMUST00000158334.1 | -7226 | Gm24947 | ENSMUSG00000088959.1 |
| chr12 | 50999494 | 50999581 | 3.49118 | 4.29083 | 1.28776 | Distal Intergenic | 12 | 51085687 | 51085803 | 117 | 1 | ENSMUST00000158341.1 | -86106 | Gm22088 | ENSMUSG00000088966.1 |
| chr9 | 62189046 | 62189149 | 3.45086 | 4.15049 | 1.22645 | Distal Intergenic | 9 | 62242763 | 62242867 | 105 | 2 | ENSMUST00000158361.1 | 53718 | Gm25603 | ENSMUSG00000088986.1 |
| chr8 | 111889146 | 111889448 | 3.51617 | 4.21252 | 1.31746 | 3' UTR | 8 | 111879304 | 111879407 | 104 | 1 | ENSMUST00000158375.1 | 9843 | Gm23853 | ENSMUSG00000089000.1 |
| chr8 | 111889910 | 111889998 | 3.74847 | 4.62562 | 1.56926 | Exon (ENSMUST00000034430.4/ENSMUSG00000031952.4, exon 2 of 2) | 8 | 111879304 | 111879407 | 104 | 1 | ENSMUST00000158375.1 | 10607 | Gm23853 | ENSMUSG00000089000.1 |
| chr8 | 111890863 | 111891287 | 3.77156 | 4.66143 | 1.59017 | 5' UTR | 8 | 111879304 | 111879407 | 104 | 1 | ENSMUST00000158375.1 | 11560 | Gm23853 | ENSMUSG00000089000.1 |
| chr19 | 48337593 | 48337671 | 3.523 | 4.16753 | 1.20859 | Intron (ENSMUST00000078880.5/ENSMUSG00000063434.5, intron 1 of 26) | 19 | 48240535 | 48240645 | 111 | 2 | ENSMUST00000158379.1 | -96949 | Gm23857 | ENSMUSG00000089004.1 |
| chr1 | 186470415 | 186470506 | 3.36041 | 3.89912 | 1.12247 | Distal Intergenic | 1 | 186676742 | 186676876 | 135 | 1 | ENSMUST00000158420.1 | -206236 | Gm26169 | ENSMUSG00000089045.1 |
| chr5 | 75518488 | 75518563 | 3.42072 | 4.00081 | 1.16564 | Distal Intergenic | 5 | 75561046 | 75561150 | 105 | 2 | ENSMUST00000158453.1 | 42587 | Gm24502 | ENSMUSG00000089078.1 |
| chr3 | 50887229 | 50887583 | 3.5077 | 4.18174 | 1.29654 | Distal Intergenic | 3 | 50838581 | 50838715 | 135 | 2 | ENSMUST00000158454.1 | -48515 | Gm24503 | ENSMUSG00000089079.1 |
| chr13 | 43960440 | 43960663 | 3.52044 | 4.15982 | 1.19334 | Distal Intergenic | 13 | 43872802 | 43872909 | 108 | 1 | ENSMUST00000158482.1 | 87639 | Gm23117 | ENSMUSG00000089107.1 |
| chr12 | 33029032 | 33029328 | 3.5388 | 4.22101 | 1.31482 | Distal Intergenic | 12 | 32999909 | 33000010 | 102 | 2 | ENSMUST00000158511.1 | -29023 | Gm24922 | ENSMUSG00000089136.1 |
| chr19 | 37232364 | 37232496 | 3.56004 | 4.26483 | 1.33632 | Promoter | 19 | 37233382 | 37233484 | 103 | 2 | ENSMUST00000158518.2 | 988 | Gm25268 | ENSMUSG00000089143.2 |
| chr19 | 37233359 | 37234196 | 3.87106 | 5.25397 | 1.94806 | Promoter | 19 | 37233382 | 37233484 | 103 | 2 | ENSMUST00000158518.2 | 0 | Gm25268 | ENSMUSG00000089143.2 |
| chr19 | 37235925 | 37236097 | 4.0225 | 5.20568 | 1.93484 | Distal Intergenic | 19 | 37233382 | 37233484 | 103 | 2 | ENSMUST00000158518.2 | -2442 | Gm25268 | ENSMUSG00000089143.2 |
| chr19 | 37236751 | 37237121 | 3.60923 | 4.33683 | 1.32506 | Distal Intergenic | 19 | 37233382 | 37233484 | 103 | 2 | ENSMUST00000158518.2 | -3268 | Gm25268 | ENSMUSG00000089143.2 |
| chr19 | 37237607 | 37237934 | 3.46819 | 4.09711 | 1.1616 | Distal Intergenic | 19 | 37233382 | 37233484 | 103 | 2 | ENSMUST00000158518.2 | -4124 | Gm25268 | ENSMUSG00000089143.2 |
| chr19 | 37239400 | 37239613 | 3.45224 | 4.03698 | 1.12412 | Distal Intergenic | 19 | 37233382 | 37233484 | 103 | 2 | ENSMUST00000158518.2 | -5917 | Gm25268 | ENSMUSG00000089143.2 |
| chr8 | 91186446 | 91186708 | 3.82928 | 4.79688 | 1.68773 | Distal Intergenic | 8 | 91184919 | 91185183 | 265 | 2 | ENSMUST00000158523.1 | -1264 | Gm23408 | ENSMUSG00000089148.1 |
| chr10 | 80463493 | 80463774 | 3.66875 | 4.44785 | 1.41209 | Distal Intergenic | 10 | 80476352 | 80476455 | 104 | 1 | ENSMUST00000158603.1 | -12578 | Gm25044 | ENSMUSG00000089228.1 |
| chr10 | 80471820 | 80472348 | 3.41224 | 4.08049 | 1.16676 | Distal Intergenic | 10 | 80476352 | 80476455 | 104 | 1 | ENSMUST00000158603.1 | -4004 | Gm25044 | ENSMUSG00000089228.1 |
| chr10 | 80481492 | 80481754 | 3.81167 | 4.78777 | 1.6227 | Distal Intergenic | 10 | 80476352 | 80476455 | 104 | 1 | ENSMUST00000158603.1 | 5141 | Gm25044 | ENSMUSG00000089228.1 |
| chr7 | 38174676 | 38174806 | 3.85589 | 4.78825 | 1.59544 | Downstream (1-2kb) | 7 | 38176614 | 38176671 | 58 | 2 | ENSMUST00000158605.1 | 1865 | Gm22203 | ENSMUSG00000089230.1 |
| chr18 | 81089938 | 81090174 | 3.42097 | 4.07577 | 1.21012 | Distal Intergenic | 18 | 81026773 | 81026890 | 118 | 1 | ENSMUST00000158632.1 | 63166 | Gm25718 | ENSMUSG00000089257.1 |
| chr3 | 149446255 | 149446465 | 3.4476 | 4.04636 | 1.18344 | Distal Intergenic | 3 | 149340903 | 149341008 | 106 | 2 | ENSMUST00000158644.1 | -105248 | Gm26468 | ENSMUSG00000089269.1 |
| chr3 | 149519378 | 149519640 | 3.59648 | 4.31012 | 1.31374 | Distal Intergenic | 3 | 149340903 | 149341008 | 106 | 2 | ENSMUST00000158644.1 | -178371 | Gm26468 | ENSMUSG00000089269.1 |
| chr7 | 91262204 | 91262395 | 3.52463 | 4.2623 | 1.31694 | Intron (ENSMUST00000107196.3/ENSMUSG00000052572.10, intron 1 of 22) | 7 | 91242995 | 91243097 | 103 | 1 | ENSMUST00000158646.1 | 19210 | Gm24552 | ENSMUSG00000089271.1 |
| chr4 | 139966354 | 139966952 | 3.63556 | 4.42324 | 1.39155 | Promoter | 4 | 139967526 | 139967620 | 95 | 2 | ENSMUST00000158669.1 | 668 | Mir2139 | ENSMUSG00000089294.1 |
| chr8 | 87600554 | 87600910 | 3.67785 | 4.45537 | 1.38122 | Distal Intergenic | 8 | 87636657 | 87637013 | 357 | 1 | ENSMUST00000158676.1 | -35747 | Gm24841 | ENSMUSG00000089301.1 |
| chr8 | 87607391 | 87607490 | 3.78643 | 4.77424 | 1.65463 | Distal Intergenic | 8 | 87636657 | 87637013 | 357 | 1 | ENSMUST00000158676.1 | -29167 | Gm24841 | ENSMUSG00000089301.1 |
| chr8 | 87608531 | 87609128 | 3.71328 | 4.53746 | 1.48048 | Distal Intergenic | 8 | 87636657 | 87637013 | 357 | 1 | ENSMUST00000158676.1 | -27529 | Gm24841 | ENSMUSG00000089301.1 |
| chr8 | 87615106 | 87615328 | 3.67251 | 4.45097 | 1.40776 | Distal Intergenic | 8 | 87636657 | 87637013 | 357 | 1 | ENSMUST00000158676.1 | -21329 | Gm24841 | ENSMUSG00000089301.1 |
| chr2 | 147230961 | 147231159 | 3.34237 | 3.84441 | 1.02946 | Intron (ENSMUST00000136998.2/ENSMUSG00000086509.2, intron 1 of 5) | 2 | 147250462 | 147250785 | 324 | 2 | ENSMUST00000158691.1 | 19626 | Gm22261 | ENSMUSG00000089316.1 |
| chr1 | 77227877 | 77228026 | 3.35101 | 3.89498 | 1.14679 | Distal Intergenic | 1 | 77144753 | 77144846 | 94 | 1 | ENSMUST00000158701.1 | 83125 | Gm25974 | ENSMUSG00000089326.1 |
| chr14 | 100478525 | 100478605 | 4.43713 | 5.92019 | 2.37779 | Distal Intergenic | 14 | 100525244 | 100525367 | 124 | 2 | ENSMUST00000158706.1 | 46762 | Gm24345 | ENSMUSG00000089331.1 |
| chr10 | 14339294 | 14339523 | 3.53355 | 4.18775 | 1.22401 | Distal Intergenic | 10 | 14233513 | 14233623 | 111 | 1 | ENSMUST00000158708.1 | 105782 | Gm24343 | ENSMUSG00000089333.1 |
| chr13 | 54447299 | 54447737 | 3.714 | 4.57721 | 1.46463 | Distal Intergenic | 13 | 54429441 | 54429700 | 260 | 2 | ENSMUST00000158716.1 | -17600 | Gm22668 | ENSMUSG00000089341.1 |
| chr1 | 186347559 | 186347737 | 3.37071 | 3.88366 | 1.01733 | Distal Intergenic | 1 | 186123201 | 186123306 | 106 | 1 | ENSMUST00000158738.1 | 224359 | Gm23821 | ENSMUSG00000089363.1 |
| chr11 | 50527798 | 50527873 | 3.35127 | 3.85743 | 1.02823 | Distal Intergenic | 11 | 50510902 | 50511177 | 276 | 2 | ENSMUST00000158741.1 | -16622 | Gm23824 | ENSMUSG00000089366.1 |
| chr8 | 32293775 | 32294185 | 3.59303 | 4.3857 | 1.41263 | Distal Intergenic | 8 | 32186129 | 32186248 | 120 | 1 | ENSMUST00000158745.1 | 107647 | Gm22140 | ENSMUSG00000089370.1 |
| chr7 | 73655264 | 73655546 | 3.4166 | 4.20134 | 1.26359 | Distal Intergenic | 7 | 73662048 | 73662300 | 253 | 1 | ENSMUST00000158766.1 | -6502 | Gm23020 | ENSMUSG00000089391.1 |
| chr7 | 73657391 | 73657711 | 3.63124 | 4.41027 | 1.43858 | Distal Intergenic | 7 | 73662048 | 73662300 | 253 | 1 | ENSMUST00000158766.1 | -4337 | Gm23020 | ENSMUSG00000089391.1 |
| chr7 | 73669900 | 73670319 | 3.57987 | 4.43989 | 1.37912 | Distal Intergenic | 7 | 73662048 | 73662300 | 253 | 1 | ENSMUST00000158766.1 | 7853 | Gm23020 | ENSMUSG00000089391.1 |
| chr5 | 41050943 | 41051077 | 3.41461 | 4.21204 | 1.27089 | Distal Intergenic | 5 | 40405750 | 40405929 | 180 | 2 | ENSMUST00000158768.1 | -645015 | Gm23022 | ENSMUSG00000089393.1 |
| chr3 | 86181005 | 86181308 | 3.89669 | 4.8542 | 1.61804 | Distal Intergenic | 3 | 86191416 | 86191526 | 111 | 1 | ENSMUST00000158769.1 | -10108 | Gm23025 | ENSMUSG00000089394.1 |
| chr15 | 76035580 | 76036195 | 3.34527 | 3.94243 | 1.1224 | Promoter | 15 | 76036459 | 76036616 | 158 | 1 | ENSMUST00000158771.1 | -264 | G | |

### GREAT version 4.0.4
**Species assembly: mm10**
**Association rule: Basal+extension: 5000 bp upstream, 1000 bp downstream, 1000000 bp max extension, curated regulatory domains included**

#### GO Biological Process

| # Term Name | Binom Rank | Binom Raw P-Value | Binom FDR Q-' | Binom Fold Enr | Binom Observe | Binom Region ! | Hyper Rank | Hyper FDR Q-\ | Hyper Fold Enr | Hyper Observe | Hyper Total Ge | Hyper Gene Set Coverage |
|---|---|---|---|---|---|---|---|---|---|---|---|---|
| animal organ morphogenesis | 1 | 0 | 0 | 2.021339 | 4258 | 0.2502645 | 16 | 3.37E-68 | 1.924146 | 539 | 962 | 0.08651685 |
| embryonic morphogenesis | 1 | 0 | 0 | 2.246941 | 3068 | 0.1803221 | 30 | 4.34E-48 | 1.971783 | 360 | 627 | 0.05778491 |
| cell fate commitment | 1 | 0 | 0 | 3.275611 | 2076 | 0.1220172 | 43 | 1.59E-40 | 2.447034 | 176 | 247 | 0.0282504 |
| pattern specification process | 1 | 0 | 0 | 2.782147 | 2552 | 0.1499941 | 56 | 3.74E-34 | 1.982464 | 254 | 440 | 0.04077047 |
| central nervous system neuron differentiation | 1 | 0 | 0 | 2.891821 | 1771 | 0.1040907 | 78 | 4.39E-30 | 2.407473 | 136 | 194 | 0.02182986 |
| regionalization | 1 | 0 | 0 | 2.978037 | 2142 | 0.1258963 | 87 | 1.80E-28 | 2.016588 | 202 | 344 | 0.03242376 |
| neuron fate commitment | 1 | 0 | 0 | 4.317885 | 1061 | 0.06236041 | 91 | 3.61E-27 | 3.052613 | 72 | 81 | 0.01155698 |
| positive regulation of transcription from RNA polymerase II promoter | 1 | 0 | 0 | 2.069365 | 4375 | 0.2571412 | 92 | 4.54E-27 | 1.520641 | 507 | 1145 | 0.08138042 |
| transcription from RNA polymerase II promoter | 1 | 0 | 0 | 2.573575 | 2908 | 0.1709181 | 141 | 4.30E-21 | 1.660553 | 279 | 577 | 0.04478331 |
| negative regulation of transcription from RNA polymerase II promoter | 1 | 0 | 0 | 2.048224 | 3249 | 0.1909604 | 212 | 7.67E-15 | 1.450082 | 353 | 836 | 0.05666132 |
| embryonic organ morphogenesis | 81 | 1.67E-301 | 2.70E-299 | 2.583456 | 1918 | 0.1127307 | 55 | 8.87E-35 | 2.177514 | 201 | 317 | 0.03226324 |
| cell differentiation in spinal cord | 86 | 9.42E-281 | 1.43E-278 | 5.377673 | 726 | 0.04267074 | 229 | 4.13E-14 | 2.809791 | 45 | 55 | 0.007223114 |
| dorsal/ventral pattern formation | 91 | 1.22E-266 | 1.76E-264 | 4.240859 | 874 | 0.05136946 | 347 | 3.63E-10 | 2.137608 | 61 | 98 | 0.009791332 |
| embryonic organ development | 96 | 2.07E-253 | 2.82E-251 | 2.159522 | 2274 | 0.1336546 | 50 | 8.30E-36 | 1.952361 | 278 | 489 | 0.04462279 |
| forebrain development | 105 | 8.48E-229 | 1.06E-226 | 2.068403 | 2268 | 0.133302 | 54 | 6.33E-35 | 2.021078 | 246 | 418 | 0.03948636 |
| cell fate specification | 106 | 2.45E-227 | 3.02E-225 | 3.870097 | 828 | 0.0486658 | 152 | 6.81E-20 | 2.83893 | 62 | 75 | 0.009951846 |
| skeletal system development | 107 | 8.33E-227 | 1.02E-224 | 2.112485 | 2149 | 0.1263077 | 114 | 3.50E-24 | 1.798861 | 242 | 462 | 0.0388443 |
| dorsal spinal cord development | 108 | 1.64E-223 | 1.99E-221 | 9.863962 | 364 | 0.02139415 | 361 | 6.82E-10 | 3.434189 | 20 | 20 | 0.003210273 |
| gland development | 111 | 2.83E-214 | 3.33E-212 | 2.110619 | 2042 | 0.1200188 | 114 | 3.50E-24 | 1.798861 | 242 | 462 | 0.0388443 |
| spinal cord development | 113 | 1.53E-204 | 1.77E-202 | 3.439779 | 863 | 0.05072293 | 215 | 9.15E-15 | 2.344495 | 71 | 104 | 0.01139647 |

The test set of 17,014 genomic regions picked 6,230 (29%) of all 21,395 genes.
*GO Biological Process* has 13,090 terms covering 17,925 (84%) of all 21,395 genes, and 1,163,819 term - gene associations.
13,090 ontology terms (100%) were tested using an annotation count range of [1, Inf].

#### GO Molecular Function

| # Term Name | Binom Rank | Binom Raw P-Value | Binom FDR Q-\ | Binom Fold Enri | Binom Observe | Binom Region S | Hyper Rank | Hyper FDR Q-V | Hyper Fold Enri | Hyper Observe | Hyper Total Gei | Hyper Gene Set Coverage |
|---|---|---|---|---|---|---|---|---|---|---|---|---|
| RNA polymerase II transcription factor activity, sequence-specific DNA binding | 1 | 0 | 0 | 2.636949 | 4192 | 0.2463853 | 2 | 6.35E-41 | 1.811243 | 404 | 766 | 0.06484751 |
| sequence-specific DNA binding | 1 | 0 | 0 | 2.660948 | 5136 | 0.301869 | 3 | 6.75E-39 | 1.659215 | 516 | 1068 | 0.08282504 |
| RNA polymerase II regulatory region sequence-specific DNA binding | 1 | 0 | 0 | 2.632958 | 3354 | 0.1971318 | 5 | 8.31E-30 | 1.748505 | 334 | 656 | 0.05361156 |
| RNA polymerase II regulatory region DNA binding | 1 | 0 | 0 | 2.6286 | 3360 | 0.1974844 | 7 | 7.04E-30 | 1.745756 | 335 | 659 | 0.05377207 |
| regulatory region DNA binding | 1 | 0 | 0 | 2.326911 | 3885 | 0.2283414 | 11 | 6.63E-28 | 1.609536 | 419 | 894 | 0.06725522 |
| transcriptional activator activity, RNA polymerase II transcription regulatory region sequenc | 1 | 0 | 0 | 2.901689 | 2589 | 0.1521688 | 12 | 6.46E-28 | 1.914273 | 233 | 418 | 0.03739968 |
| transcription regulatory region DNA binding | 1 | 0 | 0 | 2.324422 | 3879 | 0.2279887 | 13 | 1.54E-27 | 1.605445 | 417 | 892 | 0.06693419 |
| transcription regulatory region sequence-specific DNA binding | 1 | 0 | 0 | 2.459737 | 3481 | 0.2045962 | 14 | 1.61E-27 | 1.668432 | 360 | 741 | 0.05778491 |
| sequence-specific double-stranded DNA binding | 1 | 0 | 0 | 2.440218 | 3551 | 0.2087105 | 16 | 3.85E-26 | 1.635117 | 369 | 775 | 0.05922953 |
| transcription factor activity, RNA polymerase II core promoter proximal region sequence-spe | 1 | 0 | 0 | 2.742431 | 2586 | 0.1519925 | 18 | 1.99E-25 | 1.862886 | 230 | 424 | 0.03691814 |
| transcription factor activity, sequence-specific DNA binding | 1 | 0 | 0 | 2.372858 | 4758 | 0.2796521 | 22 | 9.51E-23 | 1.475188 | 497 | 1157 | 0.07977528 |
| double-stranded DNA binding | 1 | 0 | 0 | 2.354211 | 3578 | 0.2102974 | 26 | 3.59E-21 | 1.541138 | 381 | 849 | 0.0611557 |
| core promoter proximal region DNA binding | 1 | 0 | 0 | 2.667299 | 2193 | 0.1288939 | 30 | 7.29E-21 | 1.792849 | 213 | 408 | 0.03418941 |
| core promoter proximal region sequence-specific DNA binding | 1 | 0 | 0 | 2.648663 | 2164 | 0.1271894 | 31 | 1.58E-20 | 1.78917 | 211 | 405 | 0.03386838 |
| transcriptional activator activity, RNA polymerase II core promoter proximal region sequenc | 1 | 0 | 0 | 2.911727 | 1872 | 0.110027 | 32 | 3.93E-20 | 1.940016 | 161 | 285 | 0.0258427 |
| RNA polymerase II core promoter proximal region sequence-specific DNA binding | 1 | 0 | 0 | 2.685939 | 2098 | 0.1233102 | 34 | 8.75E-20 | 1.78718 | 204 | 392 | 0.03274478 |
| DNA binding | 1 | 0 | 0 | 2.045372 | 6222 | 0.3656988 | 46 | 6.45E-08 | 1.199034 | 736 | 2108 | 0.118138 |
| transcriptional repressor activity, RNA polymerase II transcription regulatory region sequen | 22 | 1.25E-214 | 2.36E-212 | 2.624443 | 1350 | 0.07934642 | 40 | 1.06E-11 | 1.790475 | 122 | 234 | 0.01958266 |
| transcriptional repressor activity, RNA polymerase II core promoter proximal region sequen | 24 | 4.34E-144 | 7.46E-142 | 2.662296 | 893 | 0.05248619 | 51 | 2.55864E-06 | 1.728466 | 76 | 151 | 0.01219904 |
| transcription factor activity, RNA polymerase II distal enhancer sequence-specific binding | 29 | 1.07E-78 | 1.52E-76 | 2.377779 | 601 | 0.03532385 | 53 | 2.84527E-06 | 1.8822 | 57 | 104 | 0.009149278 |

The test set of 17,014 genomic regions picked 6,230 (29%) of all 21,395 genes.
*GO Molecular Function* has 4,131 terms covering 17,189 (80%) of all 21,395 genes, and 227,341 term - gene associations.
4,131 ontology terms (100%) were tested using an annotation count range of [1, Inf].

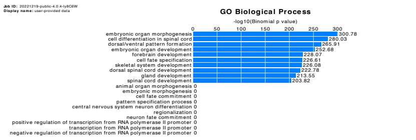

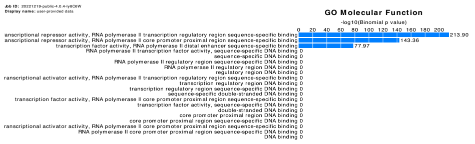

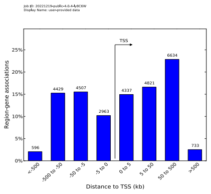

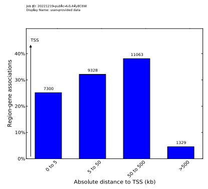

Groomed podocyte specific gene list (after Kirita et al 2020)

Thsd7a
Srgap1
Robo2
Nebl
Ptpro
Wt1
Sema3e
Synpo
Sgip1
Nphs1
Plce1
Arhgap28
Nphs2
Podxl
Ntrk3
C1qtnf7
Tmem150c
Wt1os
Ptchd4
Ntng2
Myom2
Usp13
Cdkn1c
Lmx1b
Gabra4
Chst1
Ddn
Sulf1
Tmod3
Rab3b
Kirrel
H2-Q7
Sncaip
Wipf3
St6galnac3
Cdc14a
H2-Q6
Zfp385b
Pard3b
Tdrd5
Plekhh2
Agrp
Dpp4
Gm3772
Pde3a
Zfp423
Sema3g
Zbtb7c
Il6ra
Magi2
Rasl11a
Dach1
Rhbdl3
Npr3
Tmem108
Prkg1
Tcf21
Nell2
Iqgap2
Gabrb1
Mcc
Dync1i1
Asic2
Elmo1
Clic3
St6gal1
Maml3
Plekhg1
Mertk
Cmtm7
Clic5
Ncam1
Lrrfip1
Speg
Lingo2
Flrt2
Aff3
Cttnbp2
Nsf
Cldn5
Mafb
Fam78b
St3gal1
Dock4
Zfp521
Astn2
Rhpn1
Ksr1
Myo1e
Hspa12a
Ncf2
Dock5
Plagl1
Pag1
9530026P05Rik

Amph
Mrc2
Ephb1
Fryl
Golim4
Plekho1
Vim
Sox5
Pdpn
Nck2
Nars2
Abca9
Colec12
Fyn
Galnt10
Cep85l
Rnf150
Enpep
Adcy1
Itga3
Klhl29
Dpysl2
Mtcl1
Sema5a
Tjp1
Myo1d
Lctl
Alcam
Atp8b1
Man1a2
Tmtc1
Kank1
Lrch1
Shroom3
St3gal6
Hs3st3a1
1700025G04Rik
Plat
Gdpd5
Fam222a
Qk
Elf4
Samd12
Nxn
Mgat5
Pde10a
Mtss1
St6galnac6
Fry
Arhgef18
Adcy5
Asb15
Mapt
Slco2a1
Fmnl2
Arhgef16
Arhgef26
Grk5
Fat1
Fchsd2
Dcdc2a
Opcml
Ccdc92
Coro2b
Plod2
Fnbp1l
Kcnq1ot1
Rcsd1
Card10
Parva
Cd59a
Ildr2
Ptpn14
Pde4b
St3gal3
Top1mt
Gulp1
Pknox2
Msrb3
Pdlim2
Pbx2
Srgap2
Pak1
Samd4
Col4a3
4921504A21Rik
Mcf2
Dock11
Lrrc49
Csnk1a1
Pxdc1
Inpp4a
Tm4sf1
Itgb5
Eya4
Tshz2
Cdc42se2

Daam2
Myof
Frmd6
Sh2d4a
Optn
Gas7
Unc5c
Trib2
Ust
Cblb
Rbms1
Med12l
Nedd4
Metrnl
Ctdspl
Pld1
Pkia
Mpp5
Ssbp2
Rnf144b
Ccdc73
Eml2
Vegfa
Yaf2
Gsn
Nlk
Sparc
Tgfbr3
Mgat4a
Slain2
Fam81a
Anks1b
Sirpa
Gmds
Actn4
Tpm1
Zbtb8a
Agpat4
Rbpms
Npnt
Epb41l5
Parp4
Plekha5
Anxa3
Ripor1
Syne1
Crlf3
Rab30
Rad21
Arhgap23
Slc41a2
Gpcpd1
Zdhhc14
Plekha2
Nab1
Pced1b
Plekhg2
Pcsk6
Myh9
Fgfr1
Uaca
Zfp462
Smo
Cpeb1
Itgb1
Auts2
Sh3bgrl2
Acsl4
Tns2
Creb3l2
Psd3
Xkr4
Sdc4
Ccni
Nek1
Large1
Mdfic
Rbms3
Cald1
Dag1
Setd7
Sertad2
Ncald
Dnase1l1
Mill2
Colgalt2
Micu1
Osbpl9
Plcl1
Cd2ap
Bmpr1a
Hoxc5
Laptm4a
Noct

Cables1
Atxn10
Atp8a1
Arhgap24
Calm2
Fndc3b
Anxa4
Vamp5
Per1
Actn1
Etl4
Uck2
Afap1
Tanc1
Col4a5
Vamp4
E130308A19Rik
Wtip
Tjap1
Lama5
Ankrd44
Vgll4
Cwc27
E130307A14Rik
Rbms2
Rapgef6
Gnas
Akt3
Bcar3
Cers6
Taf12
Dst
Arvcf
Lrrc1
Cyp27a1
Olfr323
Jdp2
Dlc1
Herpud2
Mob2
Pmepa1
Sh3bgrl
Rasa2
Fbxo4
Spaca6
Ctnnal1
Fermt2
Pls3
Stxbp1
Araf
Palld
Msl2
Rassf8
Ptprd
Tspan5
Spata13
Ccbe1
Phka2
Nhlrc2
Phactr4
Fam184a
Col4a4
Rras2
Unc13b
Utrn
Fa2h
Usp4
Pth1r
Cpq
Kremen1
Atp2b1
P3h2
Pbx3
Ptpn12
Usp53
Hip1
Tmem164
Dlg5
Cdyl
Macf1
Malt1
Gab2
Micu2
Sbf2
Mast4
Tenm3
Arhgef12
E2f3
Prkce
Prkar2a
Pacs2
Gabpb2
L3mbtl3
Gpc3
Klf7
Ets1

Nampt
Casz1
Tet3
Fign
Akap11
Nras
Tab2
Rai14
Ankrd6
Polb
Ext2
Lrrc4c
Ccdc163
Otud7b
Pacs1
Map3k5
Btrc
Rin2
Myo6
Smyd1
Foxj3
Tmed10
Tbc1d8b
Ppp6r2
Scel
Zhx2
Arhgap44
Spred2
Crim1
Snx12
Rap2a
Nktr
Atp6v0a1
Rbm28
Smad6
Pde4dip
Kansl3
Bmpr2
Dlgap4
Cyp4a12b
Eml4
Pcsk5
Cuedc1
Pofut1
Usp14
Exoc2
Wasf2
Ctnnd1
B4galt1
Nsd1
Map3k20
Golph3
Ftx
Ppfibp2
Pip5k1a
Zzef1
Nbeal1
Hmgxb4
S100pbp
Hivep2
Rock2
Wasl
Vezt
Rsu1
Diaph2
Scfd2
Kif5b
Kat6a
Arl8b
Epn2
Fam117b
Gm14434
Arhgap17
Rhbdd1
Pcmtd1
Lsm14a
Grb2
Tspan12
Kif13a
Tmem126a
